# Transcriptome reconstruction and functional analysis of eukaryotic marine plankton communities via high-throughput metagenomics and metatranscriptomics

**DOI:** 10.1101/812974

**Authors:** Alexey Vorobev, Marion Dupouy, Quentin Carradec, Tom O. Delmont, Anita Annamalé, Patrick Wincker, Eric Pelletier

## Abstract

Large scale metagenomic and metatranscriptomic data analyses are often restricted by their genecentric approach, limiting the ability to understand organismal and community biology. *De novo* assembly of large and mosaic eukaryotic genomes from complex meta -omics data remains a challenging task, especially in comparison with more straightforward bacterial and archaeal systems. Here we use a transcriptome reconstruction method based on clustering co-abundant genes across a series of metagenomic samples. We investigated the co-abundance patterns of ~37 million eukaryotic unigenes across 365 metagenomic samples collected during the *Tara* Oceans expeditions to assess the diversity and functional profiles of marine plankton. We identified ~12 thousand co-abundant gene groups (CAGs), encompassing ~7 million unigenes, including 924 metagenomics based transcriptomes (MGTs, CAGs larger than 500 unigenes). We demonstrated the biological validity of the MGT collection by comparing individual MGTs with available references. We identified several key eukaryotic organisms involved in dimethylsulfoniopropionate (DMSP) biosynthesis and catabolism in different oceanic provinces, thus demonstrating the potential of the MGT collection to provide functional insights on eukaryotic plankton. We established the ability of the MGT approach to capture interspecies associations through the analysis of a nitrogen-fixing haptophyte-cyanobacterial symbiotic association. This MGT collection provides a valuable resource for an exhaustive analysis of eukaryotic plankton in the open ocean by giving access to the genomic content and functional potential of many ecologically relevant eukaryotic species.

## Introduction

As an alternative to individual genome or transcriptome sequencing, environmental genomics has been used for many years to access the global genomic content of organisms from a given environment (Joly and Faure 2015). However, large scale metagenomic and metatranscriptomic data analyses are often restricted by their gene-centric approach, limiting the ability to draw an integrative functional view of sampled organisms. Nevertheless, constructing gene catalogs from environmental samples provides a useful framework for a general description of the structure and functional capabilities of microbe-dominated communities (Venter et al. 2004; Qin et al. 2010; Brum et al. 2015; Sunagawa et al. 2015; Carradec et al. 2018). Gene-centric approaches allow deep and detailed exploration of communities of organisms, but they are usually undermined by limited contextual information for different genes, apart from taxonomic affiliation based on sequence similarity.

Several methods have been developed to shift the scientific paradigm from a gene-centric to an organism-centric view of environmental genomic and transcriptomic data. These methods use direct assemblies of metagenomic reads to generate contigs that encompass several genes. High recovery of bacterial and archaeal genomes has been achieved through traditional assembly strategies from both high and low diversity environmental samples (Tully et al. 2018; Dick et al. 2009; Albertsen et al. 2013). However, when dealing with complex communities of organisms, traditional genome assembly approaches are often impaired by the large amount of sequence data and the genome heterogeneity.

To circumvent these limitations, approaches based on reference genomes have been proposed (Caron et al. 2009; Pawlowski et al. 2012). Other approaches are based on binning of assembly contigs across a series of samples to extract information (Sharon et al. 2012; Albertsen et al. 2013). Significant results for bacteria and archaea have been achieved so far (Delmont et al. 2018; Parks et al. 2017; Pasolli et al. 2019; Nayfach et al. 2019). However, eukaryotic organisms have larger, more complex genomes that require significantly higher sequence coverage than bacteria and archaea. Even in cases where significantly long contigs can be obtained, the mosaic structure of eukaryotic genes and the difficulty predicting them *de-novo* from a genome sequence leads to poor gene recovery (Boeuf et al. 2019). Only eukaryotes possessing small genomes and a high proportion of mono-exonic genes are expected to provide results similar to those achieved for bacteria or archaea. Recently a semi-supervised method based on a model trained with a set of diverse references allowed eukaryotic genome reconstruction from complex natural environments (West et al. 2018; Olm et al. 2019).

Novel reference-independent clustering approaches that produce genomes from metagenomic data have recently been developed (Albertsen et al. 2013; Imelfort et al. 2014; Delmont et al. 2018; Kang et al. 2019; Nielsen et al. 2014) and successfully applied to prokaryote-dominated communities. One of these approaches efficiently delineated co-abundant gene groups (CAGs), the largest of which were termed MetaGenomic Species (MGS), across a series of human gut microbiome samples (Nielsen et al. 2014). This method uses the metagenomic abundance profiles of a reference gene catalog, determined by stringent mapping of raw metagenomic reads onto sequences in this catalog, and defines clusters of genes showing similar variations of abundance profiles across a collection of samples.

The vast majority of the planktonic biomass in the global ocean consists of single-cell eukaryotes and multicellular zooplankton (Dortch and Packard 1989; Gasol et al. 1997). Globally, these organisms play an important role in shaping the biogeochemical cycles of the ocean, and significantly impact food webs and climate. Despite recent advances in understanding their taxonomic and gene functional compositions (Joly and Faure 2015; Carradec et al. 2018; Venter et al. 2004), little is known about the biogeographical preferences and metabolic potential of many eukaryotic plankton species from an organism-centric perspective. Several collections of reference marine eukaryote planktonic organisms sequences have been created, the largest one being the MMETSP collection (Keeling et al. 2014). However, the majority of fully sequenced marine eukaryotic genomes or transcriptomes are derived from cultured organisms. Due to the limited availability of cultured representatives of many dominant in the open ocean plankton, notably zooplankton representatives, reference sequences represent only a small fraction of the natural biological diversity (De Vargas et al. 2015; Sibbald and Archibald 2017).

Here, we used the rationale of this reference independent, gene co-abundance method (Nielsen et al. 2014) to delineate transcriptomes by mapping metagenomic sequencing data obtained from 365 metagenomic readsets generated from marine water samples collected from the global ocean during the *Tara* Oceans expedition onto the metatranscriptome-derived Marine Atlas of *Tara* Oceans Unigenes (MATOU-v1 catalog (Carradec et al. 2018)) obtained from the same set of Tara Oceans stations (Fig. S1). The samples were collected from all the major oceanic provinces except the Arctic, typically from two photic zone depths (subsurface - SRF and deep-chlorophyll maximum - DCM), and across four size fractions (0.8–5 μm, 5–20 μm, 20–180 μm, and 180– 2000 μm).

## Results

### Construction of the MGT collection

*Of the* 116,849,350 metatranscriptomic-based unigenes of the MATOU-v1 catalog, 37,381,609 (32%) were detected by metagenomic reads mapping in at least three different Tara Oceans samples, and displayed no more than 90% of their total genomic occurrence signal in a single sample. The metagenomic RPKM-based abundance matrix of these unigenes was submitted to a canopy clustering process (see Materials and Methods) that regrouped unigenes based on the covariation of their genomic abundance across the samples. 7,254,163 (19.5%) of these unigenes were clustered into 11,846 co-abundance gene groups (CAGs) with sizes varying from 2 to 226,807 unigenes. 924 CAGs consisting of at least 500 unigenes were termed MetaGenomic based Transcriptomes (MGTs) as they may constitute a significant part of an organism’s transcriptome, and which encompass 6,946,068 unigenes (Table S1). For subsequent analyses we focused on these more complete 924 MGTs since they more accurately represent organisms’ transcriptomes (Fig. S1). This MGT collection recruited a significant number of metagenomic reads across Tara Oceans stations with an average of 58.5 % (up to 94.5 % for some samples) (Fig S2).

### Taxonomic diversity of the MGT collection

We studied the distribution of taxonomically assigned unigenes for each MGT across major planktonic taxa. In several cases, we observed a homogeneous distribution of taxonomic affiliations, suggesting that the MGTs represented transcriptomes of individual organisms (Table S1 and Fig. A1). The accuracy of the taxonomic affiliations of the unigenes varied throughout the samples and depended on i) the conservation level of a given sequence across species and ii) the adequacy and robustness of a reference database in regard to a given taxonomic unit (Carradec et al. 2018).

For each MGT, global taxonomic affiliation was determined by the taxonomic node that covered at least 75% of the taxonomically assigned unigenes of that MGT (see Materials and Methods for more details). The MGT collection was mostly comprised by eukaryotic representatives (728 MGTs - 78%, 6,380,849 unigenes), followed by bacteria (148 MGTs - 16%, 454,253 unigenes), archaea (2 MGTs - 0.2%, 2,844 unigenes), and viruses (1 MGT - 0.1%, 877 unigenes). Presence of bacteria and archaea in the MATOU-v1 catalog, despite using polyadenylated RNA for the sequencing step, can be explained by (i) the true non-polyadenylated nature of these transcripts or (ii) low level of eukaryotic annotations in regard to prokaryotes in reference databases (Carradec et al. 2018). In this study we focused only on the MGTs from the domain Eukaryota.

The overall taxonomic analysis of the MGT collection revealed that most of the major eukaryotic marine planktonic kingdoms (Worden et al. 2012) were covered, with the notable exception of Amoebozoa, Cryptophyta and Rhodophyta (Fig. 1). Most of the MGTs with a low resolution global taxonomic assignment (i.e. those for which the taxonomic affiliation could be assigned only at the kingdom level or higher) were related to the Opisthokonta group (447 out of 728 MGTs) or unclassified Eukaryota (105), whereas the well-defined eukaryotic MGTs (i.e. those for which the taxonomic affiliation could be assigned at the class level or deeper) belonged to unicellular algae: Stramenopiles (62), Alveolata (48), Viridiplantae (30) and Haptophyceae (29) lineages. This low taxonomic resolution of the MGTs could be due to (i) a low number of zooplankton organisms, including representatives from the Opisthokonta group, in the reference databases or (ii) the presence of associations of several organisms in a given MGT. Overall, these observations suggest that the MGTs correspond to either organisms with available transcriptomes or those without sequenced representatives. Taxonomic diversity of the MGT collection differs significantly compared to the collections of reference transcriptomes derived from cultured organisms, including the MMETSP project (Keeling et al. 2014). Possible explanations for this include i) the essentially coastal origin of cultured strains and ii) the absence of zooplankton in the MMETSP selected organisms.

**Figure 1.**
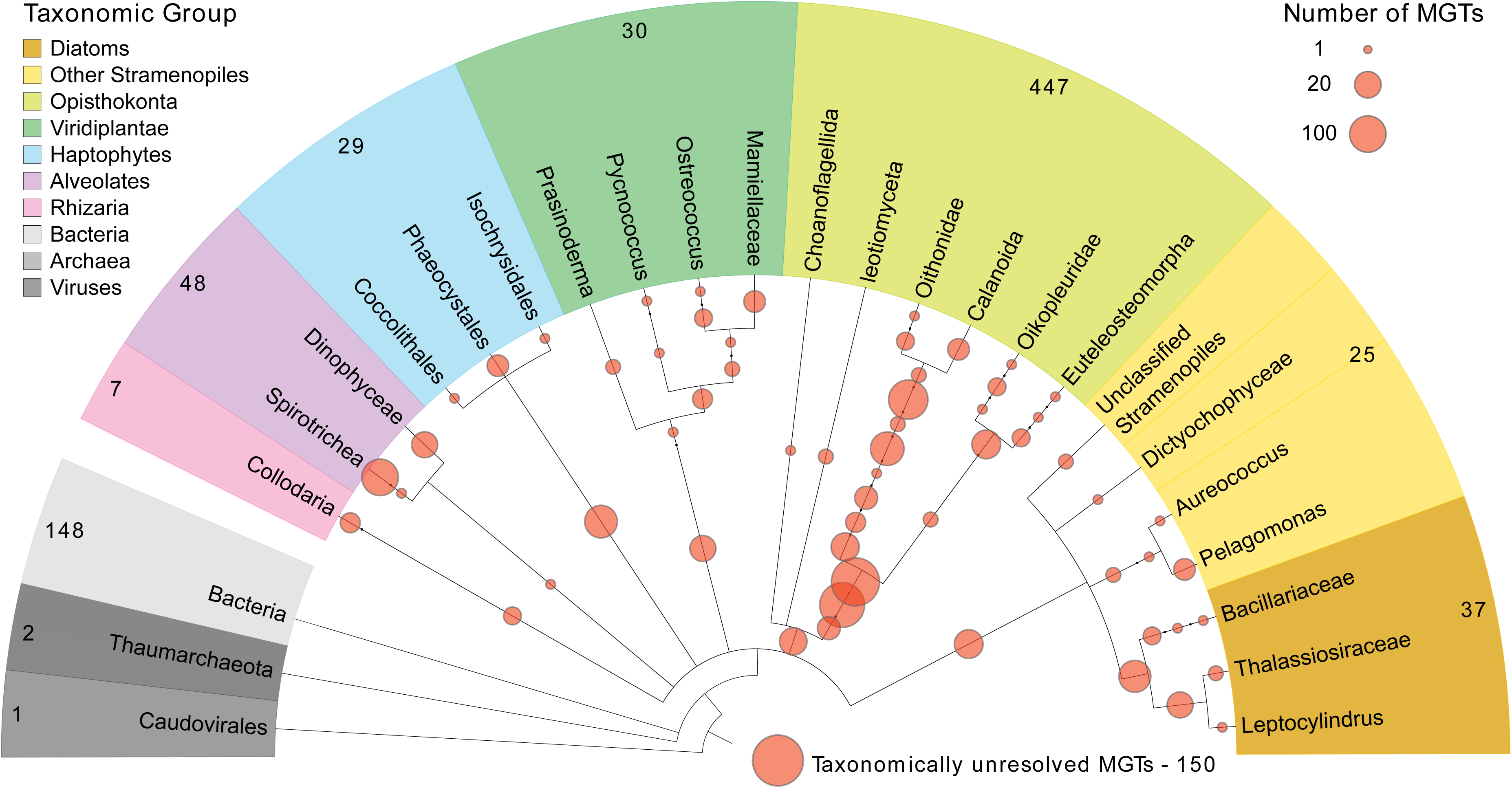
A taxonomic dendrogram representing the eukaryotic tree of life shows taxonomic positions of MGTs (orange circles) in relation to the major eukaryotic lineages. The size of the circles represents the number of MGTs positioned at a given taxonomic node. The total number of MGTs assigned to each taxonomic group is indicated on the outside of the tree.

While 48.3% of the unigenes in the MATOU-v1 catalog and 47.3% of the unigenes captured in the MGTs were taxonomically assigned, we observed an uneven bimodal distribution of taxonomically assigned unigenes among the MGTs resulting in two distinct MGT groups (Fig. 2). The first group (228 MGTs, 24.7%) consisted of well-defined MGTs (more than 75% of their unigenes had a taxonomic assignment), representing “known” organisms. Whereas the second group included 385 MGTs (41.7%) that were taxonomically poorly characterized less than 25 % of their unigenes were taxonomically assigned) which may represent currently undescribed genomes or mainly contain non-coding sequences and thus do not match with known proteins. The observed discontinuous distribution was not correlated with the MGT size or the number of samples in which an MGT was observed (Fig. S3).

**Figure 2.**
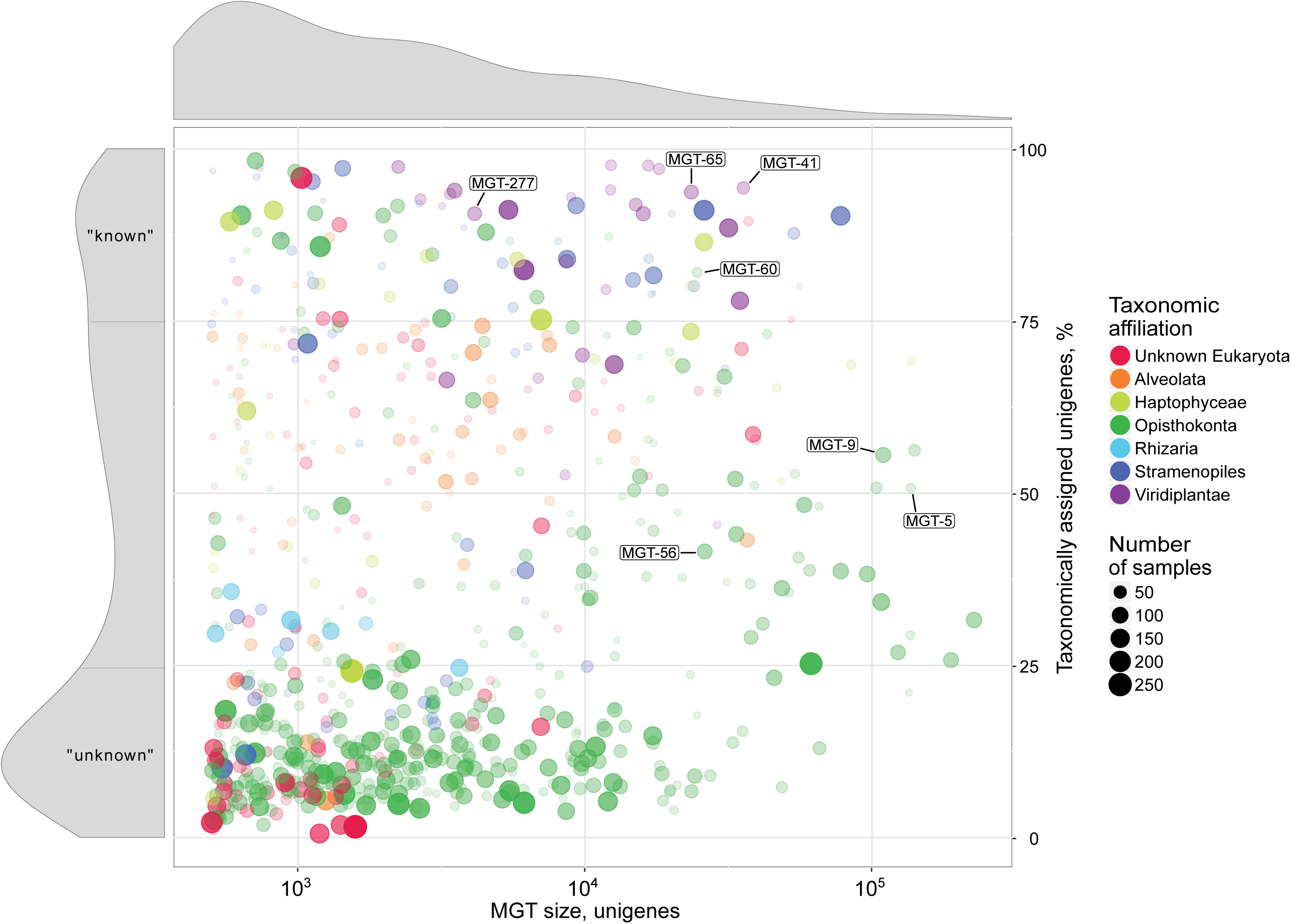
A visual representation of the major MGT statistics including the MGT size (represented by the number of unigenes, x-axis) and the fraction of taxonomically assigned unigenes (y-axis). The circle size and its opacity represent the number of samples in which a given MGT was detected. Taxonomic affiliation of the MGTs to major Eukaryotic lineages is color coded. Size distribution of MGTs based on the number of unigenes is displayed on top of the main figure. Distribution of taxonomically assigned unigenes among MGTs is presented on the left-hand side panel of the figure. “Known” and “unknown” sections of the panel indicate the MGTs comprised by more than 75% and less than 25% of taxonomically assigned unigenes, respectively. Highlighted MGTs were used for the biological validation of the MGT collection (see Results for more detail); Bathycoccus prasinos - MGT-41, MGT-65, MGT-277; Oithona nana - MGT-5, MGT-9, MGT-56, MGT-60.

### Comparison with available transcriptomes

To assess the biological validity of the obtained MGTs, we investigated the distribution of unigenes from two marine planktonic reference organisms in the MGT collection. We analyzed reference transcriptomes from a single-celled microeukaryote *Bathycoccus prasinos* and a small multicellular zooplankton *Oithona nana*. The rationale for choosing these organisms as references was as follows: (i) they play an important role in the functioning of marine ecosystems; (ii) their transcriptomes were publically available (Keeling et al. 2014; Madoui et al. 2017); (iii) they were hypothesized to be present in the datasets because of their high abundance in marine waters as demonstrated by an 18S rDNA survey from the same samples (De Vargas et al. 2015); (iv) they cover a range of organisms with substantially different transcriptome sizes (5.6-24 Mb), from phytoplankton and zooplankton groups, and from small and large size fractions of planktonic communities. We were able to recover an average of 68% (up to 77%) of the reference transcriptomes utilizing the MGT unigenes with at least 95% sequence identity over at least 50 amino acids (Fig. 2).

### Segregation of the Bathycoccus ecotypes

To assess the potential of the MGT approach to segregate closely related biological entities, we focused on the MGTs highly similar to the reference transcriptomes of *Bathycoccus prasinos*. *Bathycoccus* is a genus of green algae from the order Mamiellales which is ecologically relevant because it is widely distributed in the global ocean and contributes significantly to primary production within these ecosystems (Vannier et al. 2016; Limardo et al. 2017). Recent omics-based studies demonstrated the existence of at least two ecotypes of *Bathycoccus* (B1 and B2) which have identical 18S rRNA sequences but whose orthologous proteins share only 82 ± 6% nucleotide identity (Vannier et al. 2016; Vaulot et al. 2012).

Both of these *Bathycoccus* ecotypes were detected in the MGT collection and 99% of the total number of unigenes similar to *B. prasinos* were divided into three MGTs (MGT-41, MGT-65, MGT-277) (Table S2). We focused on MGT-41 and MGT-65 because they comprised 95.2% of the signal in this group. Pangenomic analysis demonstrated a clear segregation between the two MGTs (Fig. 3A). The average nucleotide identity (ANI) analysis indicated less than 90% sequence similarity between them, further confirming their affiliation to different ecotypes. On the other hand, the ANI values between MGT-41 and its closest reference (isolate RCC1105) and MGT-65 and its closest reference (RCC716) were 98.2% and 98.8%, respectively. The estimated completeness of the assembled unigenes (computed based on a set of 83 protistan specific single copy core genes (Simão et al. 2015) was 85.5% for MGT-41 and 73.5% for MGT-65, suggesting a high level of the transcriptome recovery. The completeness estimation method applied here is based on the identification of a set of well conserved single-copy core orthologs which usually demonstrate high levels of expression. Since not all genes from a given organism are always transcribed and thus some of them may not be captured in the MATOU-v1 catalog, this technique may overestimate the real completeness of our MGTs. However, to the best of our knowledge, this is currently the best available computational approach that estimates transcriptome completeness with a fairly high degree of confidence. MGT-41 and MGT-65 demonstrated different biogeographical preferences associated with environmental parameters, including temperature and oxygen concentrations (Fig. S4). This observation supports previous findings about the differential biogeography of the *Bathycoccus prasinos* ecotypes B1 and B2 described in (Vannier et al. 2016), leading us to assign MGT-41 to the ecotype B2, and MGT-65 to the ecotype B1. Together, these results further confirm the ability of the MGT analysis to segregate closely related eukaryotic plankton (even species with identical 18S rRNA gene sequences) in complex environmental samples.

**Figure 3.**
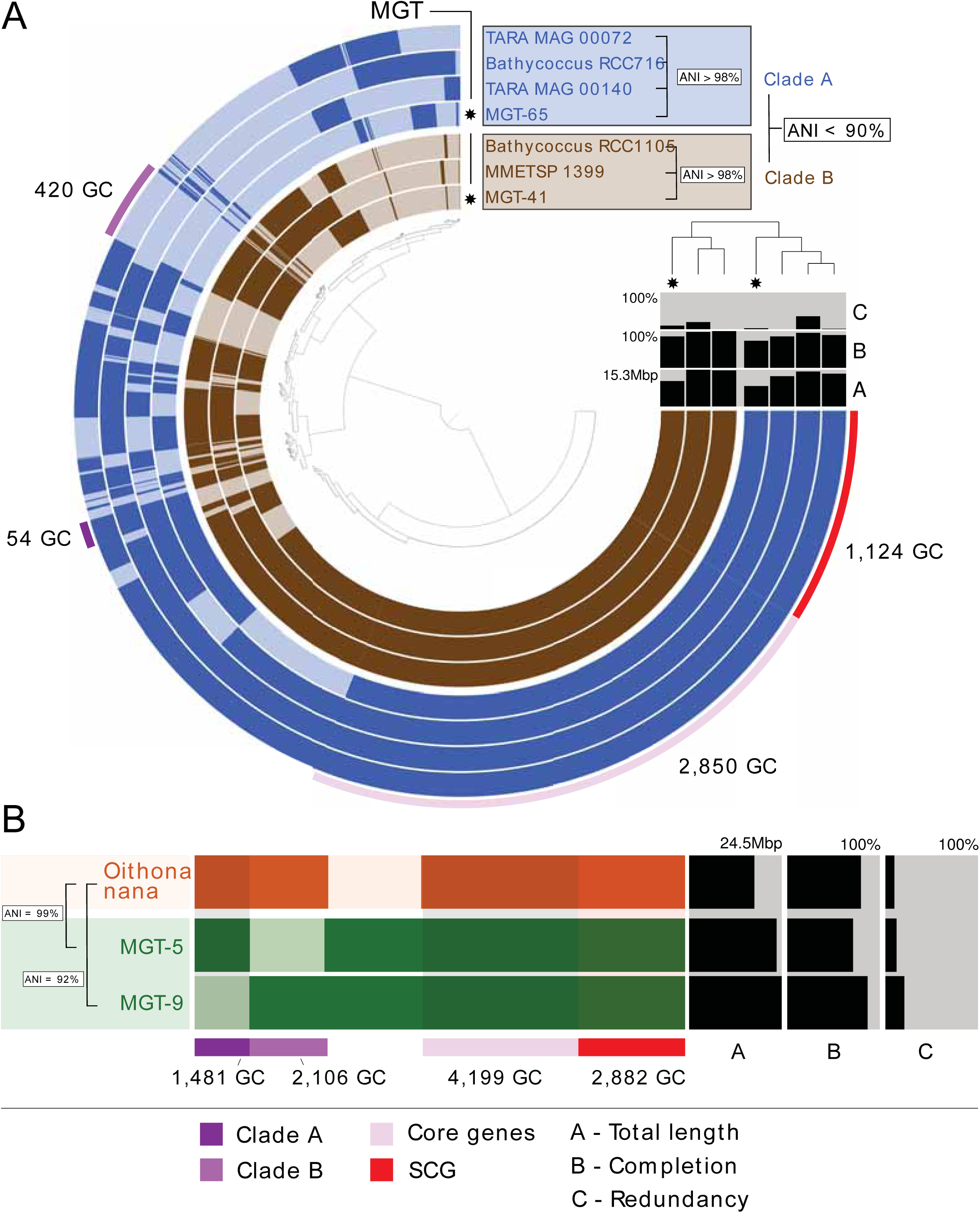
The pangenomes of A) Bathycoccus prasinos and B) Oithona nana compared to available sequenced references. Each layer represents an MGT or a reference genome or transcriptome. Gene clusters are organized based on their distribution across samples. The dendrogram in the center organizes gene clusters based on their presence or absence in the samples. The top right dendrogram represents the hierarchical clustering of the samples based on the abundance of gene clusters. ANI - Average Nucleotide Identity, SCG - Single Copy Core Genes, GC - Gene Cluster.

### Biogeography of the genus Oithona

Zooplankton, including cyclopoid copepods, also play an important role in marine ecosystems, they impact biogeochemical cycles and are key components of the oceanic food web (Roemmich and McGowan 1995; Steinberg and Landry 2017; Keister et al. 2012). However, only a few ecologically relevant references are currently available in public databases. Despite recent advances in the population genomics of the genus *Oithona*, one of the most abundant and widespread copepods in the pelagic ocean (Madoui et al. 2017; Gallienne and Robins 2001), more data are required to study the distribution of *Oithona* species and other copepods in the global ocean.

Ninety eight percent of the *Oithona-*related unigenes were detected in four MGTs (MGT-5, MGT-9, MGT-56, and MGT-60), with MGT-5 and MGT-9 alone generating 87% of the signal (Table S2). The estimated completeness of the assembled unigenes was 71% for MGT-5 and 87% for MGT-9 (Fig. 3B). MGT-5 demonstrated highly specific biogeographical preferences being observed at only twelve *Tara* Oceans stations, all from the Mediterranean Sea, which correlates well with the demonstrated biogeographic distribution of *Oithona nana* (Madoui et al. 2017). On the other hand, MGT-9 was detected at 41 stations, mostly located in the Pacific Ocean and the Mediterranean Sea, but also in the Indian and Atlantic Oceans (Fig. A2). These biogeographical preferences along with the significant difference in the ANI values relative to the reference transcriptome of *O. nana* (99% and 92% for MGT-5 and MGT-9, respectively) (Fig. 3B) suggests that MGT-9 may represent a different, yet genomically undescribed, species within the genus *Oithona*.

## Functional insights from MGTs

After demonstrating the biological validity of the MGTs, we studied their potential to assess the functional state of the ecosystem through the analysis of ecologically relevant metabolic pathways and individual marine organisms. We analyzed the expression patterns, taxonomic affiliation, and geographical distribution of the genes coding for the key enzymes involved in the cycling of dimethylsulfoniopropionate (DMSP). We also investigated the interspecies relationship between an uncultivated unicellular cyanobacterium Candidatus *Atelocyanobacterium thalassa* (UCYN-A) and a haptophyte picoplankton alga of the class Prymnesiophyceae.

### DMSP synthesis and degradation

Eukaryotic plankton, along with bacteria, are actively involved in the cycling of DMSP, an ecologically relevant organosulfur compound that can reach high concentrations in marine waters. DMSP is the precursor of the climate-active gas dimethyl sulfide (DMS) - the largest natural source of sulfur to the atmosphere. This compound is also important from an organismal point of view because of its ability to act as an osmolyte, antioxidant, predator deterrent, and cryoprotectant in phytoplankton (Bullock et al. 2017; Stefels et al. 2007).

We investigated the expression patterns and geographical distribution of the genes involved in the production and degradation of DMSP across *Tara* Oceans stations. While the role of bacteria in DMSP cycling is well described (Bullock et al. 2017; Curson et al. 2017; Johnston et al. 2016; Reisch et al. 2011), recent identification of eukaryote counterpart genes (i) *DSYB*, coding for a methyl-thiohydroxybutyrate methyltransferase, a key enzyme involved in the DMSP synthesis in eukaryotes (Curson et al. 2018) and (ii) *Alma1* coding for dimethylsulfoniopropionate lyase 1, an enzyme responsible for the cleavage of DMSP into dimethyl sulfide (DMS) and acrylate (Alcolombri et al. 2015; Johnson et al. 2016) allows to investigate the role of marine phytoplankton in the global DMSP cycle. Both of these genes were detected in the MGT collection, which allowed us to study their expression and biogeography from a genome-centric point of view (Table S3).

Out of 1220 *DSYB*-related unigenes detected in the MATOU-v1 catalog, 1214 were taxonomically assigned to eukaryotes (Fig. S5A). They were detected at all of the 66 sampling stations of the *Tara* Oceans cruise analyzed in this study and were mainly attributed to the pico-eukaryote size fraction (0.8 - 5 µm) (Fig. S6). Forty six *DSYB*-related unigenes were detected in 20 different MGTs and their expression in 10 out of 20 MGTs represented more than 10% of the total *DSYB* expression in at least one sample. This further confirms the importance of the organisms related to these MGTs in the DMSP production (Fig. 4A). Two MGTs affiliated to the genus *Phaeocystis* (MGT-4 and MGT-13) were prevalent (up to 45% of the total *DSYB* expression signal) at the Southern Ocean stations (82 to 85), whereas a Haptophyte-related organism (MGT-29) was prevalent (from 10% to 35%, depending on the size fraction) at the station 80 in Southern Atlantic. Interestingly, *Phaeocystis* affiliated MGTs demonstrated high levels of *DSYB* expression both in the small size fraction (0.8-5 µm, corresponding to individual cells) and in large size fractions (20-180 and 180-2000 µm, corresponding to multicellular colonies which may form during *P. antarctica* blooms (Carlson et al. 1998; Jr et al. 1998; Wang et al. 2018)). A large portion of the total *DSYB* expression (from 10% to 70%) was also attributed to the Prasinophytes clade VII group (Lopes Dos Santos et al. 2017): MGT-44 at the Pacific Ocean stations (93, 102, 109, 110, 122-125, 128, 136, 137, 139, 144) and MGT-166 and MGT-179 at the Indian Ocean stations (36, 38, 39 and 41). Additionally to these differences in their geographical distribution, MGT-166 and MGT-179 shared more genomic similarity between them (ANI = 95.23%), whereas MGT-44 was more distantly related (ANI = 86.86 and ANI = 85.97% with MGT-166 and MGT-179, respectively), suggesting that the former two may represent closely related organisms and the third one corresponds to a different transcriptome. However, given the small number of available Prasinophyte-related references in DNA databases, this hypothesis requires further investigation.

**Figure 4.**
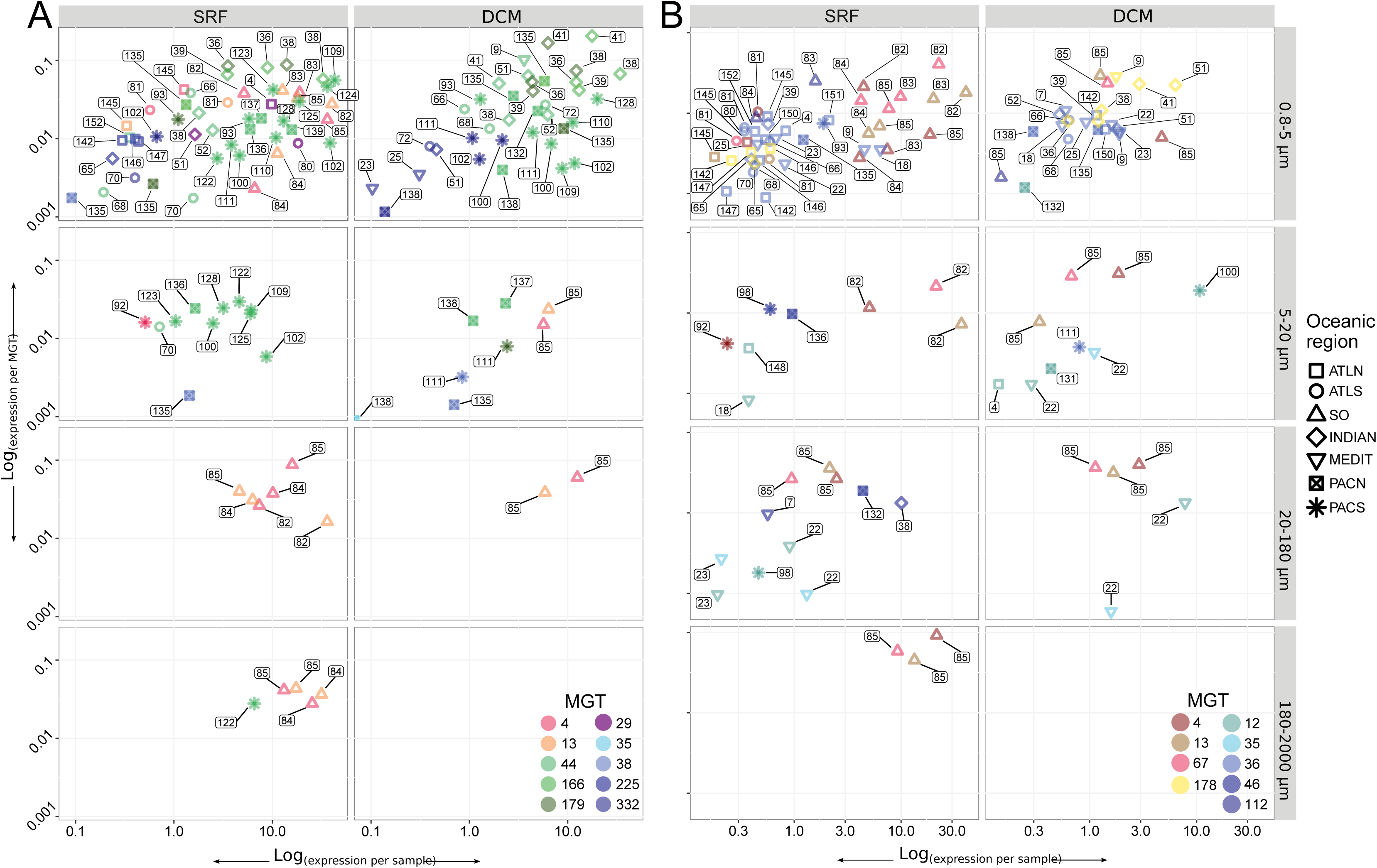
Comparison of the DSYB (A) and Alma1 (B) relative gene expression per sample (x axis) and per MGT (y axis) across samples. Numbers near each point represent a Tara Oceans station. SRF - surface, DCM - deep chlorophyll maximum. Y axis (to the left of Figure A) and size fractionation (to the right of Figure B) are common for both figures. Tara Oceans provinces are applicable to both graphs and are specified on the right side of Figure B. ATLN - North Atlantic Ocean, ATLS - South Atlantic Ocean, SO - Southern Ocean, INDIAN - Indian Ocean, MEDIT - Mediterranean Sea, PACN - North Pacific Ocean, and PACS - South Pacific Ocean. A - DSYB expression profiles across Tara Oceans stations and size fractions for ten MGTs contributing significantly to the overall DSYB expression (at least 10% of the total DSYB expression in at least one sample). Red circles (MGT-4 and MGT-13) represent MGTs taxonomically assigned to the genus Phaeocystis, green circles (MGT-44, MGT-166, and MGT-179) represent Prasinophytes clade-VII affiliated MGTs, the rest of the circles represent other organisms. B - Alma1 expression profiles across Tara Oceans stations and size fractions for nine MGTs contributing significantly to the overall Alma1 expression (at least 10% of the total Alma1 expression in at least one sample). Red circles (MGT-4, MGT-13, and MGT-67) represent MGTs taxonomically assigned to the genus Phaeocystis, yellow circle (MGT-178) represents Pelagomonas spp. affiliated MGTs, and the rest of the circles represent other organisms.

It is well established that organisms from the classes Dinophyceae (dinoflagellates) and Prymnesiophyceae (coccolithophores) are major producers of DMSP, but only few studies are available that investigate the involvement of Prasinophytes in DMSP production (Keller et al. 1989; Keller 1989; Kiene et al. 1997). Our data suggest that there are at least two distinct representatives of the Prasinophytes clade VII involved in DMSP biosynthesis and provide preliminary results on a possible pathway for DMSP production in this group.

We also investigated the expression of the *Alma1* gene coding for a key enzyme of the DMSP degradation pathway (Alcolombri et al. 2015). We detected 1059 *Alma1*-related unigenes in the MATOU-v1 catalog (Fig. S5B). The expression of these unigenes was detected mostly in the smallest size fraction (0.8 - 5 µm) at both depths (surface and DCM) at 66 sampled stations (Fig. S7). No taxonomic affiliation was found for 153 *Alma1* unigenes. The highest levels of *Alma1* abundance and expression were detected at the Southern Ocean stations (Fig. S7). Thirty six *Alma1-* related unigenes were detected in 13 MGTs. Most of these *Alma1* unigenes were taxonomically affiliated to the clade Alveolates (78%) followed by the family Haptophyceae (5%). However, 30 out of the 36 *Alma1*-related unigenes present in the MGT collection were concentrated in 6 MGTs taxonomically assigned to Haptophytes. Even though 48 MGTs possessed unigenes assigned to Alveolates, we did not detect any *Alma1*-containing MGTs affiliated to this group. In 9 out of 13 MGTs containing *Alma1* unigenes, *Alma1* expression contributed more than 10% (in some cases up to 40%) to the total *Alma1* expression detected across 43 samples (Fig. 4B). MGTs demonstrating the highest levels of *Alma1* expression were taxonomically assigned to *Phaeocystis* (MGT-4, MGT-13, and MGT-67) and *Pelagomonas* spp. (MGT-178).

### Identification of interspecies associations

We investigated the interspecies relationship between an uncultivated diazotrophic unicellular cyanobacterium Candidatus *Atelocyanobacterium thalassa* (UCYN-A) (Zehr et al. 2008; Thompson et al. 2012) and a haptophyte picoplankton alga *Braarudosphaera bigelowii* (*B. bigelowii*) from the class Prymnesiophyceae. Both members of this association are abundant and widely distributed in the ocean and are ecologically relevant because of their ability to fix N_2_ (Zehr and Kudela 2011; Farnelid et al. 2016). Several UCYN-A genomes have been previously sequenced (Tripp et al. 2010; Bombar et al. 2014), whereas no genomic information is currently available for the algal host.

2,616 UCYN-A affiliated unigenes were detected in the MATOU-v1 catalog (see Materials and Methods). They were distributed among 41 *Tara* Oceans stations and were mostly present in the small size fraction (0.8 - 5 µm). The majority of the UCYN-A affiliated unigenes (96%) were detected in two MGTs: 1,742 unigenes in MGT-29 and 771 unigenes in MGT-176 (Fig. S8). In addition to the unigenes affiliated with the diazotrophic cyanobacterium, MGT-29 also possessed ~20 thousand unigenes taxonomically assigned to the Haptophyte clade and possibly representing the eukaryotic host of this symbiosis, a Prymnesiophyte closely related to *B. bigelowii*. Together with the observation that the host’s 18S rDNA V4 region was identified in the same samples as MGT-29, this suggests that the non-UCYN-A affiliated genes of MGT-29 could be a part of the transcriptome of the host. Comparison of the MGTs comprising UCYN-A-related unigenes with reference genomes (Tripp et al. 2010; Bombar et al. 2014) and metagenome-assembled genomes (MAGs) (Delmont et al. 2018; Parks et al. 2017) demonstrated that MGT-29 unigenes covered 90.3% of the UCYN-A1 genome (Fig. 5). The estimated completeness of the UCYN-A genome computed based on a set of 139 bacterial specific single copy core genes (Campbell et al. 2013) 82.7% for MGT-29 and 42.4% for MGT-176. The ANI value between MGT-29 and UCYN-A1 isolate ALOHA was 99.7% which indicates a remarkably high genomic similarity. We hypothesize that MGT-176 may represent UCYN-A2 or another UCYN-A sublineage because the ANI analysis demonstrated its higher genomic similarity with isolate SIO64986 (UCYN-A2) than isolate ALOHA (UCYN-A1) (97.1% and 94.3%, respectively).

**Figure 5.**
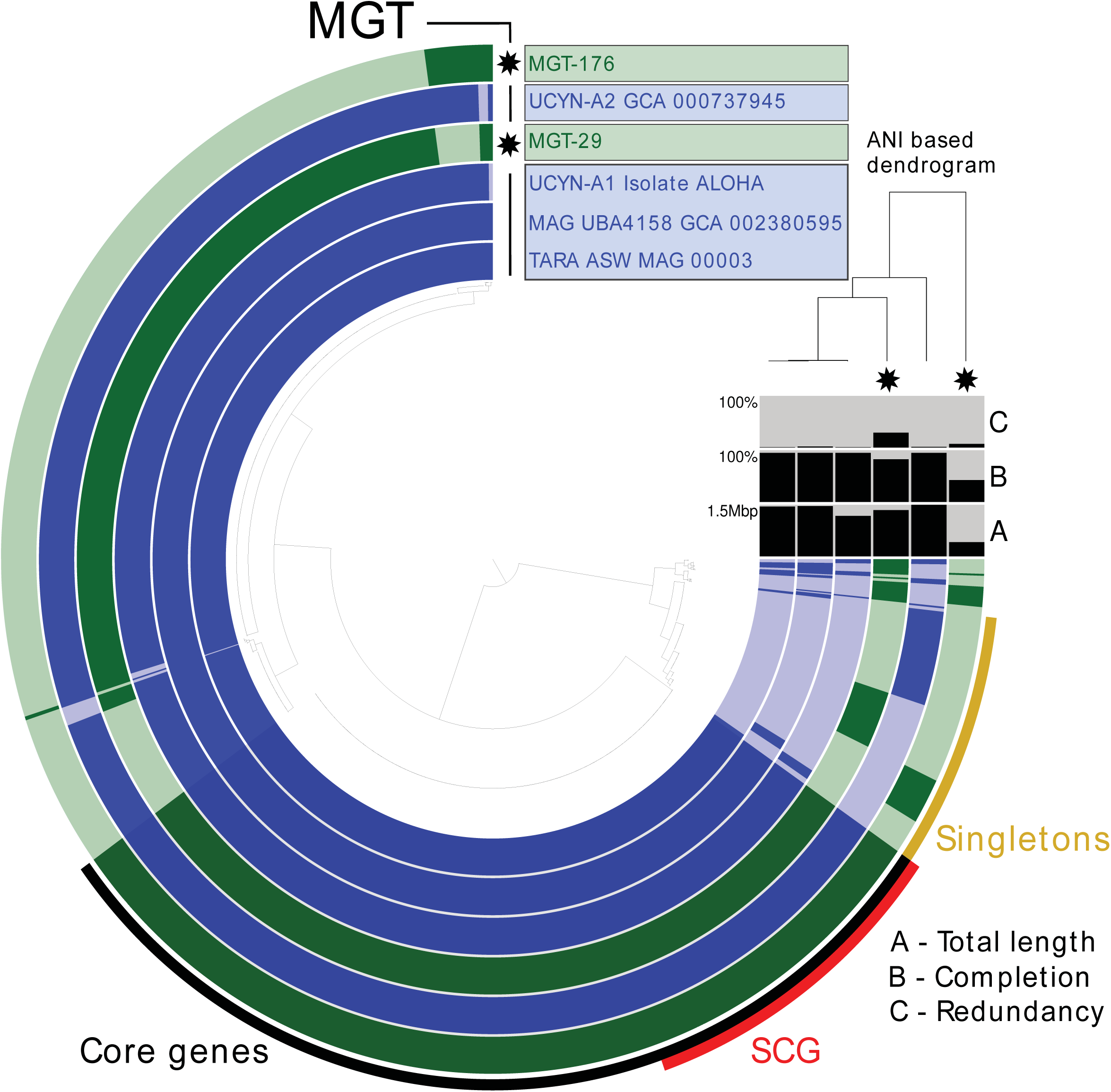
The pangenomes of MGT-29 and MGT-176 compared to available reference sequences of UCYN-A. Each layer represents an MGT, a reference genome, or a MAG. Gene clusters are organized based on their distribution across samples. The dendrogram in the center organizes gene clusters based on their presence or absence in the samples. The top right dendrogram represents the hierarchical clustering of the samples based on the ANI values. ANI - Average Nucleotide Identity, SCG - Single Copy Core Genes.

In addition to the UCYN-A-related genes, MGT-29 also contained multiple core metabolism genes taxonomically assigned to the Haptophyte clade which may belong to the eukaryotic host of this symbiotic association, *B. bigelowii* (Table A2). More specifically, we detected genes coding for enzymes driving major metabolic pathways in the haptophyte algae including glycolysis, tricarboxylic acid (TCA) cycle, pentose phosphate pathway, GS-GOGAT cycle (ammonium assimilation through sequential actions of glutamine synthetase (GS) and glutamate synthase (GOGAT)), as well as multiple genes affiliated with the metabolism of fatty acids and amino acids. We also observed the presence of the gene *bacA*, coding for the ABC transporter involved in the transport of vitamin B12.

In addition to the UCYN-A symbiosis with a single-celled haptophyte, we detected other important known microbial associations in the MGT collection (Table S4). For example, MGT-738 partially captured an association between a diatom from the class Coscinodiscophyceae (47 unigenes) and the nitrogen-fixing heterocystous cyanobacteria, *Richelia intracellularis* (132 unigenes), an abundant organism in tropical and subtropical waters (Janson et al. 1999; Lyimo 2011). This MGT collection also detected an association between a pennate diatom from the family Bacillariaceae and a tintinnid ciliate, genes from both of which were detected in MGT-136 (243 and 192 unigenes affiliated to the diatom and tintinnid, respectively) (Vincent et al. 2018).

## Discussion

Many eukaryotic lineages of ocean plankton remain largely undersampled, as a result there are few sequenced representatives for many ecologically important marine eukaryotic organisms. The MGT collection reported here represents a valuable resource for studying a range of eukaryotic planktonic organisms including those that are largely unexplored using traditional omics techniques.

The gene clustering approach applied here provides an organism-centric view of the most abundant plankton populations. This approach allowed us to focus on the diversity and functional potential of marine eukaryotic organisms across major taxonomic lineages. The 924 MGTs generated from the *Tara* Oceans datasets contain an impressive diversity of taxa allowing for comparative genomic studies in major eukaryotic groups including Opisthokonta, Haptophytes, Stramenopiles, Alveolates, Archaeplastida, and Rhizaria (Fig. 1). Additionally, this collection of MGTs provides the first glimpse of the genomic content of a variety of organisms currently not available in culture, including copepods, one of the most prevalent zooplankton of the photic open ocean. Only a small number of MGTs have closely related cultured references, suggesting that many MGTs represent organisms which have only distant relatives within the publicly available collections of sequenced genomes or transcriptomes. These MGTs provide access to valuable genomic information currently not accessible through other DNA-based resources. For example, limited availability of full genomes or transcriptomes representing heterotrophic organisms prevents advances in studying their distribution, population structure, and functional potential. Access to the zooplankton transcriptomes through the MGT collection will increase our knowledge on their biogeography and complement the general lack of references for the copepods group. Alternatively, the analysis of the MGTs closely related to organisms which have sequenced representatives may lead to a re-evaluation of their genomic potential and provide additional information on their ecology.

### DMSP biosynthesis and degradation by eukaryotes

We demonstrated the ability of the MGT approach to assess the contribution of eukaryotic plankton to ecologically important processes by focusing on the MGTs expressing genes coding for key enzymes involved in DMSP cycling. These genes included *DSYB*, coding for a methyl-thiohydroxybutyrate methyltransferase, a key enzyme of the eukaryotic DMSP synthesis pathway (Curson et al. 2018) and *Alma1* coding for dimethylsulfoniopropionate lyase 1, an algal enzyme that cleaves DMSP into DMS and acrylate (Alcolombri et al. 2015). We revealed the importance of *Phaeocystis* spp. in the Southern Ocean as potential DMSP producers and degraders. Our data also indicated the involvement of at least two representatives from the Prasinophyte clade VII in the biosynthesis of DMSP. One of them was primarily active at the oligotrophic equatorial Pacific stations, while the other one appeared to be restricted to the low oxygen stations (the Arabian basin and upwelling stations in the East Pacific). However, involvement of Prasinophytes in DMSP production is currently supported only by circumstantial evidence mostly in early studies (Keller et al. 1989; Keller 1989; Kiene et al. 1997) which demonstrates a clear need for more sequenced references of this group of organisms. Our results also support recent findings stating that picoeukaryotes should be considered as important contributors to DMSP production through the *DSYB* pathway (Curson et al. 2018). These organisms may represent interesting targets for the experimental validation of their role in the global biogeochemical sulfur cycle and their impact on climate change as proposed in the CLAW hypothesis (Charlson et al. 1987; Ayers and Cainey 2008). Thus, the MGT approach allowed us to identify candidate organisms responsible for a large part of the eukaryotic DMSP biosynthesis and catabolism in the different regions of the open ocean.

Presence of different dominating groups of organisms involved in the DMSP cycling across oceanic regions suggests the importance of environmental conditions in shaping microbial community composition that defines the DMSP fate in the ocean. If environmental conditions change in a given ecological niche, we may expect that DMSP production and degradation rates would also change because of the transformations in the microbial community structure which may lead to significant effects on climate change.

### Interspecies associations

The importance of the MGT collection as a resource for studying marine interspecies interactions was demonstrated through detection of the ecologically relevant symbiosis between the metabolically streamlined nitrogen-fixing cyanobacterium UCYN-A and a single-celled haptophyte picoplankton alga. Initially, this symbiosis was discovered using a targeted approach that involved several culture-dependent and molecular techniques proving it to be a challenging task (Zehr et al. 2017). Several UCYN-A sublineages have been defined but limited information is currently available regarding their global distribution and for some, the identity of the host (Farnelid et al. 2016; Thompson et al. 2014; Turk‐Kubo et al. 2017). MGT-29 from our collection encompasses genes similar to those from UCYN-A1 strain ALOHA and genes taxonomically affiliated with the Haptophyte clade potentially representing the eukaryotic host. This suggests that MGT-29 may represent UCYN-A1 specifically associated with a closely related to *B. bigelowii* prymnesiophyte. More information is needed about the genomic content of the host cells for different UCYN-A sublineages to confidently state which symbiosis was detected.

No genomic information is available about these symbiotic hosts beyond 18S rRNA sequences (Hagino et al. 2013). As a result, many questions remain unanswered regarding the evolution of this symbiosis and the exact nature of the relationship between the two organisms. Through partial reconstruction of the host transcriptome we provide the first glimpse of its genomic content which will dramatically change the way in which this interspecies association can be studied. Better methods are needed for the accurate targeting of distinct UCYN-A/host associations, which will improve the understanding of the evolution and ecological characteristics of this symbiosis. Access to the genomic content of the host through the MGT collection, used in conjunction with the two closed UCYN-A genomes currently available in the databases (Tripp et al. 2010; Bombar et al. 2014), will undoubtedly provide a much needed push in this direction.

Other ecologically important microbial associations were detected in the MGT collection. The diatom-cyanobacteria symbiotic populations captured in the MGT-738 which were previously observed in all major ocean basins (Foster and O’Mullan 2008) may encompass diatoms from several genera, including *Hemiaulus*, *Rhizosolenia* and *Chaetoceros*. These associations may contribute as much new nitrogen (N) as the free-living diazotroph *Trichodesmium* which is widely regarded as the most important player responsible for N_2_ fixation in the open ocean (Capone et al. 2008). It was reported that the contribution of the diatom symbioses to the global pool of N had been underestimated and that they should be included in global N models (Foster et al. 2011). In order to accurately do so, additional information on their genomic potential is required and can be accessed through the MGT collection.

Little is known about the nature of a diatom-tintinnid association detected in the MGT-136. One hypothesis suggests a mutualistic symbiosis, where diatoms acquire increased motility and tintinnids benefit from silicification through increased protection (Vincent et al. 2018). Other data indicate that the tintinnid can be the only beneficiary of the association, whereas the diatom would play the role of the “victim” (Armbrecht et al. 2017). Recent studies suggest that diatom–tintinnid associations may be more common in the ocean than previously thought. However, their global ecological and biogeographical patterns remain poorly characterized (references within Vincent et al., 2018).

The MGT collection provides a valuable resource for the evaluation of these ecologically relevant associations by studying their distribution in major oceanic provinces and by exploring the expression patterns of key genes. These findings illustrate the ability of the MGT collection to depict more interspecies relationships in the ocean, thus potentially discovering previously unknown microbial associations (Table S4), as well as study their gene expression patterns.

### Fragmentation of the MGTs

In addition to the MGTs, comprised of tens of thousands of unigenes, which cover eukaryotic reference transcriptomes with a high level of completeness, we also detected a number of smaller gene clusters which cannot reliably cover a full eukaryotic transcriptome. Several reasons may lead to the presence of these smaller MGTs representing partial eukaryotic transcriptomes. In some cases not all genes from a certain organism can be detected in all the samples where this organism is present; some genes may be missing or be present at levels below the achieved sequence coverage. This may lead to the fragmentation of the MGTs, i.e. to the fact that genes from the same organism may be allocated to multiple CAGs and MGTs of various sizes (comprised of a different number of unigenes). Several possible scenarios exist: (i) some accessory genes may be present and expressed in some subpopulations and missing in others; (ii) a sufficient sequencing depth was not achieved for some of the samples resulting in only a partial genomic coverage. Thus, for organisms with sequencing depths below or near the limit of detection, some genes may lack corresponding reads which would lead to incomplete coverage of the transcriptome by metagenomic reads. The situation when several CAGs of various sizes represent the same organism has been observed for the prokaryotic compartment of a human gut microbiome (Nielsen et al. 2014).

### Limitations and advantages of the MGT method

General limitations relevant to interpreting the gene co-abundance data obtained using the MGT approach described here include its inability to incorporate the accessory genes in the analysis due to their inherent nature of not being present in all strains and its intrinsic inability to segregate organisms that form obligate symbioses because of their identical gene co-abundance profiles.

A recently developed computational tool may solve the former problem (Plaza Oñate et al. 2019), although further analyses on environmental datasets are needed to confirm its accuracy and efficiency. Alternatively, post-processing of the MGT collection using methods based on differential sequence coverage of genes may be effective in cases where a significant bias in genome copy number of the associated organisms exists. Another caveat specific to our datasets is that genes expressed below the level of detection may be overlooked because the gene reconstruction has been performed using the metatranscriptomics data. However, the MGT approach has a number of advantages compared to other metagenome and metatranscriptome assembly methods. These include: (i) access to genomic content of organisms not available in culture (including zooplankton species) because of the culture-independent assembly and clustering of sequence data; (ii) de-novo definition of gene clusters which allows for the reconstruction of transcriptomes with no need for references.

## Conclusions

In this study we applied a gene co-abundance clustering approach on a series of samples provided by the *Tara* Oceans expedition and demonstrated its efficiency for reconstructing high quality eukaryotic transcriptomes. The resulting MGT collection provides a valuable resource for a comprehensive analysis of the eukaryotic plankton in the open sunlit ocean by providing access to biogeography, genomic content, and functional potential of many ecologically relevant eukaryotic species. This universal methodological framework can be implemented for transcriptome reconstruction of microscopic eukaryotic organisms in any environment provided that both metagenomic and metatranscriptomic data are available.

## Materials and methods

### Sampling of eukaryotic plankton communities

The samples were collected during the 2009-2013 *Tara* Oceans expeditions from all the major oceanic provinces except the Arctic. For the majority of stations samples were collected from two depths in the photic zone: subsurface (SRF) and deep-chlorophyll maximum (DCM). Planktonic eukaryotic communities were collected in the 0.8–2000 µm range and divided into four size fractions (0.8–5 μm, 5–20 μm, 20–180 μm, and 180–2000 μm). A detailed description of the sampling strategies and protocols is available in the Supplementary Materials and in (Pesant et al. 2015). Biogeochemical data measured during the expedition are available in the Supplementary Materials and in the Pangaea database (https://www.pangaea.de/).

DNA and RNA libraries were constructed and sequenced as detailed in Alberti et al., 2017 (Alberti et al. 2017), and processed as described in Carradec et al., 2018 (Carradec et al. 2018). Briefly, the raw data were filtered and cleaned to remove low quality reads, adapters, primers and ribosomal RNA-like reads. Resulting metatranscriptomic reads were assembled using velvet v.1.2.07 (Zerbino et al. 2009) with a kmer size of 63. Isoform detection was performed using oases 0.2.08 (Schulz et al. 2012). Contigs smaller than 150 bp were removed from further analysis. The longest sequence from each cluster of contigs was kept as a reference for the gene catalog. The MATOU-v1 unigene catalog is accessible at https://www.genoscope.cns.fr/tara/.

### Abundance computing and canopy clustering

The raw metagenomic (metaG) reads from 365 samples were mapped against the MATOU-v1 catalog as described in Carradec et al, 2018. Briefly, raw metagenomic reads from each sample were compared with the MATOU-v1 unigenes using the bwa tool (version 0.7.4) (Li and Durbin 2009) and those covering at least 80% of the read length with at least 95% of identity were retained for further analysis. In the case of several possible best matches, a random one was picked. For each unigene in each sample the metagenomic abundance was determined in RPKM (reads per kilo base per million of mapped reads). To improve the clustering efficiency we selected unigenes detected with metagenomic reads in at least three different samples, and which had no more than 90% of their total genomic occurrence signal in a single sample. These 2 criteria are the default parameters of the canopy clustering tool (--filter_min_obs 3 and --filter_max_top3_sample_contribution=0.9). The metagenomic RPKM-based abundance matrix of these unigenes was submitted to the canopy clustering algorithm described in (Nielsen et al. 2014) [the original code is available at https://www.genoscope.cns.fr/tara/] which is a density based clustering that does not taken into account the sequence composition, as opposed to the most binning tools. We used a max pearson correlation difference of 0.1 to define clusters and then clusters were merged if canopy centroids’ distances were smaller than 0.05 (250k iterations, default parameters).

A total of 7,254,163 unigenes were clustered into 11,846 co-abundant gene groups (CAGs) of at least 2 unigenes. CAGs with more than 500 unigenes are hereafter termed MetaGenomic based Transcriptomes (MGTs). 924 MGTs were generated which encompassed 6,946,068 unigenes (95.8%). Since this method has never been applied to eukaryotic data, a smaller cutoff of 500 unigenes was used (compared to the original method applied to prokaryote-dominated communities (Nielsen et al. 2014)) to increase the number of resulting MGTs potentially representing individual organisms. For each sampling filter we determined the fraction of metagenomics reads captured by the unigenes that compose the MGTs (Fig S2).

### Taxonomic assignment

Taxonomic assignment of the unigenes is described in Carradec *et al*., 2018. Briefly, to determine a taxonomic affiliation for each of the unigenes, a reference database was built from UniRef90 (release of 2014–09–04) (Suzek et al. 2015), the MMETSP project (release of 2014–07– 30) (Keeling et al. 2014), and *Tara* Oceans Single-cell Amplified Genomes (PRJEB6603)). The database was supplemented with three Rhizaria transcriptomes (*Collozoum*, *Phaeodaea* and *Eucyrtidium*, available through the European Nucleotide Archive under the reference PRJEB21821 (https://www.ebi.ac.uk/ena/data/view/PRJEB21821) and transcriptomes of *Oithona nana* (Madoui et al. 2017) available through the European Nucleotide Archive under the reference PRJEB18938 (https://www.ebi.ac.uk/ena/data/view/PRJEB18938). Sequence similarities between the gene catalog and the reference database were computed in protein space using Diamond (version 0.7.9) (Buchfink et al. 2015) with the following parameters: -e 1e-5 -k 500 -a 8 --more-sensitive. Taxonomic affiliation was performed using a weighted Lowest Common Ancestor approach. Subsequently, for each MGT, representative taxonomic level was determined by computing the deepest taxonomic node covering at least 75% of the taxonomically assigned unigenes of that MGT.

### Completeness and contamination assessment

For each MGT, unigenes were further assembled using CAP3 (version date: 02/10/15) (Huang and Madan 1999). Assembled contigs and singletons were pooled and completeness and contamination were computed using the Anvi’o package (ver 5.2) (Eren et al. 2015) with default parameters and a set of 83 protistan specific single copy core genes (Simão et al. 2015) for eukaryotes or a set of 139 bacterial specific single copy core genes (Campbell et al. 2013) for bacteria (Table S1). Average Nucleotide Identity (ANI) was computed using the dnadiff tool from the MUMmer package (ver 3.23) (Kurtz et al. 2004).

### Functional characterization

*DSYB-*related unigenes were identified using Hidden Markov Models (HMMs) generated from 135 sequences extracted from Curson et al., 2018. These sequences were clustered using Mmseqs2 (Steinegger and Söding 2017), and for each of the resulting 24 clusters sequences were aligned using MUSCLE (Edgar 2004). HMM construction and unigenes catalog scanning were performed using HMMer (Wheeler and Eddy 2013). The *DSYB* hmm profile had significant matches (e-value ≤ 10^−50^) with 1220 unigenes in the MATOU-v1 catalog, 46 of which were found in the MGT collection (Table S3).

*Alma1-*related unigenes were identified using HMMs generated from 5 sequences with demonstrated DMSP lyase activity, extracted from (Alcolombri et al. 2015). These sequences were clustered using Mmseqs2 (Steinegger and Söding 2017), and for each of the 2 resulting clusters sequences were aligned using Mafft v7.407 (Katoh and Standley 2013). HMM construction and unigenes catalog scanning were performed using HMMer (Wheeler and Eddy 2013). We identified 1069 positive unigenes (e ≤ 10^−50^) from the MATOU-v1 catalog, 36 of them were found in the MGT collection (Table S3).

Unigene expression values were computed in RPKM (reads per kilo base per million of mapped reads). The expression of DSYB- and Alma1-related unigenes was normalized to the total number of reads mapped to a given MGT and to the total number of reads in a given sample (Figure 4).

### Comparison with reference transcriptomes

To assess the biological validity of the resulting MGT the reference transcriptomes of *Bathycoccus prasinos* from the MMETSP database (release of 2014-07-30, (Keeling et al. 2014)) and the reference transcriptome of *Oithona nana* from Genoscope (Madoui et al. 2017) were used. Sequence similarities between the unigenes and the reference transcriptomes were computed in protein space using DIAMOND (version 0.7.9) (Buchfink et al. 2015), with the following parameters: -e 1e-5 -k 500 -a 8, and positive matches were defined as ≥ 95% identity over at least 50 amino acids.

#### Identification of potential interspecies interactions

We screened the MGT collection for potential interspecies associations by focusing on the MGTs that meet two criteria: (i) these MGTs must contain at least 10 unigenes from two different sub-kingdom taxonomic units; (ii) the number of unigenes associated with one of these taxonomic units must account for at least 5% of the number of unigenes associated with the other taxonomic unit.

For example, MGT-29 contains 19,652 unigenes assigned to Haptophyceae and 1,940 unigenes (9.9%) assigned to cyanobacteria. All the MGTs that met these criteria are listed in Table S4.

### Statistical analysis

All statistical analyses and graphical representations were conducted in R (v 3.3.2) with the R package ggplot2 (v 2.2.1). The taxonomic dendrogram shown in Fig. 1 was built using the PhyloT and NCBITaxonomy toolkits of the Python ETE3 package and visualized using iTol (Letunic and Bork 2016). The world maps were obtained using the R packages grid (v 3.3.2) and maps (v 3.2.0). Inkscape 0.92.3 was used to finalize the figures.

### Data availability

Sequencing data are archived at the European Nucleotide Archive (ENA) under the accession number PRJEB4352 for the metagenomics data and PRJEB6609 for the metatranscriptomics data. The unigene catalog is available at the ENA under the accession number ERZ480625. The MGT collection data and environmental data are available in the Supplemental Material in Table A1, at https://www.genoscope.fr/tara/, and in the Pangaea database (https://www.pangaea.de/). Additional supplemental Figures A1 and A2, MGT nucleic sequences in FASTA format, and MGT post-assemblies generated through CAP3 are available at https://www.genoscope.fr/tara/. See the Supplemental Material for more detail.

## Acknowledgments

We thank the commitment of the following people and sponsors who made this singular expedition possible: CNRS (in particular Groupement de Recherche GDR3280), European Molecular Biology Laboratory (EMBL), Genoscope/CEA, the French Government ‘Investissement d’Avenir’ programs Oceanomics (ANR-11-BTBR-0008), and FRANCE GENOMIQUE (ANR-10-INBS-09), Fund for Scientific Research – Flanders, VIB, Stazione Zoologica Anton Dohrn, UNIMIB, agnès b., the Veolia Environment Foundation, Region Bretagne, World Courier, Illumina, Cap L’Orient, the EDF Foundation EDF Diversiterre, FRB, the Prince Albert II de Monaco Foundation, Etienne Bourgois, the Tara schooner and its captain and crew. *Tara* Oceans would not exist without continuous support from 23 institutes (https://oceans.taraexpeditions.org). We also acknowledge C. Scarpelli for support in high-performance computing. Computations were performed using the platine, titane and curie HPC machine provided through GENCI grants (t2011076389, t2012076389, t2013036389, t2014036389, t2015036389 and t2016036389). All authors approved the final manuscript. This article is contribution number XX of *Tara* Oceans.

## Authors contributions

A. V., P. W., & E. P. designed research, A. V., Q. C., & E. P. generated data, A. V., M. D., Q. C., A. A., T. O. D., & E. P. analysed data, A. V. & E. P. wrote the paper, with assistance from all authors.

