## Supplemental Information for "Transcriptome reconstruction and functional analysis of eukaryotic marine plankton communities via high-throughput metagenomics and metatranscriptomics"

**This PDF file includes:**

- Materials and Methods
- Supplementary information figures S1 to S8
- Supplementary information additional figures A1 and A2 (separate files)
- Supplementary information tables S1 to S4 (separate files)
- Supplementary information datasets A1 and A2 (separate files)

### Materials and methods

*Sampling of eukaryotic plankton communities*

The samples were collected during the 2009-2013 *Tara* Oceans expeditions from all the major oceanic provinces except the Arctic. For the majority of stations samples were collected from two depths in the photic zone: subsurface (SRF) and deep-chlorophyll maximum (DCM). Planktonic eukaryotic communities were collected in the 0.8–2000 μm range and divided into four size fractions (0.8–5 μm, 5–20 μm, 20–180 μm, and 180–2000 μm).

A low-shear and non-intrusive industrial peristaltic pump was used for the 0.8–5 μm fraction and plankton nets for the others. The volumes of filtered seawater were scaled according to known organismal concentrations within each size fraction, from 0.1 $m^3$ for the most concentrated pico-plankton to 148 ± 136 $m^3$ for the most-diluted meso-plankton, in order to get near-exhaustive recovery of total eukaryotic biodiversity in each sample. Water was filtered immediately after sampling. Whole-plankton communities were subsequently filtered on polycarbonate membranes, rapidly flash-frozen, and preserved in liquid nitrogen.

DNA and RNA were extracted simultaneously by cryogenic grinding of cryopreserved membranes followed by nucleic acid extraction. Metagenomic libraries were prepared manually or in a semi-automatic manner depending on the available DNA quantity. For RNA samples, a poly(A)+ RNA selection strategy was used to limit the presence of rRNA sequences. Libraries were sequenced on the Illumina HiSeq2000 instruments with a read length of 101 bp in the paired-end mode. On average, 160 million reads per sample were obtained.

Resulting reads from each metatranscriptomic sample were assembled using velvet v.1.2.07 (Zerbino et al. 2009) with a kmer size of 63. Isoform detection was performed using oases 0.2.08 (Schulz et al. 2012). Contigs smaller than 150 bp were removed from further analysis. Similar sequences from more than one sample were removed using Cdhit-est v 4.6.1, with the following parameters: -id 95 -aS 90 (95% of nucleic identity over 90% of the length of the smallest sequence). For each cluster of contigs, the longest sequence was kept as a reference for the gene catalog. The resulting set of representative sequences was termed the MATOU-v1 catalog. For a more detailed workflow, see (Carradec et al. 2018).

*Abundance computing and canopy clustering*

The raw metagenomic (metaG) reads from 365 samples were mapped against the MATOU-v1 catalog using the bwa tool (version 0.7.4) (Li and Durbin 2009). The following parameters were used: bwa aln -l 30 -O 11 -R 1; bwa sampe -a 20000 -n 1 –N; samtools; rmdup. Low complexity reads were removed. Reads covering at least 80% of read length with at least 95% of identity were retained for further analysis. In the case of several possible best matches, a random one was picked. Unigene expression values and genomic occurrences were computed in RPKM (reads per kilobase covered per million mapped reads). RPKM values for each unigenes in each sample are accessible at https://www.genoscope.cns.fr/tara/.

To improve the clustering efficiency we selected unigenes detected with metagenomic reads in at least 3 different samples, and which had no more than 90% of their total genomic occurrence signal in a single sample. These 2 criteria are these are the default parameters of the canopy clustering tool (--filter_min_obs 3 and --

filter_max_top3_sample_contribution=0.9). The metagenomic abundance matrix of these unigenes was submitted to the canopy clustering algorithm described in (Nielsen et al. 2014) [the original code is available at http://www.genoscope.cns.fr/tara/] which is a density based clustering that does not taken into account the sequence composition, as opposed to the most binning tools. We used a max pearson correlation difference of 0.1 to define clusters and then clusters were merged if canopy centroids' distances were smaller than 0.05 (250k iterations, default parameters).

A total of 7,254,163 unigenes were clustered into 11,846 co-abundant gene groups (CAGs) of at least 2 unigenes. CAGs with more than 500 unigenes are hereafter termed MetaGenomic based Transcriptomes (MGTs). 924 MGTs were generated which encompassed 6,946,068 unigenes. Since this method has never been applied to eukaryotic data, a smaller cutoff of 500 unigenes was used (compared to the original method applied to prokaryote-dominated communities (Nielsen et al. 2014a)) to increase the number of resulting MGTs potentially representing individual organisms. For each sampling filter we determined the fraction of metagenomics reads captured by the unigenes that compose the MGTs (Fig S2).

*Taxonomic assignment*

Taxonomic assignment of the unigenes is described in (Carradec et al. 2018). Briefly, to determine a taxonomic affiliation for each of the unigenes, a reference database was built from UniRef90 (release of 2014–09–04) (Suzek et al. 2014), the MMETSP project (release of 2014–07–30) (Keeling et al. 2014), and *Tara* Oceans Single-cell Amplified Genomes (PRJEB6603)). The database was supplemented with three Rhizaria transcriptomes (Collozoum, Phaeodaea and Eucyrtidium, available through the European Nucleotide Archive under the reference PRJEB21821 (https://www.ebi.ac.uk/ena/data/view/PRJEB21821) and transcriptomes of *Oithona nana* (Madoui et al. 2017). Sequence similarities between the gene catalog and the reference database were computed in protein space using Diamond (version 0.7.9) (Buchfink et al. 2014) with the following parameters: -e 1e-5 -k 500 -a 8. Taxonomic affiliation was performed using a weighted Lowest Common Ancestor (wLCA ) approach and defined as the taxonomic node that covers at least 67 % of all the bitscores of the top matches (those having a bitscore equal or greater than 90% of the bitscore of the best match). Then a correction of the deepest possible taxonomic rank of the computed wLCA was performed based on the percentage of identity of the best score (95% identity → species, 80% identity → genus, 65% identity → family and 50% identity → order).

*Identification of potential interspecies interactions*

We have screened the MGT collection for potential interspecies associations by focusing on the MGTs that meet two criteria. (i) These MGTs must contain at least 10 unigenes from two different sub-kingdom taxonomic units. (ii) The number of unigenes associated with one of these taxonomic units must account for at least 5% of the number of unigenes associated with the other one. For example MGT-29 contains 19652 unigenes assigned to Haptophyceae and 1940 unigenes assigned to cyanobacteria. All the MGTs that met these criteria are listed in Table S4.

*Statistical analysis*.

All statistical analyses and graphical representations were conducted in R (v 3.3.2) with the R package ggplot2 (v 2.2.1). The phylogenetic tree of life shown in Fig. 1 was built using iTol (Letunic and Bork 2016). The world maps were obtained using the R packages grid (v 3.3.2) and maps (v 3.2.0). Inkscape 0.92.3 was used to finalize the figures.

*Data availability*

Sequencing data are archived at the European Nucleotide Archive (ENA) under the accession number PRJEB4352 for the metagenomics data and PRJEB6609 for the metatranscriptomics data. The unigene catalog is available at the ENA under the accession number ERZ480625. The MGT collection data and environmental data are available in the Supplemental Material in Table A1, at https://www.genoscope.fr/tara/, and in the Pangaea database (https://www.pangaea.de/). Additional supplemental Figures A1 and A2, MGT nucleic sequences in FASTA format, and MGT post-assemblies generated through CAP3 are available at https://www.genoscope.fr/tara/.

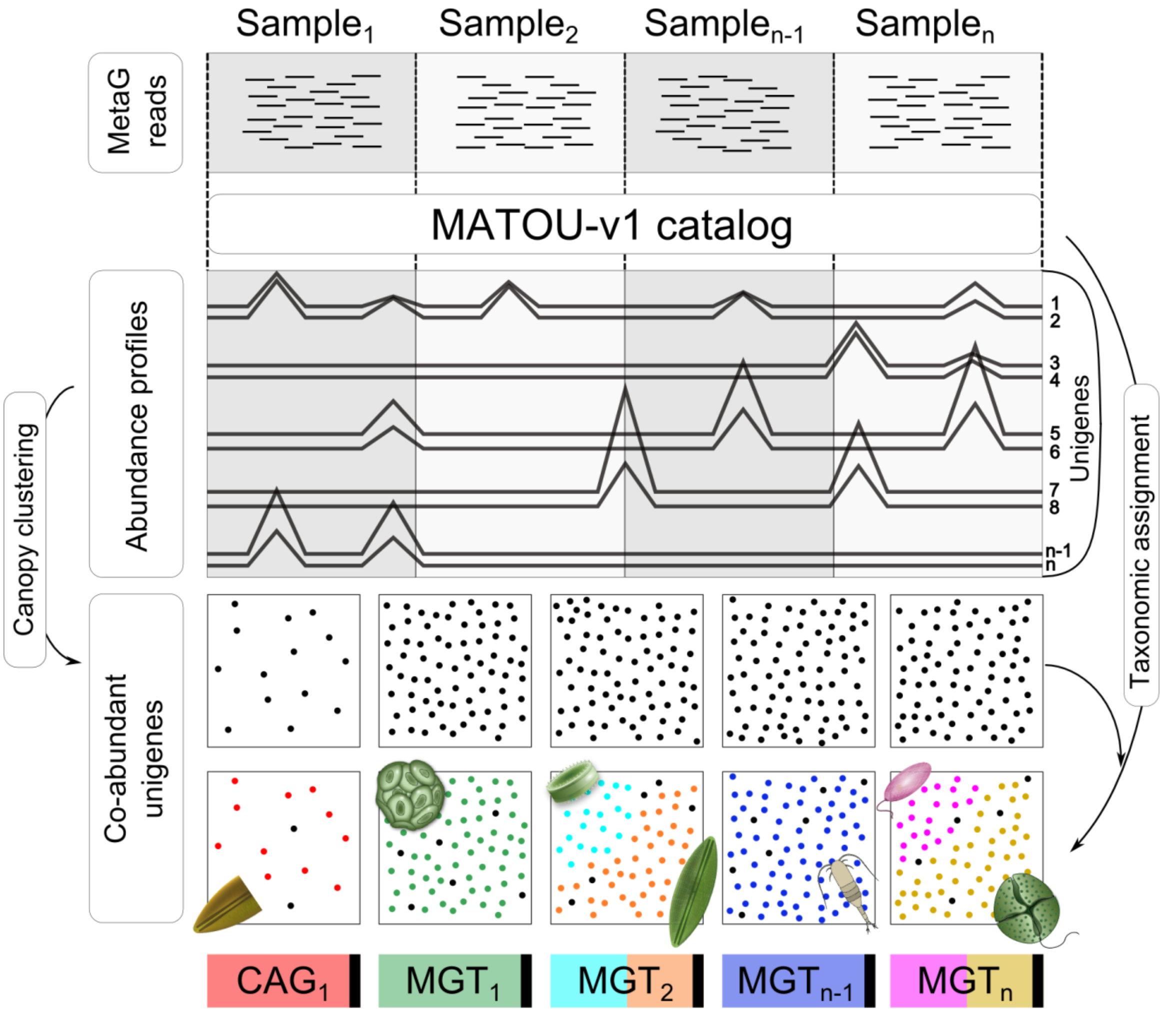


**Fig. S1.** Conceptual basis of the MGT approach.
Metagenomic reads from a series of *Tara* Oceans samples were mapped against the MATOU-v1 catalog of unigenes constructed as described in Carradec et al., 2018. The canopy clustering method is adapted from Nielsen et al., 2014. The resulting gene clusters (black squares) are called co-abundance gene groups (CAGs); those CAGs containing more than 500 unigenes are referred to as metagenomics based transcriptomes (MGTs). Black dots in the bottom row of the black squares represent taxonomically unassigned unigenes. The possibility of having more than one taxon in an MGT is demonstrated by different colors of the dots in the same black square. 11,846

CAGs are composed of 7,254,163 unigenes,  924 MGTs are composed of 6,946,068 unigenes.

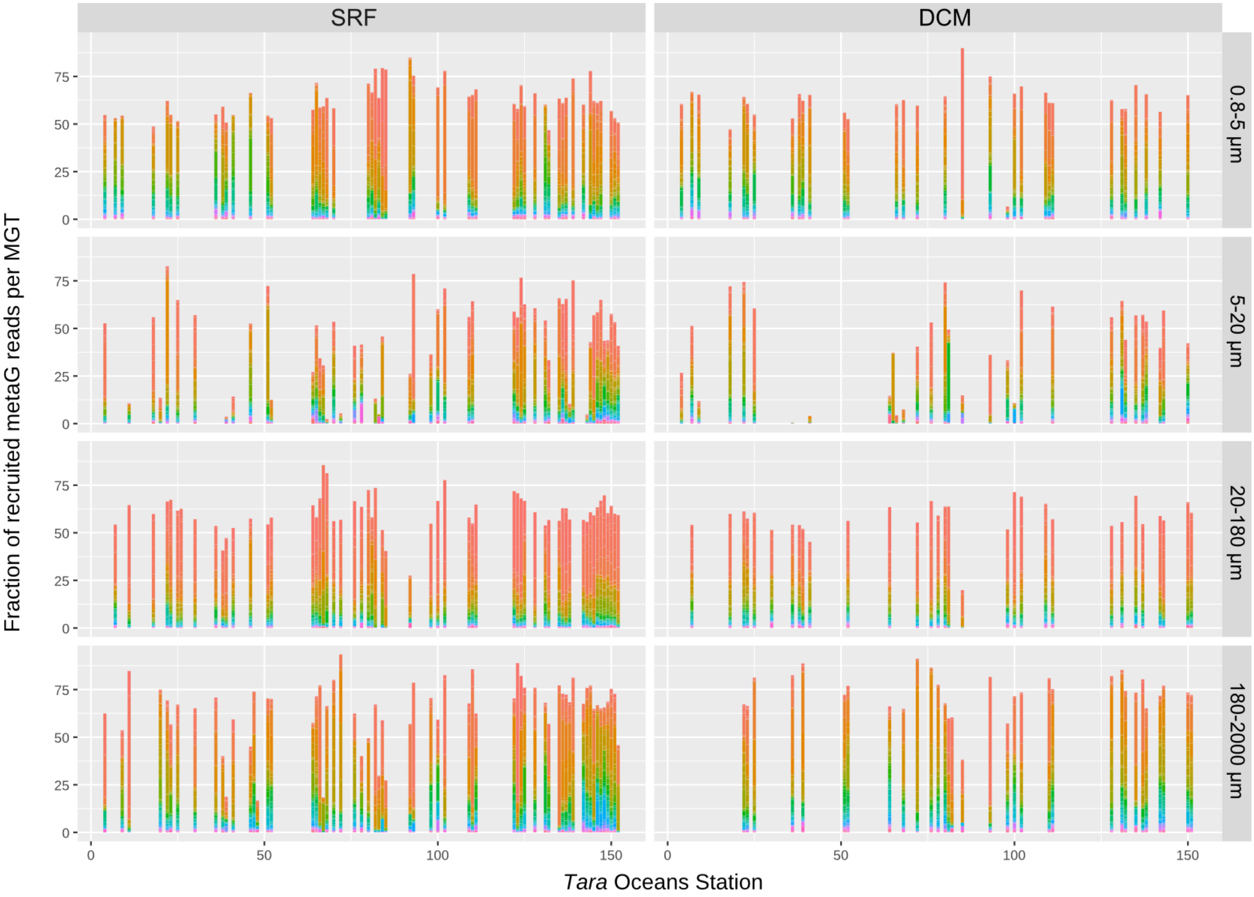


**Fig. S2.** Distribution of fractions of metagenomic reads recruited by MGTs across *Tara* Oceans stations. Metagenomic readsets were mapped onto the MGT unigenes with at least 95% of identity over at least 80% of the read length. Each column represents a total proportion of metagenomic reads mapped onto the MATOU-v1 catalog and assigned to an MGT. Color legend - from MGT-1 in red to MGT-924 in purple. SRF - surface, DCM - deep-chlorophyll maximum.

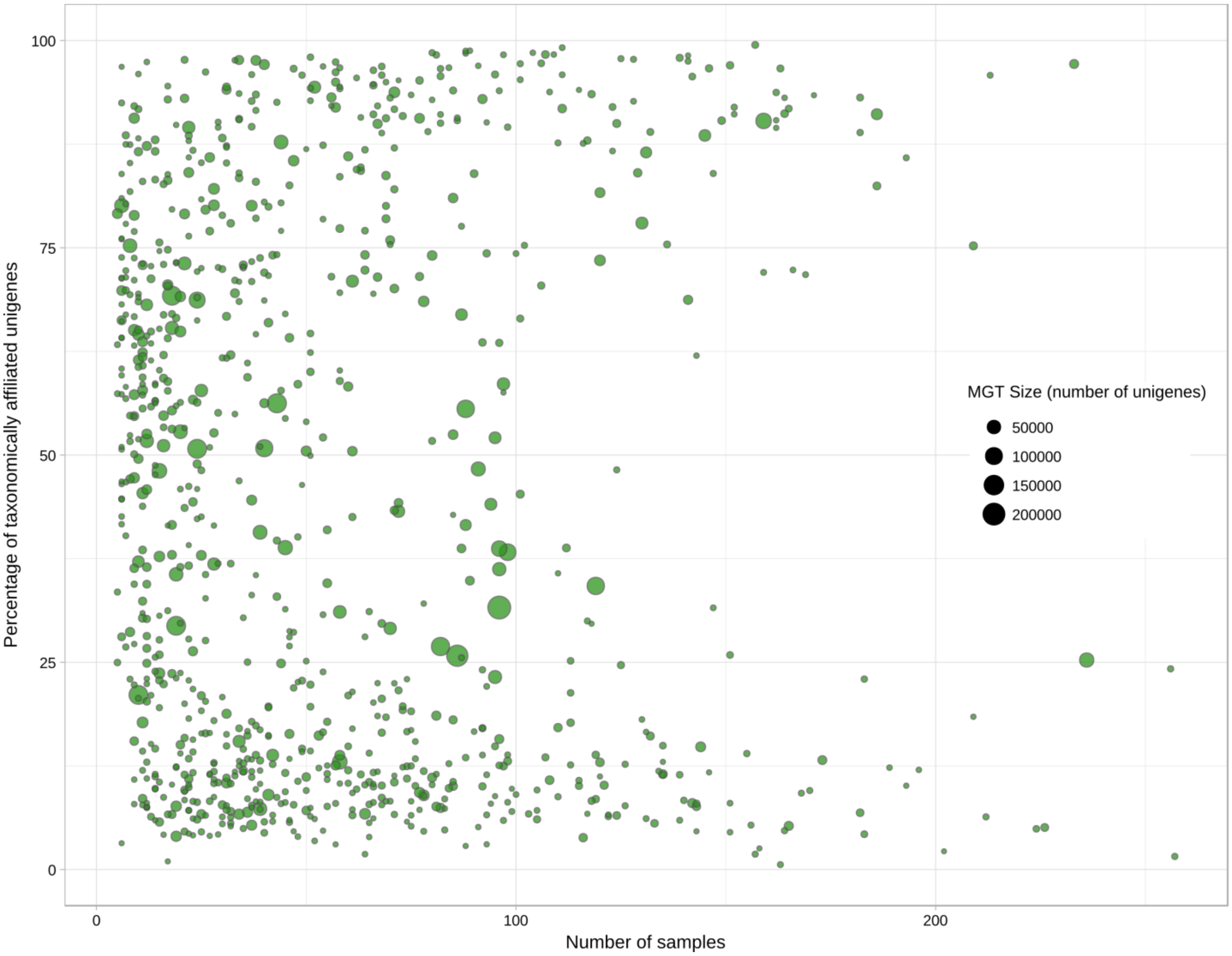


**Fig. S3.** Correlation between the number of samples recruiting an MGT (x axis), the number of taxonomically assigned unigenes in that MGT (y axis), and the MGT size (circle size).

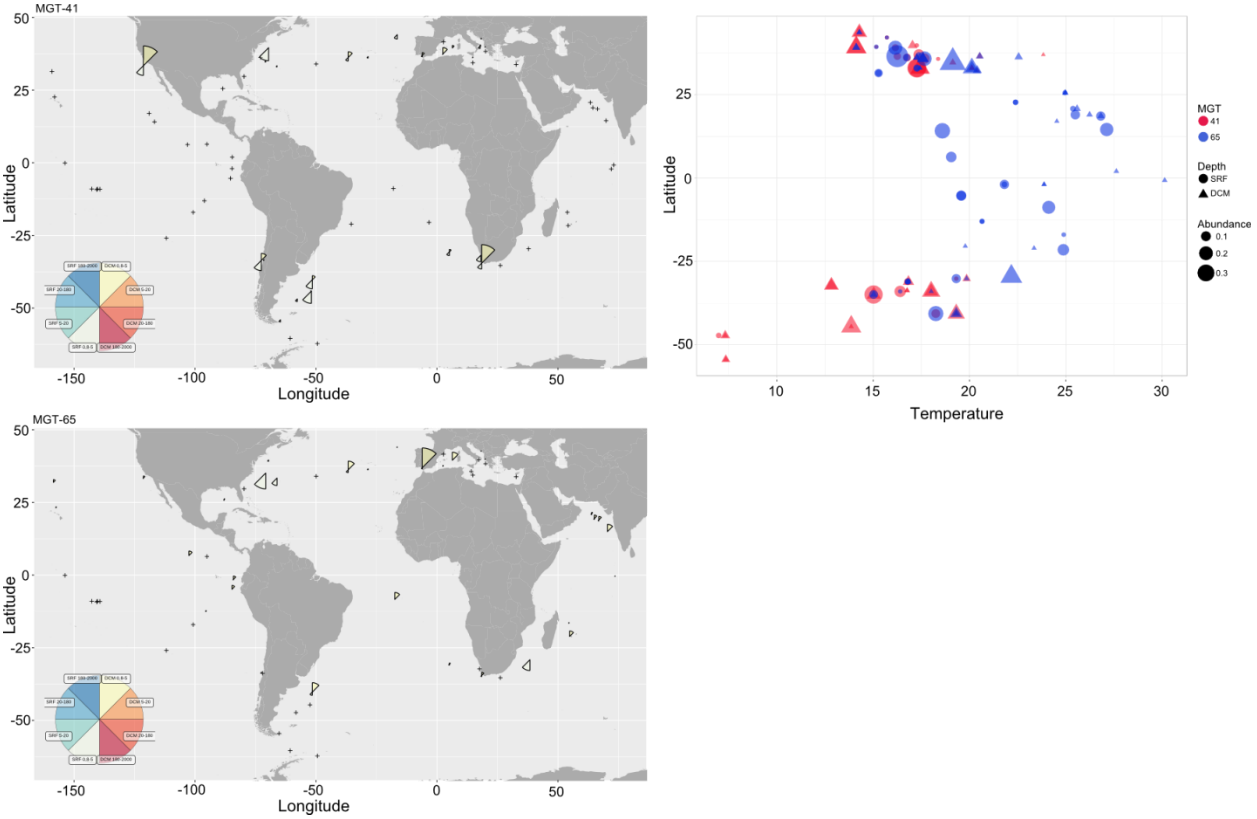


**Fig. S4.** Geographical distribution of MGT-41 (A) and MGT-65 (B) representing the B2 and B1 ecotypes of *Bathycoccus prasinos*, respectively. Panel C shows differential environmental preferences of MGT-41 and MGT-65 in relation to latitude (y axis), sea water temperature (x axis), and sampling depth (the symbol shape).

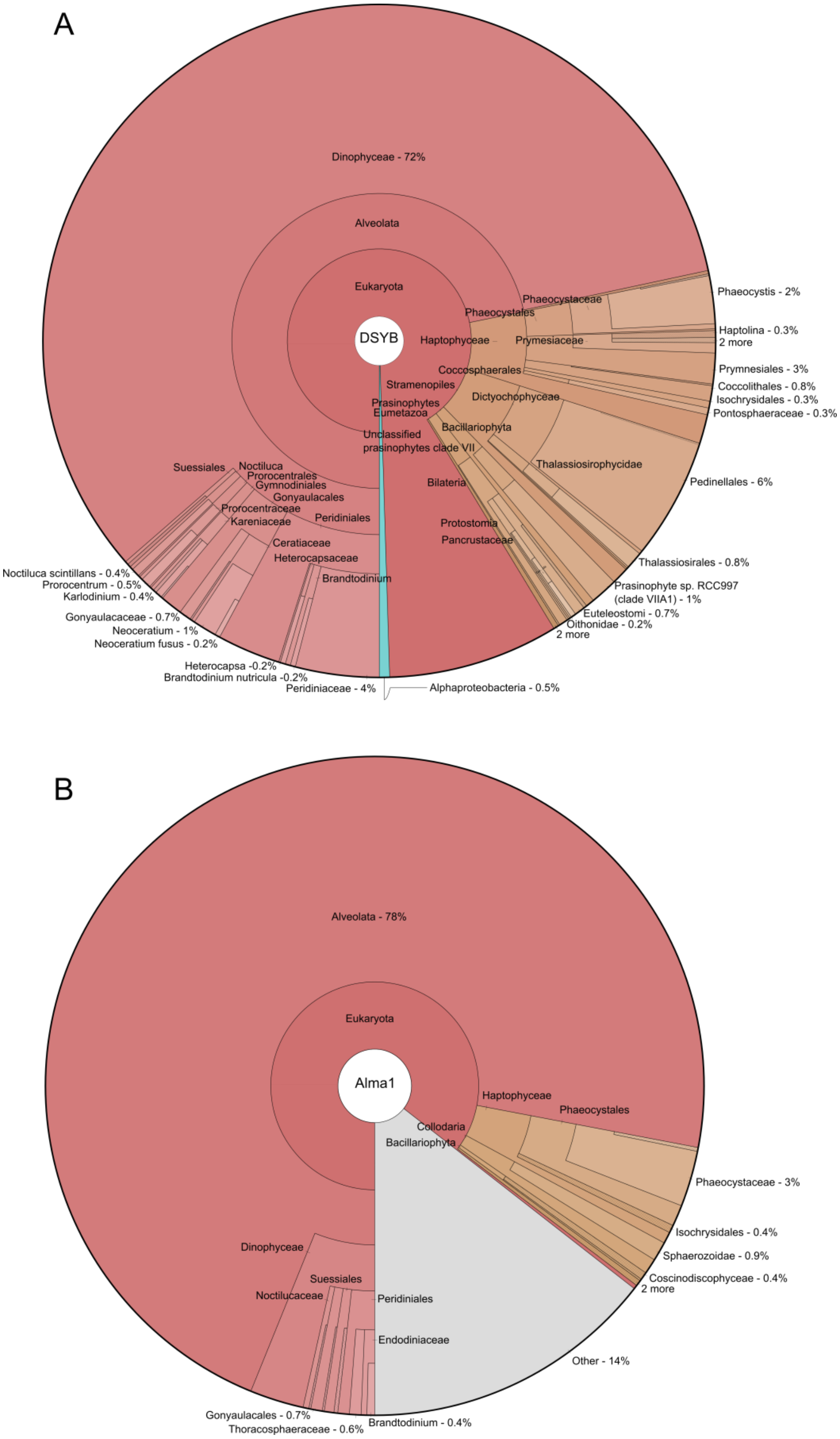


**Fig. S5.** Krona maps demonstrating the taxonomic diversity of *DSYB*-related (A) and *Alma1*-related (B) unigenes in the MATOU-v1 catalog.

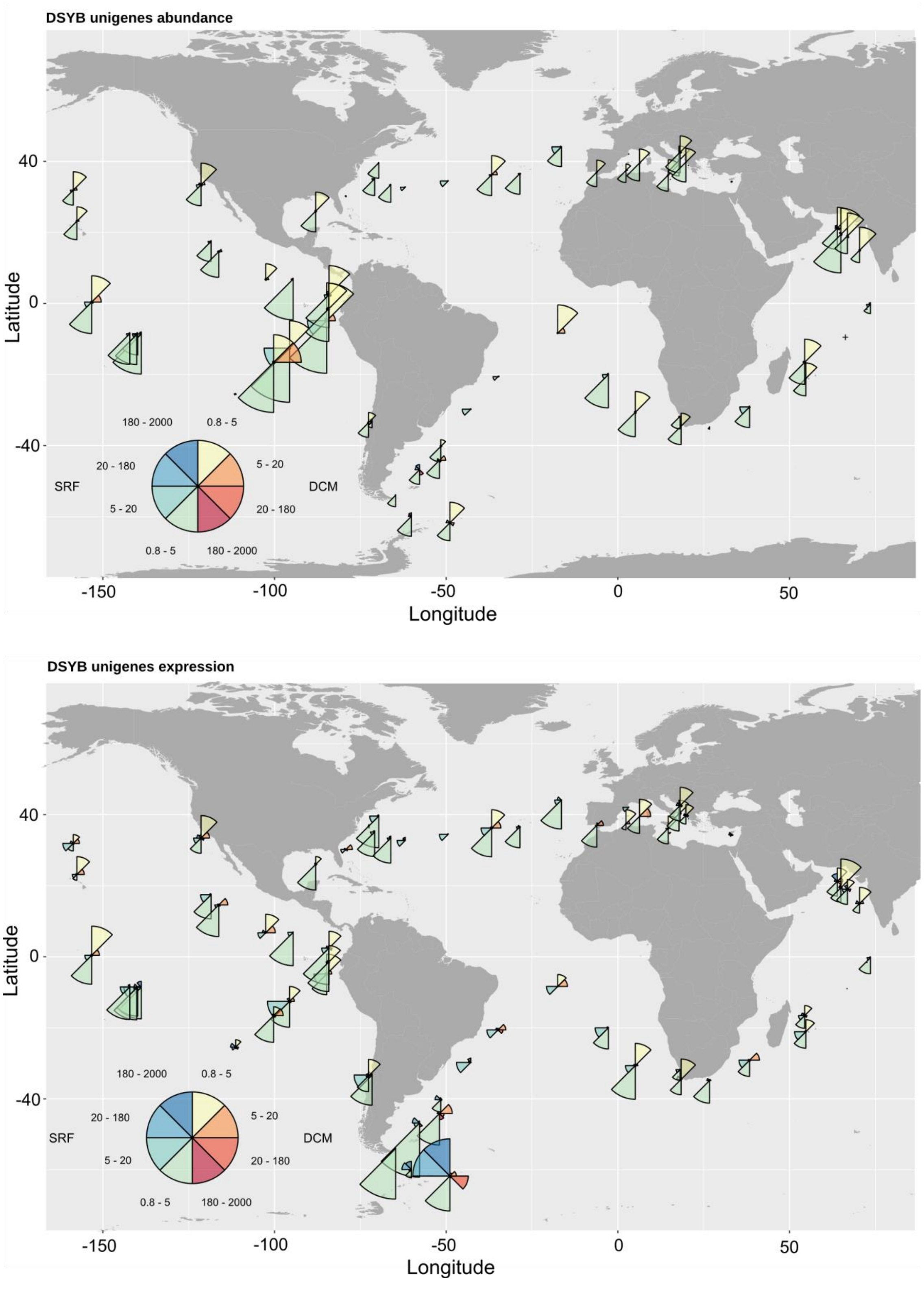


**Fig. S6.** Geographical distribution of *DSYB*-related unigenes across *Tara* Oceans stations based on their abundance (top) and expression (bottom).

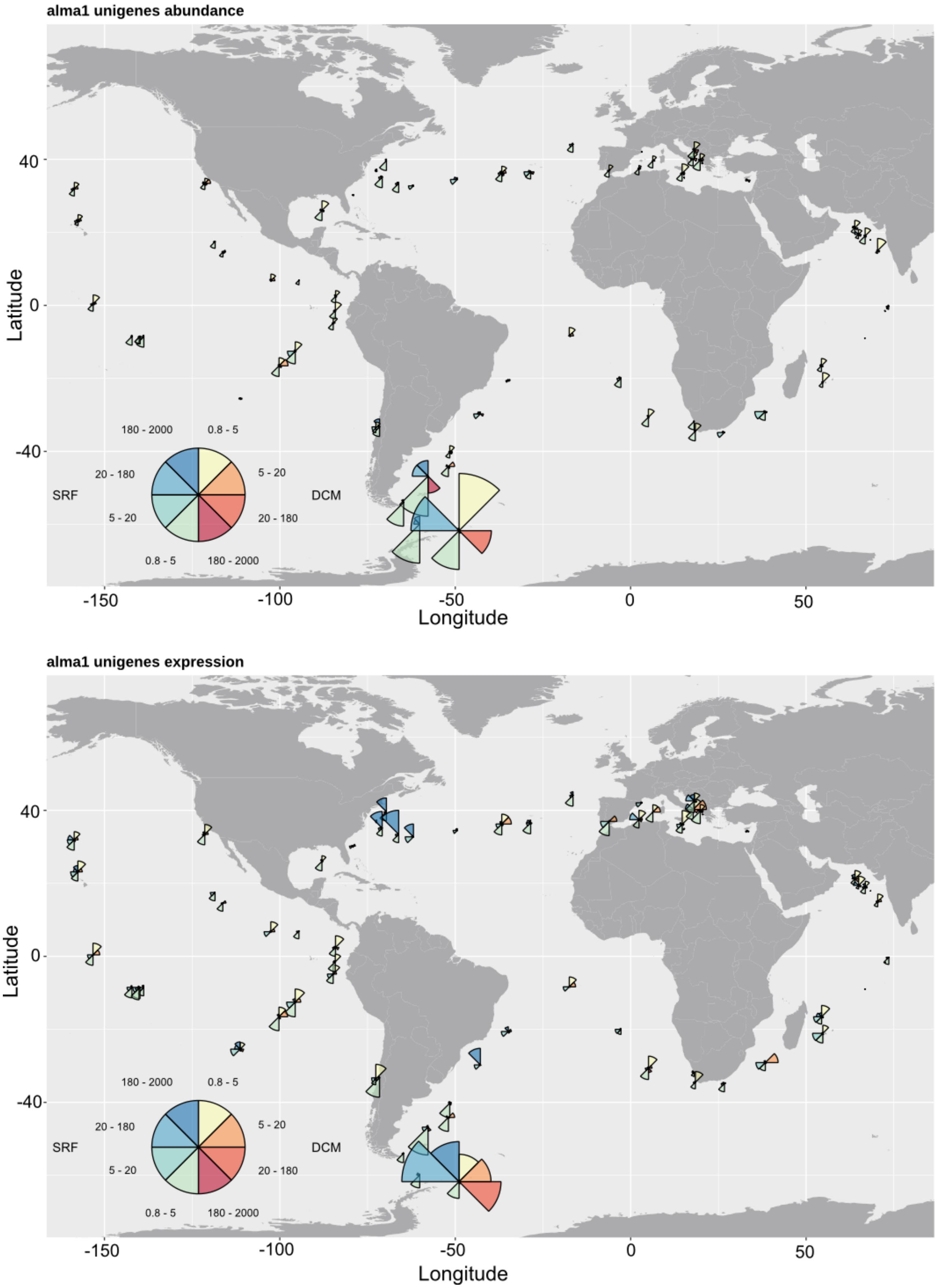


**Fig. S7.** Geographical distribution of *Alma1*-related unigenes across *Tara* Oceans stations based on their abundance (top) and expression (bottom).

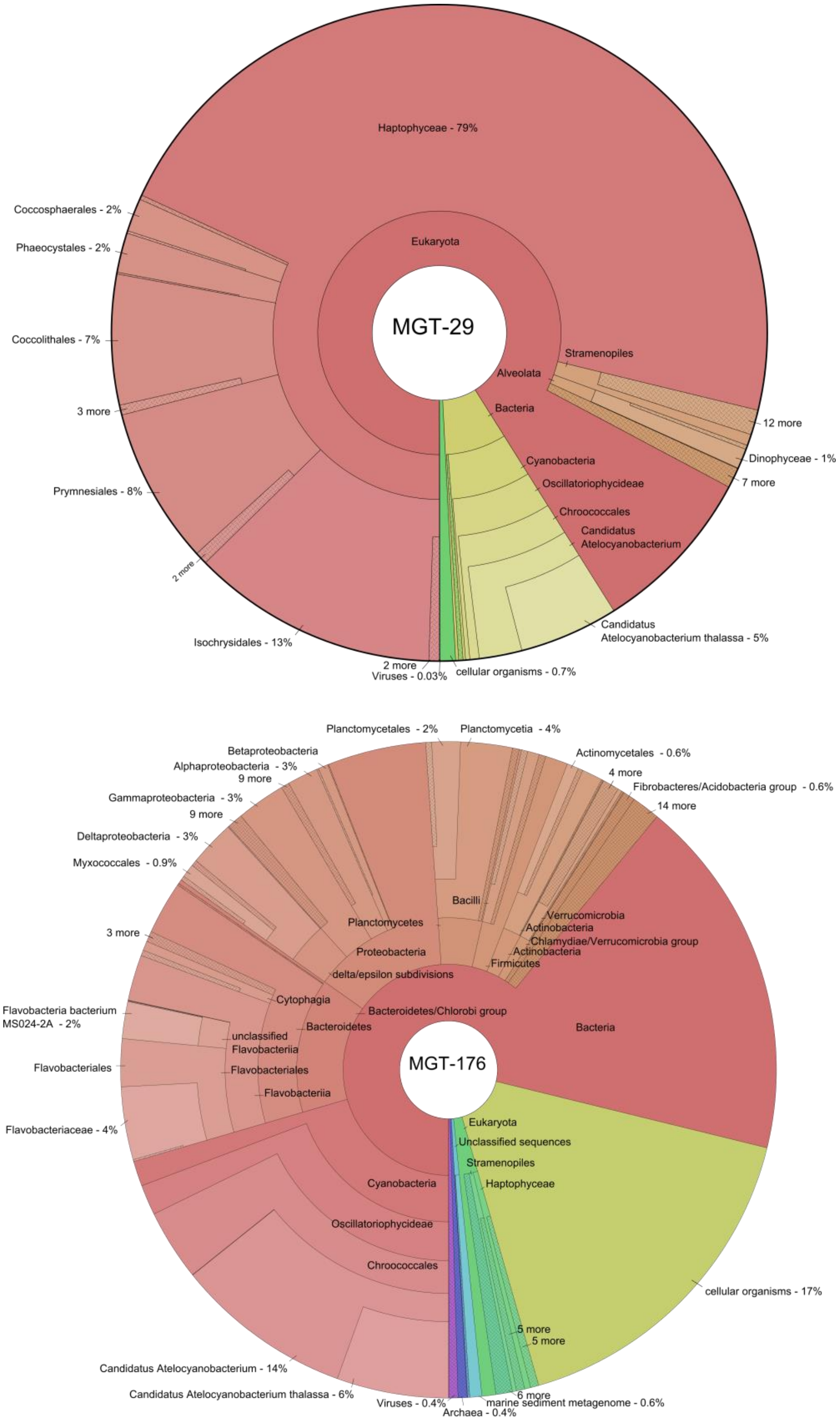


**Fig. S8.** Krona maps demonstrating the taxonomic diversity of unigenes within MGT-29 (A) and MGT-176 (B).

Additional data table S1 (separate file). Major characteristics of the individual MGTs, including their size, percentage of the taxonomically affiliated unigenes, global taxonomic assignment, completeness, and contamination.

Additional data table S2 (separate file). Sequence coverage of the reference organisms (*Bathycoccus prasinos* and *Oithona nana*) by the MGTs.

Additional data table S3 (separate file). Abundance and expression of the *DSYB*- and *Alma1*-related genes across *Tara* Oceans stations.

Additional data table S4 (separate file). Potential interspecies associations detected in the MGT collection.

Supplementary additional Figure A1 (separate file) – Graphical representation of taxonomic affiliation of unigenes in MGTs. O/U - other unclassified (a separate file at http://www.genoscope.cns.fr/tara/ ).

Supplementary additional Figure A2 (separate file) – Geographical distribution of MGTs across *Tara* Oceans stations separated by depth (SRF and DCM) and size fractions (0.8 - 5 μm; 5 - 20 μm; 20 - 180 μm; 180 - 2000 μm). For each sample, the MGTs' abundance is defined as the third quartile of the abundance of their unigenes. Each map shows relative values. SRF - surface, DCM - deep chlorophyll maximum (a separate file at http://www.genoscope.cns.fr/tara/).

Supplementary information dataset A1 (separate file) – Major statistical characteristics of the MGT collection and distribution of the MGTs across samples collected from the *Tara* Oceans expedition (http://www.genoscope.cns.fr/tara/).

Supplementary information dataset A2 (separate file) – Functional annotations of the host-related unigenes in MGT-29 based on the Kyoto Encyclopedia of Genes and Genomes (KEGG) database (http://www.genoscope.cns.fr/tara/).
