## Additional supplemental Fig A1 for "Transcriptome reconstruction and functional analysis of eukaryotic marine plankton communities via high-throughput metagenomics and metatranscriptomics"

MGT−v1_1 : 226807 unigenes, 71674 (31.60%) taxonomically assigned

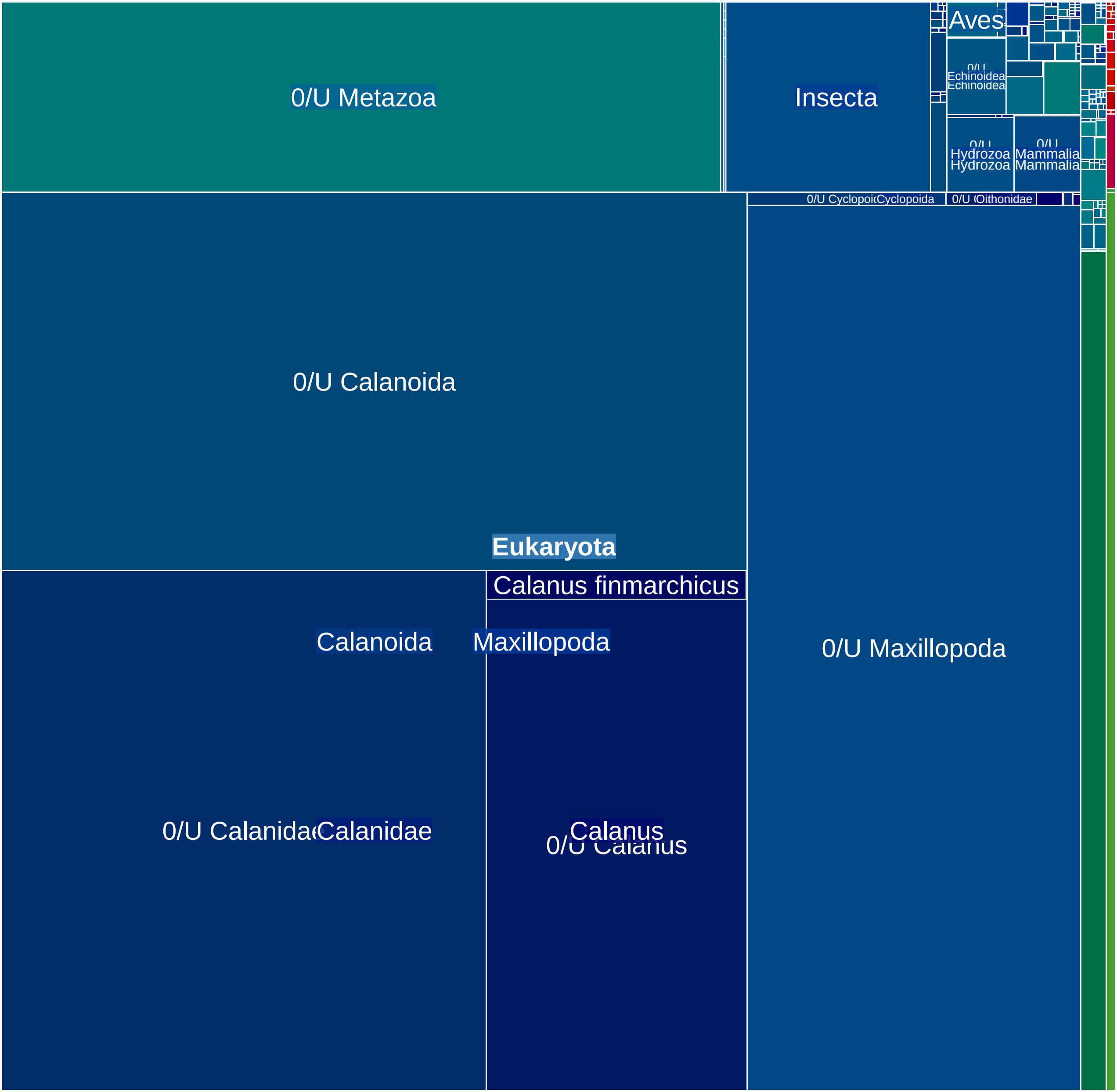

MGT−v1_2 : 188464 unigenes, 48605 (25.79%) taxonomically assigned

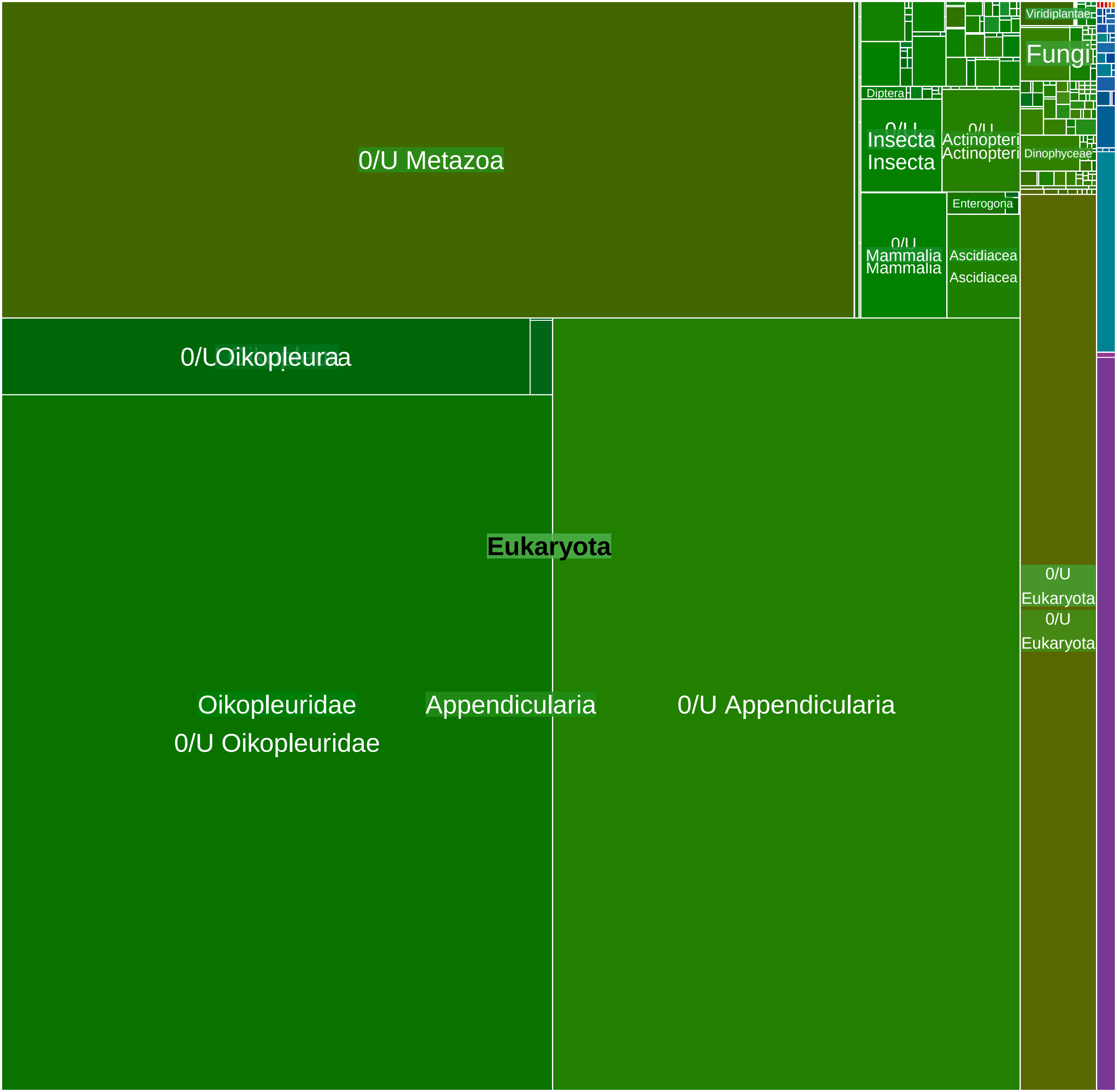

MGT−v1_3 : 140612 unigenes, 79081 (56.24%) taxonomically assigned

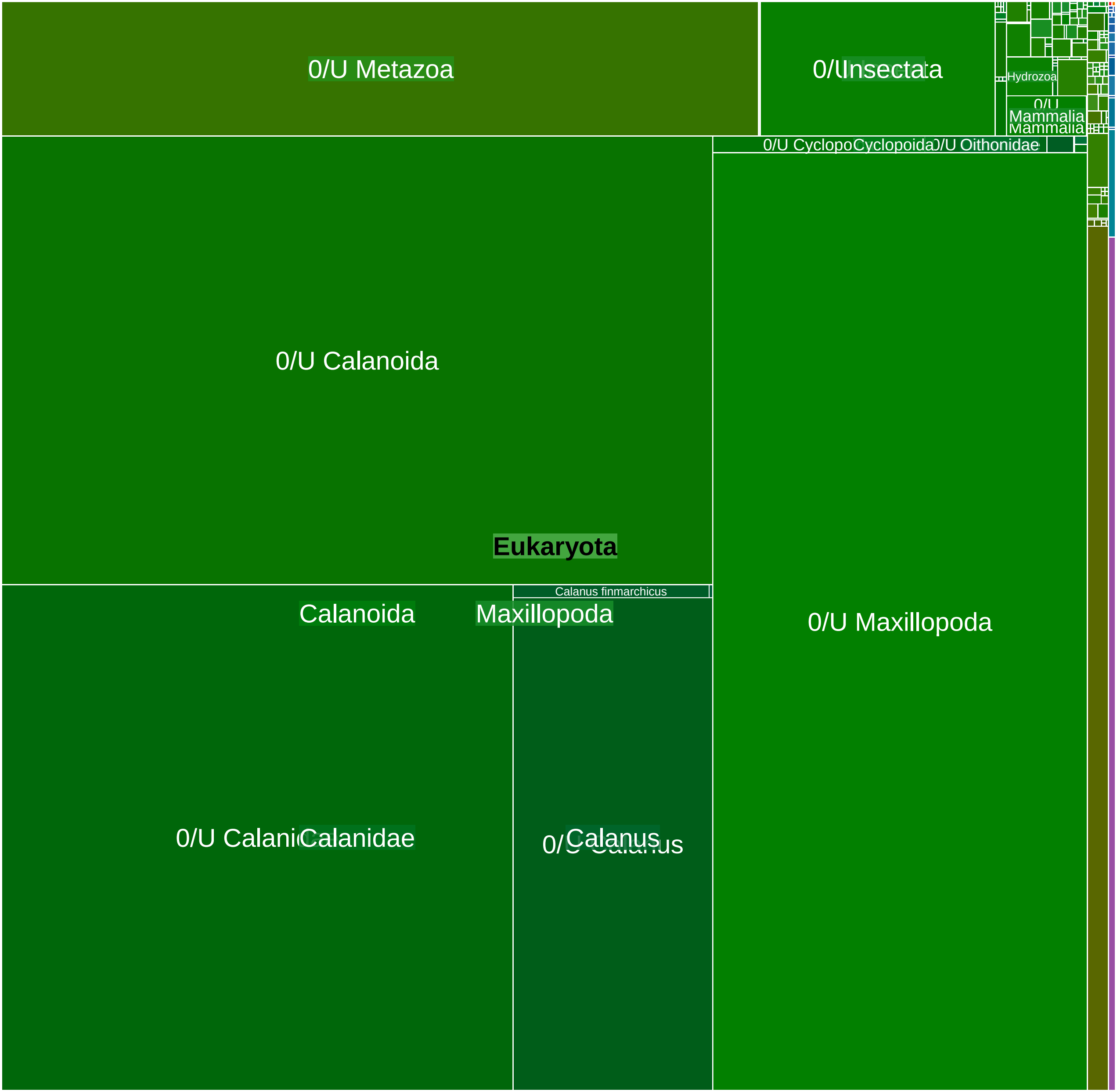

MGT−v1_4 : 137550 unigenes, 95215 (69.22%) taxonomically assigned

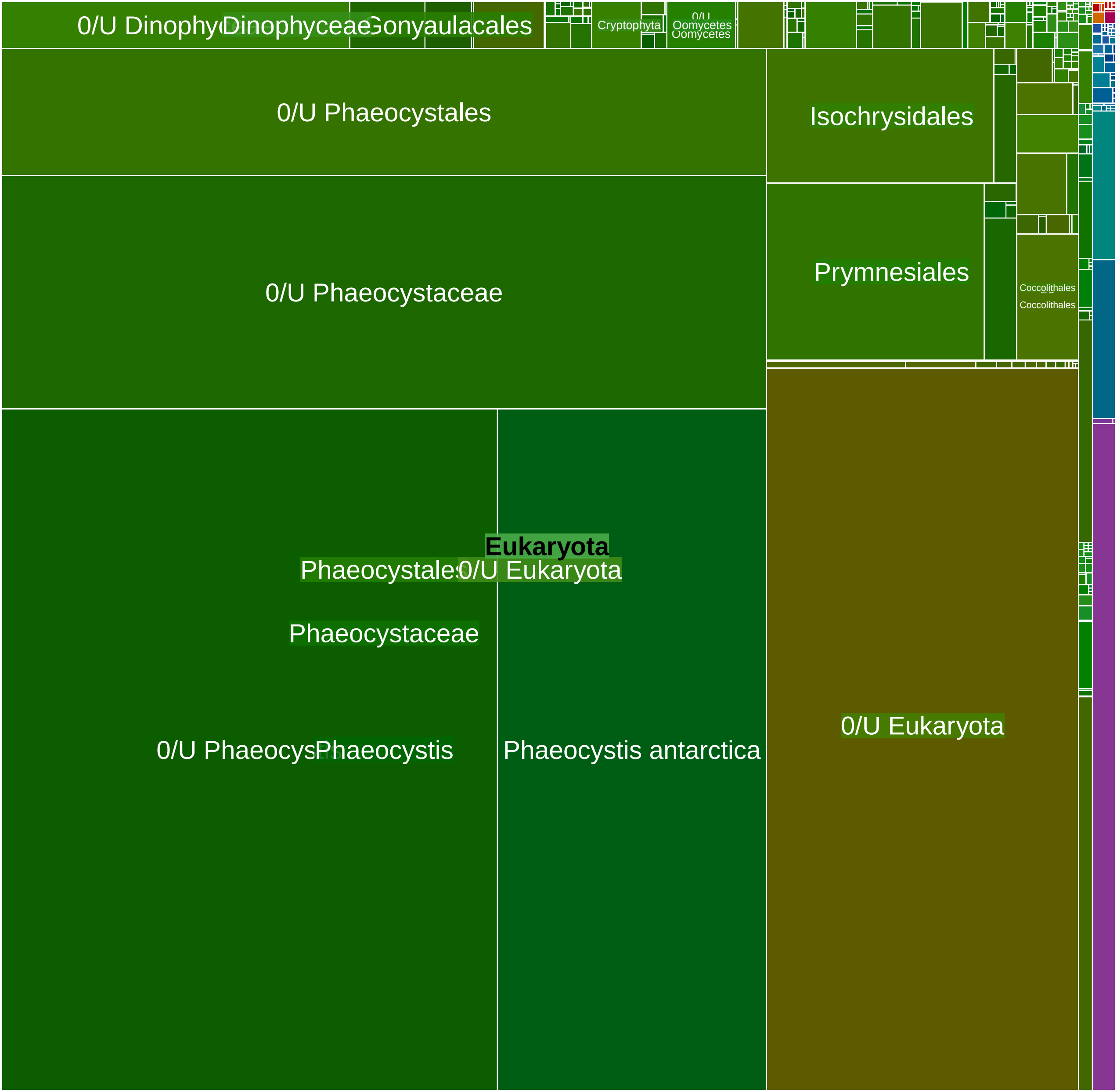

MGT−v1_5 : 136601 unigenes, 69325 (50.75%) taxonomically assigned

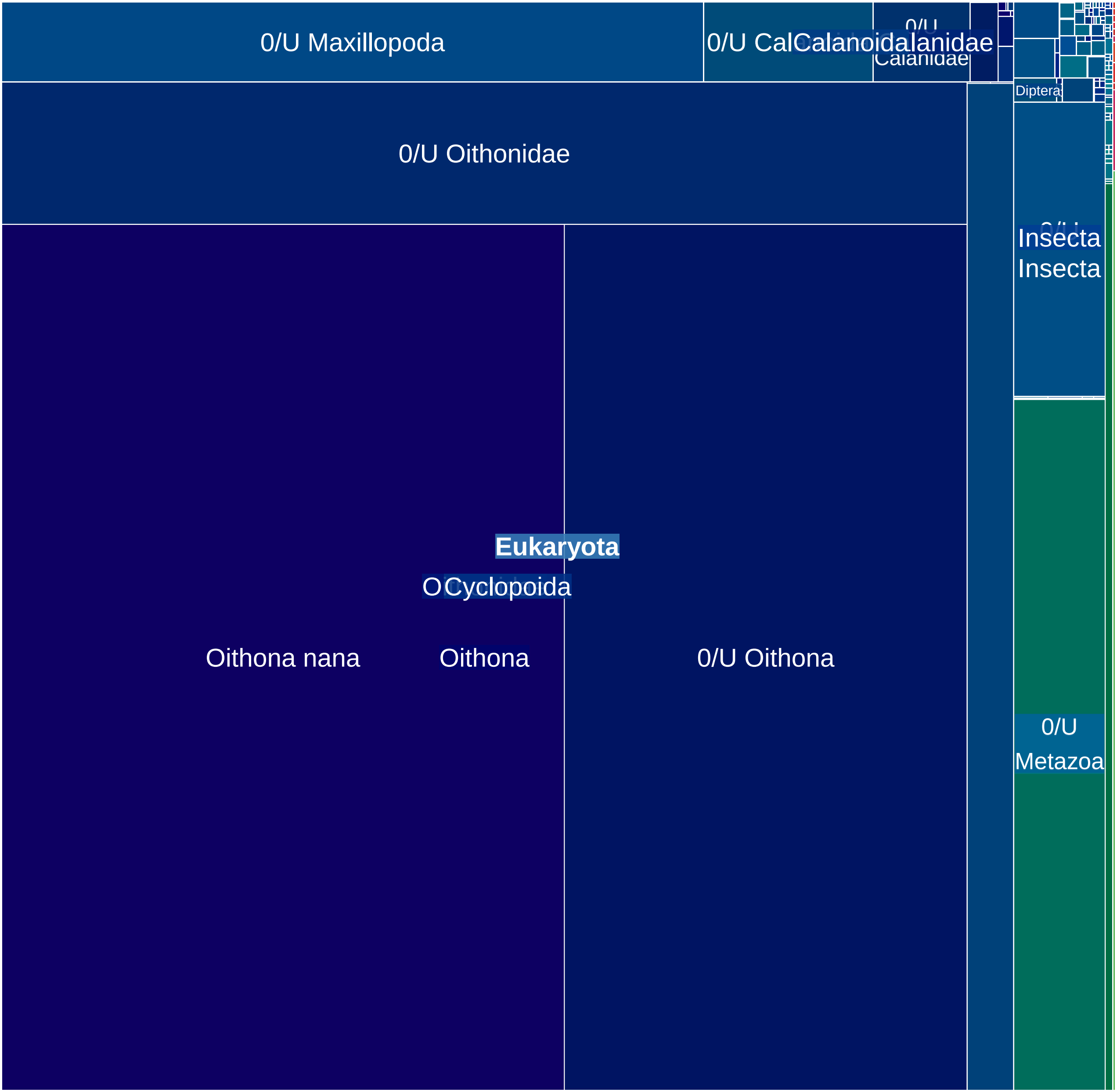

MGT−v1_6 : 135892 unigenes, 39950 (29.40%) taxonomically assigned

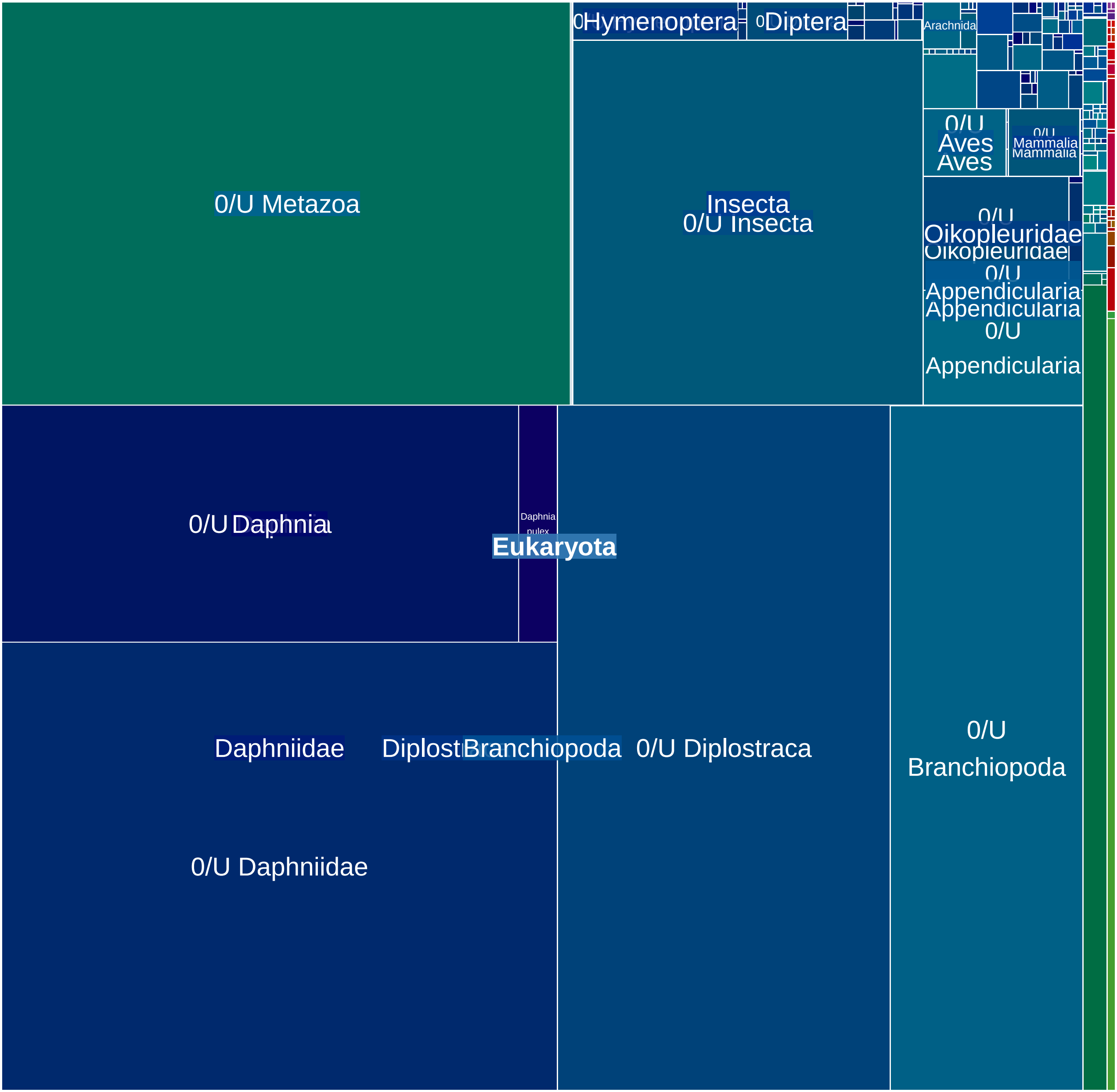

MGT−v1_7 : 134359 unigenes, 28296 (21.06%) taxonomically assigned

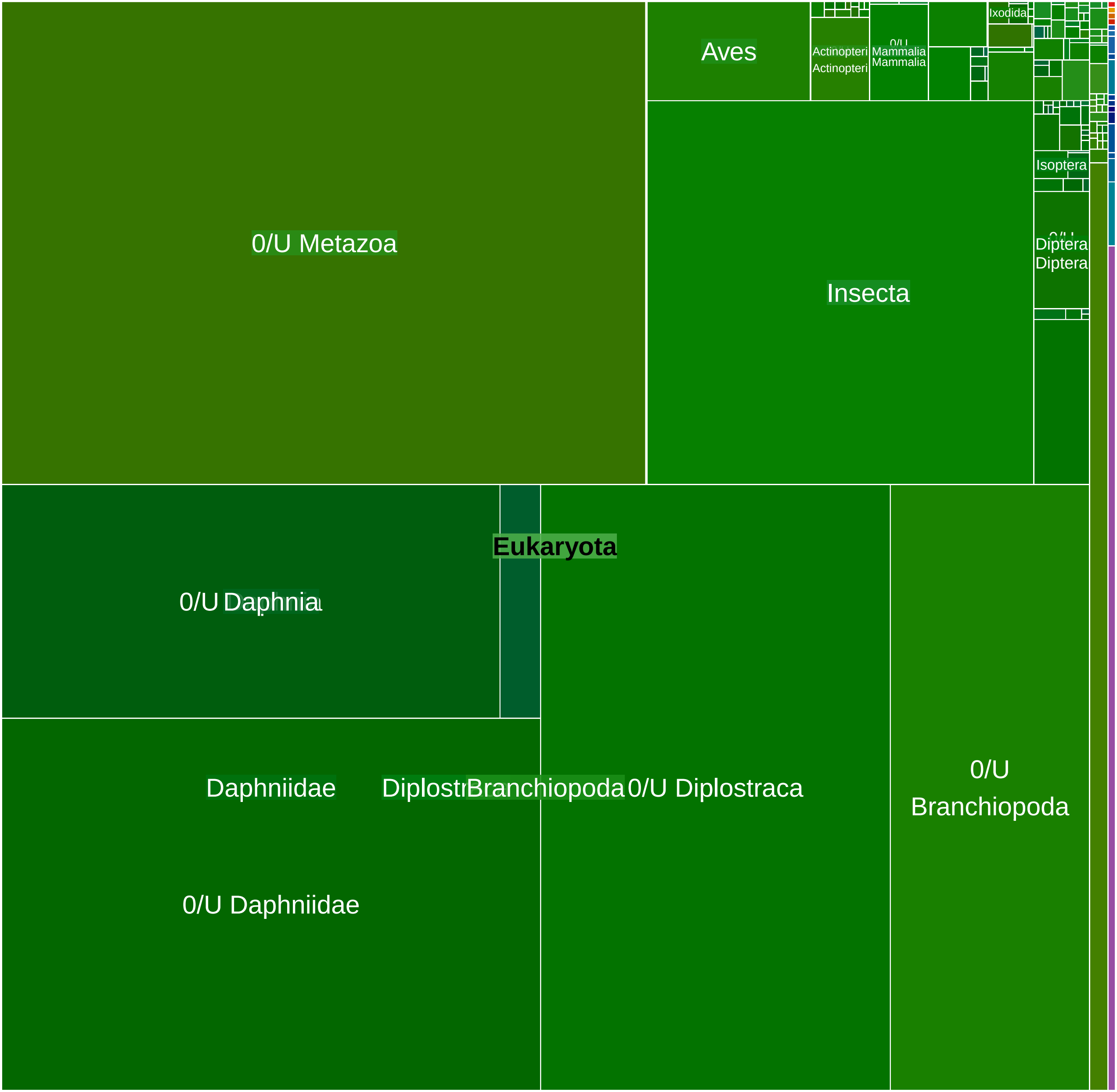

MGT−v1_8 : 123134 unigenes, 33117 (26.90%) taxonomically assigned

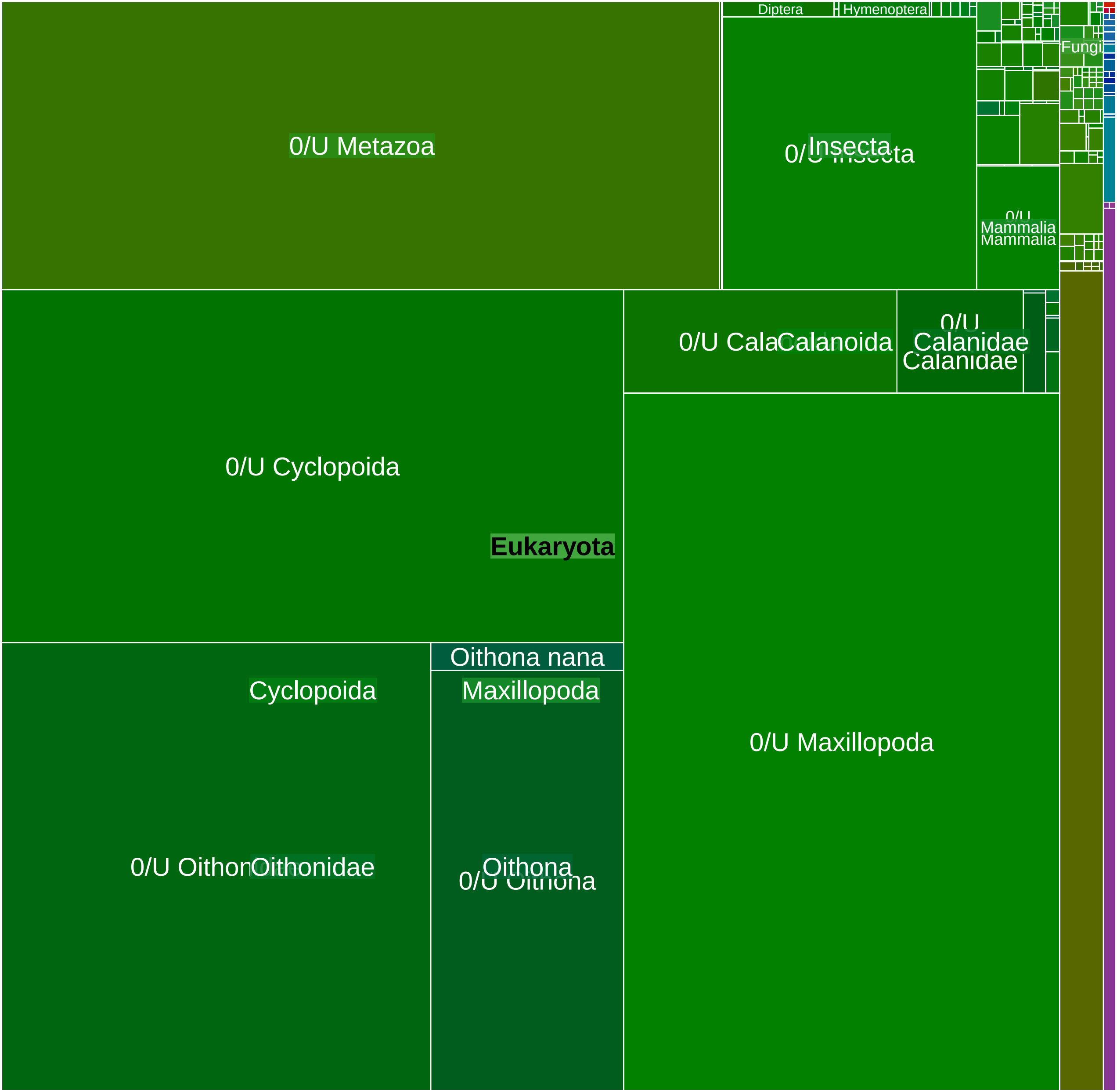

MGT−v1_9 : 109368 unigenes, 60778 (55.57%) taxonomically assigned

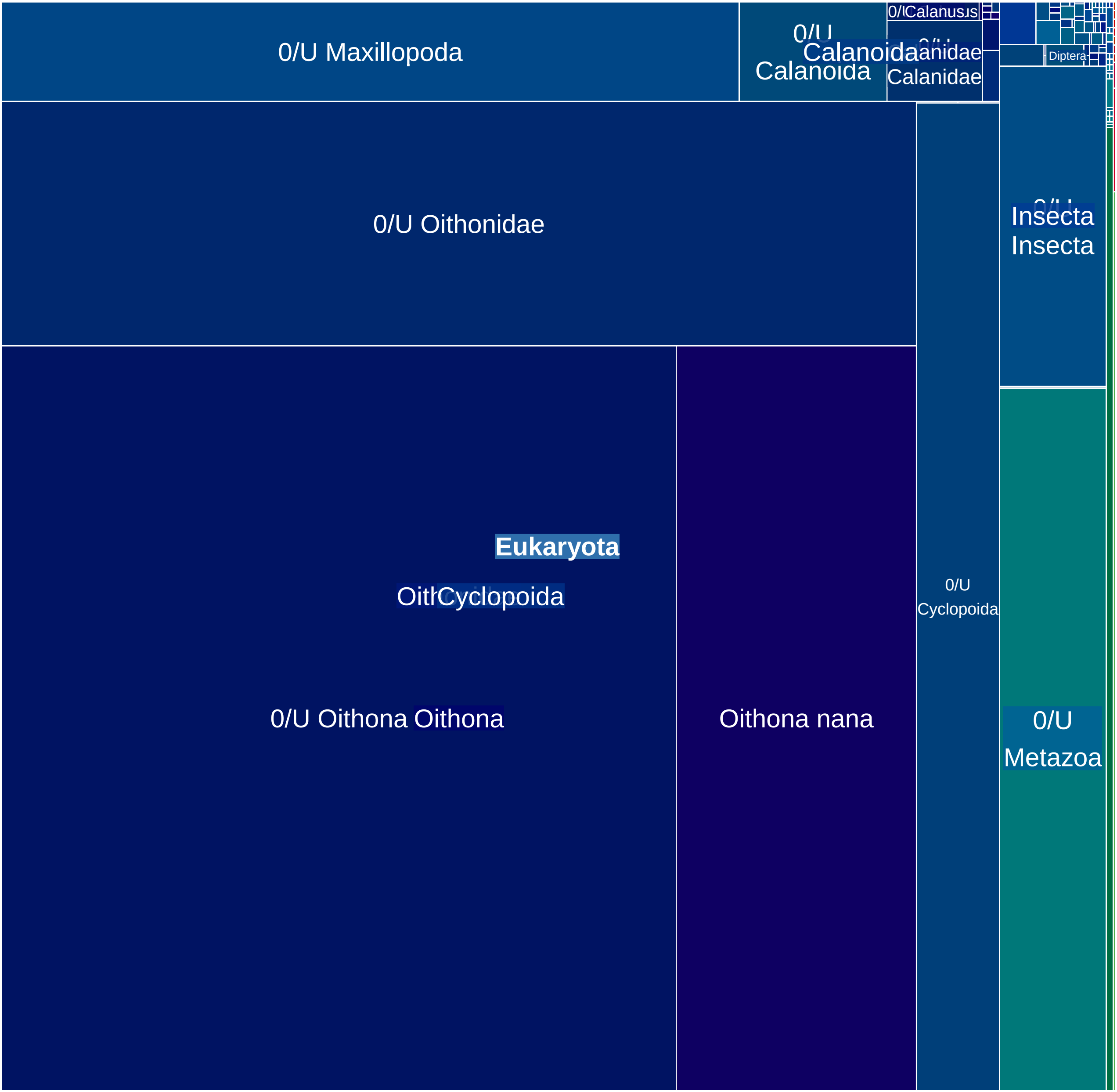

MGT−v1_10 : 107664 unigenes, 36834 (34.21%) taxonomically assigned

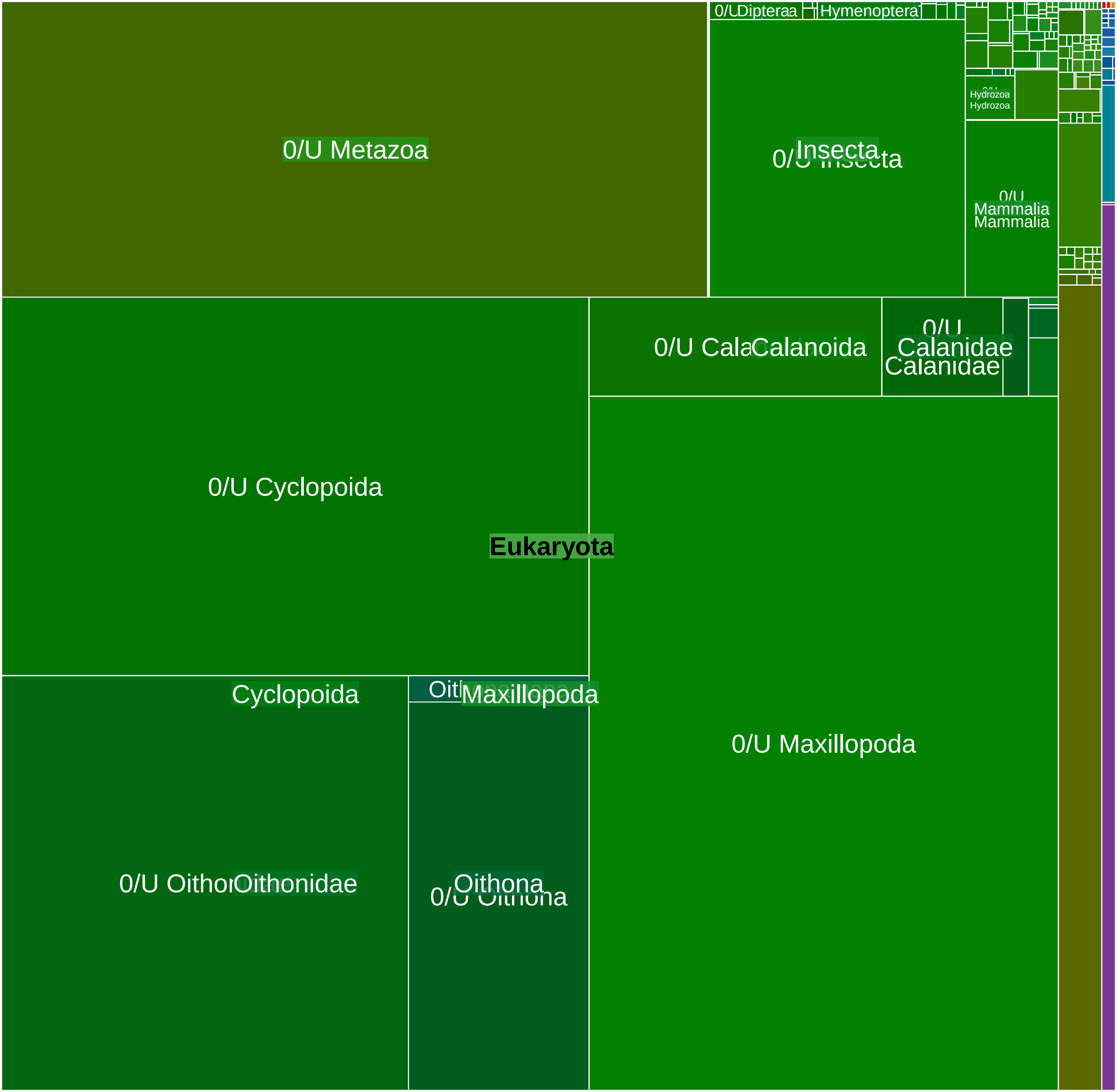

MGT−v1_11 : 103565 unigenes, 52631 (50.82%) taxonomically assigned

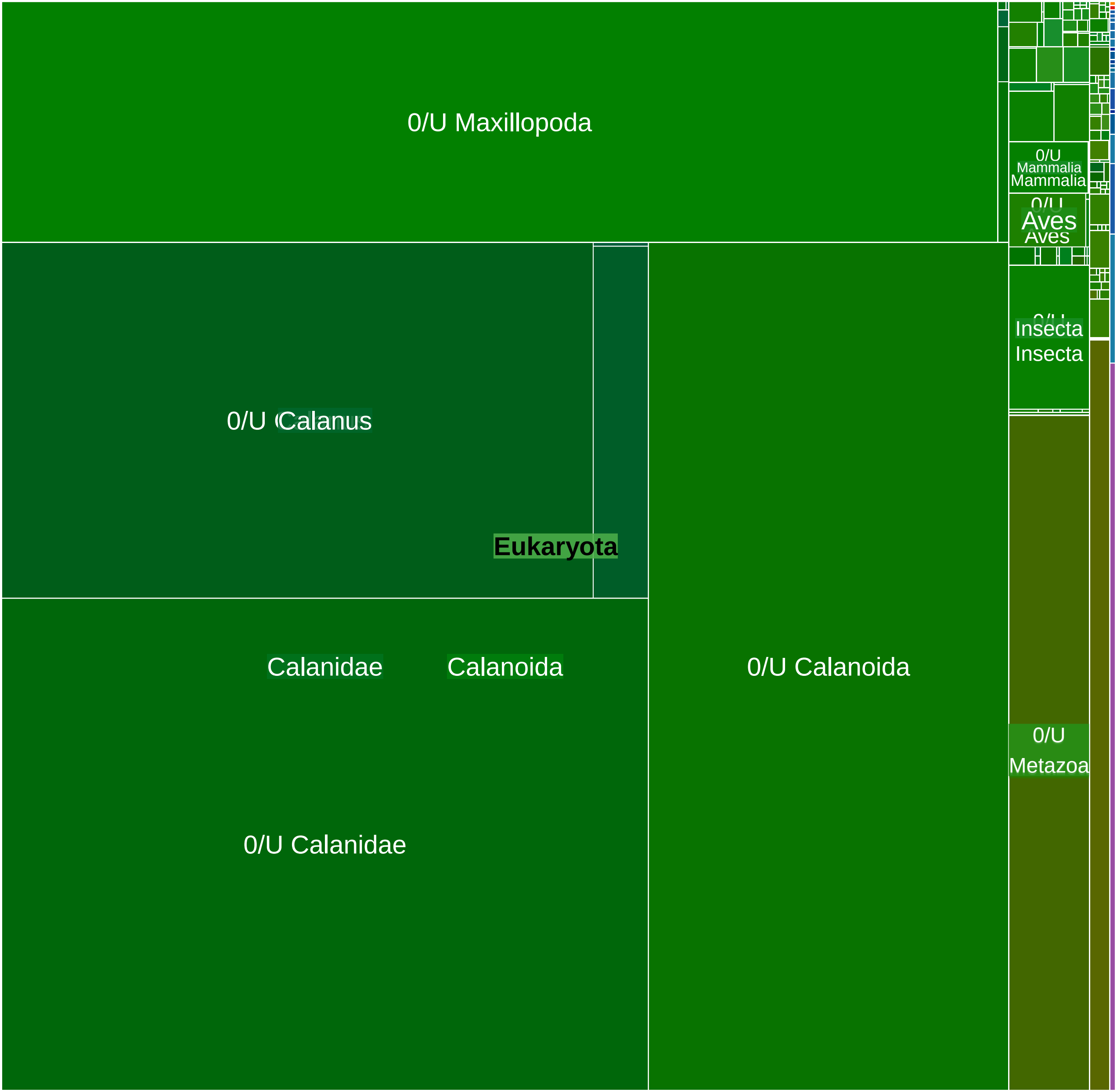

MGT−v1_12 : 96216 unigenes, 36827 (38.28%) taxonomically assigned

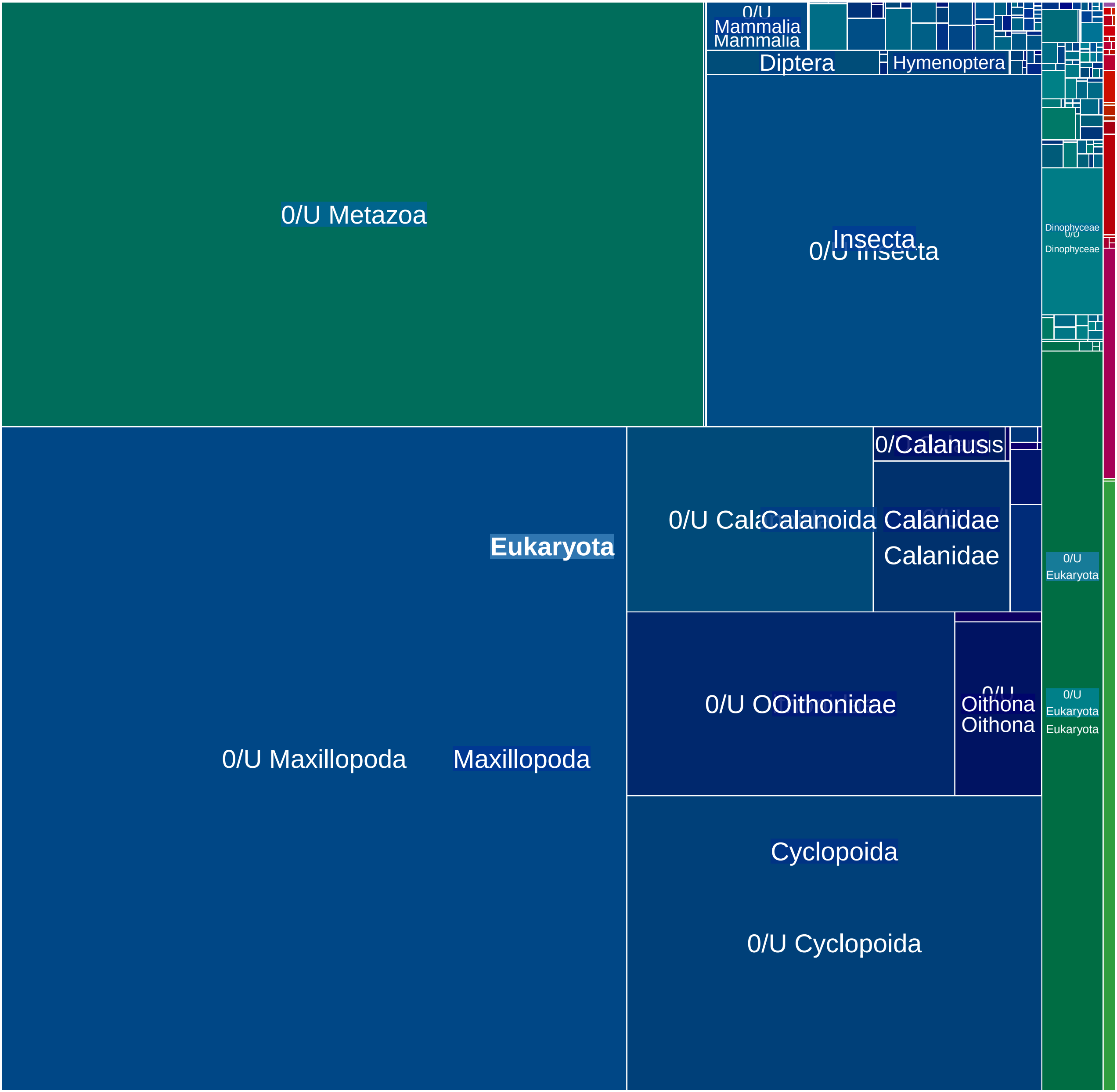

MGT−v1_13 : 84384 unigenes, 57966 (68.69%) taxonomically assigned

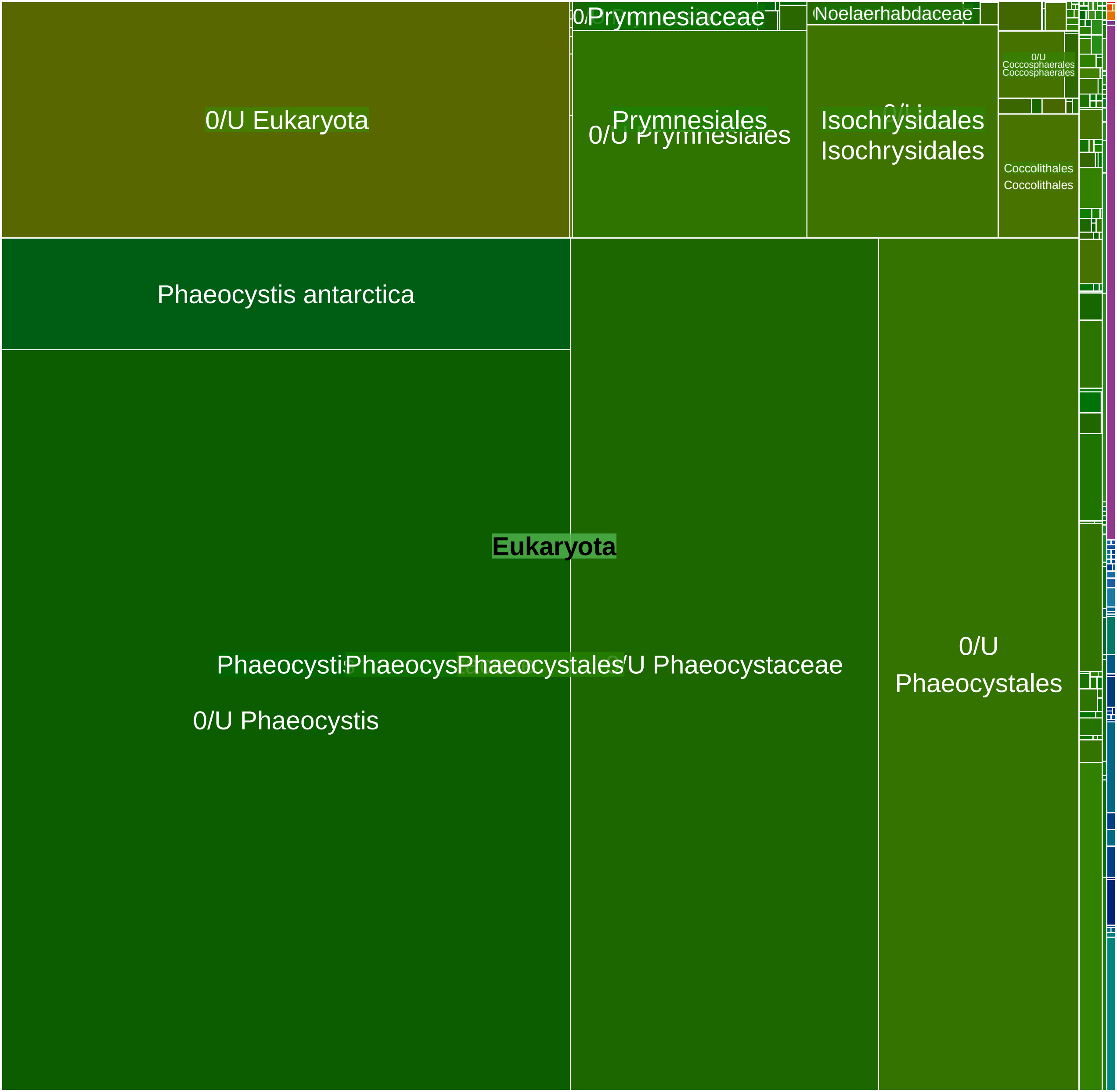

MGT−v1_14 : 77802 unigenes, 30120 (38.71%) taxonomically assigned

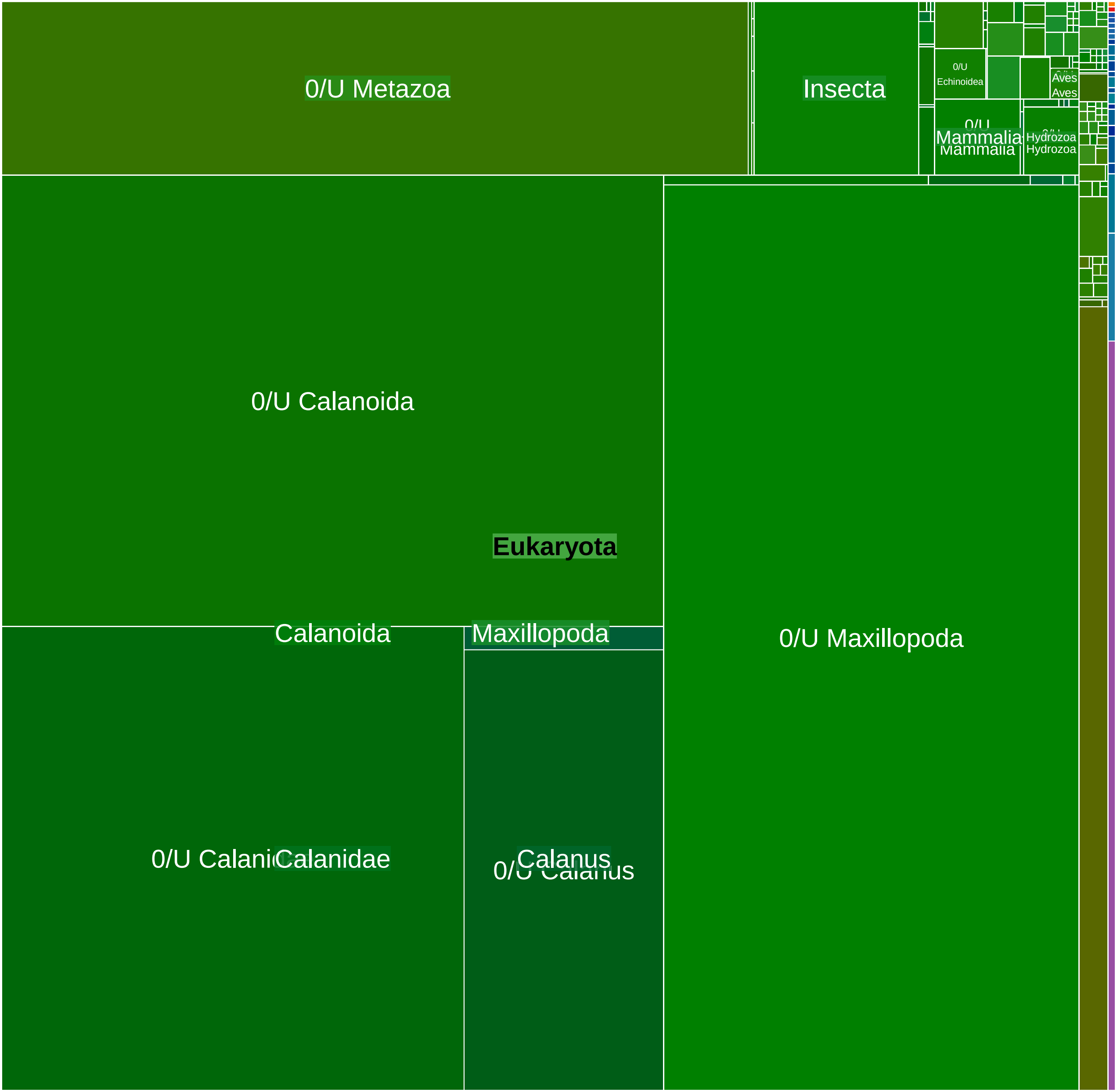

MGT−v1_15 : 77705 unigenes, 70164 (90.30%) taxonomically assigned

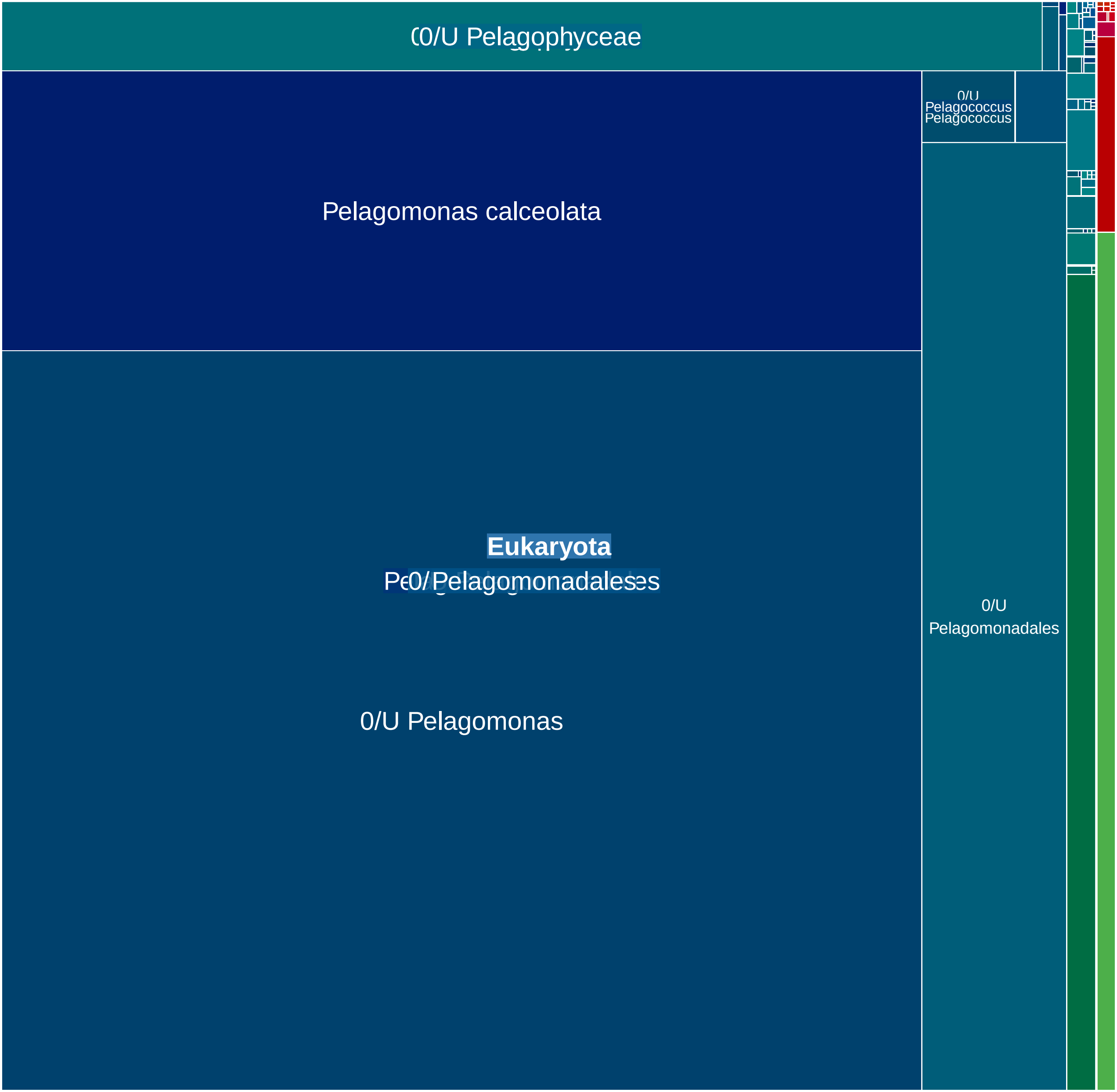

MGT−v1_16 : 65566 unigenes, 8518 (12.99%) taxonomically assigned

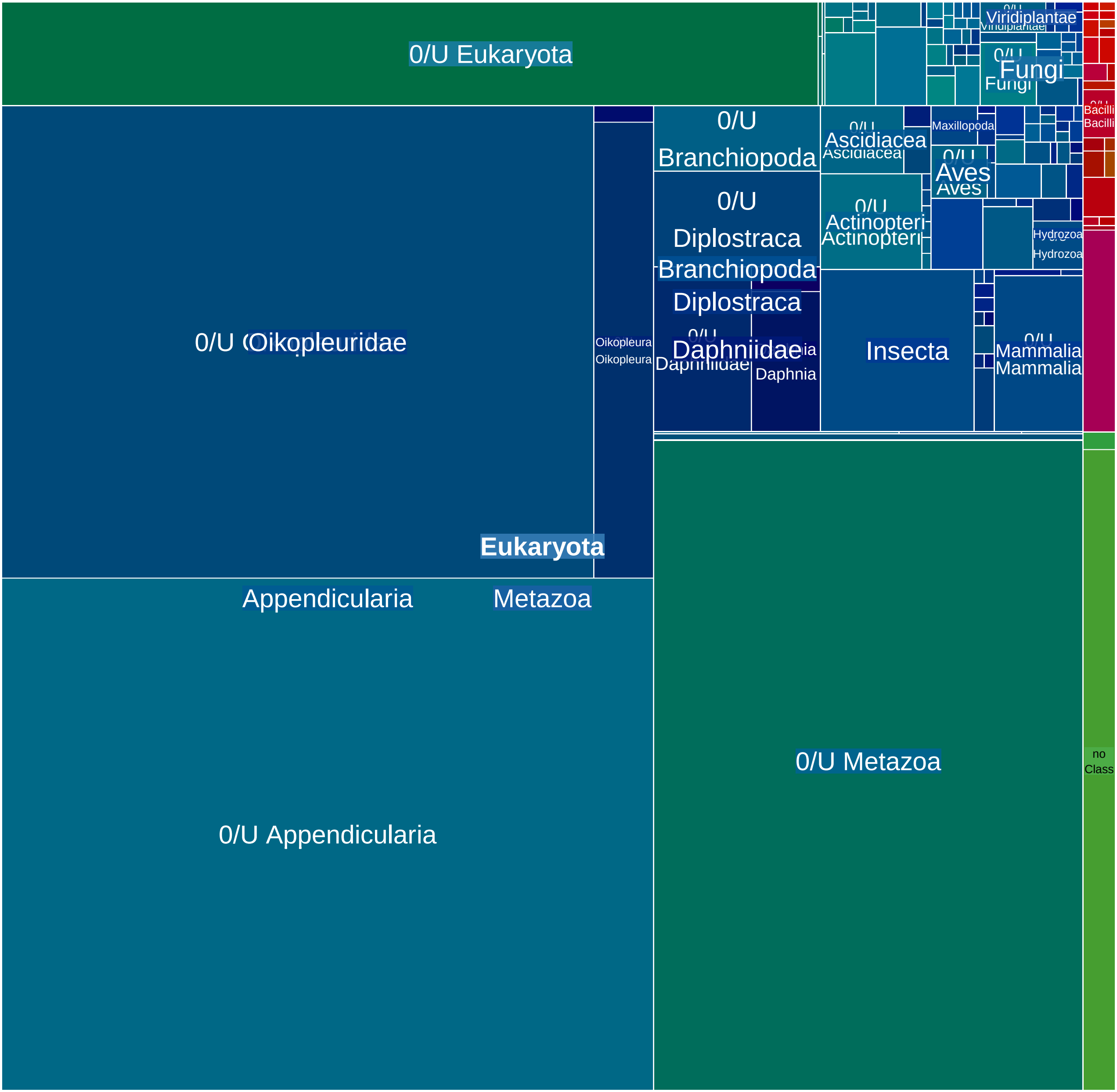

MGT−v1_17 : 65404 unigenes, 31450 (48.09%) taxonomically assigned

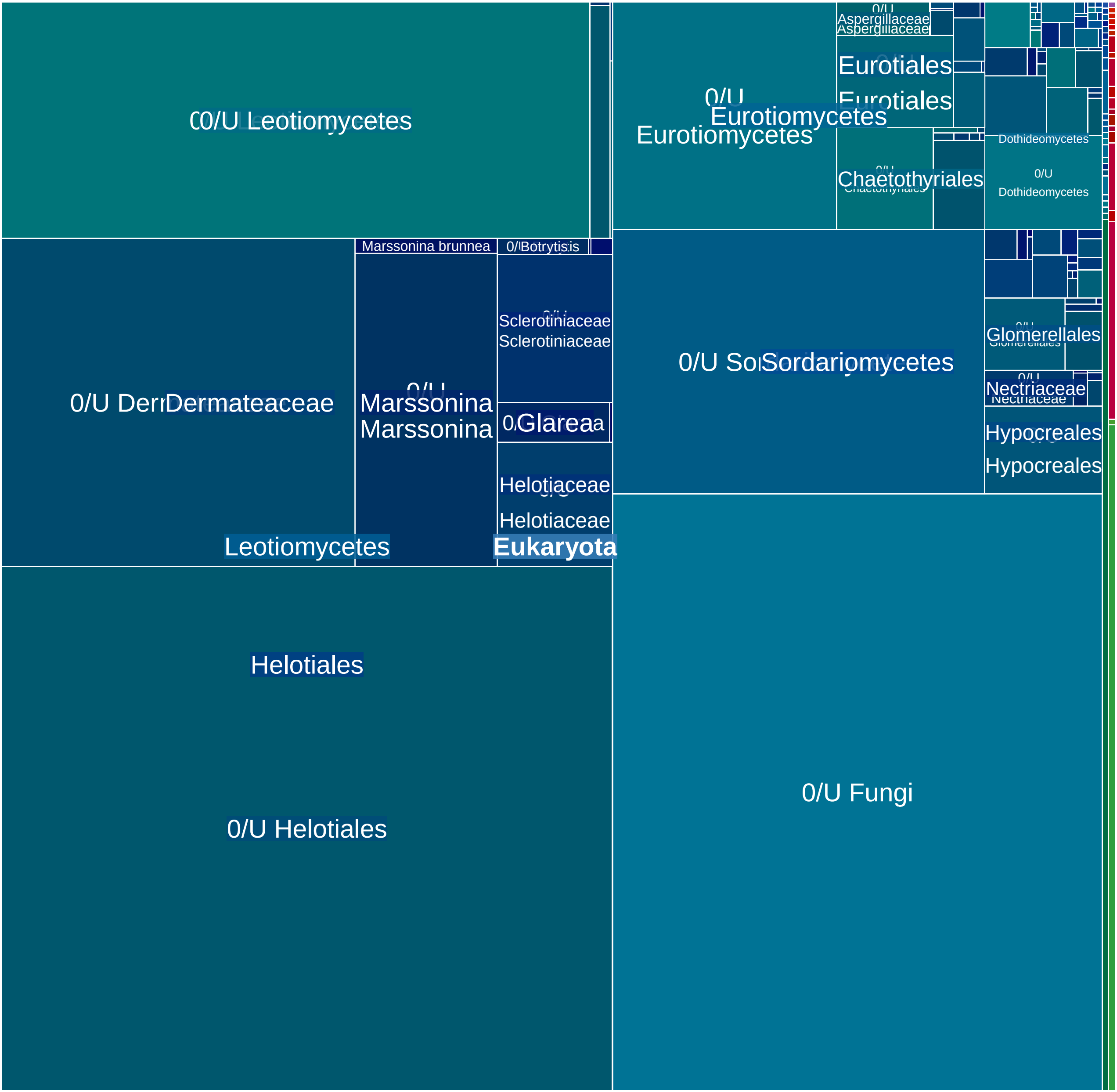

MGT−v1_18 : 61296 unigenes, 15487 (25.27%) taxonomically assigned

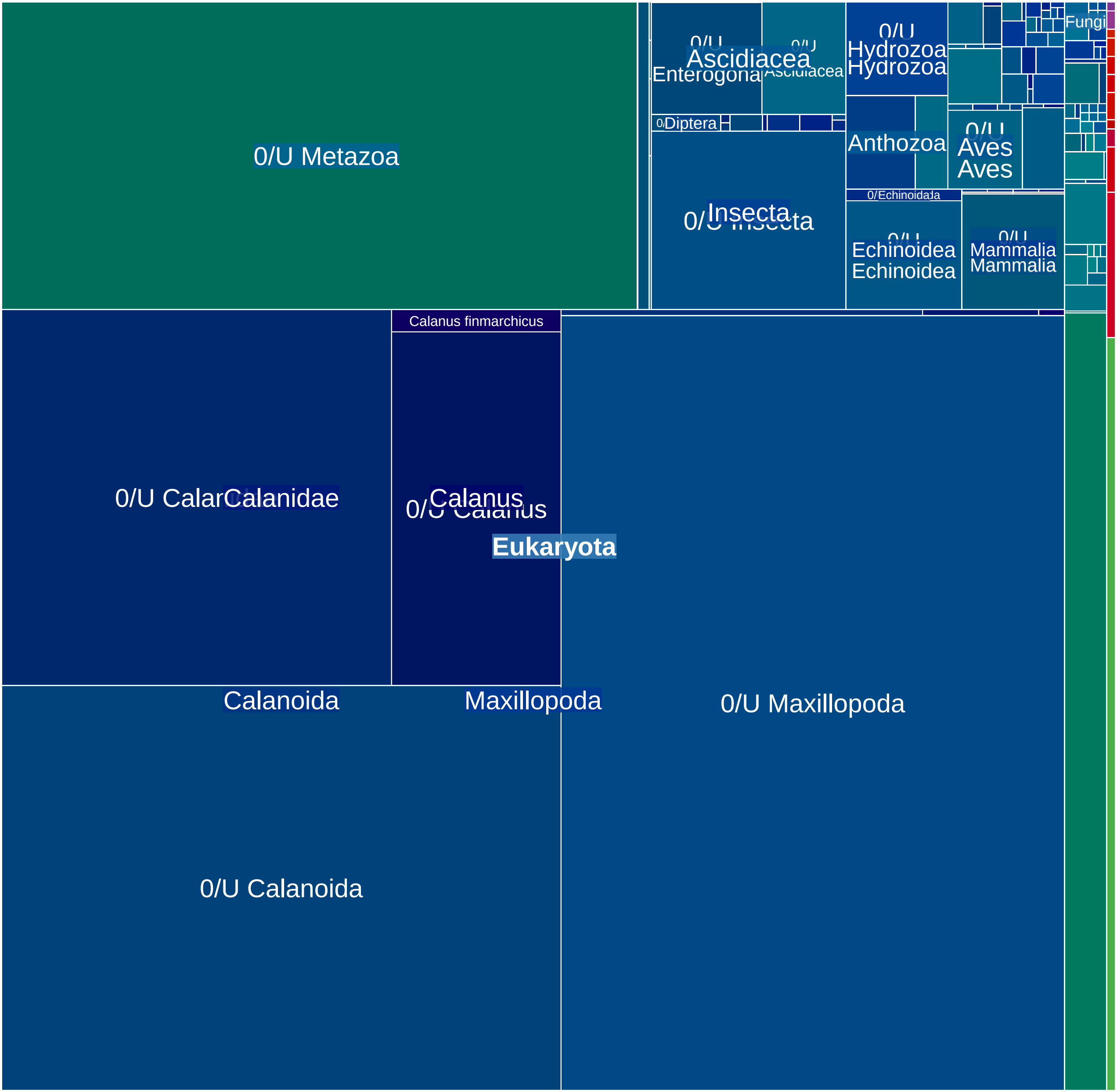

MGT−v1_19 : 60584 unigenes, 23526 (38.83%) taxonomically assigned

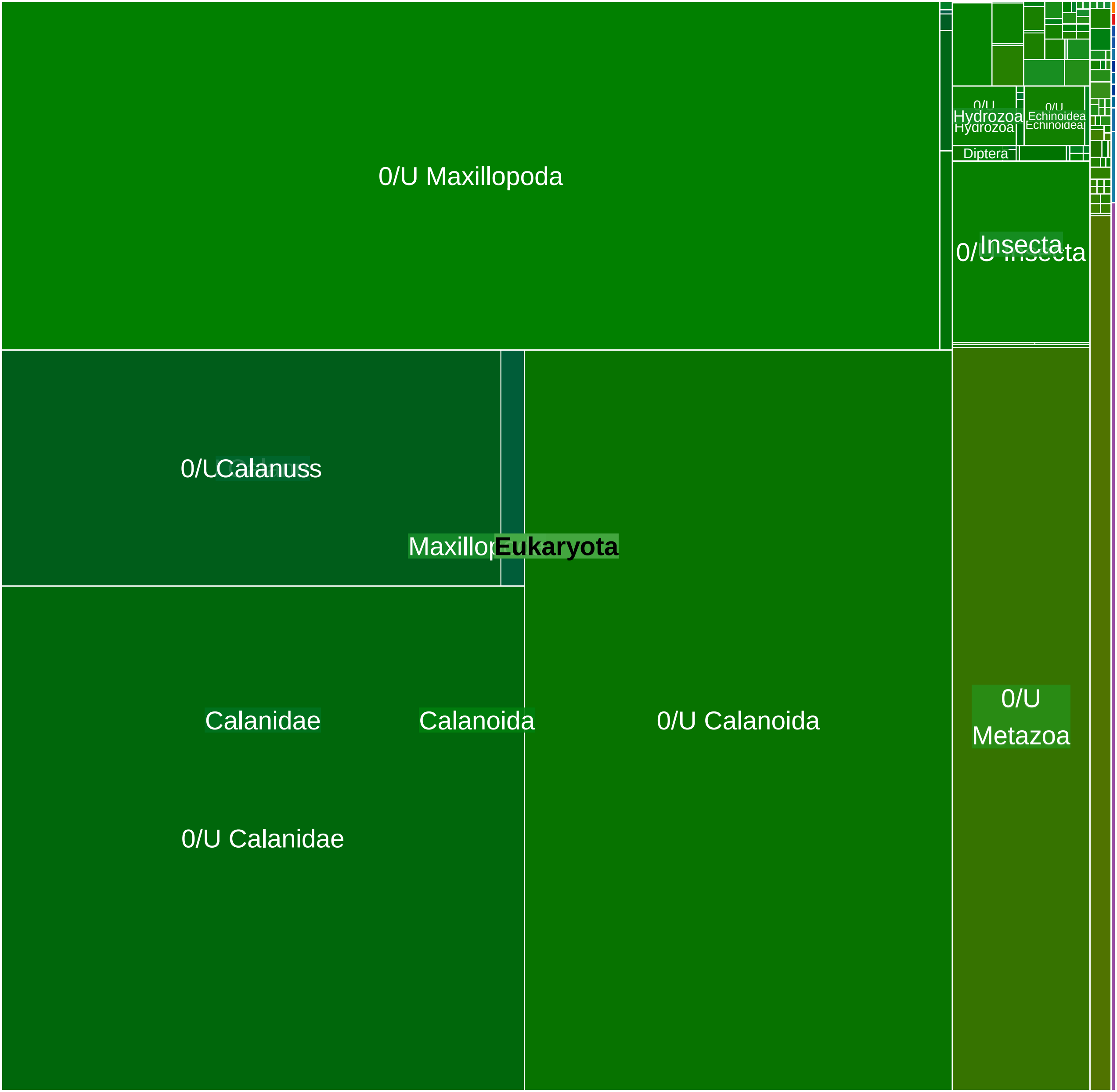

MGT−v1_20 : 58041 unigenes, 28047 (48.32%) taxonomically assigned

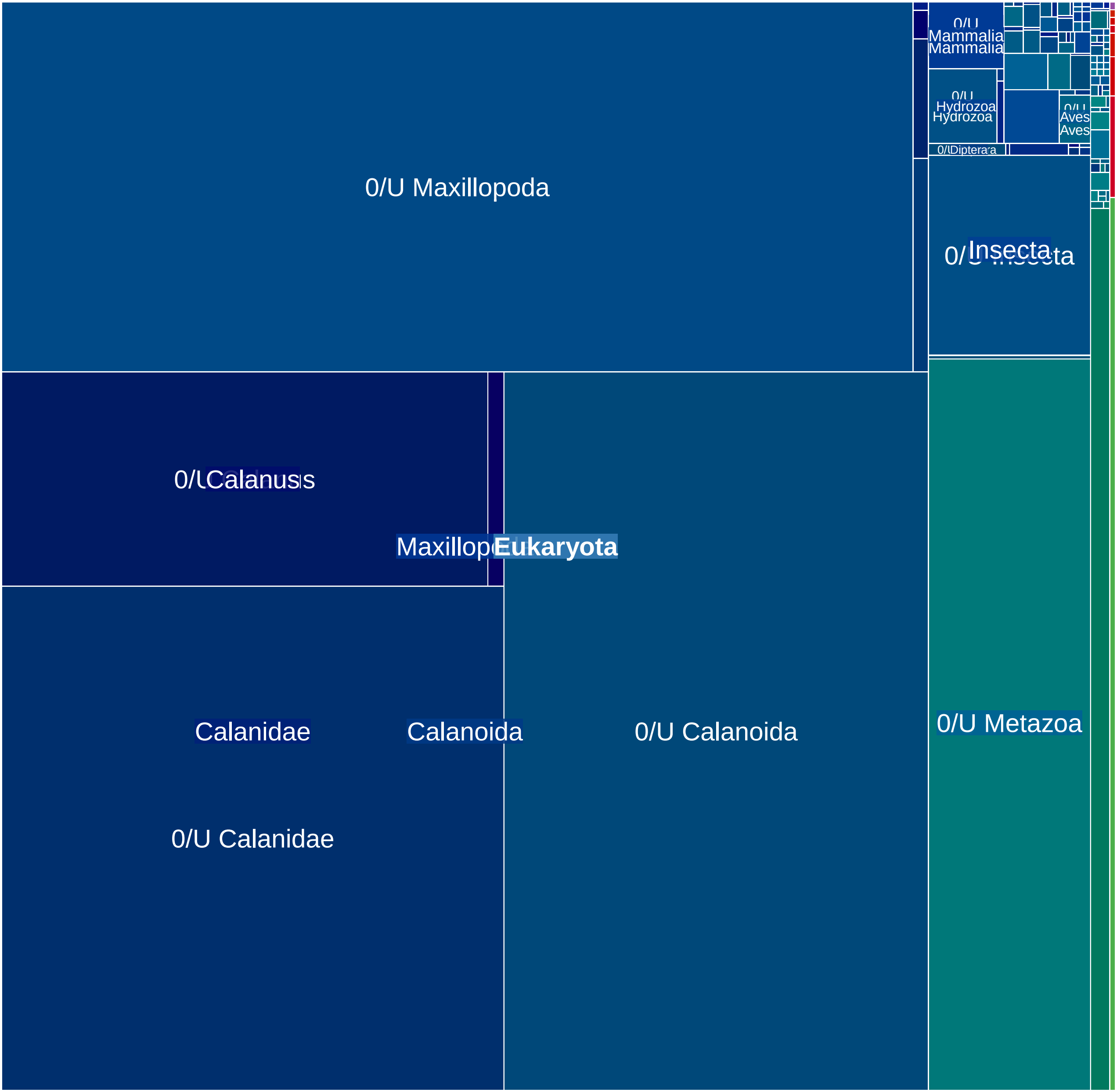

MGT−v1_21 : 55924 unigenes, 44790 (80.09%) taxonomically assigned

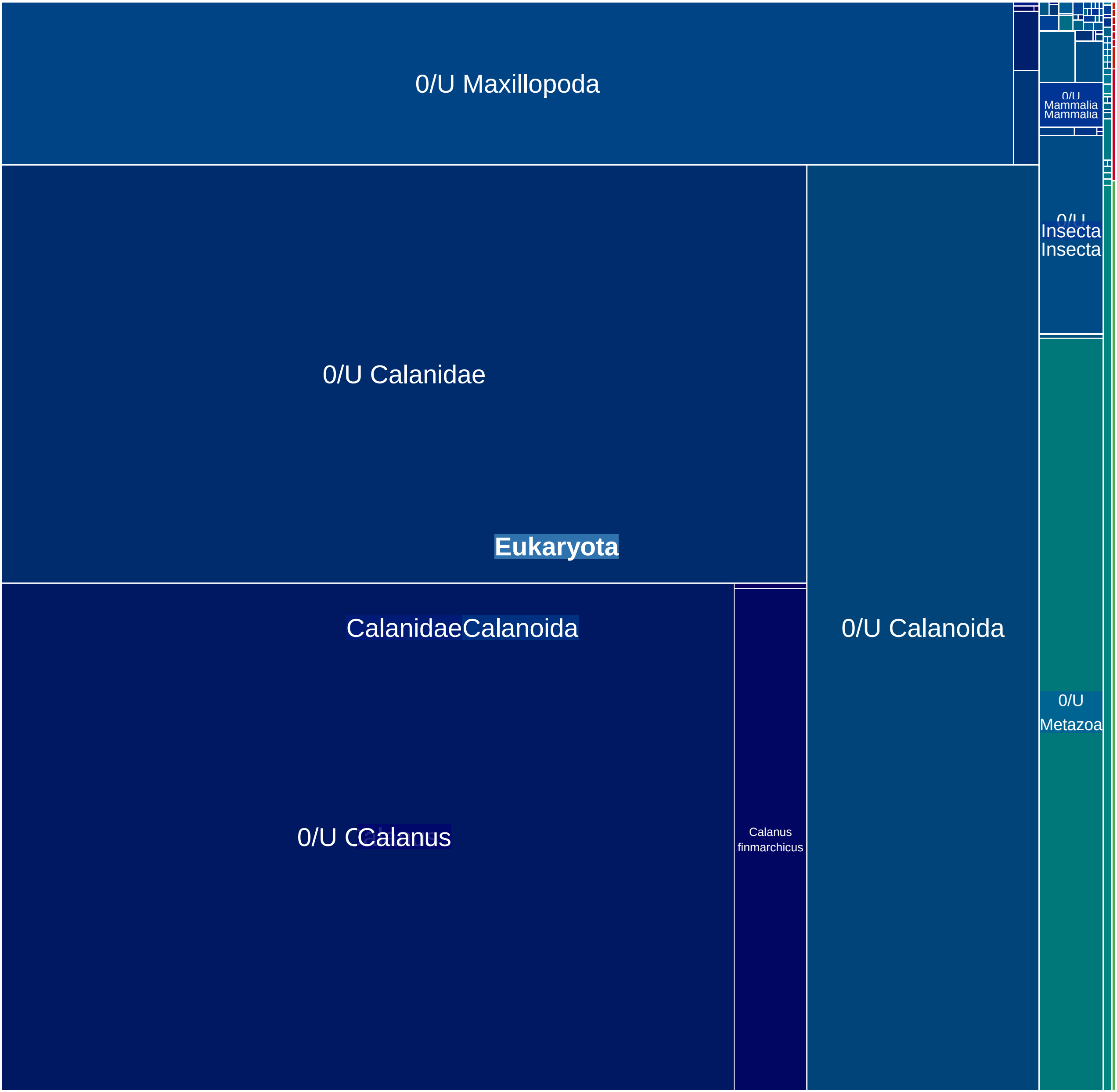

MGT−v1_22 : 55444 unigenes, 22551 (40.67%) taxonomically assigned

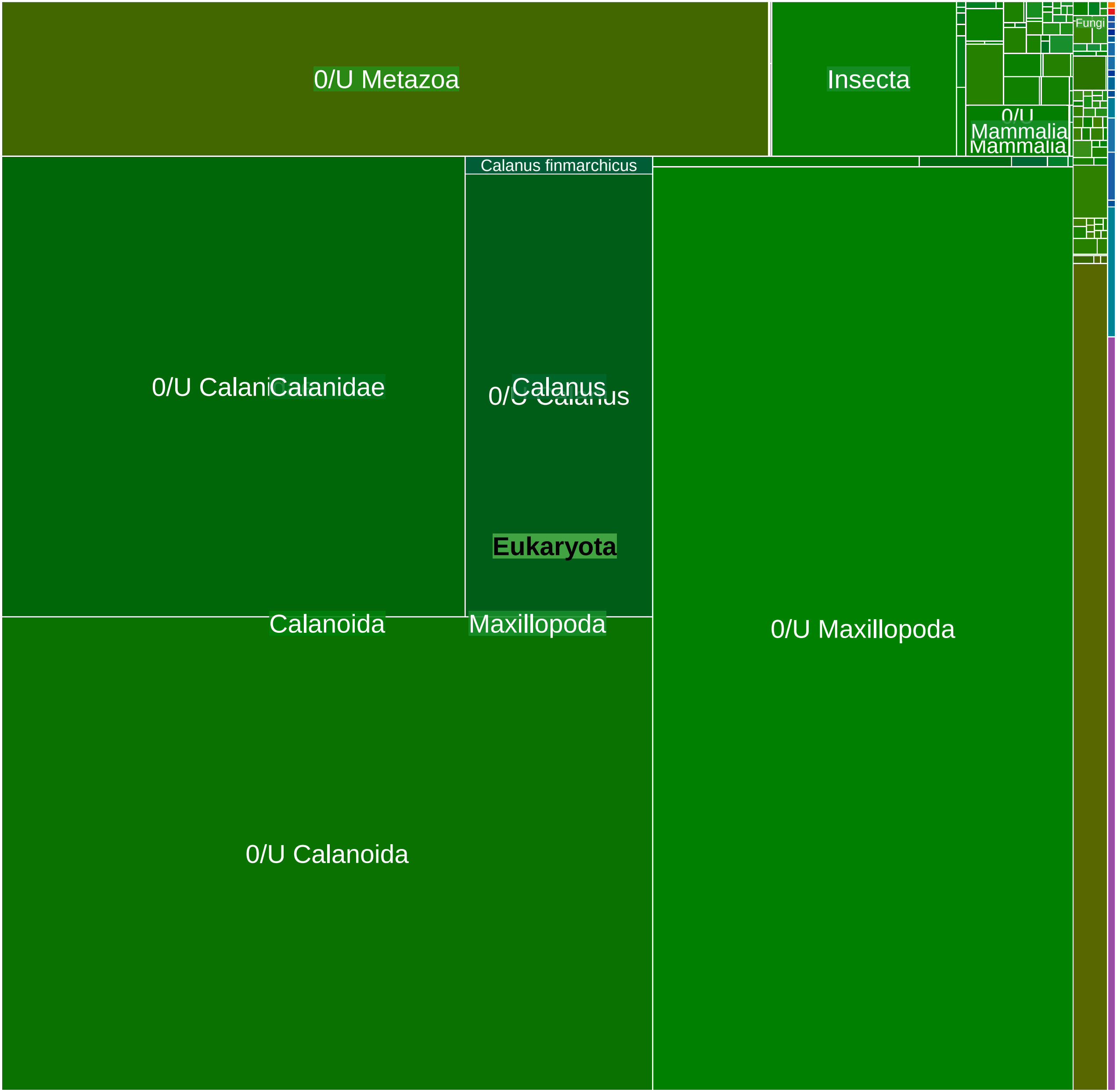

MGT−v1_23 : 53910 unigenes, 28470 (52.81%) taxonomically assigned

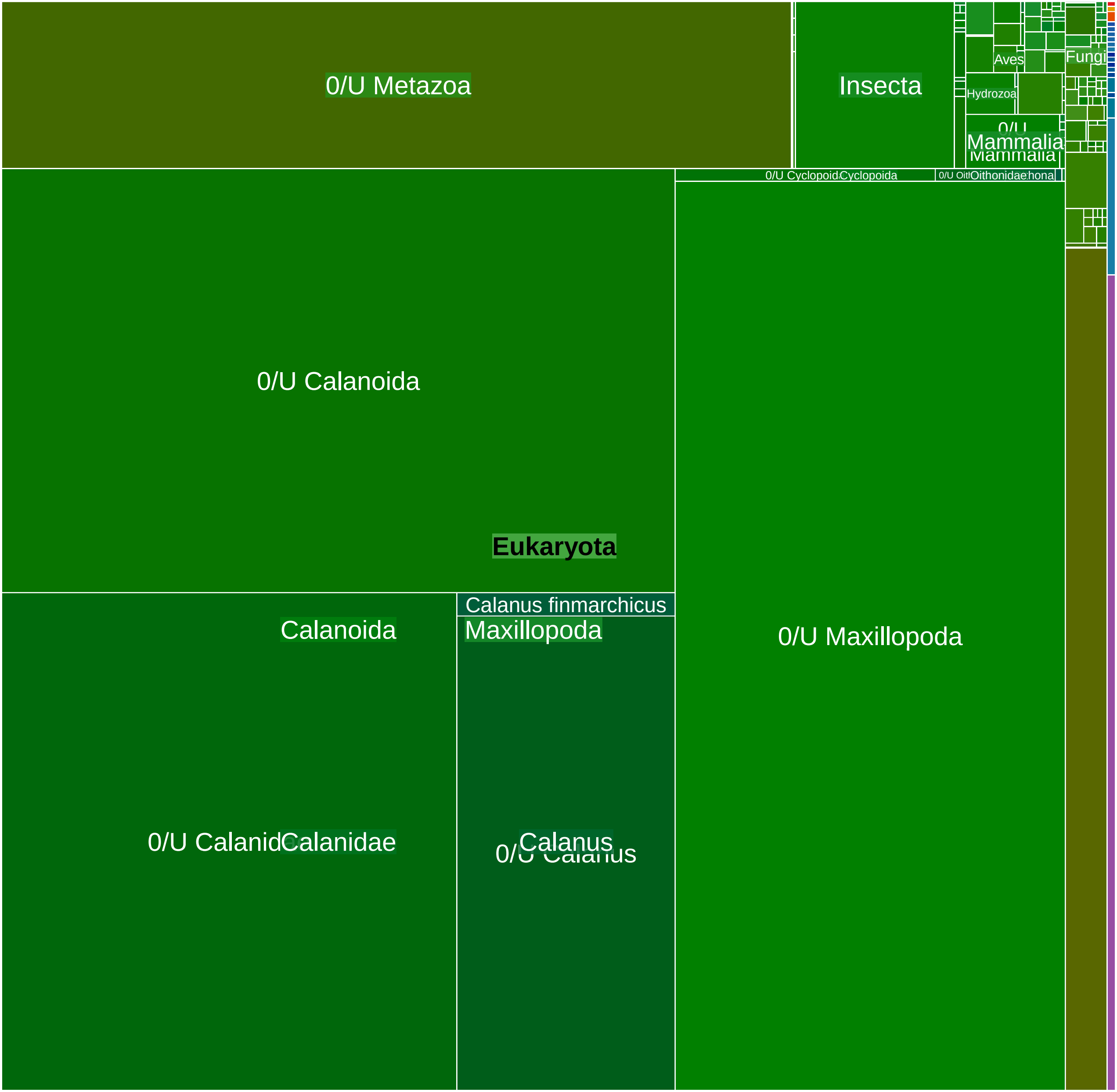

MGT−v1_24 : 53408 unigenes, 46865 (87.75%) taxonomically assigned

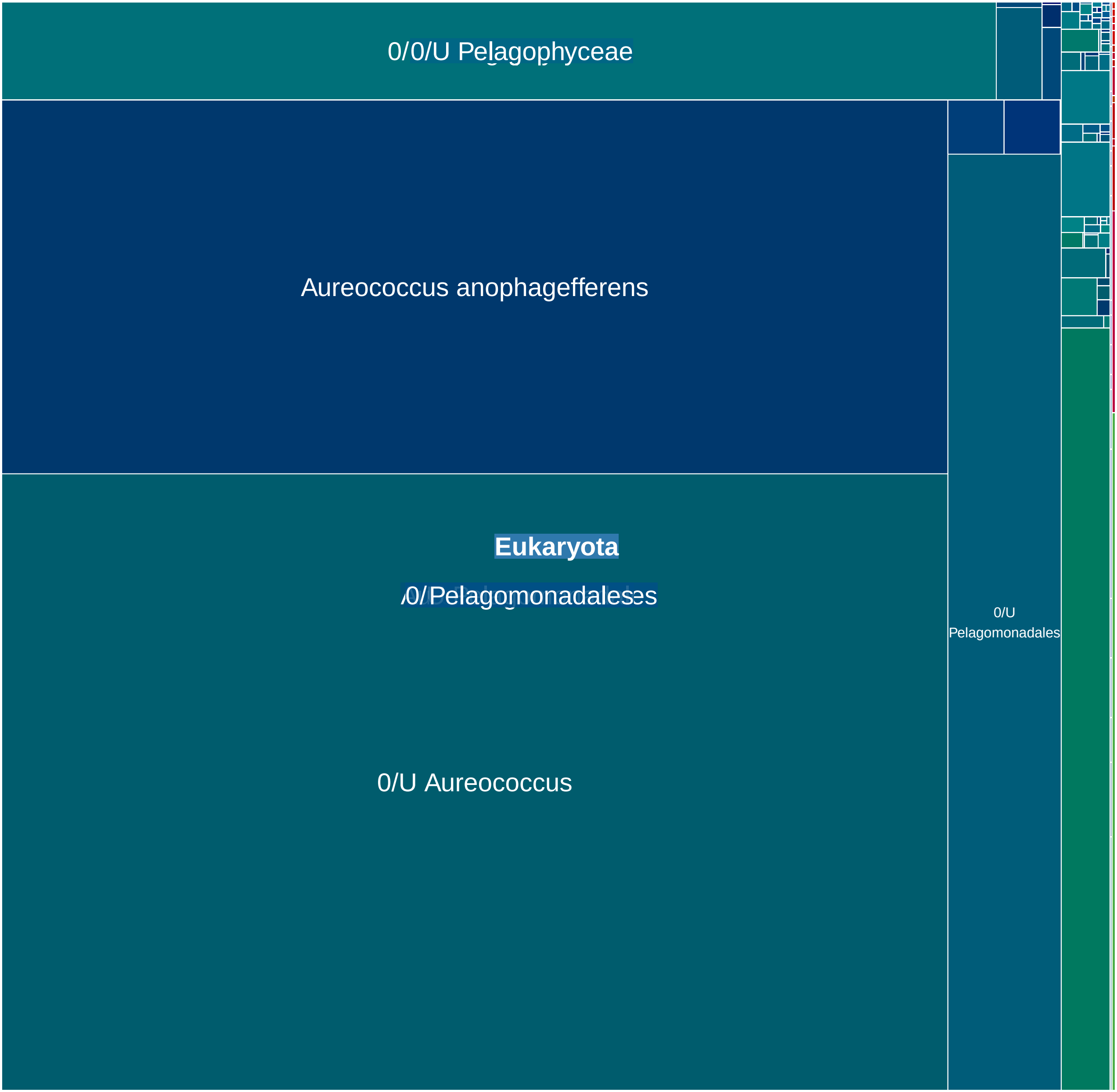

MGT−v1_25 : 52253 unigenes, 39318 (75.25%) taxonomically assigned

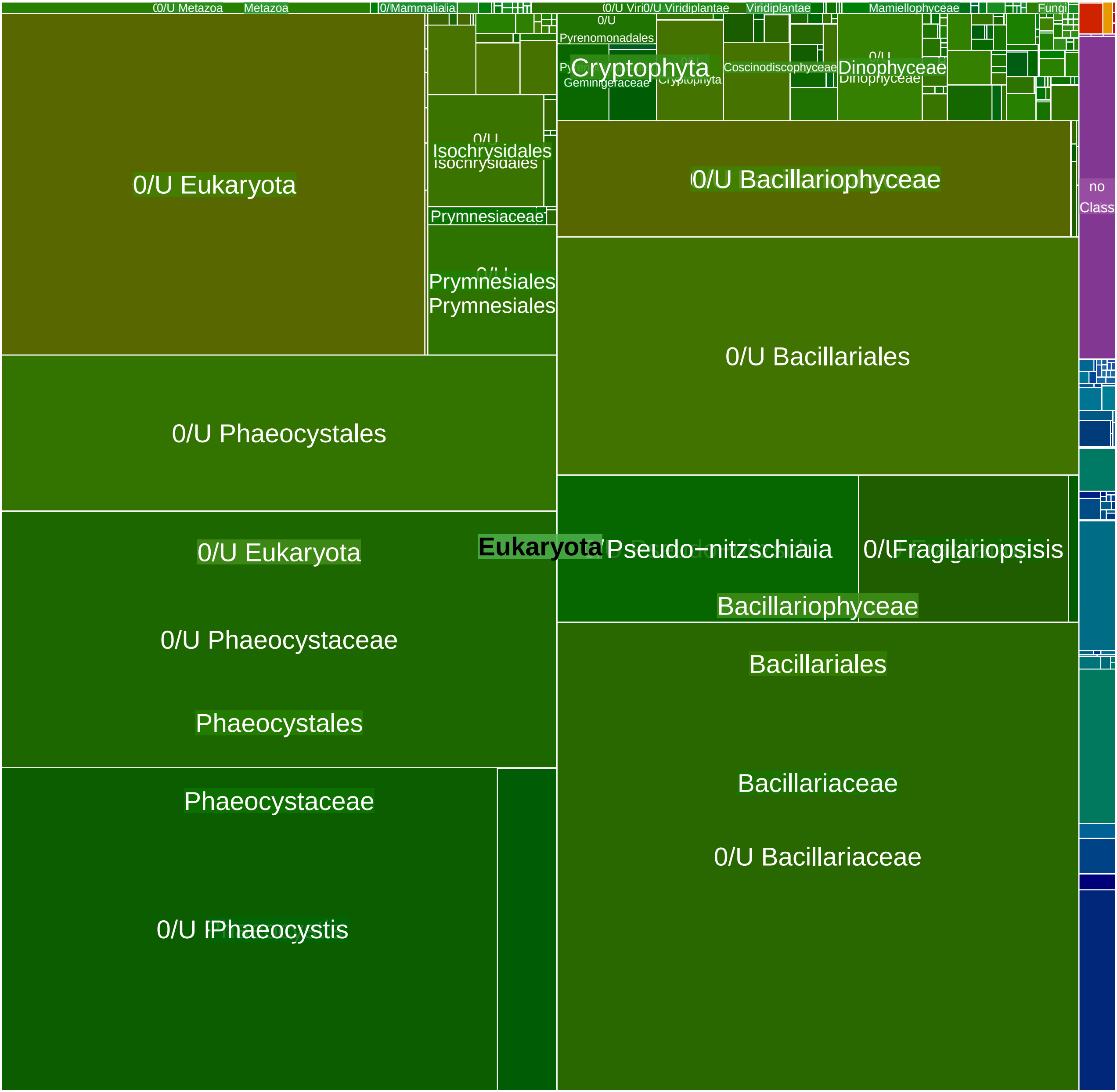

MGT−v1_26 : 51411 unigenes, 18308 (35.61%) taxonomically assigned

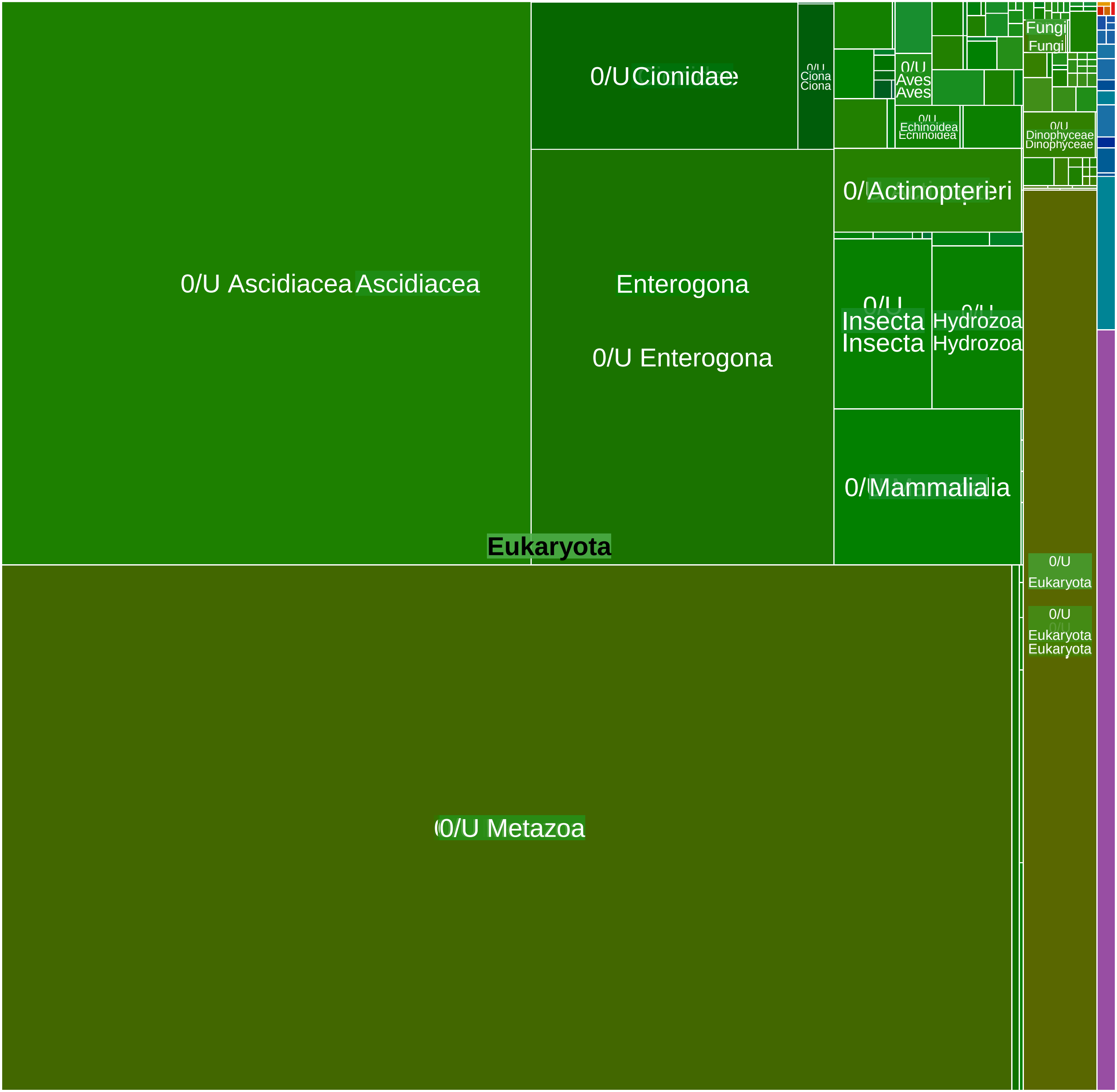

MGT−v1_27 : 48515 unigenes, 17576 (36.23%) taxonomically assigned

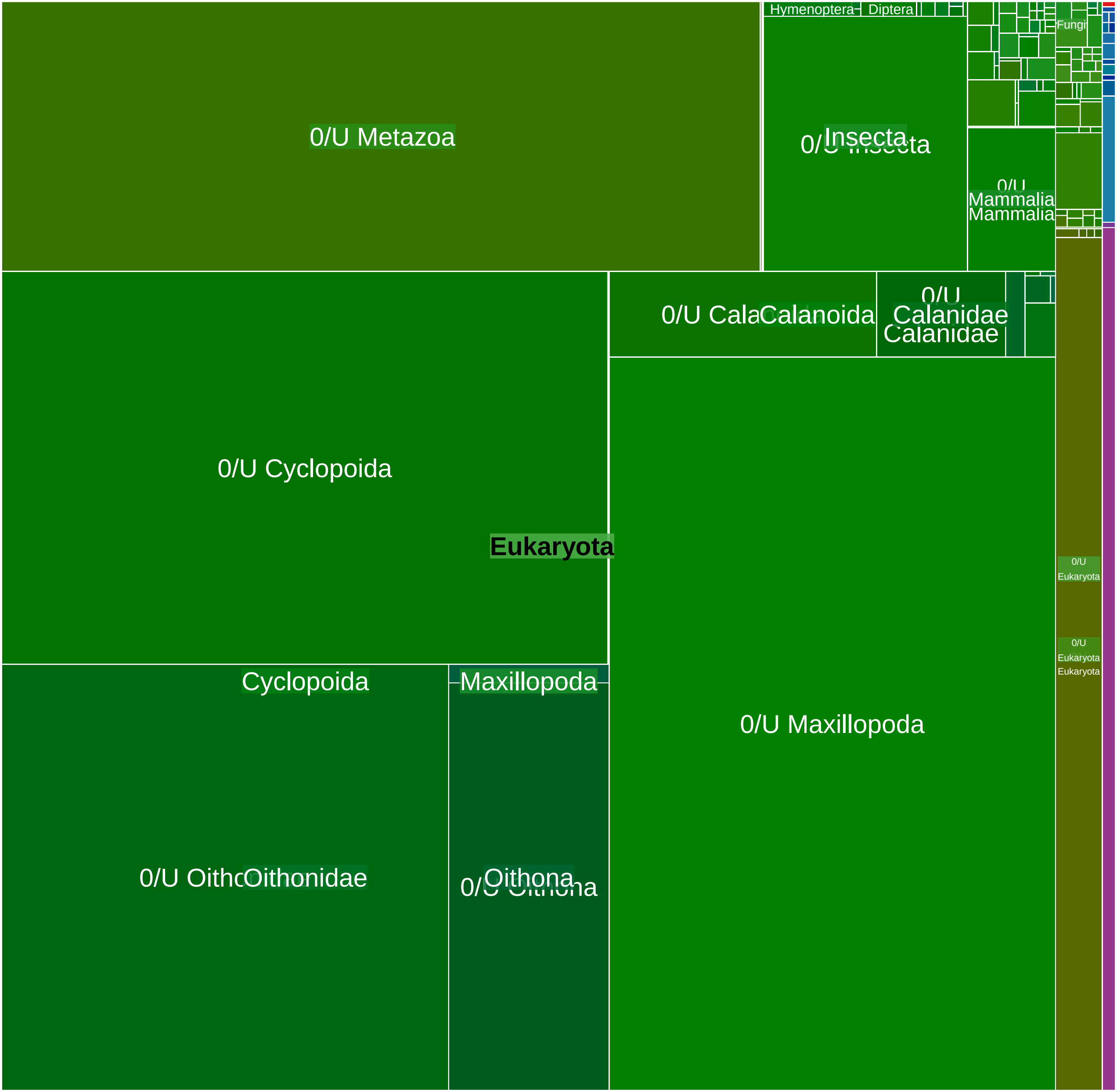

MGT−v1_28 : 48509 unigenes, 3551 (7.32%) taxonomically assigned

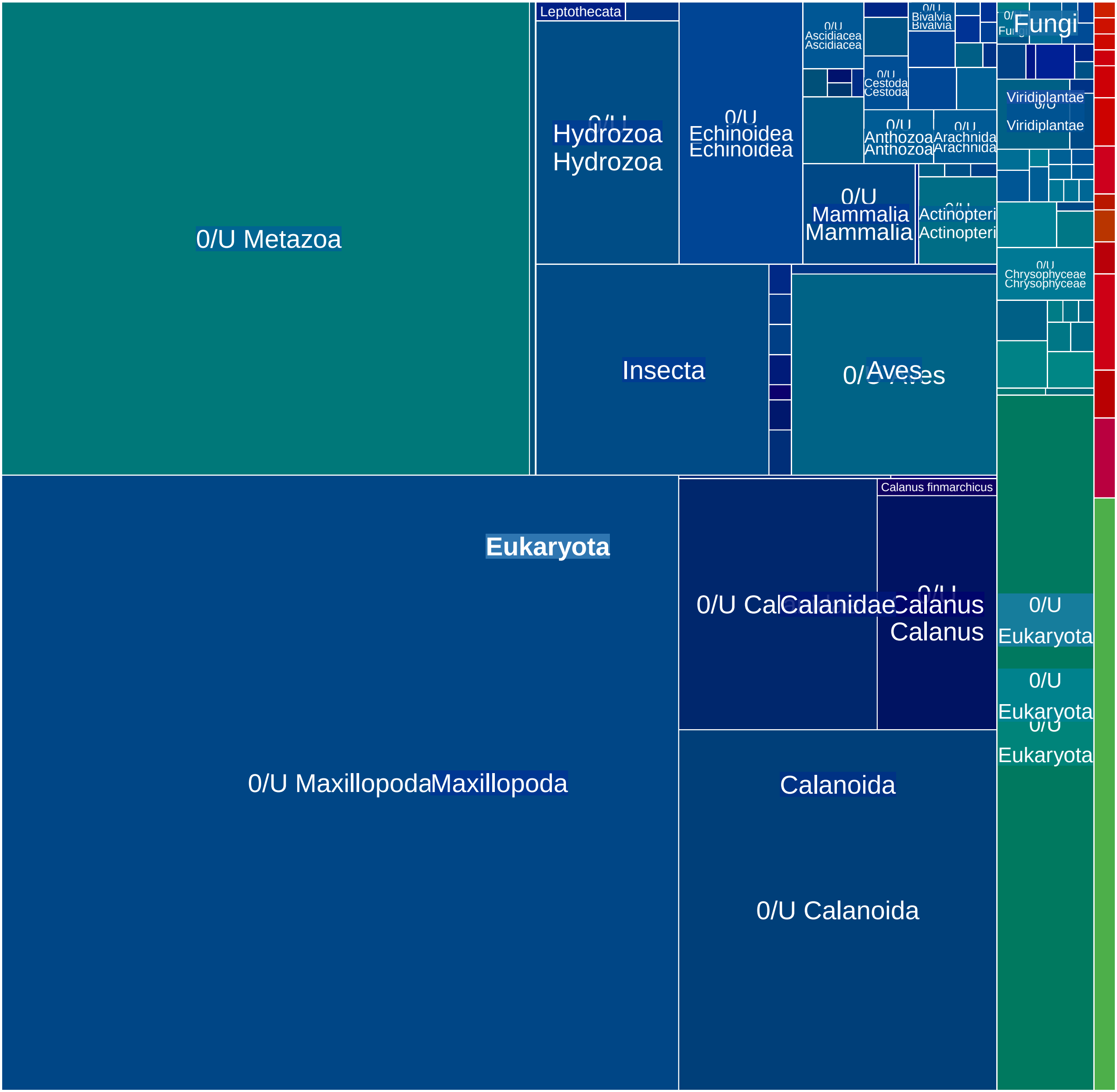

MGT−v1_29 : 48249 unigenes, 24942 (51.69%) taxonomically assigned

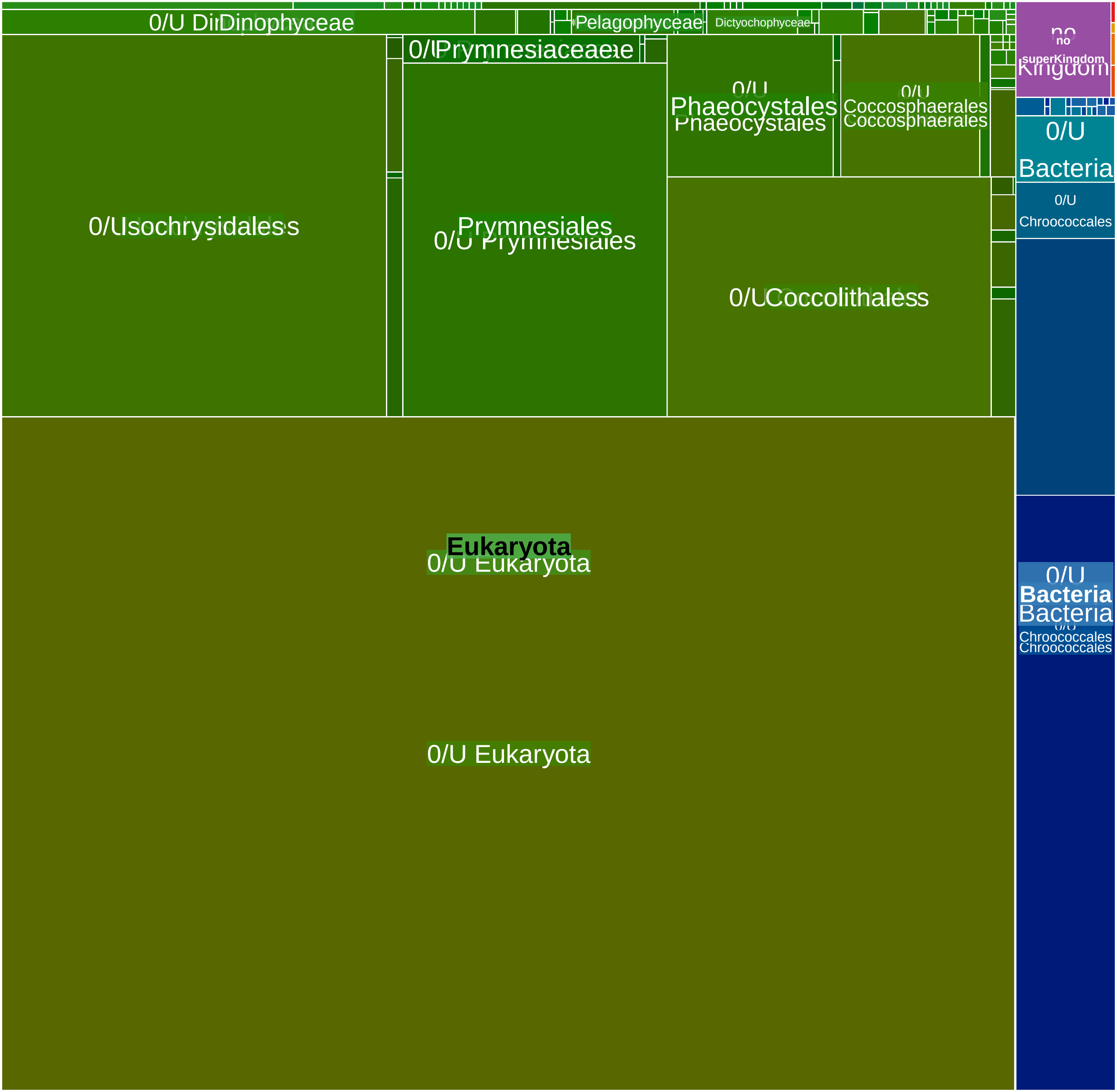

MGT−v1_30 : 46533 unigenes, 30399 (65.33%) taxonomically assigned

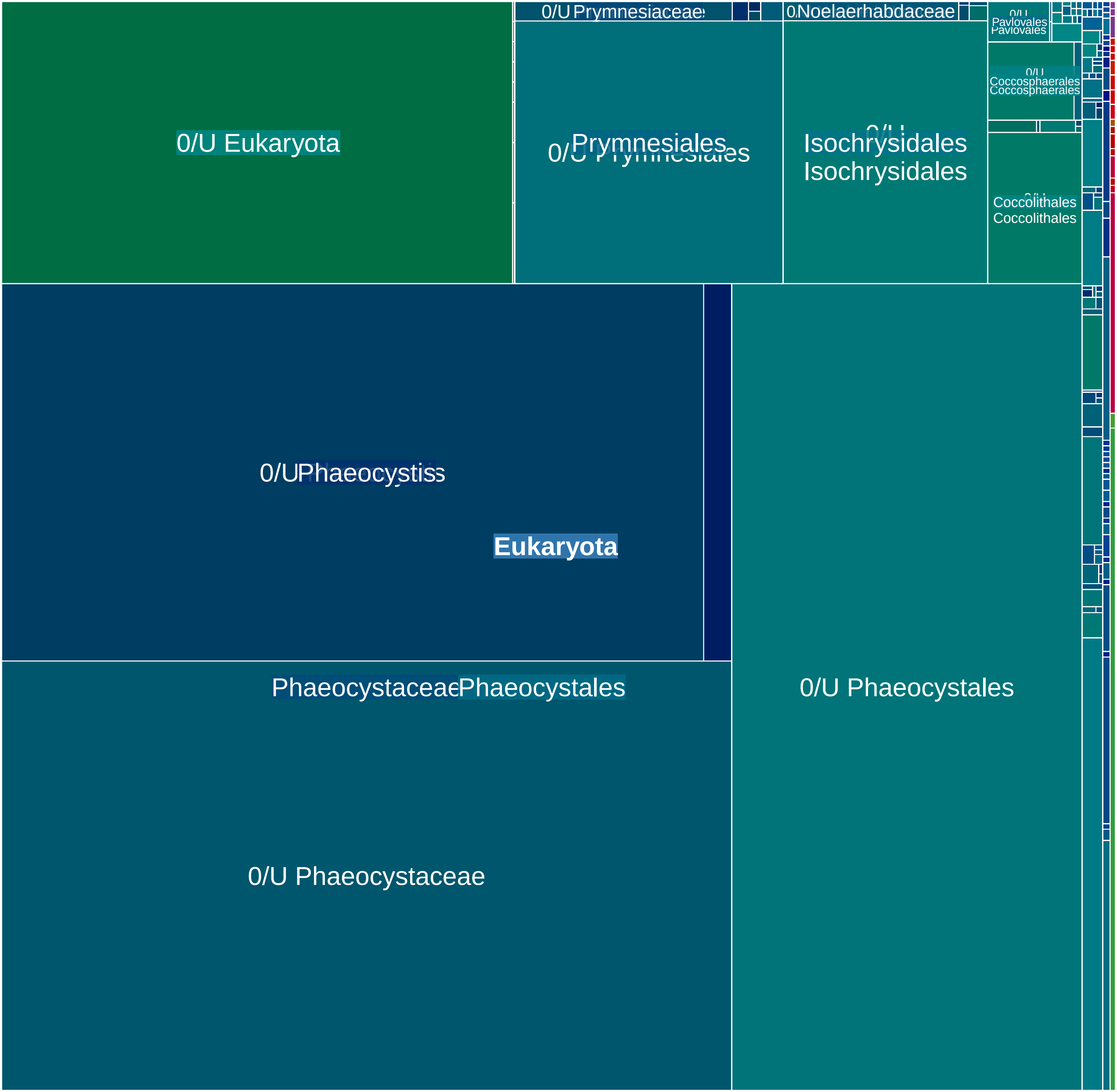

MGT−v1_31 : 45593 unigenes, 10587 (23.22%) taxonomically assigned

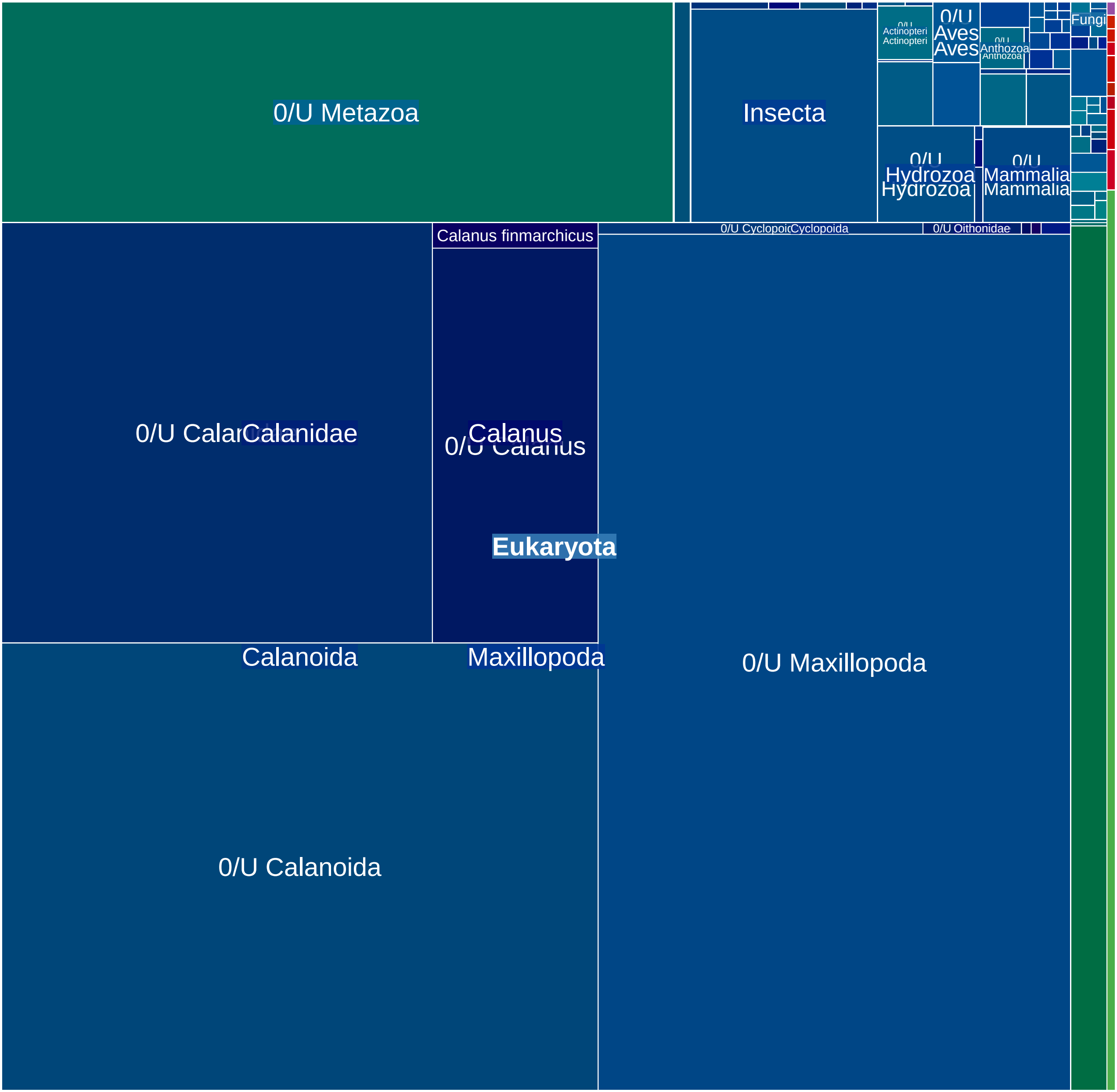

MGT−v1_32 : 43096 unigenes, 31508 (73.11%) taxonomically assigned

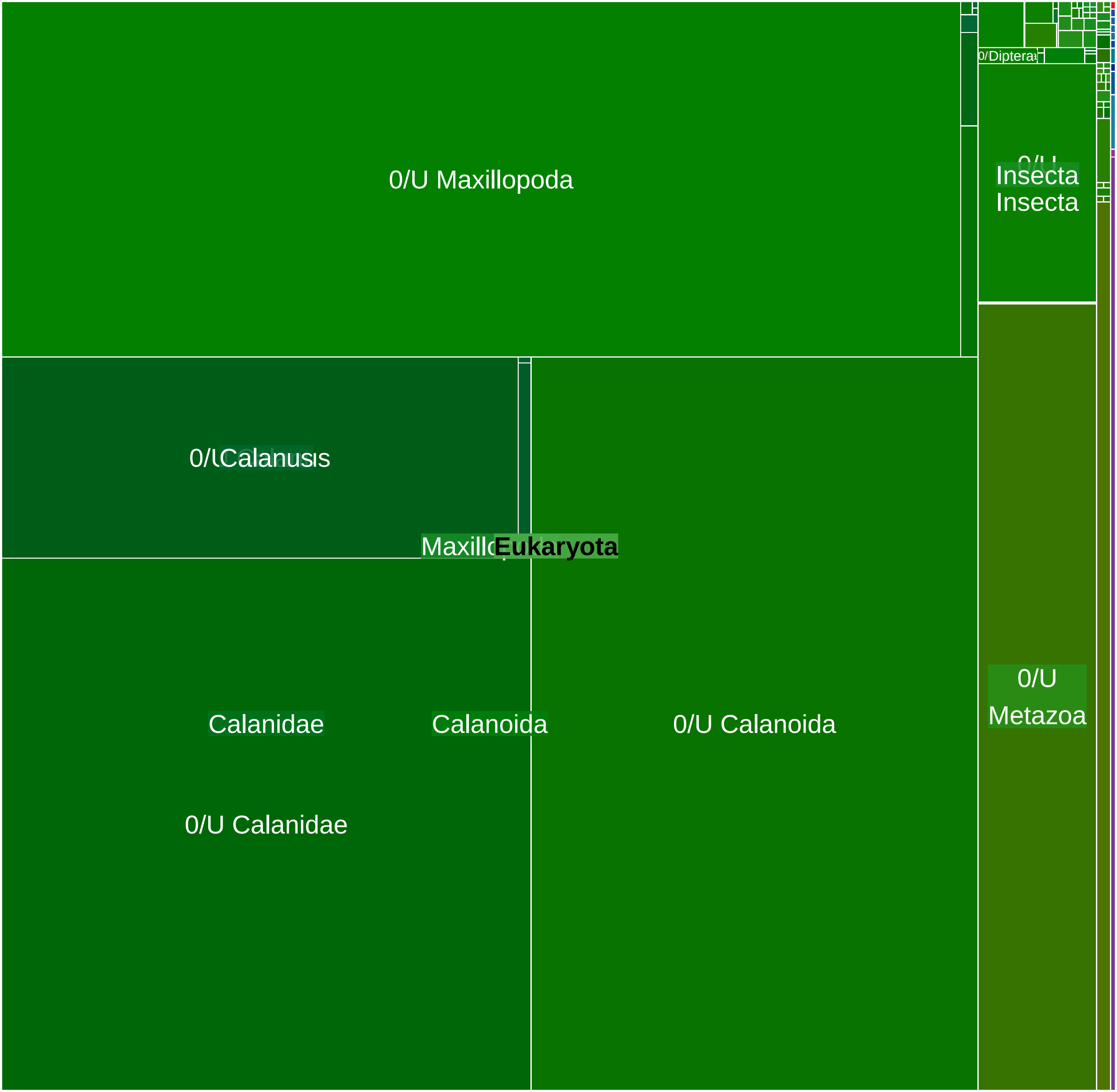

MGT−v1_33 : 41640 unigenes, 12939 (31.07%) taxonomically assigned

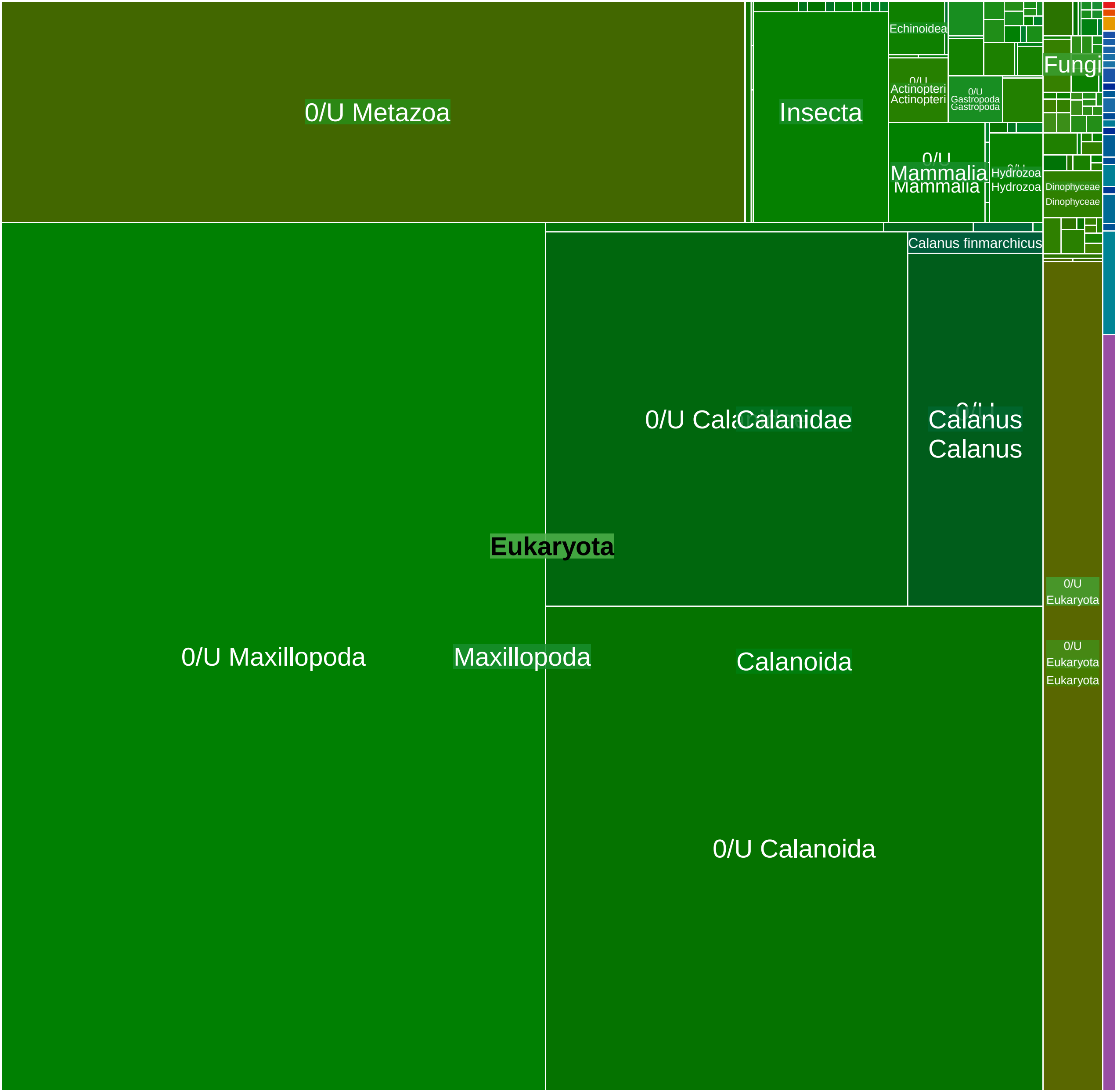

MGT−v1_34 : 39353 unigenes, 22740 (57.78%) taxonomically assigned

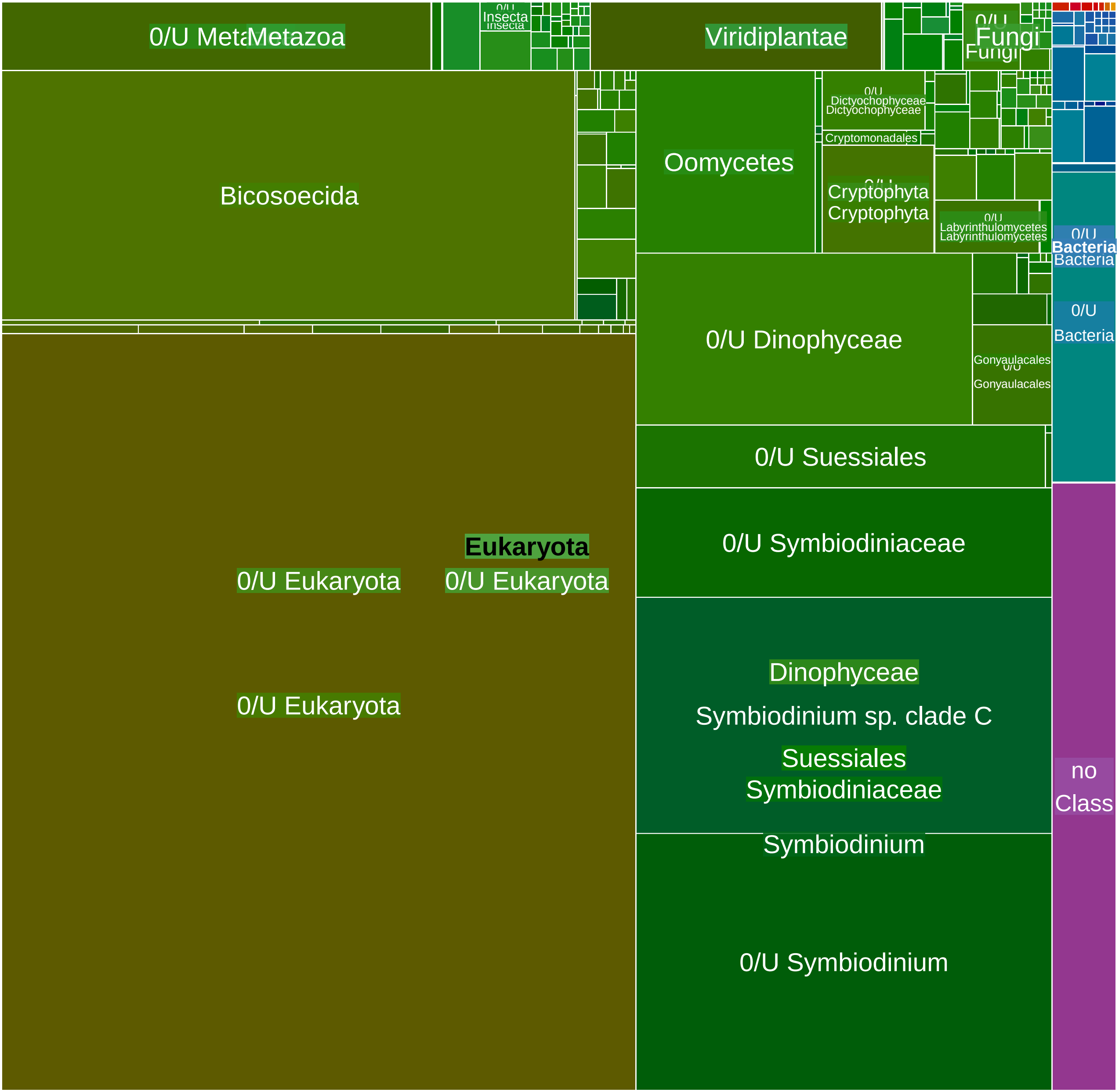

MGT−v1_35 : 38940 unigenes, 19910 (51.13%) taxonomically assigned

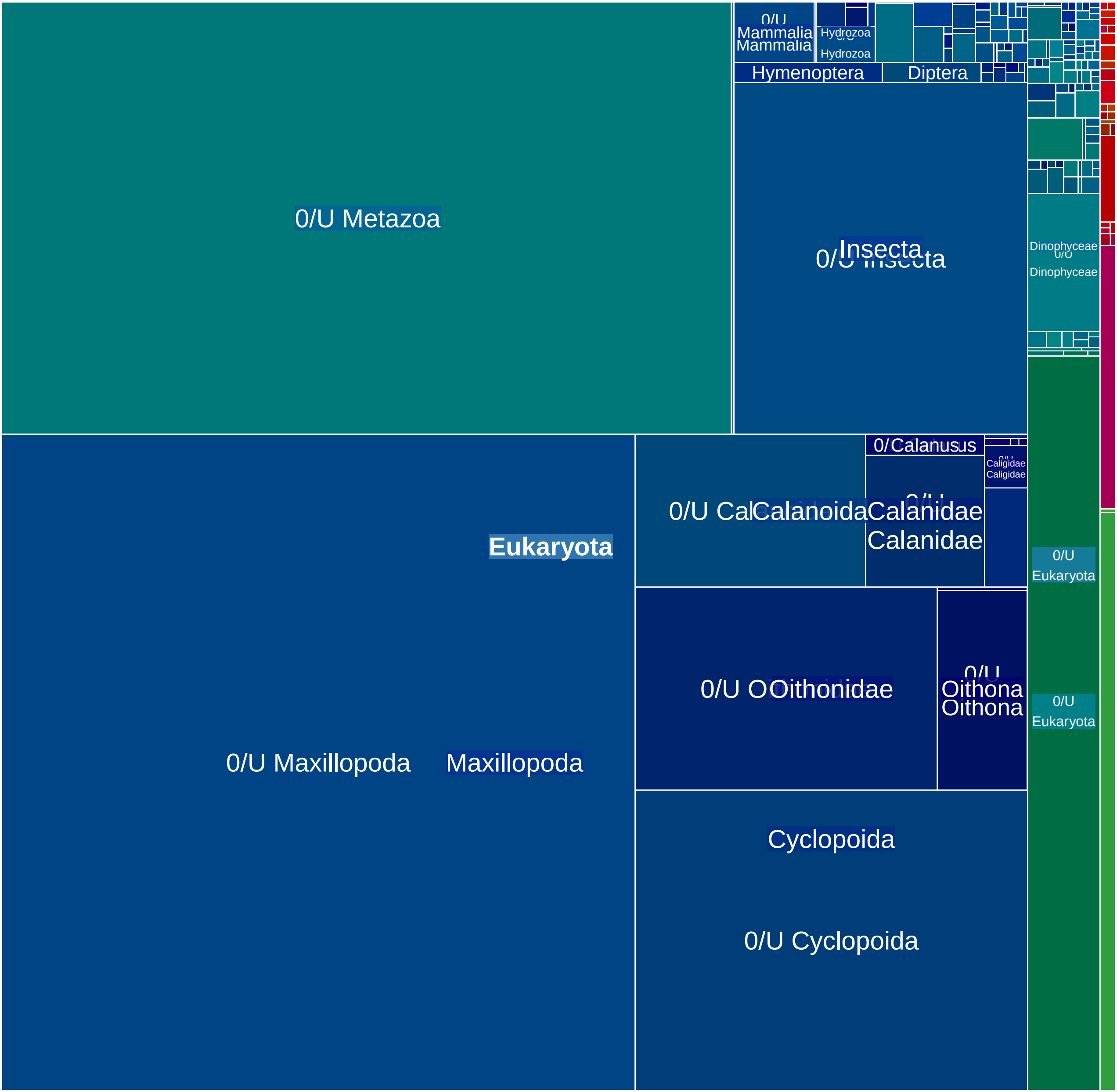

### MGT−v1_36 : 38502 unigenes, 22549 (58.57%) taxonomically assigned

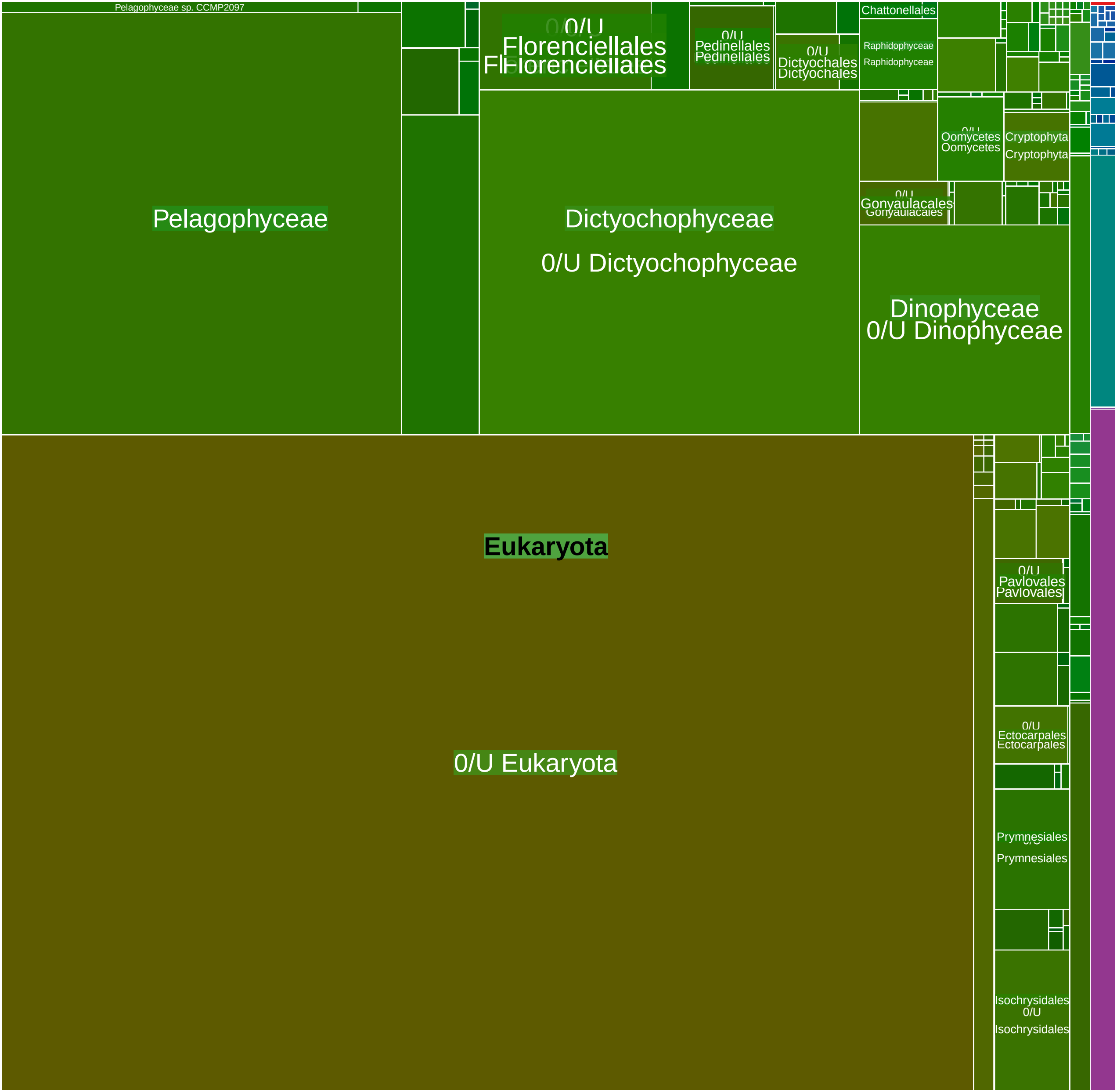

MGT−v1_37 : 37876 unigenes, 11017 (29.09%) taxonomically assigned

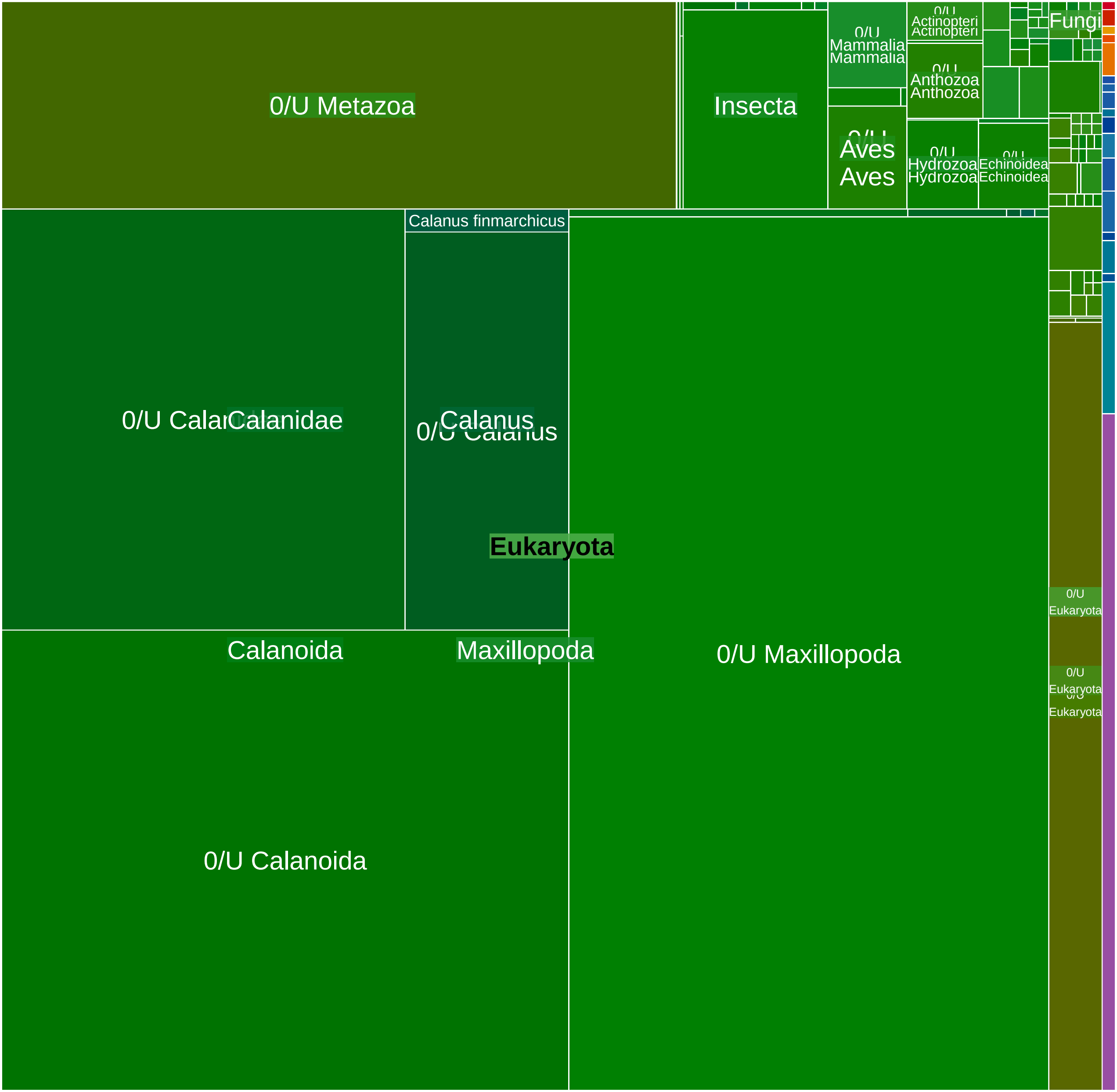

MGT−v1_38 : 37557 unigenes, 13839 (36.85%) taxonomically assigned

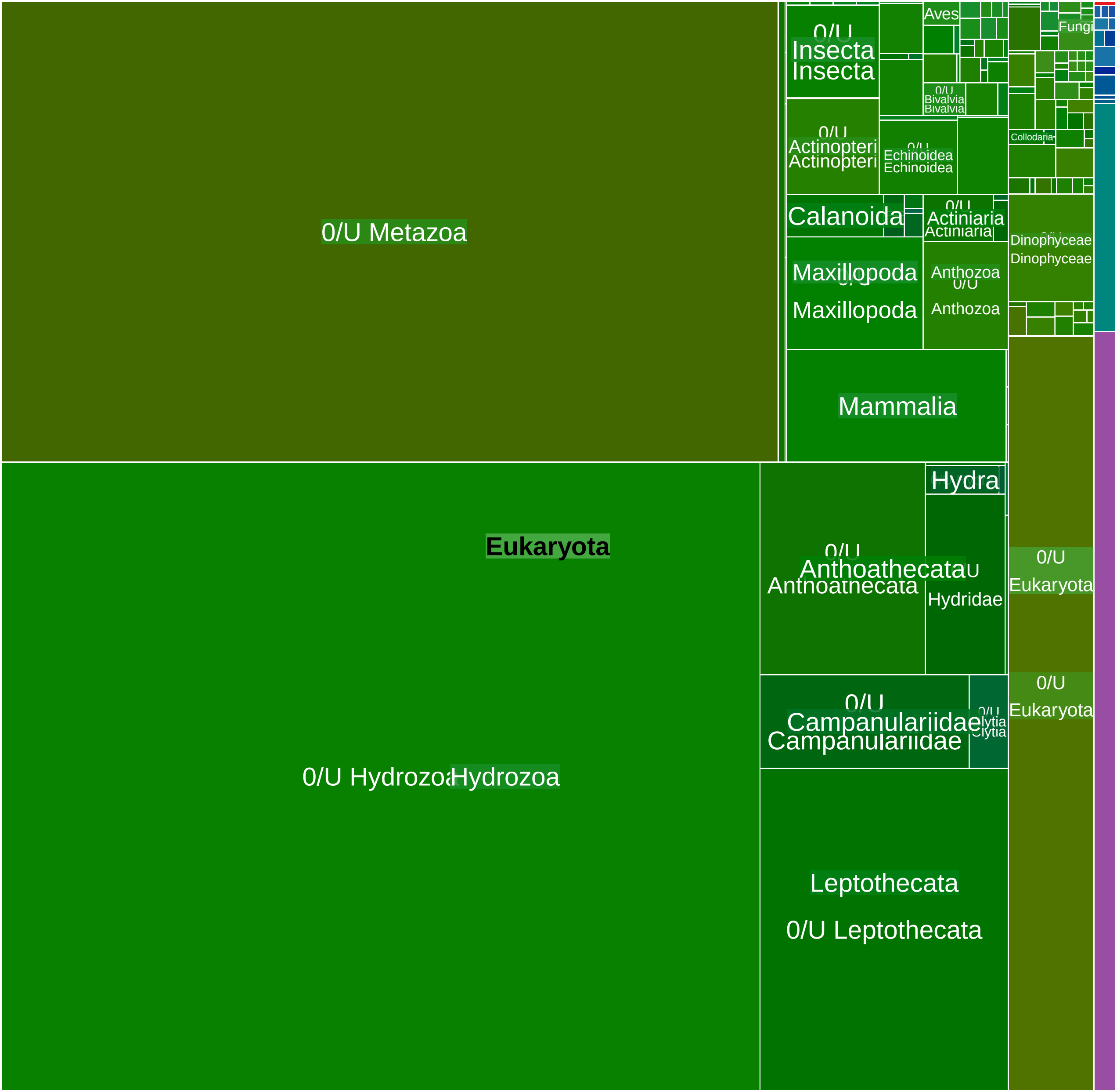

MGT−v1_39 : 37116 unigenes, 33228 (89.52%) taxonomically assigned

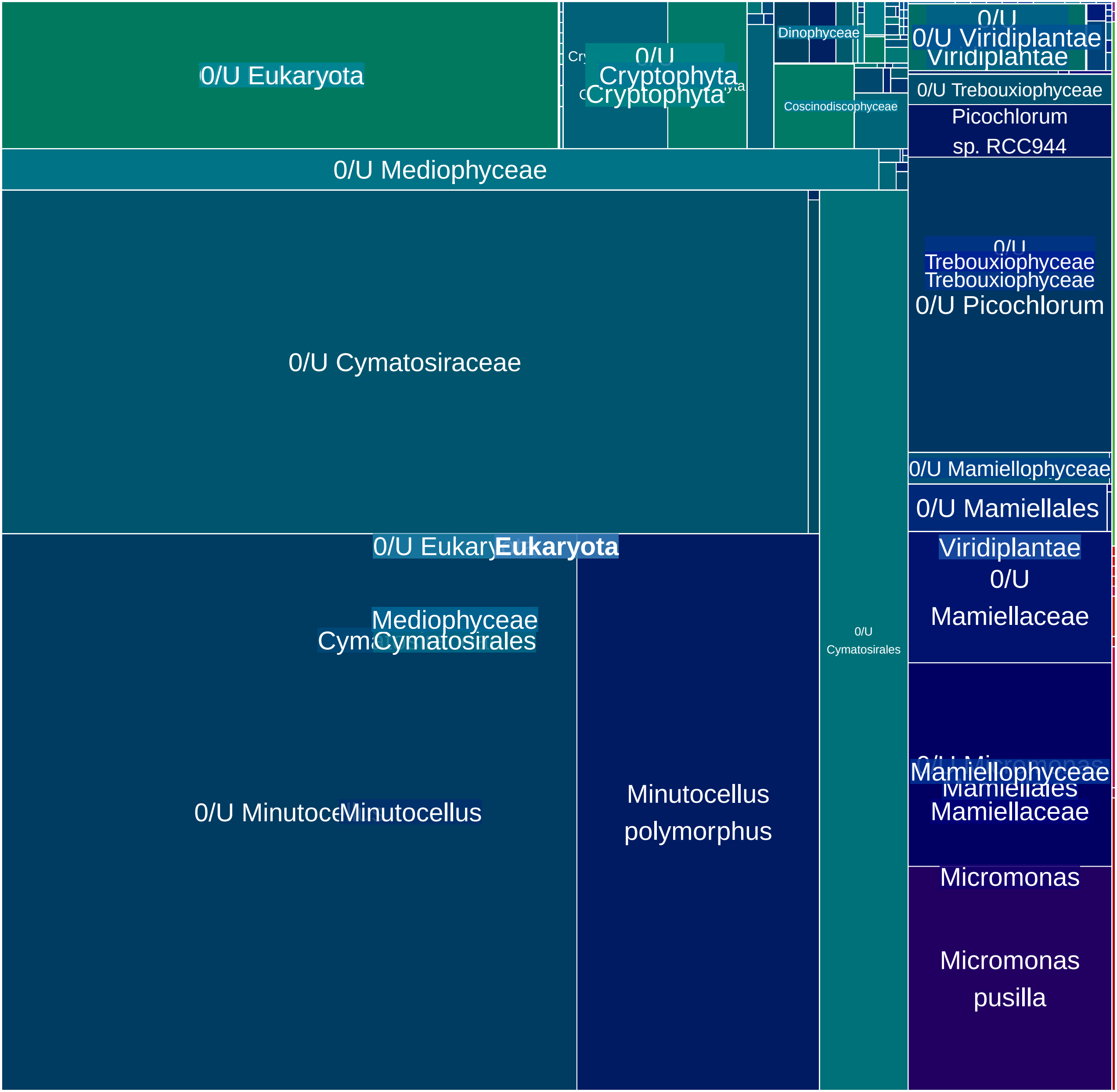

MGT−v1_40 : 36721 unigenes, 15868 (43.21%) taxonomically assigned

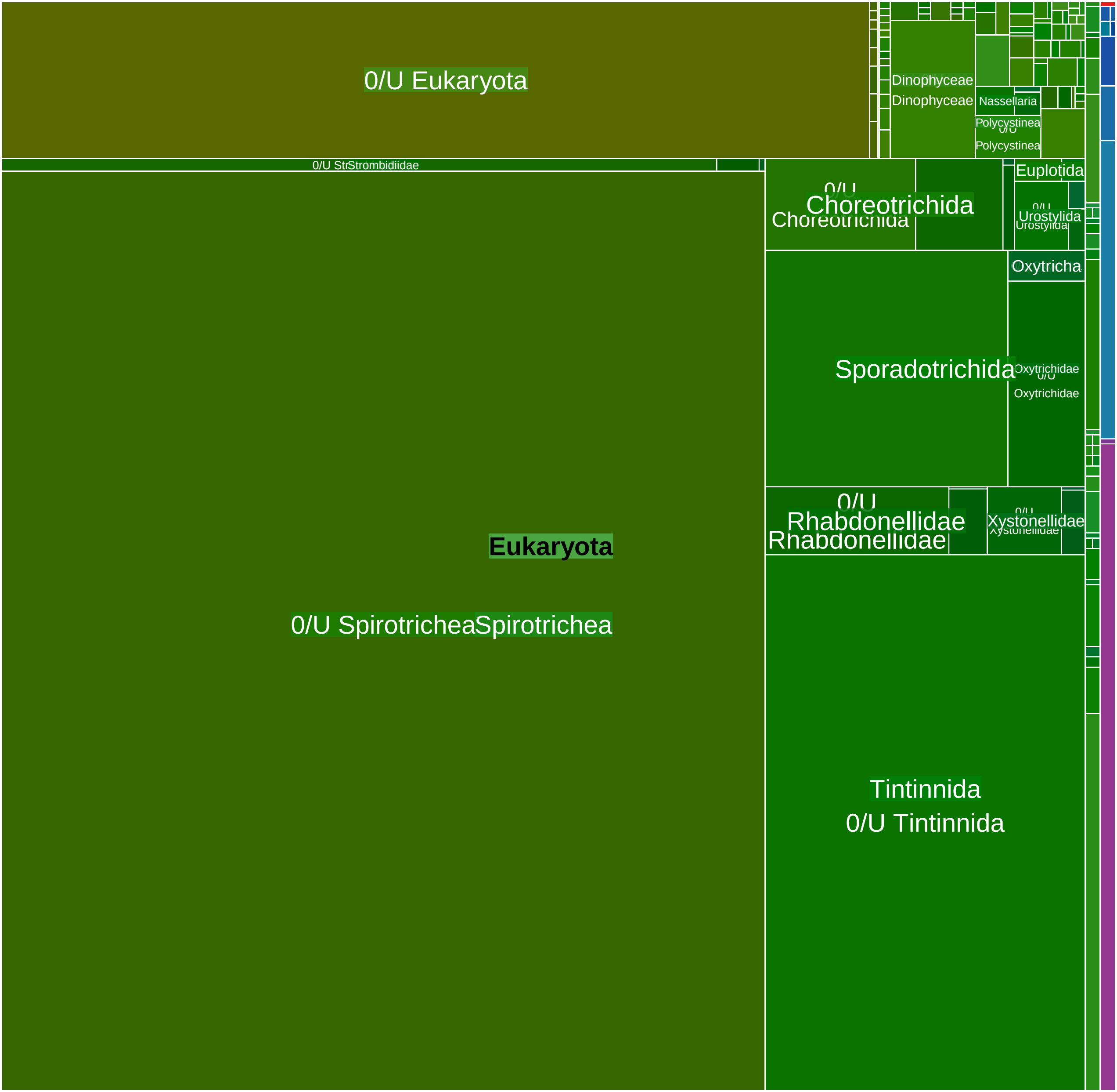

MGT−v1_41 : 35610 unigenes, 33605 (94.37%) taxonomically assigned

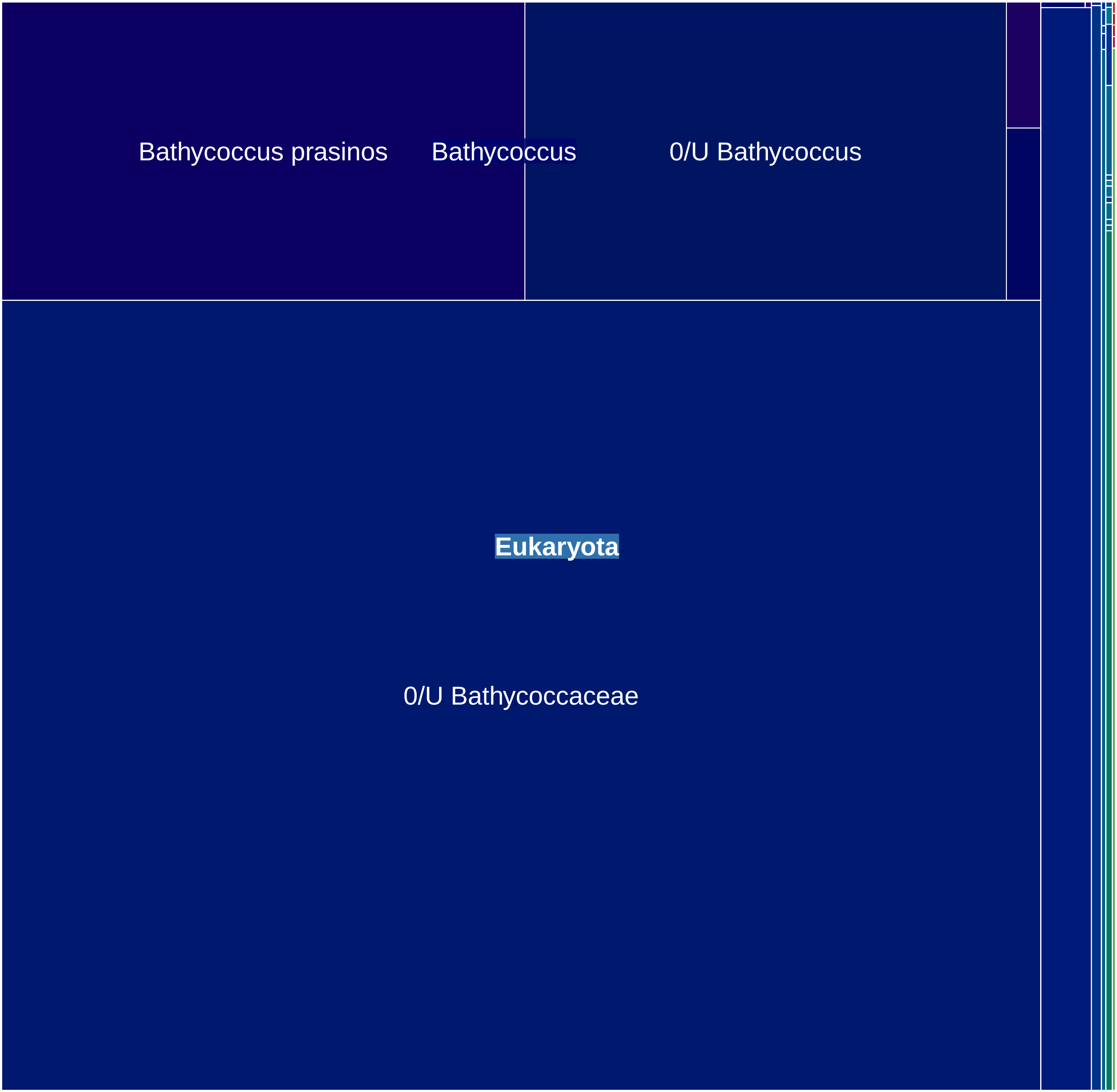

MGT−v1_42 : 35355 unigenes, 5467 (15.46%) taxonomically assigned

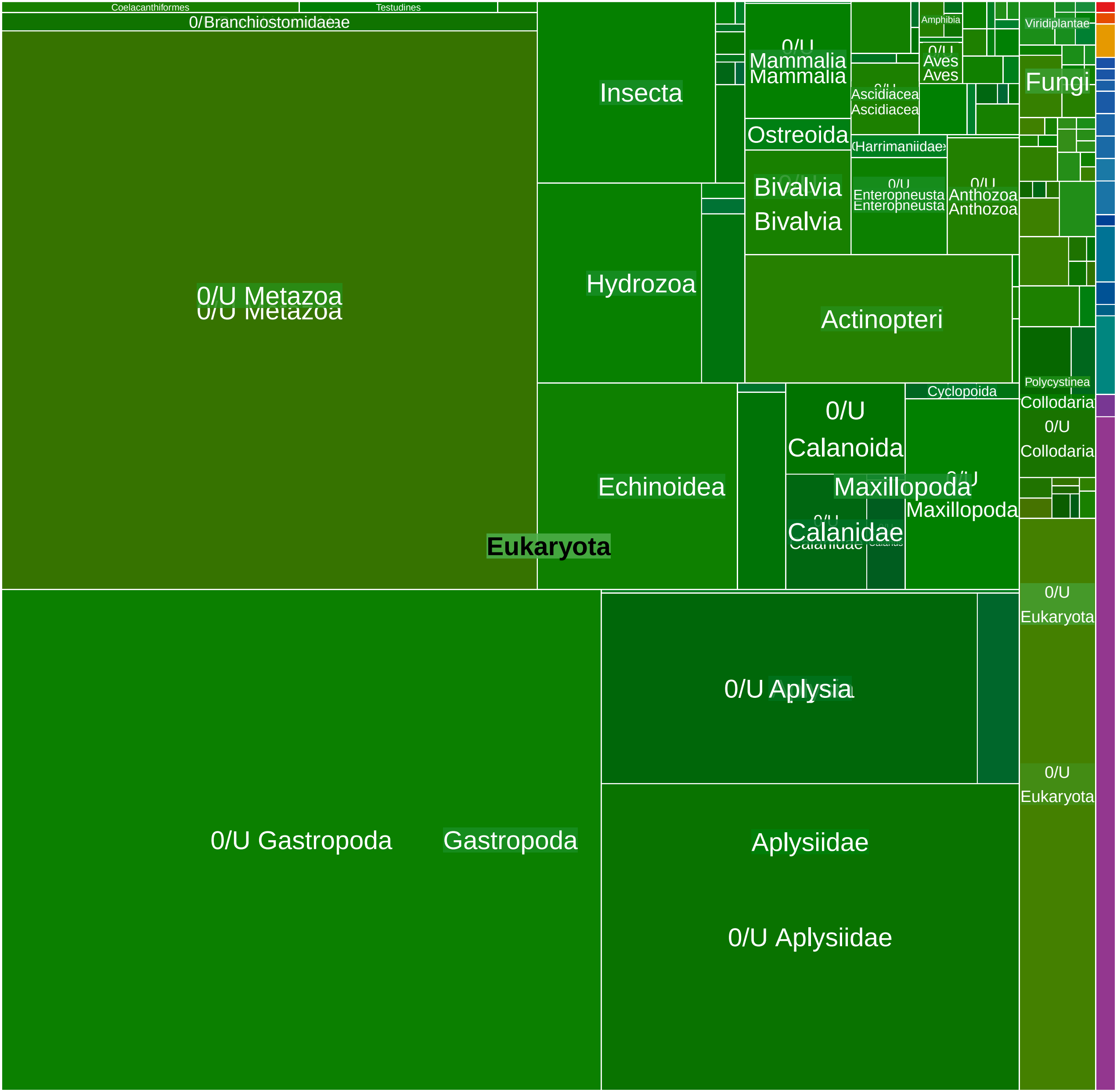

### MGT−v1_43 : 35097 unigenes, 24909 (70.97%) taxonomically assigned

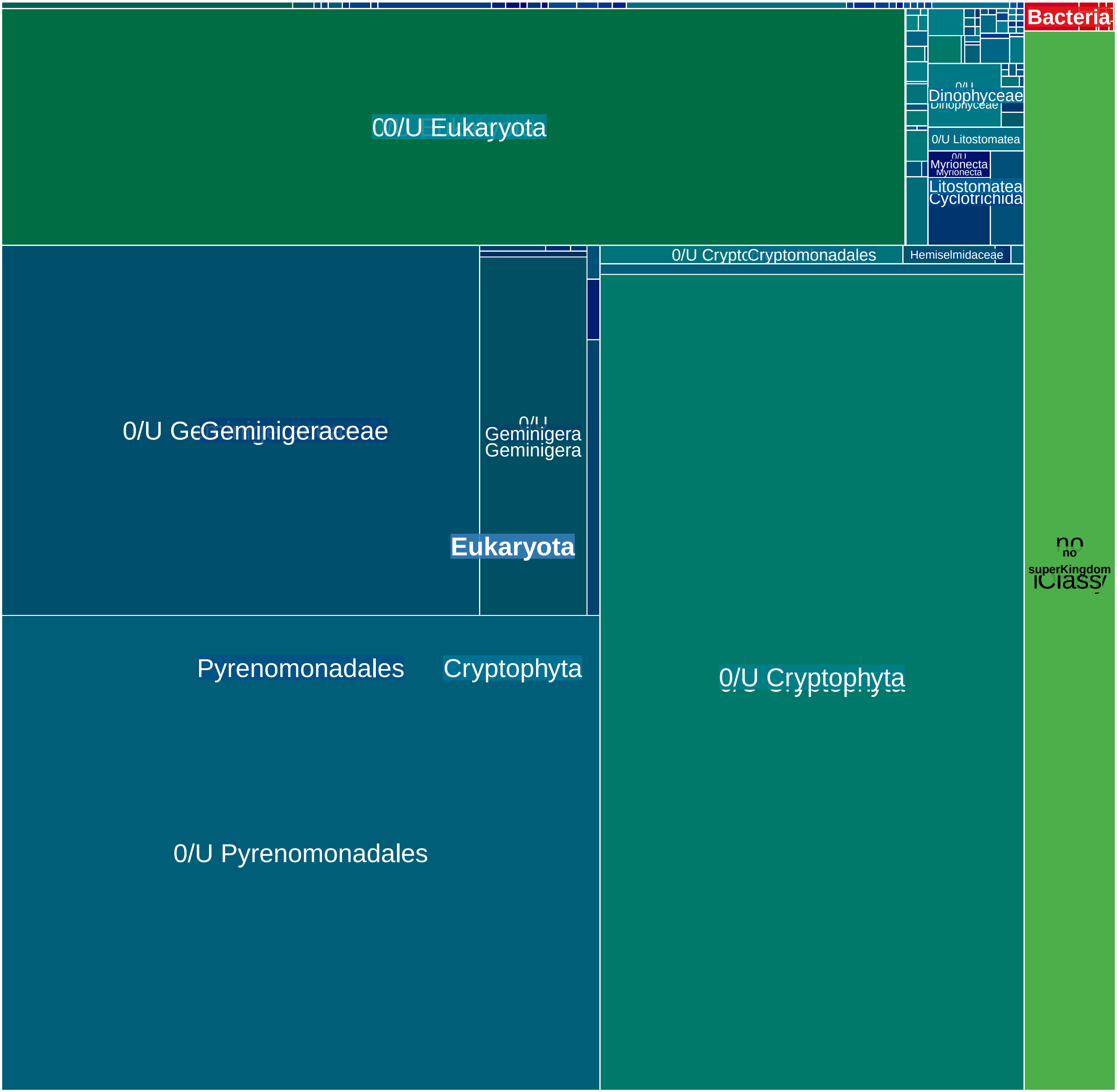

MGT−v1_44 : 34643 unigenes, 27014 (77.98%) taxonomically assigned

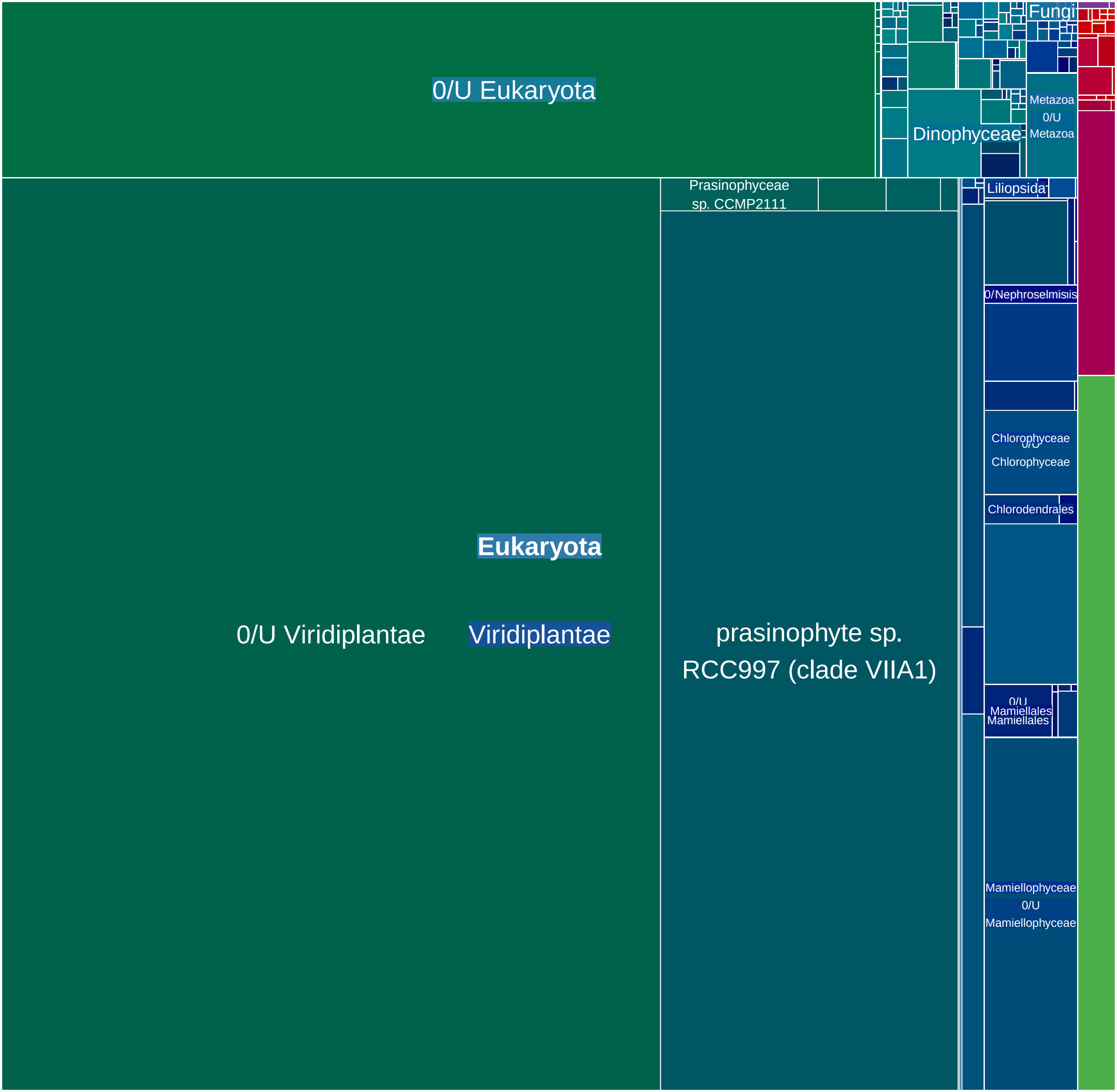

MGT−v1_45 : 33662 unigenes, 14831 (44.06%) taxonomically assigned

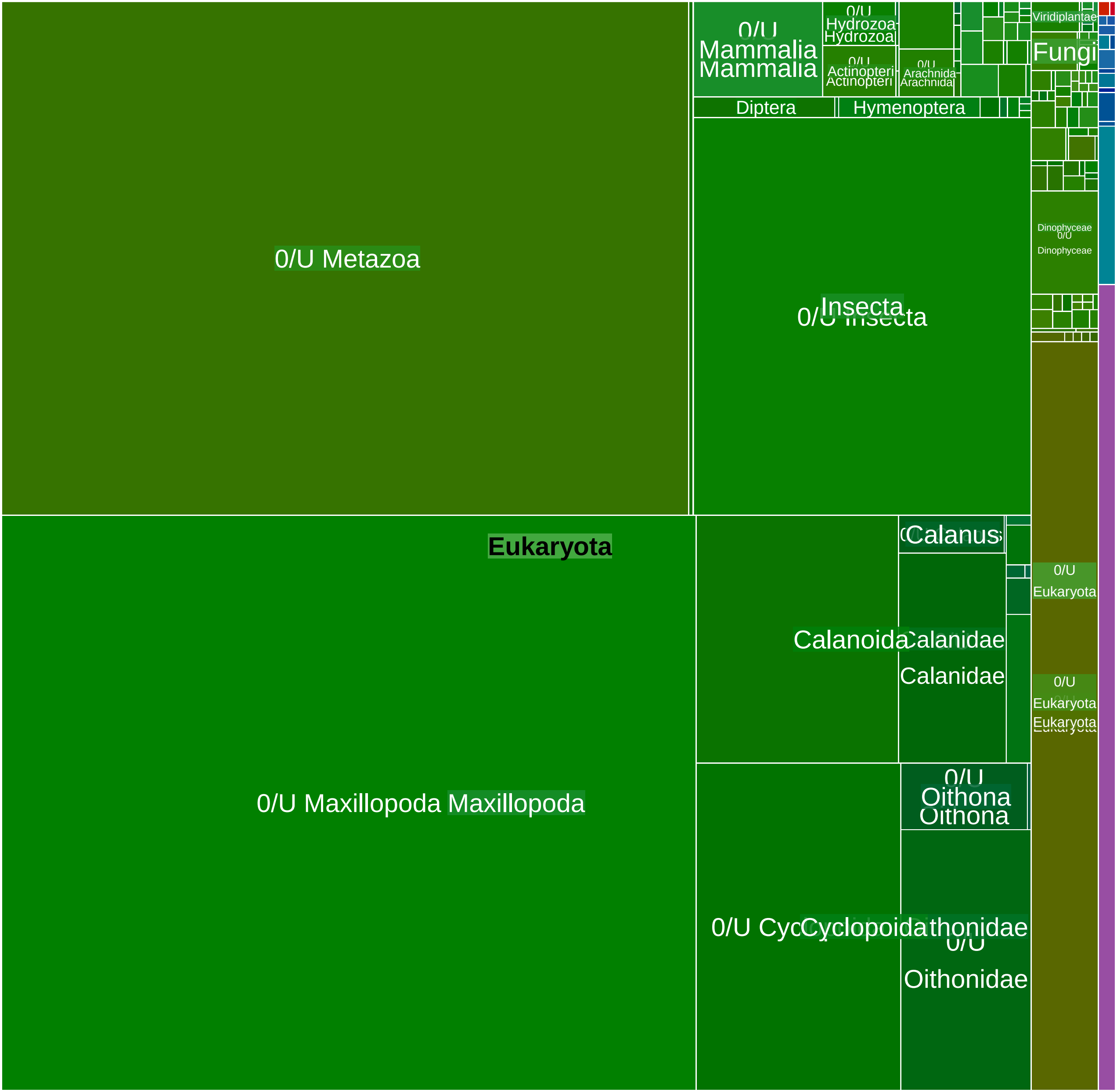

MGT−v1_46 : 33378 unigenes, 17382 (52.08%) taxonomically assigned

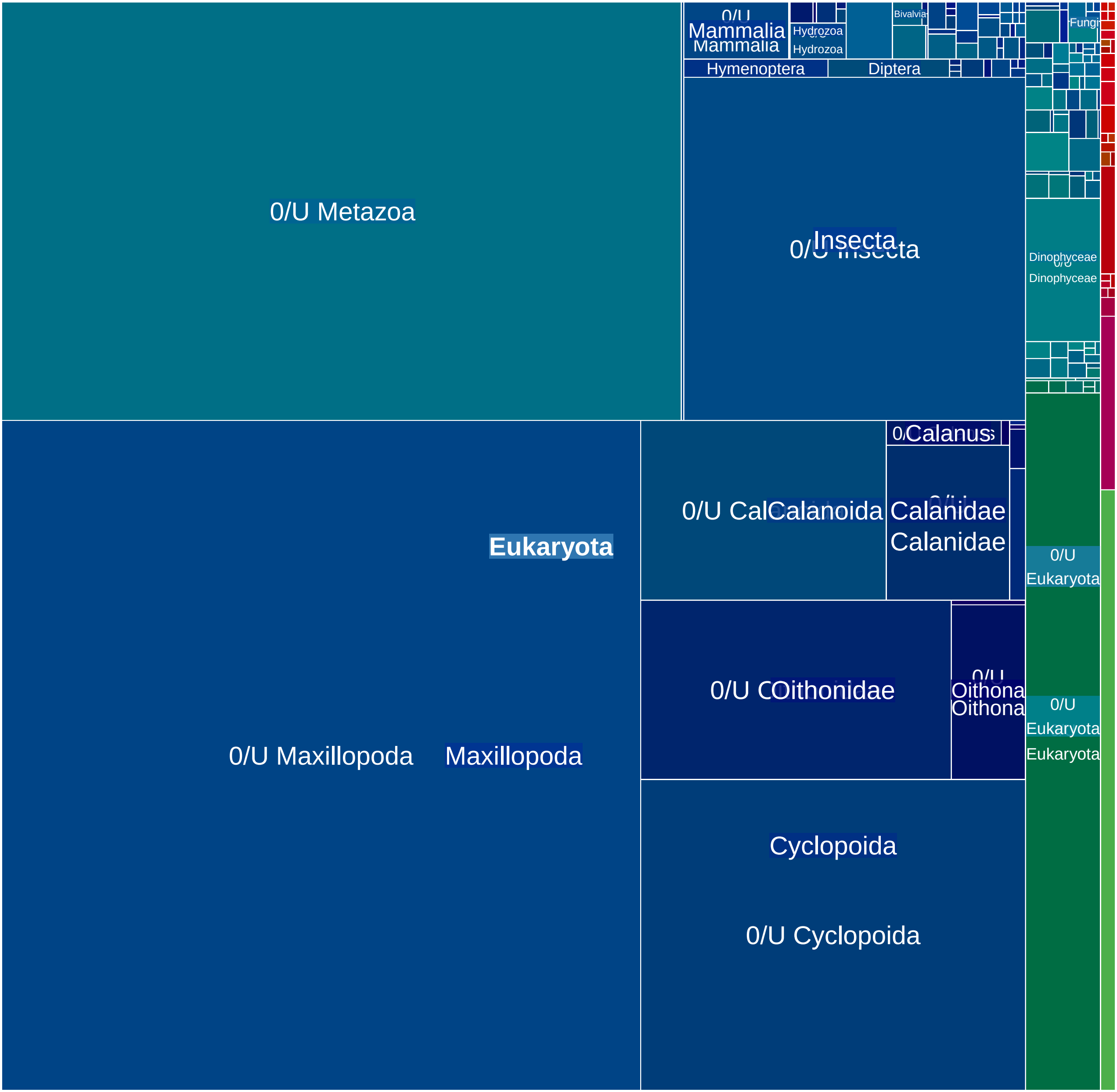

MGT−v1_47 : 32740 unigenes, 4518 (13.80%) taxonomically assigned

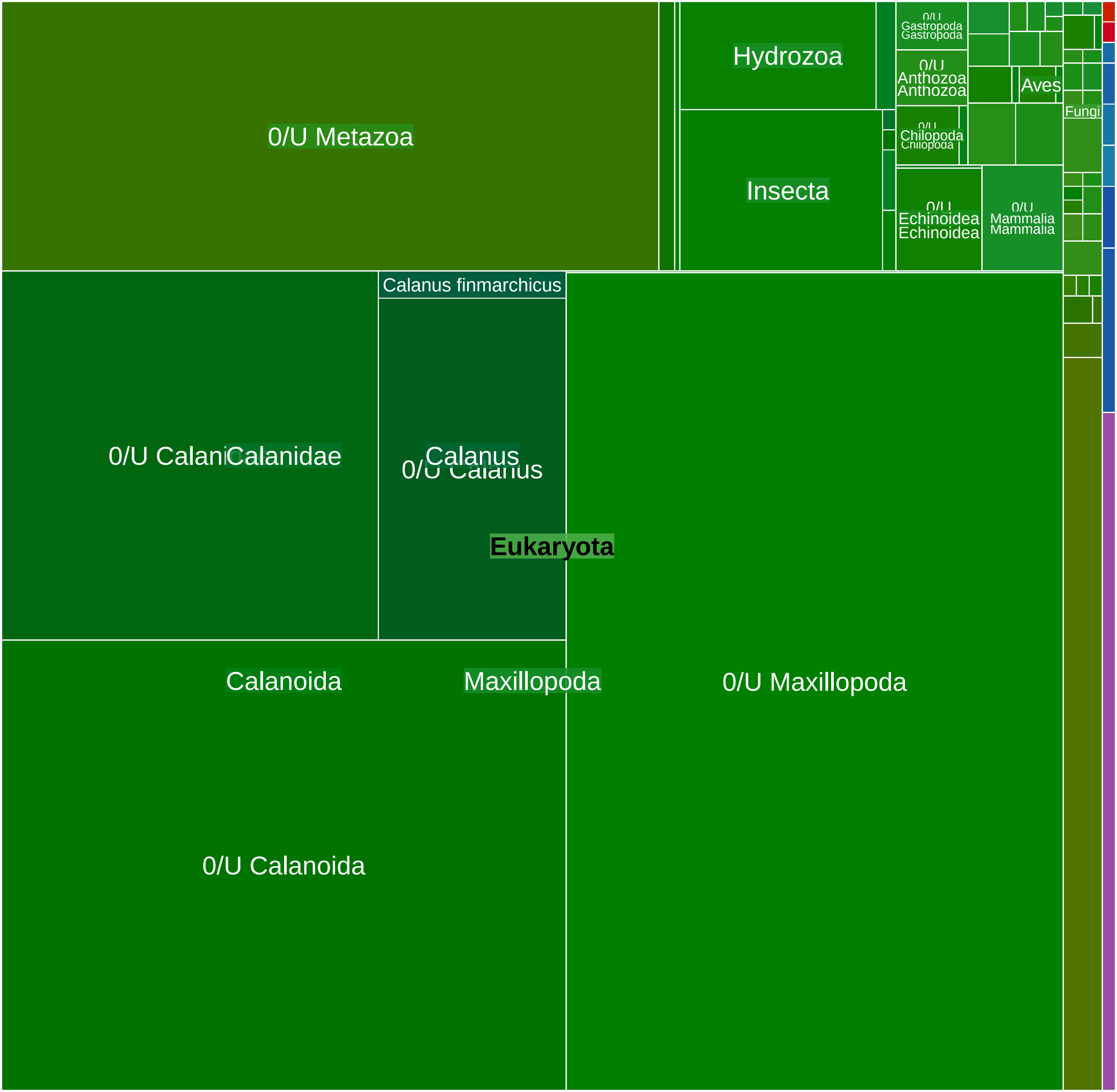

MGT−v1_48 : 31633 unigenes, 28011 (88.55%) taxonomically assigned

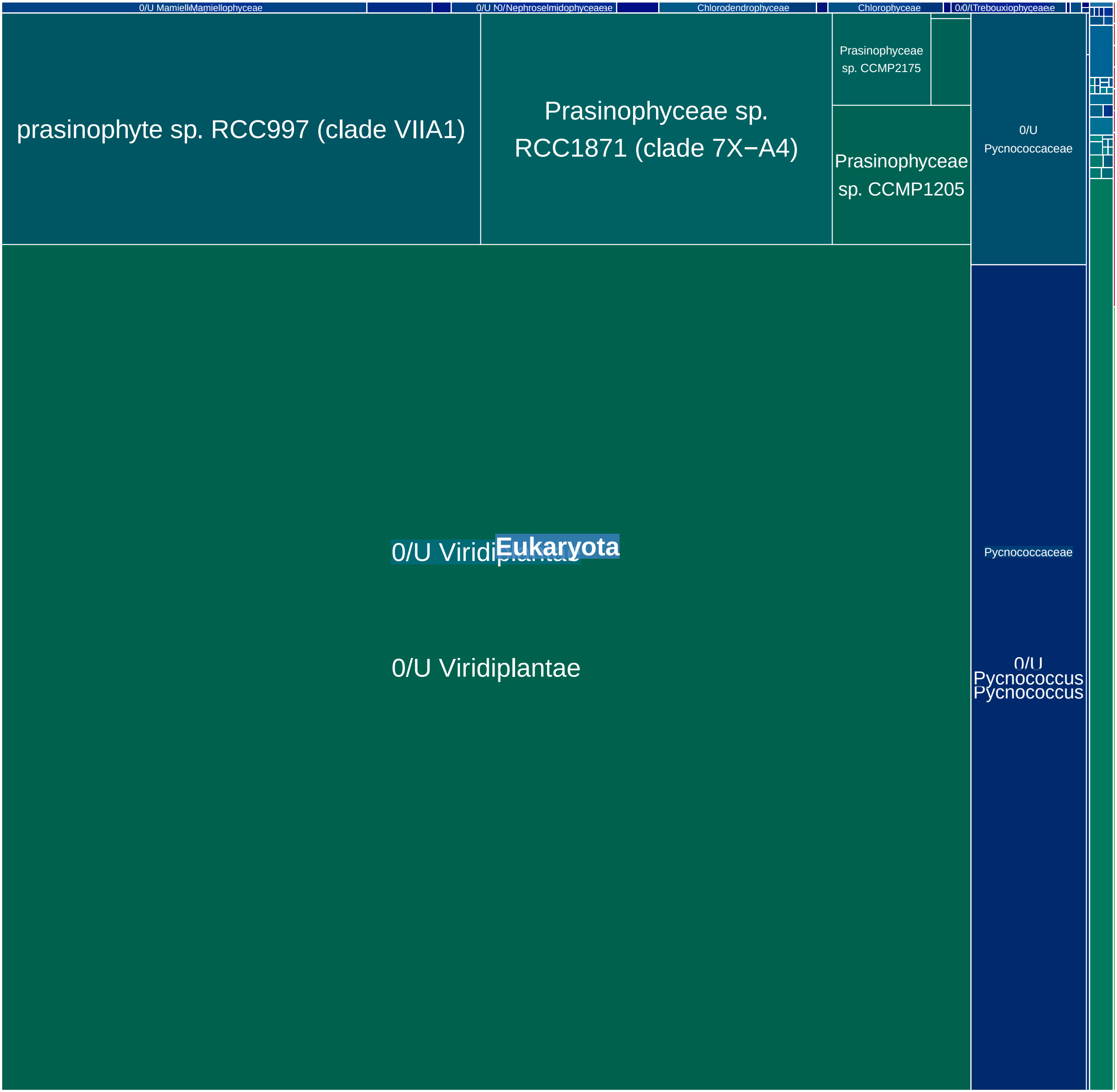

MGT−v1_49 : 31155 unigenes, 11572 (37.14%) taxonomically assigned

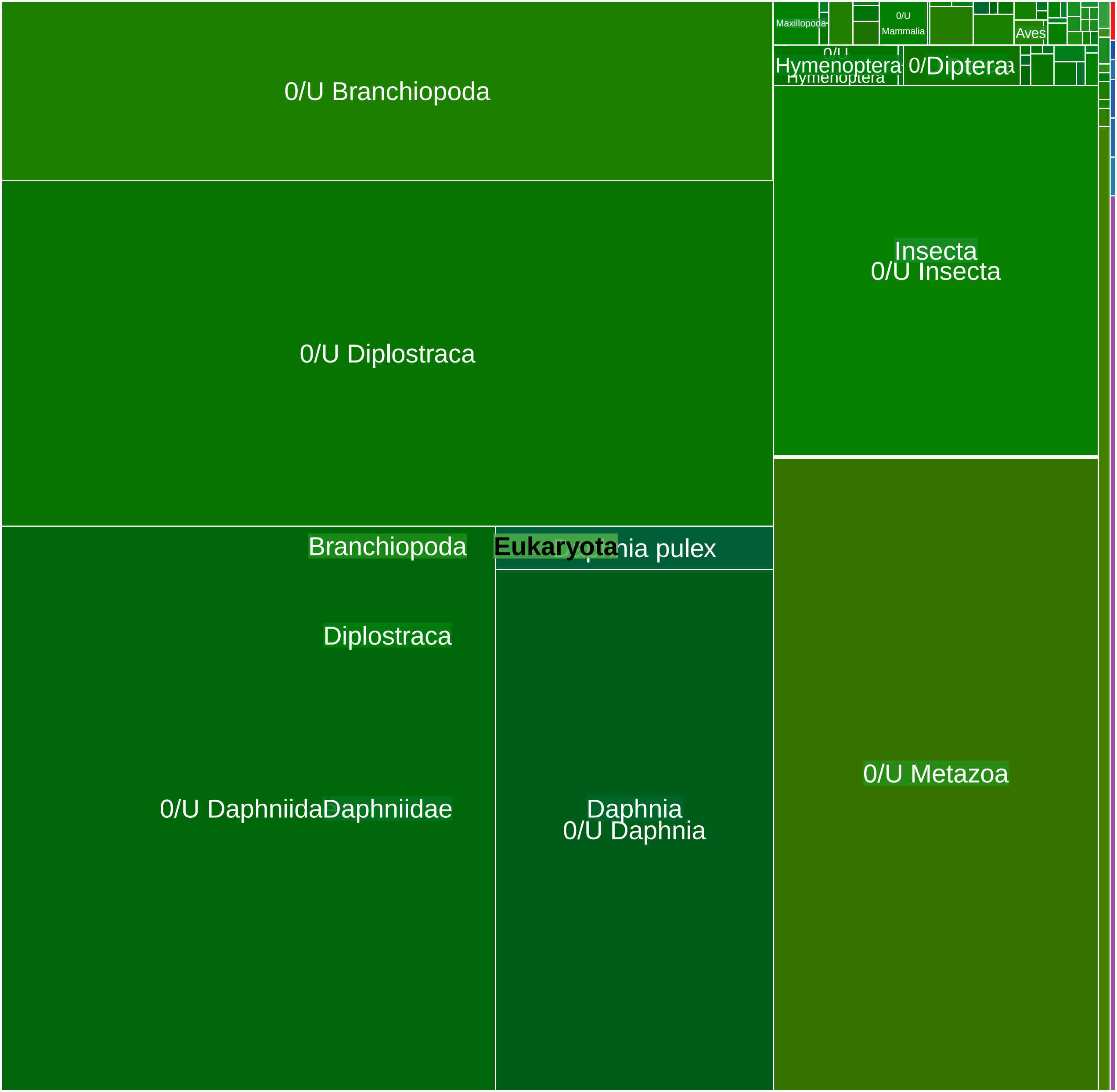

MGT−v1_50 : 30764 unigenes, 20955 (68.12%) taxonomically assigned

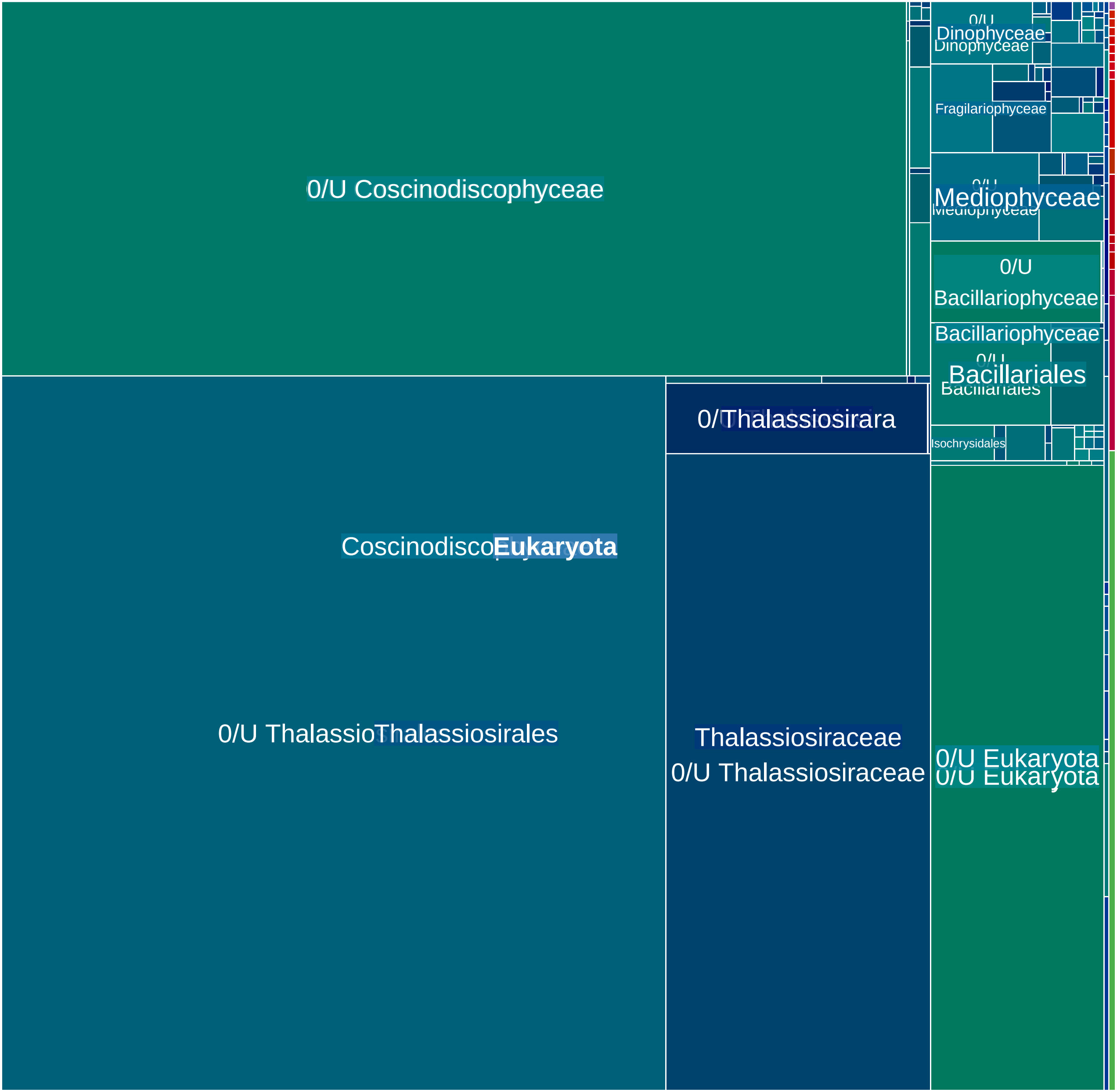

MGT−v1_51 : 30594 unigenes, 20480 (66.94%) taxonomically assigned

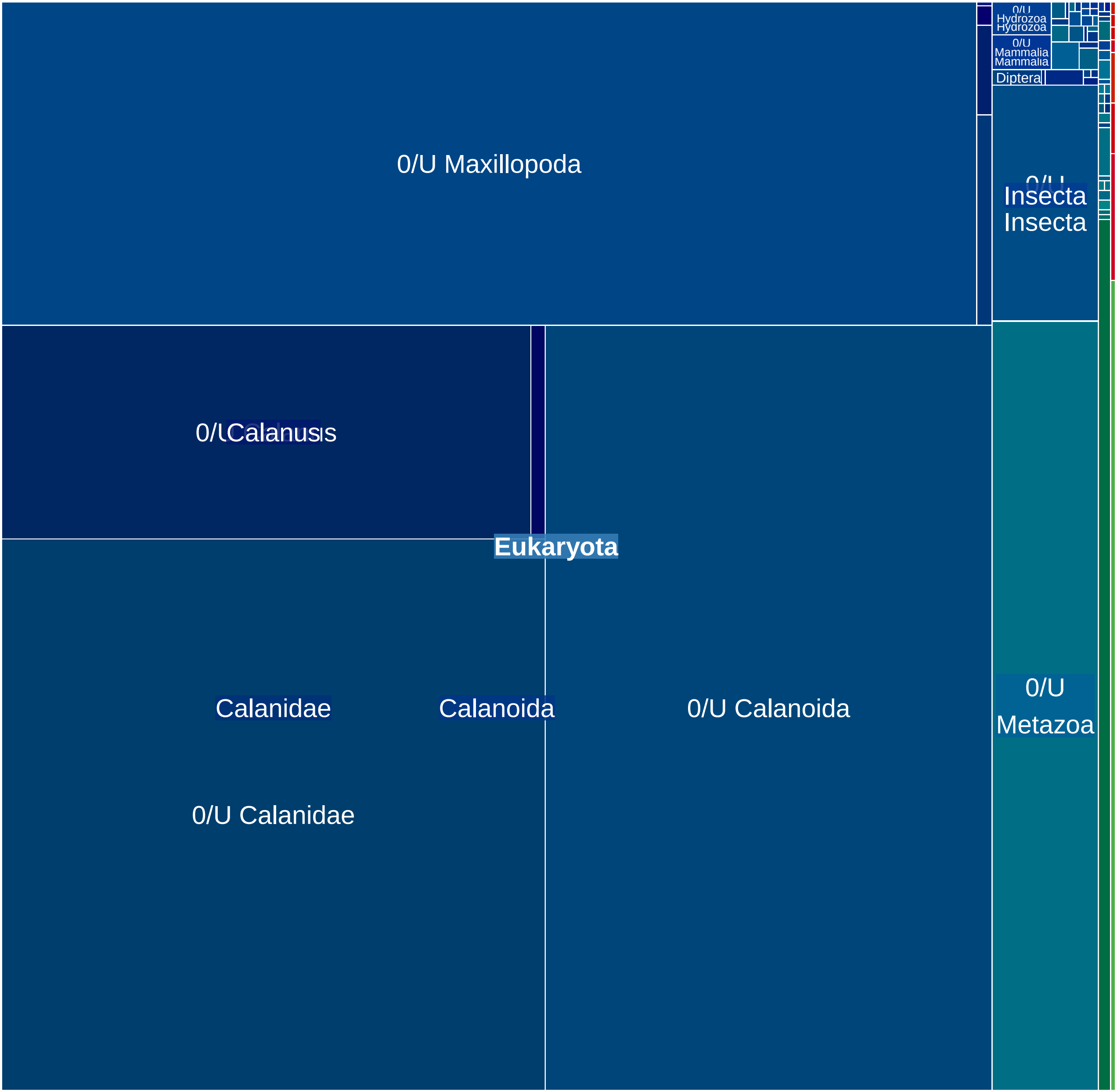

MGT−v1_52 : 29110 unigenes, 18939 (65.06%) taxonomically assigned

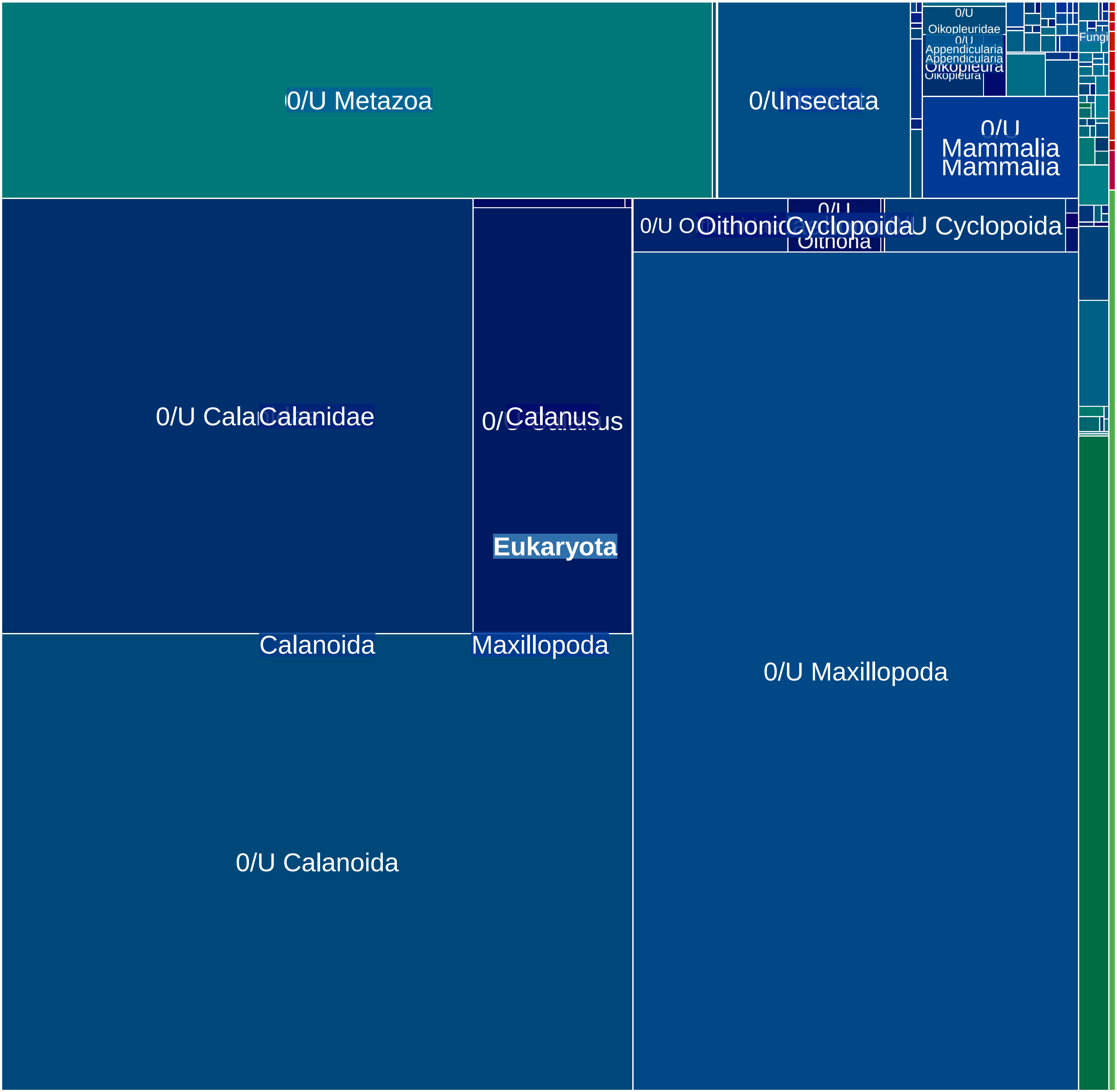

MGT−v1_53 : 28858 unigenes, 18621 (64.53%) taxonomically assigned

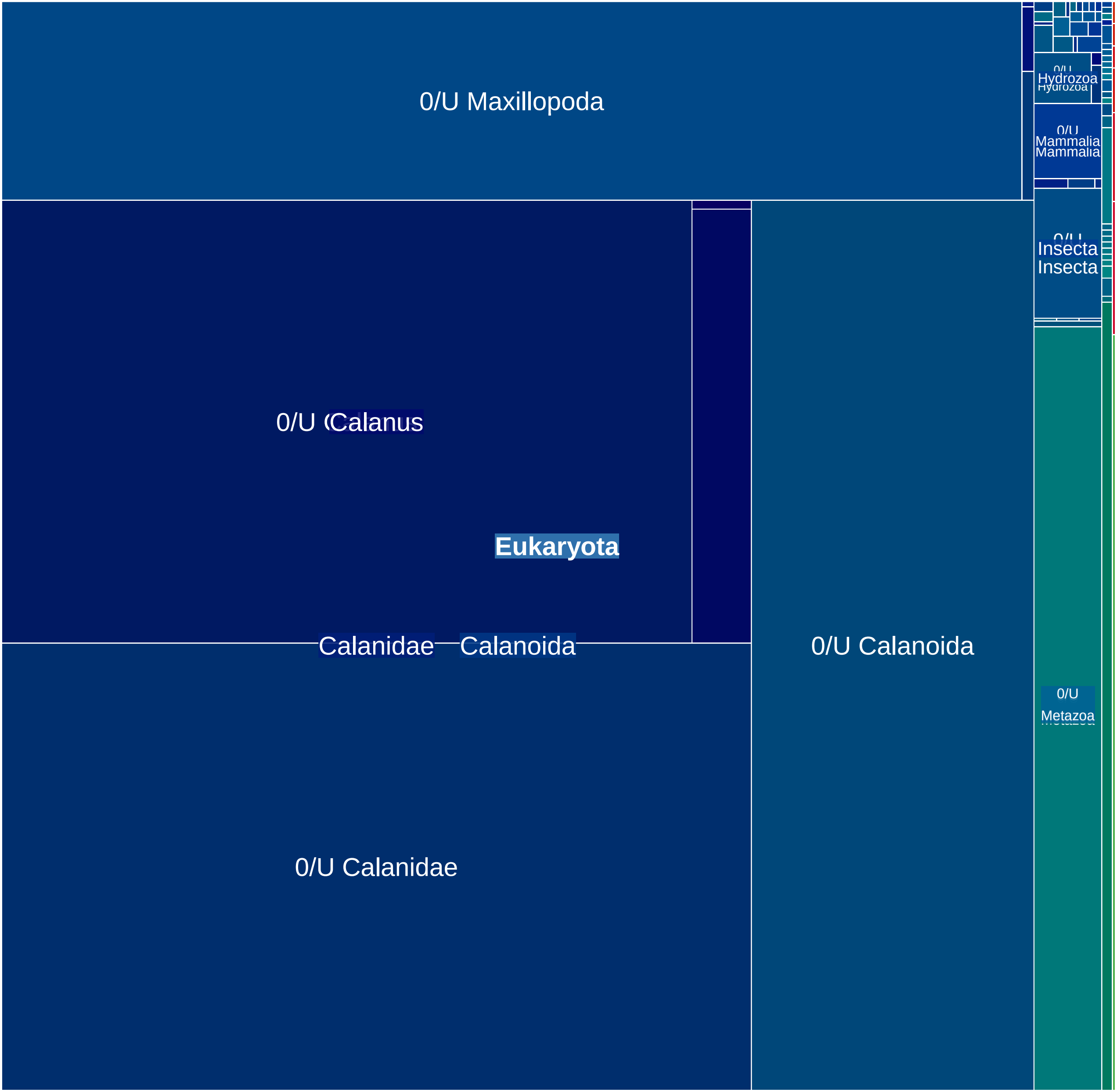

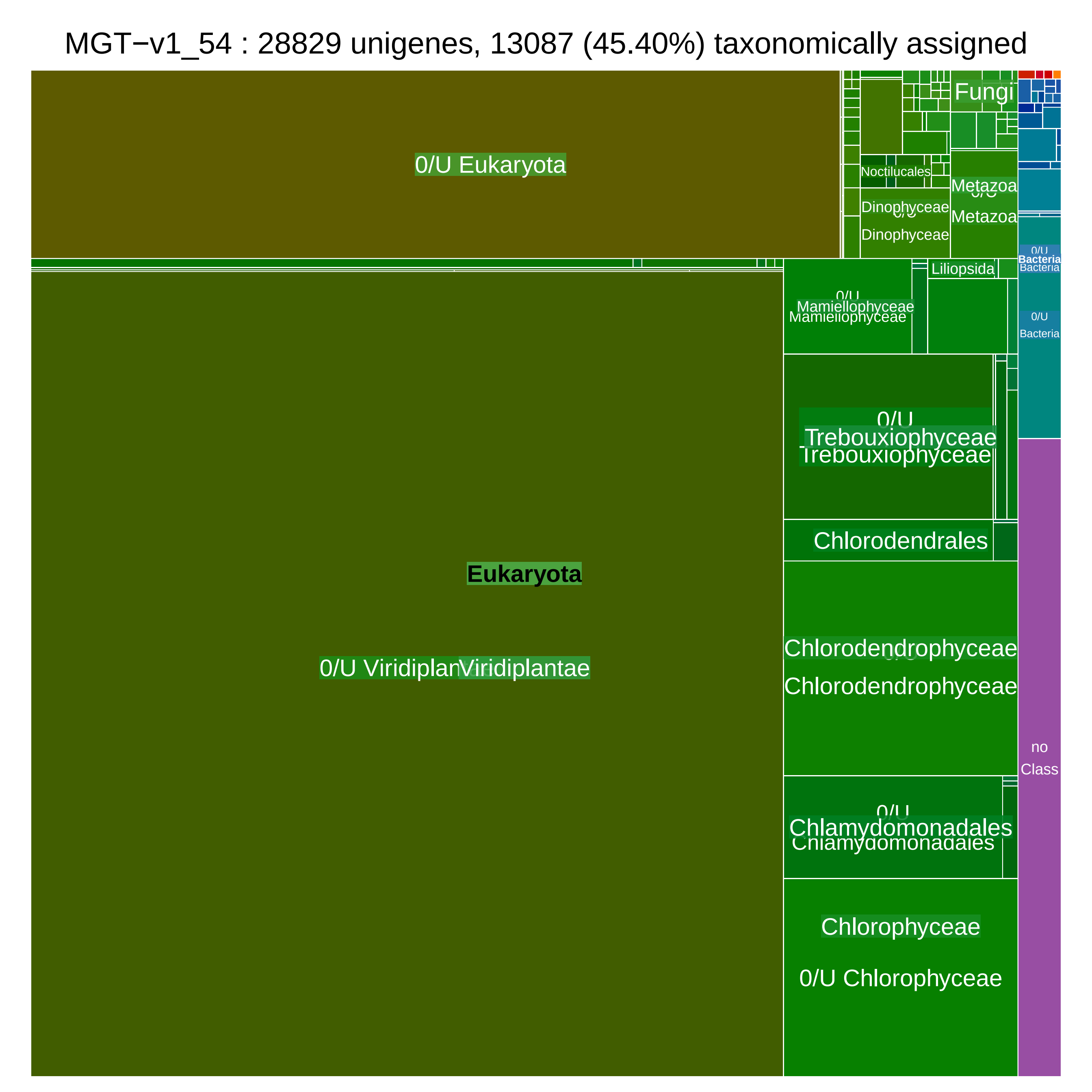
MGT−v1_54 : 28829 unigenes, 13087 (45.40%) taxonomically assigned
0/U Eukaryota
Fungi
Noctilucales
Metazoa
0/U Metazoa
Dinophyceae
0/U Dinophyceae
Liliopsida
0/U Mamiellophyceae
Mamiellophyceae
0/U Trebouxiophyceae
Trebouxiophyceae
Chlorodendrales
Eukaryota
0/U Viridiplantae
Viridiplantae
Chlorodendrophyceae
0/U Chlorodendrophyceae
0/U Chlamydomonadales
Chlamydomonadales
Chlorophyceae
0/U Chlorophyceae
0/U Bacteria
Bacteria
no Class

MGT−v1_55 : 27304 unigenes, 2461 (9.01%) taxonomically assigned

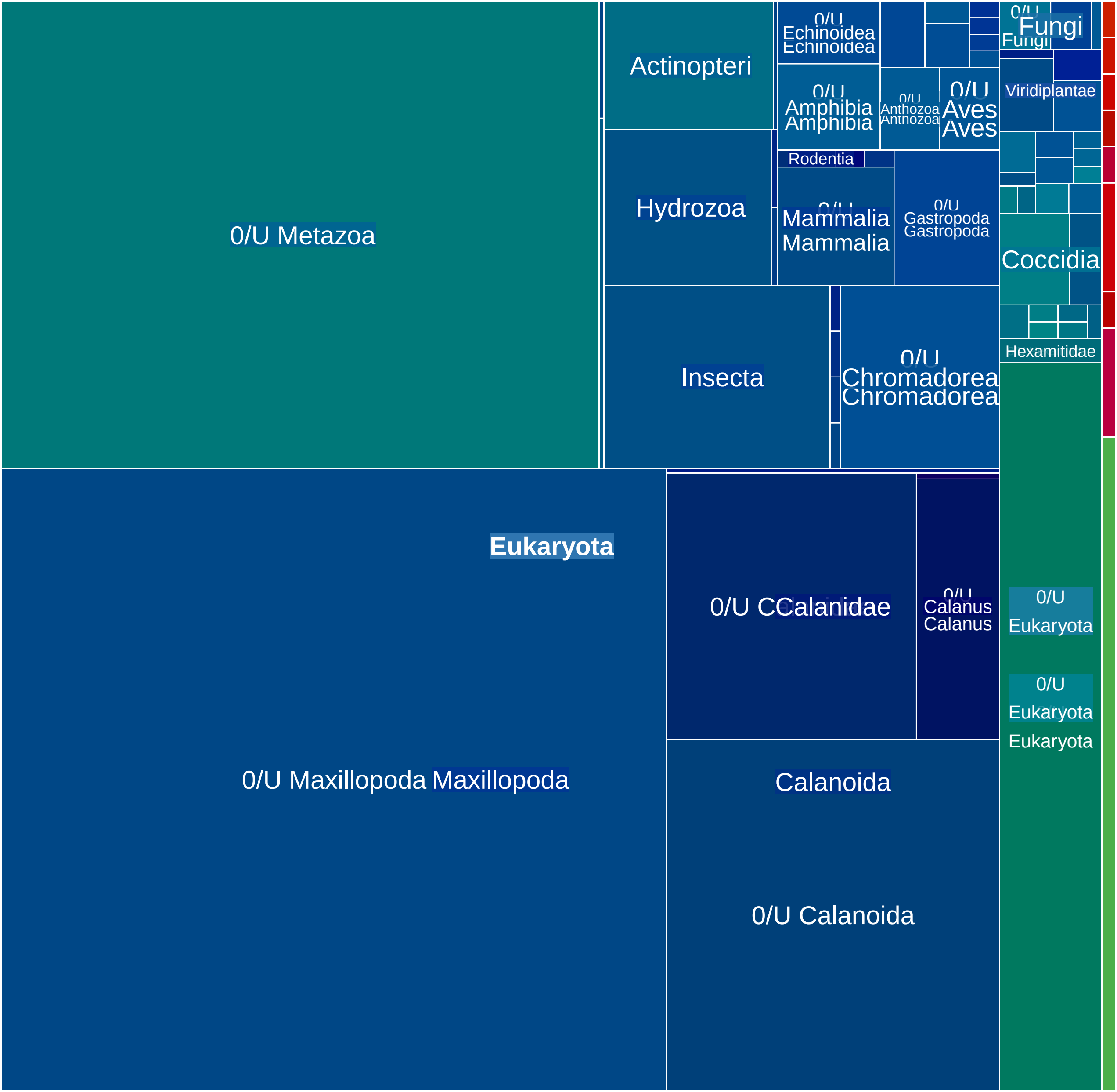

MGT−v1_56 : 26118 unigenes, 10854 (41.56%) taxonomically assigned

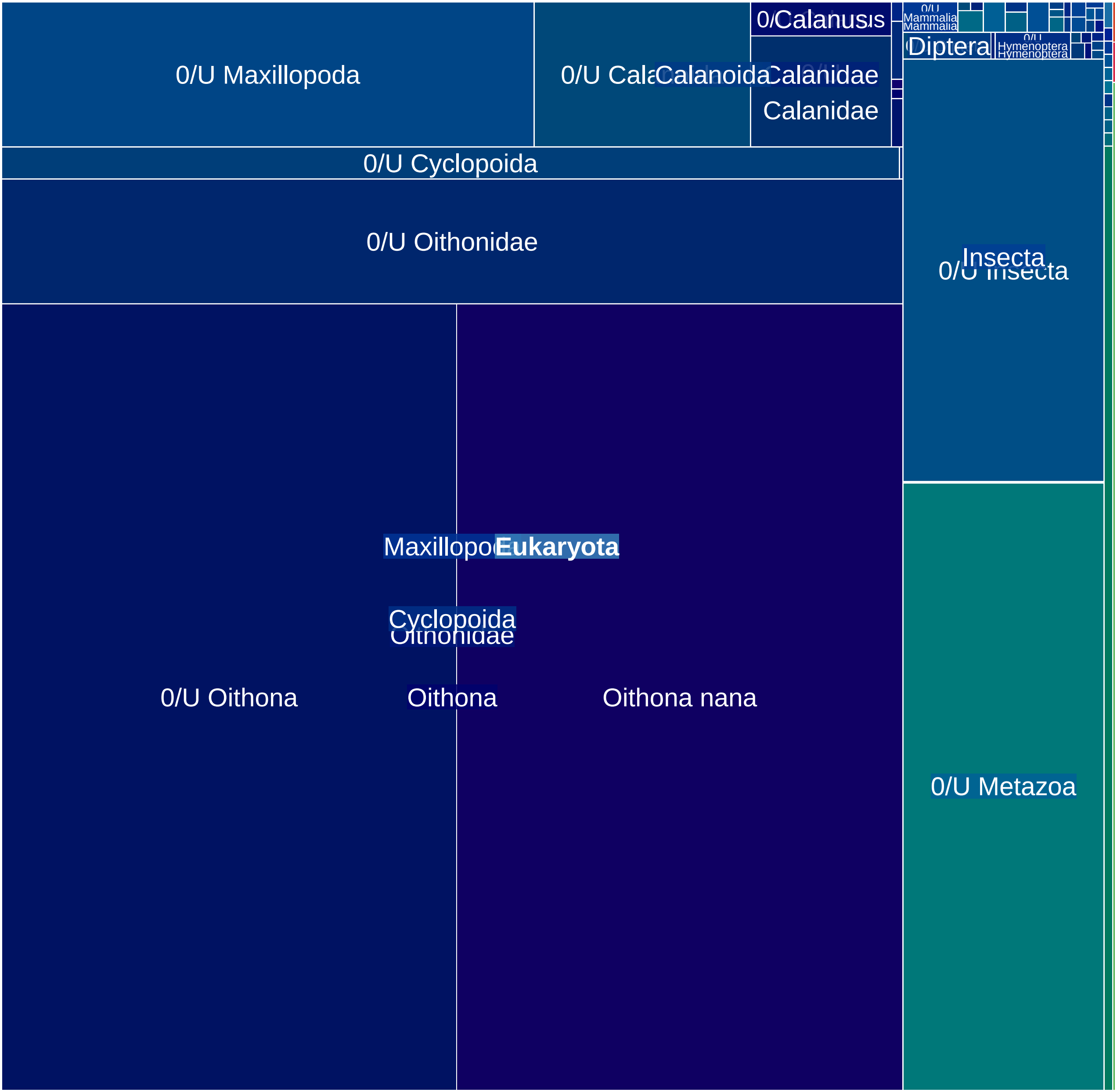

MGT−v1_57 : 25971 unigenes, 23662 (91.11%) taxonomically assigned

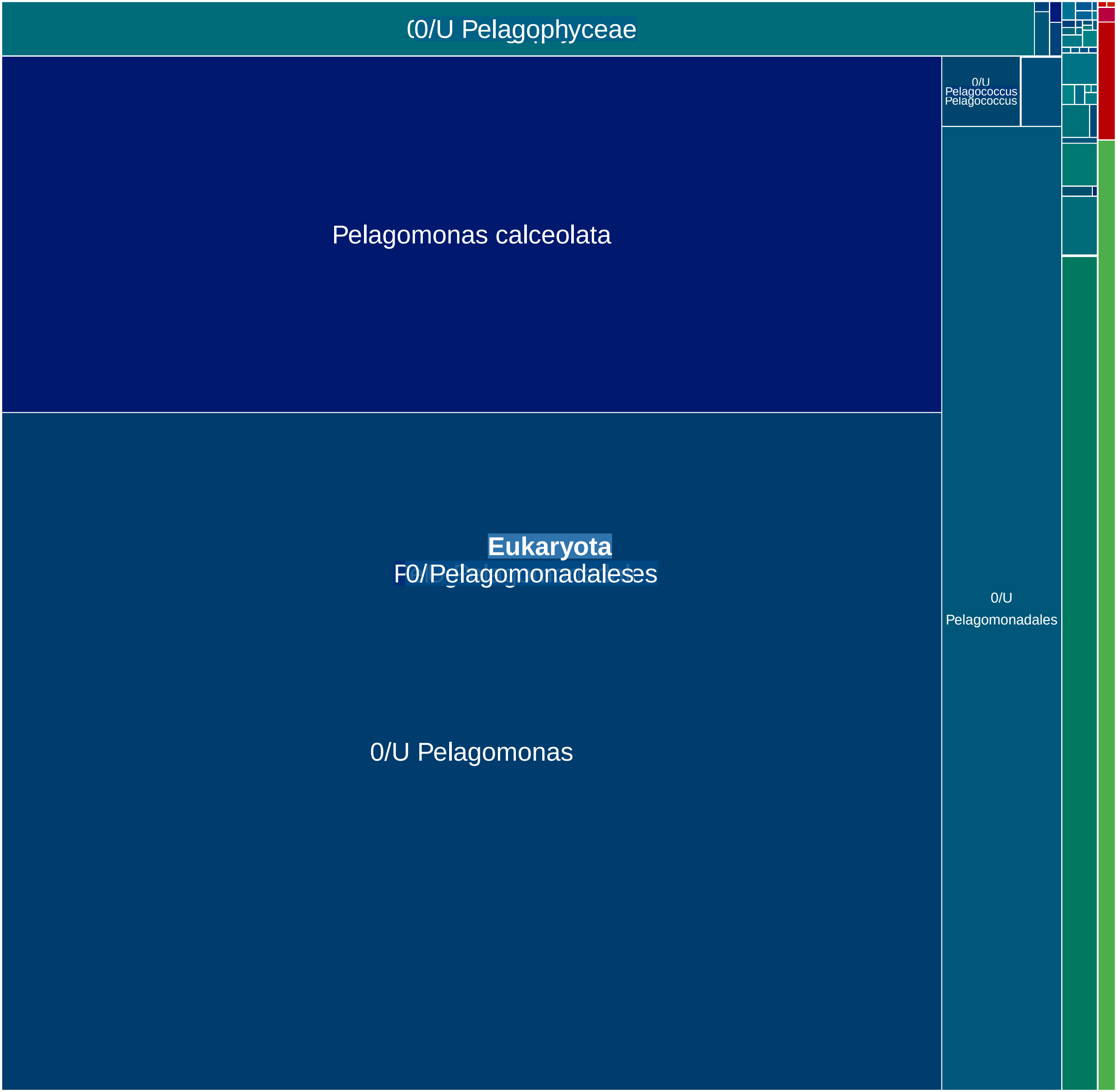

### MGT−v1_58 : 25964 unigenes, 22457 (86.49%) taxonomically assigned

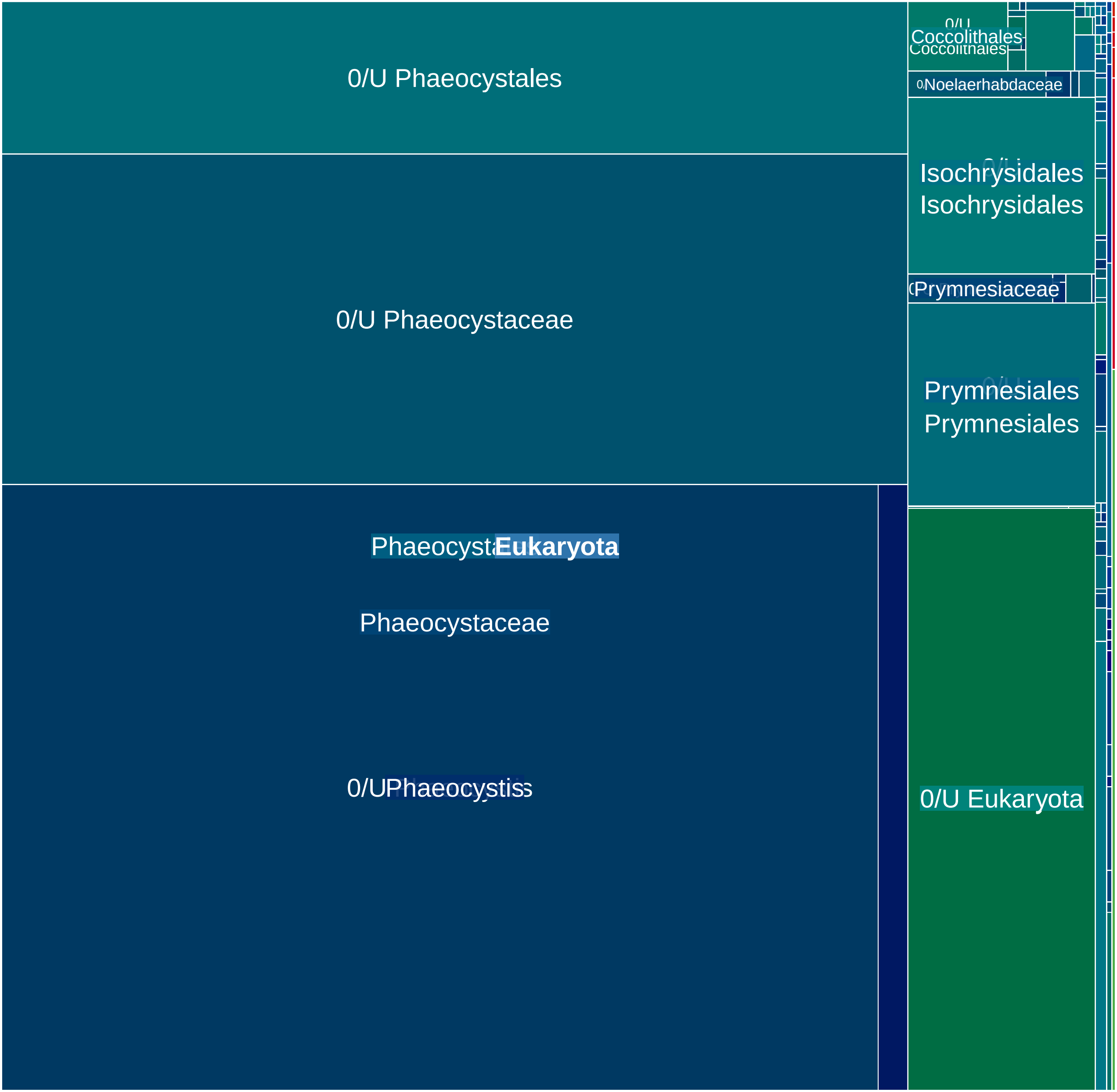

MGT−v1_59 : 24885 unigenes, 16150 (64.90%) taxonomically assigned

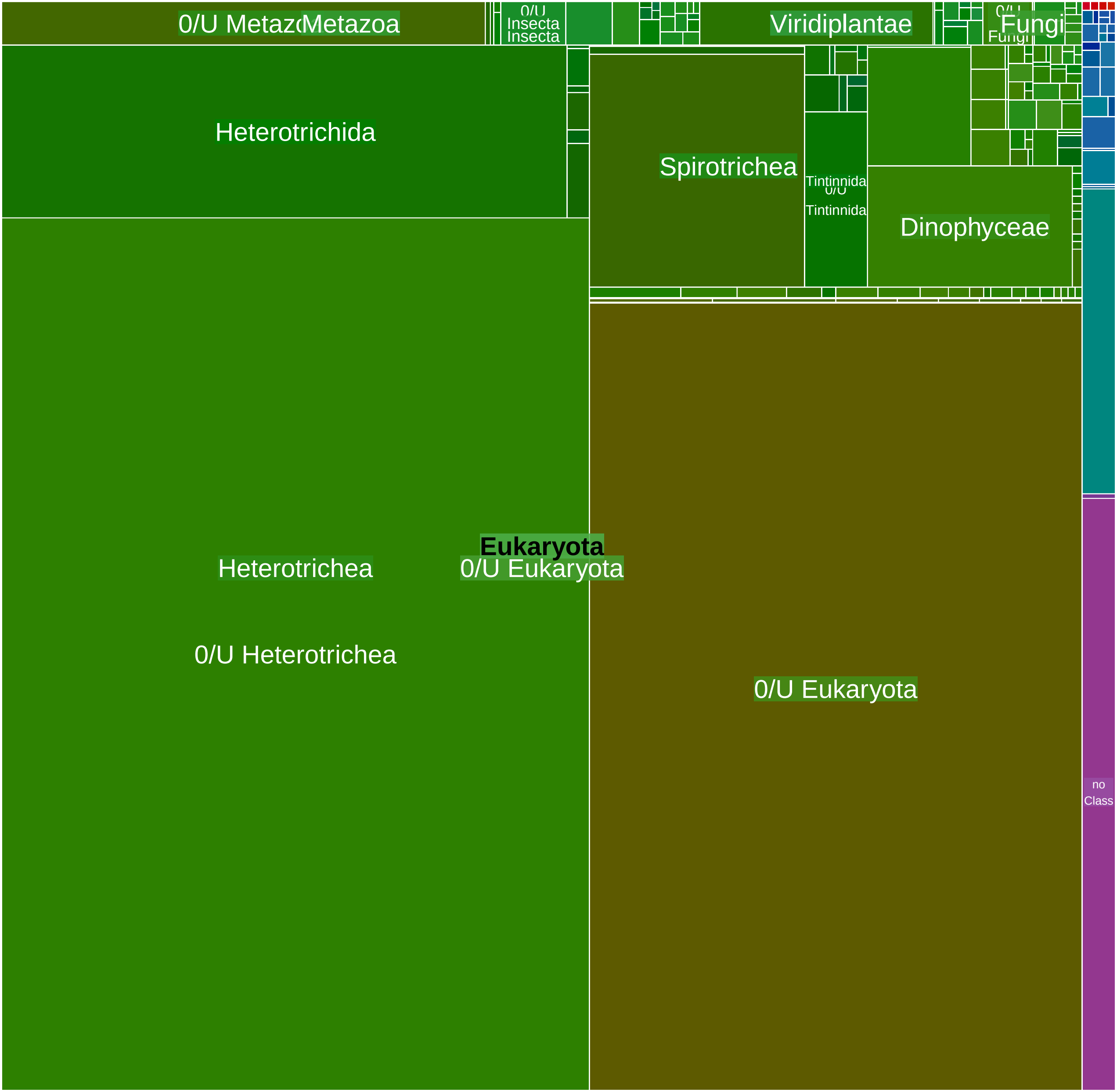

MGT−v1_60 : 24590 unigenes, 20191 (82.11%) taxonomically assigned

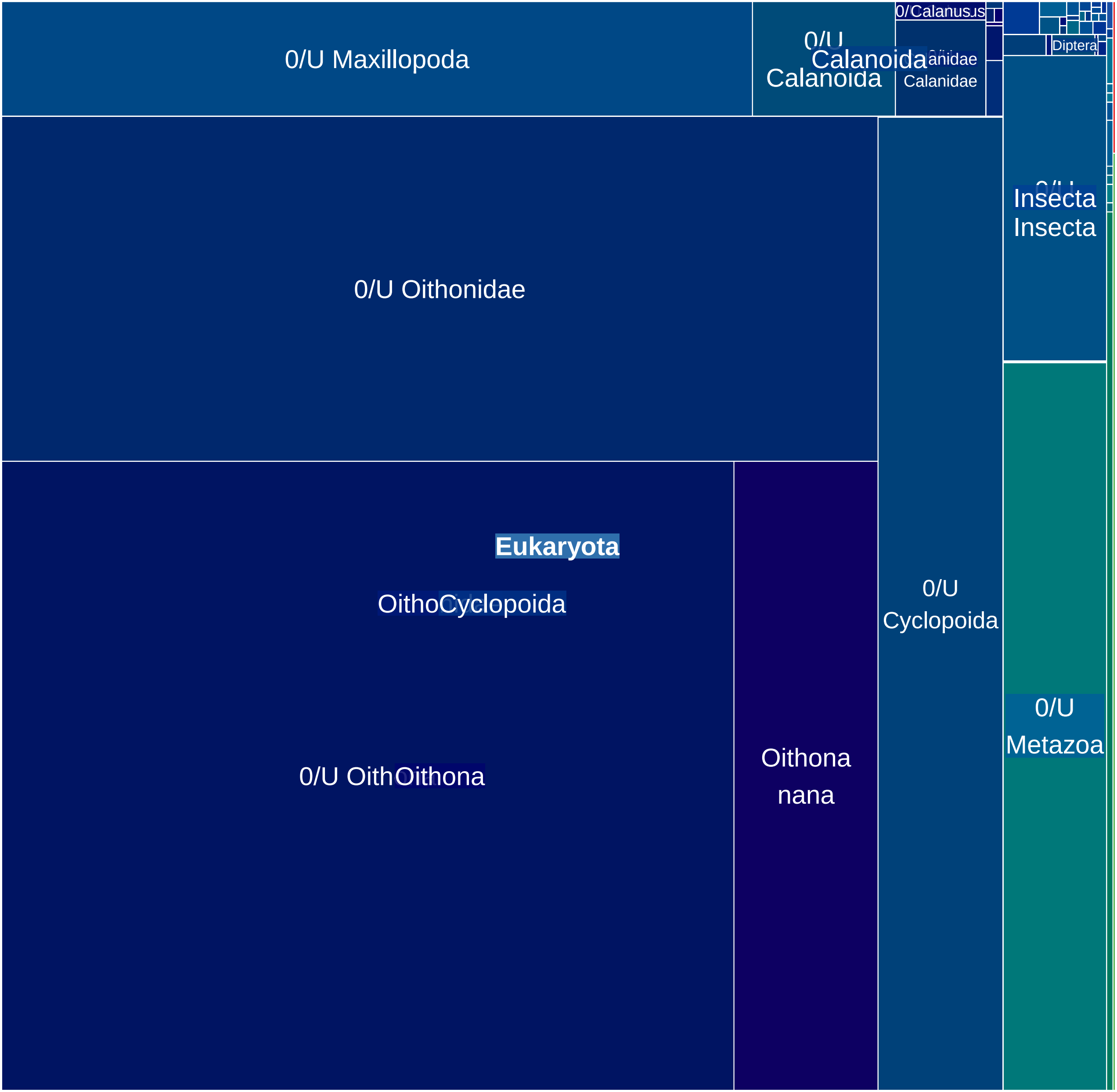

MGT−v1_61 : 24539 unigenes, 4353 (17.74%) taxonomically assigned

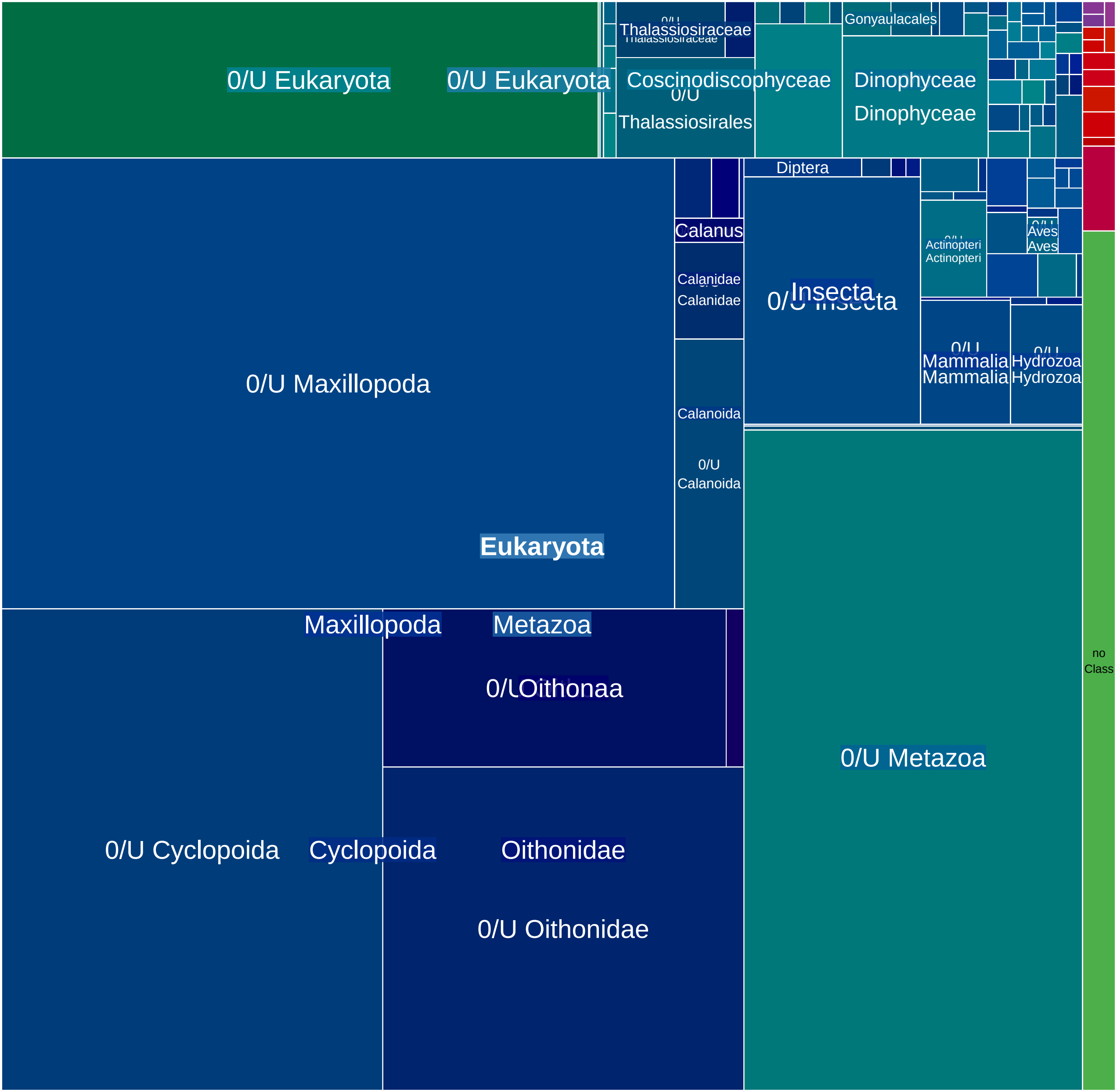

MGT−v1_62 : 24118 unigenes, 2159 (8.95%) taxonomically assigned

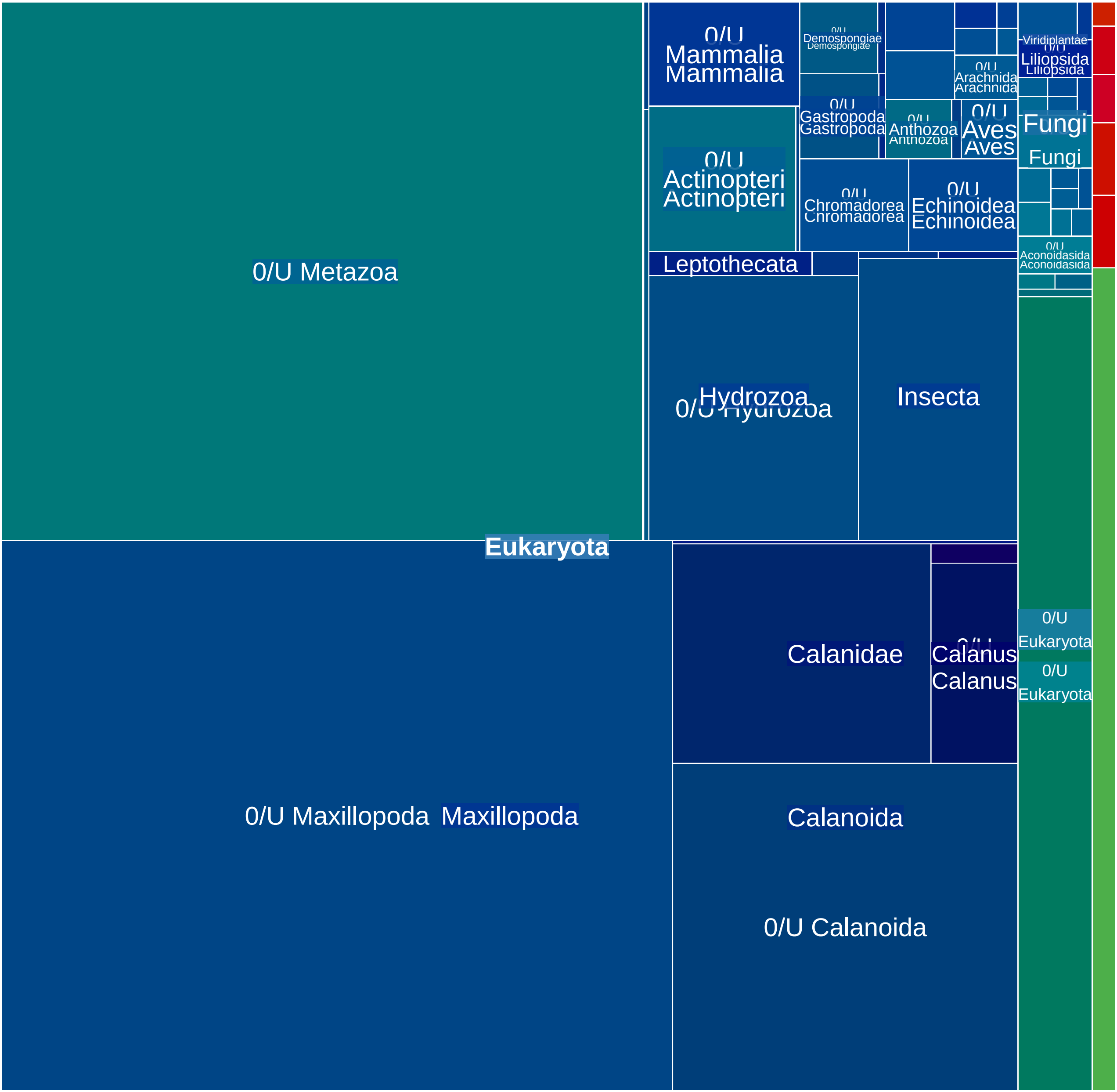

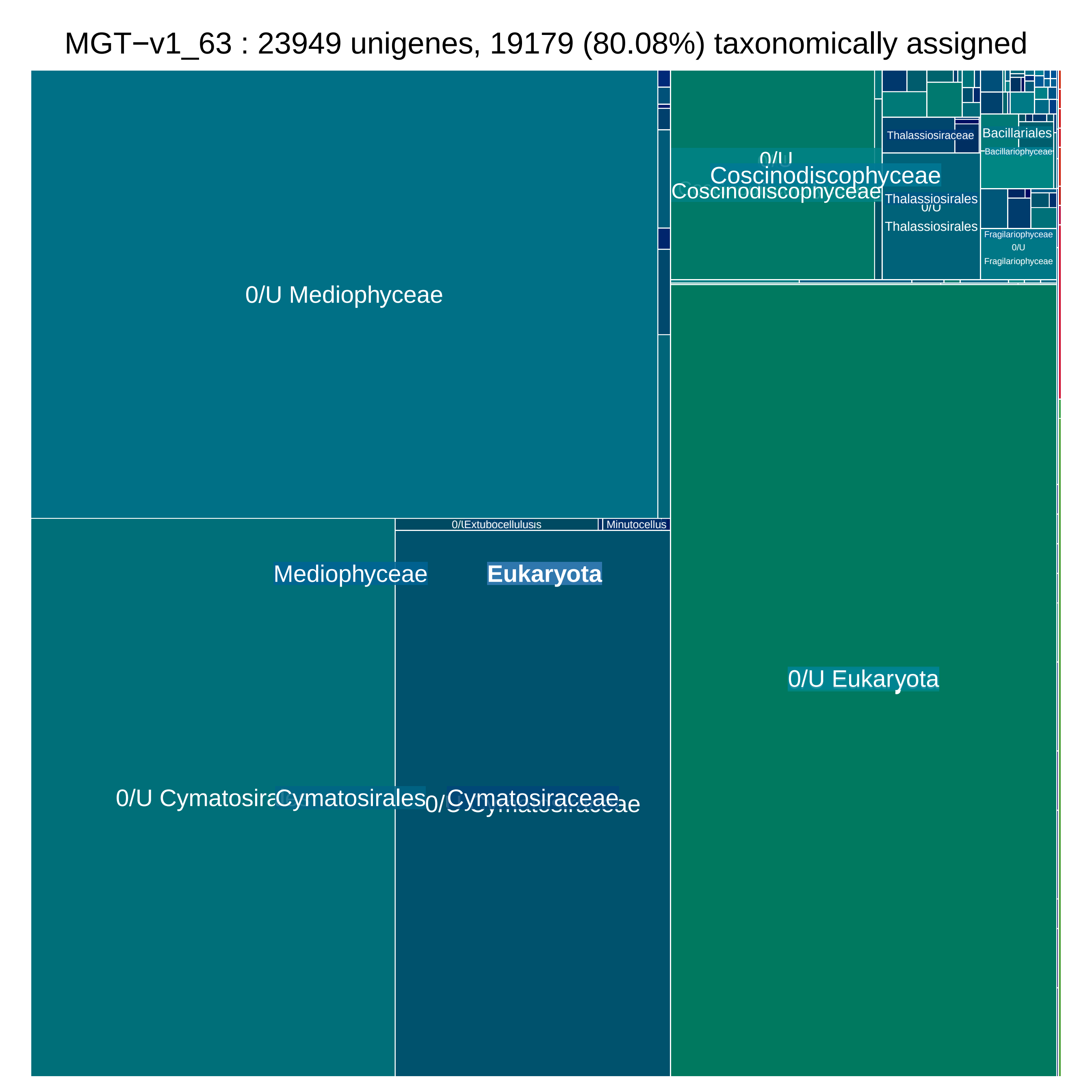

MGT−v1_63 : 23949 unigenes, 19179 (80.08%) taxonomically assigned
0/U Mediophyceae
Mediophyceae
Eukaryota
0/U Cymatosira
Cymatosirales
Cymatosiraceae
0/U Cymatosiraceae
Extubocellulus
Minutocellus
0/U Eukaryota
Coscinodiscophyceae
0/U Coscinodiscophyceae
Thalassiosiraceae
Thalassiosirales
0/U Thalassiosirales
Bacillariales
Bacillariophyceae
Fragilariophyceae
0/U Fragilariophyceae

### MGT−v1_64 : 23483 unigenes, 1577 (6.72%) taxonomically assigned

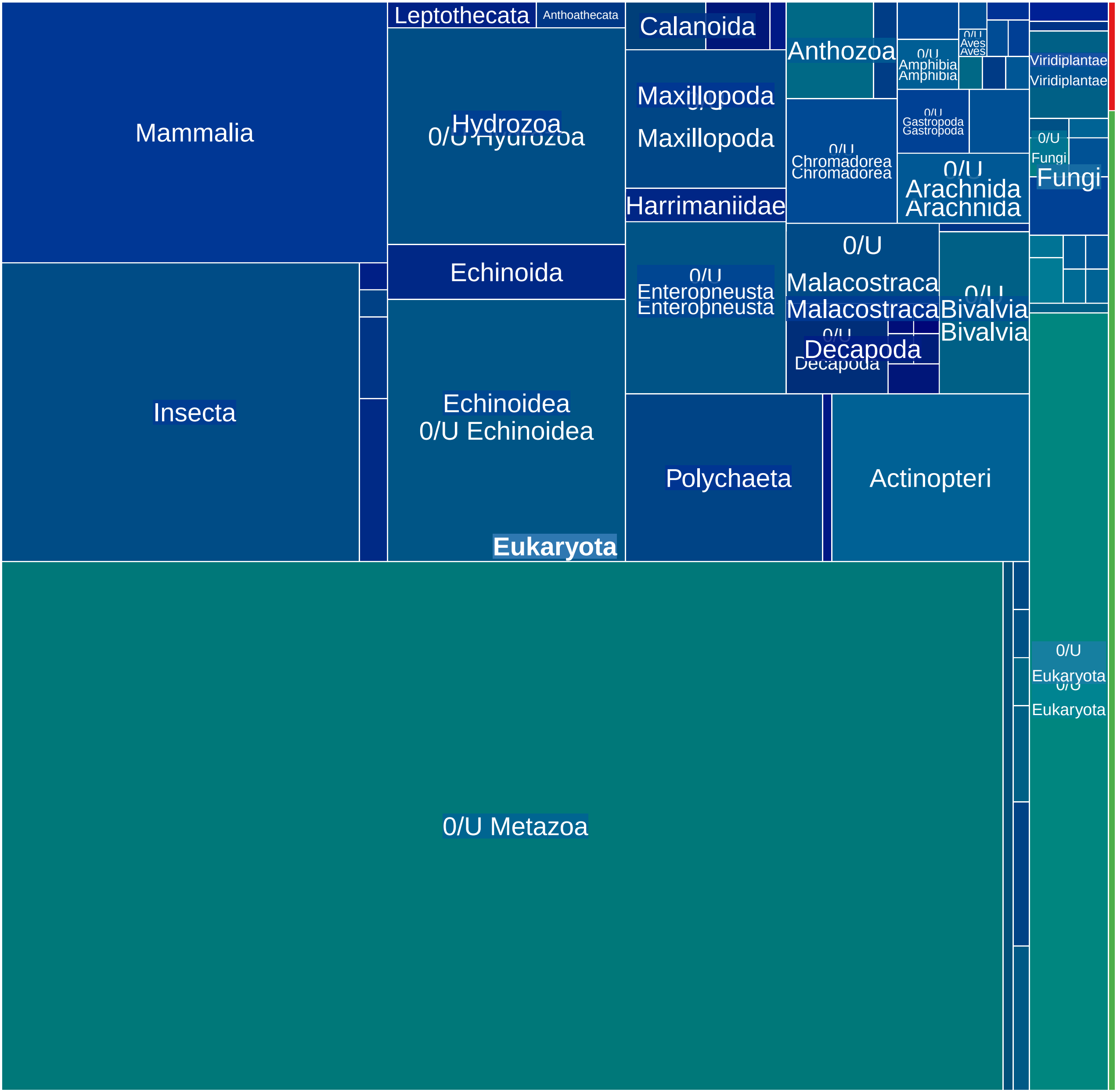

MGT−v1_65 : 23454 unigenes, 21987 (93.75%) taxonomically assigned

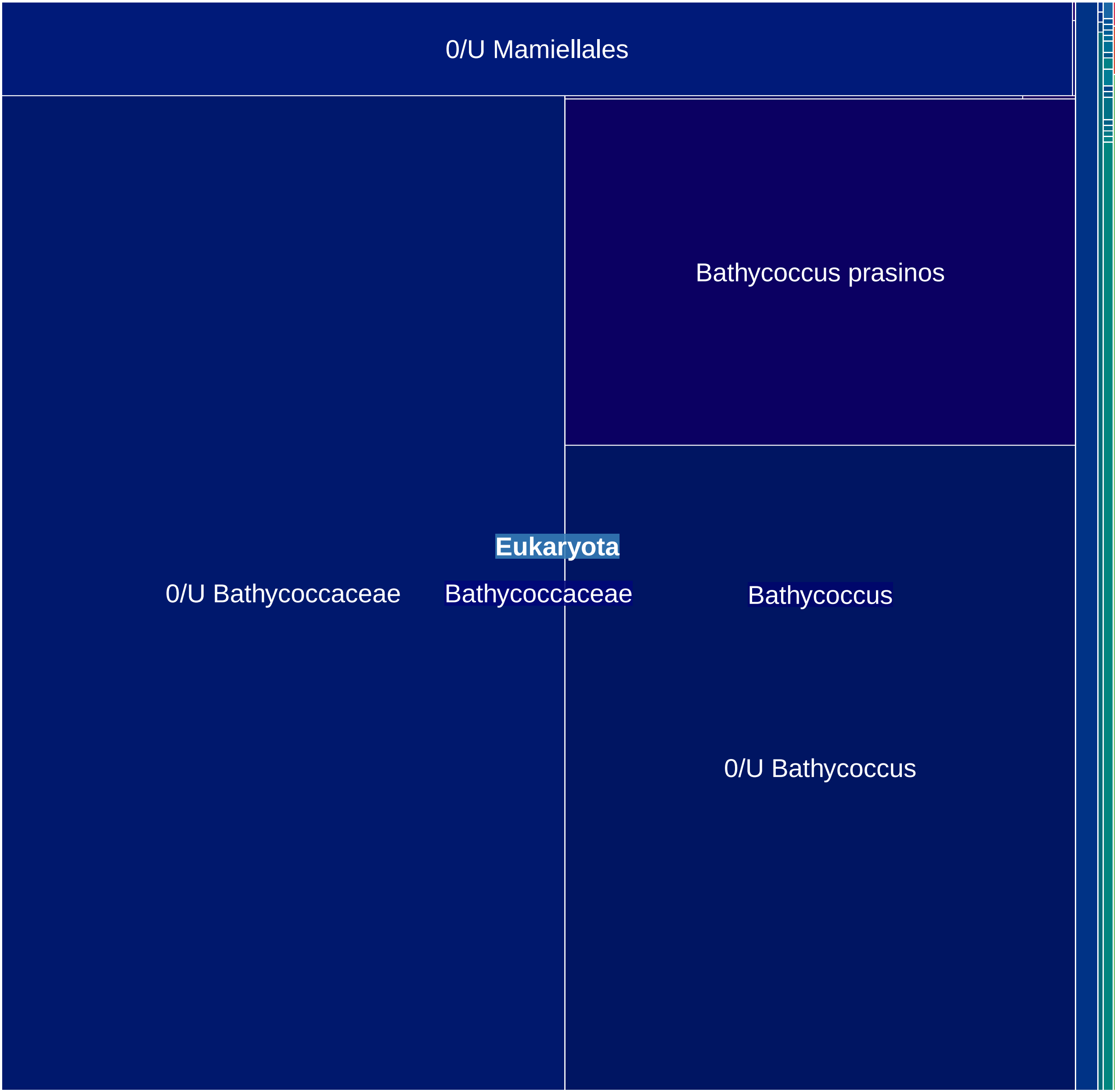

MGT−v1_66 : 23371 unigenes, 17172 (73.48%) taxonomically assigned

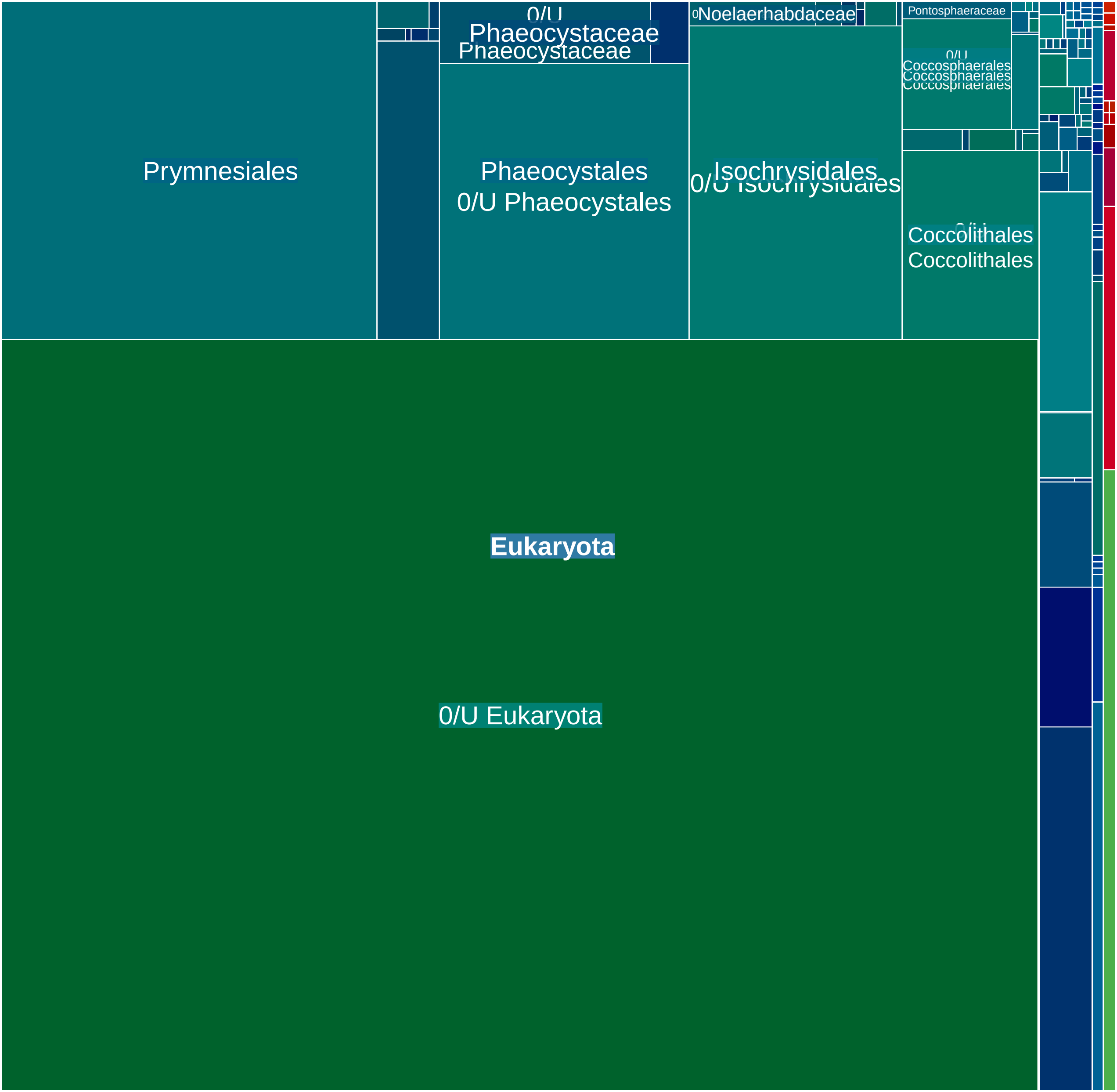

MGT−v1_67 : 23357 unigenes, 18717 (80.13%) taxonomically assigned

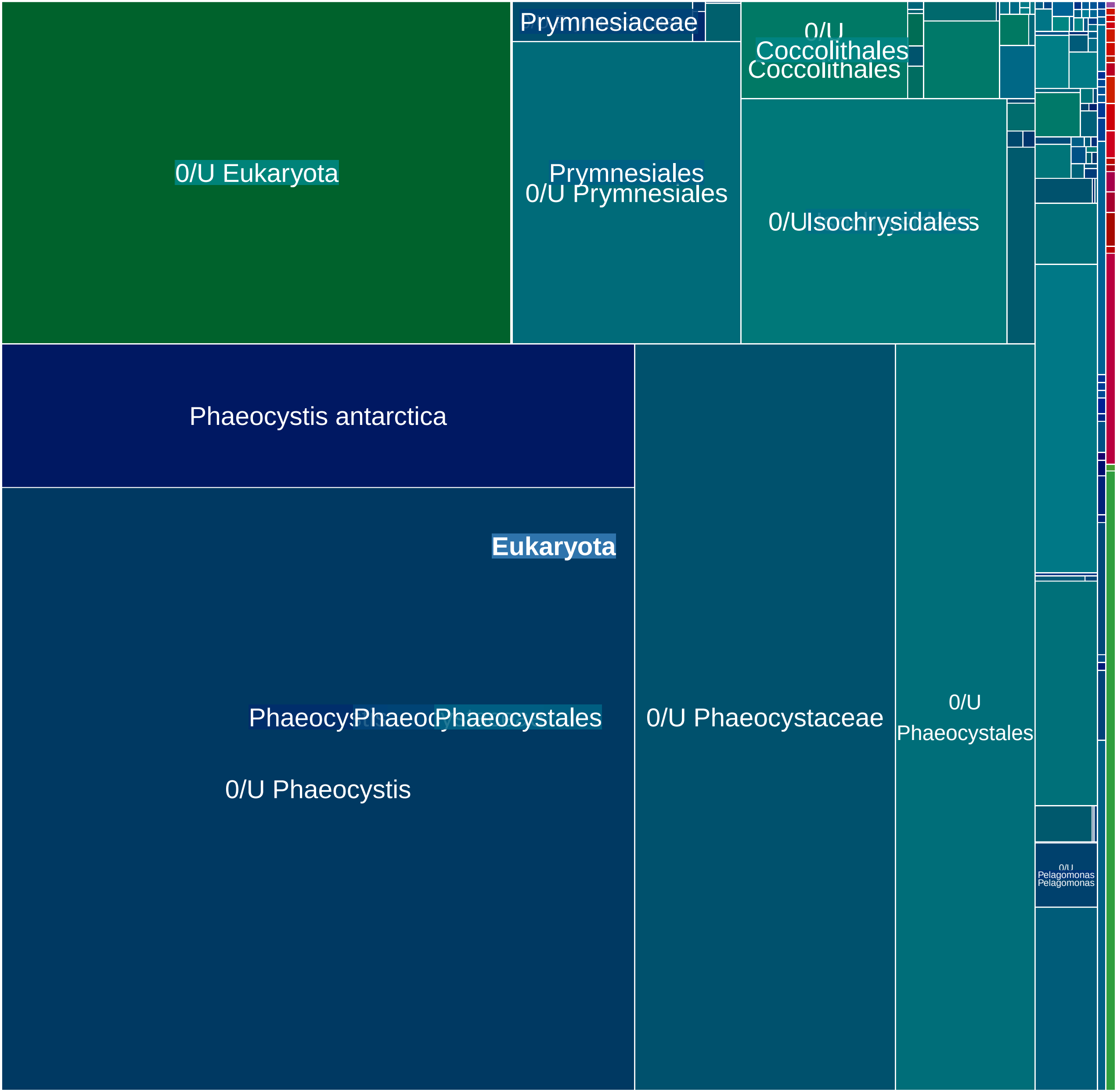

### MGT−v1_68 : 23241 unigenes, 5500 (23.67%) taxonomically assigned

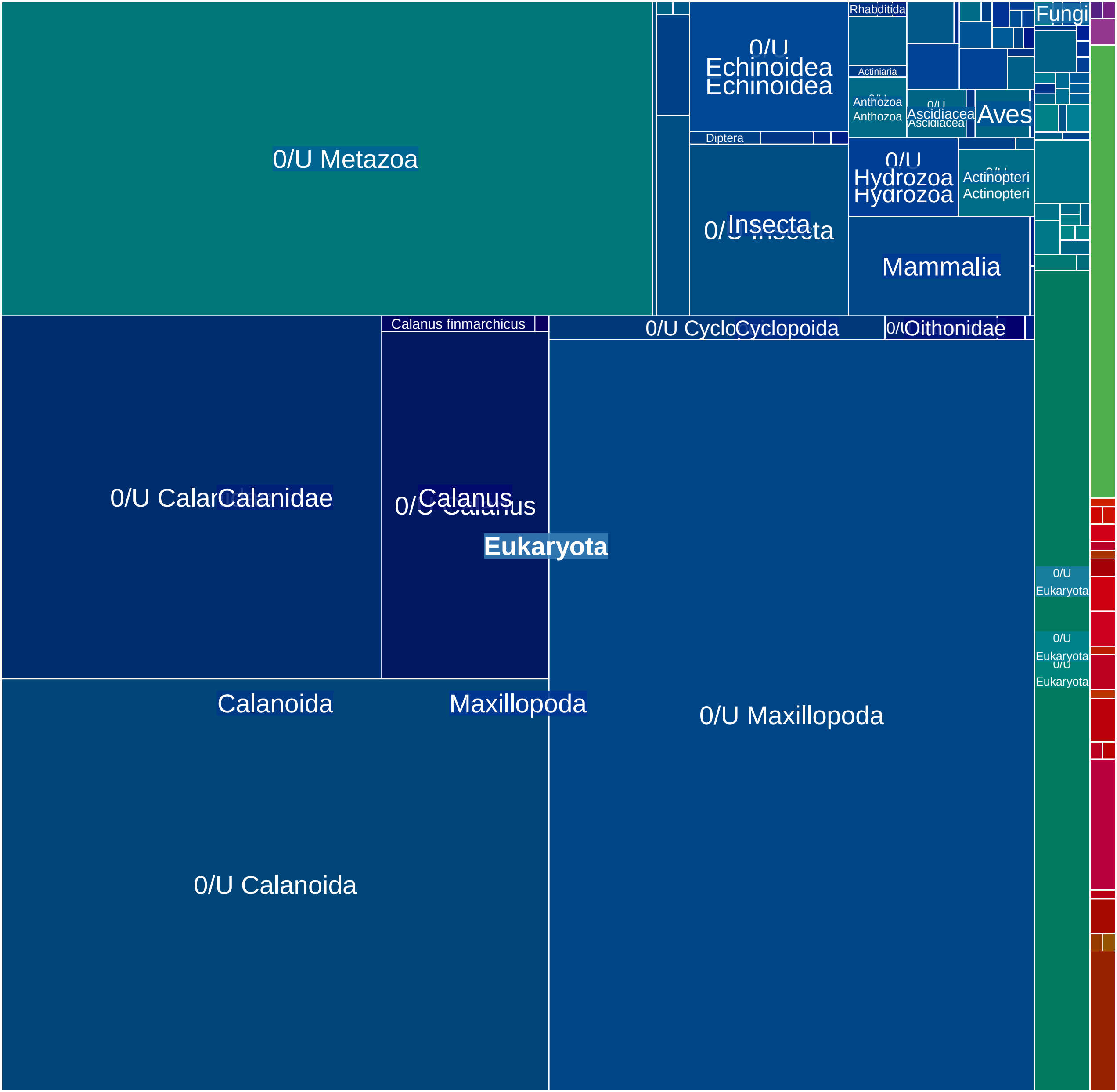

MGT−v1_69 : 21901 unigenes, 15008 (68.53%) taxonomically assigned

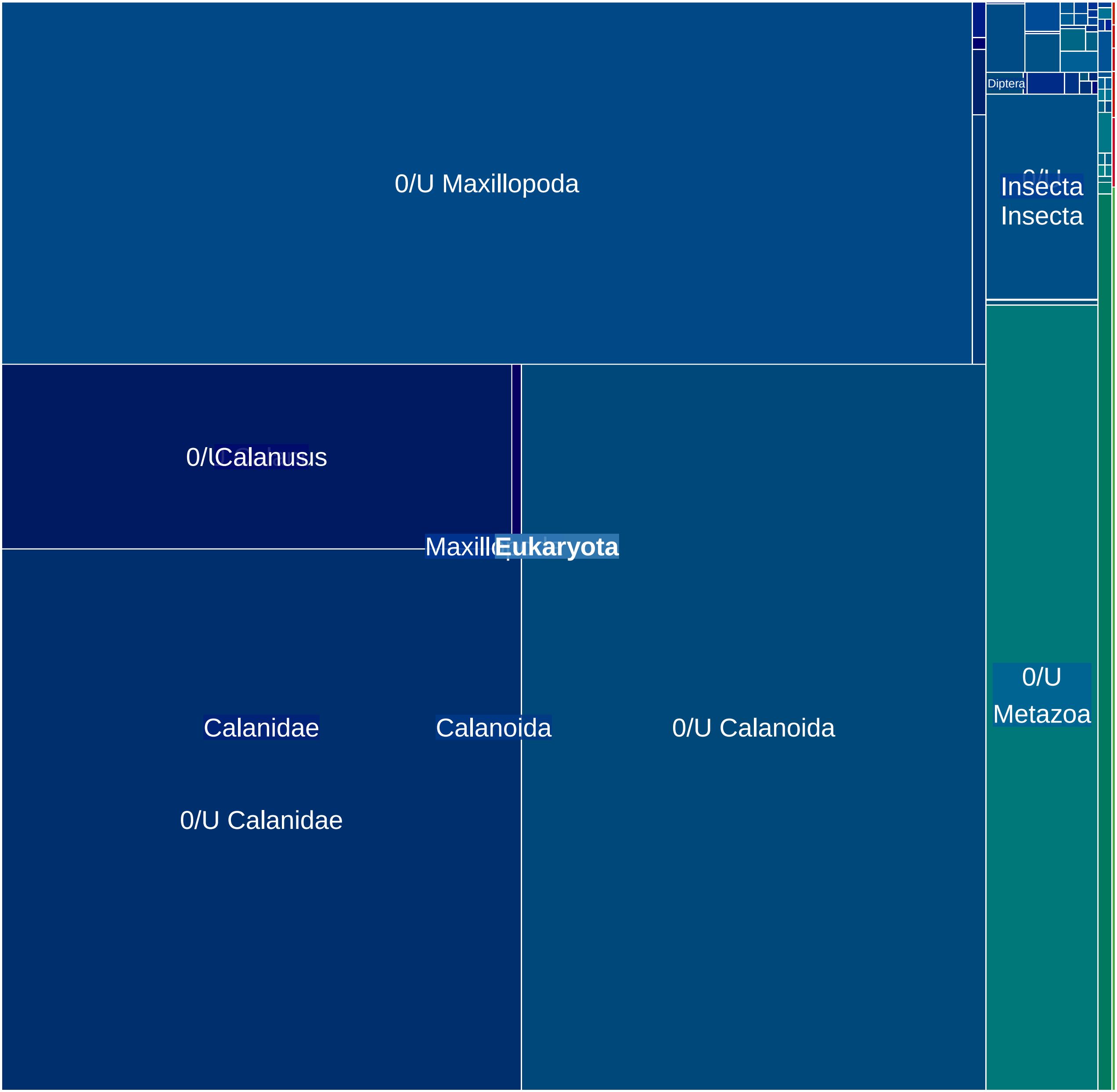

MGT−v1_70 : 21806 unigenes, 8234 (37.76%) taxonomically assigned

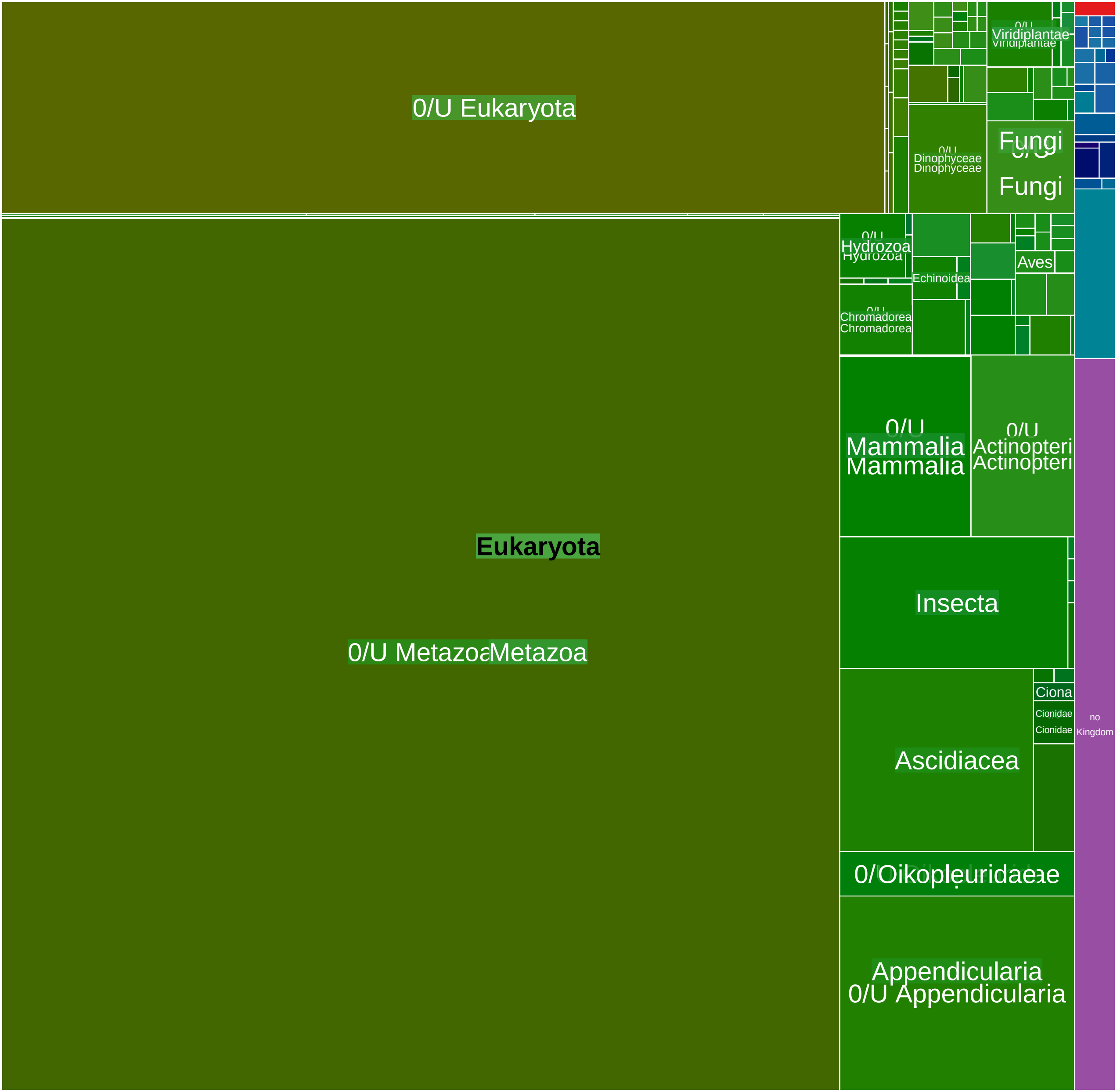

MGT−v1_71 : 21792 unigenes, 18591 (85.31%) taxonomically assigned

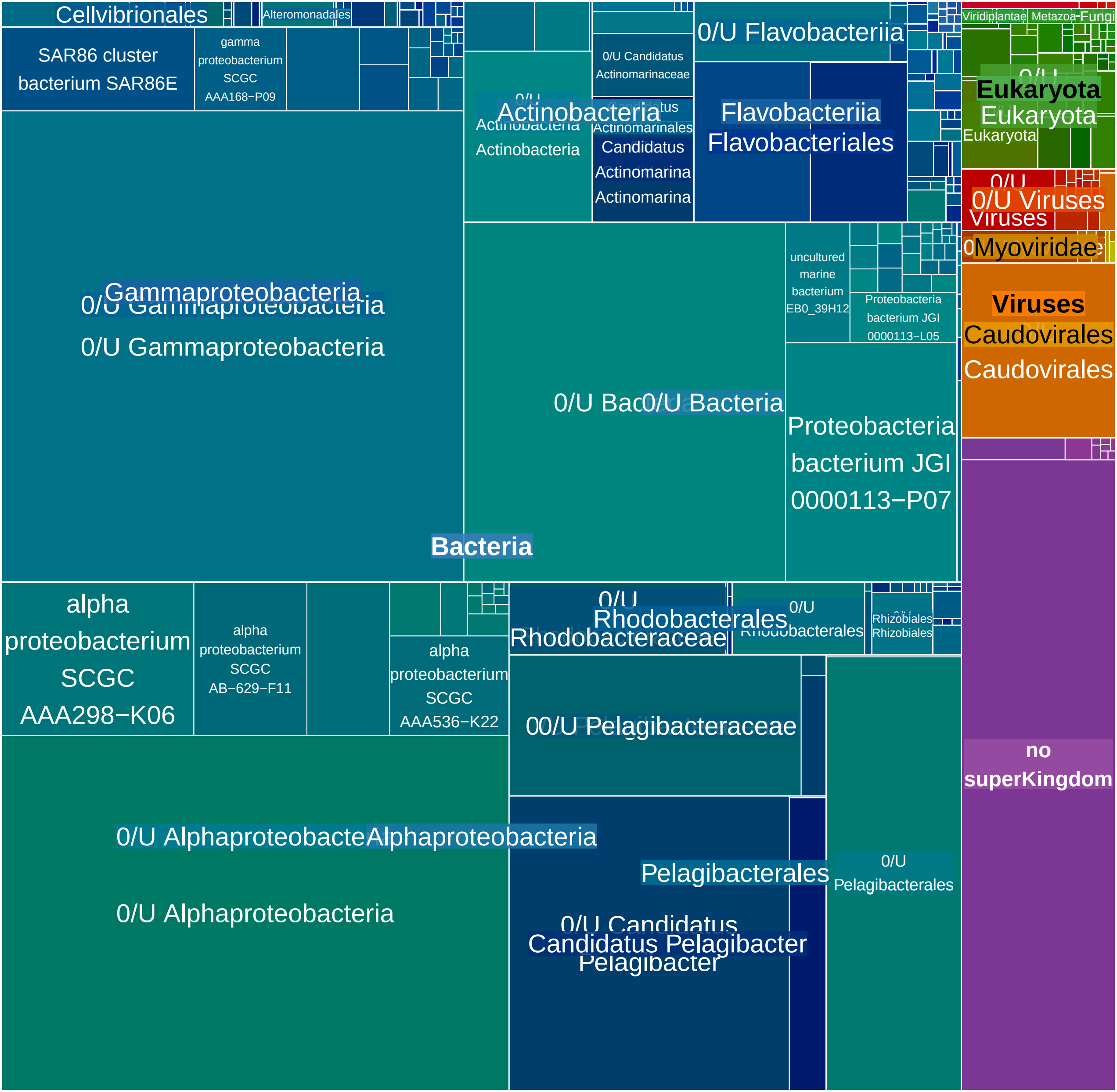

MGT−v1_72 : 21014 unigenes, 9930 (47.25%) taxonomically assigned

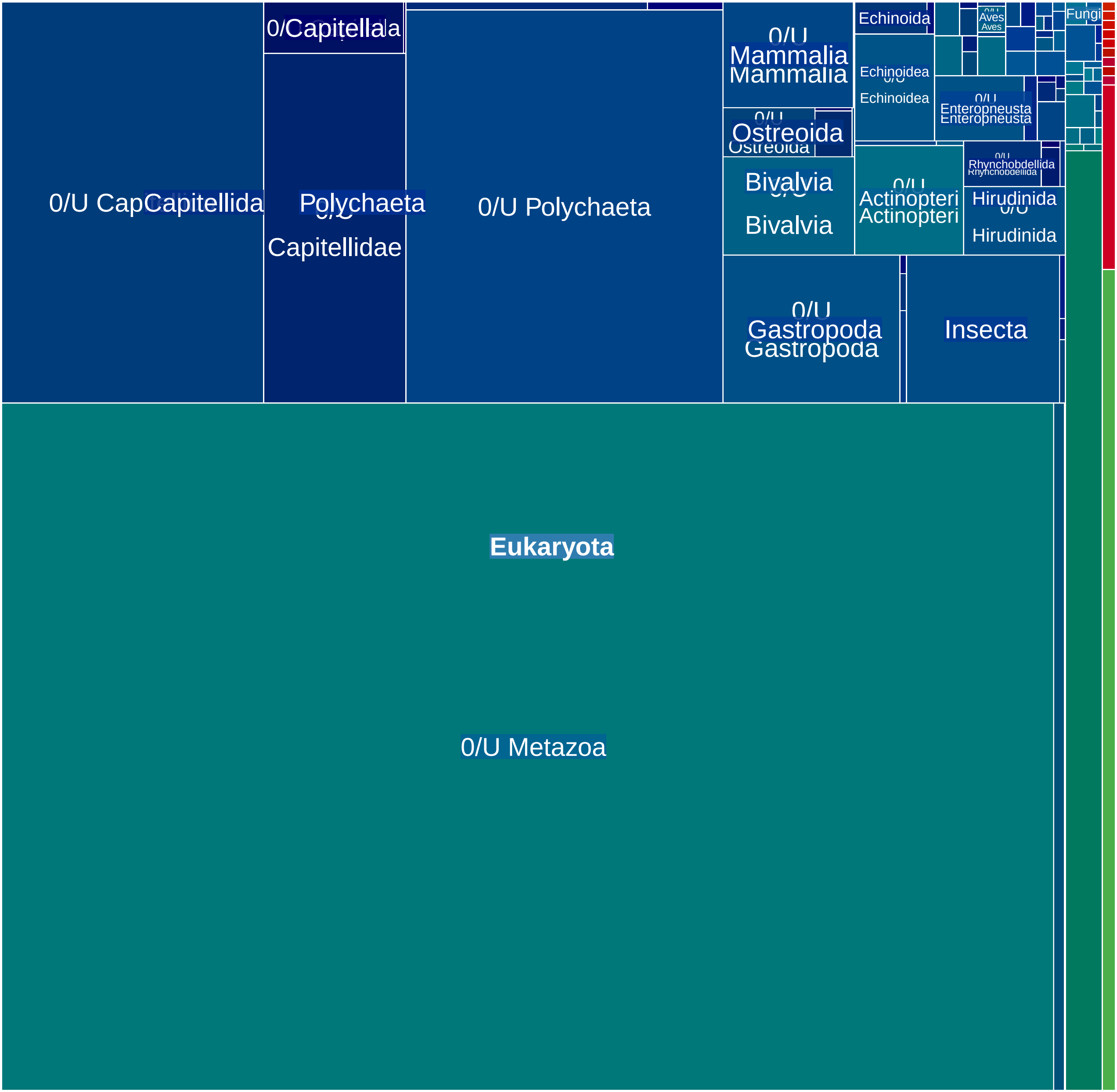

MGT−v1_73 : 20936 unigenes, 1600 (7.64%) taxonomically assigned

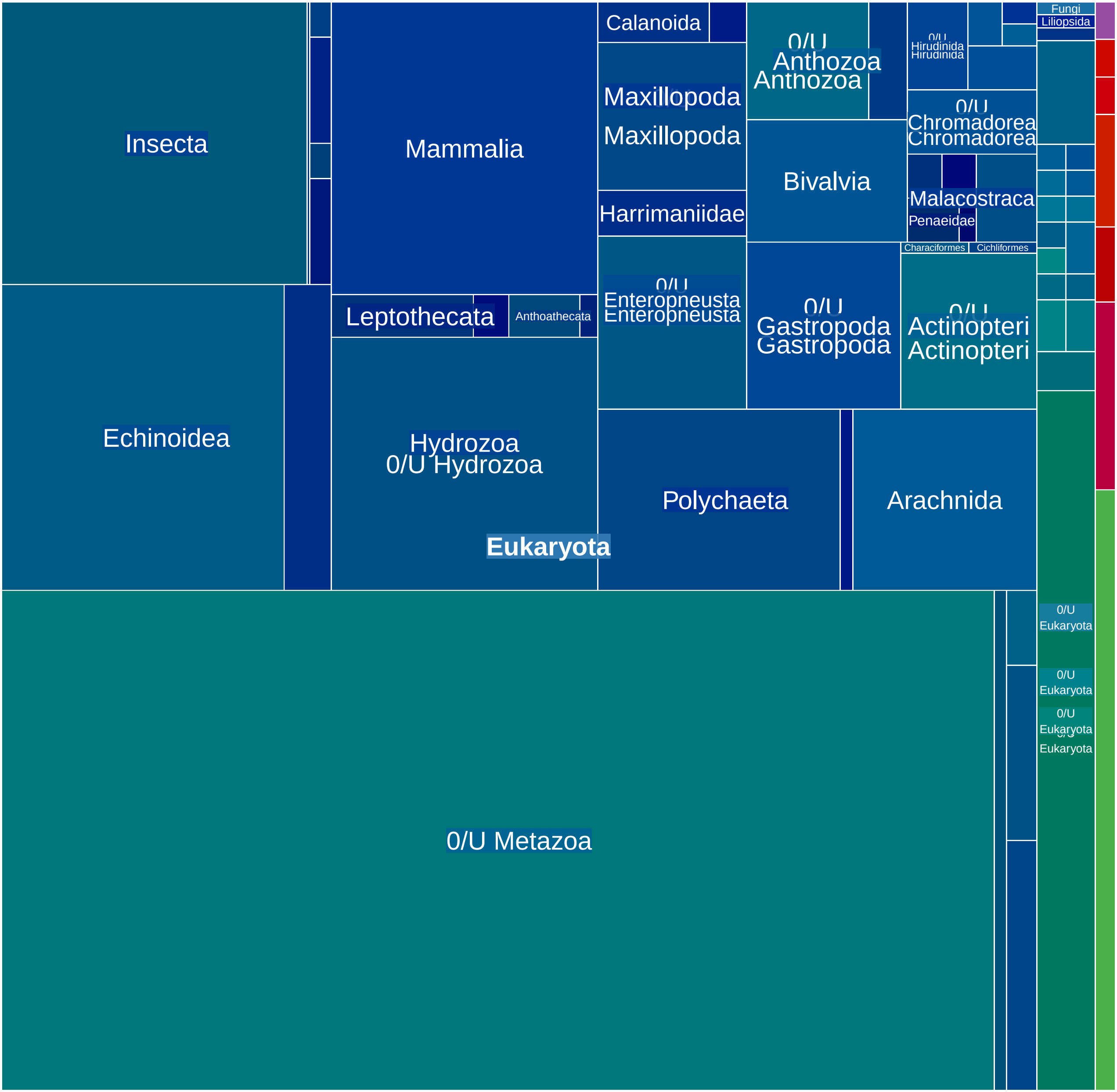

MGT−v1_74 : 20616 unigenes, 14249 (69.12%) taxonomically assigned

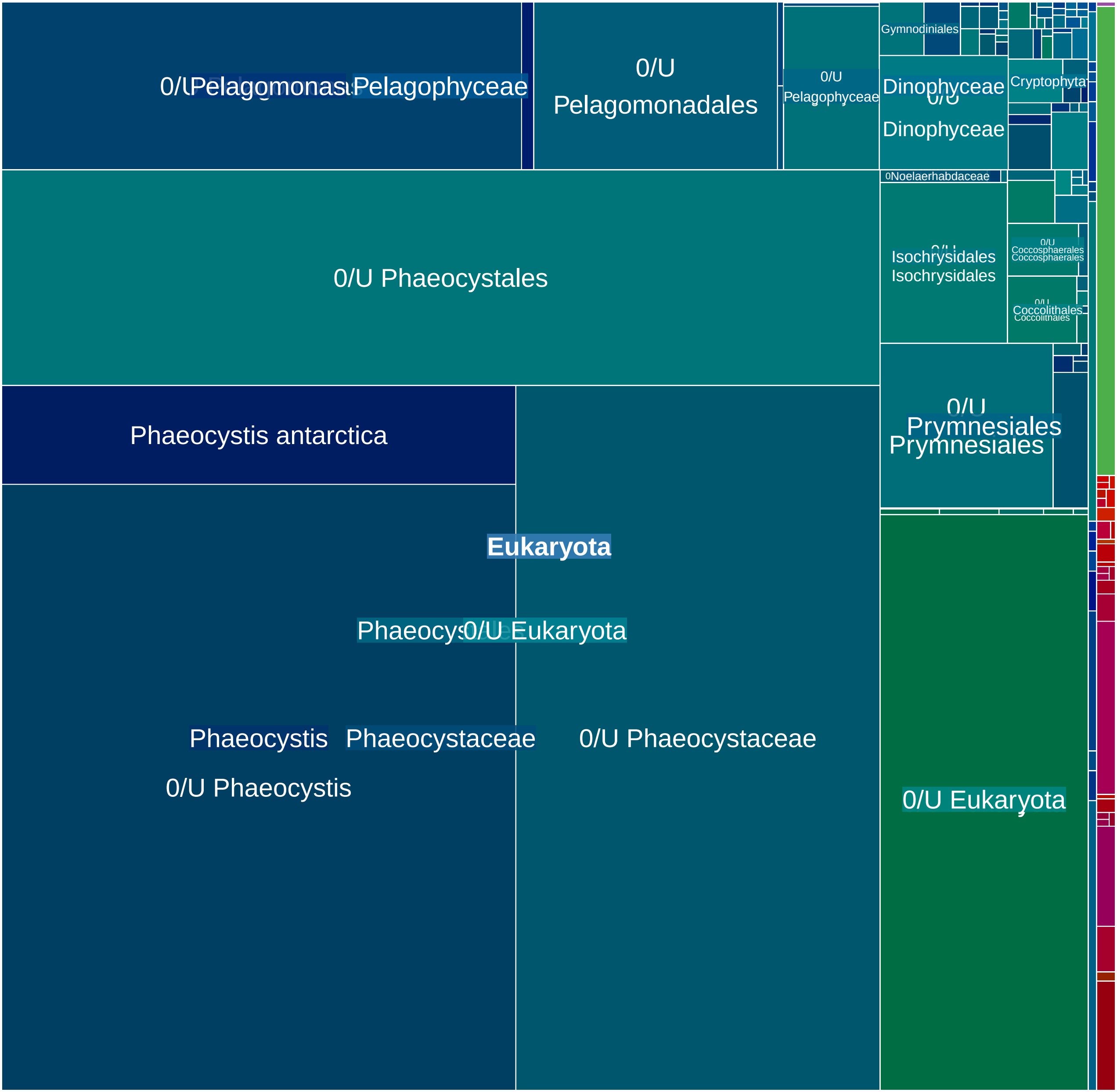

MGT−v1_75 : 20421 unigenes, 819 (4.01%) taxonomically assigned

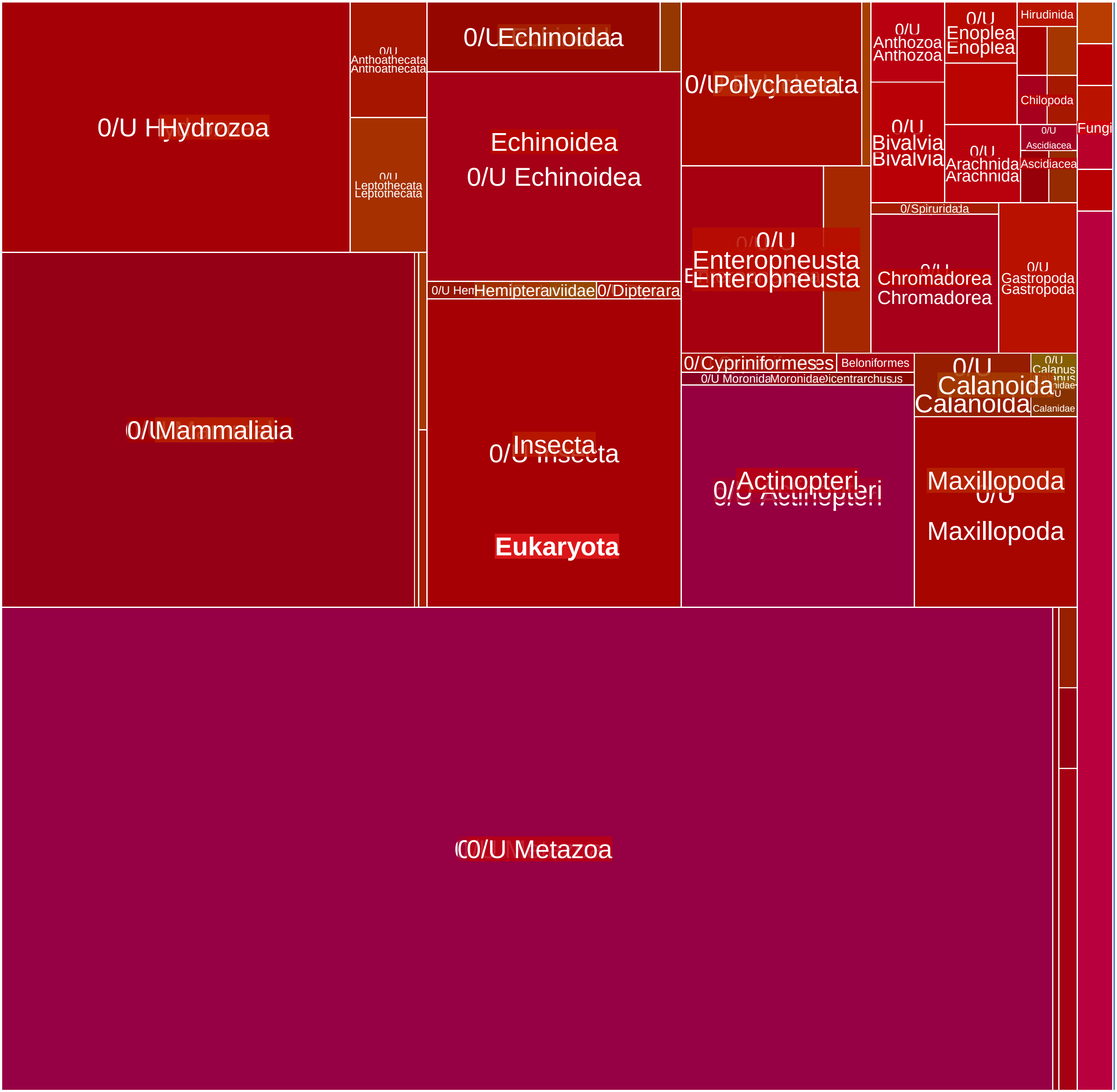

MGT−v1_76 : 20019 unigenes, 18143 (90.63%) taxonomically assigned

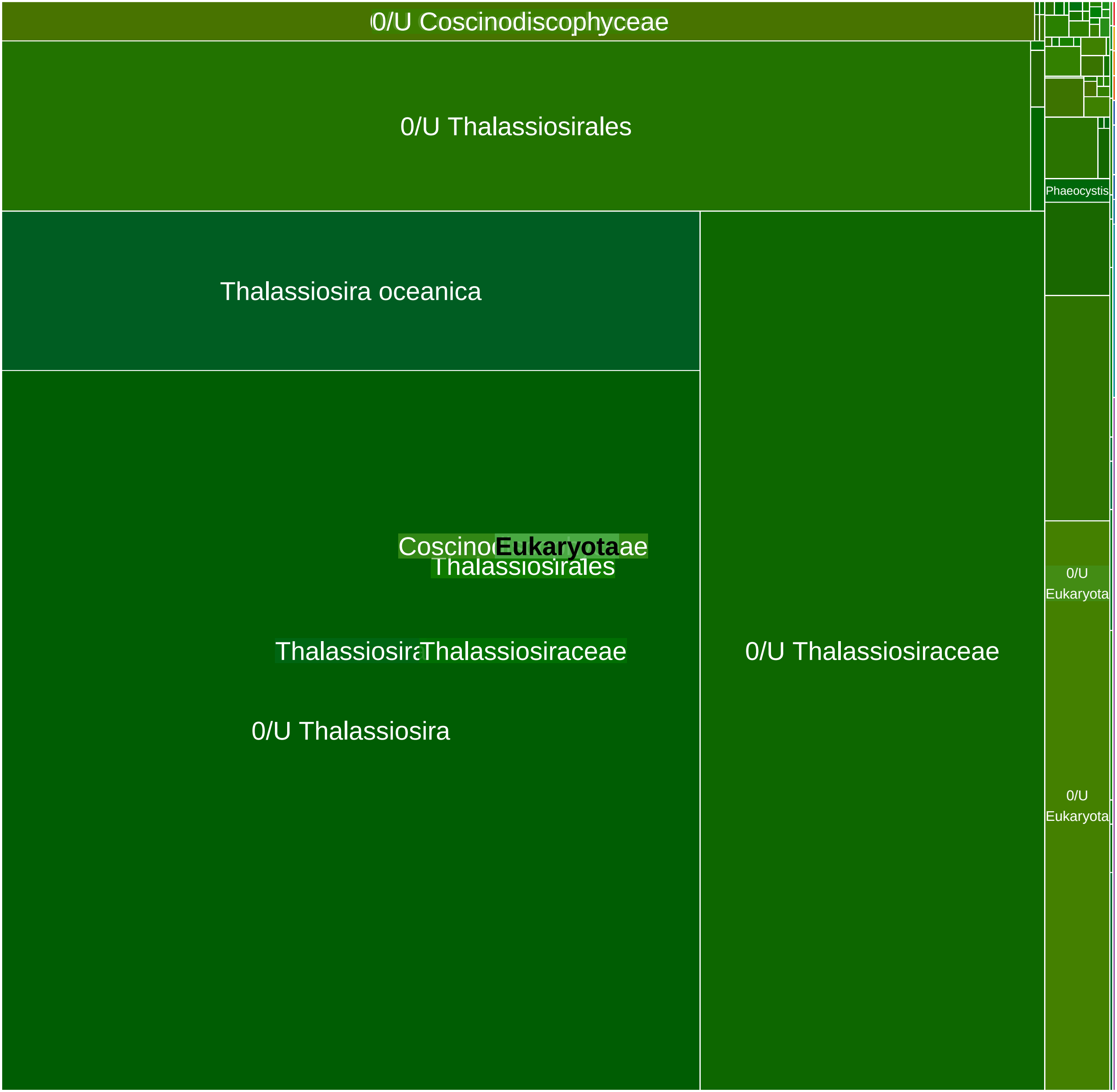

MGT−v1_77 : 19847 unigenes, 1372 (6.91%) taxonomically assigned

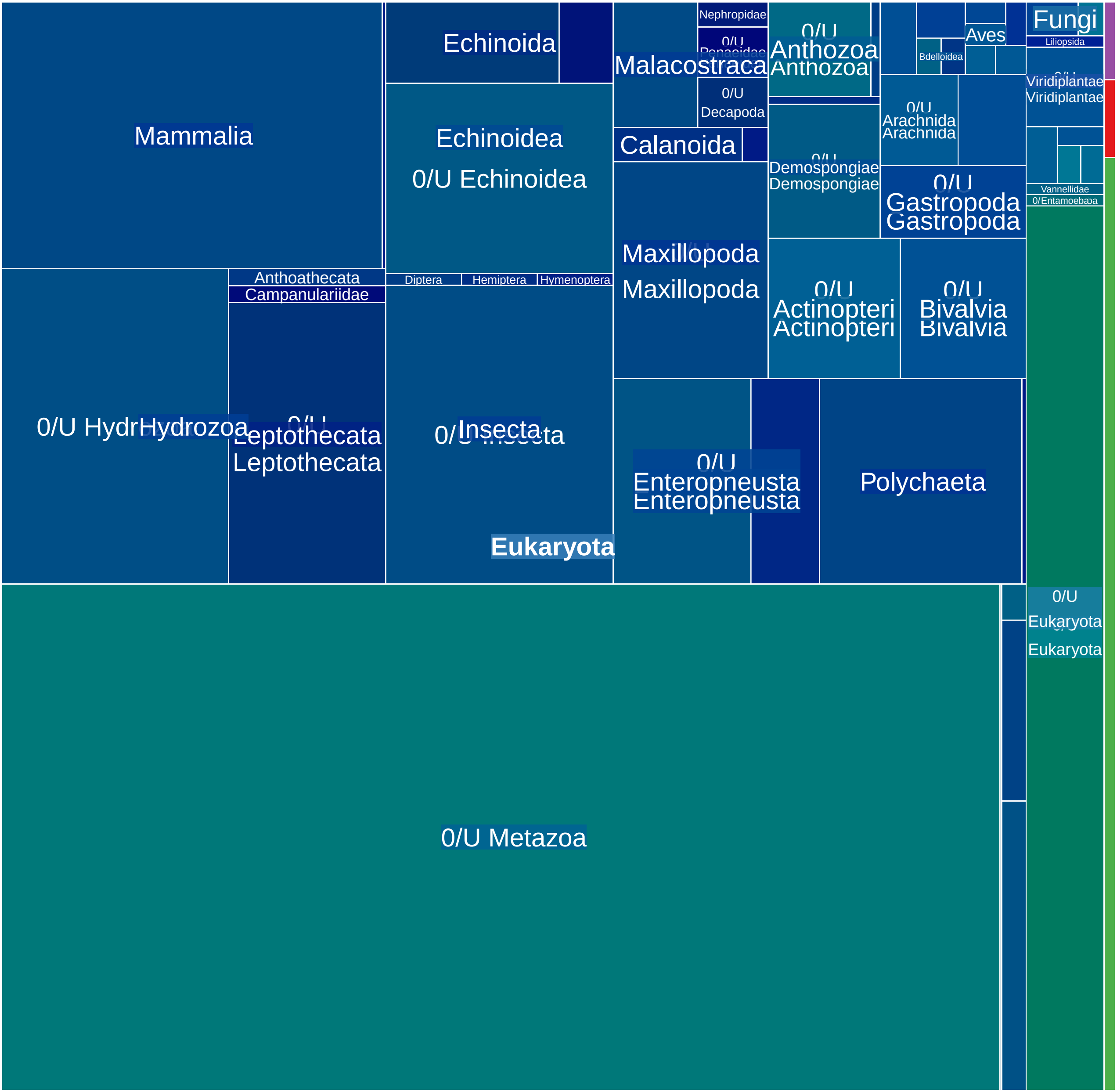

MGT−v1_78 : 19718 unigenes, 13905 (70.52%) taxonomically assigned

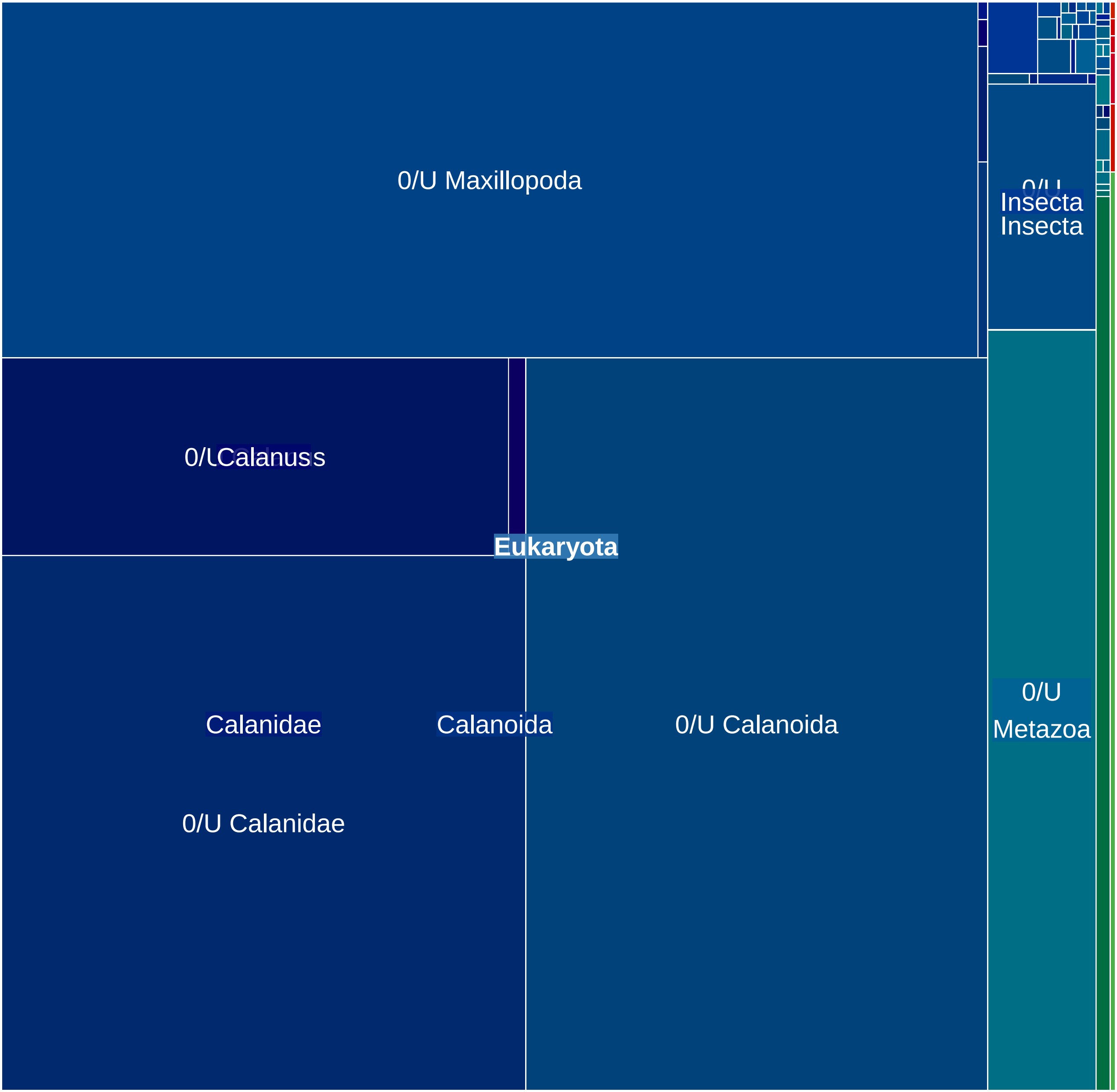

### MGT−v1_79 : 19705 unigenes, 1317 (6.68%) taxonomically assigned

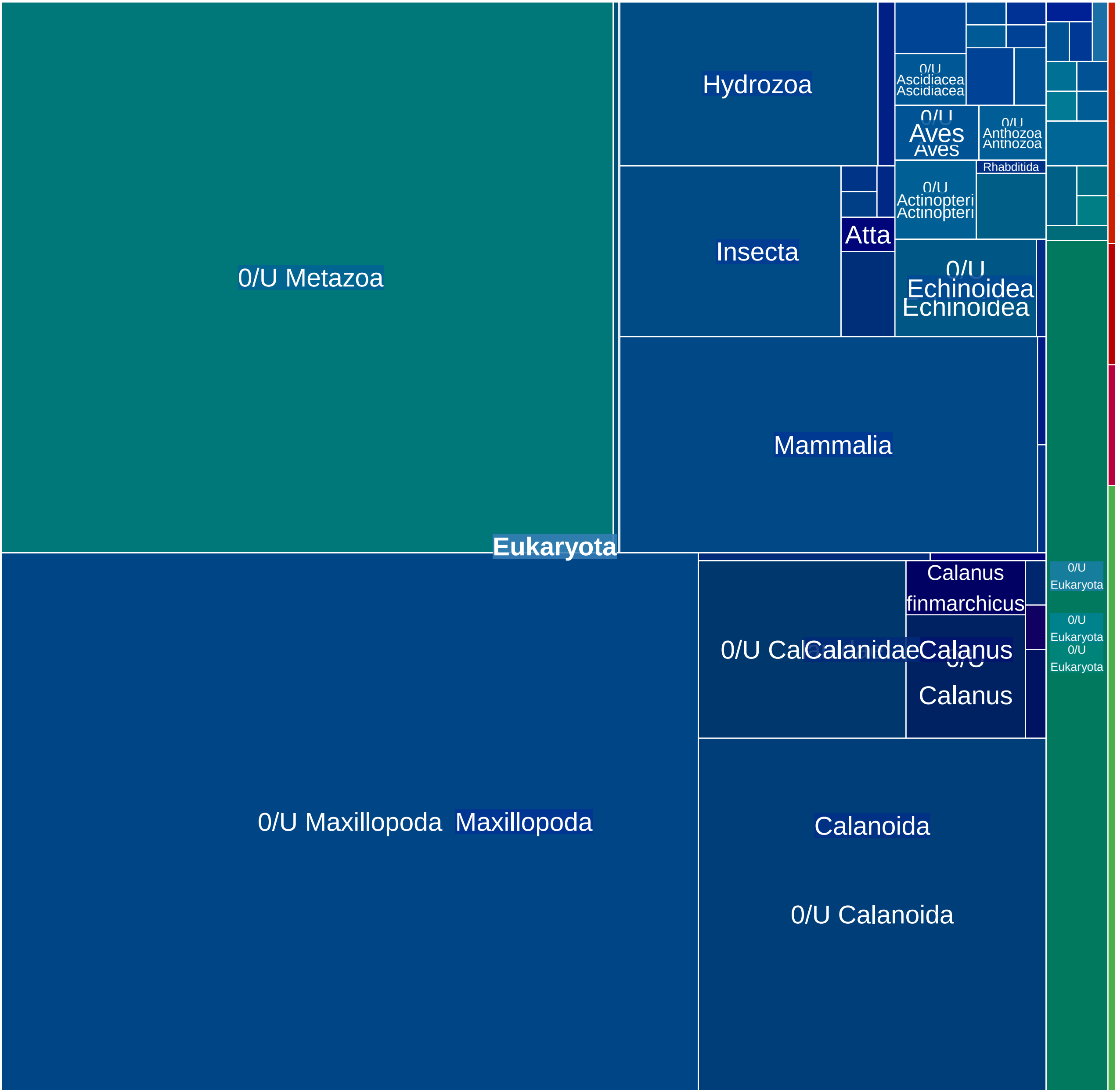

MGT−v1_80 : 18684 unigenes, 999 (5.35%) taxonomically assigned

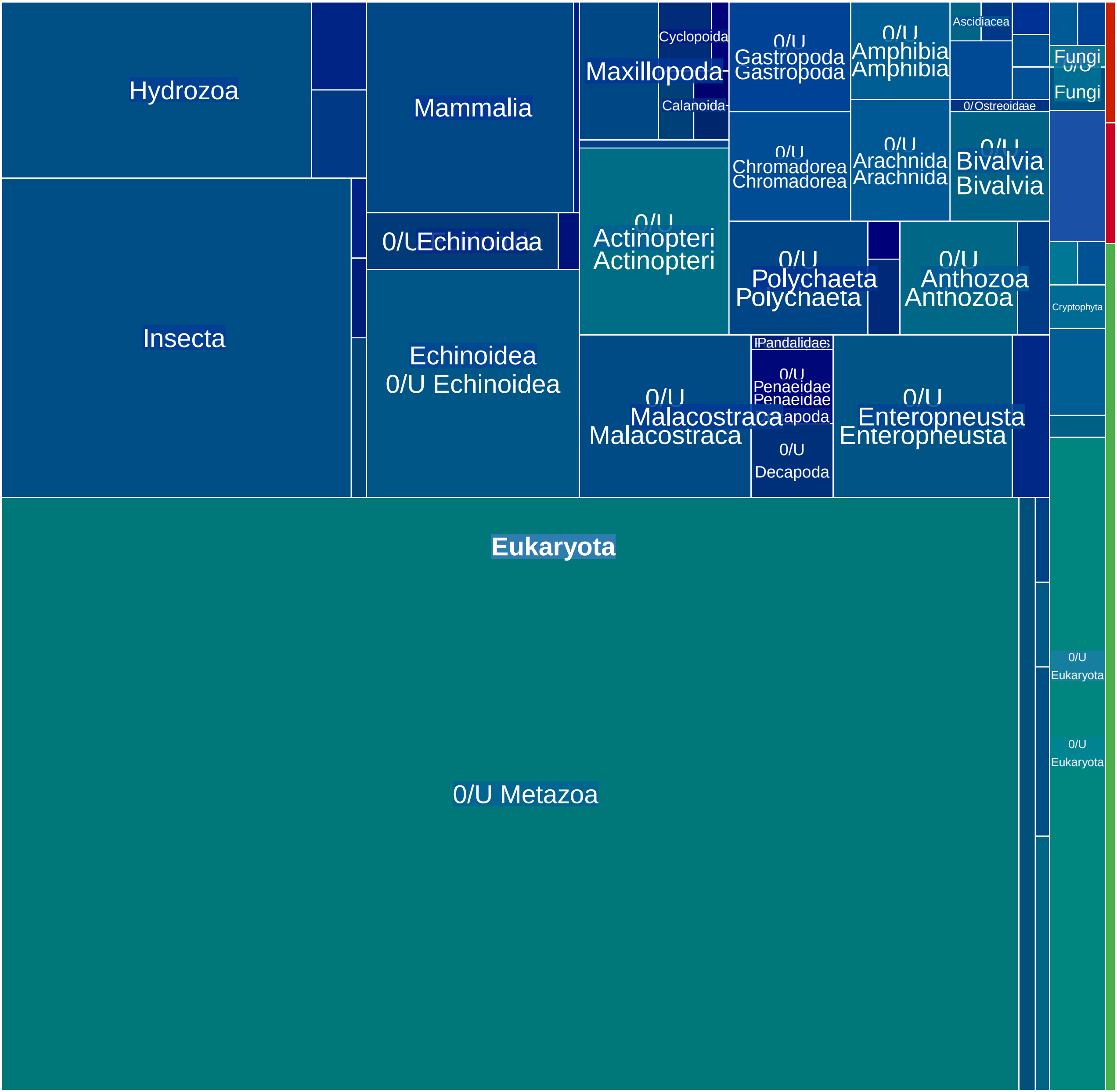

### MGT−v1_81 : 18570 unigenes, 9375 (50.48%) taxonomically assigned

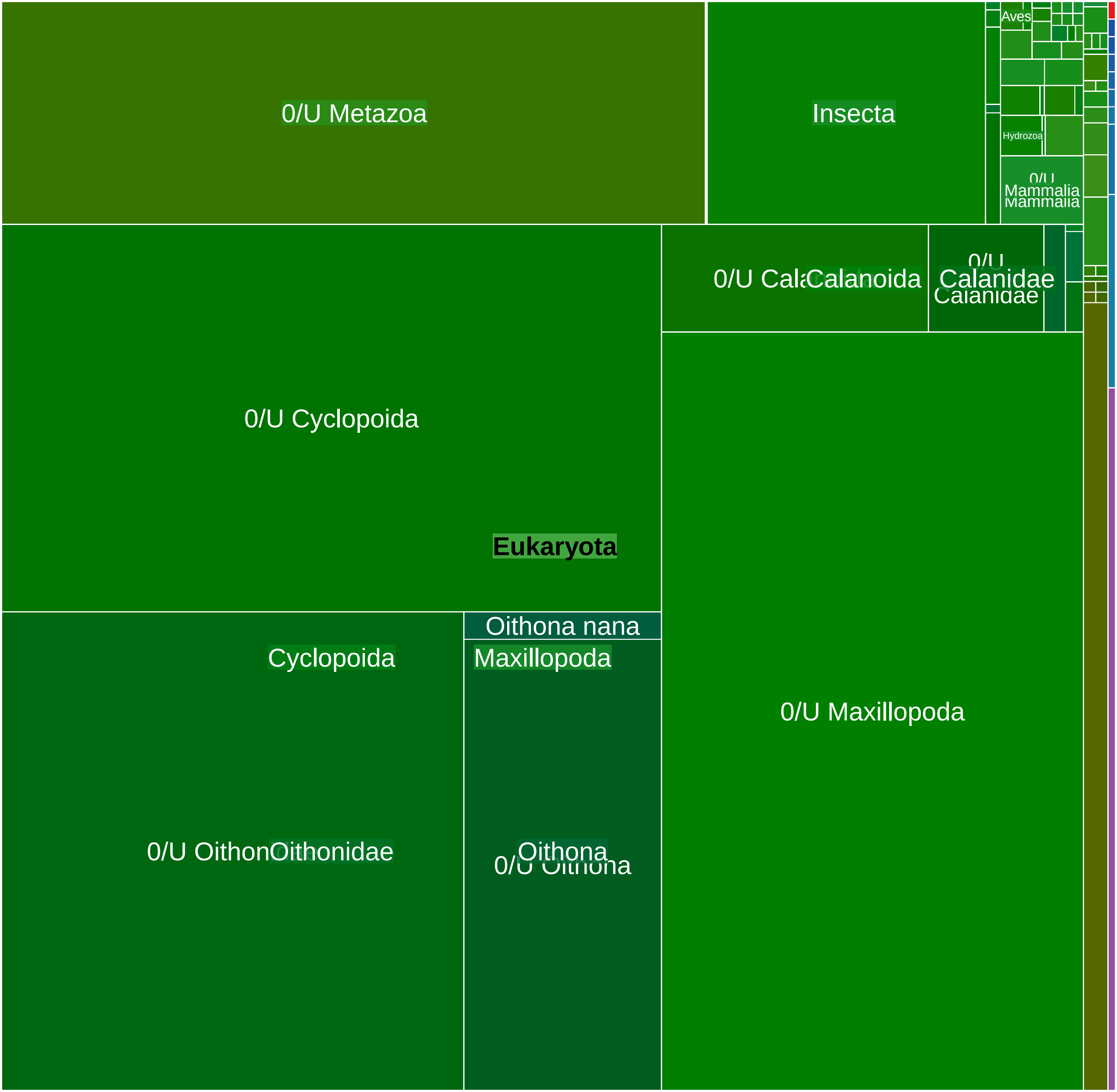

### MGT−v1_82 : 18520 unigenes, 1722 (9.30%) taxonomically assigned

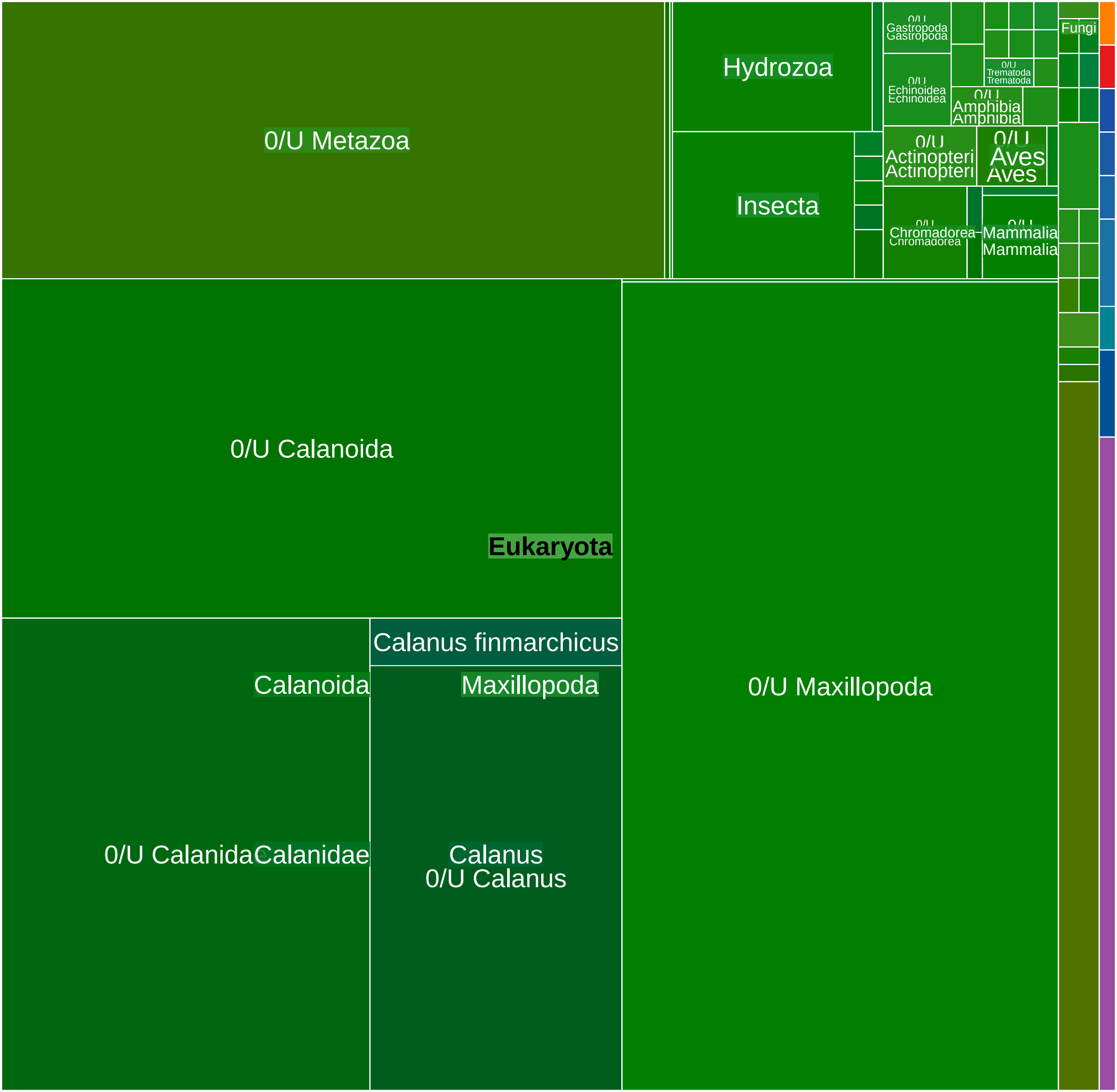

MGT−v1_83 : 18203 unigenes, 1900 (10.44%) taxonomically assigned

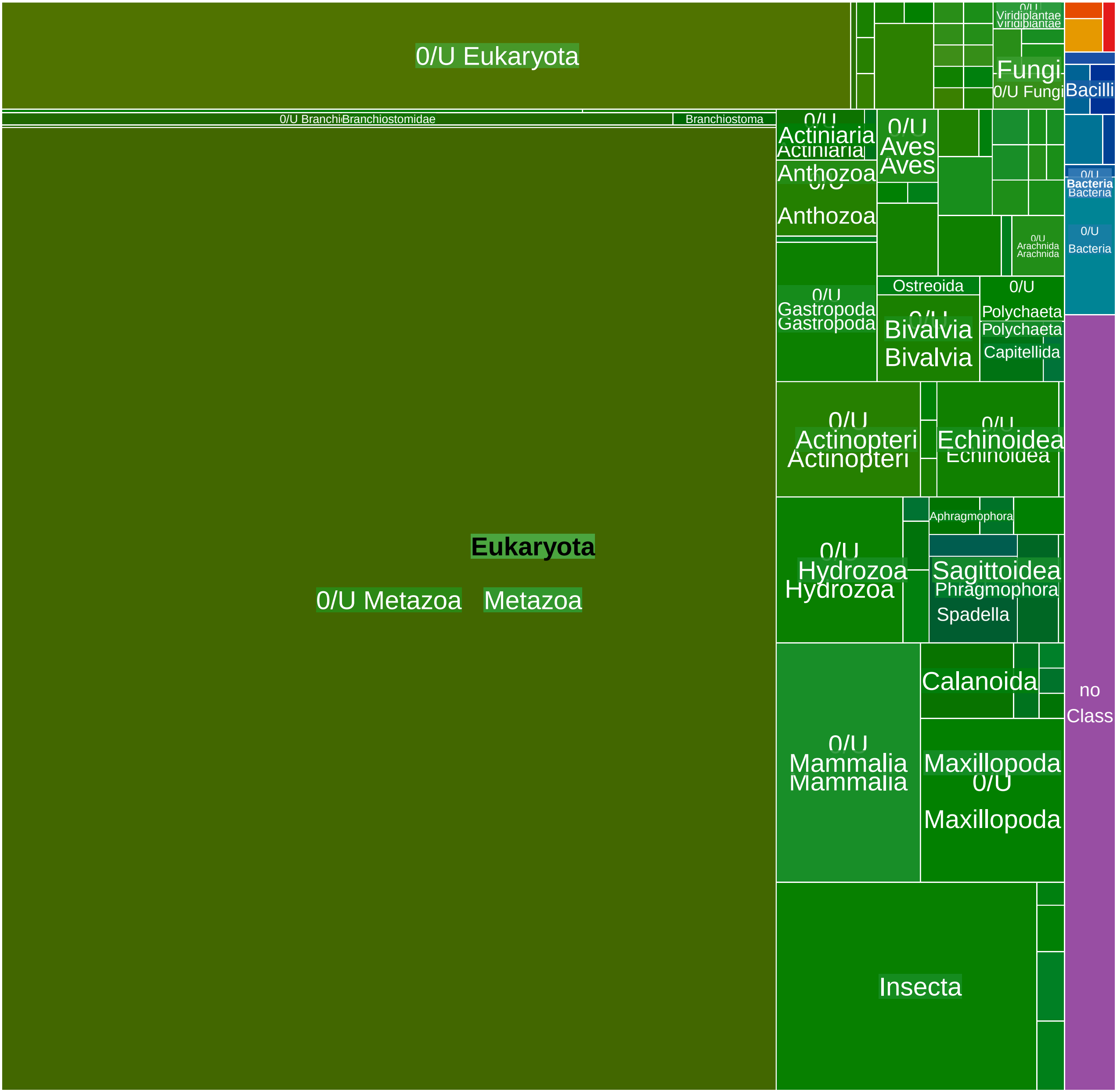

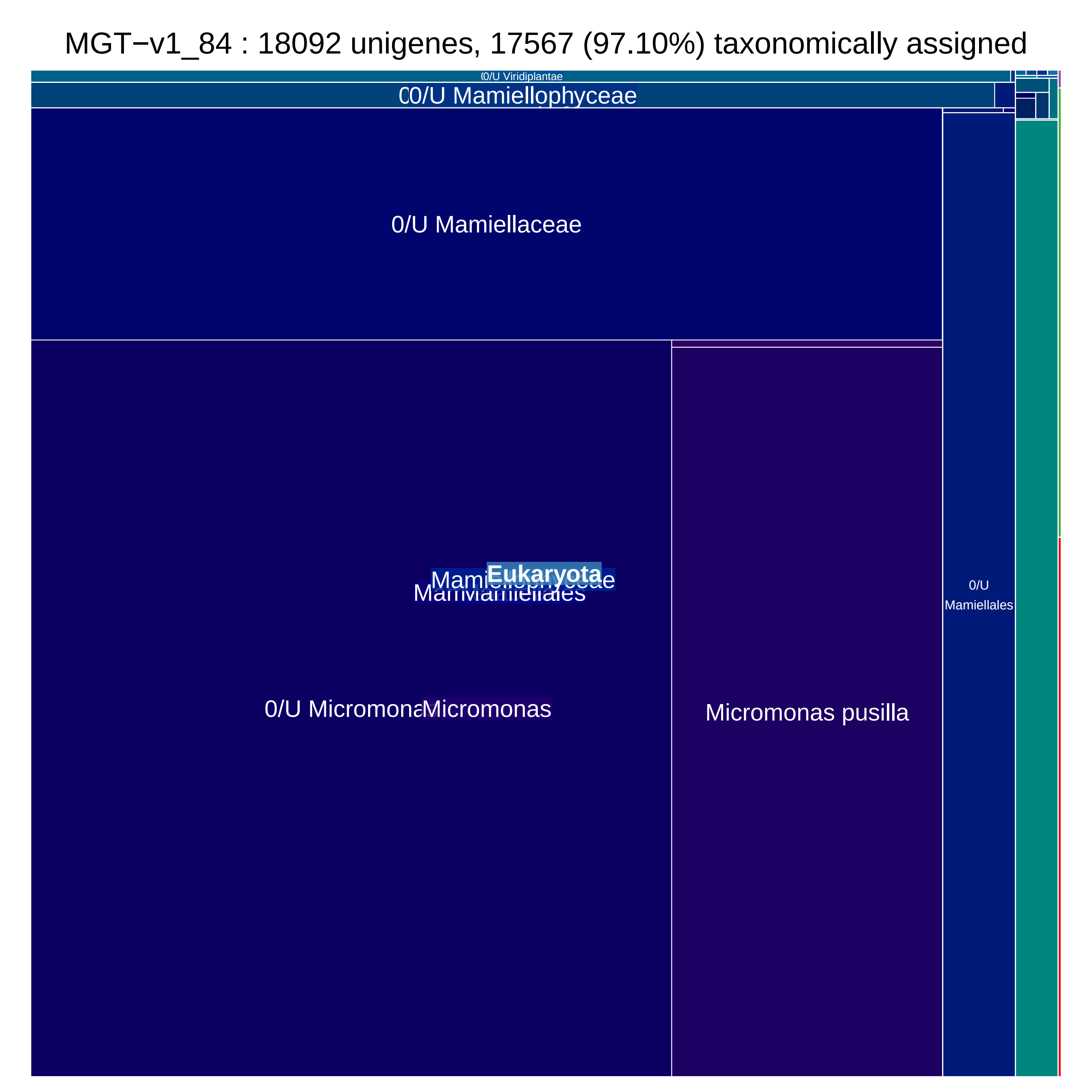
MGT−v1_84 : 18092 unigenes, 17567 (97.10%) taxonomically assigned
0/U Viridiplantae
0/U Mamiellophyceae
0/U Mamiellaceae
Eukaryota
Mamiellophyceae
Mamiellales
0/U Micromonas
Micromonas
Micromonas pusilla
0/U Mamiellales

MGT−v1_85 : 17568 unigenes, 10797 (61.46%) taxonomically assigned

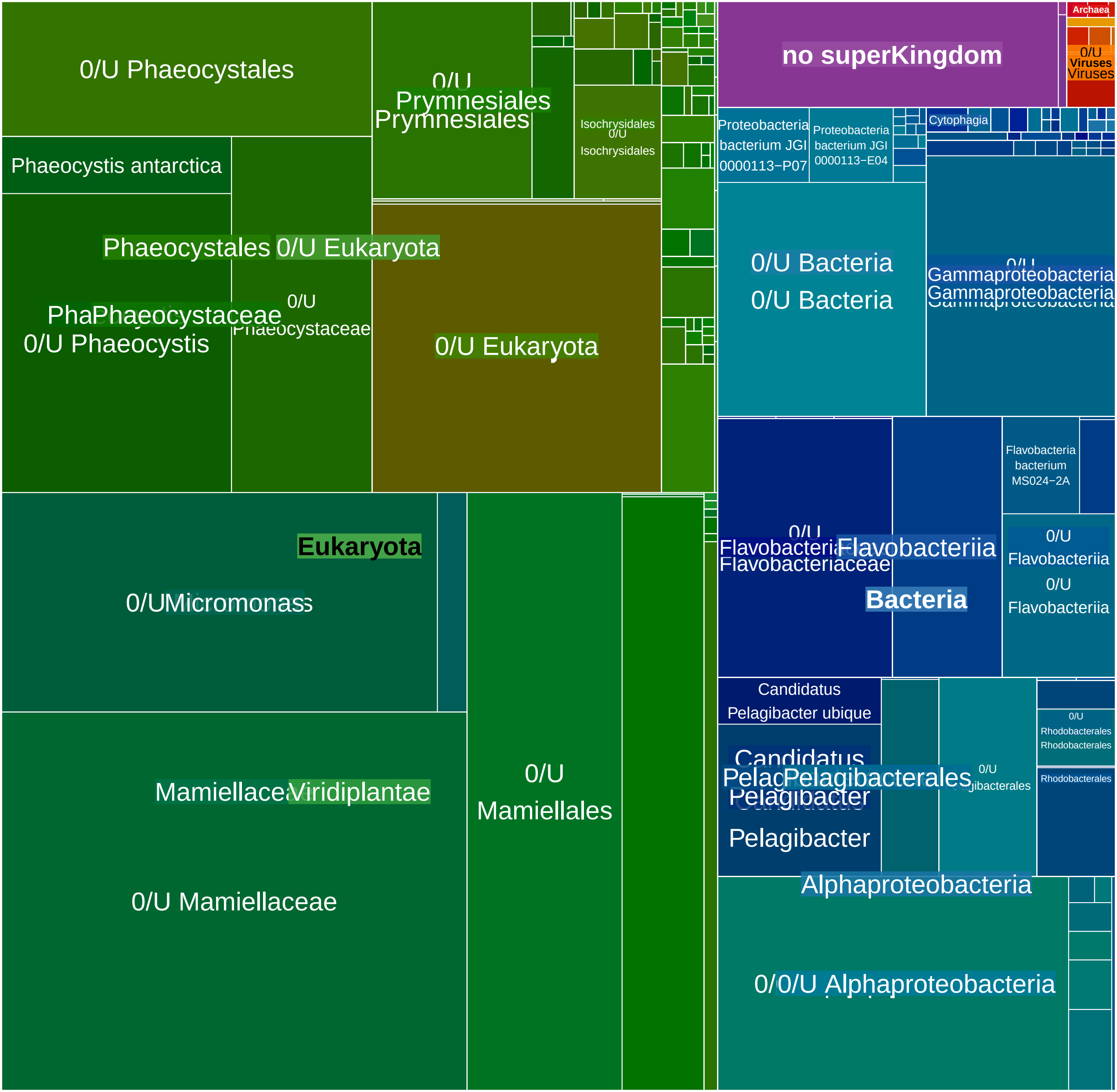

### MGT−v1_86 : 17321 unigenes, 13696 (79.07%) taxonomically assigned

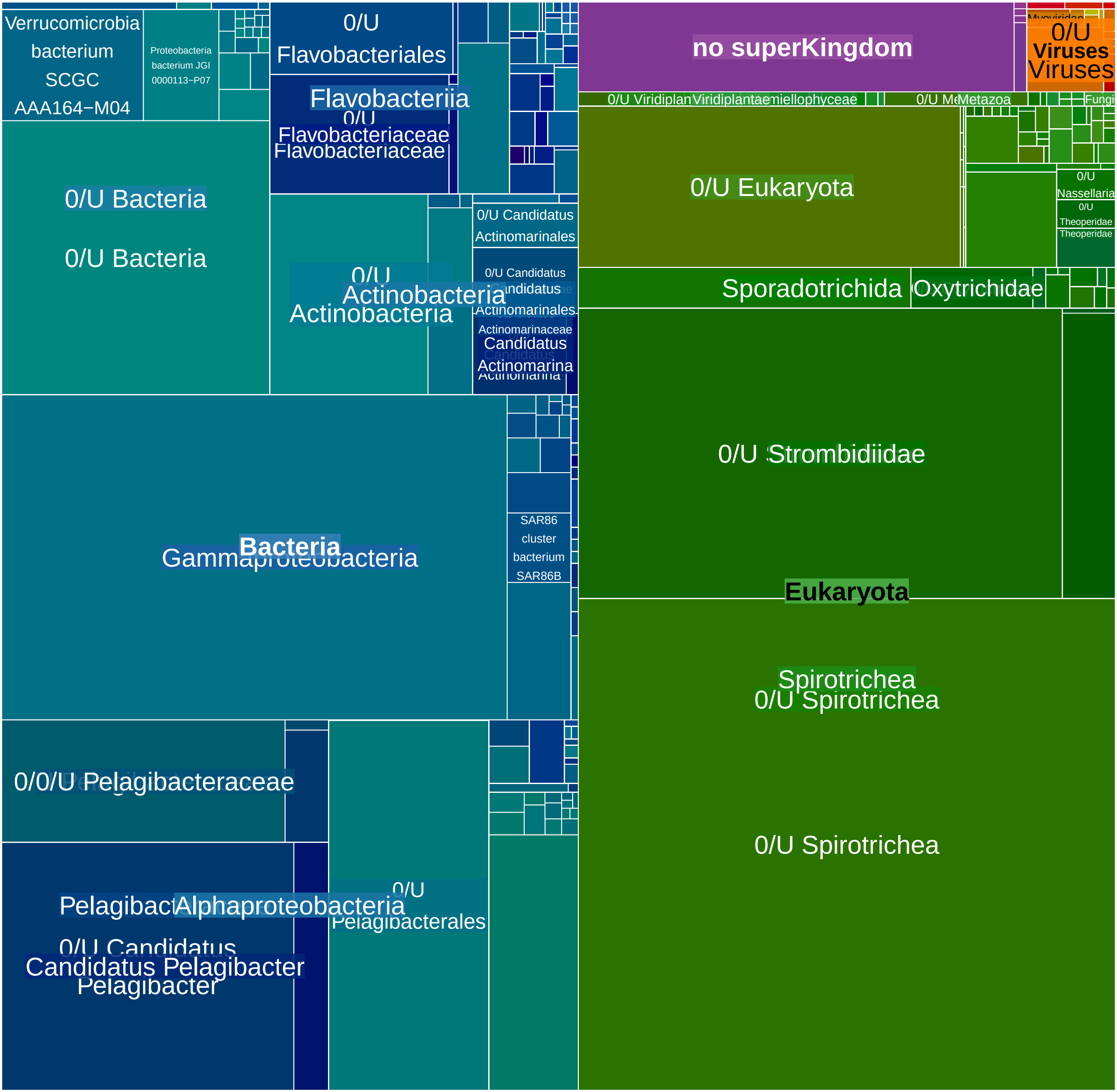

MGT−v1_87 : 17319 unigenes, 14141 (81.65%) taxonomically assigned

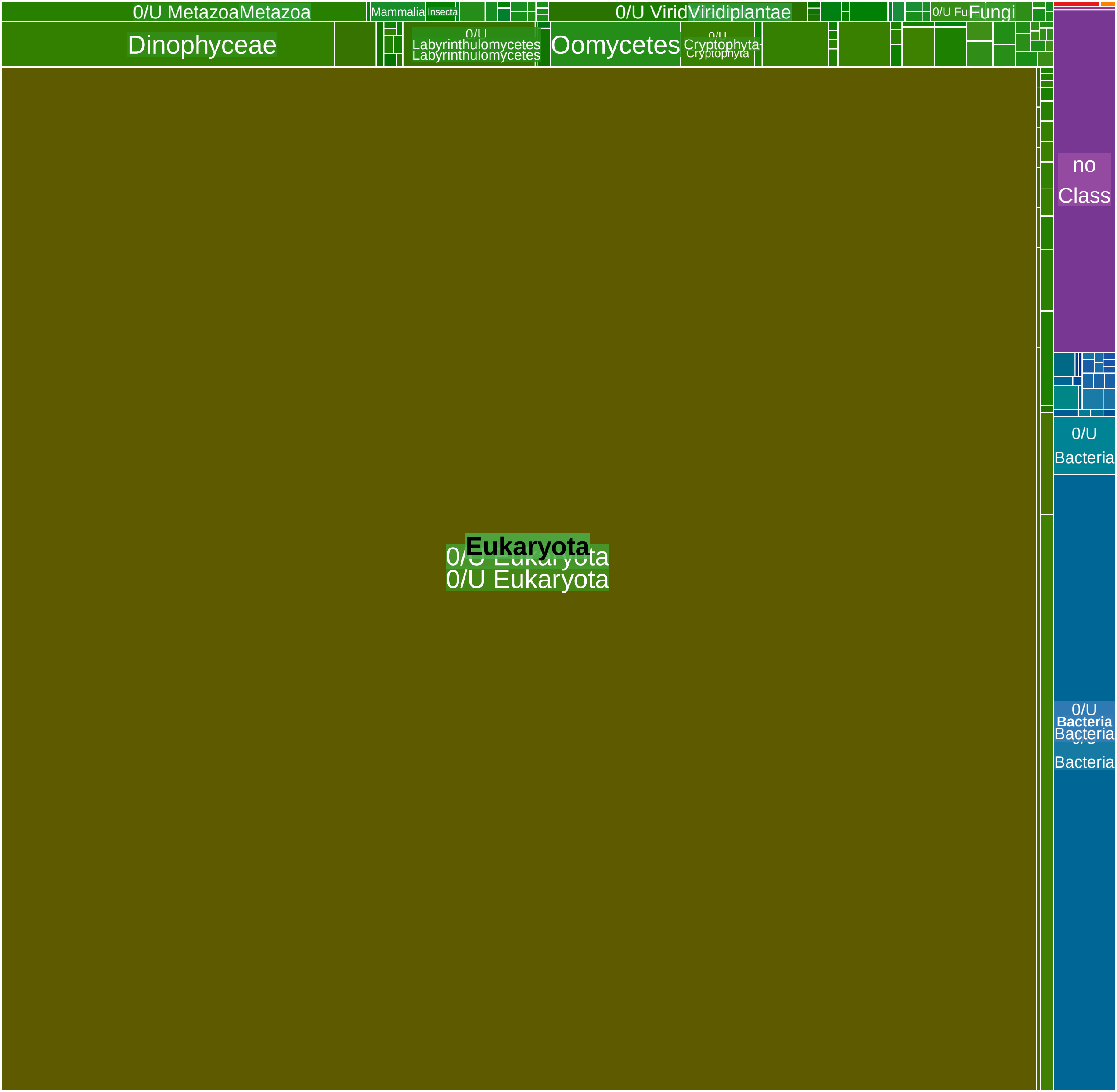

MGT−v1_88 : 17312 unigenes, 13697 (79.12%) taxonomically assigned

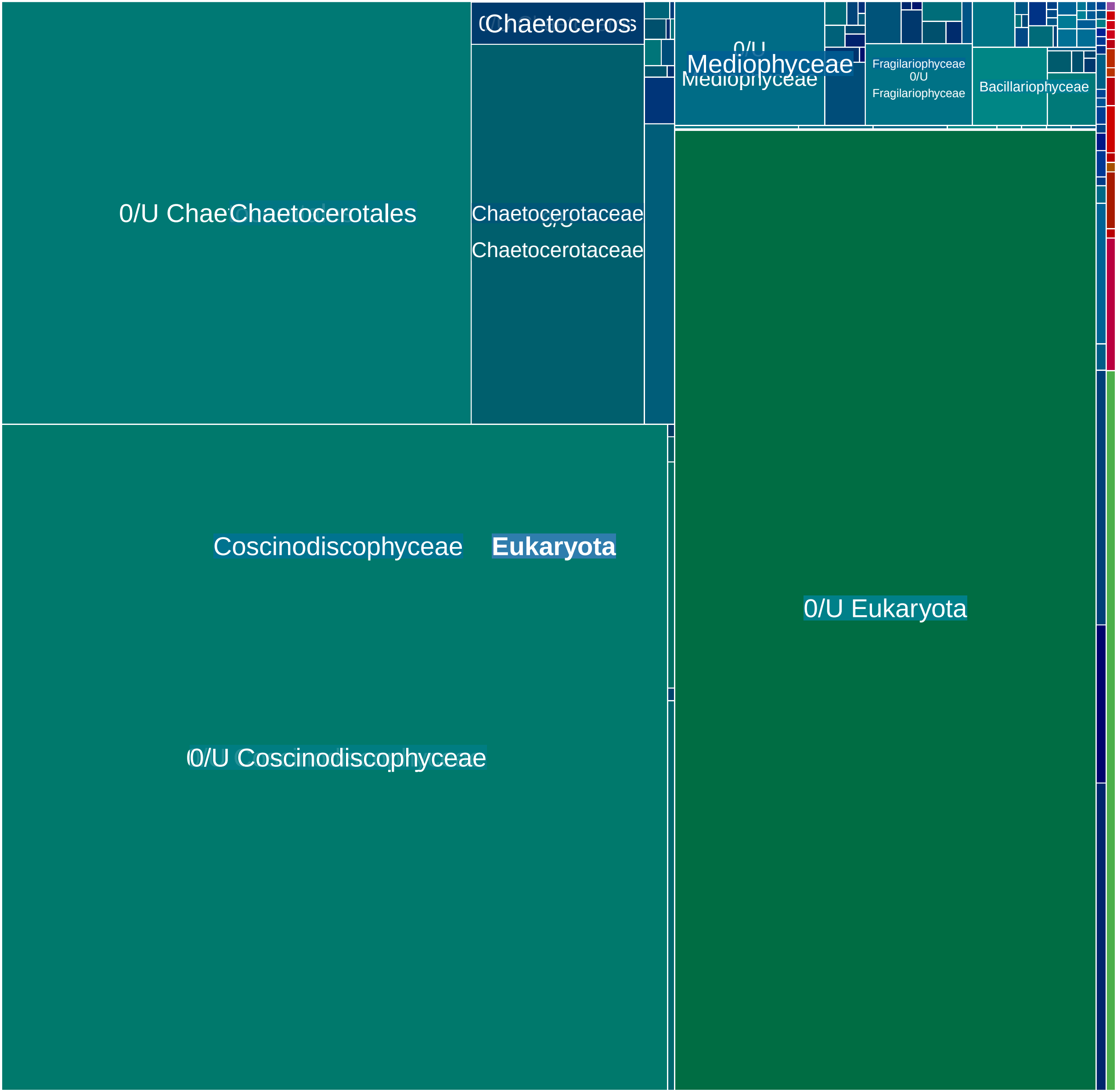

### MGT−v1_89 : 17308 unigenes, 9914 (57.28%) taxonomically assigned

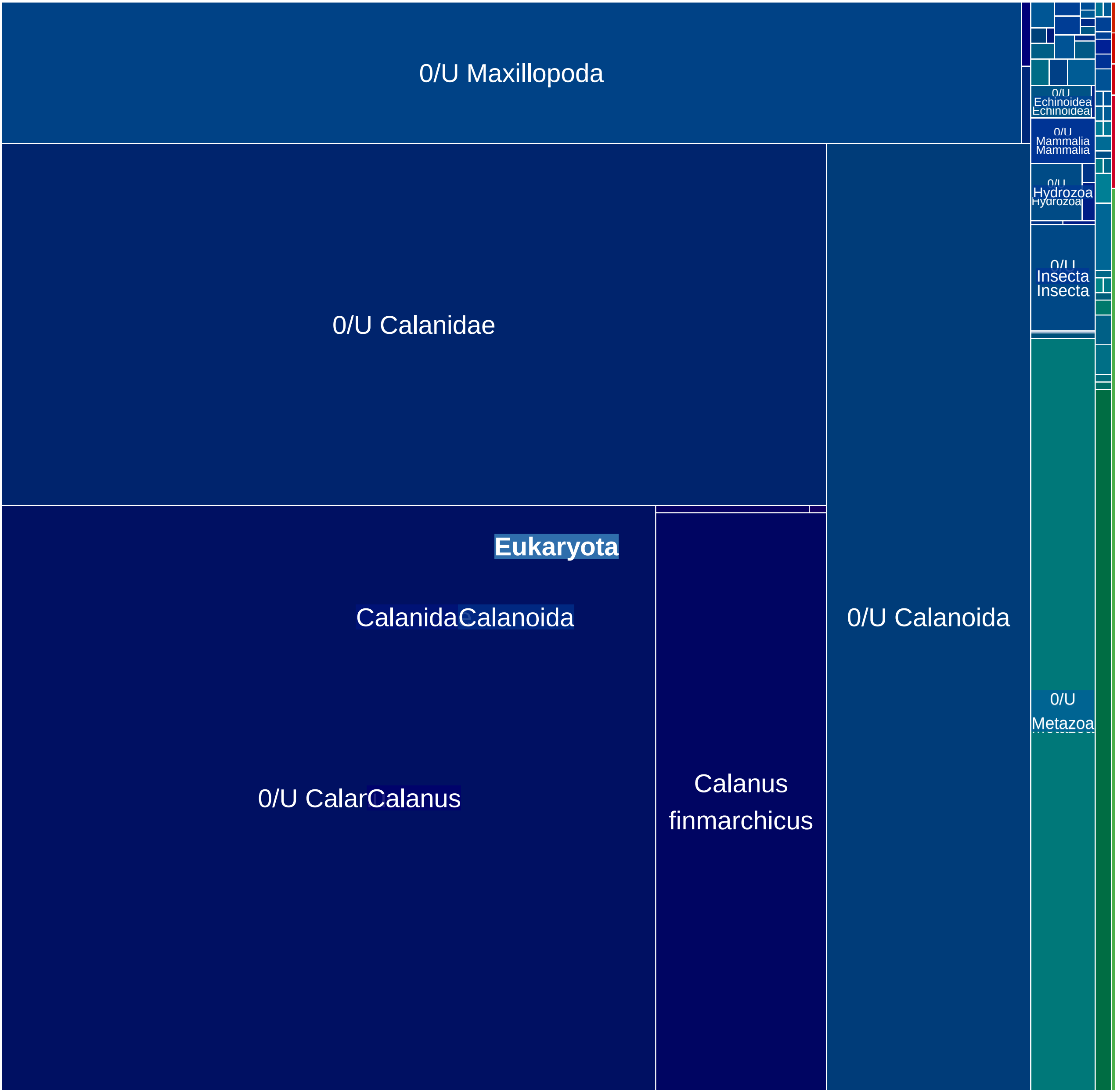

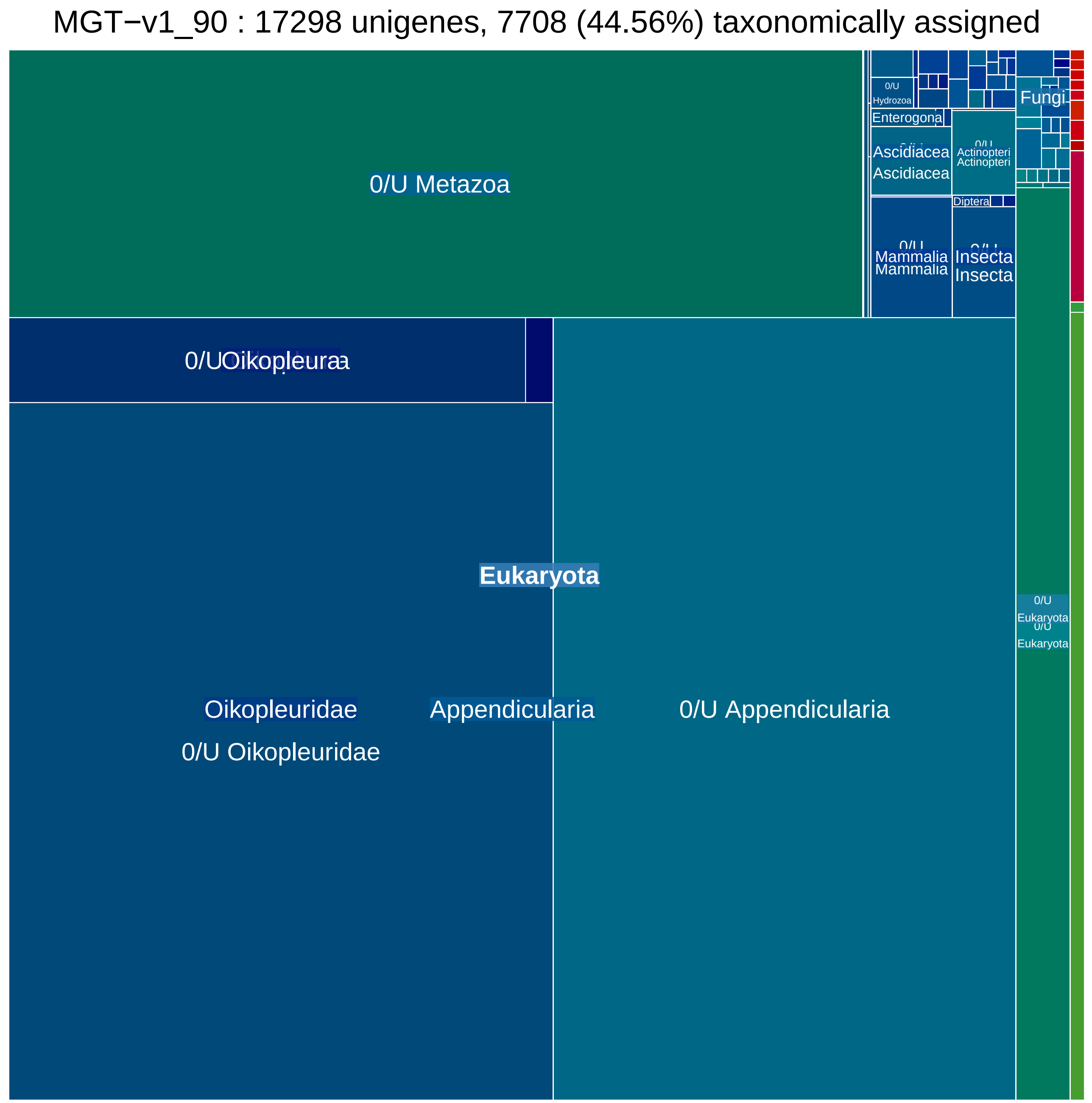
MGT−v1_90 : 17298 unigenes, 7708 (44.56%) taxonomically assigned
0/U Metazoa
0/U Oikopleura
Oikopleura
Eukaryota
Oikopleuridae
0/U Oikopleuridae
Appendicularia
0/U Appendicularia
0/U Hydrozoa
Enterogona
Ascidiacea
Ascidiacea
0/U Actinopteri
Actinopteri
Diptera
0/U Mammalia
Mammalia
0/U Insecta
Insecta
Fungi
0/U Eukaryota
0/U Eukaryota

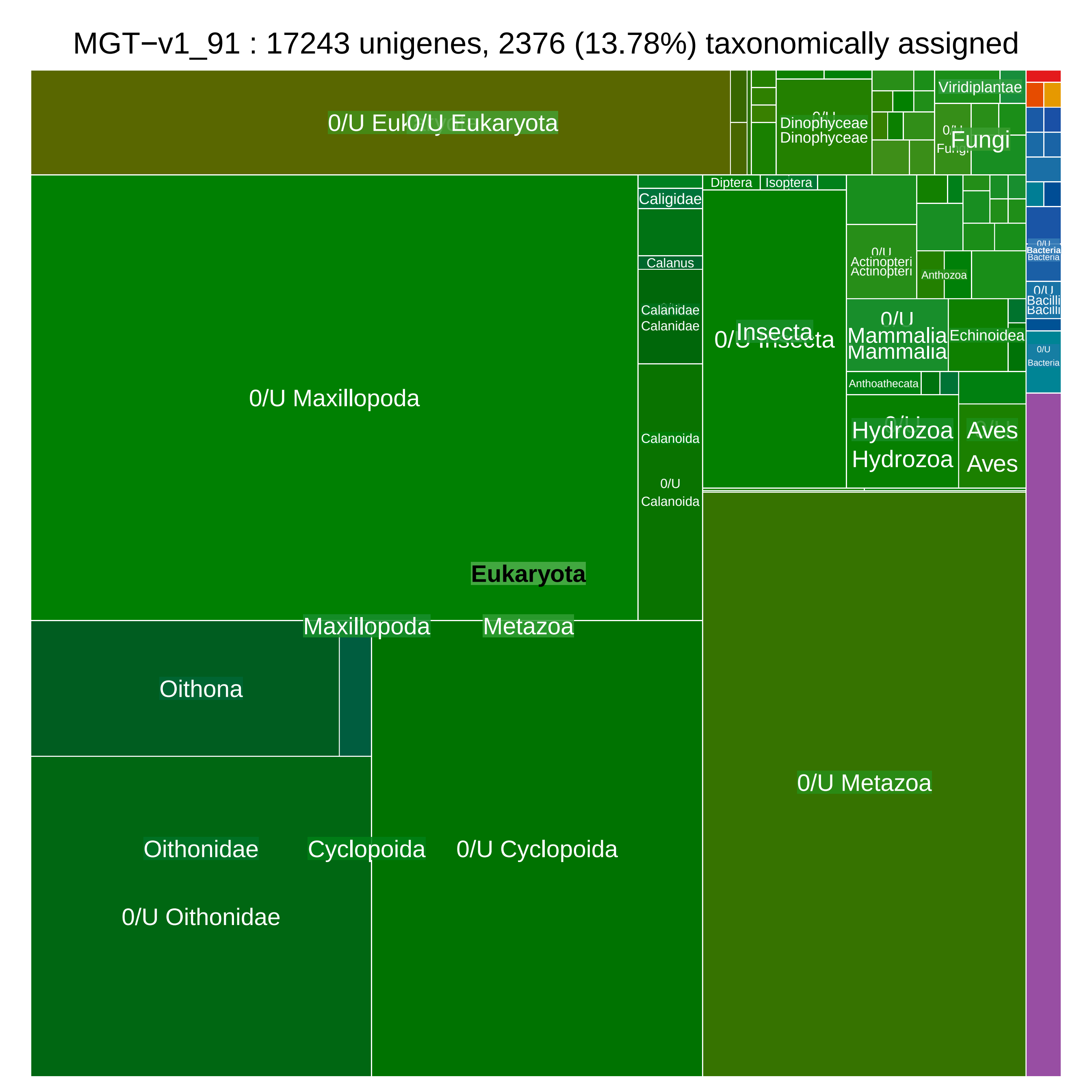
MGT−v1_91 : 17243 unigenes, 2376 (13.78%) taxonomically assigned
0/U Eukaryota
Viridiplantae
Dinophyceae
Fungi
0/U Maxillopoda
Caligidae
Calanus
Calanidae
Calanoida
0/U Calanoida
Diptera
Isoptera
0/U Insecta
Insecta
Actinopteri
Anthozoa
0/U Mammalia
Mammalia
Echinoidea
Anthoathecata
Hydrozoa
Aves
Bacteria
Bacilli
Eukaryota
Maxillopoda
Metazoa
Oithona
Oithonidae
Cyclopoida
0/U Cyclopoida
0/U Oithonidae
0/U Metazoa

### MGT−v1_92 : 17212 unigenes, 2547 (14.80%) taxonomically assigned

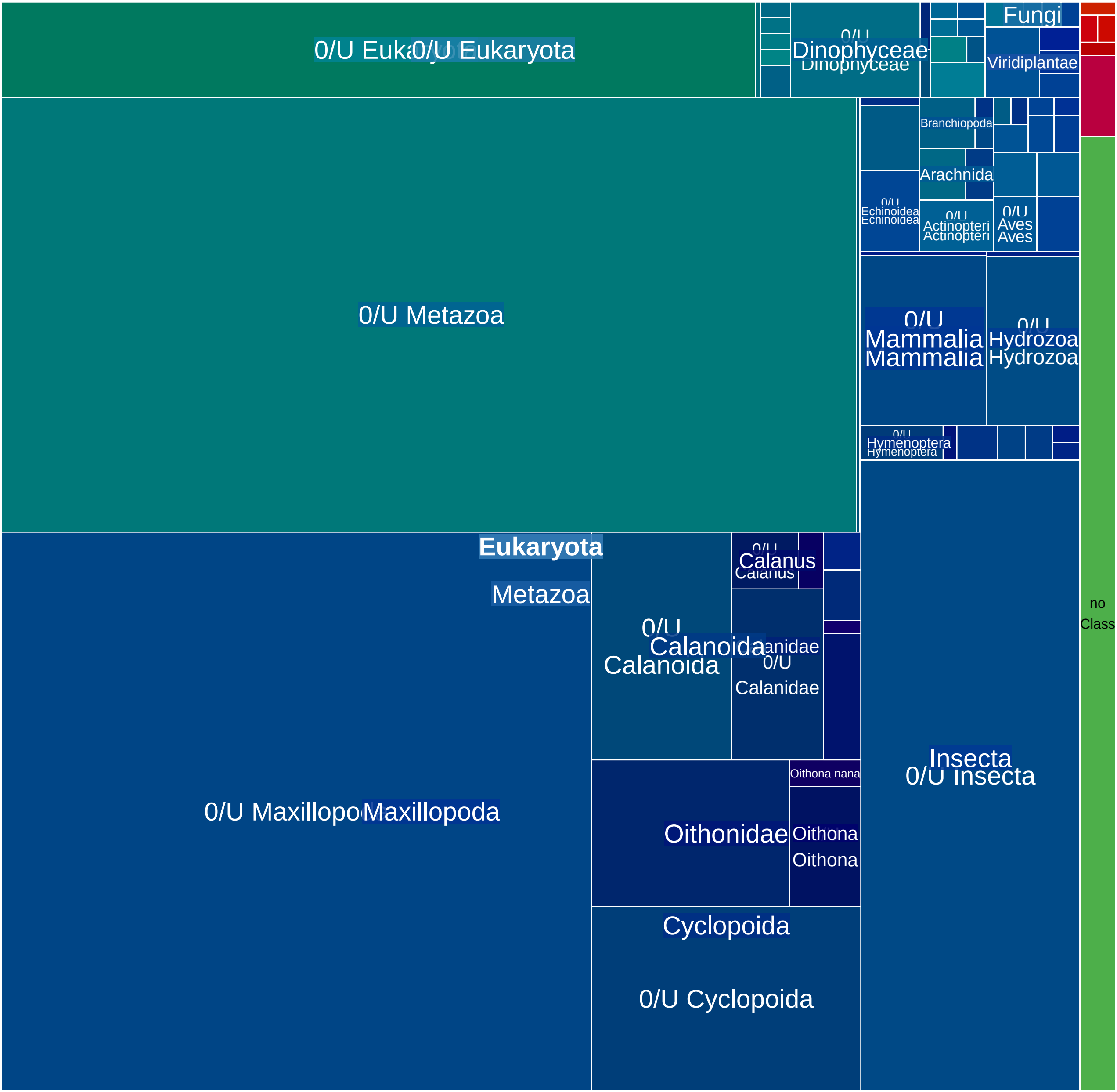

MGT−v1_93 : 17030 unigenes, 13439 (78.91%) taxonomically assigned

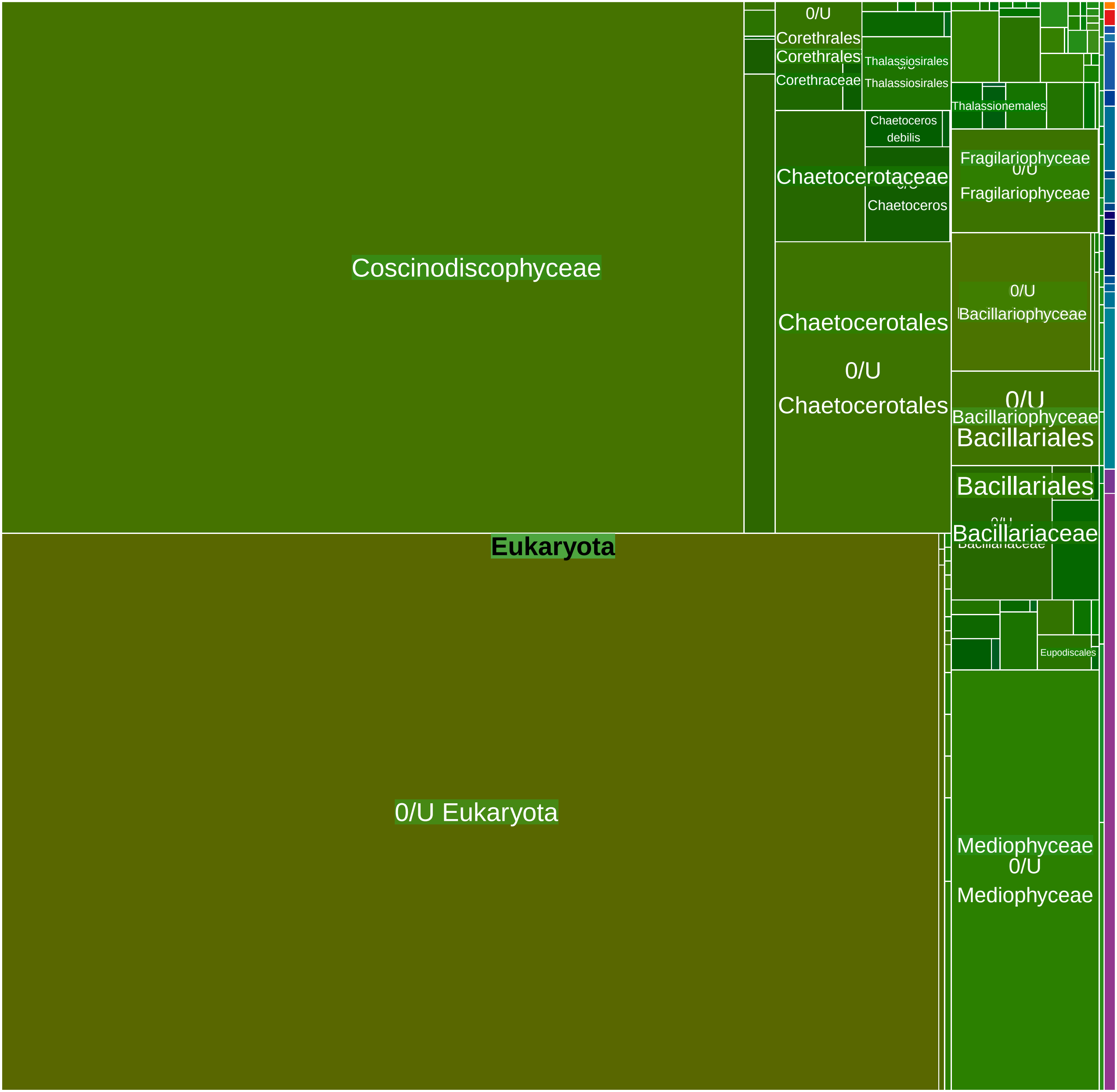

MGT−v1_94 : 16921 unigenes, 10774 (63.67%) taxonomically assigned

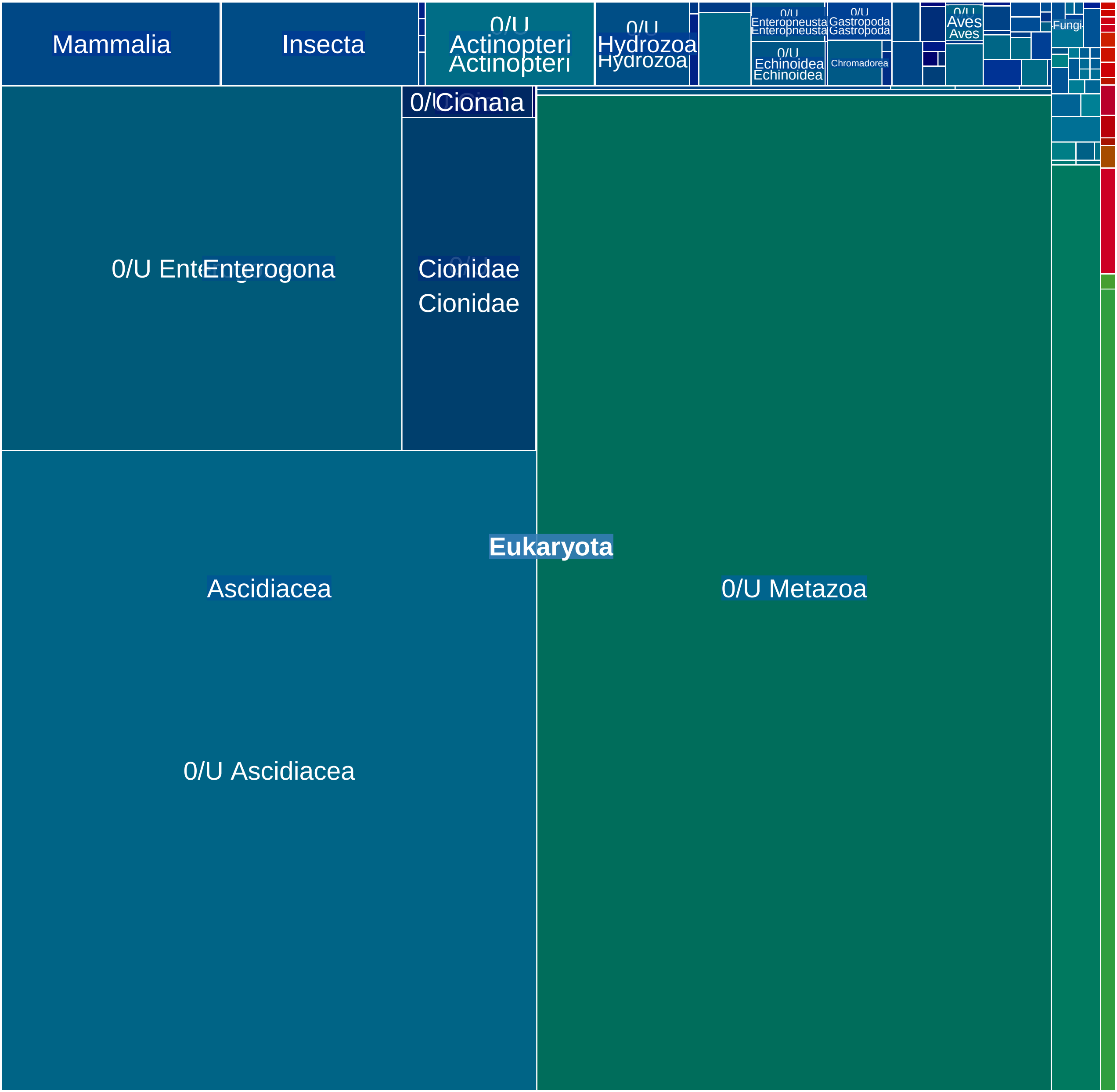

MGT−v1_95 : 16663 unigenes, 14010 (84.08%) taxonomically assigned

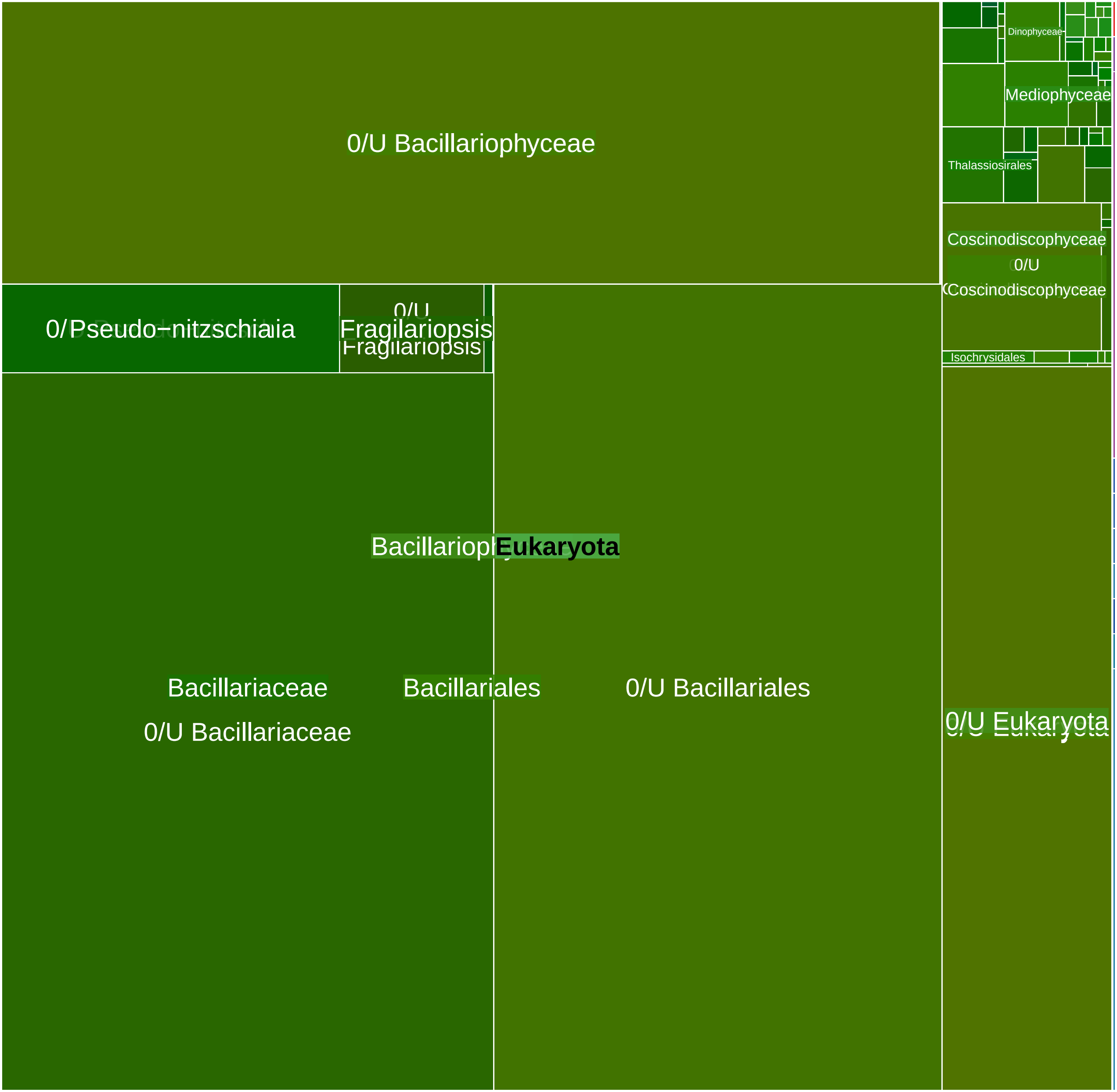

MGT−v1_96 : 16663 unigenes, 8754 (52.54%) taxonomically assigned

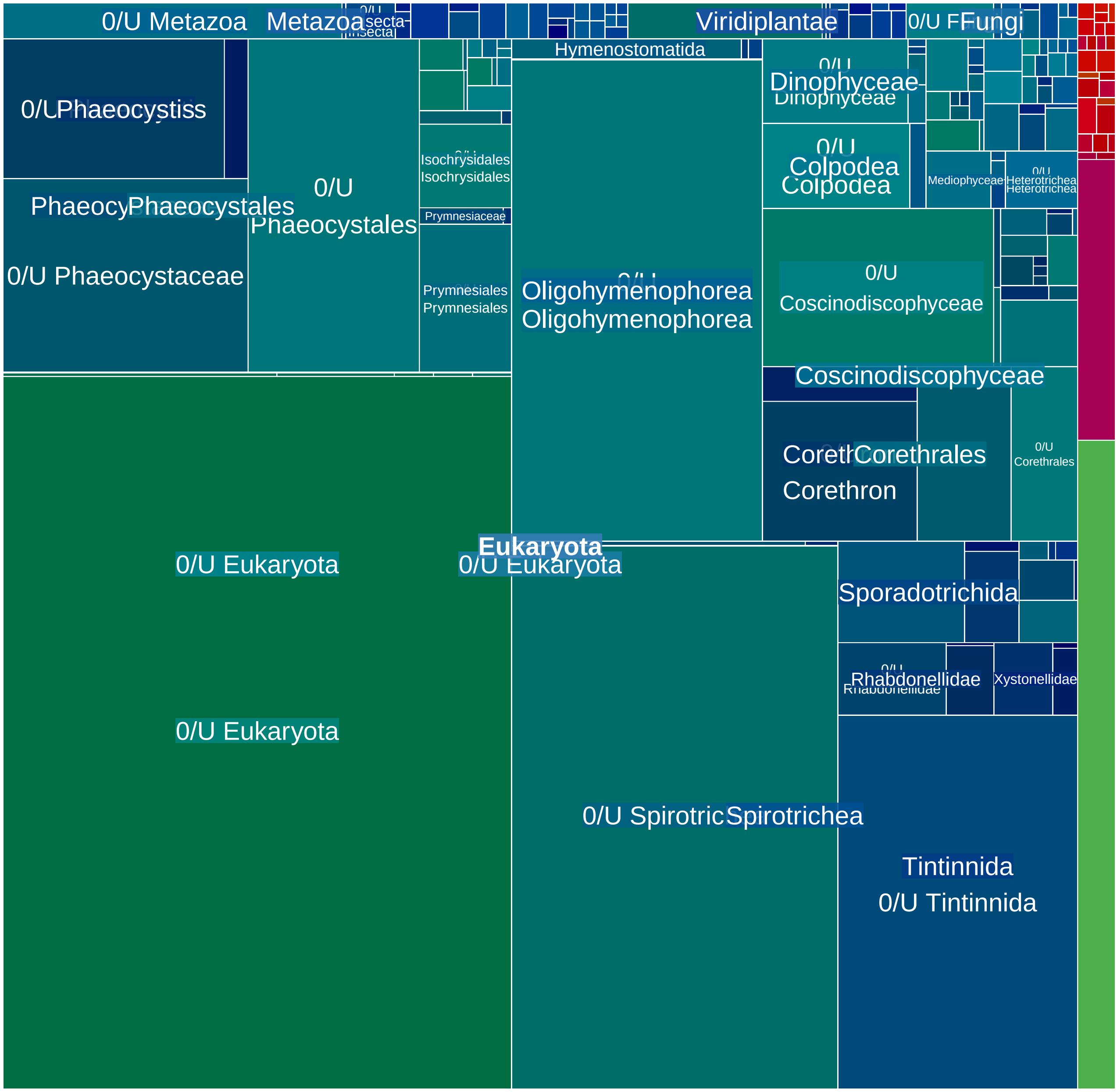

MGT−v1_97 : 16619 unigenes, 9608 (57.81%) taxonomically assigned

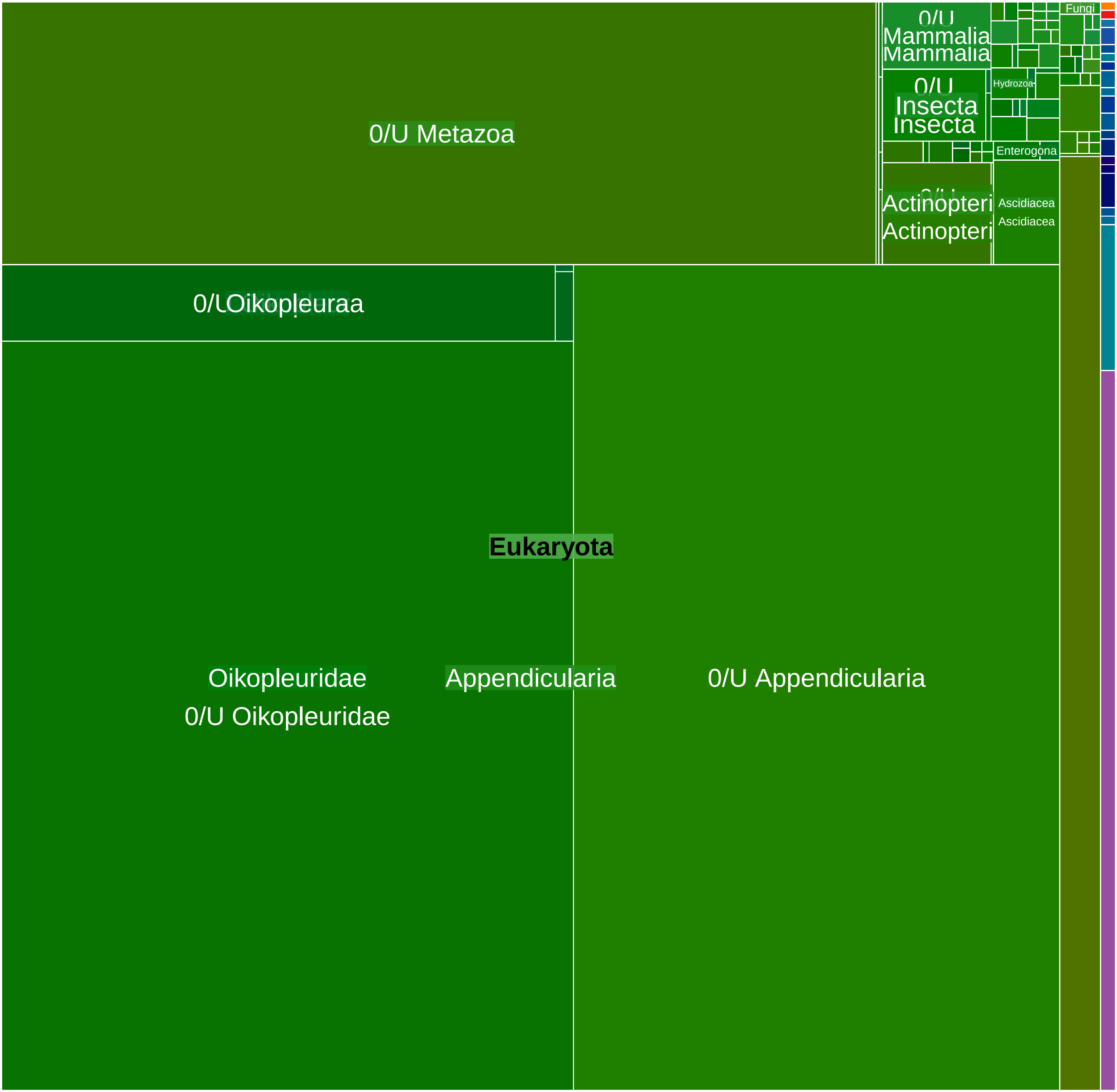

MGT−v1_98 : 16604 unigenes, 16205 (97.60%) taxonomically assigned

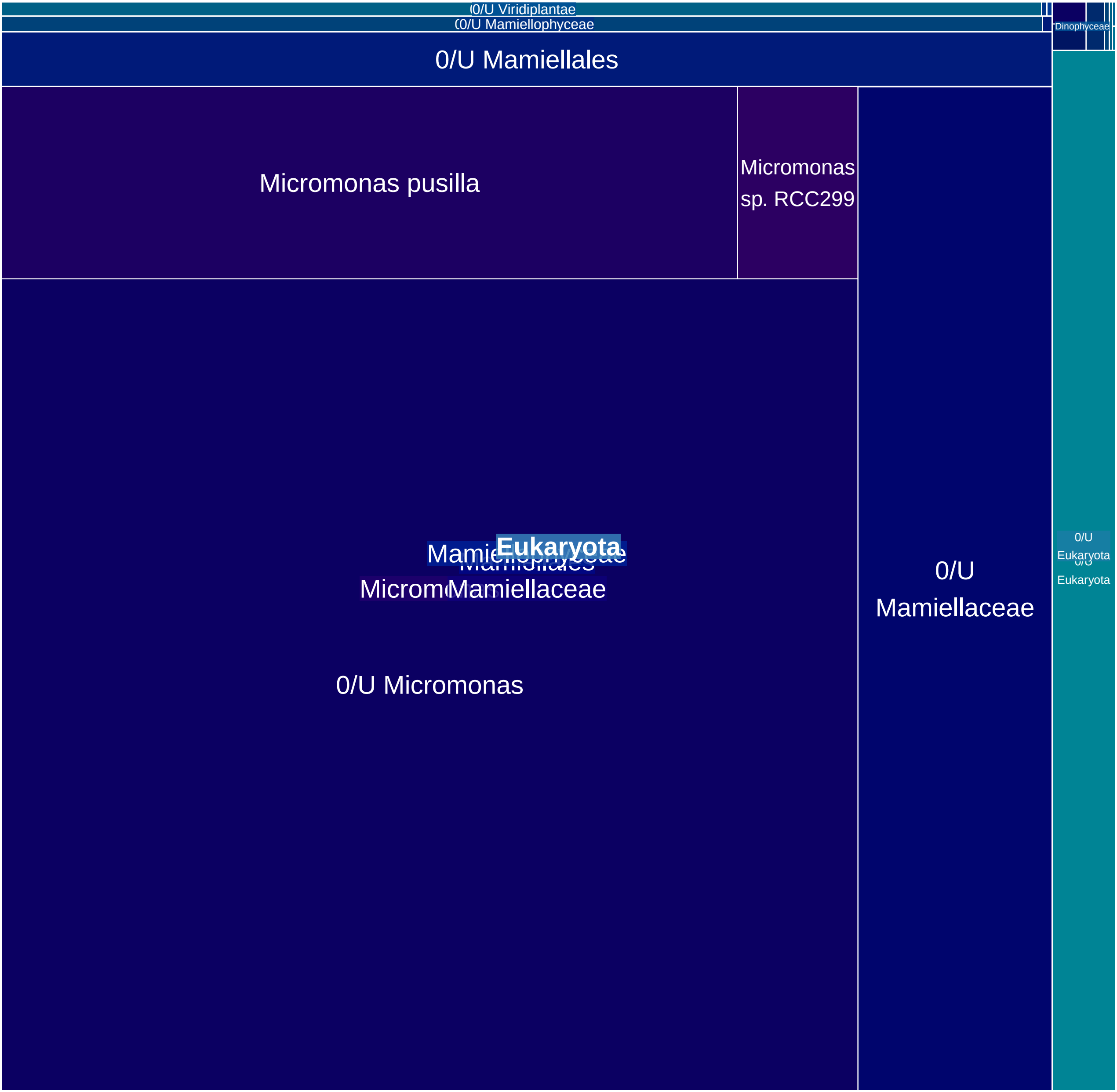

MGT−v1_99 : 16366 unigenes, 6202 (37.90%) taxonomically assigned

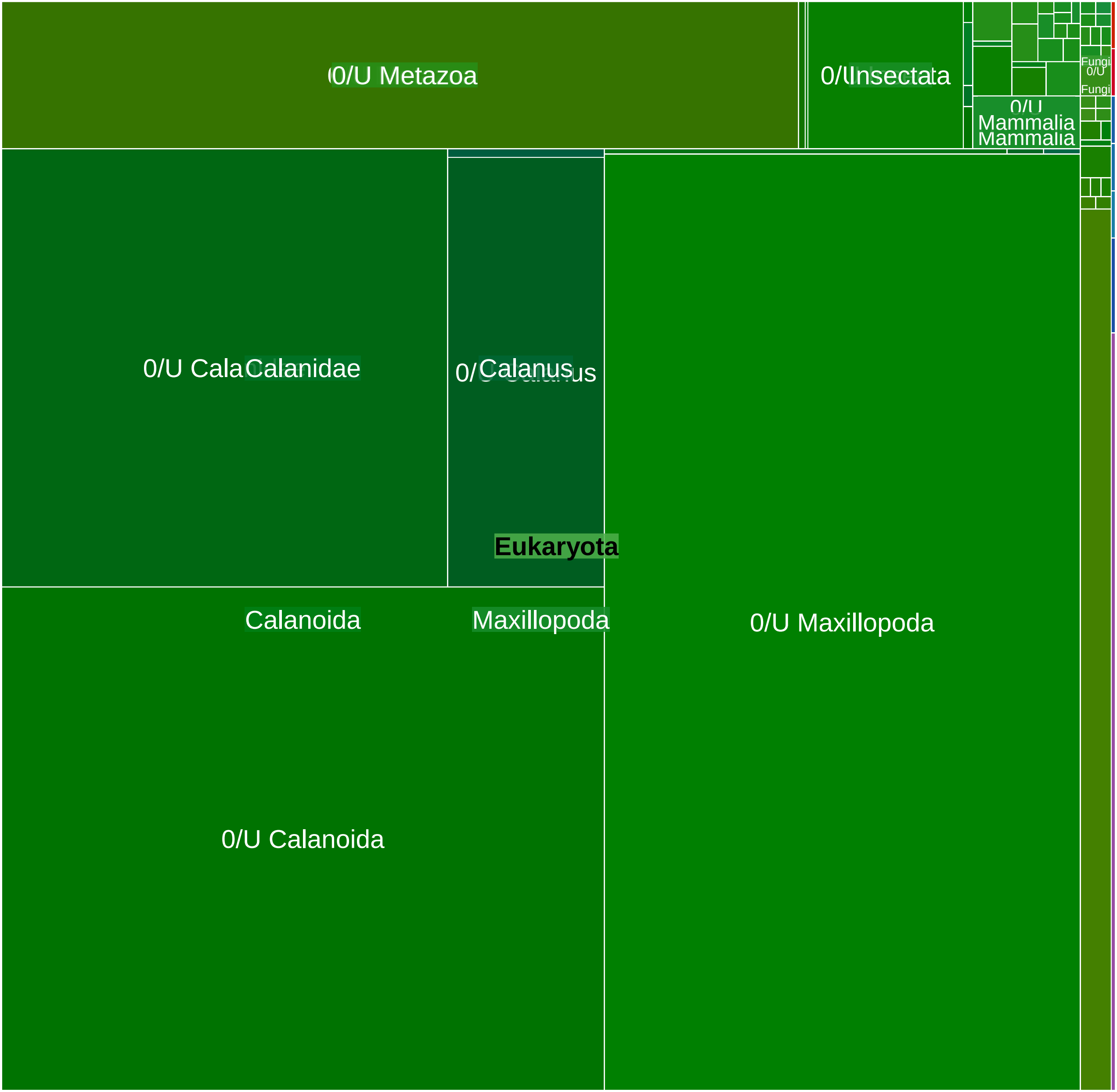

MGT−v1_100 : 15951 unigenes, 14454 (90.62%) taxonomically assigned

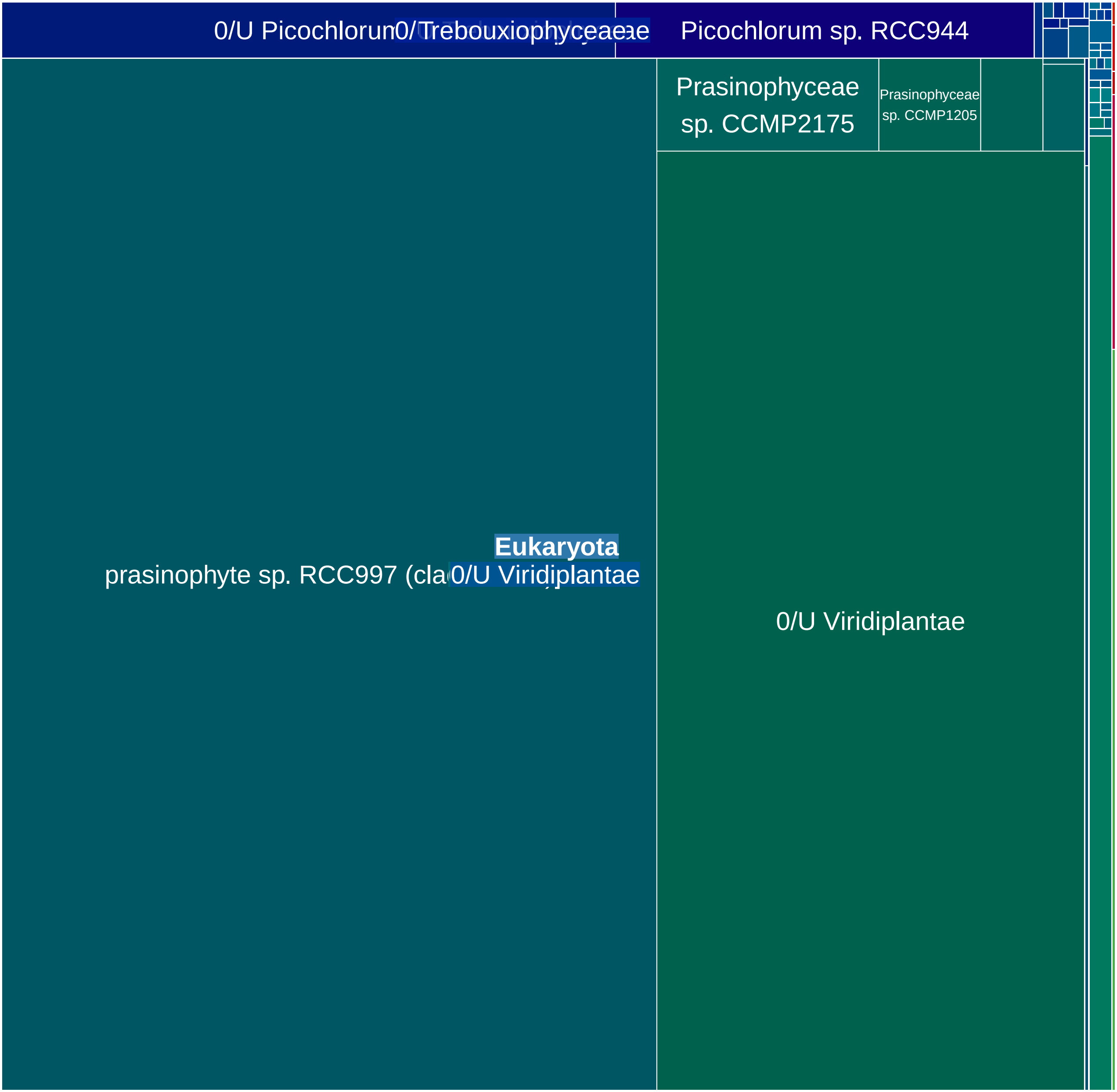

MGT−v1_101 : 15622 unigenes, 7159 (45.83%) taxonomically assigned

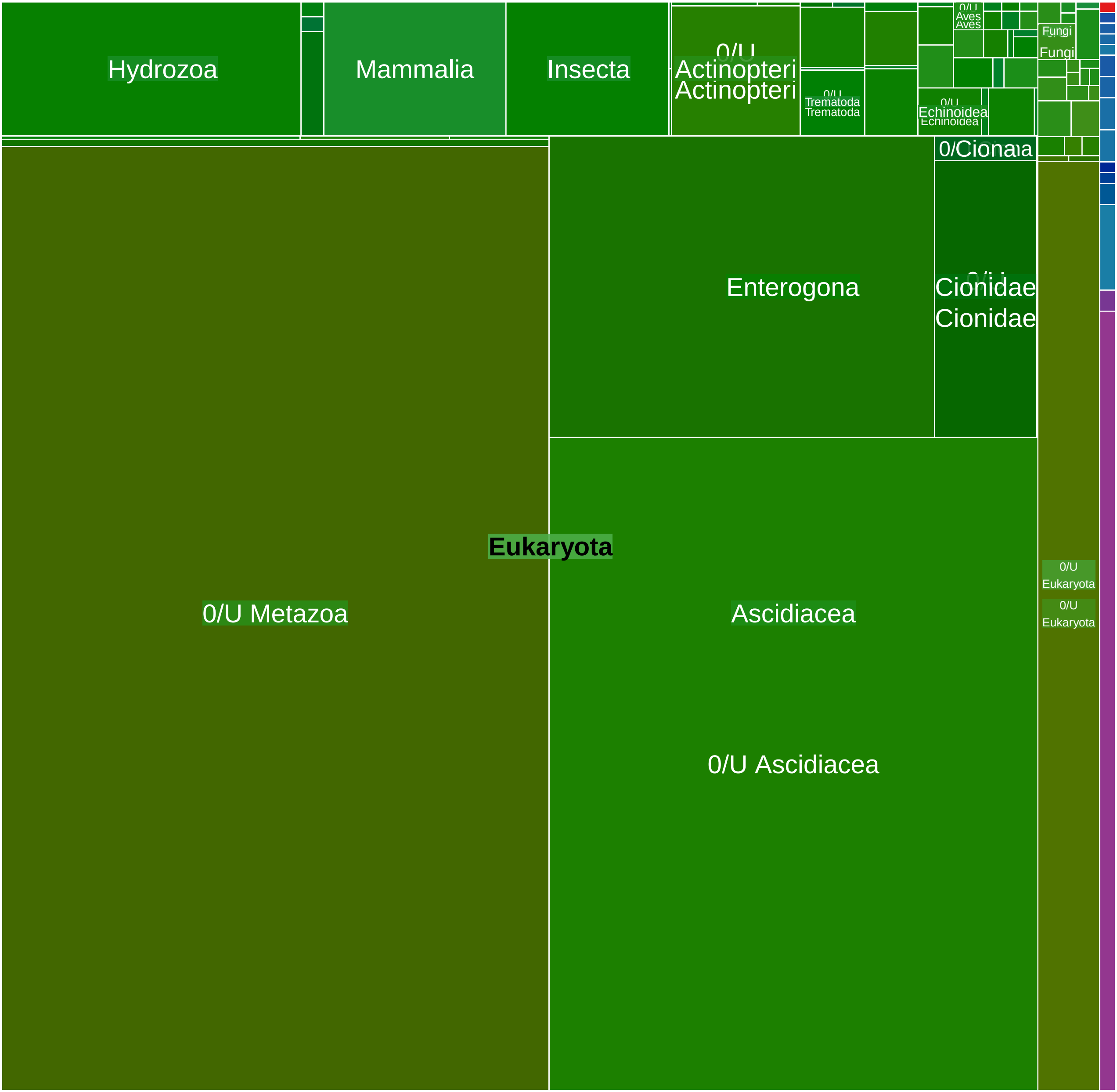

### MGT−v1_102 : 15538 unigenes, 10851 (69.84%) taxonomically assigned

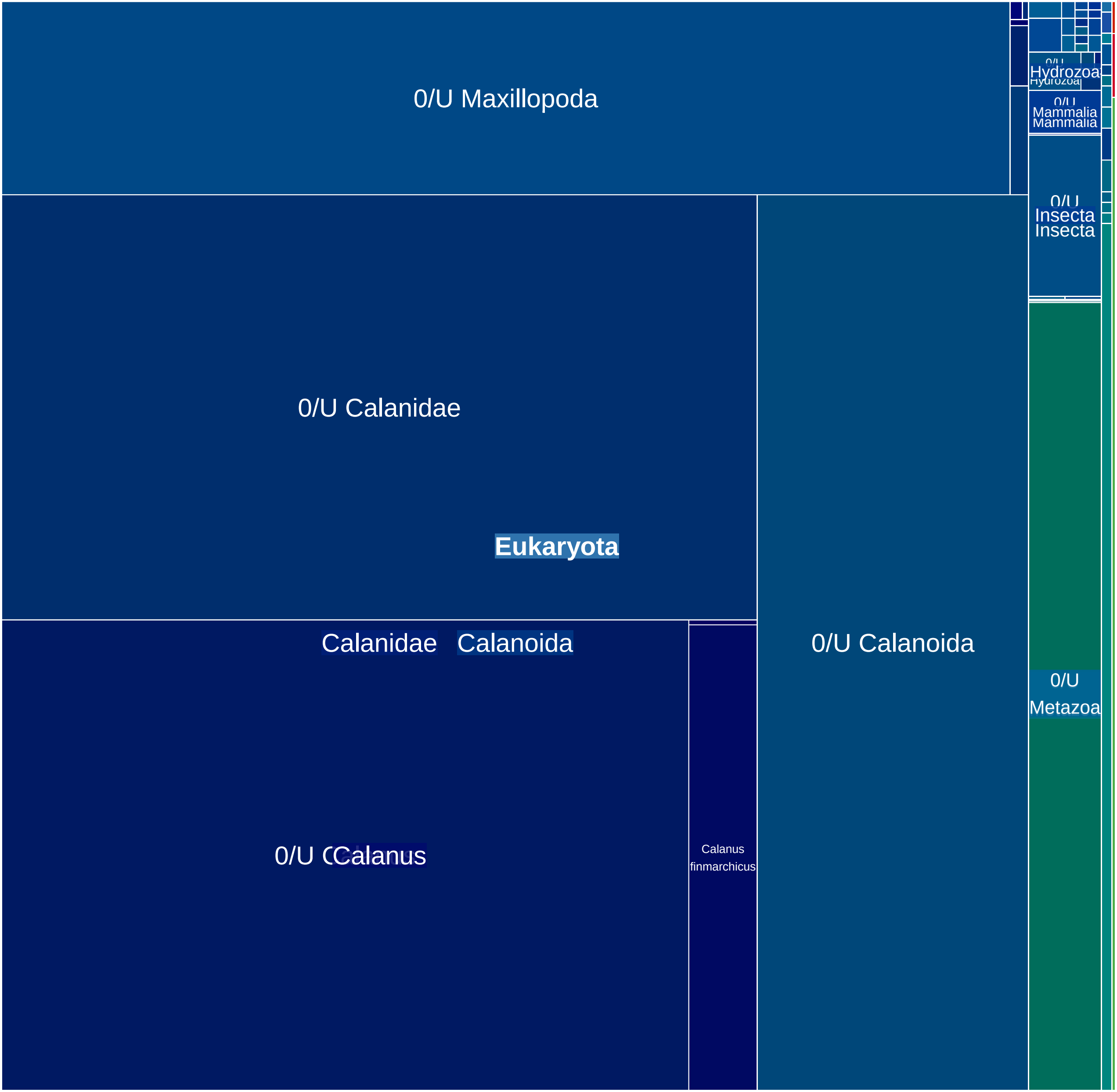

MGT−v1_103 : 15538 unigenes, 8152 (52.46%) taxonomically assigned

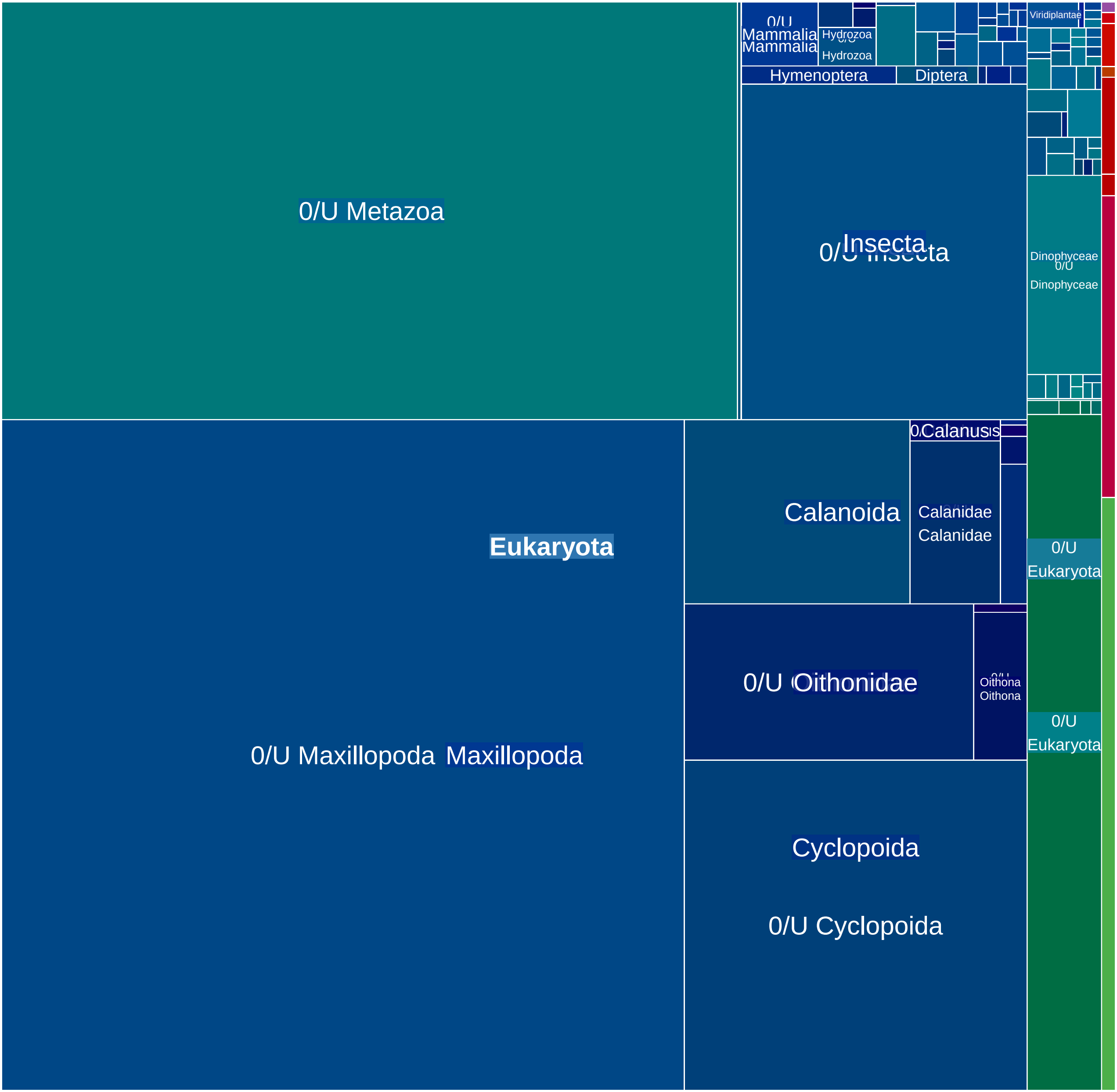

MGT−v1_104 : 15010 unigenes, 13799 (91.93%) taxonomically assigned

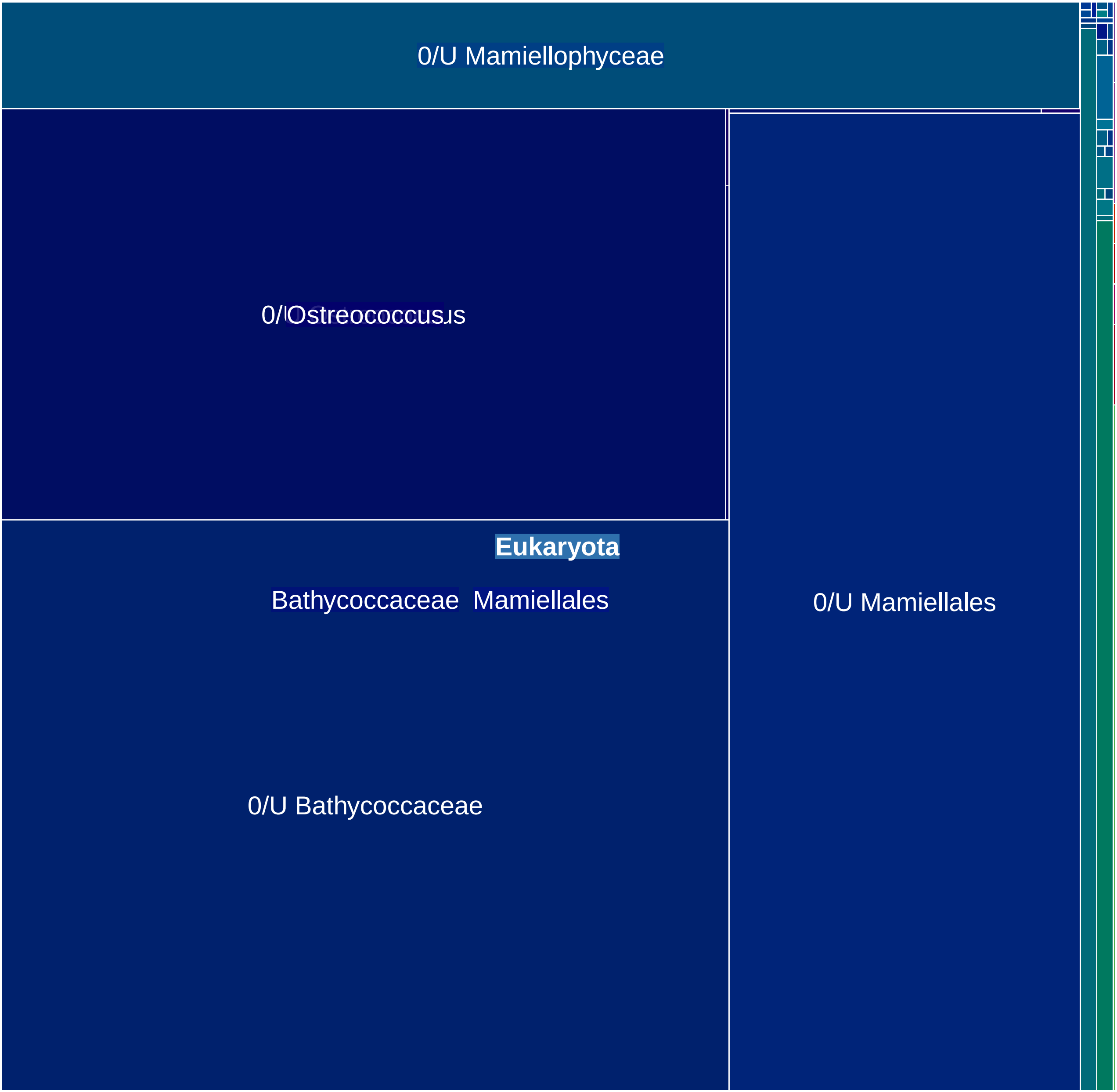

MGT−v1_105 : 14939 unigenes, 12833 (85.90%) taxonomically assigned

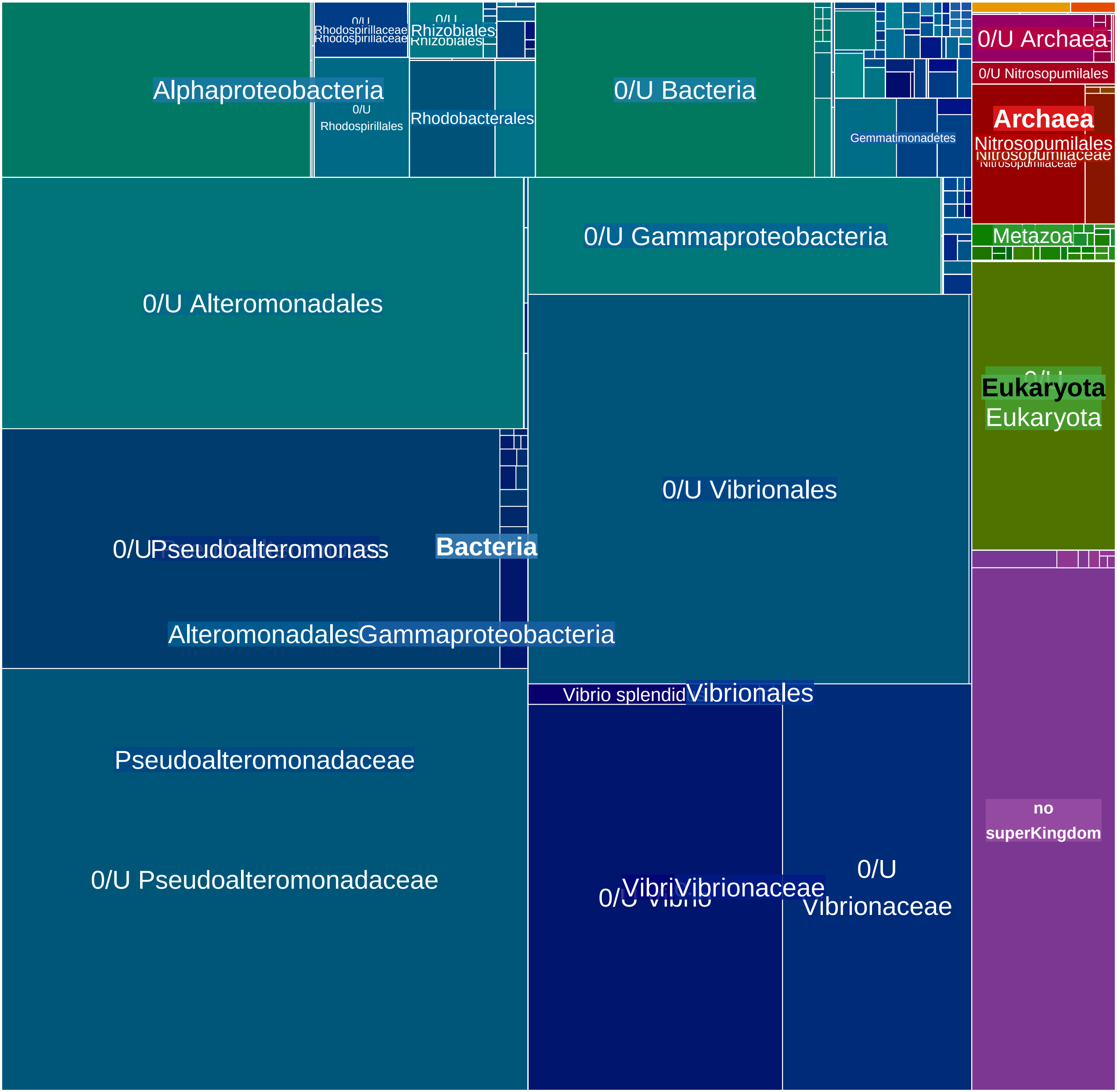

MGT−v1_106 : 14931 unigenes, 8174 (54.75%) taxonomically assigned

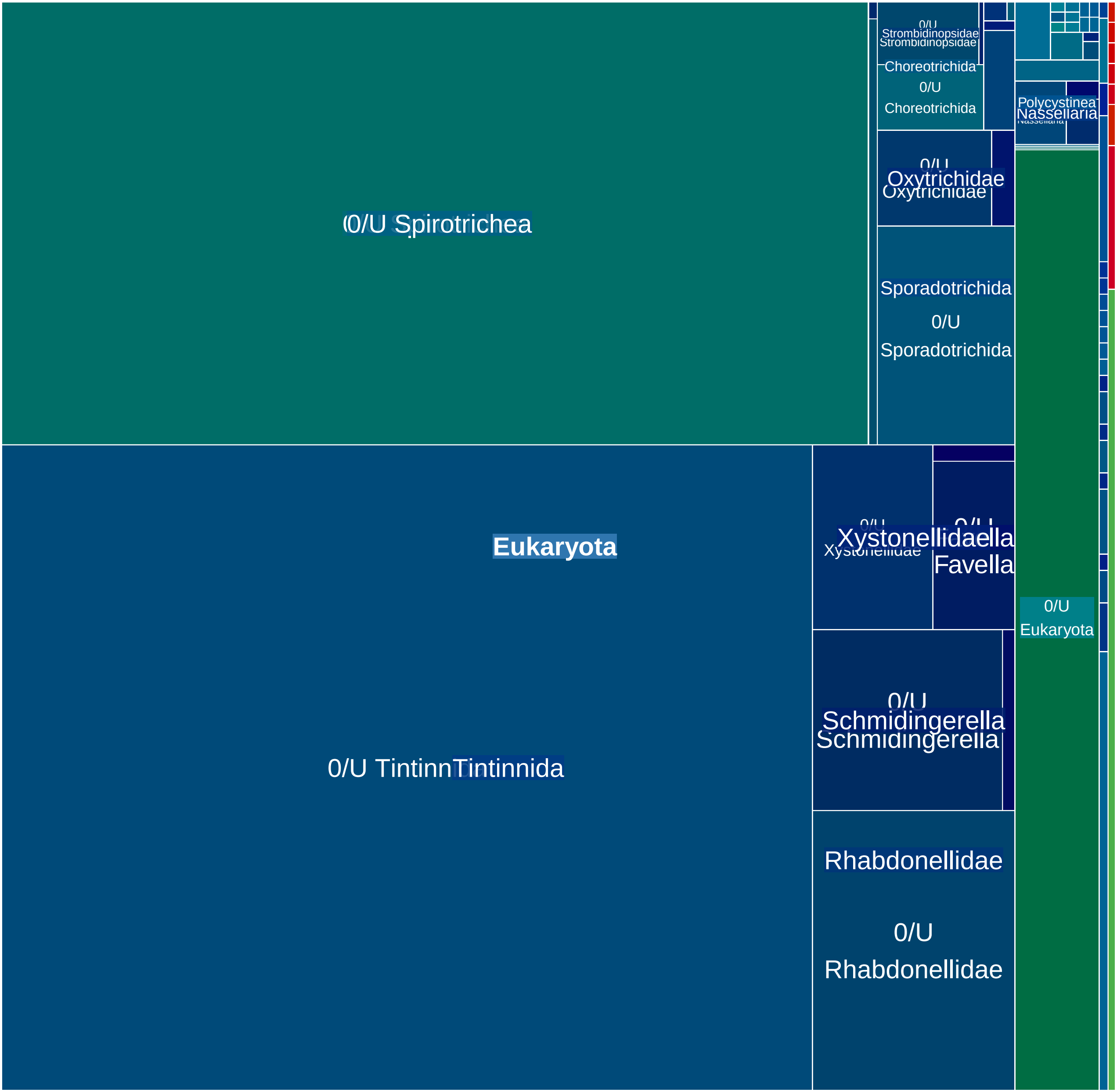

MGT−v1_107 : 14820 unigenes, 7478 (50.46%) taxonomically assigned

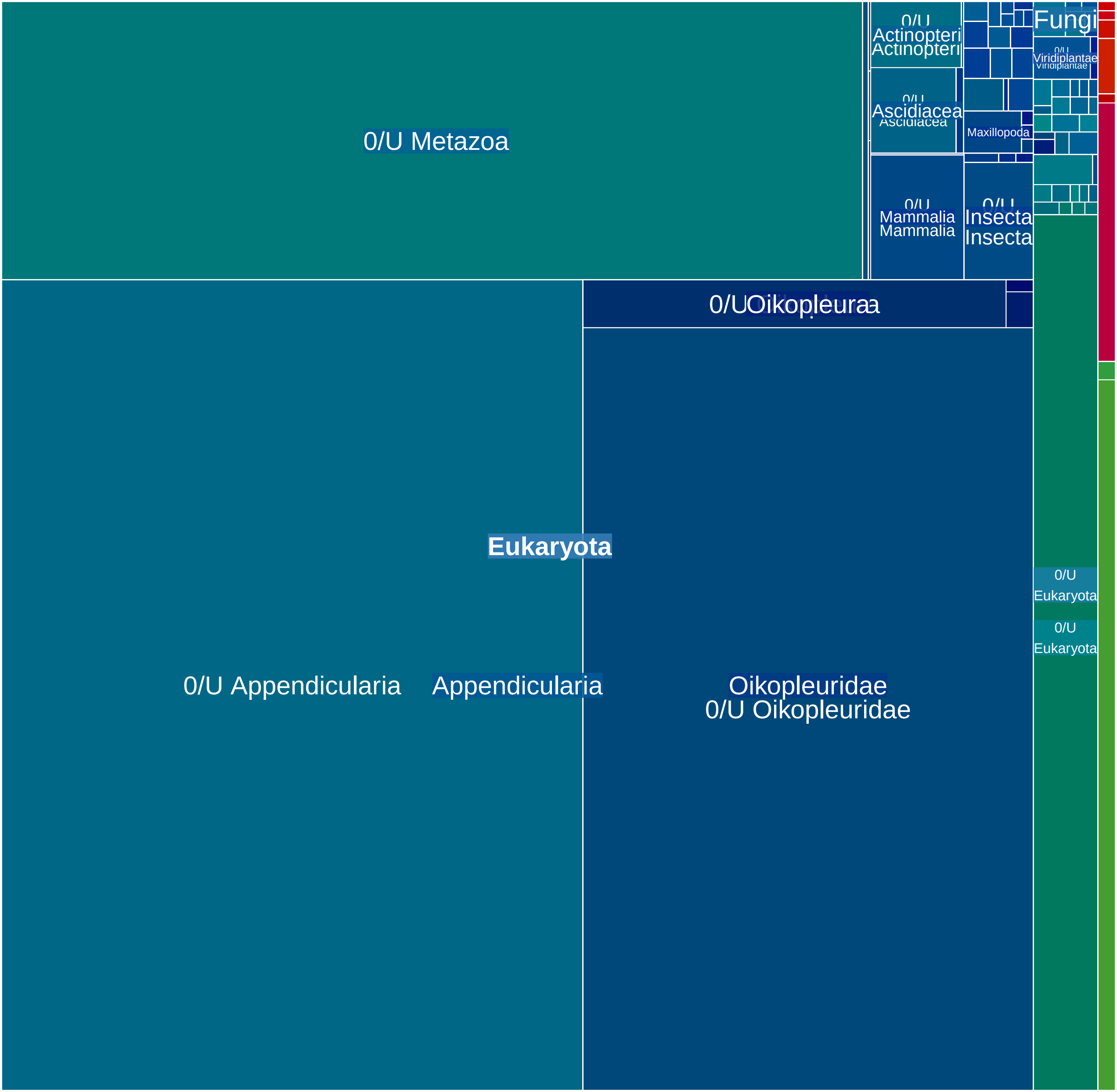

MGT−v1_108 : 14792 unigenes, 10957 (74.07%) taxonomically assigned

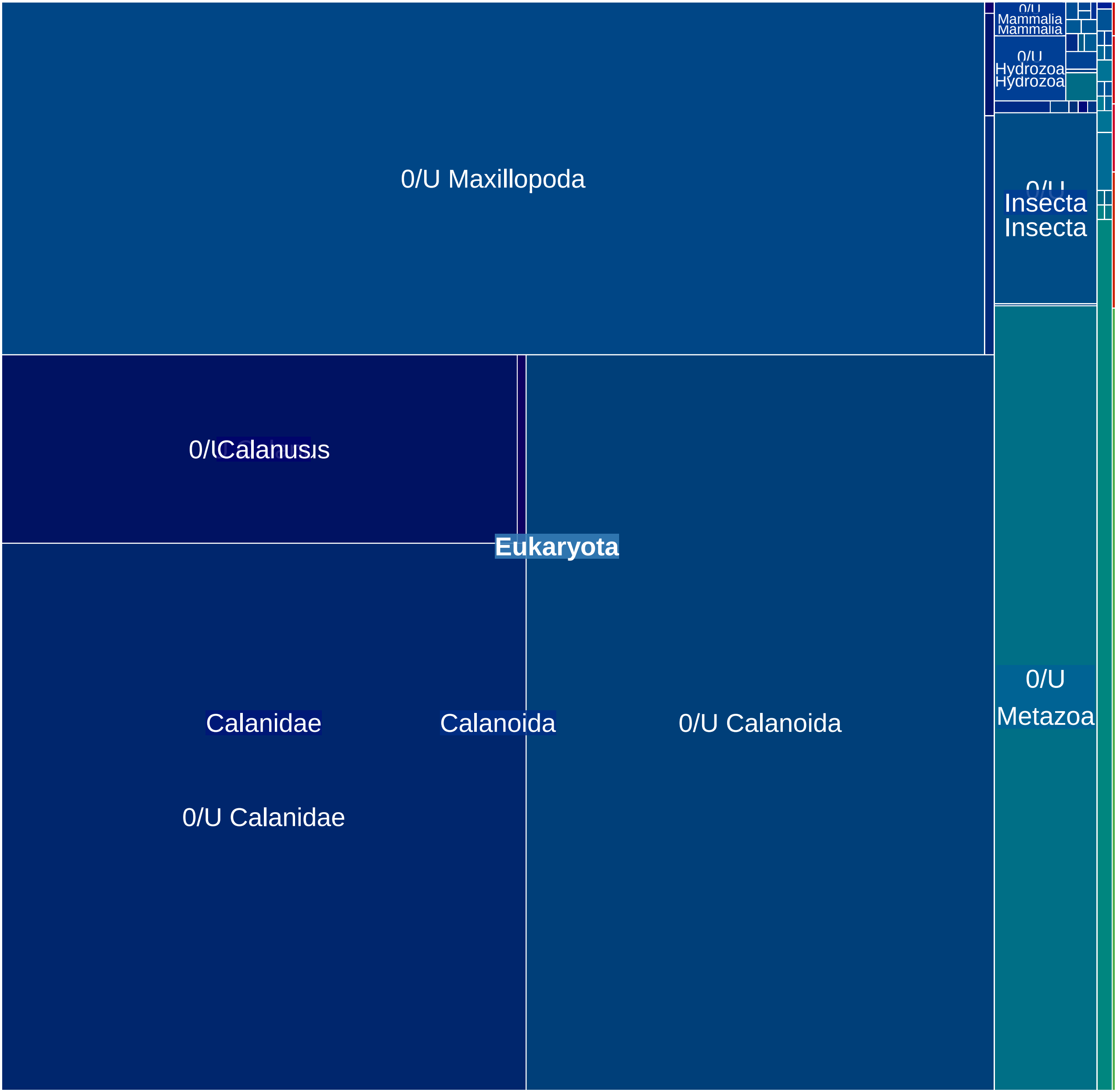

MGT−v1_109 : 14689 unigenes, 11897 (80.99%) taxonomically assigned

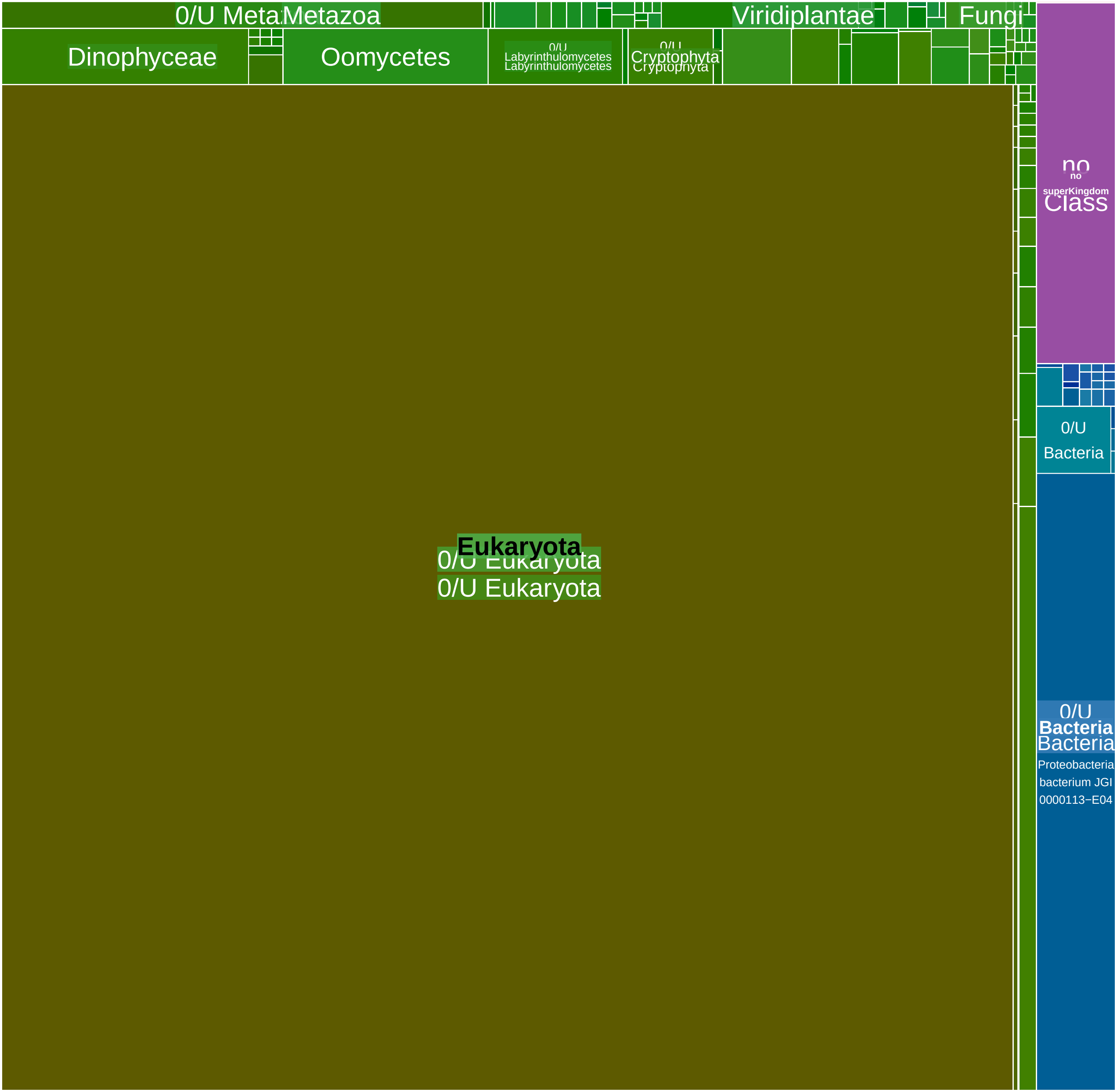

MGT−v1_110 : 13810 unigenes, 2232 (16.16%) taxonomically assigned

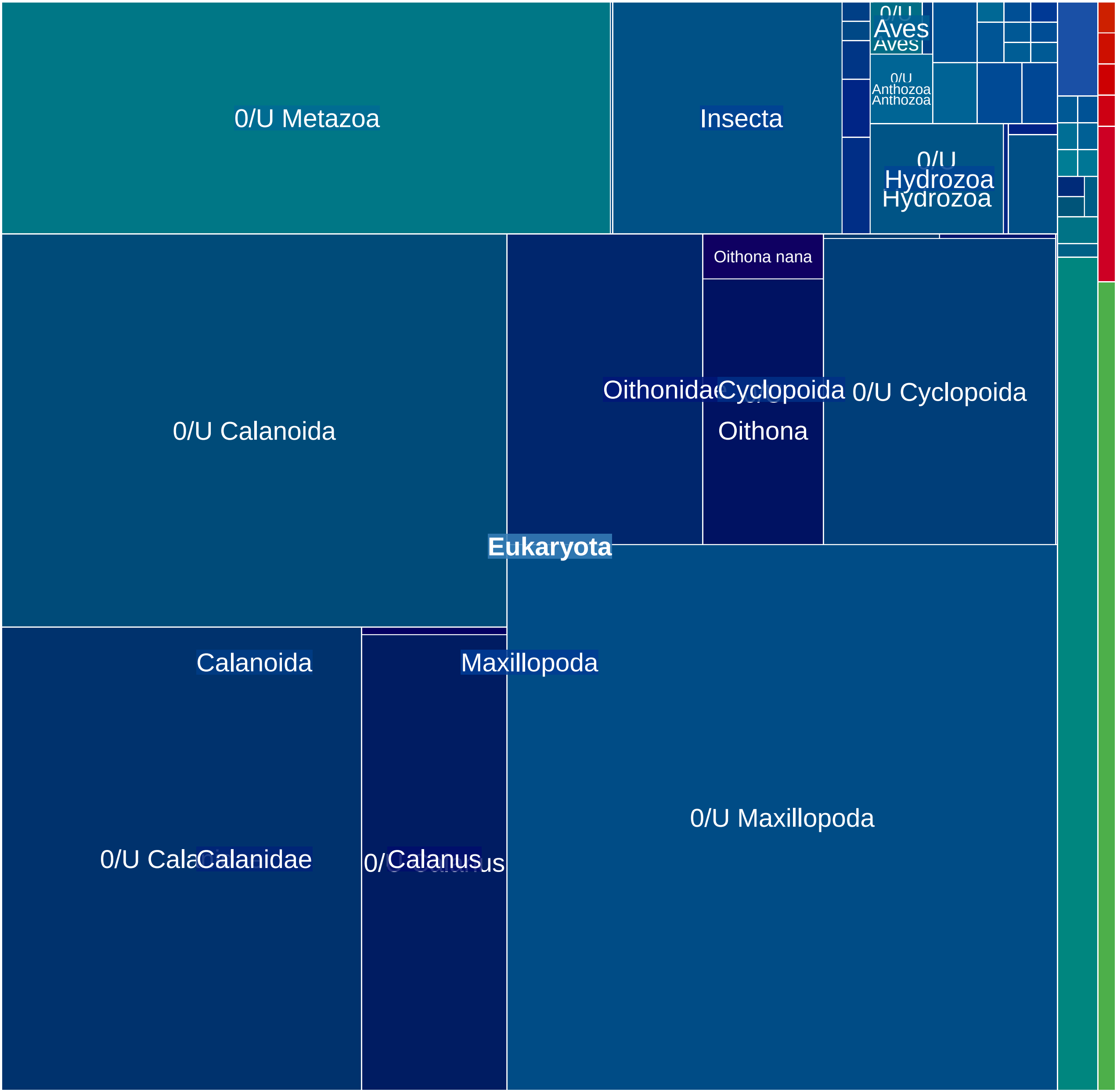

MGT−v1_111 : 13474 unigenes, 6676 (49.55%) taxonomically assigned

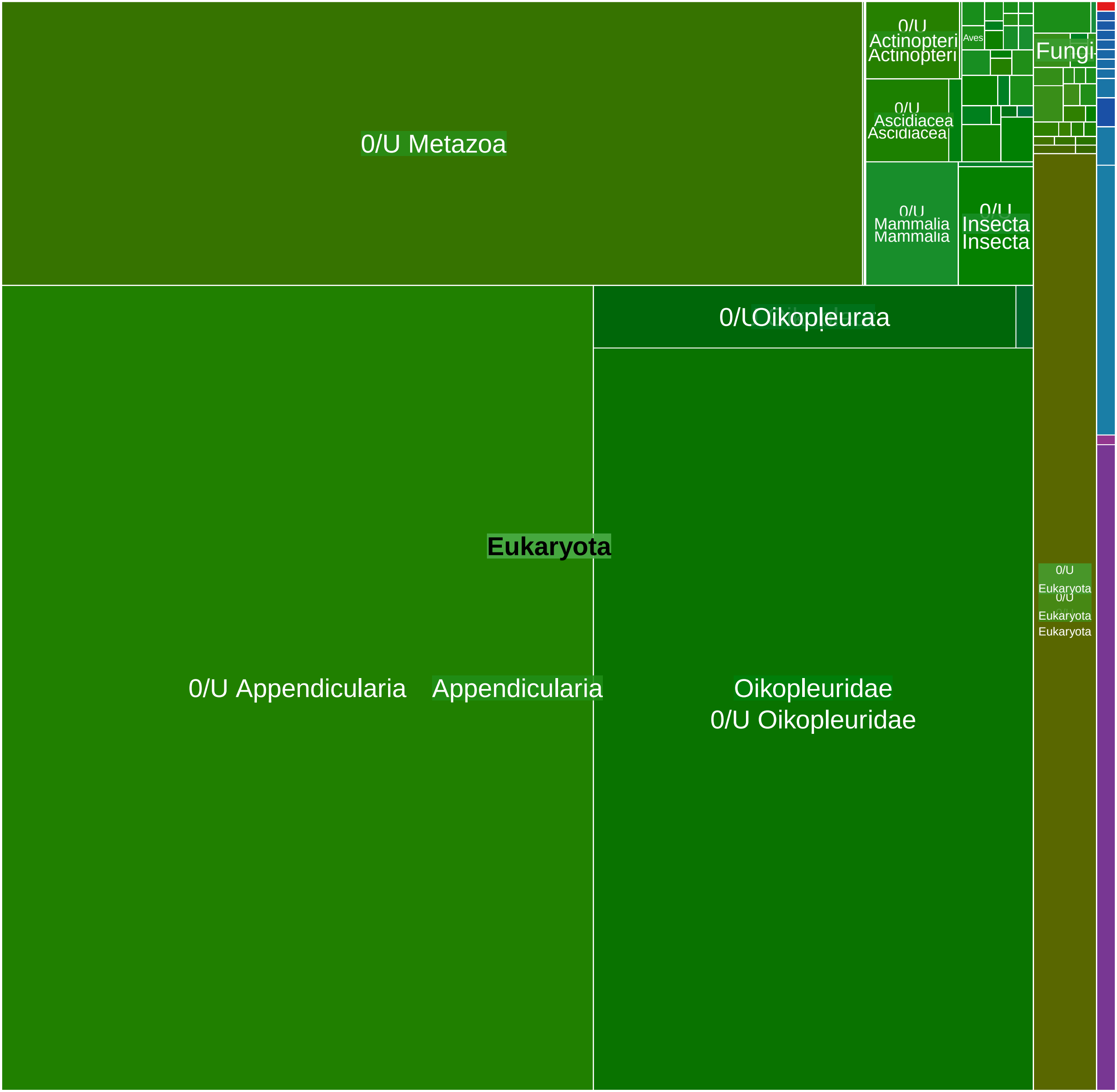

MGT−v1_112 : 13318 unigenes, 8306 (62.37%) taxonomically assigned

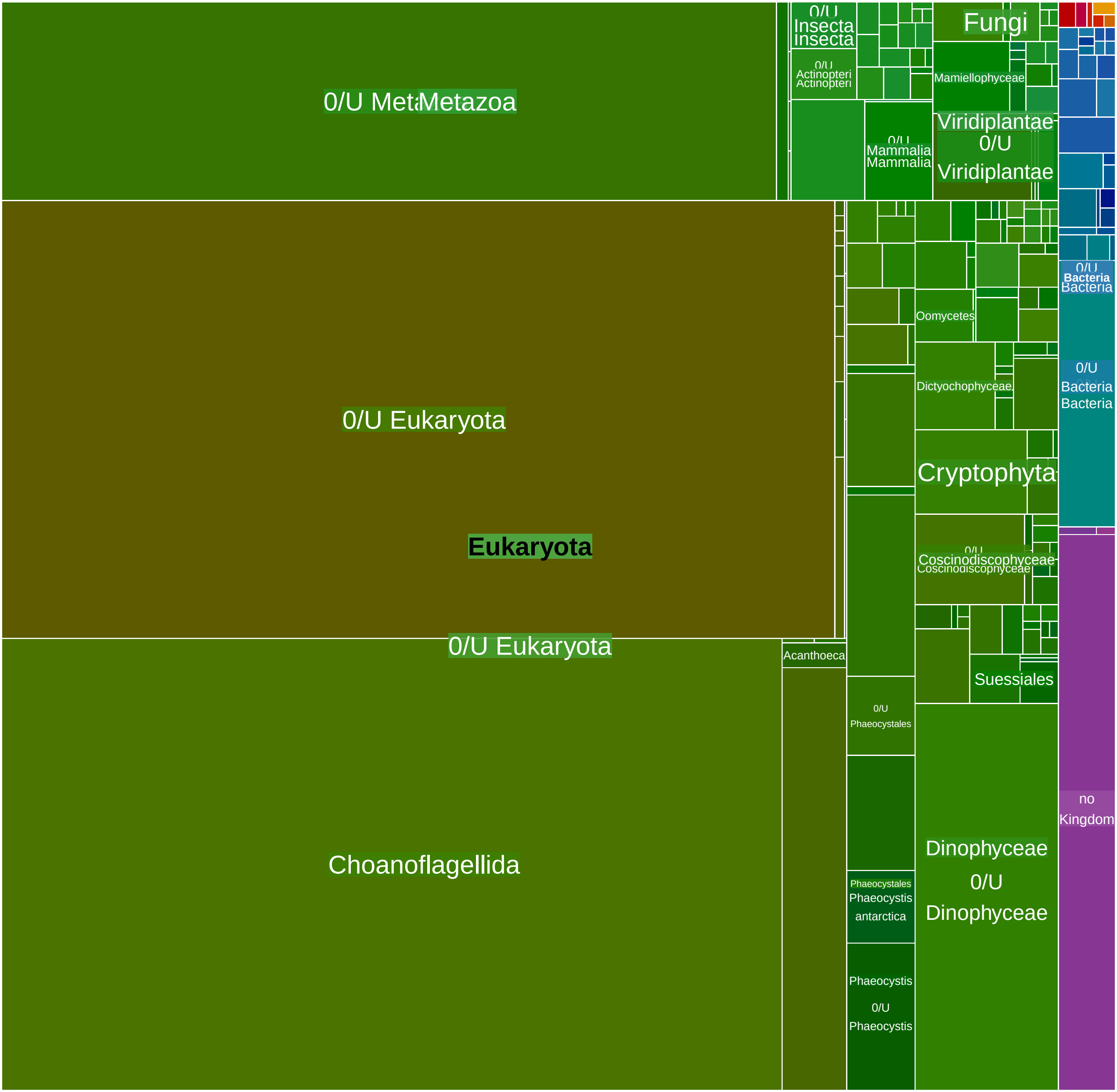

MGT−v1_113 : 13271 unigenes, 1667 (12.56%) taxonomically assigned

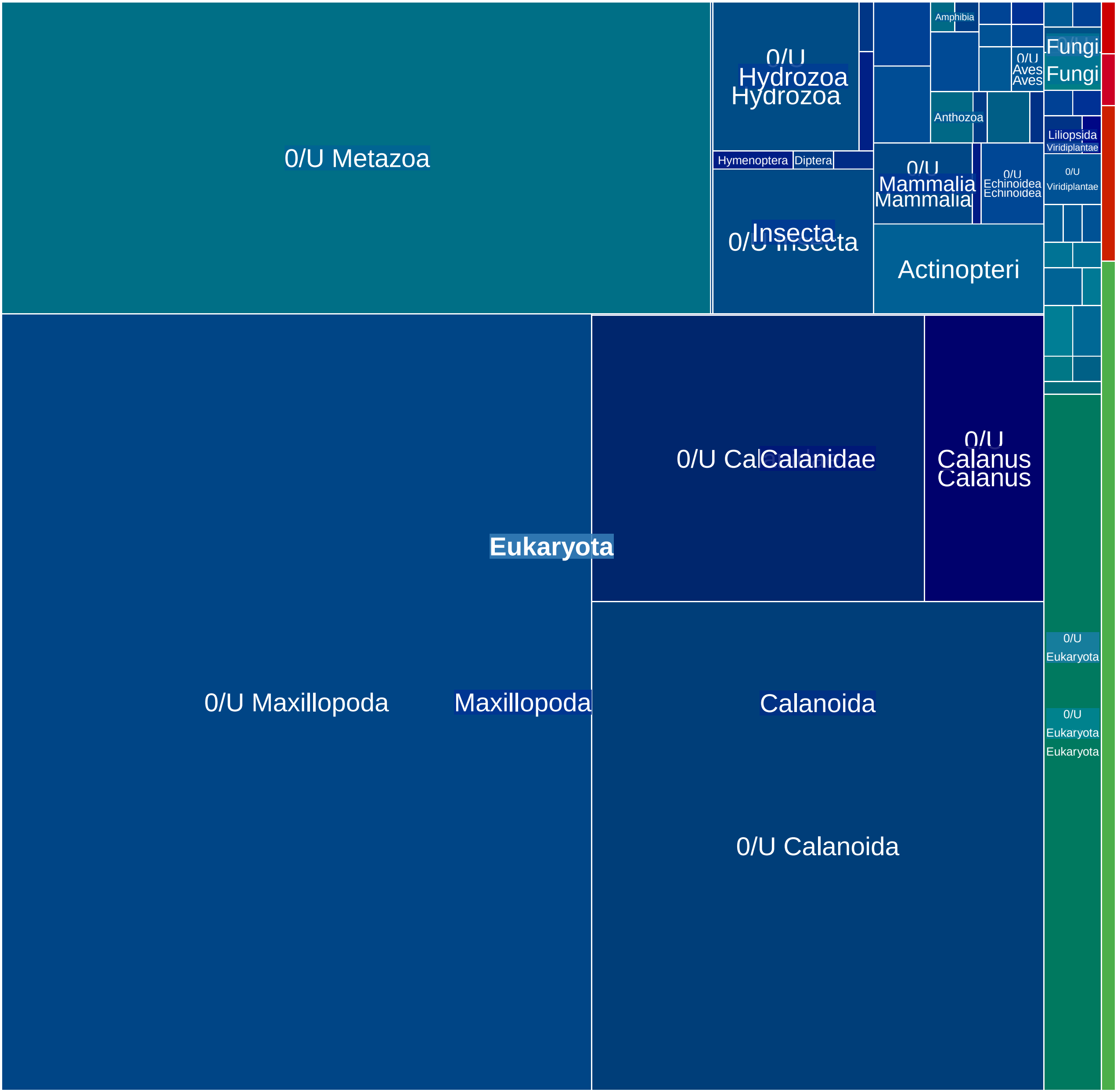

MGT−v1_114 : 13255 unigenes, 985 (7.43%) taxonomically assigned

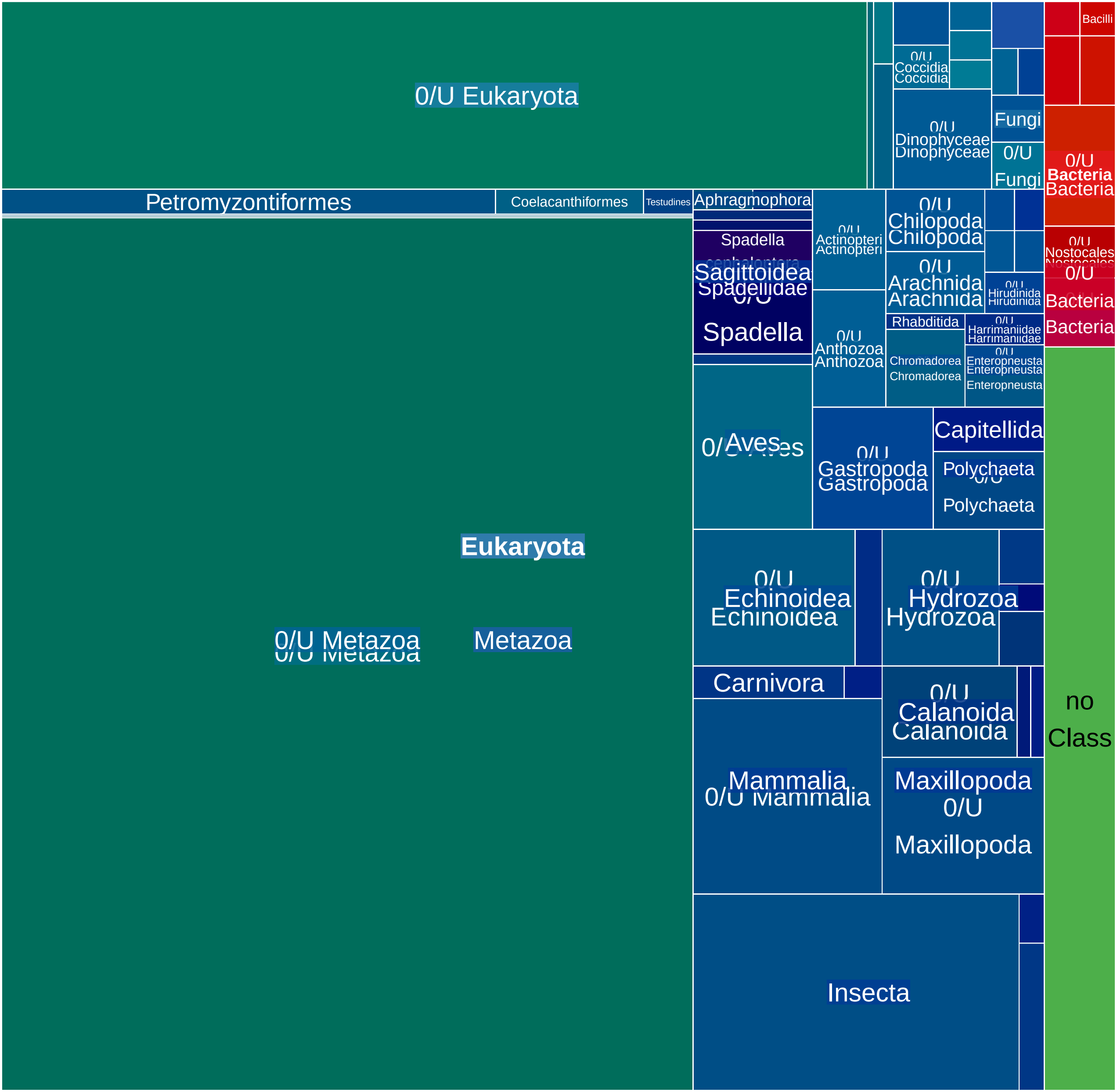

MGT−v1_115 : 12923 unigenes, 11604 (89.79%) taxonomically assigned

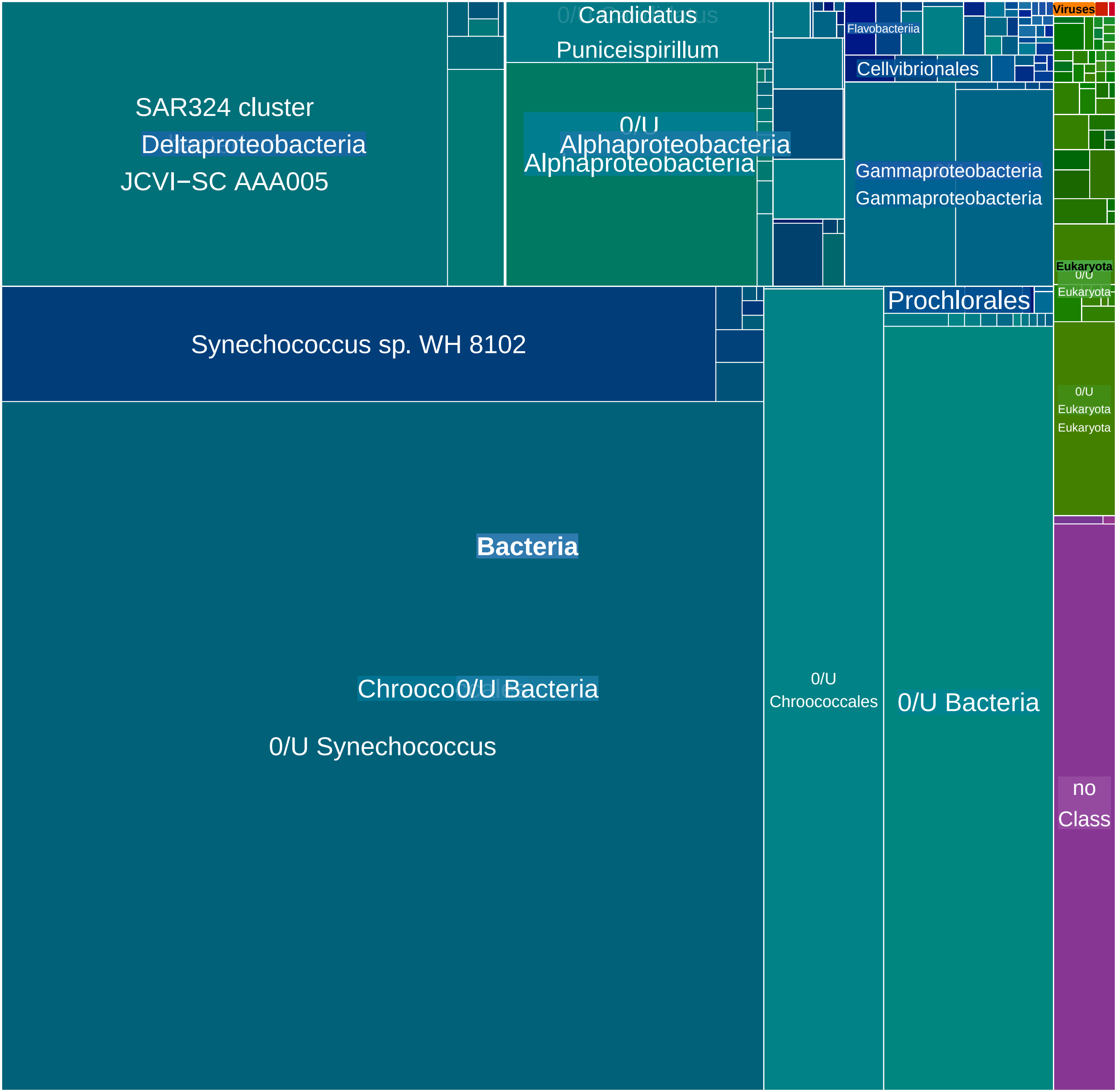

MGT−v1_116 : 12873 unigenes, 3688 (28.65%) taxonomically assigned

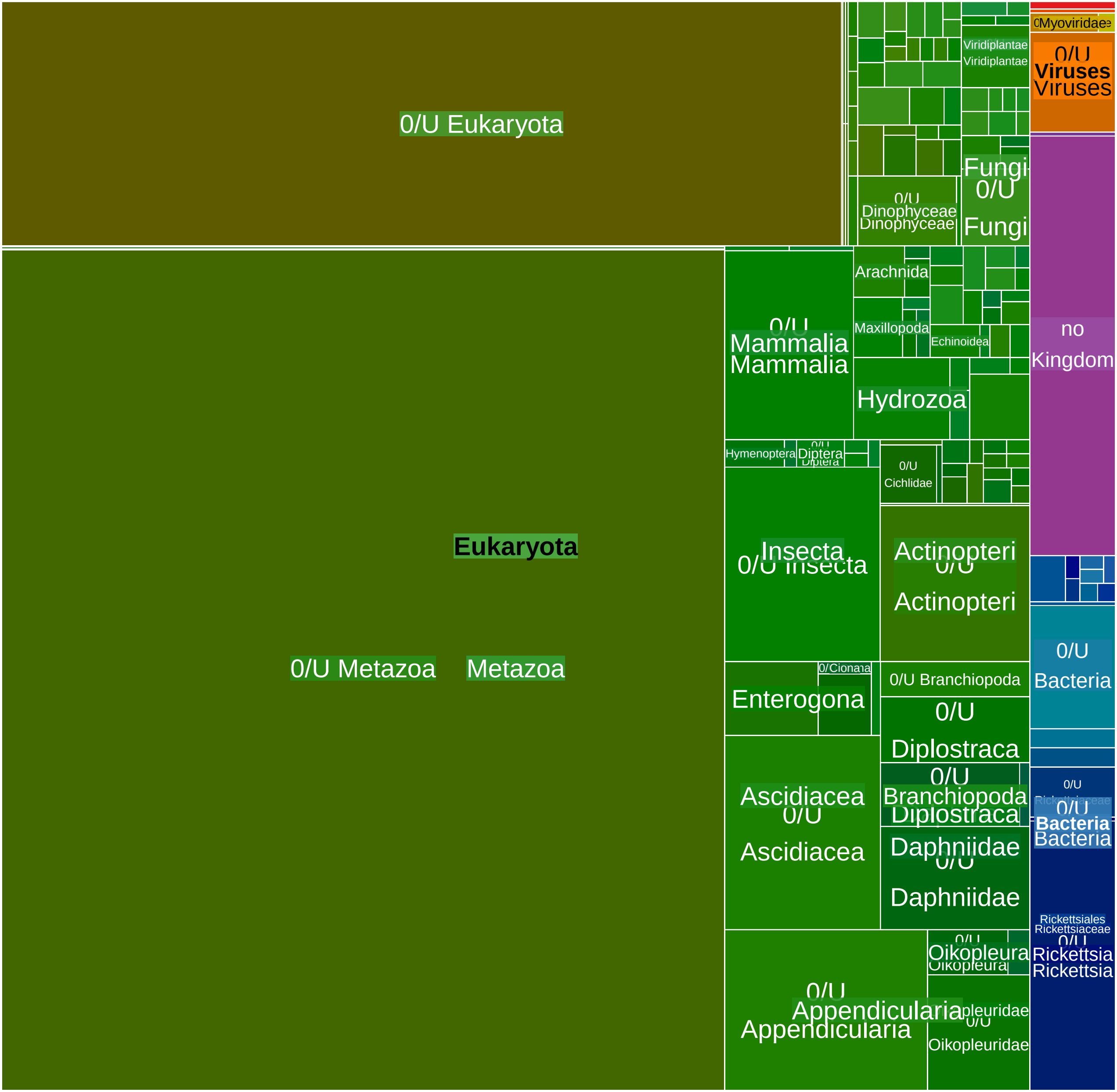

MGT−v1_117 : 12859 unigenes, 11970 (93.09%) taxonomically assigned

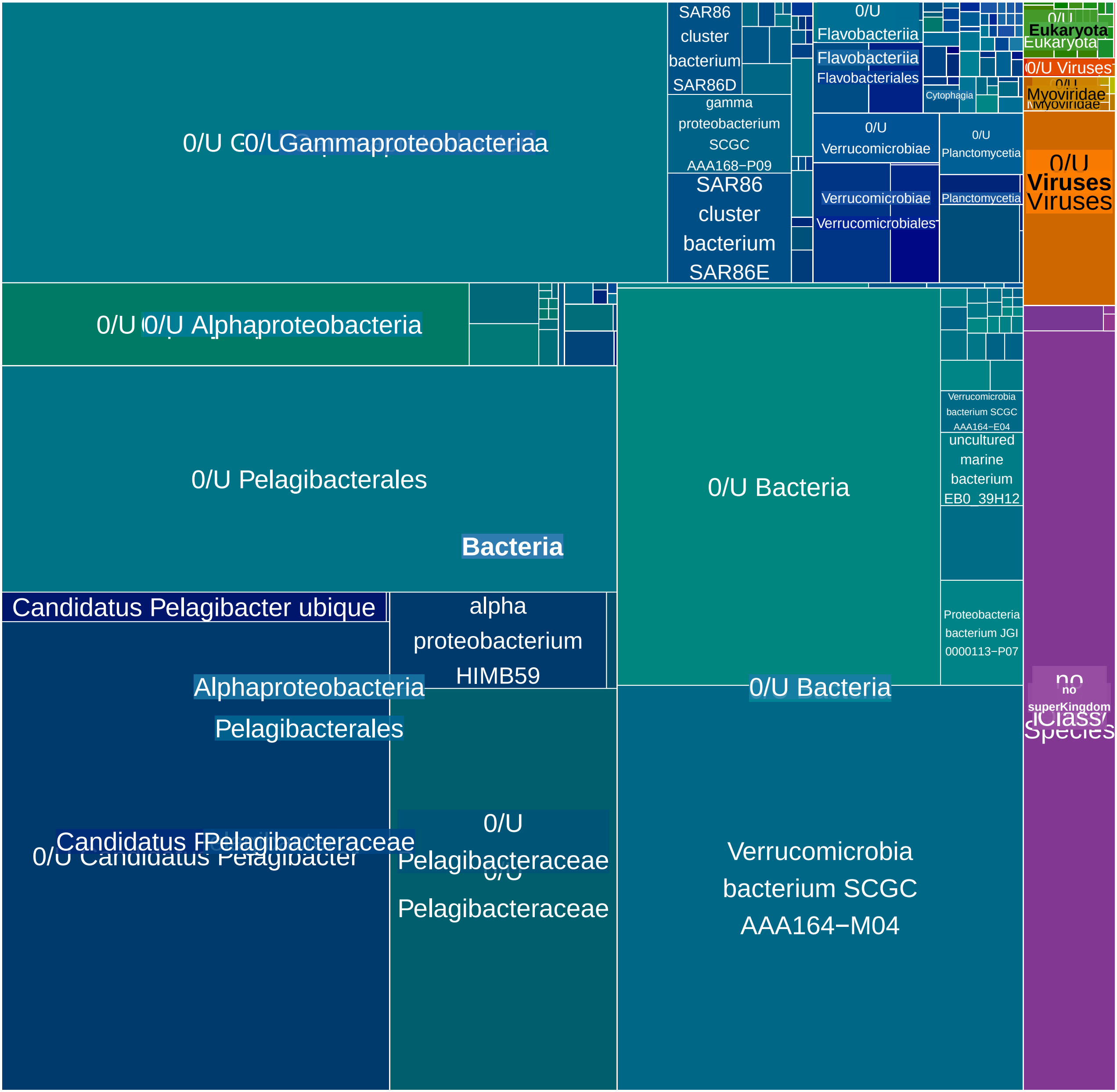

### MGT−v1_118 : 12698 unigenes, 3344 (26.33%) taxonomically assigned

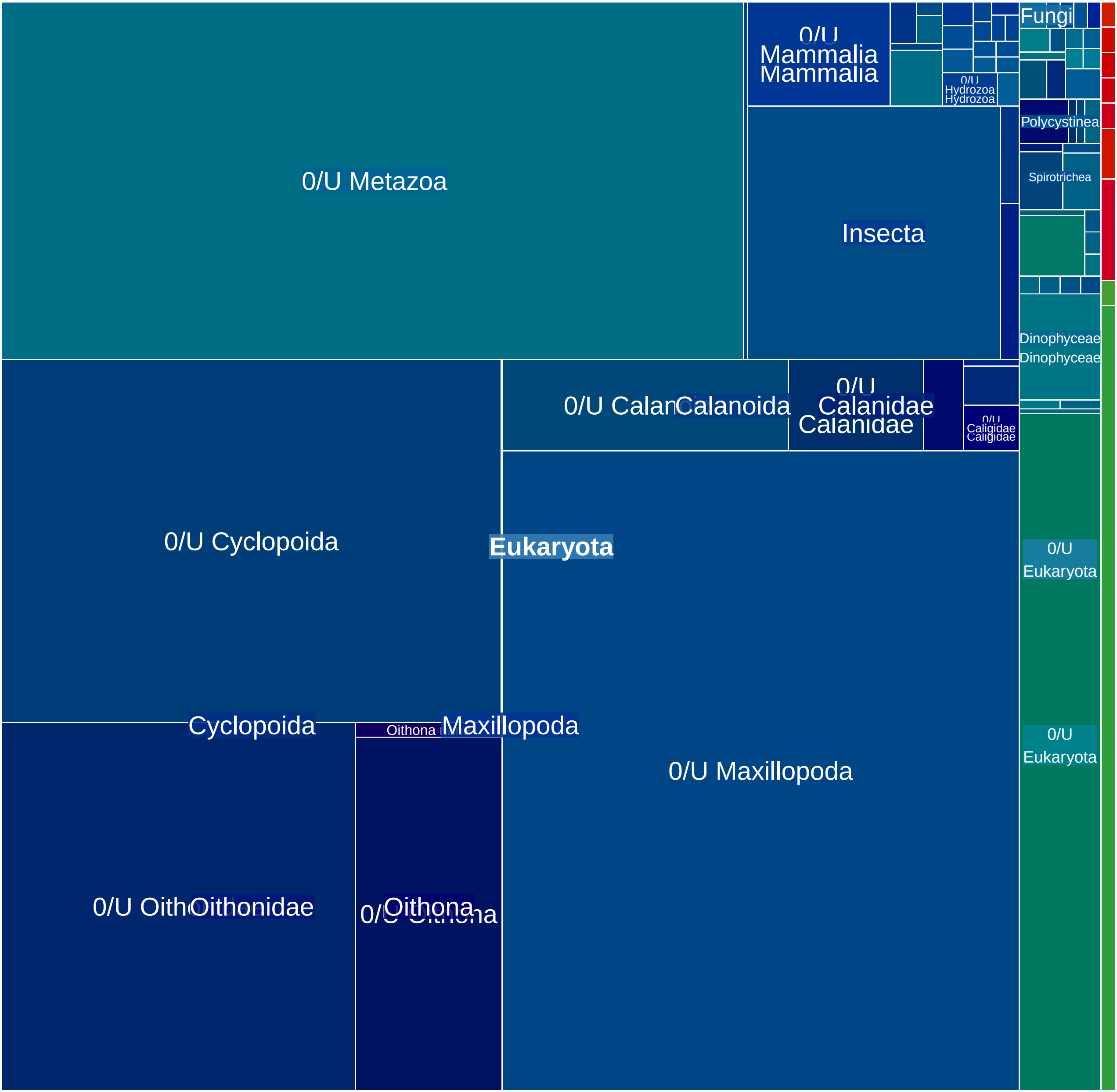

MGT−v1_119 : 12683 unigenes, 2354 (18.56%) taxonomically assigned

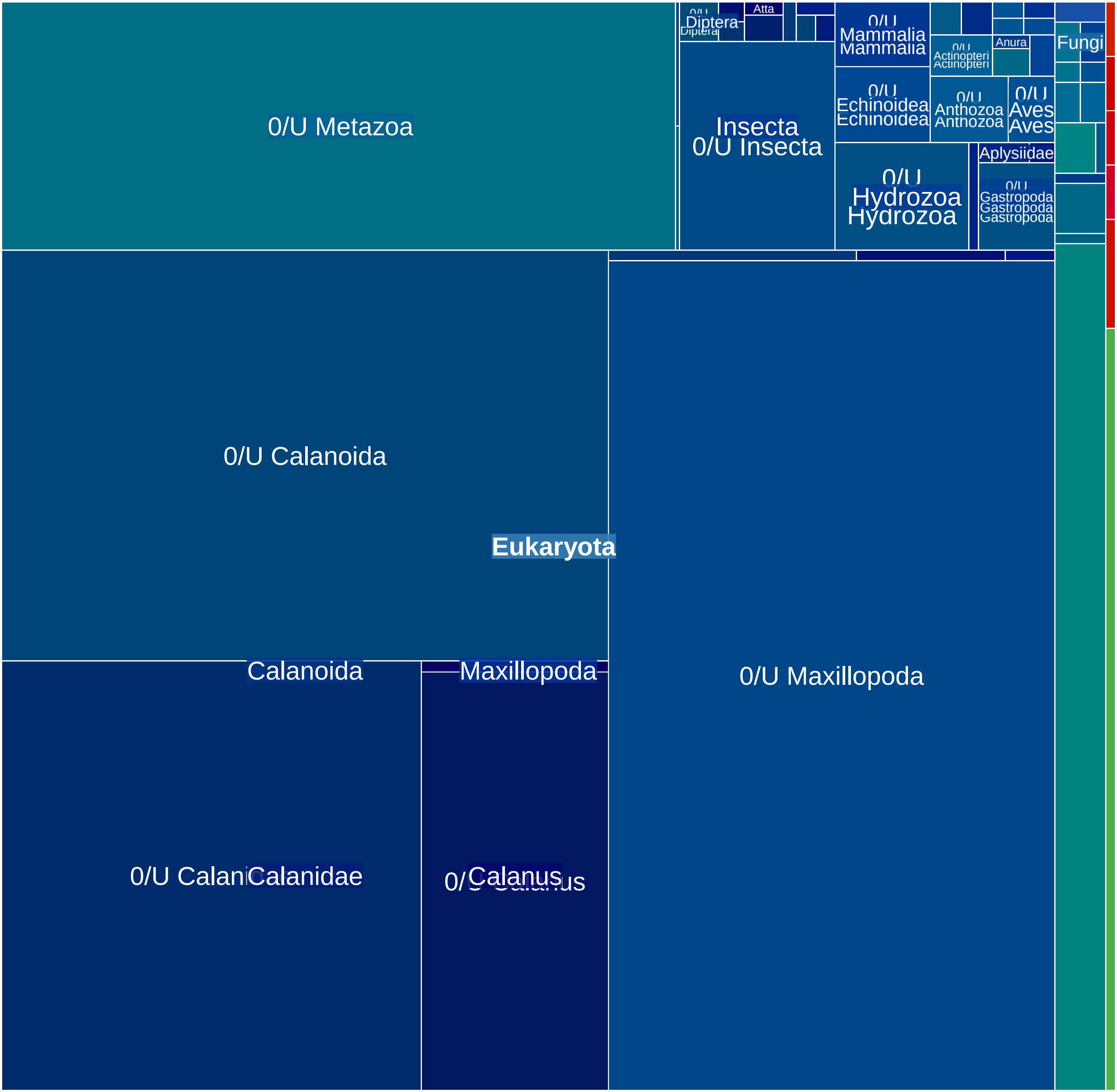

MGT−v1_120 : 12668 unigenes, 7381 (58.26%) taxonomically assigned

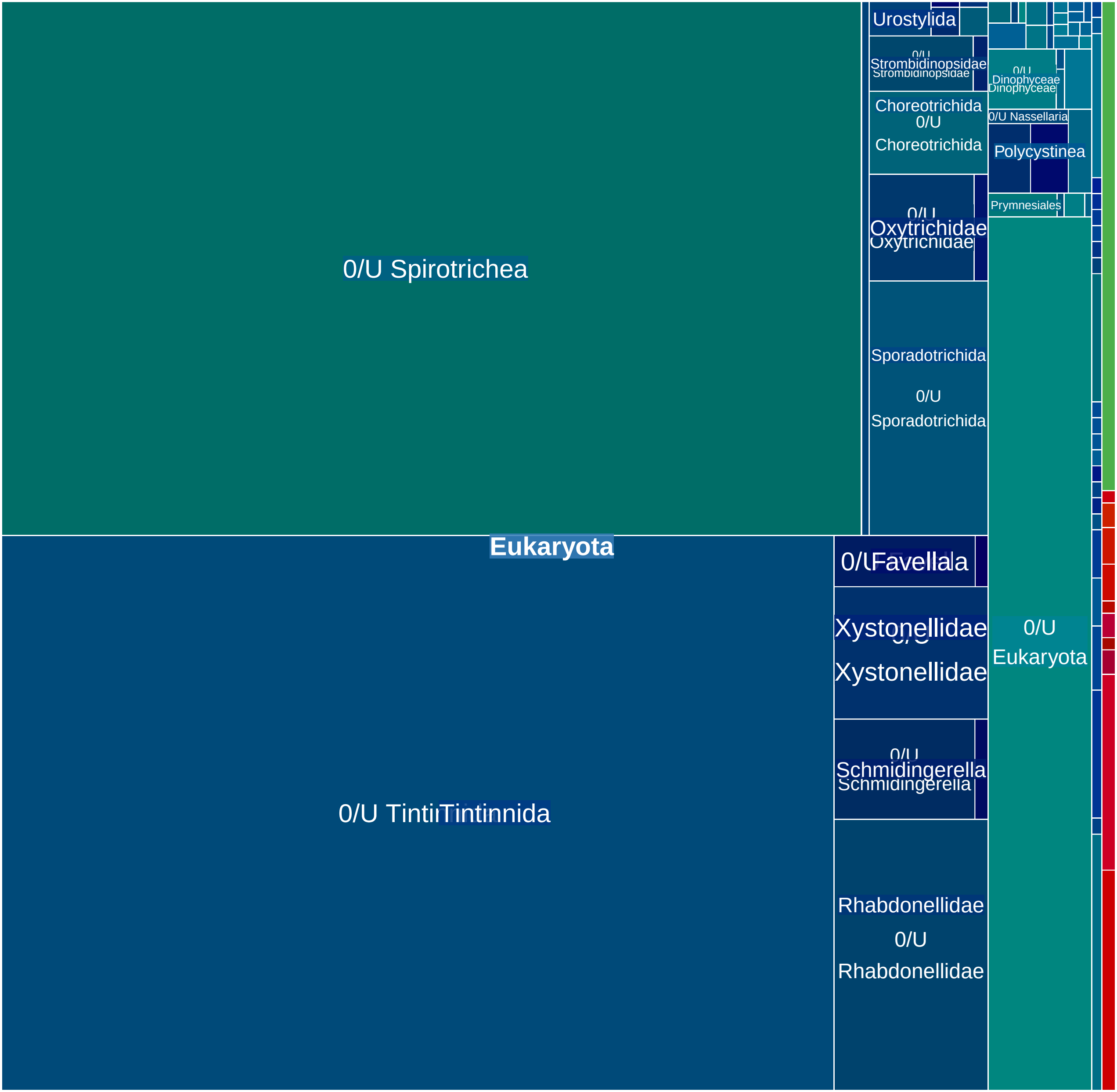

### MGT−v1_121 : 12643 unigenes, 2376 (18.79%) taxonomically assigned

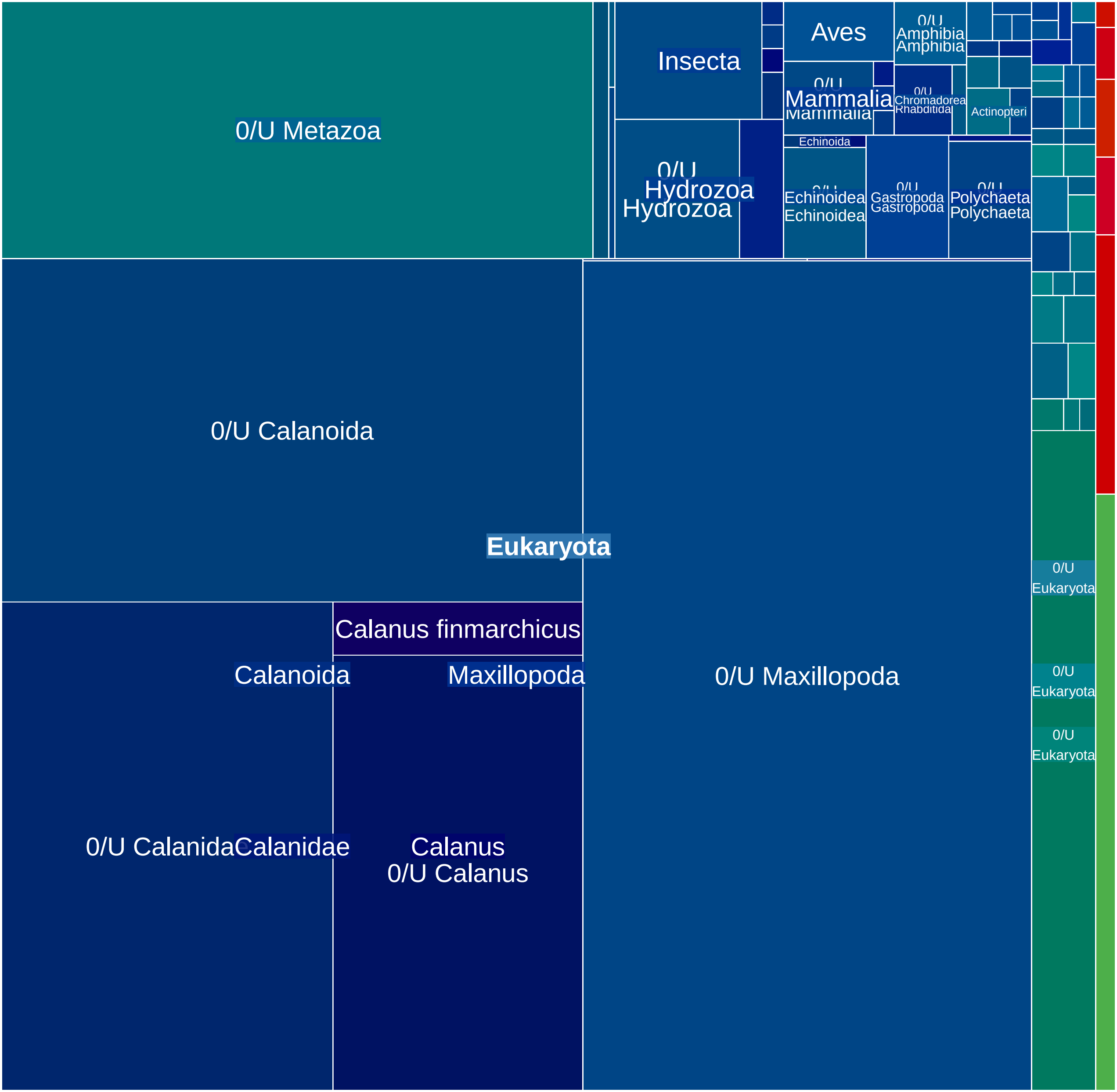

MGT−v1_122 : 12629 unigenes, 8679 (68.72%) taxonomically assigned

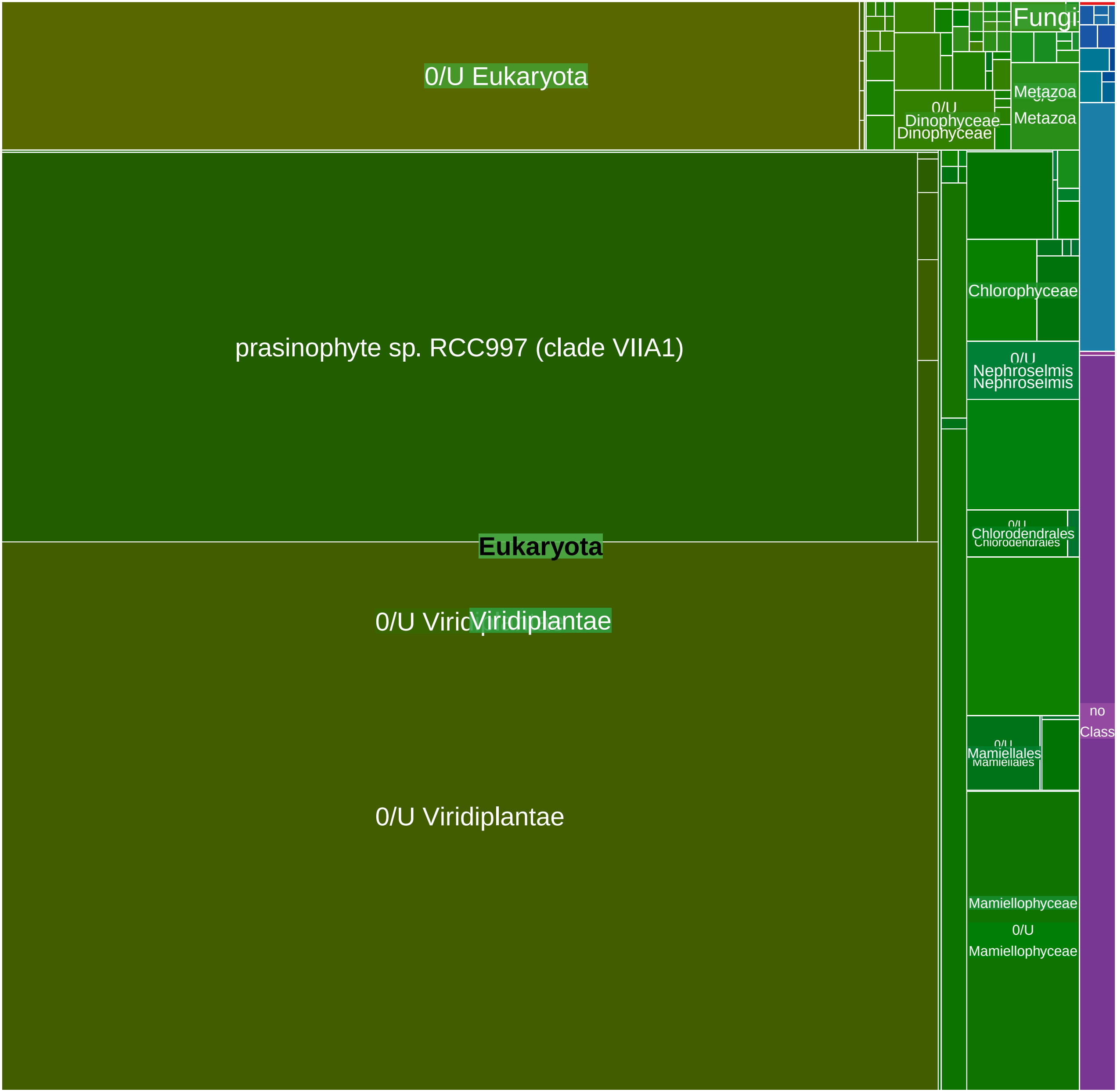

MGT−v1_123 : 12584 unigenes, 11695 (92.94%) taxonomically assigned

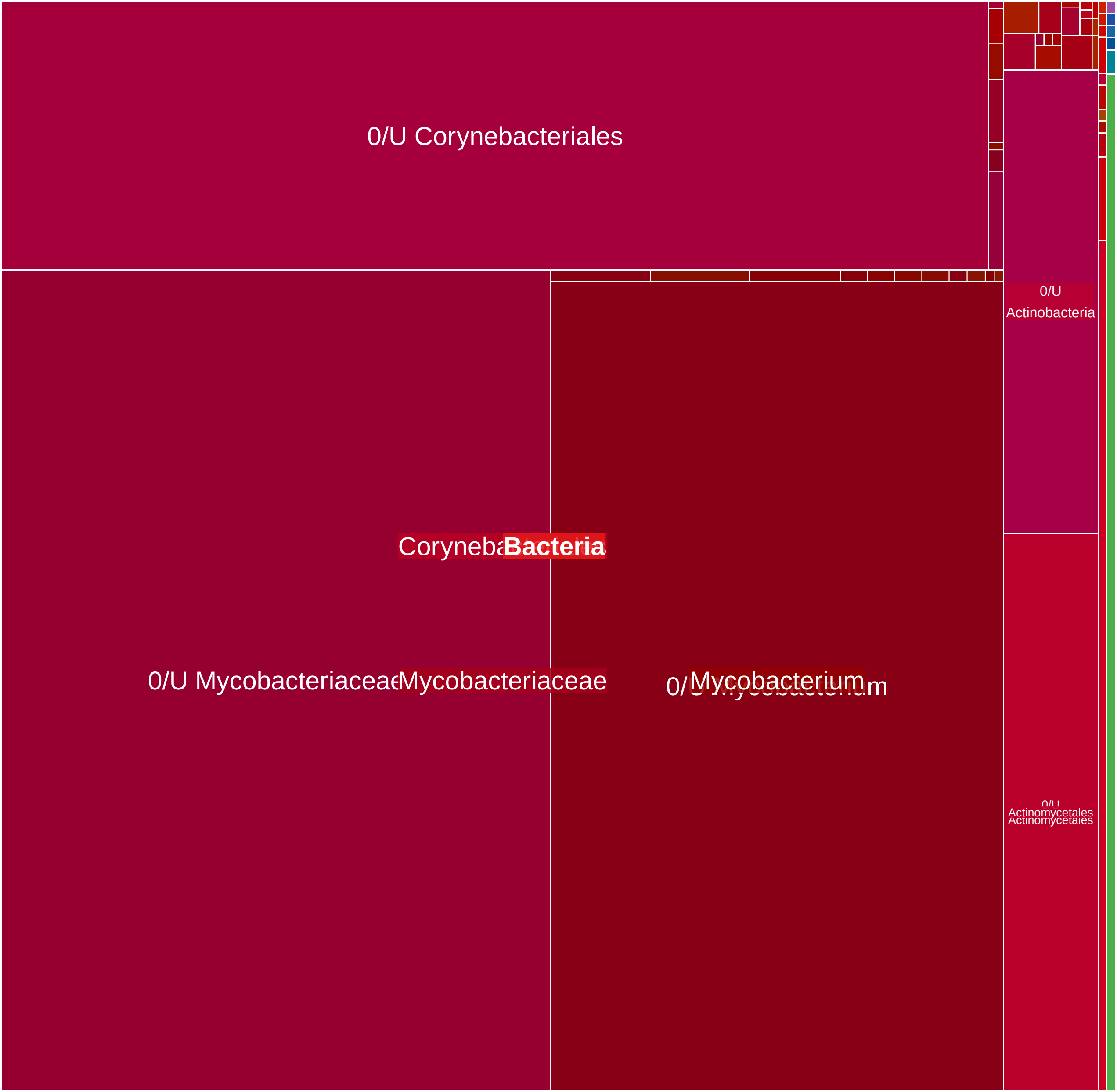

MGT−v1_124 : 12520 unigenes, 1003 (8.01%) taxonomically assigned

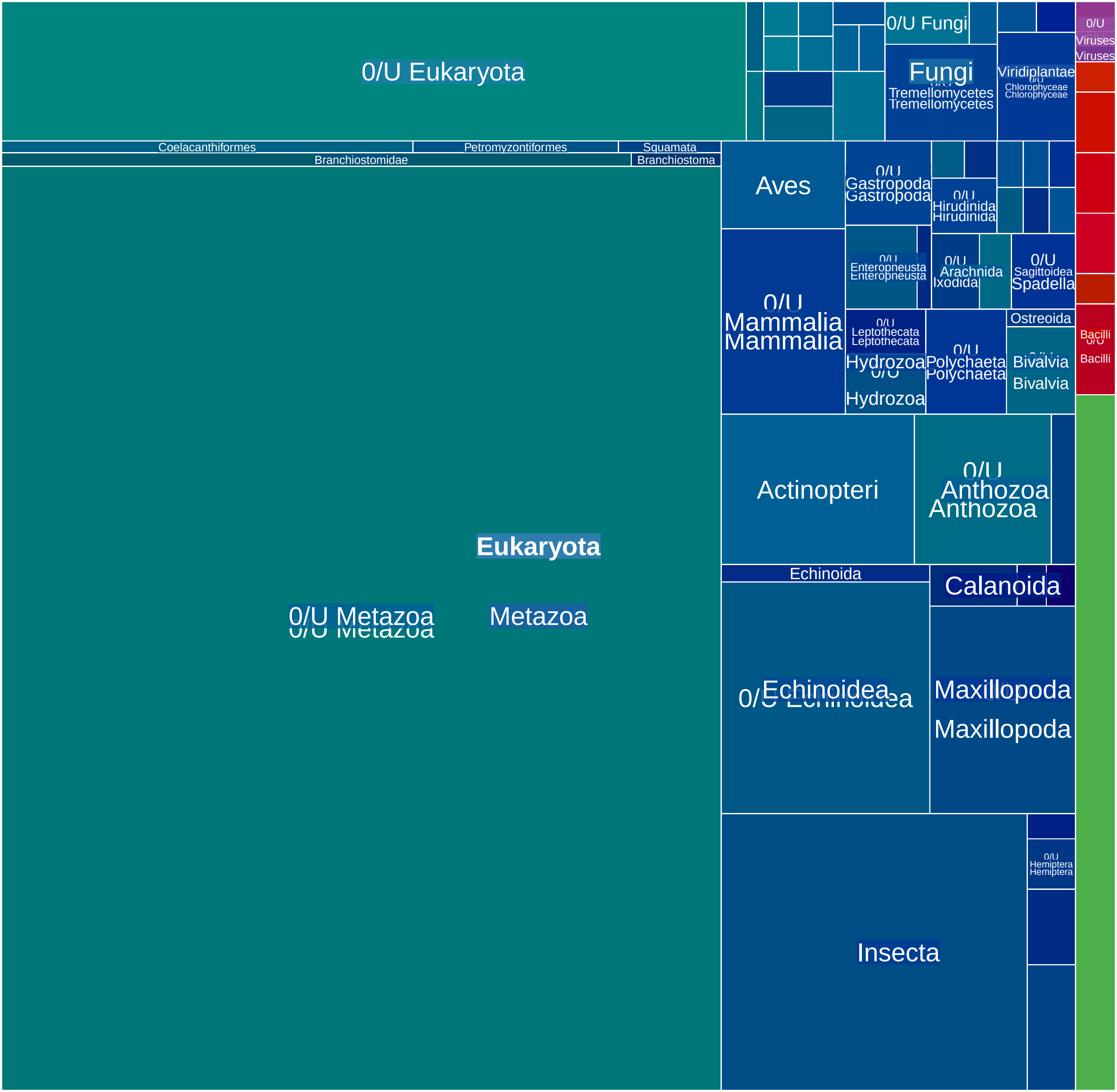

MGT−v1_125 : 12316 unigenes, 12026 (97.65%) taxonomically assigned

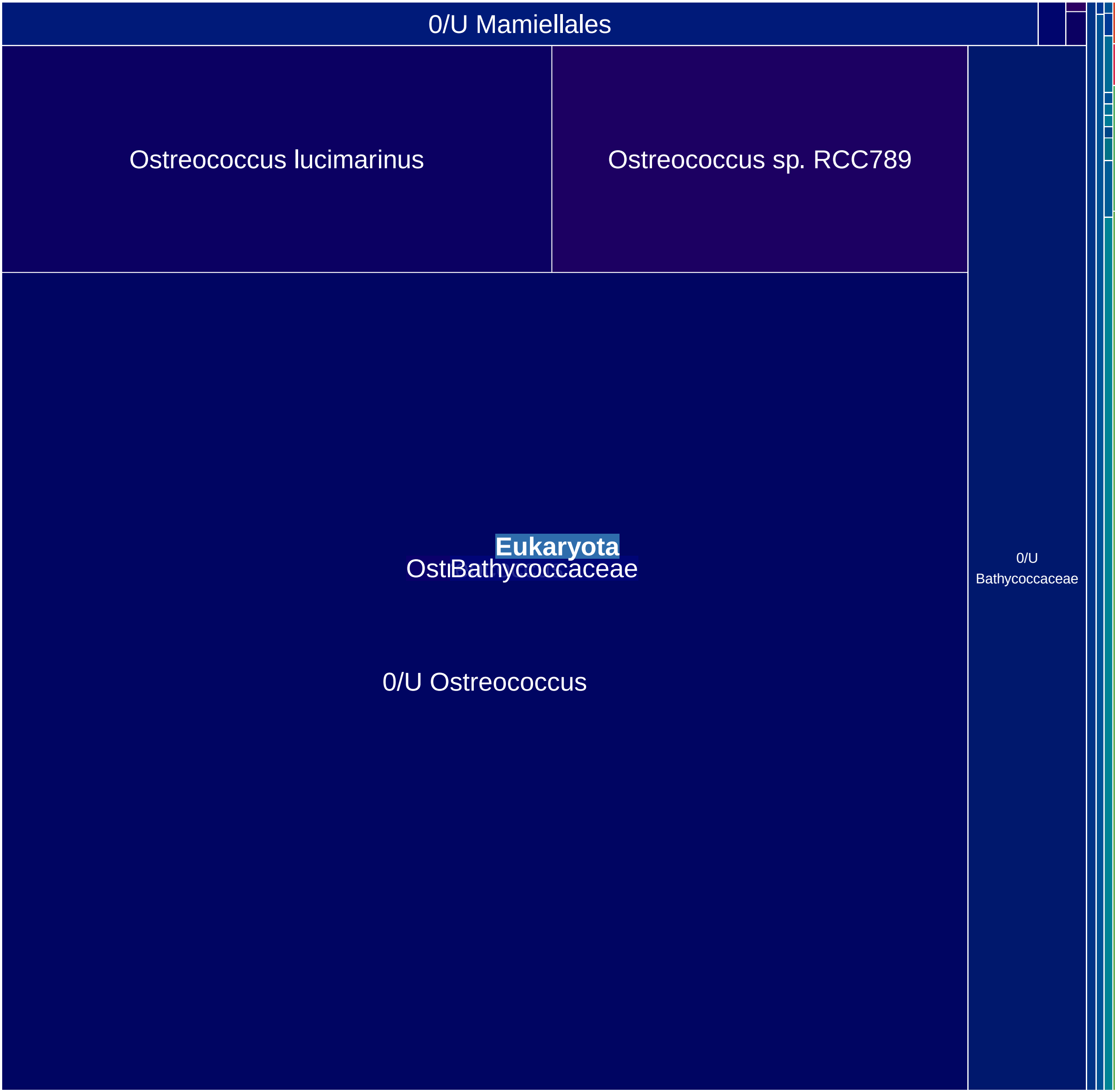

MGT−v1_126 : 12306 unigenes, 822 (6.68%) taxonomically assigned

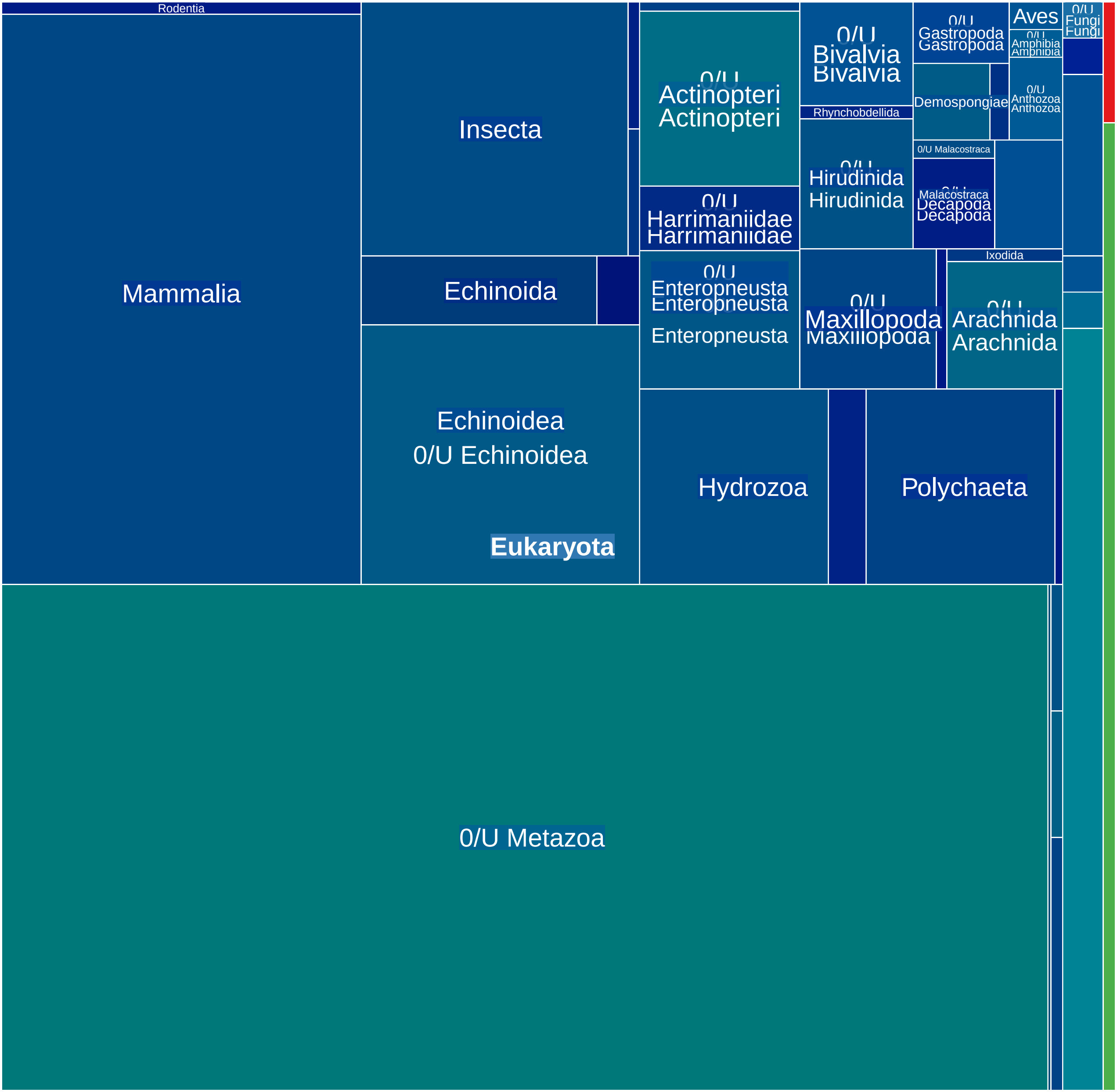

MGT−v1_127 : 12269 unigenes, 11540 (94.06%) taxonomically assigned

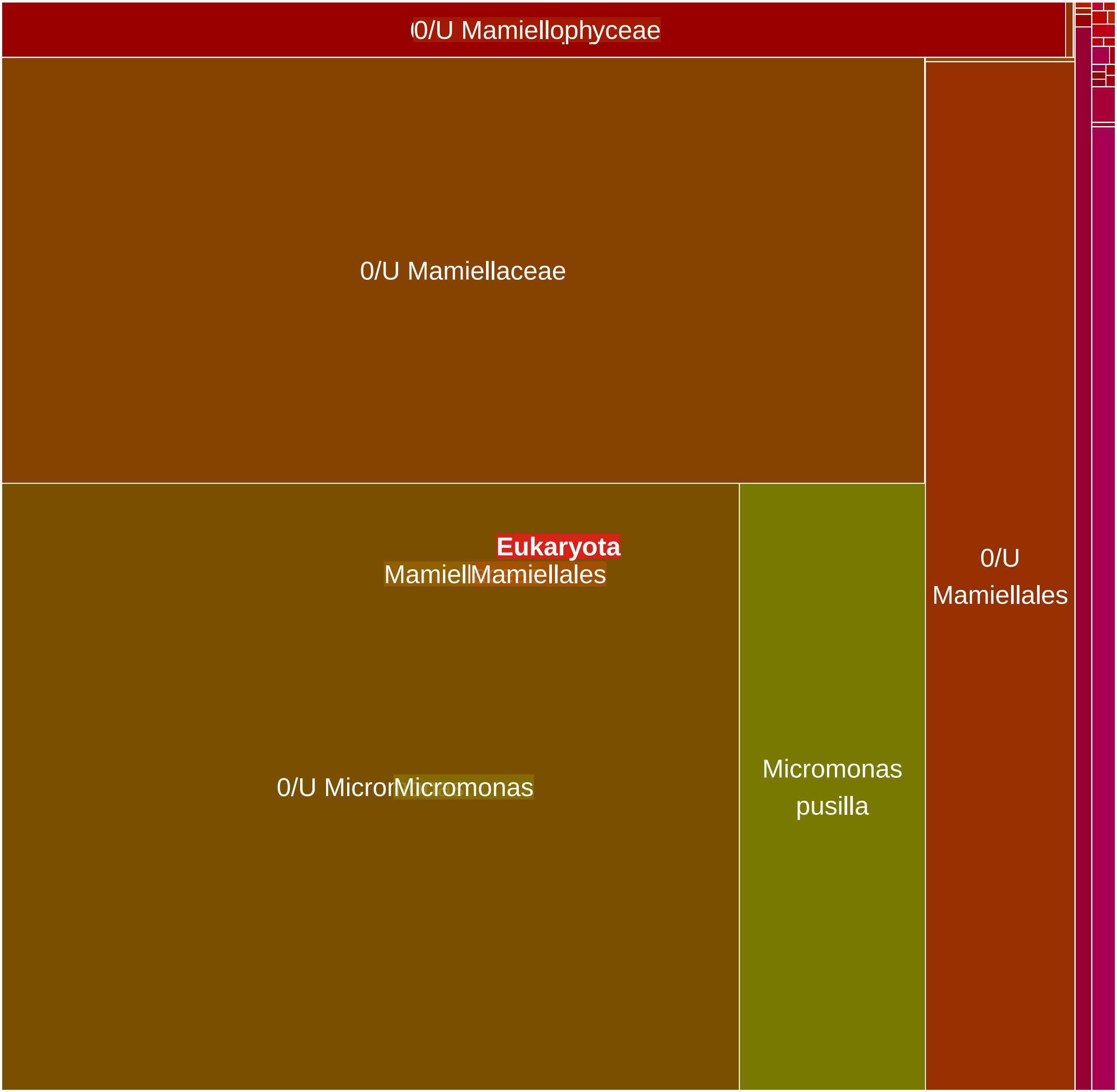

MGT−v1_128 : 12153 unigenes, 6805 (55.99%) taxonomically assigned

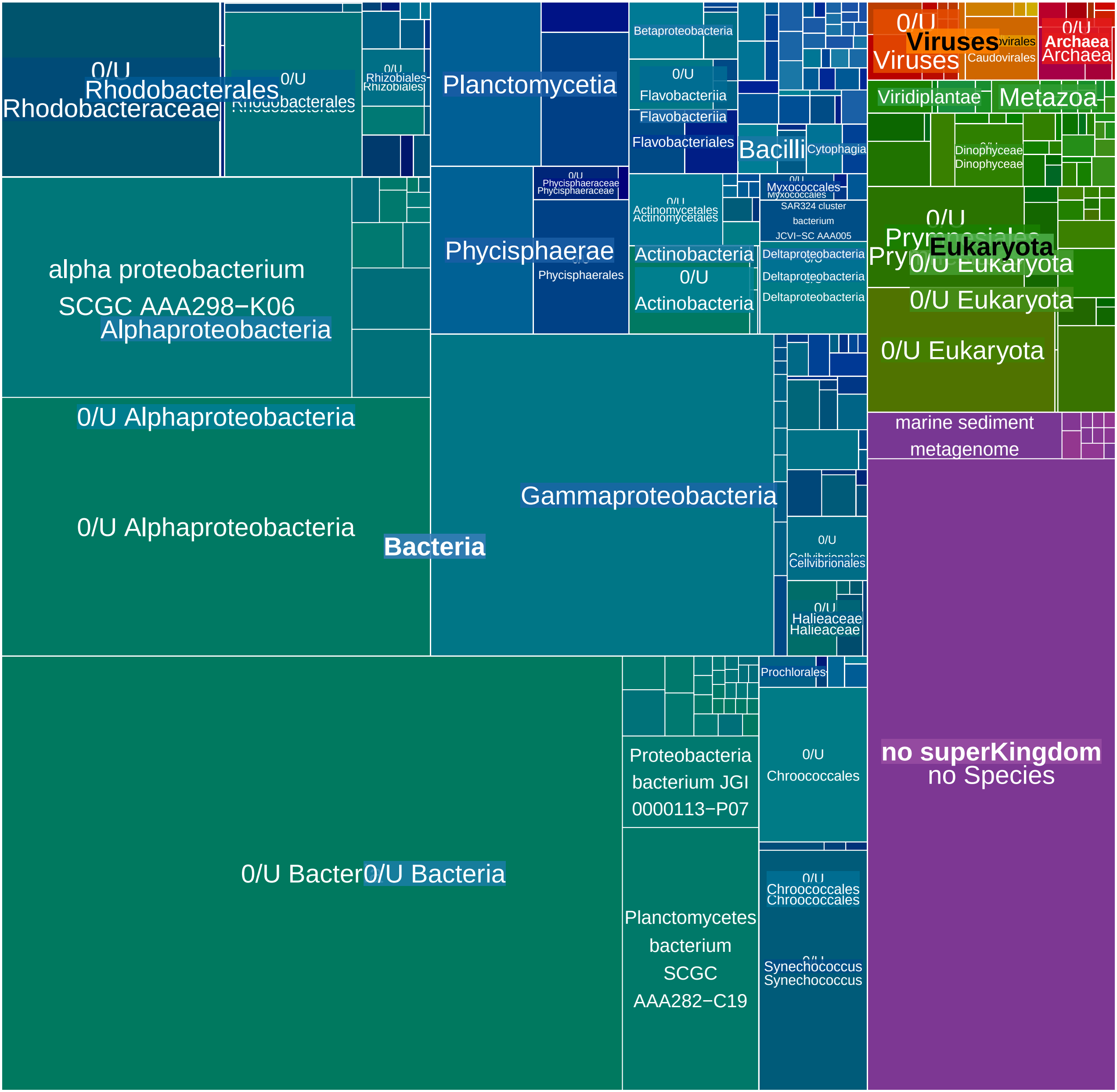

MGT−v1_129 : 12093 unigenes, 10555 (87.28%) taxonomically assigned

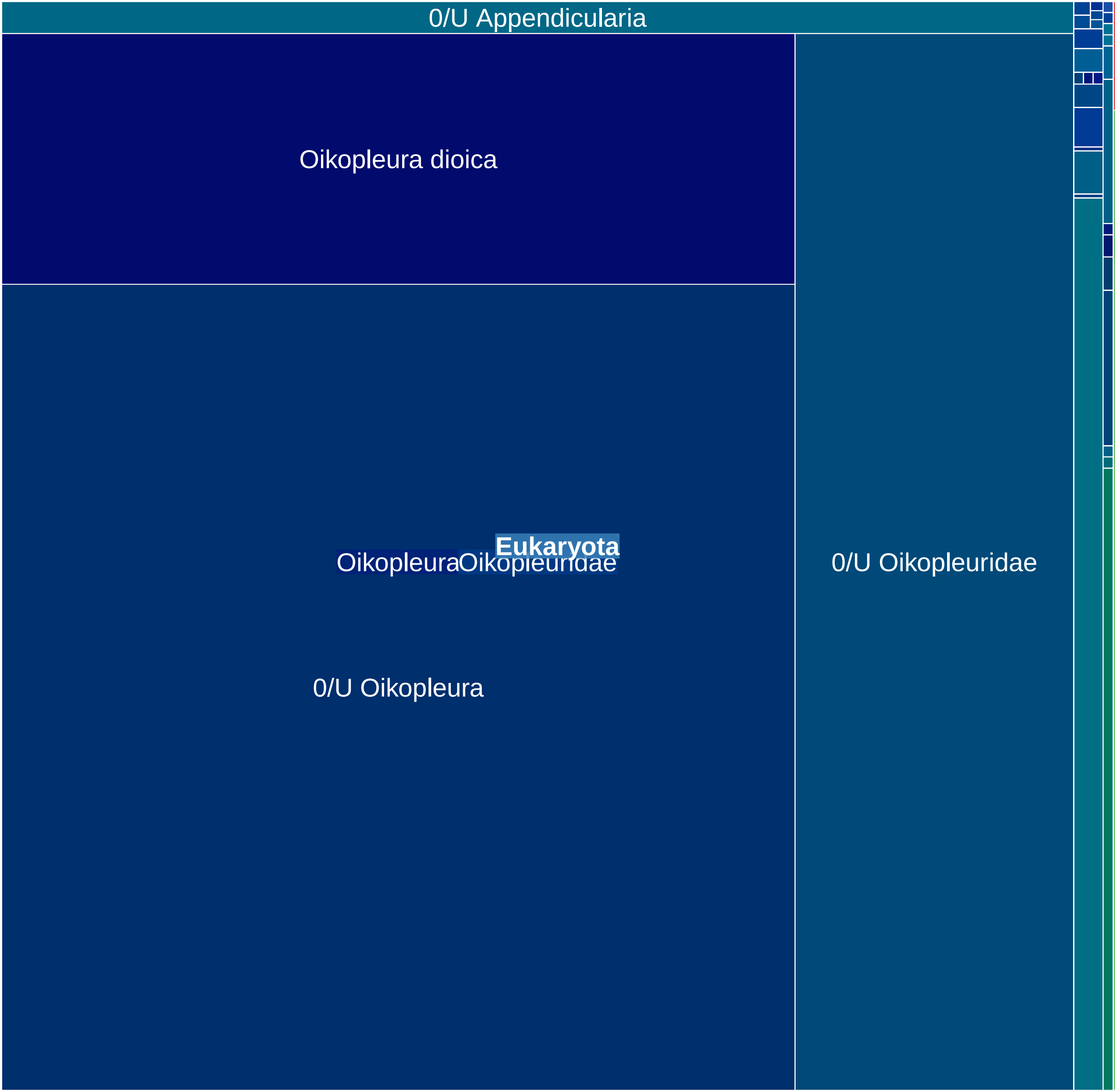

MGT−v1_130 : 12014 unigenes, 632 (5.26%) taxonomically assigned

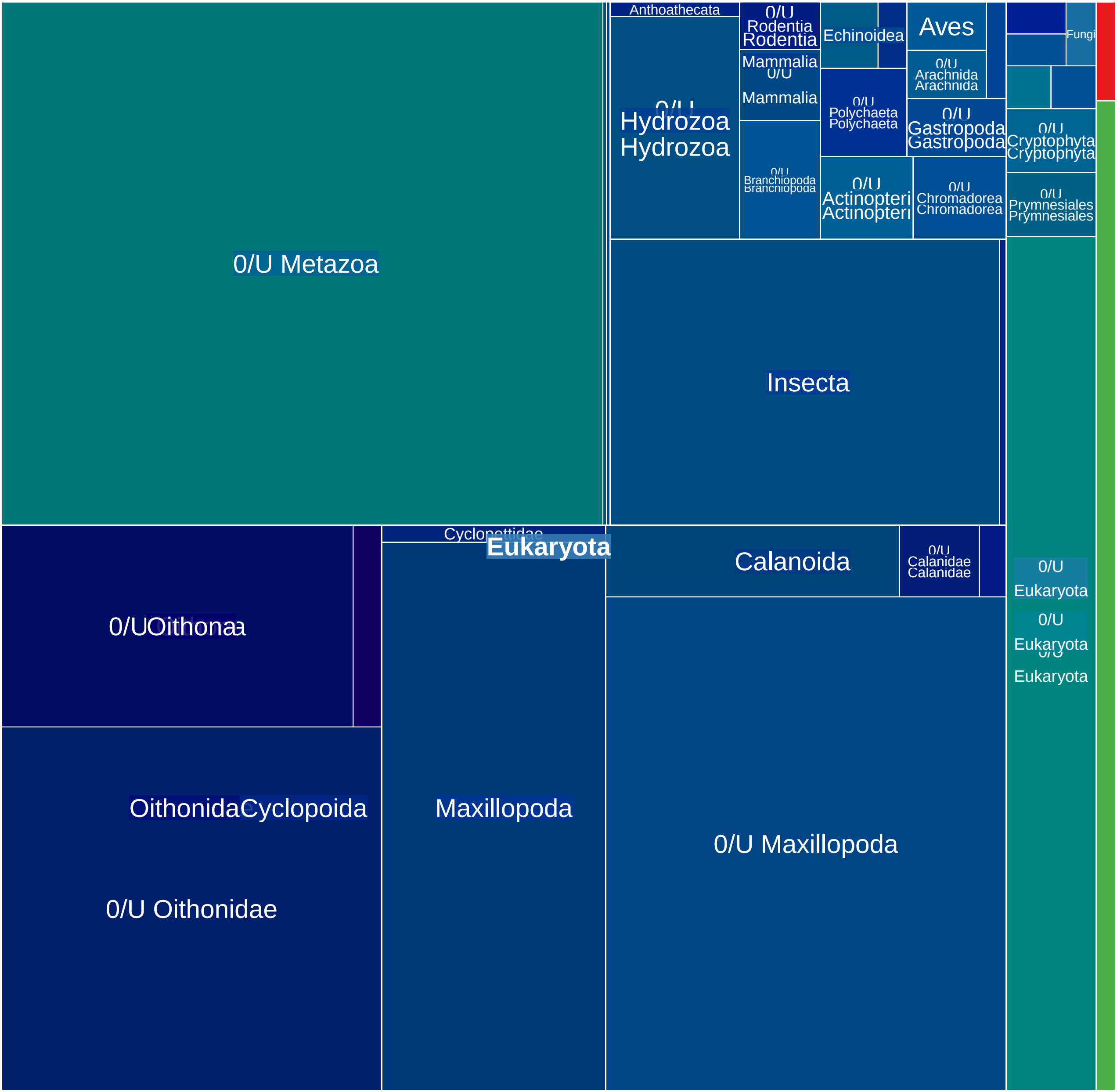

MGT−v1_131 : 11922 unigenes, 8692 (72.91%) taxonomically assigned

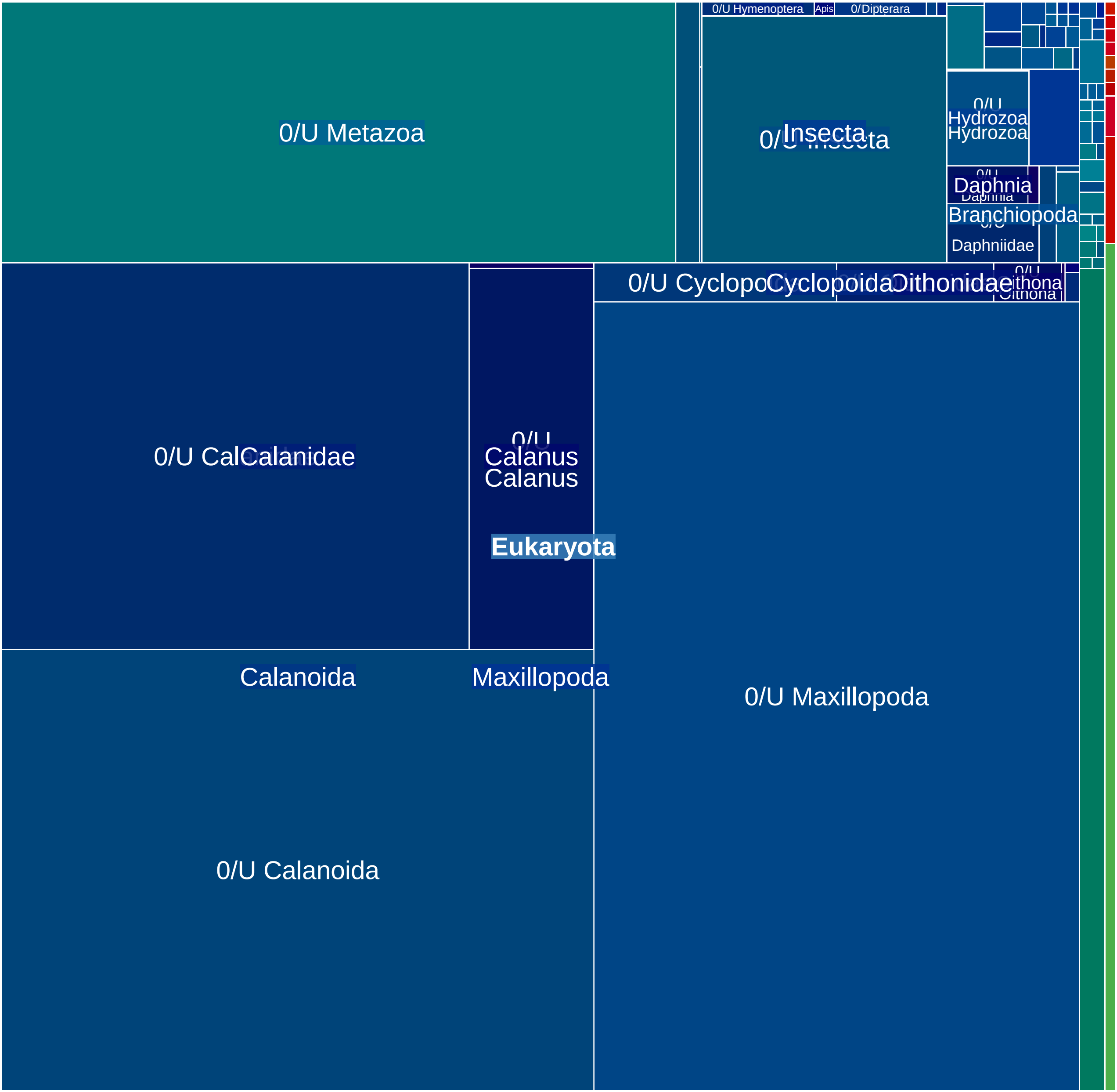

### MGT−v1_132 : 11894 unigenes, 10226 (85.98%) taxonomically assigned

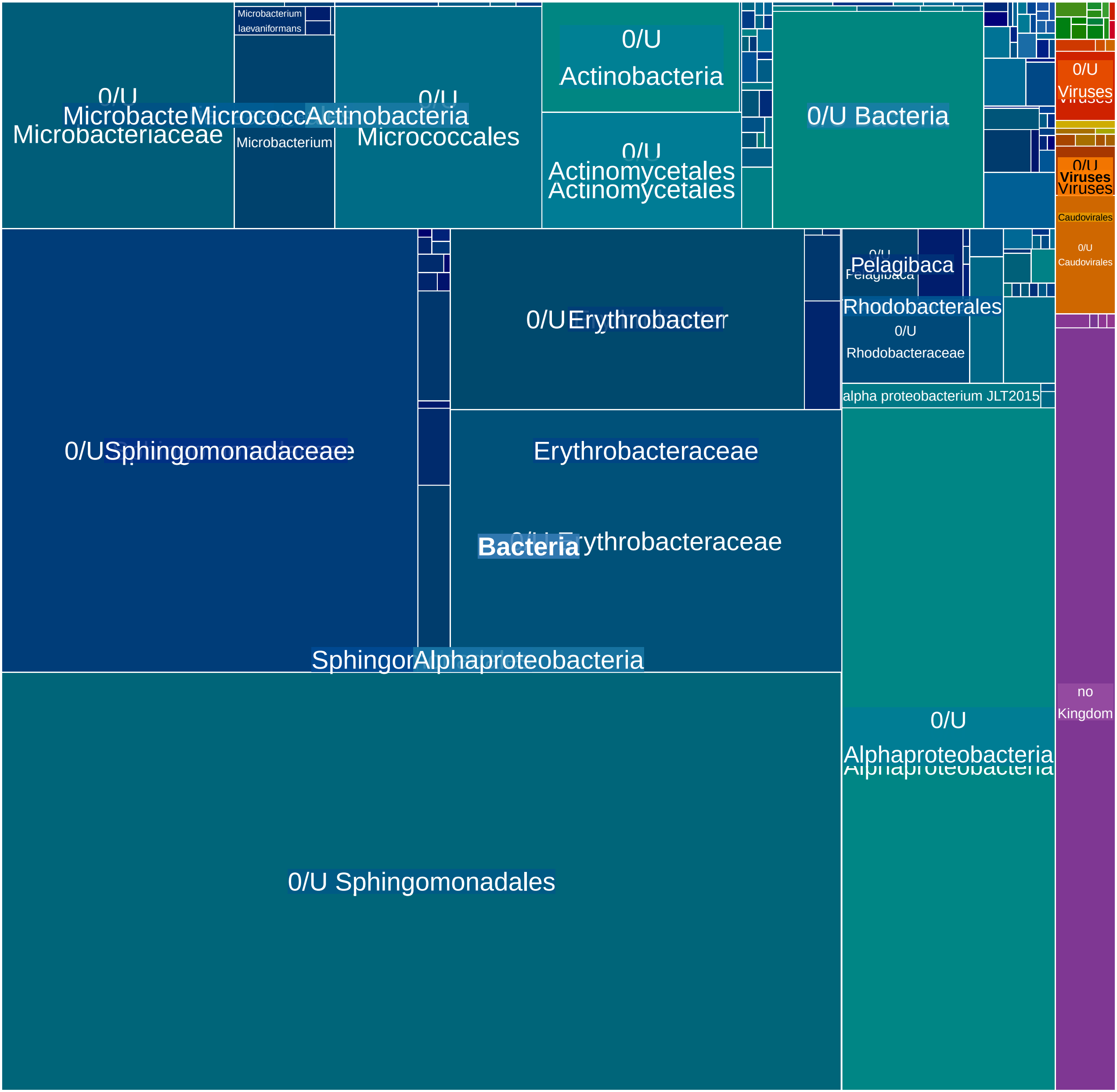

MGT−v1_133 : 11827 unigenes, 9413 (79.59%) taxonomically assigned

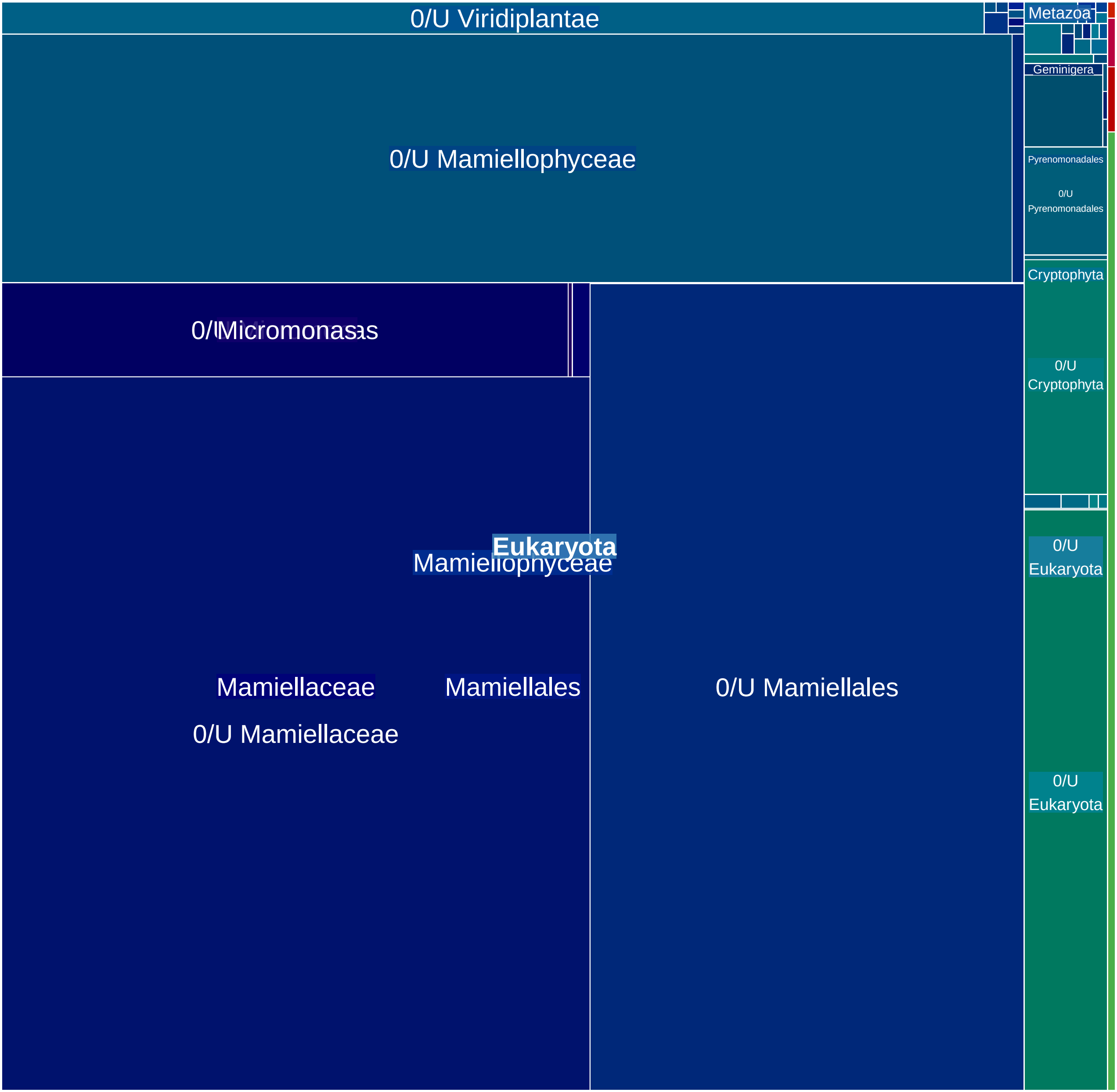

MGT−v1_134 : 11696 unigenes, 1840 (15.73%) taxonomically assigned

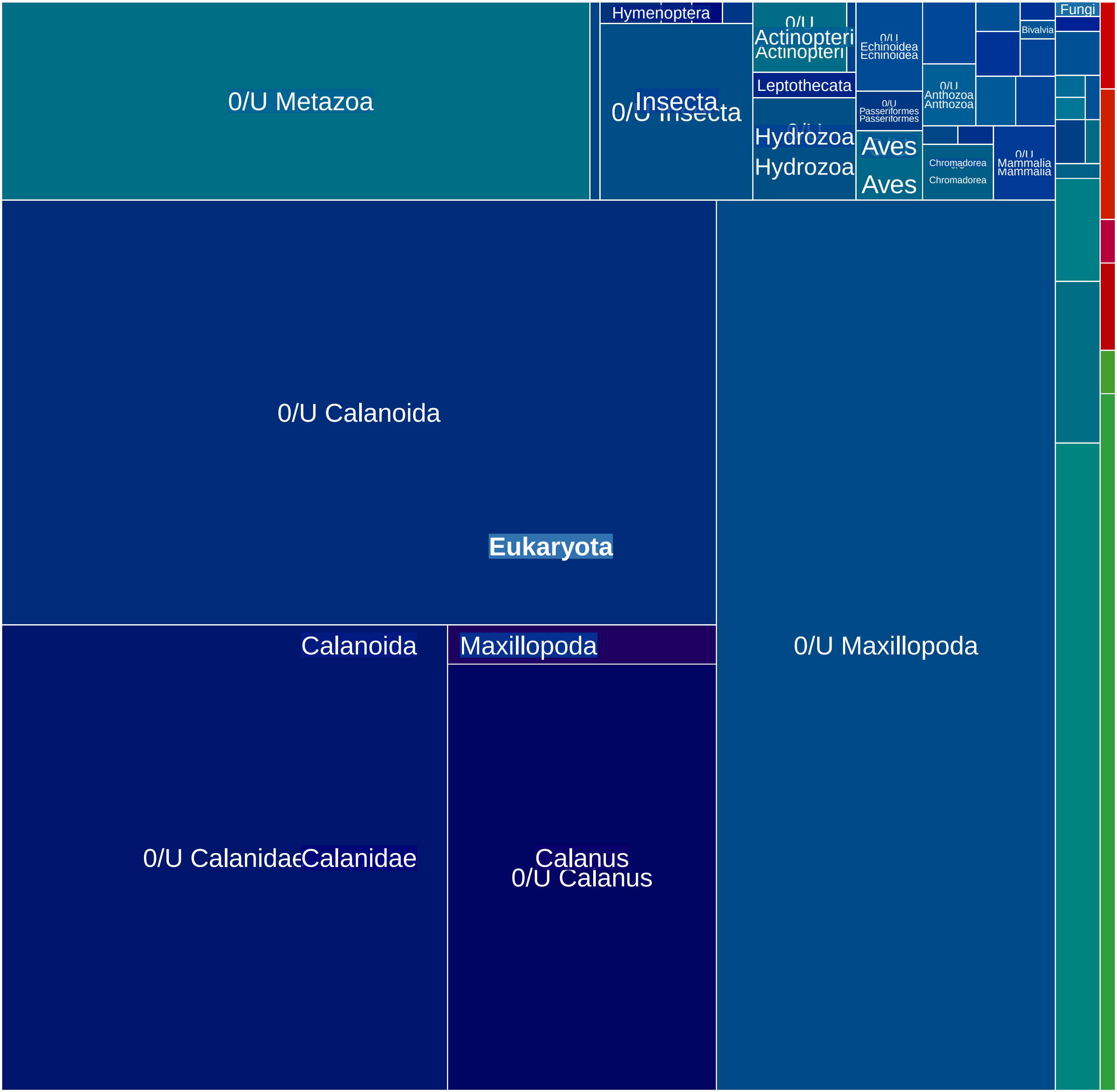

MGT−v1_135 : 11672 unigenes, 6558 (56.19%) taxonomically assigned

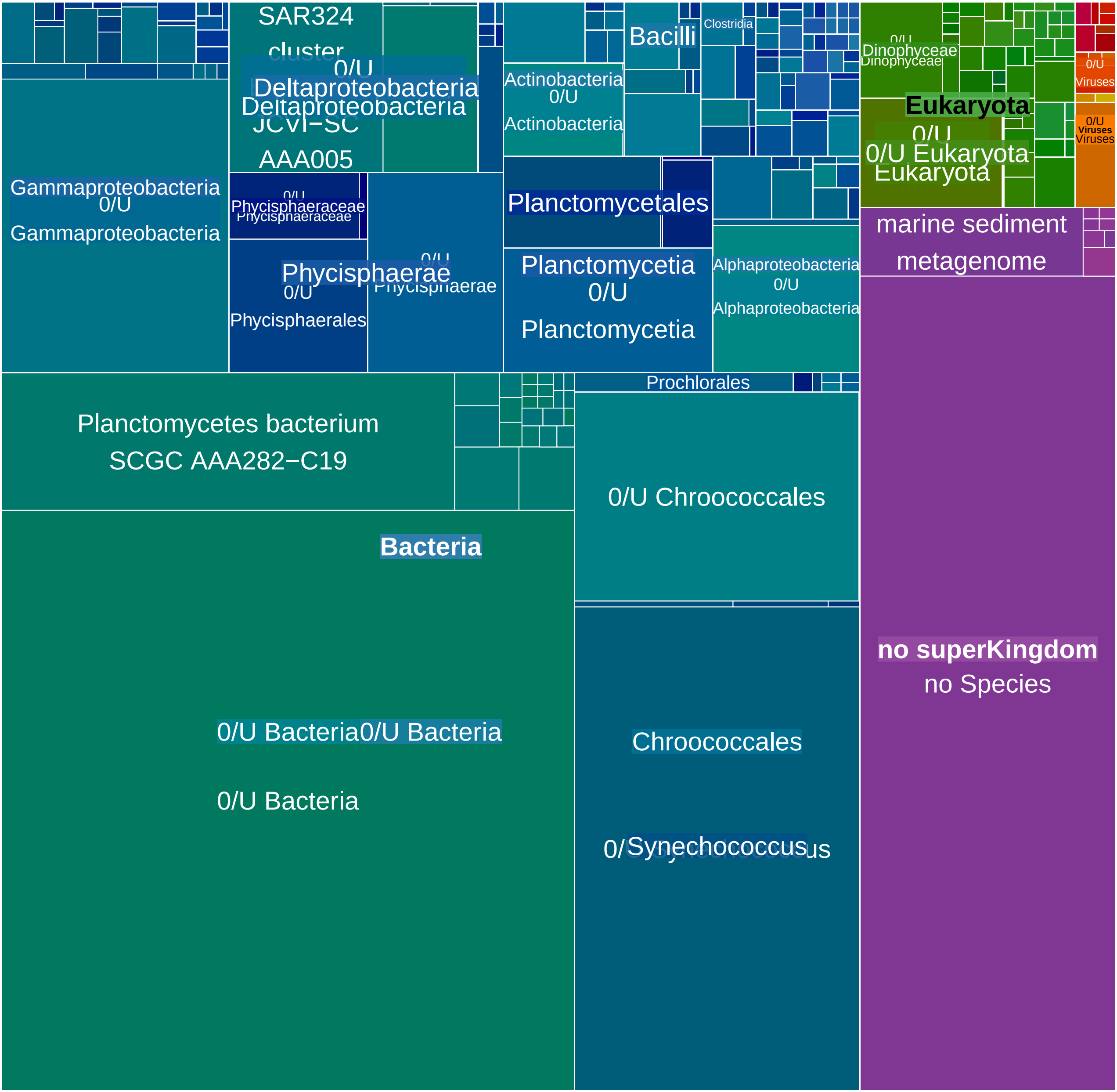

MGT−v1_136 : 11628 unigenes, 7703 (66.25%) taxonomically assigned

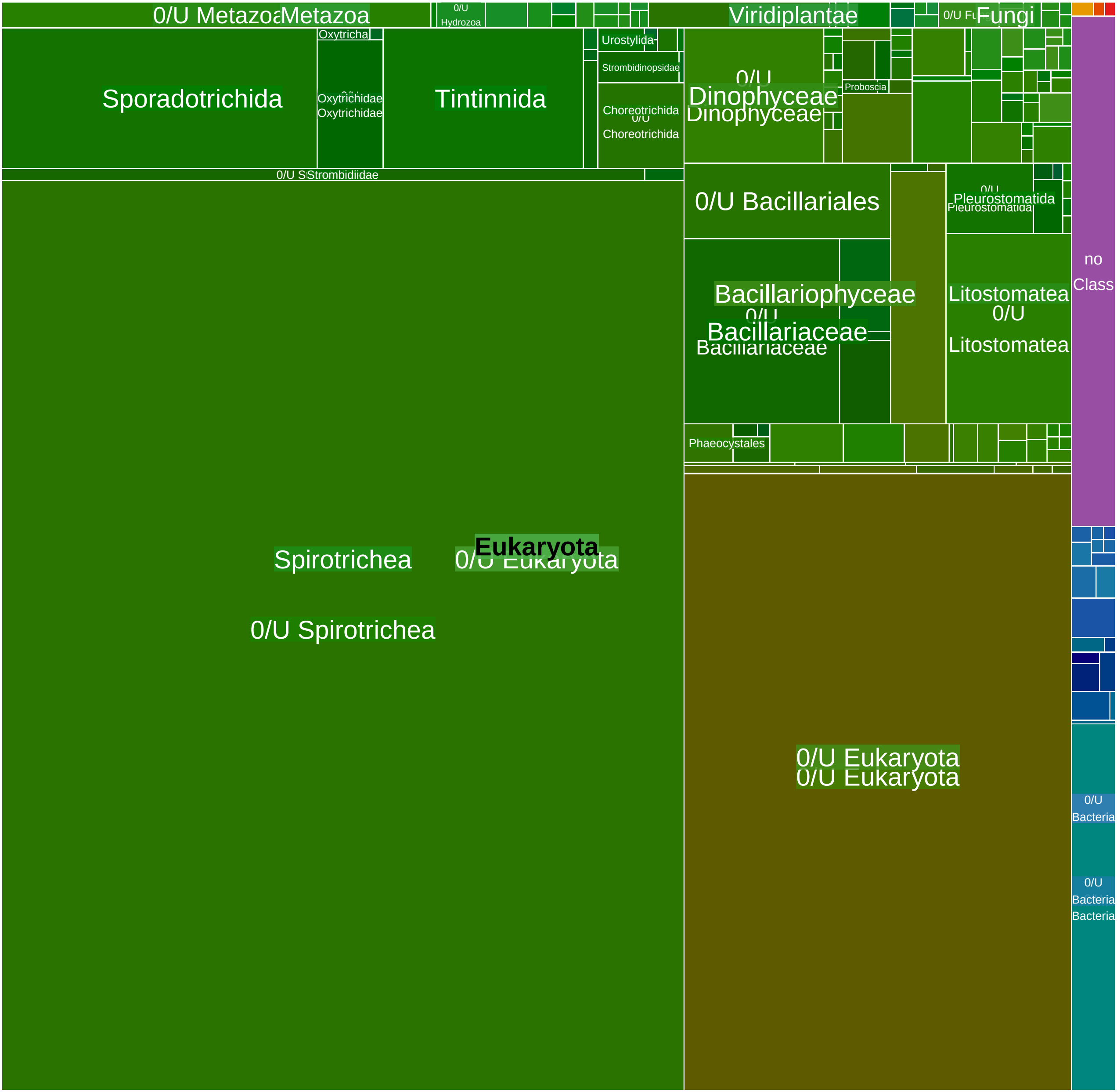

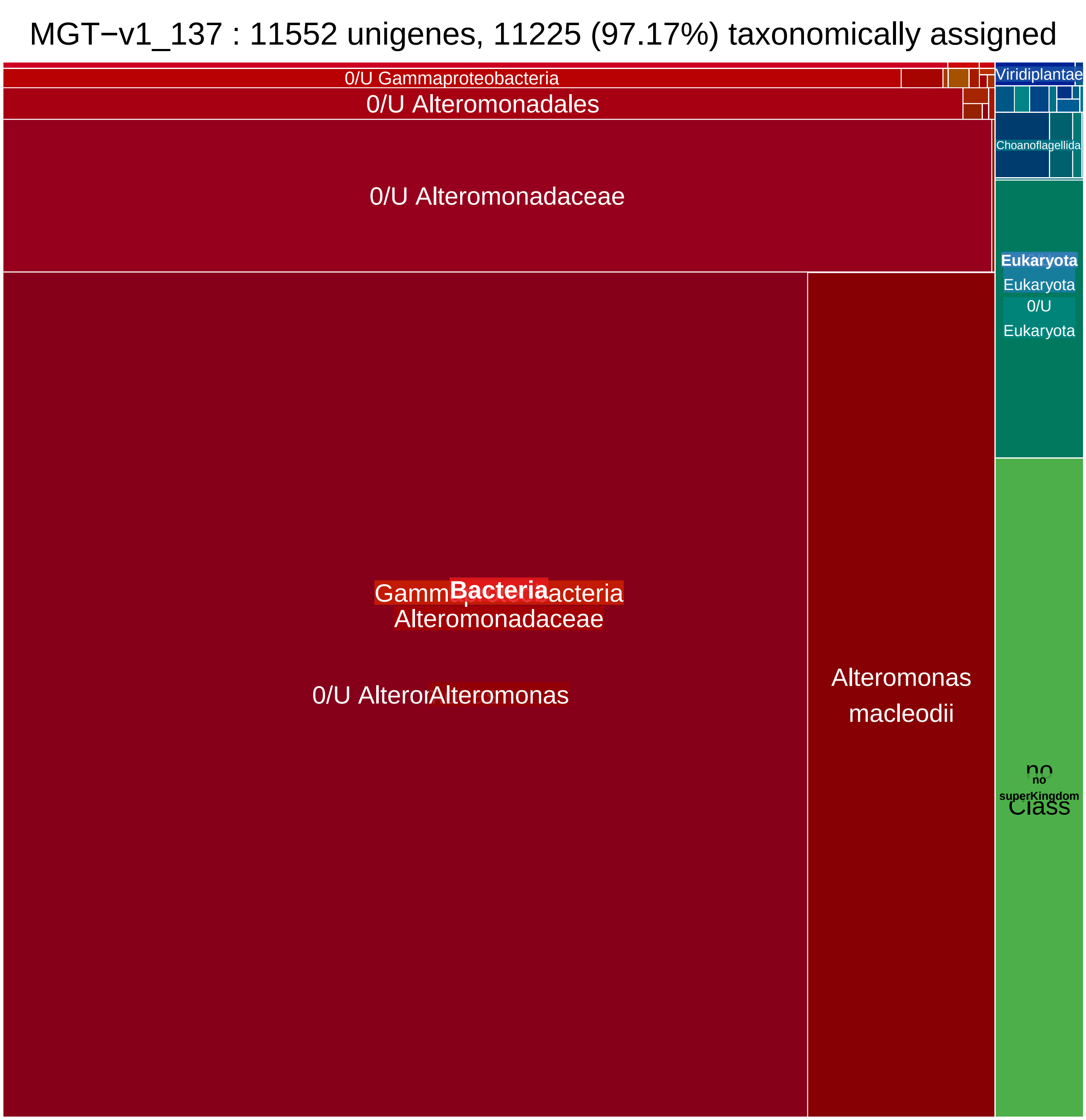
MGT−v1_137 : 11552 unigenes, 11225 (97.17%) taxonomically assigned
0/U Gammaproteobacteria
0/U Alteromonadales
0/U Alteromonadaceae
Bacteria
Gammaproteobacteria
Alteromonadaceae
Alteromonas
Alteromonas macleodii
Viridiplantae
Choanoflagellida
Eukaryota
Eukaryota
0/U Eukaryota
no Class
no superKingdom

MGT−v1_138 : 11537 unigenes, 1887 (16.36%) taxonomically assigned

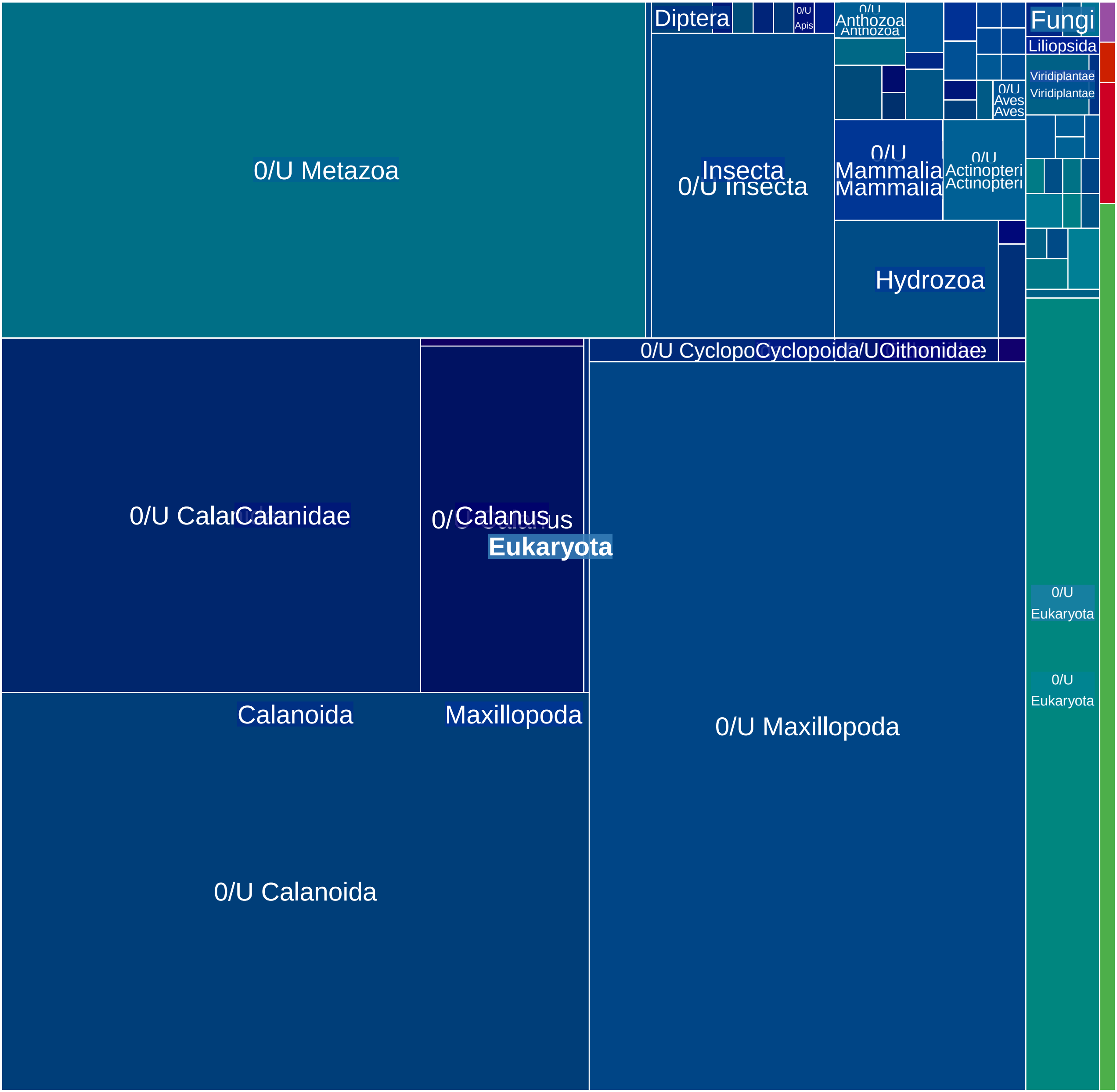

MGT−v1_139 : 11413 unigenes, 6319 (55.37%) taxonomically assigned

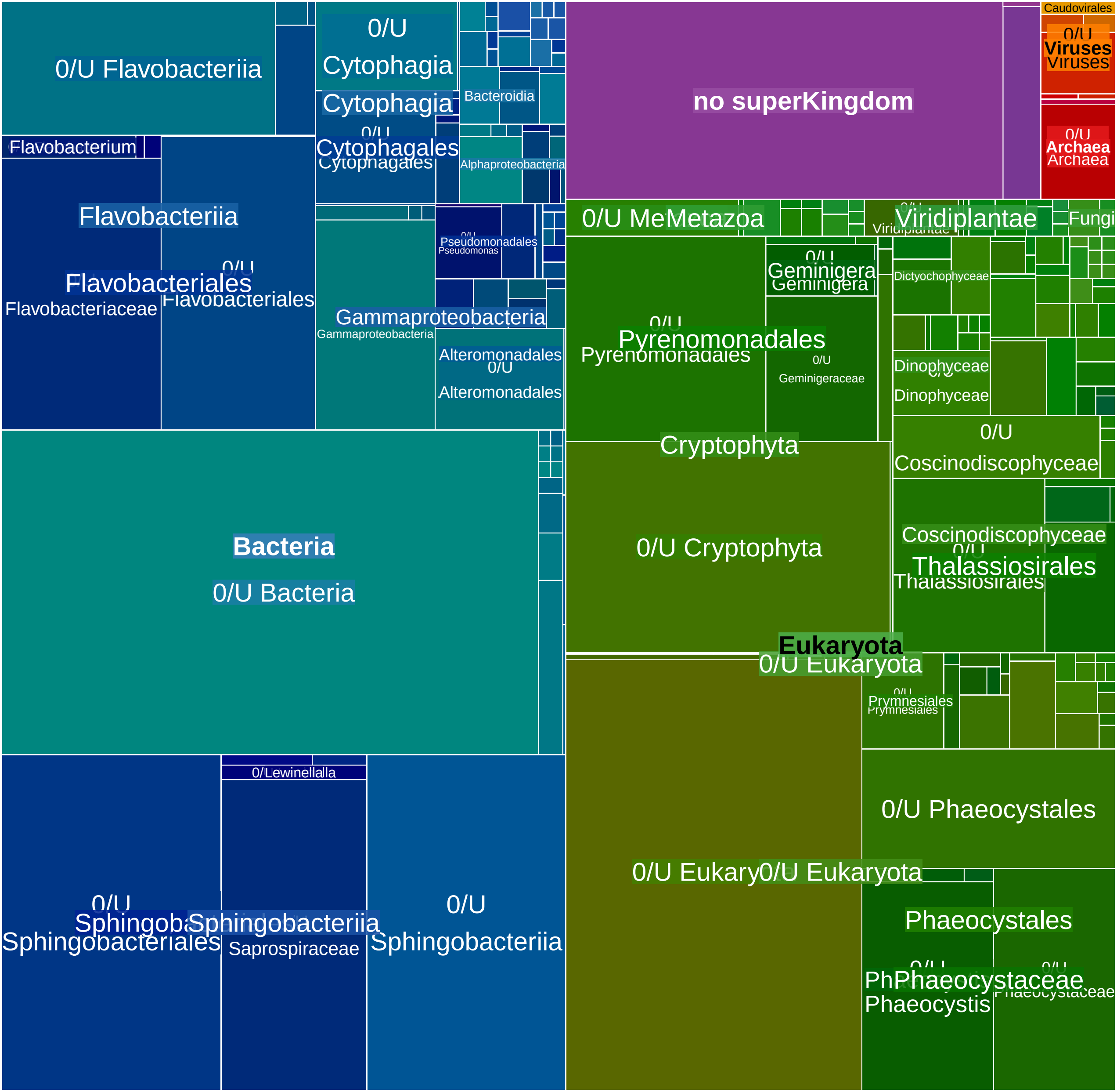

MGT−v1_140 : 11221 unigenes, 8518 (75.91%) taxonomically assigned

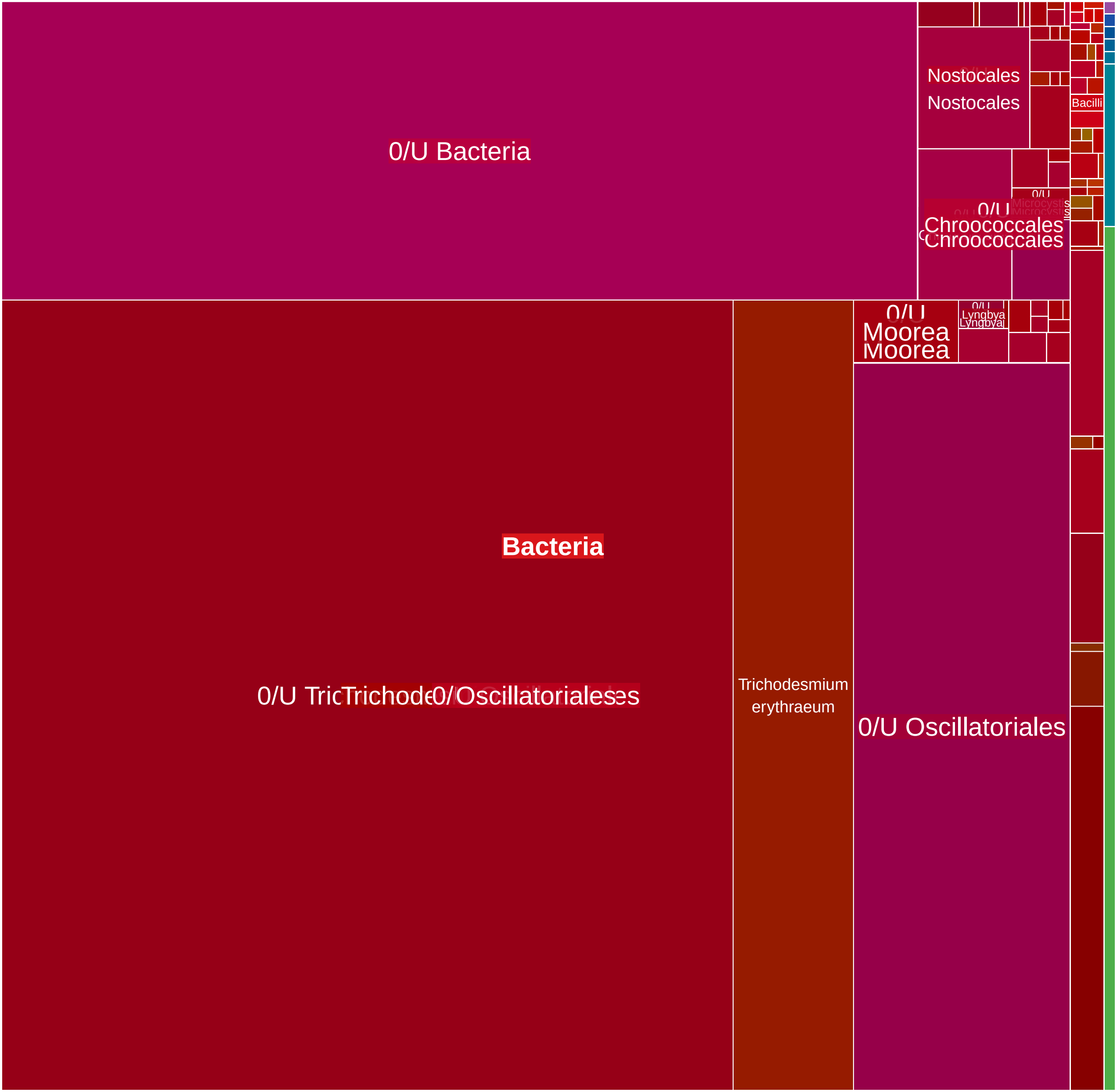

MGT−v1_141 : 11184 unigenes, 6116 (54.69%) taxonomically assigned

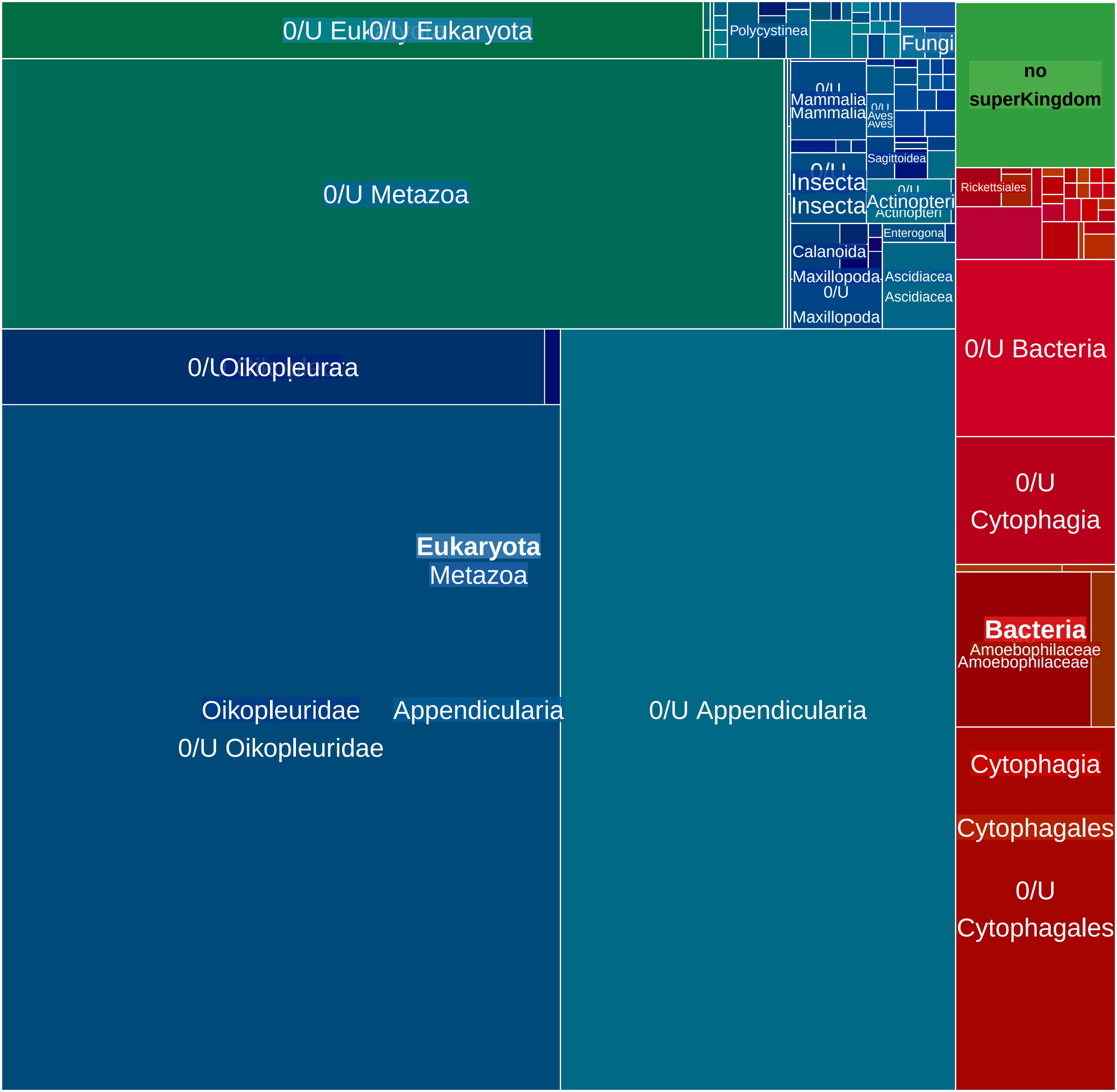

MGT−v1_142 : 11167 unigenes, 1238 (11.09%) taxonomically assigned

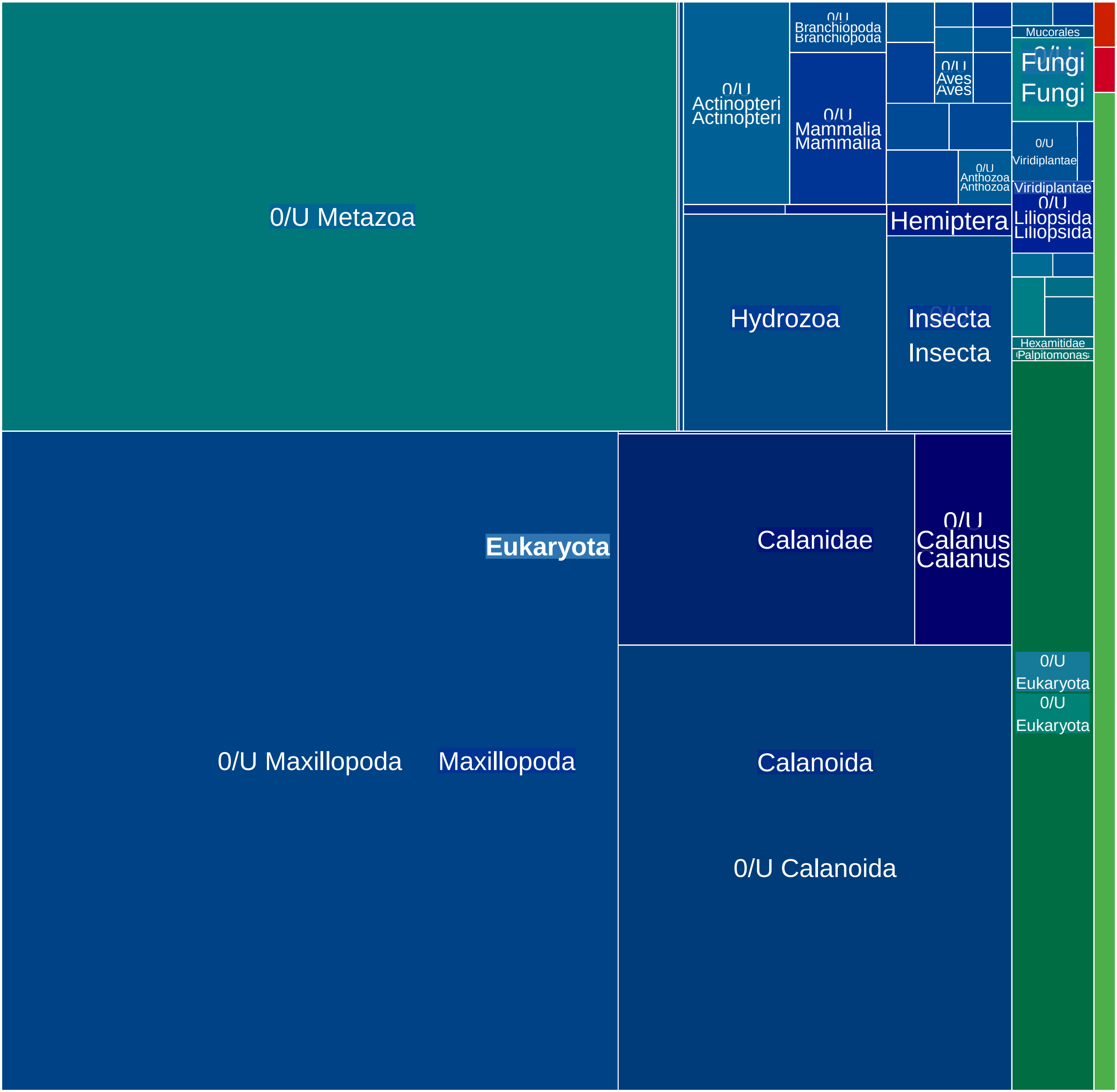

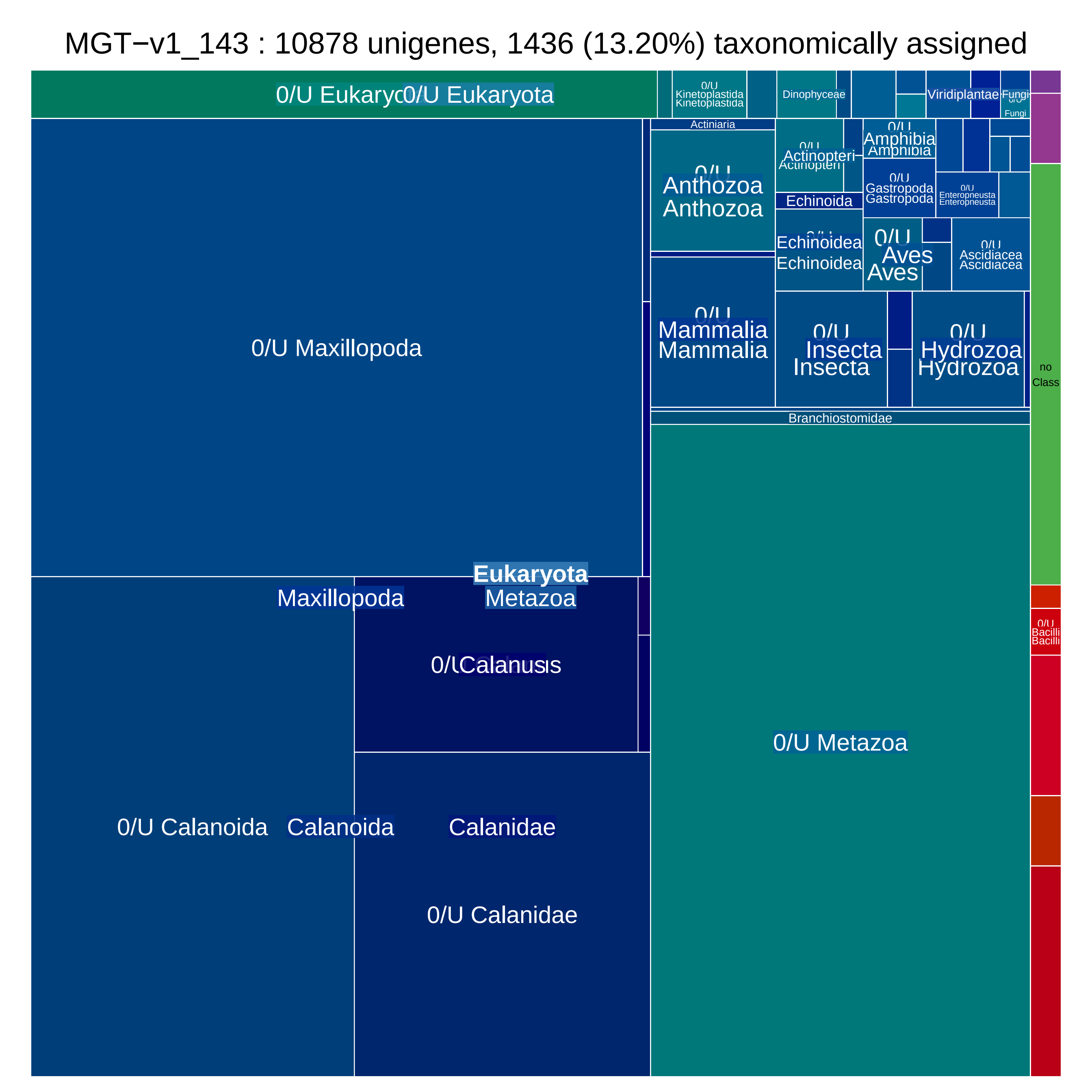
MGT−v1_143 : 10878 unigenes, 1436 (13.20%) taxonomically assigned
0/U Eukaryota
0/U Kinetoplastida
Kinetoplastida
Dinophyceae
Viridiplantae
Fungi
0/U Fungi
0/U Maxillopoda
Maxillopoda
Eukaryota
Metazoa
0/U Calanoida
Calanoida
Calanus
Calanidae
0/U Calanidae
Actiniaria
0/U Anthozoa
Anthozoa
0/U Actinopteri
Actinopteri
0/U Amphibia
Amphibia
0/U Gastropoda
Gastropoda
Echinoida
0/U Enteropneusta
Enteropneusta
0/U Echinoidea
Echinoidea
0/U Aves
Aves
0/U Ascidiacea
Ascidiacea
0/U Mammalia
Mammalia
0/U Insecta
Insecta
0/U Hydrozoa
Hydrozoa
Branchiostomidae
0/U Metazoa
no Class
0/U Bacilli
Bacilli

MGT−v1_144 : 10820 unigenes, 7522 (69.52%) taxonomically assigned

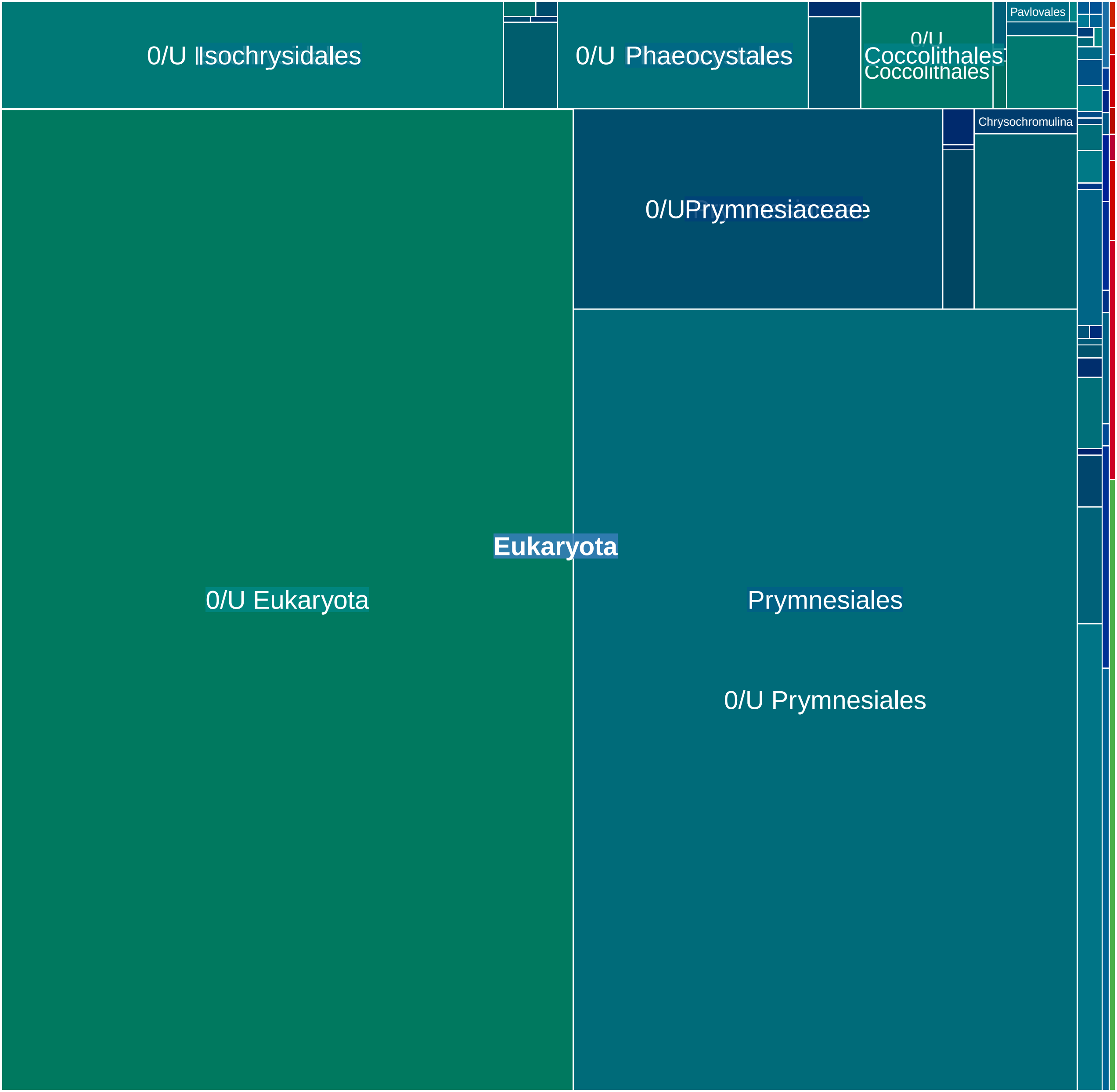

MGT−v1_145 : 10810 unigenes, 6684 (61.83%) taxonomically assigned

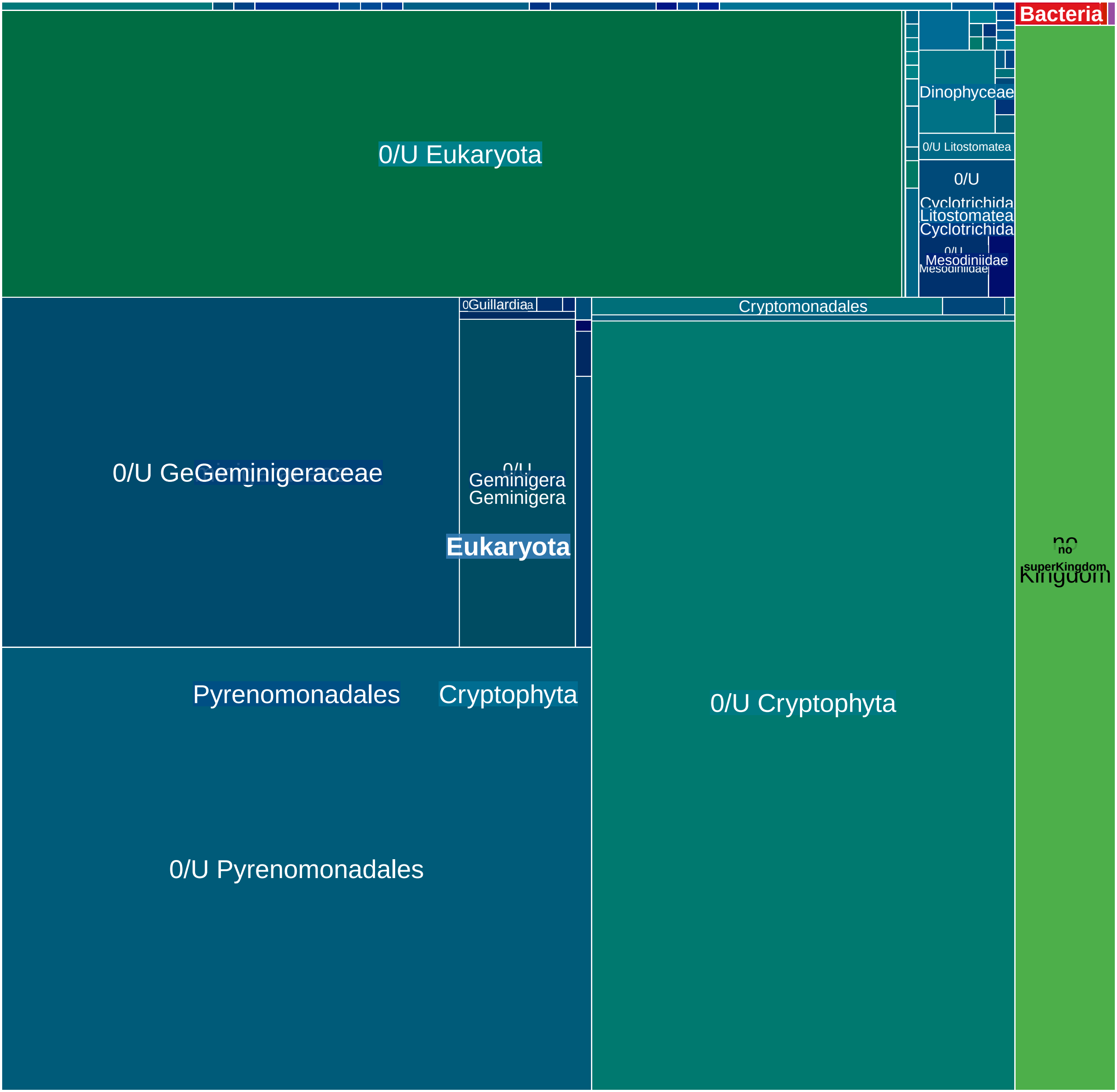

MGT−v1_146 : 10714 unigenes, 4454 (41.57%) taxonomically assigned

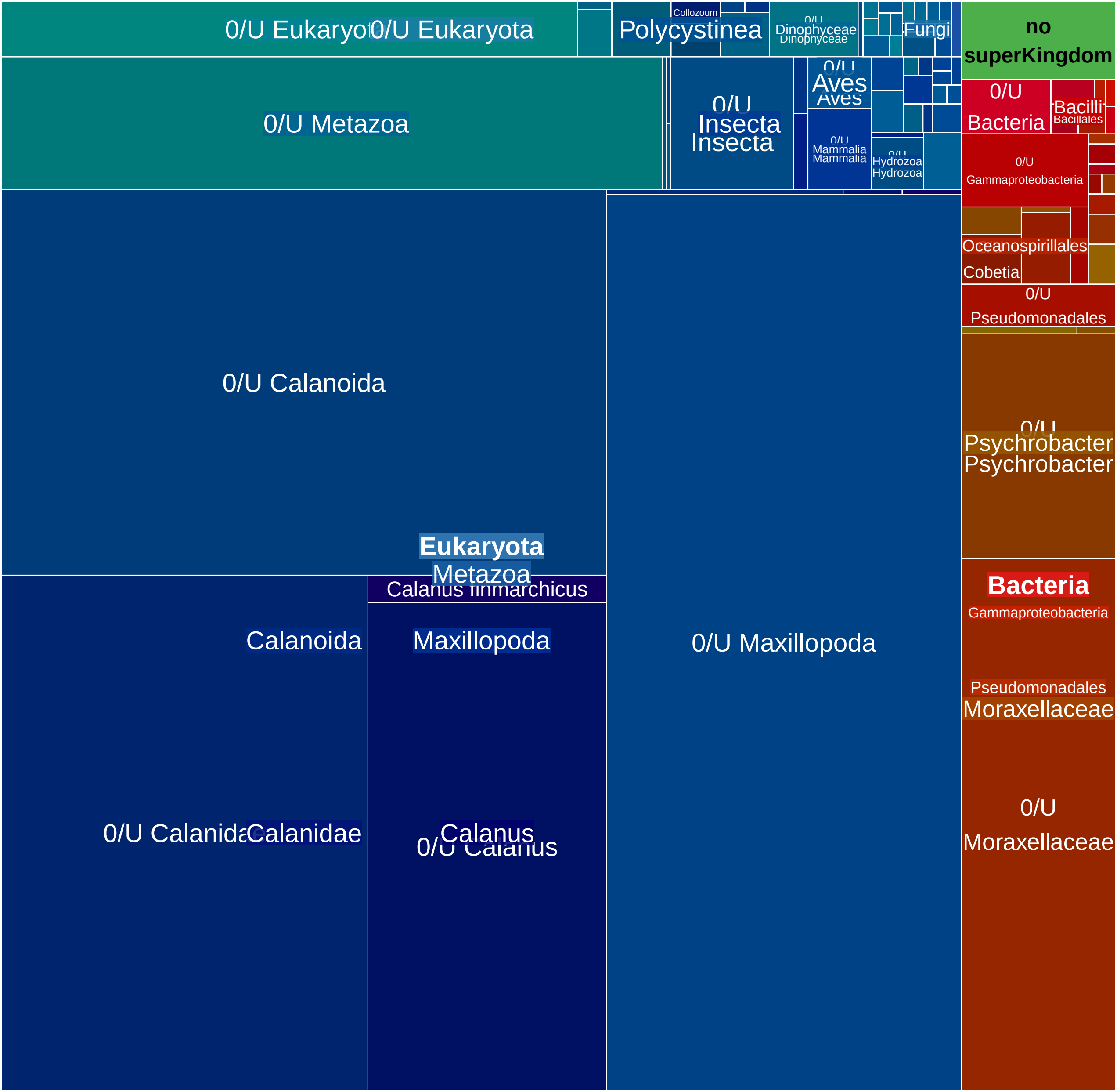

MGT−v1_147 : 10706 unigenes, 814 (7.60%) taxonomically assigned

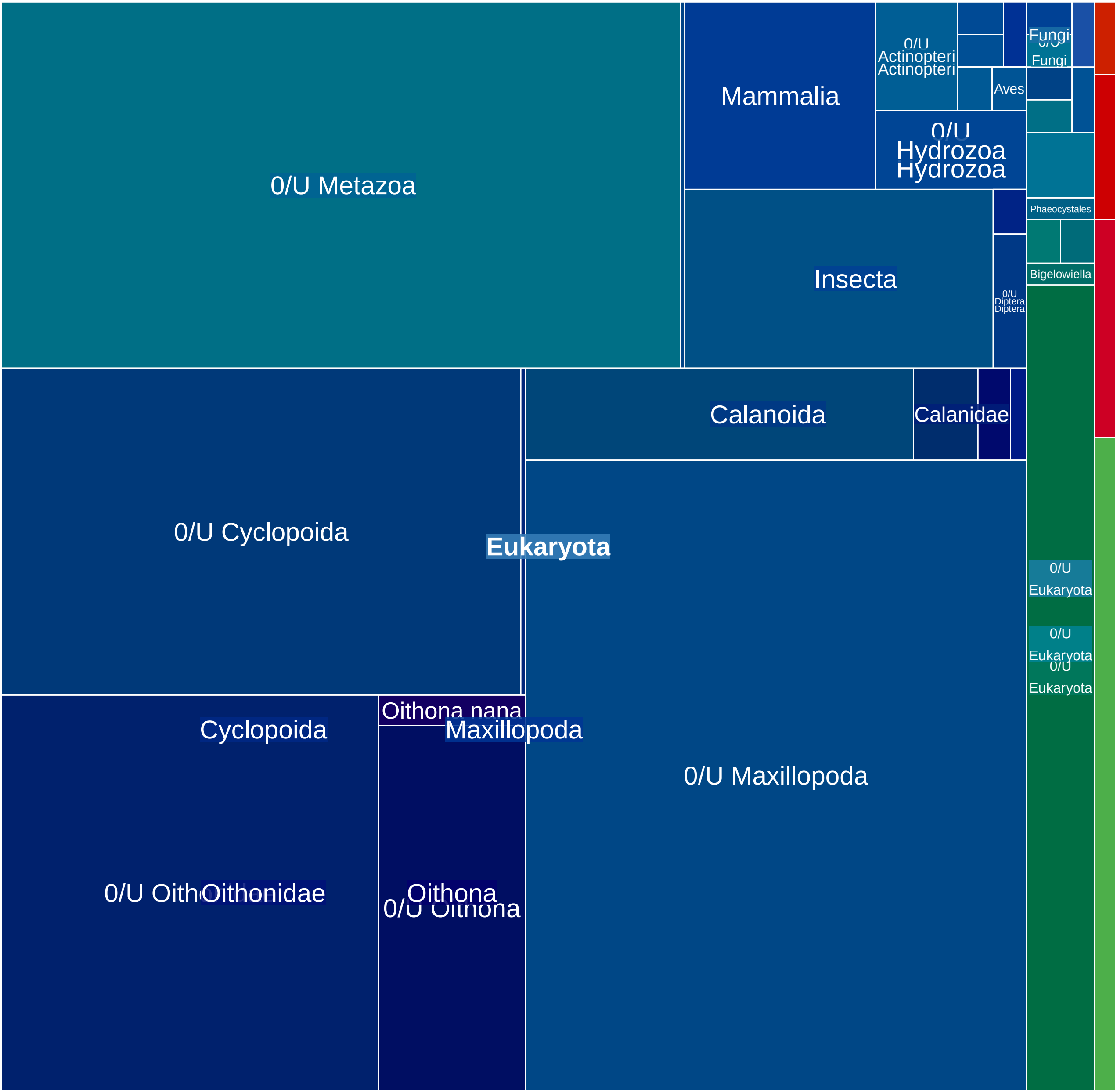

MGT−v1_148 : 10675 unigenes, 4052 (37.96%) taxonomically assigned

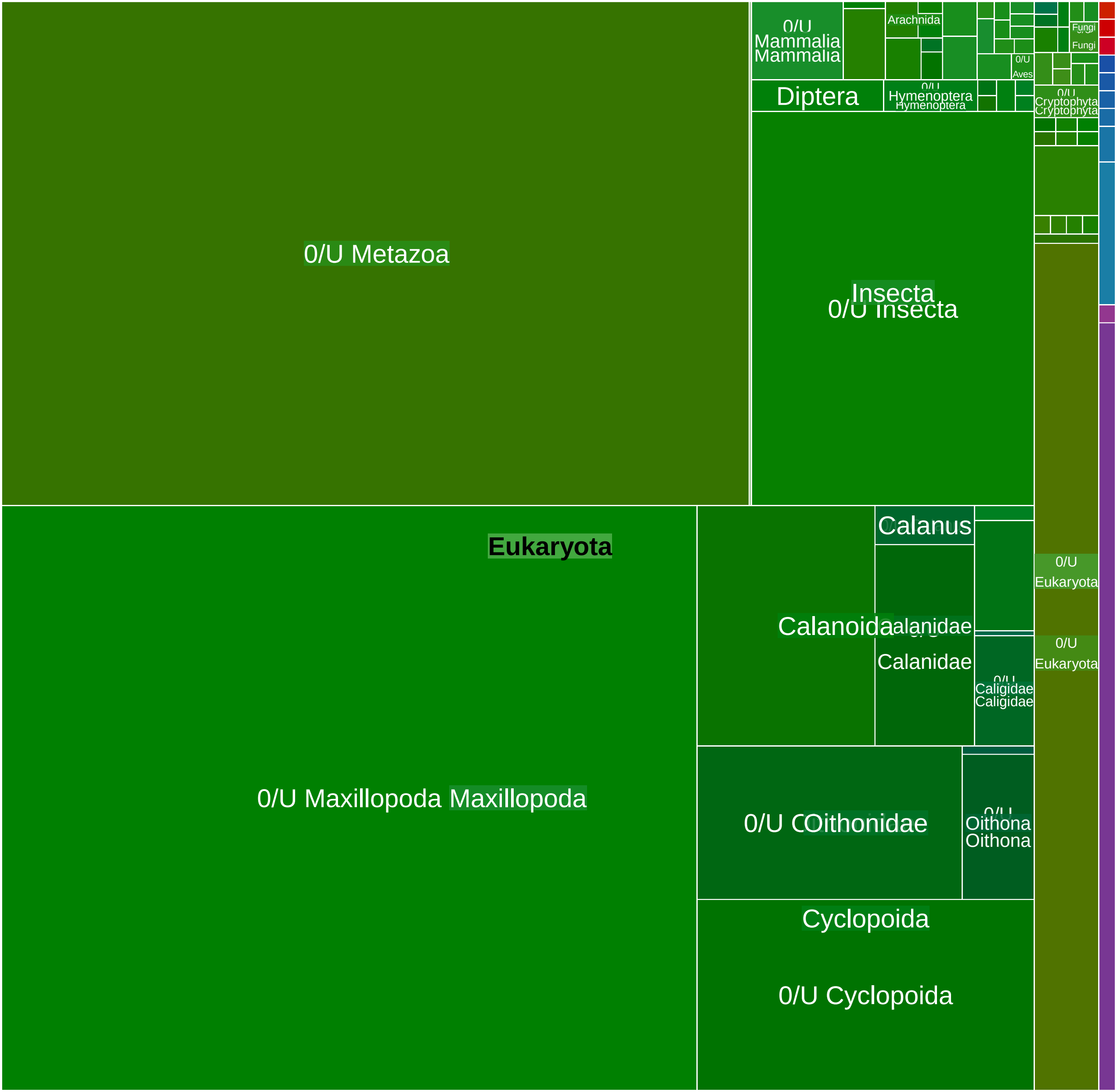

MGT−v1_149 : 10436 unigenes, 3636 (34.84%) taxonomically assigned

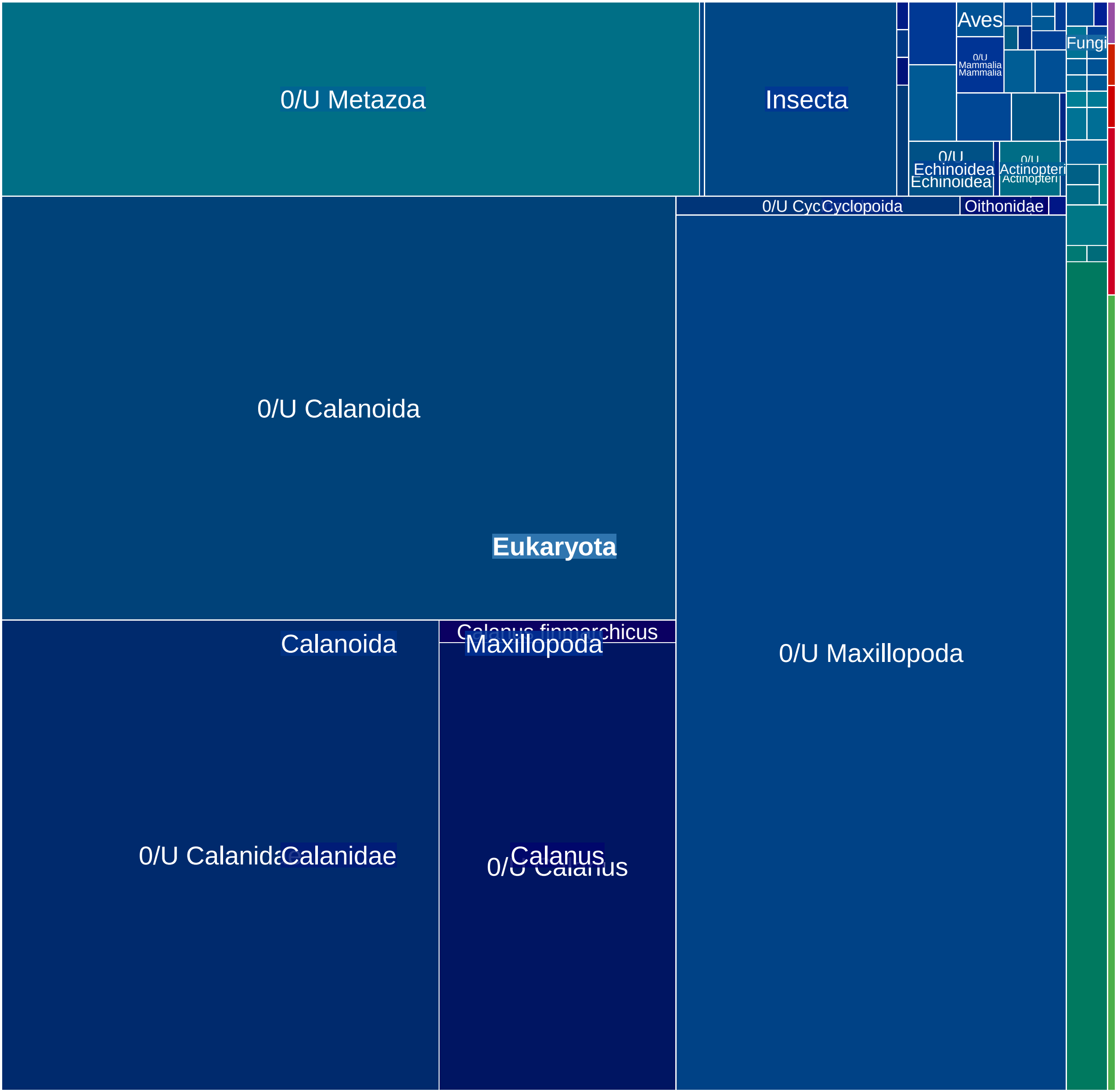

MGT−v1_150 : 10255 unigenes, 3542 (34.54%) taxonomically assigned

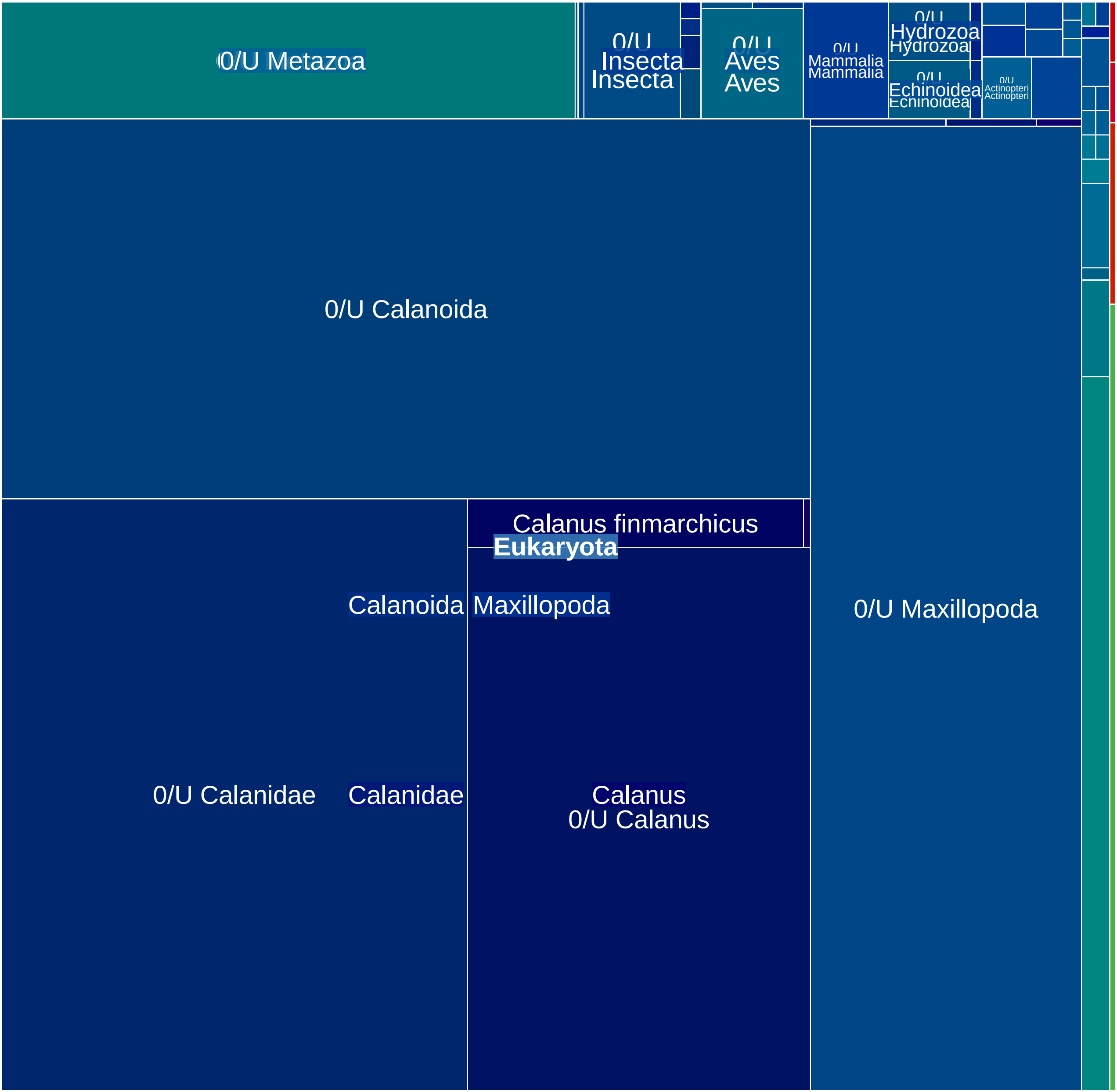

MGT−v1_151 : 10224 unigenes, 1177 (11.51%) taxonomically assigned

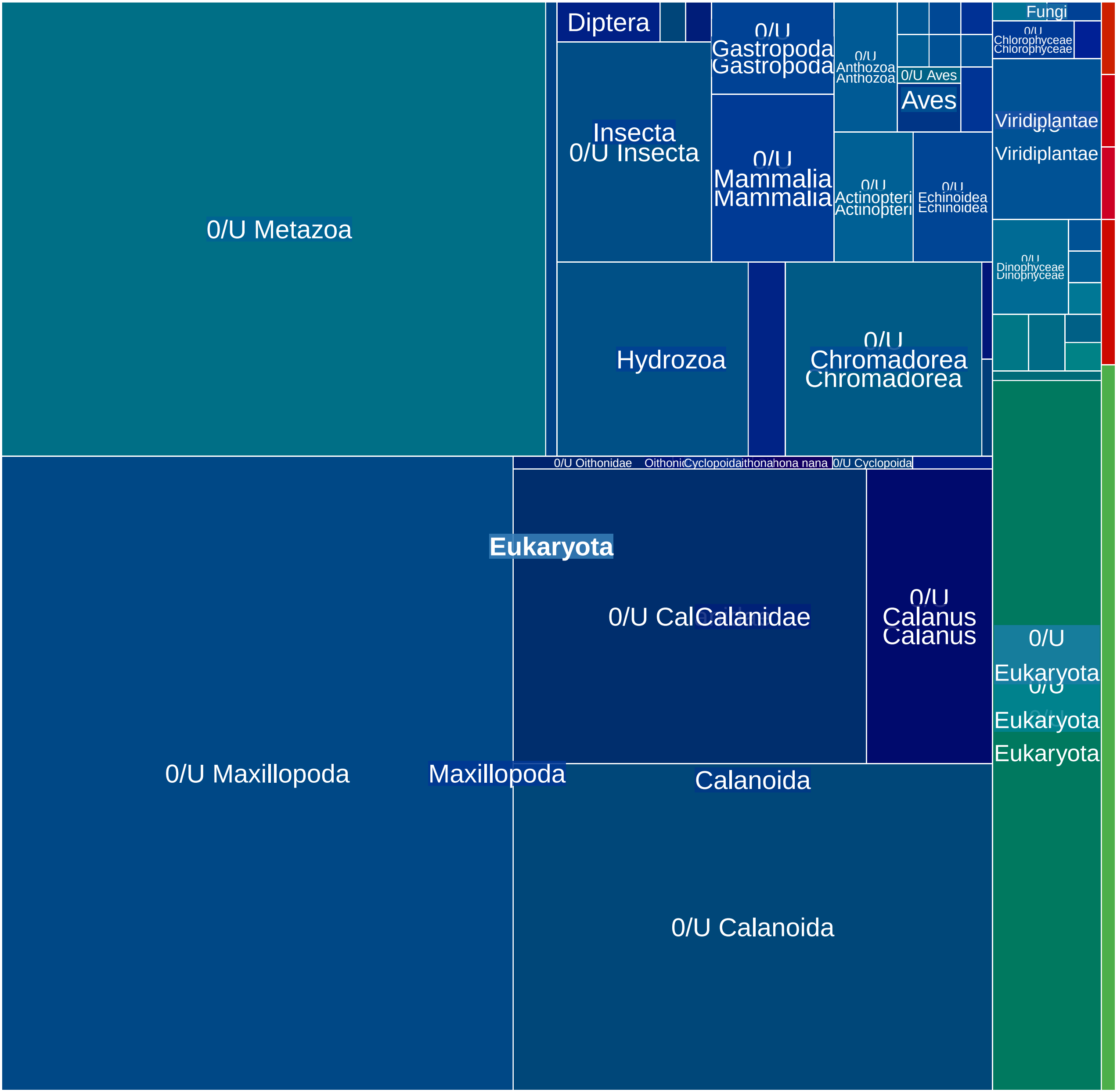

### MGT−v1_152 : 10177 unigenes, 2530 (24.86%) taxonomically assigned

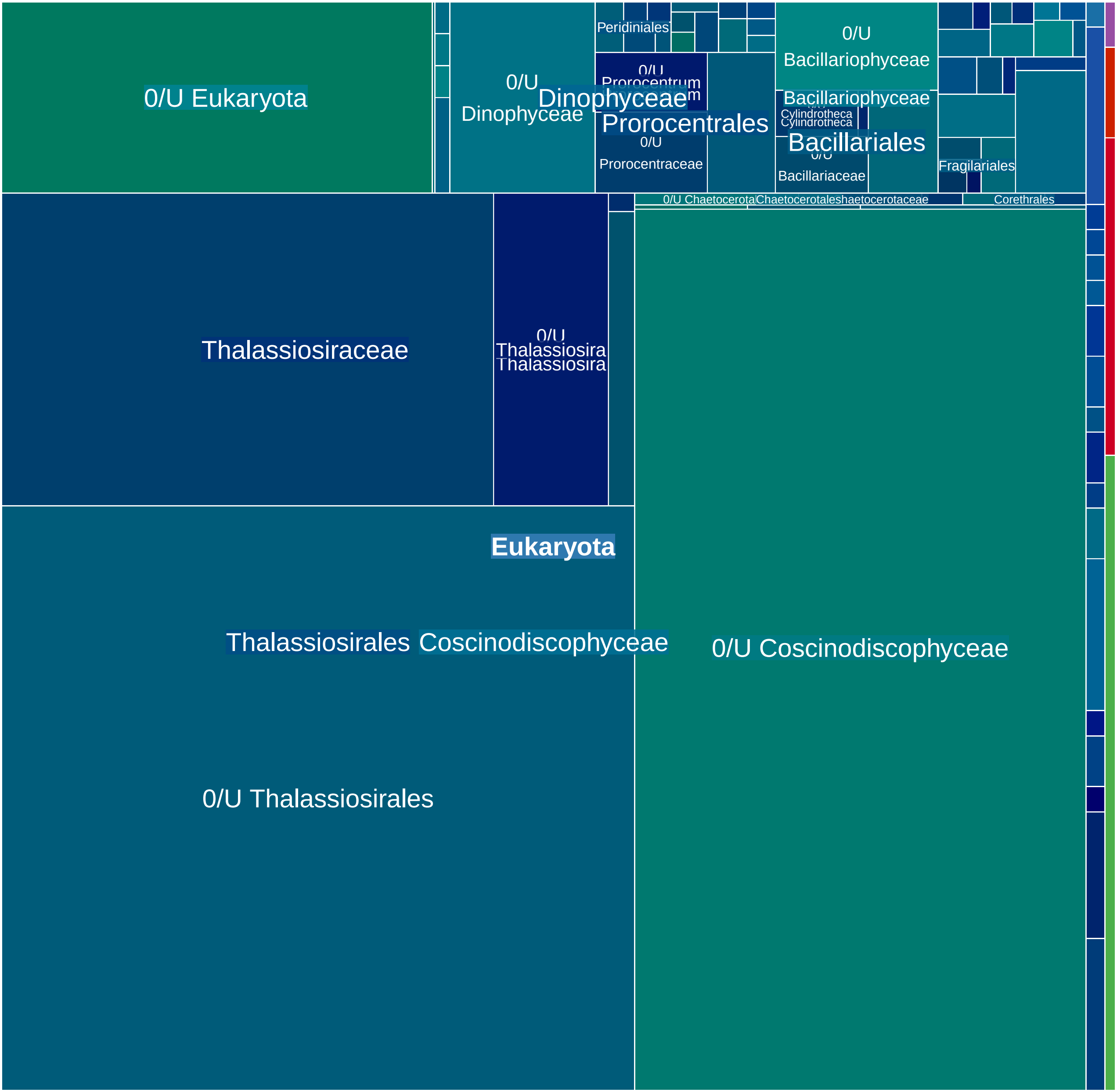

MGT−v1_153 : 9985 unigenes, 3643 (36.48%) taxonomically assigned

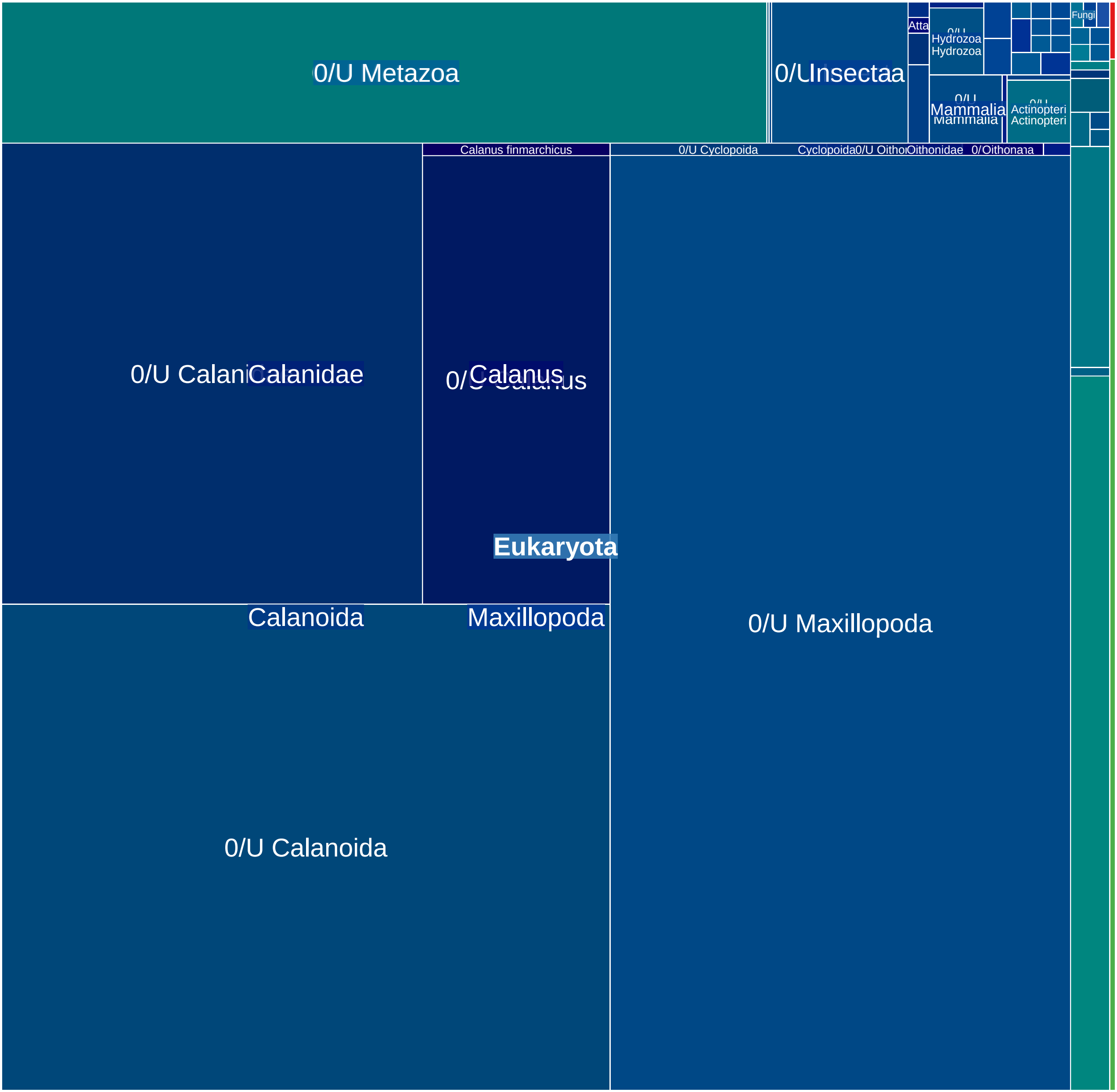

### MGT−v1_154 : 9894 unigenes, 3832 (38.73%) taxonomically assigned

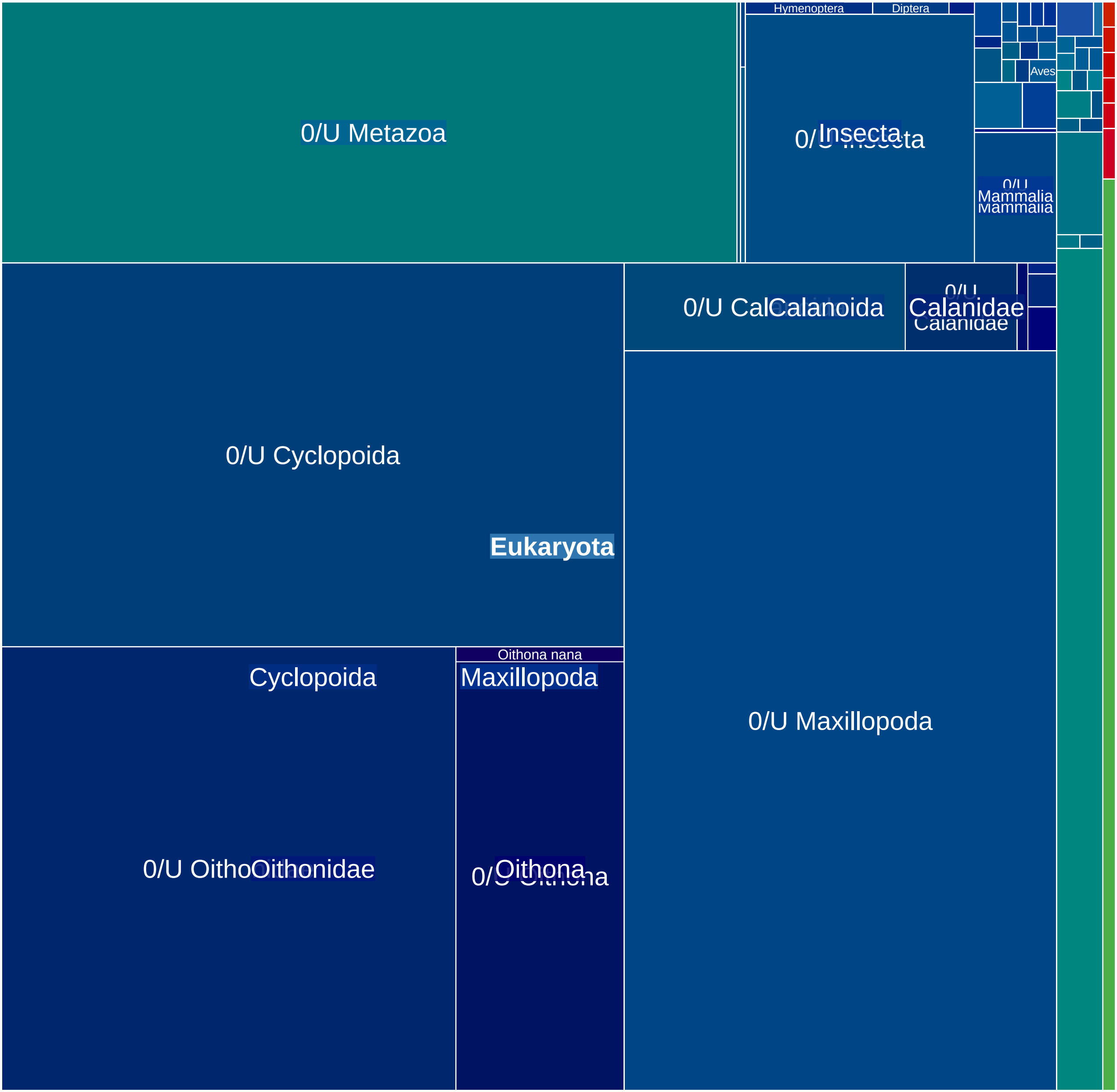

MGT−v1_155 : 9887 unigenes, 4373 (44.23%) taxonomically assigned

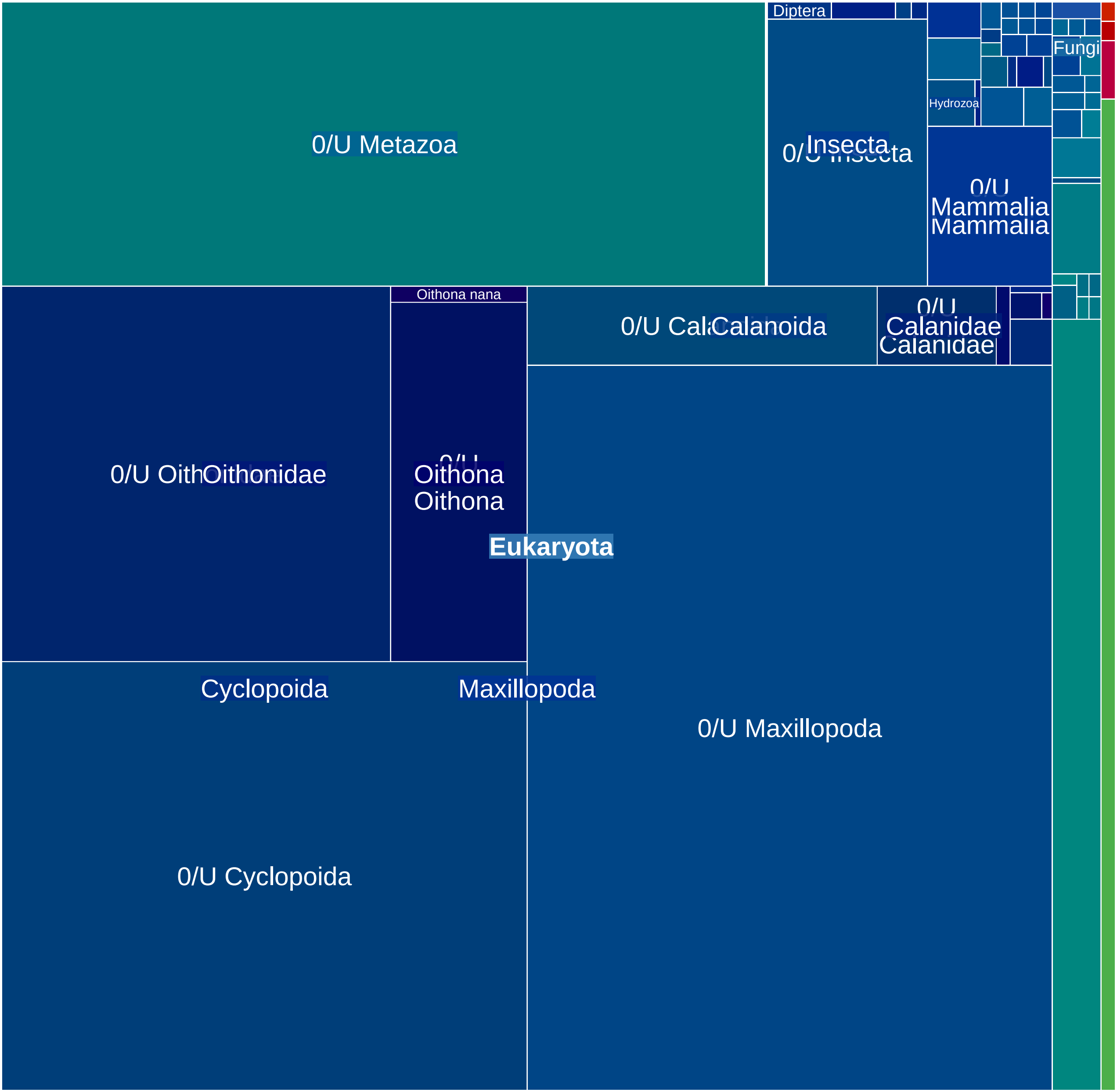

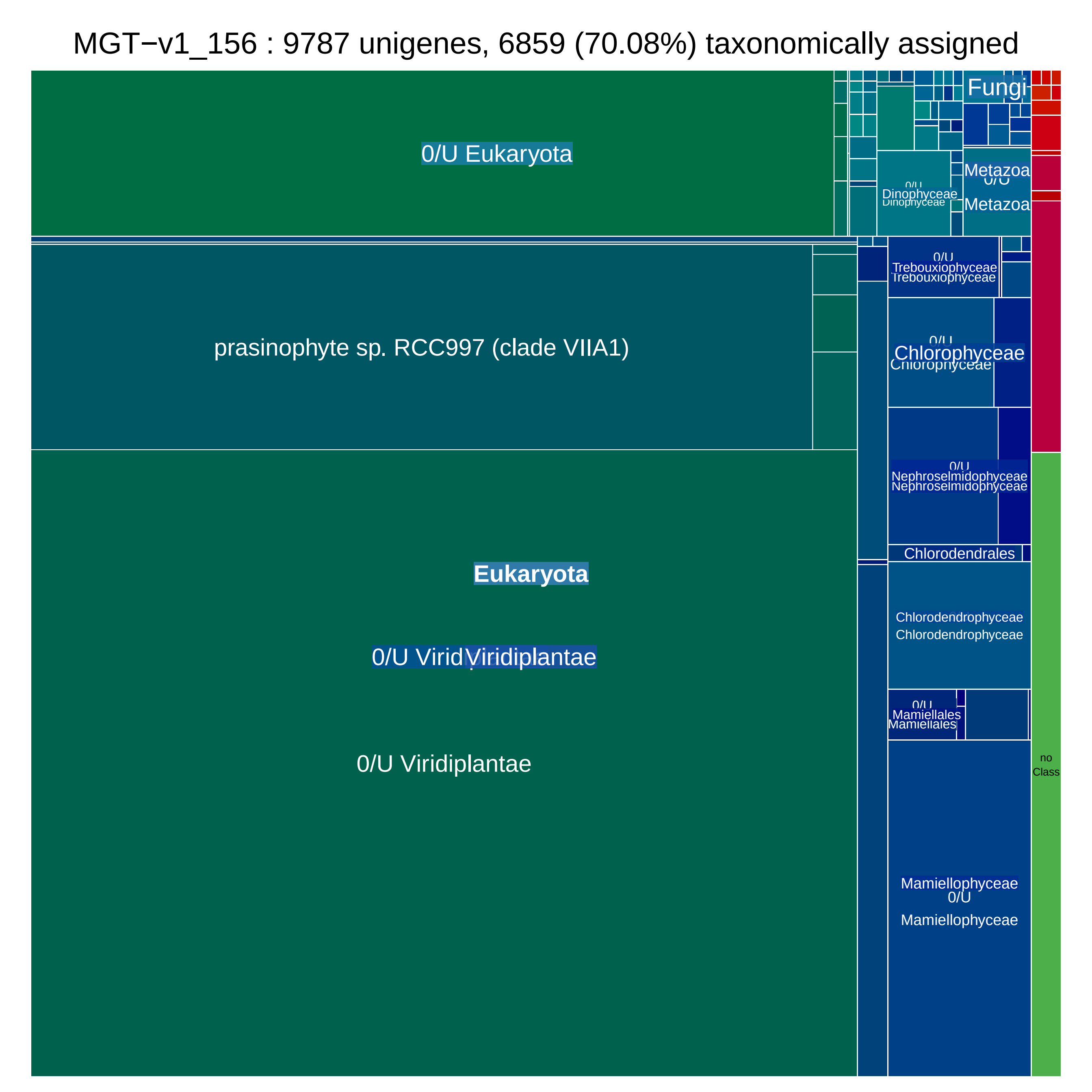
MGT−v1_156 : 9787 unigenes, 6859 (70.08%) taxonomically assigned
0/U Eukaryota
Fungi
Metazoa
0/U
Metazoa
0/U
Dinophyceae
Dinophyceae
prasinophyte sp. RCC997 (clade VIIA1)
0/U
Trebouxiophyceae
Trebouxiophyceae
0/U
Chlorophyceae
Chlorophyceae
0/U
Nephroselmidophyceae
Nephroselmidophyceae
Chlorodendrales
Chlorodendrophyceae
Chlorodendrophyceae
Eukaryota
0/U Viridiplantae
0/U Viridiplantae
0/U
Mamiellales
Mamiellales
Mamiellophyceae
0/U
Mamiellophyceae
no Class

### MGT−v1_157 : 9740 unigenes, 1050 (10.78%) taxonomically assigned

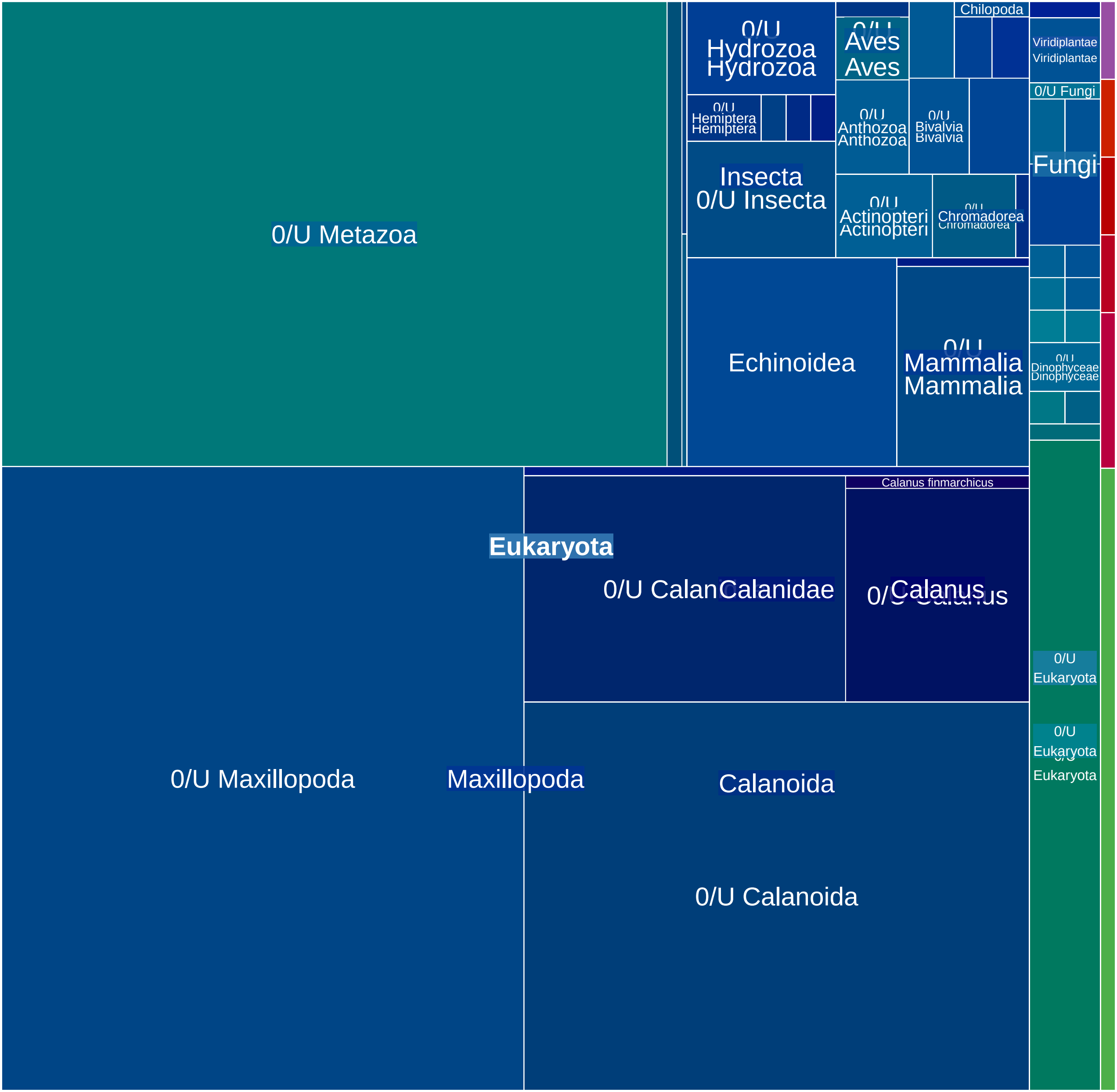

MGT−v1_158 : 9693 unigenes, 975 (10.06%) taxonomically assigned

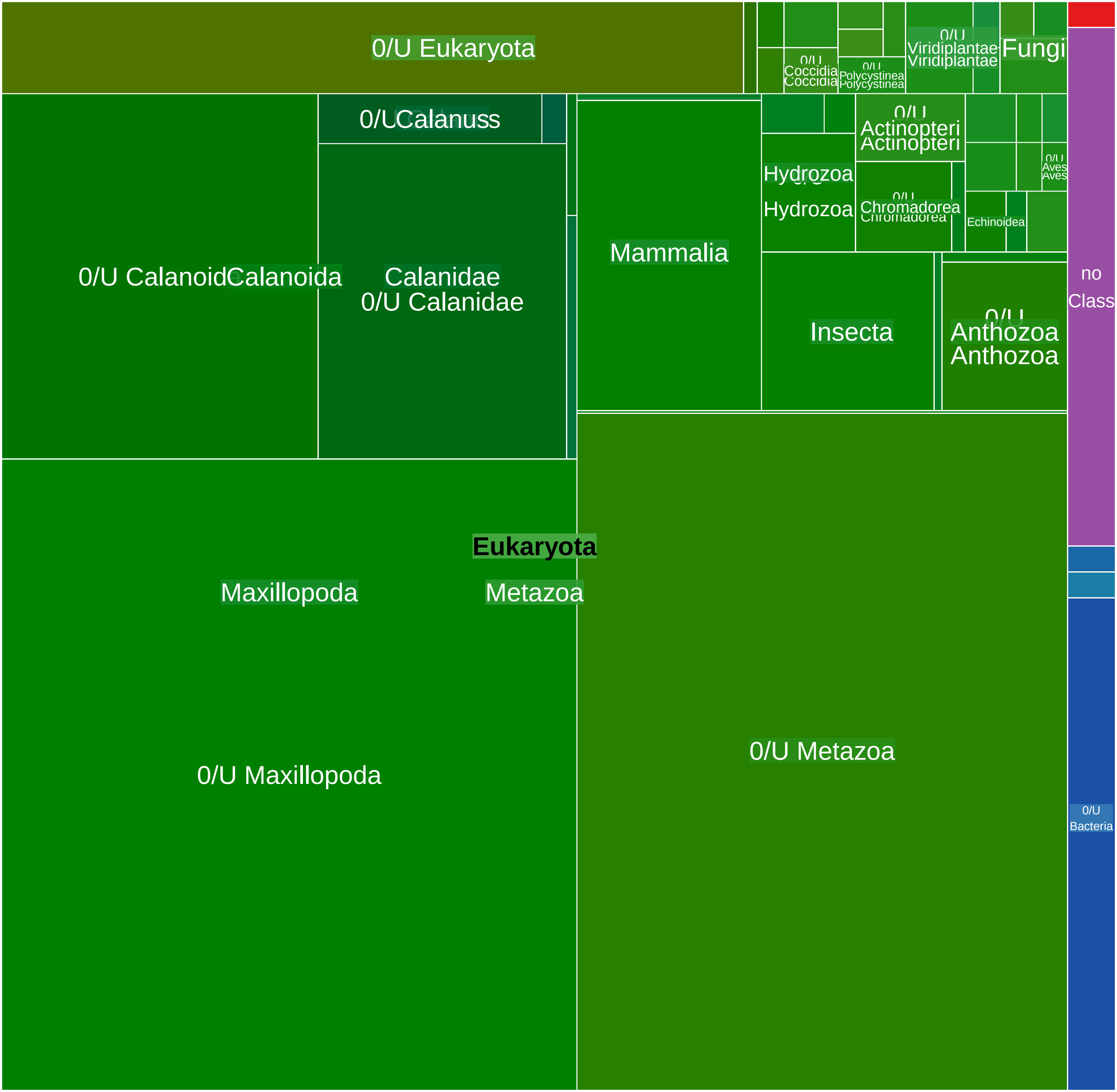

MGT−v1_159 : 9623 unigenes, 685 (7.12%) taxonomically assigned

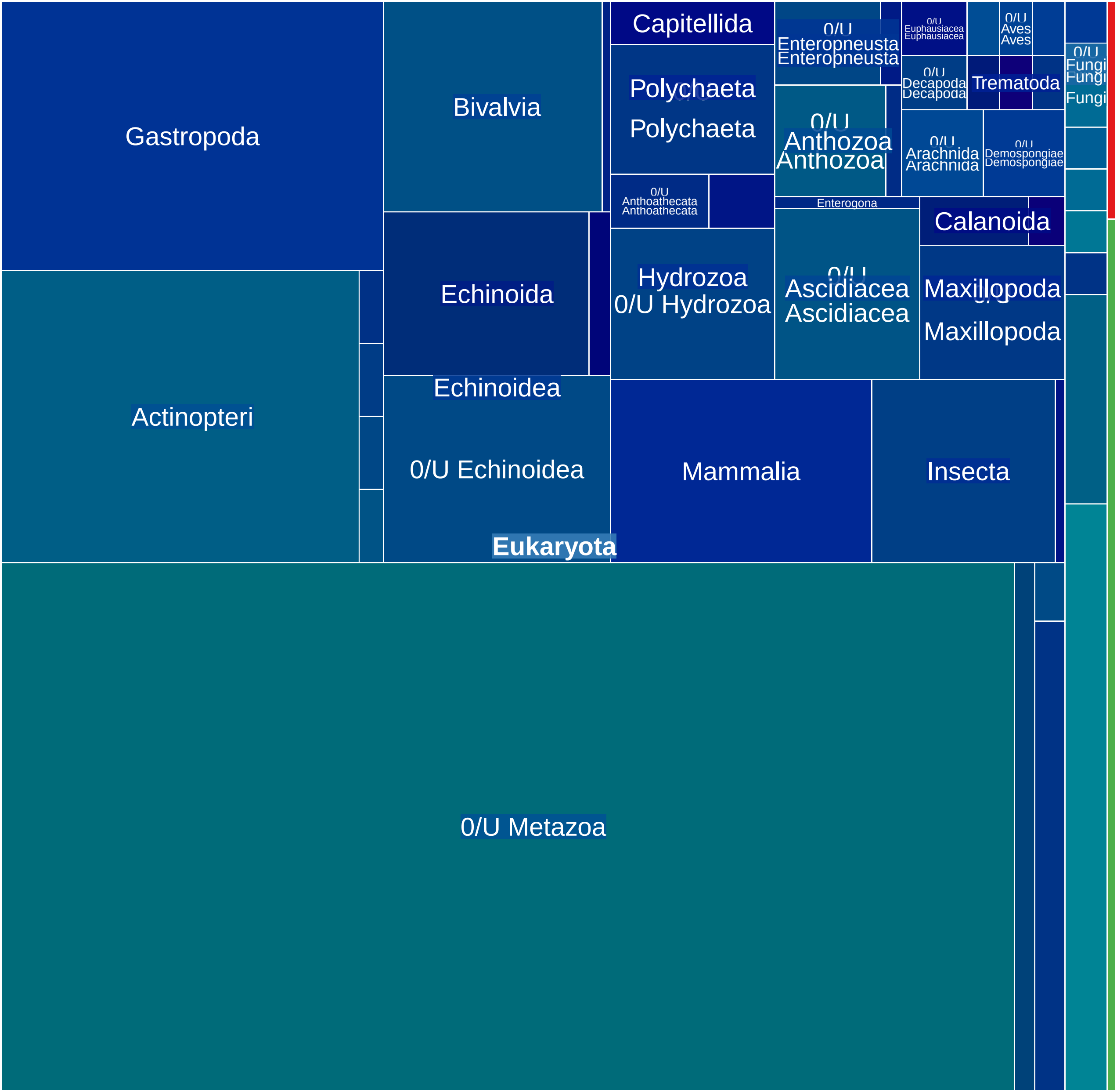

MGT−v1_160 : 9494 unigenes, 3450 (36.34%) taxonomically assigned

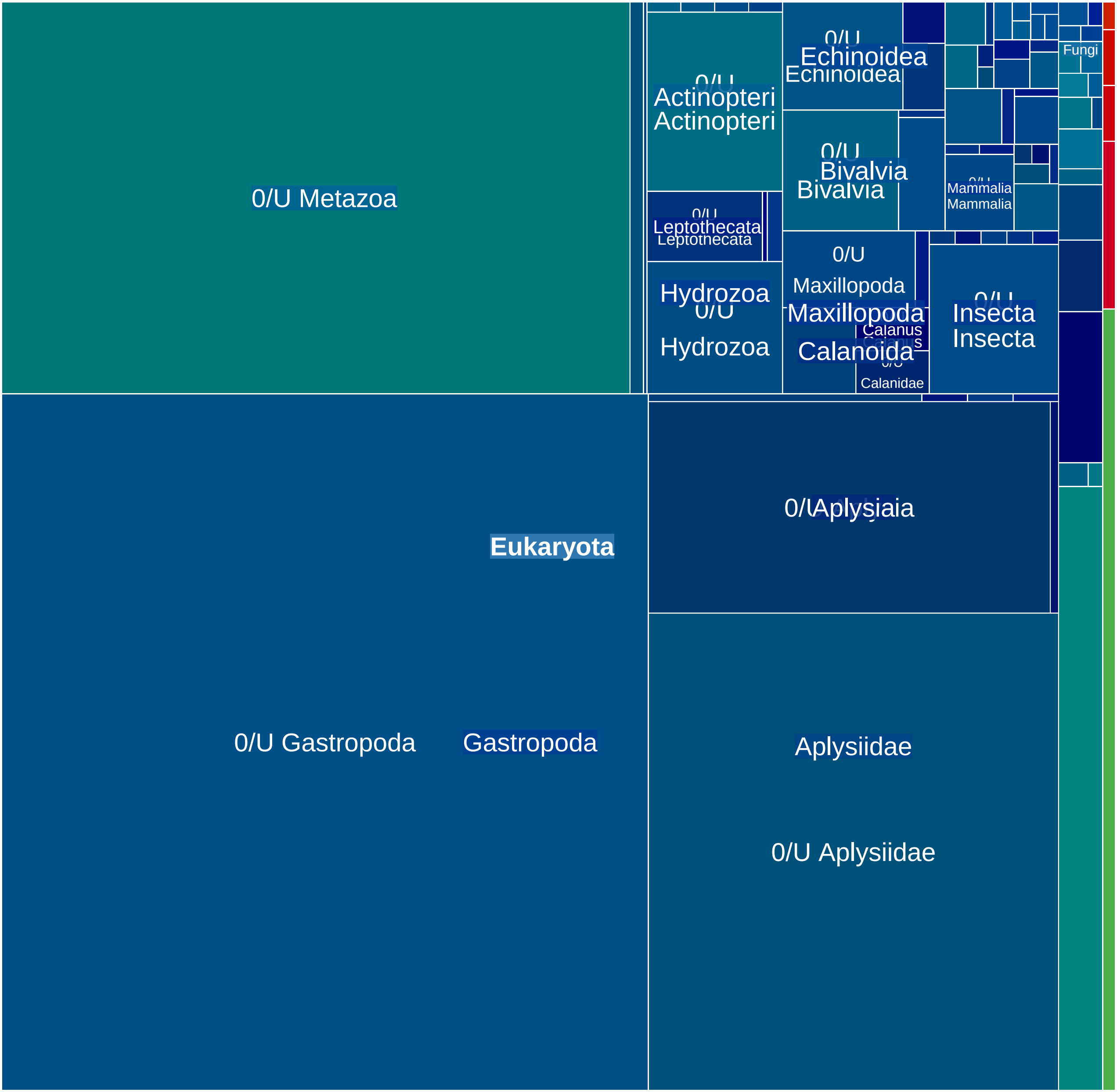

MGT−v1_161 : 9466 unigenes, 6245 (65.97%) taxonomically assigned

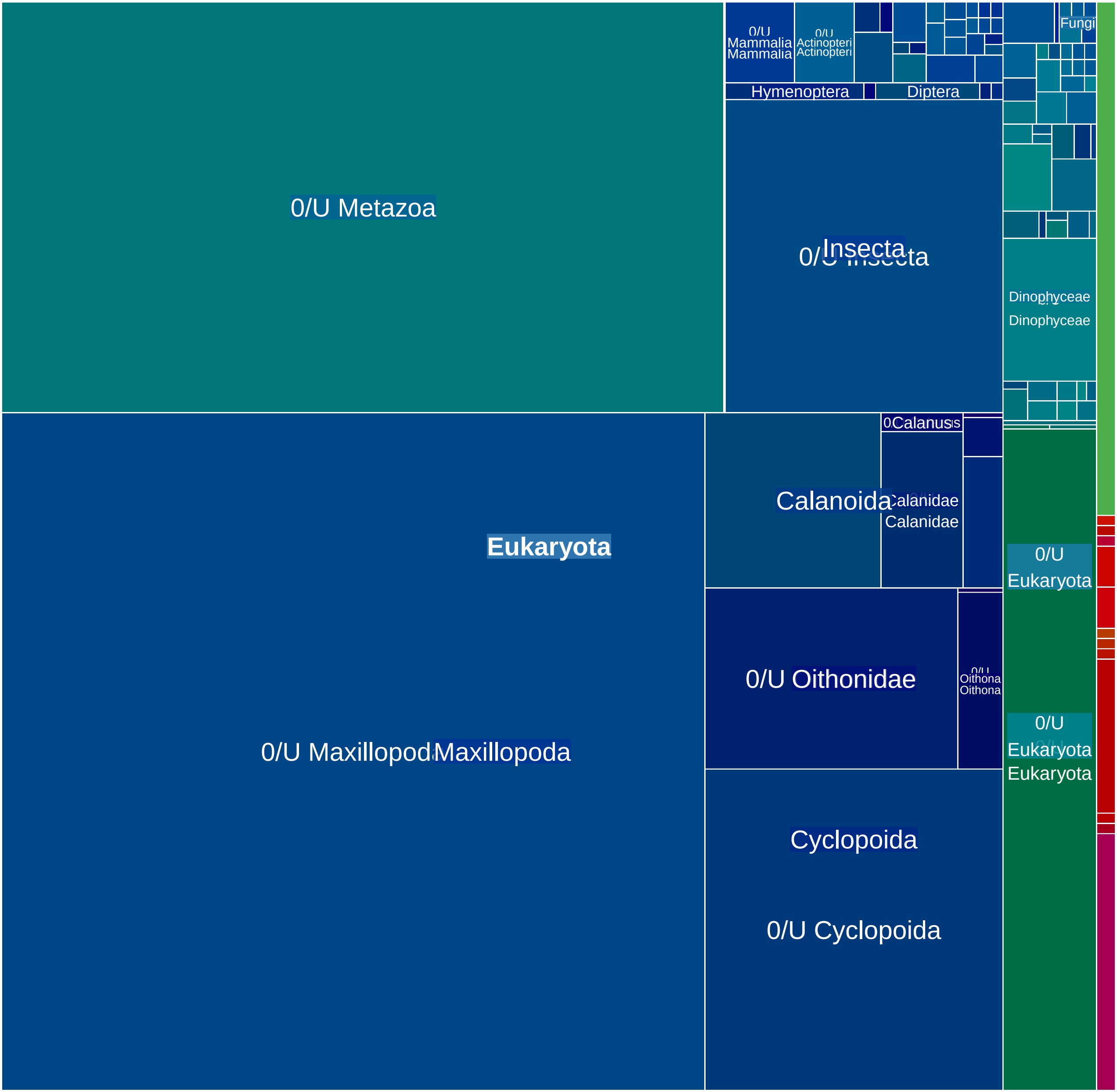

MGT−v1_162 : 9459 unigenes, 1223 (12.93%) taxonomically assigned

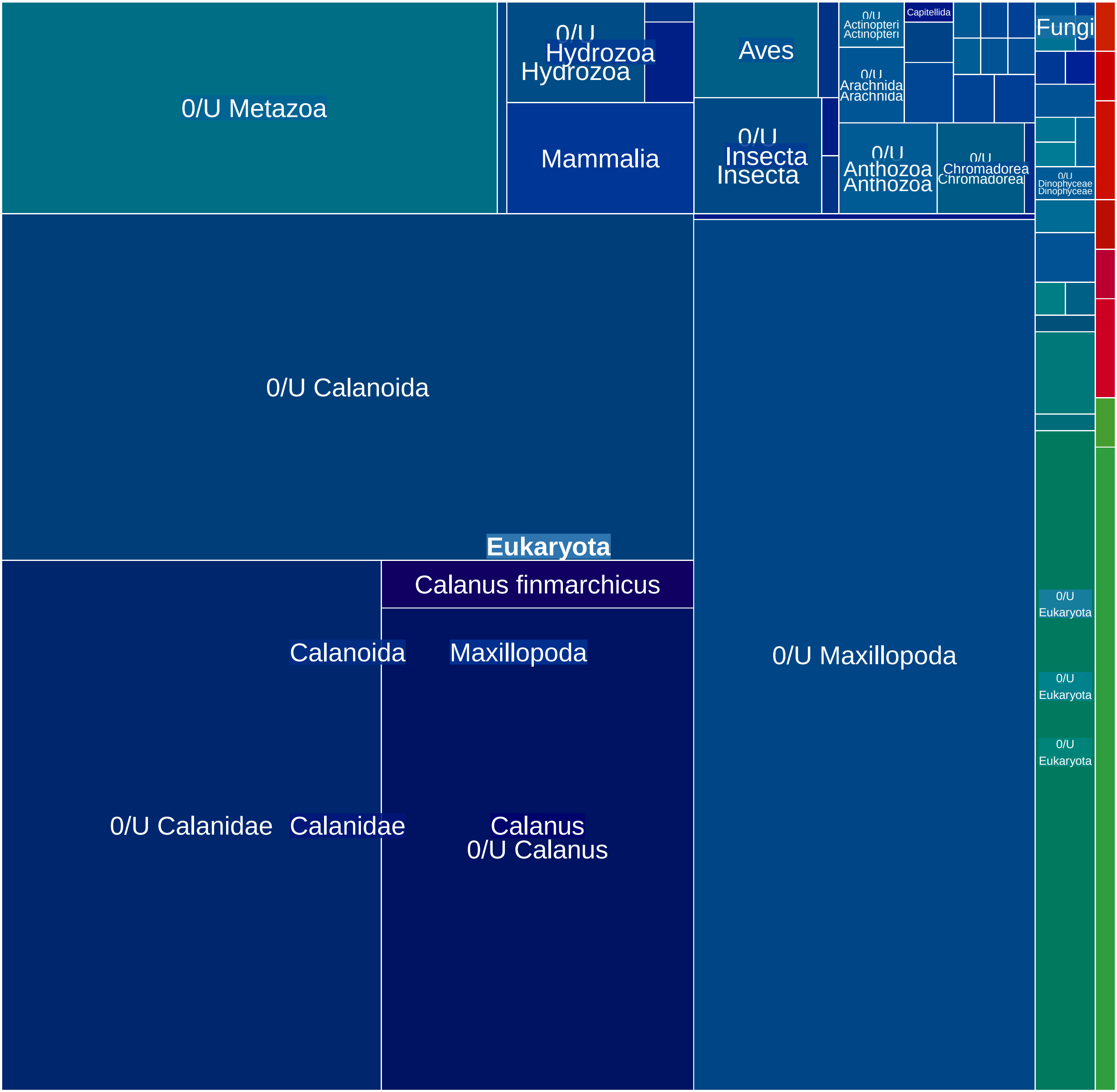

MGT−v1_163 : 9400 unigenes, 1456 (15.49%) taxonomically assigned

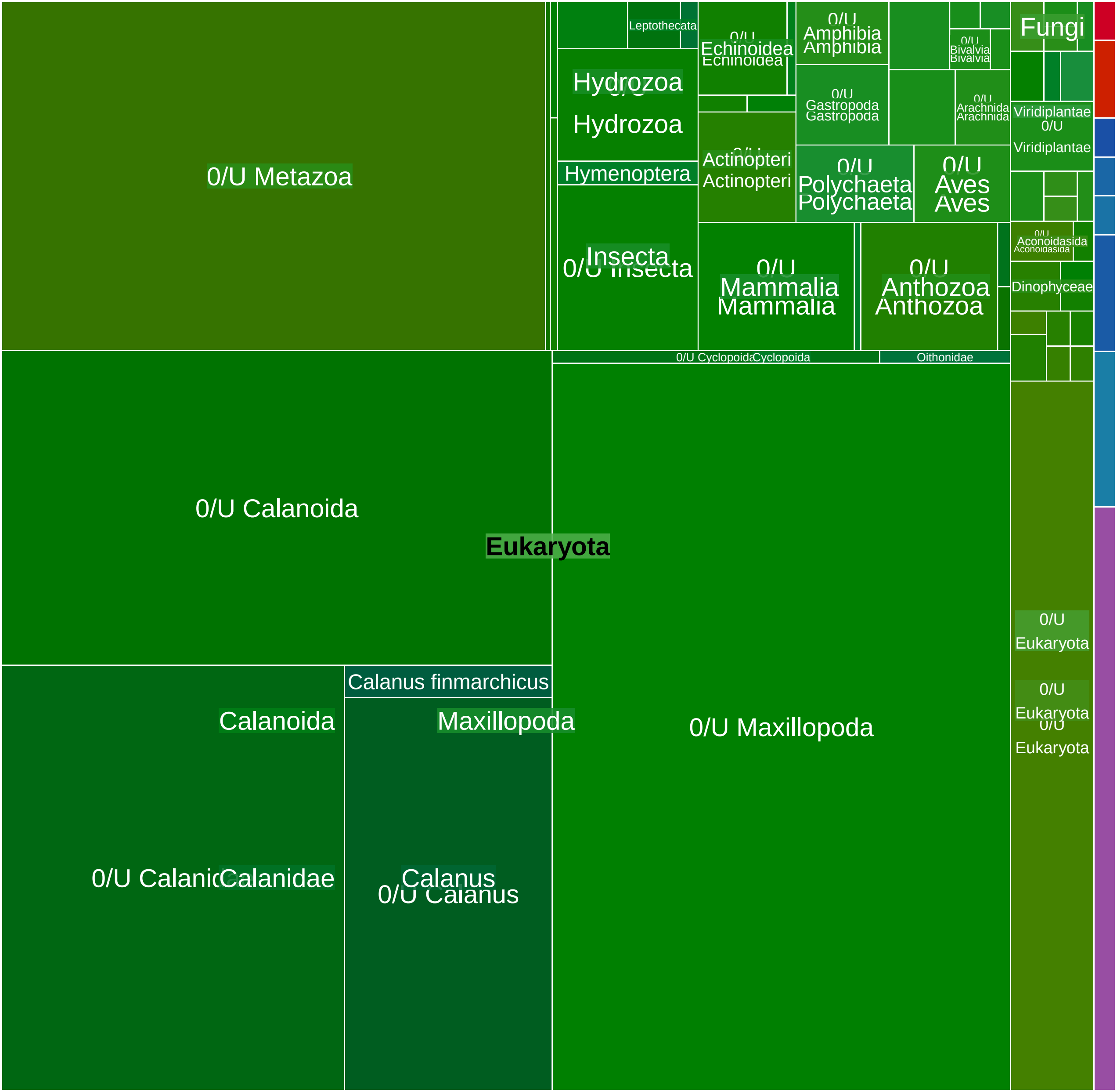

MGT−v1_164 : 9309 unigenes, 8545 (91.79%) taxonomically assigned

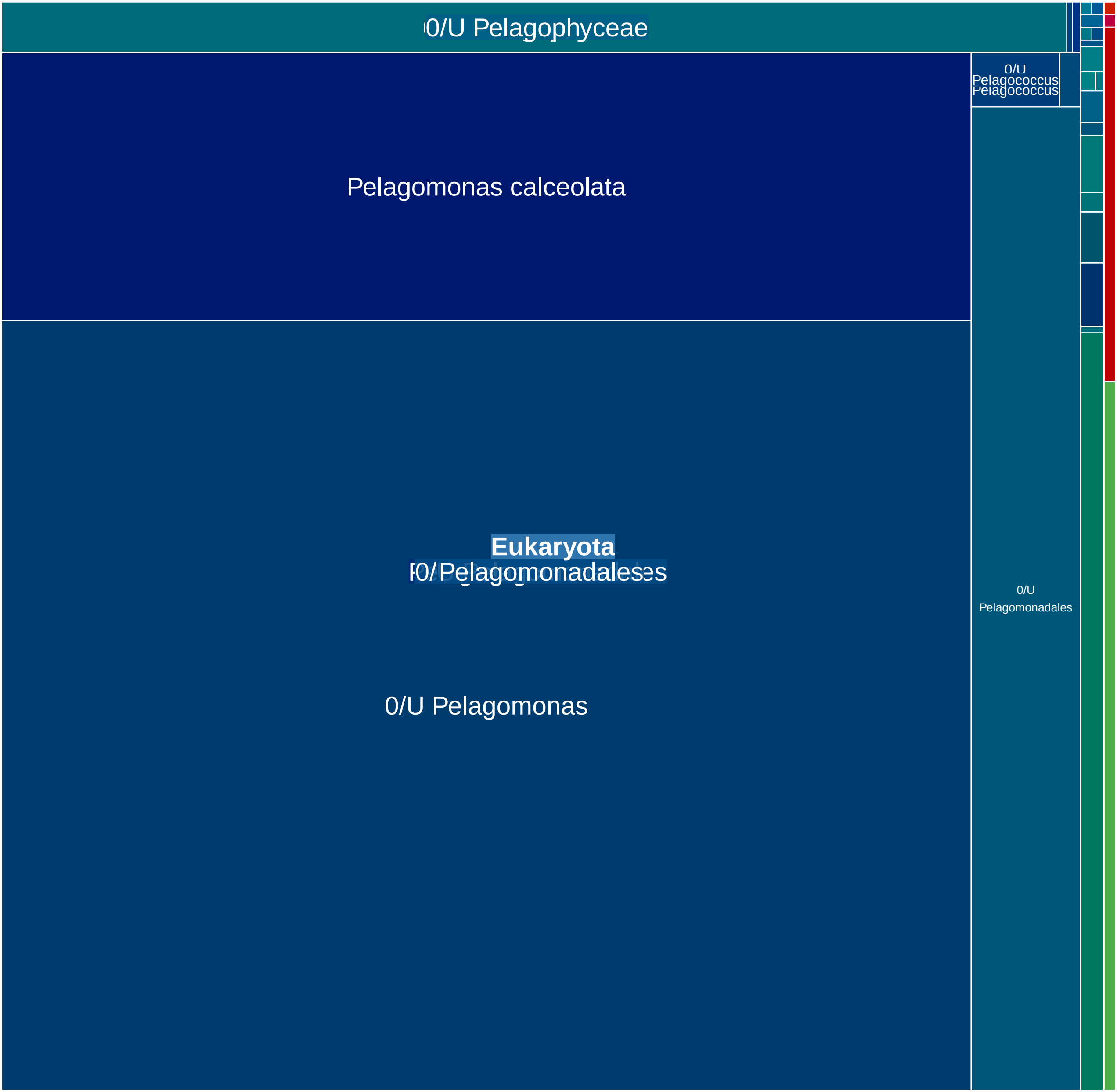

MGT−v1_165 : 9304 unigenes, 1400 (15.05%) taxonomically assigned

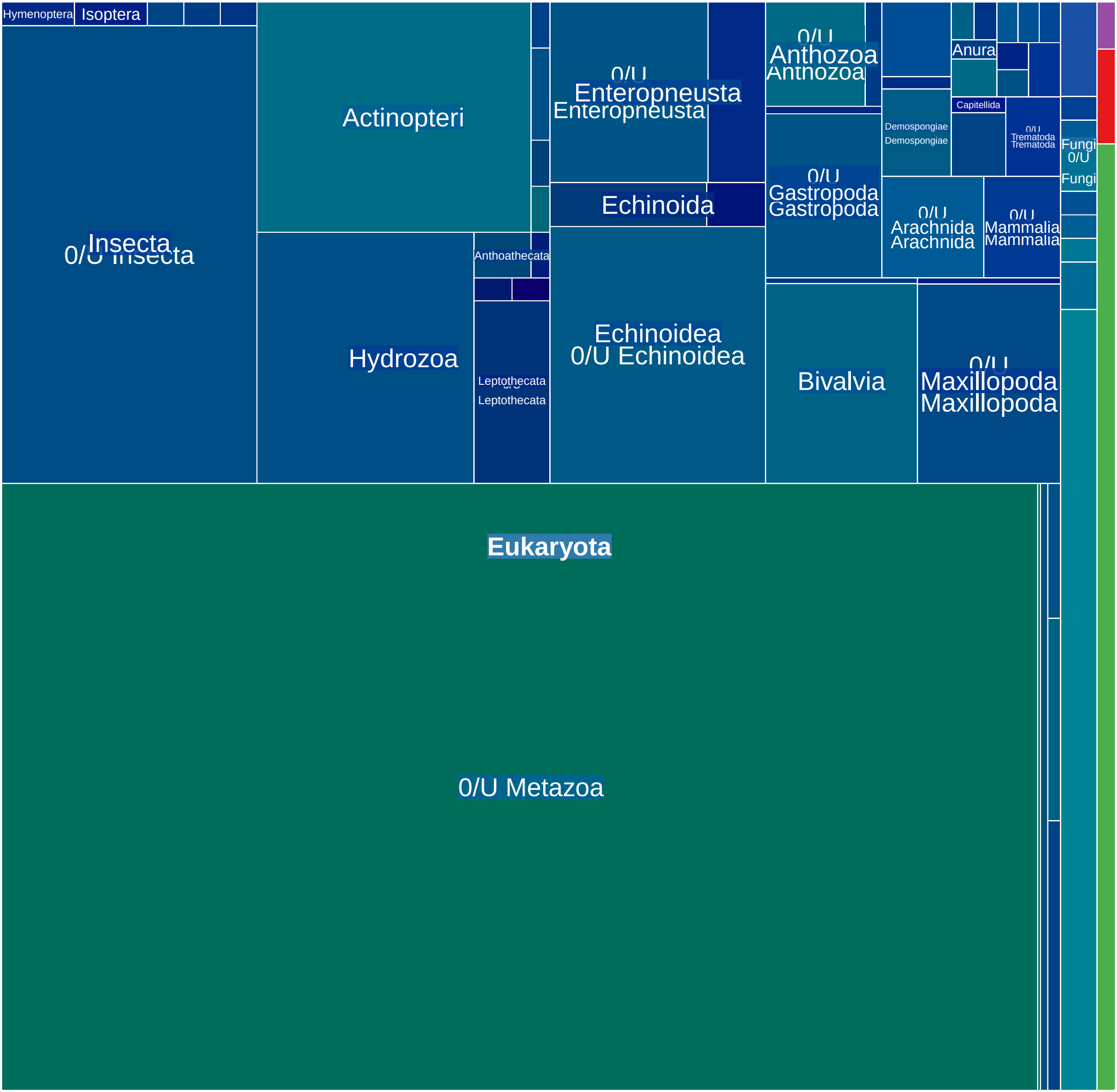

### MGT−v1_166 : 9247 unigenes, 5931 (64.14%) taxonomically assigned

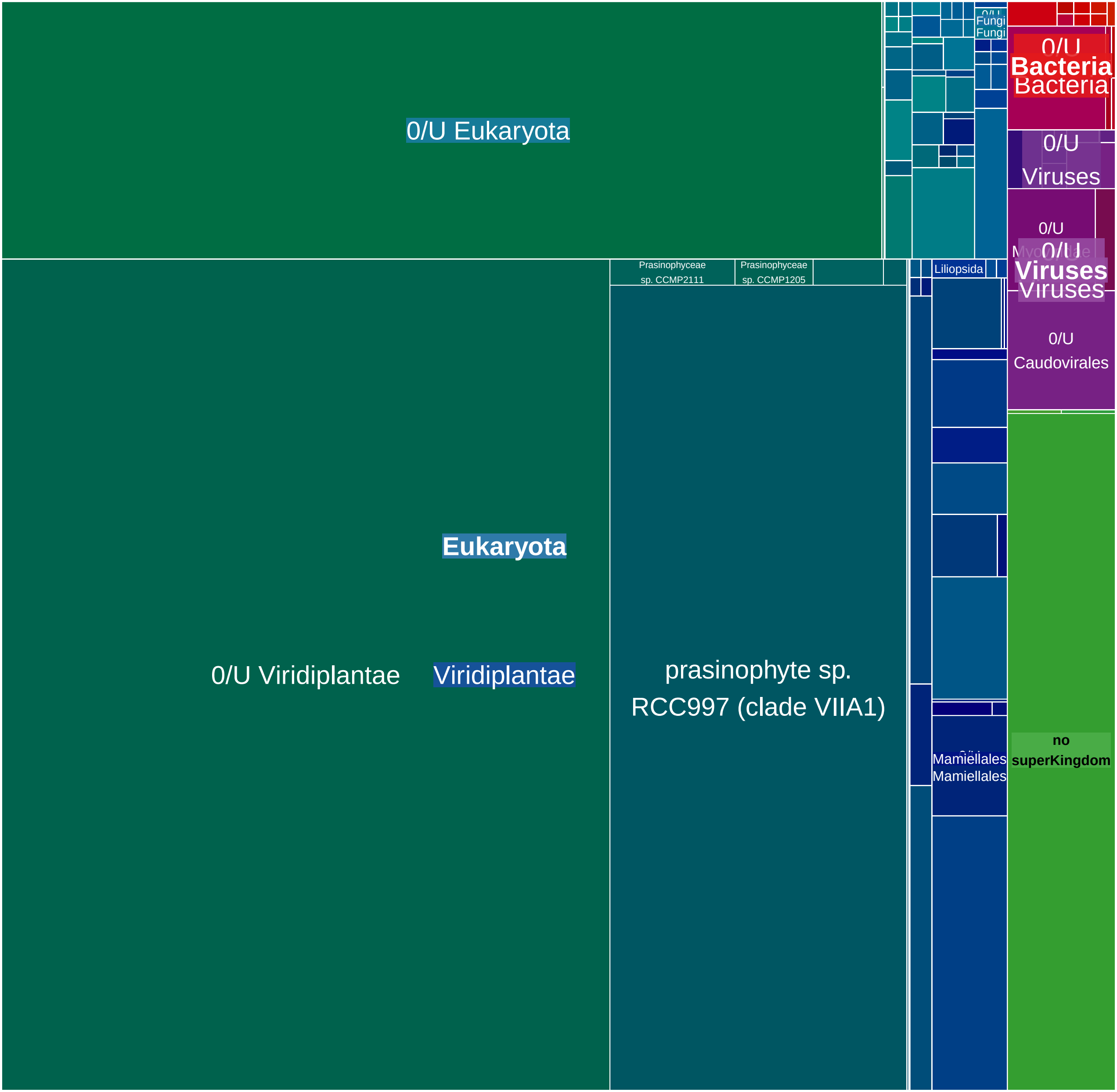

MGT−v1_167 : 9138 unigenes, 1020 (11.16%) taxonomically assigned

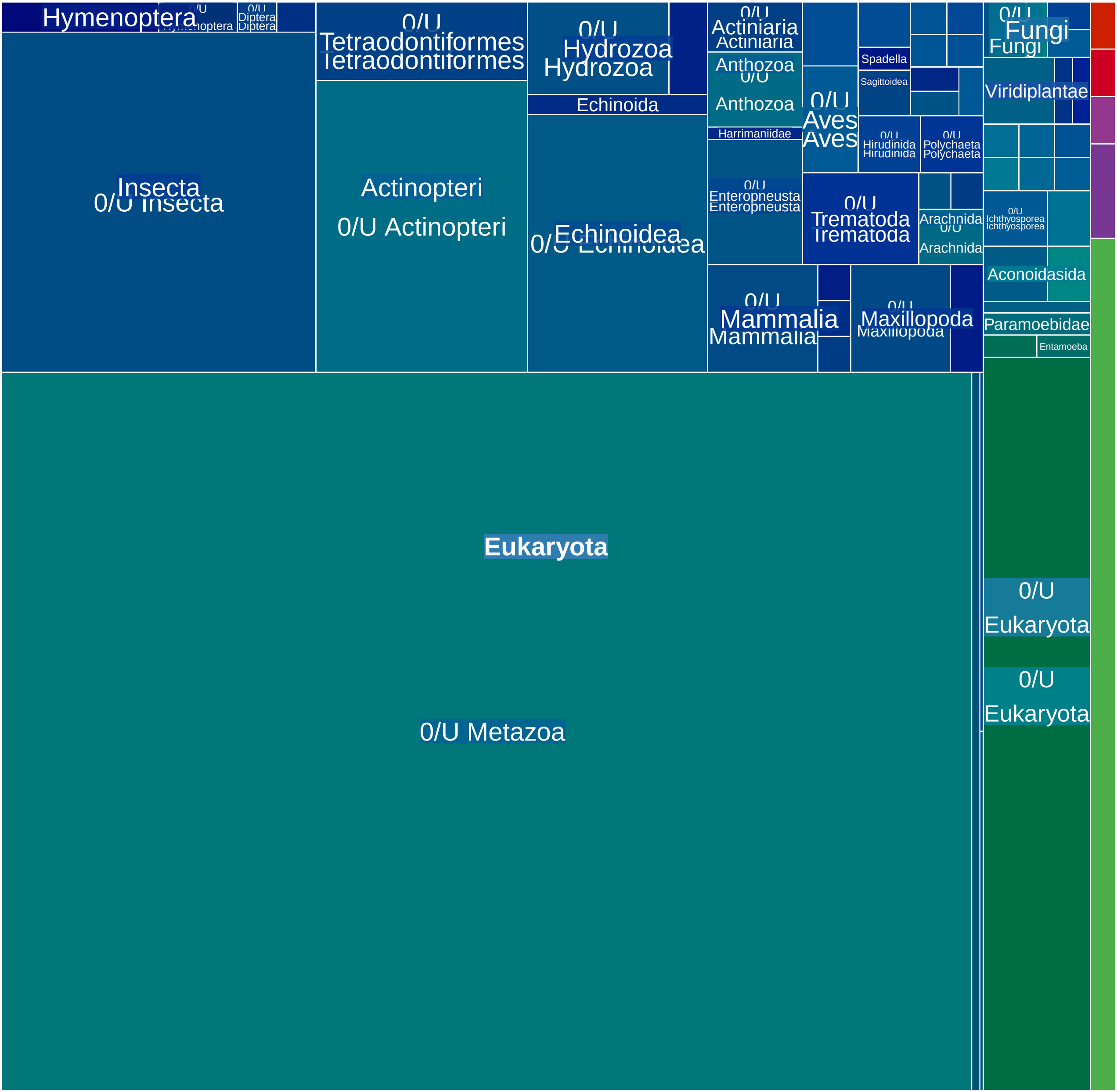

MGT−v1_168 : 9069 unigenes, 6479 (71.44%) taxonomically assigned

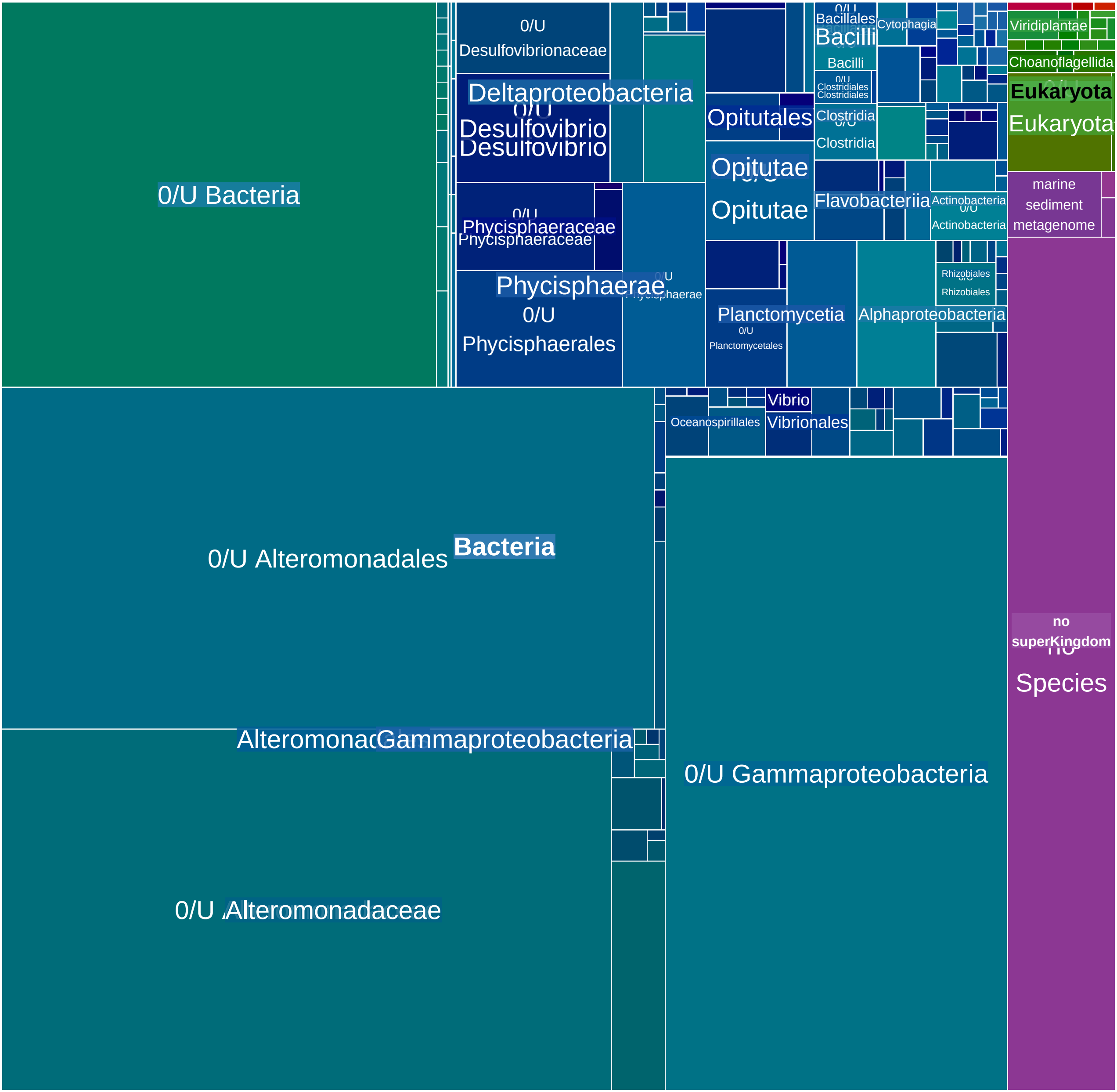

MGT−v1_169 : 9056 unigenes, 990 (10.93%) taxonomically assigned

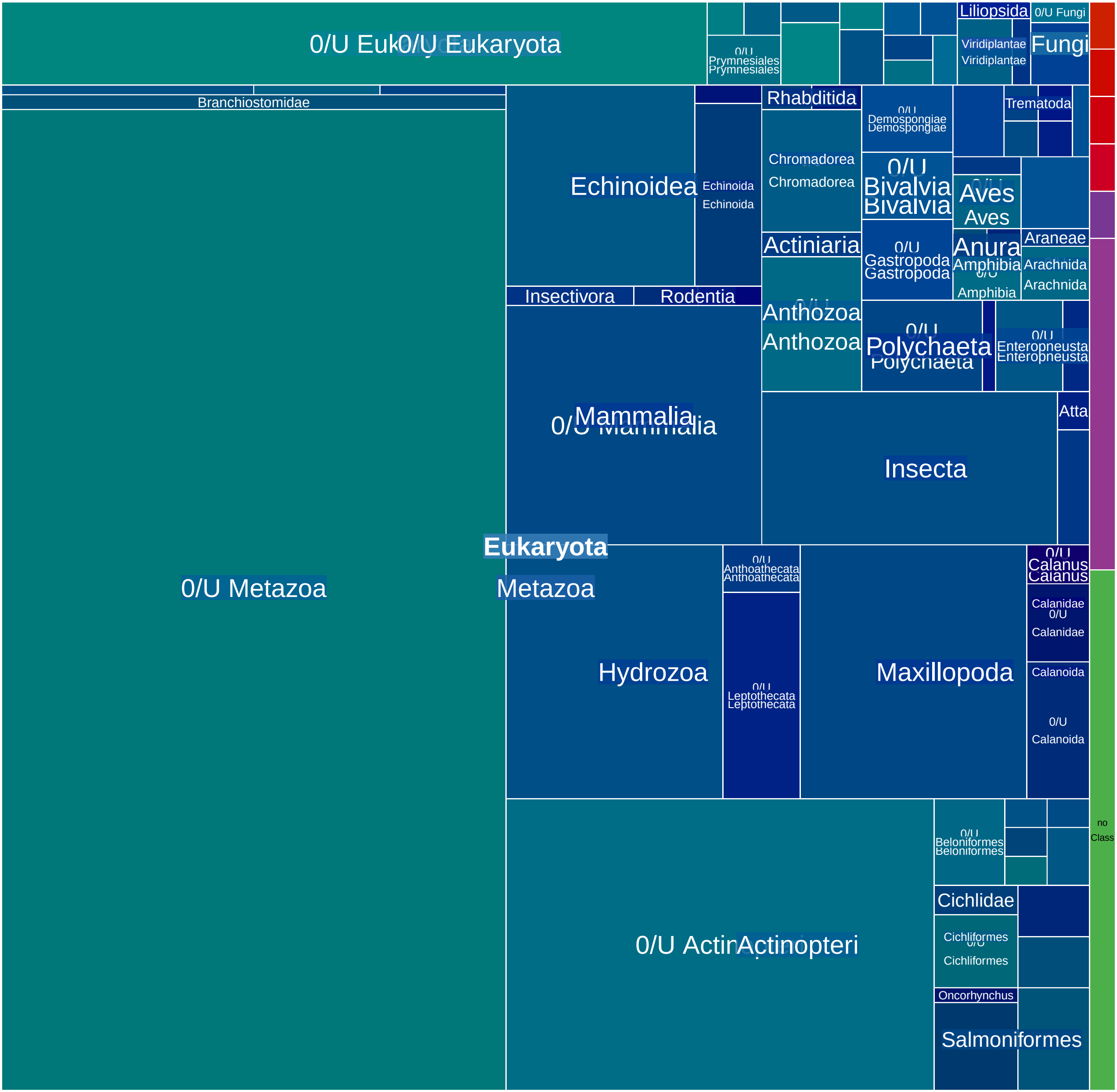

MGT−v1_170 : 9053 unigenes, 6712 (74.14%) taxonomically assigned

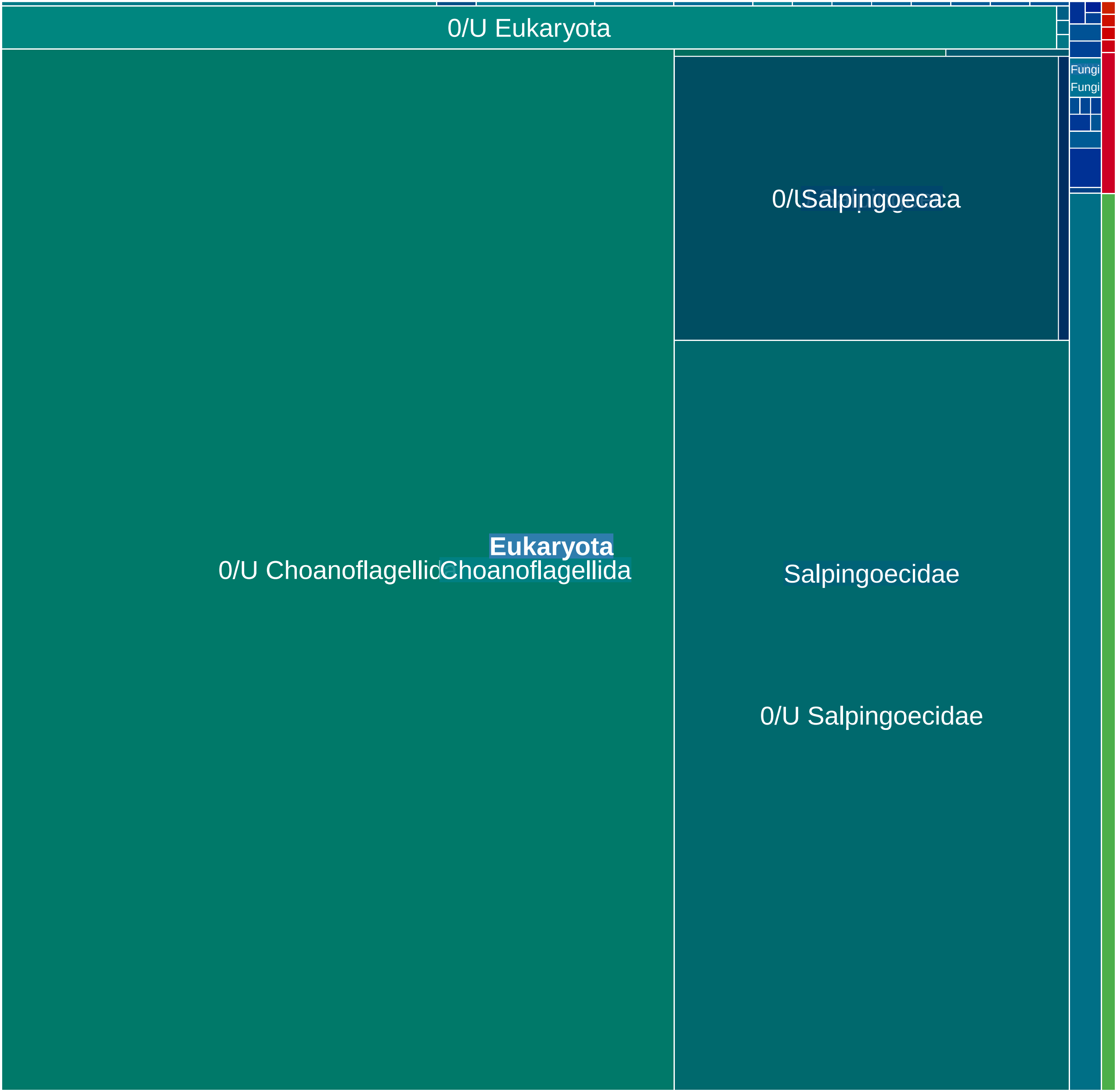

MGT−v1_171 : 8950 unigenes, 3967 (44.32%) taxonomically assigned

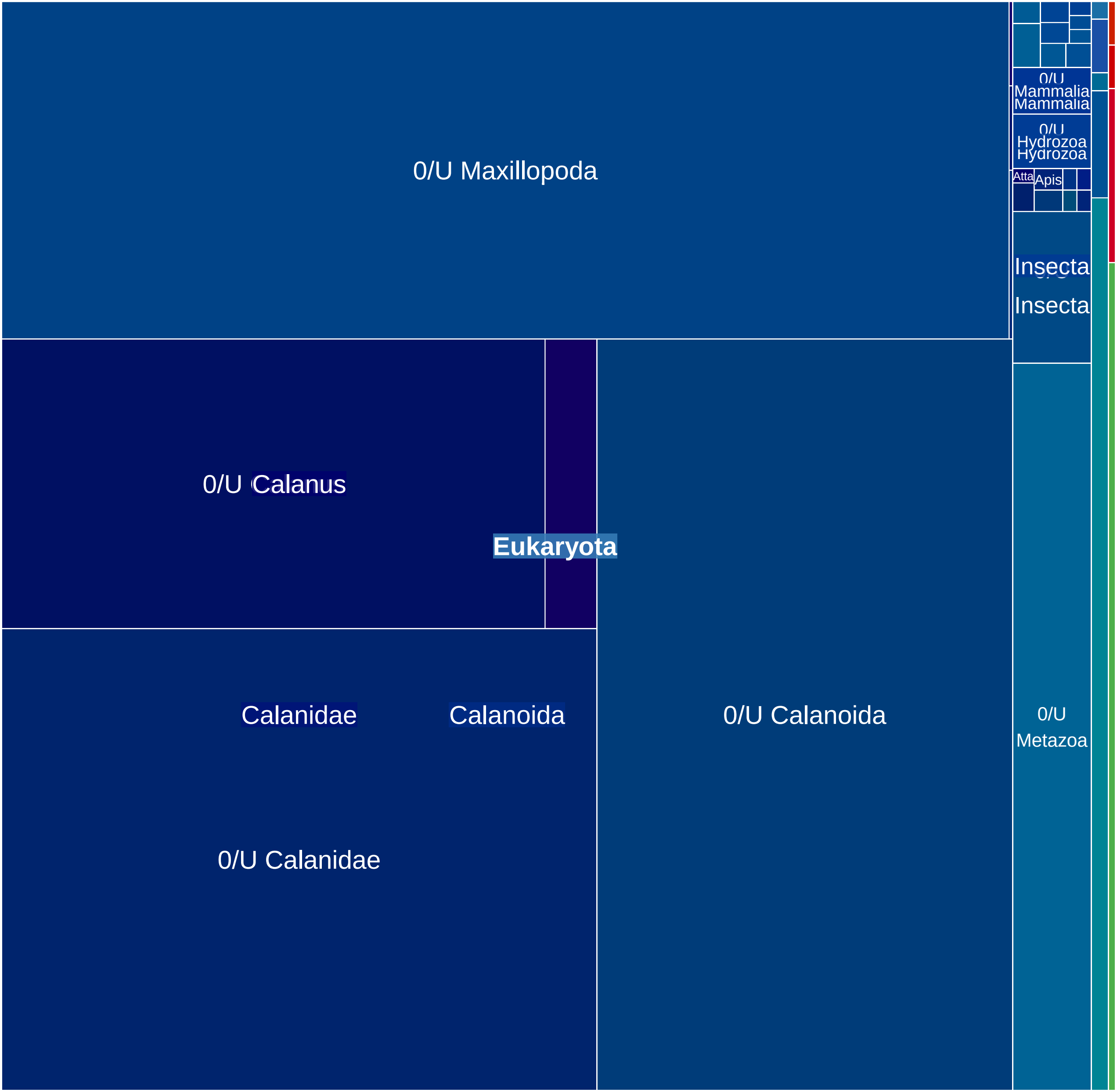

MGT−v1_172 : 8950 unigenes, 1066 (11.91%) taxonomically assigned

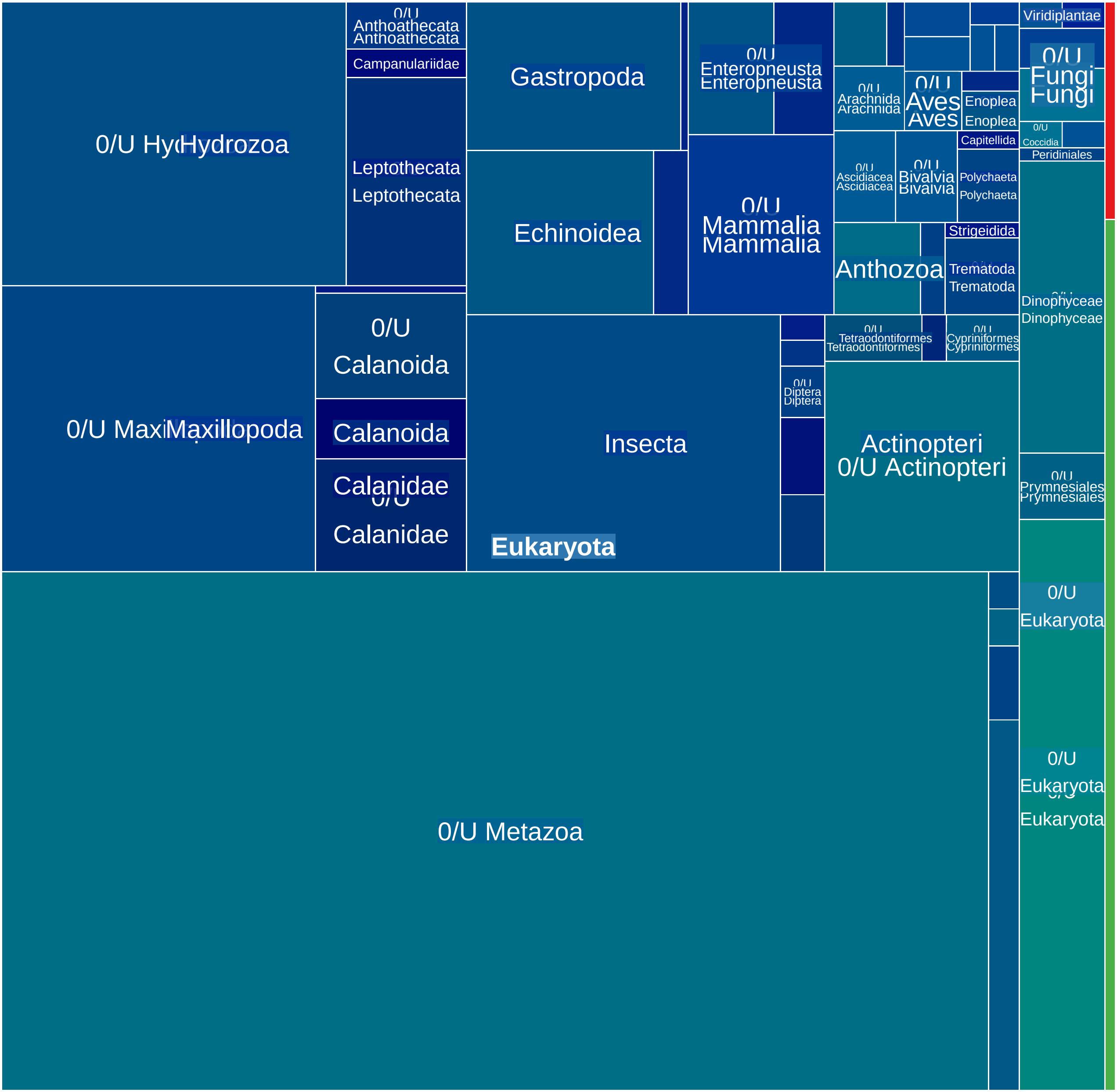

MGT−v1_173 : 8939 unigenes, 8315 (93.02%) taxonomically assigned

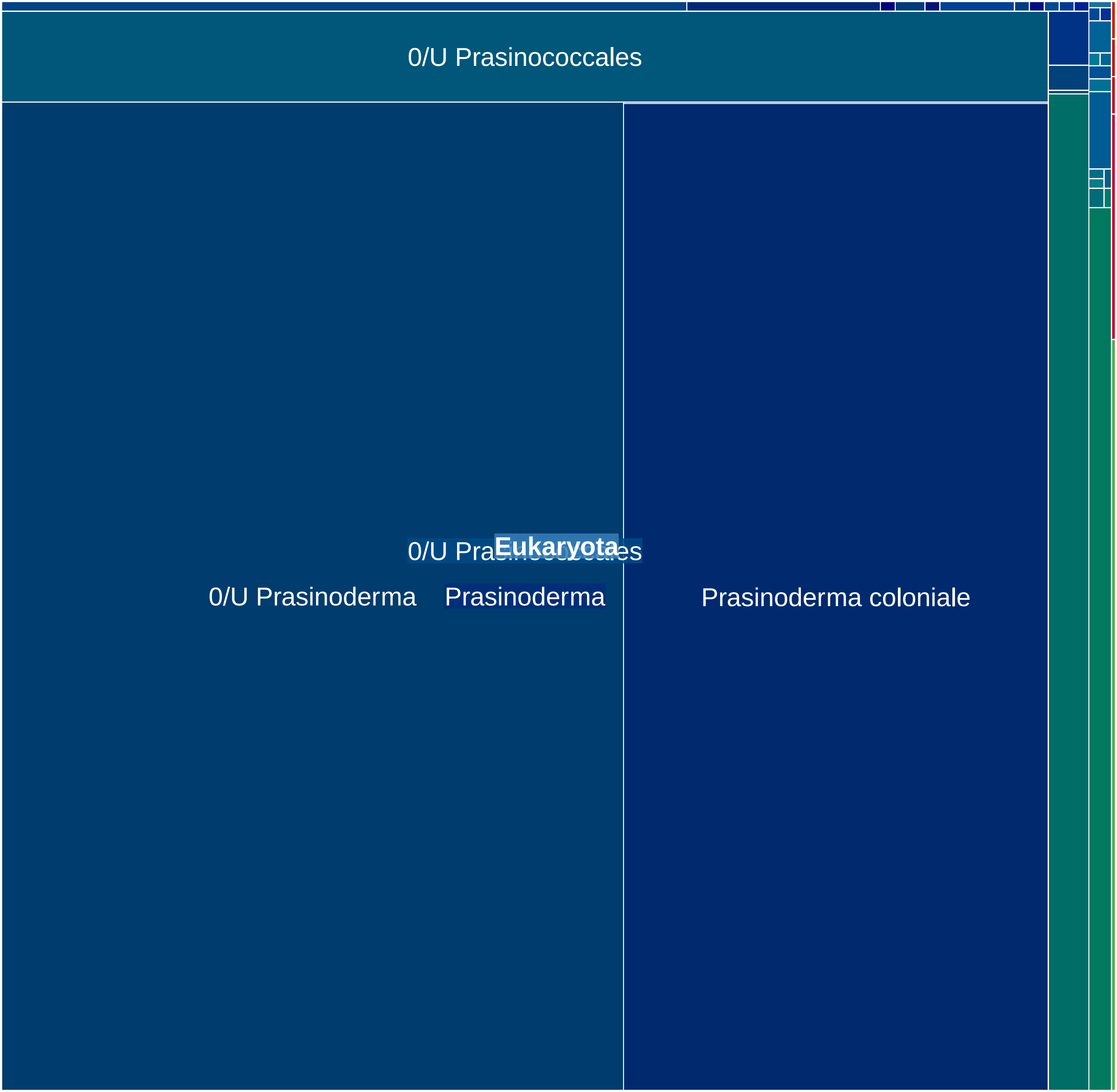

MGT−v1_174 : 8888 unigenes, 4186 (47.10%) taxonomically assigned

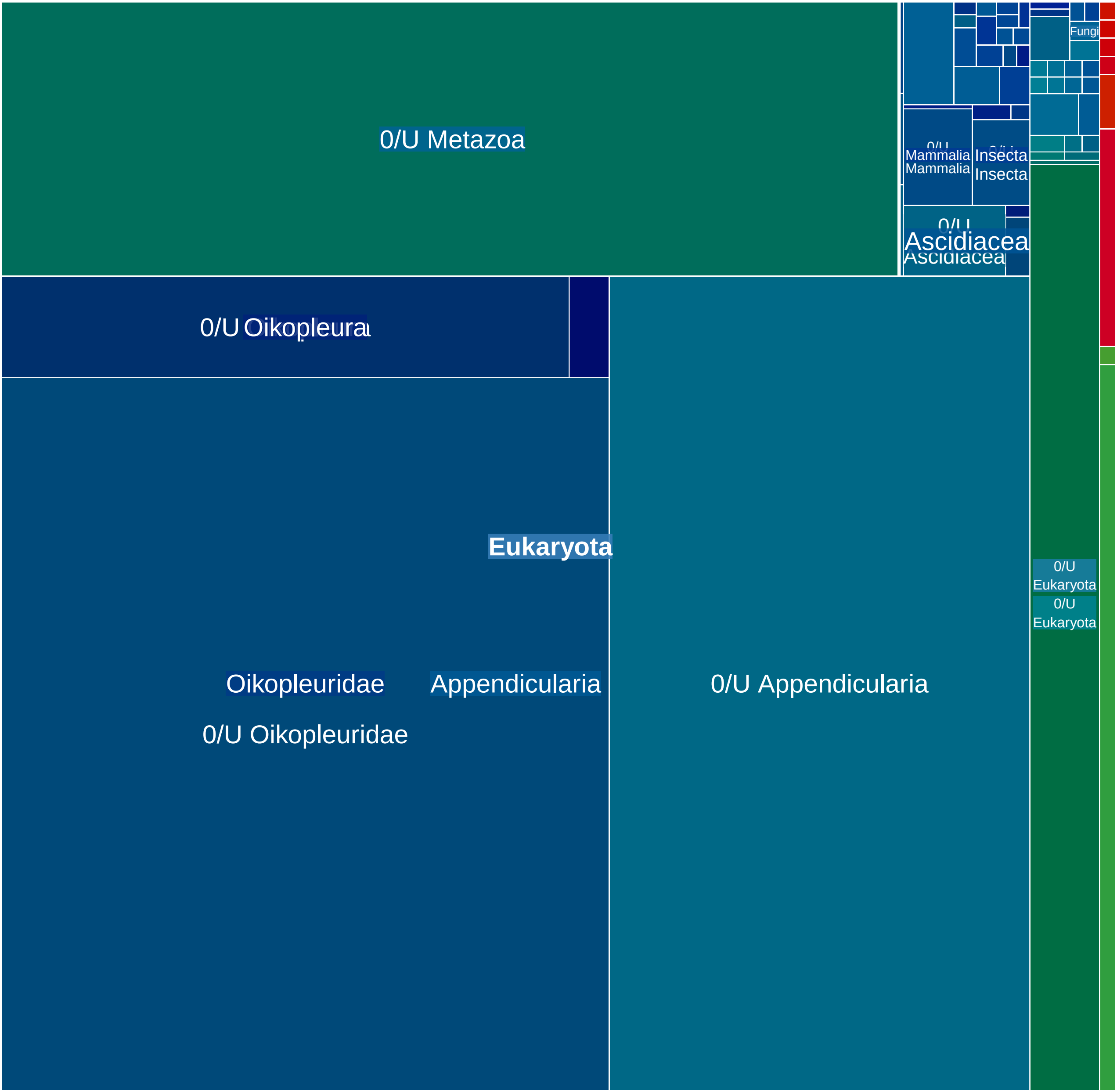

MGT−v1_175 : 8797 unigenes, 7314 (83.14%) taxonomically assigned

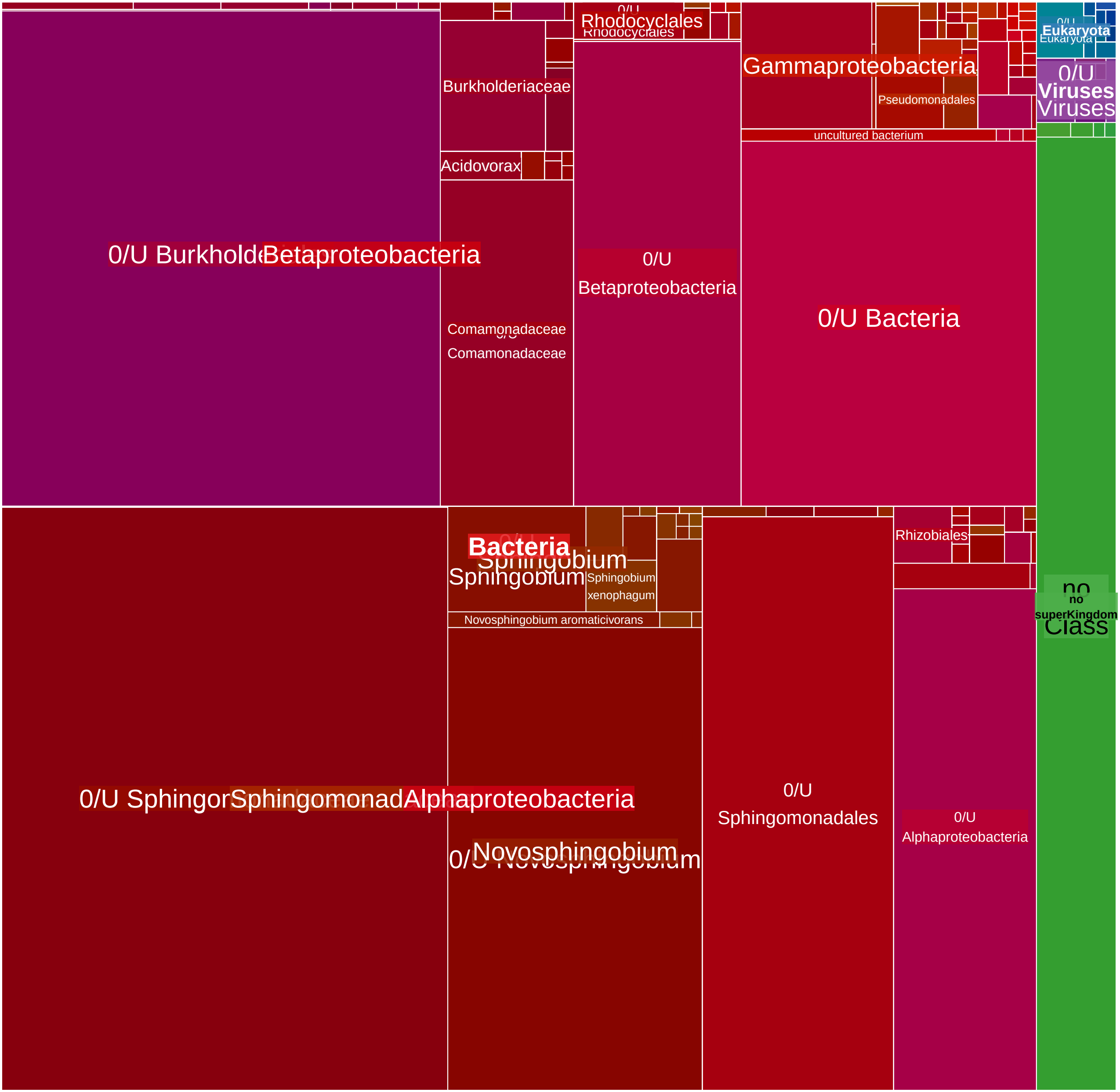

### MGT−v1_176 : 8717 unigenes, 5410 (62.06%) taxonomically assigned

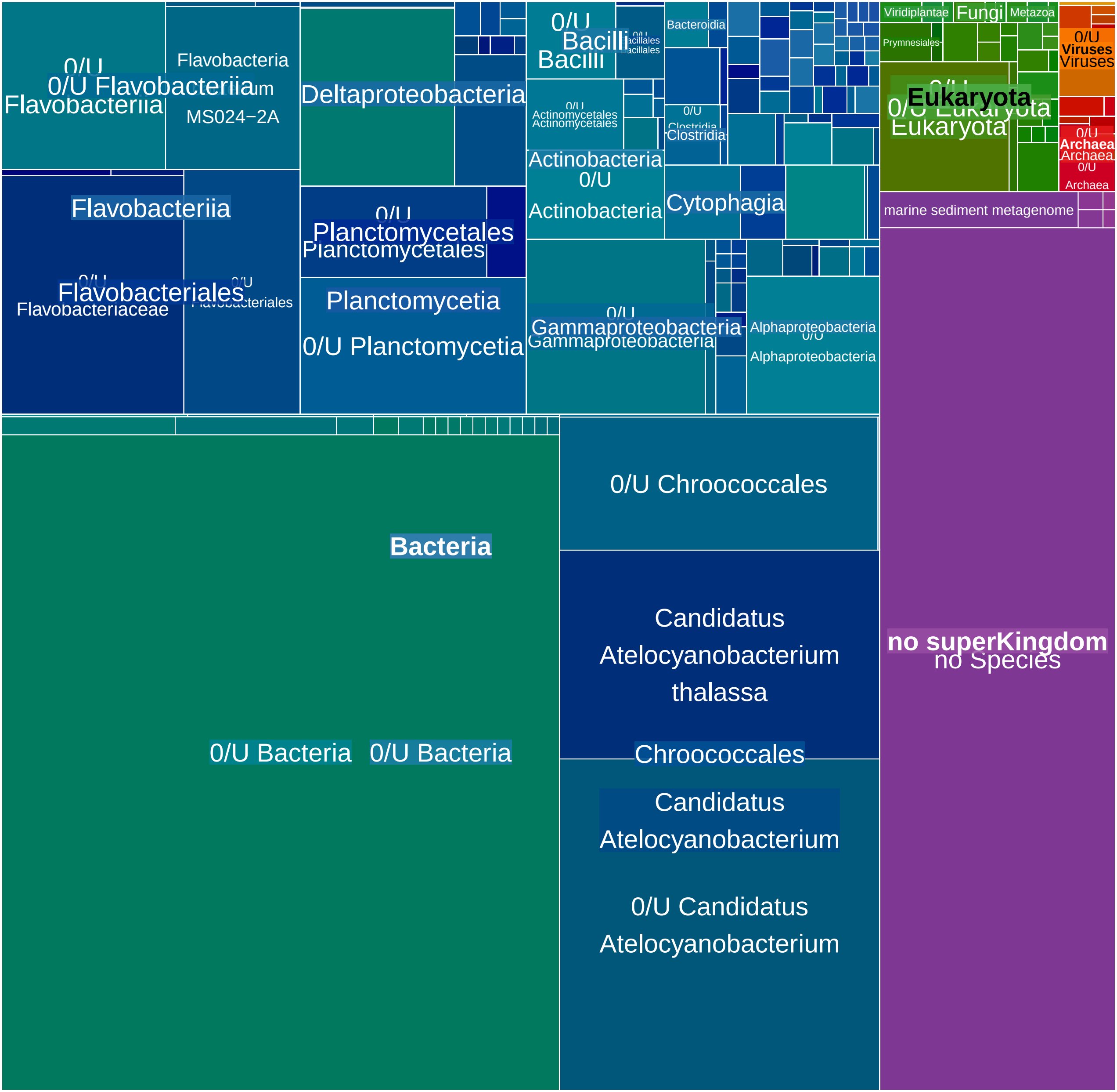

MGT−v1_177 : 8676 unigenes, 743 (8.56%) taxonomically assigned

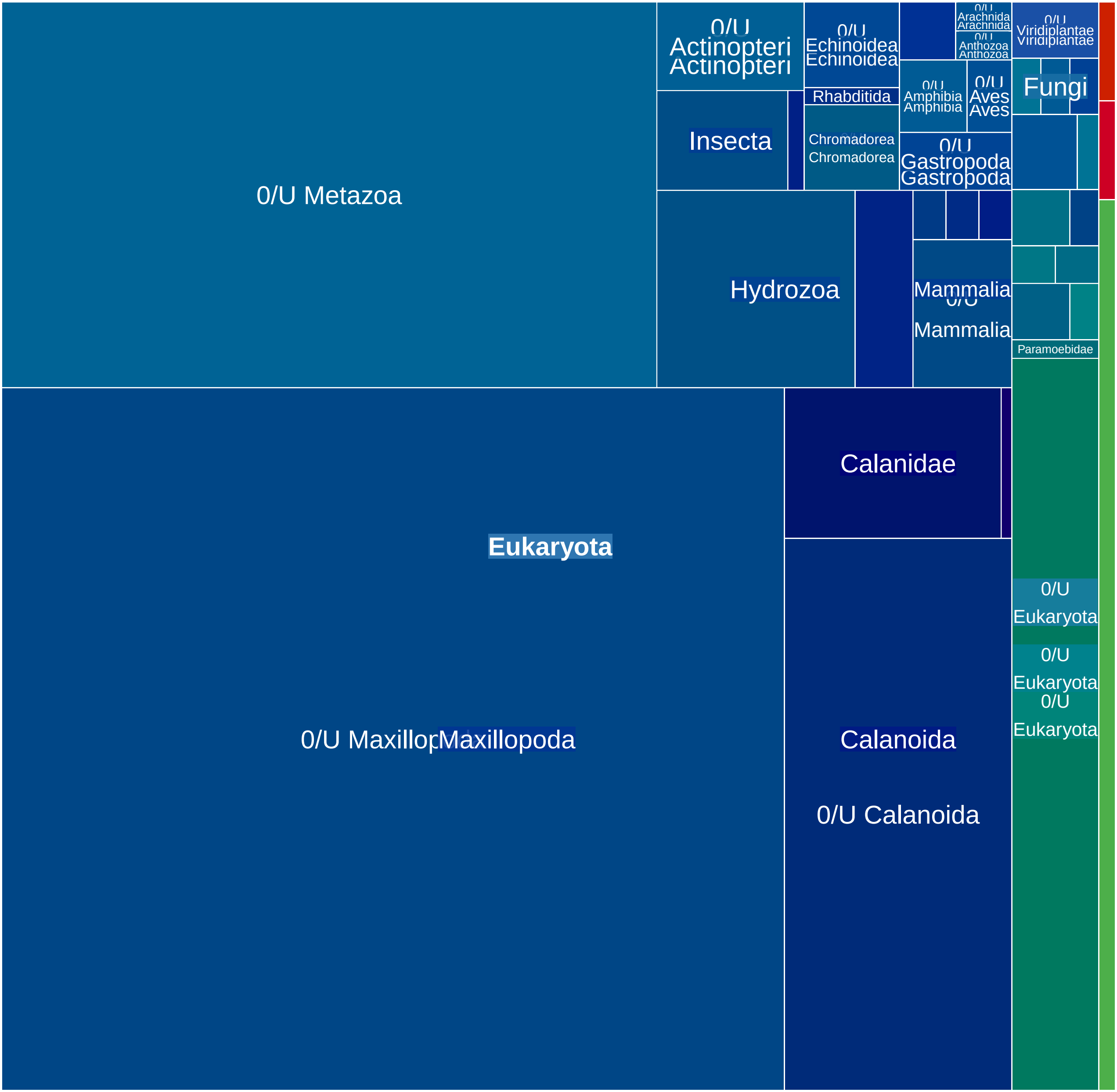

MGT−v1_178 : 8653 unigenes, 7272 (84.04%) taxonomically assigned

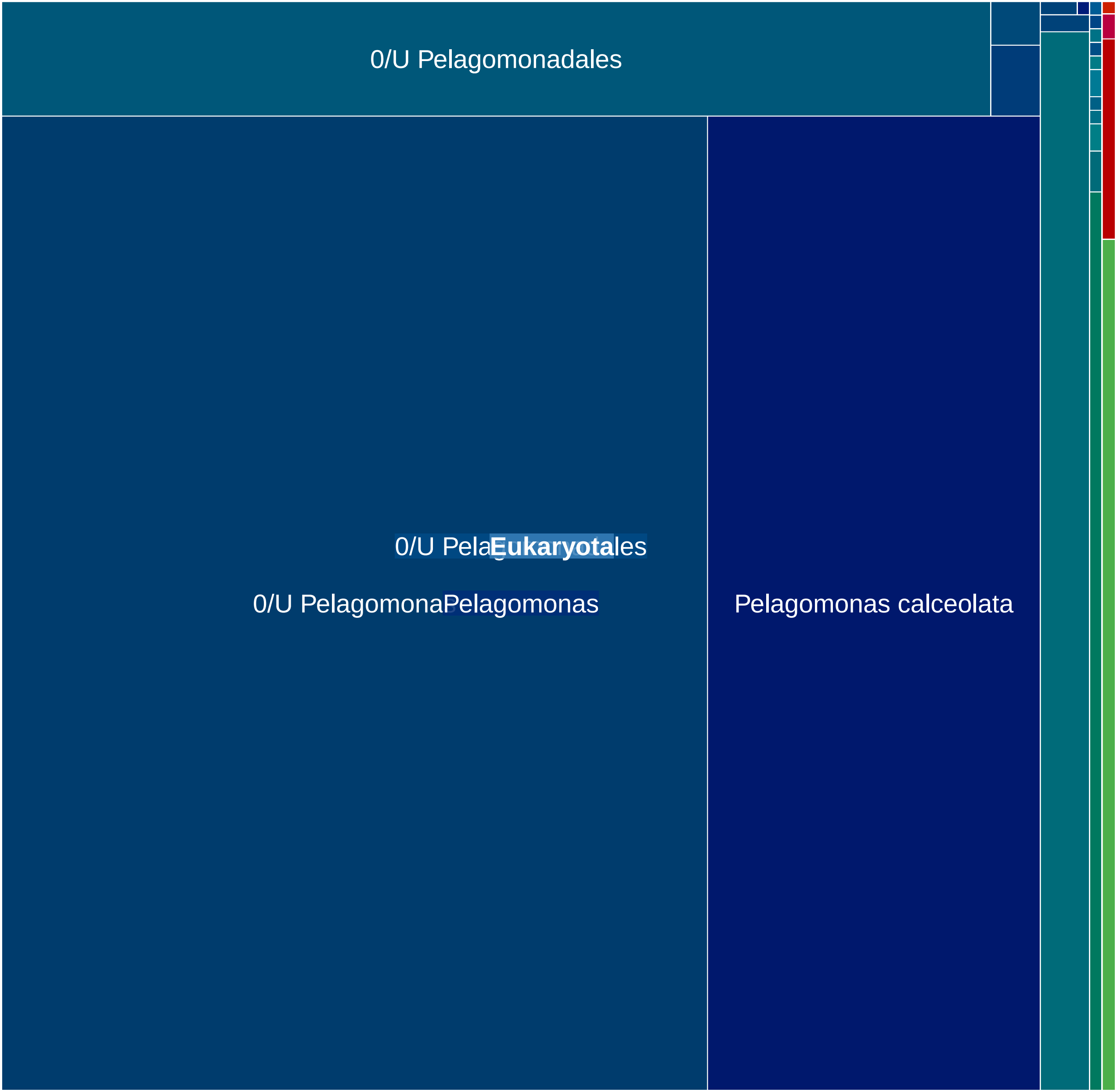

### MGT−v1_179 : 8603 unigenes, 7202 (83.71%) taxonomically assigned

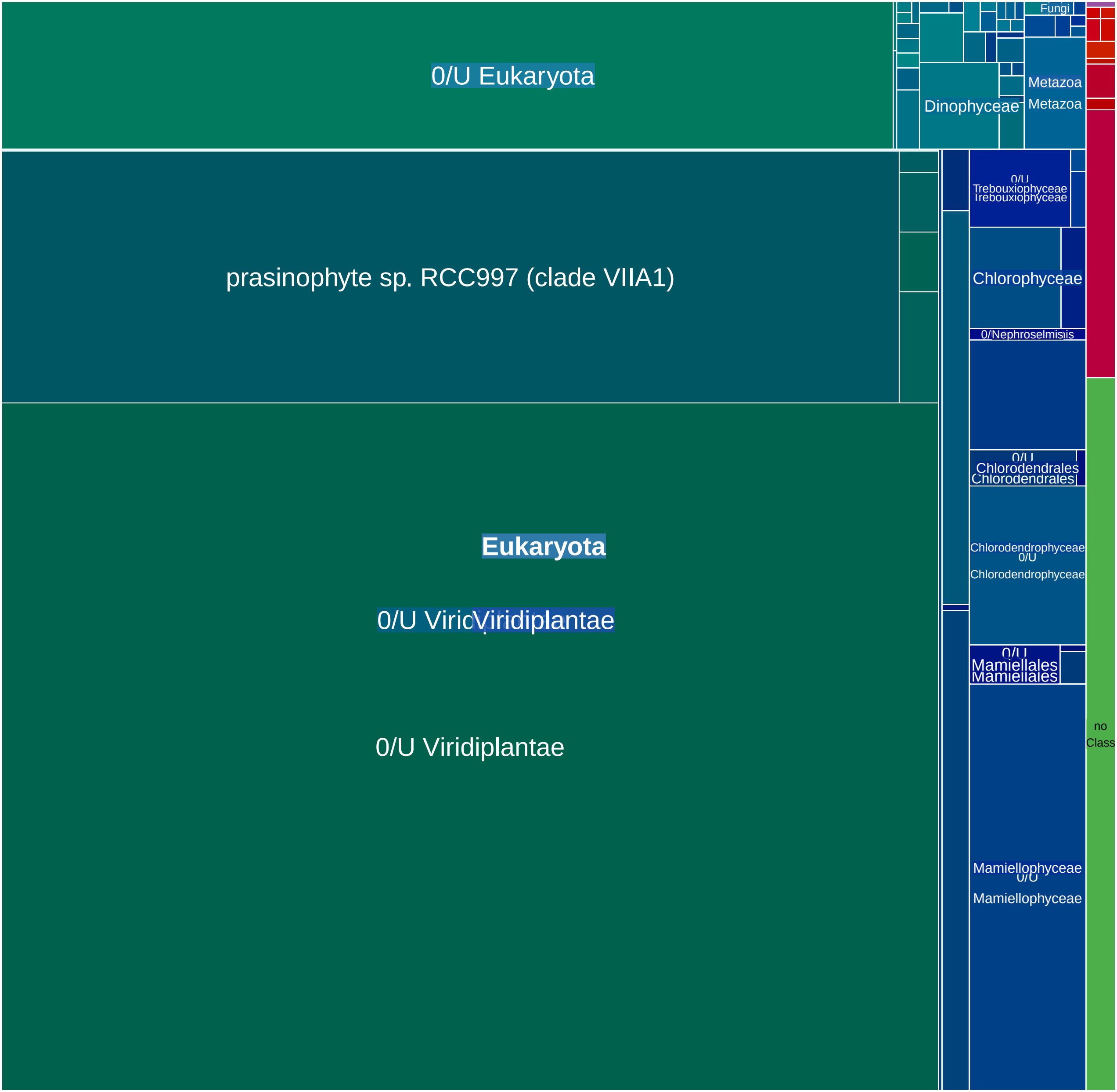

MGT−v1_180 : 8598 unigenes, 330 (3.84%) taxonomically assigned

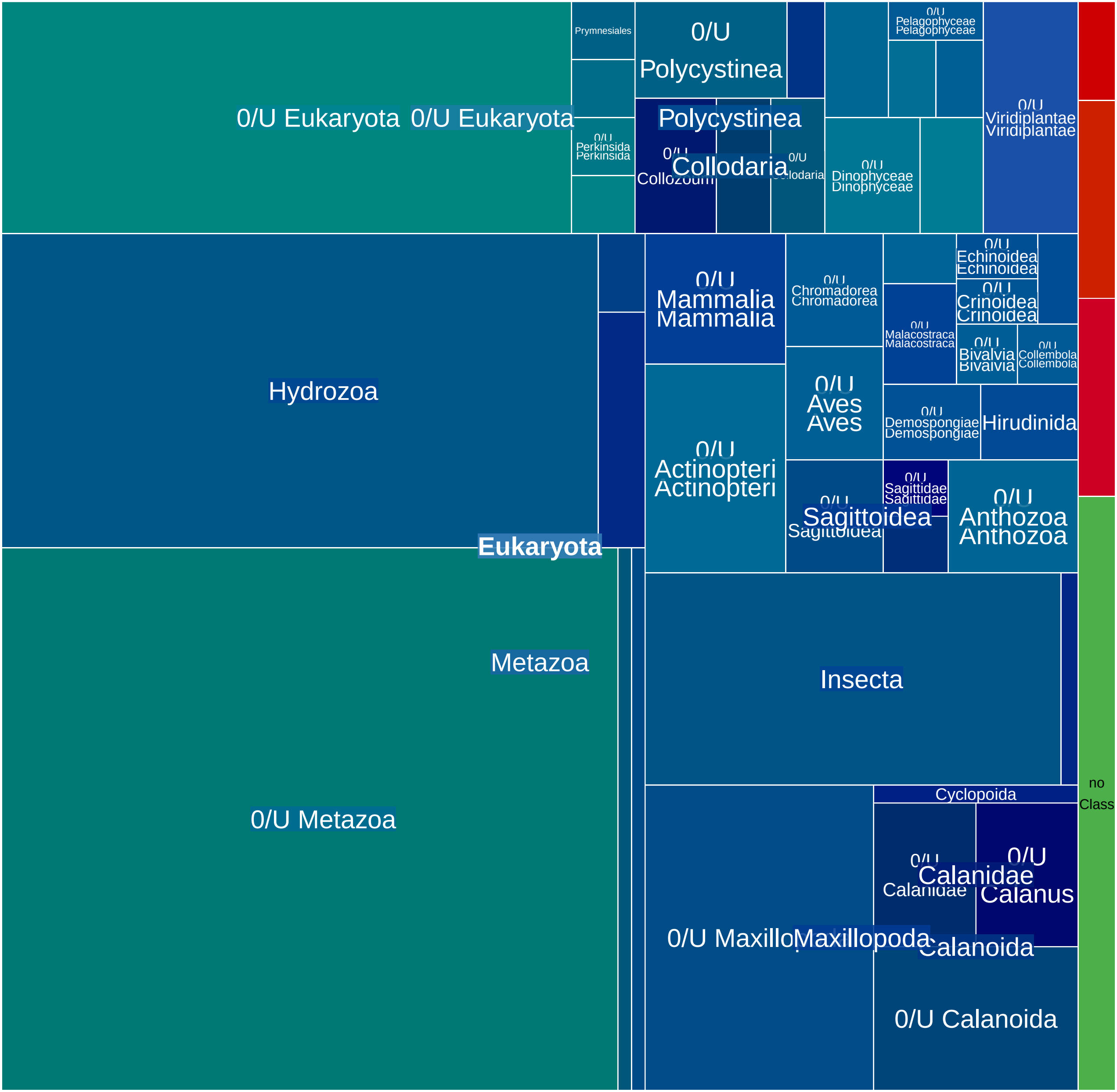

MGT−v1_181 : 8514 unigenes, 4484 (52.67%) taxonomically assigned

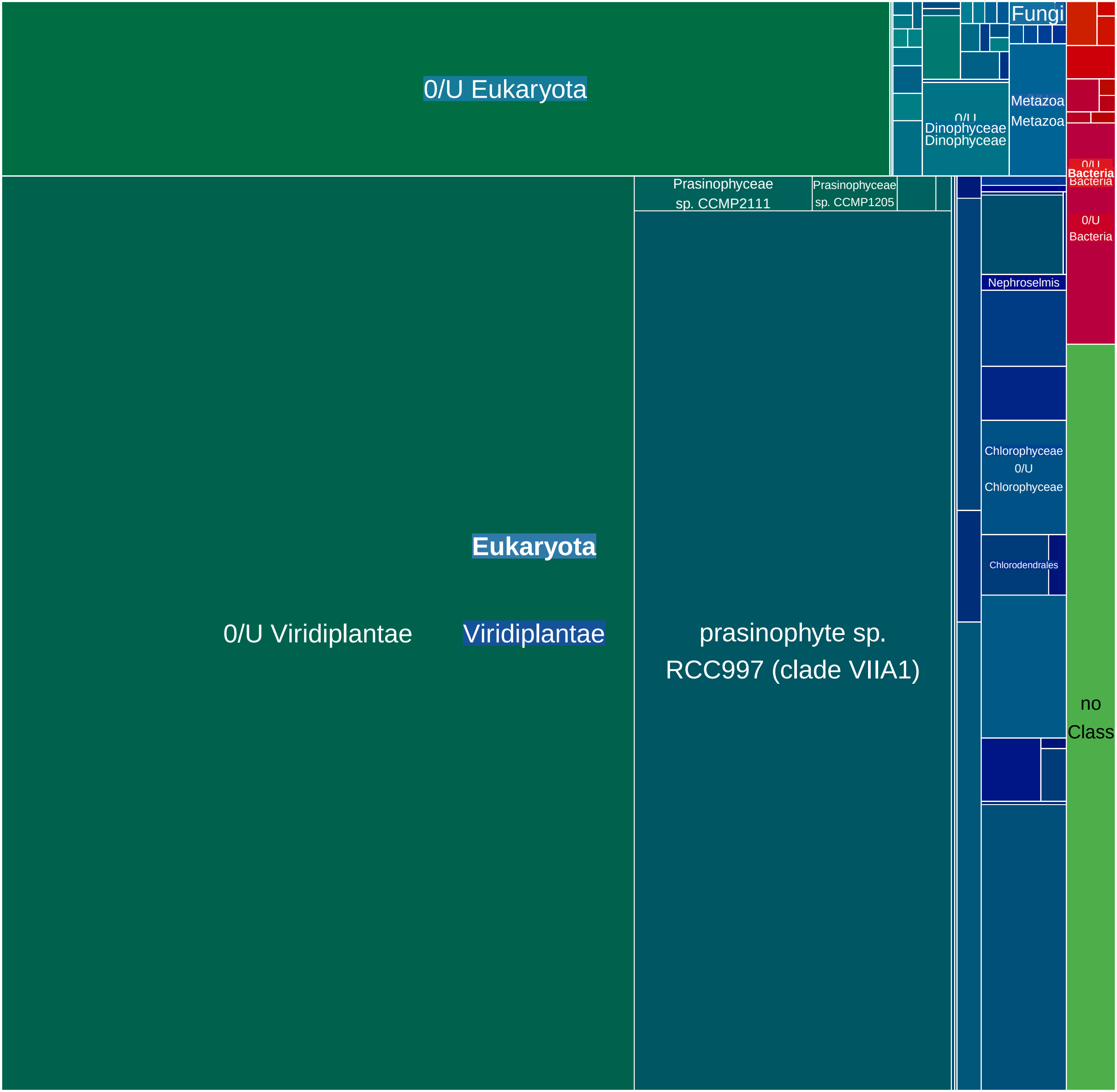

MGT−v1_182 : 8495 unigenes, 1454 (17.12%) taxonomically assigned

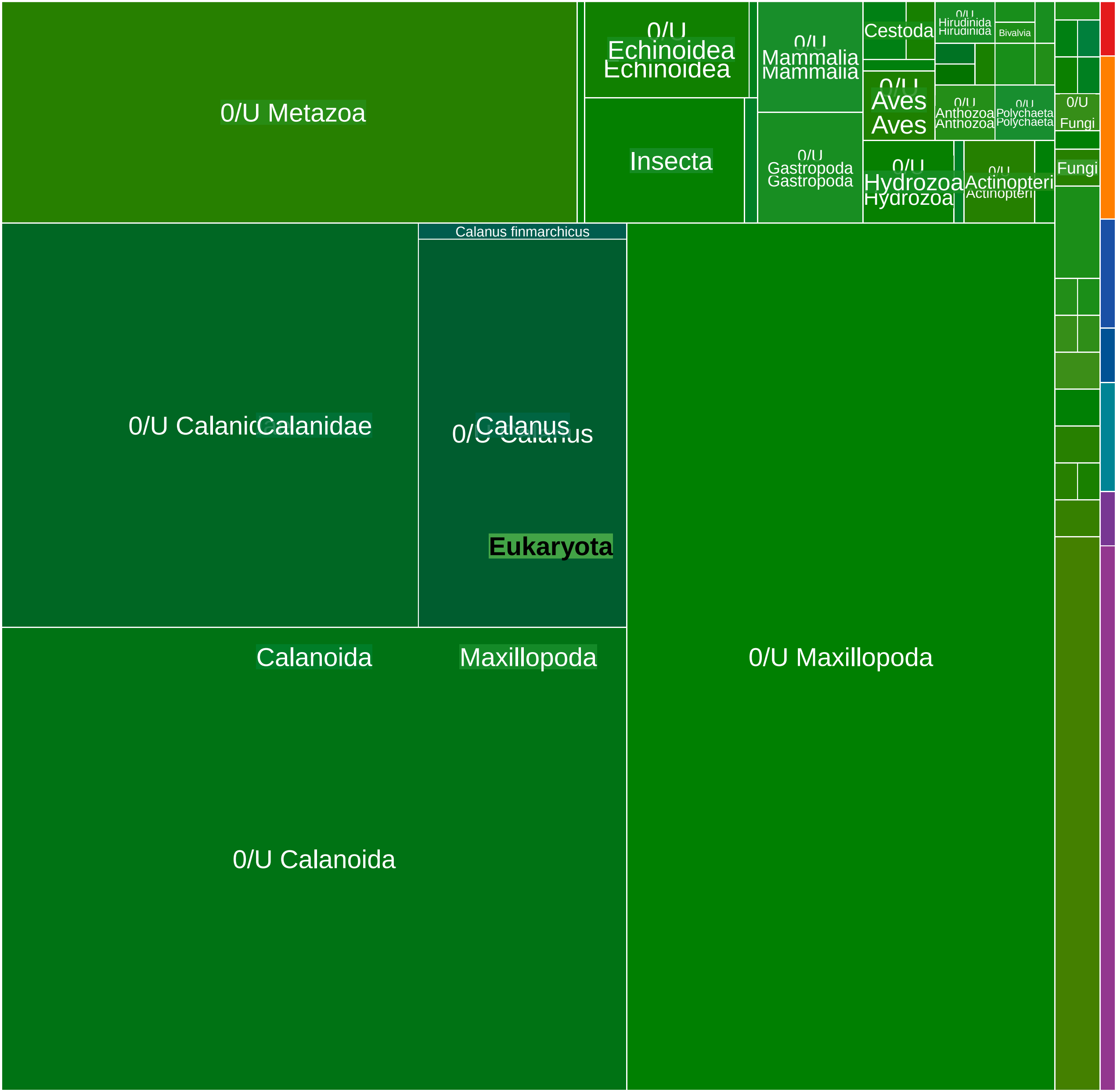

MGT−v1_183 : 8433 unigenes, 3653 (43.32%) taxonomically assigned

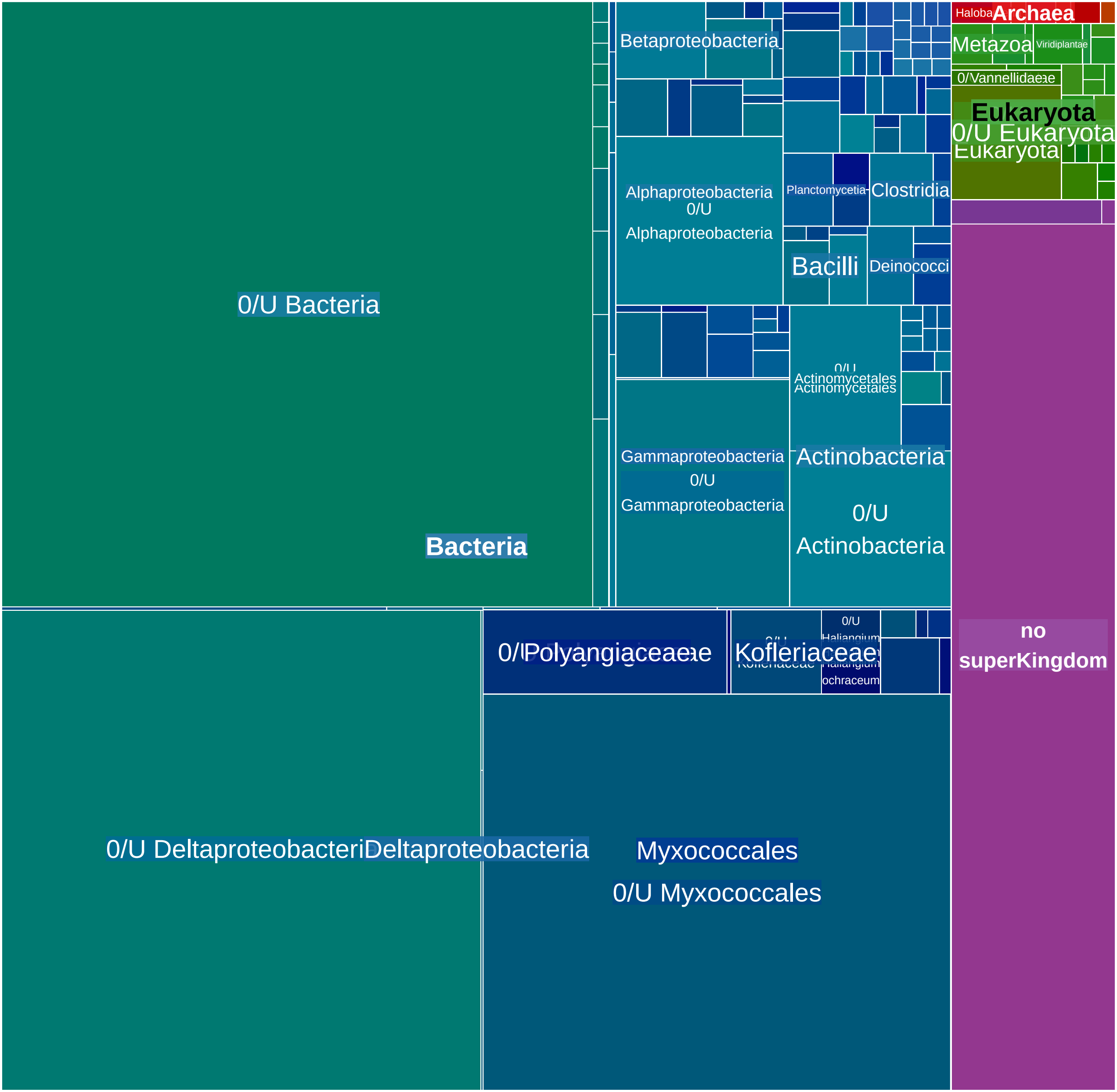

MGT−v1_184 : 8378 unigenes, 480 (5.73%) taxonomically assigned

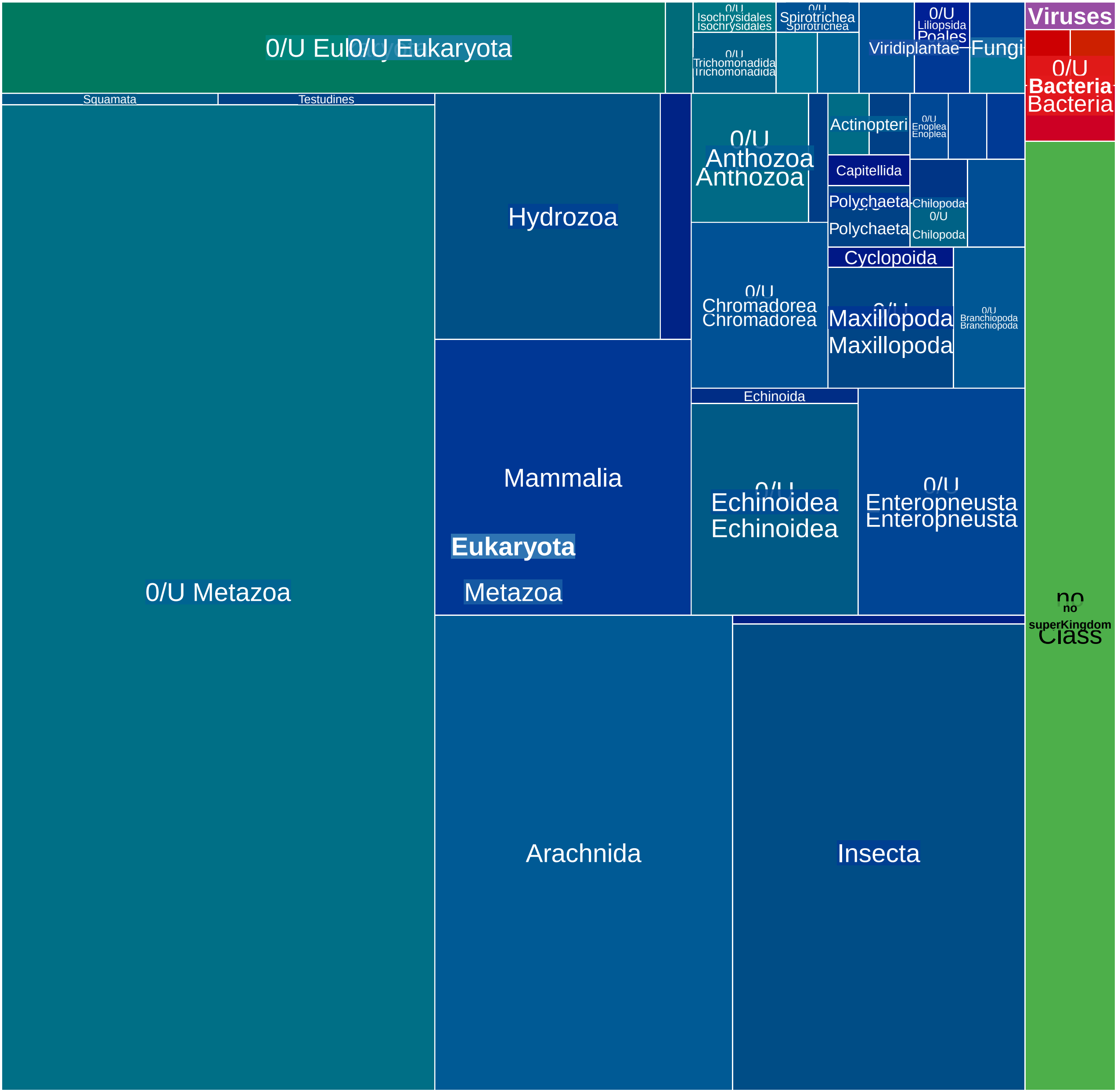

MGT−v1_185 : 8243 unigenes, 627 (7.61%) taxonomically assigned

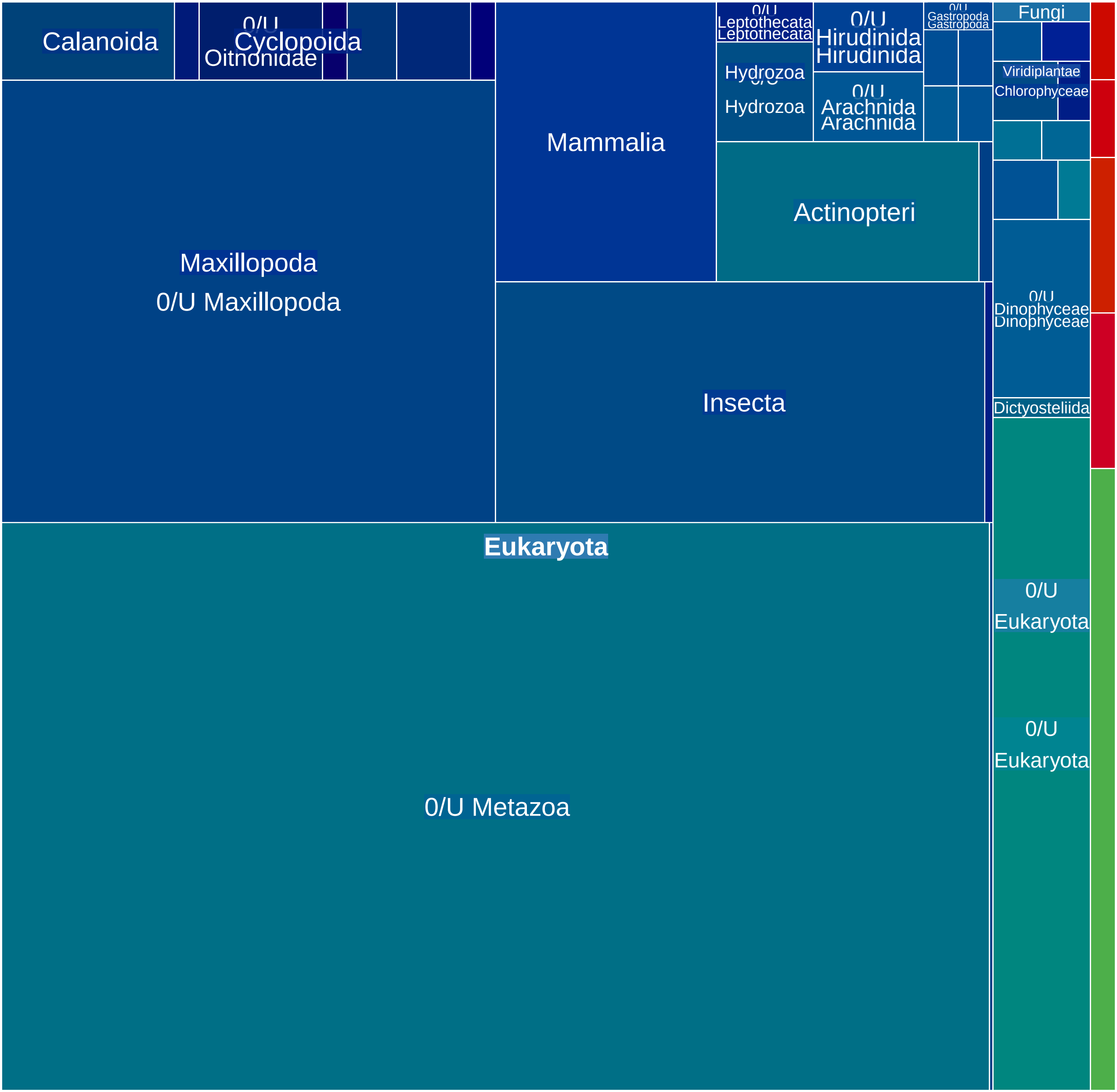

MGT−v1_186 : 8121 unigenes, 1919 (23.63%) taxonomically assigned

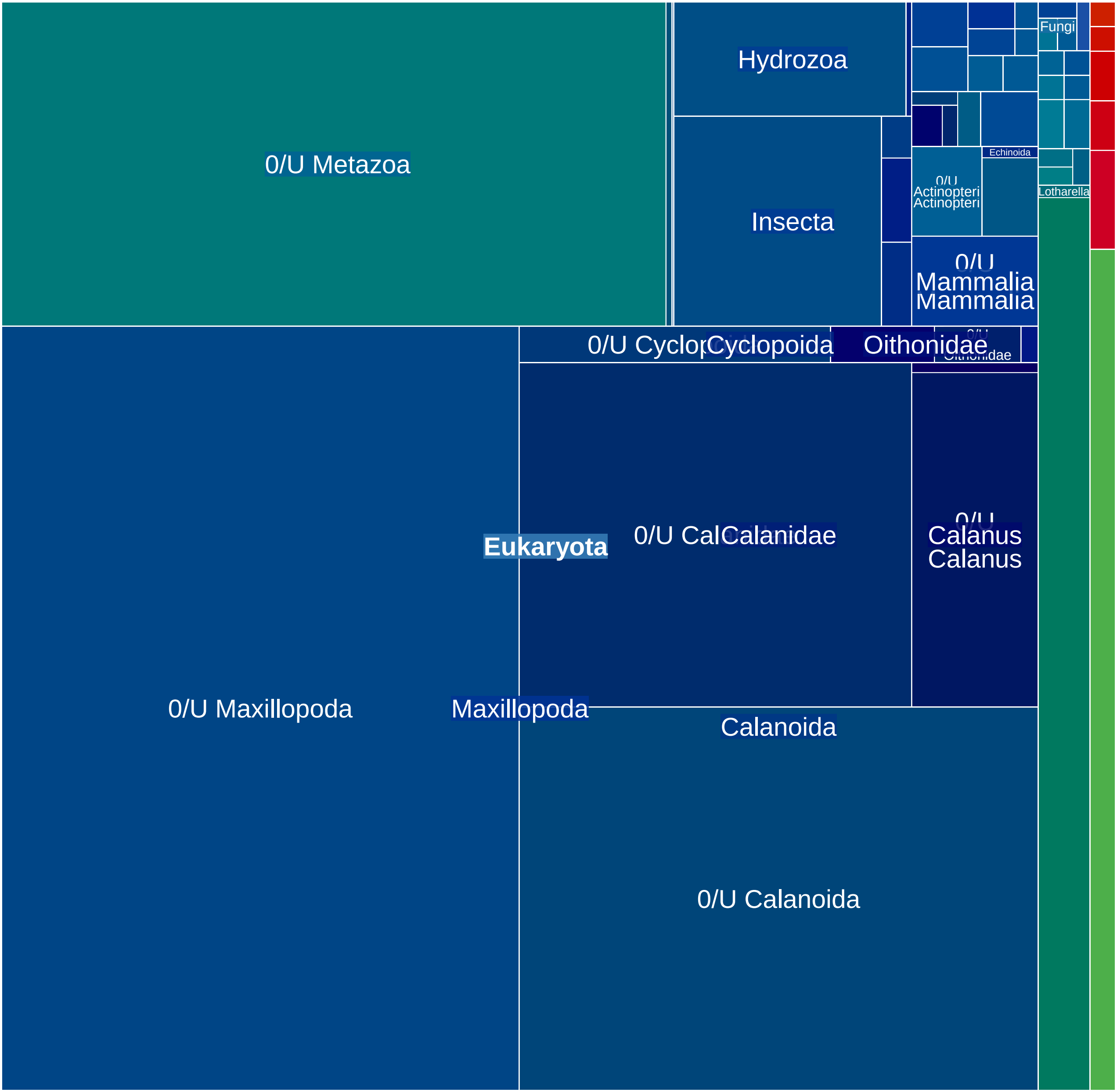

MGT−v1_187 : 8054 unigenes, 6974 (86.59%) taxonomically assigned

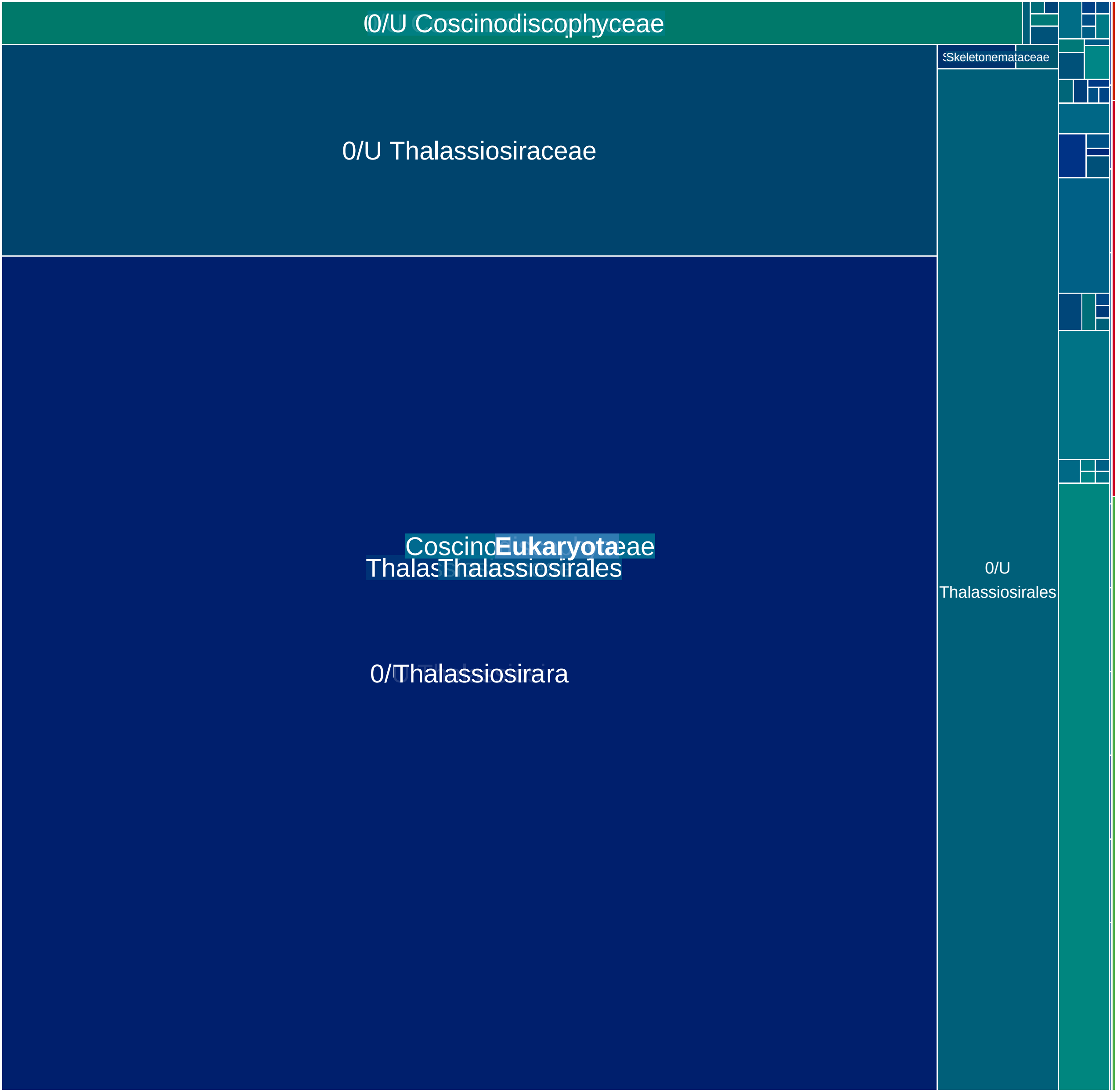

MGT−v1_188 : 8036 unigenes, 2437 (30.33%) taxonomically assigned

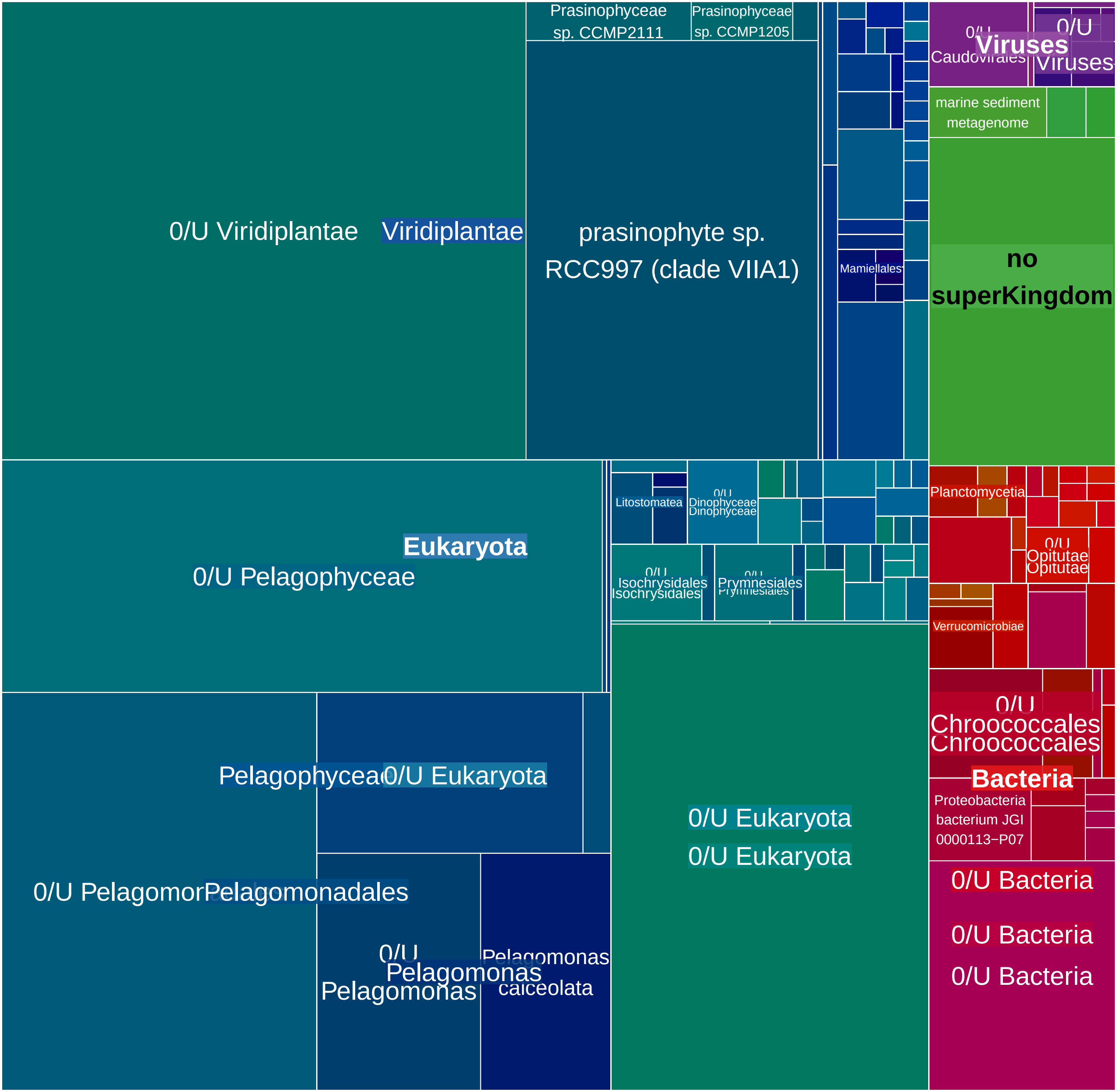

MGT−v1_189 : 7758 unigenes, 605 (7.80%) taxonomically assigned

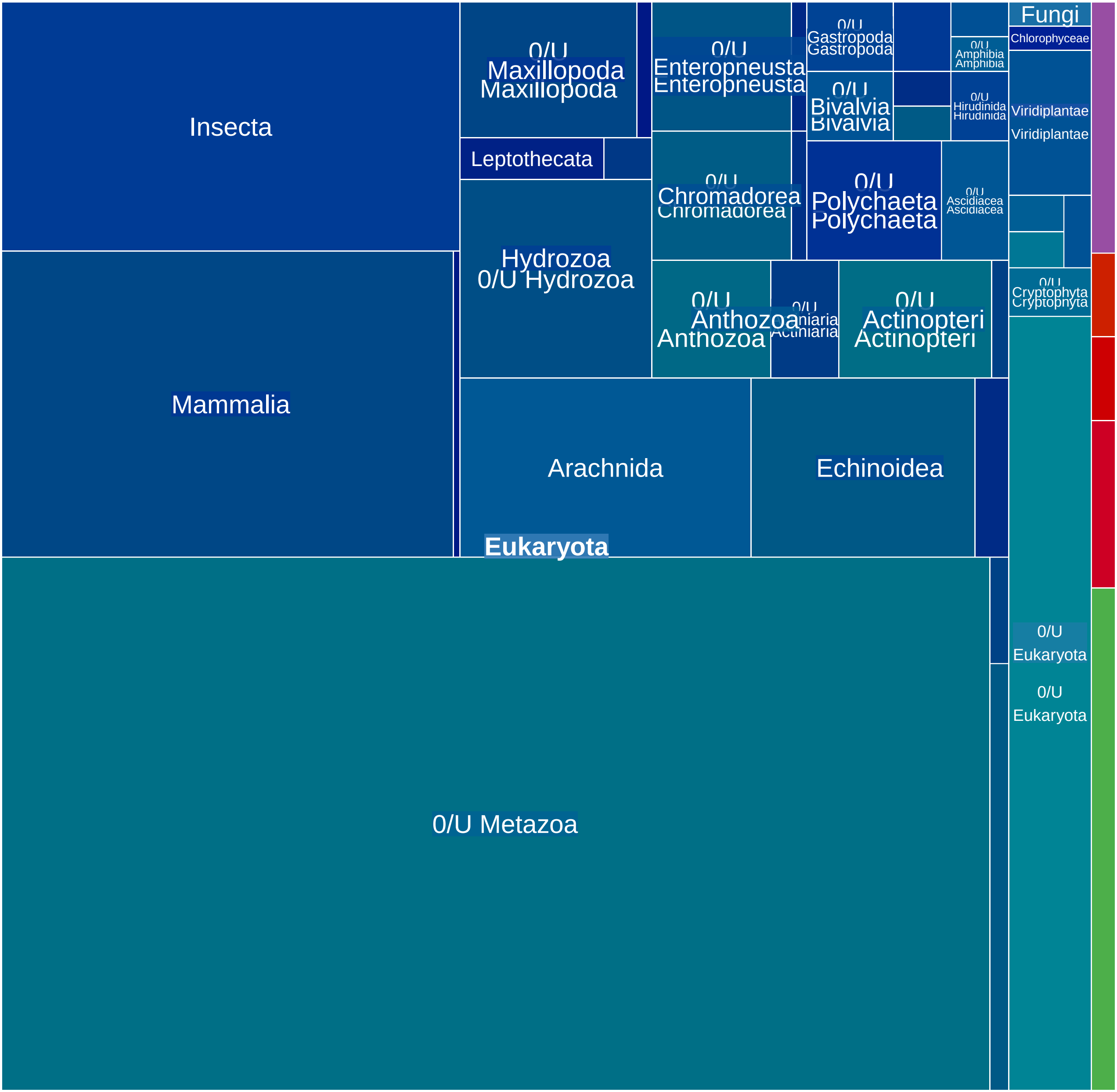

### MGT−v1_190 : 7520 unigenes, 3679 (48.92%) taxonomically assigned

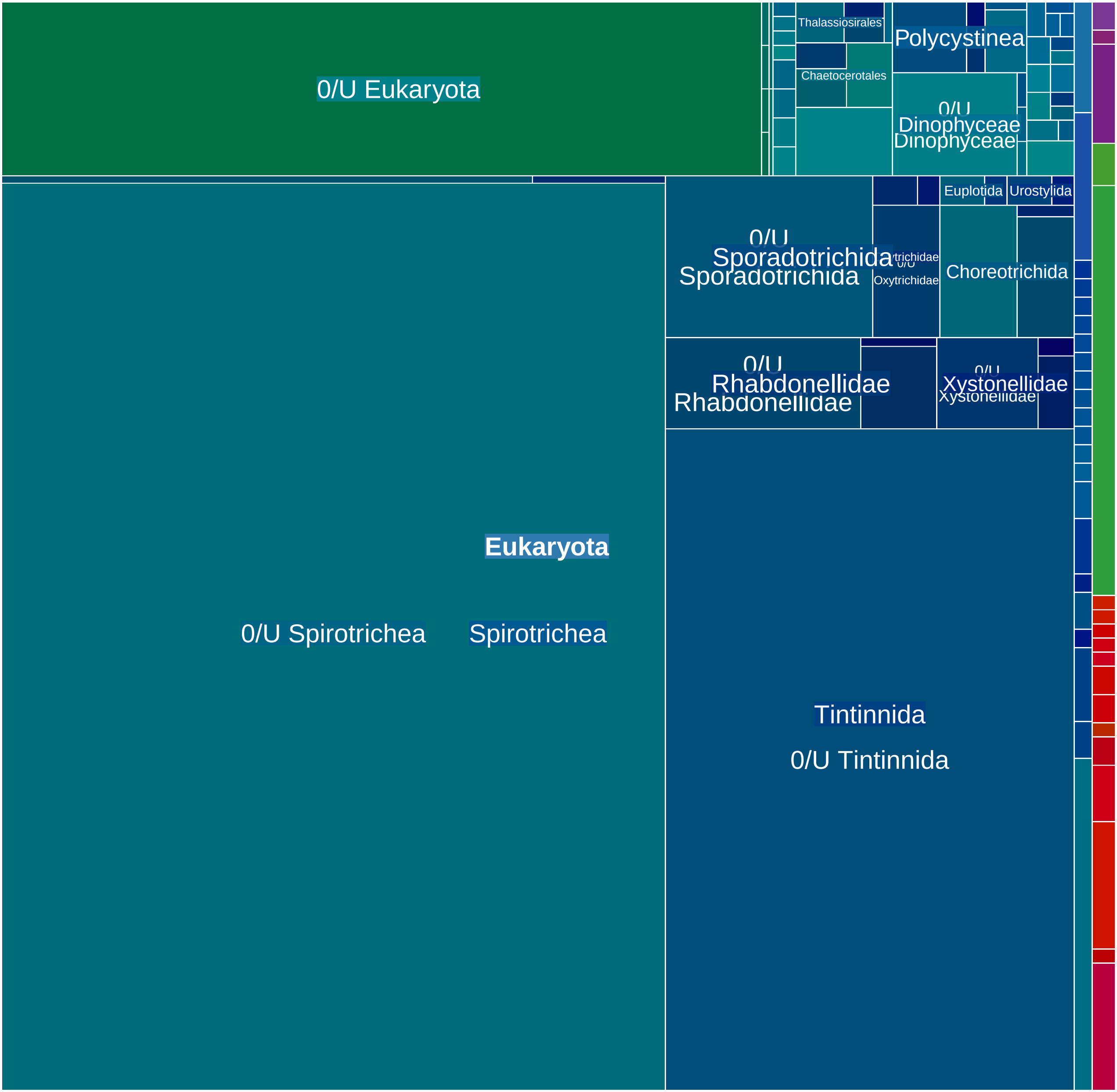

MGT−v1_191 : 7517 unigenes, 5377 (71.53%) taxonomically assigned

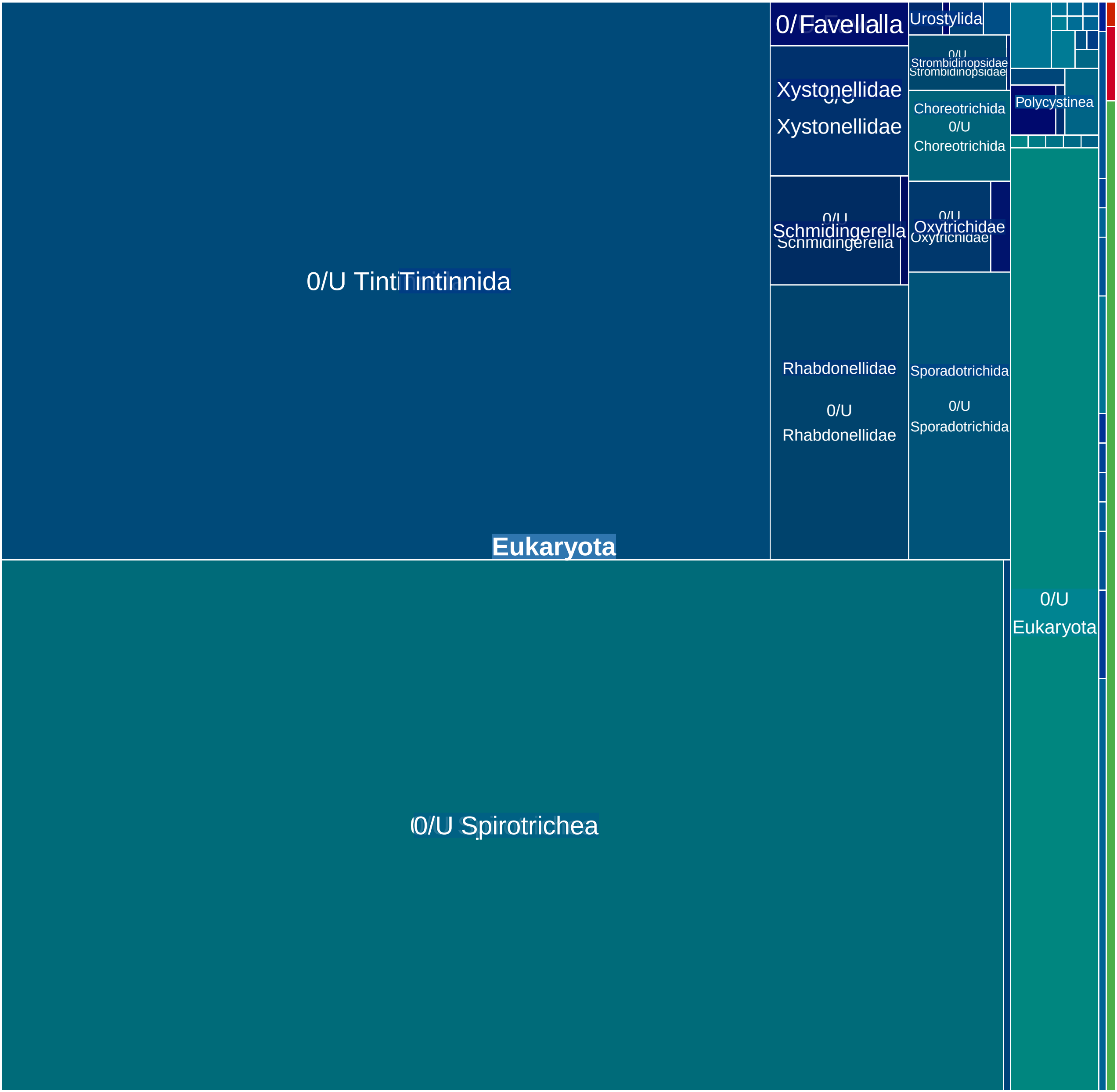

### MGT−v1_192 : 7455 unigenes, 758 (10.17%) taxonomically assigned

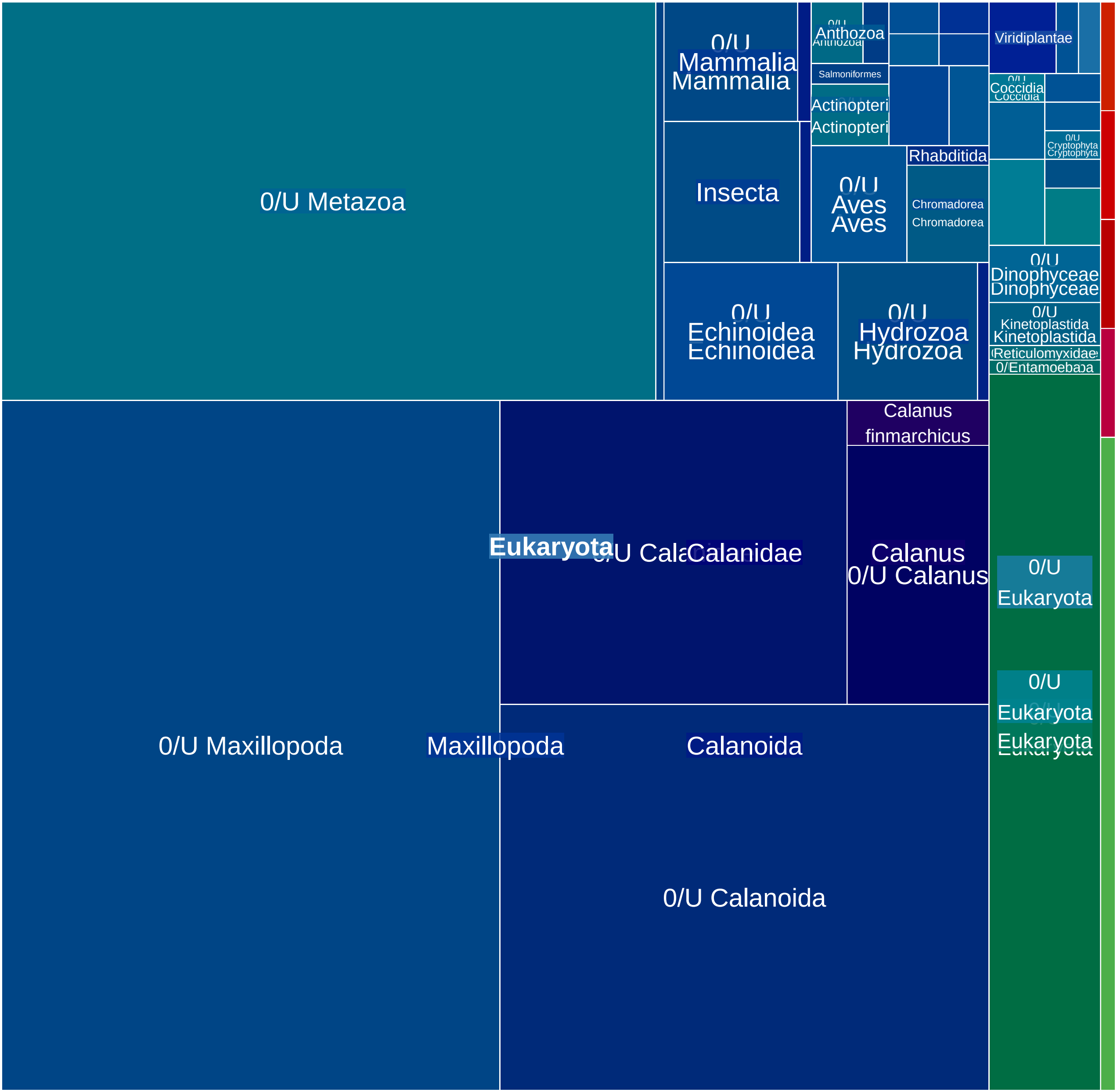

MGT−v1_193 : 7449 unigenes, 4414 (59.26%) taxonomically assigned

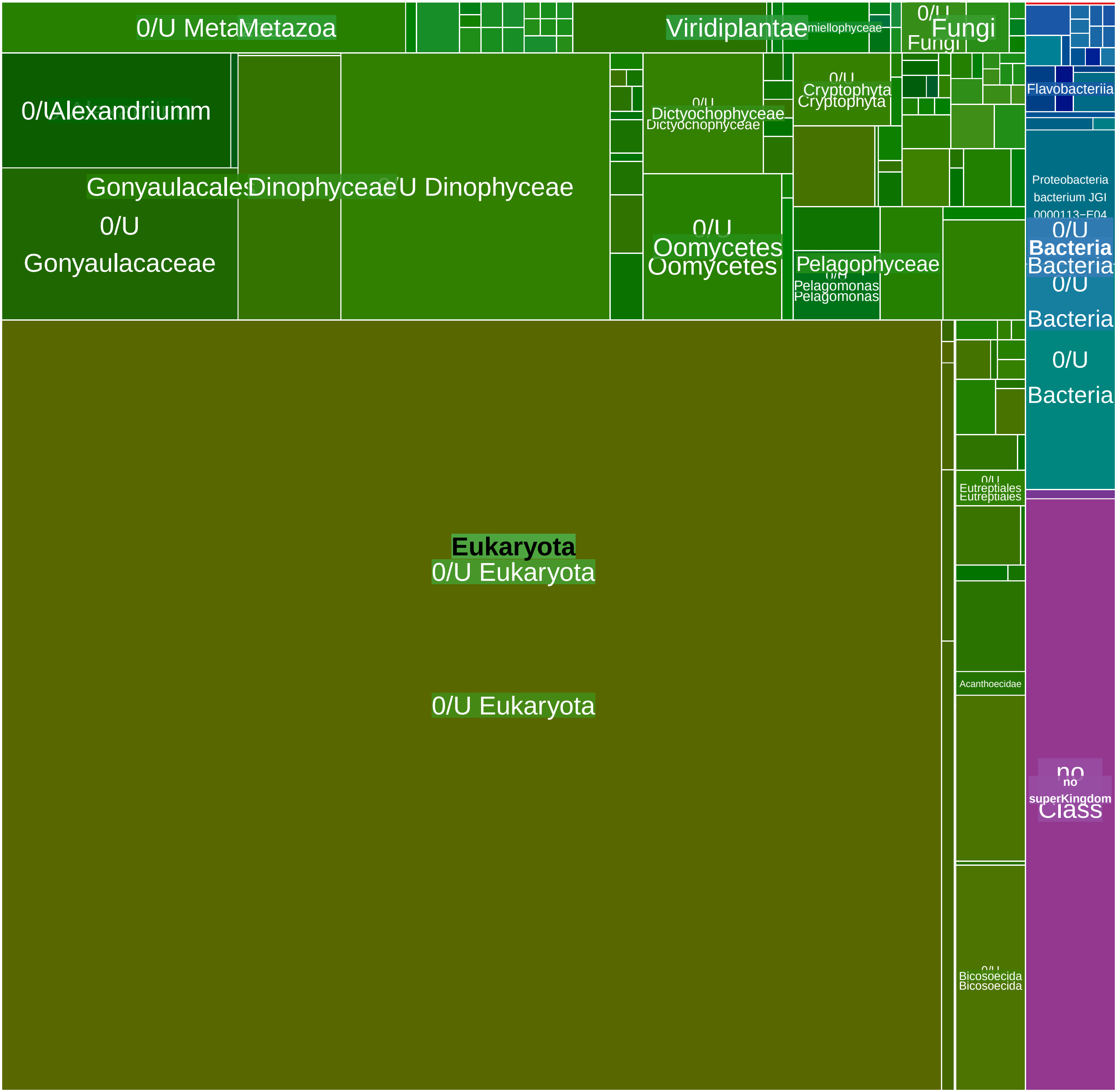

MGT−v1_194 : 7397 unigenes, 1335 (18.05%) taxonomically assigned

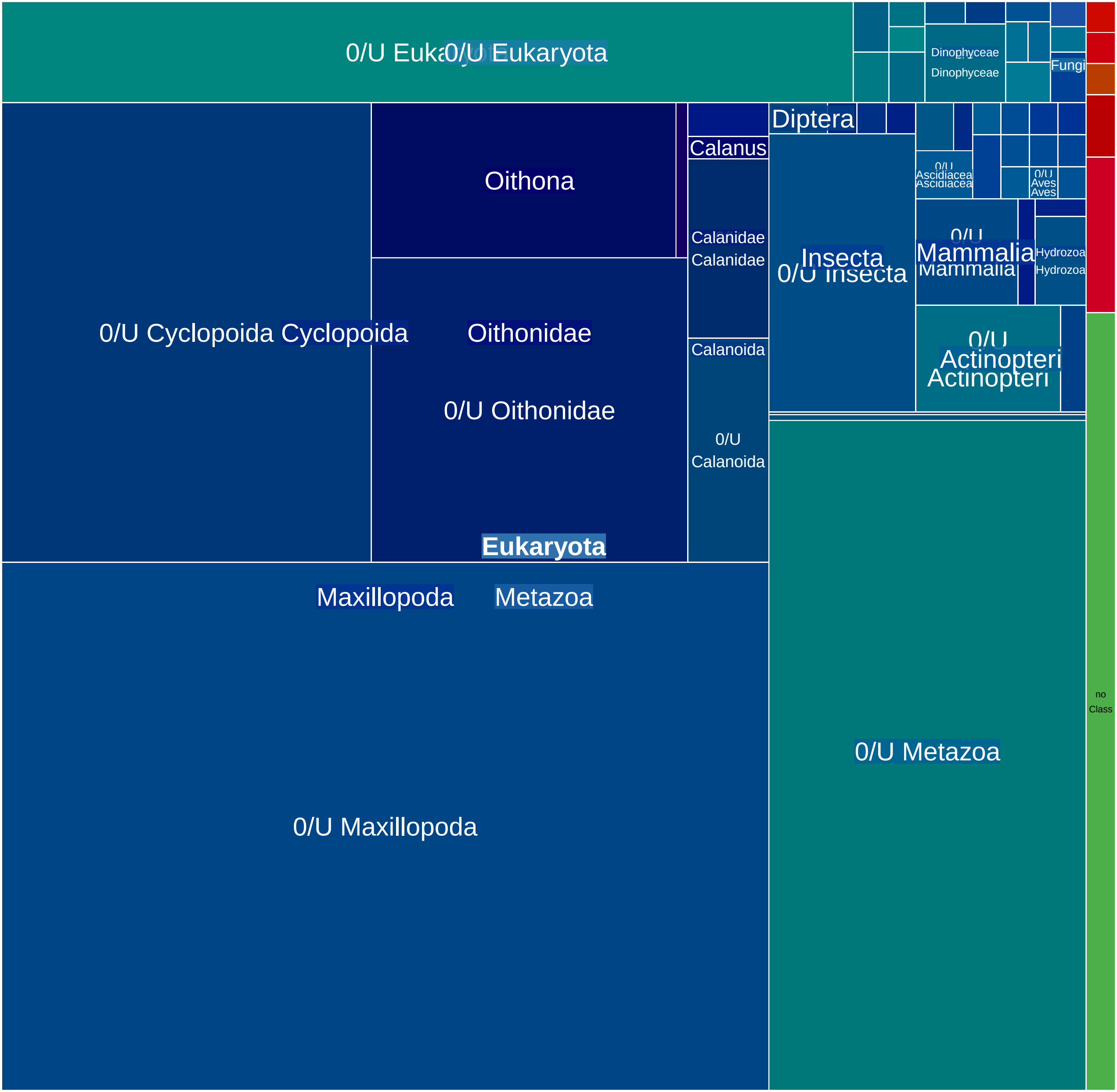

MGT−v1_195 : 7388 unigenes, 7017 (94.98%) taxonomically assigned

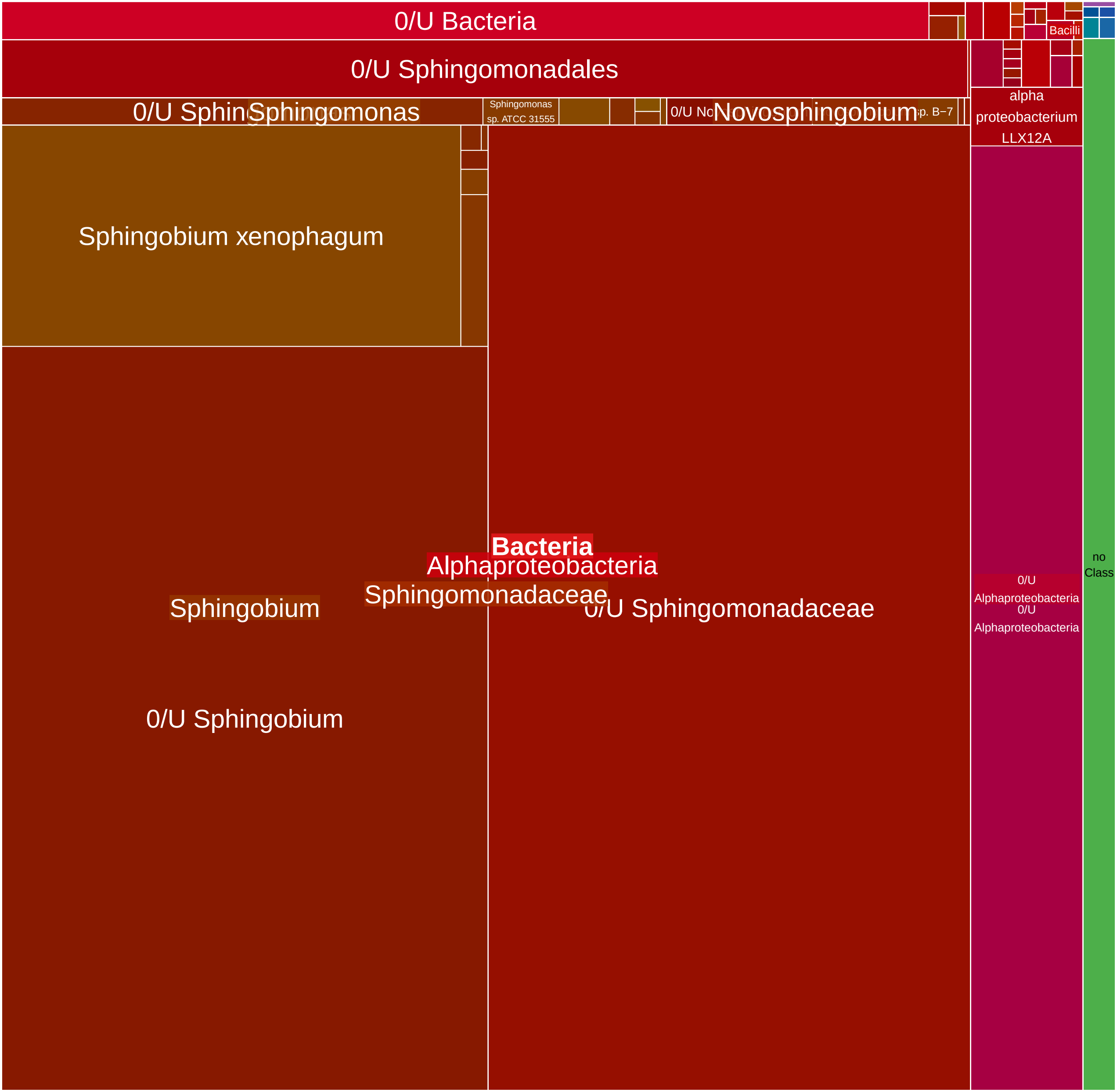

MGT−v1_196 : 7378 unigenes, 5380 (72.92%) taxonomically assigned

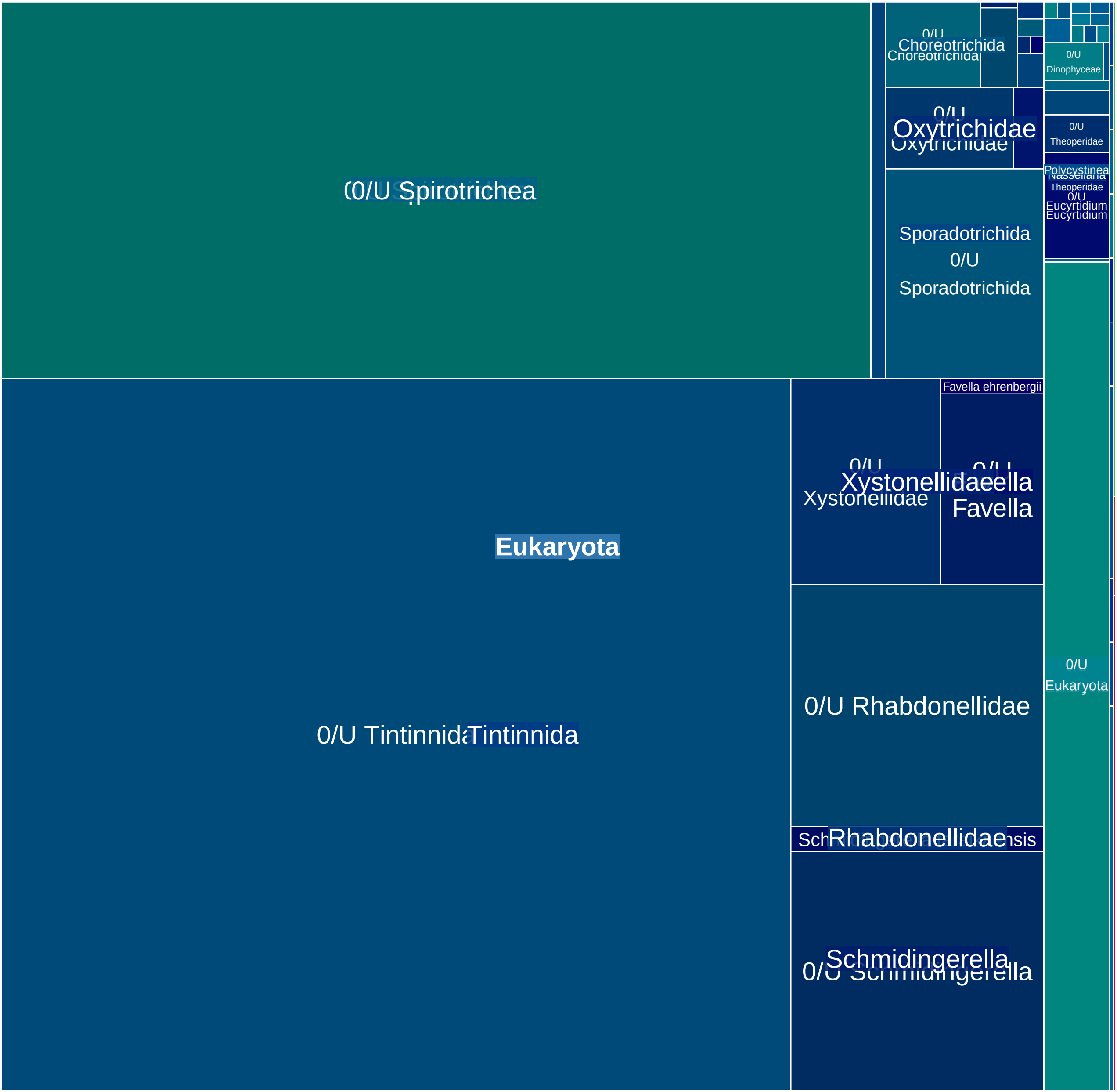

MGT−v1_197 : 7244 unigenes, 2345 (32.37%) taxonomically assigned

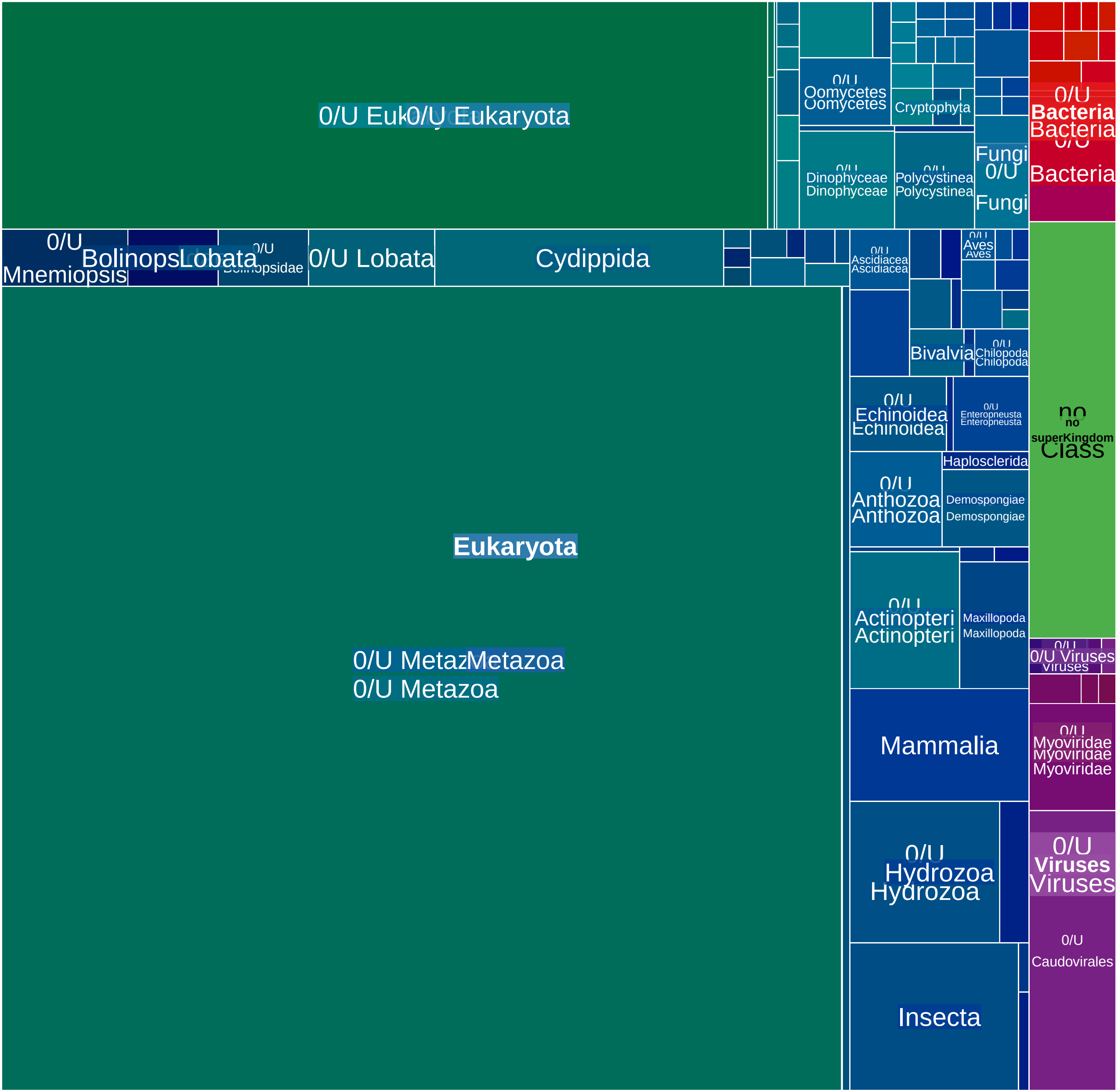

MGT−v1_198 : 7167 unigenes, 476 (6.64%) taxonomically assigned

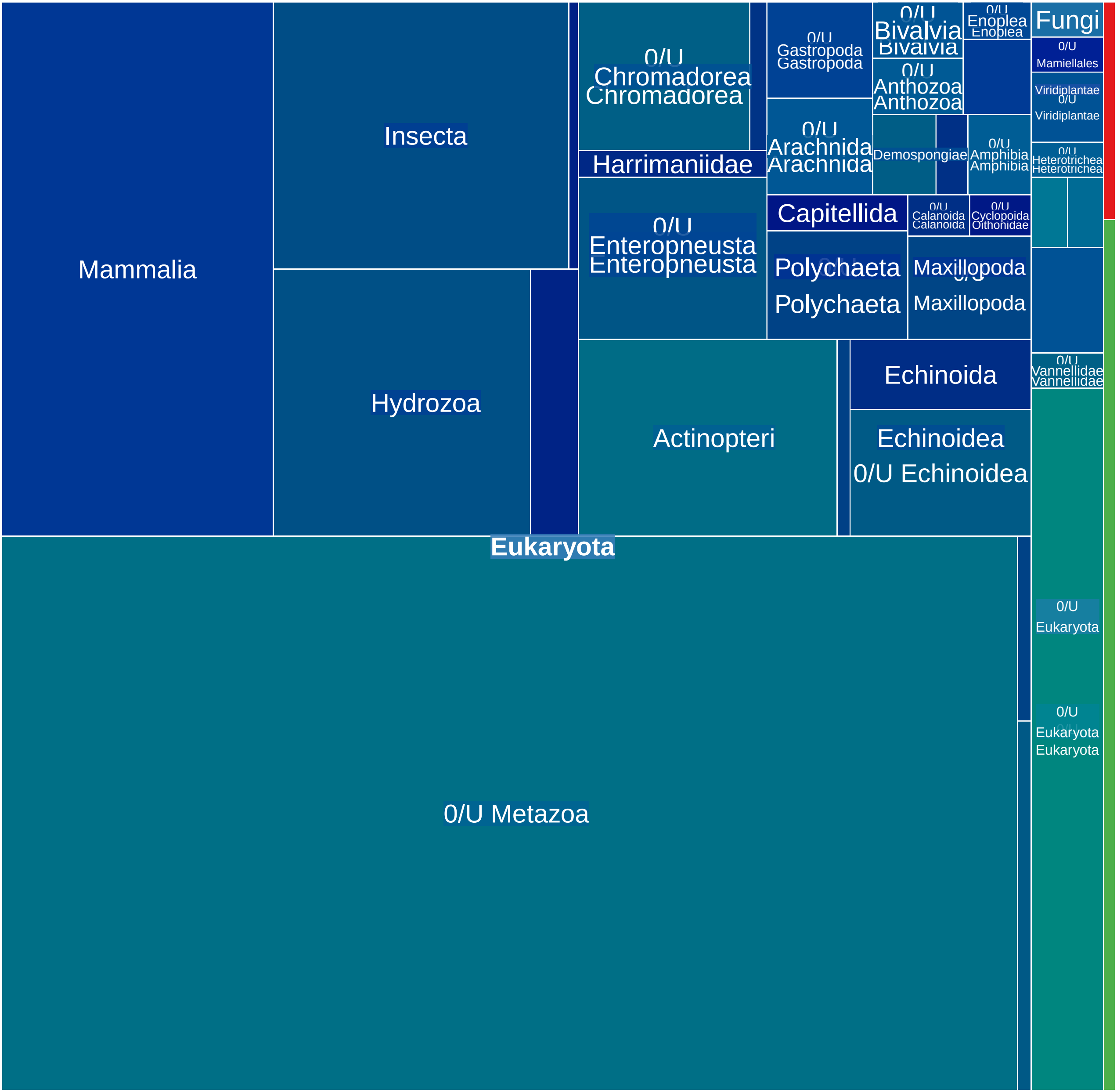

### MGT−v1_199 : 7050 unigenes, 3192 (45.28%) taxonomically assigned

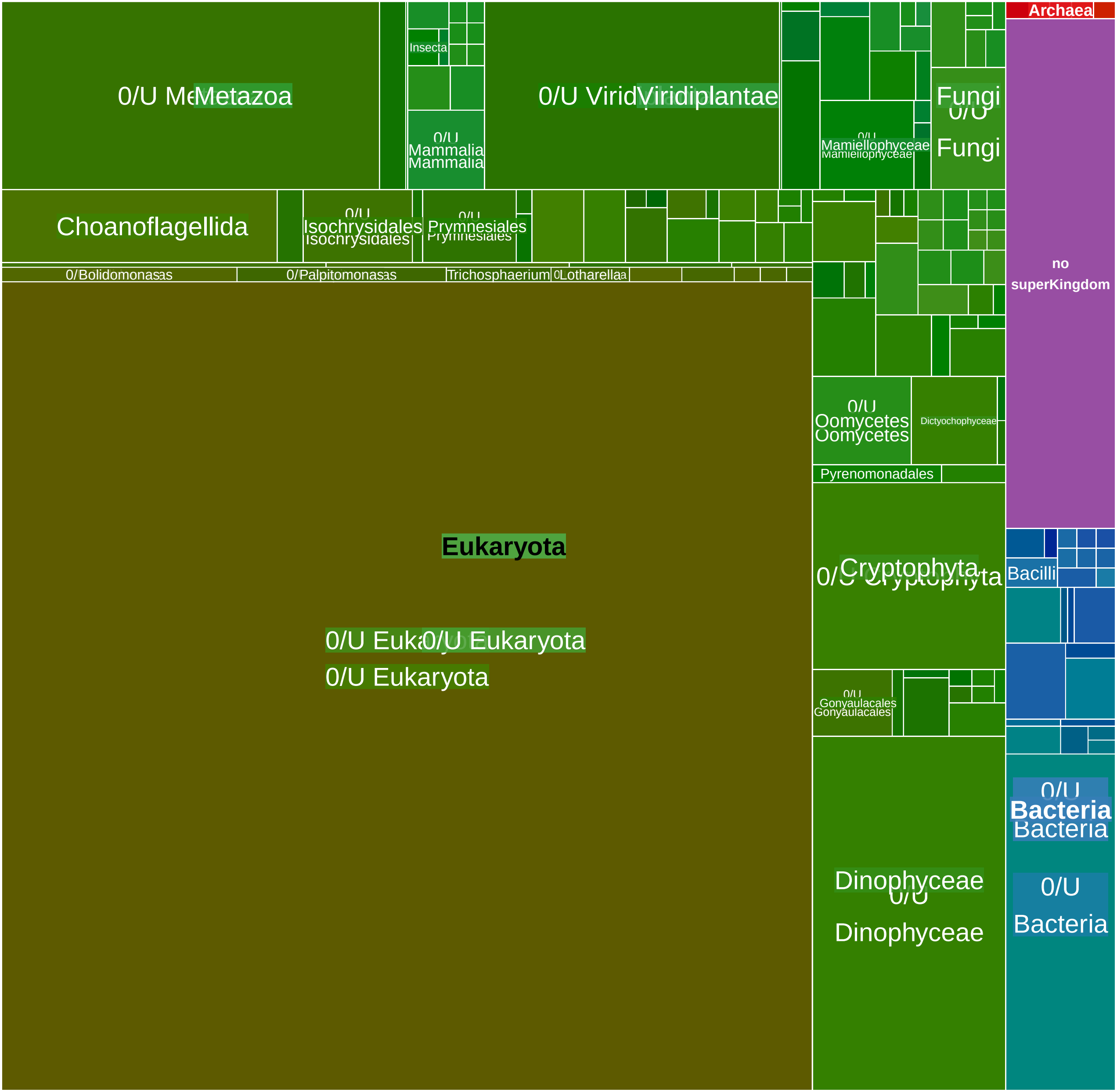

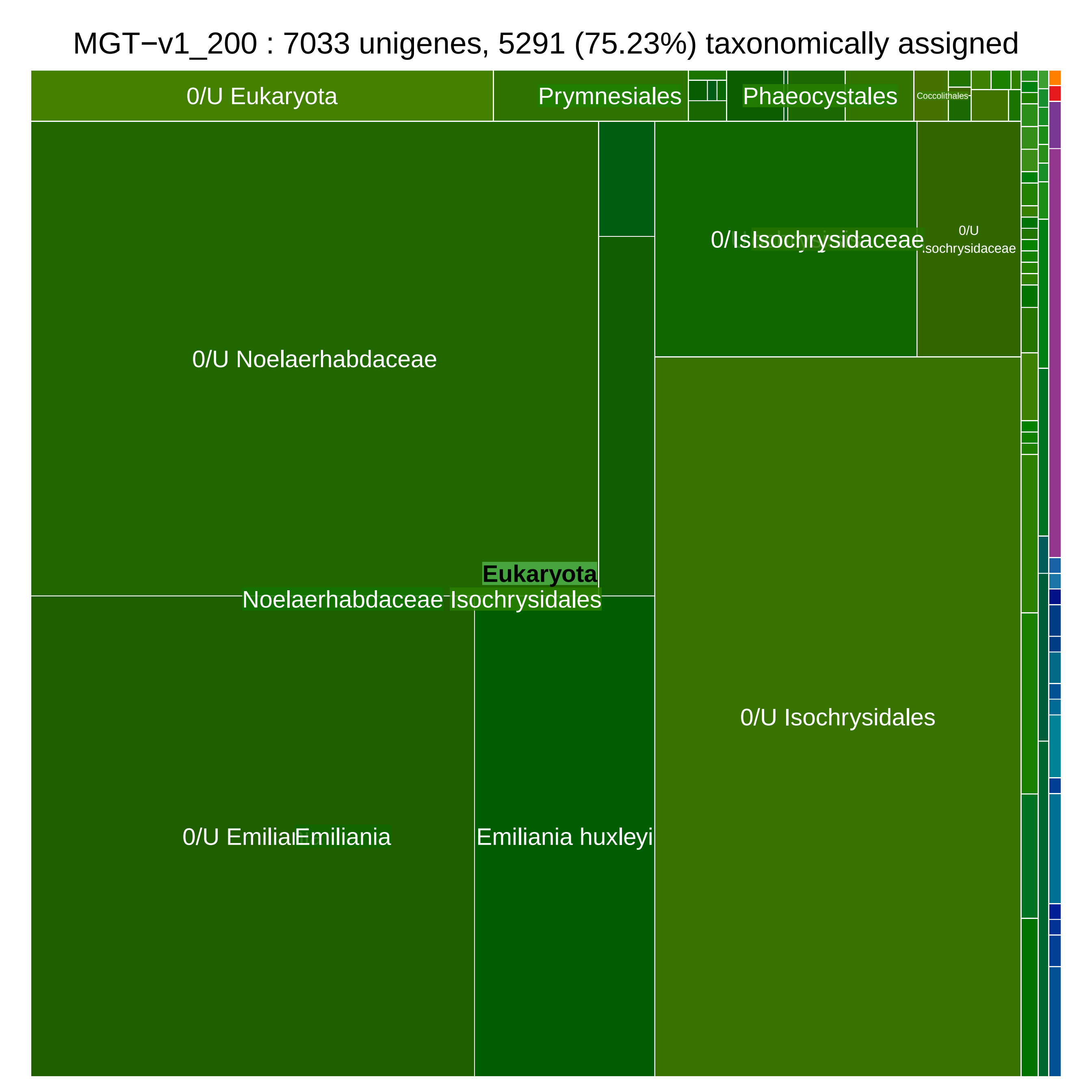

MGT−v1_200 : 7033 unigenes, 5291 (75.23%) taxonomically assigned
0/U Eukaryota
Prymnesiales
Phaeocystales
Coccolithales
0/U Noelaerhabdaceae
0/U Isochrysidaceae
Isochrysidaceae
0/U Isochrysidaceae
Eukaryota
Noelaerhabdaceae
Isochrysidales
0/U Isochrysidales
0/U Emiliania
Emiliania
Emiliania huxleyi

MGT−v1_201 : 7006 unigenes, 1128 (16.10%) taxonomically assigned

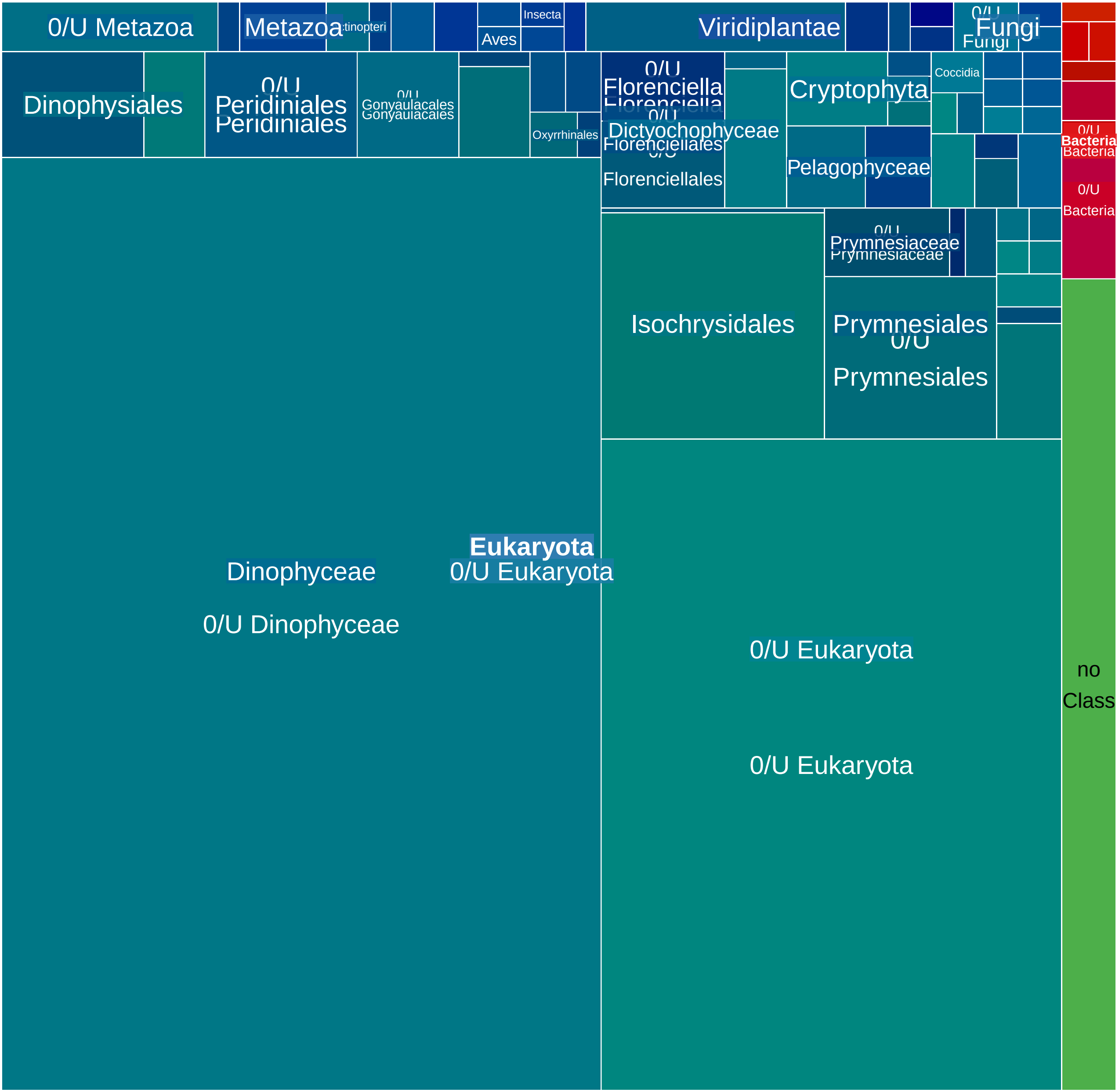

MGT−v1_202 : 6983 unigenes, 1464 (20.97%) taxonomically assigned

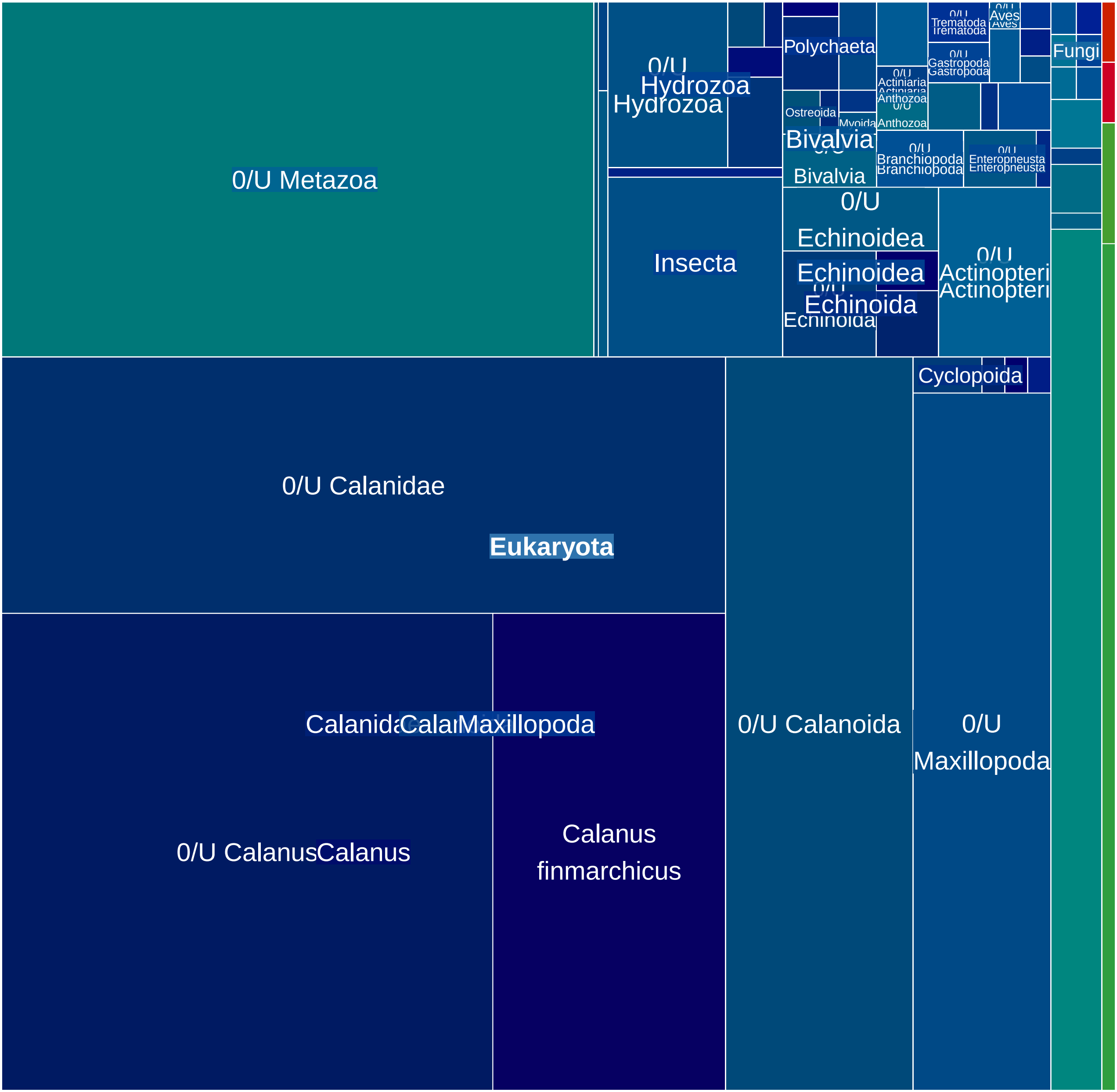

MGT−v1_203 : 6961 unigenes, 5033 (72.30%) taxonomically assigned

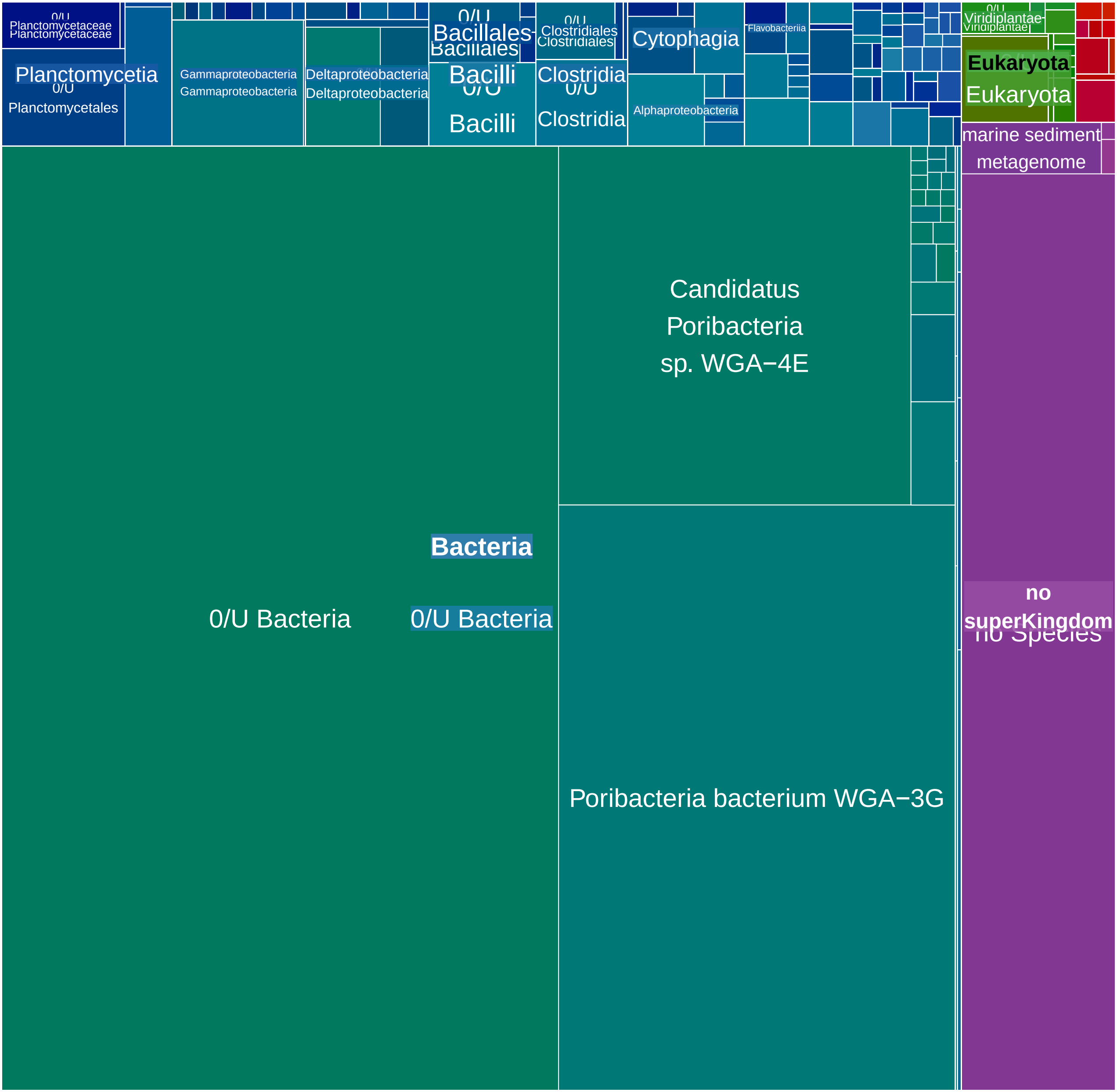

MGT−v1_204 : 6941 unigenes, 1727 (24.88%) taxonomically assigned

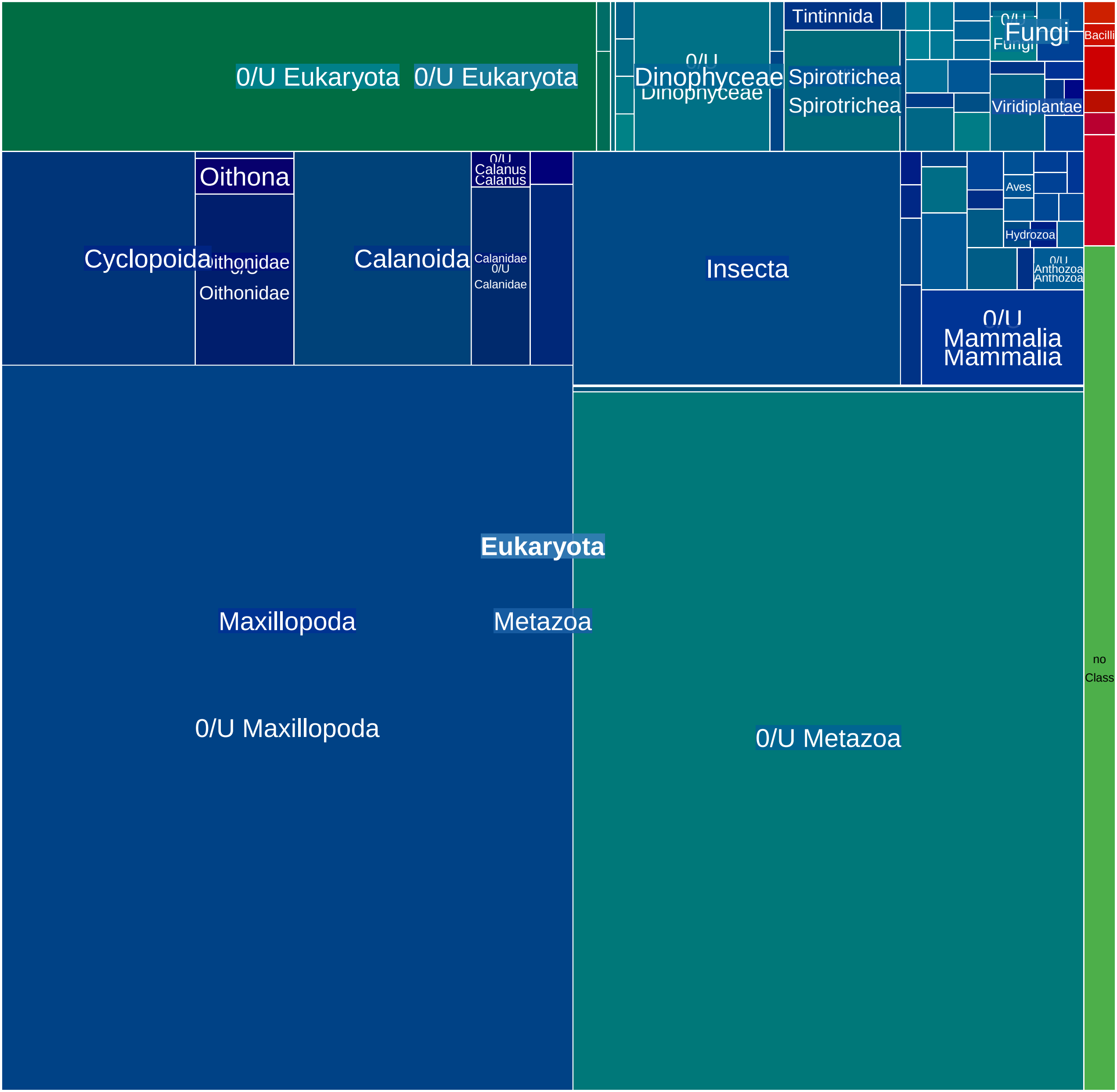

MGT−v1_205 : 6869 unigenes, 4584 (66.73%) taxonomically assigned

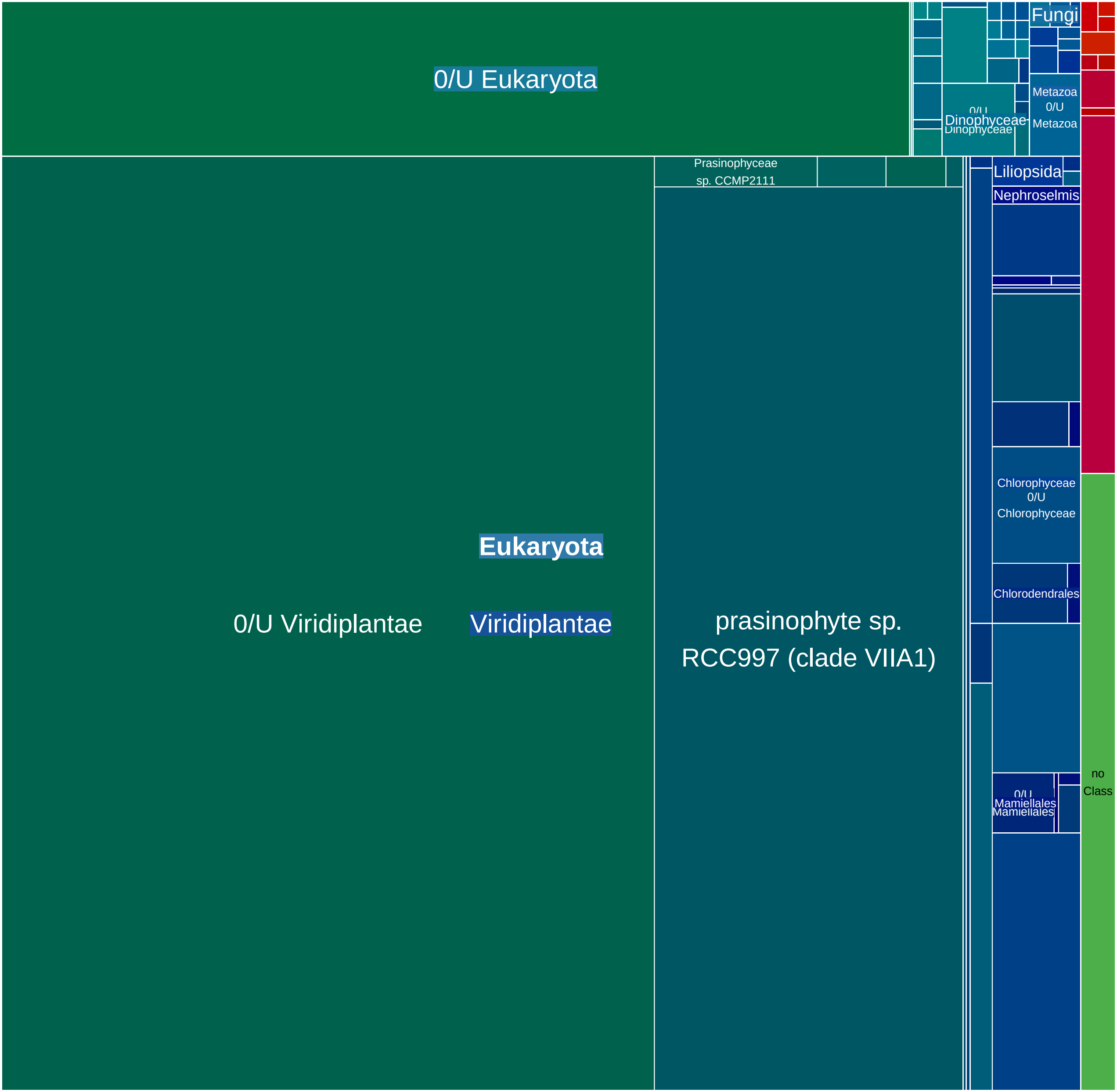

MGT−v1_206 : 6804 unigenes, 5341 (78.50%) taxonomically assigned

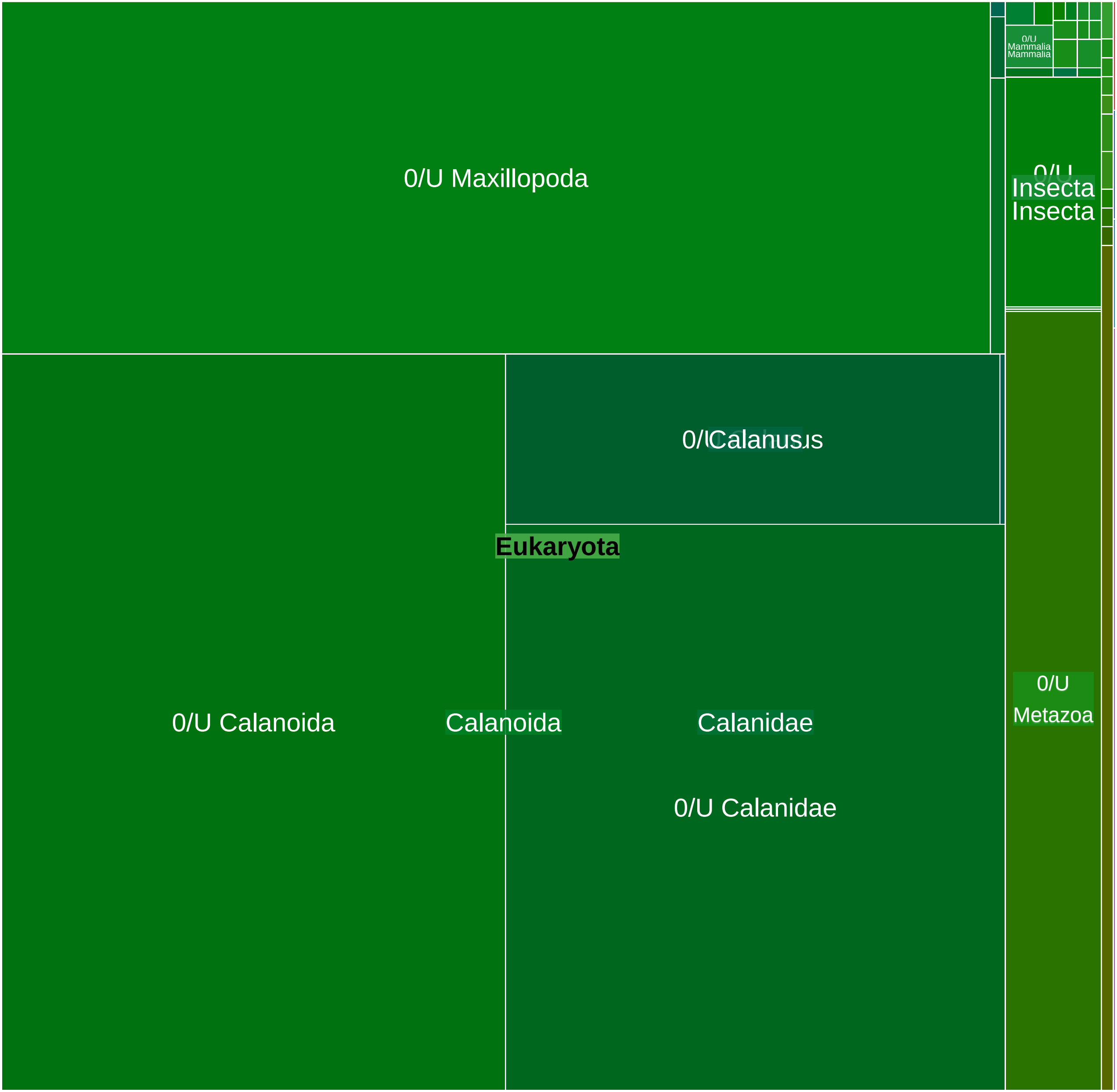

MGT−v1_207 : 6715 unigenes, 6043 (89.99%) taxonomically assigned

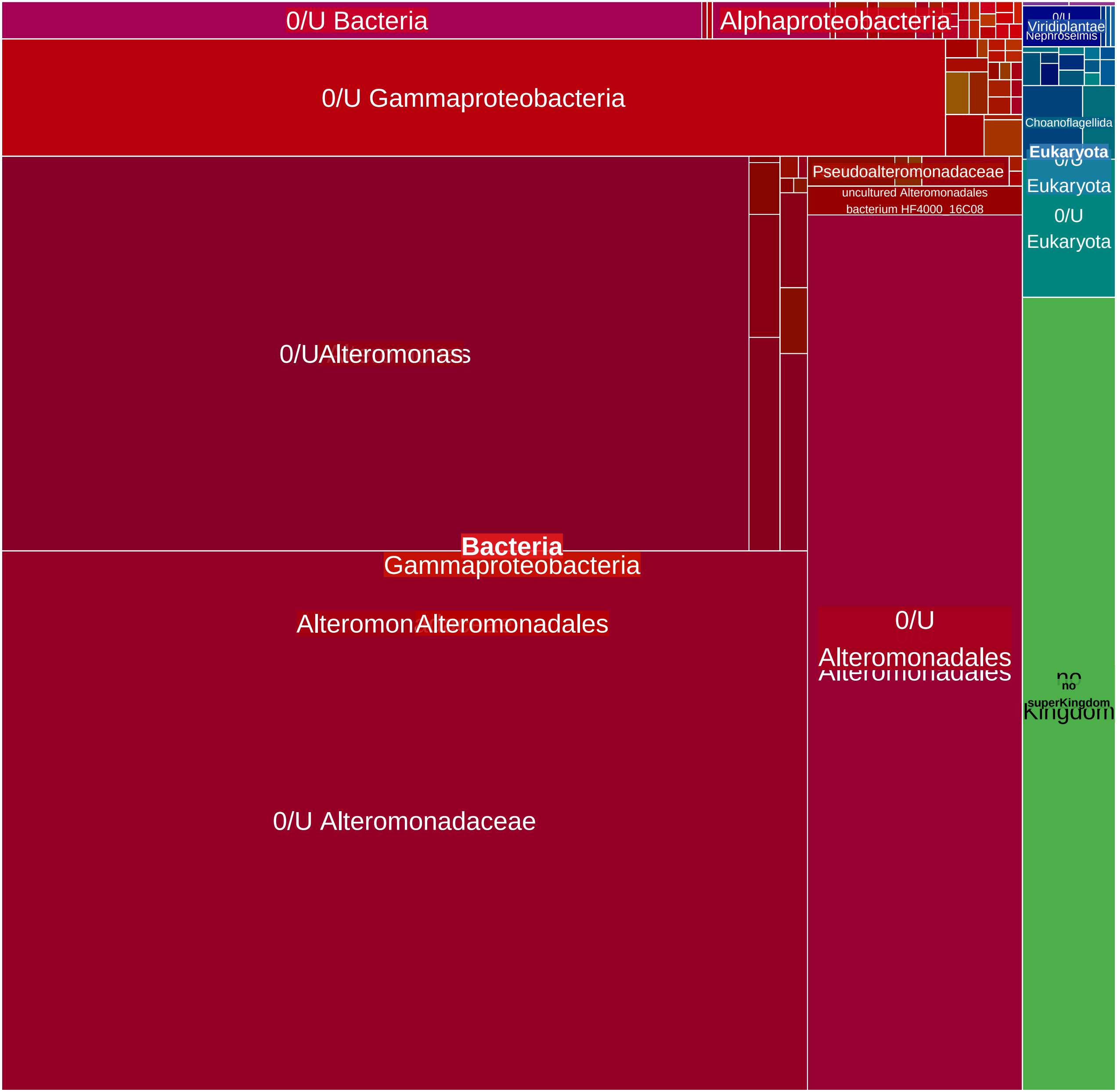

MGT−v1_208 : 6656 unigenes, 1776 (26.68%) taxonomically assigned

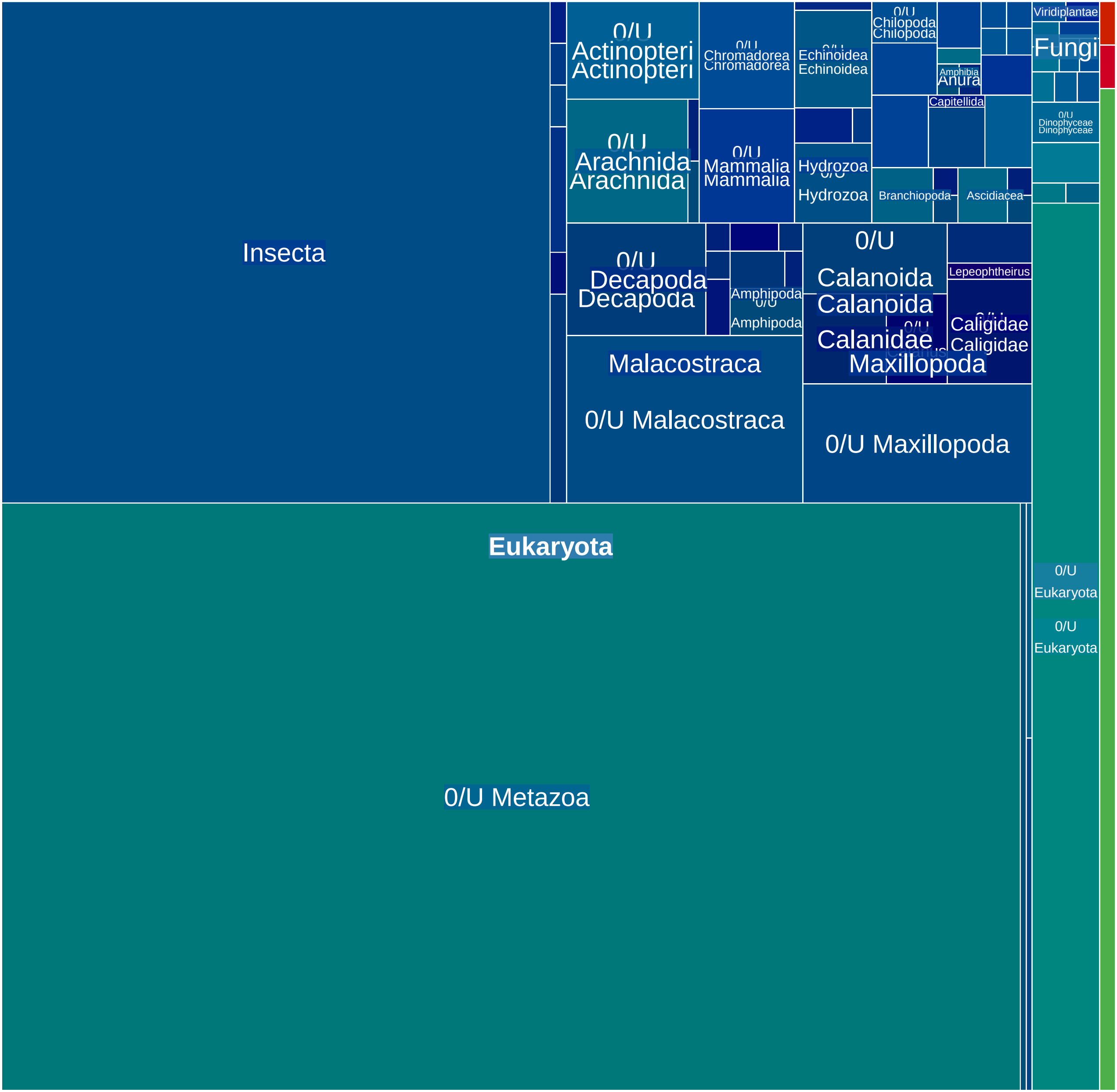

MGT−v1_209 : 6345 unigenes, 830 (13.08%) taxonomically assigned

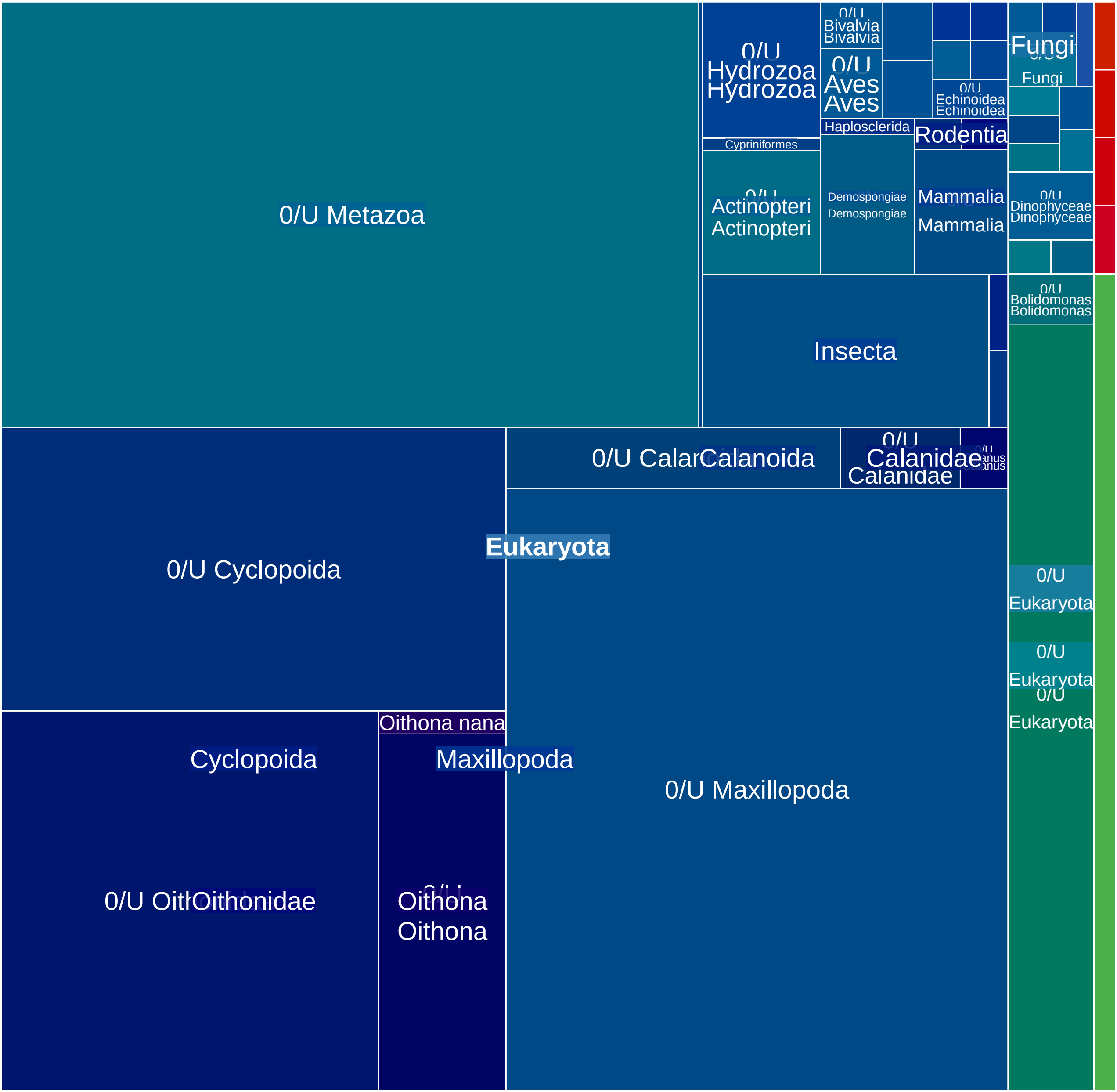

MGT−v1_210 : 6327 unigenes, 4118 (65.09%) taxonomically assigned

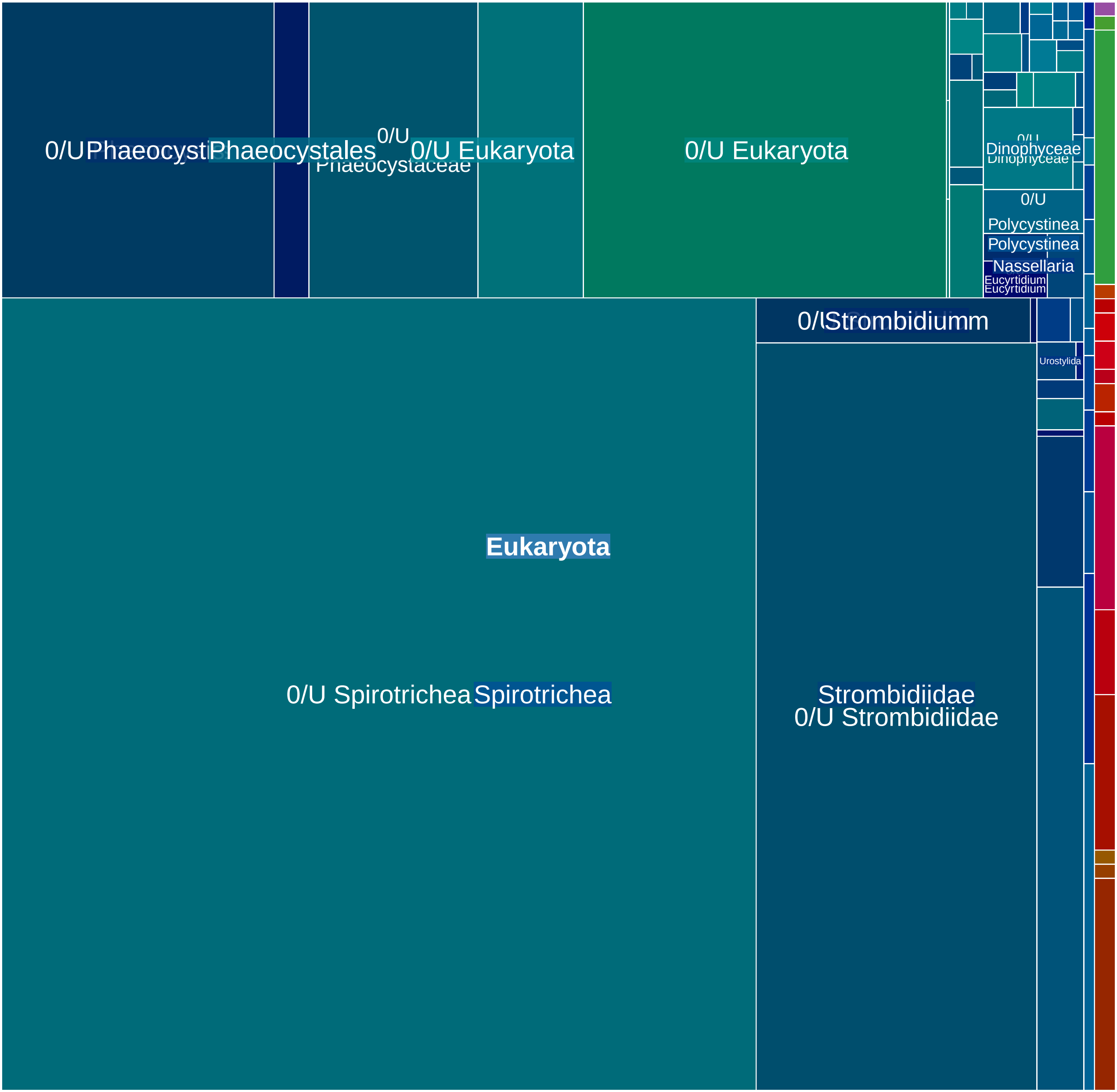

MGT−v1_211 : 6304 unigenes, 4492 (71.26%) taxonomically assigned

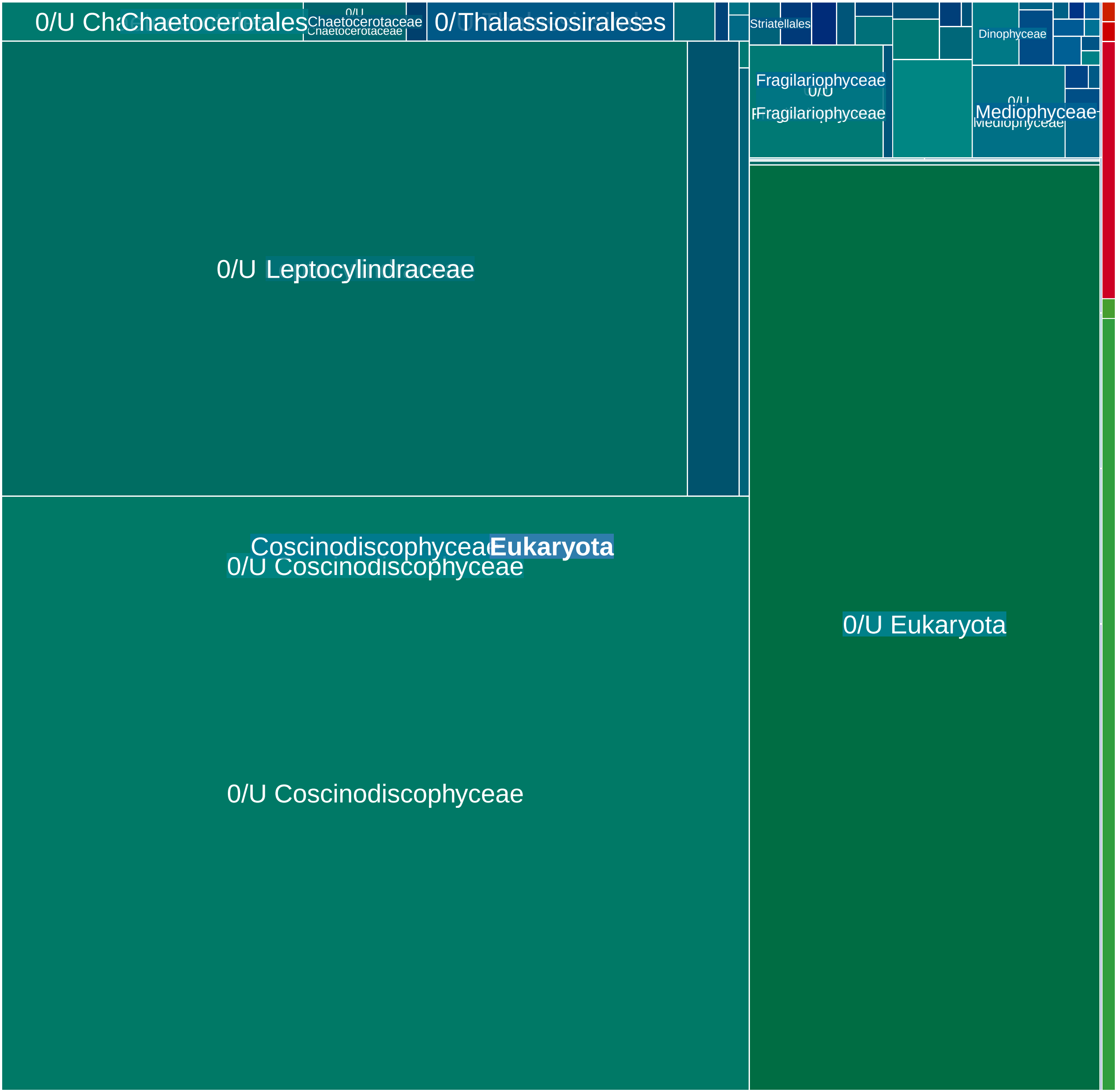

MGT−v1_212 : 6215 unigenes, 2546 (40.97%) taxonomically assigned

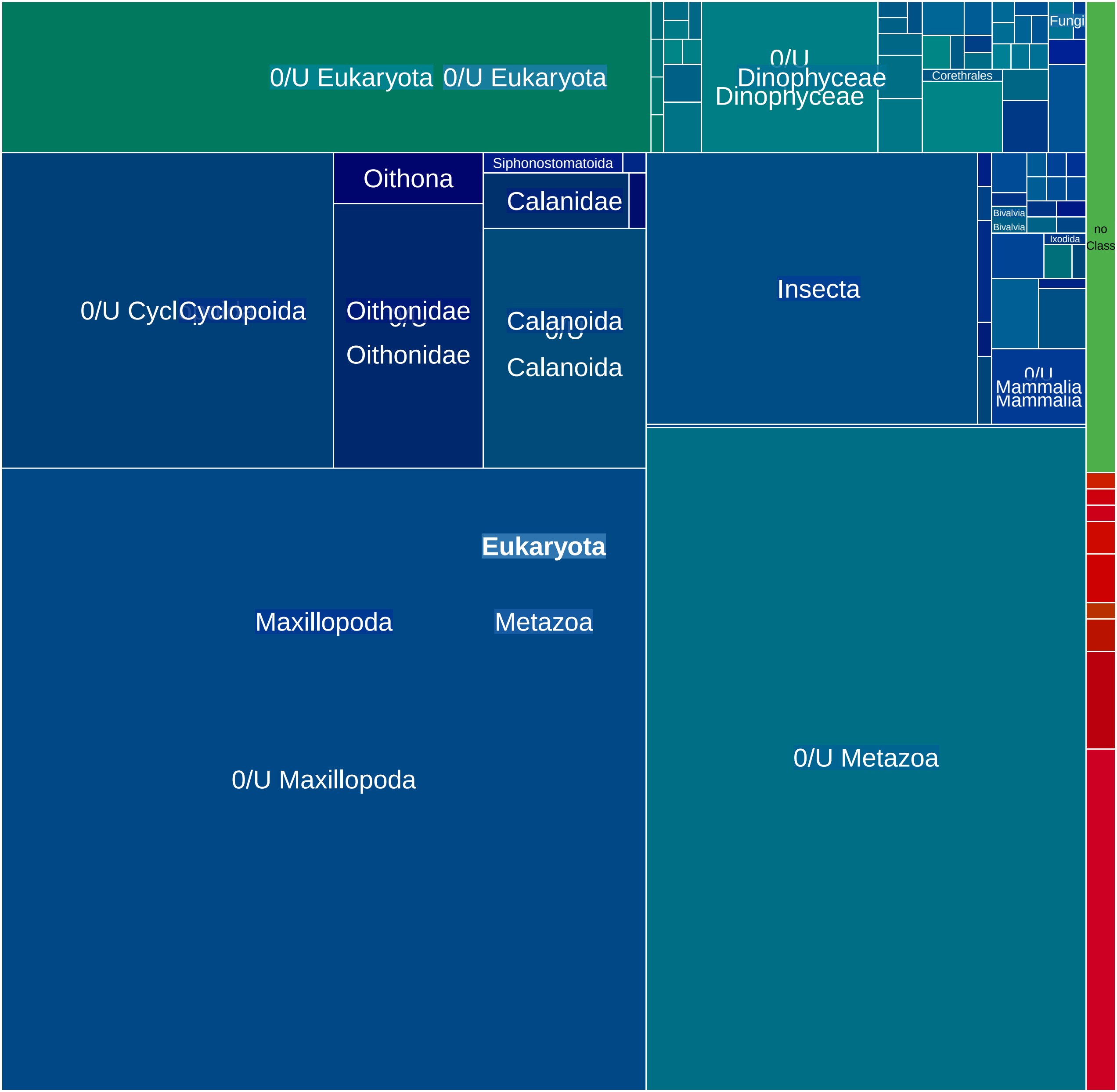

MGT−v1_213 : 6203 unigenes, 2406 (38.79%) taxonomically assigned

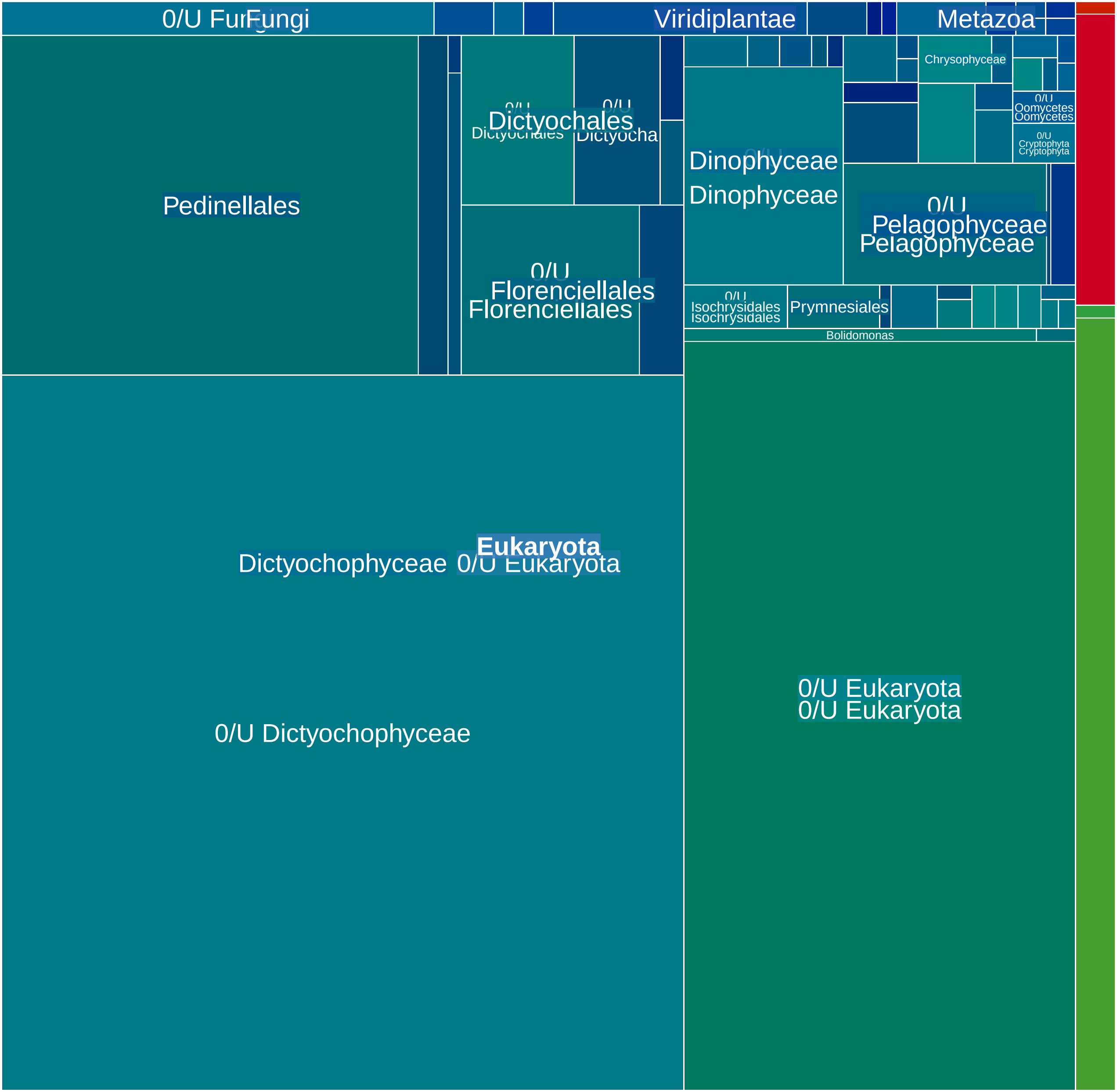

MGT−v1_214 : 6183 unigenes, 2128 (34.42%) taxonomically assigned

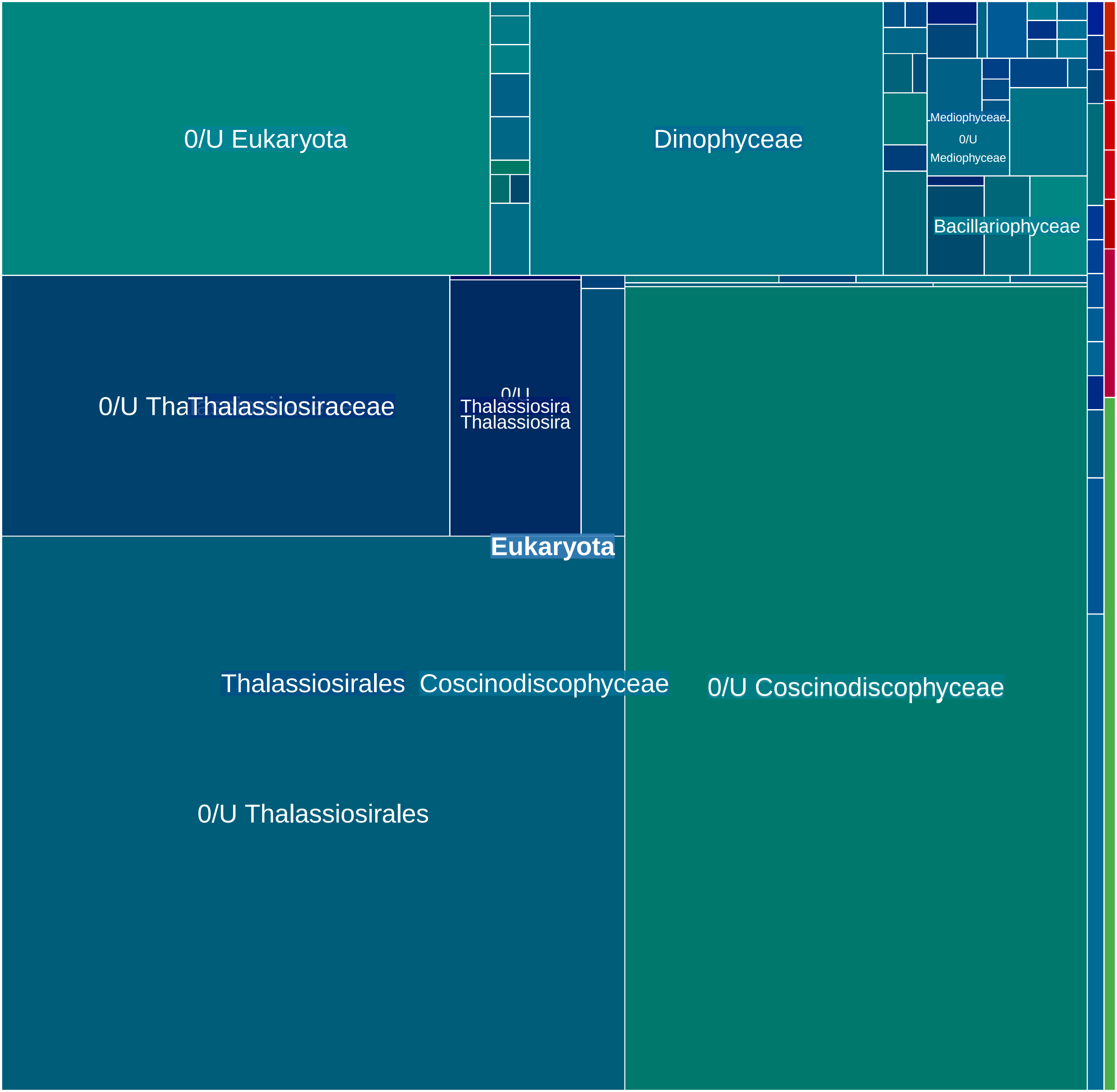

### MGT−v1_215 : 6182 unigenes, 4780 (77.32%) taxonomically assigned

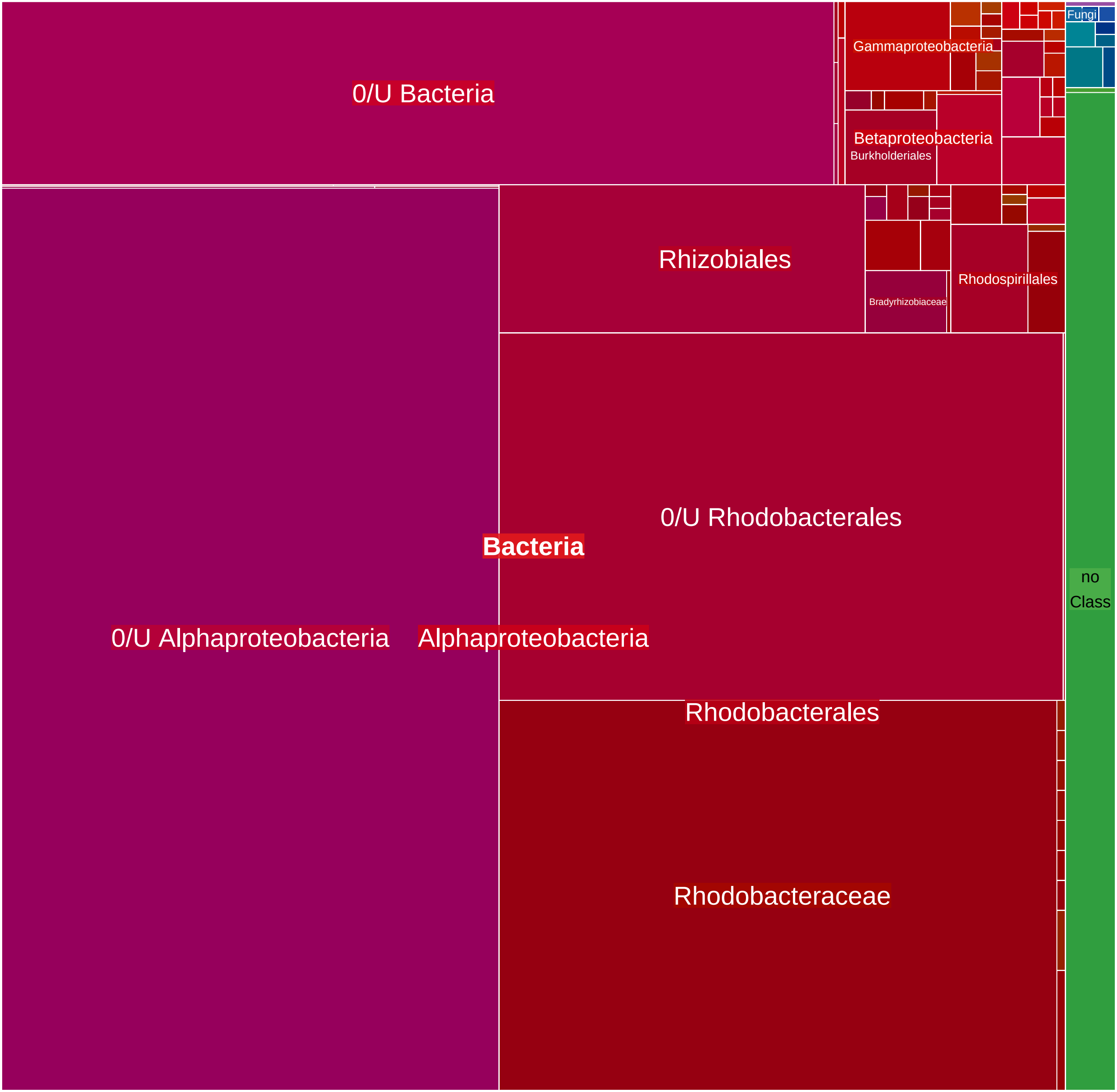

MGT−v1_216 : 6139 unigenes, 1723 (28.07%) taxonomically assigned

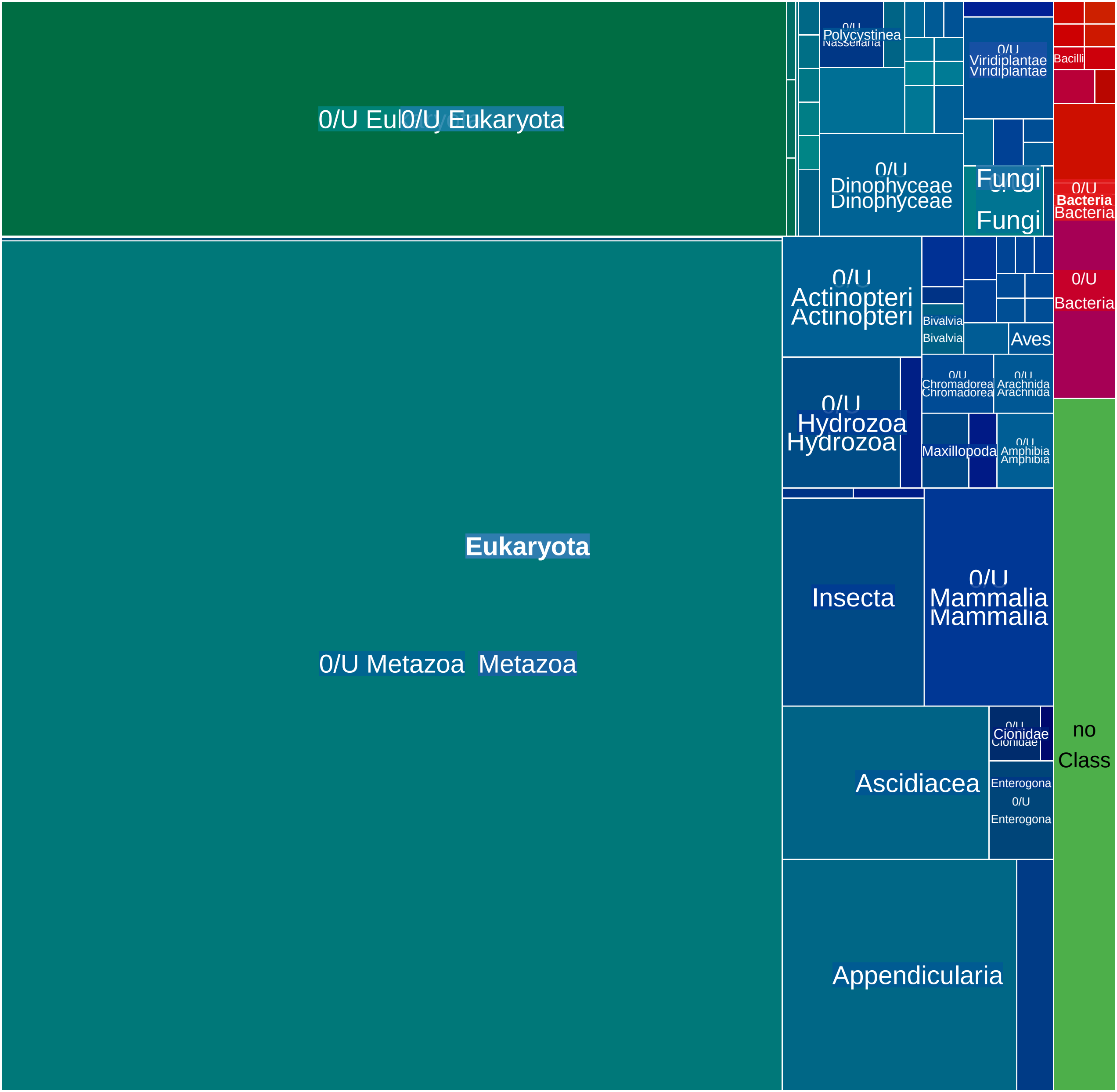

MGT−v1_217 : 6129 unigenes, 311 (5.07%) taxonomically assigned

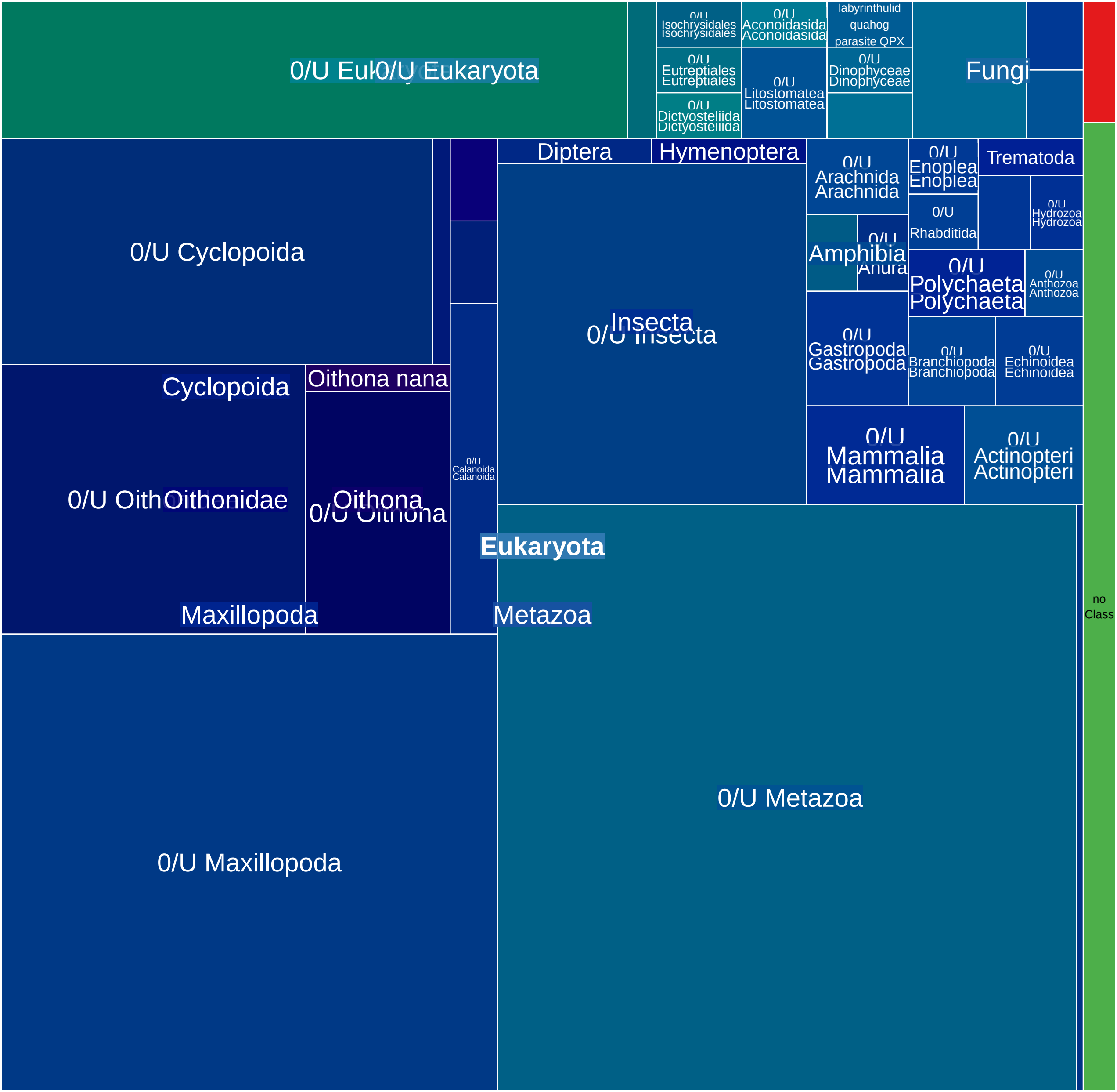

MGT−v1_218 : 6124 unigenes, 5050 (82.46%) taxonomically assigned

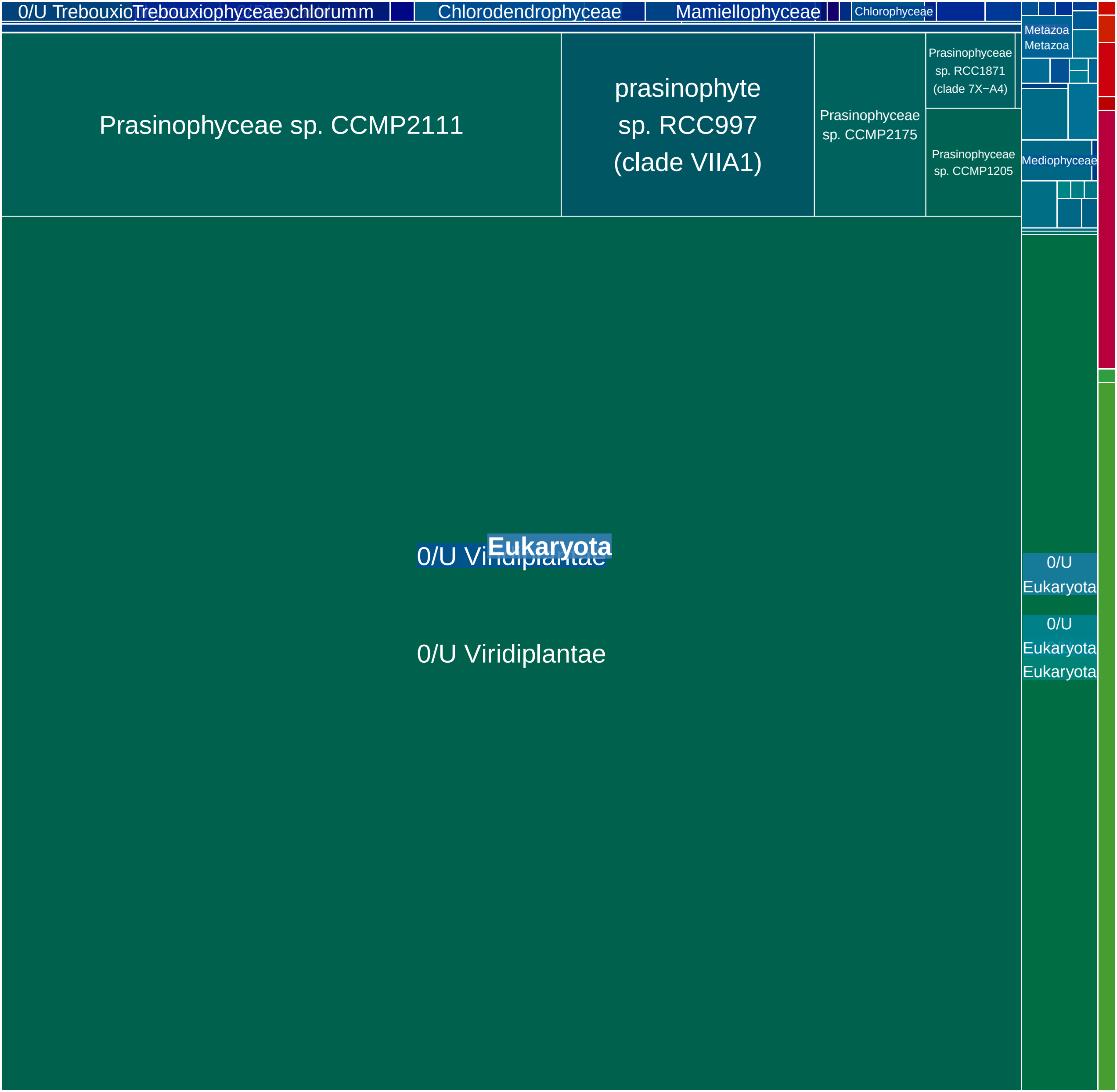

MGT−v1_219 : 6115 unigenes, 3600 (58.87%) taxonomically assigned

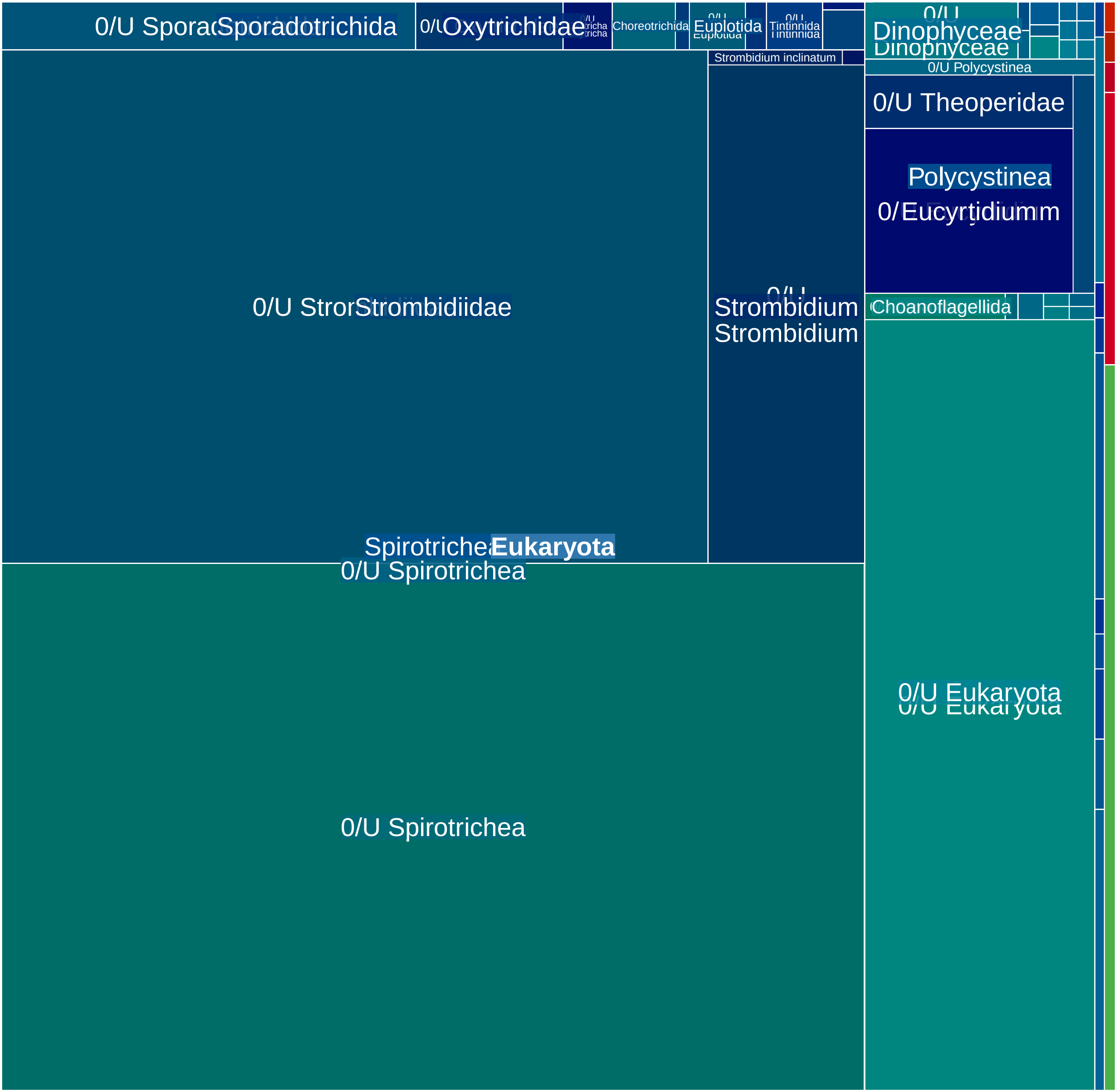

MGT−v1_220 : 6109 unigenes, 428 (7.01%) taxonomically assigned

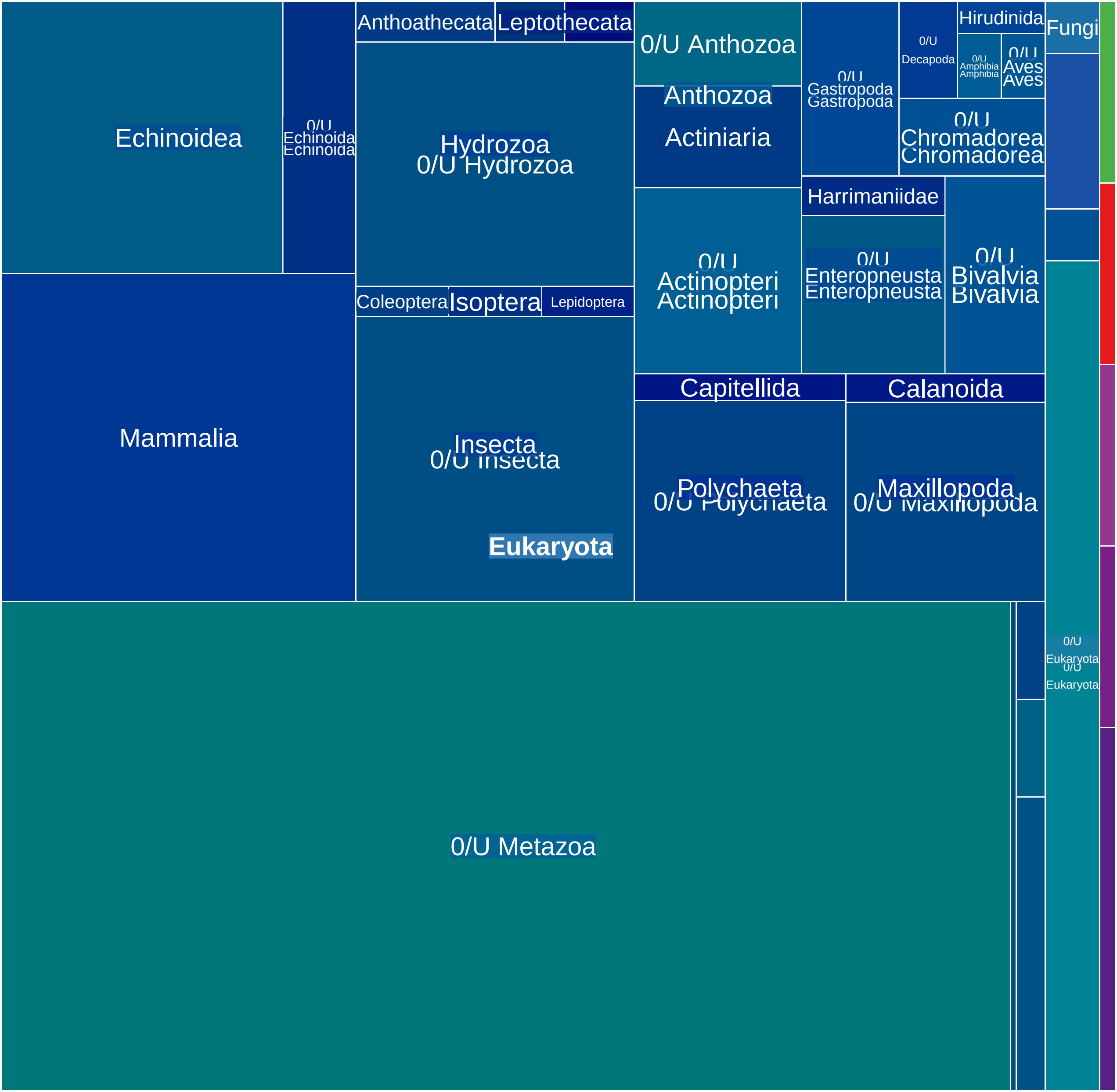

### MGT−v1_221 : 6033 unigenes, 753 (12.48%) taxonomically assigned

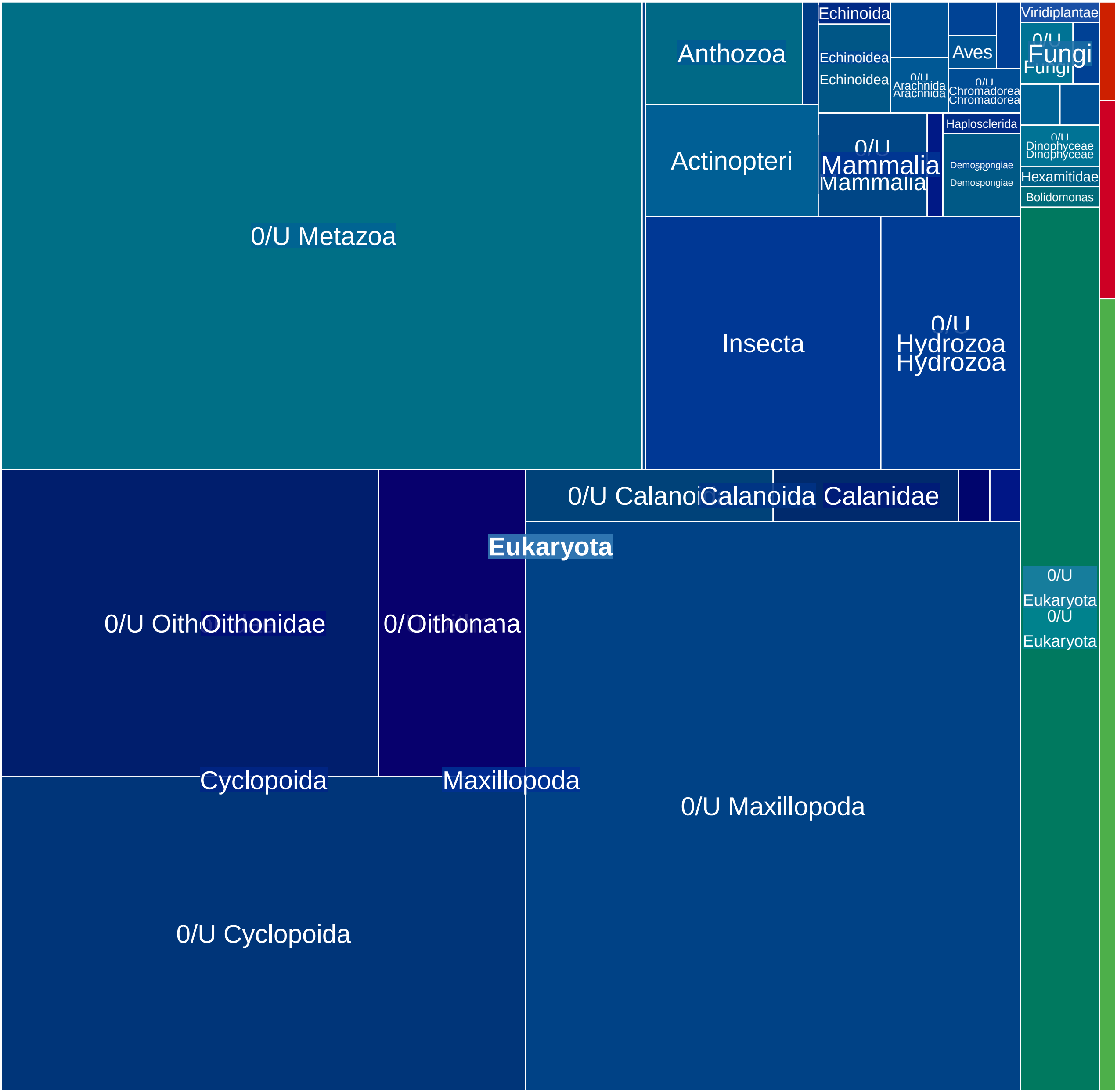

### MGT−v1_222 : 5923 unigenes, 3147 (53.13%) taxonomically assigned

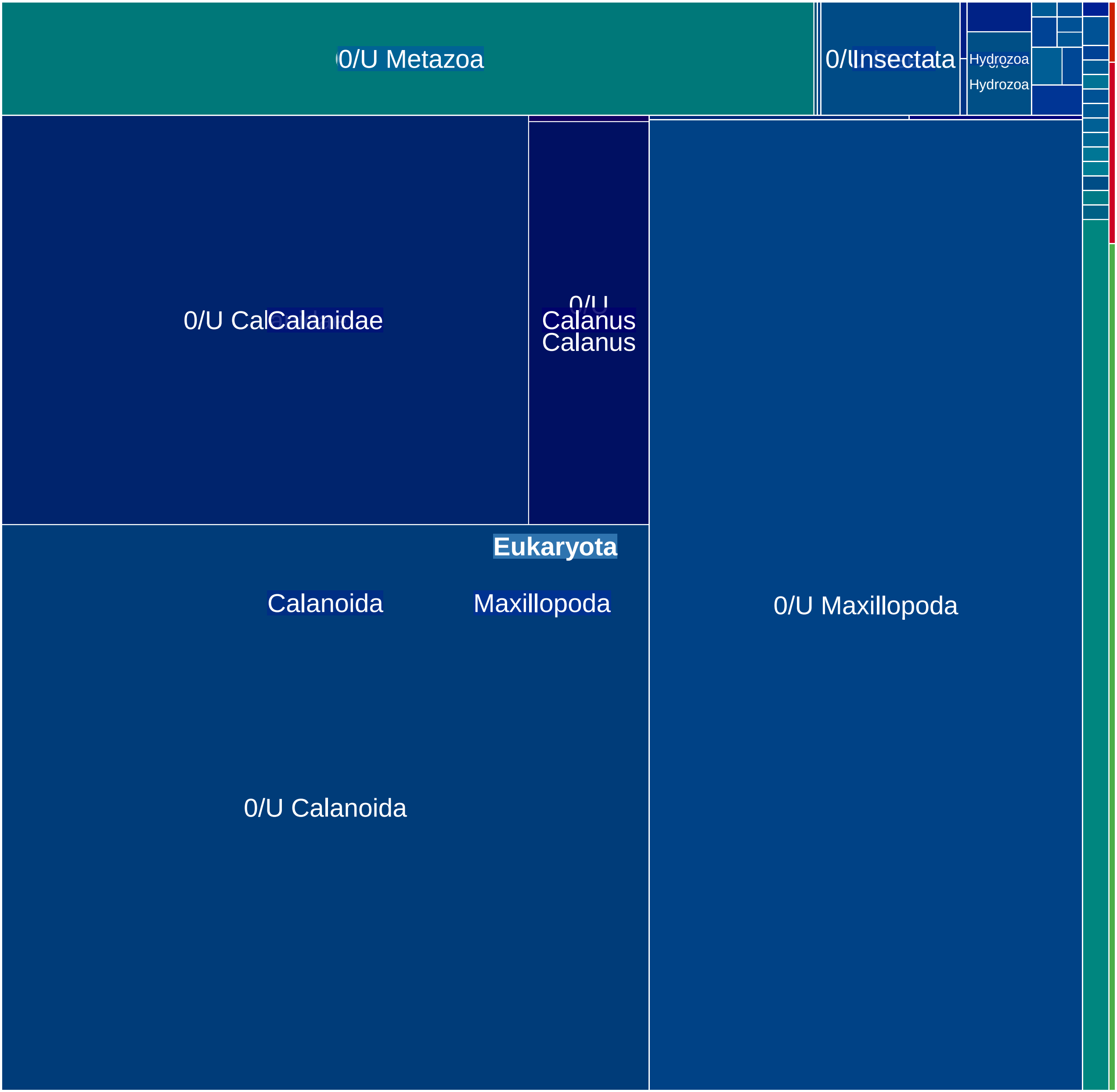

### MGT−v1_223 : 5916 unigenes, 5207 (88.02%) taxonomically assigned

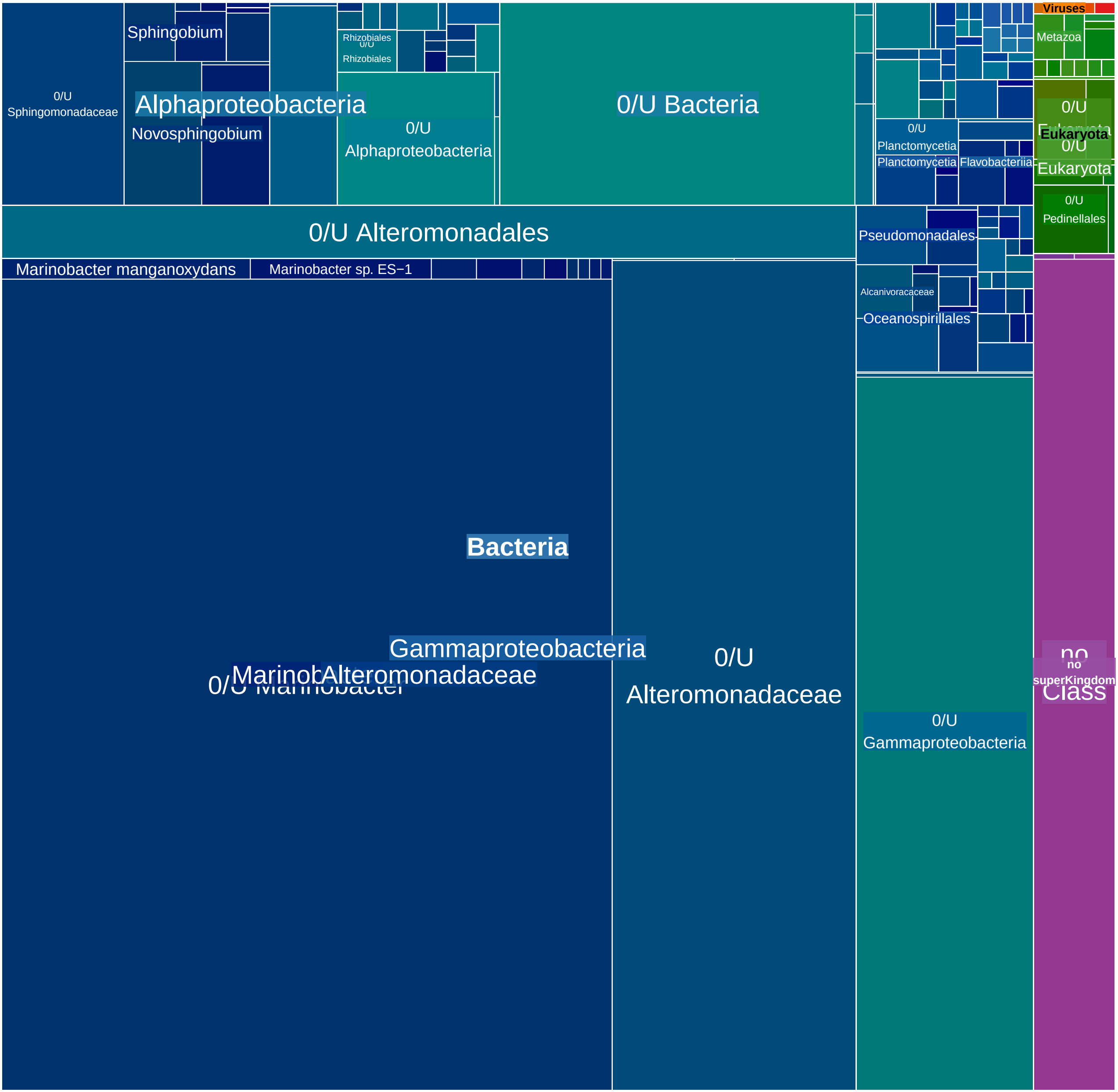

MGT−v1_224 : 5900 unigenes, 3454 (58.54%) taxonomically assigned

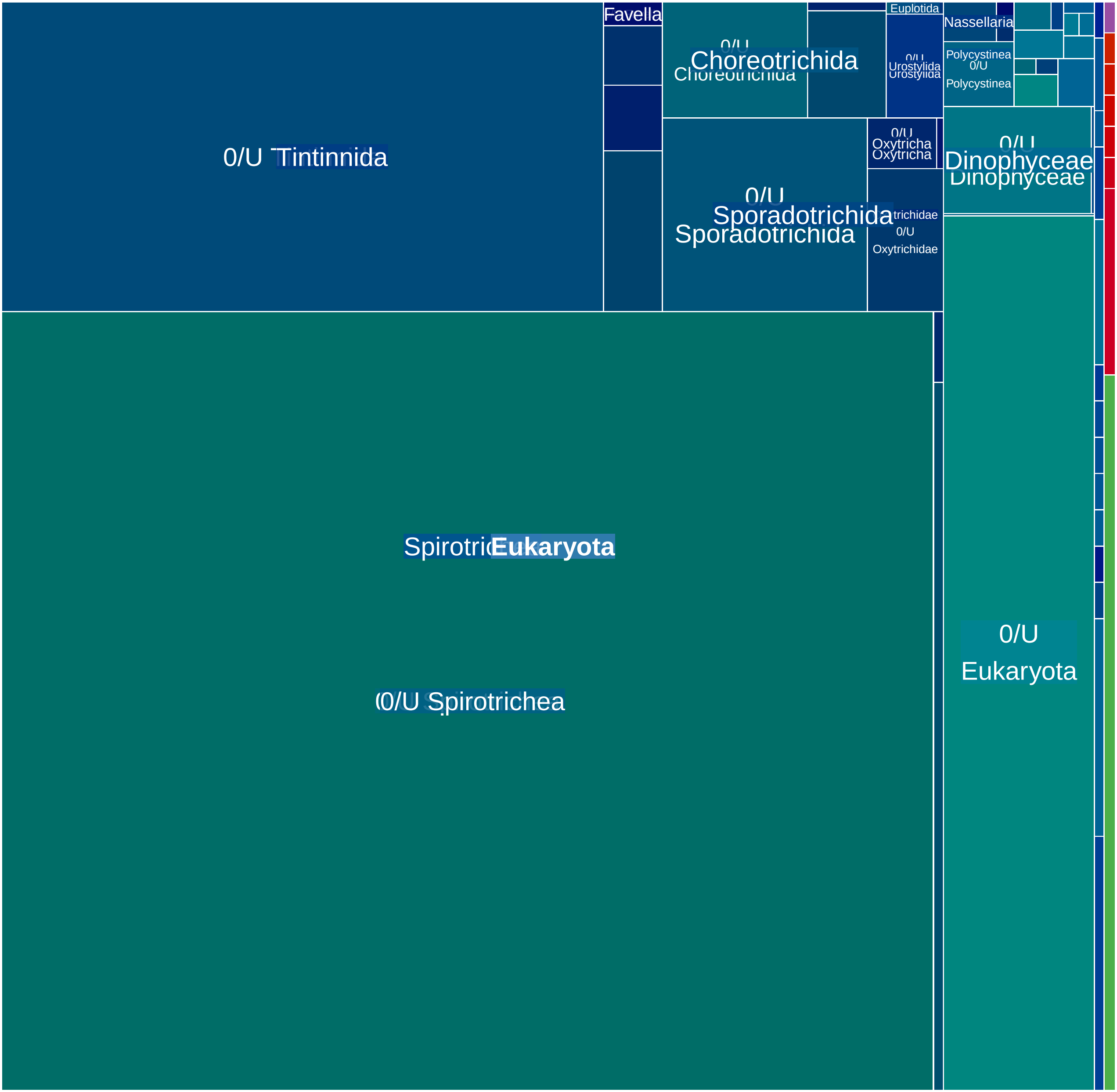

MGT−v1_225 : 5805 unigenes, 4873 (83.94%) taxonomically assigned

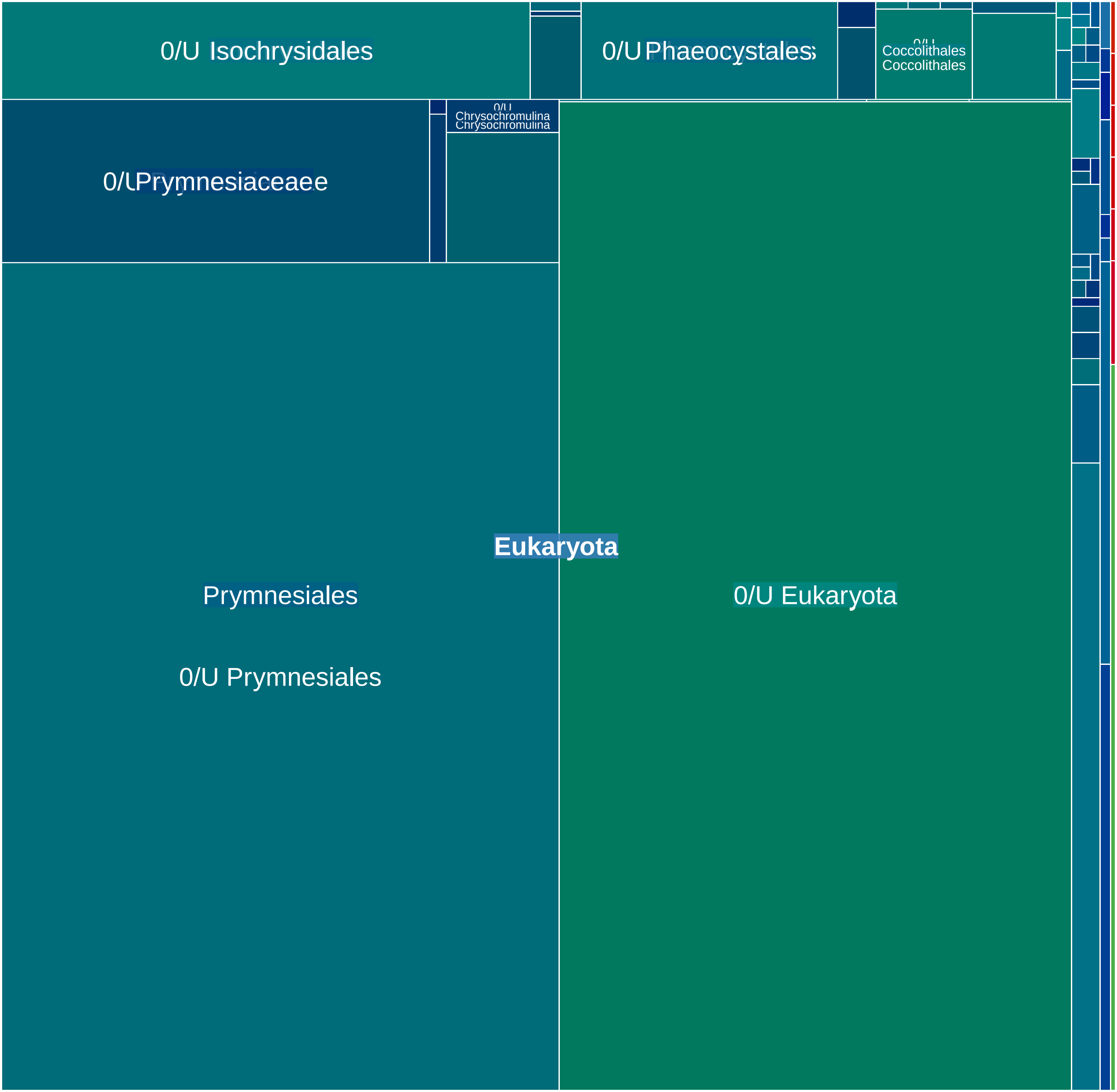

MGT−v1_226 : 5748 unigenes, 1706 (29.68%) taxonomically assigned

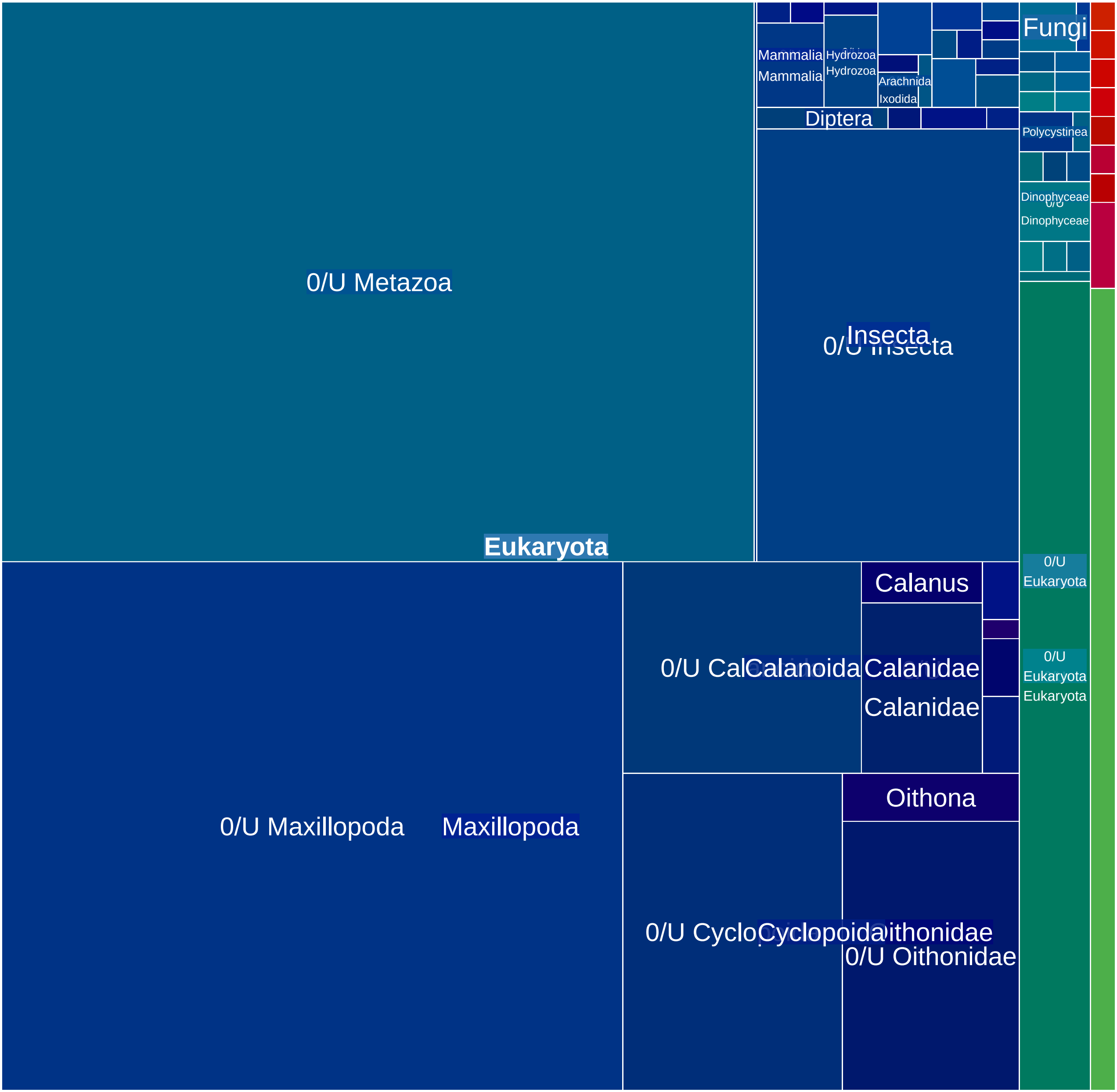

### MGT−v1_227 : 5738 unigenes, 5641 (98.31%) taxonomically assigned

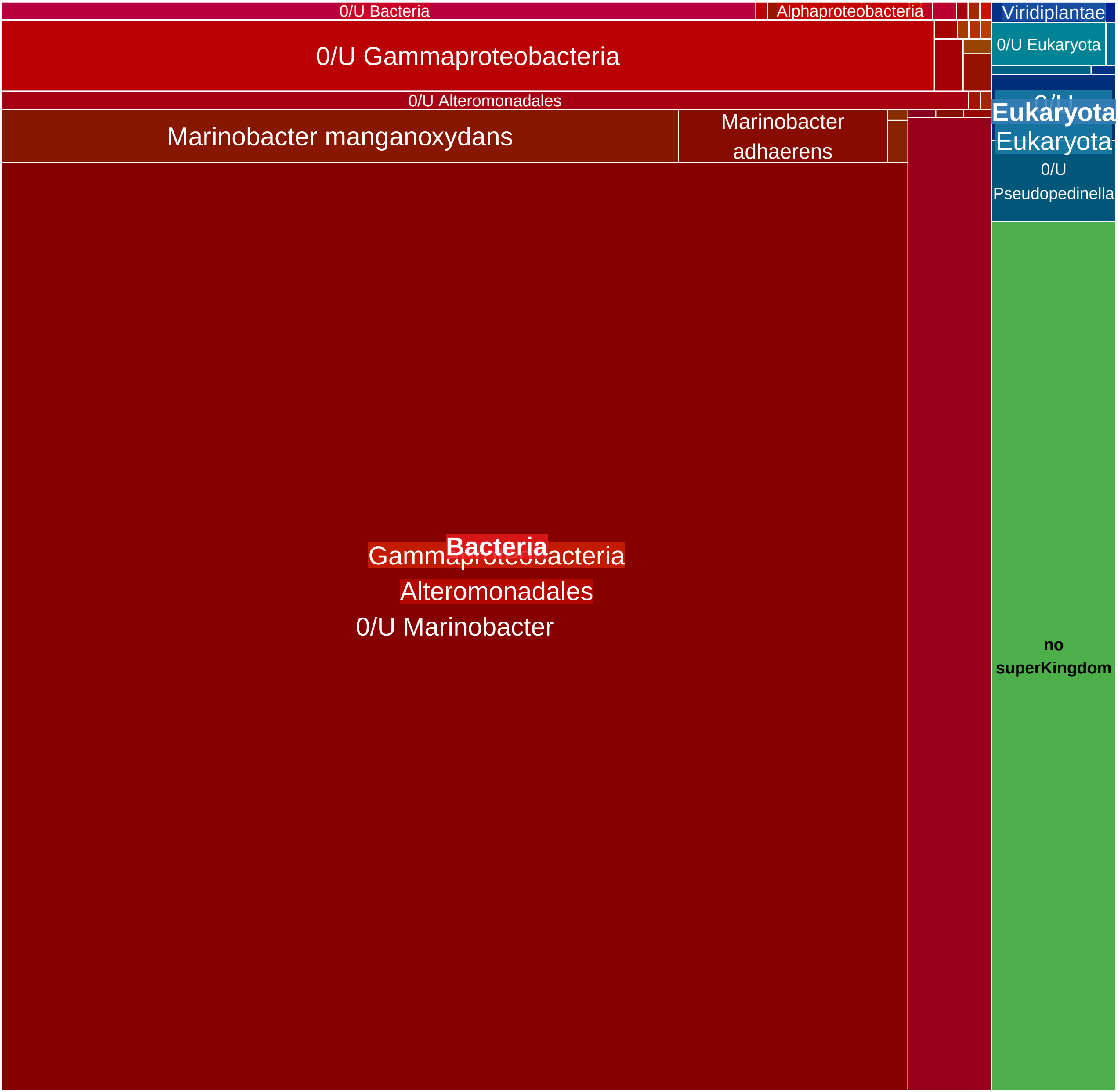

MGT−v1_228 : 5716 unigenes, 4025 (70.42%) taxonomically assigned

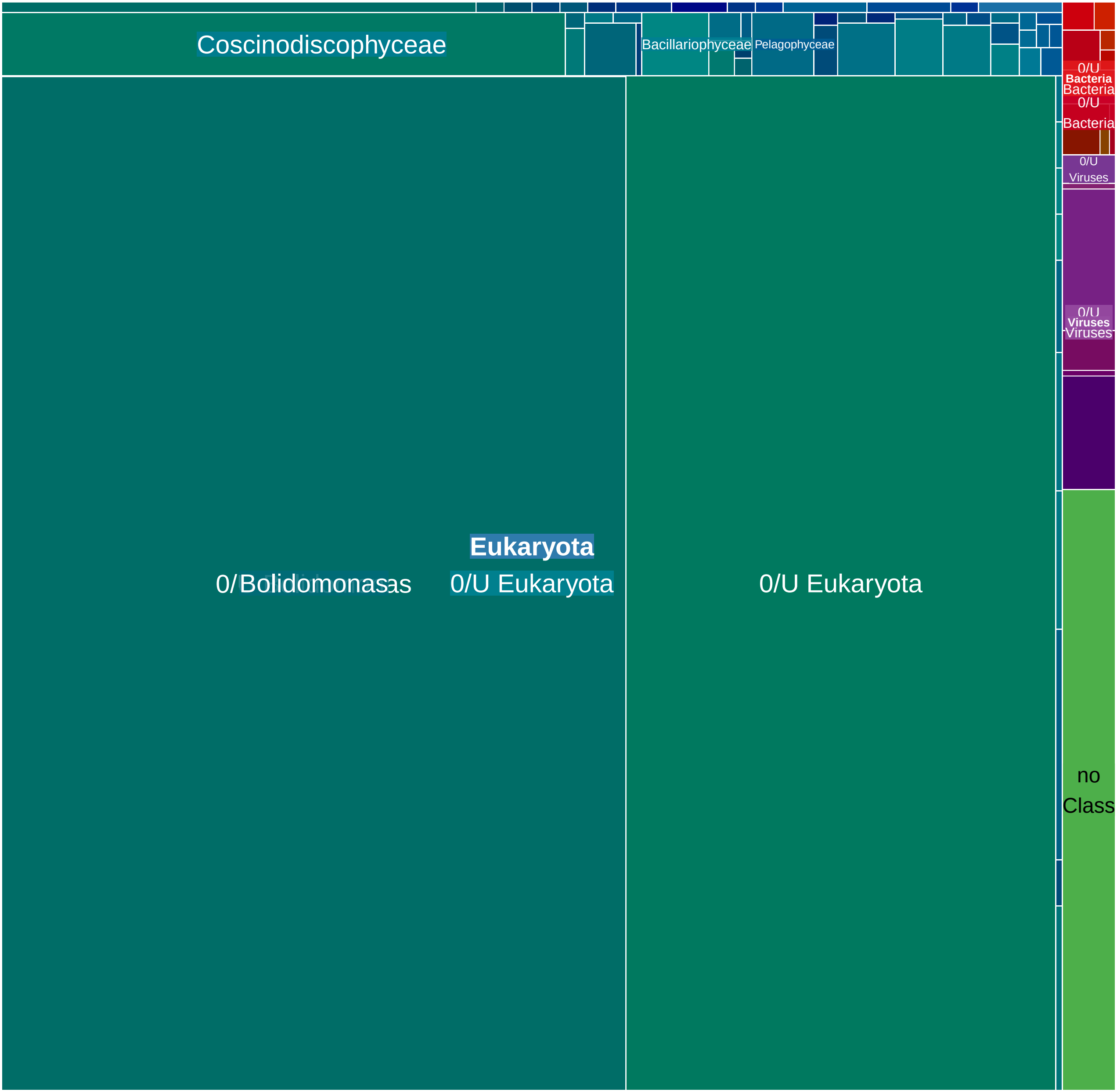

MGT−v1_229 : 5666 unigenes, 2184 (38.55%) taxonomically assigned

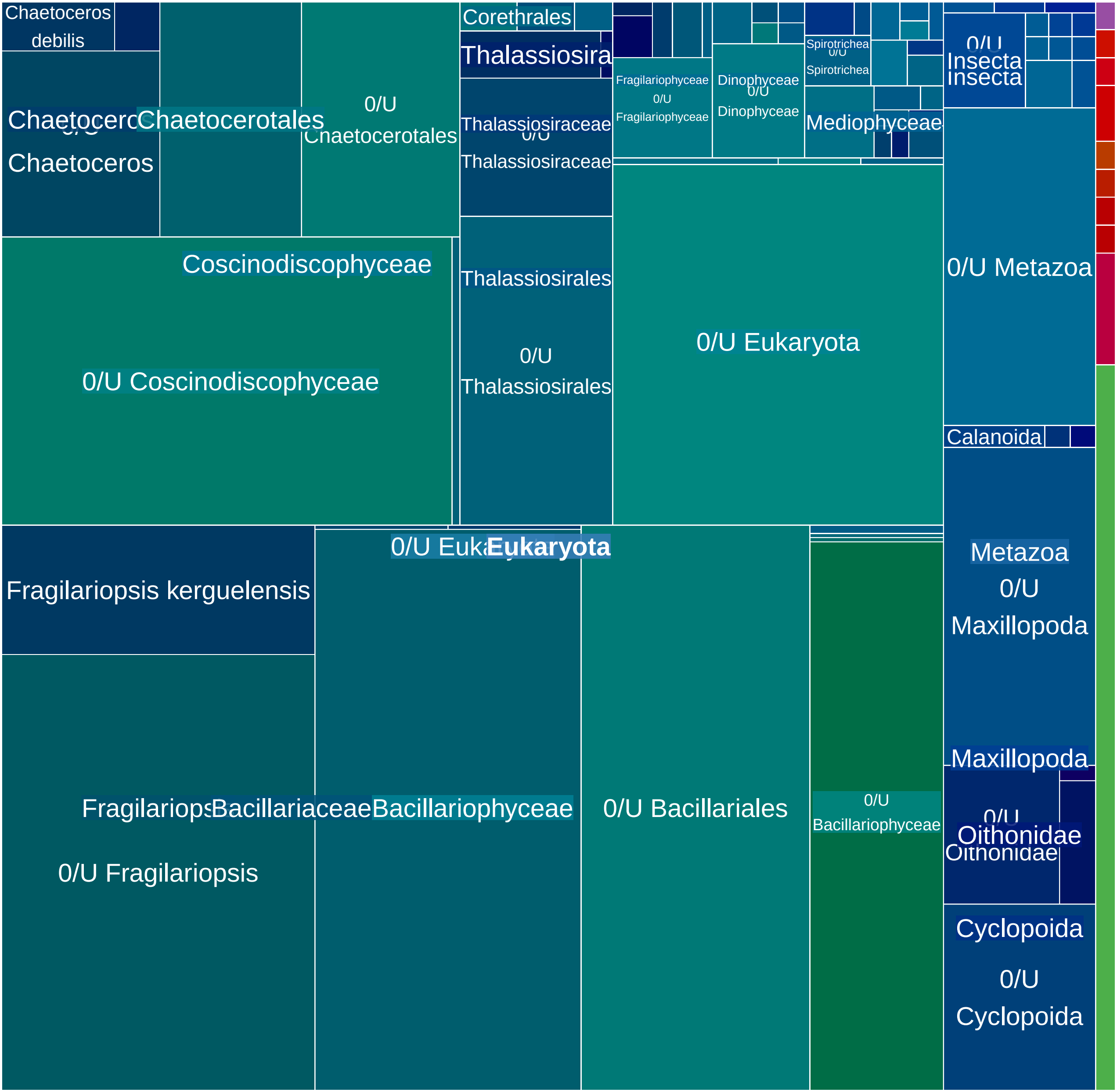

MGT−v1_230 : 5654 unigenes, 4190 (74.11%) taxonomically assigned

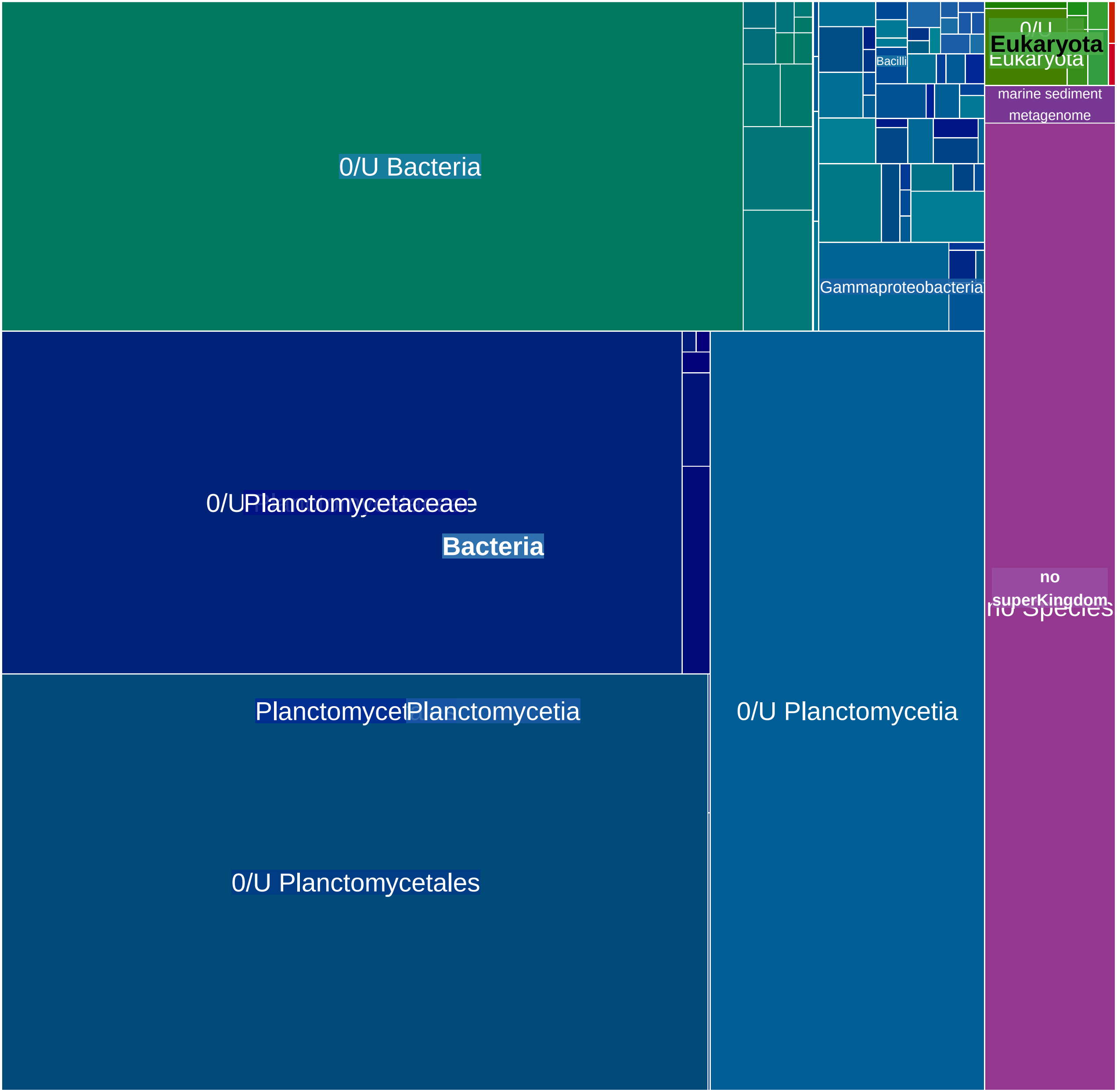

### MGT−v1_231 : 5611 unigenes, 4321 (77.01%) taxonomically assigned

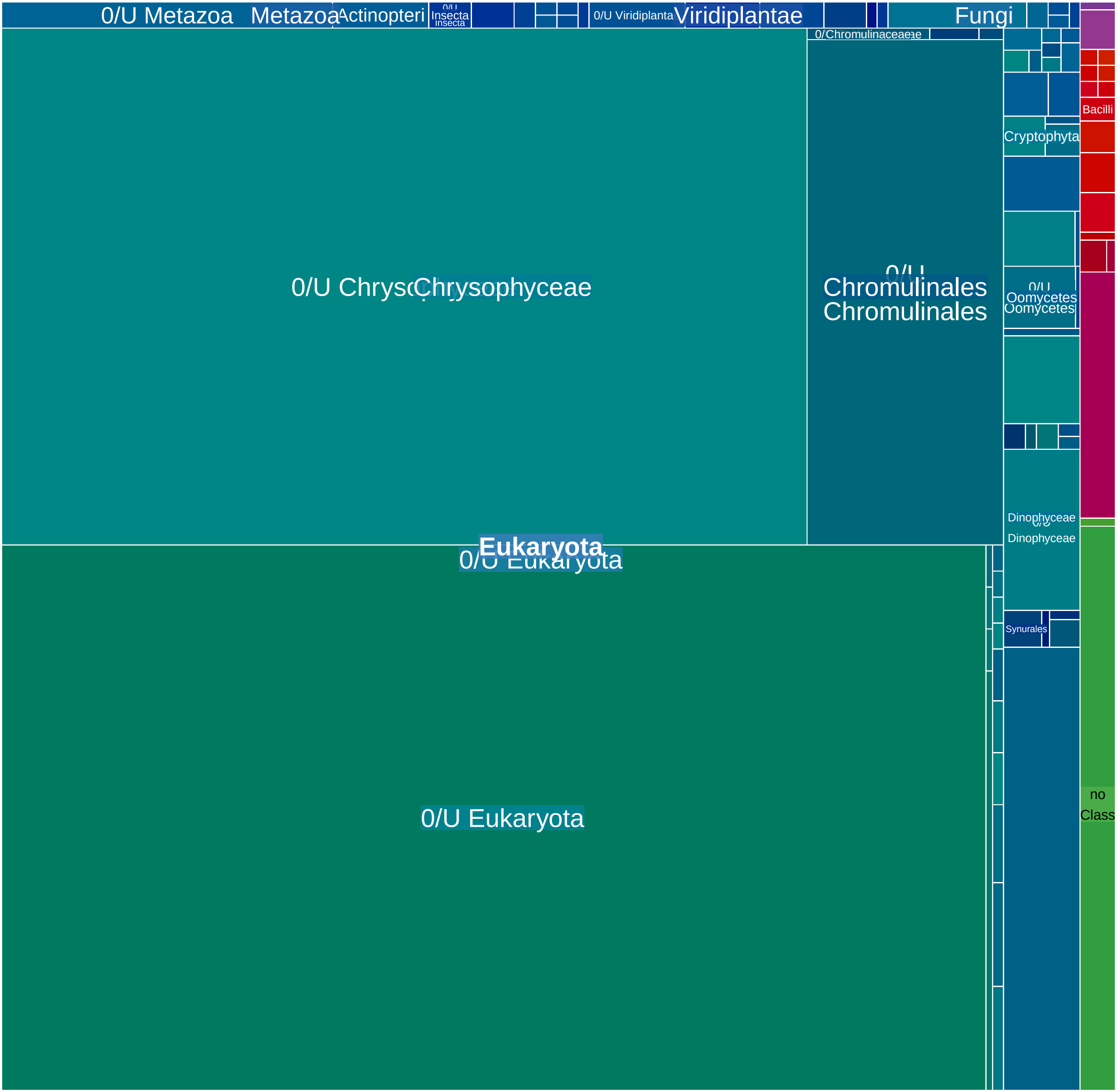

### MGT−v1_232 : 5587 unigenes, 536 (9.59%) taxonomically assigned

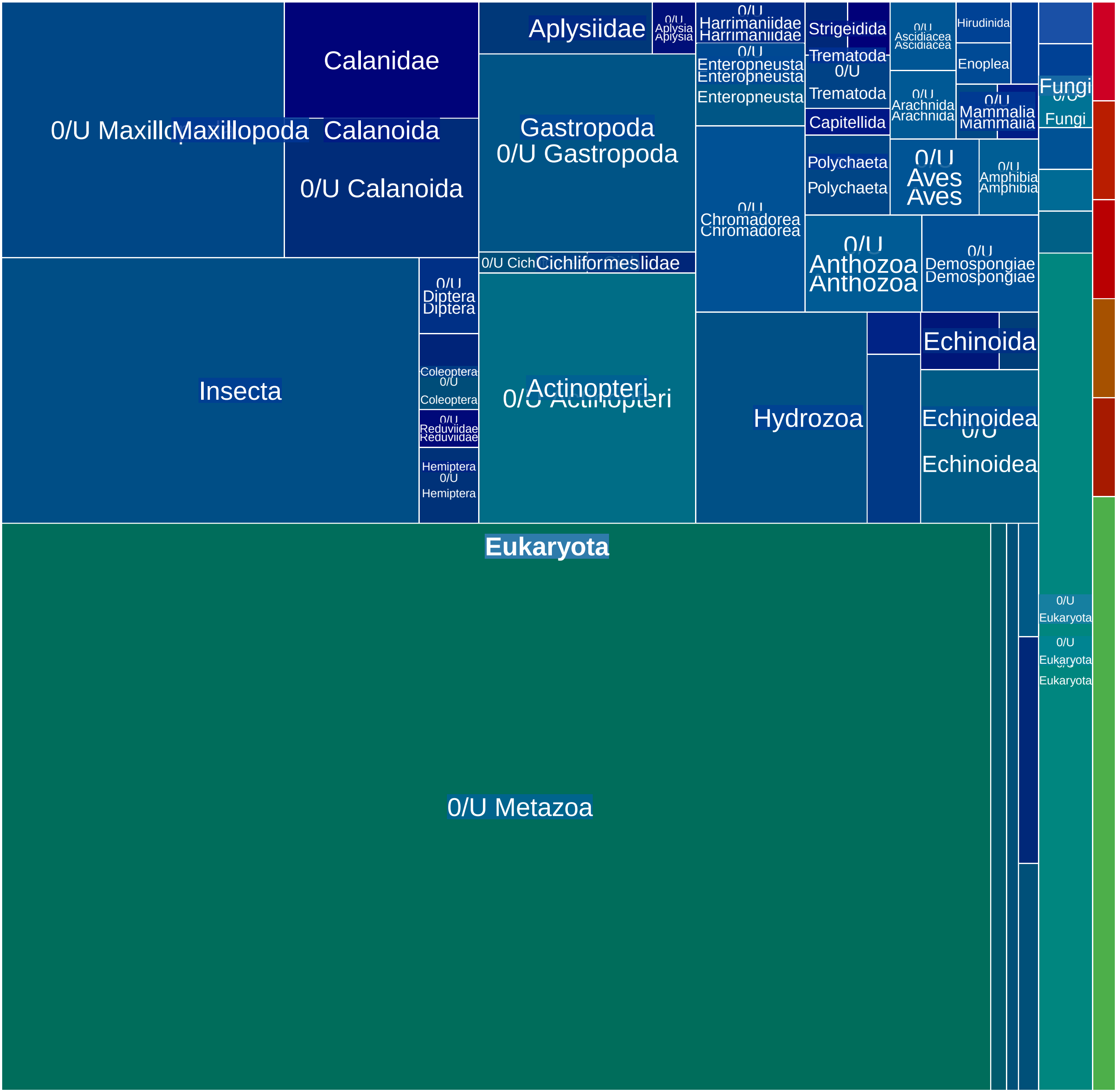

MGT−v1_233 : 5494 unigenes, 5187 (94.41%) taxonomically assigned

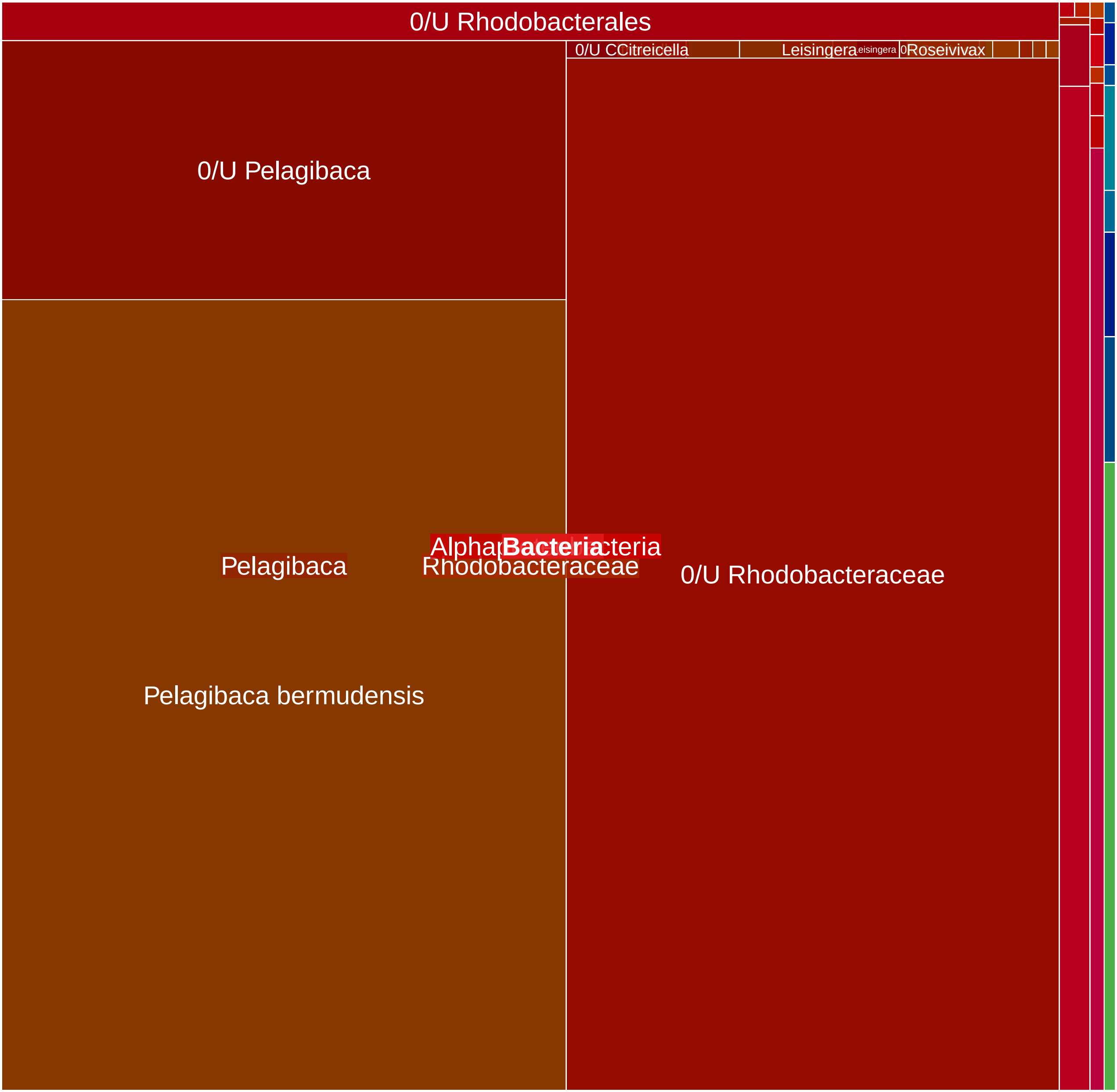

MGT−v1_234 : 5480 unigenes, 577 (10.53%) taxonomically assigned

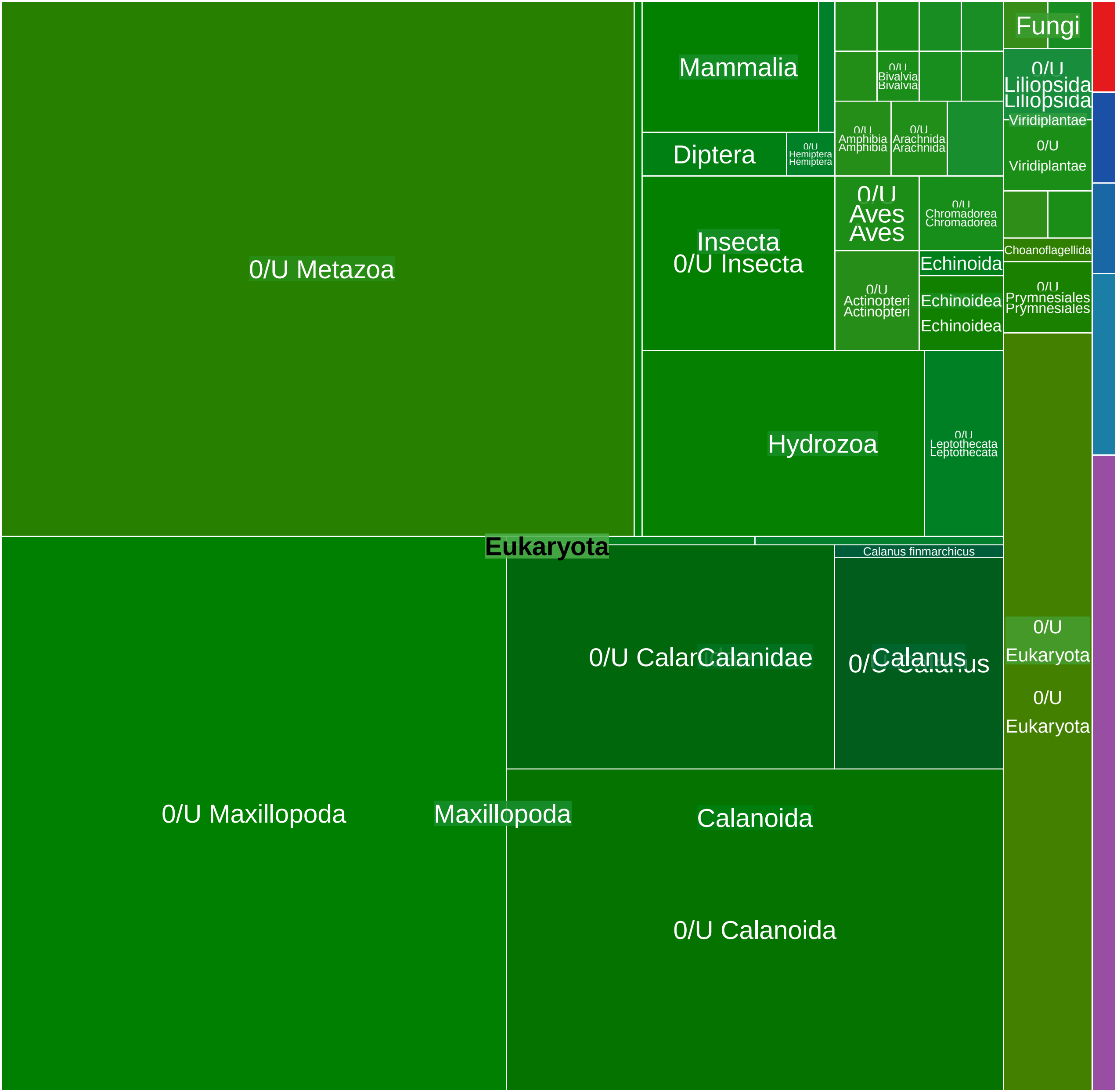

MGT−v1_235 : 5473 unigenes, 4740 (86.61%) taxonomically assigned

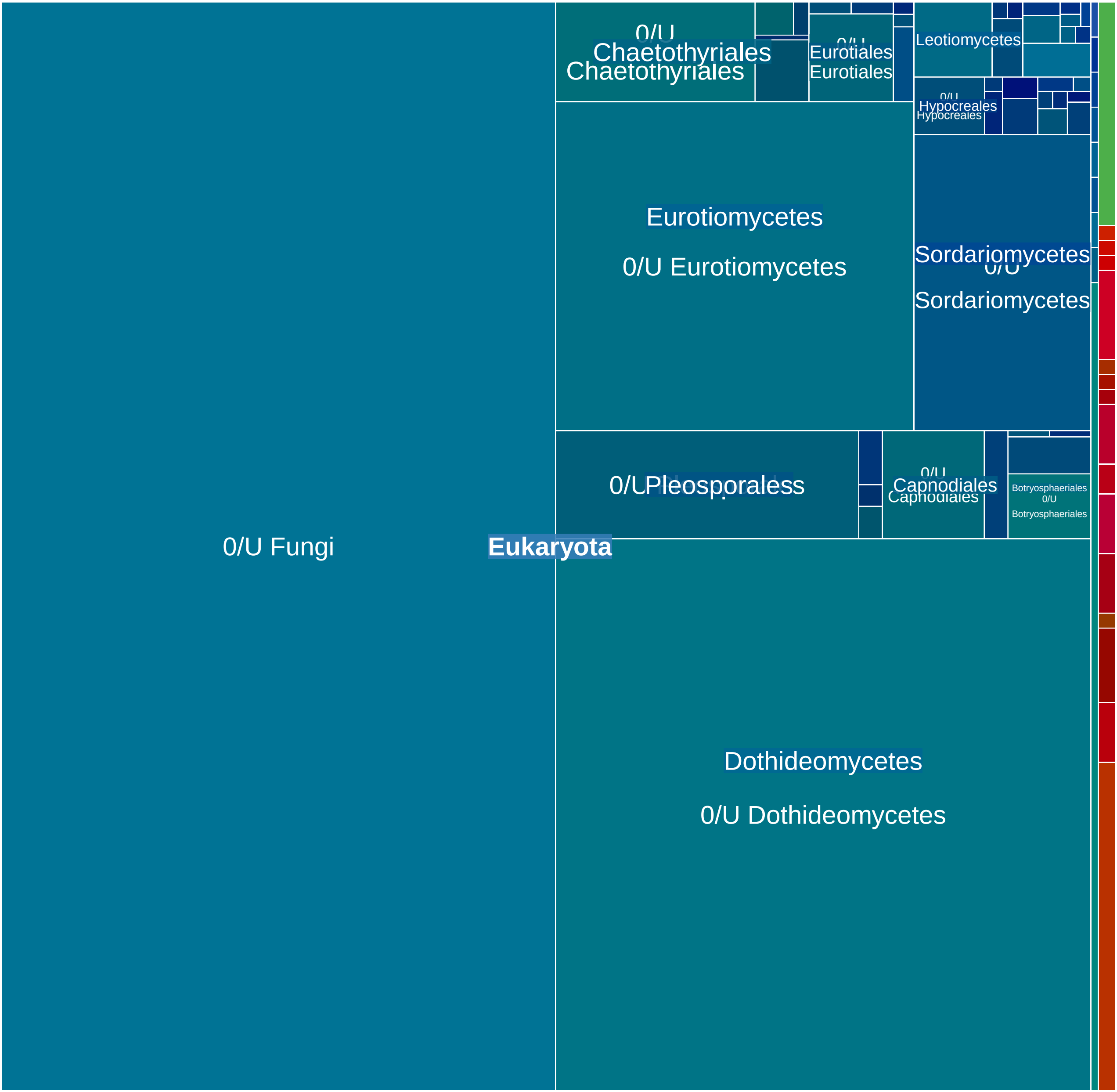

MGT−v1_236 : 5456 unigenes, 356 (6.52%) taxonomically assigned

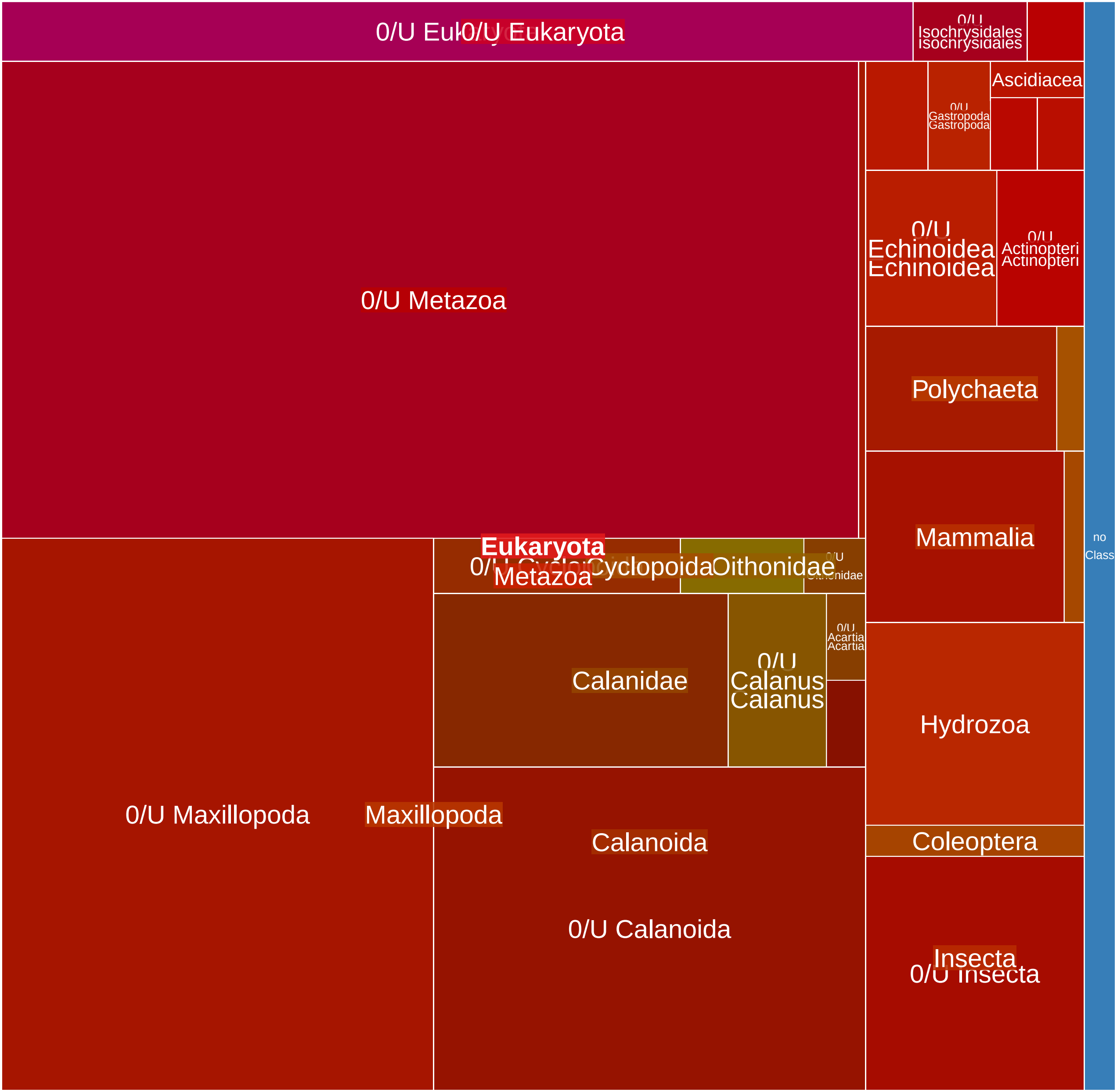

MGT−v1_237 : 5449 unigenes, 373 (6.85%) taxonomically assigned

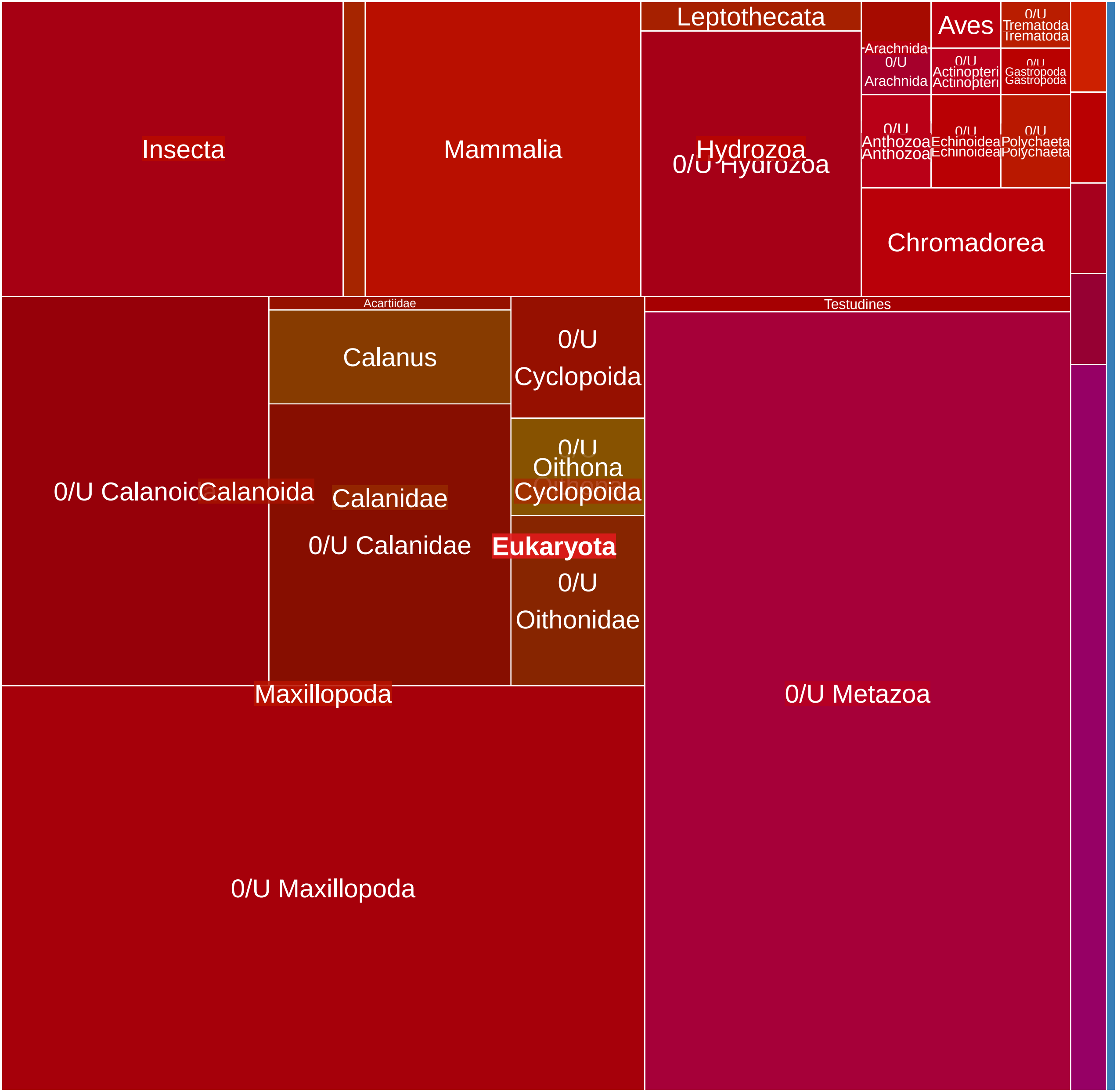

MGT−v1_238 : 5409 unigenes, 4757 (87.95%) taxonomically assigned

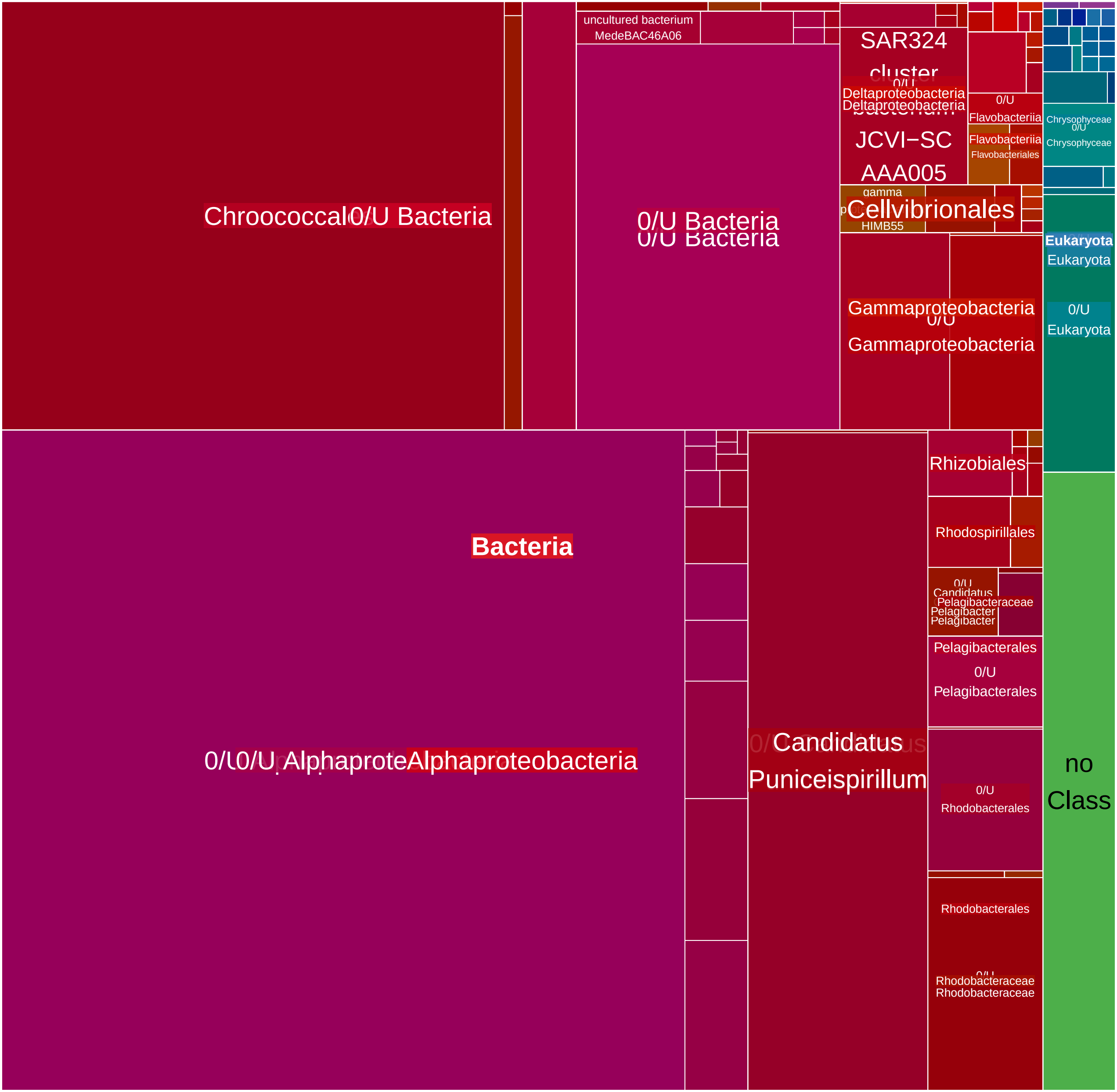

### MGT−v1_239 : 5408 unigenes, 4931 (91.18%) taxonomically assigned

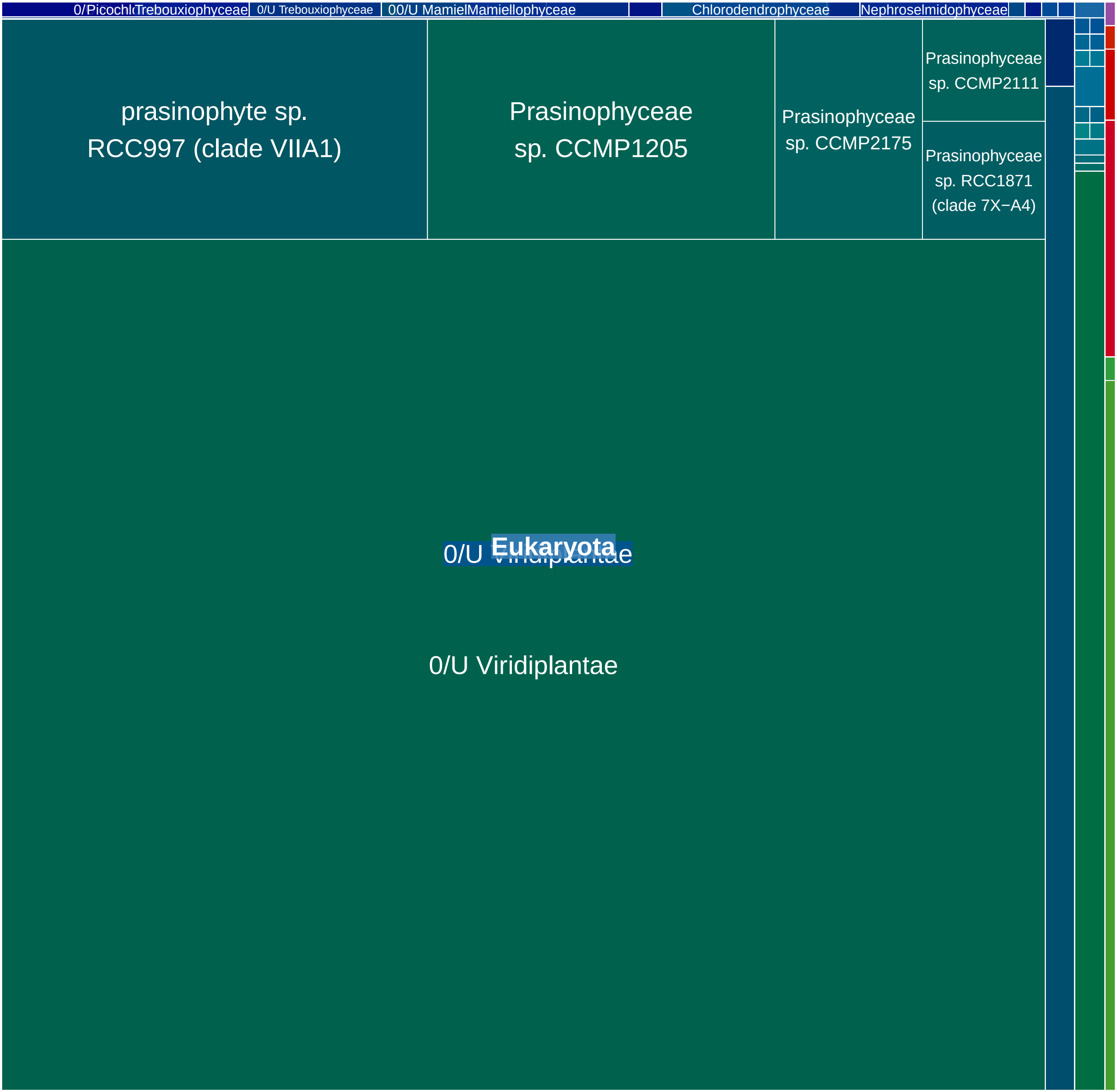

MGT−v1_240 : 5292 unigenes, 4781 (90.34%) taxonomically assigned

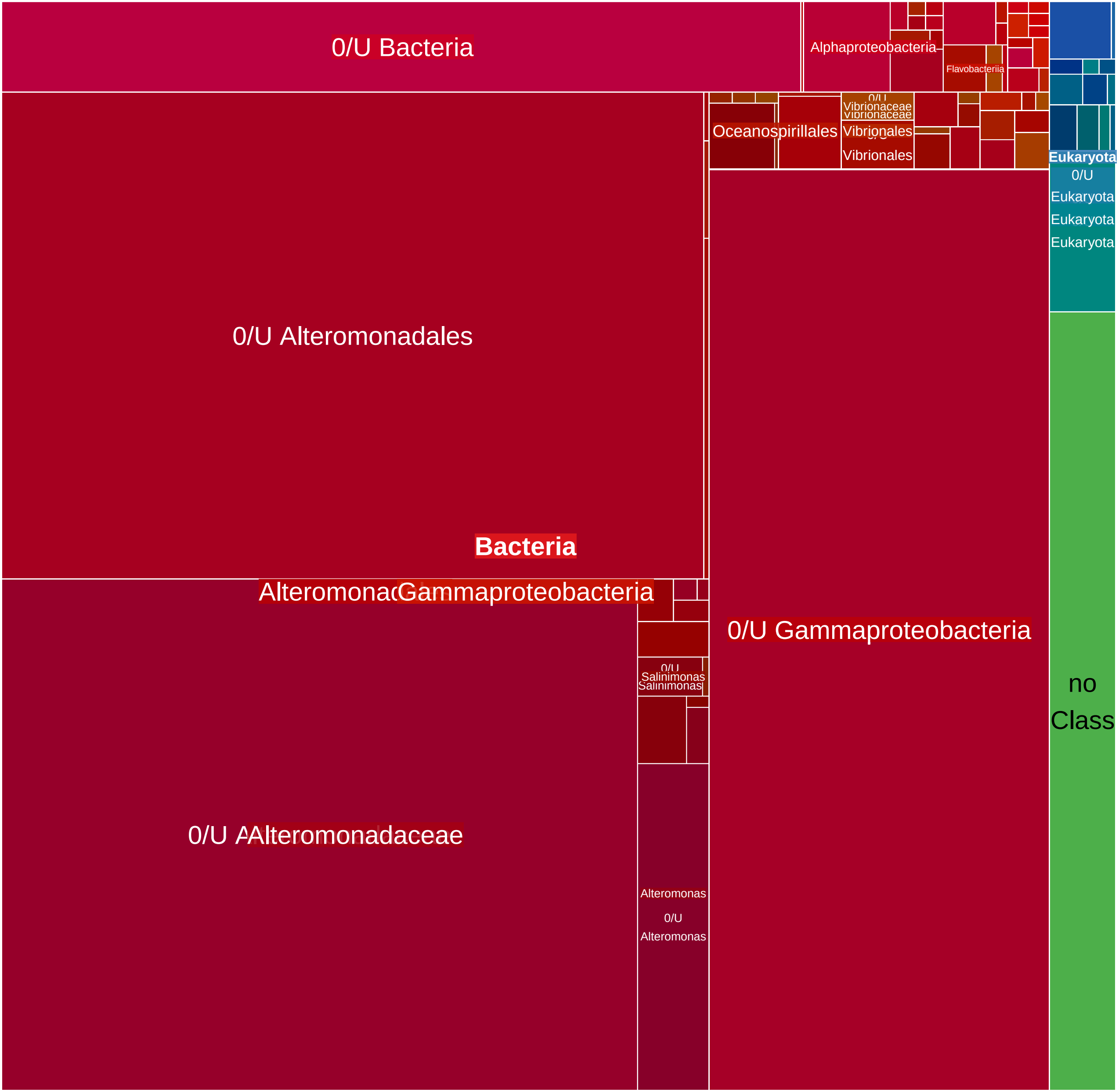

MGT−v1_241 : 5274 unigenes, 1736 (32.92%) taxonomically assigned

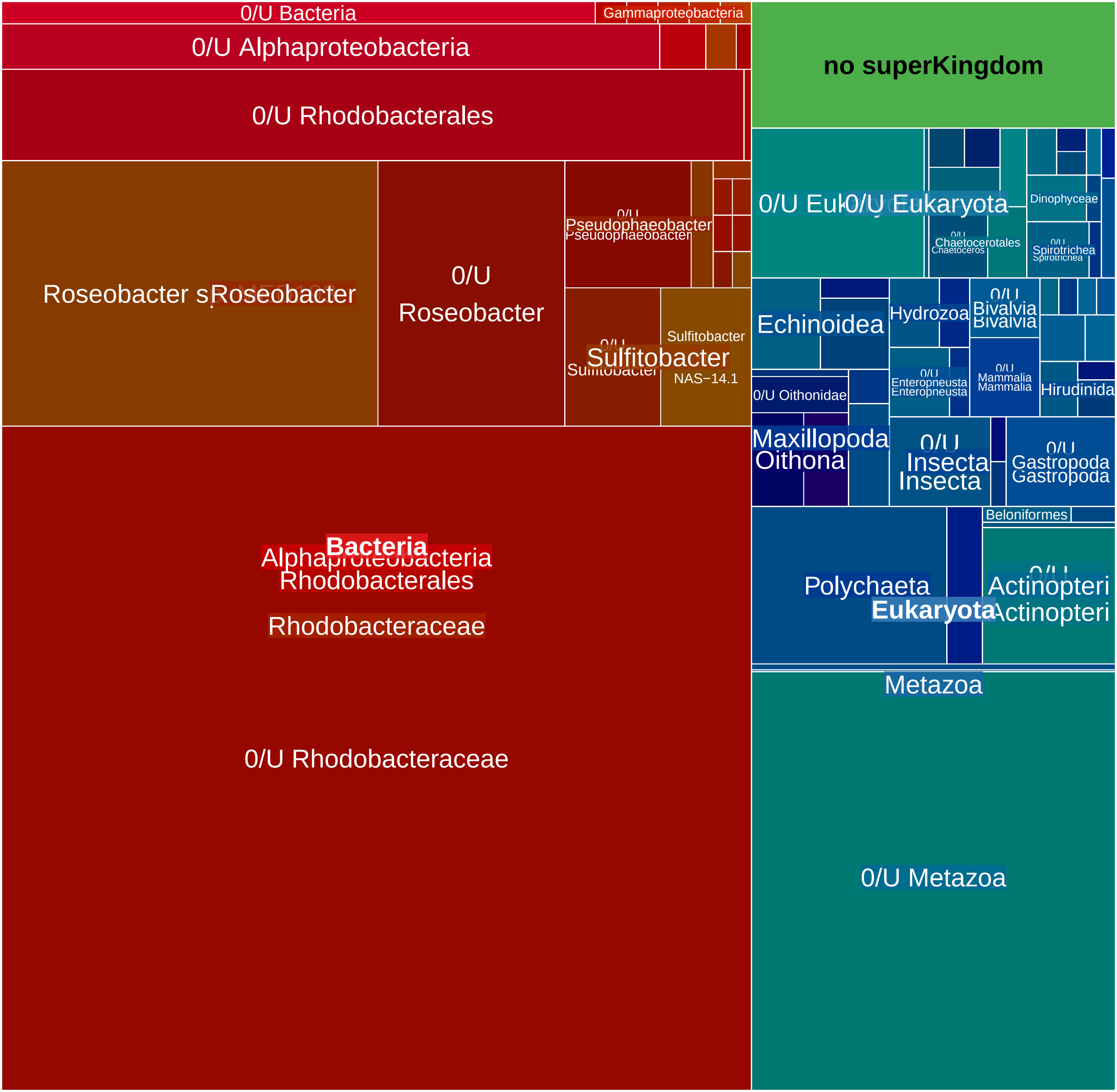

### MGT−v1_242 : 5257 unigenes, 1361 (25.89%) taxonomically assigned

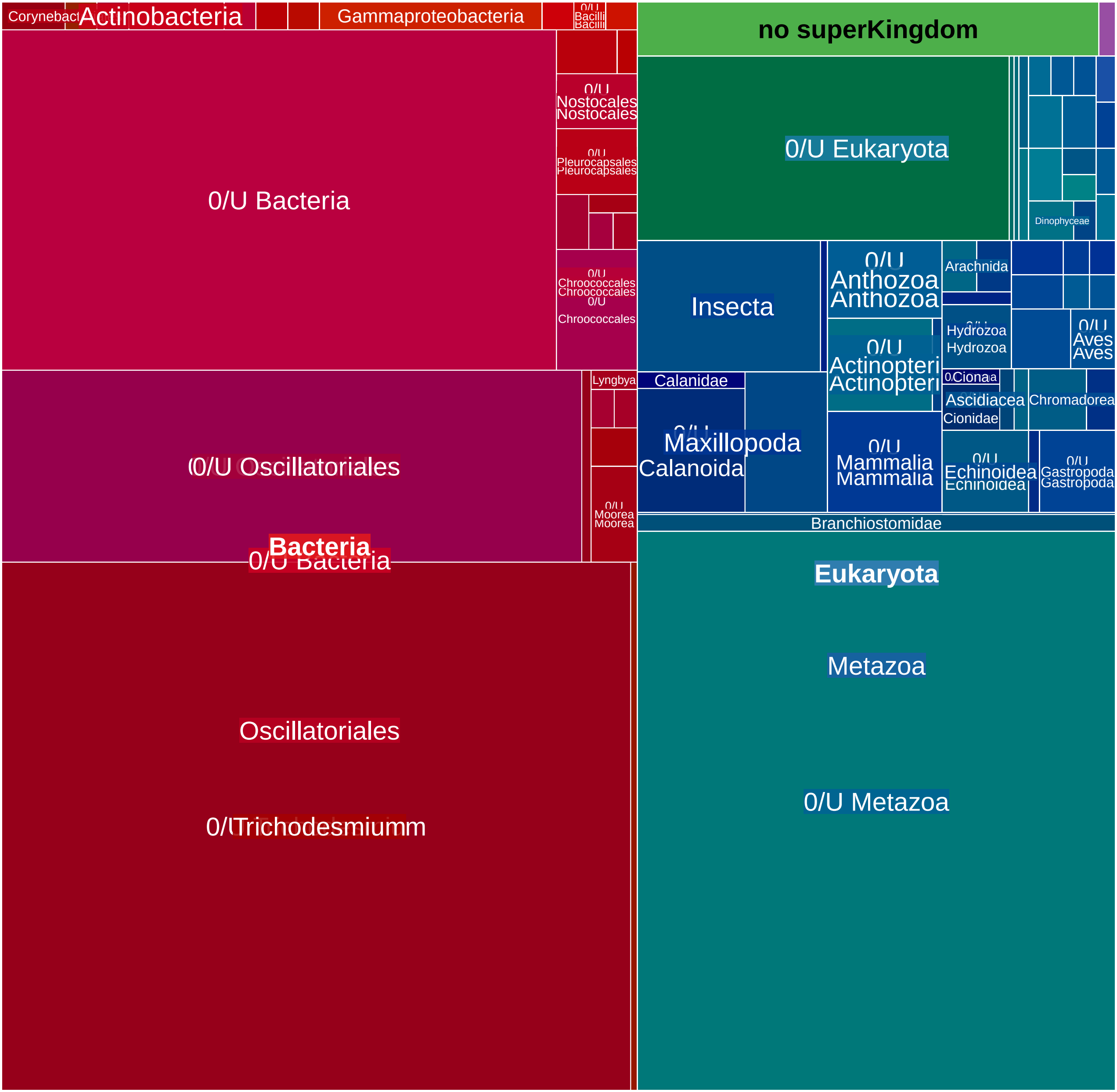

MGT−v1_243 : 5158 unigenes, 4303 (83.42%) taxonomically assigned

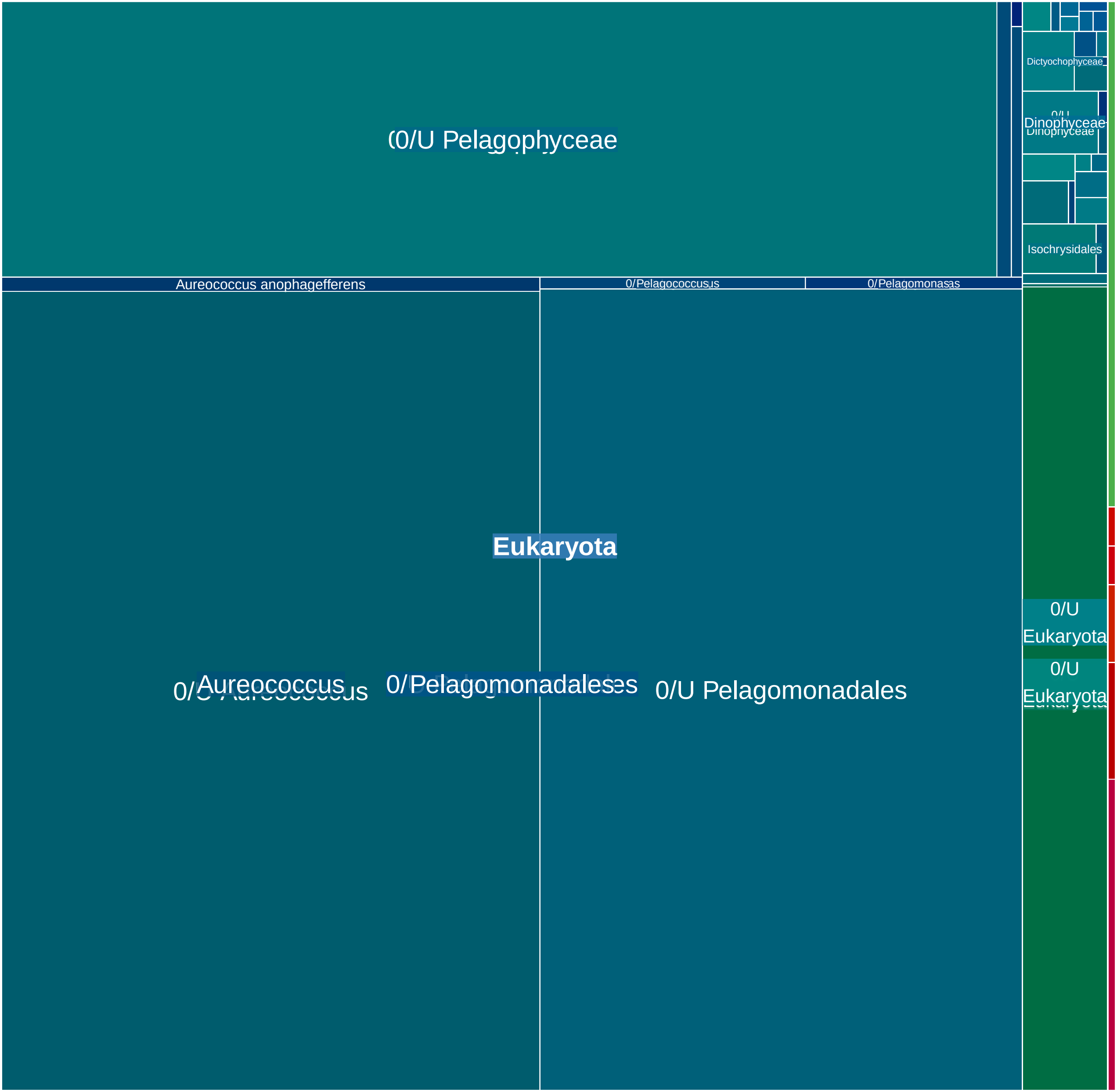

MGT−v1_244 : 5150 unigenes, 287 (5.57%) taxonomically assigned

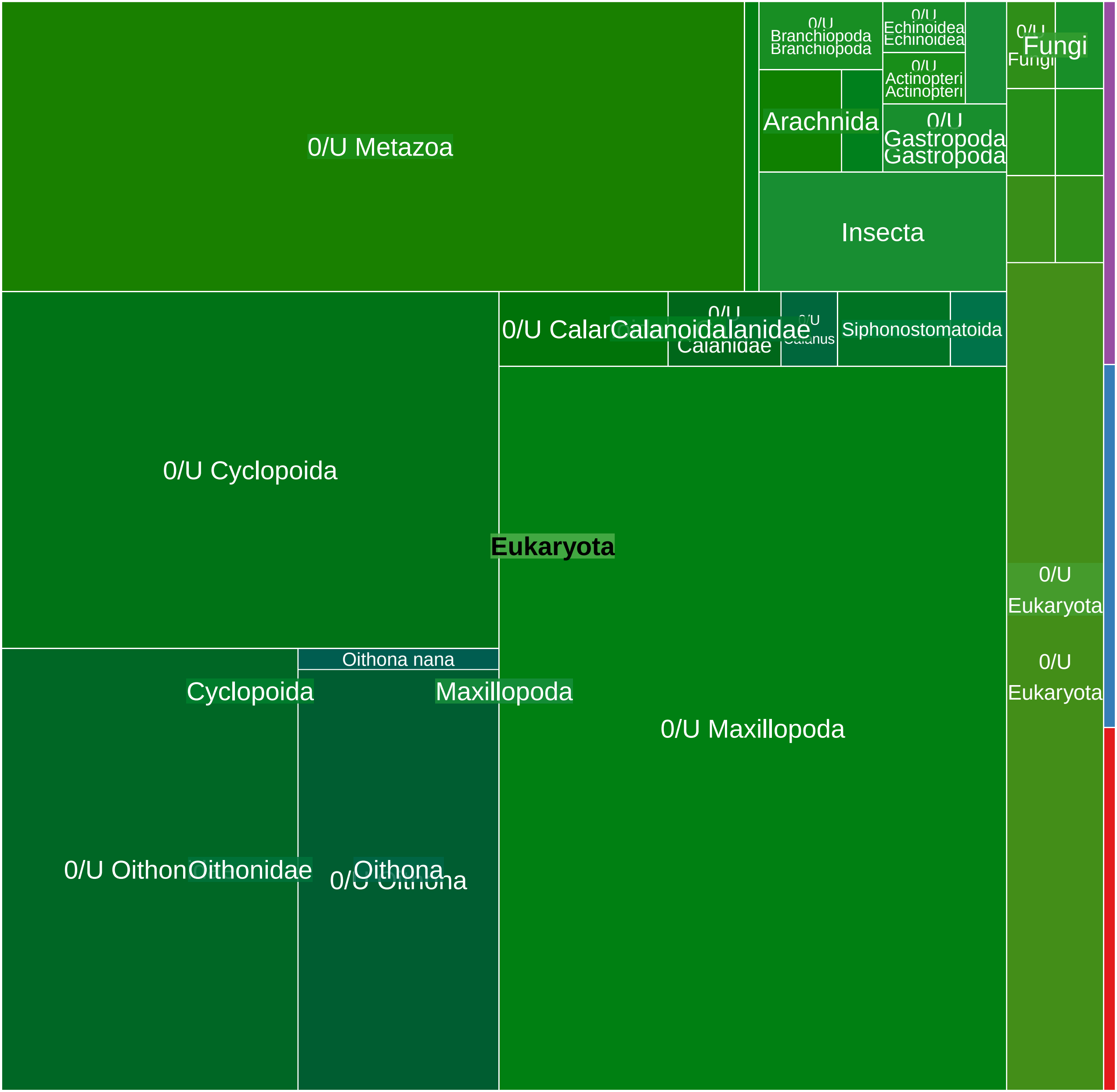

MGT−v1_245 : 5092 unigenes, 2869 (56.34%) taxonomically assigned

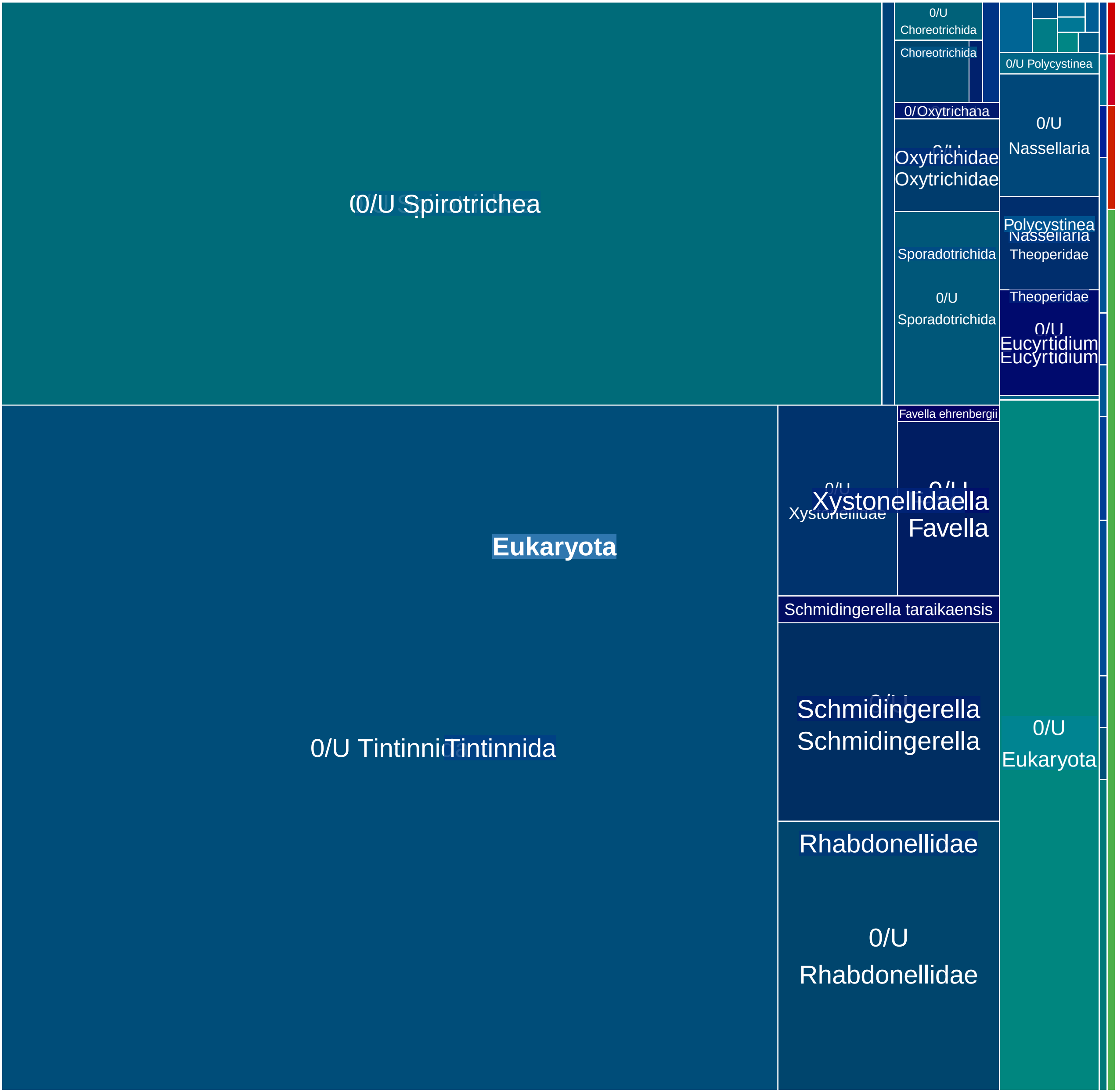

### MGT−v1_246 : 5043 unigenes, 4800 (95.18%) taxonomically assigned

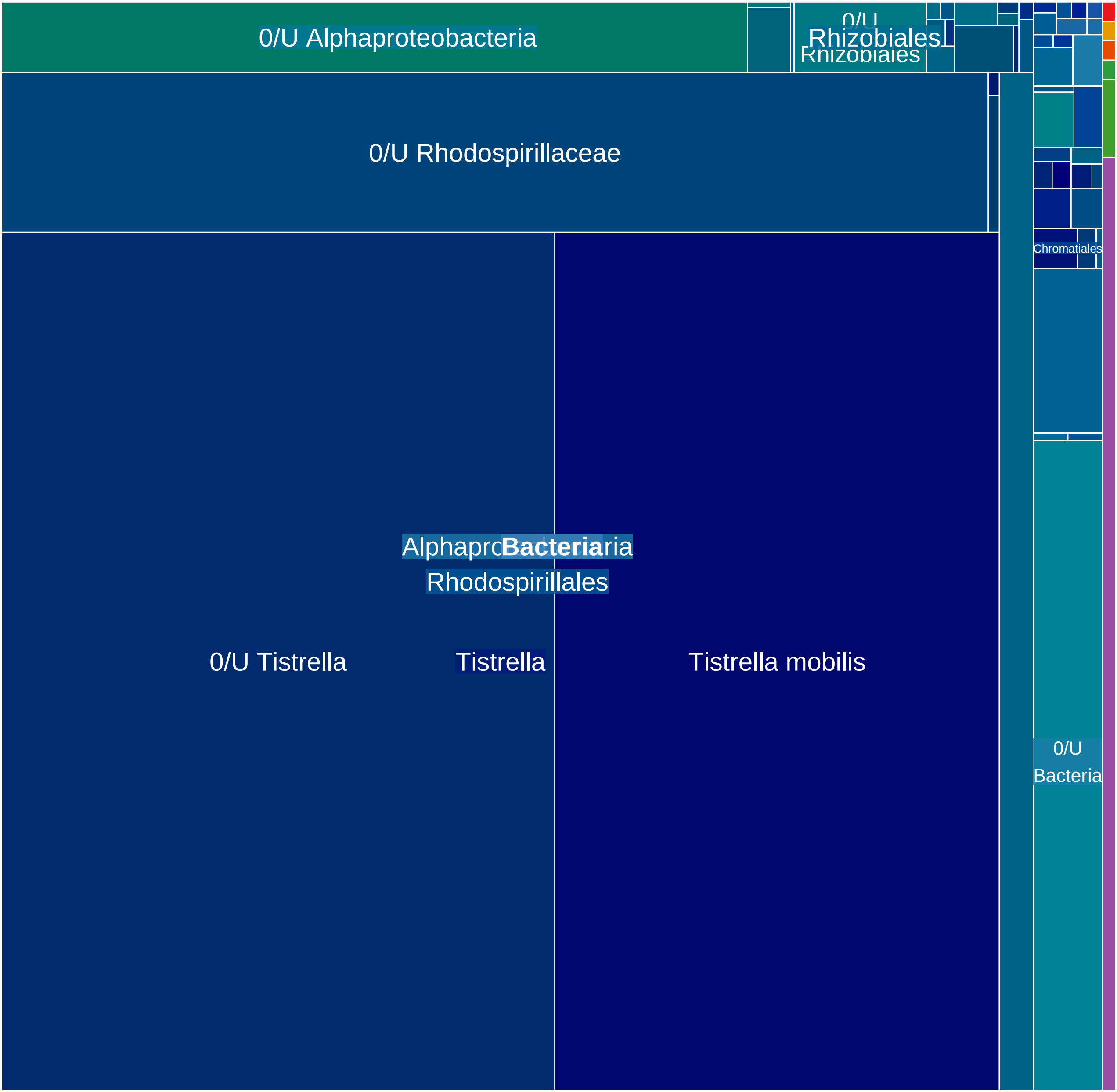

MGT−v1_247 : 5039 unigenes, 2958 (58.70%) taxonomically assigned

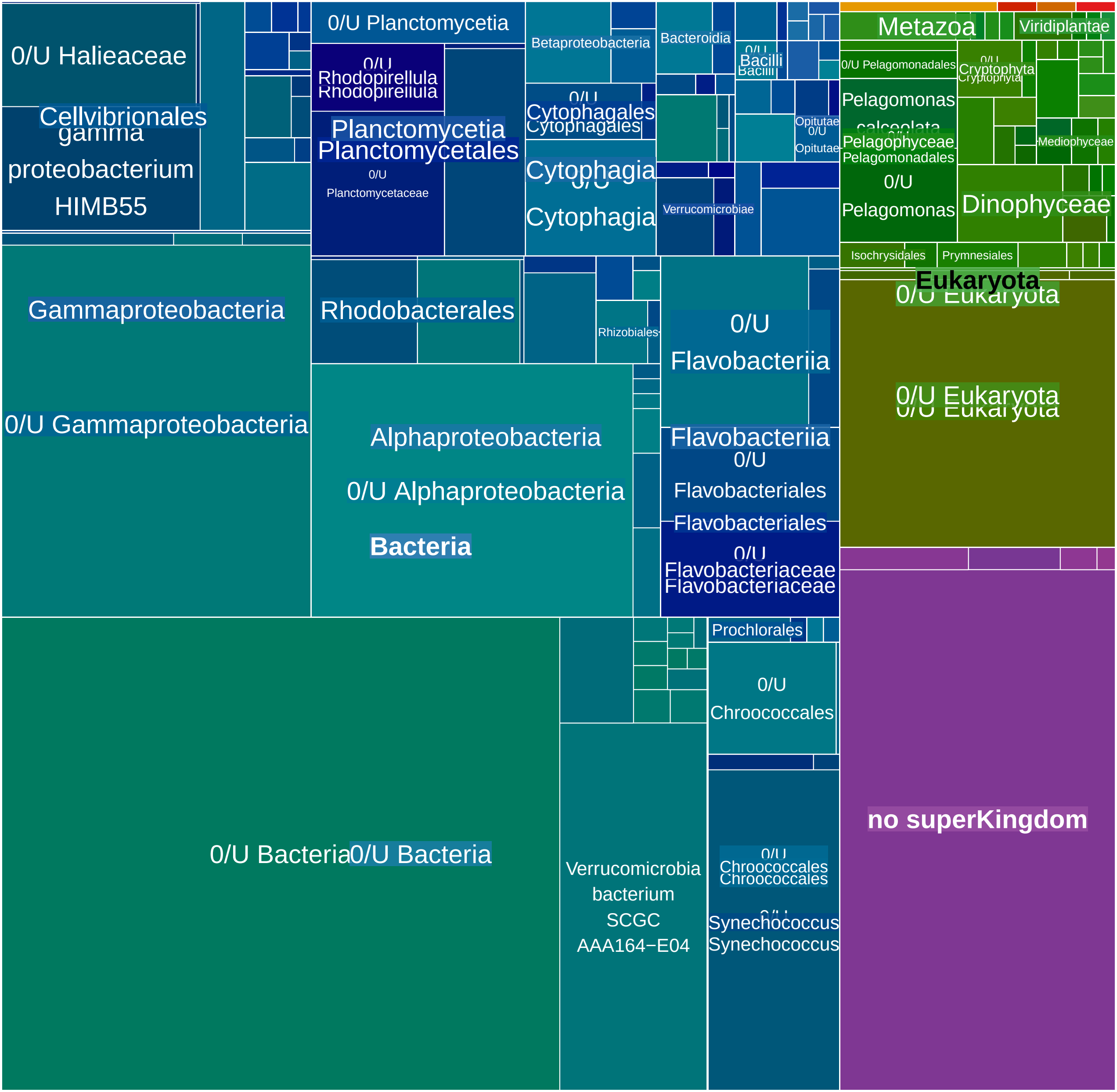

MGT−v1_248 : 4981 unigenes, 488 (9.80%) taxonomically assigned

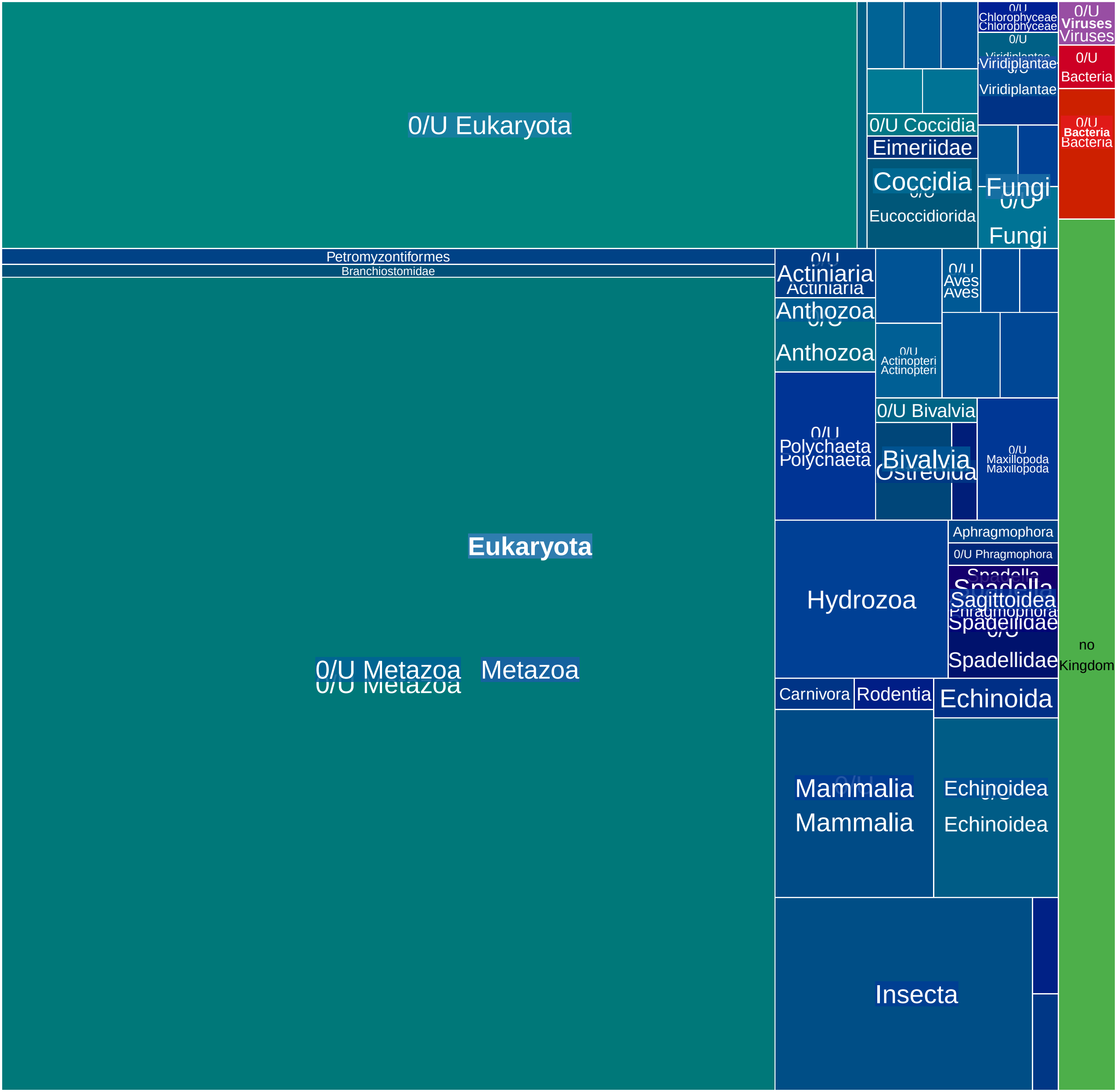

MGT−v1_249 : 4980 unigenes, 3882 (77.95%) taxonomically assigned

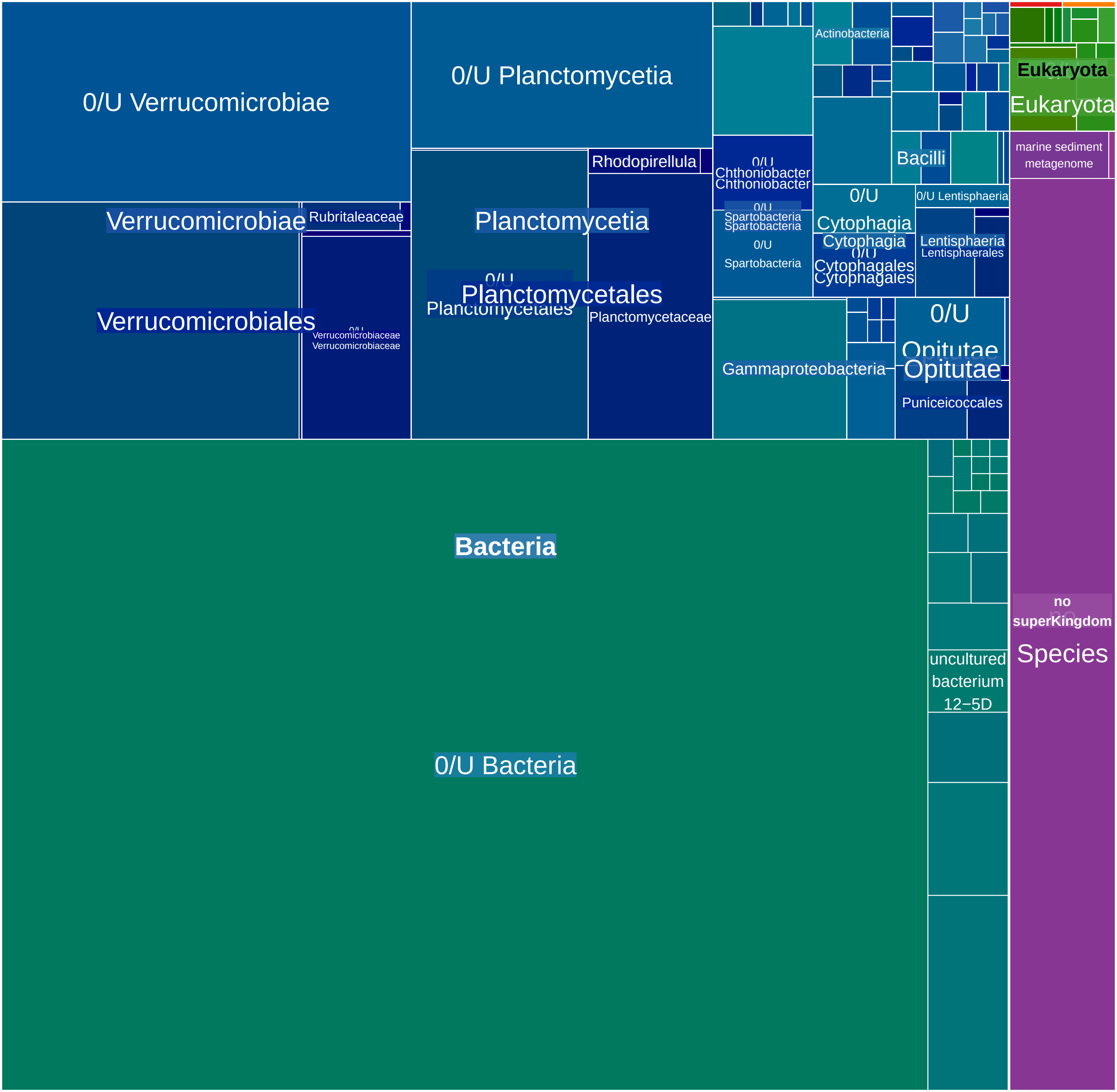

MGT−v1_250 : 4893 unigenes, 541 (11.06%) taxonomically assigned

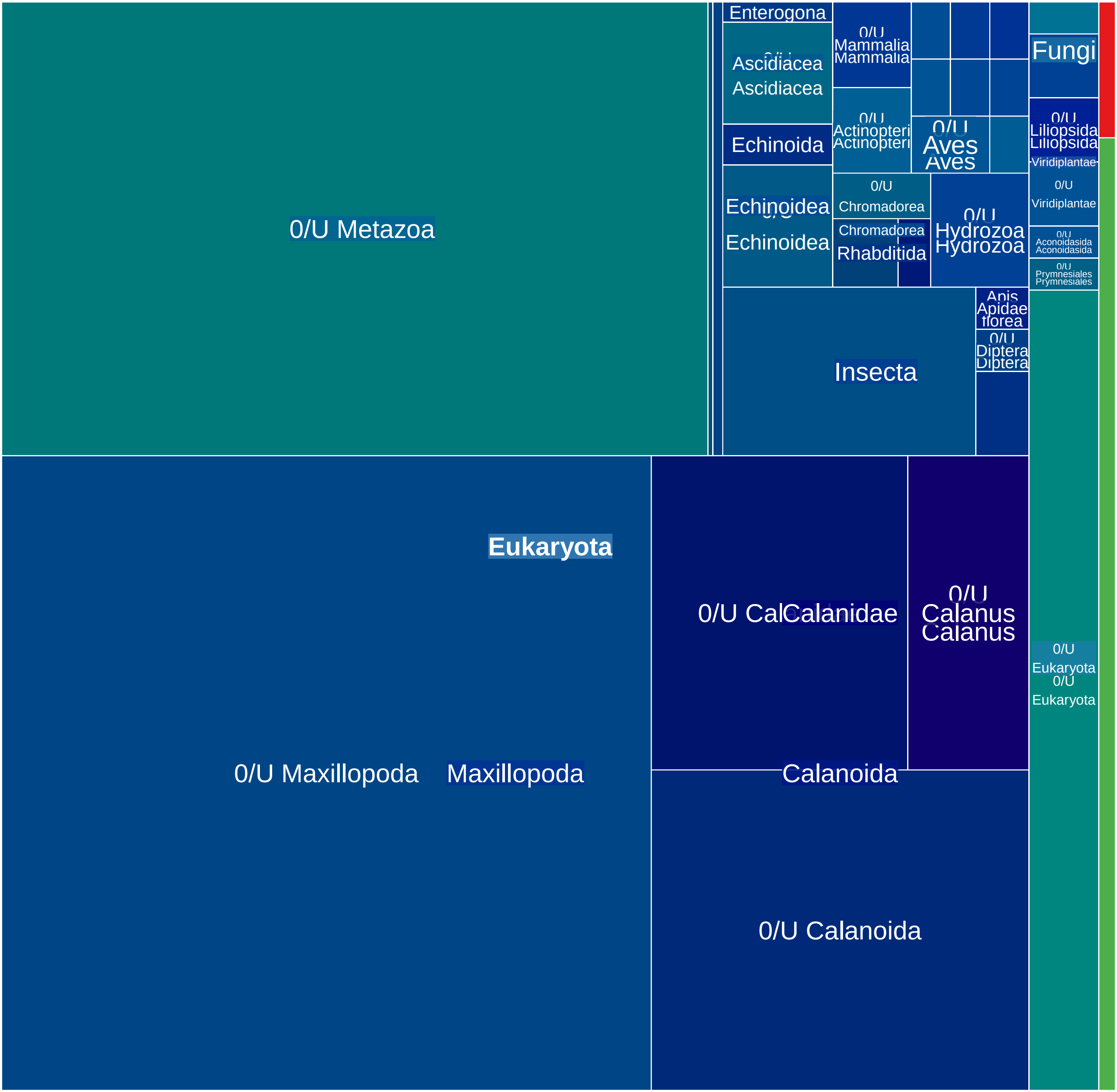

MGT−v1_251 : 4889 unigenes, 866 (17.71%) taxonomically assigned

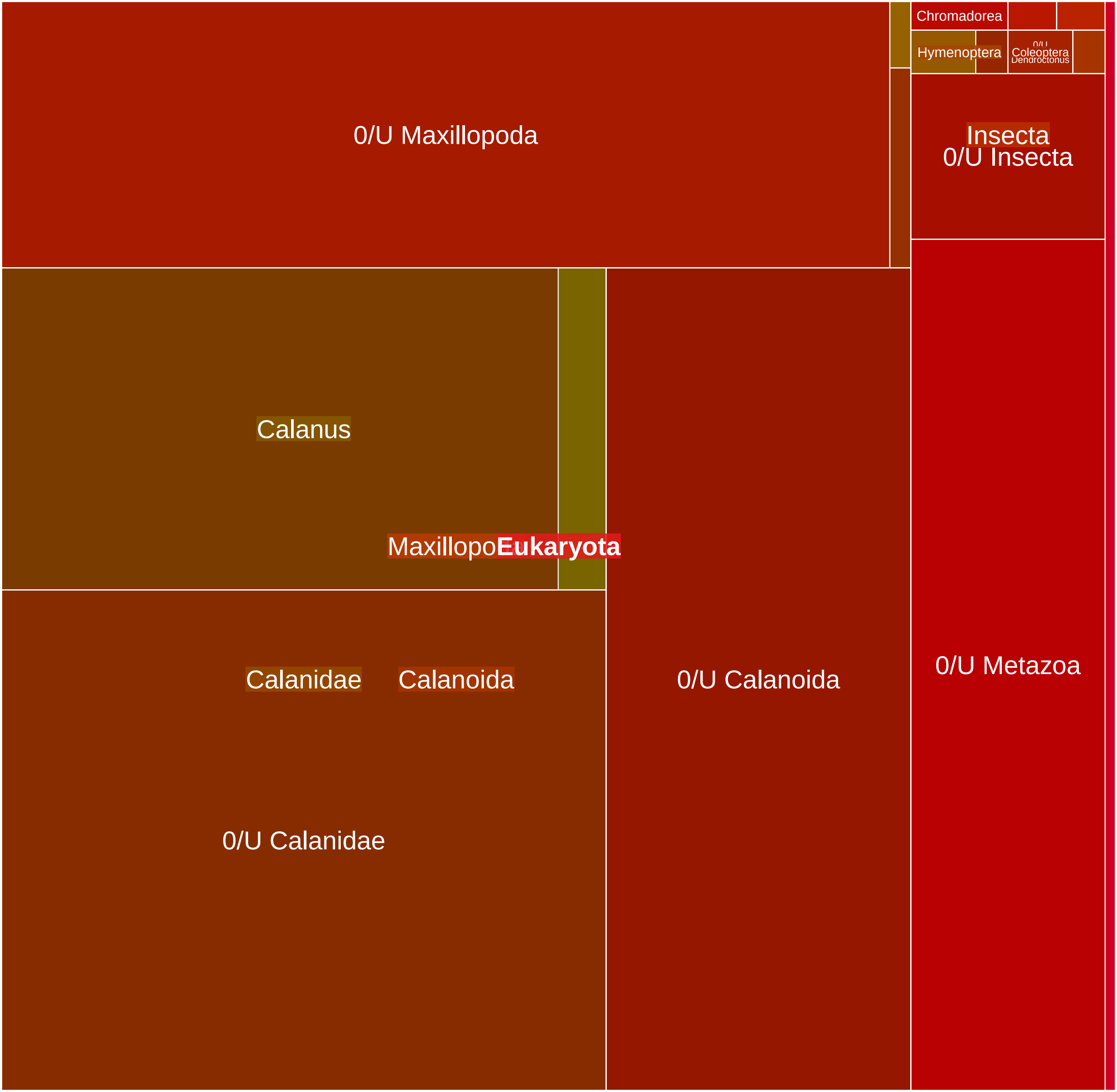

MGT−v1_252 : 4881 unigenes, 1092 (22.37%) taxonomically assigned

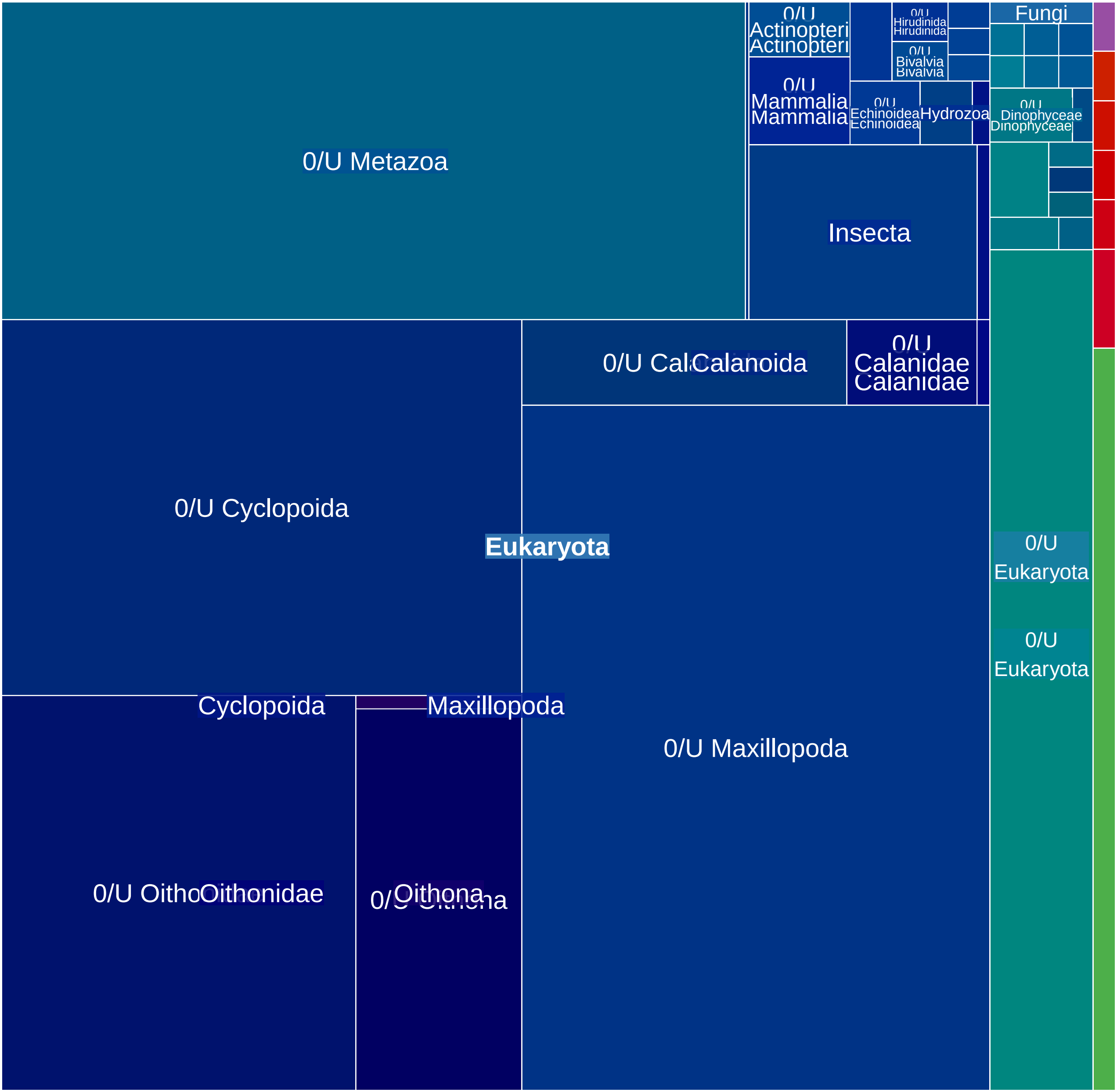

MGT−v1_253 : 4857 unigenes, 1780 (36.65%) taxonomically assigned

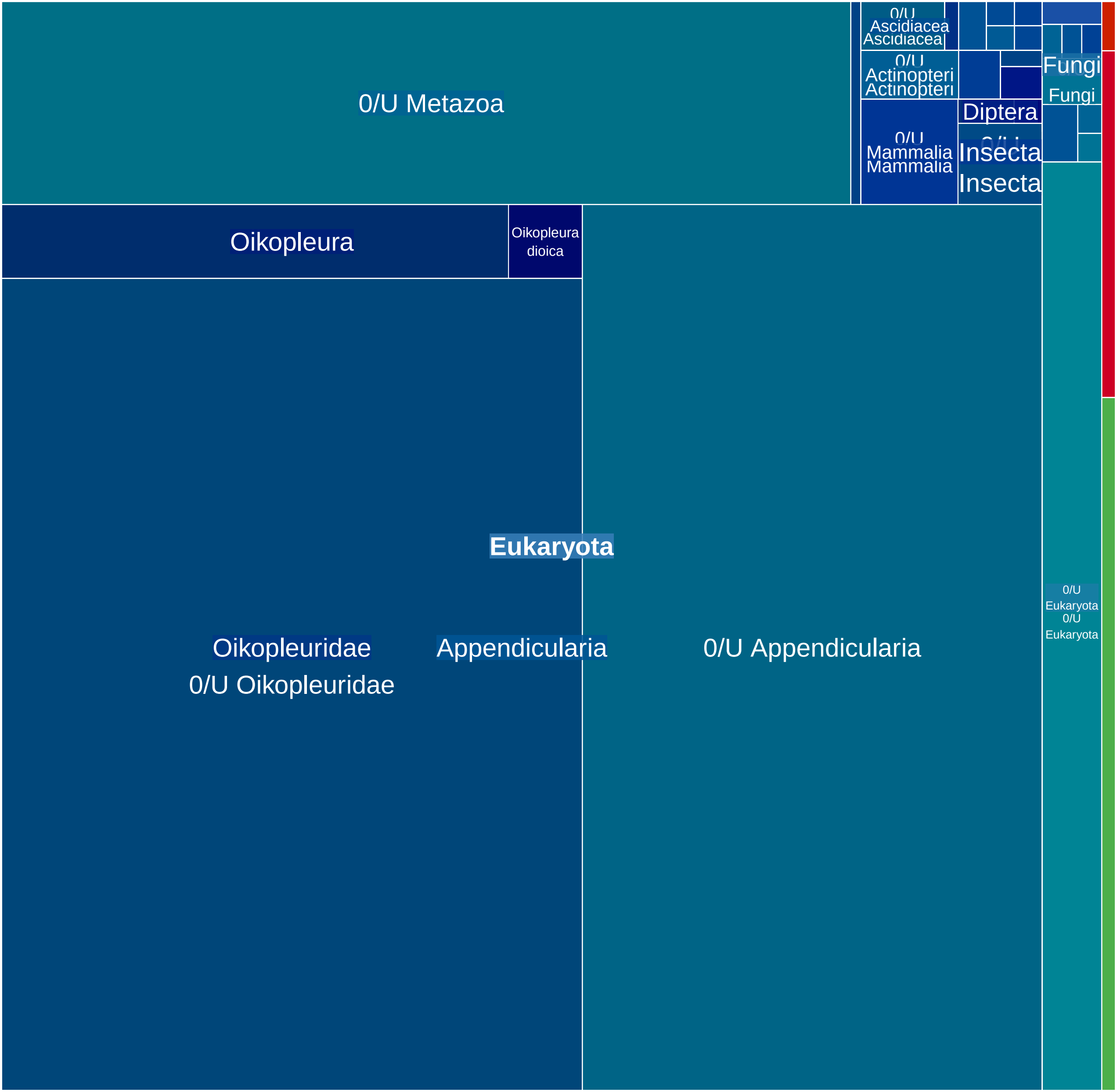

### MGT−v1_254 : 4787 unigenes, 4035 (84.29%) taxonomically assigned

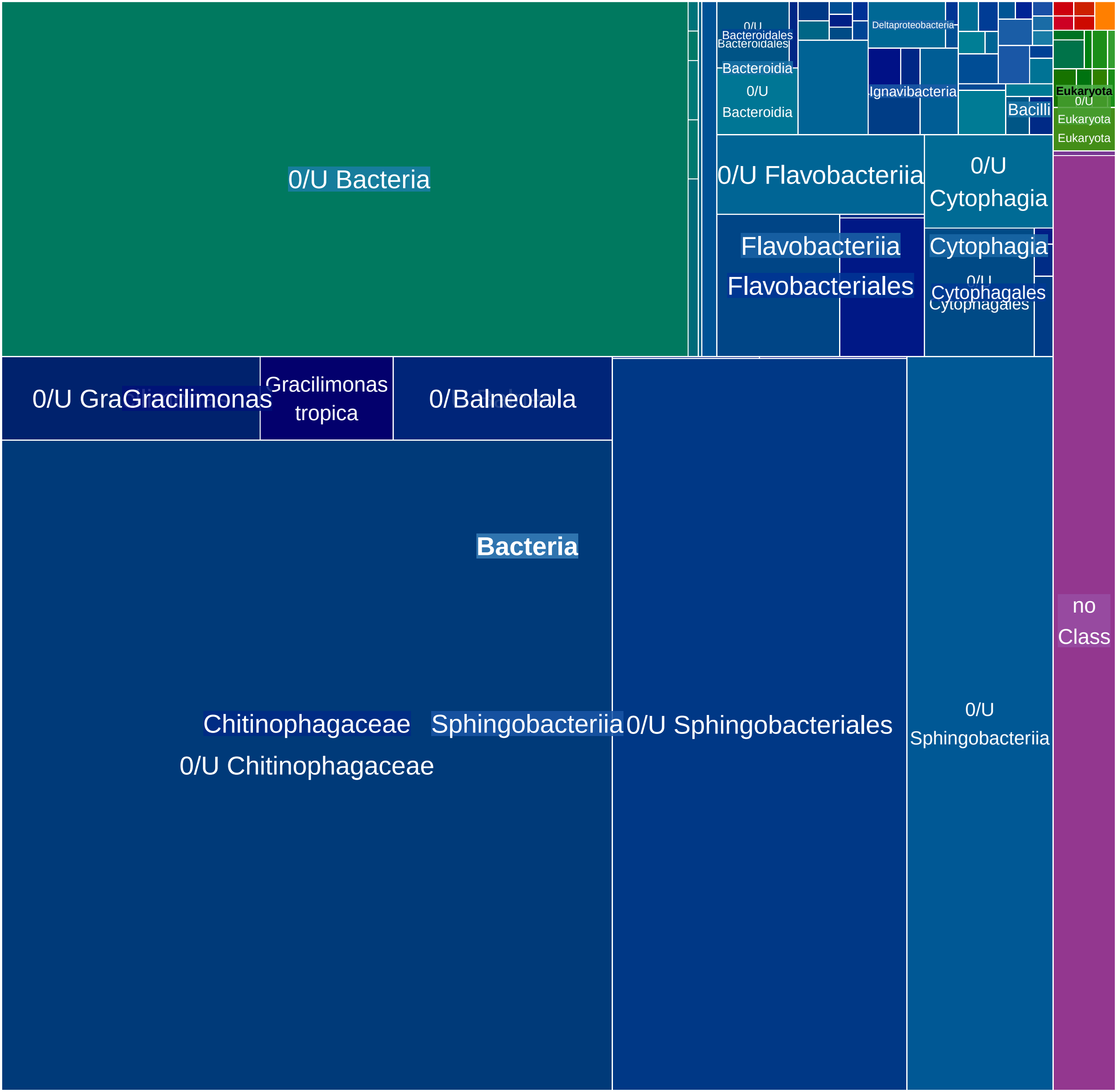

MGT−v1_255 : 4743 unigenes, 962 (20.28%) taxonomically assigned

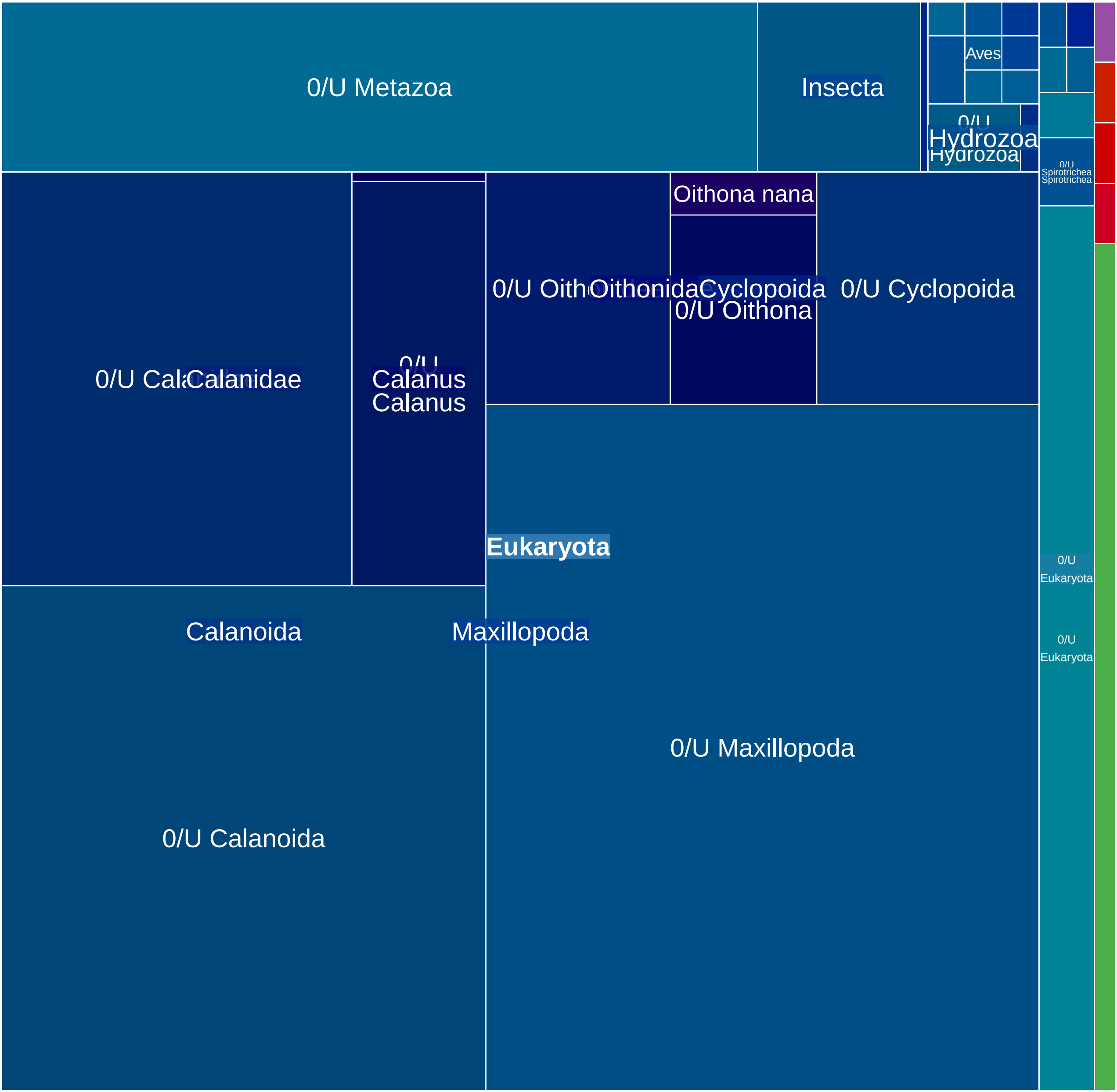

### MGT−v1_256 : 4737 unigenes, 2940 (62.06%) taxonomically assigned

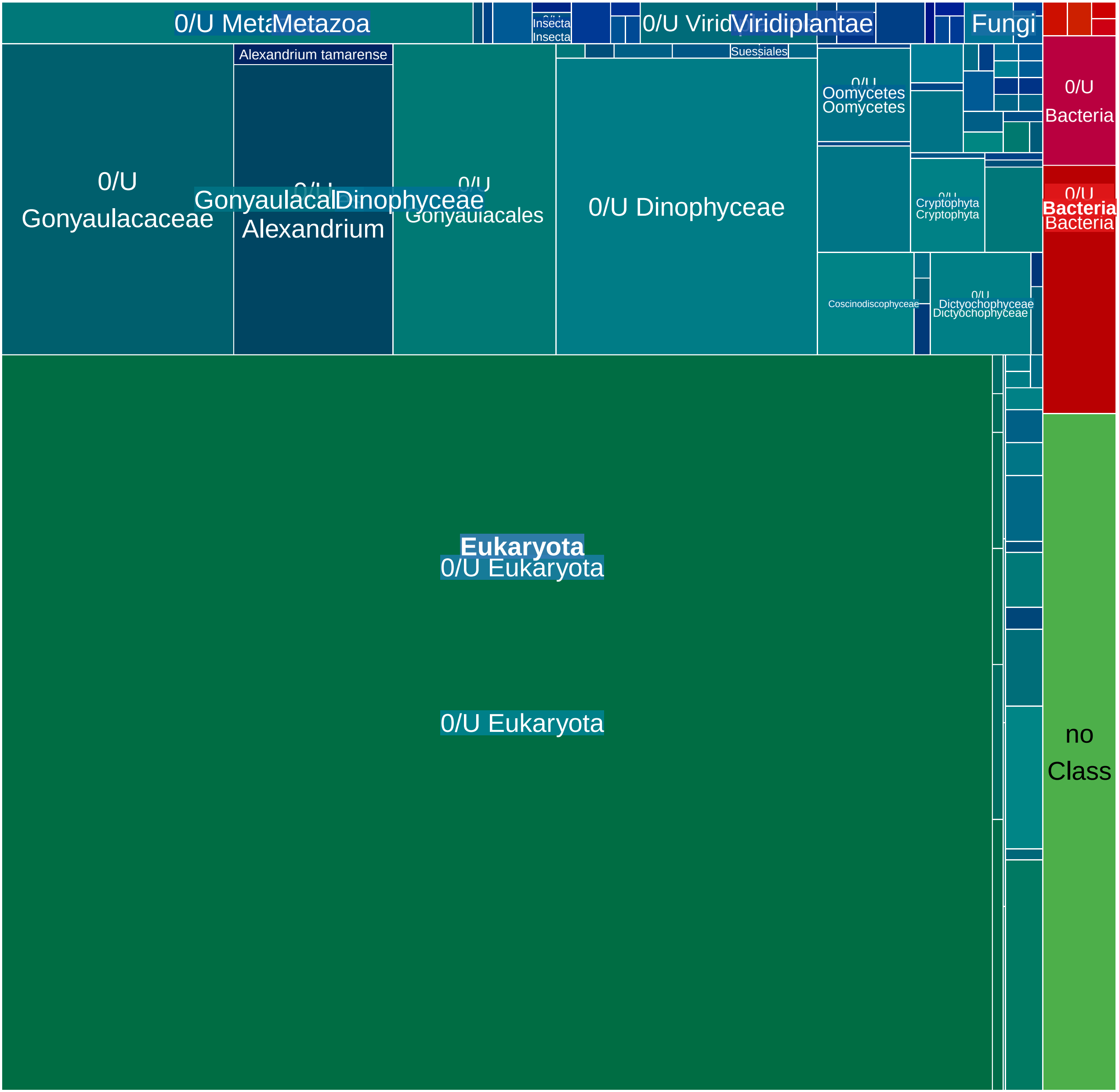

### MGT−v1_257 : 4729 unigenes, 2839 (60.03%) taxonomically assigned

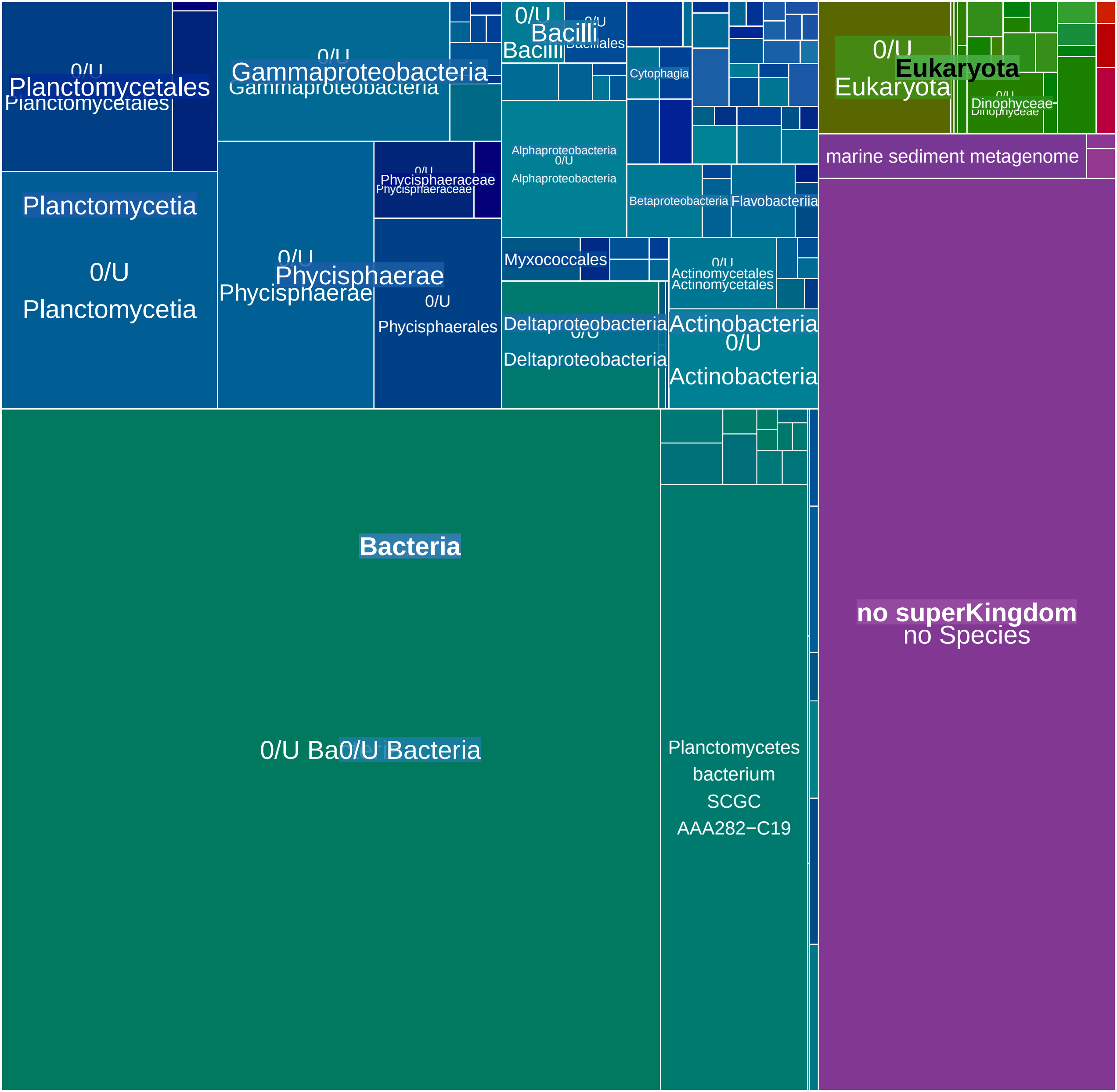

MGT−v1_258 : 4724 unigenes, 3143 (66.53%) taxonomically assigned

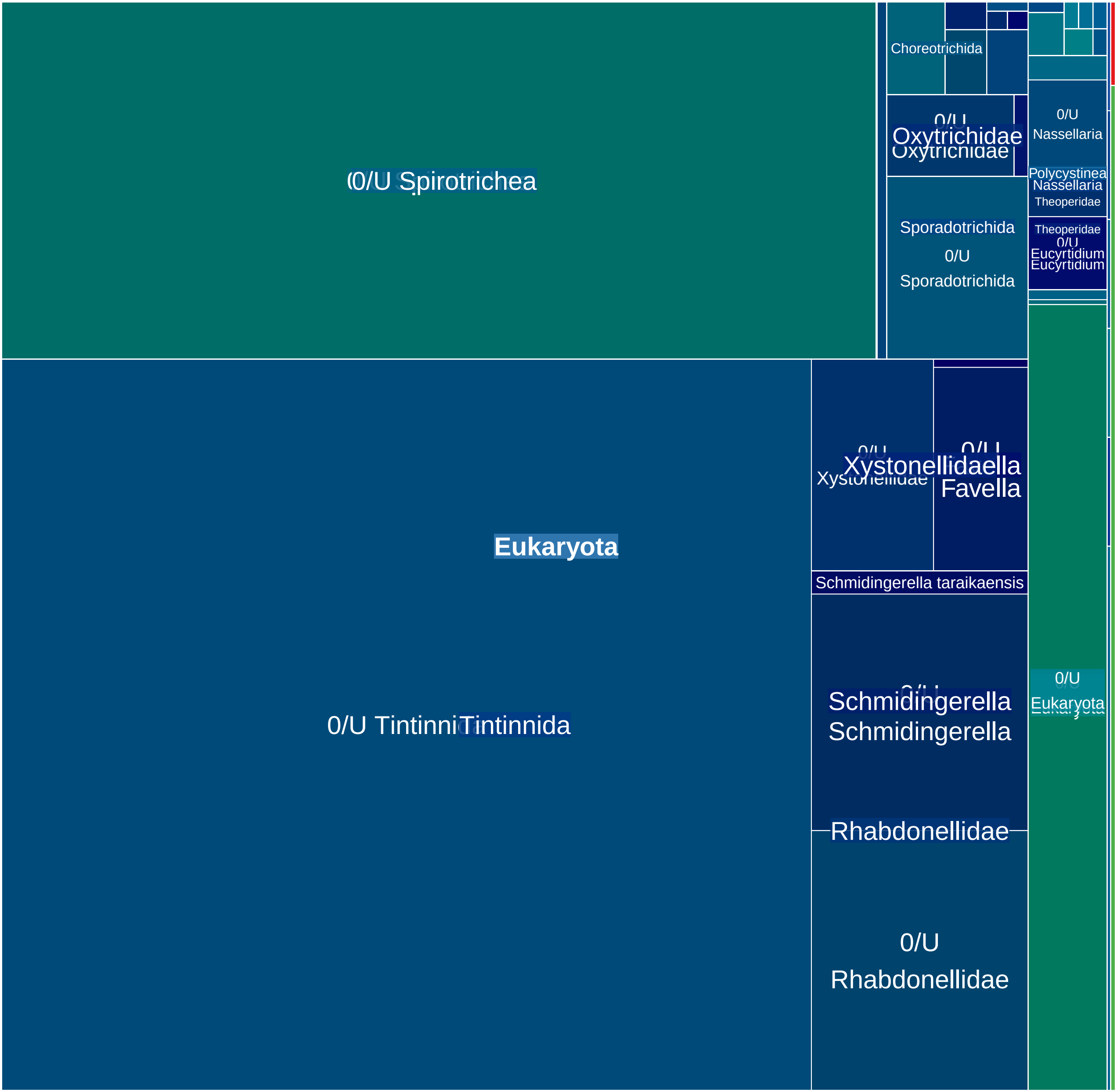

MGT−v1_259 : 4707 unigenes, 651 (13.83%) taxonomically assigned

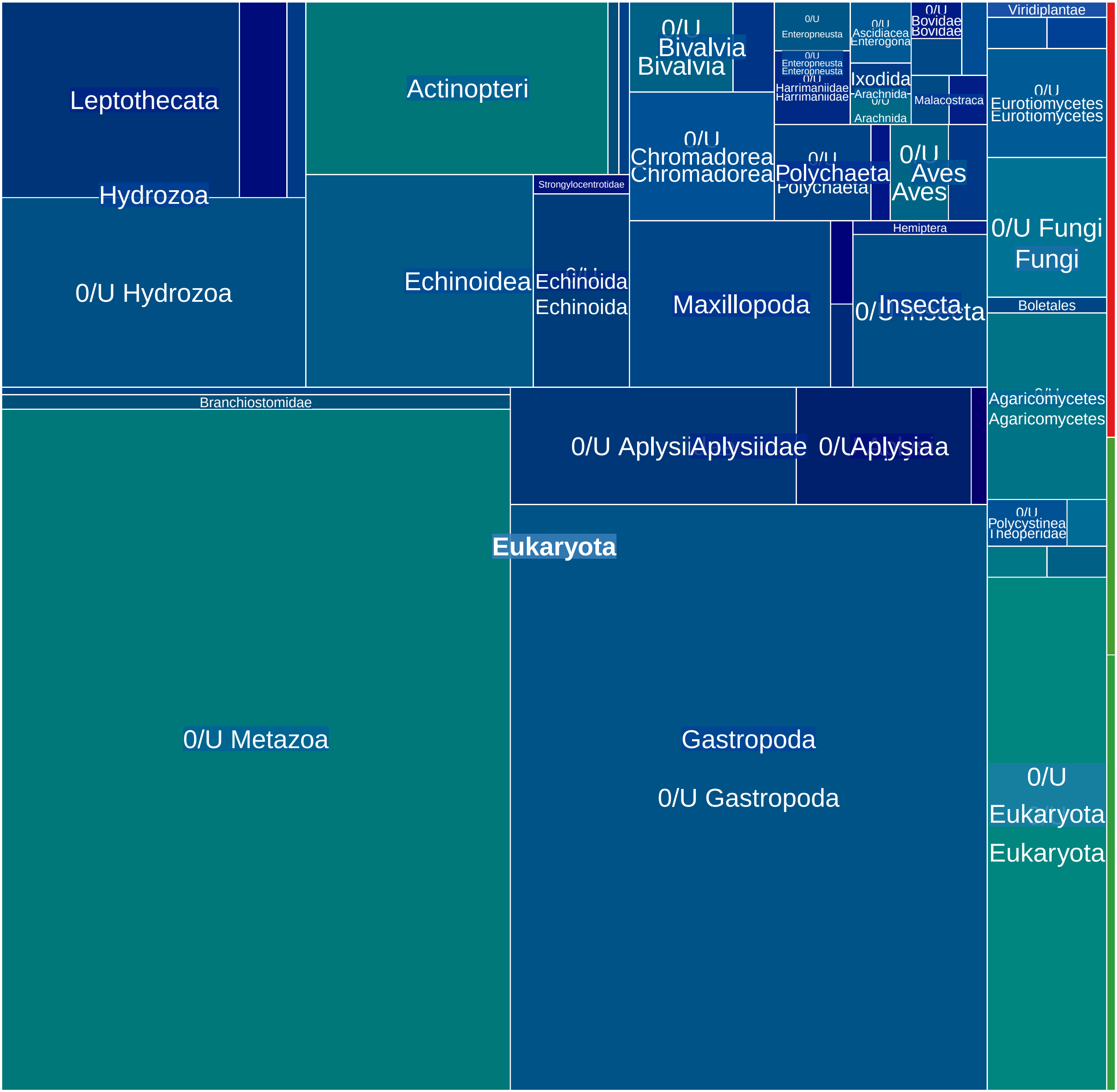

MGT−v1_260 : 4688 unigenes, 1070 (22.82%) taxonomically assigned

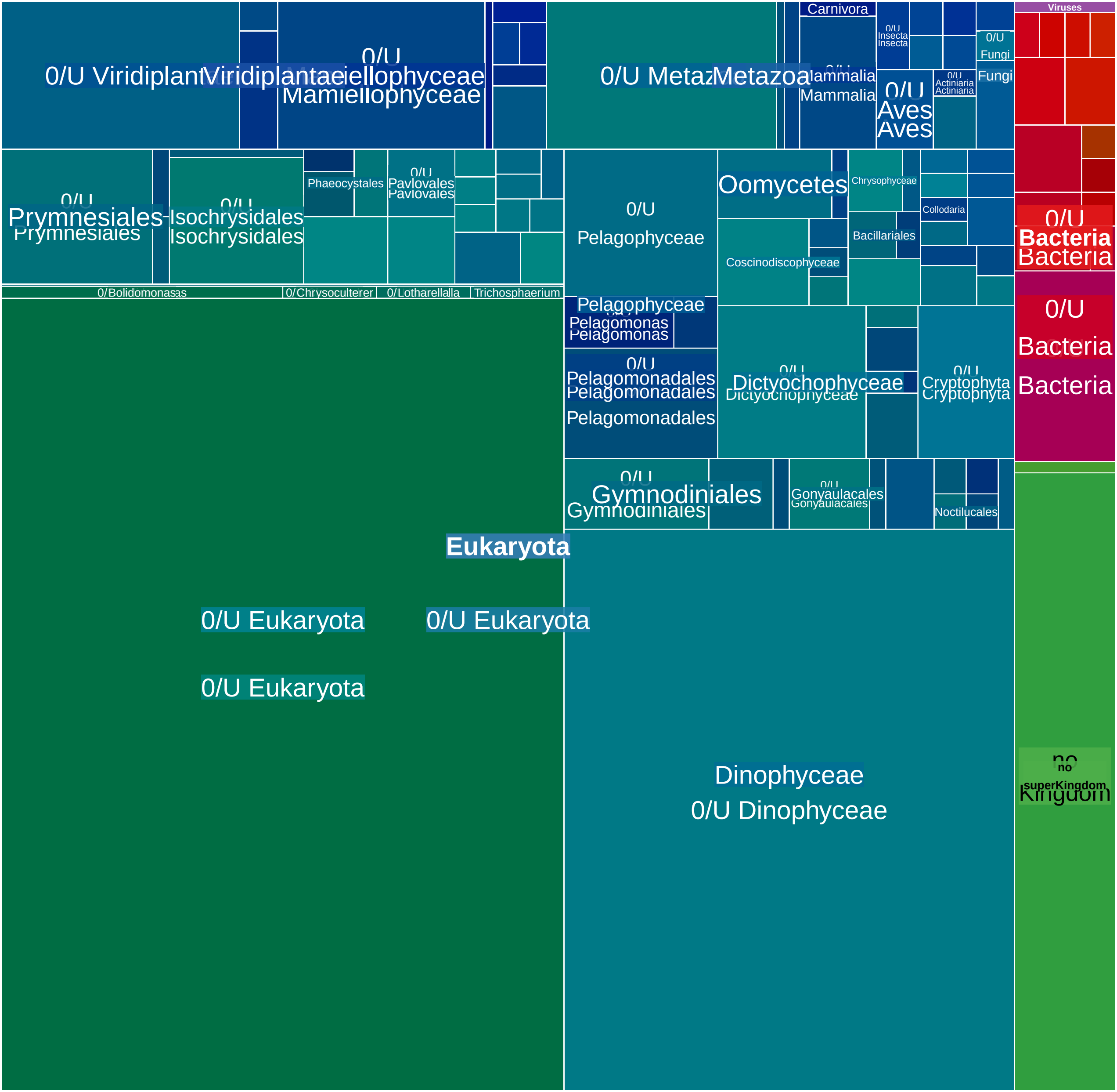

MGT−v1_261 : 4671 unigenes, 2970 (63.58%) taxonomically assigned

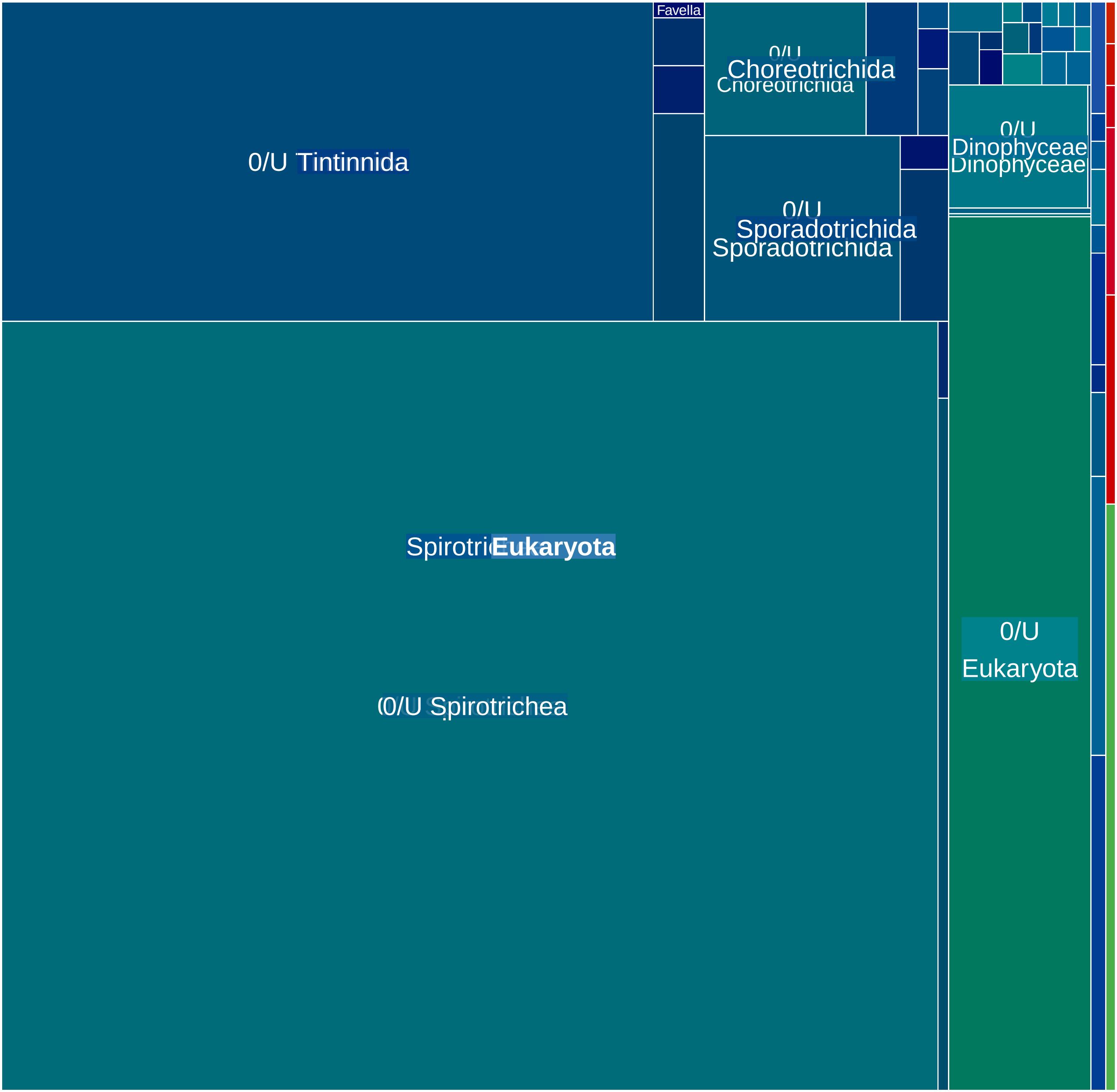

MGT−v1_262 : 4631 unigenes, 4381 (94.60%) taxonomically assigned

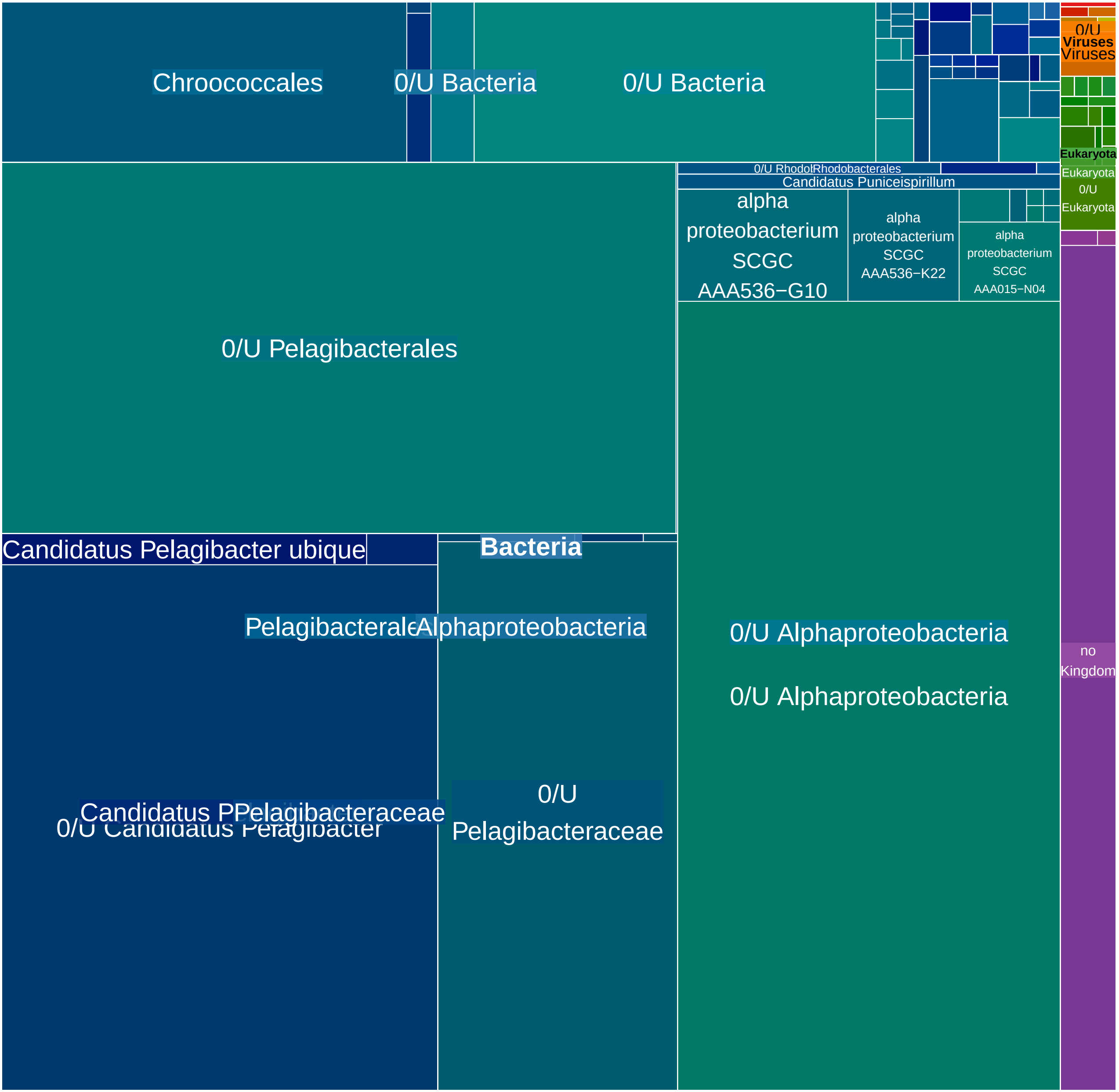

MGT−v1_263 : 4539 unigenes, 274 (6.04%) taxonomically assigned

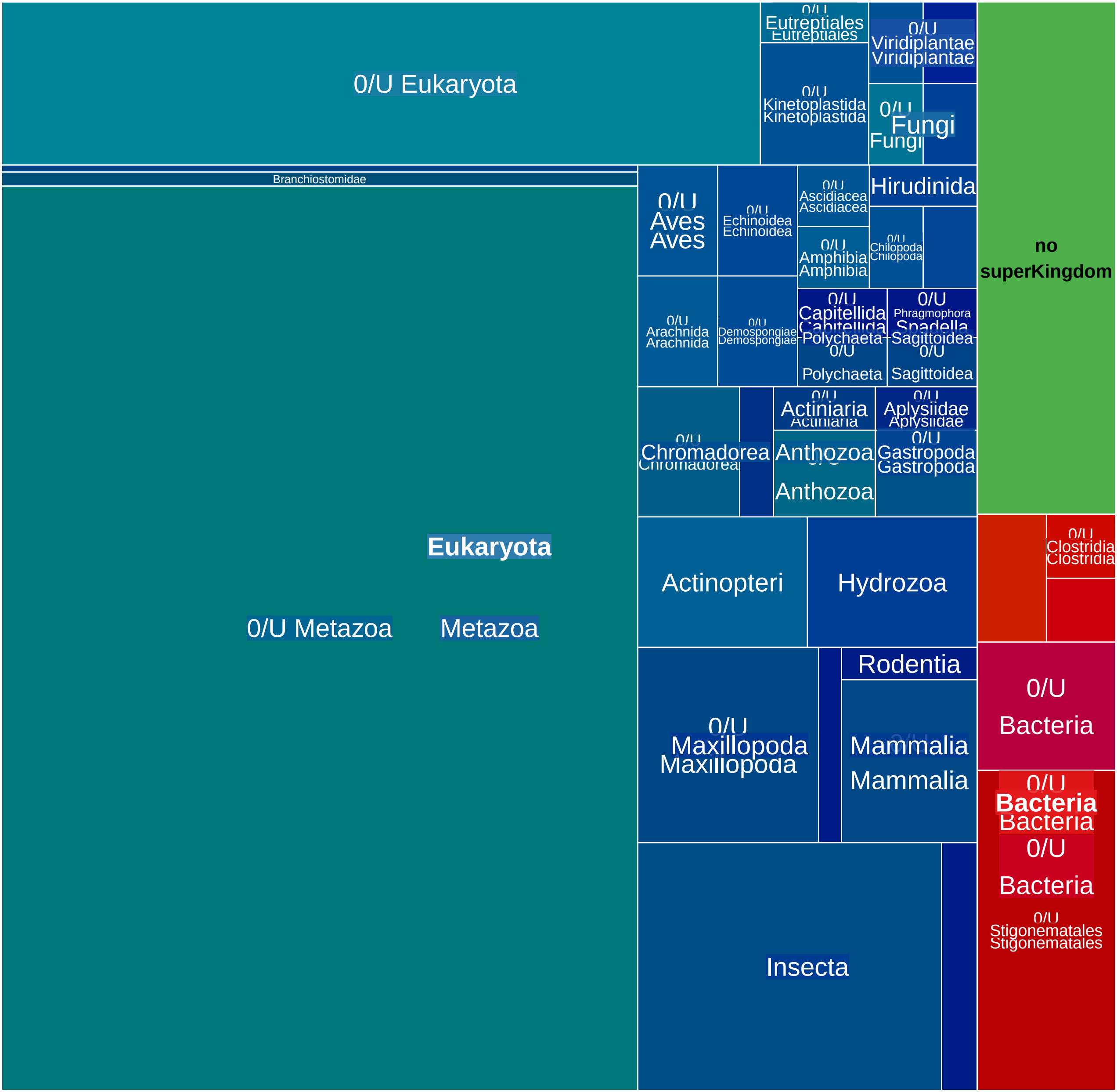

MGT−v1_264 : 4519 unigenes, 3974 (87.94%) taxonomically assigned

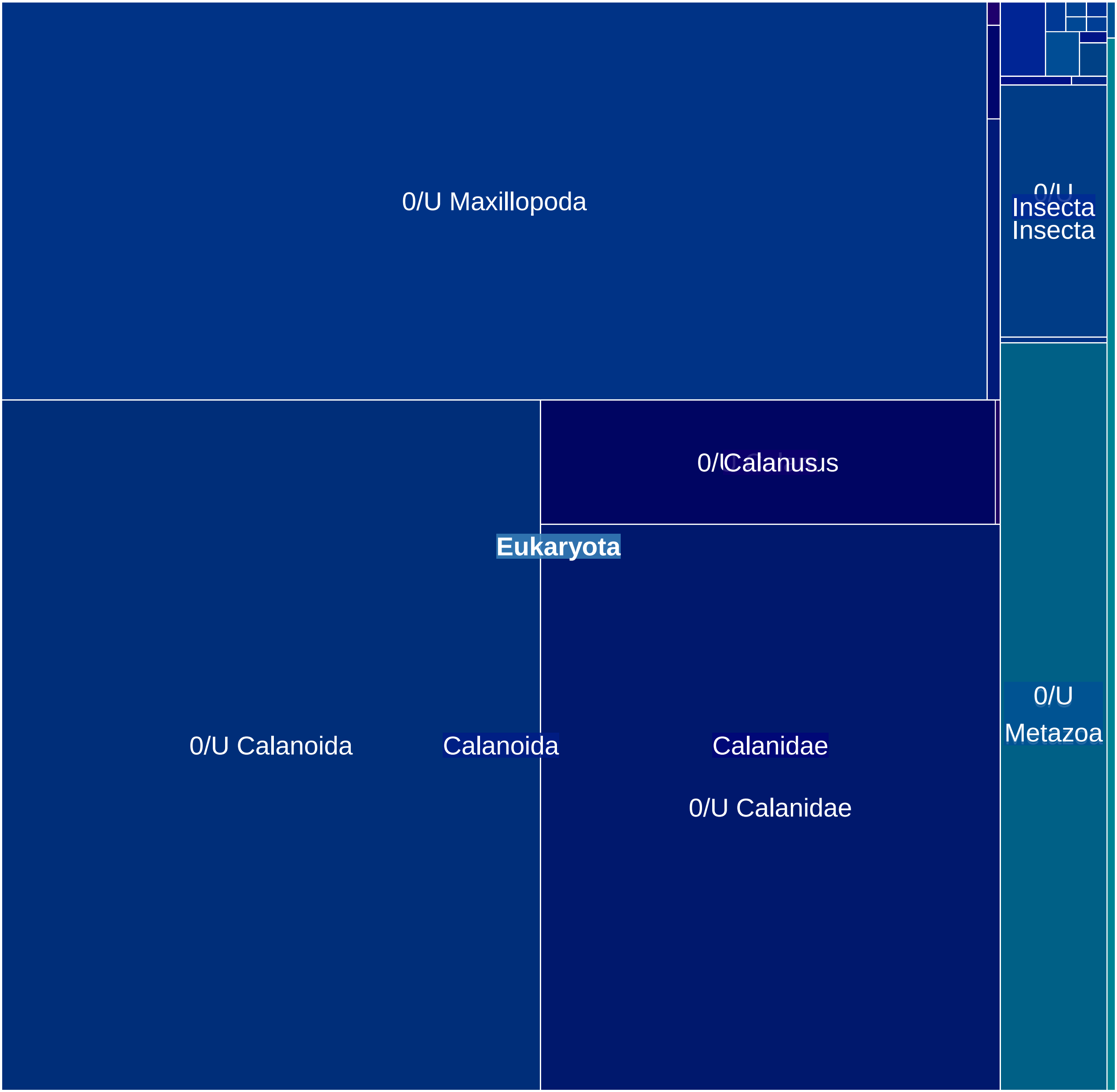

### MGT−v1_265 : 4491 unigenes, 620 (13.81%) taxonomically assigned

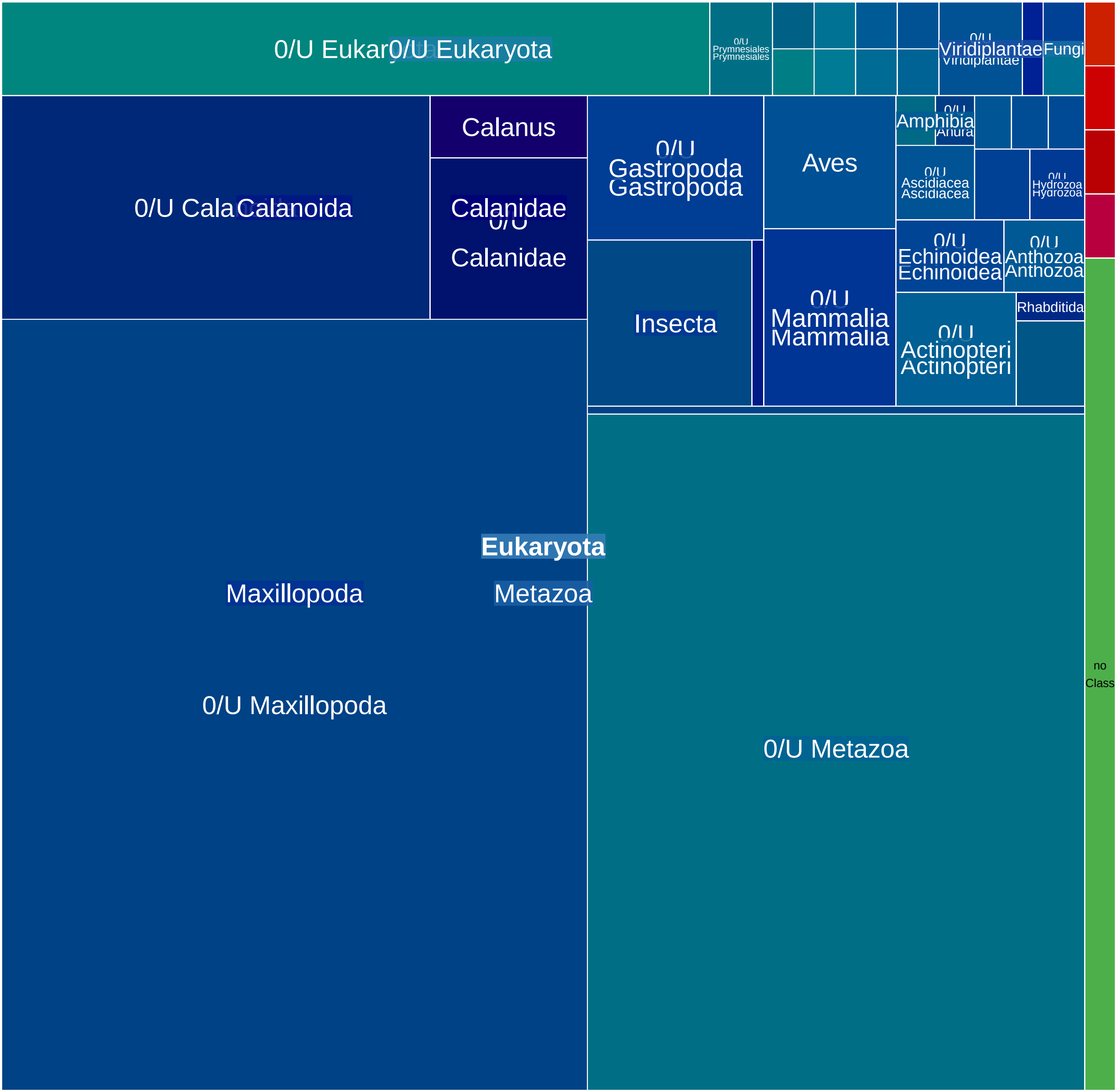

MGT−v1_266 : 4489 unigenes, 381 (8.49%) taxonomically assigned

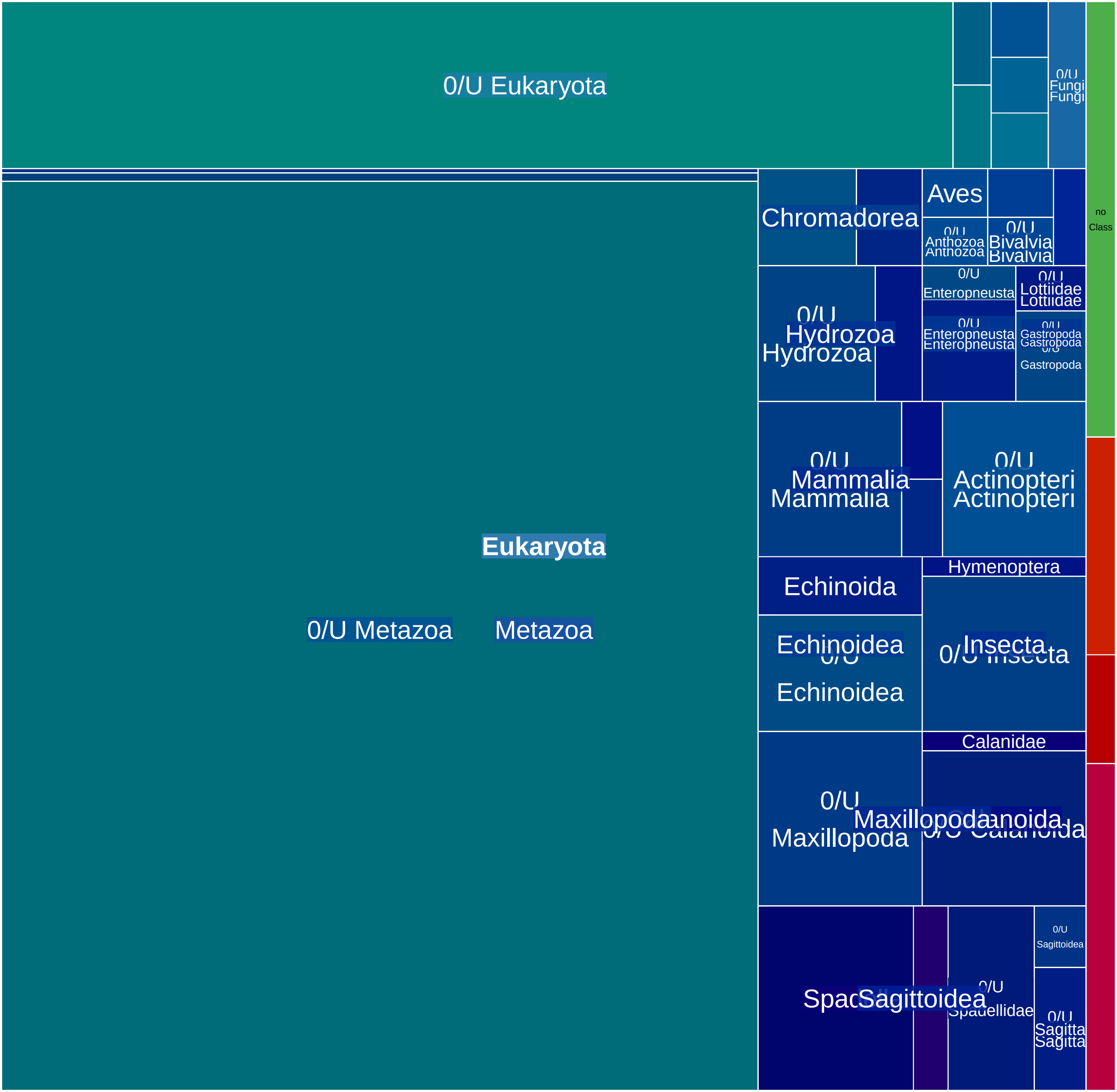

### MGT−v1_267 : 4470 unigenes, 921 (20.60%) taxonomically assigned

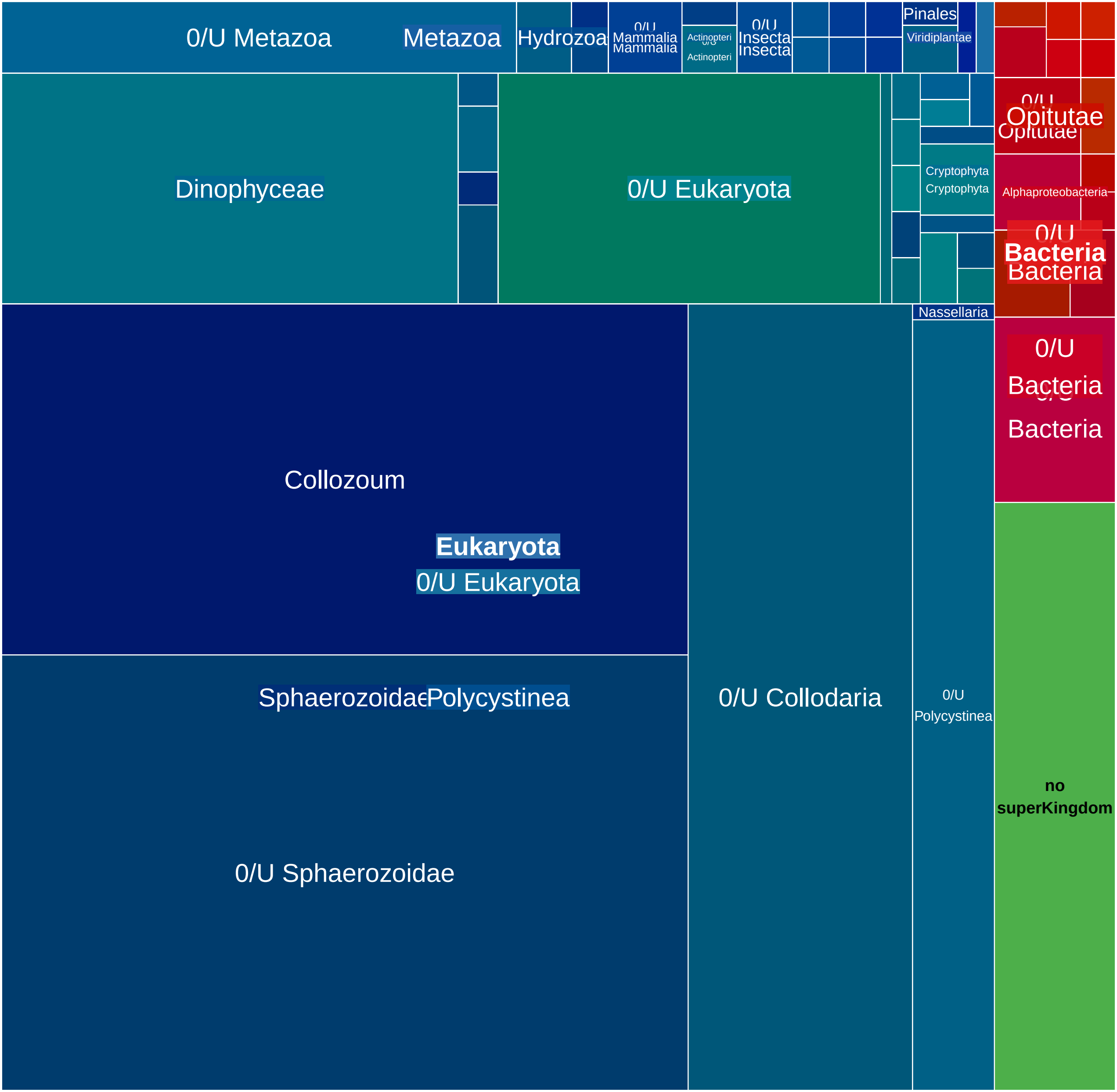

MGT−v1_268 : 4453 unigenes, 3206 (72.00%) taxonomically assigned

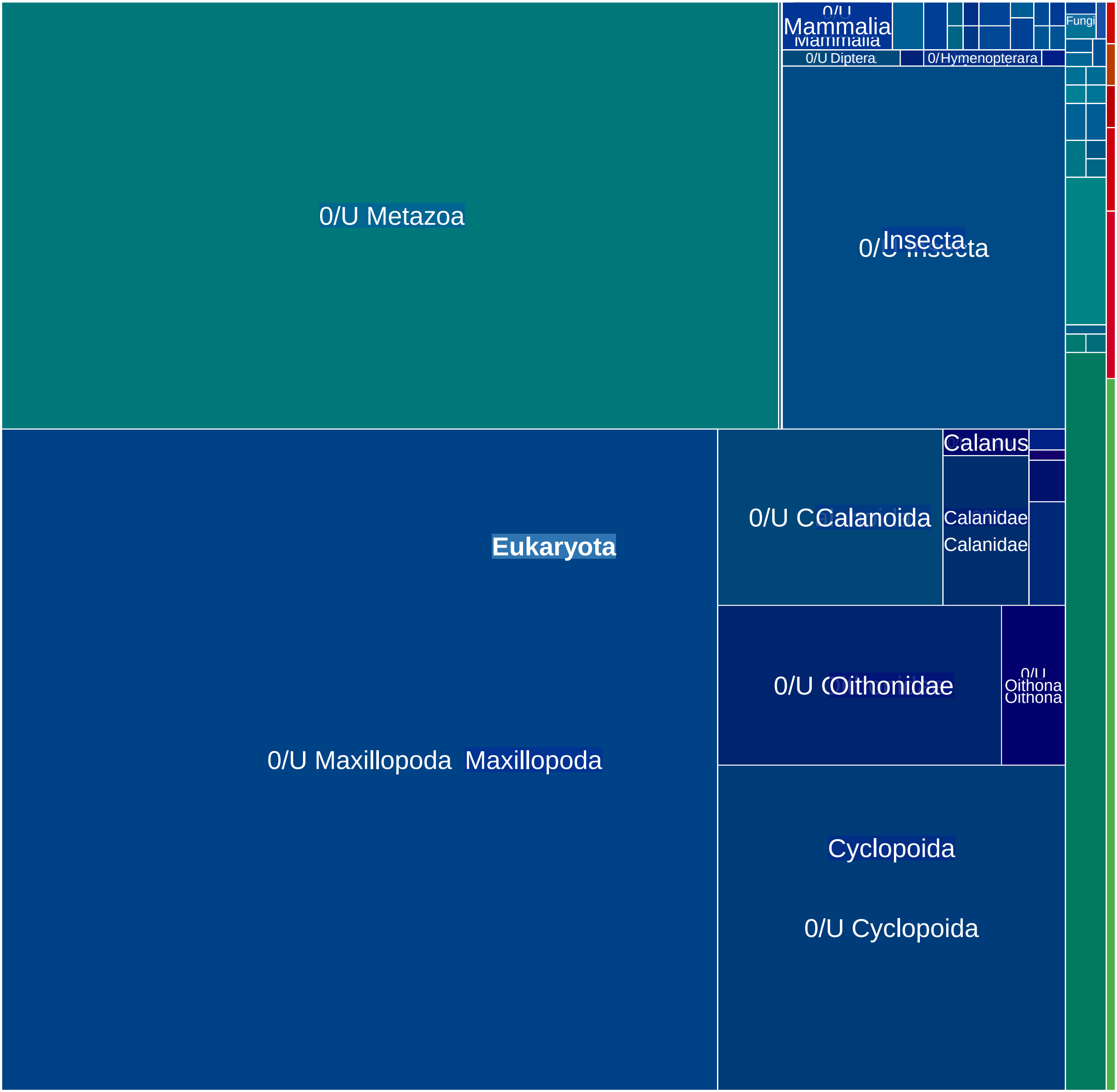

MGT−v1_269 : 4436 unigenes, 1249 (28.16%) taxonomically assigned

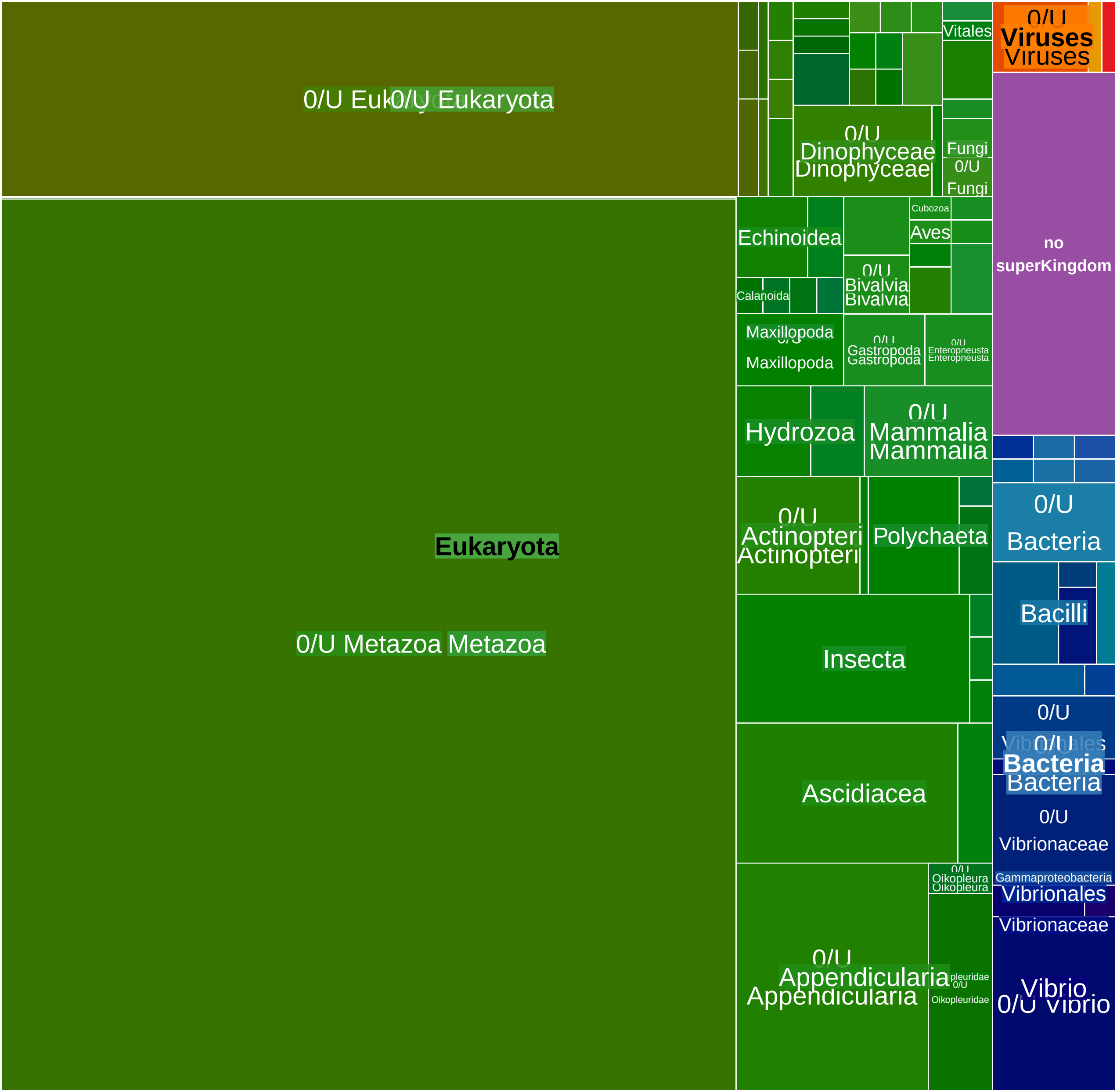

MGT−v1_270 : 4415 unigenes, 4283 (97.01%) taxonomically assigned

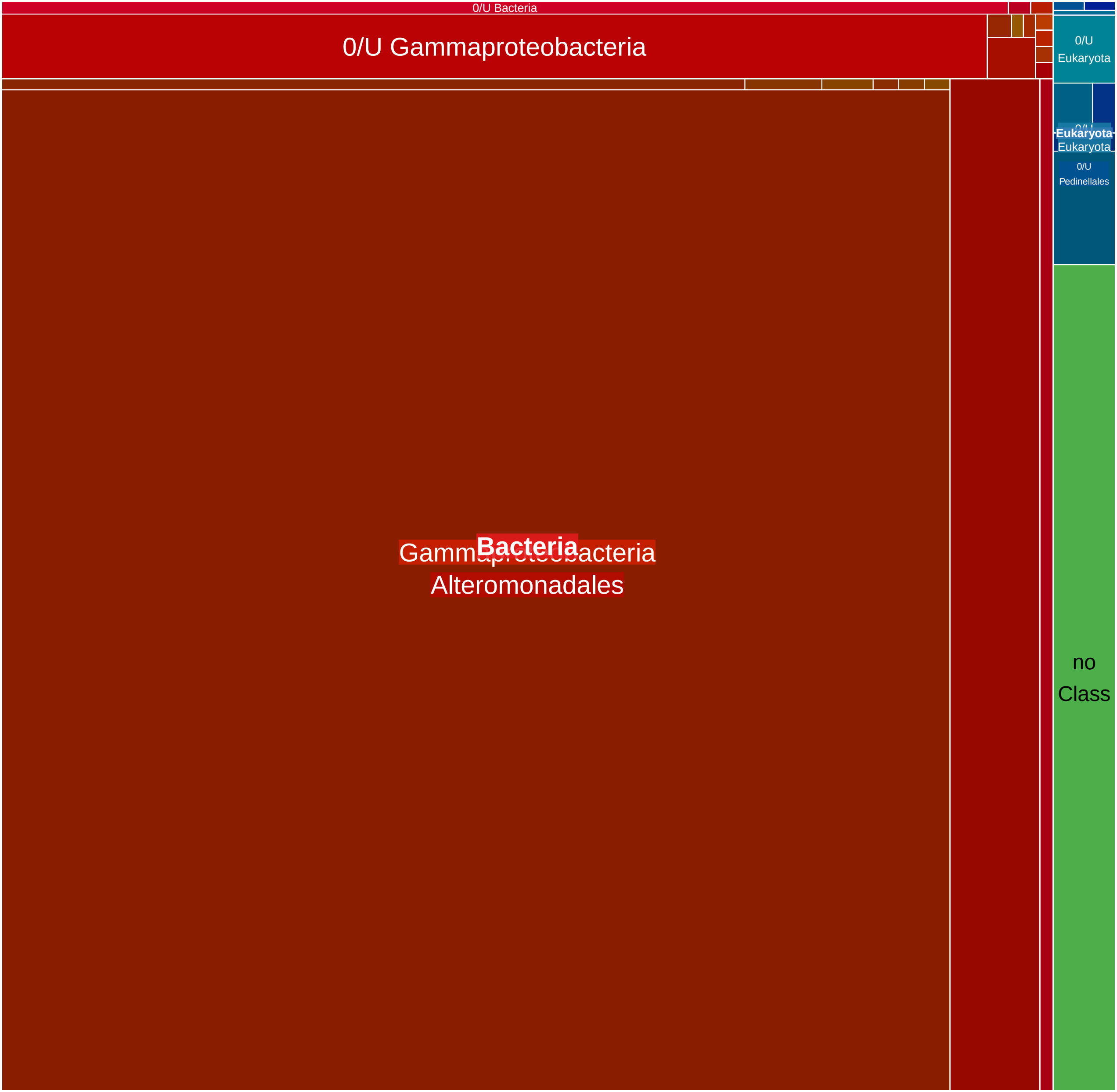

MGT−v1_271 : 4385 unigenes, 3259 (74.32%) taxonomically assigned

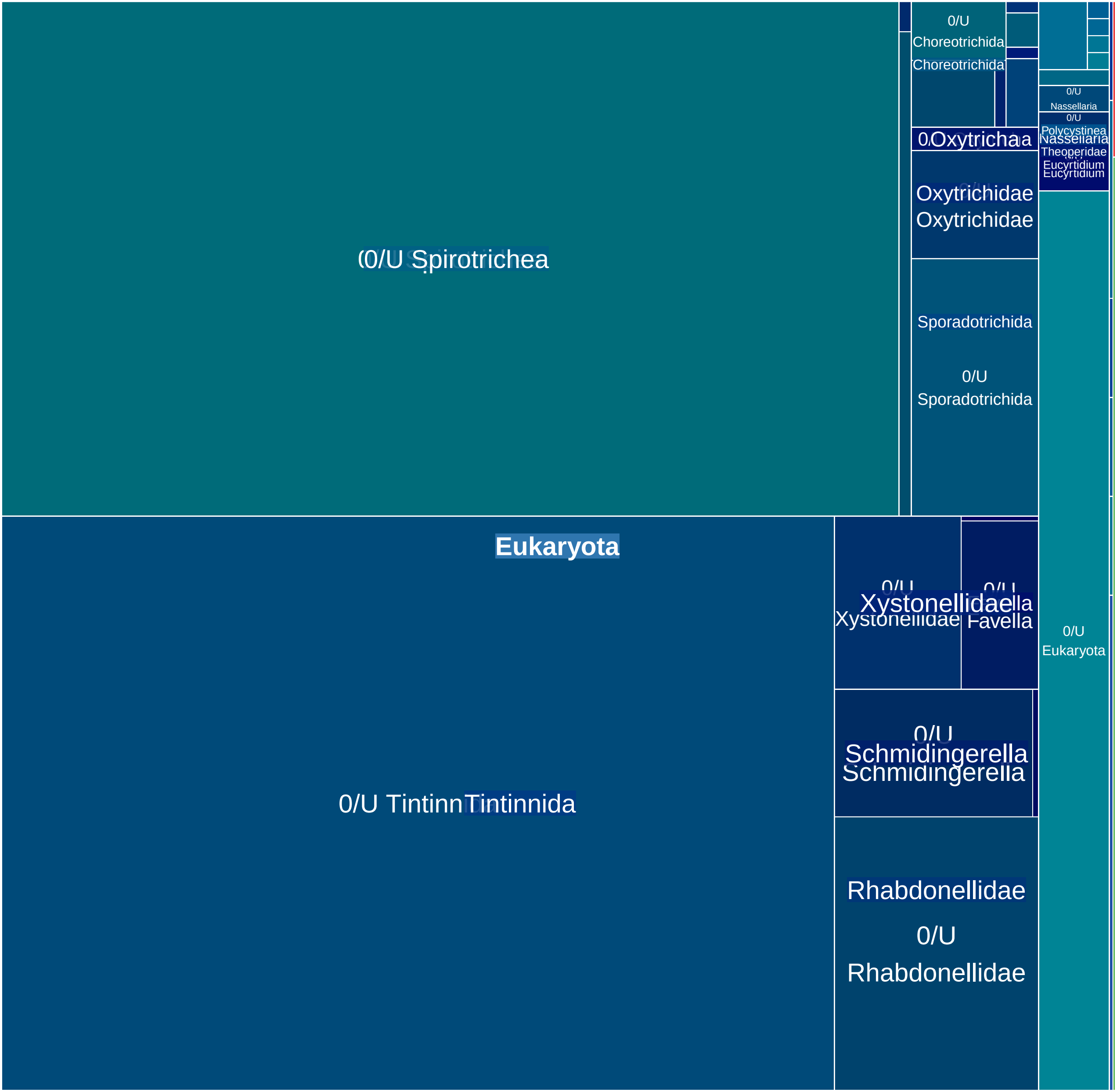

MGT−v1_272 : 4334 unigenes, 834 (19.24%) taxonomically assigned

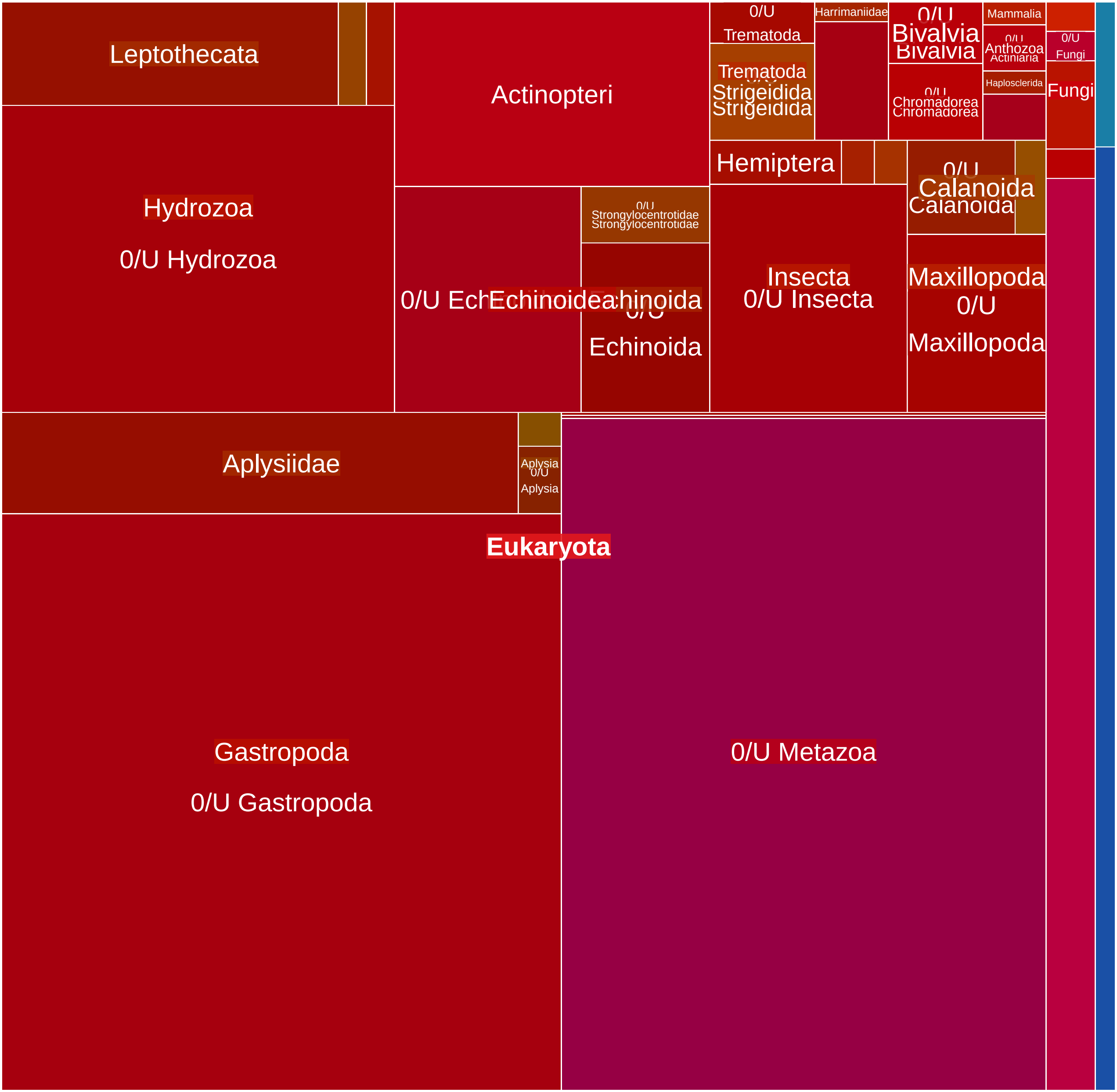

MGT−v1_273 : 4315 unigenes, 4170 (96.64%) taxonomically assigned

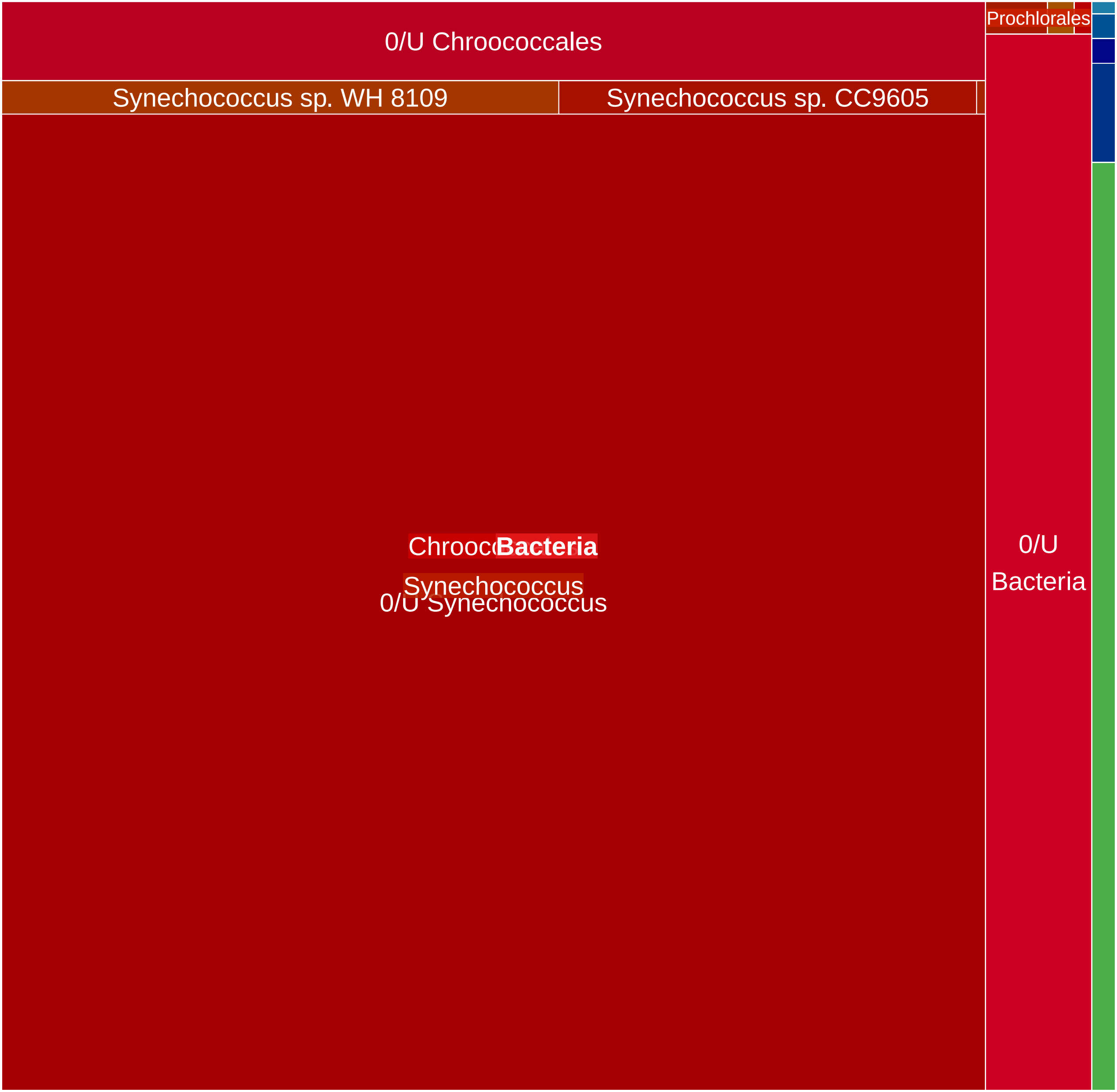

MGT−v1_274 : 4269 unigenes, 3993 (93.53%) taxonomically assigned

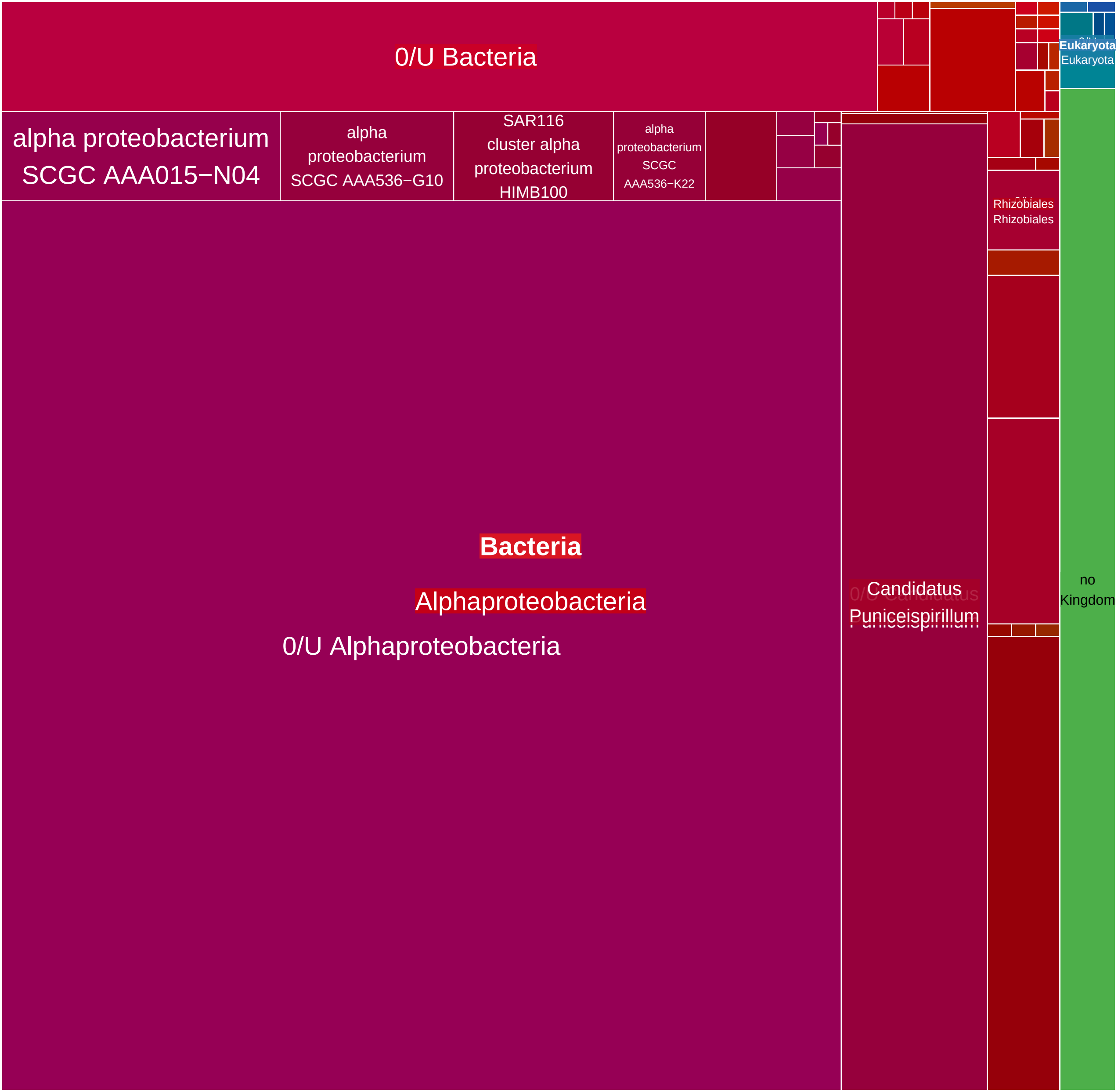

MGT−v1_275 : 4224 unigenes, 1277 (30.23%) taxonomically assigned

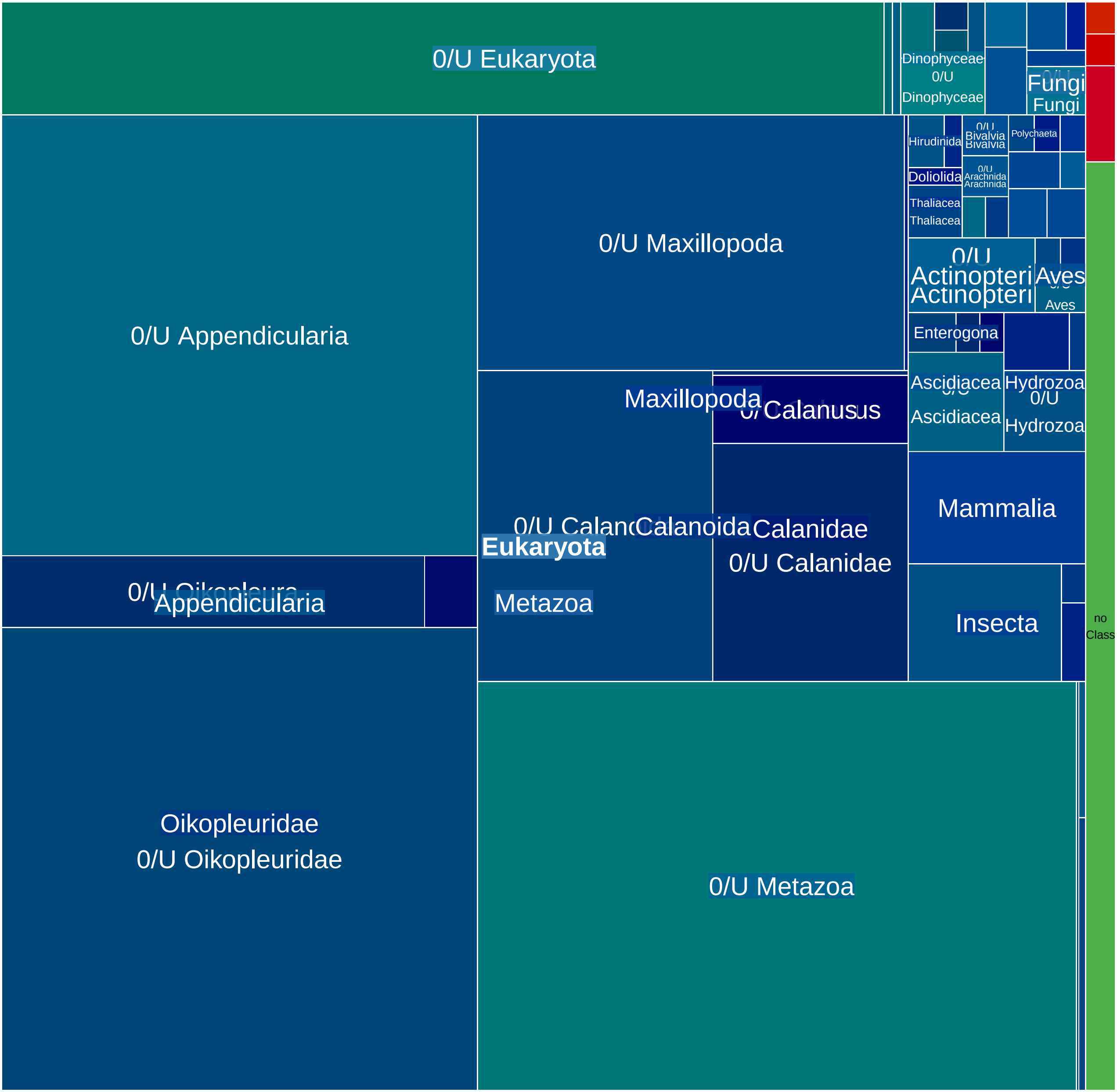

MGT−v1_276 : 4219 unigenes, 670 (15.88%) taxonomically assigned

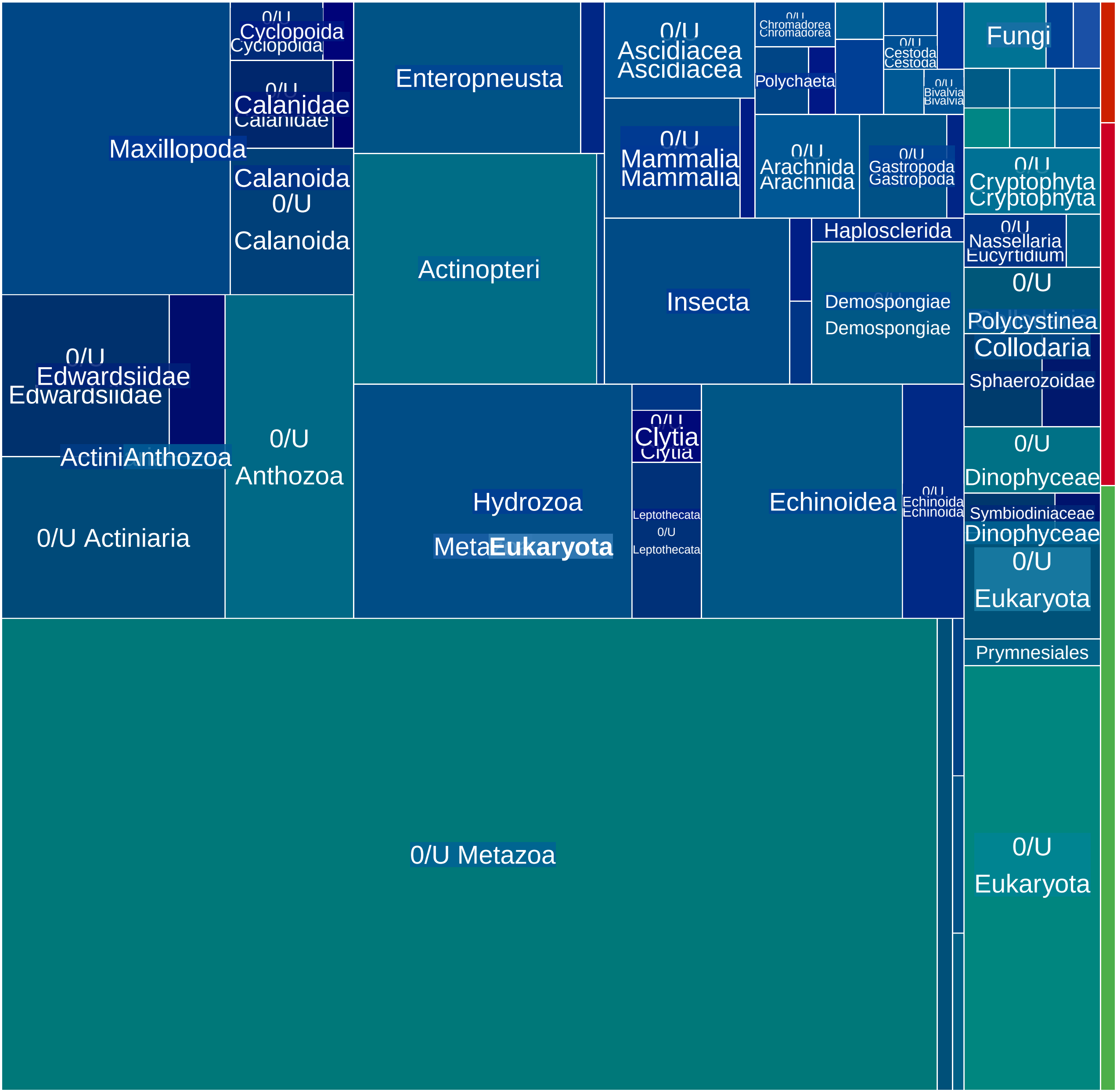

MGT−v1_277 : 4124 unigenes, 3737 (90.62%) taxonomically assigned

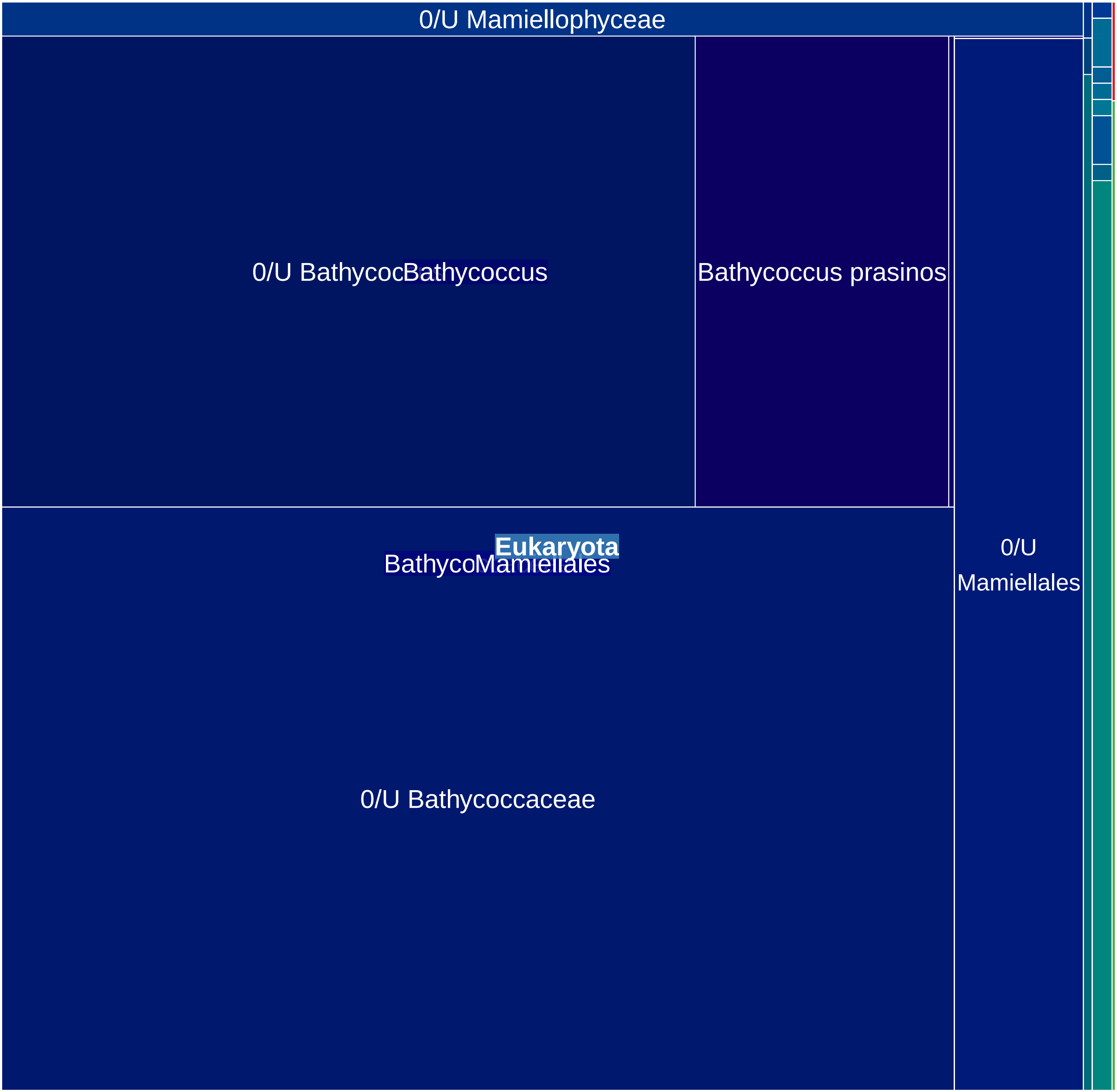

### MGT−v1_278 : 4117 unigenes, 413 (10.03%) taxonomically assigned

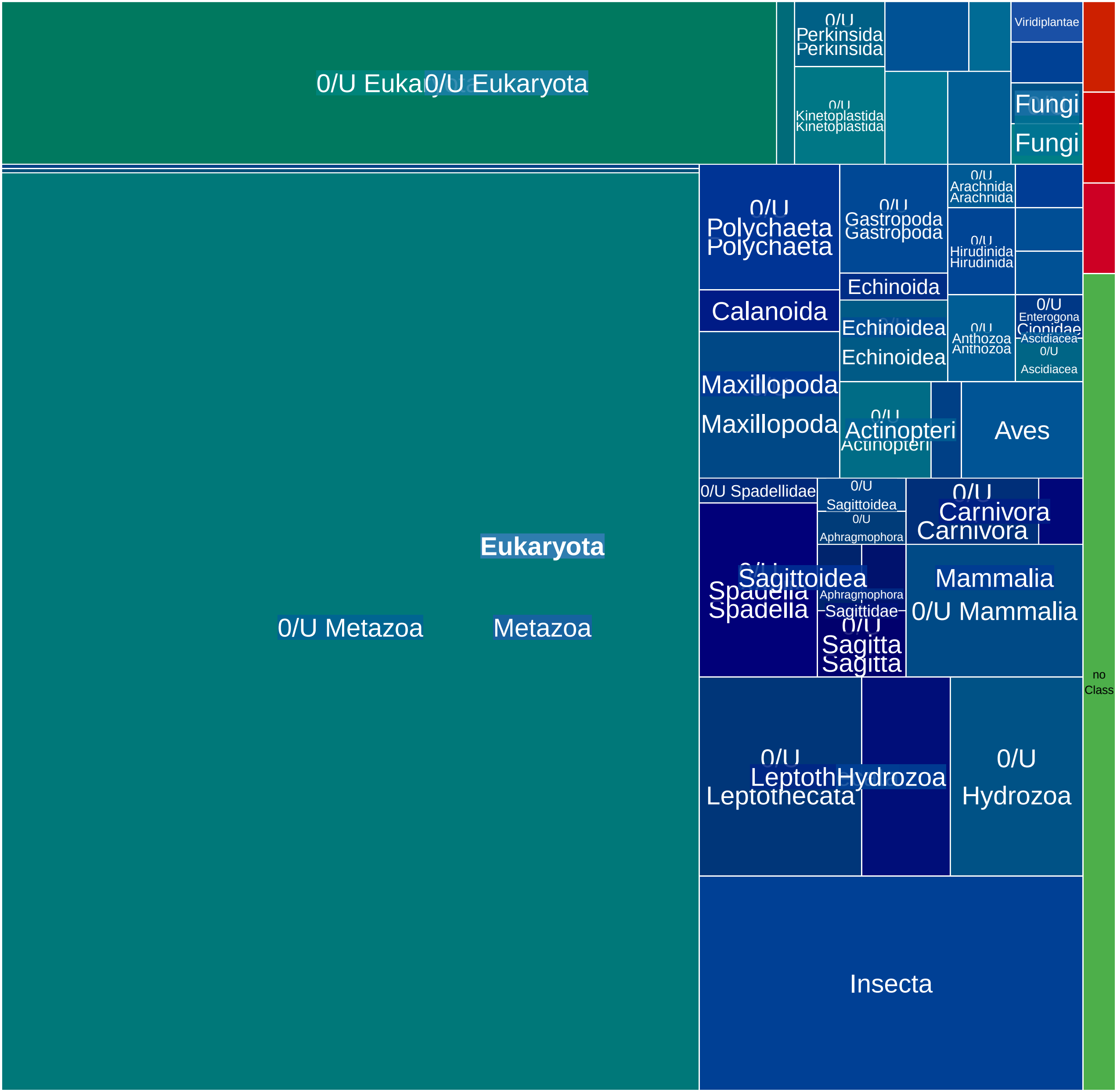

MGT−v1_279 : 4097 unigenes, 730 (17.82%) taxonomically assigned

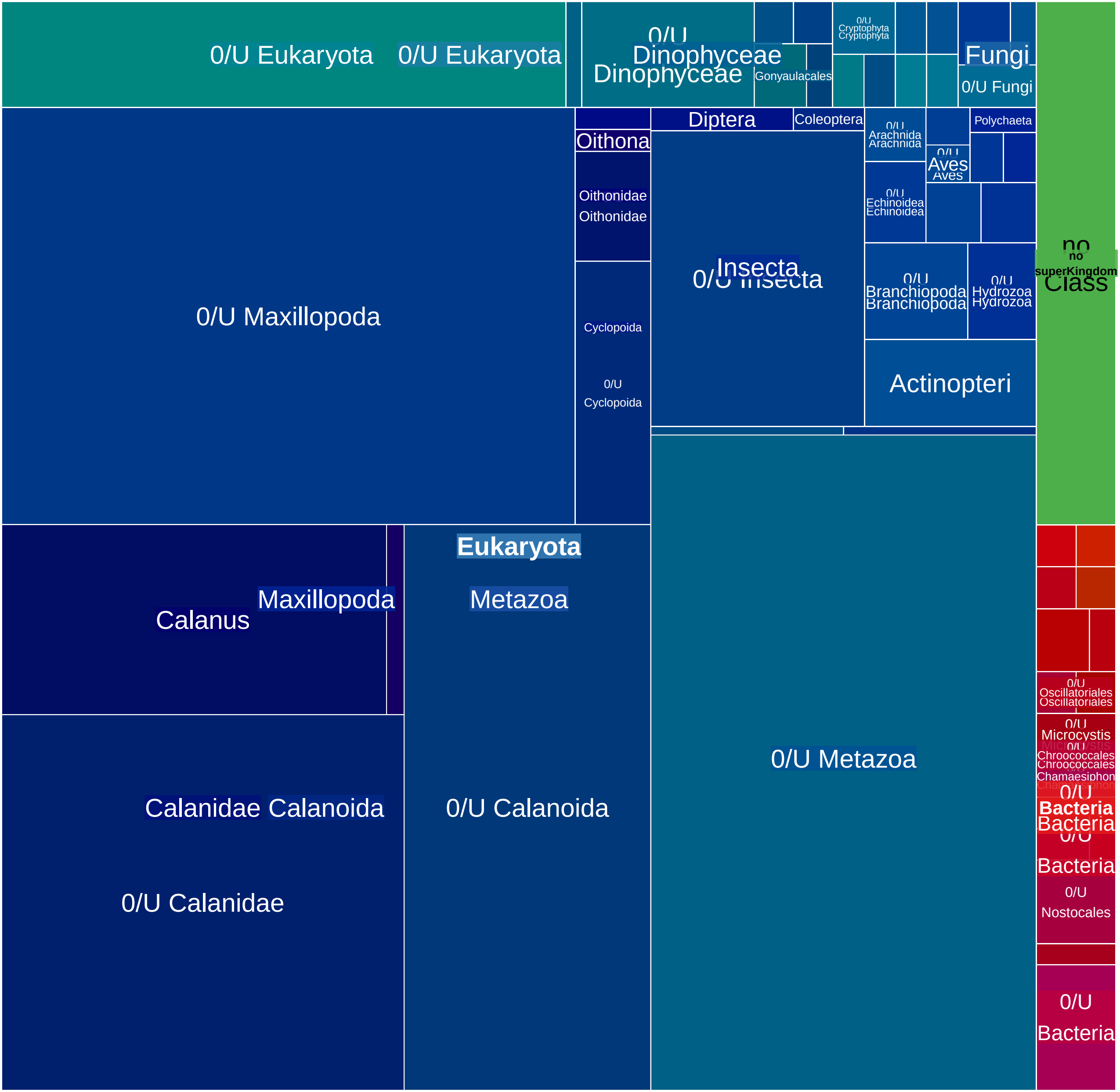

MGT−v1_280 : 4080 unigenes, 2874 (70.44%) taxonomically assigned

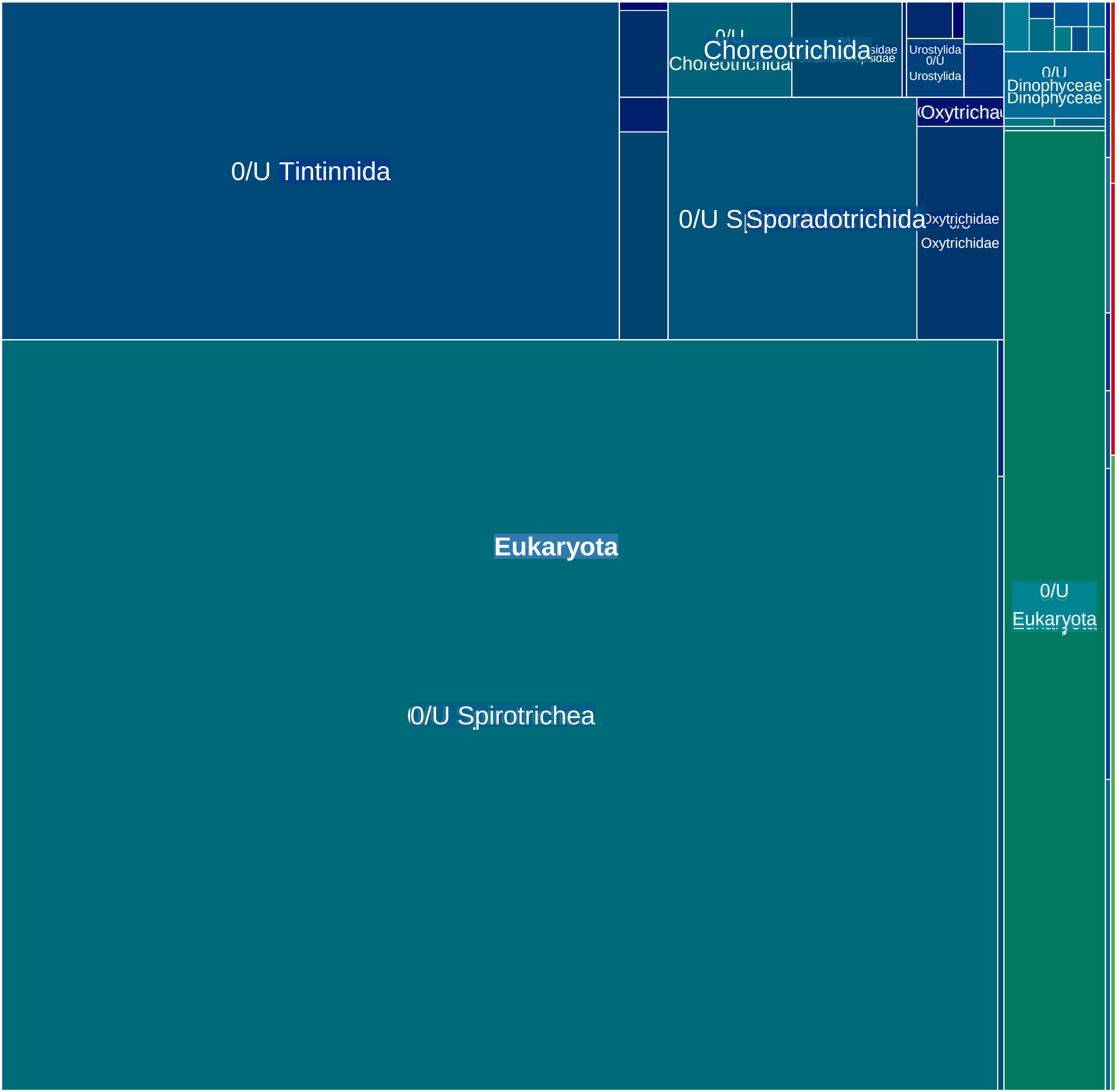

MGT−v1_281 : 4079 unigenes, 3789 (92.89%) taxonomically assigned

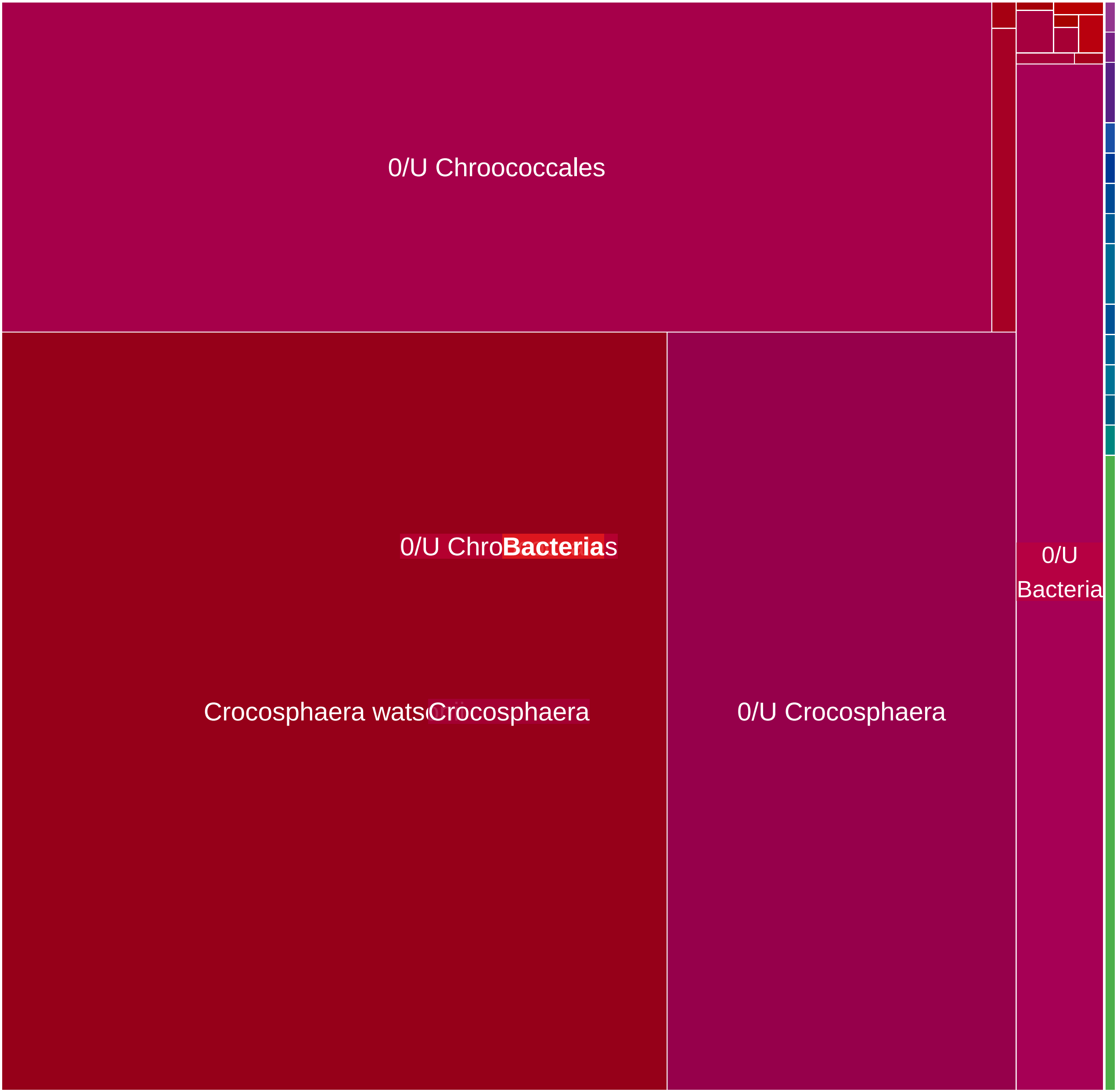

MGT−v1_282 : 4072 unigenes, 2587 (63.53%) taxonomically assigned

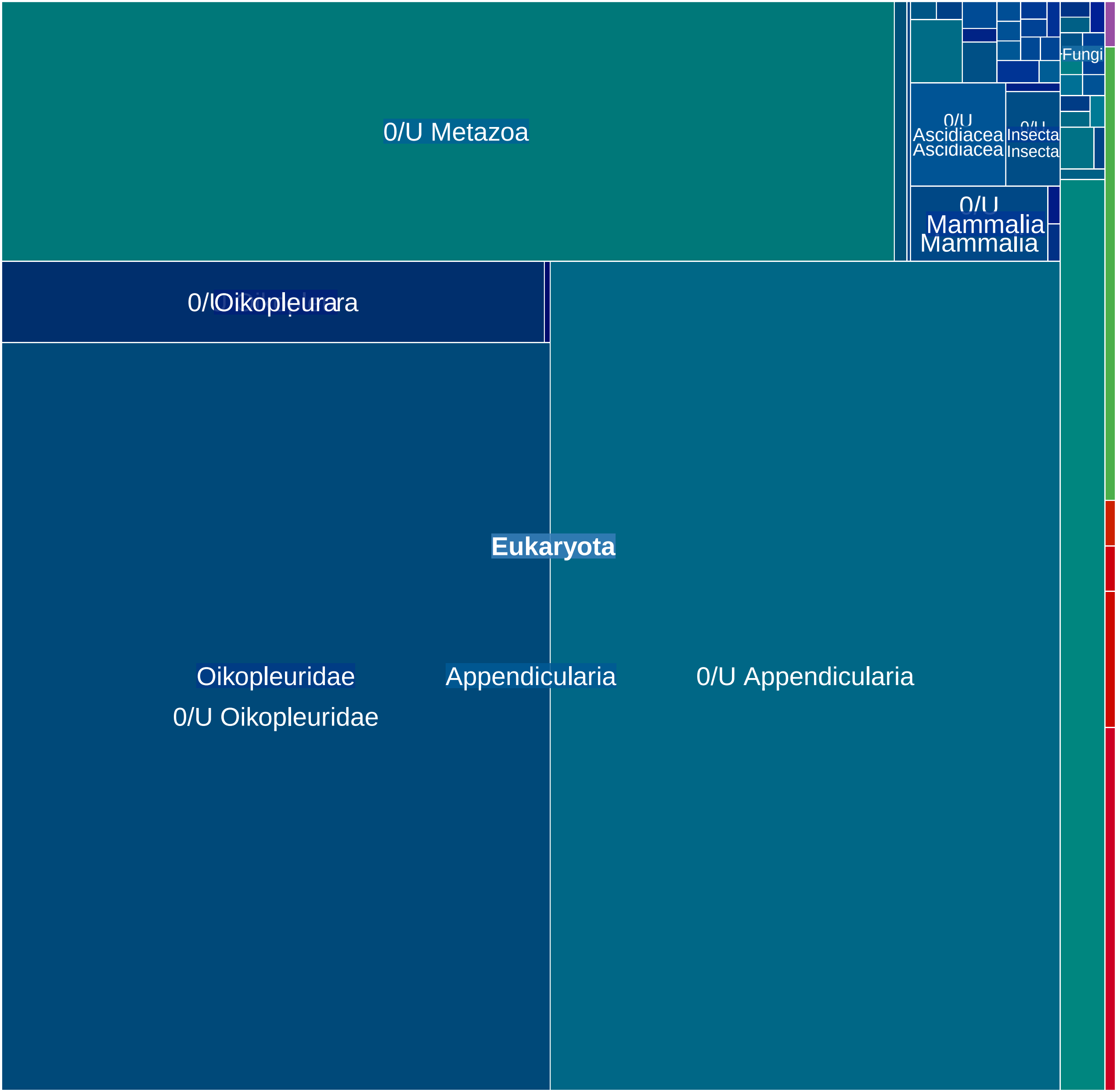

MGT−v1_283 : 4039 unigenes, 667 (16.51%) taxonomically assigned

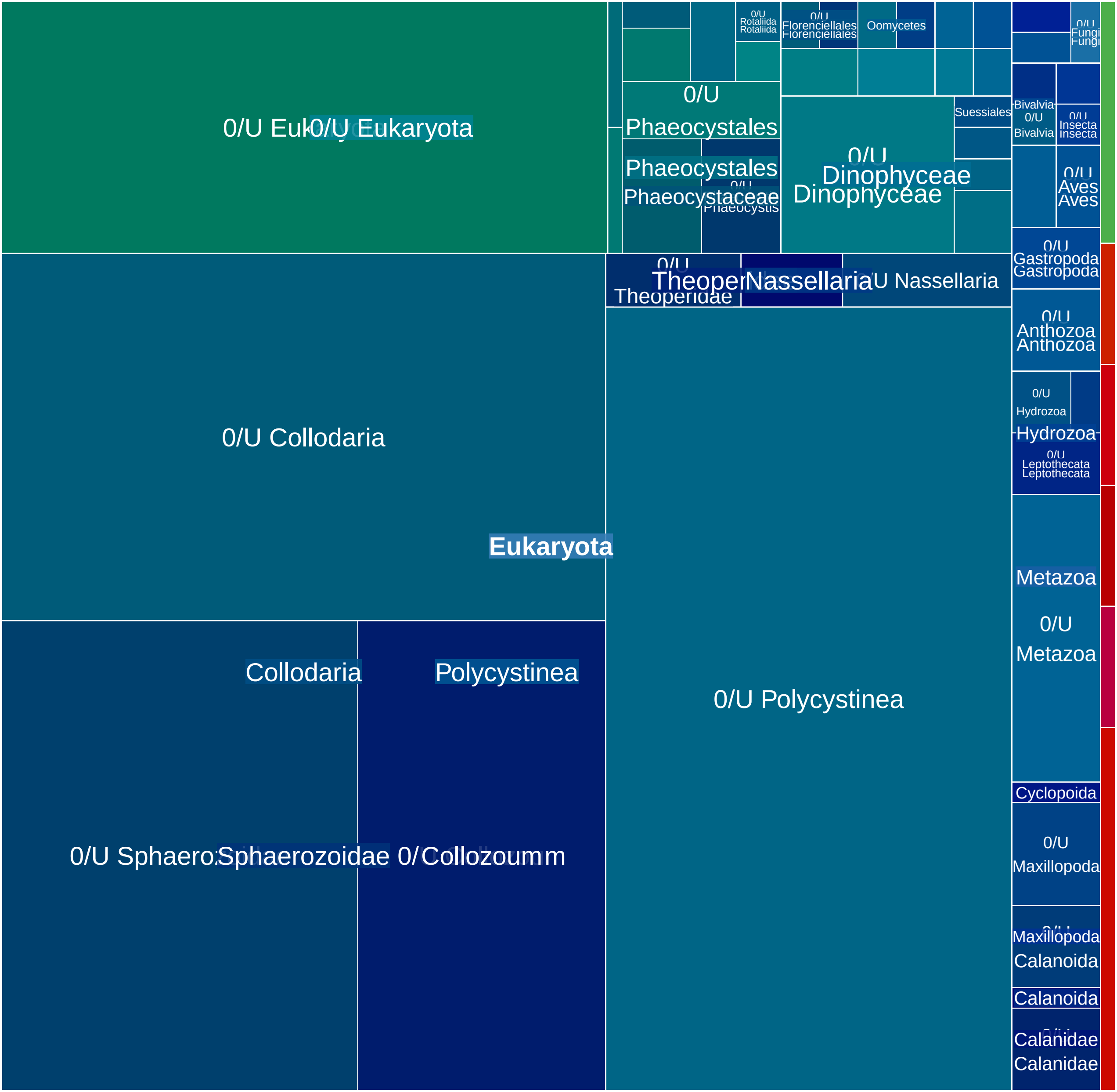

MGT−v1_284 : 4027 unigenes, 2099 (52.12%) taxonomically assigned

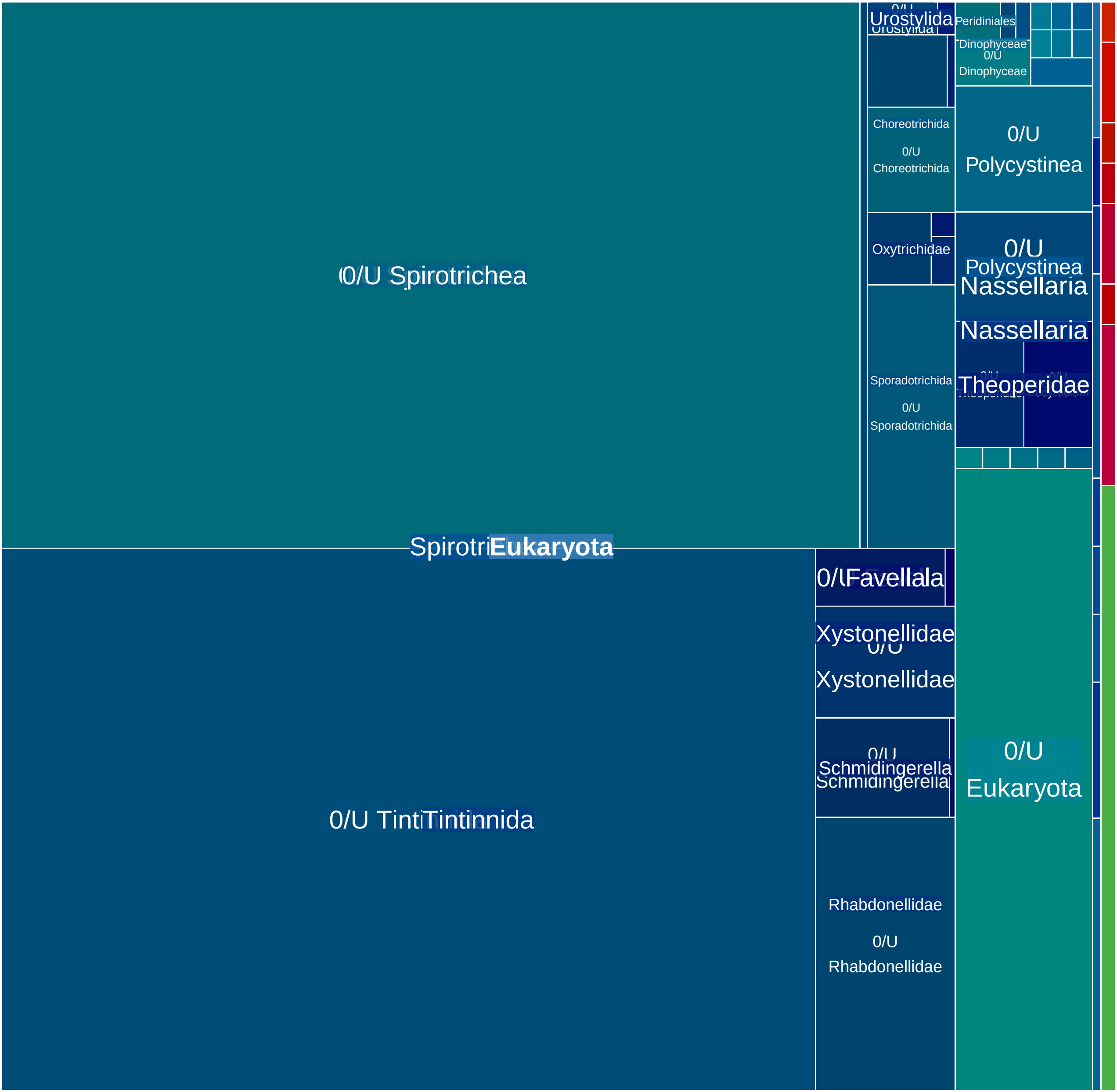

MGT−v1_285 : 3949 unigenes, 460 (11.65%) taxonomically assigned

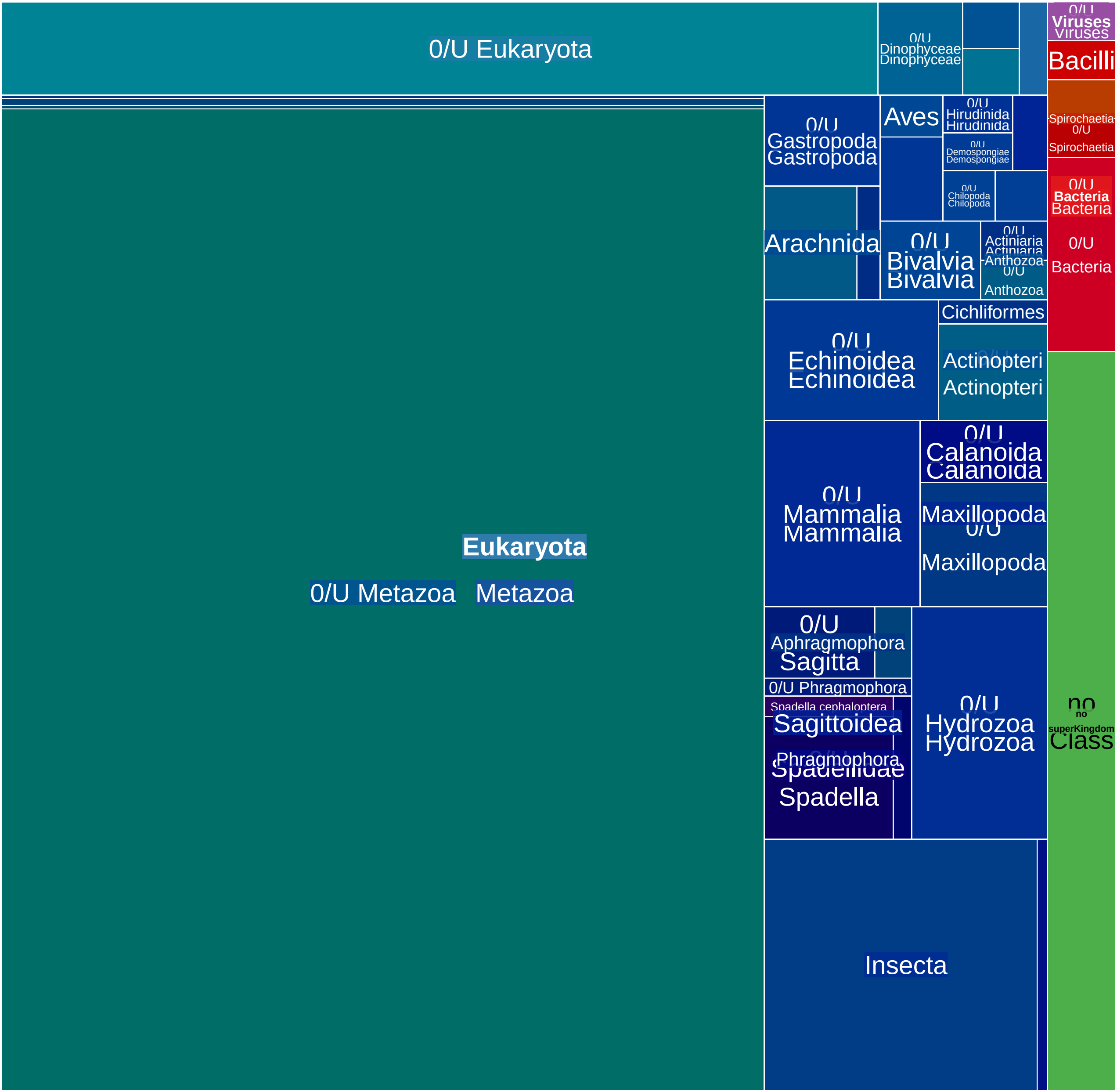

MGT−v1_286 : 3942 unigenes, 472 (11.97%) taxonomically assigned

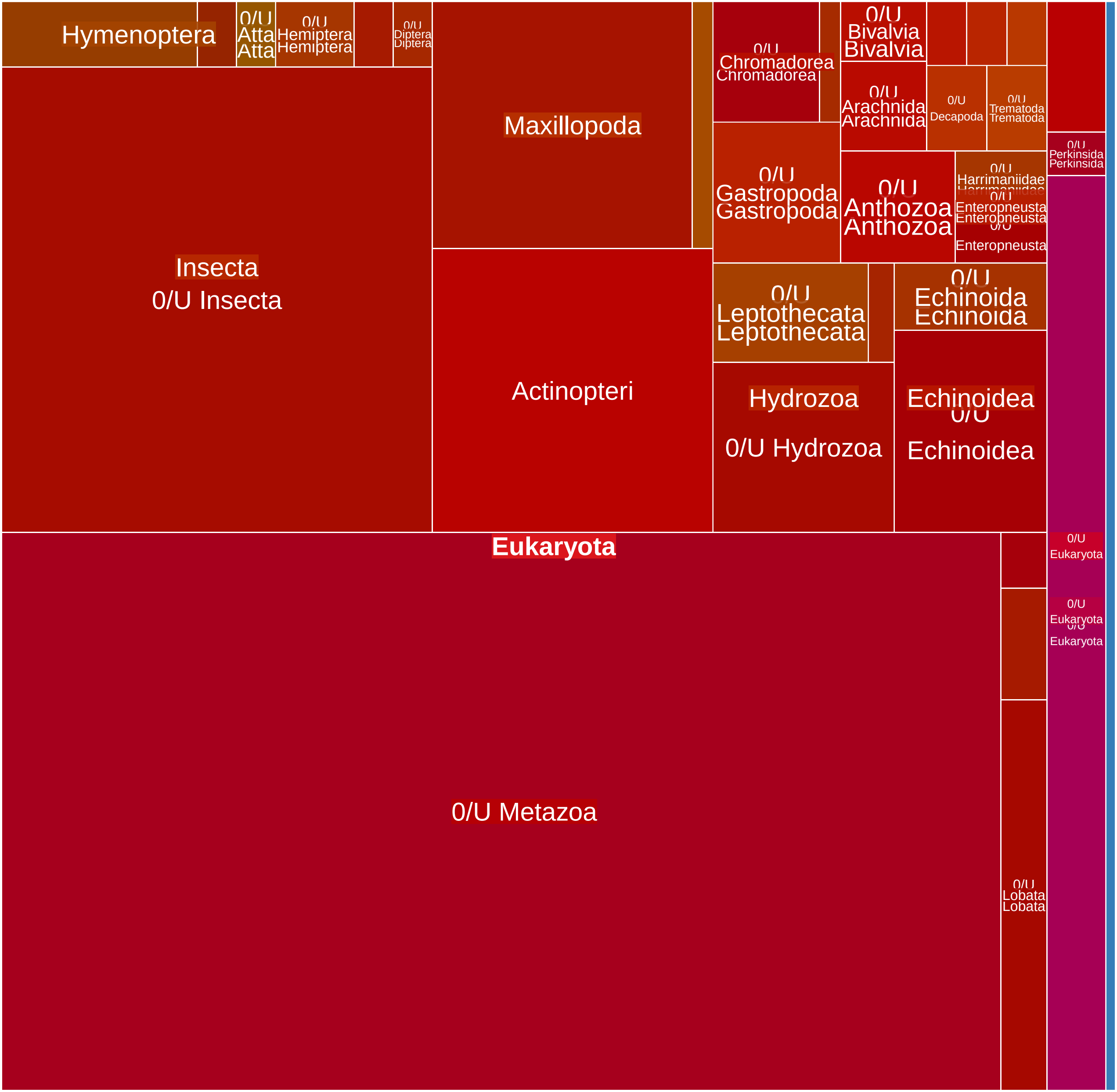

MGT−v1_287 : 3910 unigenes, 2377 (60.79%) taxonomically assigned

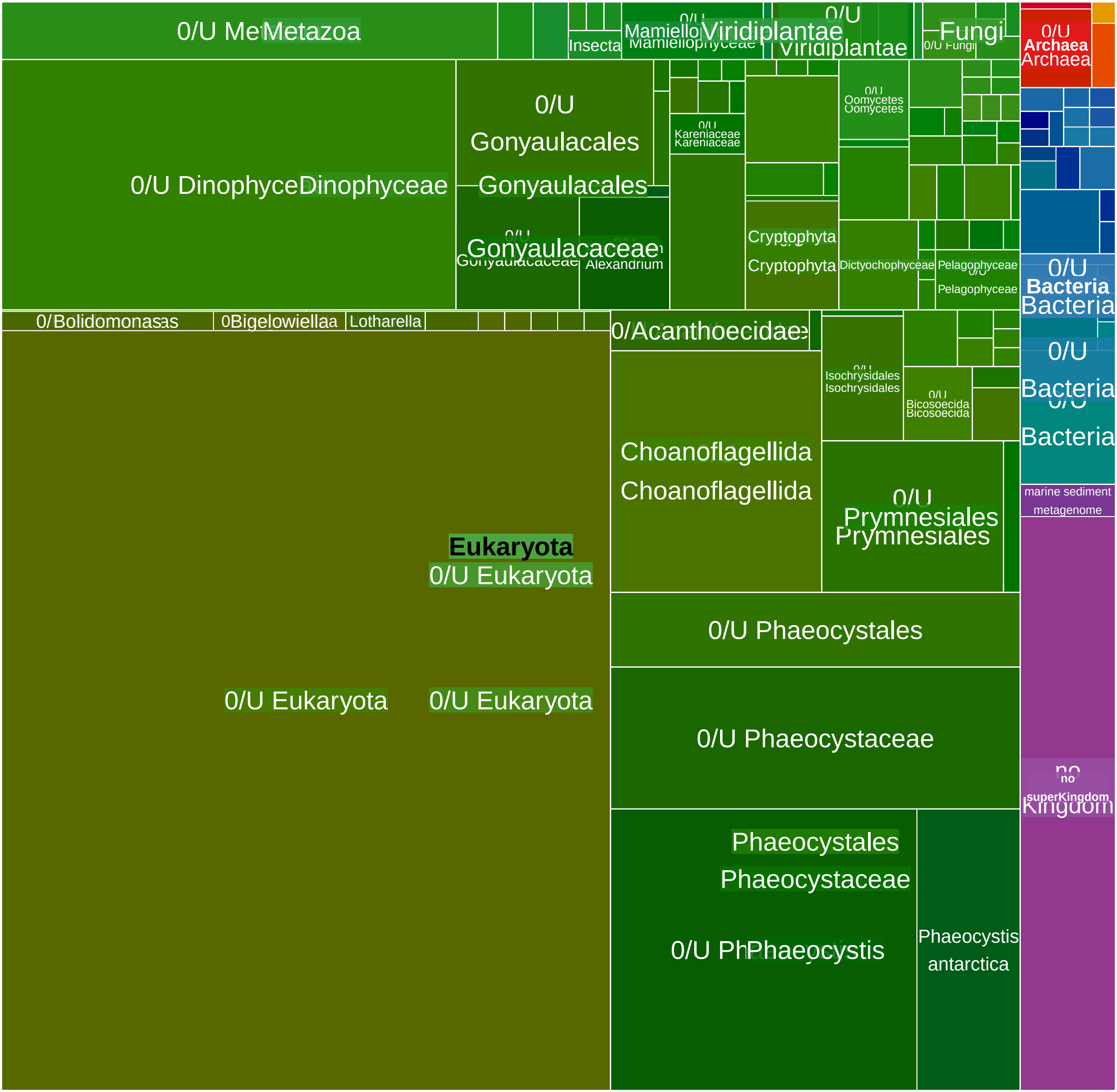

MGT−v1_288 : 3902 unigenes, 513 (13.15%) taxonomically assigned

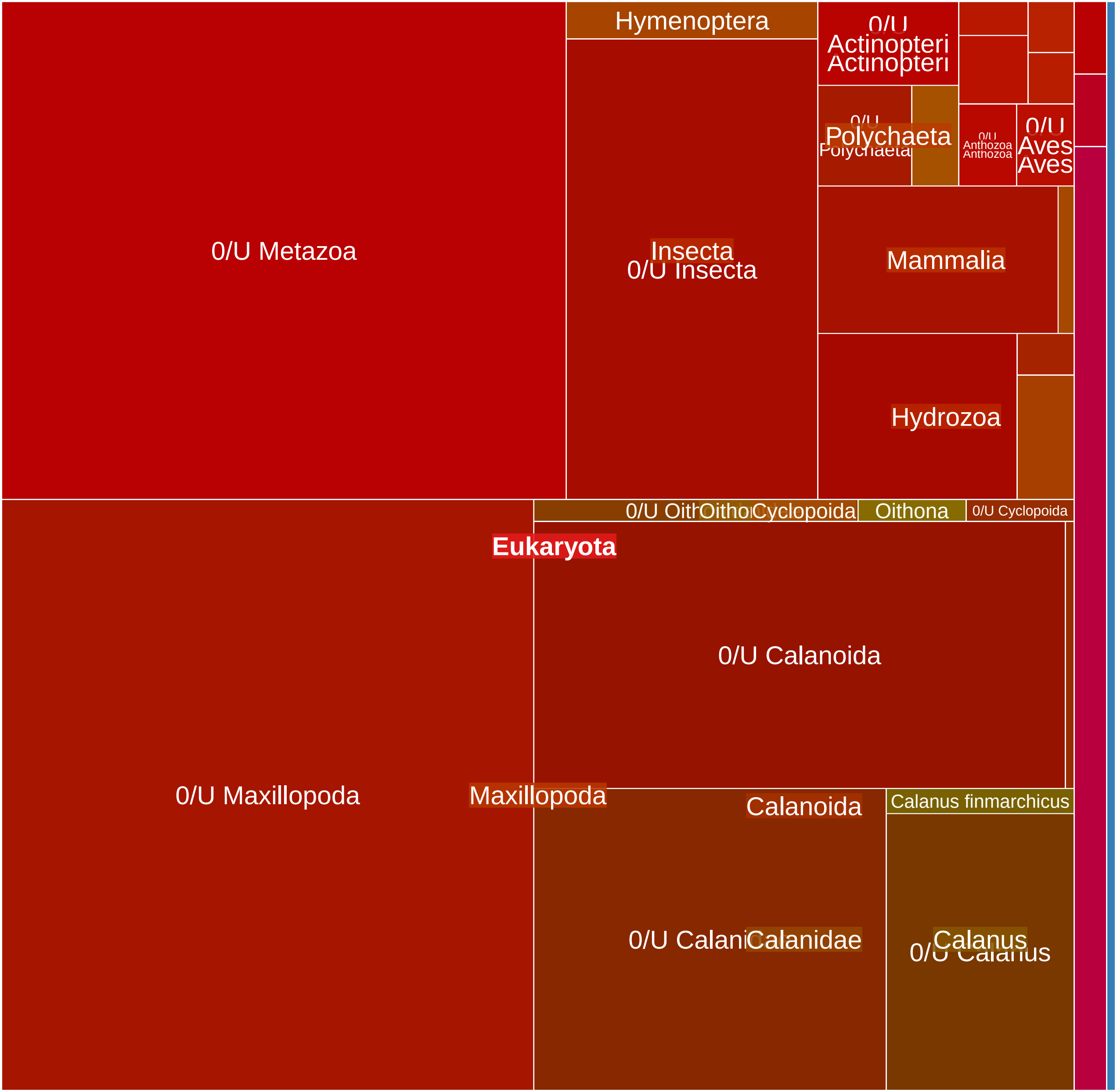

MGT−v1_289 : 3900 unigenes, 630 (16.15%) taxonomically assigned

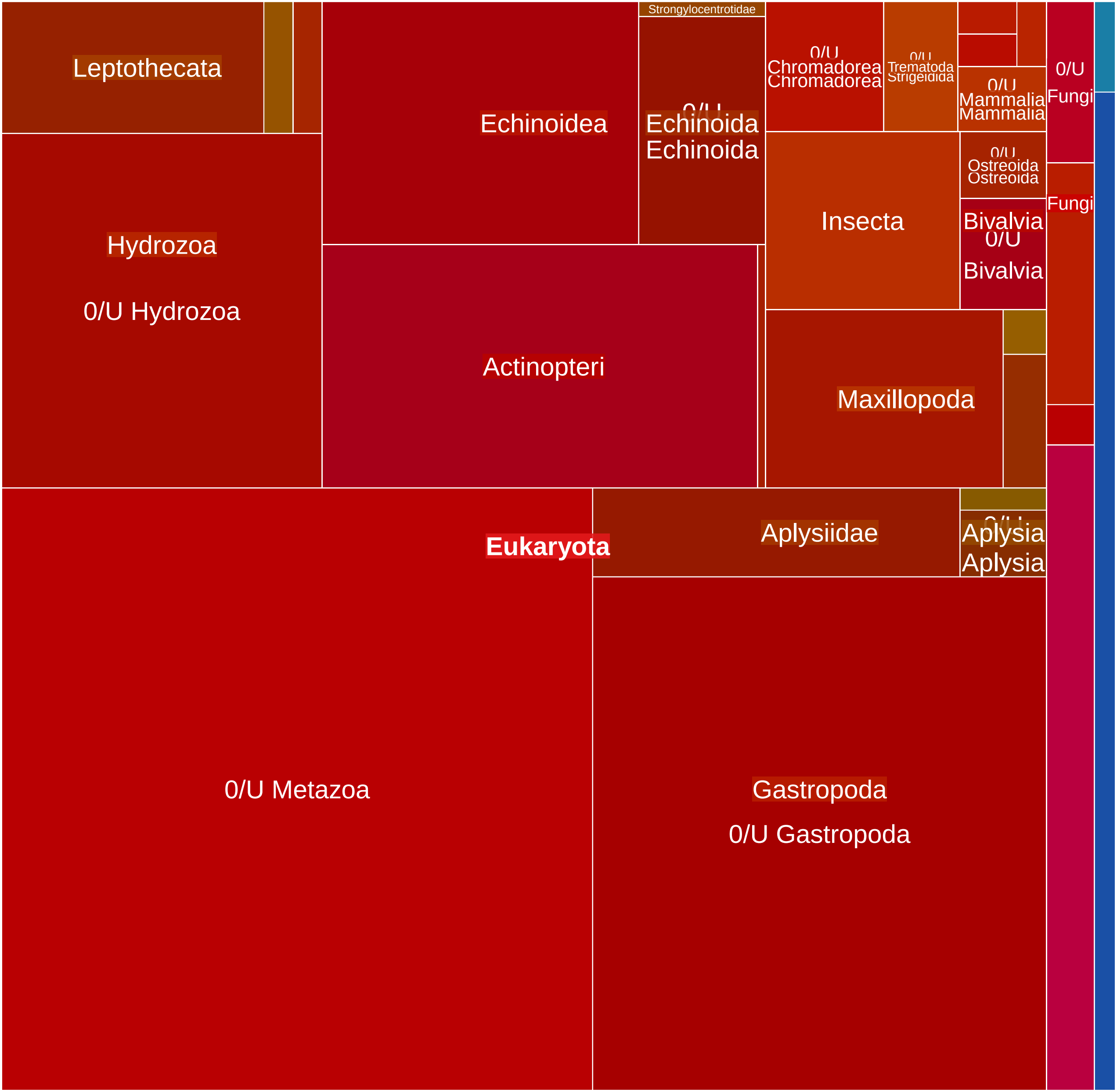

MGT−v1_290 : 3899 unigenes, 3199 (82.05%) taxonomically assigned

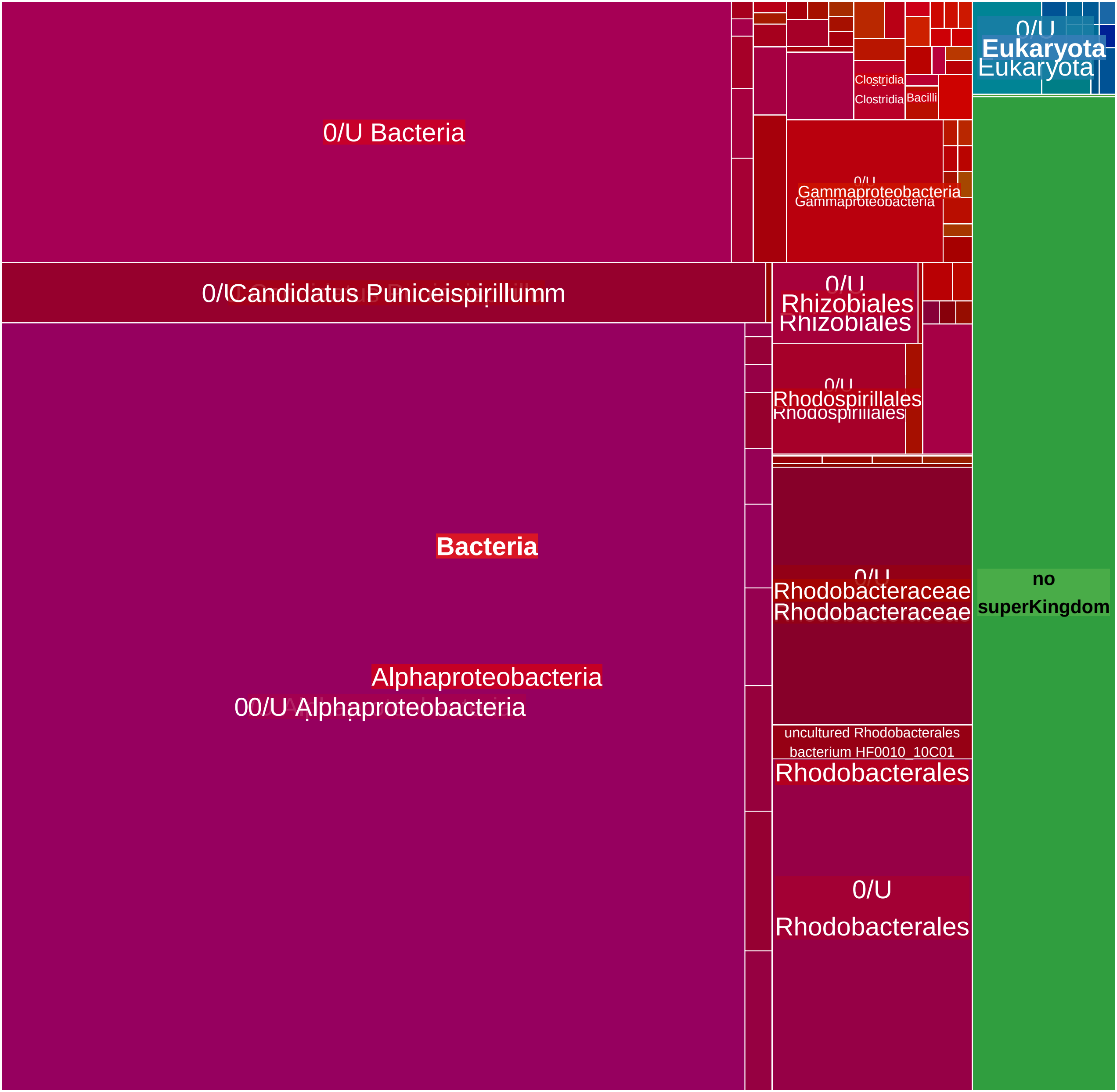

MGT−v1_291 : 3887 unigenes, 410 (10.55%) taxonomically assigned

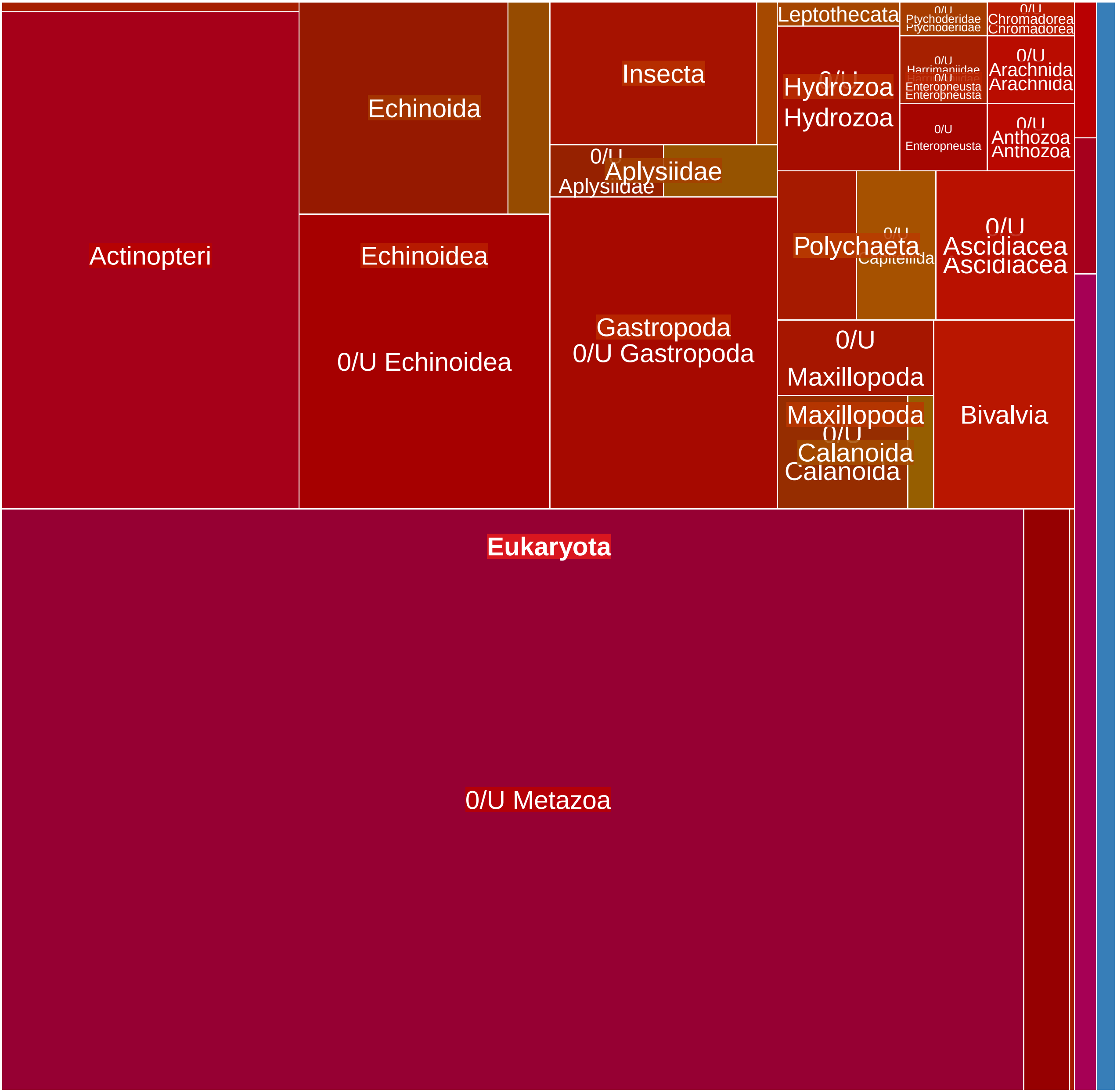

MGT−v1_292 : 3883 unigenes, 1651 (42.52%) taxonomically assigned

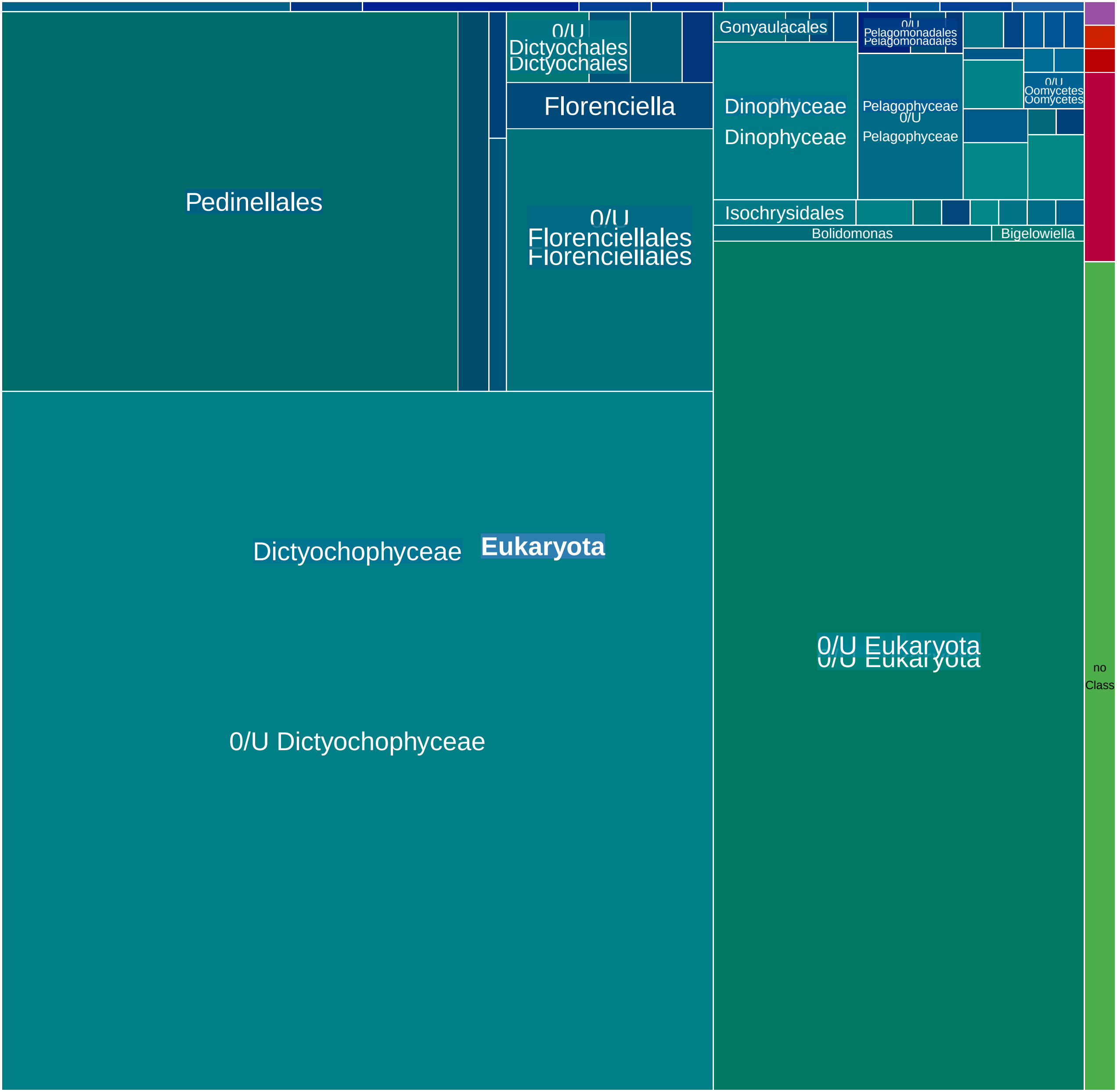

MGT−v1_293 : 3873 unigenes, 3213 (82.96%) taxonomically assigned

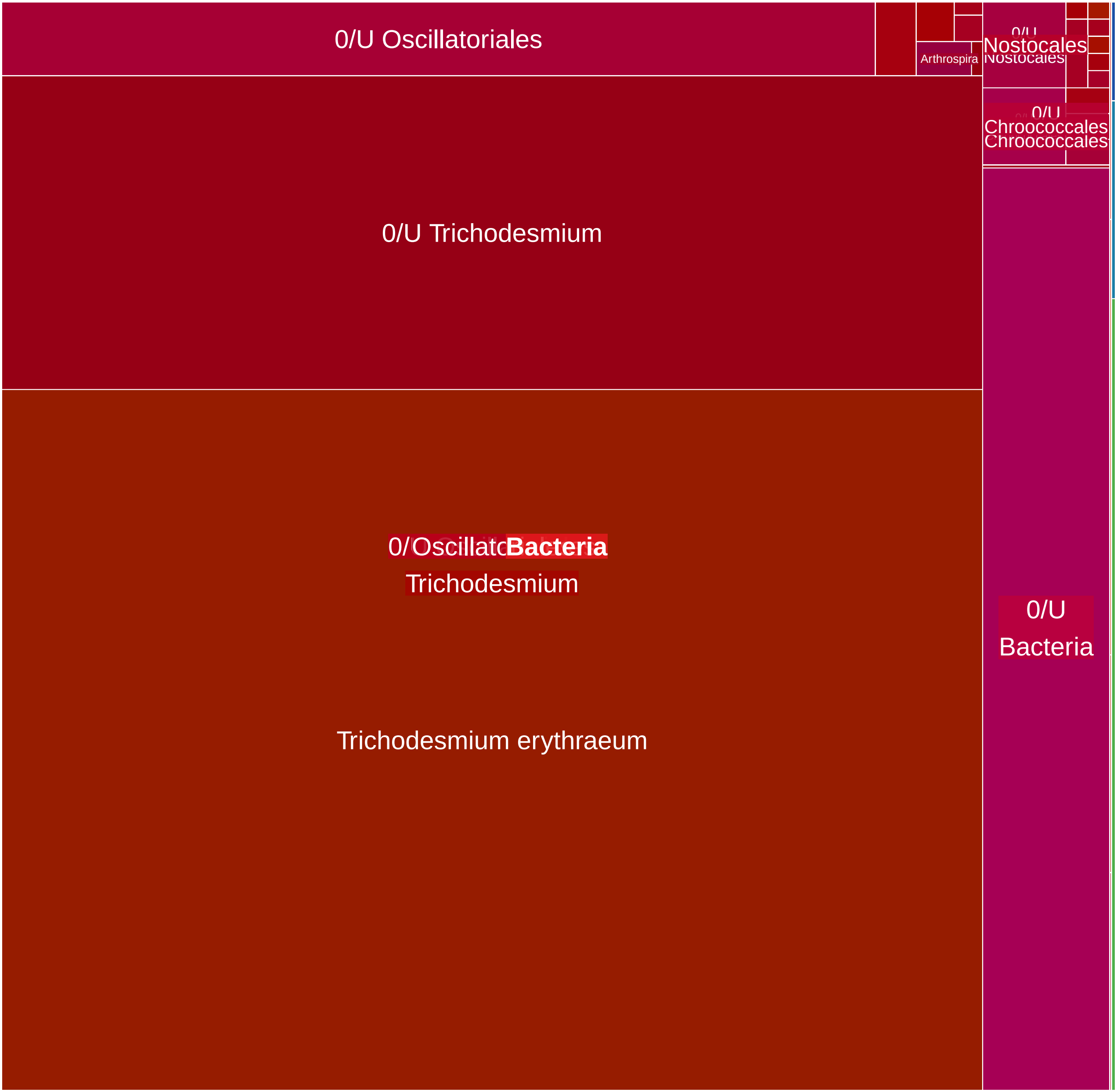

MGT−v1_294 : 3865 unigenes, 862 (22.30%) taxonomically assigned

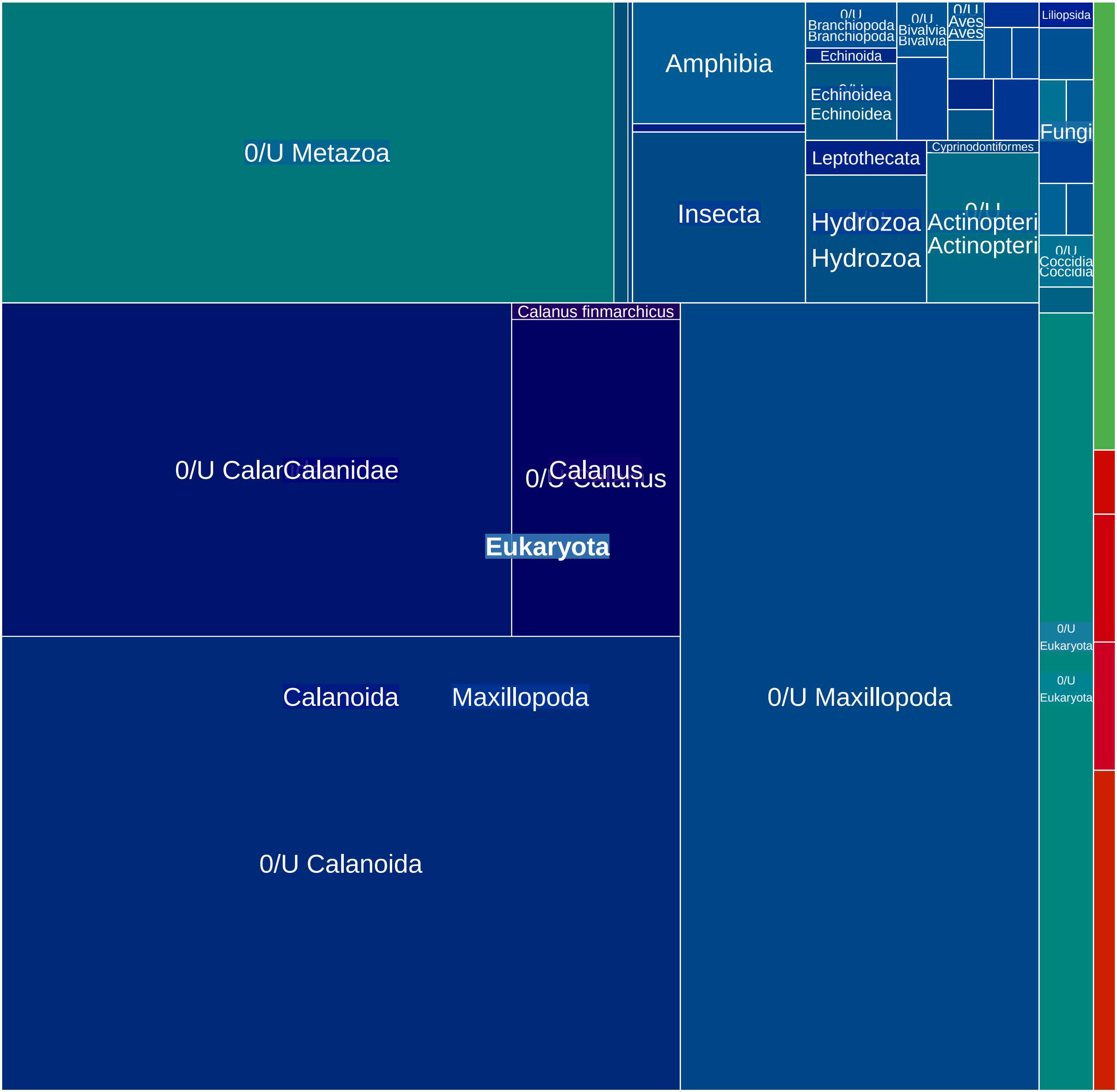

MGT−v1_295 : 3819 unigenes, 387 (10.13%) taxonomically assigned

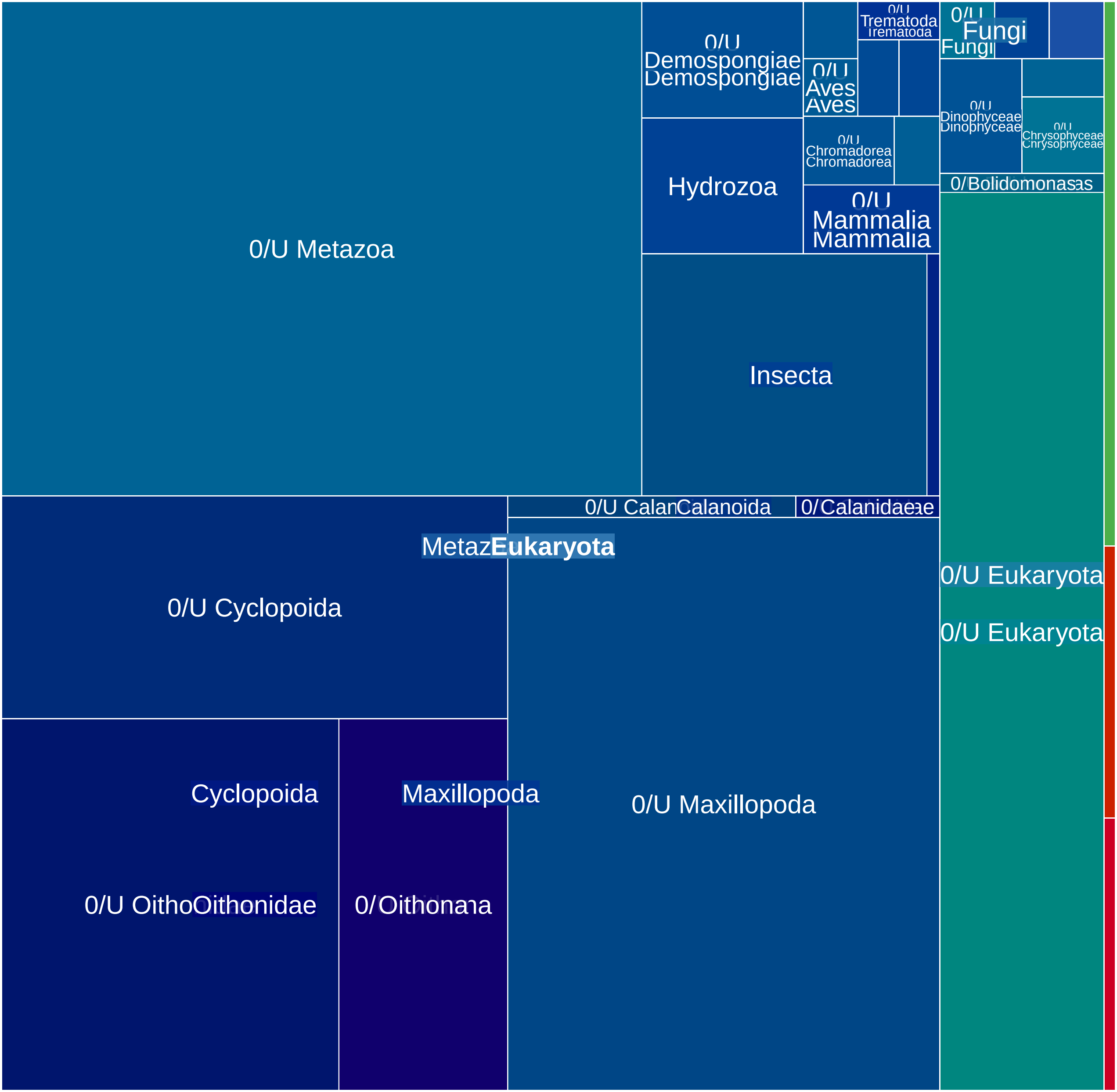

MGT−v1_296 : 3817 unigenes, 385 (10.09%) taxonomically assigned

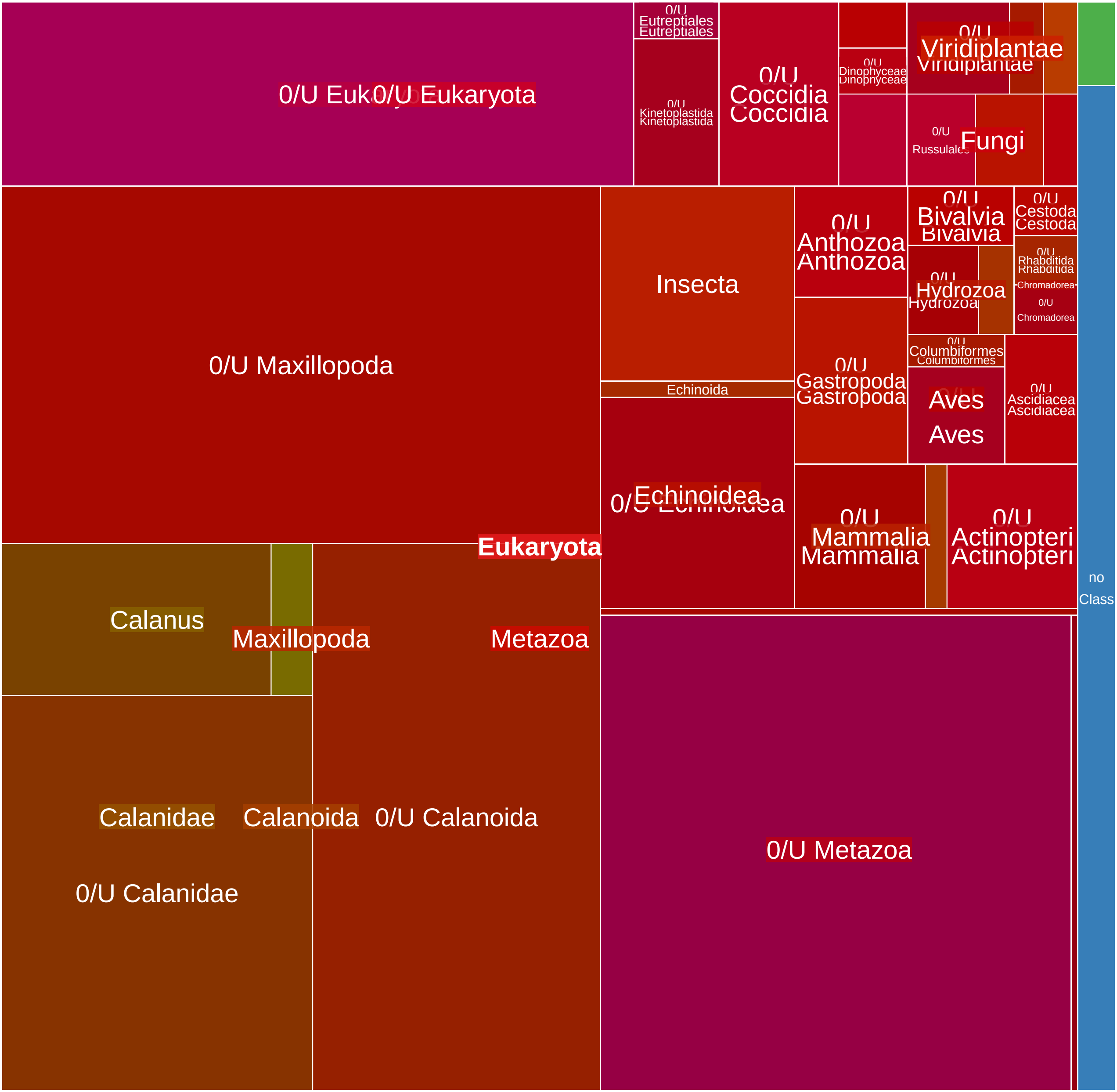

MGT−v1_297 : 3788 unigenes, 229 (6.05%) taxonomically assigned

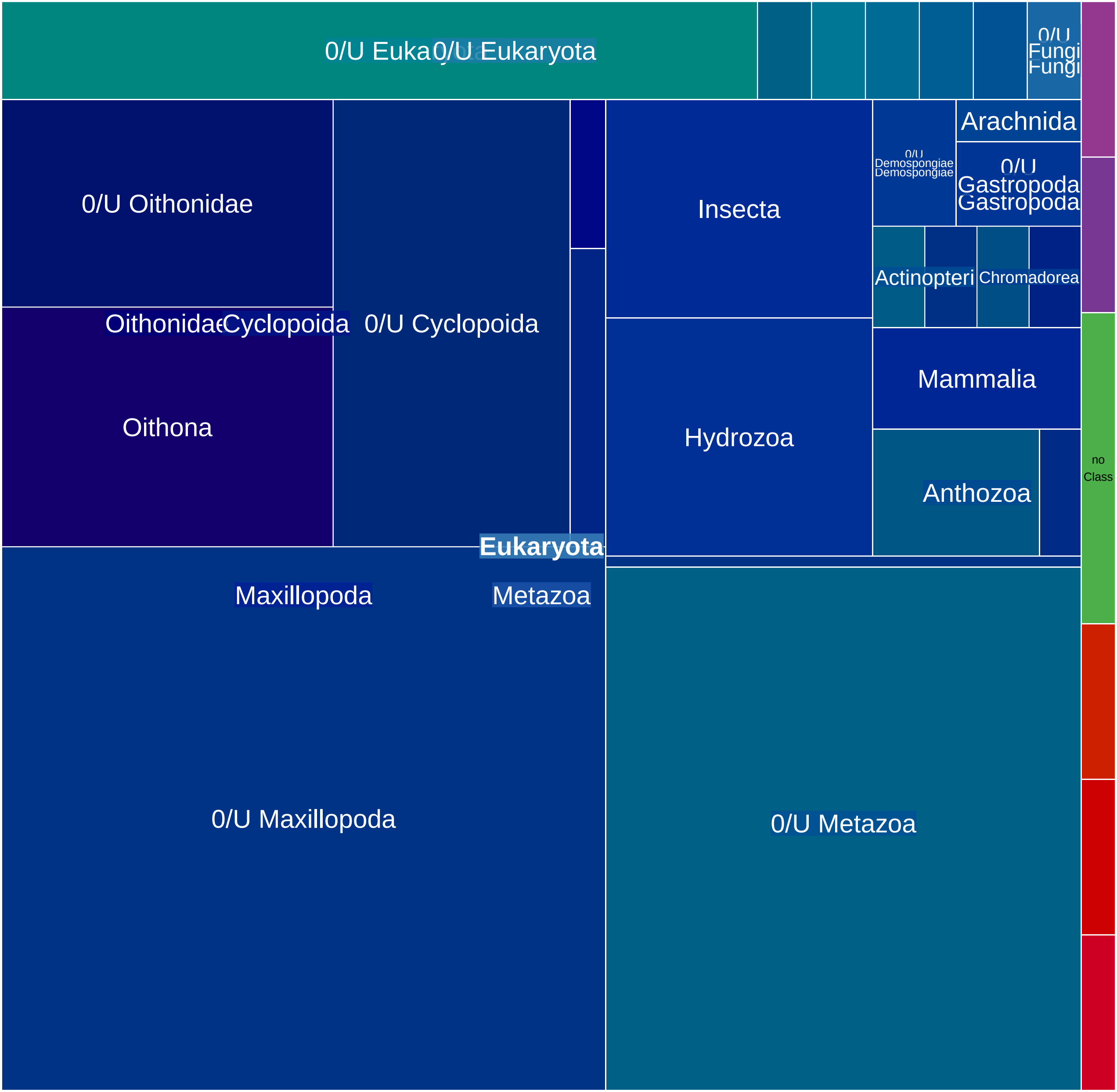

### MGT−v1_298 : 3785 unigenes, 1502 (39.68%) taxonomically assigned

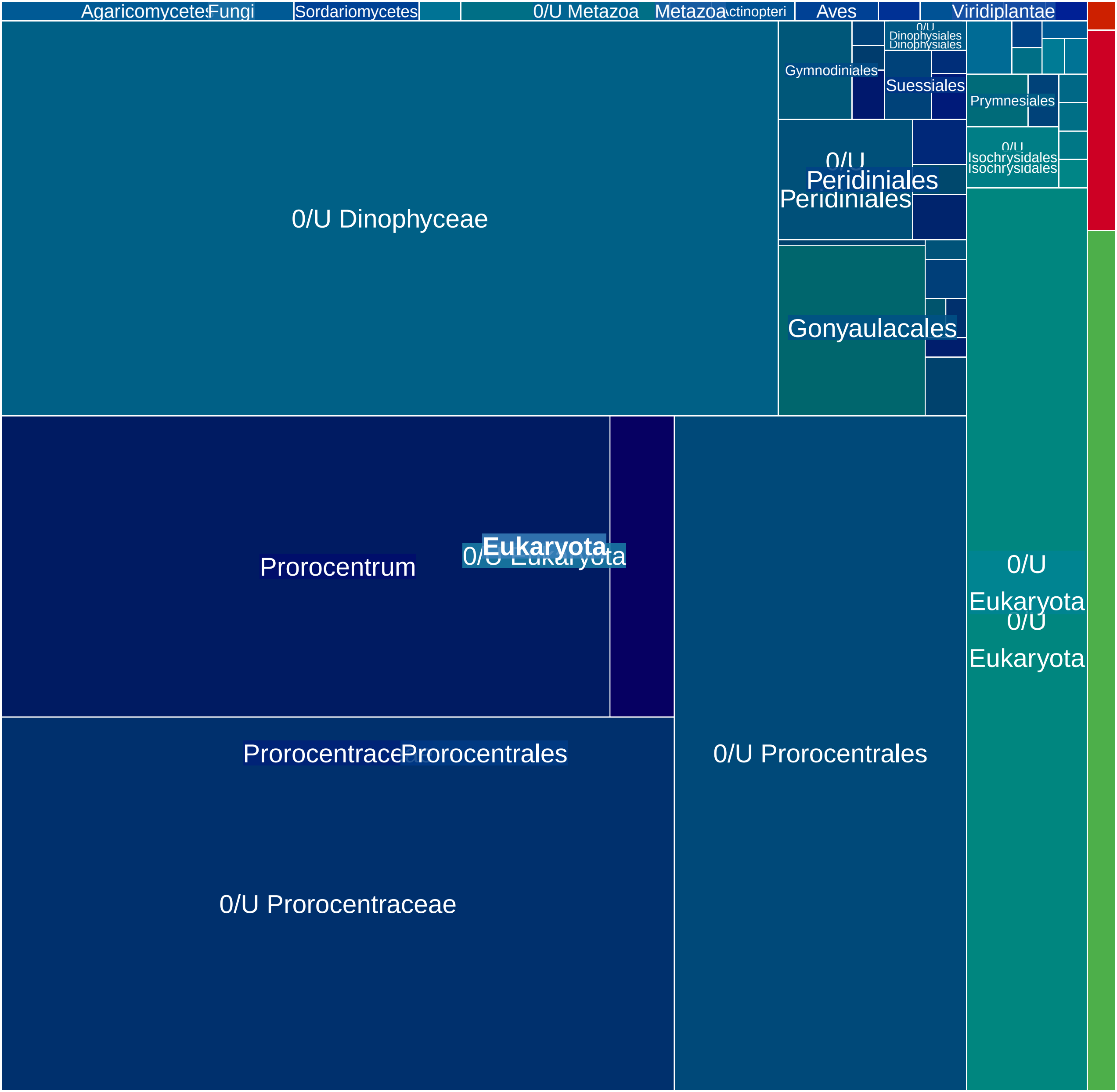

MGT−v1_299 : 3784 unigenes, 433 (11.44%) taxonomically assigned

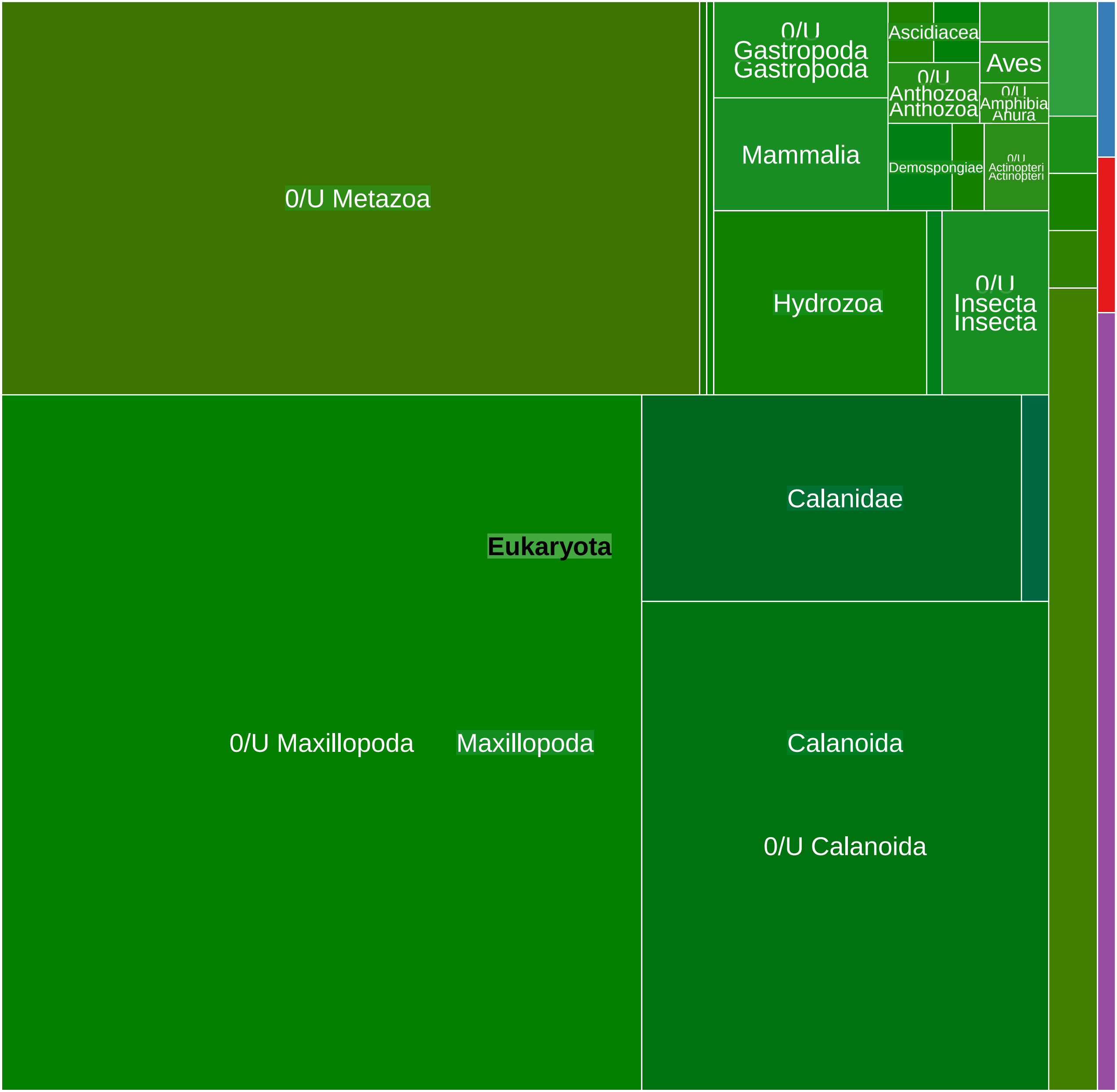

MGT−v1_300 : 3781 unigenes, 3349 (88.57%) taxonomically assigned

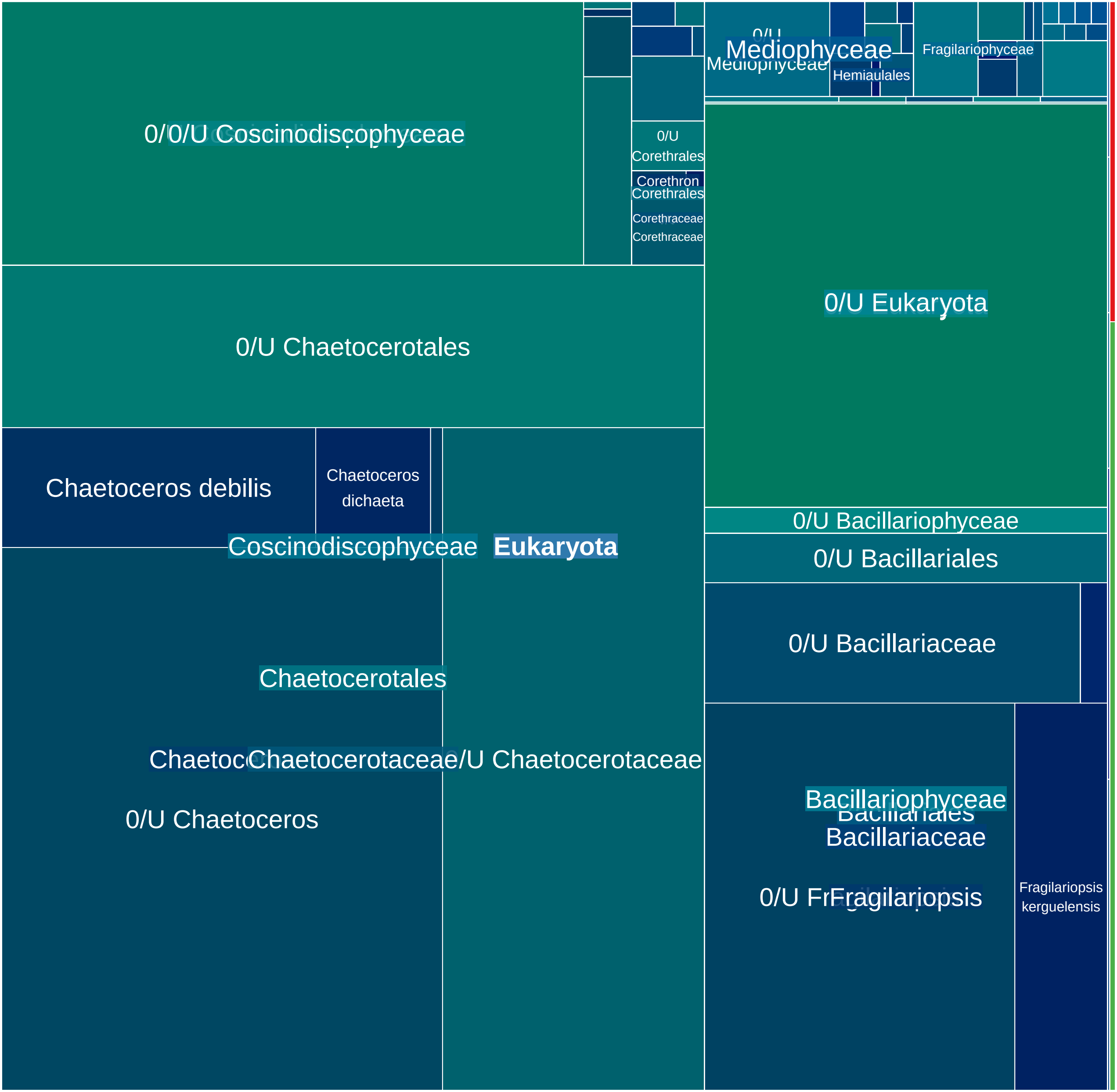

MGT−v1_301 : 3735 unigenes, 2201 (58.93%) taxonomically assigned

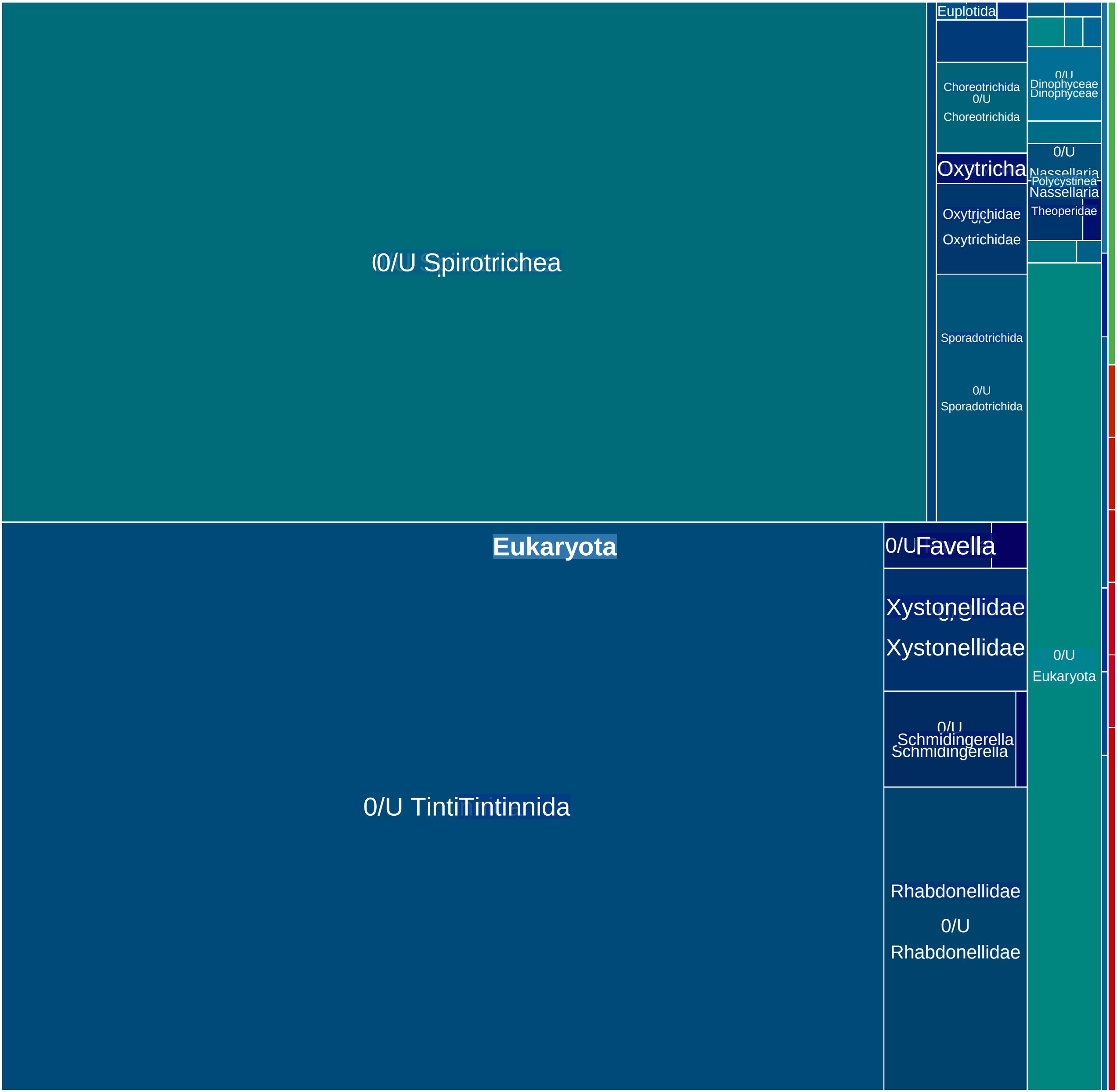

### MGT−v1_302 : 3708 unigenes, 267 (7.20%) taxonomically assigned

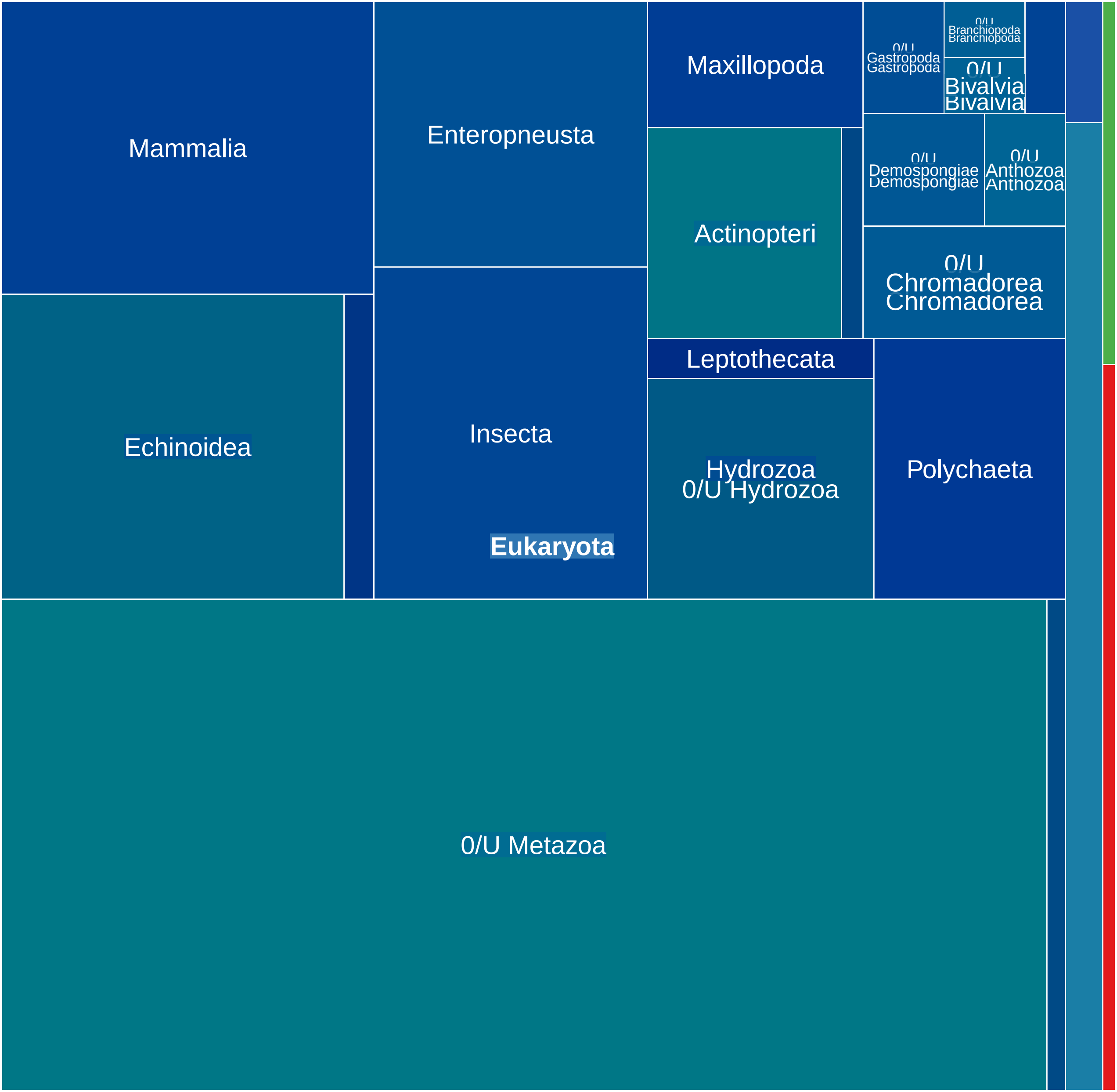

MGT−v1_303 : 3702 unigenes, 1351 (36.49%) taxonomically assigned

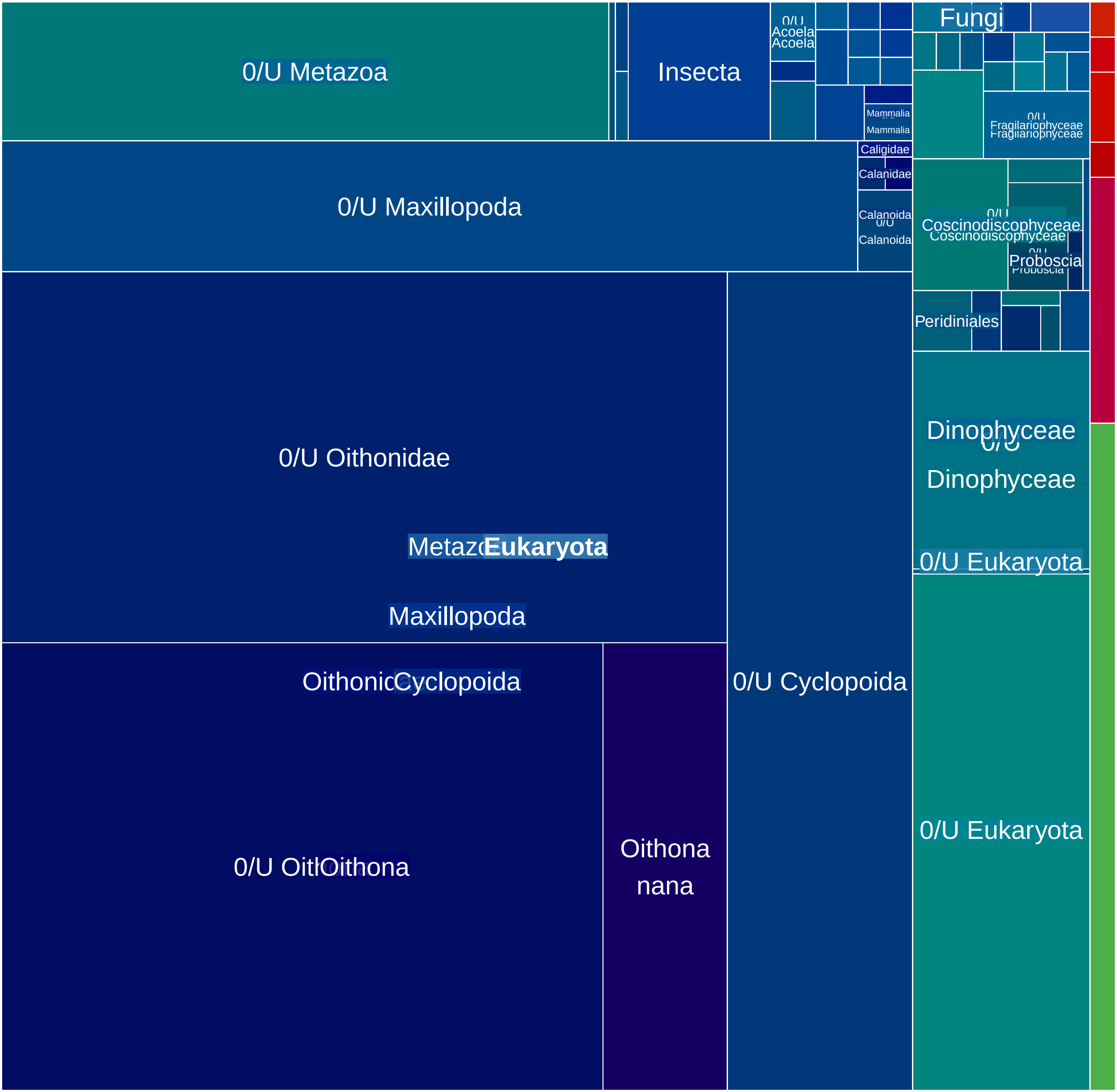

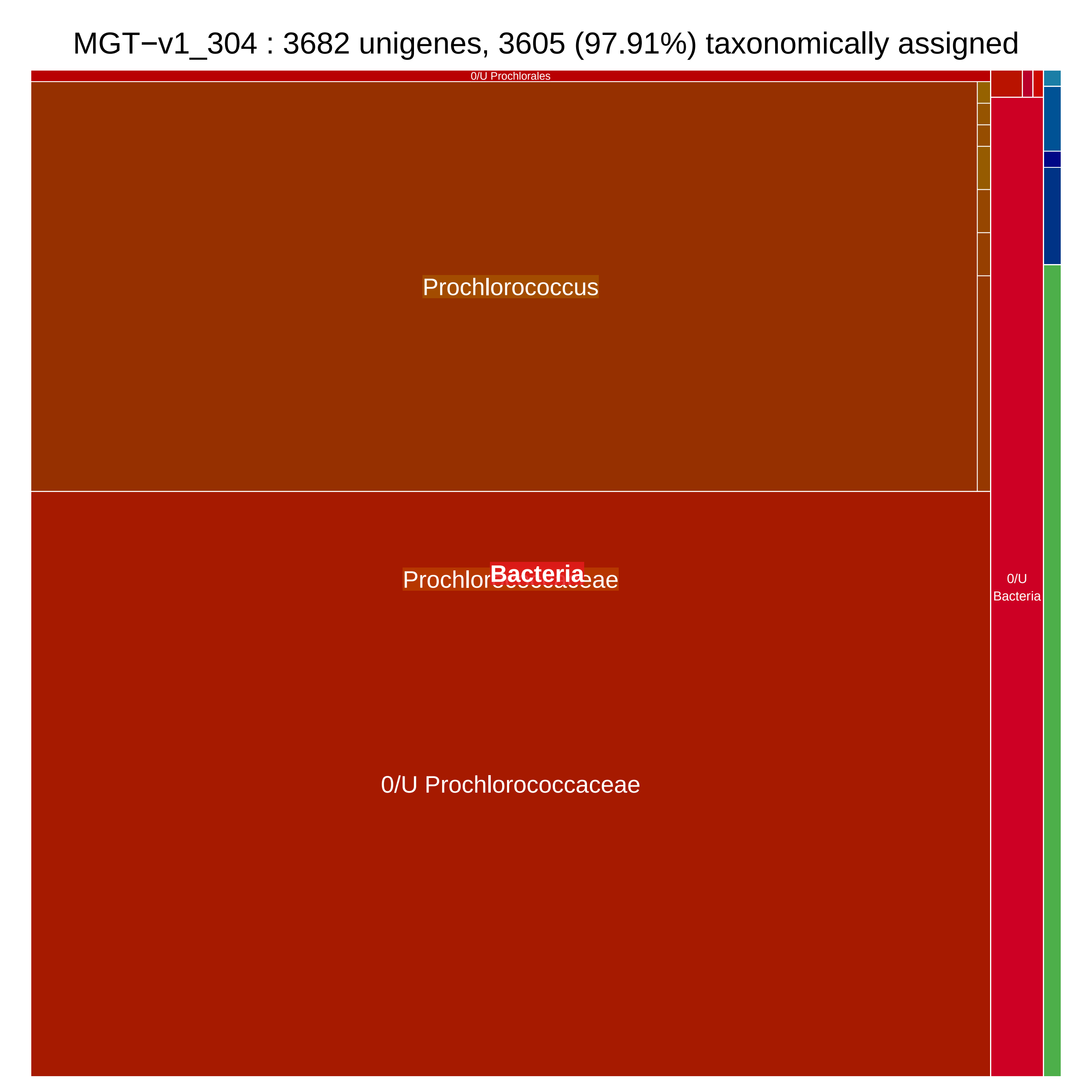
MGT−v1_304 : 3682 unigenes, 3605 (97.91%) taxonomically assigned
0/U Prochlorales
Prochlorococcus
Bacteria
Prochlorococcaceae
0/U Prochlorococcaceae
0/U
Bacteria

### MGT−v1_305 : 3661 unigenes, 3005 (82.08%) taxonomically assigned

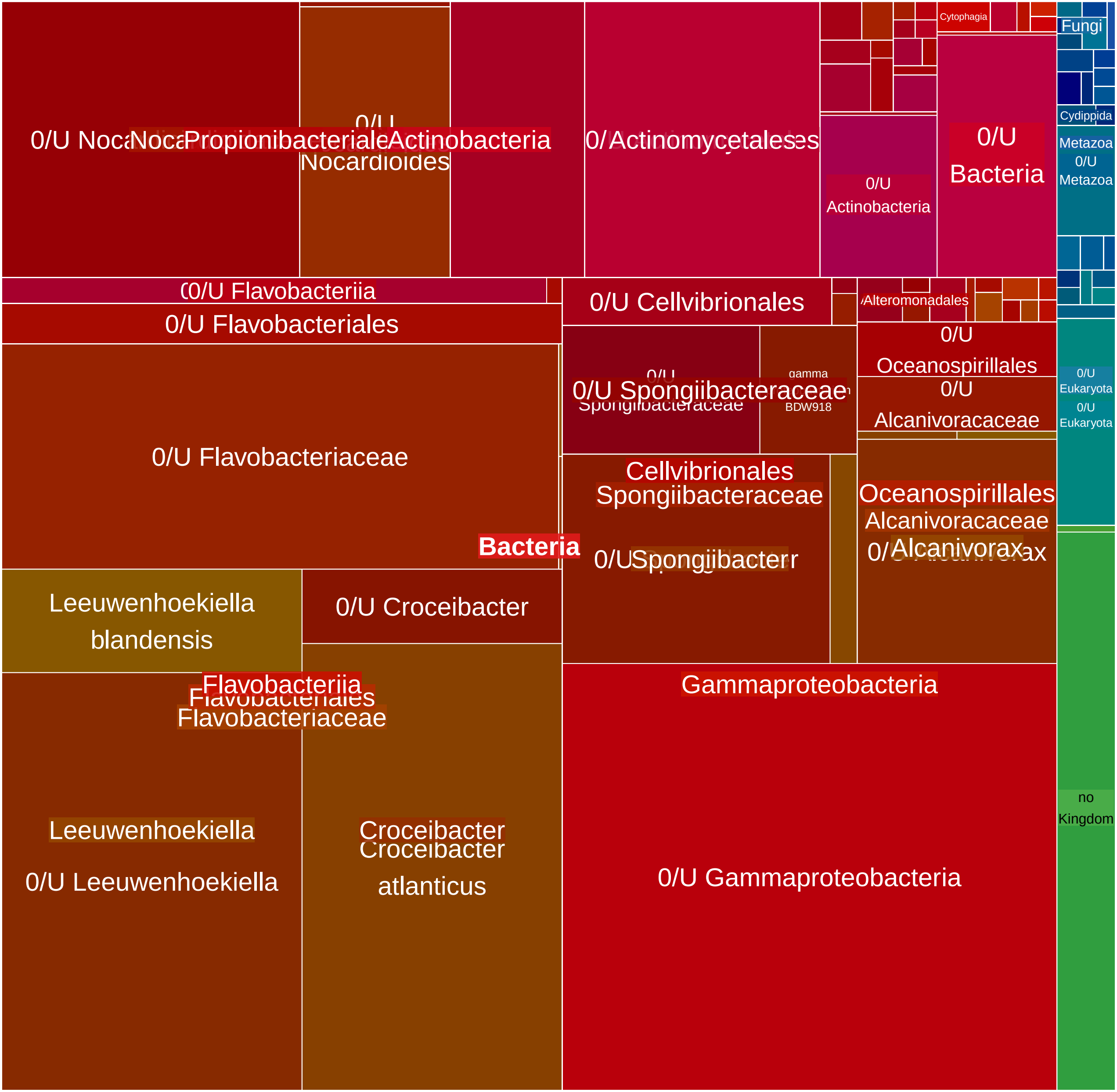

MGT−v1_306 : 3659 unigenes, 495 (13.53%) taxonomically assigned

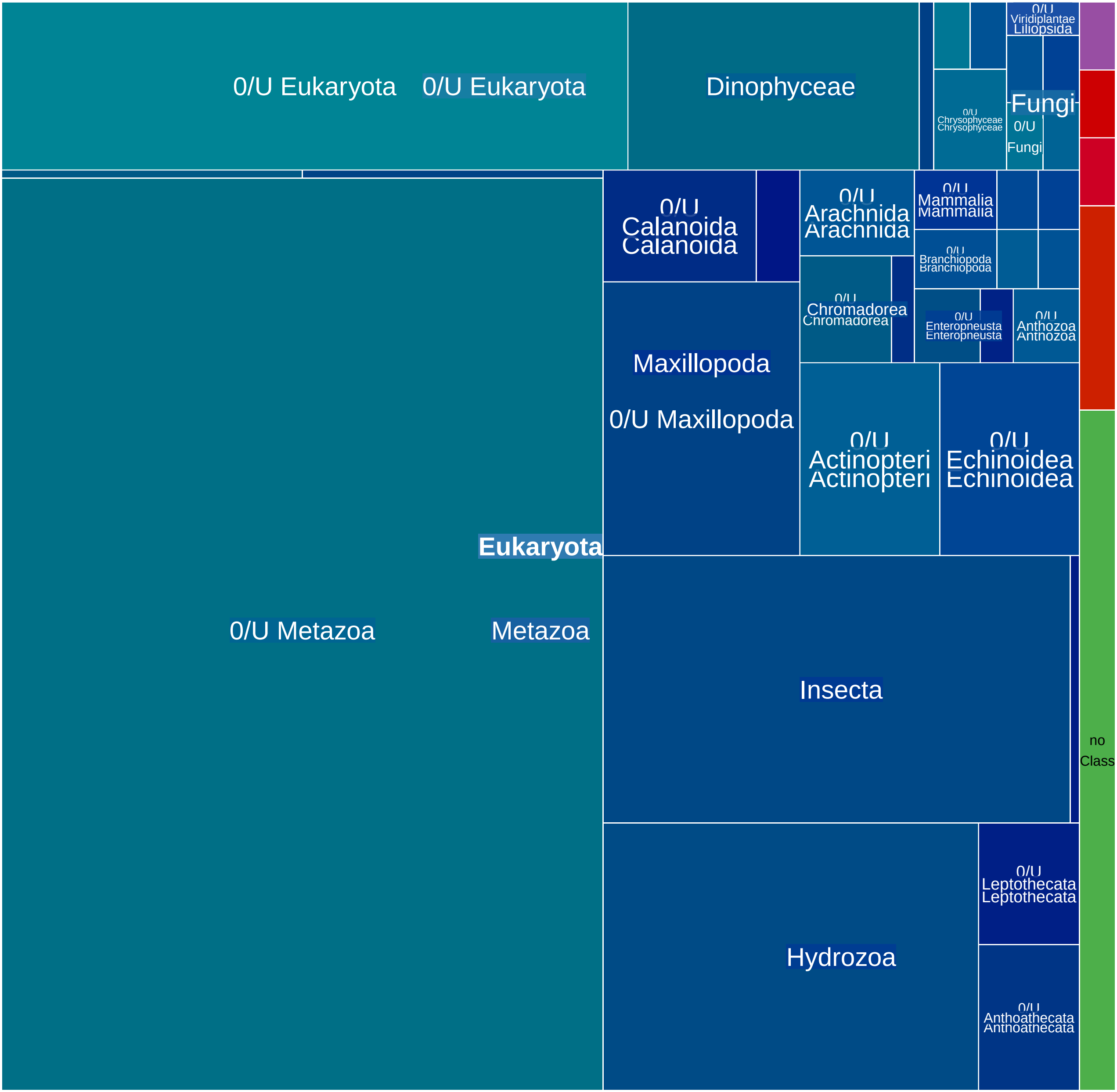

### MGT−v1_307 : 3654 unigenes, 3321 (90.89%) taxonomically assigned

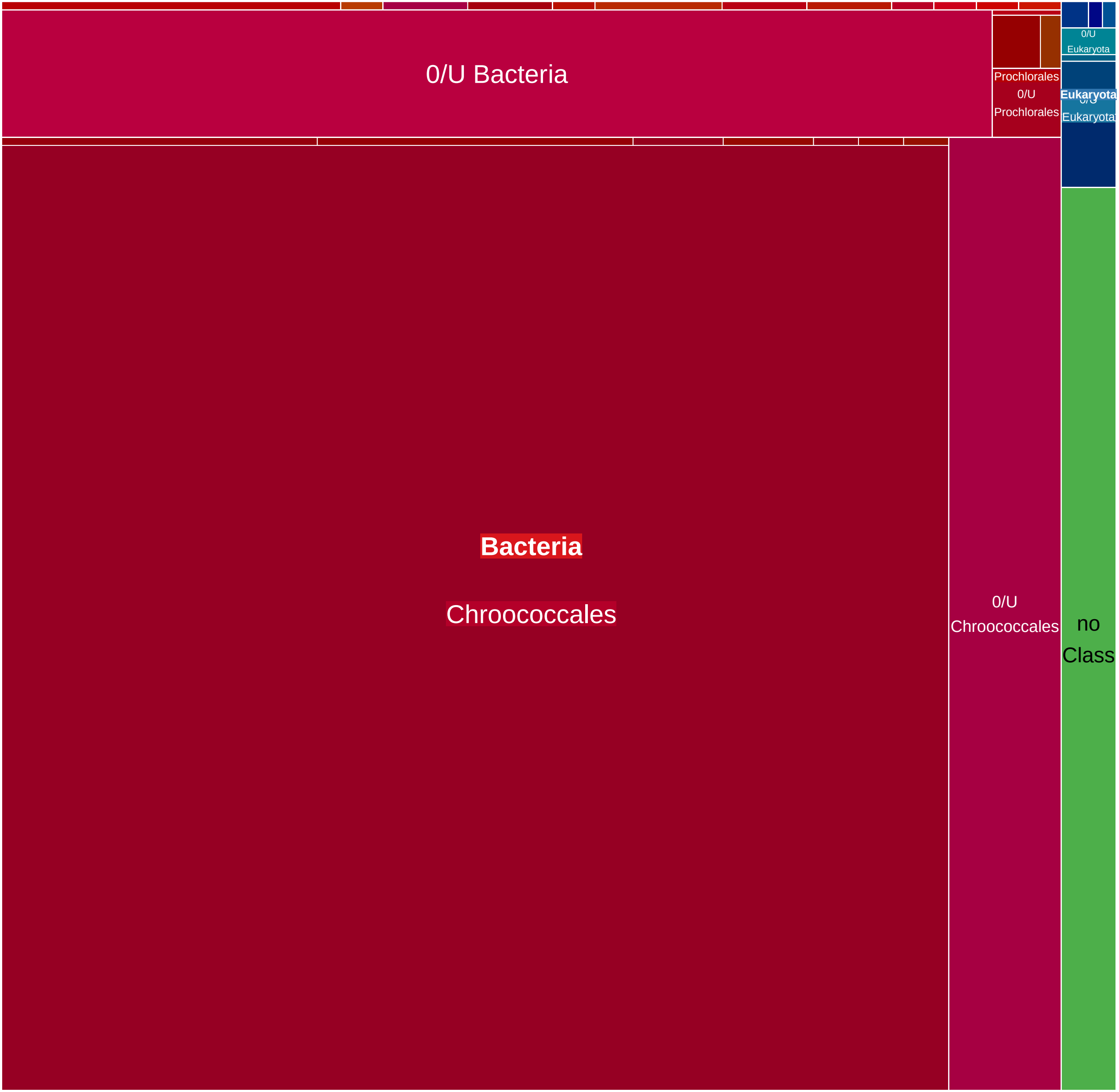

MGT−v1_308 : 3650 unigenes, 900 (24.66%) taxonomically assigned

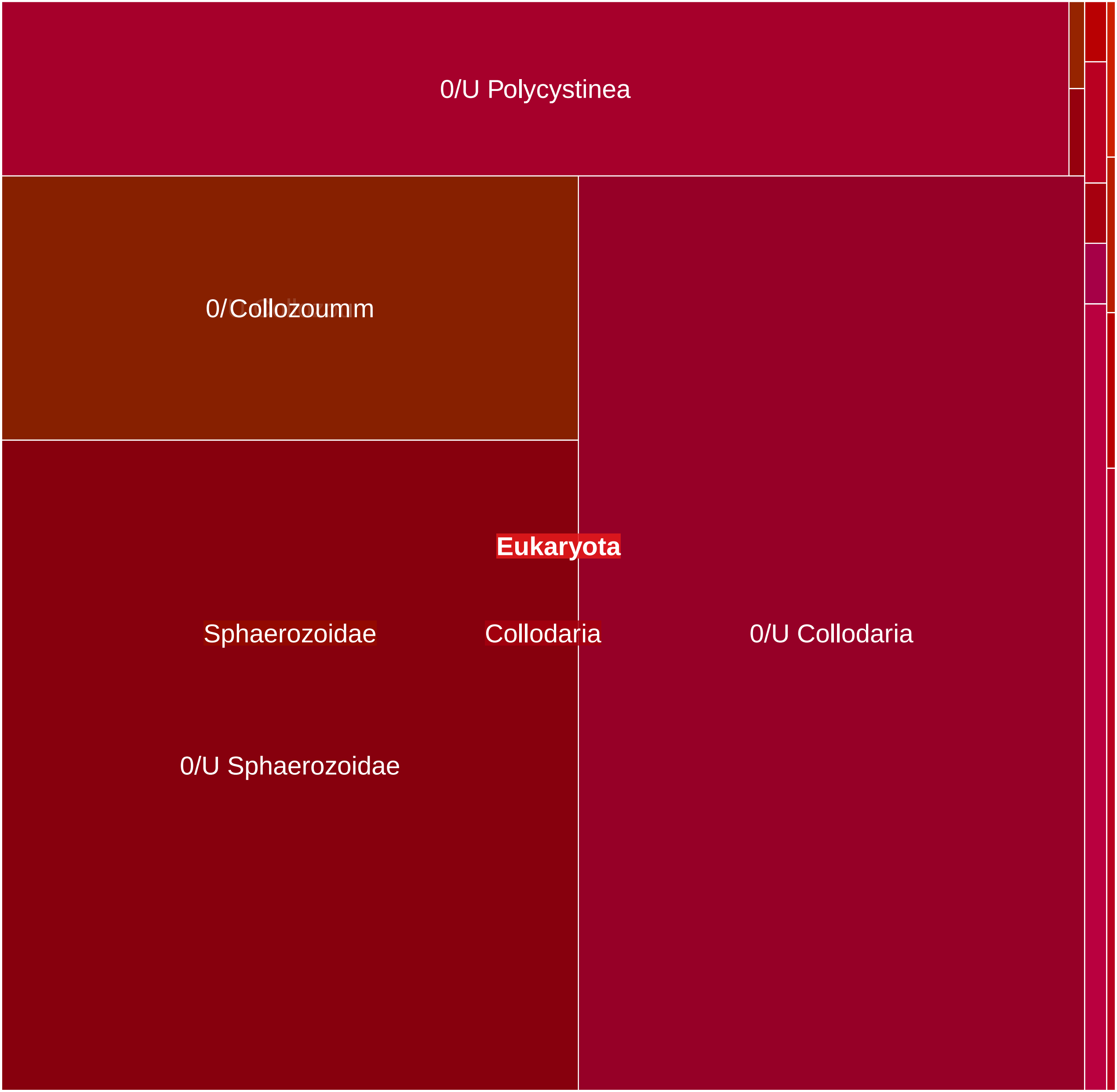

MGT−v1_309 : 3621 unigenes, 595 (16.43%) taxonomically assigned

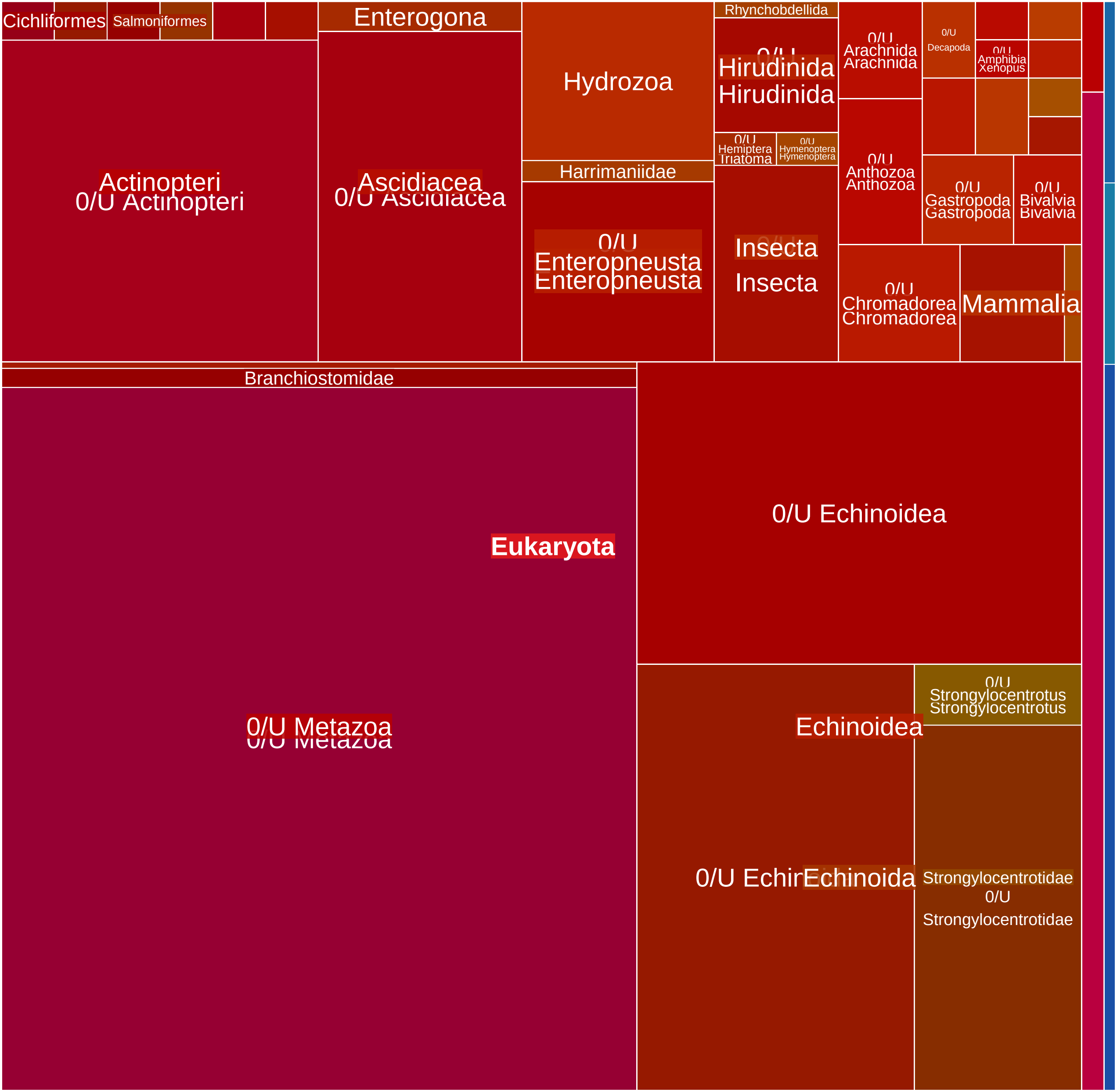

MGT−v1_310 : 3608 unigenes, 615 (17.05%) taxonomically assigned

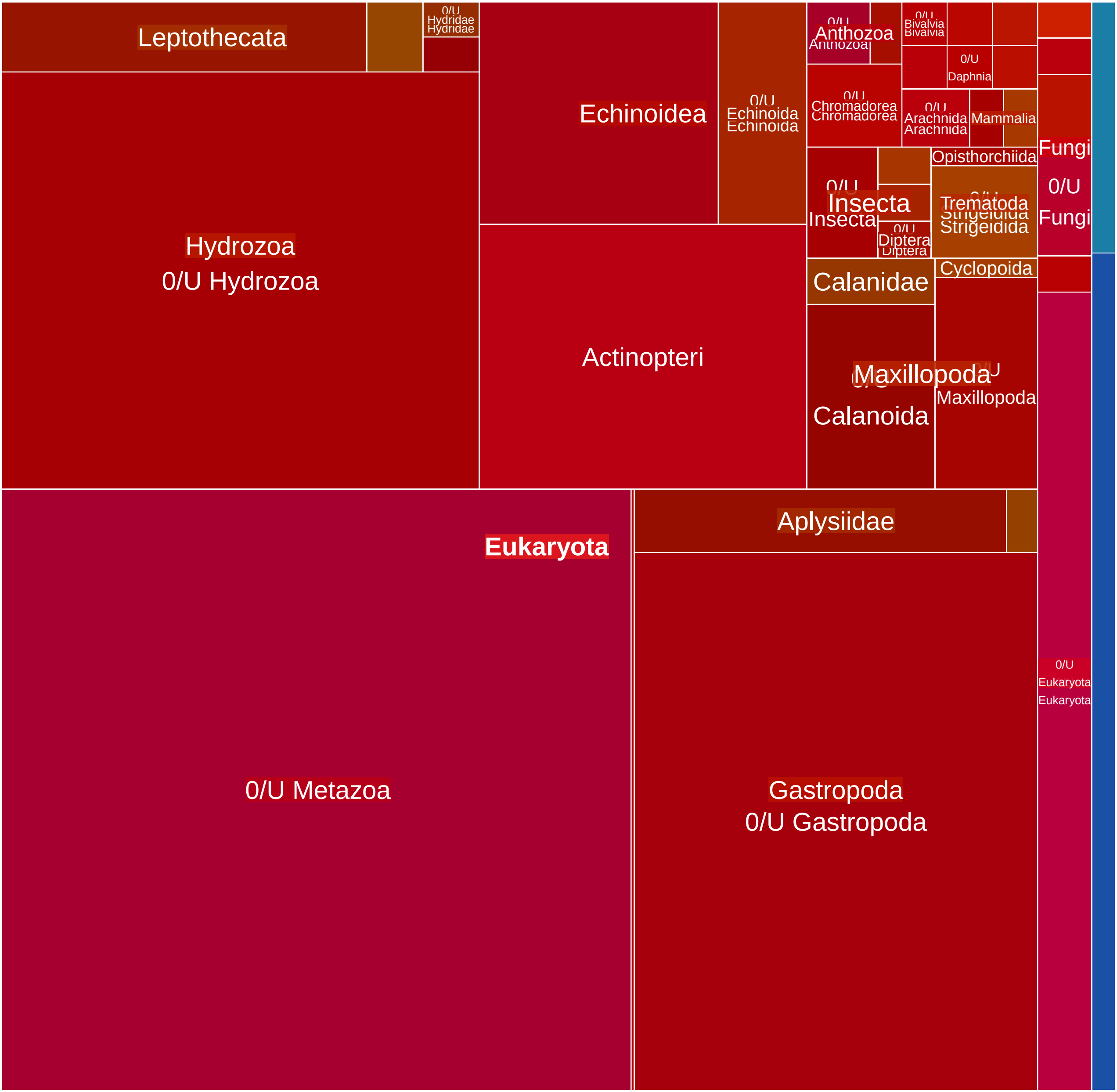

MGT−v1_311 : 3573 unigenes, 234 (6.55%) taxonomically assigned

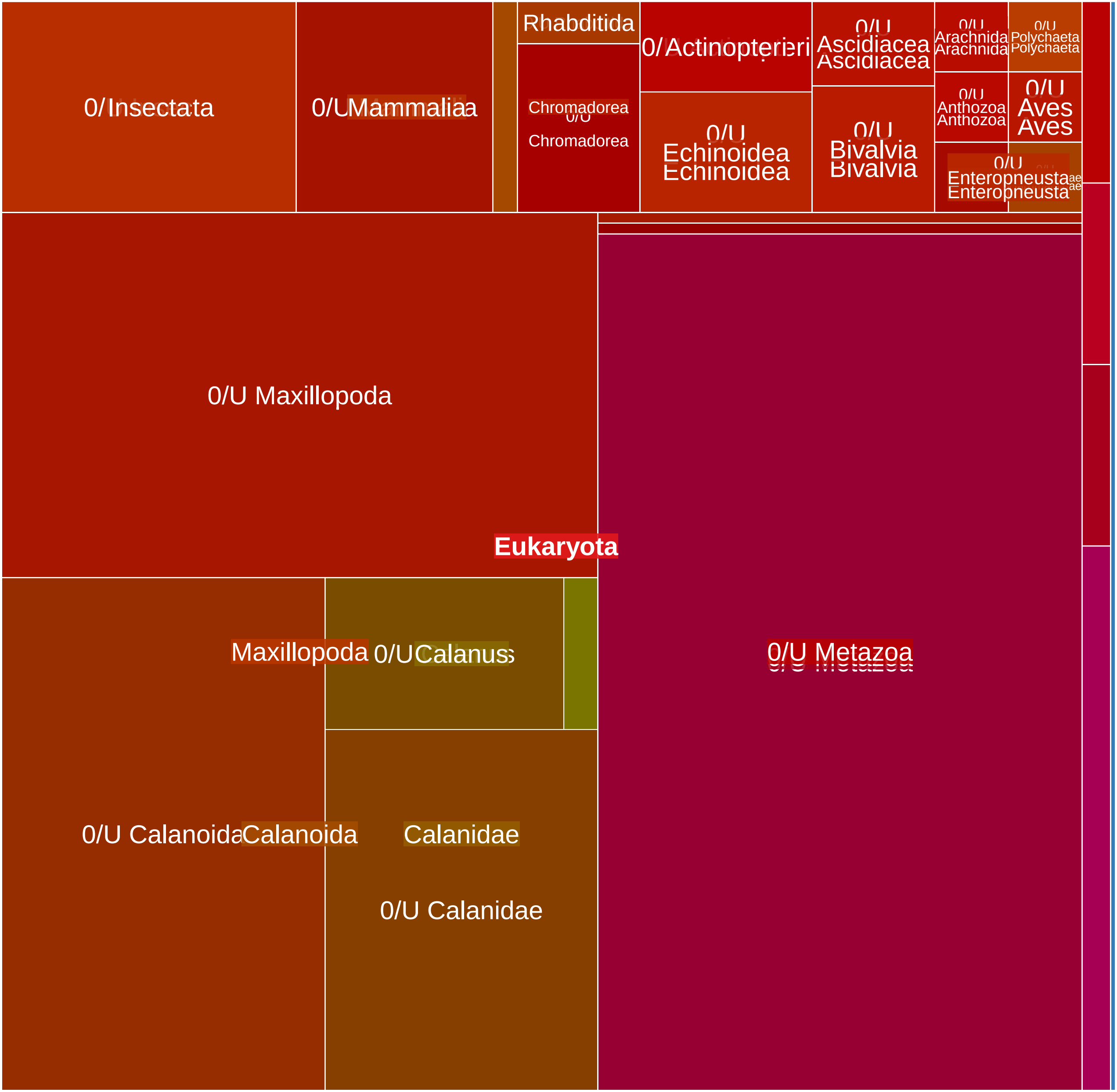

MGT−v1_312 : 3572 unigenes, 1558 (43.62%) taxonomically assigned

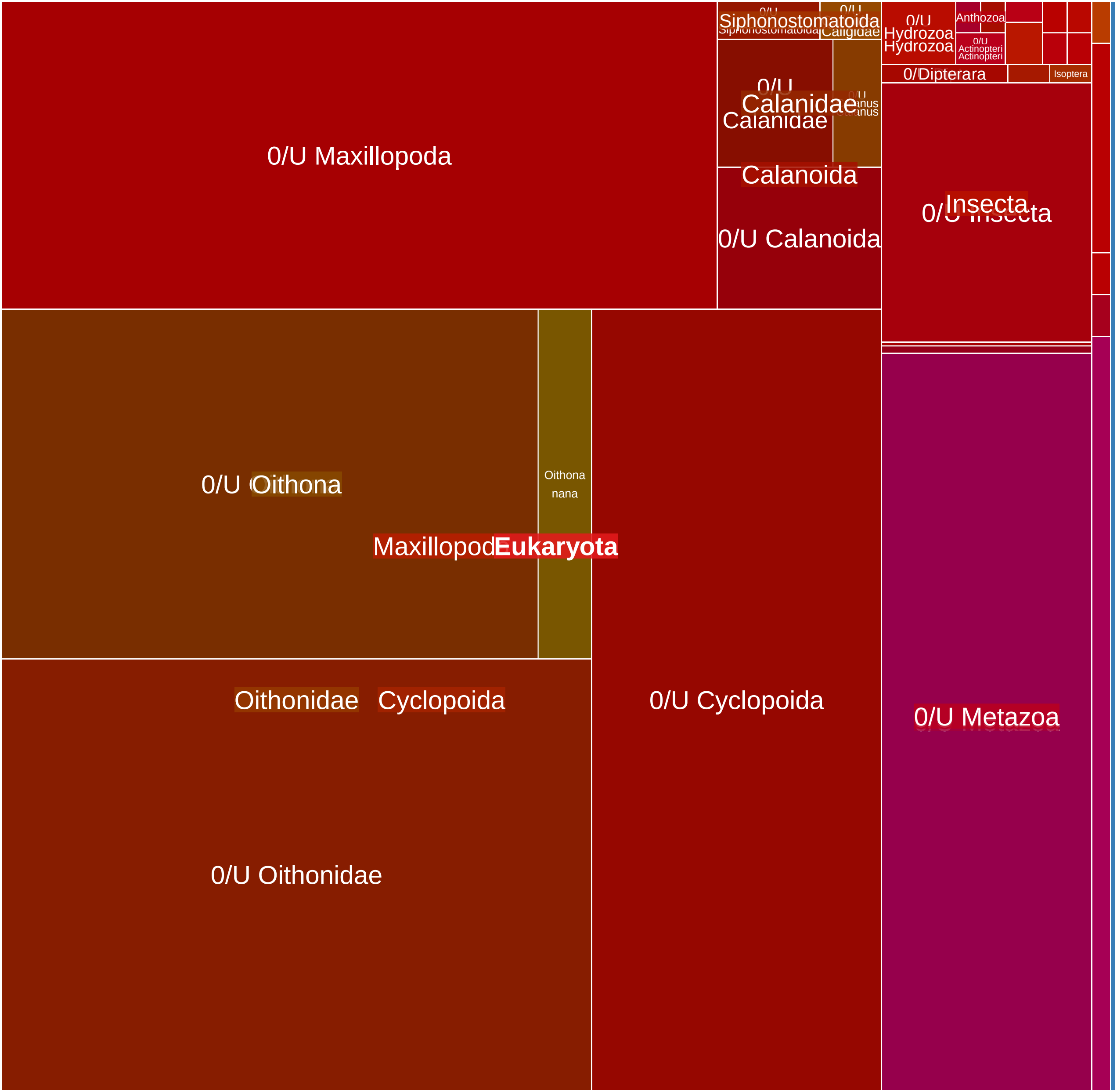

MGT−v1_313 : 3560 unigenes, 519 (14.58%) taxonomically assigned

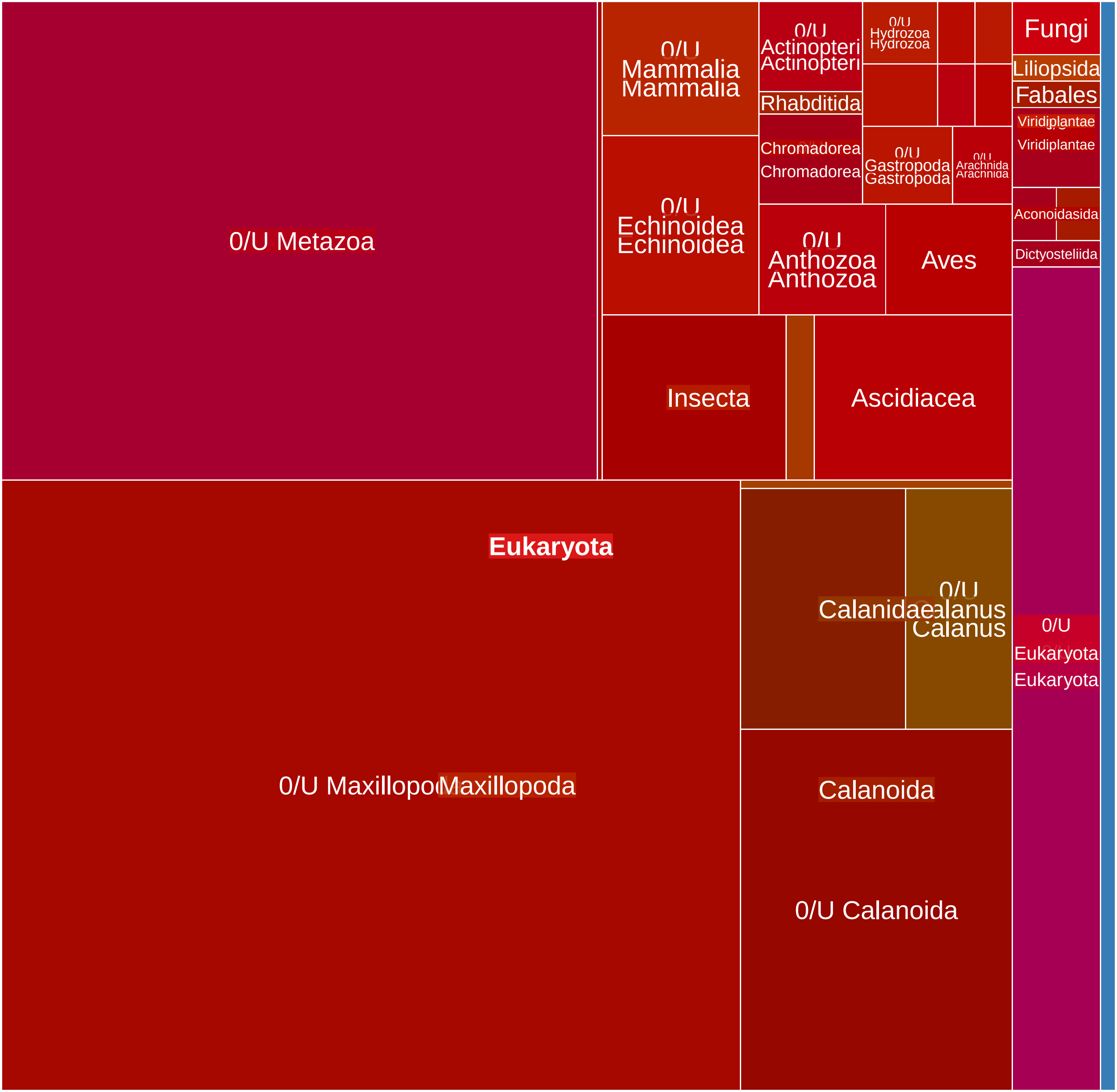

MGT−v1_314 : 3555 unigenes, 455 (12.80%) taxonomically assigned

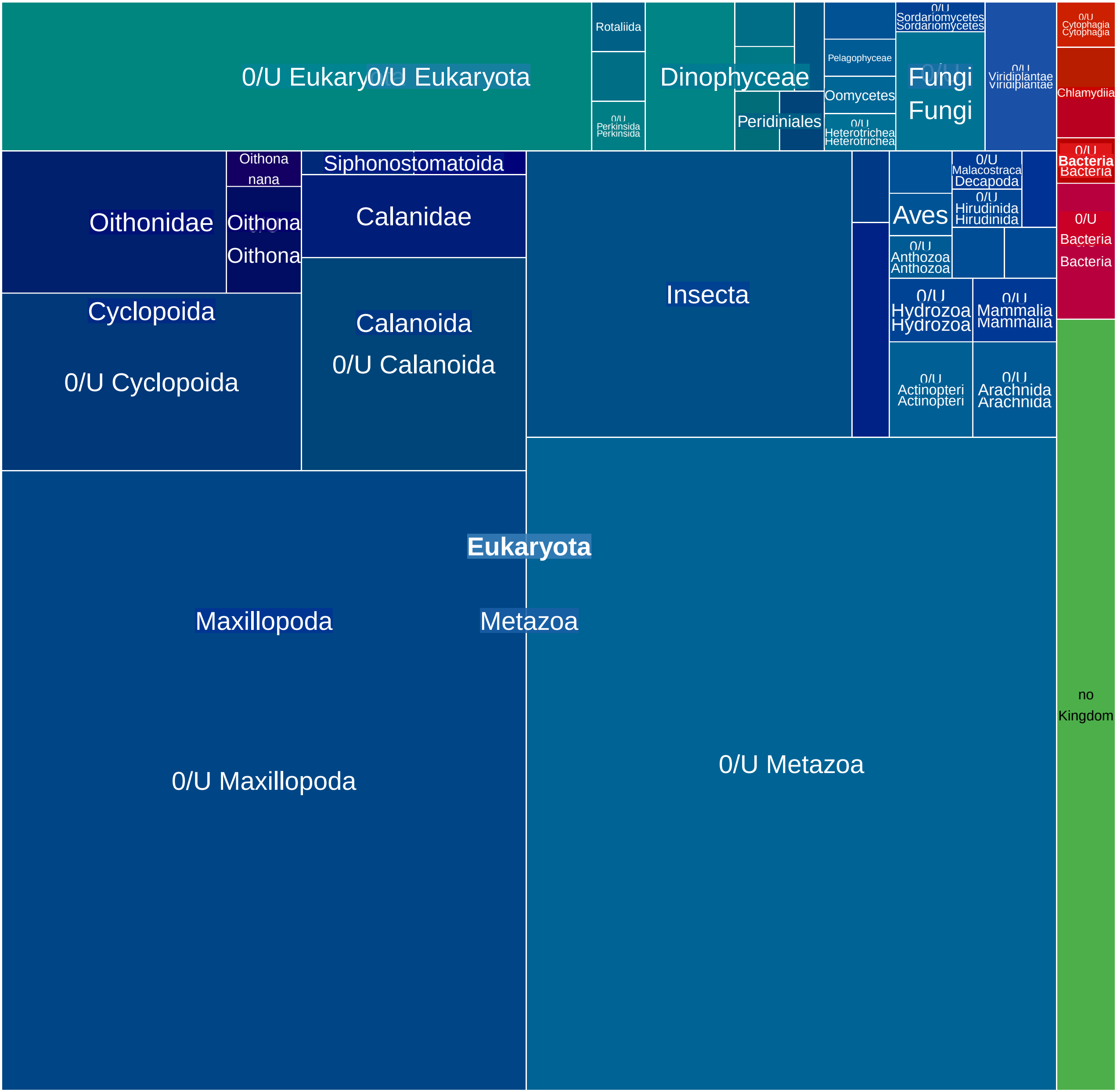

MGT−v1_315 : 3544 unigenes, 307 (8.66%) taxonomically assigned

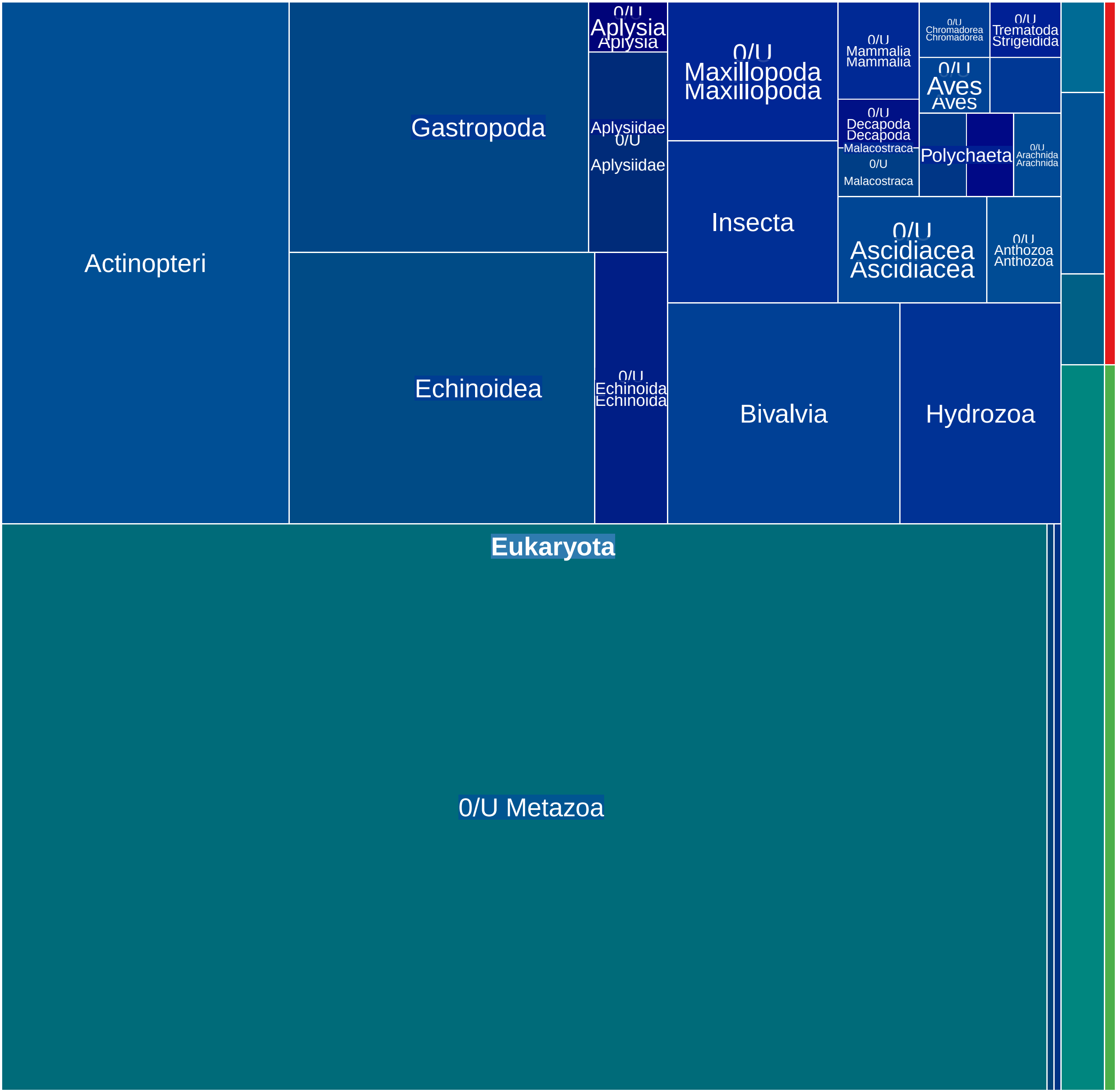

MGT−v1_316 : 3540 unigenes, 765 (21.61%) taxonomically assigned

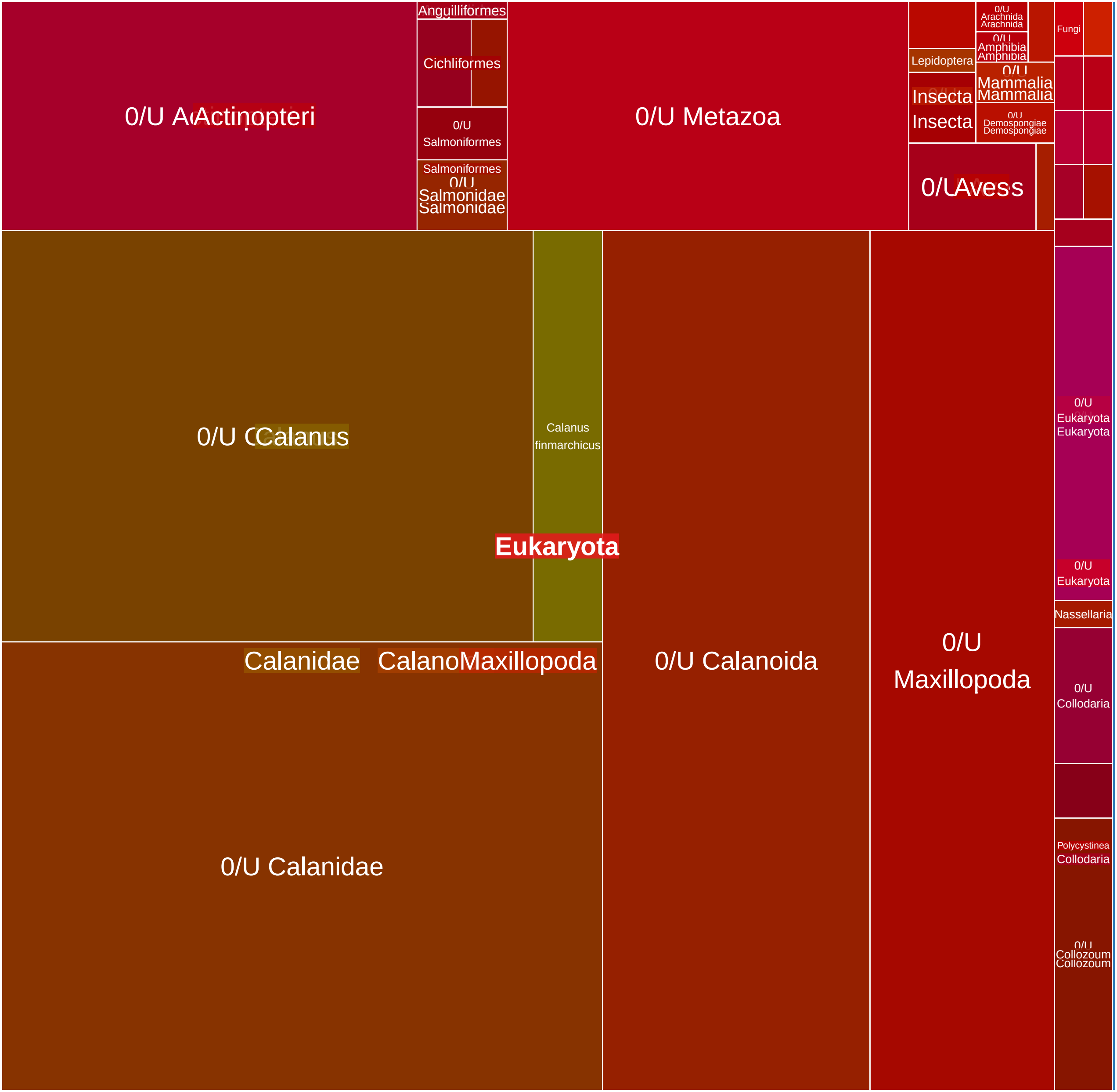

MGT−v1_317 : 3526 unigenes, 292 (8.28%) taxonomically assigned

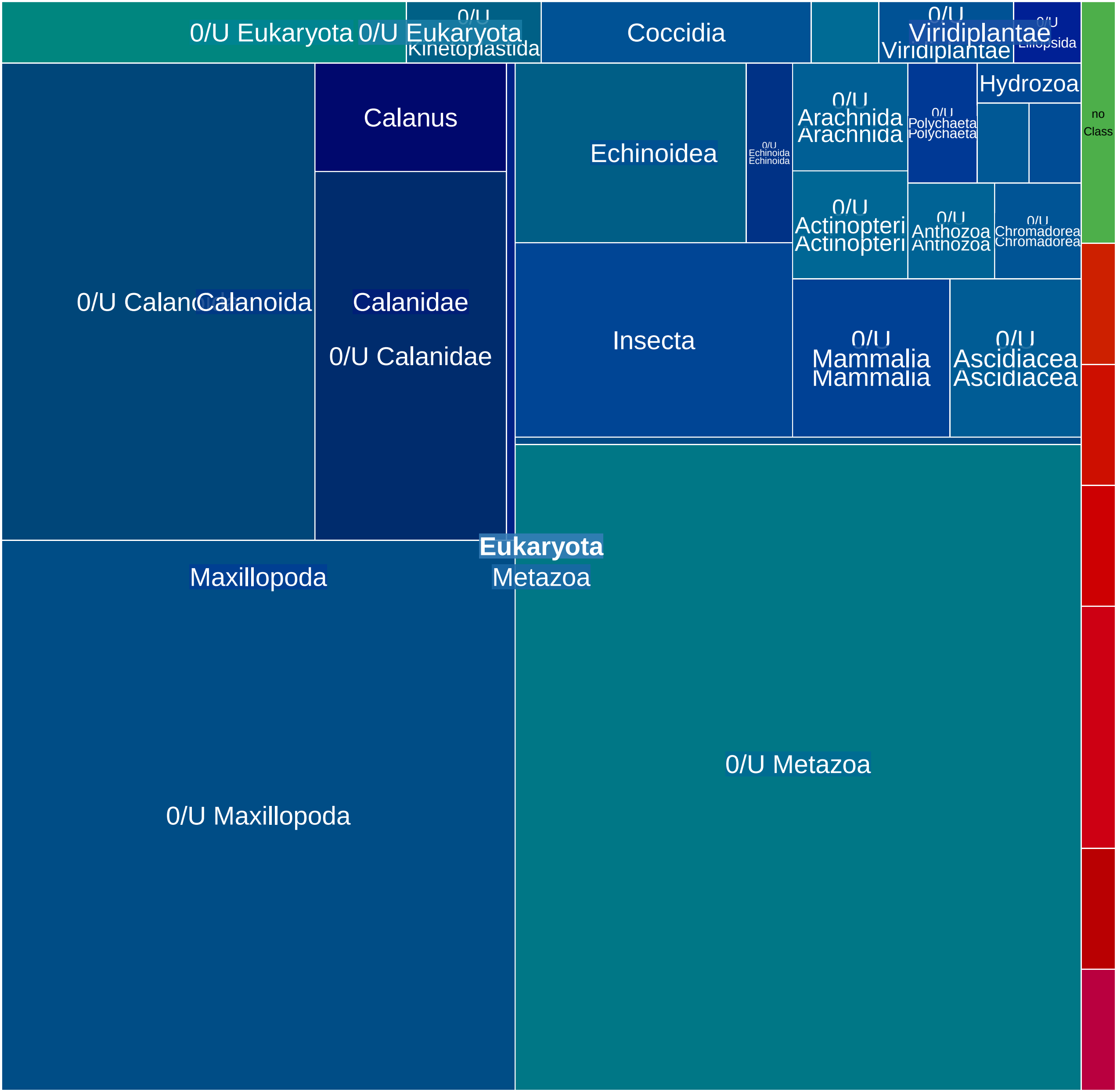

MGT−v1_318 : 3519 unigenes, 161 (4.58%) taxonomically assigned

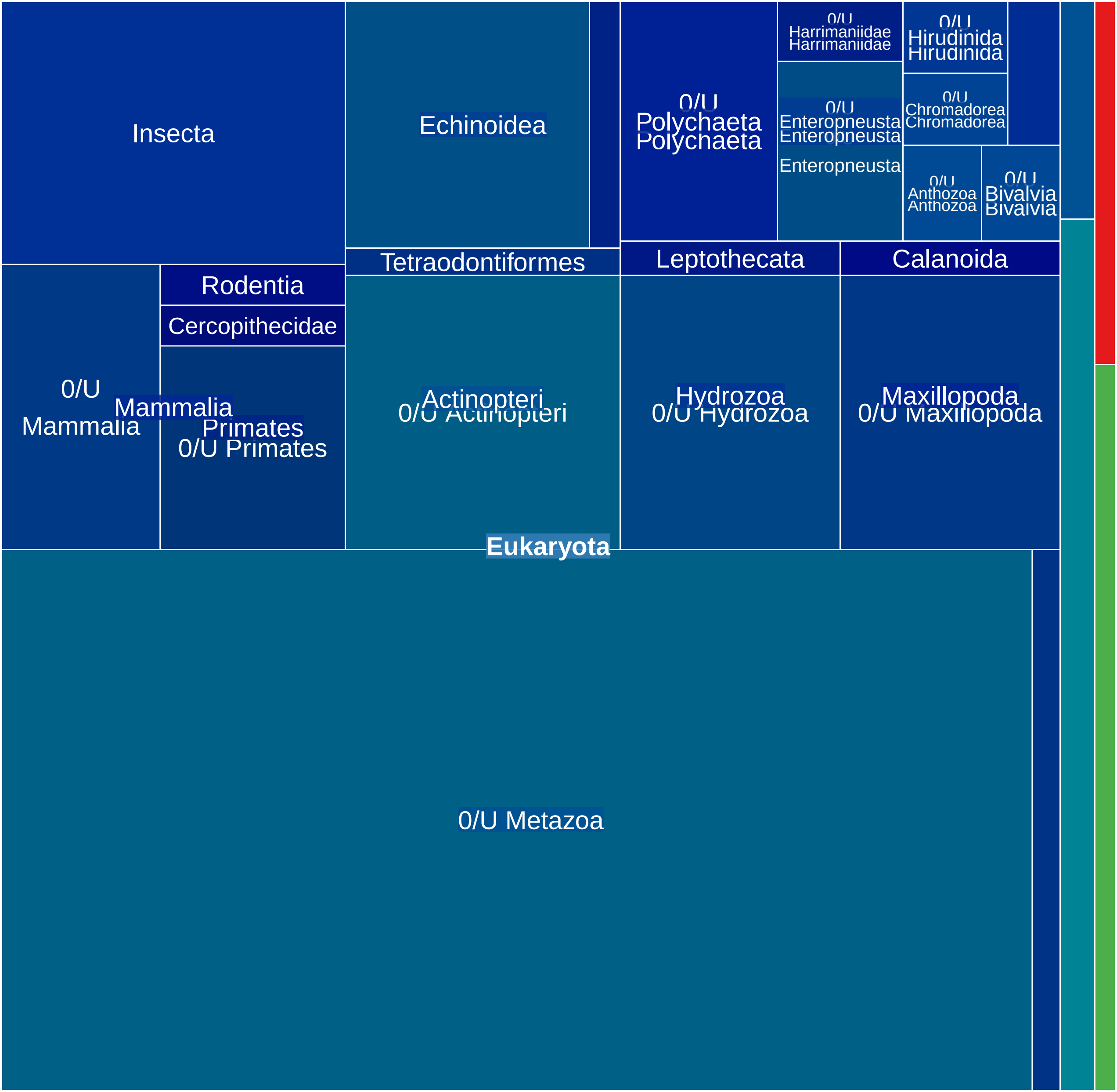

MGT−v1_319 : 3513 unigenes, 3301 (93.97%) taxonomically assigned

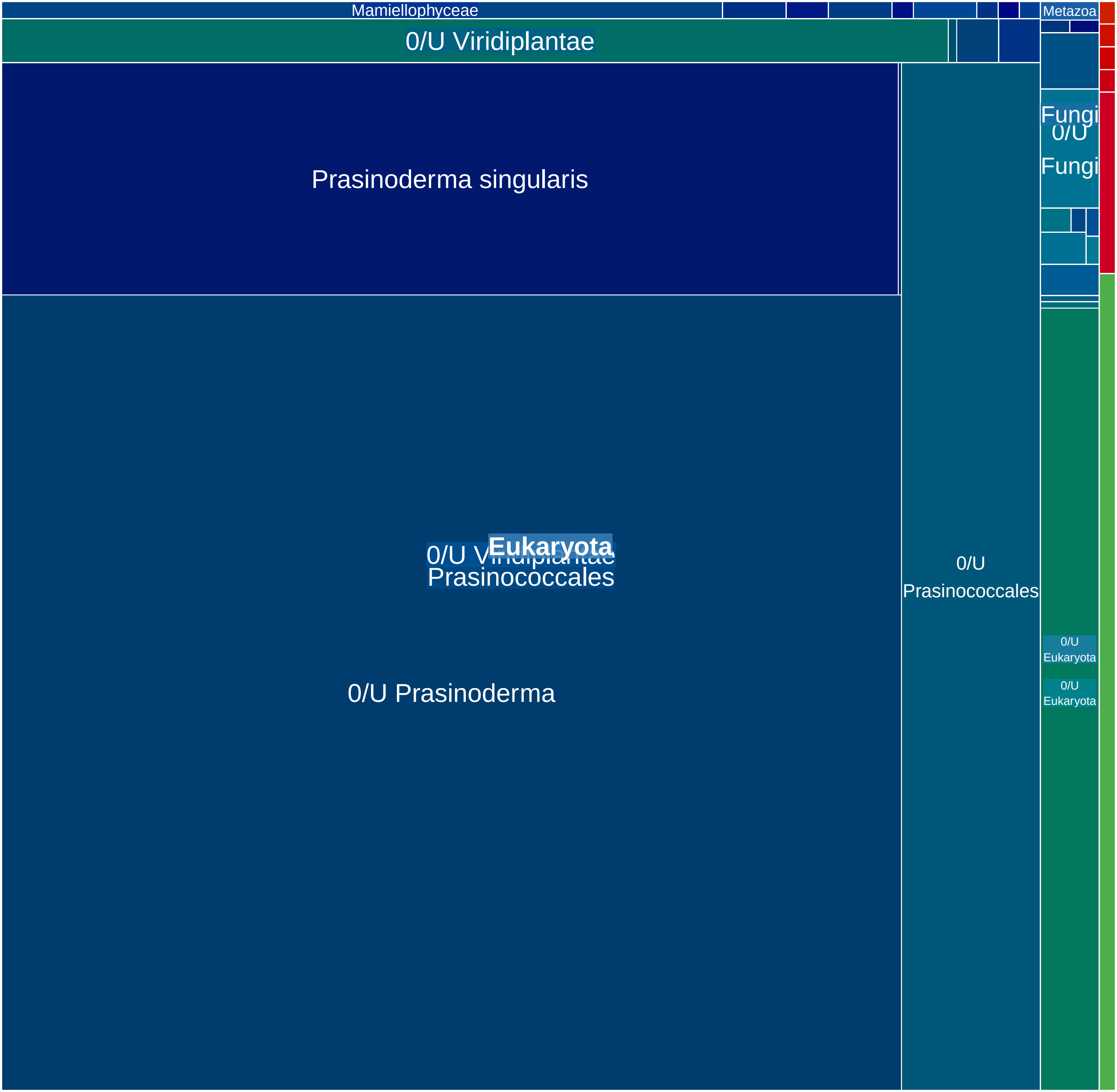

MGT−v1_320 : 3512 unigenes, 2454 (69.87%) taxonomically assigned

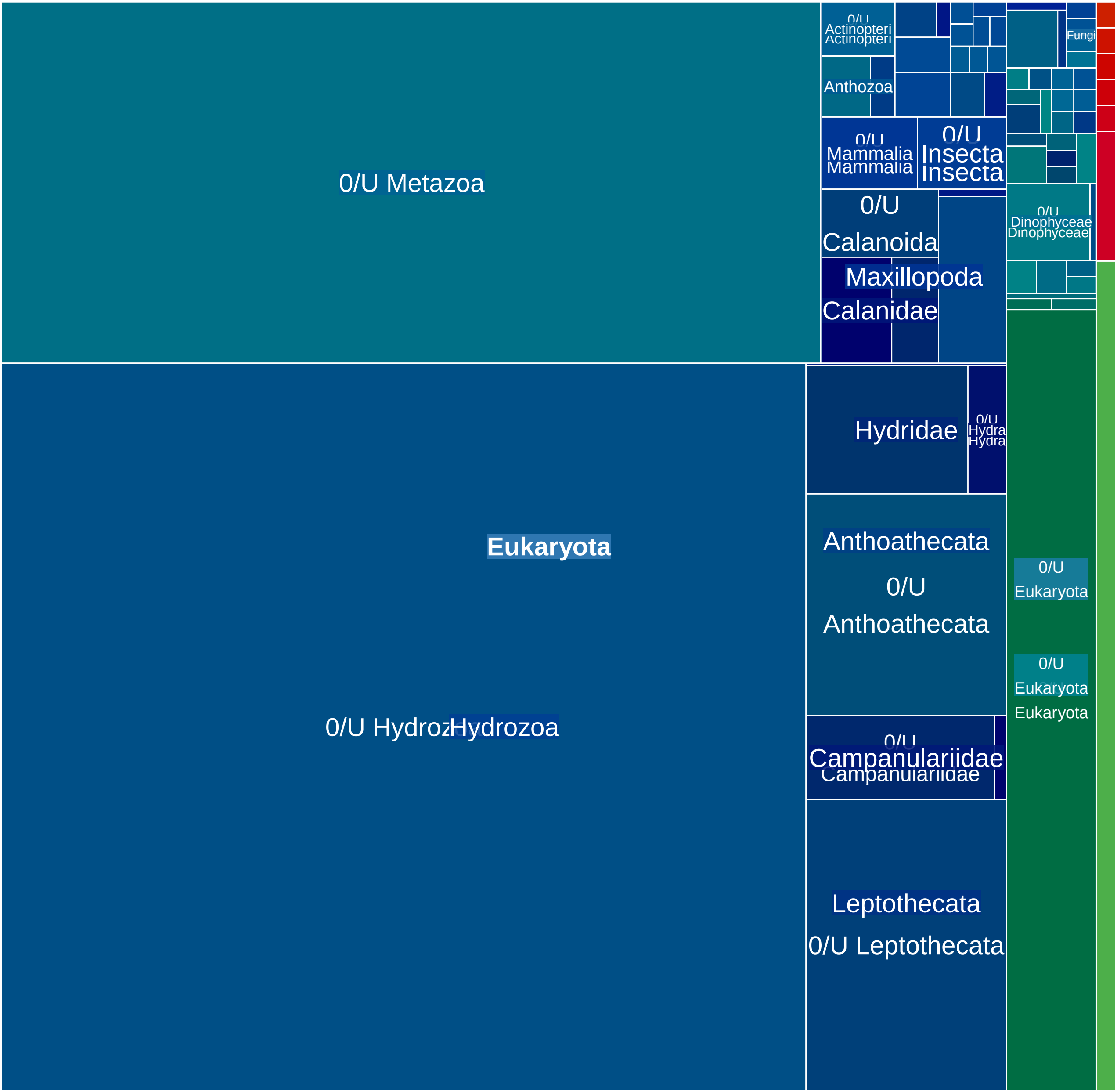

### MGT−v1_321 : 3490 unigenes, 3347 (95.90%) taxonomically assigned

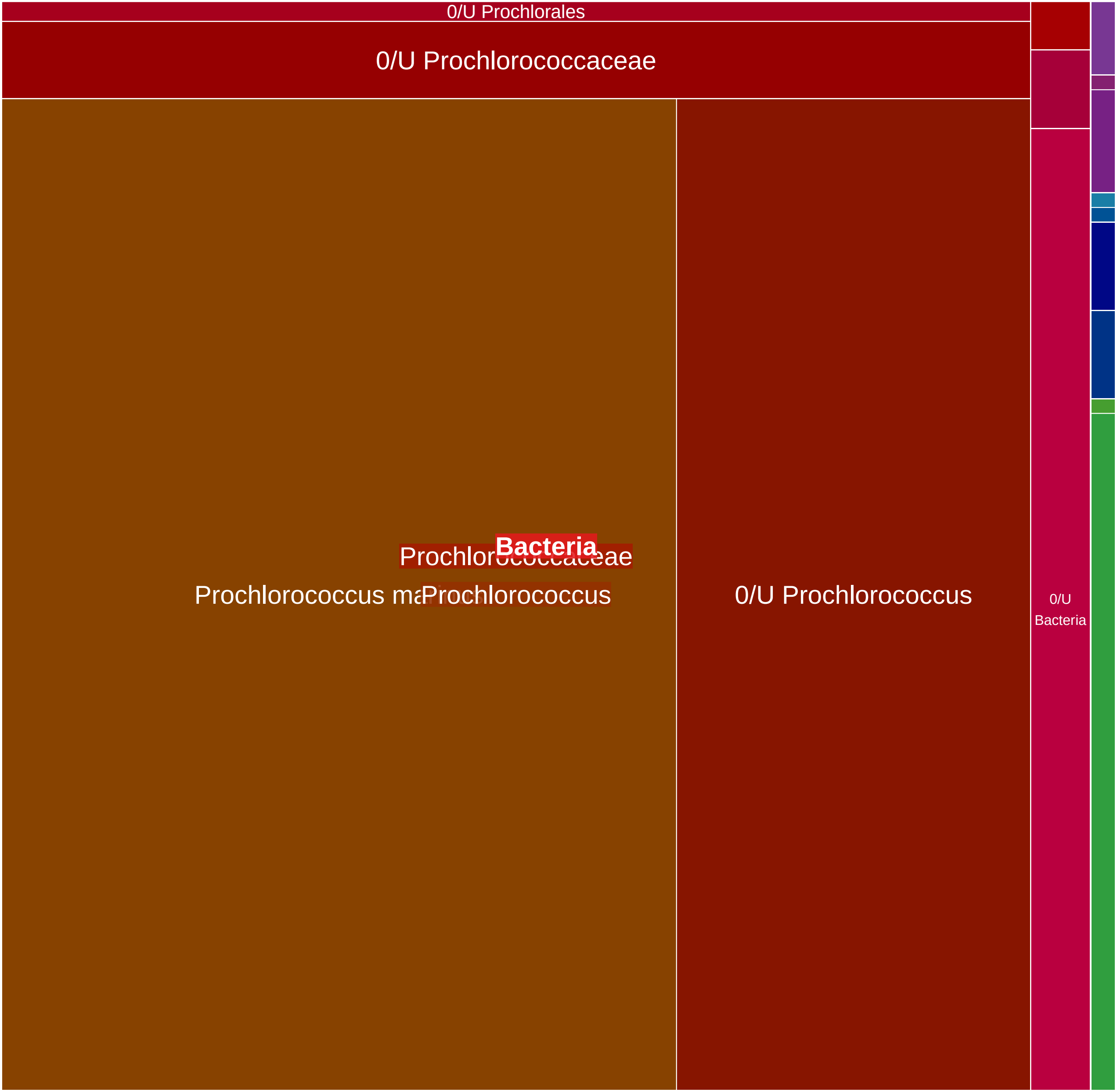

### MGT−v1_322 : 3482 unigenes, 382 (10.97%) taxonomically assigned

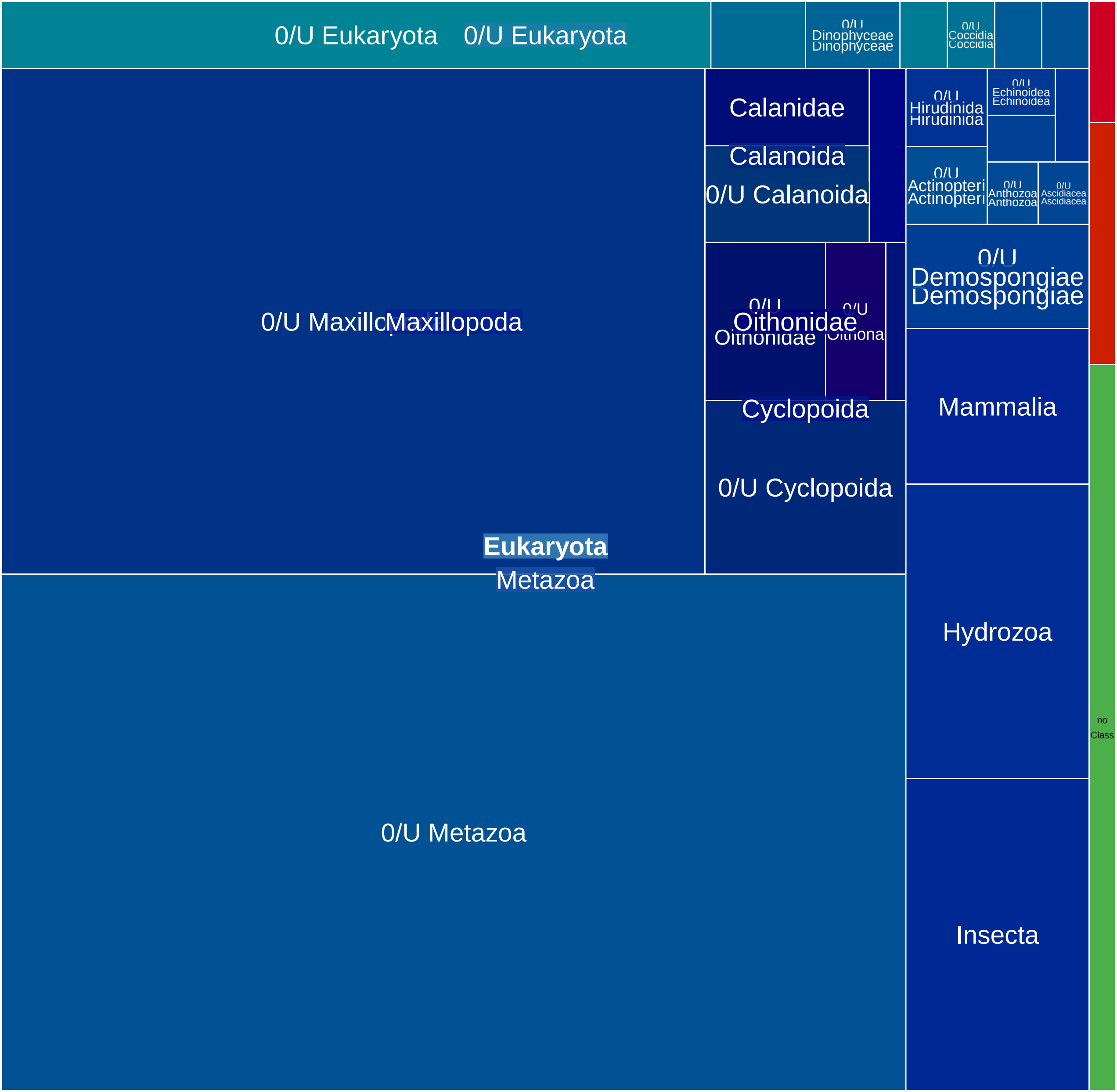

MGT−v1_323 : 3472 unigenes, 1961 (56.48%) taxonomically assigned

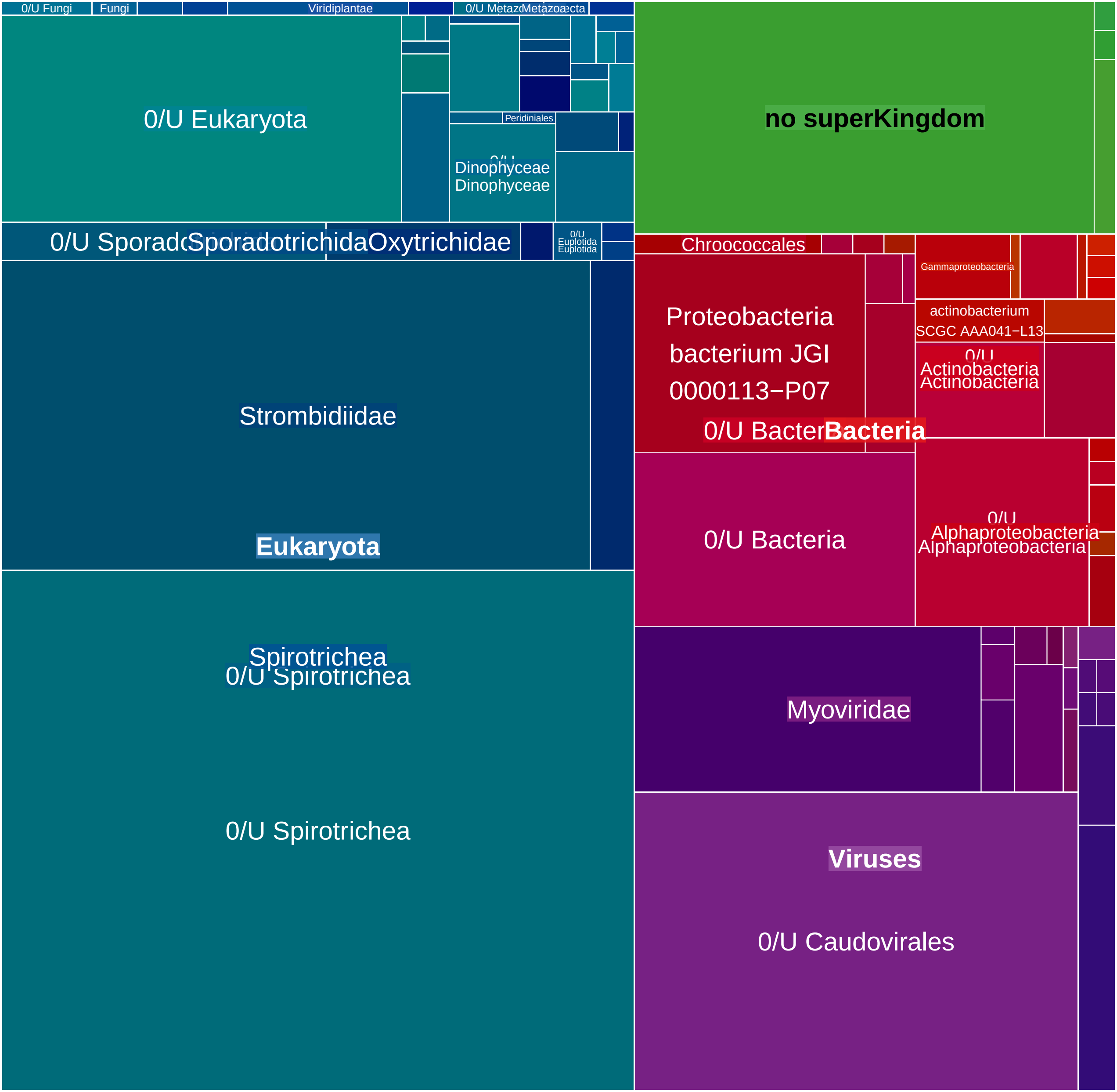

### MGT−v1_324 : 3458 unigenes, 2216 (64.08%) taxonomically assigned

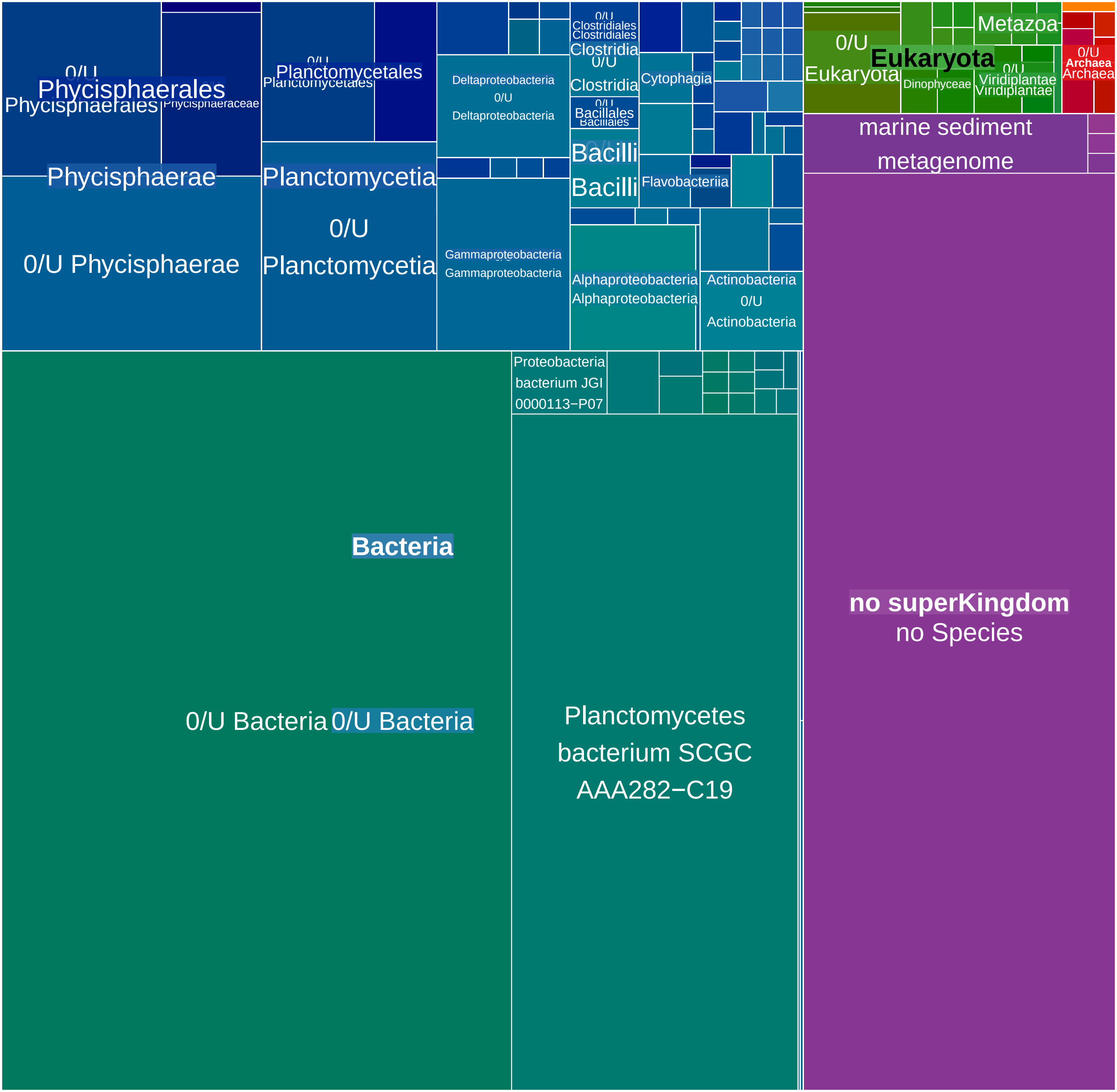

MGT−v1_325 : 3411 unigenes, 2580 (75.64%) taxonomically assigned

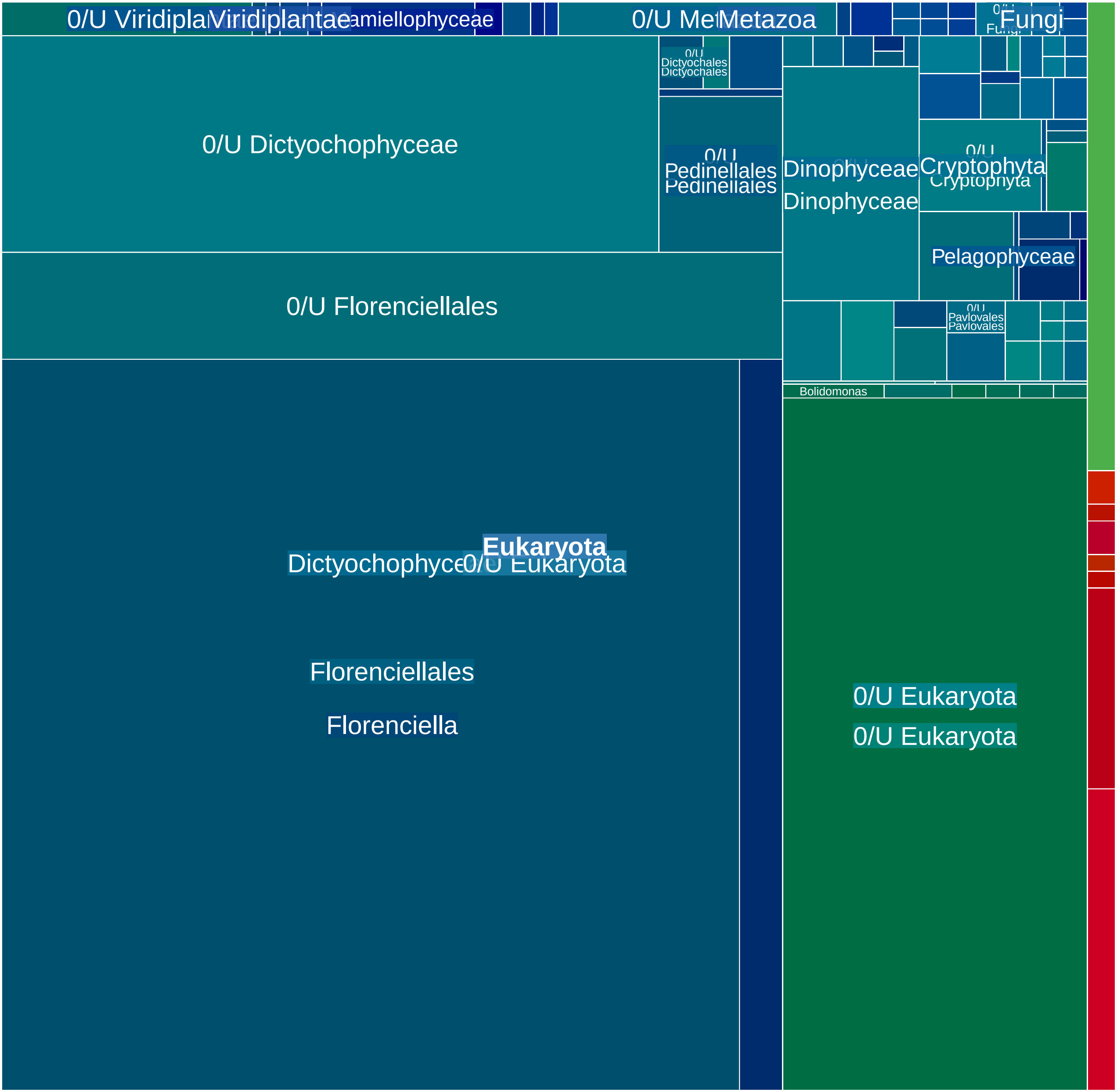

### MGT−v1_326 : 3408 unigenes, 2728 (80.05%) taxonomically assigned

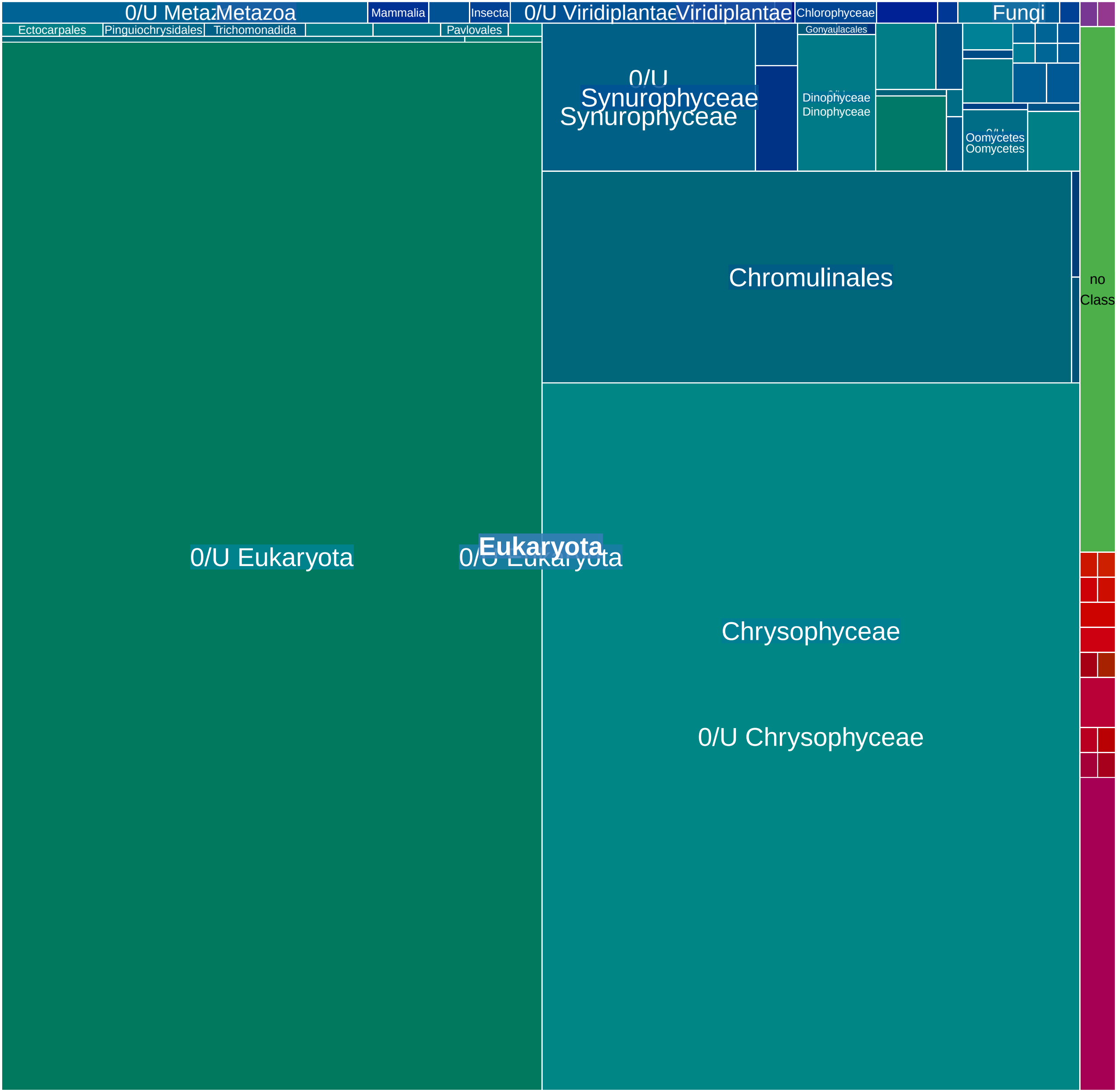

### MGT−v1_327 : 3406 unigenes, 1894 (55.61%) taxonomically assigned

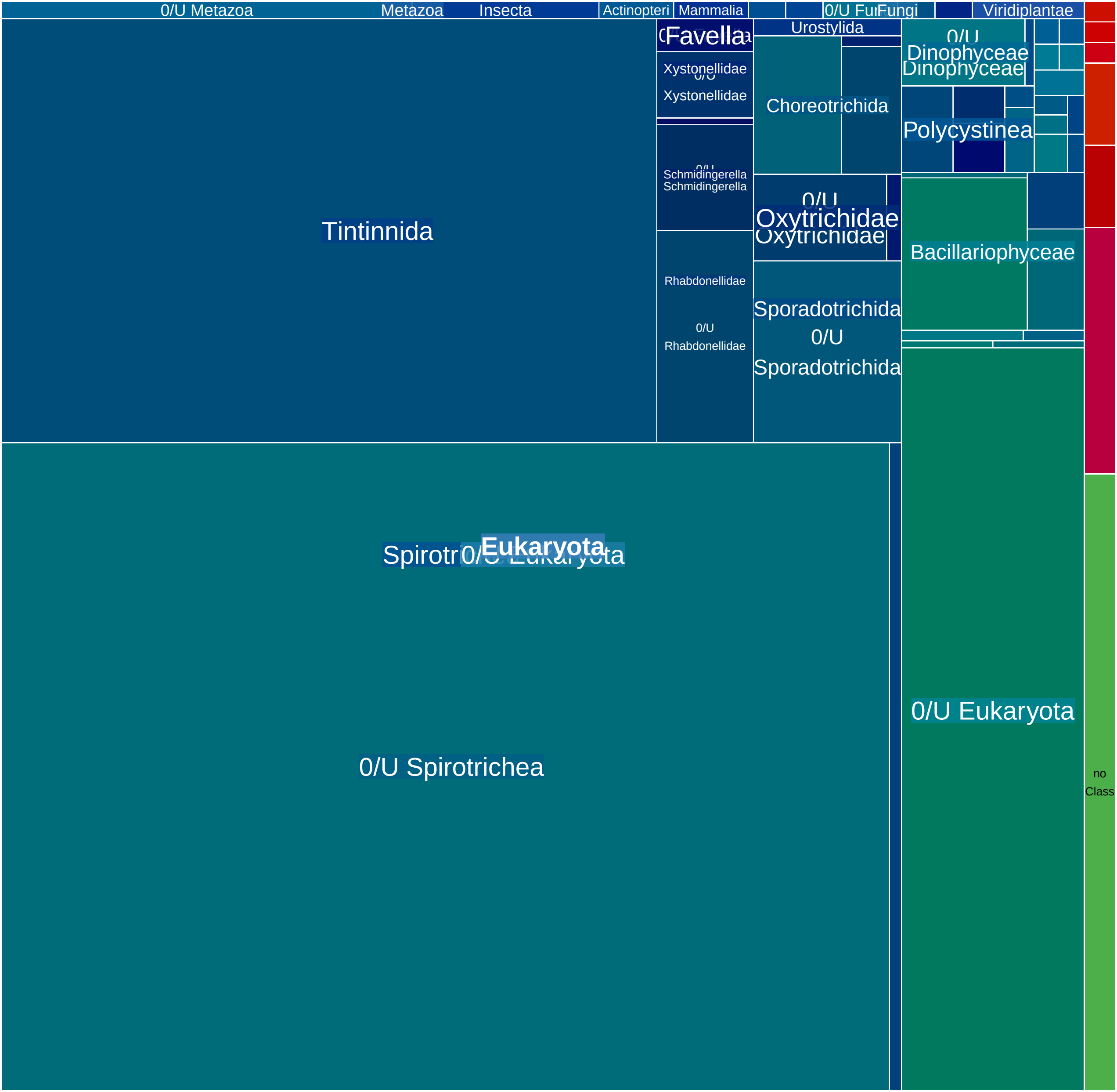

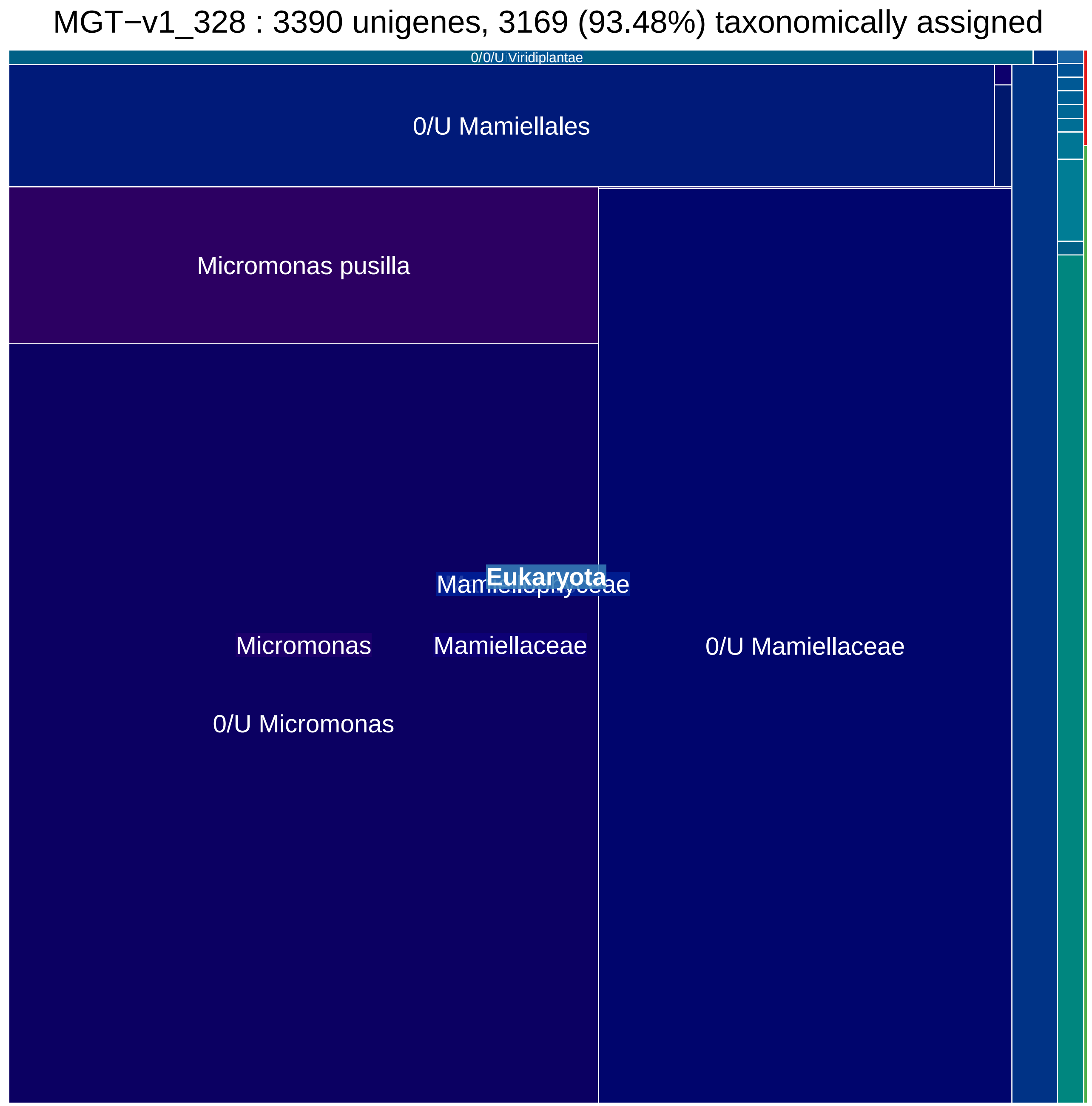

MGT−v1_328 : 3390 unigenes, 3169 (93.48%) taxonomically assigned
0/0/U Viridiplantae
0/U Mamiellales
Micromonas pusilla
Eukaryota
Mamiellophyceae
Micromonas
Mamiellaceae
0/U Mamiellaceae
0/U Micromonas

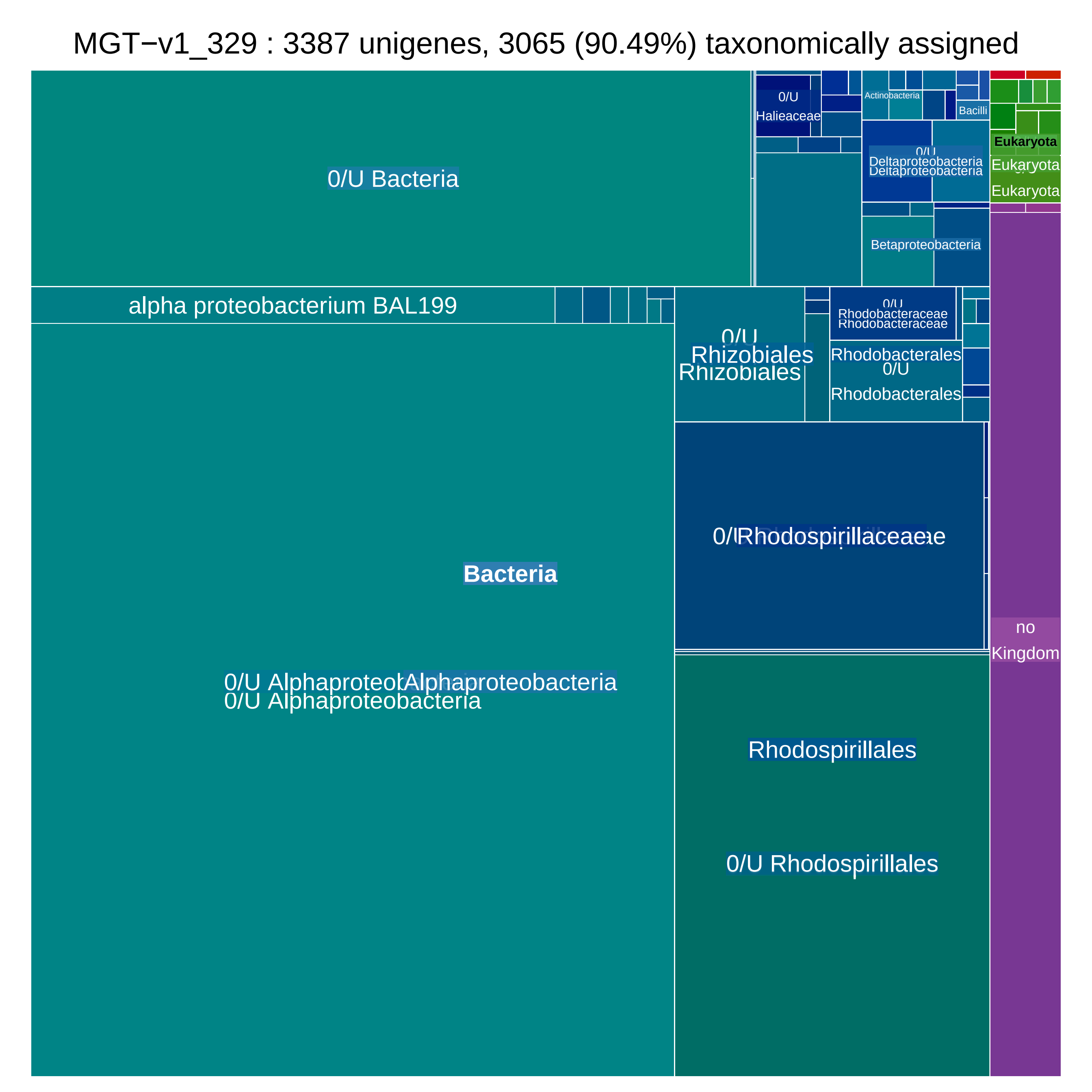
MGT−v1_329 : 3387 unigenes, 3065 (90.49%) taxonomically assigned
0/U Bacteria
alpha proteobacterium BAL199
Bacteria
0/U Alphaproteobacteria
Alphaproteobacteria
0/U Alphaproteobacteria
0/U Halieaceae
Actinobacteria
Bacilli
0/U Deltaproteobacteria
Deltaproteobacteria
Betaproteobacteria
Eukaryota
Eukaryota
Eukaryota
0/U Rhizobiales
Rhizobiales
0/U Rhodobacteraceae
Rhodobacteraceae
Rhodobacterales
0/U Rhodobacterales
Rhodospirillaceae
Rhodospirillales
0/U Rhodospirillales
no Kingdom

### MGT−v1_330 : 3376 unigenes, 1691 (50.09%) taxonomically assigned

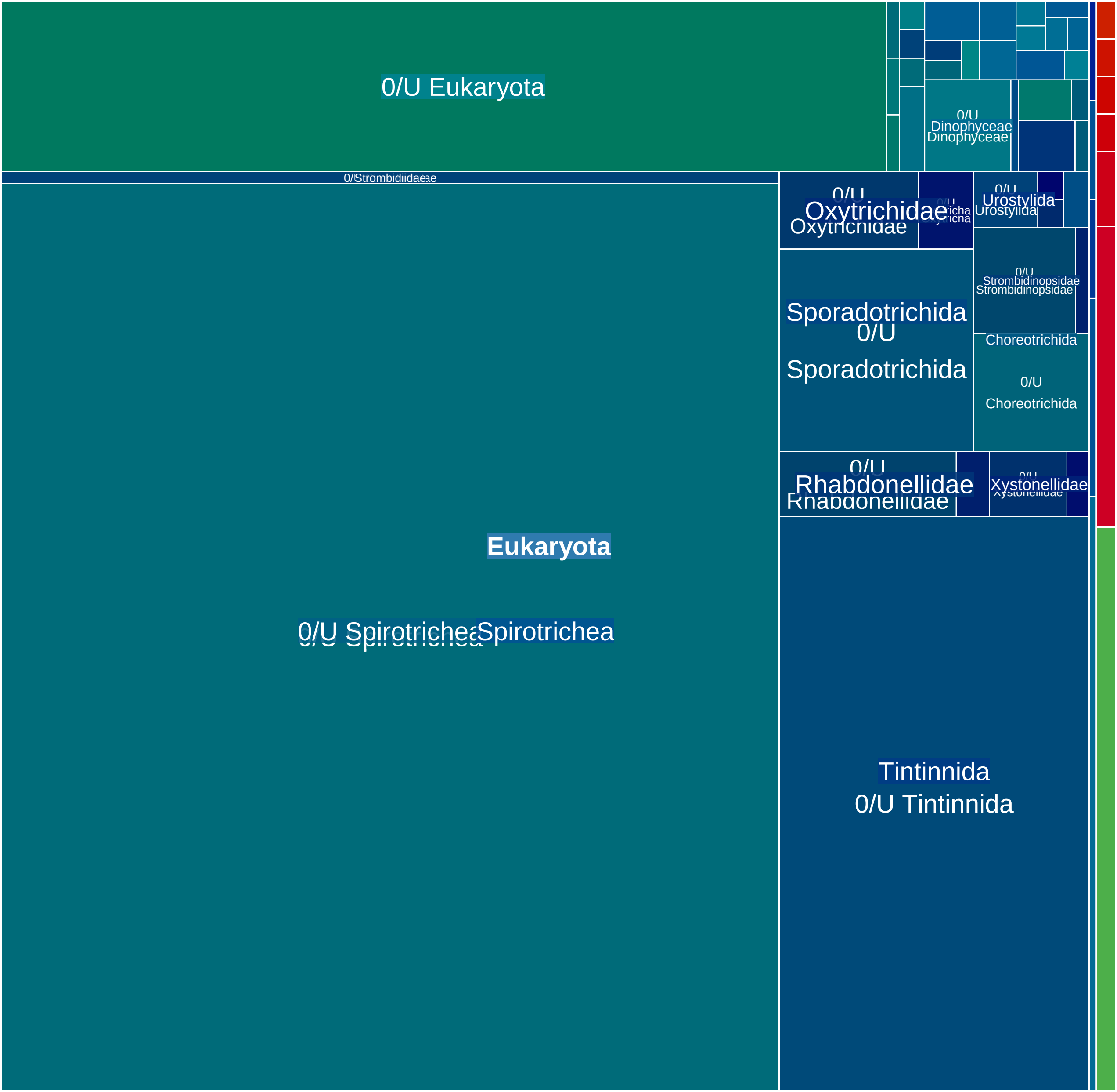

MGT−v1_331 : 3357 unigenes, 267 (7.95%) taxonomically assigned

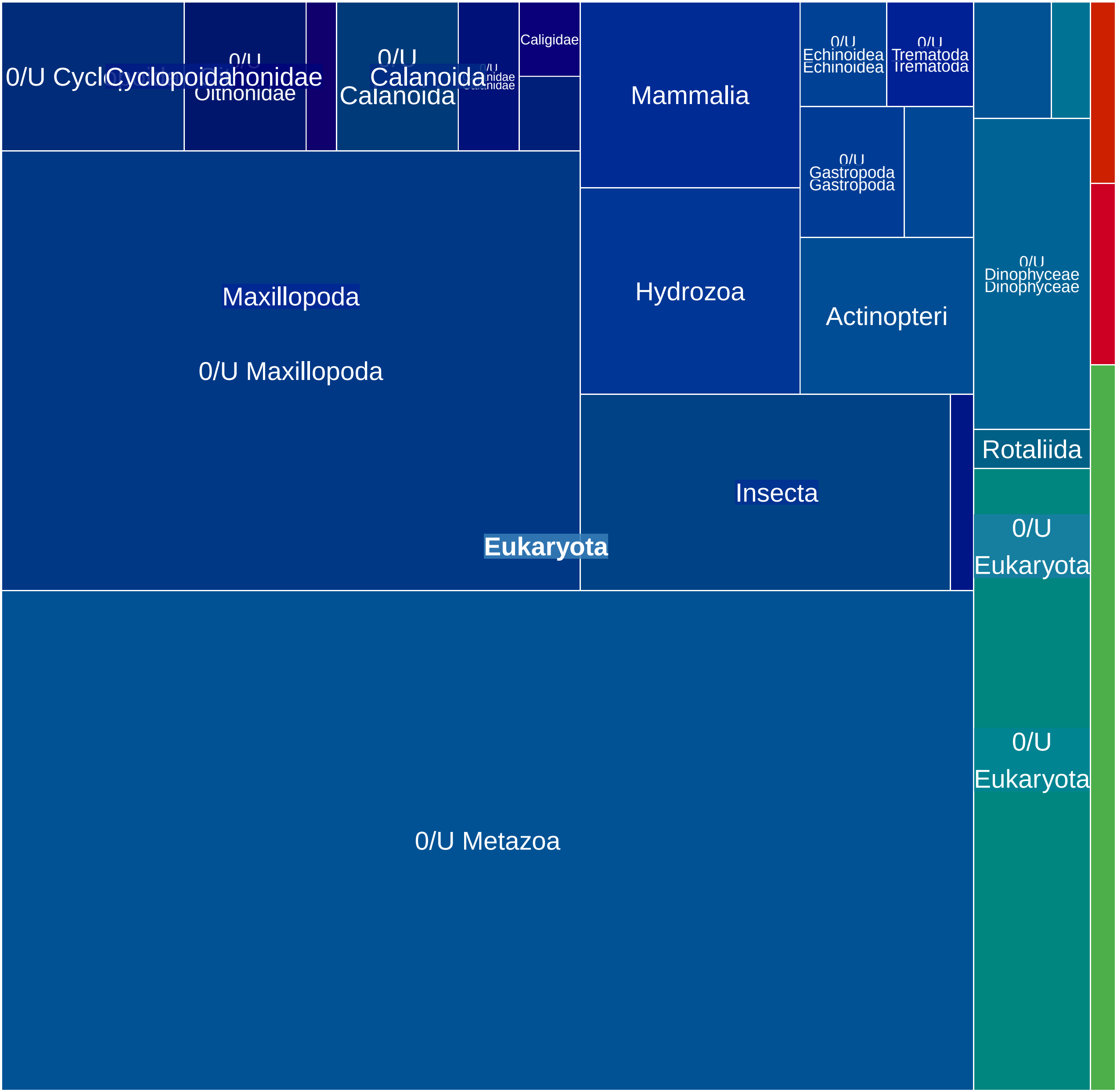

MGT−v1_332 : 3298 unigenes, 2192 (66.46%) taxonomically assigned

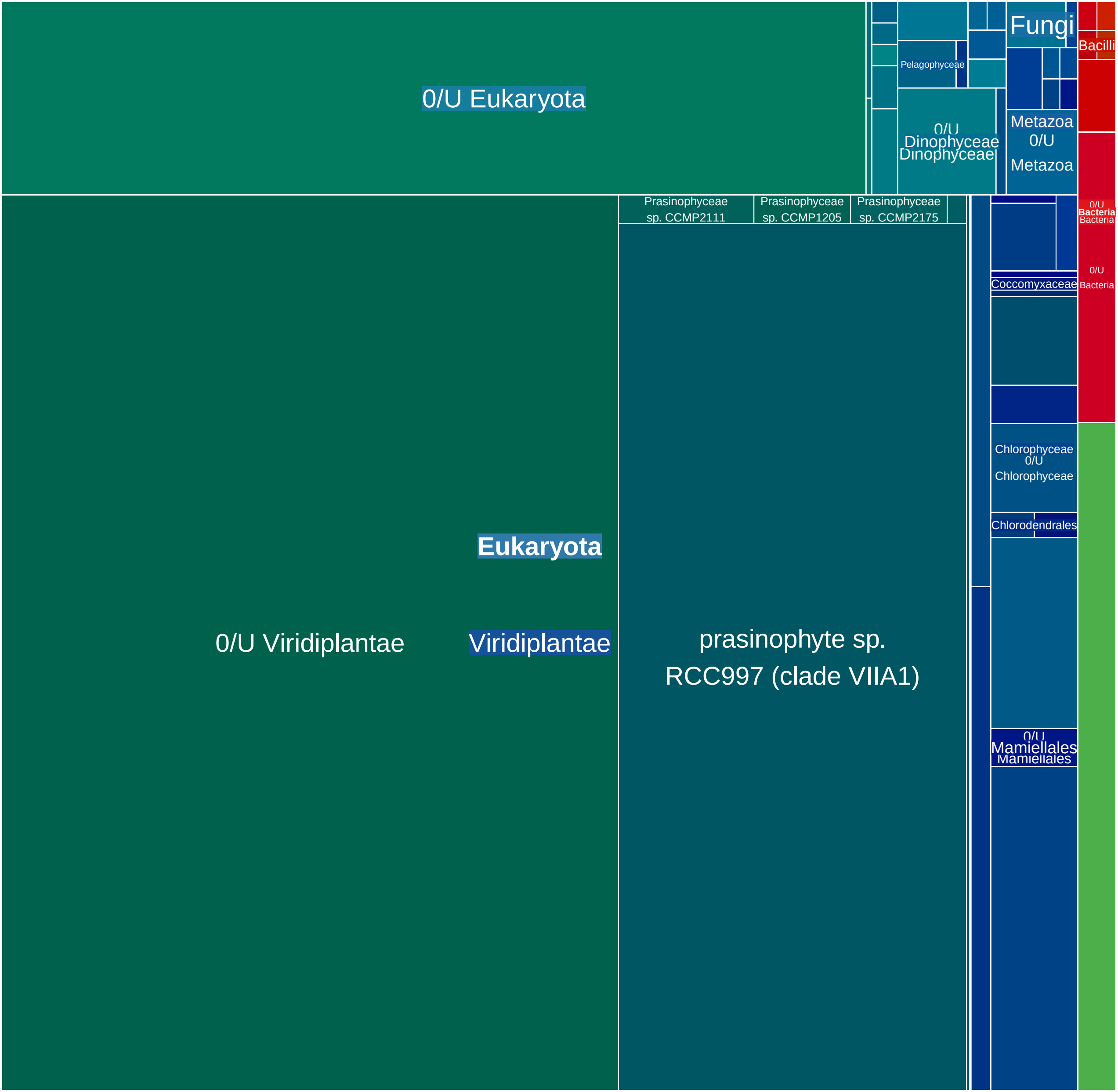

MGT−v1_333 : 3288 unigenes, 749 (22.78%) taxonomically assigned

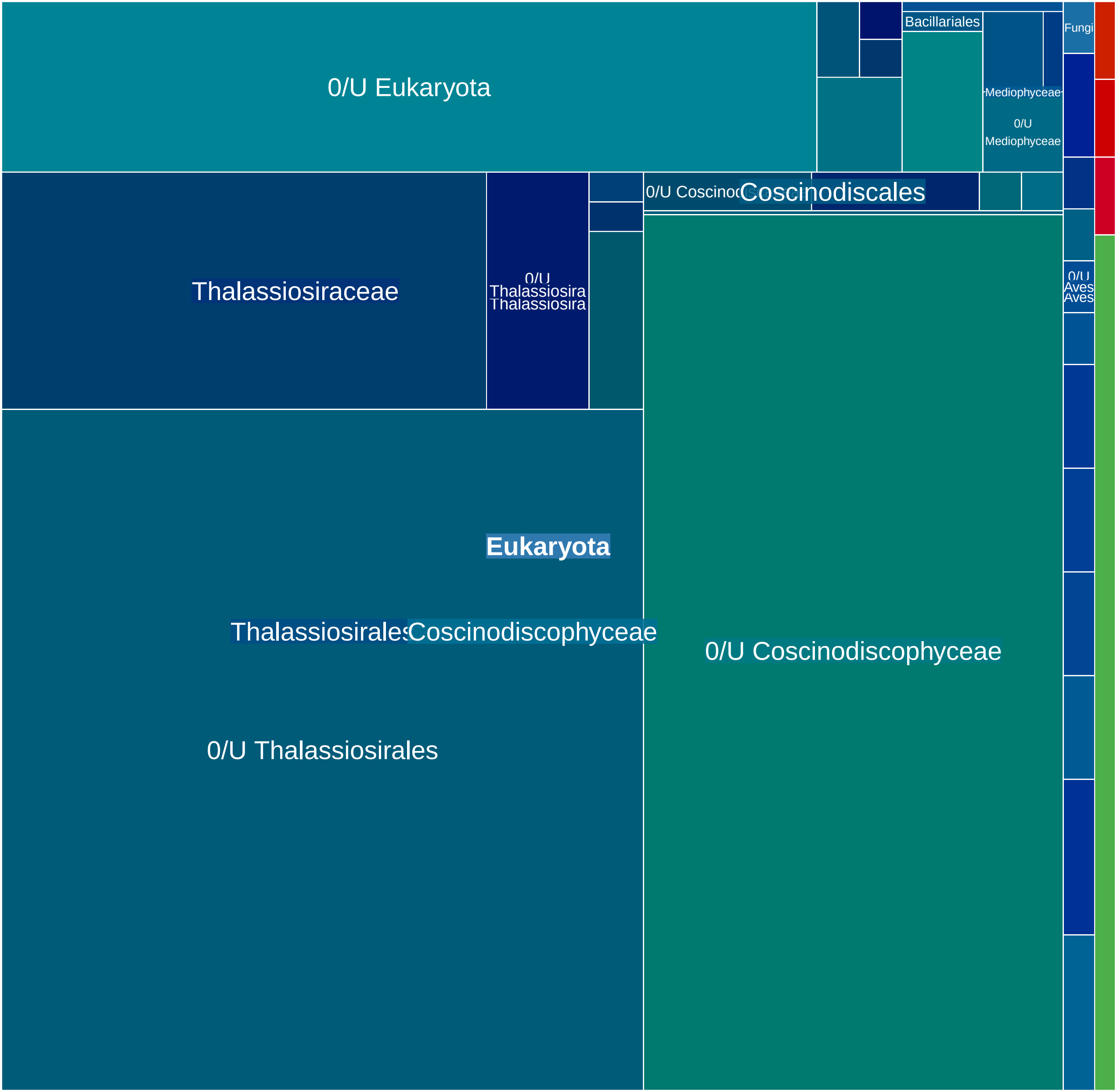

MGT−v1_334 : 3273 unigenes, 1692 (51.70%) taxonomically assigned

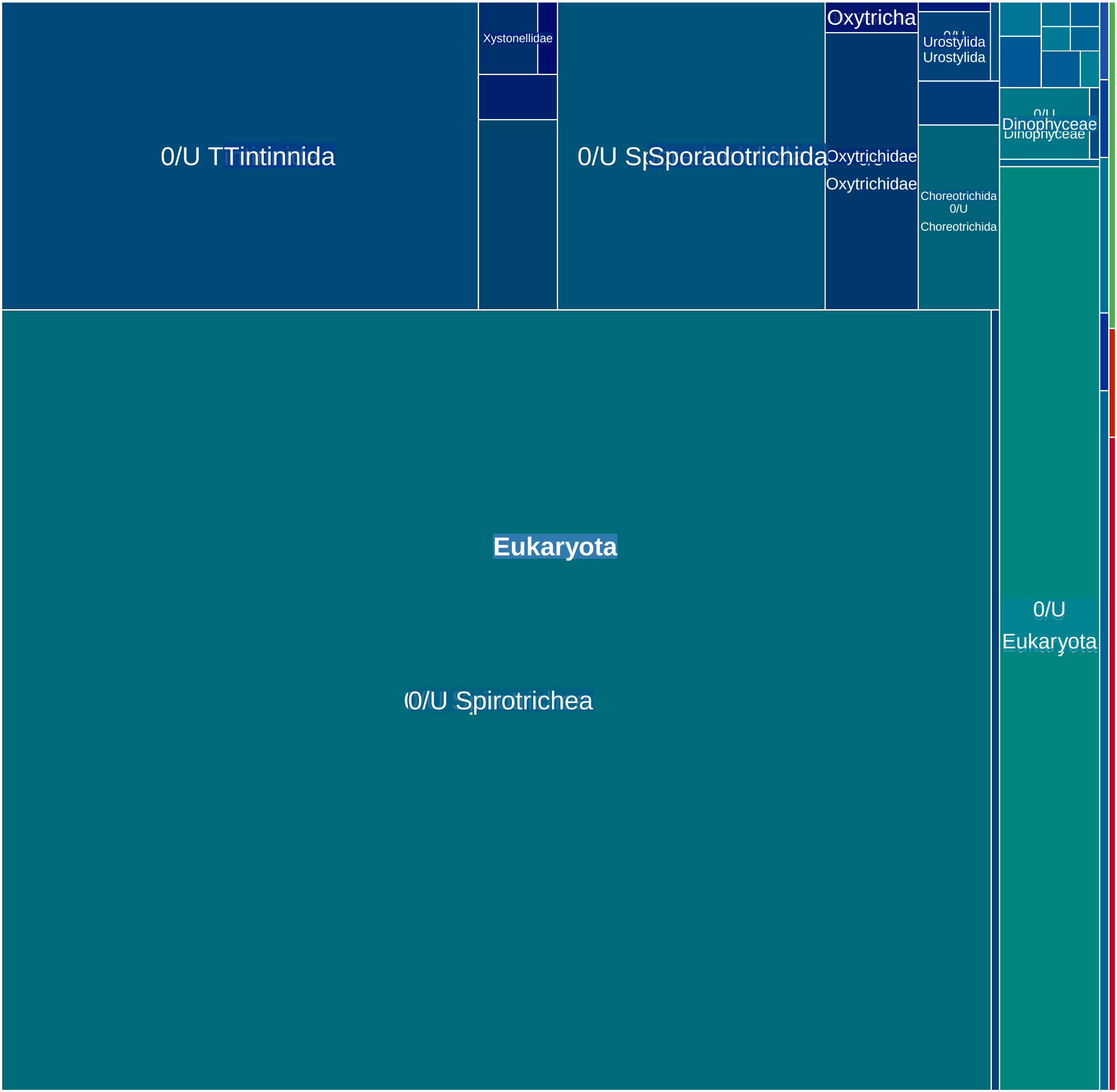

MGT−v1_335 : 3266 unigenes, 3156 (96.63%) taxonomically assigned

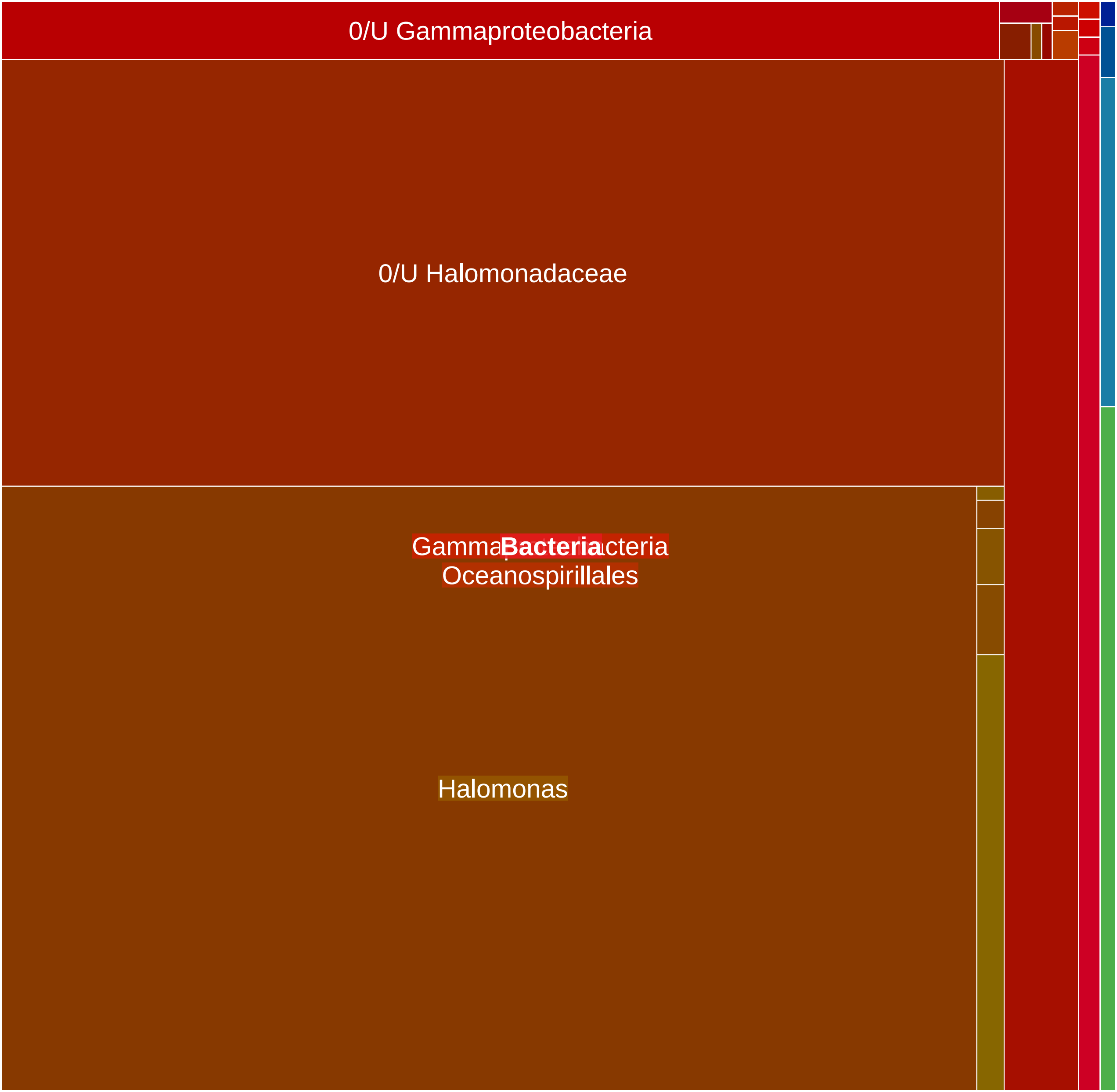

### MGT−v1_336 : 3264 unigenes, 3139 (96.17%) taxonomically assigned

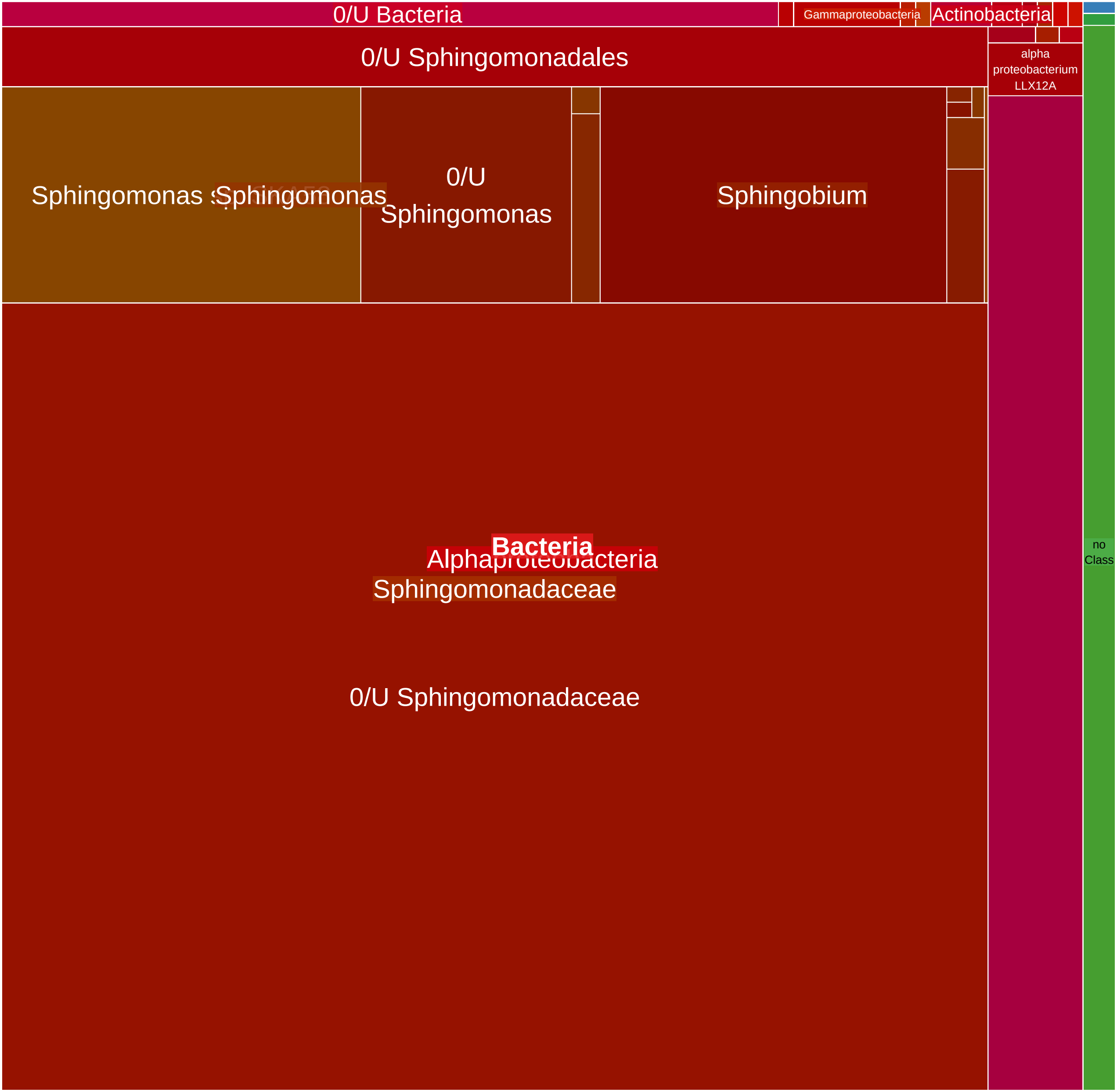

MGT−v1_337 : 3233 unigenes, 2098 (64.89%) taxonomically assigned

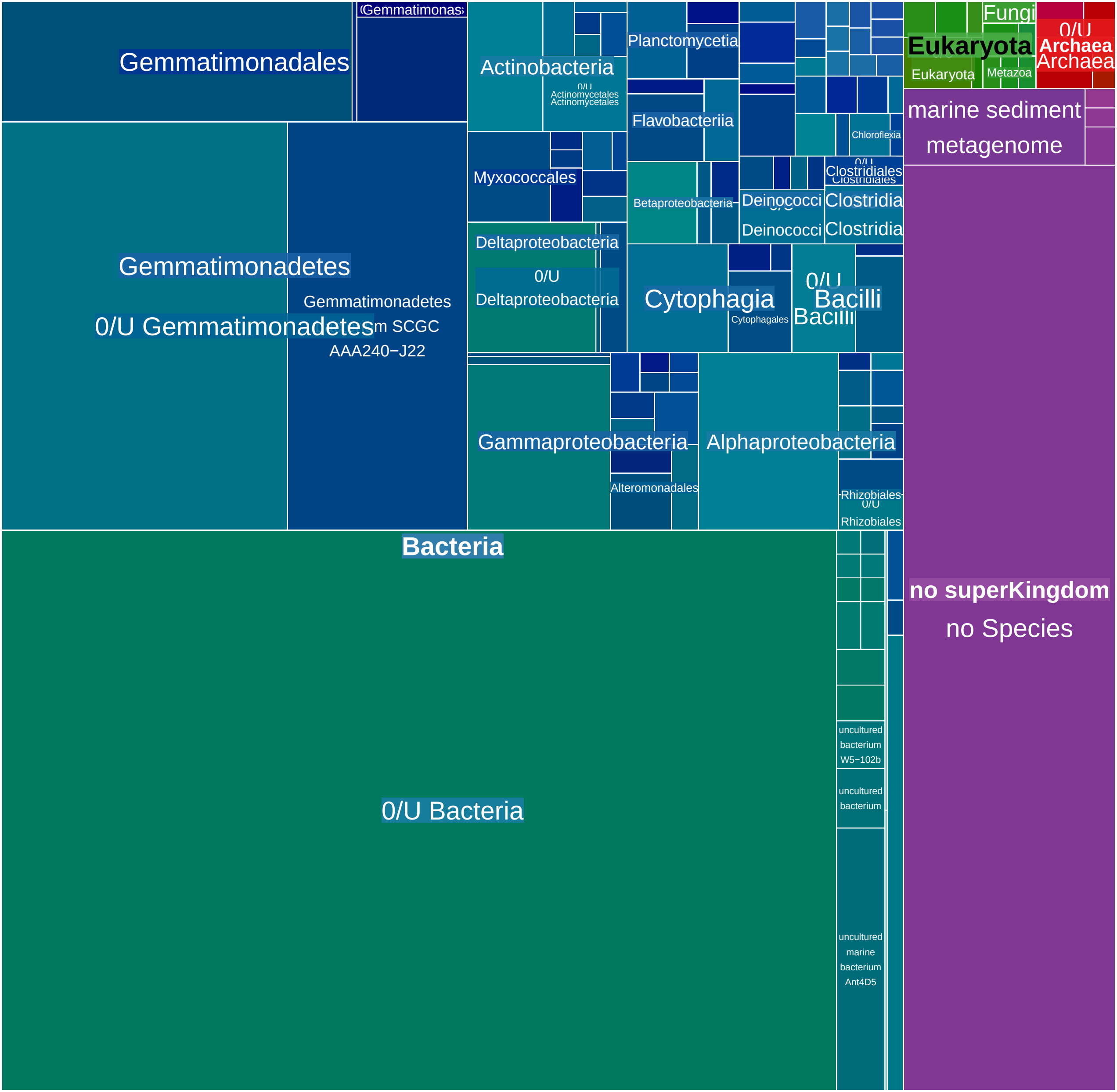

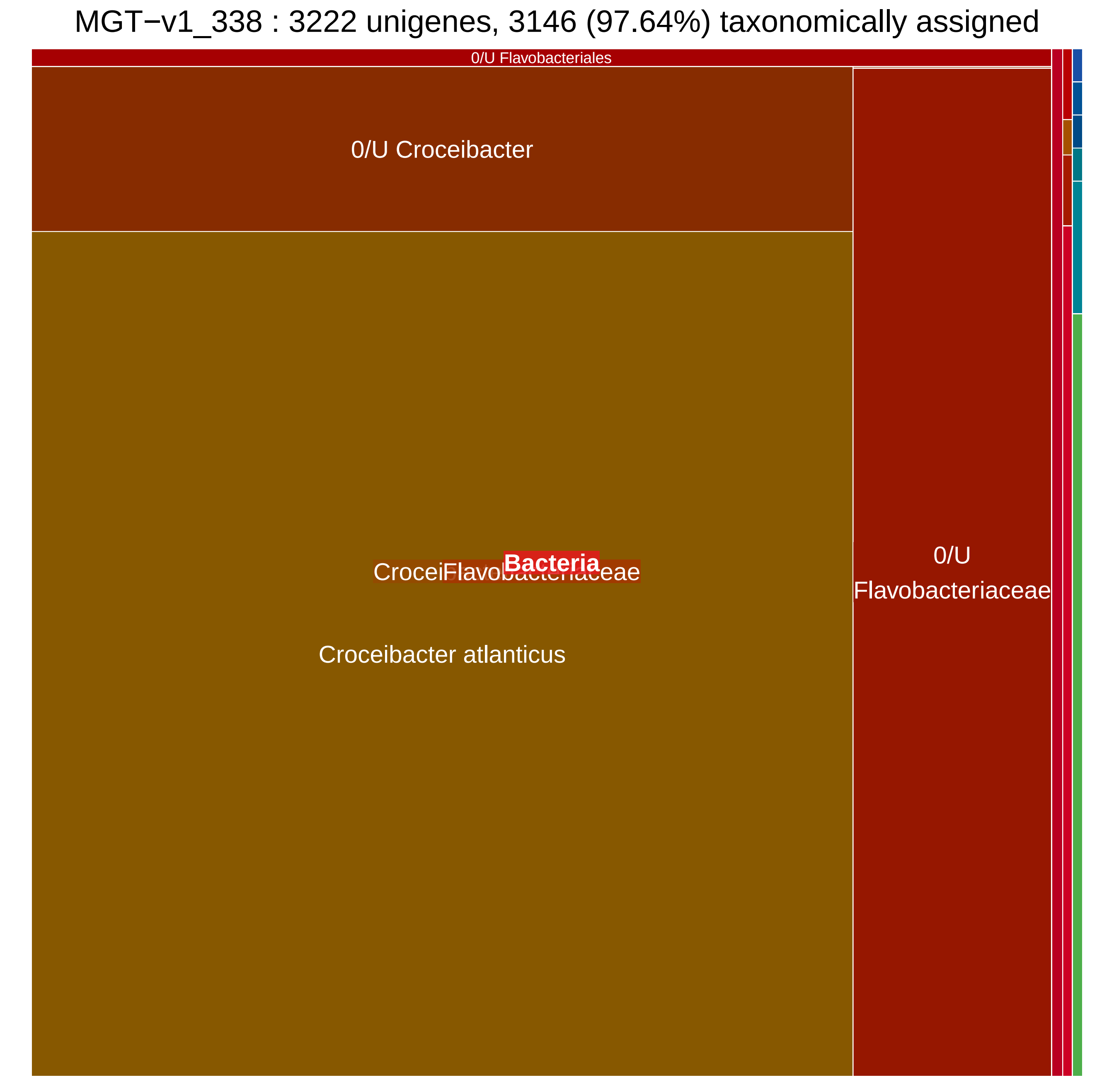
MGT−v1_338 : 3222 unigenes, 3146 (97.64%) taxonomically assigned
0/U Flavobacteriales
0/U Croceibacter
Bacteria
CroceiFlavobacteriaceae
Croceibacter atlanticus
0/U
Flavobacteriaceae

MGT−v1_339 : 3196 unigenes, 2642 (82.67%) taxonomically assigned

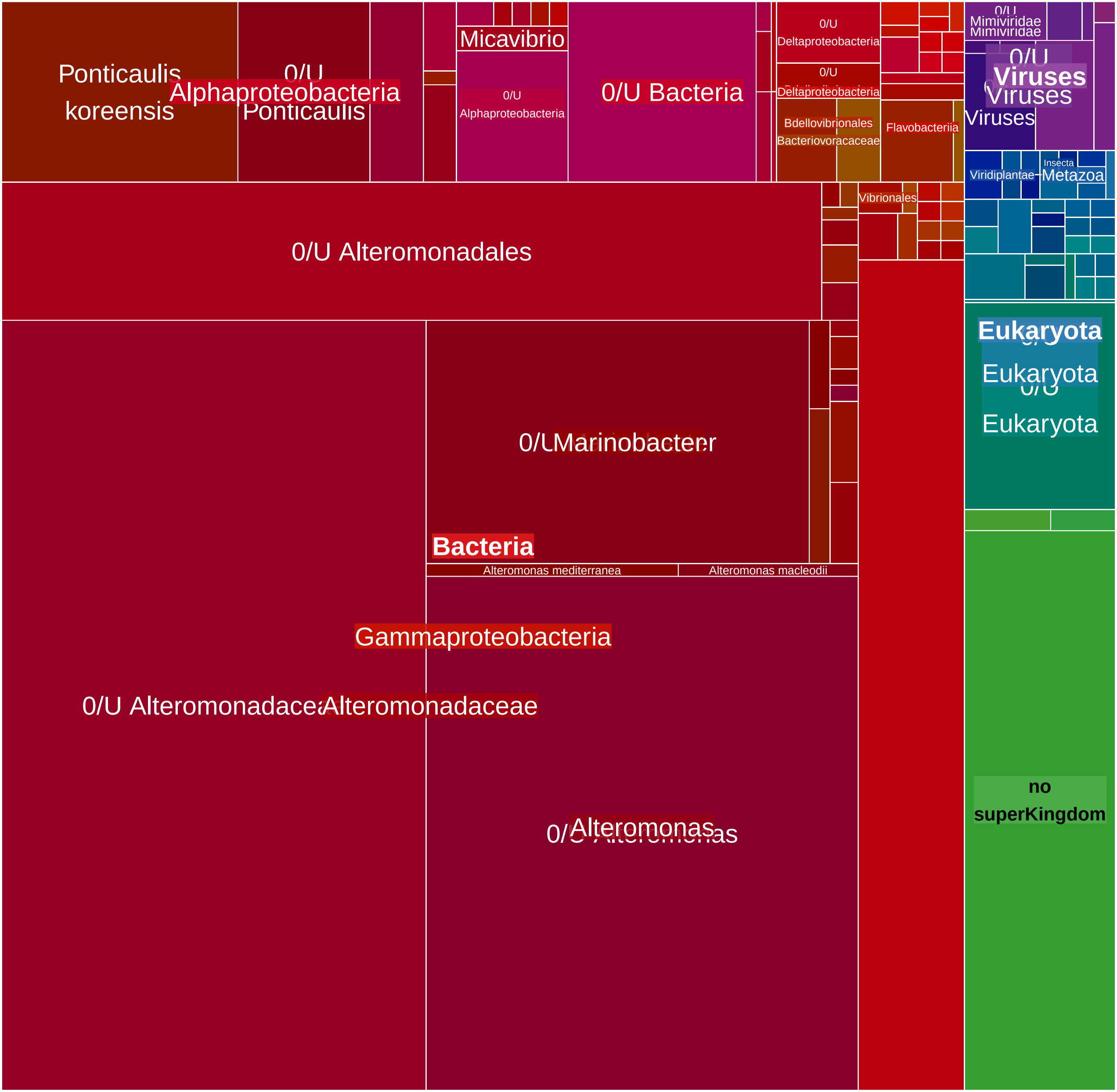

MGT−v1_340 : 3183 unigenes, 2627 (82.53%) taxonomically assigned

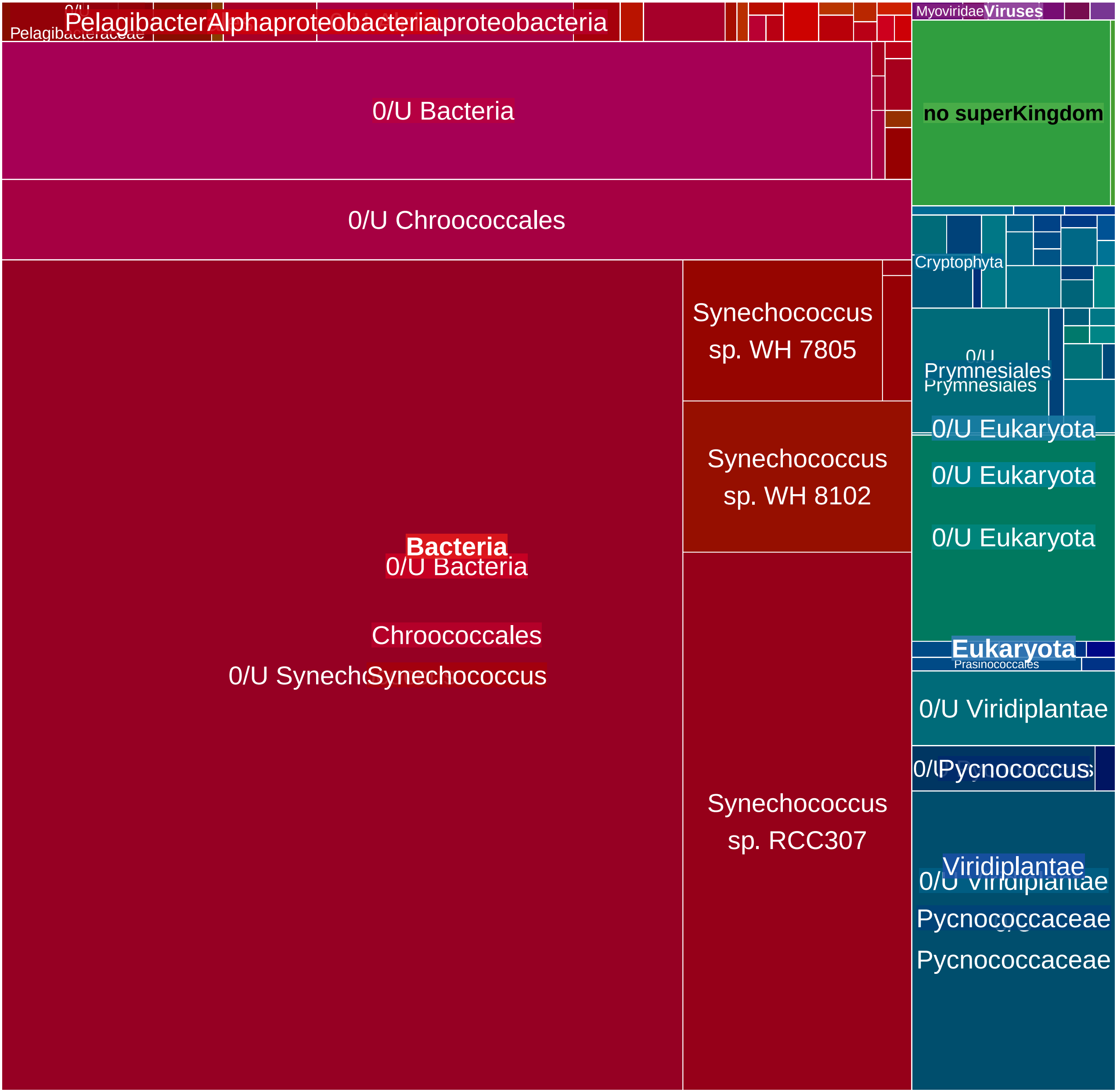

MGT−v1_341 : 3176 unigenes, 2913 (91.72%) taxonomically assigned

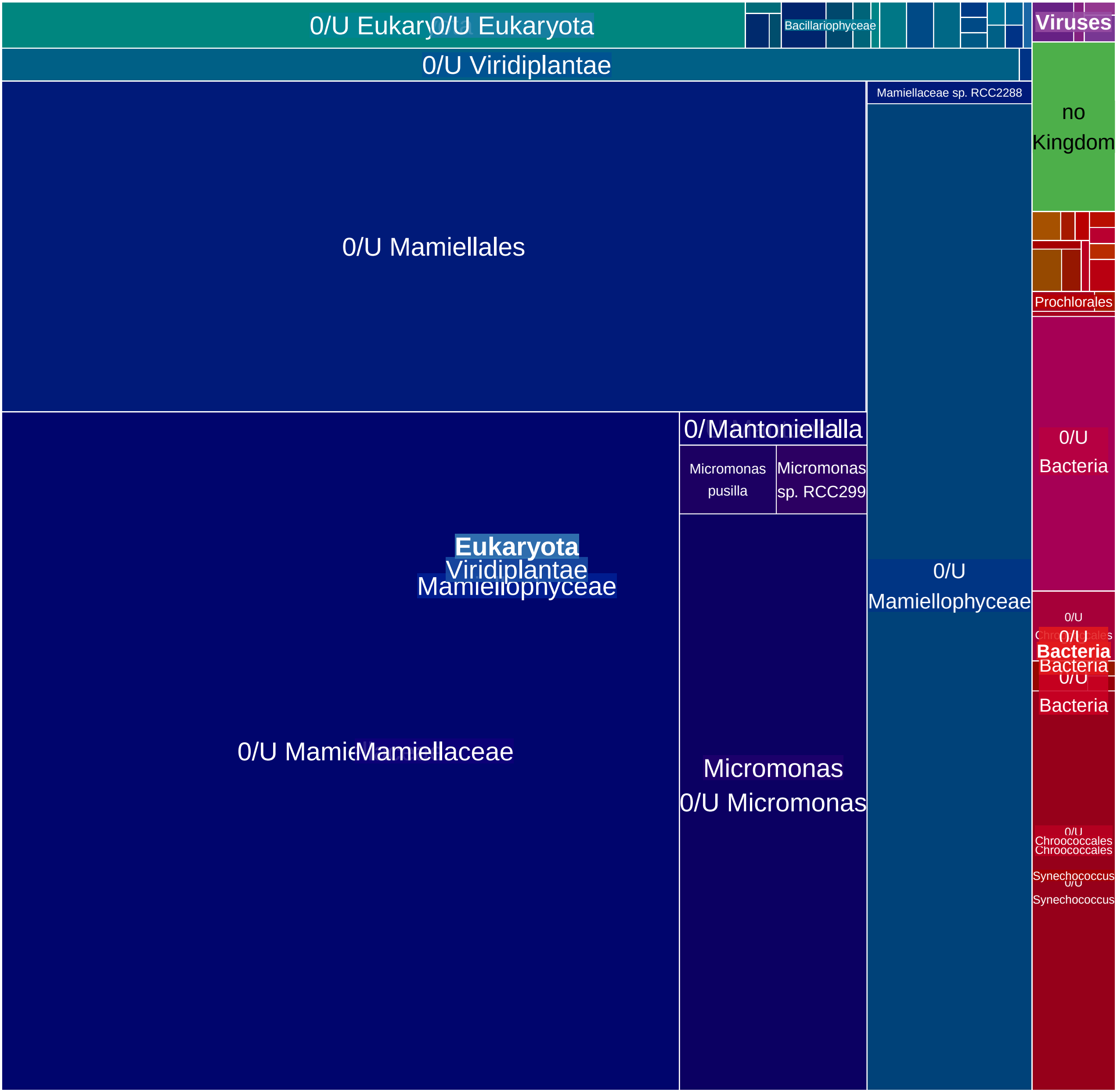

MGT−v1_342 : 3174 unigenes, 364 (11.47%) taxonomically assigned

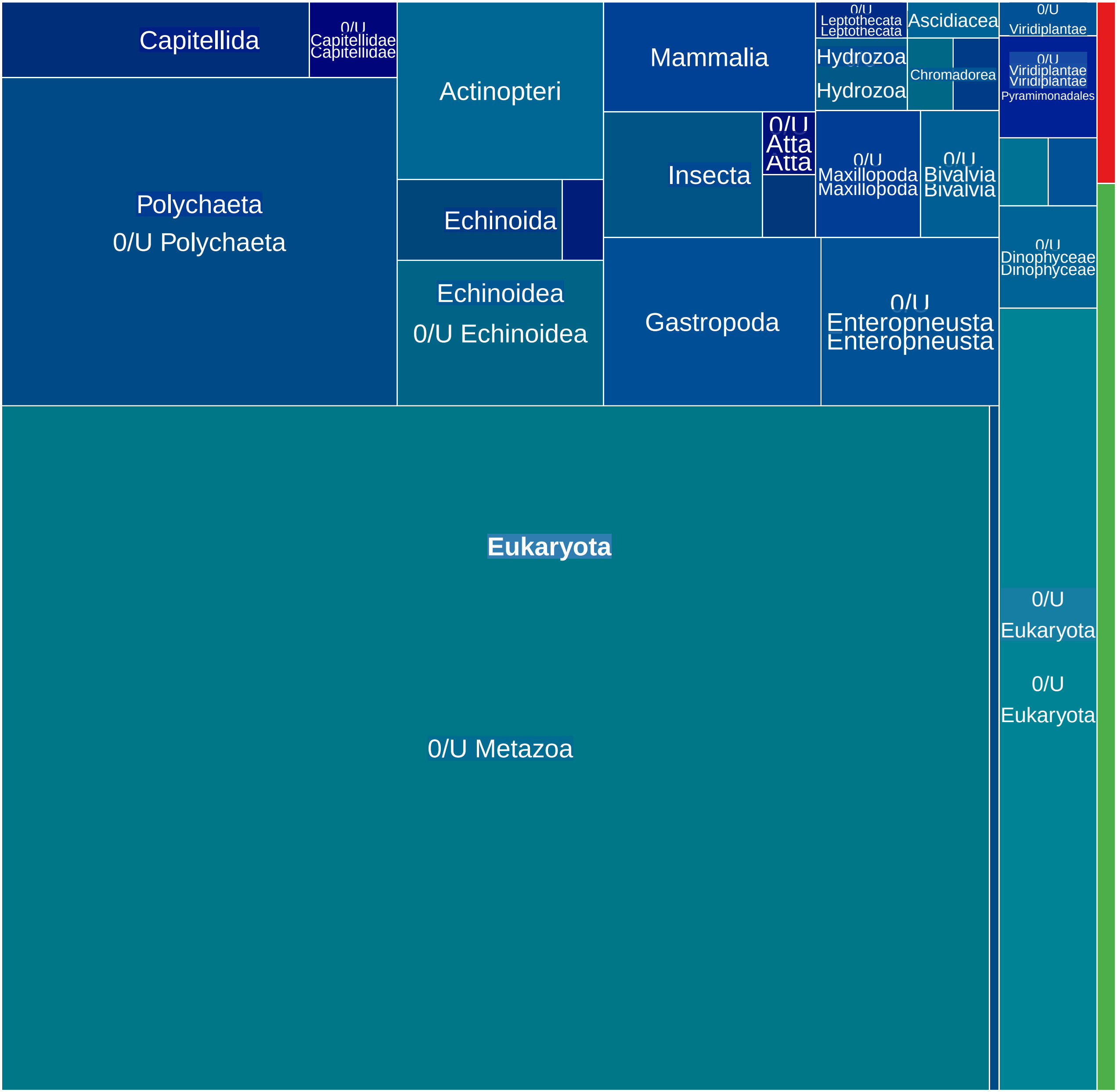

MGT−v1_343 : 3162 unigenes, 2384 (75.40%) taxonomically assigned

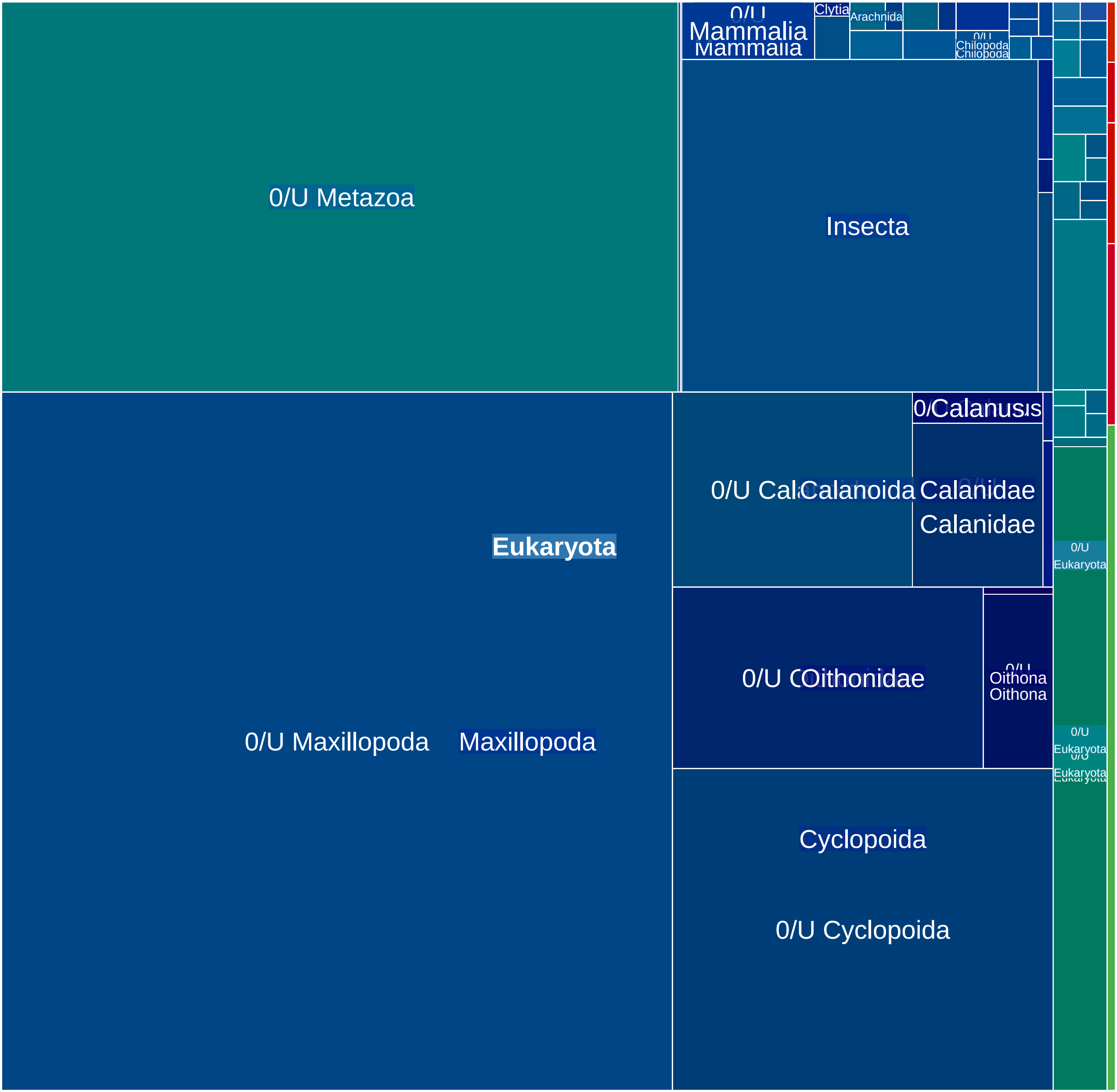

MGT−v1_344 : 3142 unigenes, 458 (14.58%) taxonomically assigned

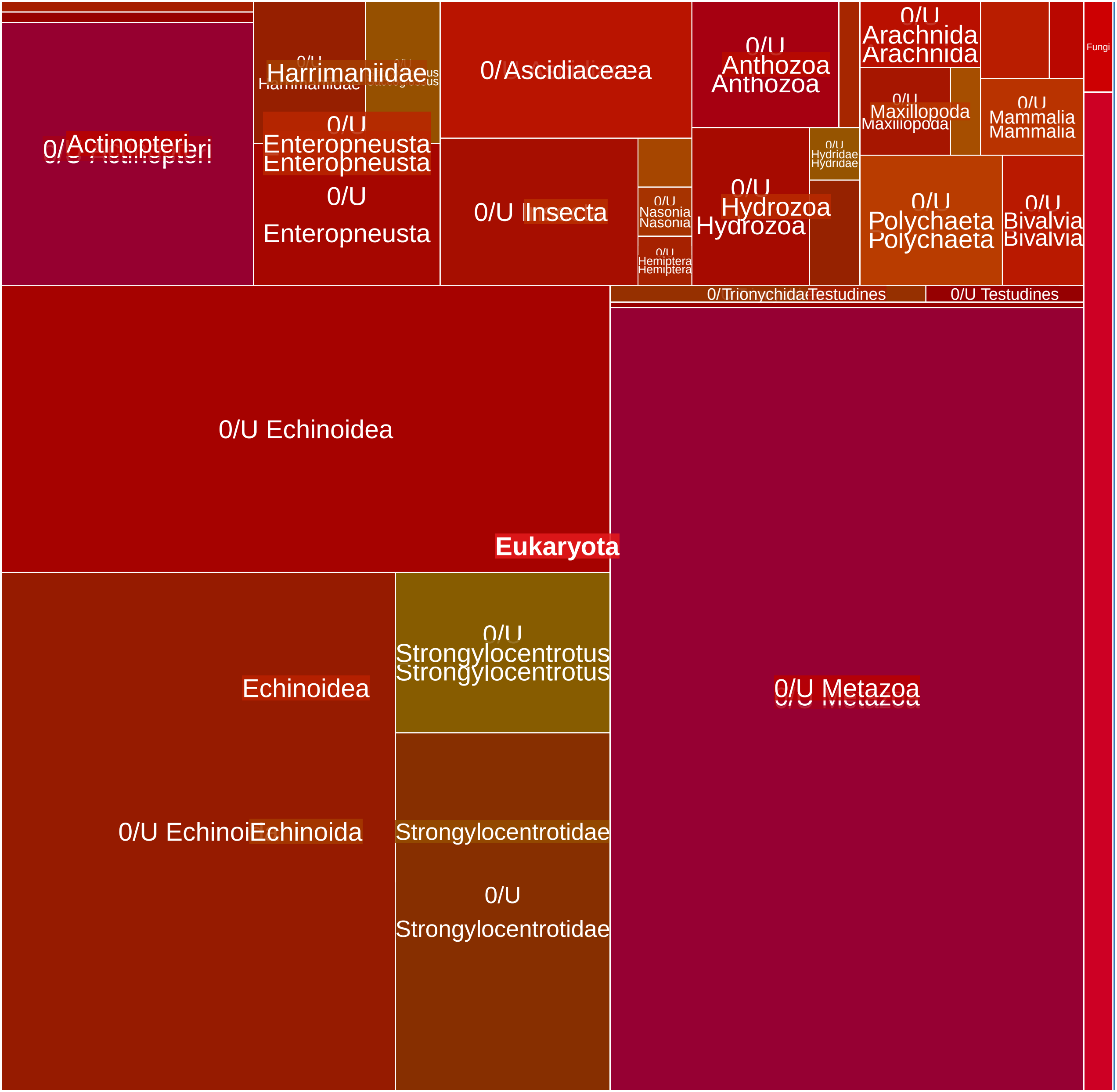

### MGT−v1_345 : 3138 unigenes, 2792 (88.97%) taxonomically assigned

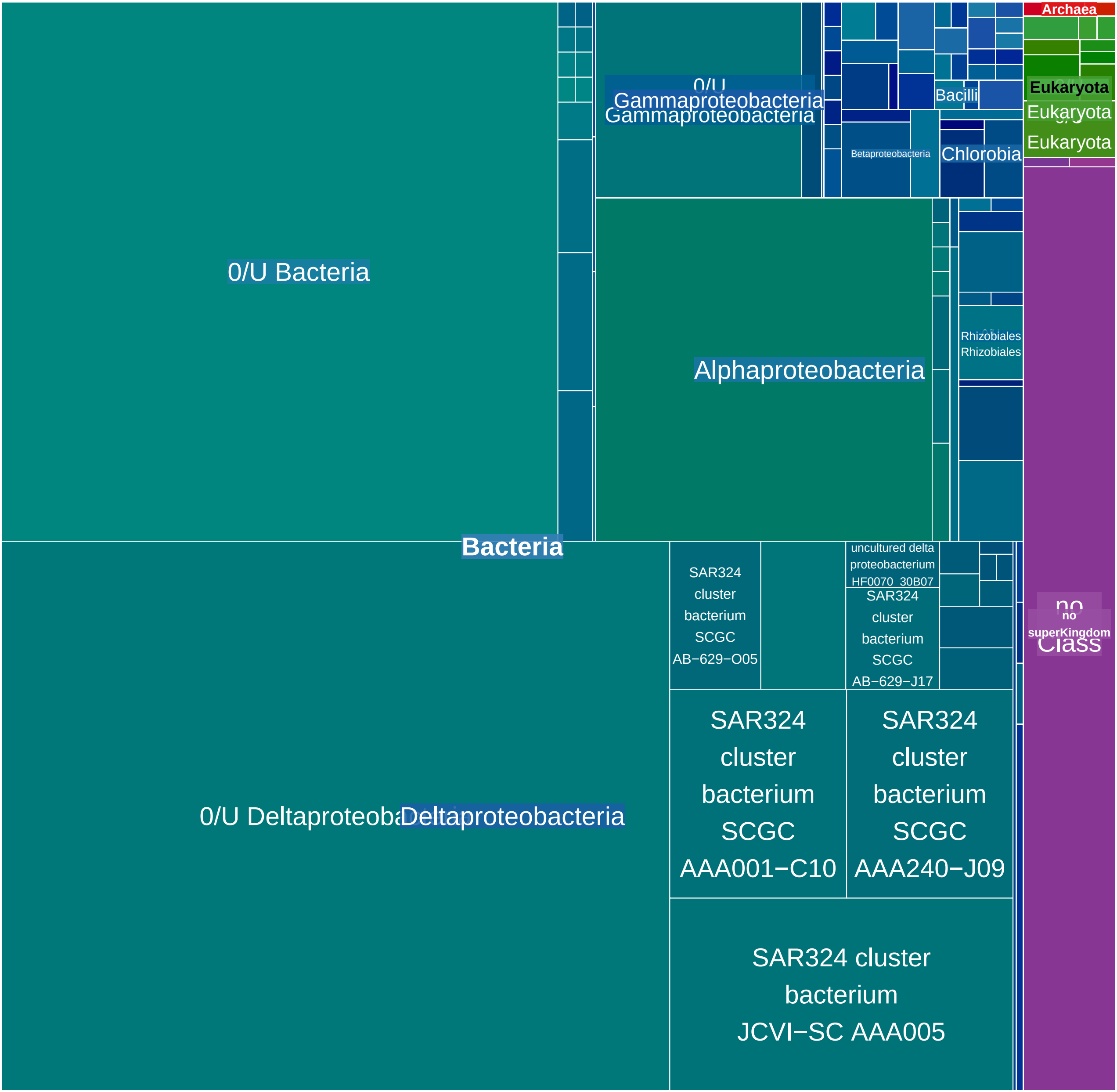

### MGT−v1_346 : 3088 unigenes, 2955 (95.69%) taxonomically assigned

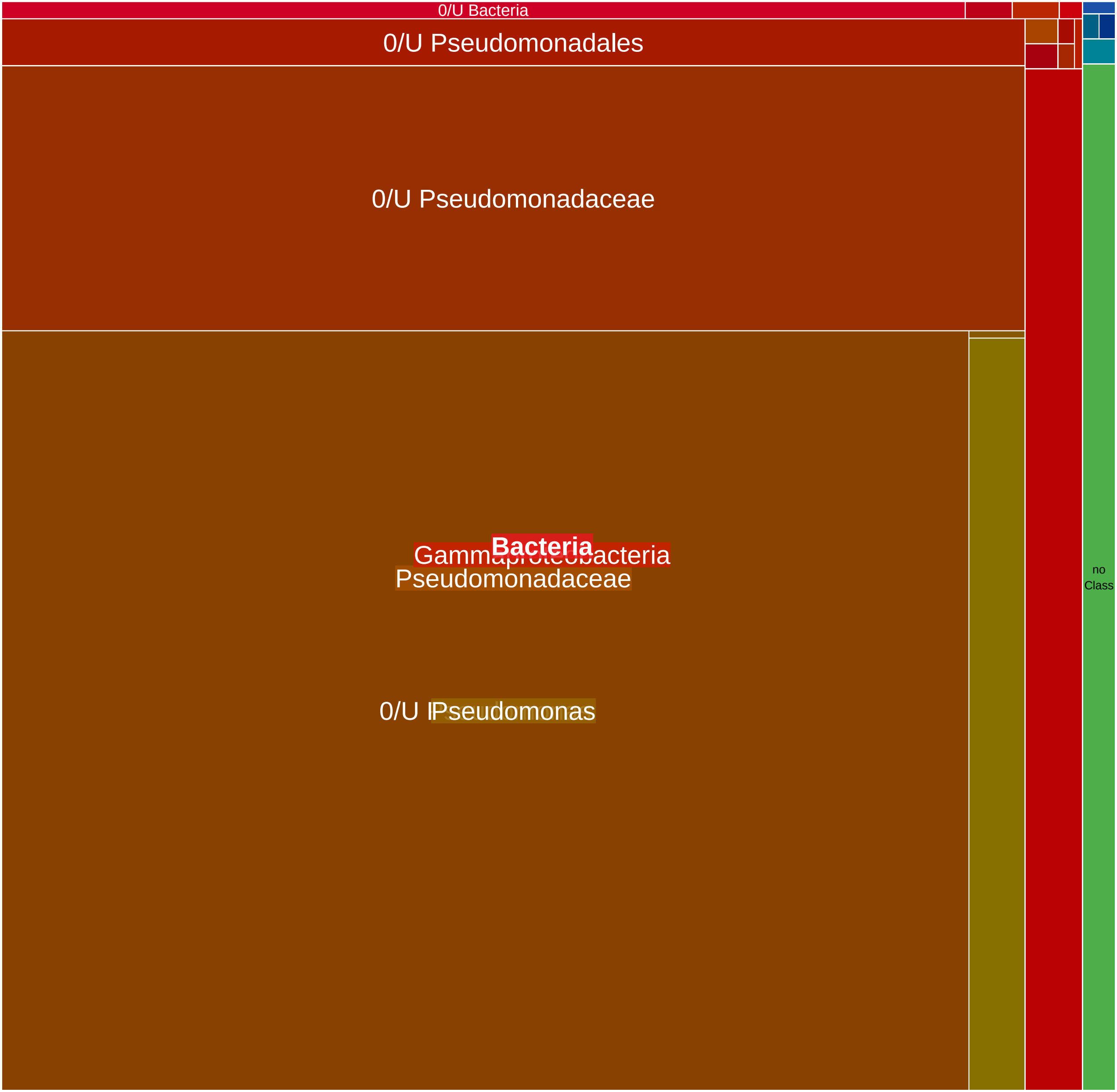

MGT−v1_347 : 3083 unigenes, 2465 (79.95%) taxonomically assigned

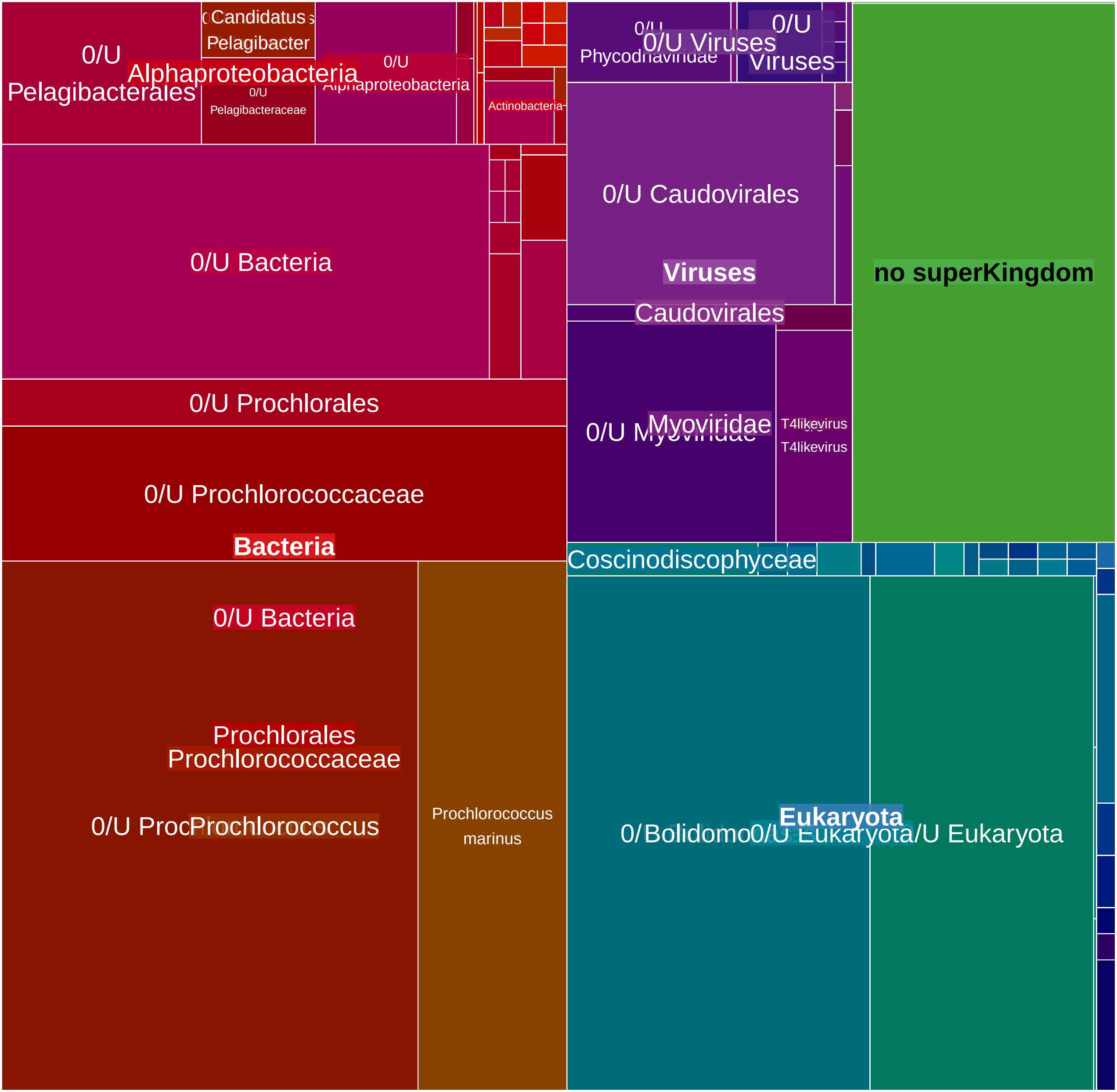

MGT−v1_348 : 3042 unigenes, 452 (14.86%) taxonomically assigned

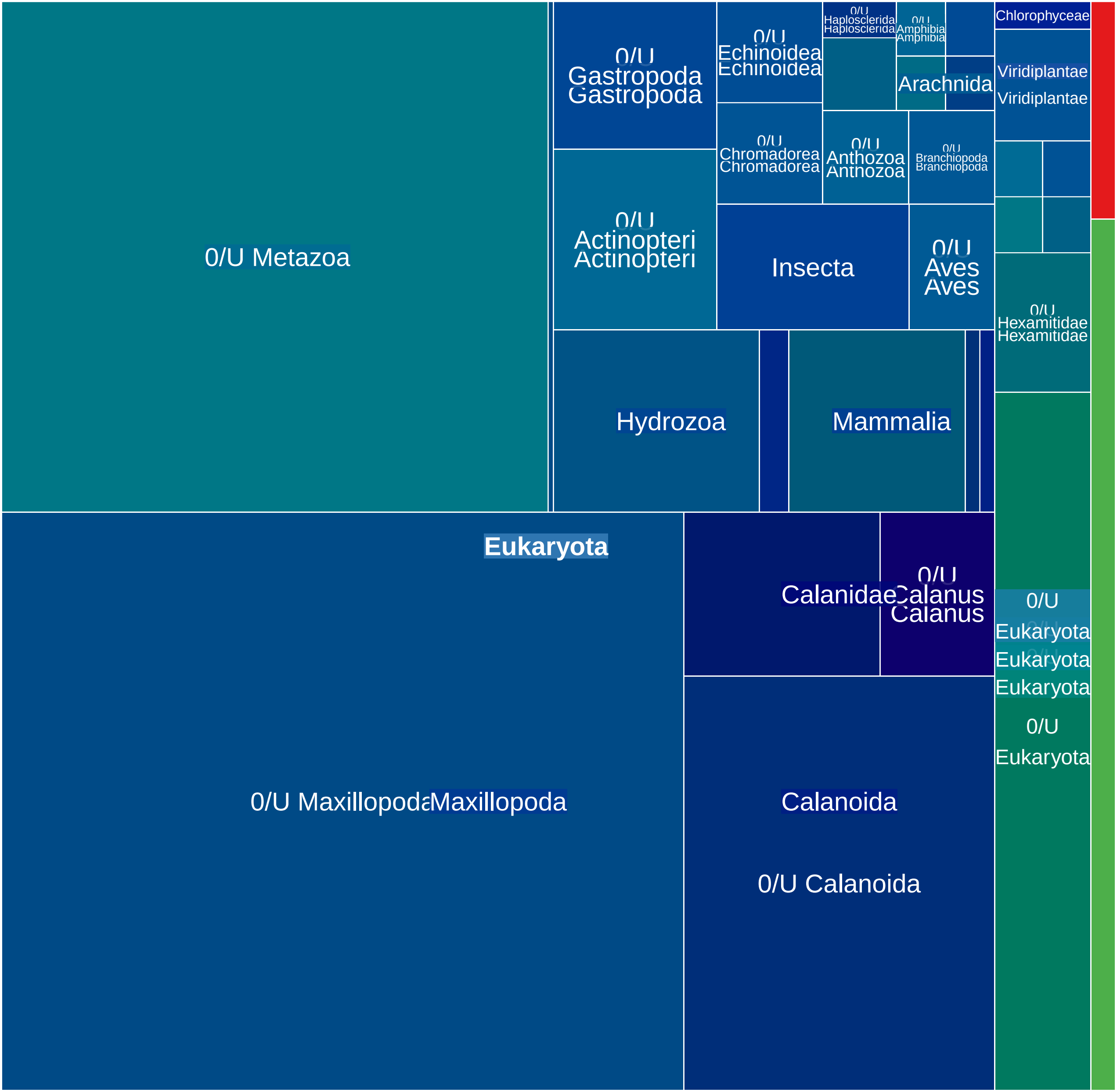

### MGT−v1_349 : 3032 unigenes, 193 (6.37%) taxonomically assigned

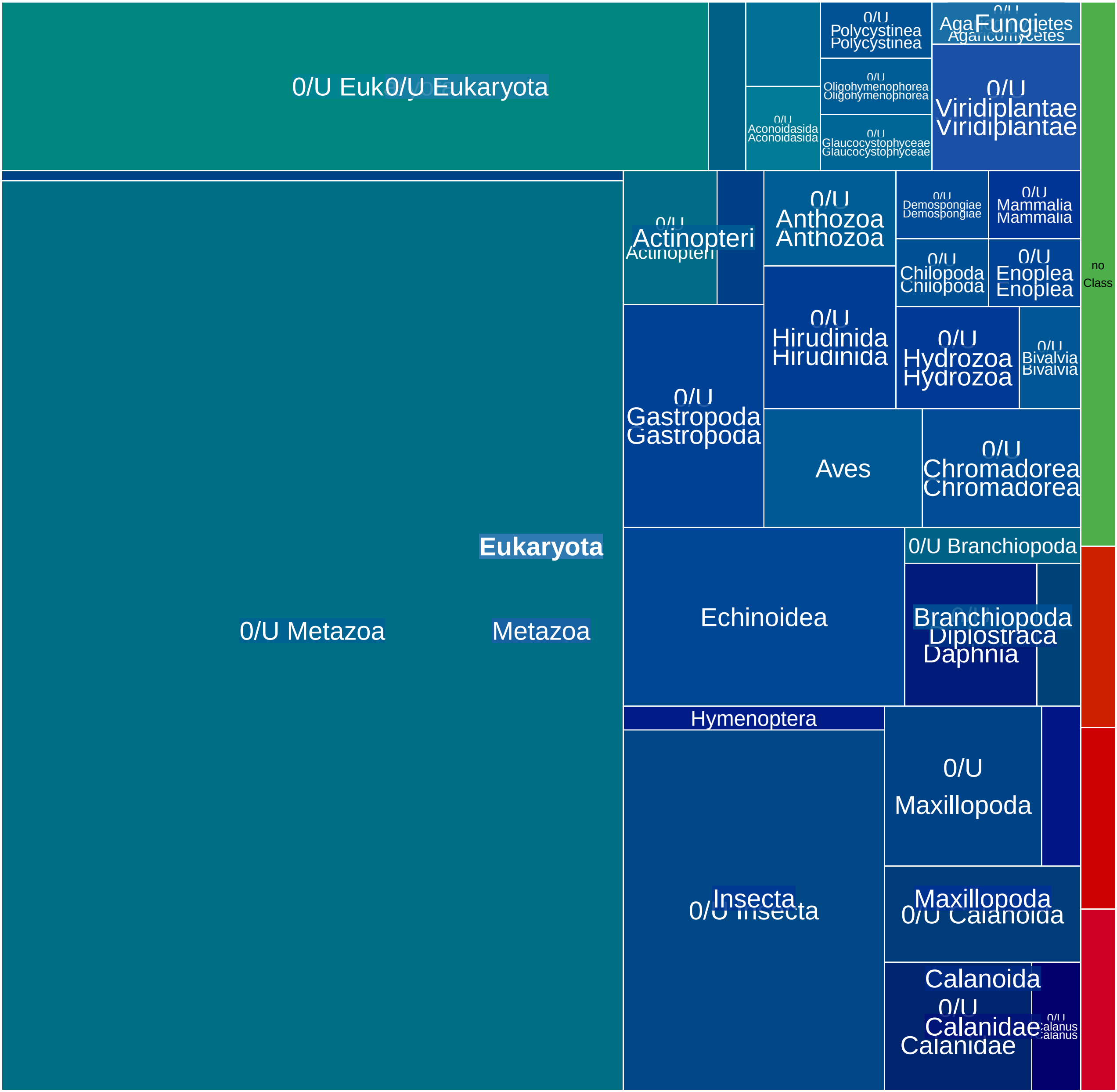

MGT−v1_350 : 3031 unigenes, 2926 (96.54%) taxonomically assigned

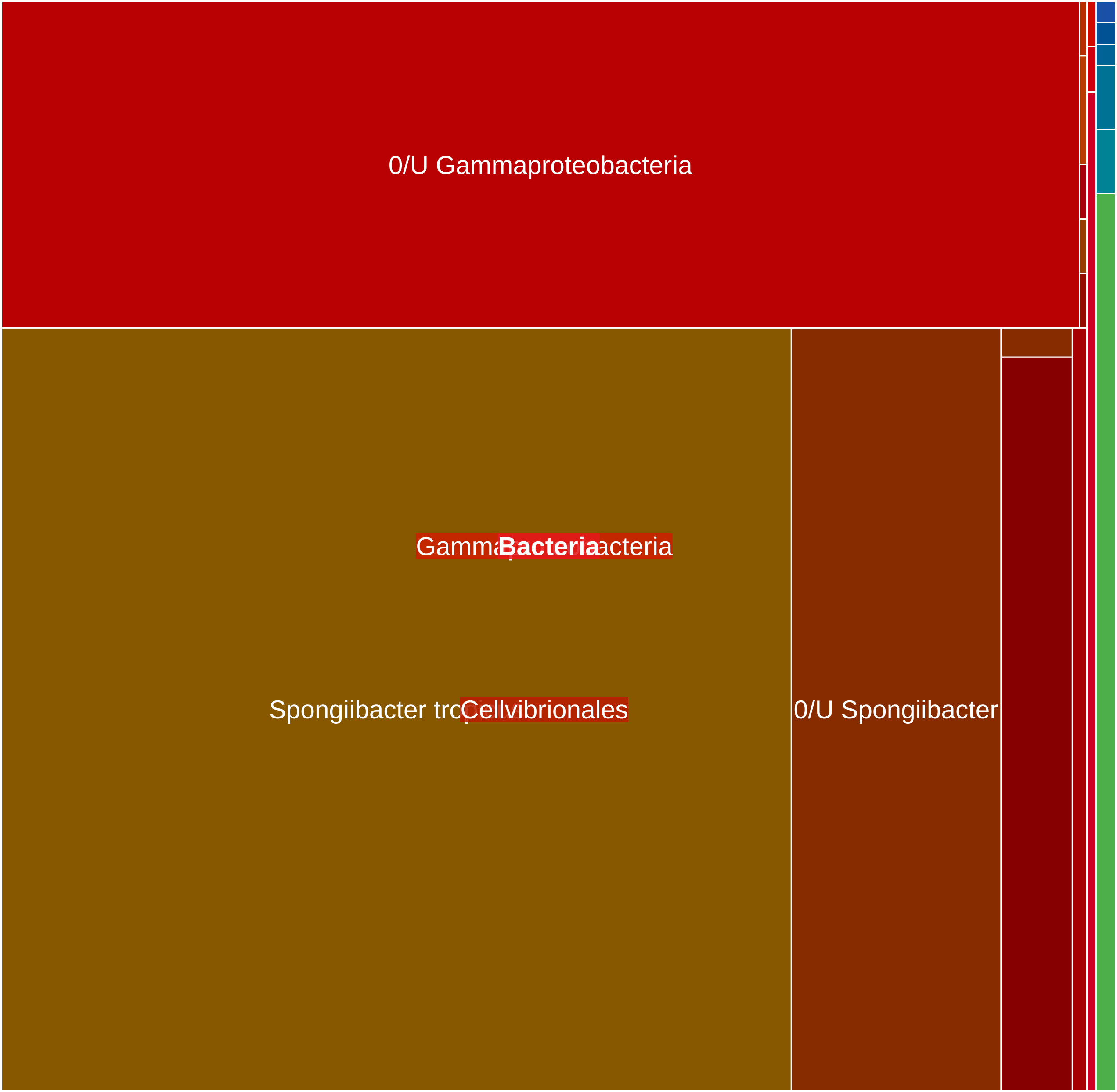

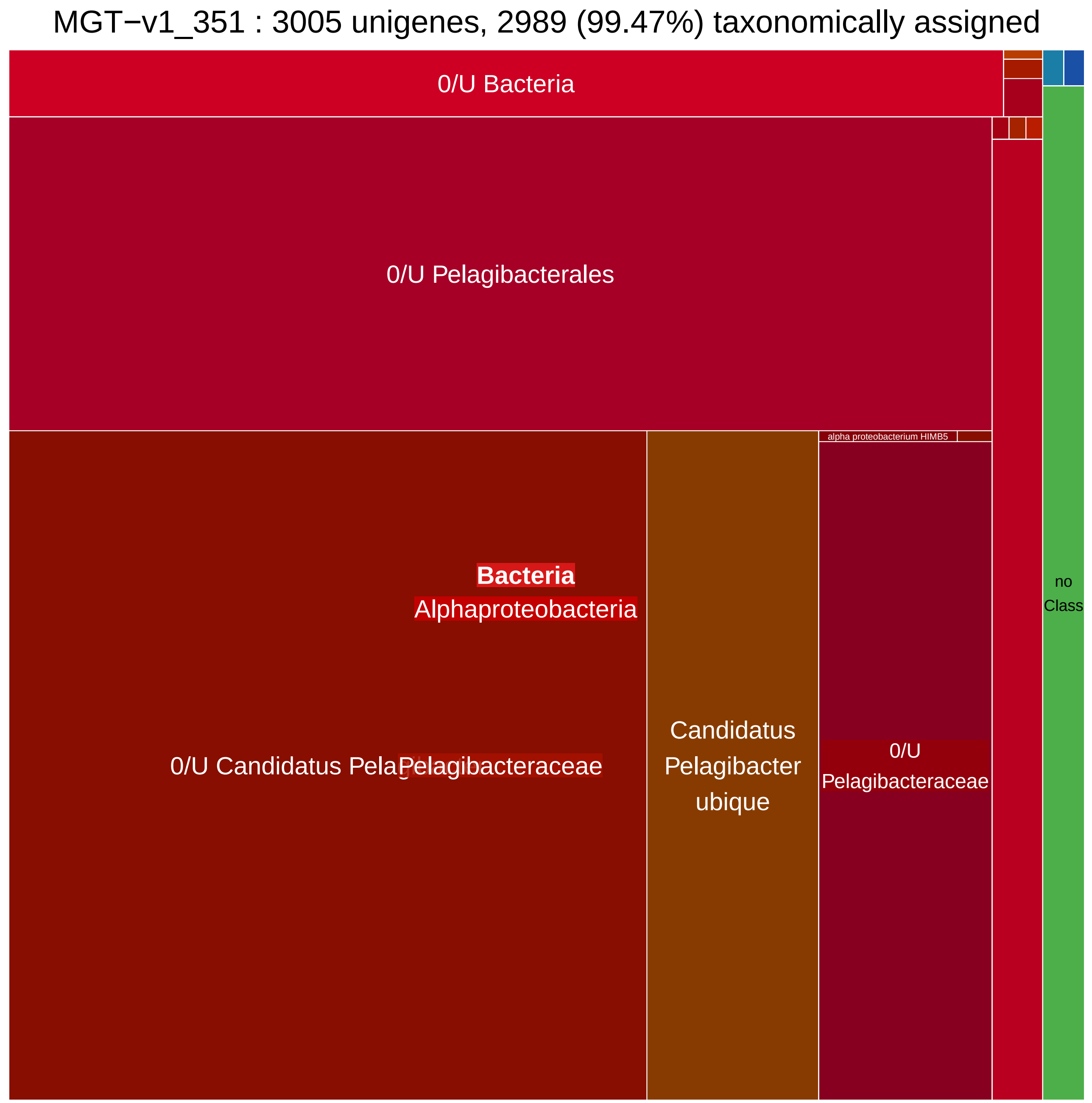

MGT−v1_351 : 3005 unigenes, 2989 (99.47%) taxonomically assigned
0/U Bacteria
0/U Pelagibacterales
Bacteria
Alphaproteobacteria
0/U Candidatus Pelagibacter
Pelagibacteraceae
Candidatus Pelagibacter ubique
alpha proteobacterium HIMB5
0/U Pelagibacteraceae
no Class

MGT−v1_352 : 3005 unigenes, 374 (12.45%) taxonomically assigned

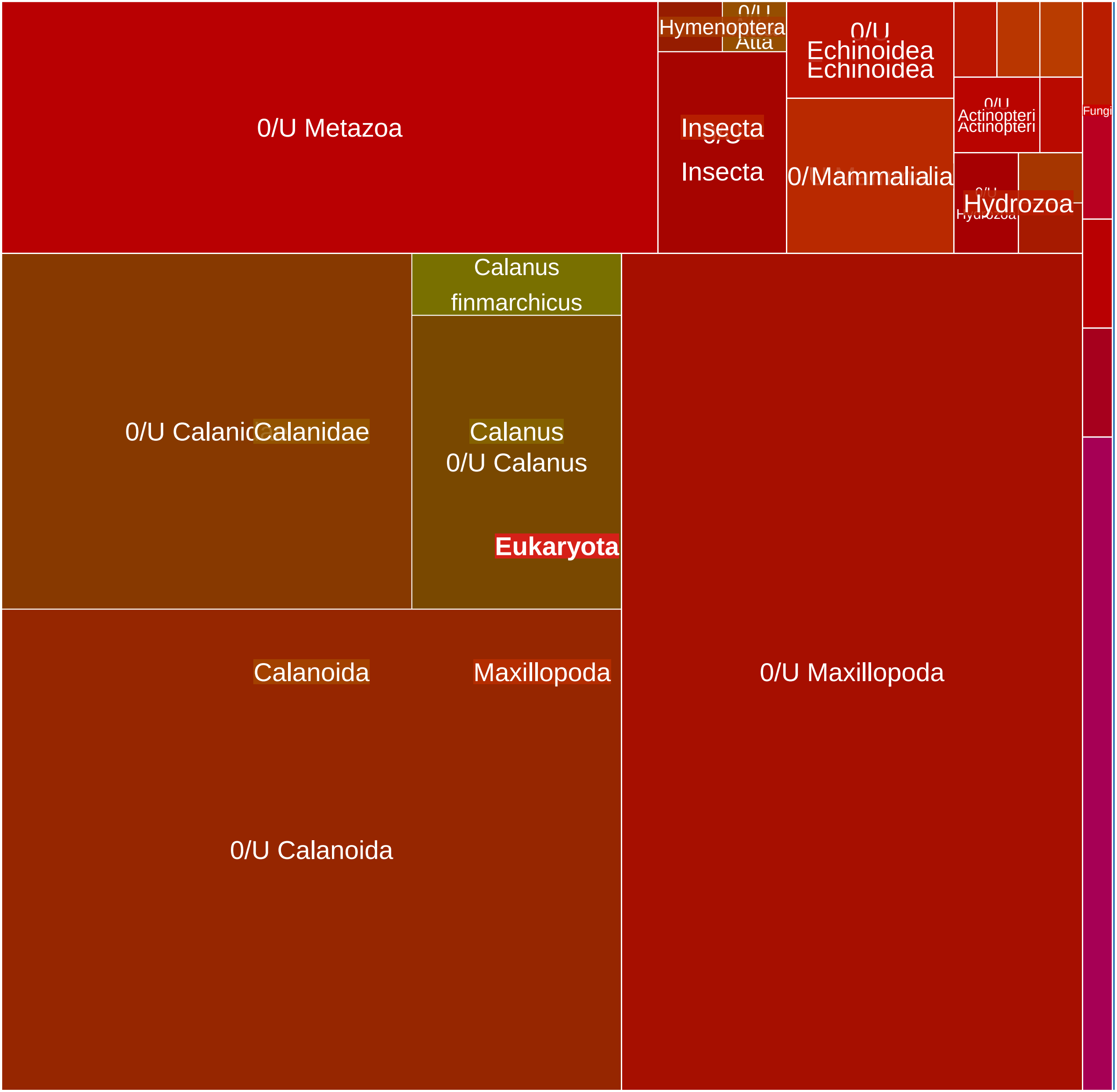

MGT−v1_353 : 2995 unigenes, 2789 (93.12%) taxonomically assigned

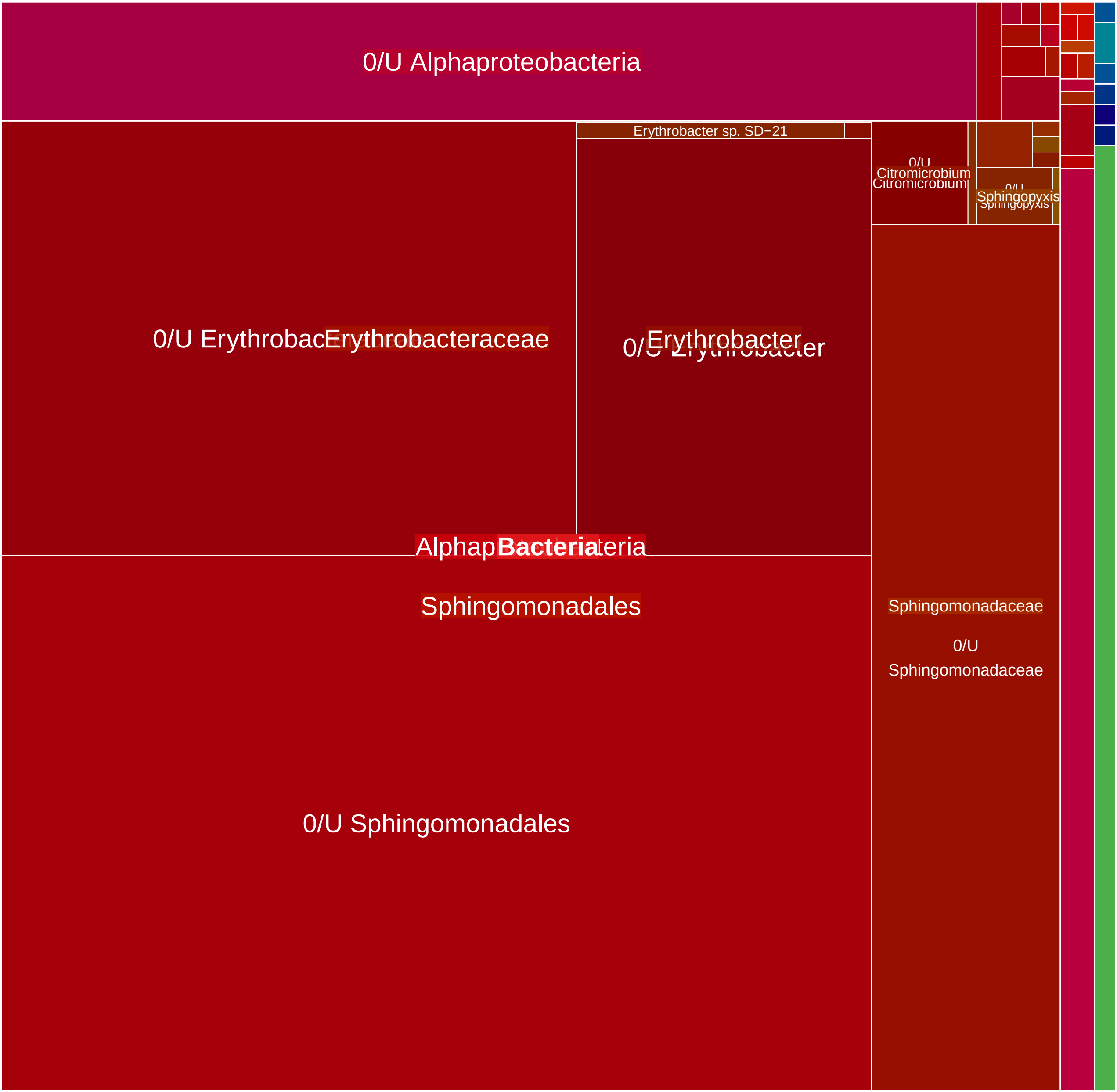

MGT−v1_354 : 2993 unigenes, 1777 (59.37%) taxonomically assigned

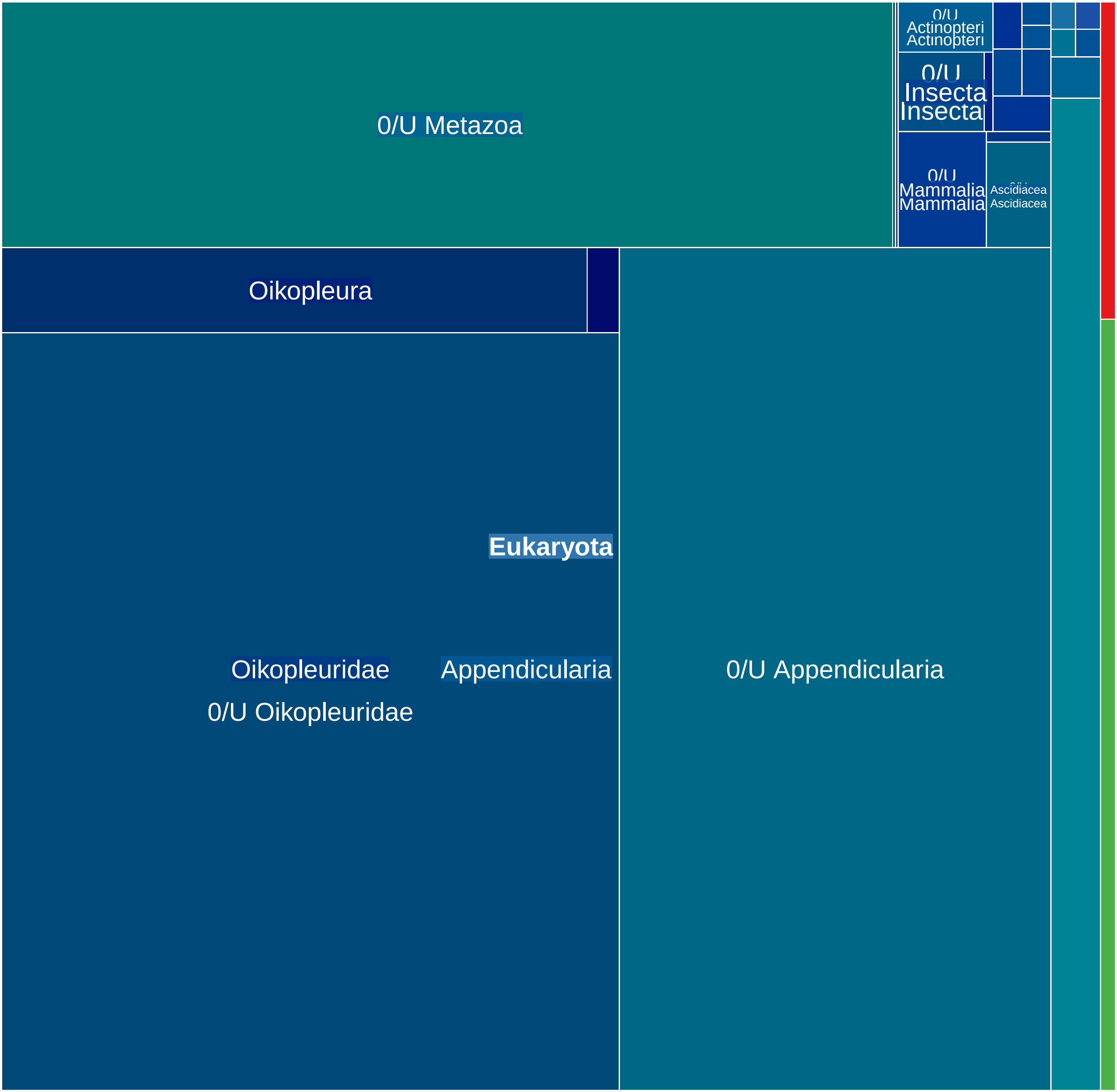

MGT−v1_355 : 2990 unigenes, 495 (16.56%) taxonomically assigned

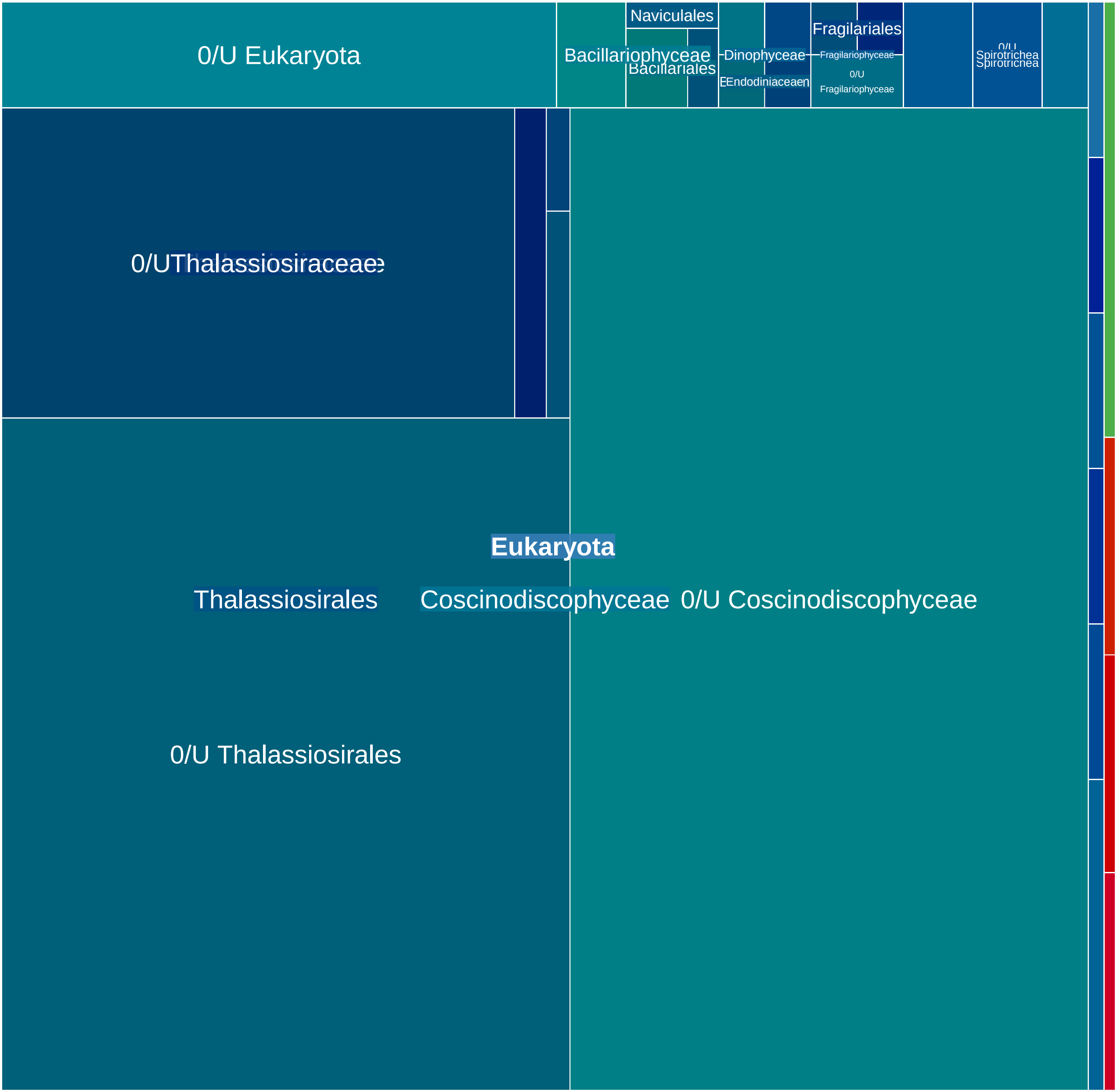

MGT−v1_356 : 2968 unigenes, 584 (19.68%) taxonomically assigned

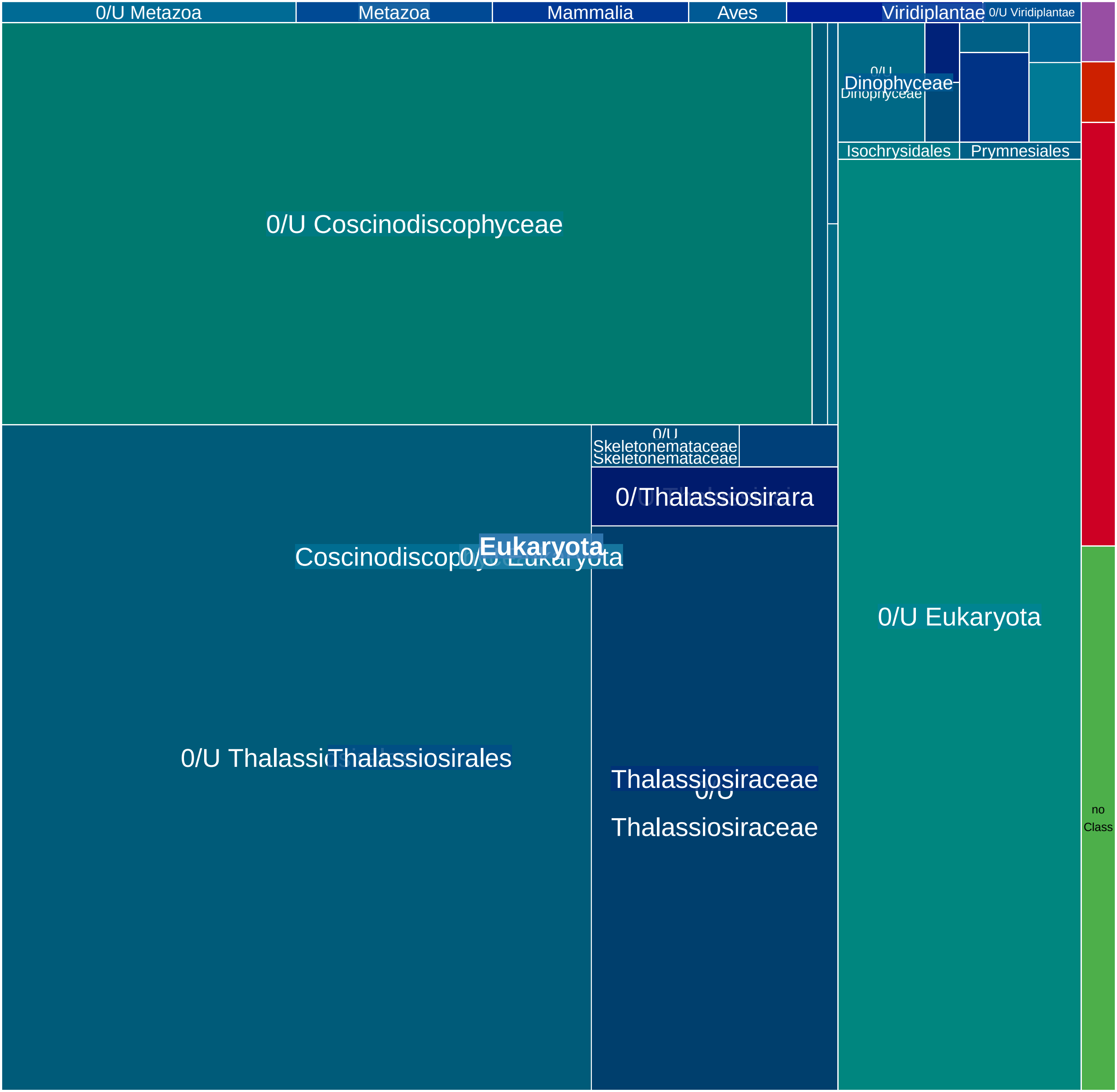

MGT−v1_357 : 2960 unigenes, 583 (19.70%) taxonomically assigned

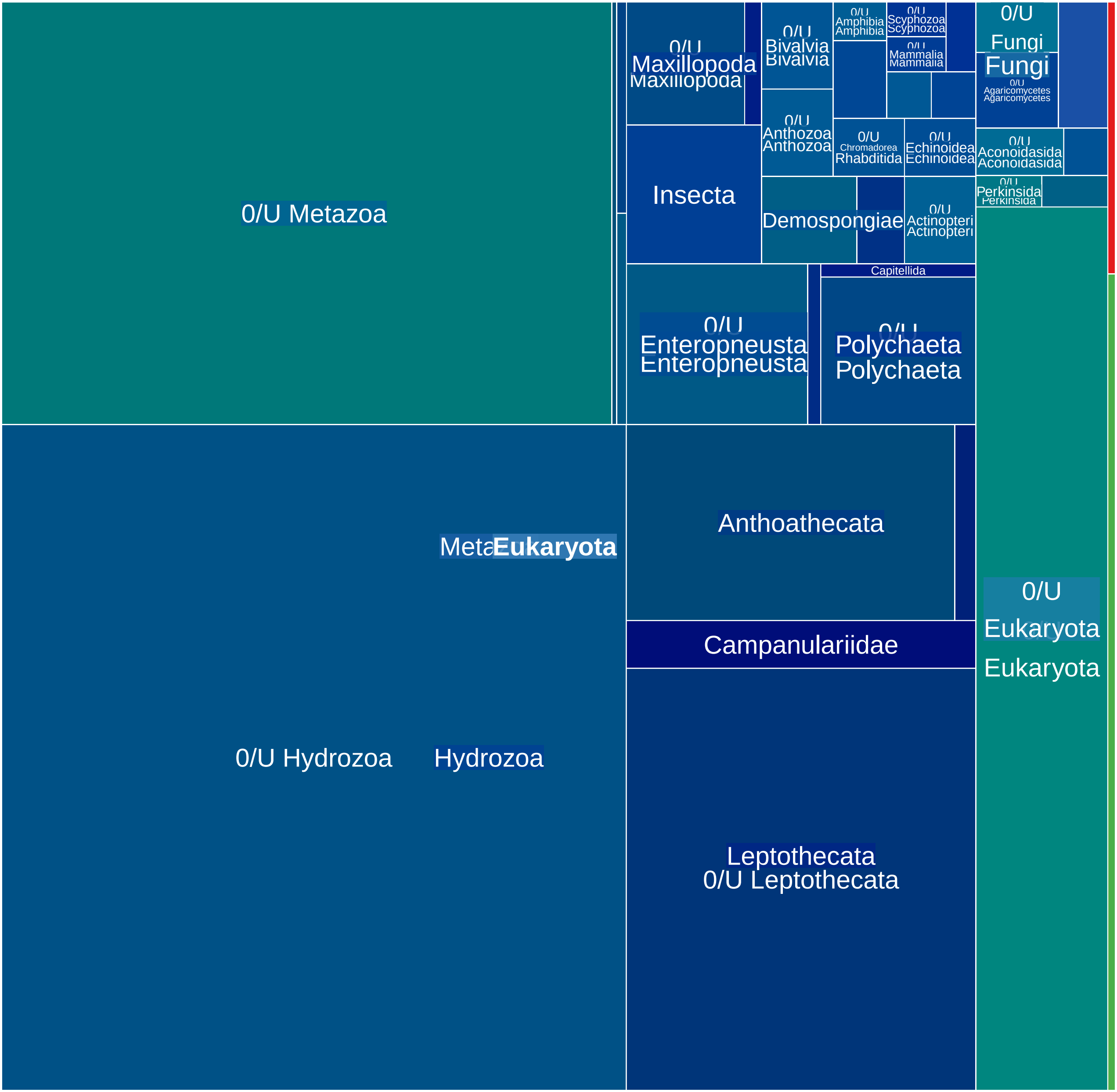

MGT−v1_358 : 2959 unigenes, 338 (11.42%) taxonomically assigned

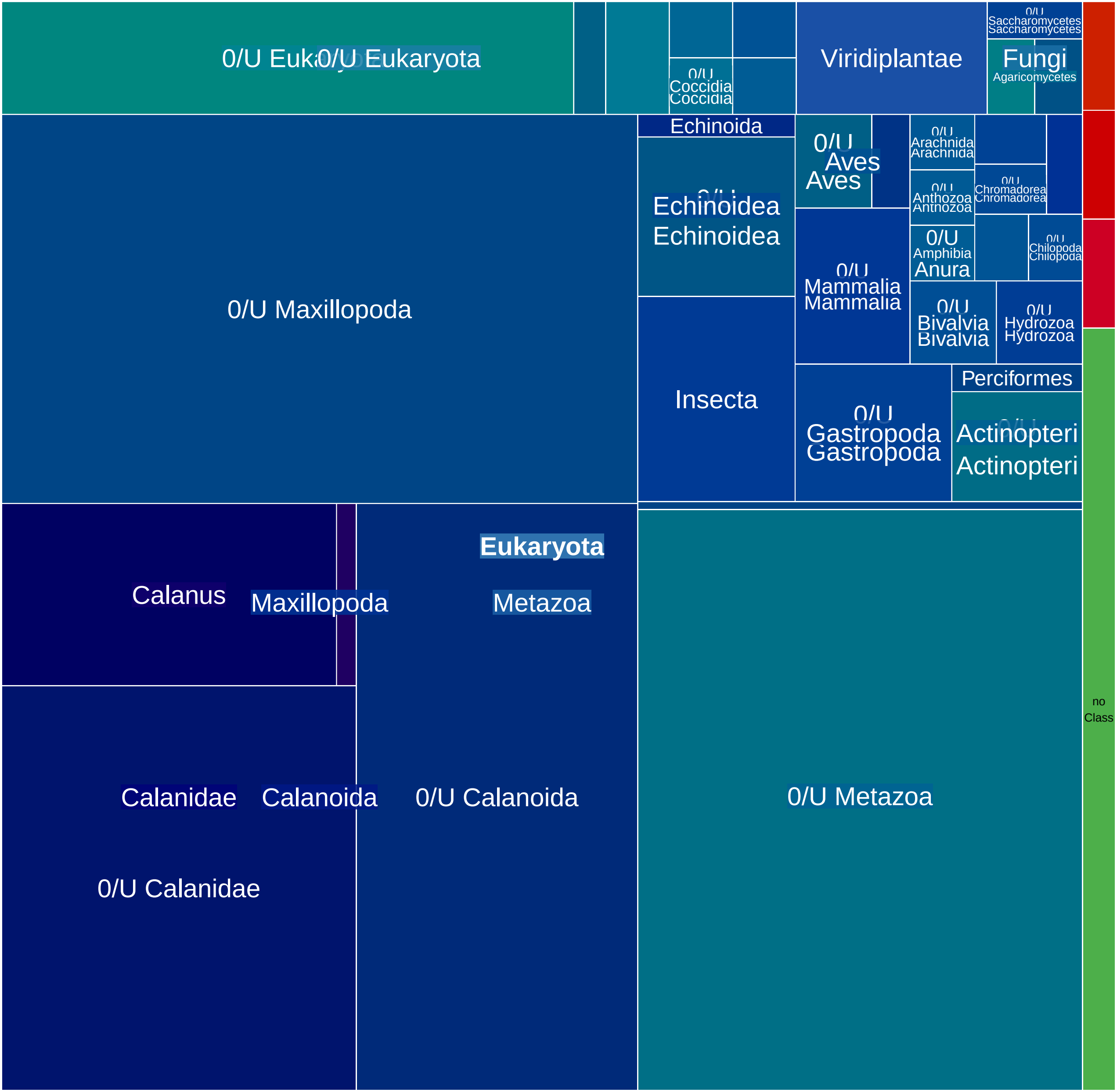

MGT−v1_359 : 2953 unigenes, 2690 (91.09%) taxonomically assigned

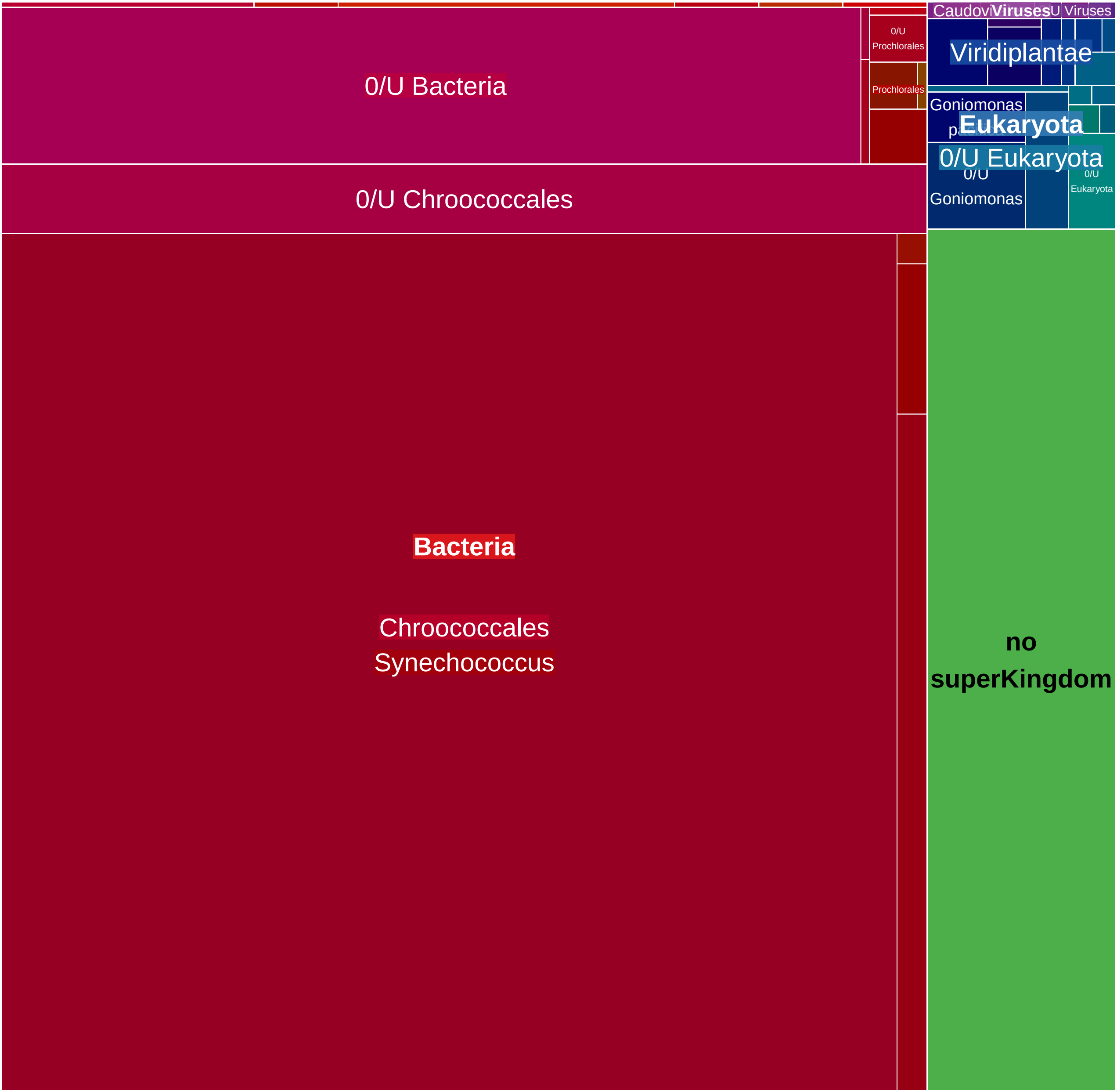

MGT−v1_360 : 2940 unigenes, 561 (19.08%) taxonomically assigned

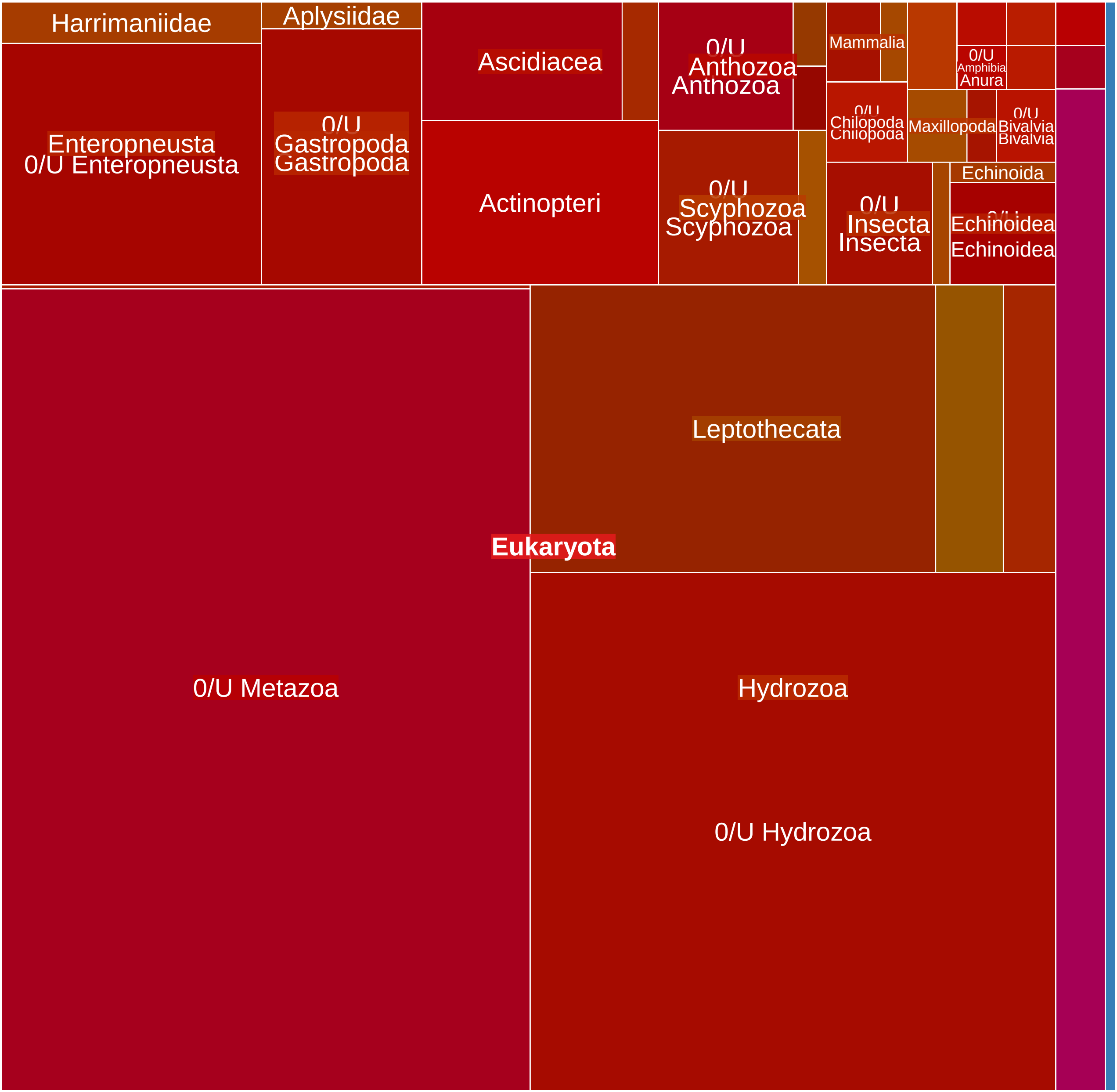

MGT−v1_361 : 2938 unigenes, 510 (17.36%) taxonomically assigned

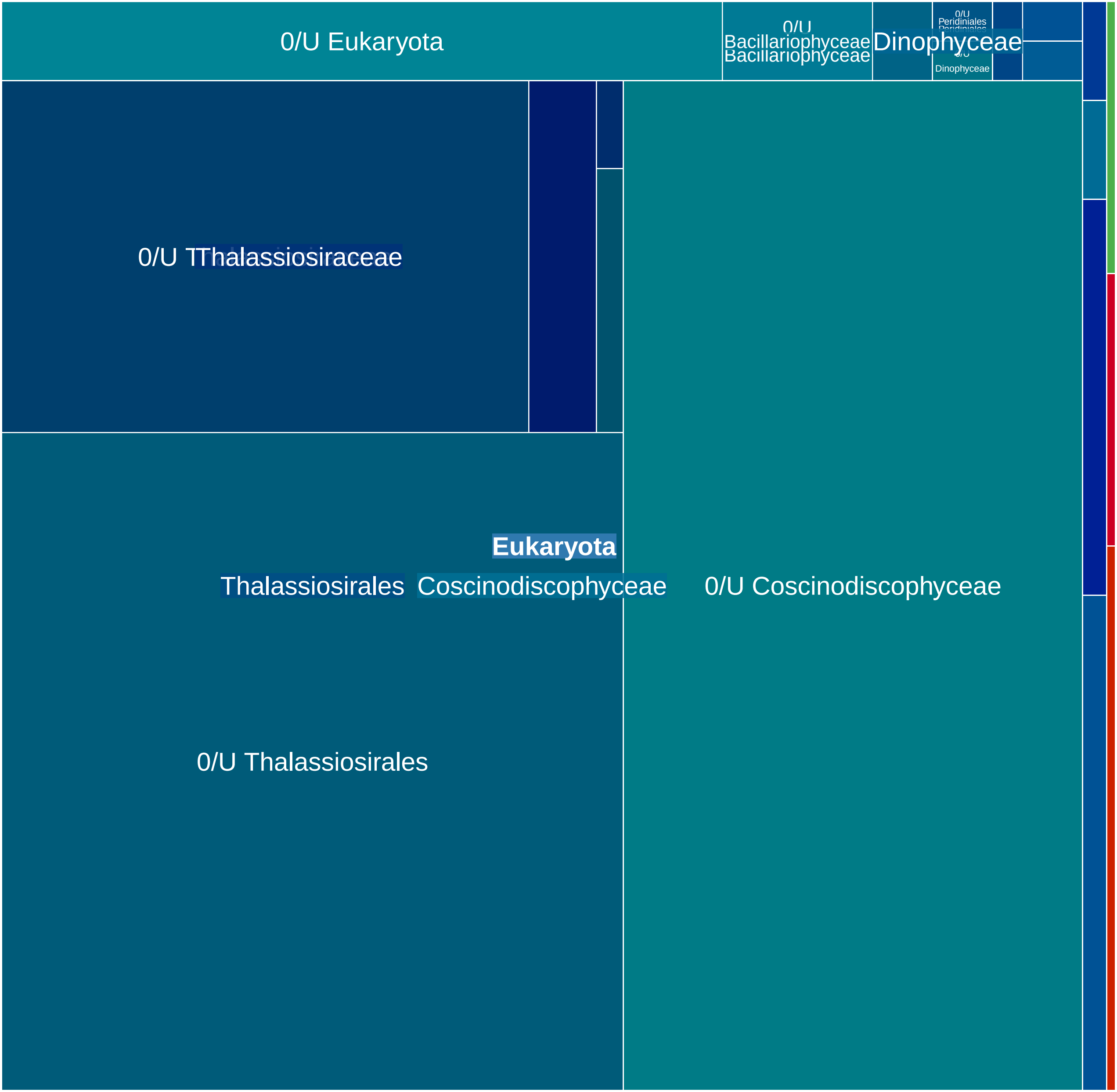

MGT−v1_362 : 2934 unigenes, 2485 (84.70%) taxonomically assigned

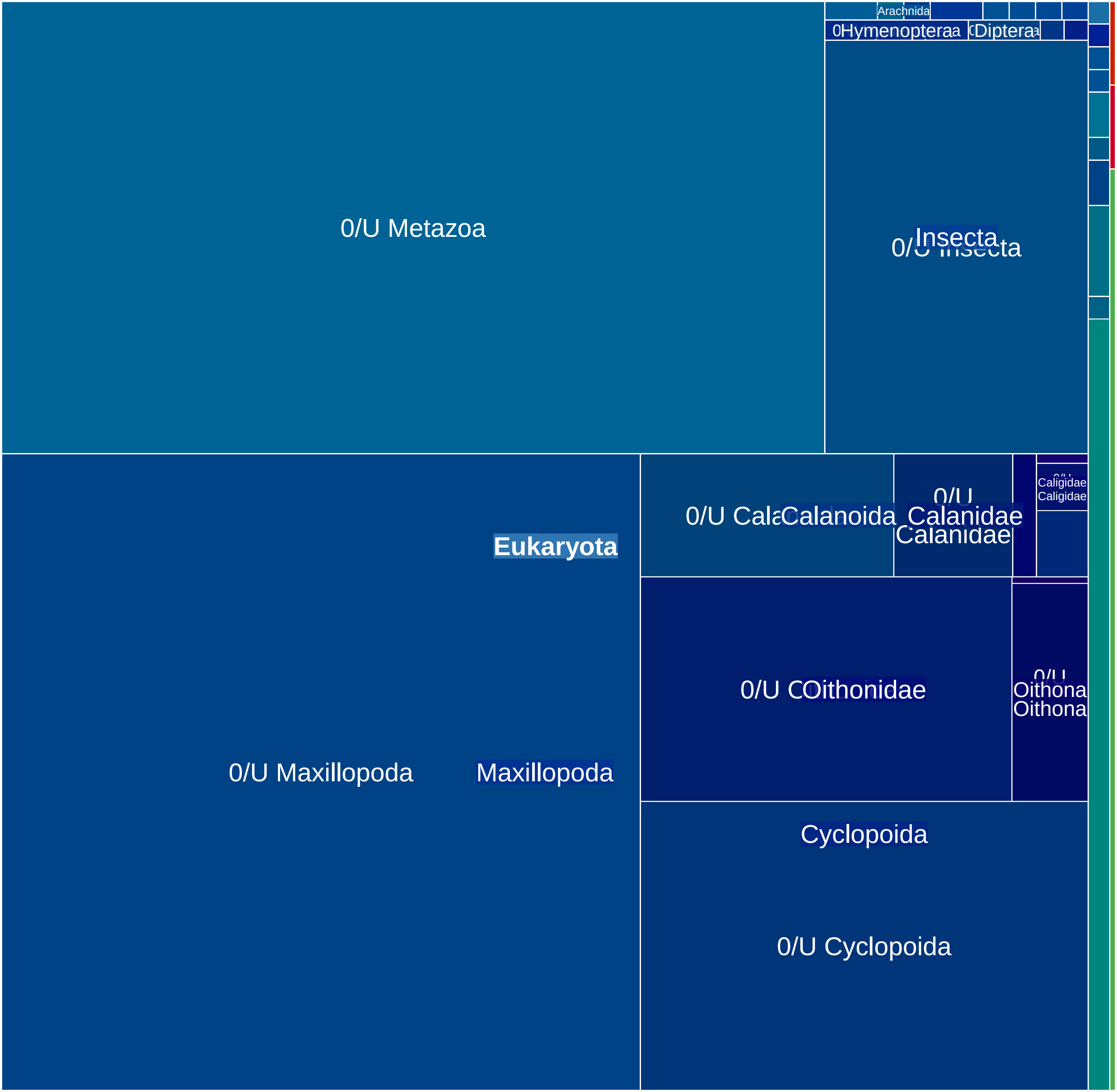

MGT−v1_363 : 2932 unigenes, 573 (19.54%) taxonomically assigned

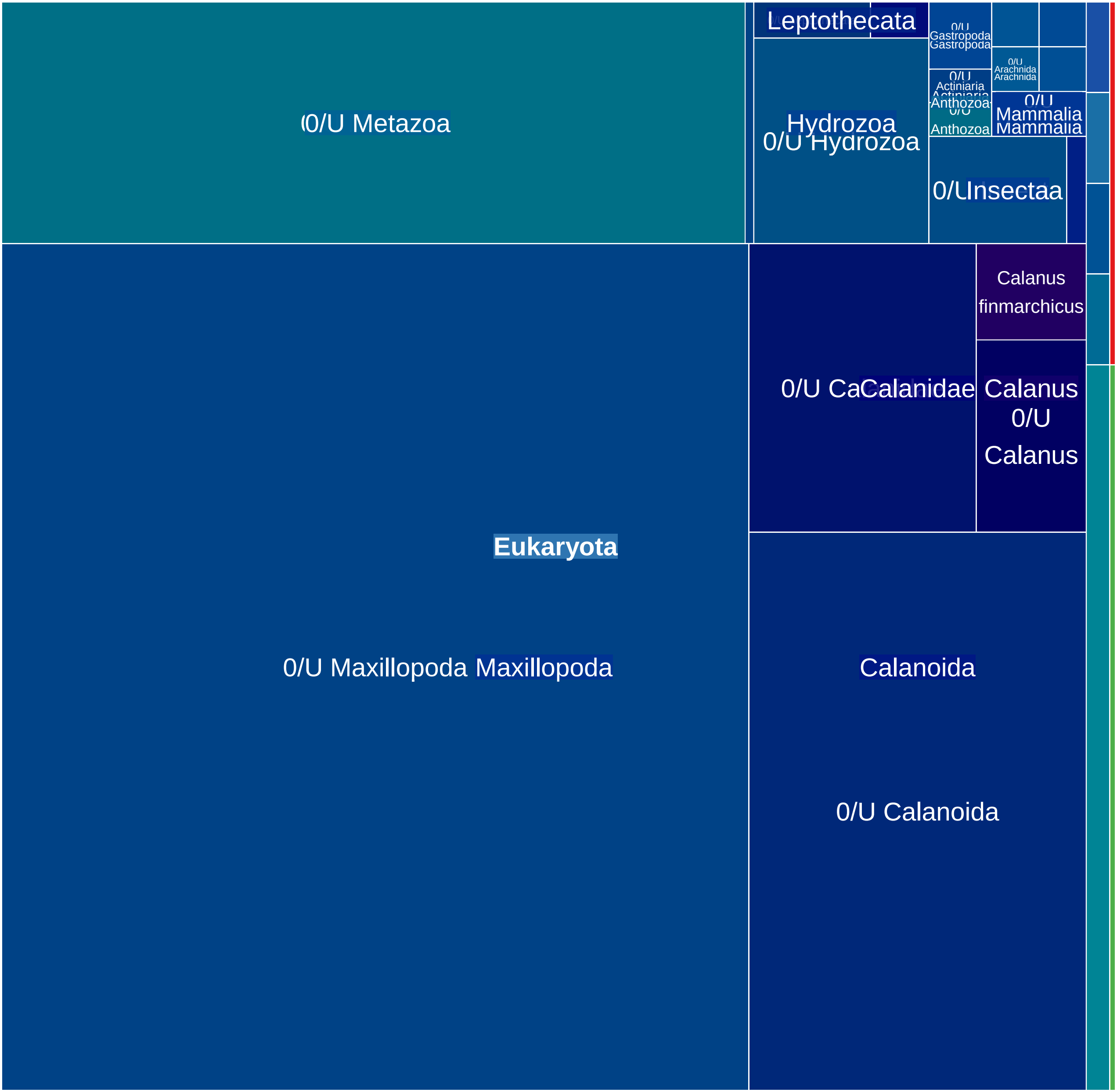

MGT−v1_364 : 2923 unigenes, 2117 (72.43%) taxonomically assigned

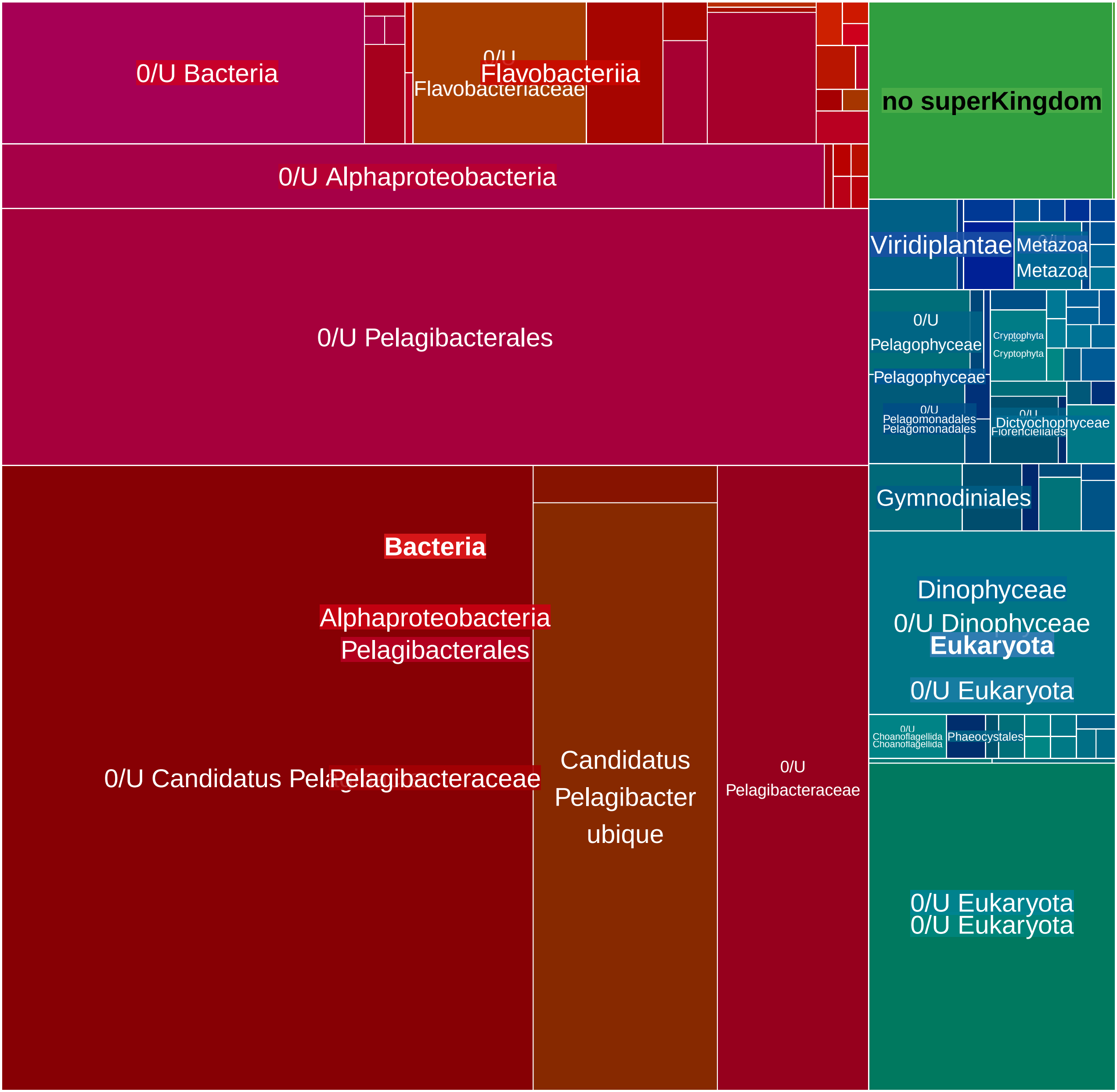

MGT−v1_365 : 2916 unigenes, 612 (20.99%) taxonomically assigned

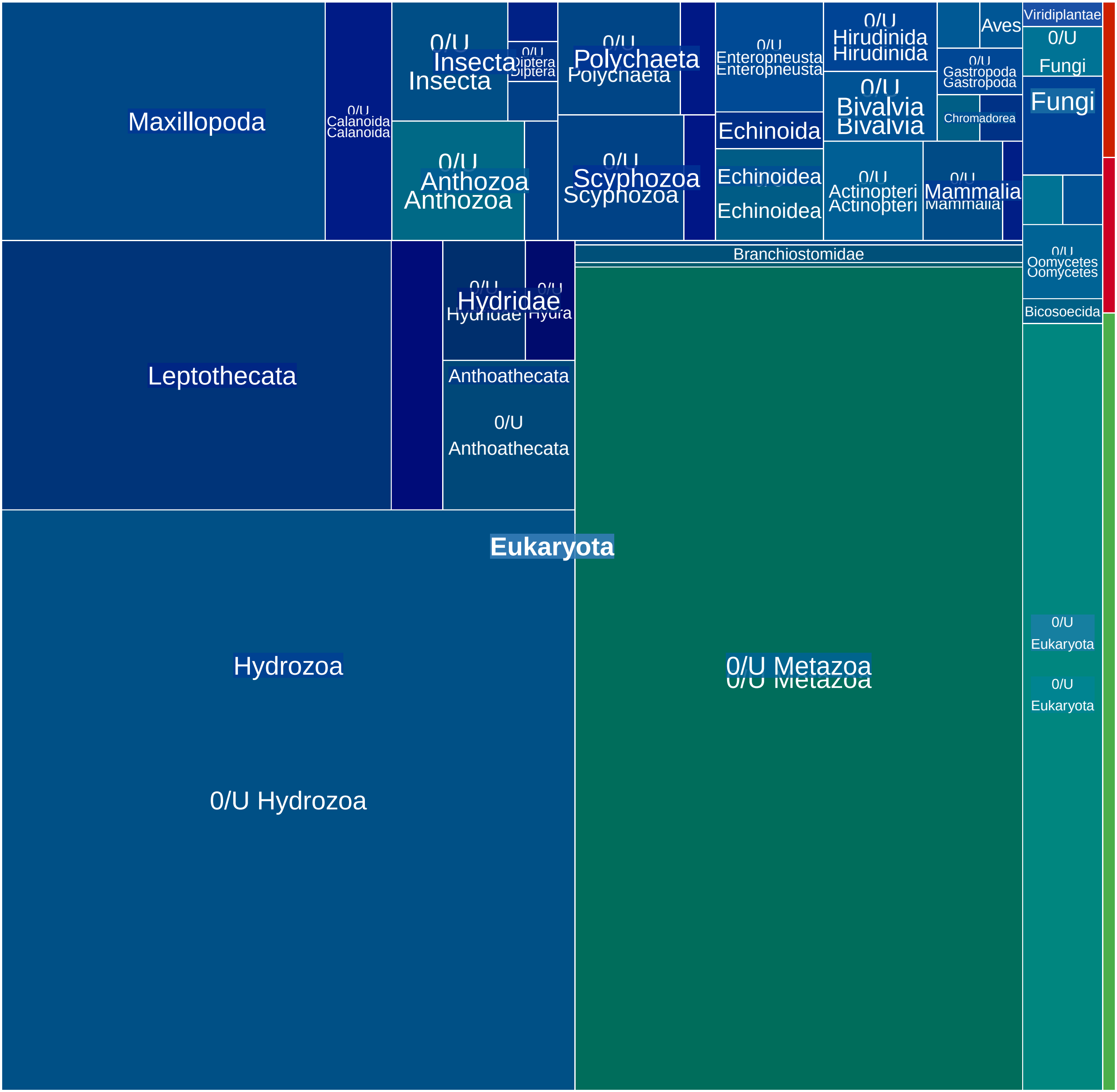

### MGT−v1_366 : 2910 unigenes, 1950 (67.01%) taxonomically assigned

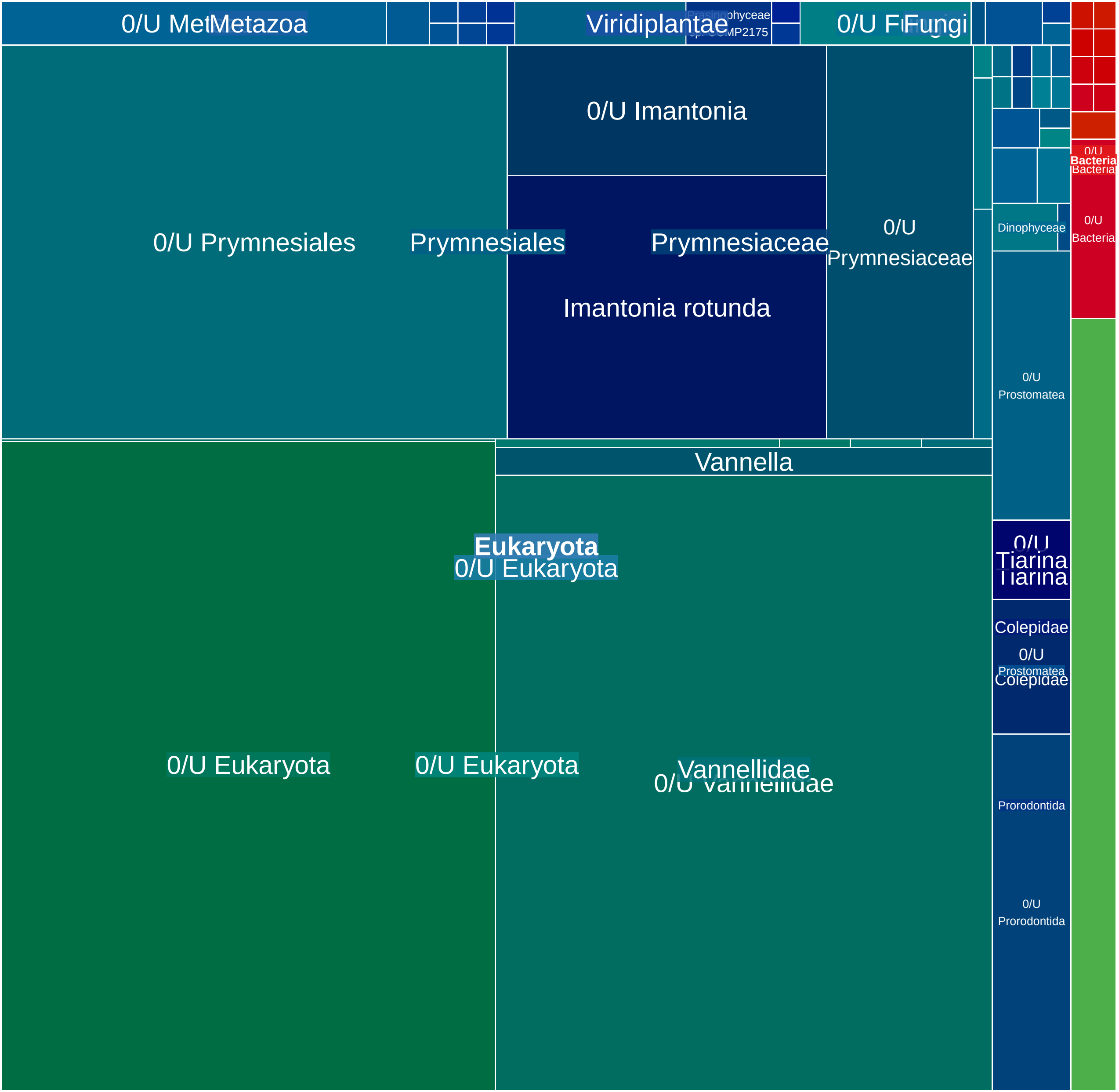

MGT−v1_367 : 2901 unigenes, 2002 (69.01%) taxonomically assigned

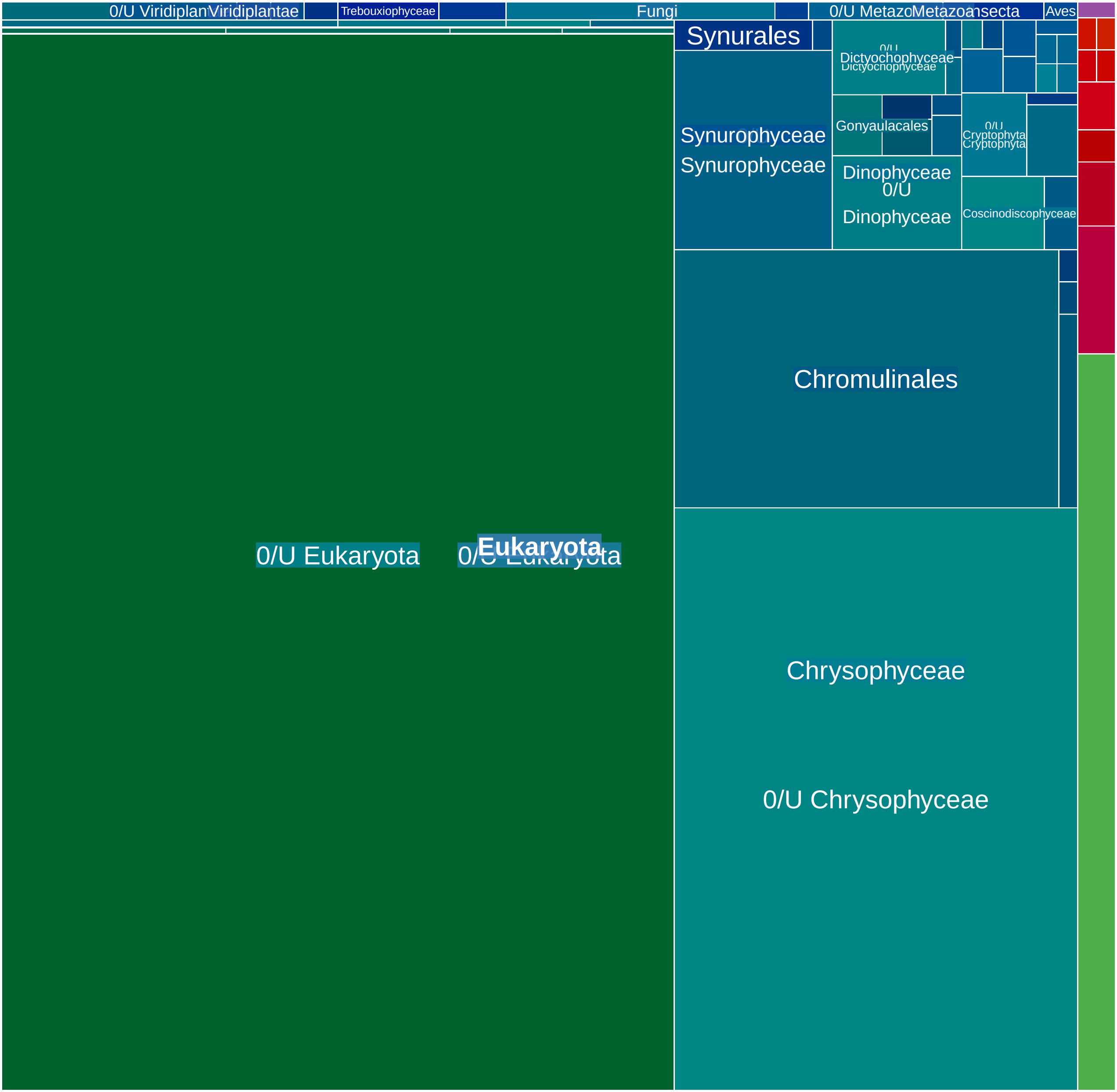

MGT−v1_368 : 2899 unigenes, 344 (11.87%) taxonomically assigned

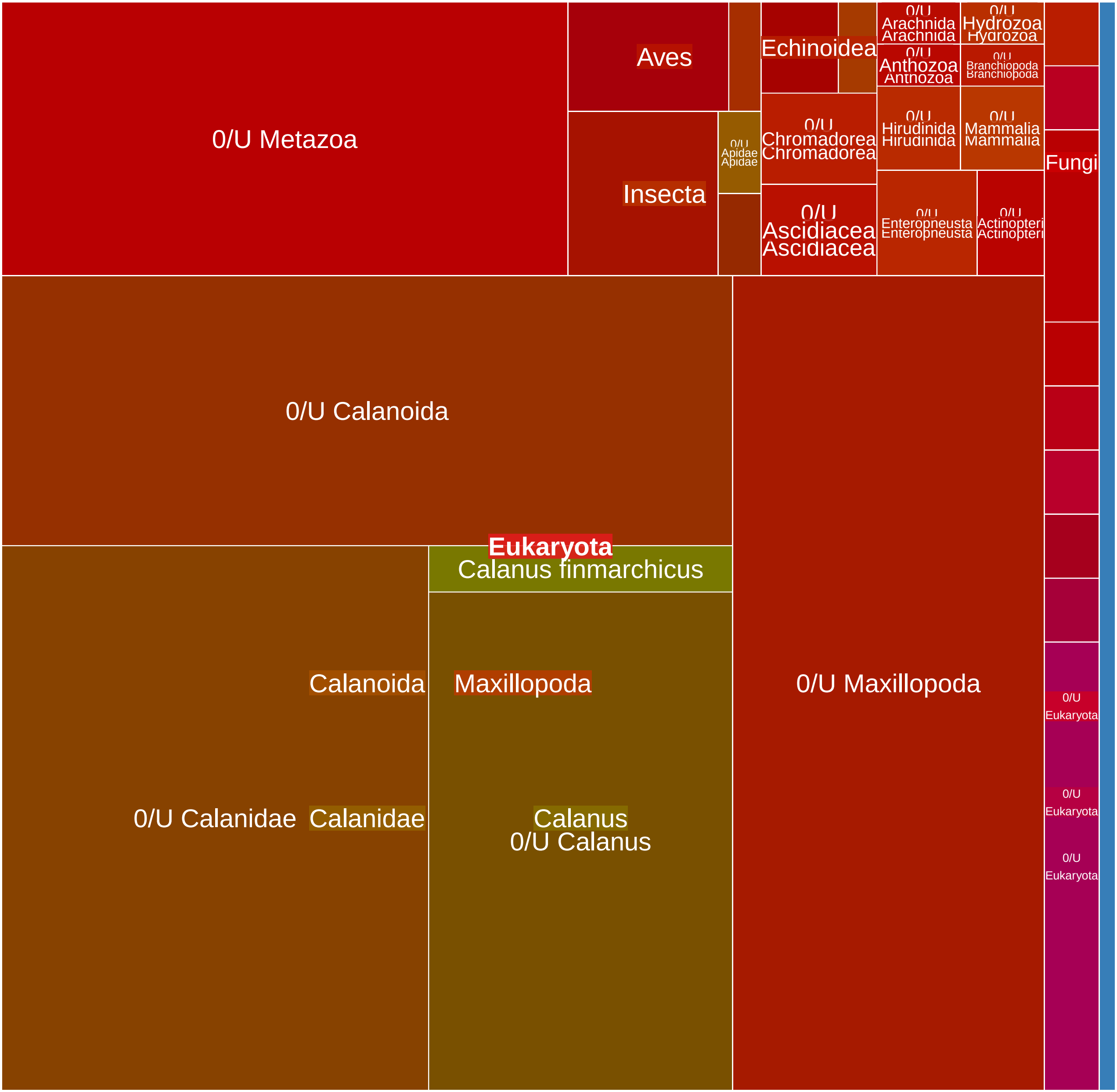

### MGT−v1_369 : 2876 unigenes, 1606 (55.84%) taxonomically assigned

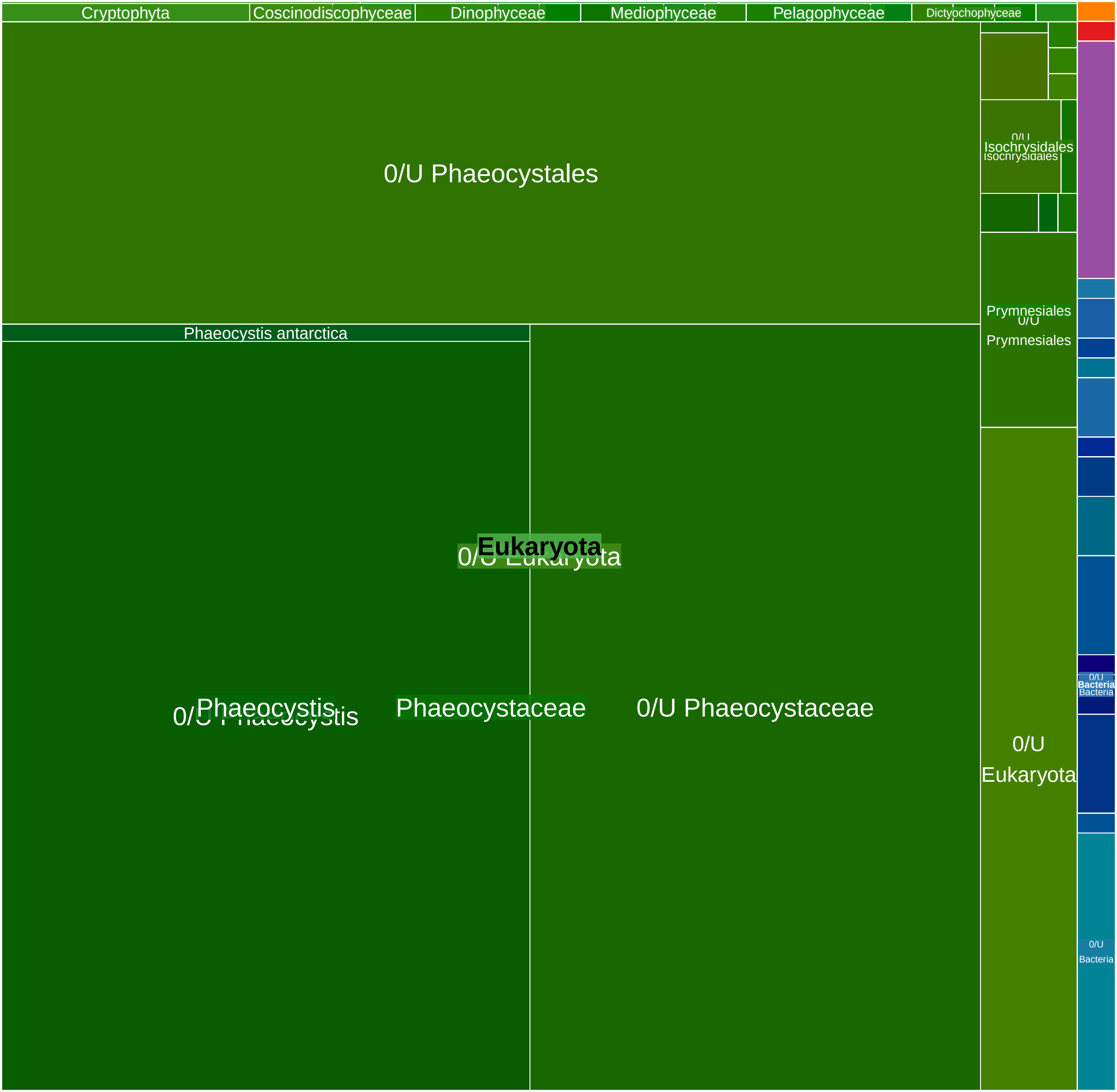

### MGT−v1_370 : 2875 unigenes, 2639 (91.79%) taxonomically assigned

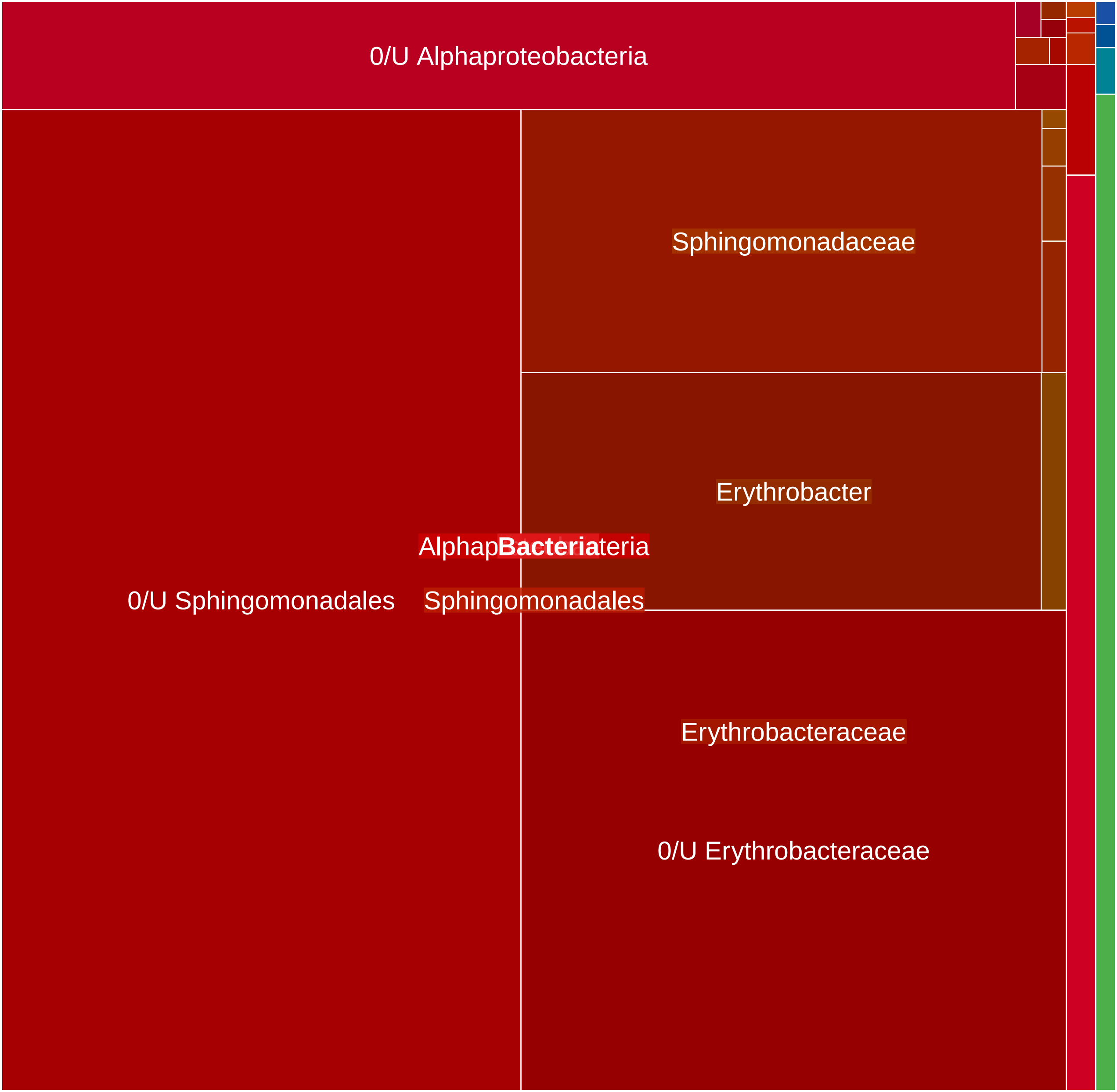

MGT−v1_371 : 2840 unigenes, 1555 (54.75%) taxonomically assigned

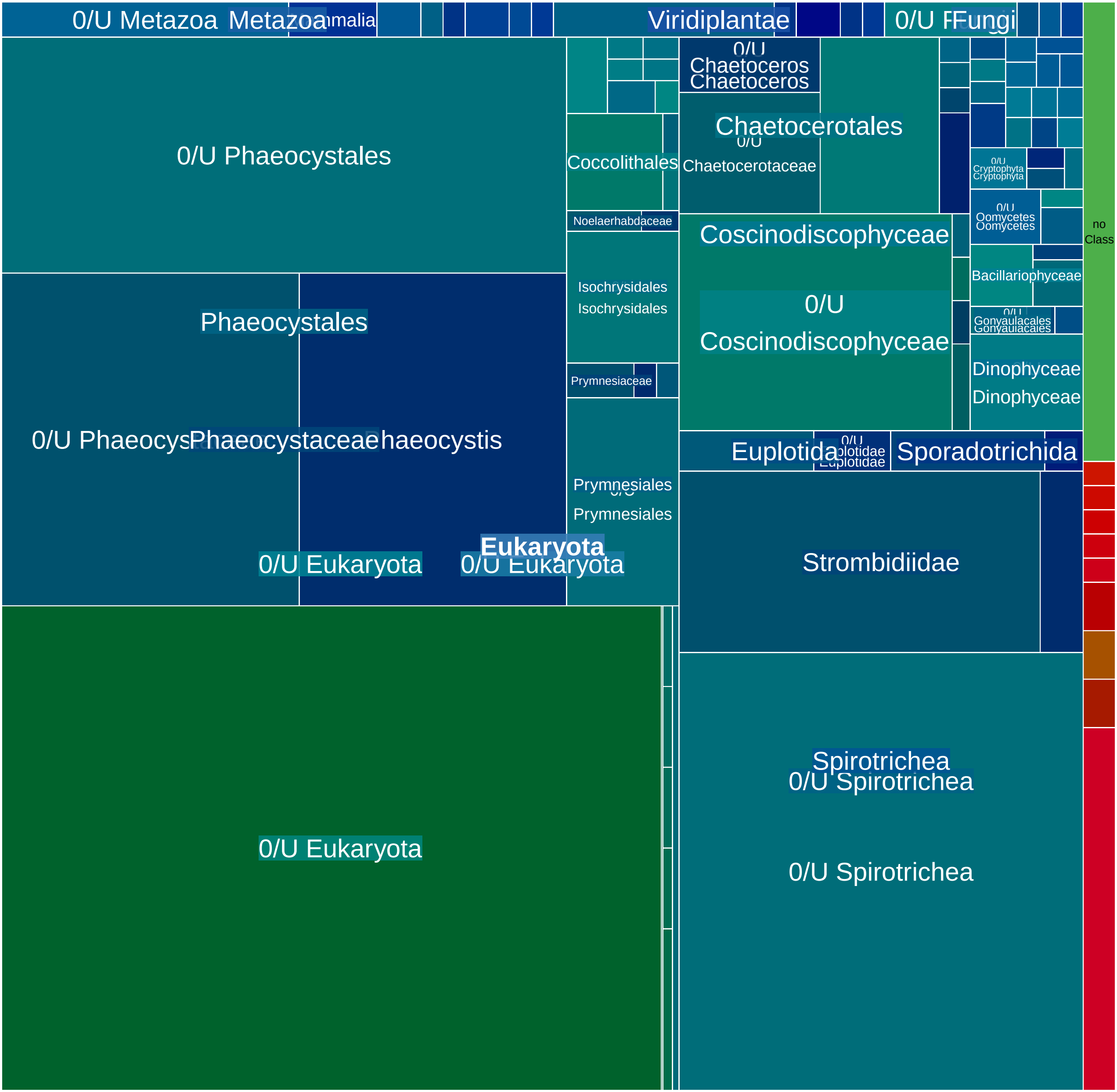

### MGT−v1_372 : 2830 unigenes, 482 (17.03%) taxonomically assigned

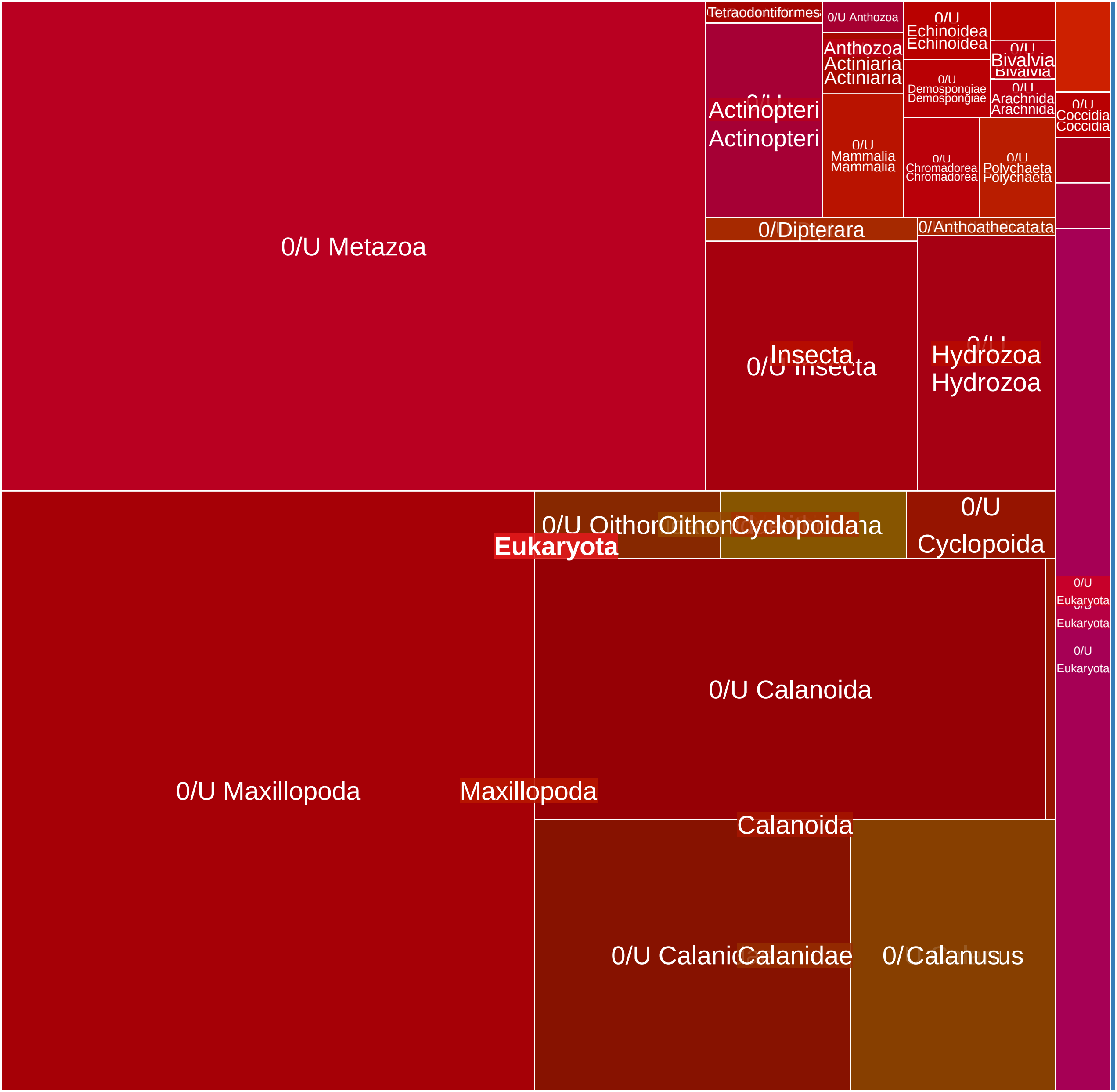

MGT−v1_373 : 2830 unigenes, 2601 (91.91%) taxonomically assigned

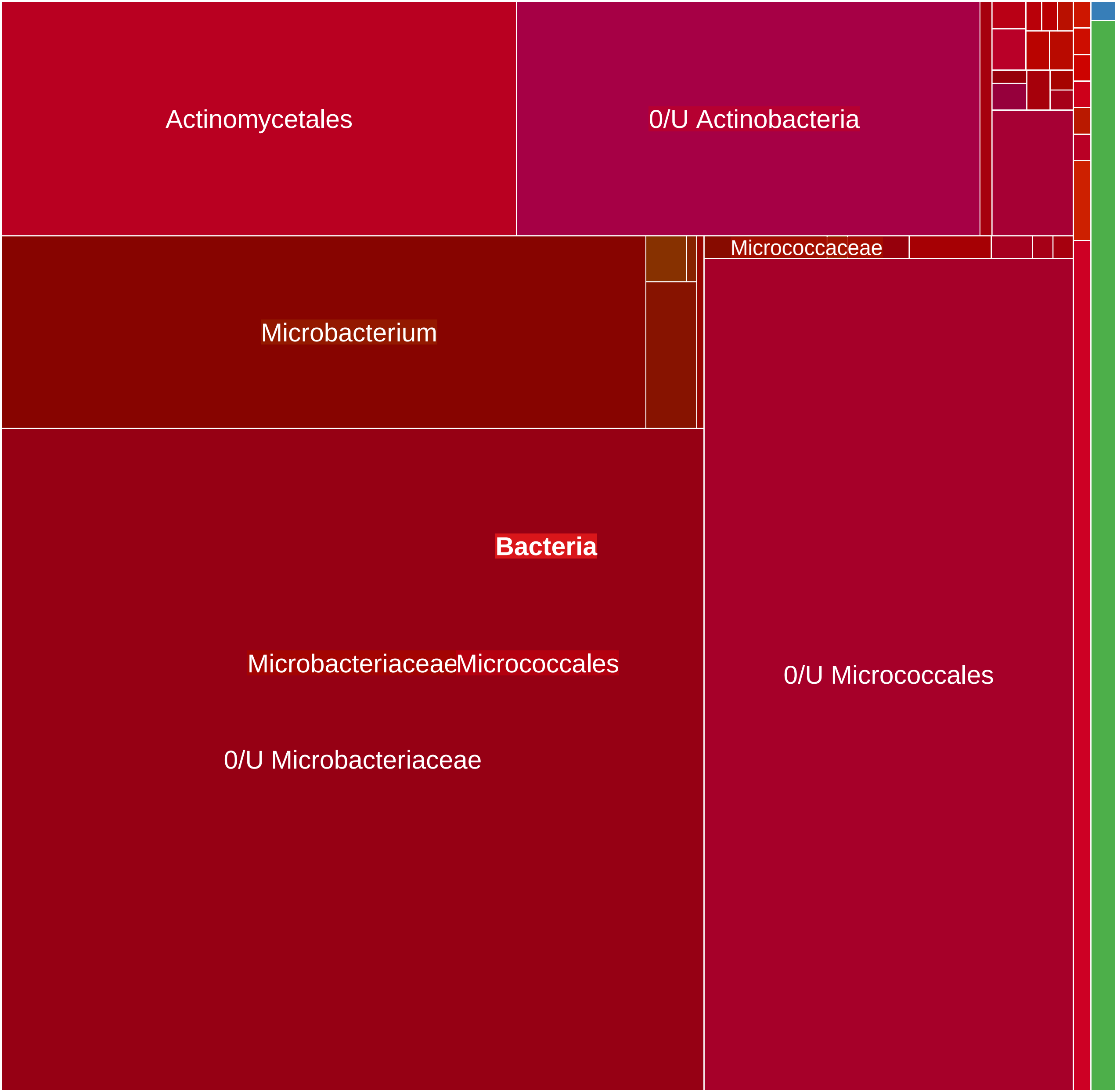

MGT−v1_374 : 2830 unigenes, 2355 (83.22%) taxonomically assigned

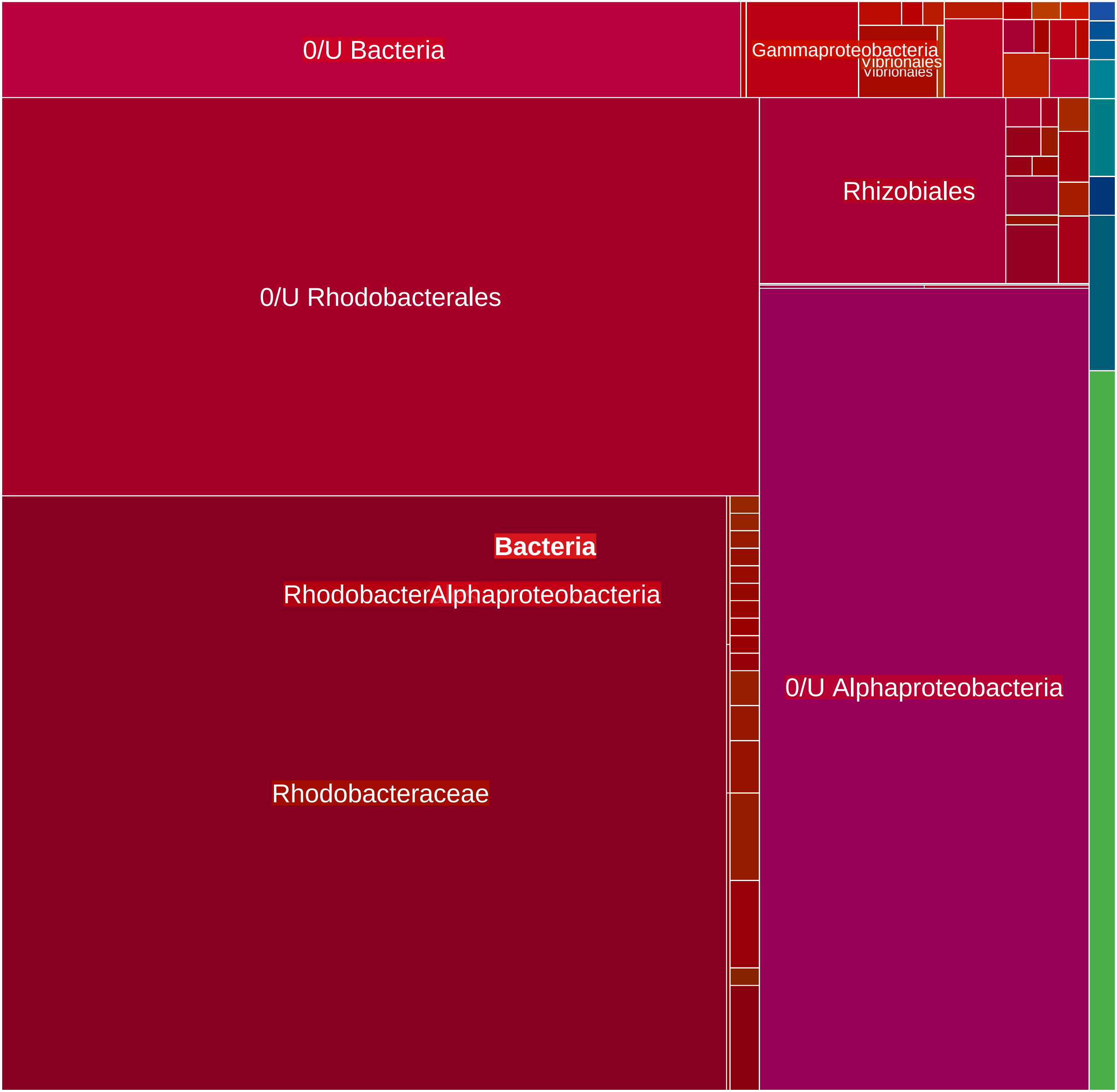

### MGT−v1_375 : 2826 unigenes, 188 (6.65%) taxonomically assigned

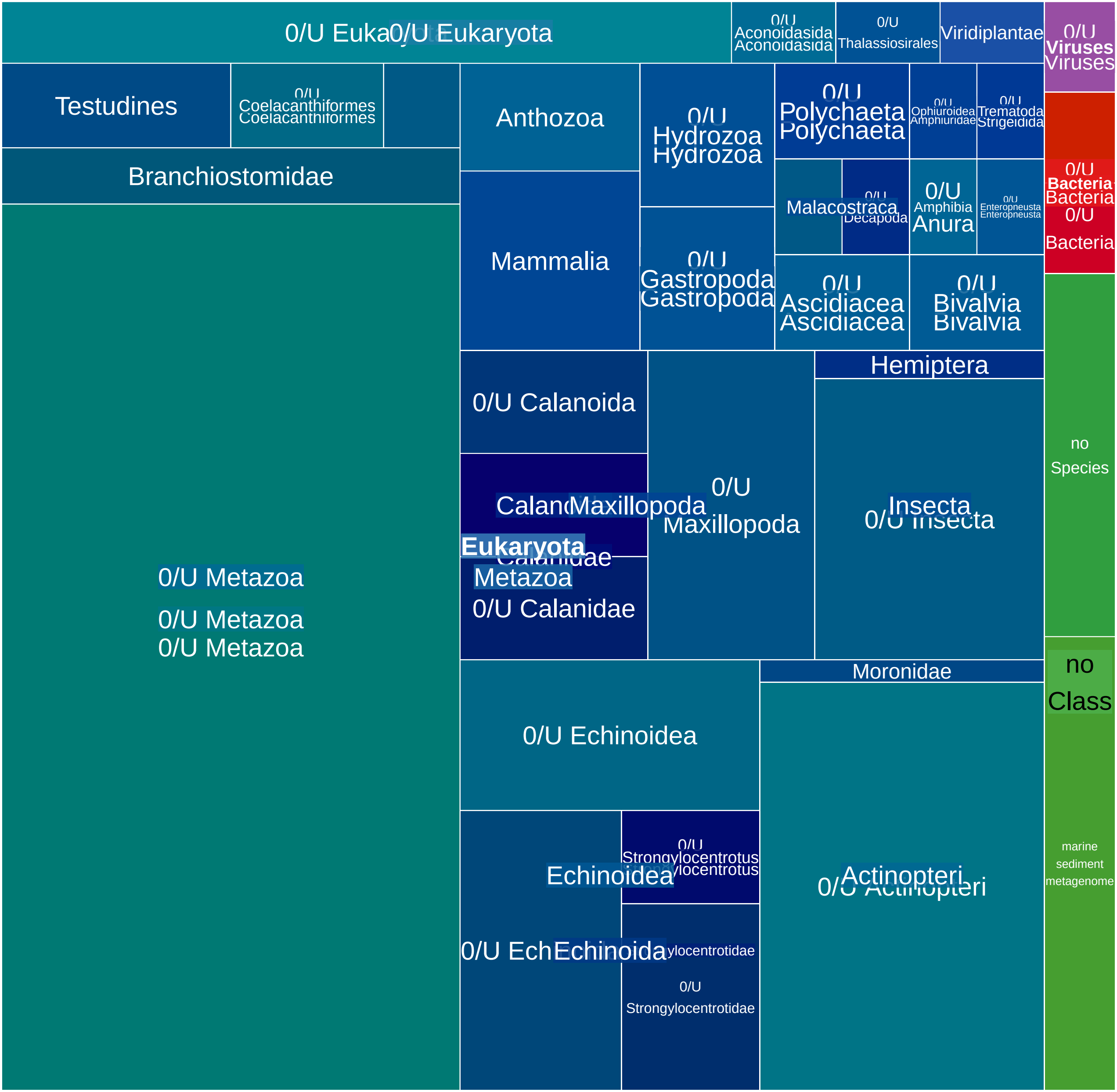

MGT−v1_376 : 2809 unigenes, 2372 (84.44%) taxonomically assigned

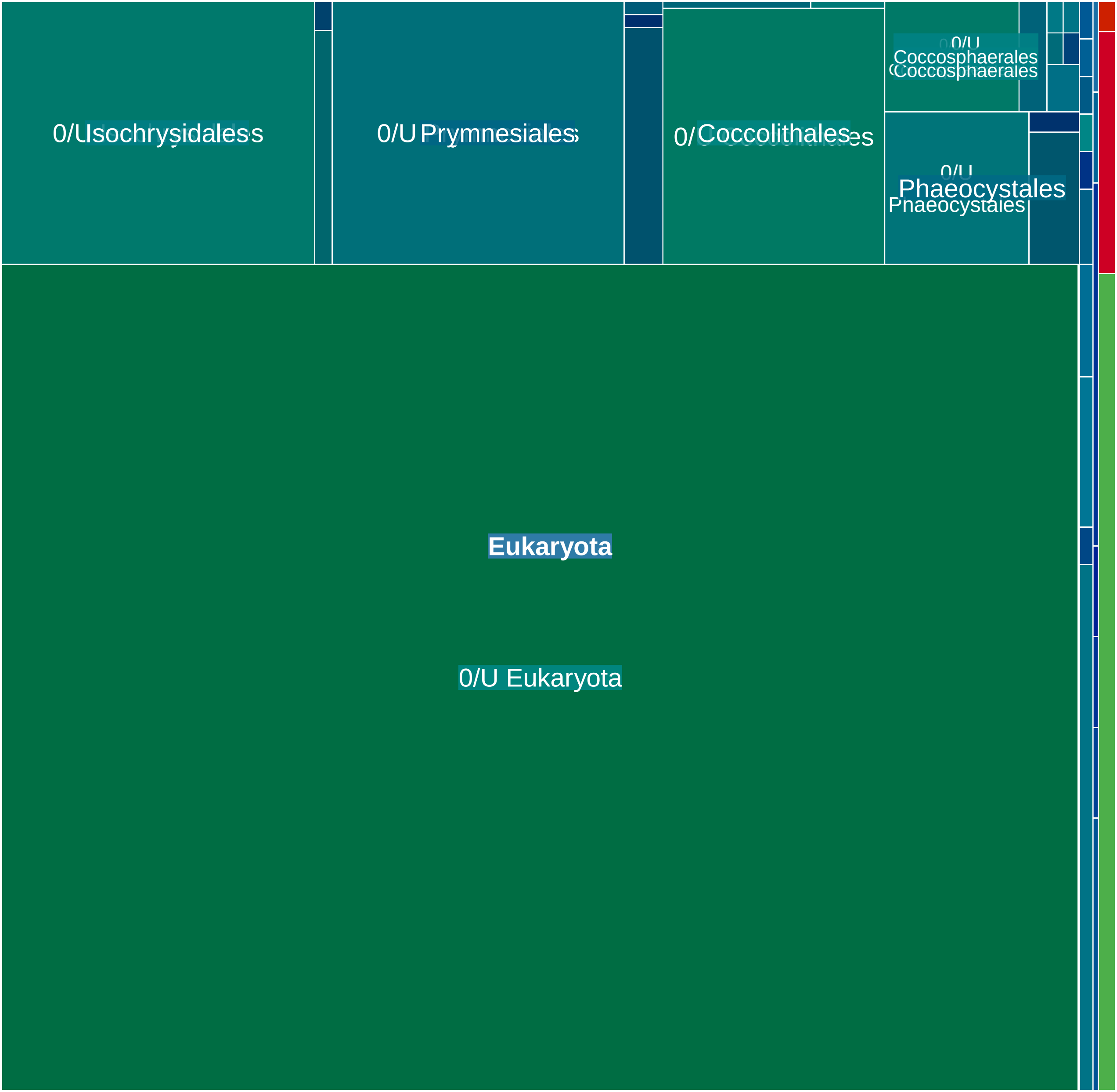

### MGT−v1_377 : 2806 unigenes, 1351 (48.15%) taxonomically assigned

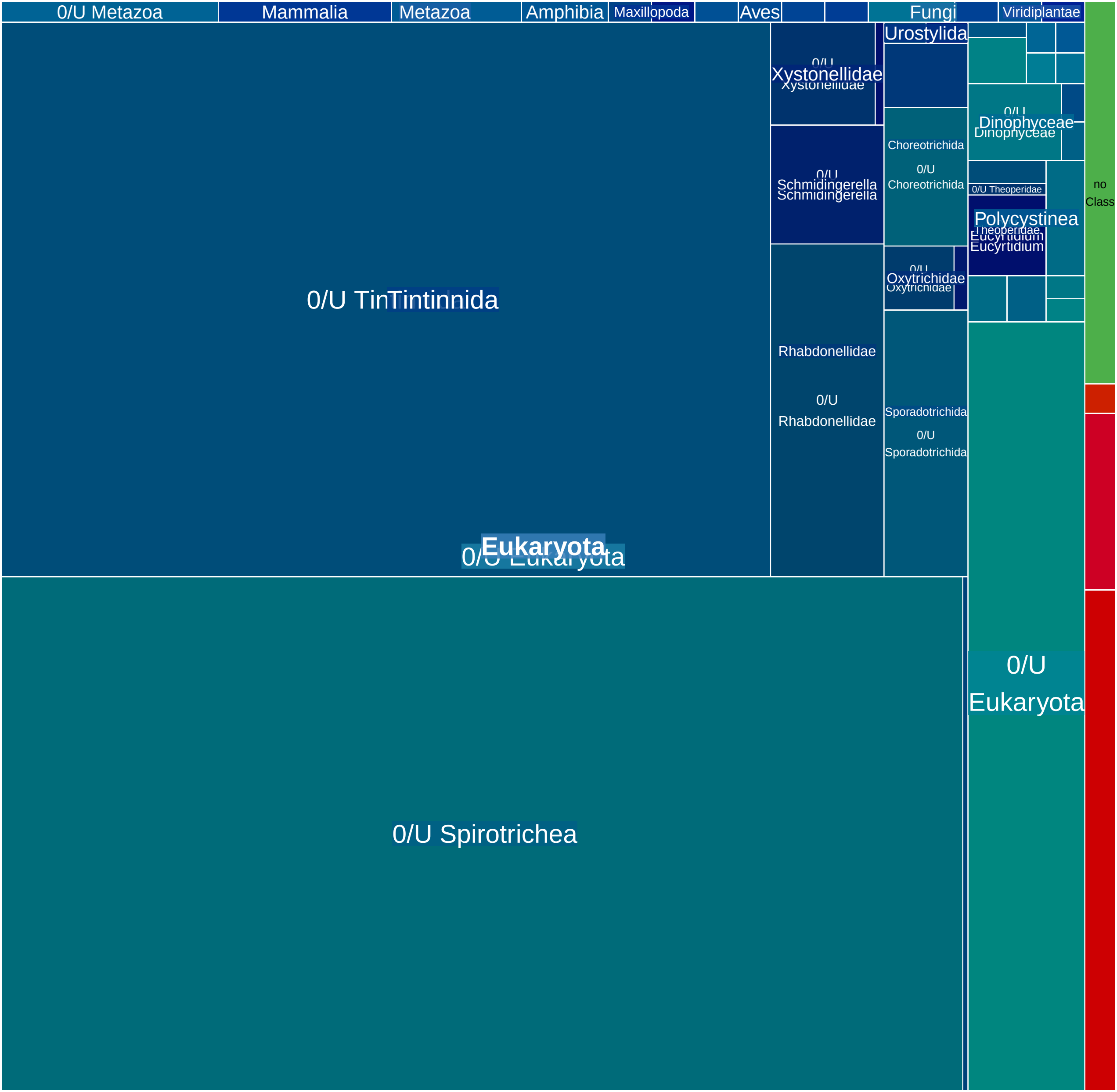

MGT−v1_378 : 2768 unigenes, 1525 (55.09%) taxonomically assigned

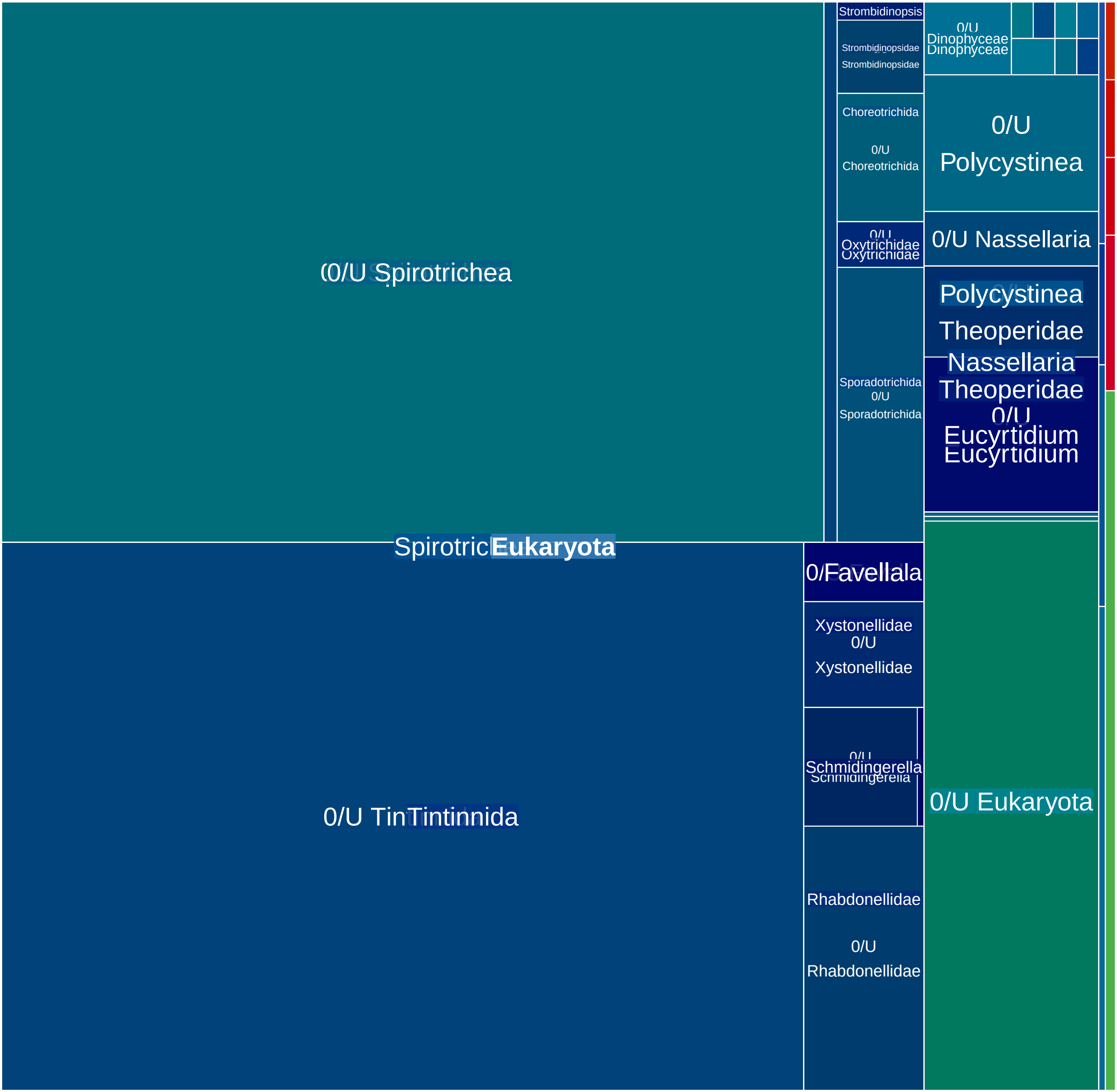

### MGT−v1_379 : 2765 unigenes, 2067 (74.76%) taxonomically assigned

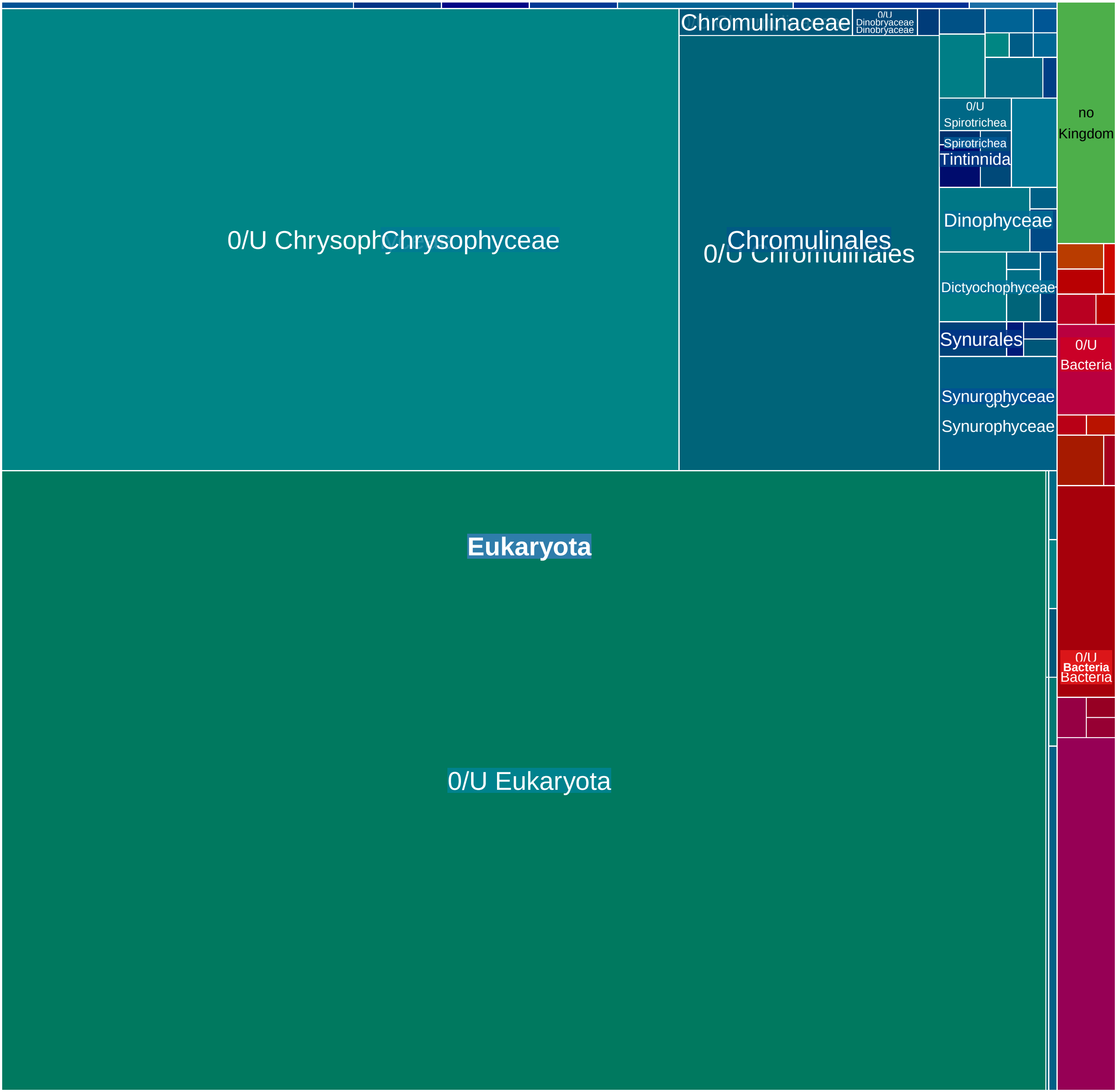

MGT−v1_380 : 2755 unigenes, 541 (19.64%) taxonomically assigned

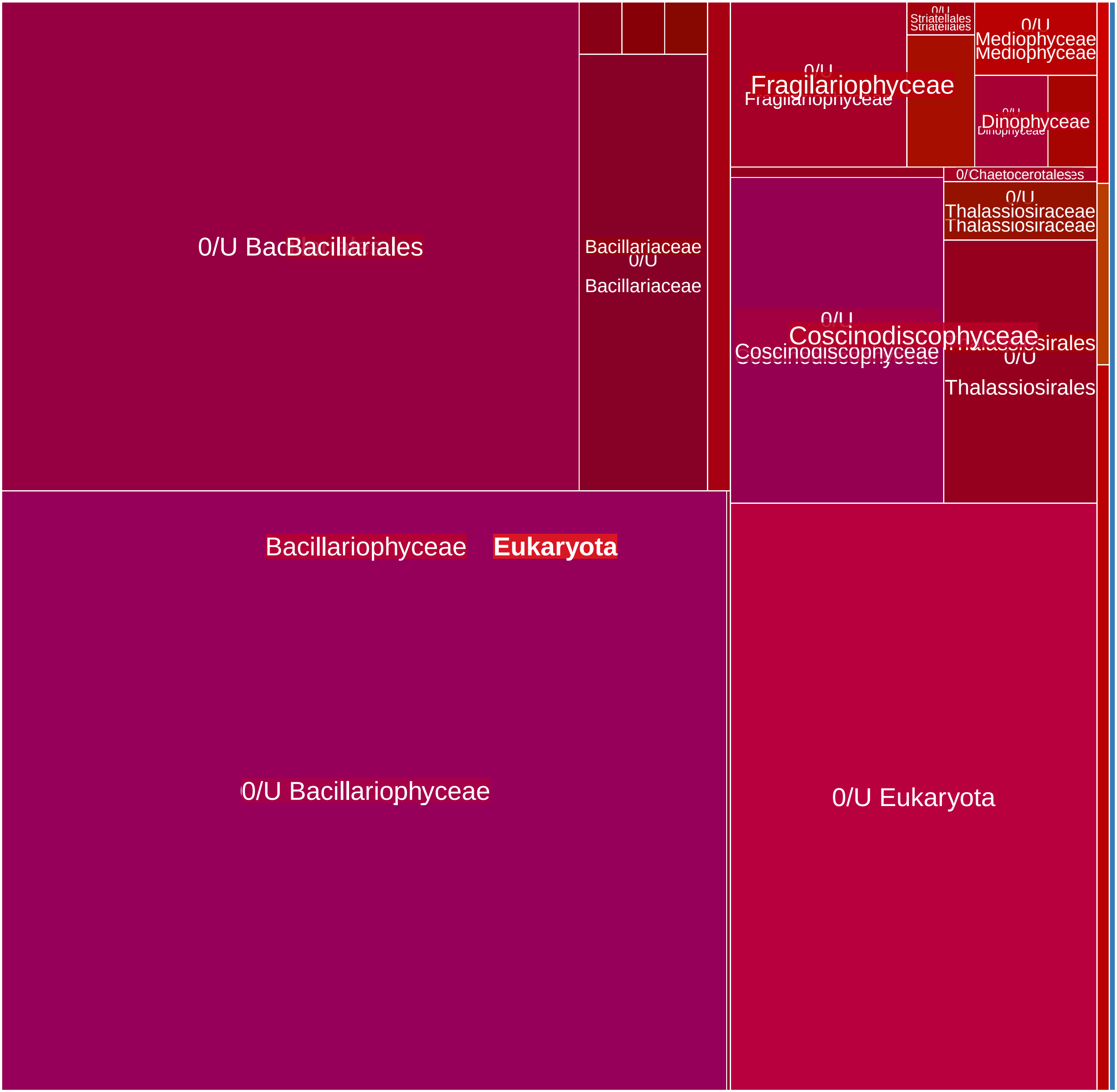

MGT−v1_381 : 2745 unigenes, 288 (10.49%) taxonomically assigned

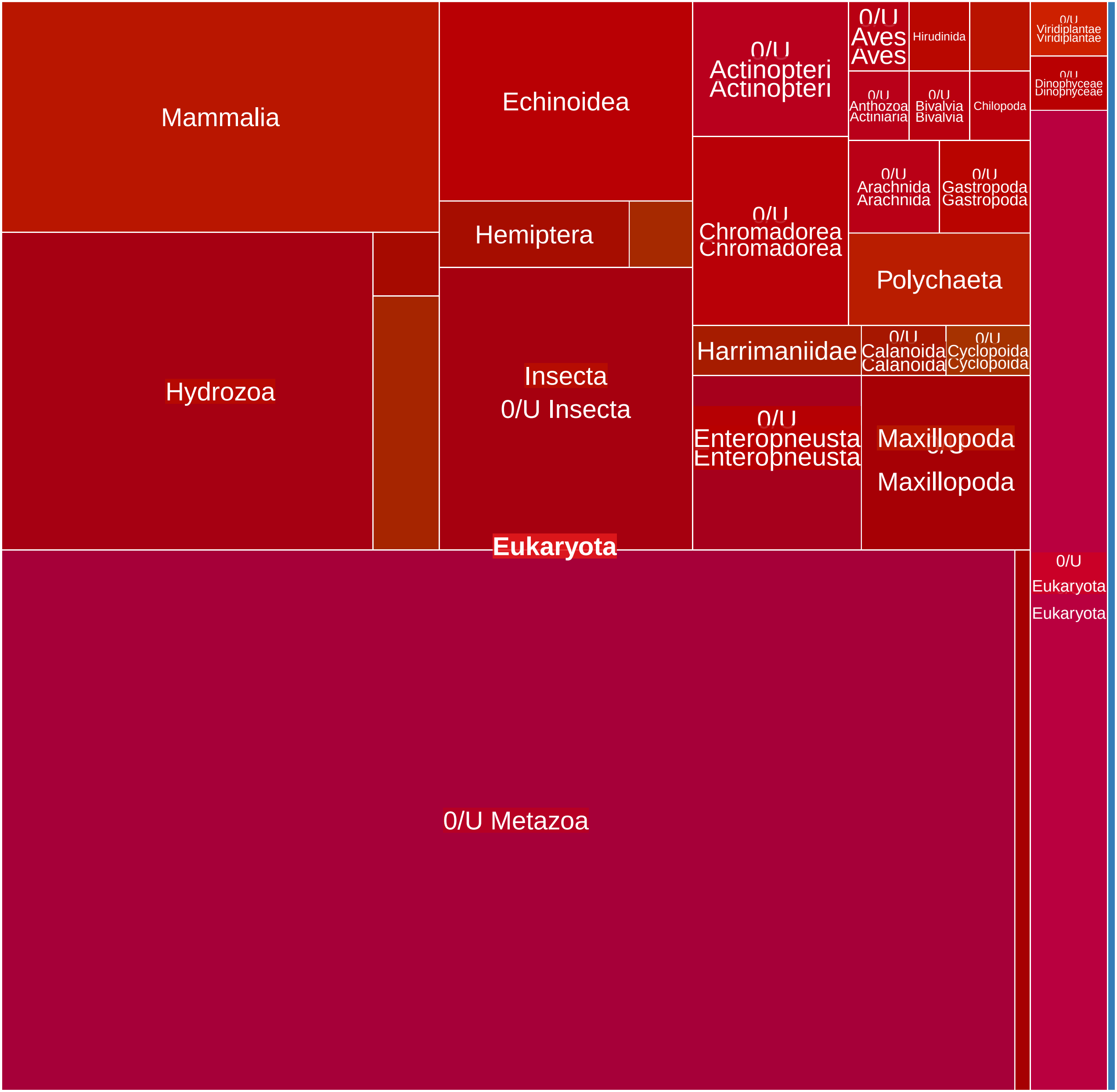

### MGT−v1_382 : 2680 unigenes, 221 (8.25%) taxonomically assigned

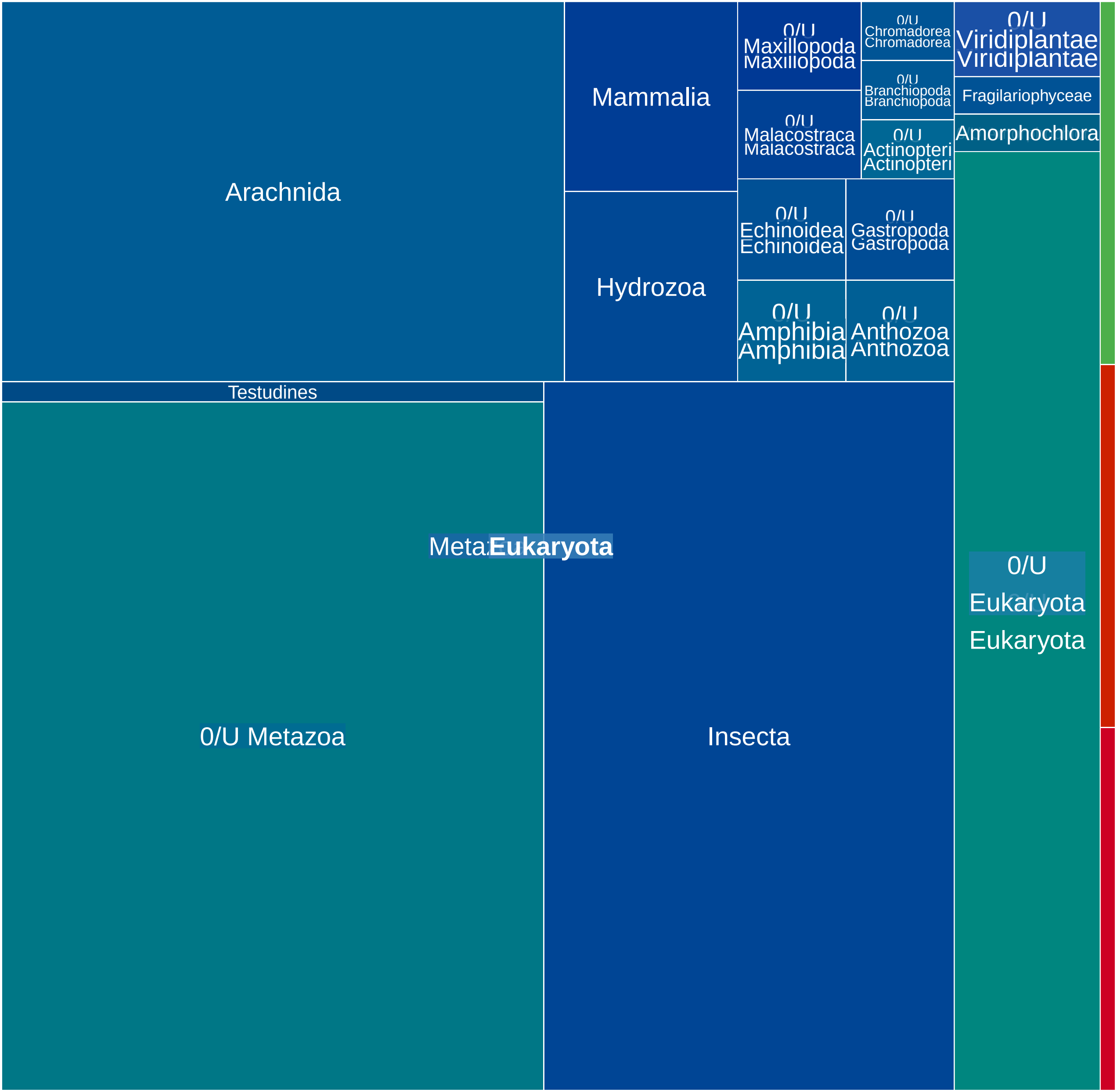

MGT−v1_383 : 2670 unigenes, 2475 (92.70%) taxonomically assigned

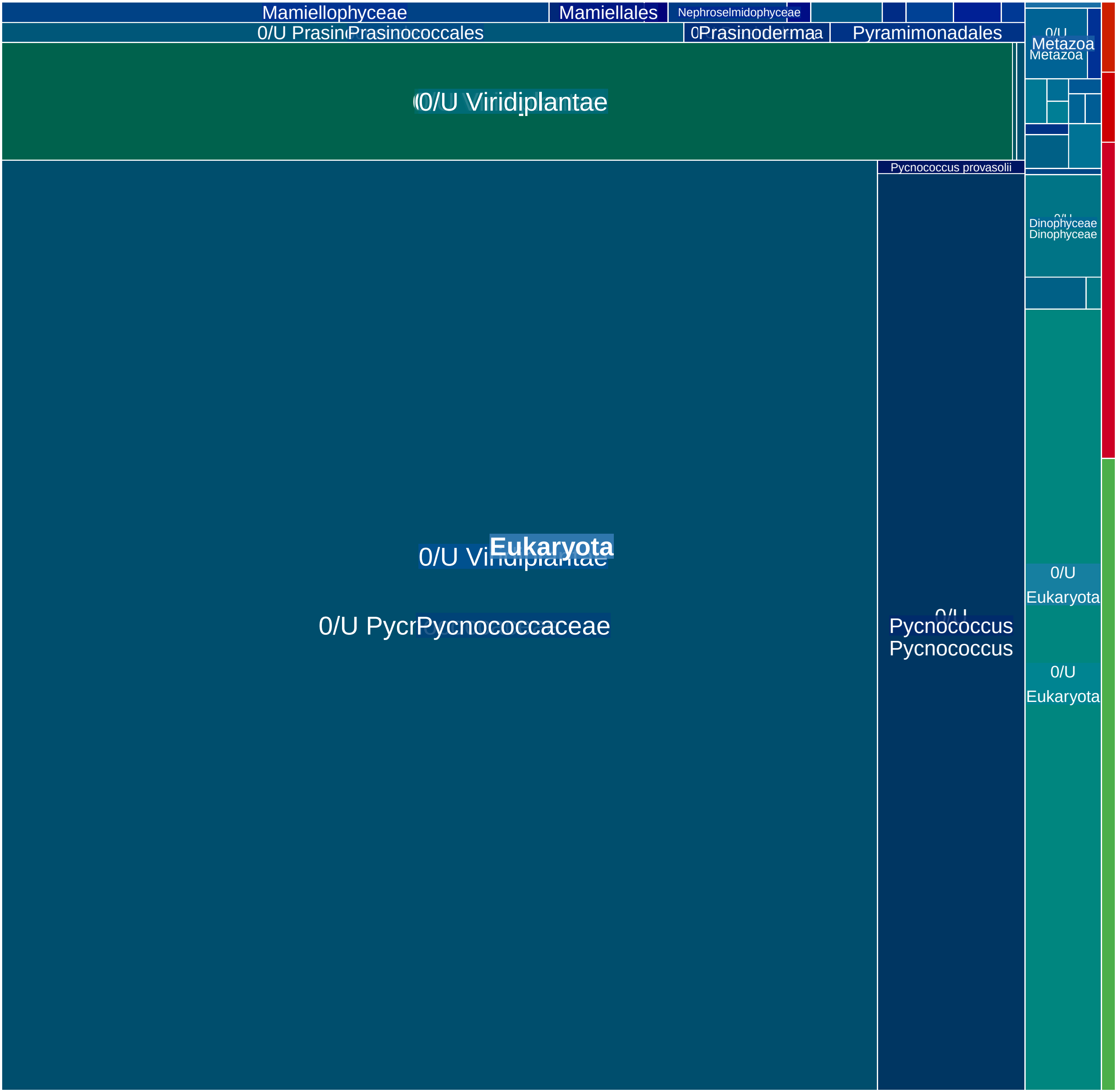

MGT−v1_384 : 2665 unigenes, 2549 (95.65%) taxonomically assigned

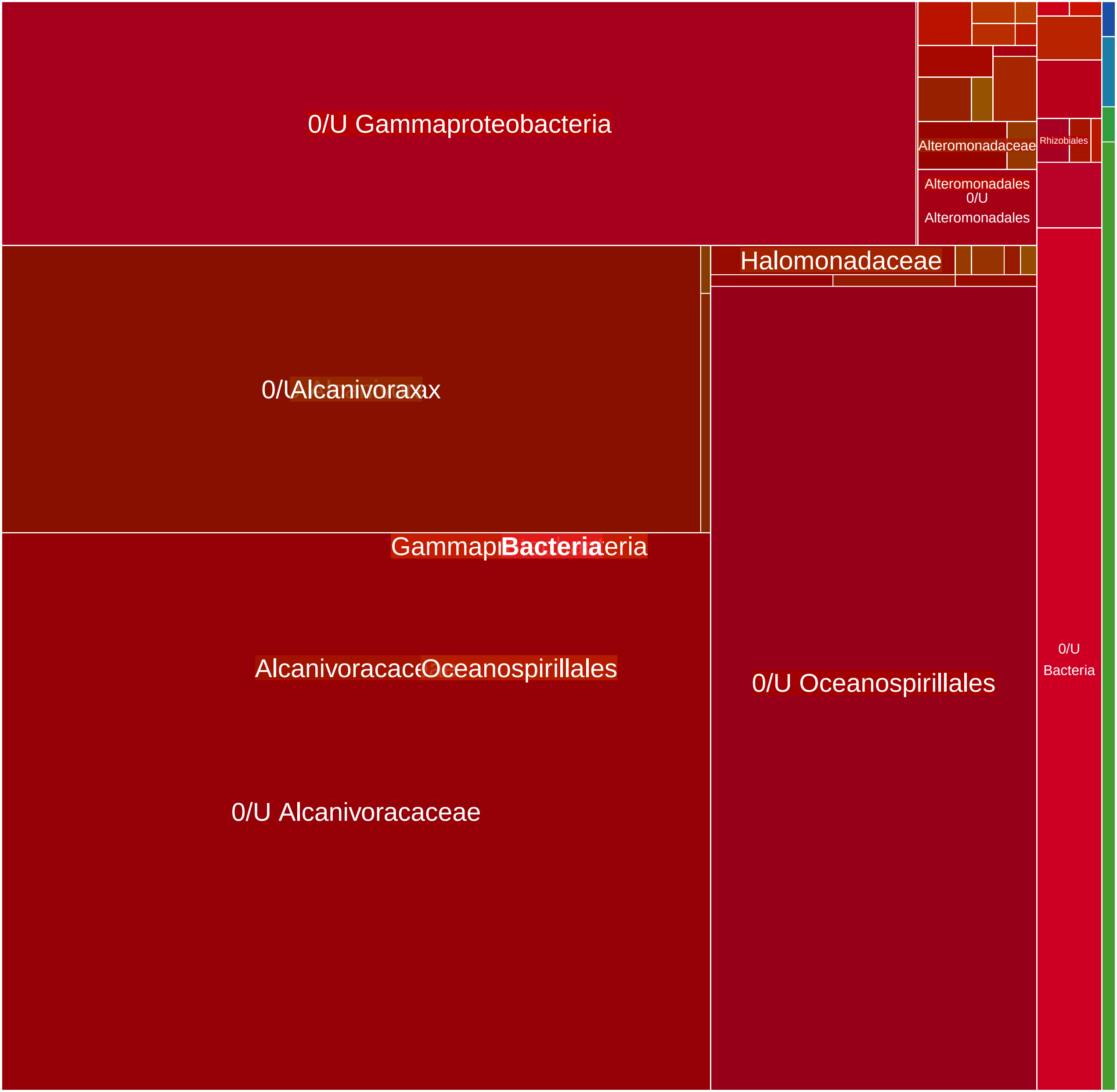

MGT−v1_385 : 2655 unigenes, 2379 (89.60%) taxonomically assigned

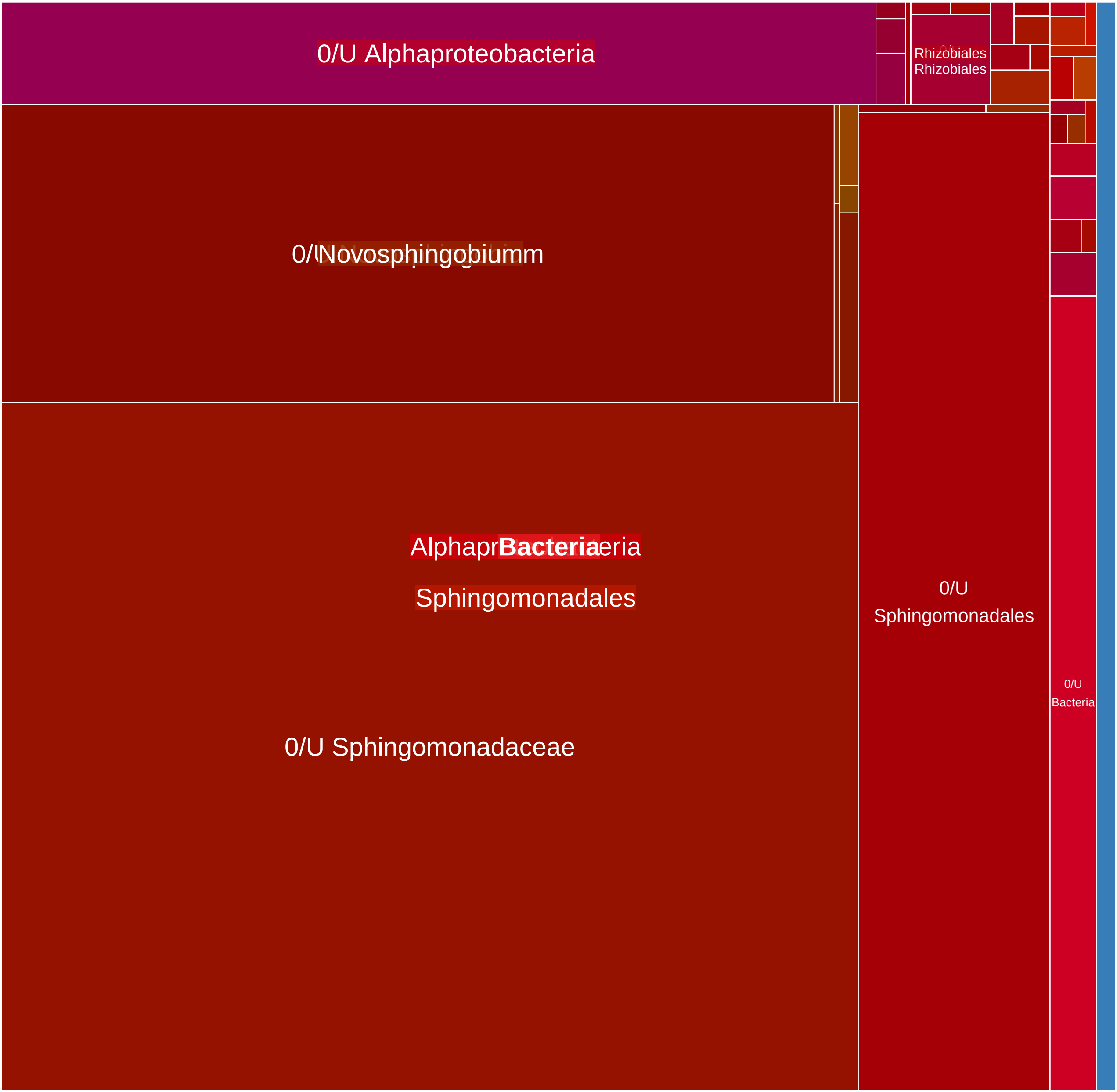

### MGT−v1_386 : 2652 unigenes, 113 (4.26%) taxonomically assigned

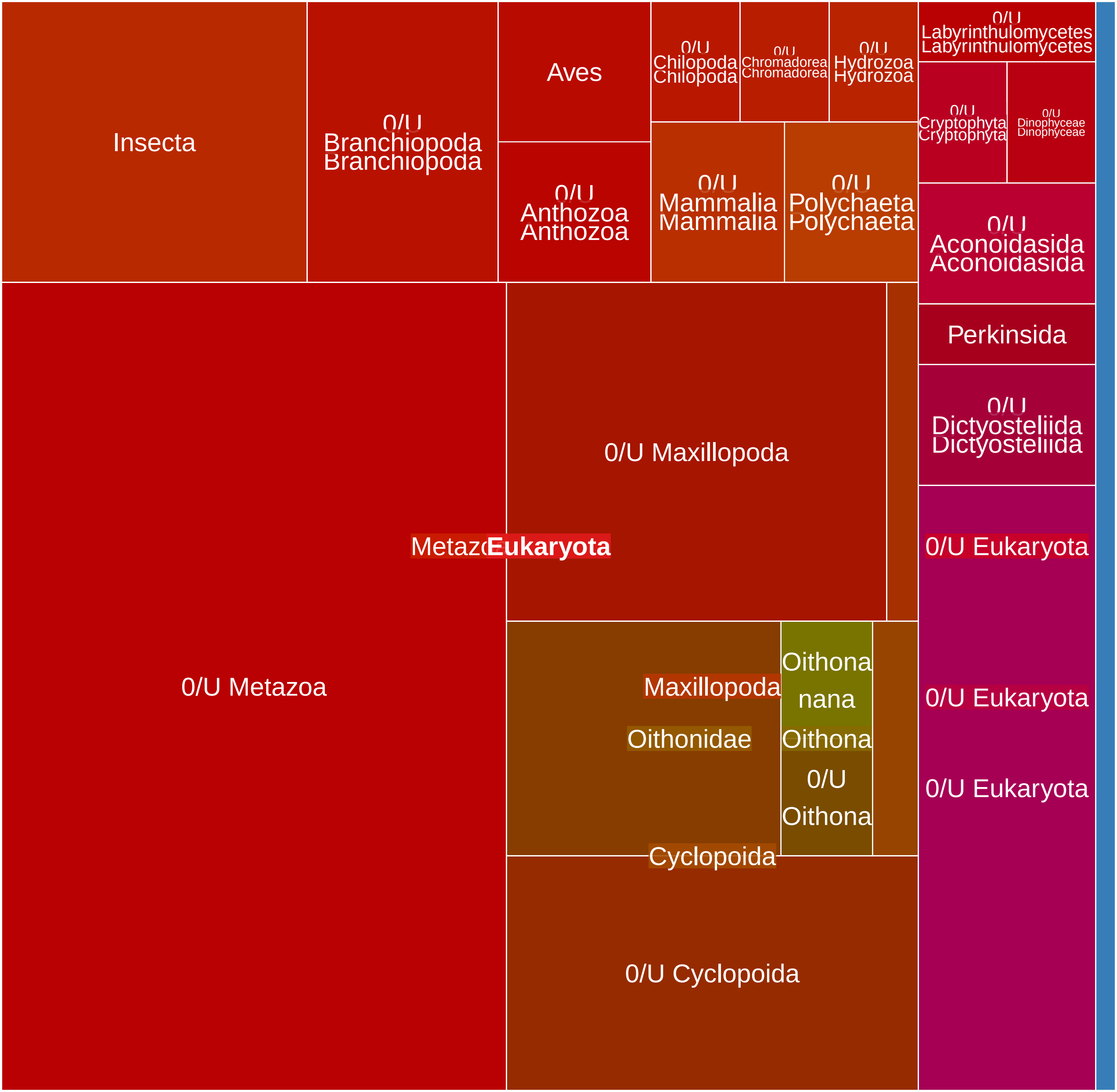

MGT−v1_387 : 2643 unigenes, 2434 (92.09%) taxonomically assigned

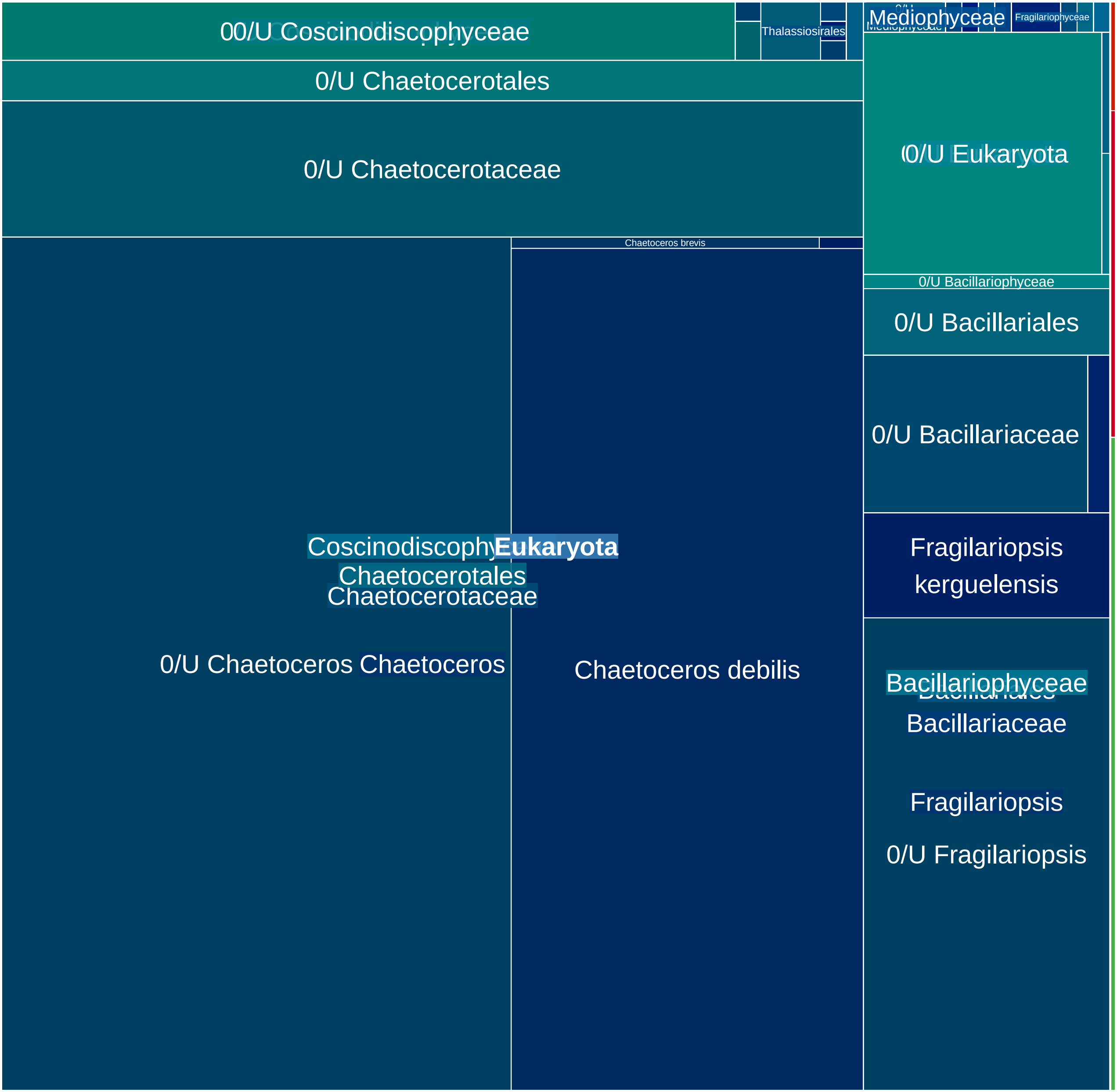

MGT−v1_388 : 2642 unigenes, 975 (36.90%) taxonomically assigned

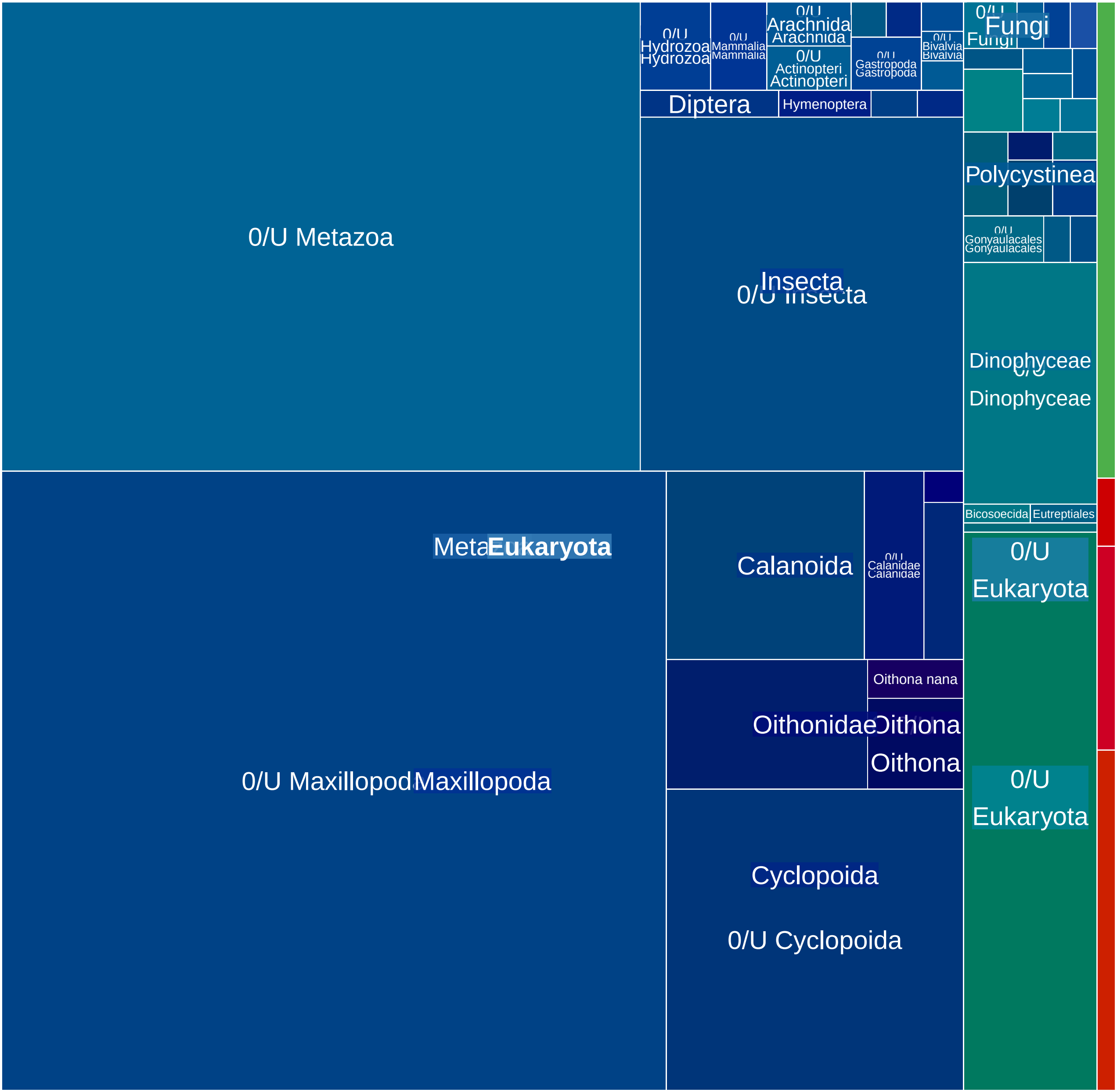

MGT−v1_389 : 2632 unigenes, 352 (13.37%) taxonomically assigned

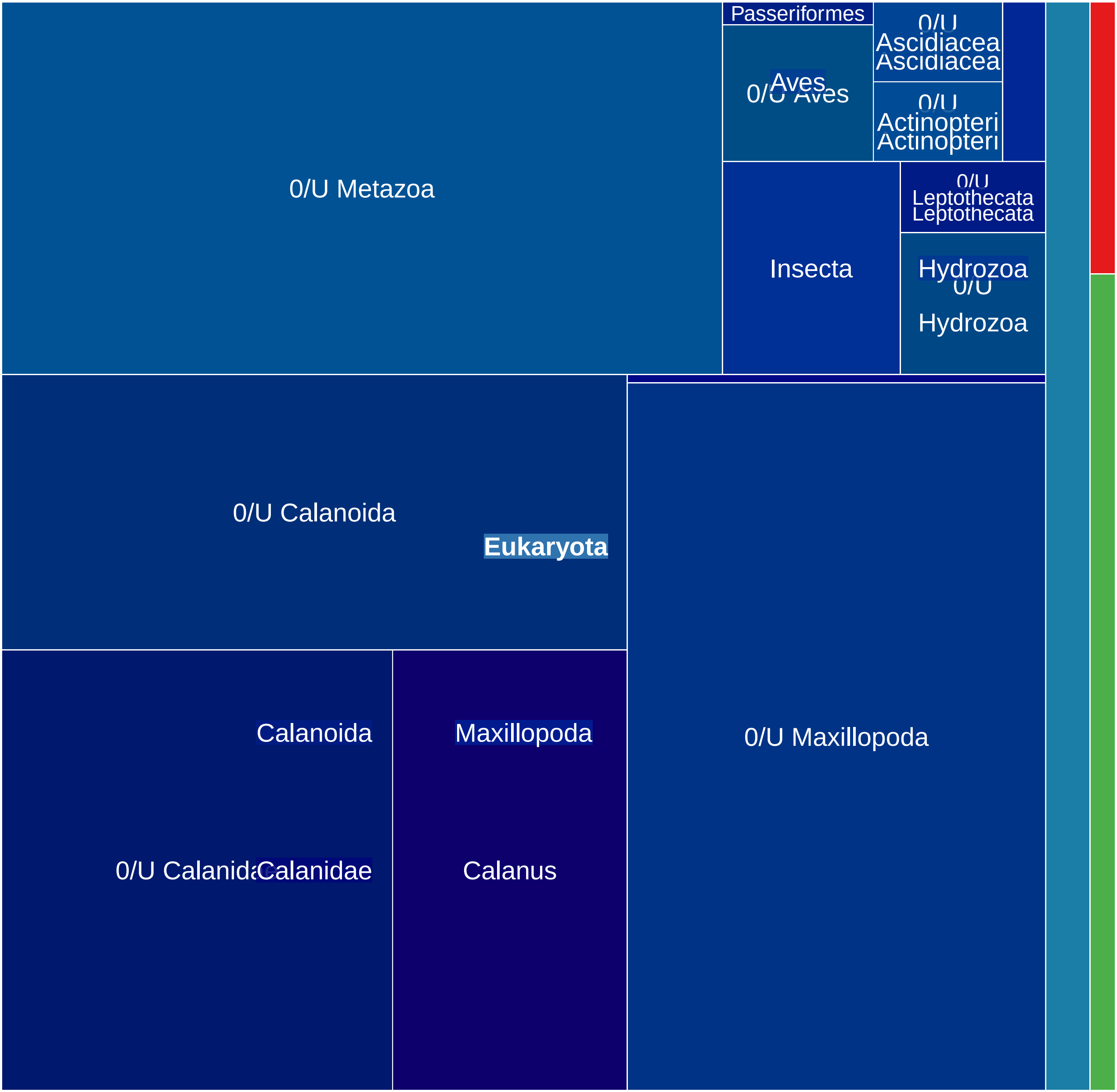

MGT−v1_390 : 2622 unigenes, 1875 (71.51%) taxonomically assigned

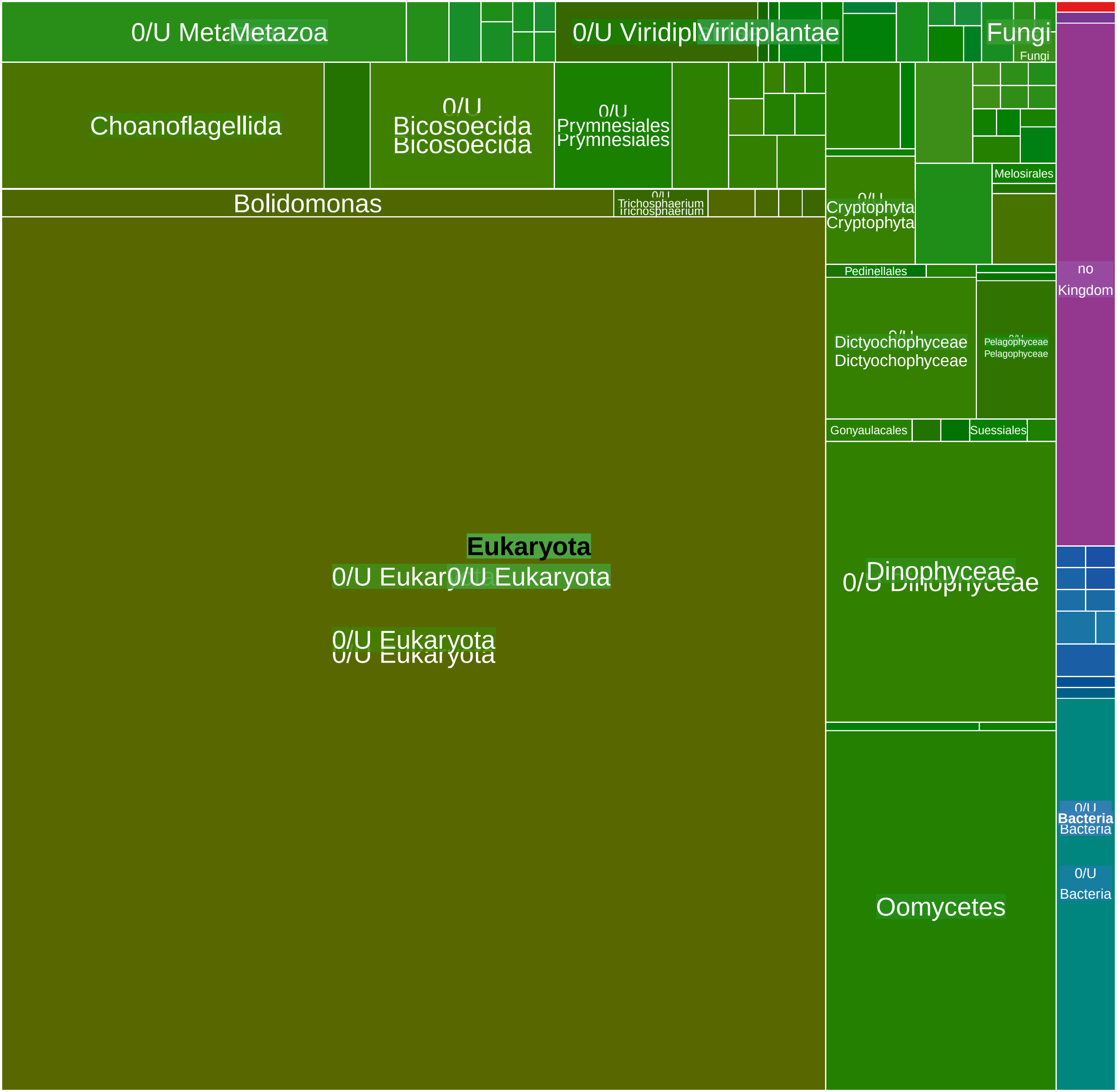

MGT−v1_391 : 2597 unigenes, 269 (10.36%) taxonomically assigned

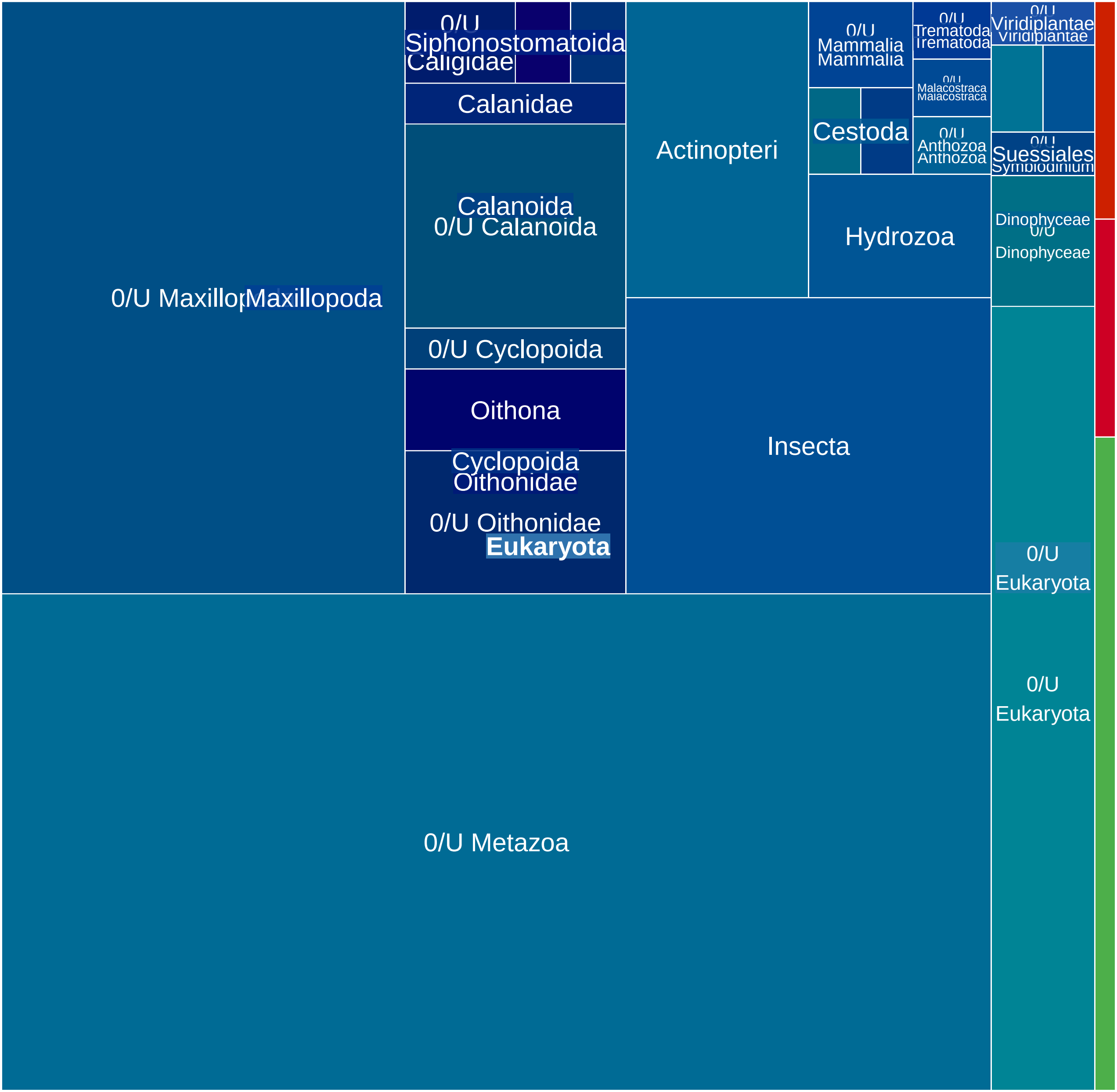

MGT−v1_392 : 2596 unigenes, 115 (4.43%) taxonomically assigned

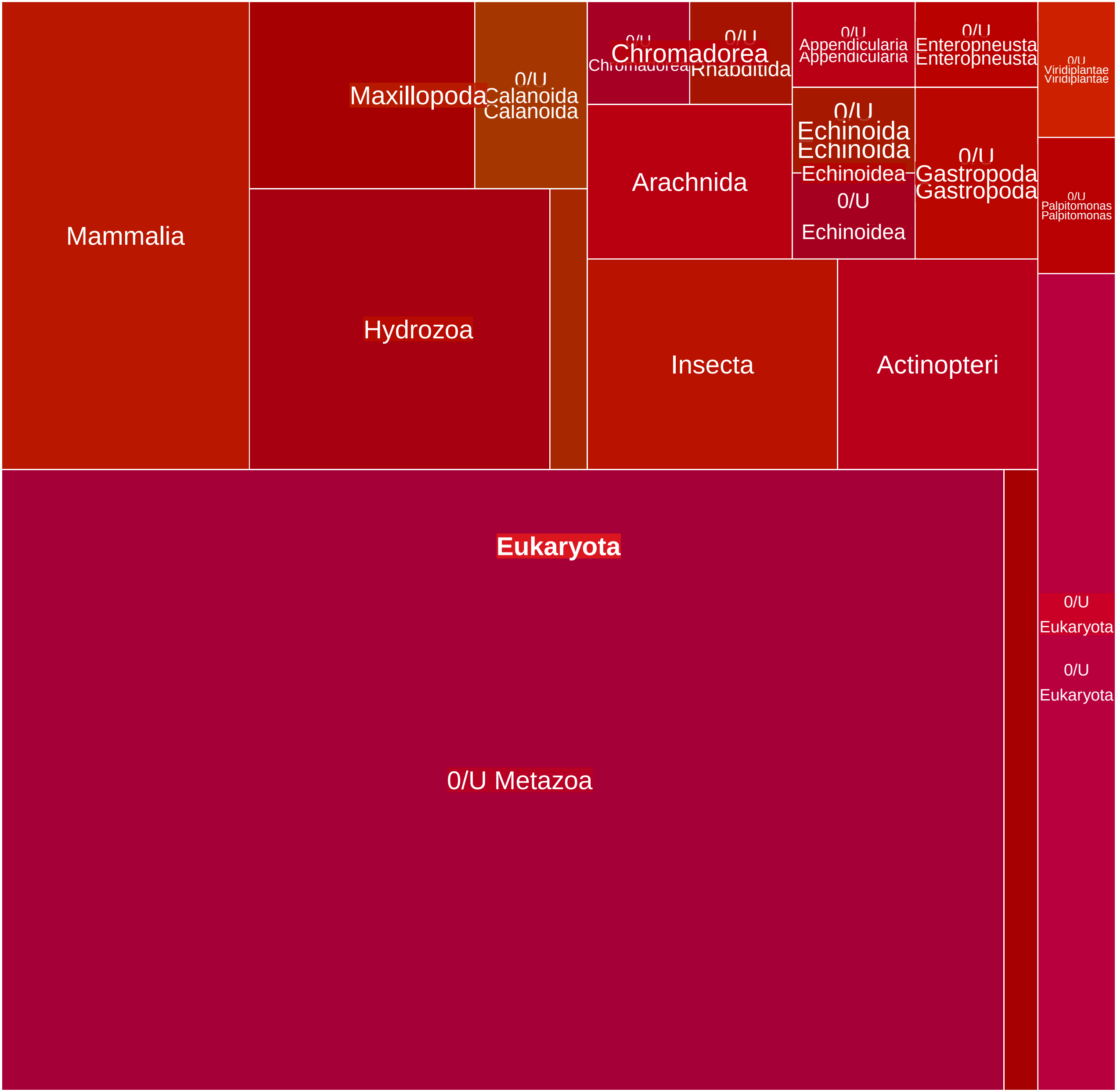

### MGT−v1_393 : 2594 unigenes, 1999 (77.06%) taxonomically assigned

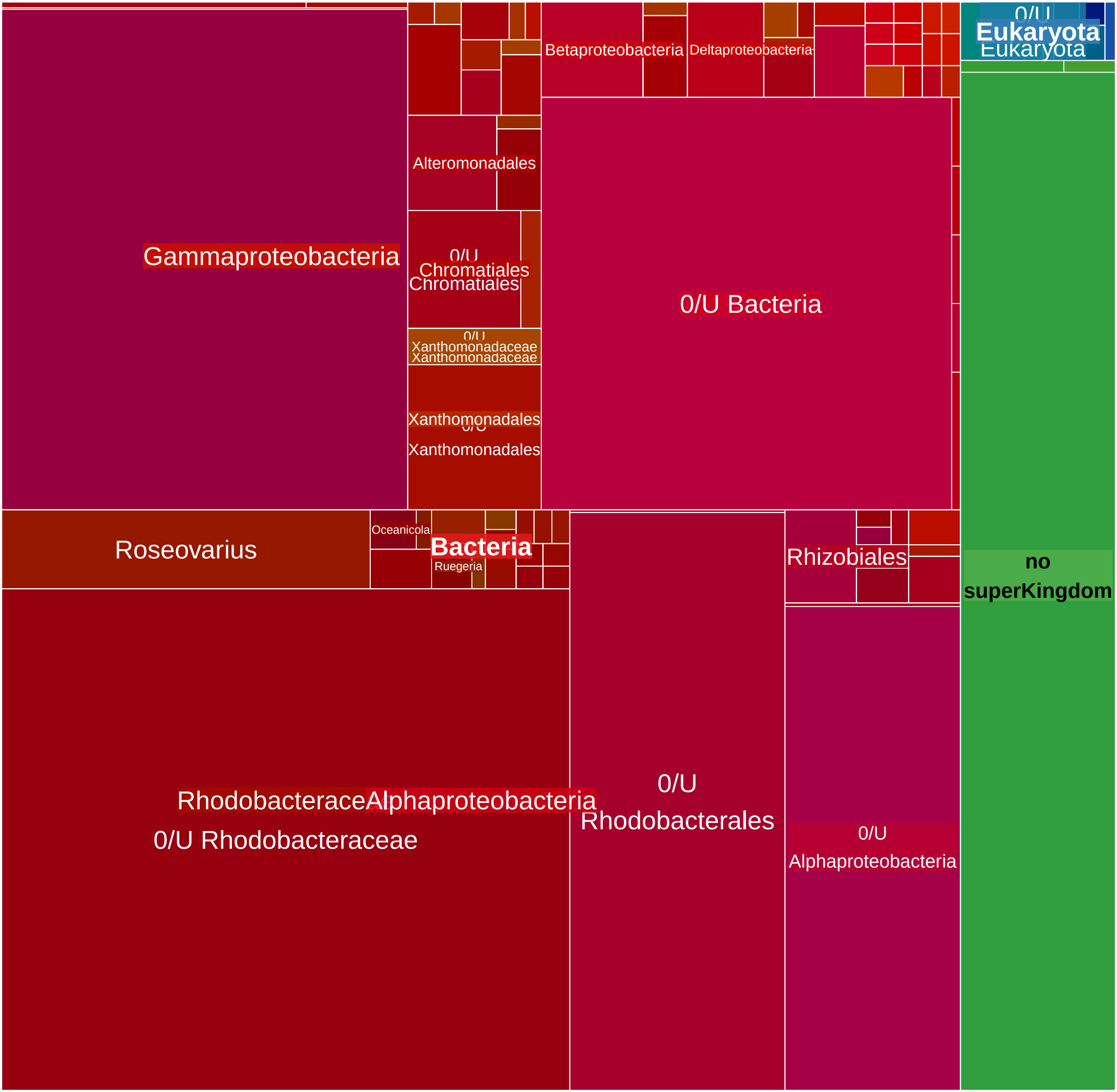

MGT−v1_394 : 2583 unigenes, 328 (12.70%) taxonomically assigned

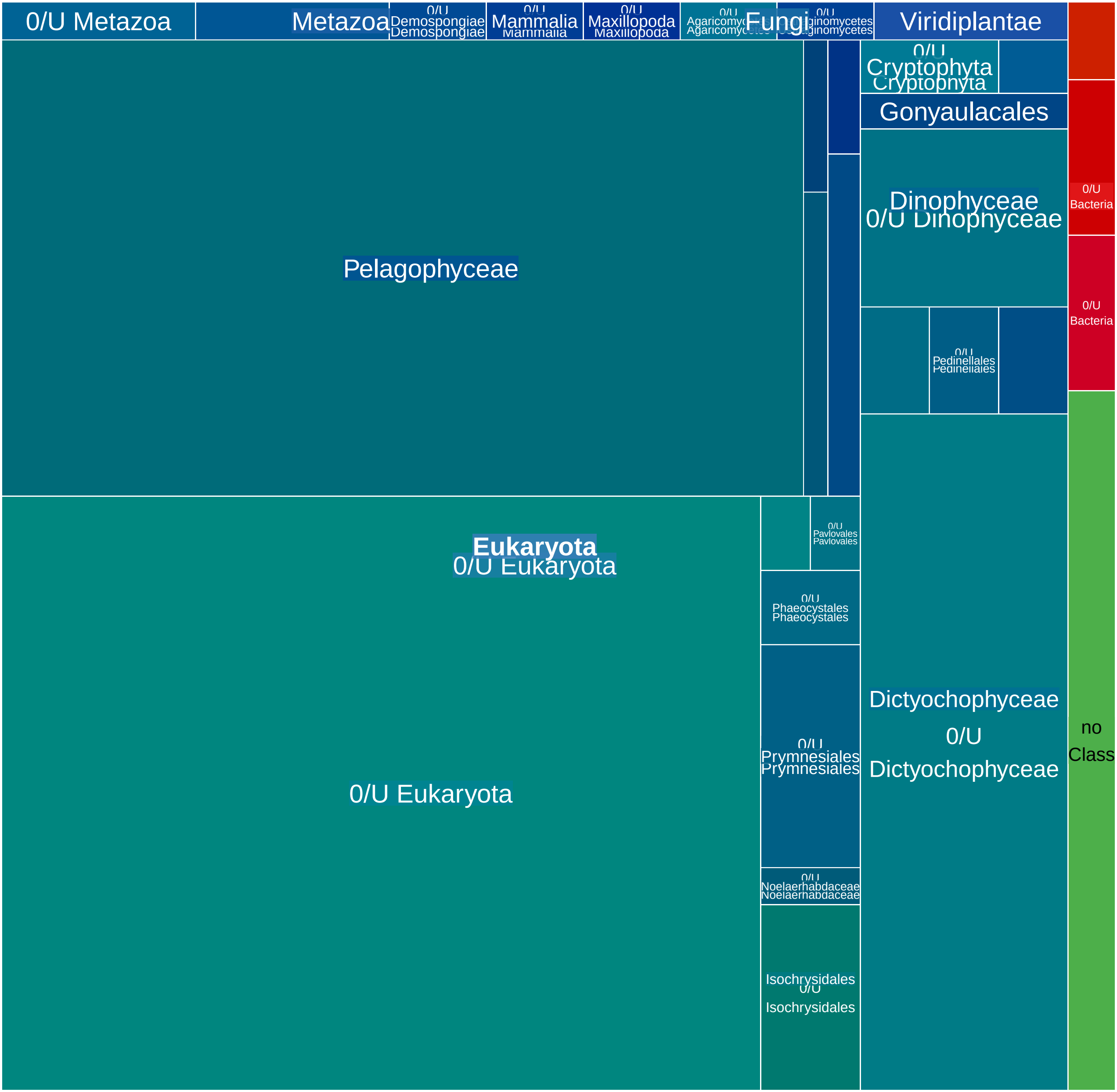

### MGT−v1_395 : 2582 unigenes, 2511 (97.25%) taxonomically assigned

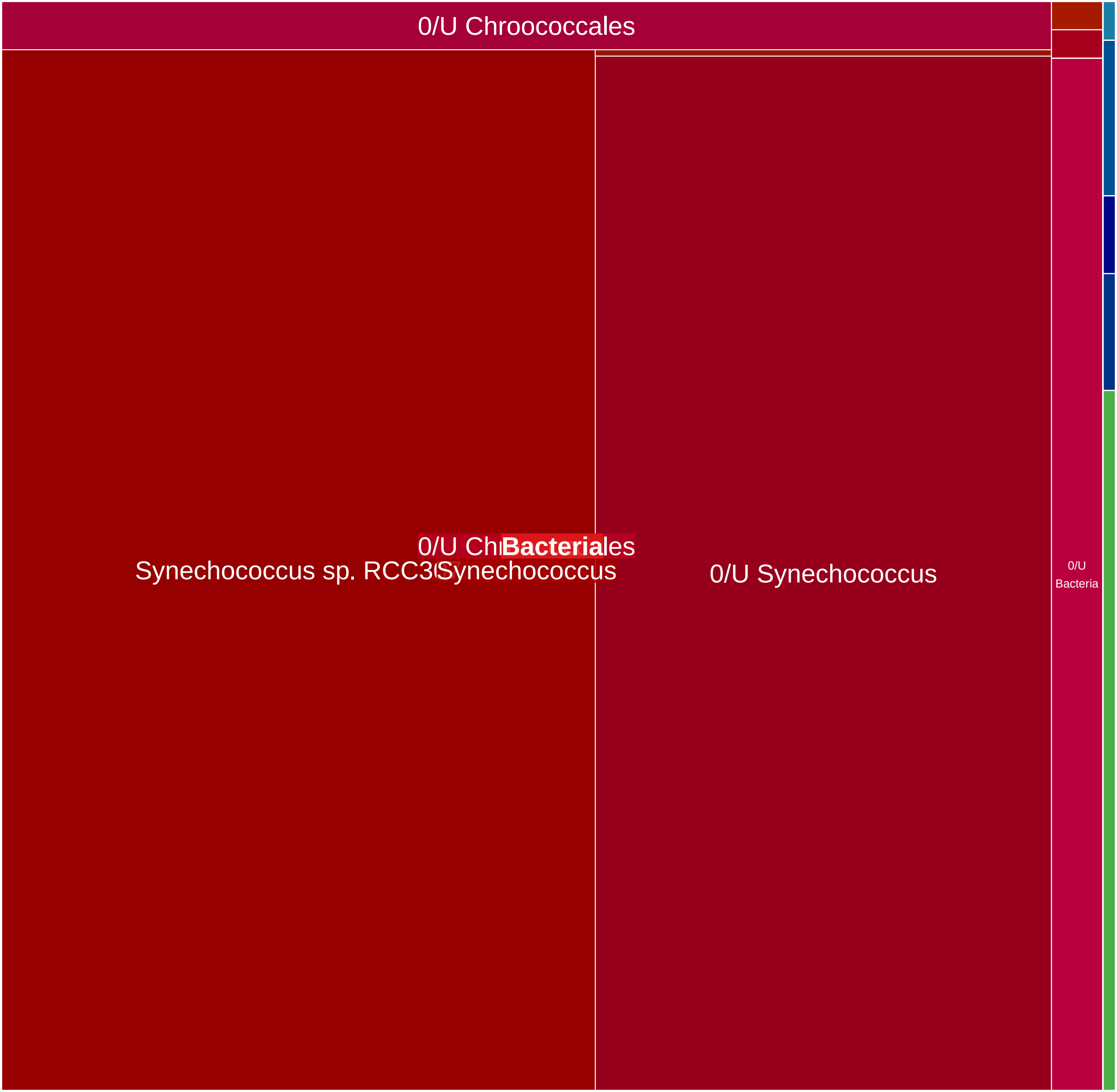

MGT−v1_396 : 2579 unigenes, 324 (12.56%) taxonomically assigned

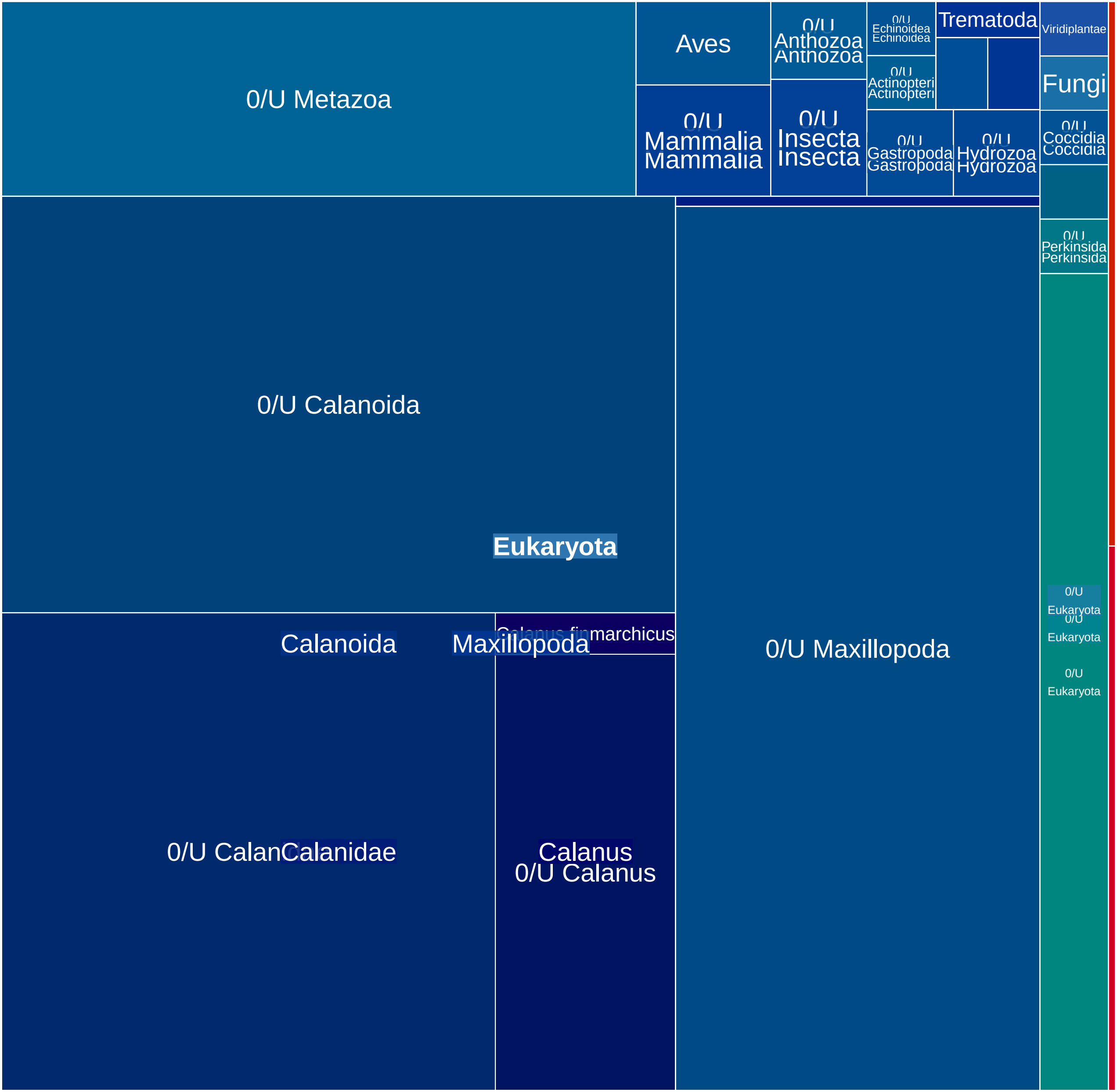

MGT−v1_397 : 2578 unigenes, 260 (10.09%) taxonomically assigned

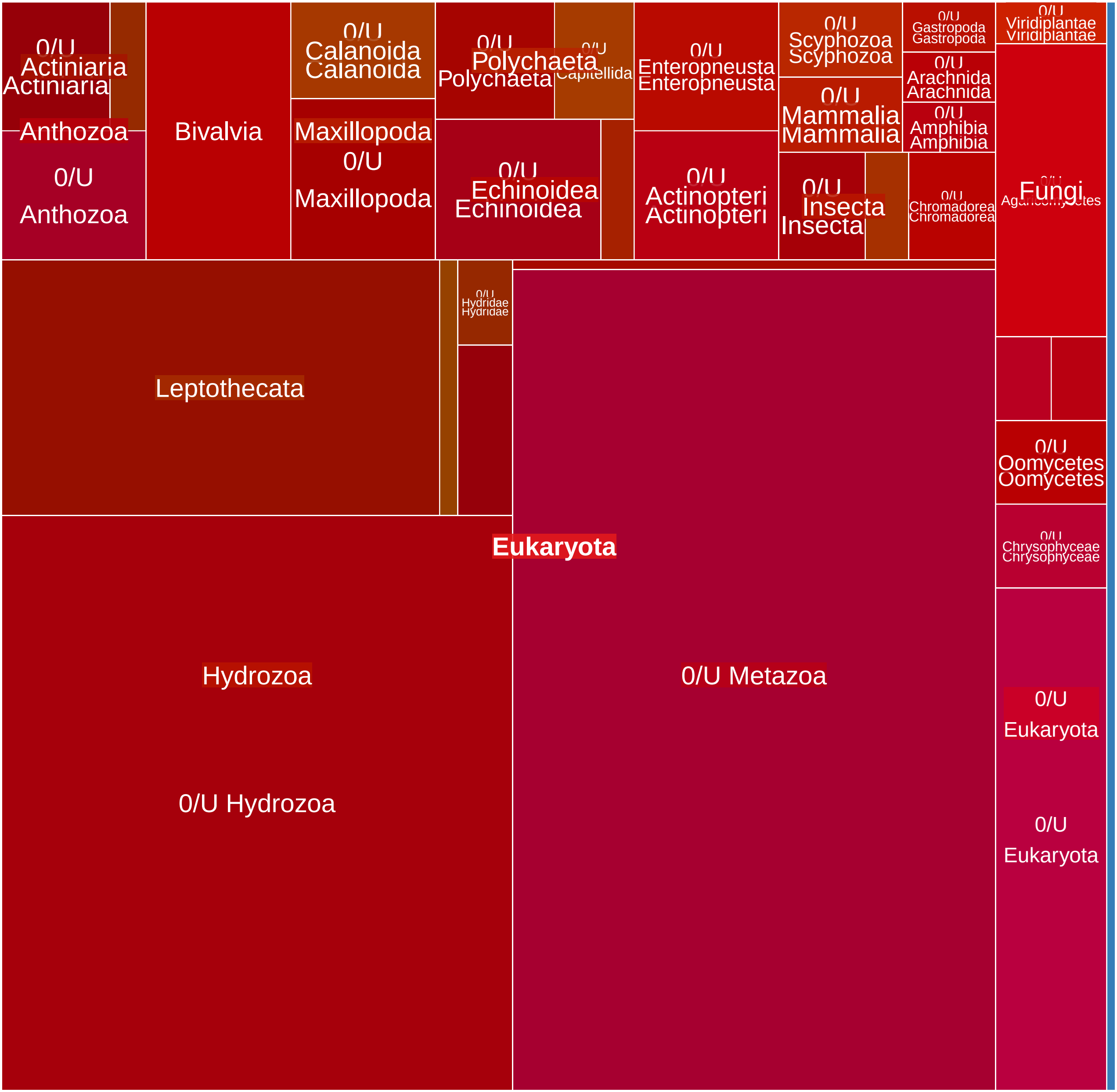

MGT−v1_398 : 2566 unigenes, 1893 (73.77%) taxonomically assigned

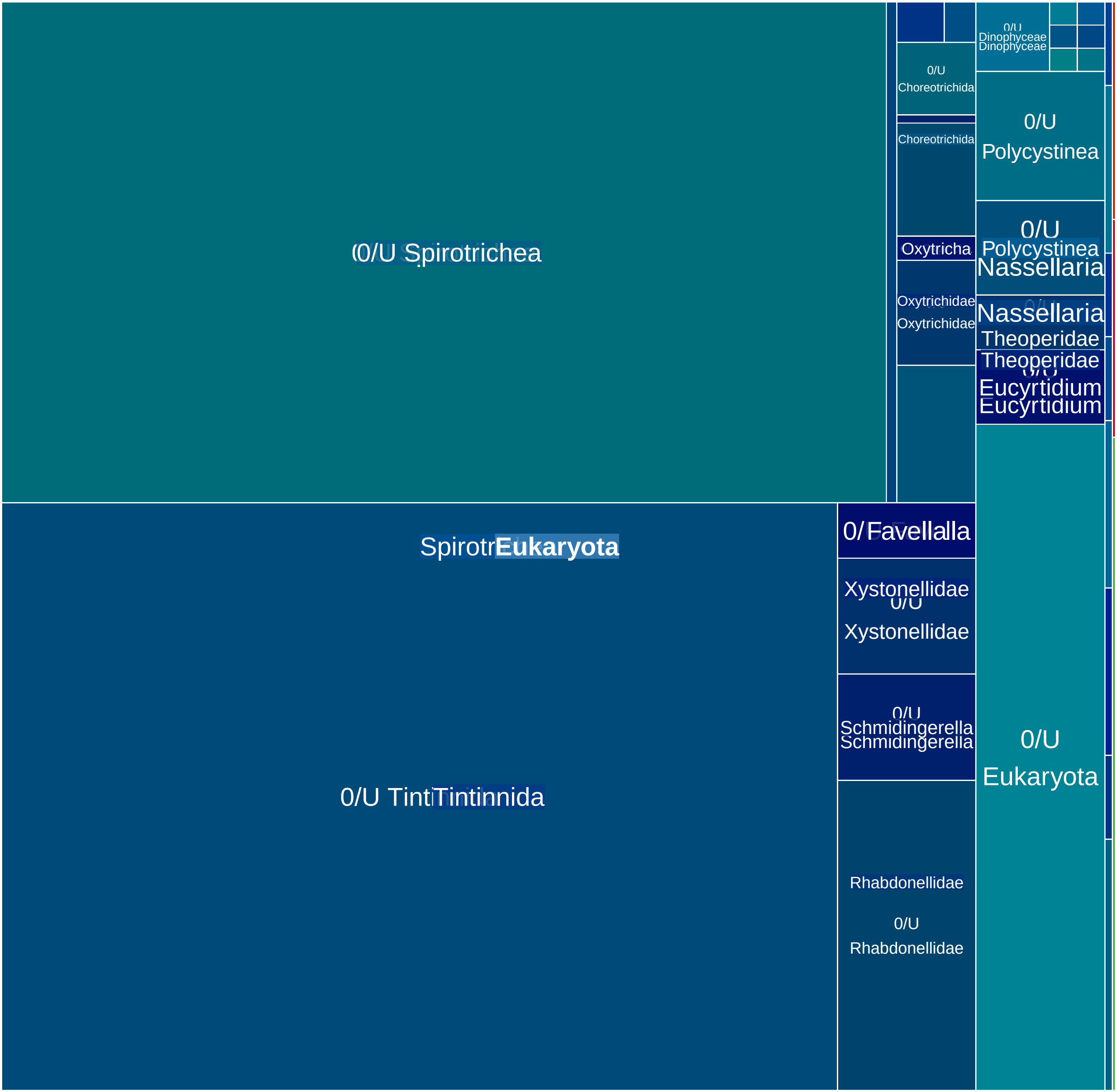

MGT−v1_399 : 2550 unigenes, 253 (9.92%) taxonomically assigned

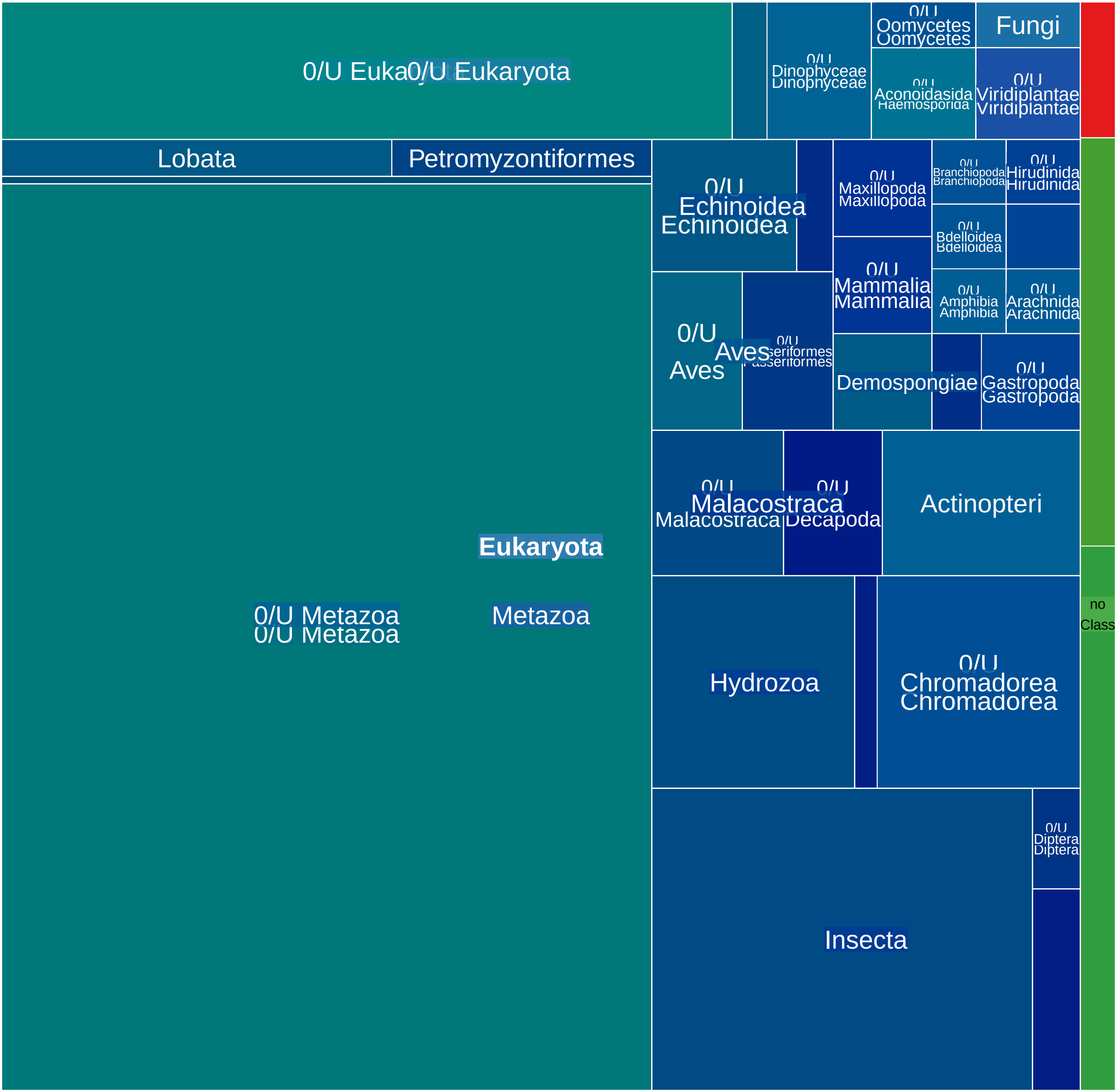

MGT−v1_400 : 2539 unigenes, 1466 (57.74%) taxonomically assigned

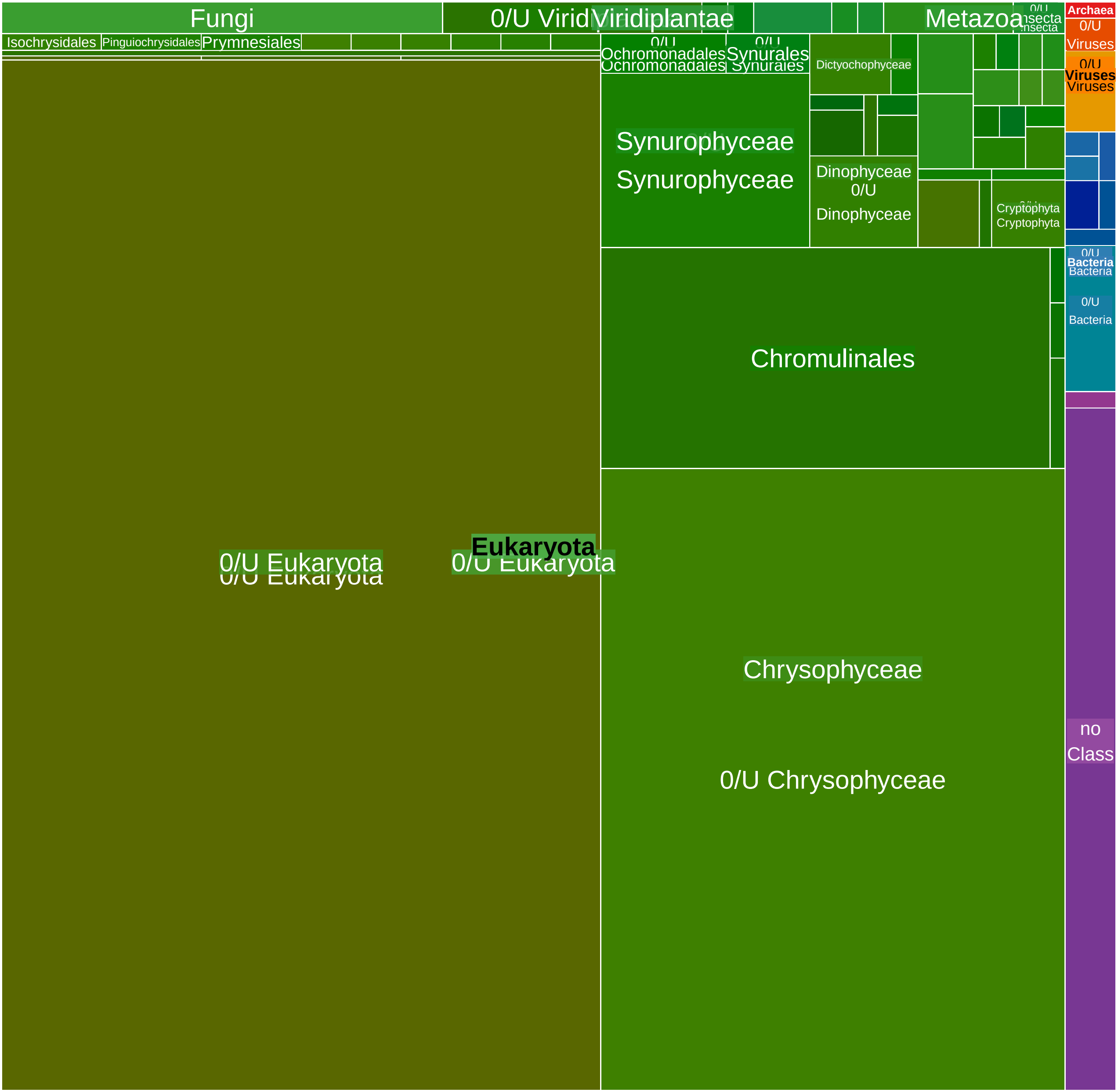

MGT−v1_401 : 2536 unigenes, 198 (7.81%) taxonomically assigned

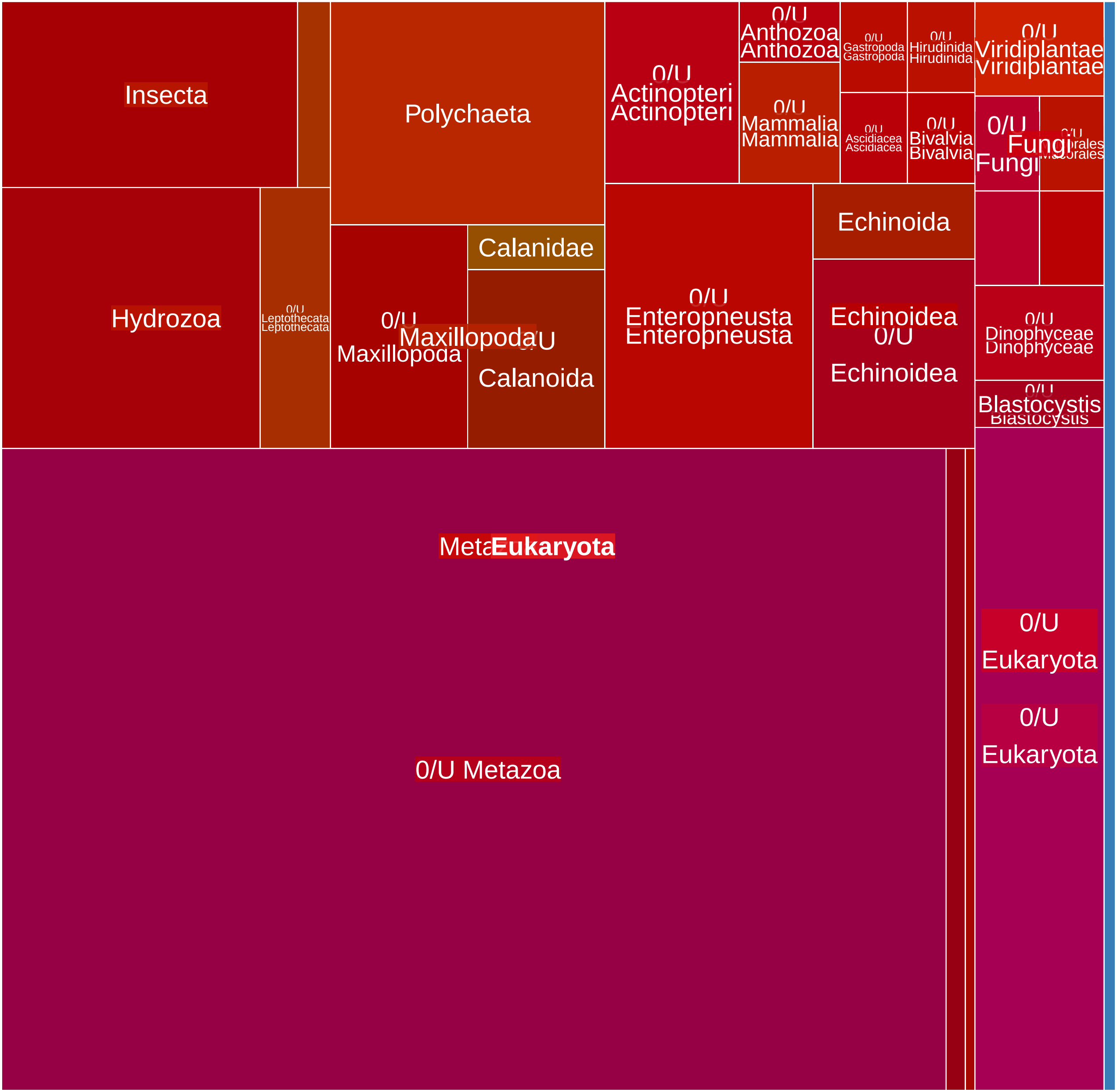

### MGT−v1_402 : 2512 unigenes, 204 (8.12%) taxonomically assigned

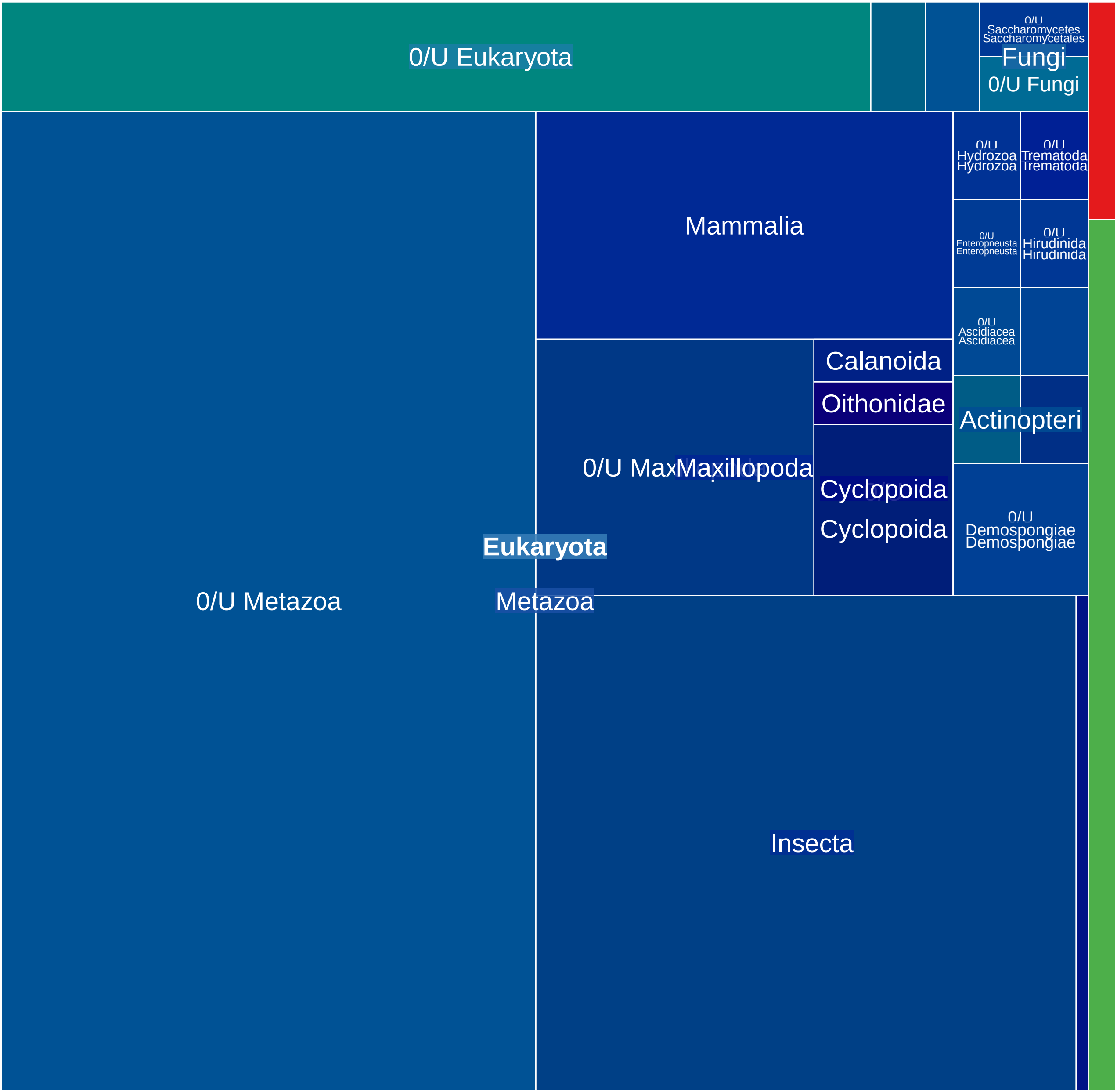

MGT−v1_403 : 2498 unigenes, 1514 (60.61%) taxonomically assigned

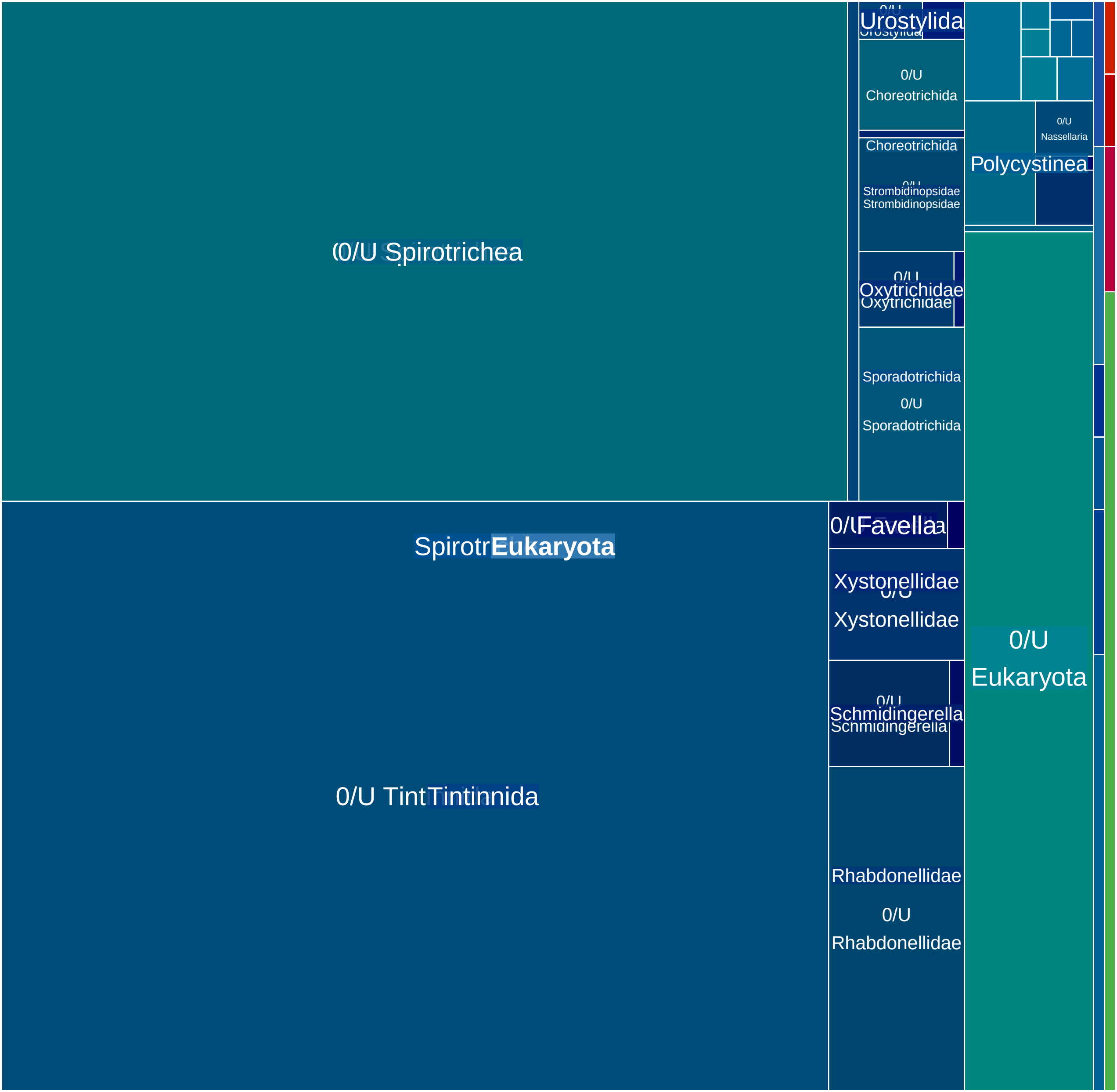

MGT−v1_404 : 2488 unigenes, 1327 (53.34%) taxonomically assigned

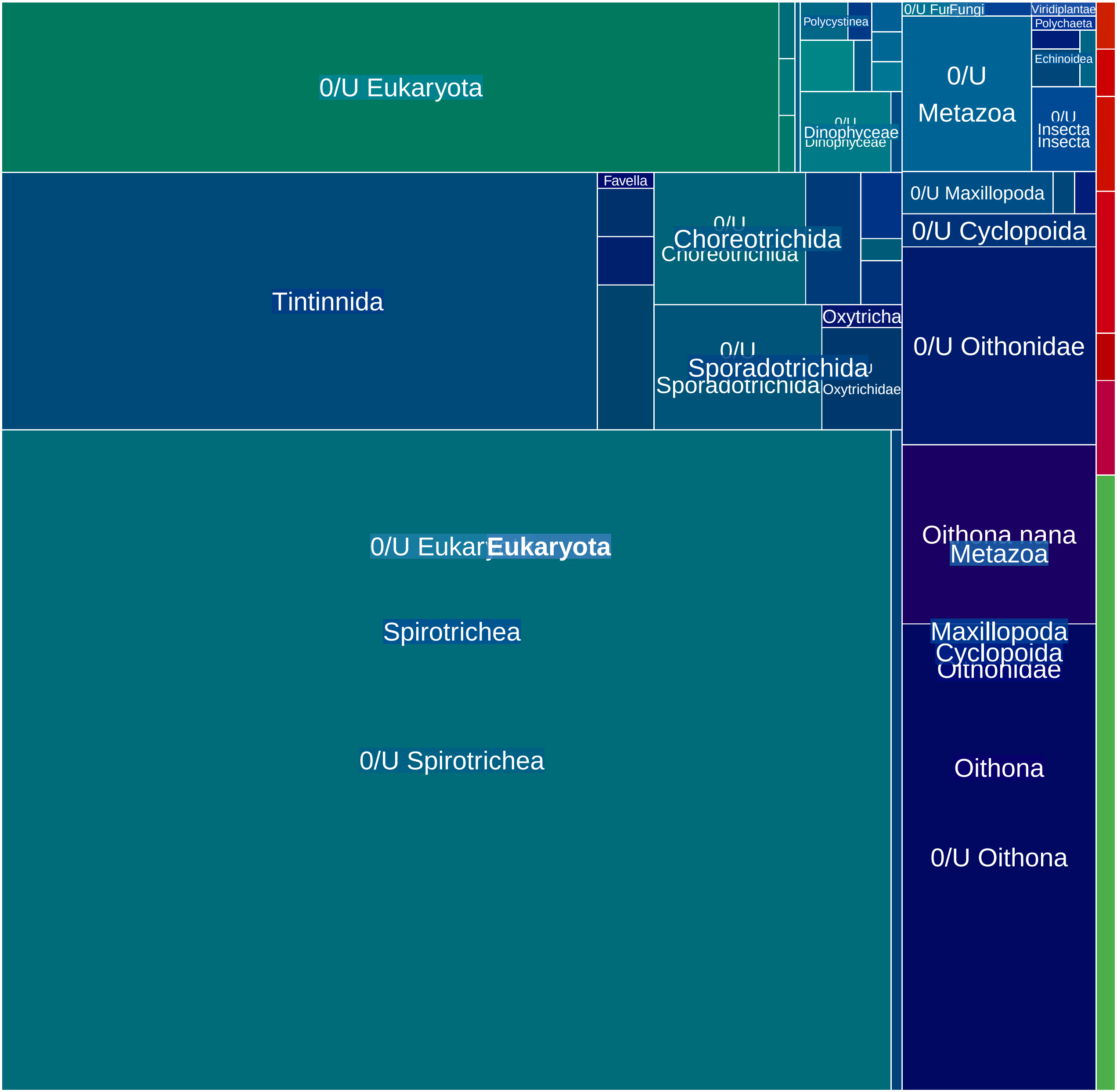

MGT−v1_405 : 2483 unigenes, 371 (14.94%) taxonomically assigned

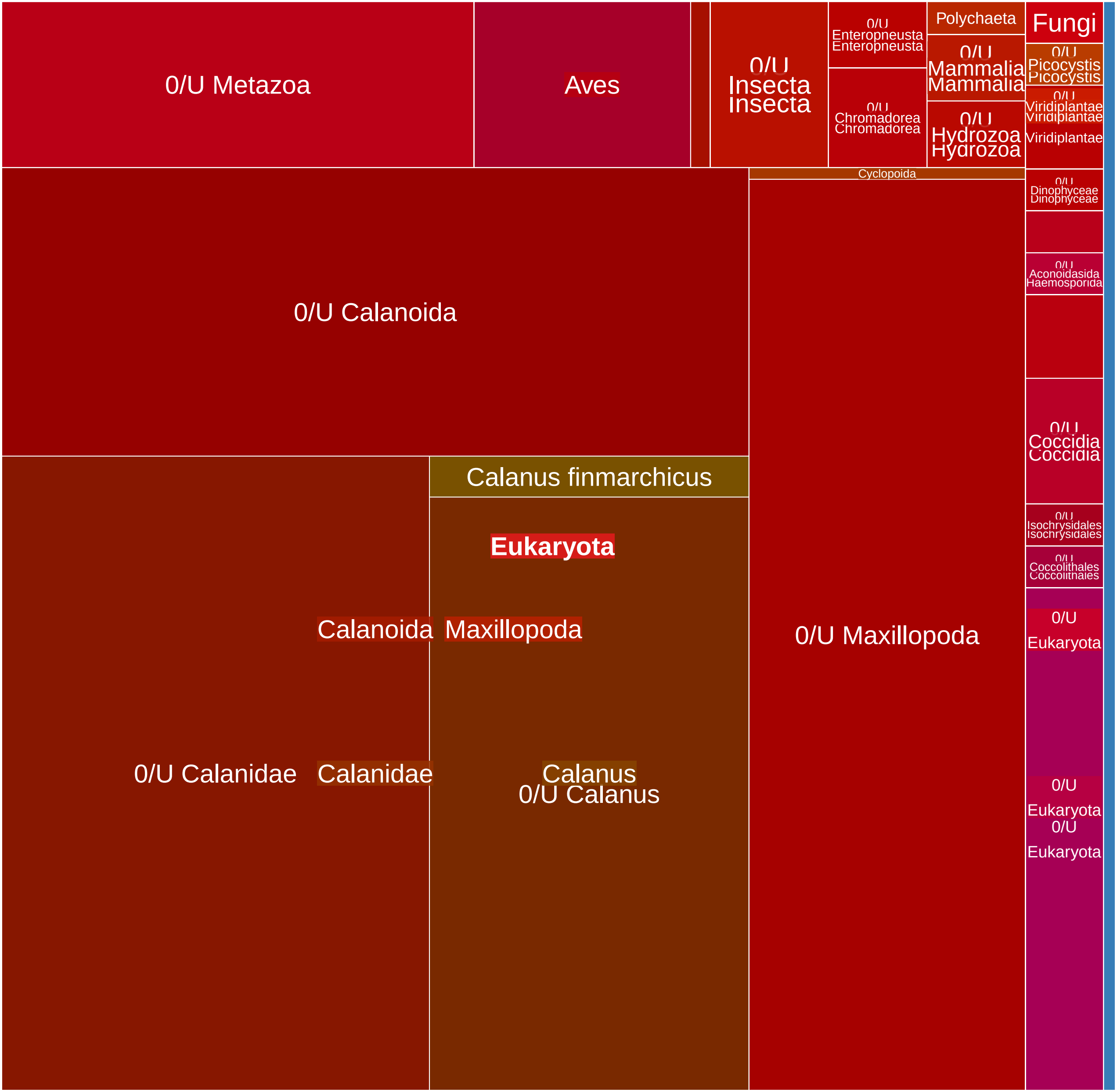

MGT−v1_406 : 2469 unigenes, 639 (25.88%) taxonomically assigned

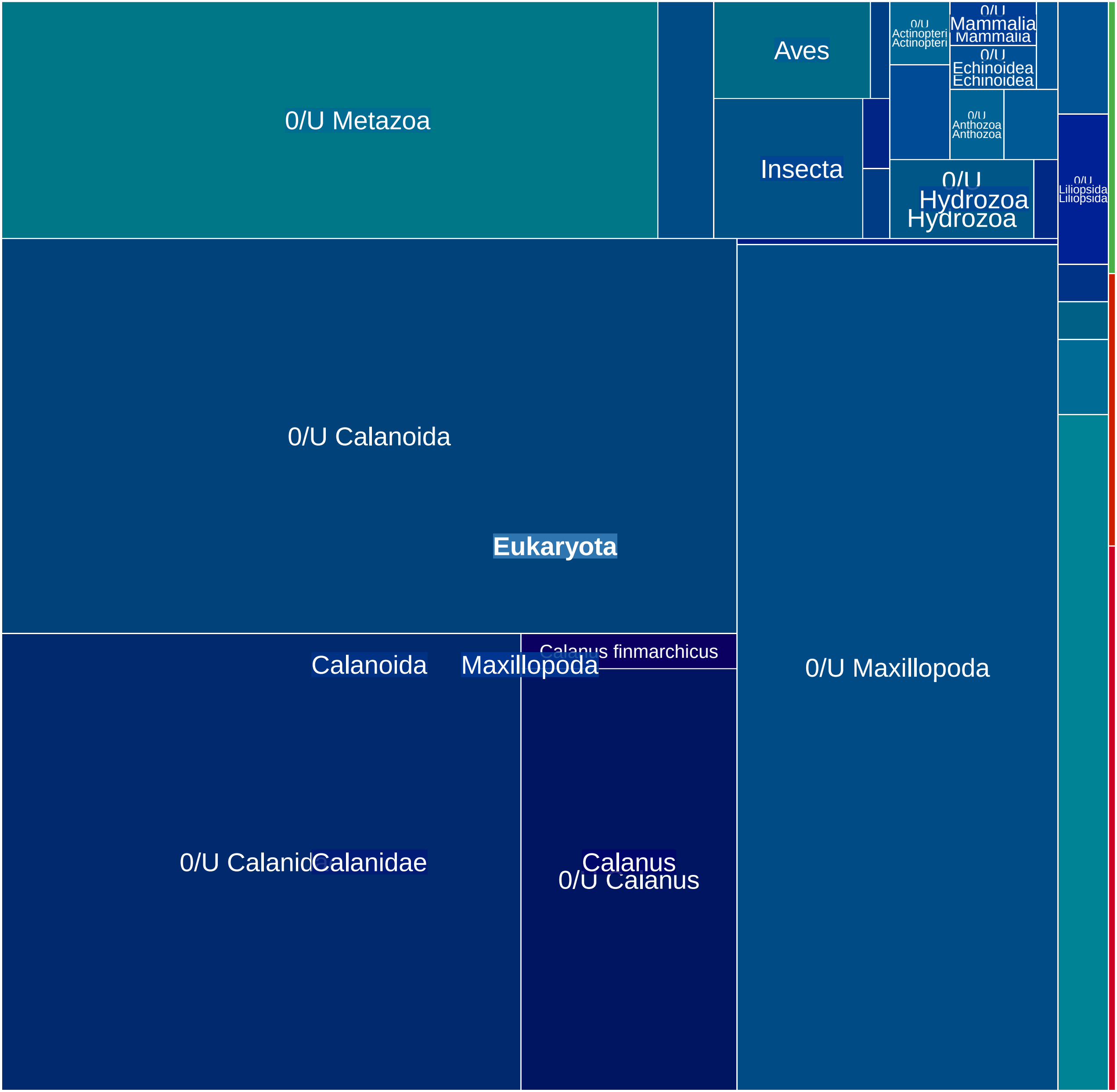

MGT−v1_407 : 2443 unigenes, 244 (9.99%) taxonomically assigned

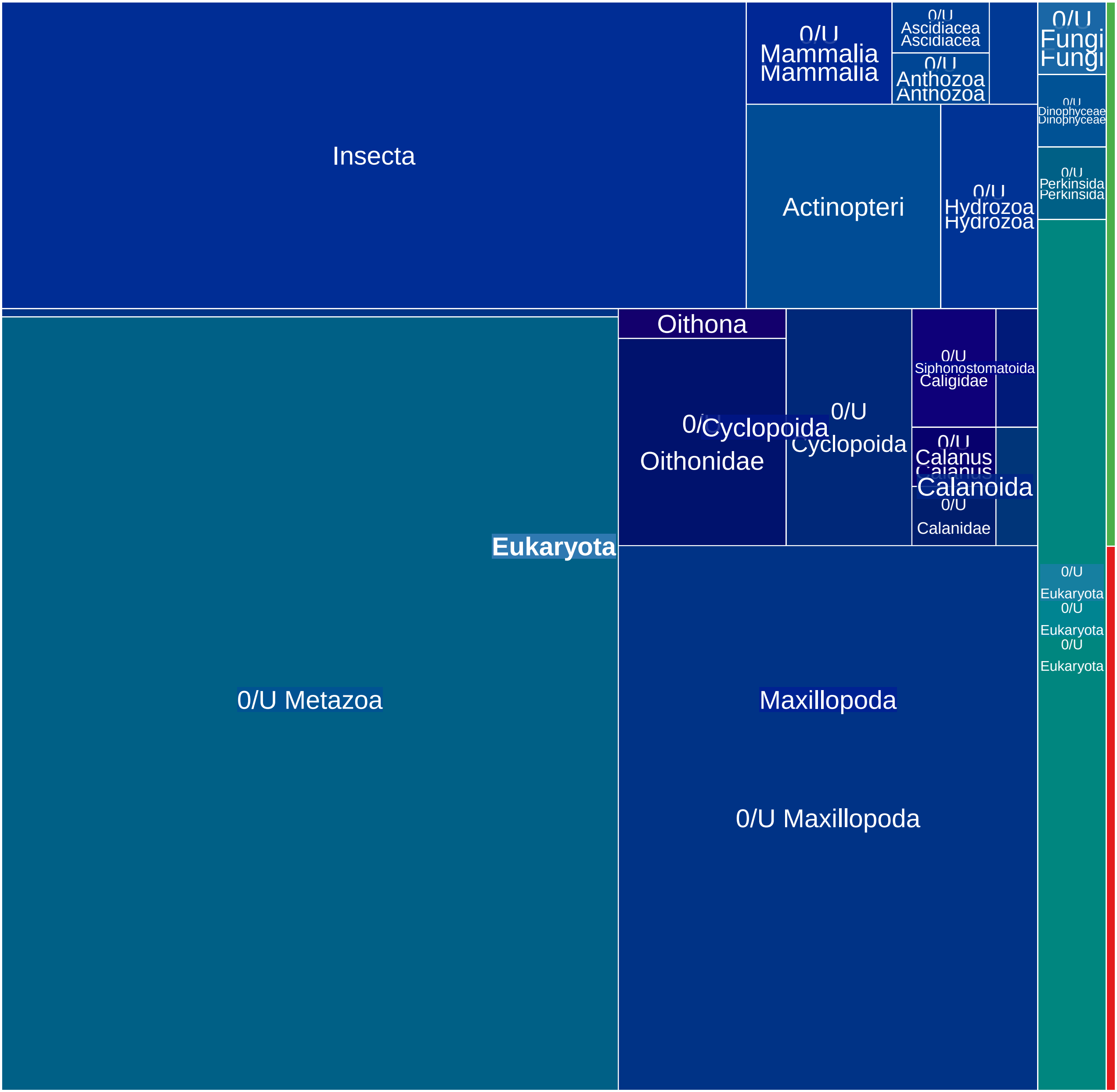

MGT−v1_408 : 2431 unigenes, 1571 (64.62%) taxonomically assigned

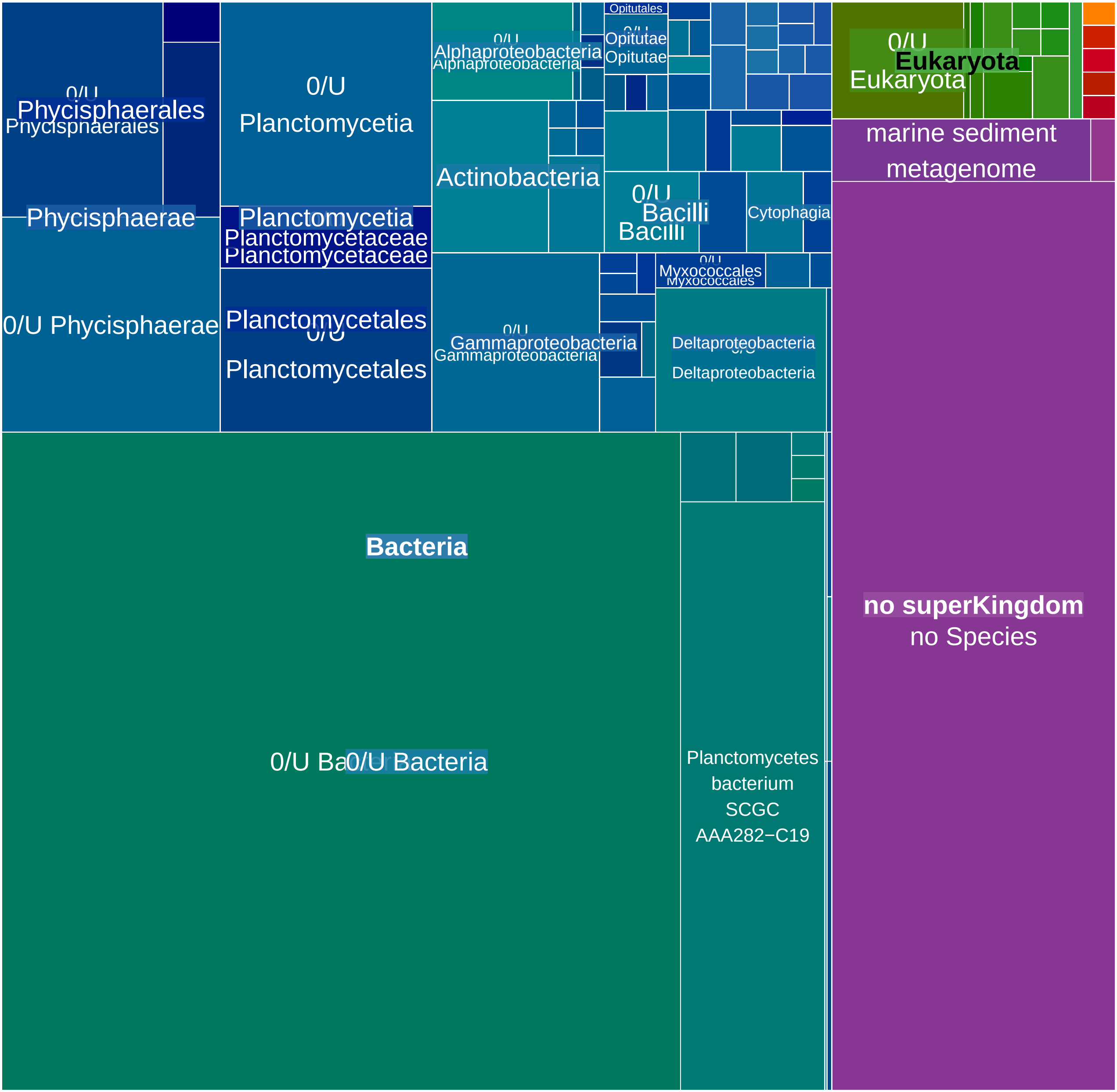

MGT−v1_409 : 2393 unigenes, 188 (7.86%) taxonomically assigned

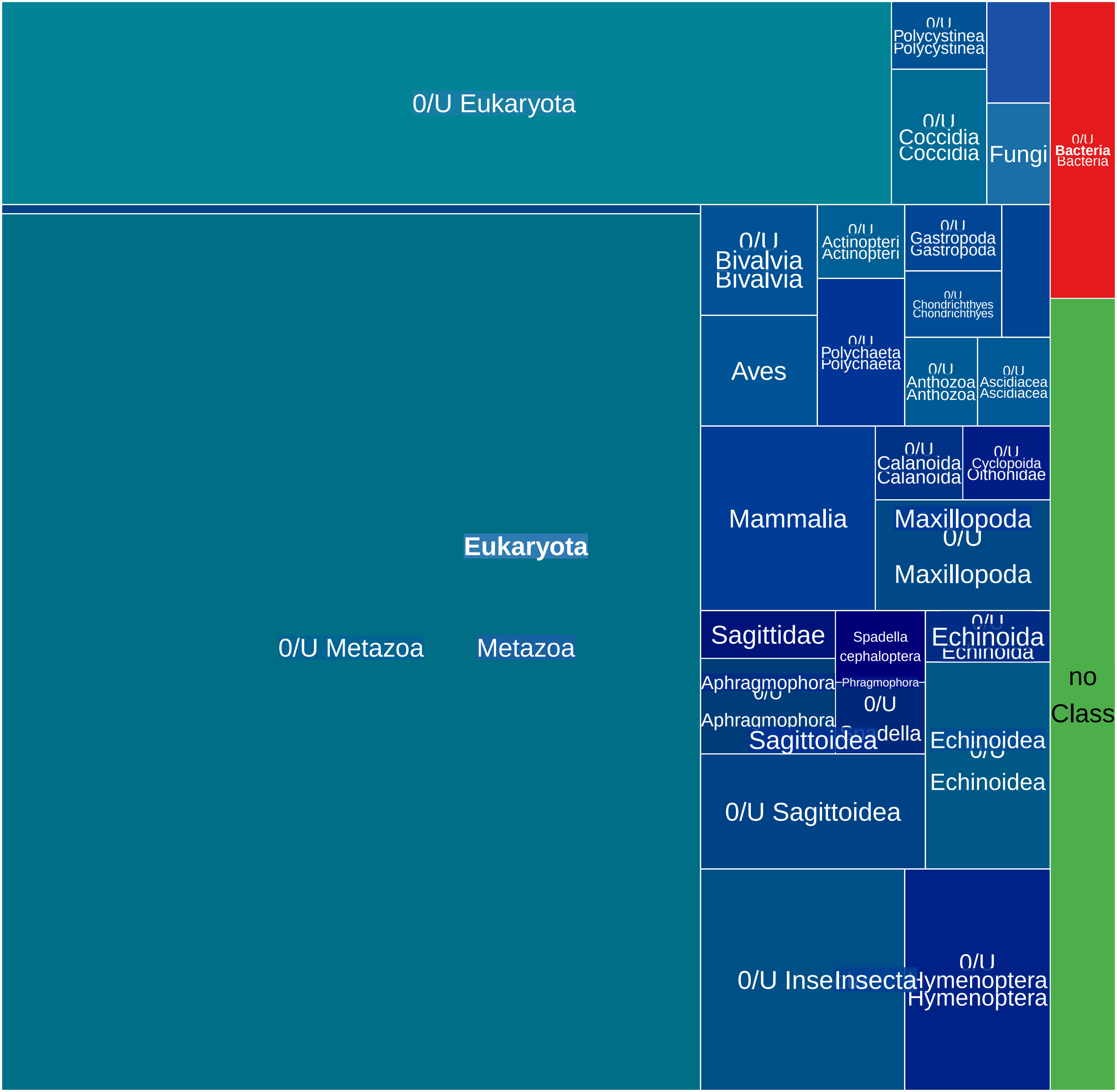

MGT−v1_410 : 2391 unigenes, 1638 (68.51%) taxonomically assigned

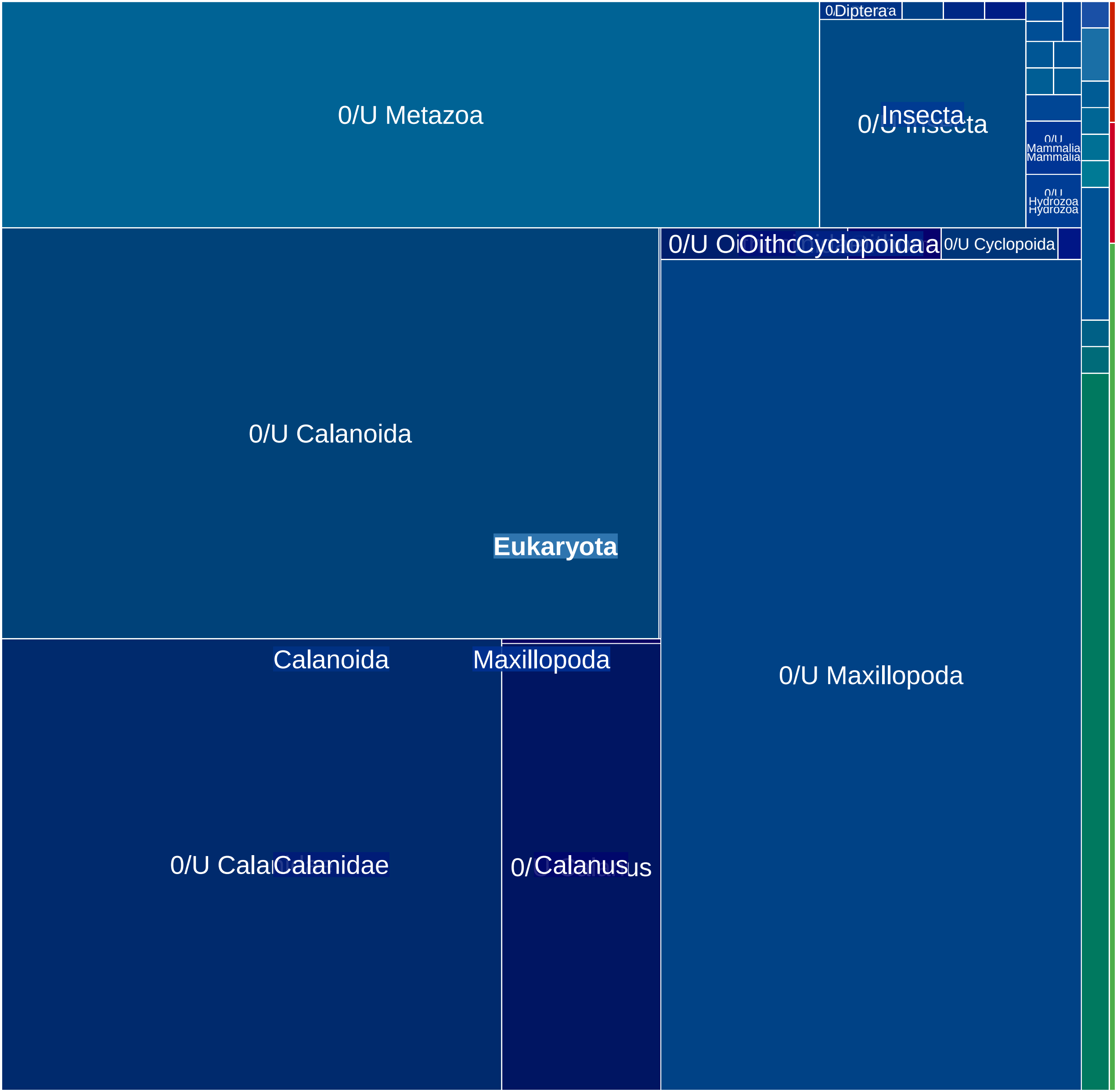

MGT−v1_411 : 2383 unigenes, 354 (14.86%) taxonomically assigned

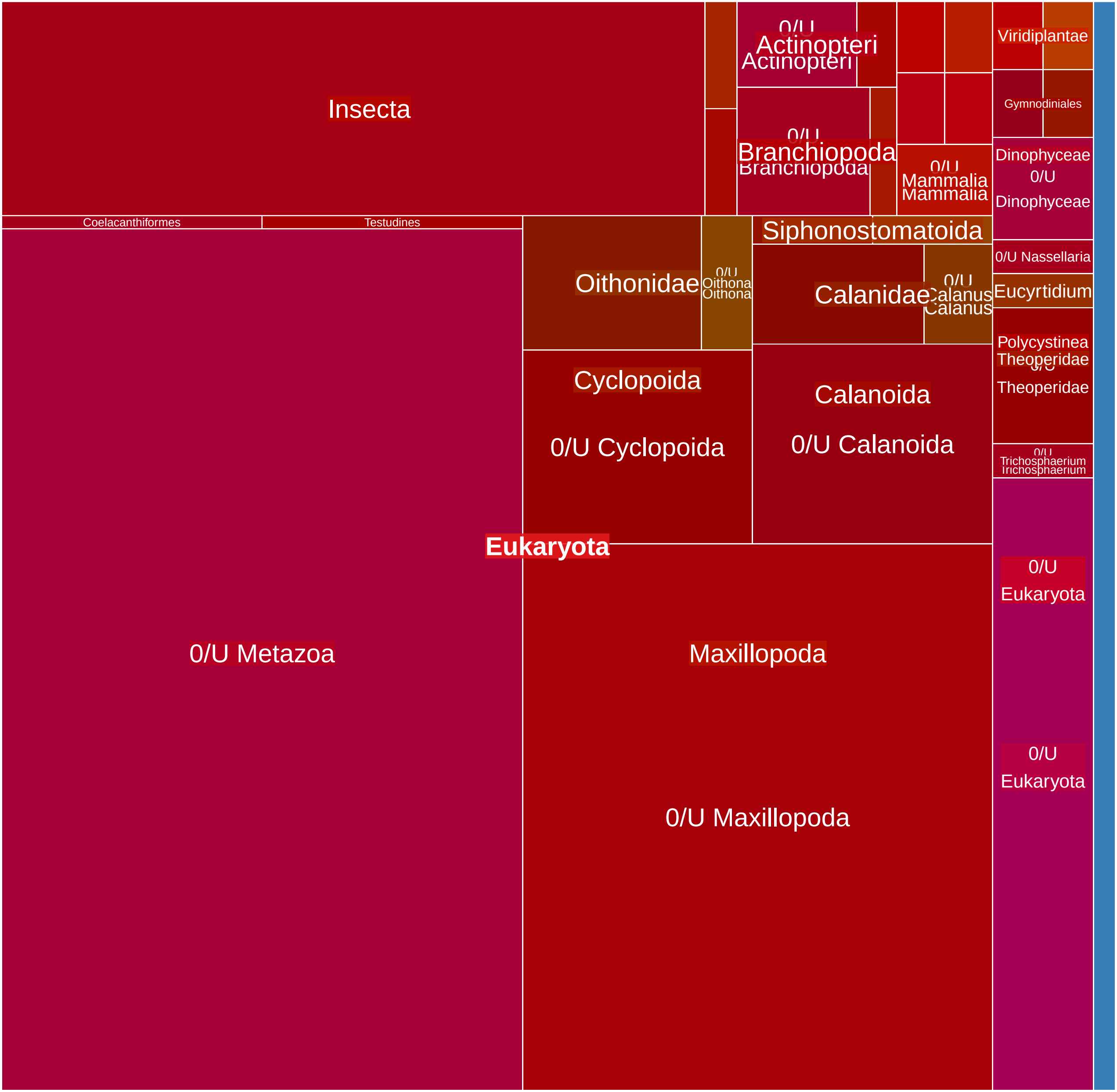

MGT−v1_412 : 2361 unigenes, 2287 (96.87%) taxonomically assigned

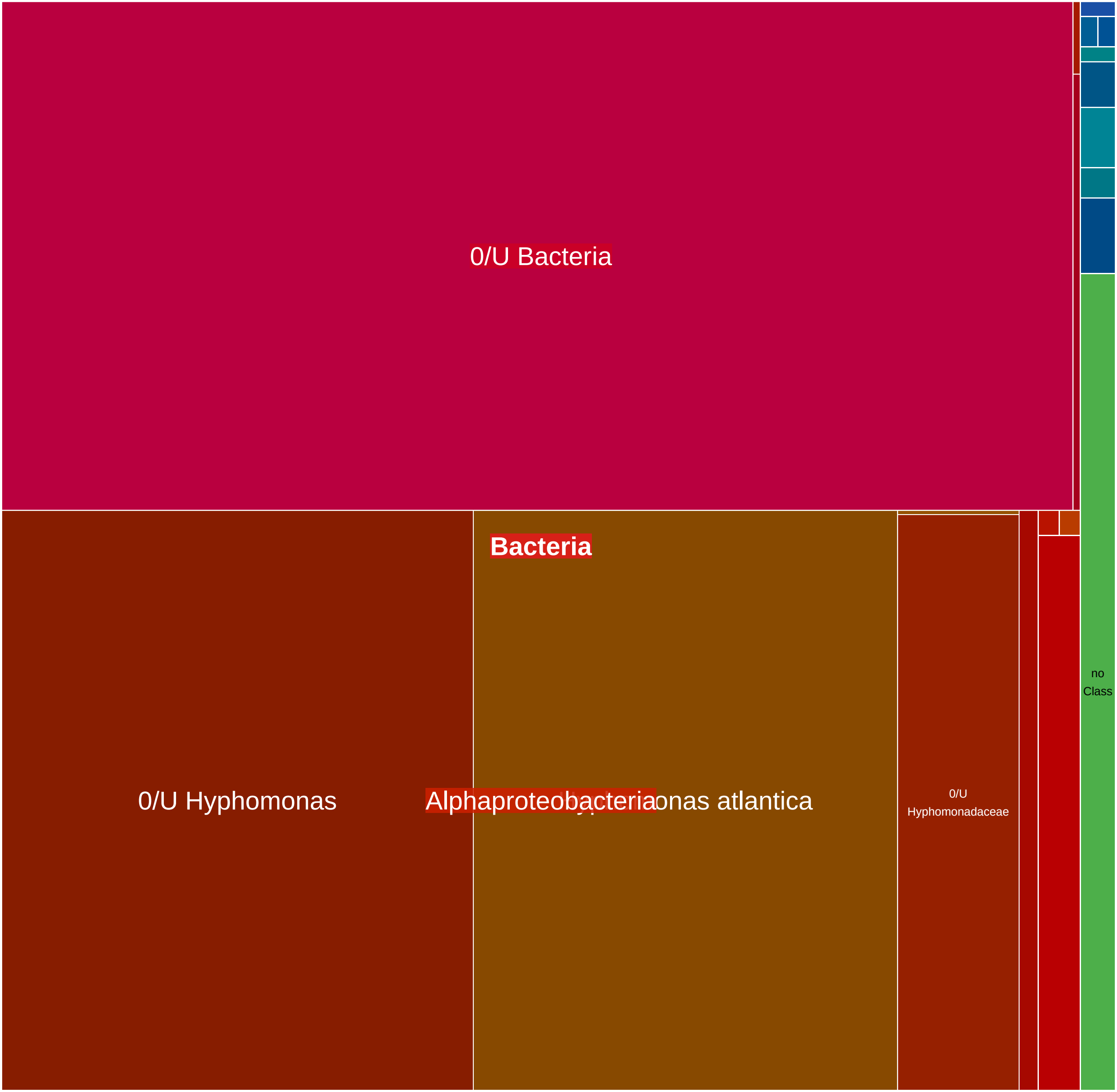

MGT−v1_413 : 2335 unigenes, 429 (18.37%) taxonomically assigned

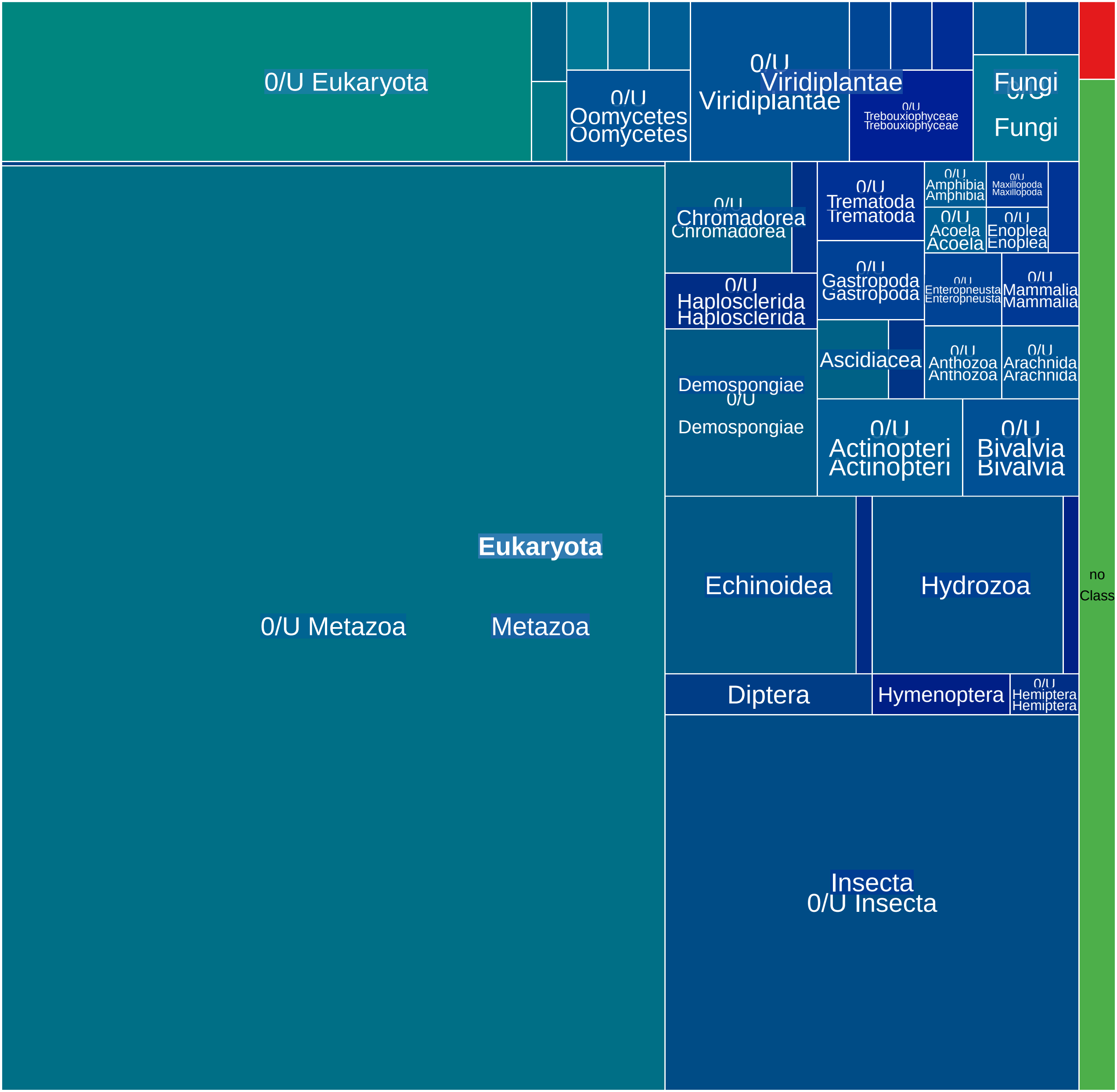

MGT−v1_414 : 2326 unigenes, 1689 (72.61%) taxonomically assigned

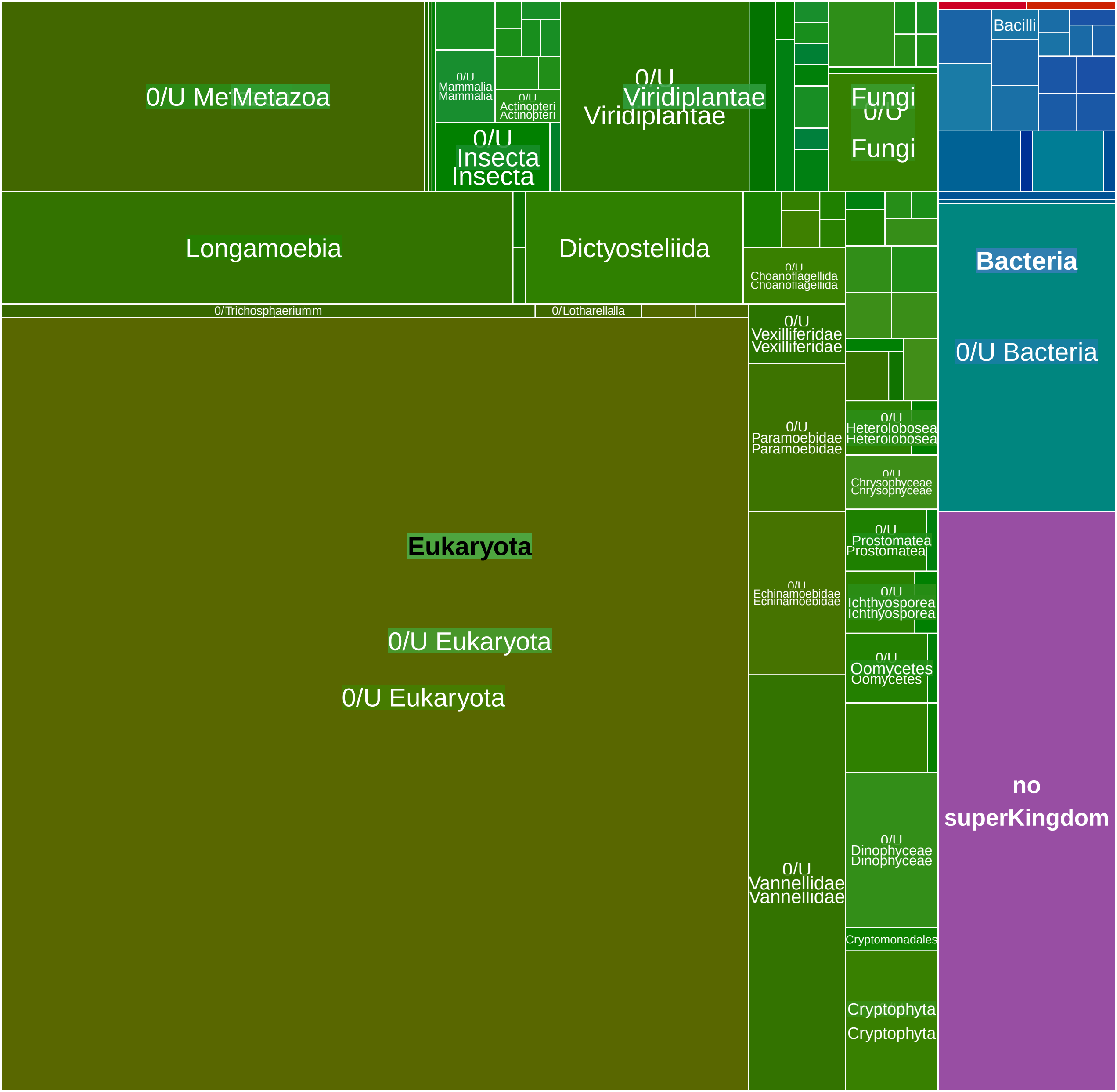

MGT−v1_415 : 2316 unigenes, 583 (25.17%) taxonomically assigned

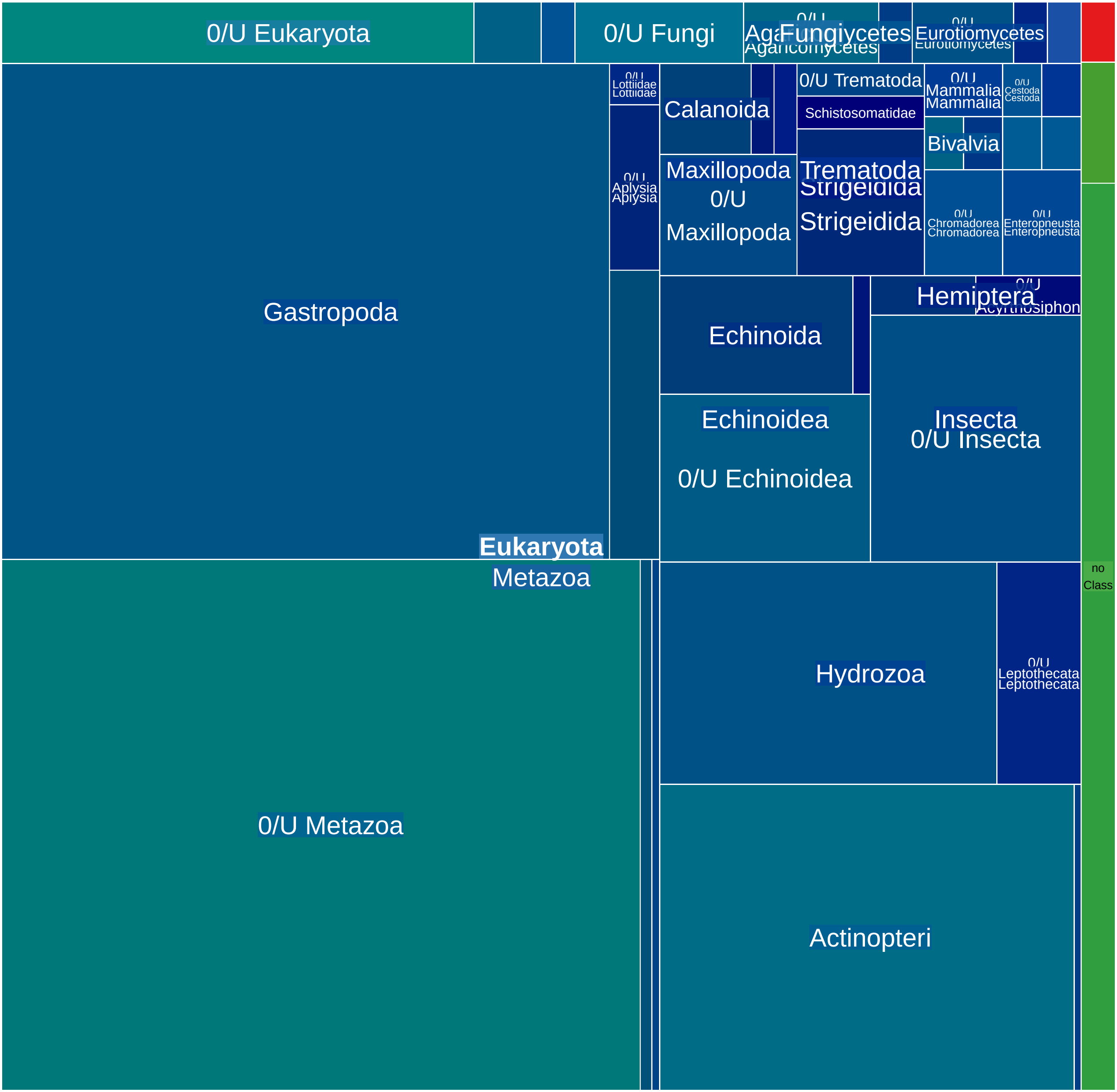

### MGT−v1_416 : 2285 unigenes, 1529 (66.91%) taxonomically assigned

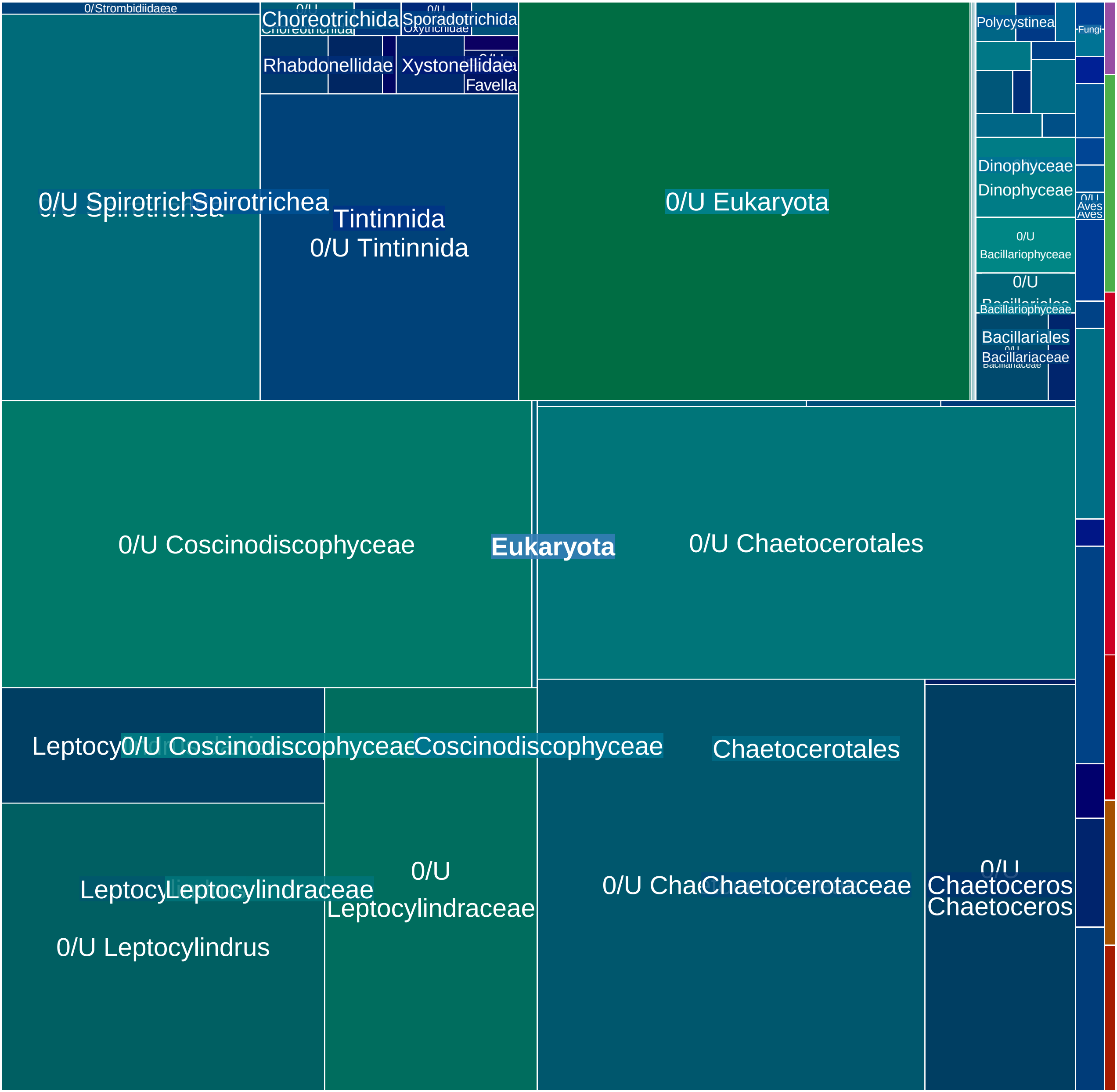

MGT−v1_417 : 2271 unigenes, 629 (27.70%) taxonomically assigned

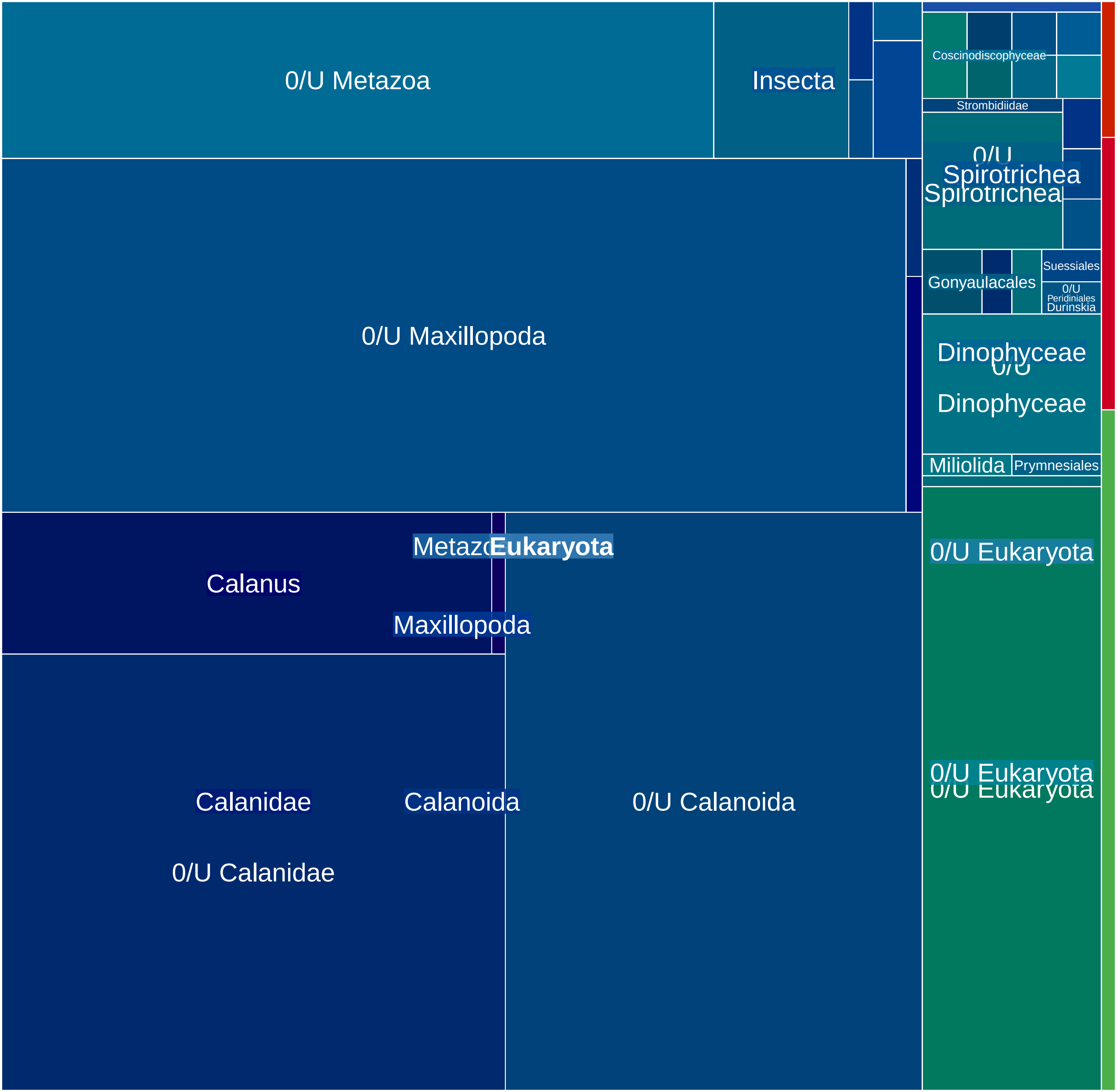

MGT−v1_418 : 2261 unigenes, 267 (11.81%) taxonomically assigned

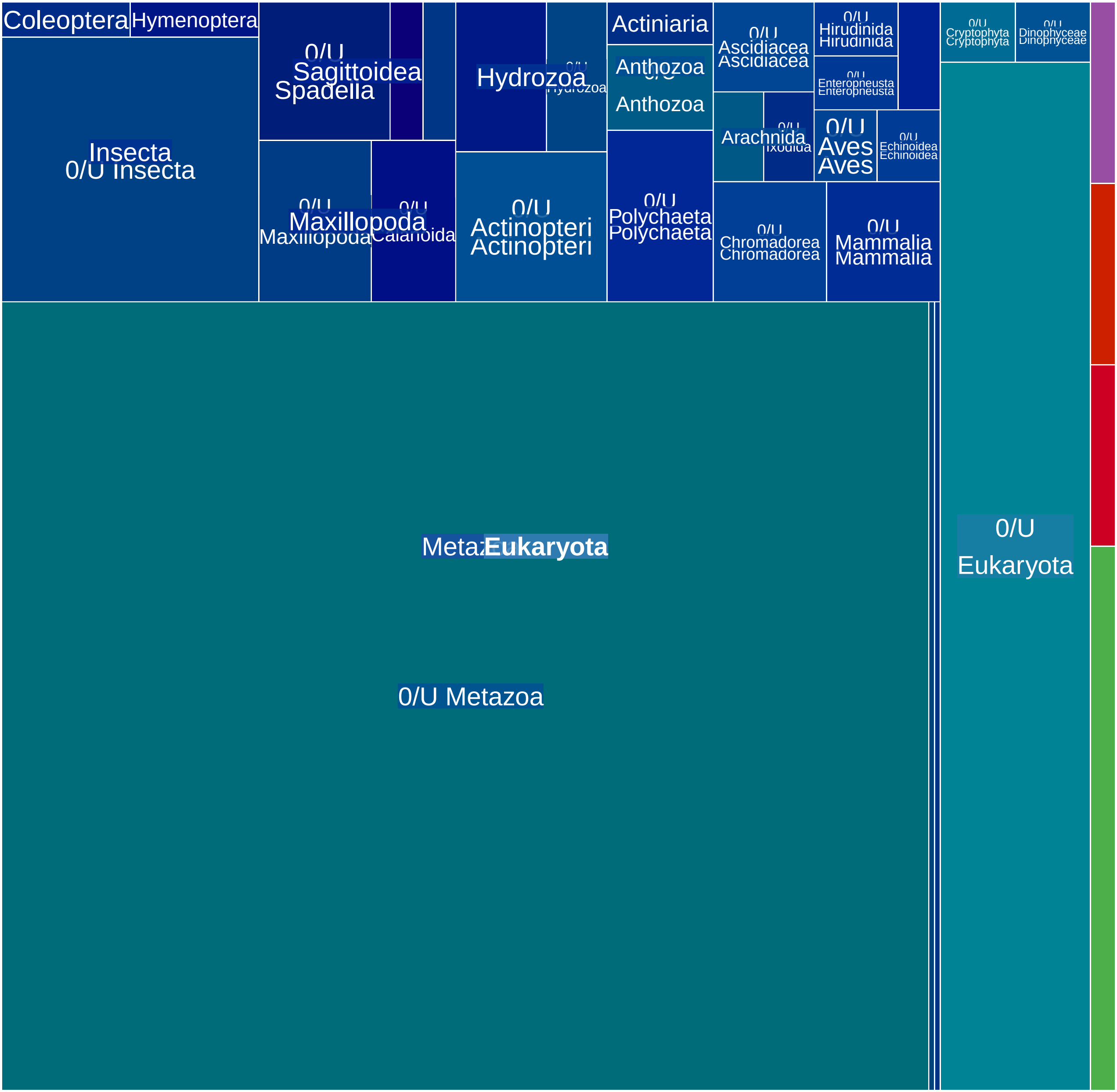

MGT−v1_419 : 2252 unigenes, 480 (21.31%) taxonomically assigned

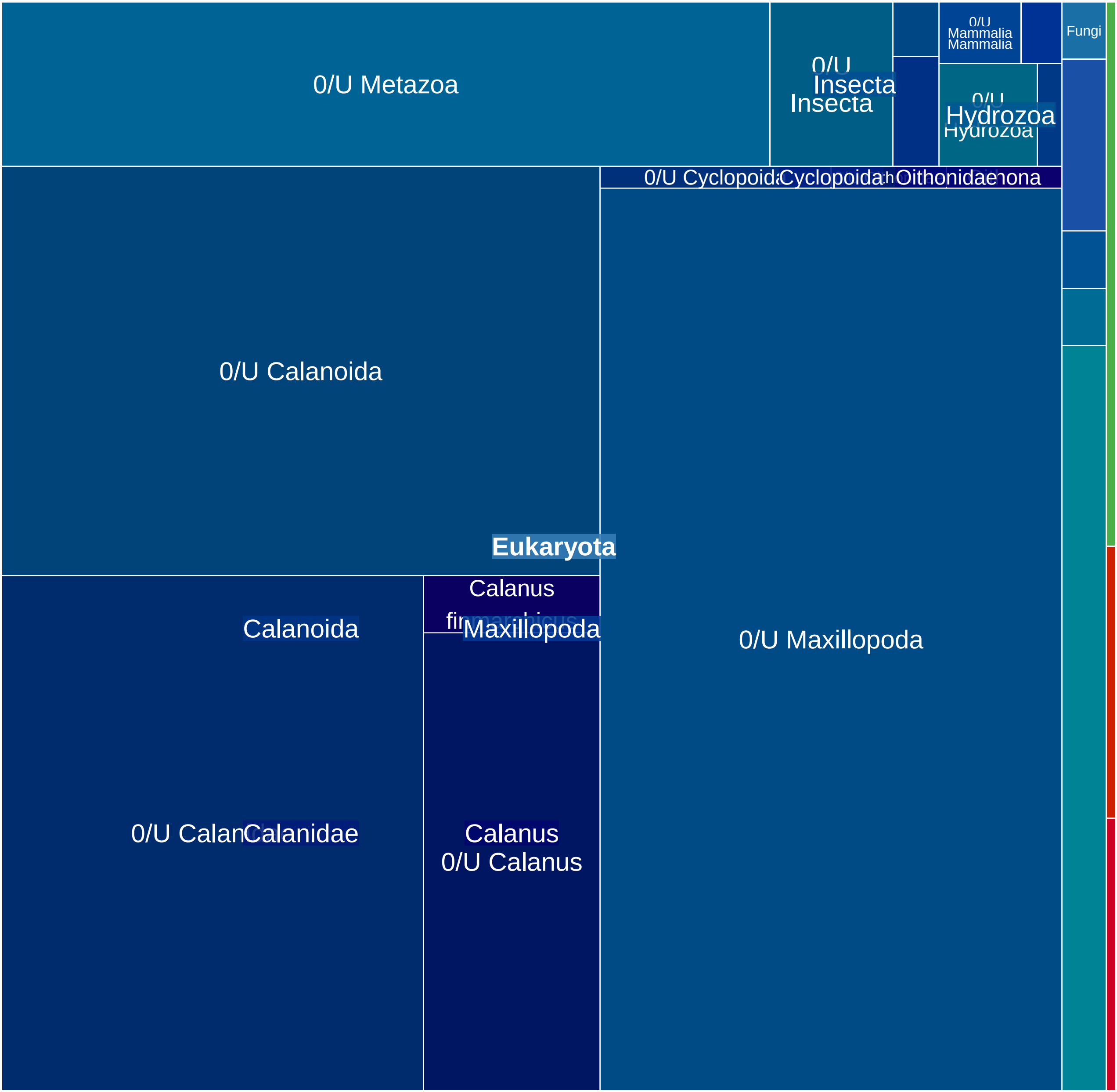

MGT−v1_420 : 2251 unigenes, 562 (24.97%) taxonomically assigned

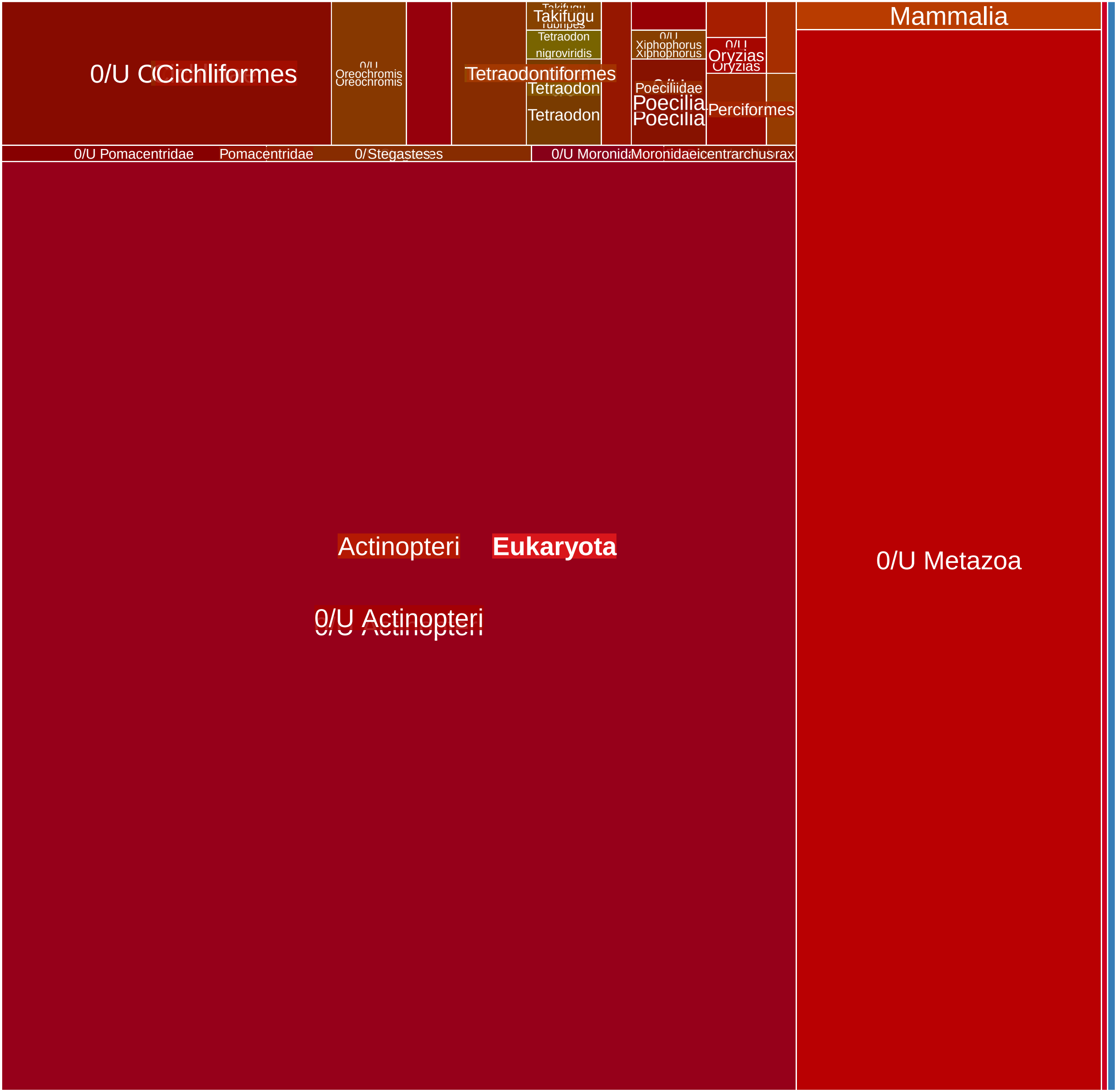

MGT−v1_421 : 2250 unigenes, 1300 (57.78%) taxonomically assigned

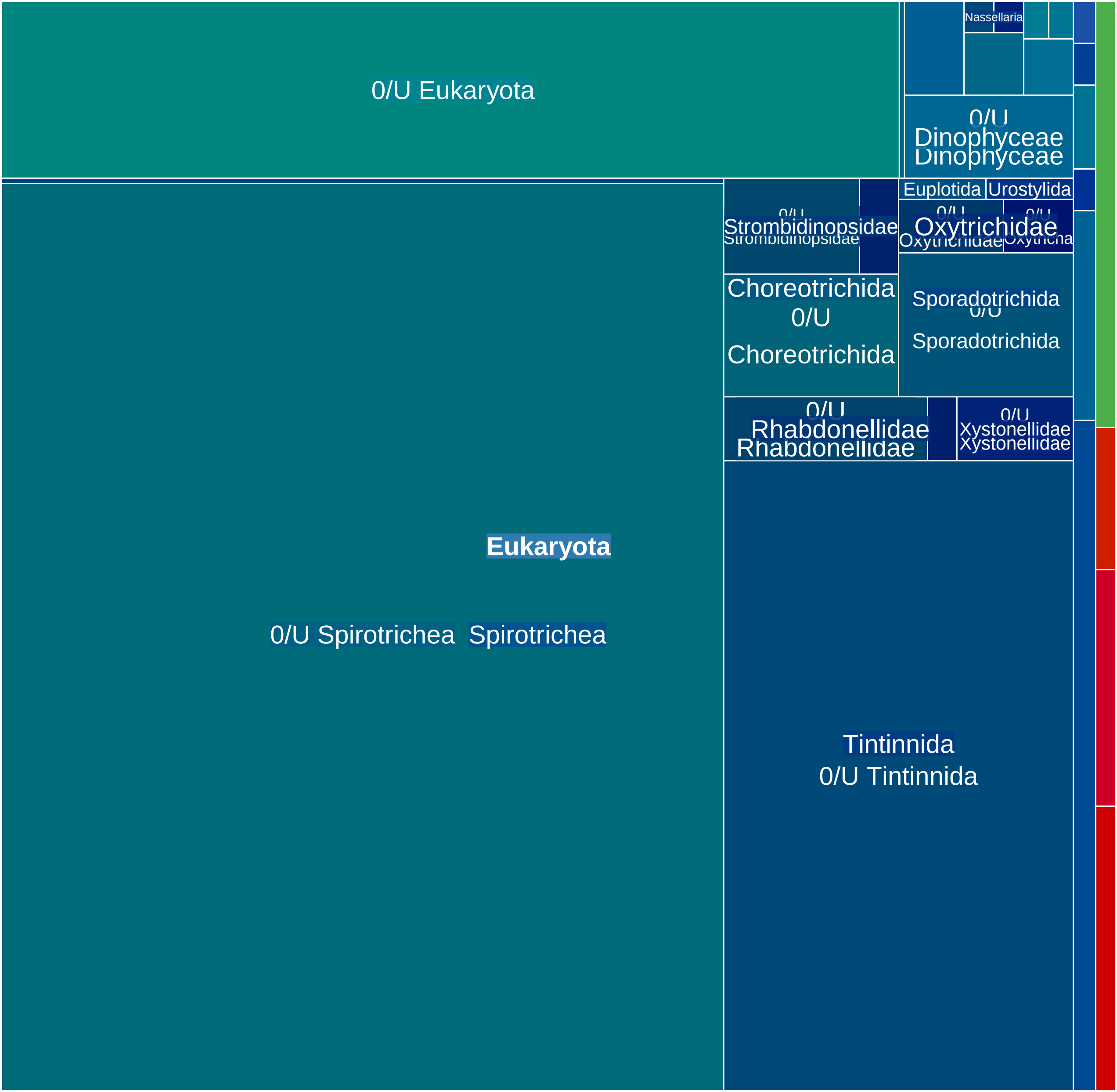

### MGT−v1_422 : 2247 unigenes, 144 (6.41%) taxonomically assigned

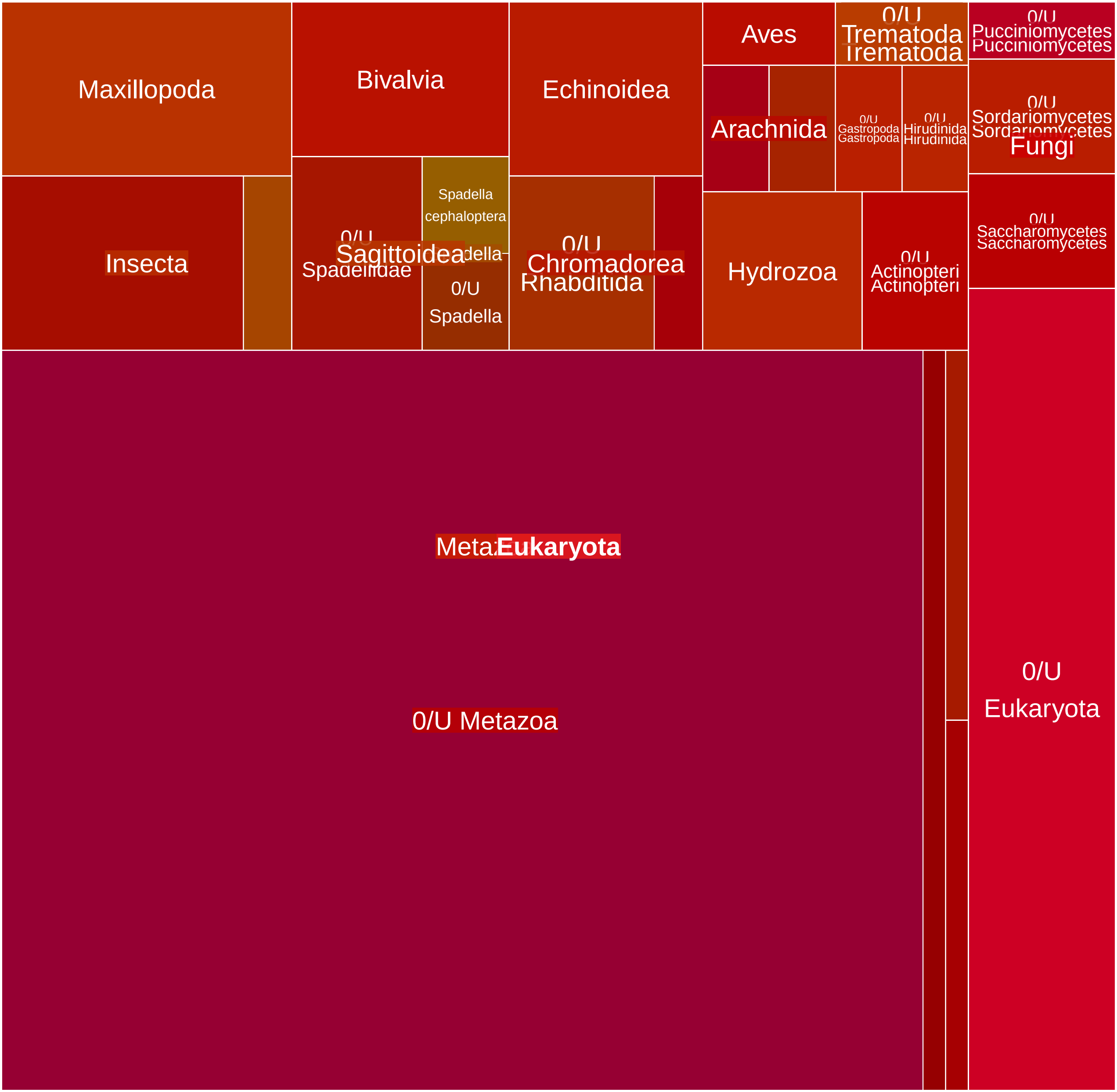

MGT−v1_423 : 2239 unigenes, 110 (4.91%) taxonomically assigned

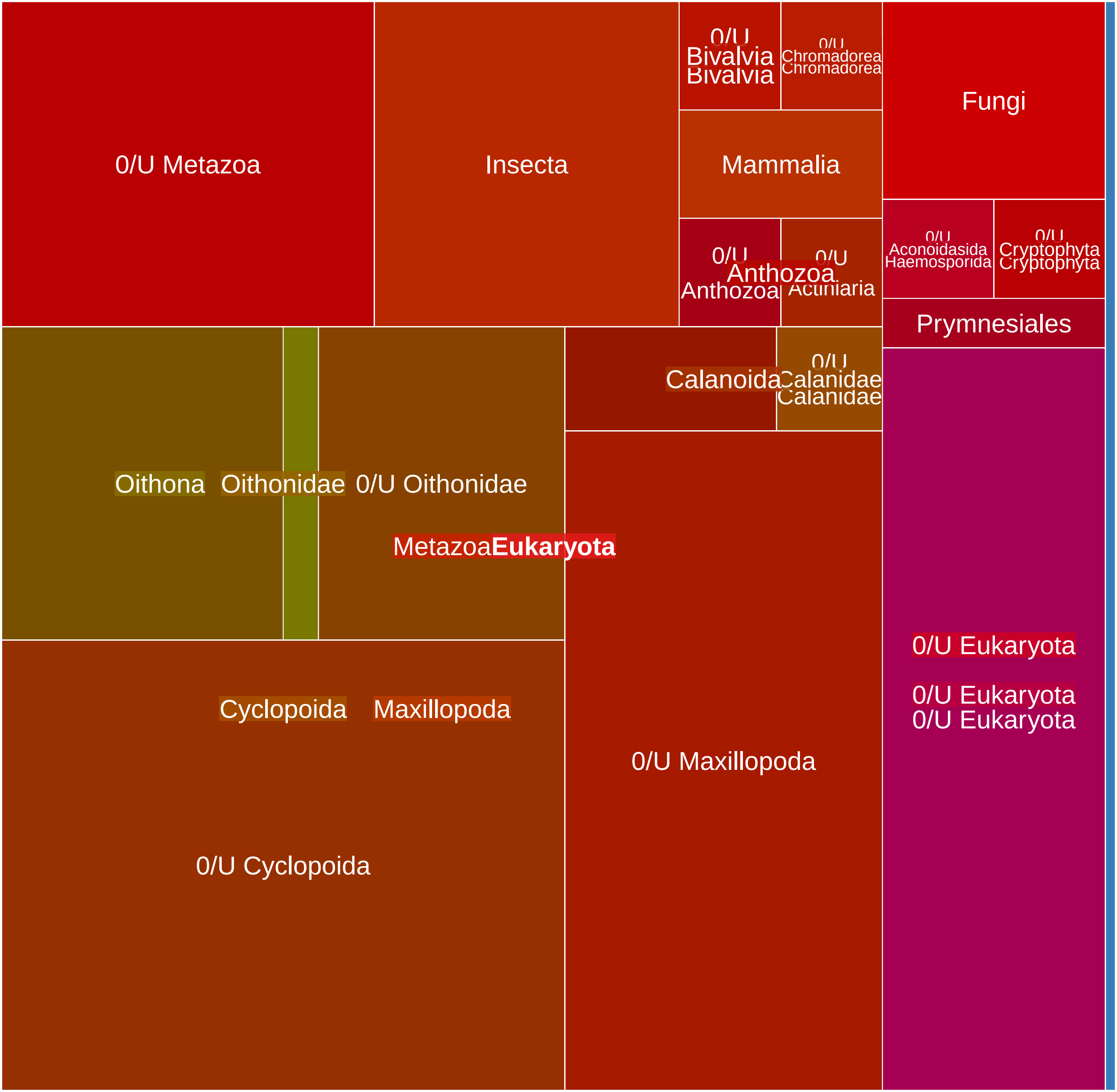

MGT−v1_424 : 2233 unigenes, 2175 (97.40%) taxonomically assigned

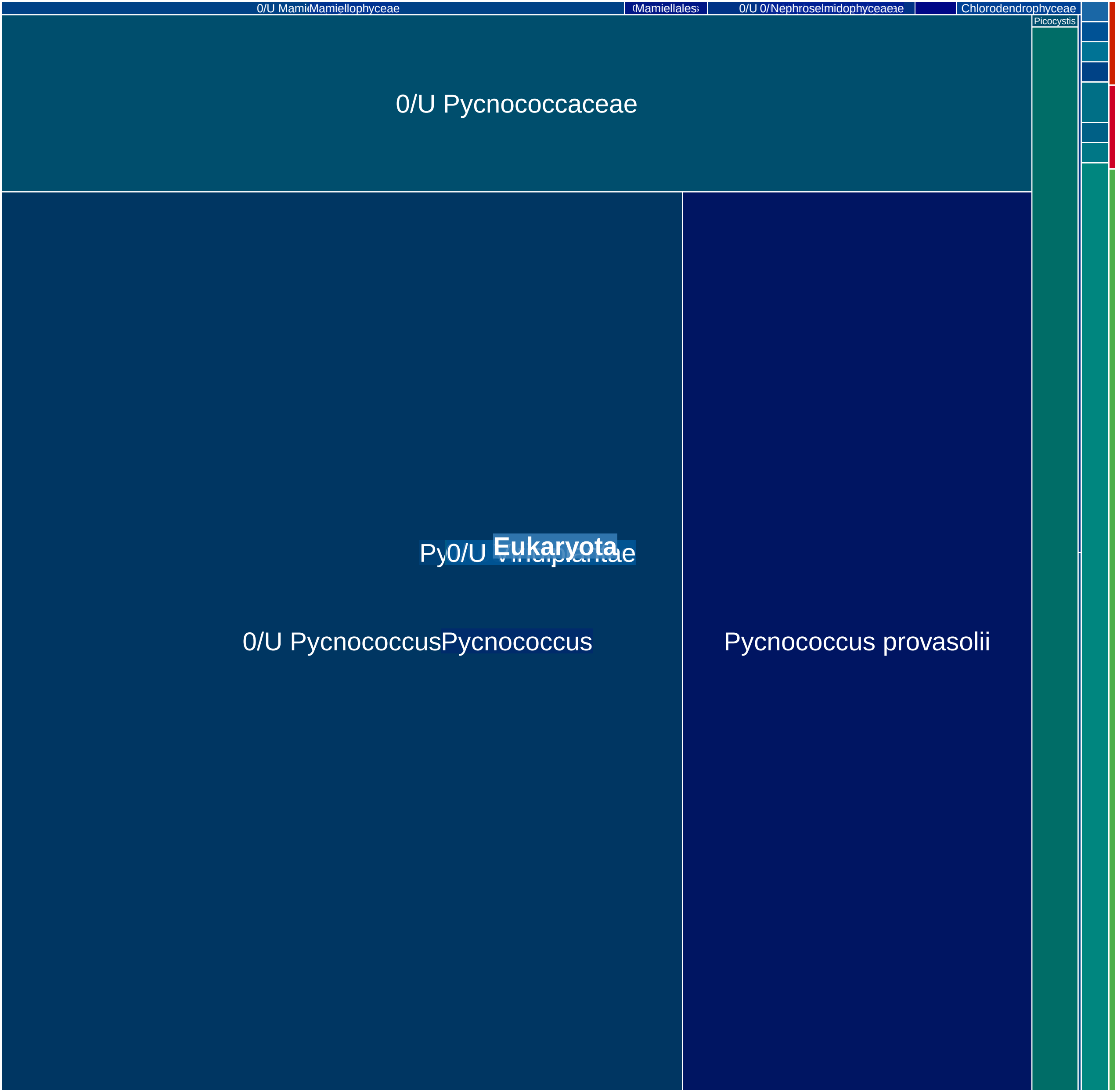

### MGT−v1_425 : 2226 unigenes, 2146 (96.41%) taxonomically assigned

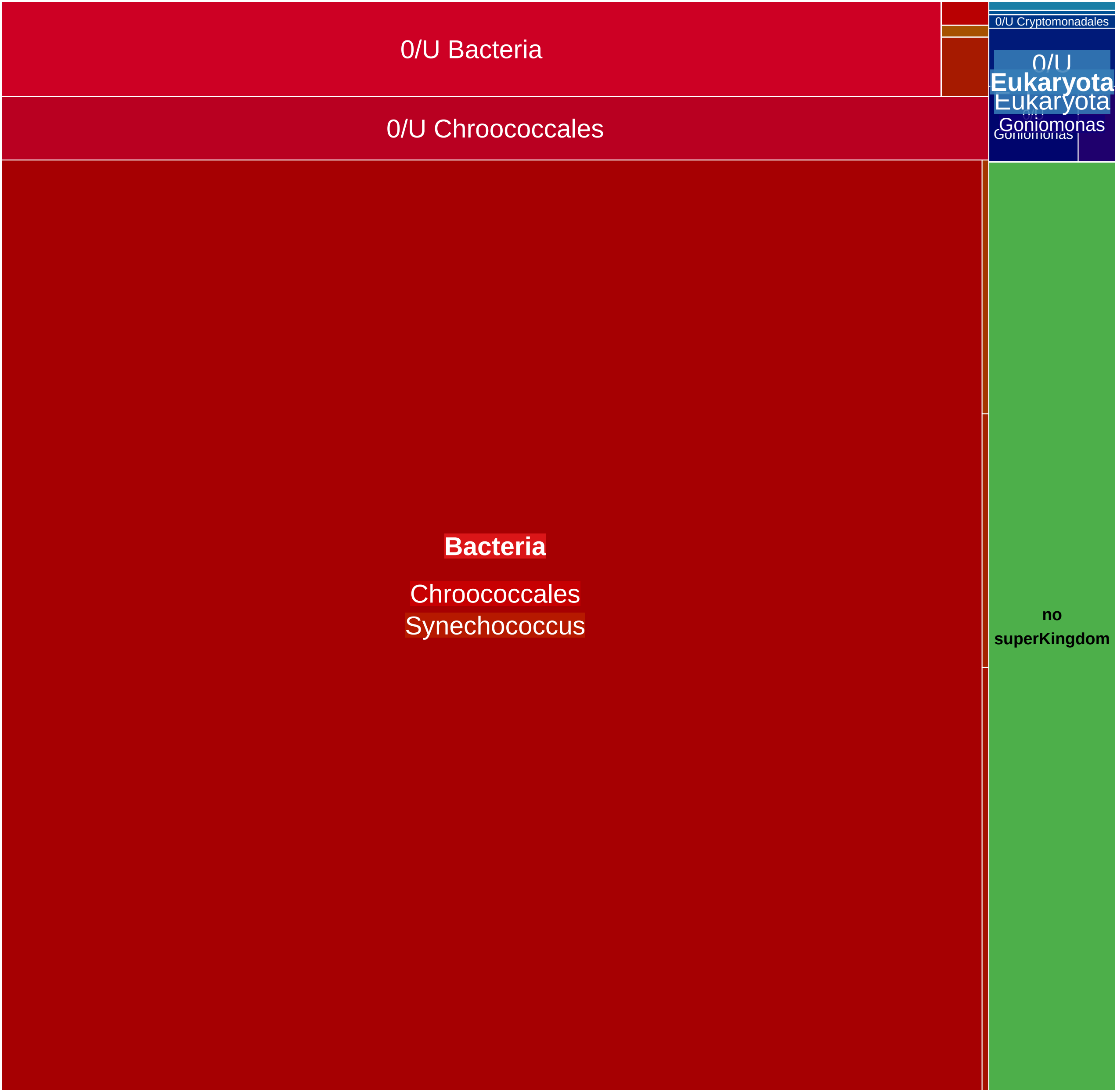

MGT−v1_426 : 2222 unigenes, 254 (11.43%) taxonomically assigned

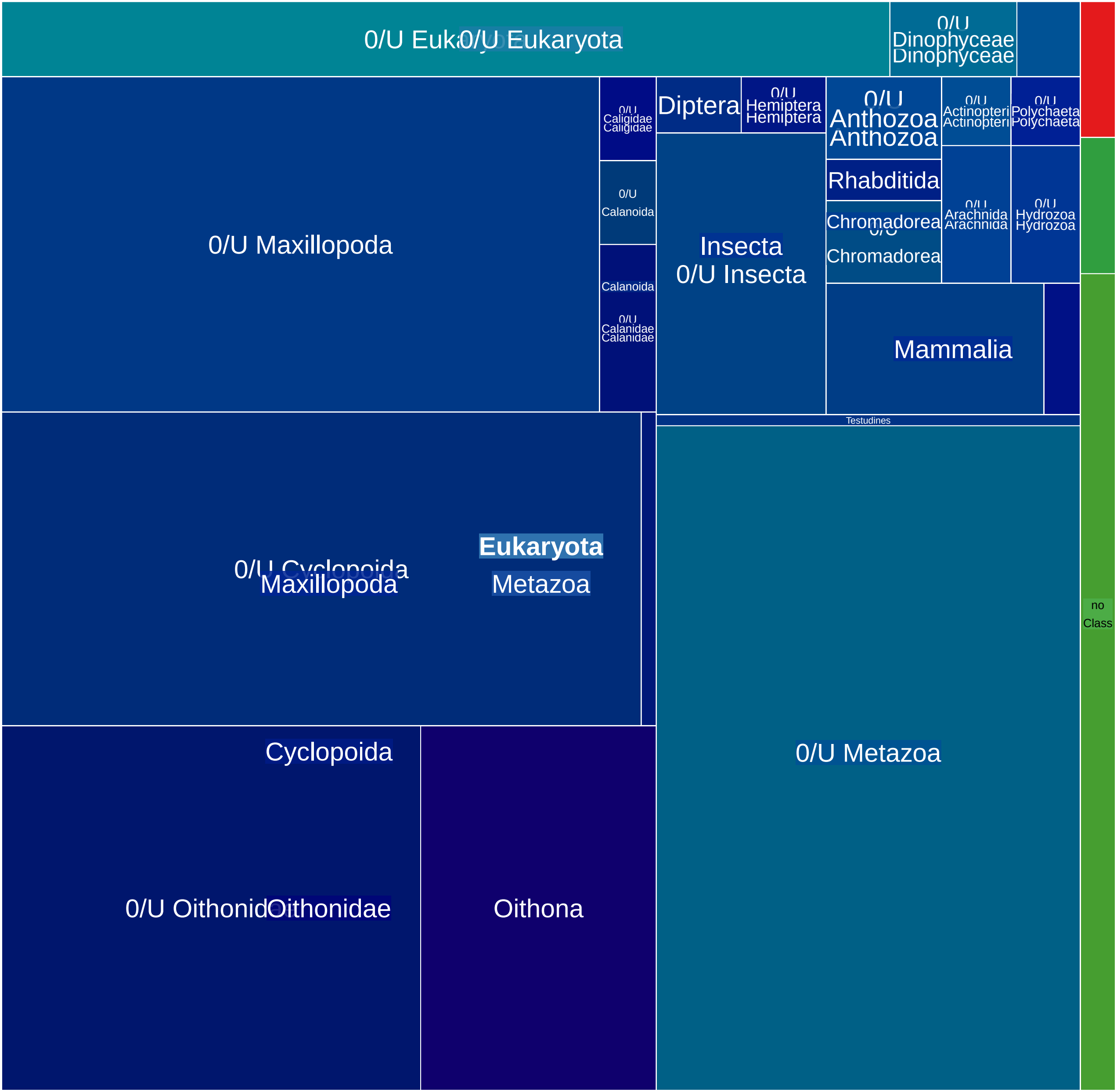

MGT−v1_427 : 2222 unigenes, 530 (23.85%) taxonomically assigned

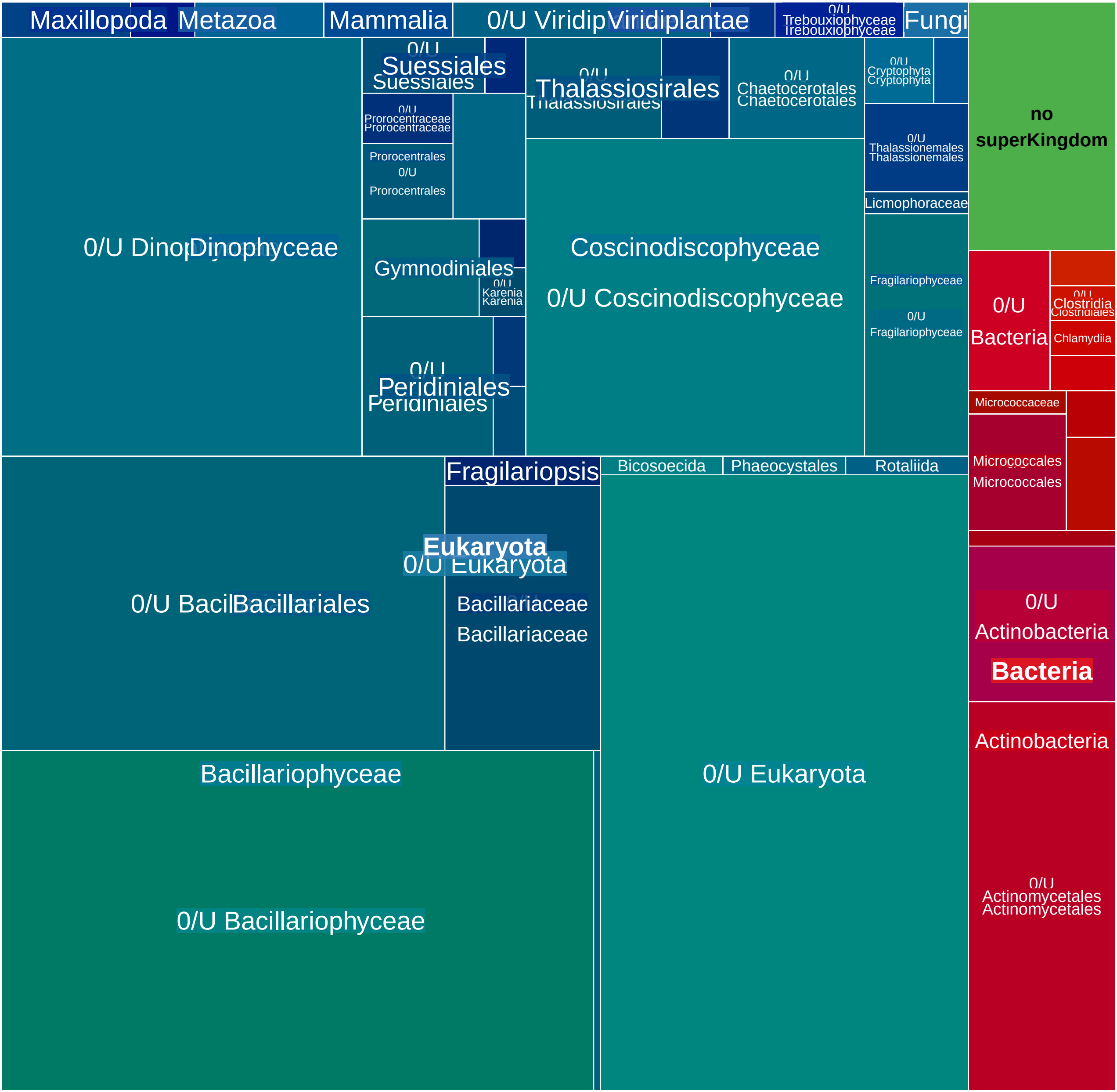

MGT−v1_428 : 2220 unigenes, 2036 (91.71%) taxonomically assigned

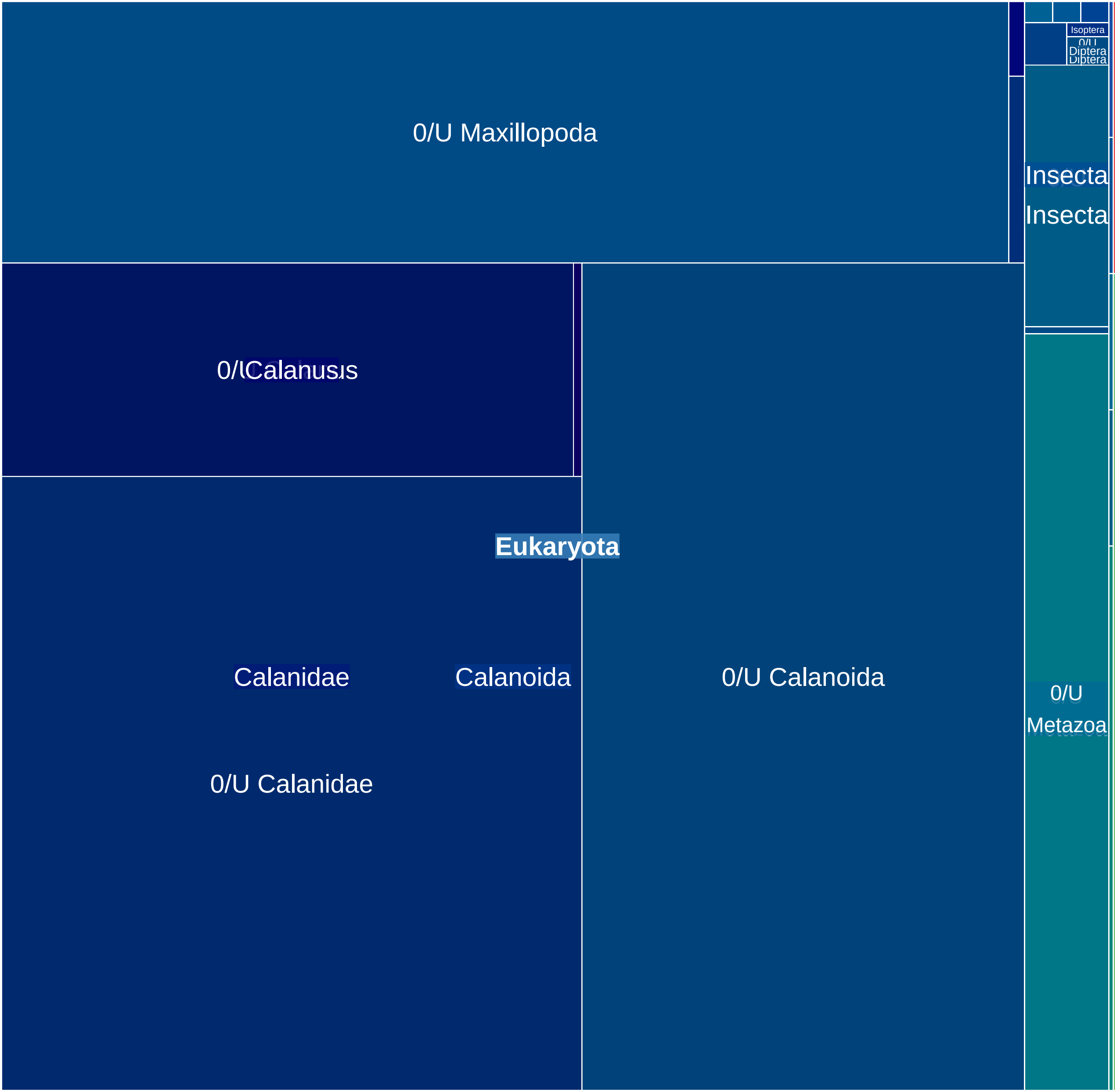

MGT−v1_429 : 2208 unigenes, 255 (11.55%) taxonomically assigned

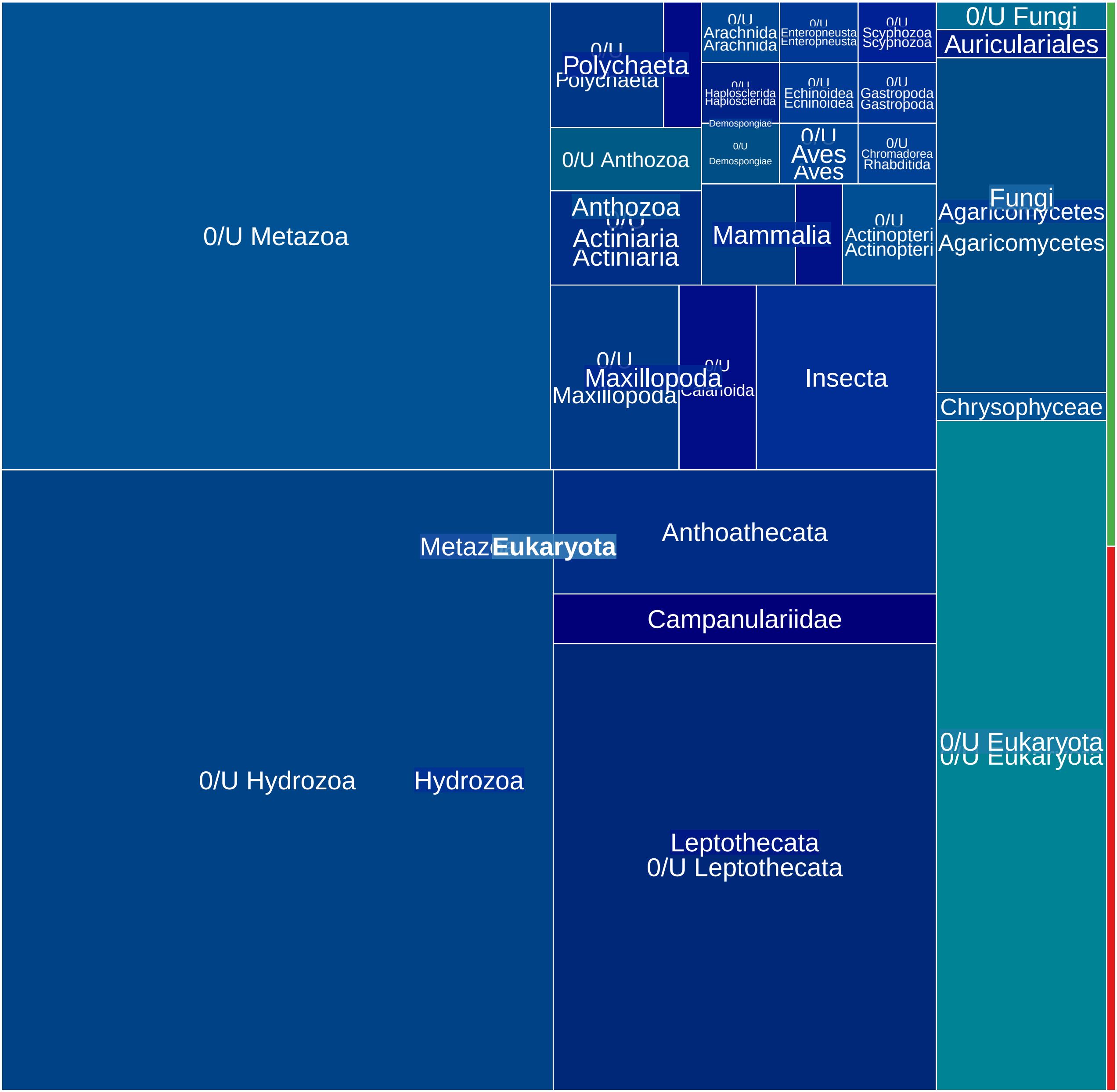

MGT−v1_430 : 2203 unigenes, 1841 (83.57%) taxonomically assigned

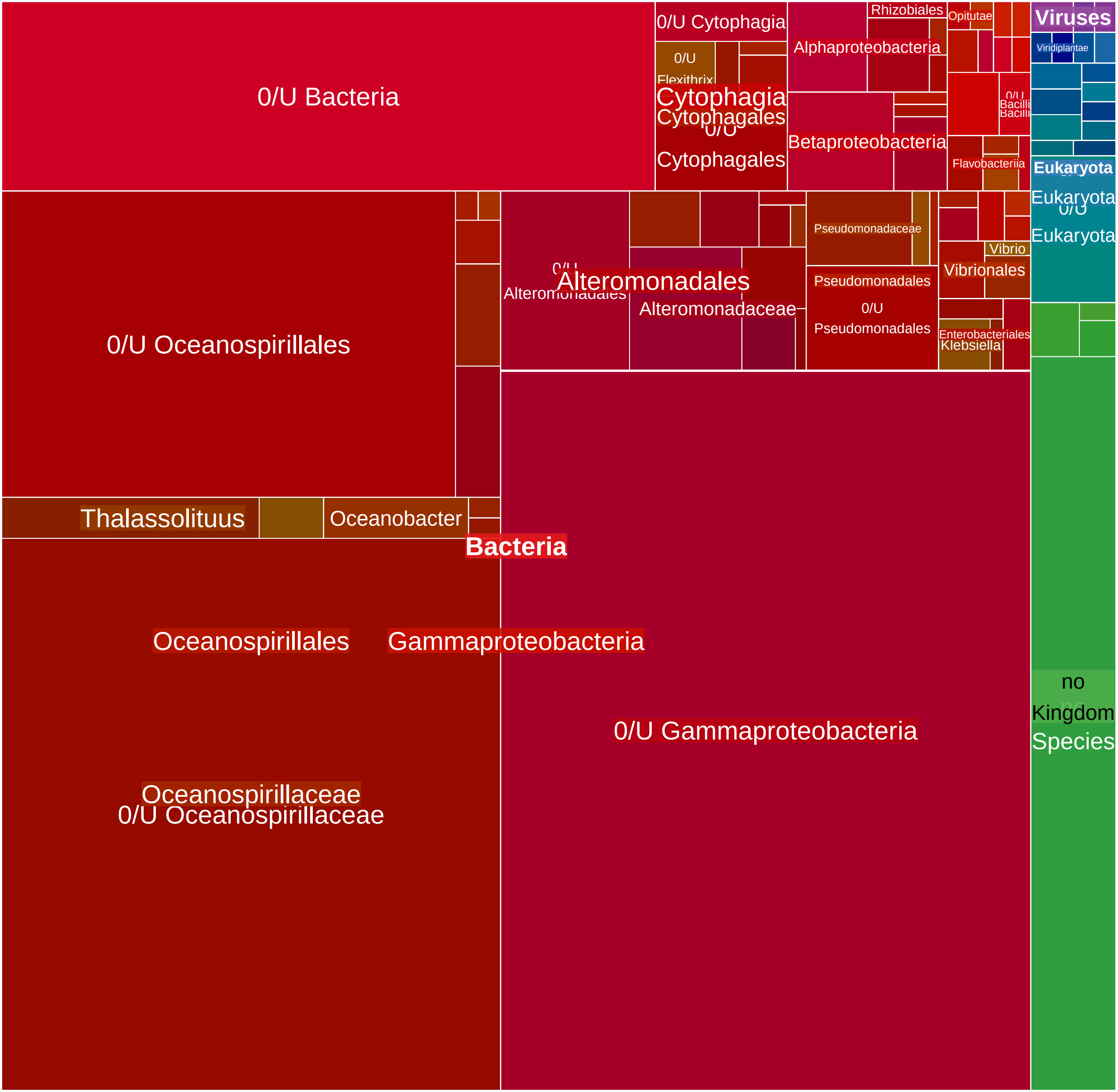

MGT−v1_431 : 2196 unigenes, 257 (11.70%) taxonomically assigned

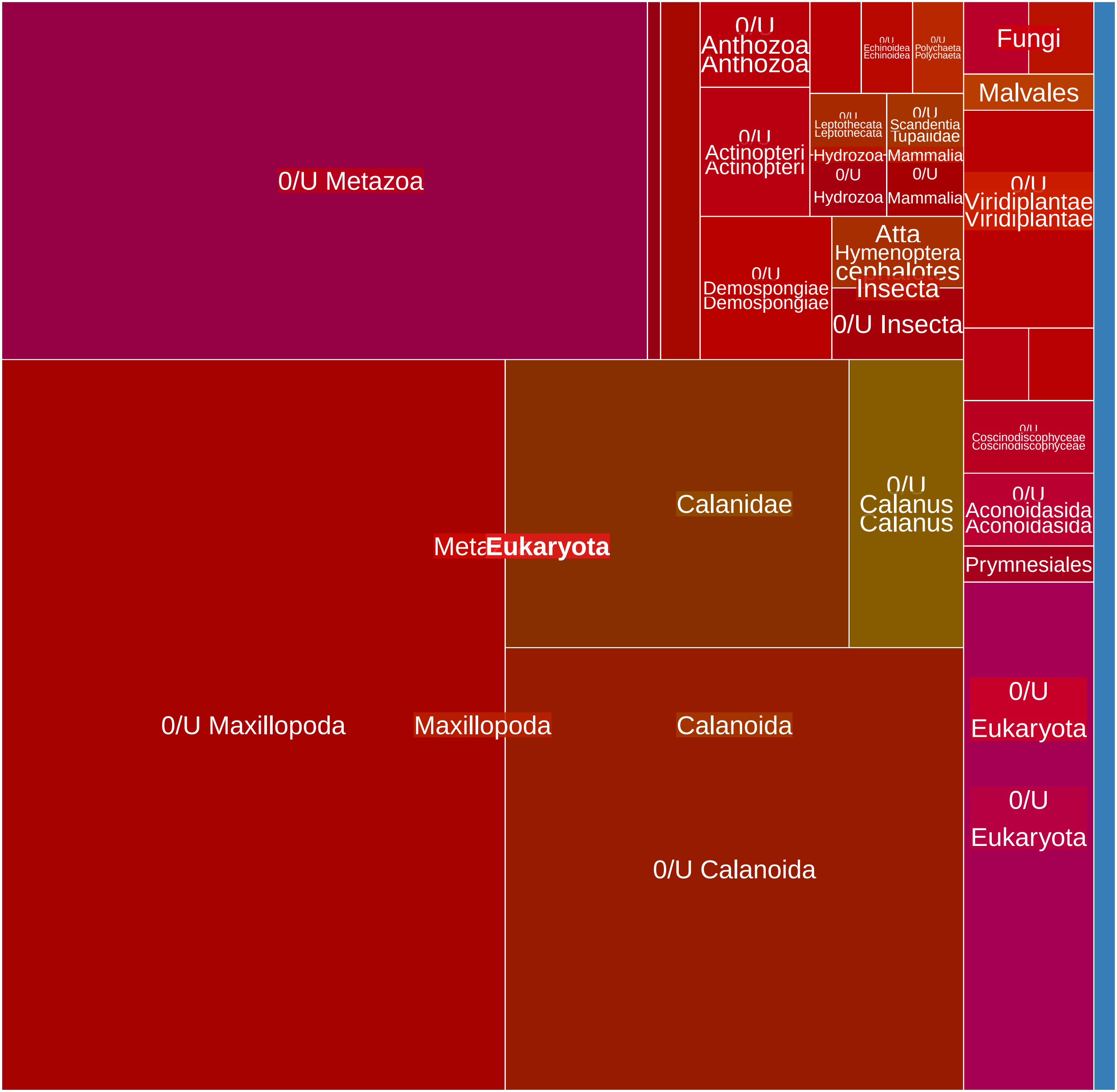

MGT−v1_432 : 2190 unigenes, 195 (8.90%) taxonomically assigned

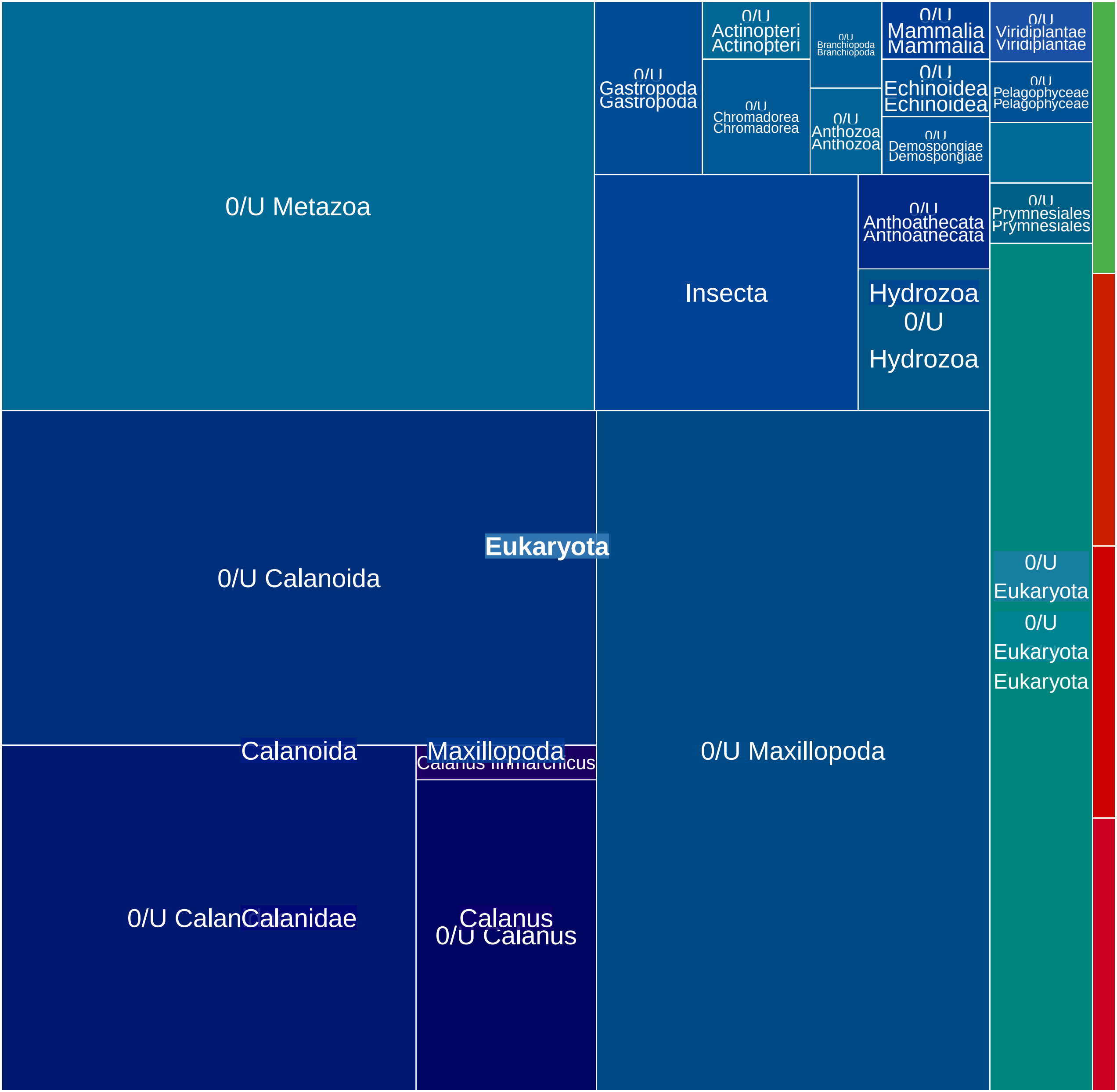

MGT−v1_433 : 2185 unigenes, 1897 (86.82%) taxonomically assigned

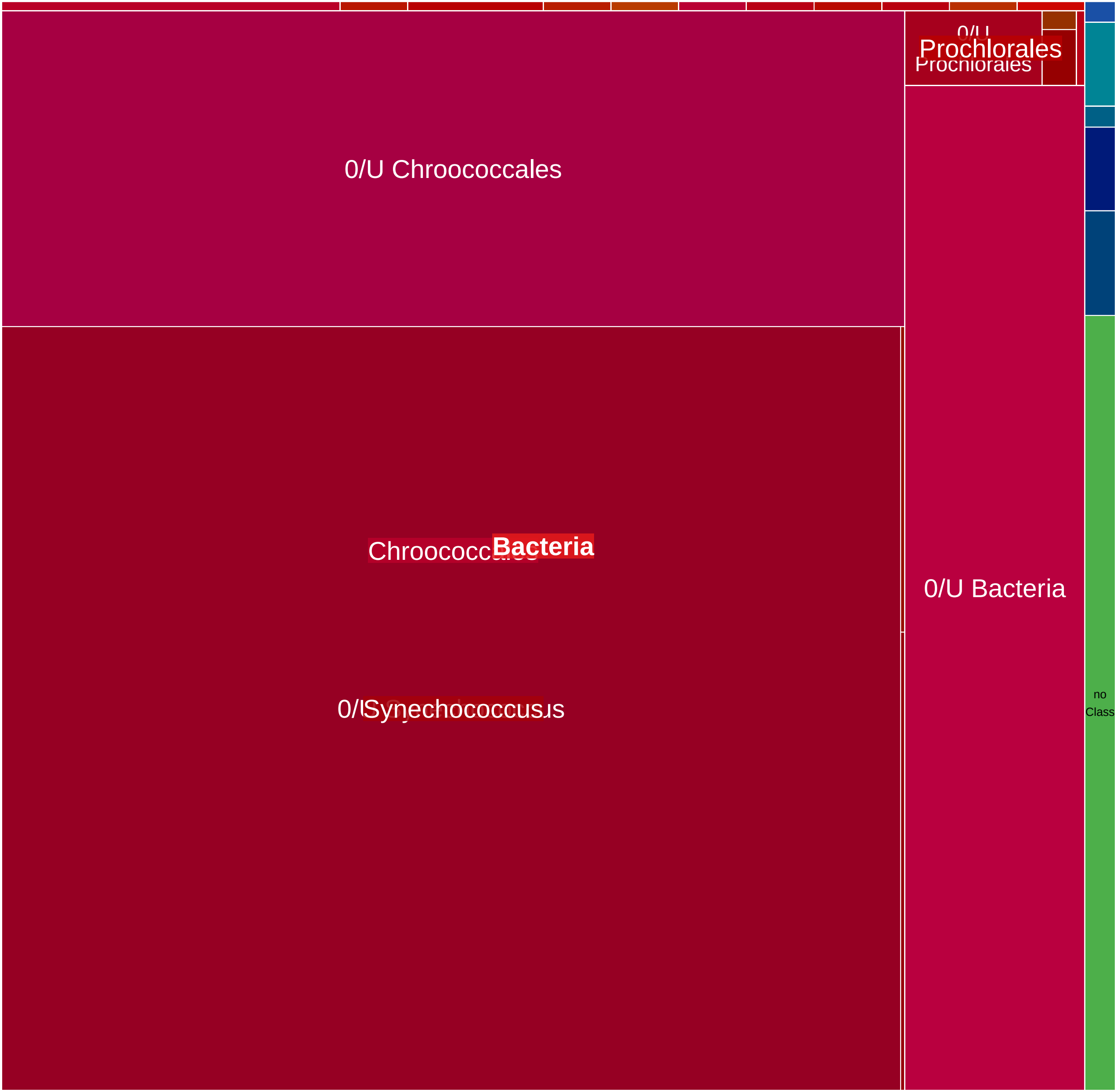

MGT−v1_434 : 2149 unigenes, 241 (11.21%) taxonomically assigned

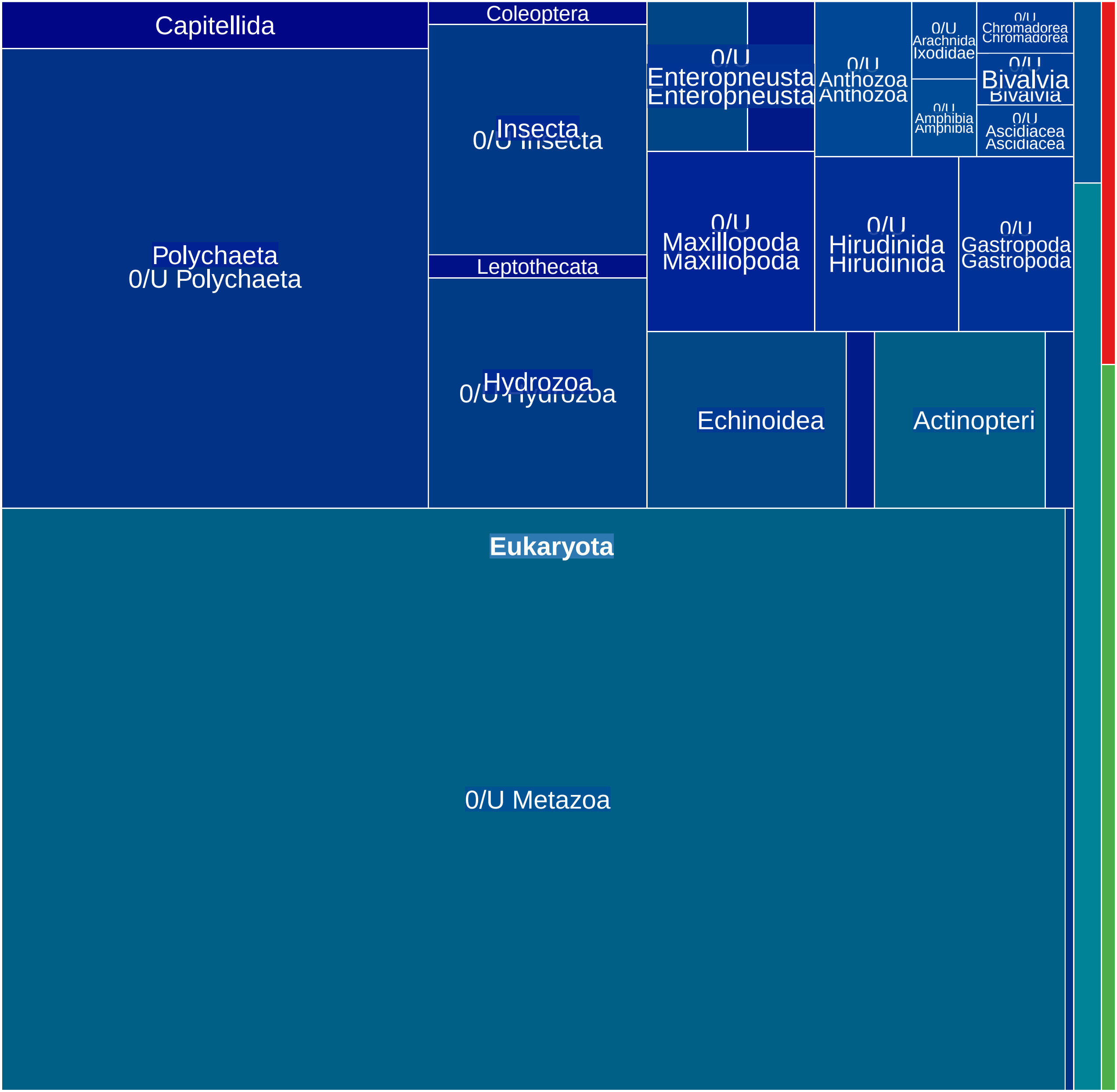

MGT−v1_435 : 2144 unigenes, 276 (12.87%) taxonomically assigned

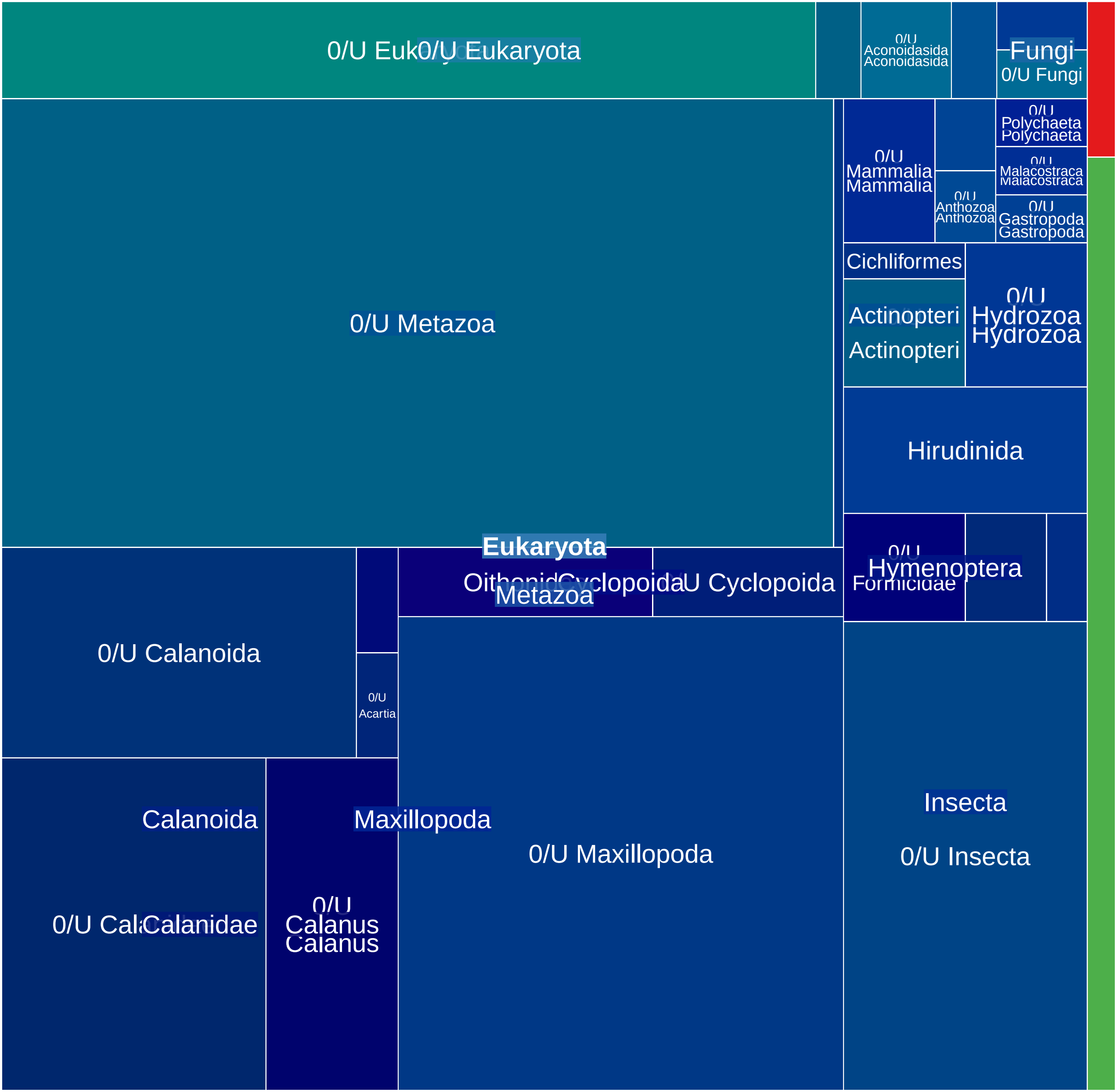

MGT−v1_436 : 2140 unigenes, 591 (27.62%) taxonomically assigned

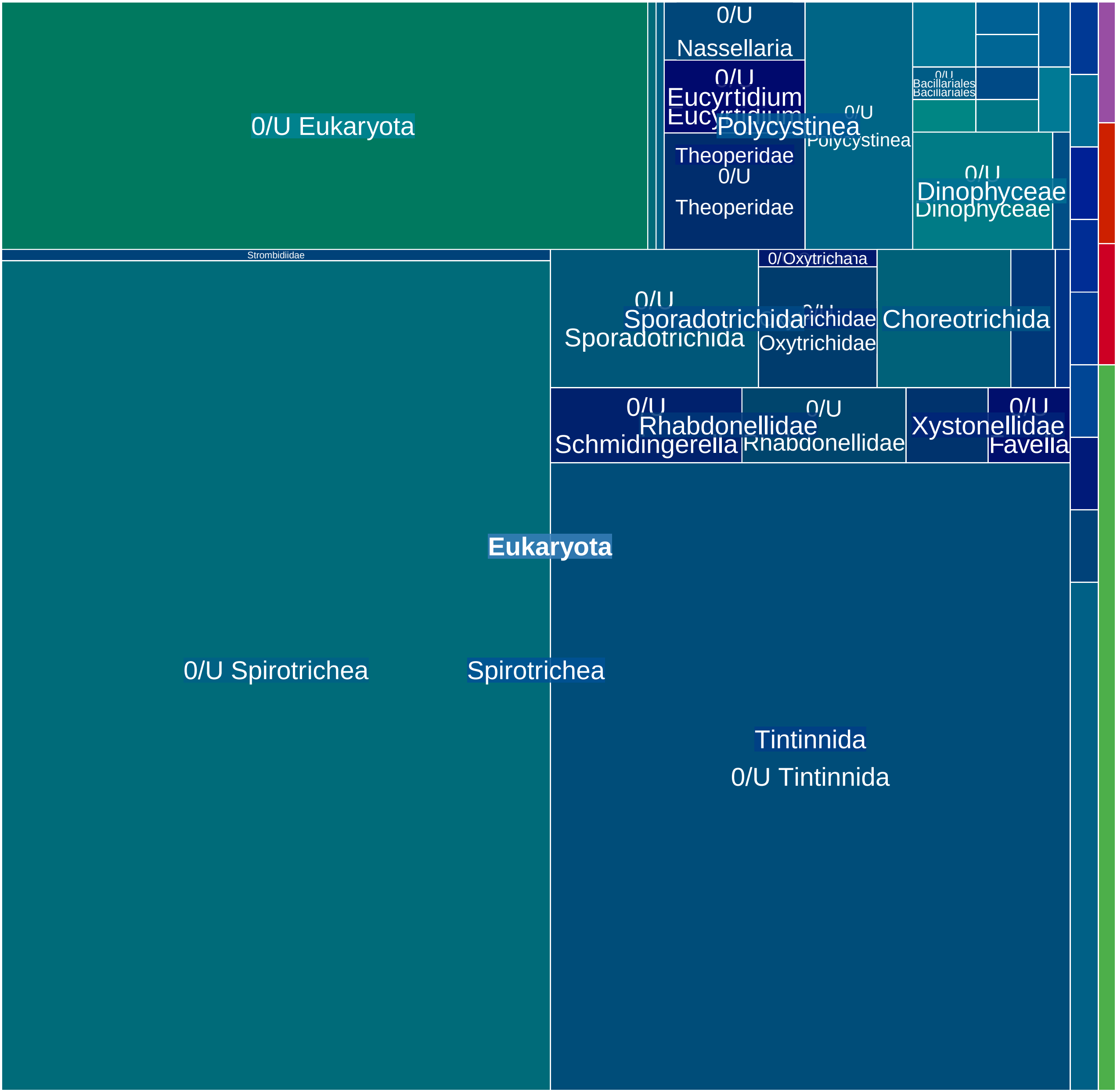

MGT−v1_437 : 2136 unigenes, 468 (21.91%) taxonomically assigned

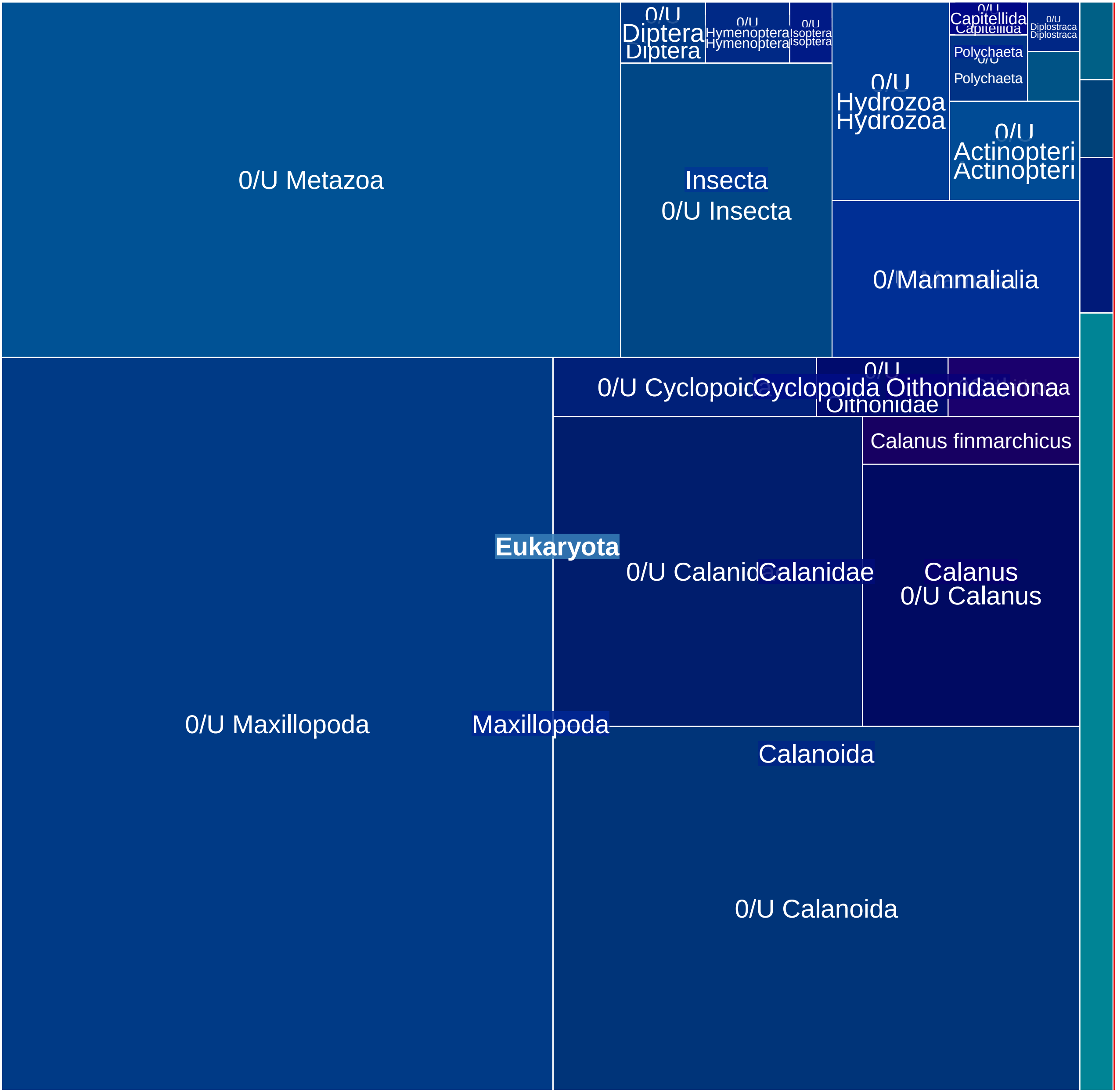

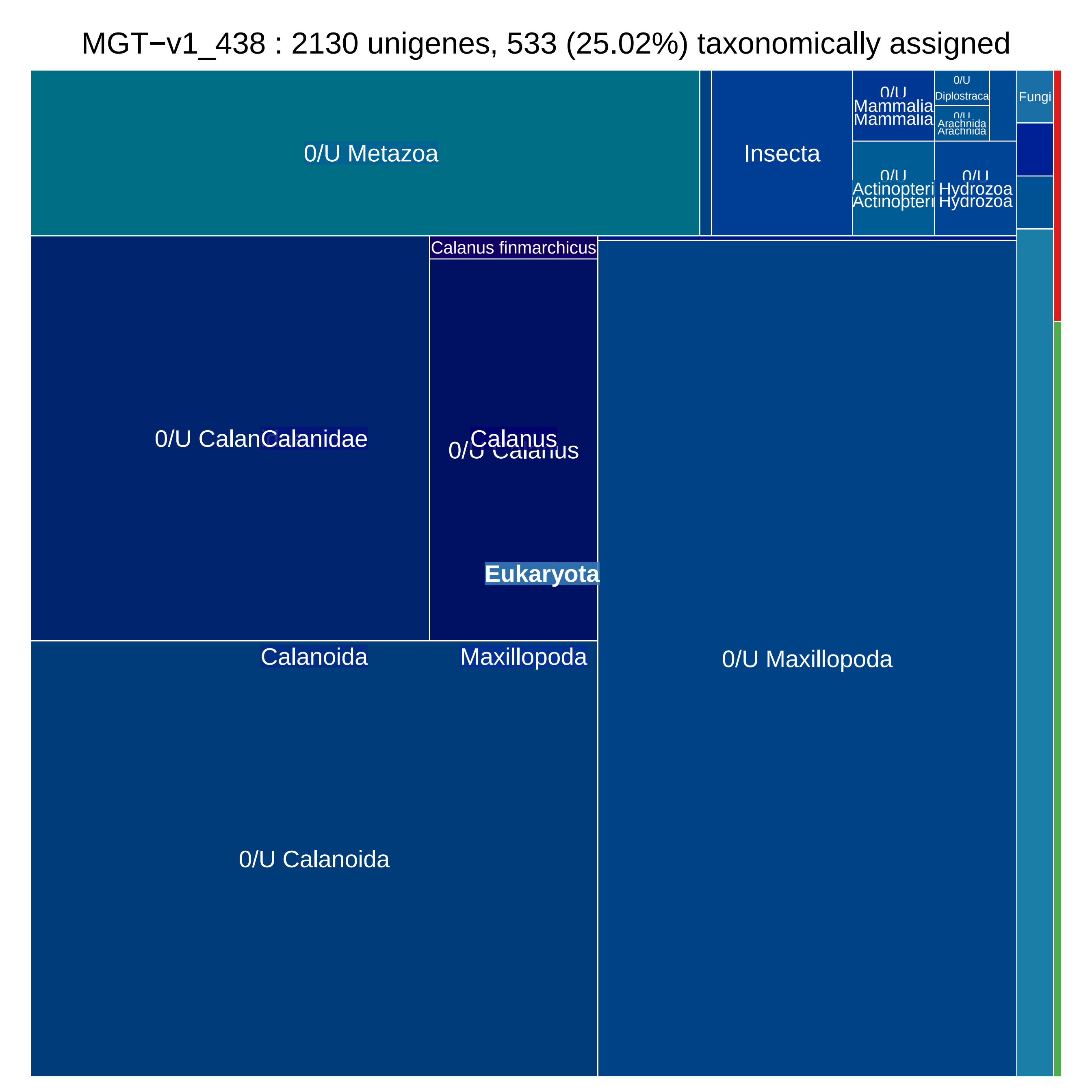
MGT−v1_438 : 2130 unigenes, 533 (25.02%) taxonomically assigned
0/U Metazoa
Insecta
0/U Mammalia
Mammalia
0/U Diplostraca
0/U Arachnida
Arachnida
Fungi
0/U Actinopteri
Actinopteri
0/U Hydrozoa
Hydrozoa
Calanus finmarchicus
0/U Calanidae
Calanidae
Calanus
0/U Calanus
Eukaryota
Calanoida
Maxillopoda
0/U Maxillopoda
0/U Calanoida

MGT−v1_439 : 2126 unigenes, 2089 (98.26%) taxonomically assigned

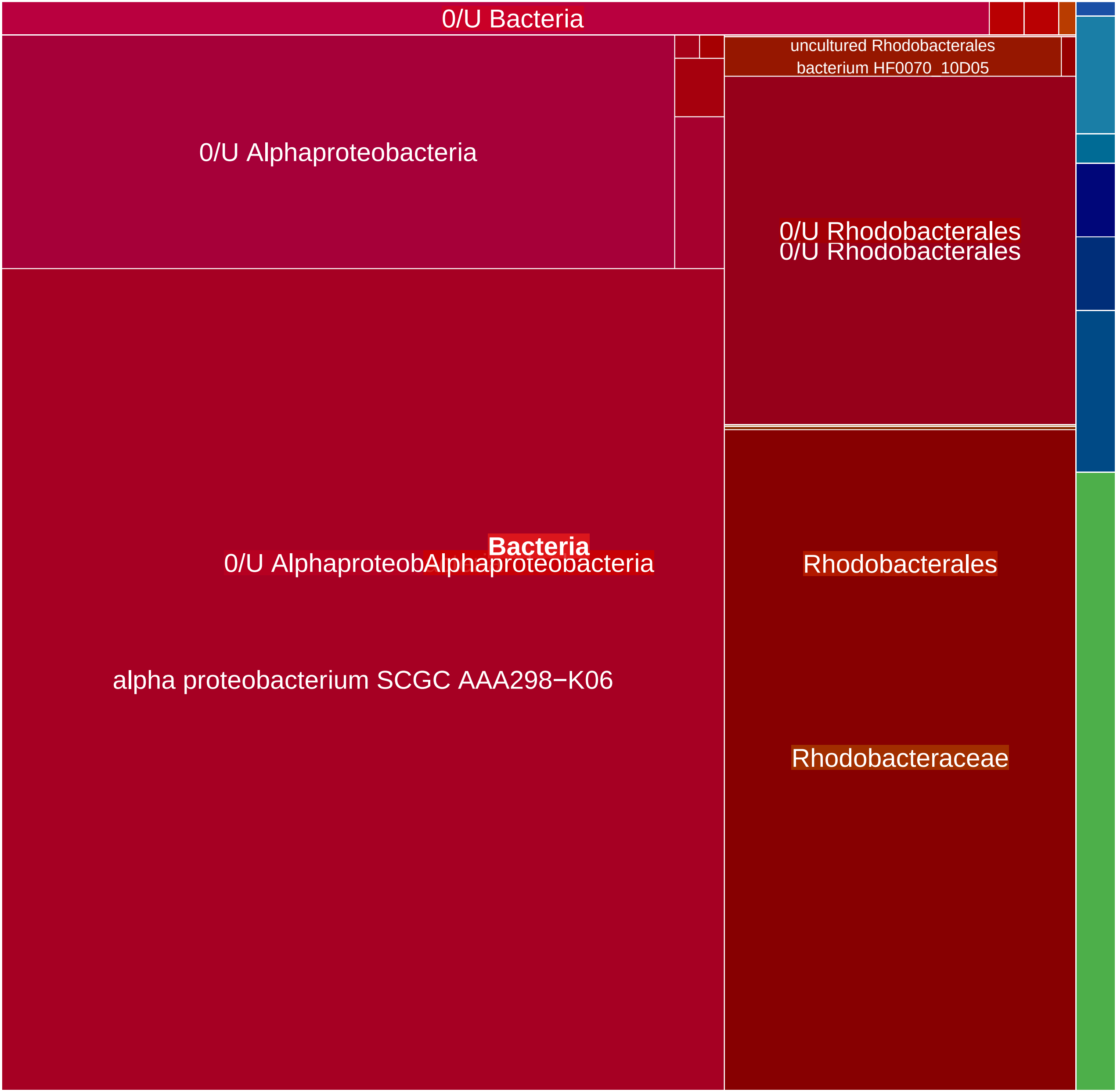

### MGT−v1_440 : 2123 unigenes, 353 (16.63%) taxonomically assigned

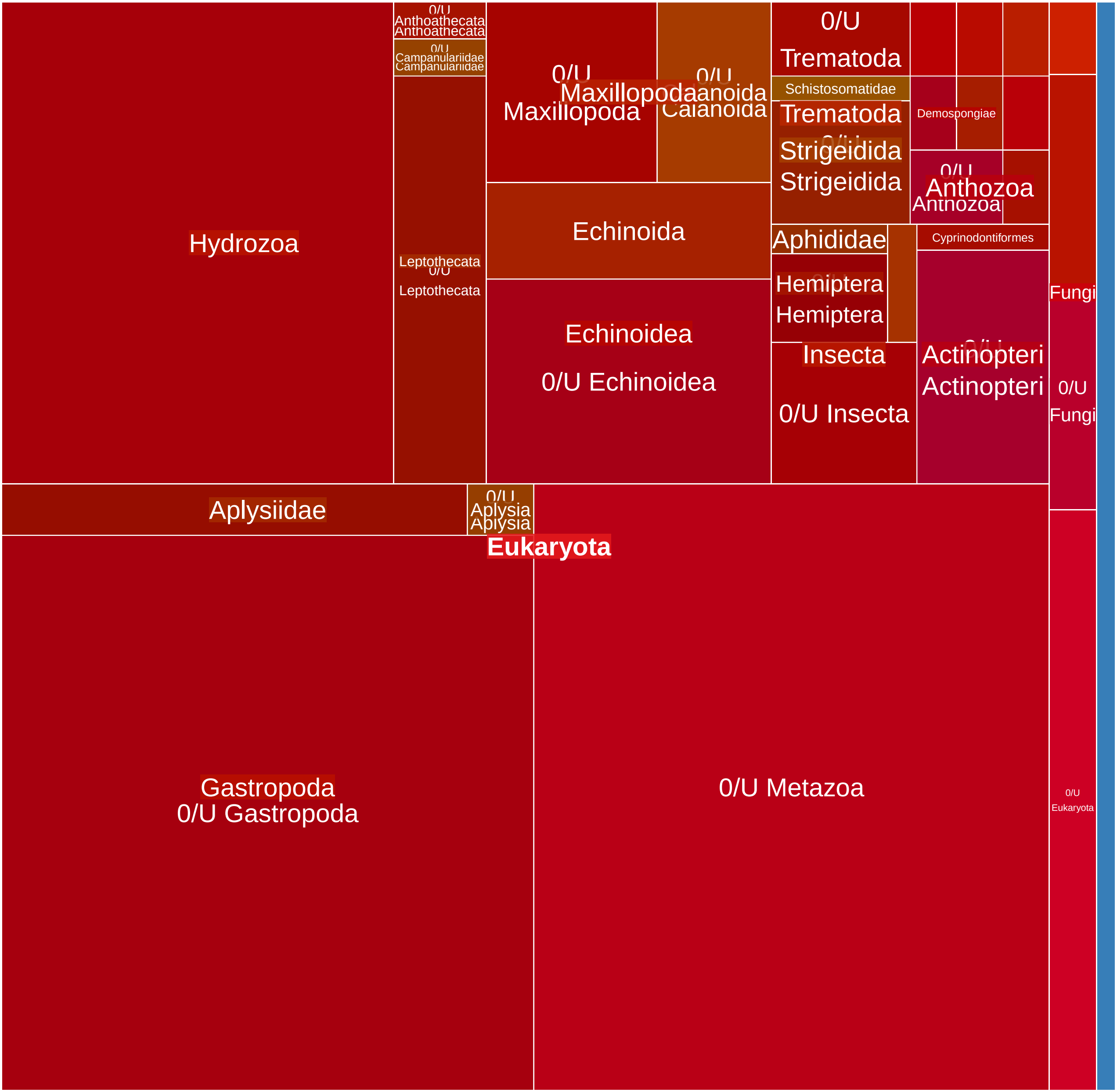

MGT−v1_441 : 2119 unigenes, 286 (13.50%) taxonomically assigned

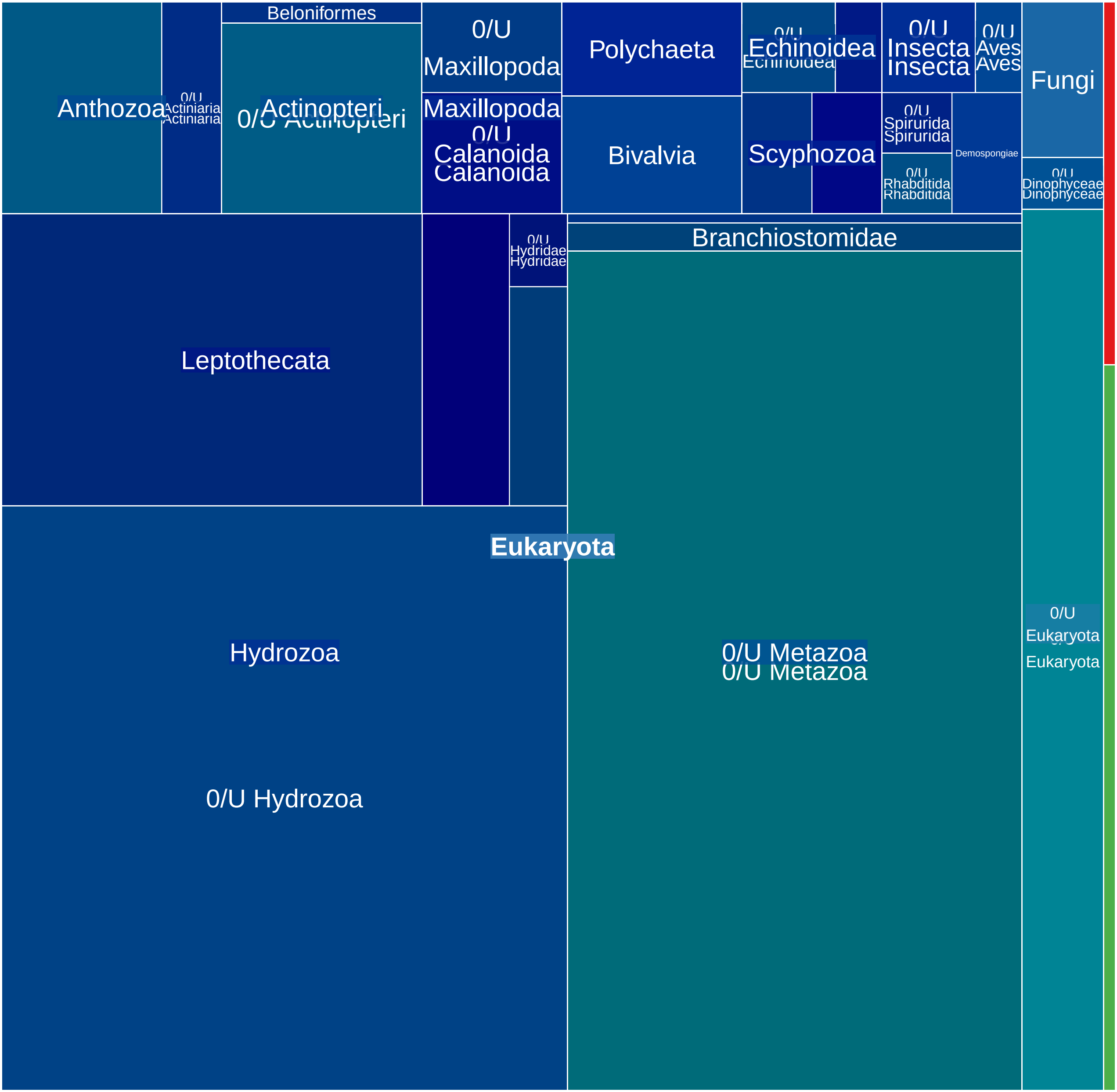

MGT−v1_442 : 2098 unigenes, 1833 (87.37%) taxonomically assigned

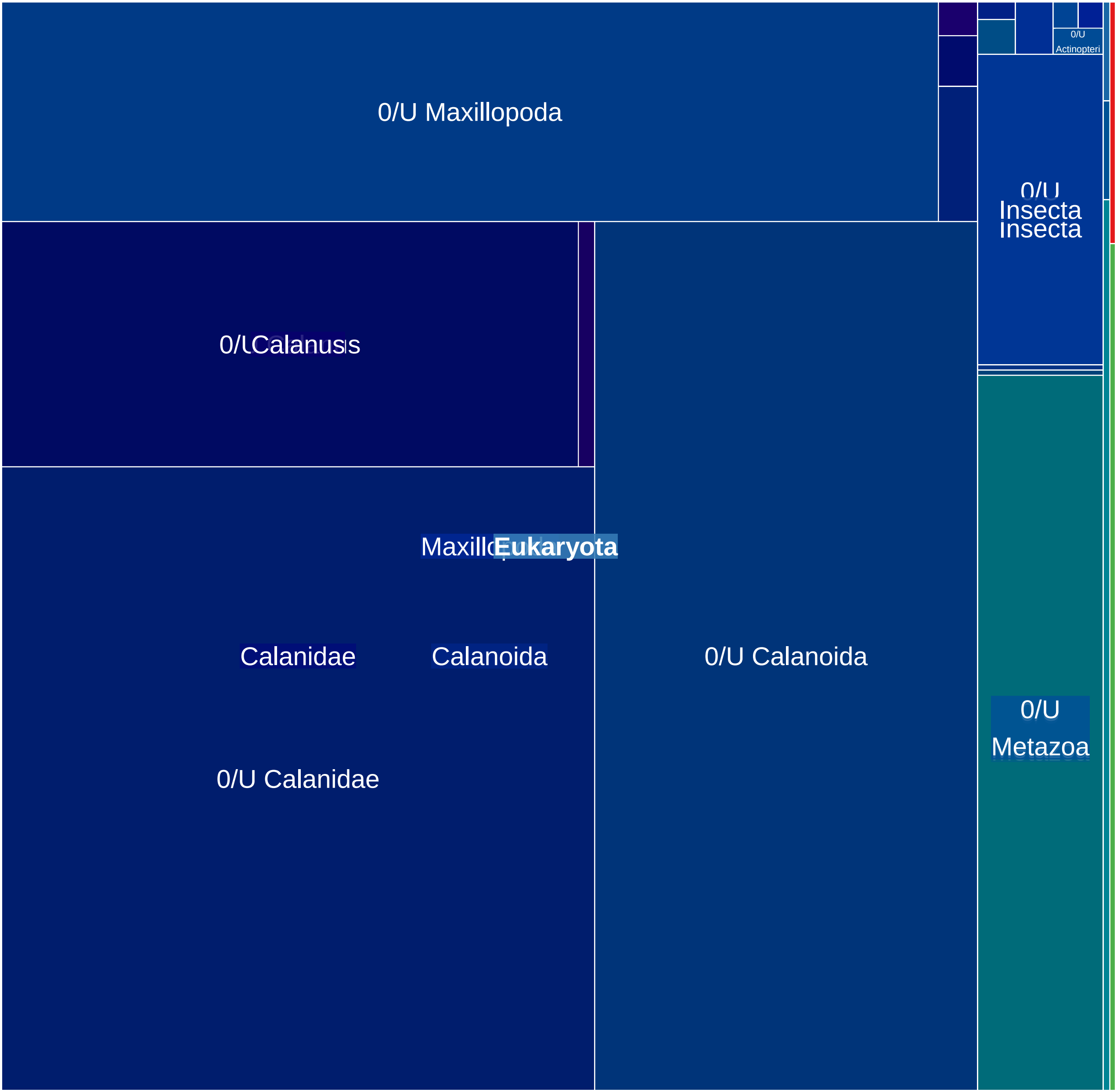

### MGT−v1_443 : 2097 unigenes, 374 (17.84%) taxonomically assigned

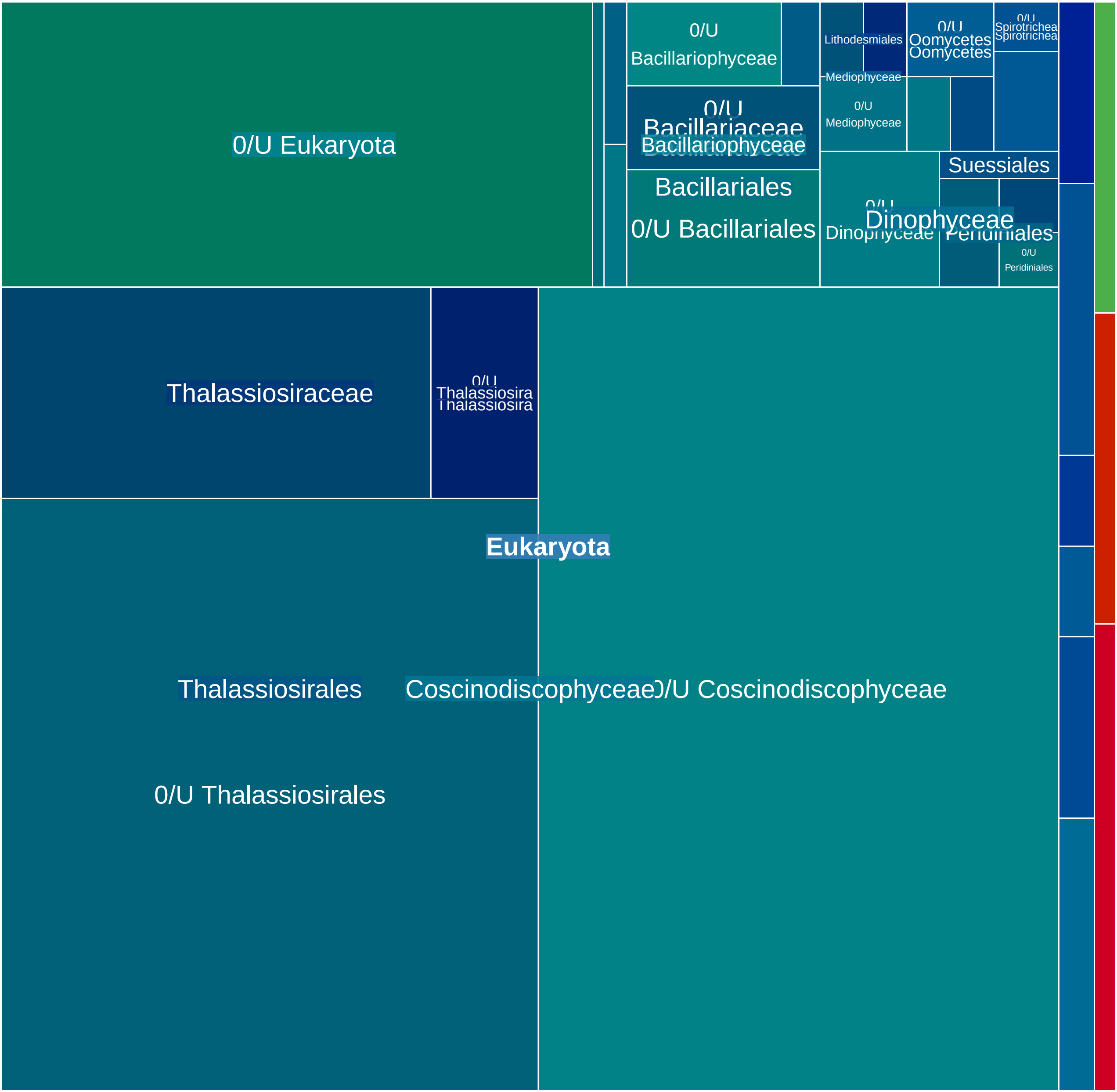

MGT−v1_444 : 2085 unigenes, 1638 (78.56%) taxonomically assigned

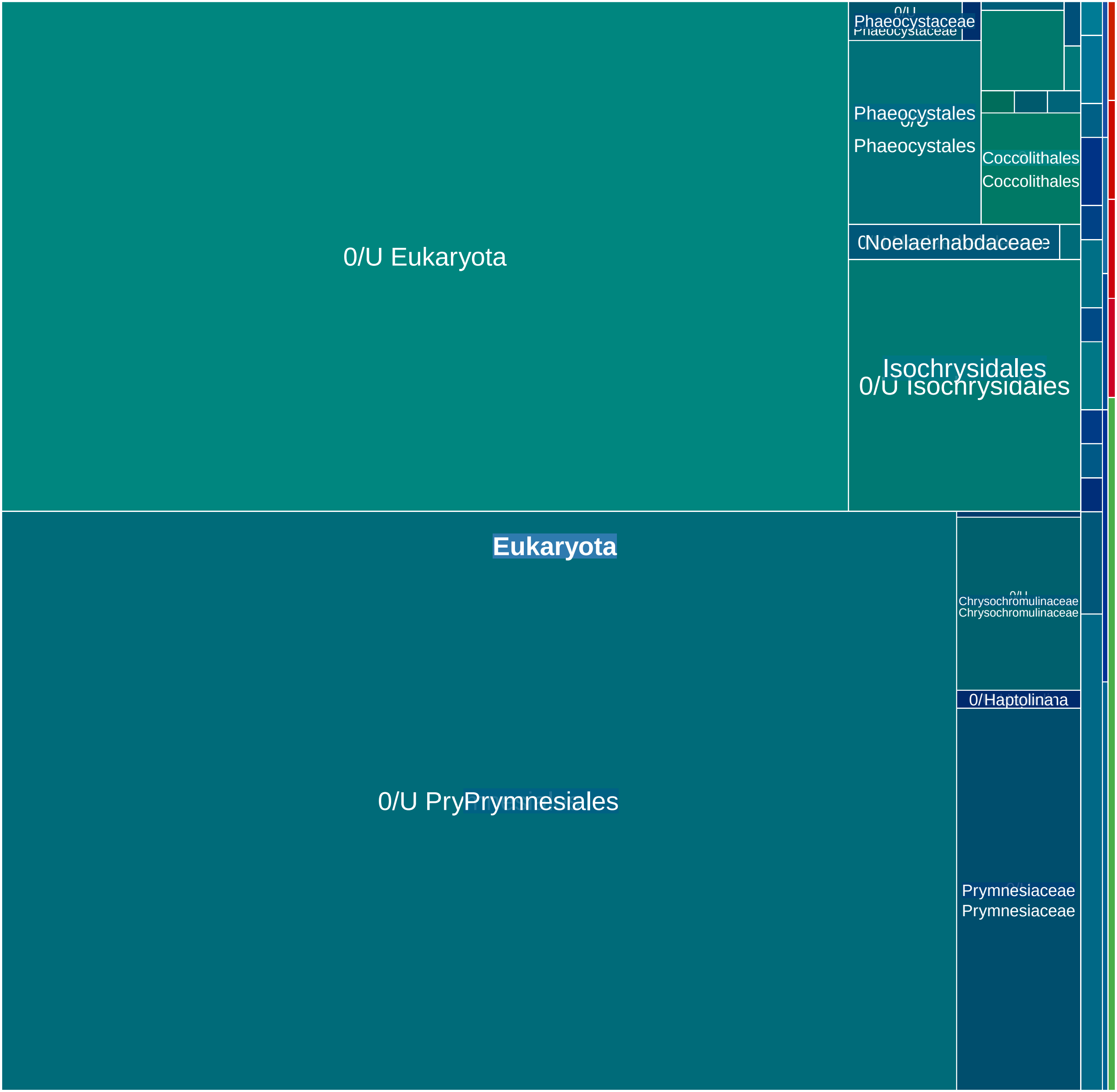

MGT−v1_445 : 2079 unigenes, 216 (10.39%) taxonomically assigned

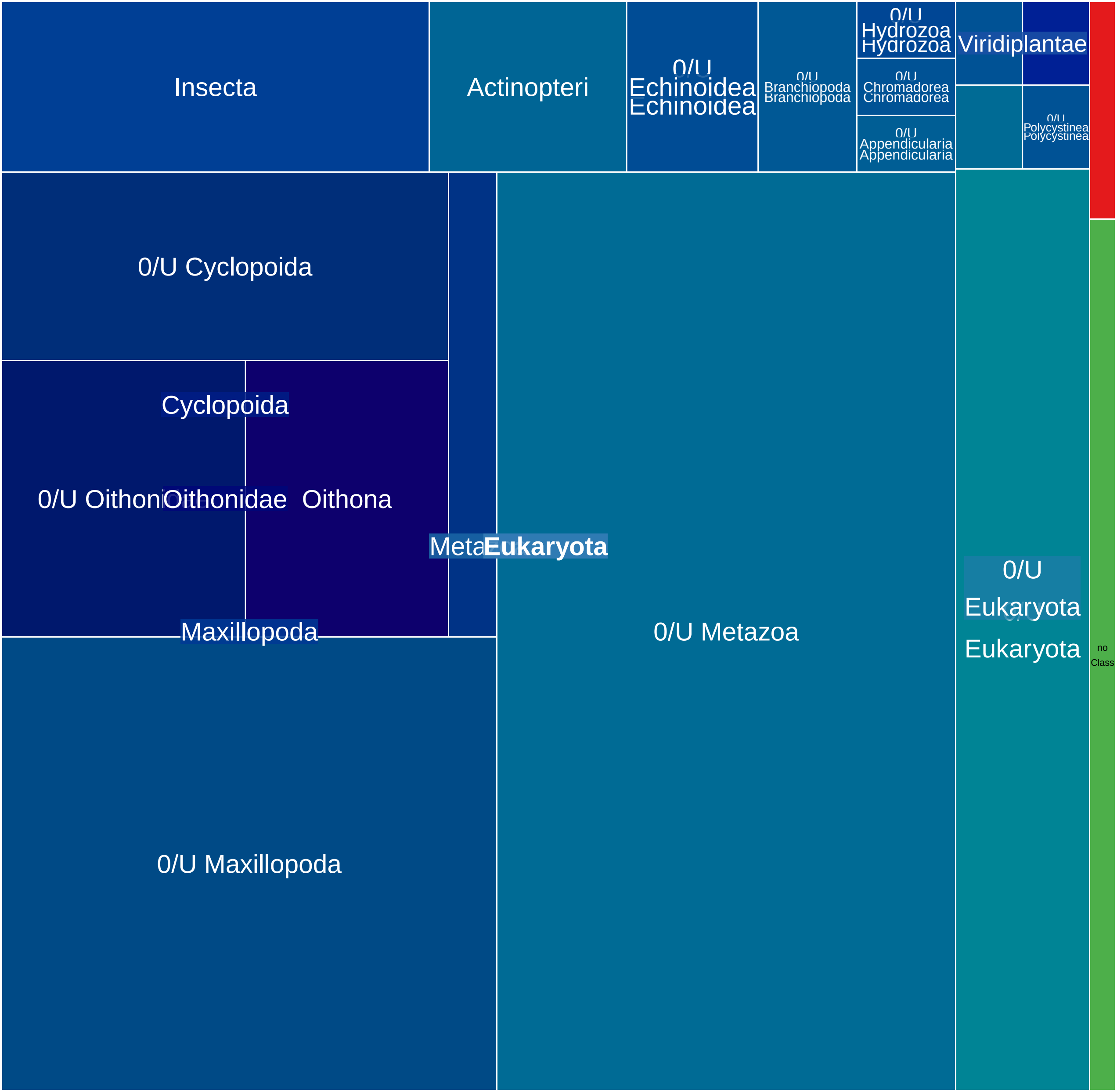

MGT−v1_446 : 2065 unigenes, 1274 (61.69%) taxonomically assigned

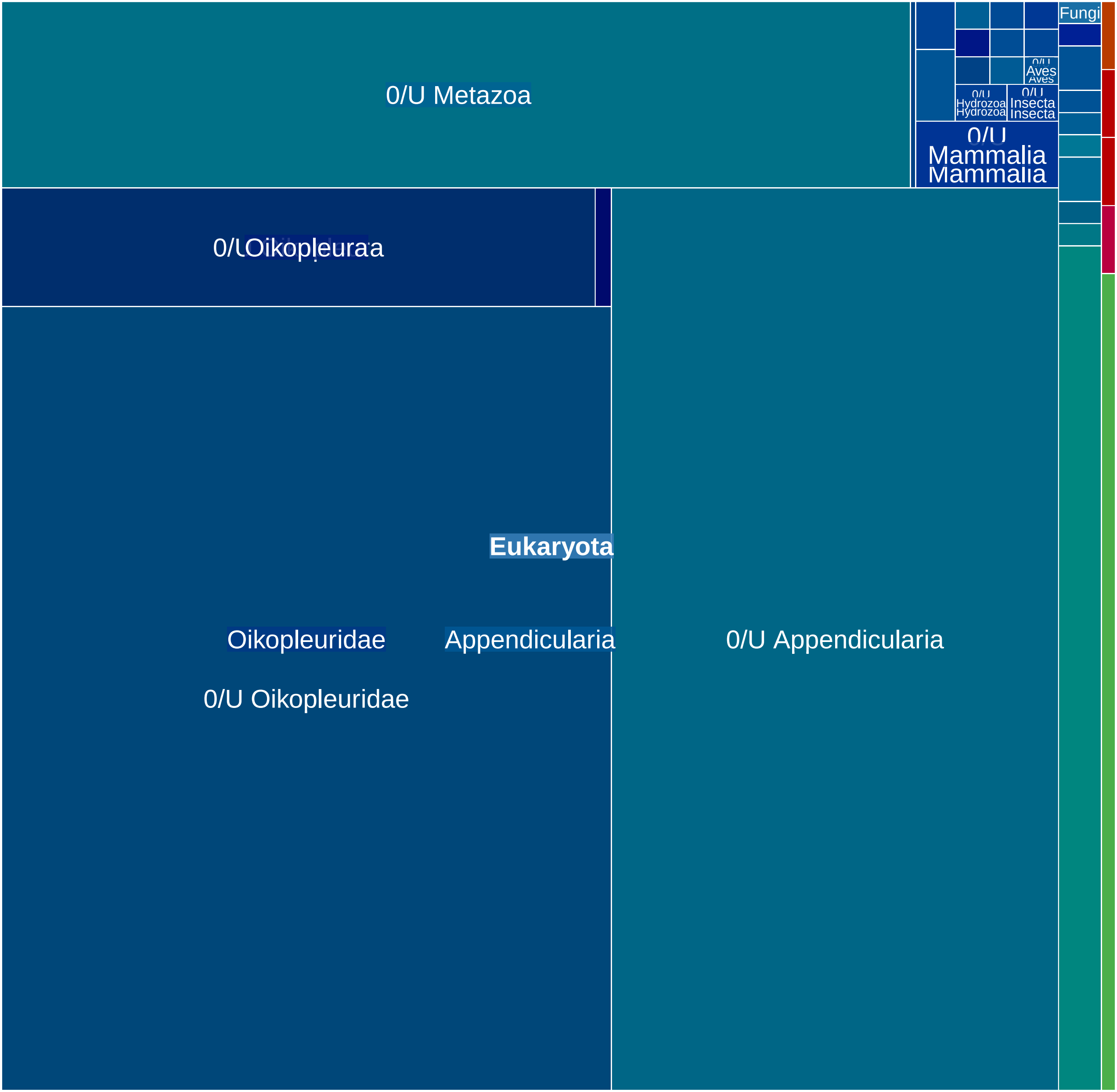

MGT−v1_447 : 2058 unigenes, 1063 (51.65%) taxonomically assigned

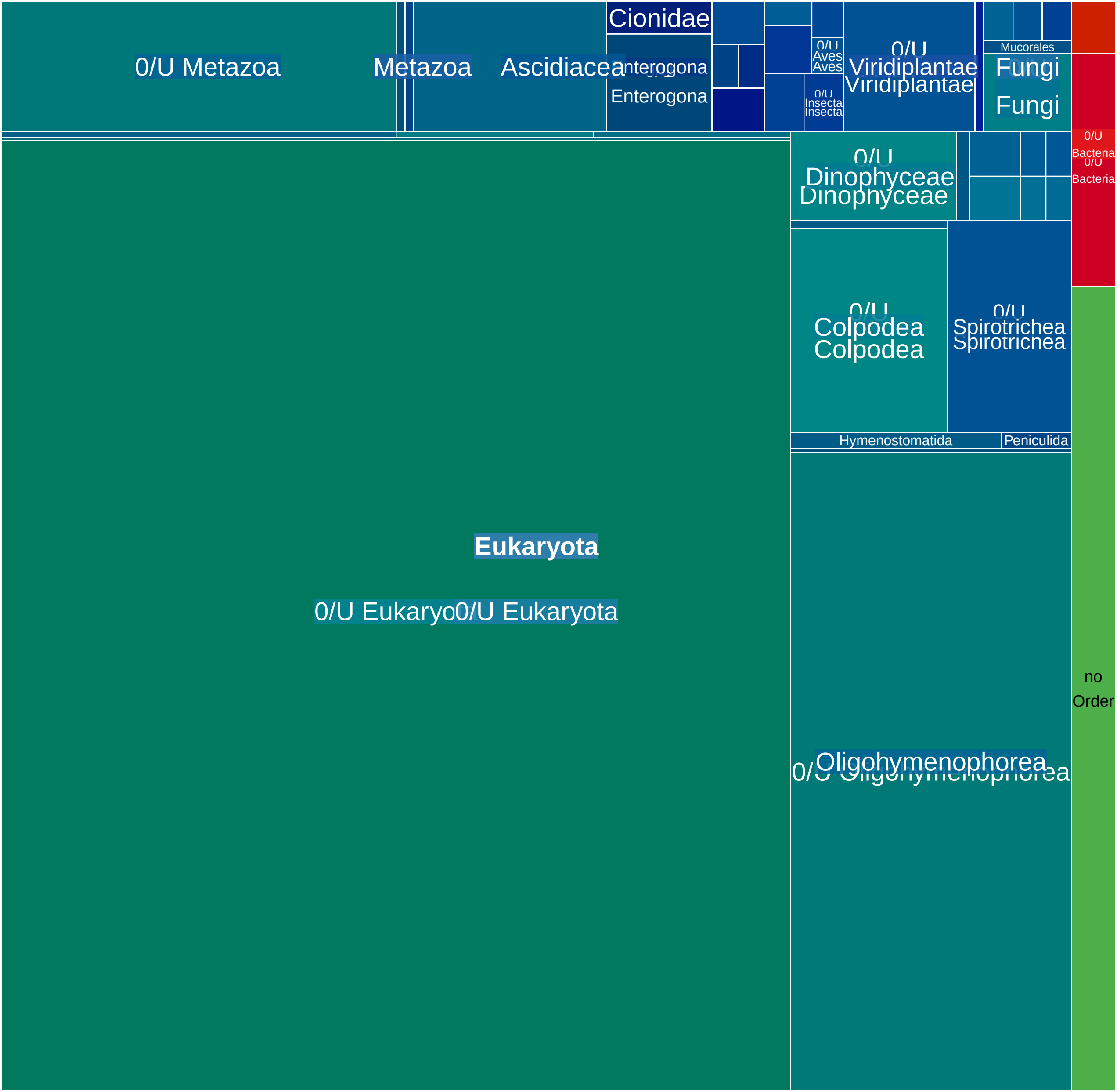

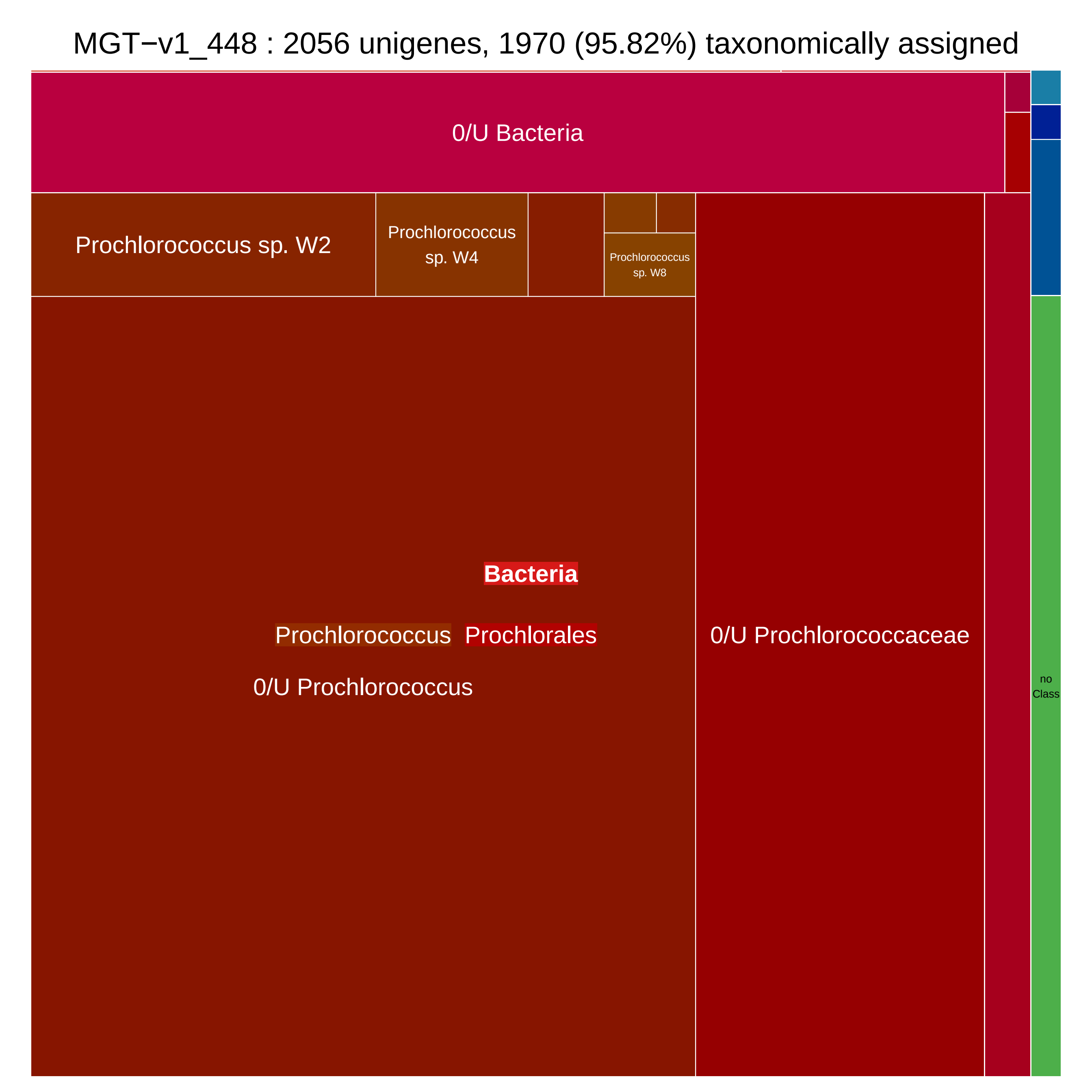
MGT−v1_448 : 2056 unigenes, 1970 (95.82%) taxonomically assigned
0/U Bacteria
Prochlorococcus sp. W2
Prochlorococcus sp. W4
Prochlorococcus sp. W8
Bacteria
Prochlorococcus
Prochlorales
0/U Prochlorococcus
0/U Prochlorococcaceae
no Class

MGT−v1_449 : 2040 unigenes, 1912 (93.73%) taxonomically assigned

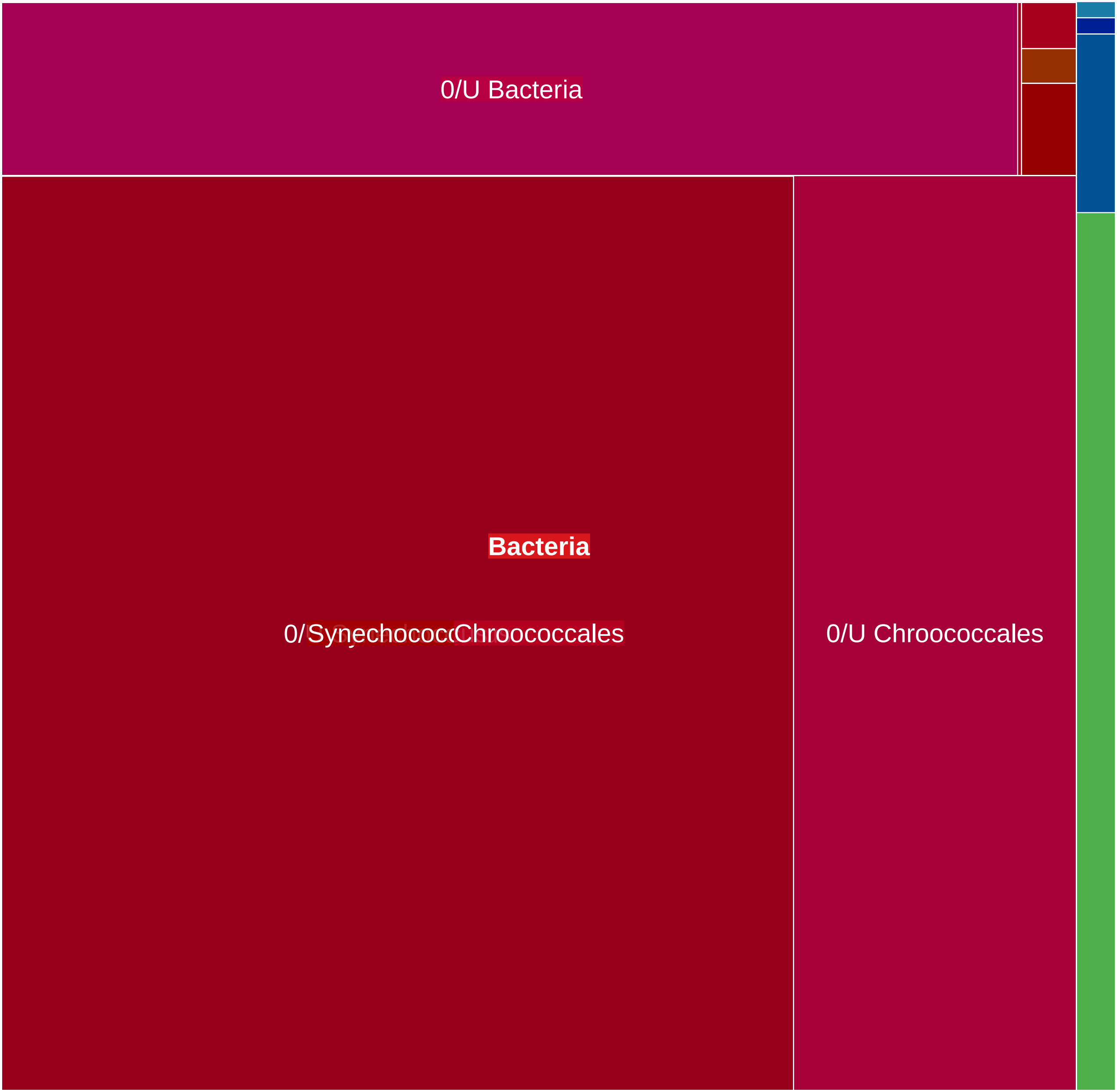

MGT−v1_450 : 2026 unigenes, 169 (8.34%) taxonomically assigned

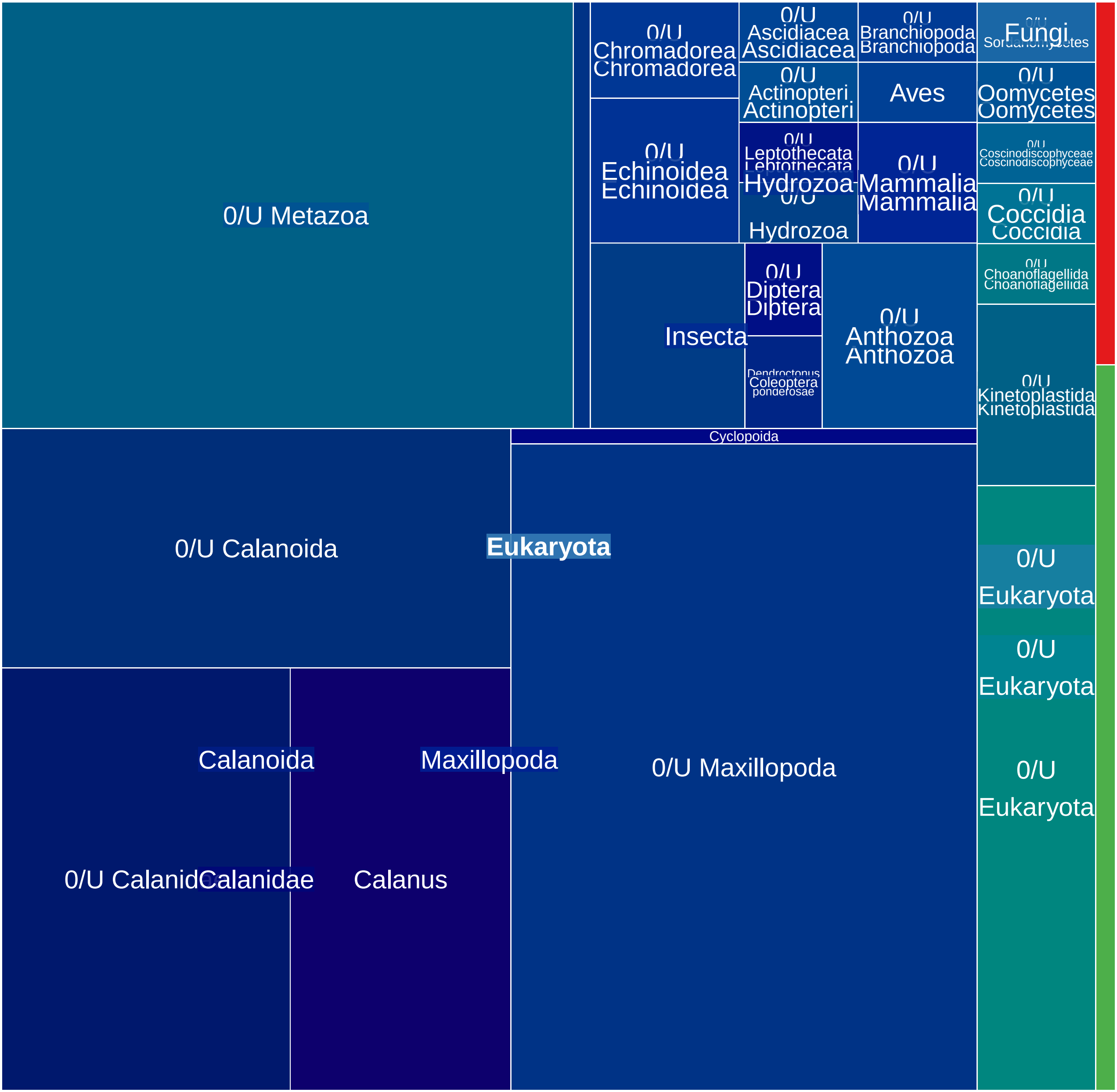

### MGT−v1_451 : 2021 unigenes, 1595 (78.92%) taxonomically assigned

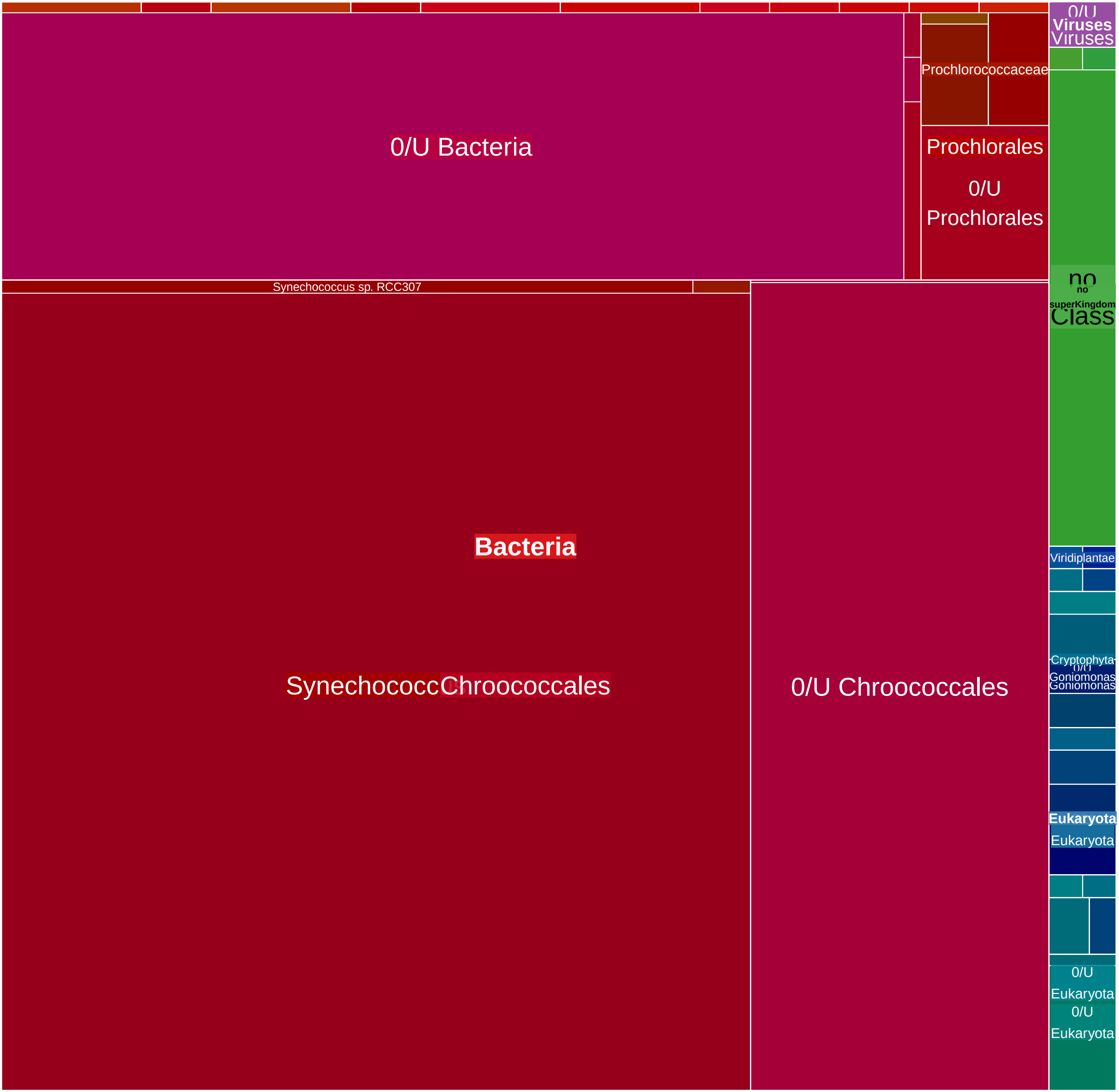

MGT−v1_452 : 2013 unigenes, 195 (9.69%) taxonomically assigned

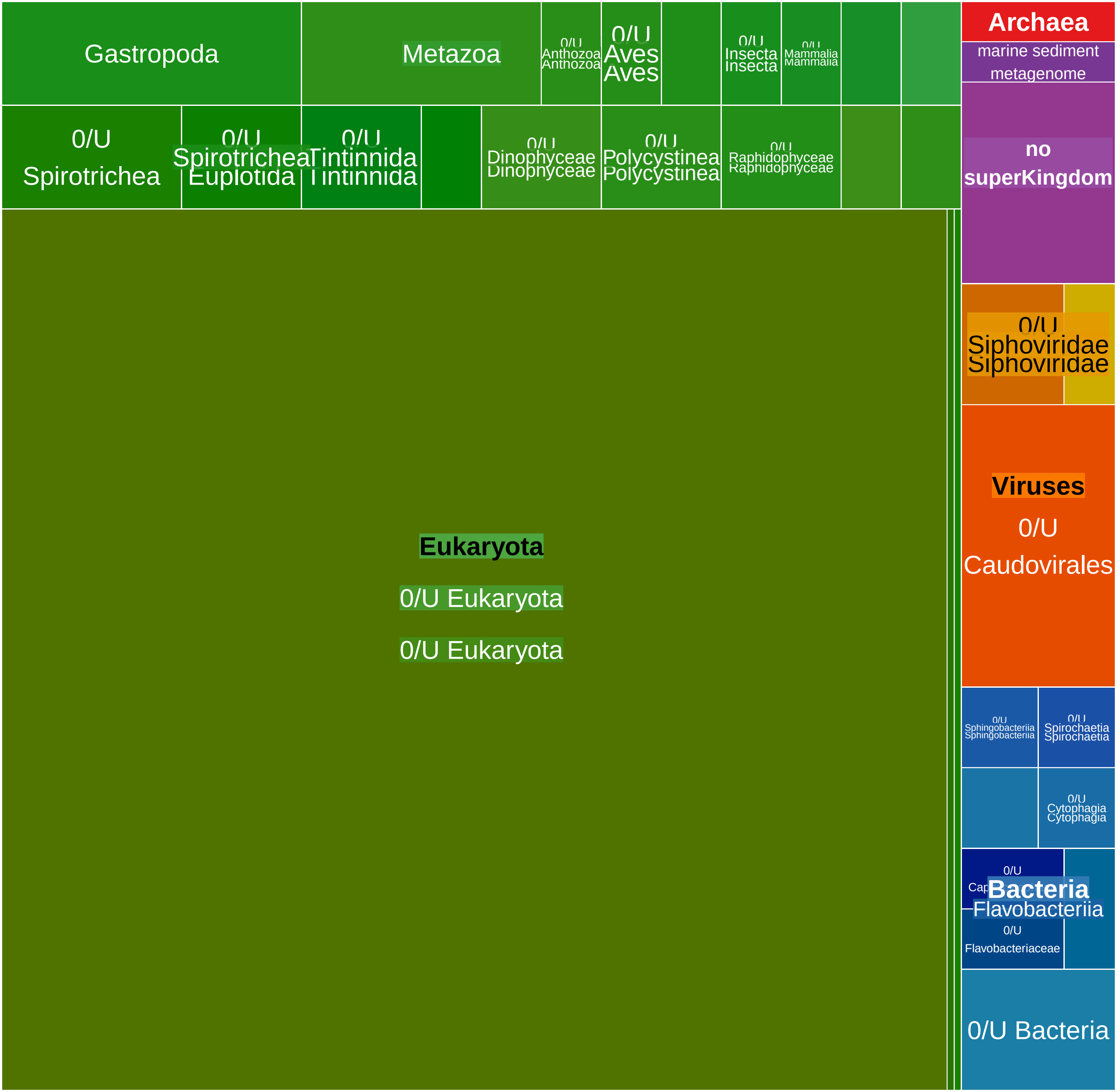

### MGT−v1_453 : 1991 unigenes, 211 (10.60%) taxonomically assigned

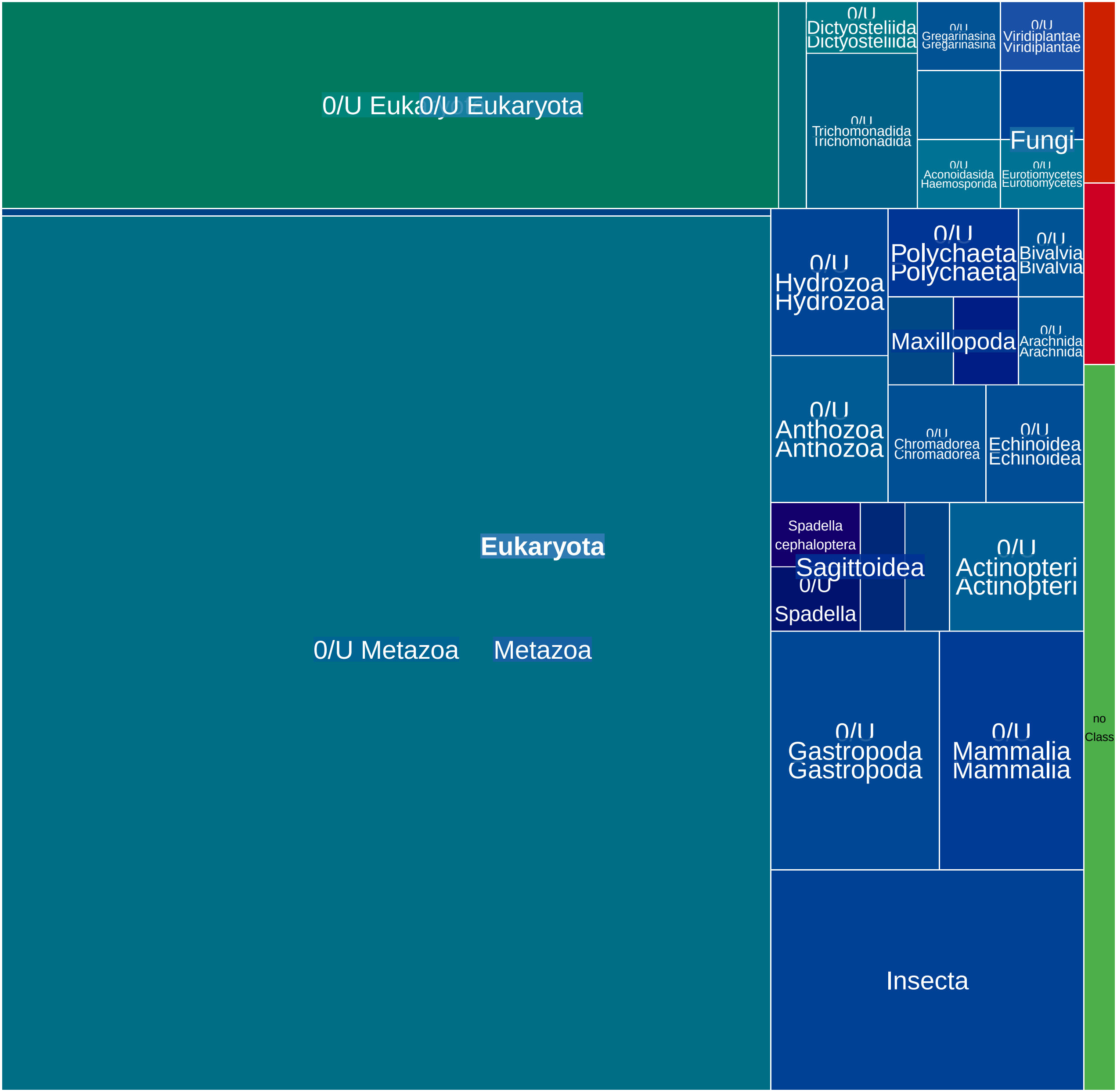

MGT−v1_454 : 1989 unigenes, 1949 (97.99%) taxonomically assigned

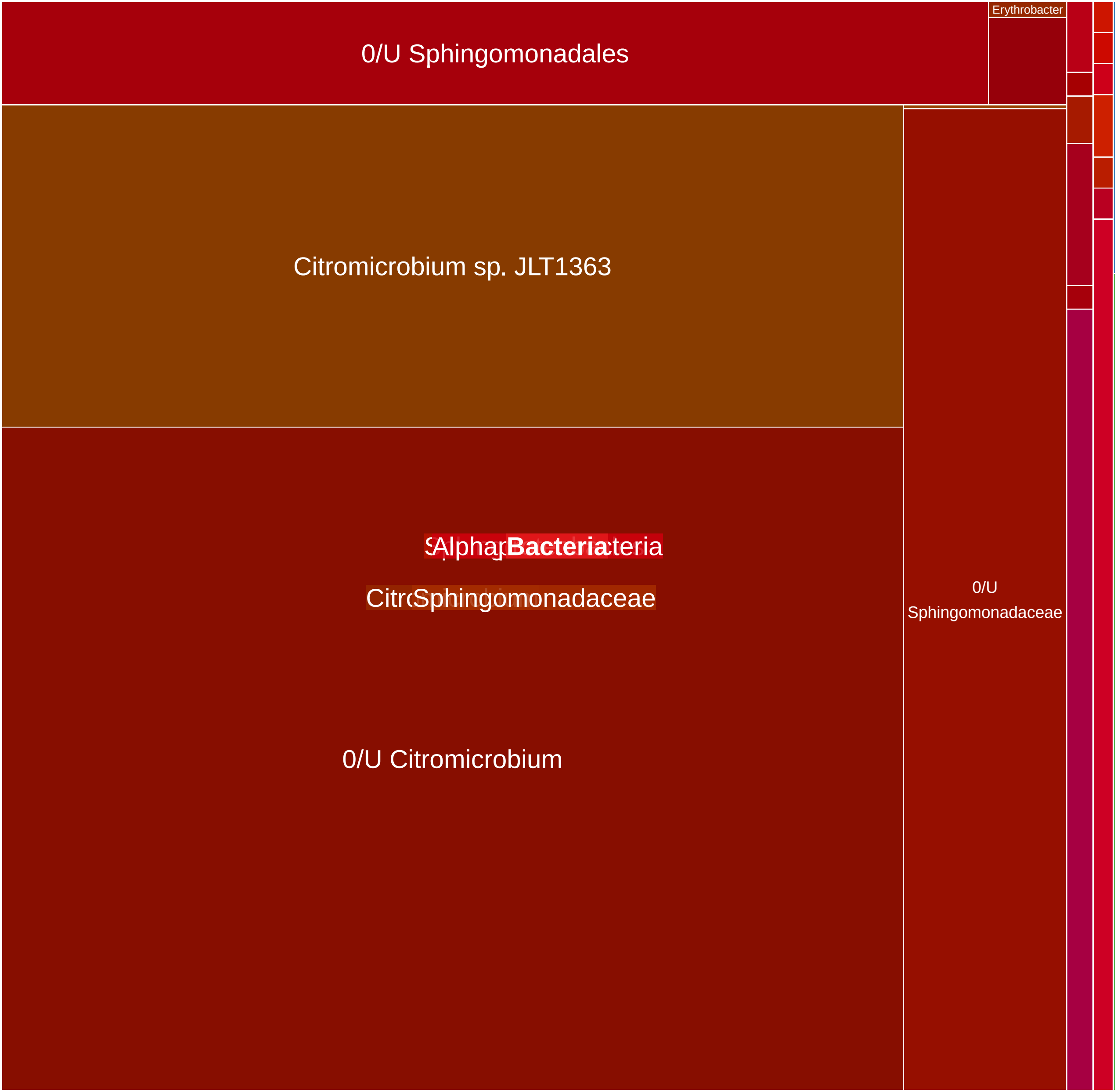

MGT−v1_455 : 1986 unigenes, 296 (14.90%) taxonomically assigned

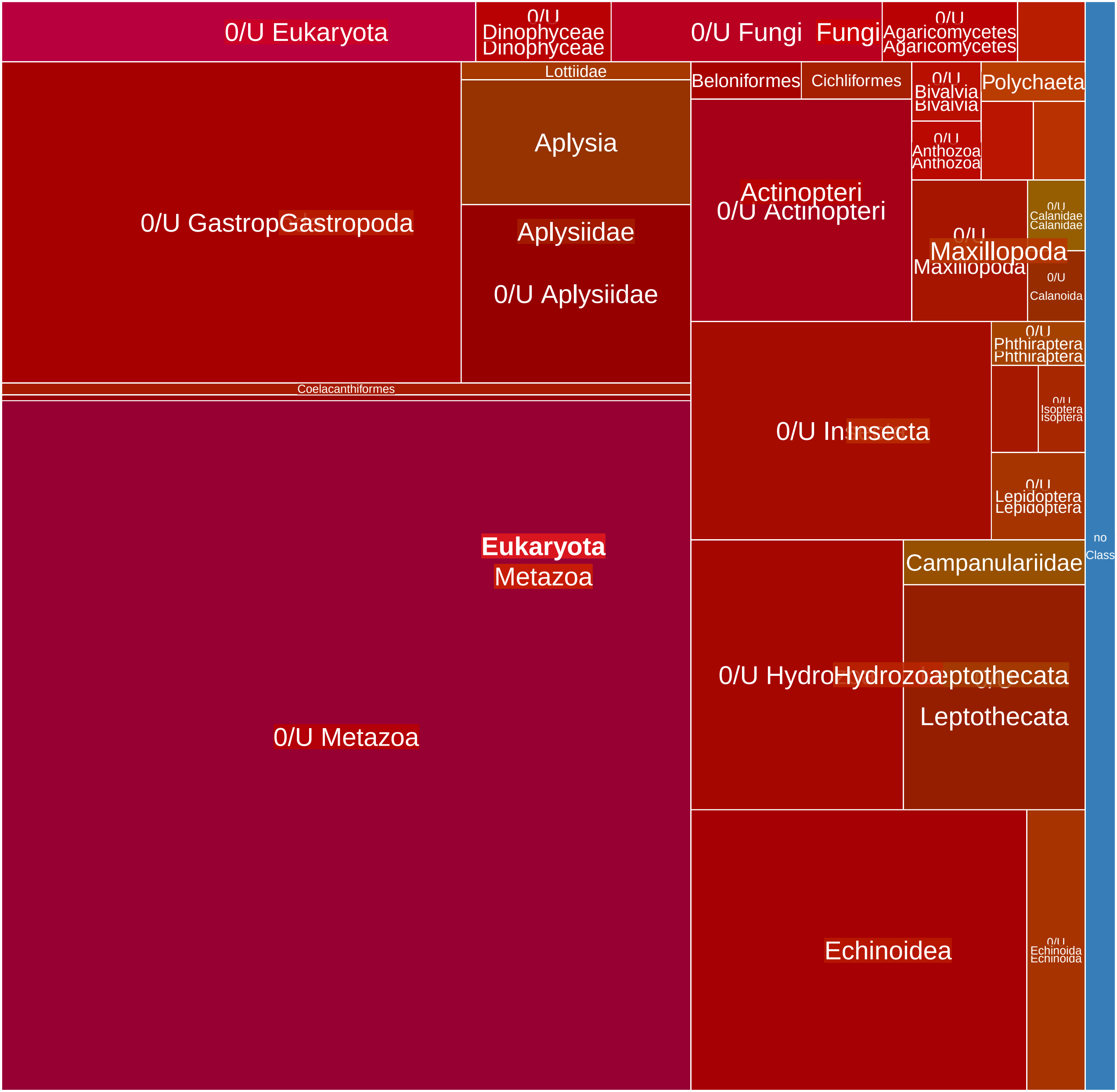

### MGT−v1_456 : 1984 unigenes, 114 (5.75%) taxonomically assigned

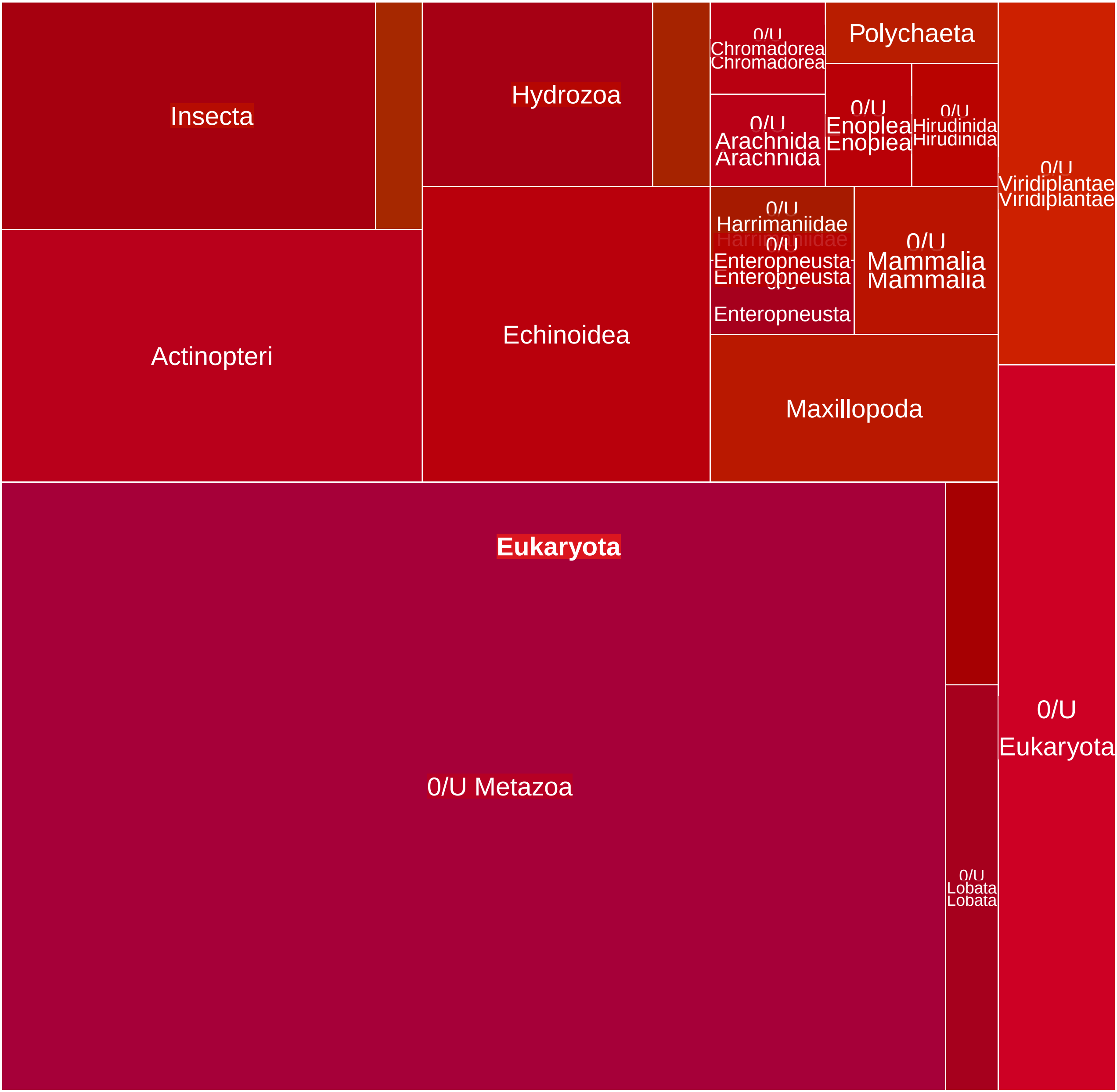

MGT−v1_457 : 1983 unigenes, 151 (7.61%) taxonomically assigned

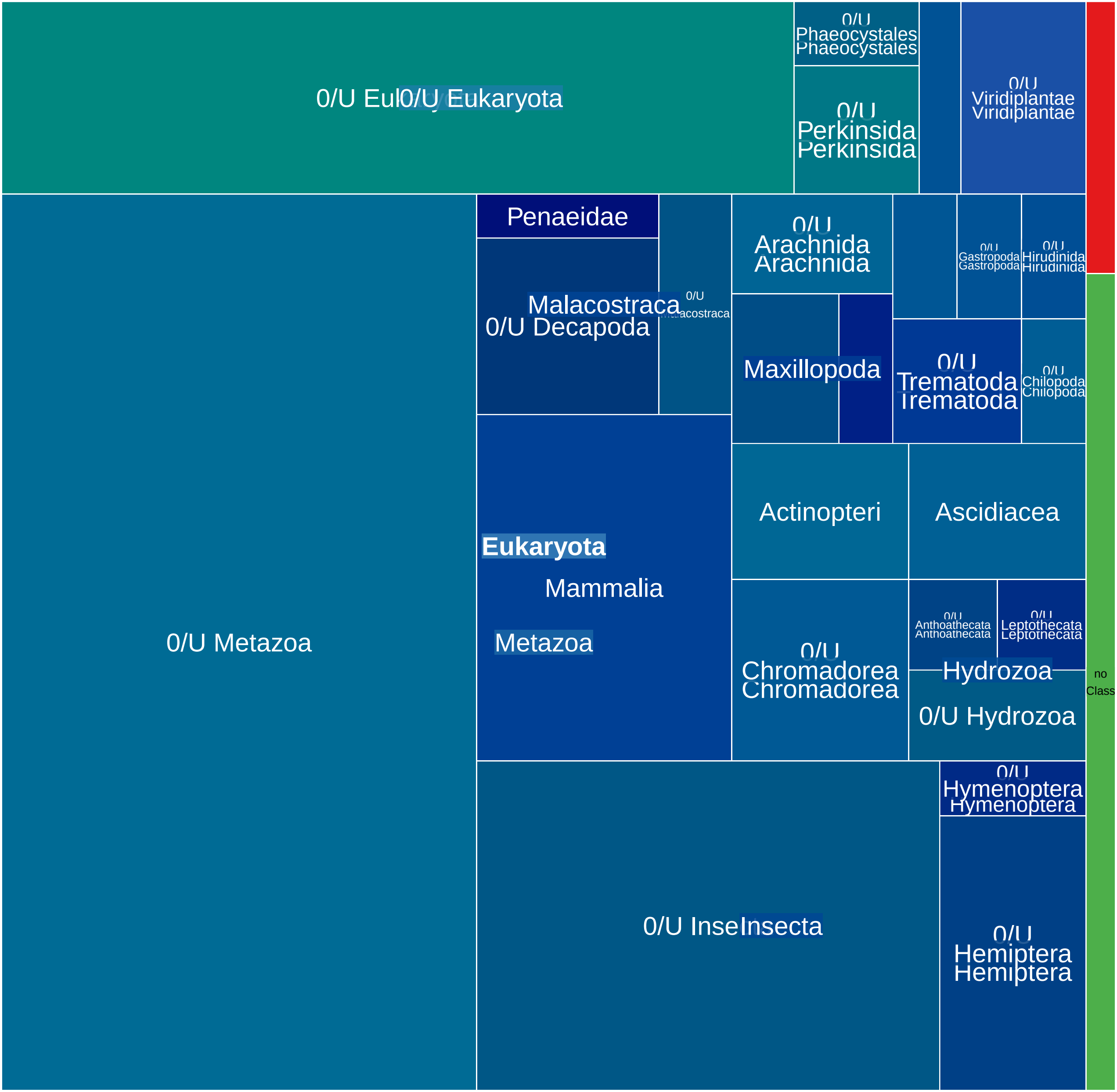

MGT−v1_458 : 1981 unigenes, 133 (6.71%) taxonomically assigned

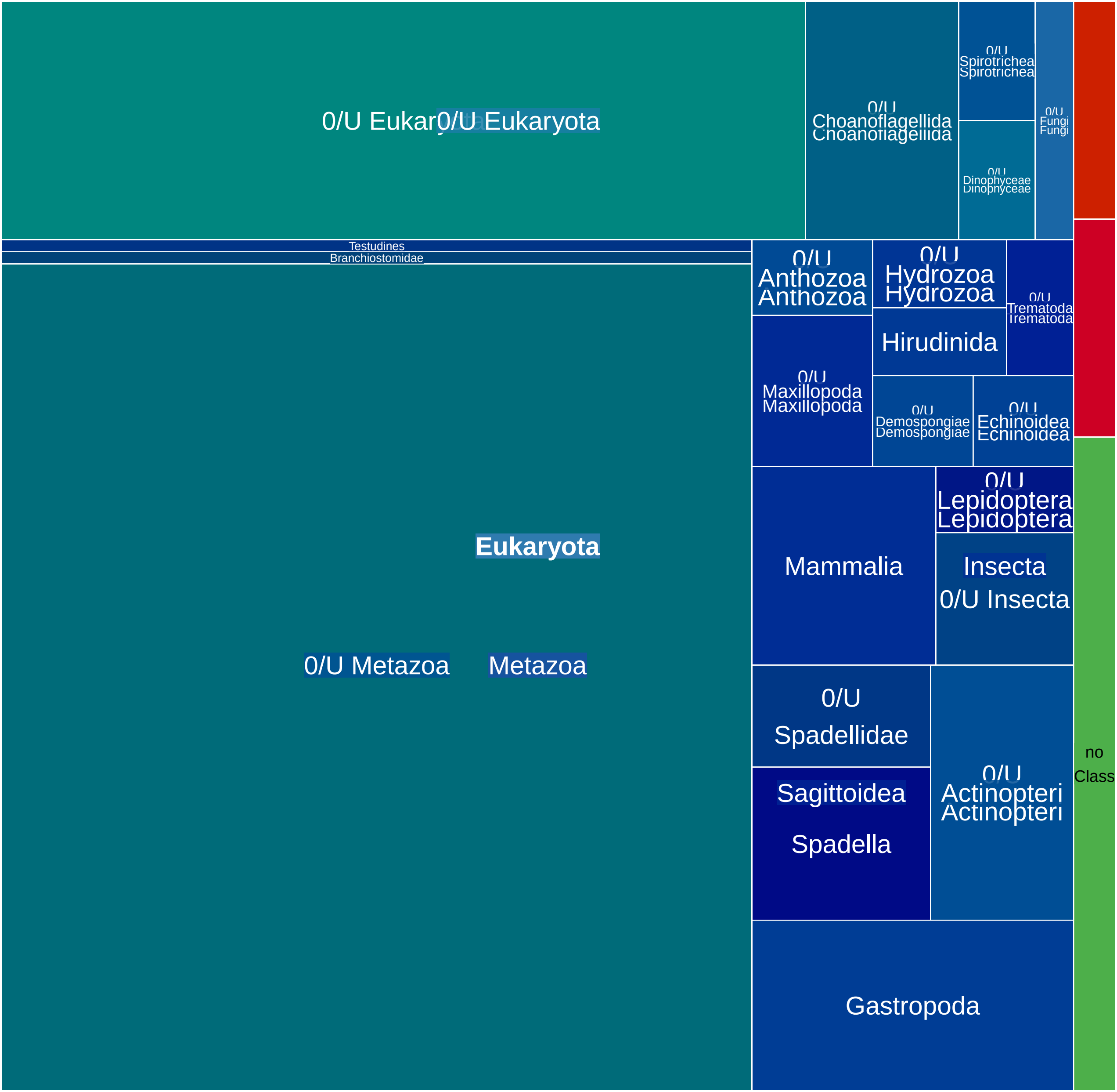

MGT−v1_459 : 1980 unigenes, 1789 (90.35%) taxonomically assigned

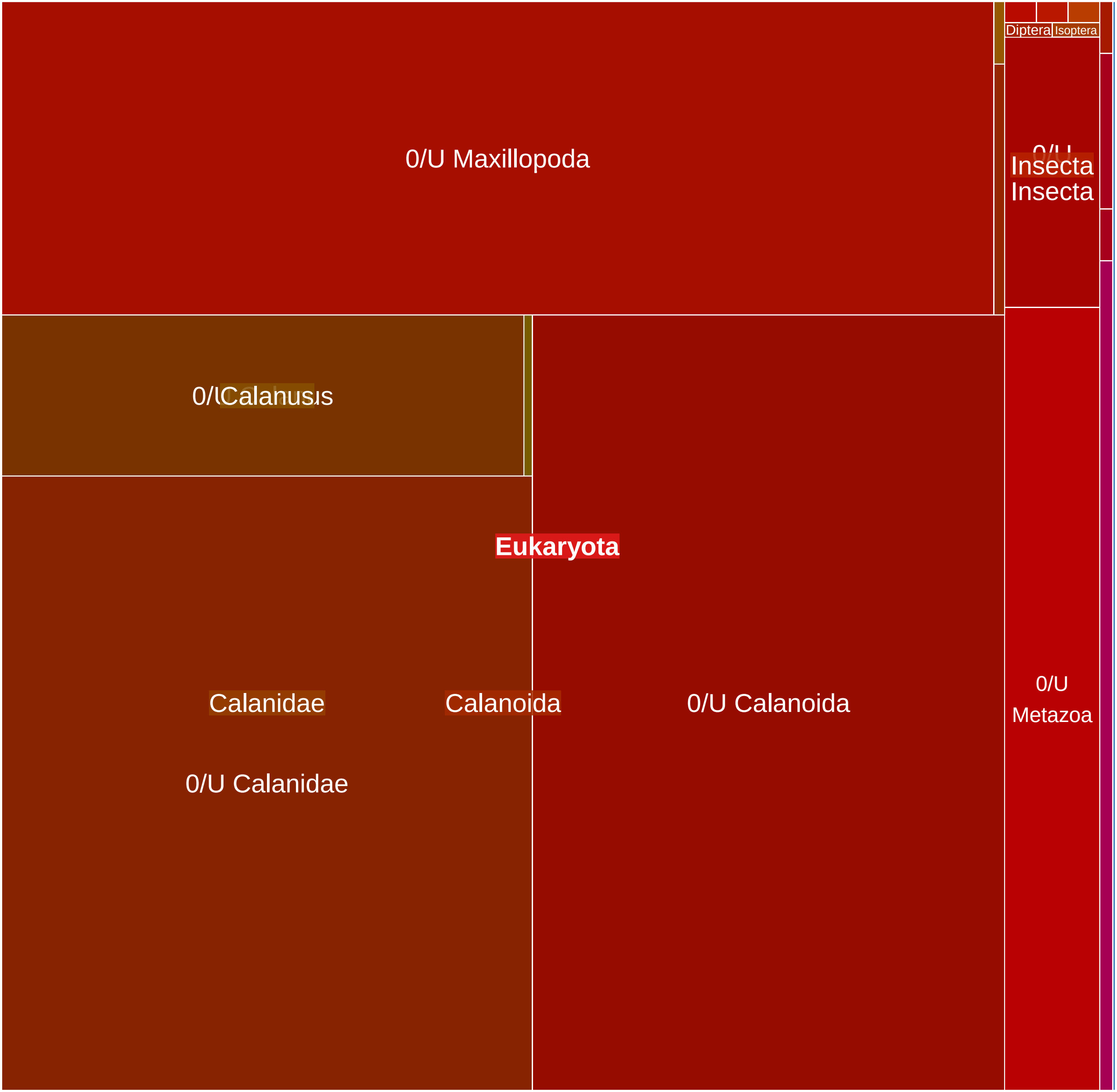

MGT−v1_460 : 1979 unigenes, 1842 (93.08%) taxonomically assigned

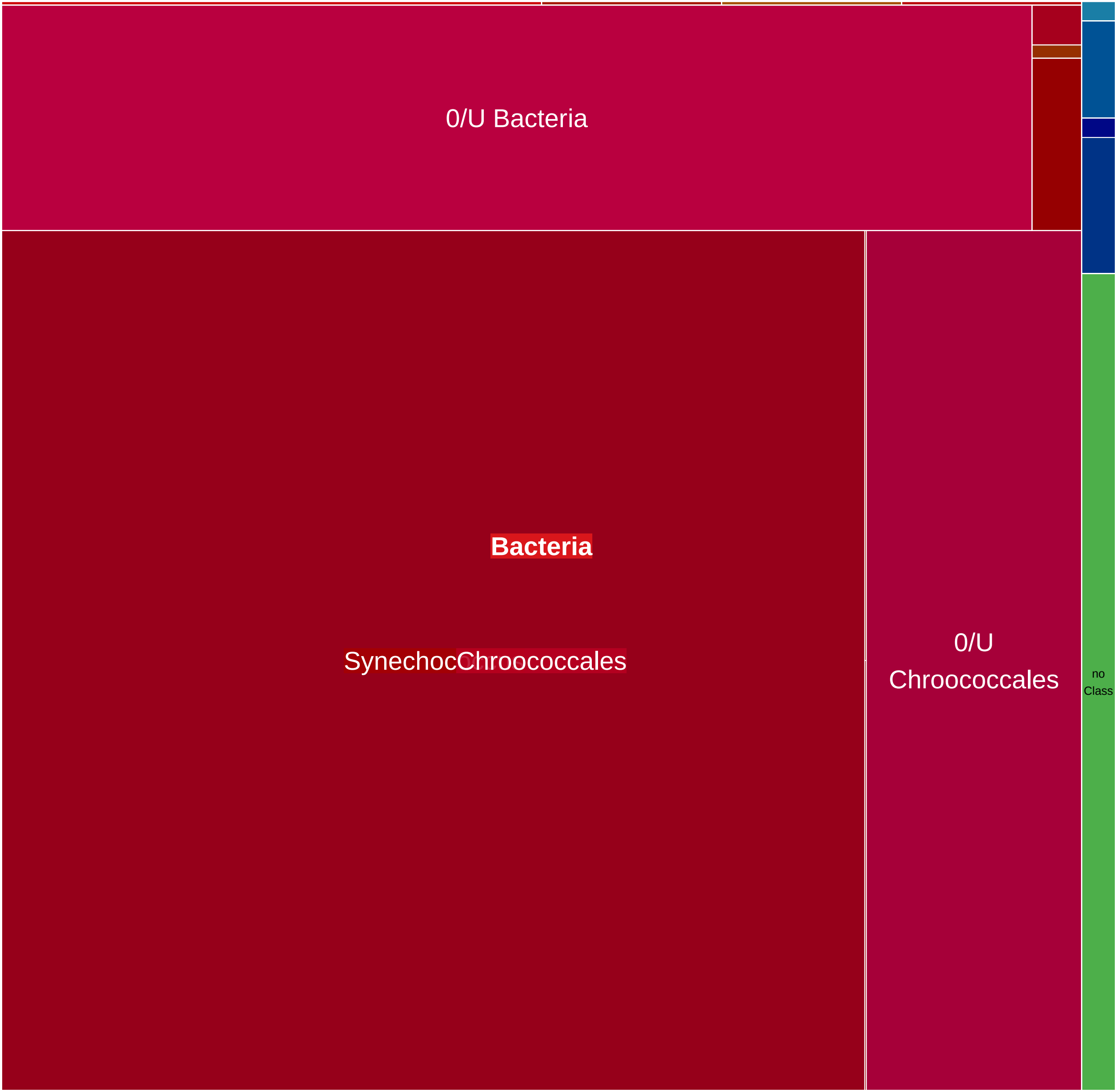

MGT−v1_461 : 1965 unigenes, 1767 (89.92%) taxonomically assigned

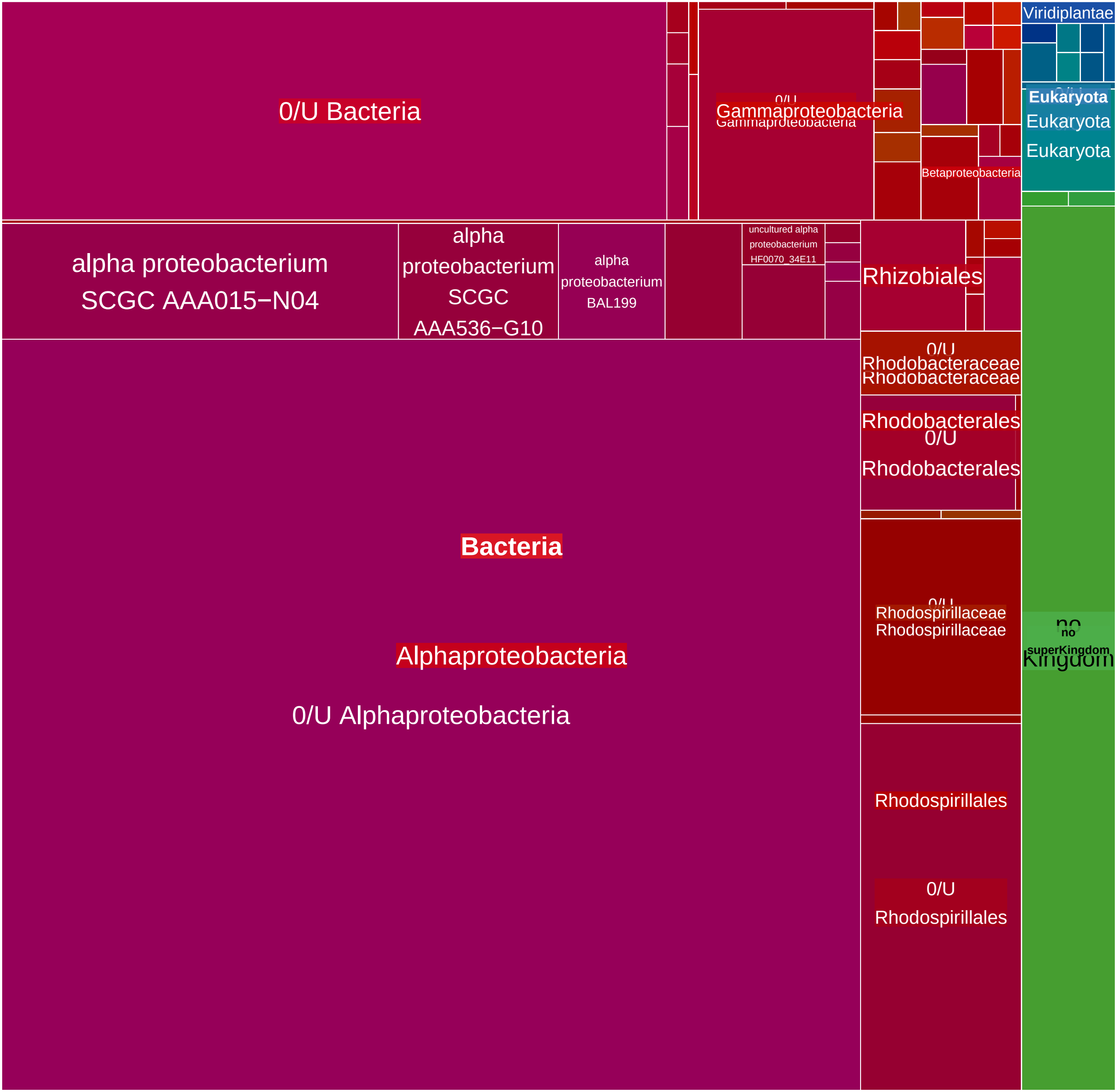

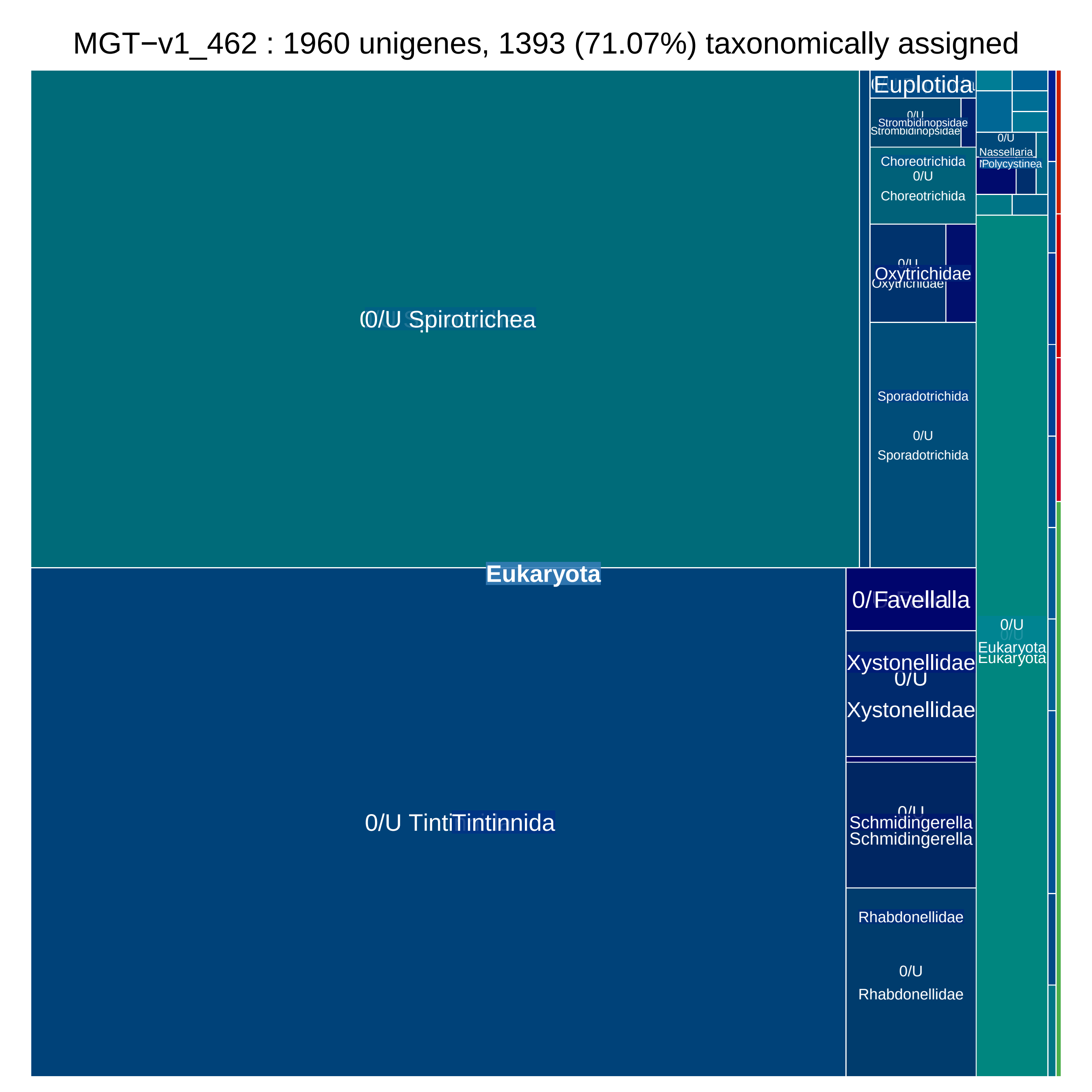
MGT−v1_462 : 1960 unigenes, 1393 (71.07%) taxonomically assigned
0/U Spirotrichea
Euplotida
0/U
Strombidinopsidae
Strombidinopsidae
Choreotrichida
0/U
Choreotrichida
0/U
Oxytrichidae
Oxytrichidae
Sporadotrichida
0/U
Sporadotrichida
0/U
Nassellaria
Polycystinea
Eukaryota
0/U Tintinnida
0/Favellalla
Xystonellidae
0/U
Xystonellidae
0/U
Schmidingerella
Schmidingerella
Rhabdonellidae
0/U
Rhabdonellidae
0/U
Eukaryota

### MGT−v1_463 : 1959 unigenes, 1626 (83.00%) taxonomically assigned

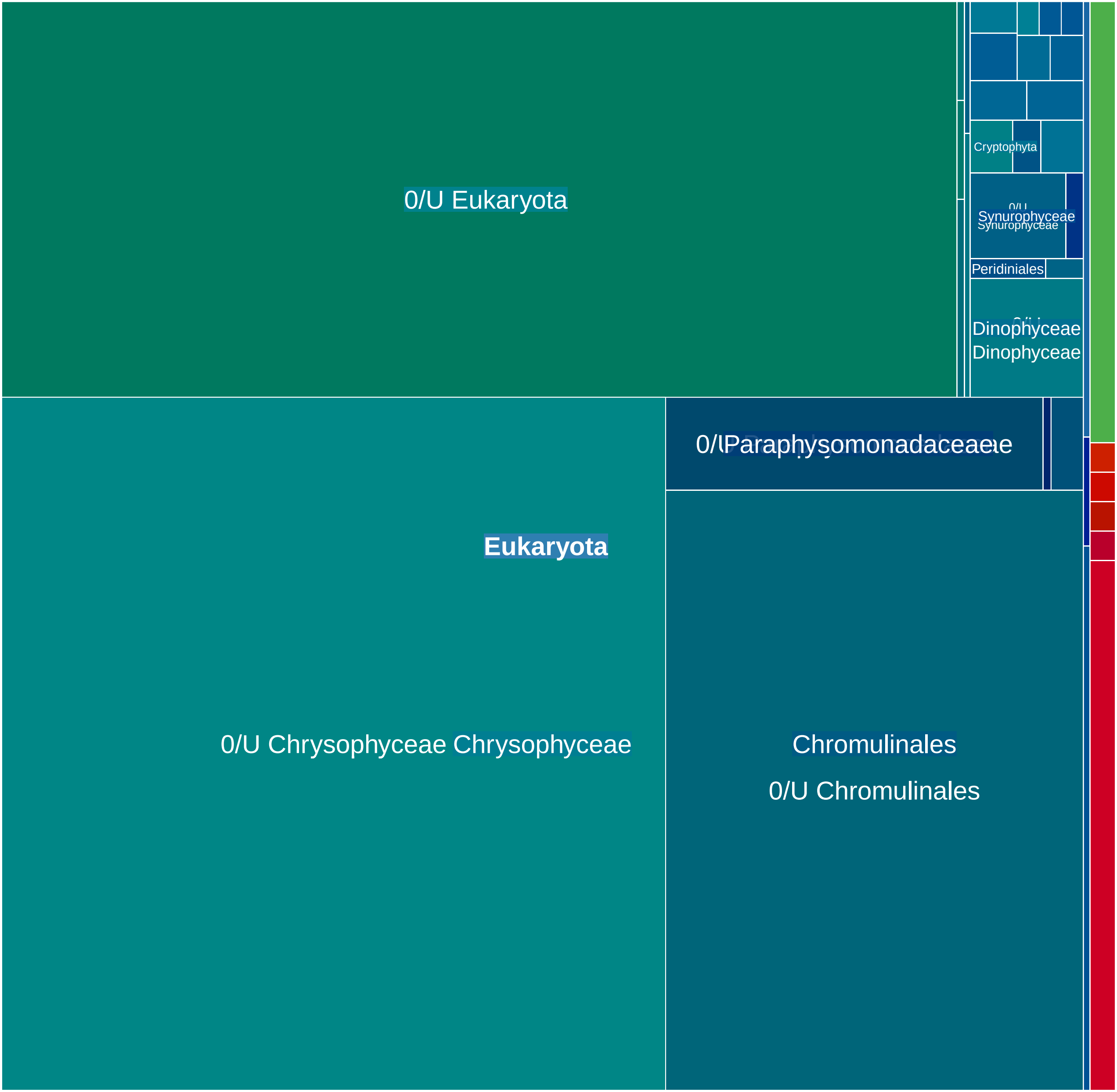

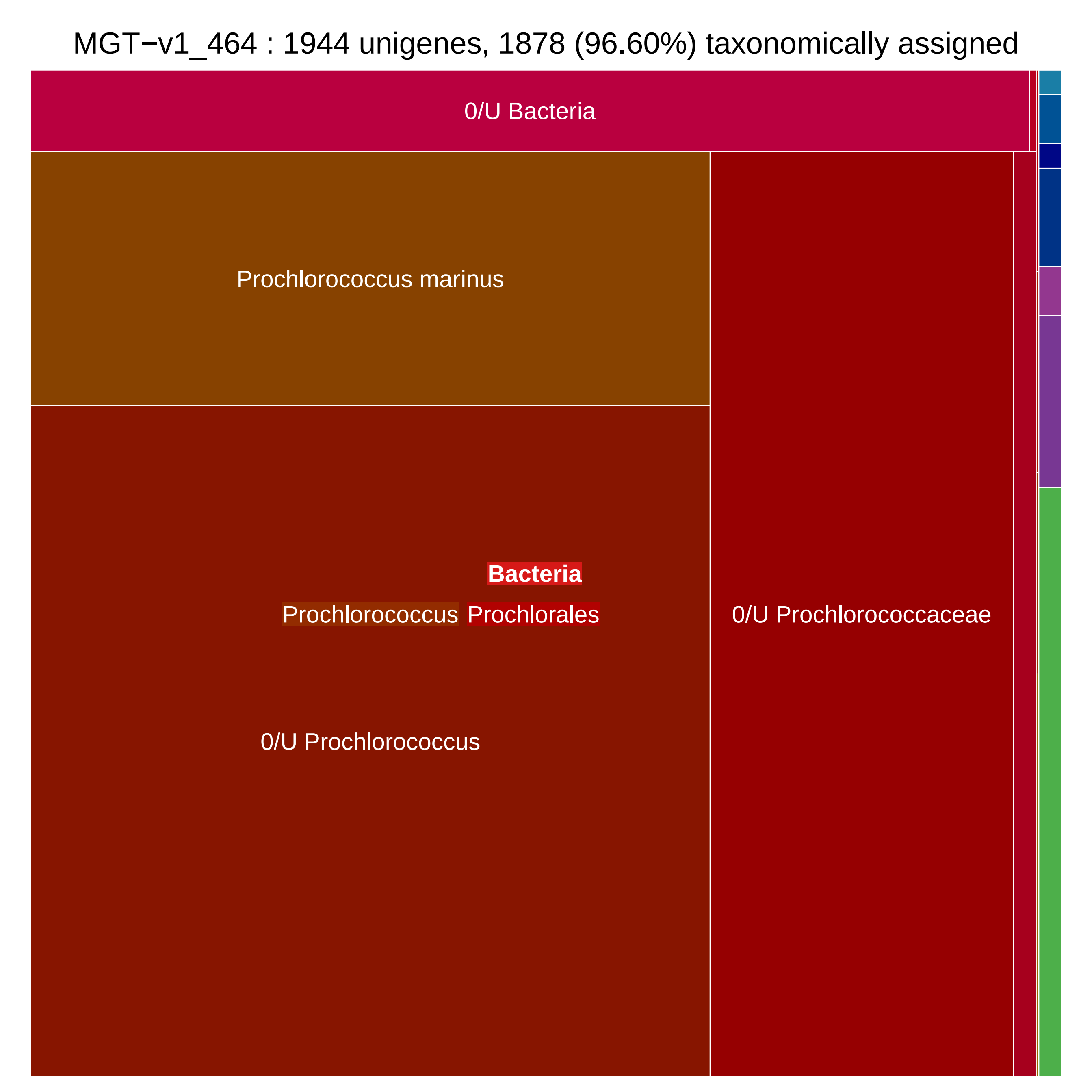
MGT−v1_464 : 1944 unigenes, 1878 (96.60%) taxonomically assigned
0/U Bacteria
Prochlorococcus marinus
Bacteria
Prochlorococcus
Prochlorales
0/U Prochlorococcaceae
0/U Prochlorococcus

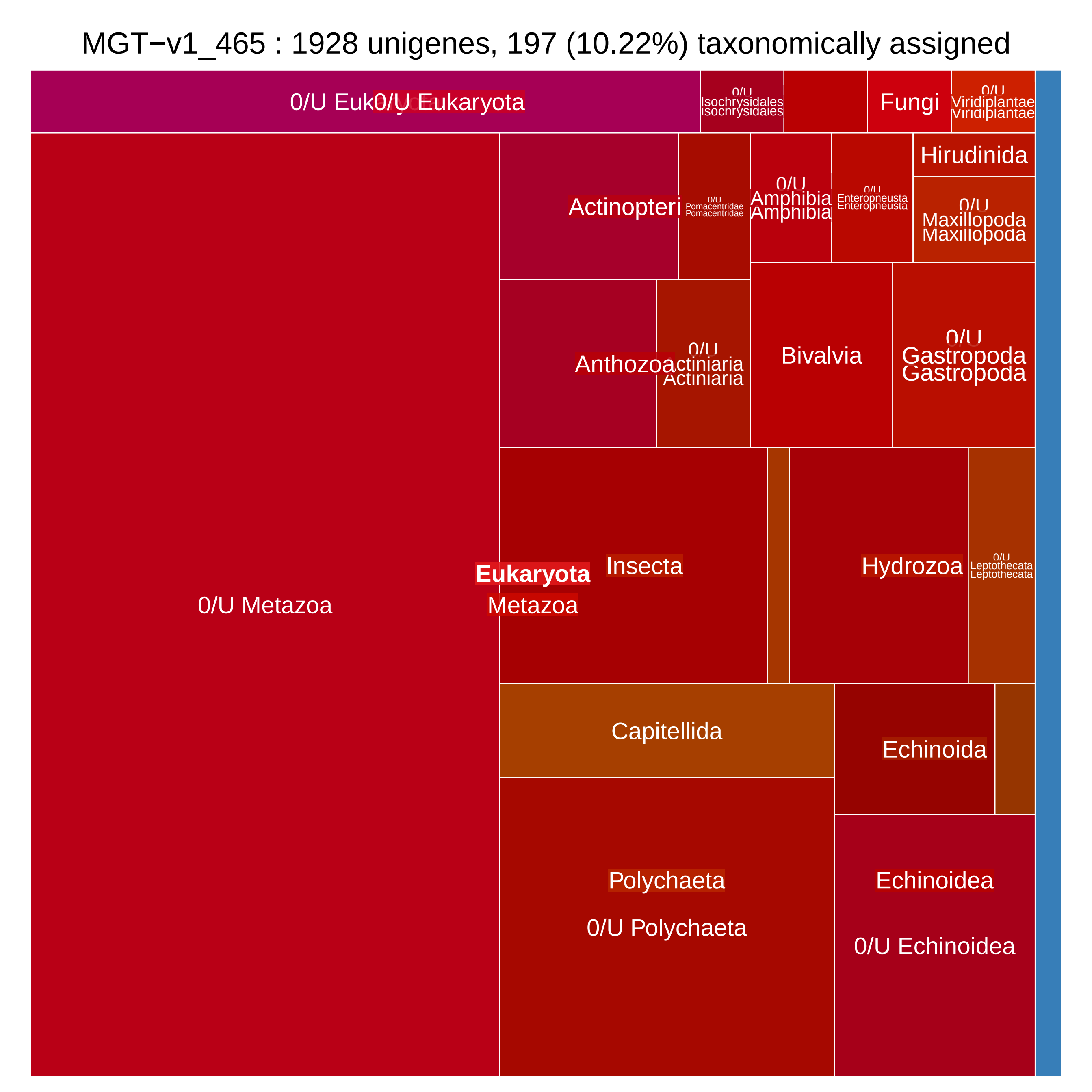
MGT−v1_465 : 1928 unigenes, 197 (10.22%) taxonomically assigned
0/U Eukaryota
0/U Isochrysidales
Isochrysidales
Fungi
0/U Viridiplantae
Viridiplantae
0/U Metazoa
Actinopteri
0/U Pomacentridae
Pomacentridae
0/U Amphibia
Amphibia
0/U Enteropneusta
Enteropneusta
Hirudinida
0/U Maxillopoda
Maxillopoda
Anthozoa
0/U Actiniaria
Actiniaria
Bivalvia
0/U Gastropoda
Gastropoda
Eukaryota
Metazoa
Insecta
Hydrozoa
0/U Leptothecata
Leptothecata
Capitellida
Echinoida
Polychaeta
0/U Polychaeta
Echinoidea
0/U Echinoidea

MGT−v1_466 : 1926 unigenes, 225 (11.68%) taxonomically assigned

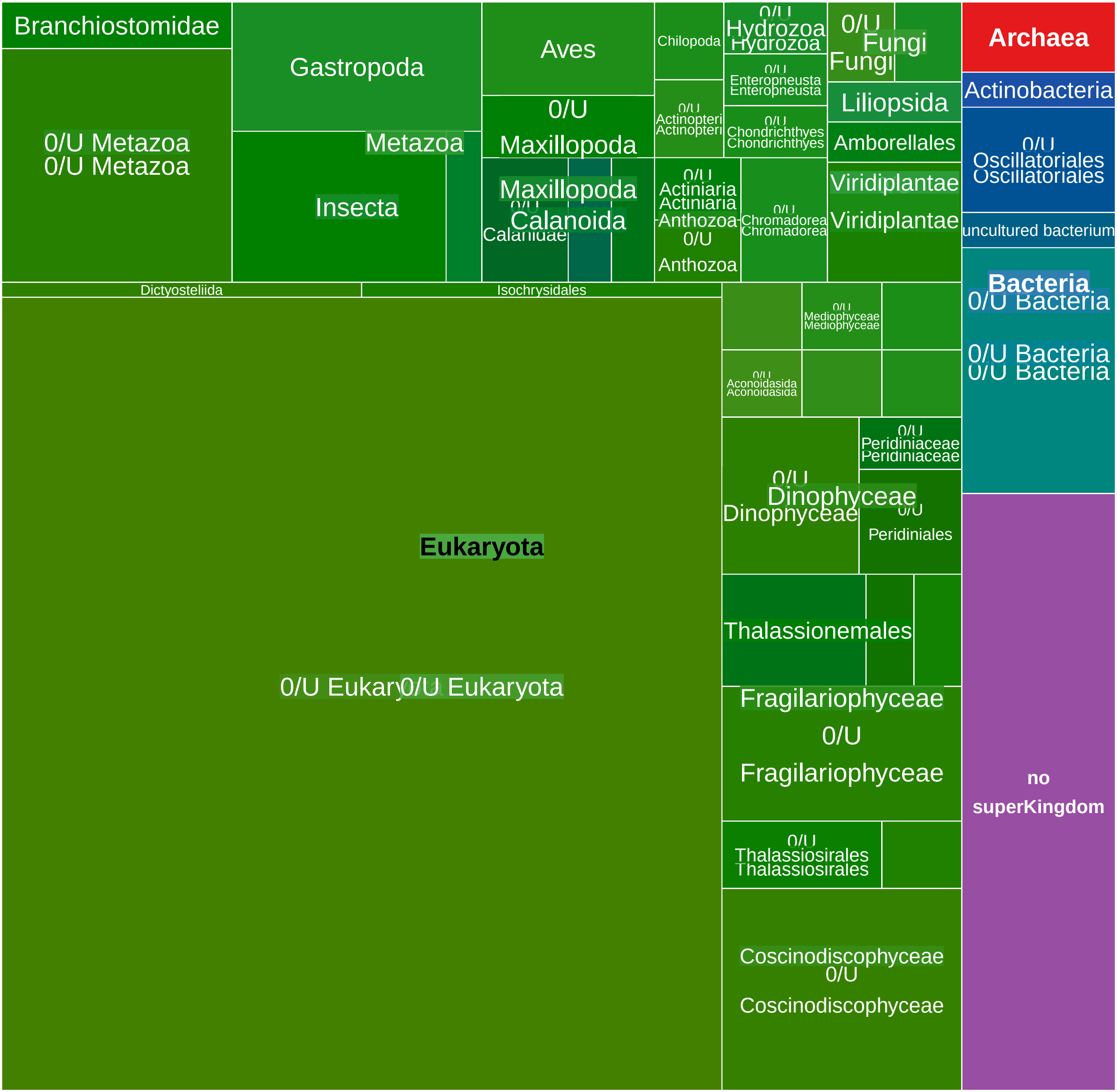

MGT−v1_467 : 1917 unigenes, 1734 (90.45%) taxonomically assigned

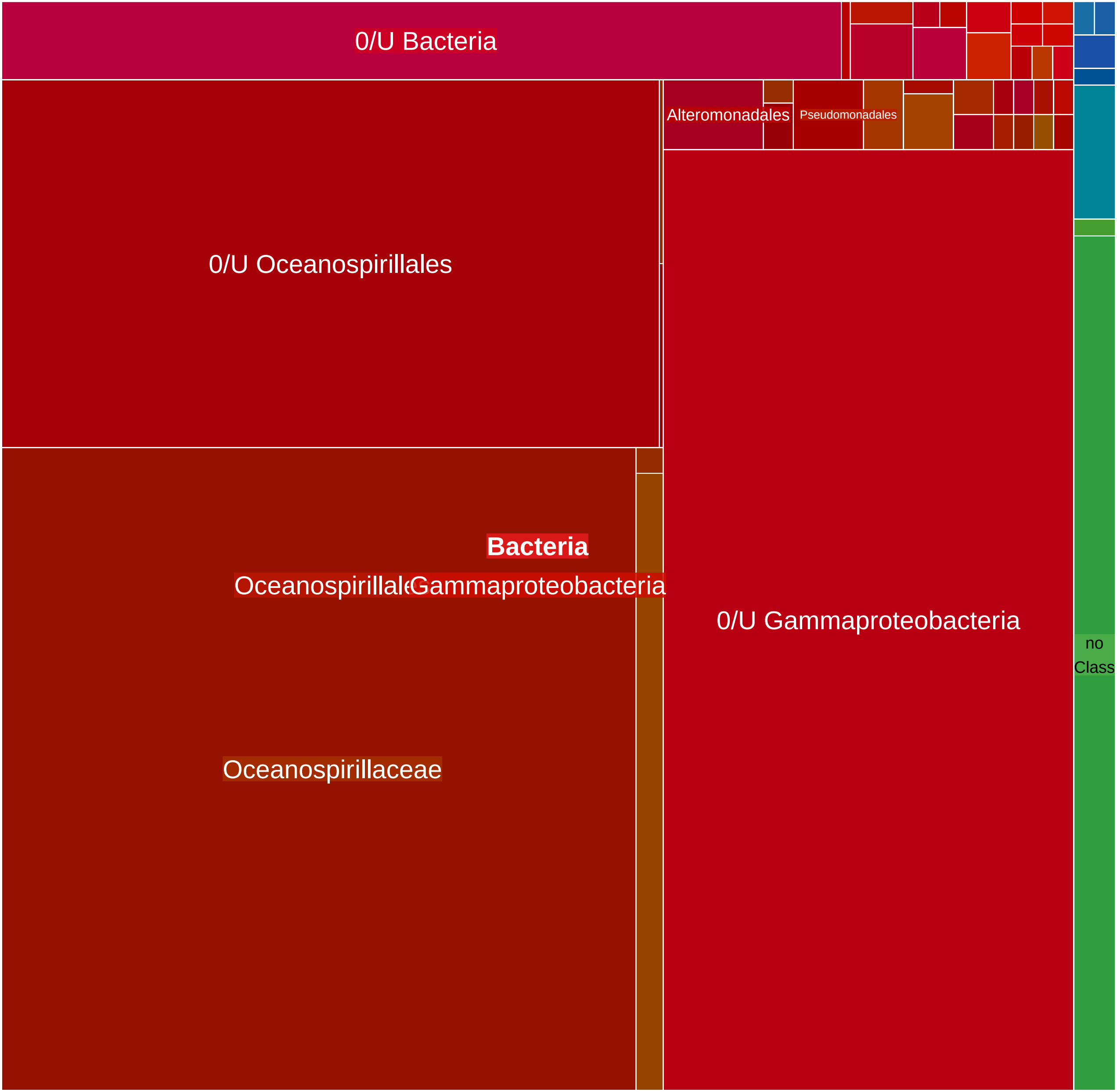

MGT−v1_468 : 1908 unigenes, 277 (14.52%) taxonomically assigned

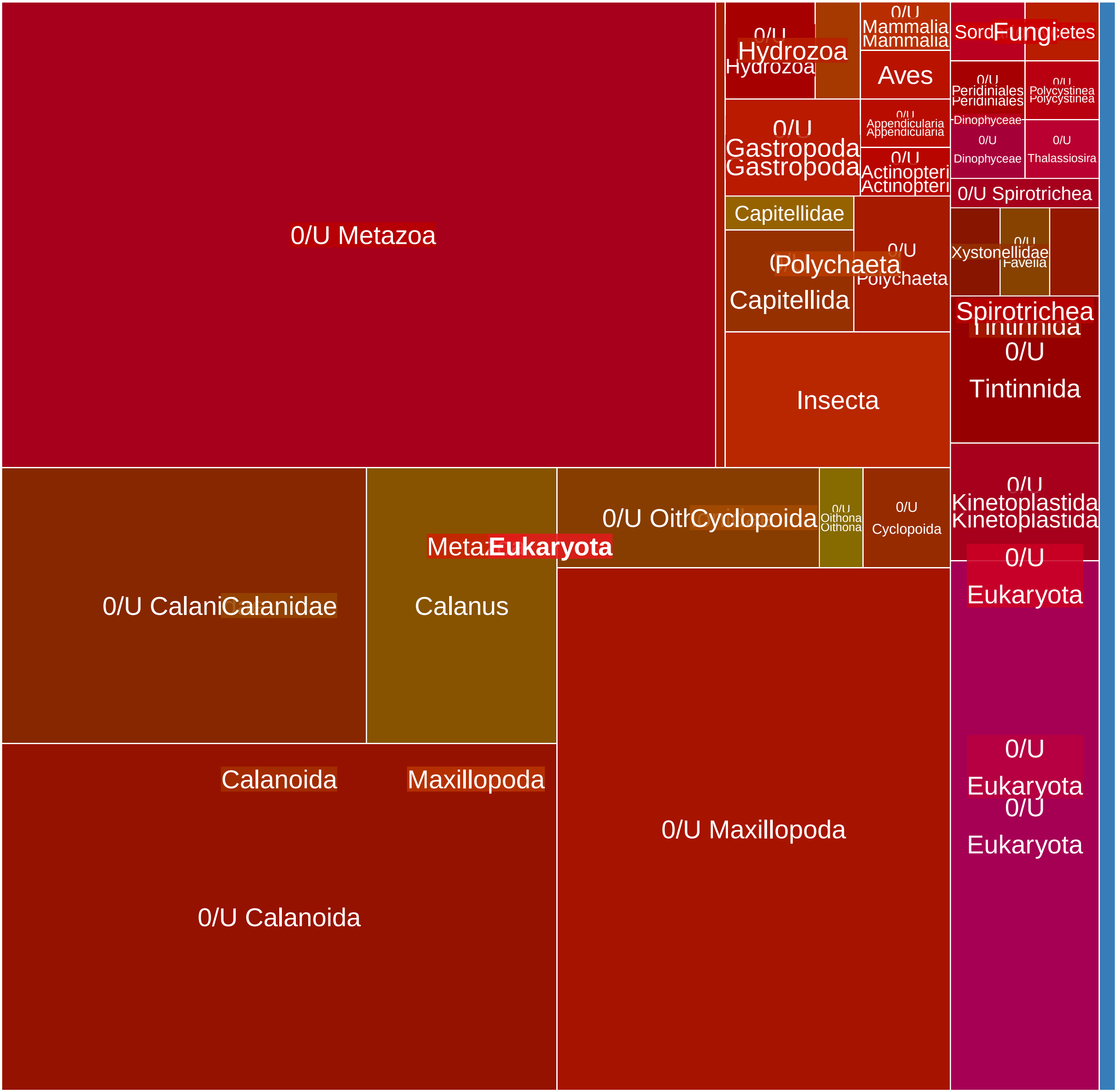

### MGT−v1_469 : 1883 unigenes, 1674 (88.90%) taxonomically assigned

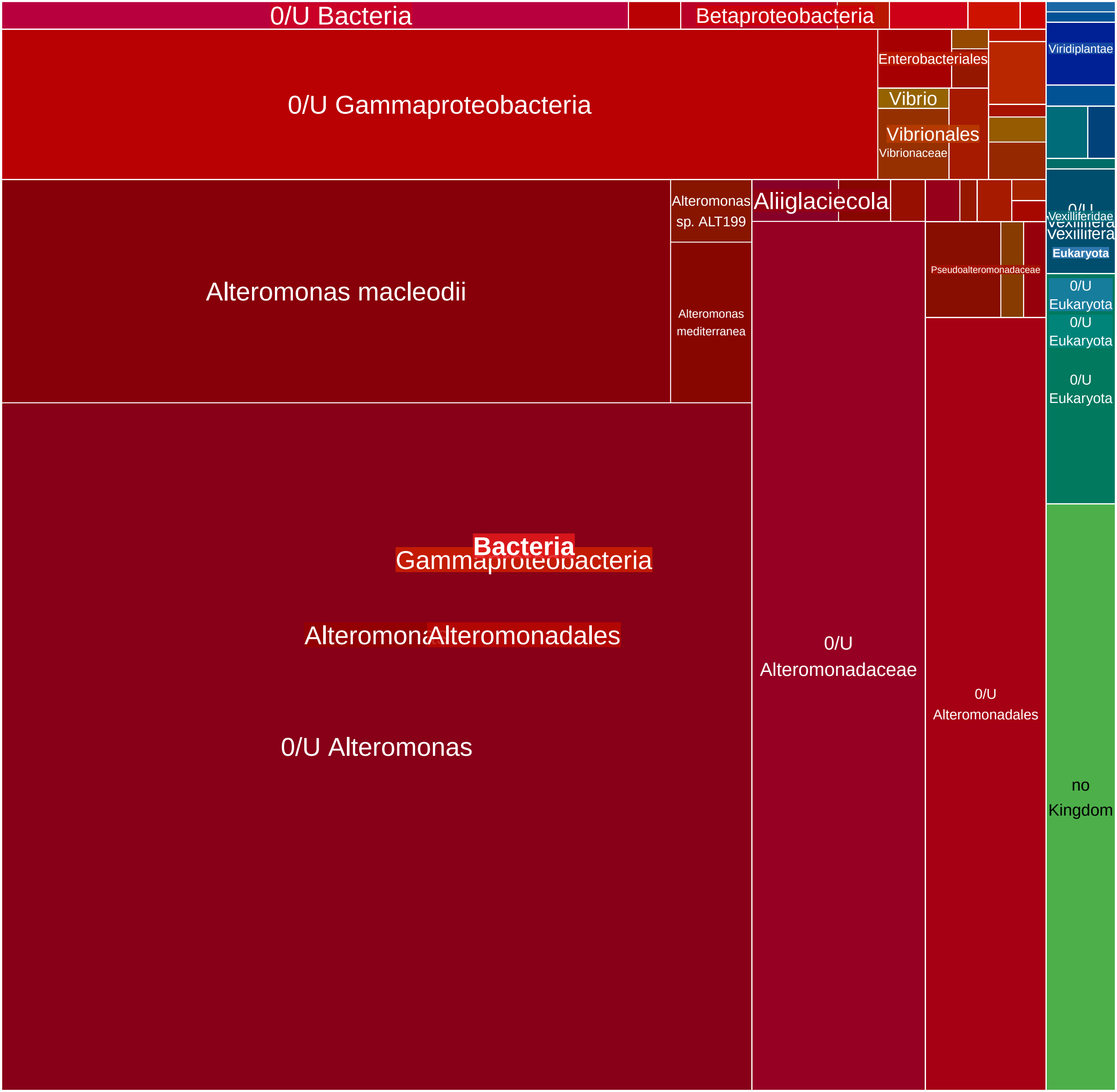

MGT−v1_470 : 1865 unigenes, 1787 (95.82%) taxonomically assigned

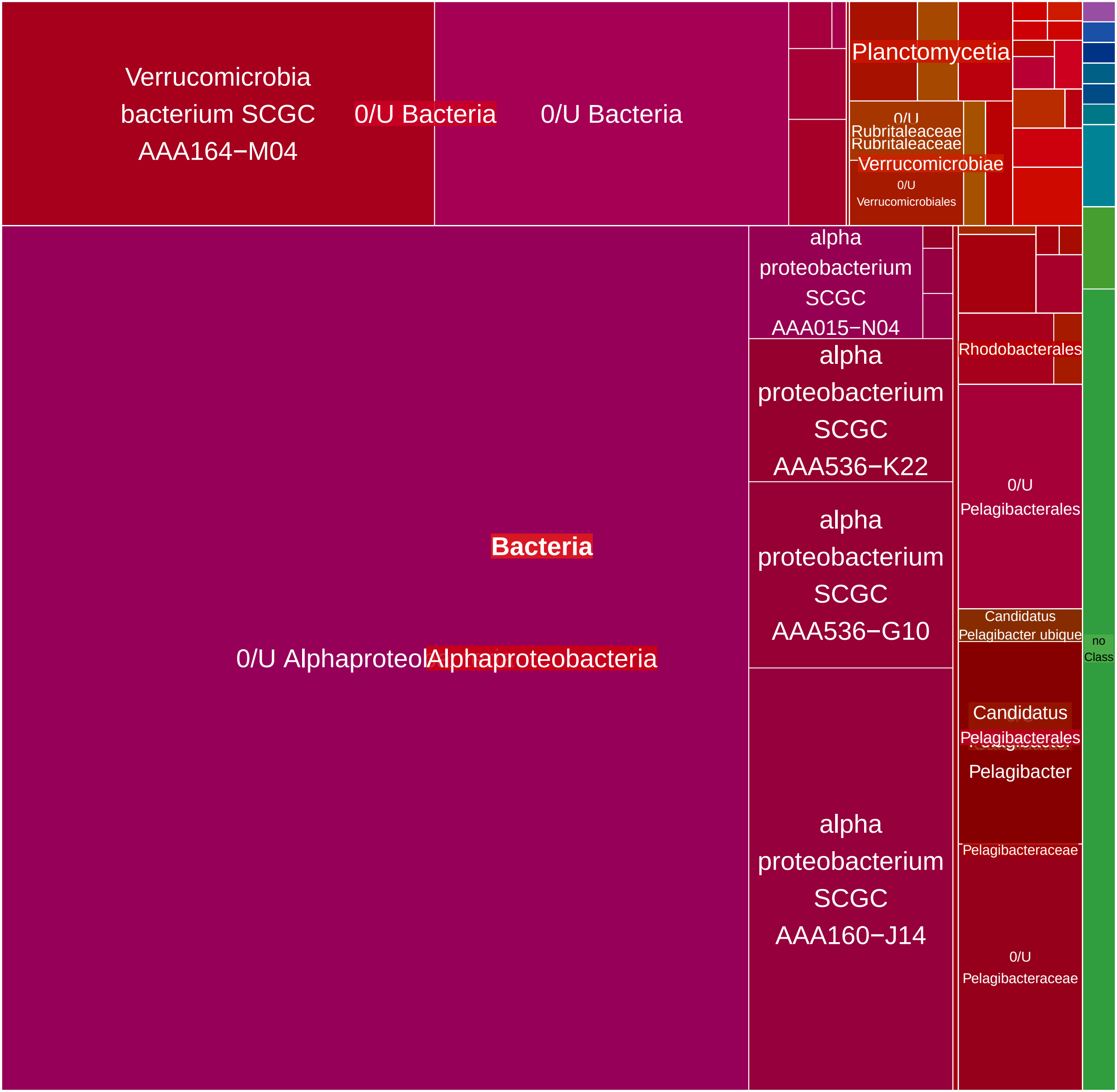

MGT−v1_471 : 1820 unigenes, 418 (22.97%) taxonomically assigned

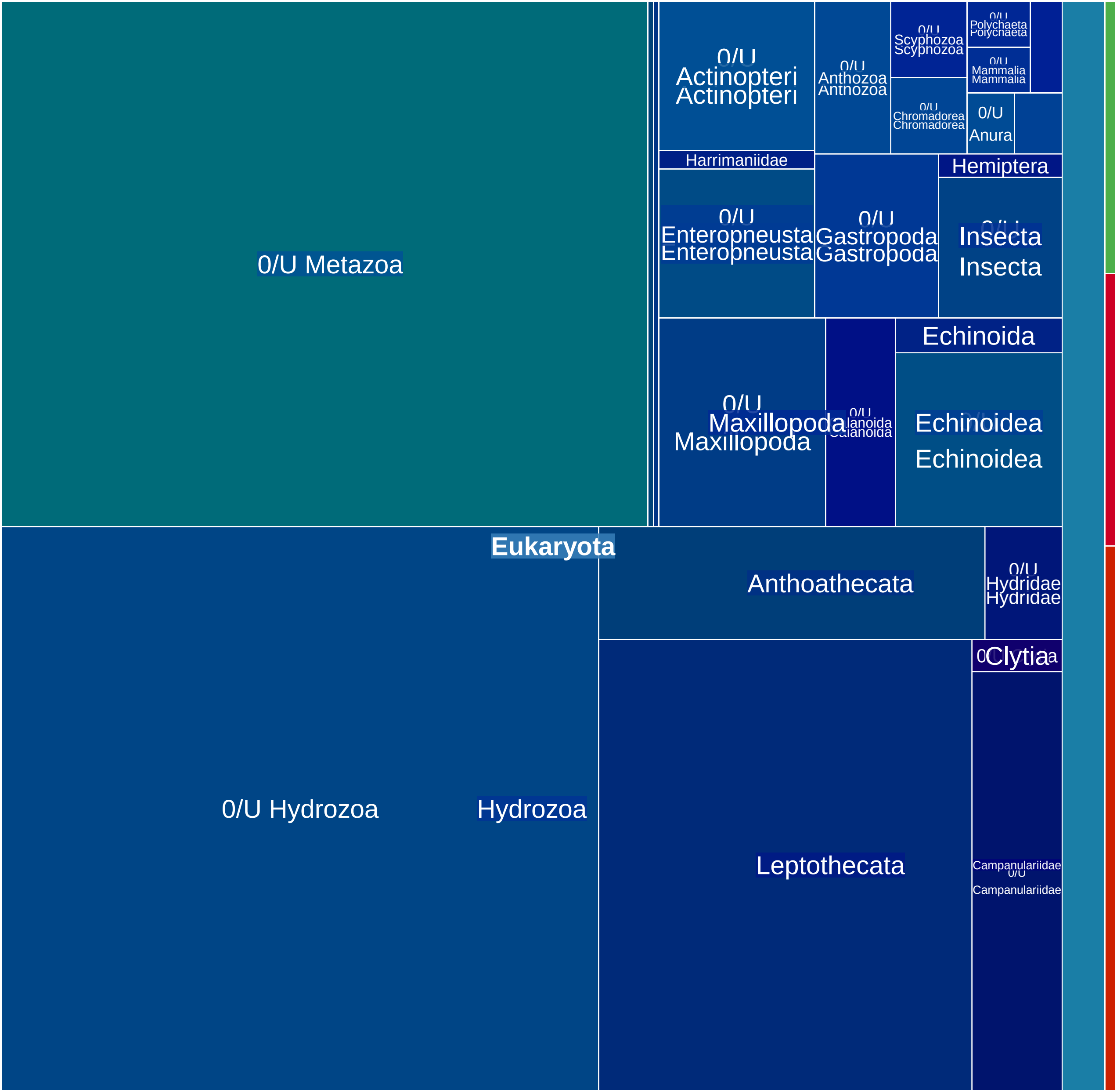

MGT−v1_472 : 1818 unigenes, 438 (24.09%) taxonomically assigned

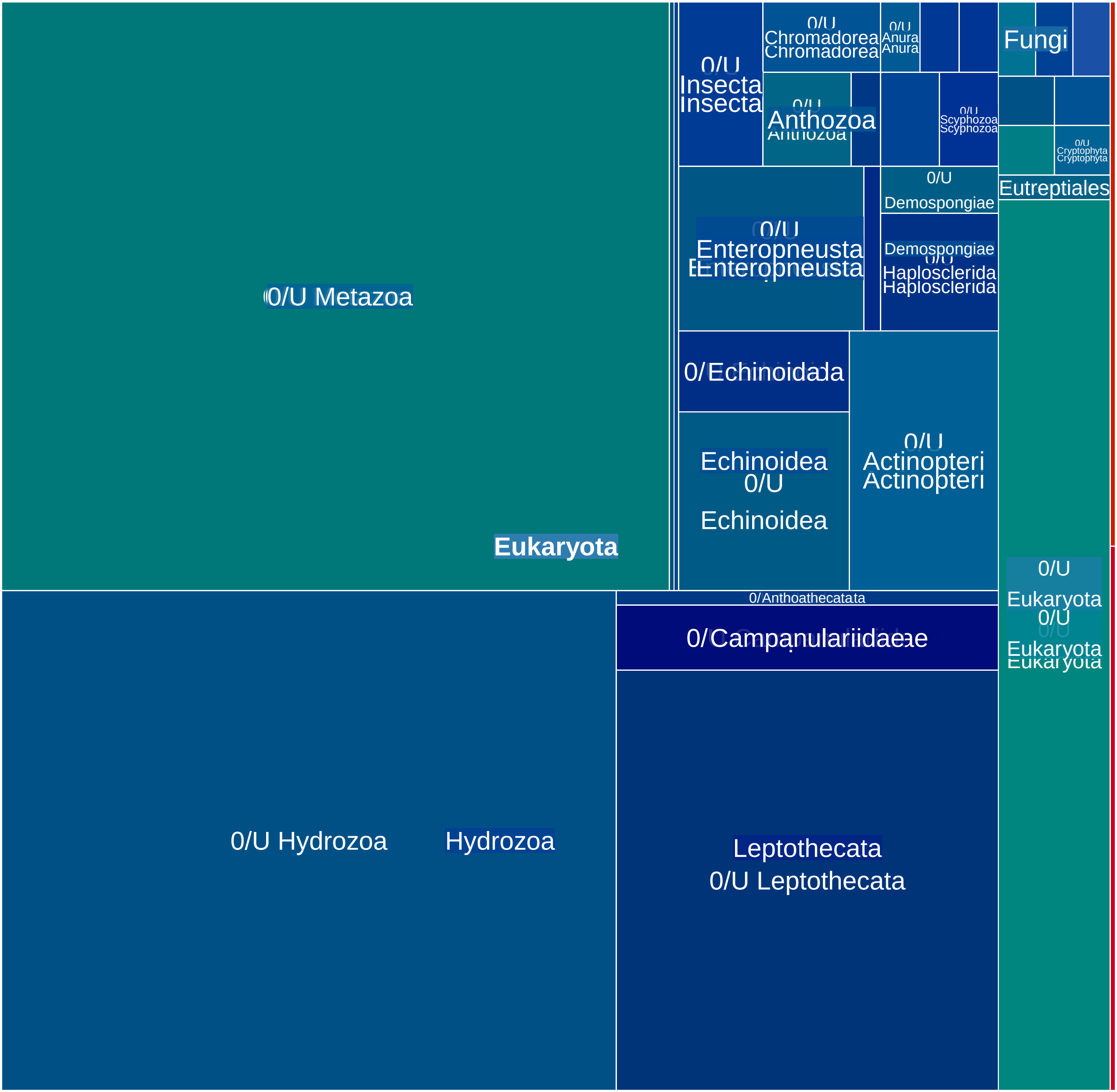

MGT−v1_473 : 1813 unigenes, 202 (11.14%) taxonomically assigned

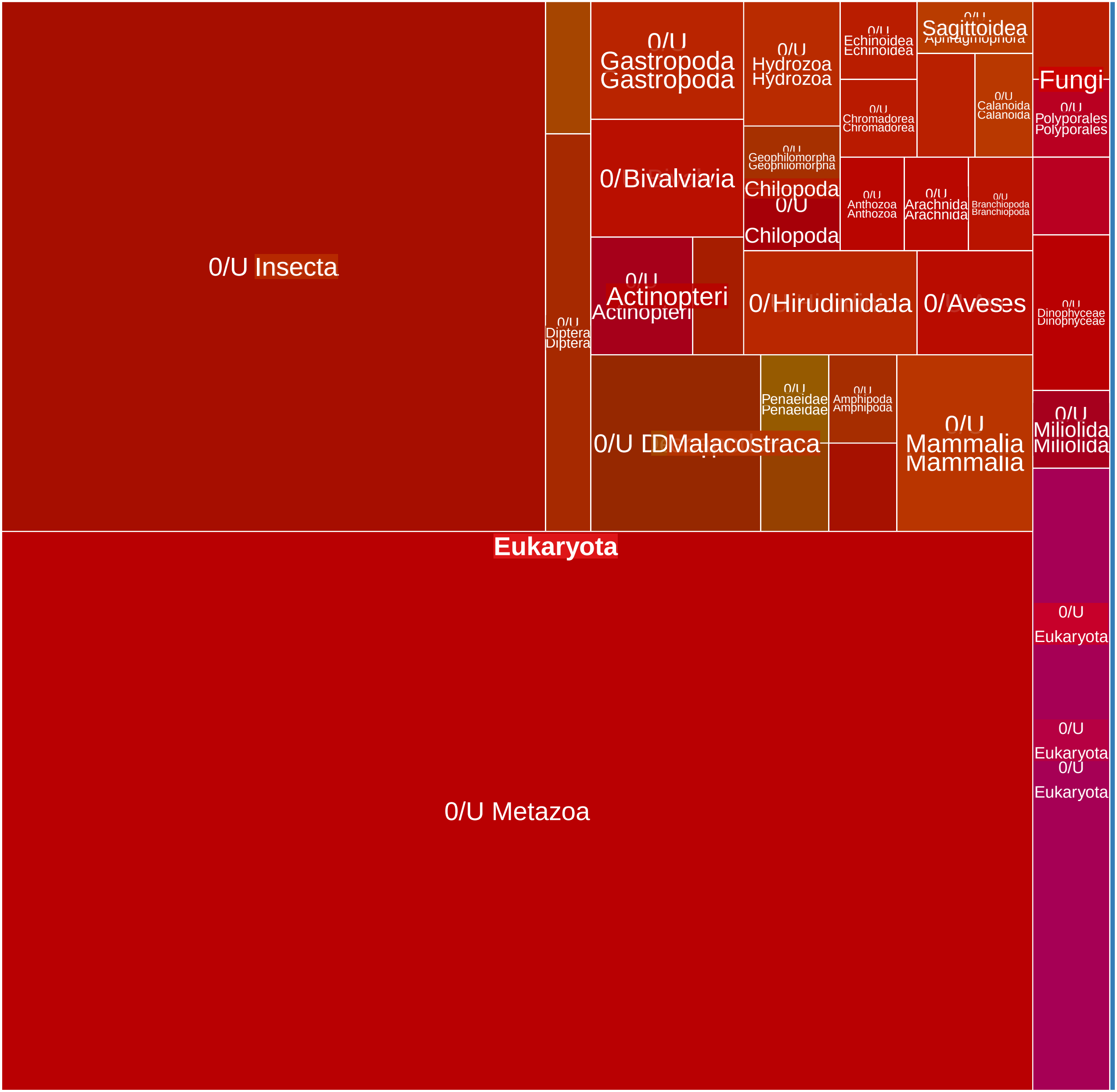

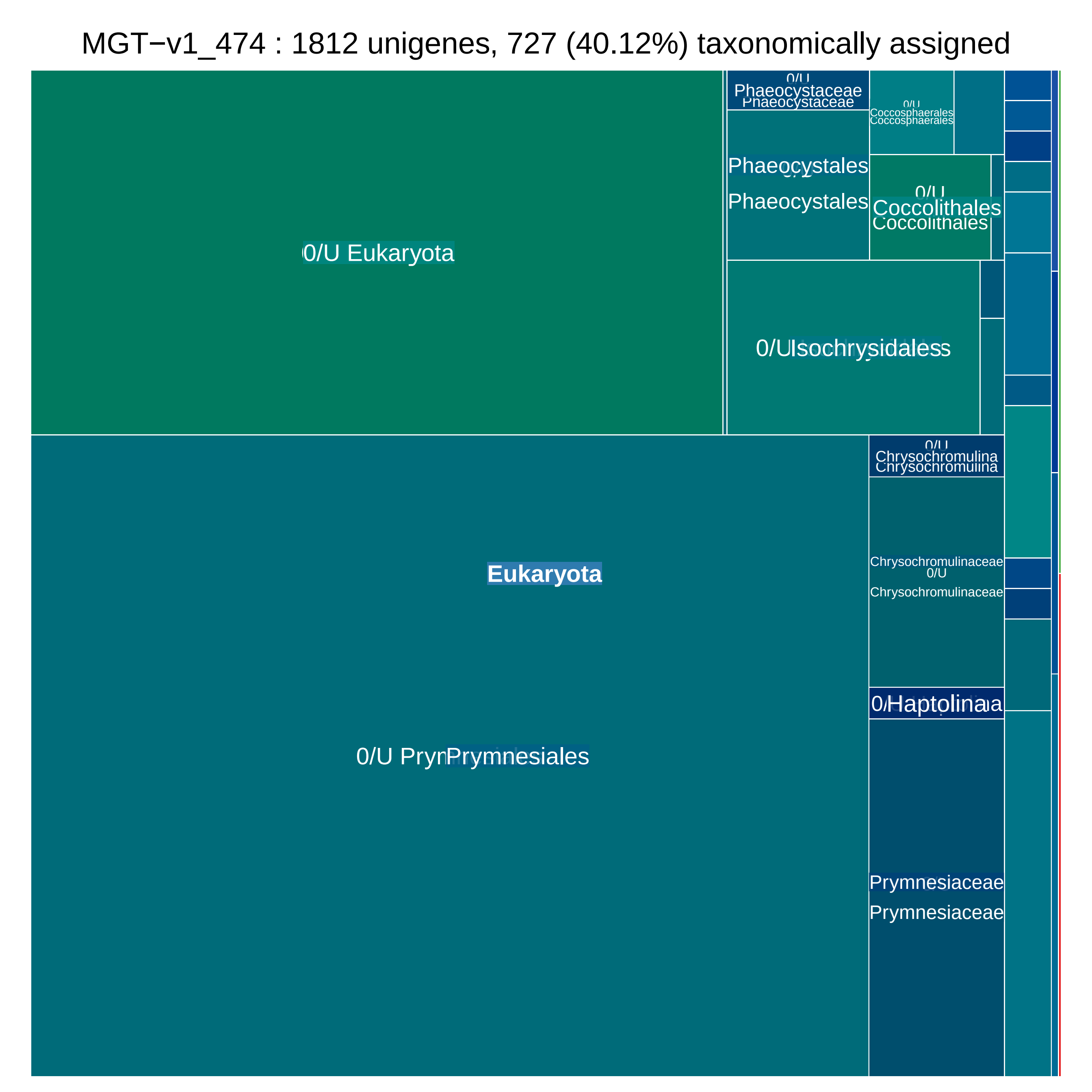
MGT−v1_474 : 1812 unigenes, 727 (40.12%) taxonomically assigned
0/U Eukaryota
0/U
Phaeocystaceae
Phaeocystaceae
Phaeocystales
Phaeocystales
0/U
Coccosphaerales
Coccosphaerales
0/U
Coccolithales
Coccolithales
0/U Isochrysidales
Eukaryota
0/U Prymnesiales
0/U
Chrysochromulina
Chrysochromulina
Chrysochromulinaceae
0/U
Chrysochromulinaceae
0/Haptolina
Prymnesiaceae
Prymnesiaceae

MGT−v1_475 : 1807 unigenes, 622 (34.42%) taxonomically assigned

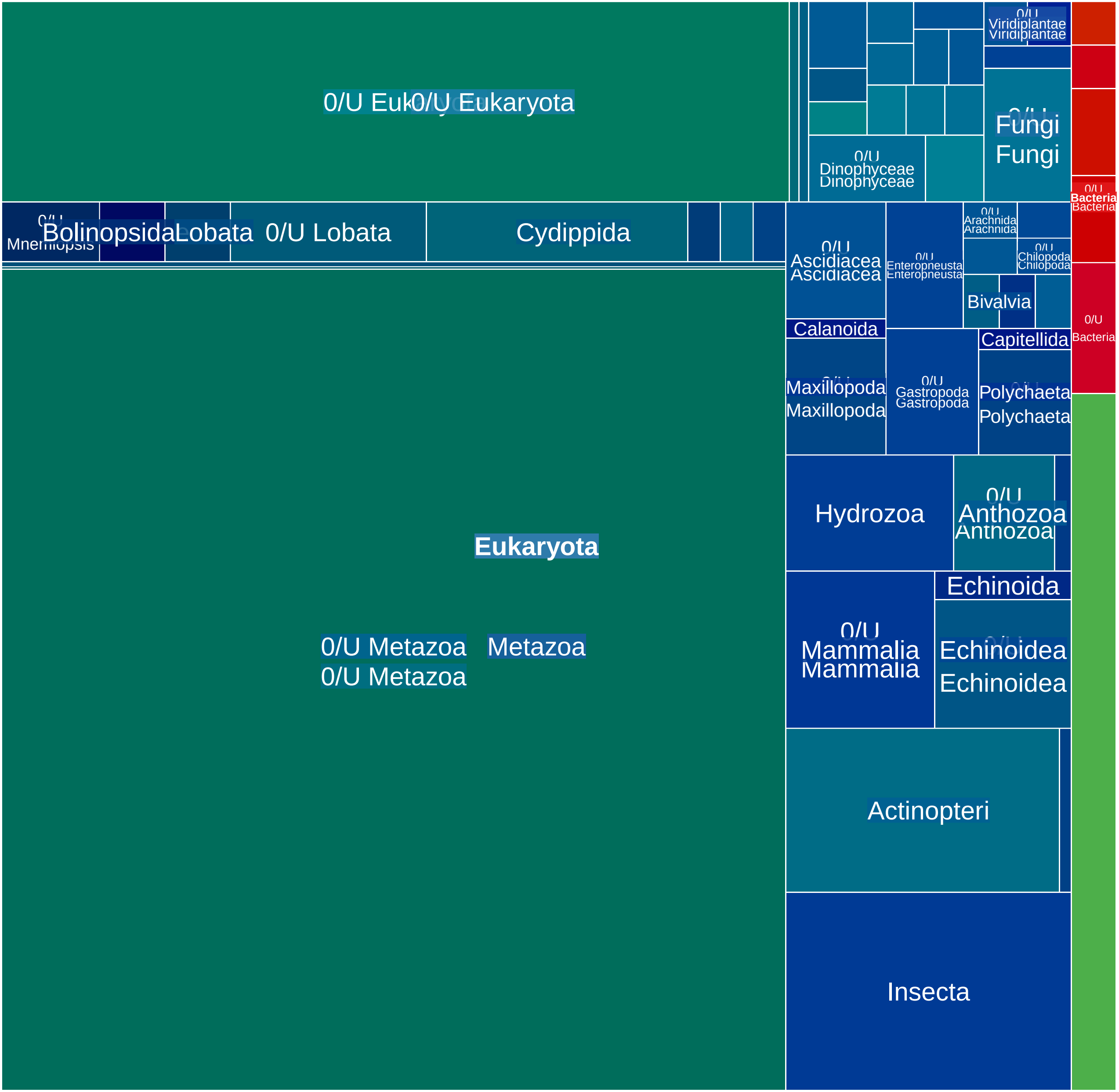

MGT−v1_476 : 1794 unigenes, 251 (13.99%) taxonomically assigned

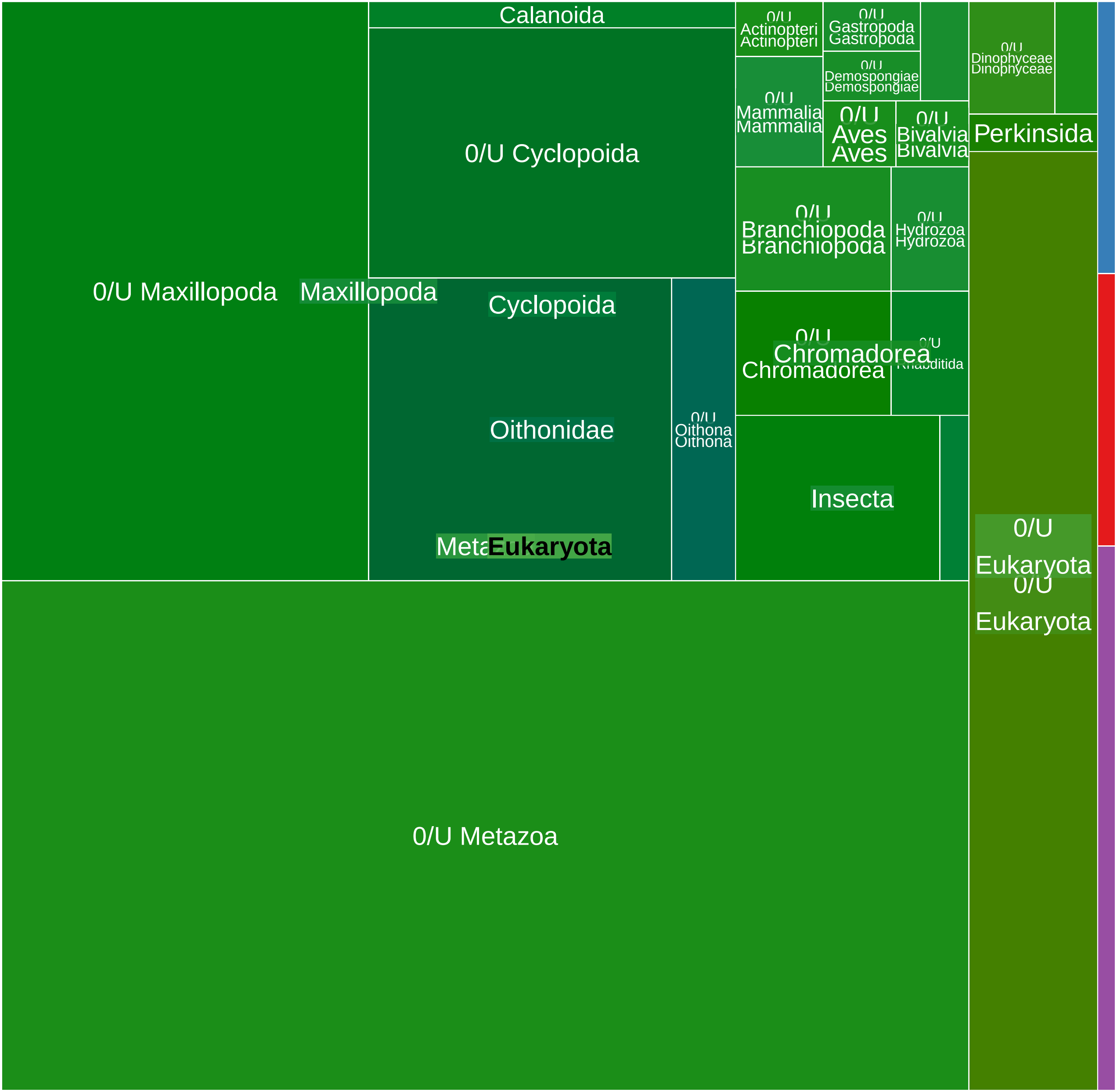

MGT−v1_477 : 1794 unigenes, 191 (10.65%) taxonomically assigned

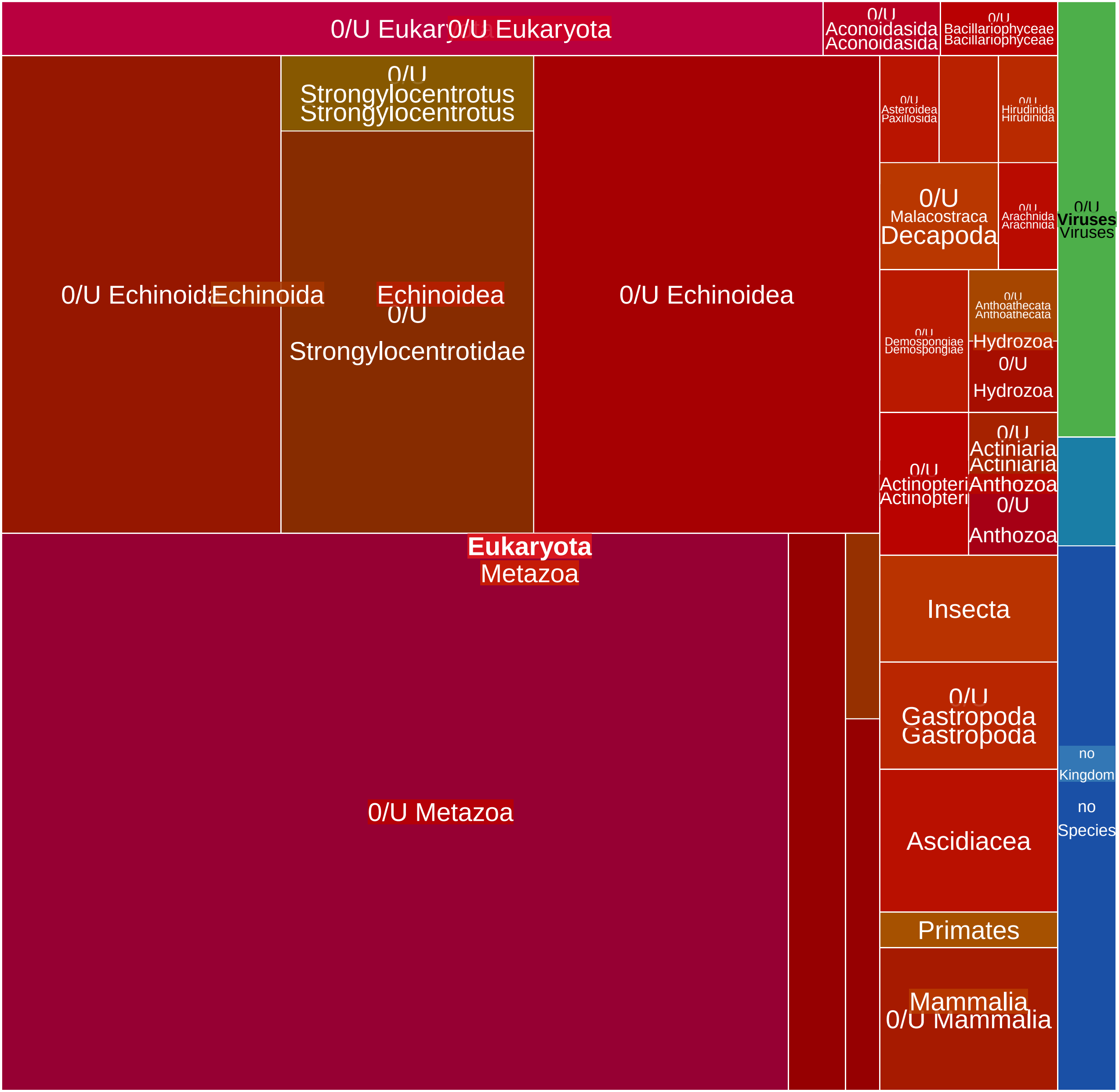

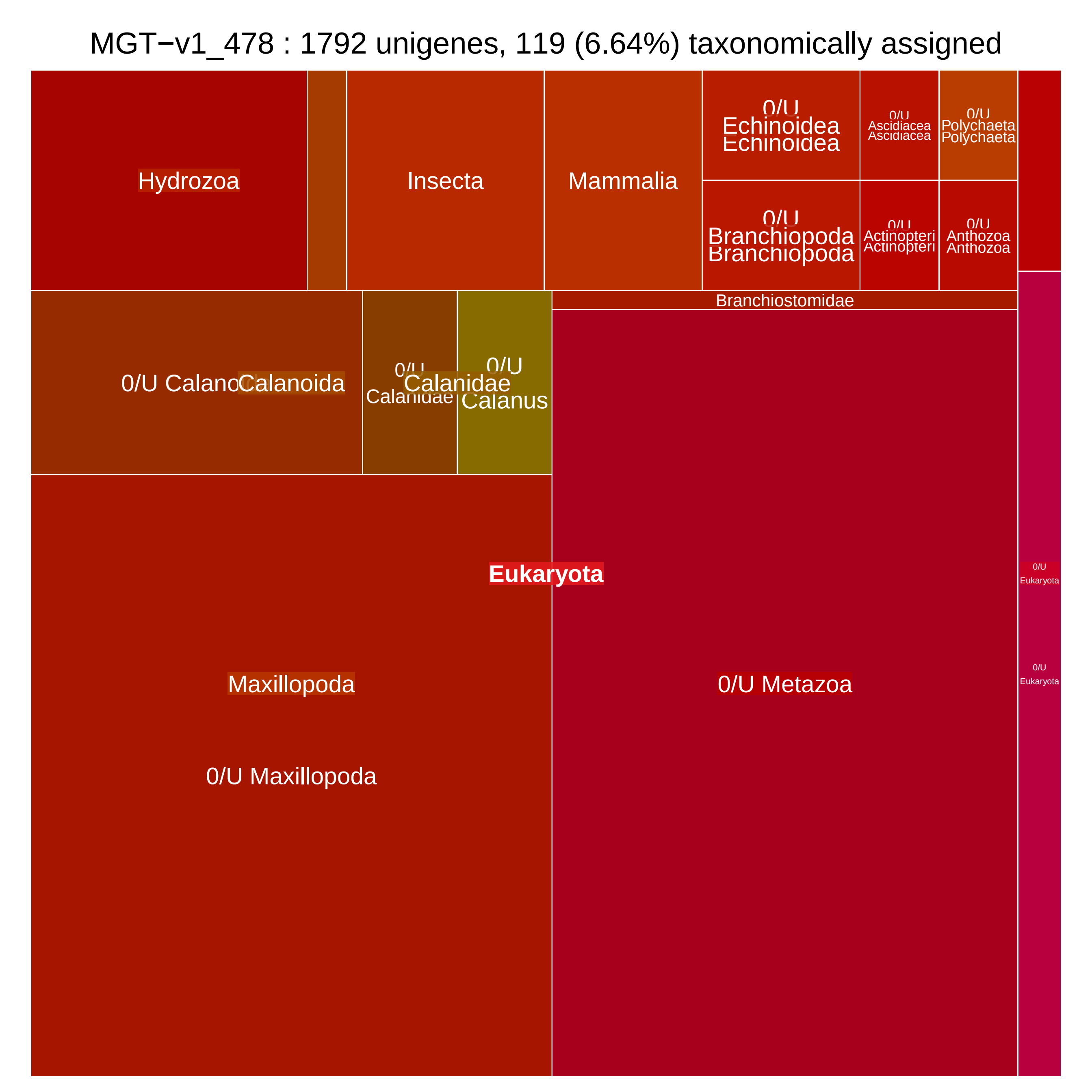
MGT−v1_478 : 1792 unigenes, 119 (6.64%) taxonomically assigned
Hydrozoa
Insecta
Mammalia
0/U Echinoidea Echinoidea
0/U Branchiopoda Branchiopoda
0/U Ascidiacea Ascidiacea
0/U Actinopteri Actinopteri
0/U Polychaeta Polychaeta
0/U Anthozoa Anthozoa
Branchiostomidae
0/U Calanoida
Calanoida
0/U Calanidae
Calanidae
0/U Calanus
Eukaryota
Maxillopoda
0/U Maxillopoda
0/U Metazoa
0/U Eukaryota
0/U Eukaryota

MGT−v1_479 : 1780 unigenes, 1594 (89.55%) taxonomically assigned

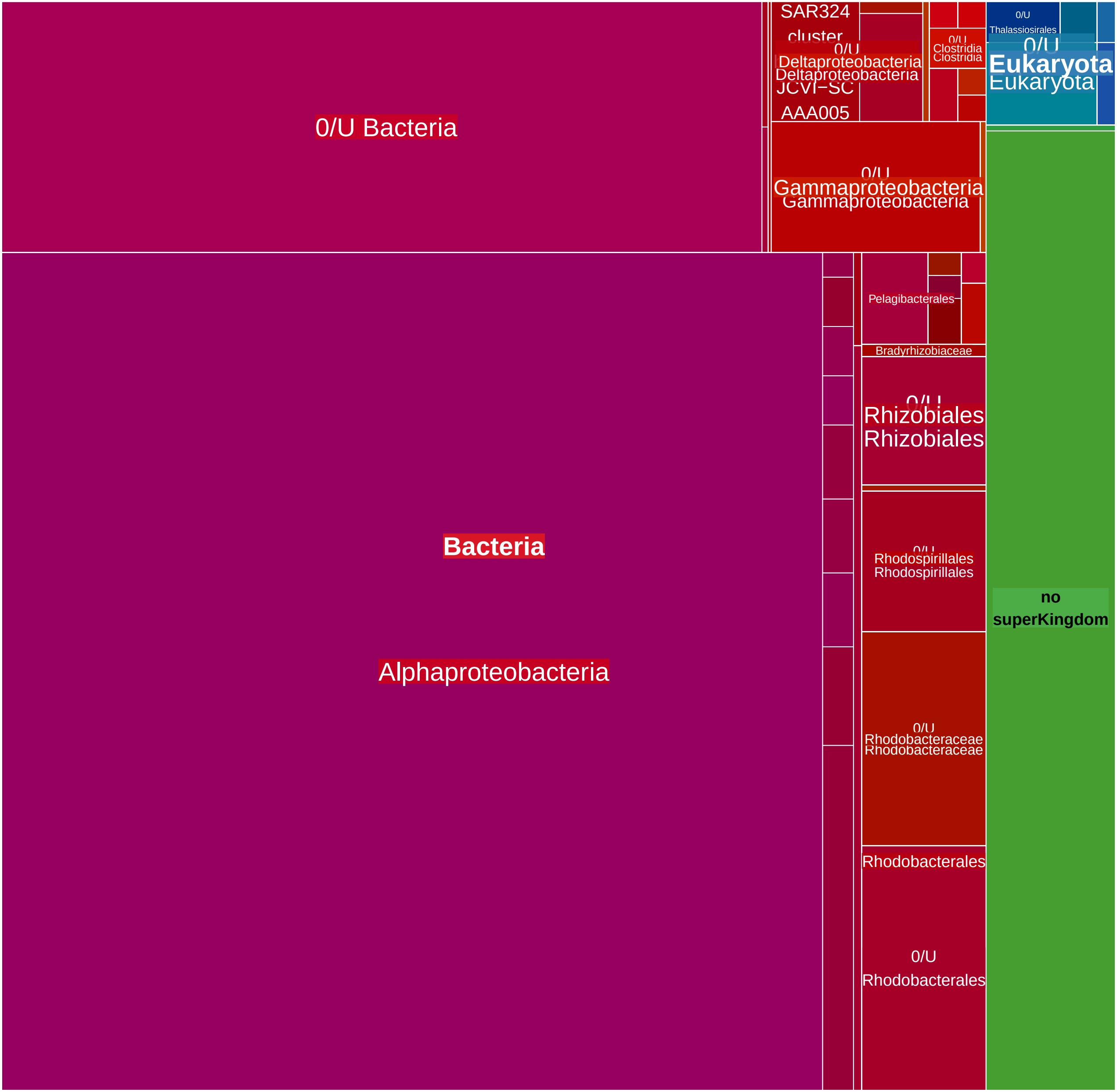

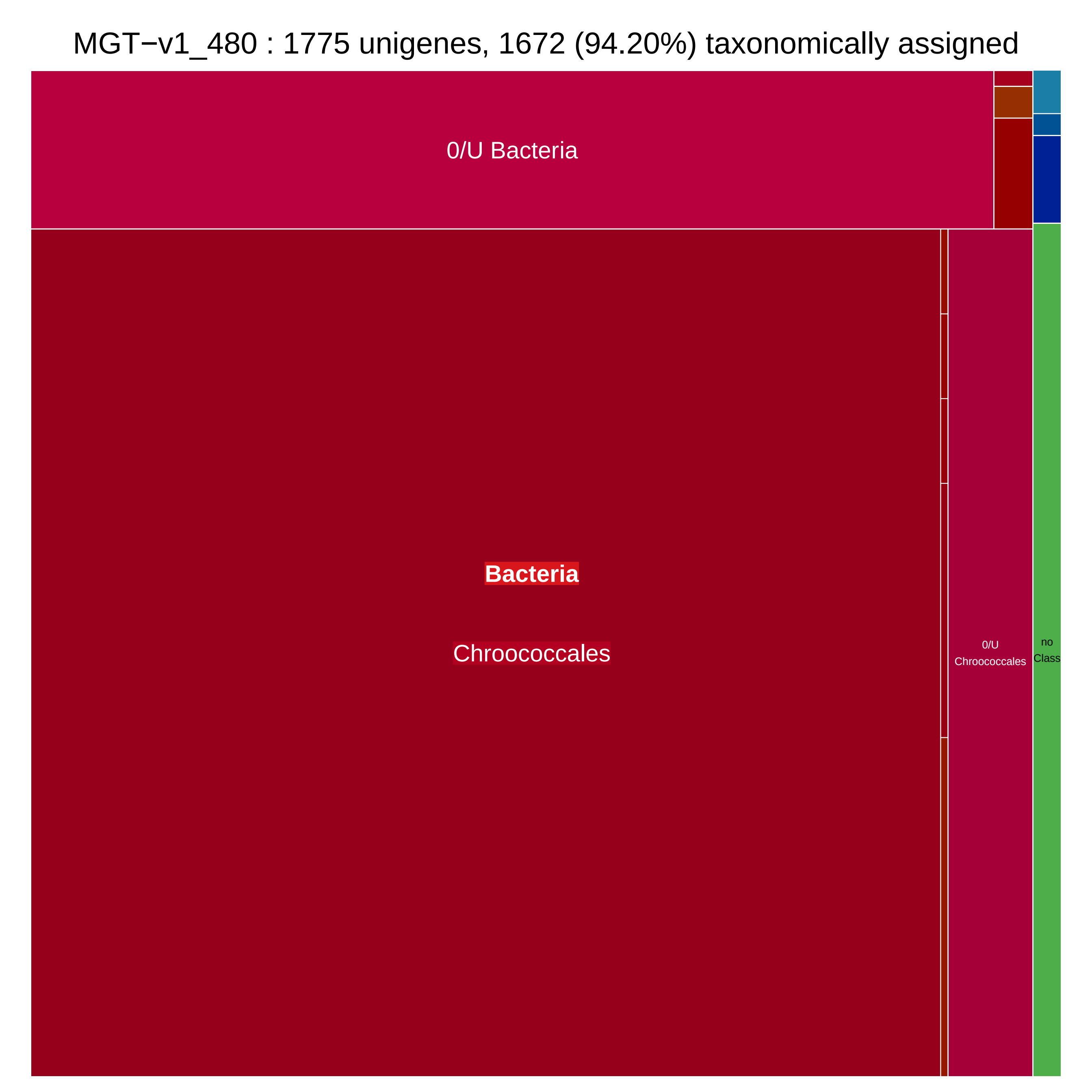
MGT−v1_480 : 1775 unigenes, 1672 (94.20%) taxonomically assigned
0/U Bacteria
Bacteria
Chroococcales
0/U
Chroococcales
no
Class

MGT−v1_481 : 1767 unigenes, 137 (7.75%) taxonomically assigned

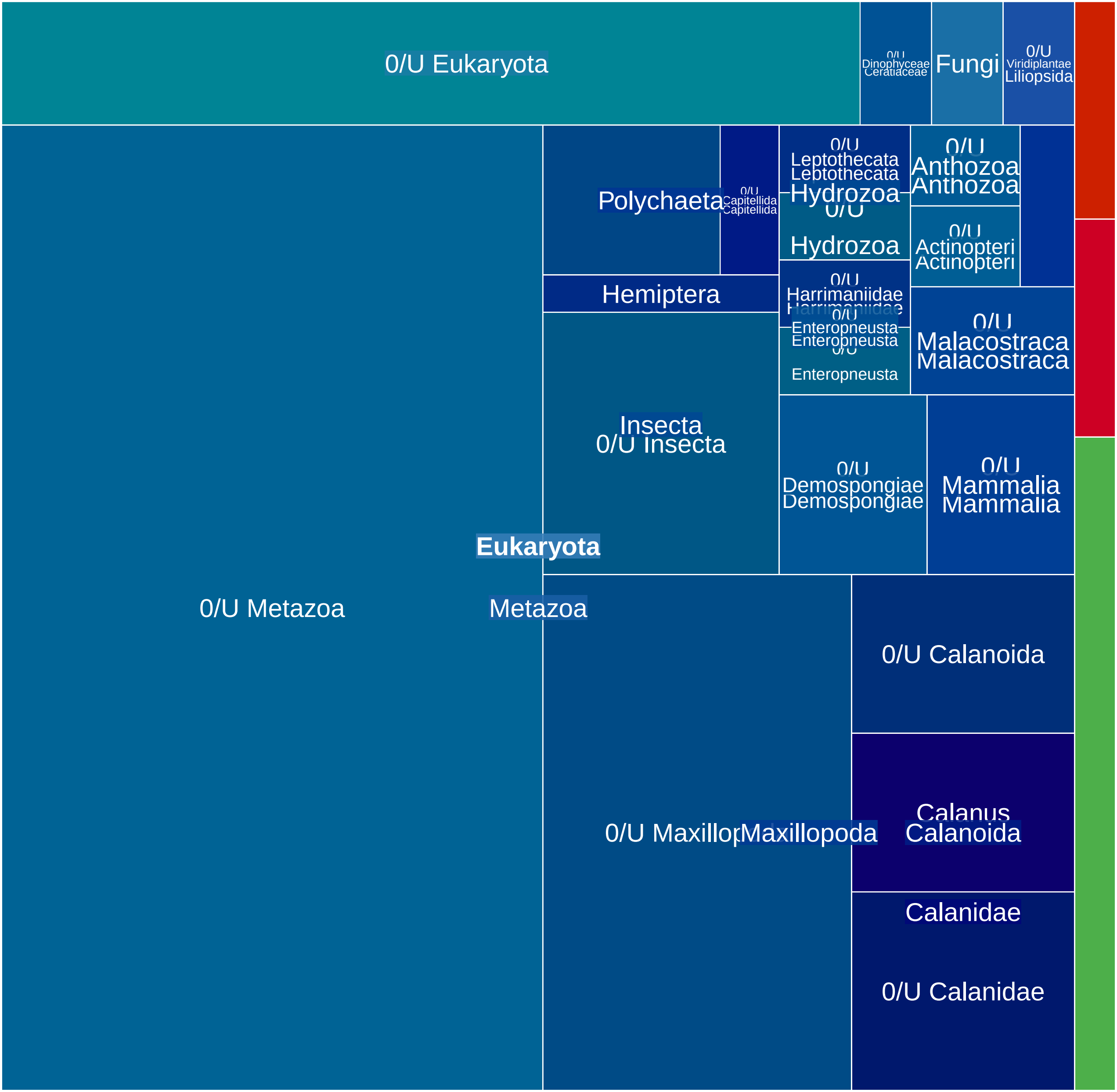

MGT−v1_482 : 1767 unigenes, 1538 (87.04%) taxonomically assigned

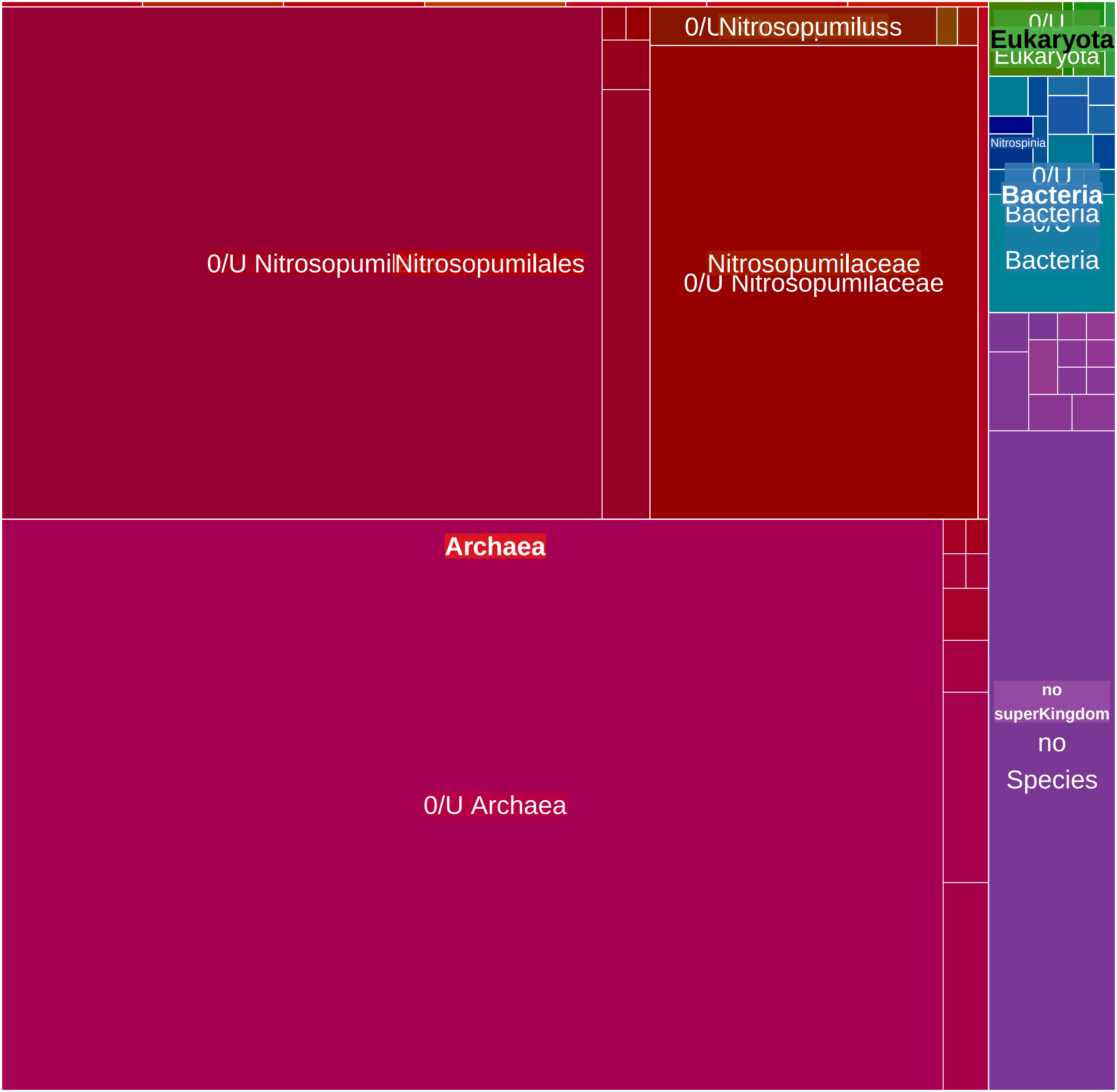

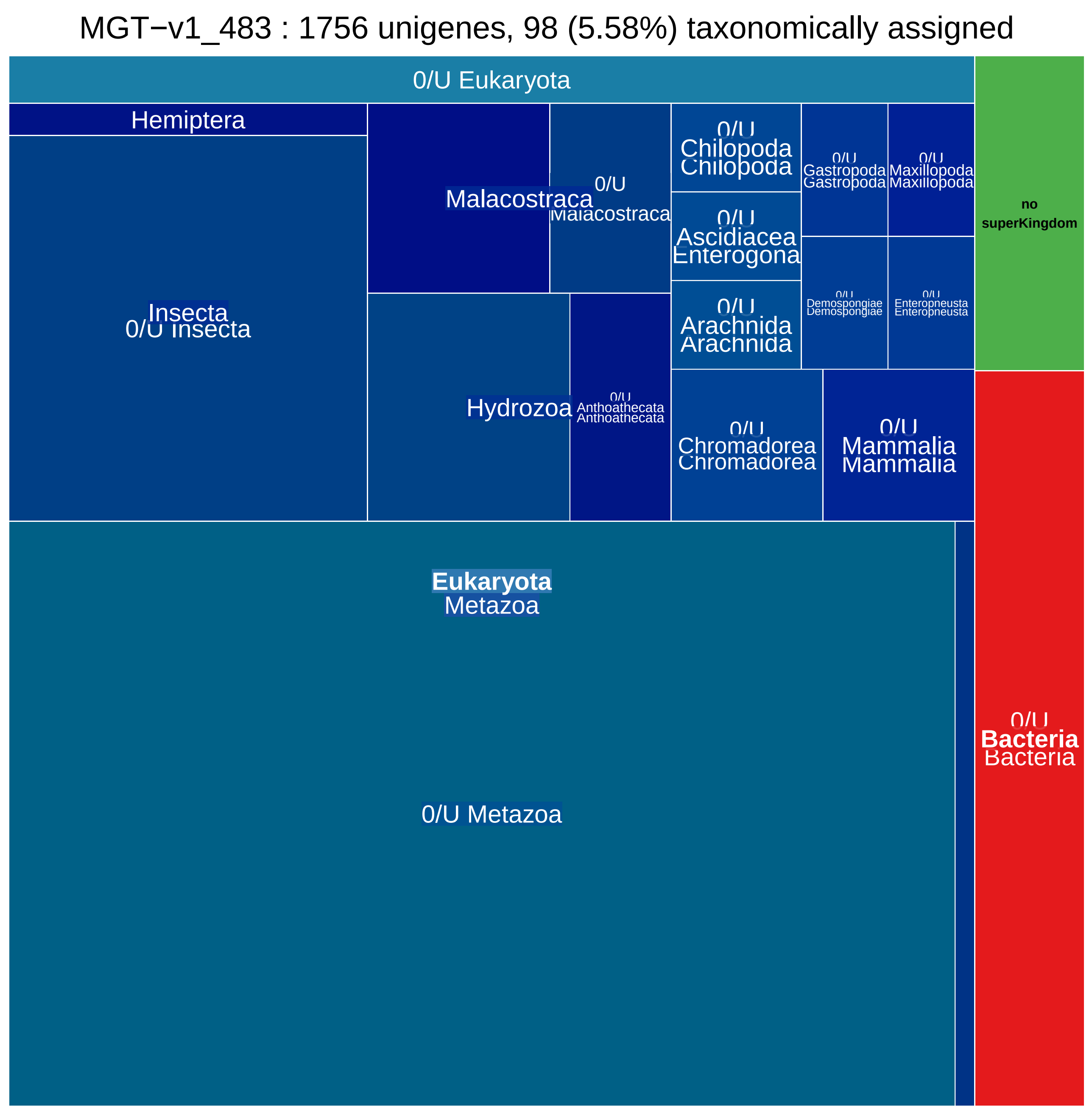
MGT−v1_483 : 1756 unigenes, 98 (5.58%) taxonomically assigned
0/U Eukaryota
Hemiptera
Insecta
0/U Insecta
Malacostraca
0/U
Malacostraca
Hydrozoa
0/U
Anthoathecata
Anthoathecata
0/U
Chilopoda
Chilopoda
0/U
Ascidiacea
Enterogona
0/U
Arachnida
Arachnida
0/U
Gastropoda
Gastropoda
0/U
Maxillopoda
Maxillopoda
0/U
Demospongiae
Demospongiae
0/U
Enteropneusta
Enteropneusta
0/U
Chromadorea
Chromadorea
0/U
Mammalia
Mammalia
Eukaryota
Metazoa
0/U Metazoa
no
superKingdom
0/U
Bacteria
Bacteria

MGT−v1_484 : 1742 unigenes, 79 (4.54%) taxonomically assigned

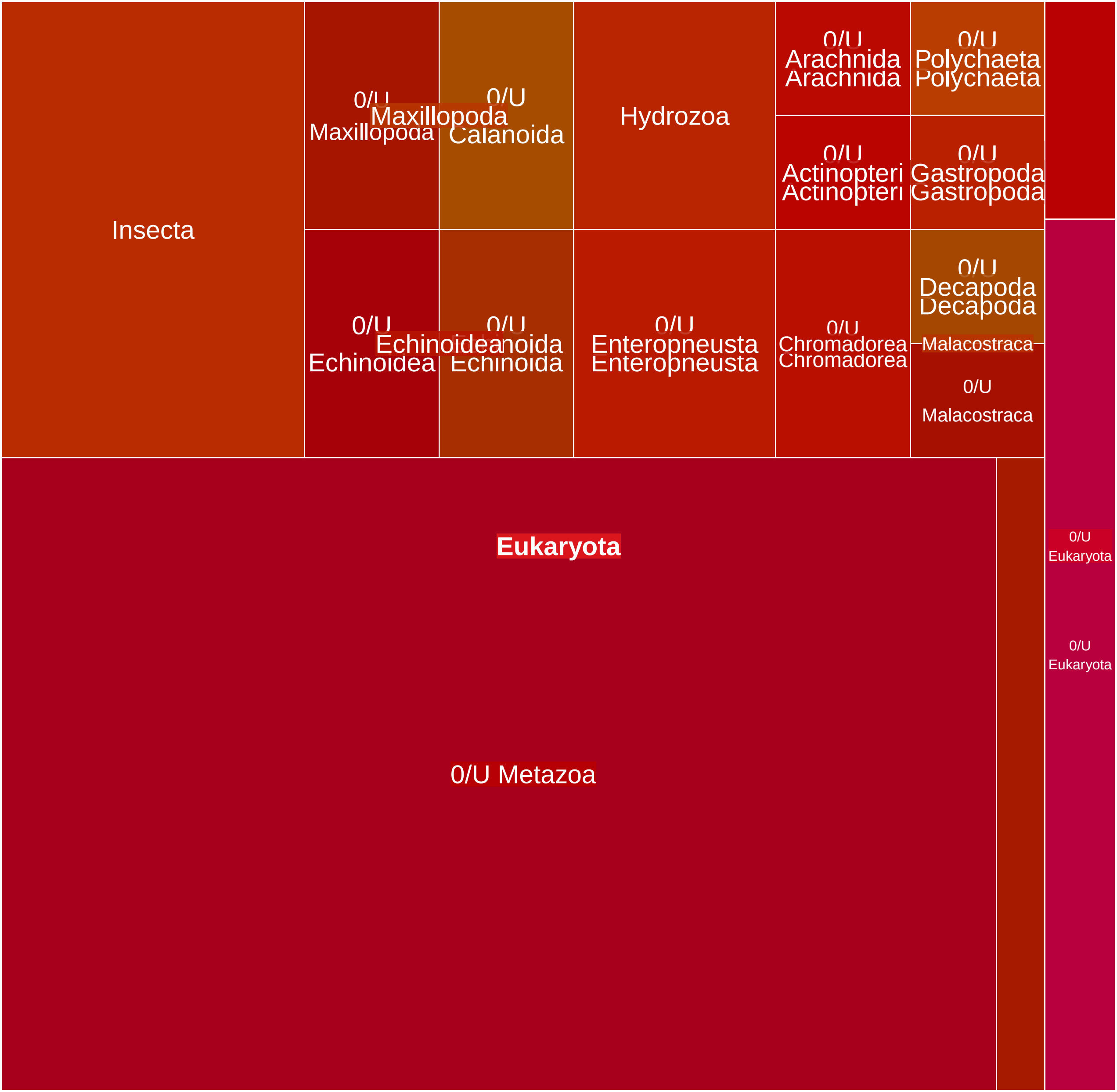

#### MGT−v1_485 : 1739 unigenes, 762 (43.82%) taxonomically assigned

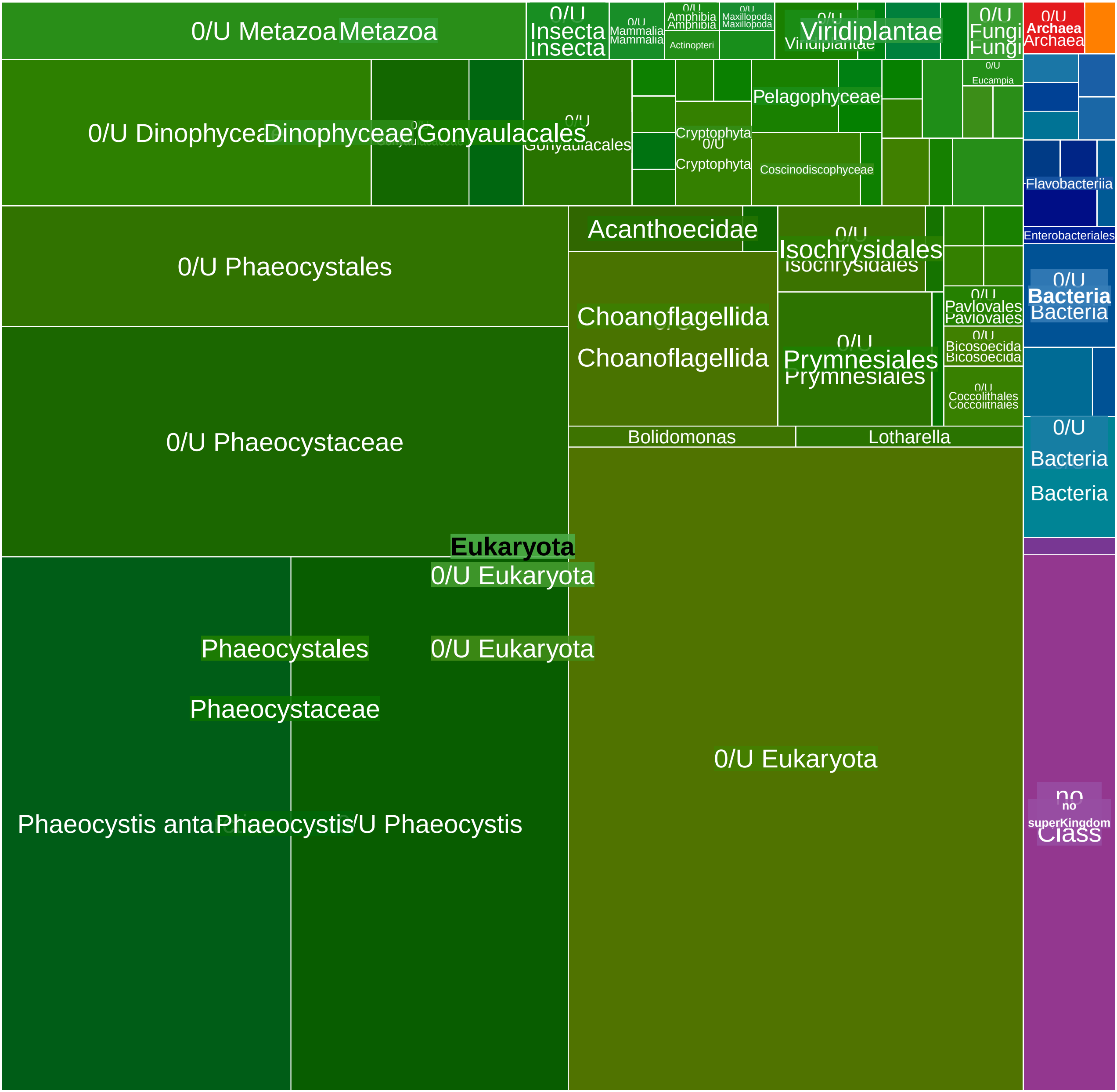

### MGT−v1_486 : 1736 unigenes, 1670 (96.20%) taxonomically assigned

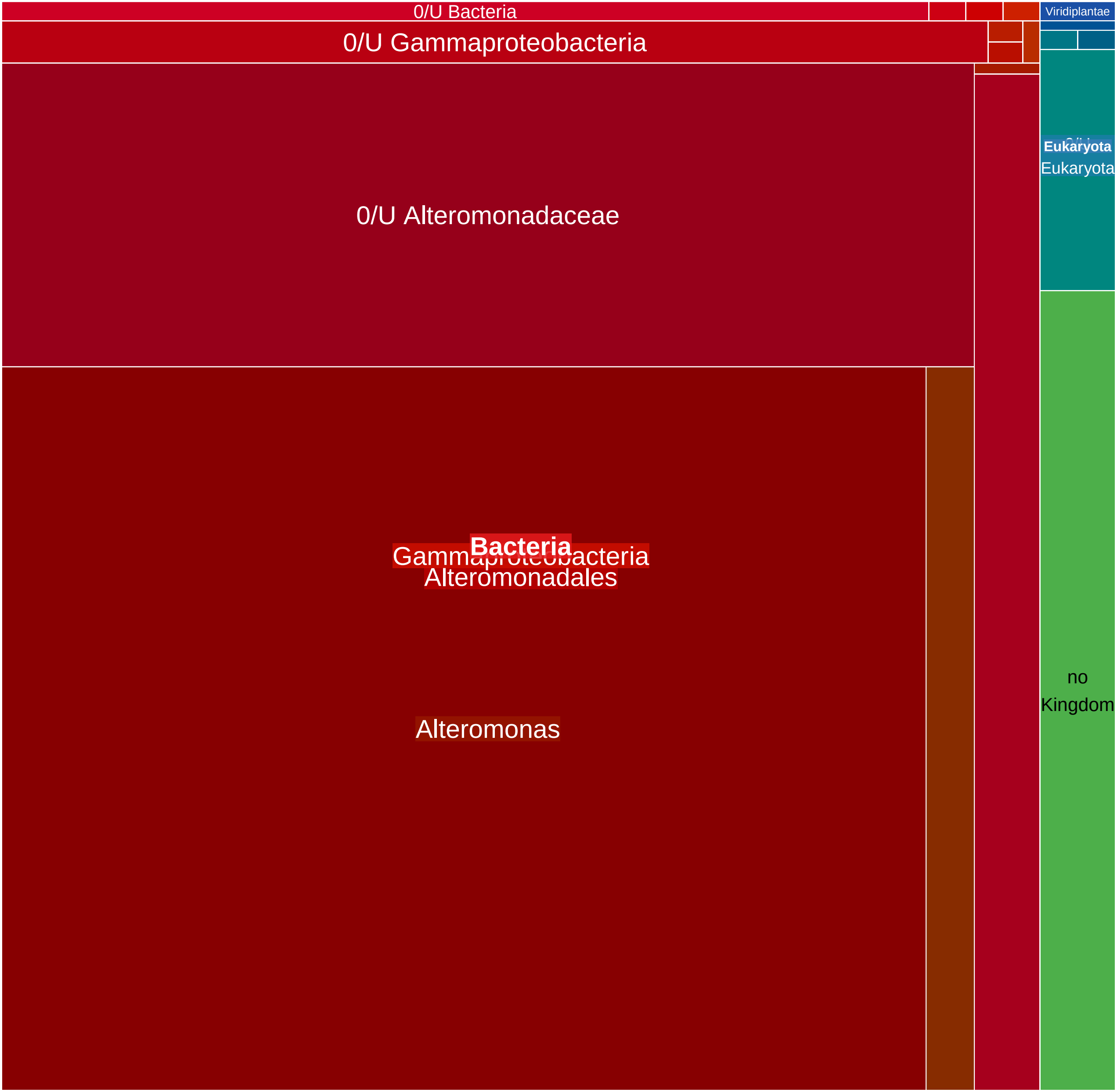

MGT−v1_487 : 1727 unigenes, 81 (4.69%) taxonomically assigned

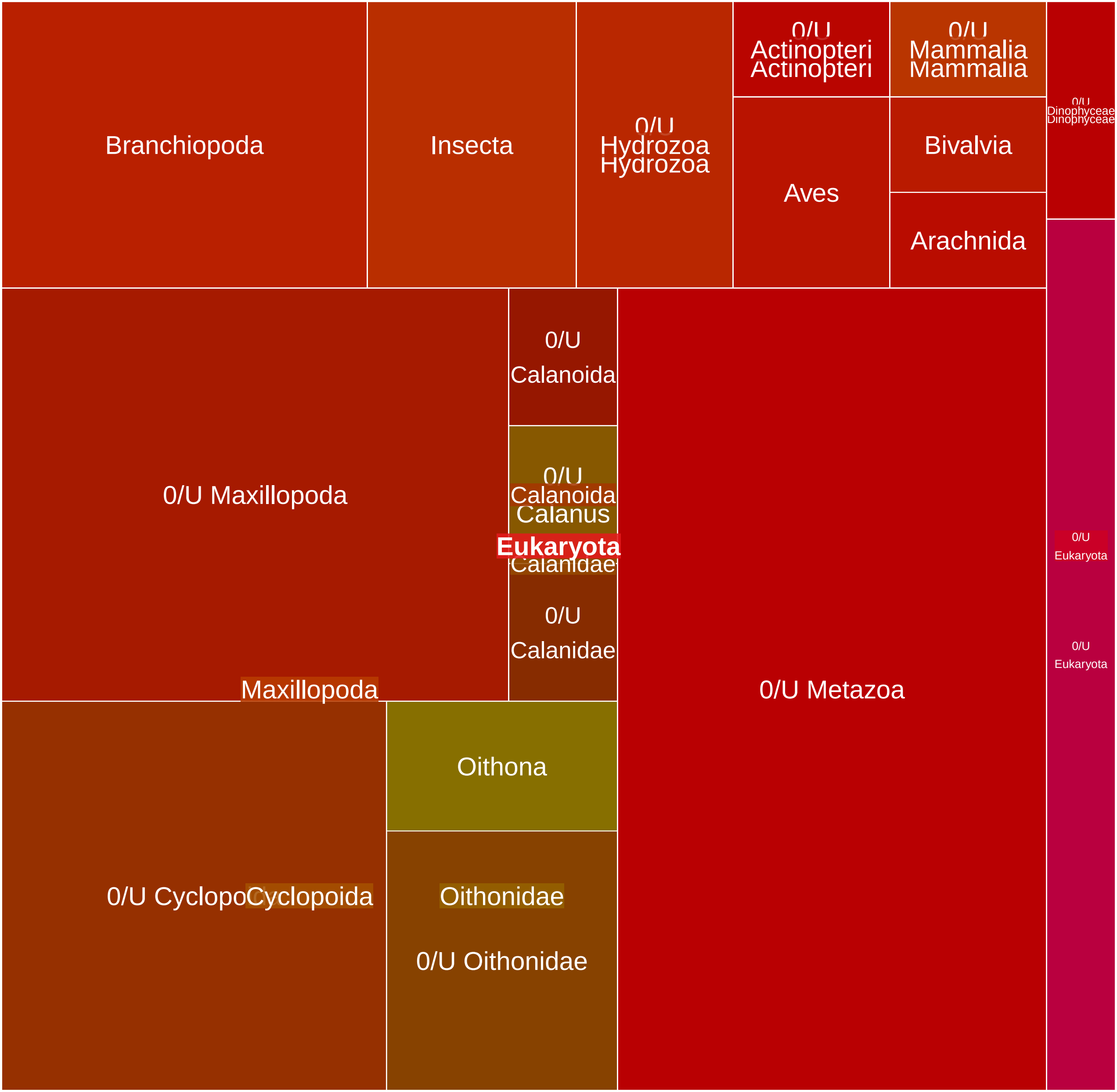

MGT−v1_488 : 1723 unigenes, 536 (31.11%) taxonomically assigned

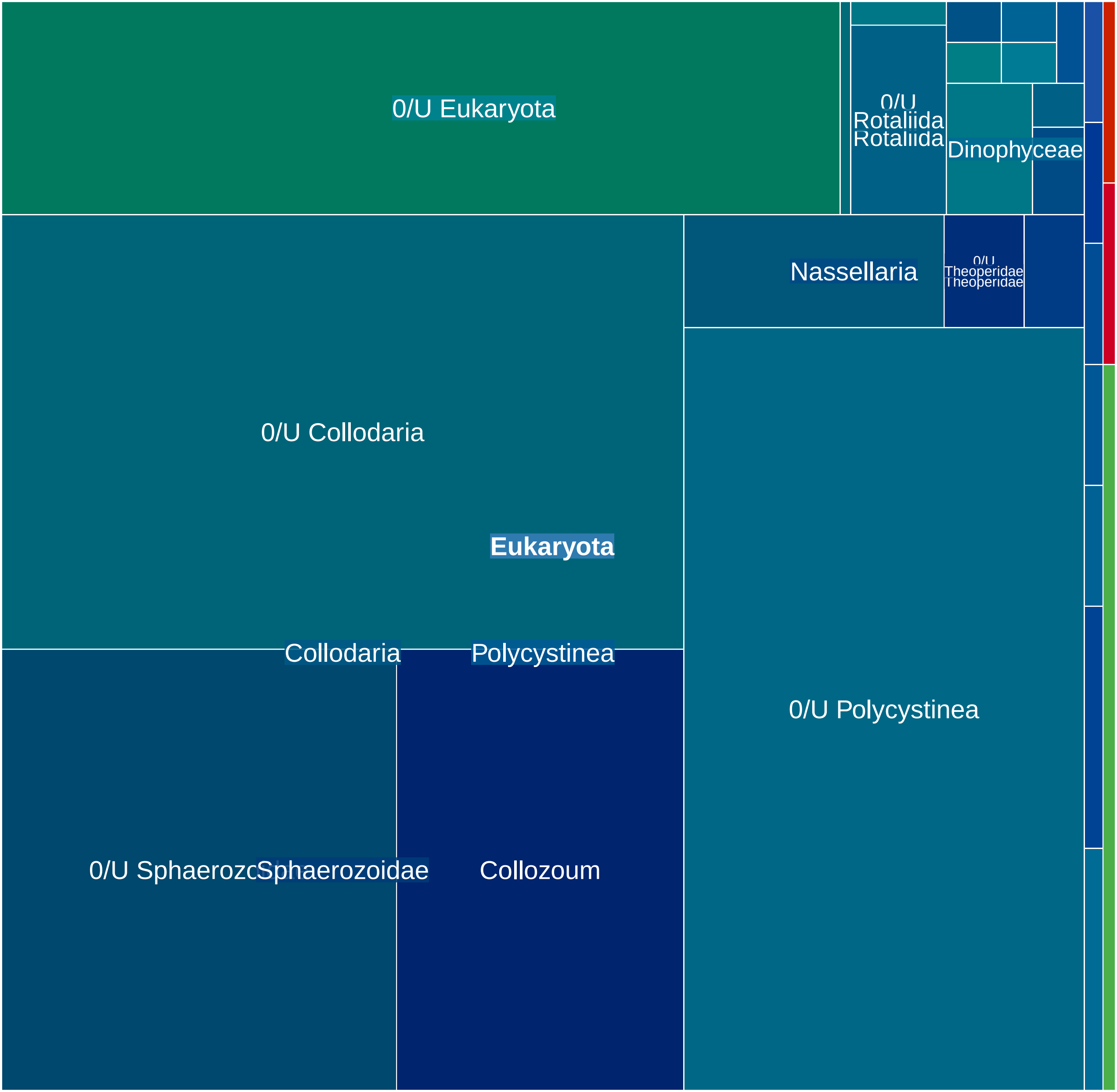

MGT−v1_489 : 1695 unigenes, 182 (10.74%) taxonomically assigned

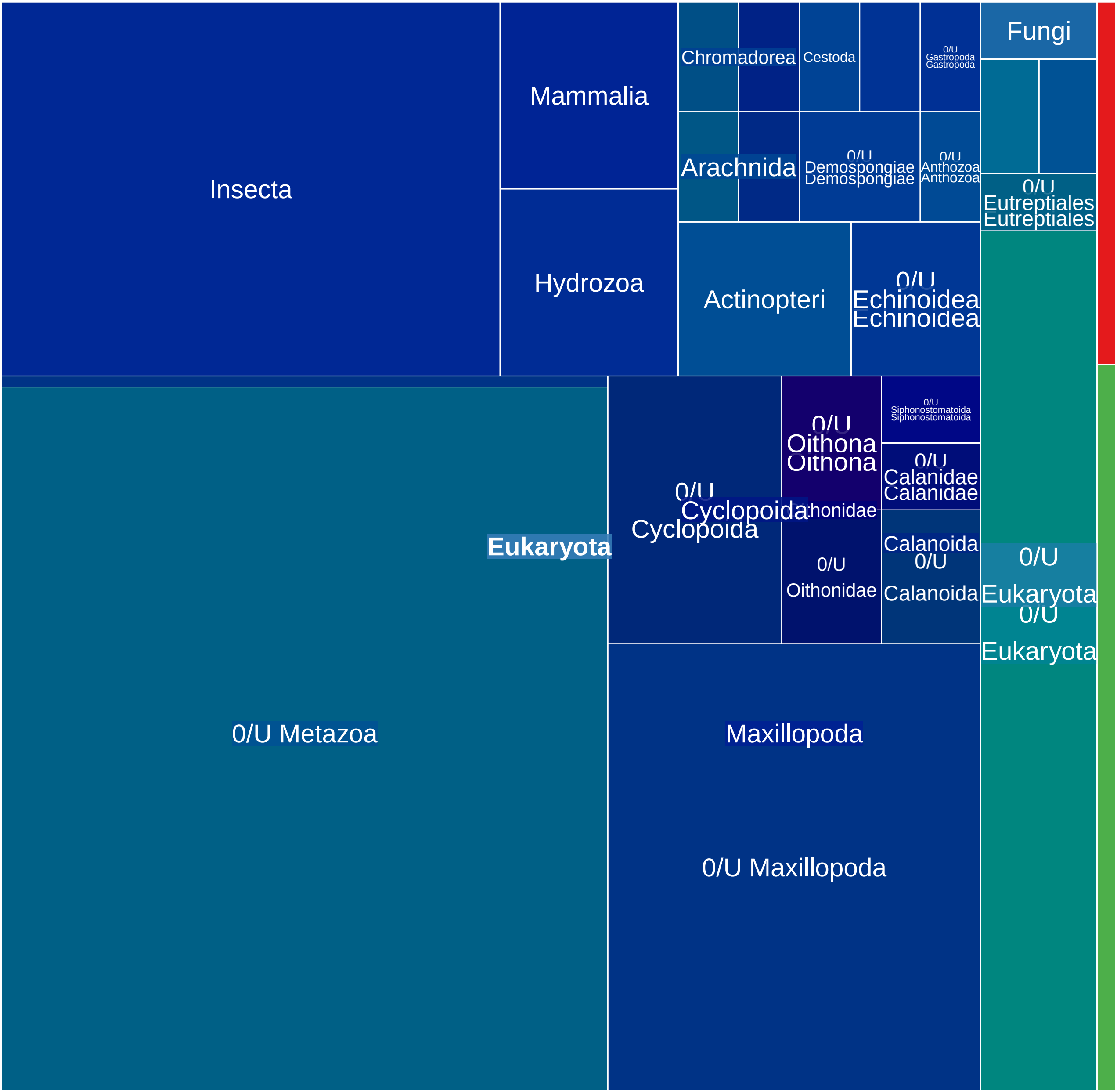

MGT−v1_490 : 1692 unigenes, 1566 (92.55%) taxonomically assigned

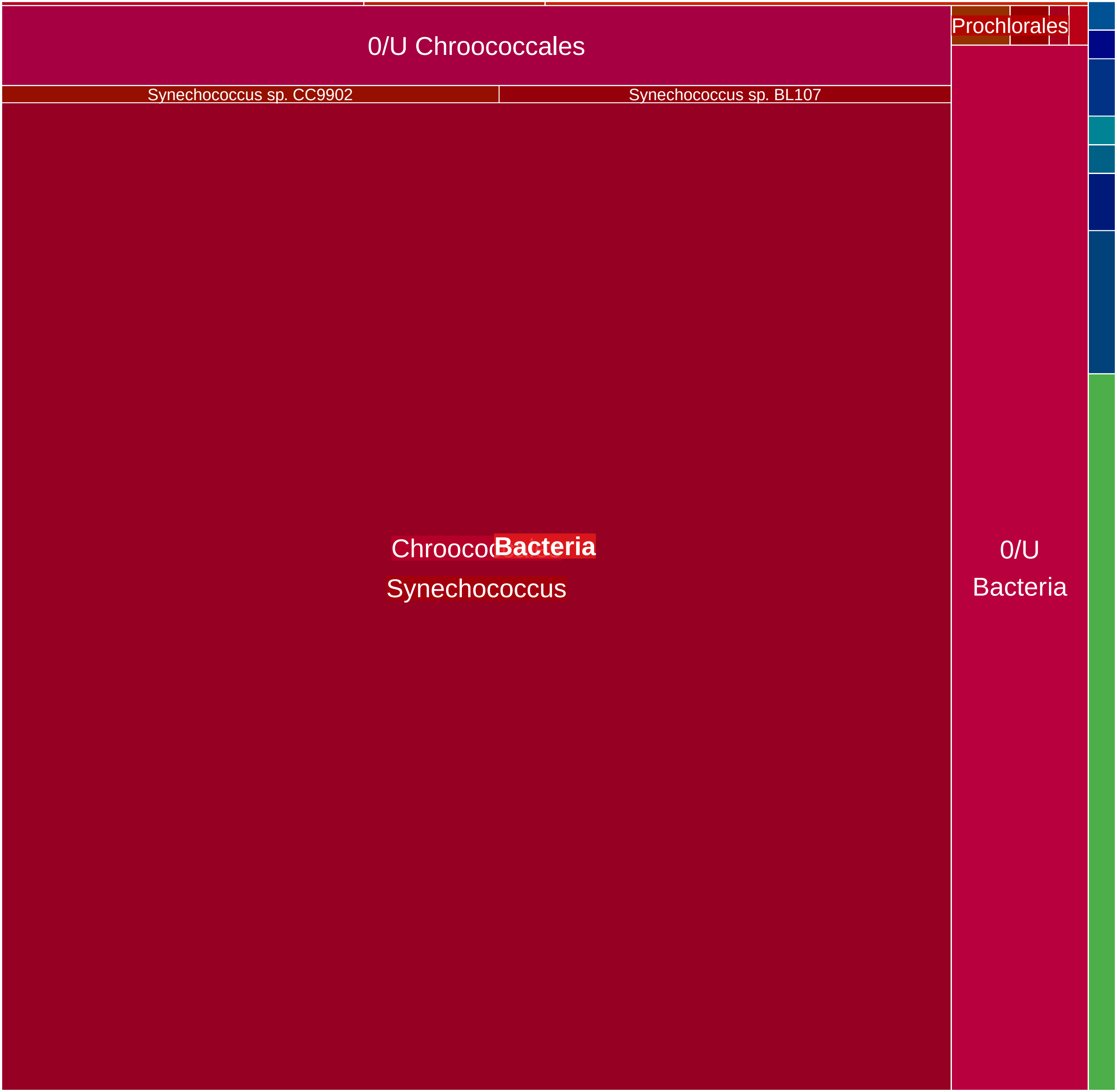

MGT−v1_491 : 1691 unigenes, 1594 (94.26%) taxonomically assigned

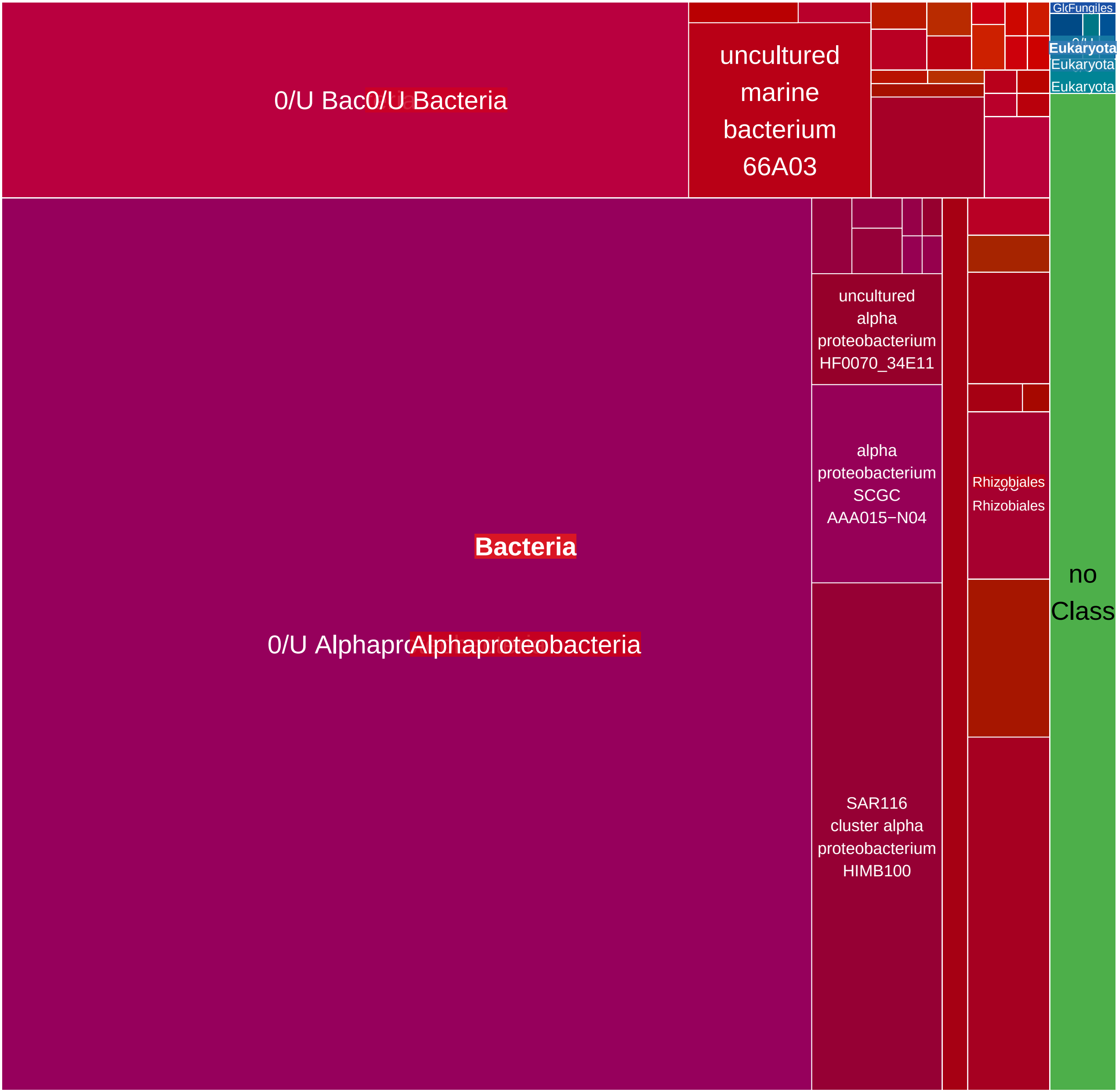

MGT−v1_492 : 1689 unigenes, 211 (12.49%) taxonomically assigned

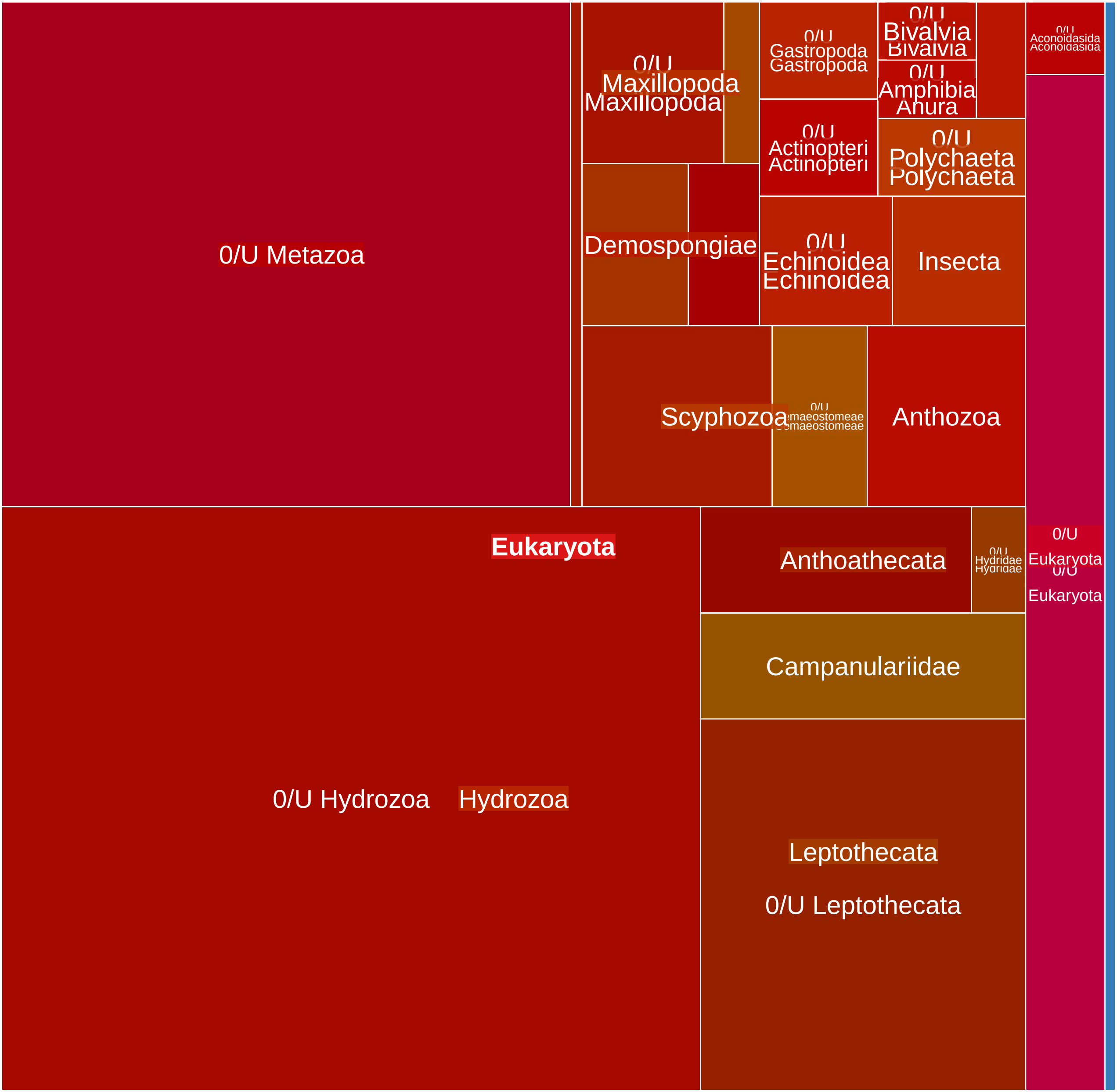

MGT−v1_493 : 1681 unigenes, 1227 (72.99%) taxonomically assigned

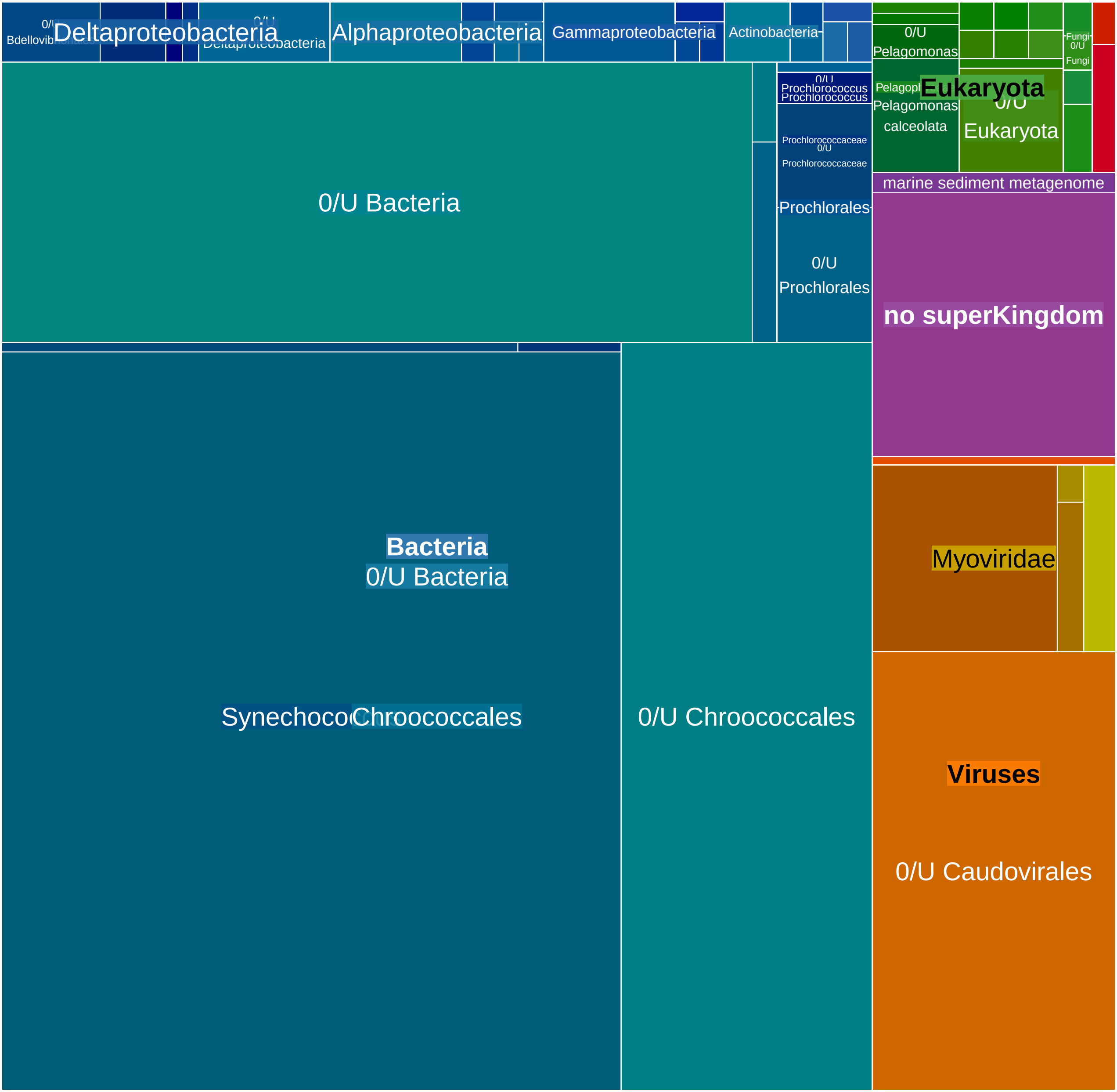

MGT−v1_494 : 1672 unigenes, 1186 (70.93%) taxonomically assigned

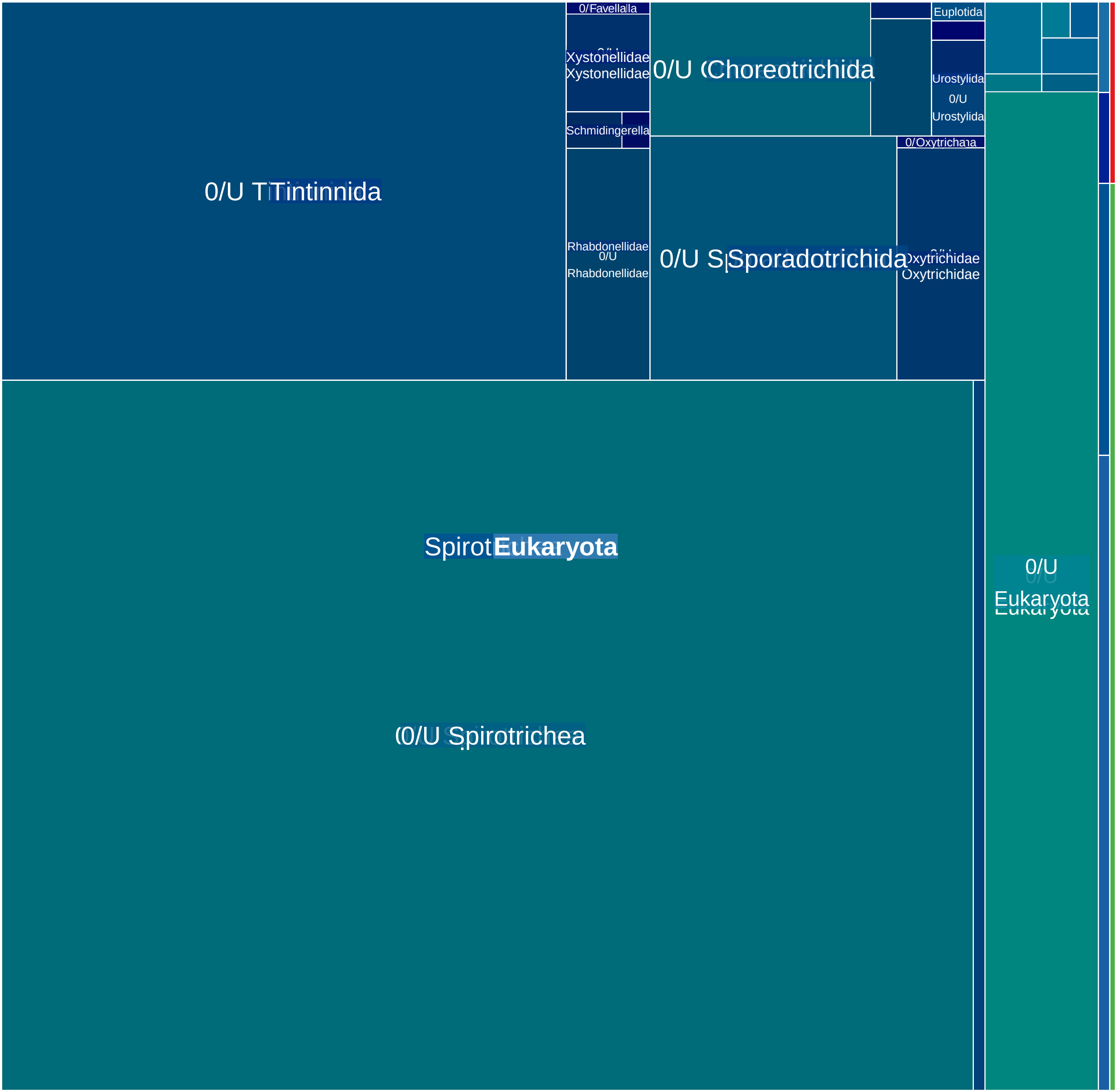

### MGT−v1_495 : 1664 unigenes, 592 (35.58%) taxonomically assigned

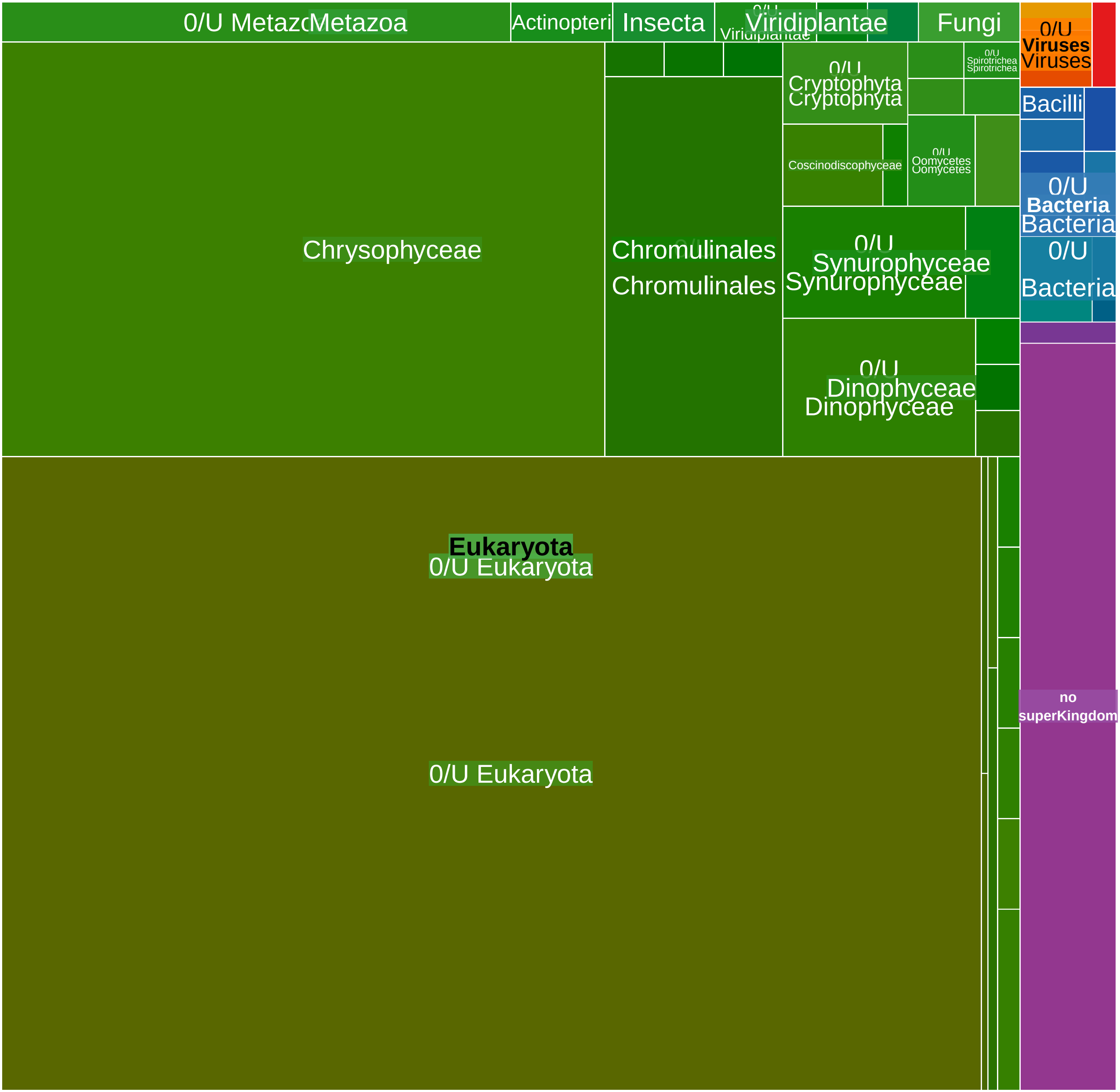

MGT−v1_496 : 1653 unigenes, 214 (12.95%) taxonomically assigned

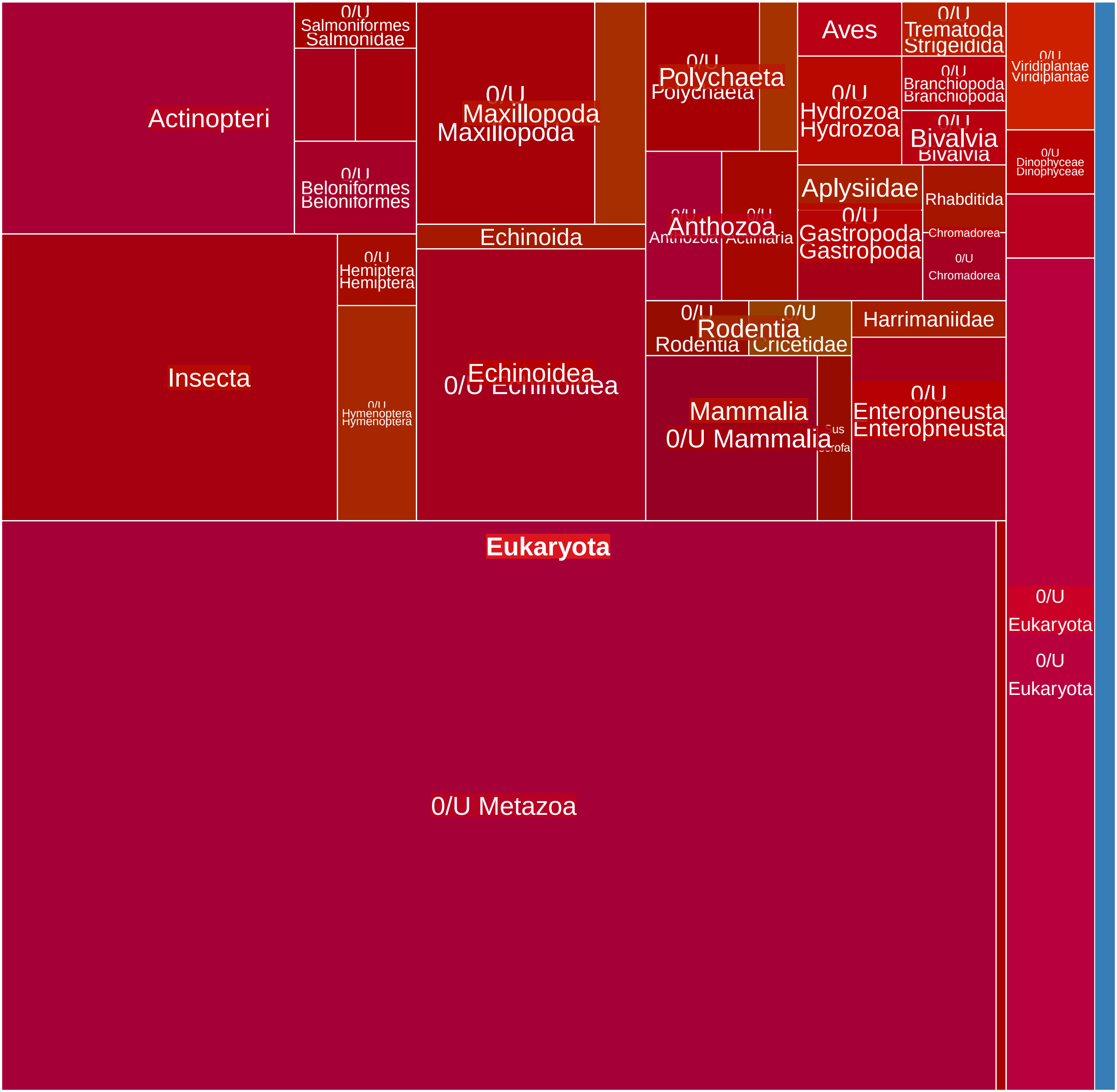

MGT−v1_497 : 1652 unigenes, 1406 (85.11%) taxonomically assigned

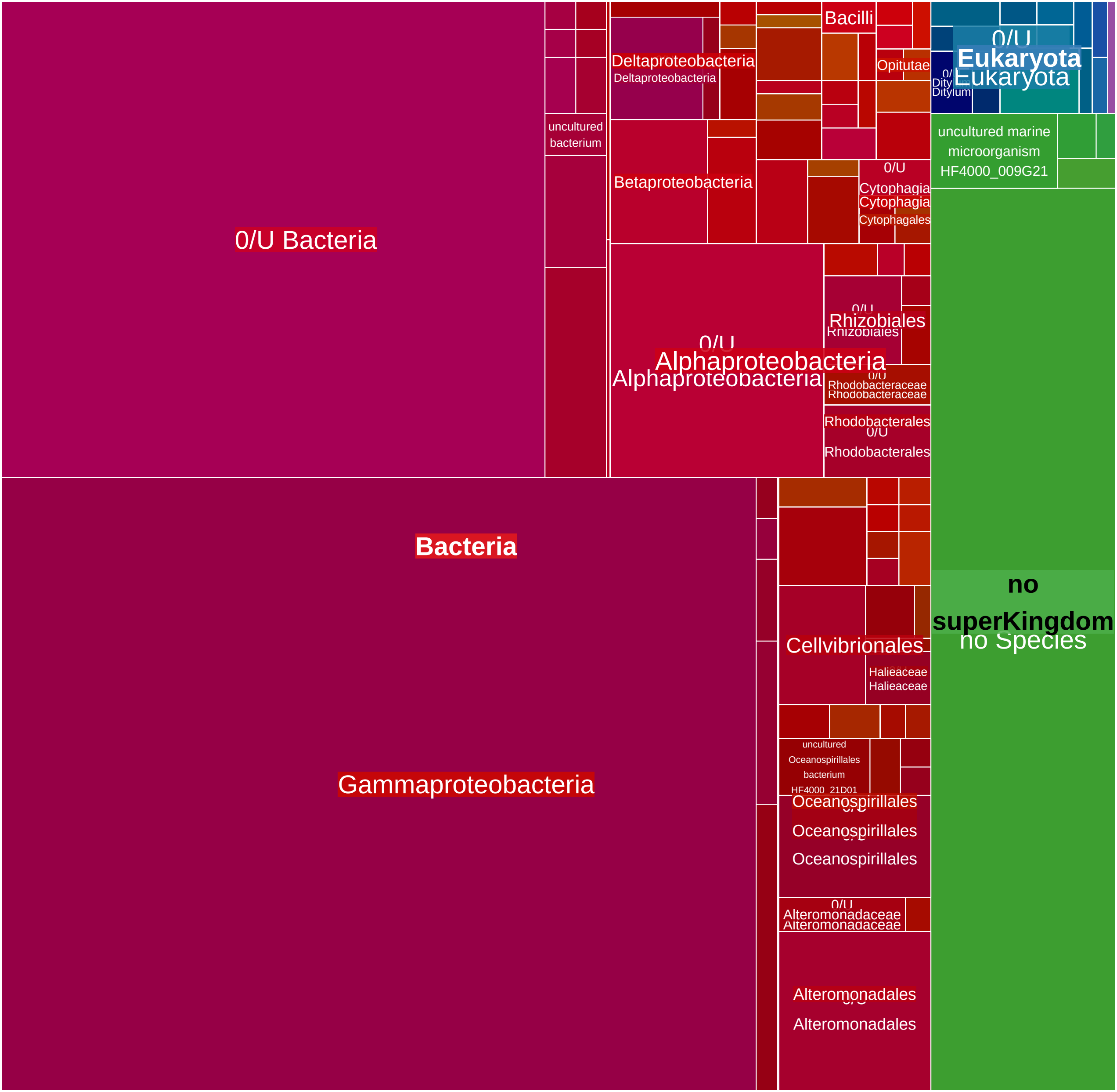

MGT−v1_498 : 1646 unigenes, 1383 (84.02%) taxonomically assigned

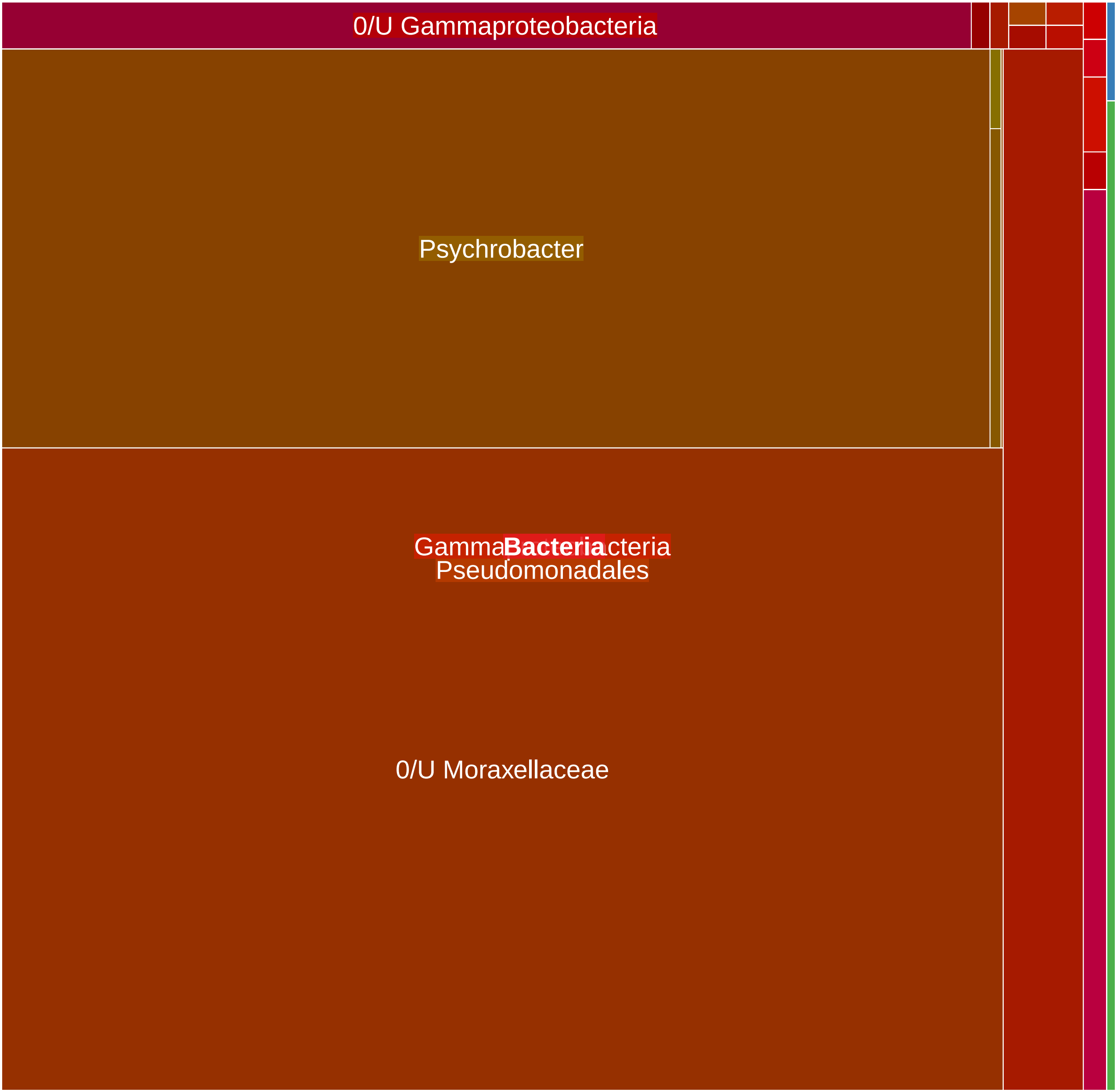

MGT−v1_499 : 1637 unigenes, 937 (57.24%) taxonomically assigned

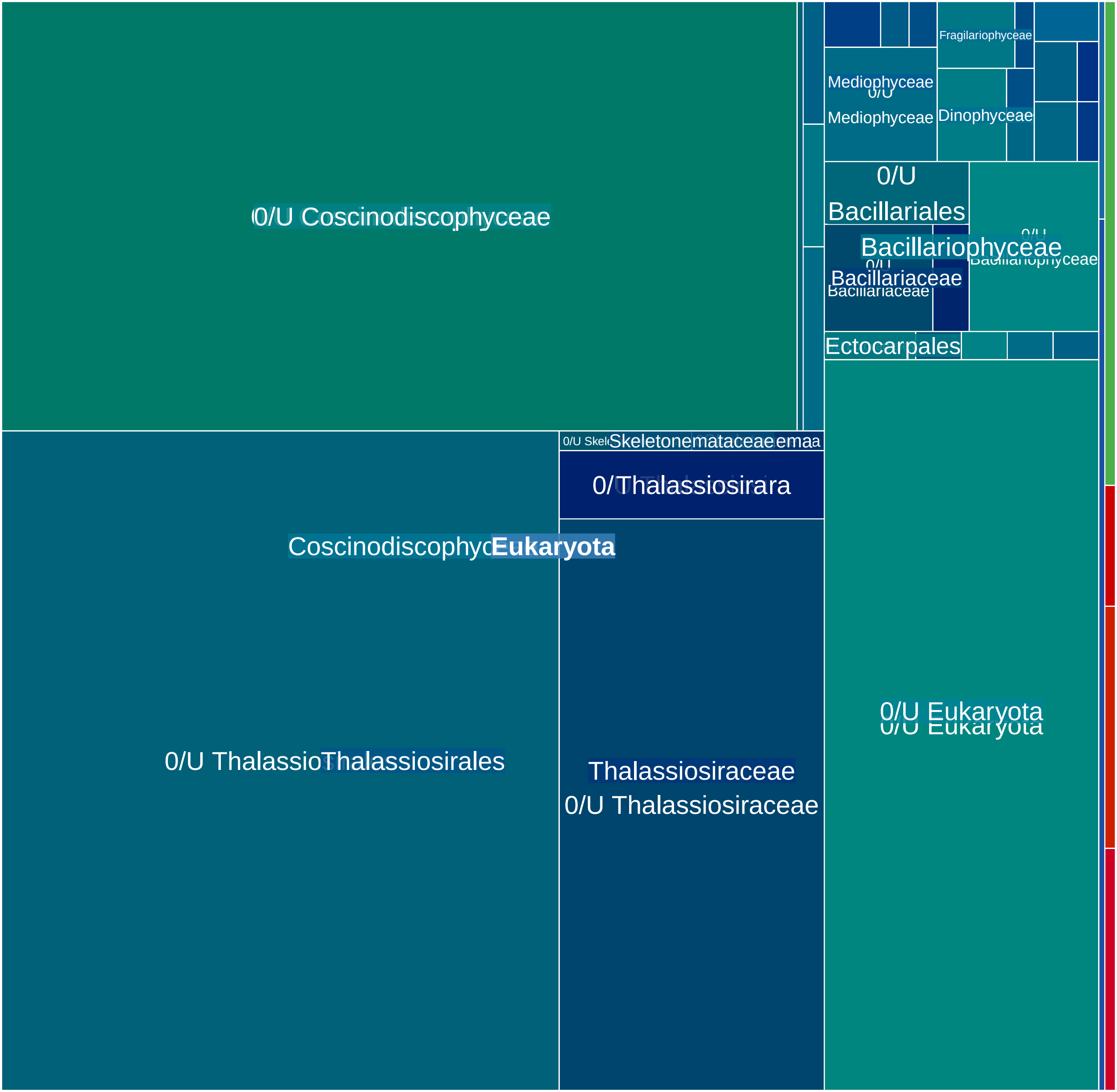

MGT−v1_500 : 1637 unigenes, 119 (7.27%) taxonomically assigned

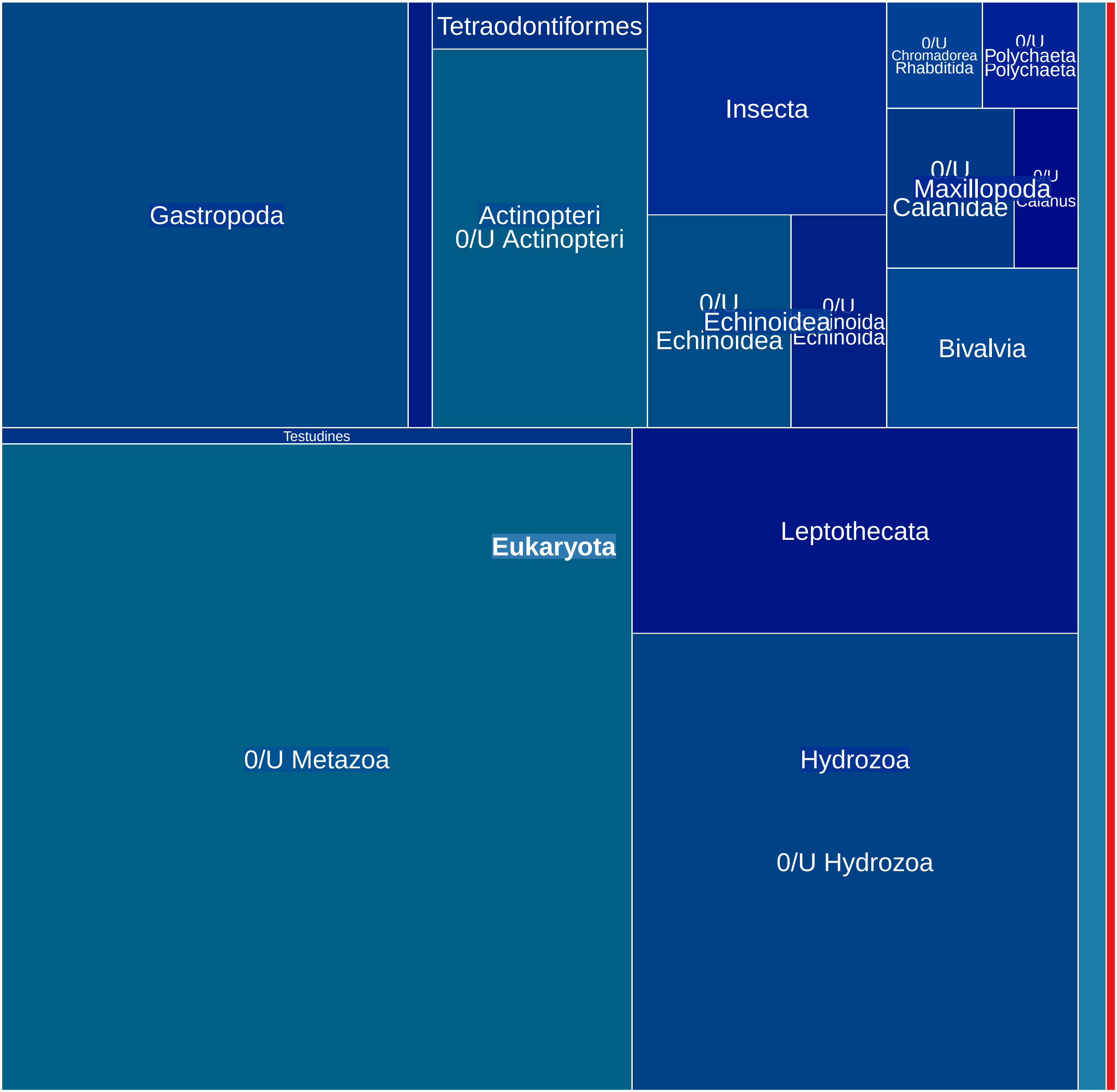

MGT−v1_501 : 1636 unigenes, 232 (14.18%) taxonomically assigned

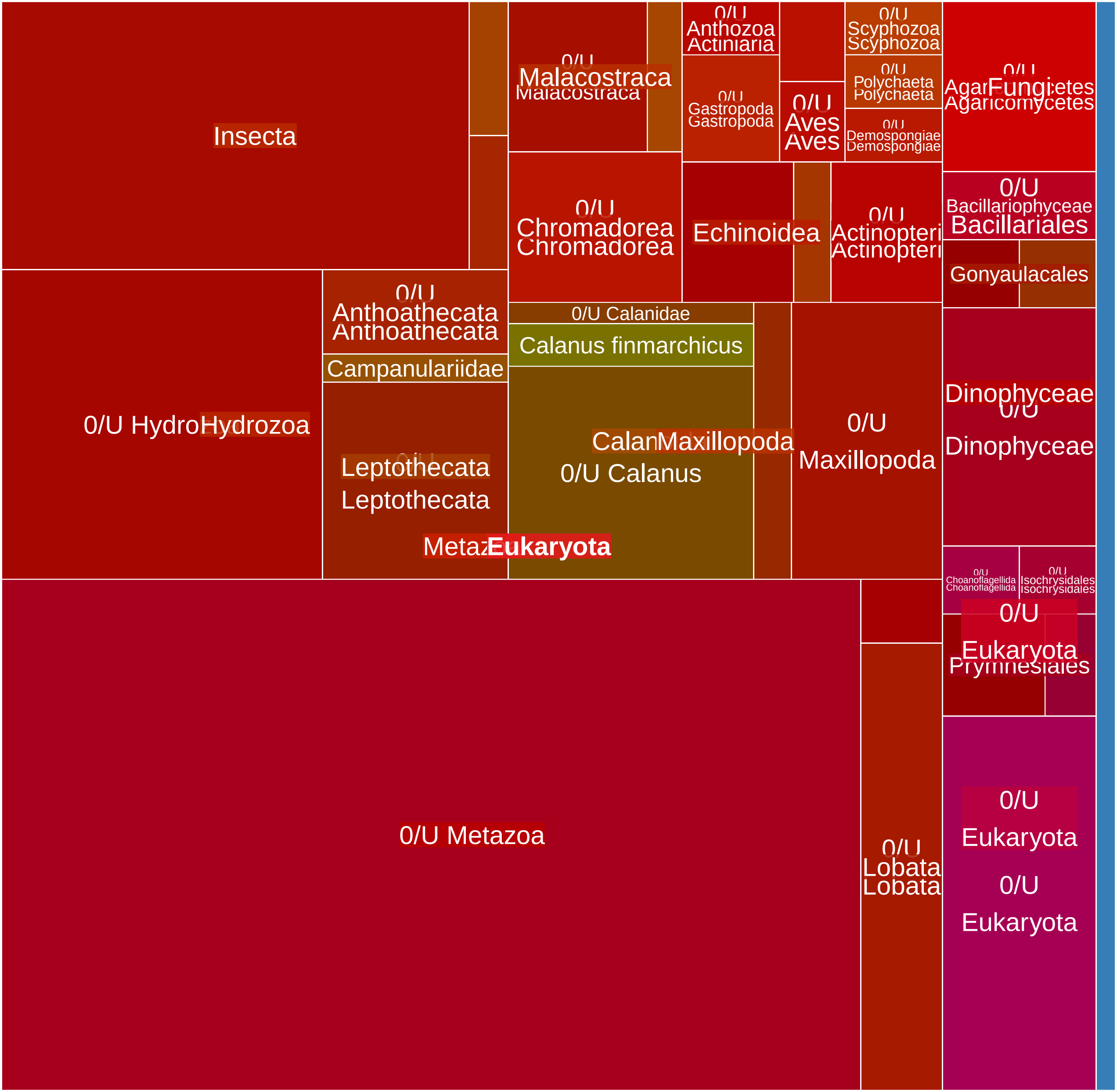

MGT−v1_502 : 1633 unigenes, 93 (5.70%) taxonomically assigned

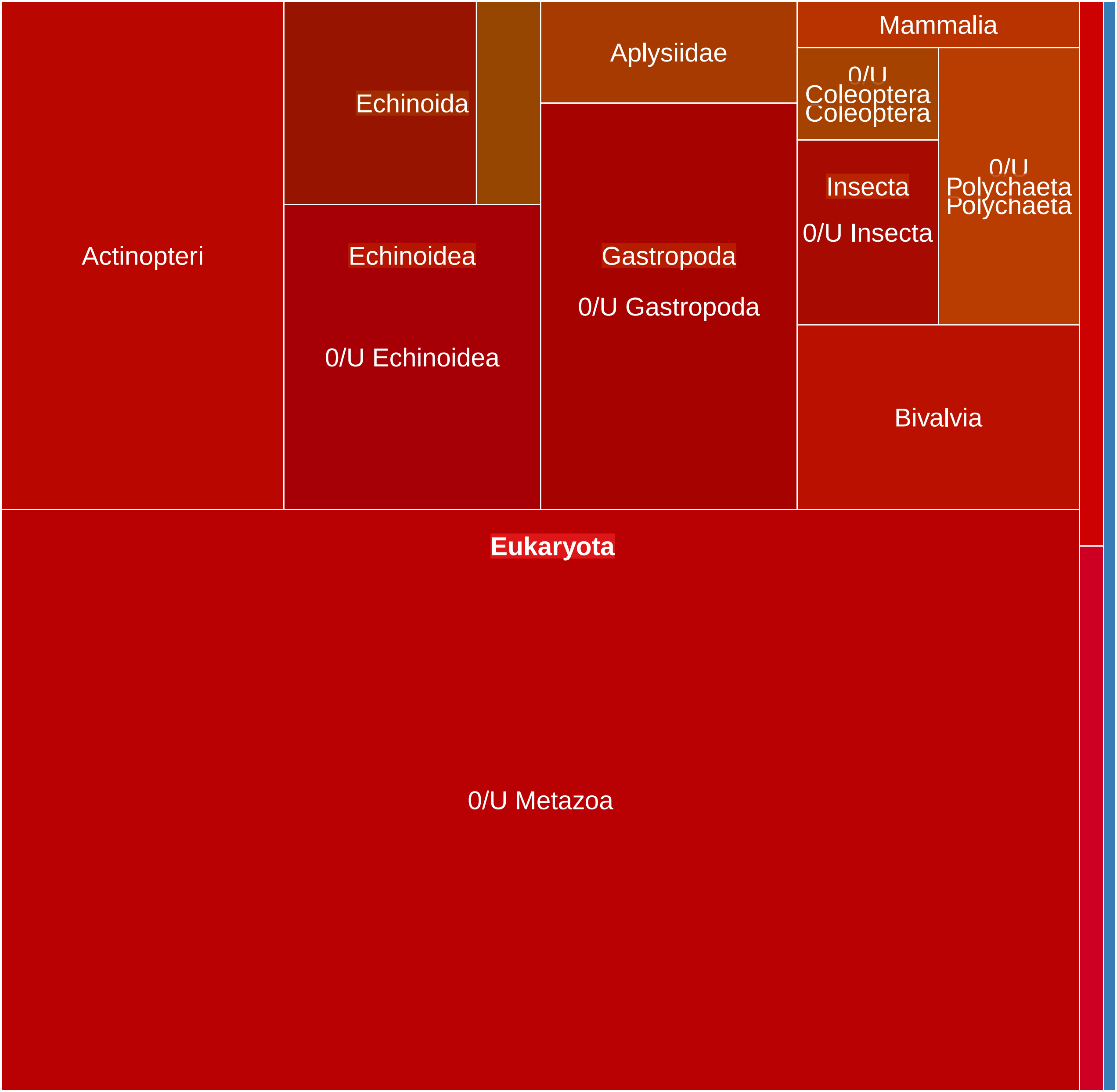

MGT−v1_503 : 1628 unigenes, 436 (26.78%) taxonomically assigned

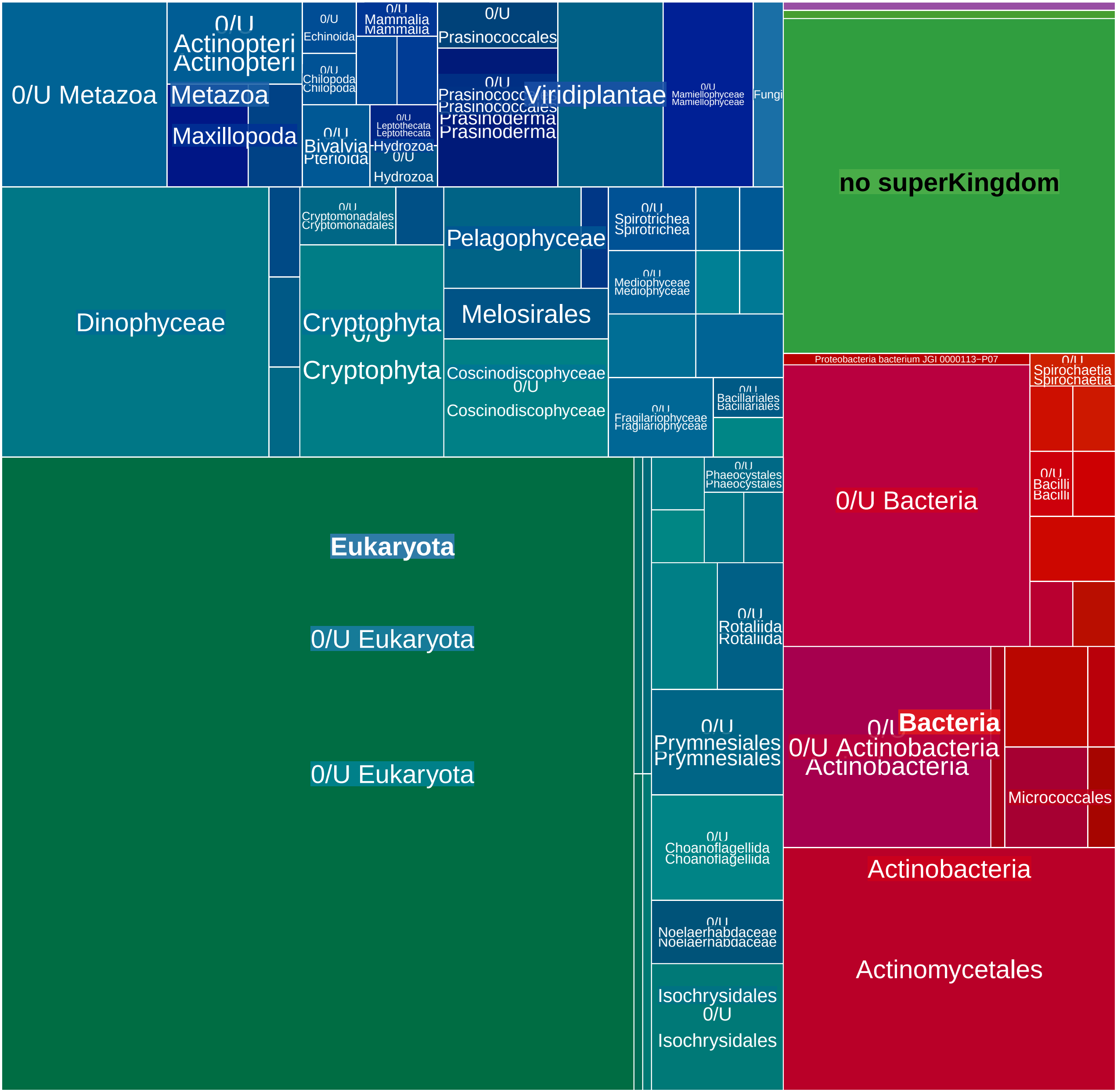

MGT−v1_504 : 1627 unigenes, 184 (11.31%) taxonomically assigned

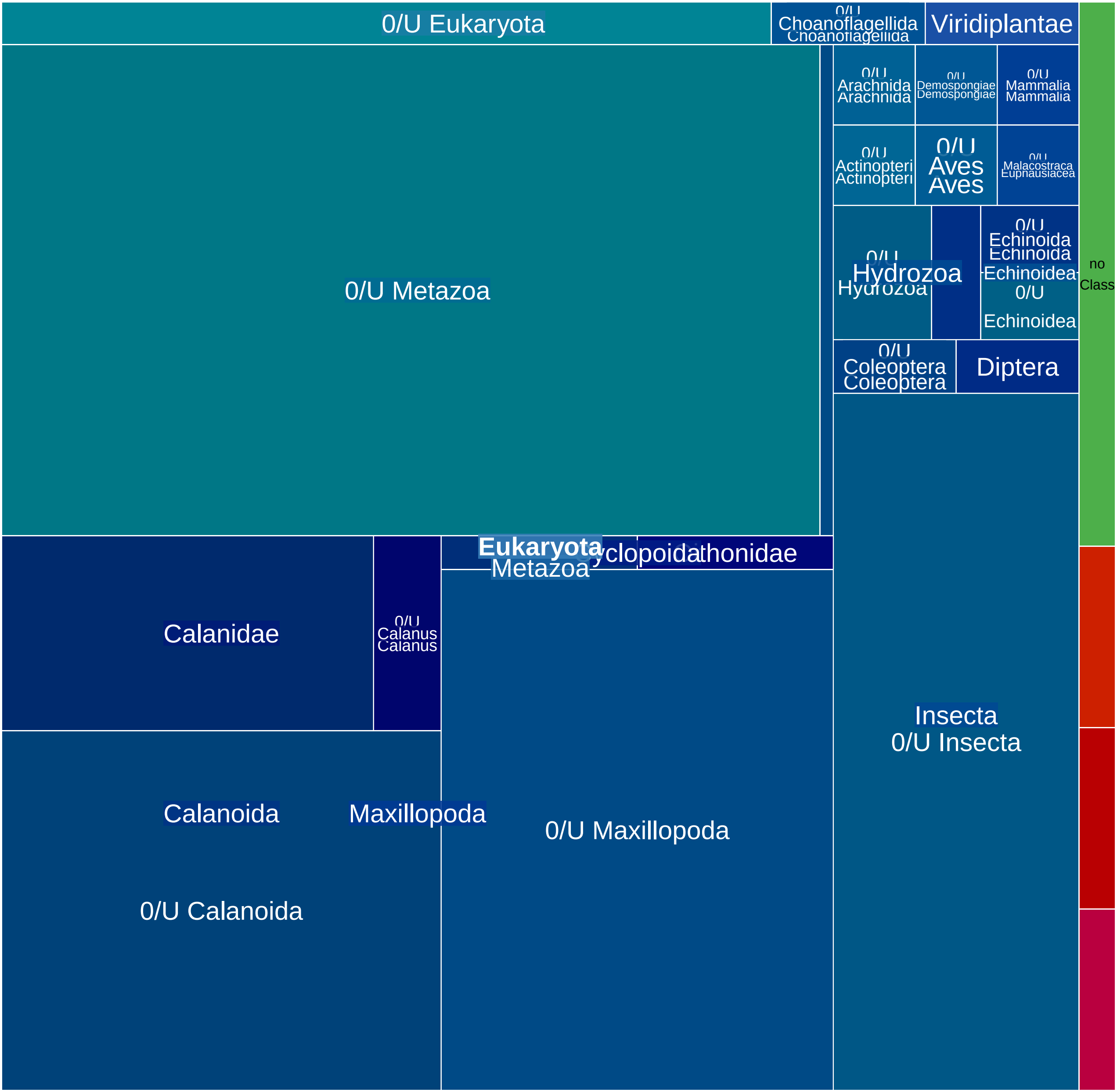

MGT−v1_505 : 1627 unigenes, 1603 (98.52%) taxonomically assigned

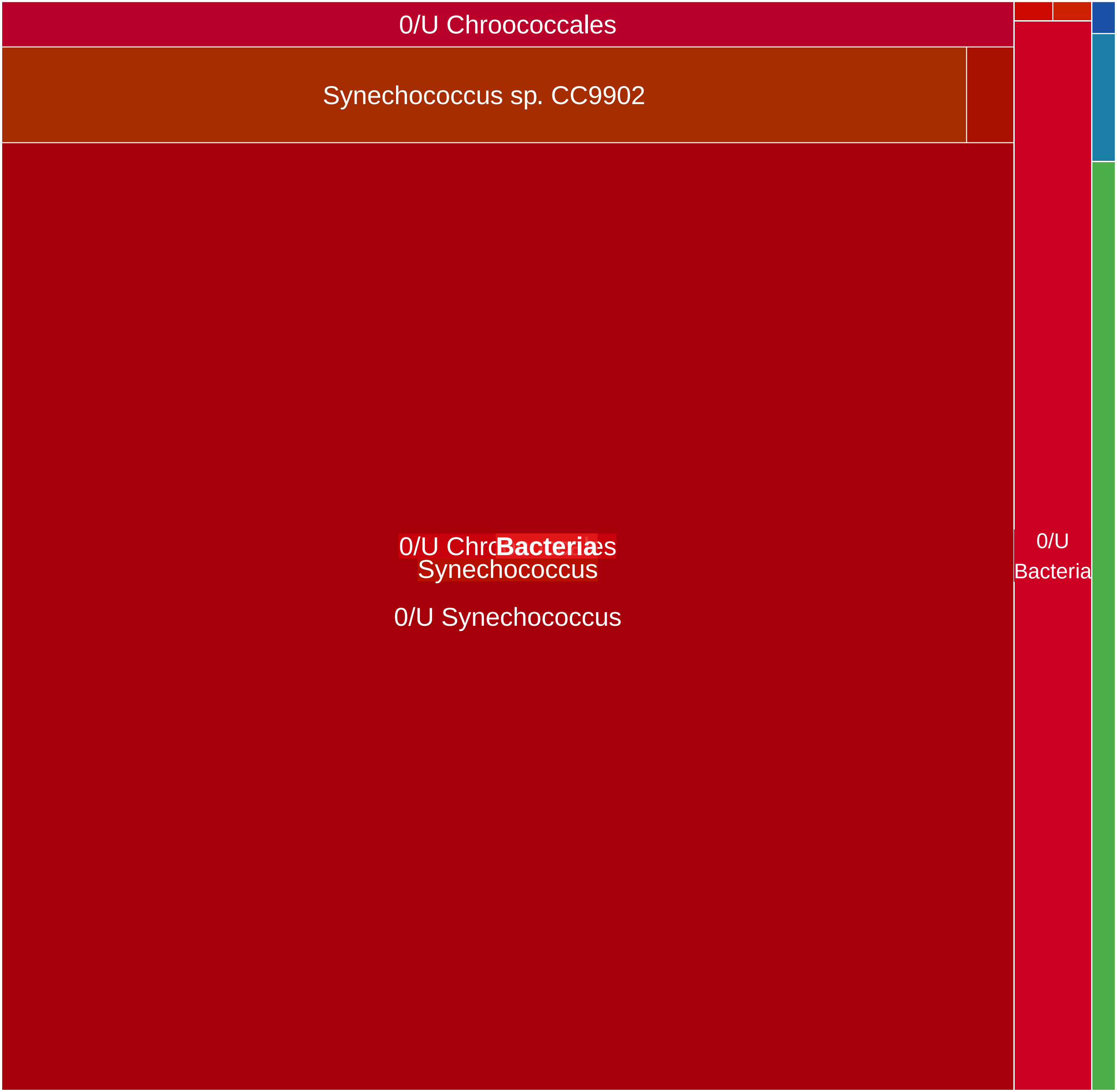

MGT−v1_506 : 1618 unigenes, 158 (9.77%) taxonomically assigned

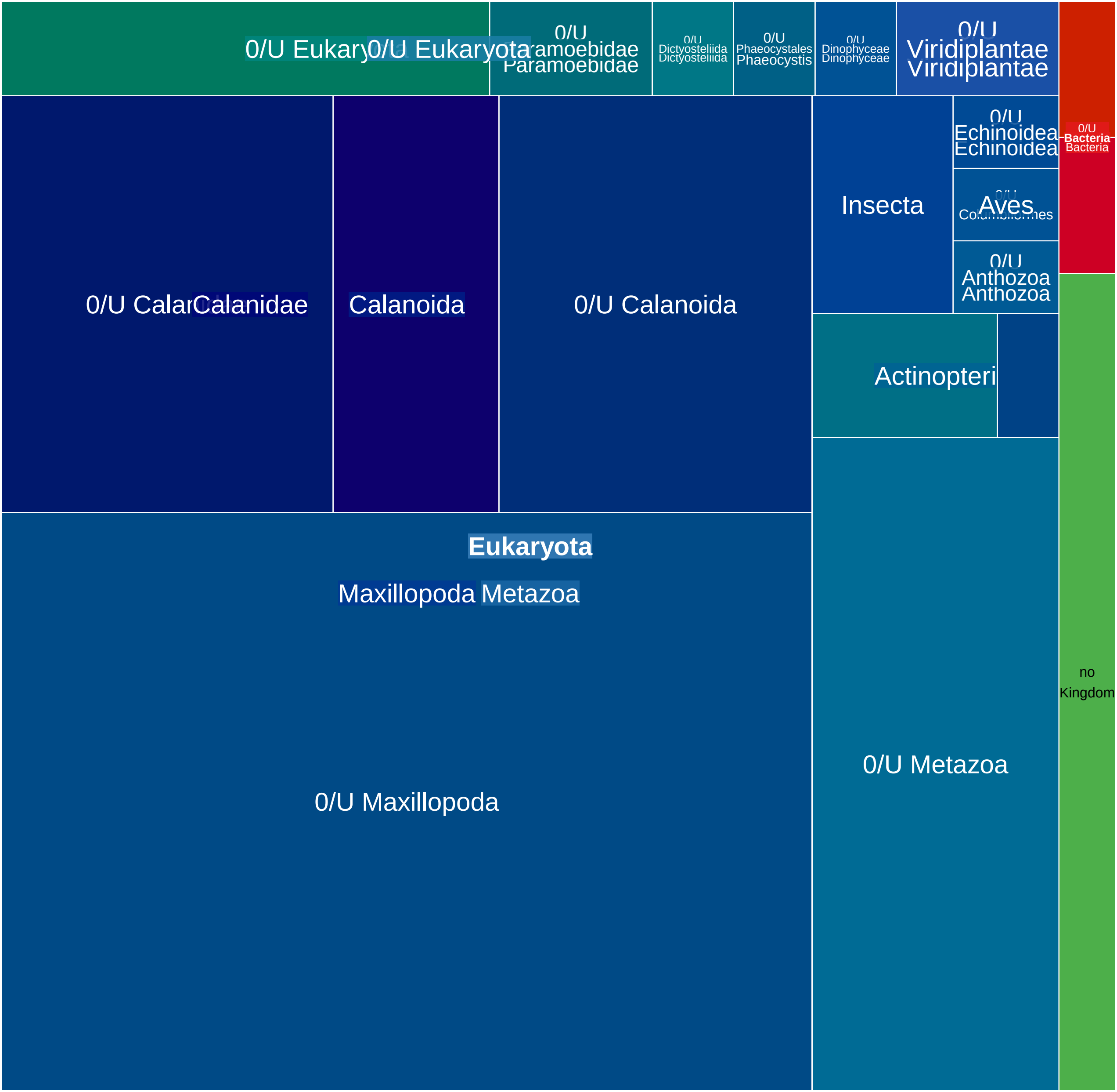

MGT−v1_507 : 1601 unigenes, 202 (12.62%) taxonomically assigned

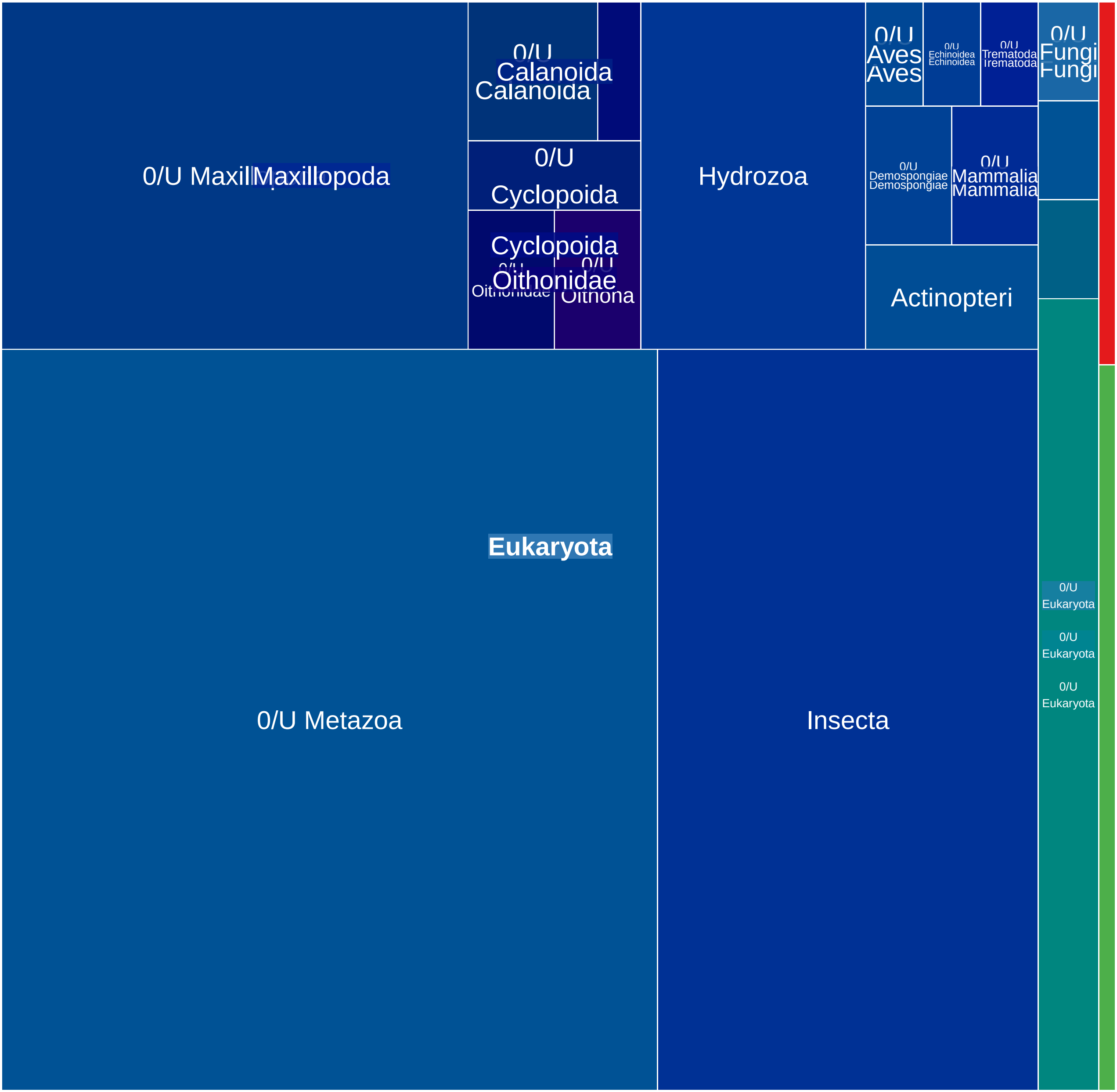

MGT−v1_508 : 1582 unigenes, 25 (1.58%) taxonomically assigned

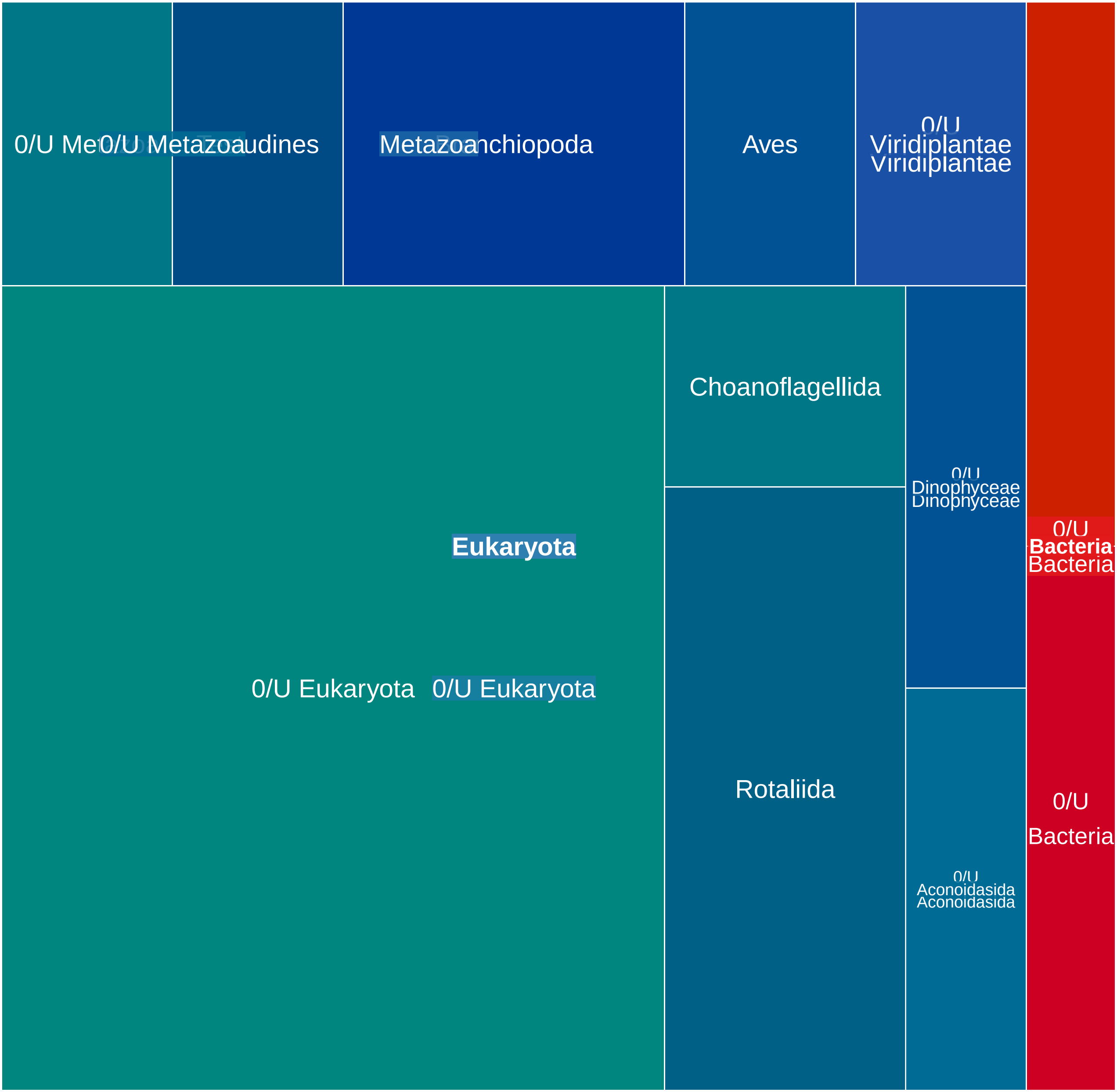

MGT−v1_509 : 1578 unigenes, 974 (61.72%) taxonomically assigned

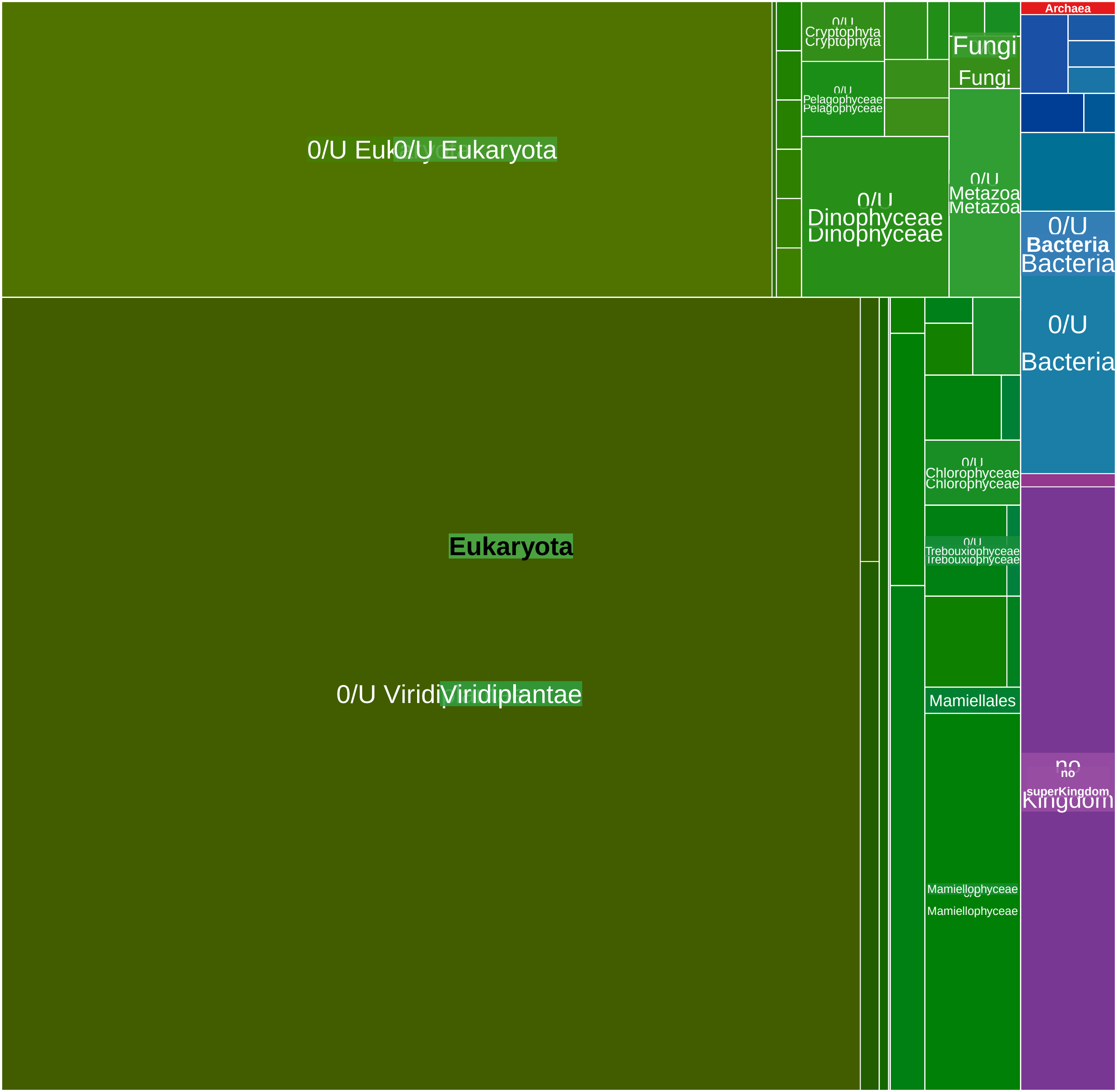

MGT−v1_510 : 1575 unigenes, 728 (46.22%) taxonomically assigned

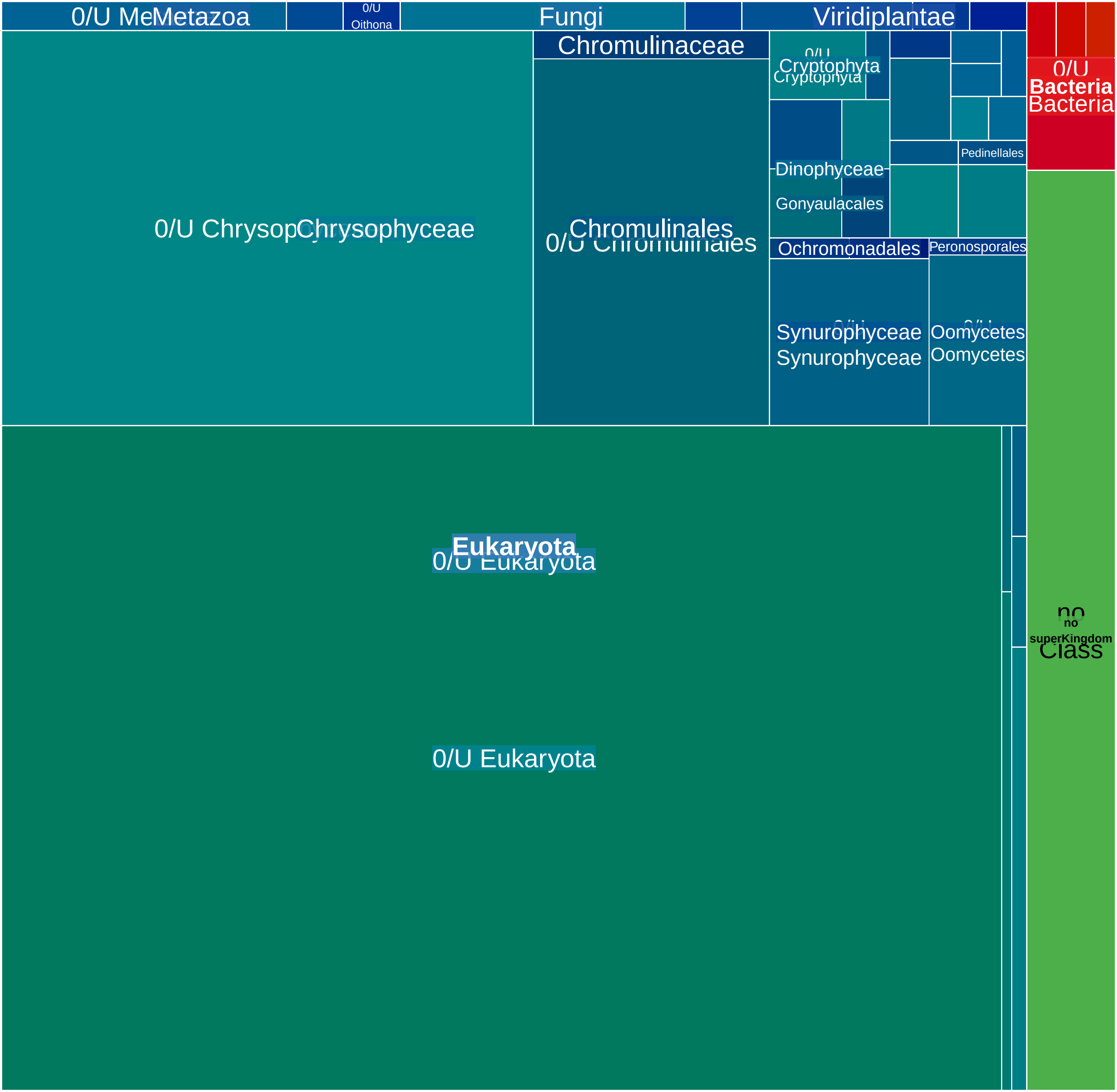

MGT−v1_511 : 1552 unigenes, 1427 (91.95%) taxonomically assigned

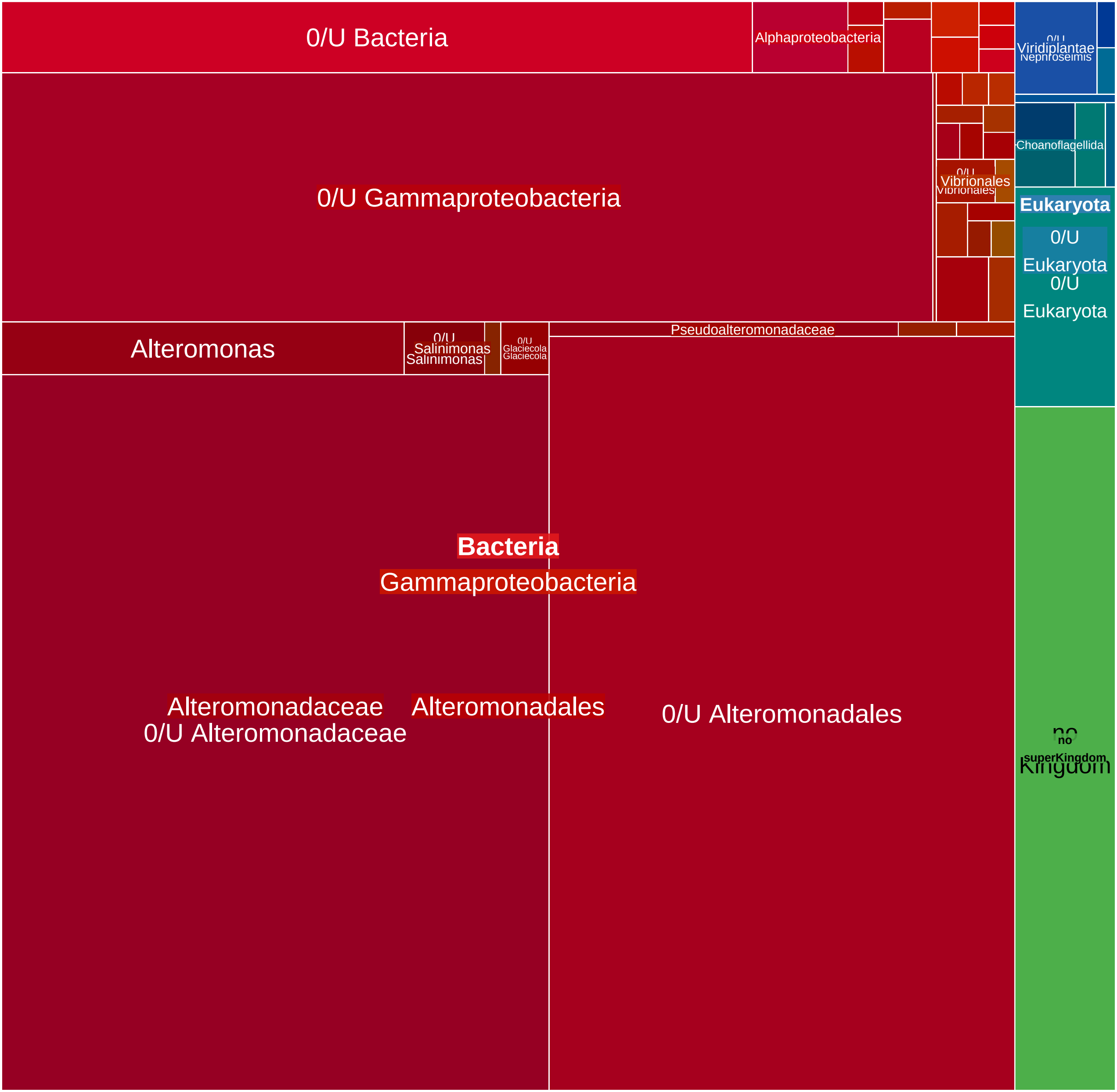

MGT−v1_512 : 1549 unigenes, 136 (8.78%) taxonomically assigned

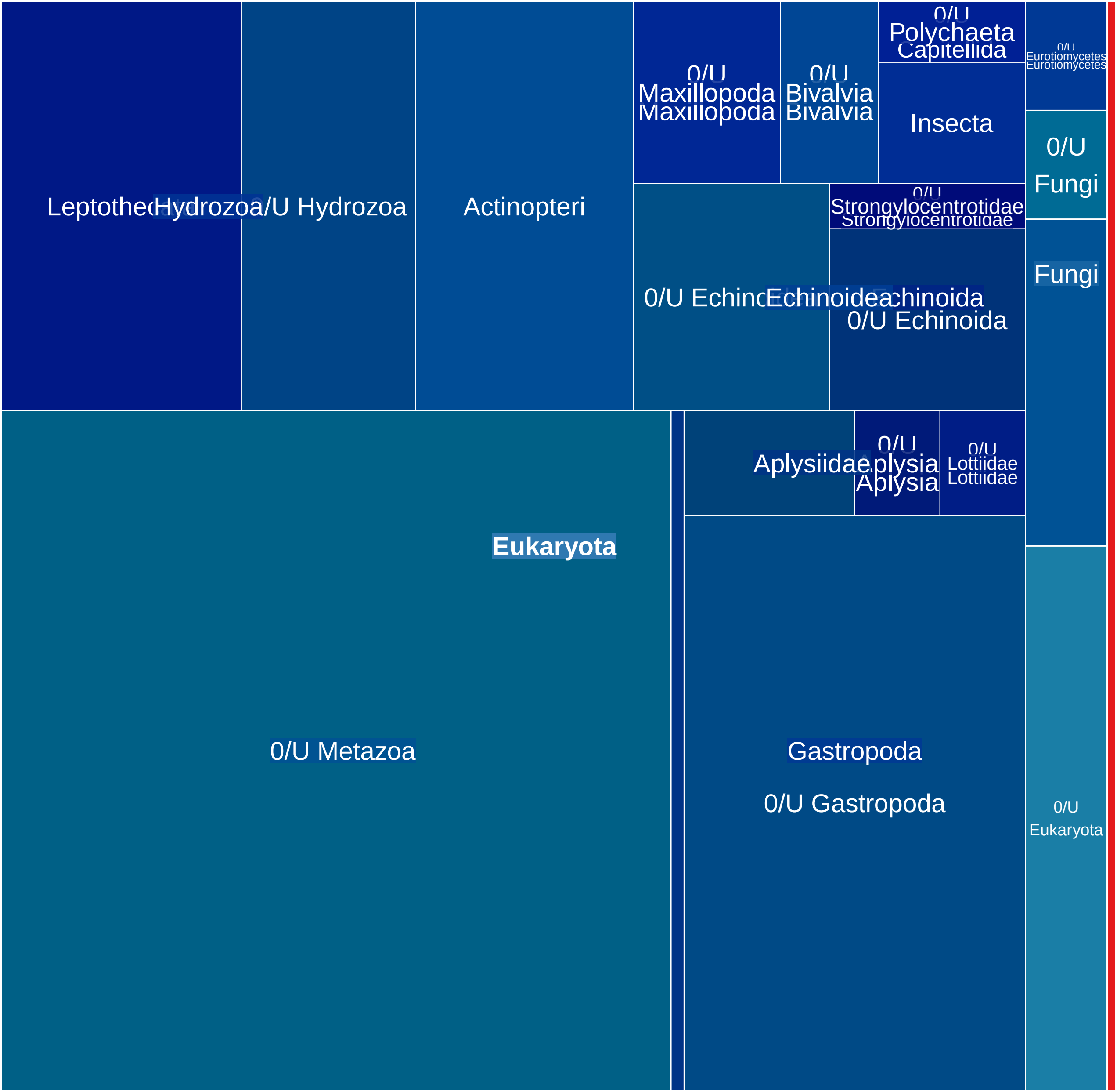

MGT−v1_513 : 1546 unigenes, 161 (10.41%) taxonomically assigned

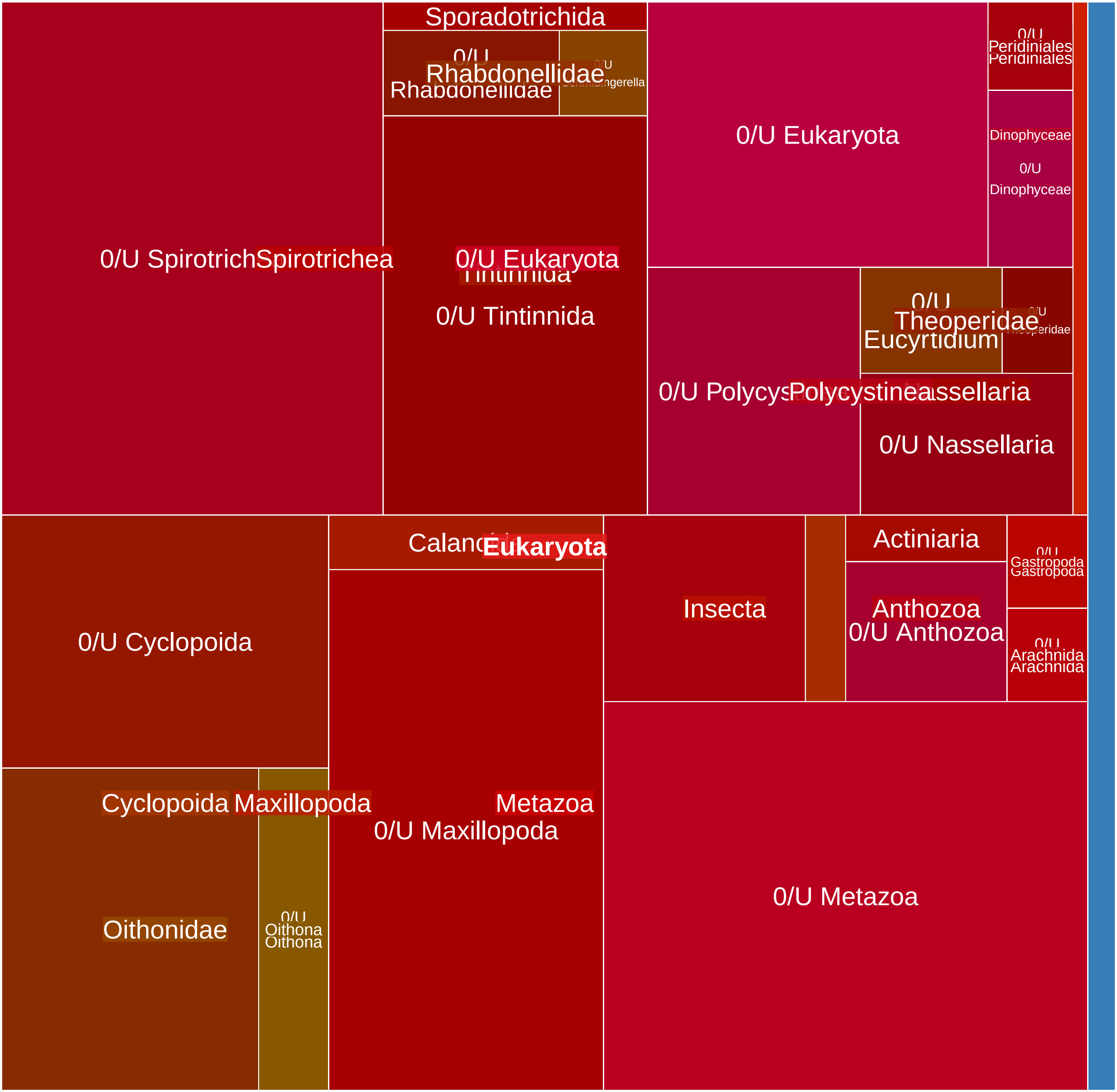

MGT−v1_514 : 1541 unigenes, 373 (24.21%) taxonomically assigned

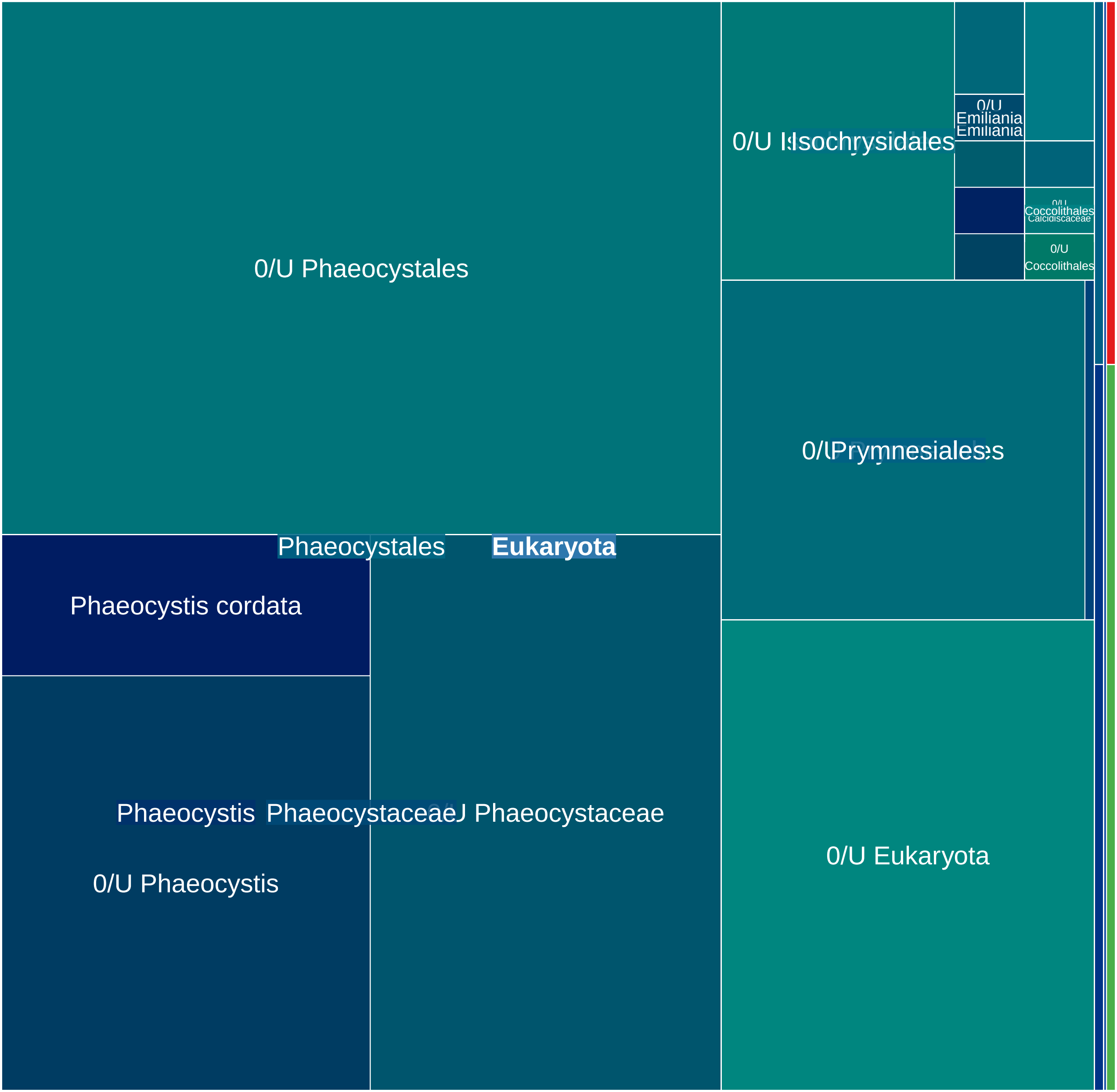

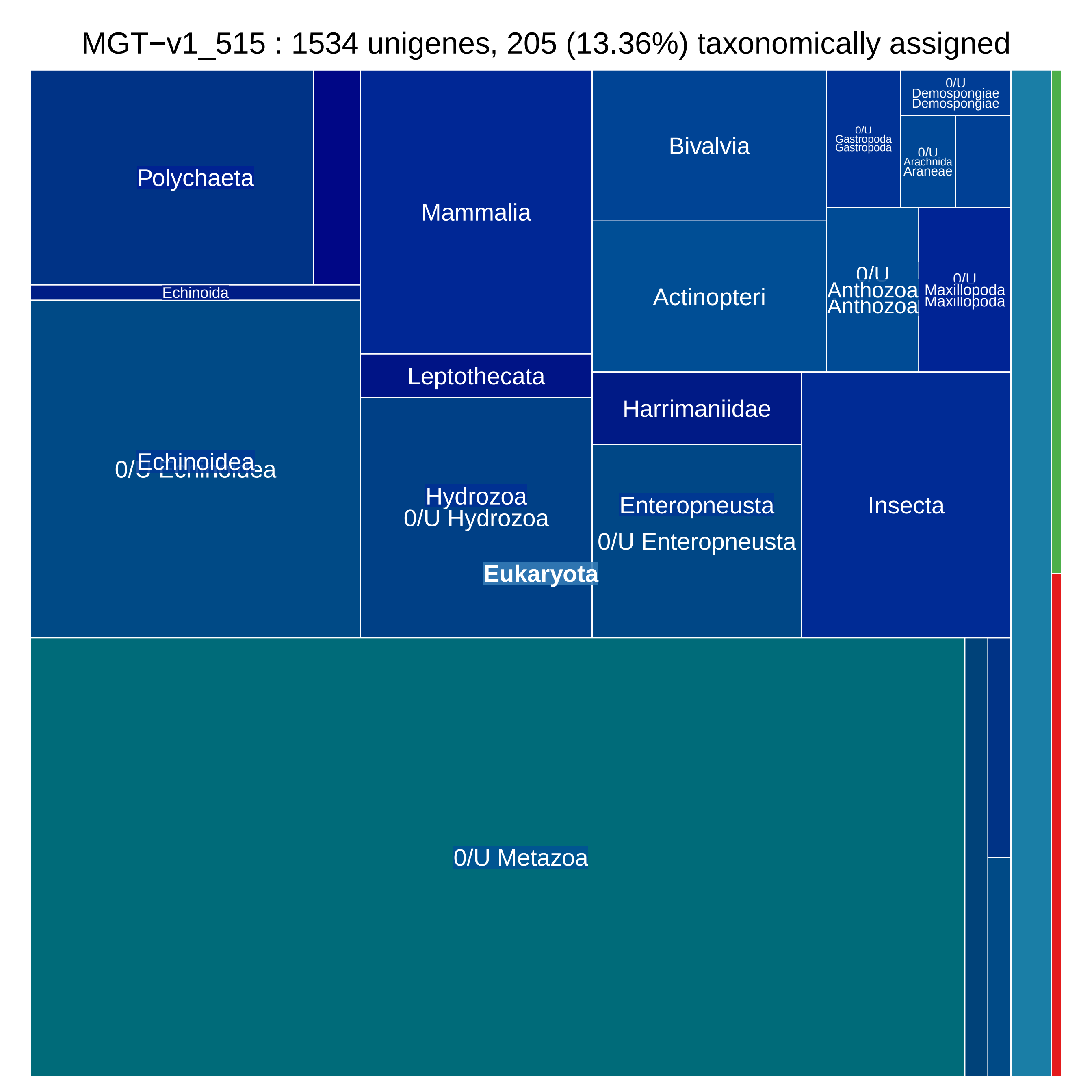
MGT−v1_515 : 1534 unigenes, 205 (13.36%) taxonomically assigned
Polychaeta
Echinoida
0/U Echinoidea
Echinoidea
Mammalia
Leptothecata
Hydrozoa
0/U Hydrozoa
Eukaryota
Bivalvia
Actinopteri
Harrimaniidae
Enteropneusta
0/U Enteropneusta
0/U Gastropoda Gastropoda
0/U Demospongiae Demospongiae
0/U Arachnida Araneae
0/U Anthozoa Anthozoa
0/U Maxillopoda Maxillopoda
Insecta
0/U Metazoa

MGT−v1_516 : 1514 unigenes, 108 (7.13%) taxonomically assigned

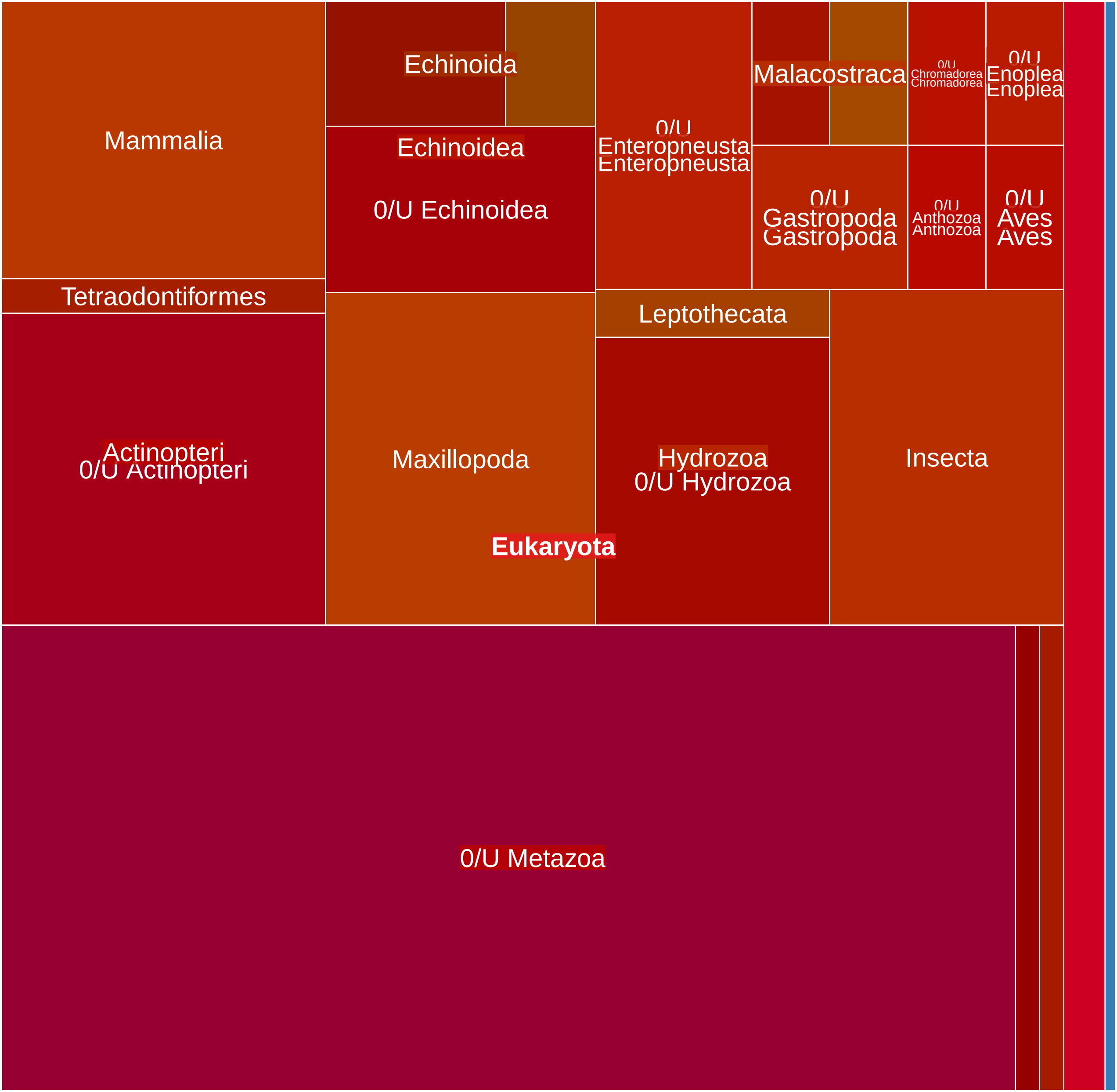

MGT−v1_517 : 1512 unigenes, 192 (12.70%) taxonomically assigned

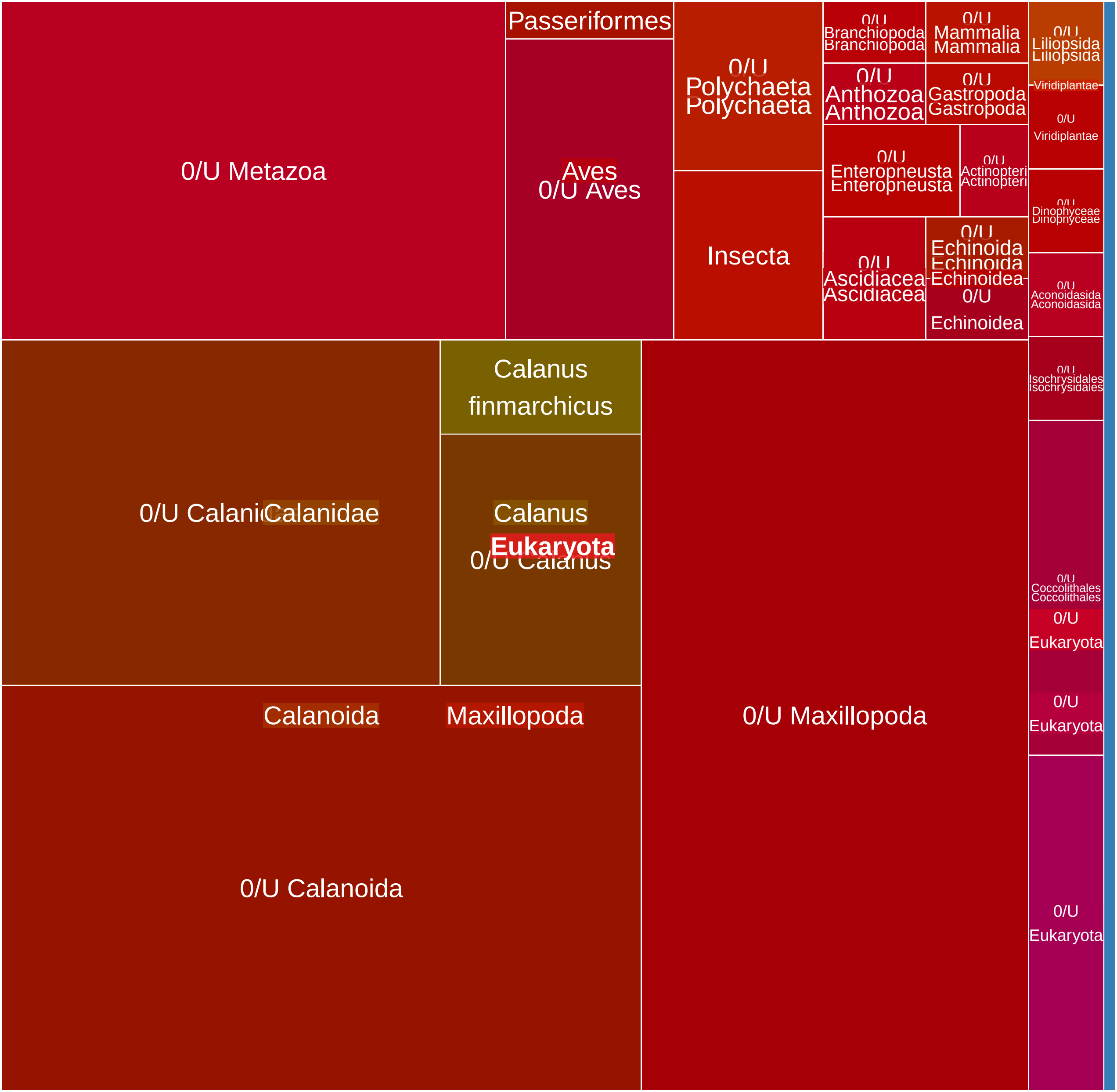

MGT−v1_518 : 1506 unigenes, 294 (19.52%) taxonomically assigned

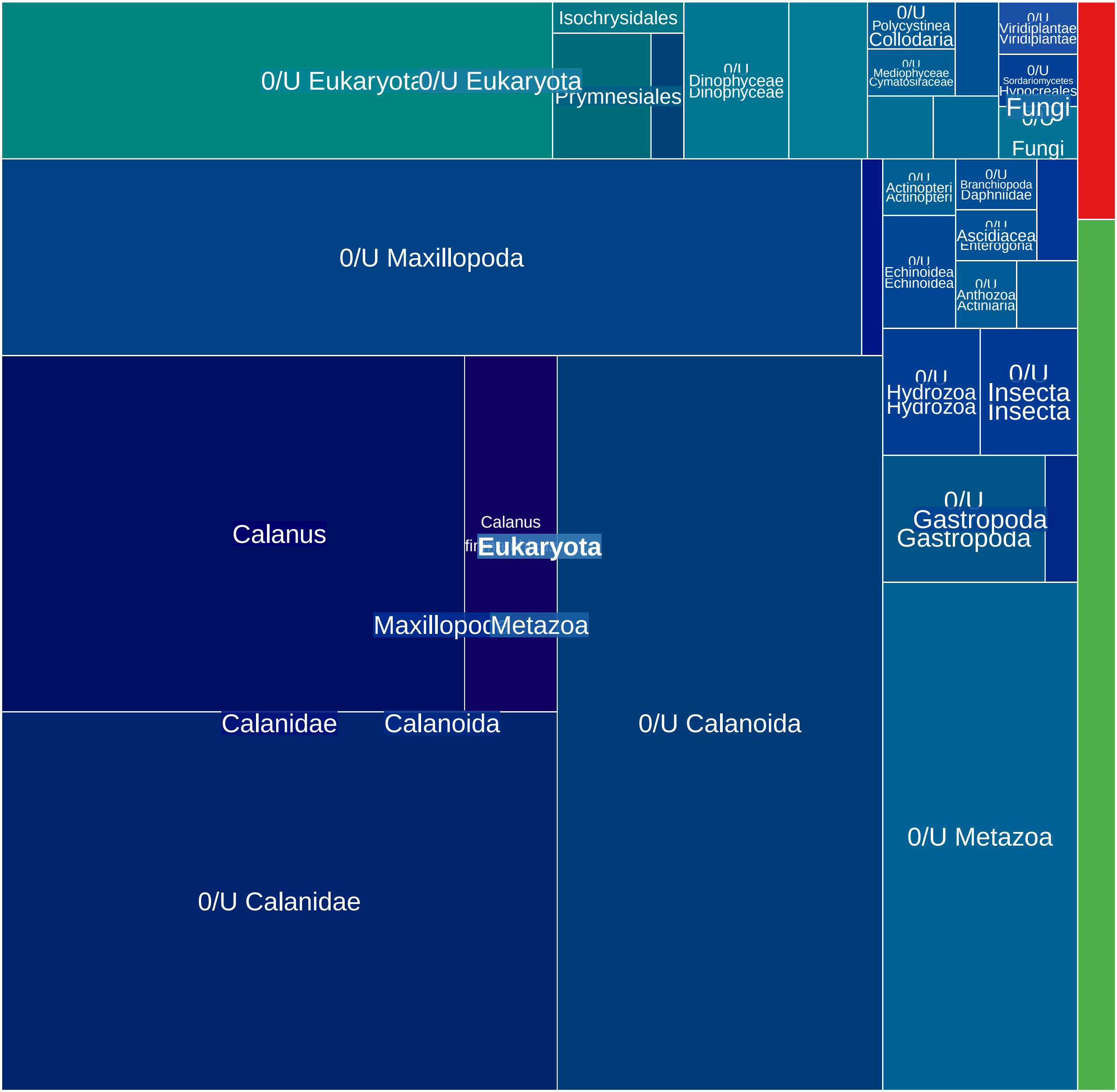

### MGT−v1_519 : 1496 unigenes, 112 (7.49%) taxonomically assigned

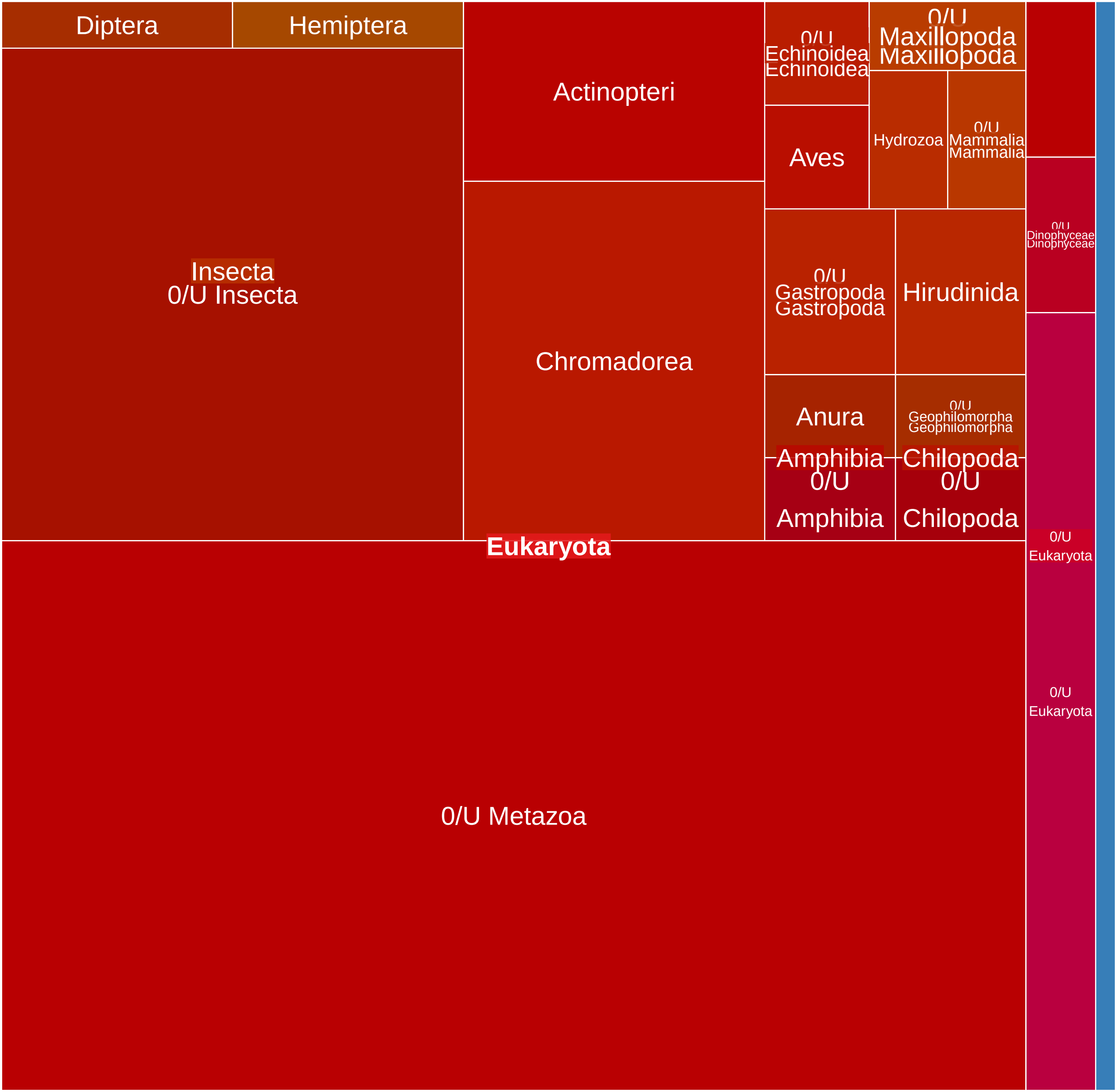

MGT−v1_520 : 1495 unigenes, 74 (4.95%) taxonomically assigned

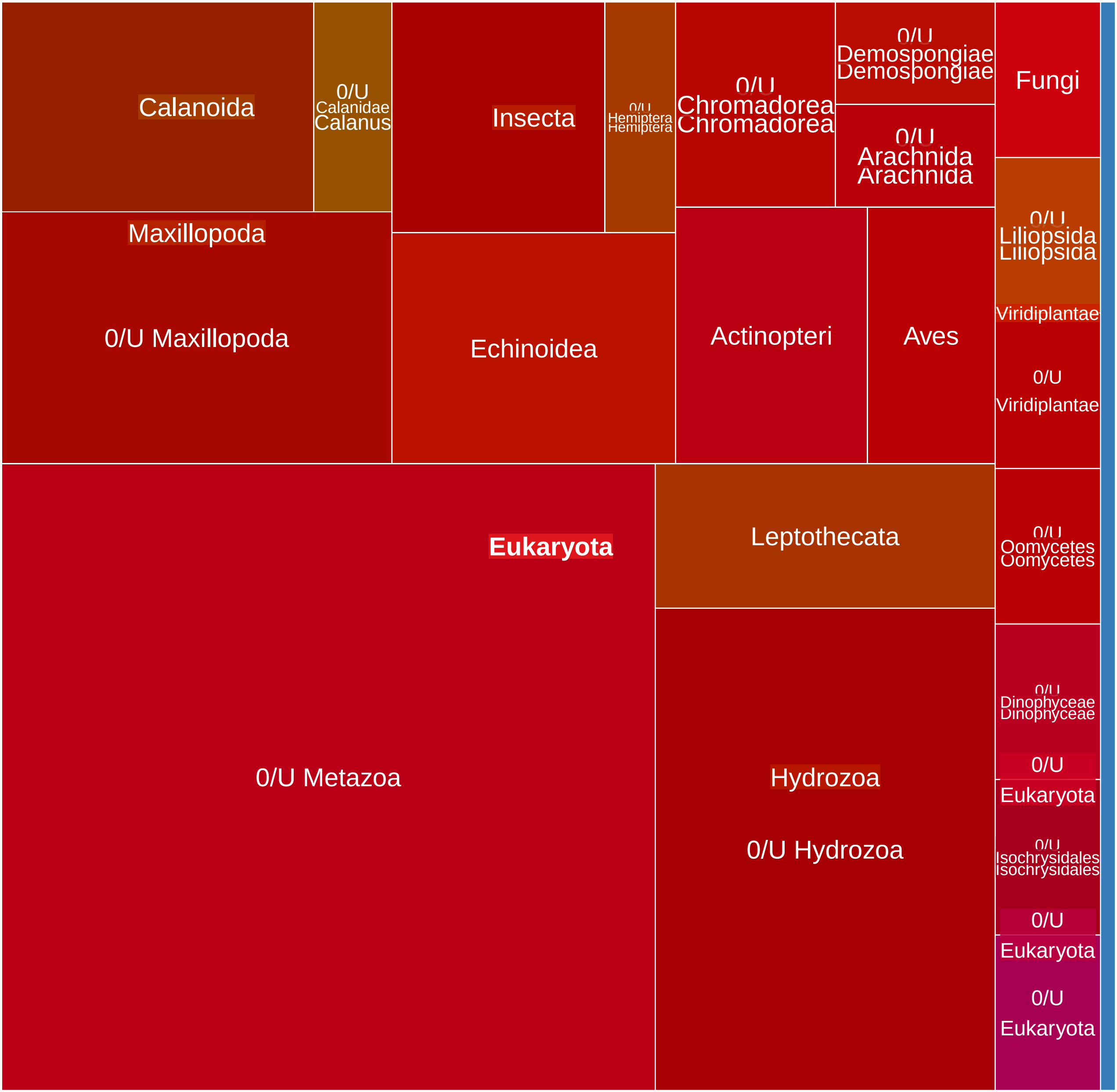

MGT−v1_521 : 1493 unigenes, 184 (12.32%) taxonomically assigned

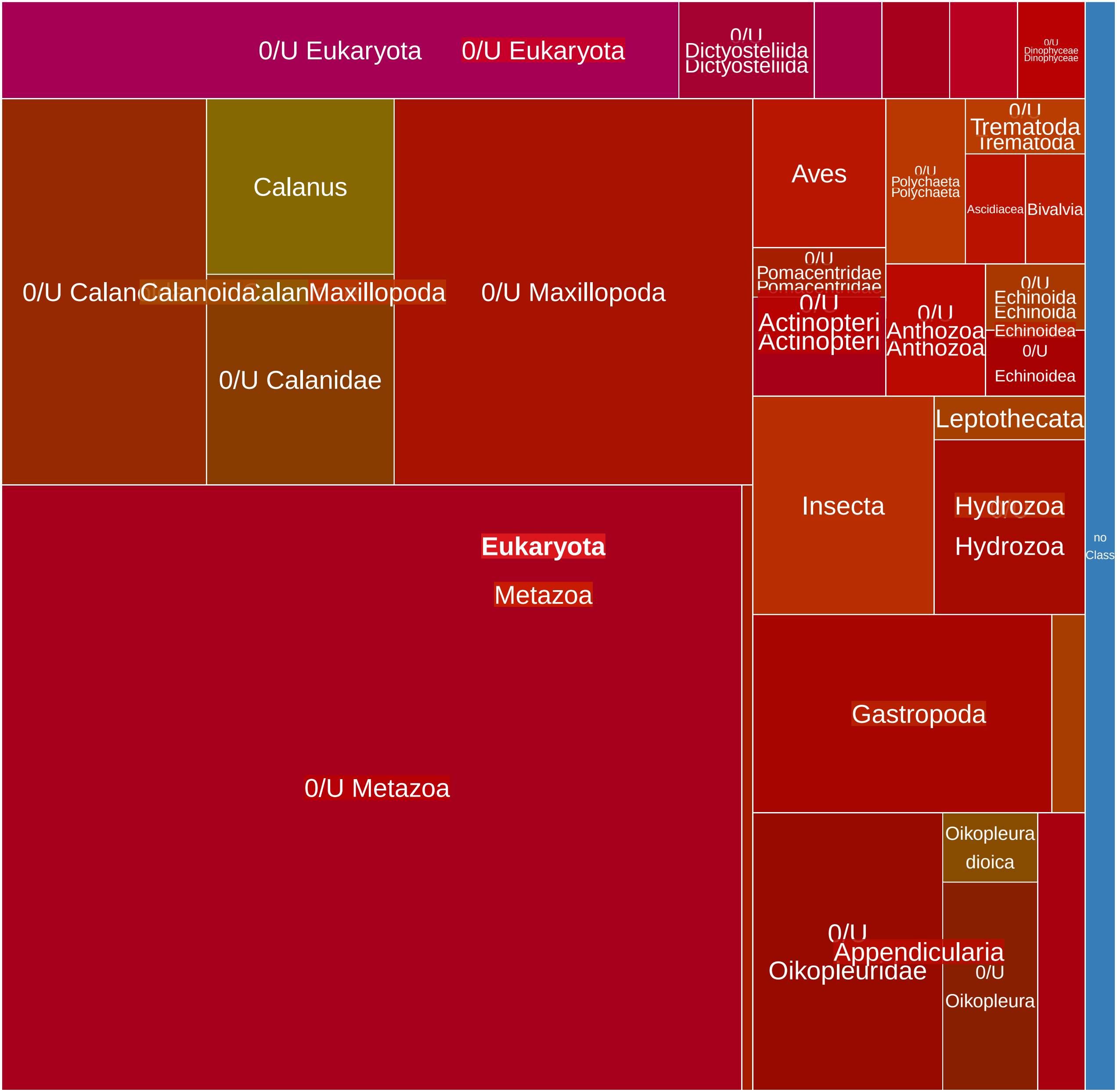

MGT−v1_522 : 1489 unigenes, 1300 (87.31%) taxonomically assigned

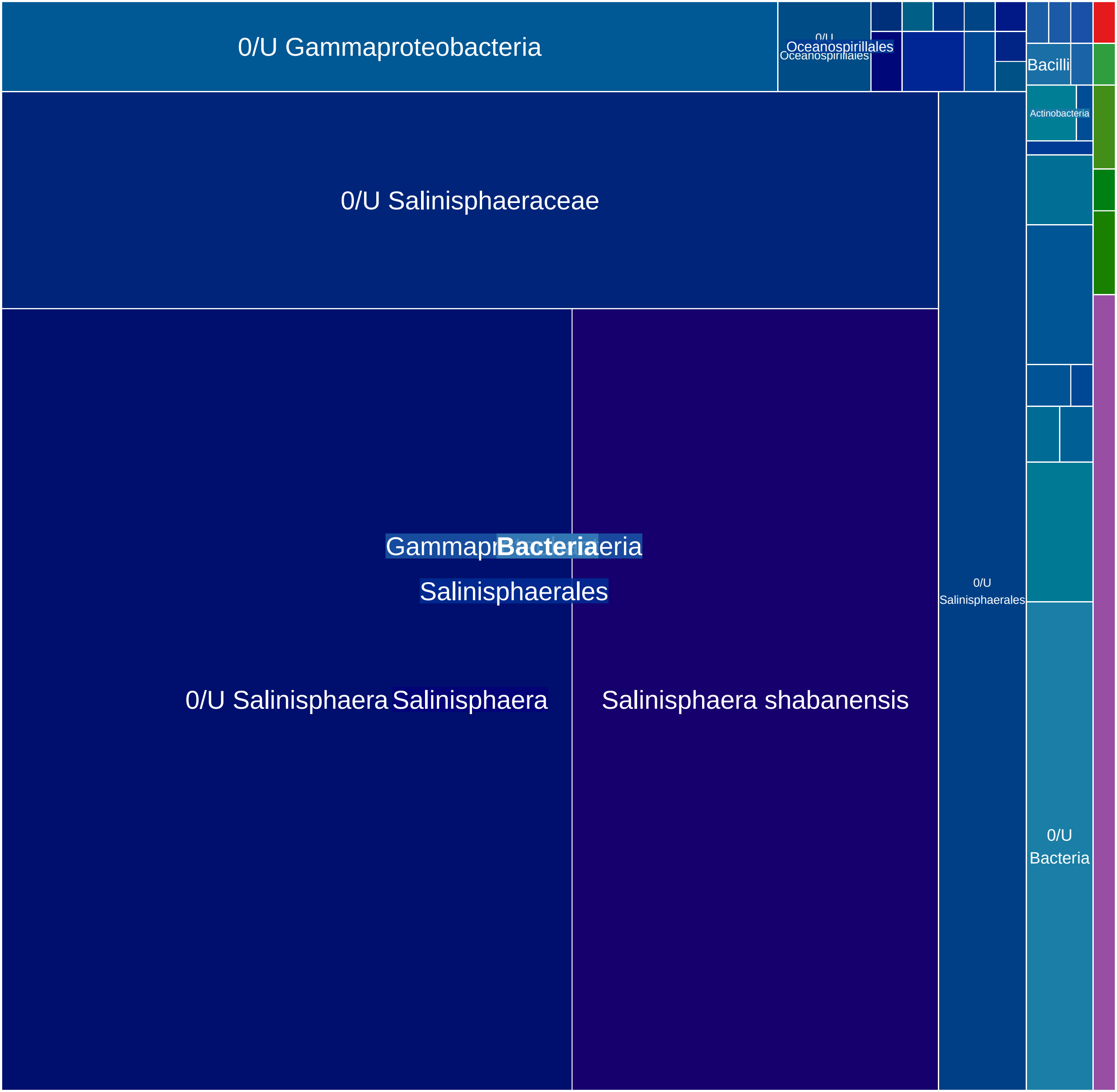

MGT−v1_523 : 1484 unigenes, 122 (8.22%) taxonomically assigned

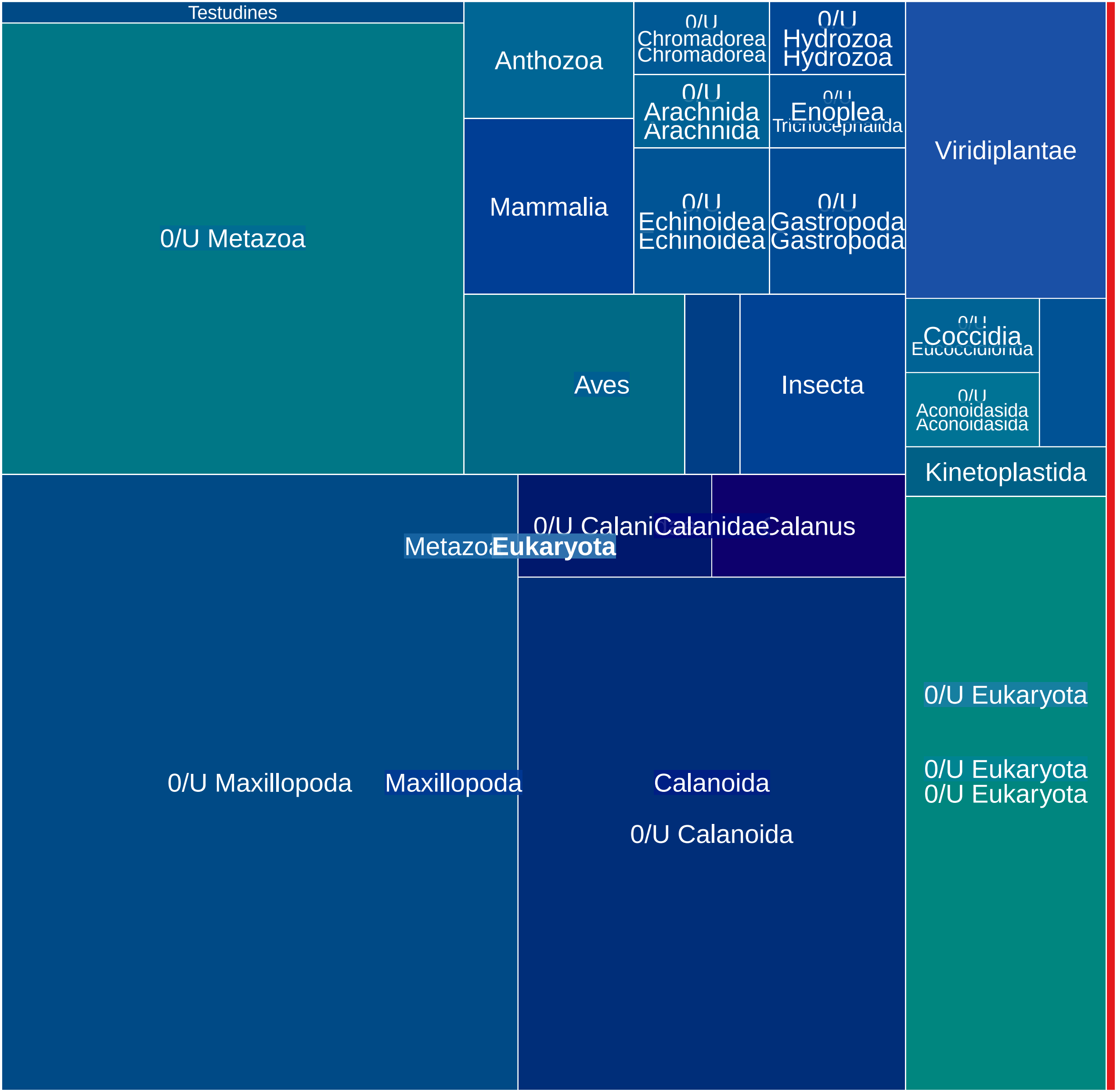

MGT−v1_524 : 1482 unigenes, 1357 (91.57%) taxonomically assigned

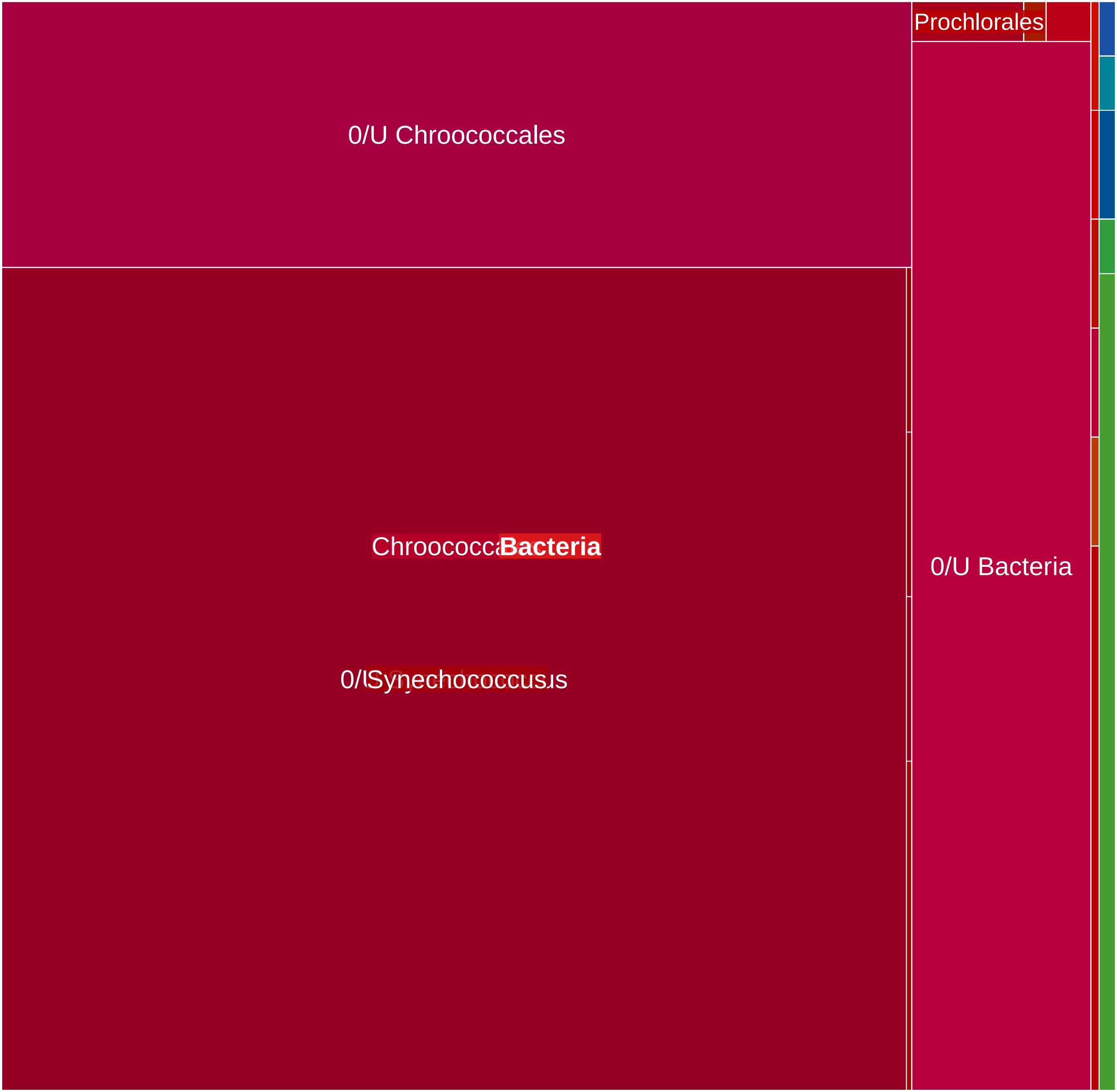

MGT−v1_525 : 1449 unigenes, 92 (6.35%) taxonomically assigned

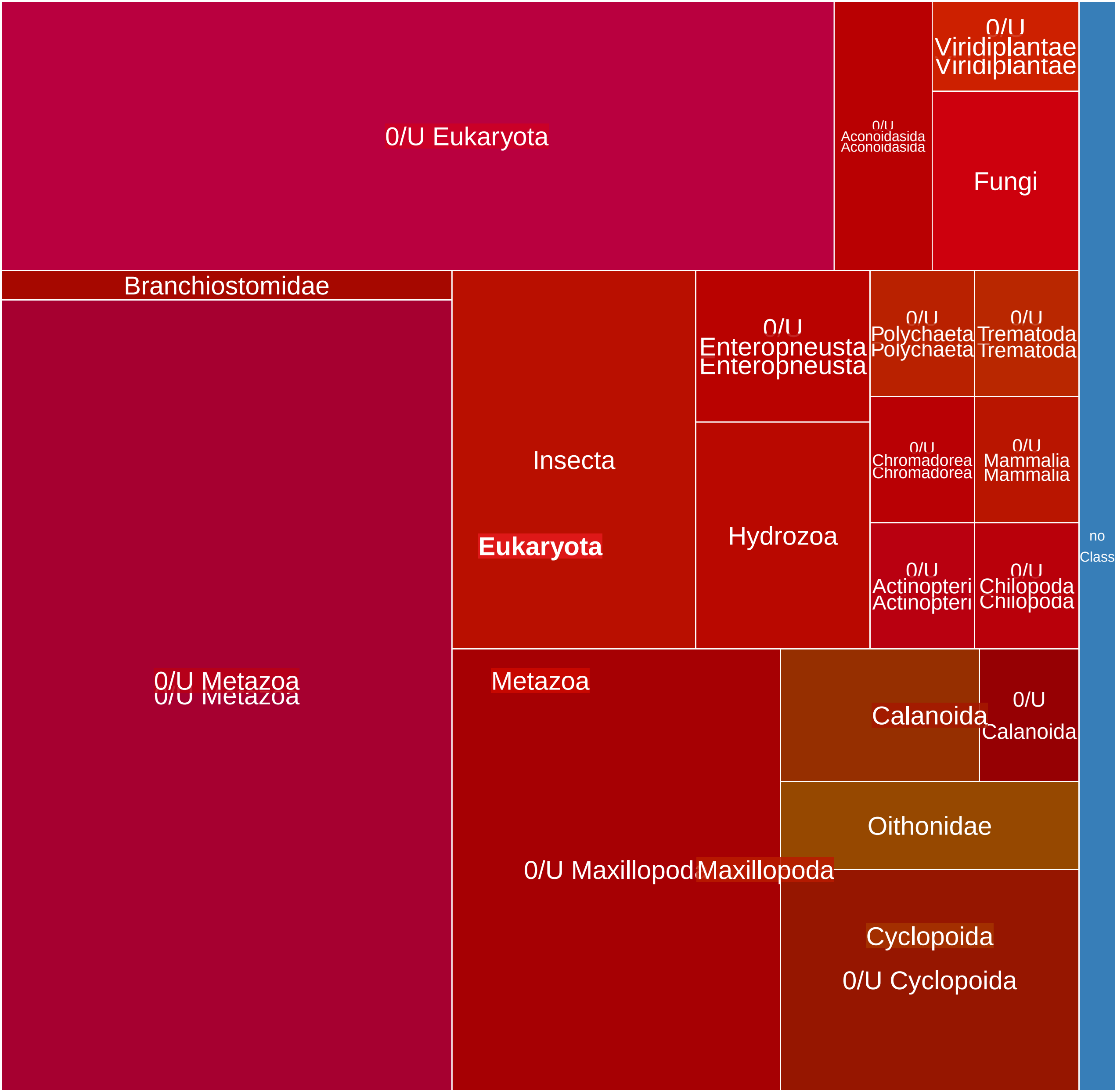

MGT−v1_526 : 1448 unigenes, 370 (25.55%) taxonomically assigned

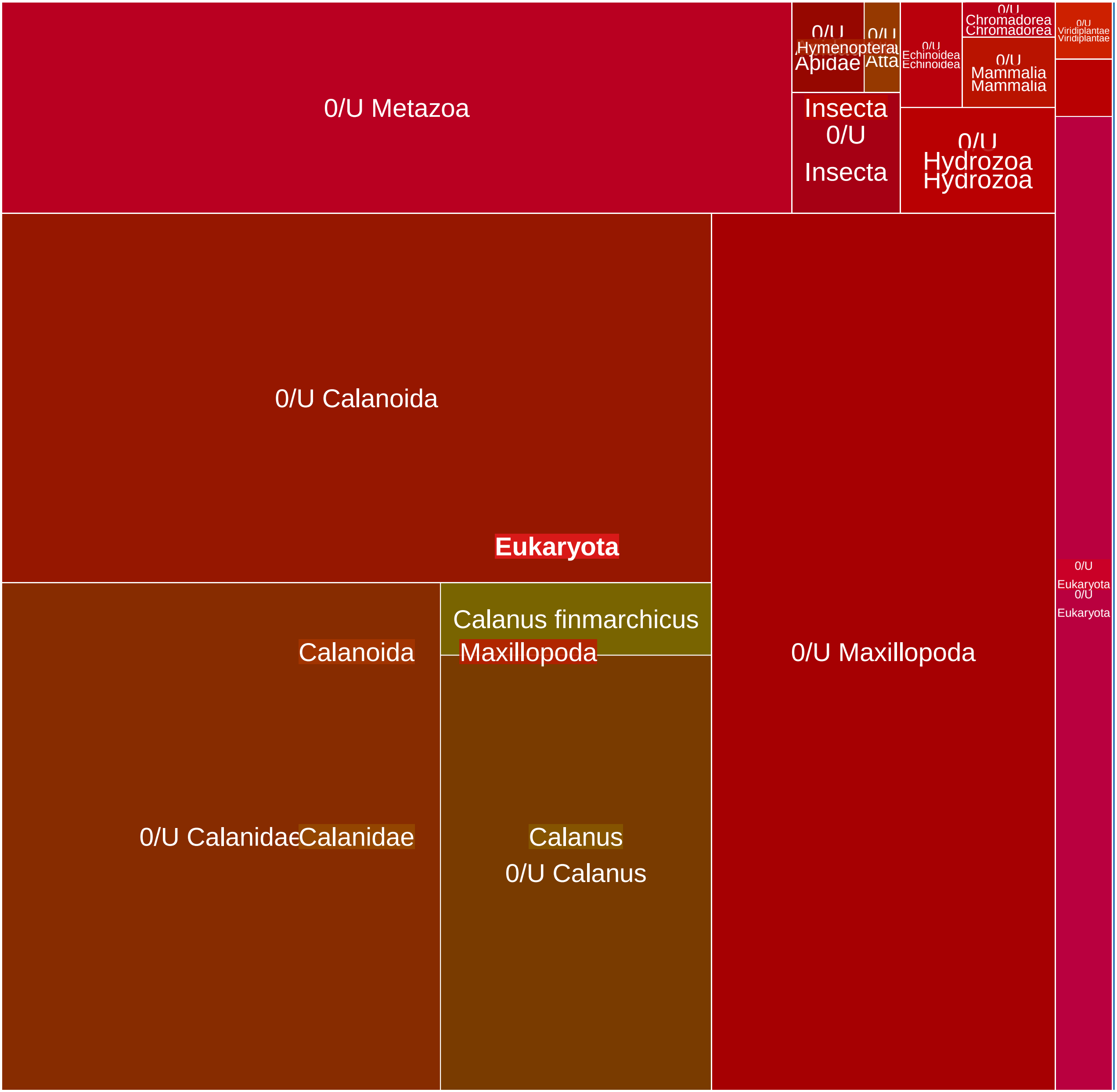

MGT−v1_527 : 1431 unigenes, 1391 (97.20%) taxonomically assigned

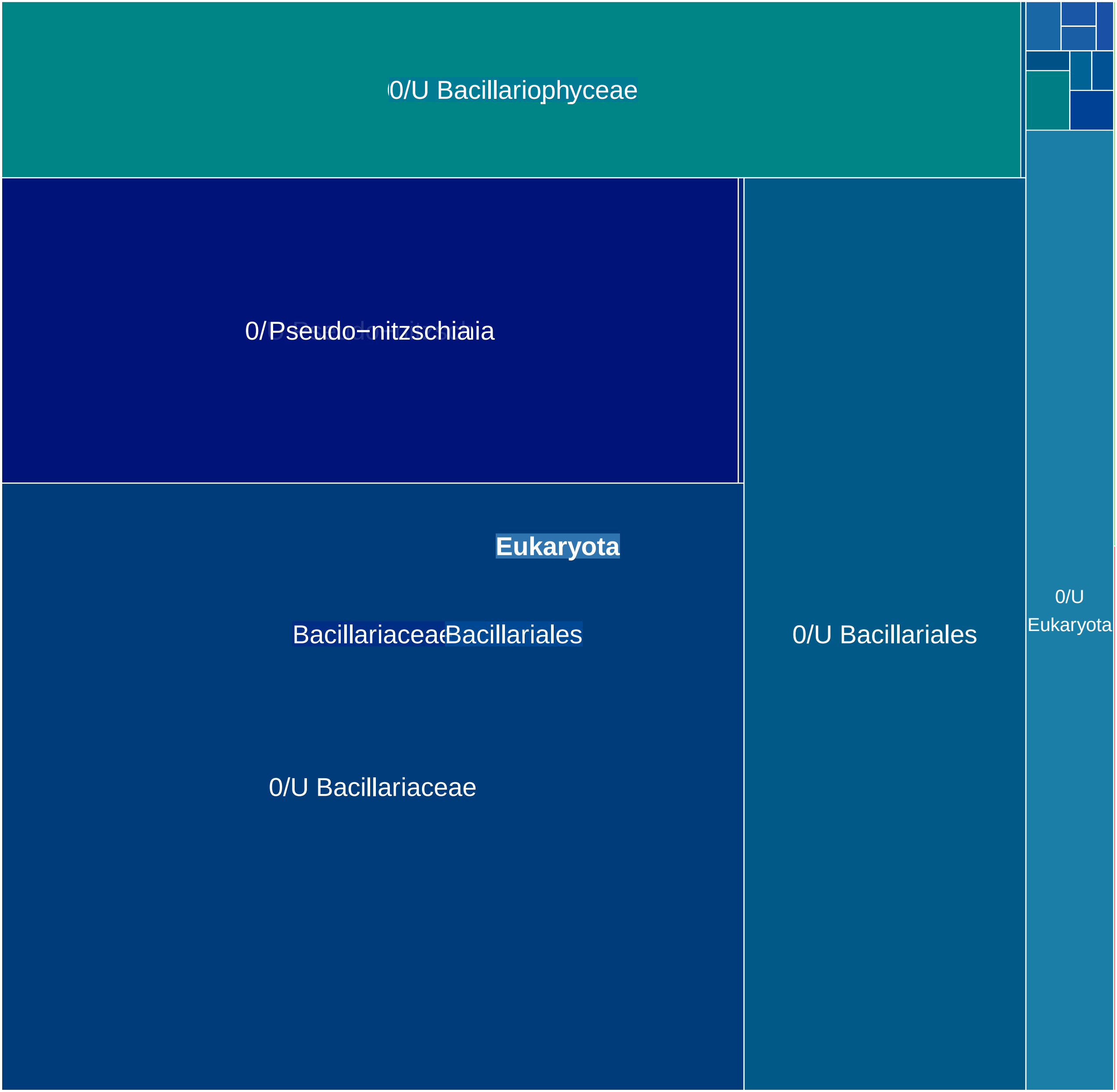

MGT−v1_528 : 1430 unigenes, 90 (6.29%) taxonomically assigned

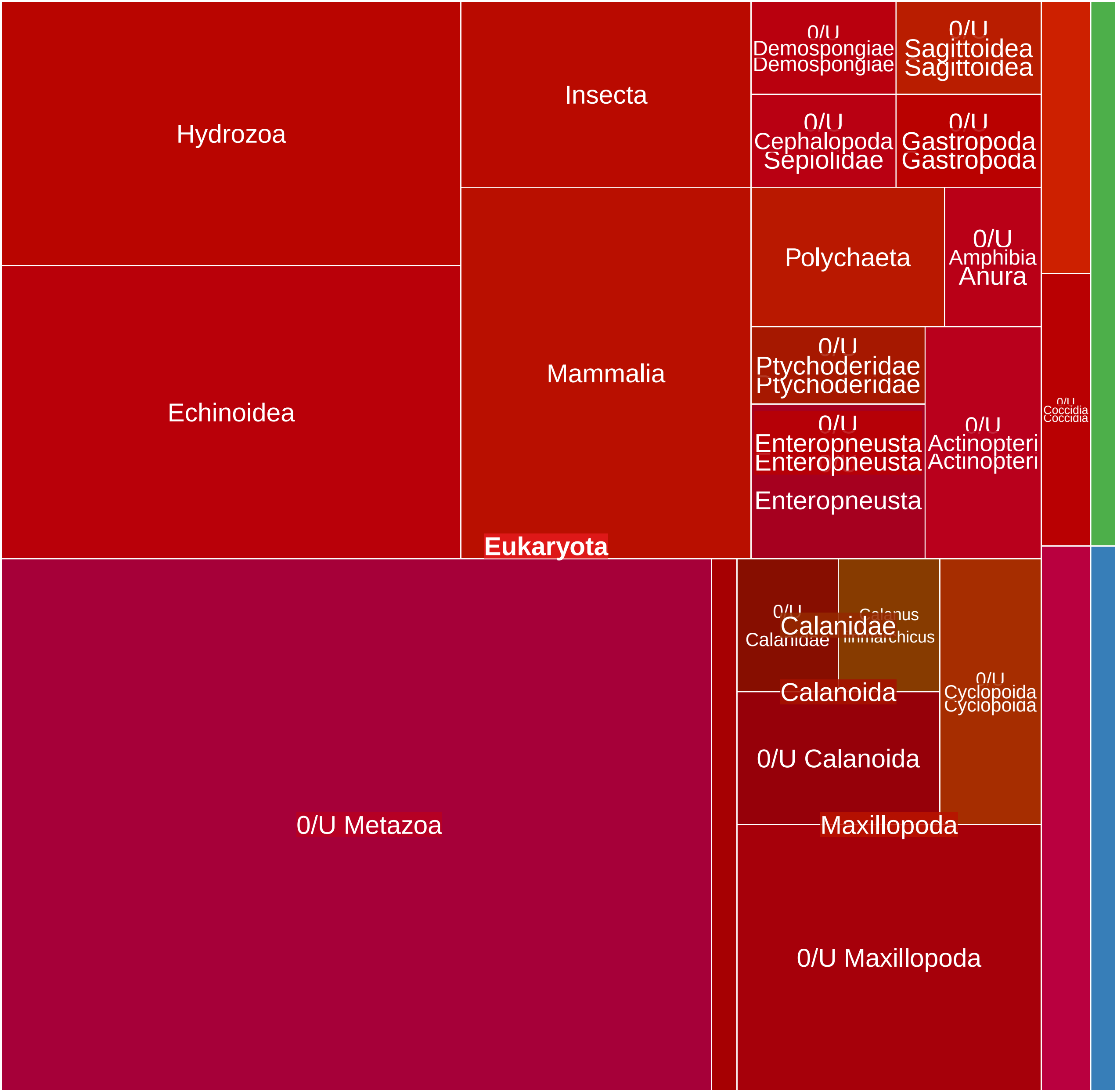

MGT−v1_529 : 1424 unigenes, 294 (20.65%) taxonomically assigned

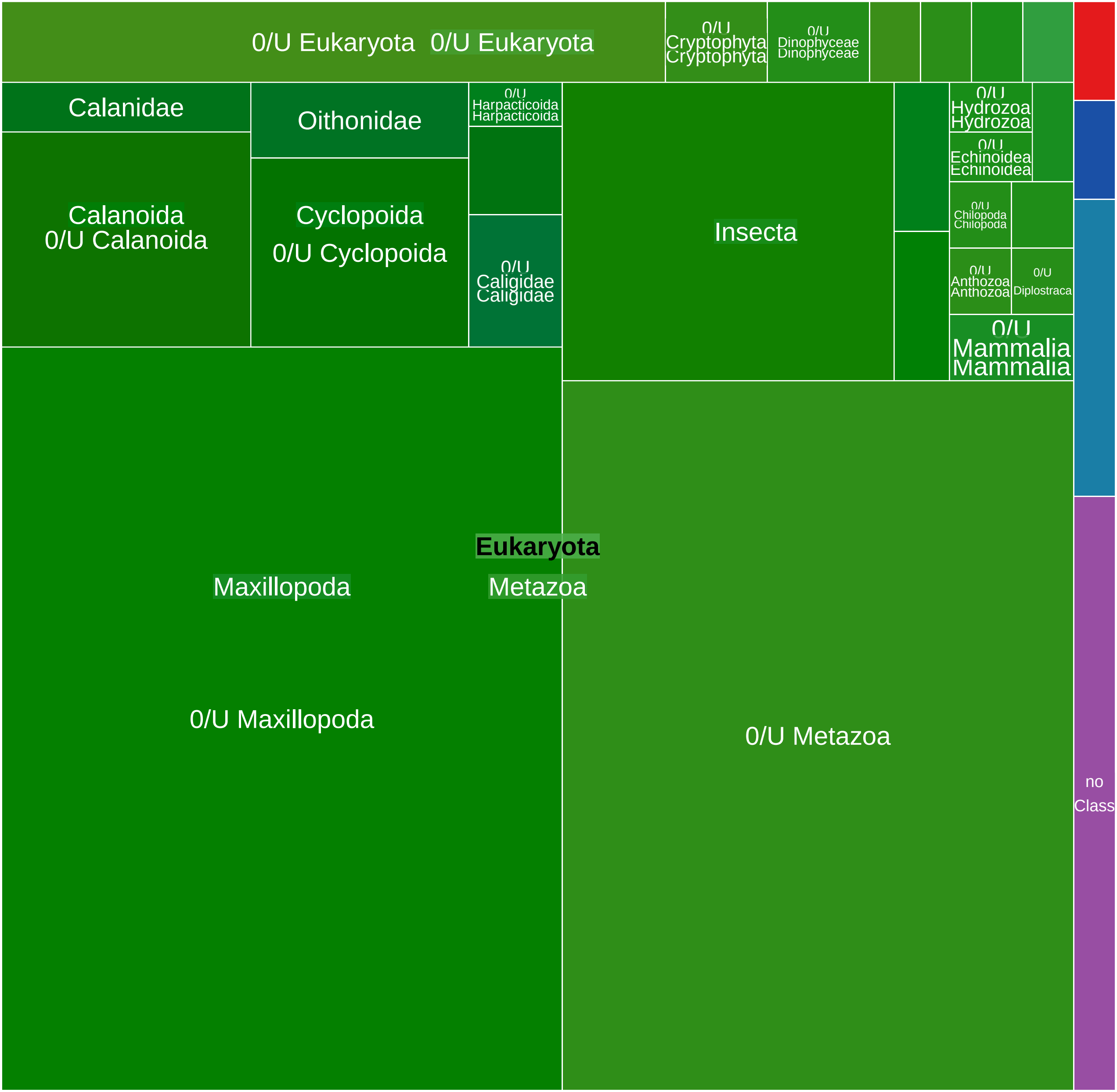

MGT−v1_530 : 1423 unigenes, 686 (48.21%) taxonomically assigned

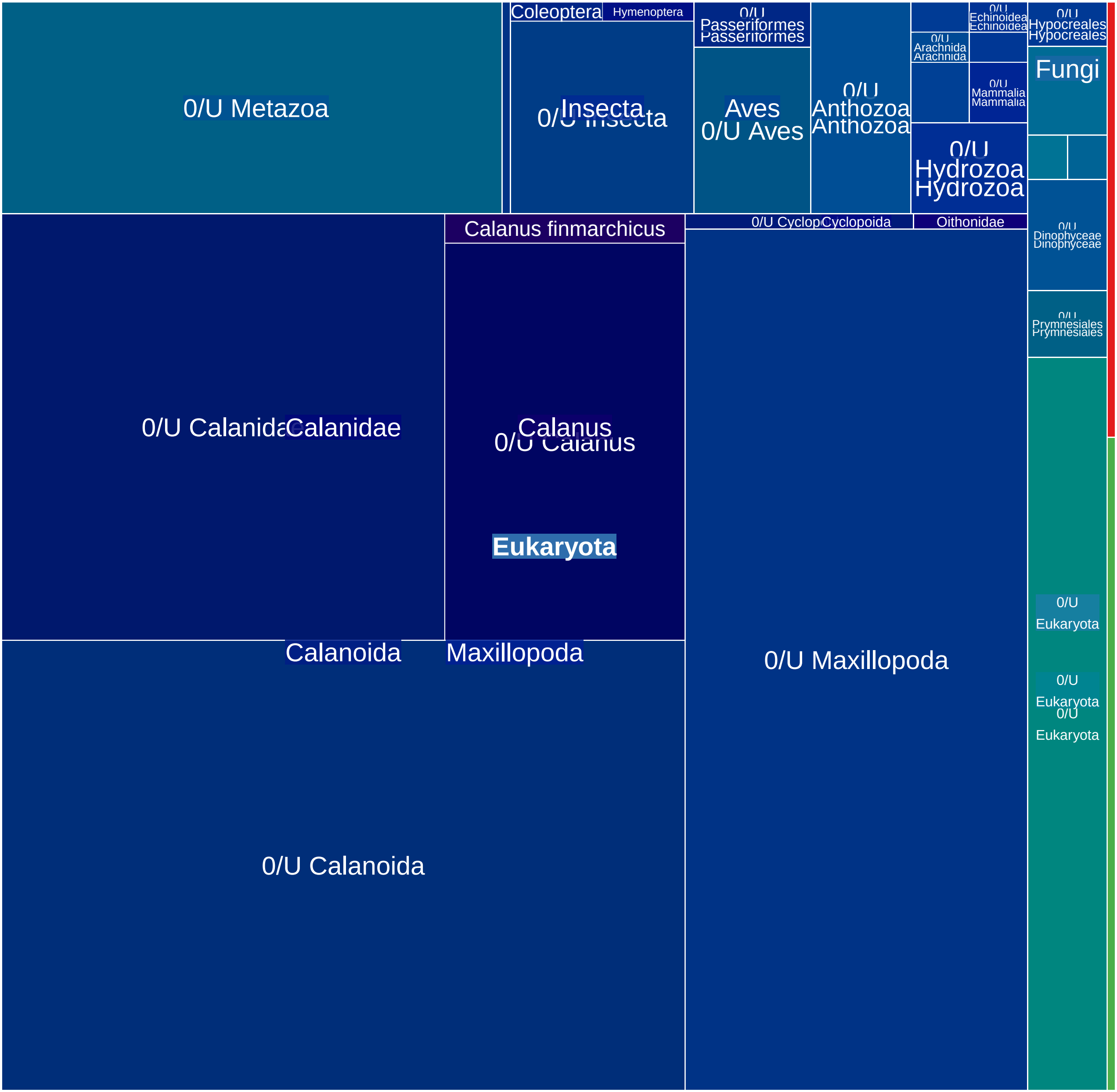

MGT−v1_531 : 1419 unigenes, 475 (33.47%) taxonomically assigned

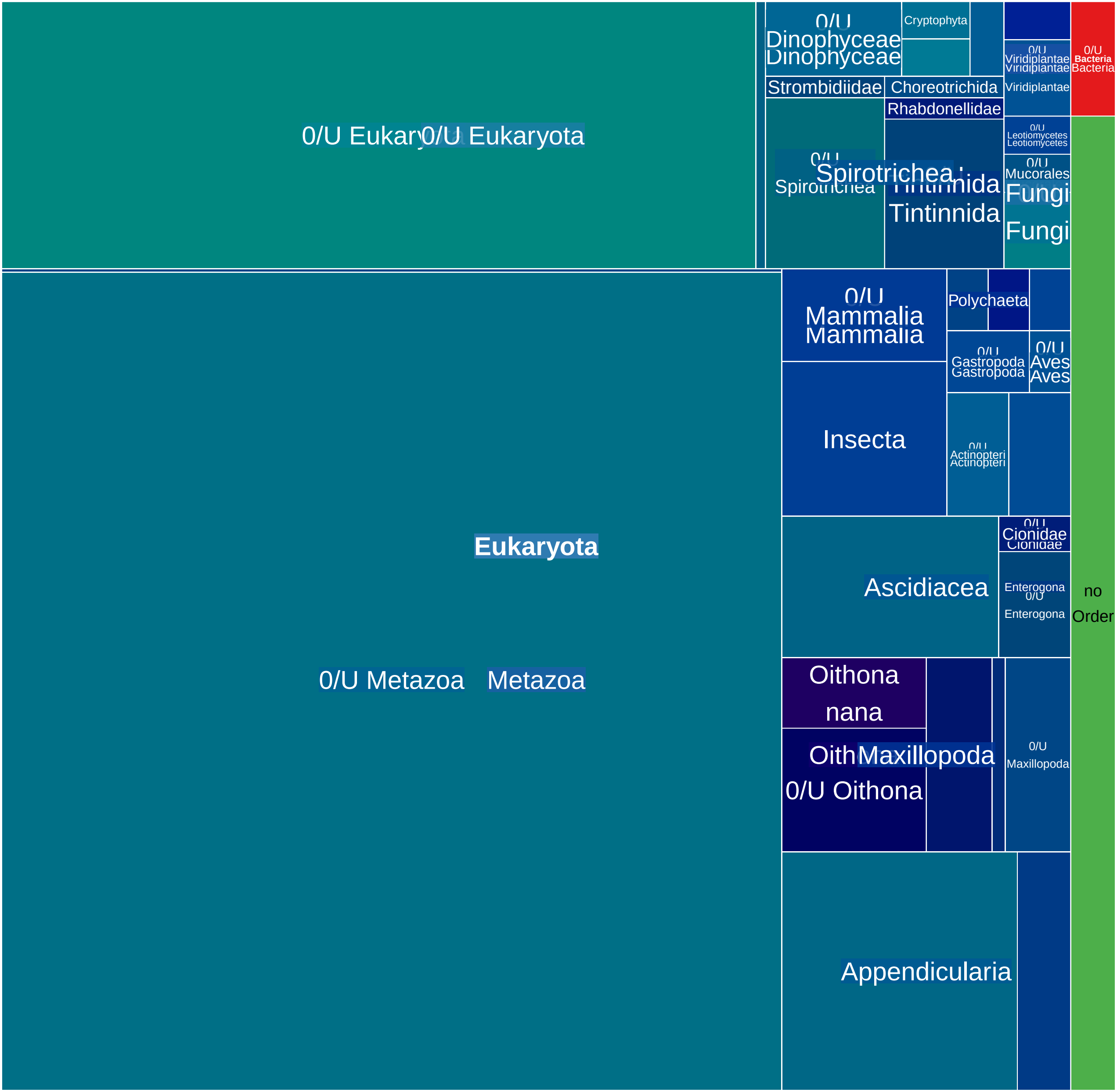

MGT−v1_532 : 1417 unigenes, 109 (7.69%) taxonomically assigned

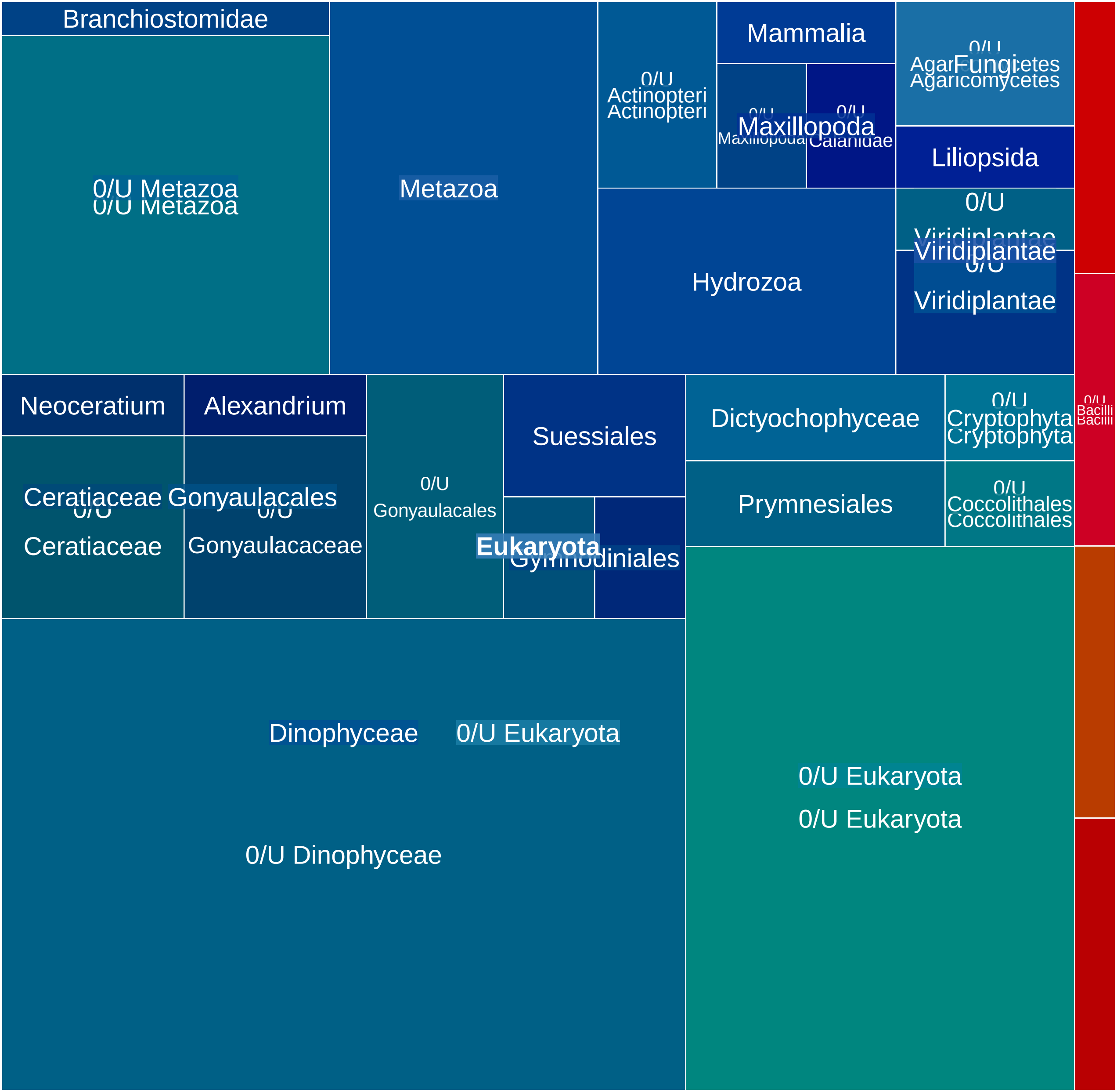

MGT−v1_533 : 1410 unigenes, 198 (14.04%) taxonomically assigned

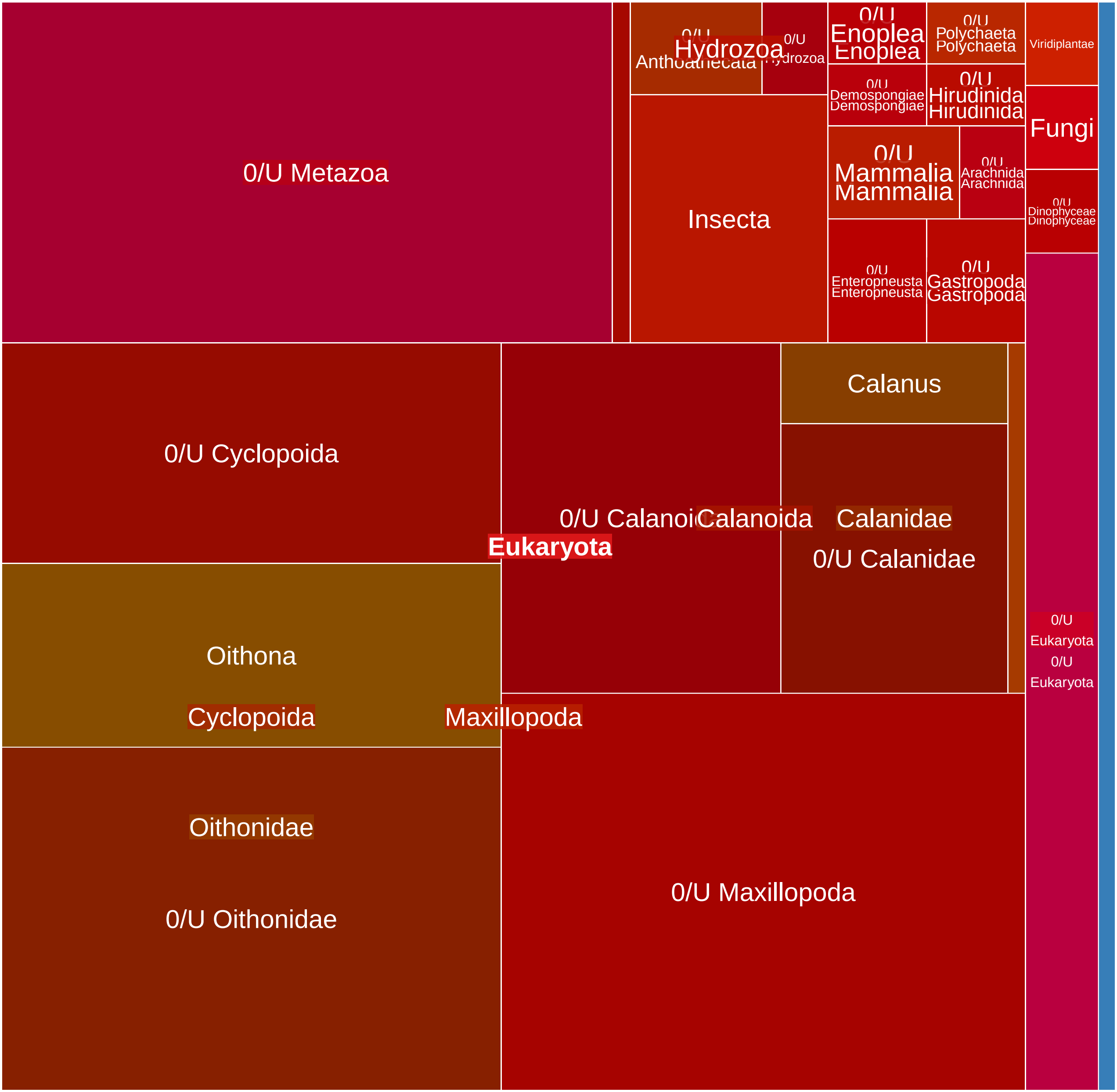

MGT−v1_534 : 1405 unigenes, 67 (4.77%) taxonomically assigned

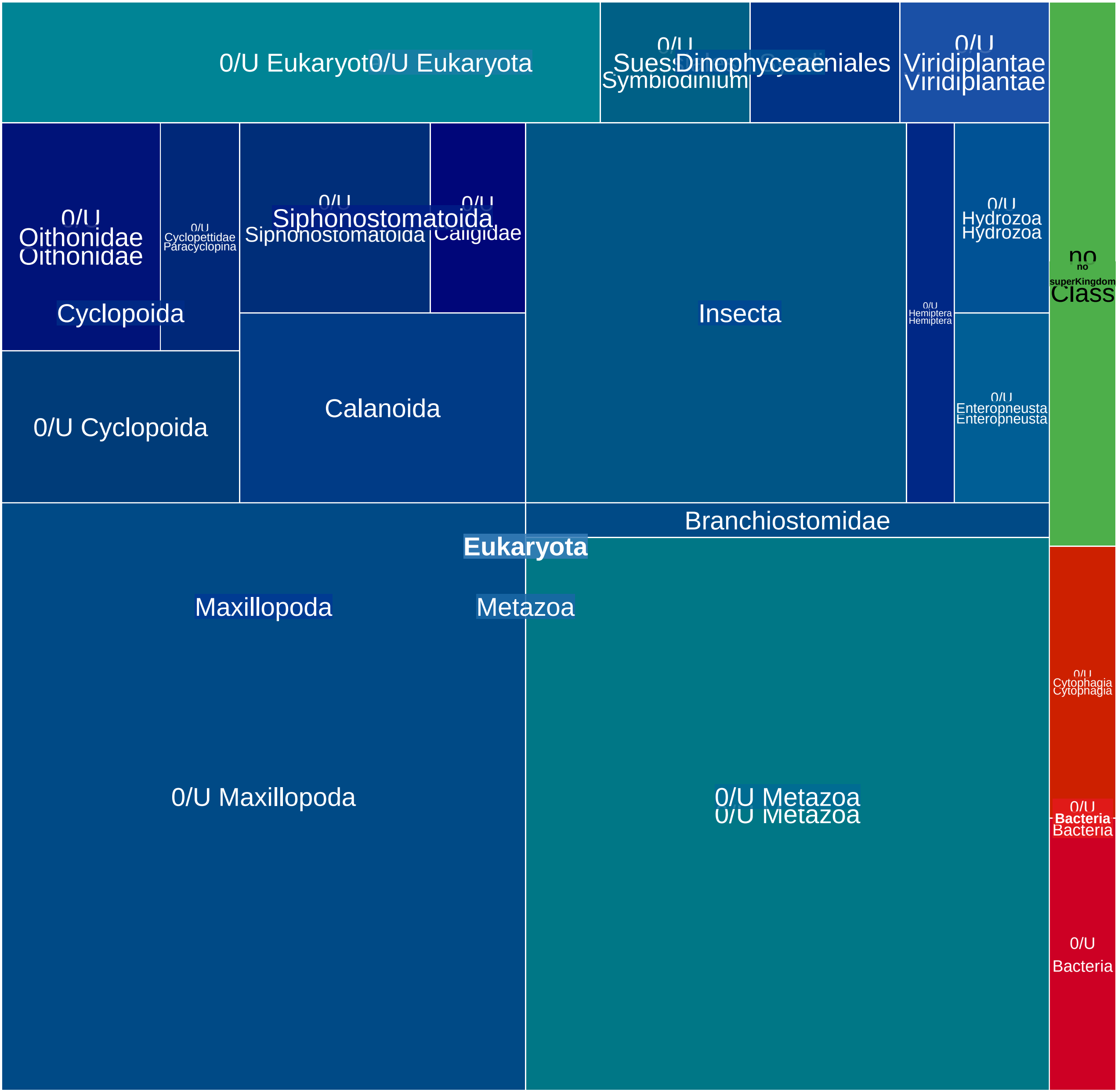

MGT−v1_535 : 1405 unigenes, 26 (1.85%) taxonomically assigned

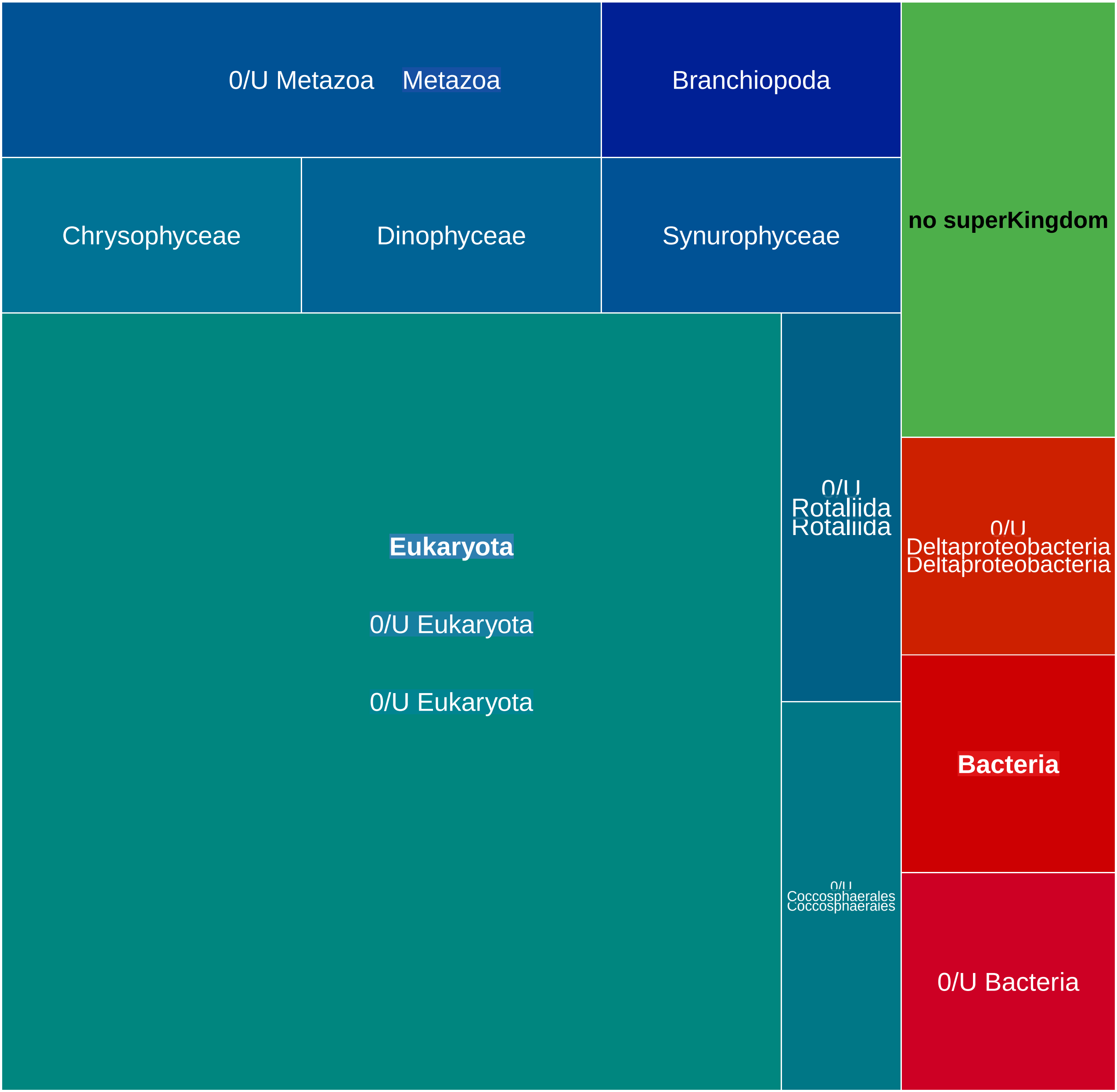

MGT−v1_536 : 1404 unigenes, 1218 (86.75%) taxonomically assigned

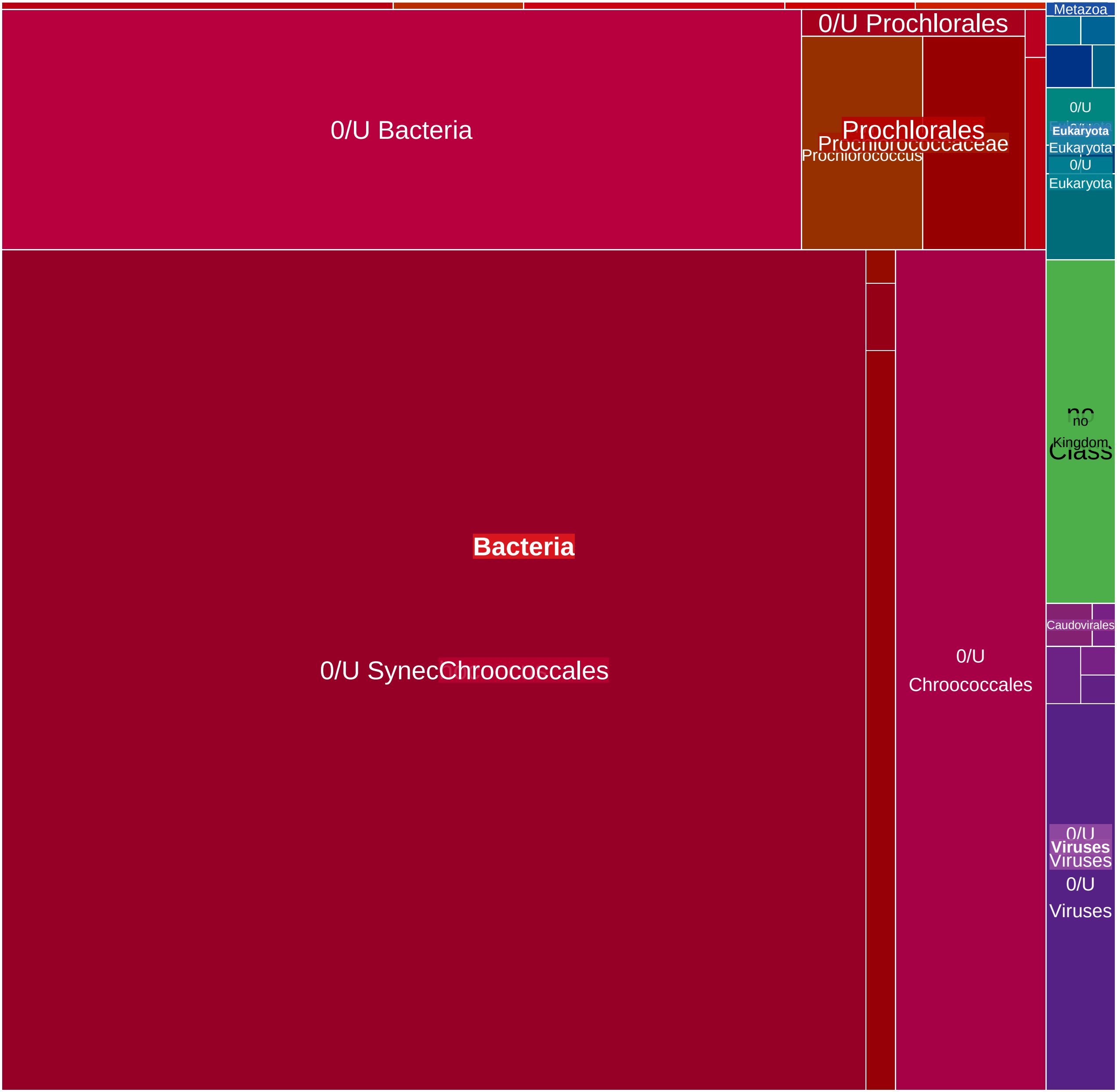

MGT−v1_537 : 1404 unigenes, 791 (56.34%) taxonomically assigned

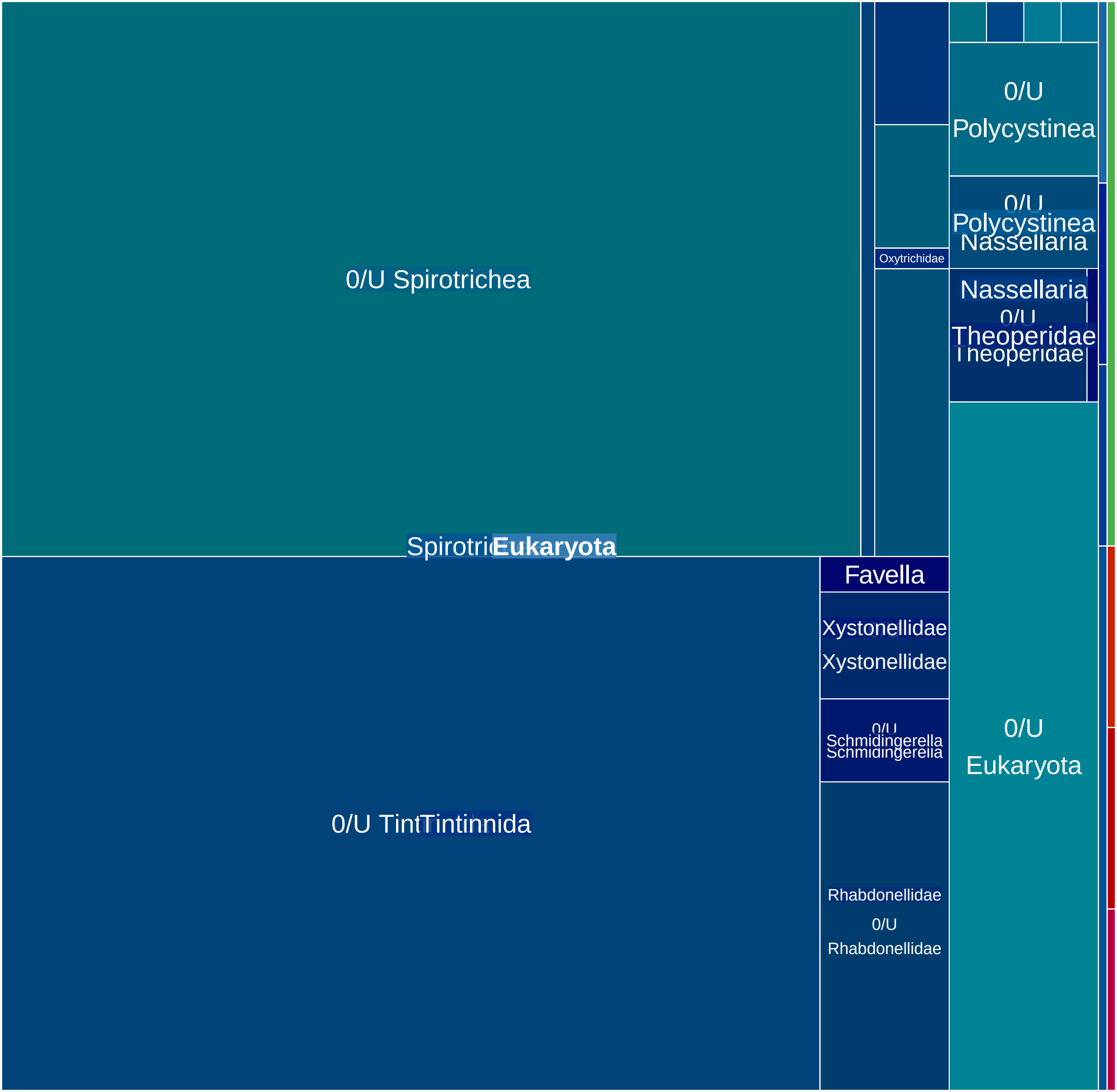

MGT−v1_538 : 1403 unigenes, 1178 (83.96%) taxonomically assigned

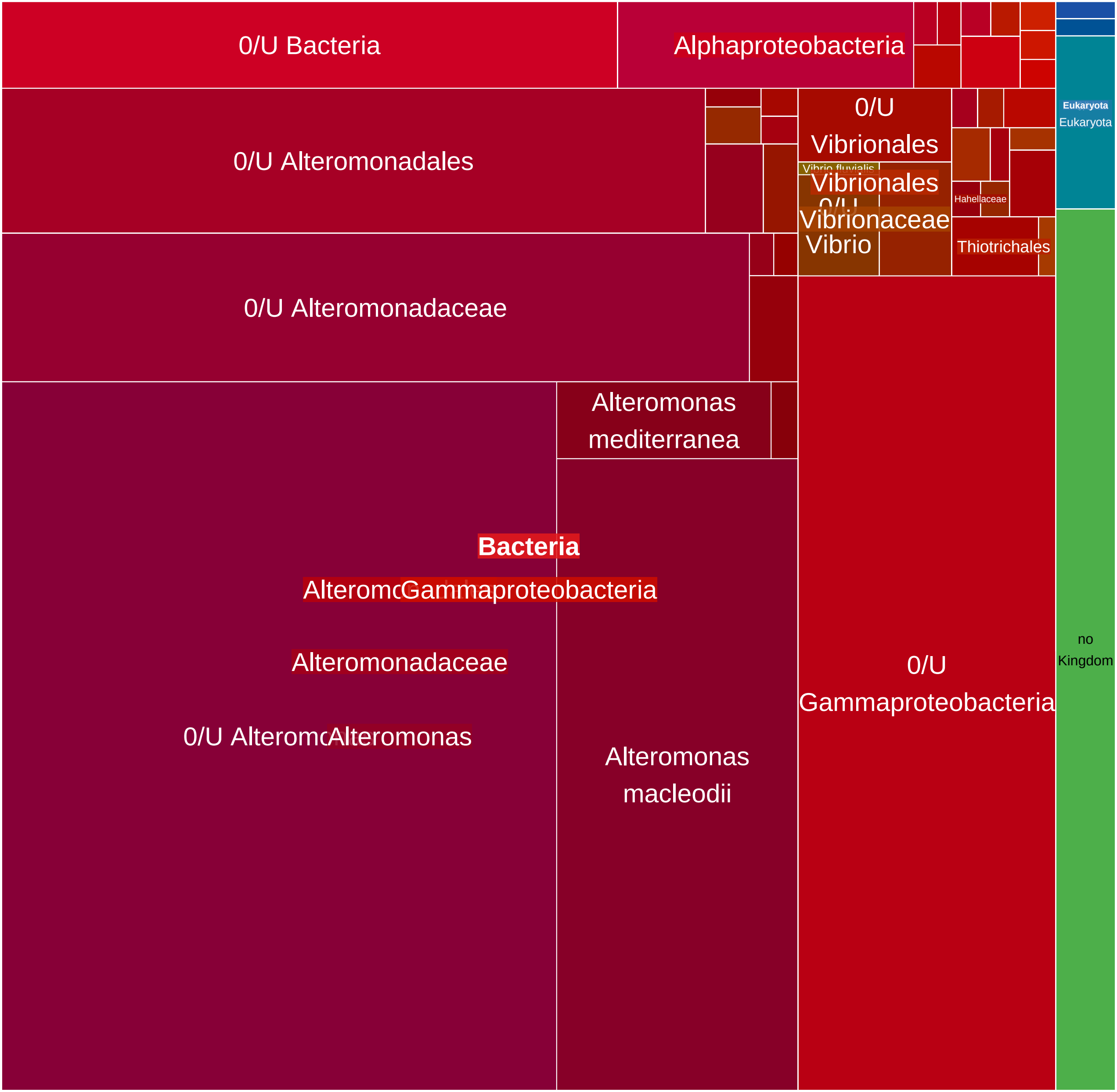

### MGT−v1_539 : 1400 unigenes, 1054 (75.29%) taxonomically assigned

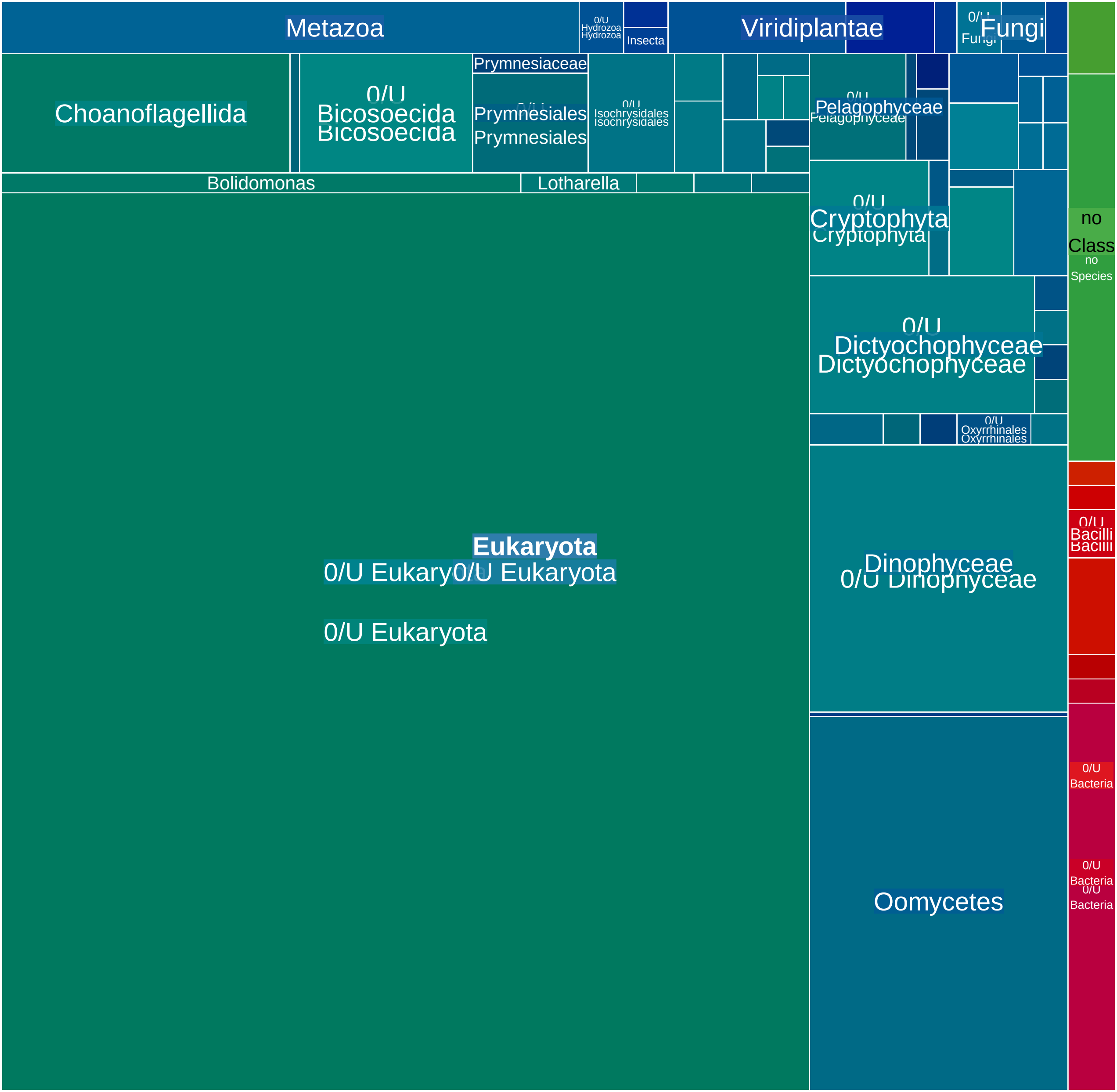

MGT−v1_540 : 1392 unigenes, 1239 (89.01%) taxonomically assigned

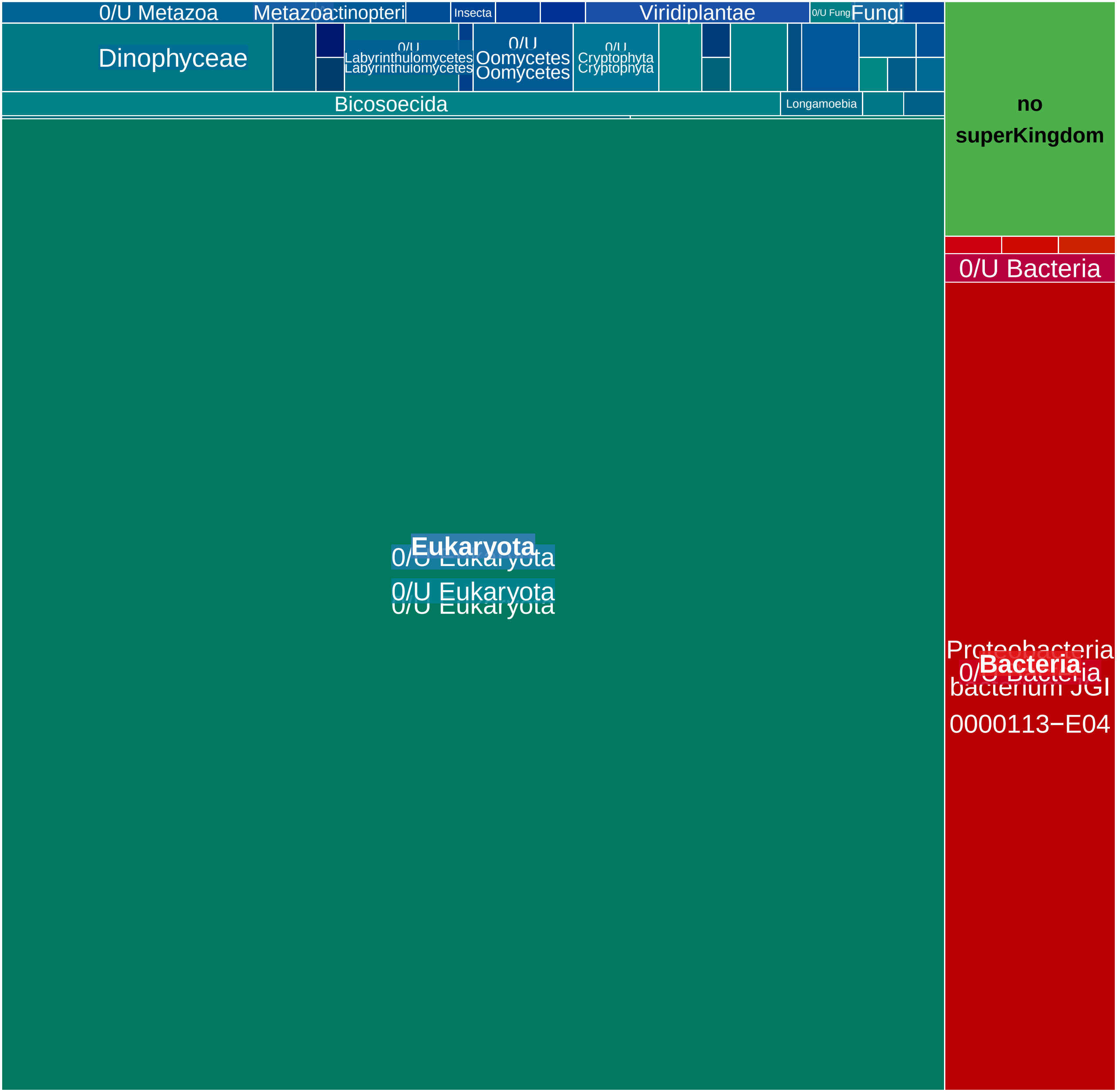

MGT−v1_541 : 1389 unigenes, 237 (17.06%) taxonomically assigned

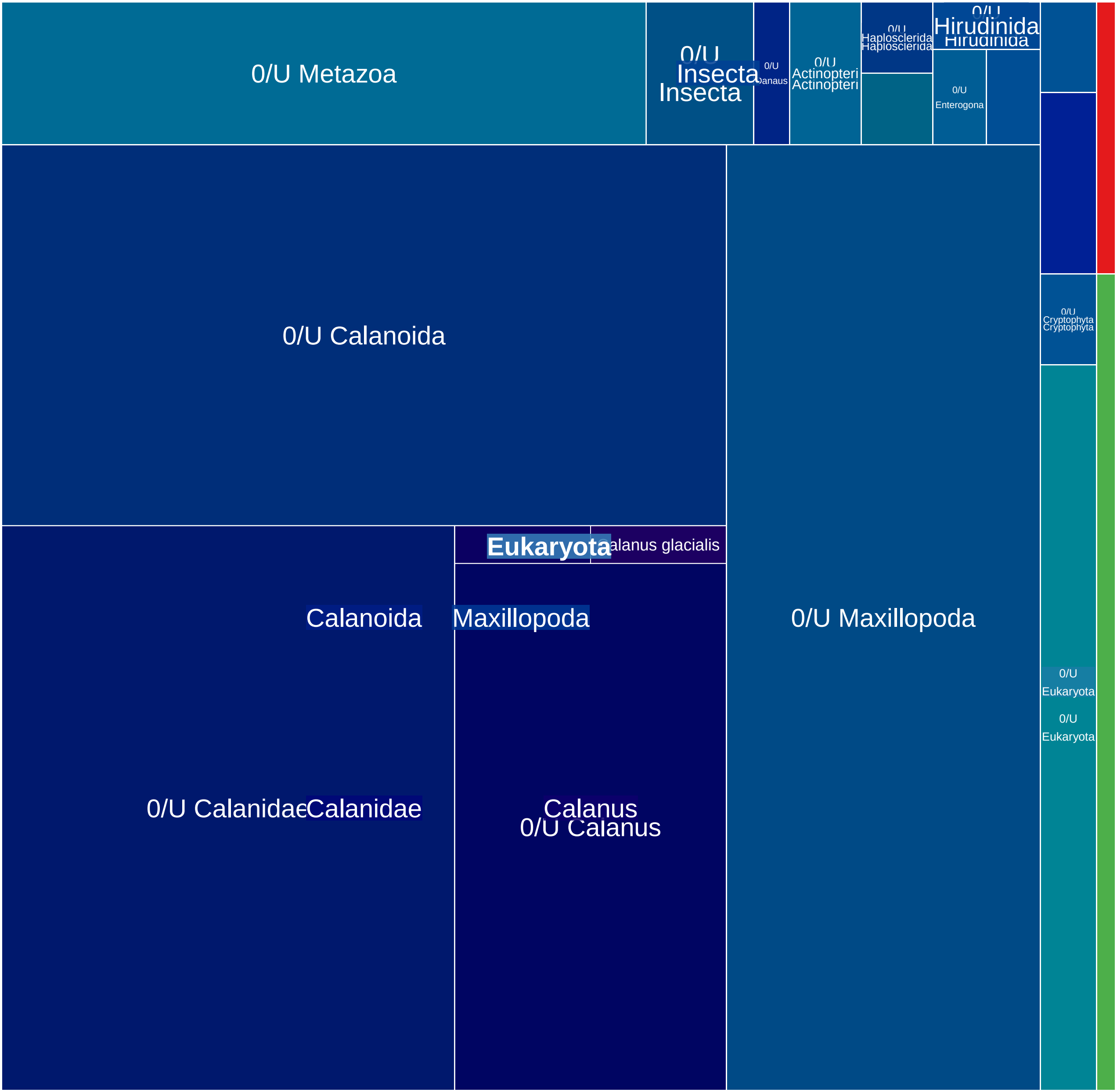

MGT−v1_542 : 1378 unigenes, 1022 (74.17%) taxonomically assigned

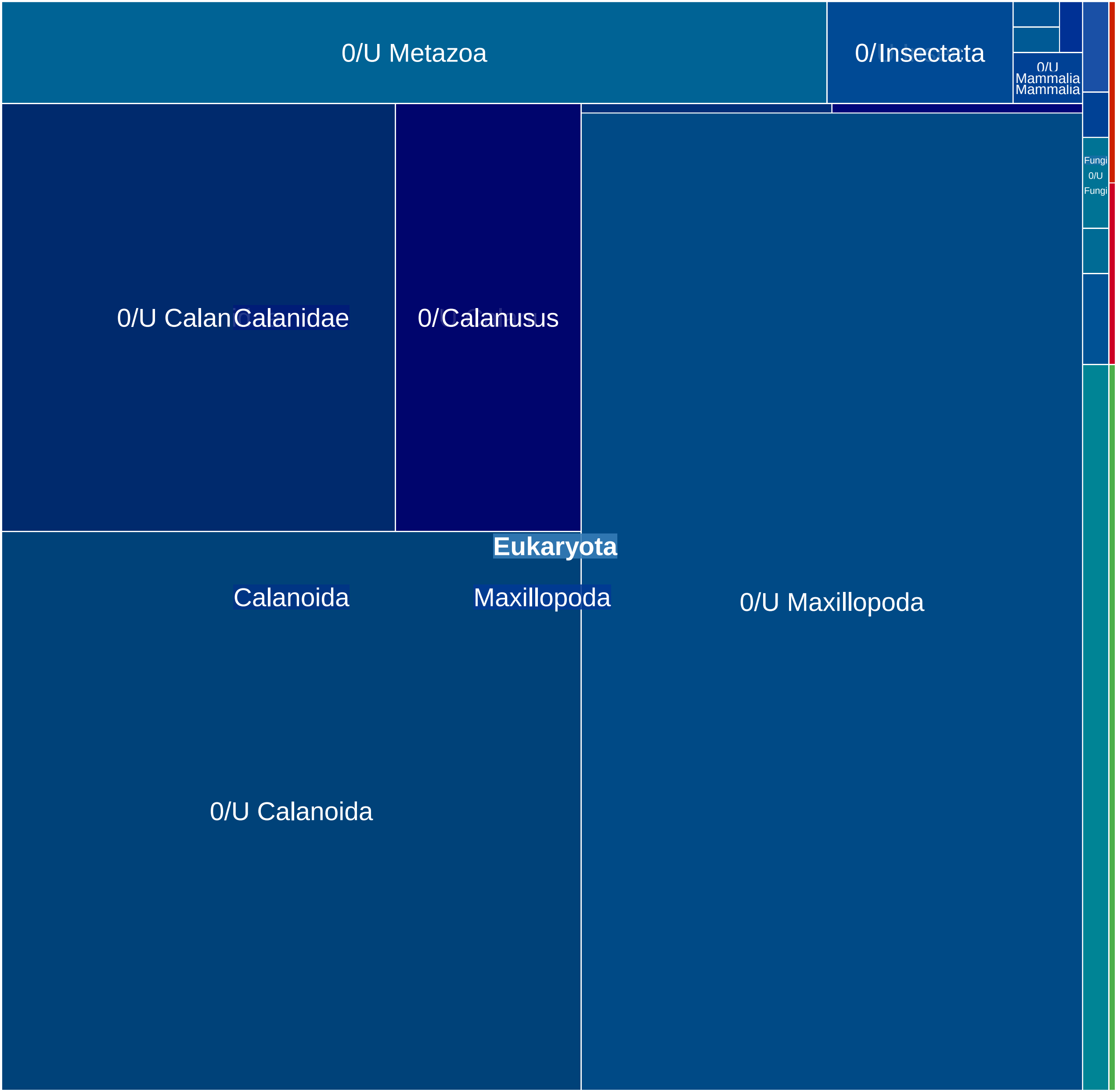

MGT−v1_543 : 1371 unigenes, 613 (44.71%) taxonomically assigned

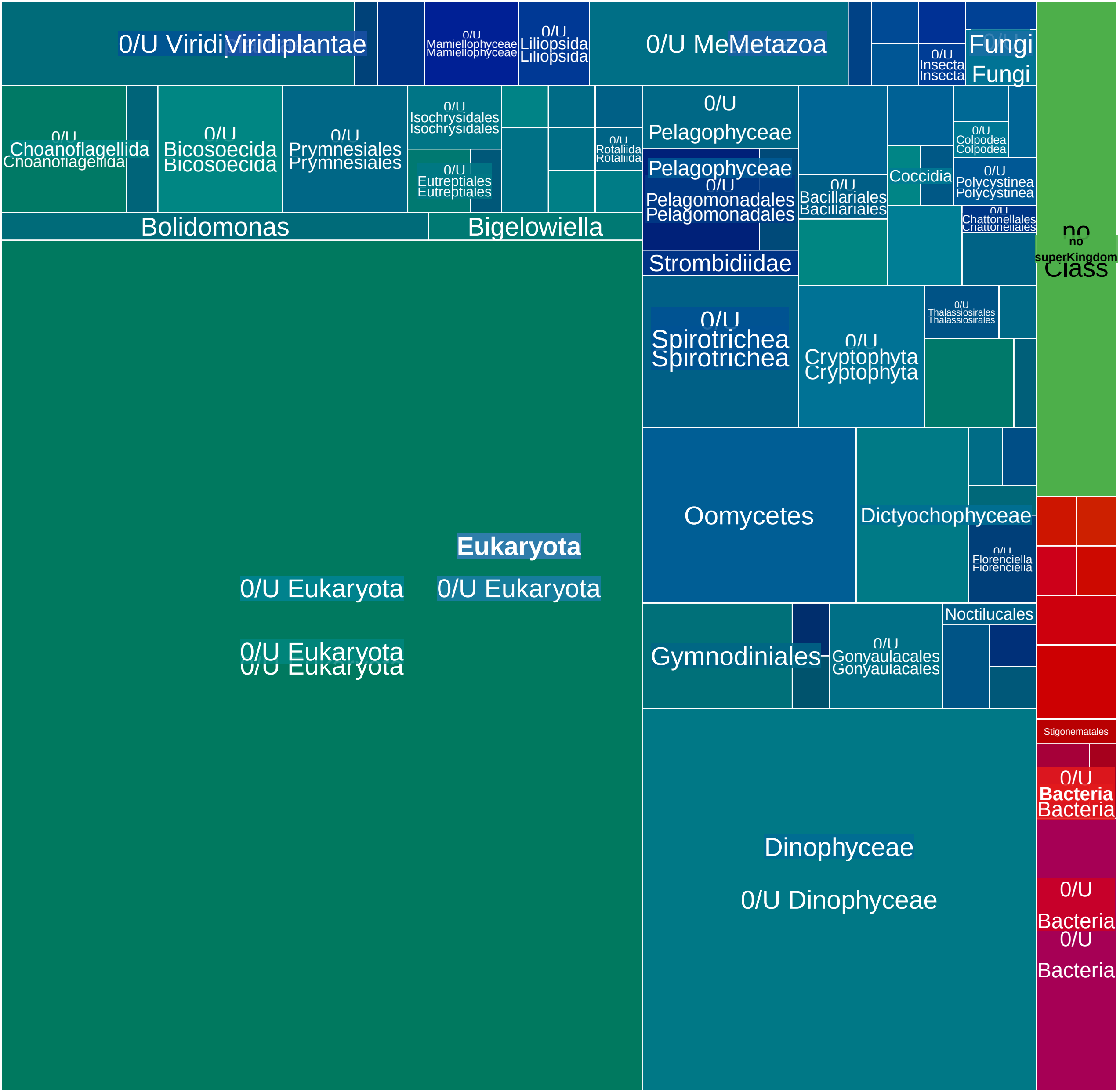

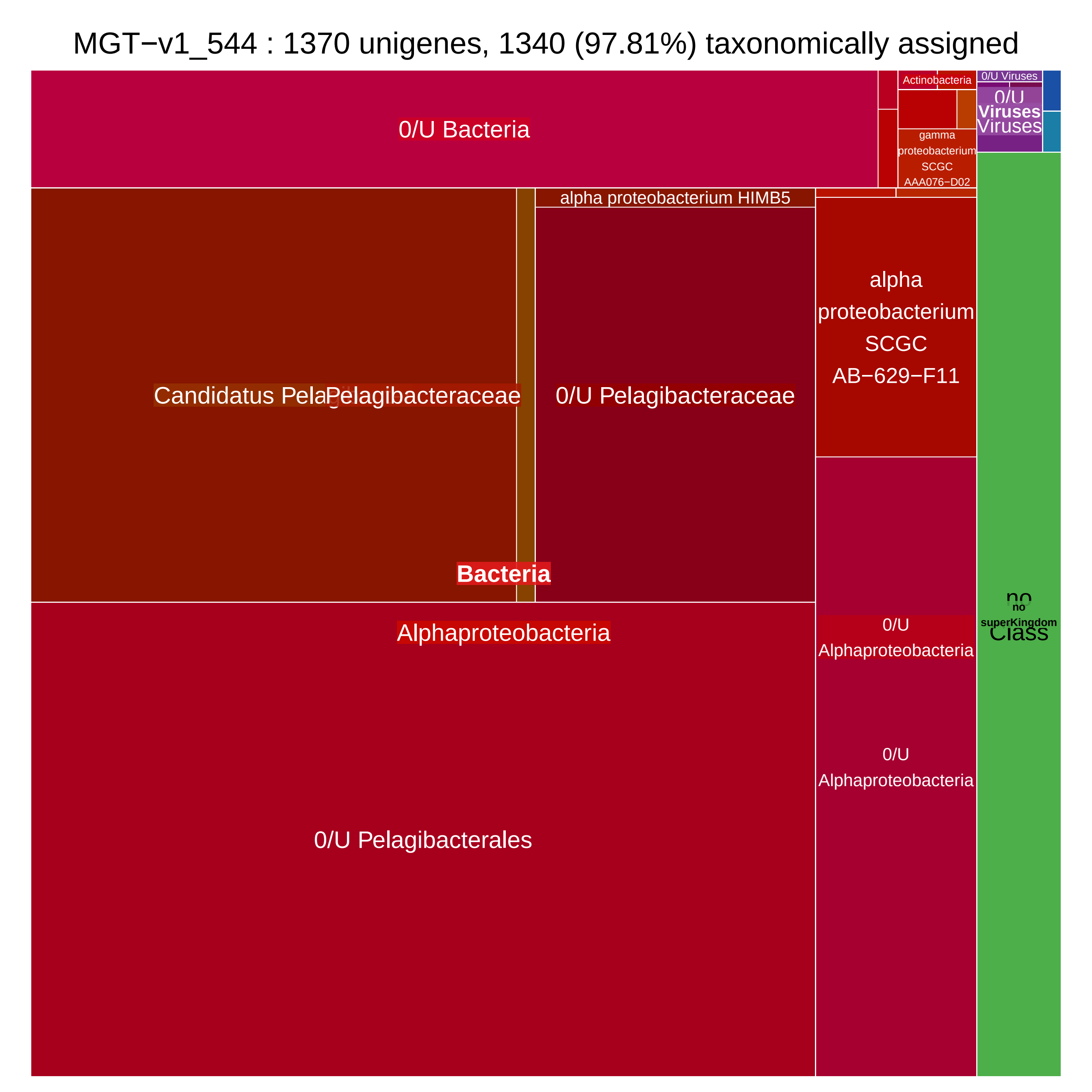
MGT−v1_544 : 1370 unigenes, 1340 (97.81%) taxonomically assigned
0/U Bacteria
Actinobacteria
0/U Viruses
0/U Viruses
Viruses
gamma proteobacterium SCGC AAA076−D02
alpha proteobacterium HIMB5
Candidatus Pelagibacteraceae
0/U Pelagibacteraceae
alpha proteobacterium SCGC AB−629−F11
Bacteria
Alphaproteobacteria
0/U Alphaproteobacteria
0/U Alphaproteobacteria
0/U Pelagibacterales
no superKingdom
no Class

MGT−v1_545 : 1366 unigenes, 1321 (96.71%) taxonomically assigned

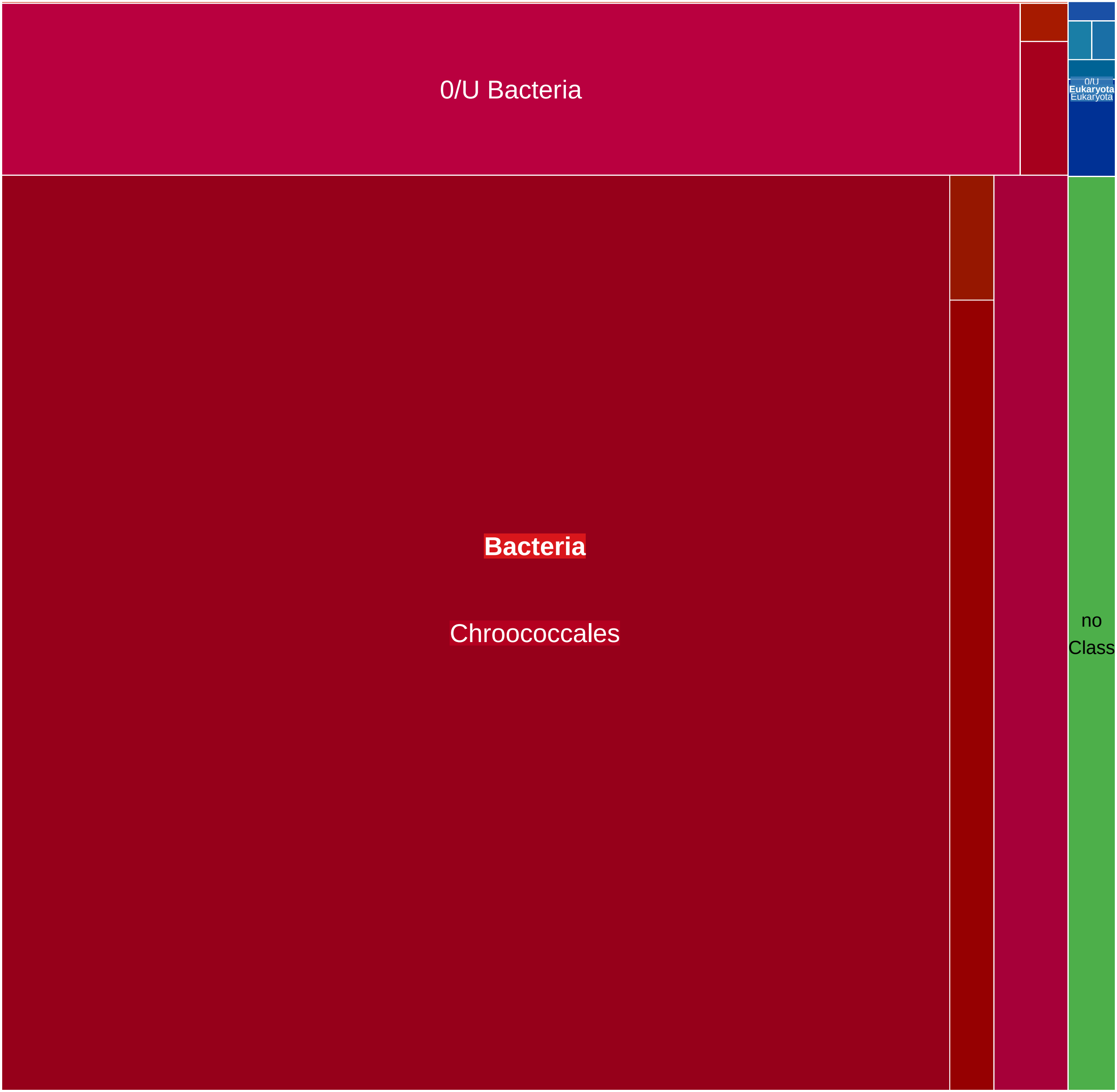

MGT−v1_546 : 1359 unigenes, 194 (14.28%) taxonomically assigned

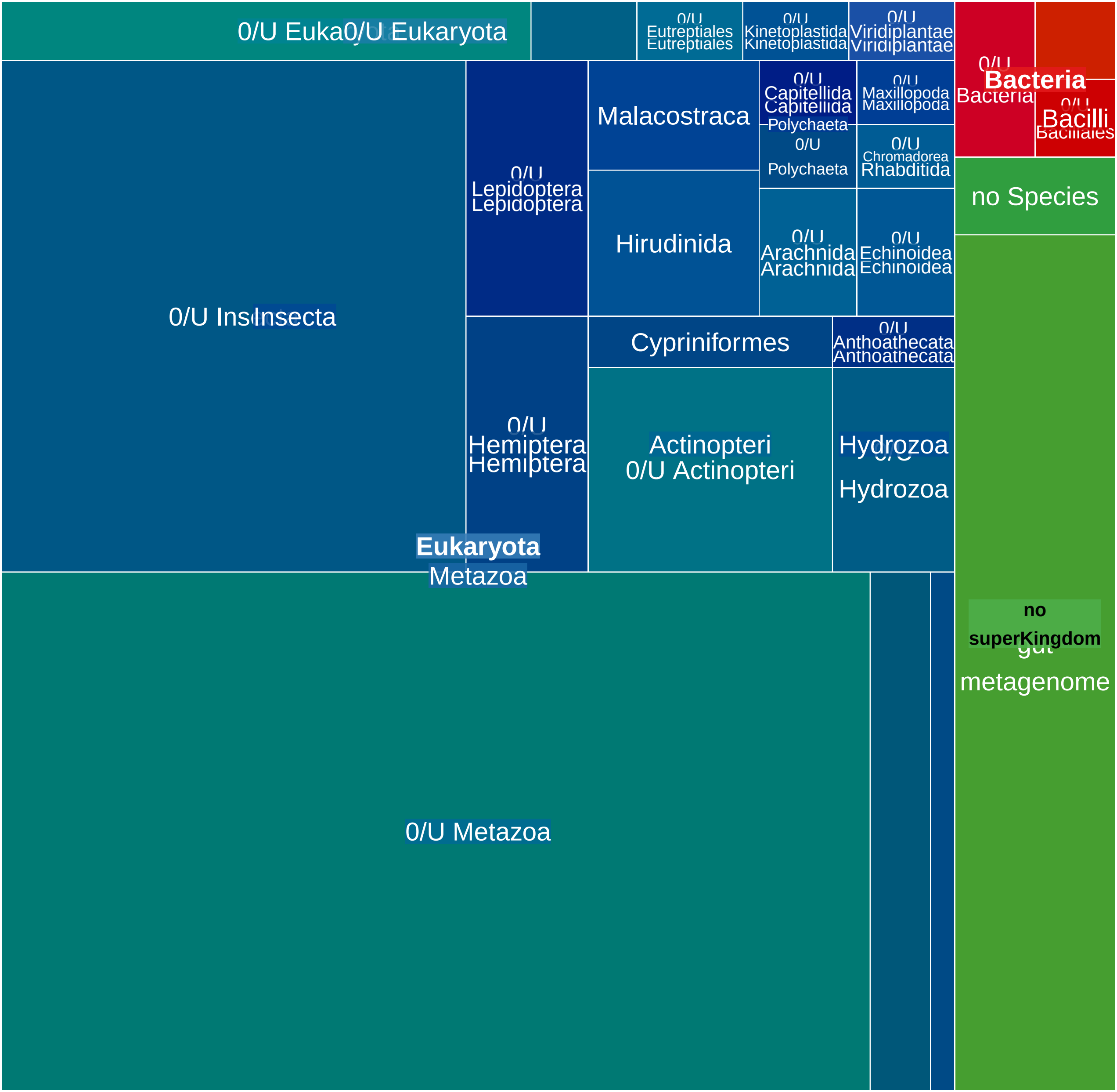

MGT−v1_547 : 1357 unigenes, 941 (69.34%) taxonomically assigned

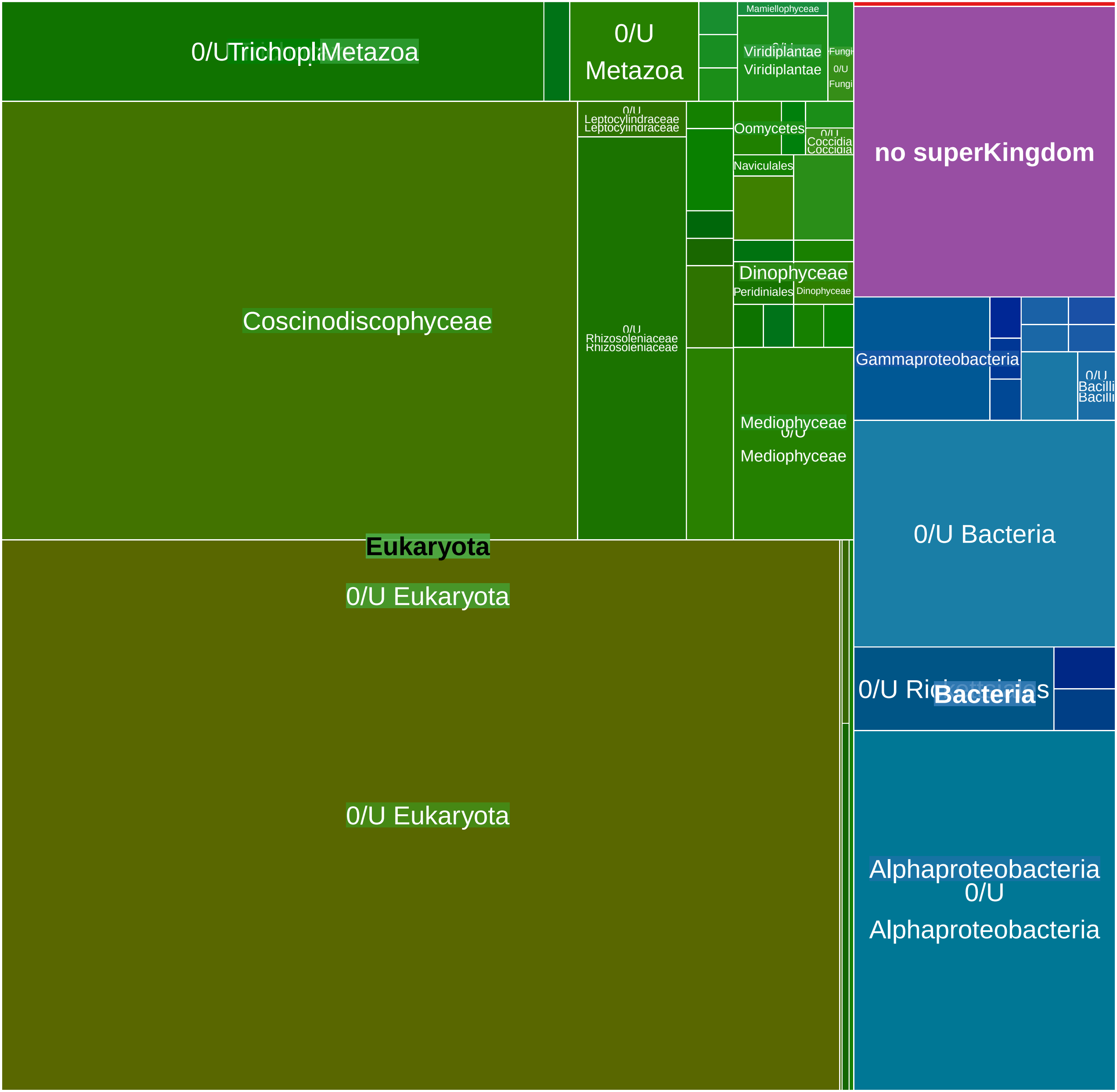

#### MGT−v1_548 : 1354 unigenes, 129 (9.53%) taxonomically assigned

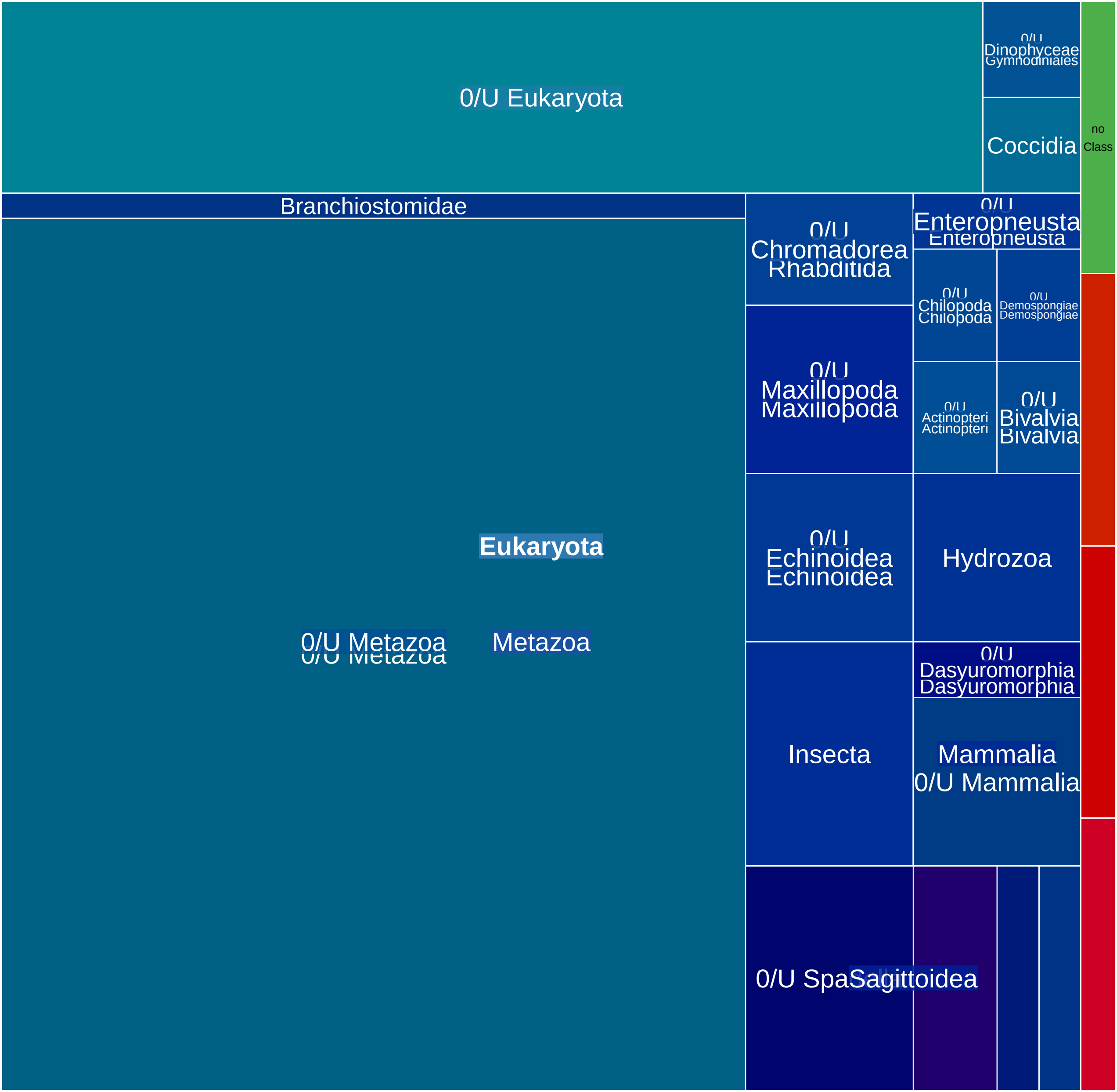

MGT−v1_549 : 1353 unigenes, 402 (29.71%) taxonomically assigned

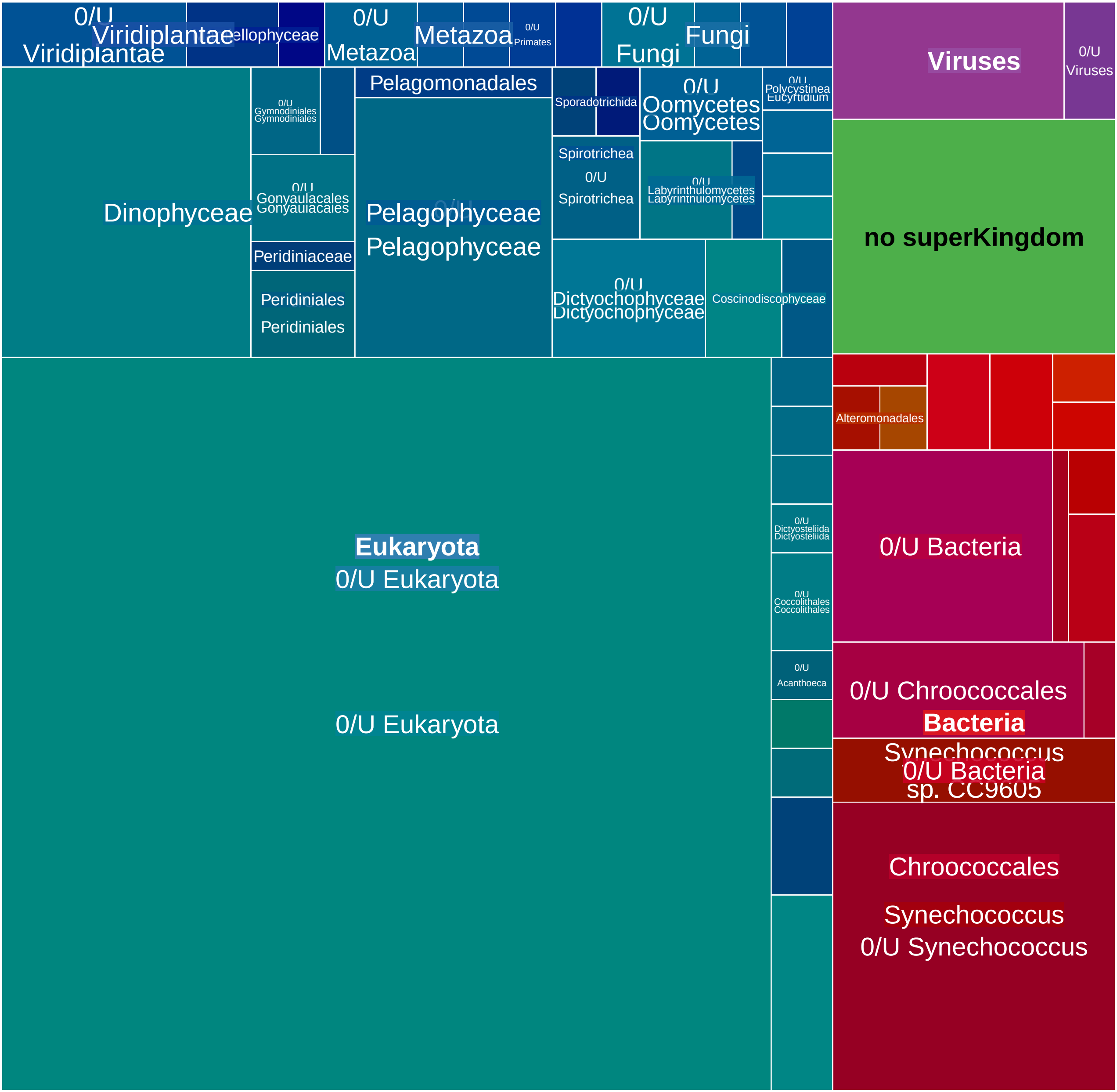

MGT−v1_550 : 1352 unigenes, 80 (5.92%) taxonomically assigned

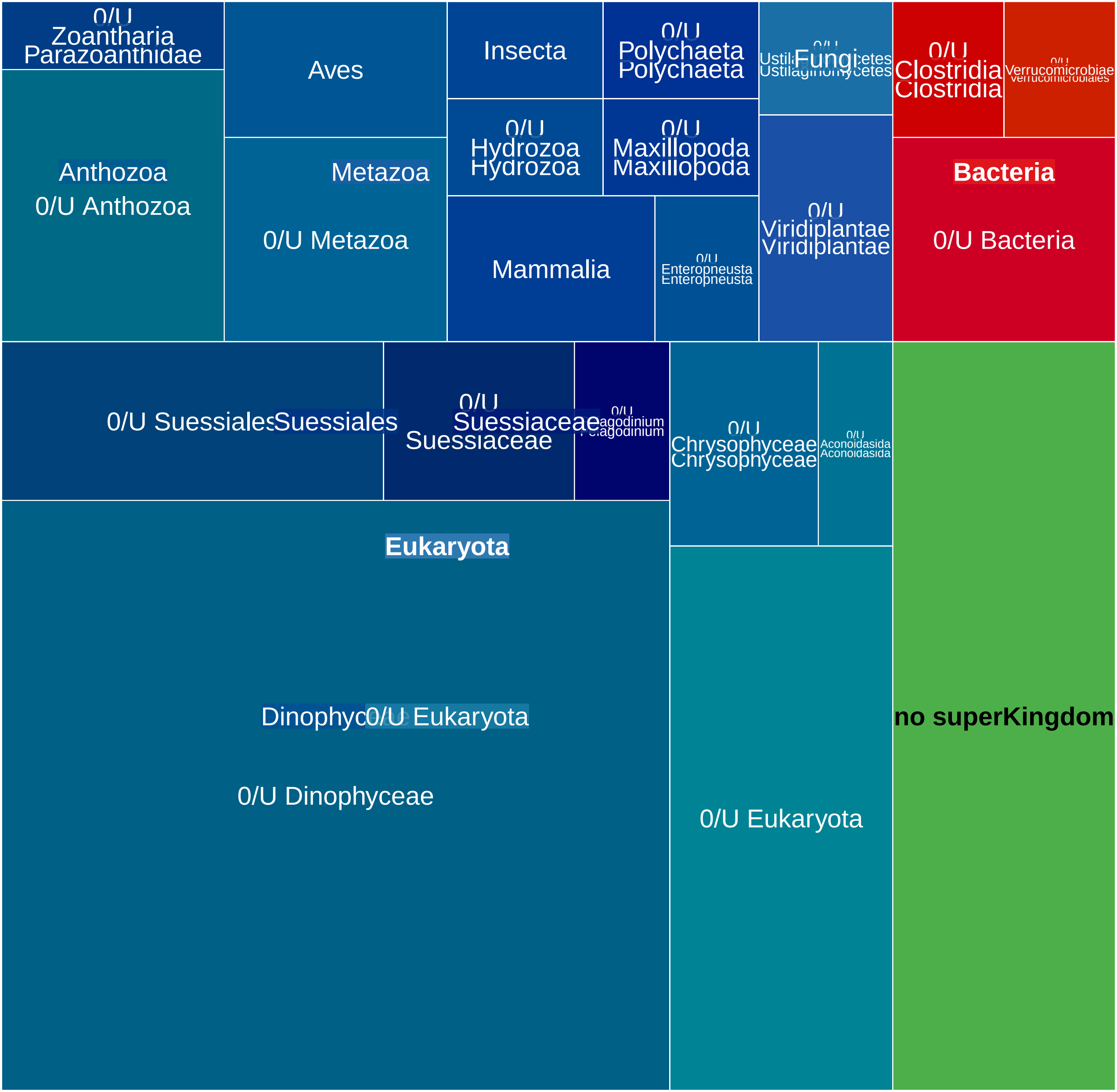

MGT−v1_551 : 1331 unigenes, 912 (68.52%) taxonomically assigned

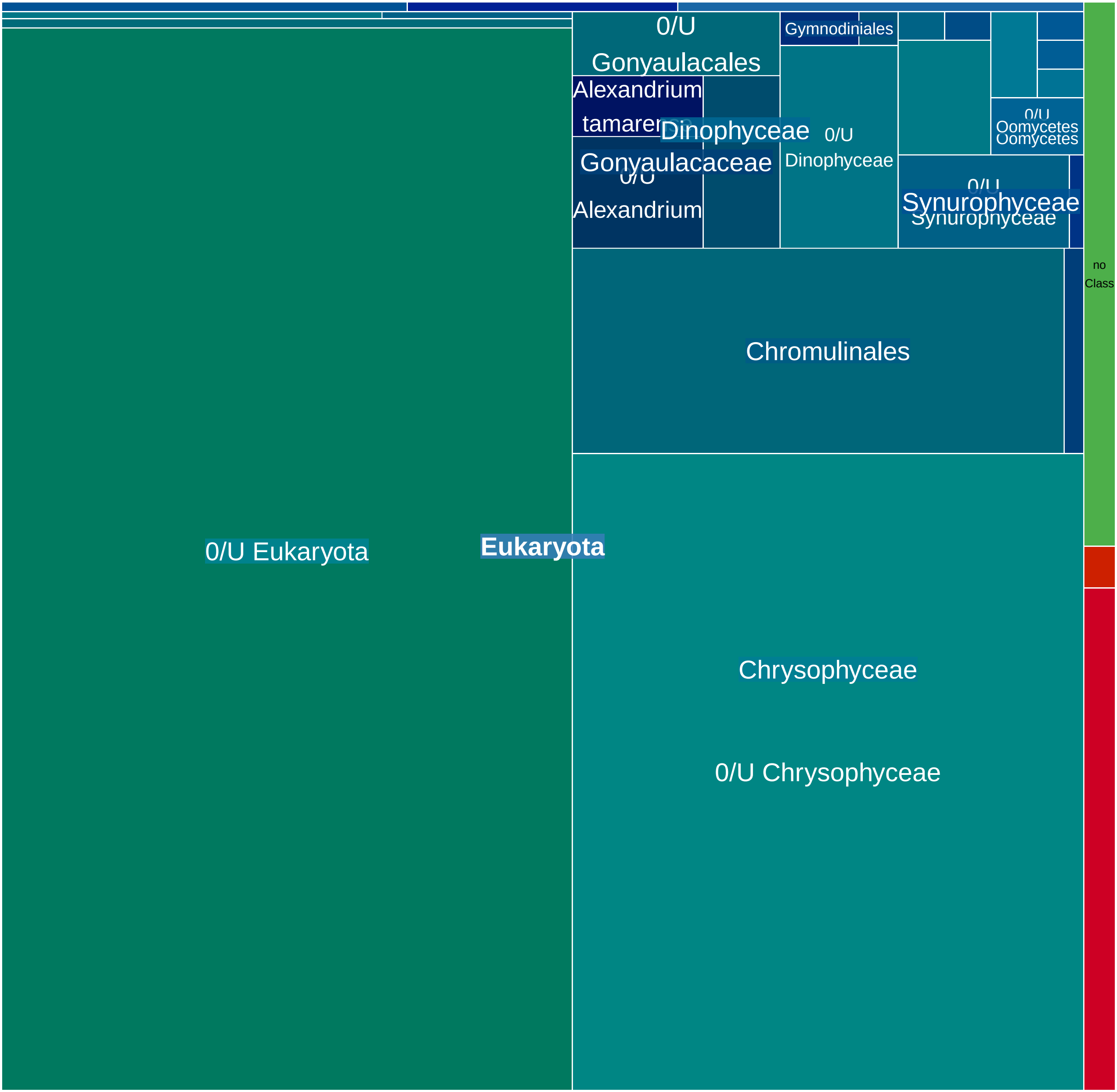

MGT−v1_552 : 1331 unigenes, 239 (17.96%) taxonomically assigned

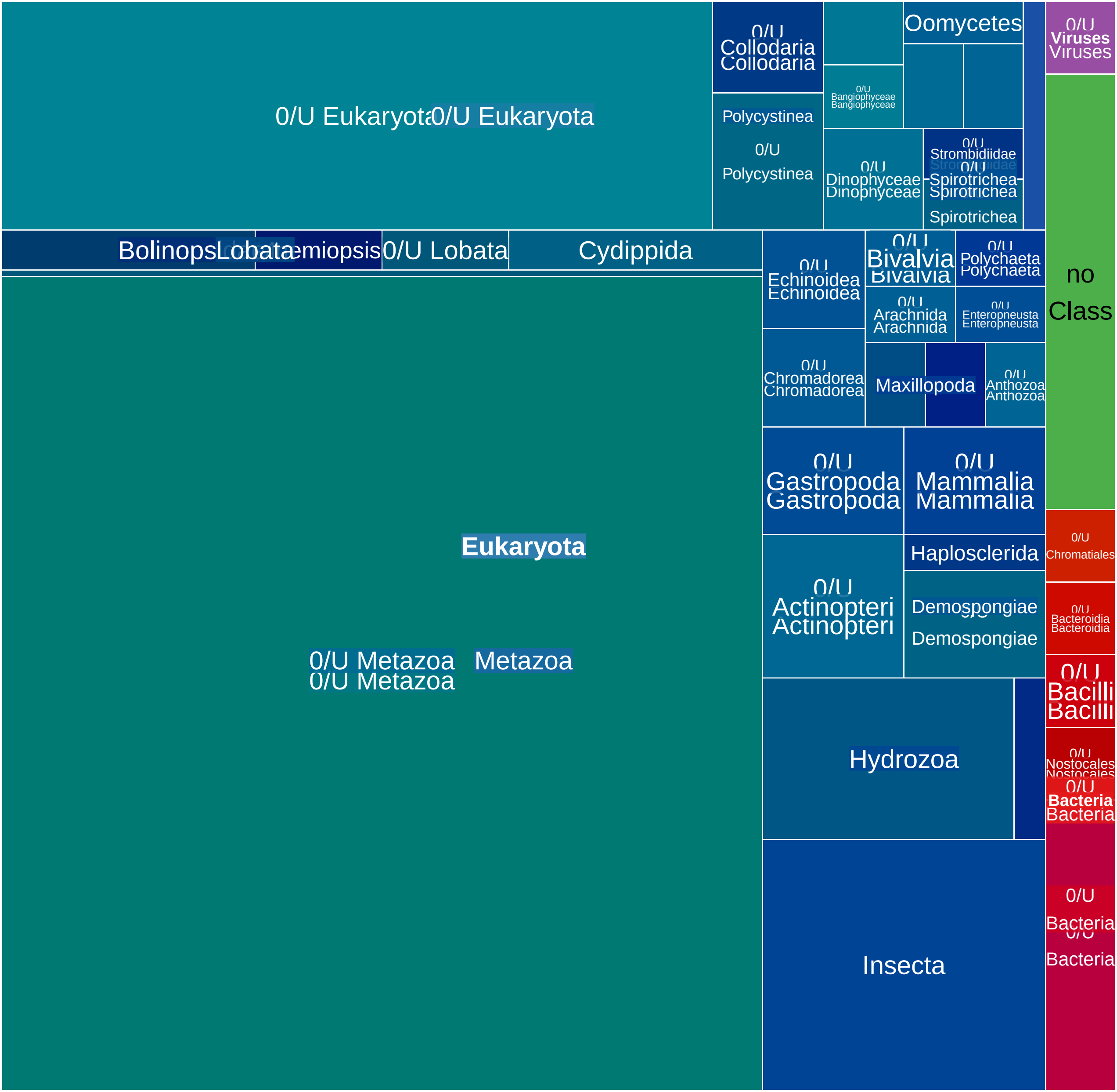

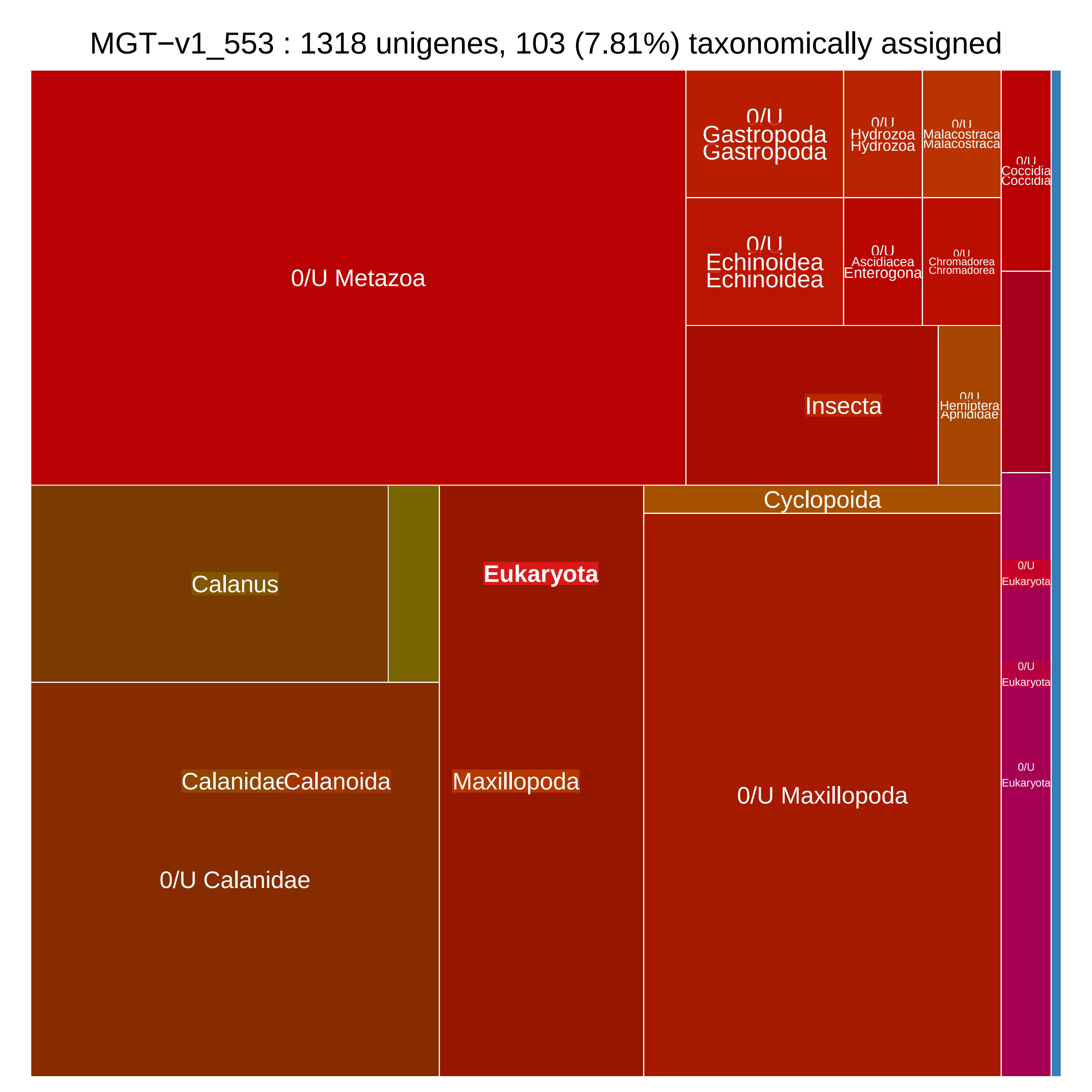

MGT−v1_553 : 1318 unigenes, 103 (7.81%) taxonomically assigned
0/U Metazoa
0/U
Gastropoda
Gastropoda
0/U
Hydrozoa
Hydrozoa
0/U
Malacostraca
Malacostraca
0/U
Coccidia
Coccidia
0/U
Echinoidea
Echinoidea
0/U
Ascidiacea
Enterogona
0/U
Chromadorea
Chromadorea
Insecta
0/U
Hemiptera
Aphididae
Cyclopoida
Calanus
Eukaryota
0/U
Eukaryota
0/U
Eukaryota
Calanidae
Calanoida
Maxillopoda
0/U Maxillopoda
0/U
Eukaryota
0/U Calanidae

### MGT−v1_554 : 1316 unigenes, 963 (73.18%) taxonomically assigned

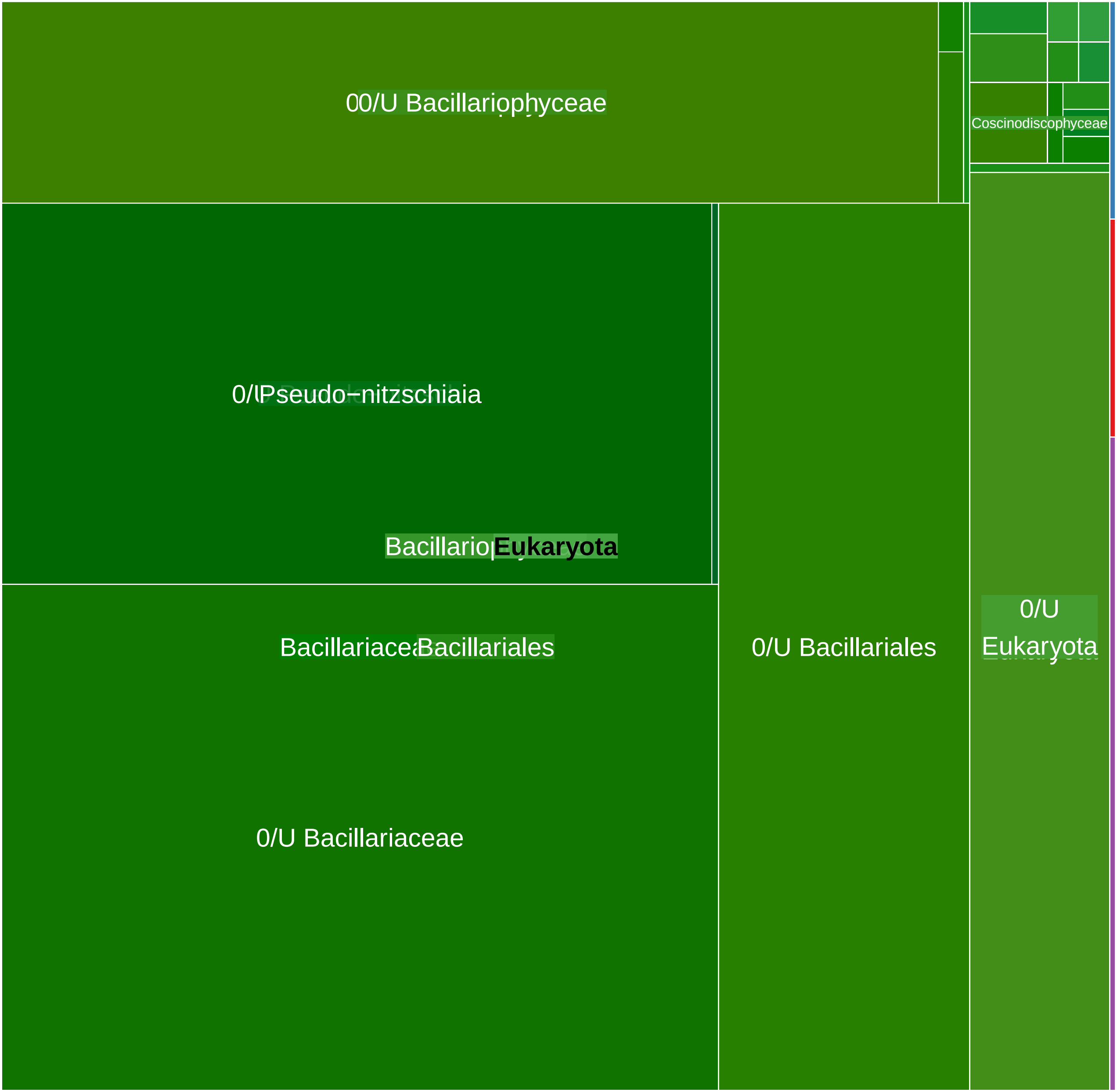

### MGT−v1_555 : 1311 unigenes, 292 (22.27%) taxonomically assigned

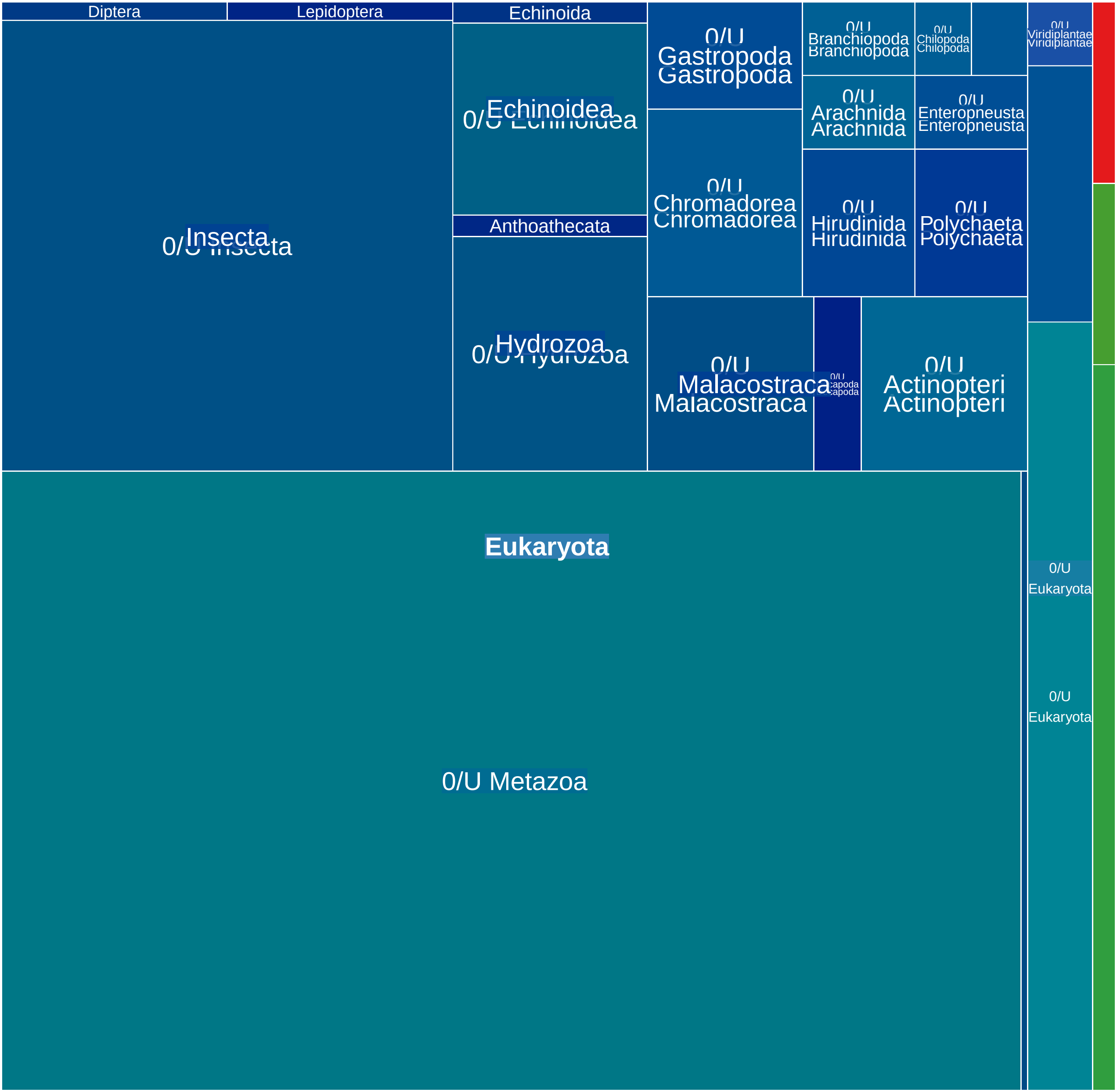

MGT−v1_556 : 1311 unigenes, 114 (8.70%) taxonomically assigned

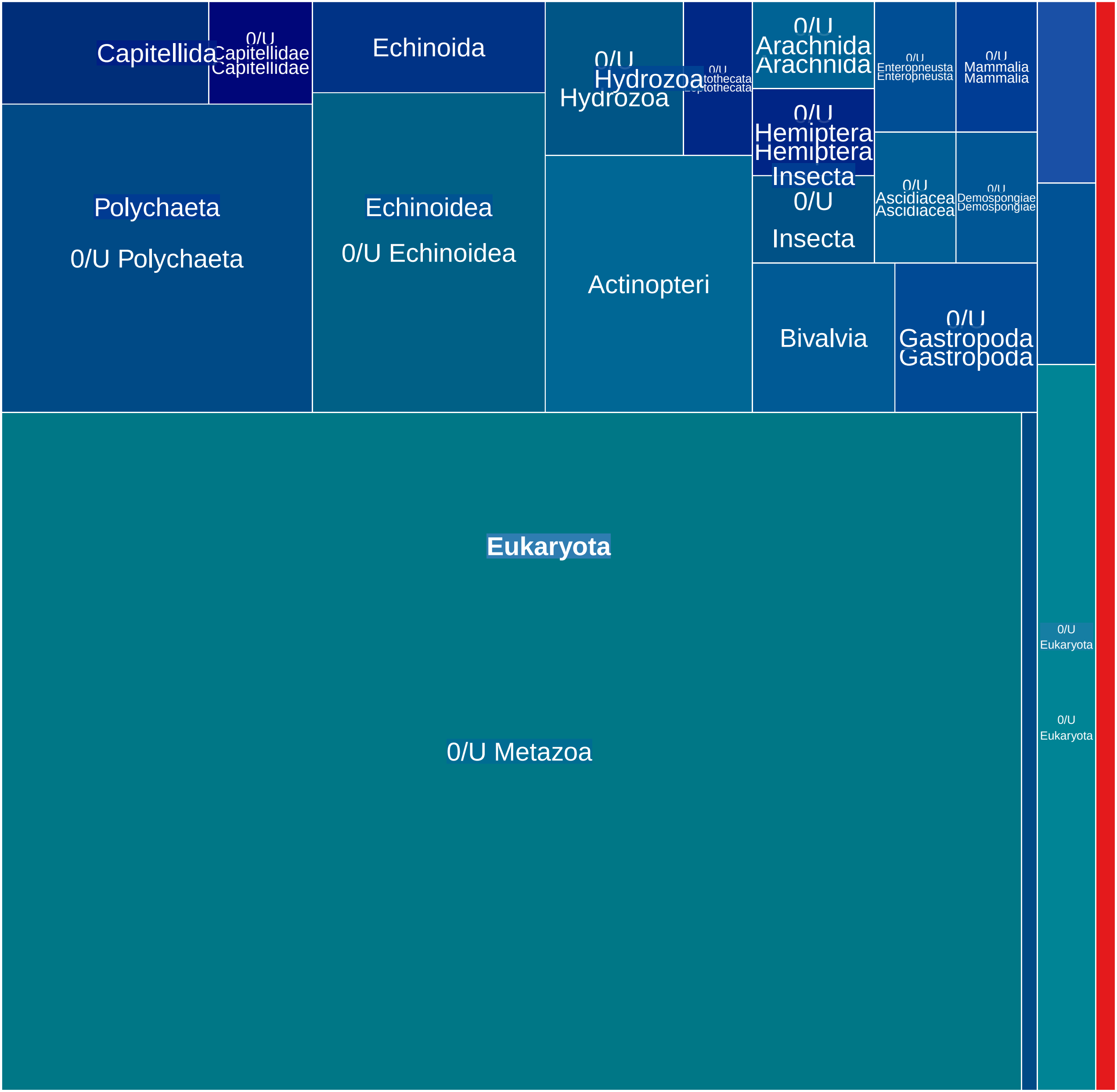

### MGT−v1_557 : 1305 unigenes, 840 (64.37%) taxonomically assigned

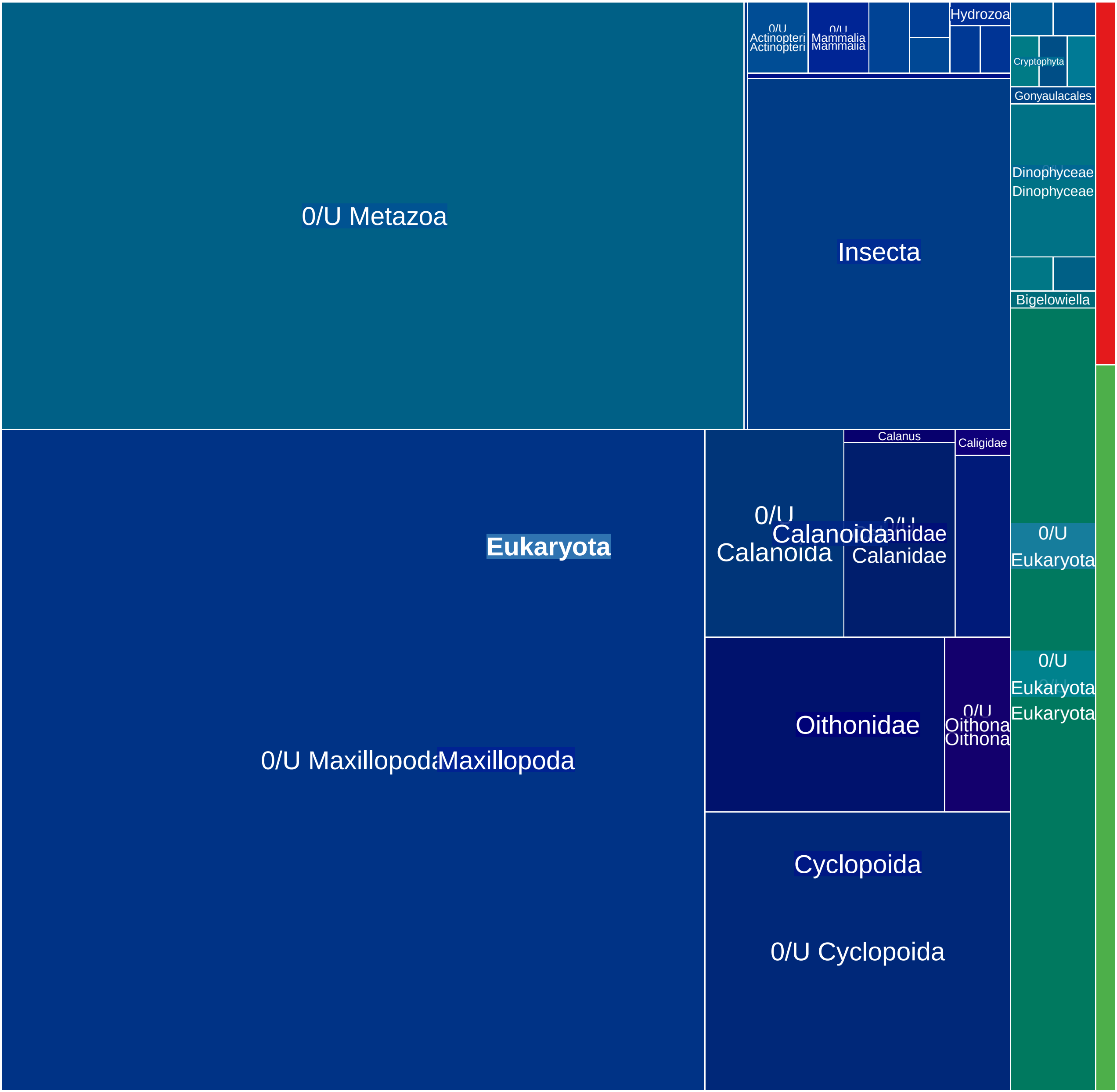

MGT−v1_558 : 1301 unigenes, 390 (29.98%) taxonomically assigned

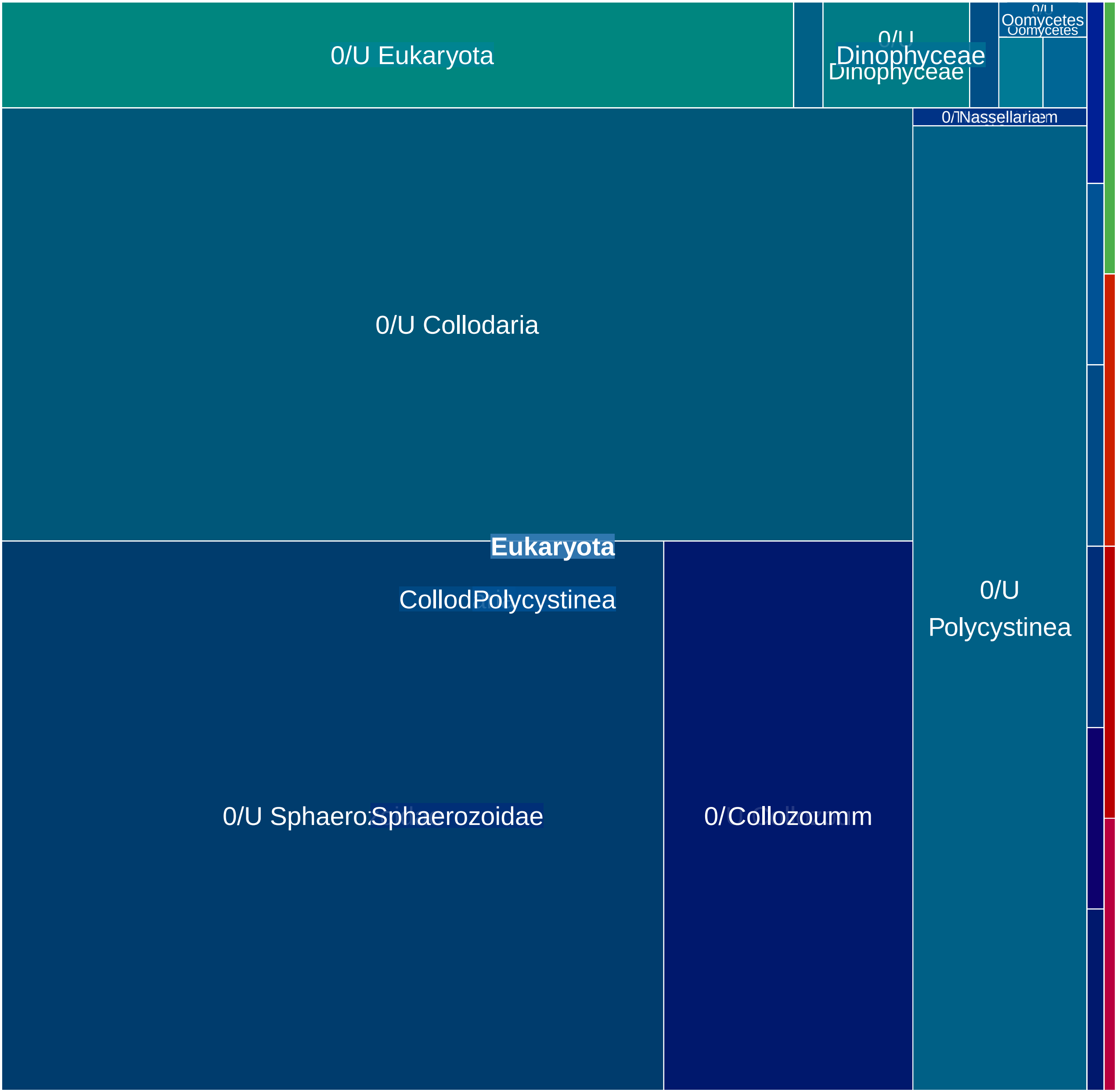

MGT−v1_559 : 1289 unigenes, 1123 (87.12%) taxonomically assigned

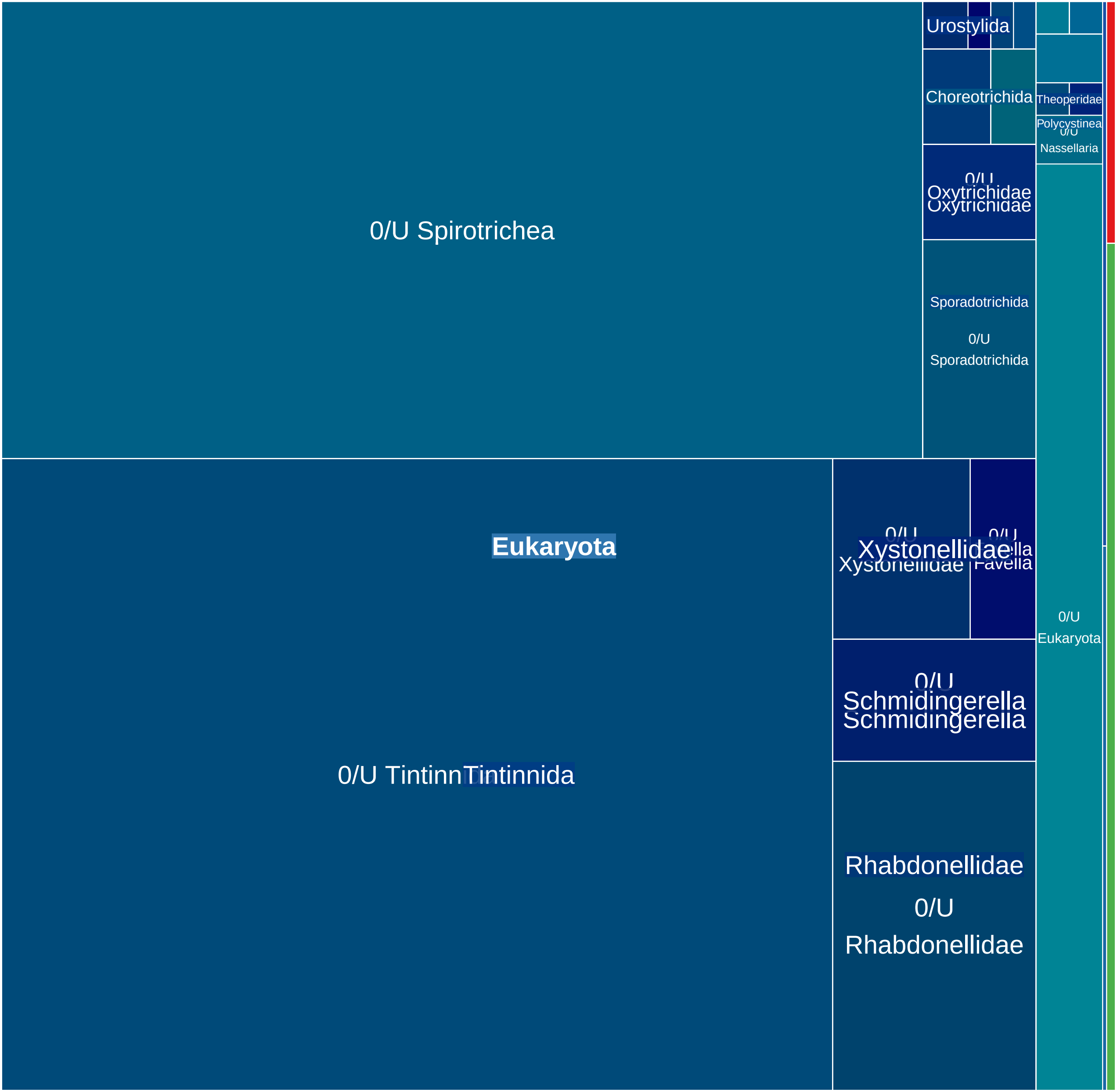

MGT−v1_560 : 1285 unigenes, 151 (11.75%) taxonomically assigned

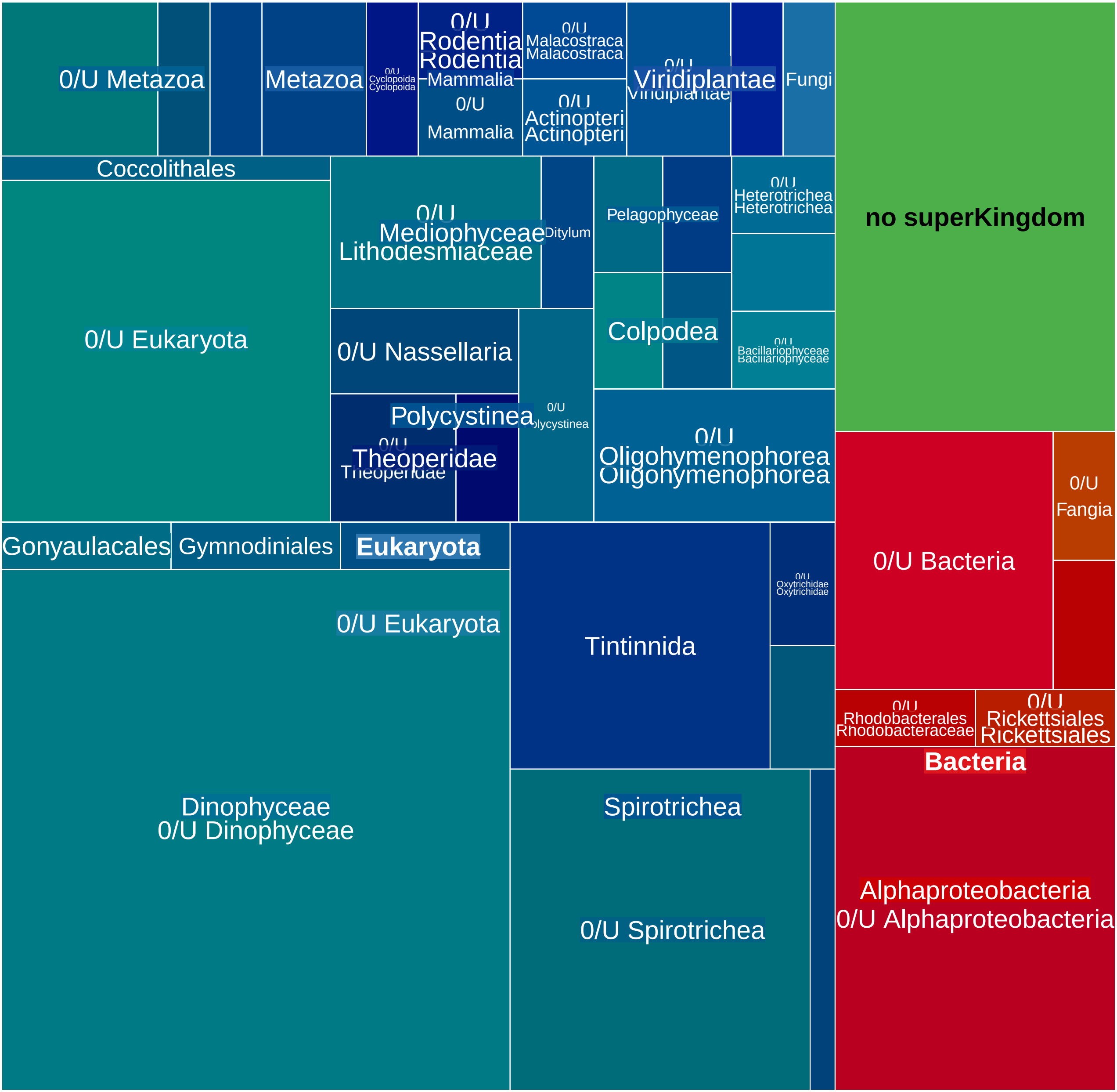

MGT−v1_561 : 1280 unigenes, 1251 (97.73%) taxonomically assigned

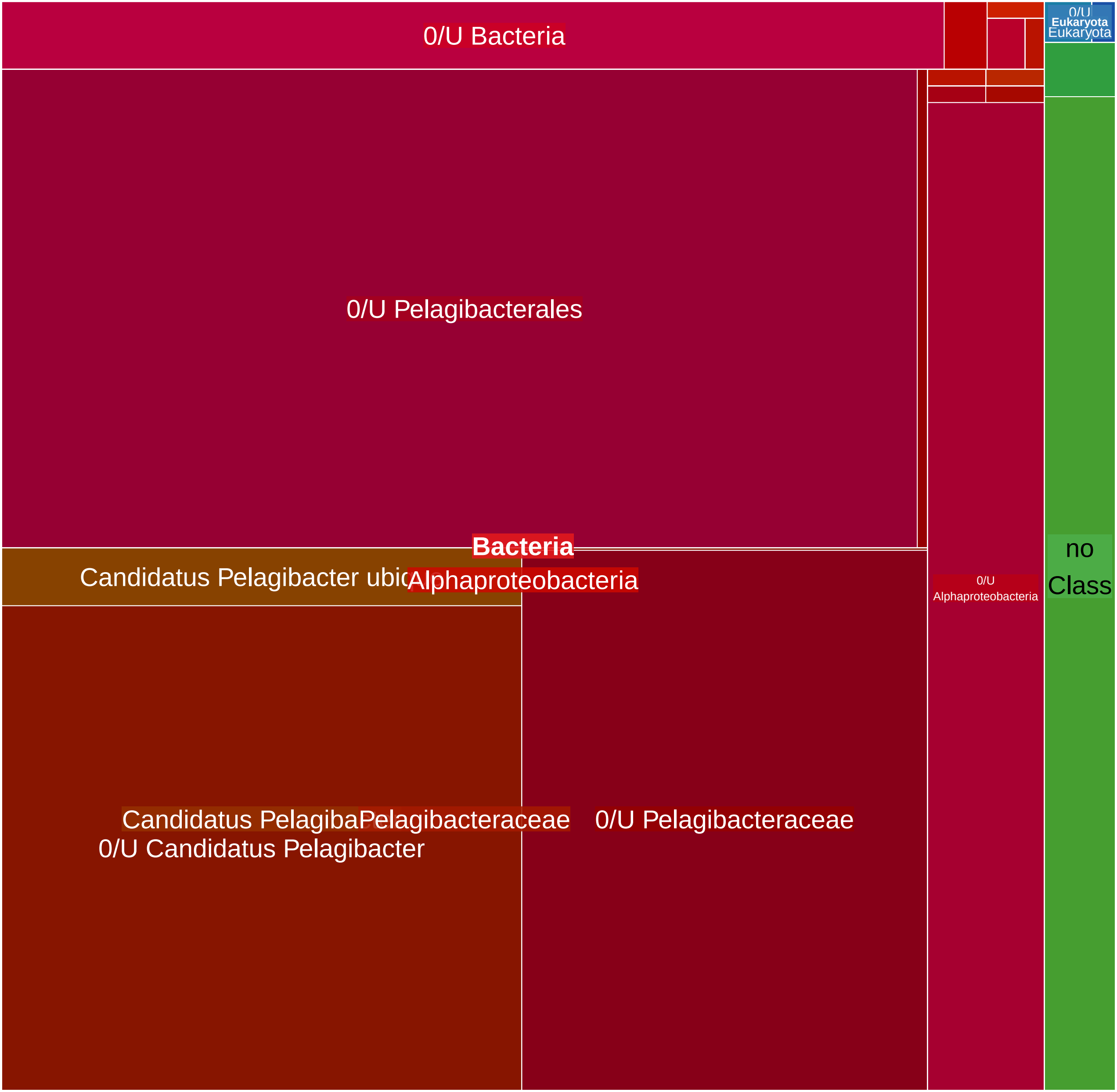

MGT−v1_562 : 1268 unigenes, 101 (7.97%) taxonomically assigned

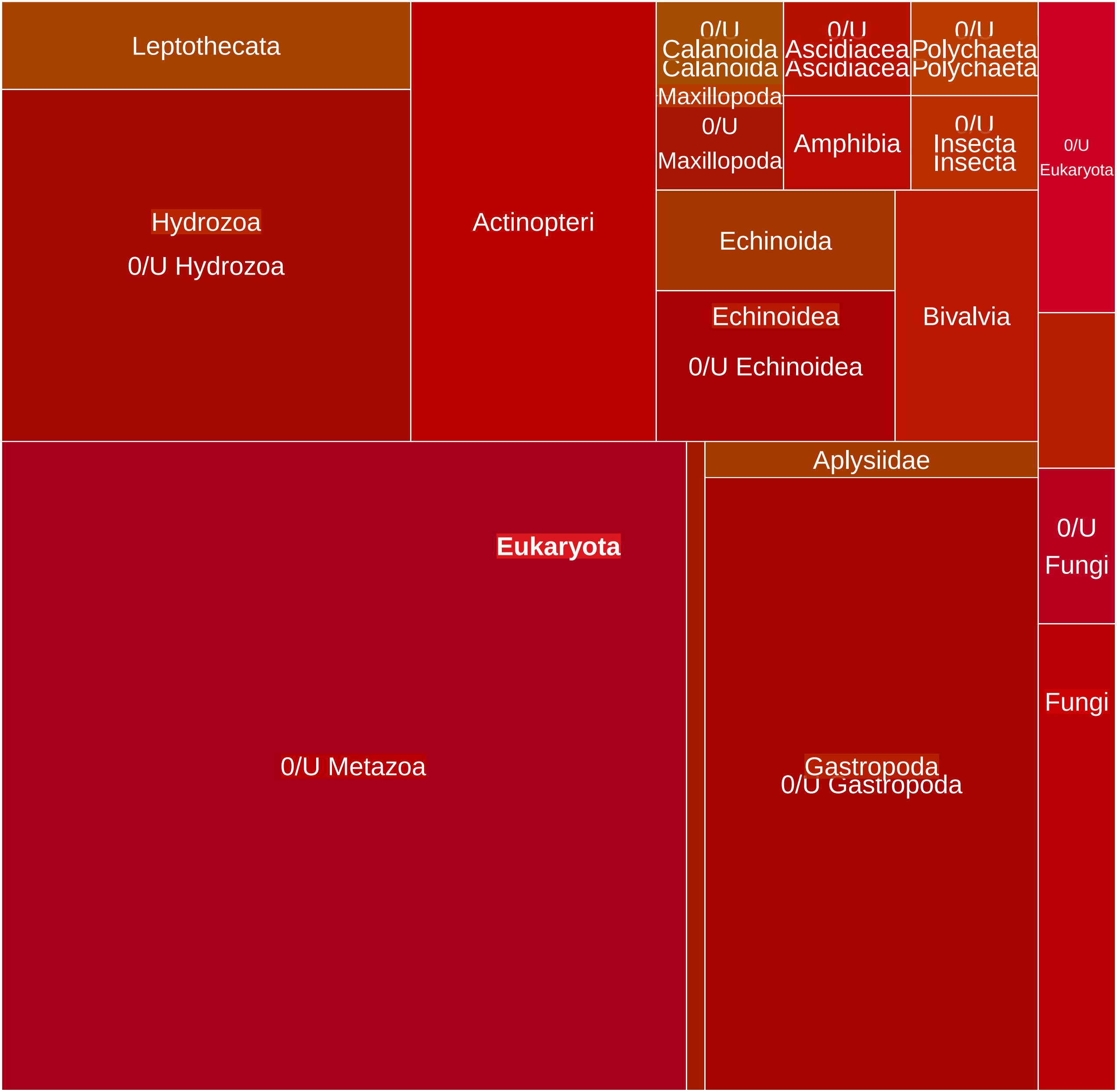

MGT−v1_563 : 1267 unigenes, 1167 (92.11%) taxonomically assigned

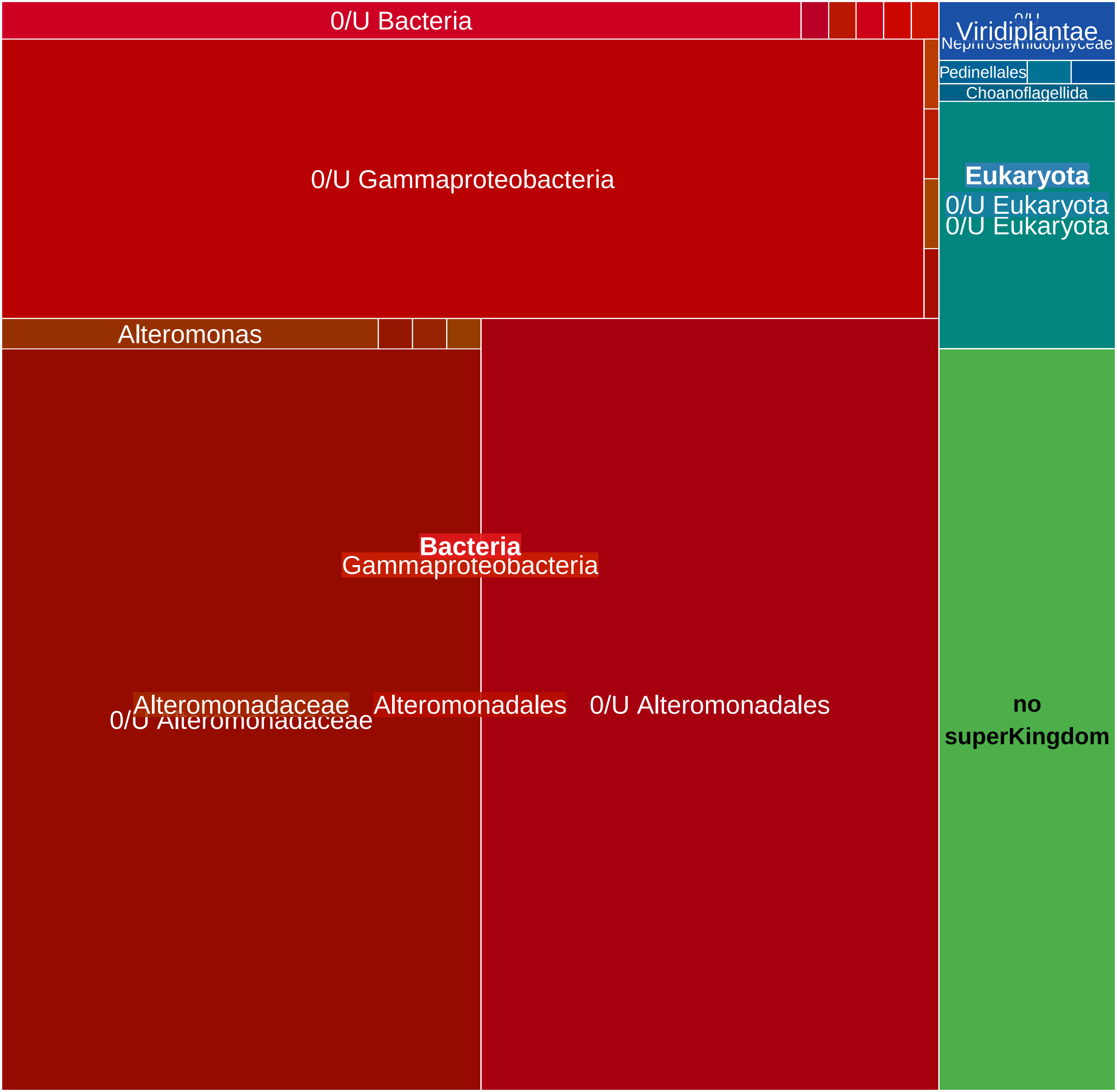

MGT−v1_564 : 1261 unigenes, 323 (25.61%) taxonomically assigned

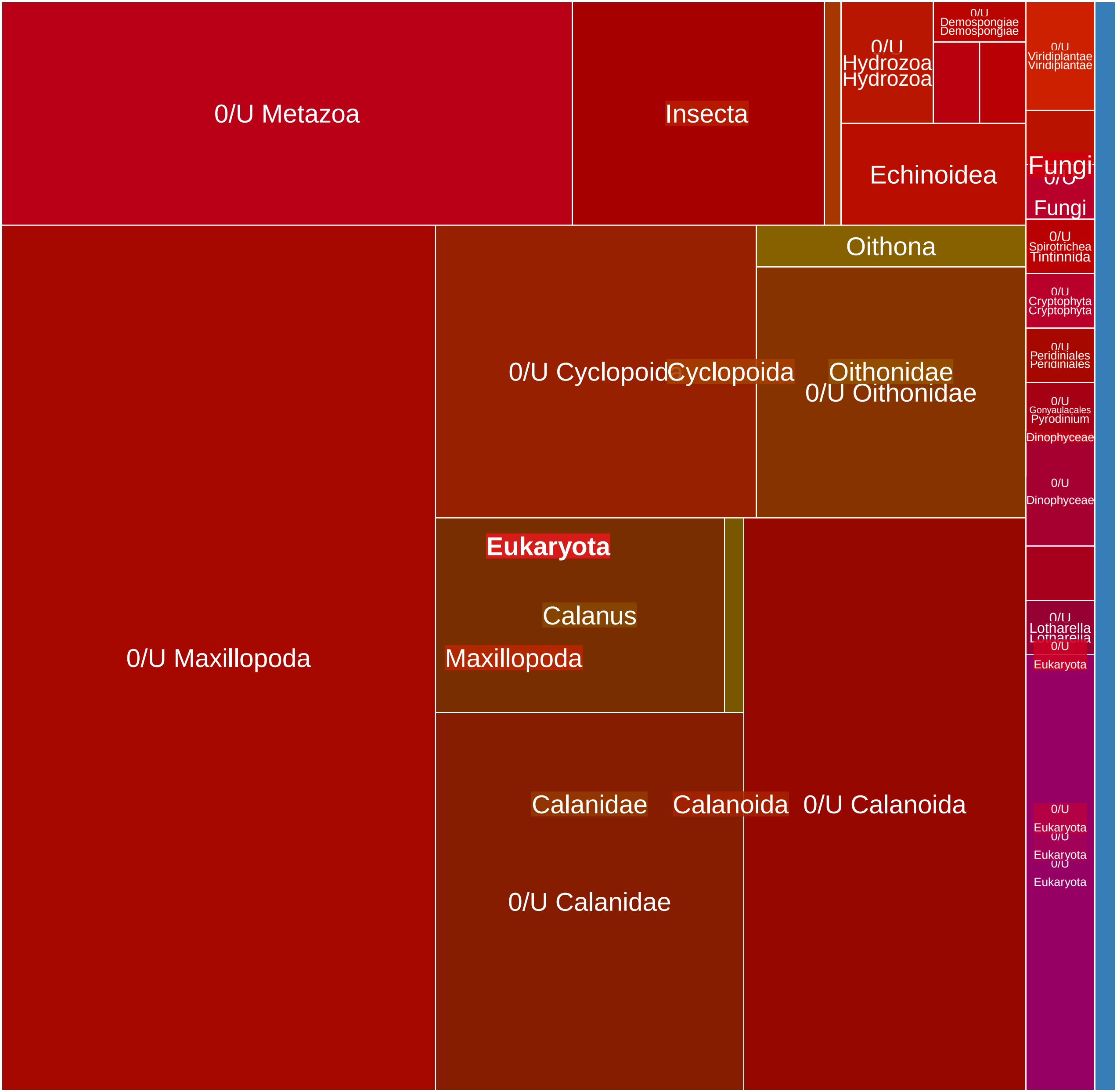

MGT−v1_565 : 1259 unigenes, 393 (31.22%) taxonomically assigned

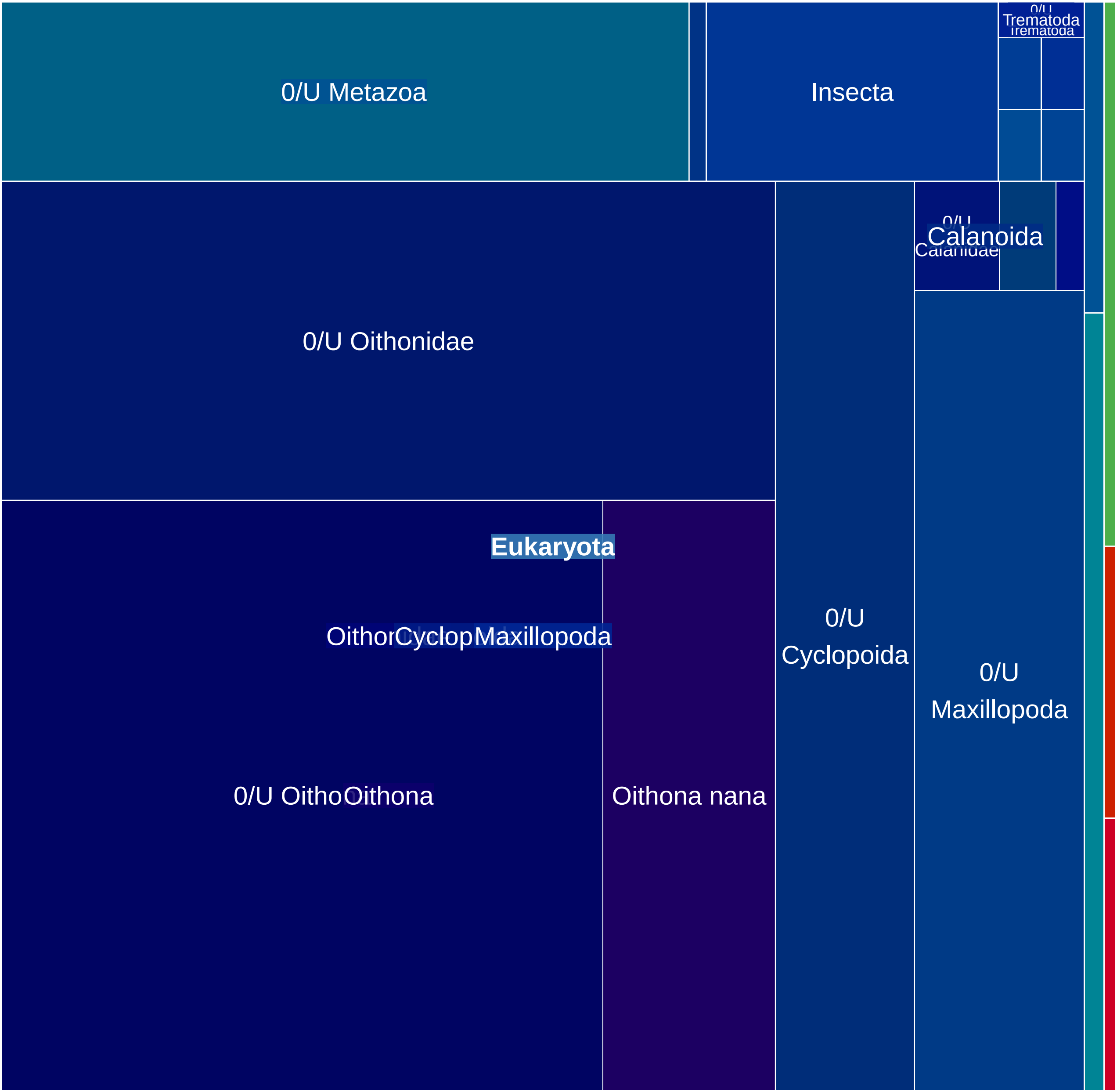

MGT−v1_566 : 1258 unigenes, 212 (16.85%) taxonomically assigned

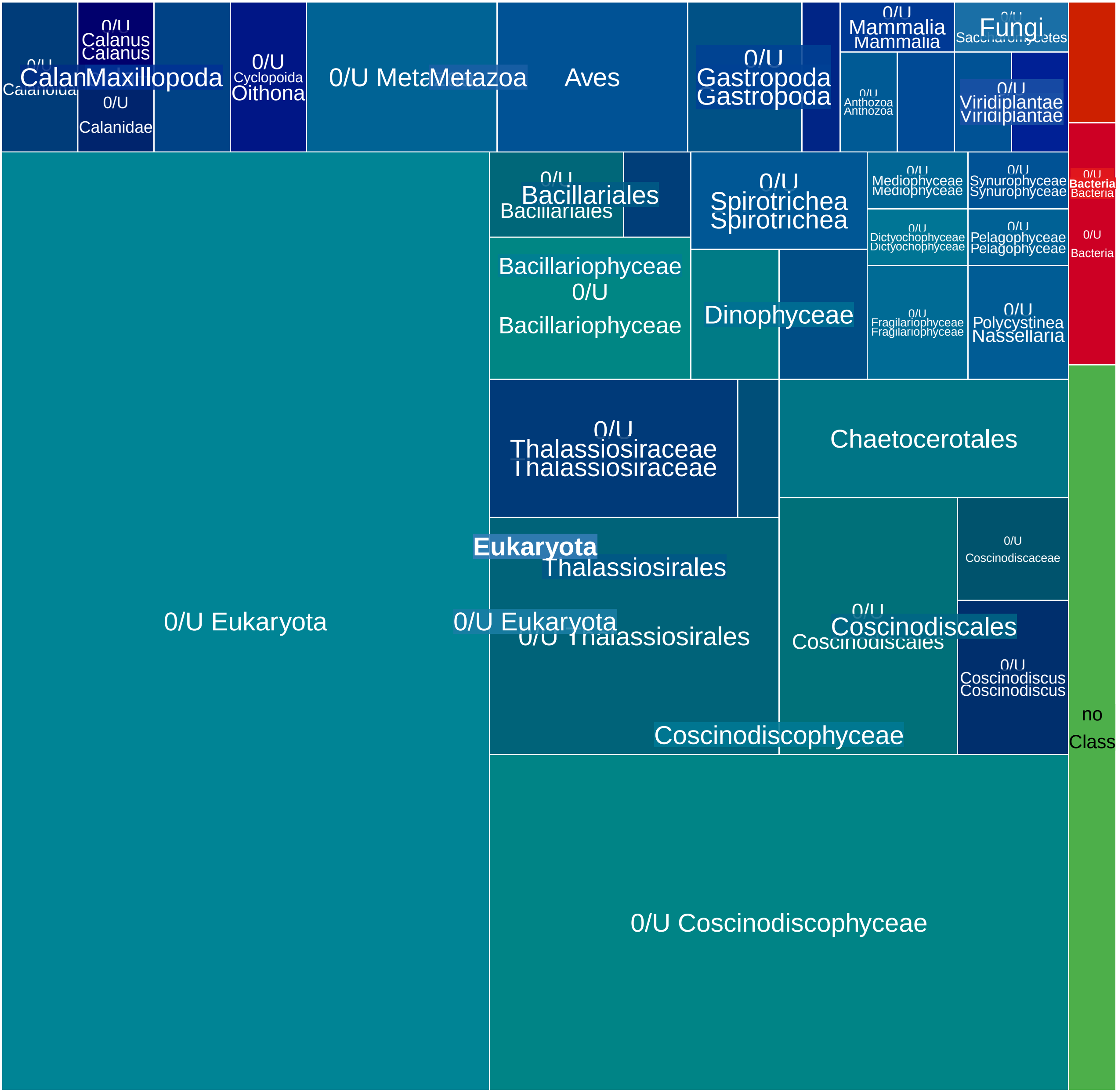

### MGT−v1_567 : 1258 unigenes, 202 (16.06%) taxonomically assigned

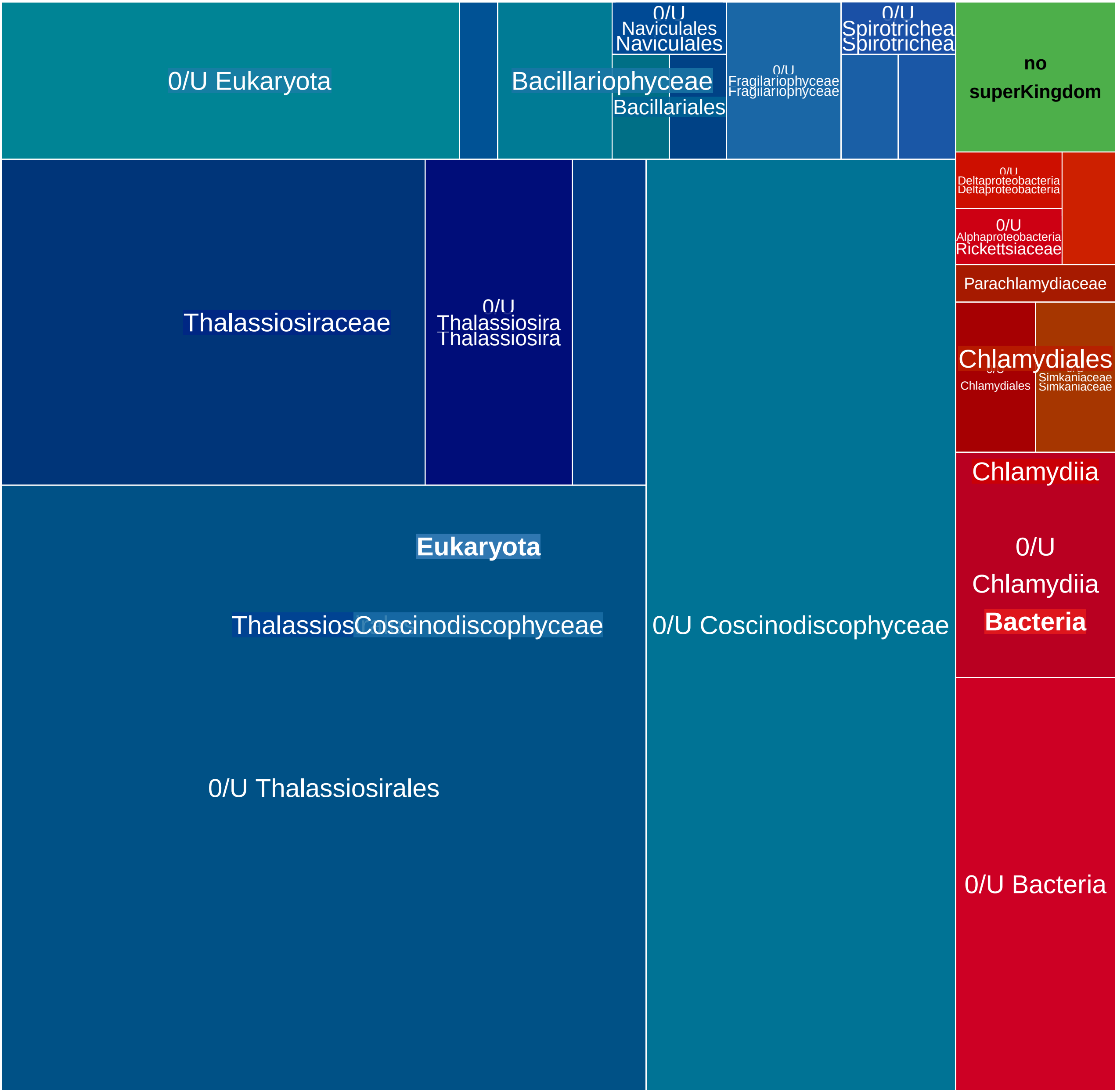

### MGT−v1_568 : 1250 unigenes, 67 (5.36%) taxonomically assigned

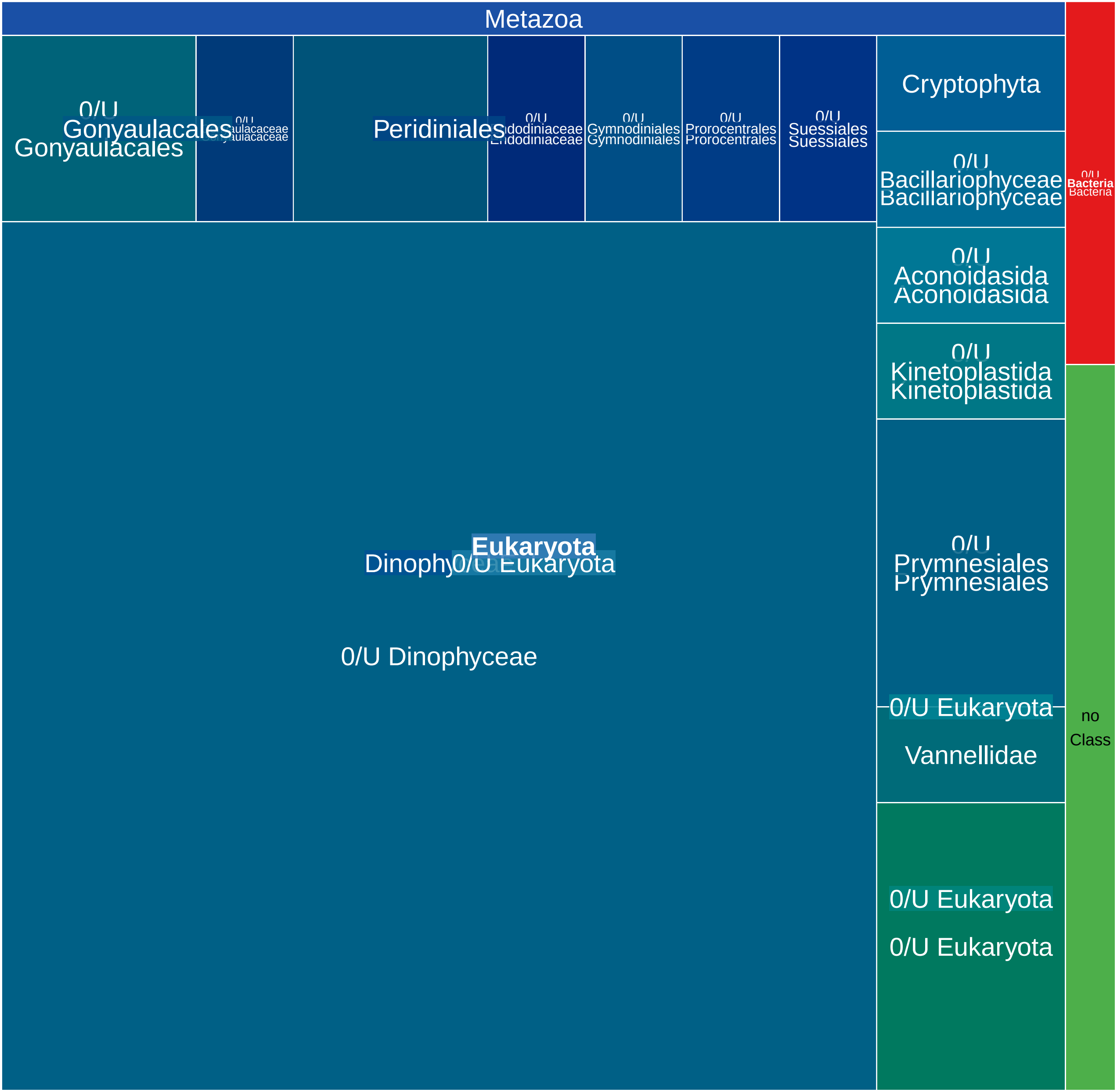

MGT−v1_569 : 1237 unigenes, 116 (9.38%) taxonomically assigned

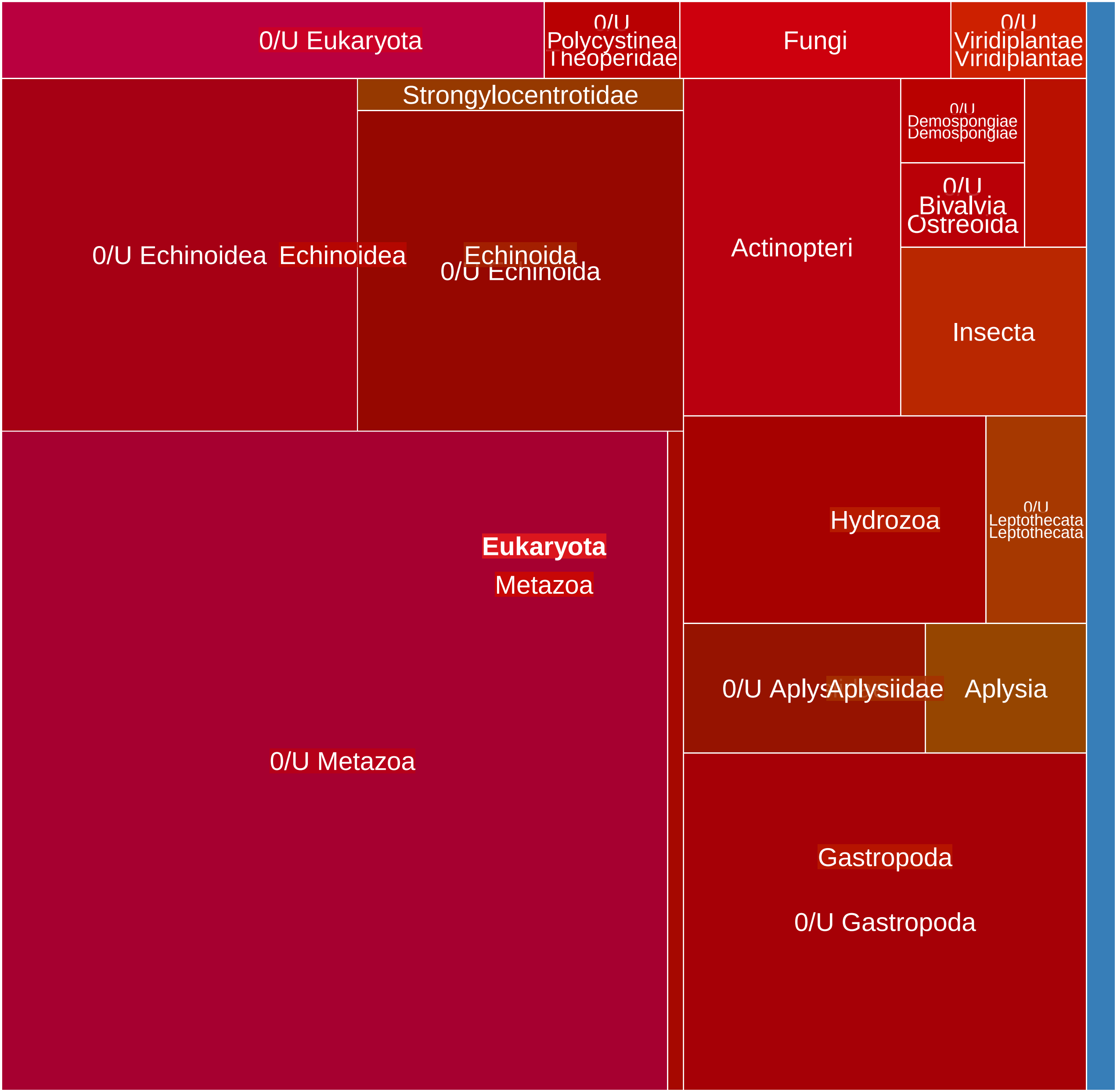

MGT−v1_570 : 1232 unigenes, 130 (10.55%) taxonomically assigned

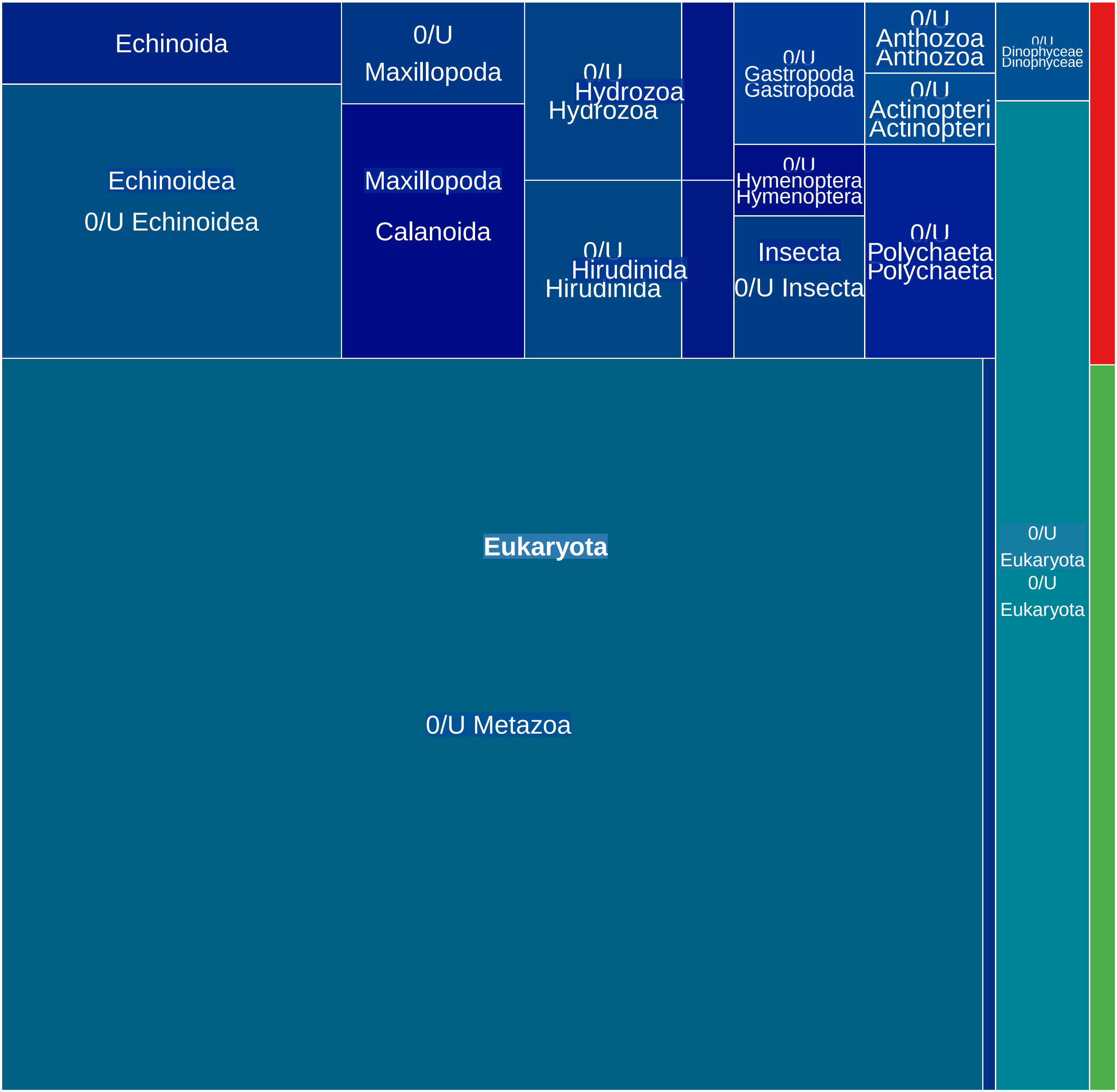

MGT−v1_571 : 1227 unigenes, 113 (9.21%) taxonomically assigned

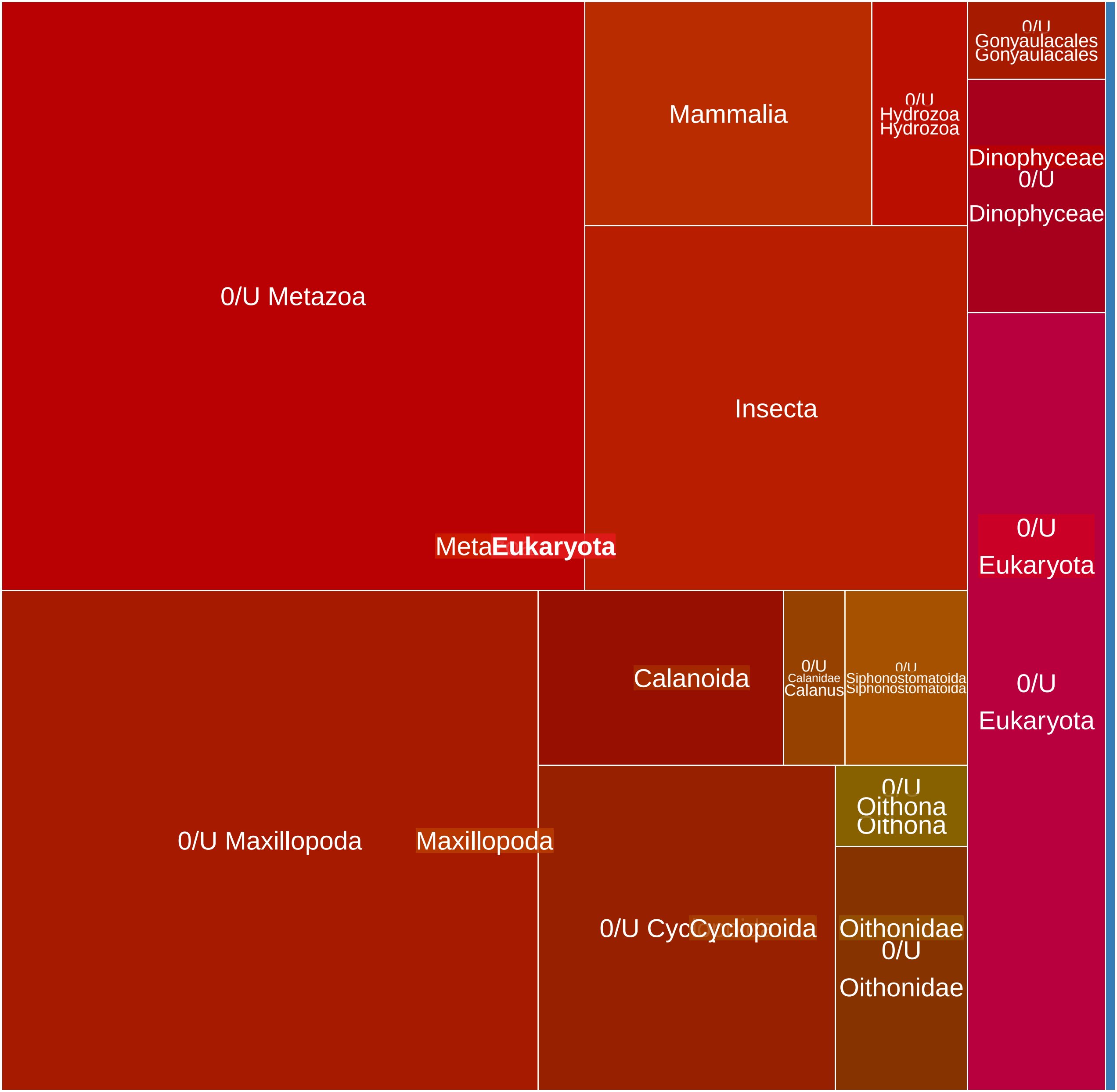

MGT−v1_572 : 1223 unigenes, 922 (75.39%) taxonomically assigned

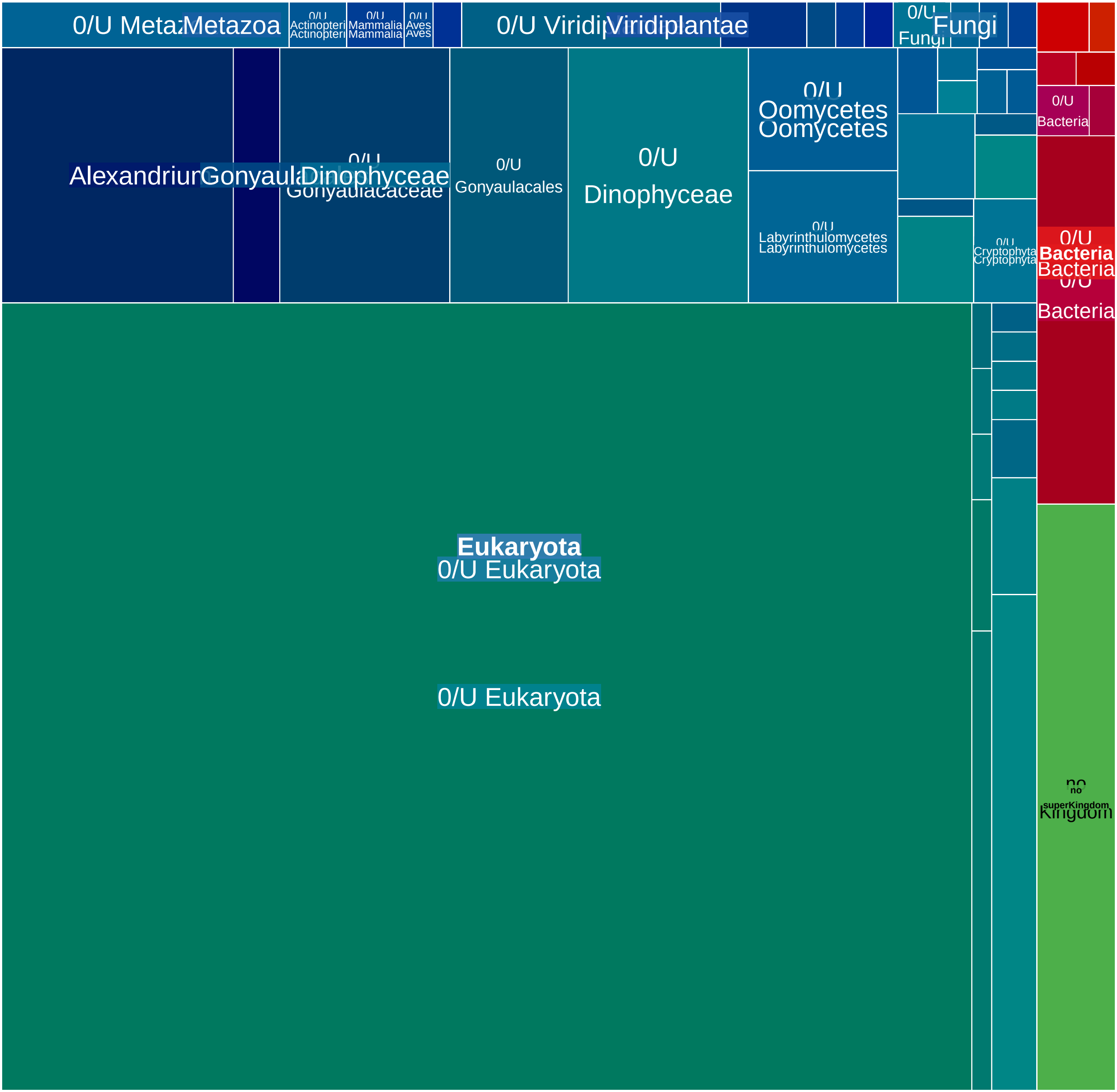

MGT−v1_573 : 1223 unigenes, 248 (20.28%) taxonomically assigned

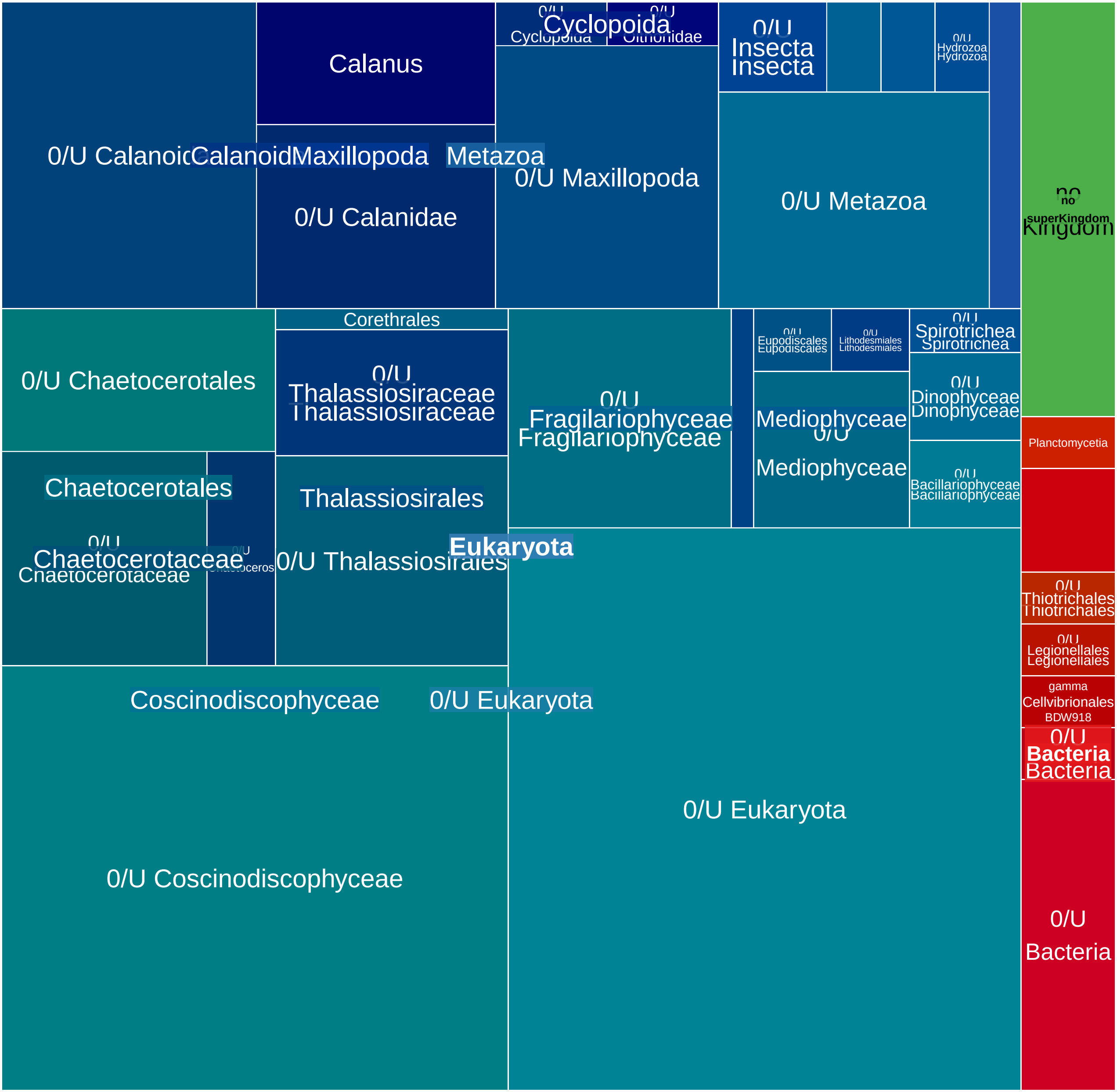

MGT−v1_574 : 1222 unigenes, 1130 (92.47%) taxonomically assigned

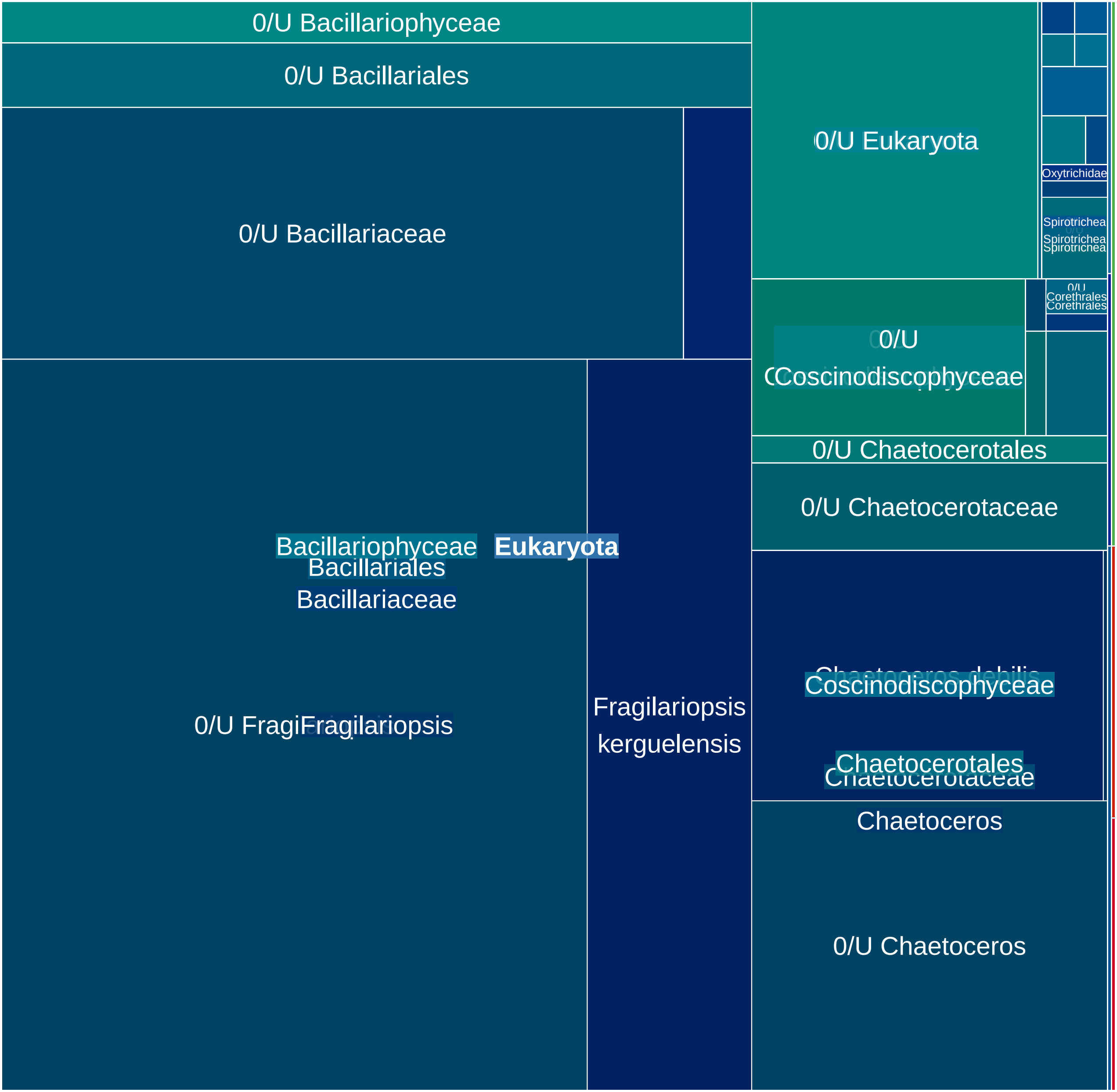

### MGT−v1_575 : 1219 unigenes, 813 (66.69%) taxonomically assigned

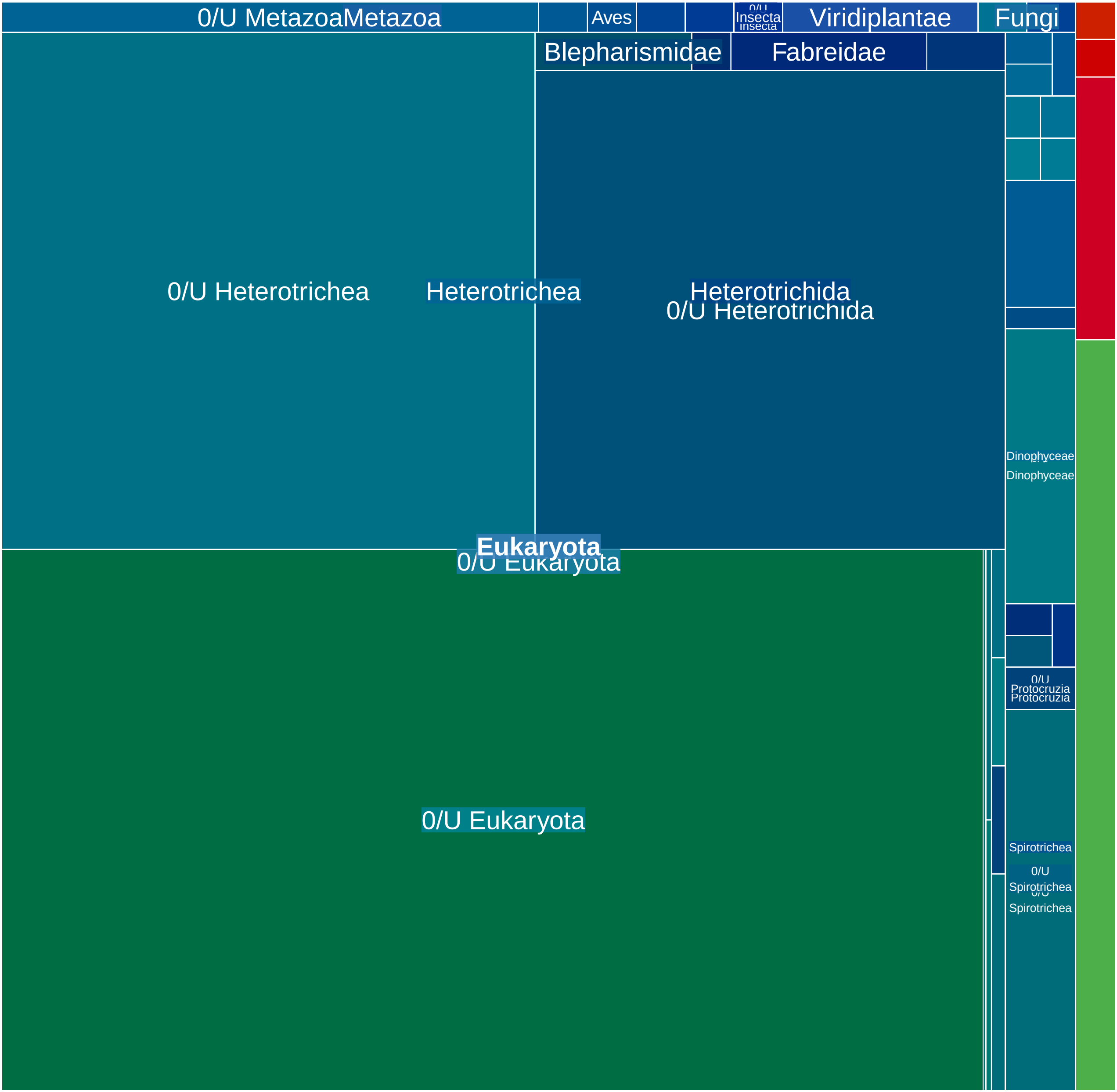

MGT−v1_576 : 1215 unigenes, 1065 (87.65%) taxonomically assigned

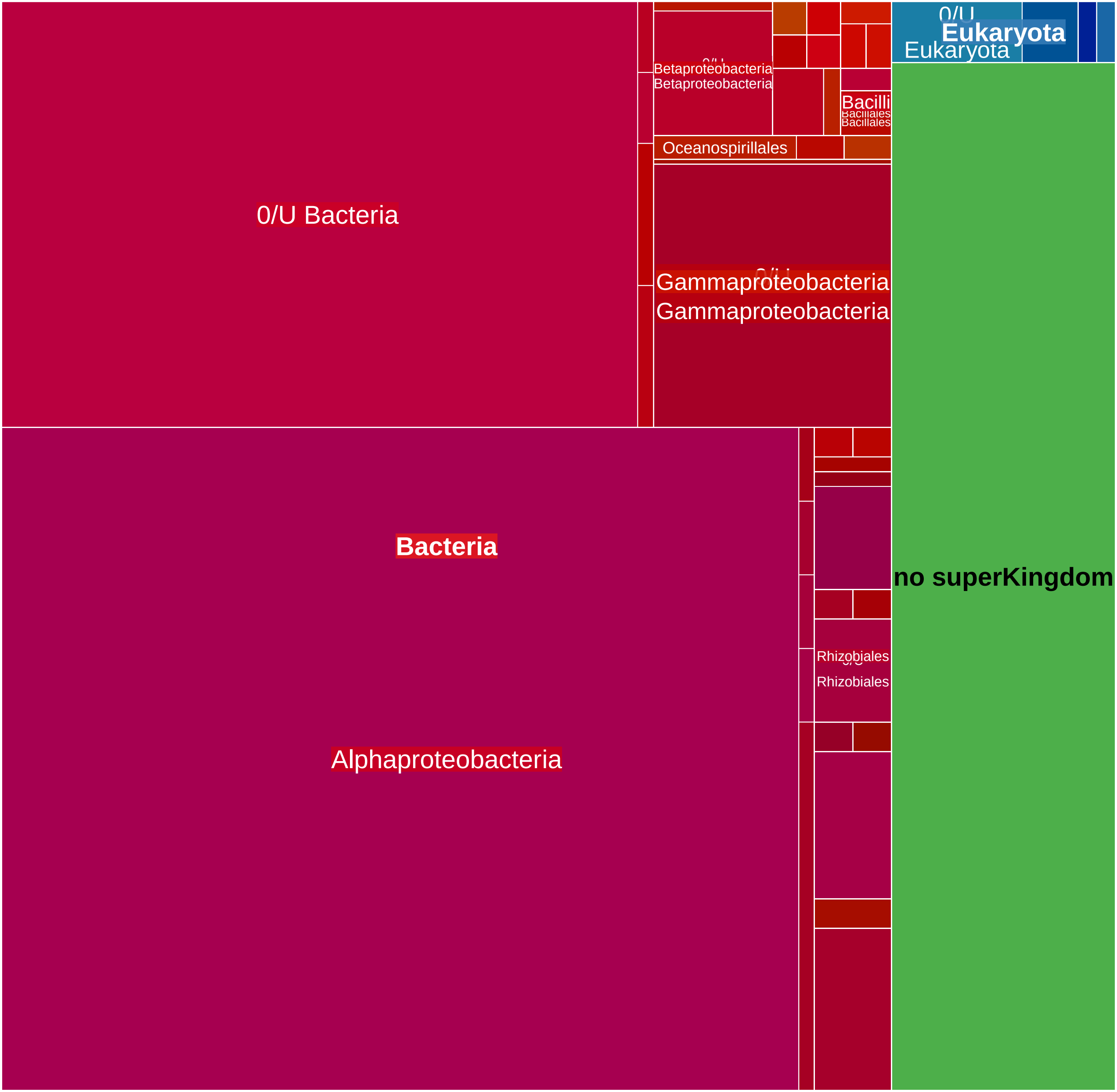

MGT−v1_577 : 1207 unigenes, 115 (9.53%) taxonomically assigned

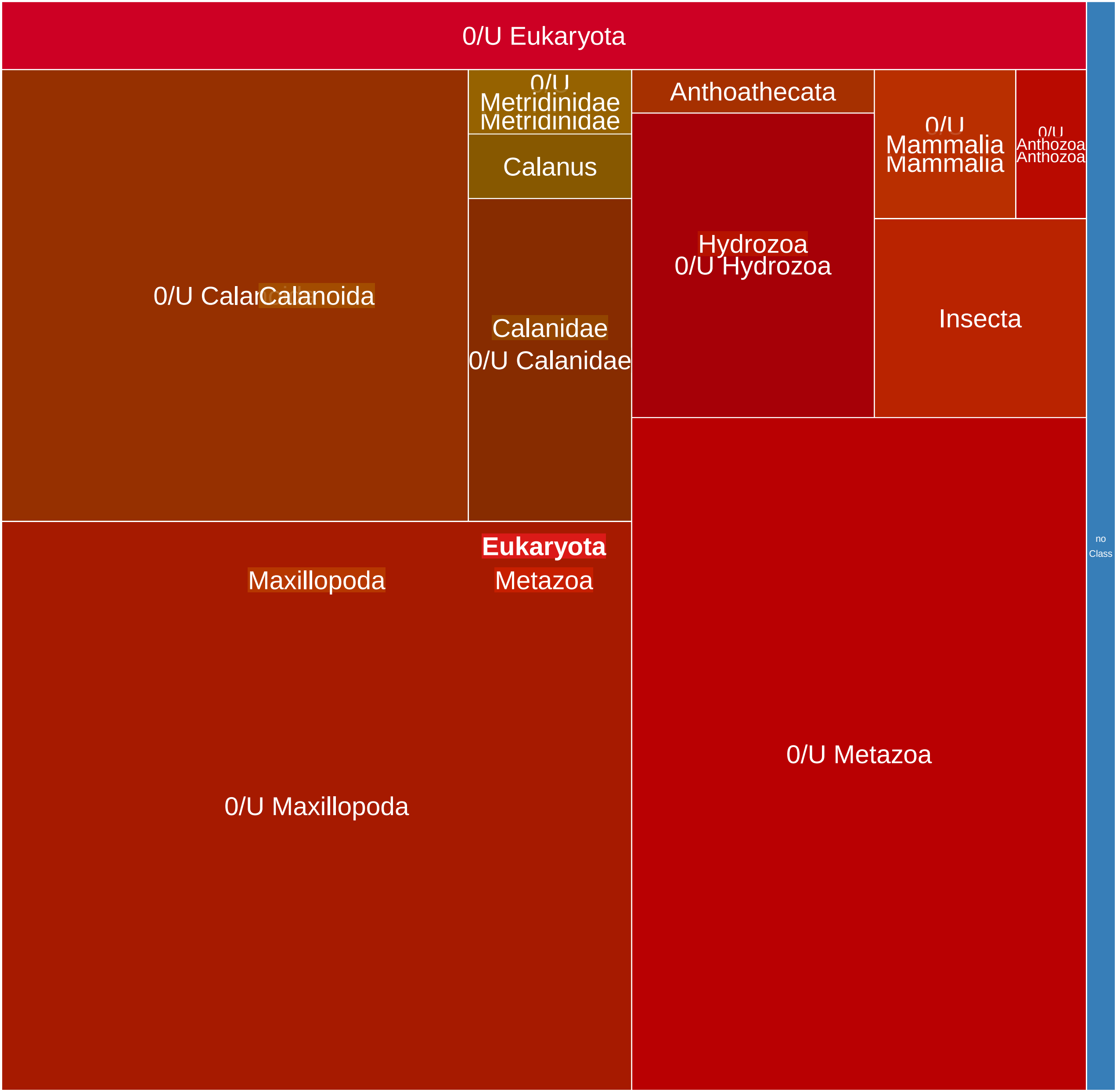

MGT−v1_578 : 1197 unigenes, 275 (22.97%) taxonomically assigned

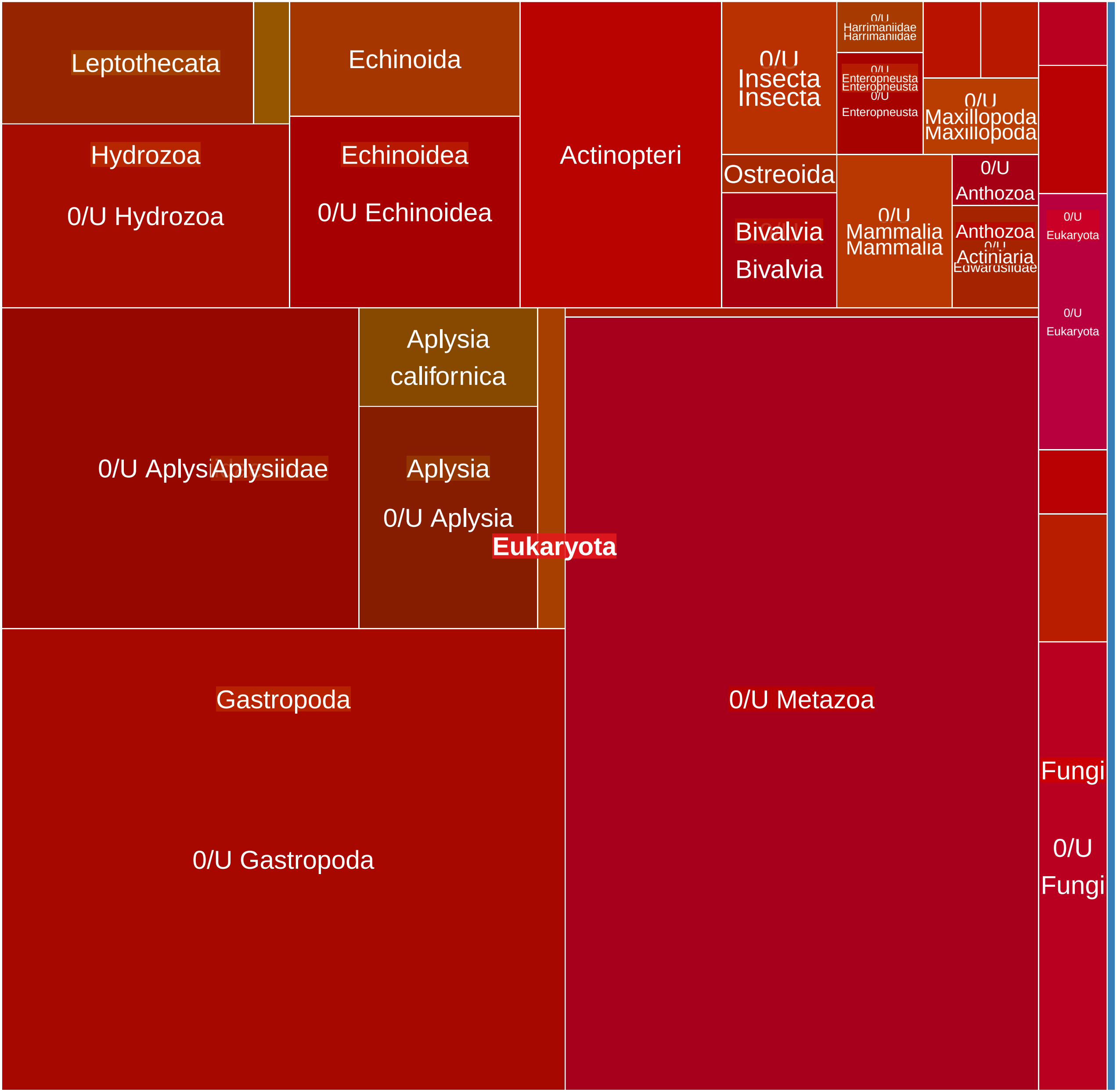

MGT−v1_579 : 1194 unigenes, 1025 (85.85%) taxonomically assigned

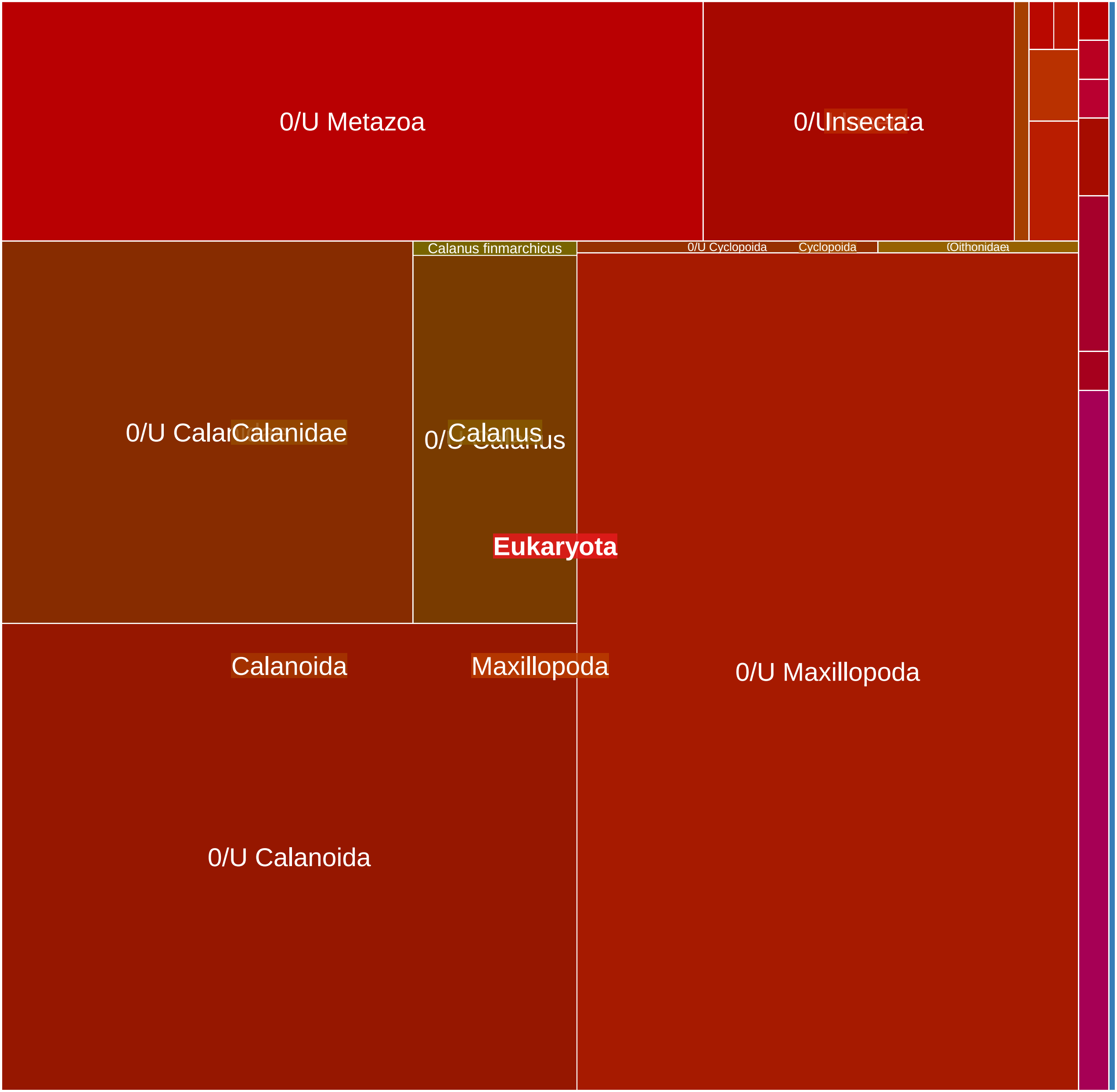

MGT−v1_580 : 1189 unigenes, 1117 (93.94%) taxonomically assigned

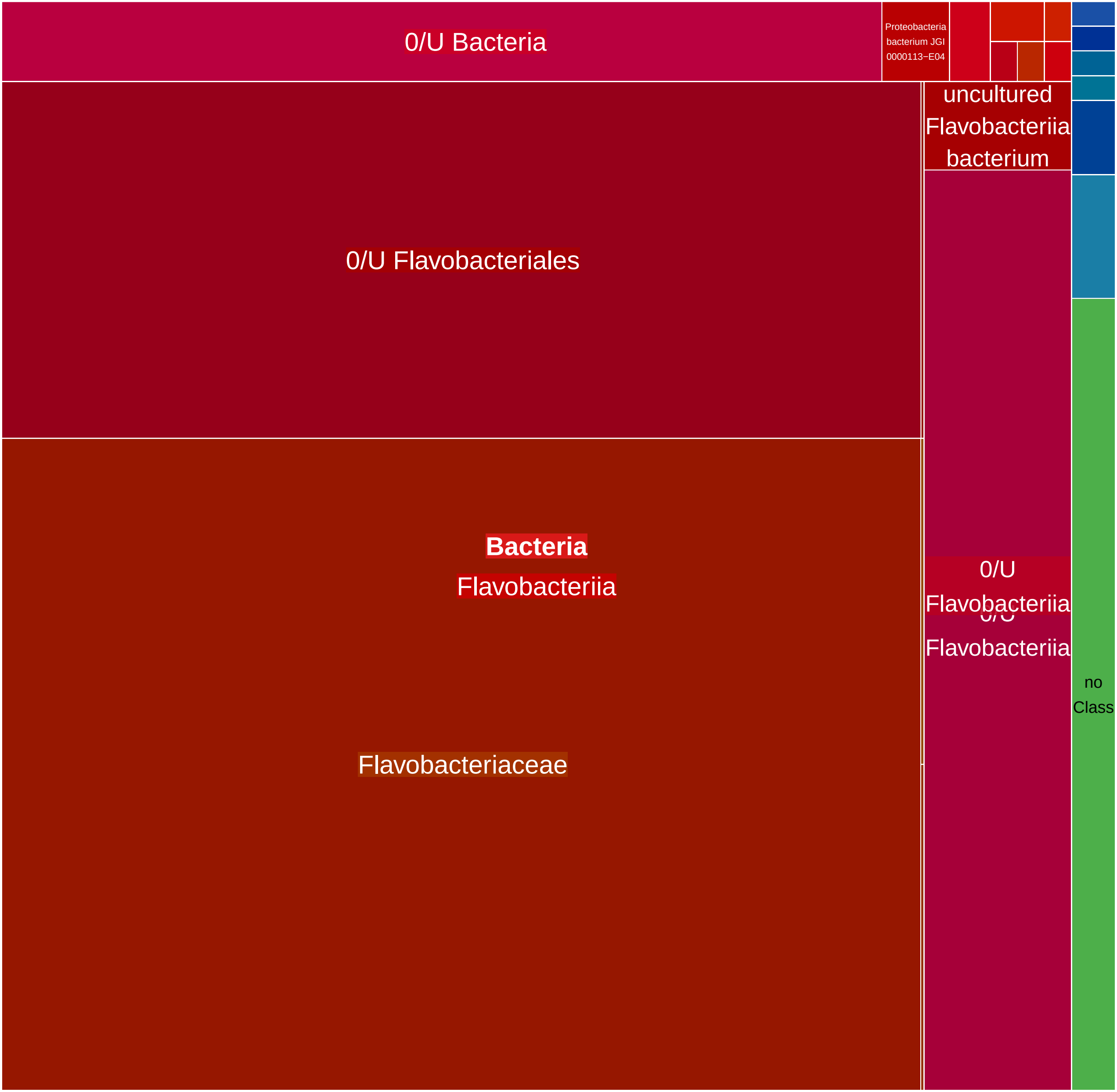

MGT−v1_581 : 1188 unigenes, 7 (0.59%) taxonomically assigned

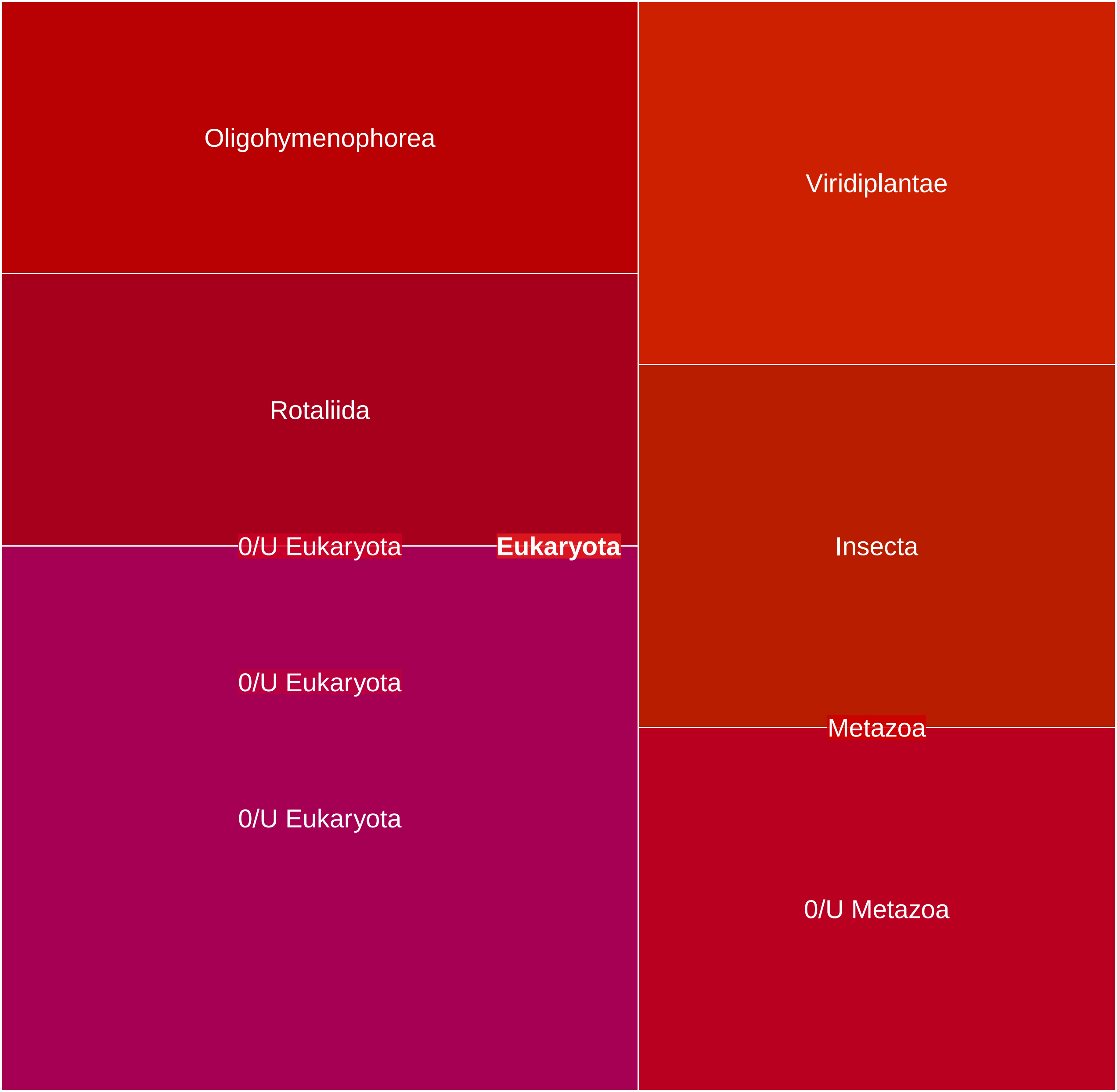

MGT−v1_582 : 1187 unigenes, 921 (77.59%) taxonomically assigned

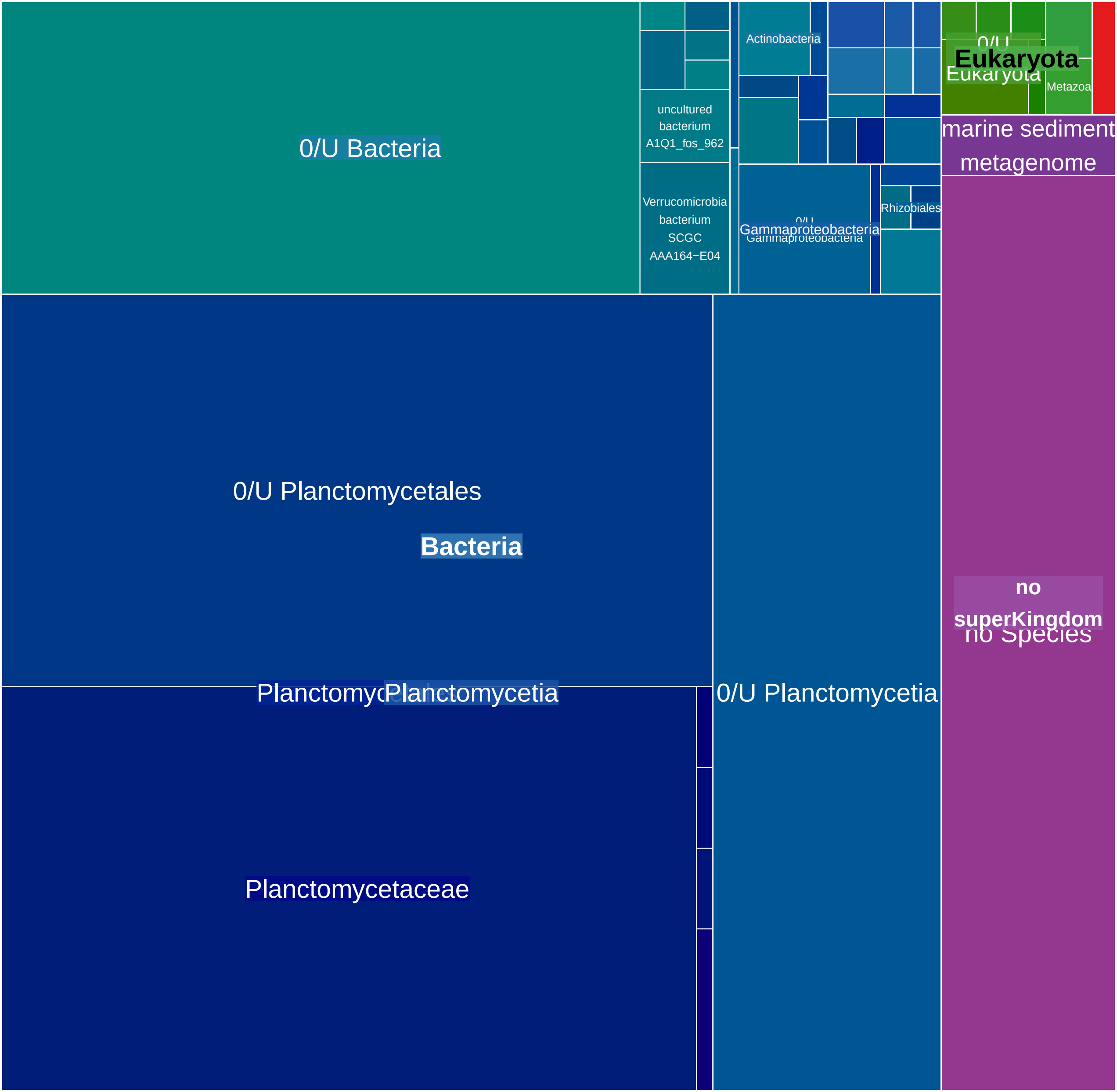

MGT−v1_583 : 1185 unigenes, 953 (80.42%) taxonomically assigned

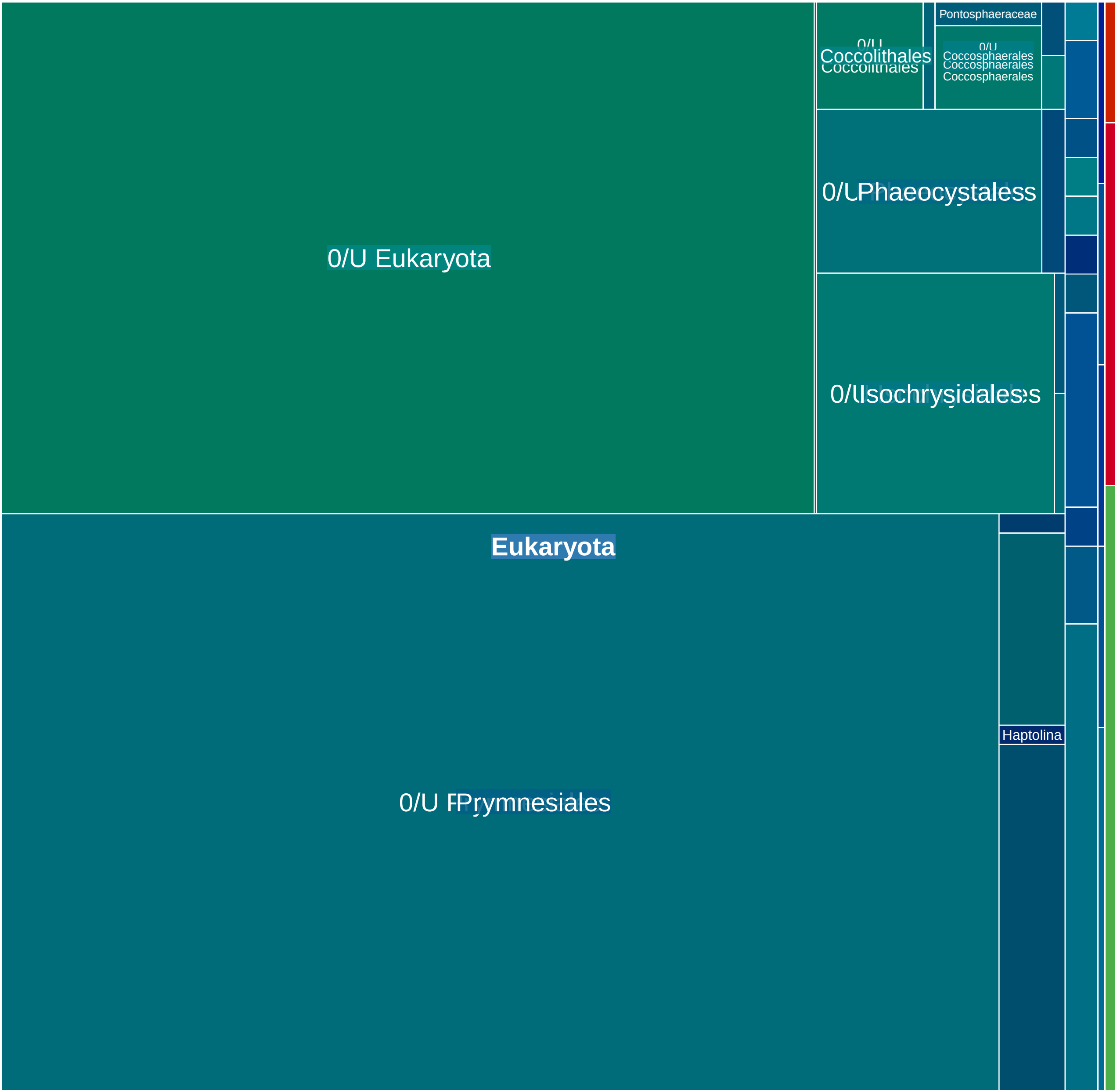

MGT−v1_584 : 1175 unigenes, 150 (12.77%) taxonomically assigned

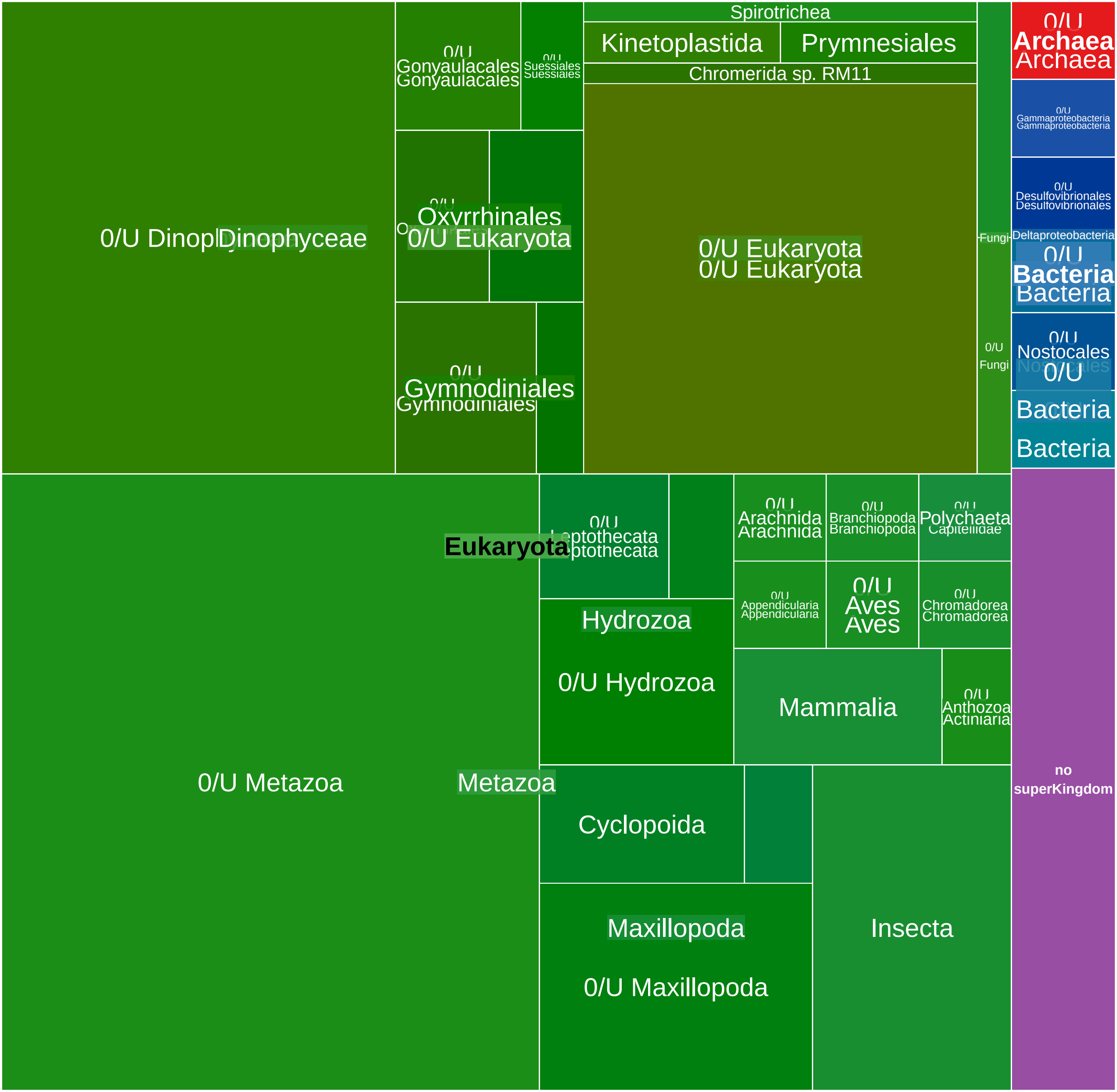

MGT−v1_585 : 1175 unigenes, 157 (13.36%) taxonomically assigned

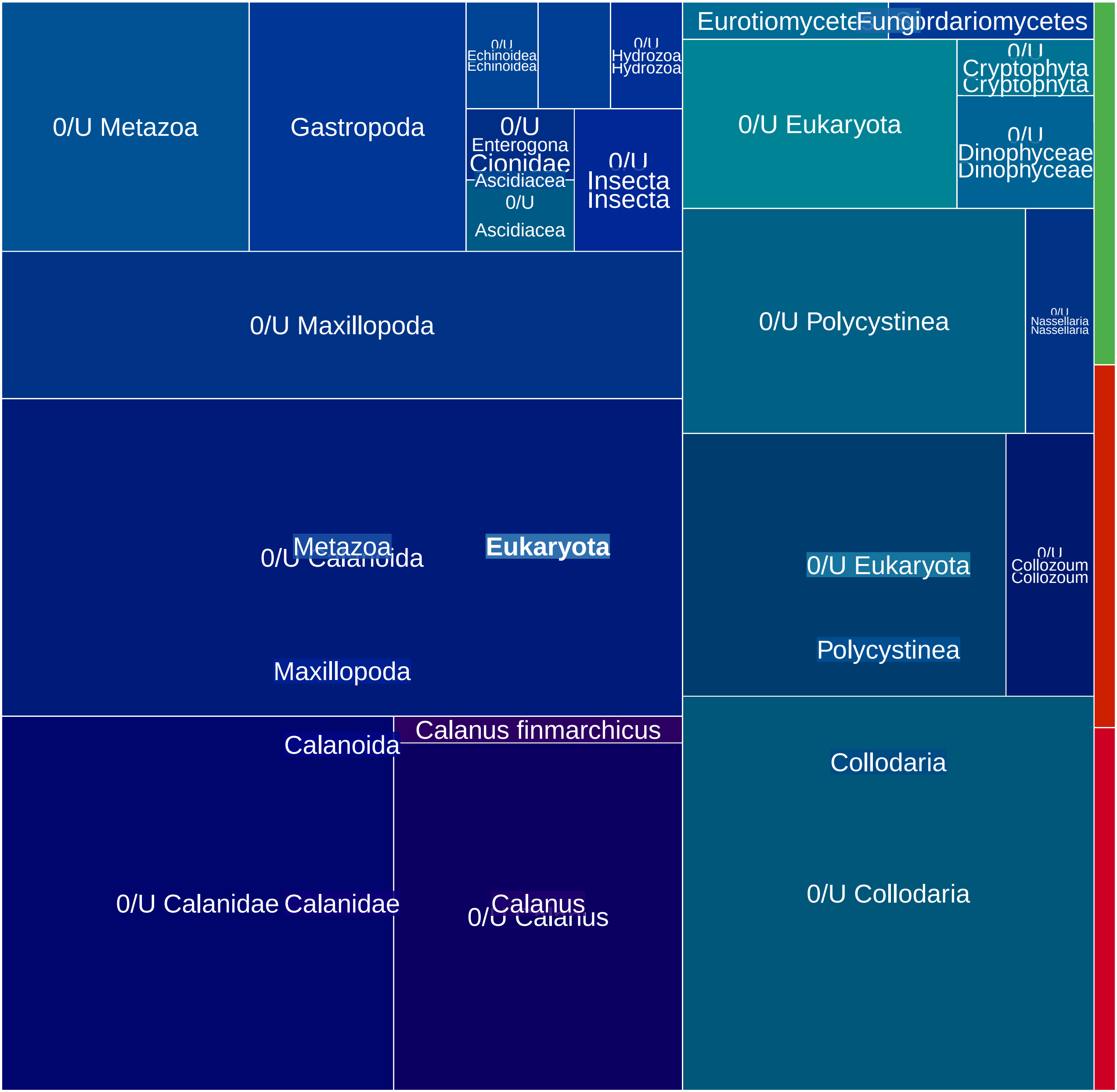

MGT−v1_586 : 1169 unigenes, 325 (27.80%) taxonomically assigned

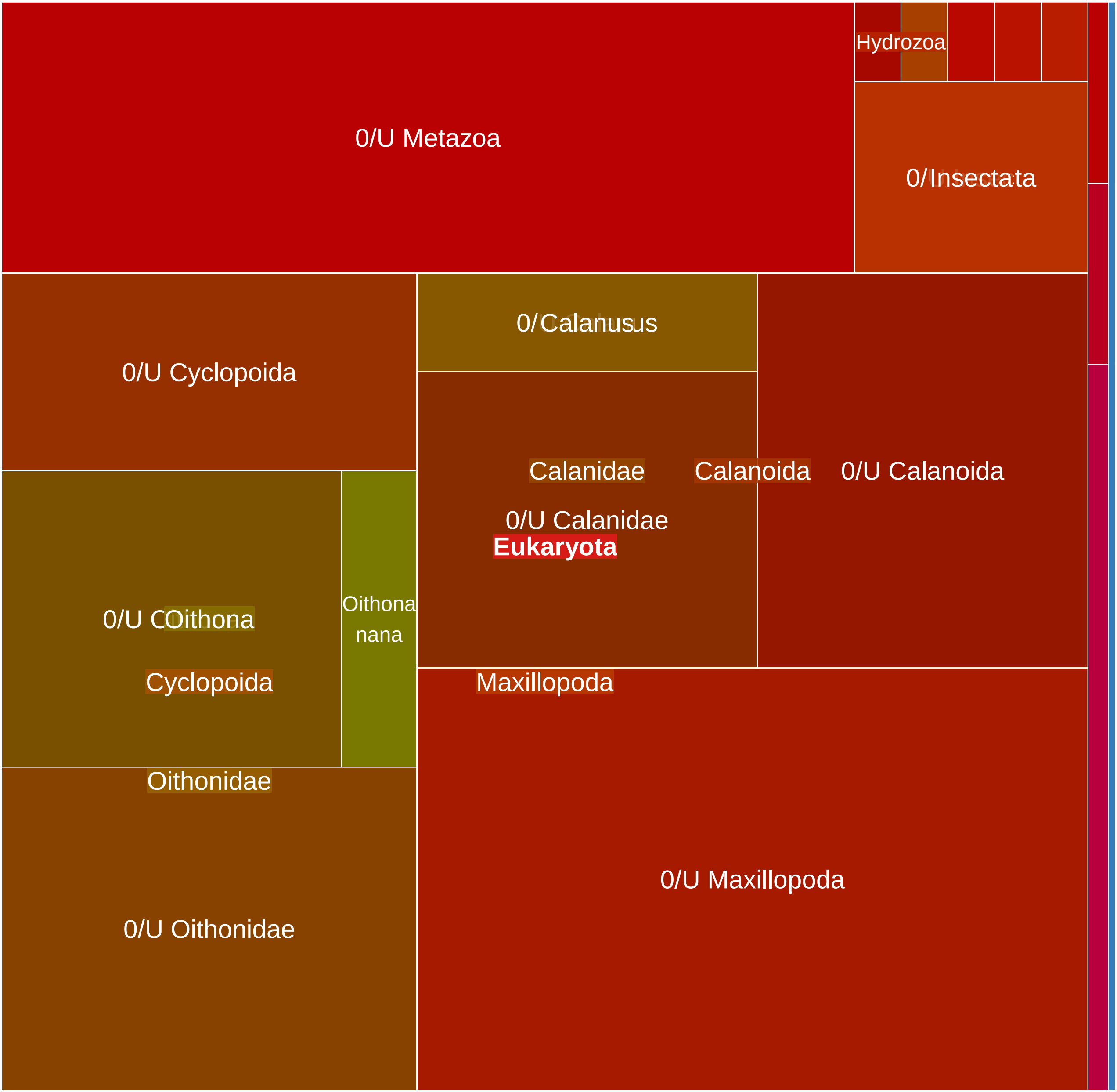

MGT−v1_587 : 1168 unigenes, 1120 (95.89%) taxonomically assigned

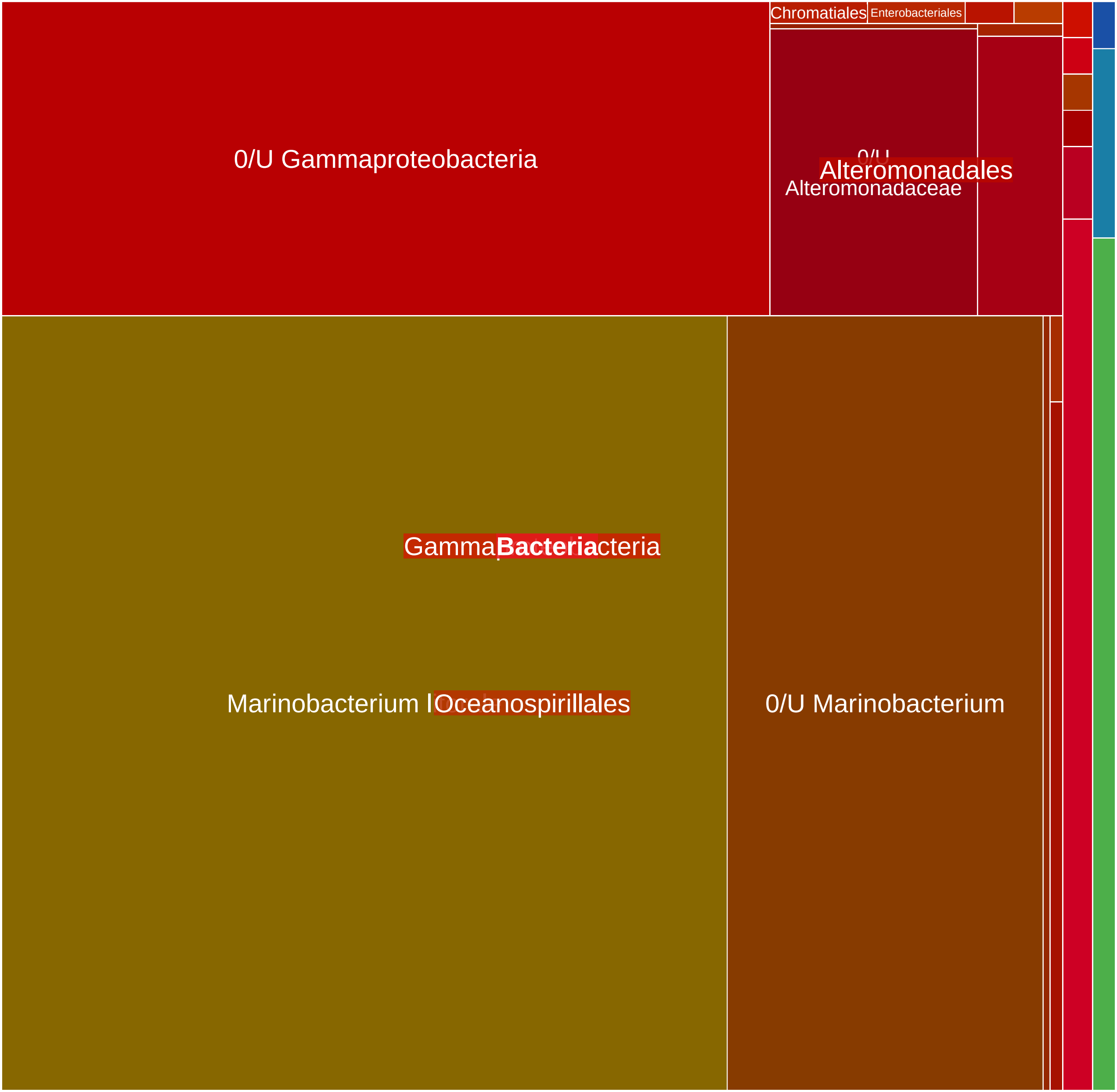

MGT−v1_588 : 1161 unigenes, 69 (5.94%) taxonomically assigned

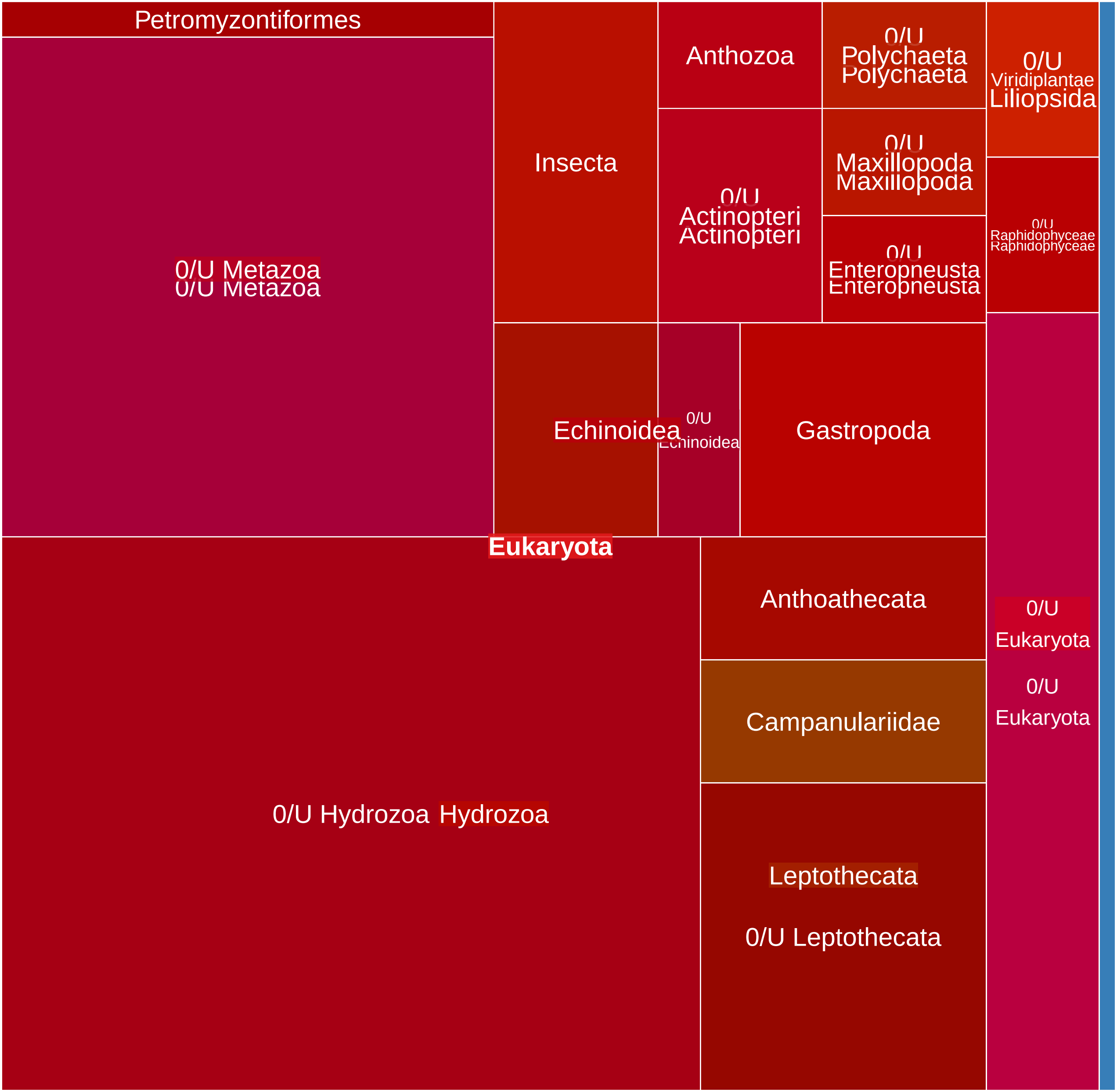

MGT−v1_589 : 1158 unigenes, 133 (11.49%) taxonomically assigned

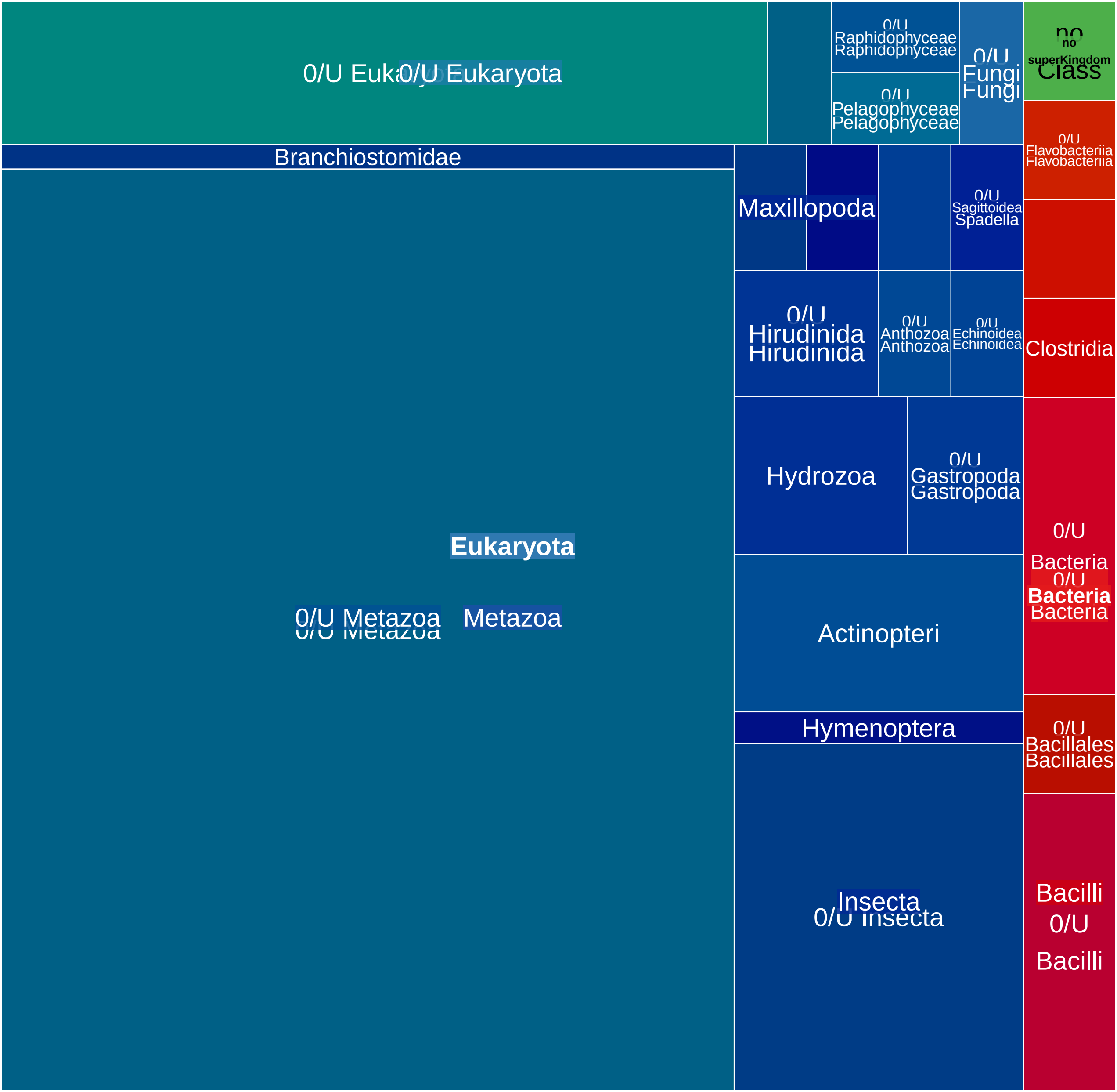

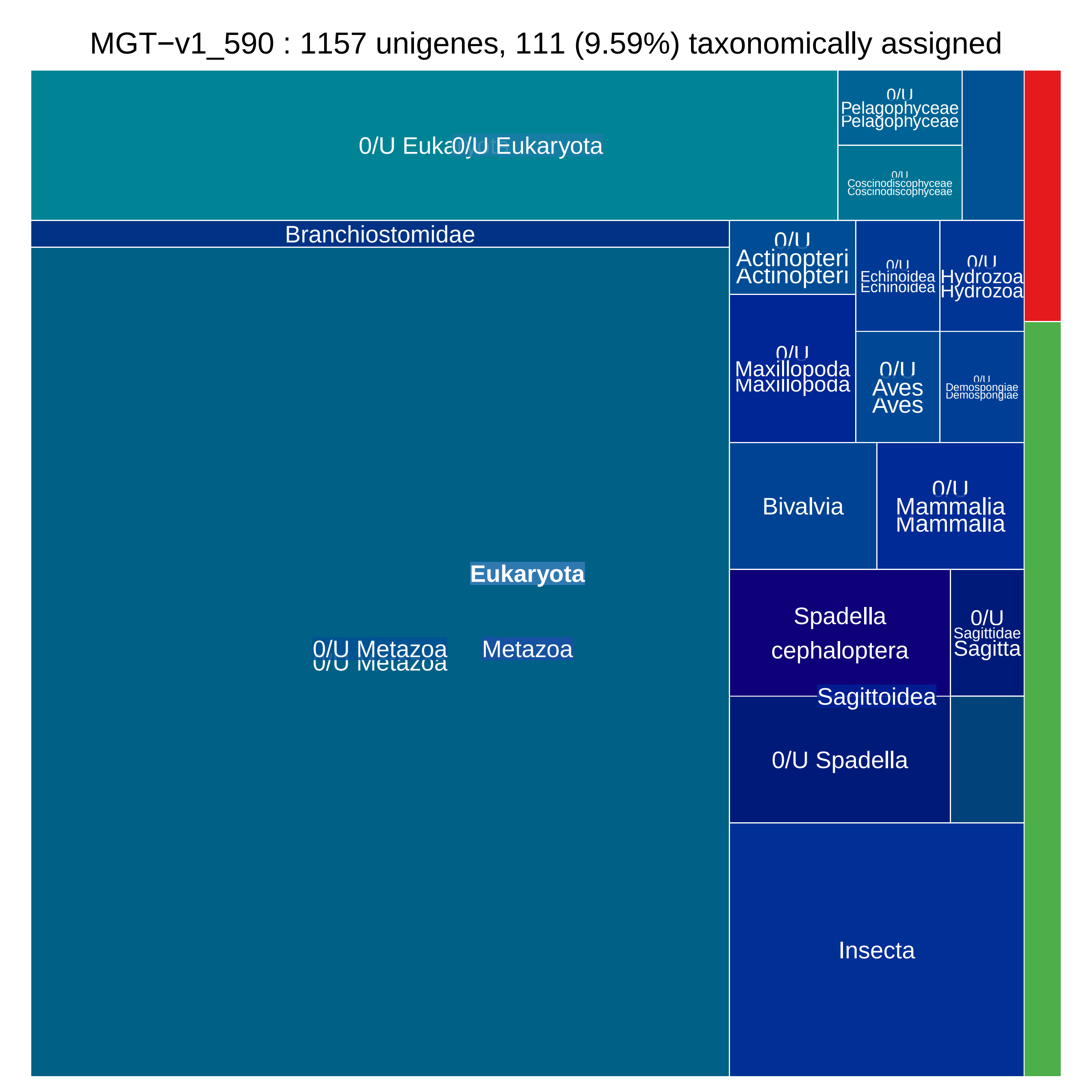

MGT−v1_590 : 1157 unigenes, 111 (9.59%) taxonomically assigned
0/U Eukaryota
0/U Eukaryota
0/U Pelagophyceae Pelagophyceae
0/U Coscinodiscophyceae Coscinodiscophyceae
Branchiostomidae
0/U Actinopteri Actinopteri
0/U Echinoidea Echinoidea
0/U Hydrozoa Hydrozoa
0/U Maxillopoda Maxillopoda
0/U Aves Aves
0/U Demospongiae Demospongiae
Bivalvia
0/U Mammalia Mammalia
Eukaryota
Spadella cephaloptera
0/U Sagittidae Sagitta
0/U Metazoa
0/U Metazoa
Metazoa
Sagittoidea
0/U Spadella
Insecta

MGT−v1_591 : 1154 unigenes, 76 (6.59%) taxonomically assigned

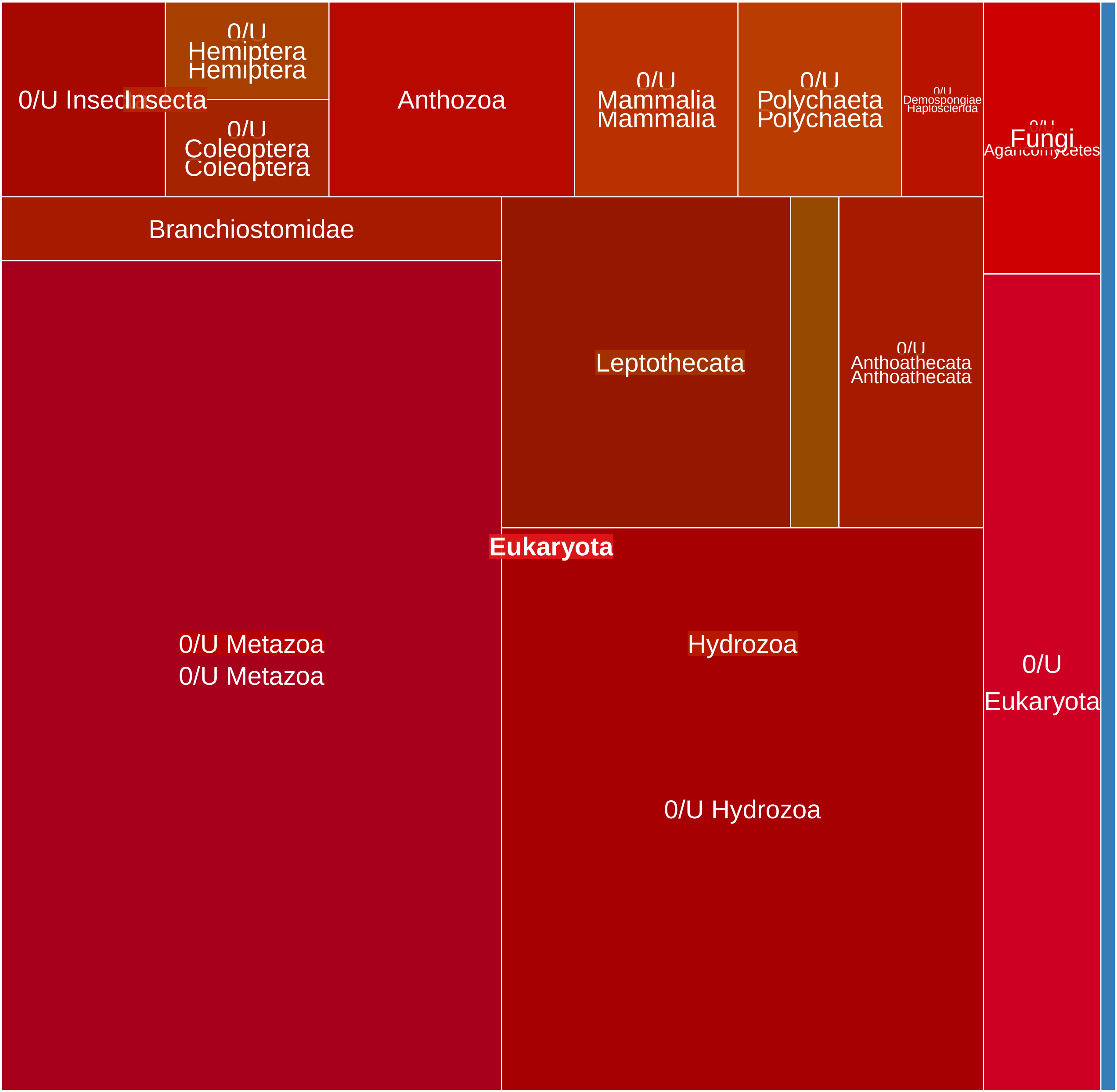

MGT−v1_592 : 1153 unigenes, 132 (11.45%) taxonomically assigned

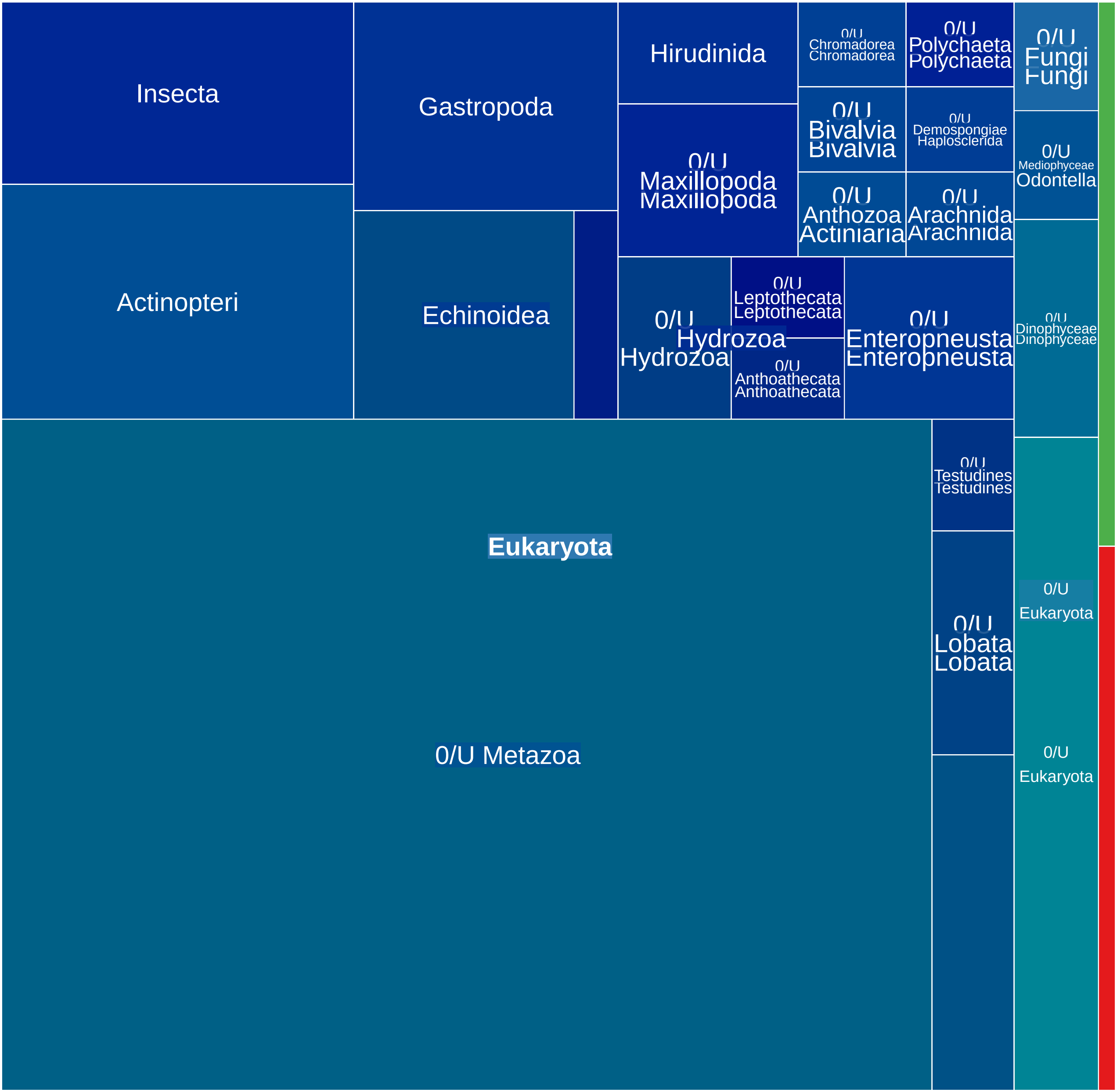

MGT−v1_593 : 1147 unigenes, 1040 (90.67%) taxonomically assigned

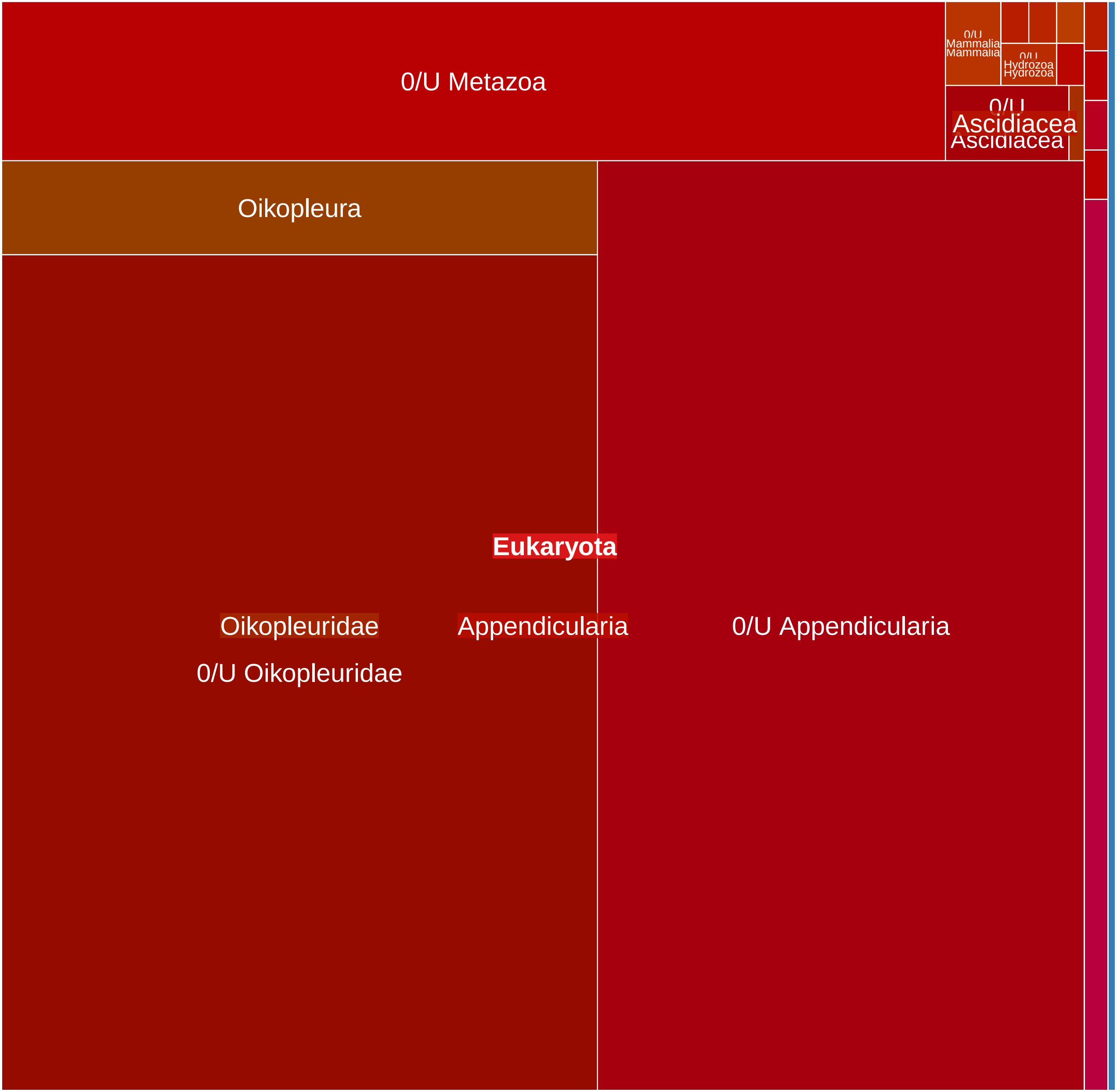

MGT−v1_594 : 1146 unigenes, 423 (36.91%) taxonomically assigned

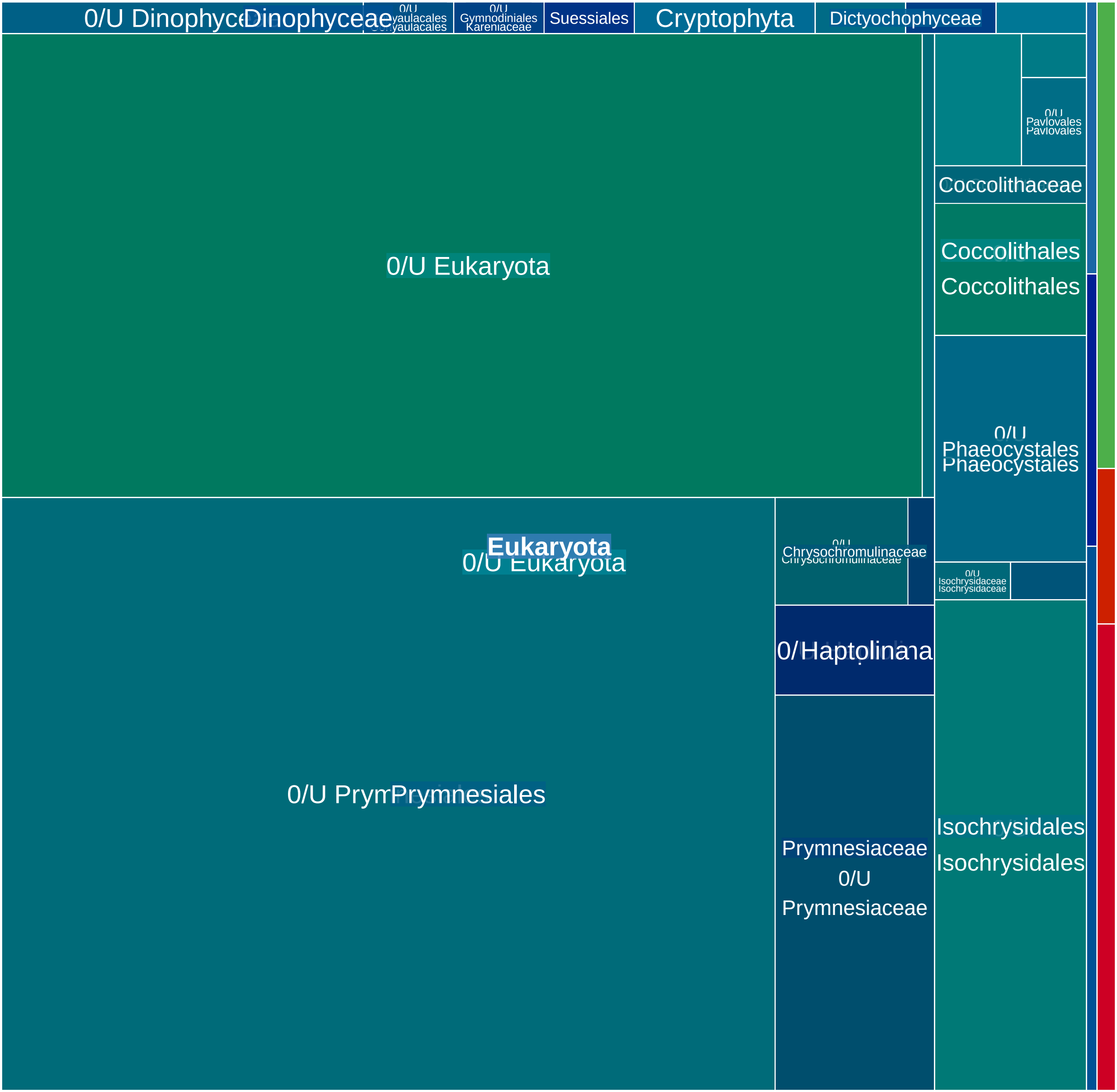

MGT−v1_595 : 1141 unigenes, 88 (7.71%) taxonomically assigned

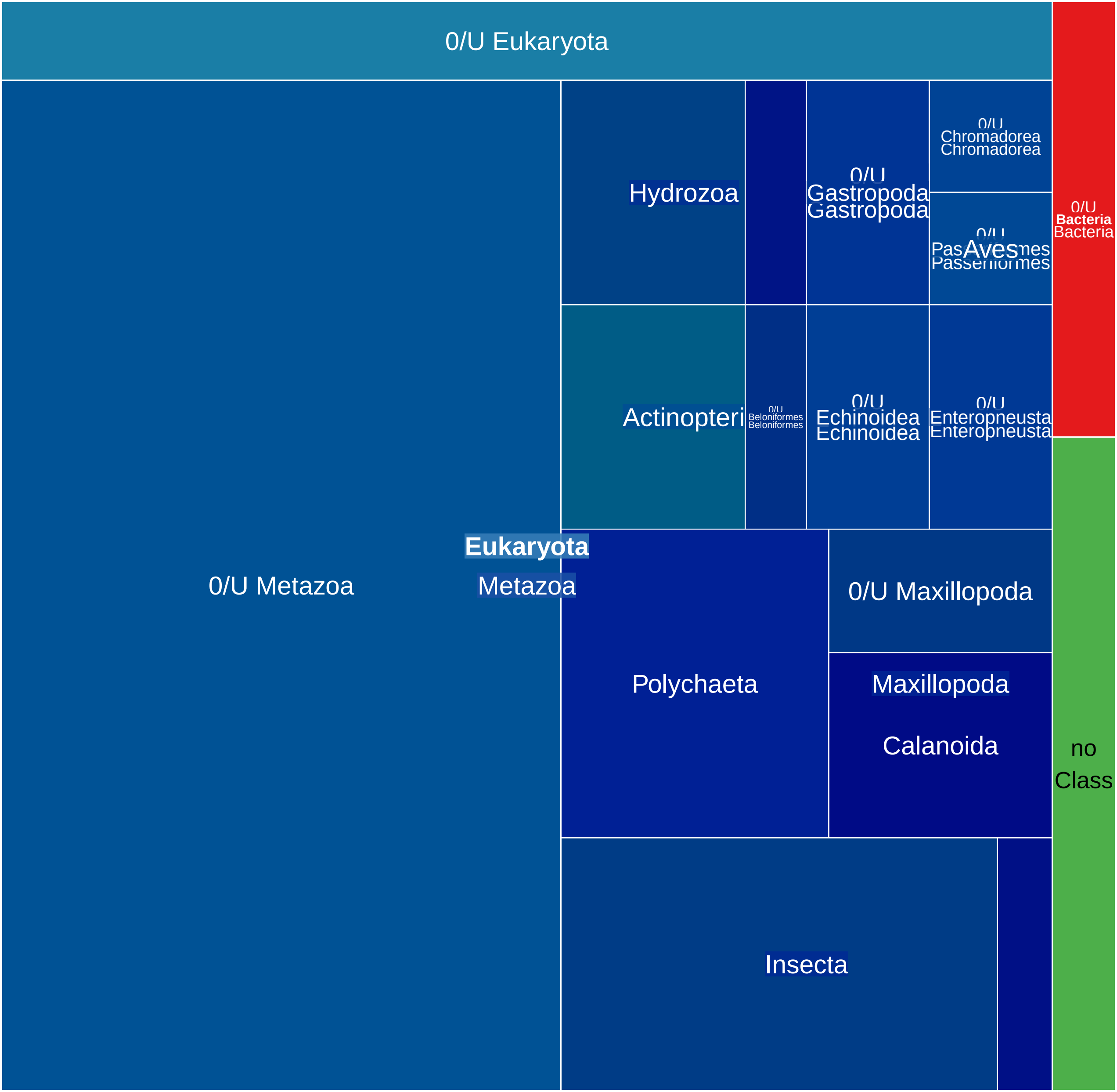

MGT−v1_596 : 1137 unigenes, 533 (46.88%) taxonomically assigned

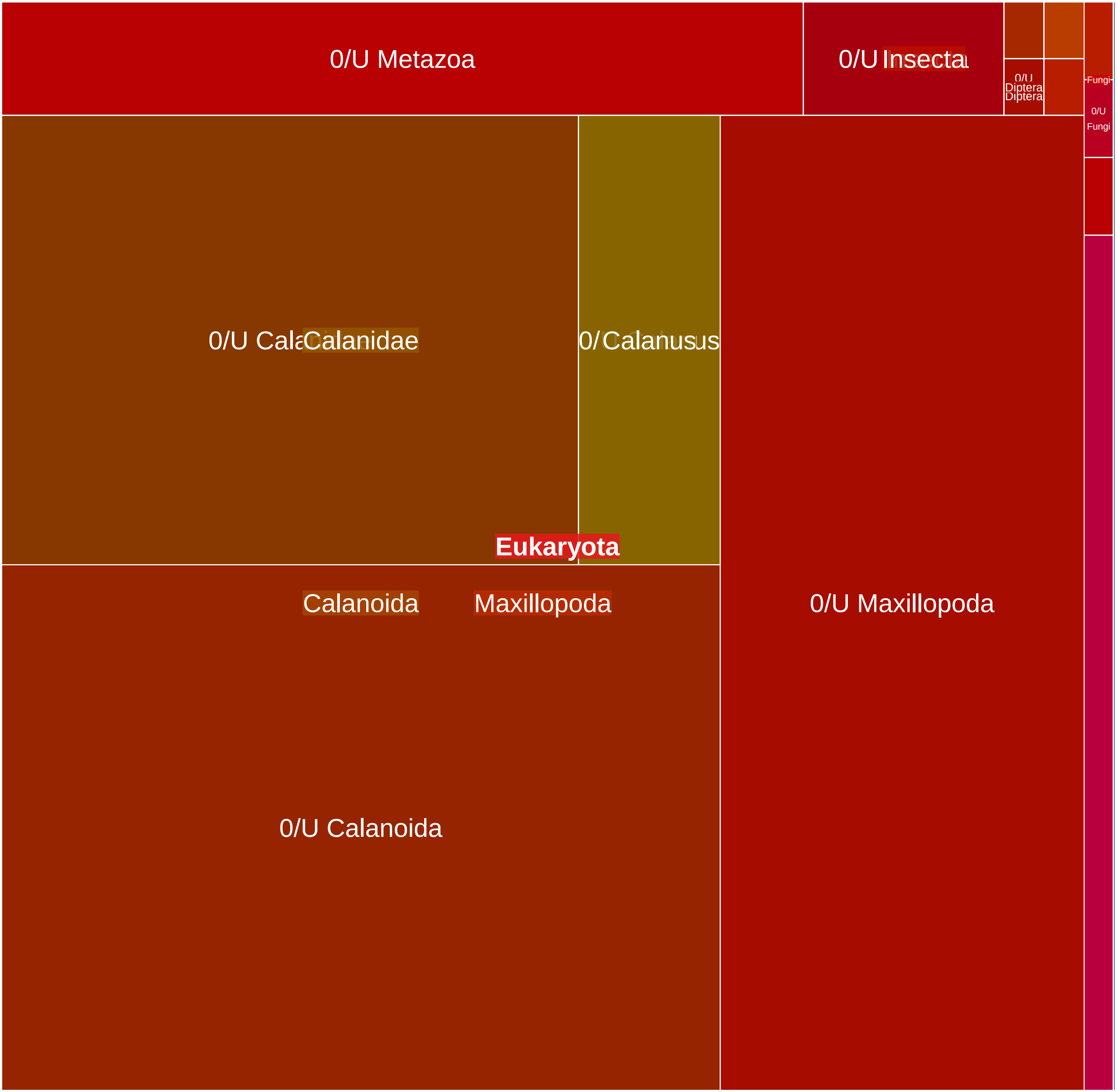

MGT−v1_597 : 1133 unigenes, 162 (14.30%) taxonomically assigned

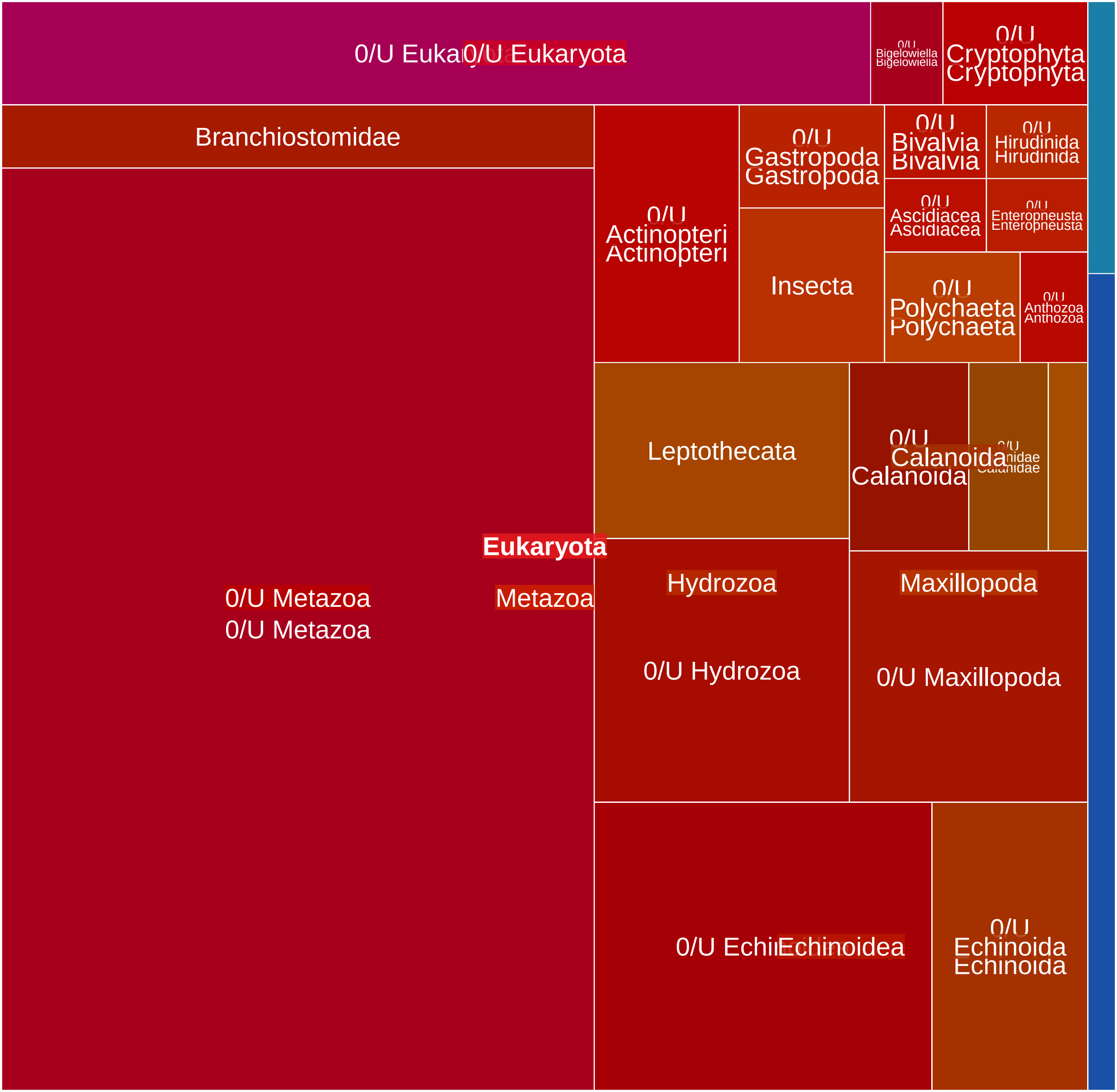

MGT−v1_598 : 1132 unigenes, 324 (28.62%) taxonomically assigned

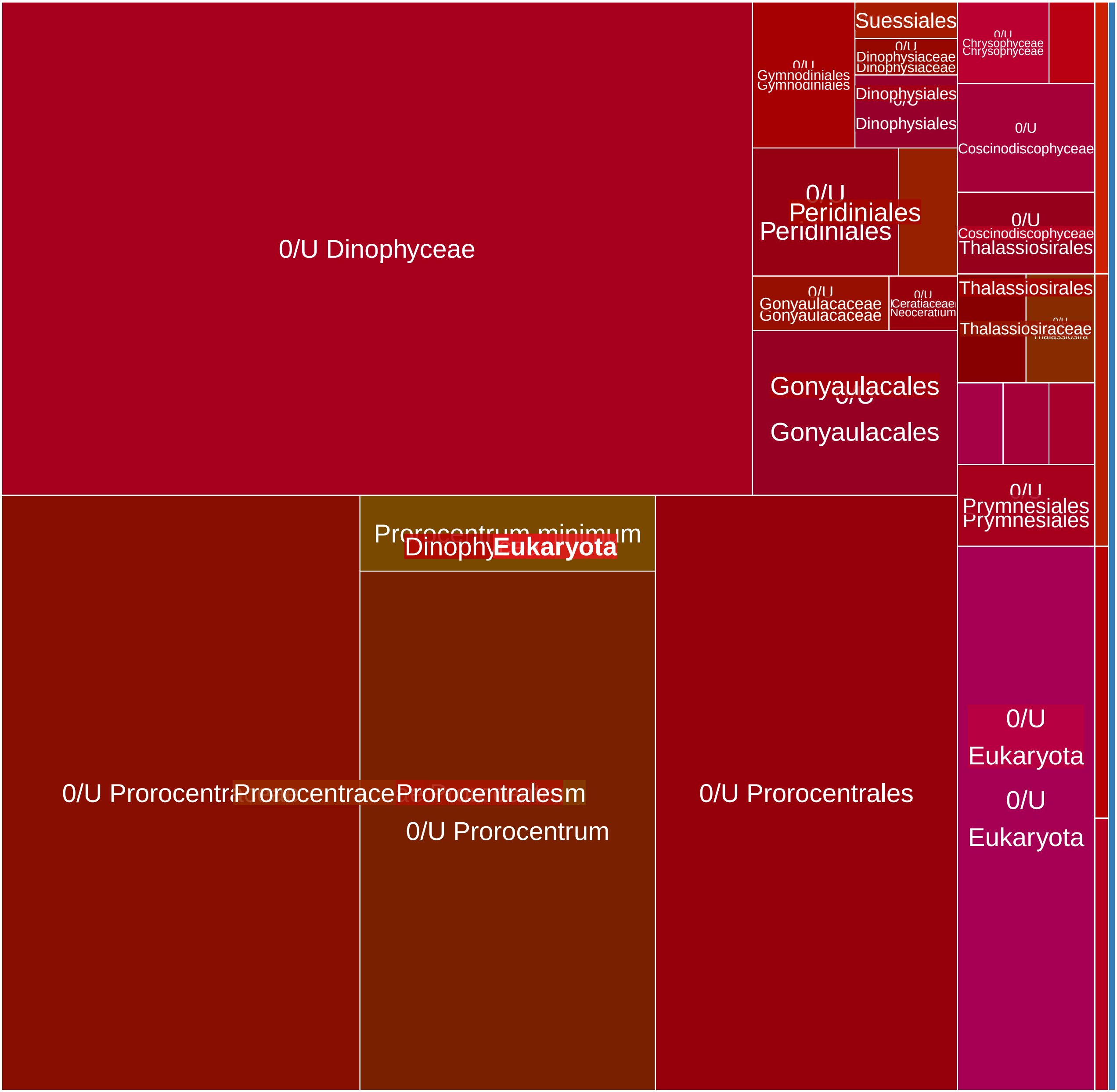

MGT−v1_599 : 1129 unigenes, 648 (57.40%) taxonomically assigned

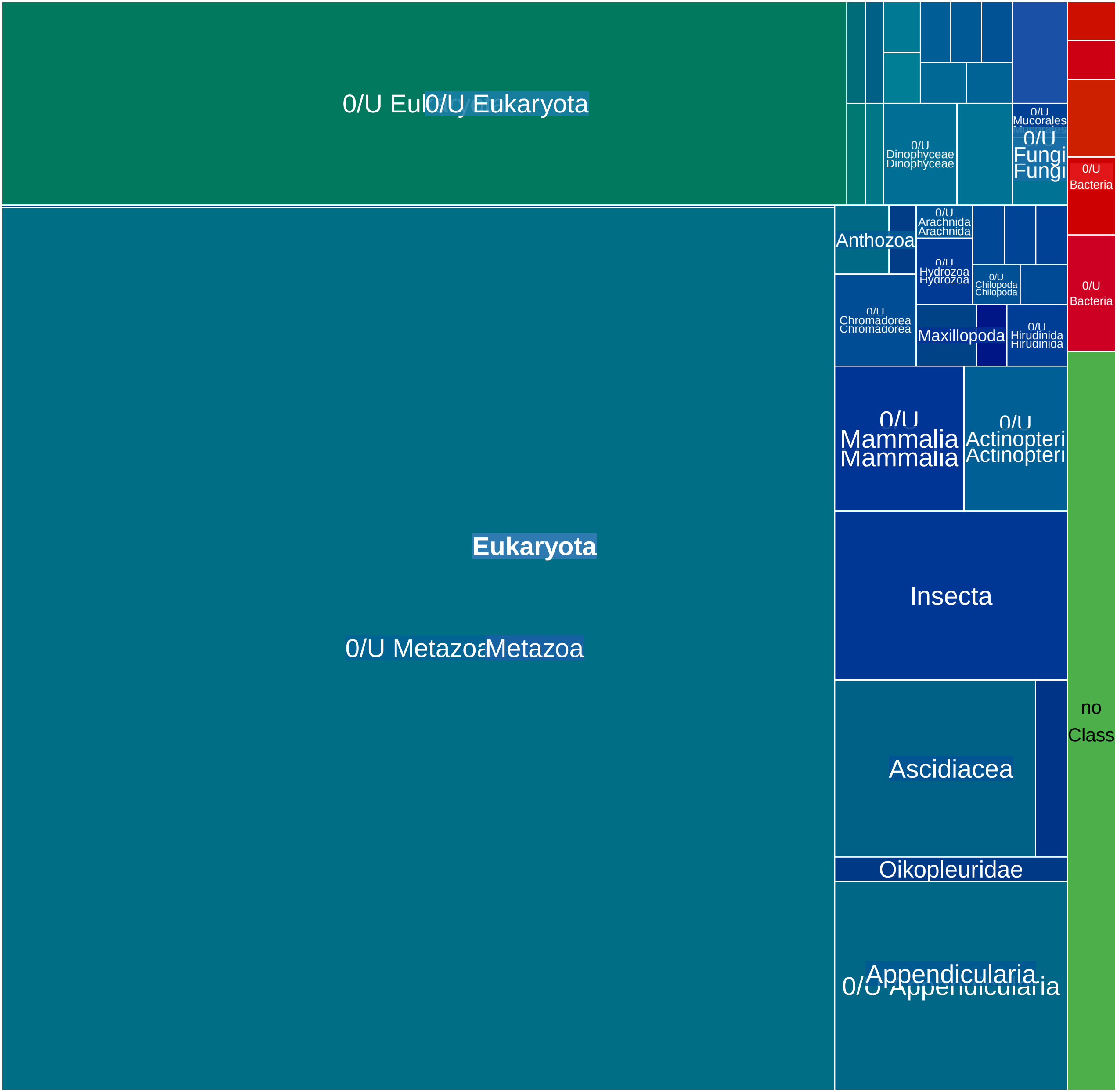

### MGT−v1_600 : 1129 unigenes, 909 (80.51%) taxonomically assigned

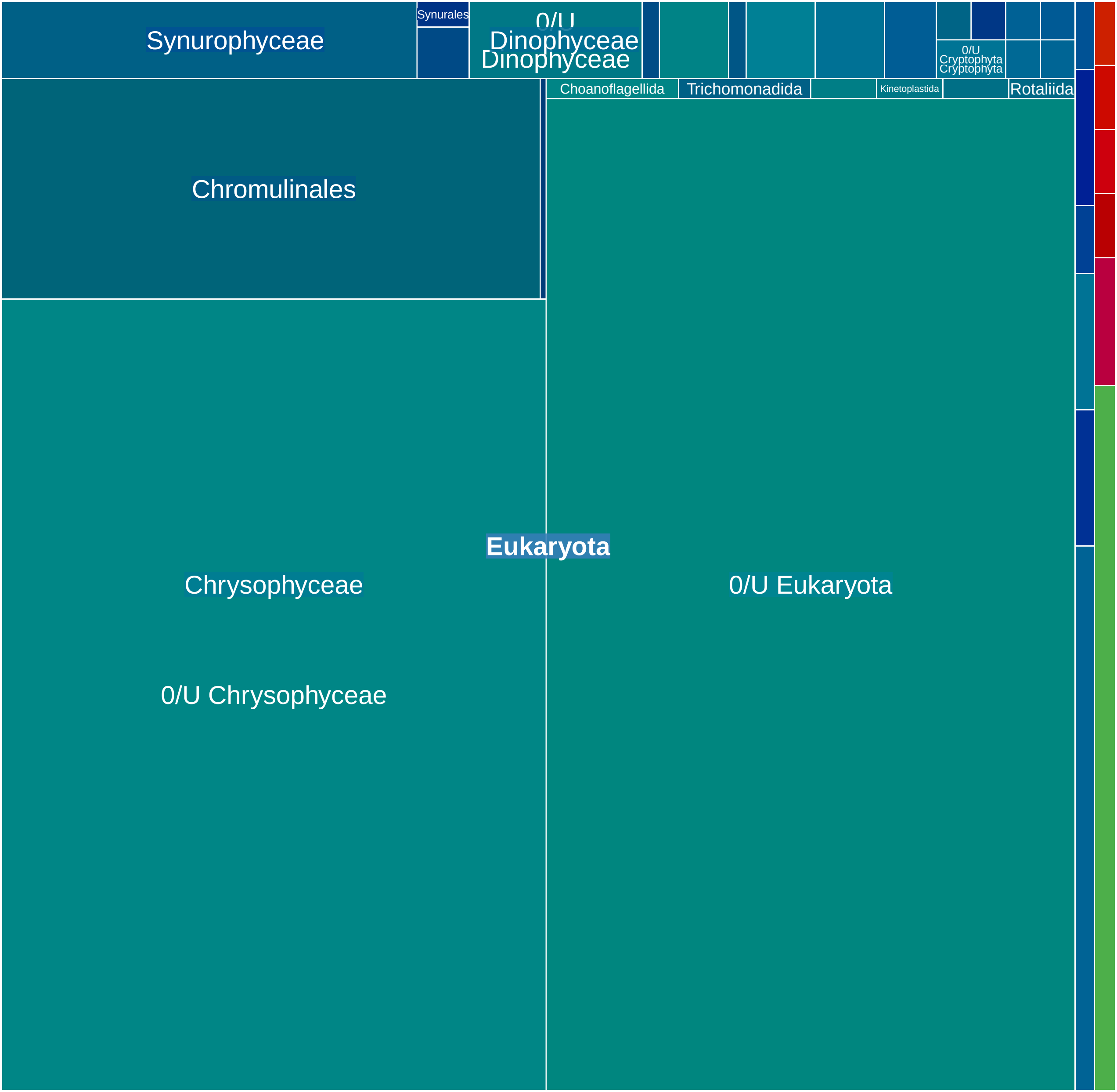

#### MGT−v1_601 : 1128 unigenes, 93 (8.24%) taxonomically assigned

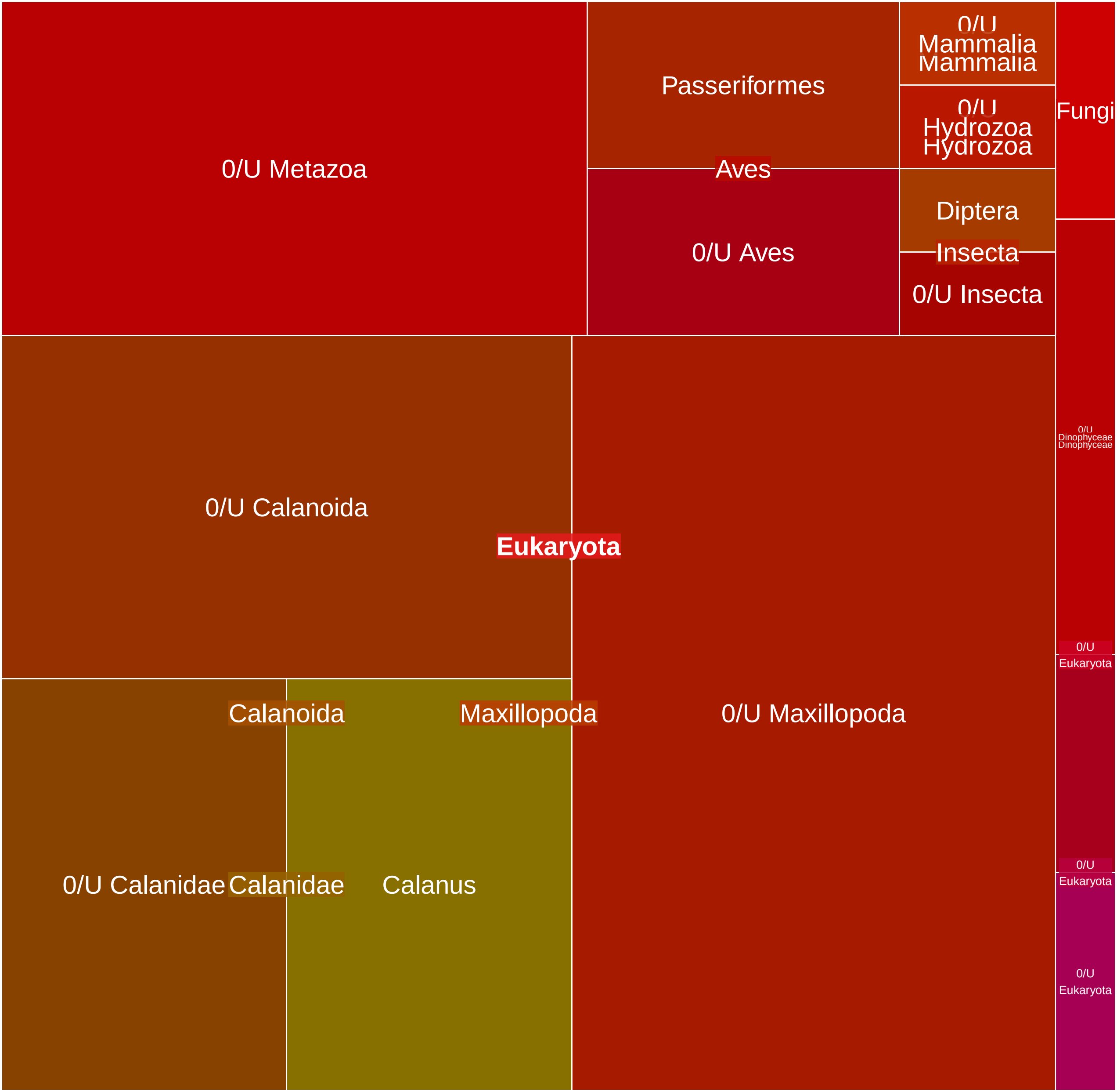

MGT−v1_602 : 1125 unigenes, 98 (8.71%) taxonomically assigned

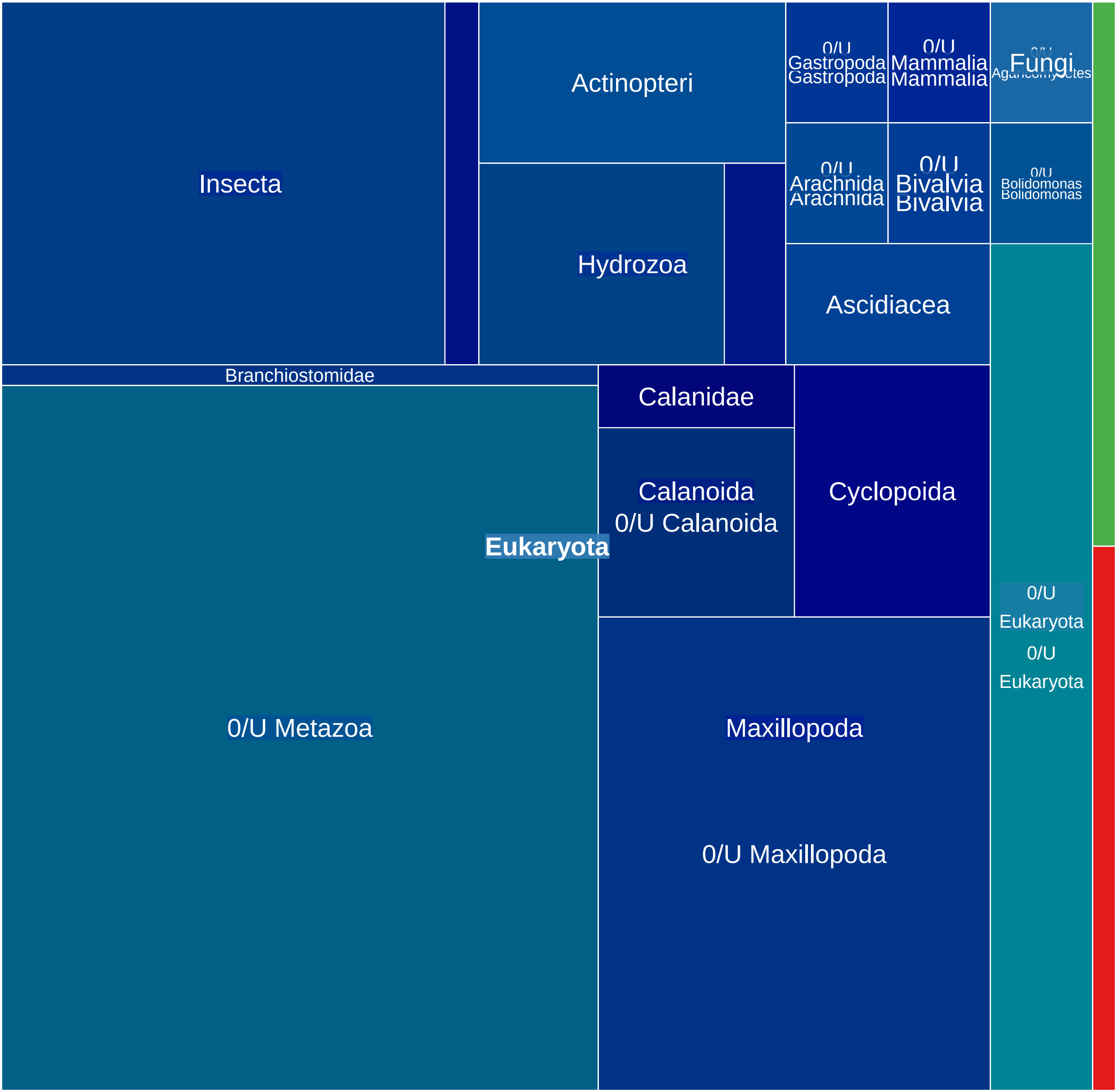

MGT−v1_603 : 1123 unigenes, 69 (6.14%) taxonomically assigned

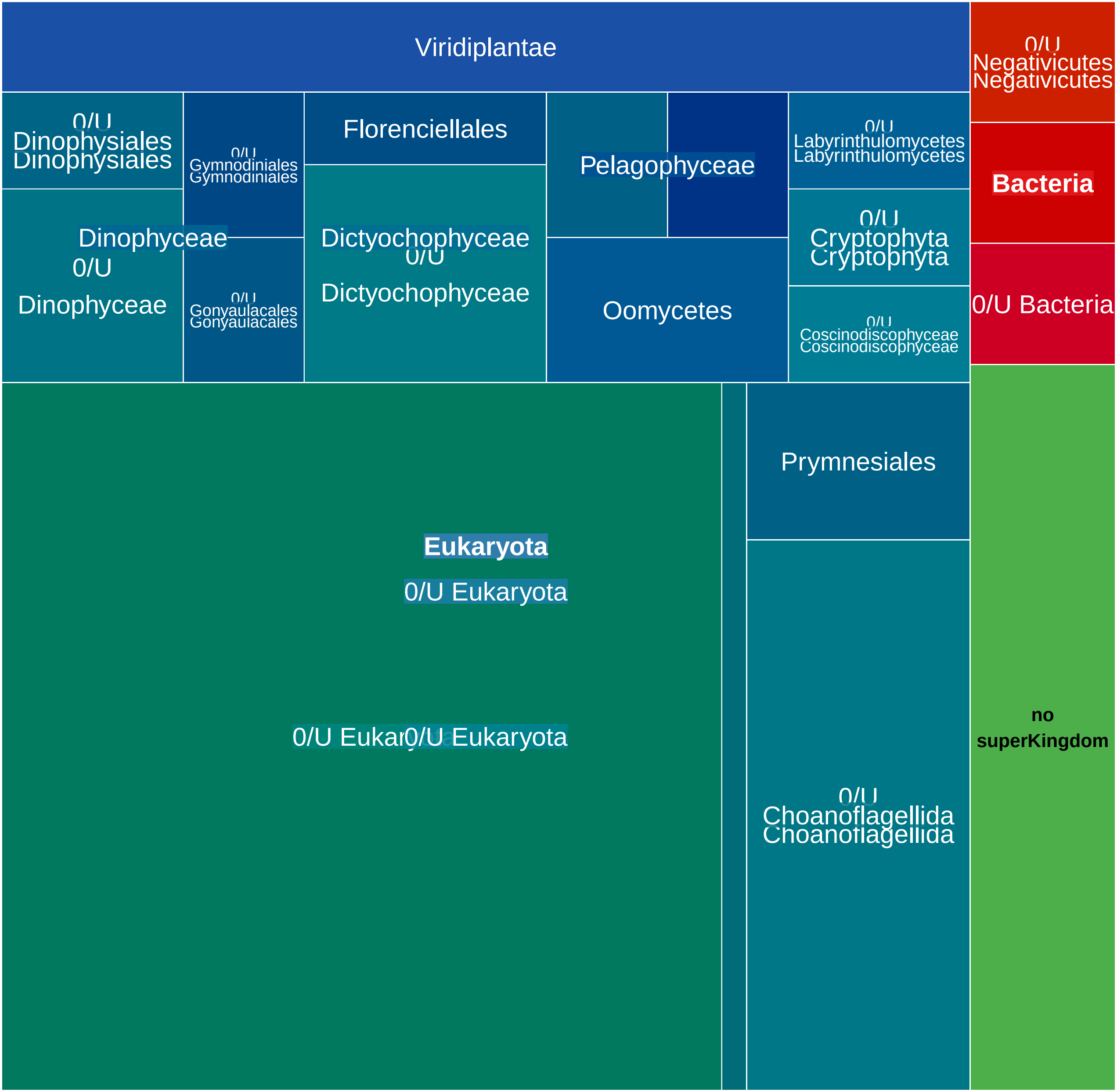

MGT−v1_604 : 1121 unigenes, 1068 (95.27%) taxonomically assigned

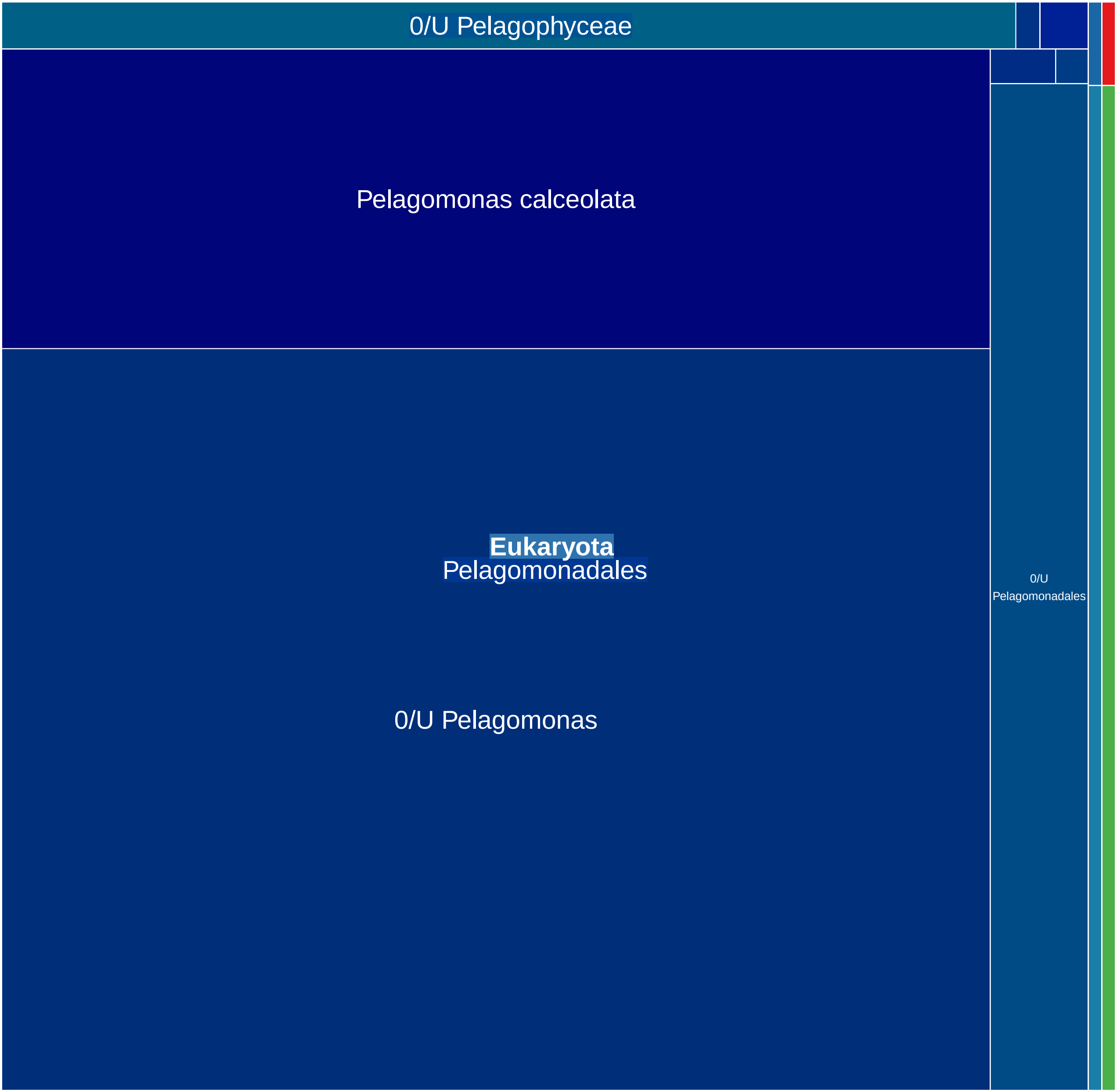

MGT−v1_605 : 1121 unigenes, 1093 (97.50%) taxonomically assigned

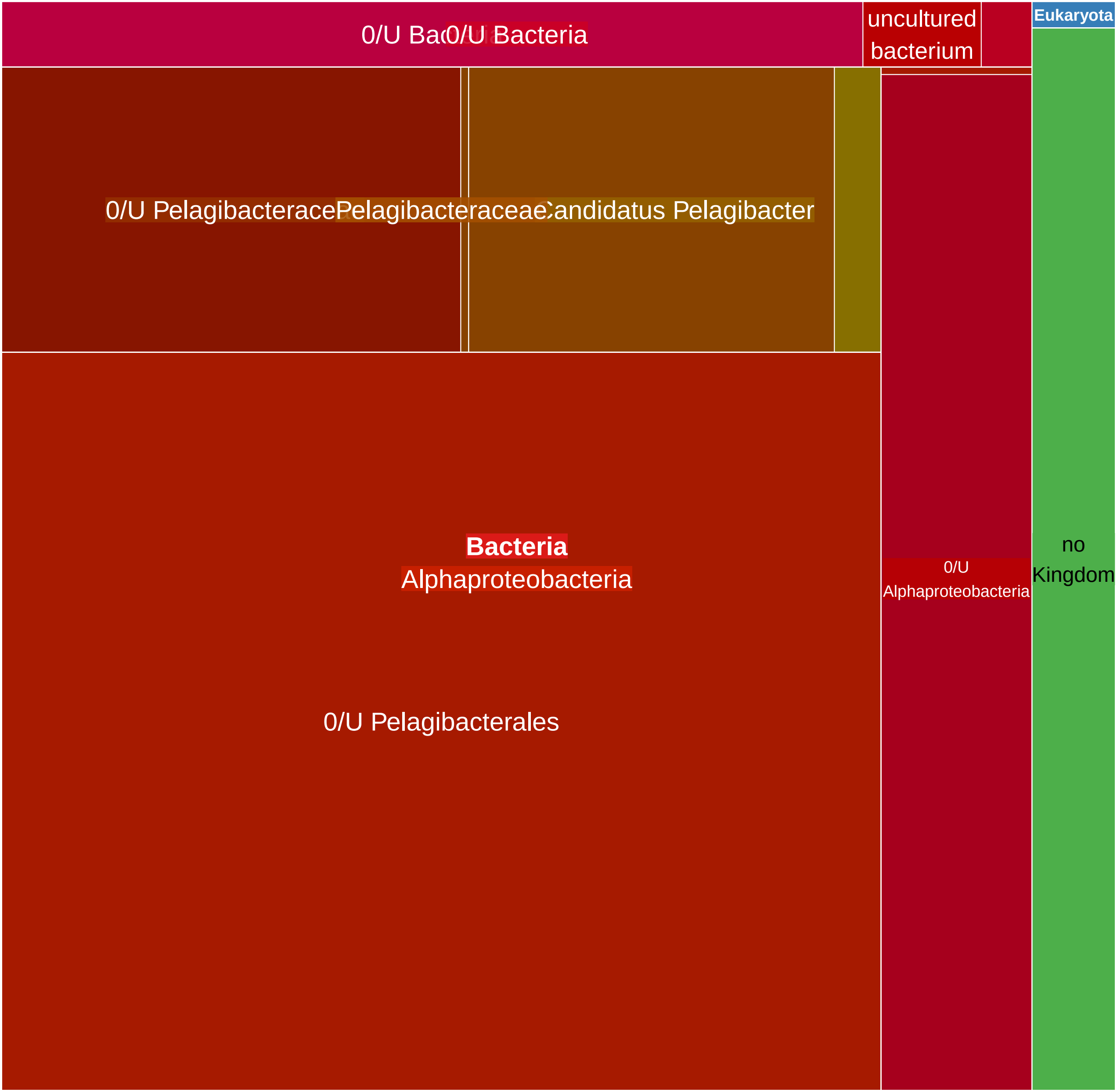

MGT−v1_606 : 1112 unigenes, 90 (8.09%) taxonomically assigned

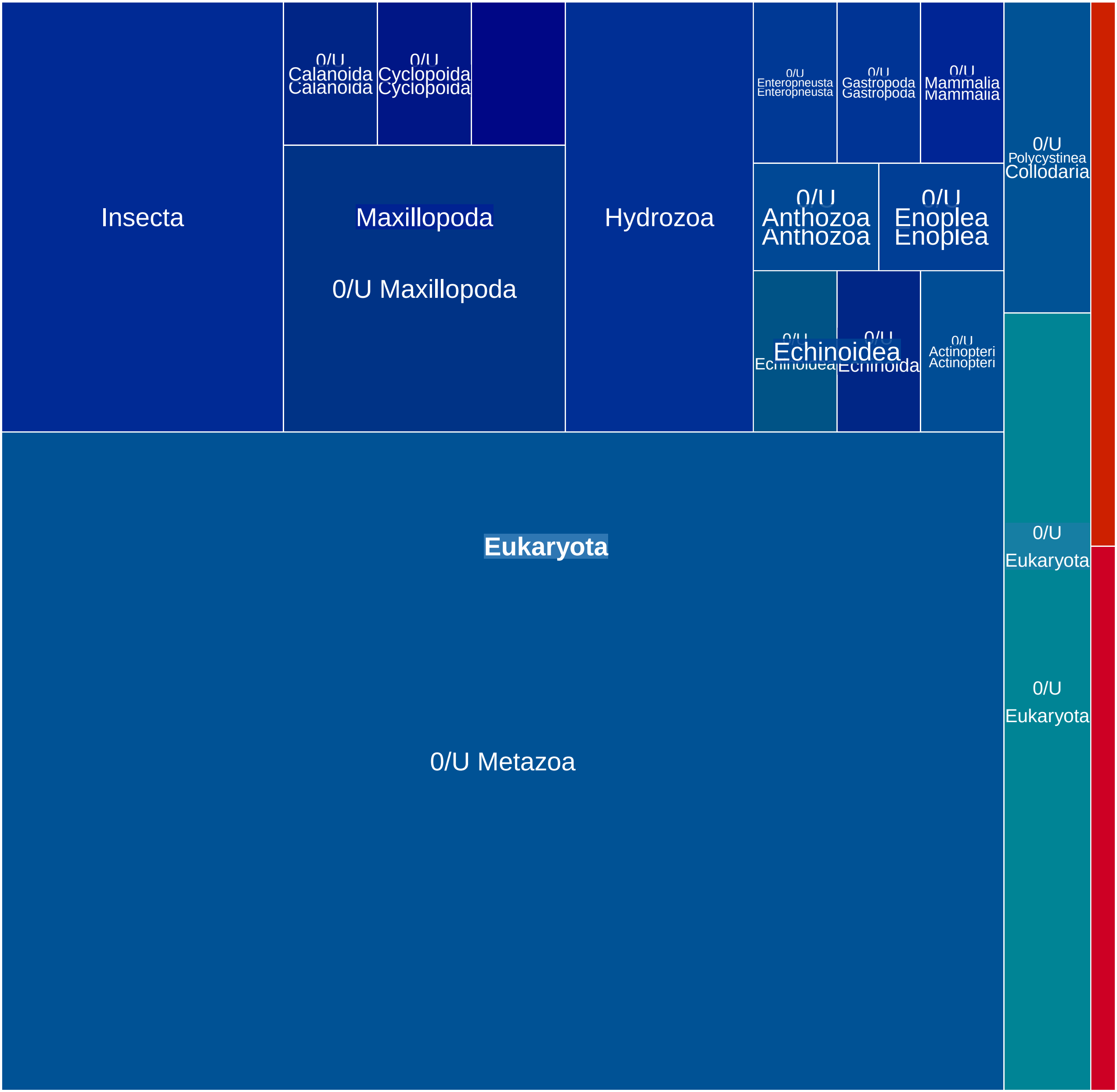

MGT−v1_607 : 1109 unigenes, 88 (7.94%) taxonomically assigned

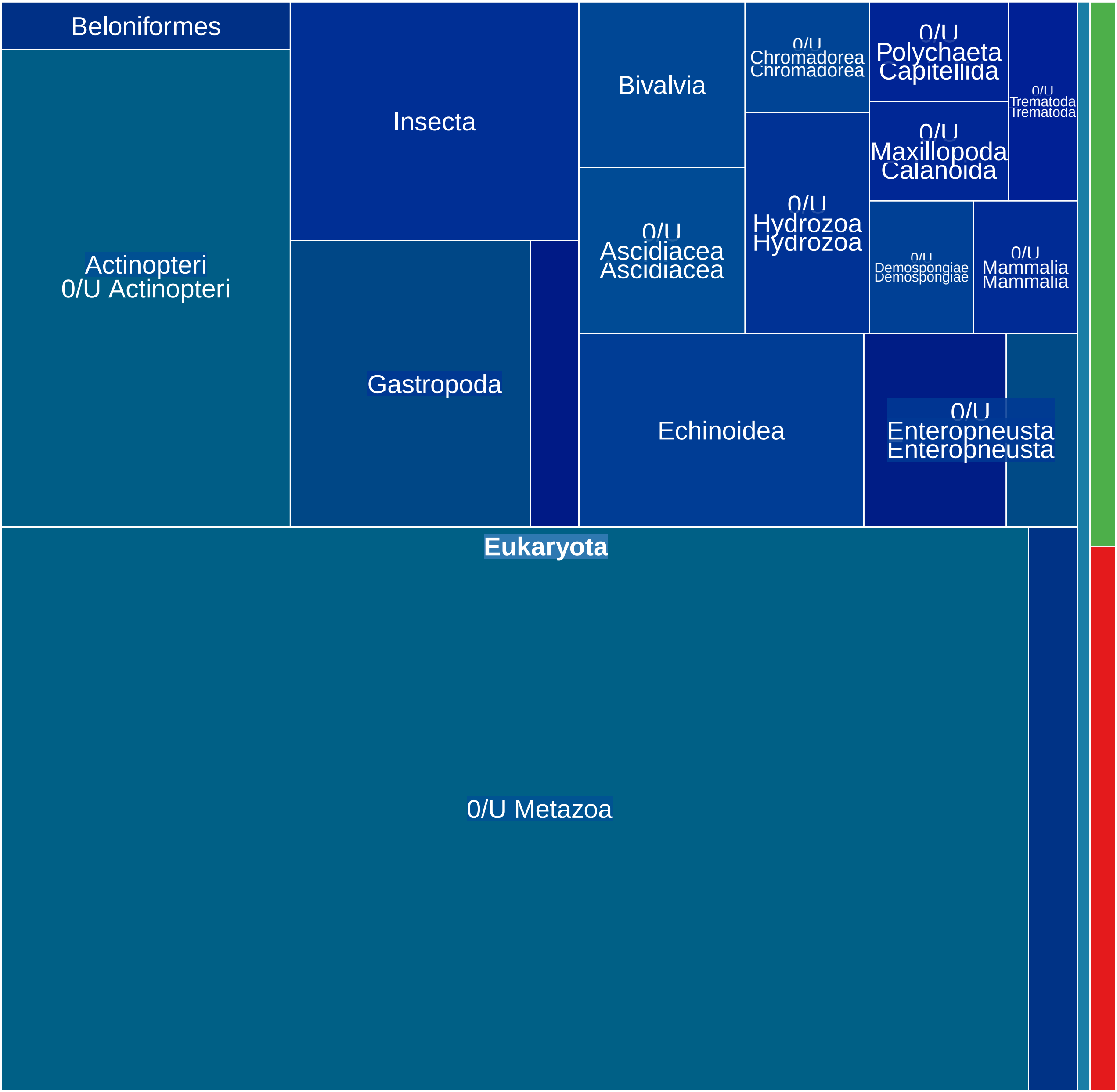

MGT−v1_608 : 1107 unigenes, 852 (76.96%) taxonomically assigned

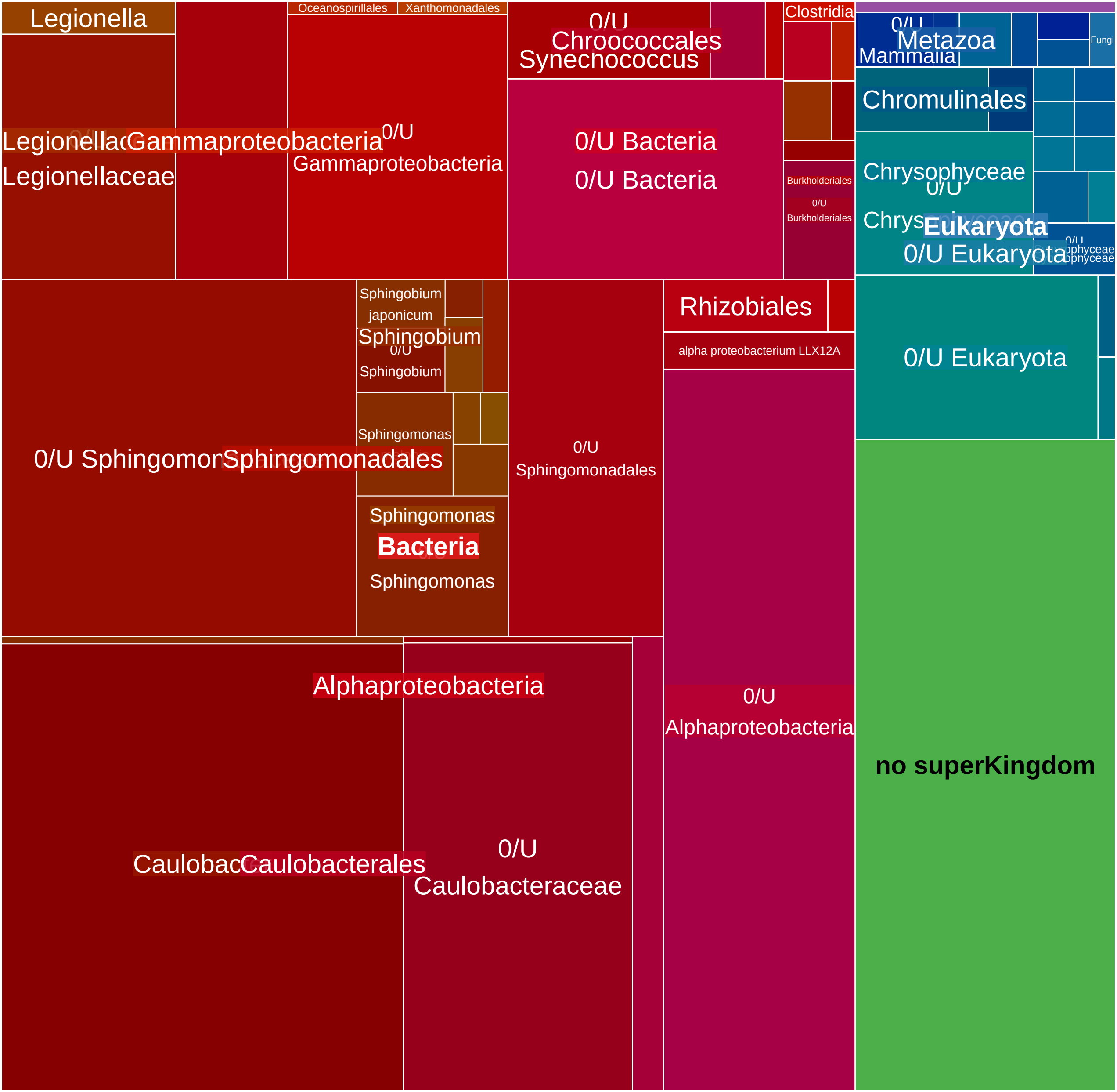

MGT−v1_609 : 1106 unigenes, 799 (72.24%) taxonomically assigned

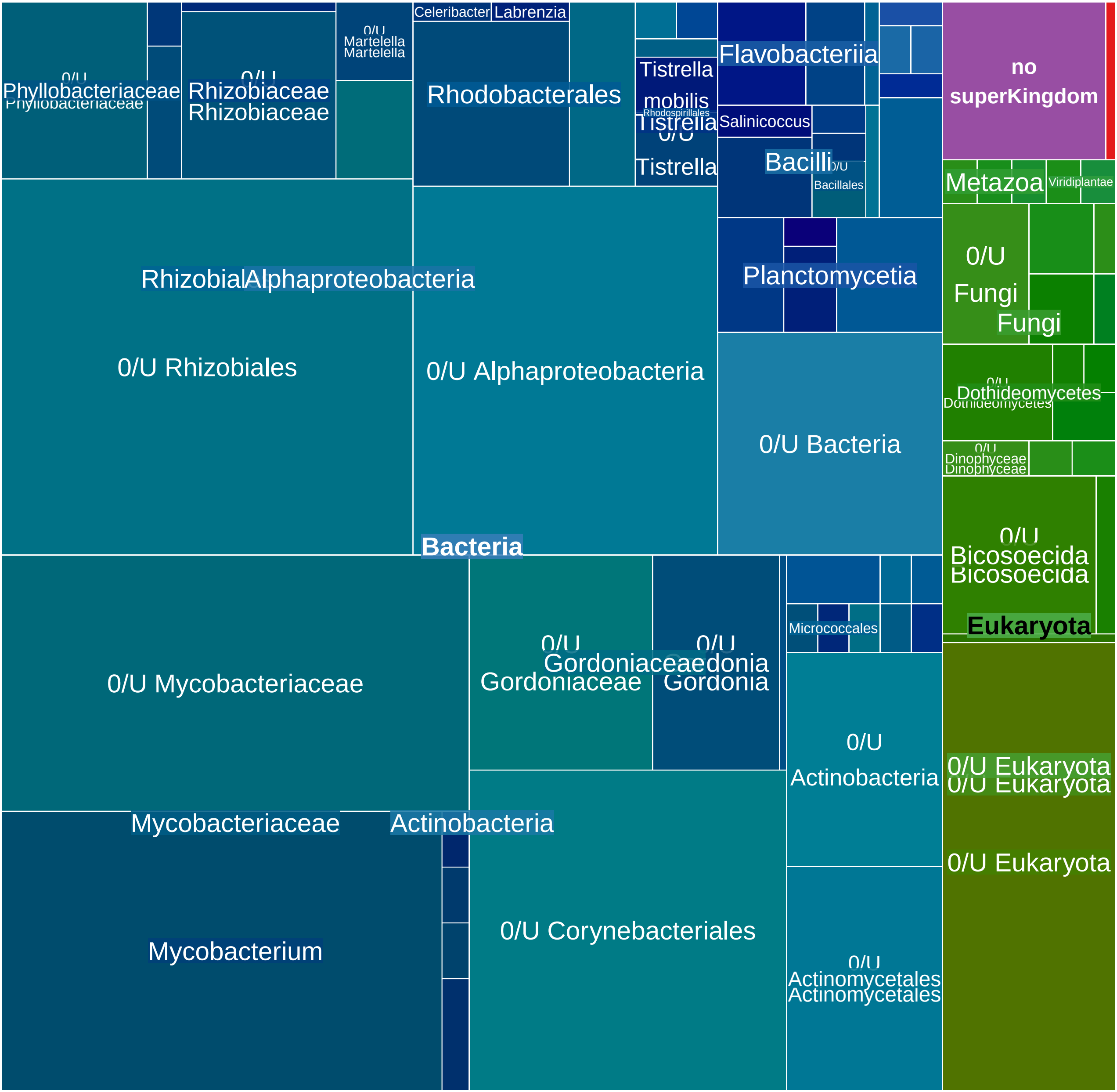

MGT−v1_610 : 1104 unigenes, 94 (8.51%) taxonomically assigned

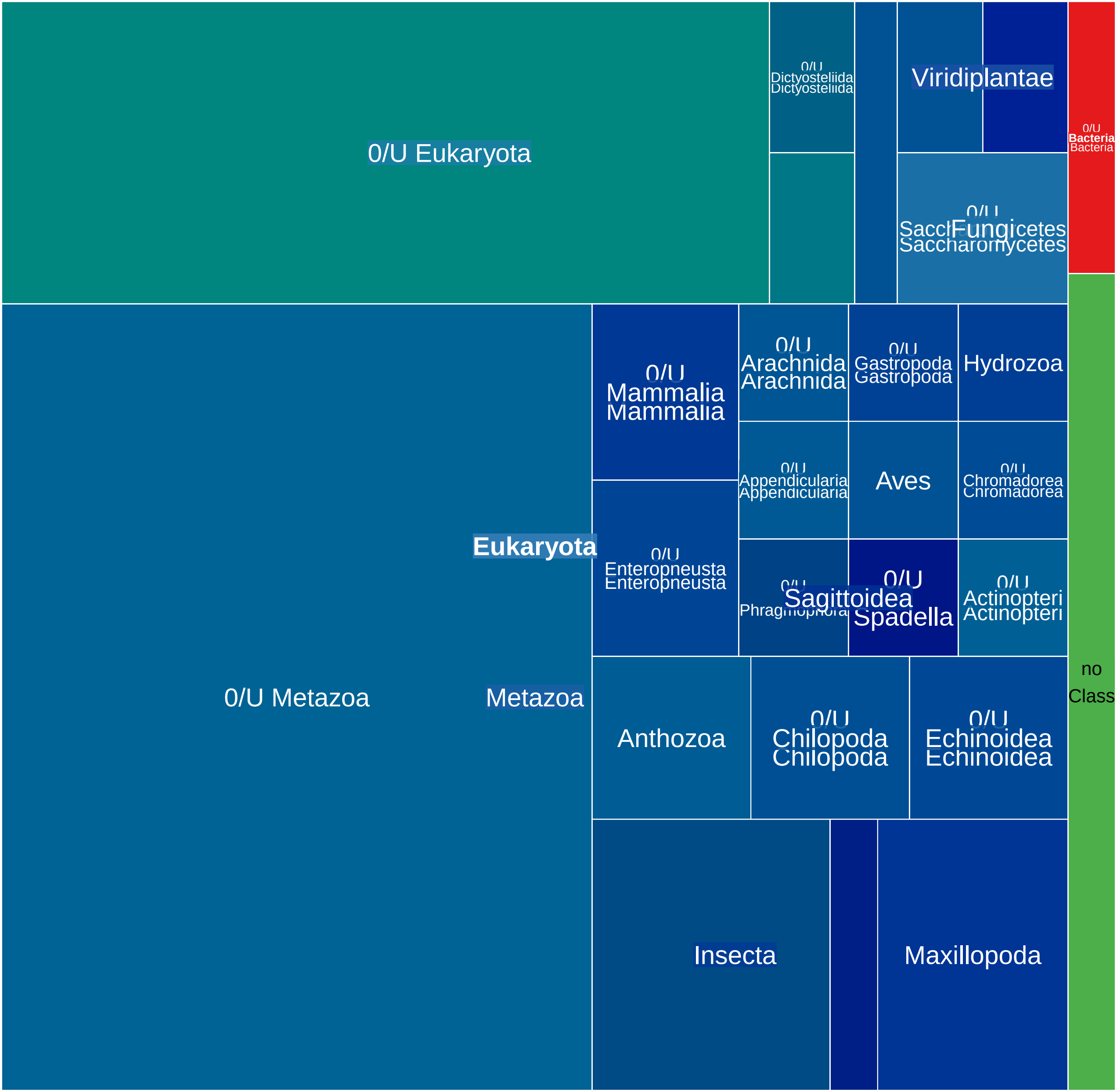

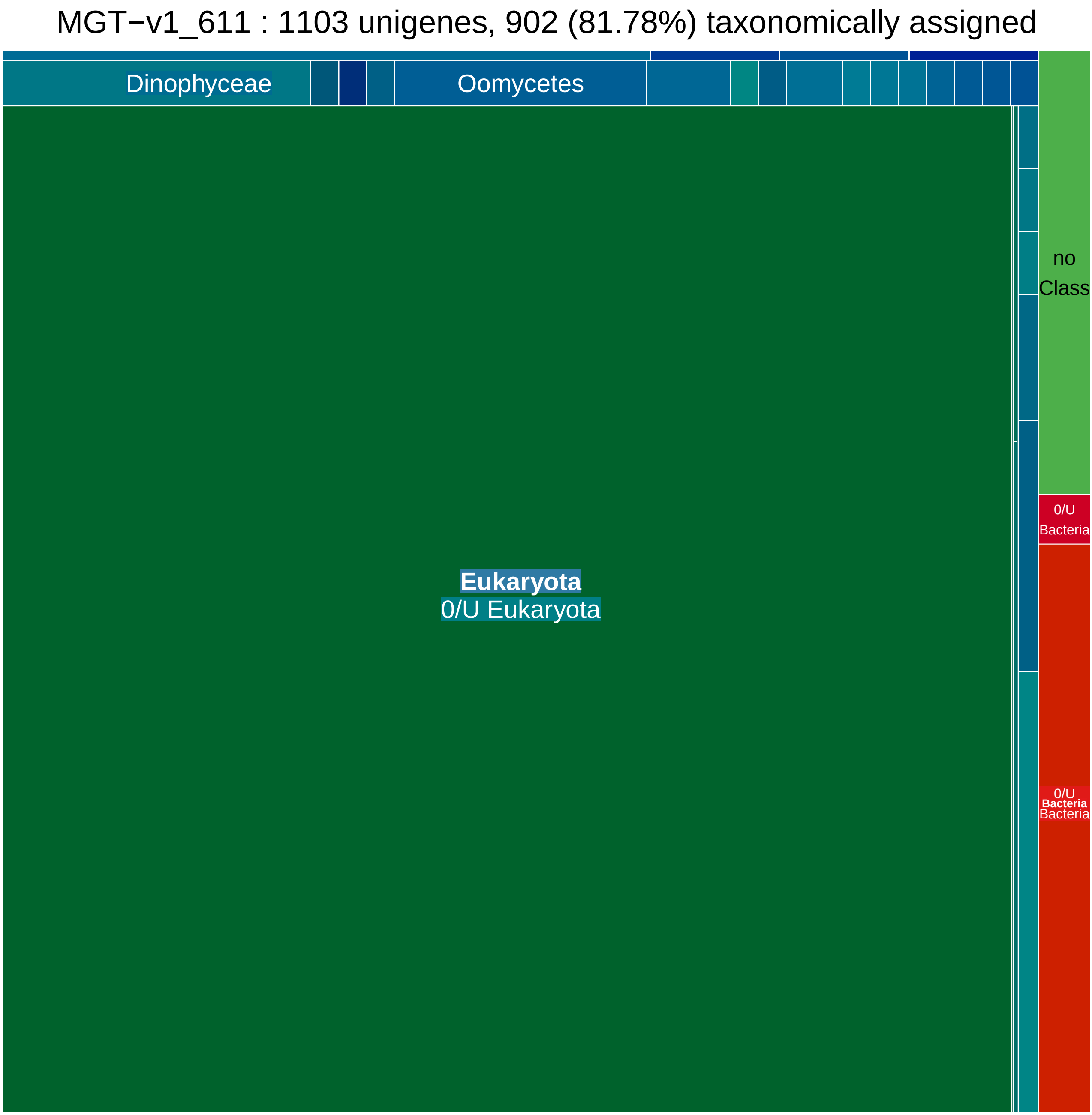
MGT−v1_611 : 1103 unigenes, 902 (81.78%) taxonomically assigned
Dinophyceae
Oomycetes
no Class
0/U Bacteria
Eukaryota
0/U Eukaryota
0/U Bacteria
Bacteria
Bacteria

MGT−v1_612 : 1086 unigenes, 111 (10.22%) taxonomically assigned

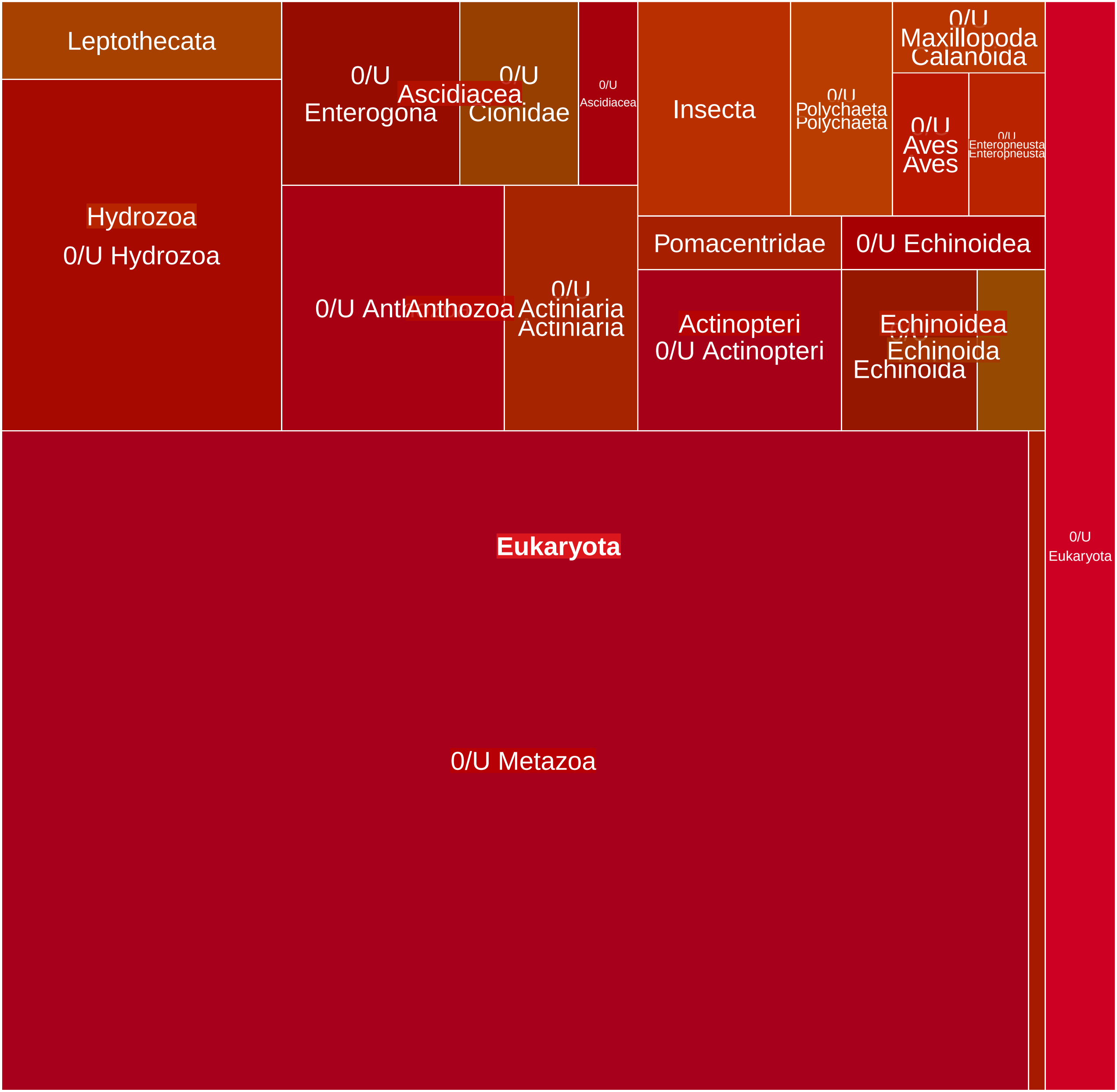

### MGT−v1_613 : 1085 unigenes, 50 (4.61%) taxonomically assigned

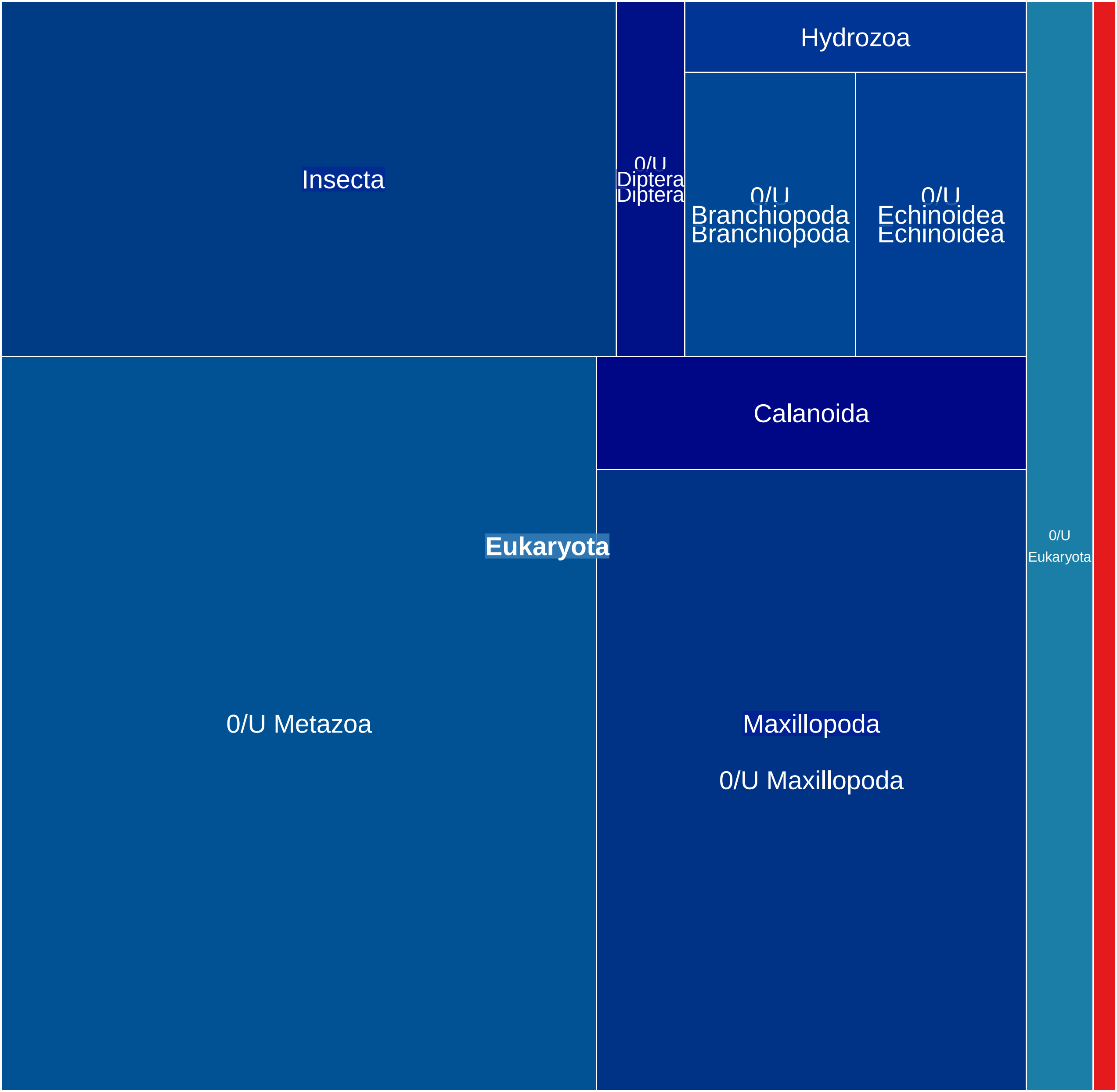

MGT−v1_614 : 1082 unigenes, 128 (11.83%) taxonomically assigned

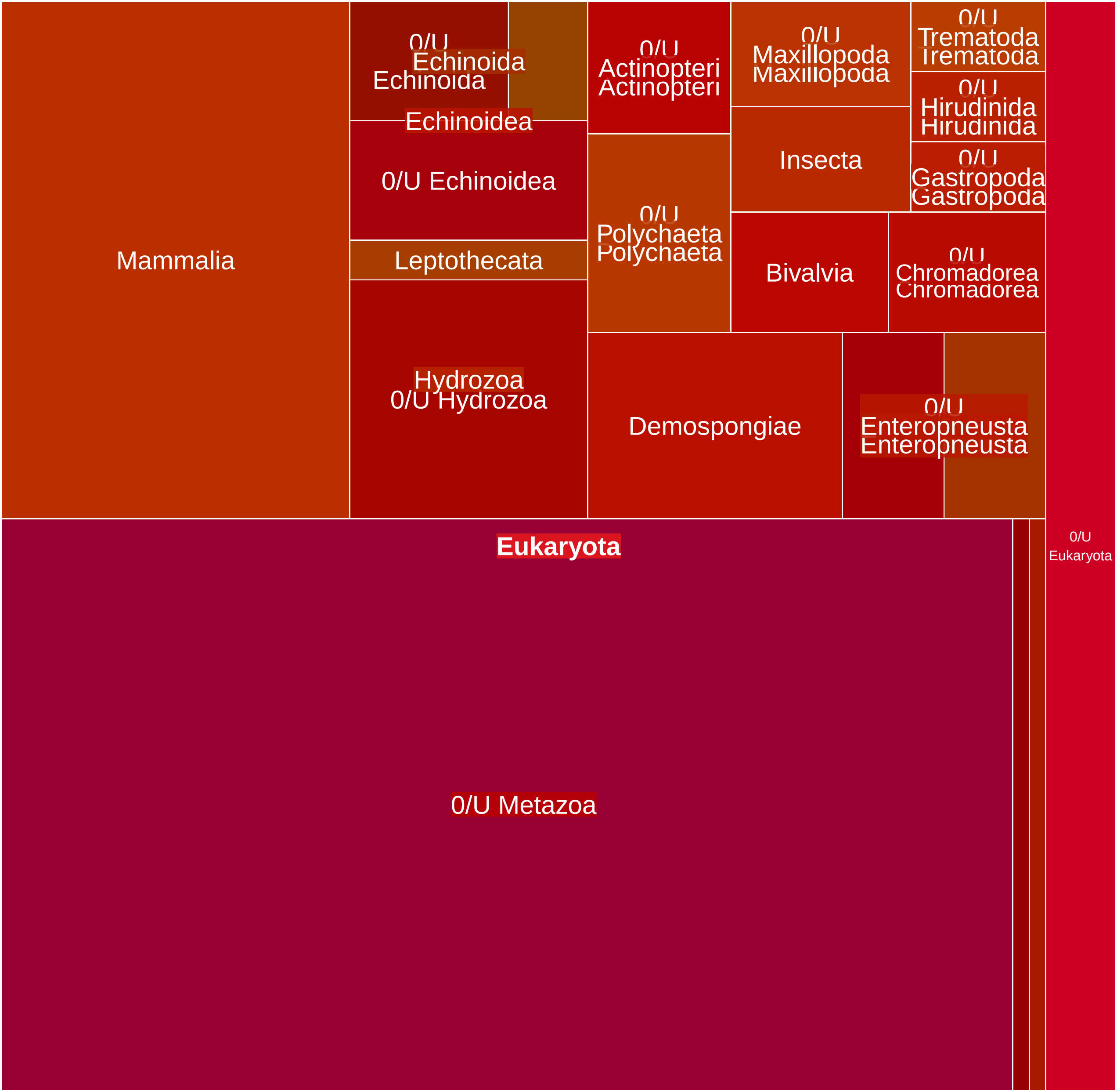

MGT−v1_615 : 1080 unigenes, 775 (71.76%) taxonomically assigned

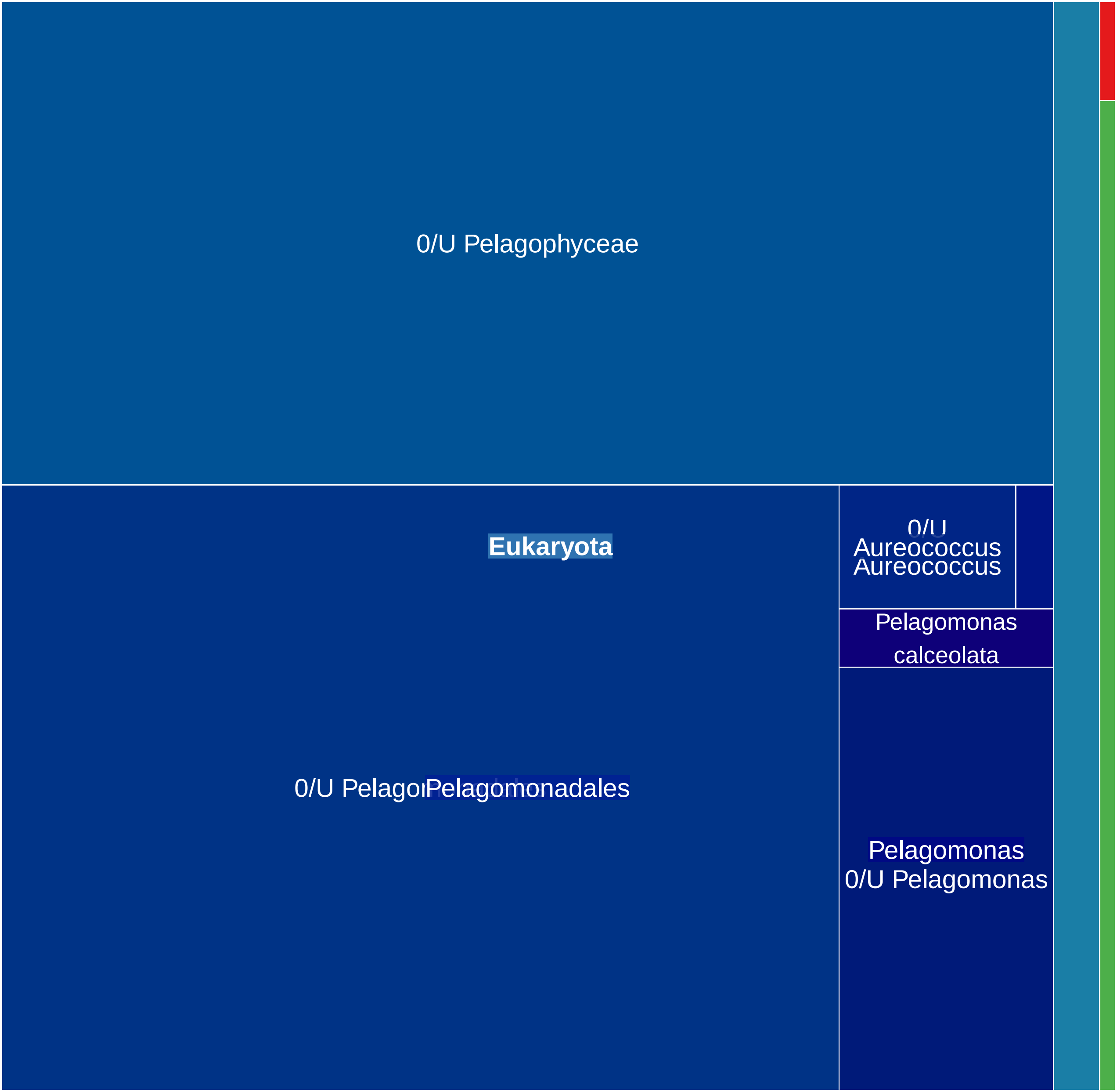

MGT−v1_616 : 1077 unigenes, 149 (13.83%) taxonomically assigned

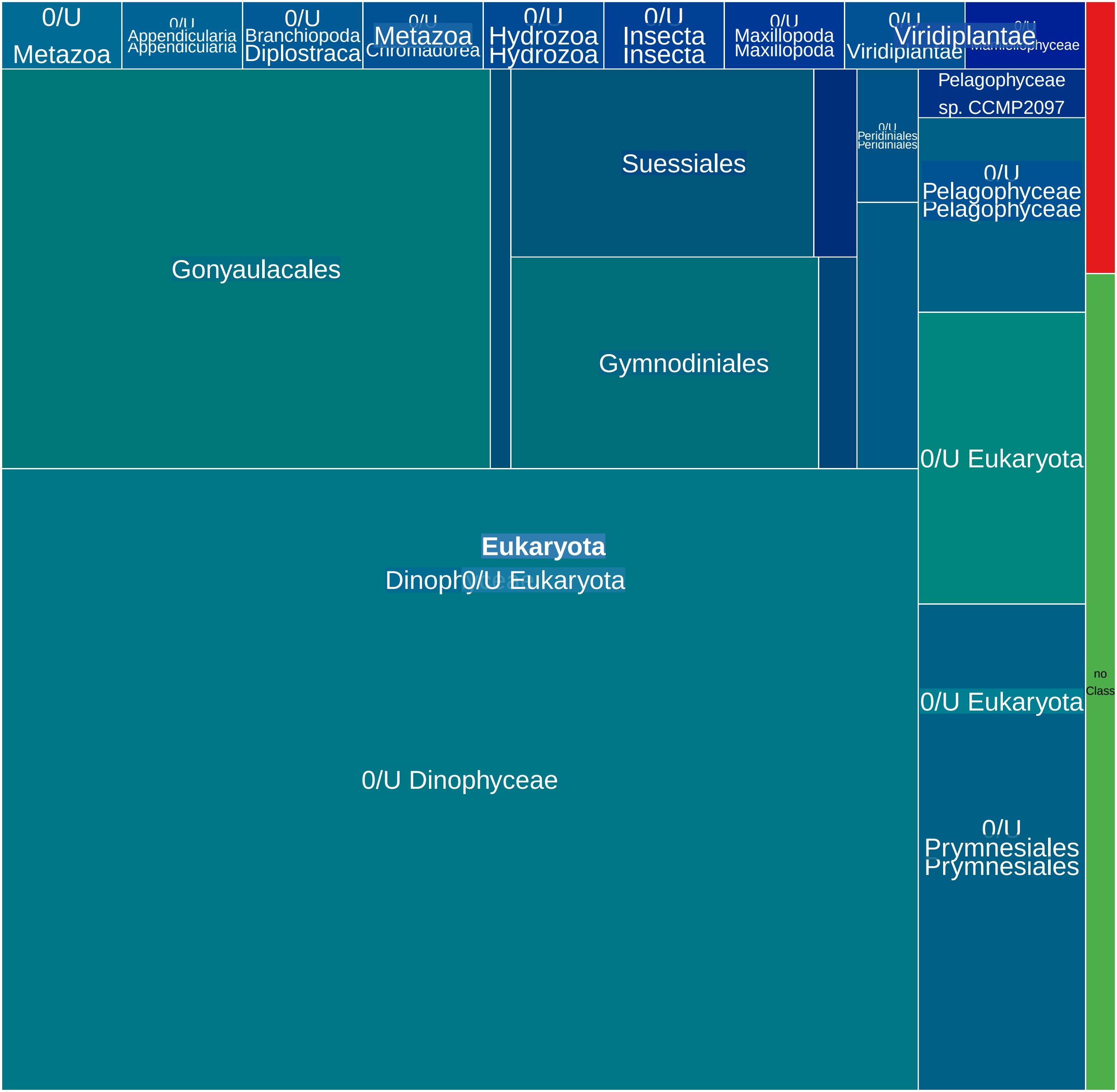

MGT−v1_617 : 1077 unigenes, 658 (61.10%) taxonomically assigned

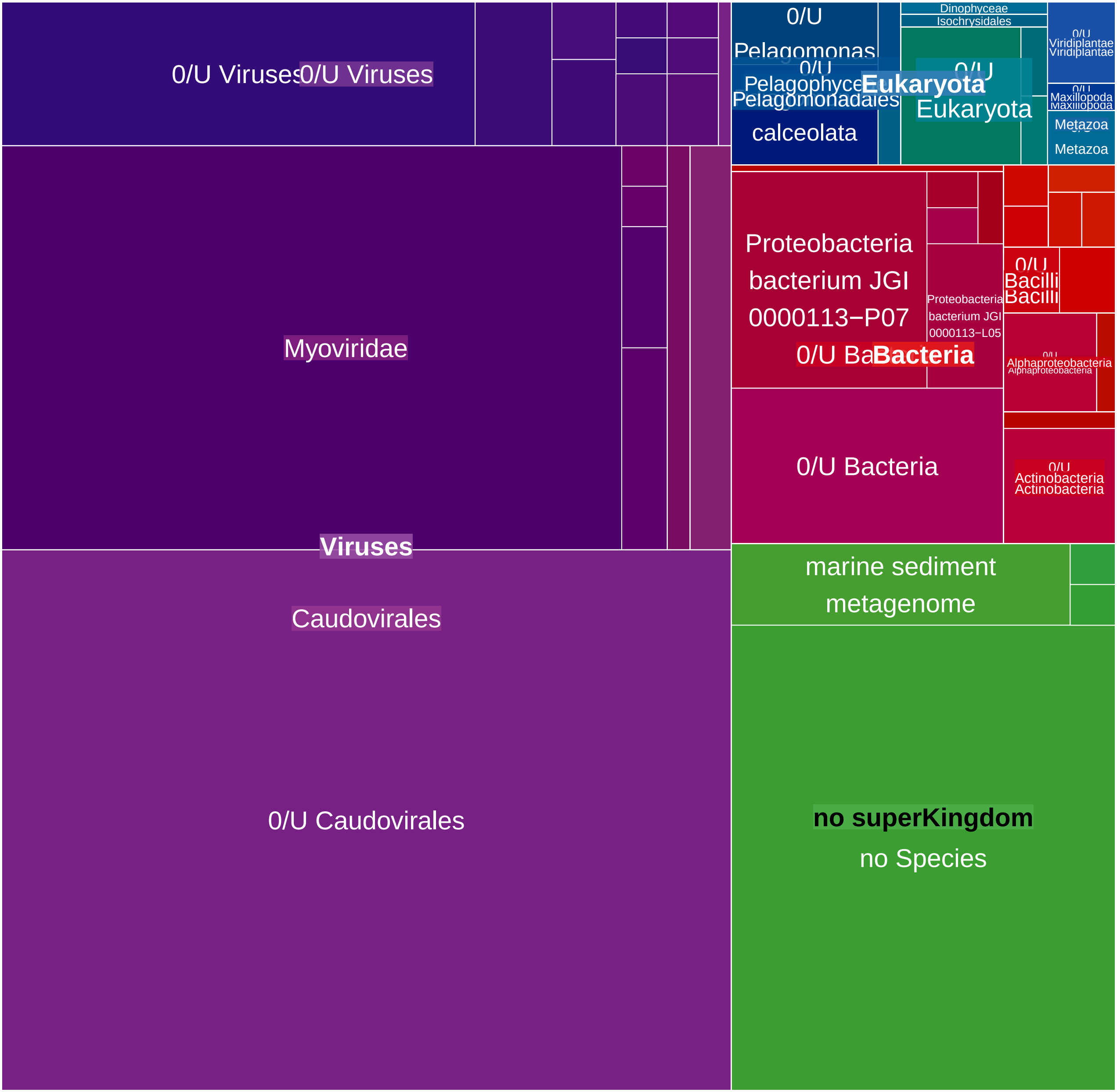

MGT−v1_618 : 1077 unigenes, 971 (90.16%) taxonomically assigned

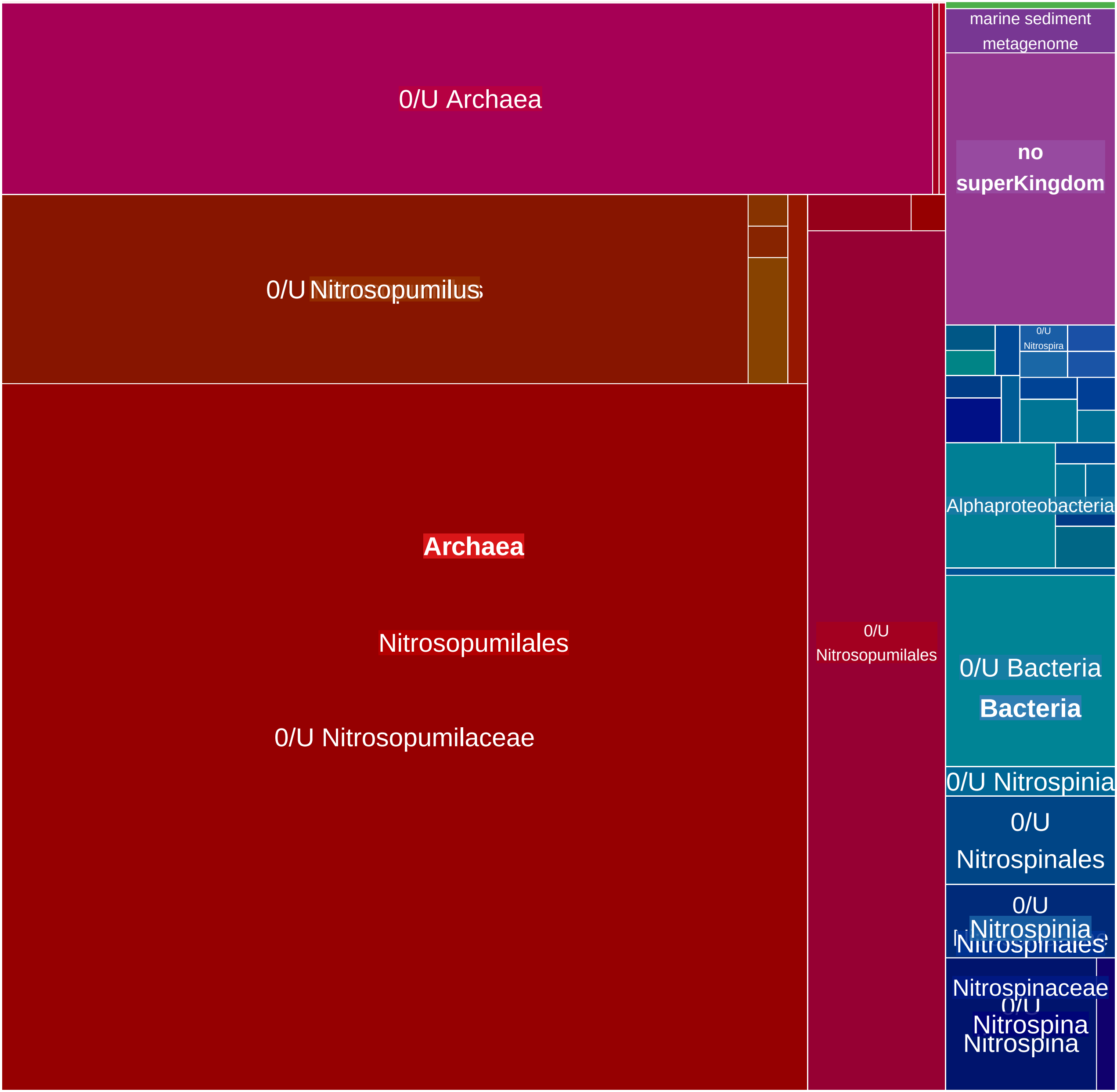

MGT−v1_619 : 1072 unigenes, 104 (9.70%) taxonomically assigned

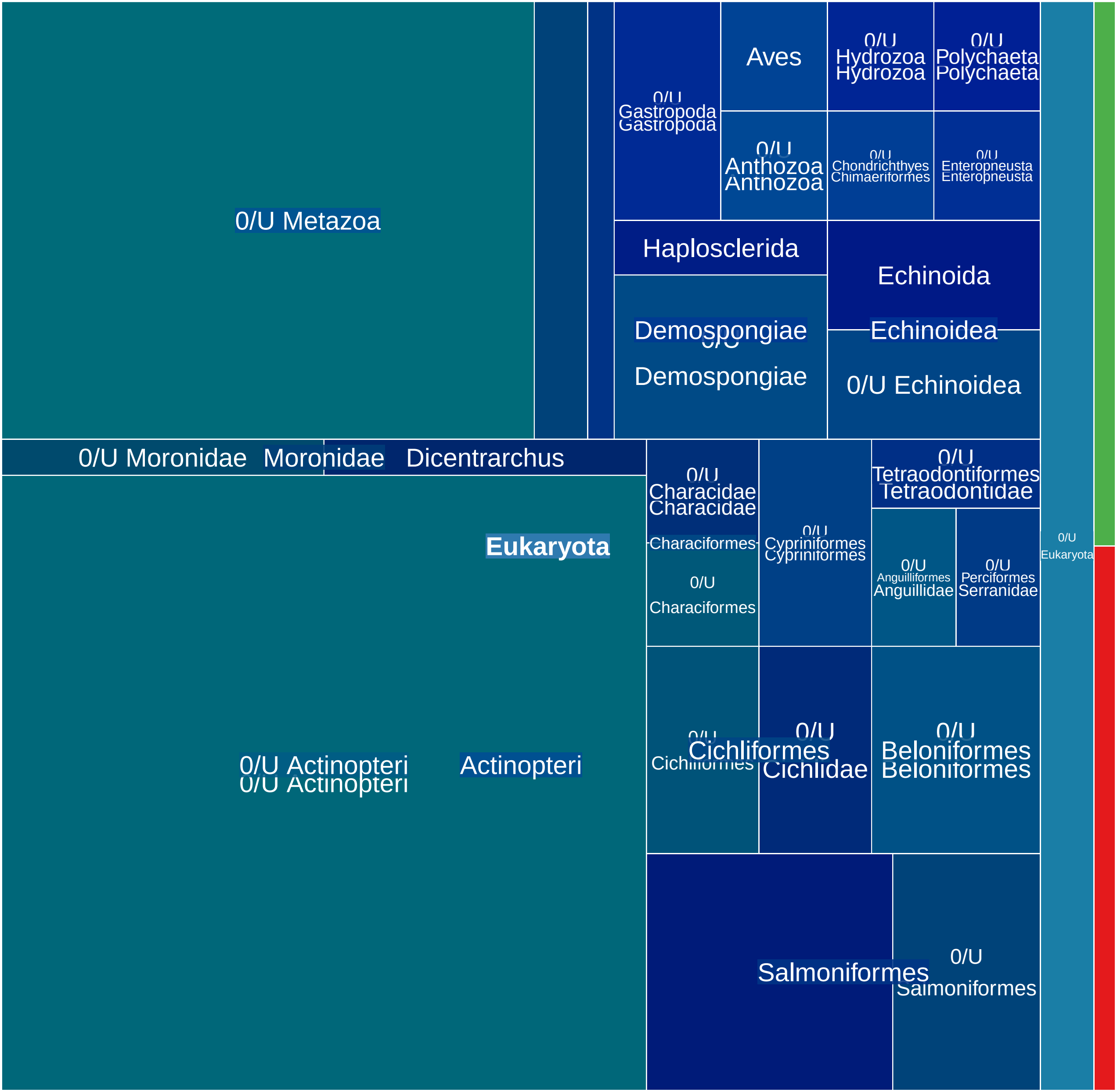

MGT−v1_620 : 1071 unigenes, 510 (47.62%) taxonomically assigned

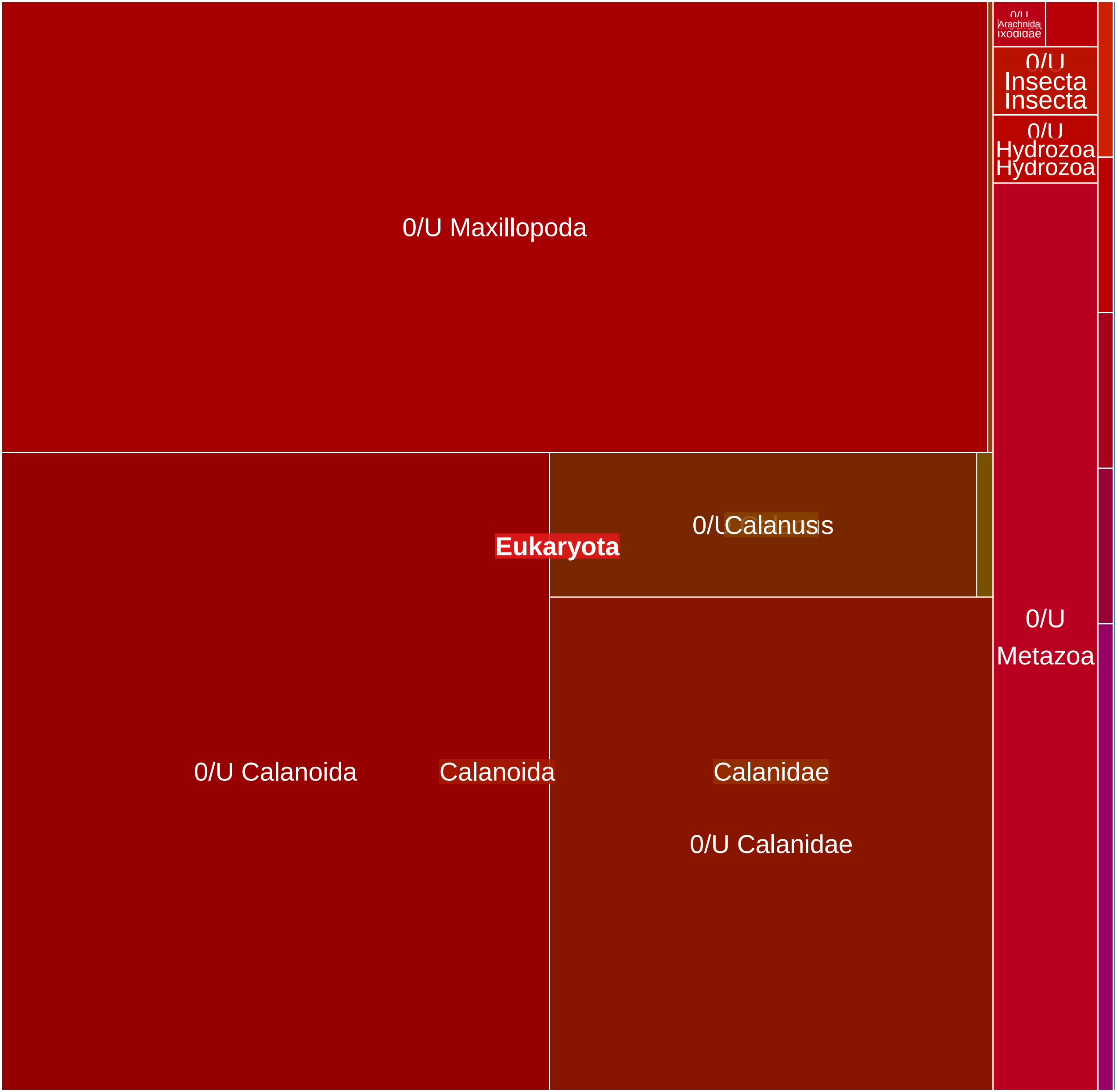

MGT−v1_621 : 1071 unigenes, 678 (63.31%) taxonomically assigned

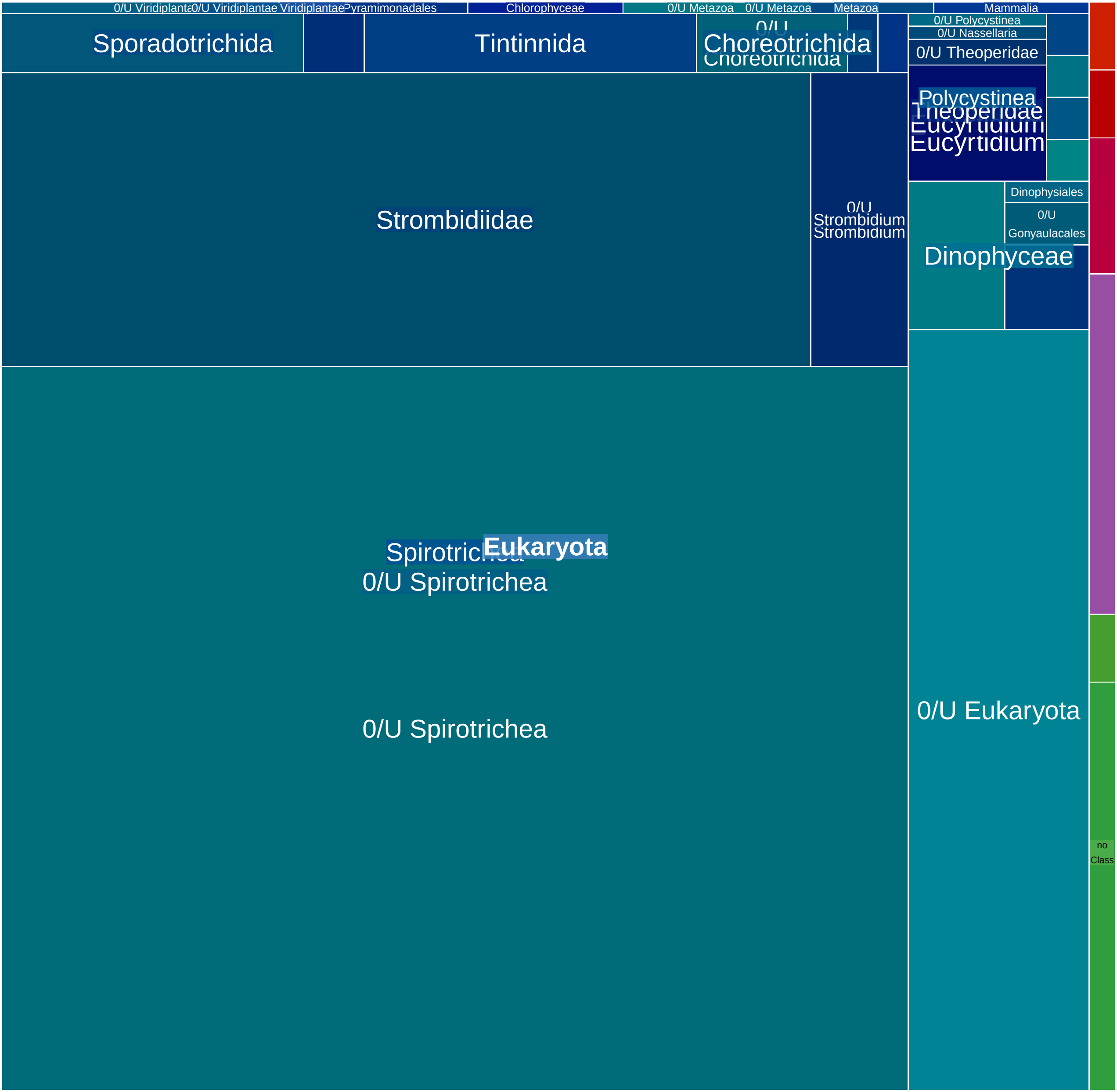

MGT−v1_622 : 1070 unigenes, 128 (11.96%) taxonomically assigned

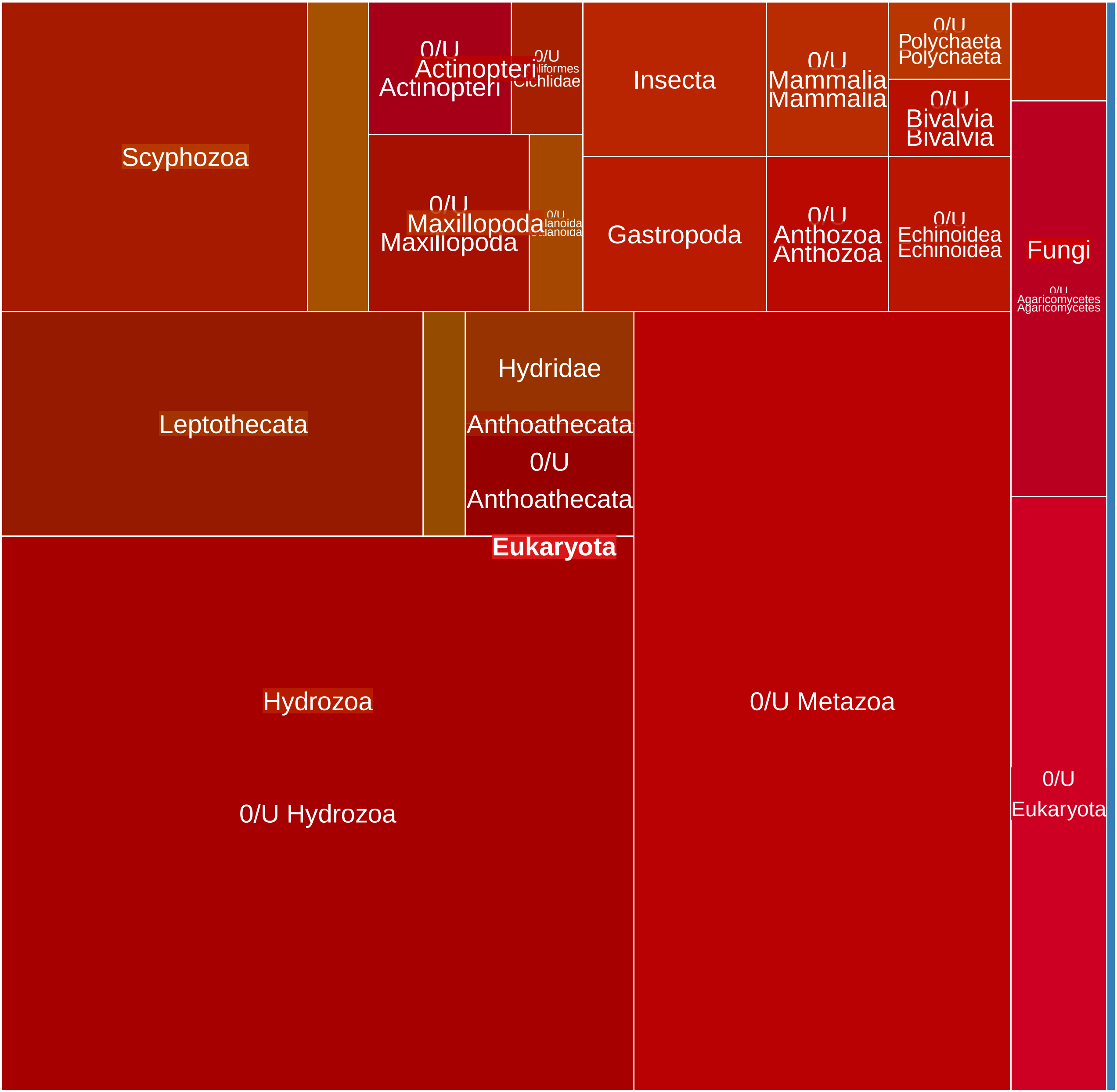

MGT−v1_623 : 1066 unigenes, 580 (54.41%) taxonomically assigned

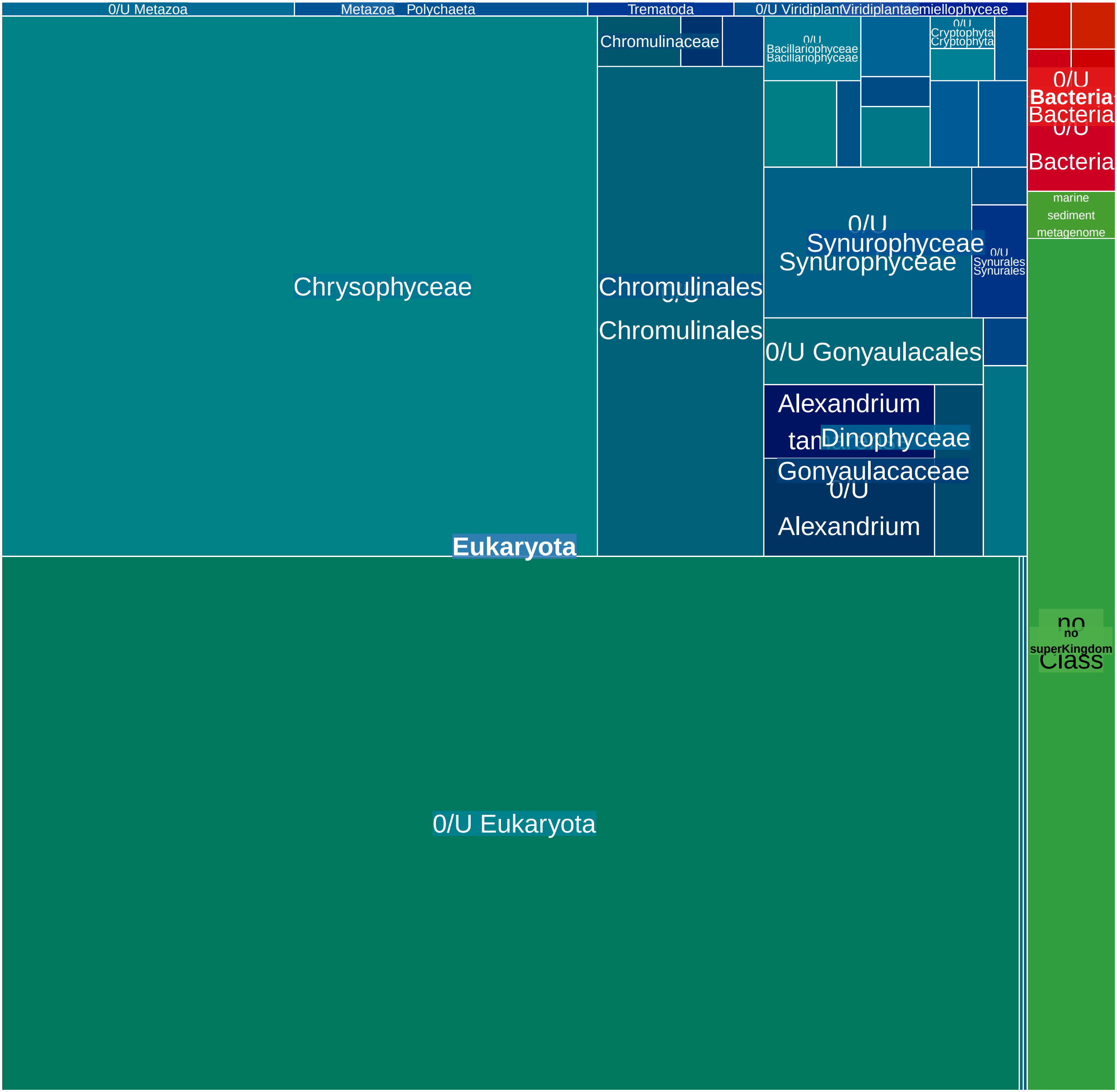

MGT−v1_624 : 1058 unigenes, 426 (40.26%) taxonomically assigned

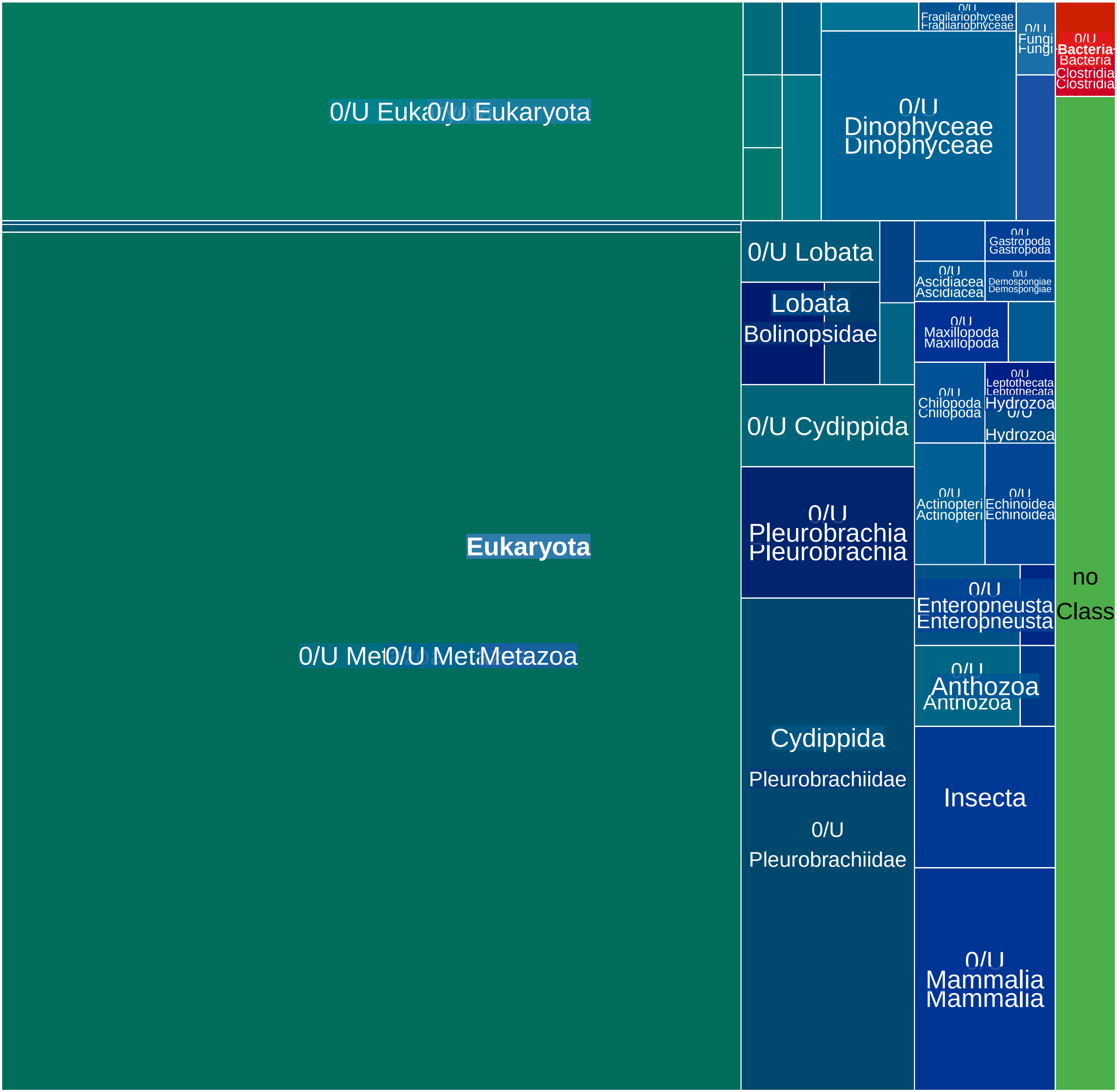

MGT−v1_625 : 1050 unigenes, 770 (73.33%) taxonomically assigned

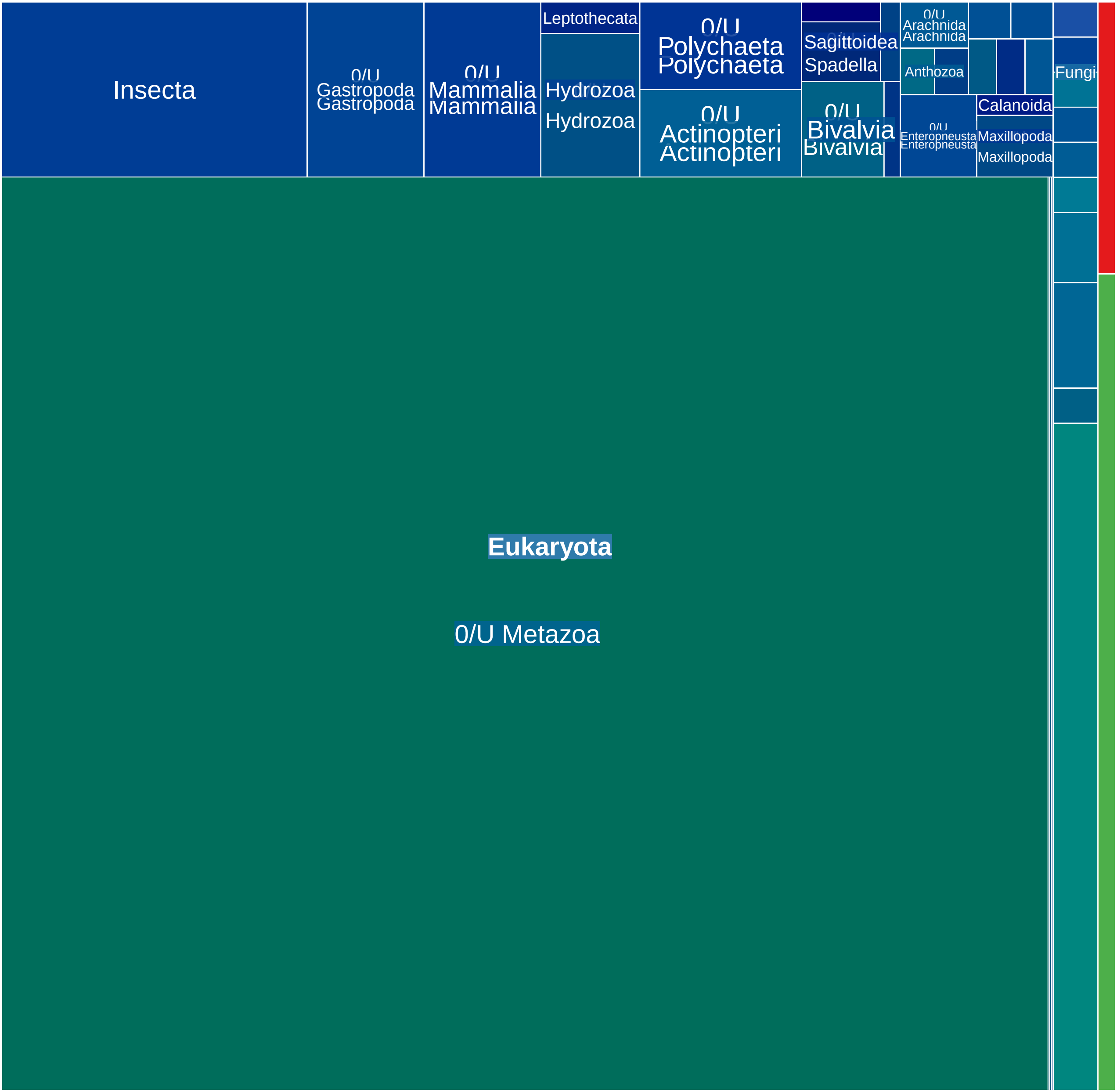

MGT−v1_626 : 1045 unigenes, 85 (8.13%) taxonomically assigned

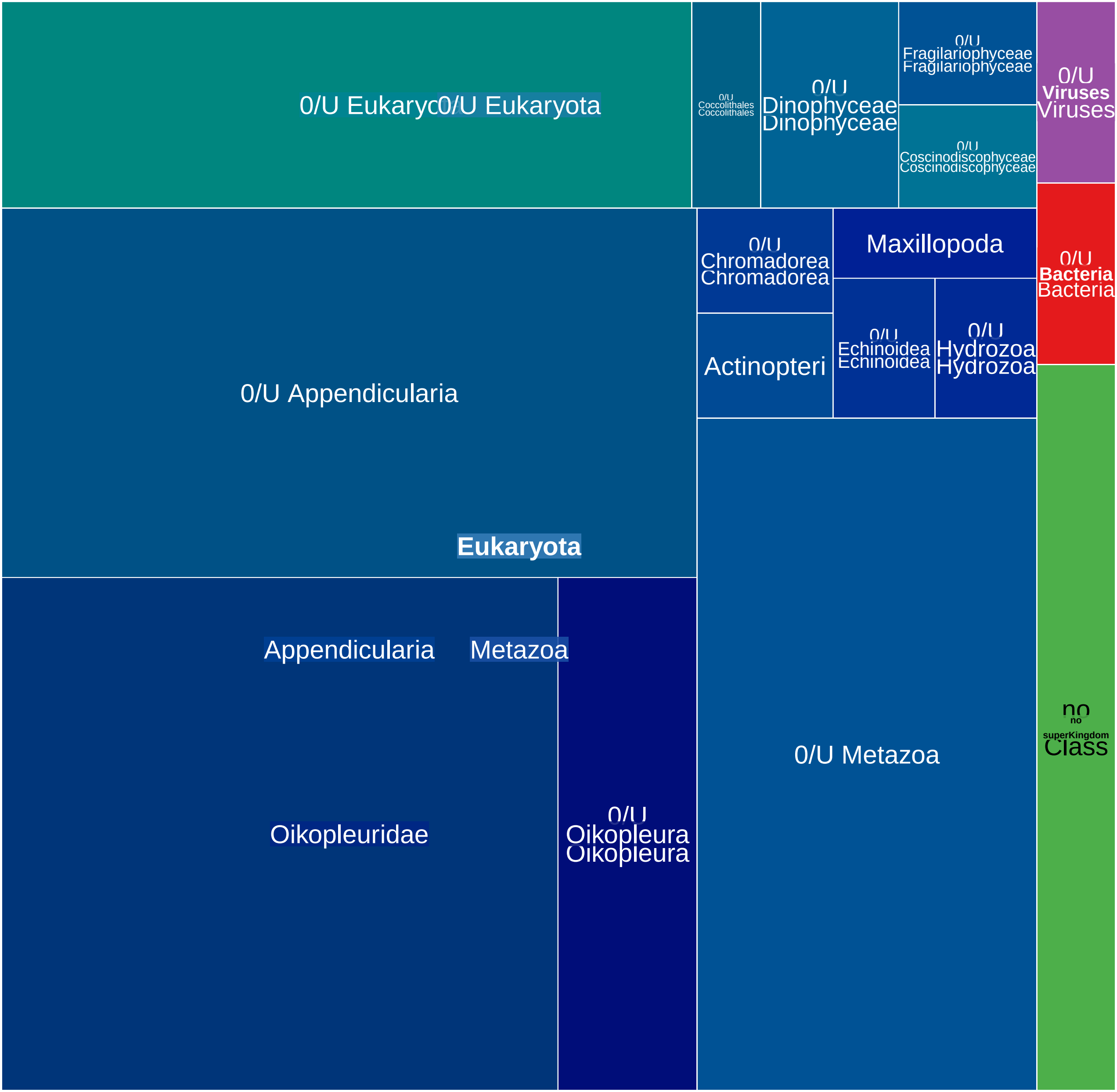

### MGT−v1_627 : 1044 unigenes, 752 (72.03%) taxonomically assigned

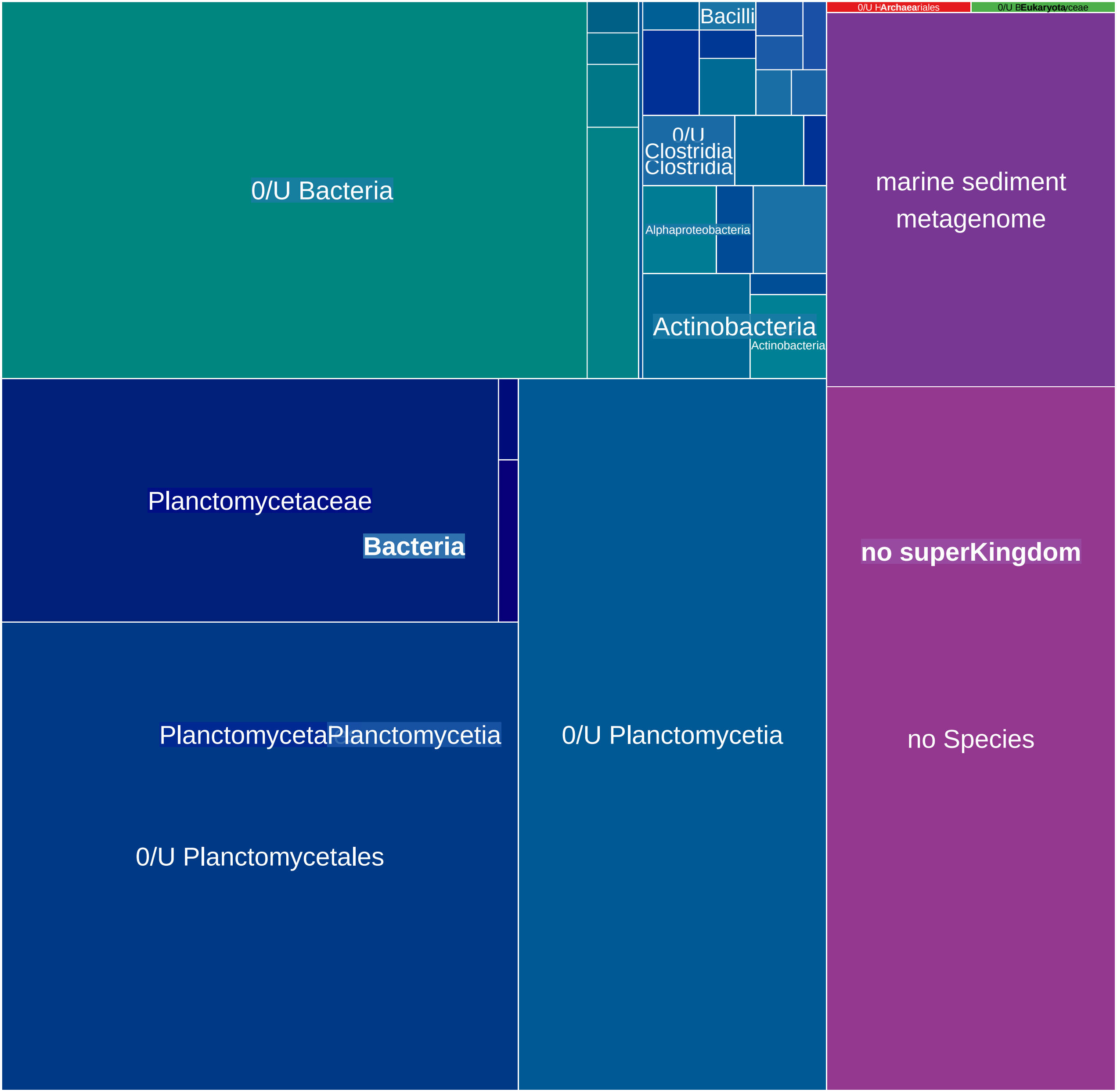

MGT−v1_628 : 1037 unigenes, 589 (56.80%) taxonomically assigned

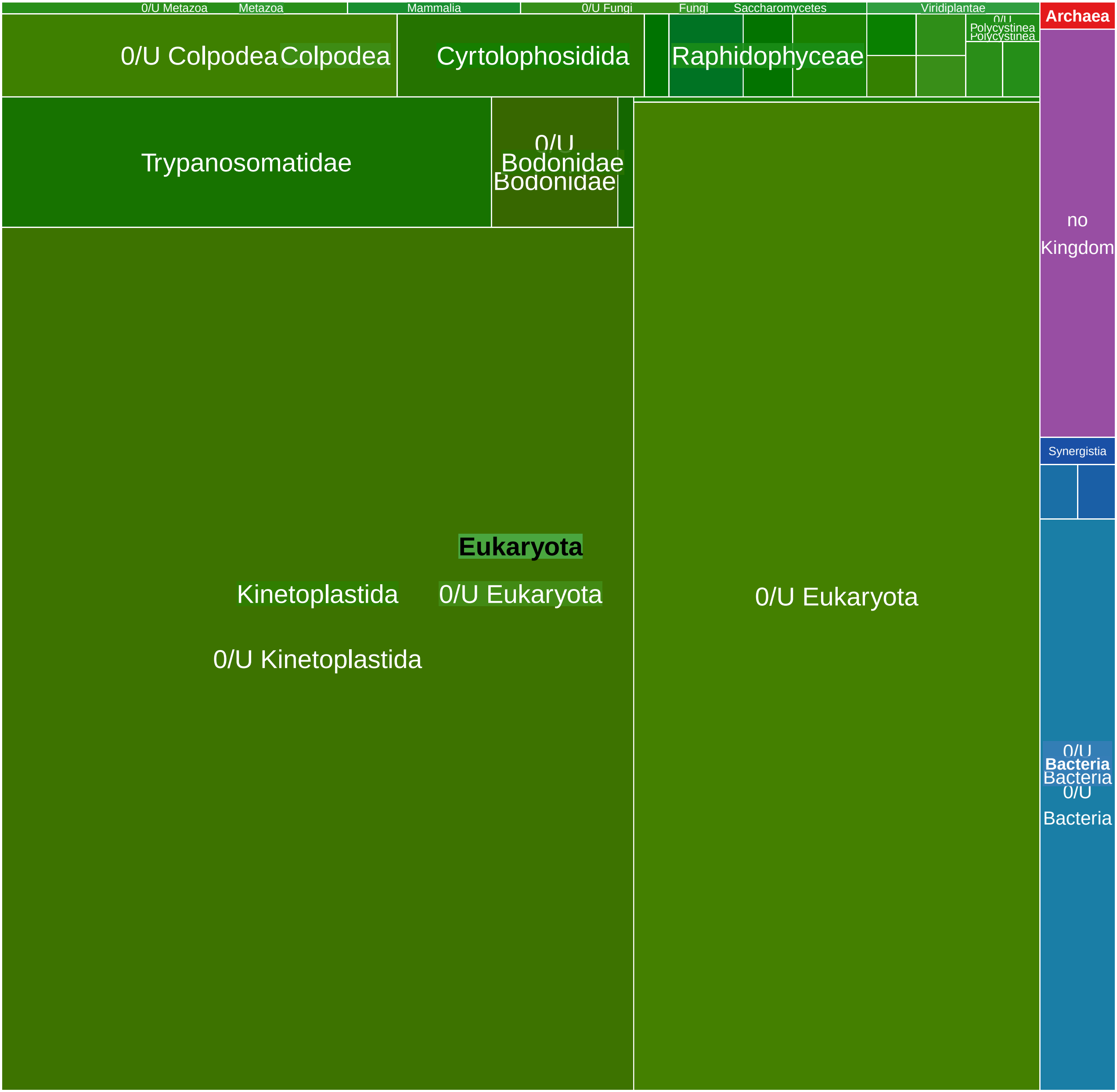

### MGT−v1_629 : 1027 unigenes, 984 (95.81%) taxonomically assigned

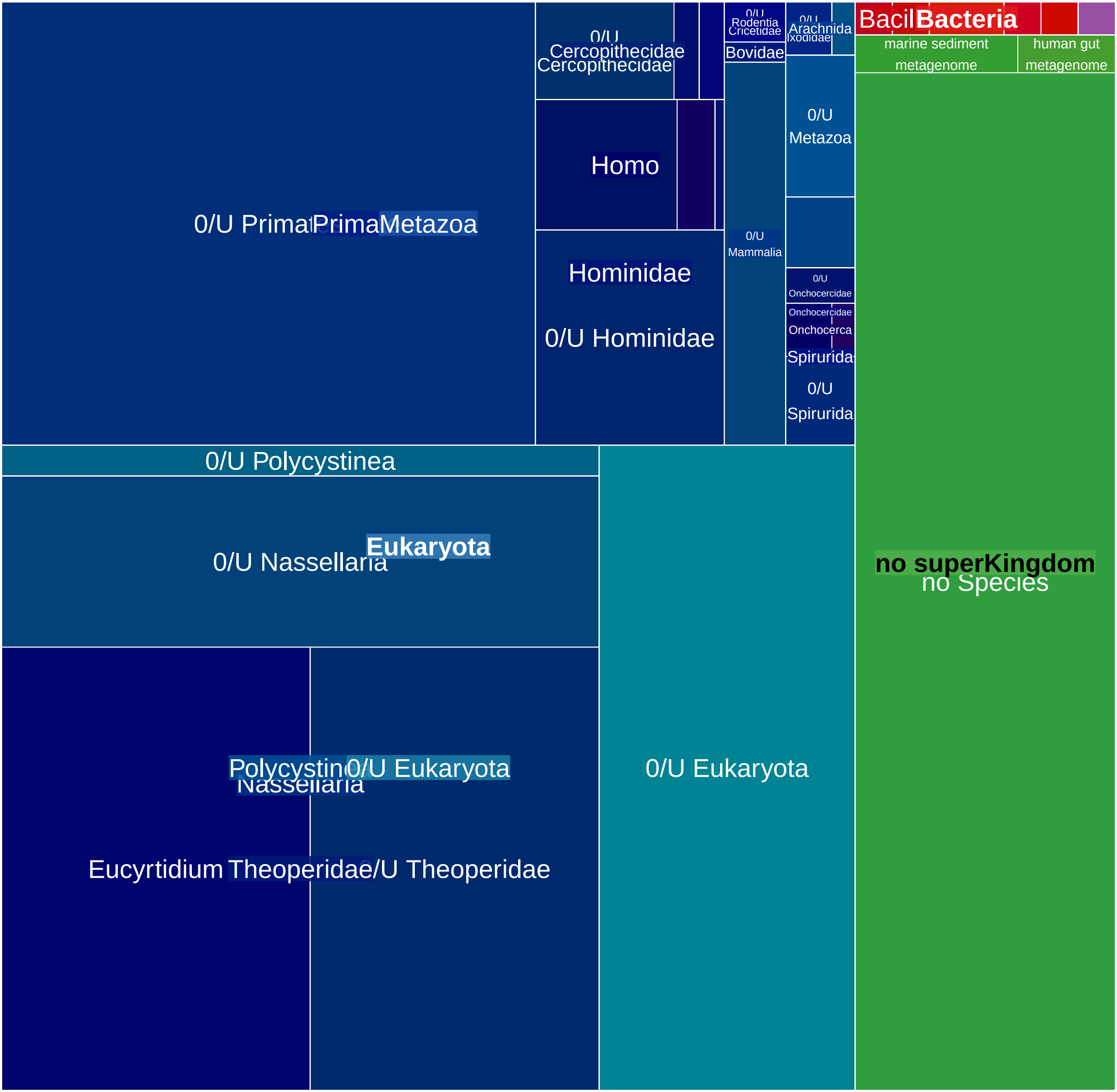

MGT−v1_630 : 1027 unigenes, 480 (46.74%) taxonomically assigned

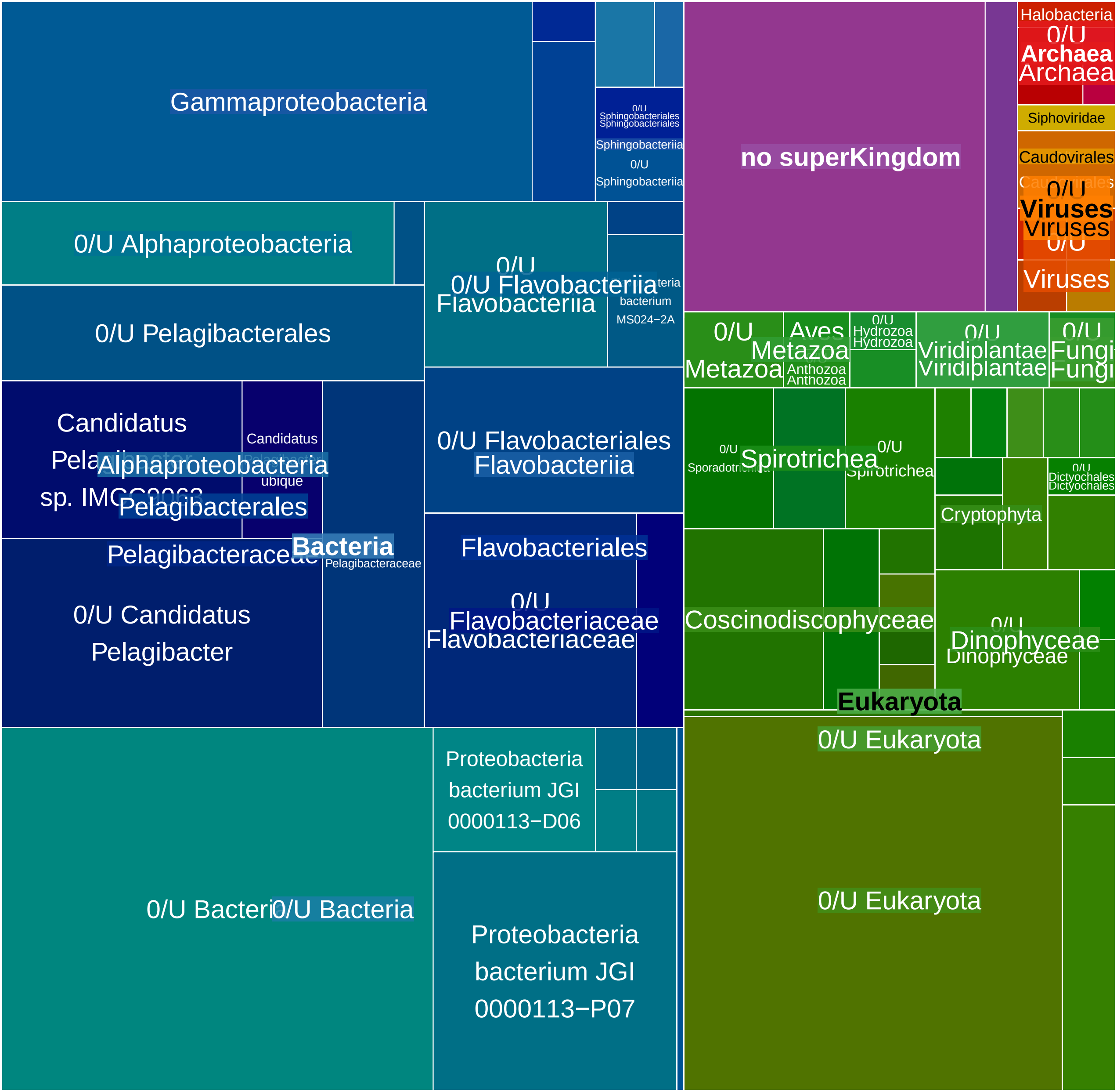

### MGT−v1_631 : 1023 unigenes, 743 (72.63%) taxonomically assigned

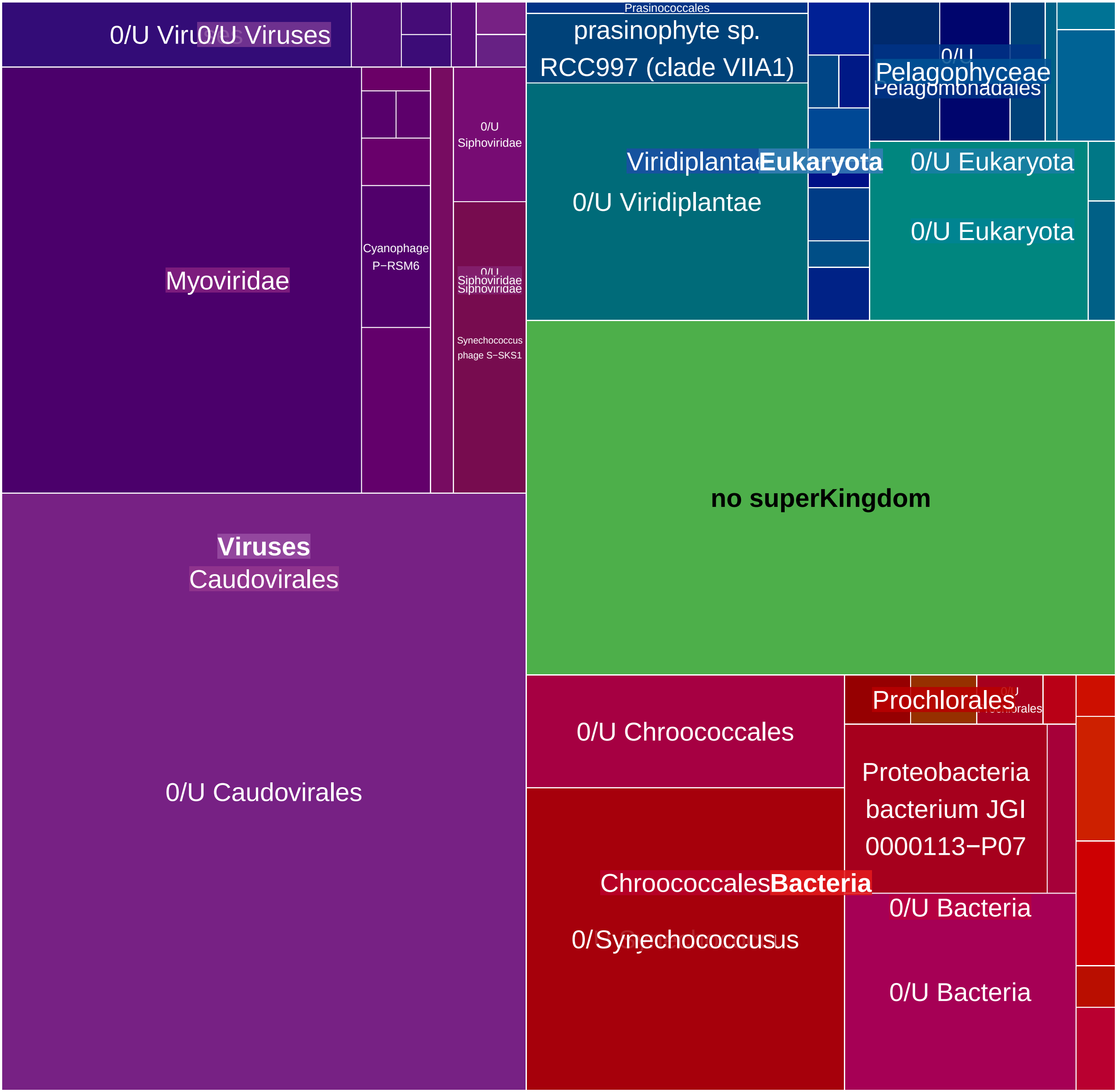

MGT−v1_632 : 1021 unigenes, 125 (12.24%) taxonomically assigned

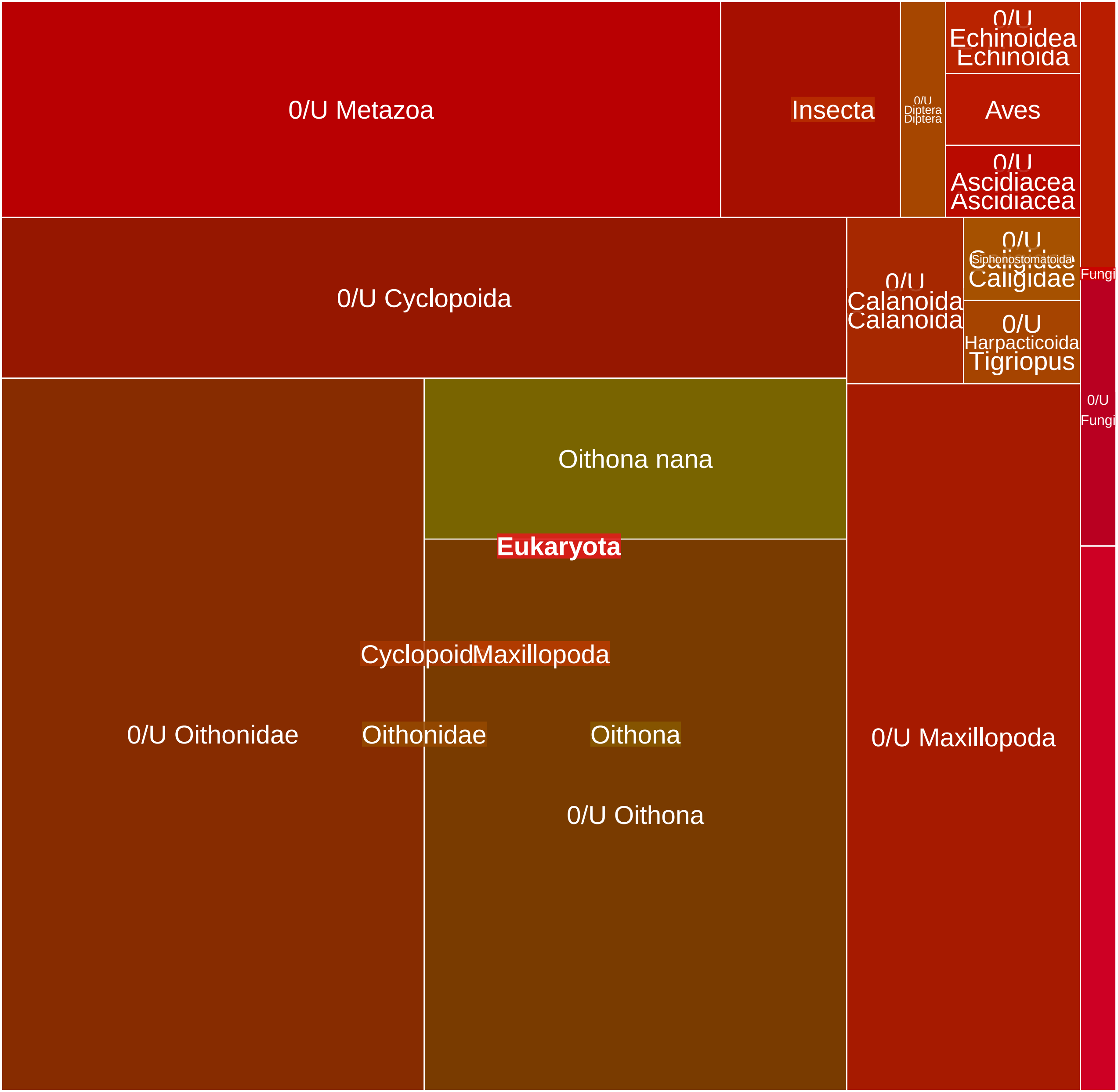

MGT−v1_633 : 1021 unigenes, 780 (76.40%) taxonomically assigned

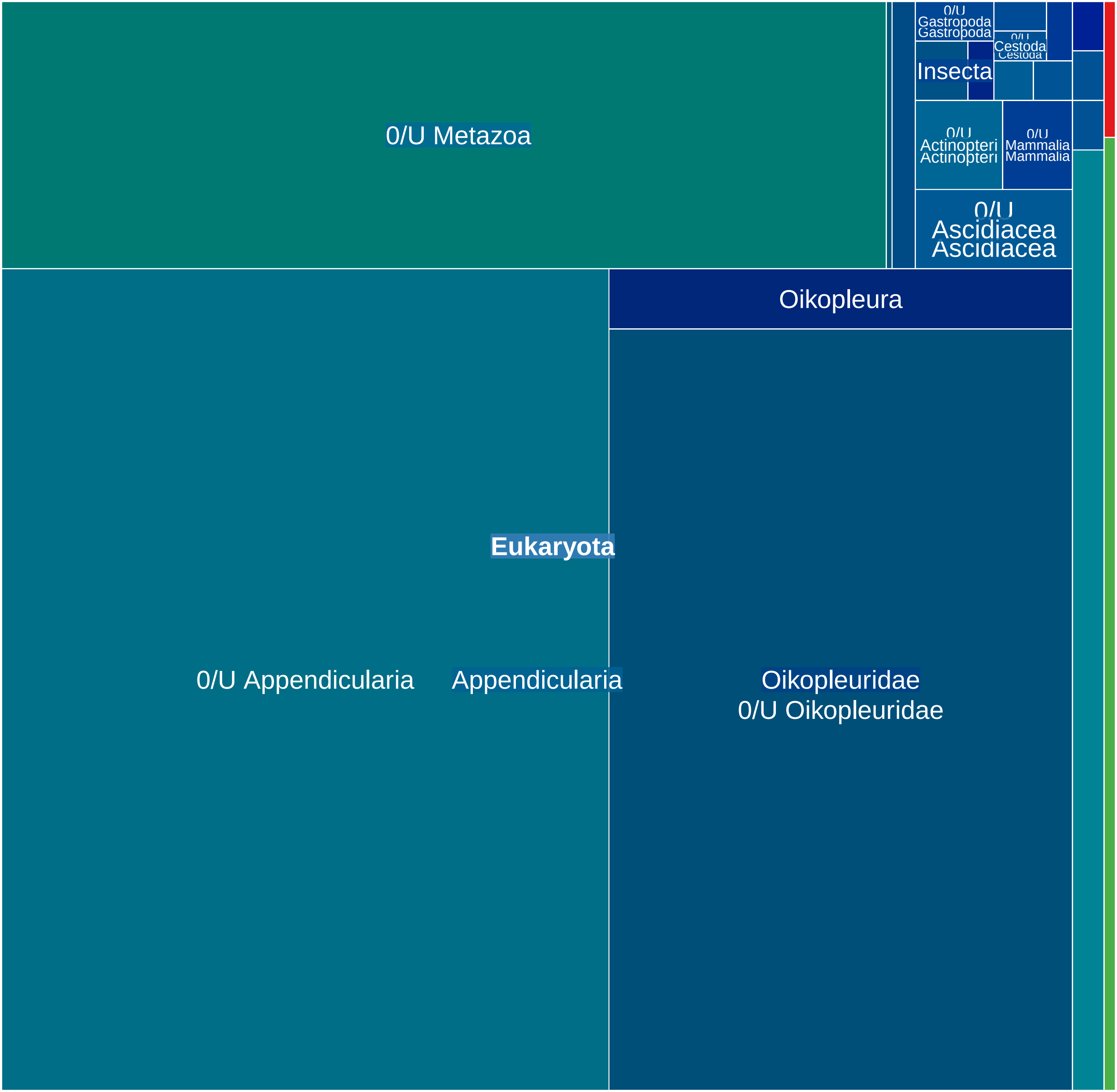

MGT−v1_634 : 1018 unigenes, 59 (5.80%) taxonomically assigned

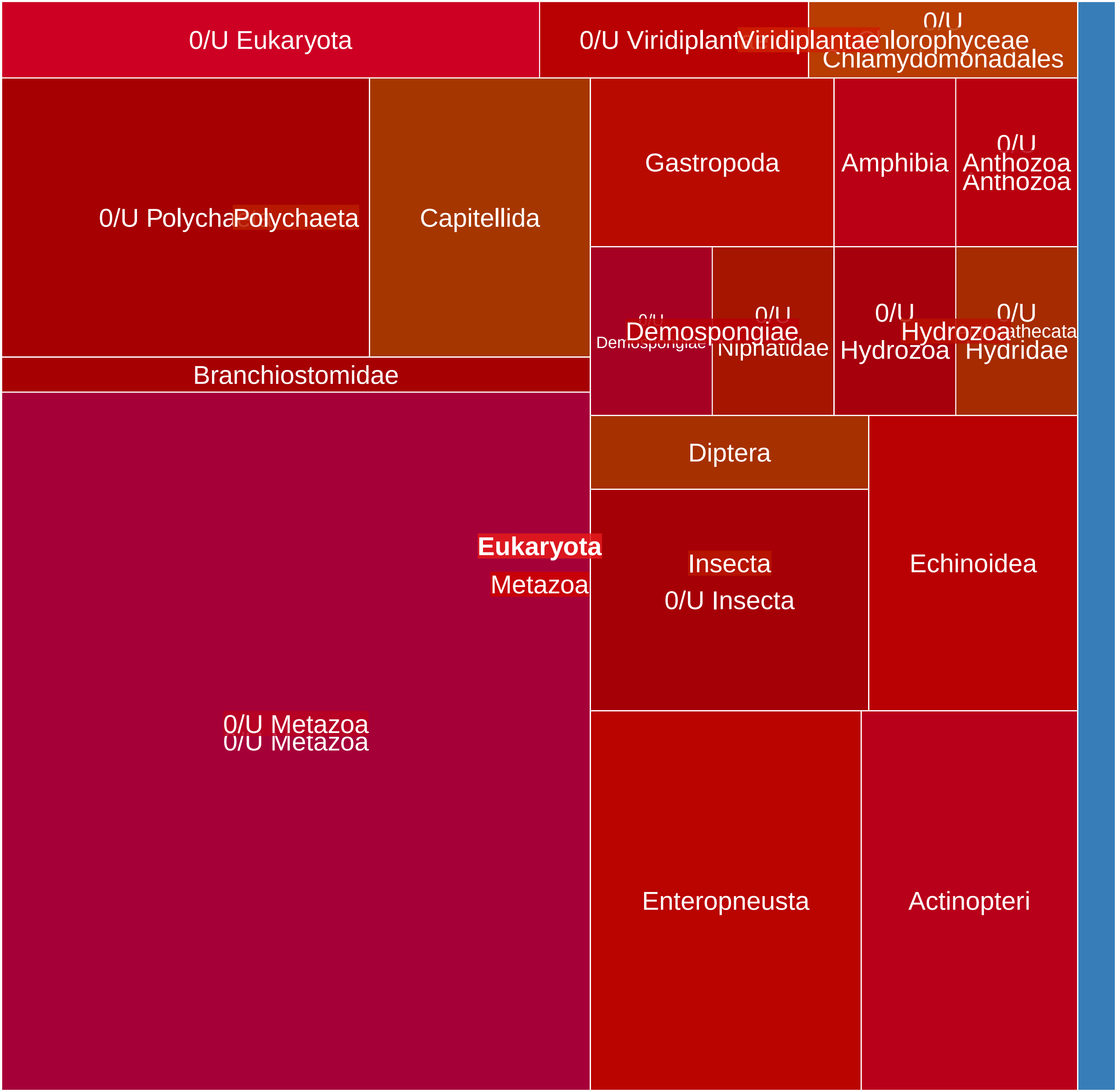

MGT−v1_635 : 1015 unigenes, 71 (7.00%) taxonomically assigned

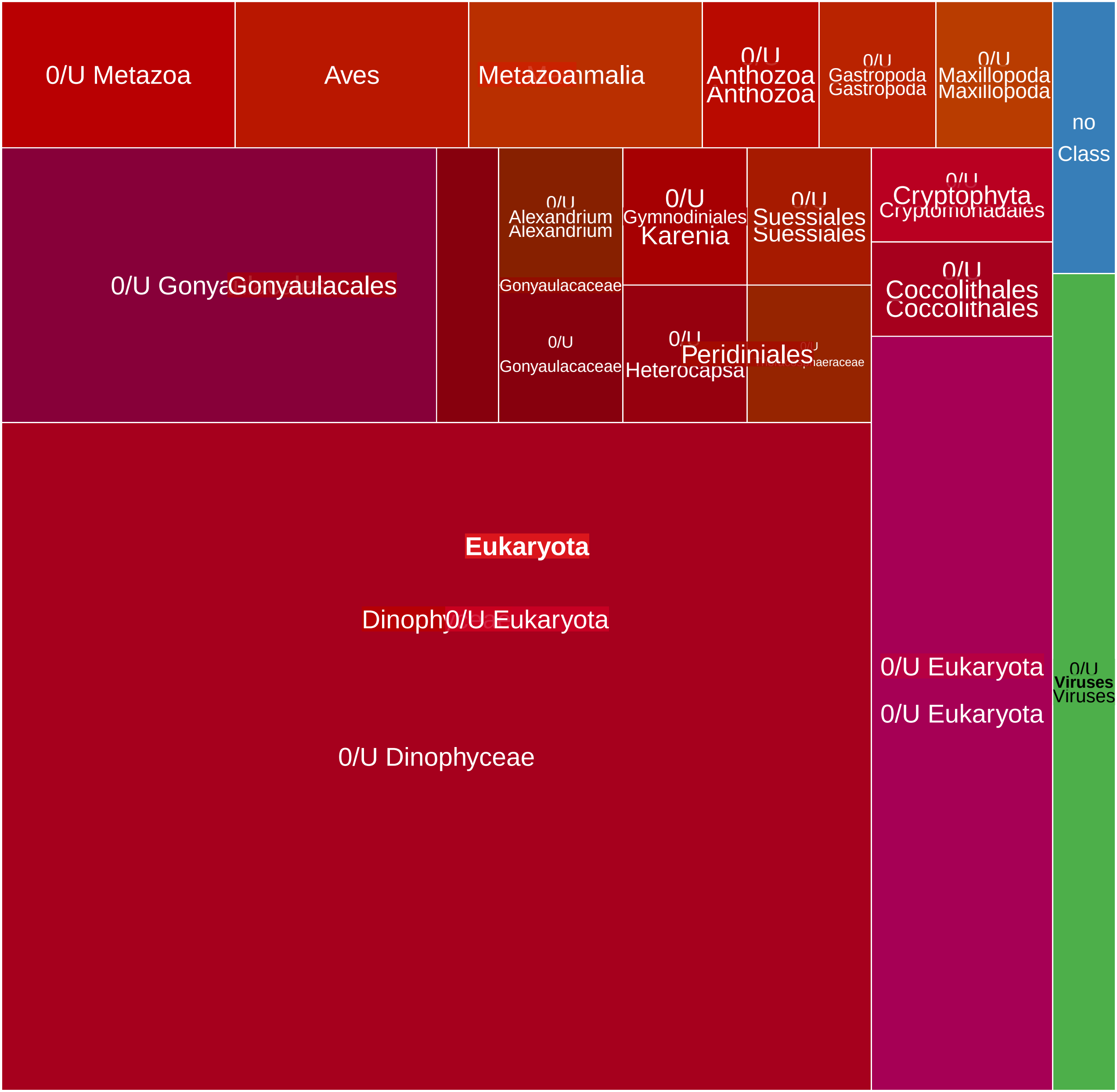

MGT−v1_636 : 1008 unigenes, 193 (19.15%) taxonomically assigned

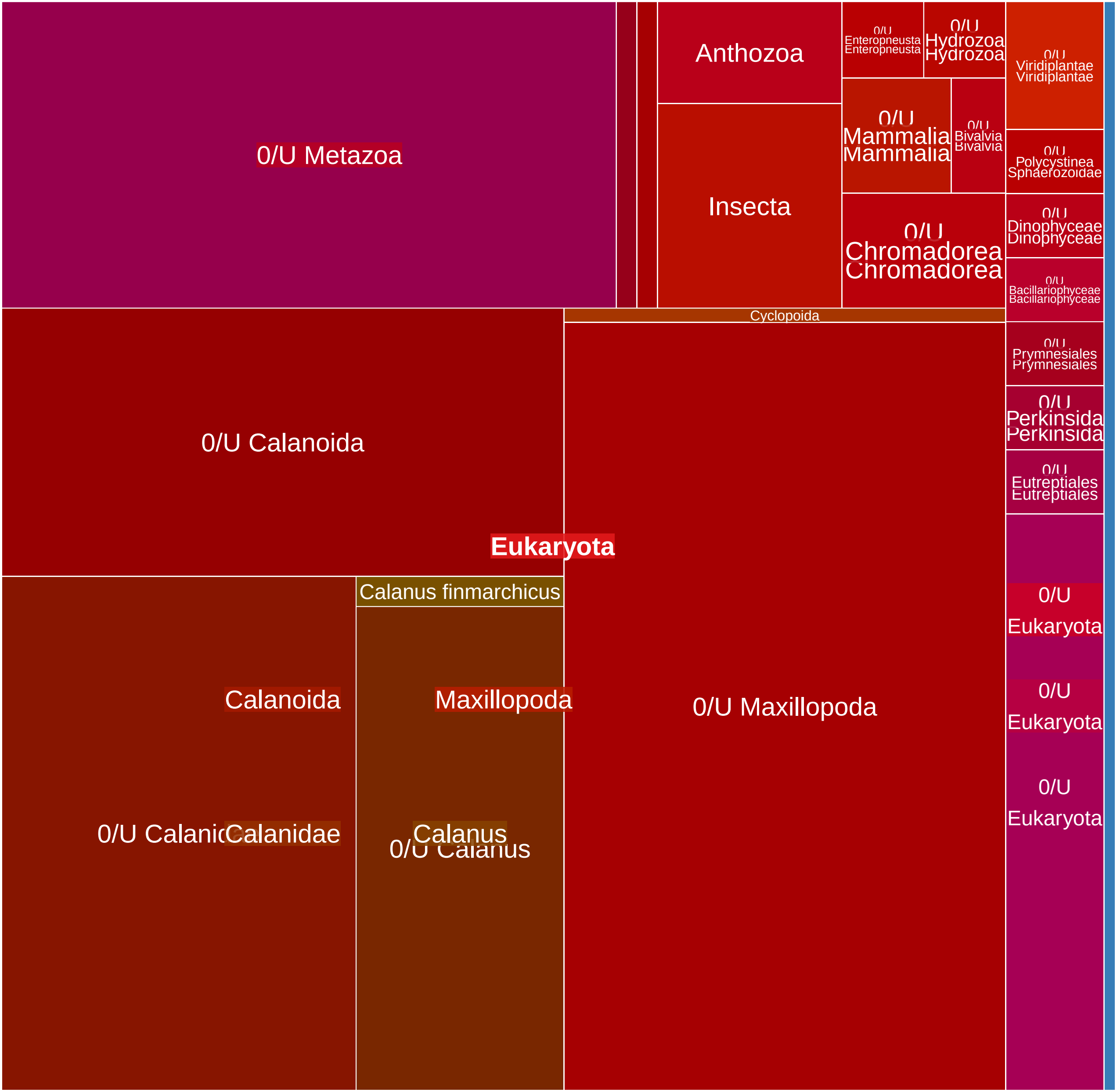

MGT−v1_637 : 1007 unigenes, 570 (56.60%) taxonomically assigned

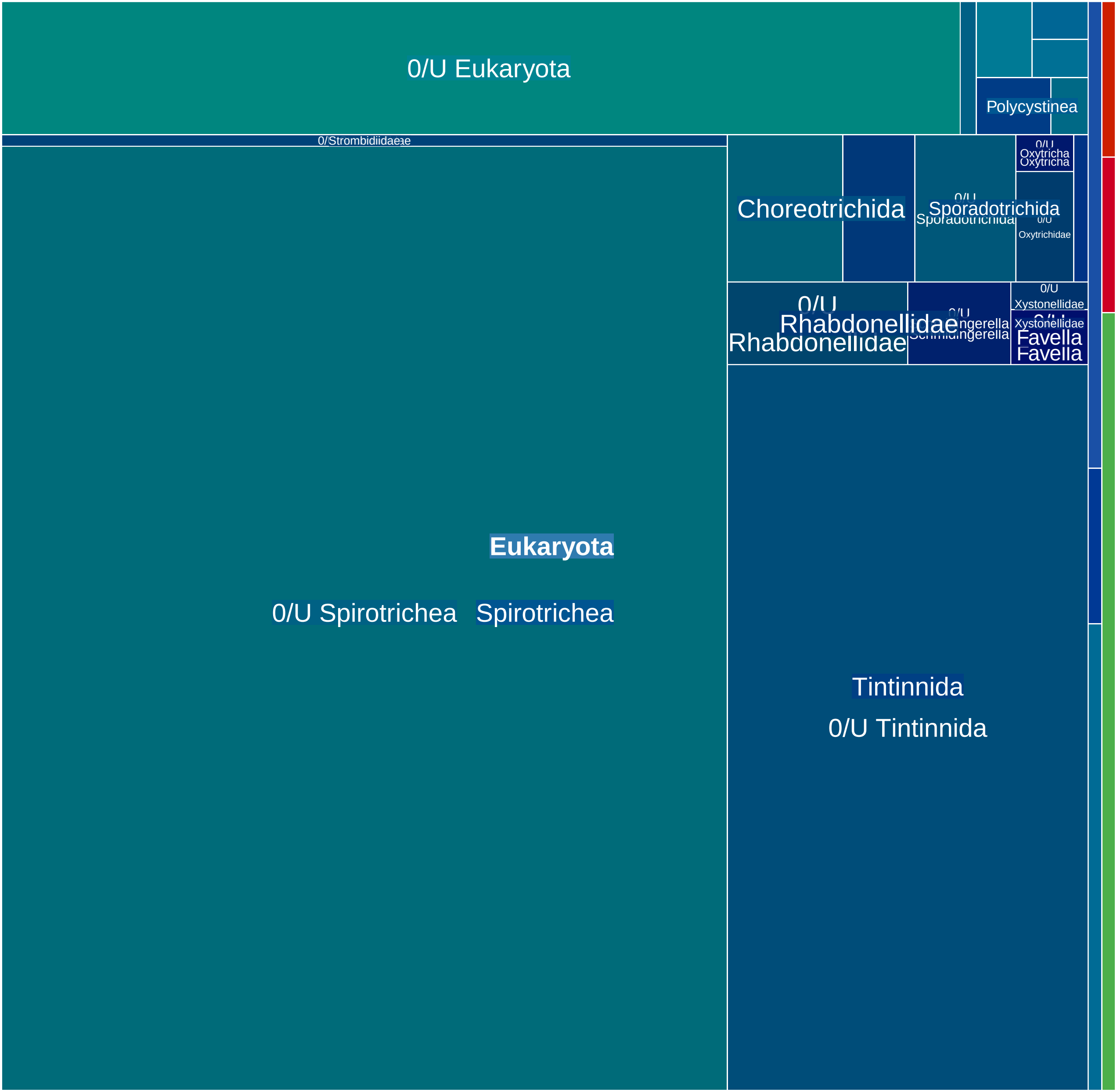

MGT−v1_638 : 995 unigenes, 108 (10.85%) taxonomically assigned

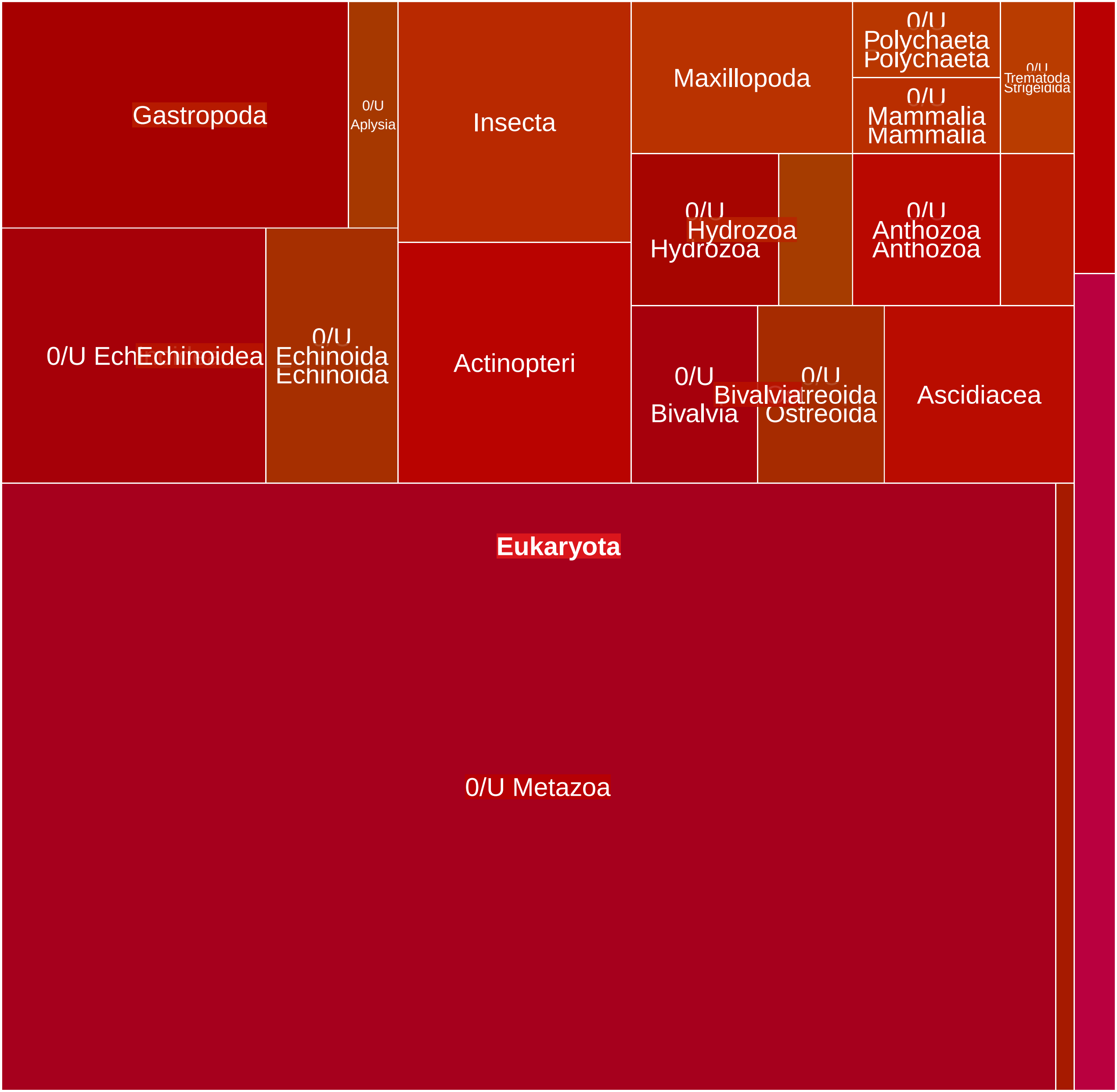

MGT−v1_639 : 993 unigenes, 141 (14.20%) taxonomically assigned

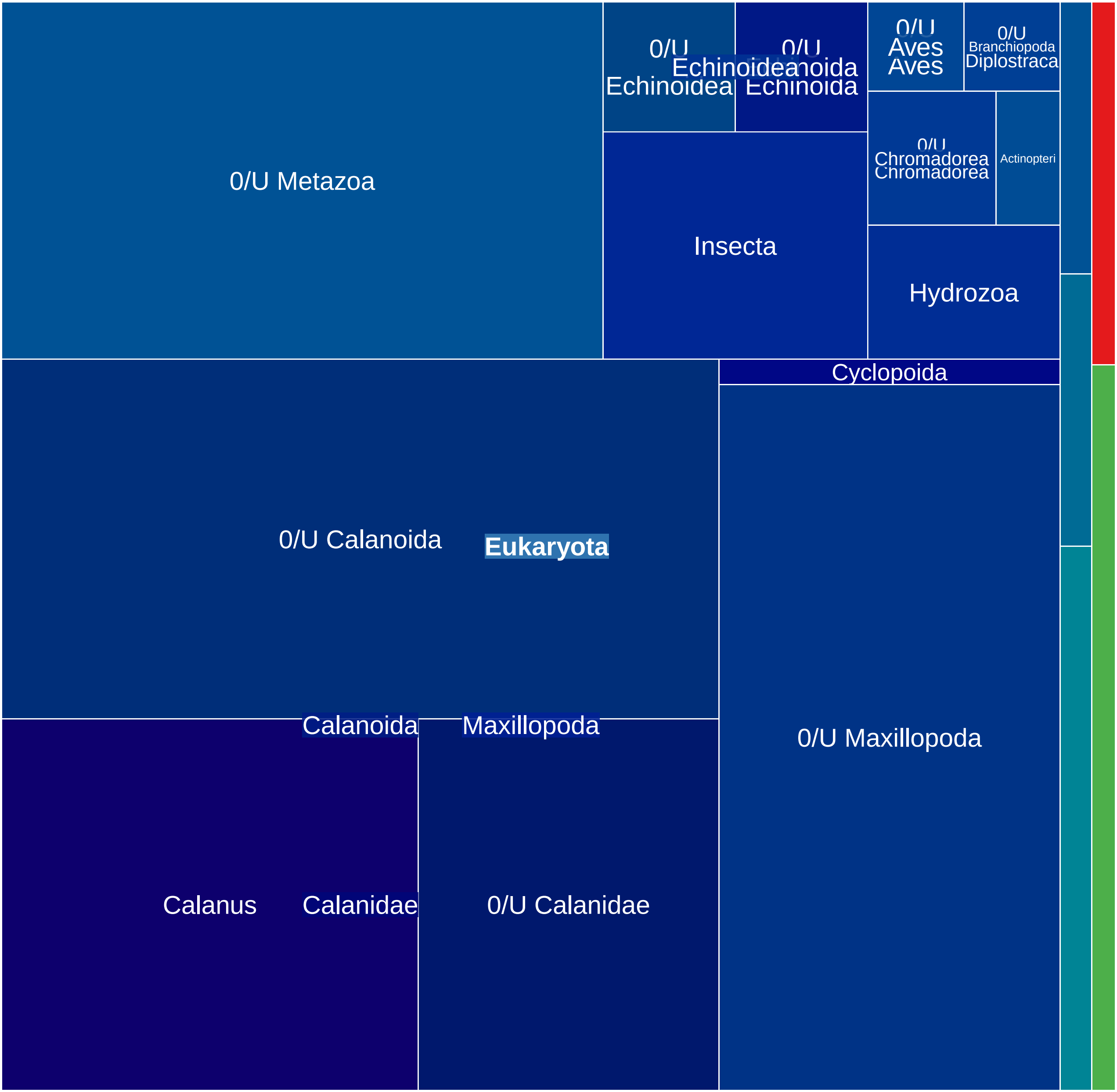

MGT−v1_640 : 988 unigenes, 971 (98.28%) taxonomically assigned

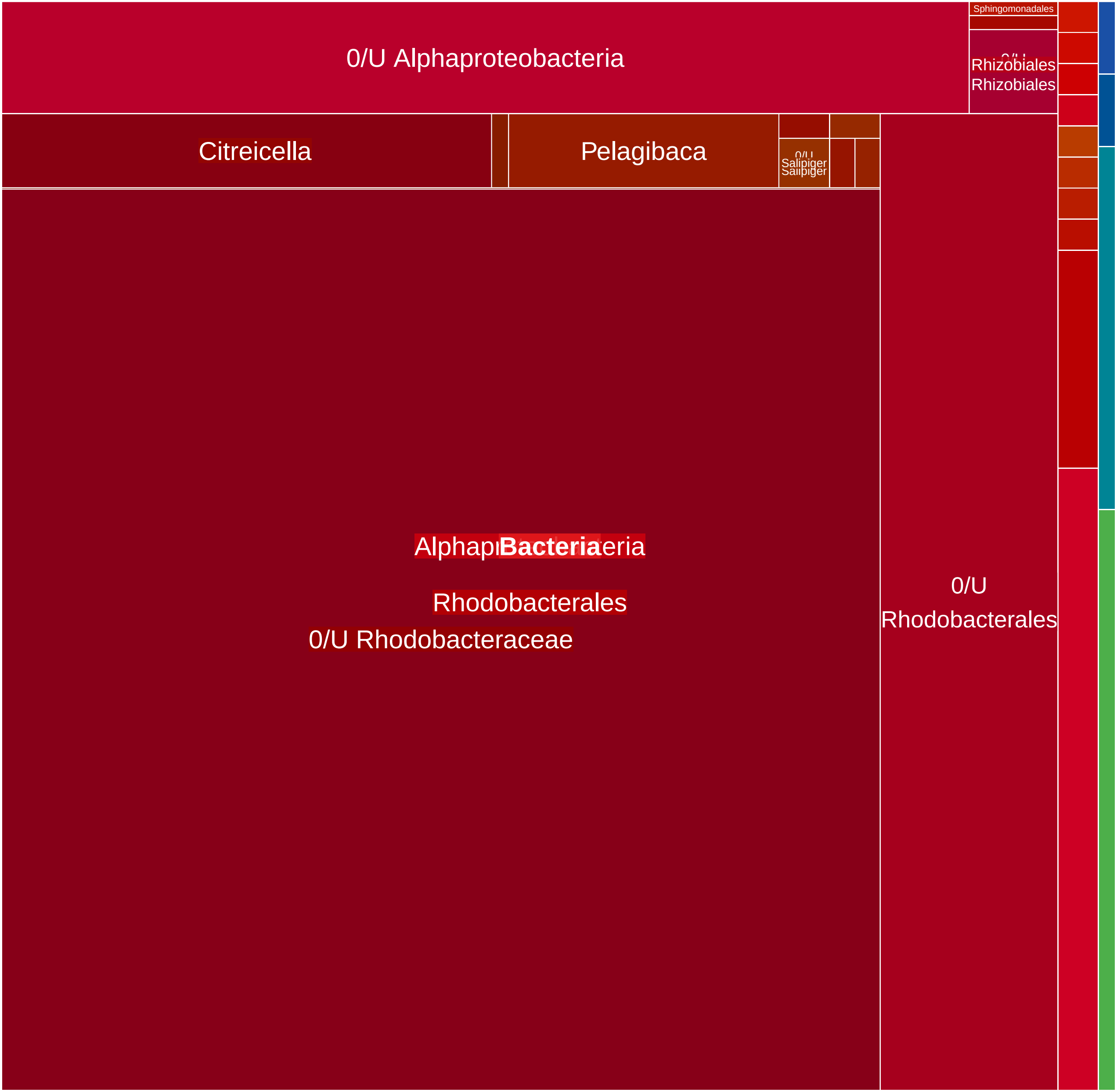

MGT−v1_641 : 980 unigenes, 64 (6.53%) taxonomically assigned

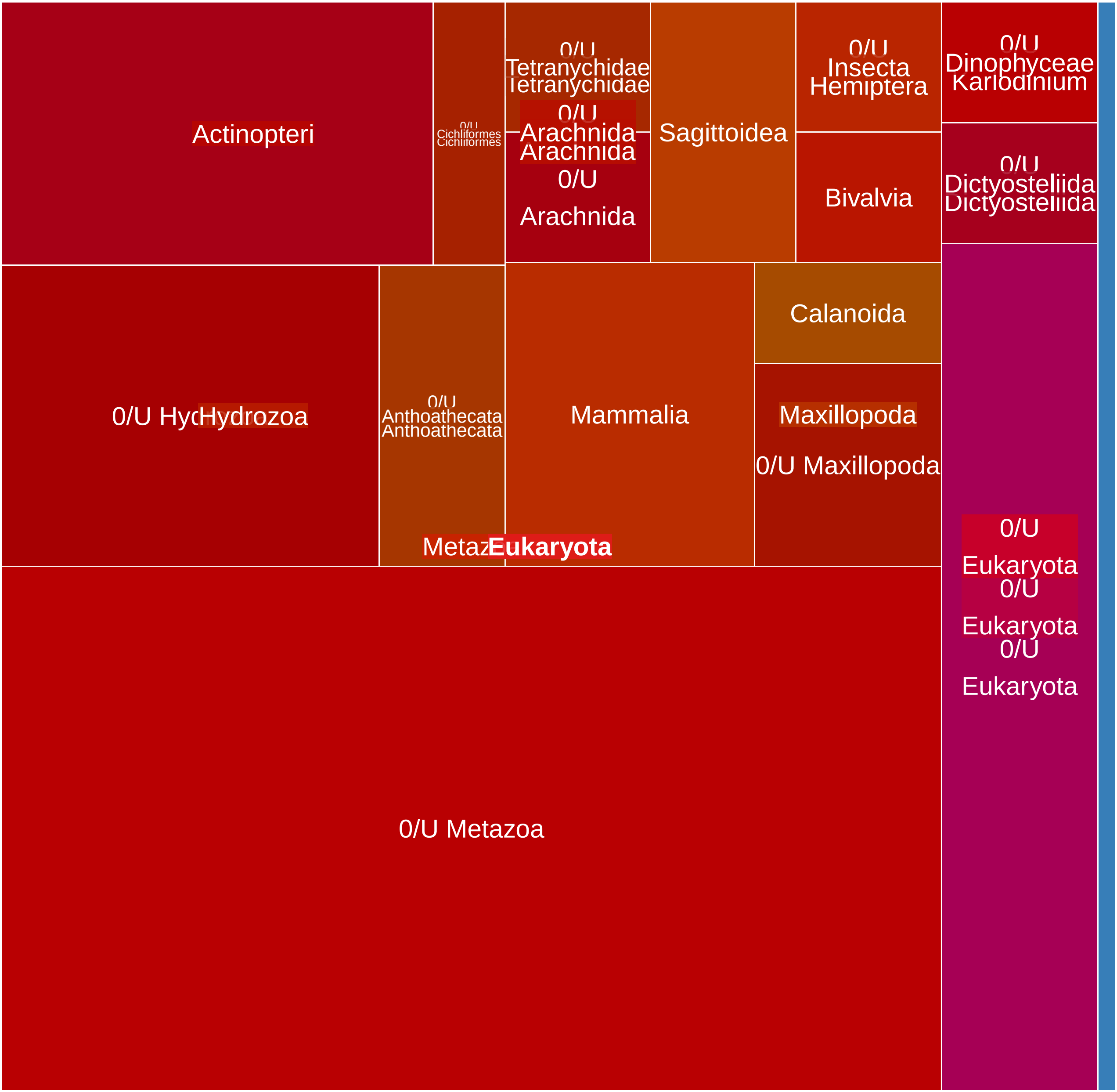

MGT−v1_642 : 980 unigenes, 112 (11.43%) taxonomically assigned

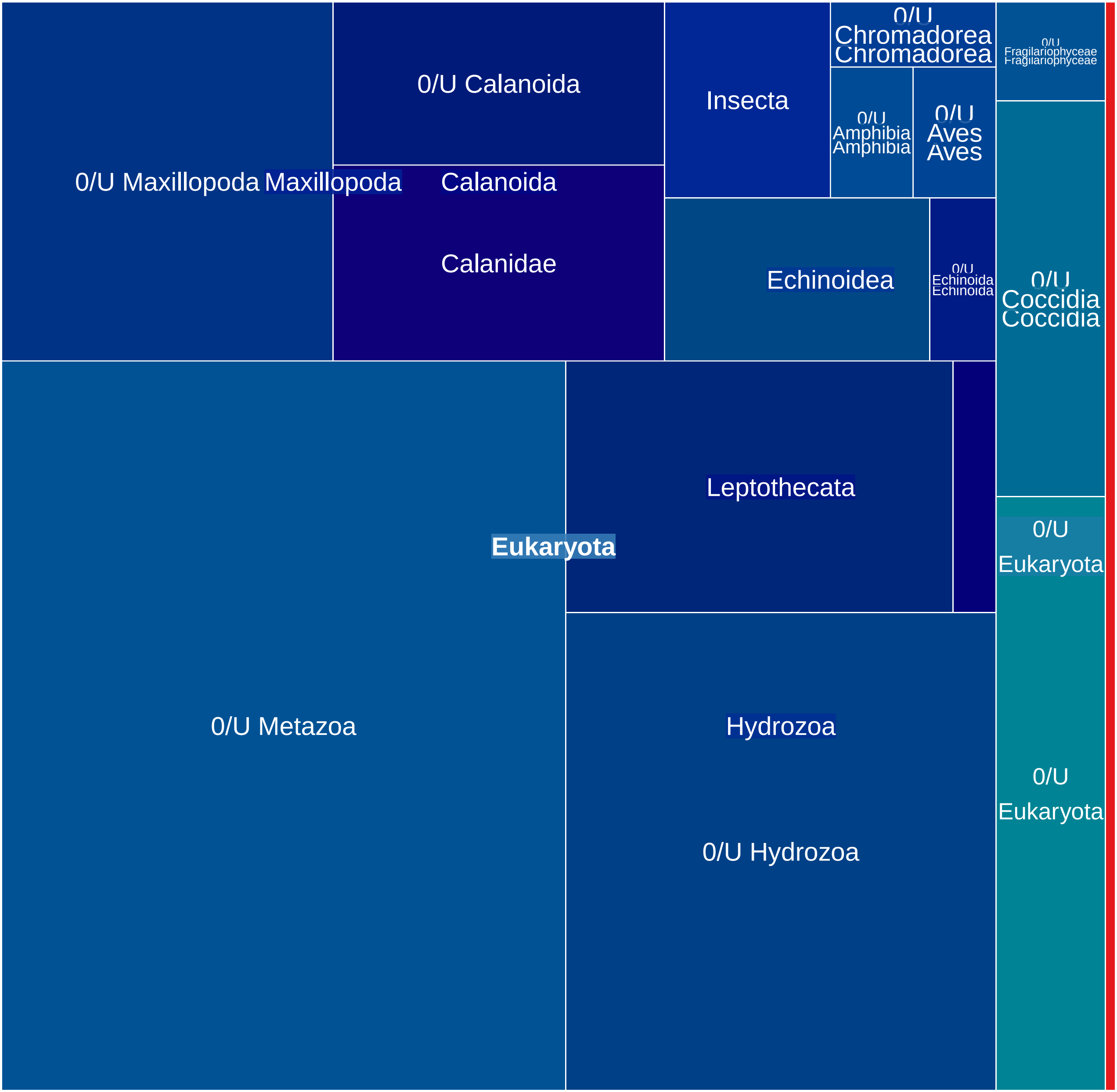

MGT−v1_643 : 978 unigenes, 907 (92.74%) taxonomically assigned

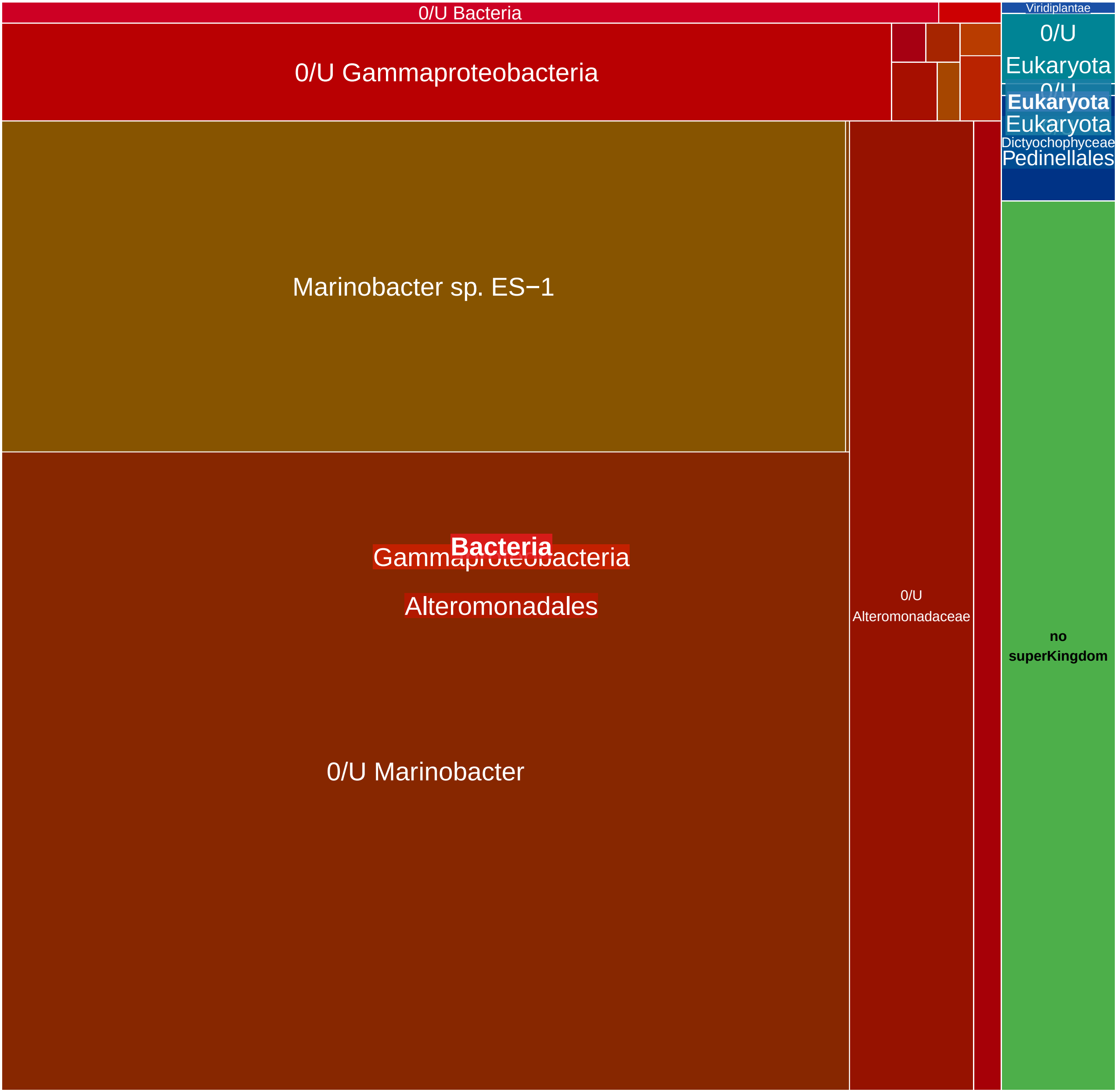

MGT−v1_644 : 978 unigenes, 679 (69.43%) taxonomically assigned

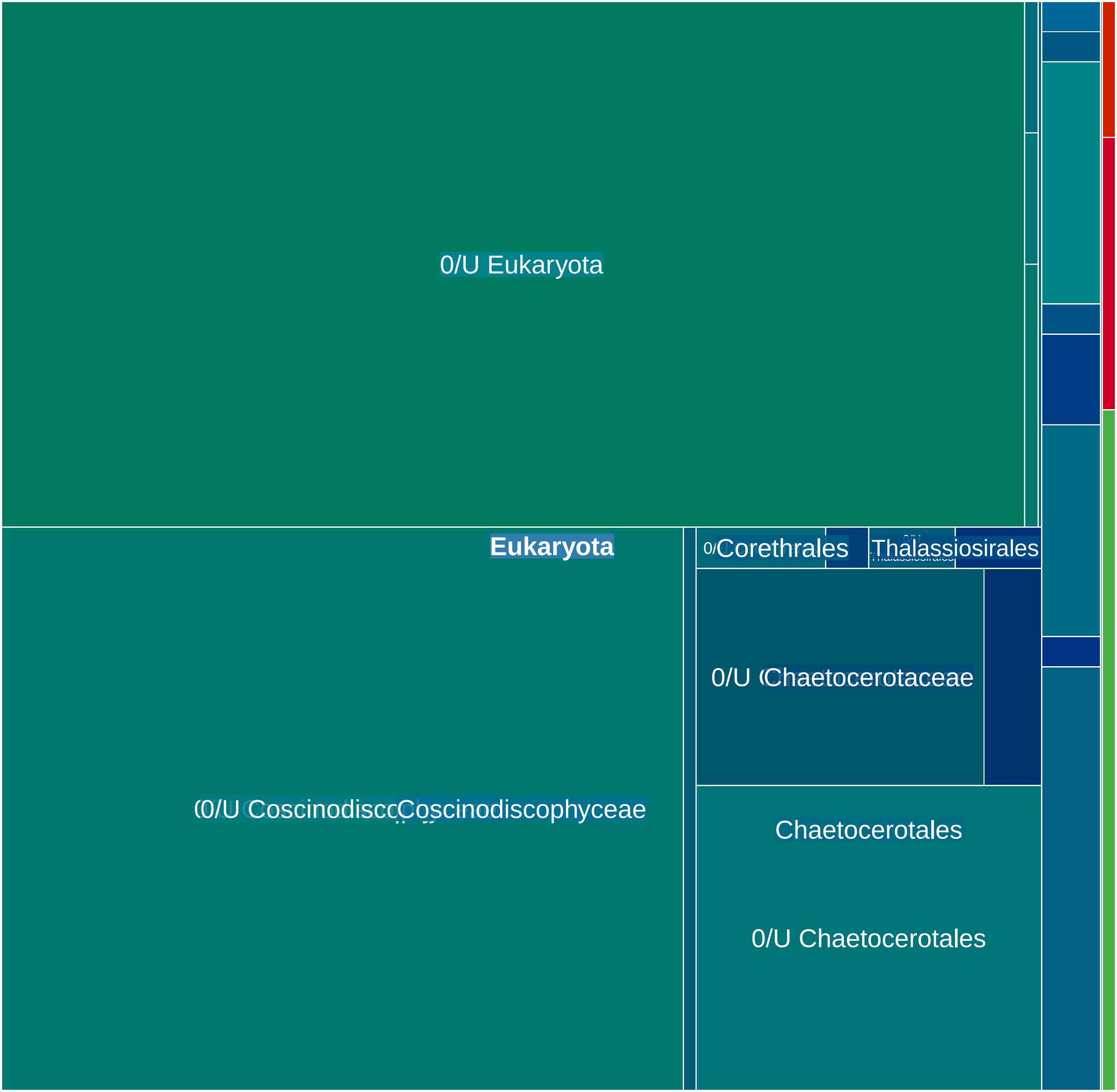

MGT−v1_645 : 978 unigenes, 297 (30.37%) taxonomically assigned

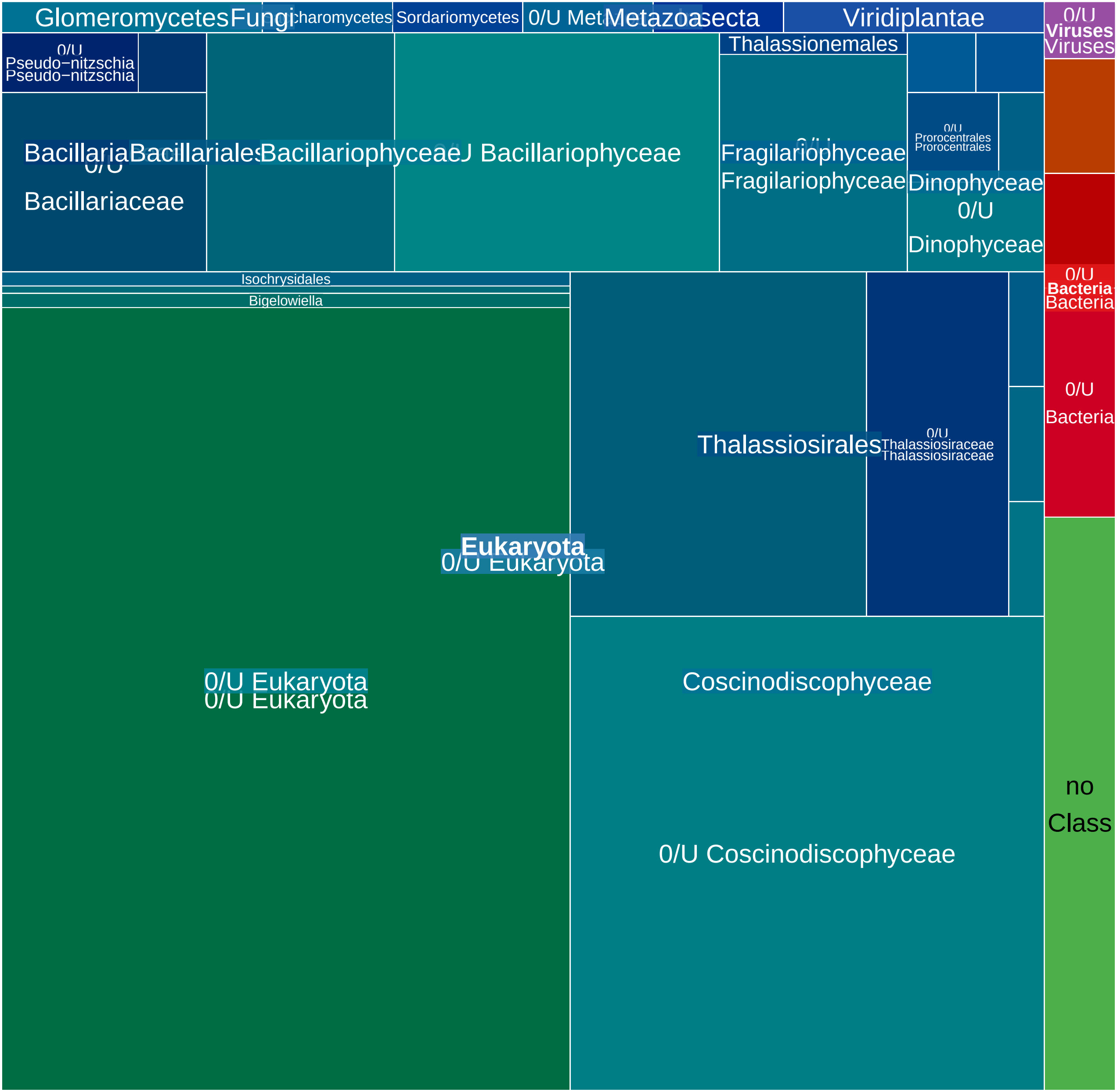

MGT−v1_646 : 978 unigenes, 216 (22.09%) taxonomically assigned

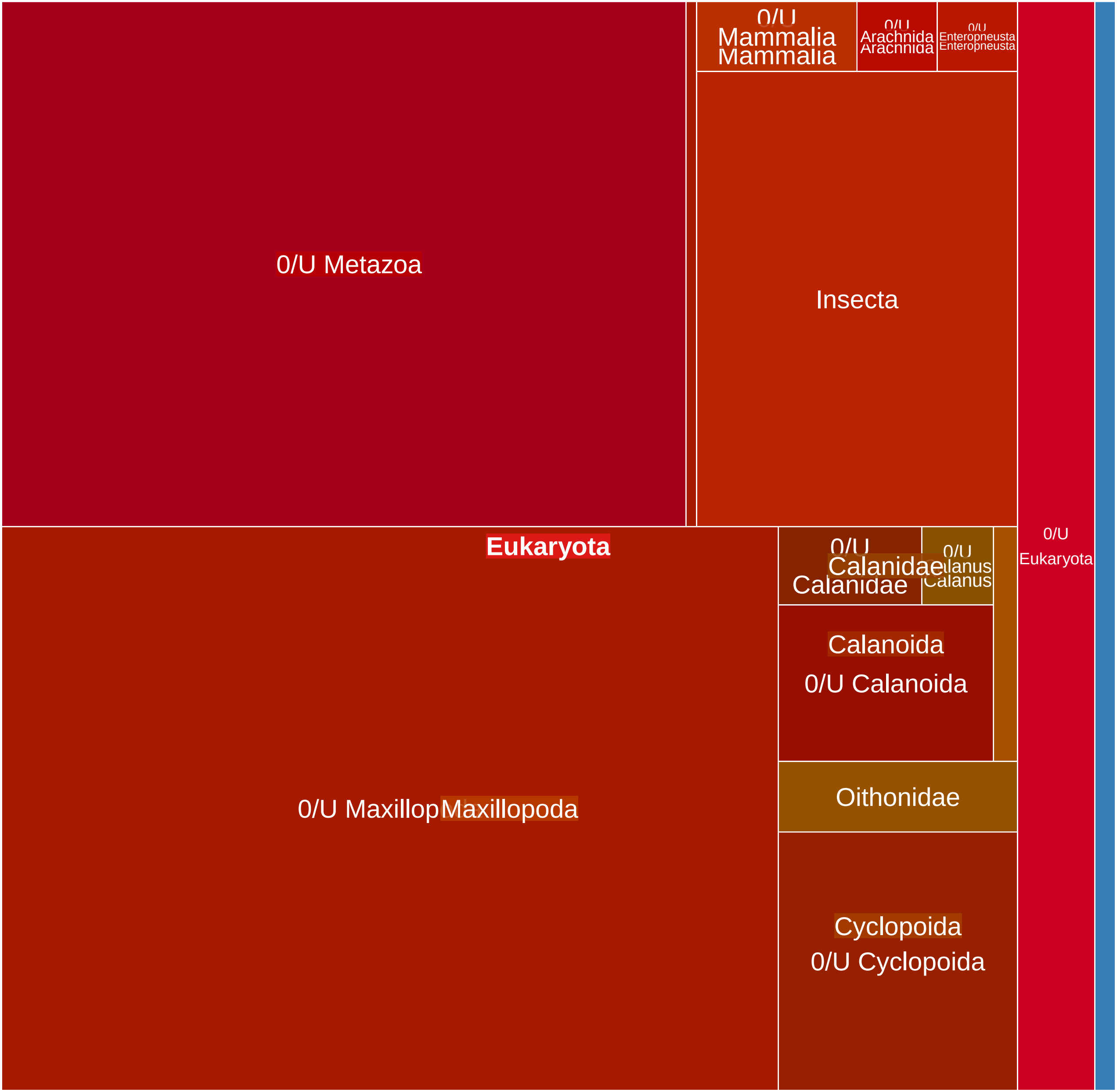

### MGT−v1_647 : 977 unigenes, 962 (98.46%) taxonomically assigned

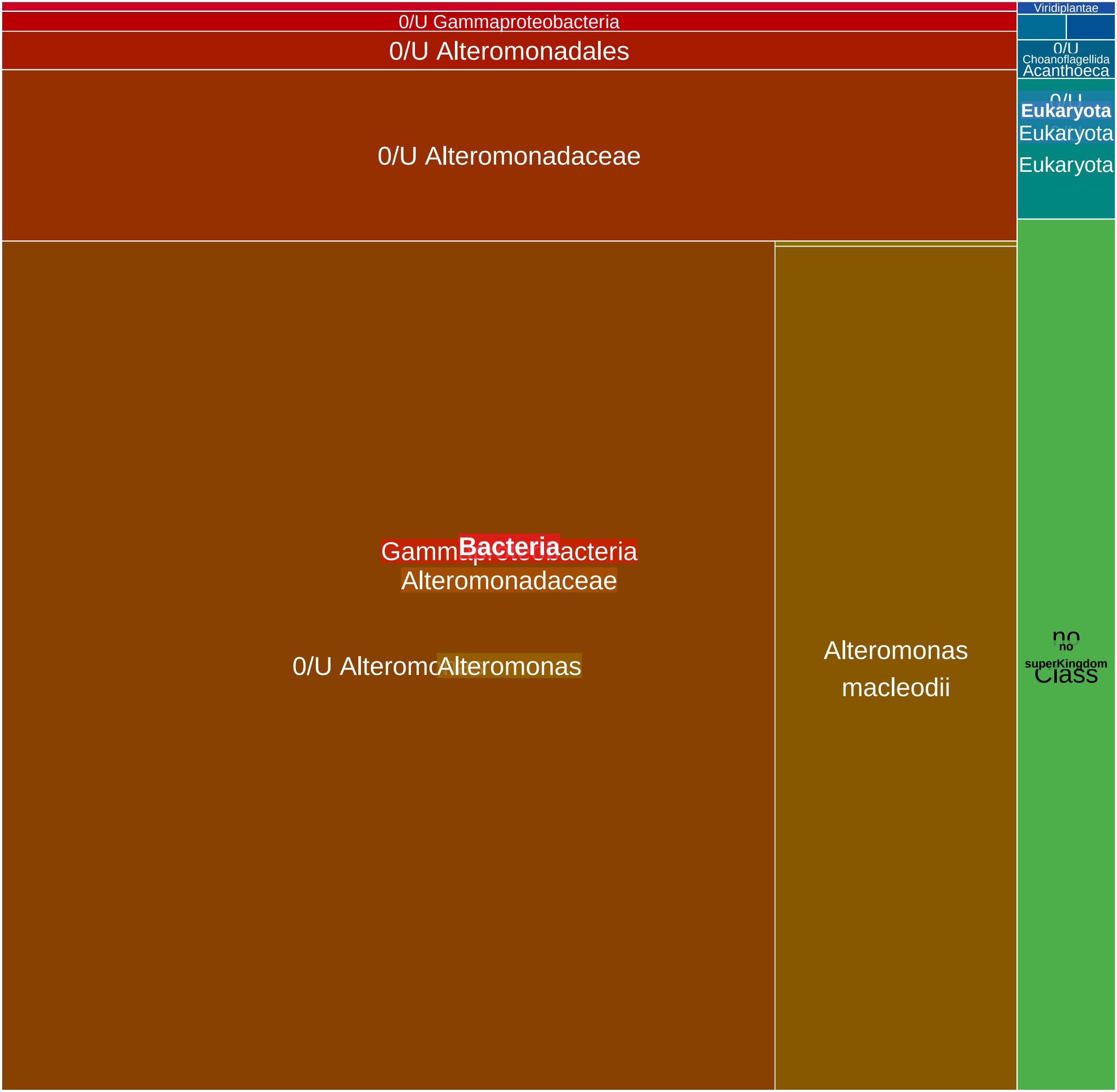

MGT−v1_648 : 977 unigenes, 79 (8.09%) taxonomically assigned

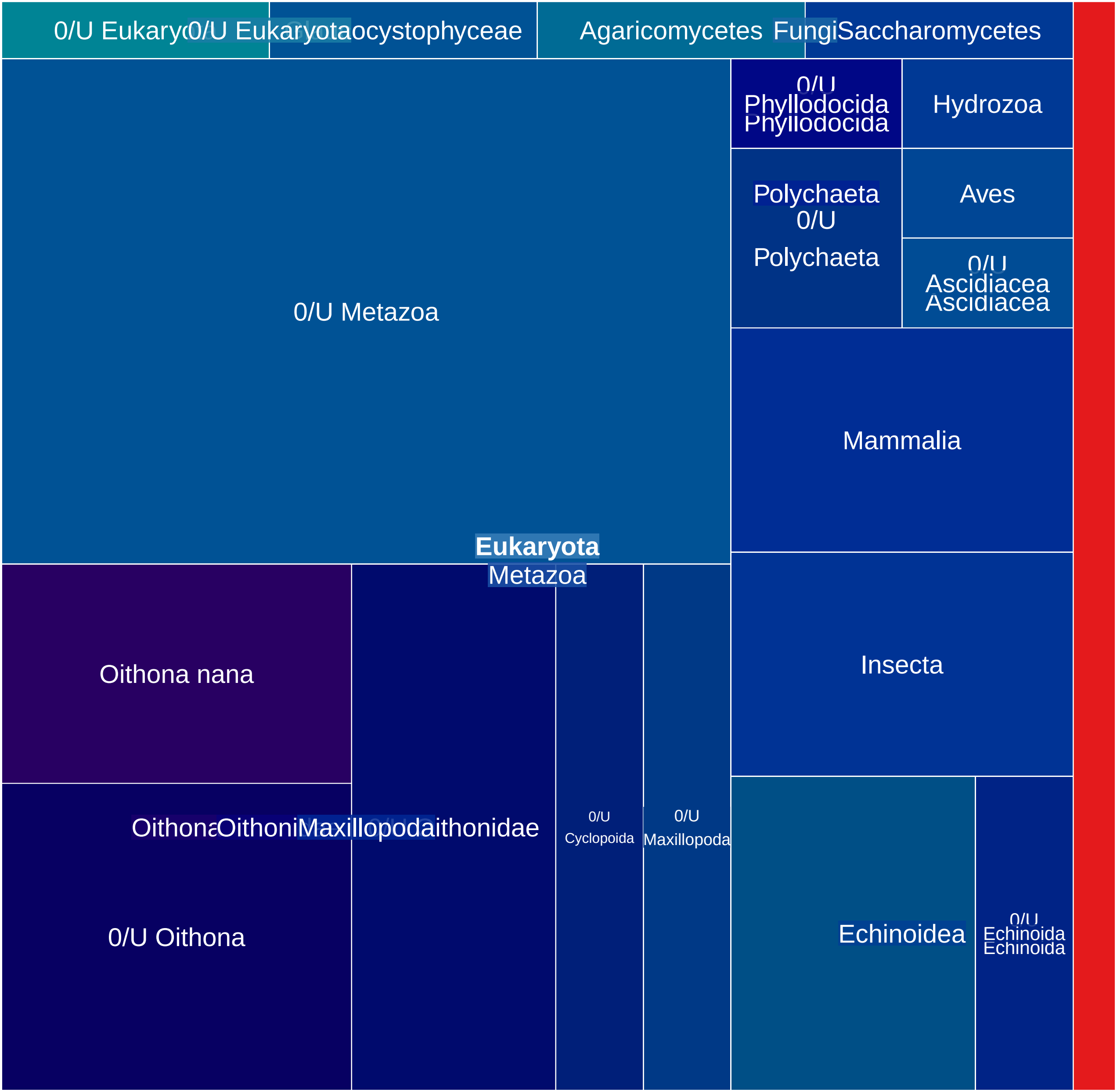

MGT−v1_649 : 976 unigenes, 944 (96.72%) taxonomically assigned

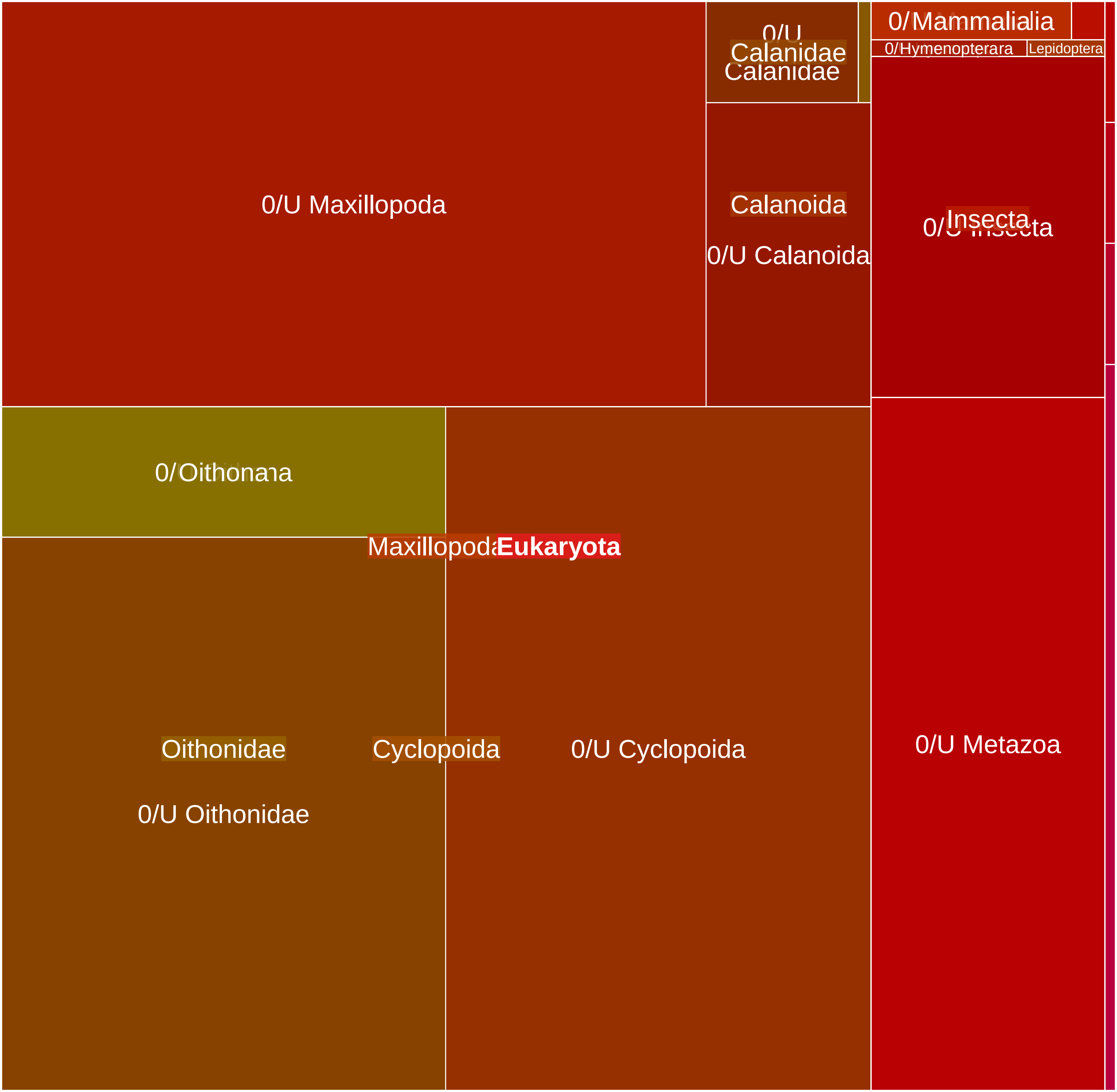

MGT−v1_650 : 976 unigenes, 300 (30.74%) taxonomically assigned

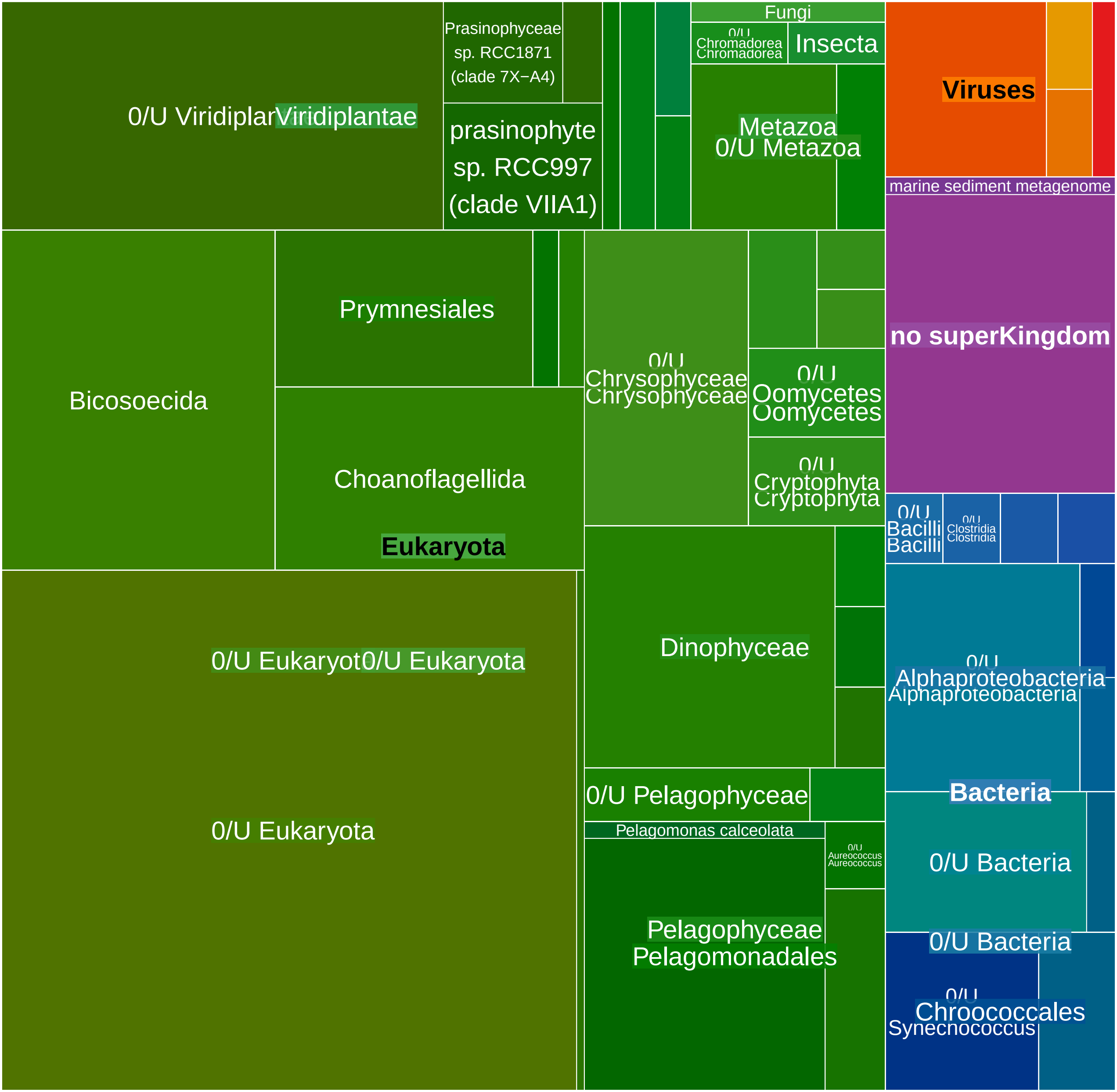

MGT−v1_651 : 975 unigenes, 854 (87.59%) taxonomically assigned

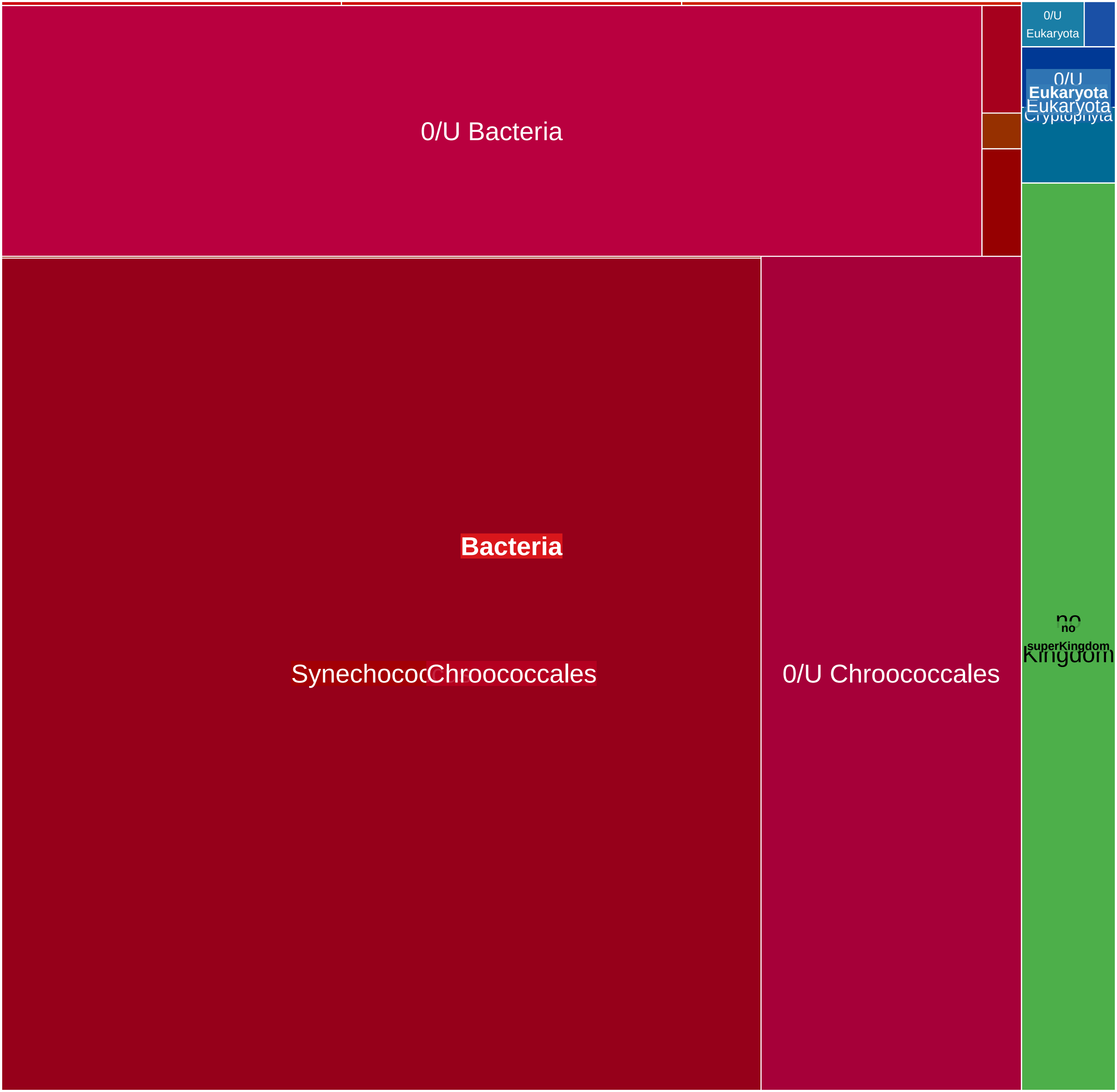

MGT−v1_652 : 974 unigenes, 782 (80.29%) taxonomically assigned

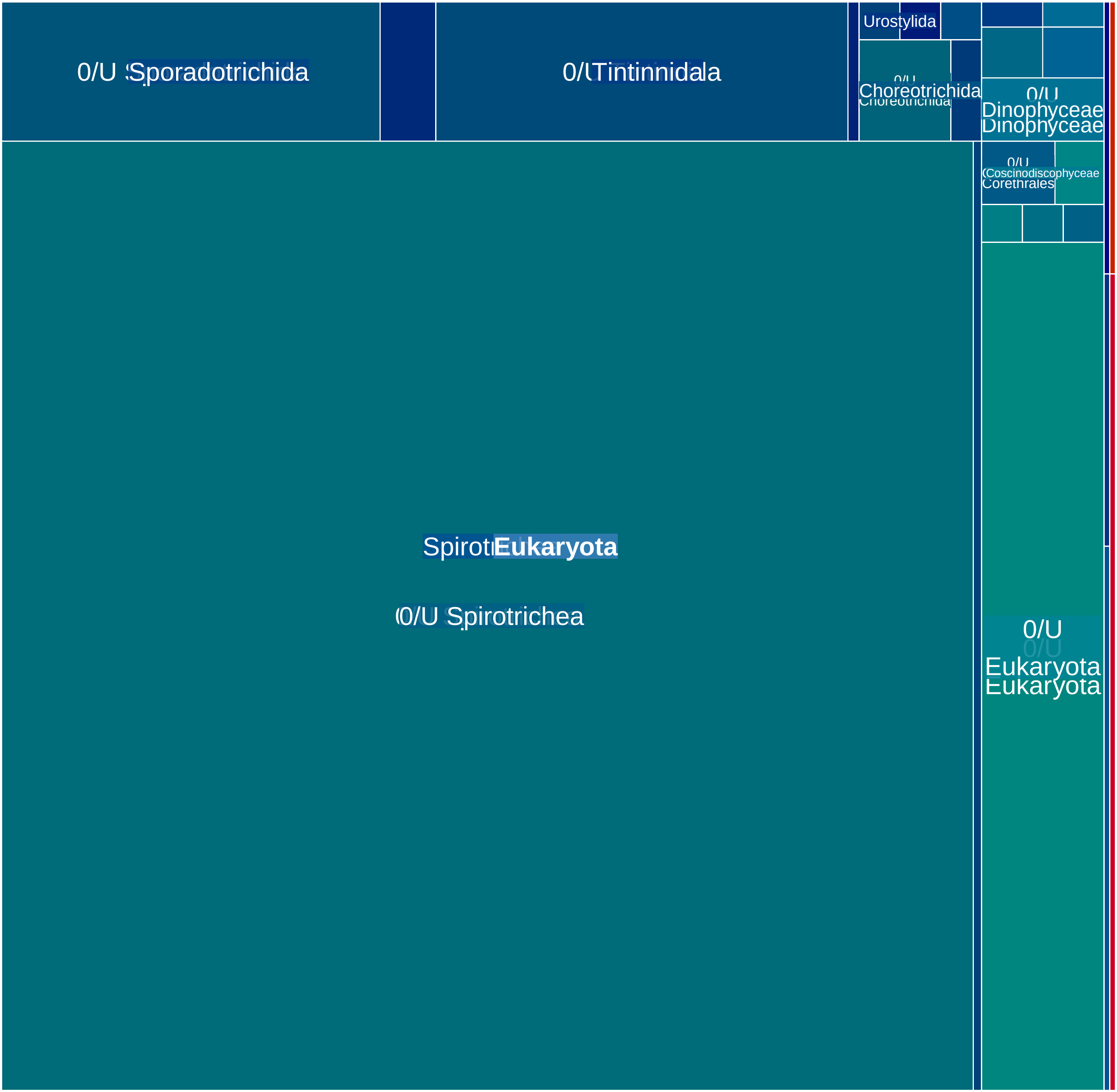

MGT−v1_653 : 973 unigenes, 232 (23.84%) taxonomically assigned

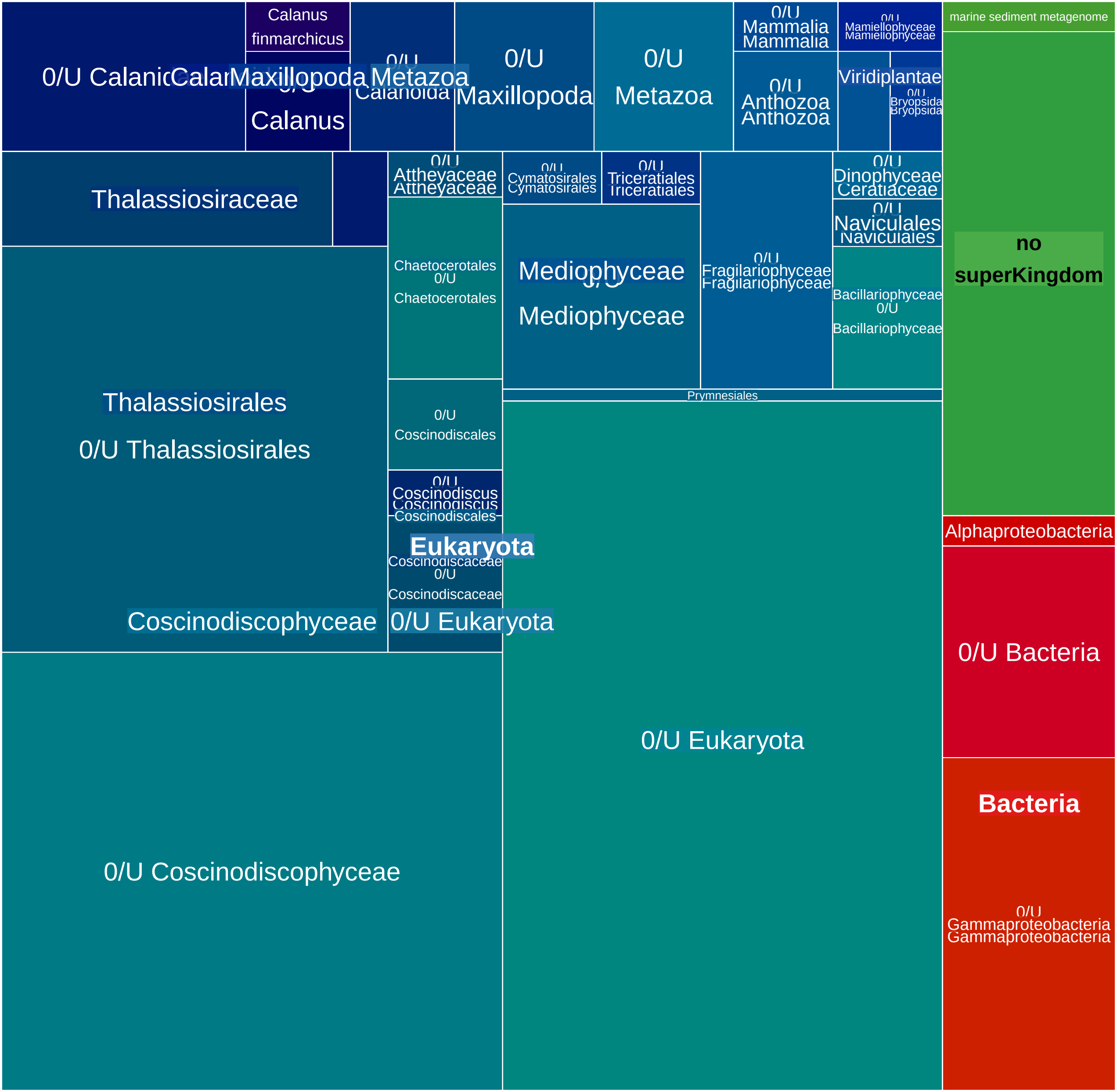

MGT−v1_654 : 970 unigenes, 134 (13.81%) taxonomically assigned

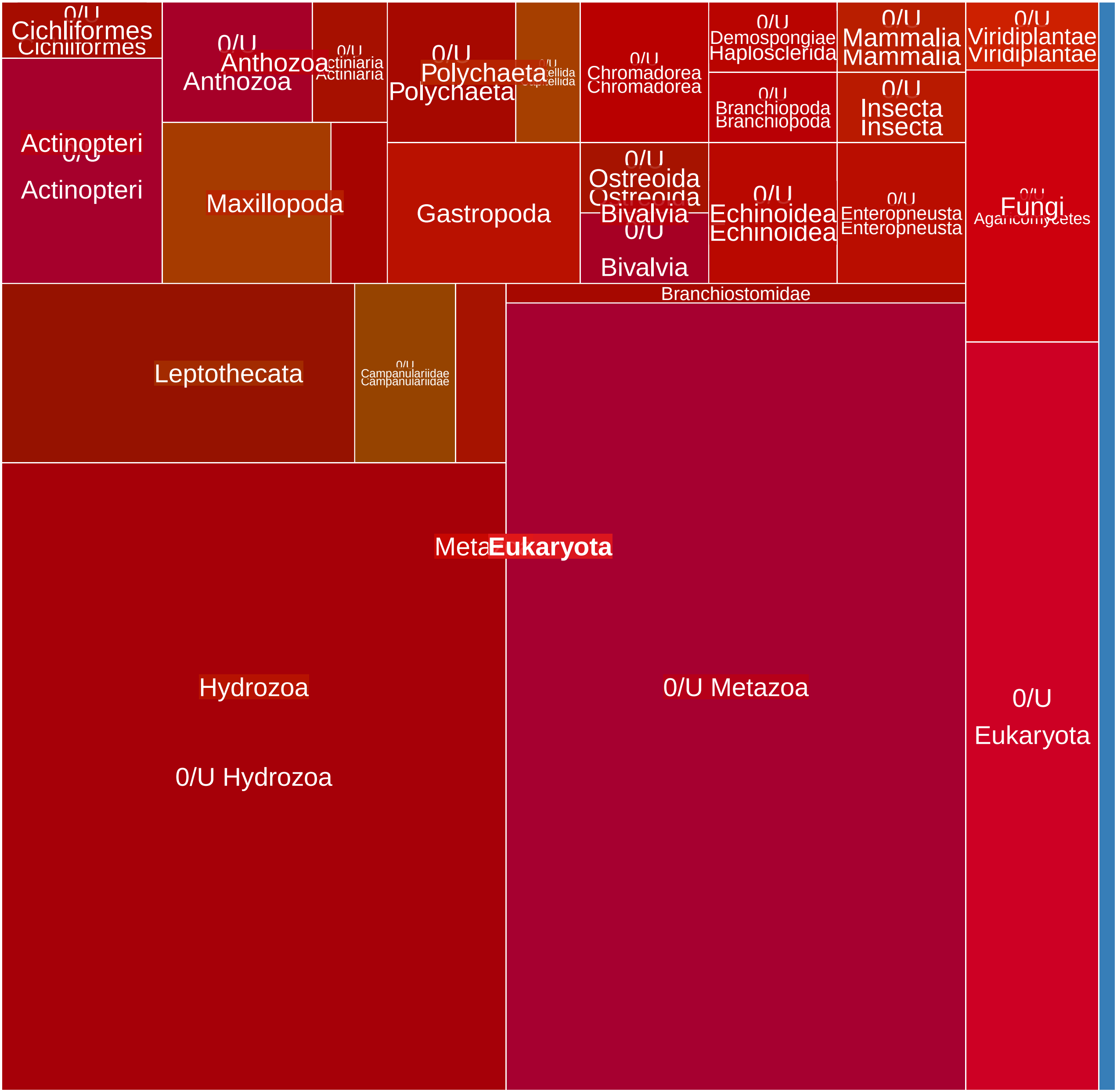

MGT−v1_655 : 969 unigenes, 115 (11.87%) taxonomically assigned

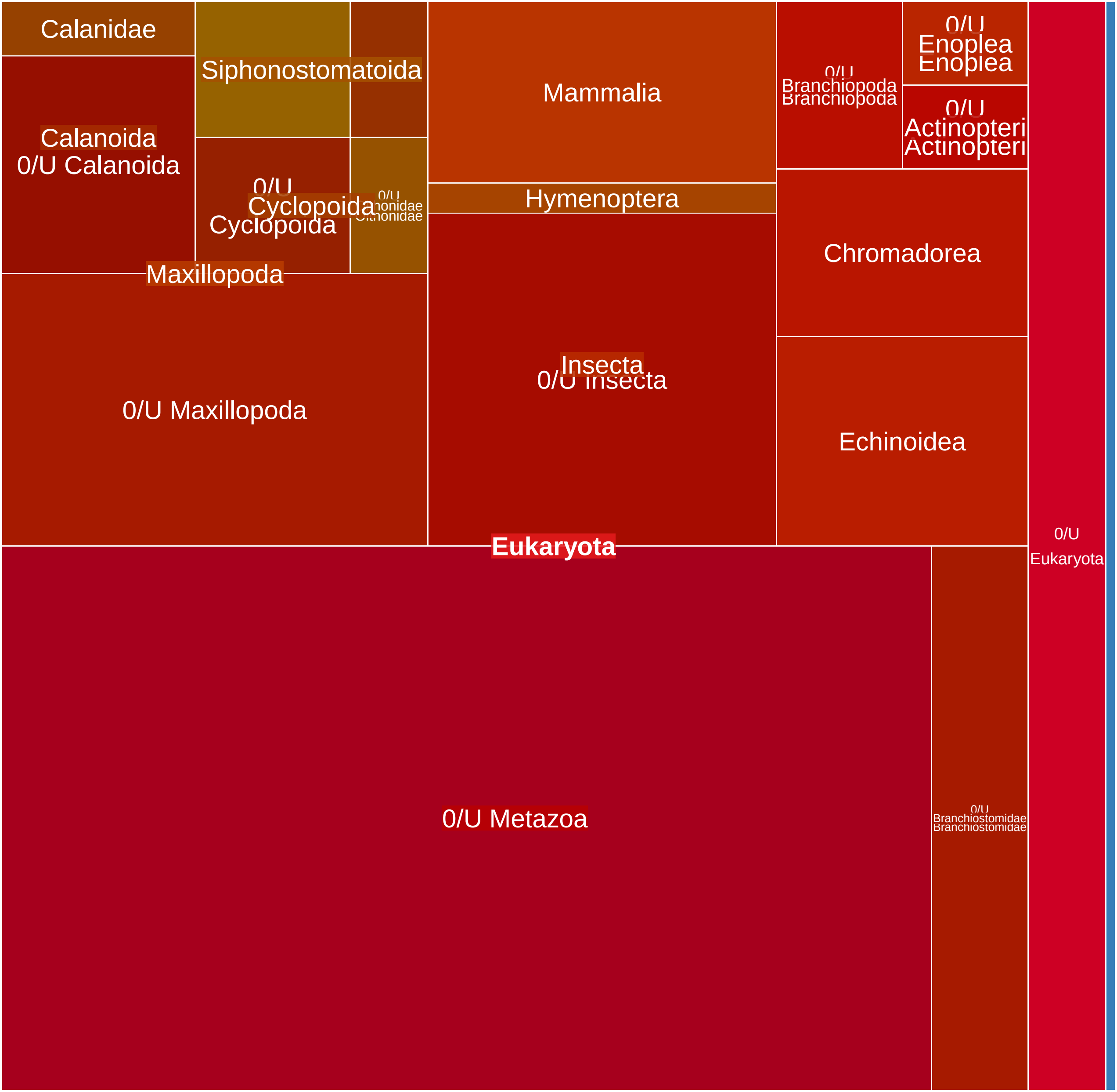

MGT−v1_656 : 968 unigenes, 714 (73.76%) taxonomically assigned

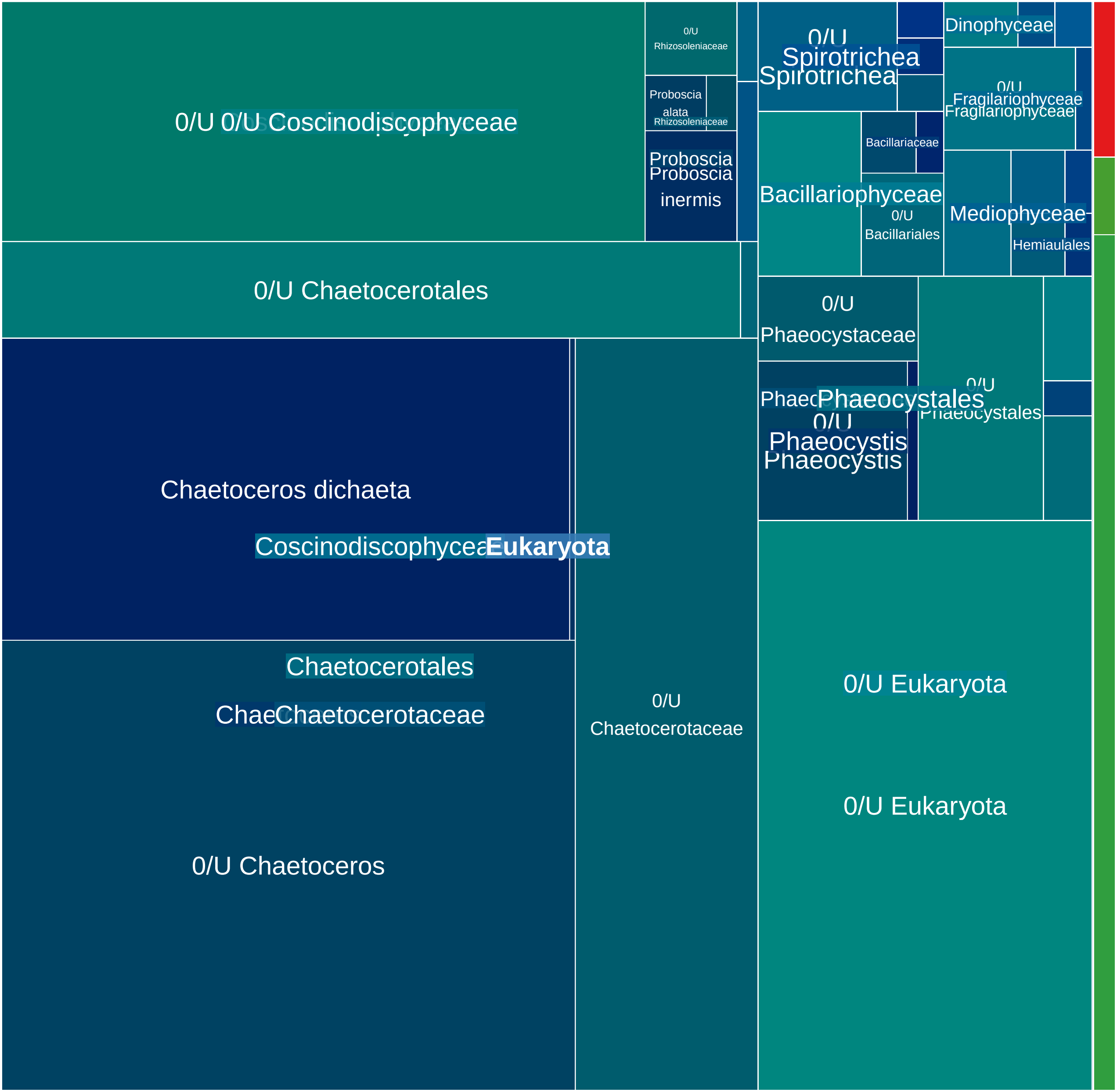

MGT−v1_657 : 967 unigenes, 603 (62.36%) taxonomically assigned

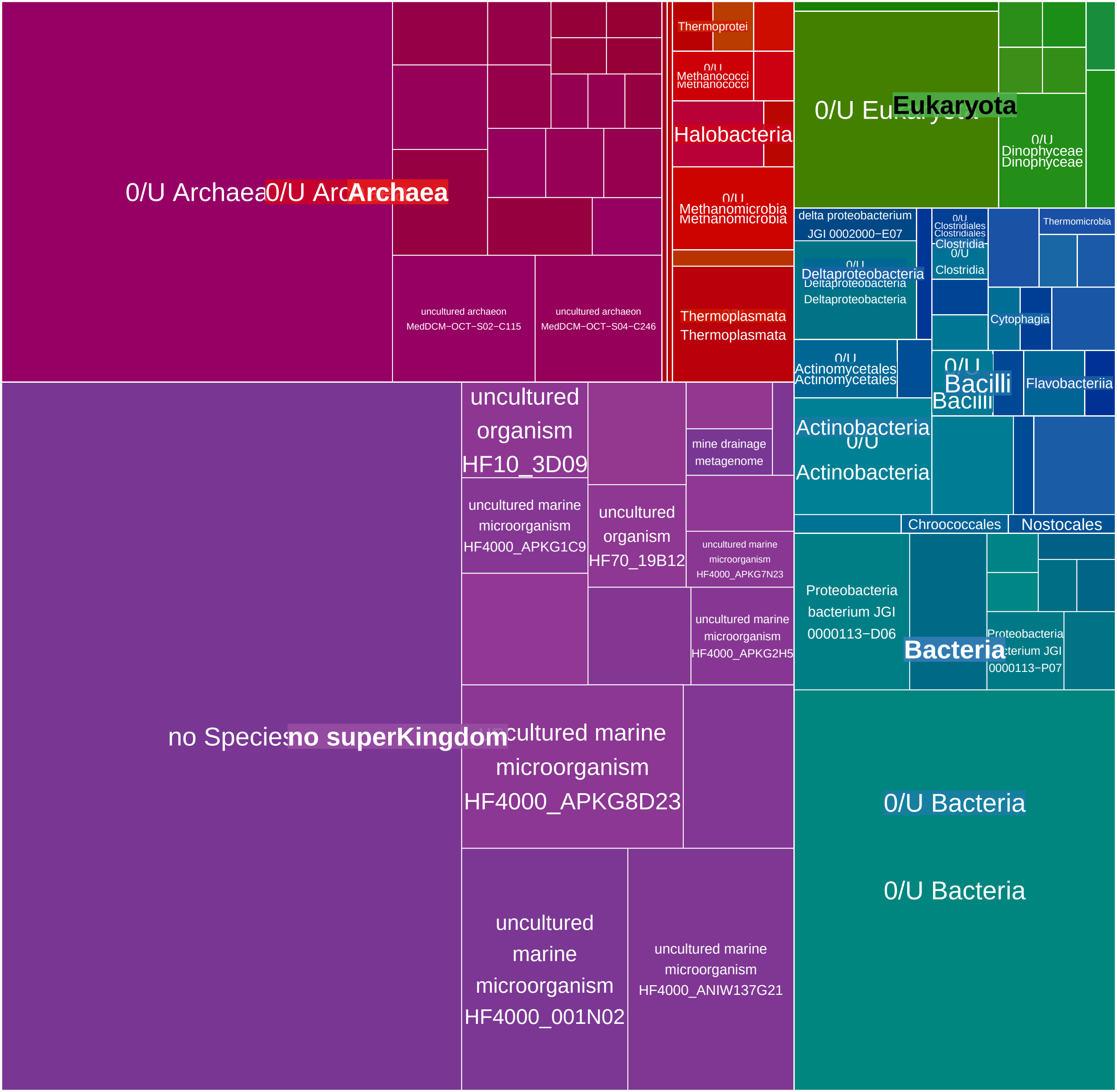

MGT−v1_658 : 966 unigenes, 692 (71.64%) taxonomically assigned

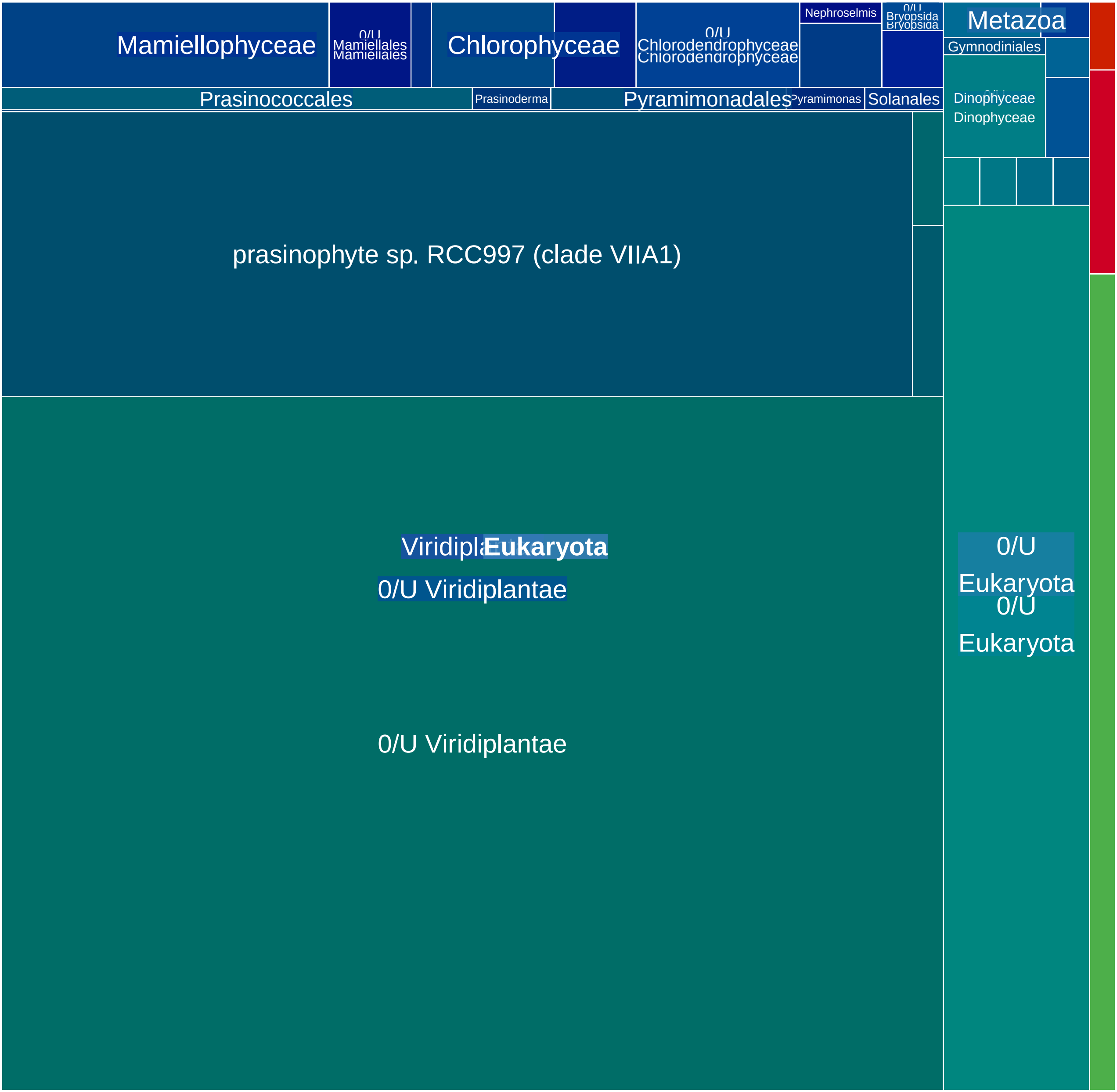

MGT−v1_659 : 964 unigenes, 878 (91.08%) taxonomically assigned

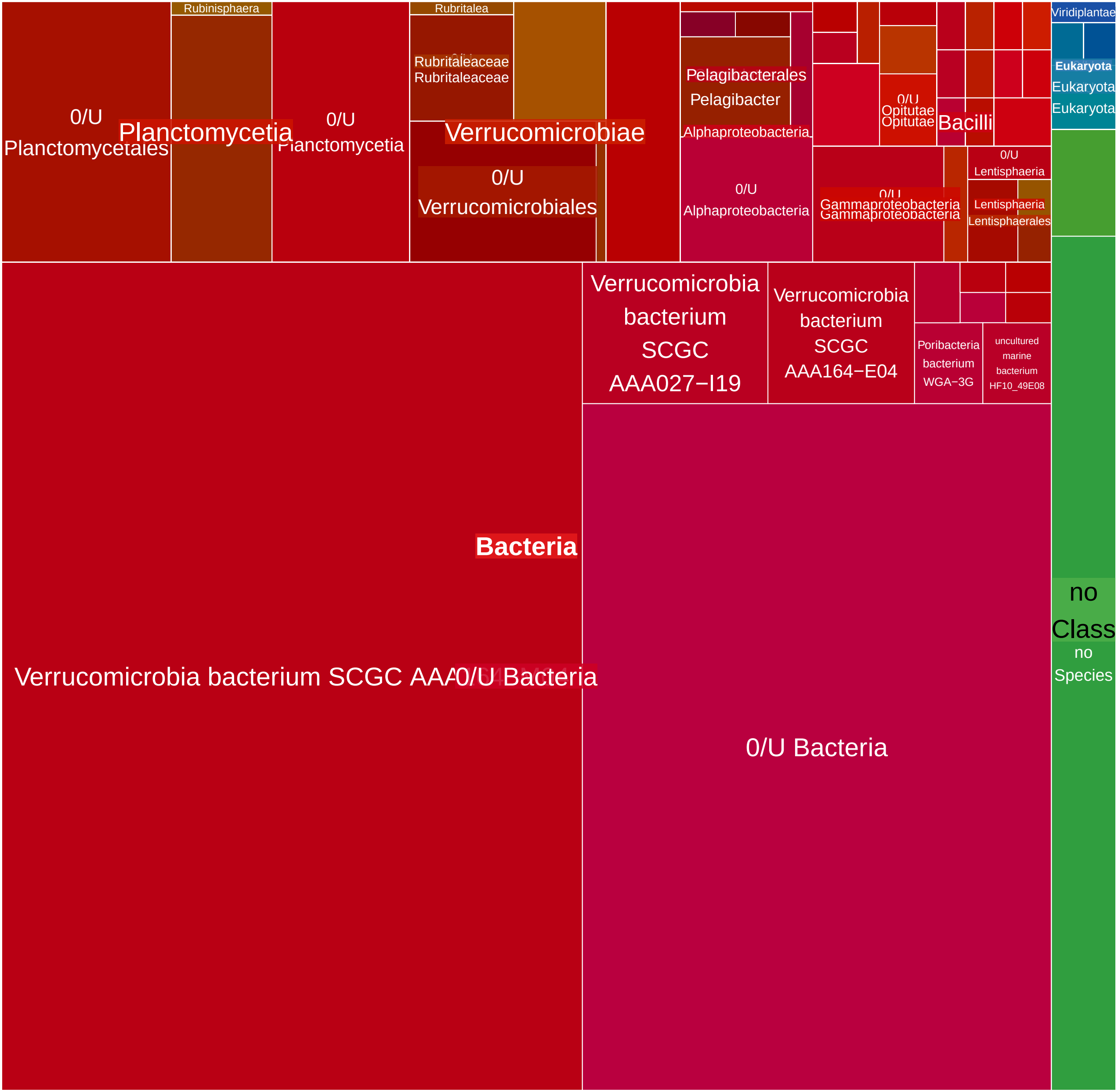

### MGT−v1_660 : 962 unigenes, 538 (55.93%) taxonomically assigned

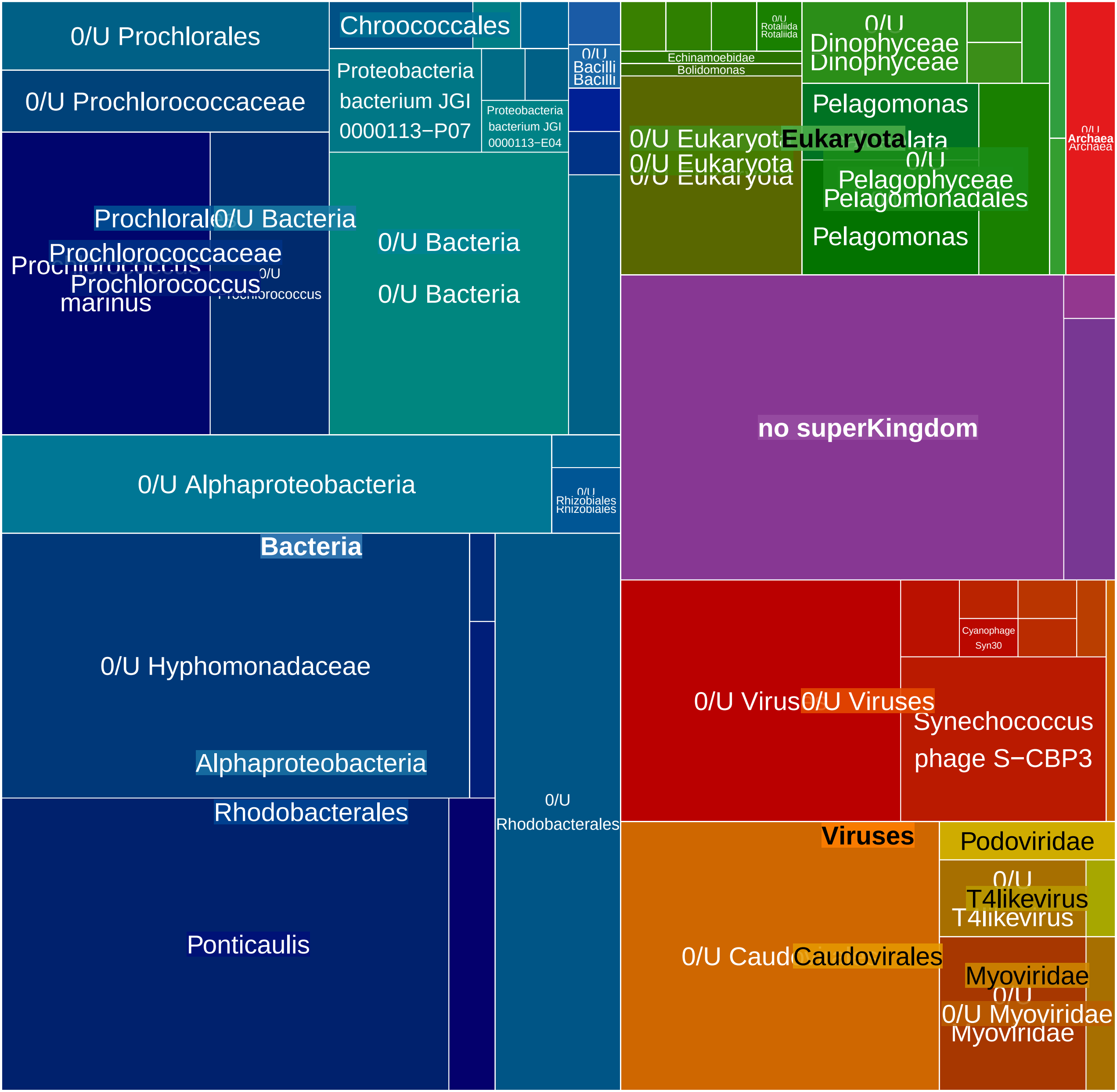

MGT−v1_661 : 959 unigenes, 63 (6.57%) taxonomically assigned

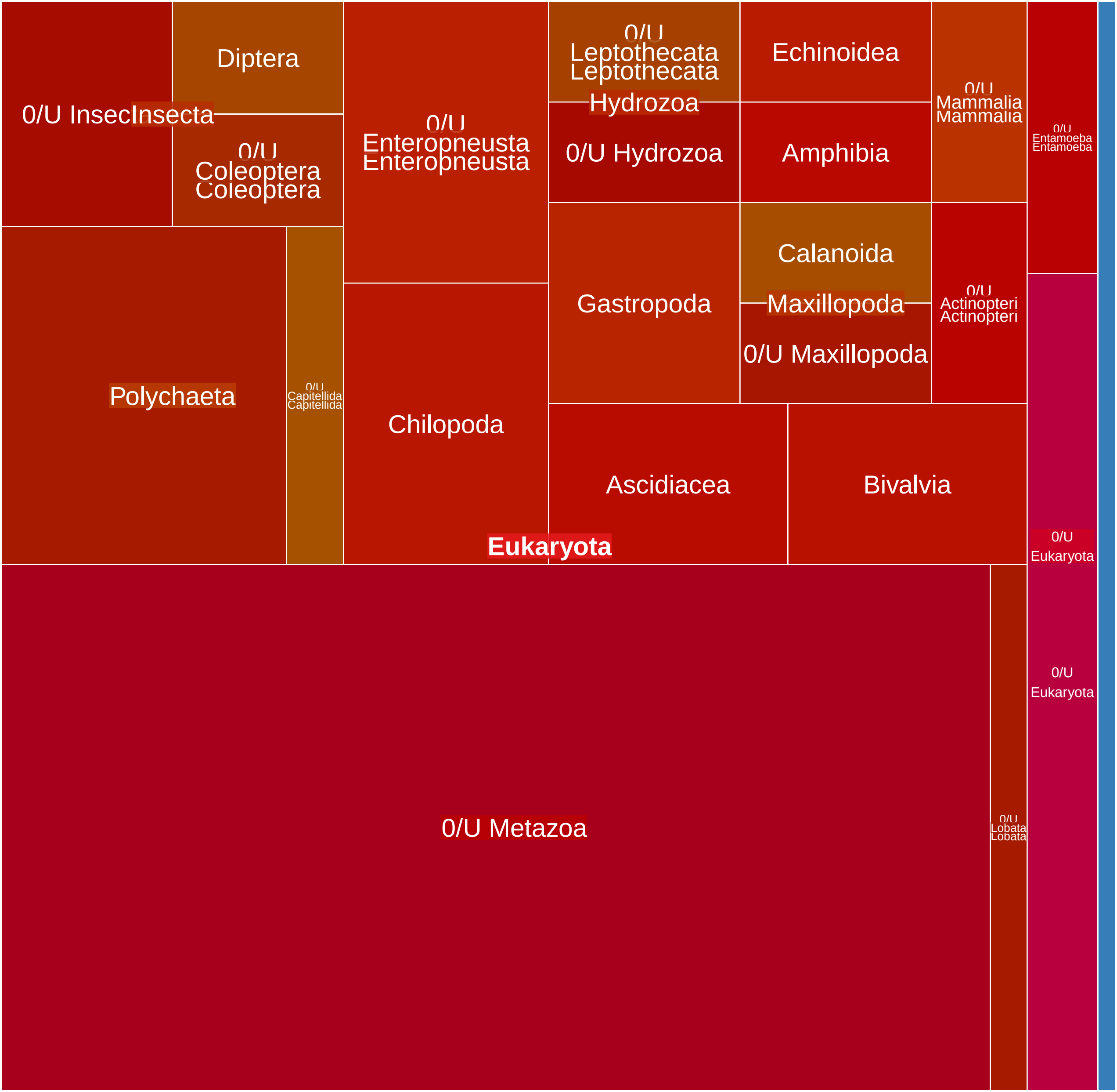

MGT−v1_662 : 958 unigenes, 104 (10.86%) taxonomically assigned

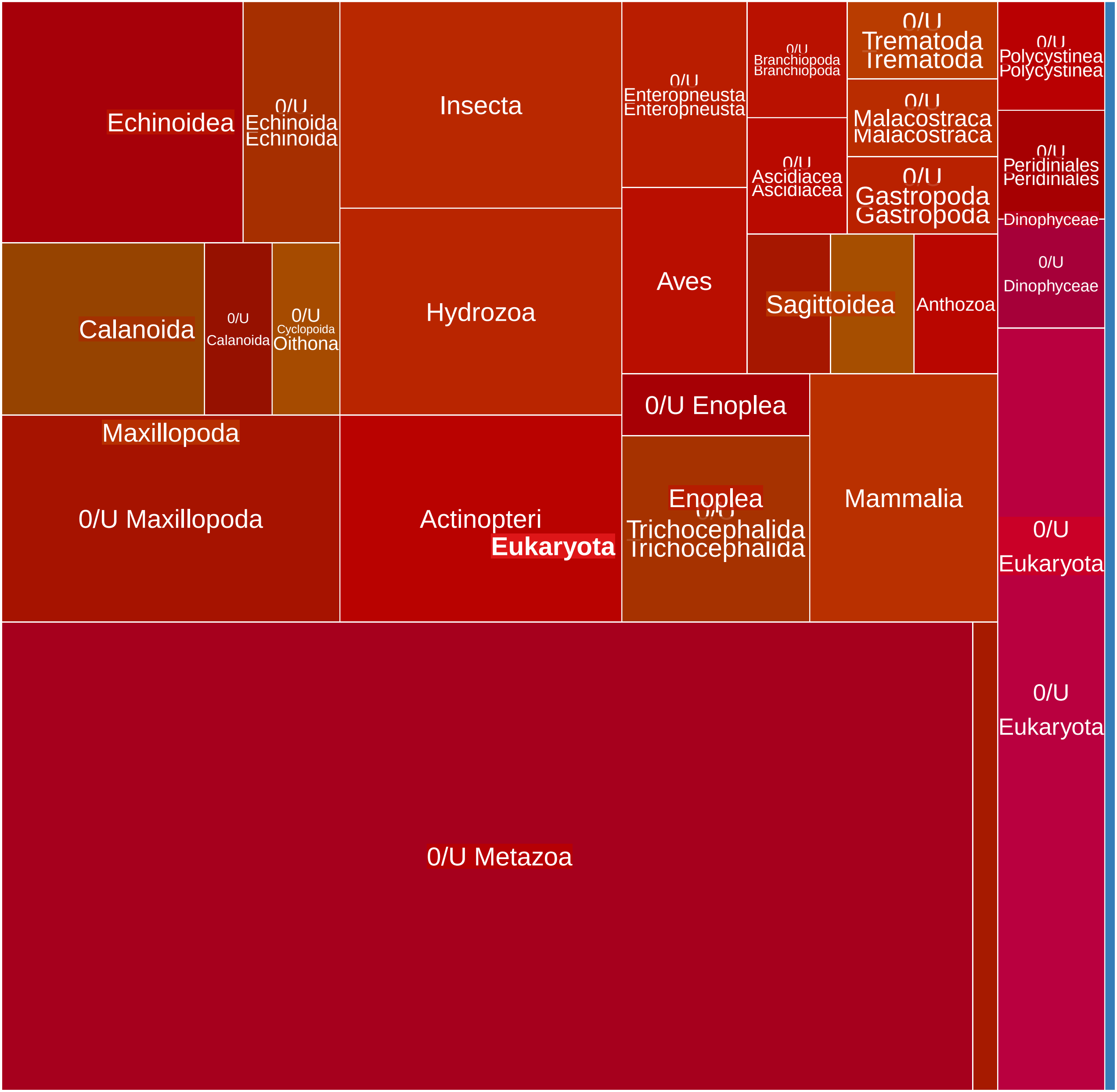

MGT−v1_663 : 953 unigenes, 910 (95.49%) taxonomically assigned

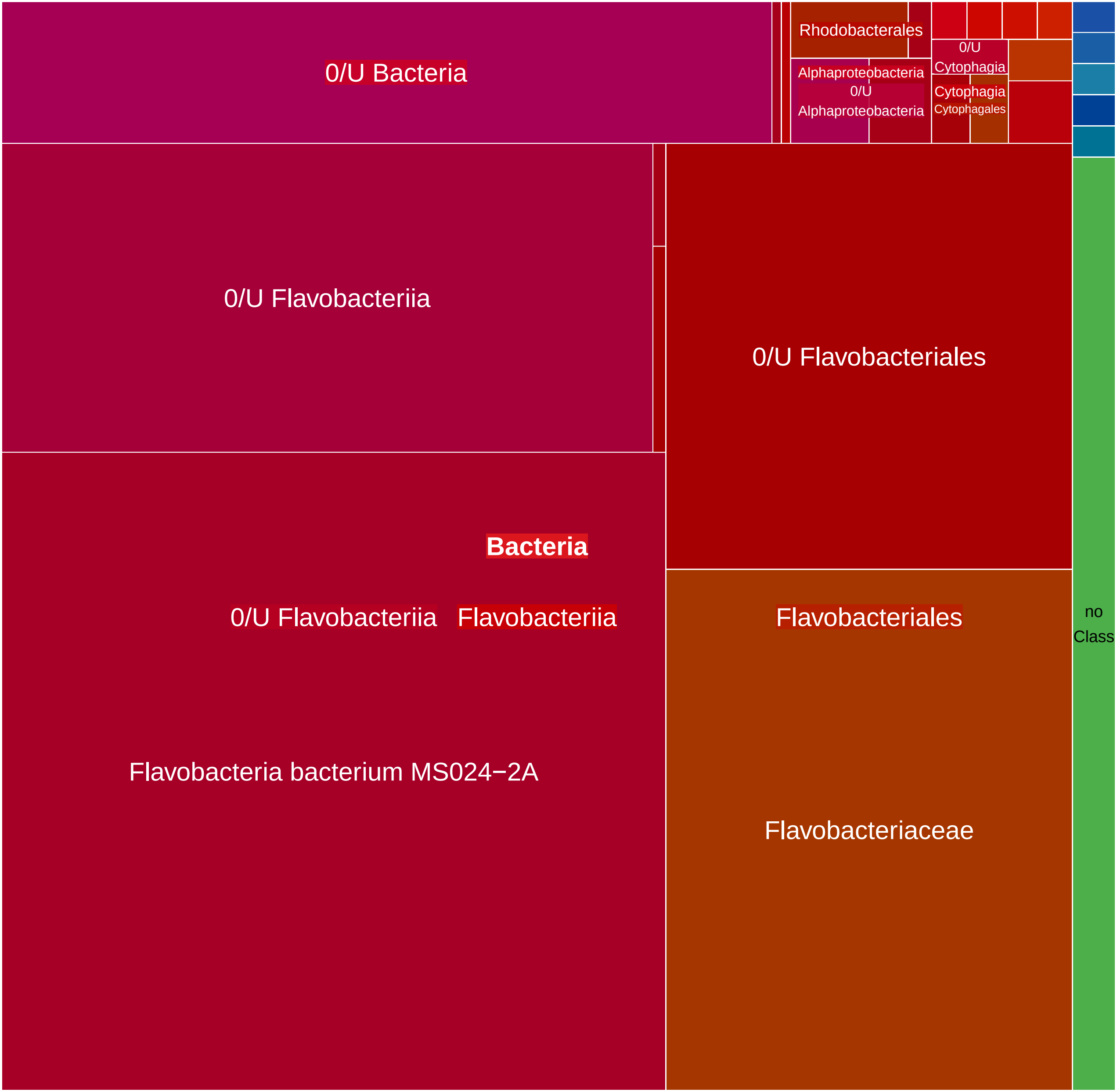

MGT−v1_664 : 950 unigenes, 436 (45.89%) taxonomically assigned

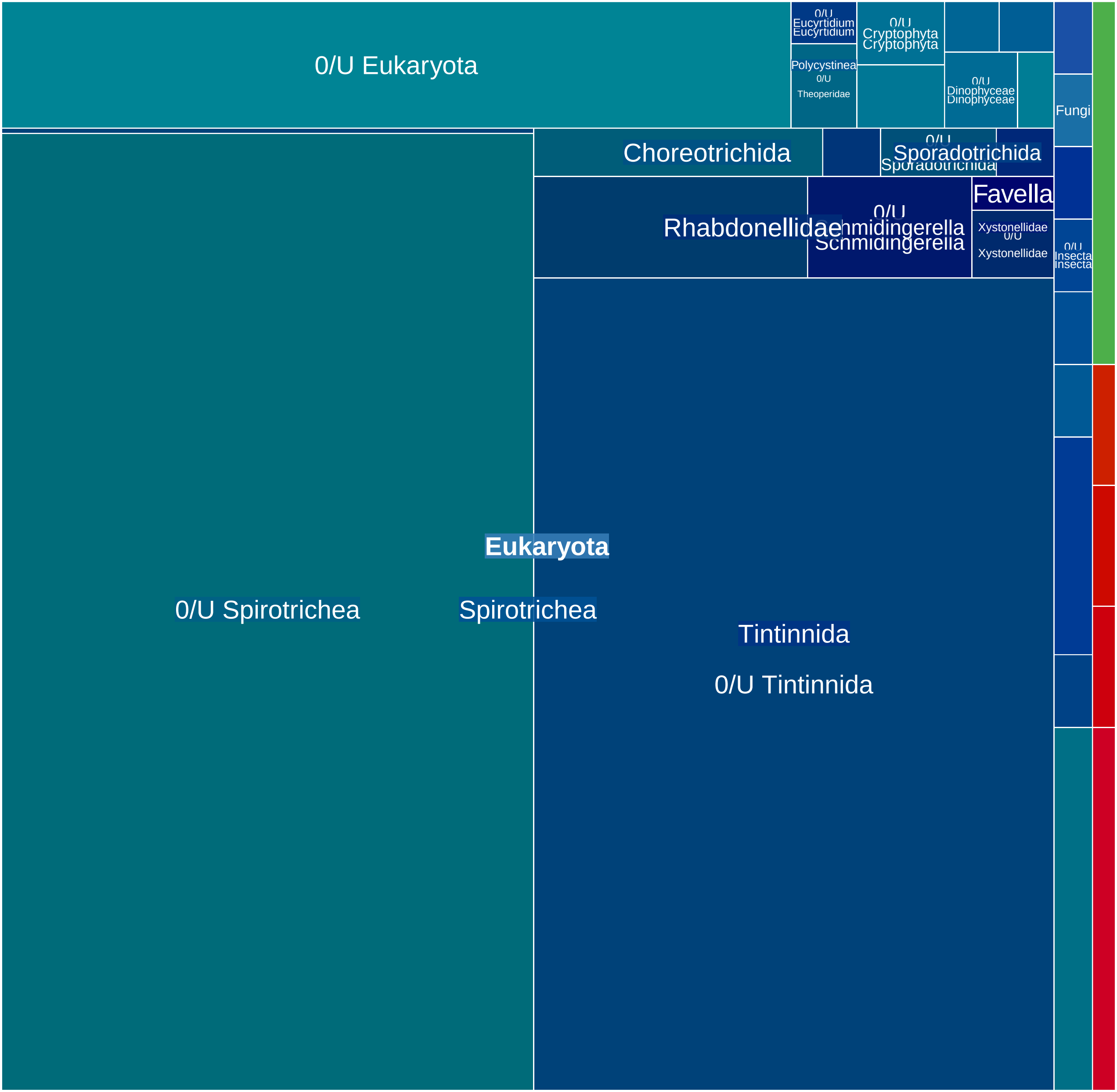

MGT−v1_665 : 950 unigenes, 810 (85.26%) taxonomically assigned

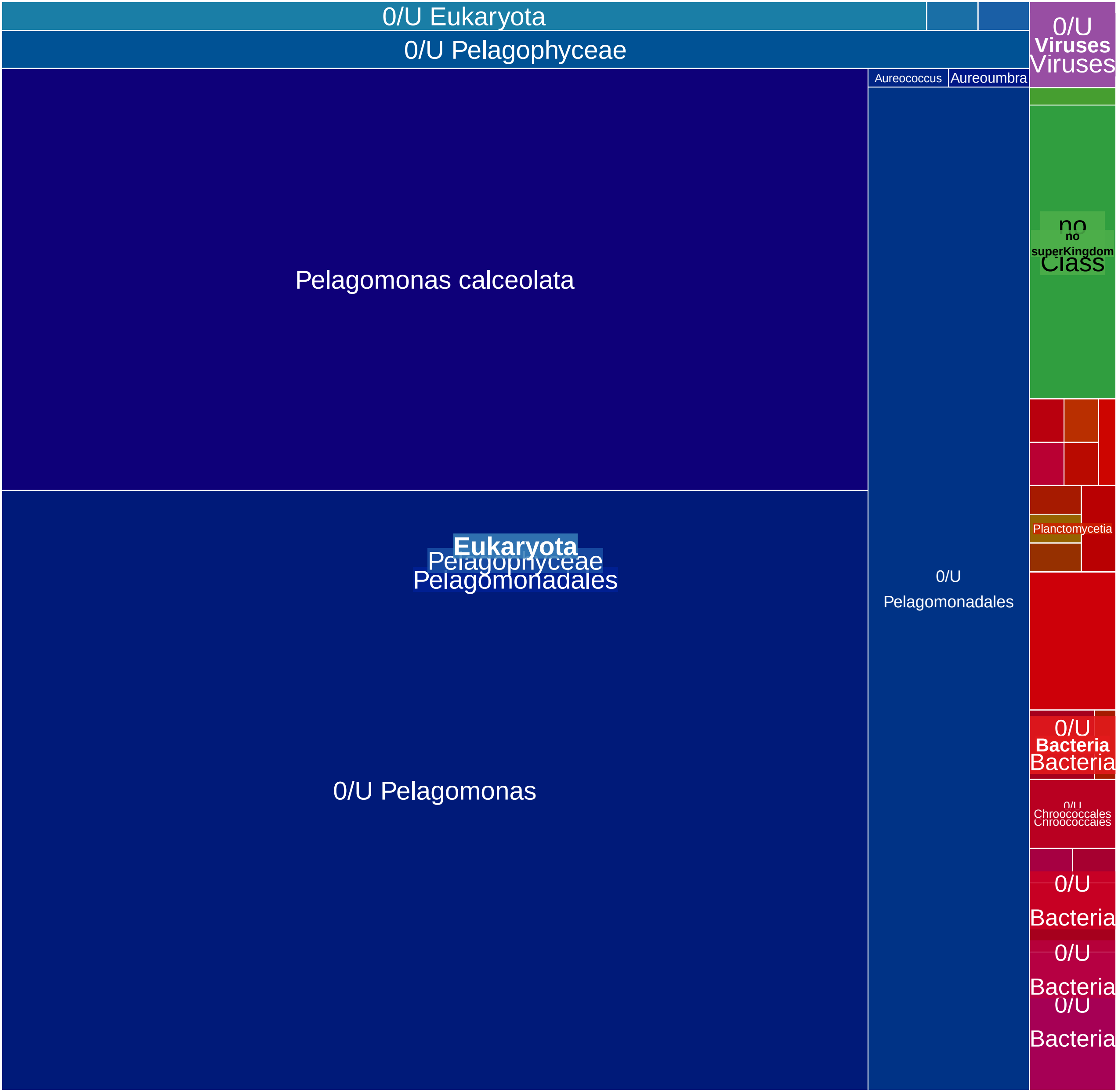

MGT−v1_666 : 946 unigenes, 894 (94.50%) taxonomically assigned

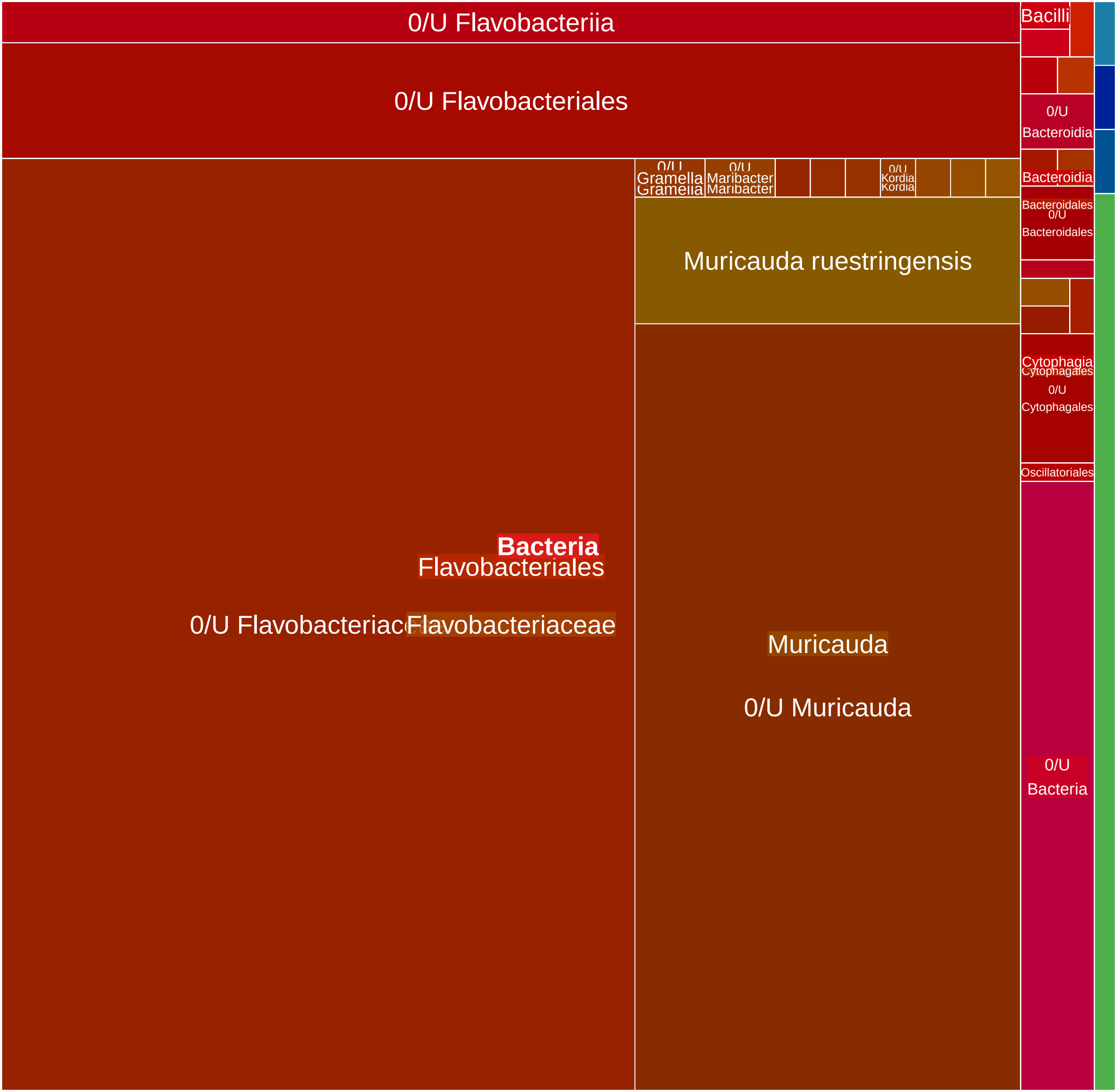

### MGT−v1_667 : 945 unigenes, 906 (95.87%) taxonomically assigned

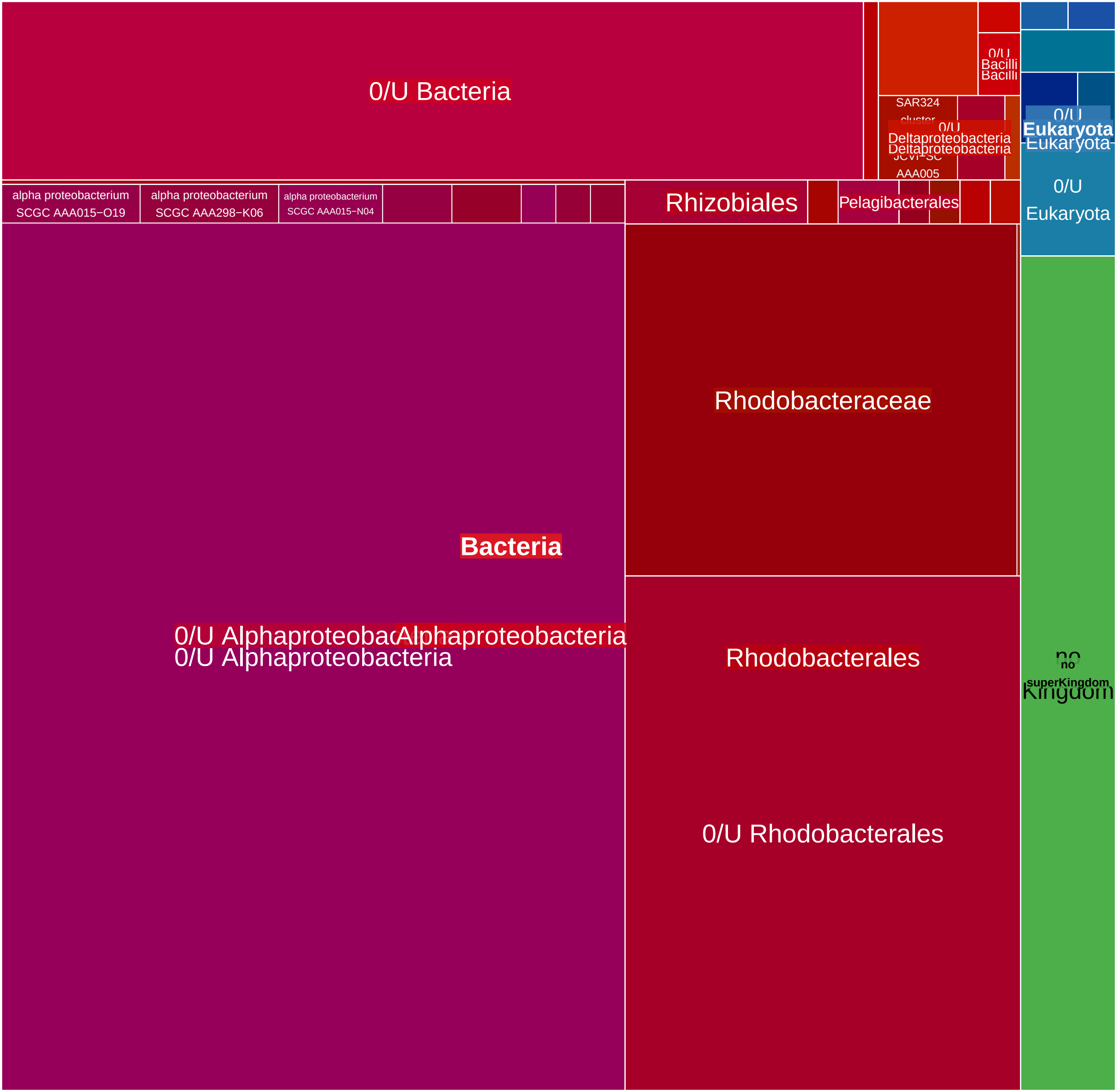

MGT−v1_668 : 944 unigenes, 298 (31.57%) taxonomically assigned

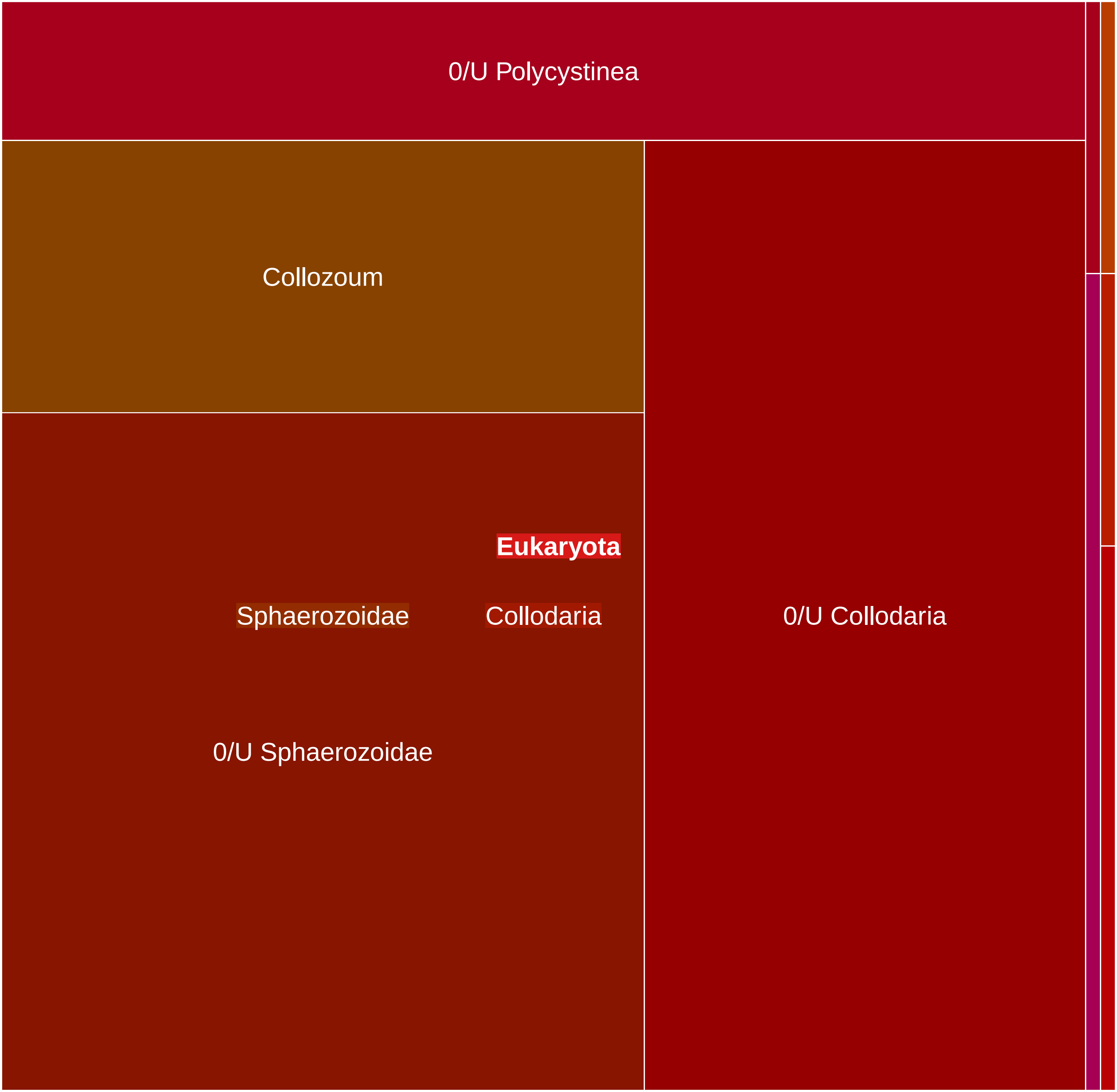

MGT−v1_669 : 941 unigenes, 872 (92.67%) taxonomically assigned

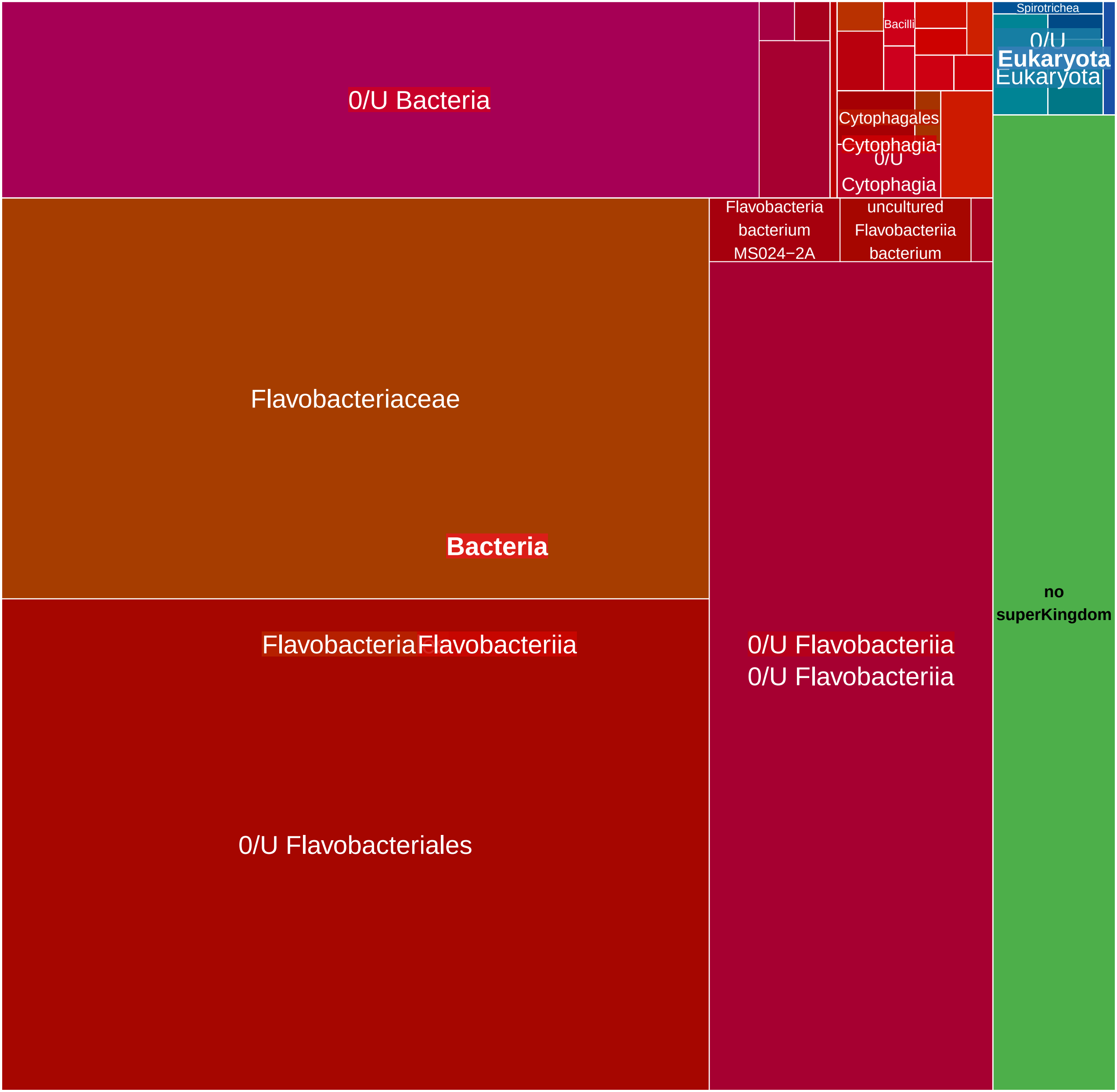

MGT−v1_670 : 939 unigenes, 108 (11.50%) taxonomically assigned

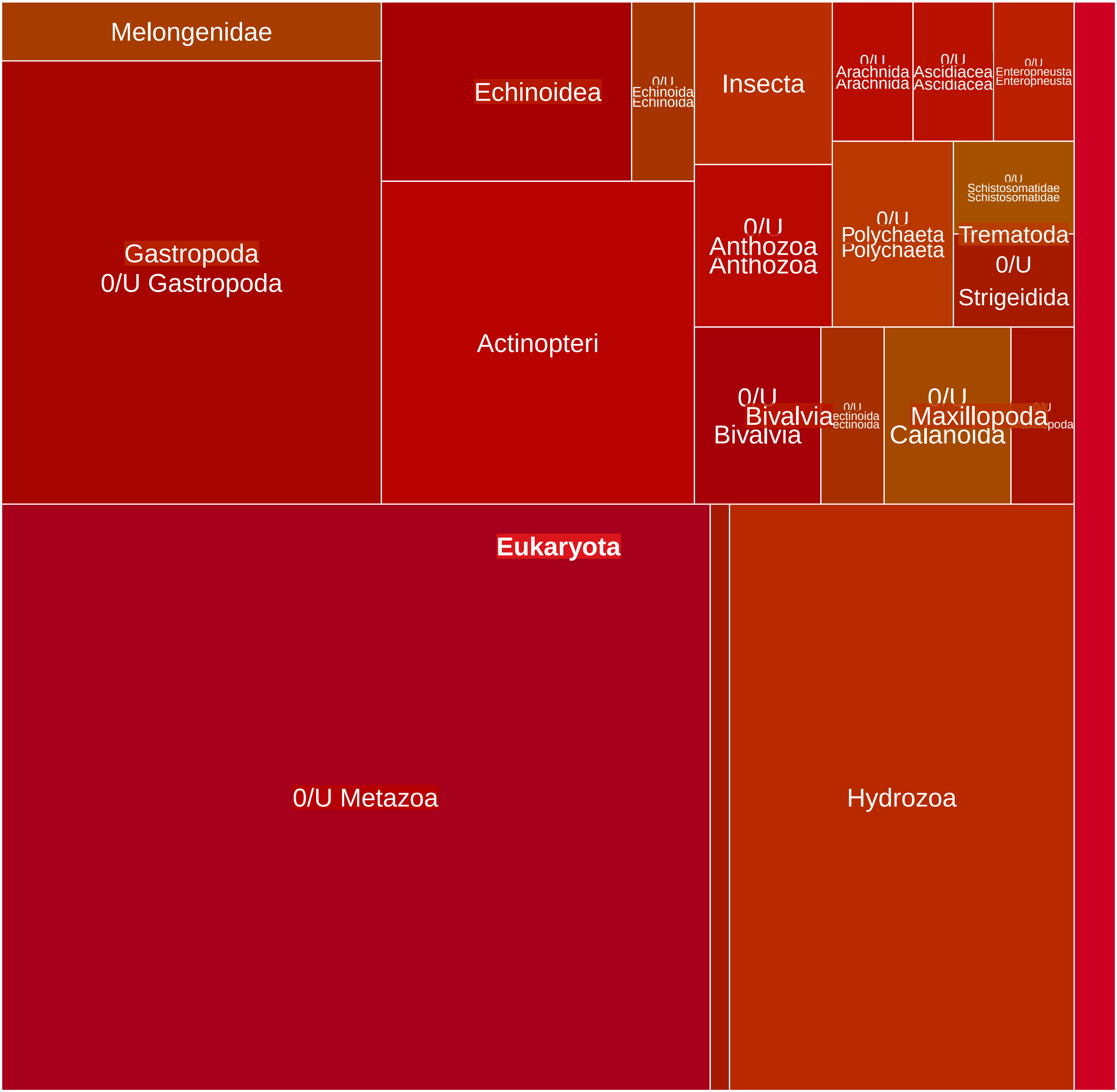

MGT−v1_671 : 937 unigenes, 80 (8.54%) taxonomically assigned

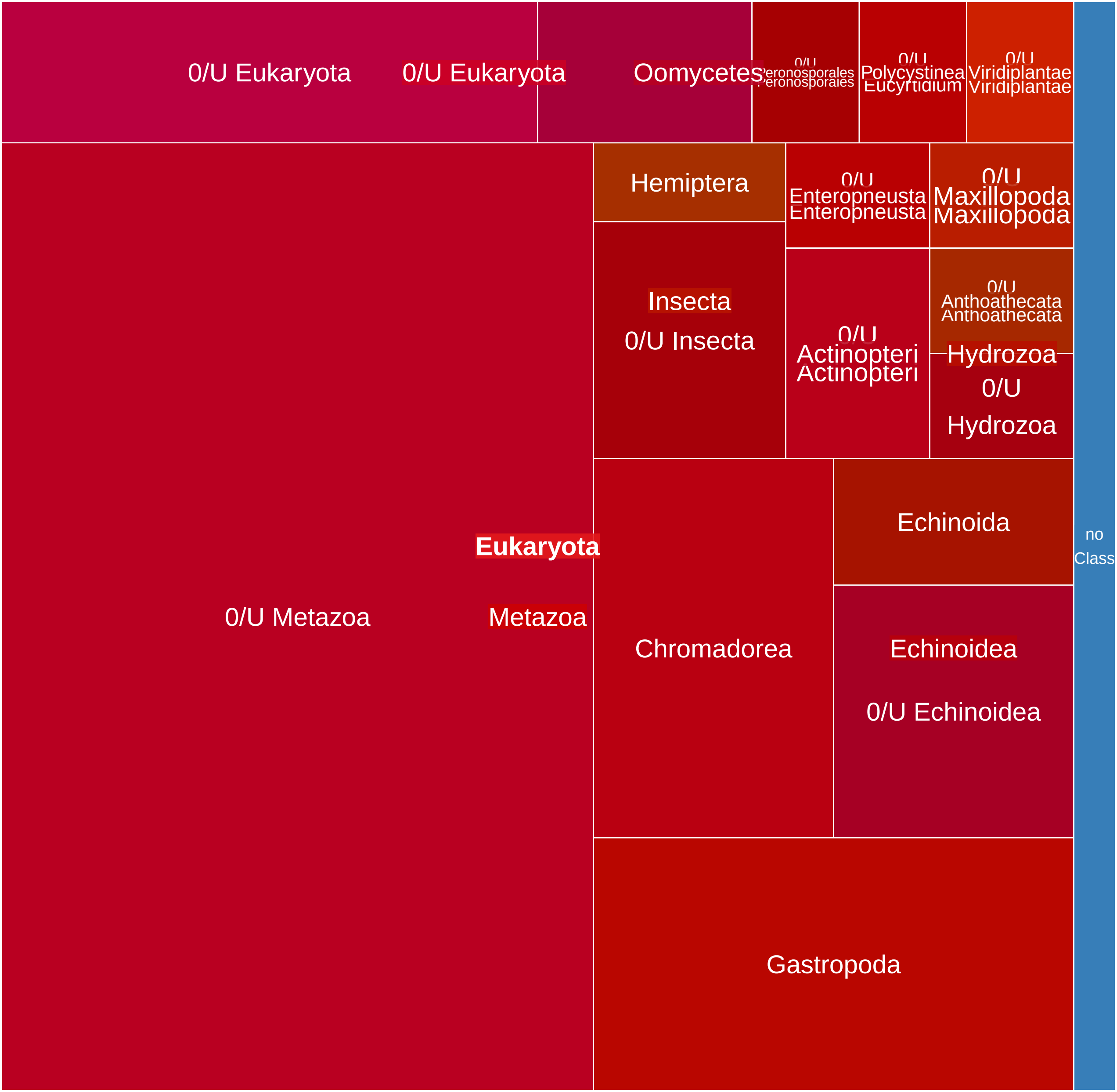

### MGT−v1_672 : 932 unigenes, 828 (88.84%) taxonomically assigned

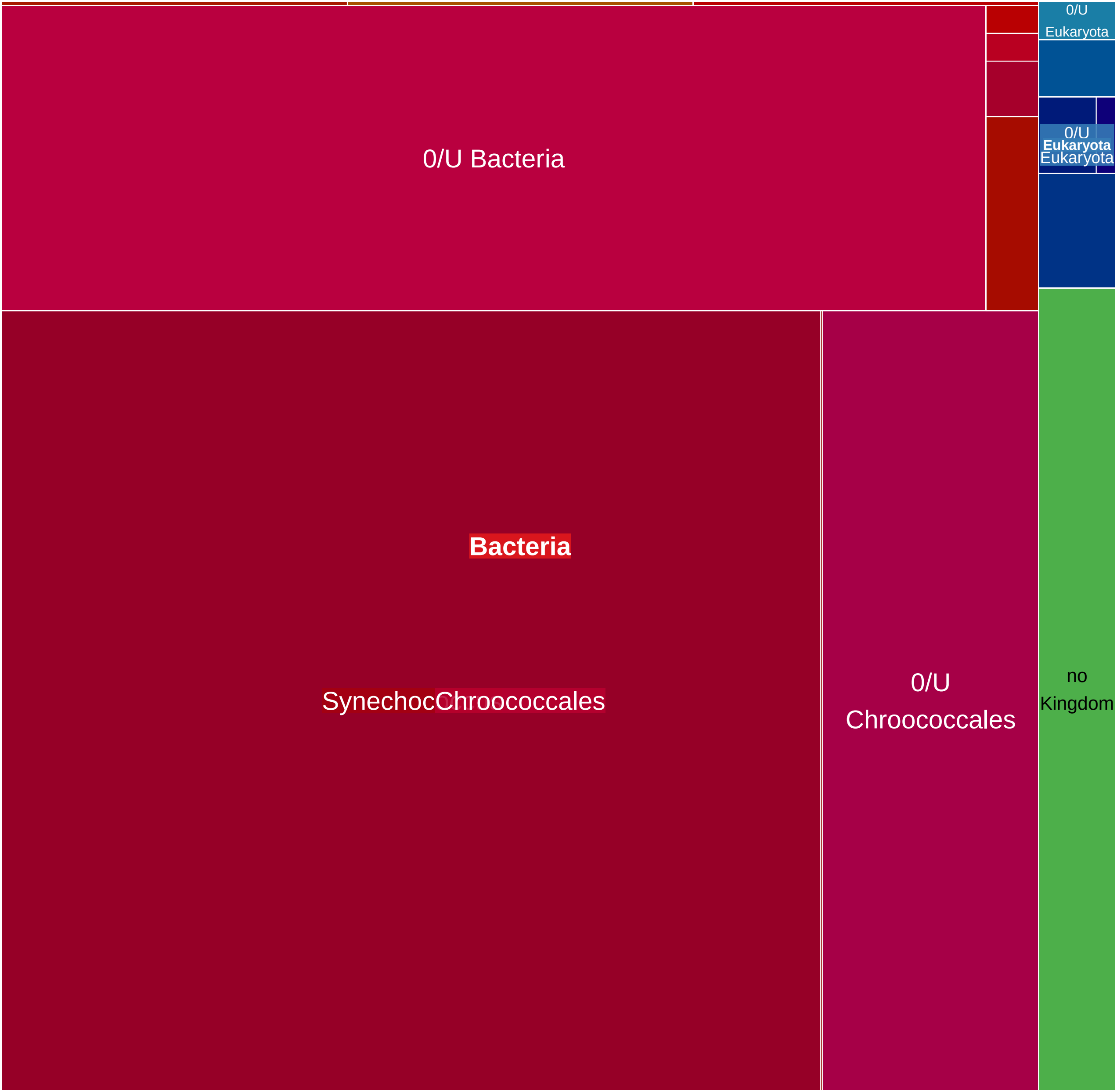

MGT−v1_673 : 930 unigenes, 691 (74.30%) taxonomically assigned

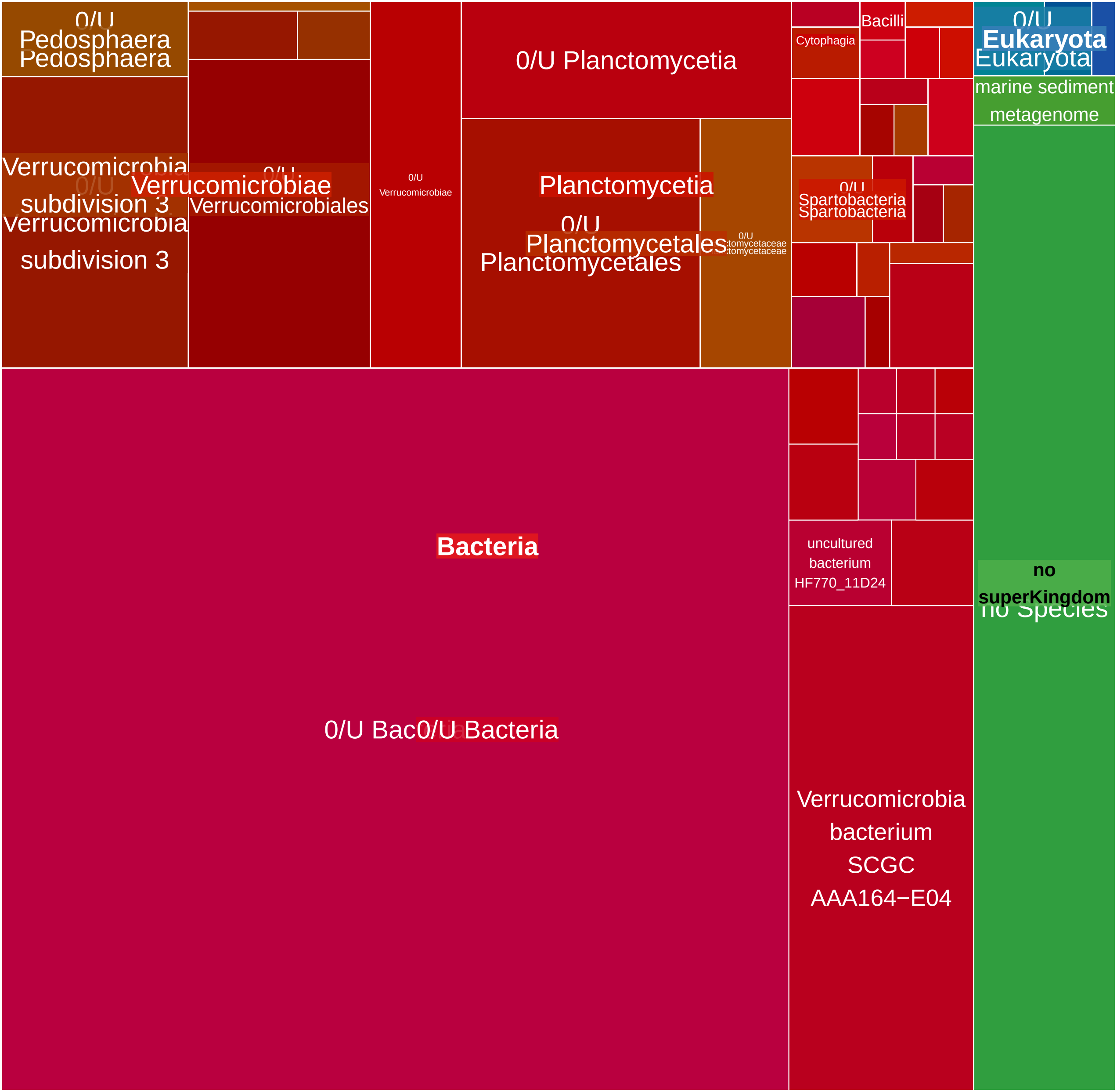

MGT−v1_674 : 928 unigenes, 920 (99.14%) taxonomically assigned

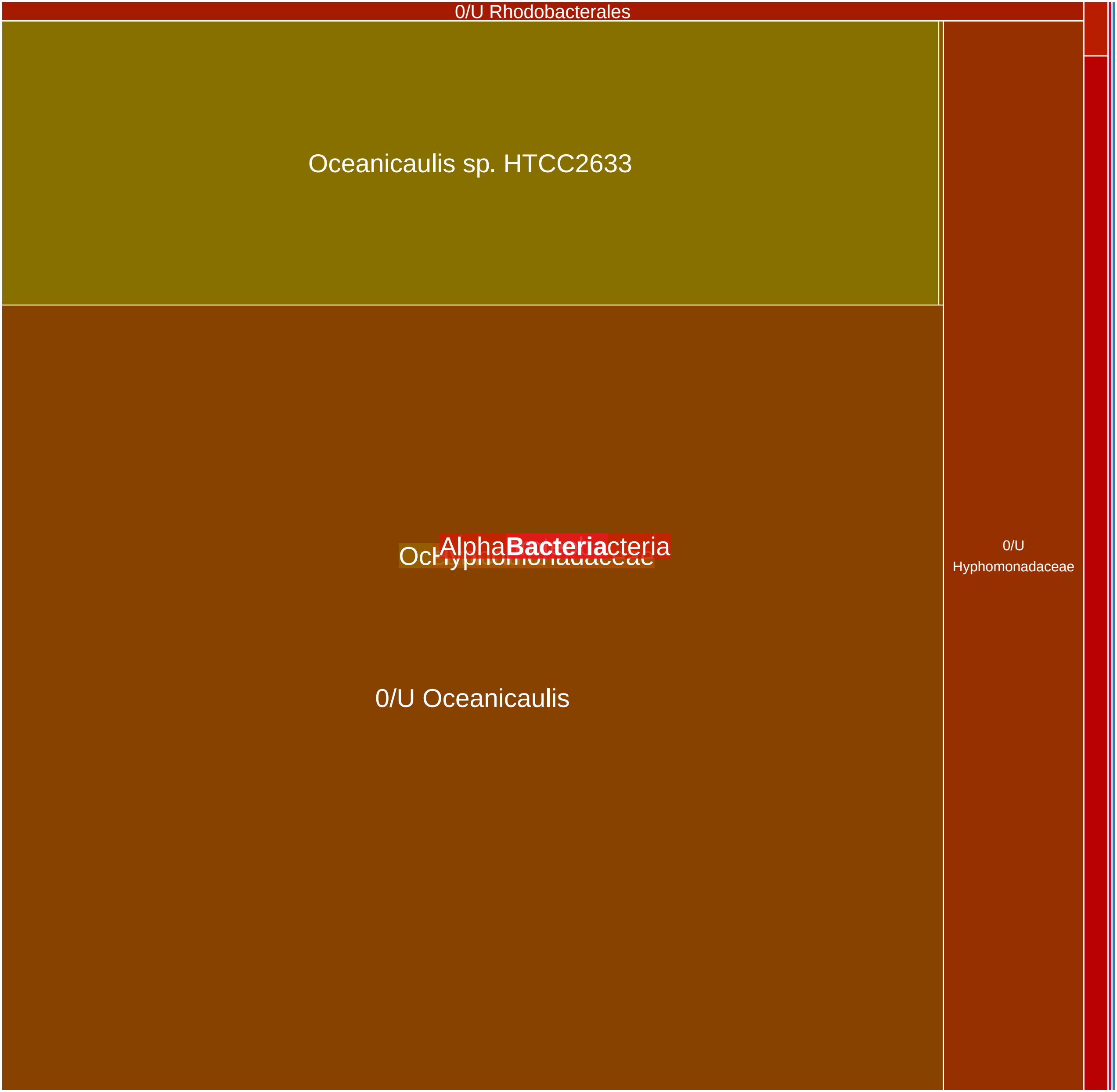

MGT−v1_675 : 926 unigenes, 66 (7.13%) taxonomically assigned

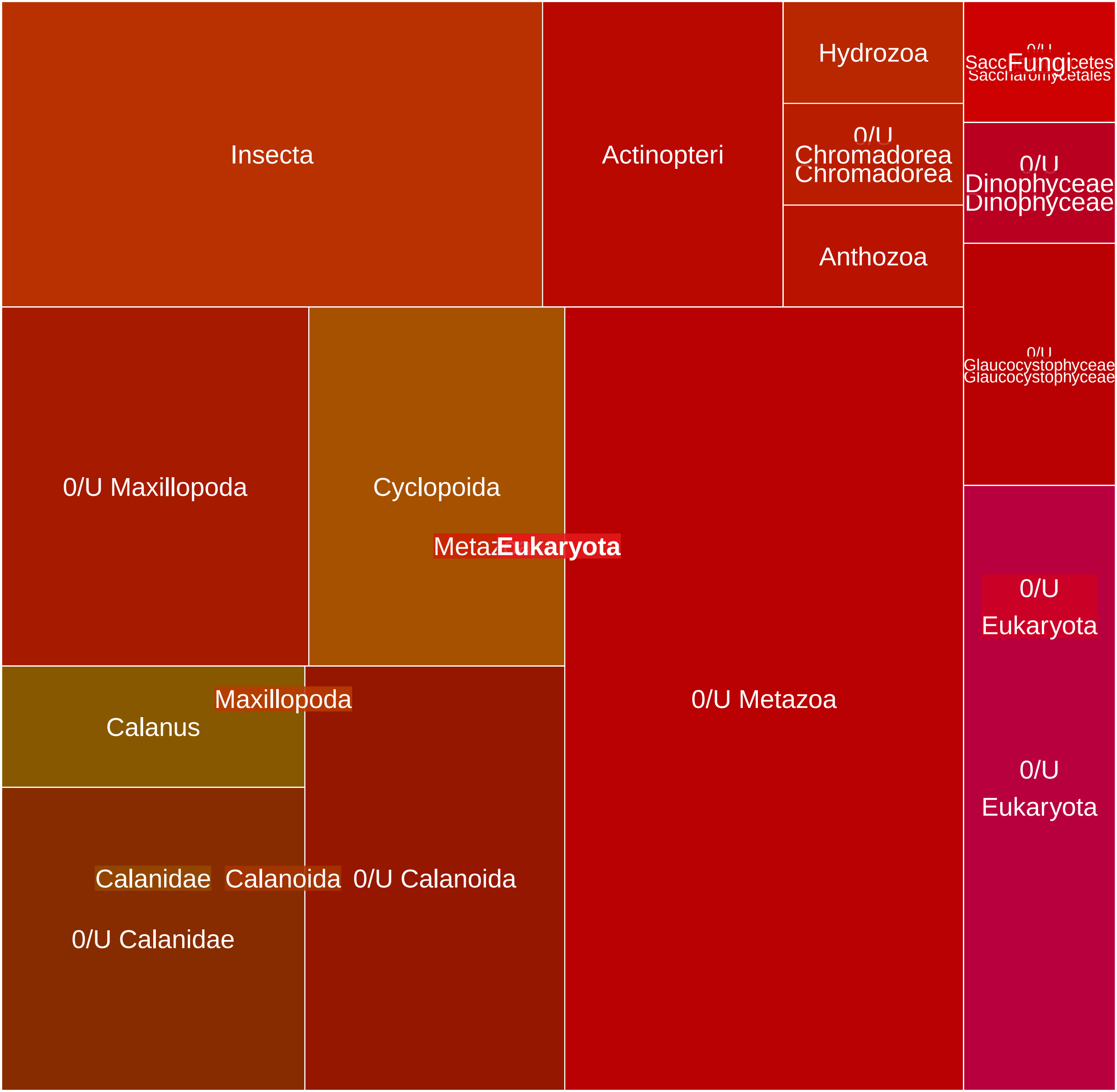

MGT−v1_676 : 923 unigenes, 194 (21.02%) taxonomically assigned

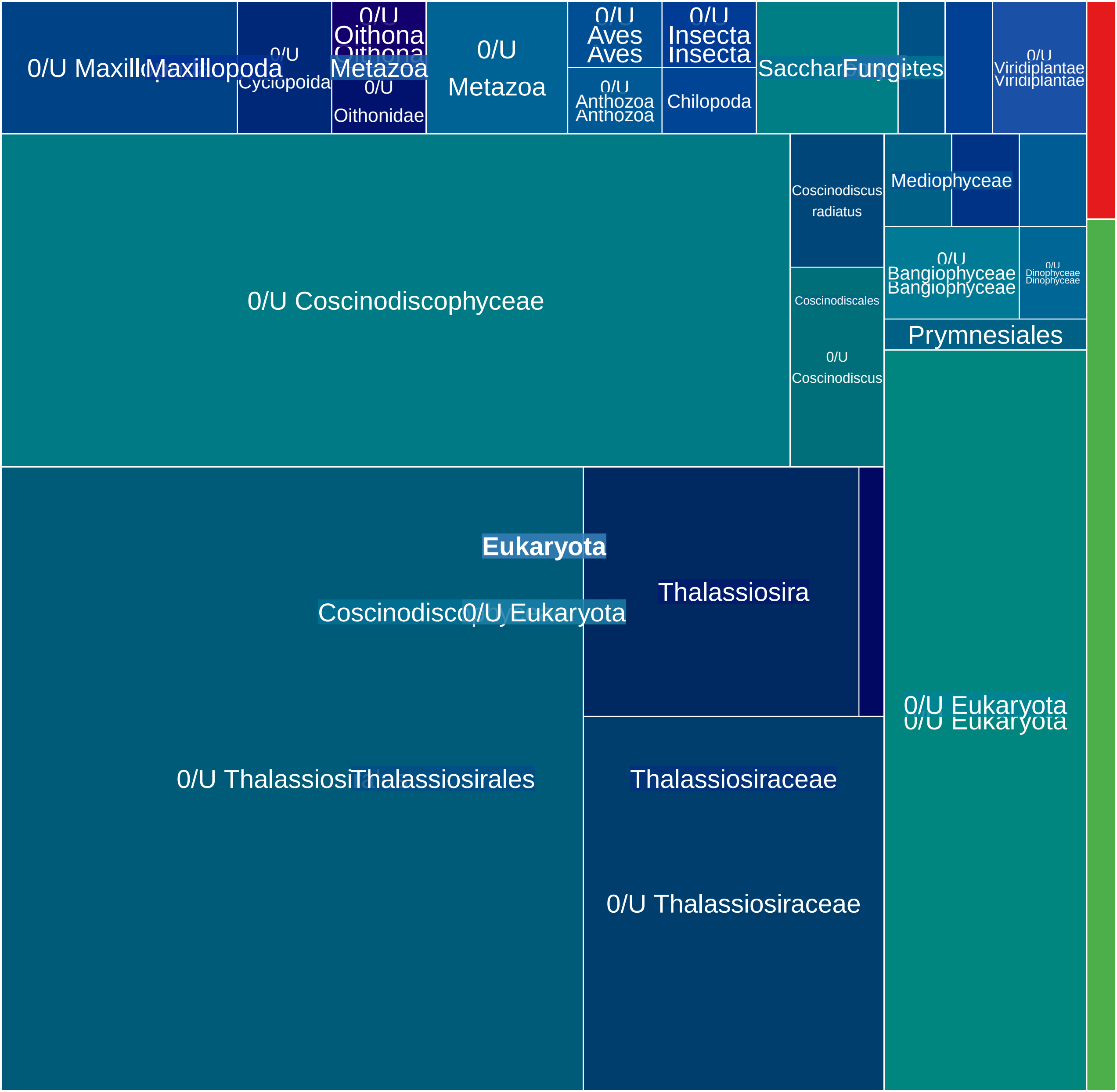

MGT−v1_677 : 922 unigenes, 863 (93.60%) taxonomically assigned

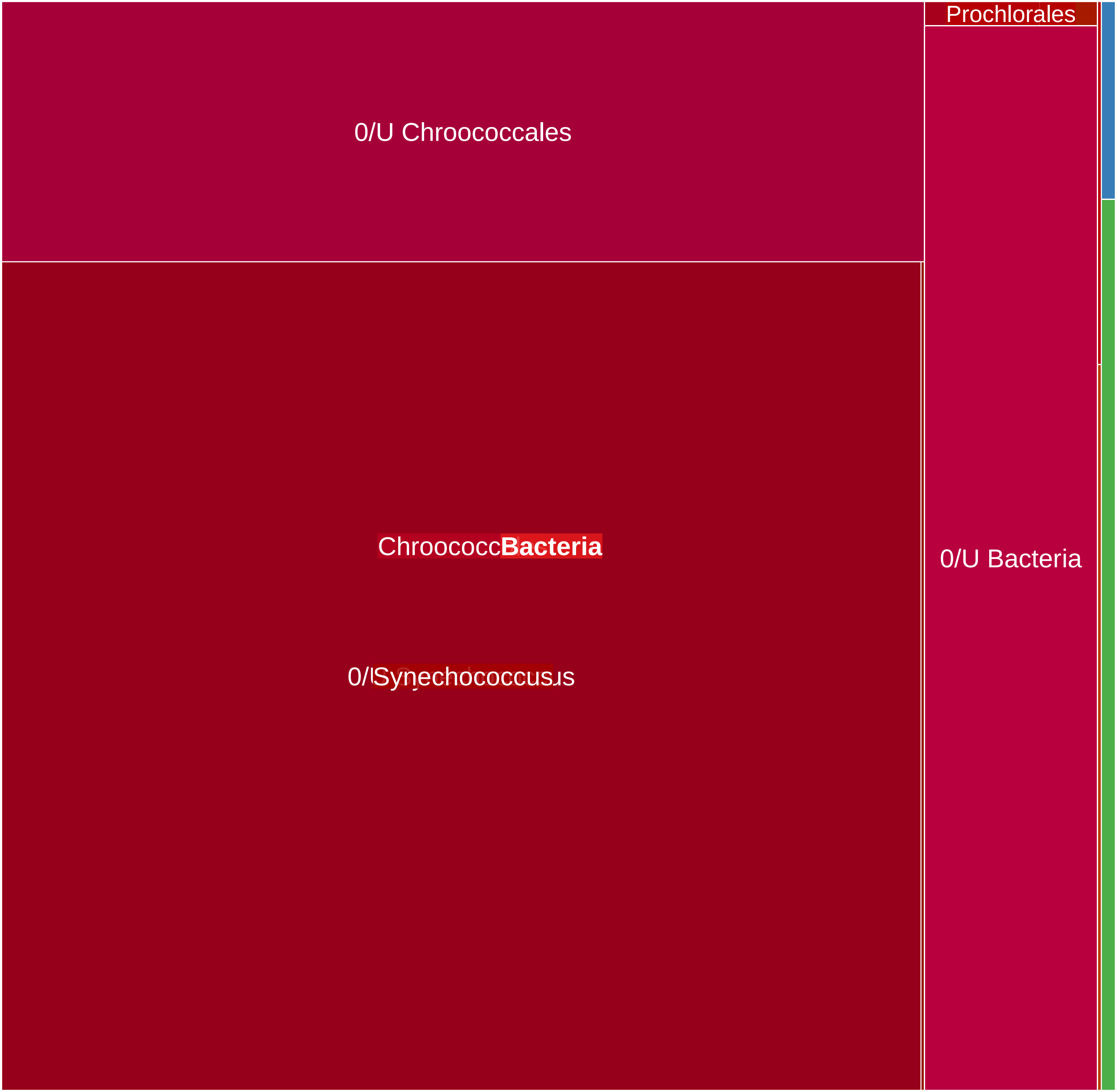

MGT−v1_678 : 922 unigenes, 390 (42.30%) taxonomically assigned

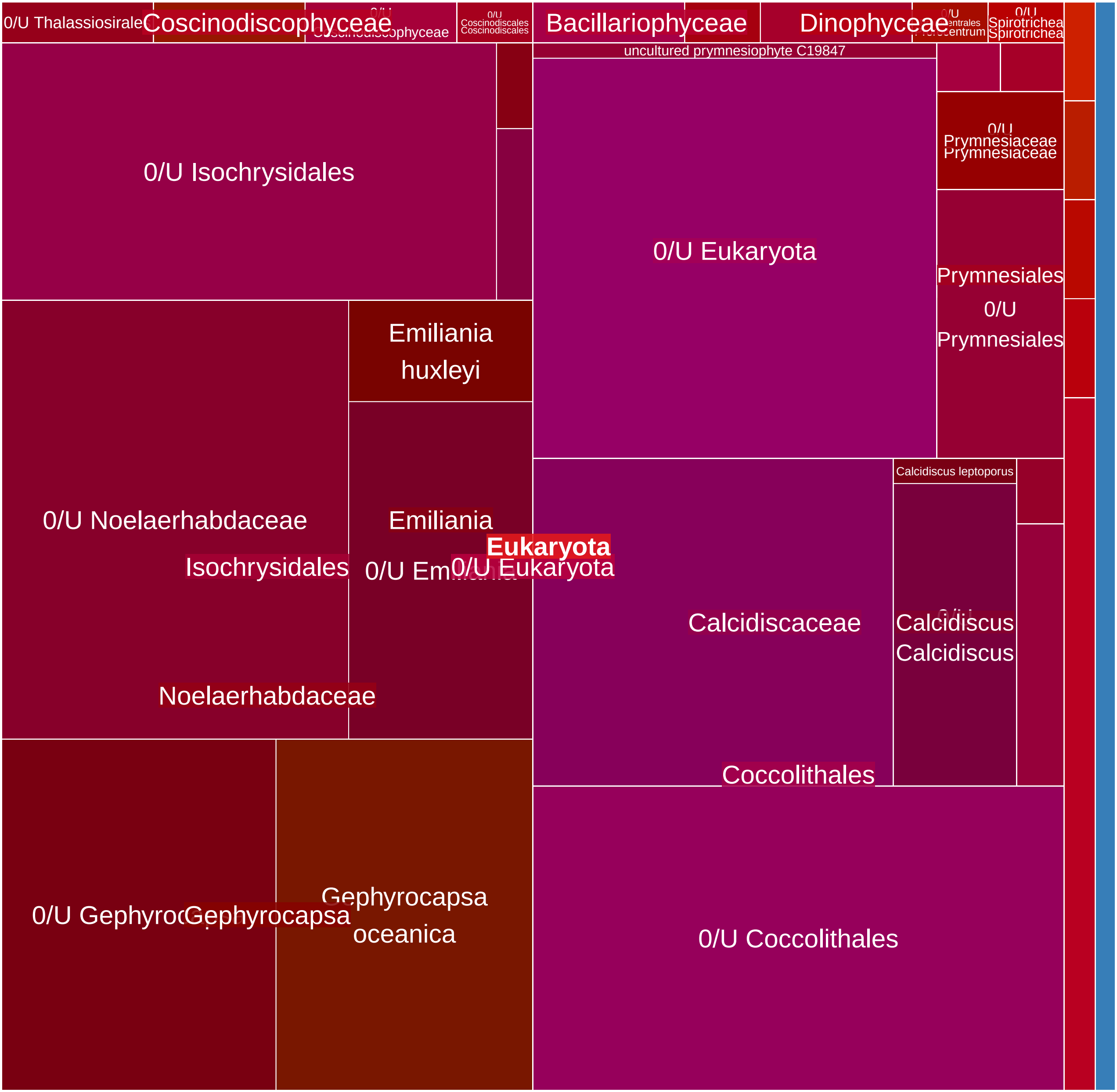

MGT−v1_679 : 914 unigenes, 535 (58.53%) taxonomically assigned

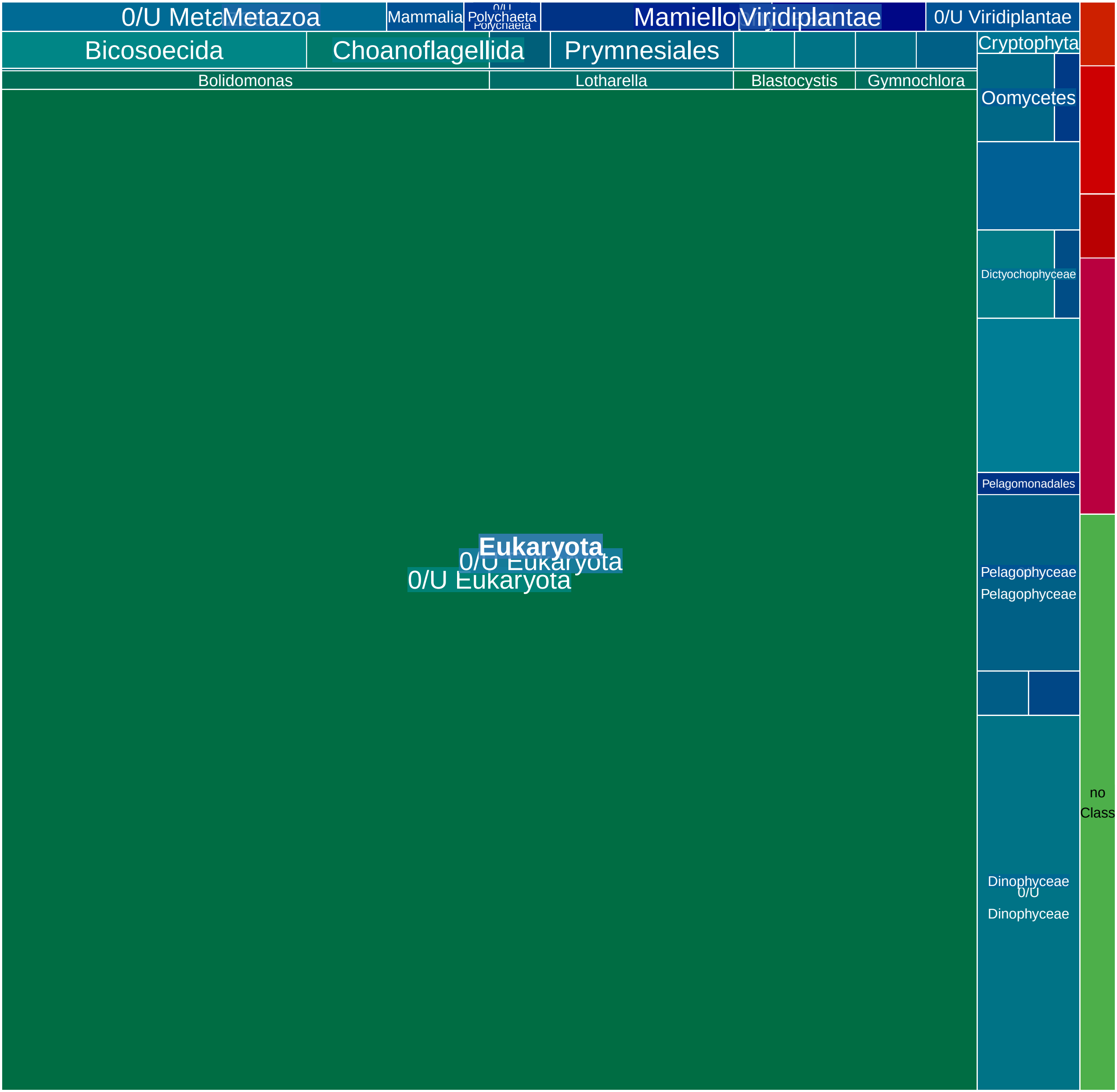

MGT−v1_680 : 913 unigenes, 58 (6.35%) taxonomically assigned

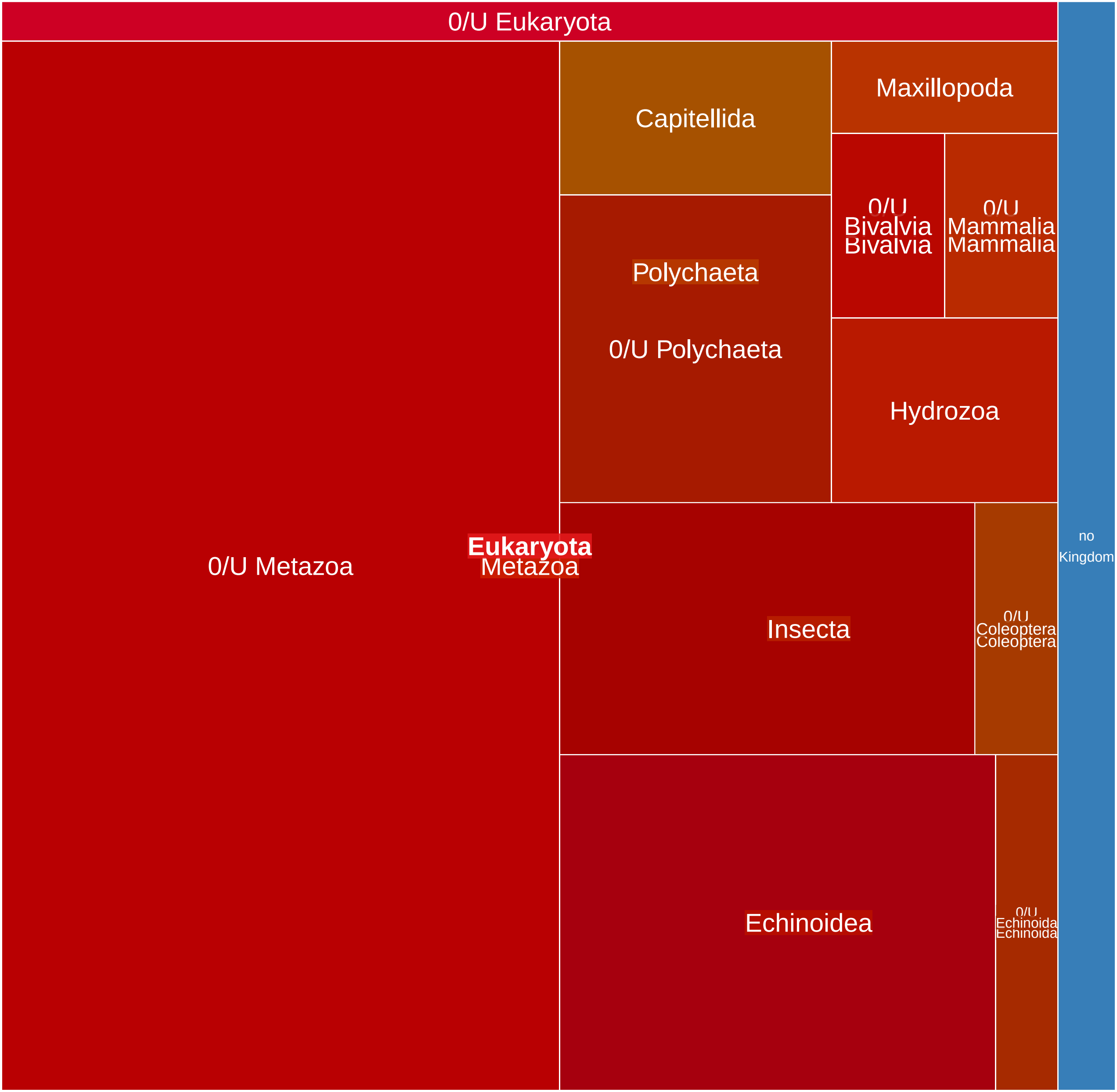

MGT−v1_681 : 913 unigenes, 74 (8.11%) taxonomically assigned

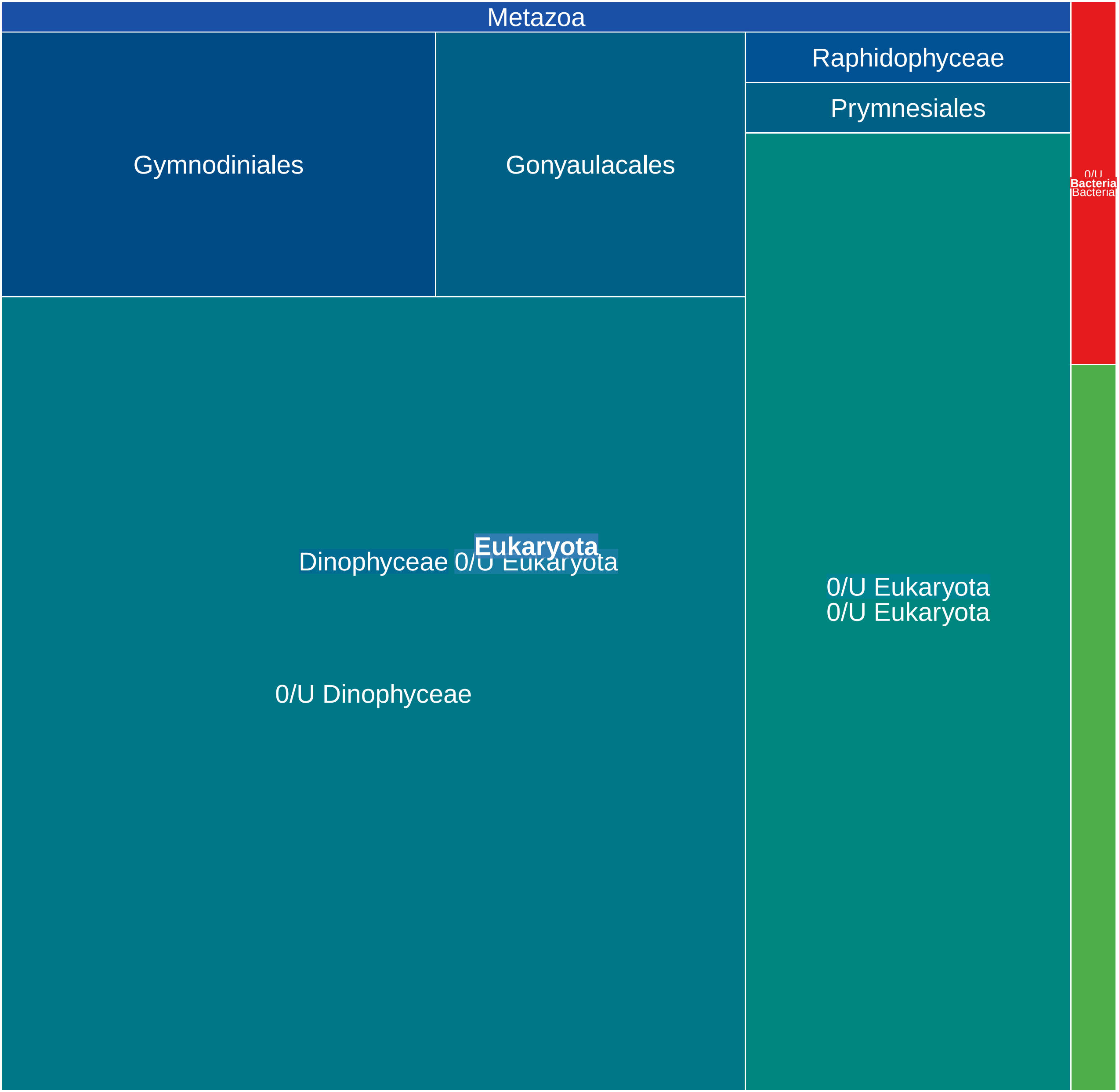

### MGT−v1_682 : 912 unigenes, 256 (28.07%) taxonomically assigned

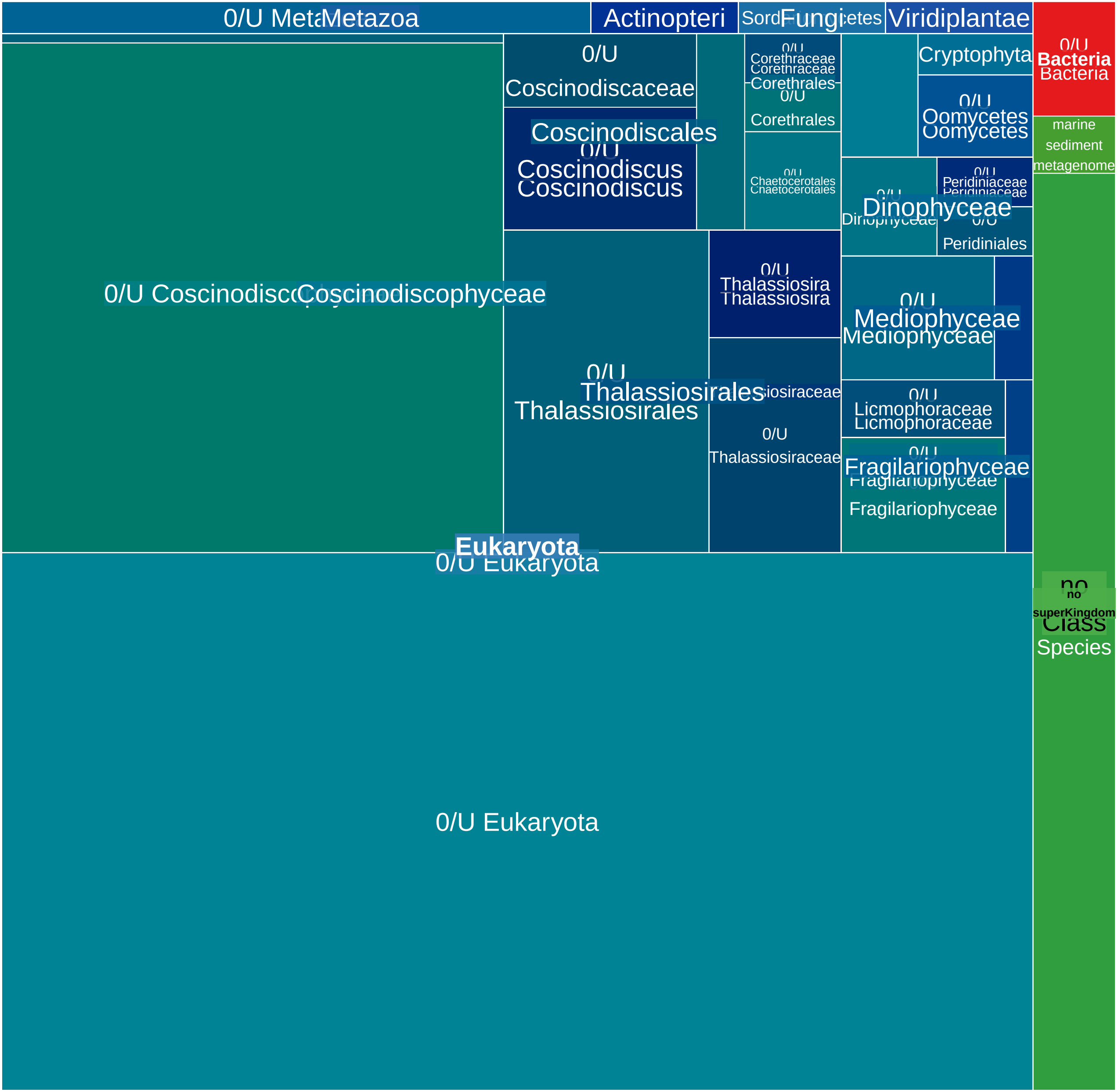

MGT−v1_683 : 910 unigenes, 75 (8.24%) taxonomically assigned

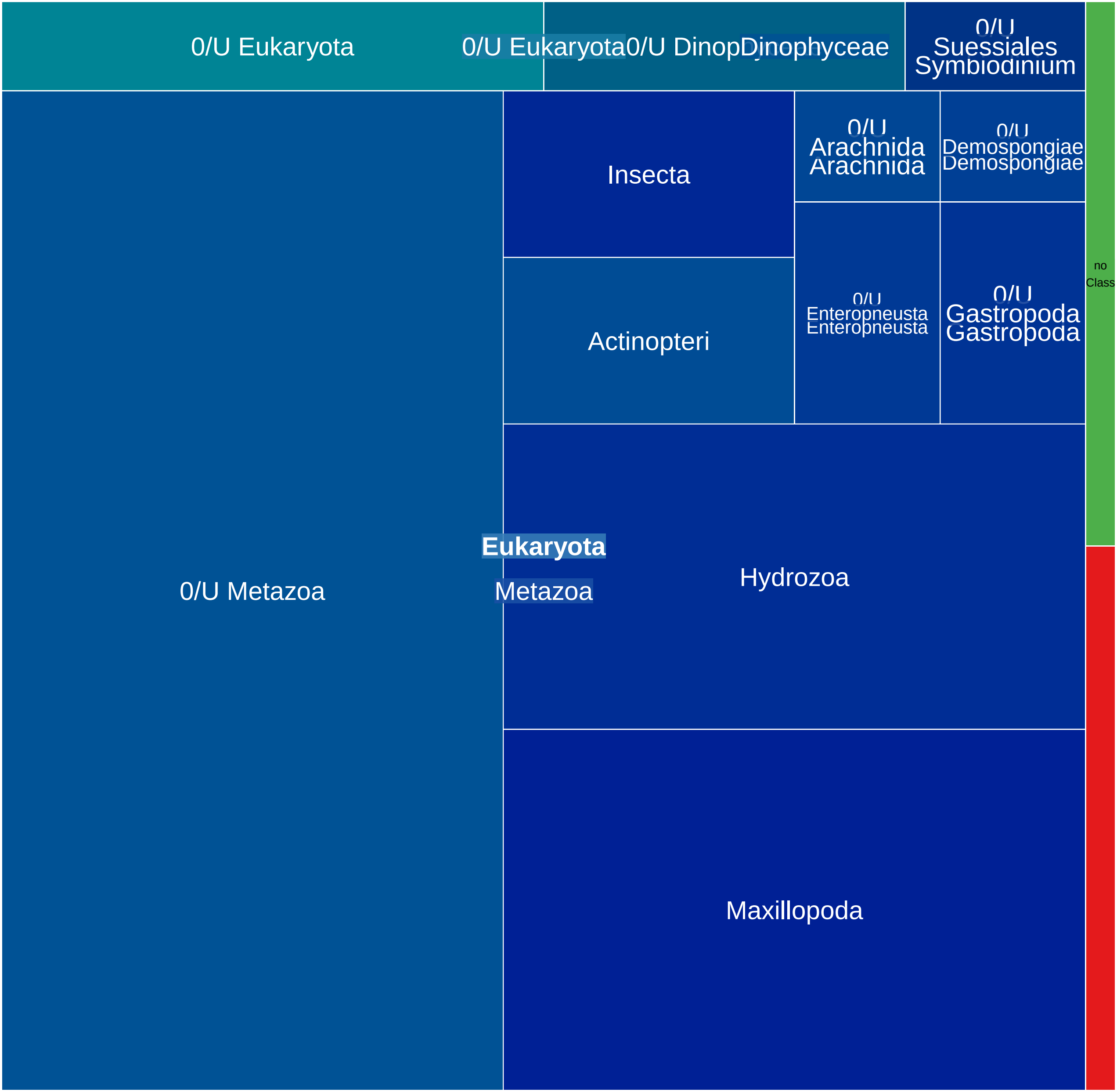

MGT−v1_684 : 909 unigenes, 50 (5.50%) taxonomically assigned

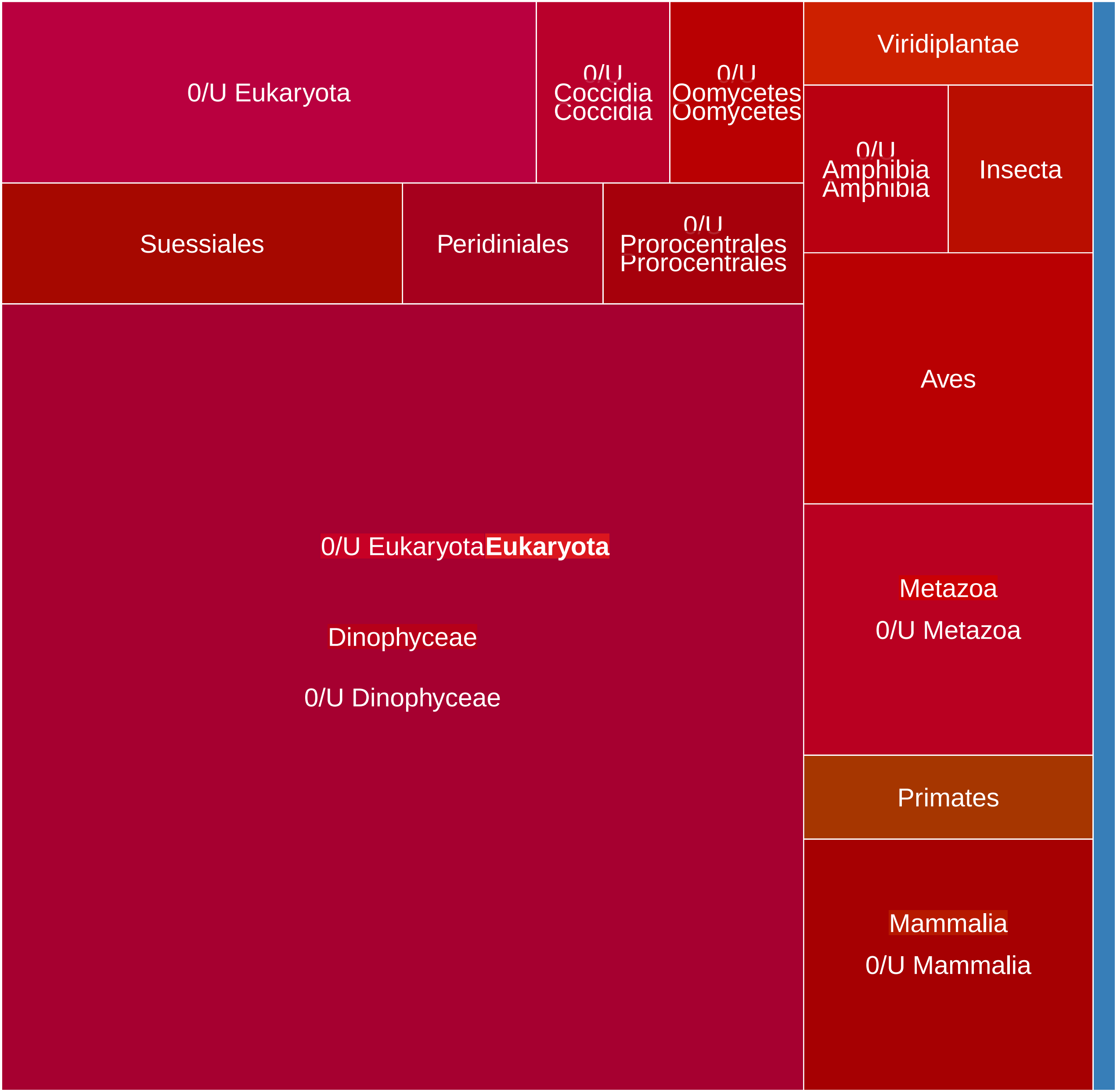

MGT−v1_685 : 907 unigenes, 140 (15.44%) taxonomically assigned

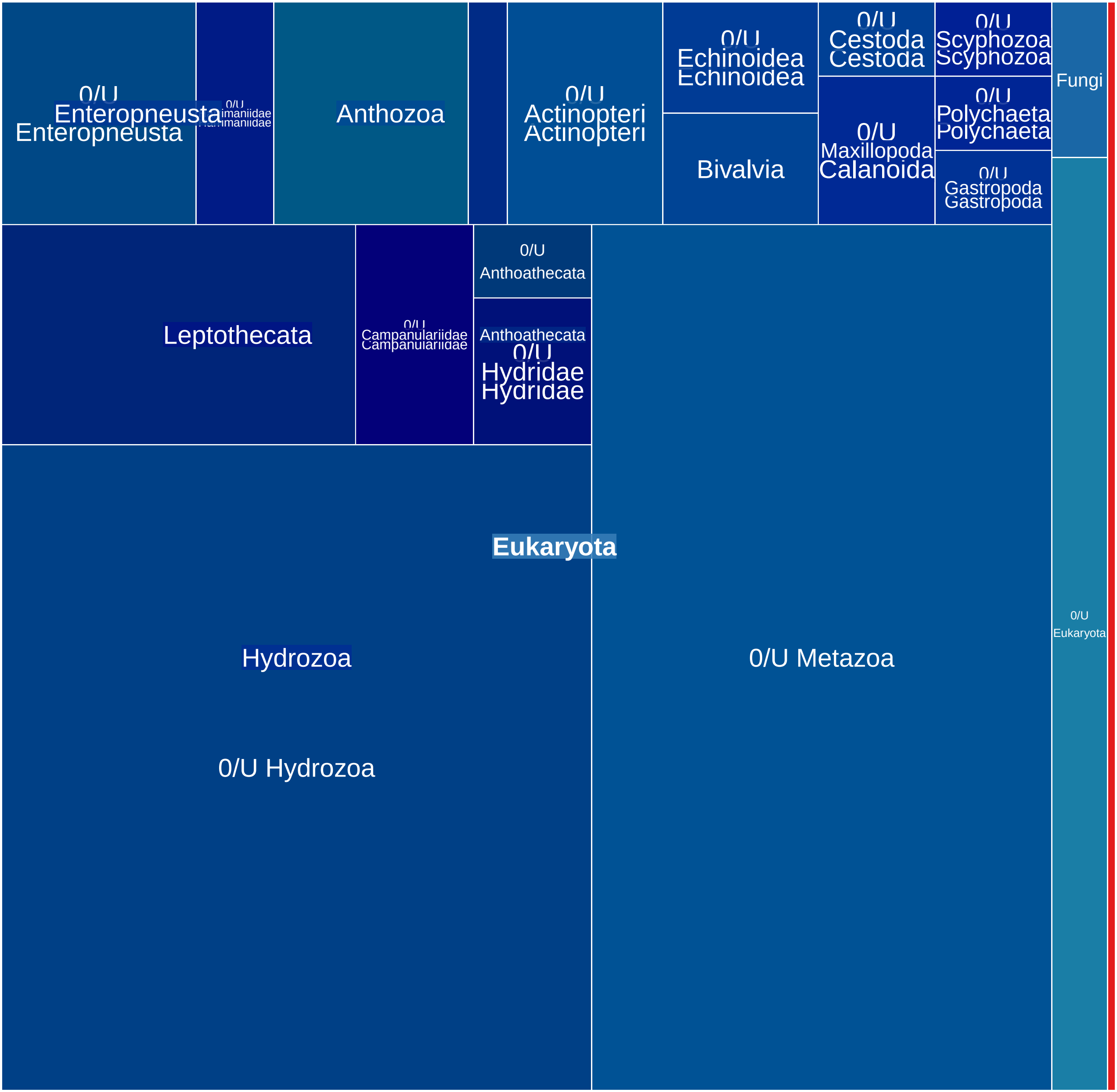

MGT−v1_686 : 901 unigenes, 47 (5.22%) taxonomically assigned

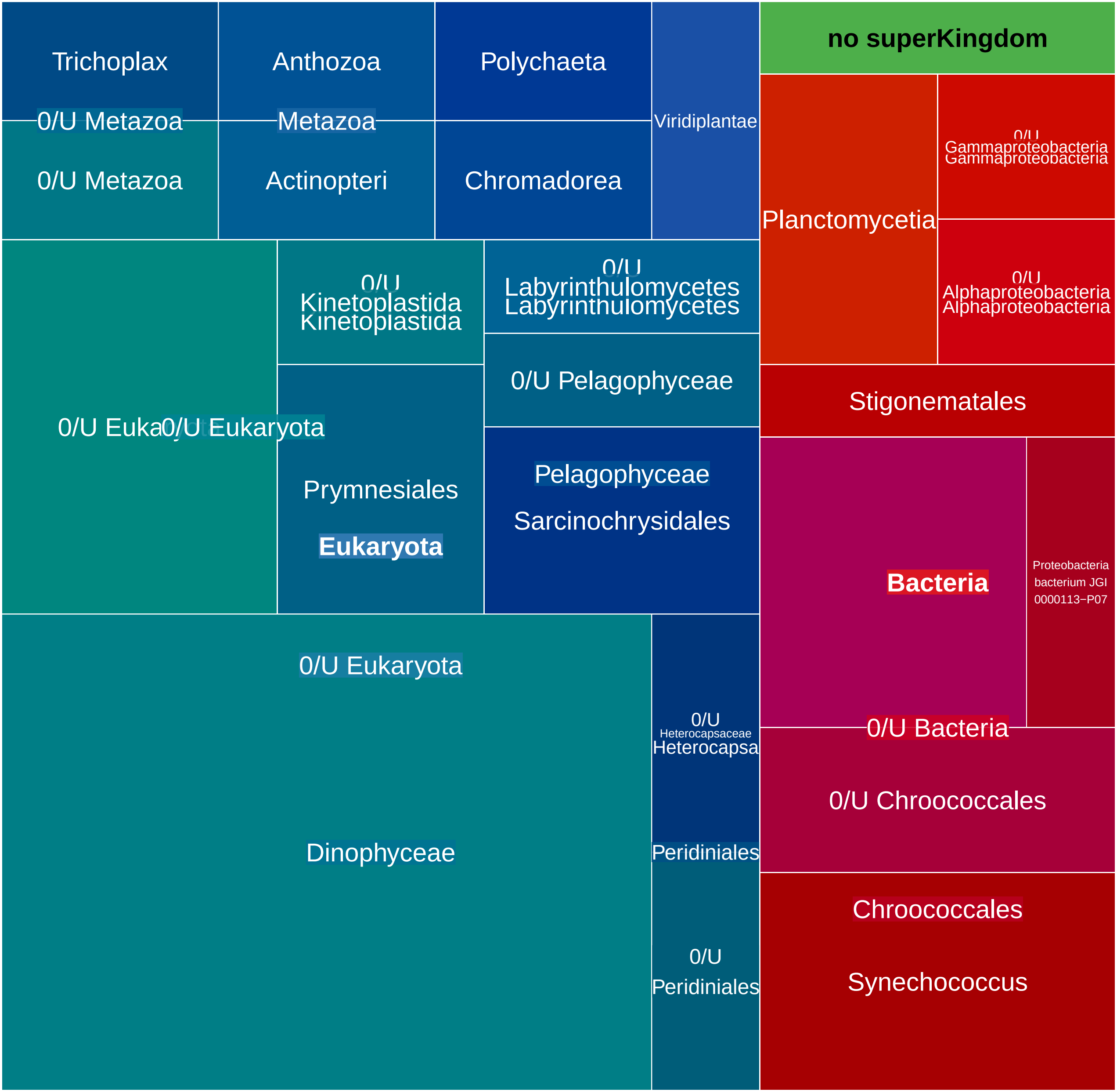

MGT−v1_687 : 900 unigenes, 72 (8.00%) taxonomically assigned

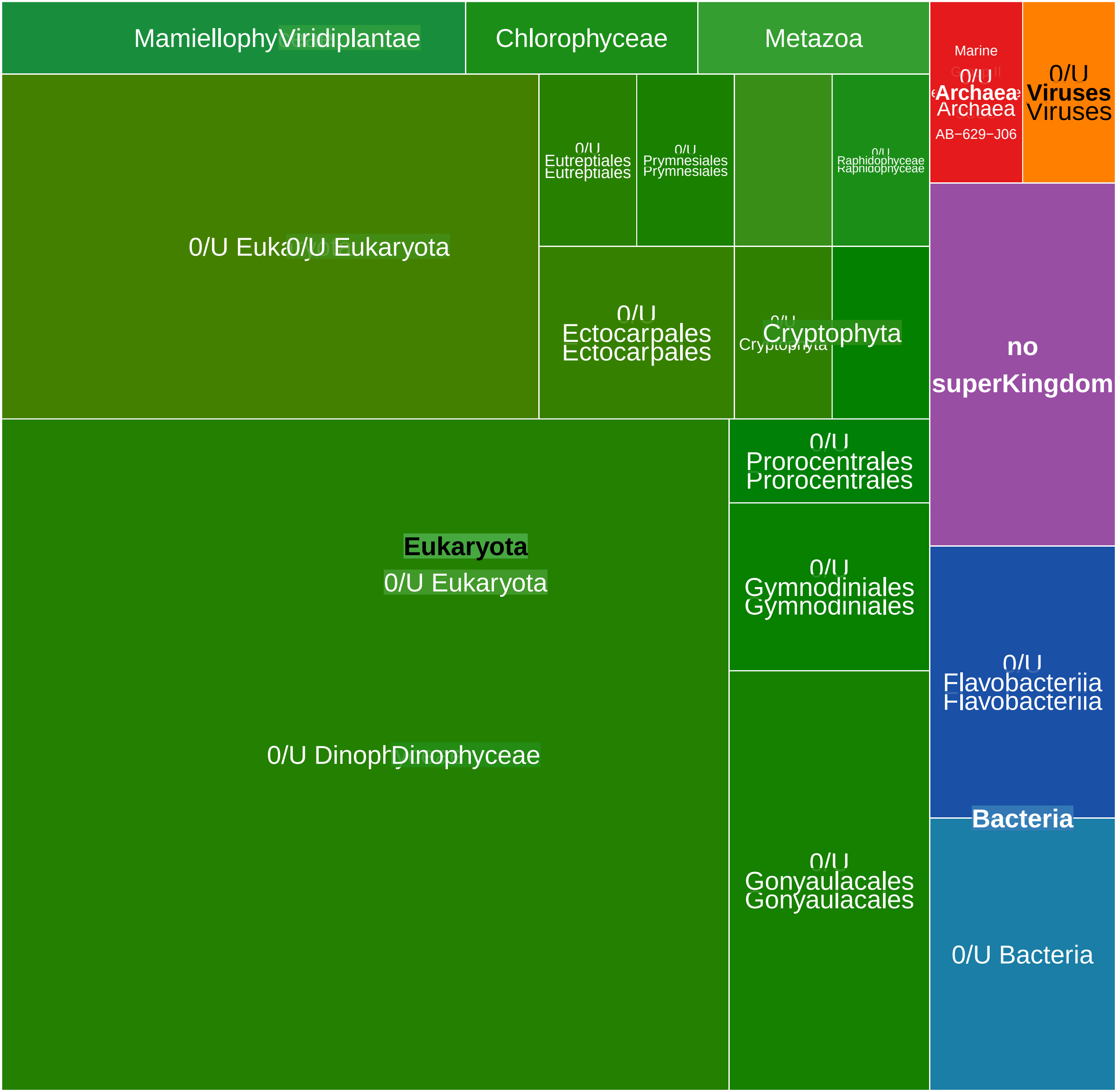

### MGT−v1_688 : 895 unigenes, 146 (16.31%) taxonomically assigned

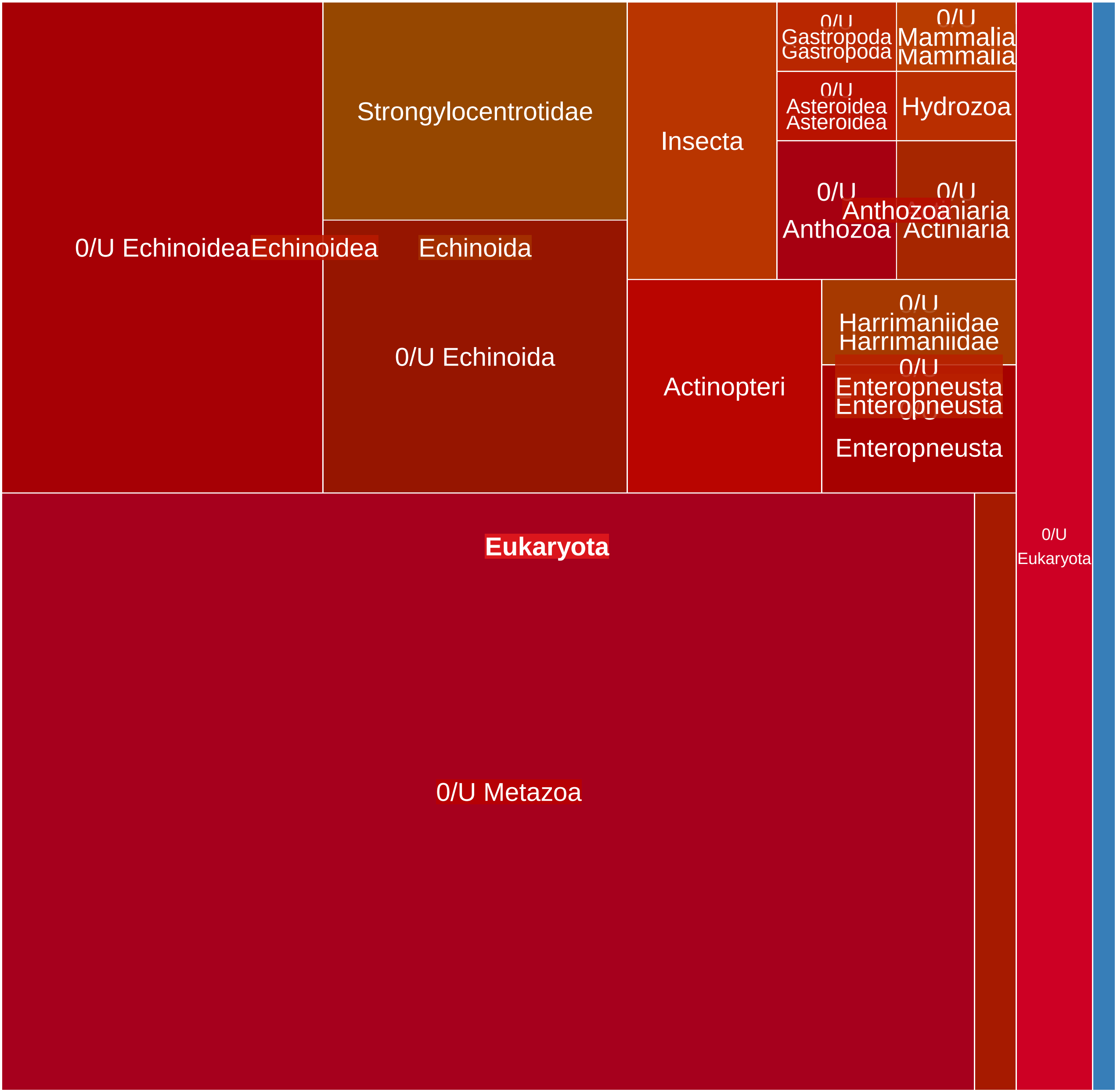

MGT−v1_689 : 892 unigenes, 780 (87.44%) taxonomically assigned

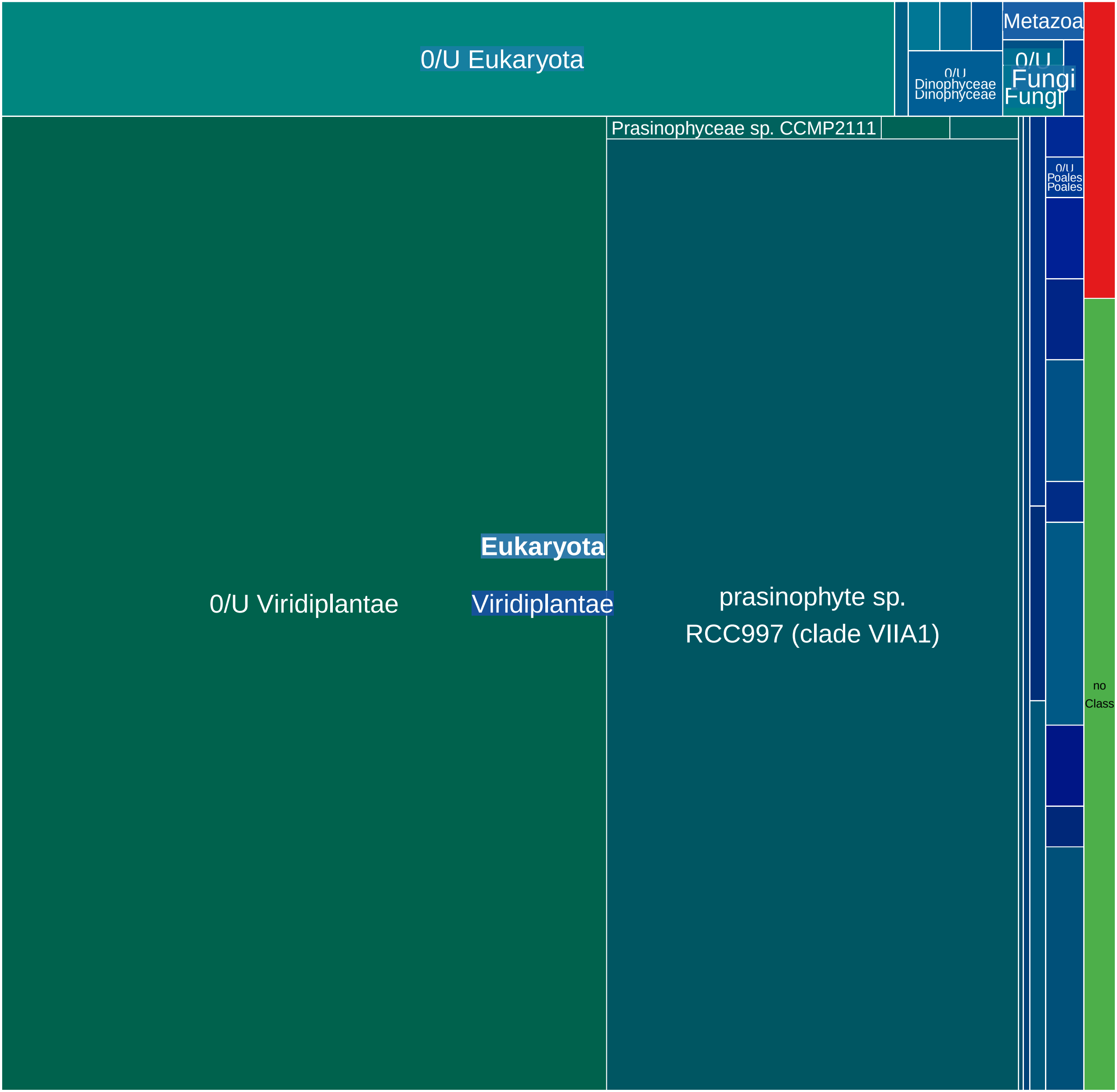

MGT−v1_690 : 887 unigenes, 69 (7.78%) taxonomically assigned

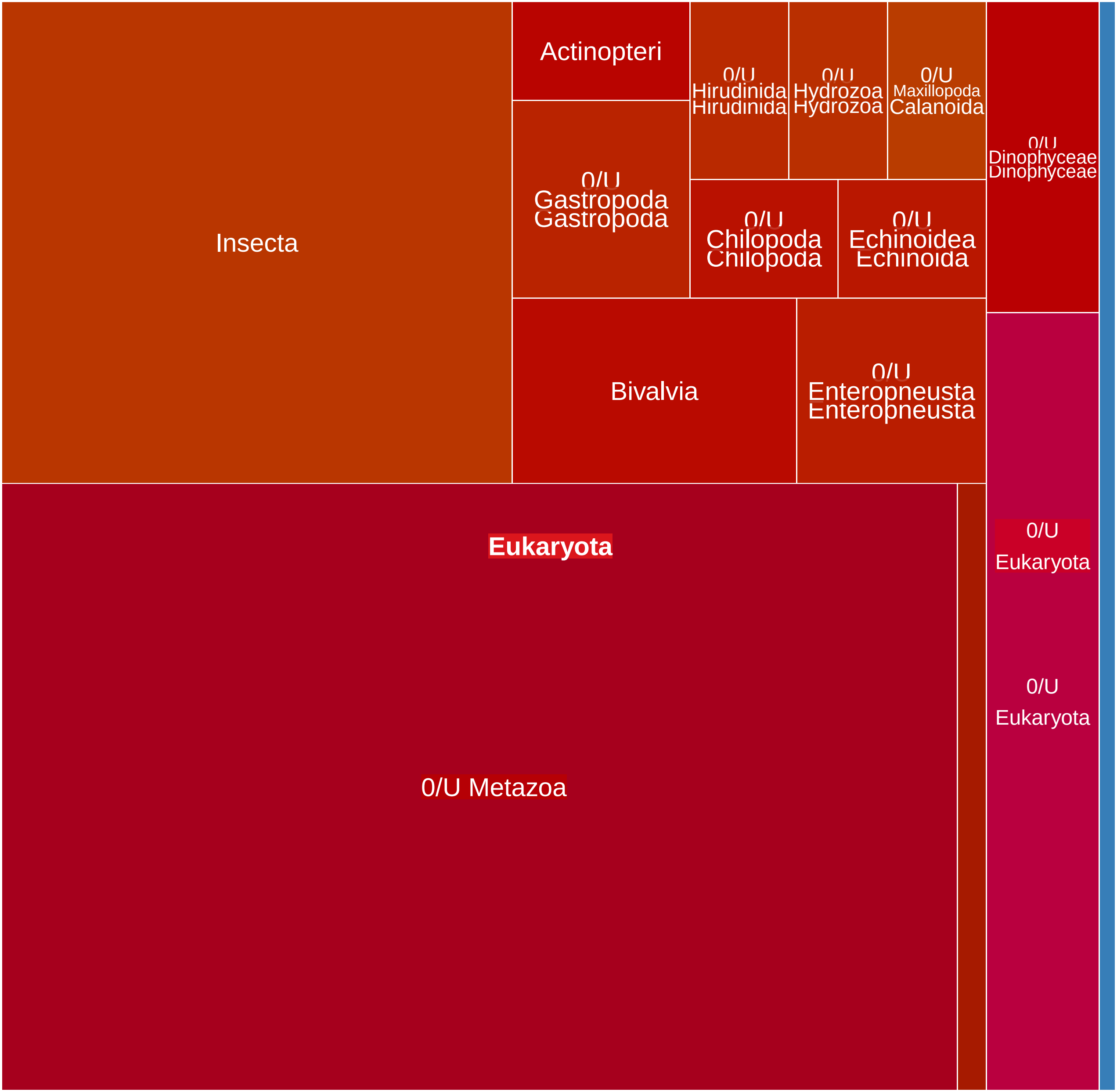

MGT−v1_691 : 886 unigenes, 377 (42.55%) taxonomically assigned

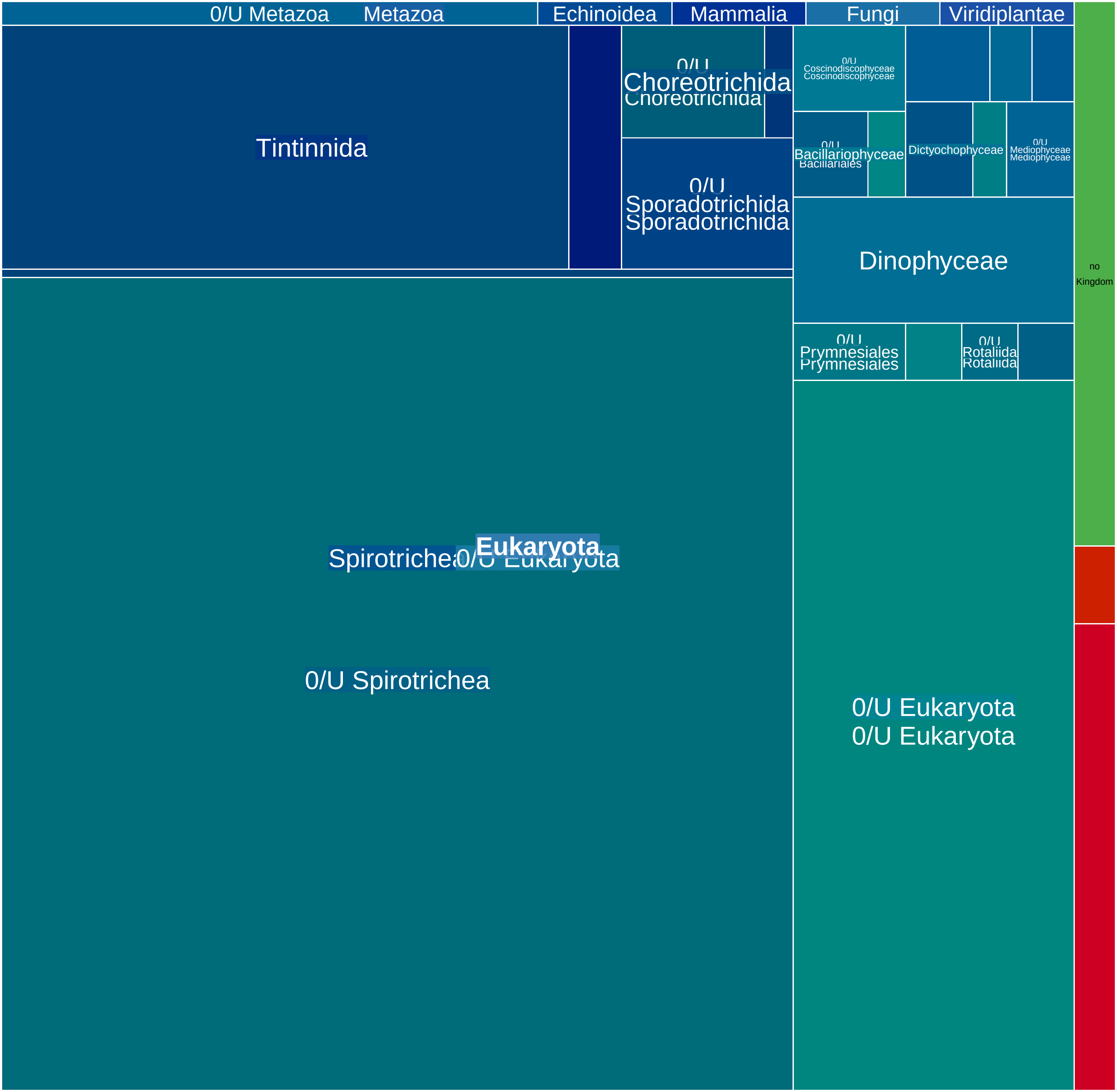

MGT−v1_692 : 885 unigenes, 830 (93.79%) taxonomically assigned

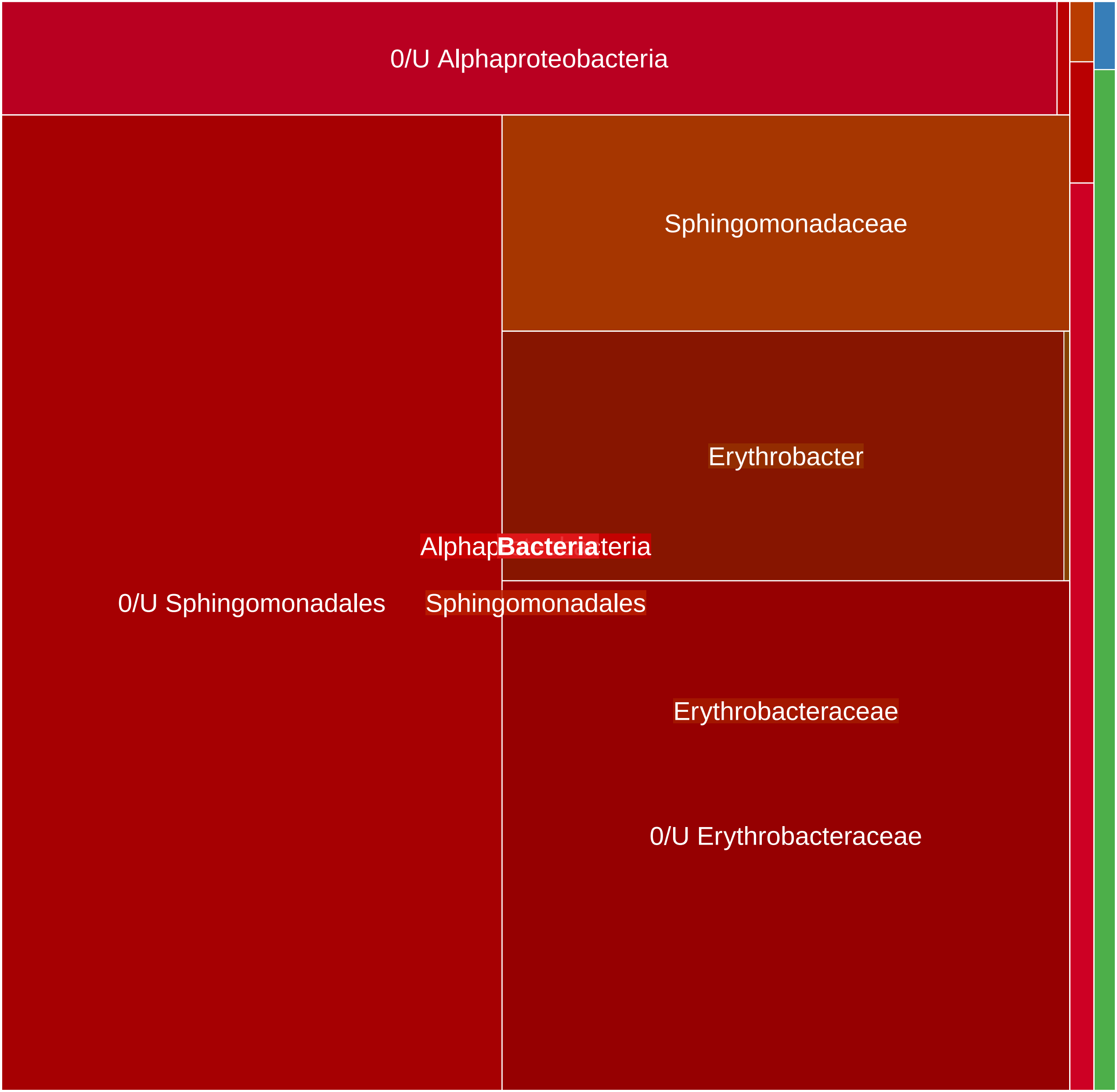

MGT−v1_693 : 885 unigenes, 533 (60.23%) taxonomically assigned

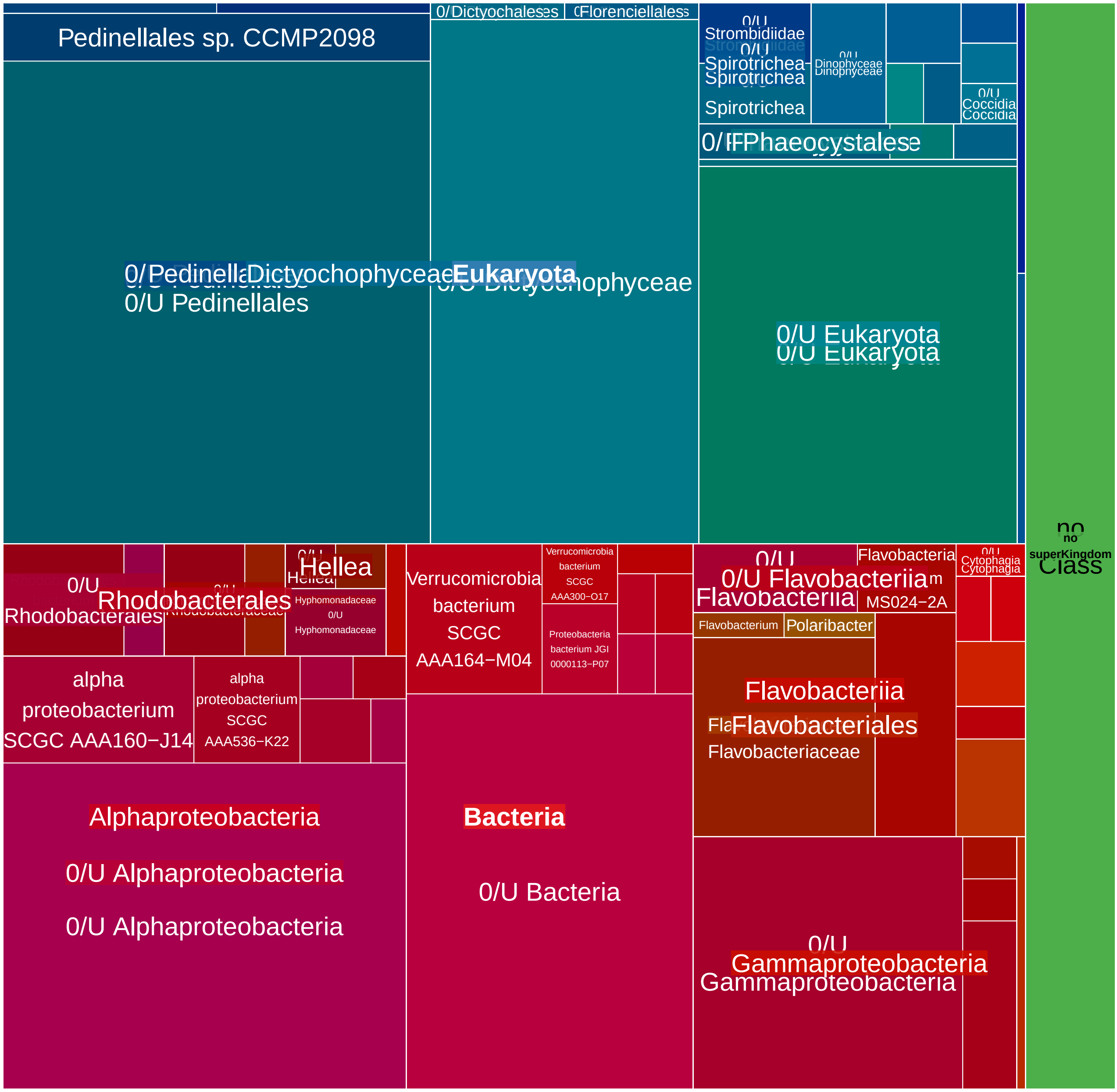

### MGT−v1_694 : 884 unigenes, 863 (97.62%) taxonomically assigned

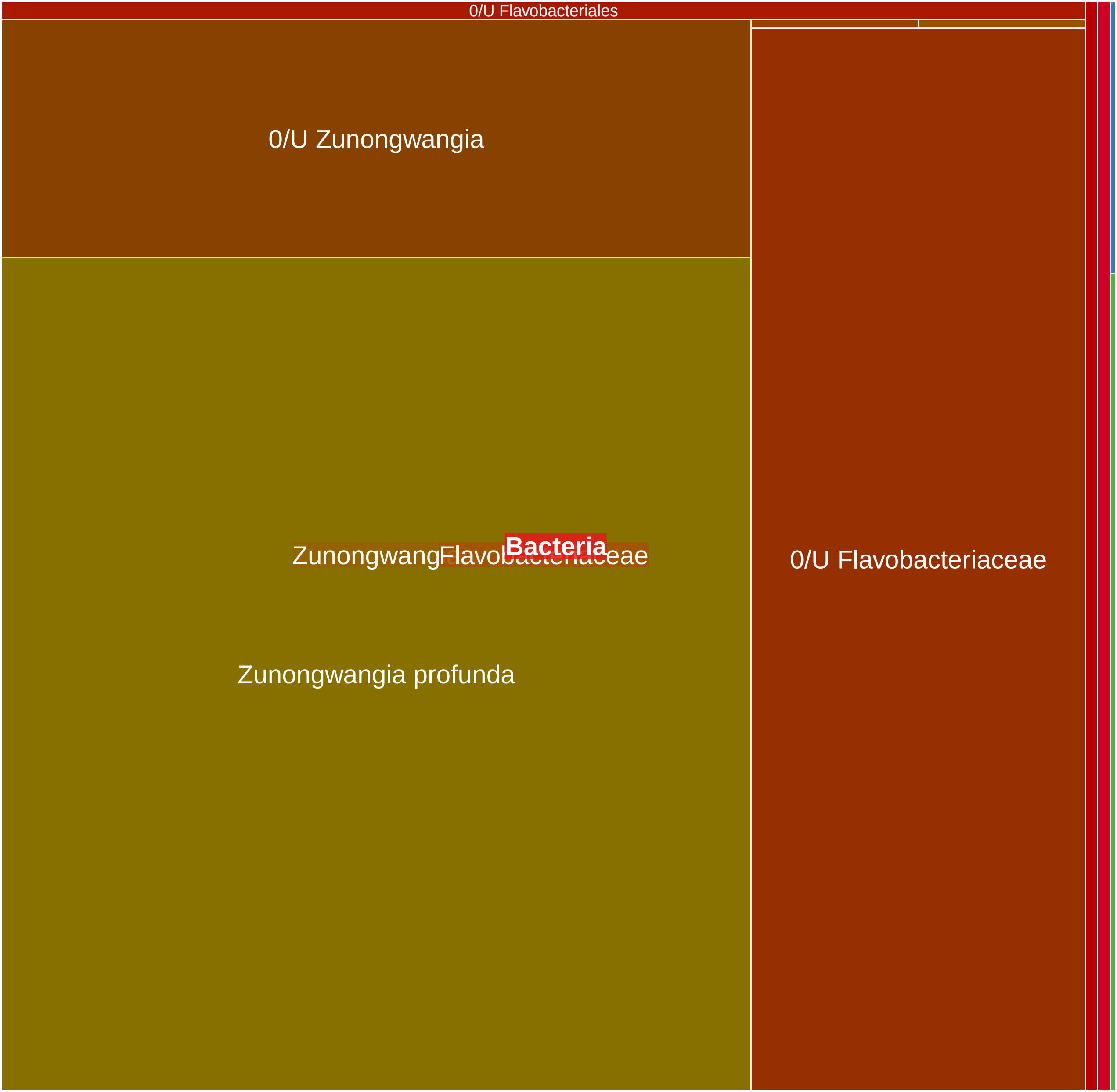

MGT−v1_695 : 883 unigenes, 63 (7.13%) taxonomically assigned

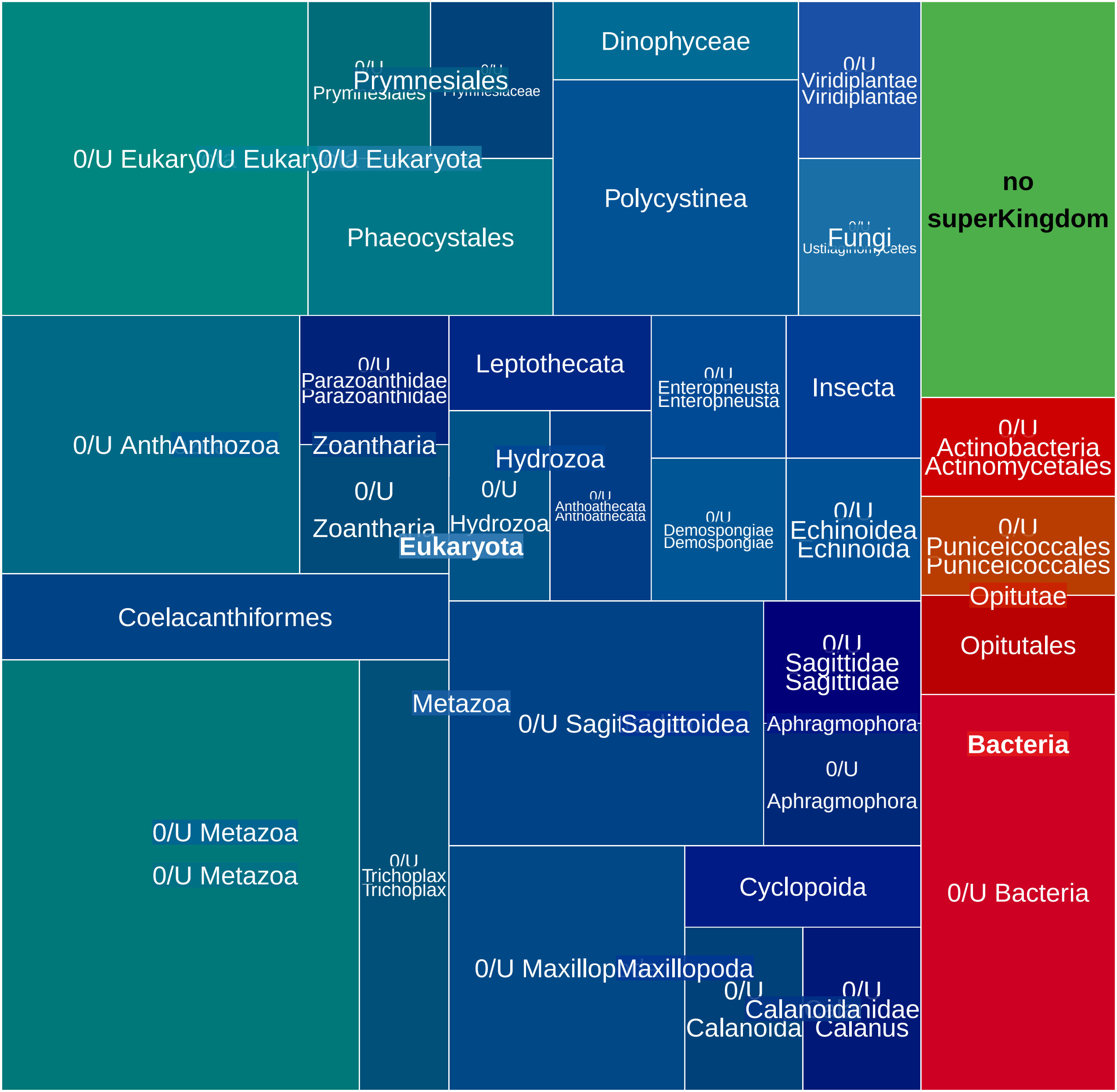

MGT−v1_696 : 883 unigenes, 652 (73.84%) taxonomically assigned

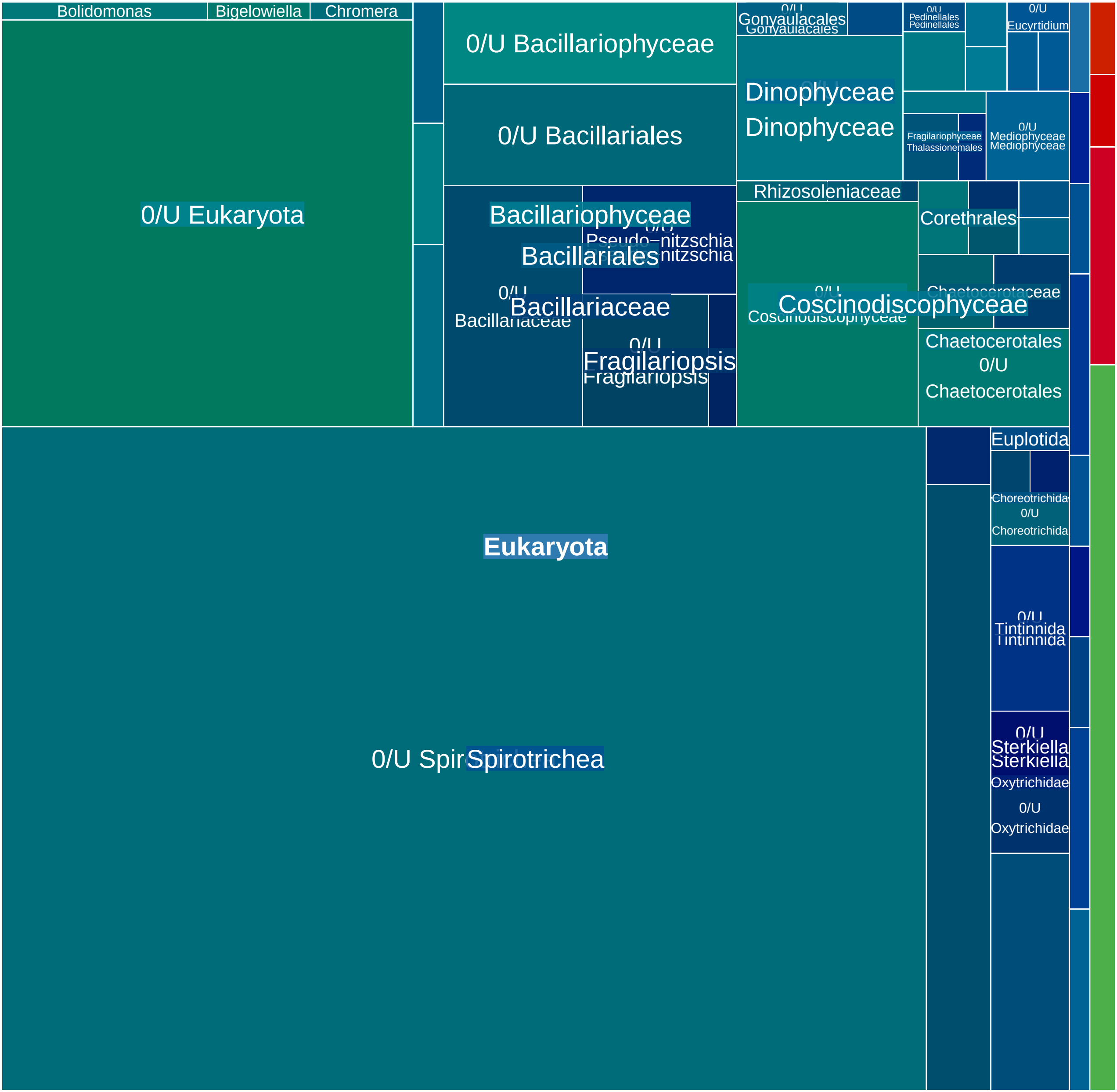

MGT−v1_697 : 877 unigenes, 688 (78.45%) taxonomically assigned

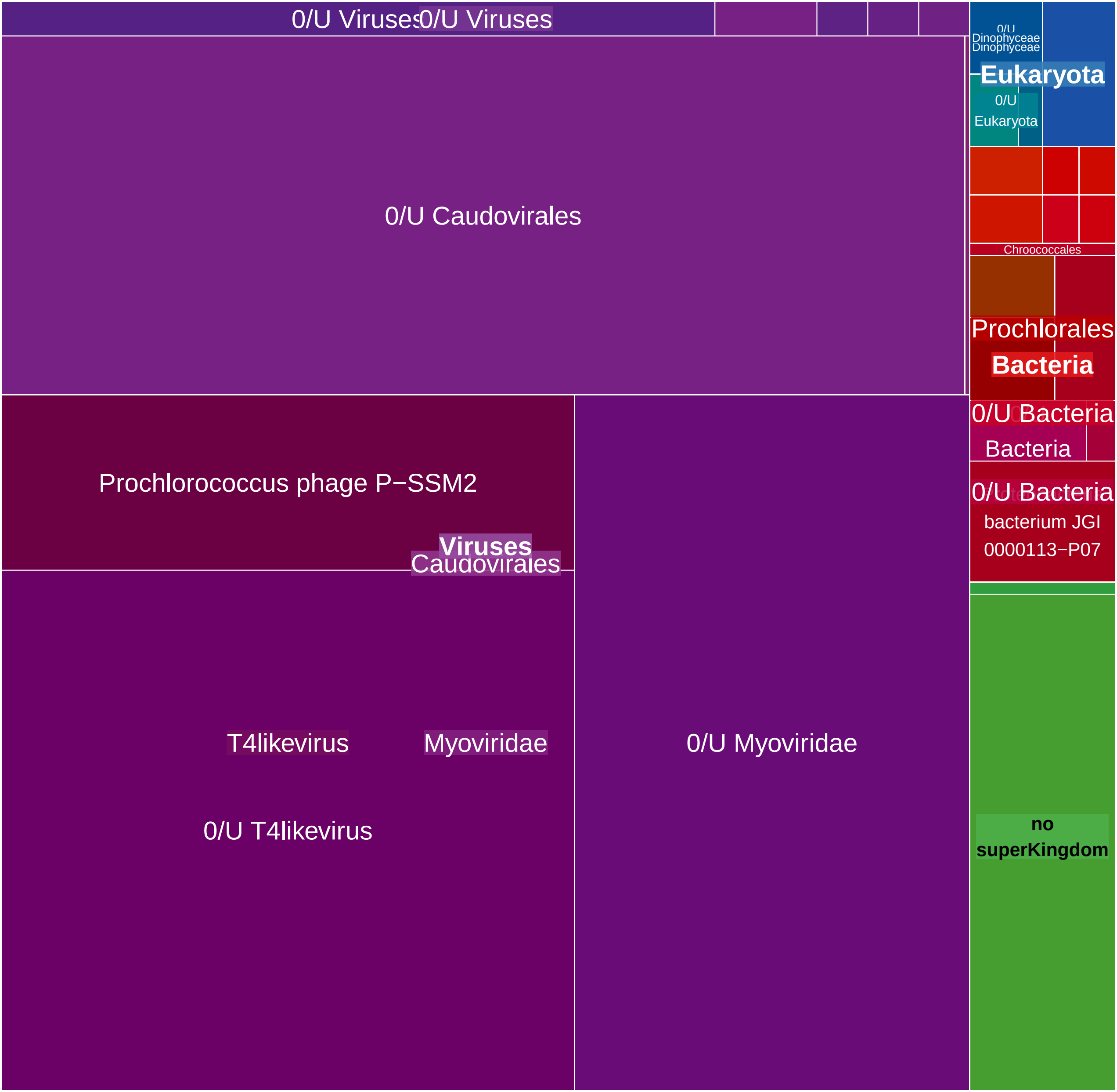

MGT−v1_698 : 873 unigenes, 792 (90.72%) taxonomically assigned

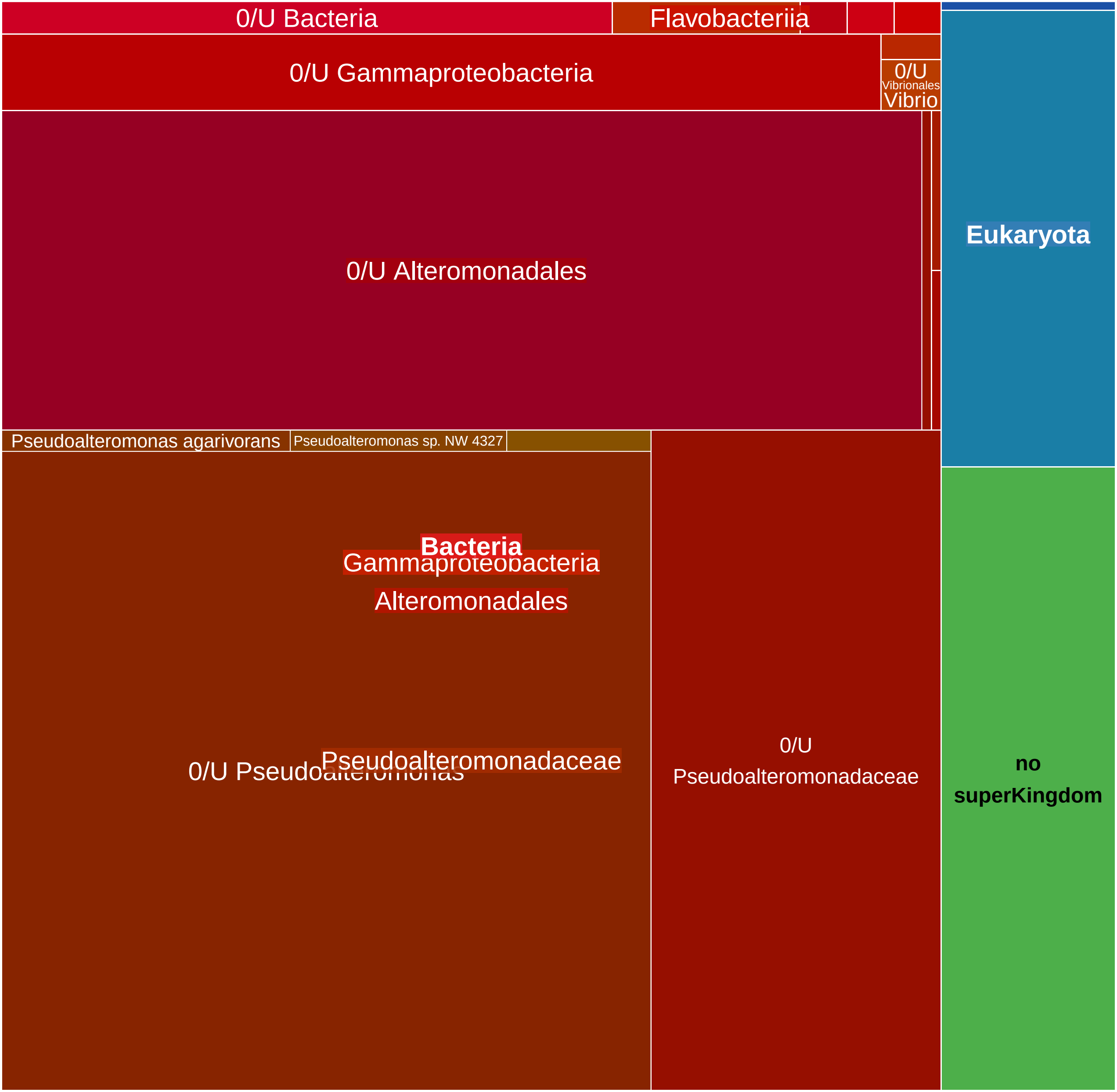

MGT−v1_699 : 873 unigenes, 79 (9.05%) taxonomically assigned

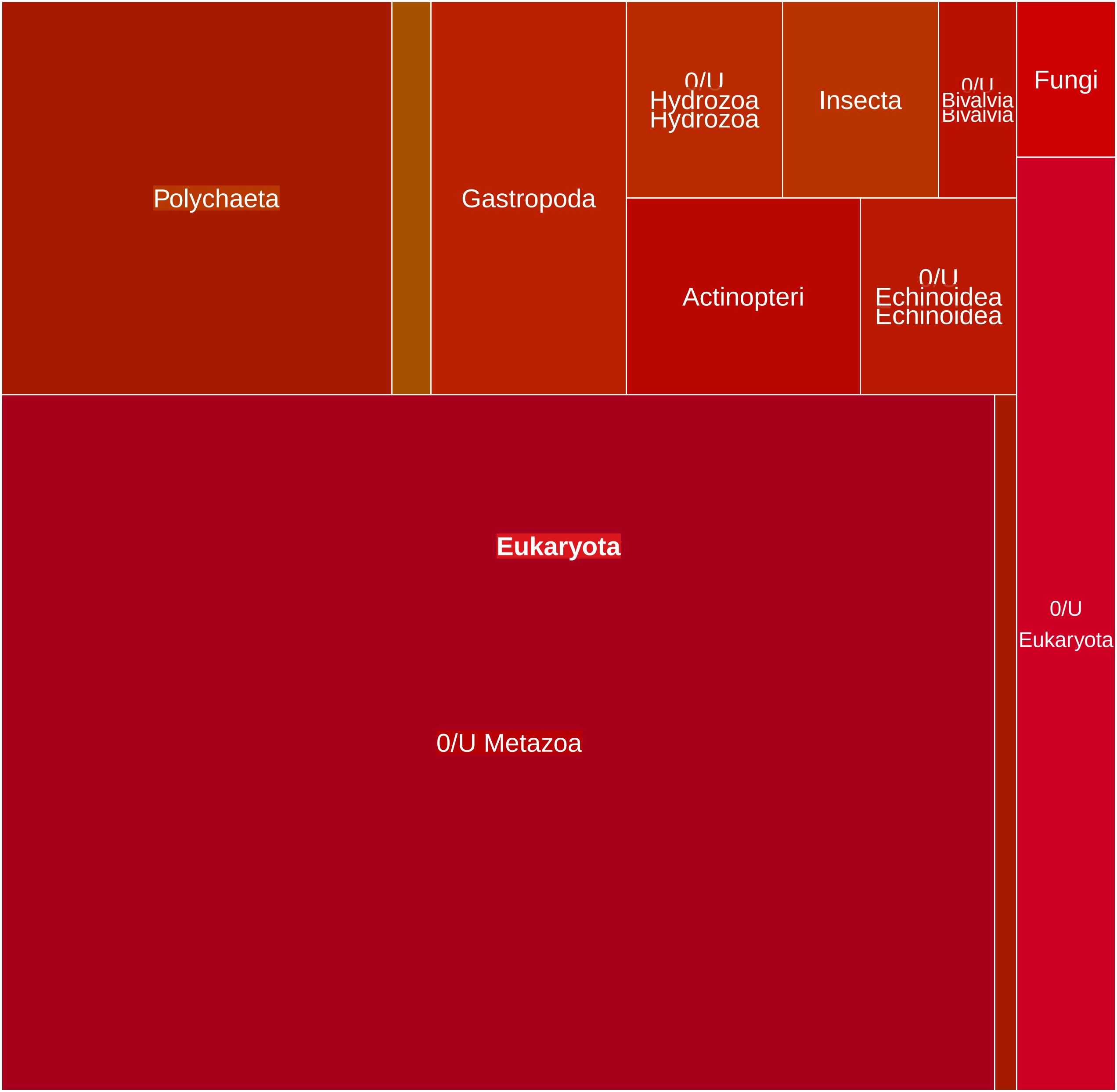

MGT−v1_700 : 871 unigenes, 606 (69.58%) taxonomically assigned

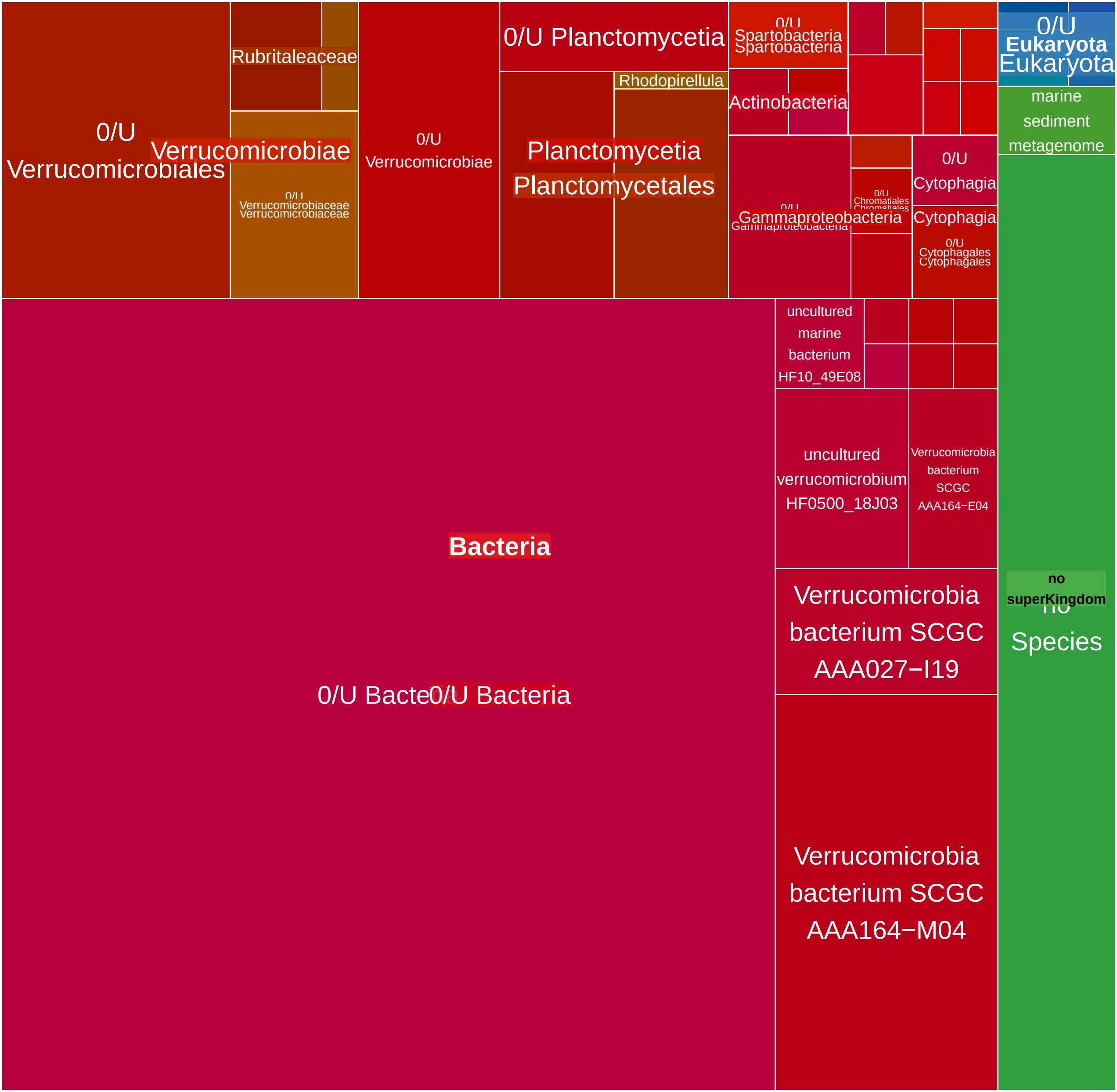

MGT−v1_701 : 871 unigenes, 219 (25.14%) taxonomically assigned

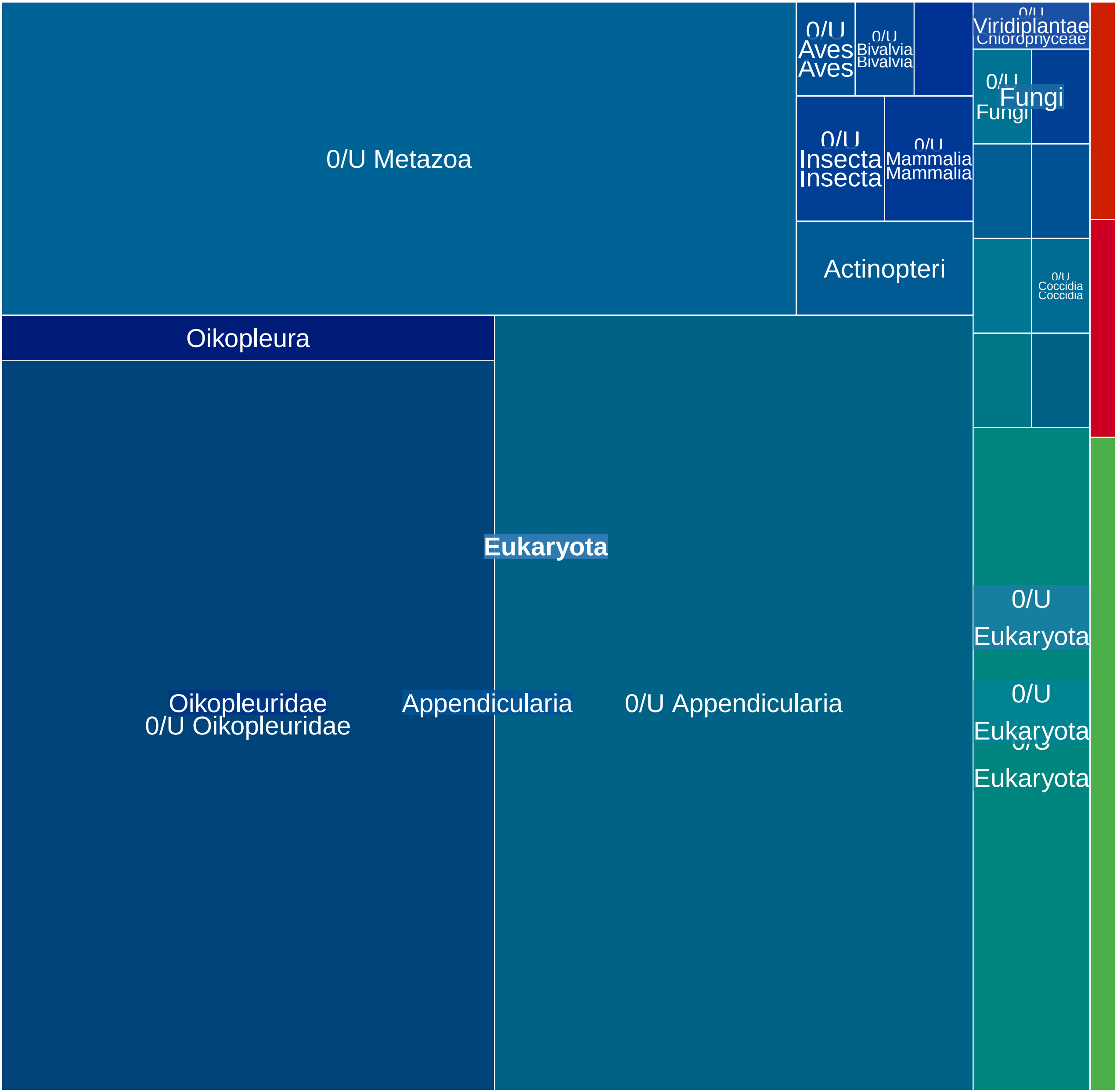

MGT−v1_702 : 870 unigenes, 754 (86.67%) taxonomically assigned

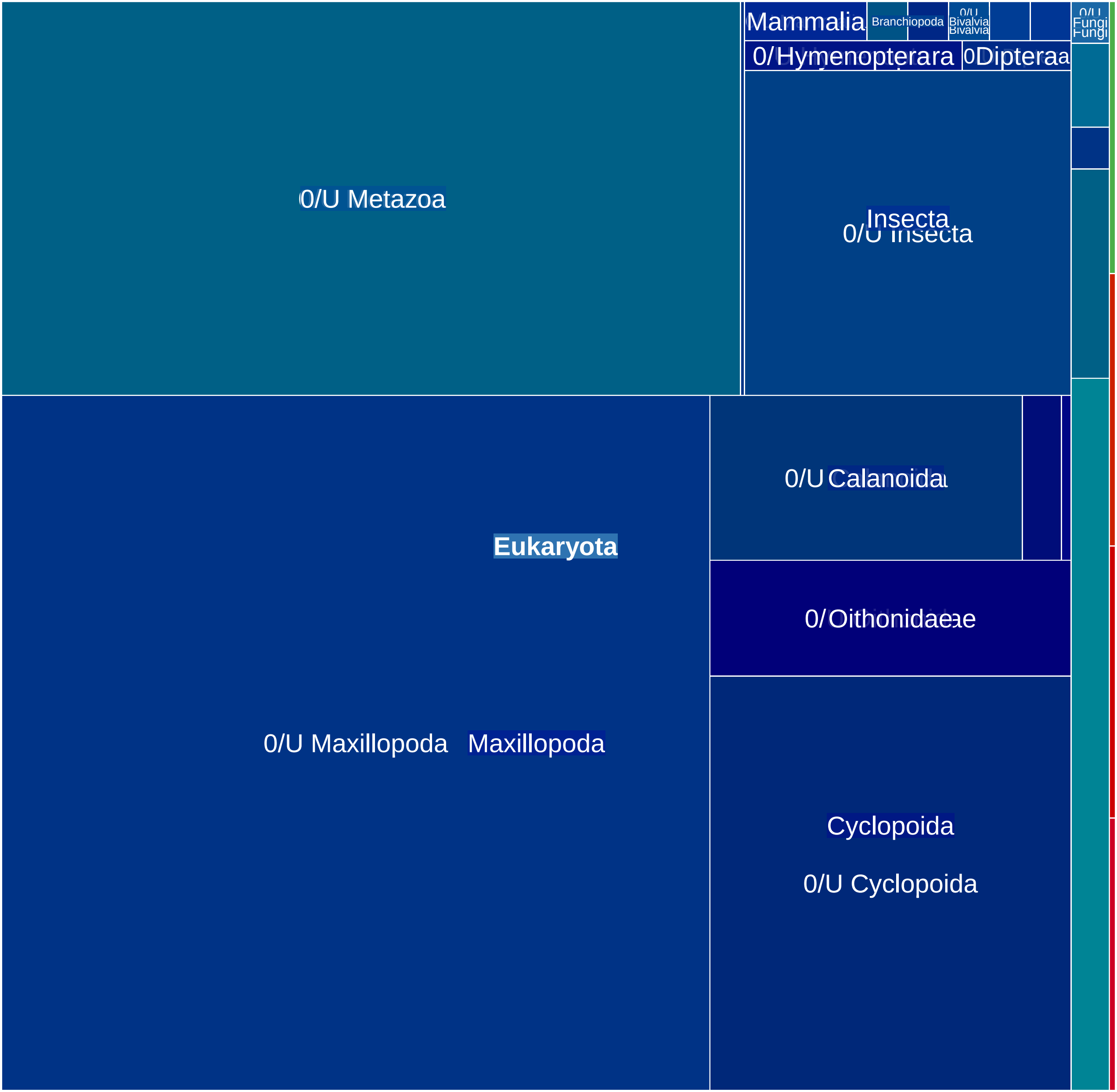

MGT−v1_703 : 870 unigenes, 761 (87.47%) taxonomically assigned

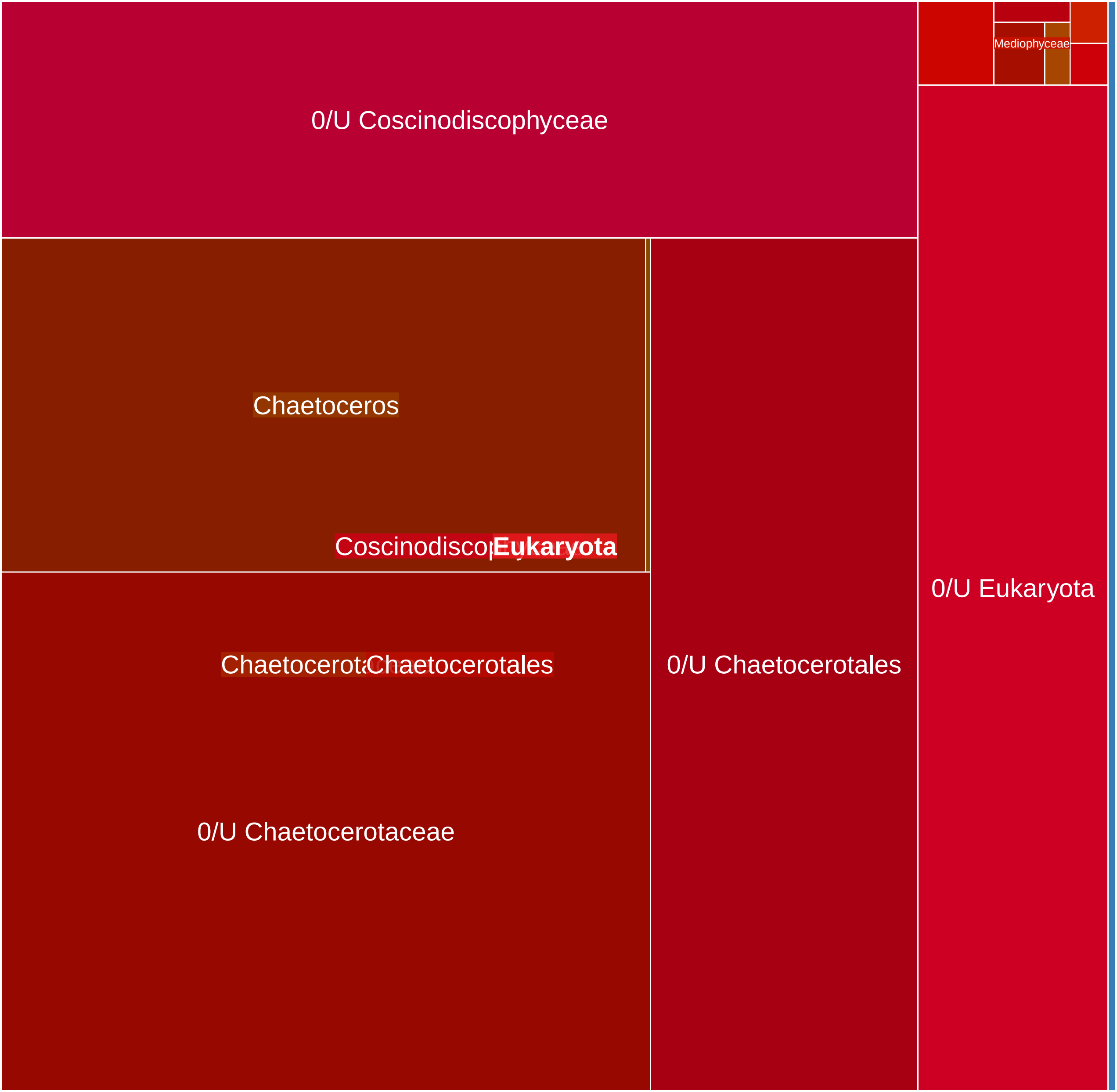

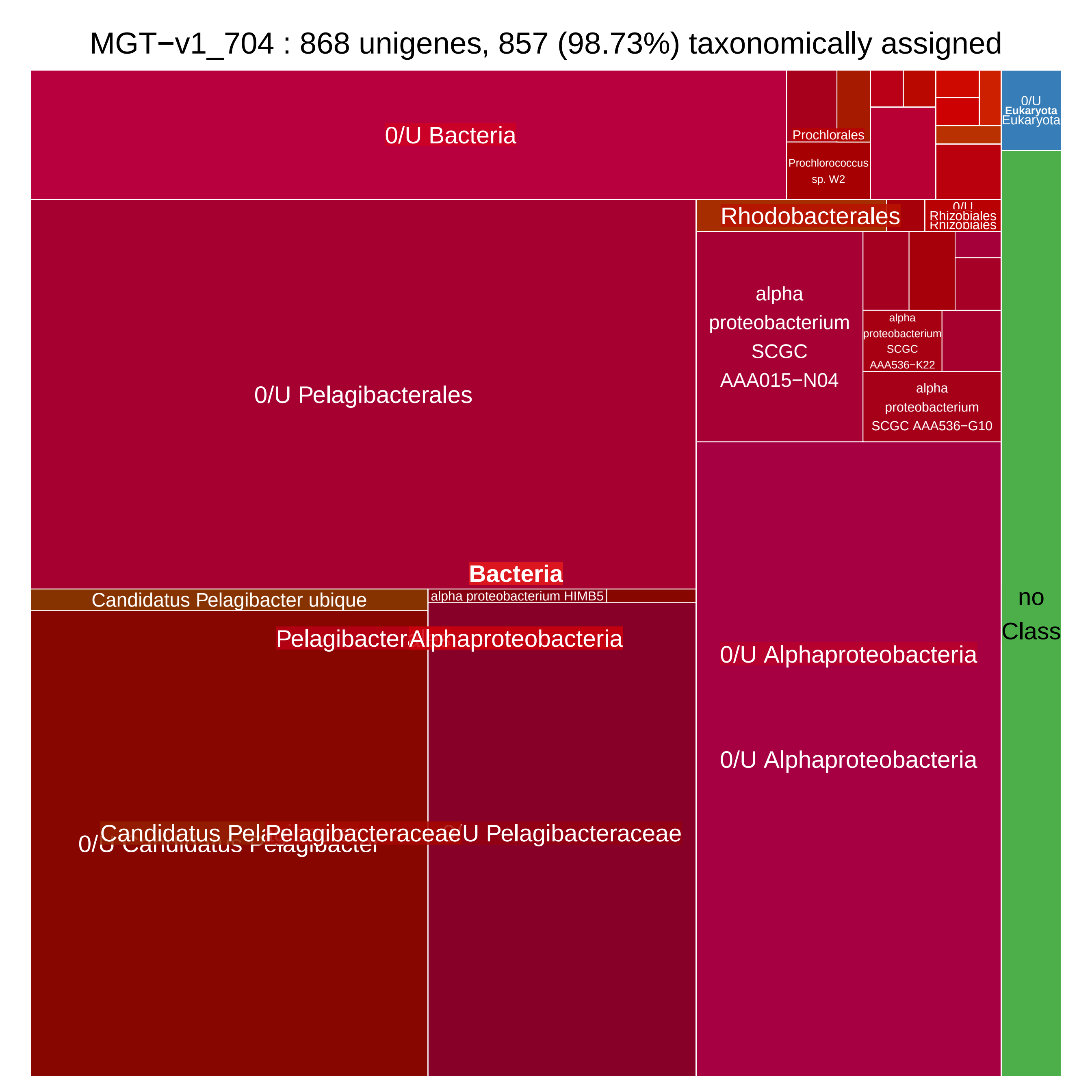
MGT−v1_704 : 868 unigenes, 857 (98.73%) taxonomically assigned
0/U Bacteria
Prochlorales
Prochlorococcus sp. W2
0/U Eukaryota
Eukaryota
Rhodobacterales
0/U Rhizobiales
Rhizobiales
0/U Pelagibacterales
alpha proteobacterium SCGC AAA015−N04
alpha proteobacterium SCGC AAA536−K22
alpha proteobacterium SCGC AAA536−G10
Bacteria
Candidatus Pelagibacter ubique
alpha proteobacterium HIMB5
Alphaproteobacteria
0/U Alphaproteobacteria
0/U Alphaproteobacteria
0/U Pelagibacteraceae
0/U Candidatus Pelagibacter
no Class

MGT−v1_705 : 867 unigenes, 107 (12.34%) taxonomically assigned

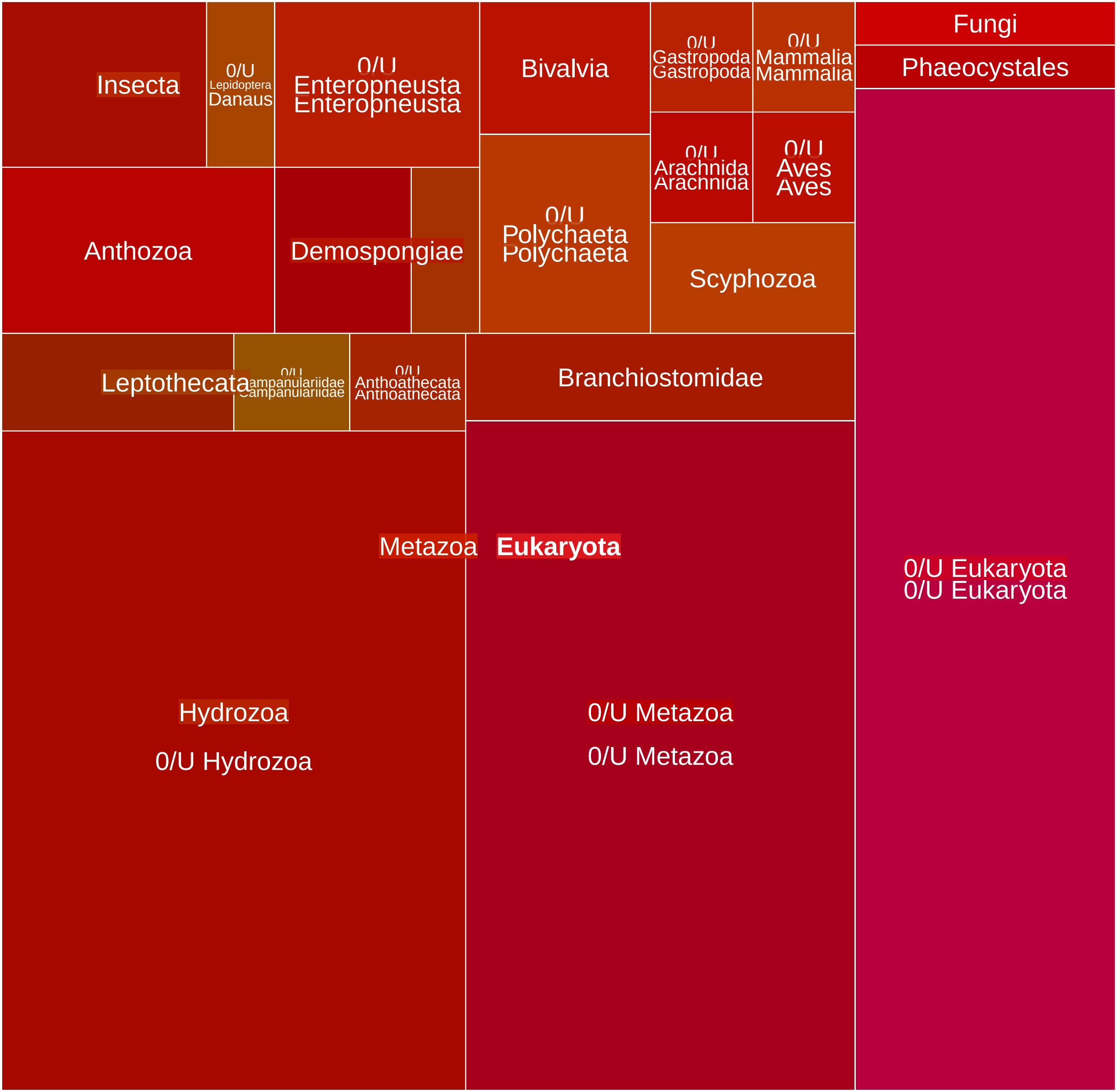

MGT−v1_706 : 853 unigenes, 230 (26.96%) taxonomically assigned

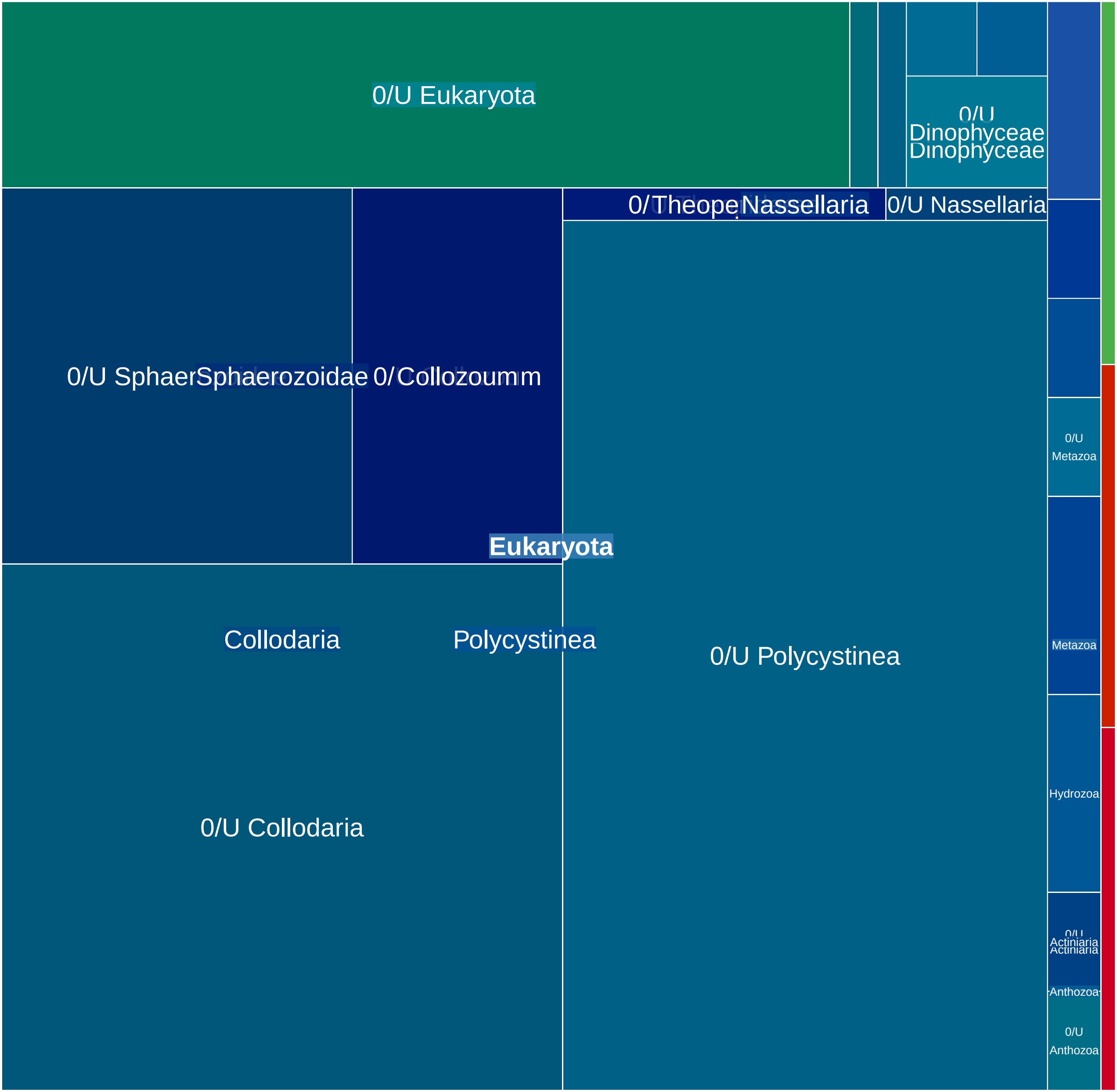

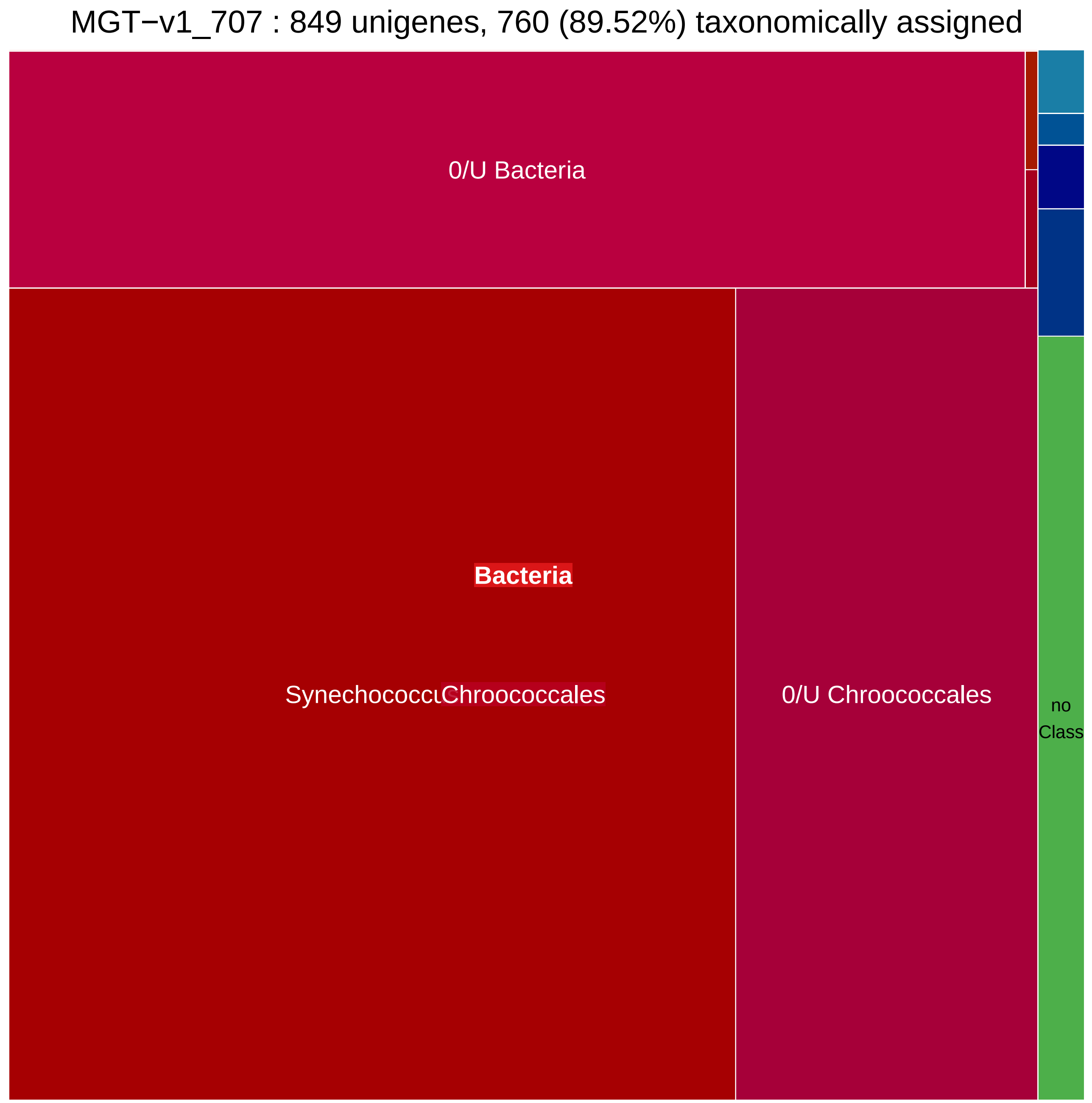
MGT−v1_707 : 849 unigenes, 760 (89.52%) taxonomically assigned
0/U Bacteria
Bacteria
Synechococcus
Chroococcales
0/U Chroococcales
no Class

MGT−v1_708 : 849 unigenes, 87 (10.25%) taxonomically assigned

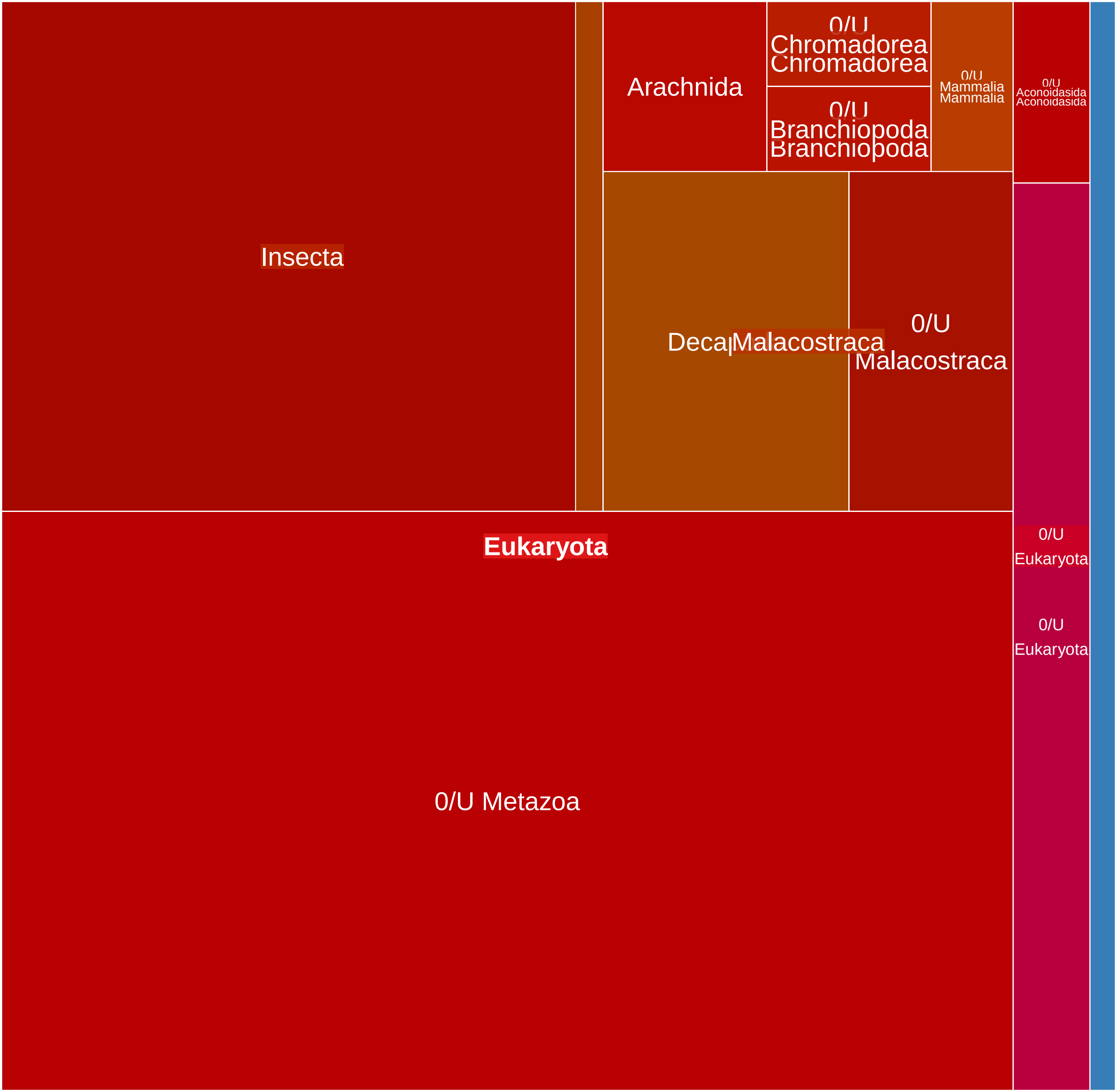

MGT−v1_709 : 845 unigenes, 431 (51.01%) taxonomically assigned

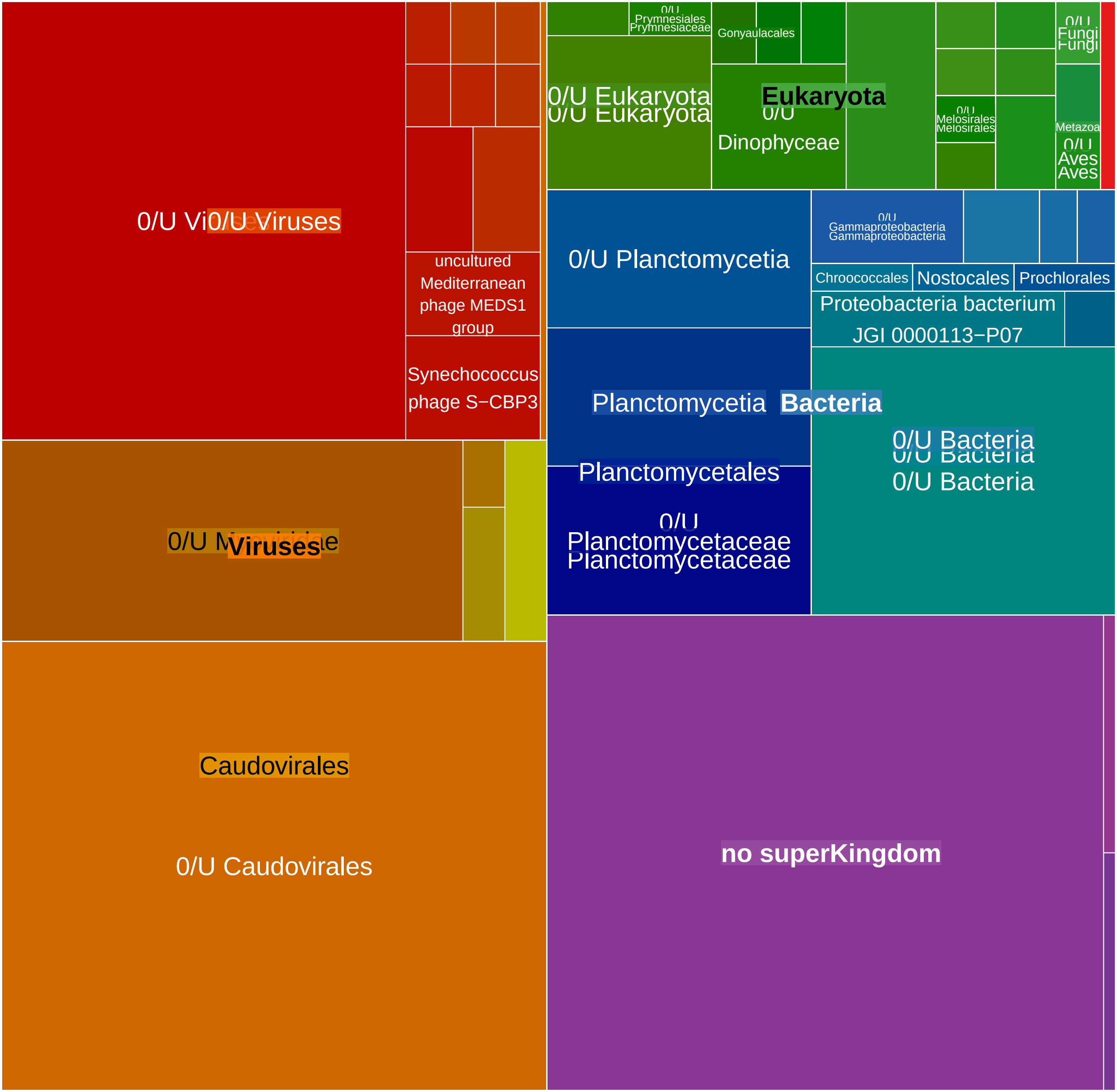

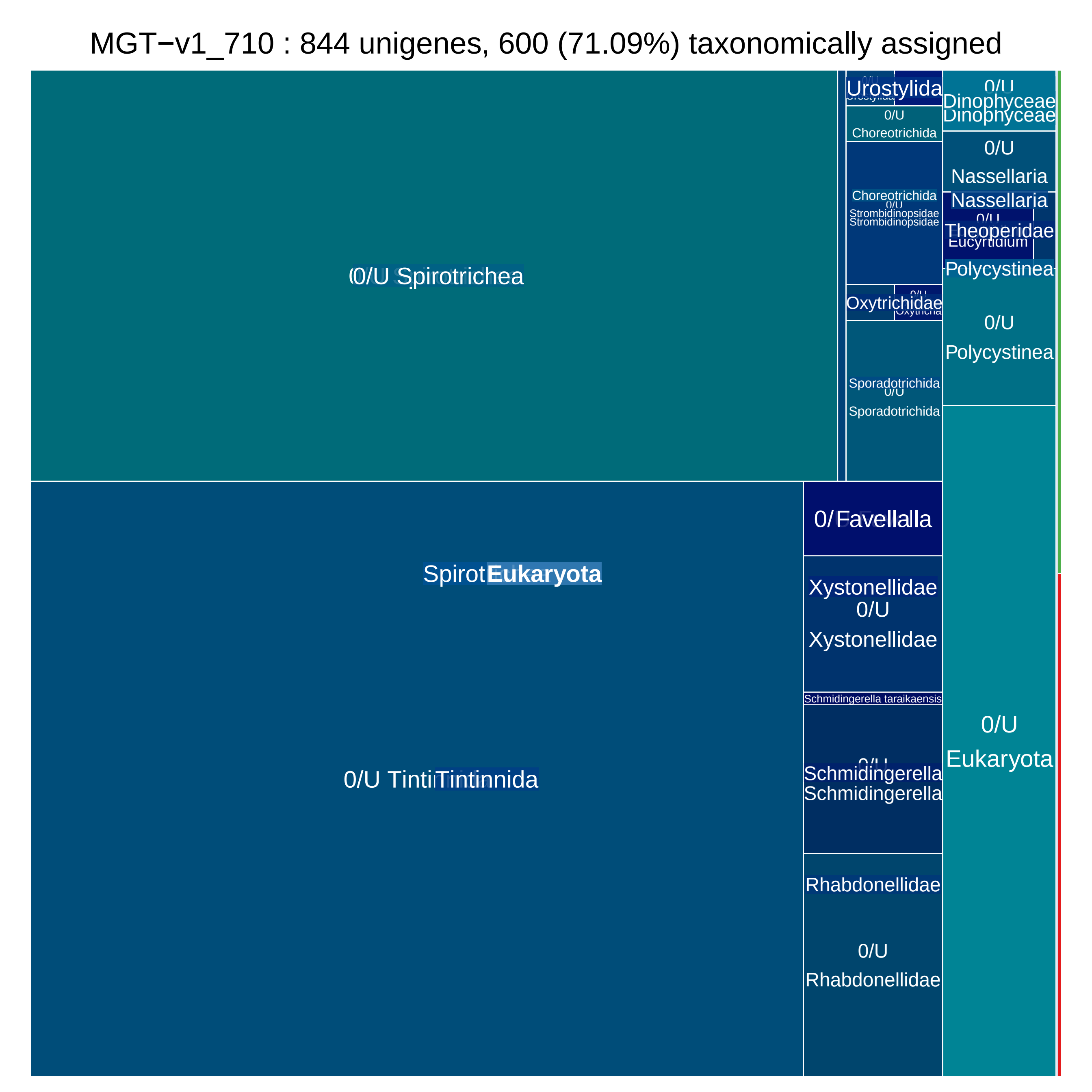
MGT−v1_710 : 844 unigenes, 600 (71.09%) taxonomically assigned
0/U Spirotrichea
Eukaryota
Spirot
0/U Tintinnida
Tintinnida
Urostylida
0/U
Choreotrichida
Choreotrichida
Strombidinopsidae
Strombidinopsidae
Oxytrichidae
Sporadotrichida
Sporadotrichida
0/U
Dinophyceae
Dinophyceae
0/U
Nassellaria
Nassellaria
Theoperidae
Eucyrtidium
Polycystinea
0/U
Polycystinea
0/U
Eukaryota
0/Favellalla
Xystonellidae
0/U
Xystonellidae
Schmidingerella taraikaensis
Schmidingerella
Schmidingerella
Rhabdonellidae
0/U
Rhabdonellidae

### MGT−v1_711 : 843 unigenes, 411 (48.75%) taxonomically assigned

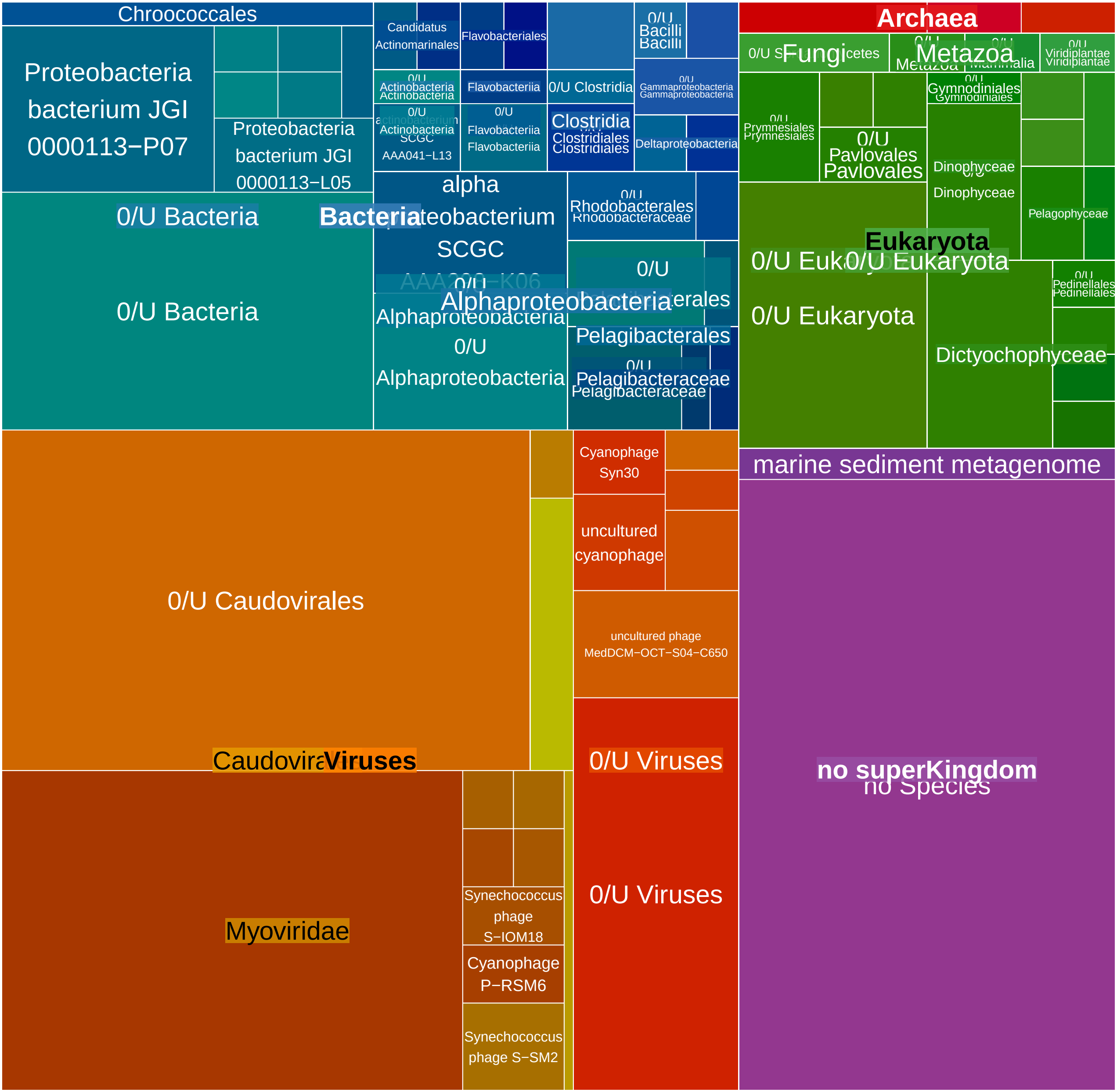

MGT−v1_712 : 841 unigenes, 493 (58.62%) taxonomically assigned

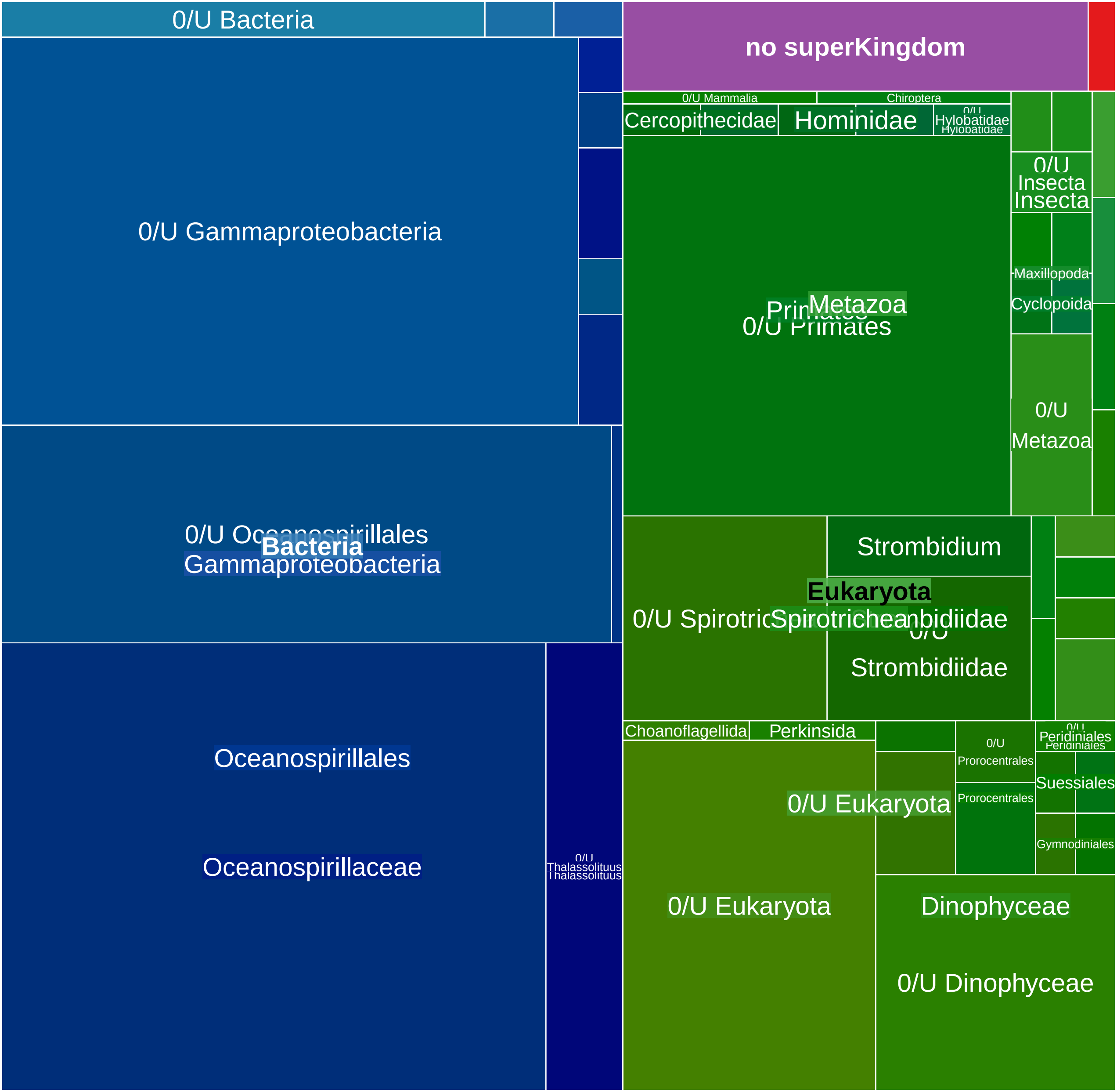

MGT−v1_713 : 840 unigenes, 29 (3.45%) taxonomically assigned

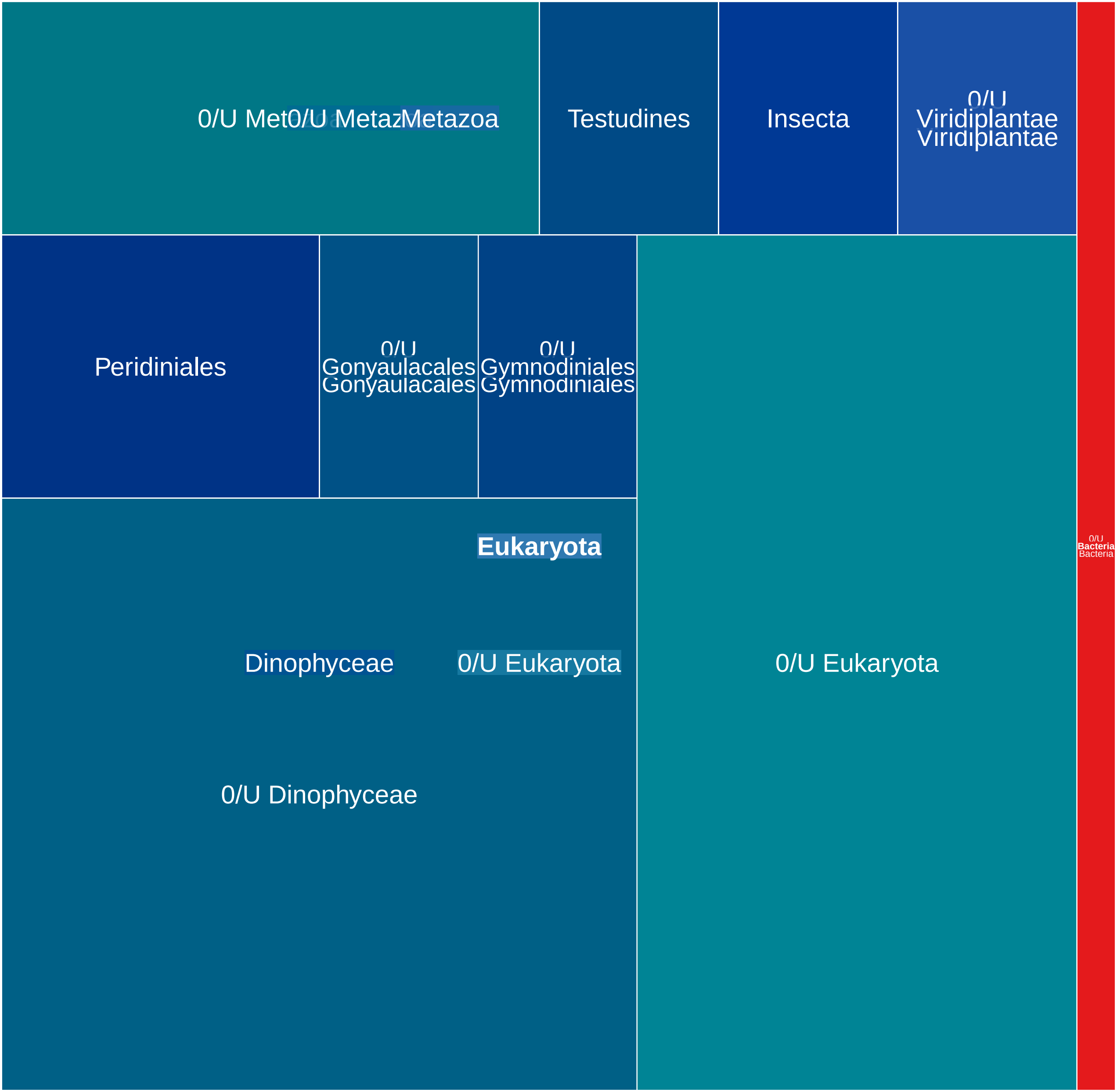

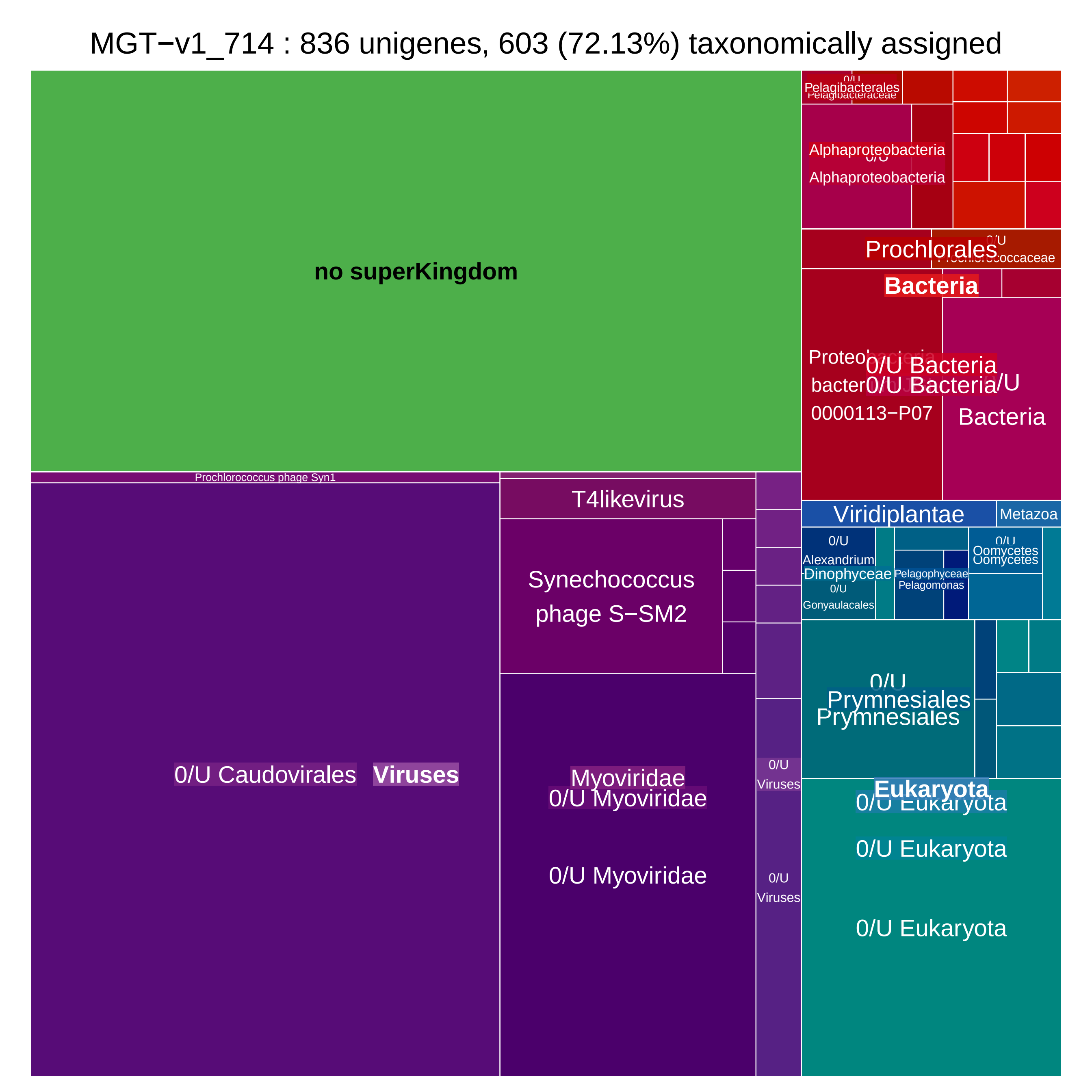
MGT−v1_714 : 836 unigenes, 603 (72.13%) taxonomically assigned
no superKingdom
Prochlorococcus phage Syn1
0/U Caudovirales
Viruses
T4likevirus
Synechococcus phage S−SM2
Myoviridae
0/U Myoviridae
0/U Myoviridae
0/U Viruses
0/U Viruses
Pelagibacterales
Alphaproteobacteria
Alphaproteobacteria
Prochlorales
Prochlorococcaceae
Bacteria
0/U Bacteria
0/U Bacteria
0000113−P07
Bacteria
Viridiplantae
Metazoa
0/U Alexandrium
Dinophyceae
0/U Gonyaulacales
Pelagophyceae
Pelagomonas
0/U Oomycetes
Oomycetes
0/U Prymnesiales
Prymnesiales
Eukaryota
0/U Eukaryota
0/U Eukaryota
0/U Eukaryota

MGT−v1_715 : 834 unigenes, 135 (16.19%) taxonomically assigned

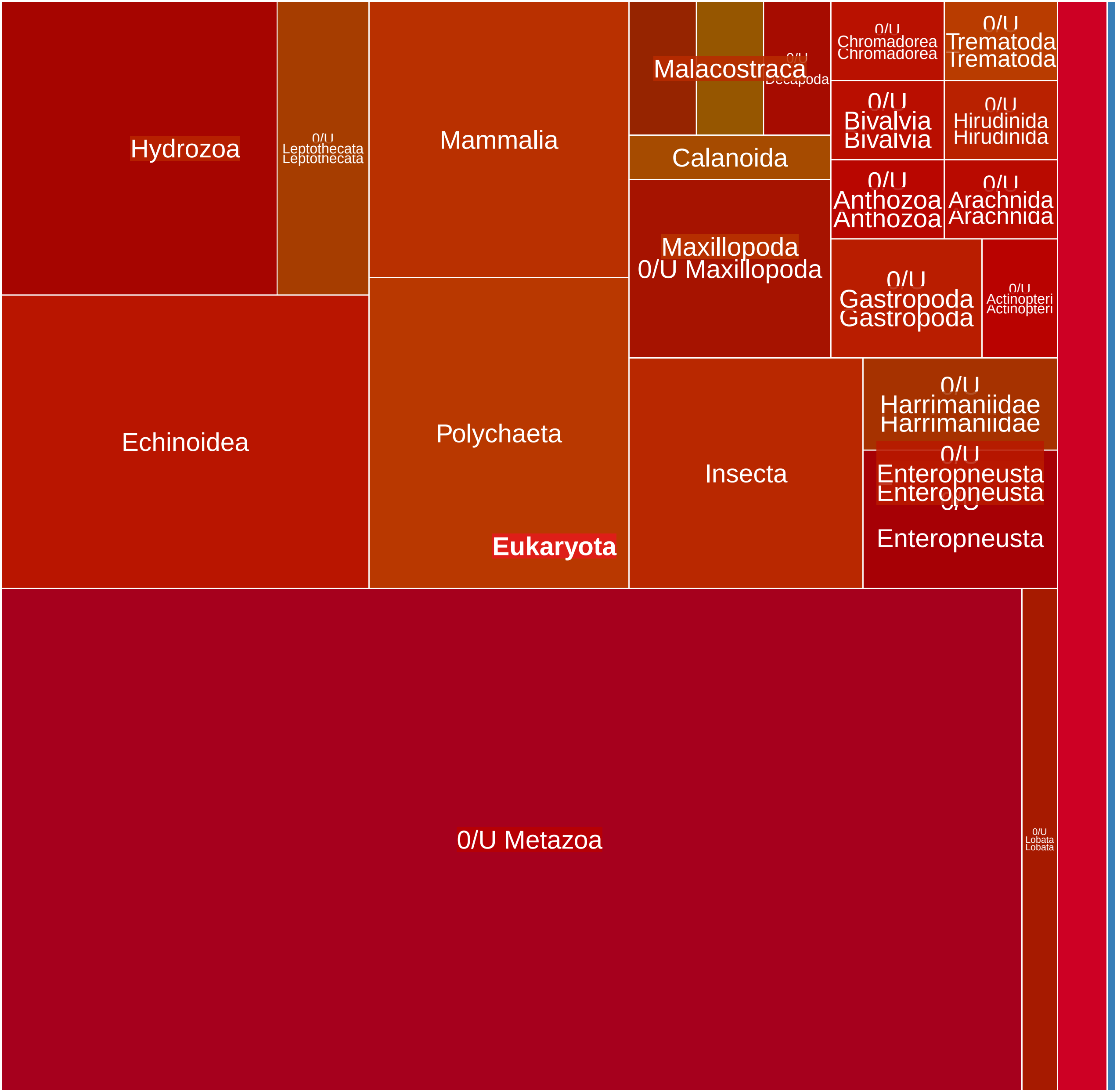

MGT−v1_716 : 831 unigenes, 601 (72.32%) taxonomically assigned

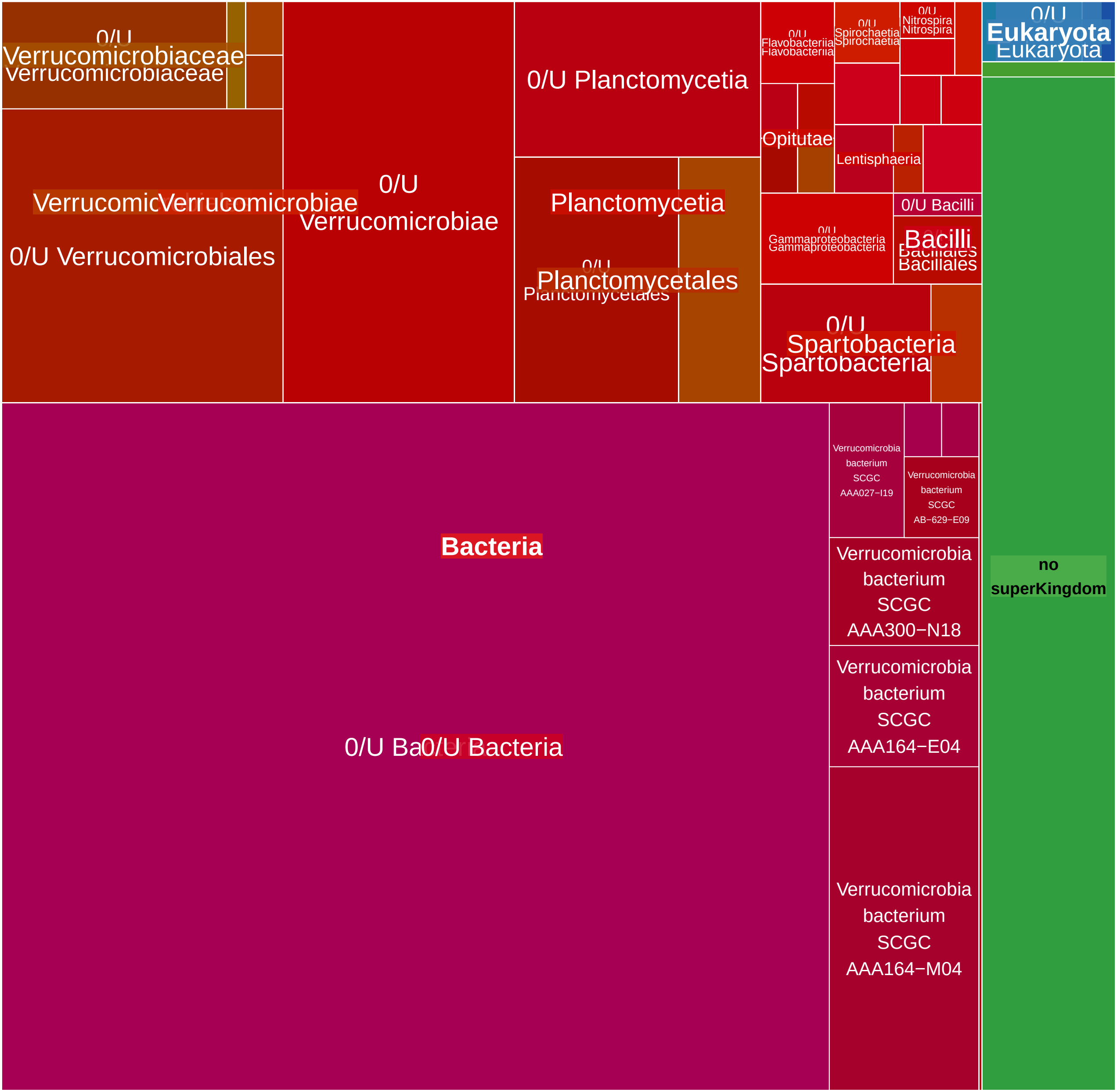

MGT−v1_717 : 822 unigenes, 444 (54.01%) taxonomically assigned

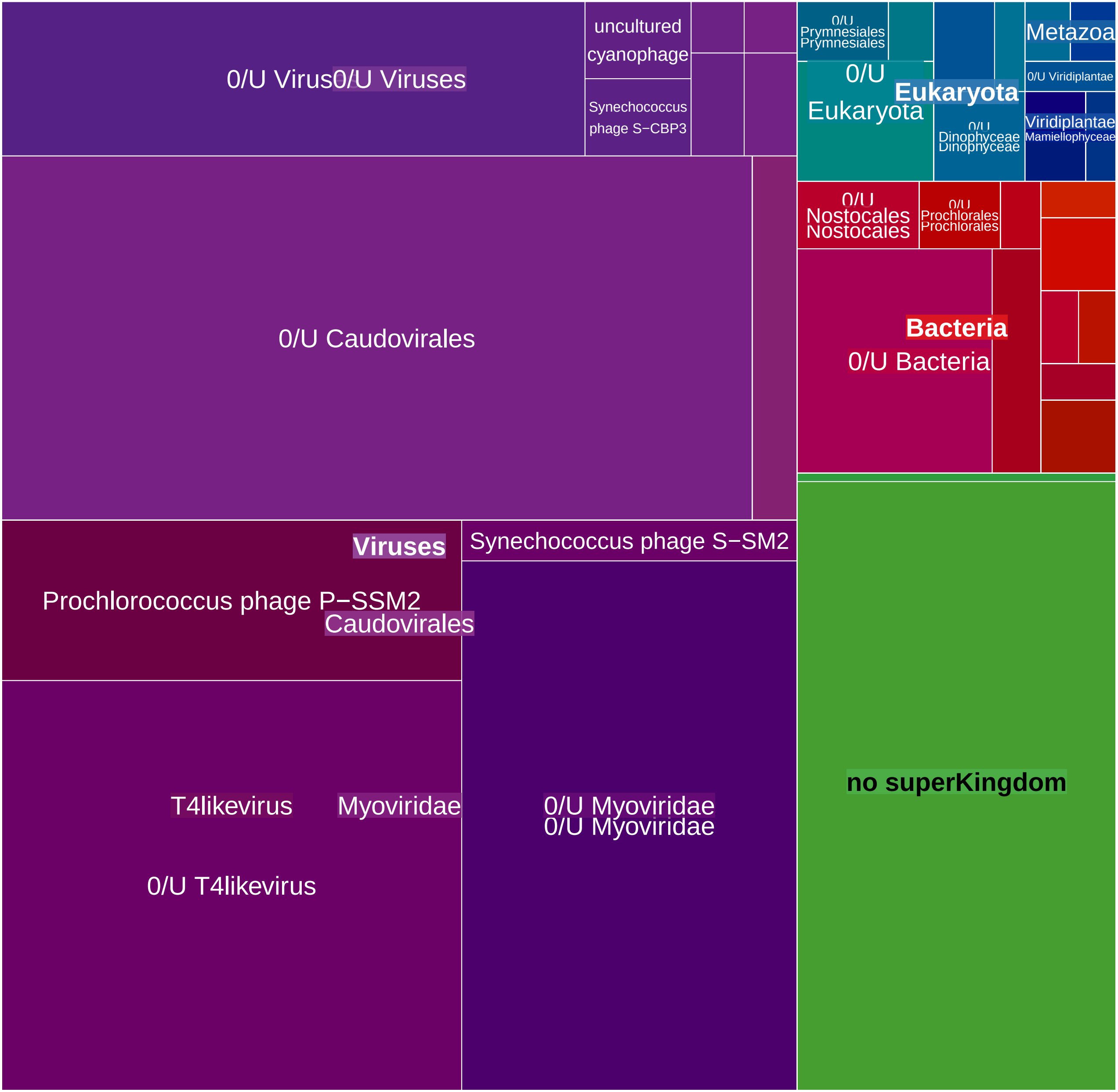

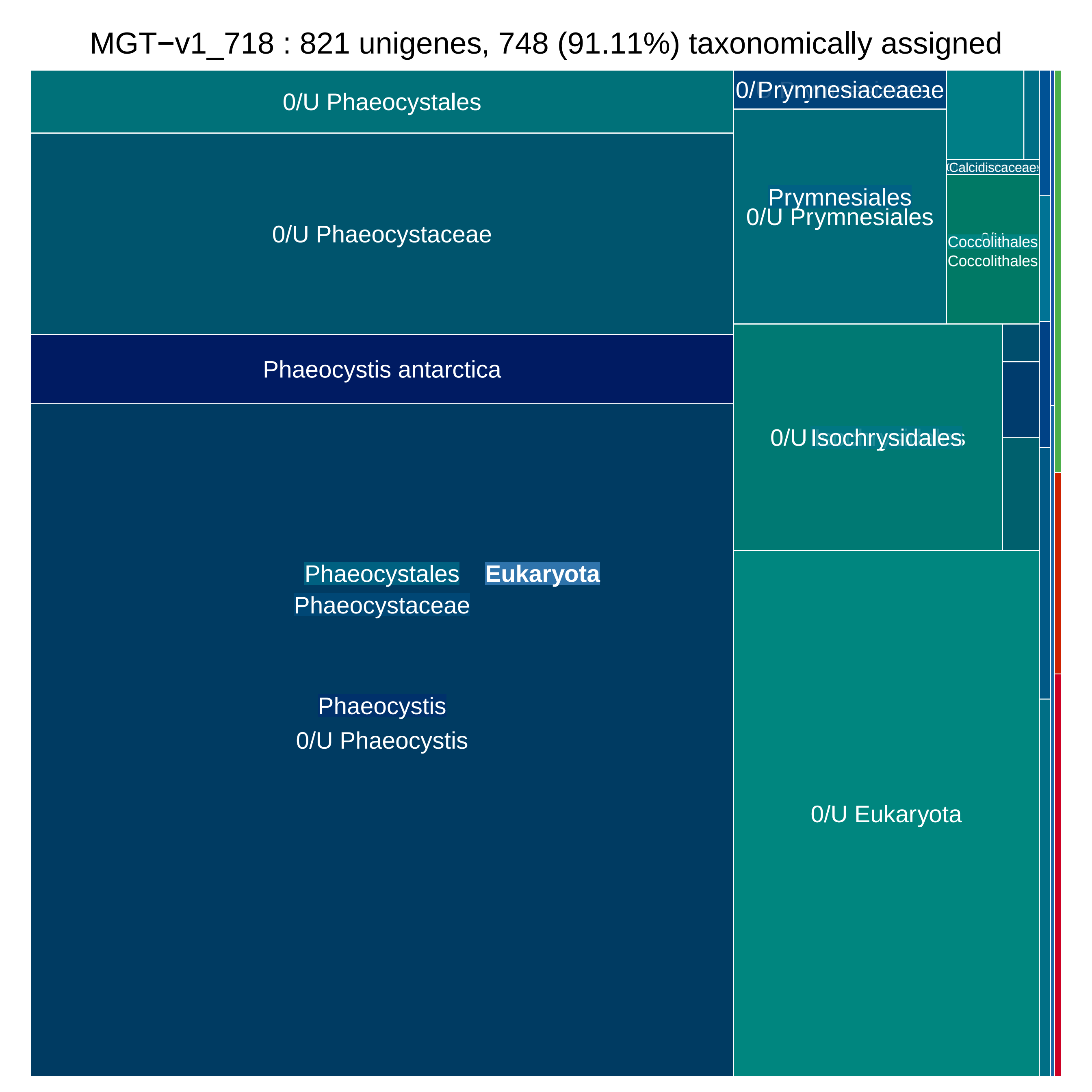

MGT−v1_718 : 821 unigenes, 748 (91.11%) taxonomically assigned
0/U Phaeocystales
0/U Phaeocystaceae
Phaeocystis antarctica
Phaeocystales
Eukaryota
Phaeocystaceae
Phaeocystis
0/U Phaeocystis
0/U Prymnesiaceae
Prymnesiales
0/U Prymnesiales
Calcidiscaceae
Coccolithales
Coccolithales
0/U Isochrysidales
0/U Eukaryota

MGT−v1_719 : 811 unigenes, 413 (50.92%) taxonomically assigned

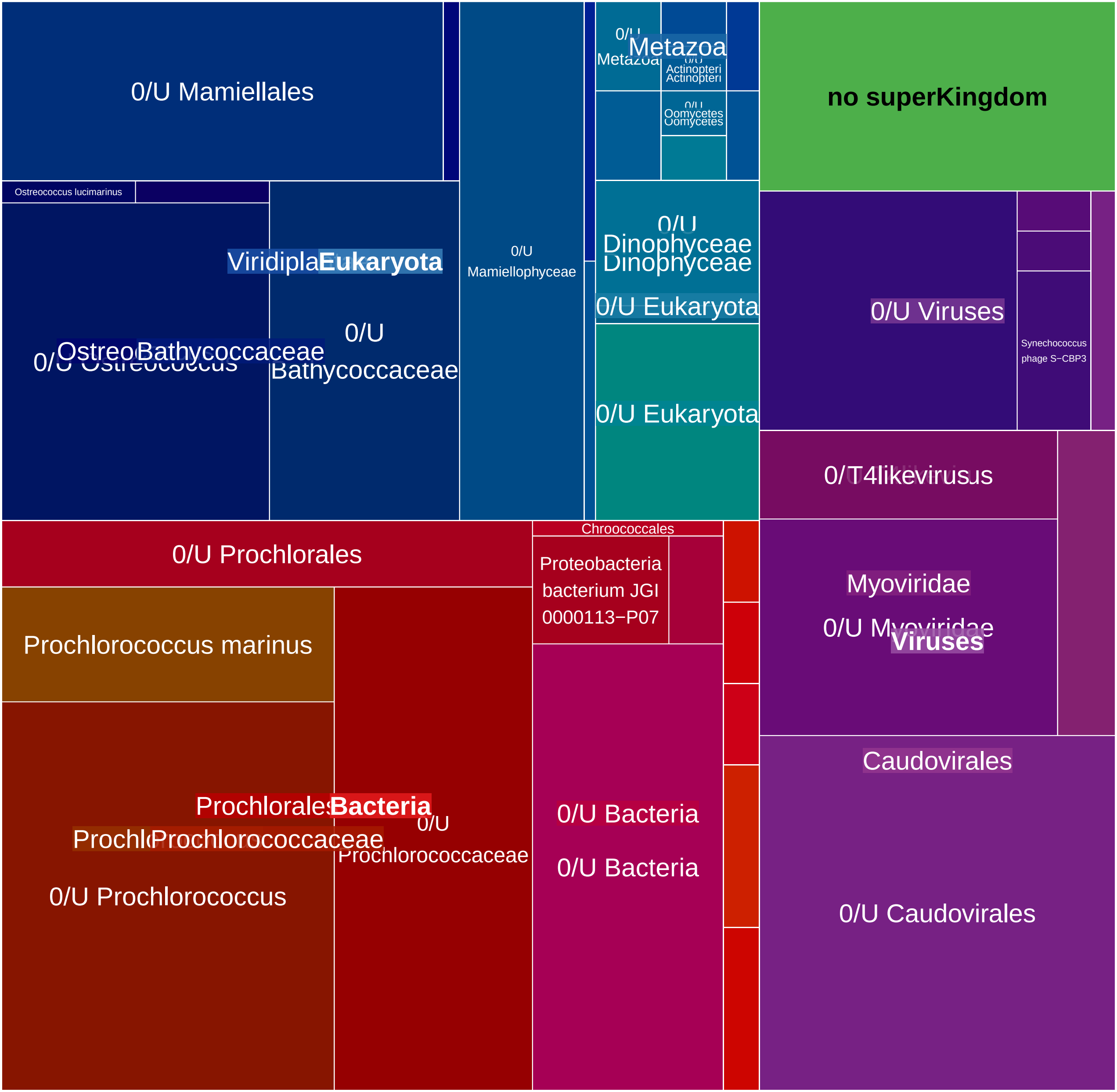

MGT−v1_720 : 806 unigenes, 32 (3.97%) taxonomically assigned

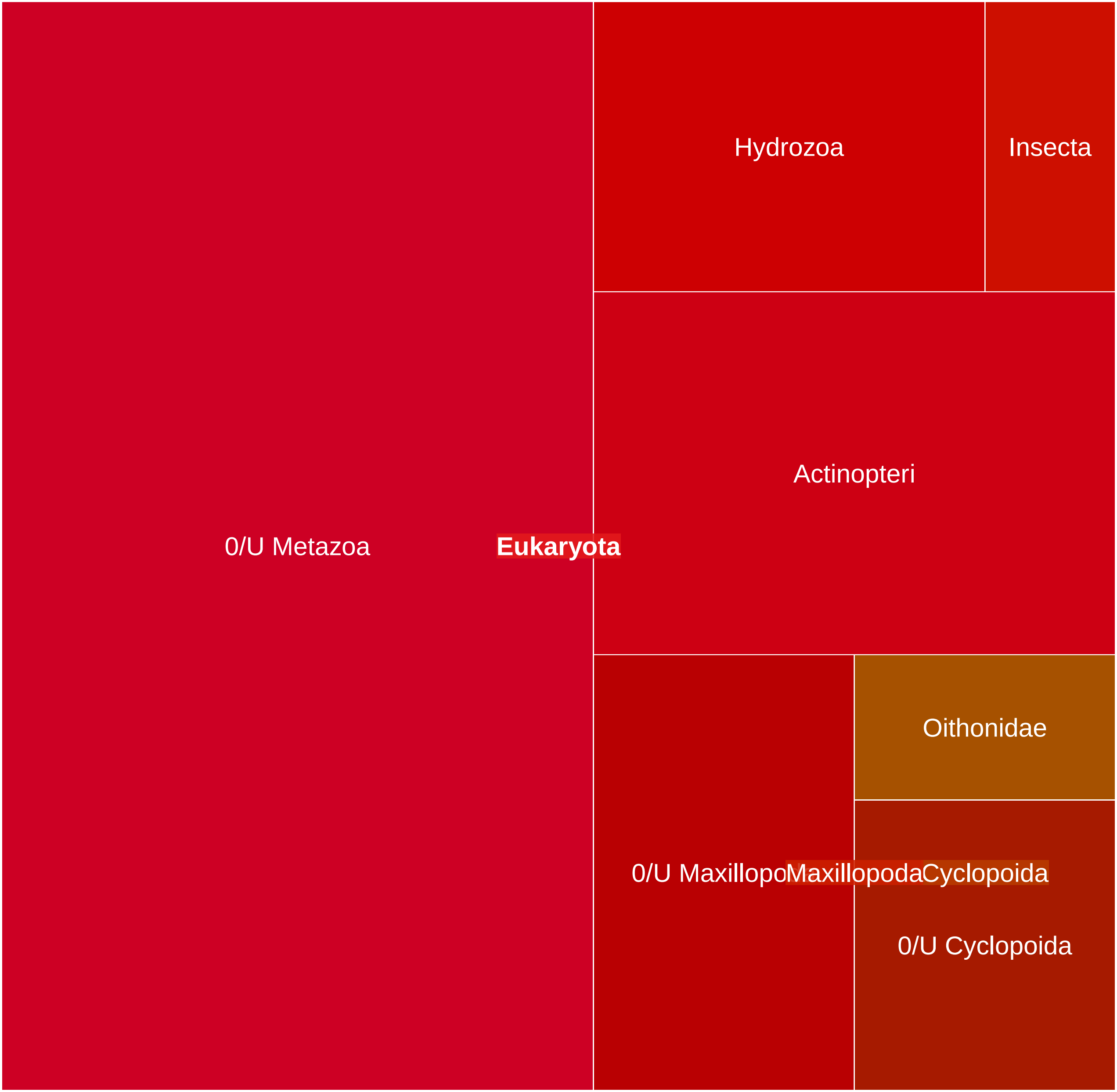

MGT−v1_721 : 805 unigenes, 675 (83.85%) taxonomically assigned

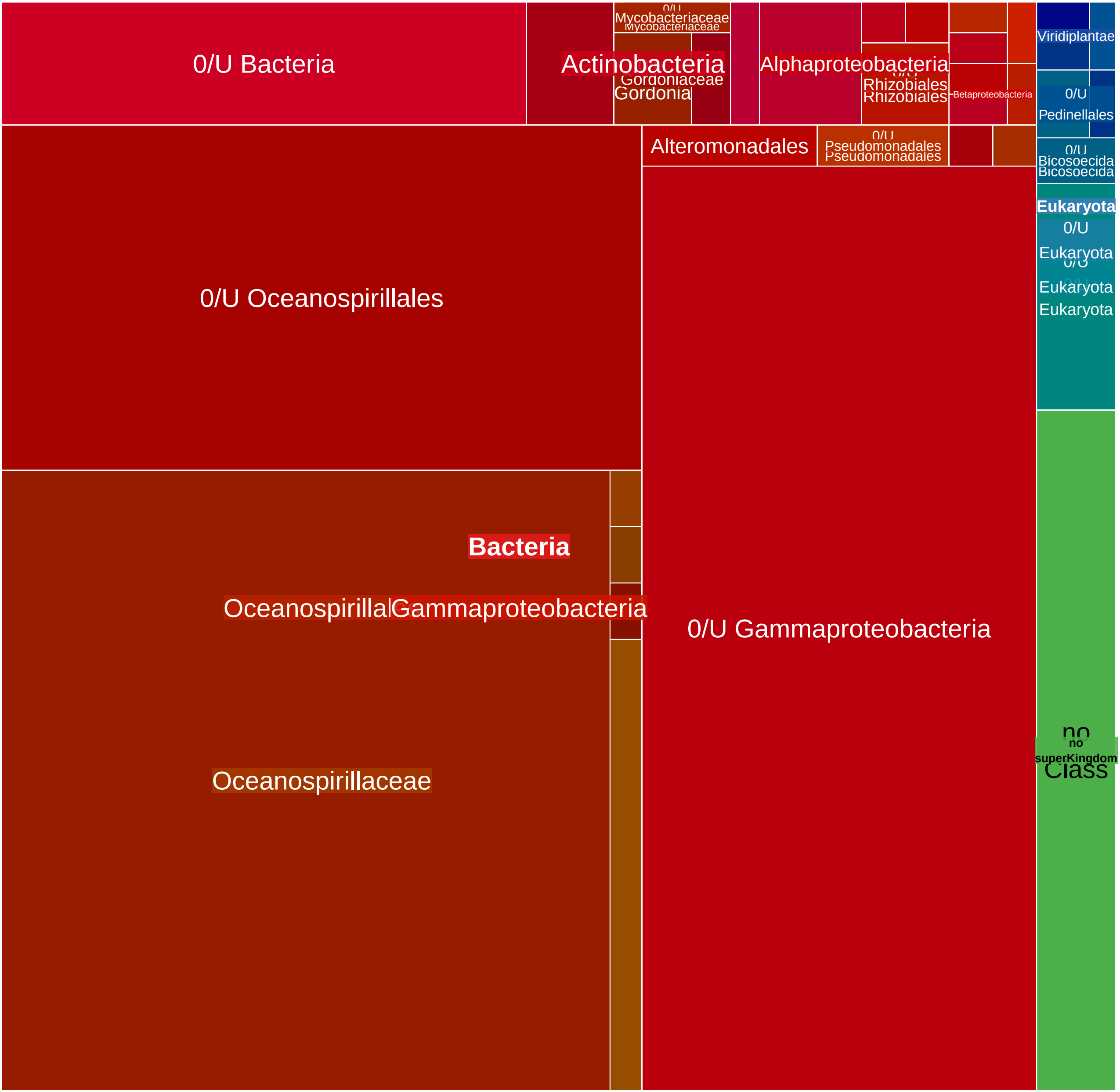

MGT−v1_722 : 798 unigenes, 171 (21.43%) taxonomically assigned

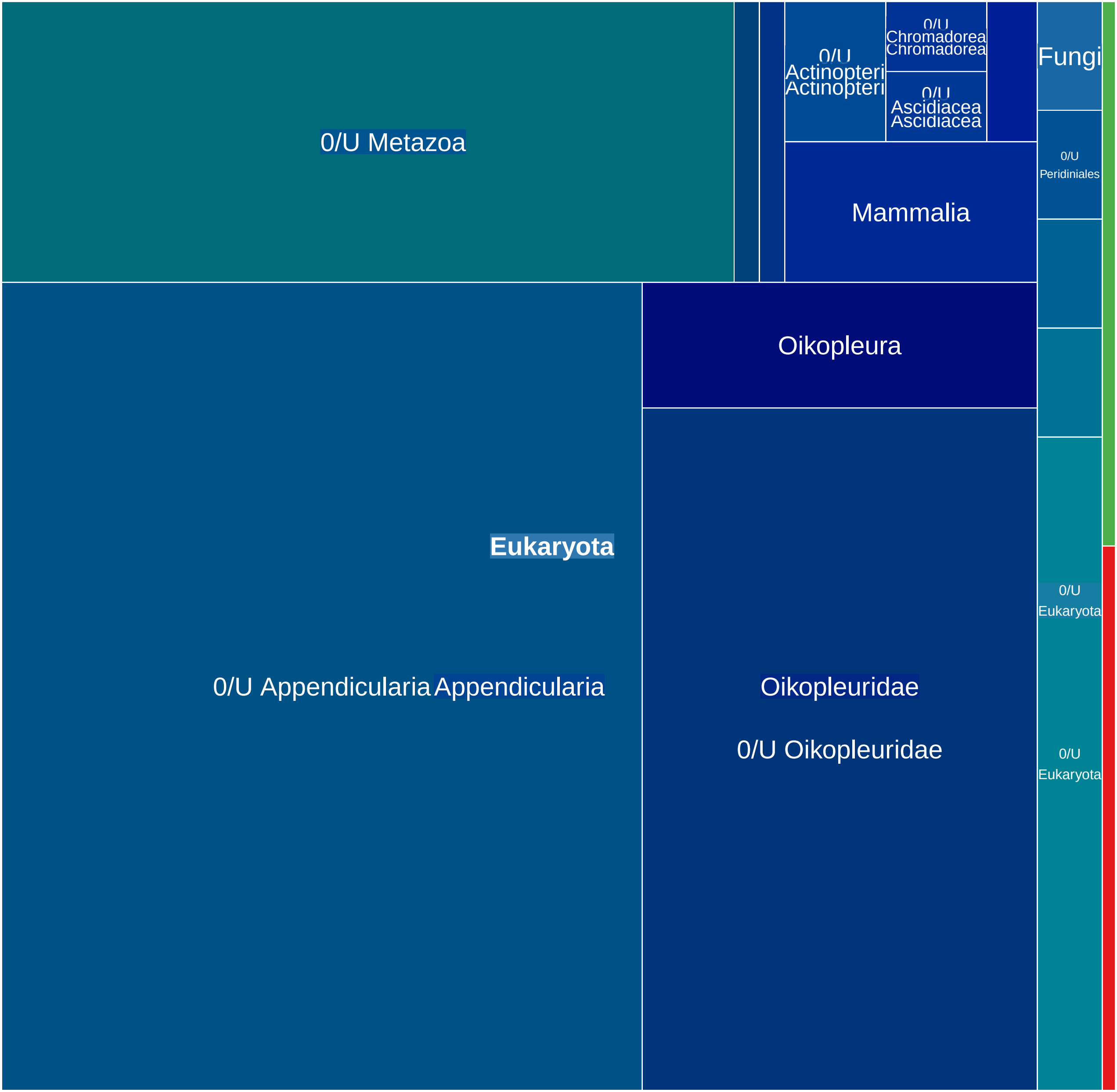

MGT−v1_723 : 797 unigenes, 757 (94.98%) taxonomically assigned

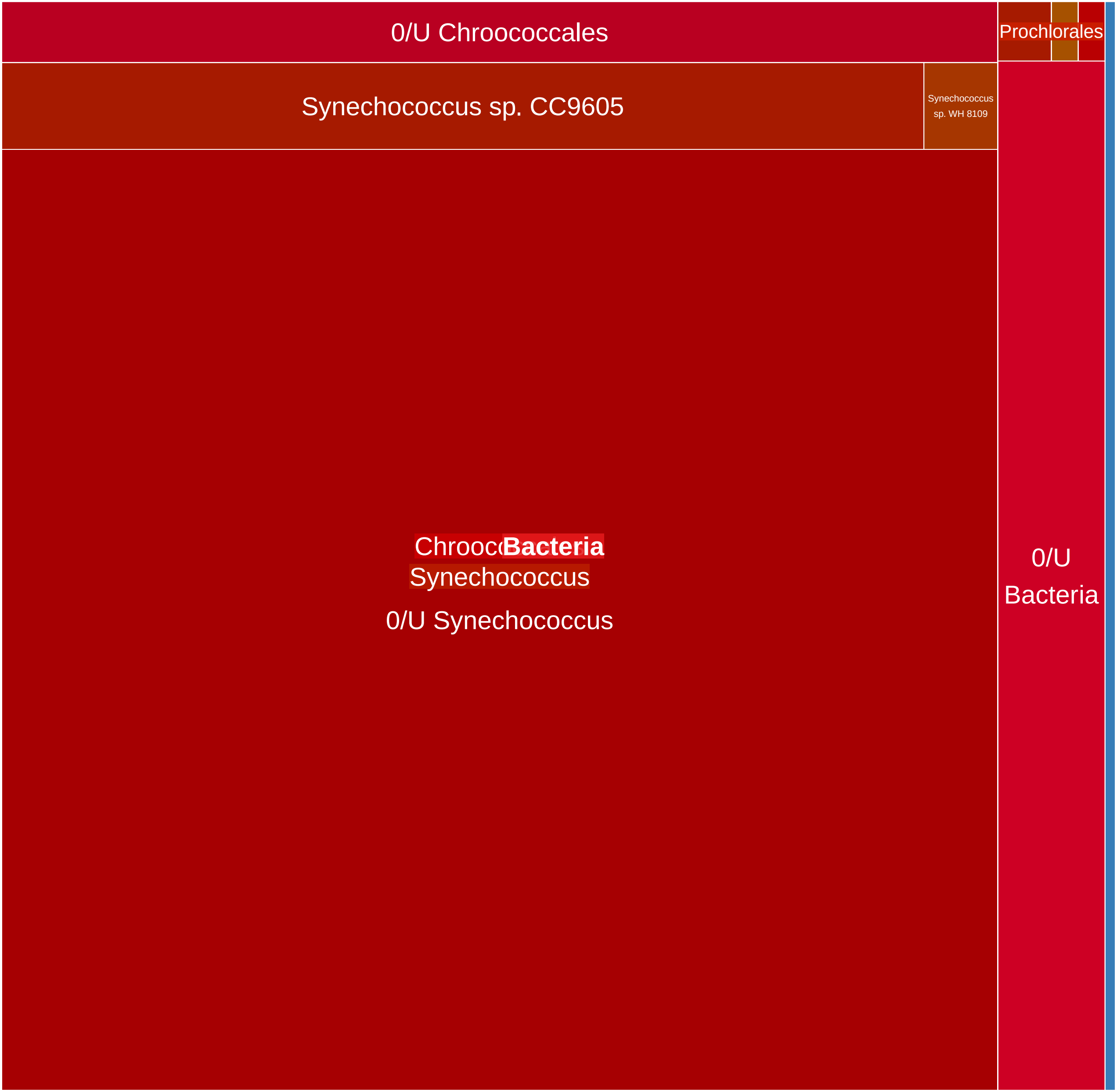

### MGT−v1_724 : 797 unigenes, 511 (64.12%) taxonomically assigned

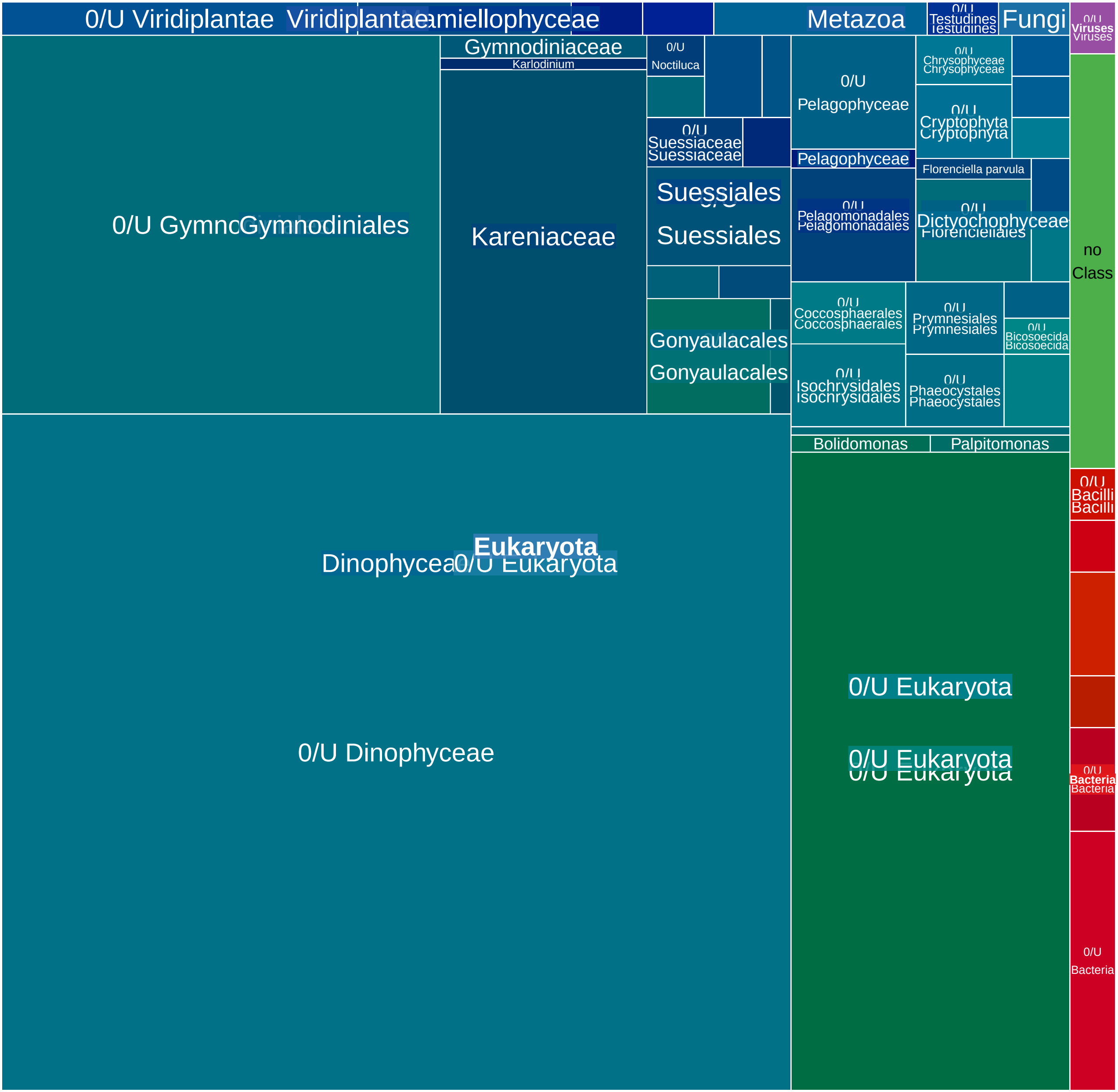

MGT−v1_725 : 797 unigenes, 404 (50.69%) taxonomically assigned

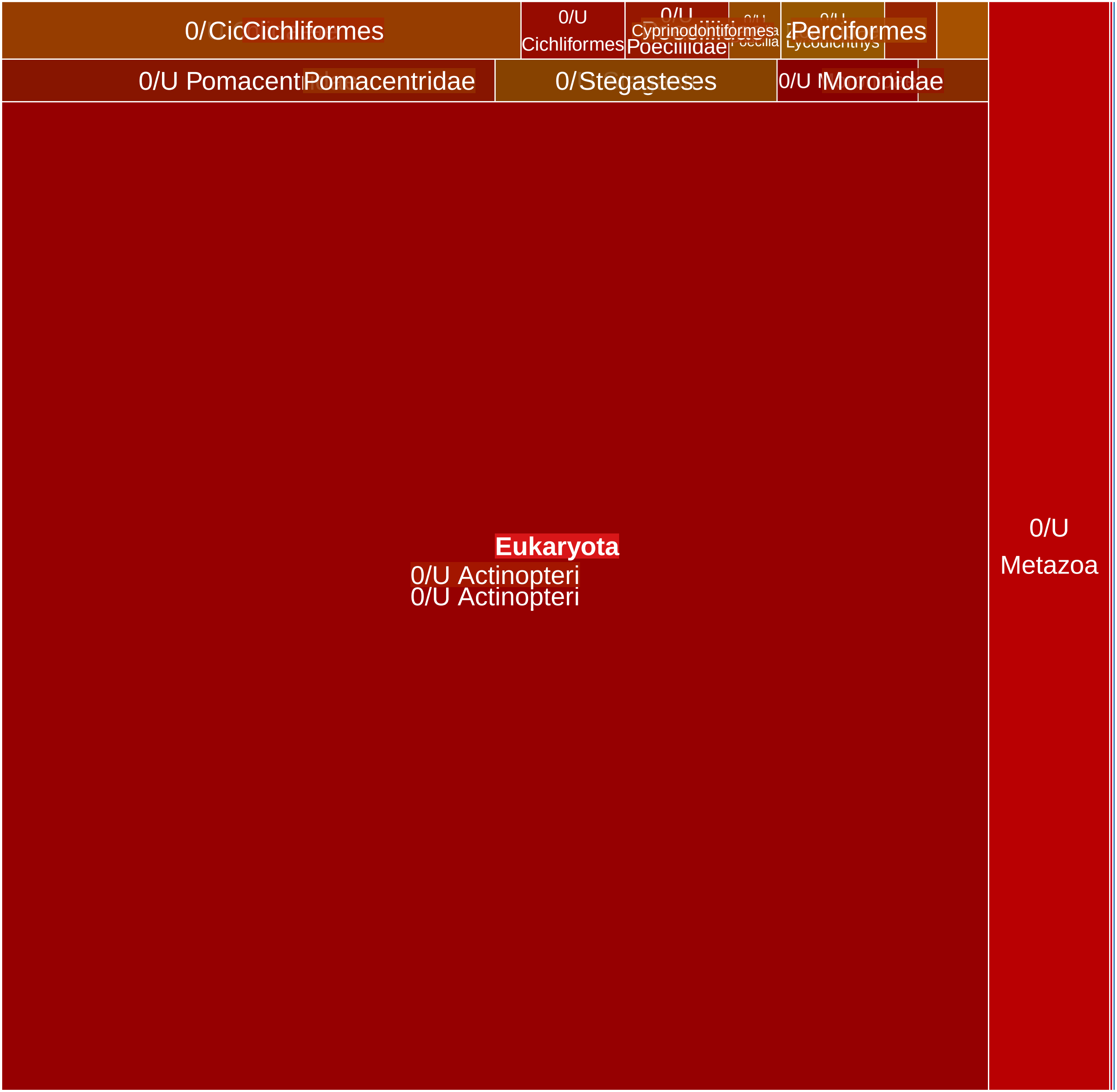

MGT−v1_726 : 796 unigenes, 465 (58.42%) taxonomically assigned

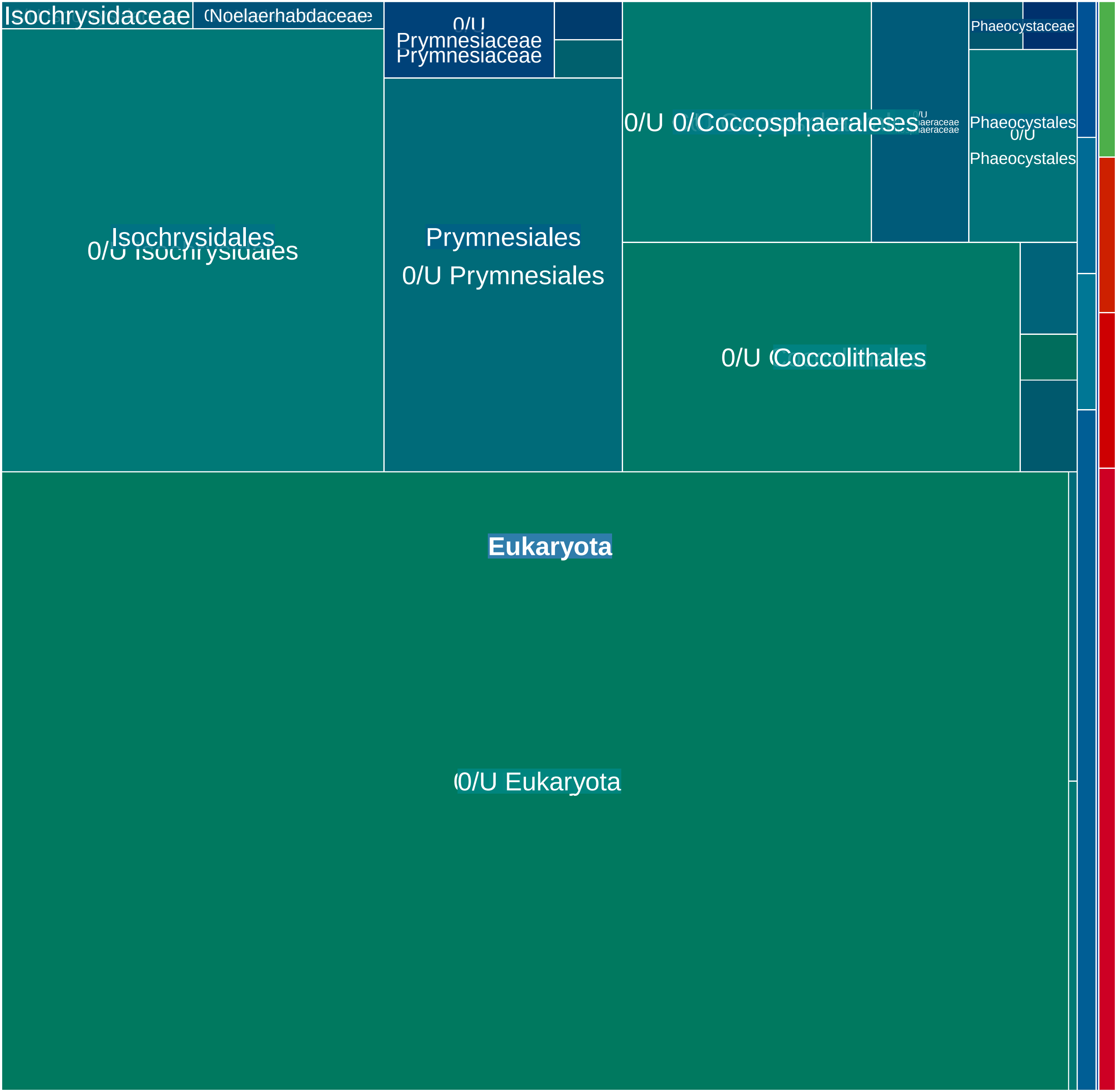

MGT−v1_727 : 793 unigenes, 183 (23.08%) taxonomically assigned

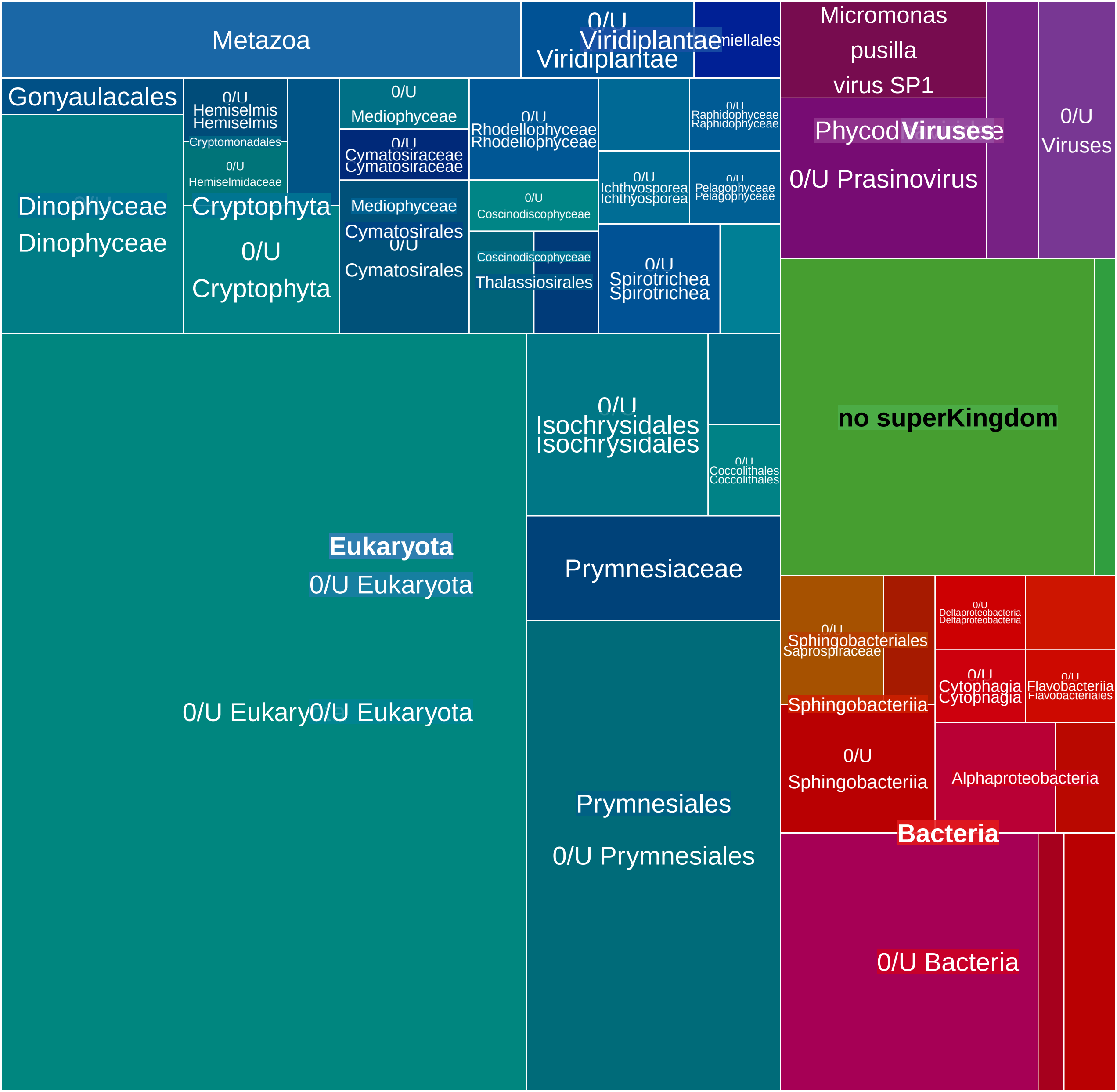

MGT−v1_728 : 789 unigenes, 563 (71.36%) taxonomically assigned

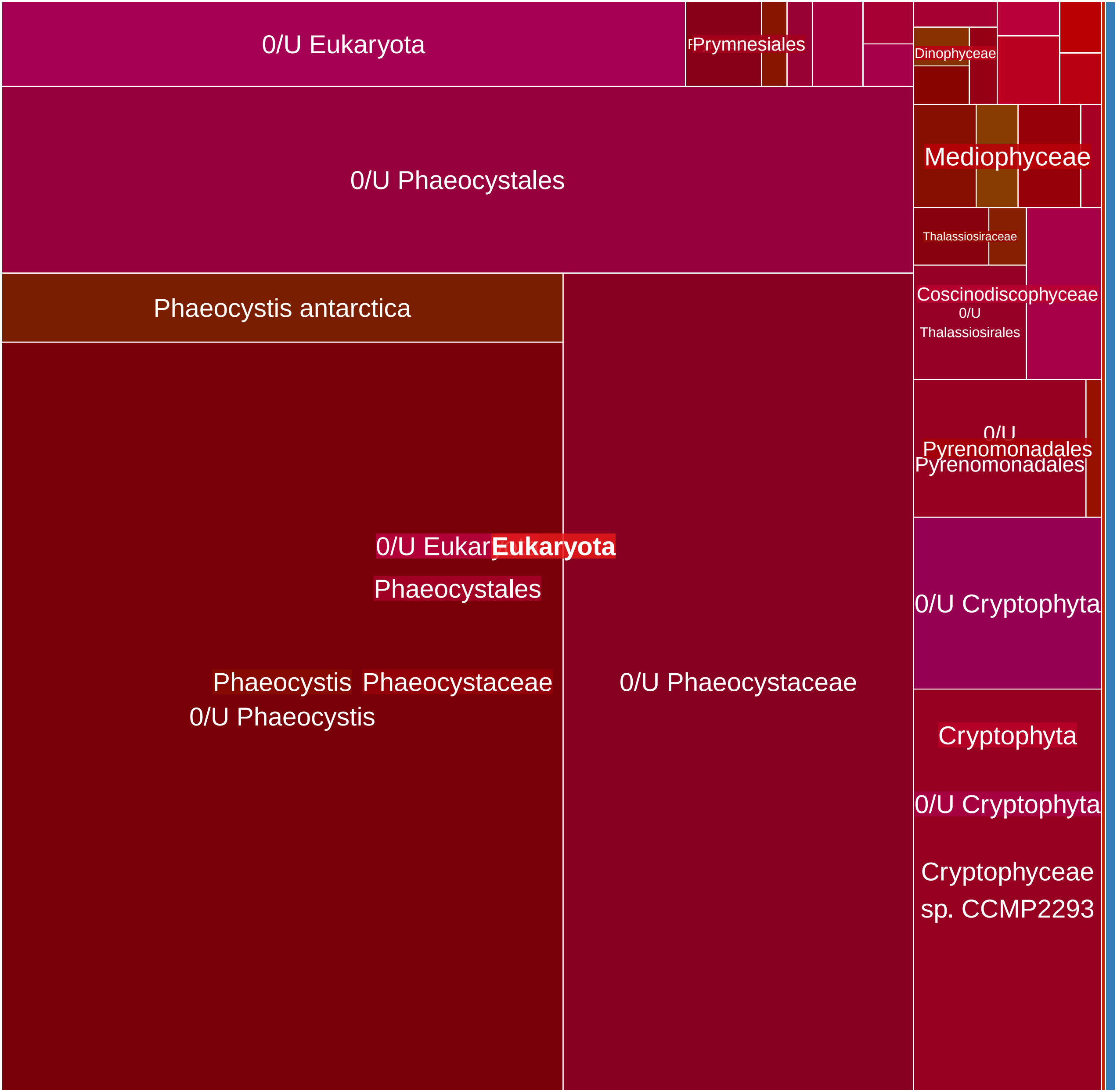

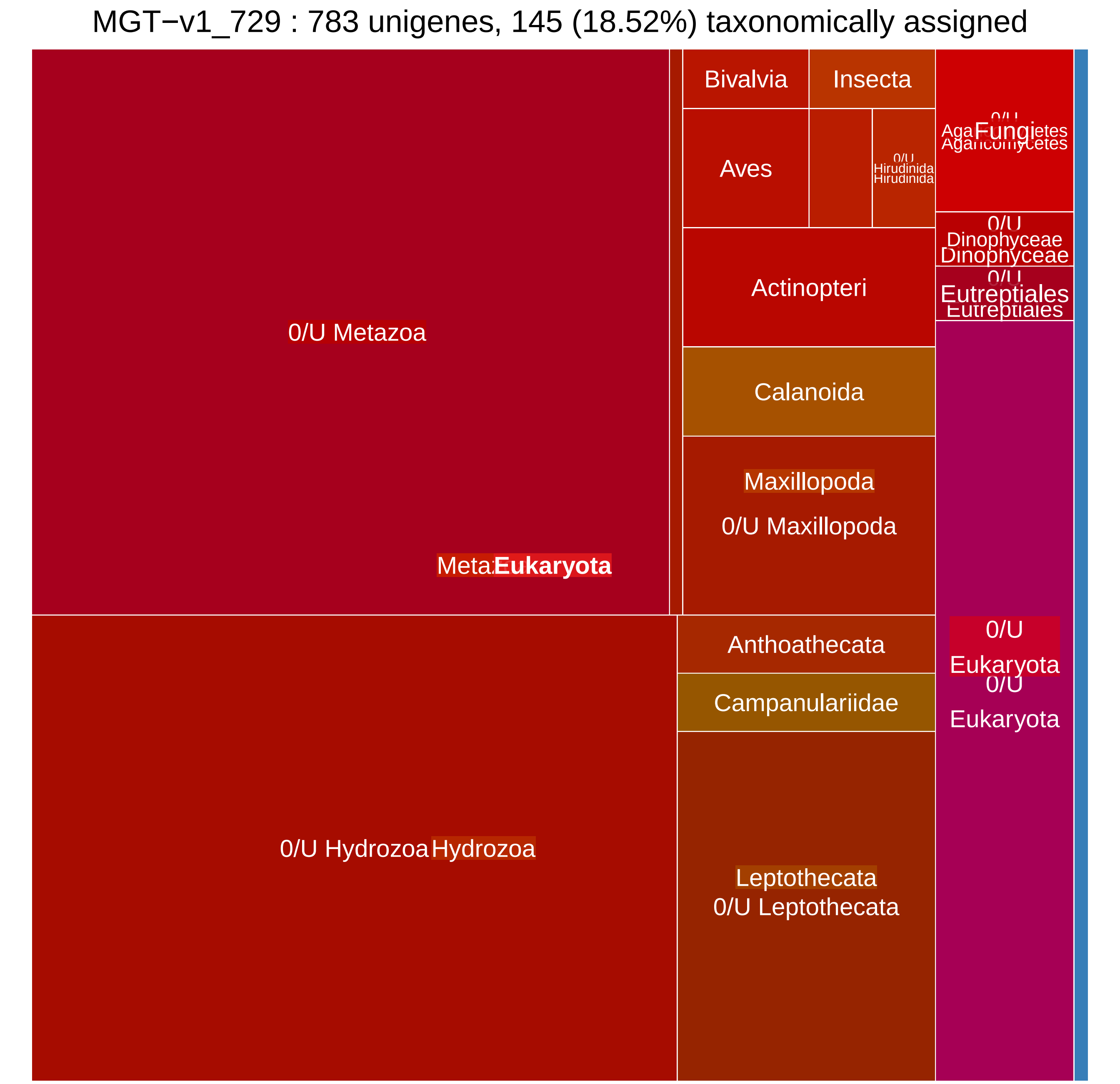
MGT−v1_729 : 783 unigenes, 145 (18.52%) taxonomically assigned
0/U Metazoa
Metazoa Eukaryota
Bivalvia
Insecta
Fungi
0/U Hirudinida
Hirudinida
Aves
0/U Dinophyceae
Dinophyceae
0/U Eutreptiales
Eutreptiales
Actinopteri
Calanoida
Maxillopoda
0/U Maxillopoda
0/U Eukaryota
Anthoathecata
Campanulariidae
0/U Hydrozoa
Hydrozoa
Leptothecata
0/U Leptothecata

MGT−v1_730 : 782 unigenes, 333 (42.58%) taxonomically assigned

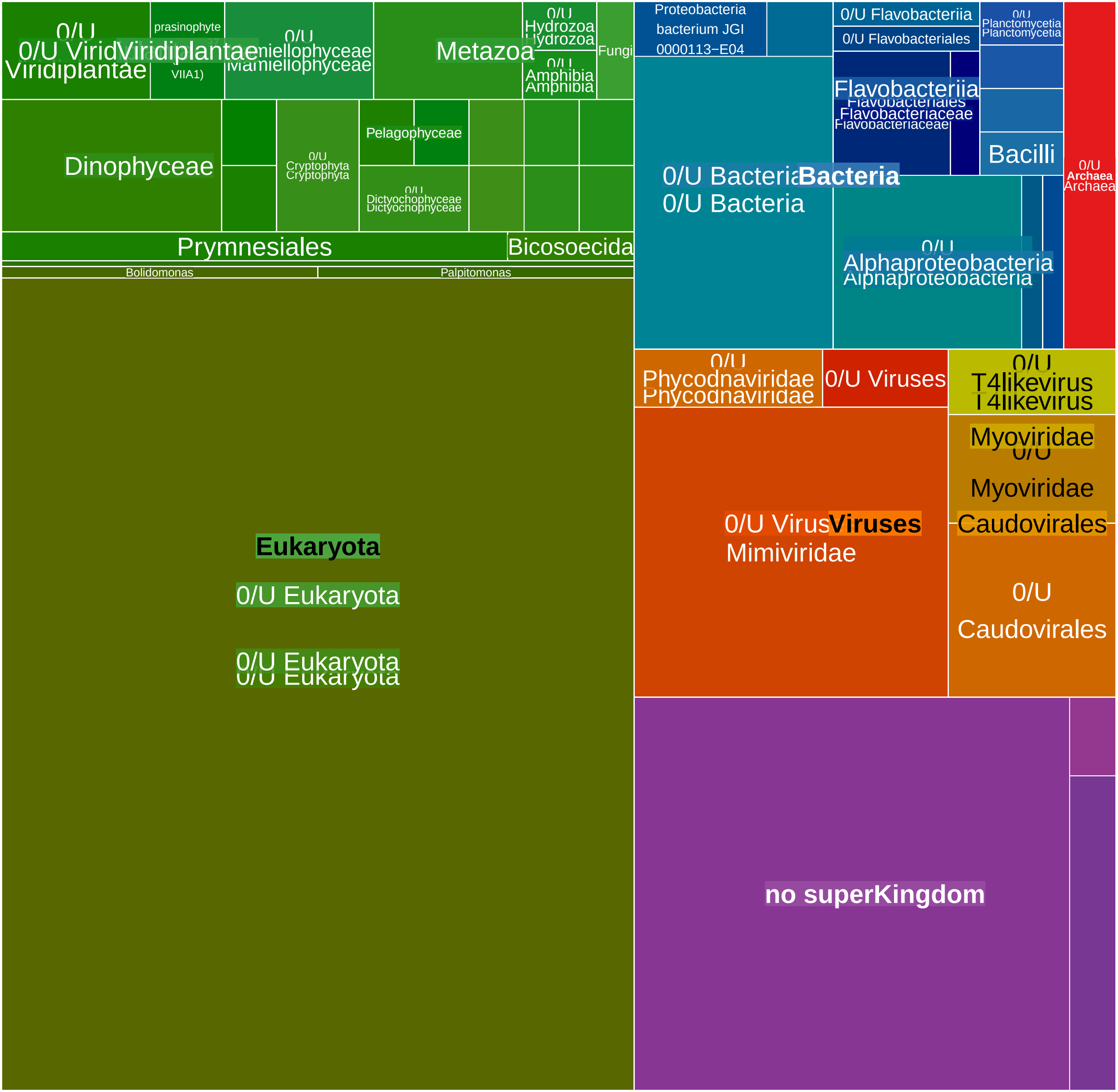

MGT−v1_731 : 782 unigenes, 510 (65.22%) taxonomically assigned

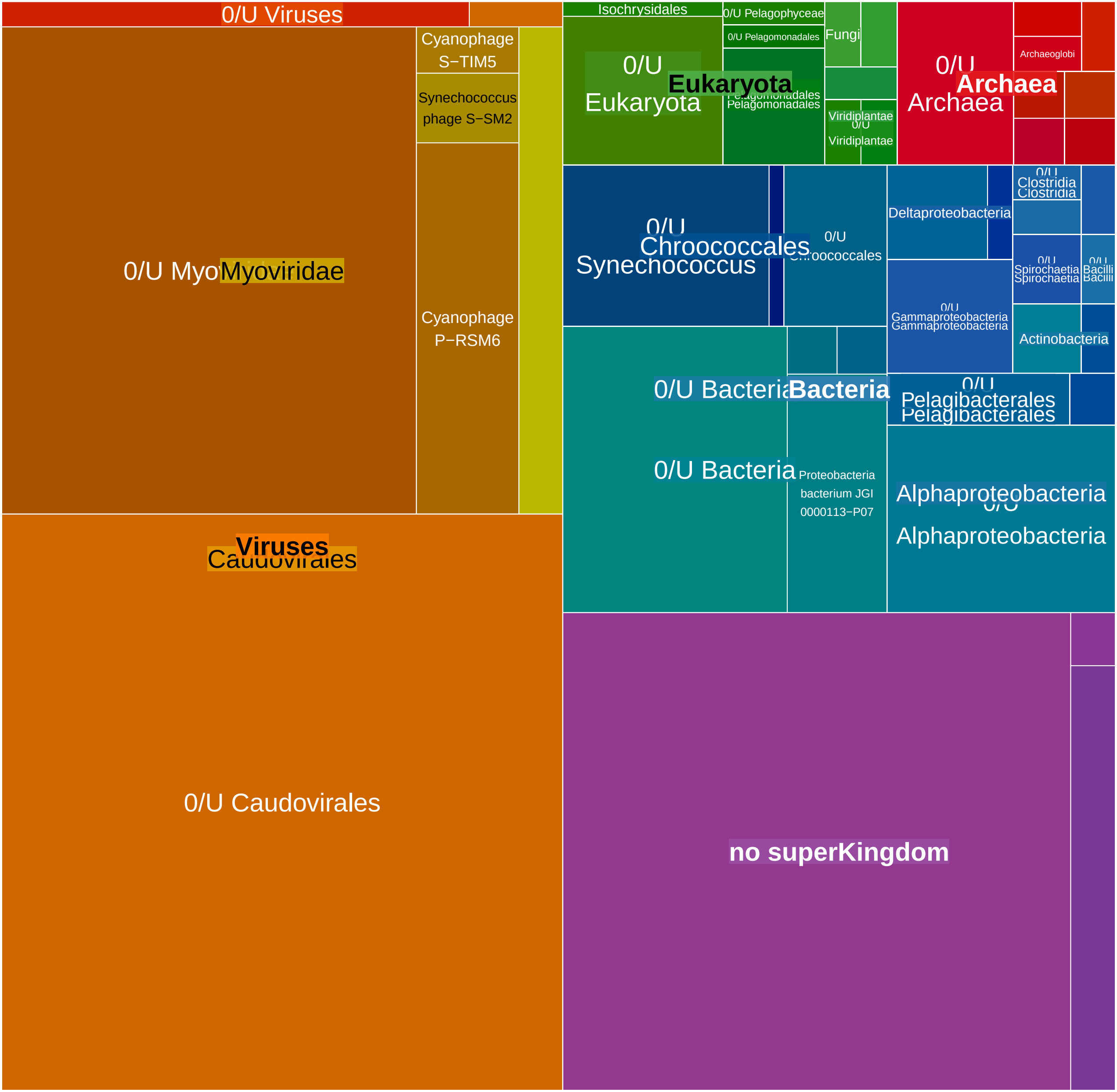

MGT−v1_732 : 781 unigenes, 97 (12.42%) taxonomically assigned

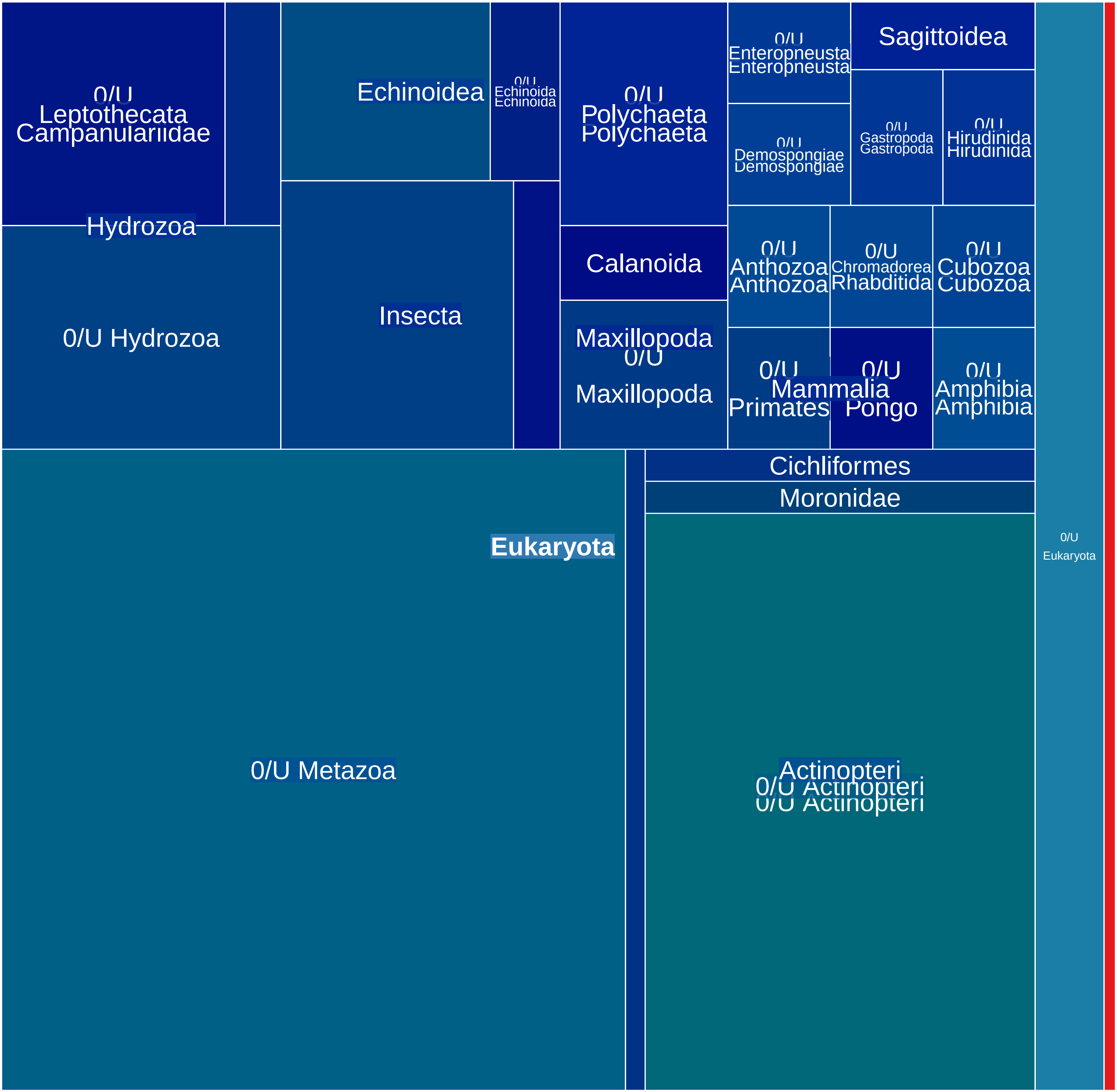

MGT−v1_733 : 780 unigenes, 51 (6.54%) taxonomically assigned

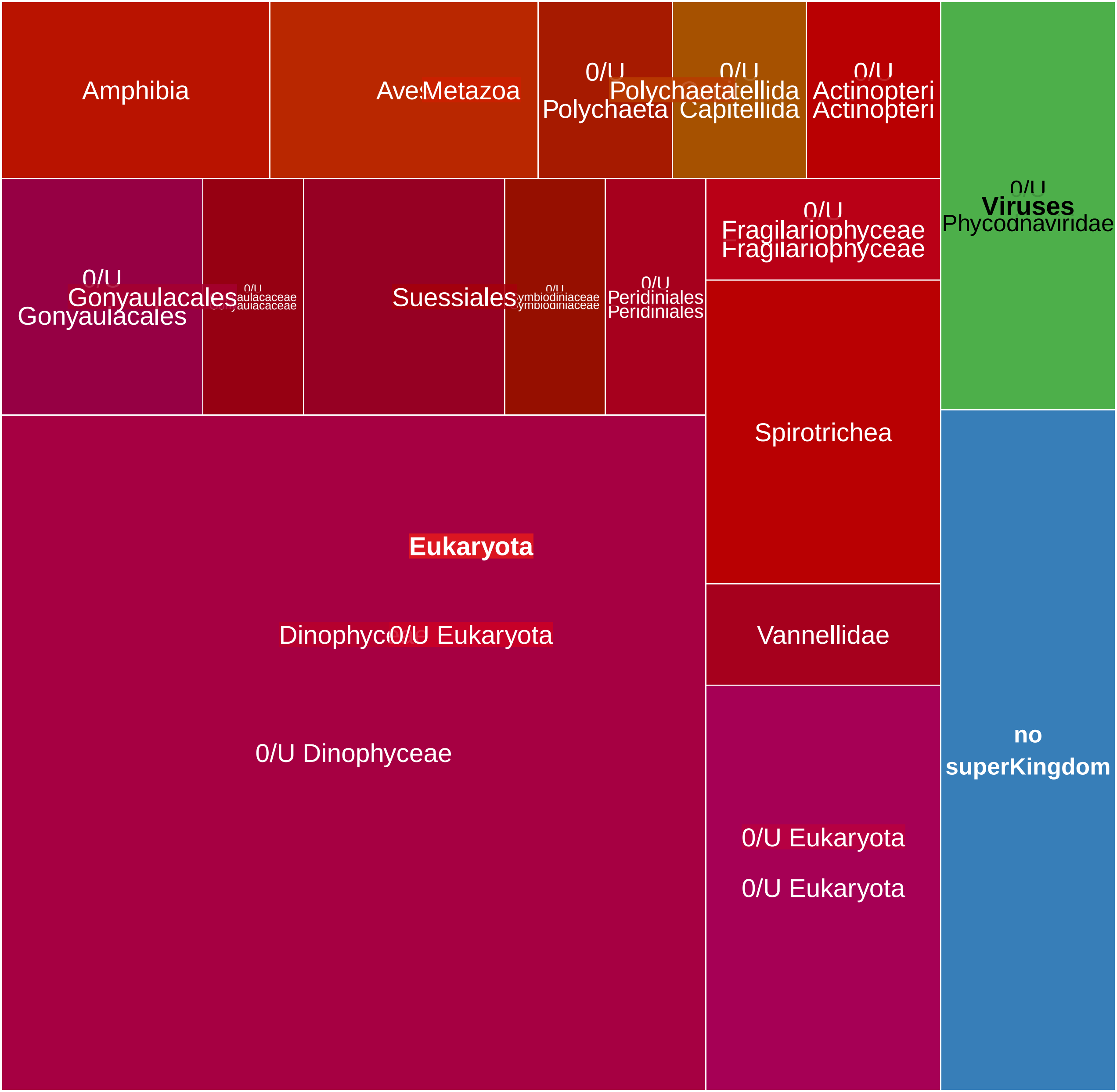

MGT−v1_734 : 780 unigenes, 142 (18.21%) taxonomically assigned

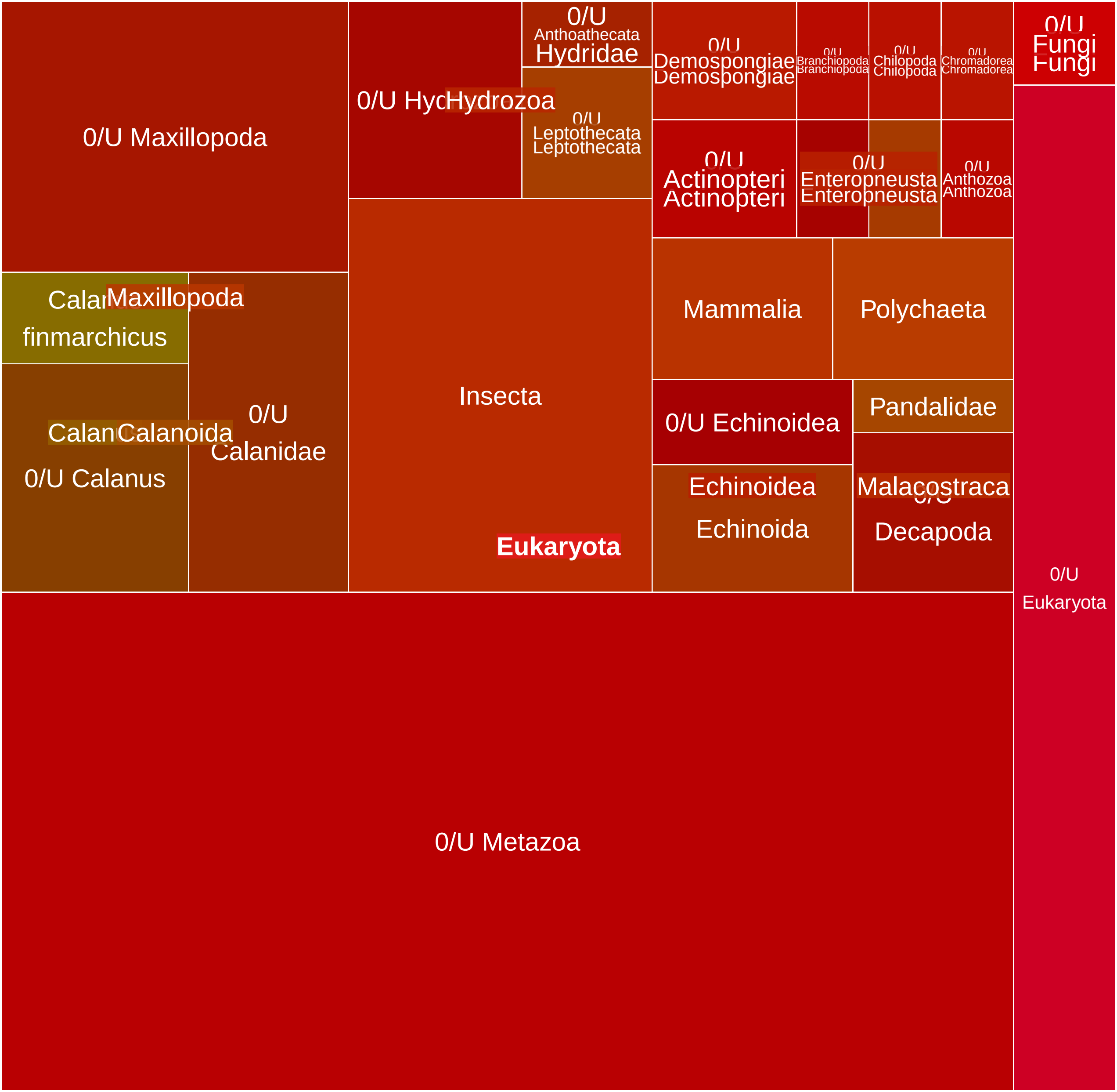

MGT−v1_735 : 777 unigenes, 97 (12.48%) taxonomically assigned

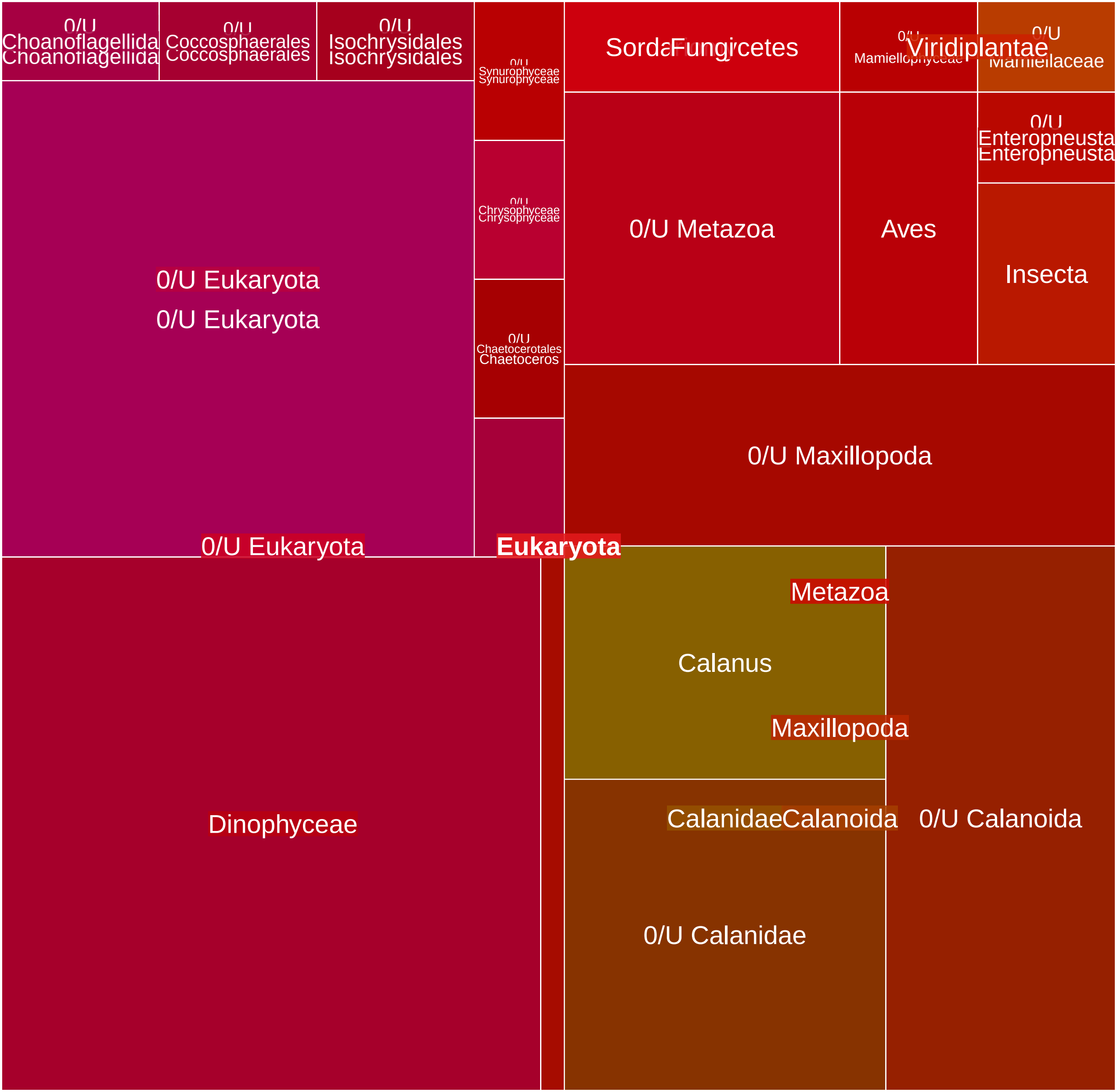

MGT−v1_736 : 777 unigenes, 725 (93.31%) taxonomically assigned

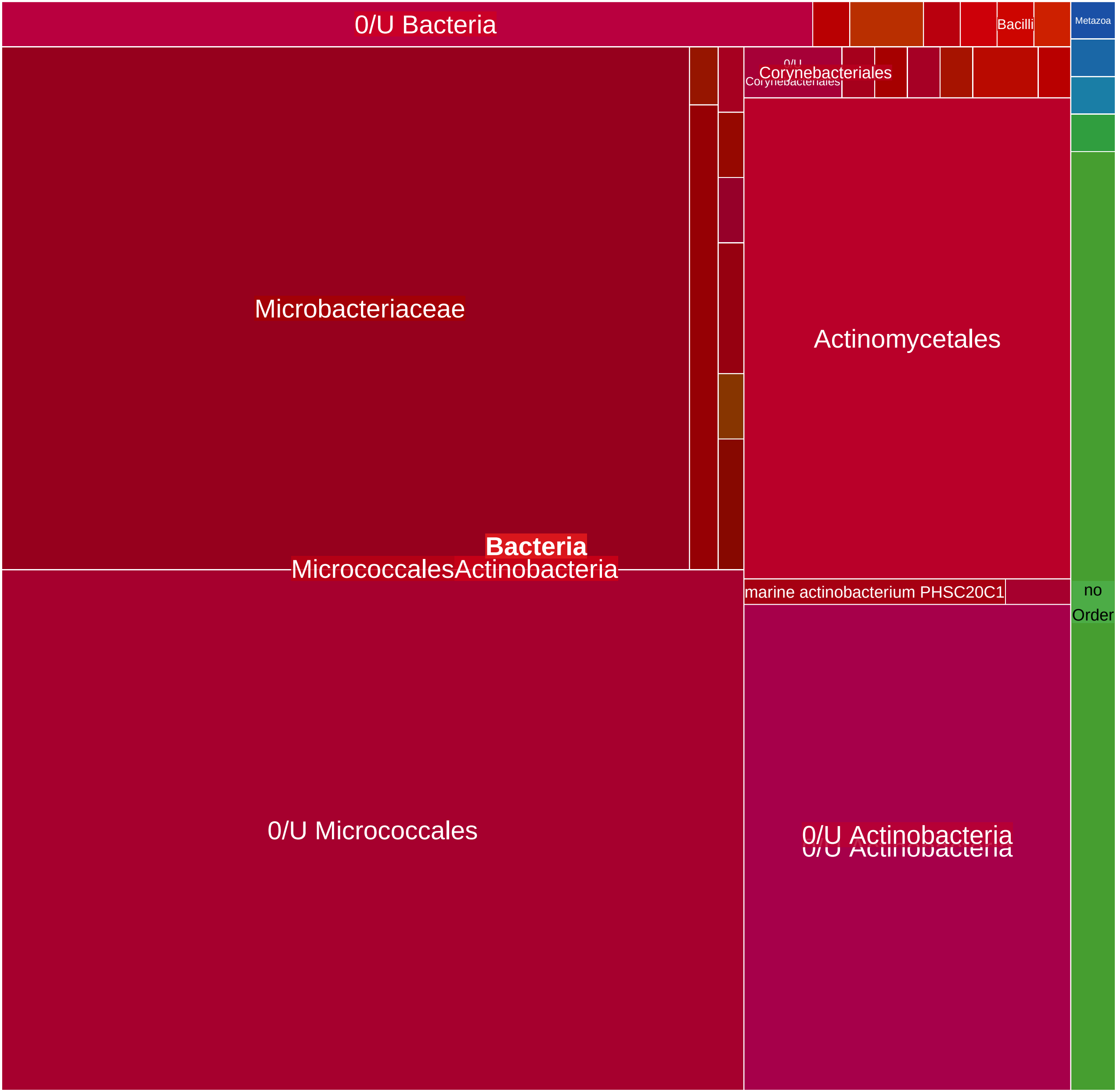

MGT−v1_737 : 774 unigenes, 56 (7.24%) taxonomically assigned

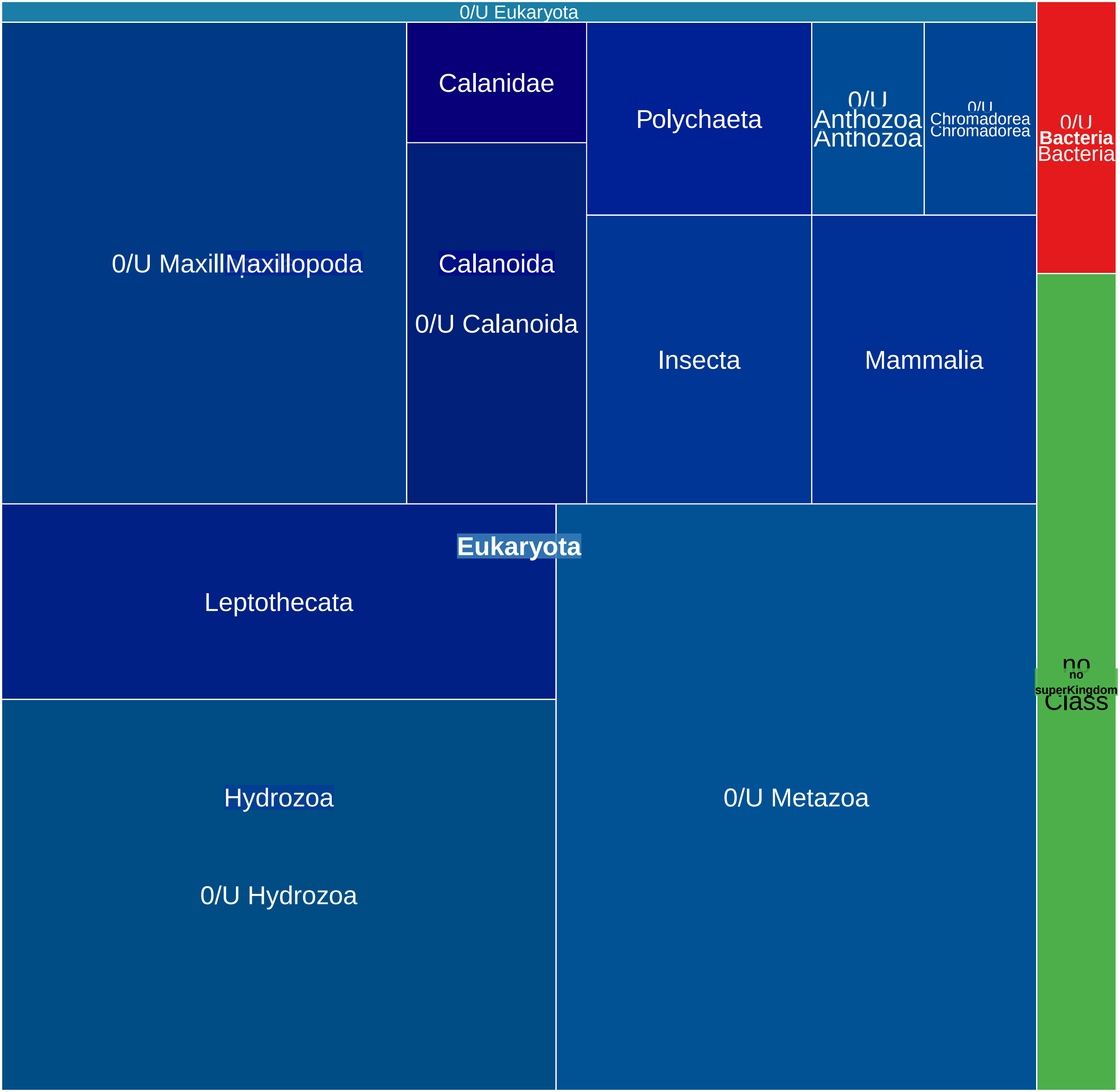

MGT−v1_738 : 770 unigenes, 423 (54.94%) taxonomically assigned

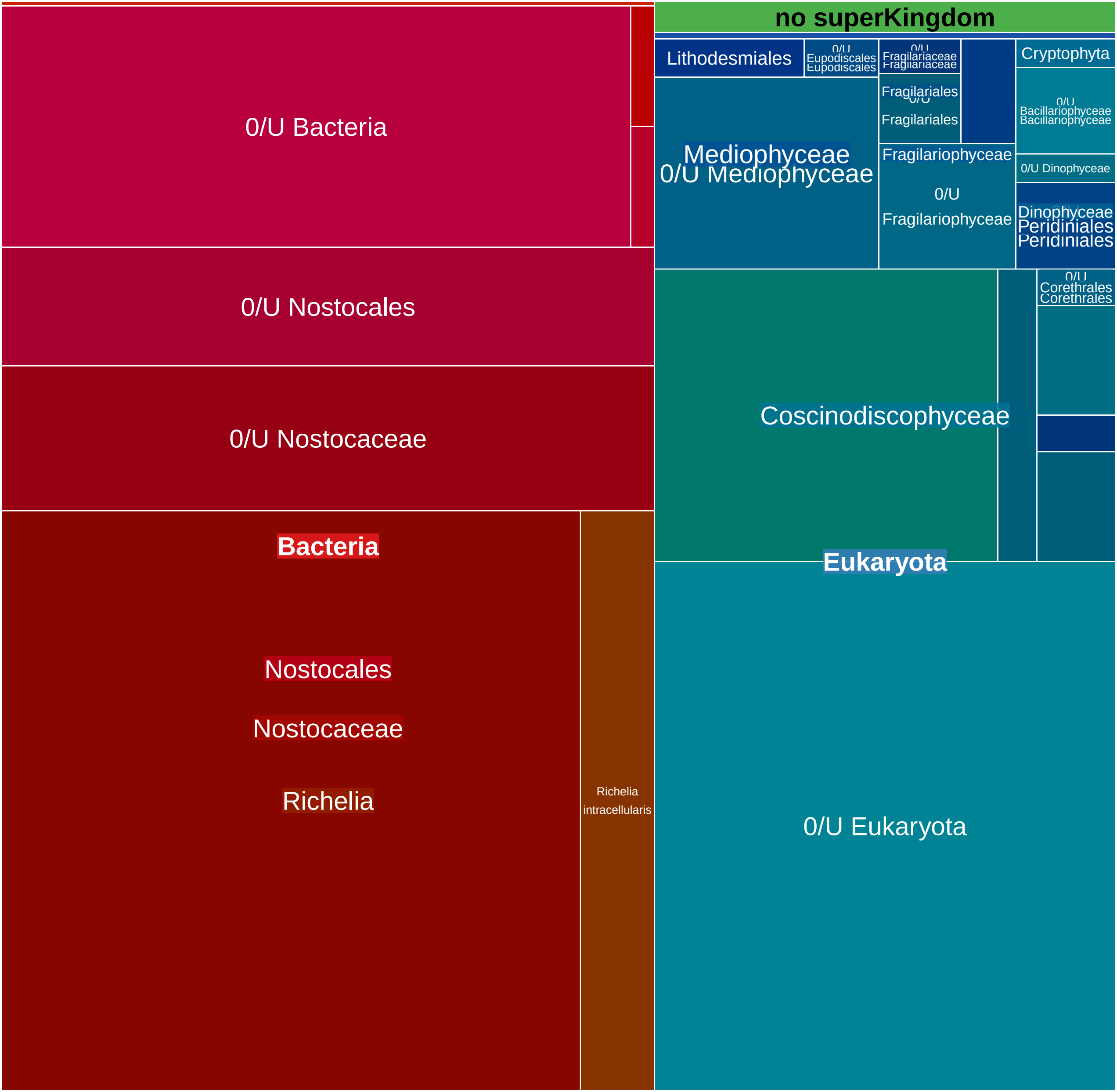

MGT−v1_739 : 769 unigenes, 128 (16.64%) taxonomically assigned

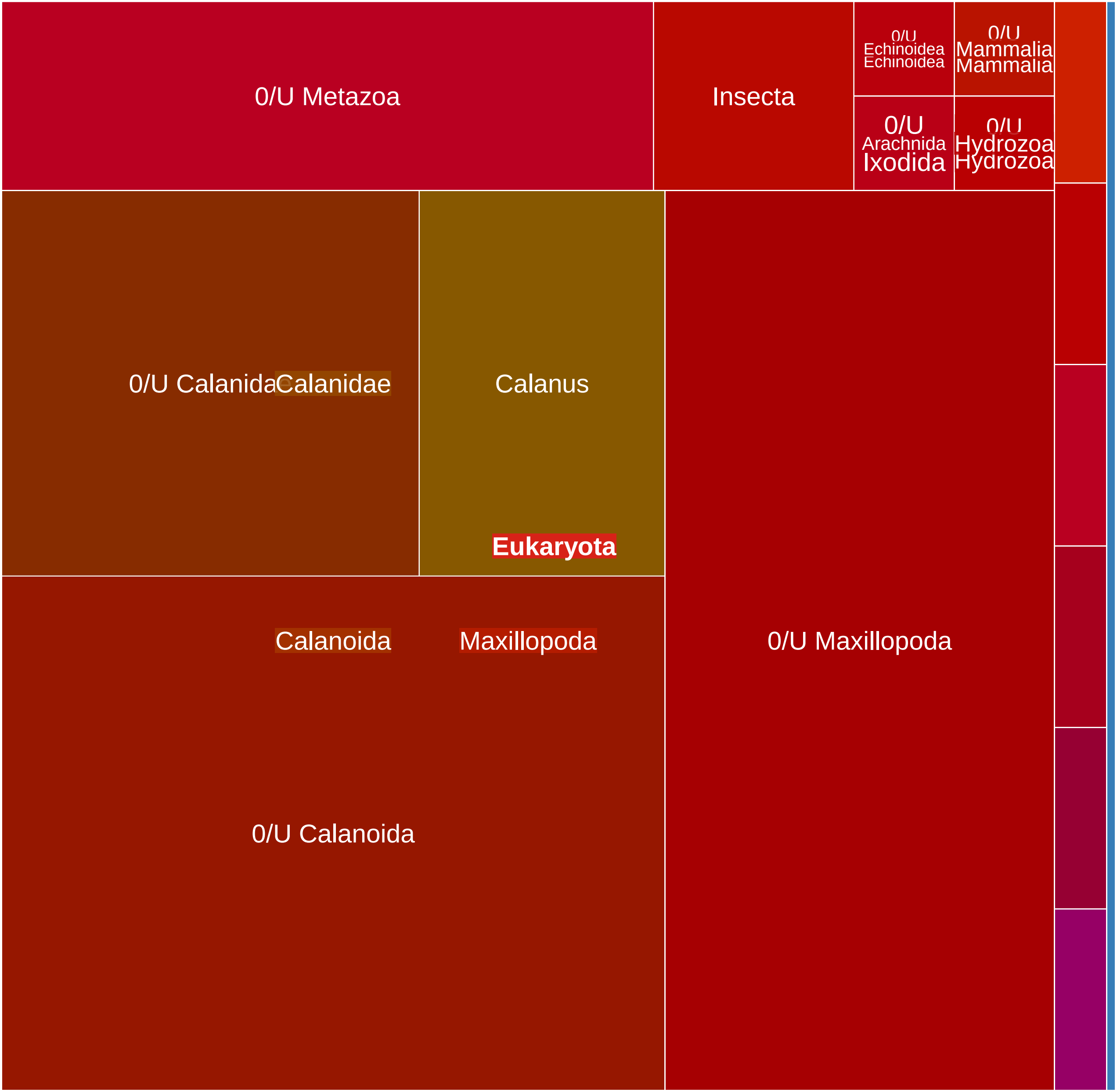

MGT−v1_740 : 768 unigenes, 139 (18.10%) taxonomically assigned

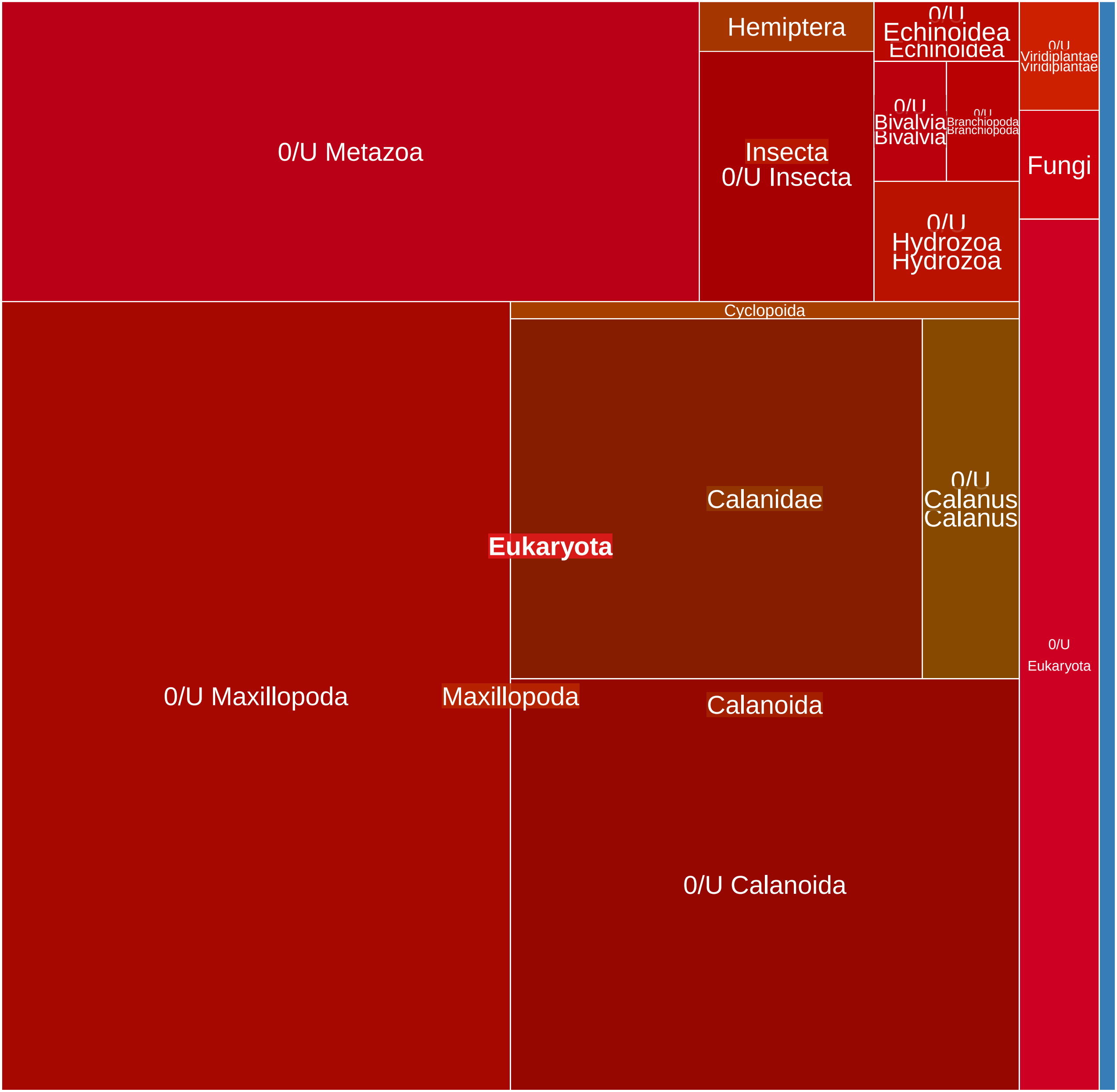

MGT−v1_741 : 766 unigenes, 120 (15.67%) taxonomically assigned

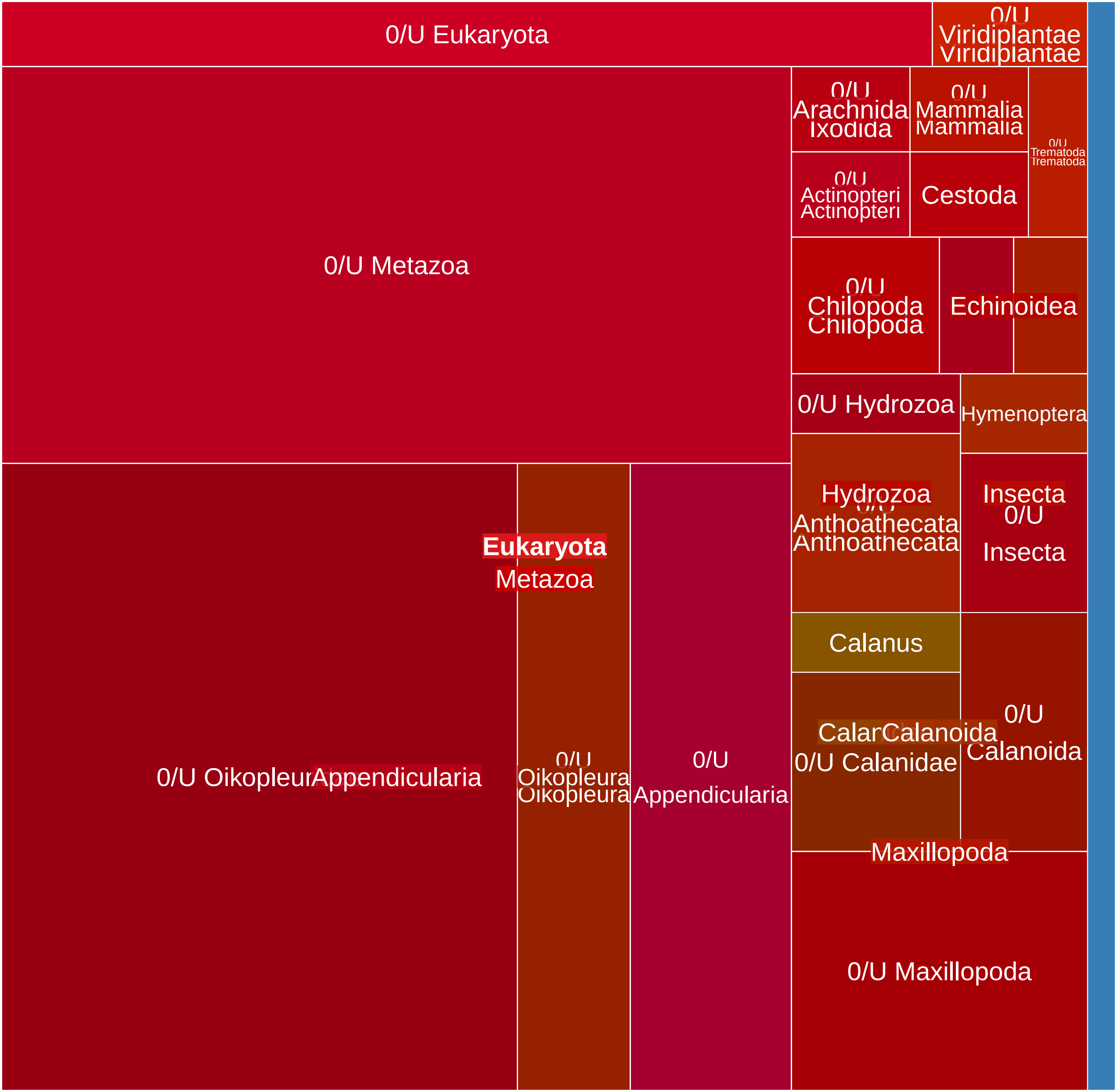

MGT−v1_742 : 765 unigenes, 49 (6.41%) taxonomically assigned

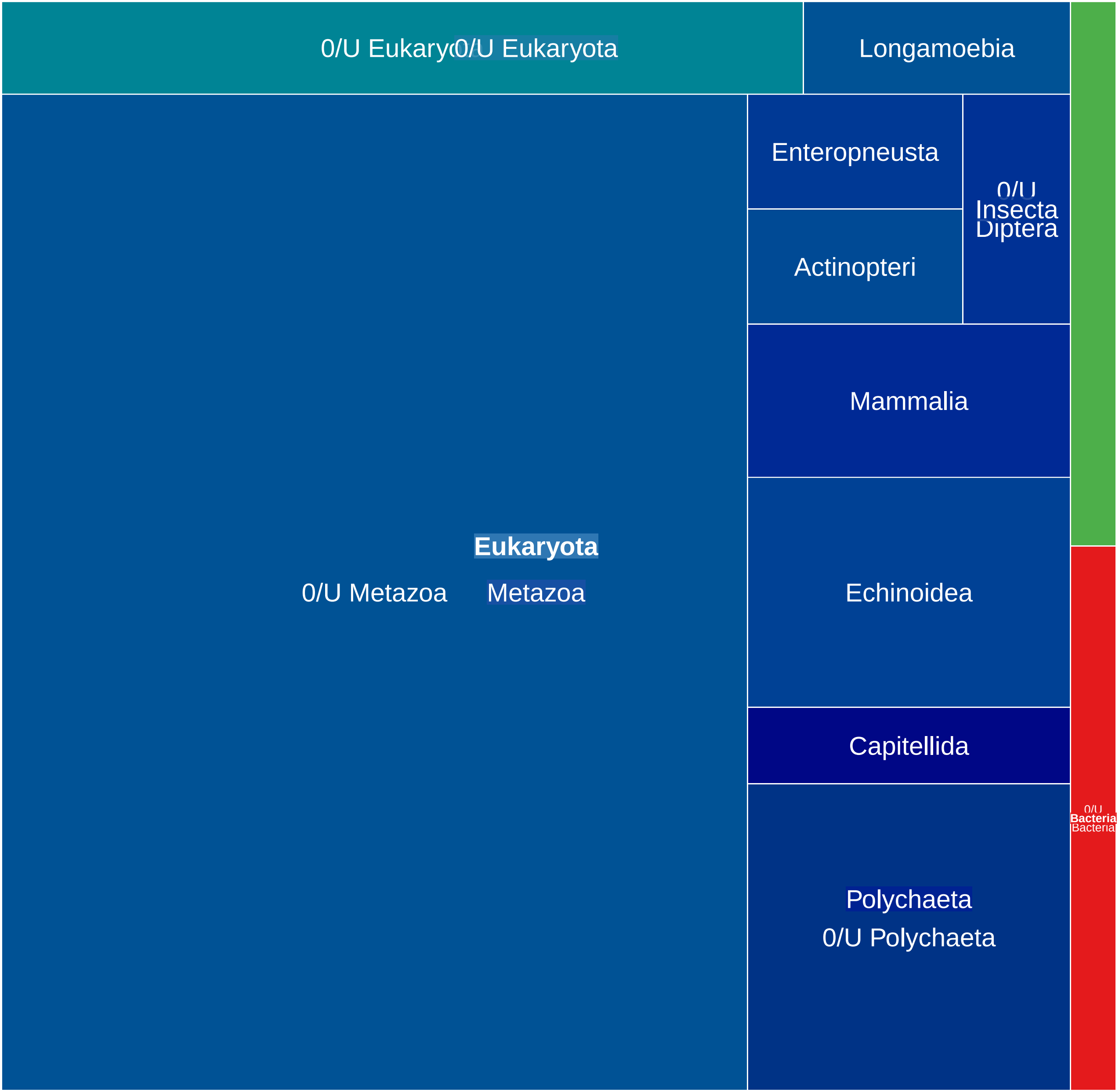

MGT−v1_743 : 764 unigenes, 512 (67.02%) taxonomically assigned

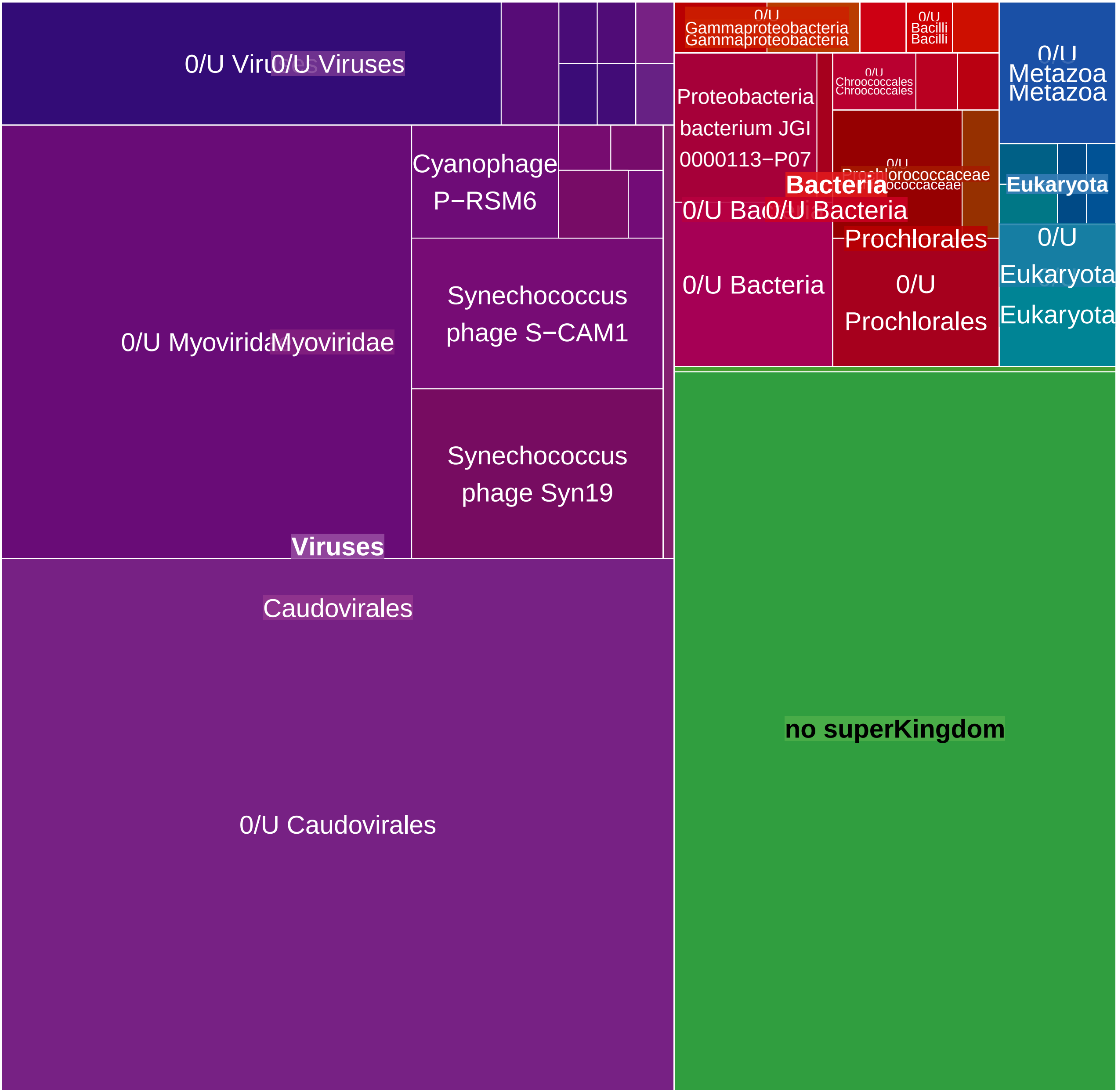

MGT−v1_744 : 764 unigenes, 64 (8.38%) taxonomically assigned

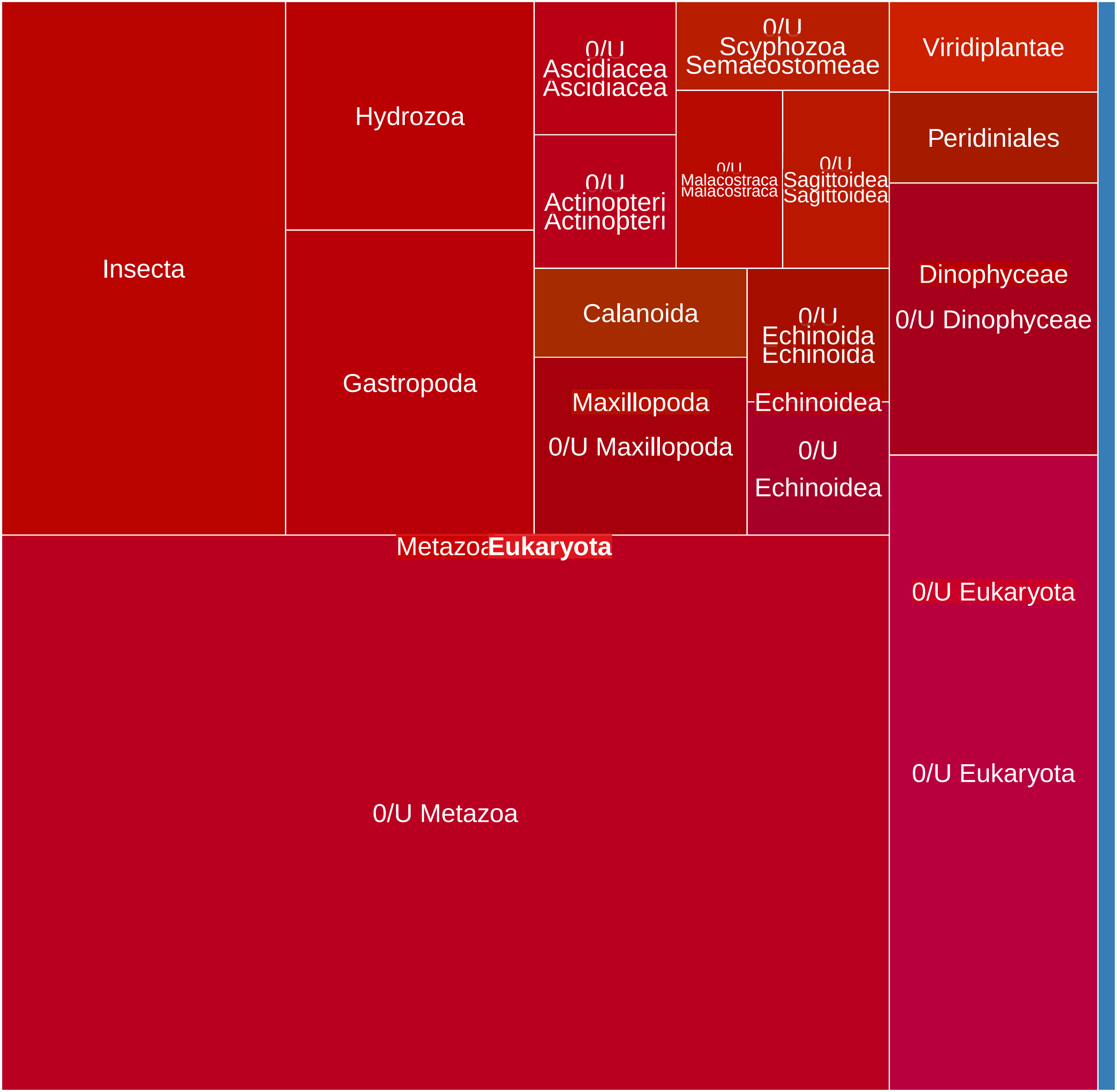

MGT−v1_745 : 758 unigenes, 165 (21.77%) taxonomically assigned

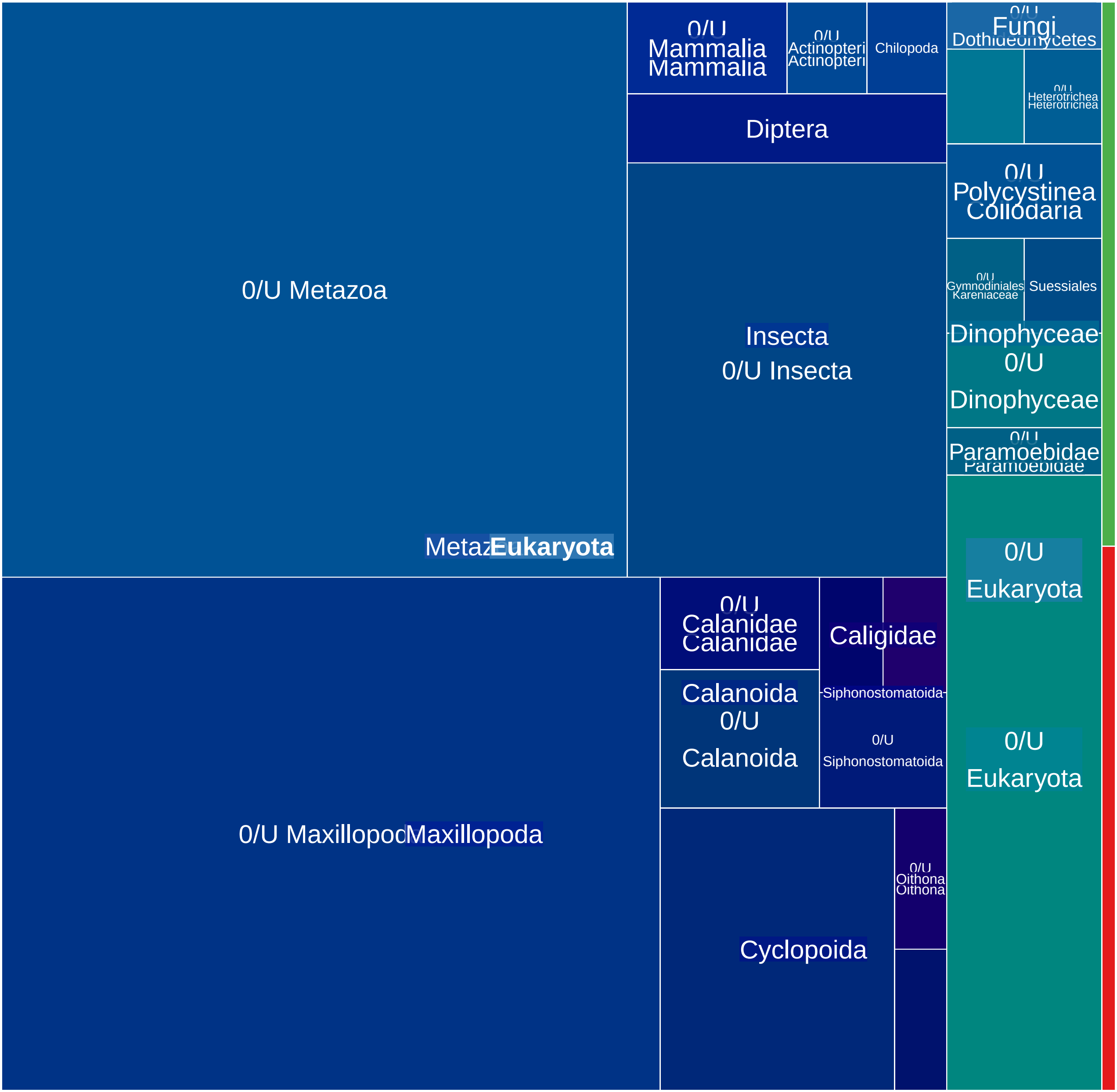

MGT−v1_746 : 757 unigenes, 14 (1.85%) taxonomically assigned

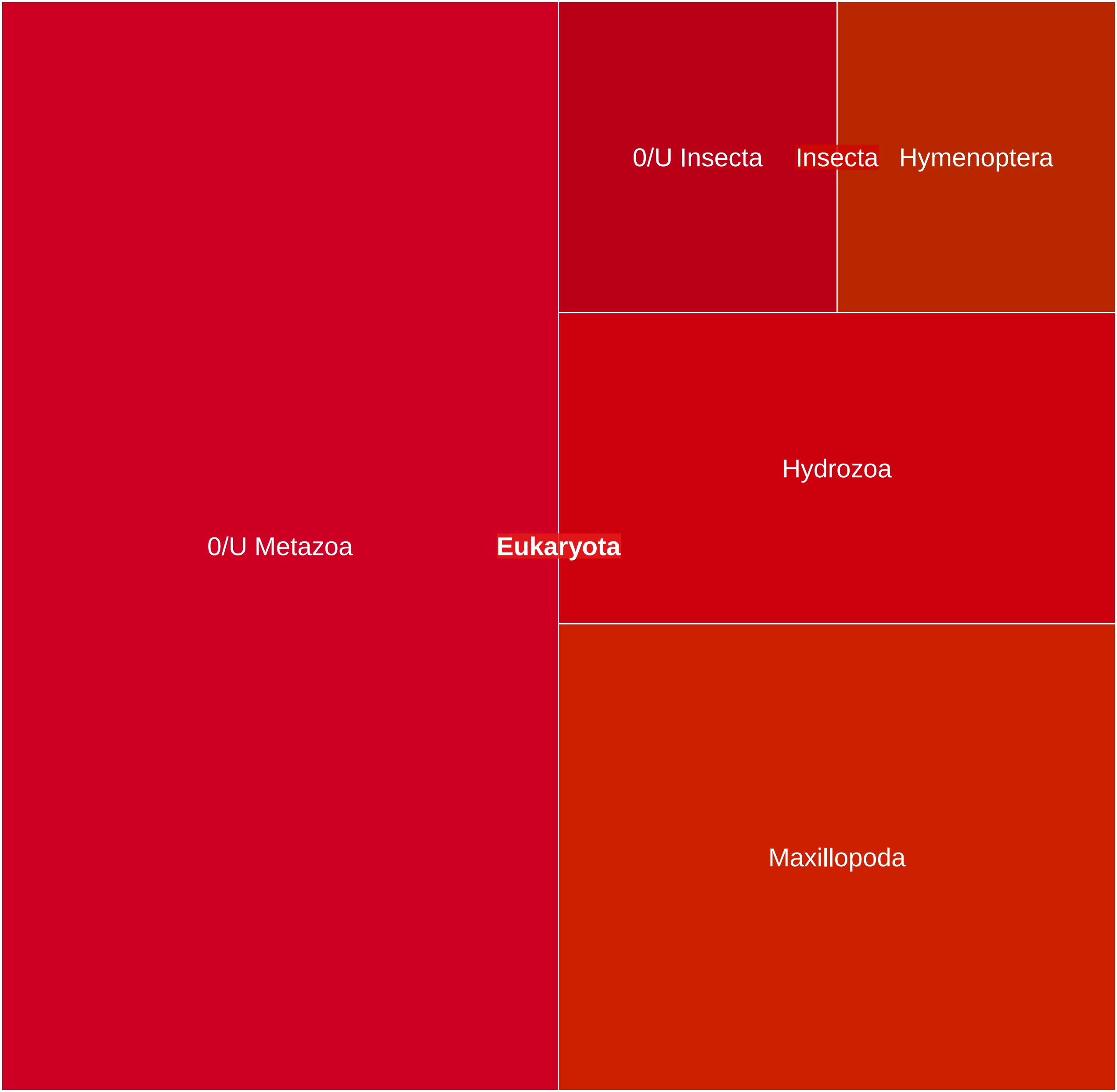

MGT−v1_747 : 756 unigenes, 540 (71.43%) taxonomically assigned

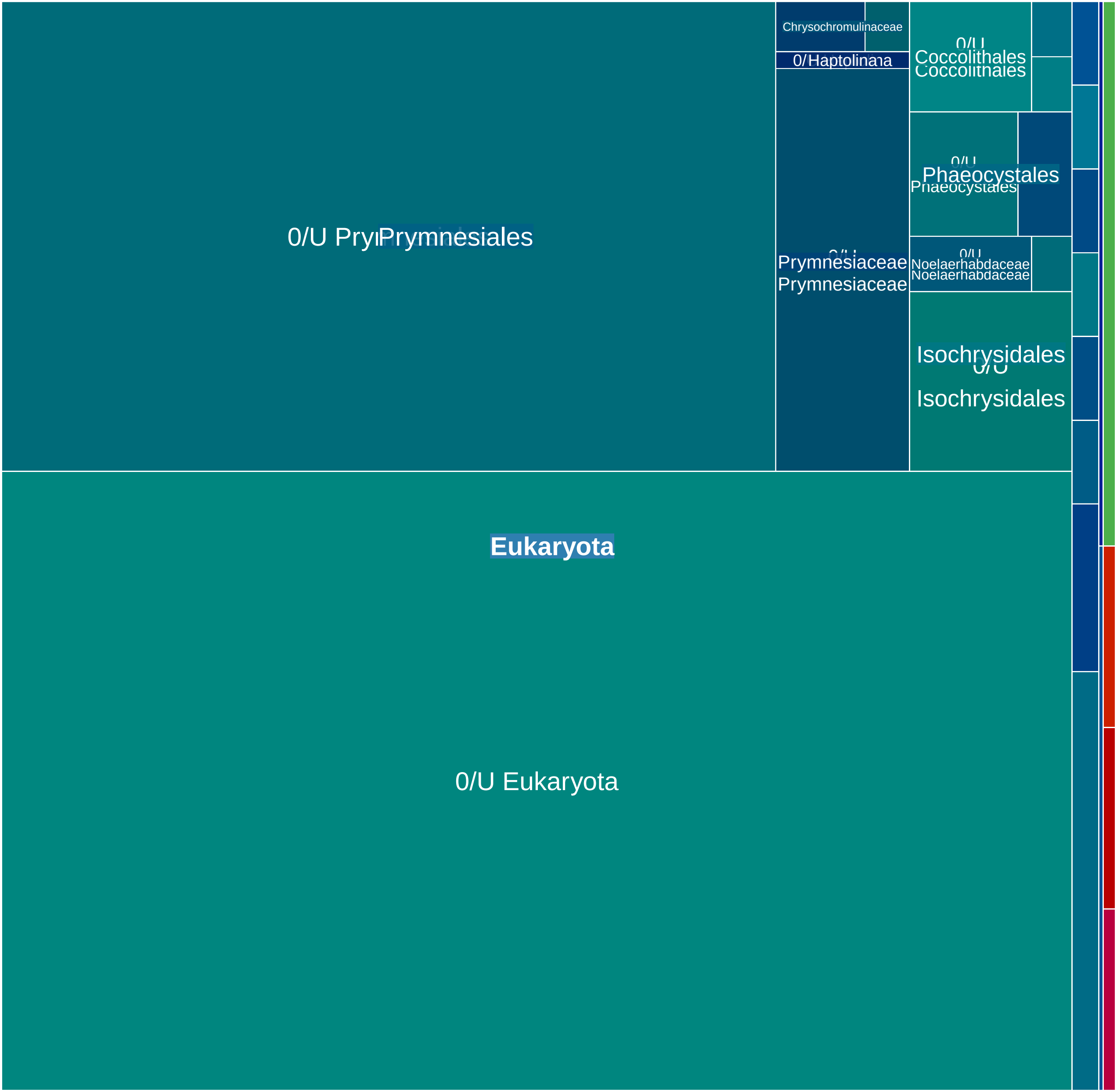

MGT−v1_748 : 750 unigenes, 150 (20.00%) taxonomically assigned

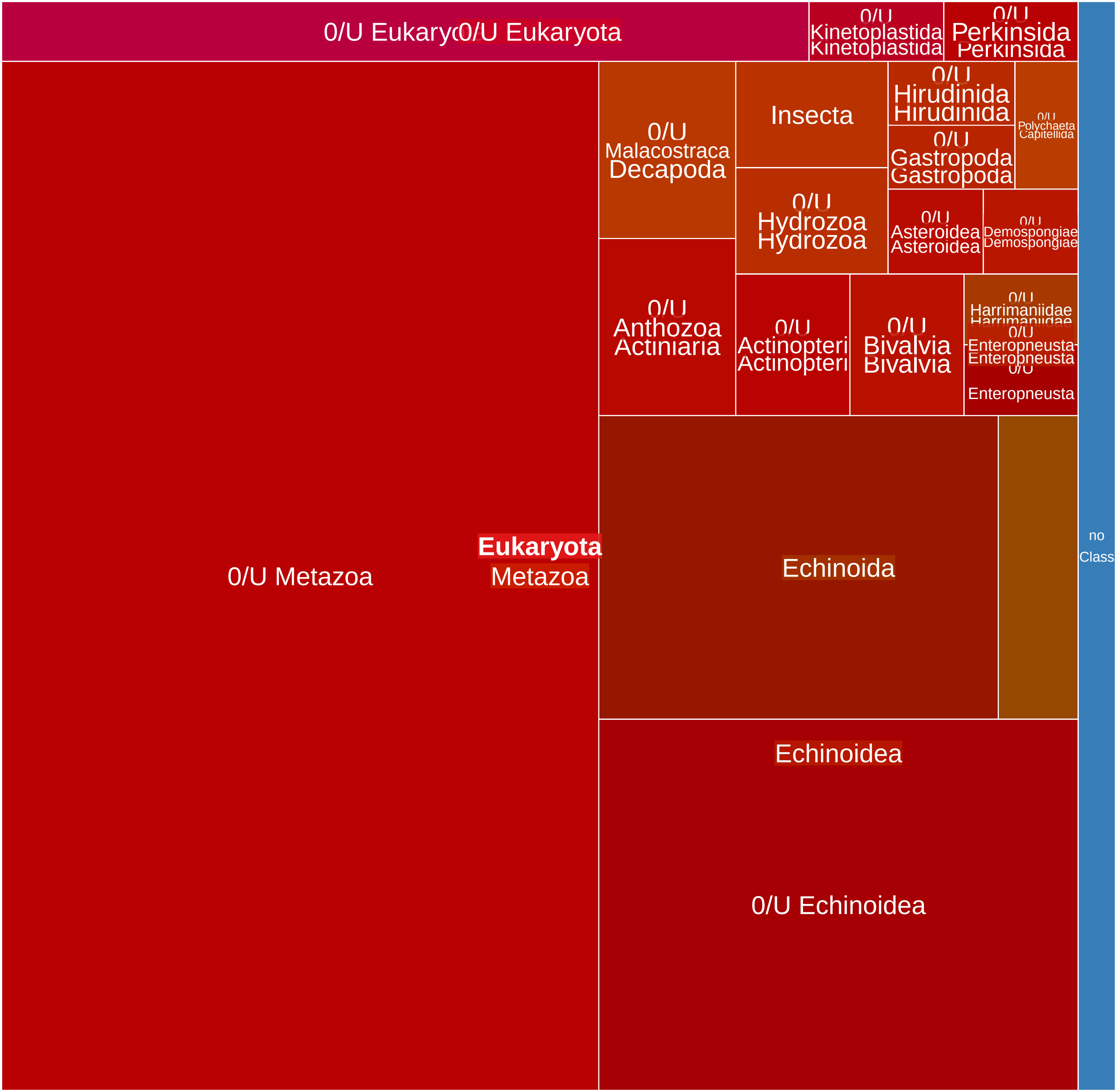

MGT−v1_749 : 746 unigenes, 167 (22.39%) taxonomically assigned

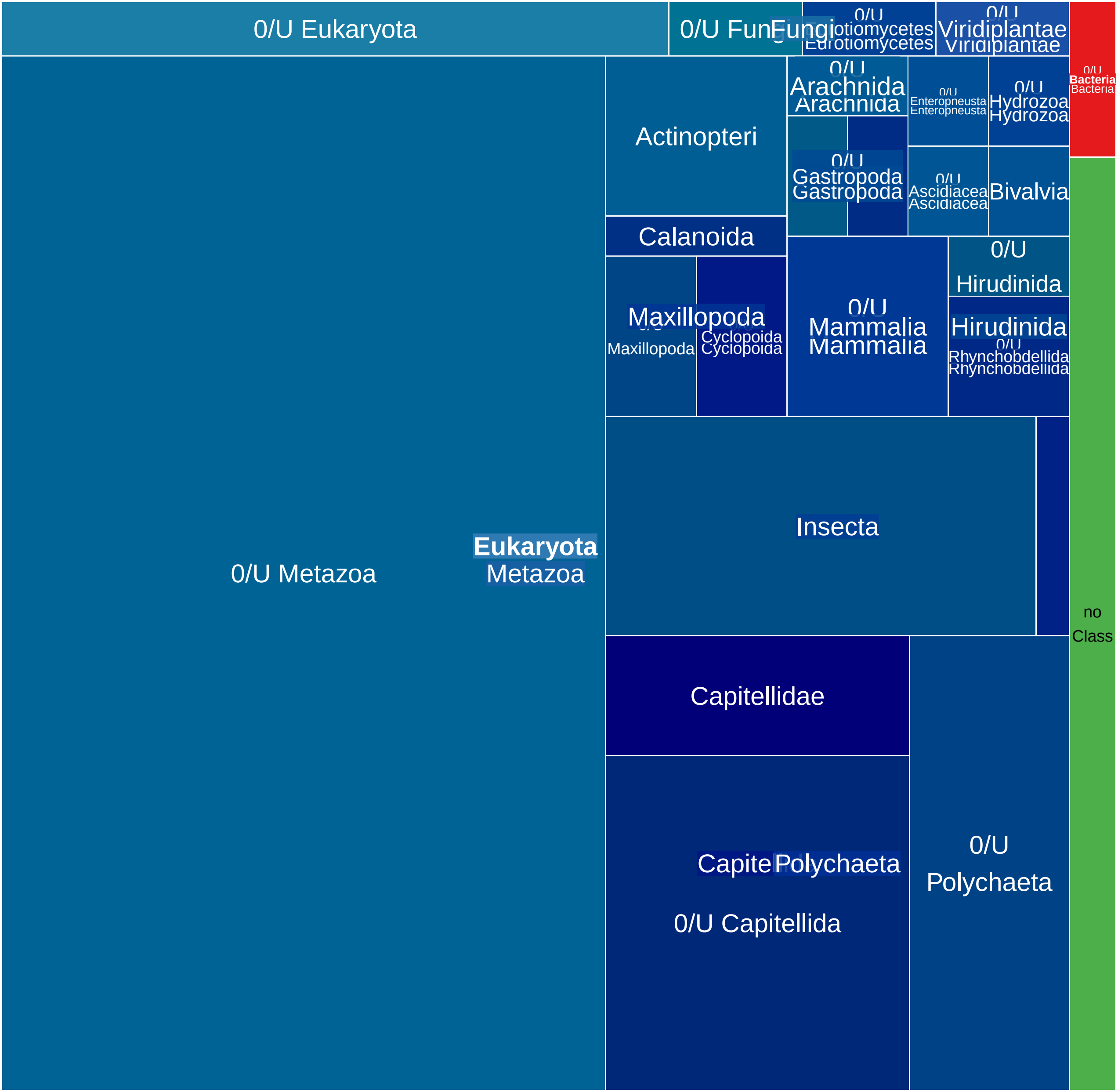

### MGT−v1_750 : 743 unigenes, 246 (33.11%) taxonomically assigned

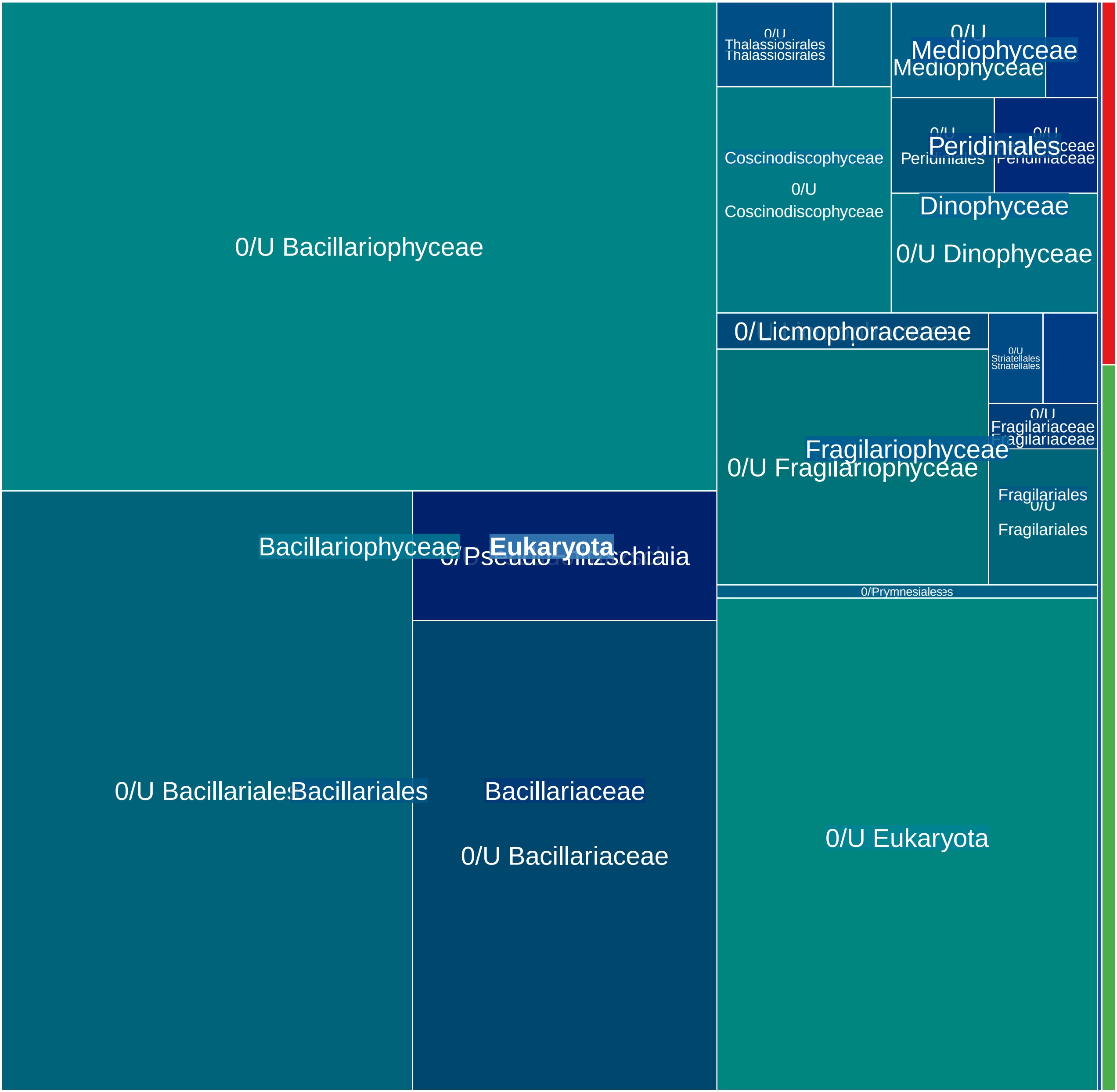

MGT−v1_751 : 737 unigenes, 55 (7.46%) taxonomically assigned

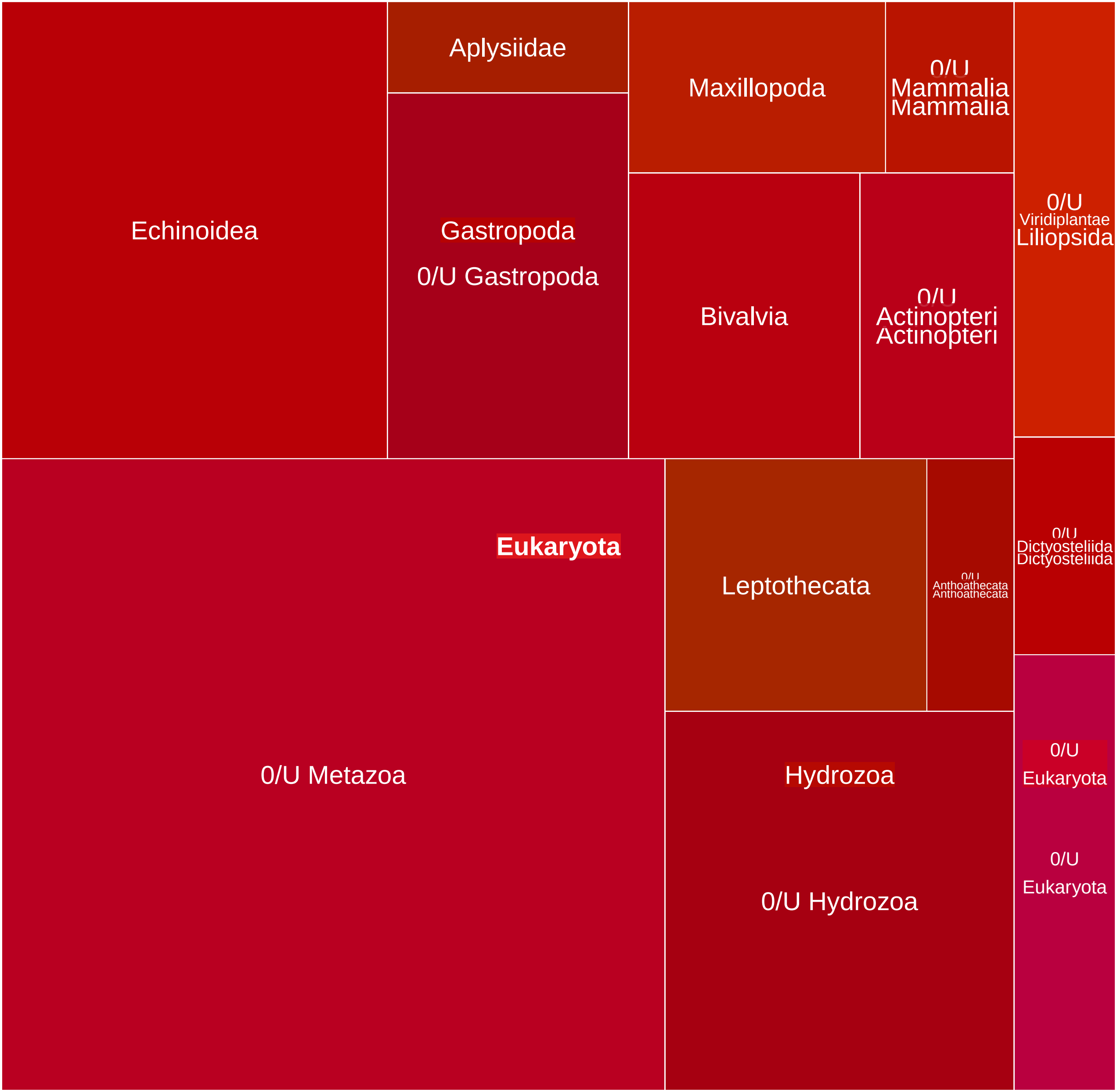

MGT−v1_752 : 734 unigenes, 33 (4.50%) taxonomically assigned

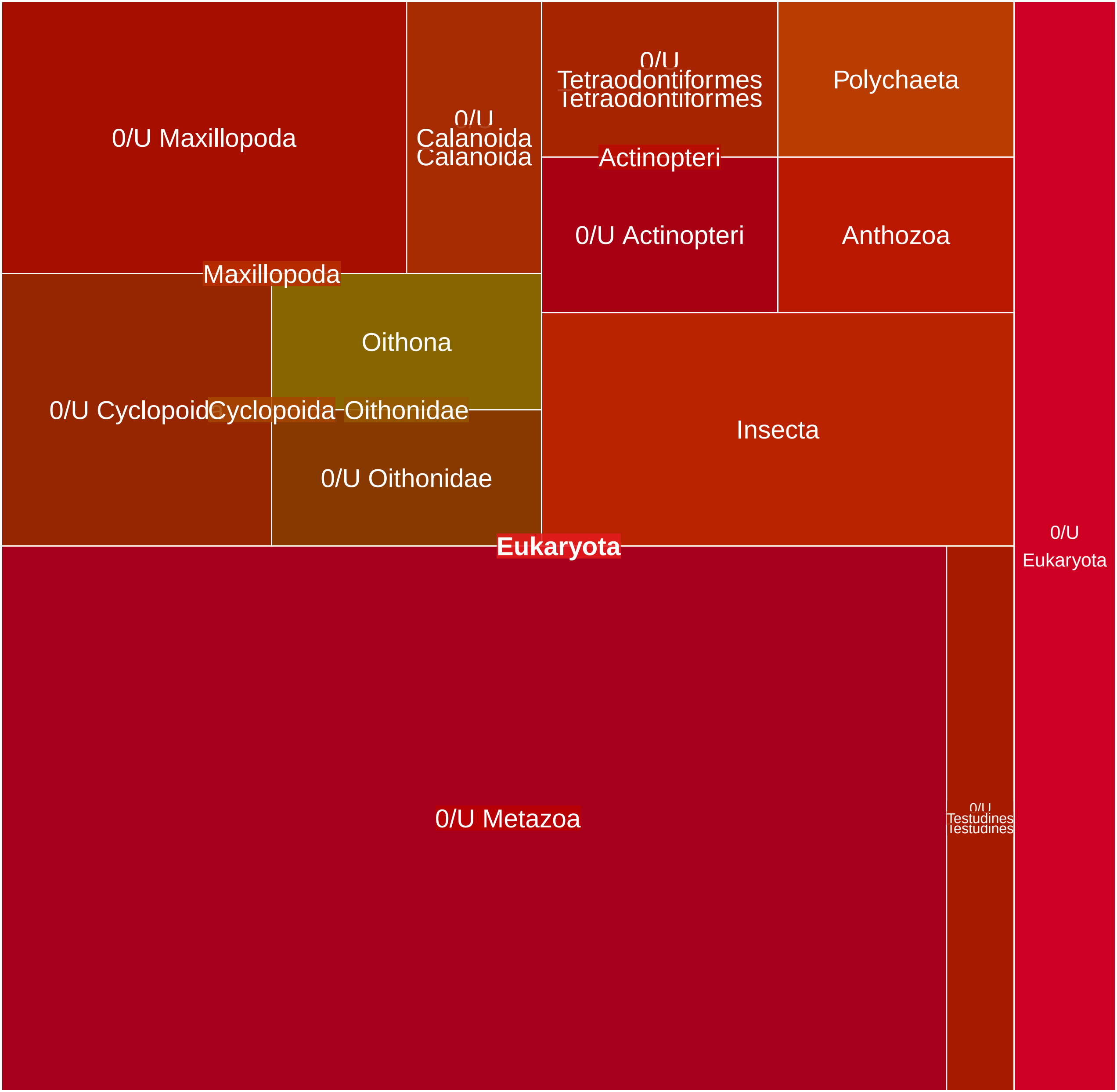

MGT−v1_753 : 734 unigenes, 715 (97.41%) taxonomically assigned

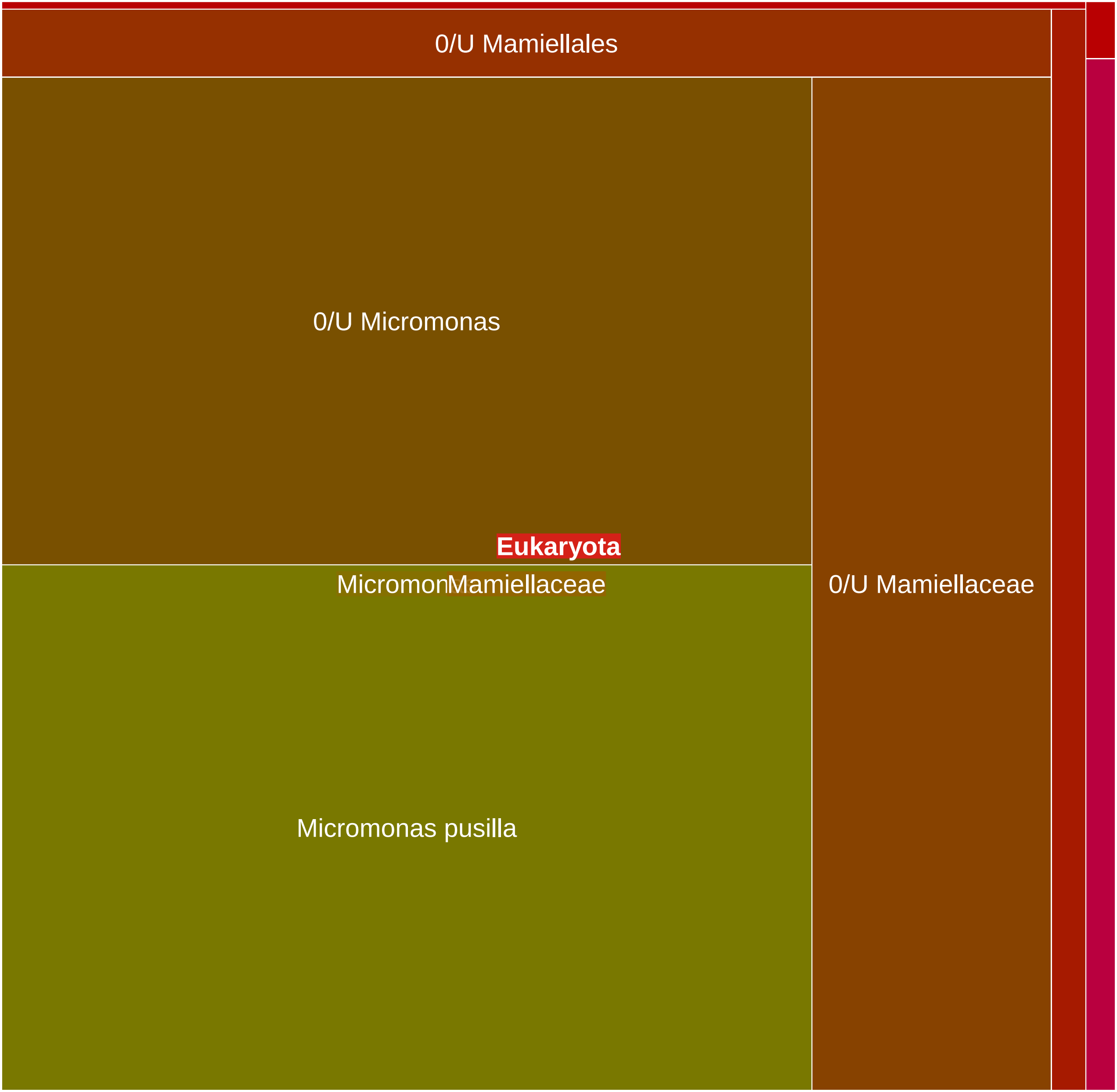

MGT−v1_754 : 733 unigenes, 724 (98.77%) taxonomically assigned

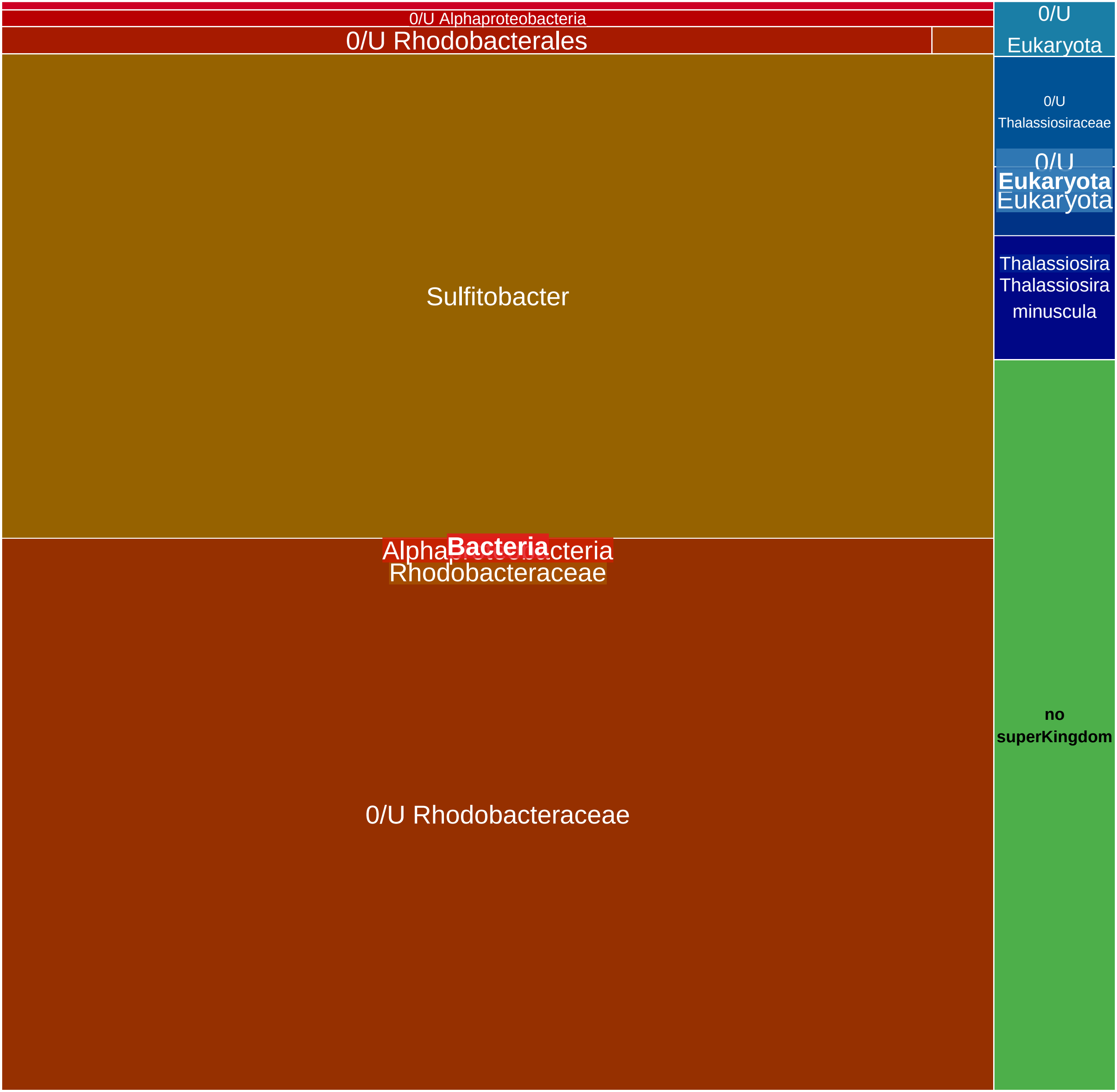

MGT−v1_755 : 726 unigenes, 674 (92.84%) taxonomically assigned

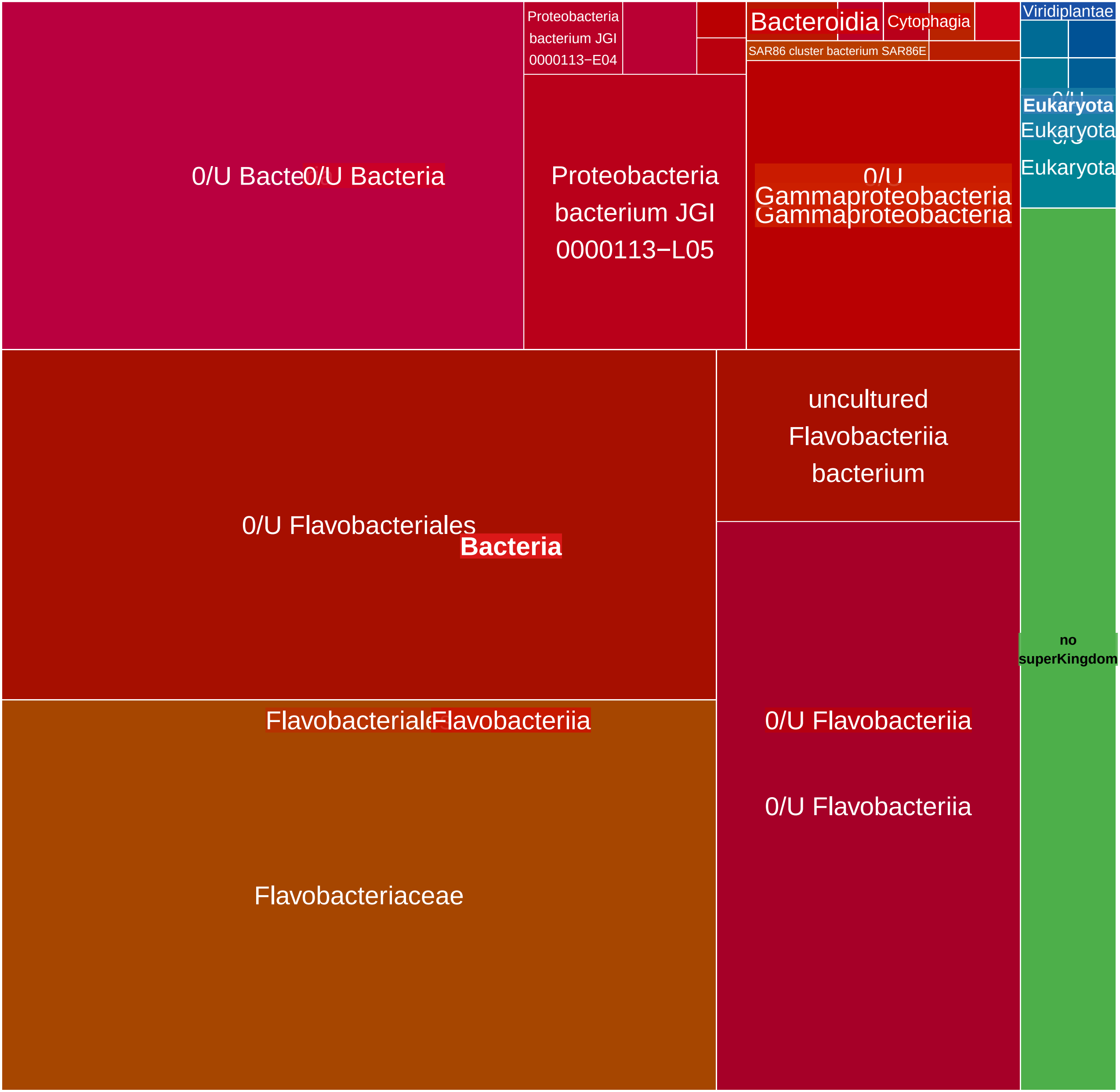

MGT−v1_756 : 726 unigenes, 48 (6.61%) taxonomically assigned

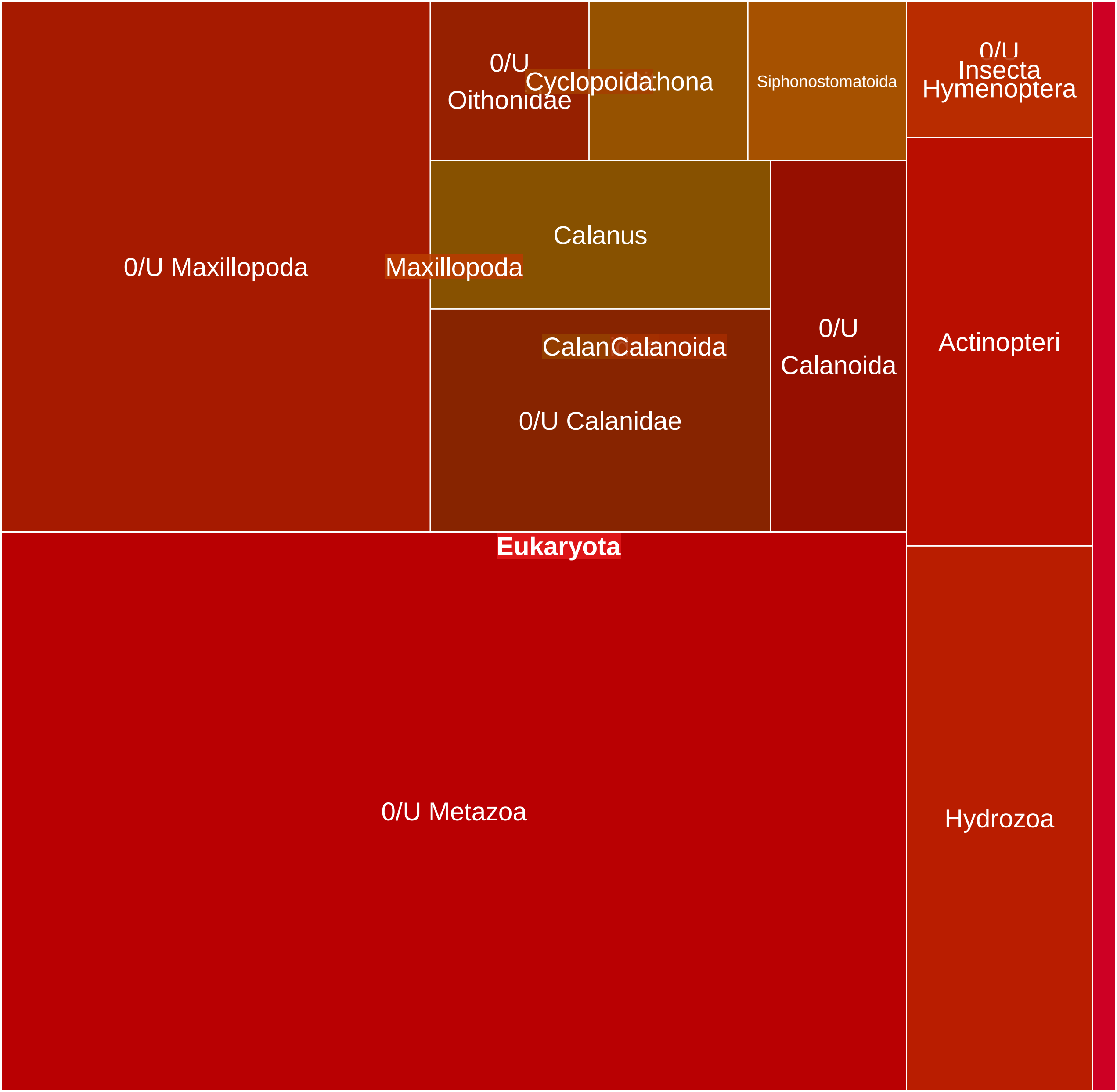

MGT−v1_757 : 725 unigenes, 67 (9.24%) taxonomically assigned

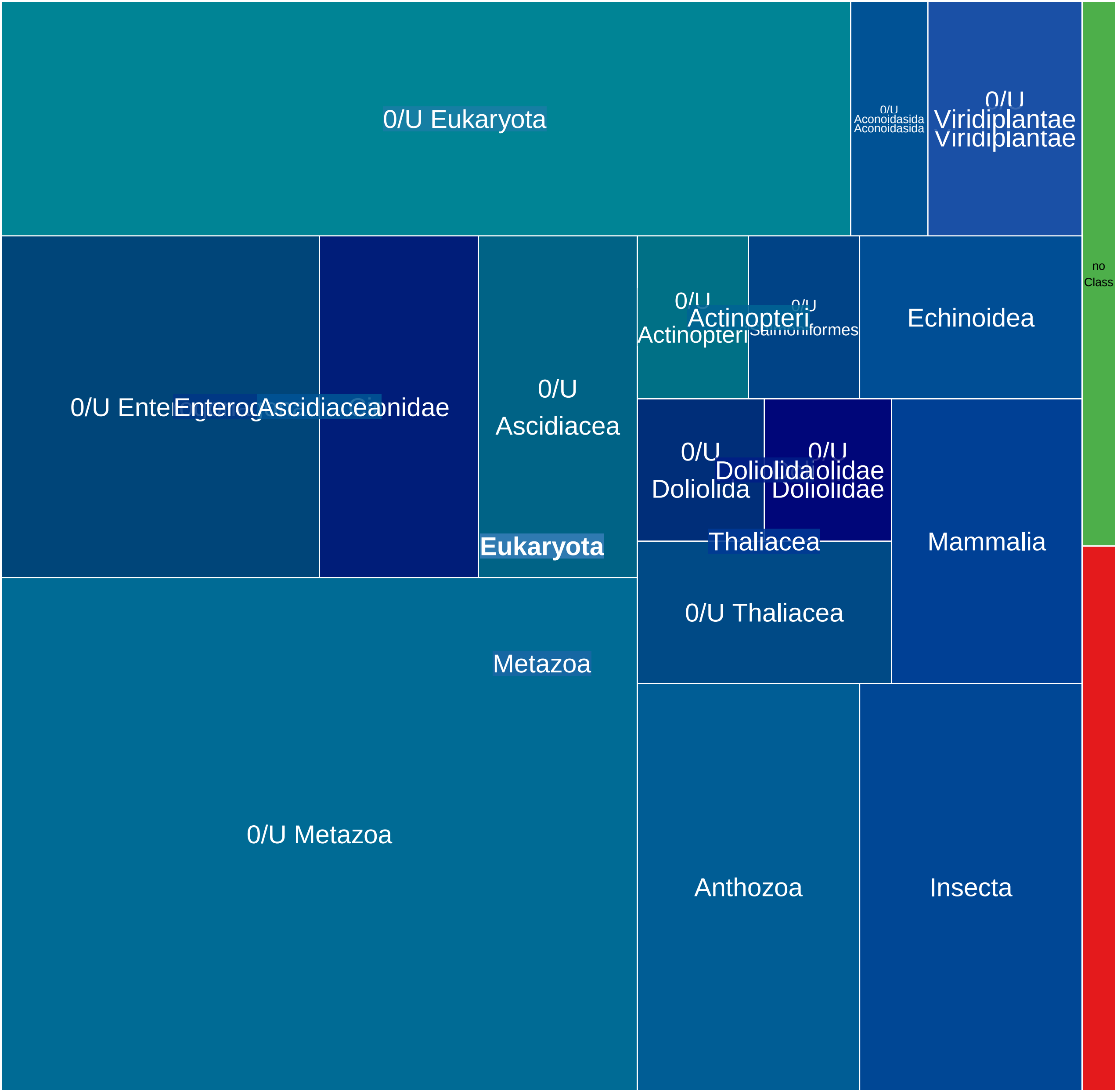

MGT−v1_758 : 724 unigenes, 123 (16.99%) taxonomically assigned

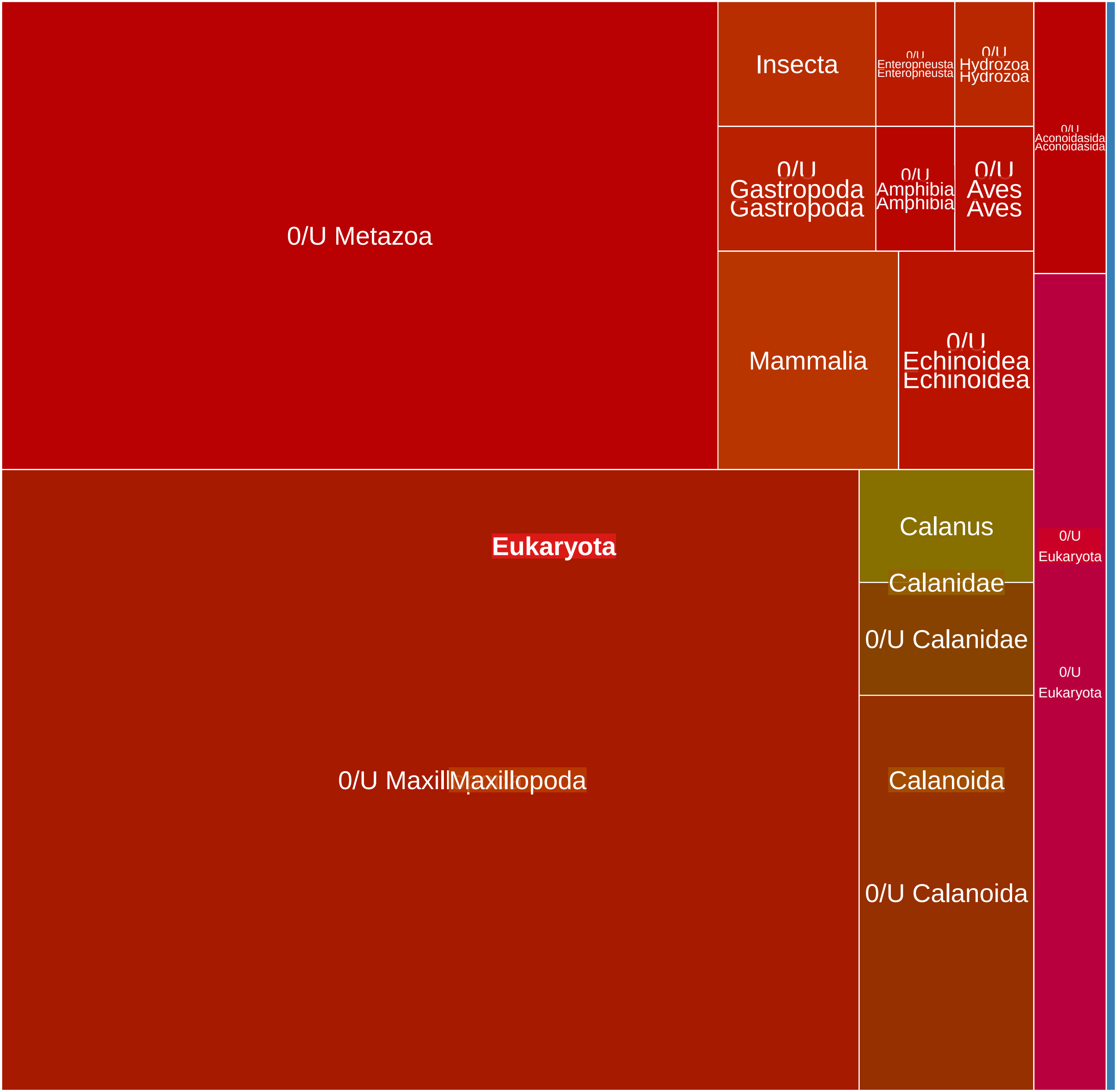

MGT−v1_759 : 721 unigenes, 150 (20.80%) taxonomically assigned

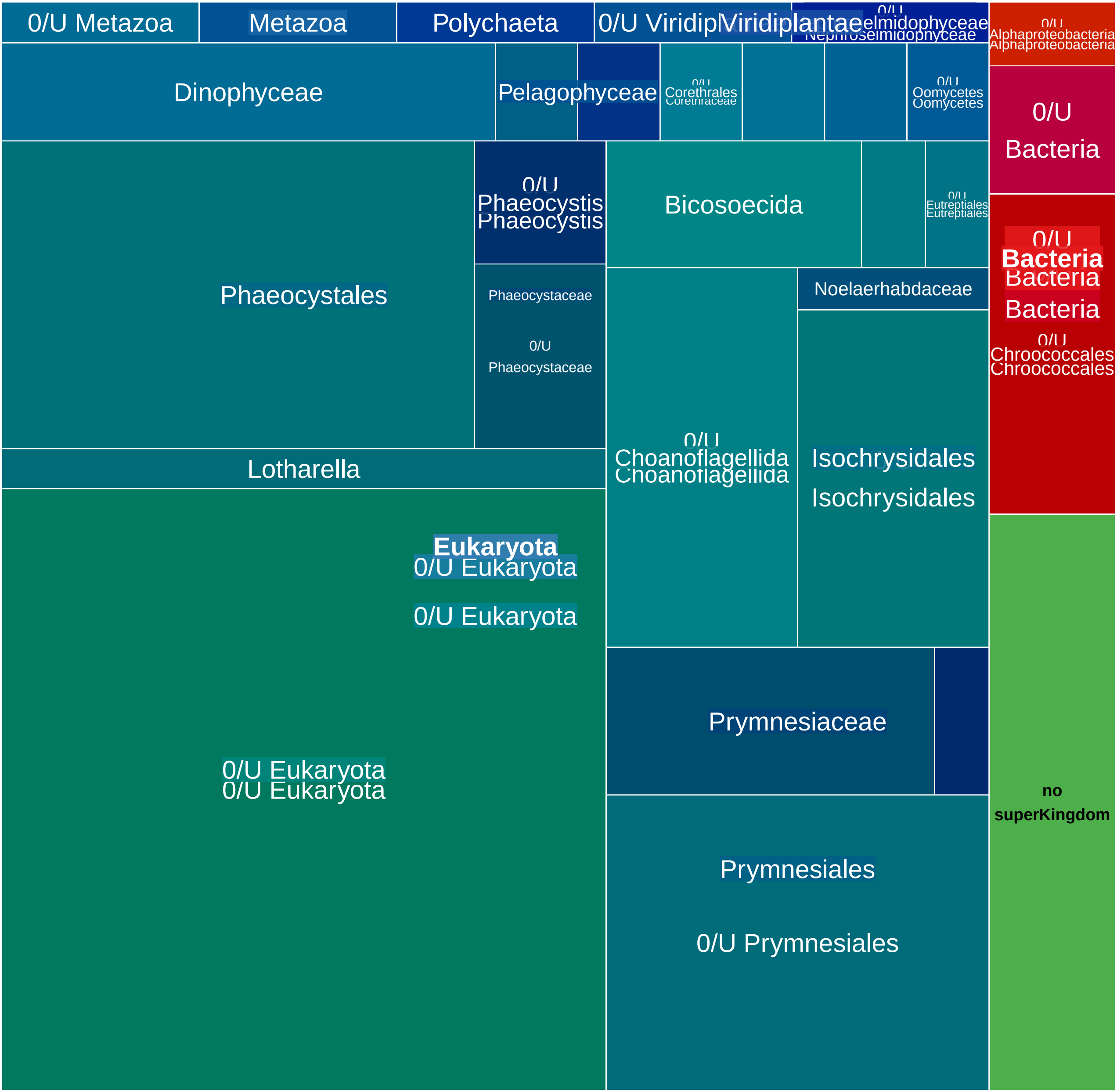

MGT−v1_760 : 720 unigenes, 524 (72.78%) taxonomically assigned

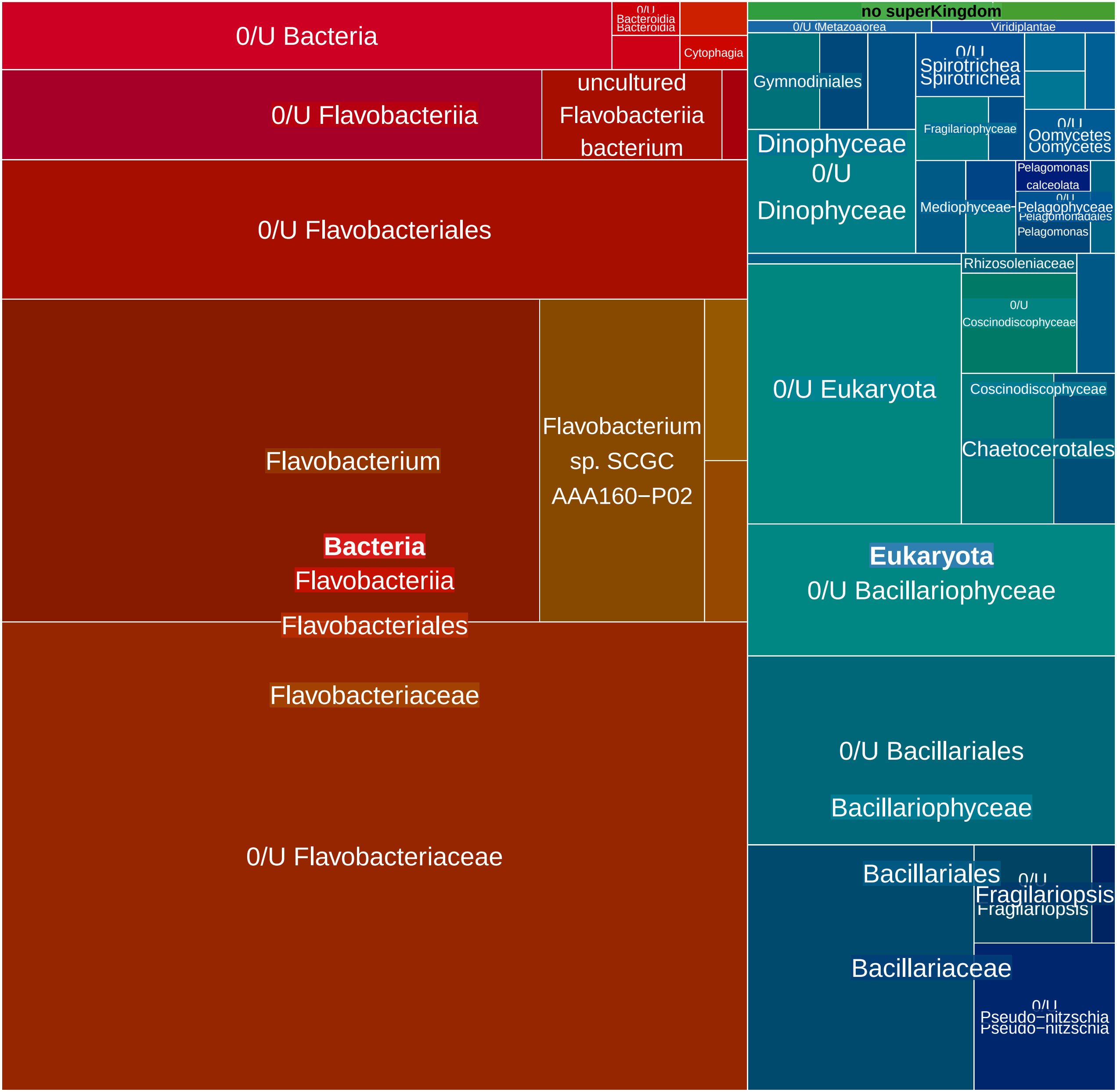

MGT−v1_761 : 719 unigenes, 632 (87.90%) taxonomically assigned

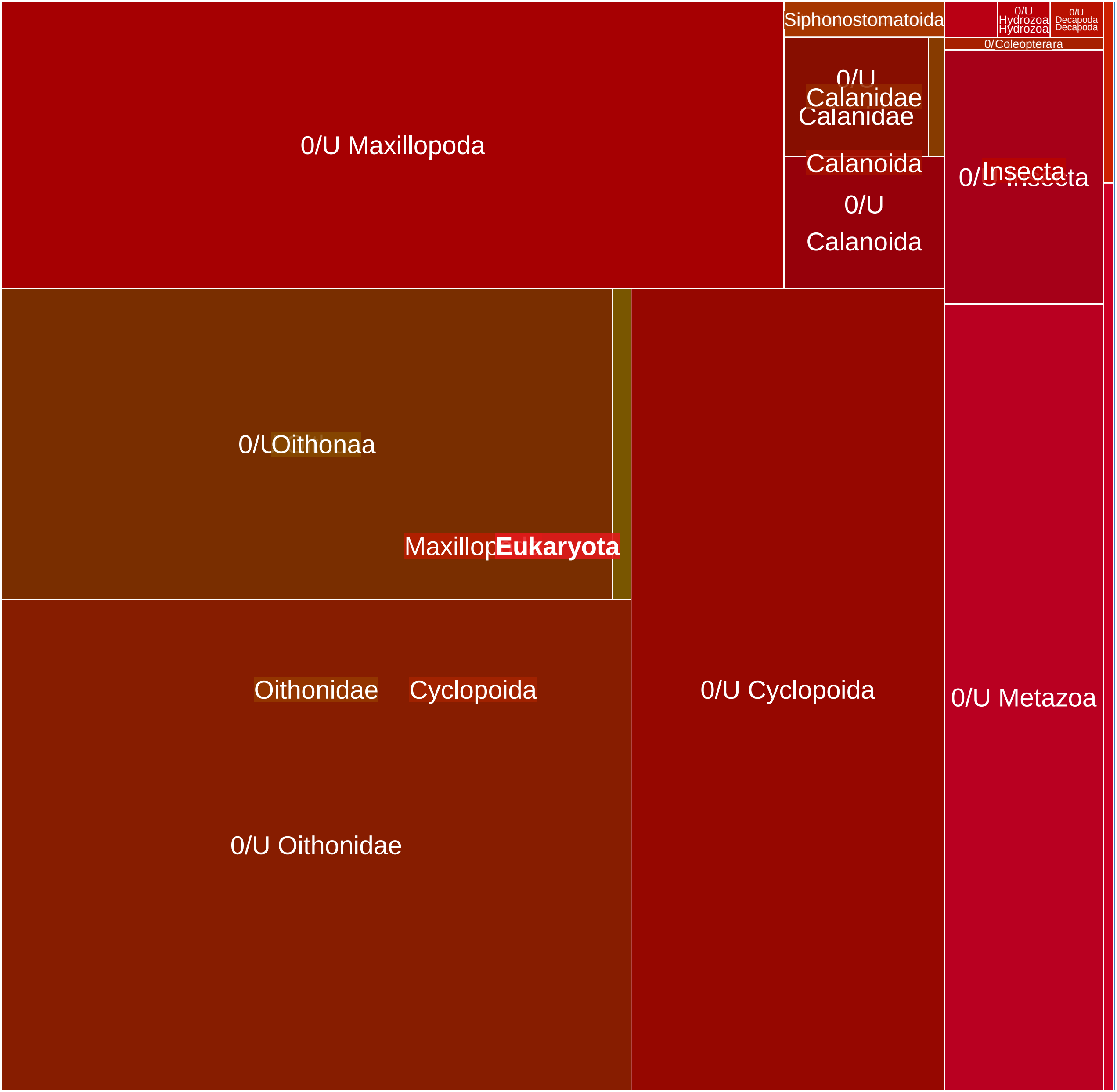

MGT−v1_762 : 717 unigenes, 688 (95.96%) taxonomically assigned

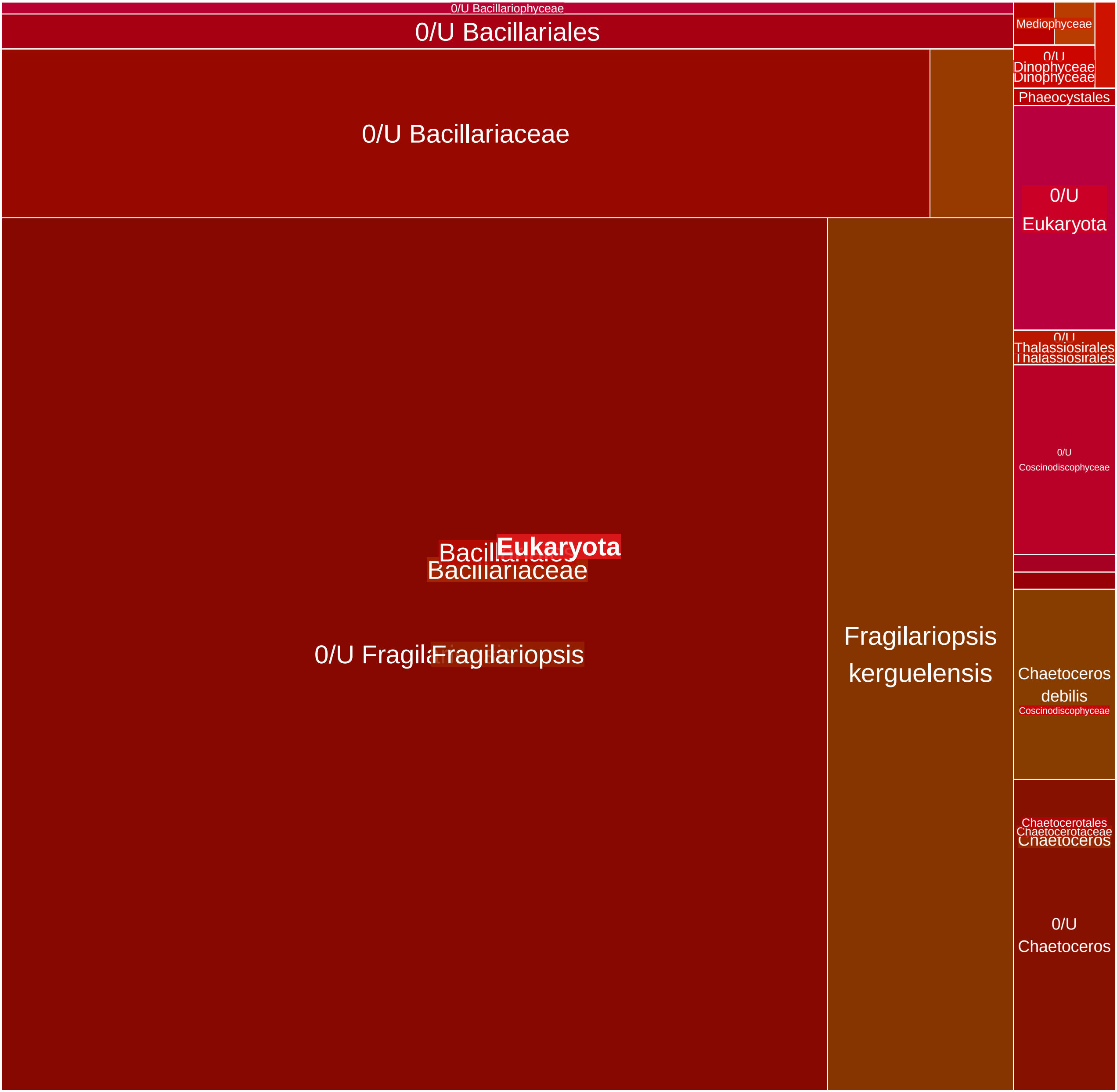

MGT−v1_763 : 712 unigenes, 567 (79.63%) taxonomically assigned

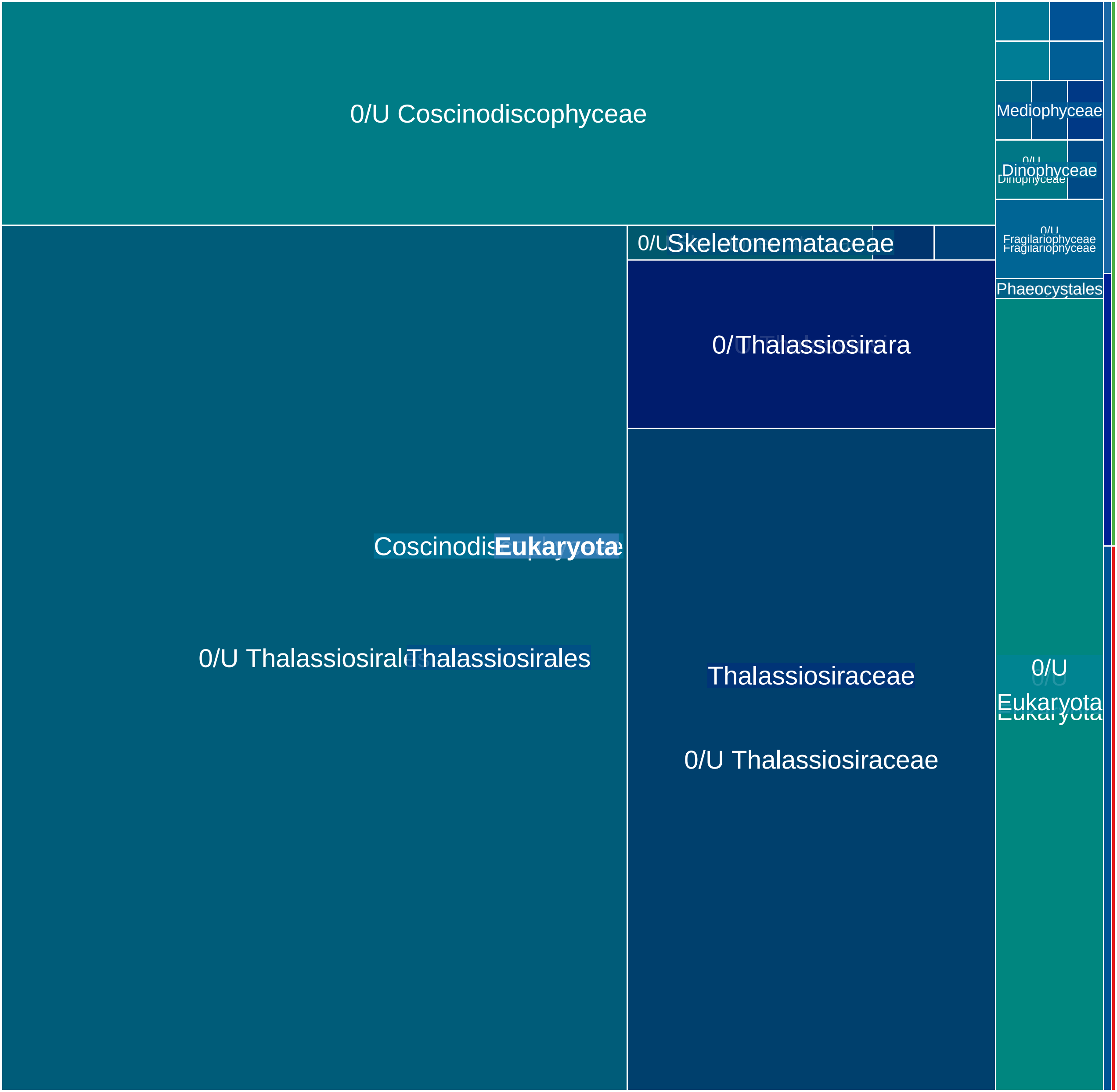

MGT−v1_764 : 711 unigenes, 54 (7.59%) taxonomically assigned

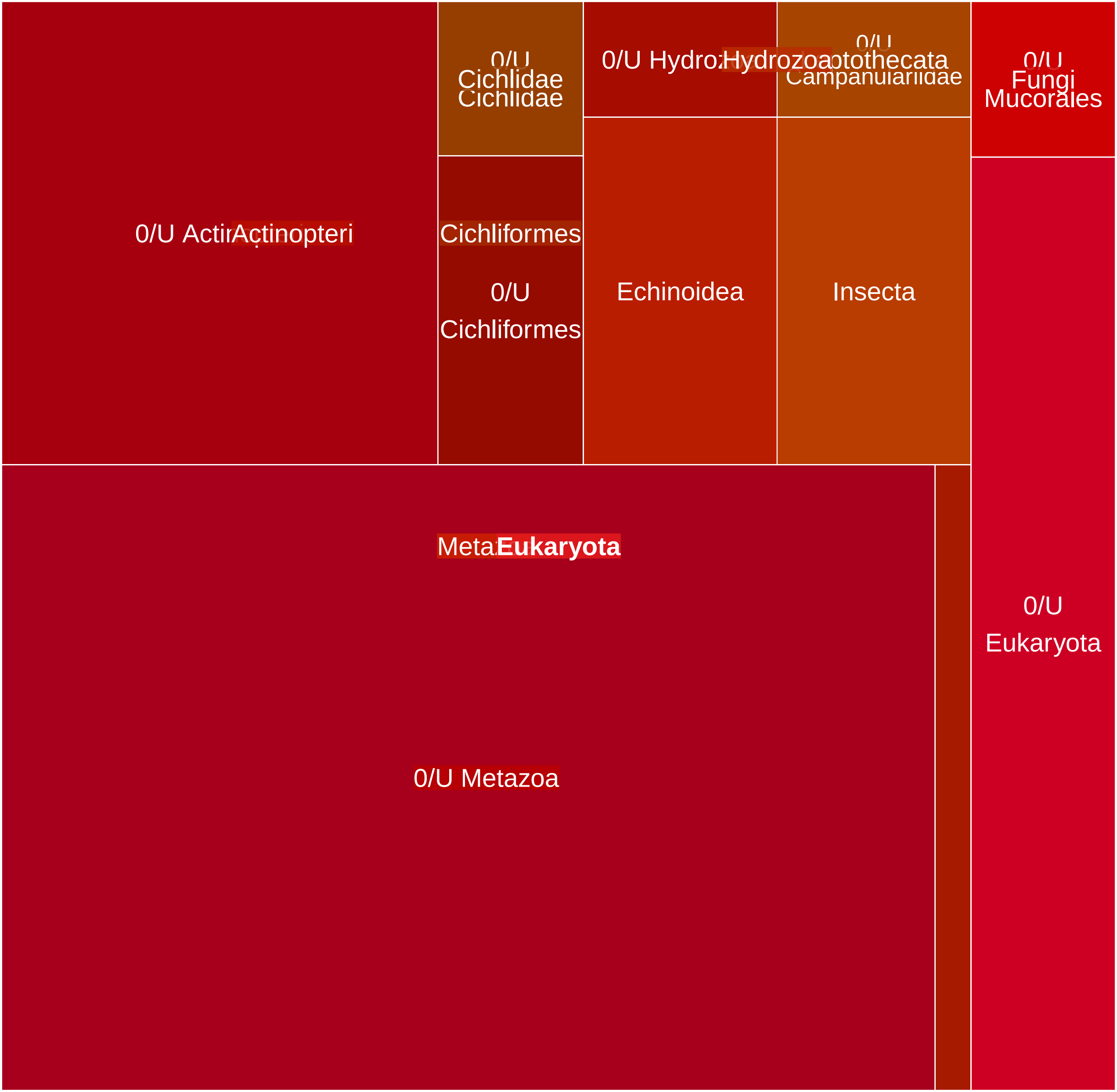

MGT−v1_765 : 711 unigenes, 40 (5.63%) taxonomically assigned

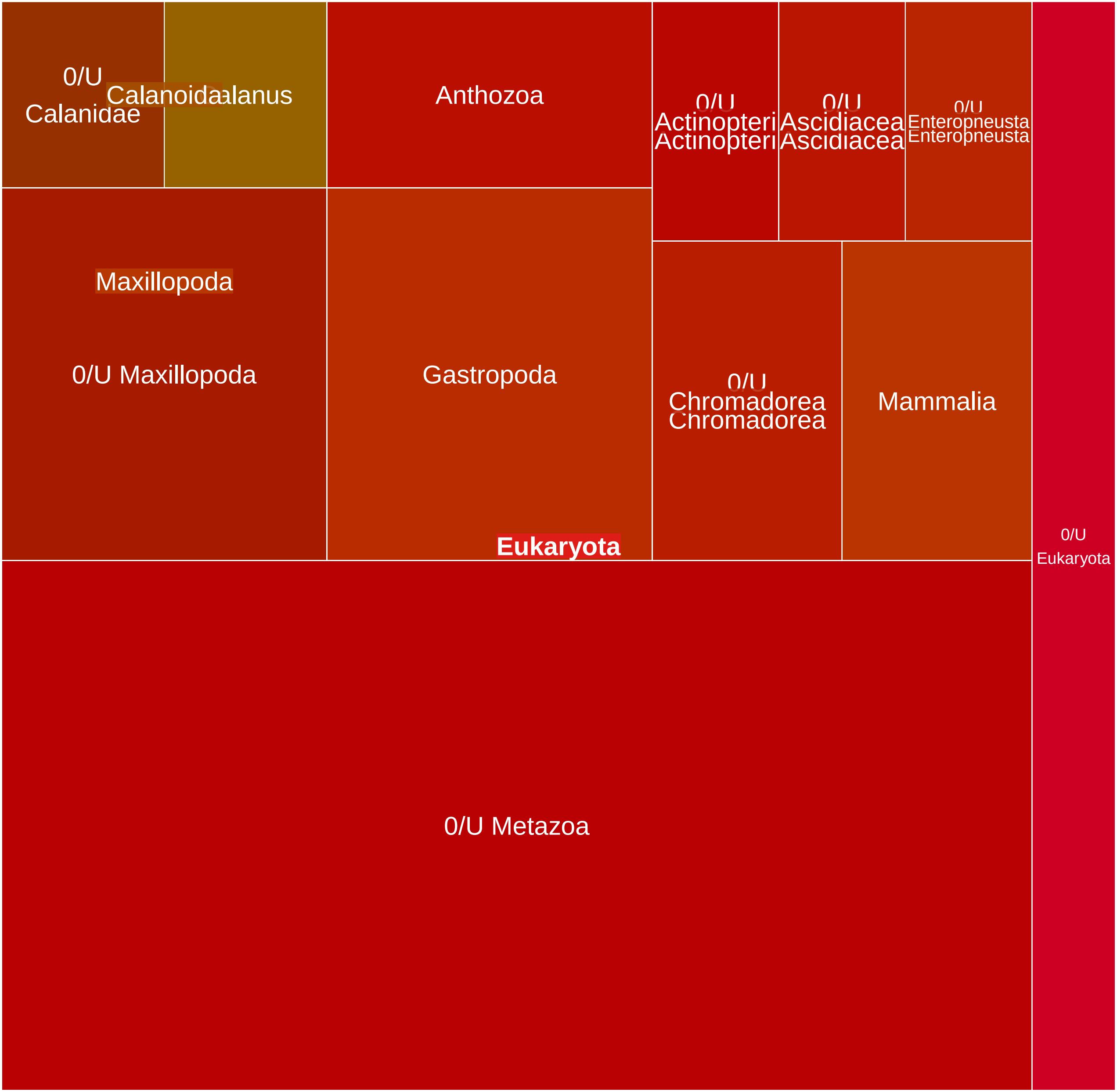

MGT−v1_766 : 710 unigenes, 698 (98.31%) taxonomically assigned

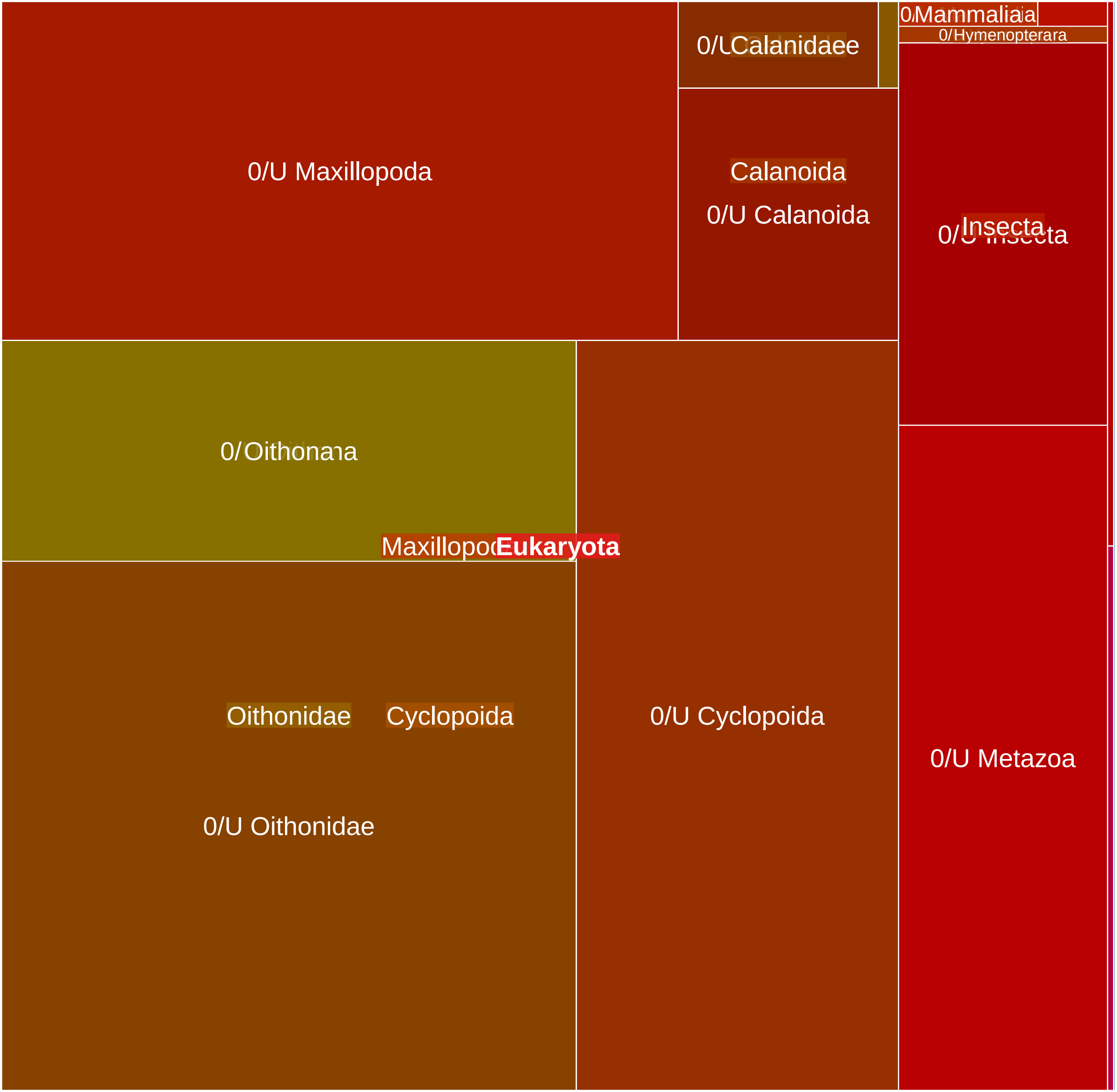

MGT−v1_767 : 710 unigenes, 56 (7.89%) taxonomically assigned

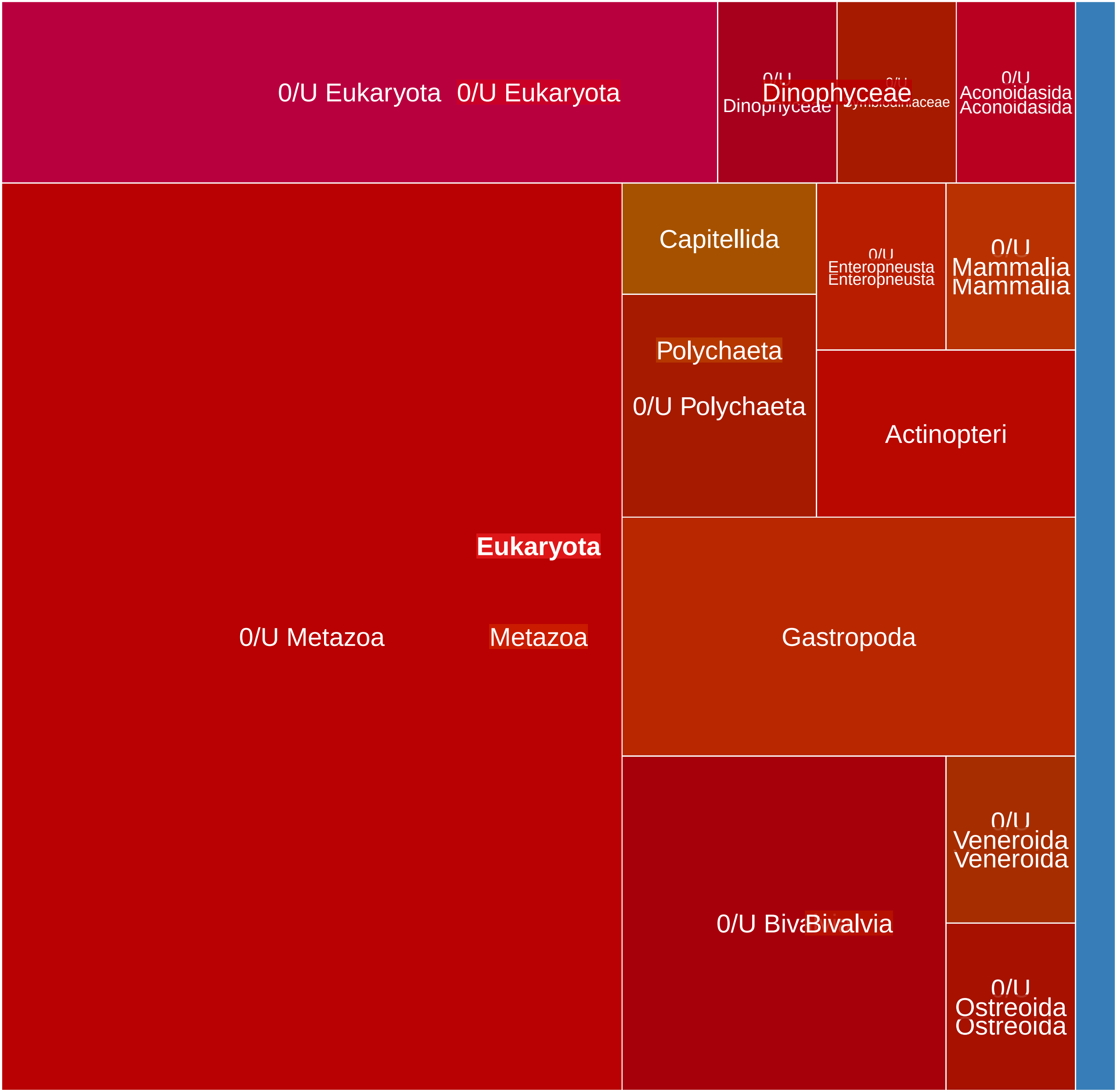

### MGT−v1_768 : 708 unigenes, 87 (12.29%) taxonomically assigned

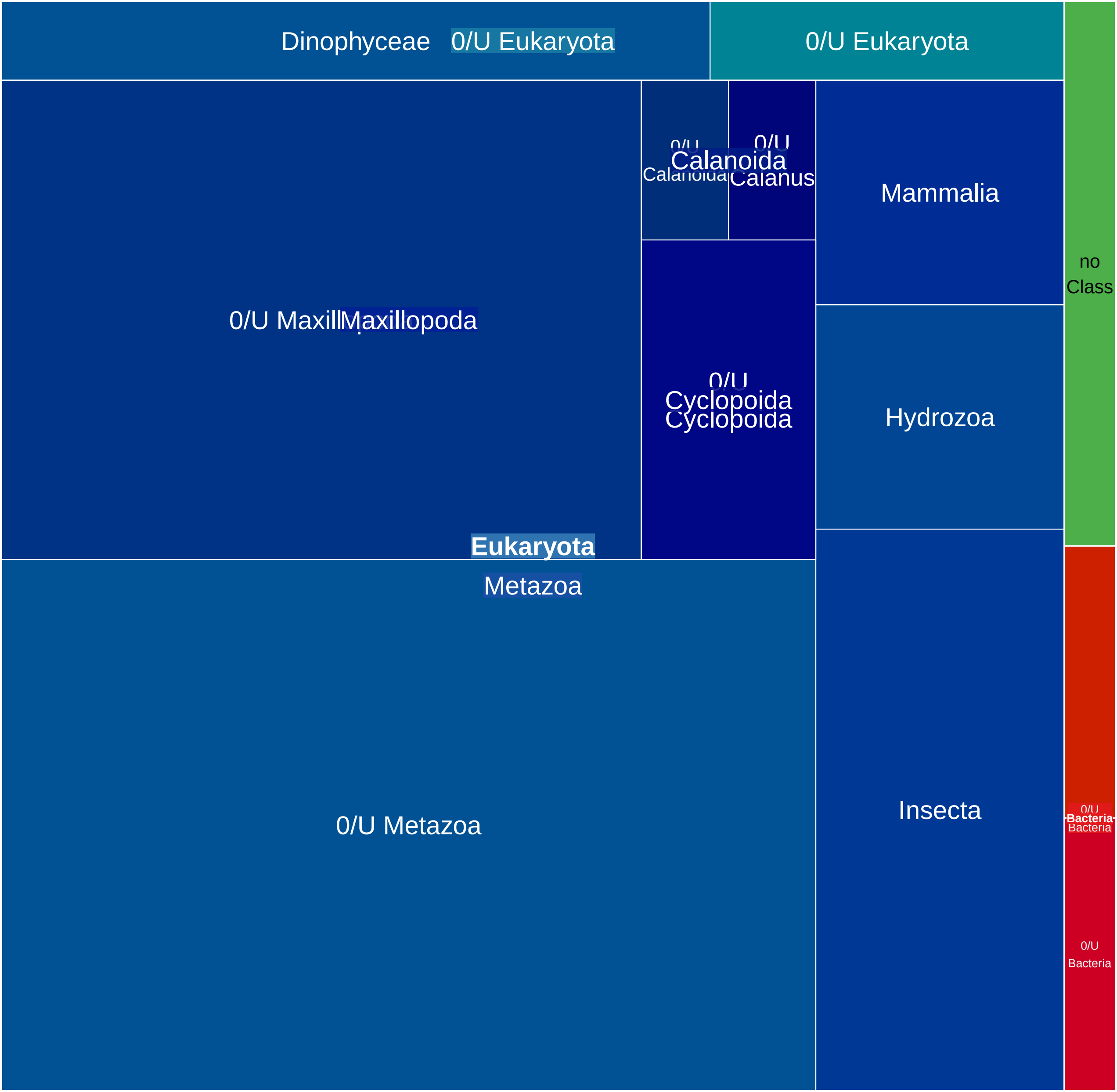

MGT−v1_769 : 705 unigenes, 142 (20.14%) taxonomically assigned

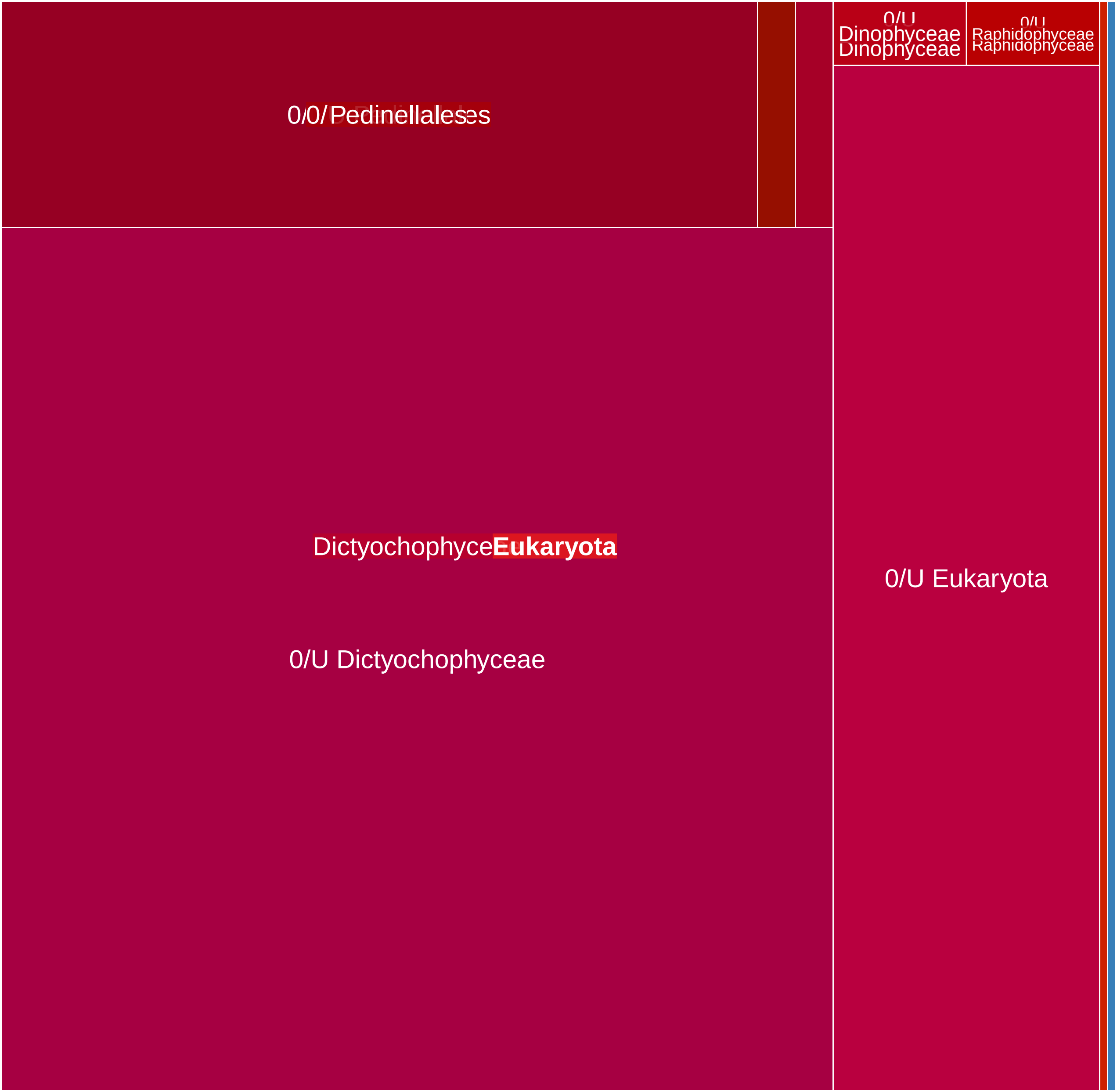

MGT−v1_770 : 705 unigenes, 365 (51.77%) taxonomically assigned

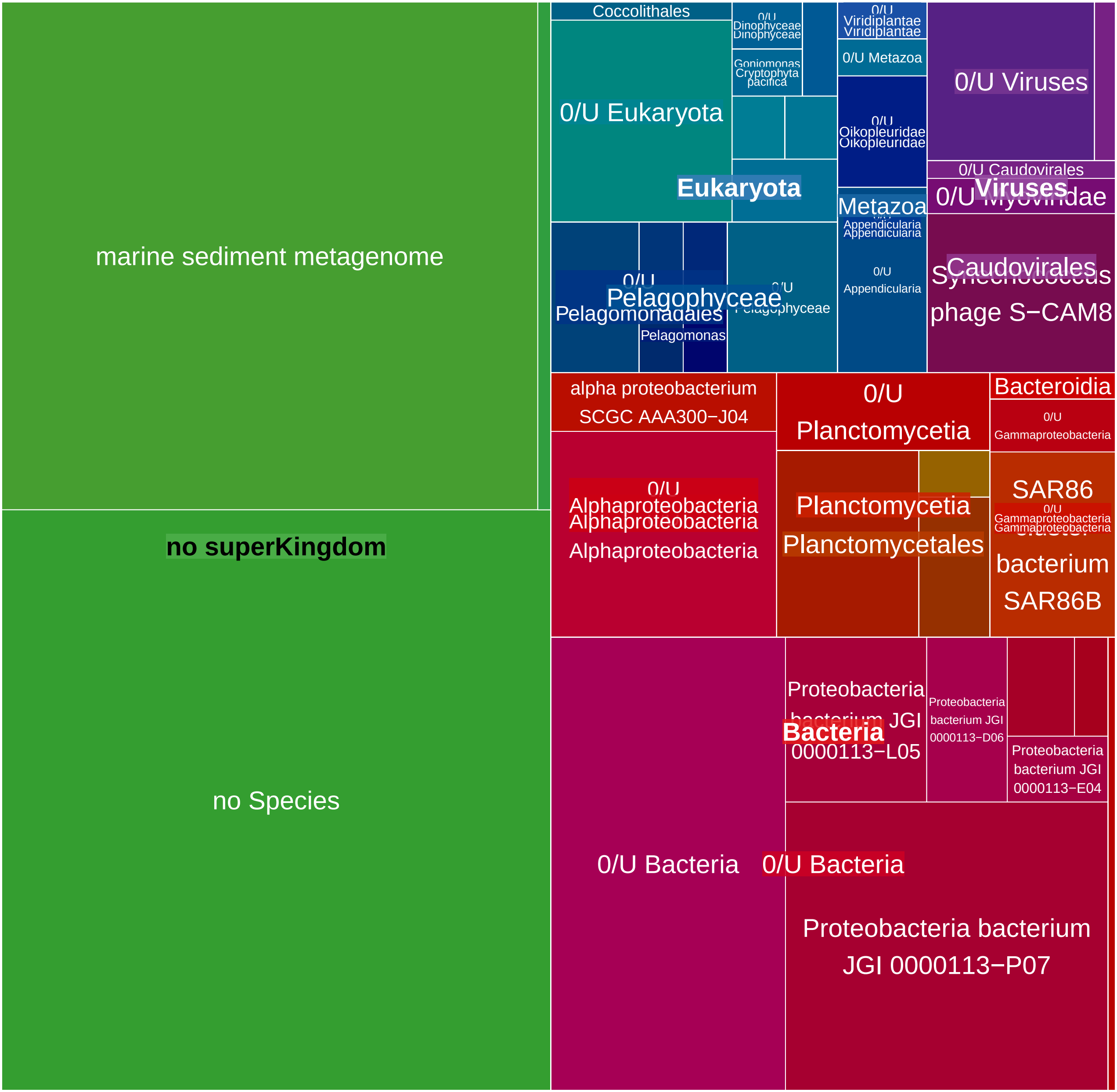

MGT−v1_771 : 702 unigenes, 88 (12.54%) taxonomically assigned

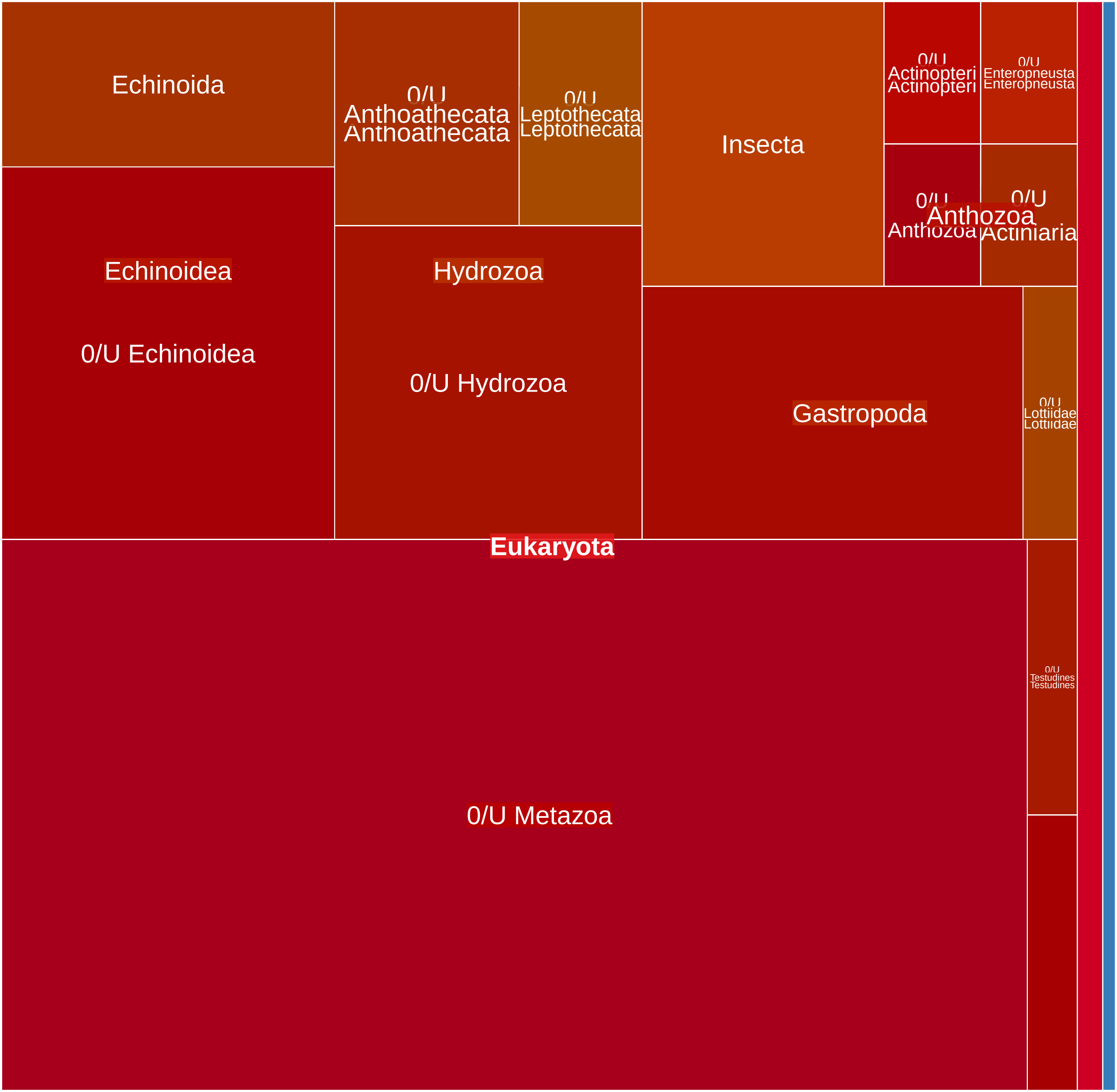

MGT−v1_772 : 700 unigenes, 229 (32.71%) taxonomically assigned

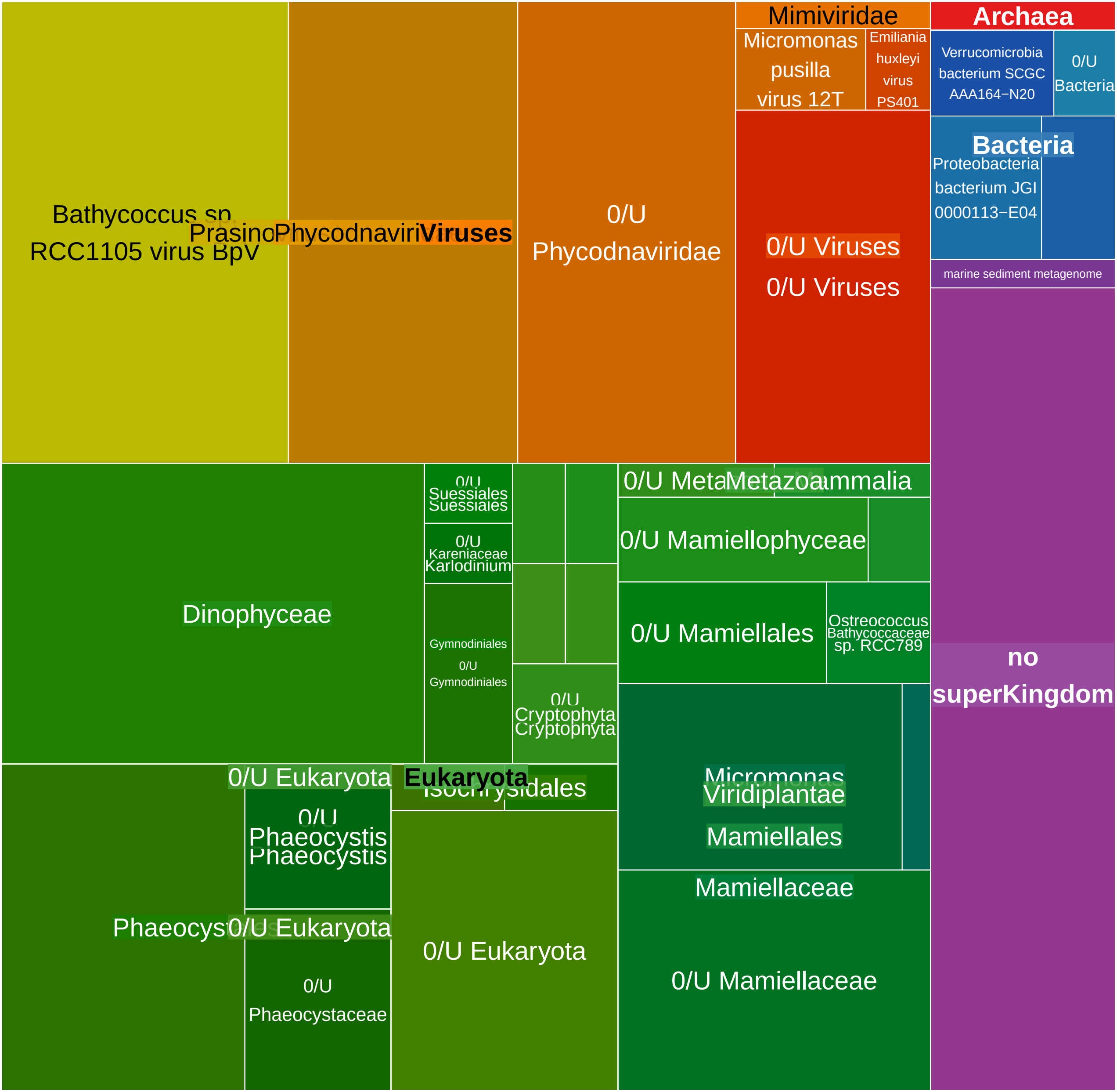

MGT−v1_773 : 699 unigenes, 312 (44.64%) taxonomically assigned

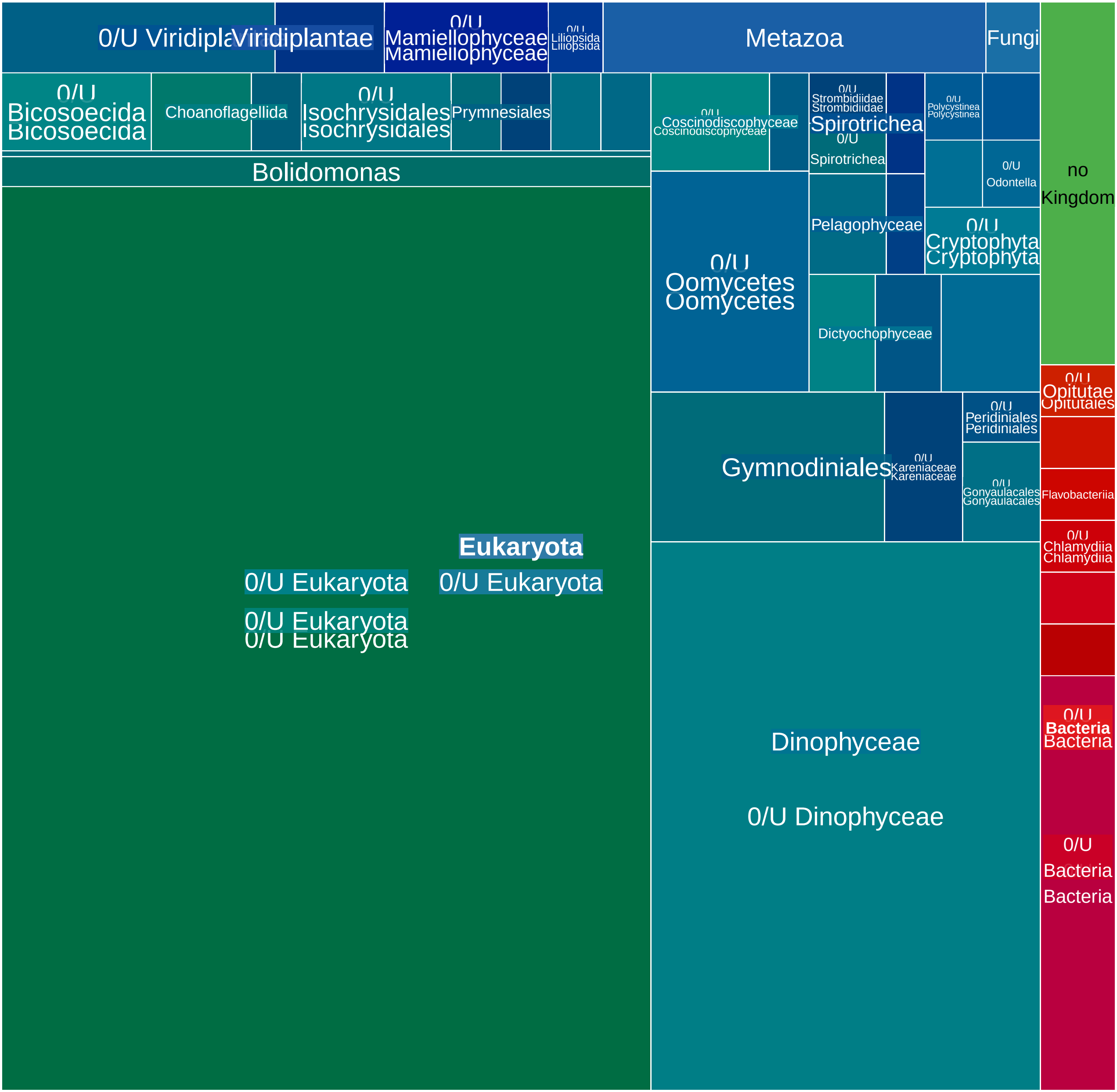

### MGT−v1_774 : 695 unigenes, 647 (93.09%) taxonomically assigned

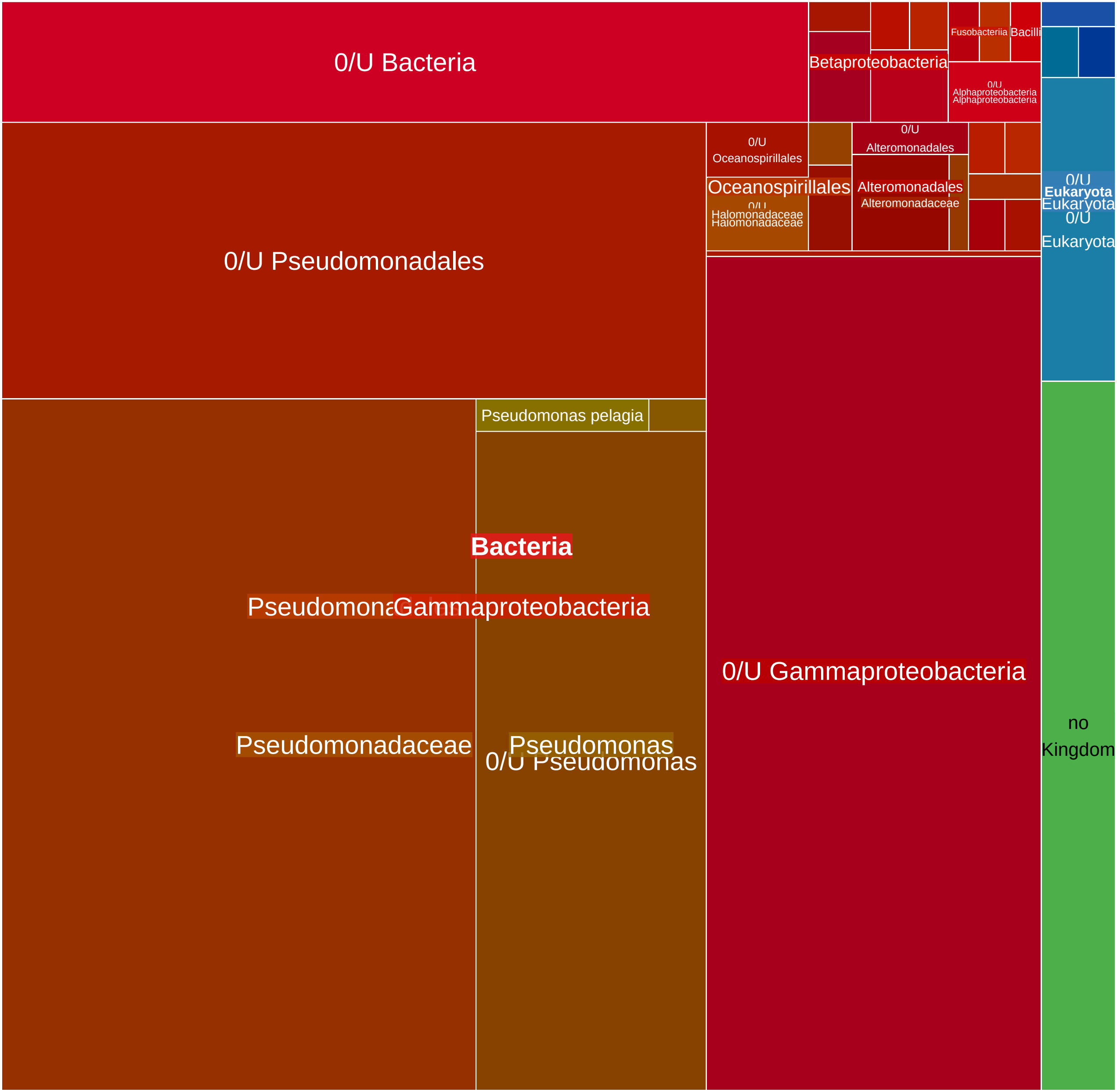

MGT−v1_775 : 694 unigenes, 51 (7.35%) taxonomically assigned

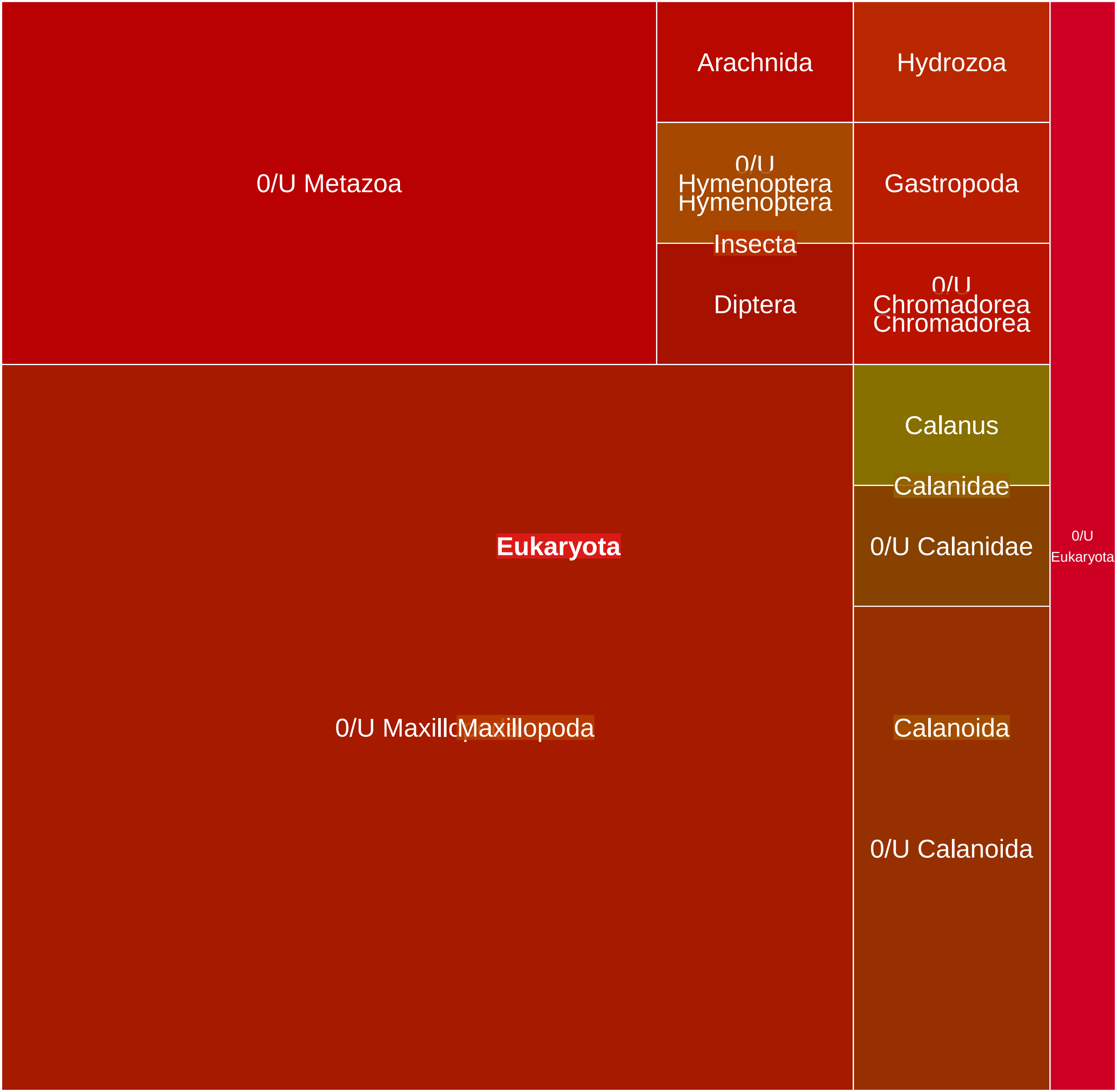

### MGT−v1_776 : 689 unigenes, 287 (41.65%) taxonomically assigned

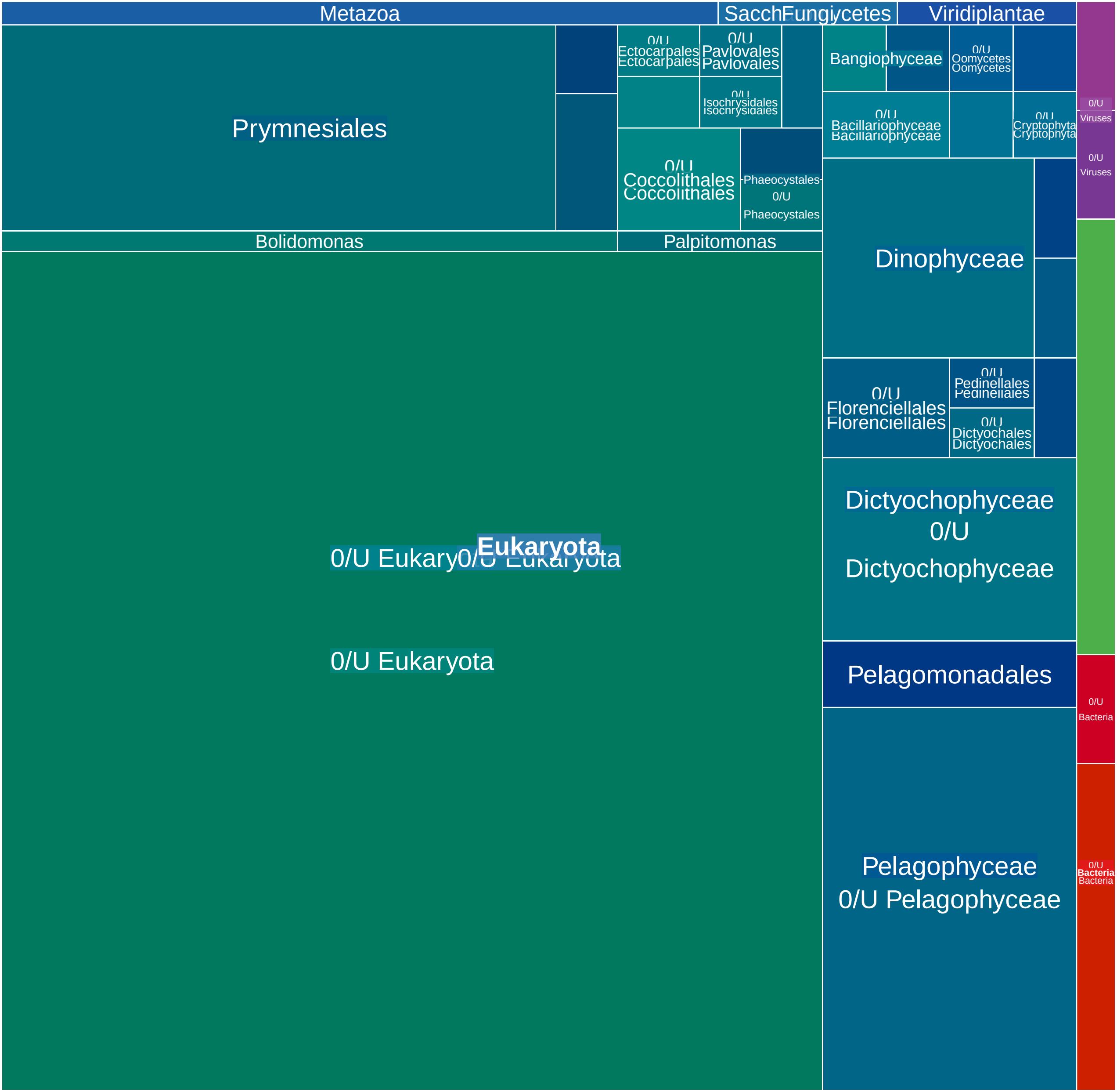

MGT−v1_777 : 689 unigenes, 668 (96.95%) taxonomically assigned

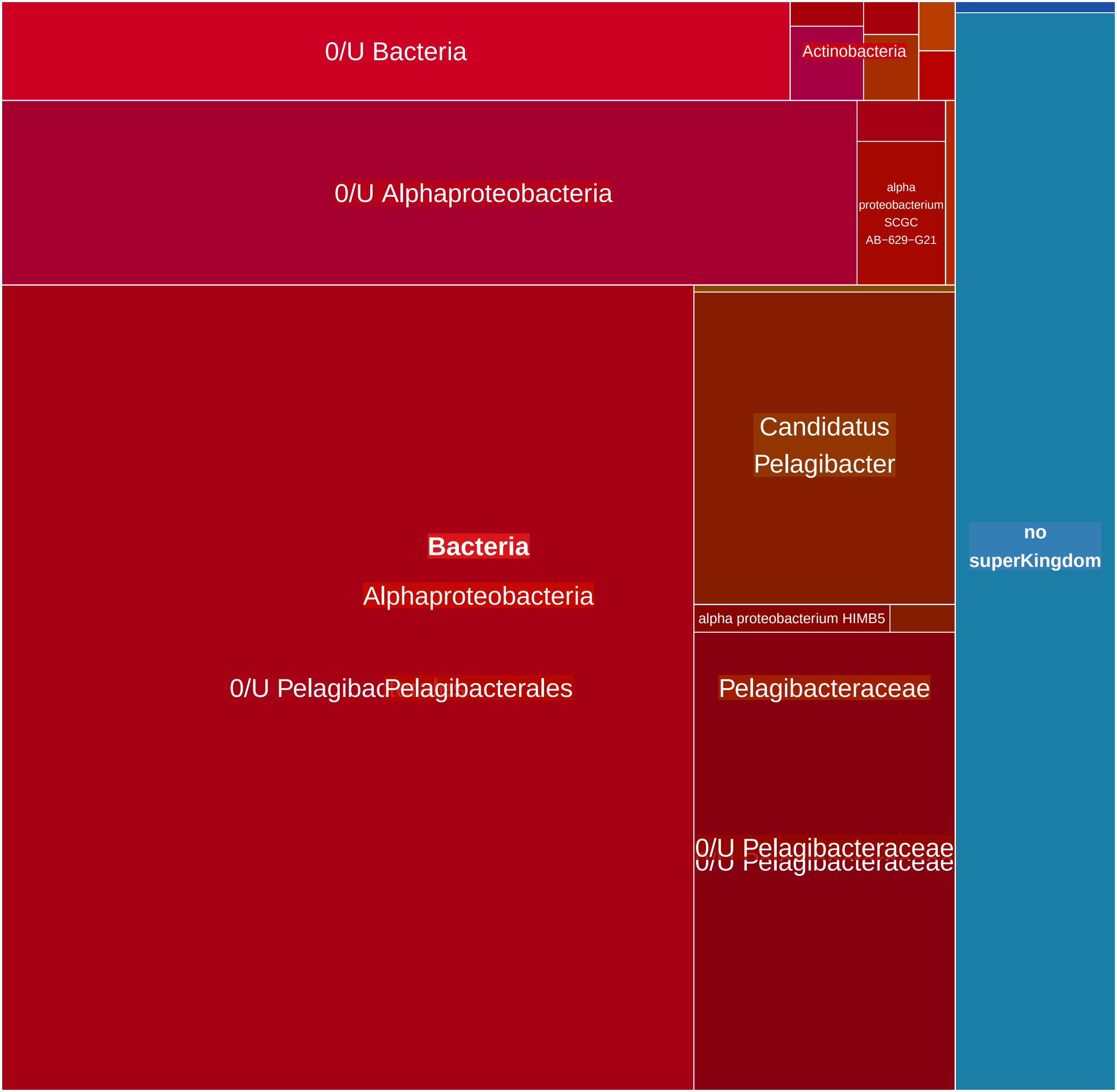

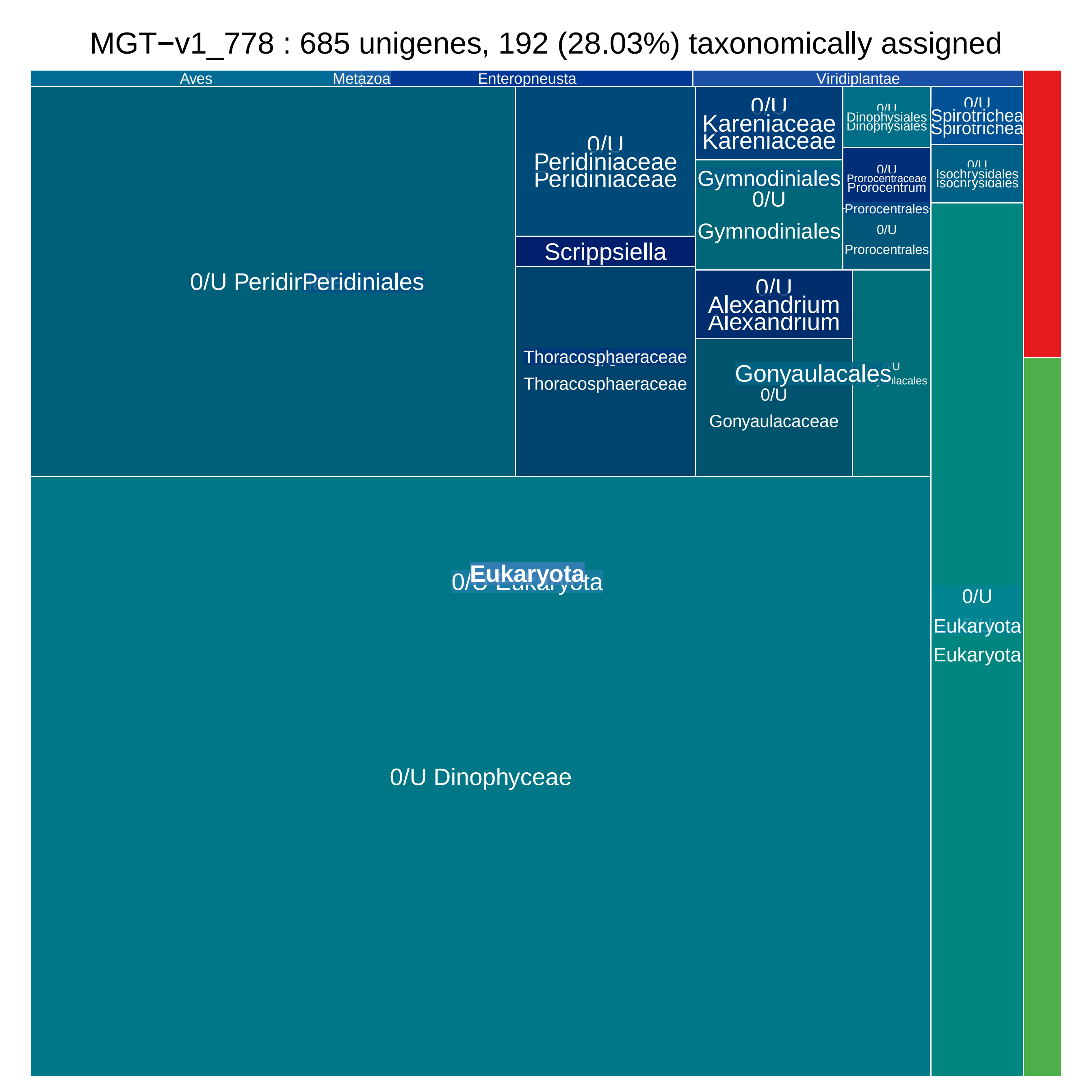
MGT−v1_778 : 685 unigenes, 192 (28.03%) taxonomically assigned
Aves
Metazoa
Enteropneusta
Viridiplantae
0/U Peridiniales
Peridiniales
0/U
Peridiniaceae
Peridiniaceae
Scrippsiella
Thoracosphaeraceae
Thoracosphaeraceae
0/U
Kareniaceae
Kareniaceae
Gymnodiniales
0/U
Gymnodiniales
0/U
Dinophysiales
Dinophysiales
0/U
Prorocentraceae
Prorocentrum
Prorocentrales
0/U
Prorocentrales
0/U
Spirotrichea
Spirotrichea
0/U
Isochrysidales
Isochrysidales
0/U
Alexandrium
Alexandrium
Gonyaulacales
0/U
Gonyaulacaceae
Eukaryota
0/U Eukaryota
0/U Dinophyceae
0/U
Eukaryota
Eukaryota

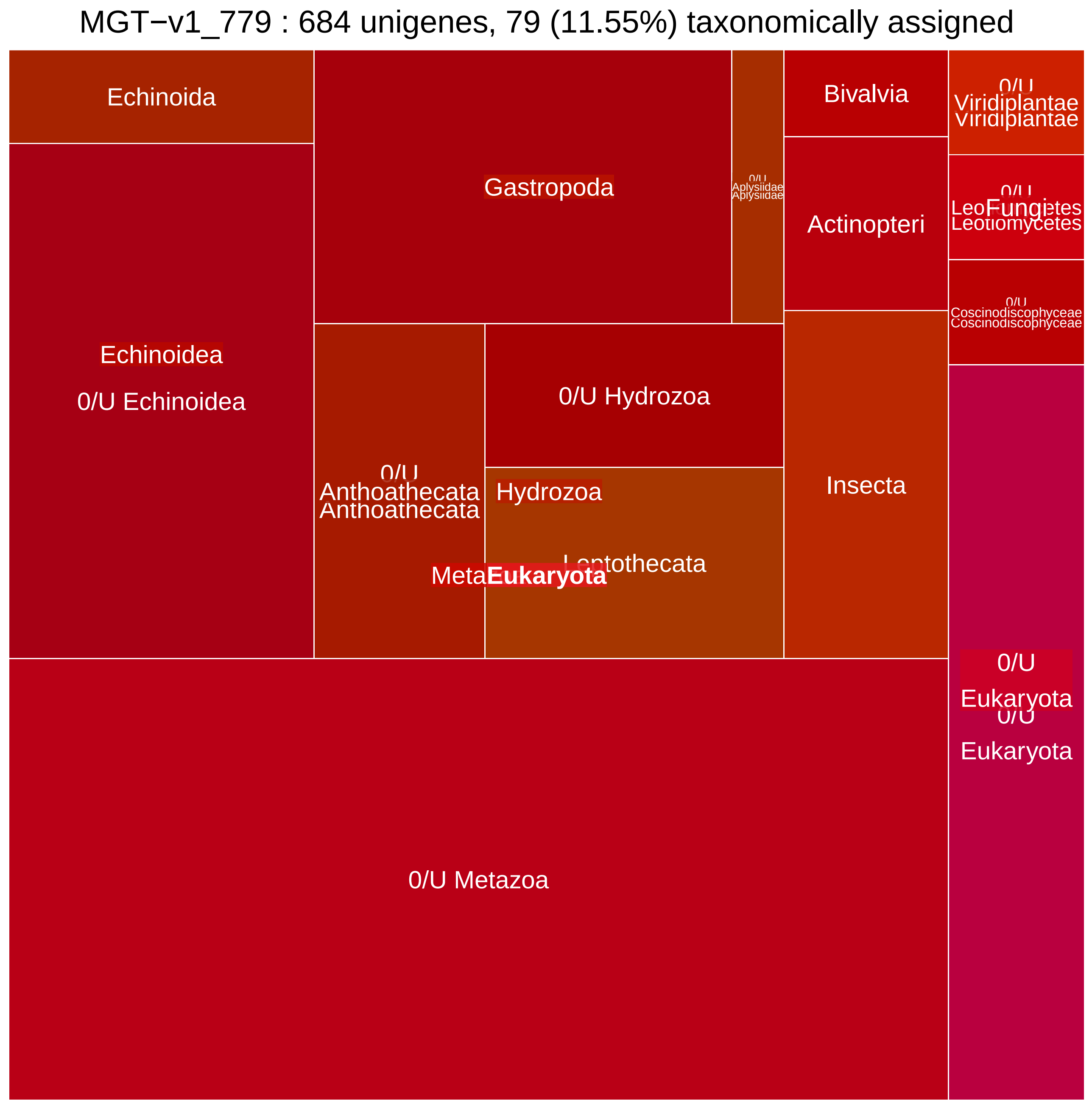
MGT−v1_779 : 684 unigenes, 79 (11.55%) taxonomically assigned
Echinoida
Echinoidea
0/U Echinoidea
Gastropoda
0/U Aplysiidae Aplysiidae
Bivalvia
Actinopteri
Insecta
0/U Viridiplantae Viridiplantae
0/U Leotiomycetes Leotiomycetes
Fungi
0/U Coscinodiscophyceae Coscinodiscophyceae
0/U Anthoathecata Anthoathecata
0/U Hydrozoa
Hydrozoa
Leptothecata
Metazoa
Eukaryota
0/U Metazoa
0/U Eukaryota
0/U Eukaryota

MGT−v1_780 : 683 unigenes, 75 (10.98%) taxonomically assigned

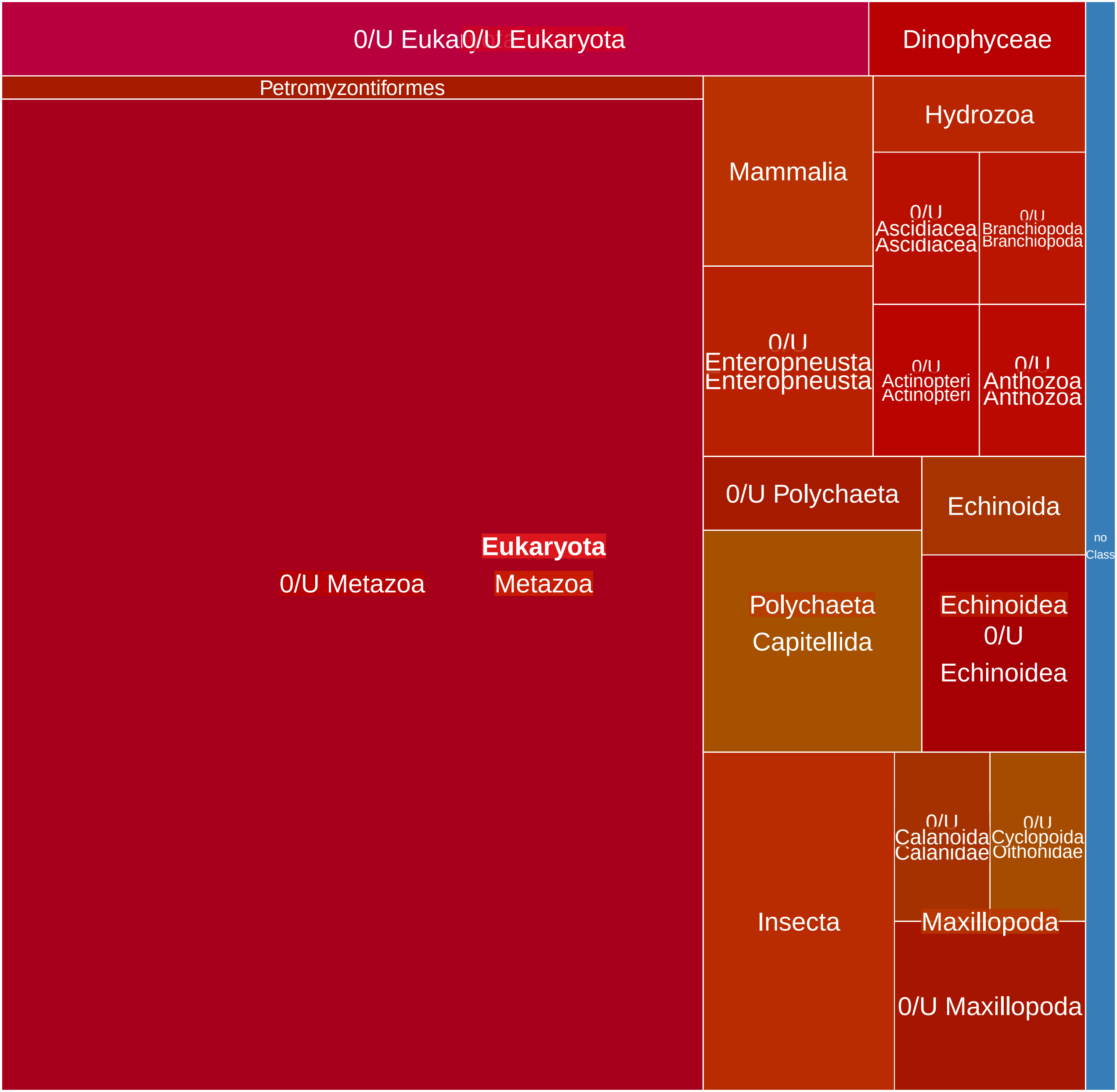

### MGT−v1_781 : 679 unigenes, 161 (23.71%) taxonomically assigned

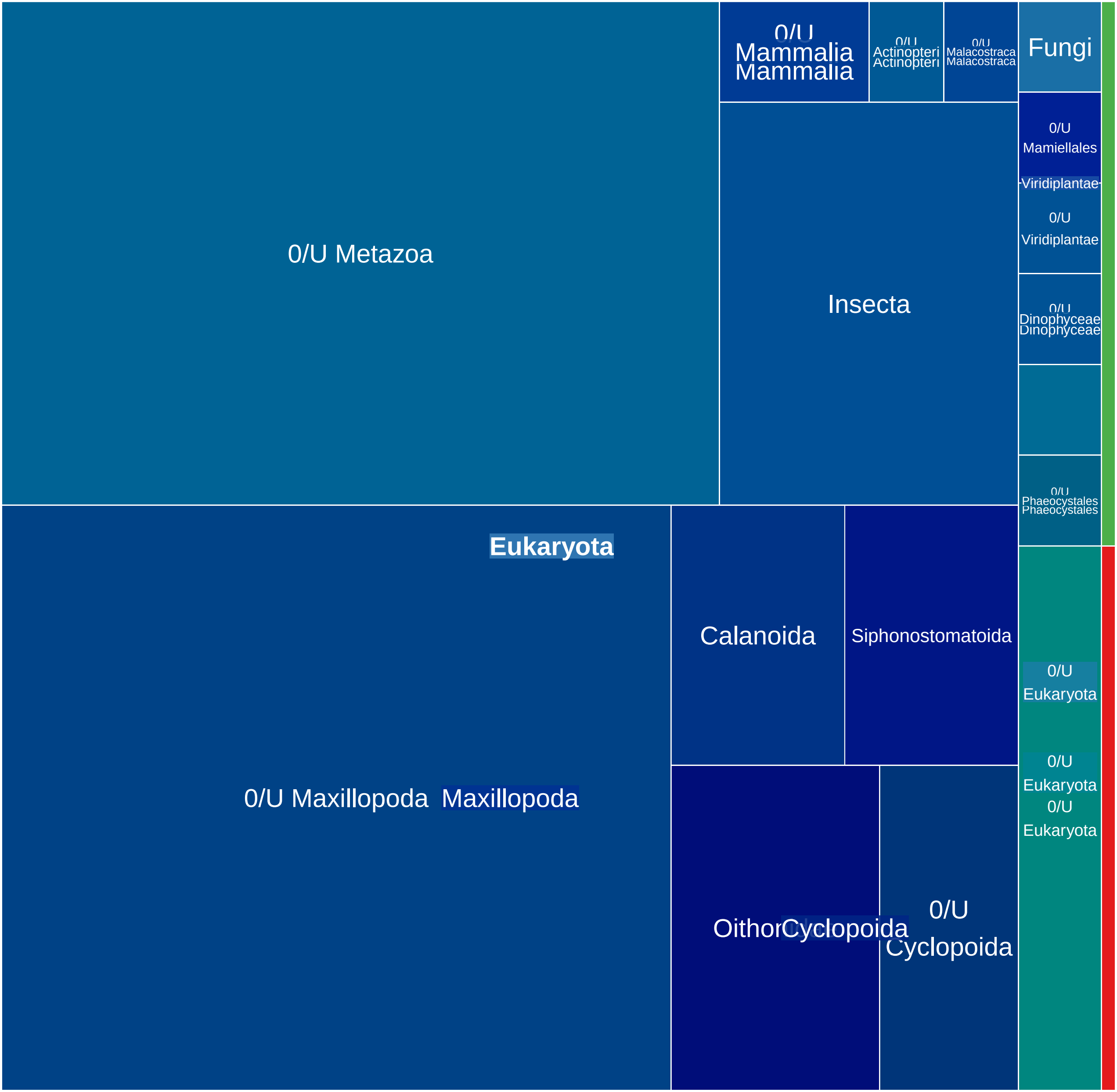

MGT−v1_782 : 675 unigenes, 665 (98.52%) taxonomically assigned

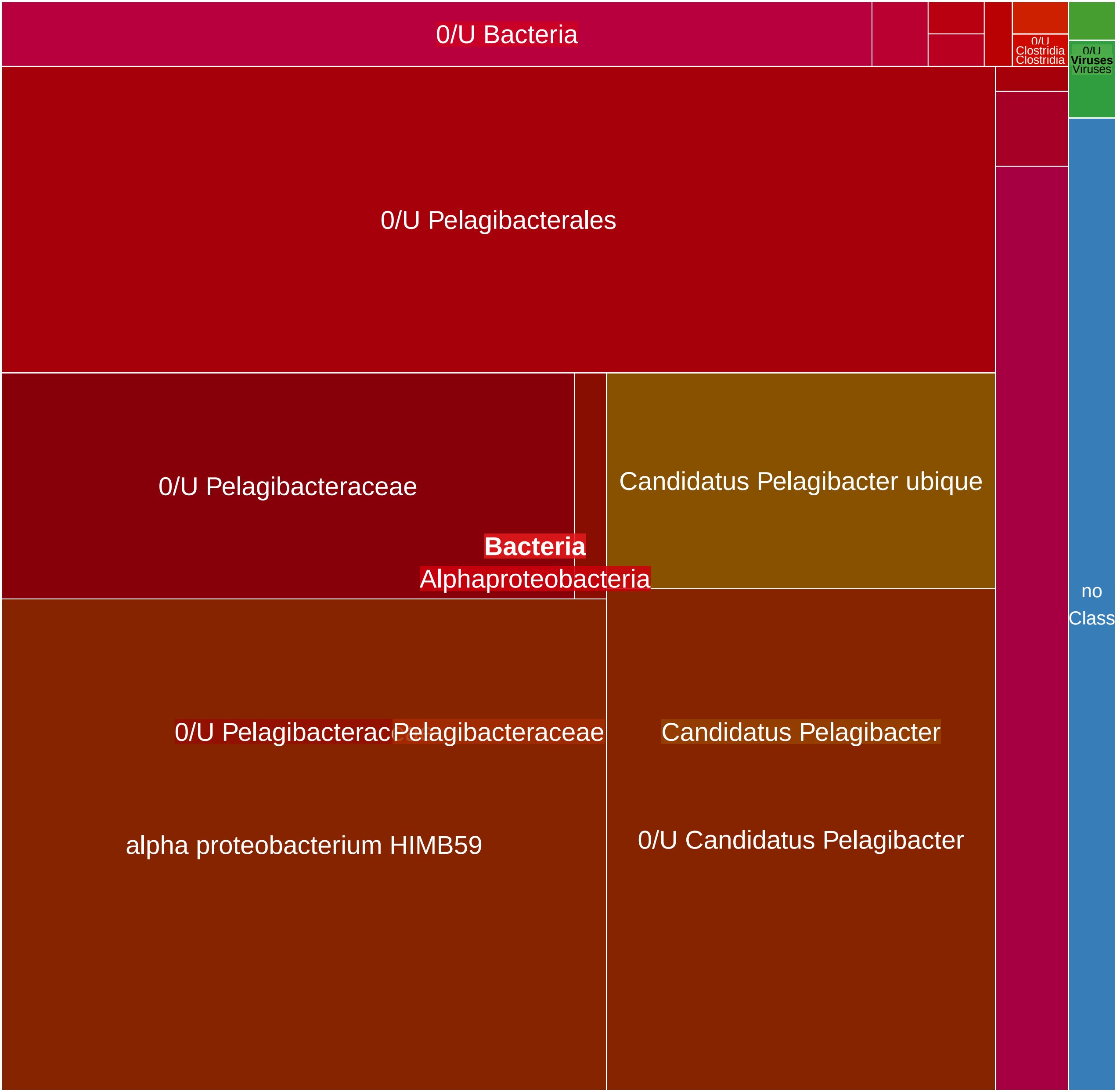

MGT−v1_783 : 675 unigenes, 112 (16.59%) taxonomically assigned

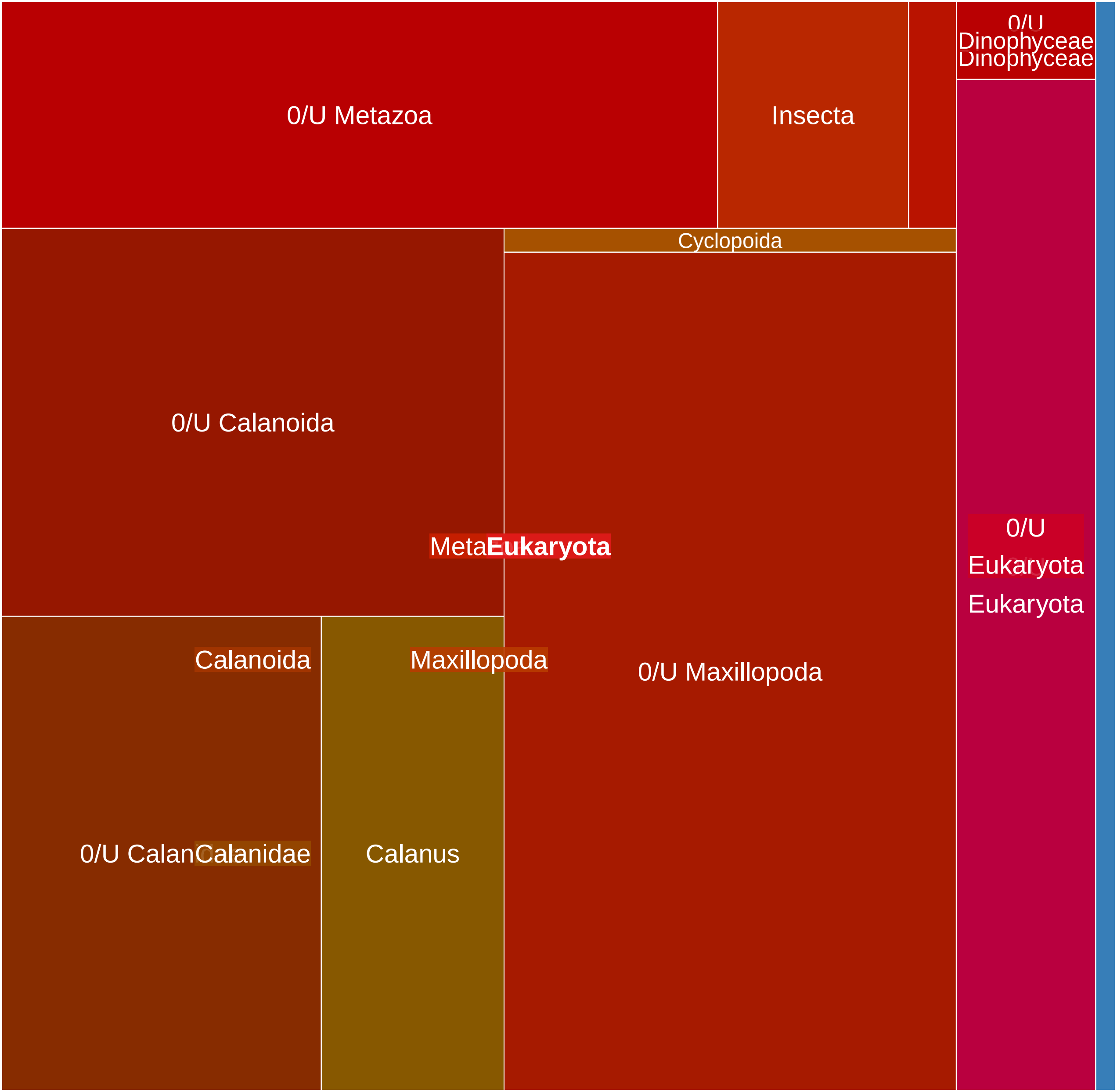

MGT−v1_784 : 675 unigenes, 59 (8.74%) taxonomically assigned

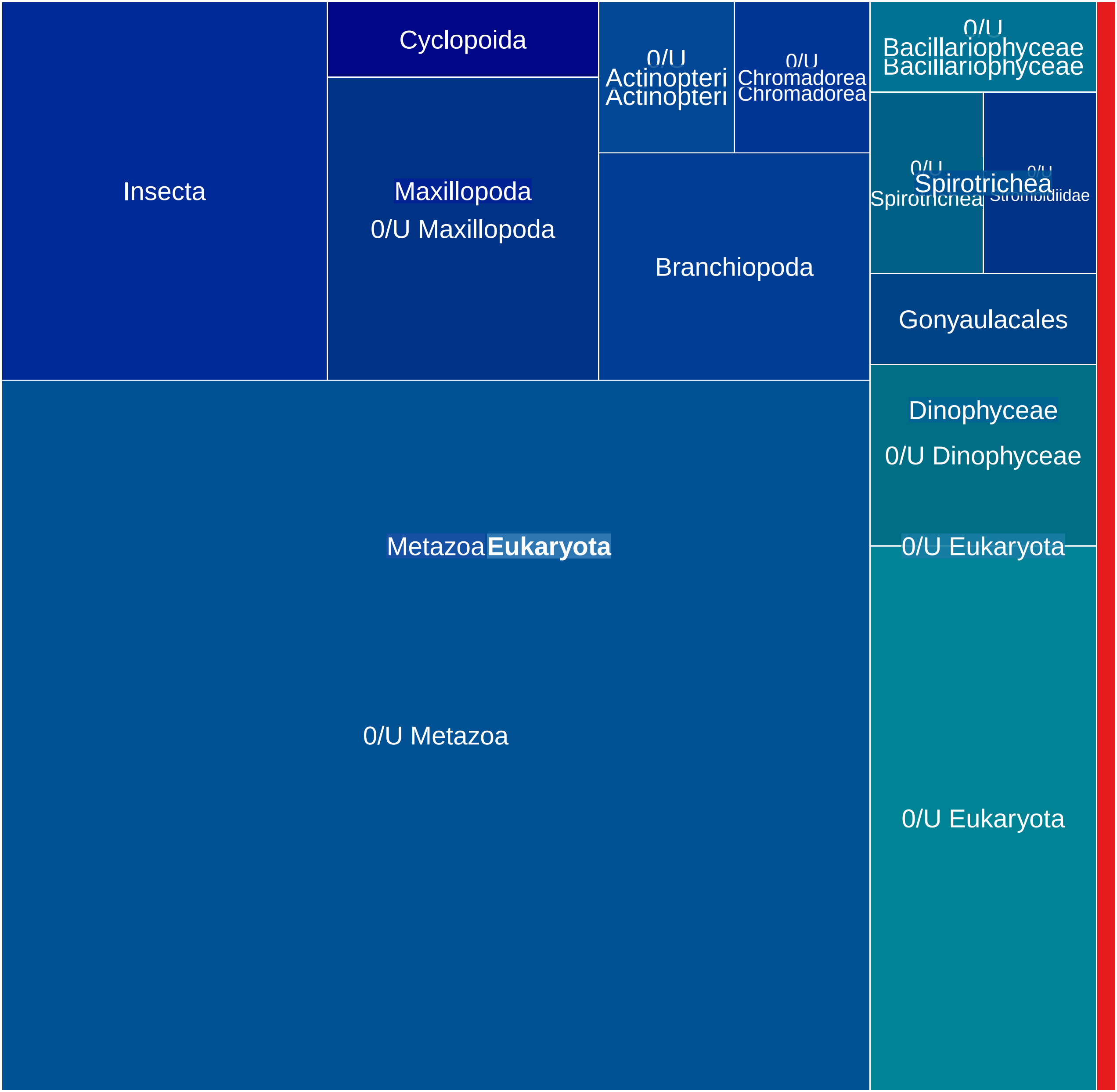

MGT−v1_785 : 673 unigenes, 368 (54.68%) taxonomically assigned

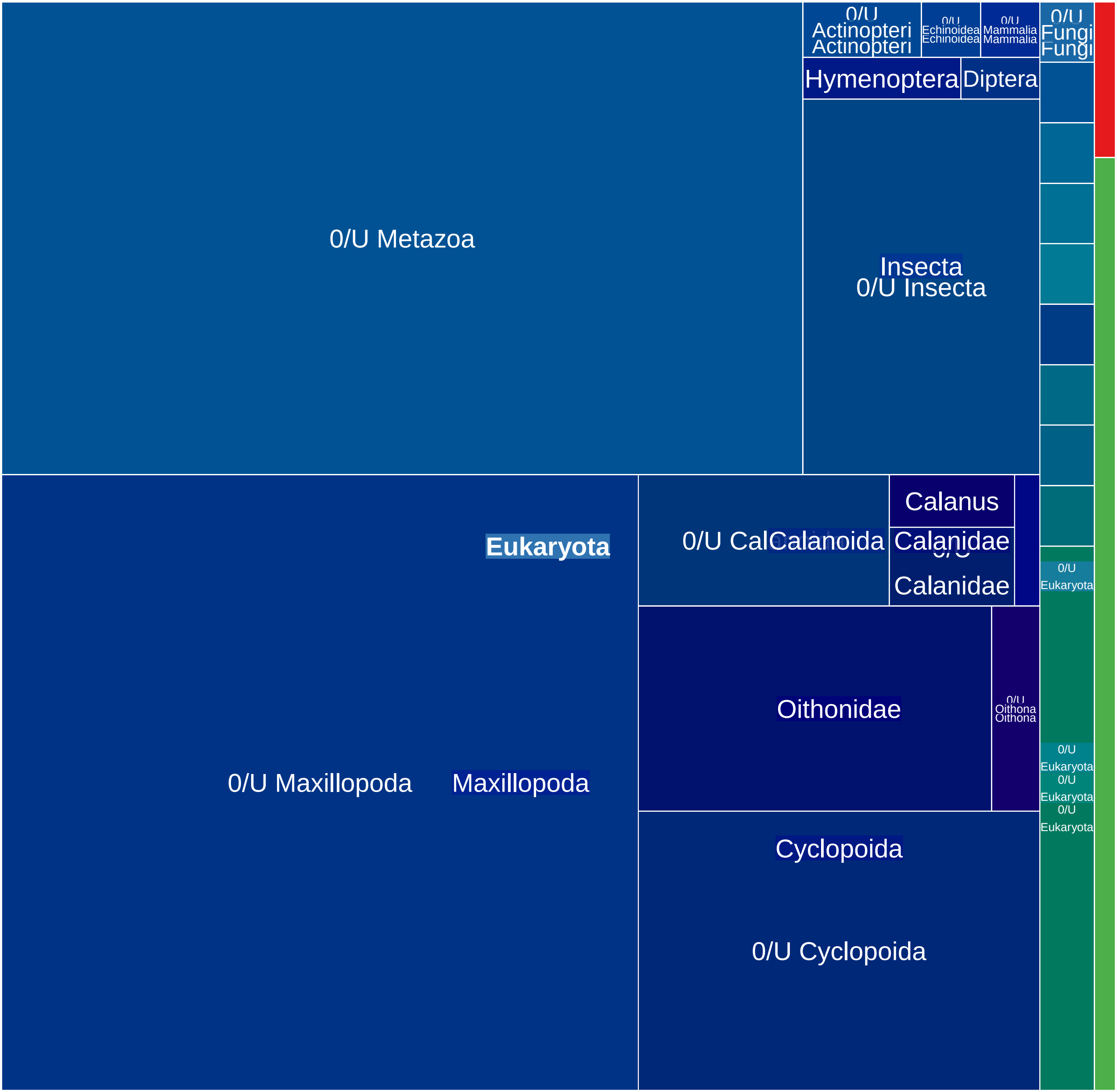

MGT−v1_786 : 672 unigenes, 211 (31.40%) taxonomically assigned

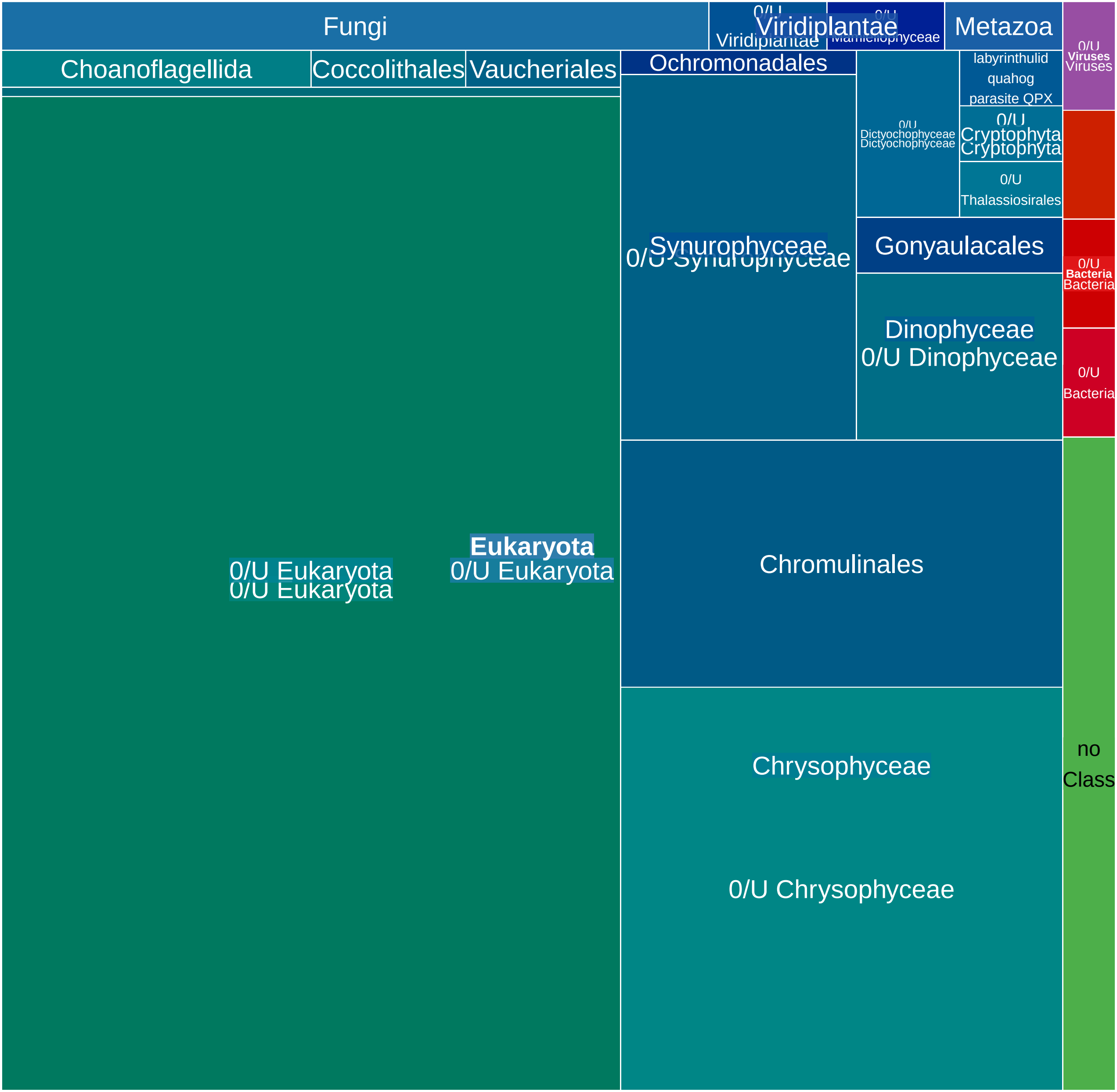

MGT−v1_787 : 671 unigenes, 650 (96.87%) taxonomically assigned

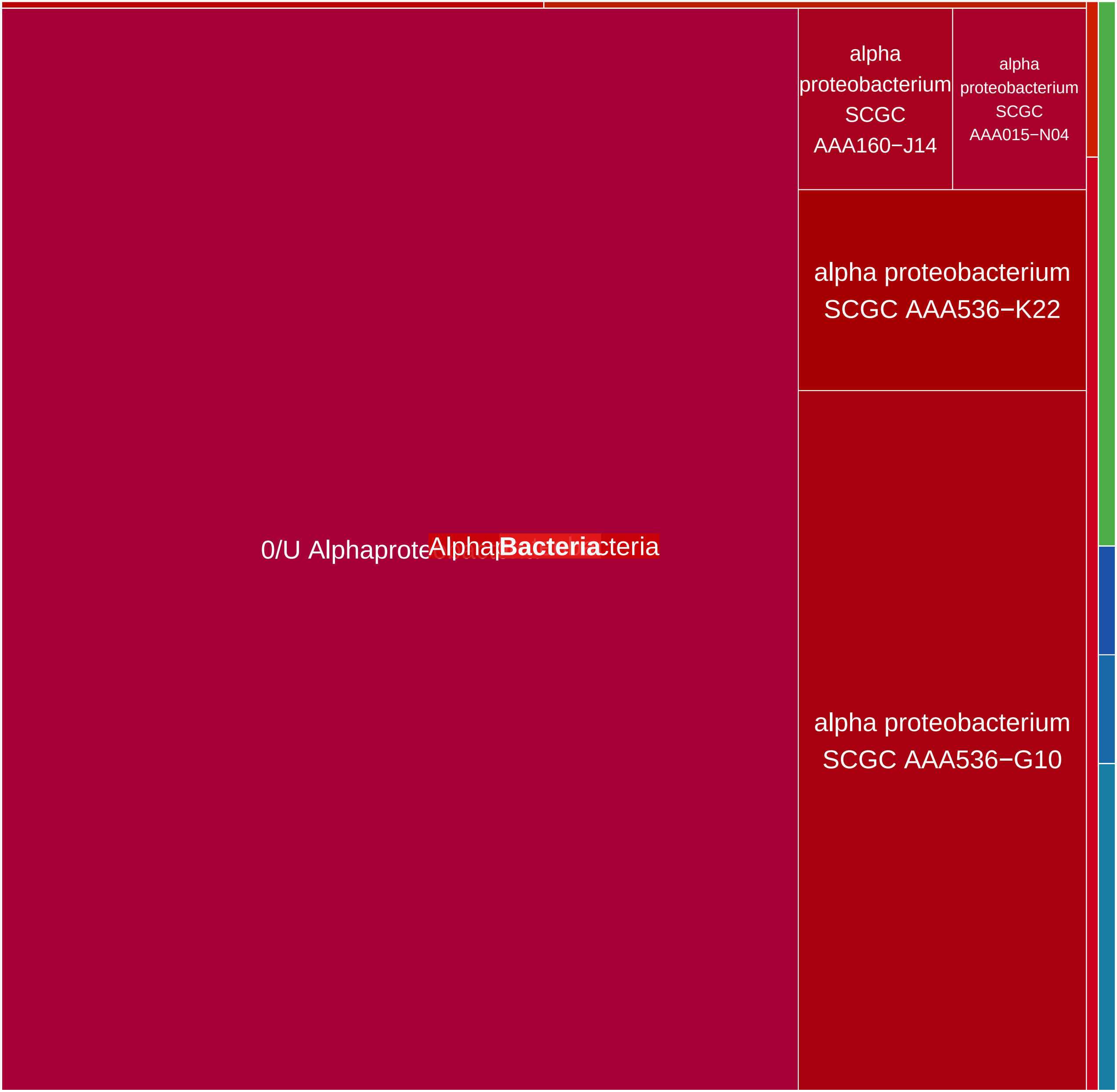

MGT−v1_788 : 670 unigenes, 80 (11.94%) taxonomically assigned

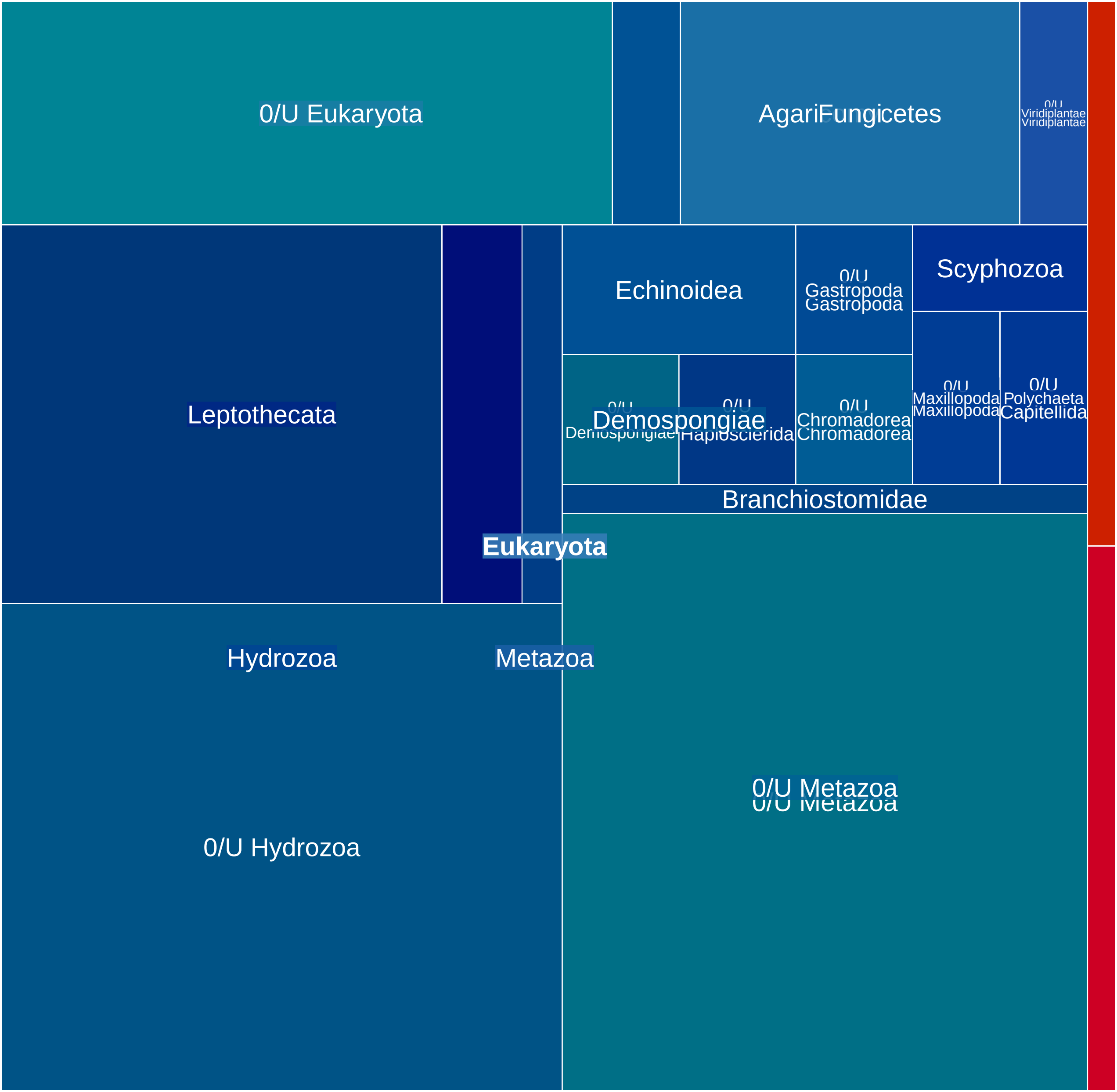

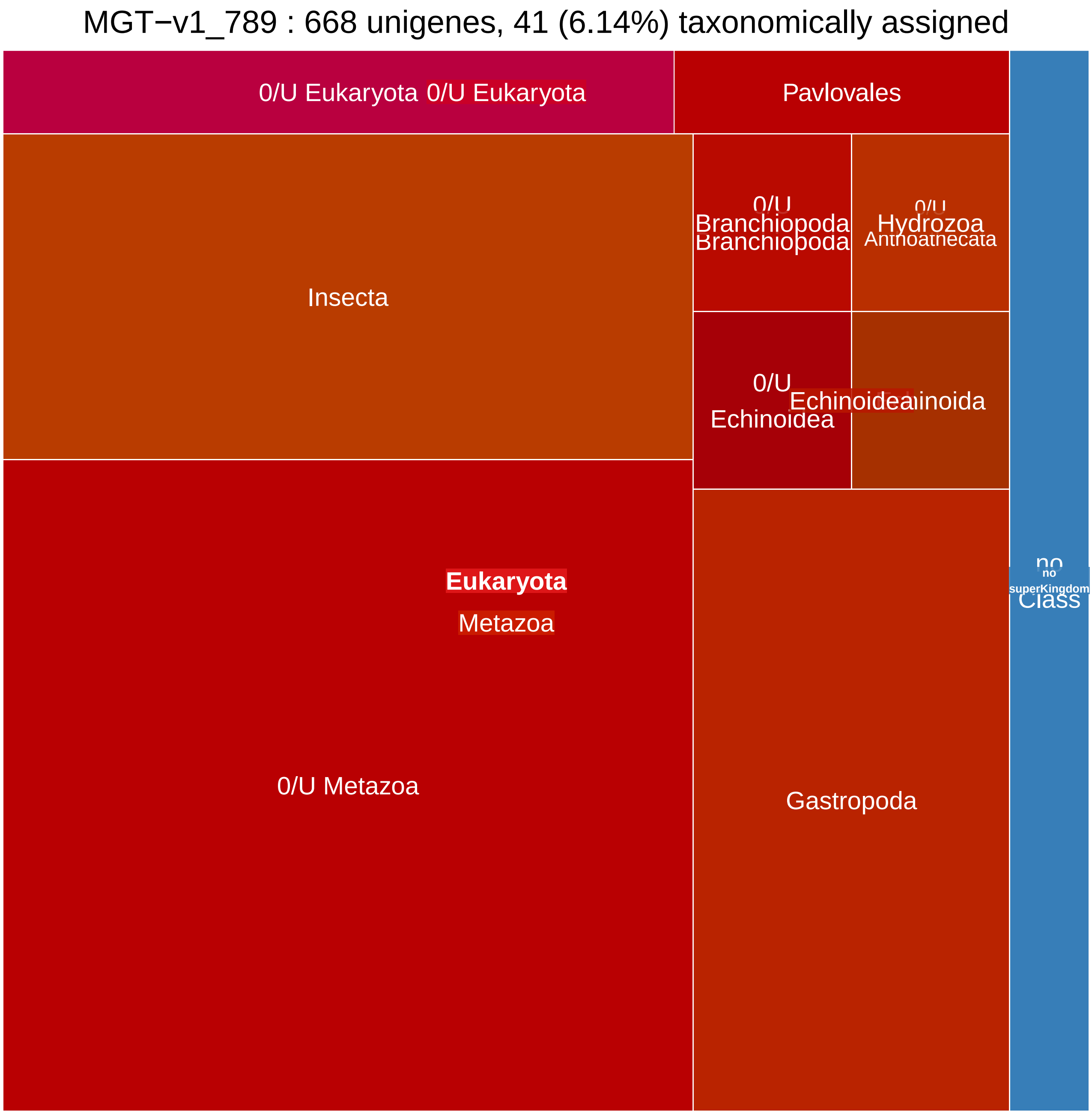
MGT−v1_789 : 668 unigenes, 41 (6.14%) taxonomically assigned
0/U Eukaryota
0/U Eukaryota
Pavlovales
Insecta
0/U Branchiopoda
Branchiopoda
0/U Hydrozoa
Anthoathecata
0/U Echinoidea
Echinoidea
Echinoida
Eukaryota
Metazoa
0/U Metazoa
Gastropoda
no Class
no superKingdom

MGT−v1_790 : 668 unigenes, 150 (22.46%) taxonomically assigned

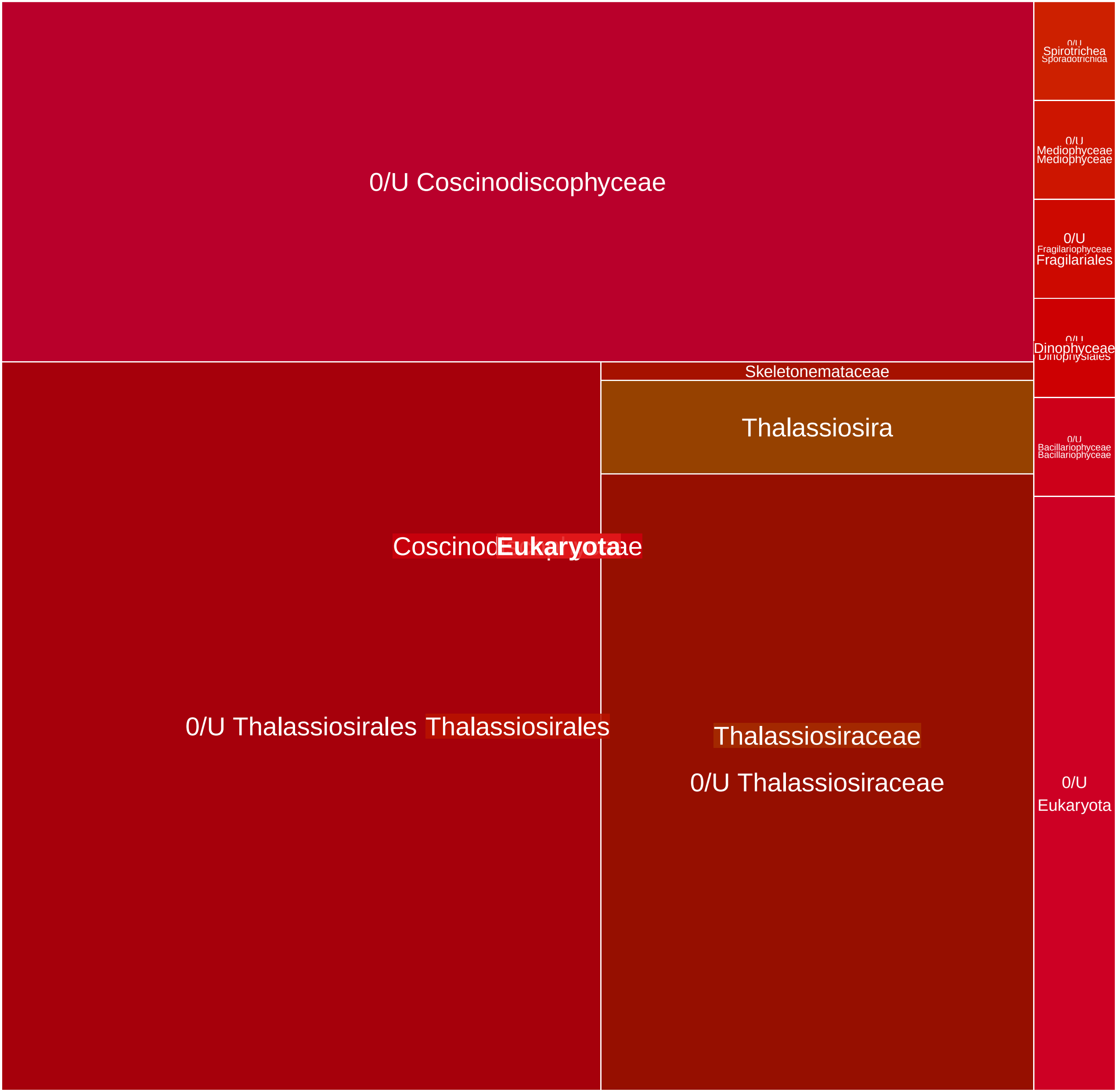

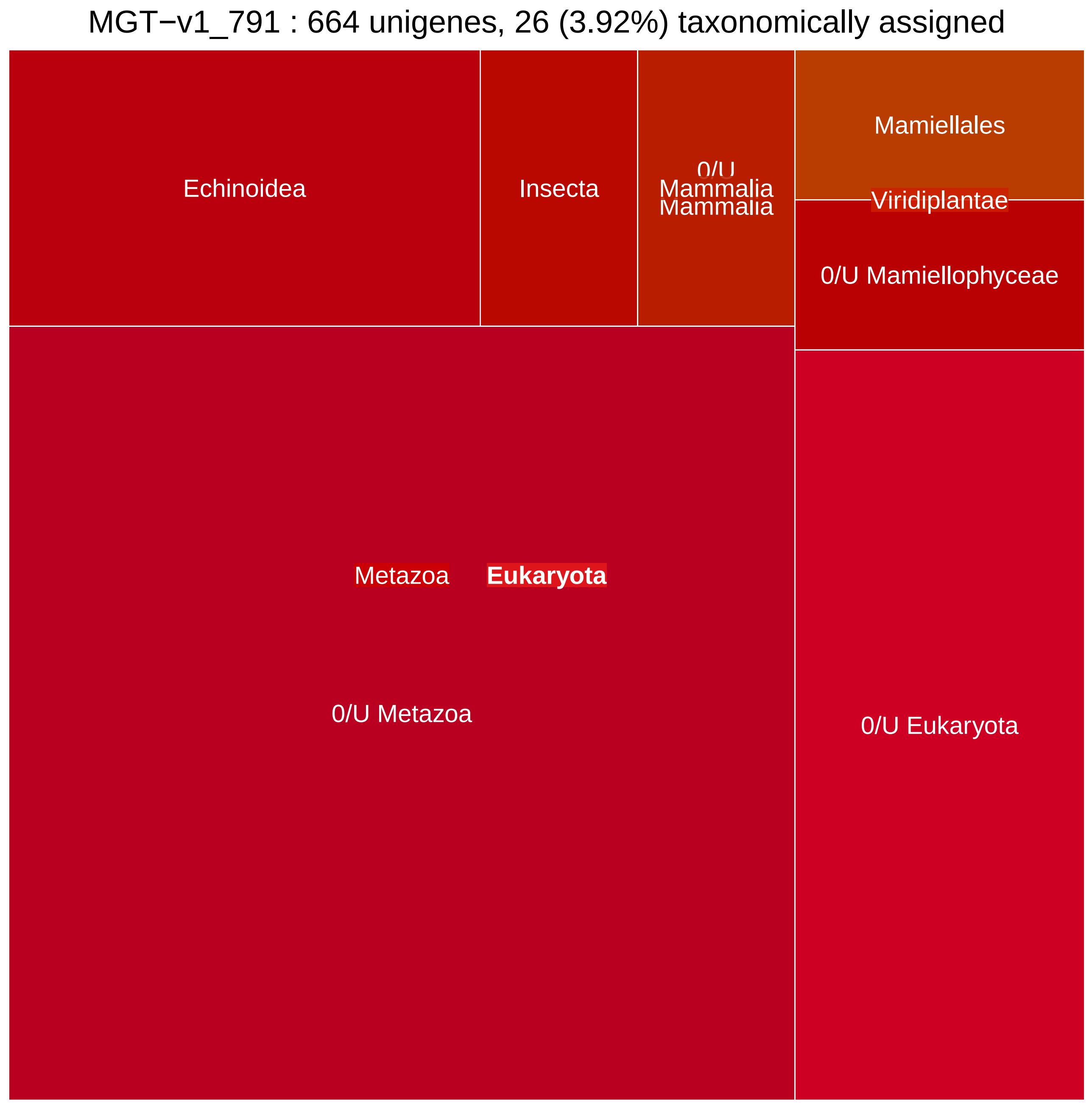
MGT−v1_791 : 664 unigenes, 26 (3.92%) taxonomically assigned
Echinoidea
Insecta
0/U Mammalia
Mammalia
Mamiellales
Viridiplantae
0/U Mamiellophyceae
Metazoa
Eukaryota
0/U Metazoa
0/U Eukaryota

MGT−v1_792 : 663 unigenes, 565 (85.22%) taxonomically assigned

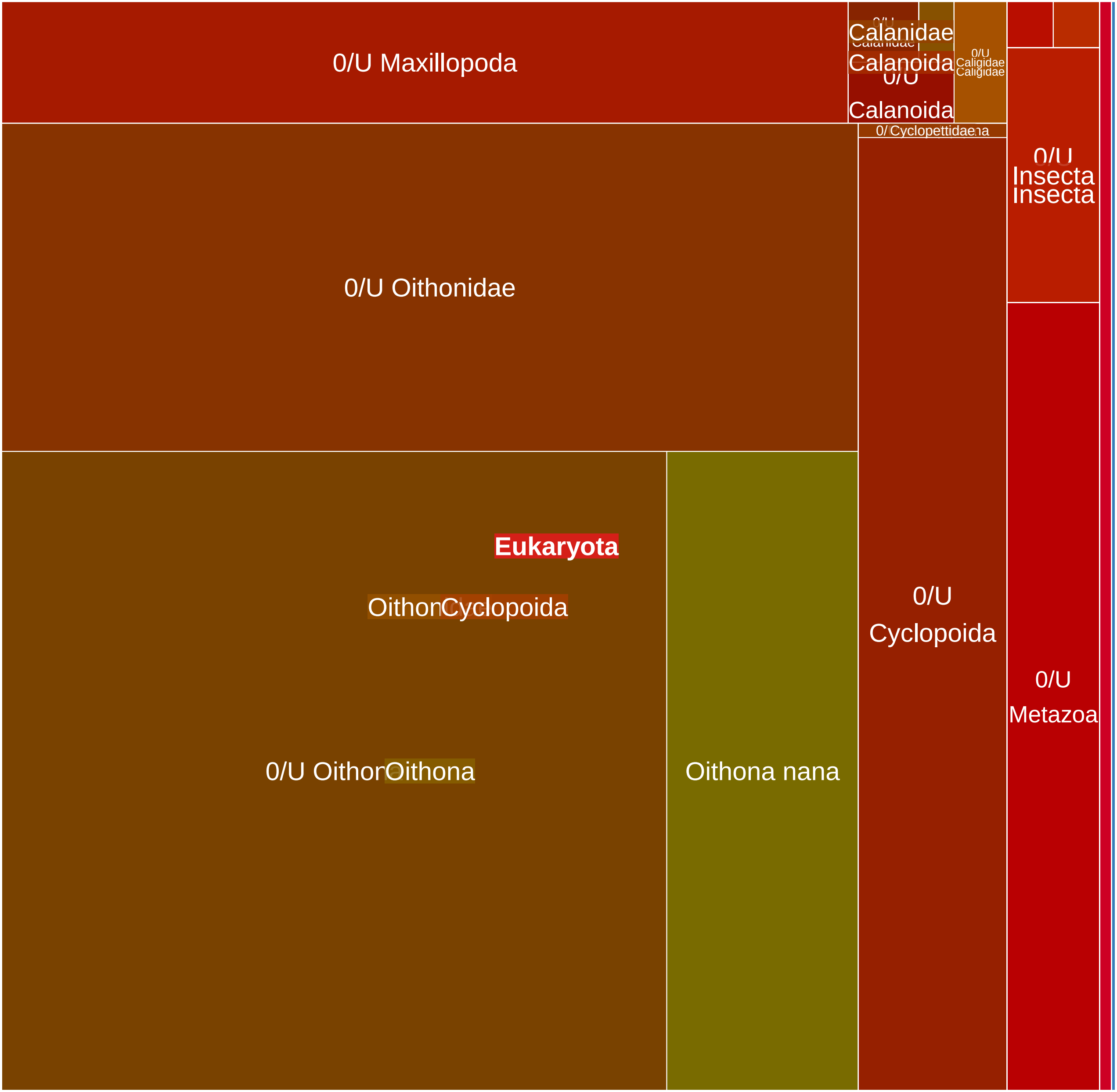

MGT−v1_793 : 663 unigenes, 411 (61.99%) taxonomically assigned

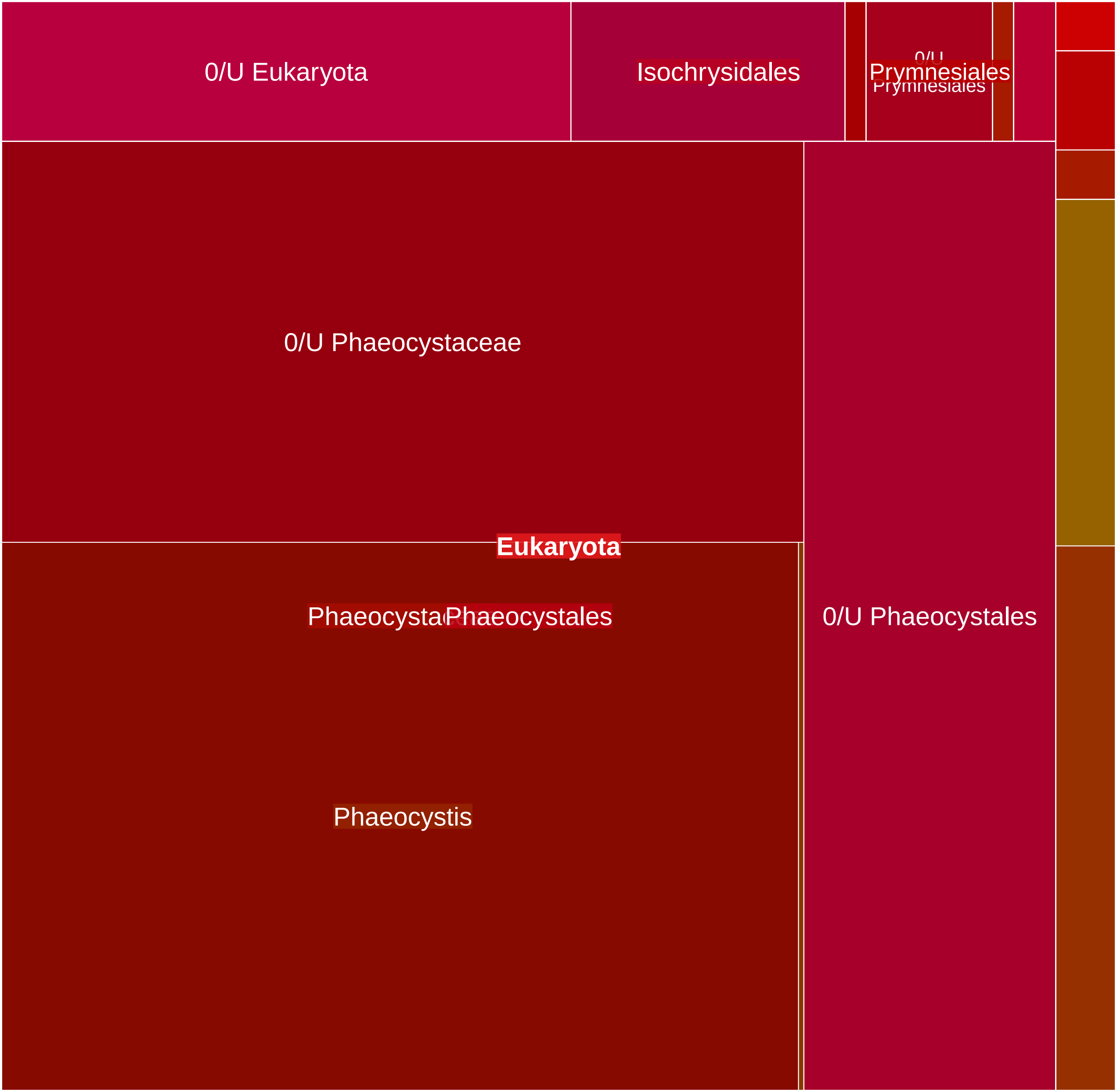

MGT−v1_794 : 663 unigenes, 380 (57.32%) taxonomically assigned

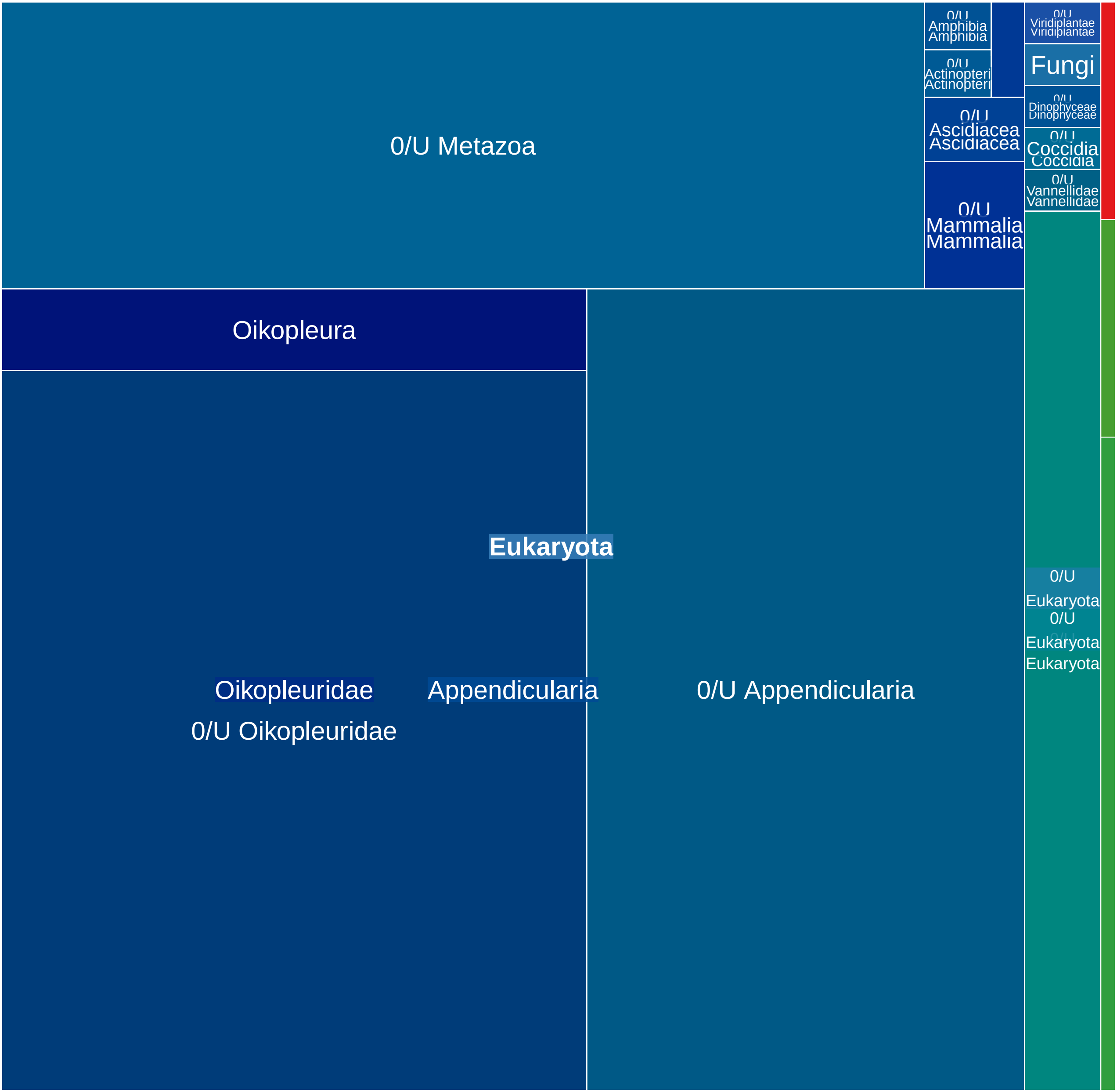

MGT−v1_795 : 661 unigenes, 64 (9.68%) taxonomically assigned

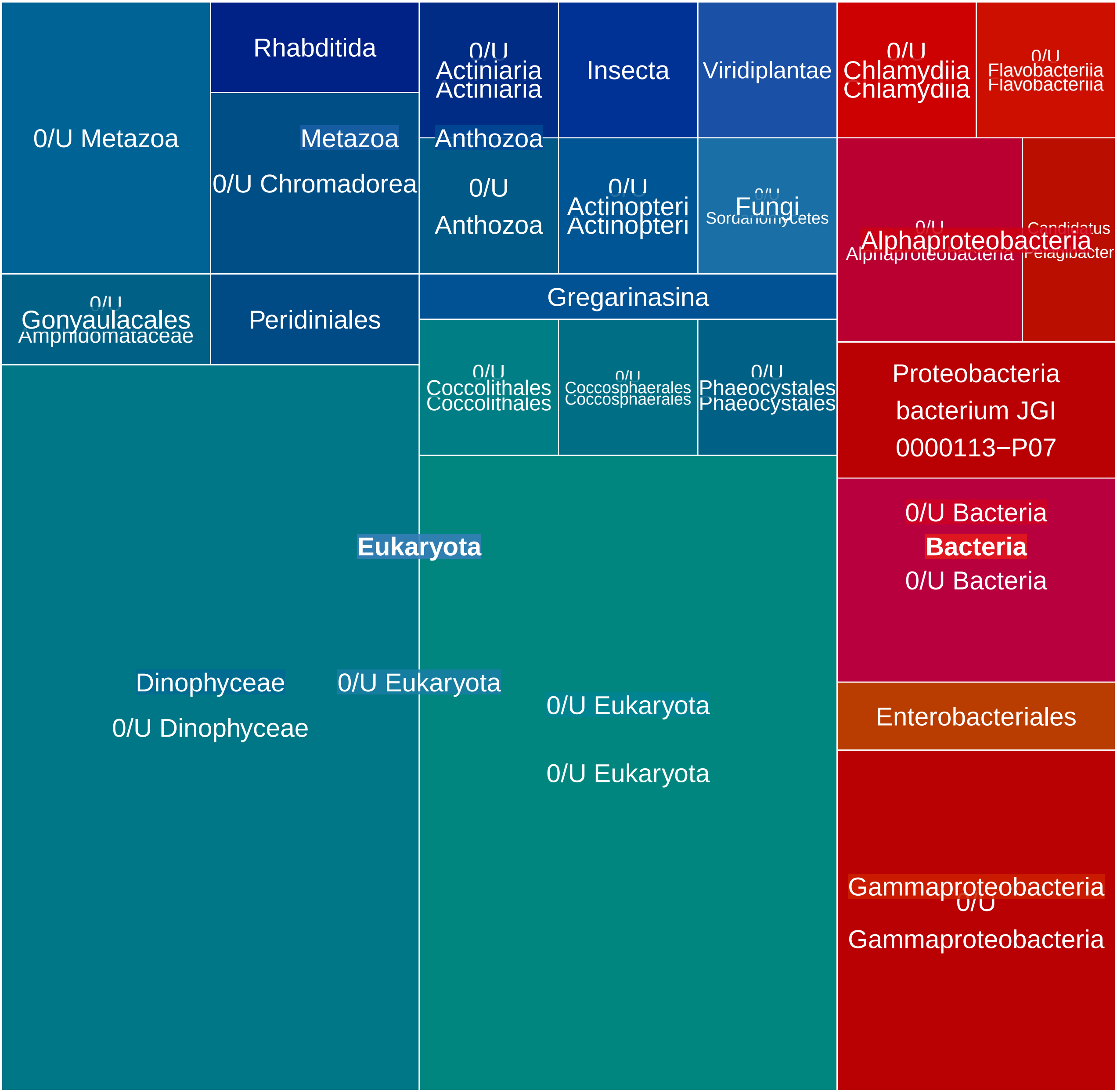

MGT−v1_796 : 660 unigenes, 62 (9.39%) taxonomically assigned

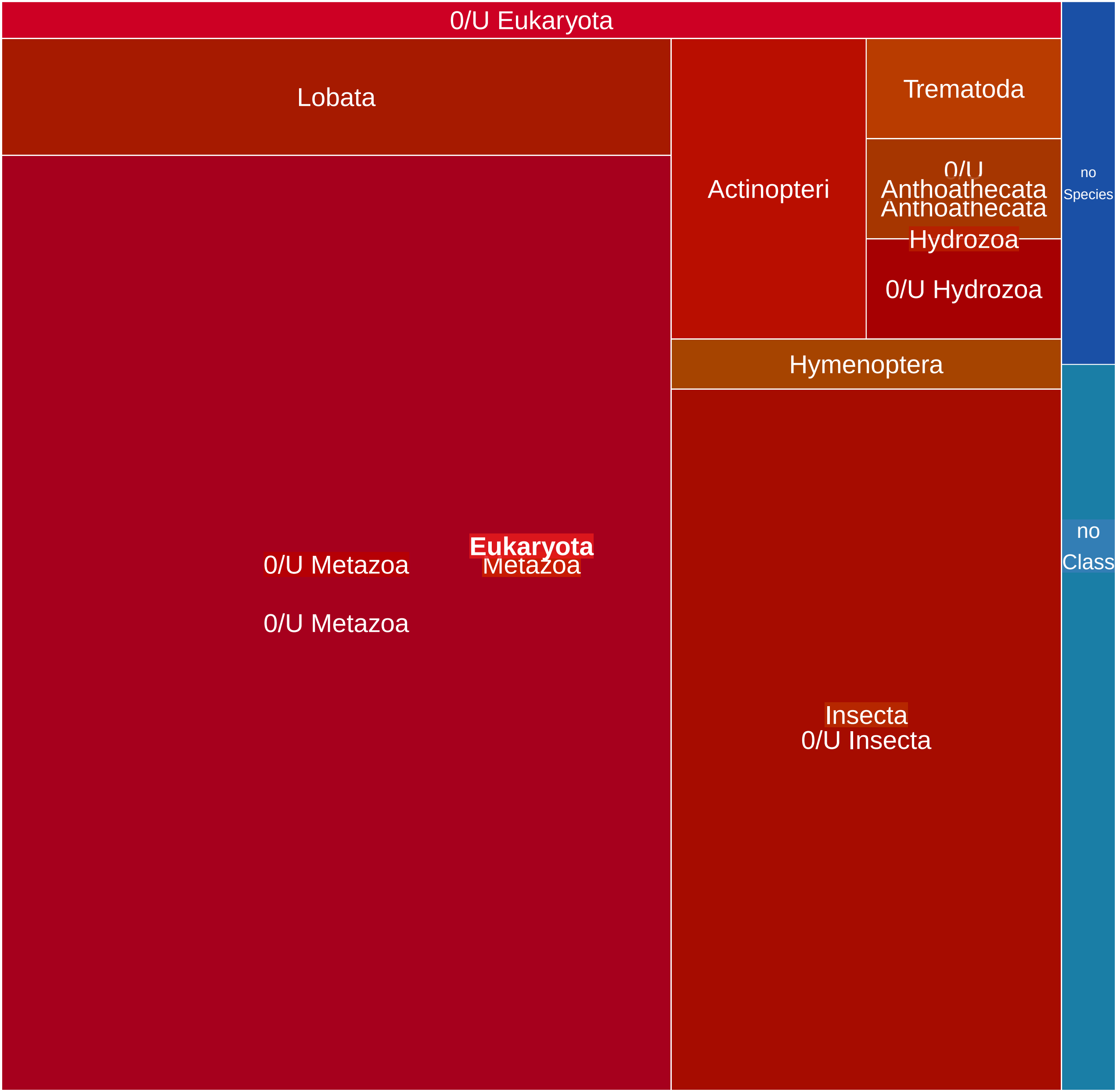

MGT−v1_797 : 659 unigenes, 149 (22.61%) taxonomically assigned

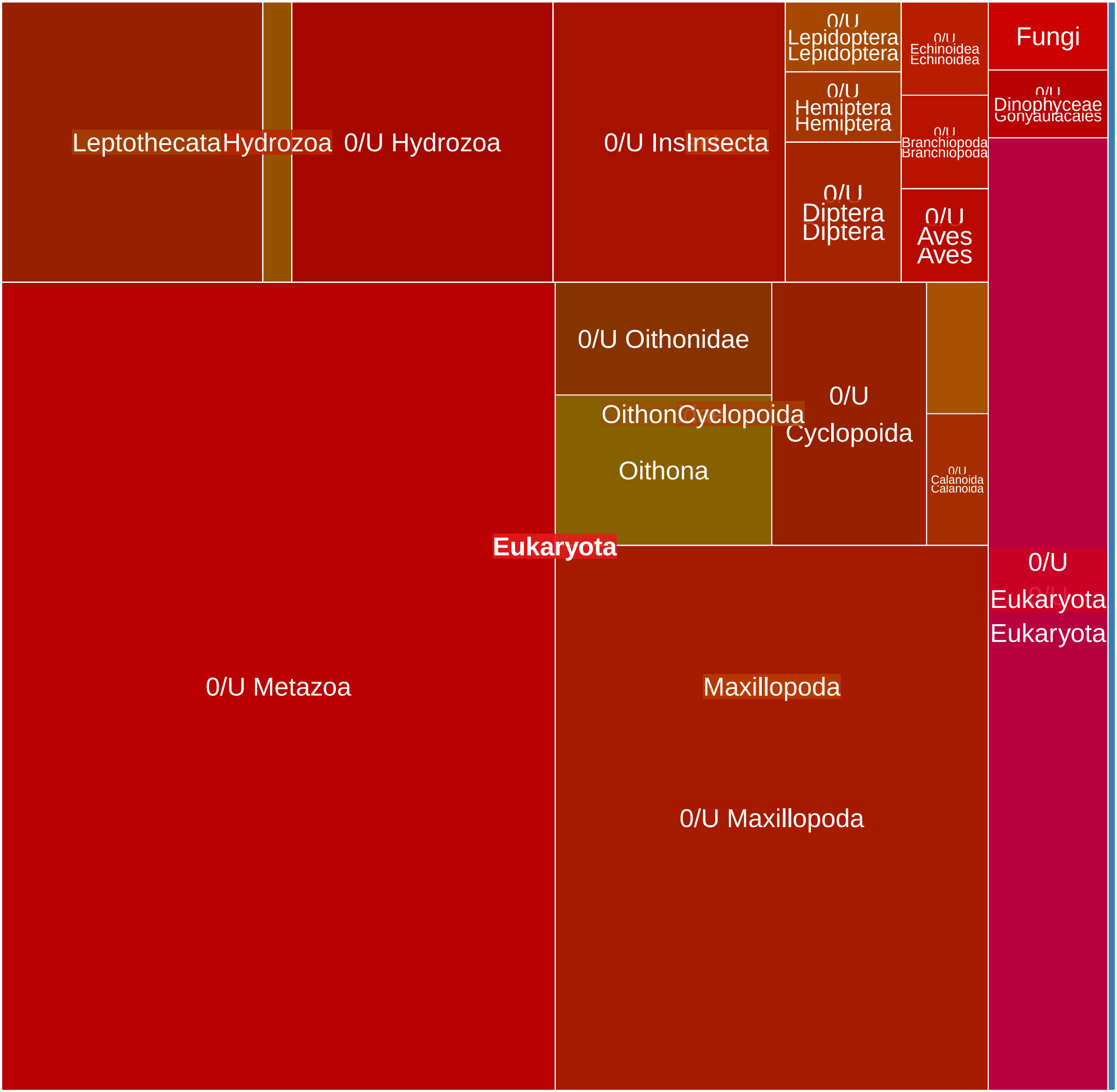

MGT−v1_798 : 657 unigenes, 79 (12.02%) taxonomically assigned

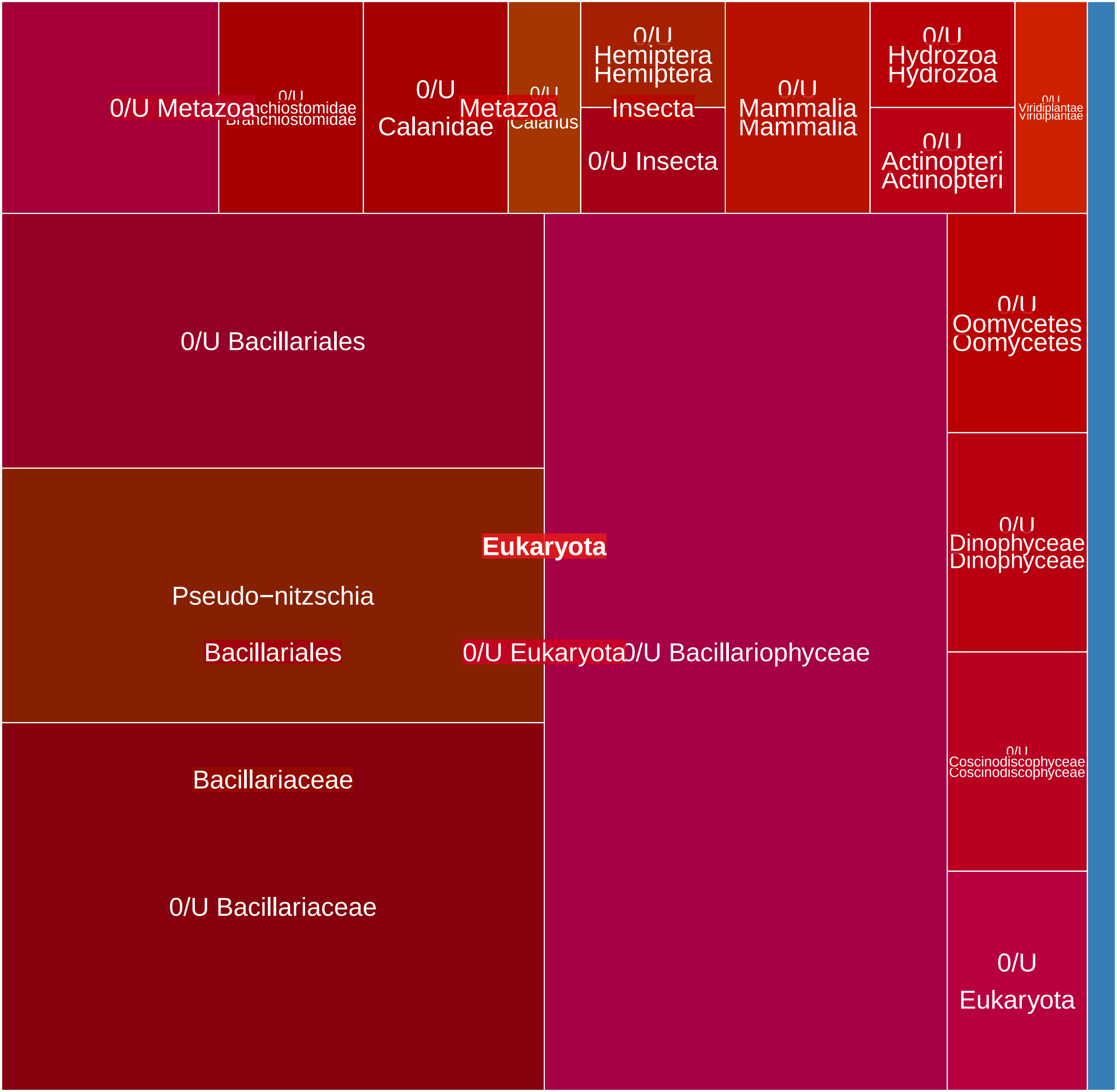

MGT−v1_799 : 656 unigenes, 54 (8.23%) taxonomically assigned

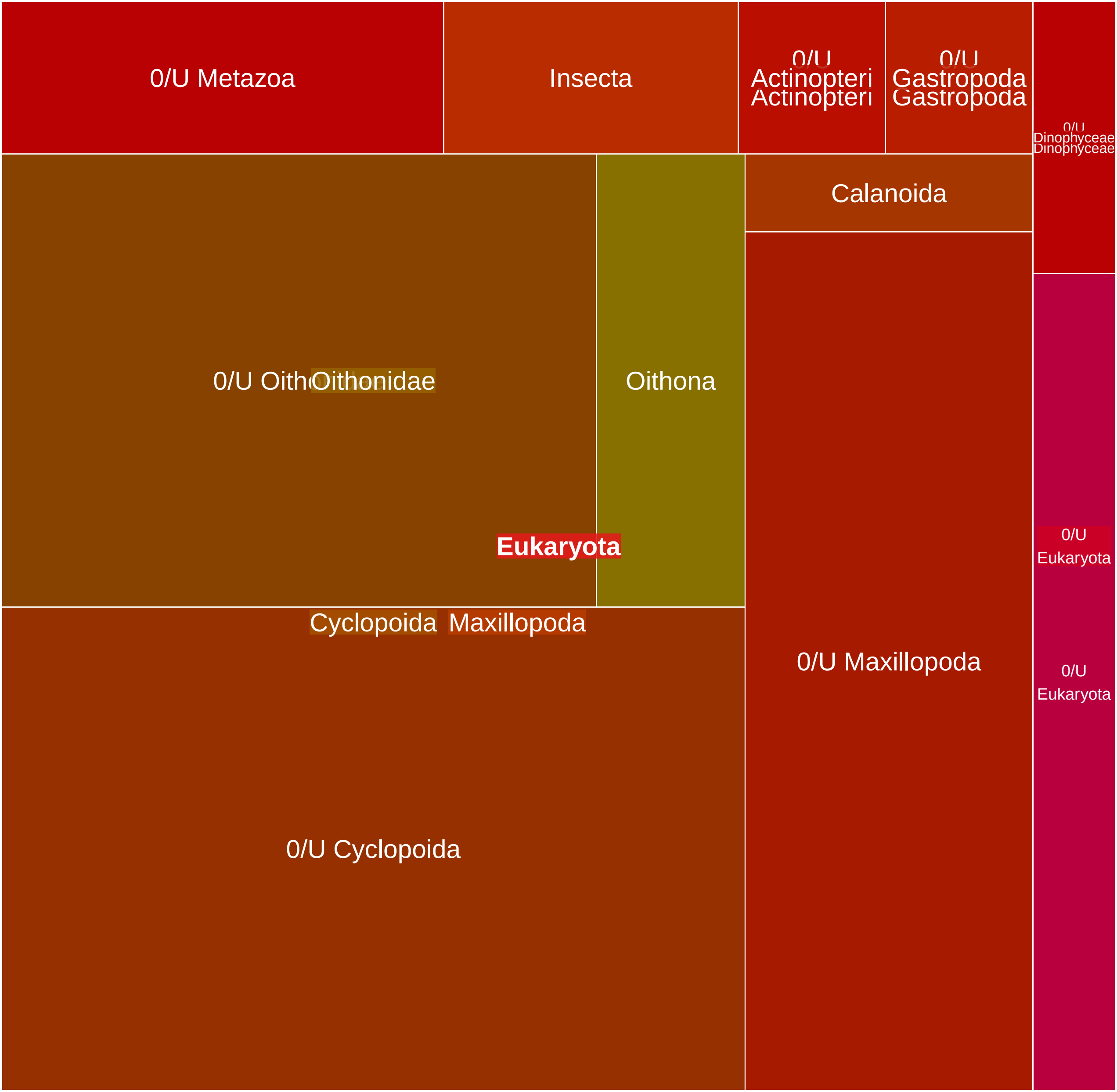

MGT−v1_800 : 652 unigenes, 450 (69.02%) taxonomically assigned

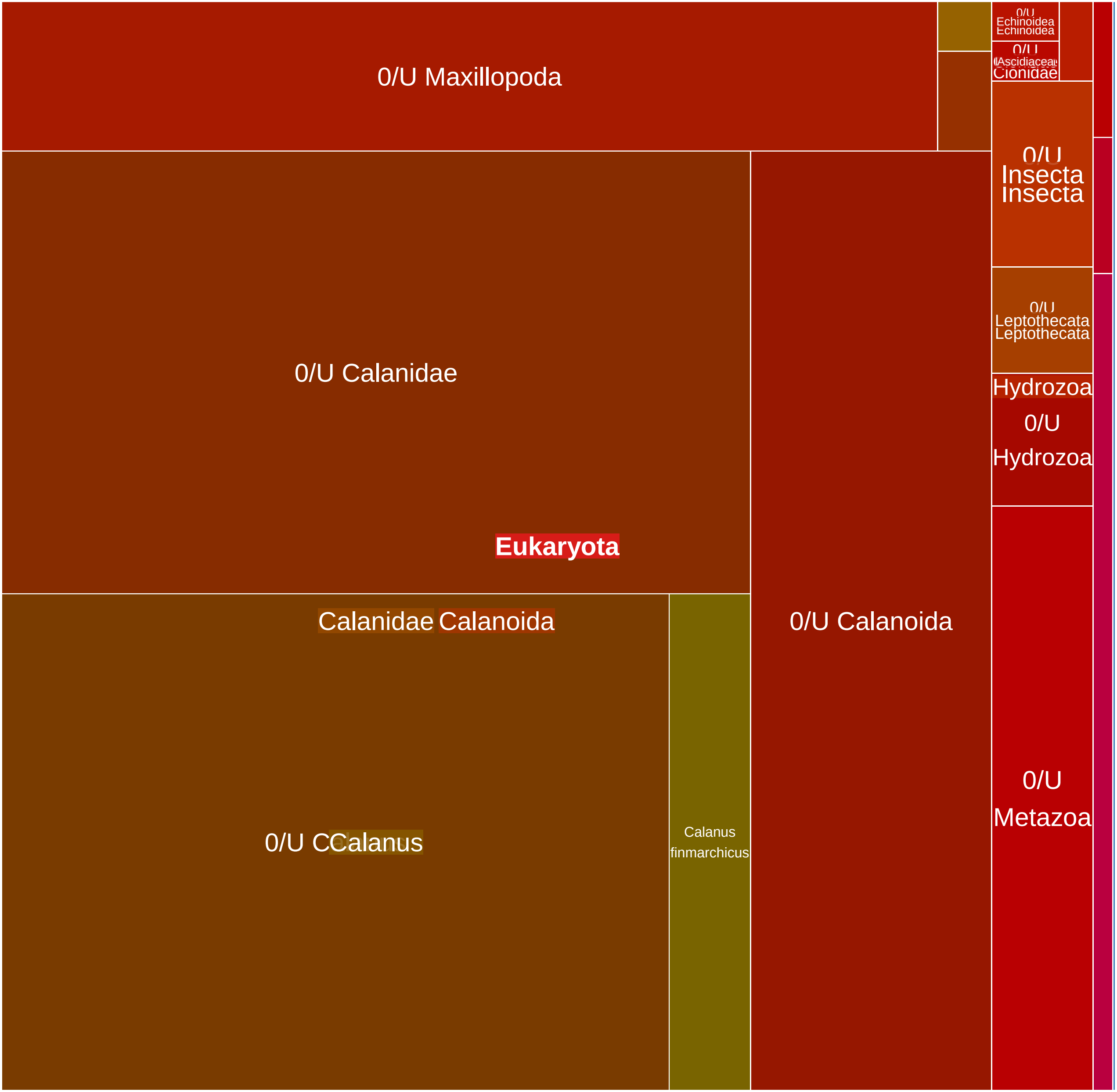

MGT−v1_801 : 651 unigenes, 341 (52.38%) taxonomically assigned

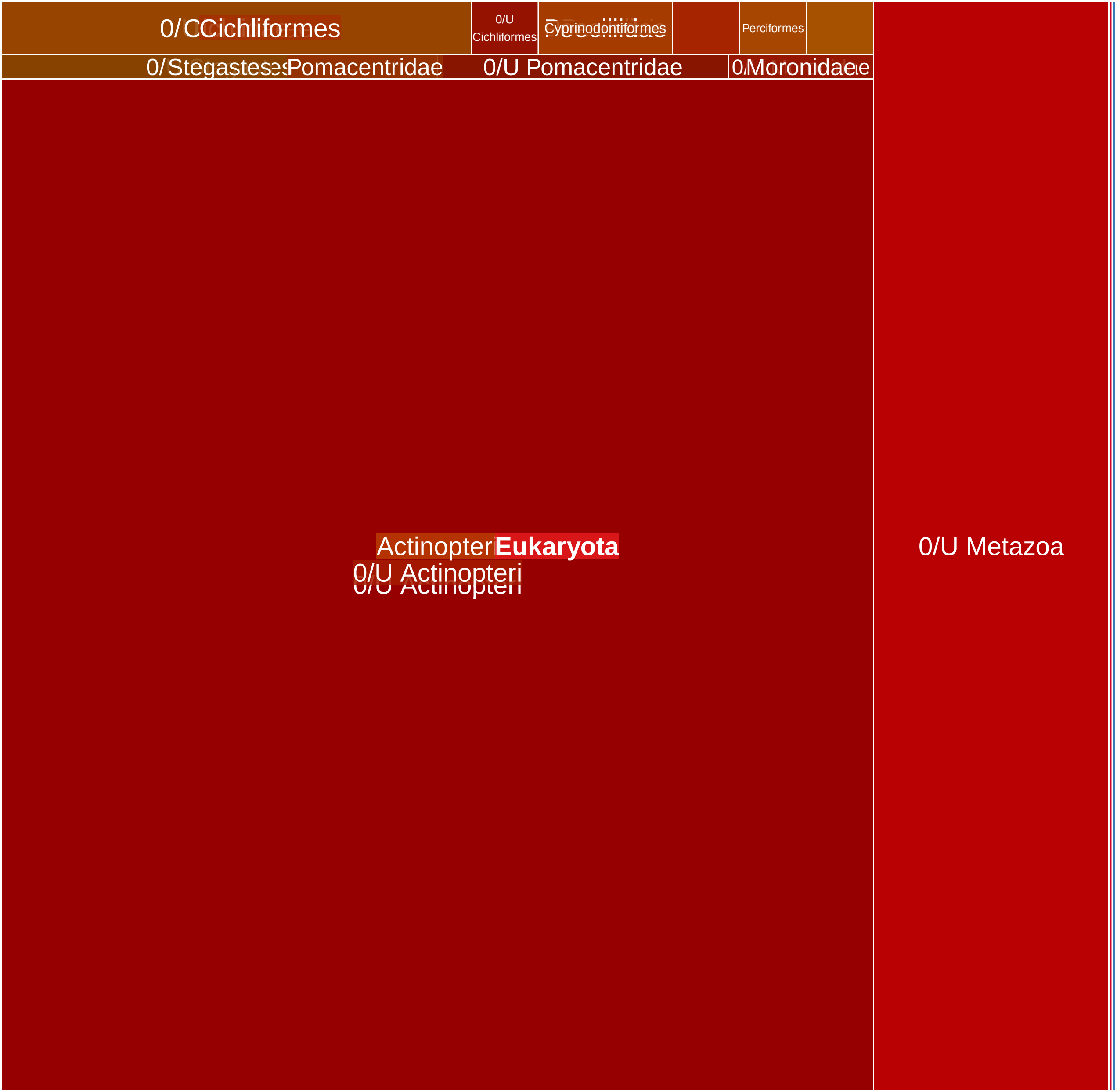

MGT−v1_802 : 649 unigenes, 365 (56.24%) taxonomically assigned

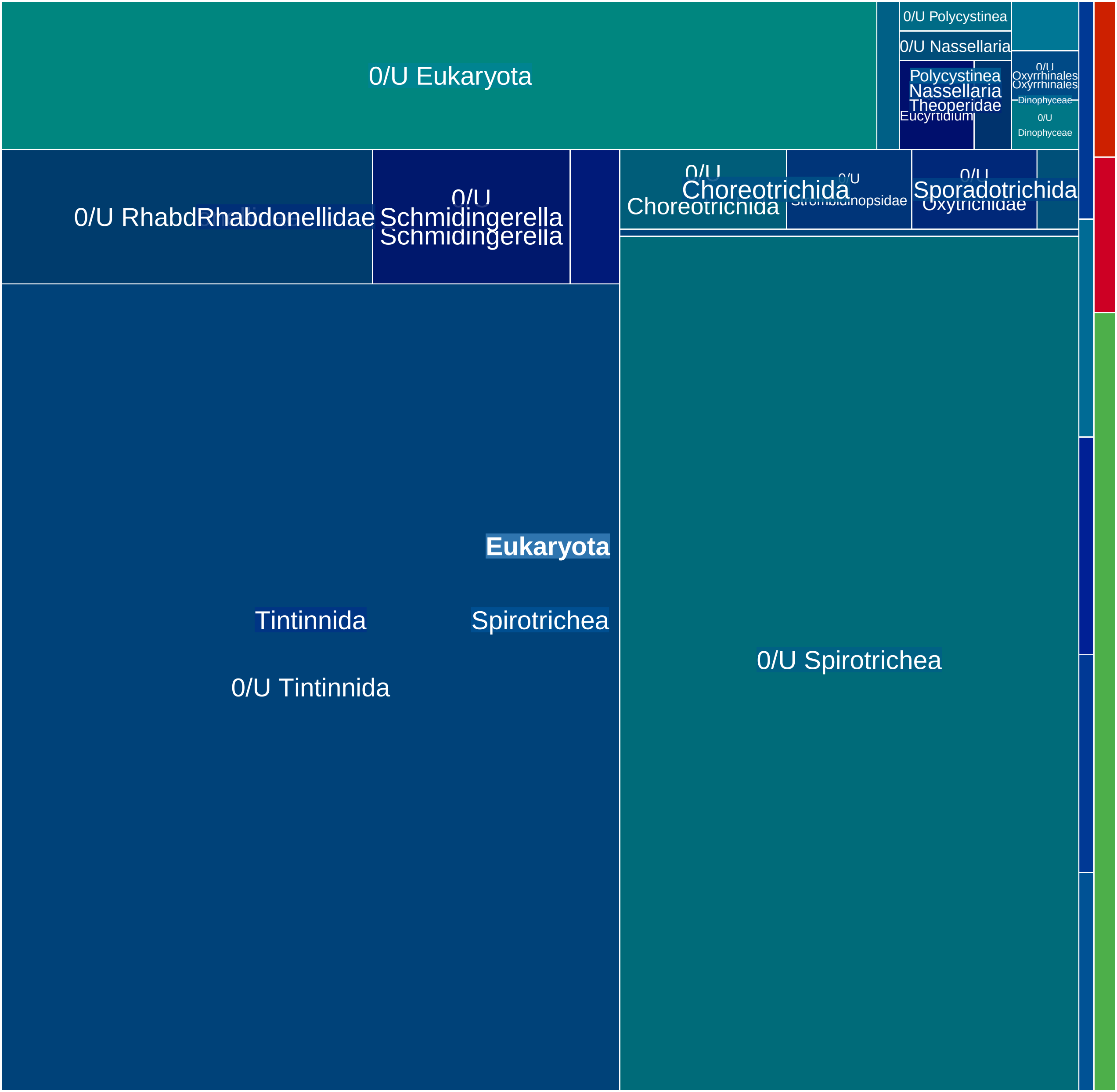

### MGT−v1_803 : 647 unigenes, 596 (92.12%) taxonomically assigned

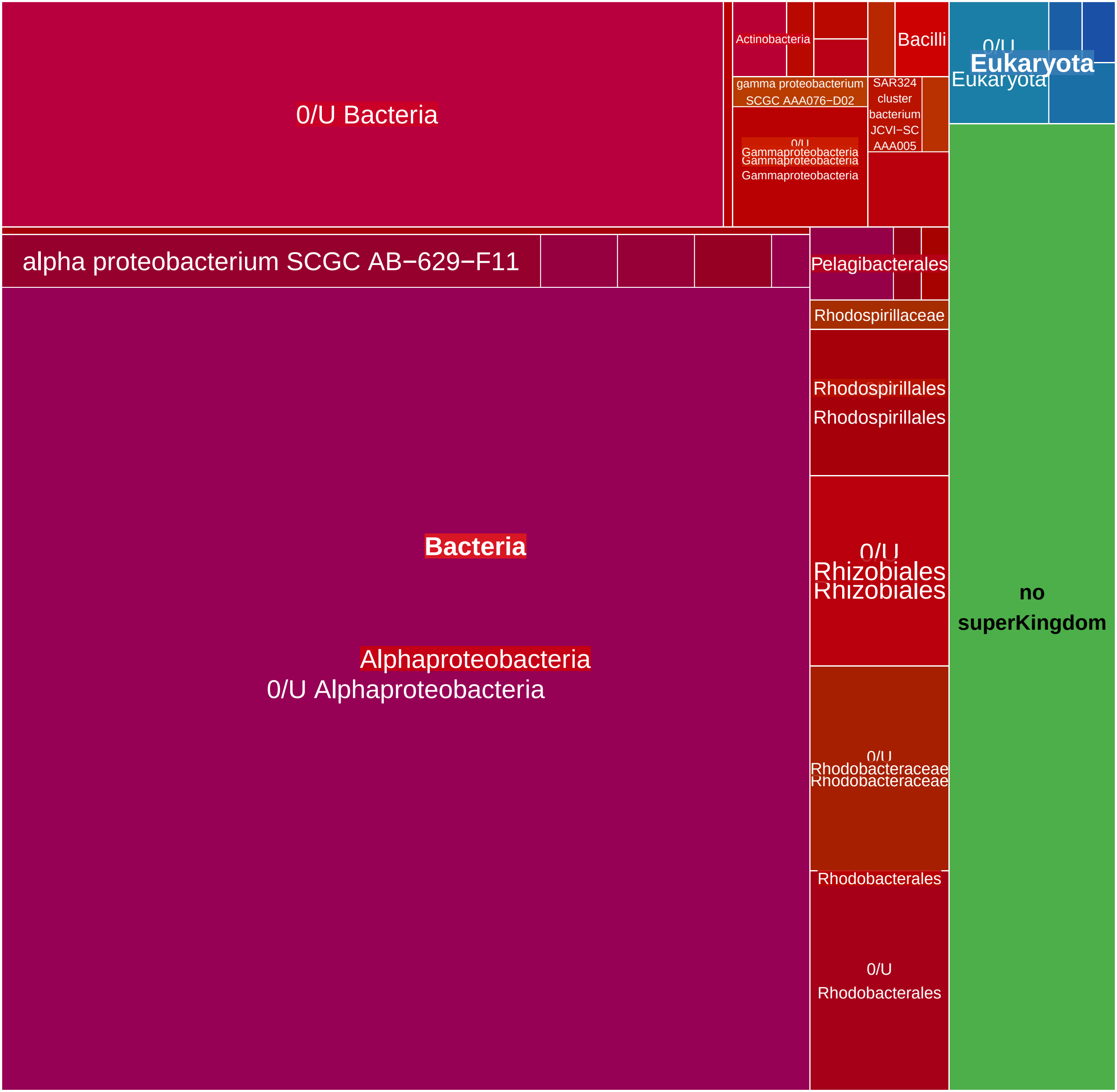

MGT−v1_804 : 644 unigenes, 76 (11.80%) taxonomically assigned

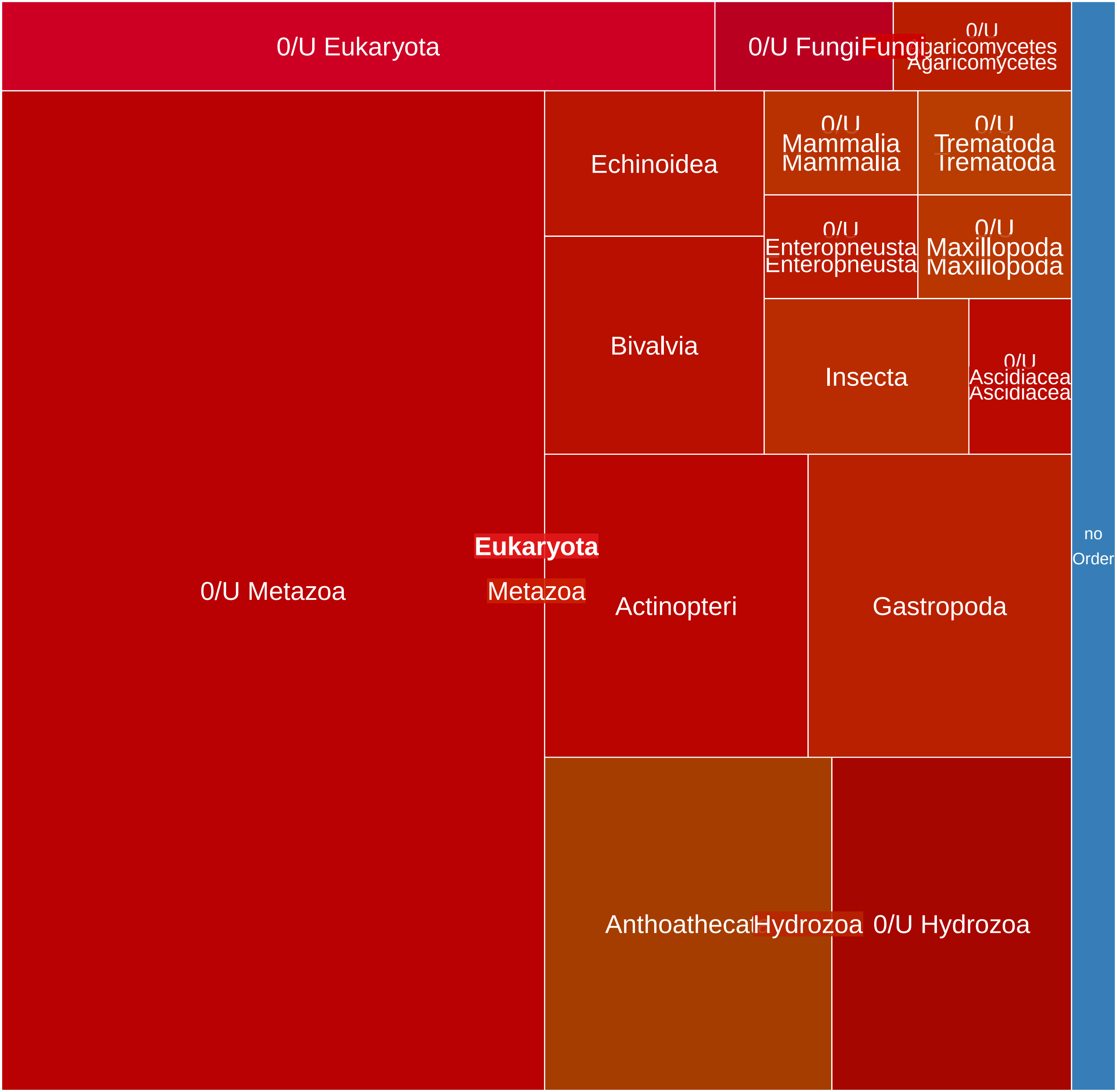

MGT−v1_805 : 641 unigenes, 465 (72.54%) taxonomically assigned

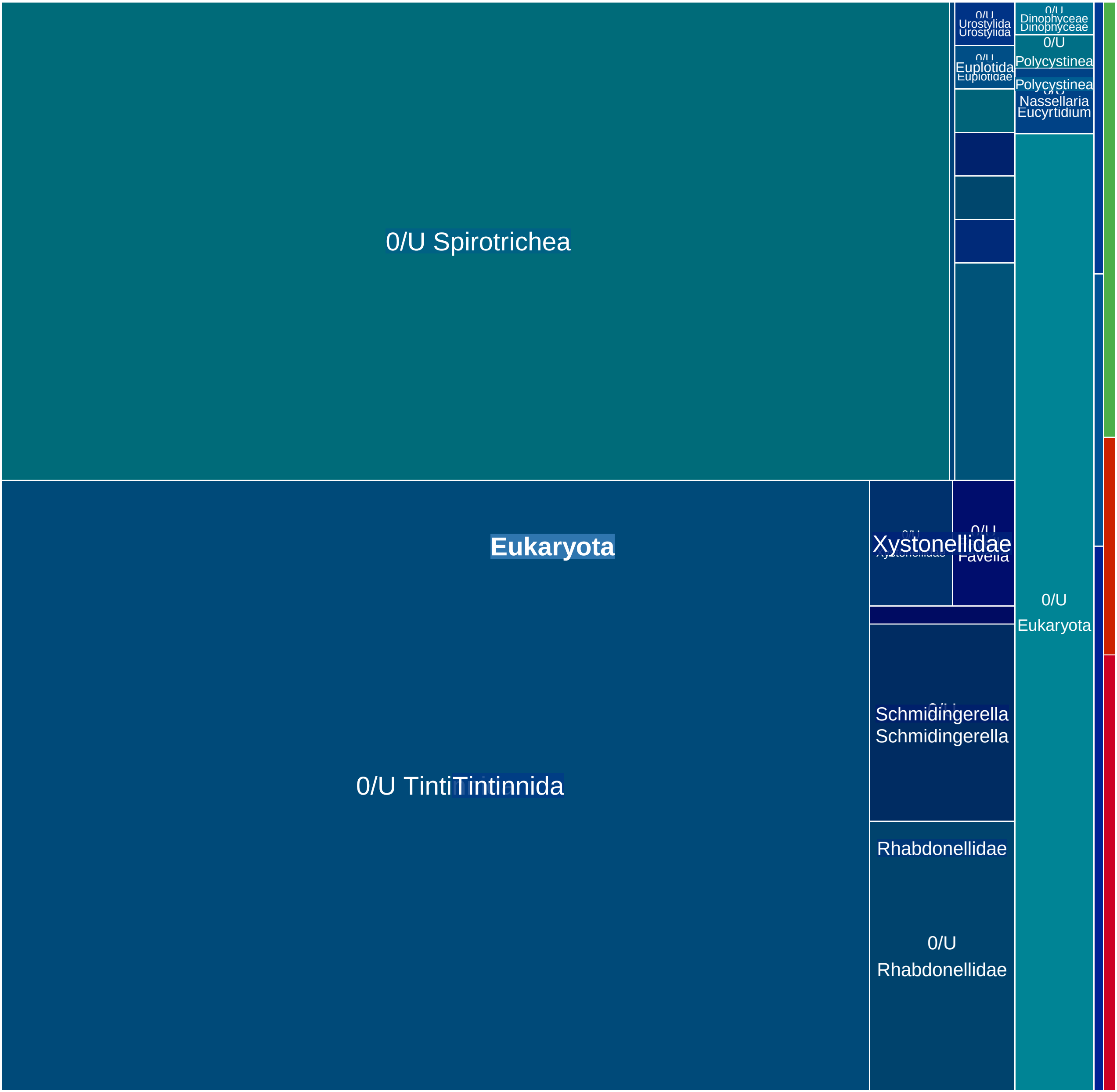

MGT−v1_806 : 641 unigenes, 440 (68.64%) taxonomically assigned

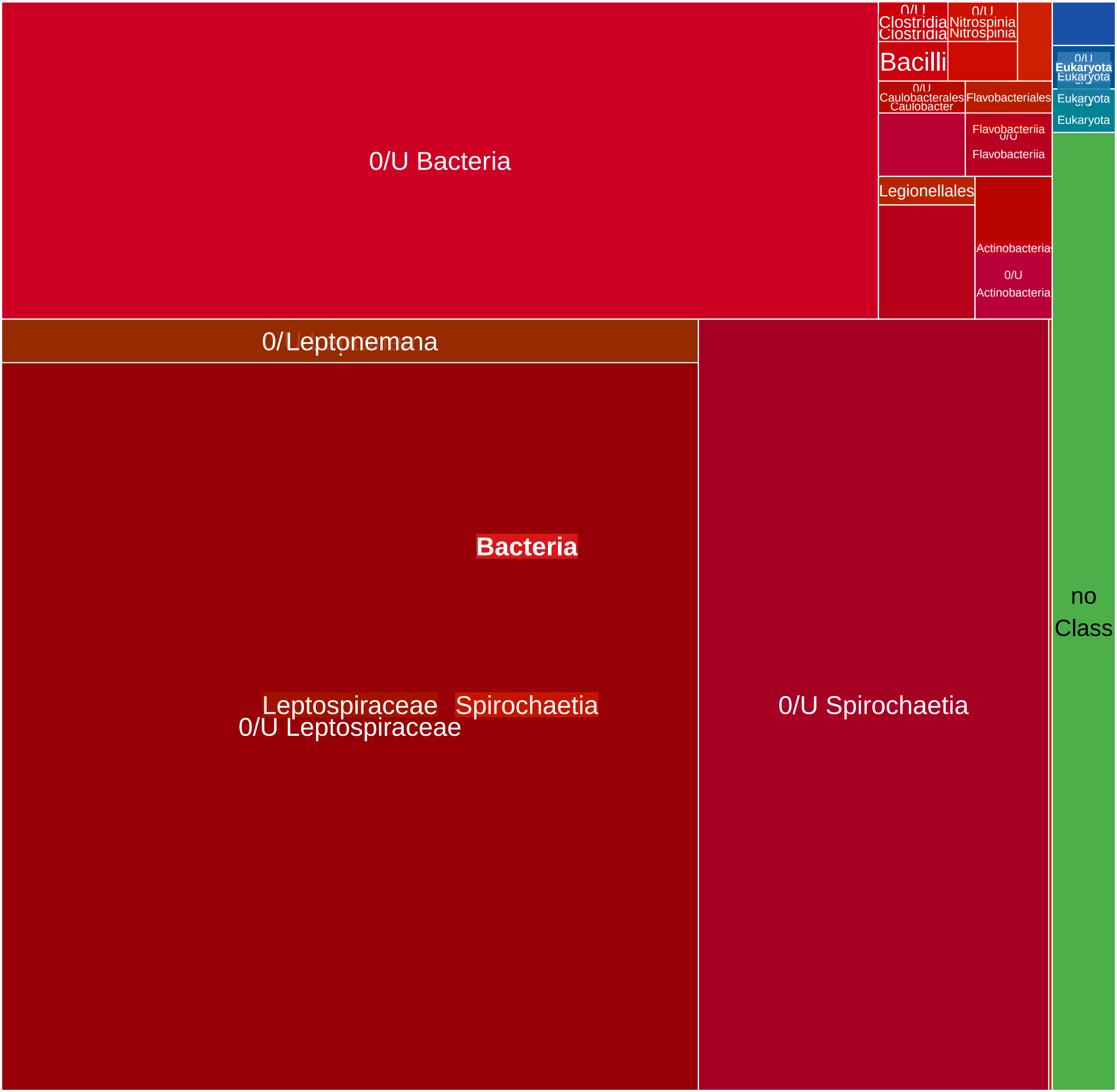

MGT−v1_807 : 640 unigenes, 85 (13.28%) taxonomically assigned

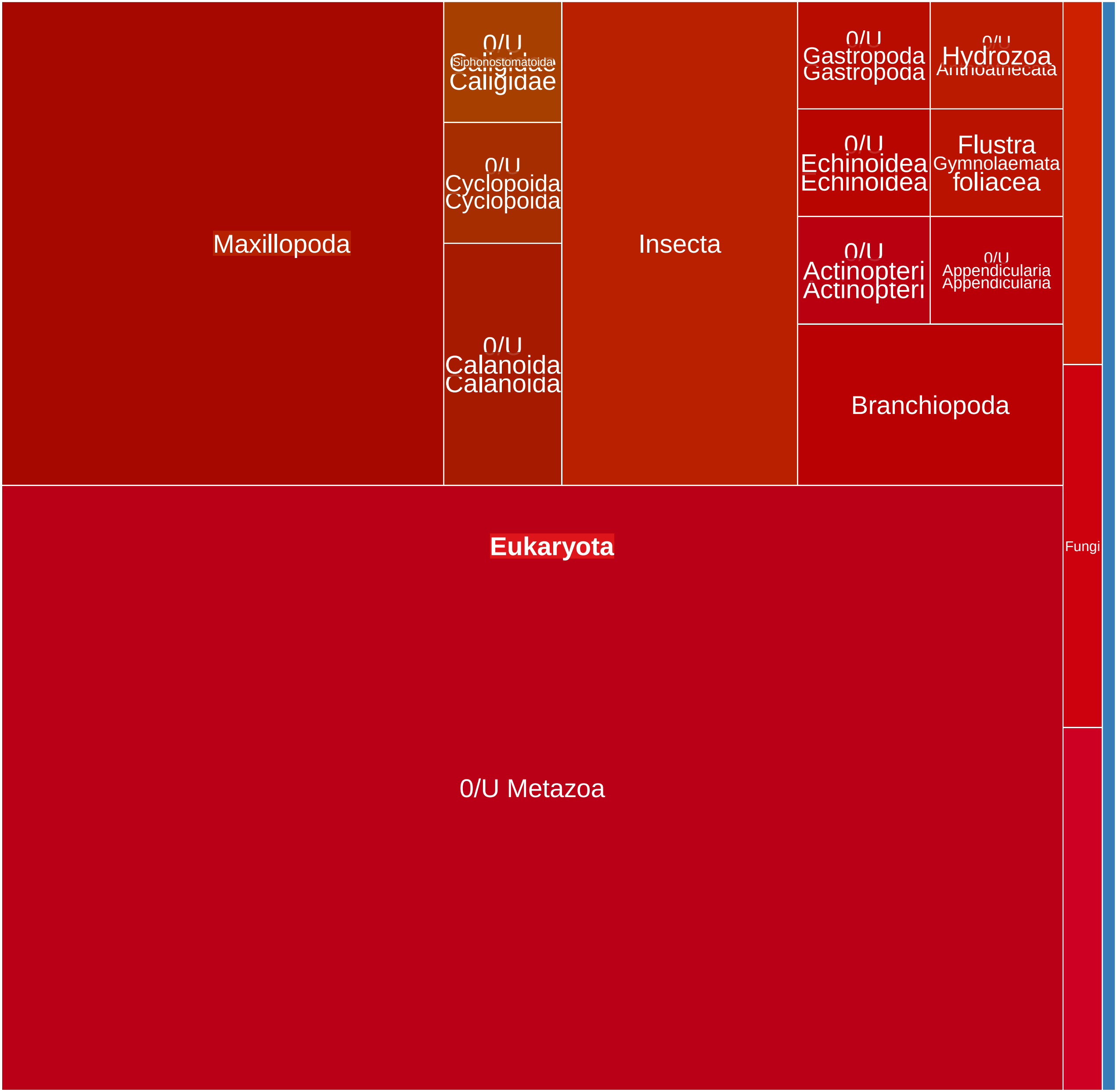

MGT−v1_808 : 638 unigenes, 27 (4.23%) taxonomically assigned

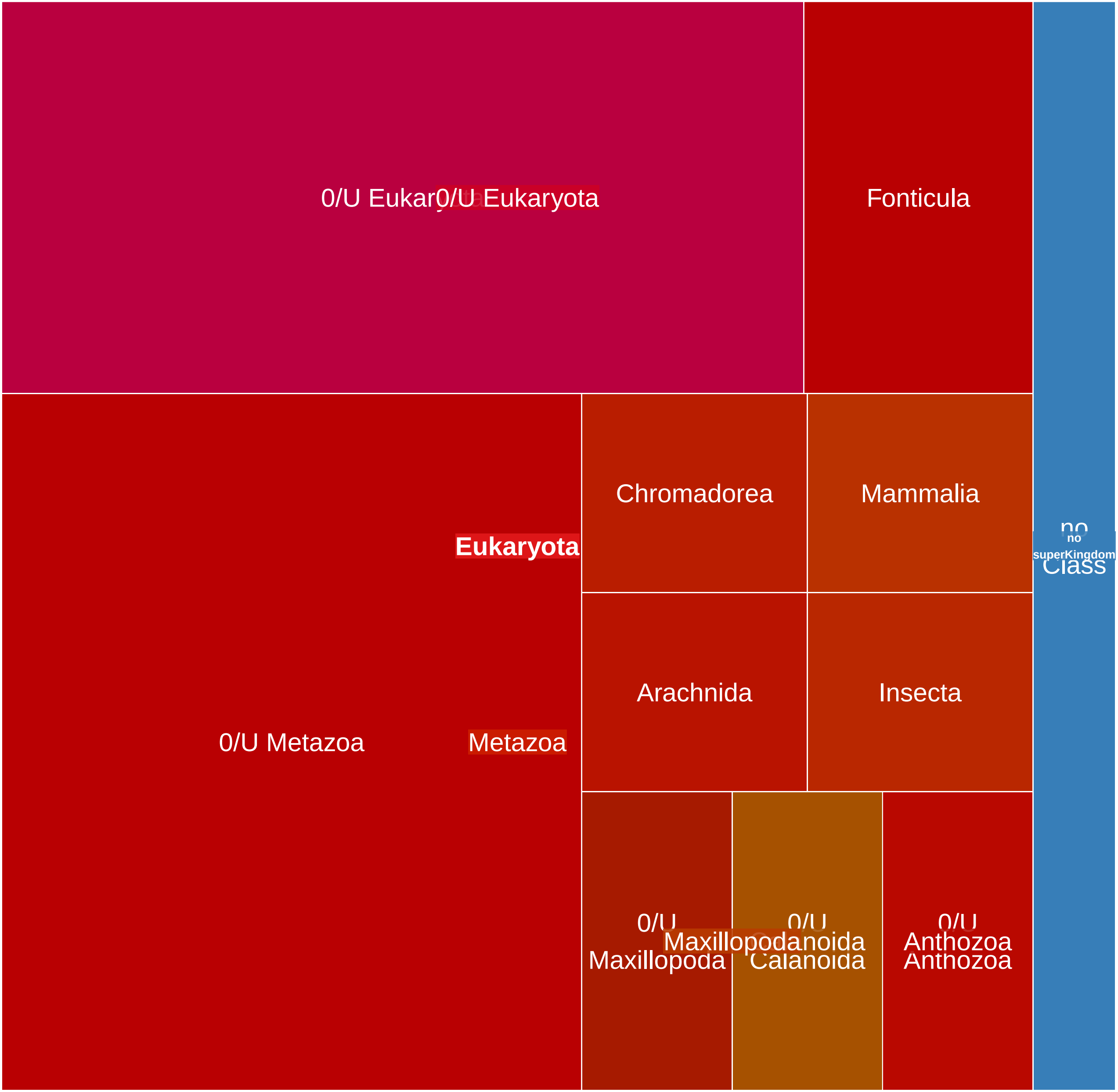

MGT−v1_809 : 636 unigenes, 69 (10.85%) taxonomically assigned

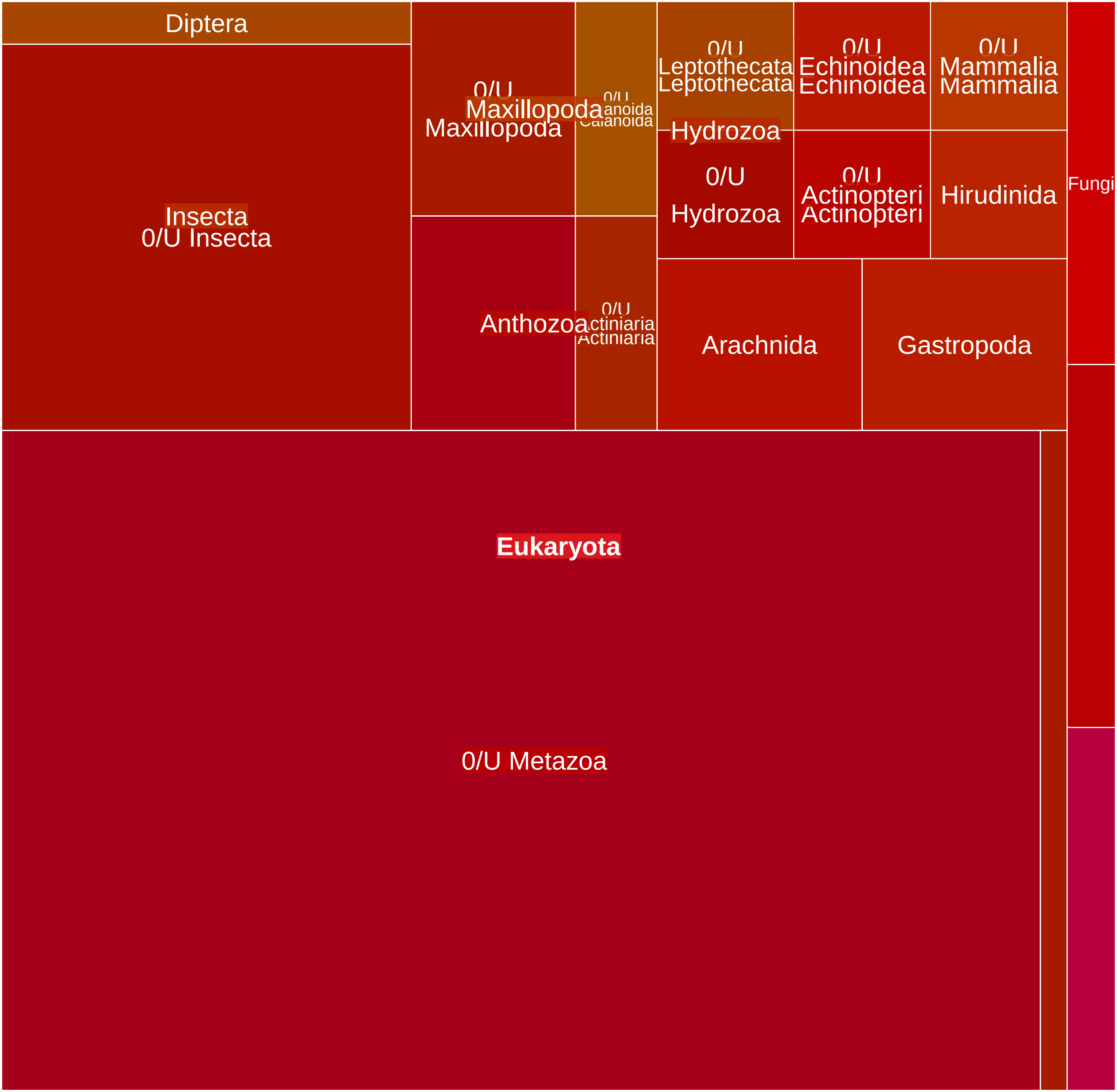

MGT−v1_810 : 636 unigenes, 515 (80.97%) taxonomically assigned

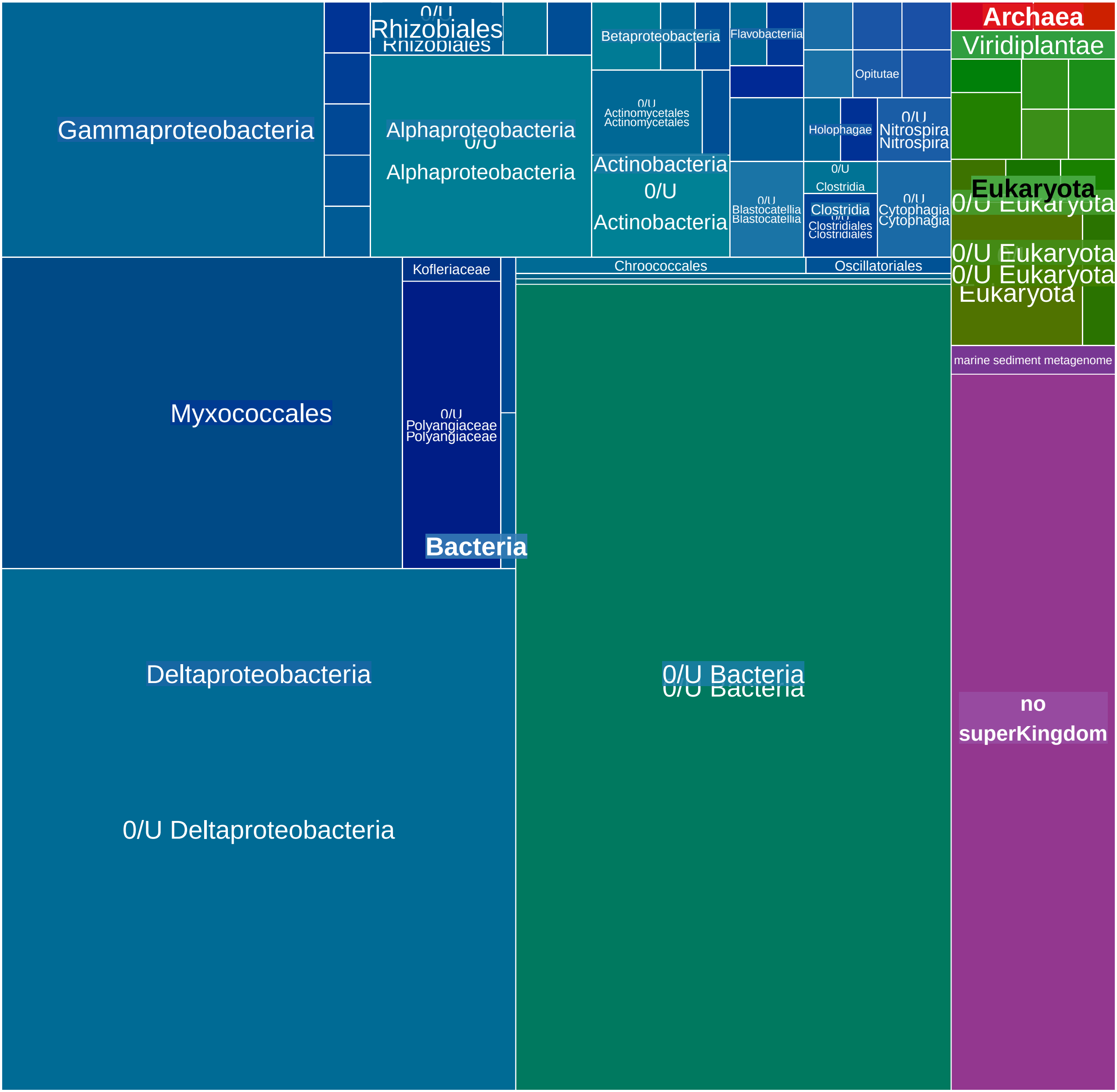

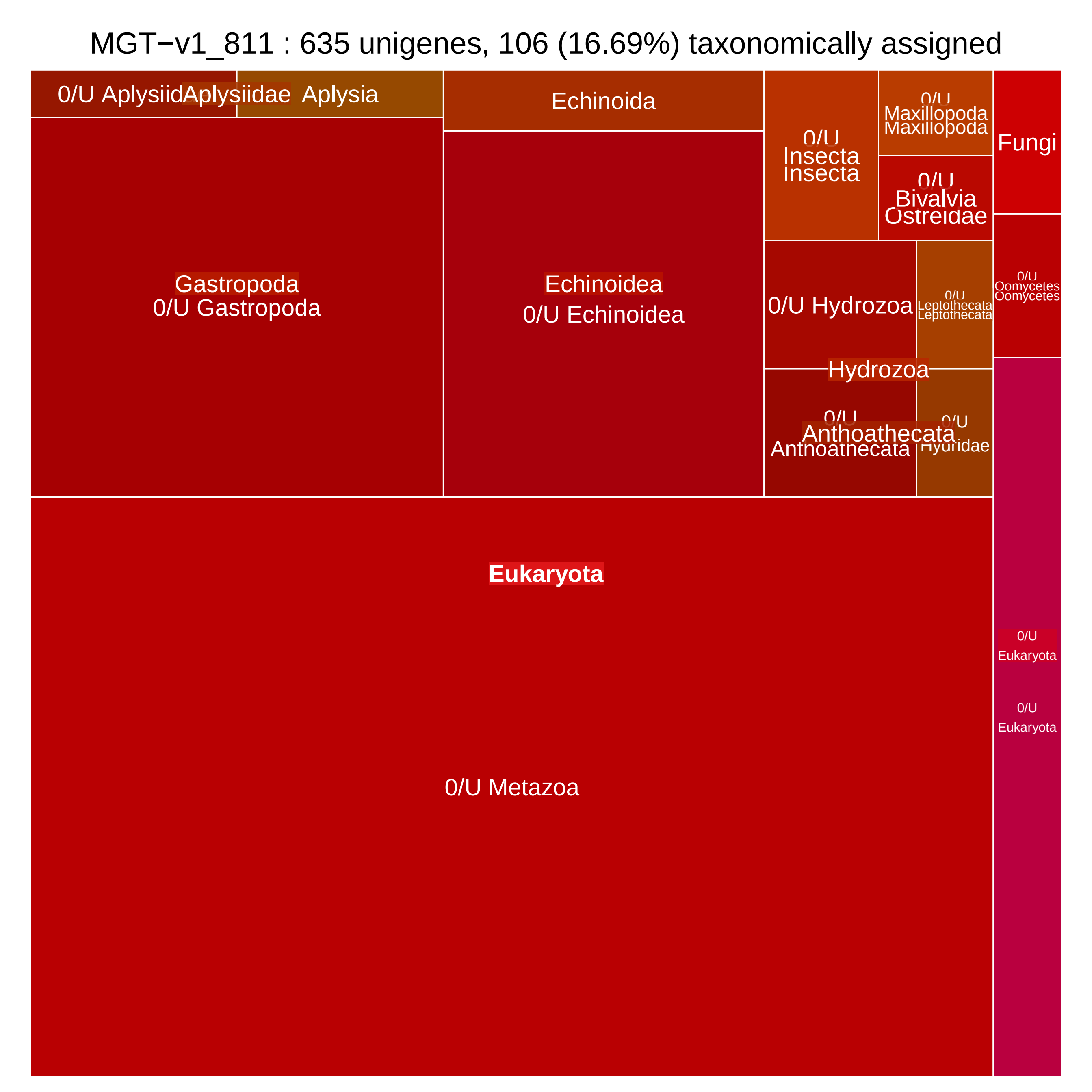
MGT−v1_811 : 635 unigenes, 106 (16.69%) taxonomically assigned
0/U Aplysiidae
Aplysiidae
Aplysia
Echinoida
Gastropoda
0/U Gastropoda
Echinoidea
0/U Echinoidea
0/U Insecta
Insecta
0/U Maxillopoda
Maxillopoda
0/U Bivalvia
Ostreidae
Fungi
0/U Oomycetes
Oomycetes
0/U Hydrozoa
0/U Leptothecata
Leptothecata
Hydrozoa
0/U Anthoathecata
Anthoathecata
0/U Hydridae
Eukaryota
0/U Metazoa
0/U Eukaryota
0/U Eukaryota

MGT−v1_812 : 634 unigenes, 573 (90.38%) taxonomically assigned

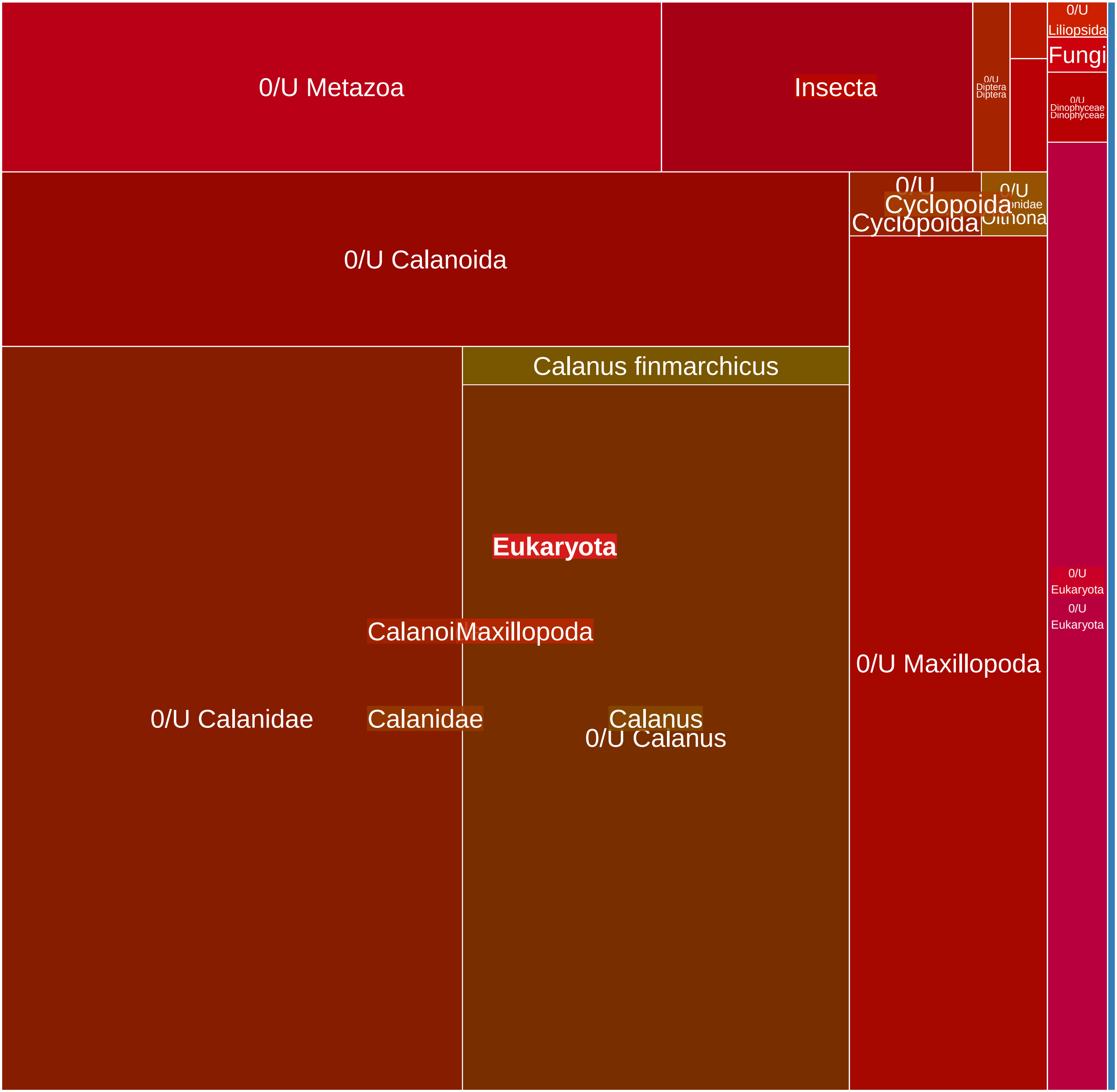

MGT−v1_813 : 634 unigenes, 83 (13.09%) taxonomically assigned

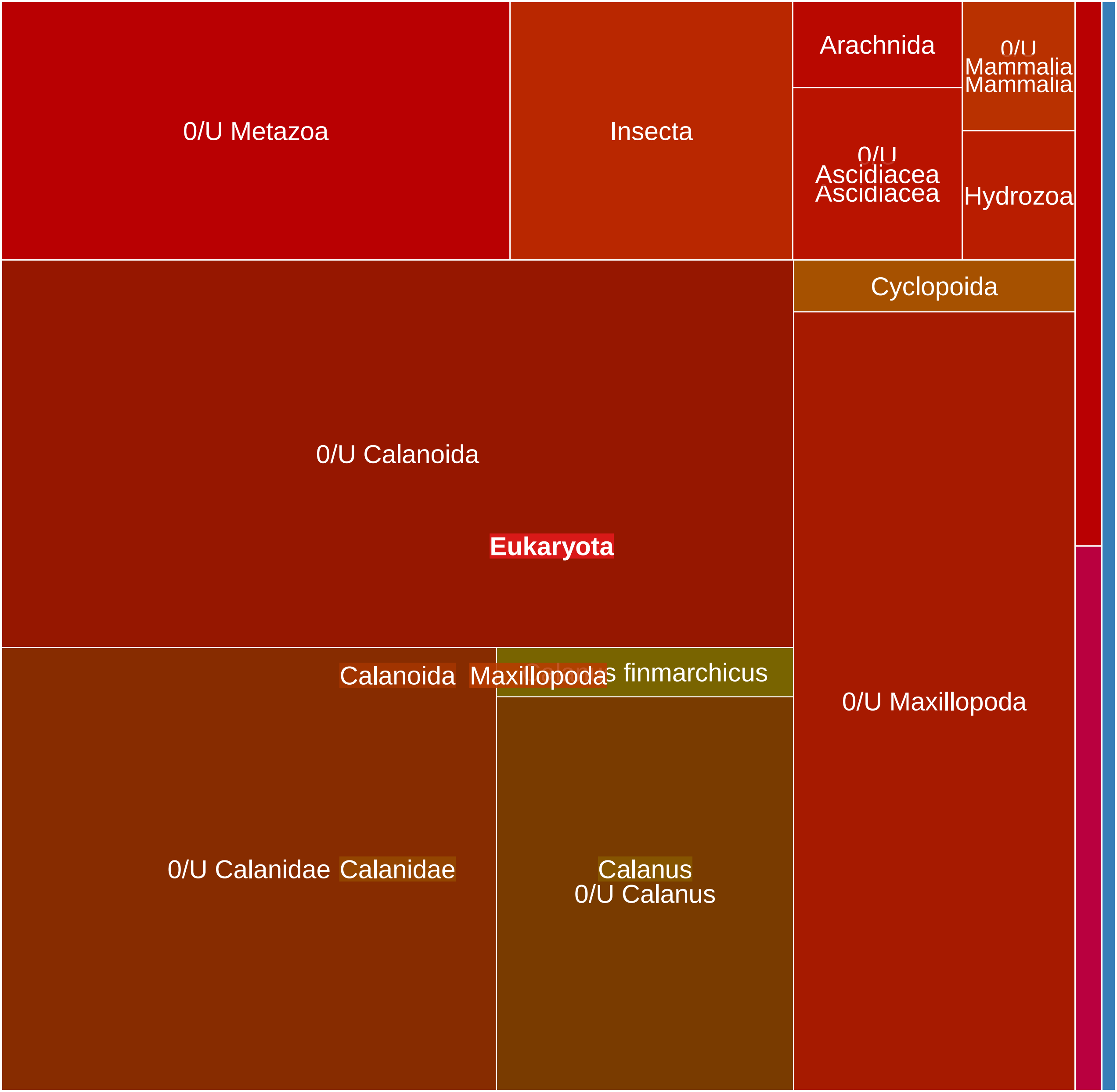

### MGT−v1_814 : 630 unigenes, 145 (23.02%) taxonomically assigned

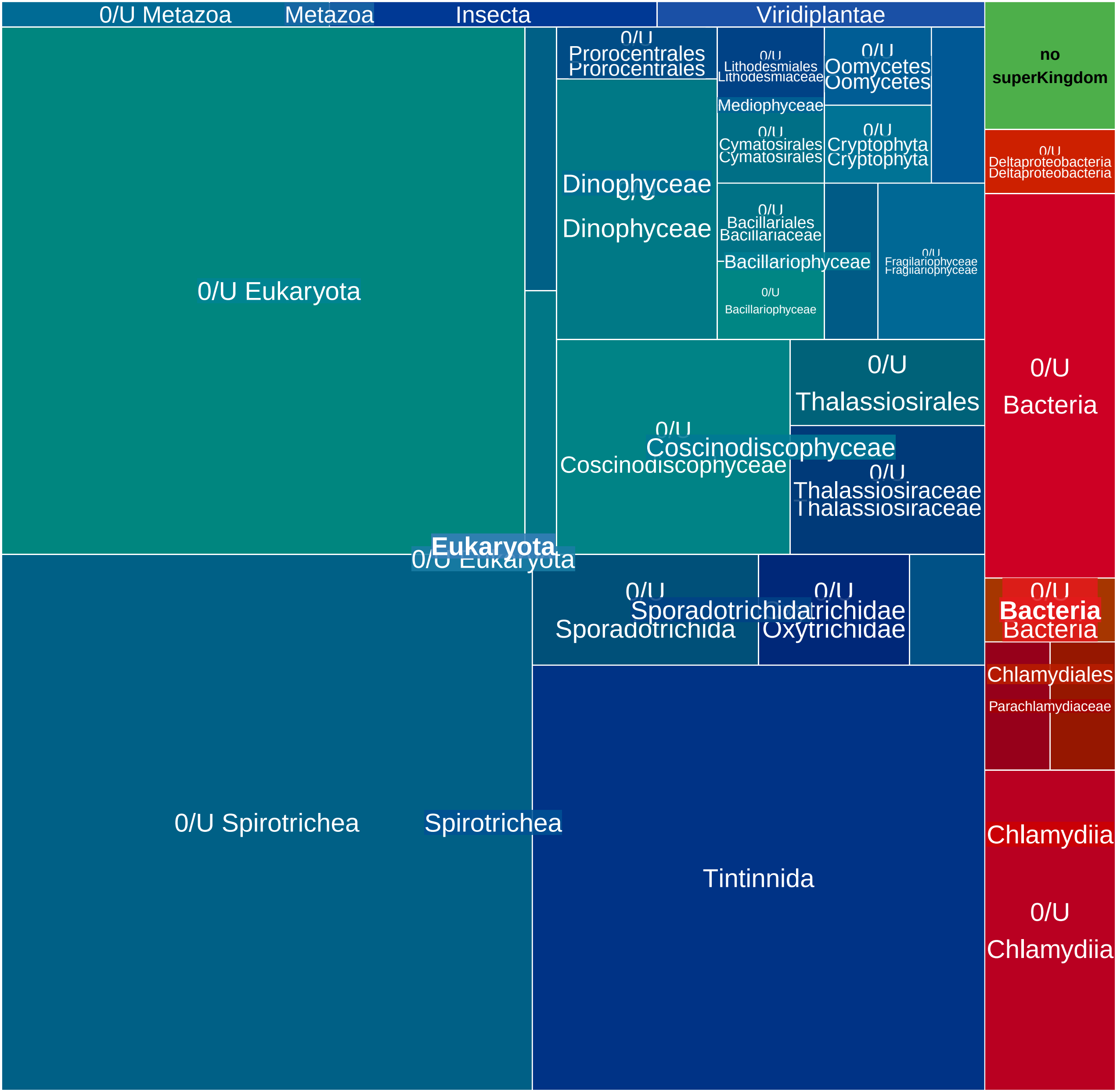

MGT−v1_815 : 629 unigenes, 335 (53.26%) taxonomically assigned

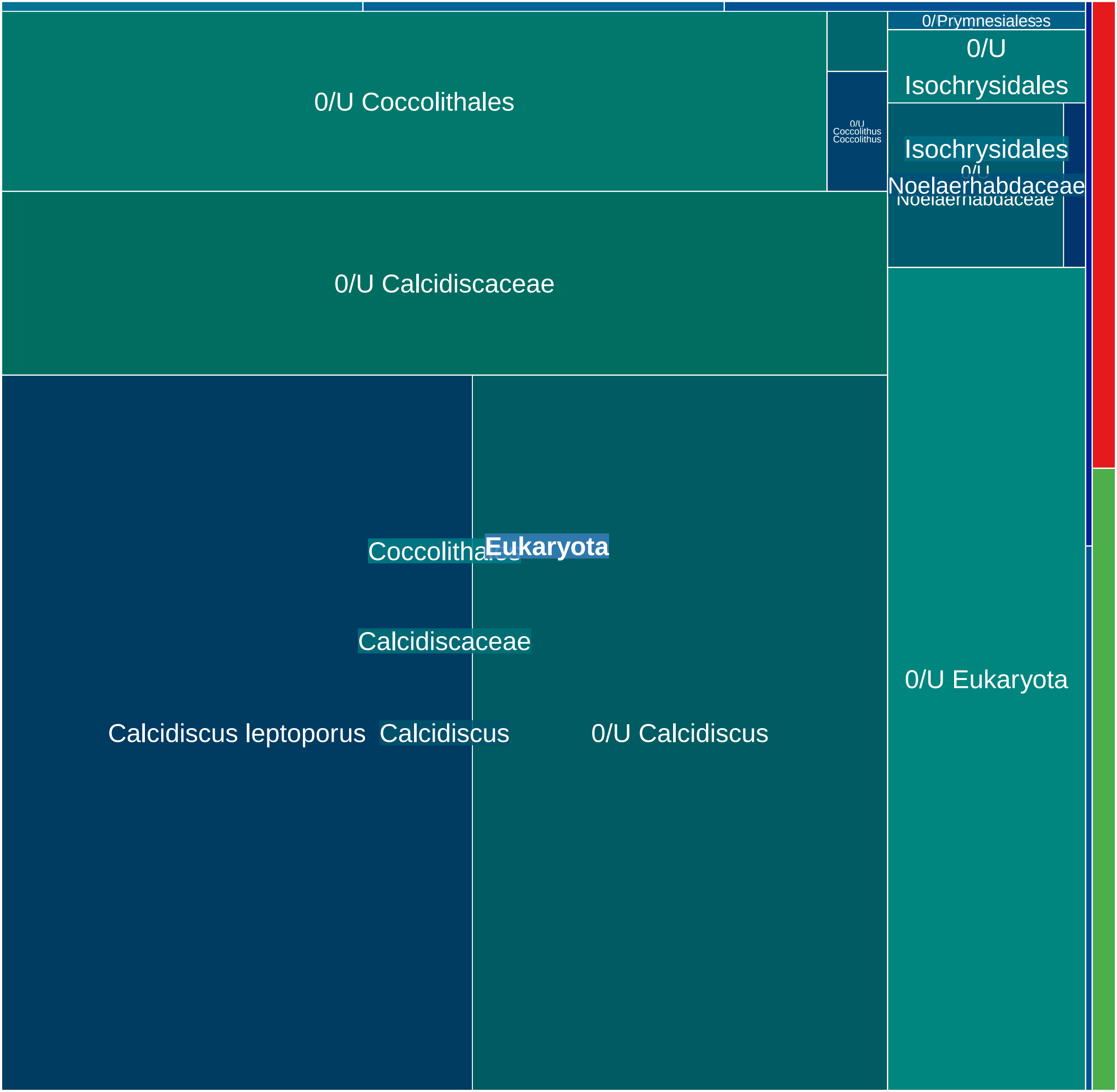

MGT−v1_816 : 628 unigenes, 593 (94.43%) taxonomically assigned

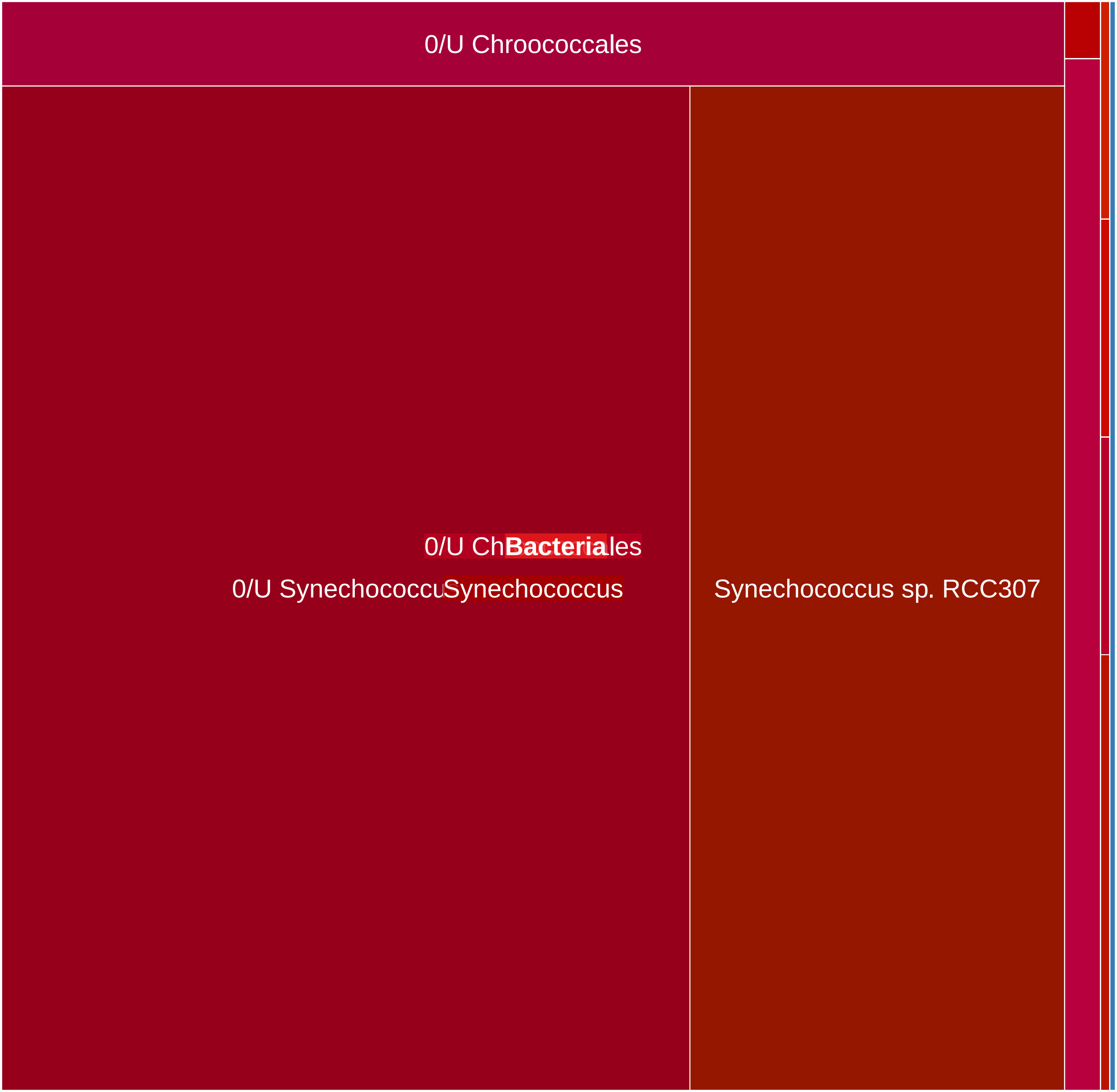

MGT−v1_817 : 628 unigenes, 292 (46.50%) taxonomically assigned

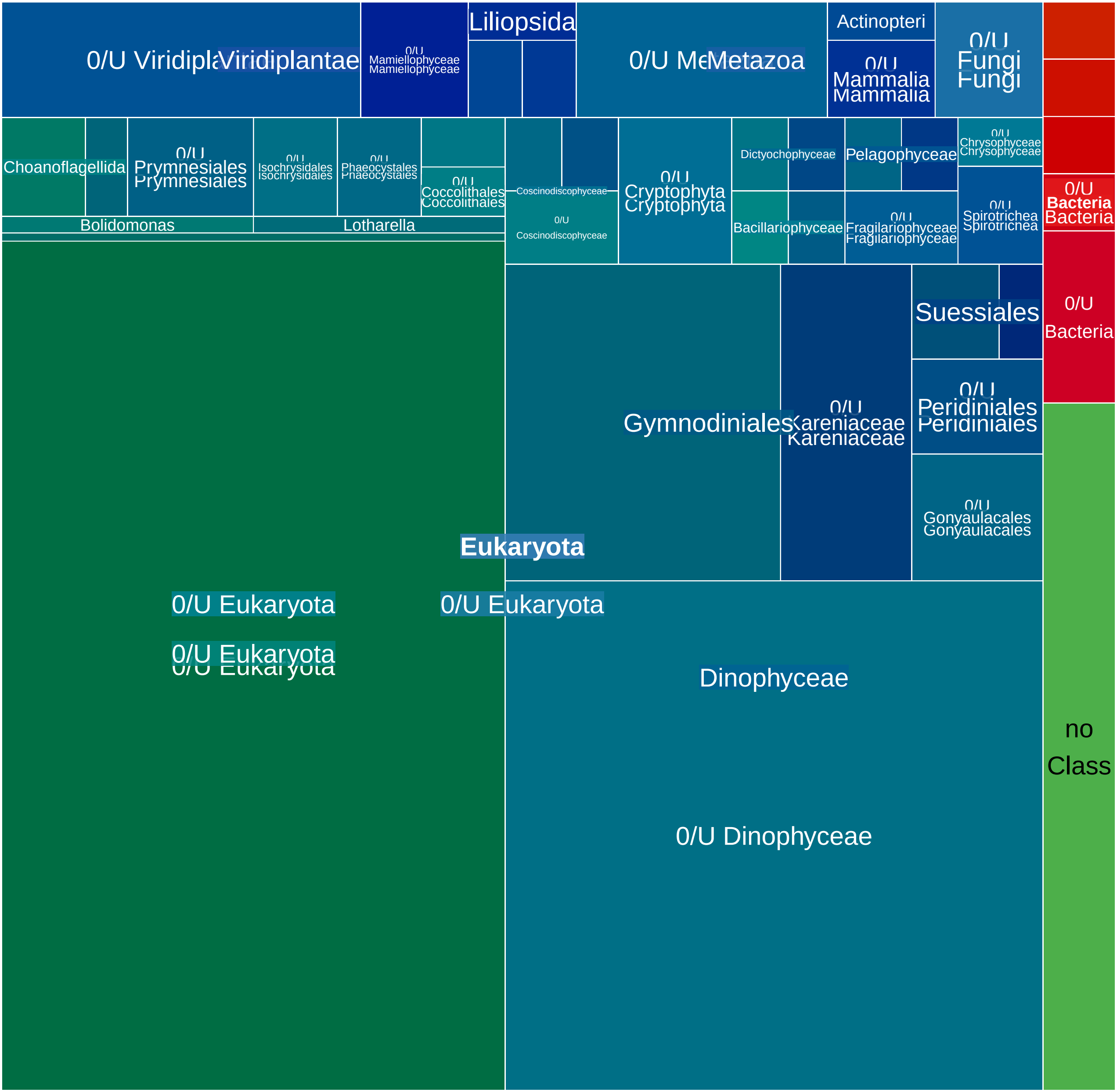

MGT−v1_818 : 627 unigenes, 65 (10.37%) taxonomically assigned

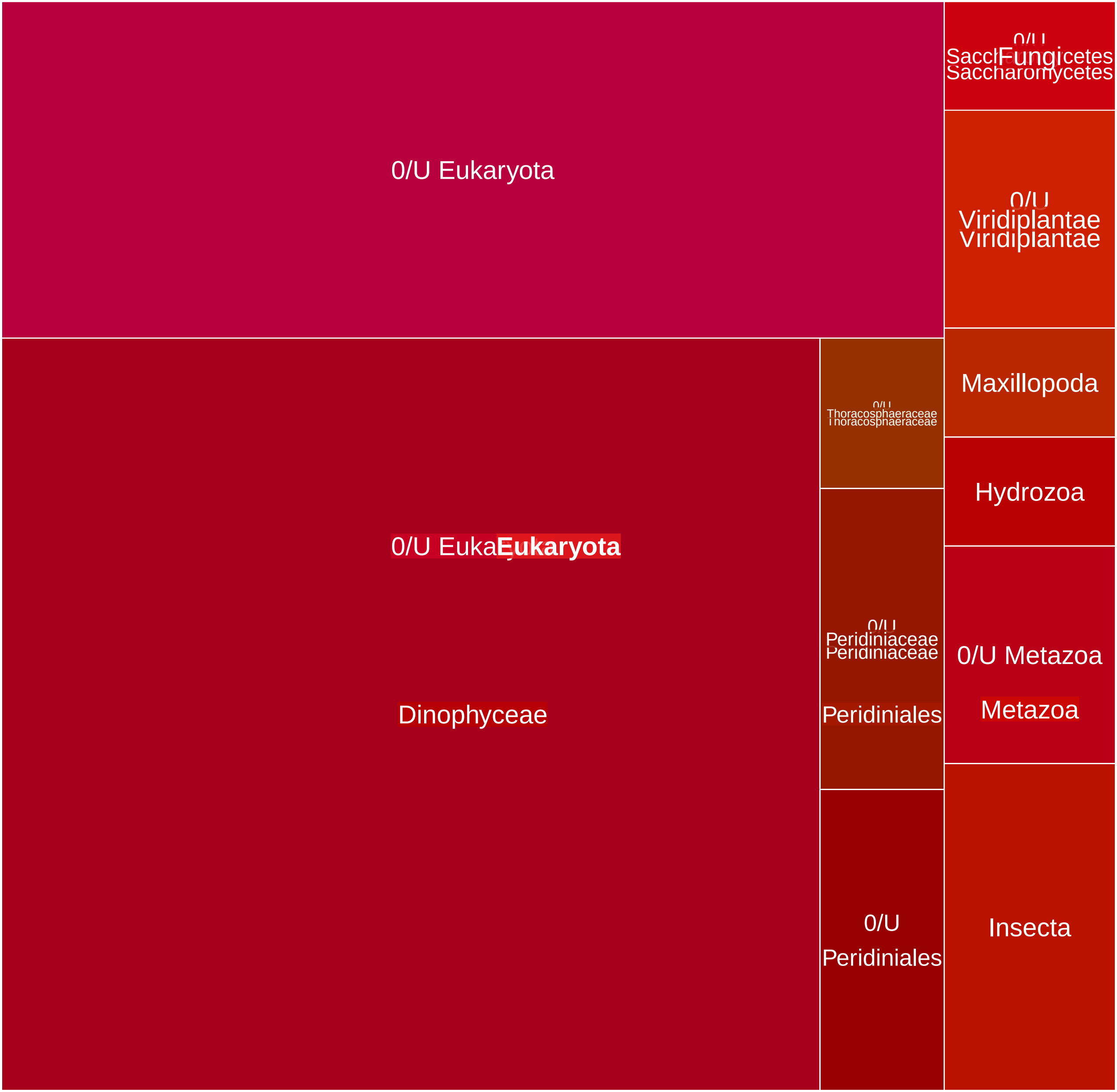

MGT−v1_819 : 627 unigenes, 143 (22.81%) taxonomically assigned

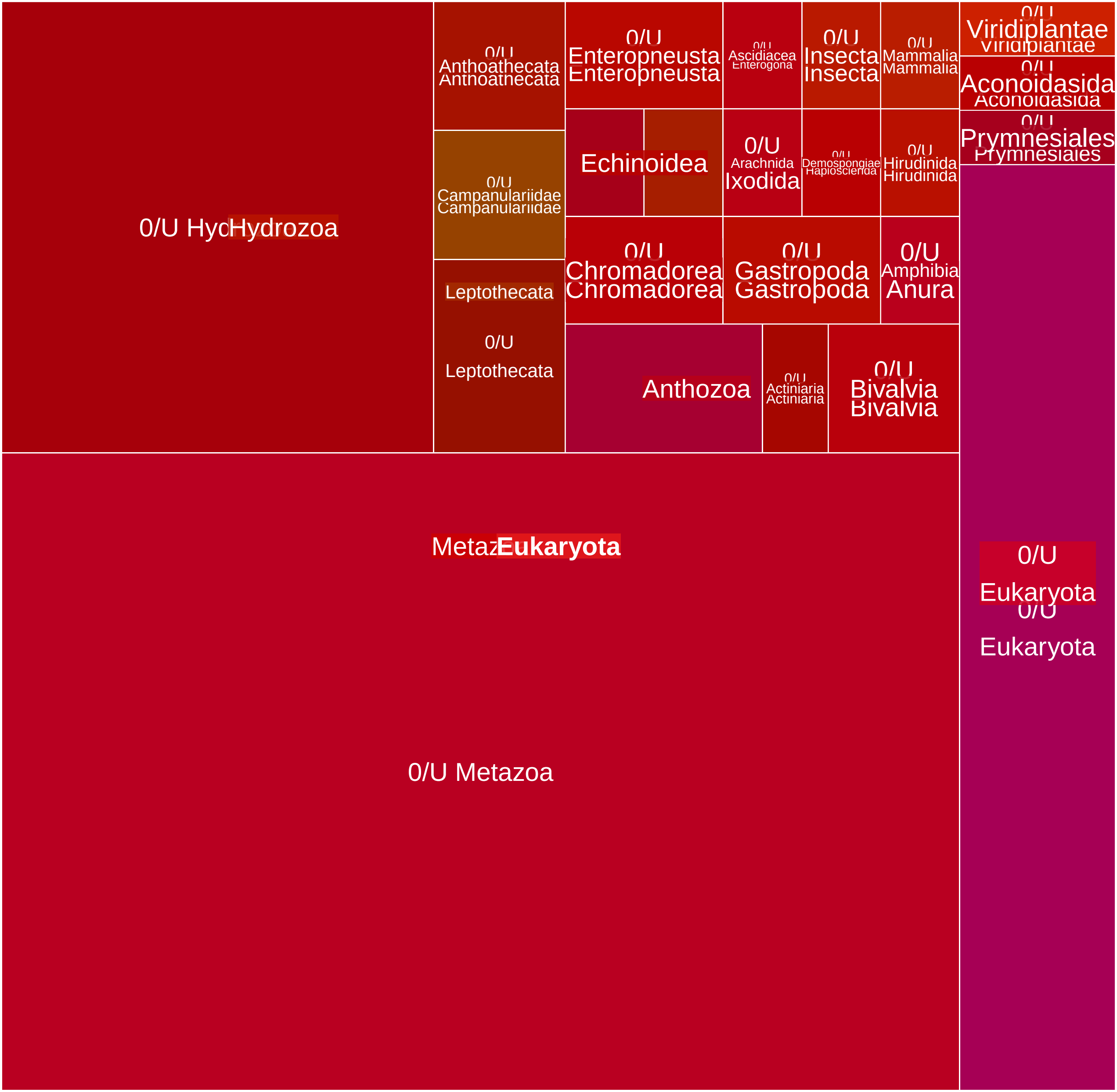

MGT−v1_820 : 627 unigenes, 447 (71.29%) taxonomically assigned

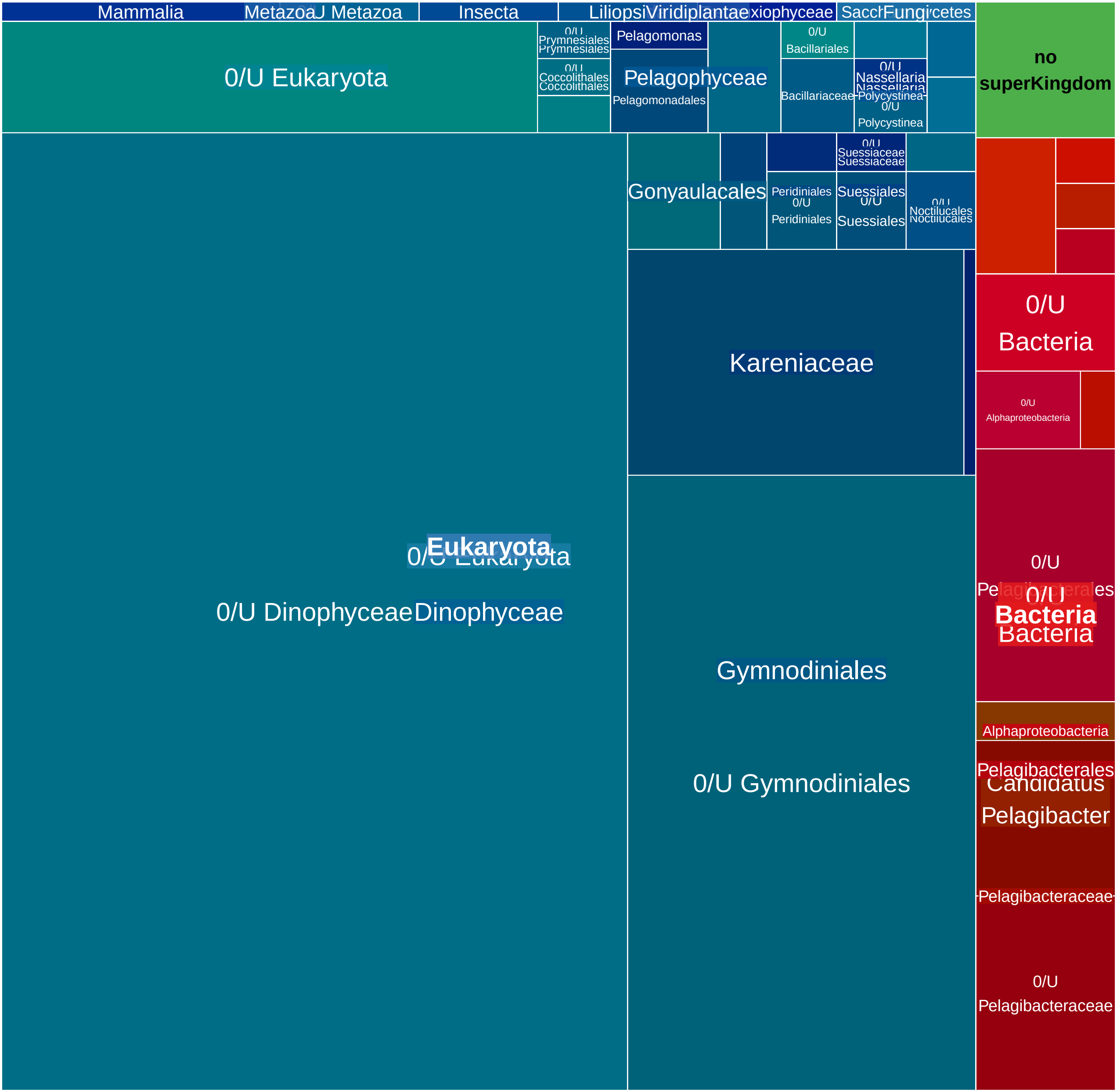

MGT−v1_821 : 626 unigenes, 419 (66.93%) taxonomically assigned

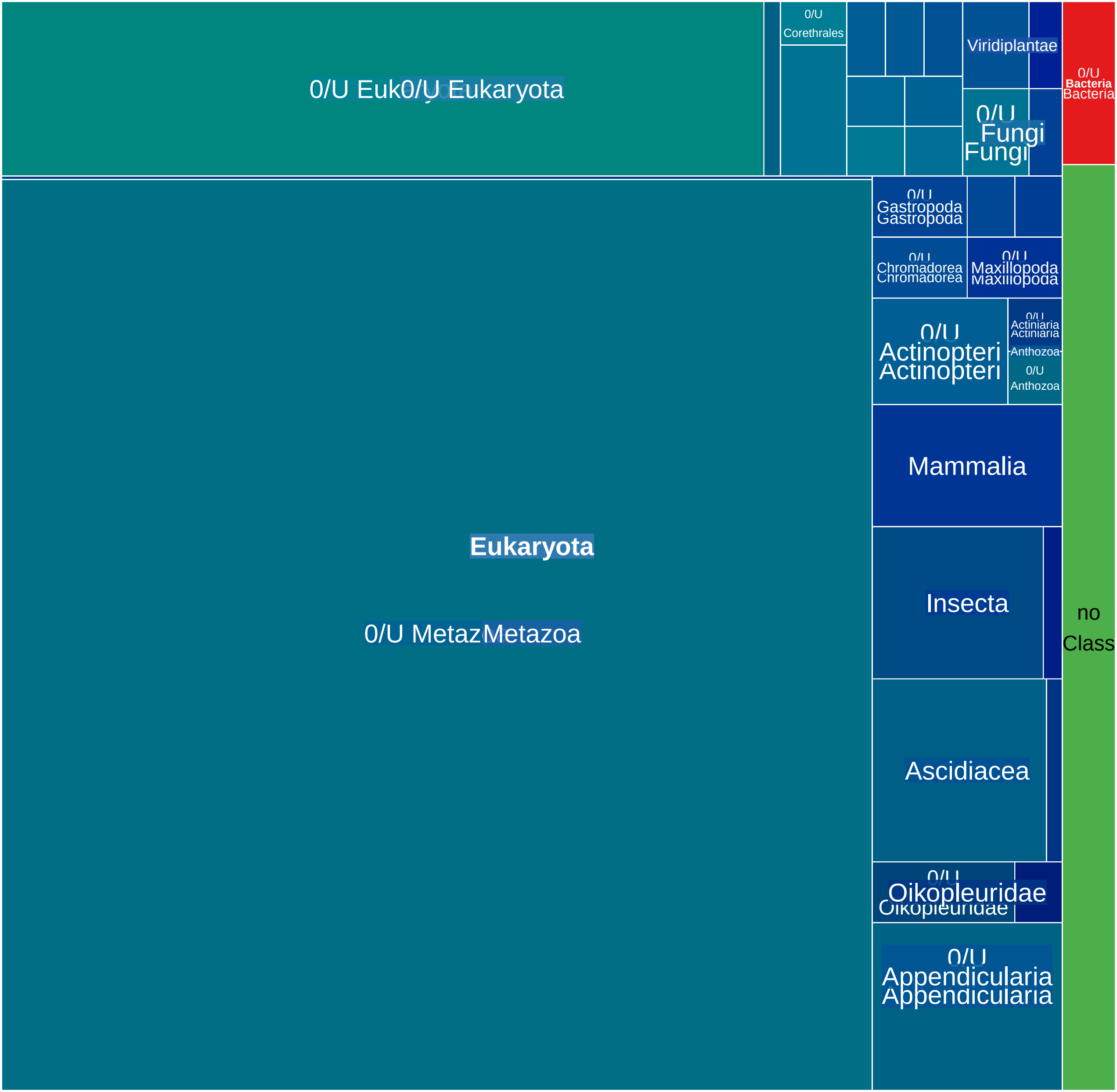

MGT−v1_822 : 626 unigenes, 476 (76.04%) taxonomically assigned

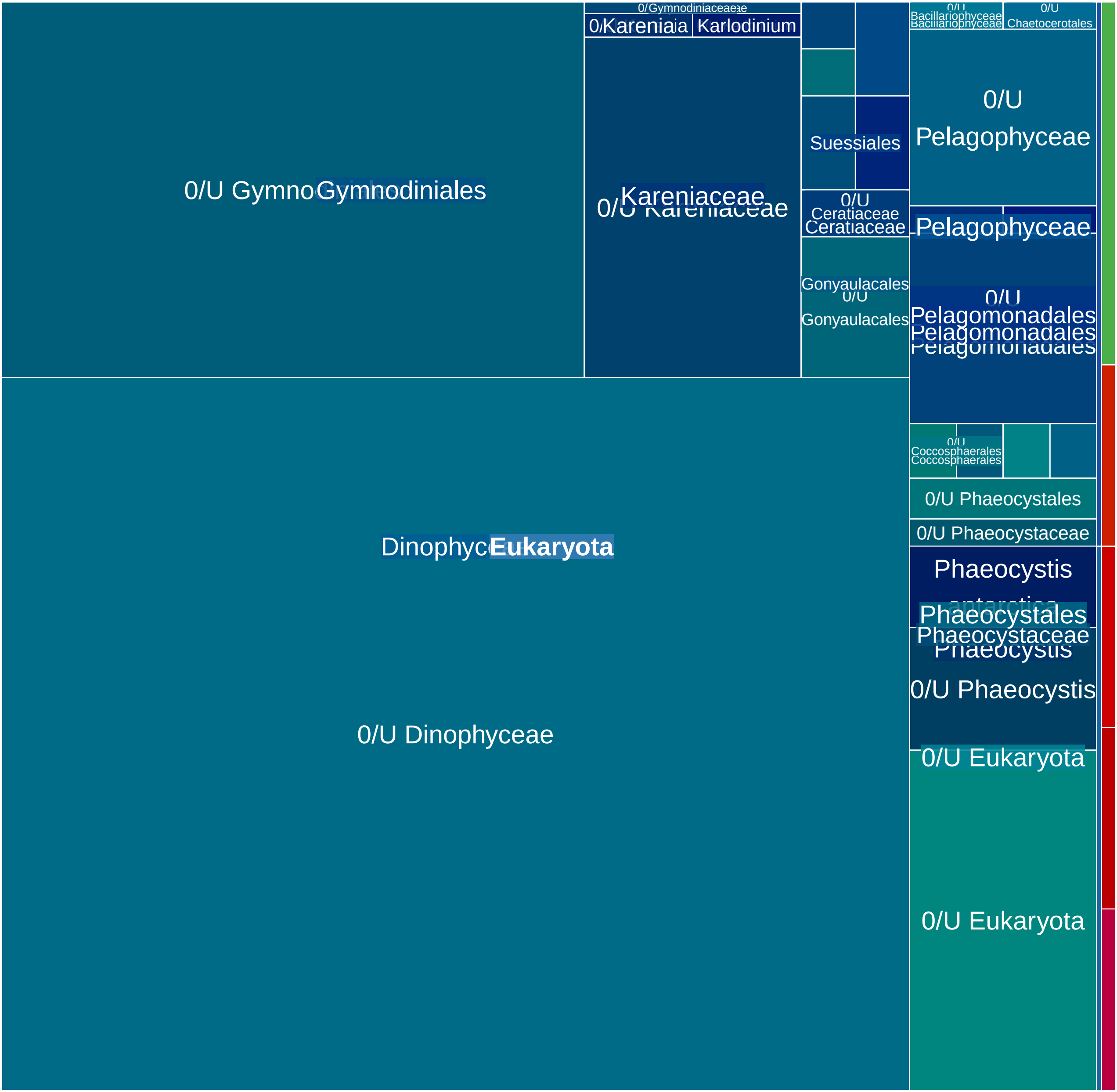

### MGT−v1_823 : 625 unigenes, 395 (63.20%) taxonomically assigned

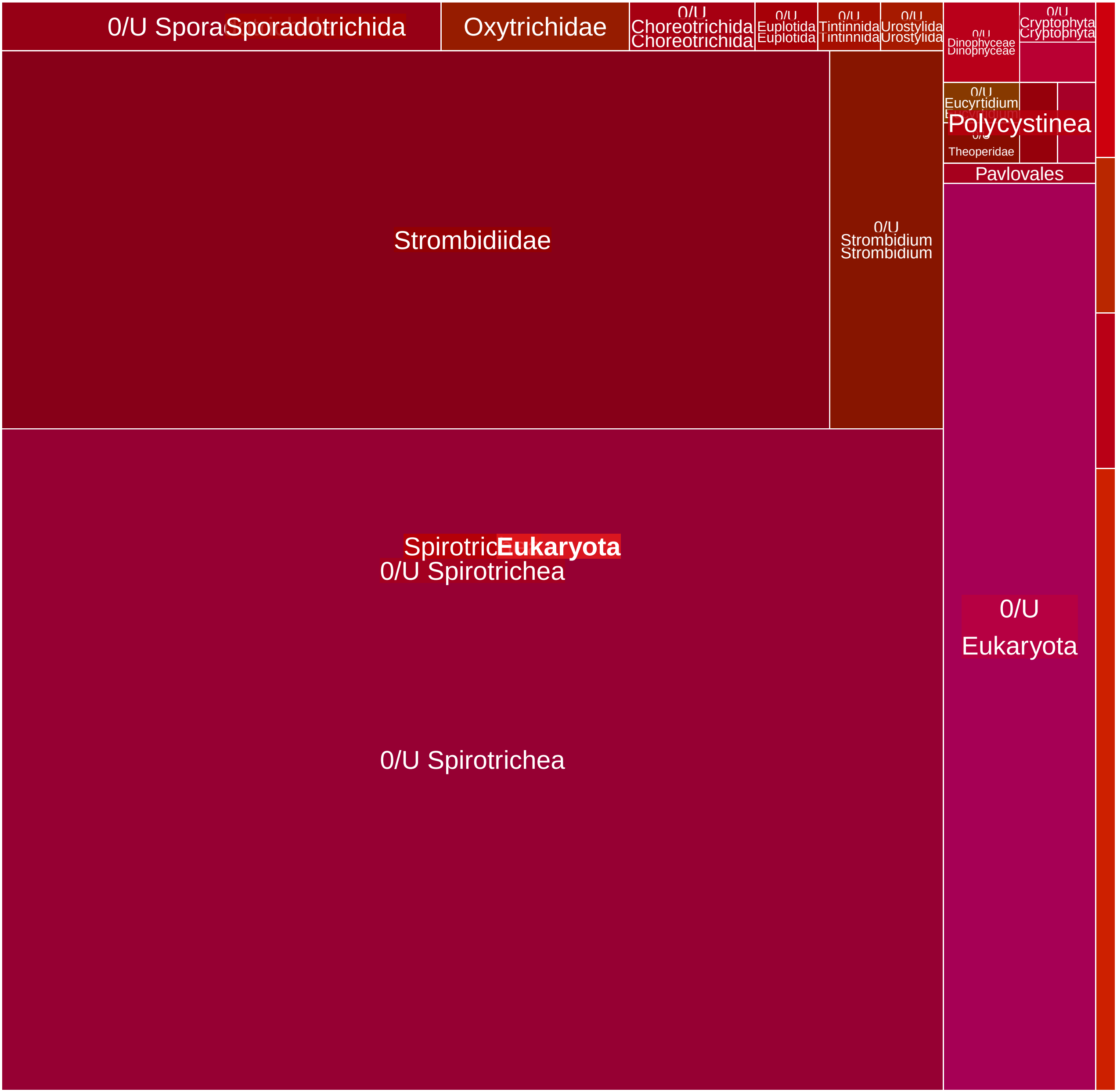

MGT−v1_824 : 625 unigenes, 170 (27.20%) taxonomically assigned

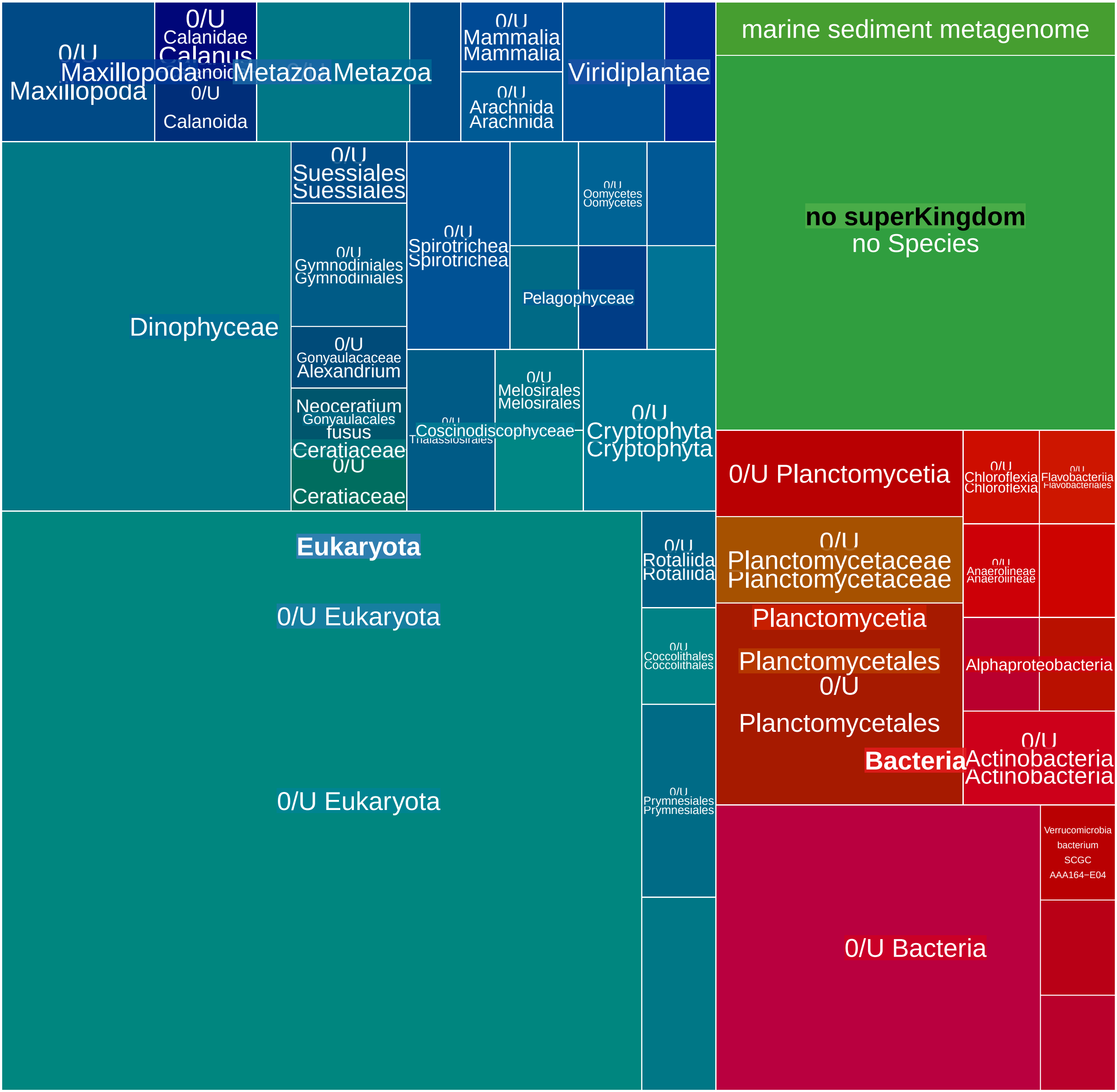

### MGT−v1_825 : 625 unigenes, 32 (5.12%) taxonomically assigned

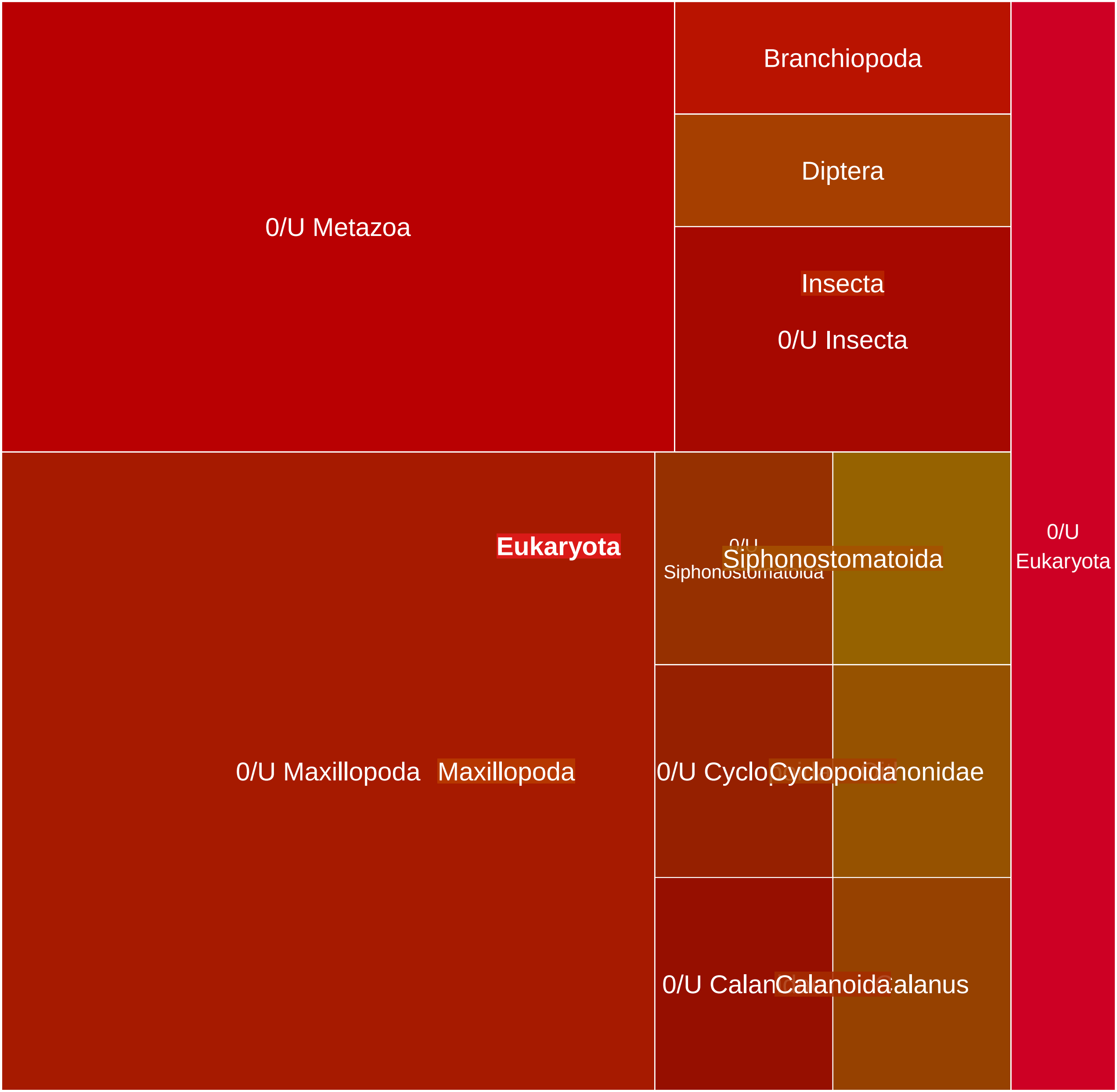

MGT−v1_826 : 624 unigenes, 396 (63.46%) taxonomically assigned

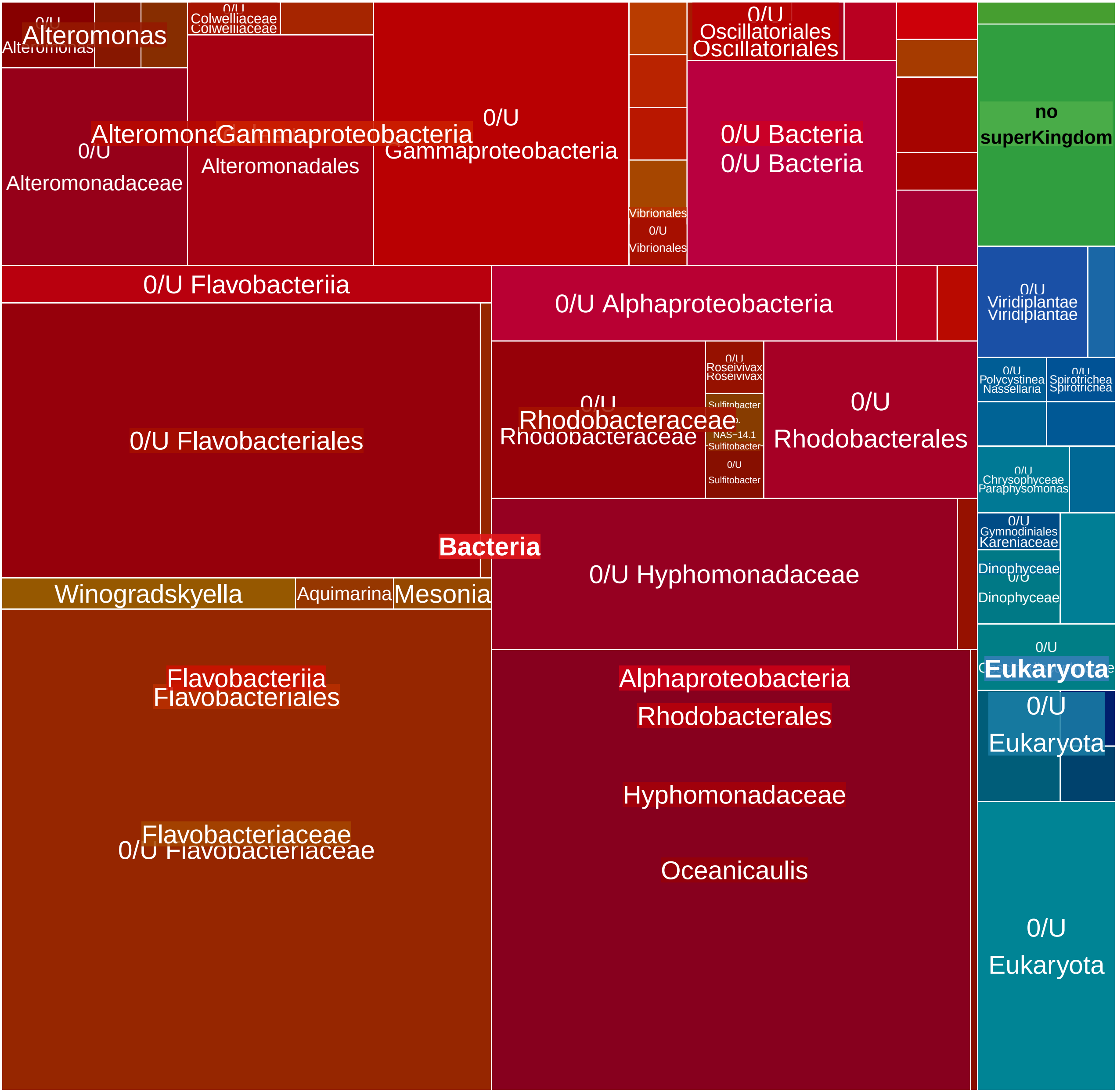

### MGT−v1_827 : 621 unigenes, 401 (64.57%) taxonomically assigned

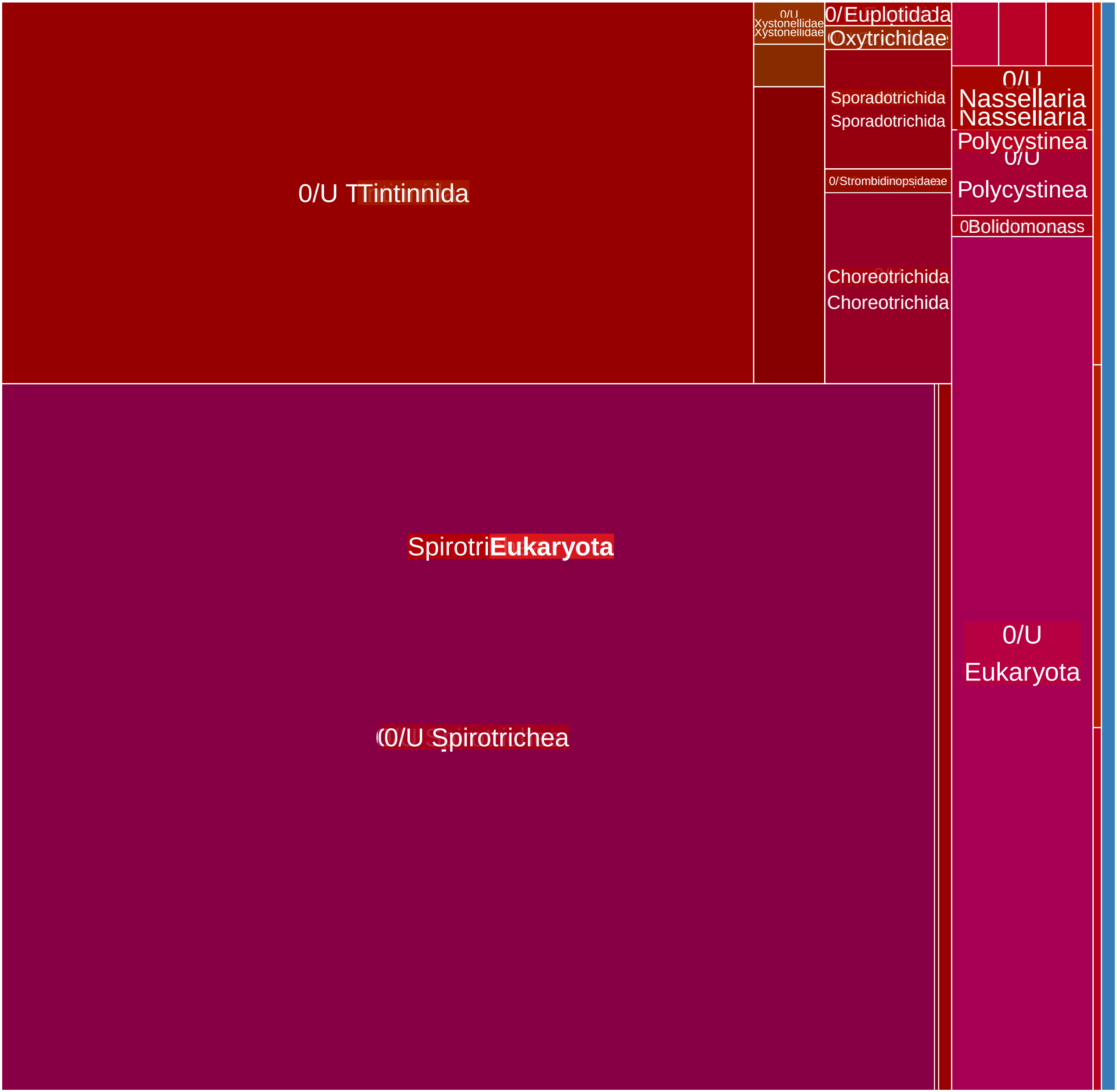

MGT−v1_828 : 619 unigenes, 369 (59.61%) taxonomically assigned

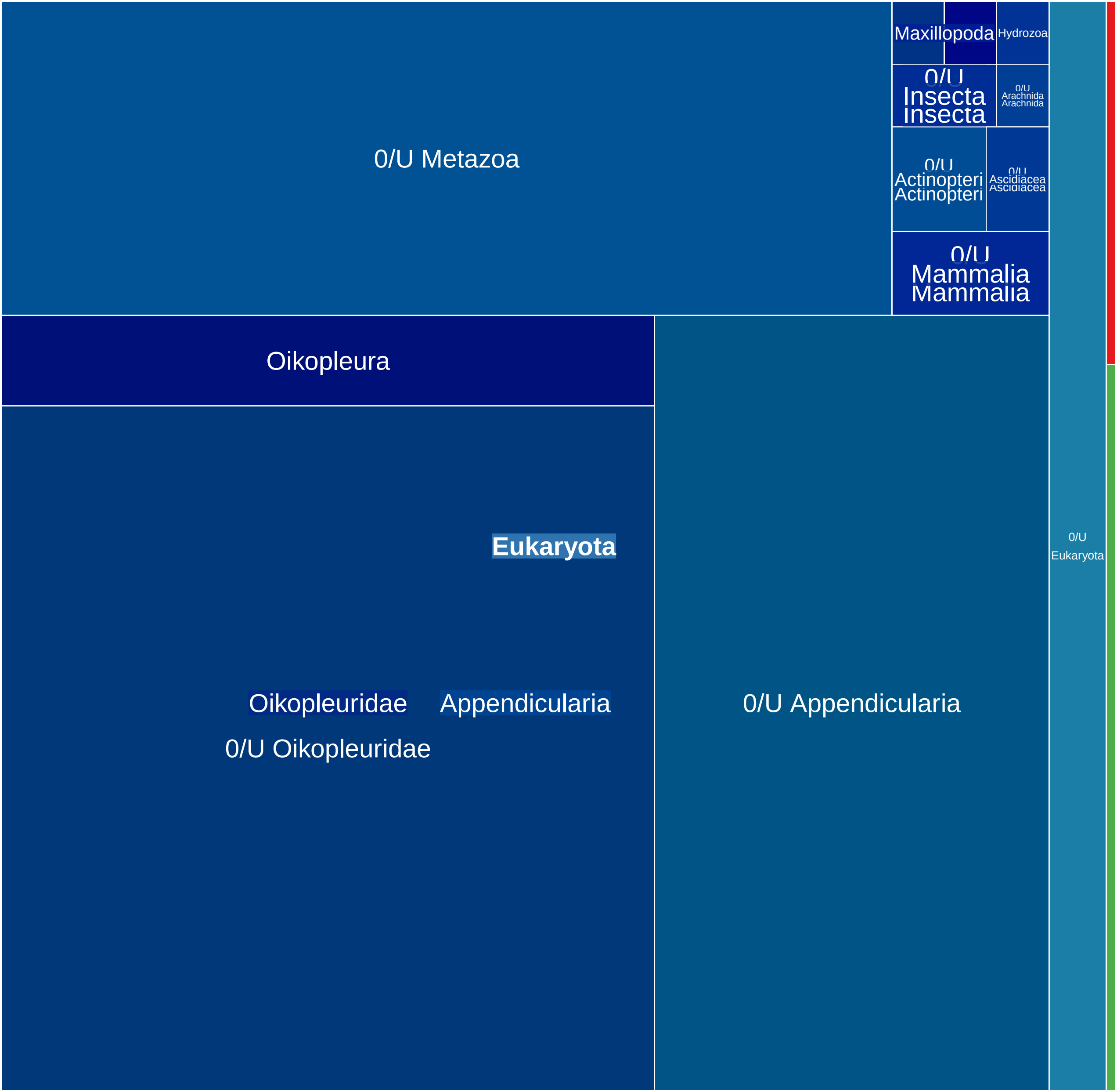

MGT−v1_829 : 619 unigenes, 556 (89.82%) taxonomically assigned

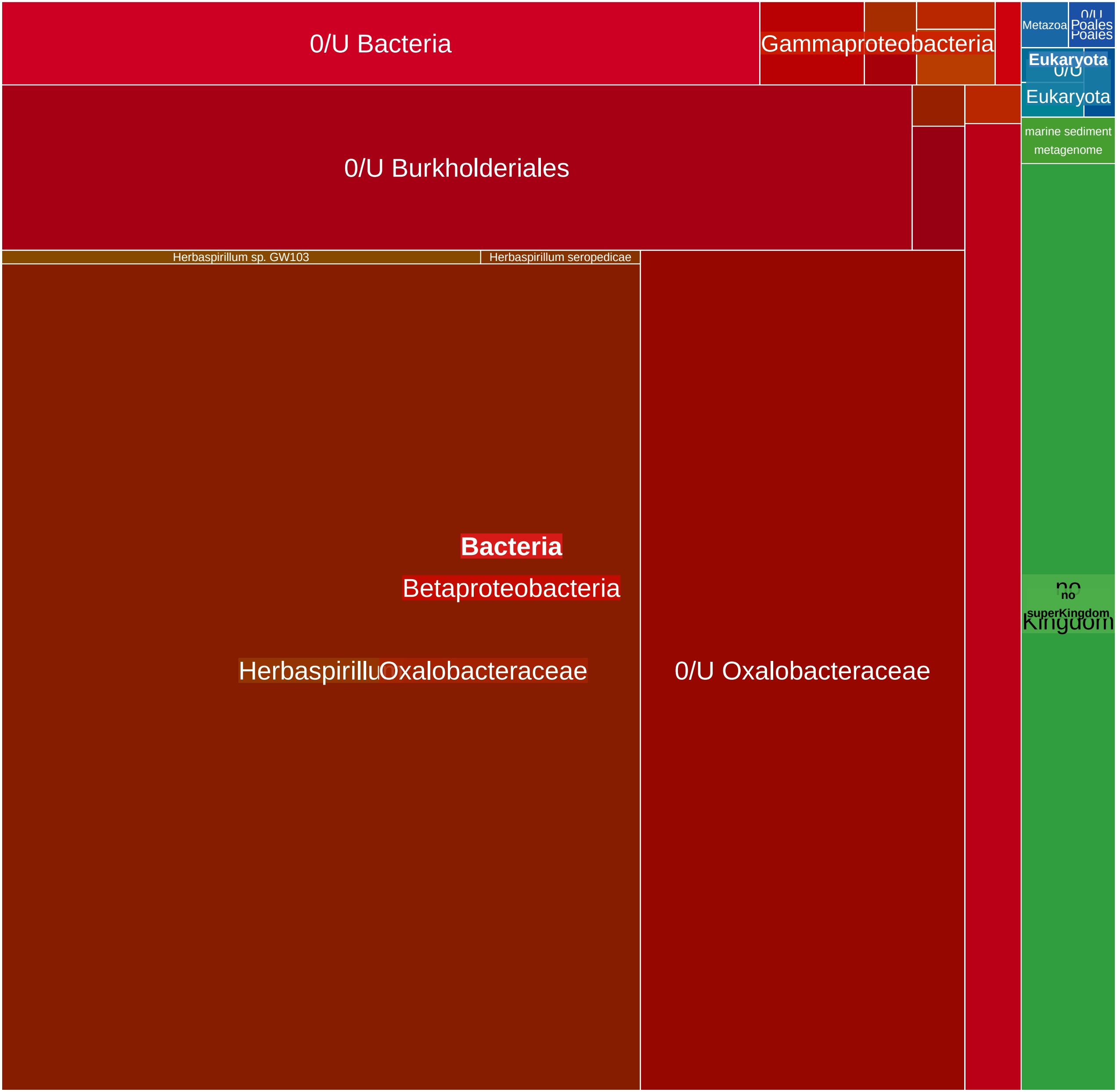

MGT−v1_830 : 616 unigenes, 498 (80.84%) taxonomically assigned

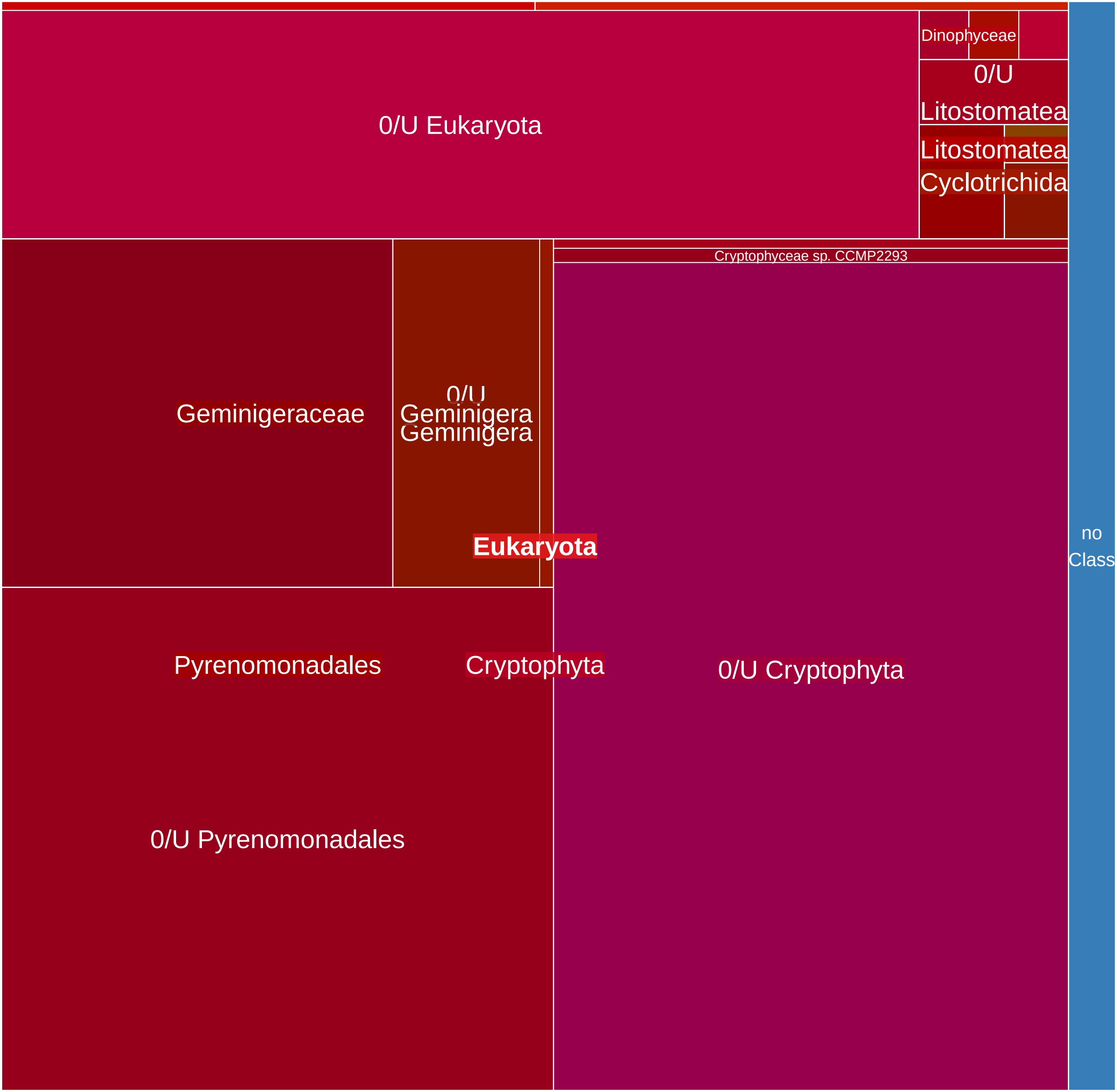

MGT−v1_831 : 615 unigenes, 46 (7.48%) taxonomically assigned

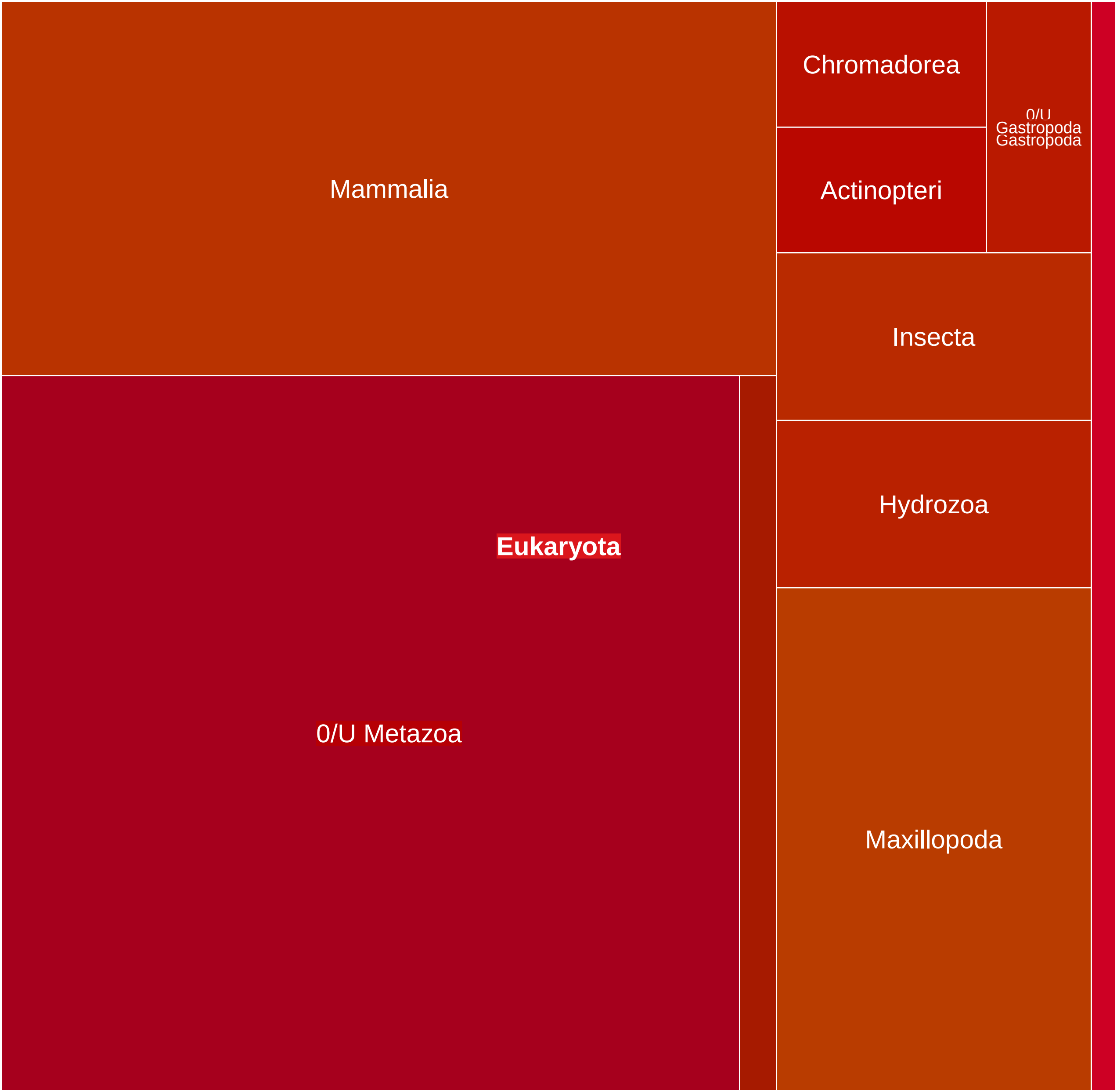

MGT−v1_832 : 614 unigenes, 197 (32.08%) taxonomically assigned

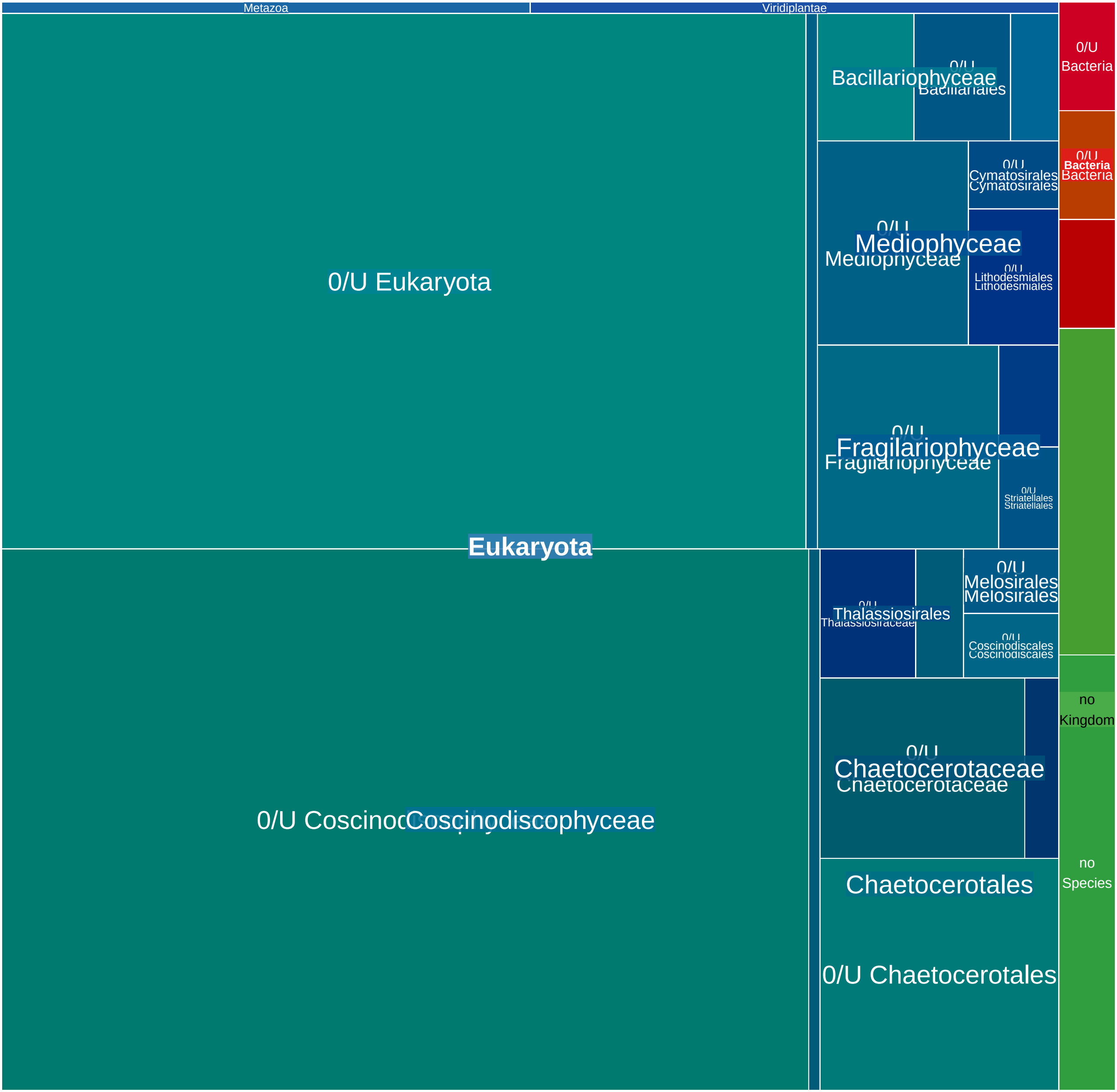

MGT−v1_833 : 614 unigenes, 141 (22.96%) taxonomically assigned

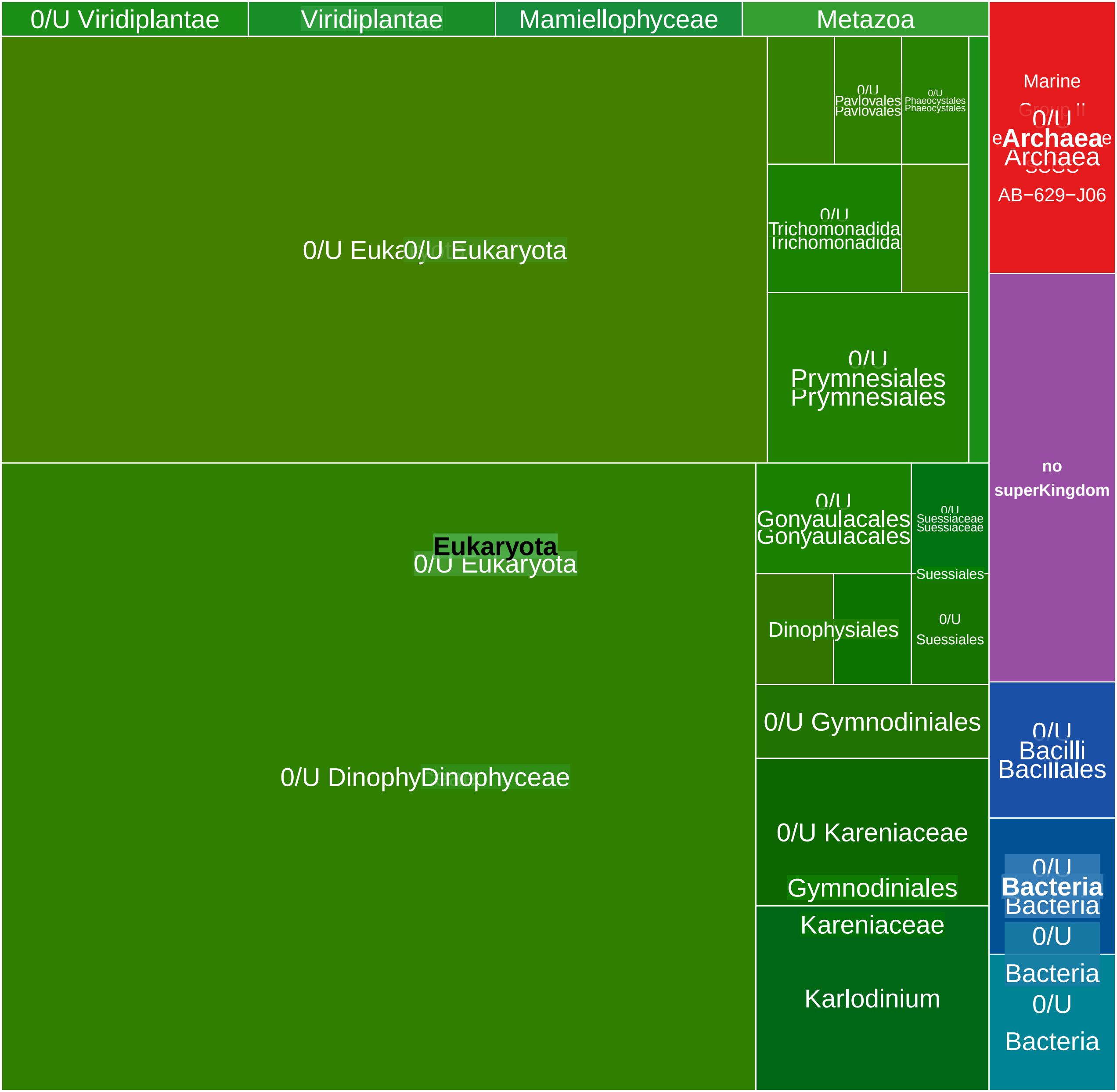

MGT−v1_834 : 614 unigenes, 371 (60.42%) taxonomically assigned

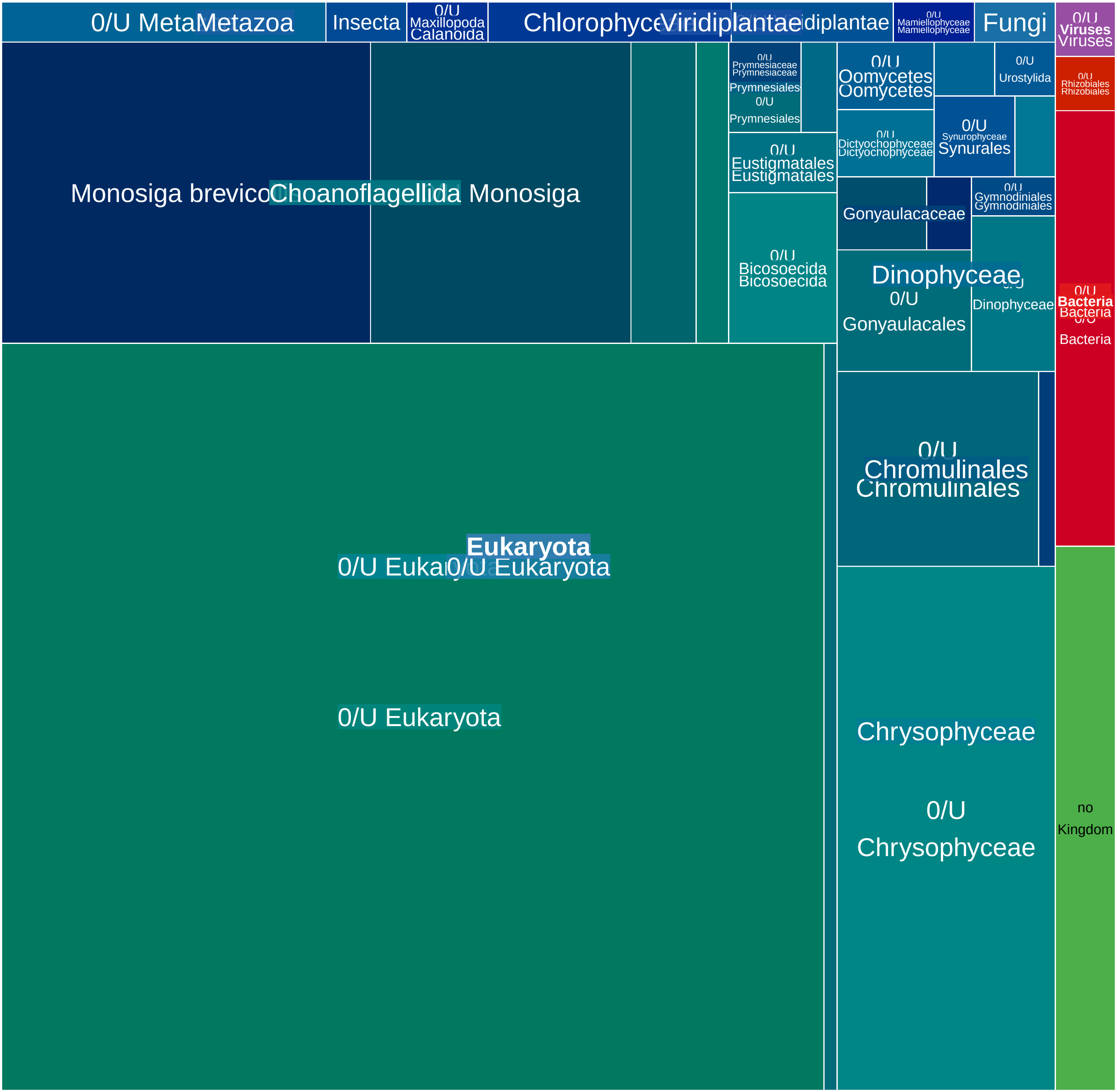

MGT−v1_835 : 611 unigenes, 44 (7.20%) taxonomically assigned

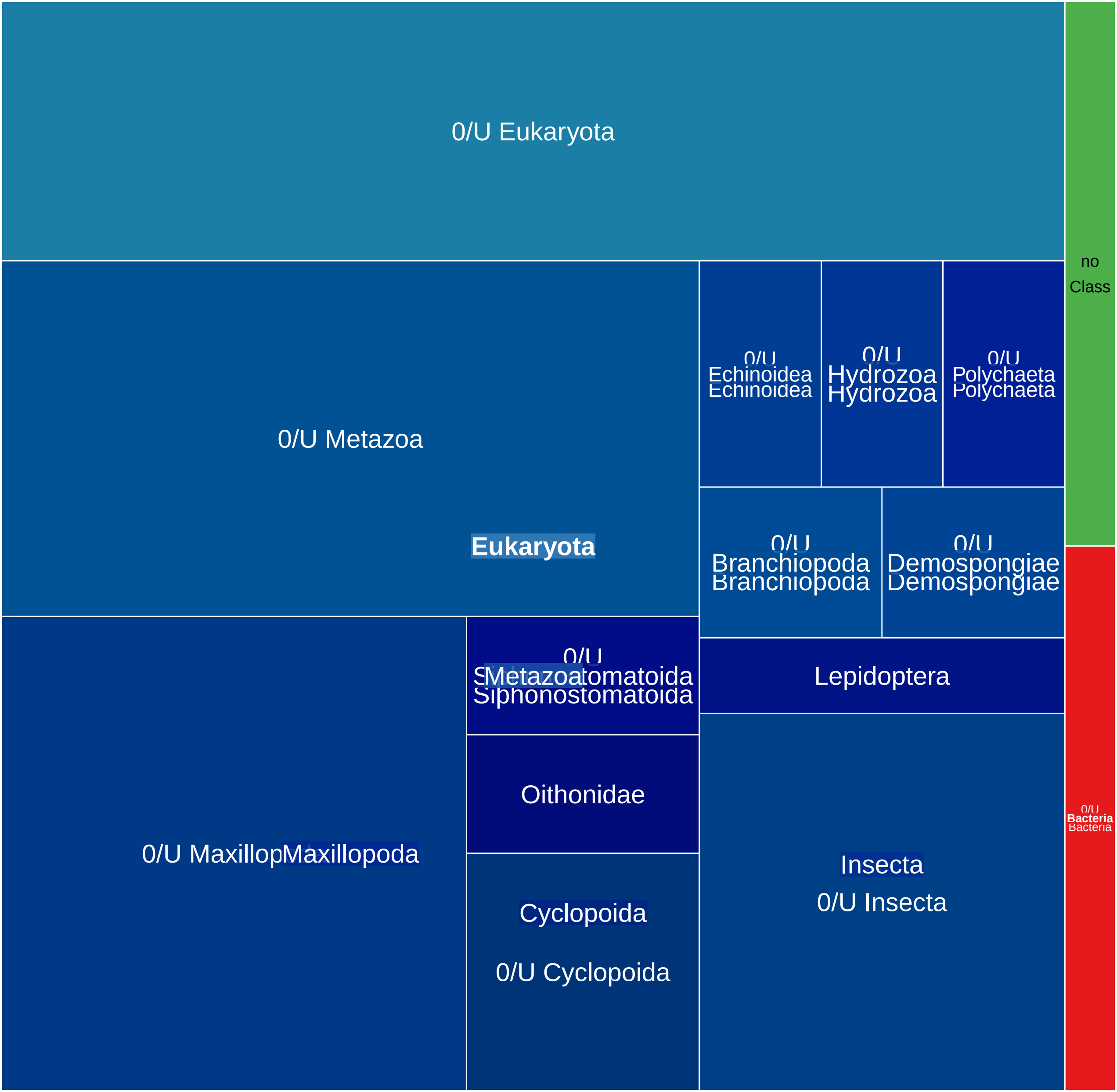

MGT−v1_836 : 610 unigenes, 404 (66.23%) taxonomically assigned

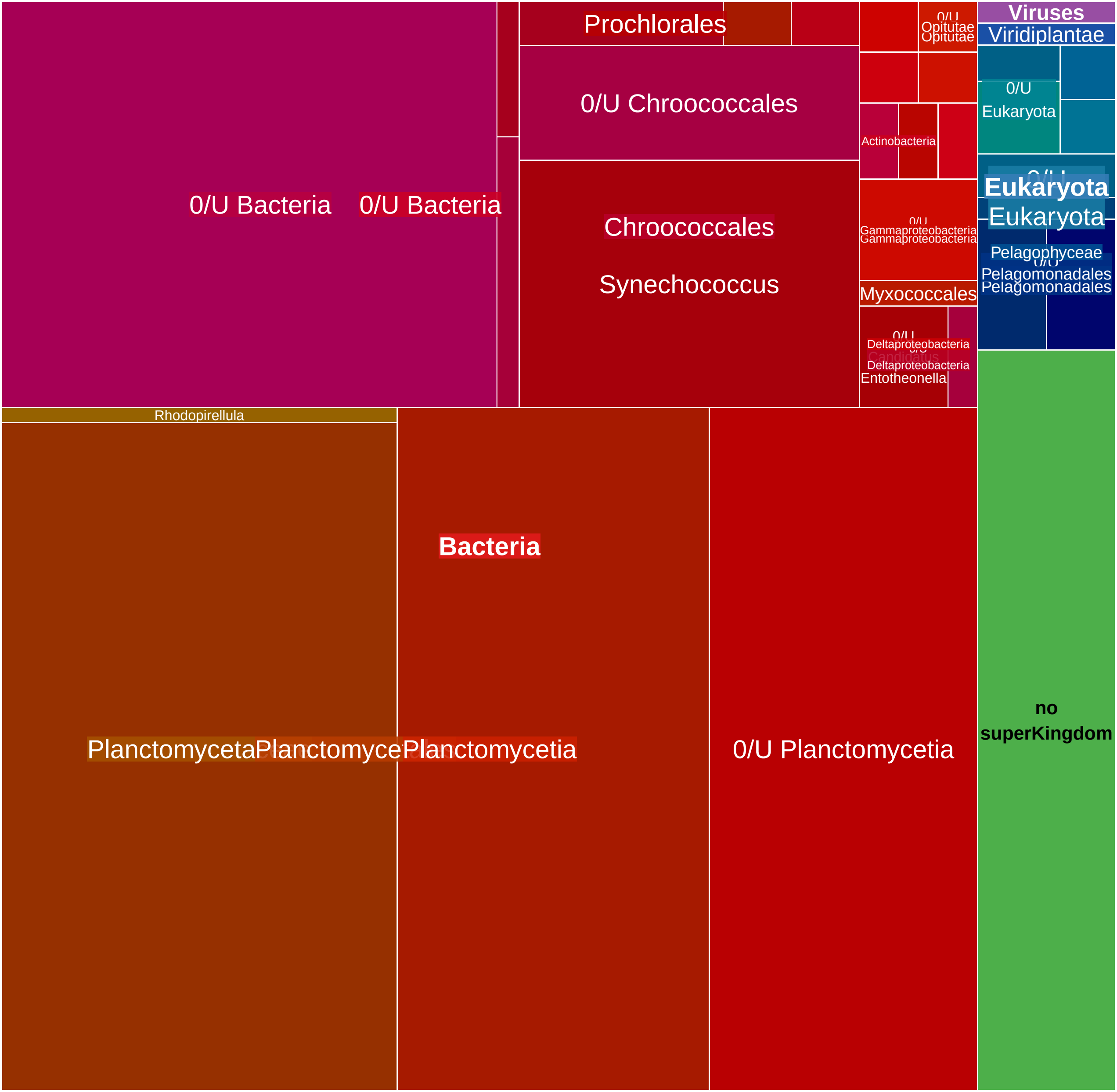

### MGT−v1_837 : 608 unigenes, 366 (60.20%) taxonomically assigned

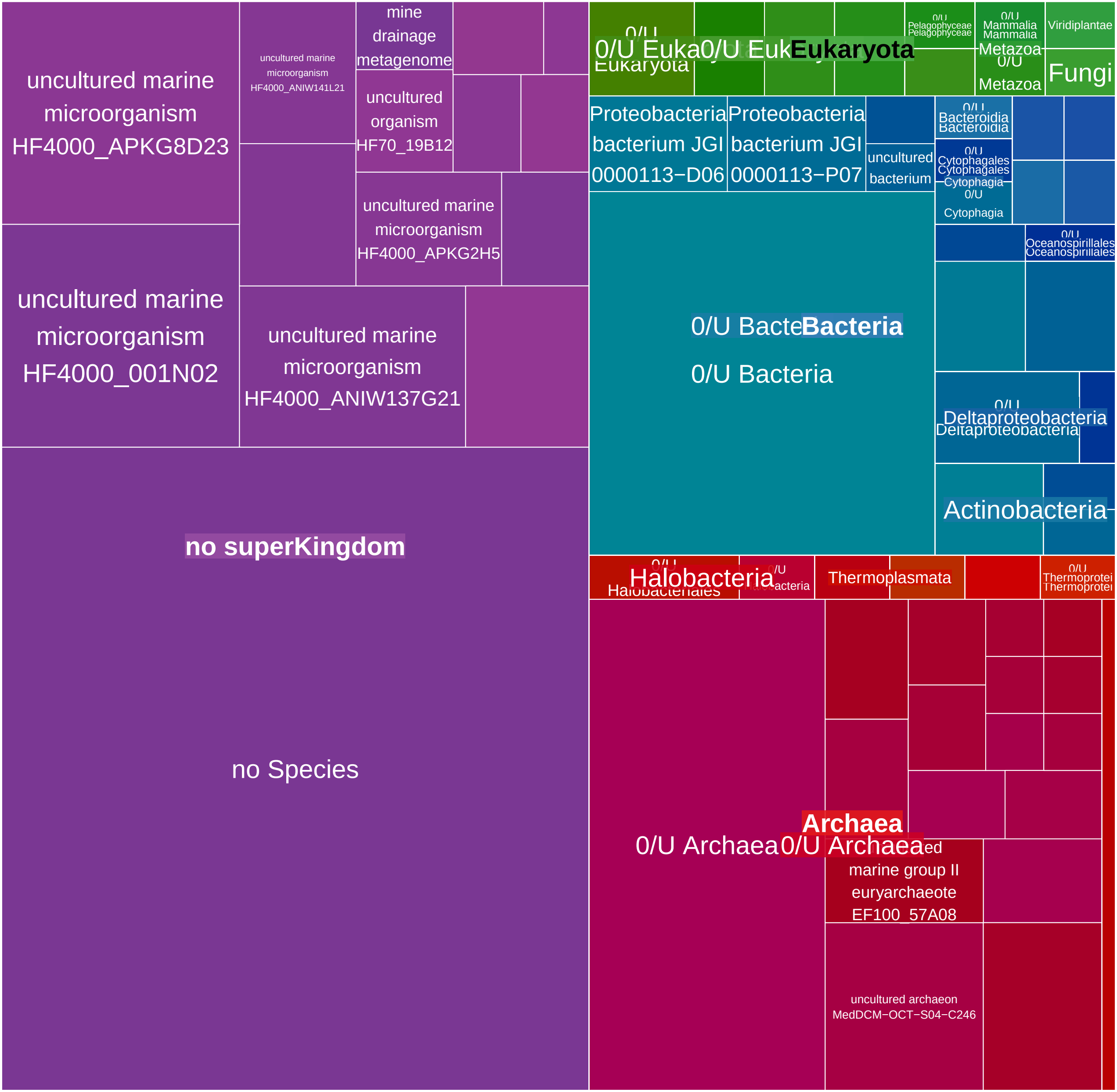

MGT−v1_838 : 607 unigenes, 37 (6.10%) taxonomically assigned

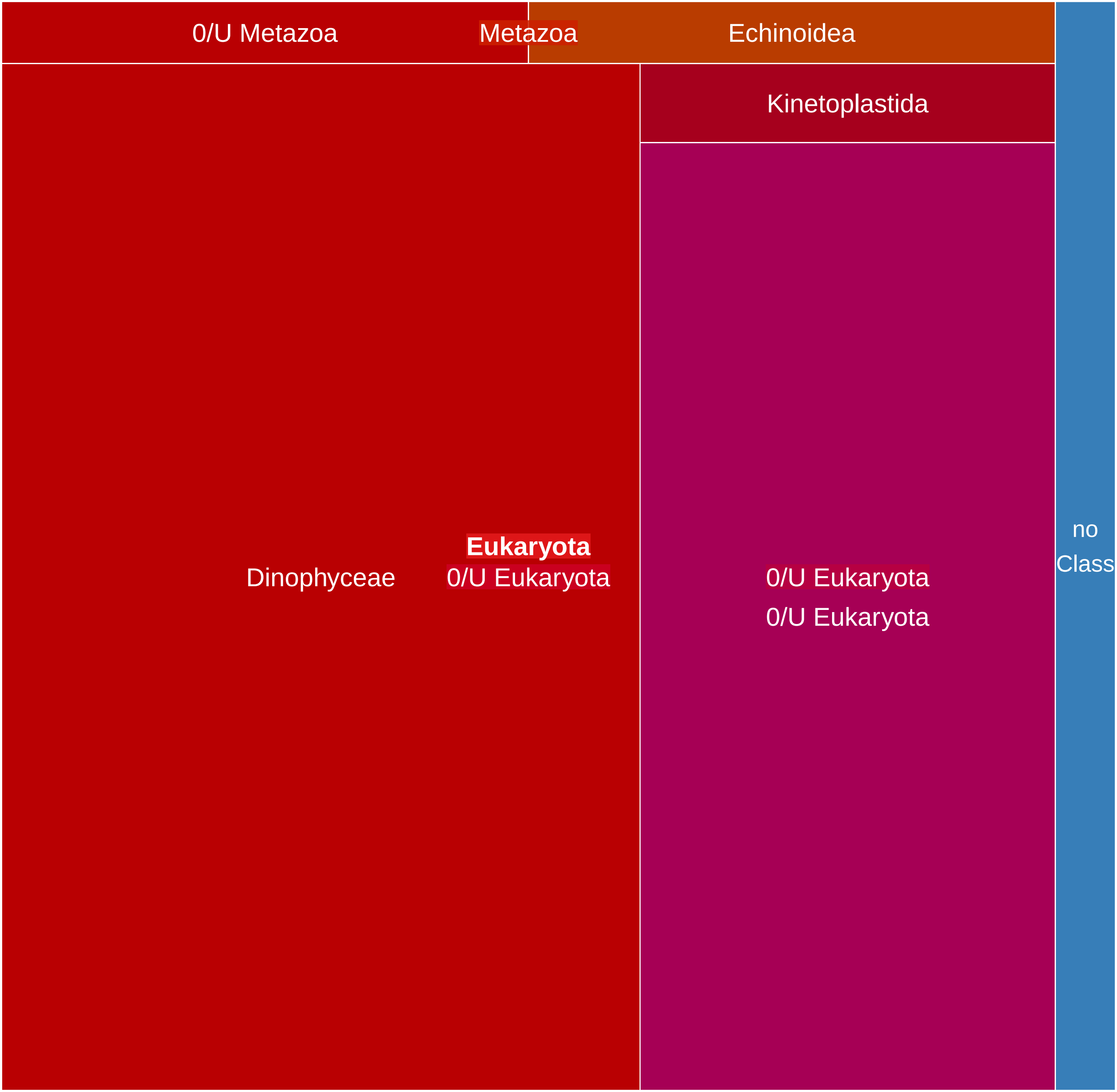

MGT−v1_839 : 607 unigenes, 85 (14.00%) taxonomically assigned

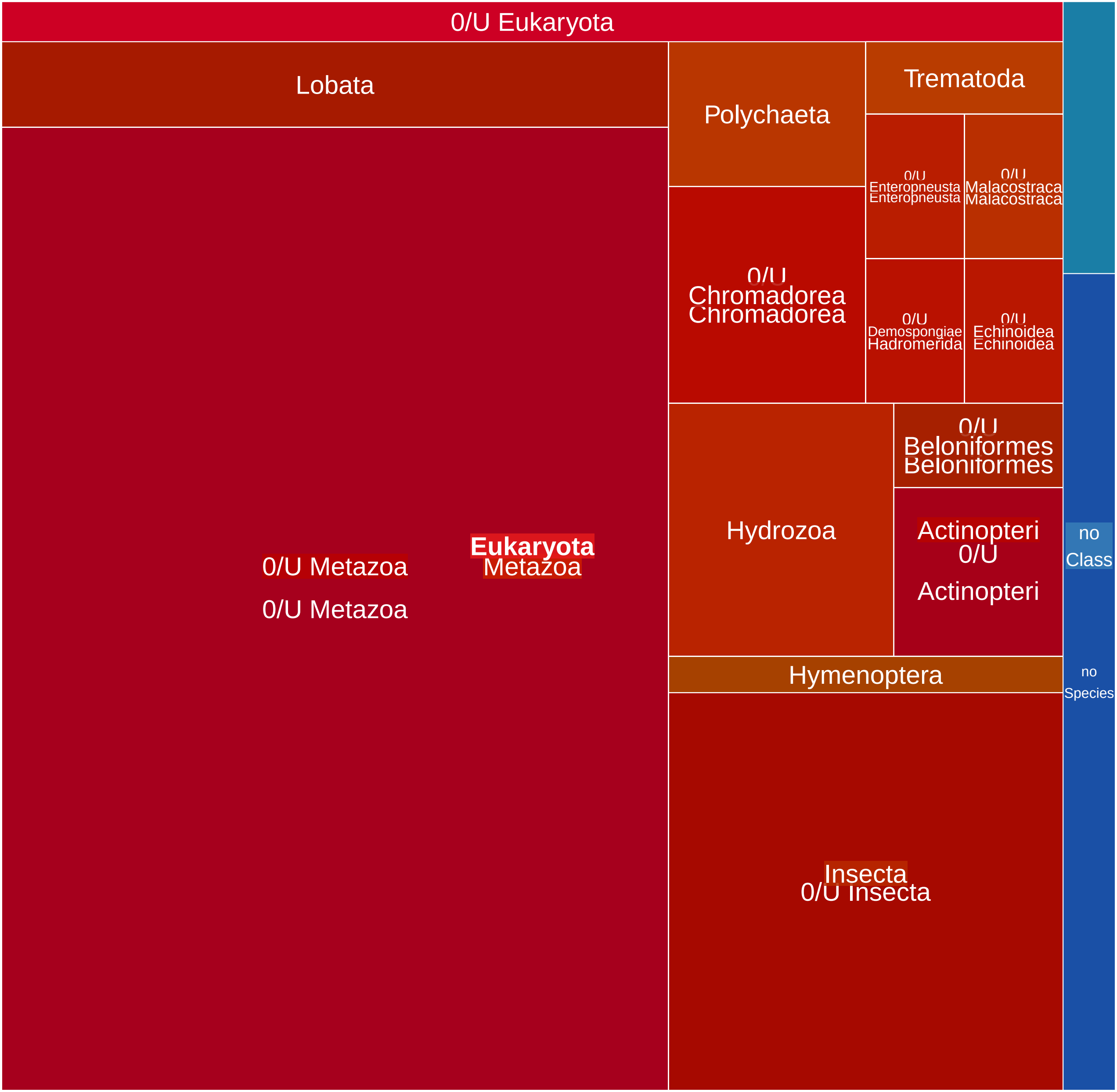

MGT−v1_840 : 605 unigenes, 251 (41.49%) taxonomically assigned

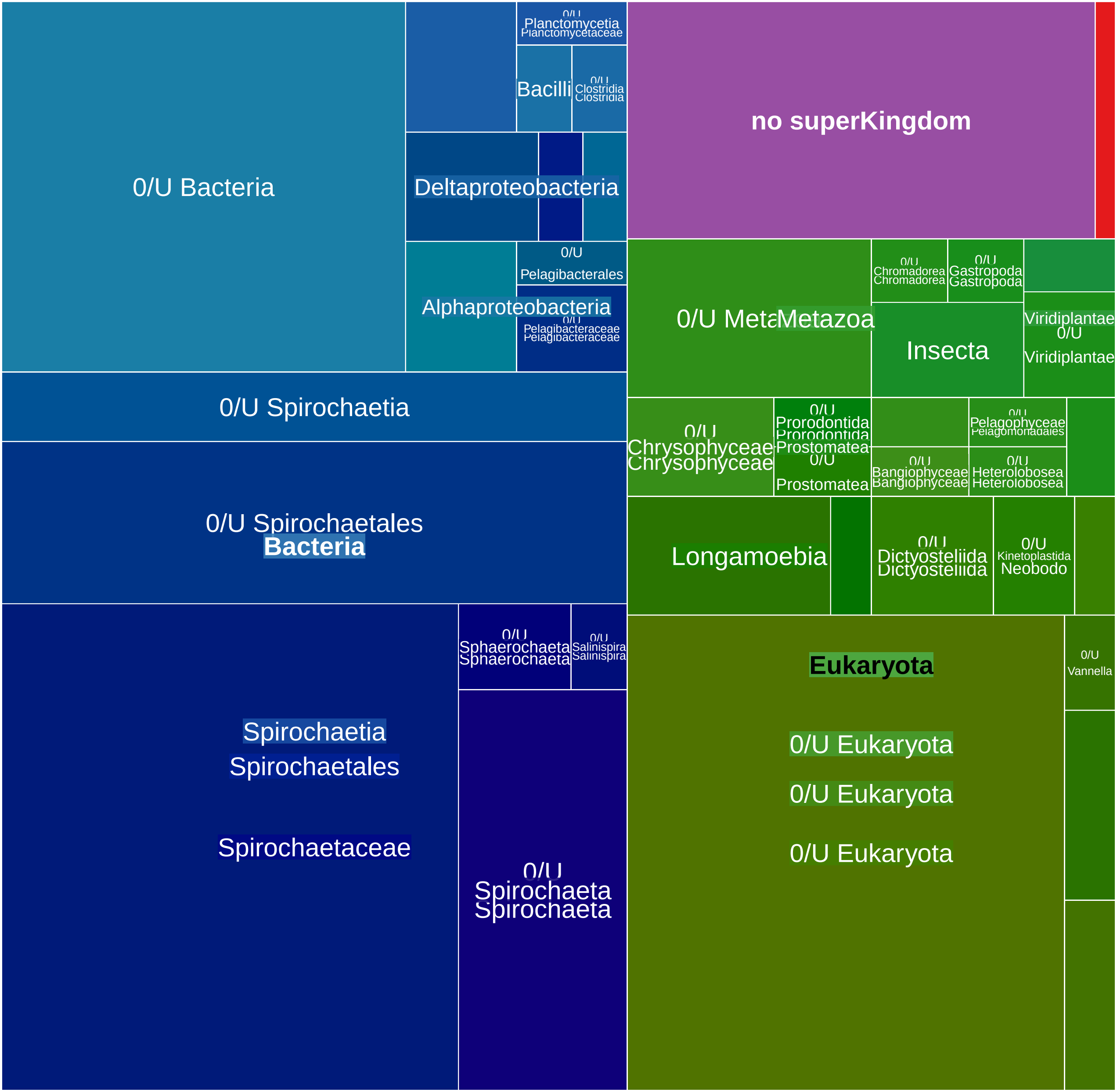

MGT−v1_841 : 604 unigenes, 77 (12.75%) taxonomically assigned

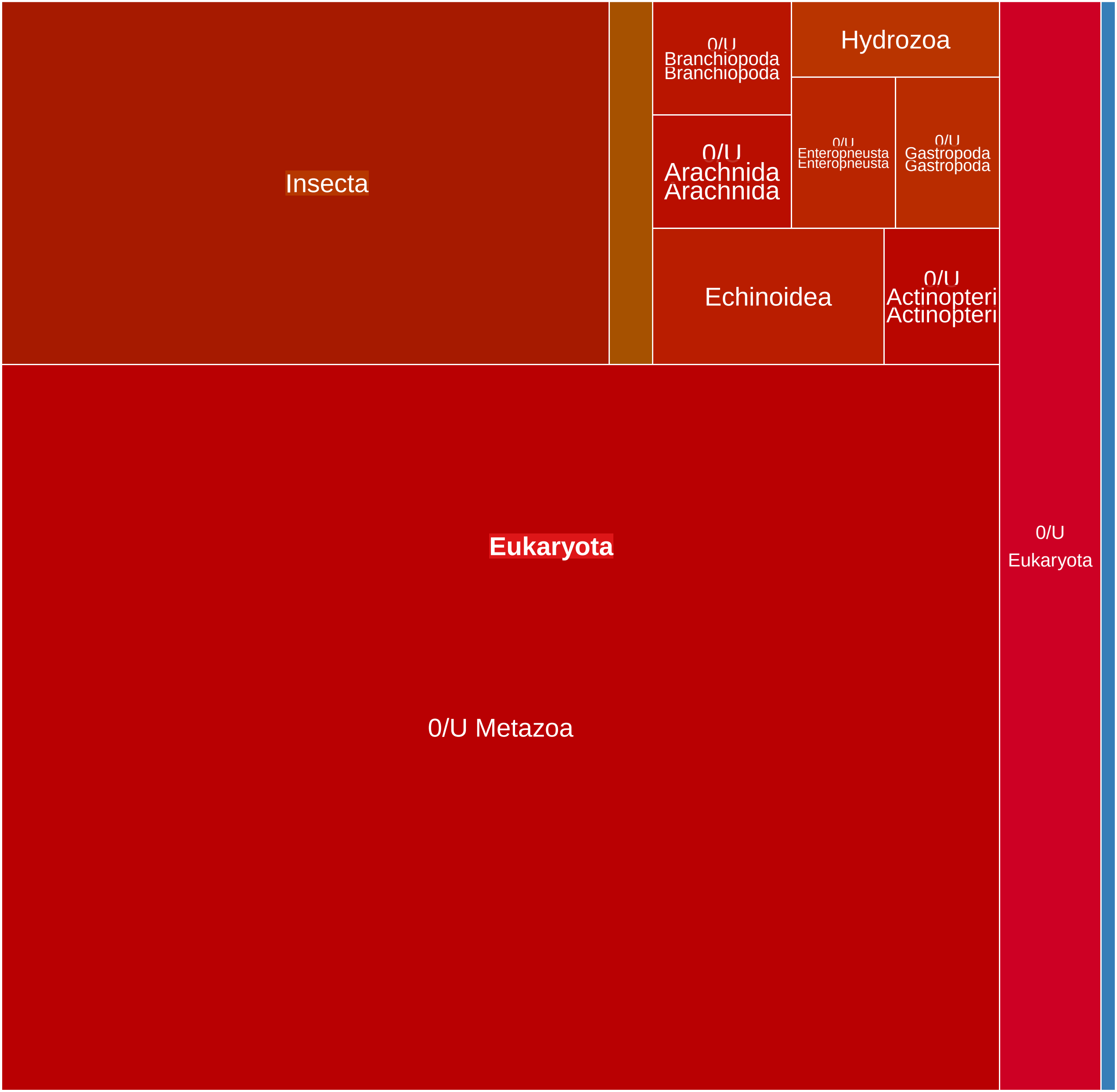

MGT−v1_842 : 599 unigenes, 416 (69.45%) taxonomically assigned

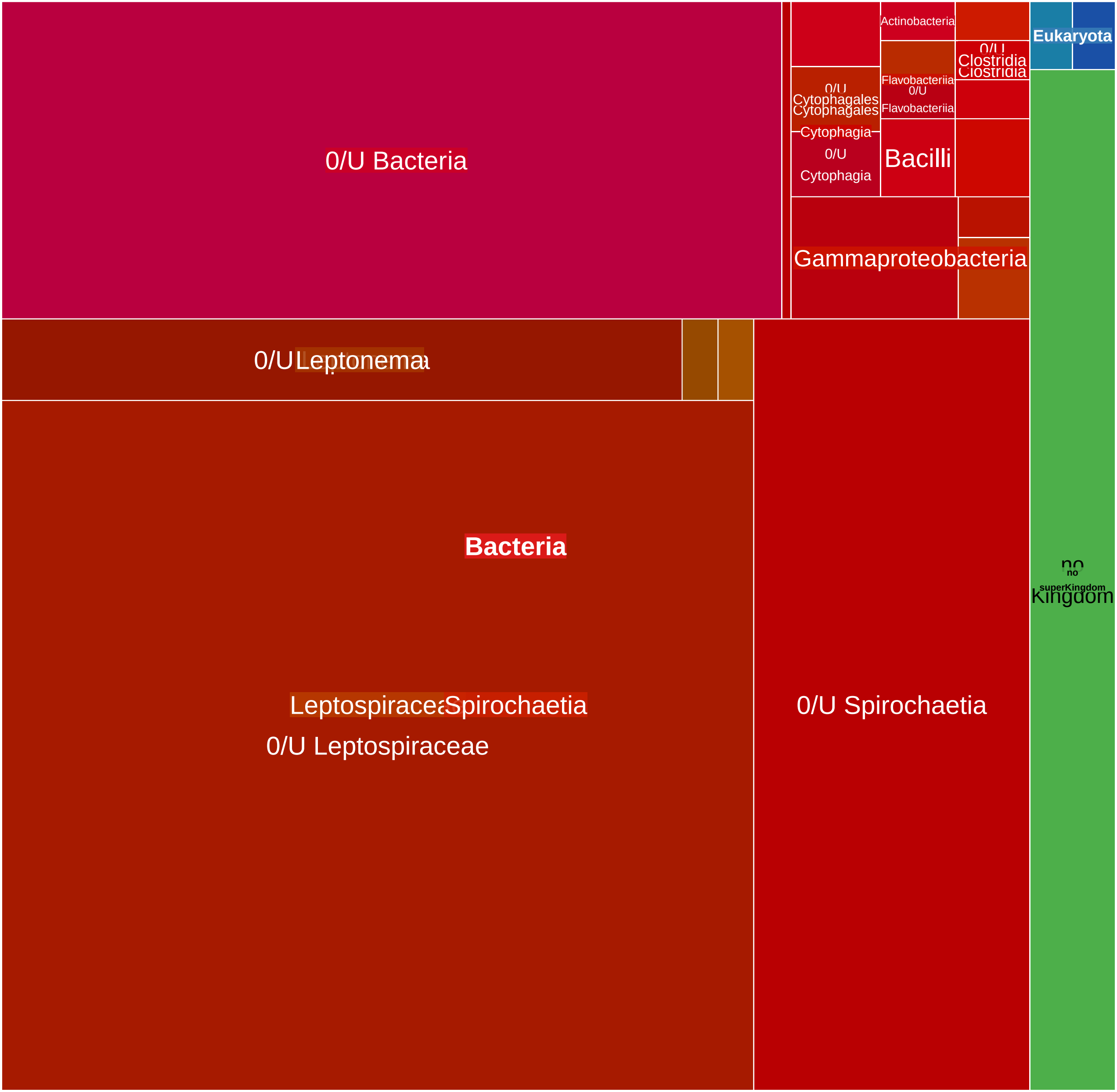

MGT−v1_843 : 596 unigenes, 134 (22.48%) taxonomically assigned

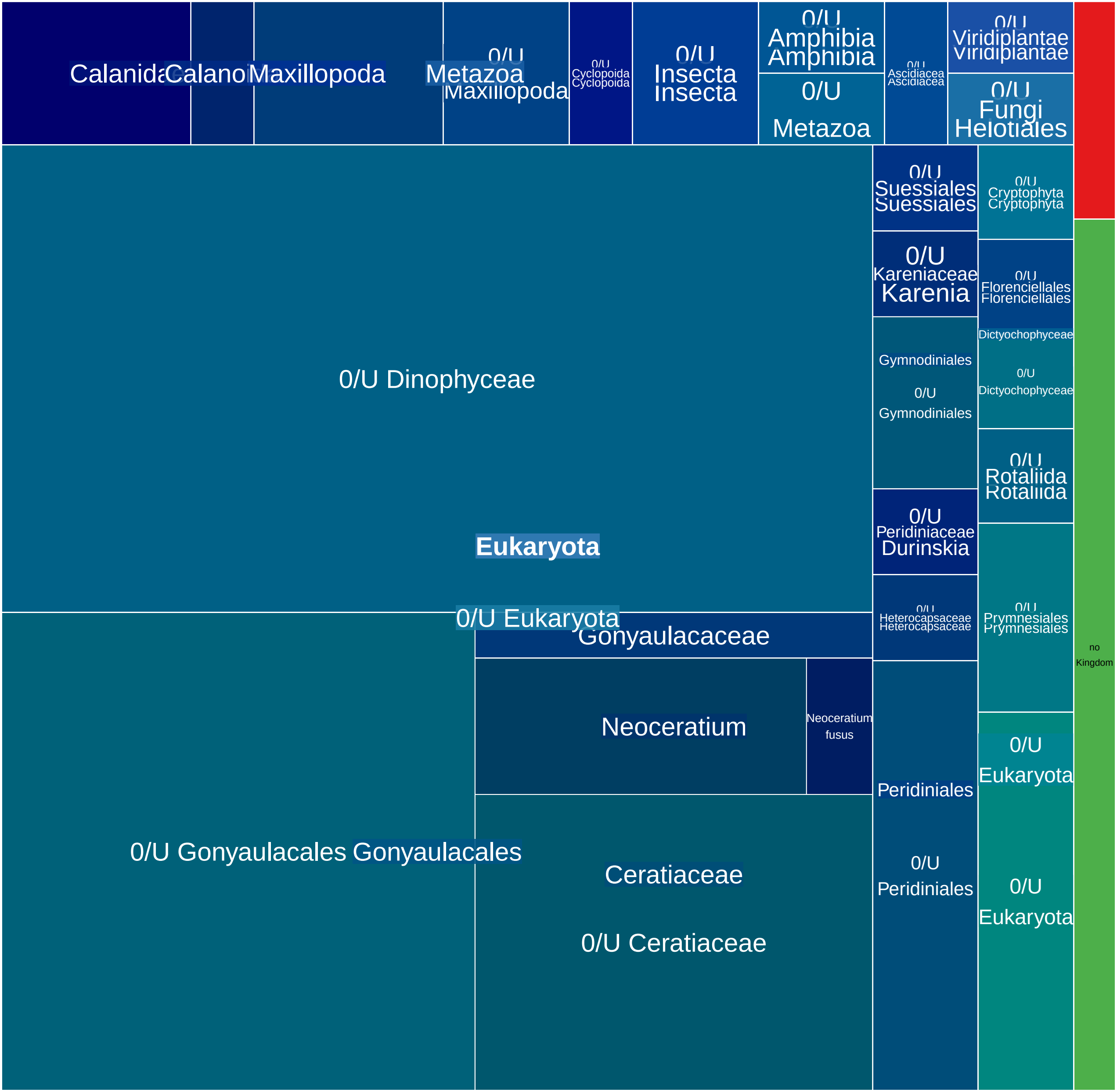

MGT−v1_844 : 595 unigenes, 25 (4.20%) taxonomically assigned

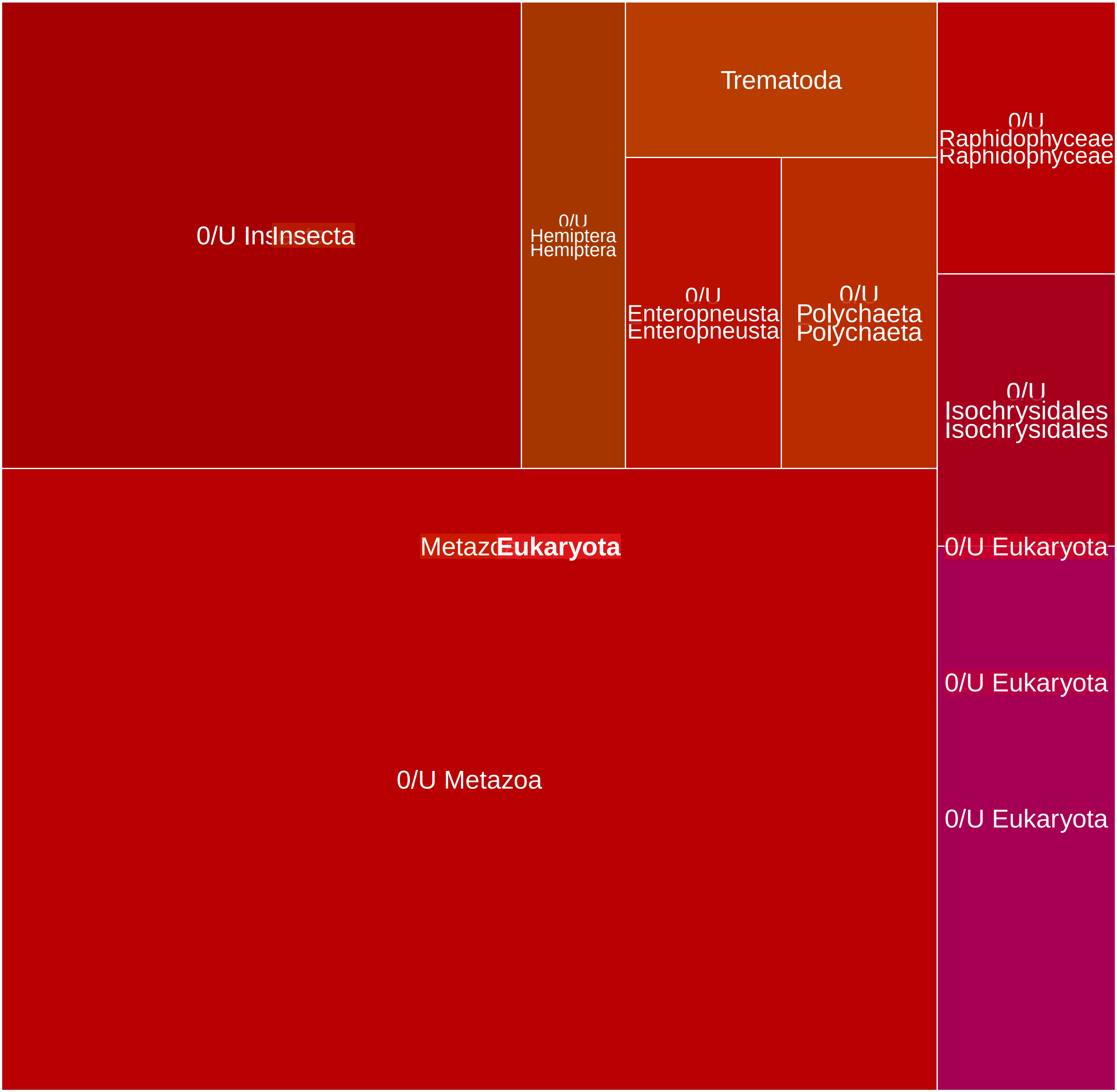

MGT−v1_845 : 594 unigenes, 31 (5.22%) taxonomically assigned

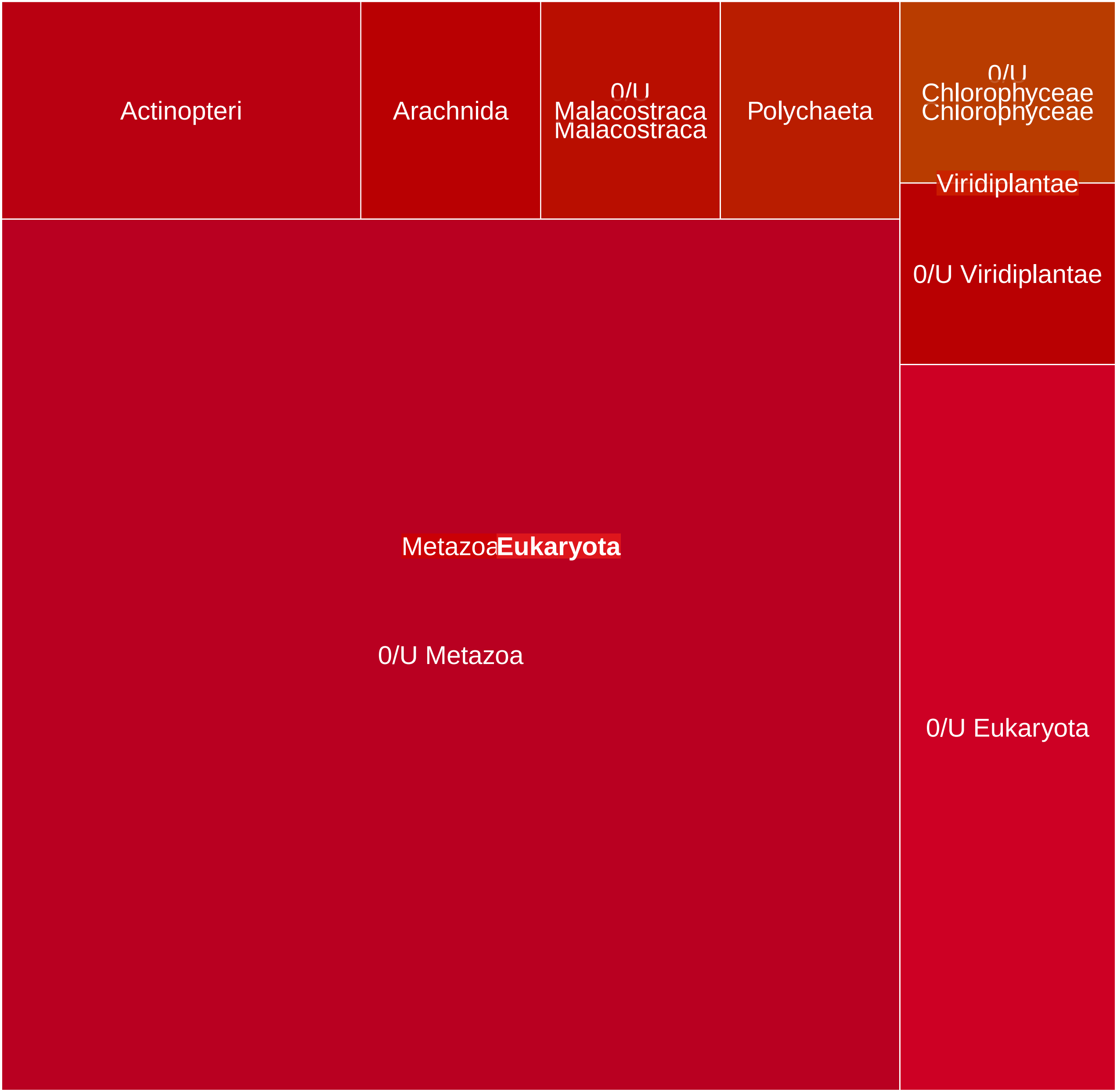

MGT−v1_846 : 591 unigenes, 295 (49.92%) taxonomically assigned

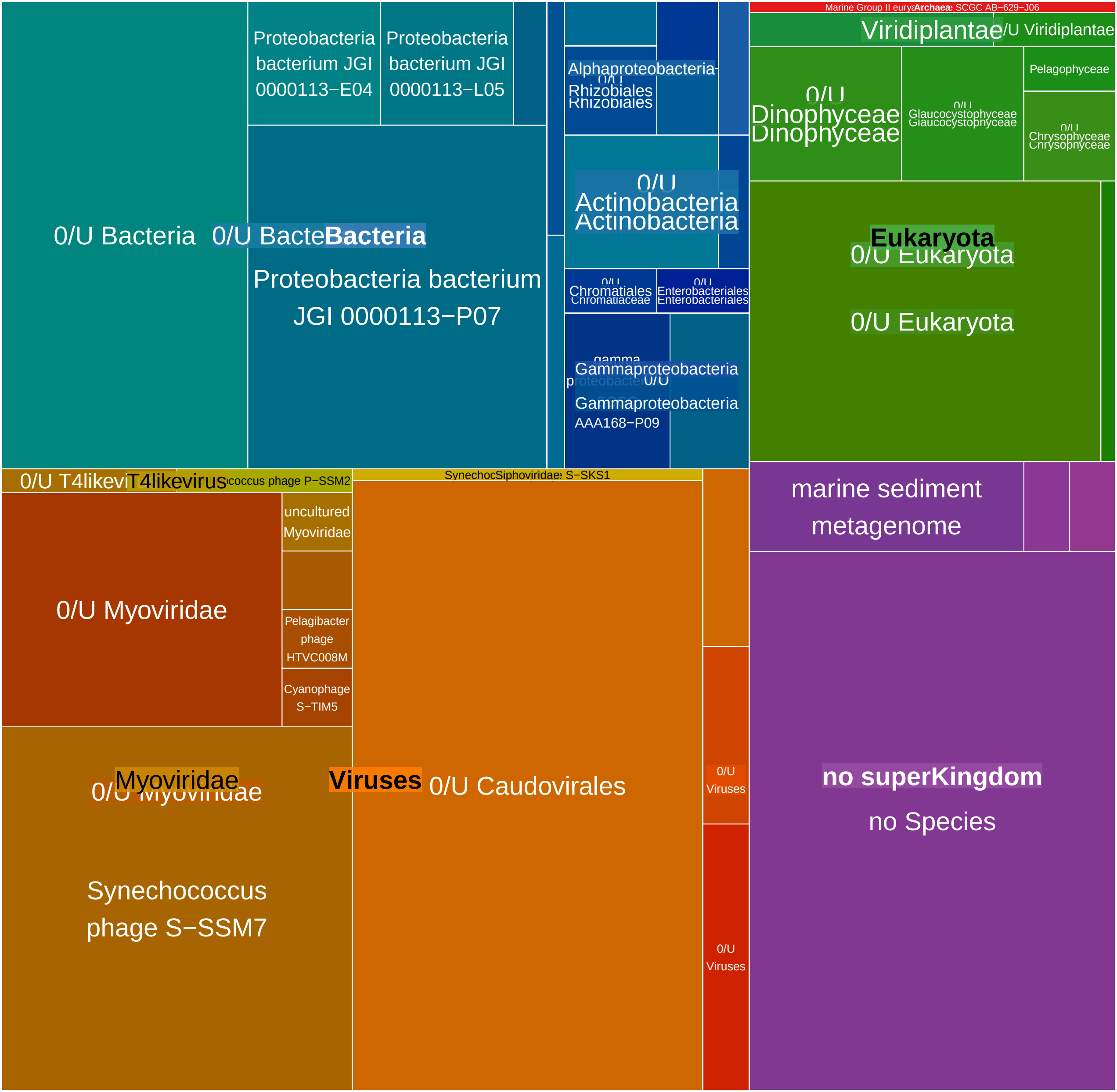

MGT−v1_847 : 590 unigenes, 495 (83.90%) taxonomically assigned

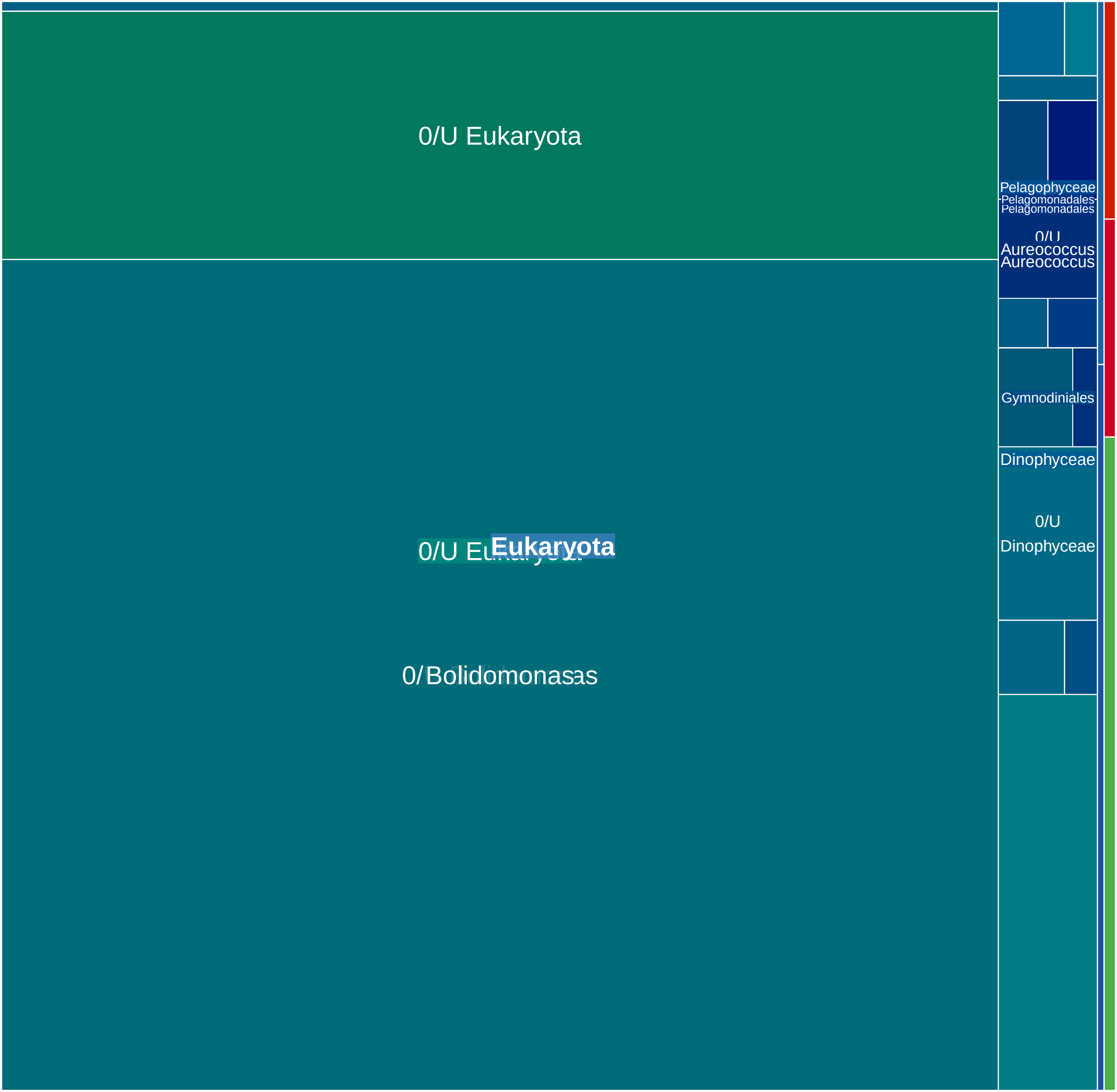

MGT−v1_848 : 589 unigenes, 35 (5.94%) taxonomically assigned

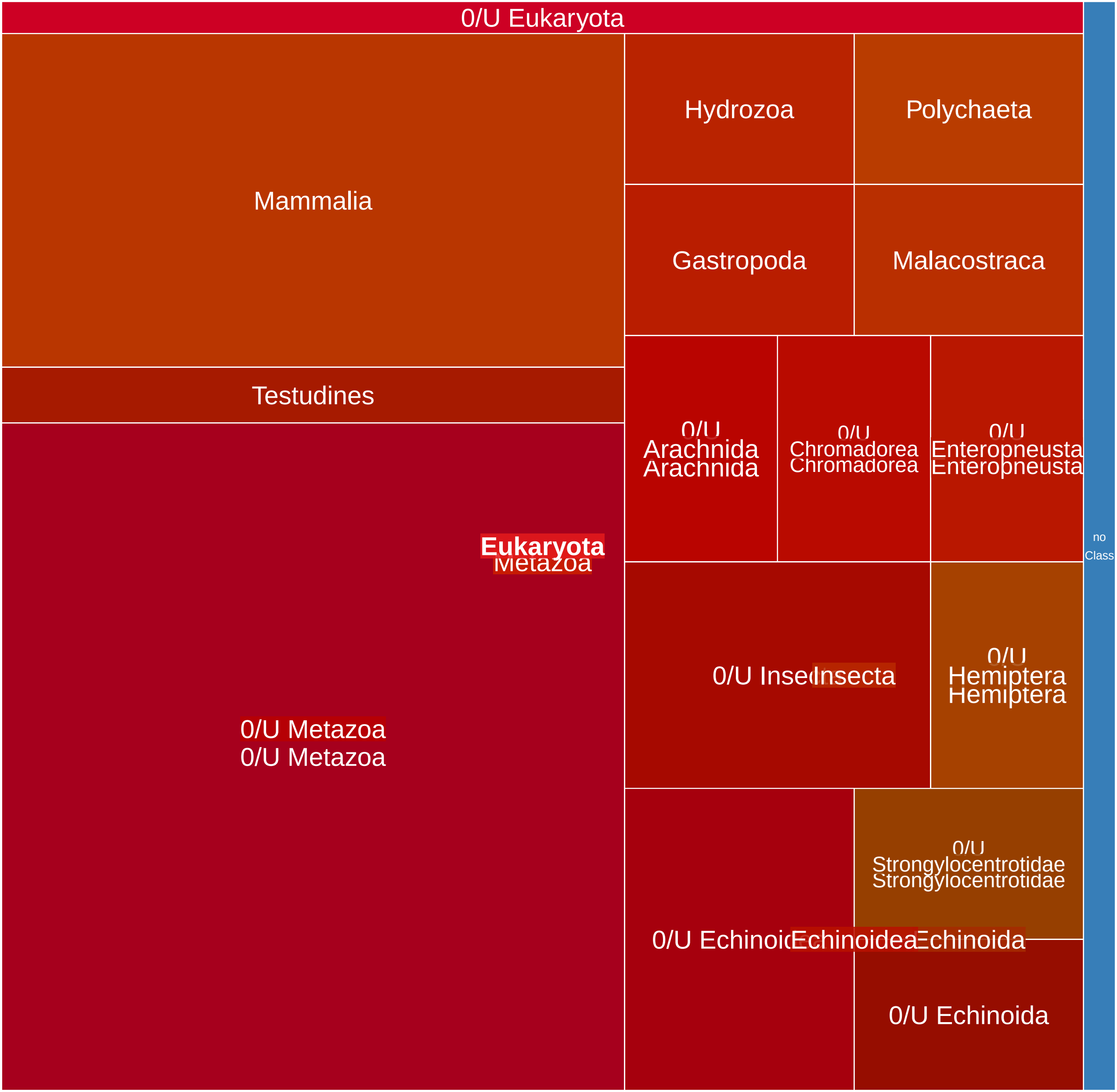

### MGT−v1_849 : 587 unigenes, 529 (90.12%) taxonomically assigned

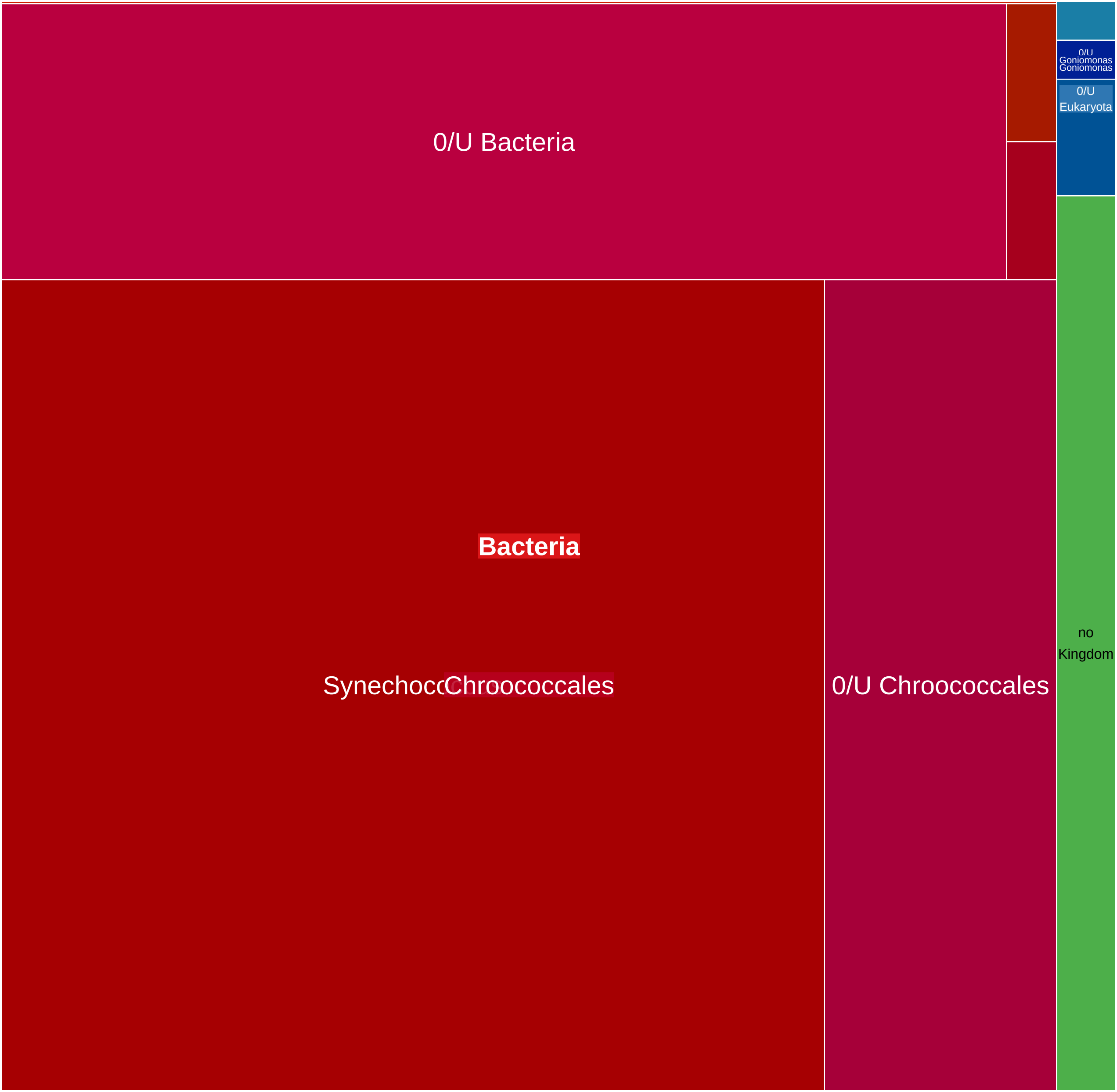

MGT−v1_850 : 586 unigenes, 35 (5.97%) taxonomically assigned

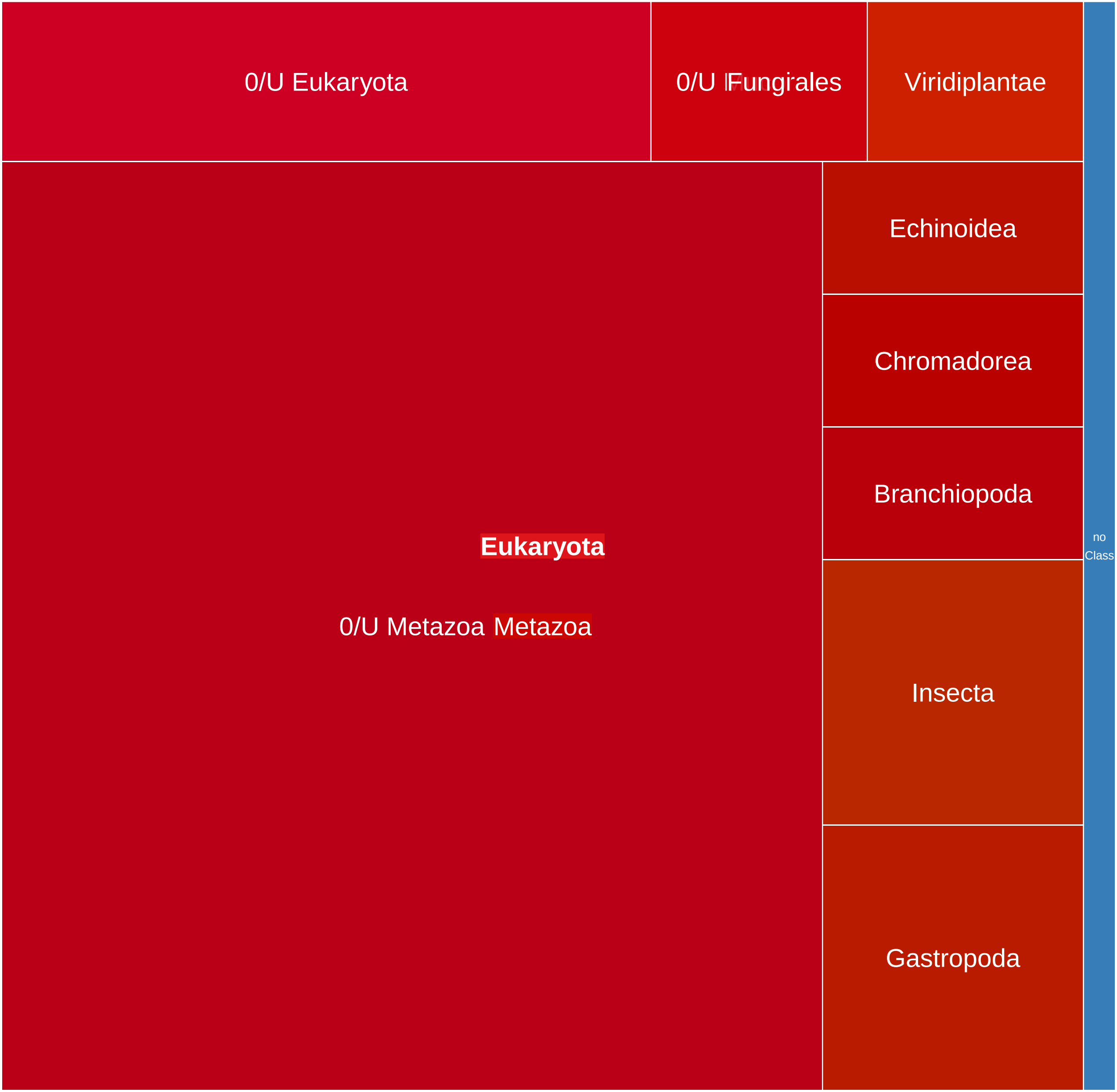

MGT−v1_851 : 586 unigenes, 376 (64.16%) taxonomically assigned

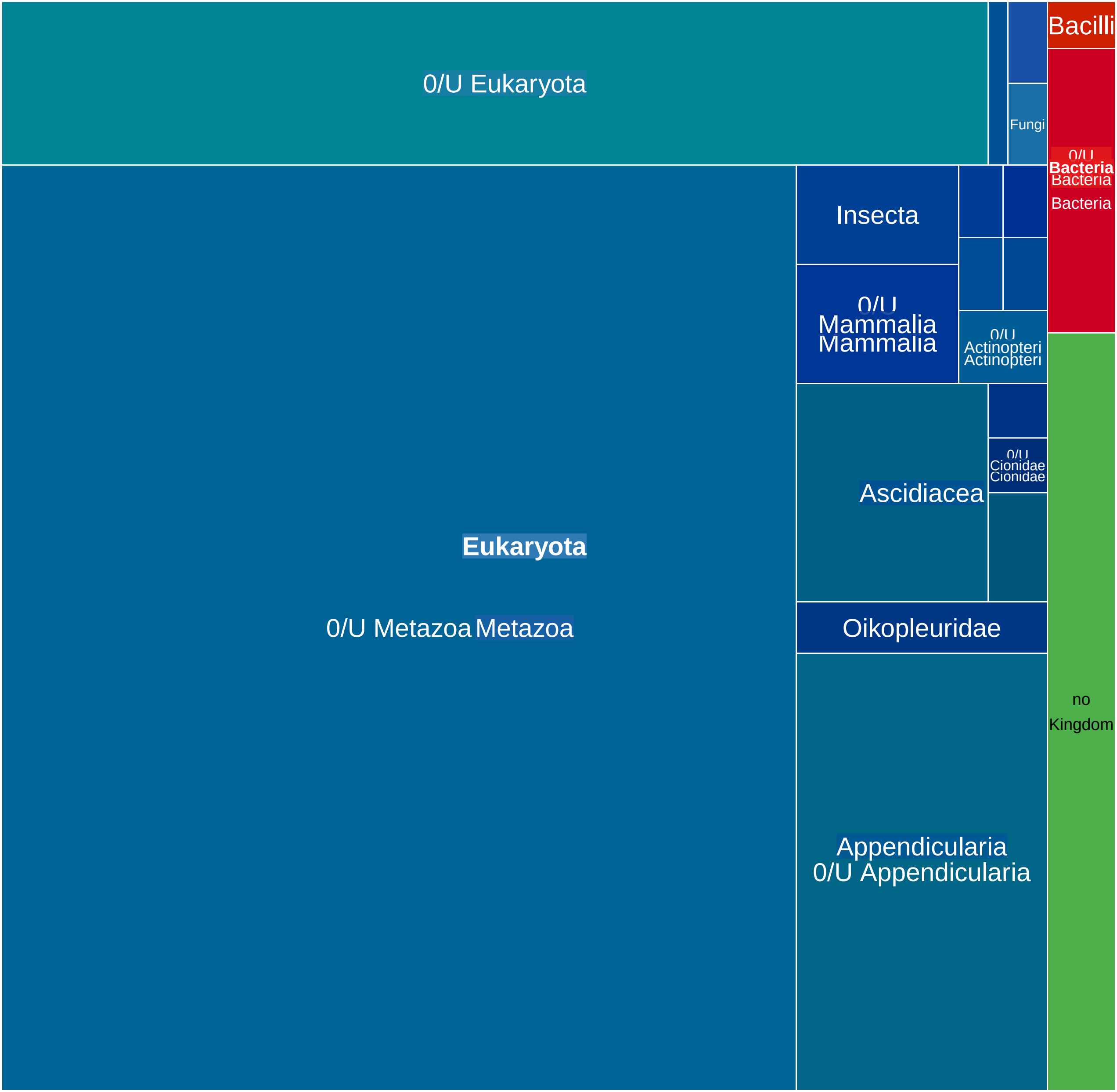

MGT−v1_852 : 585 unigenes, 209 (35.73%) taxonomically assigned

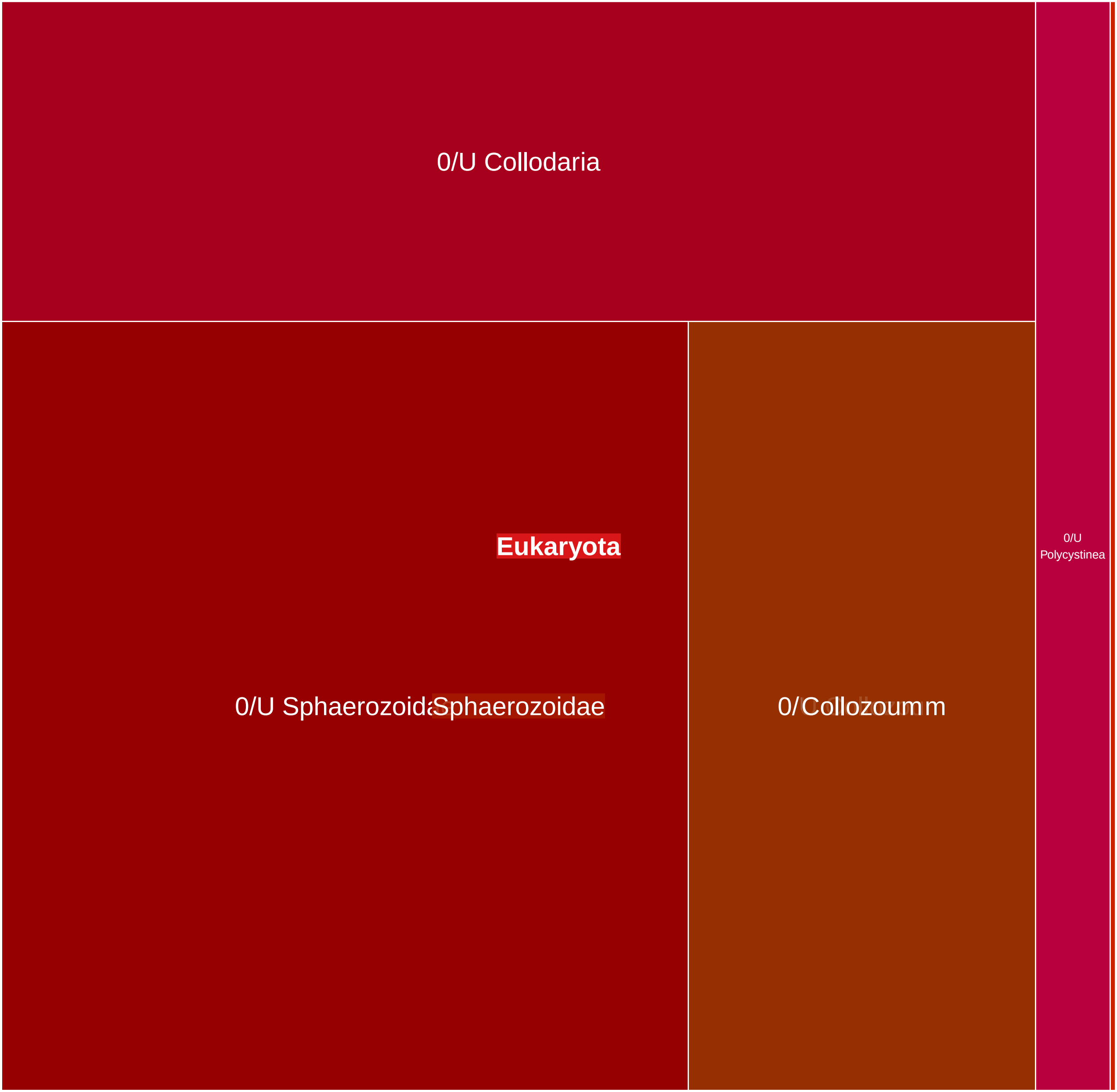

MGT−v1_853 : 583 unigenes, 96 (16.47%) taxonomically assigned

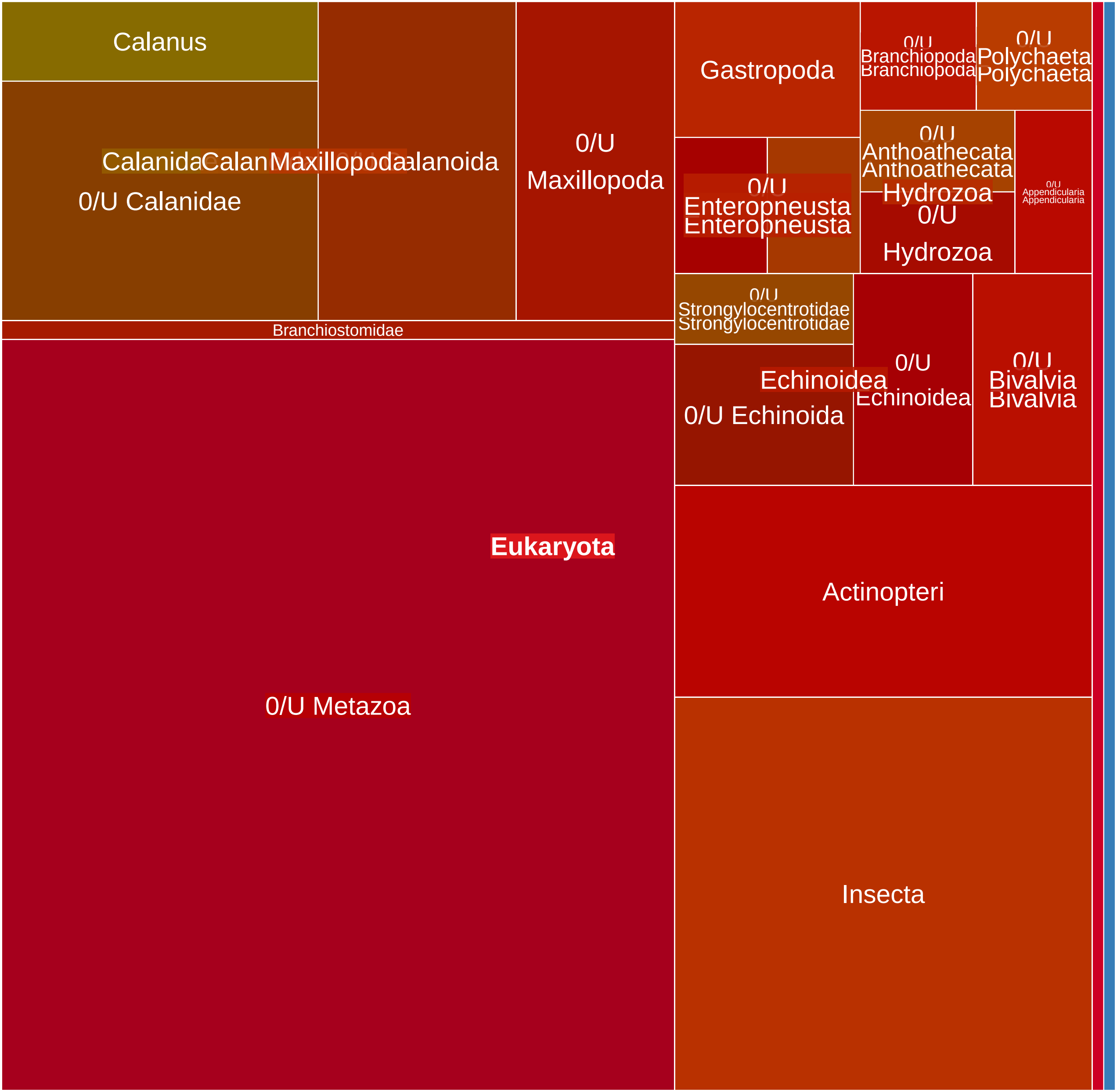

MGT−v1_854 : 581 unigenes, 24 (4.13%) taxonomically assigned

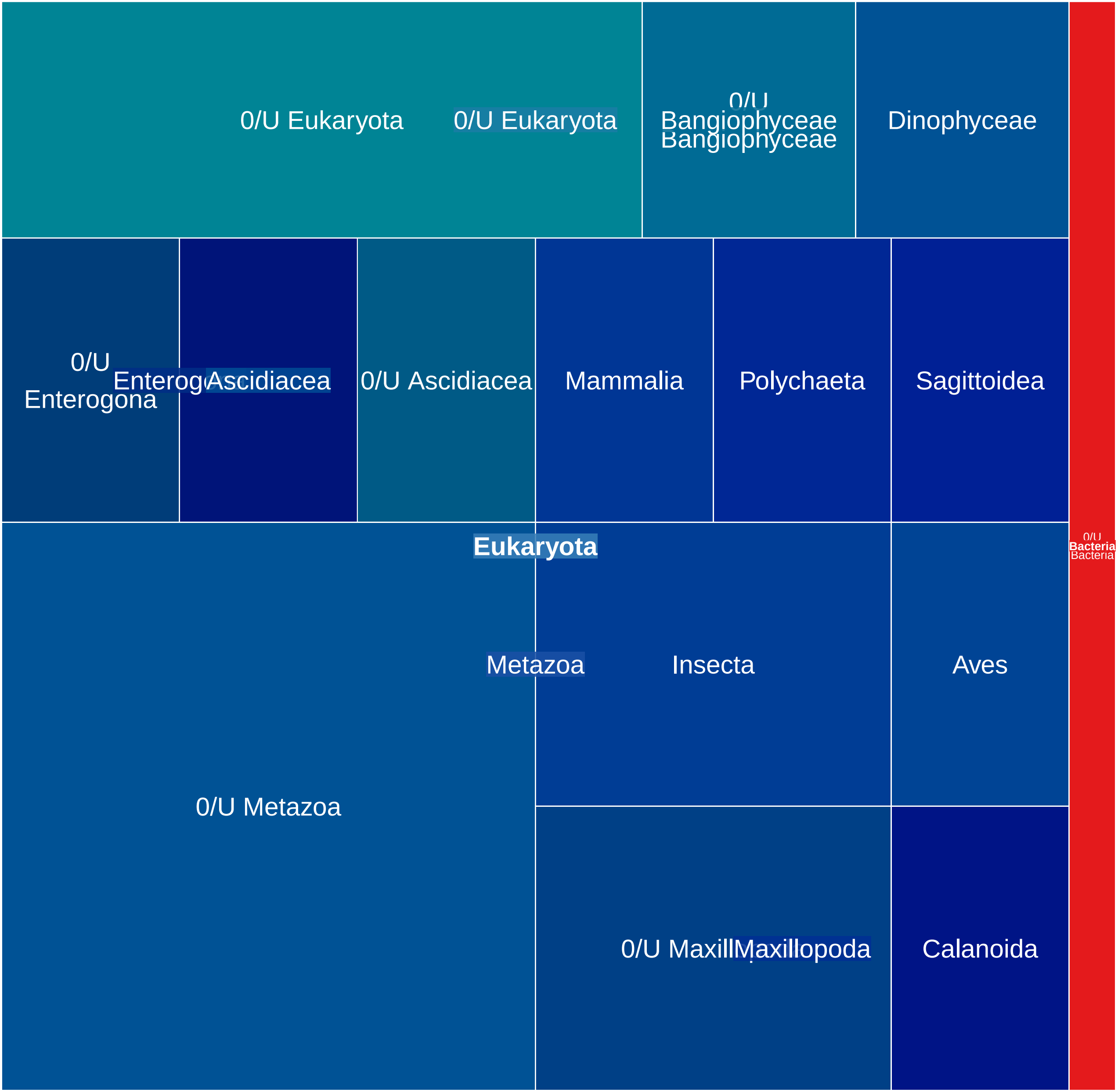

MGT−v1_855 : 579 unigenes, 43 (7.43%) taxonomically assigned

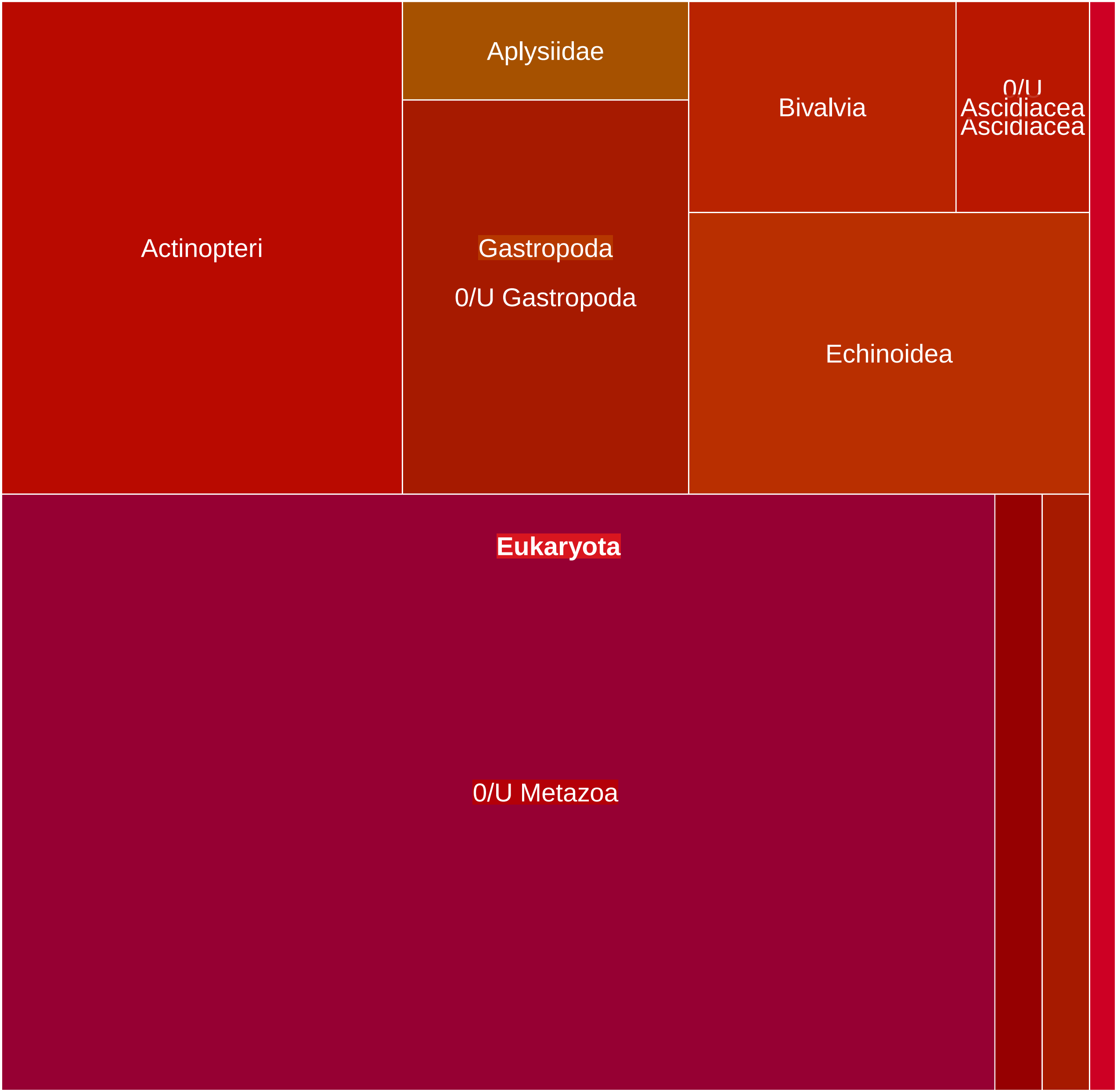

MGT−v1_856 : 579 unigenes, 518 (89.46%) taxonomically assigned

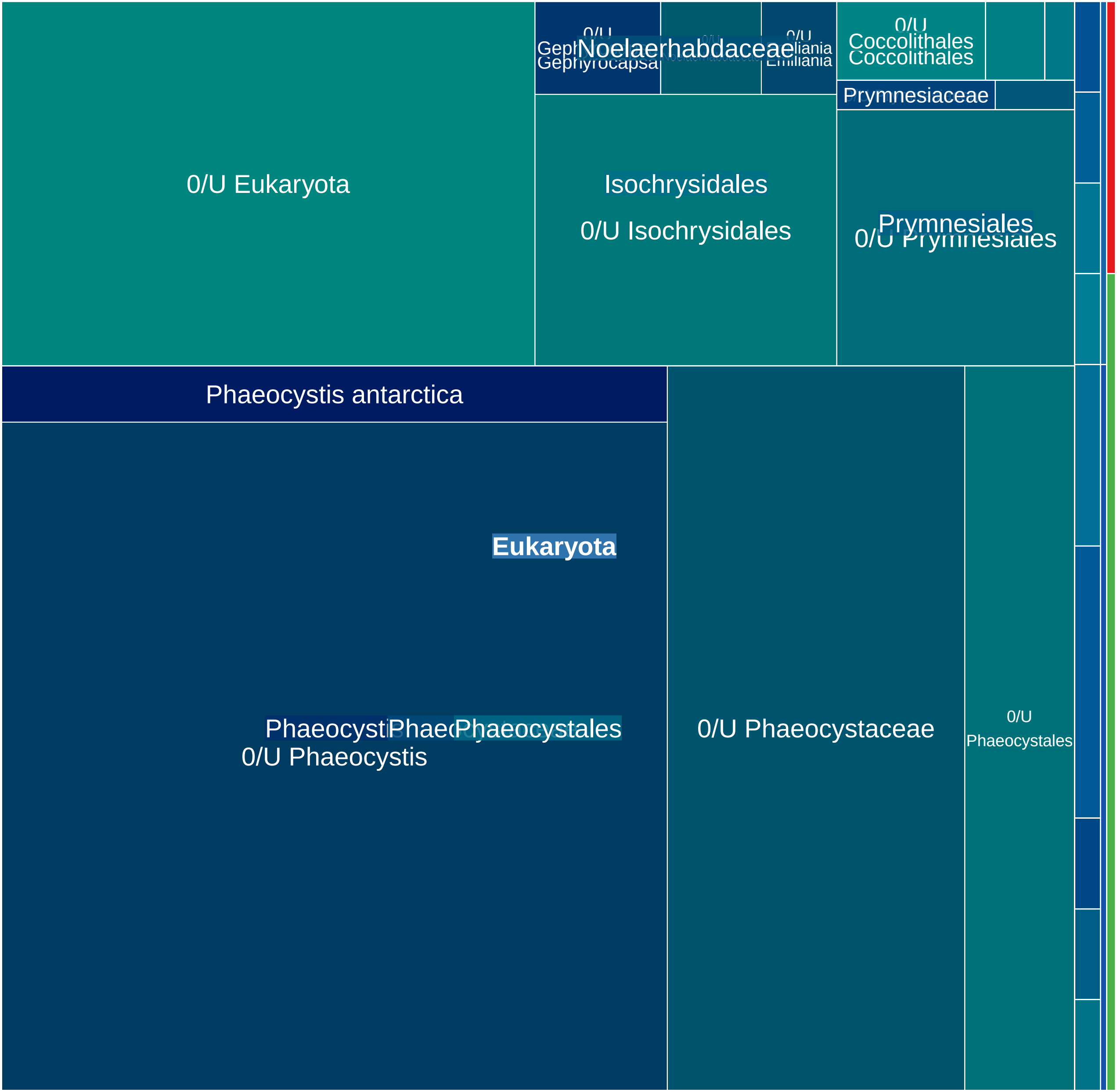

MGT−v1_857 : 578 unigenes, 394 (68.17%) taxonomically assigned

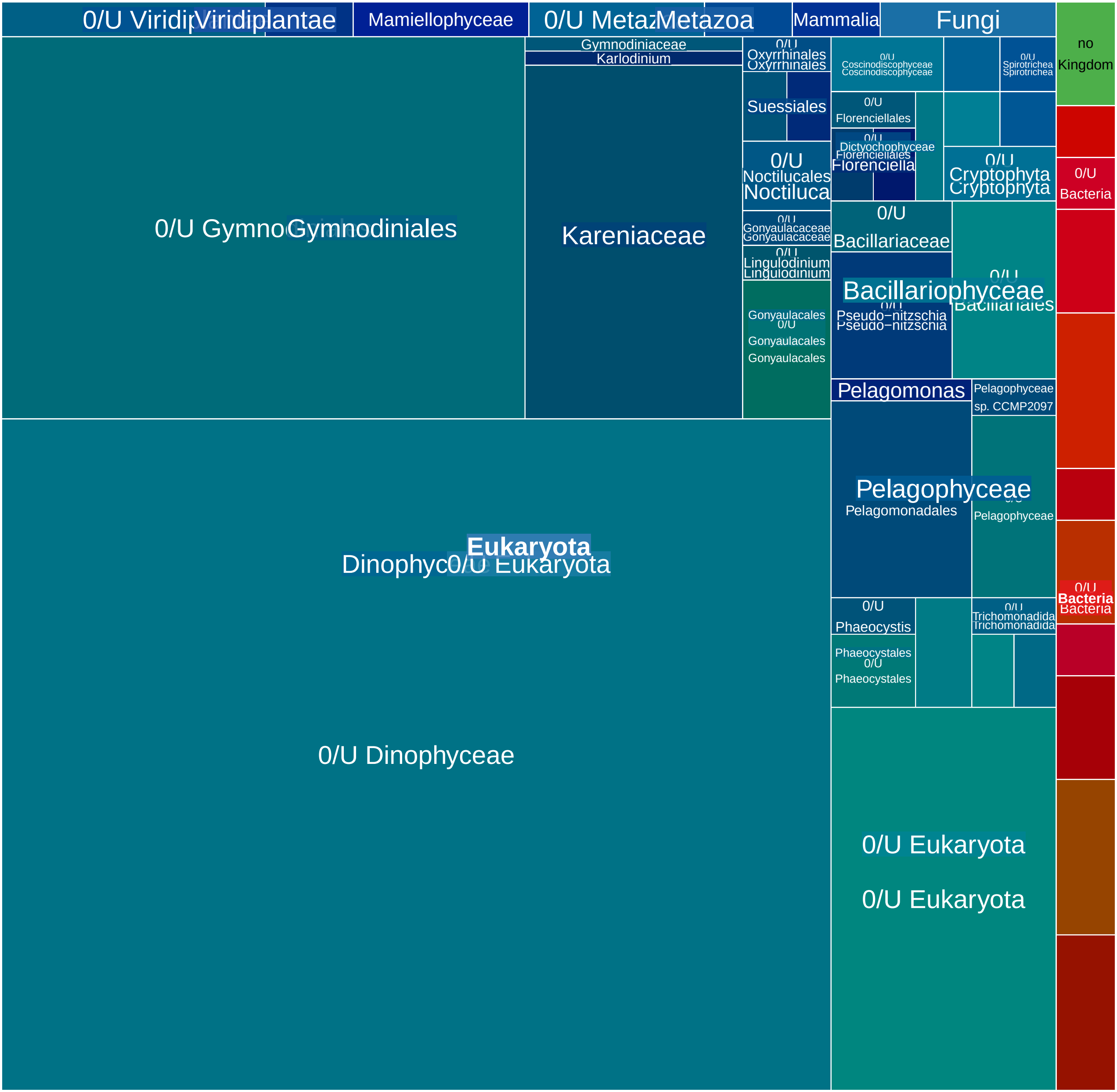

### MGT−v1_858 : 578 unigenes, 25 (4.33%) taxonomically assigned

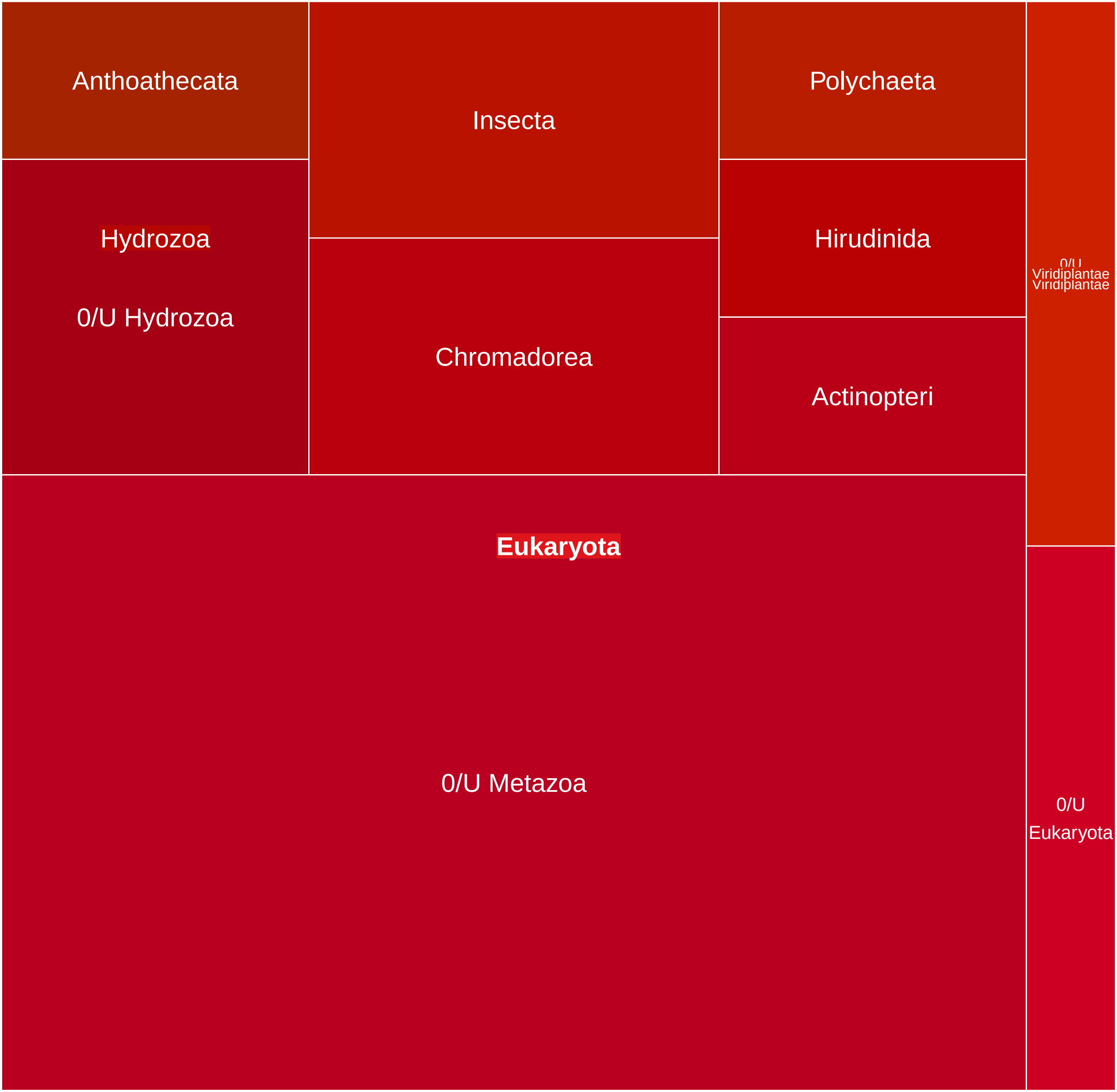

MGT−v1_859 : 575 unigenes, 176 (30.61%) taxonomically assigned

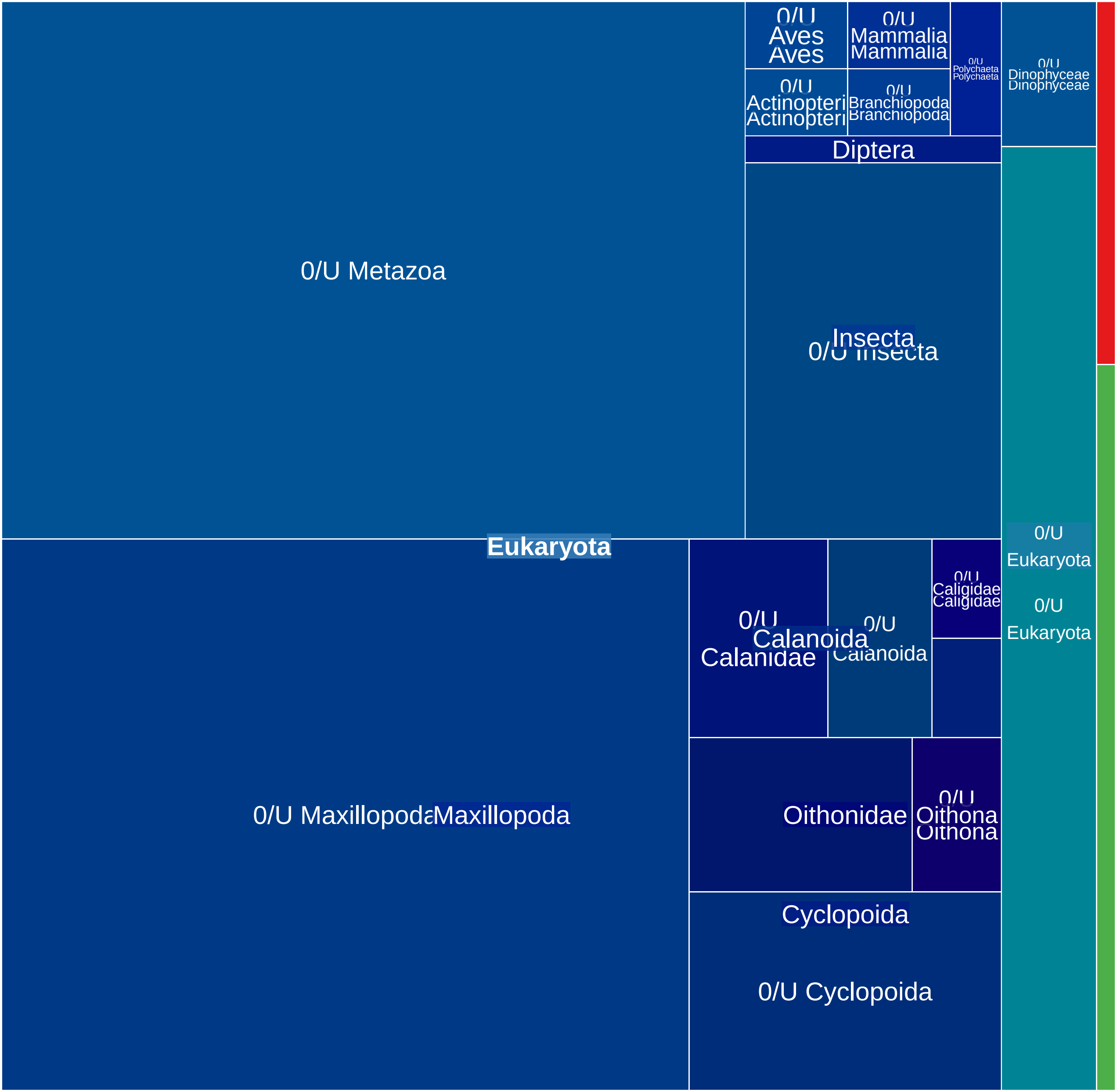

MGT−v1_860 : 568 unigenes, 501 (88.20%) taxonomically assigned

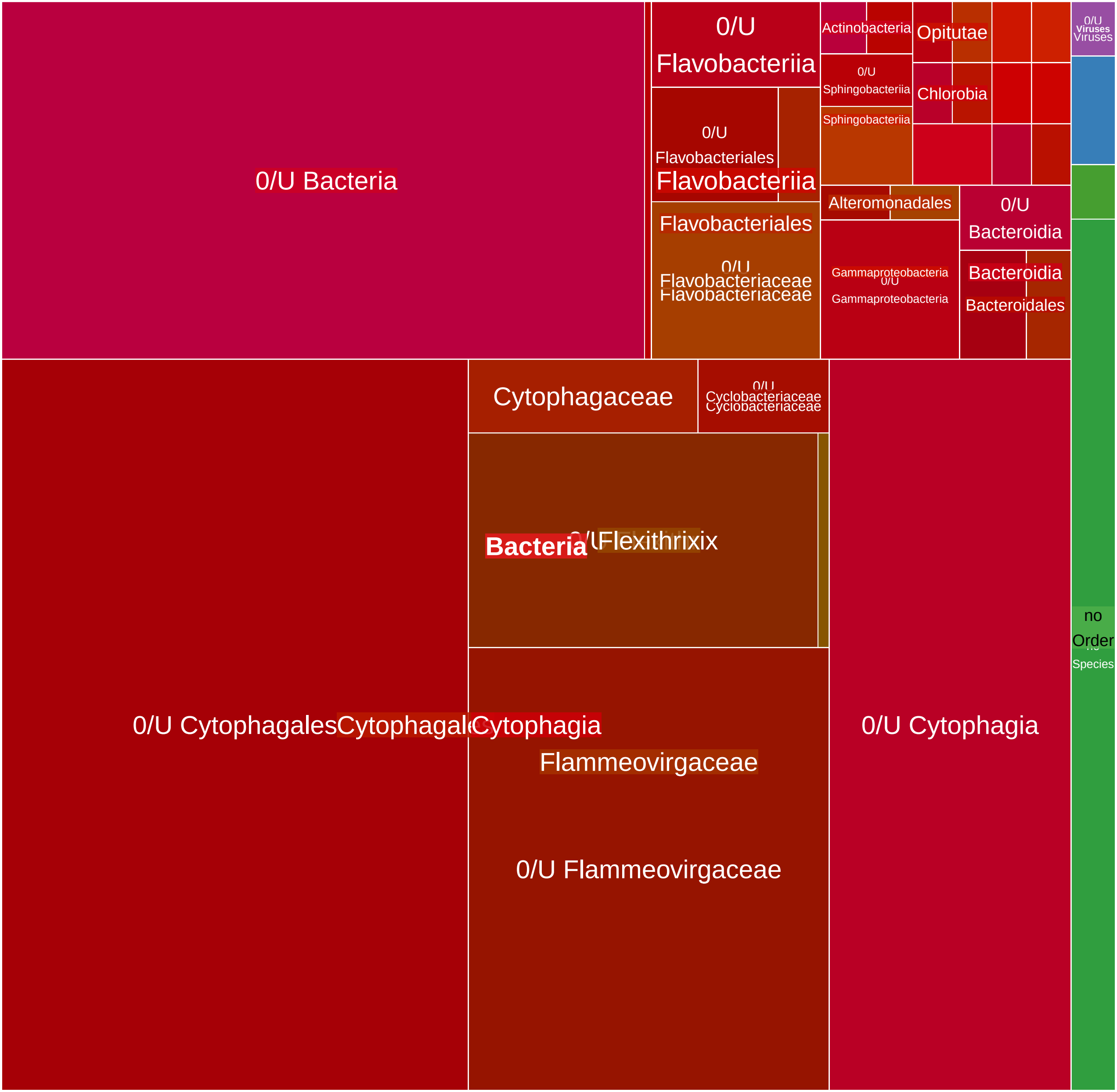

MGT−v1_861 : 568 unigenes, 415 (73.06%) taxonomically assigned

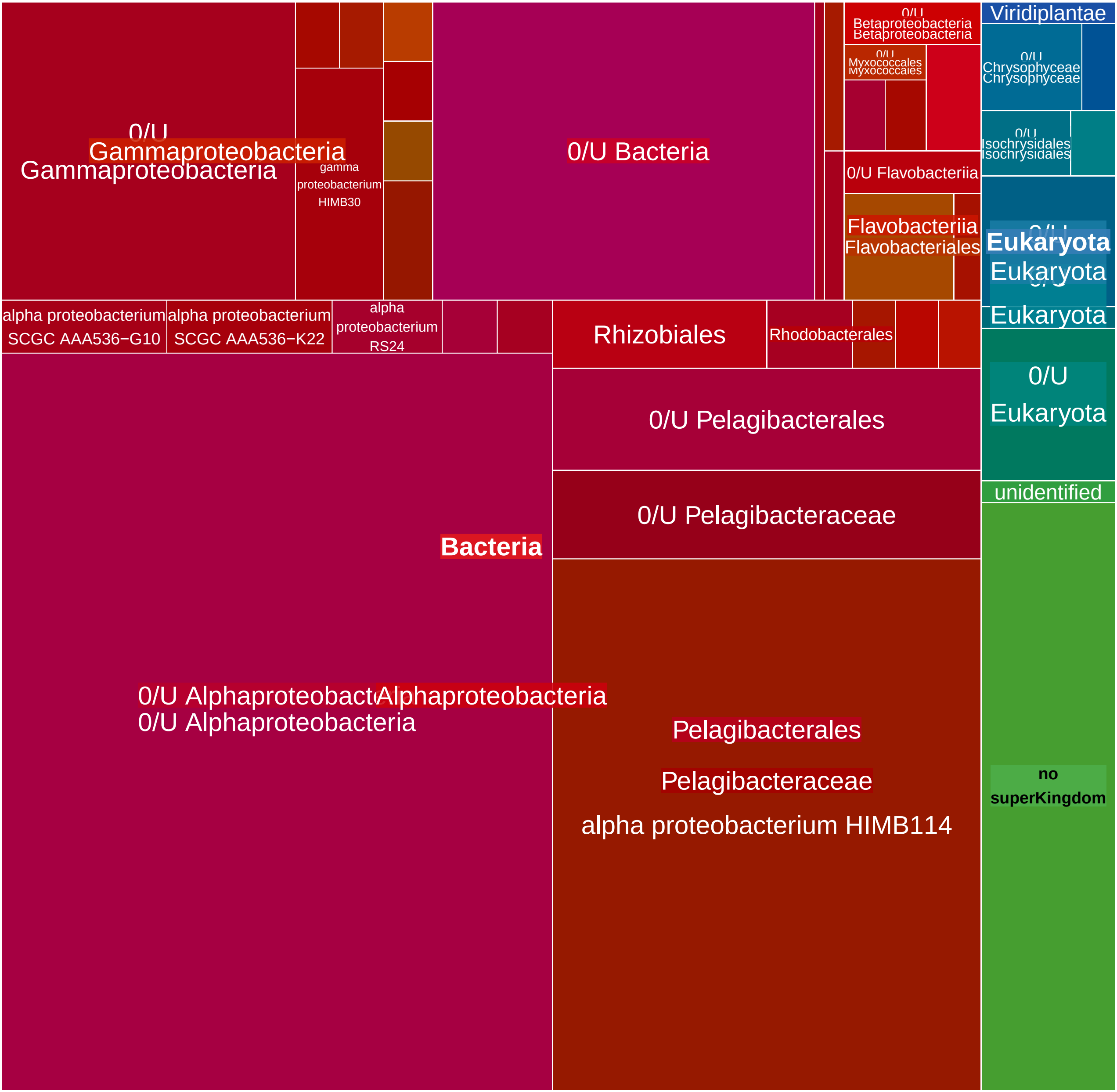

MGT−v1_862 : 562 unigenes, 58 (10.32%) taxonomically assigned

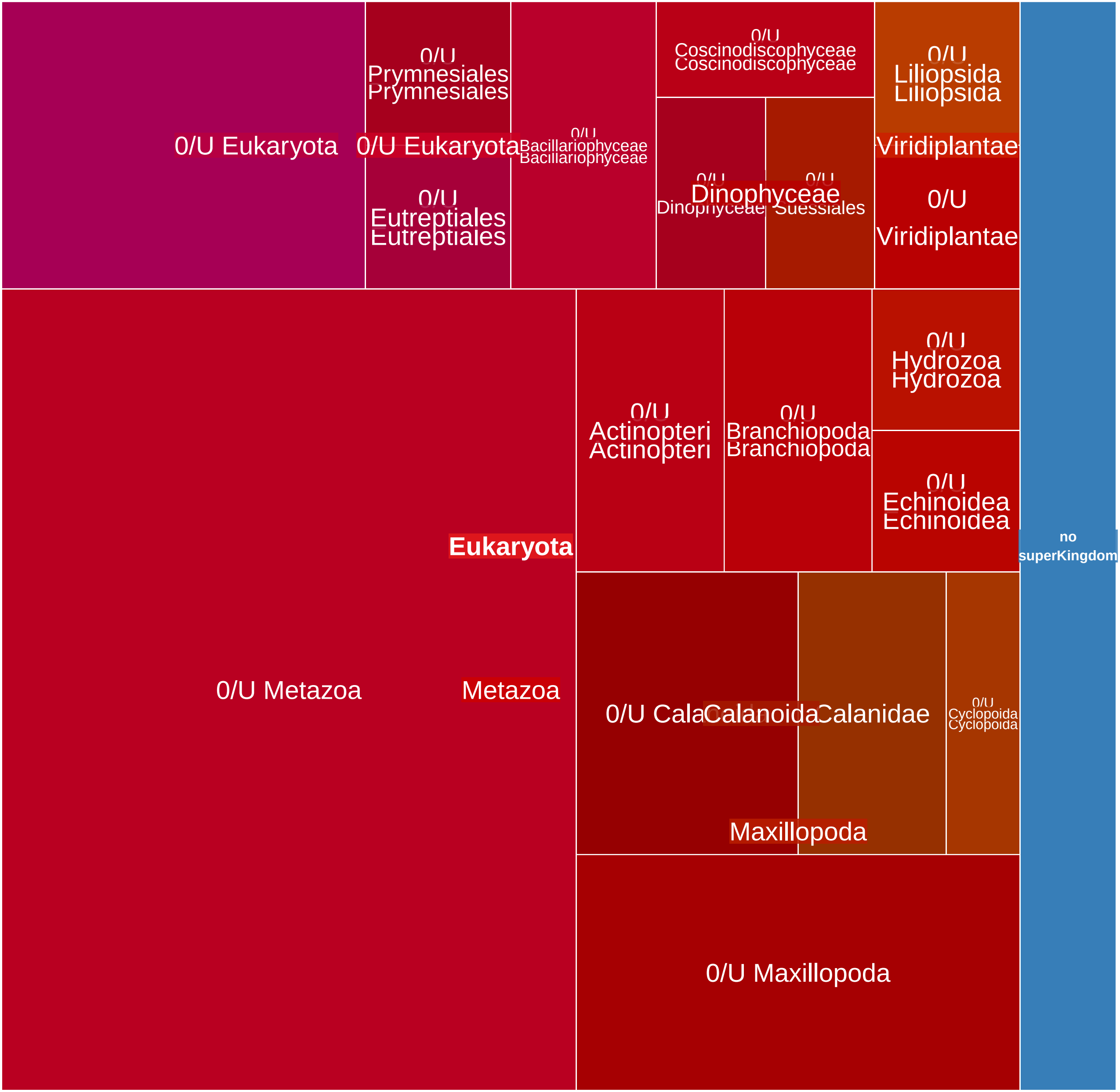

MGT−v1_863 : 560 unigenes, 46 (8.21%) taxonomically assigned

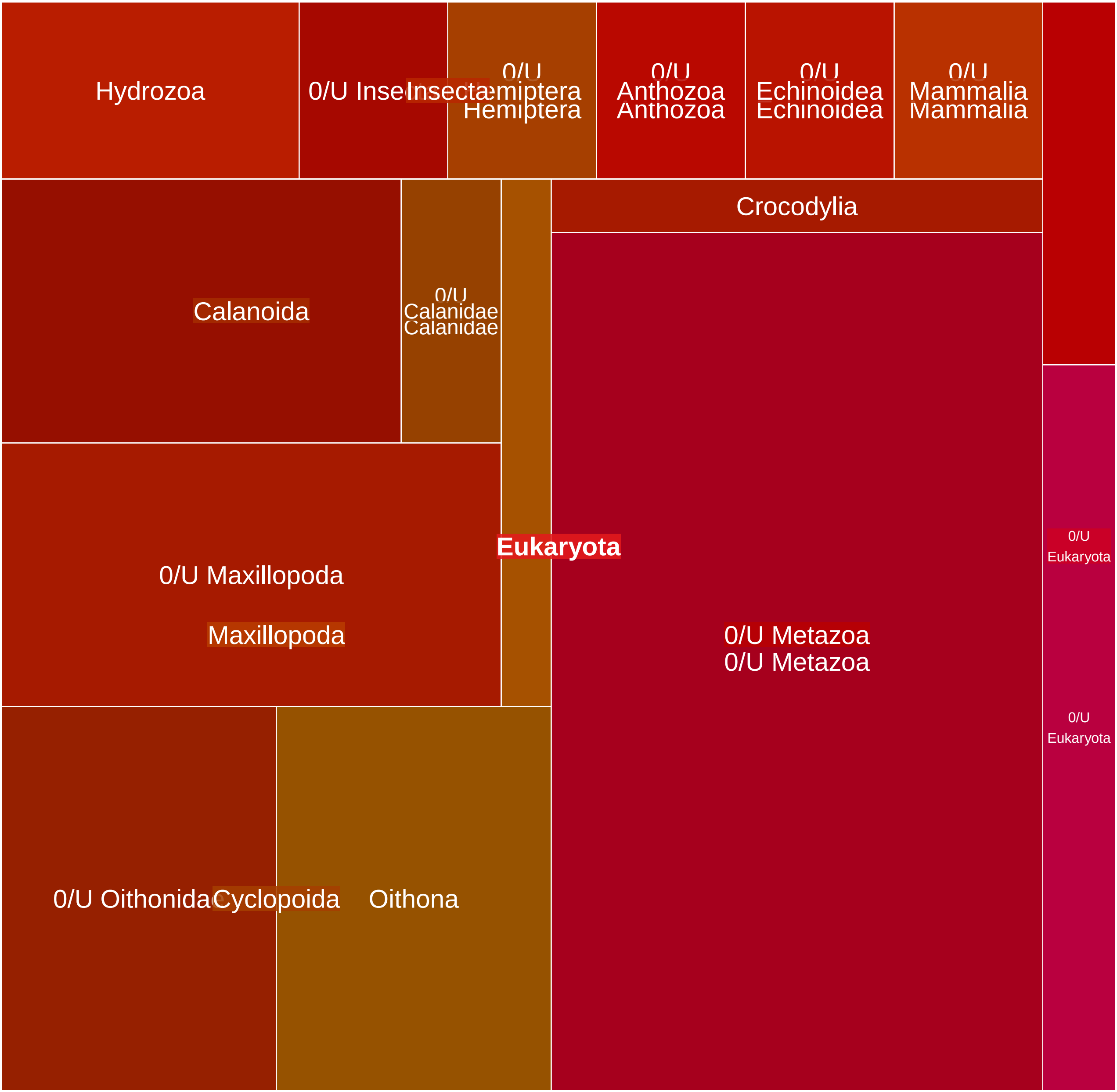

MGT−v1_864 : 560 unigenes, 17 (3.04%) taxonomically assigned

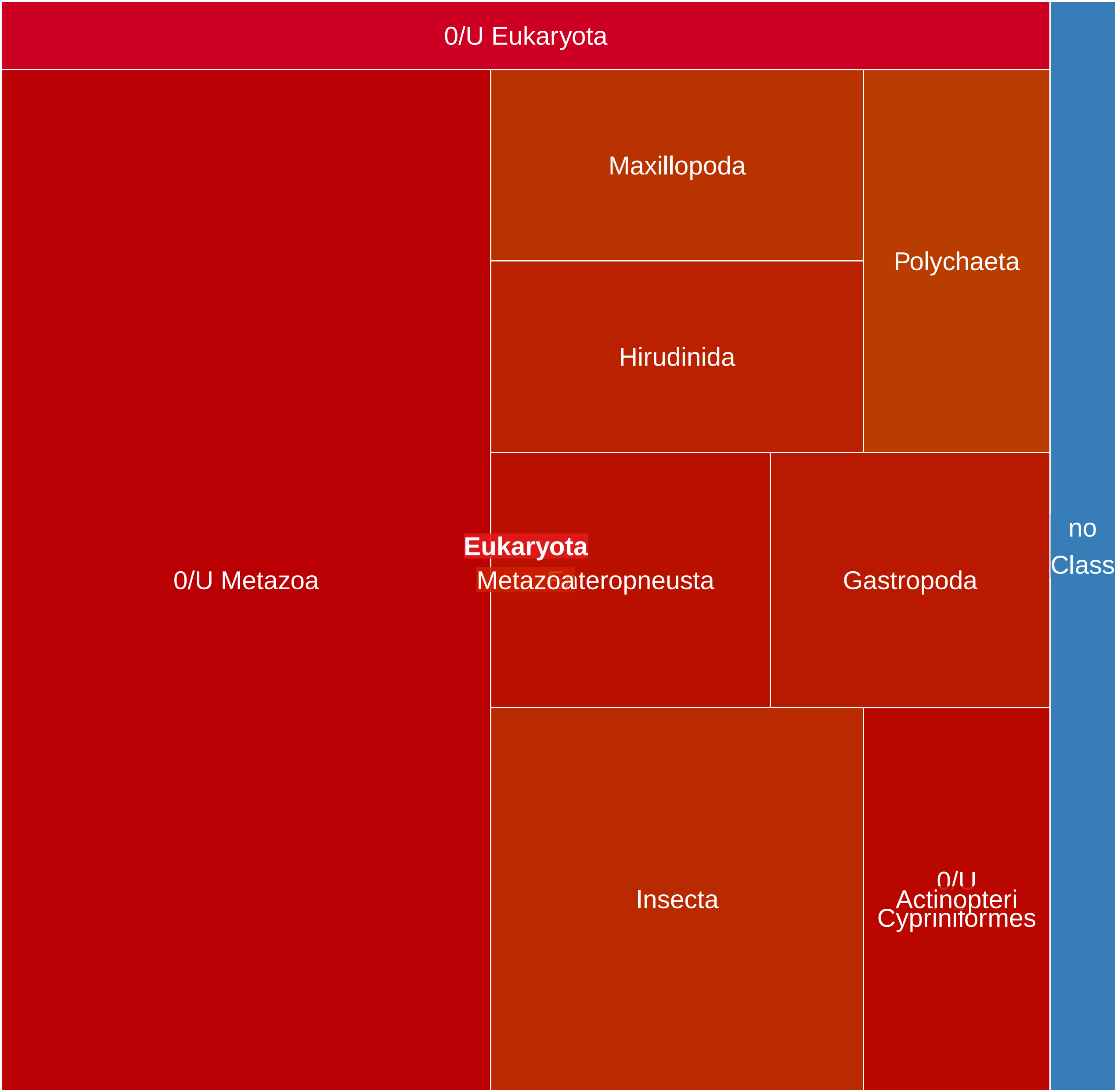

### MGT−v1_865 : 559 unigenes, 103 (18.43%) taxonomically assigned

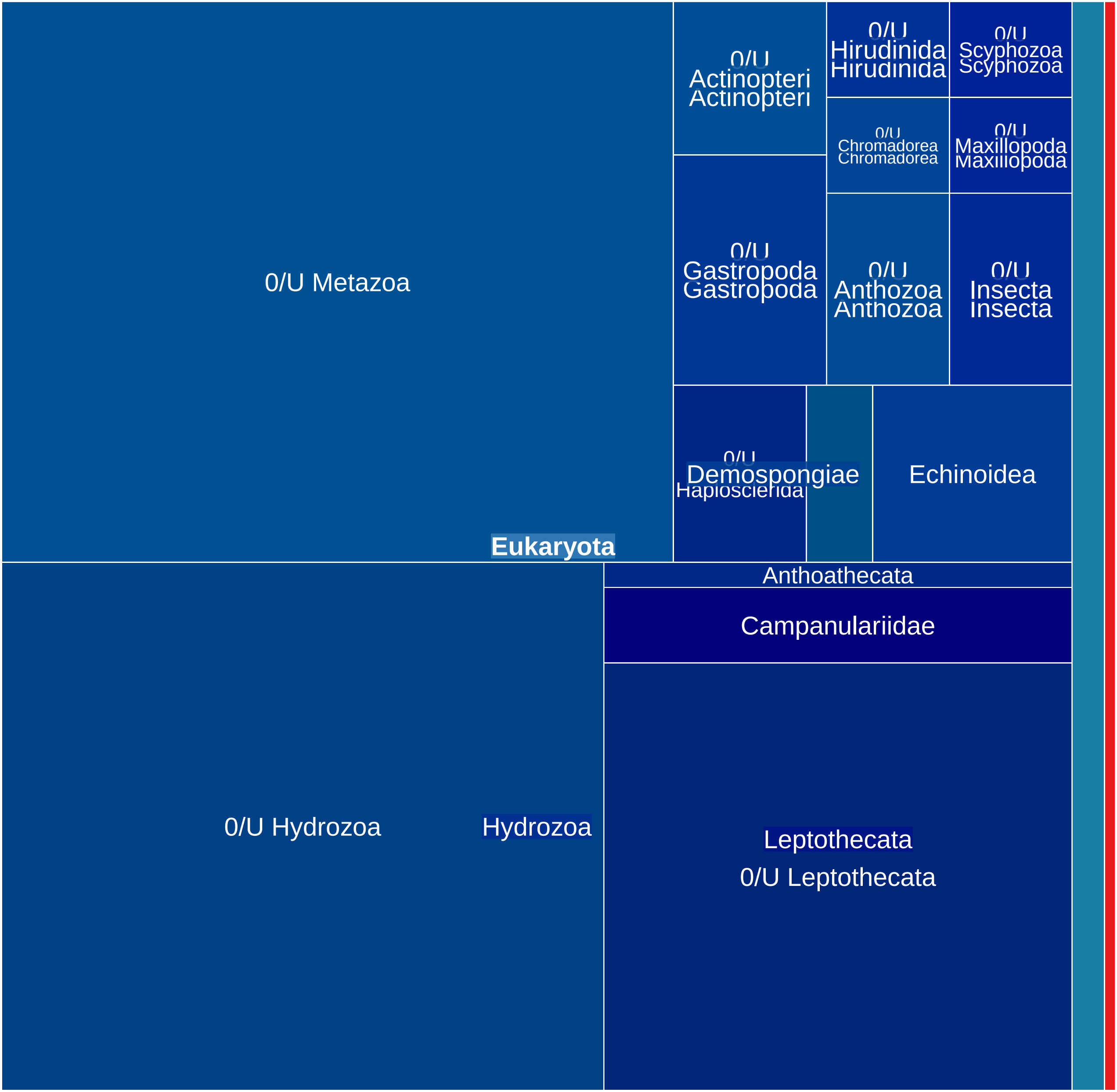

MGT−v1_866 : 556 unigenes, 433 (77.88%) taxonomically assigned

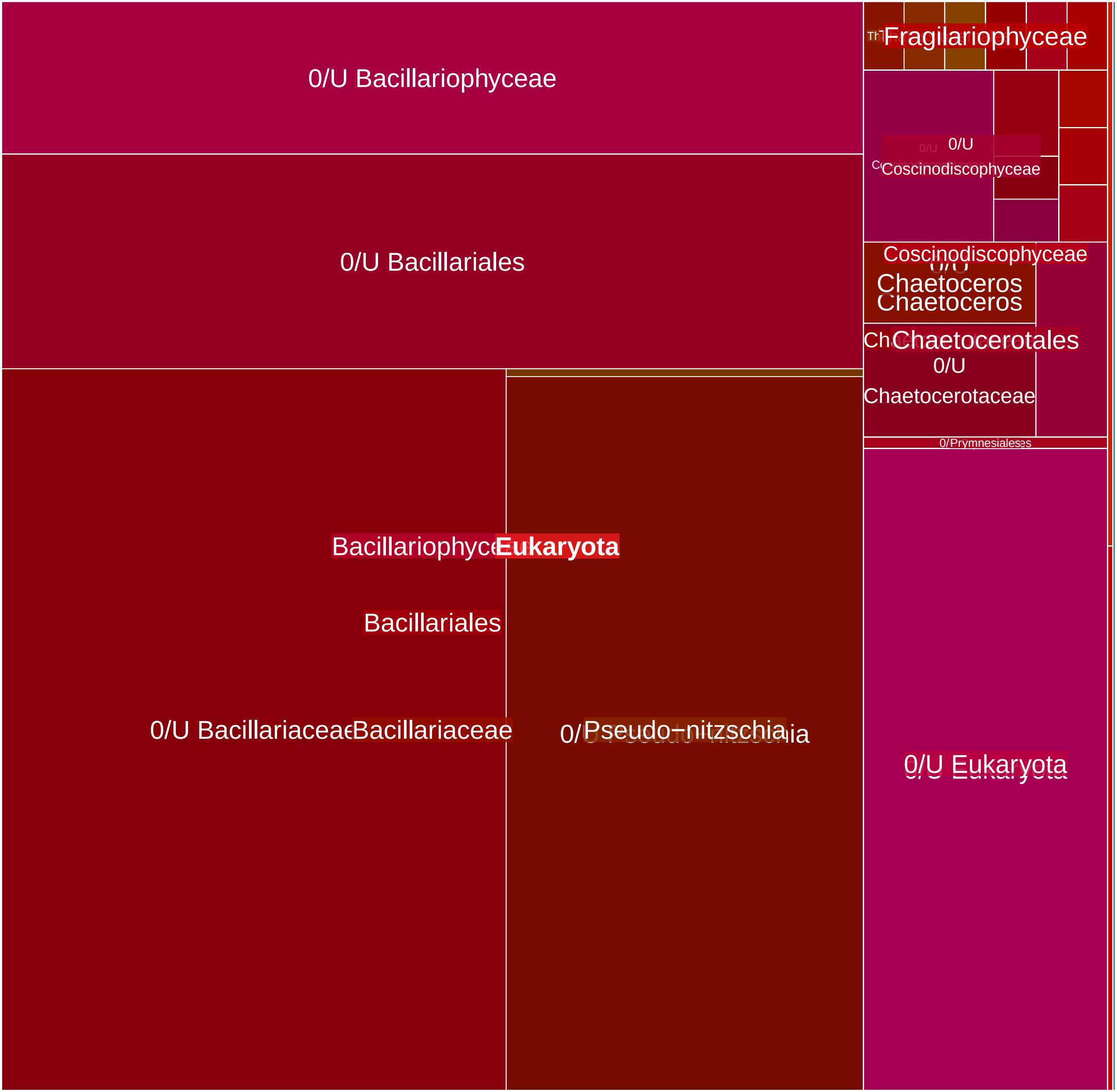

MGT−v1_867 : 555 unigenes, 26 (4.68%) taxonomically assigned

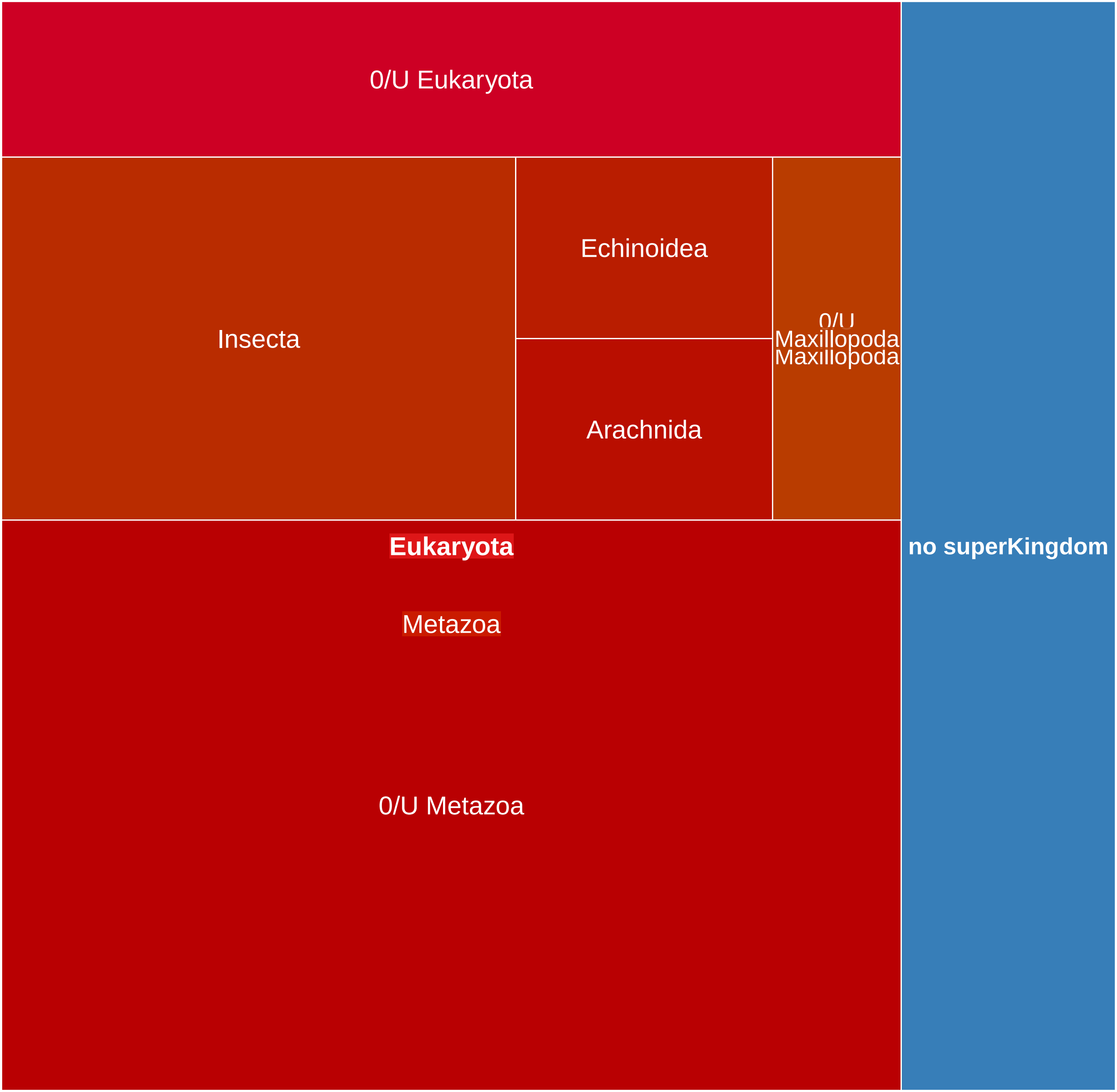

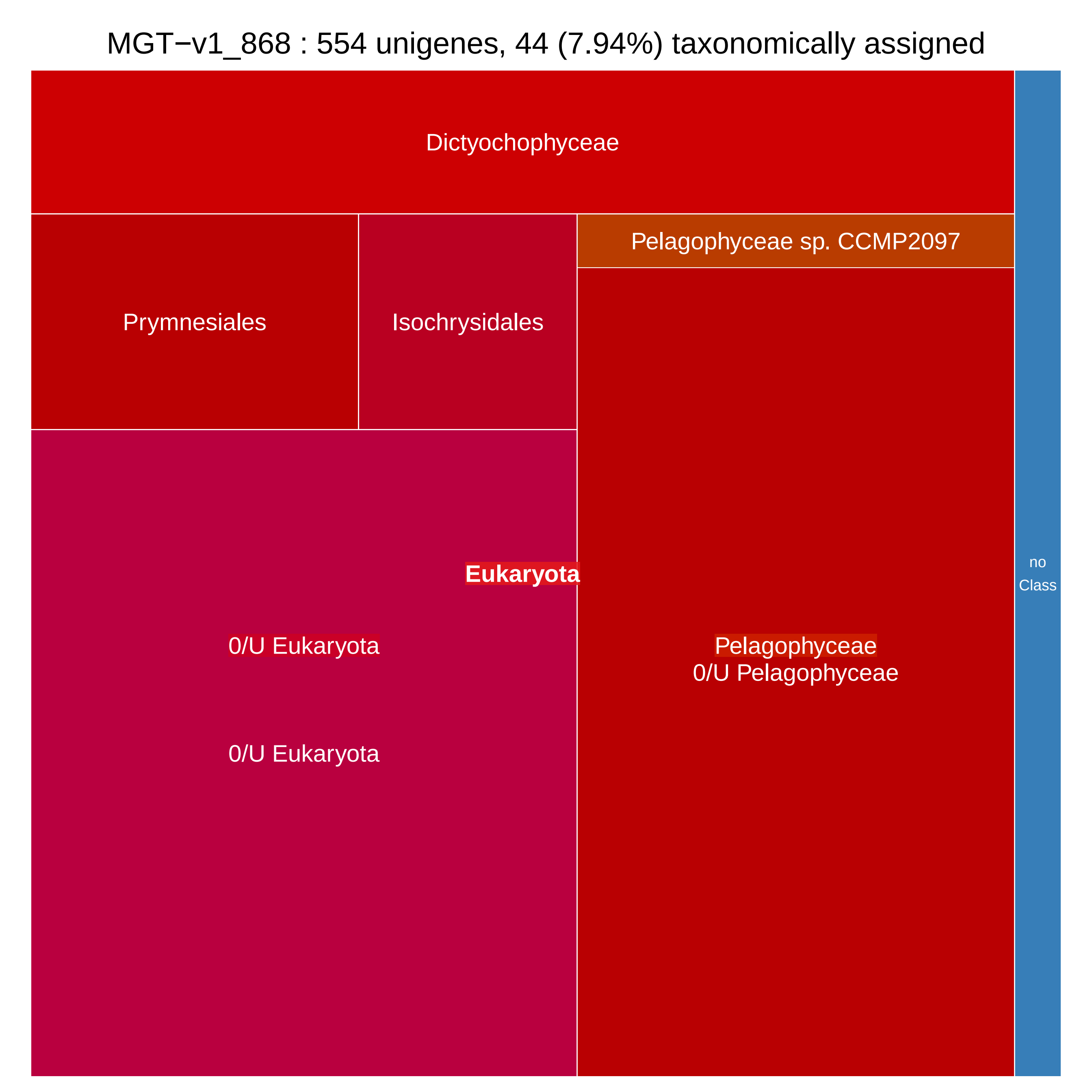
MGT−v1_868 : 554 unigenes, 44 (7.94%) taxonomically assigned
Dictyochophyceae
Prymnesiales
Isochrysidales
Pelagophyceae sp. CCMP2097
Eukaryota
0/U Eukaryota
0/U Eukaryota
Pelagophyceae
0/U Pelagophyceae
no
Class

MGT−v1_869 : 553 unigenes, 93 (16.82%) taxonomically assigned

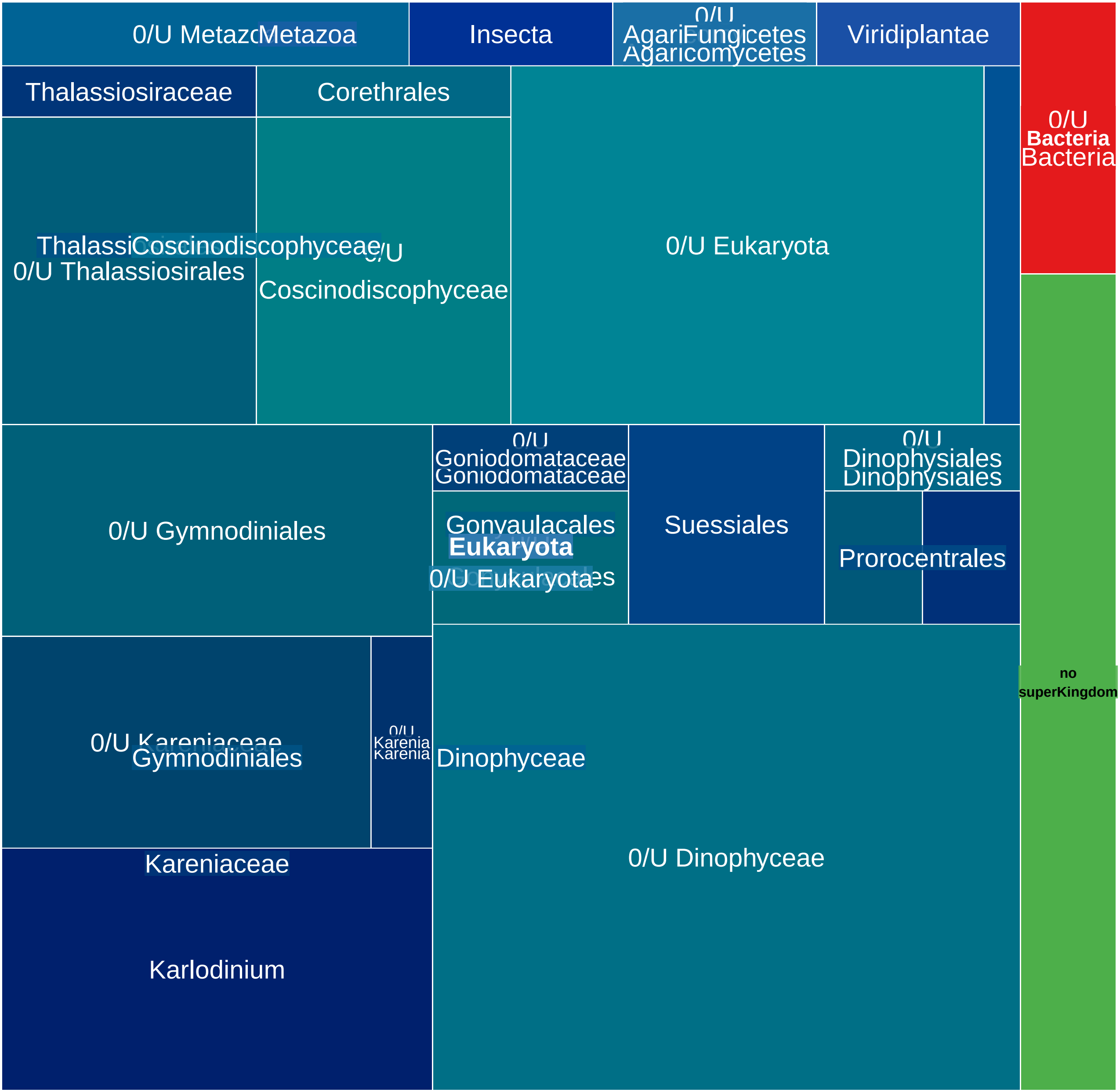

MGT−v1_870 : 552 unigenes, 542 (98.19%) taxonomically assigned

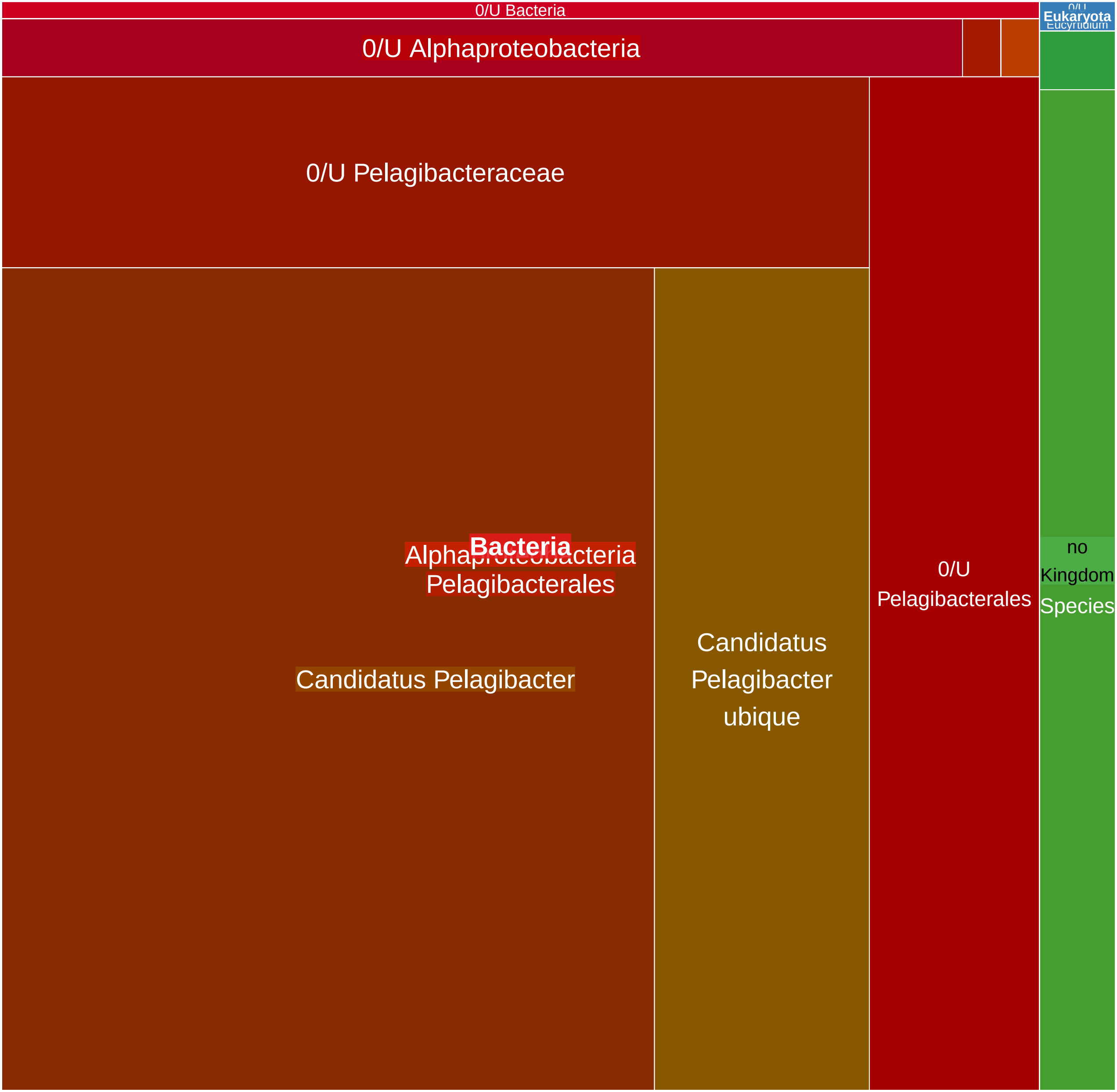

MGT−v1_871 : 551 unigenes, 37 (6.72%) taxonomically assigned

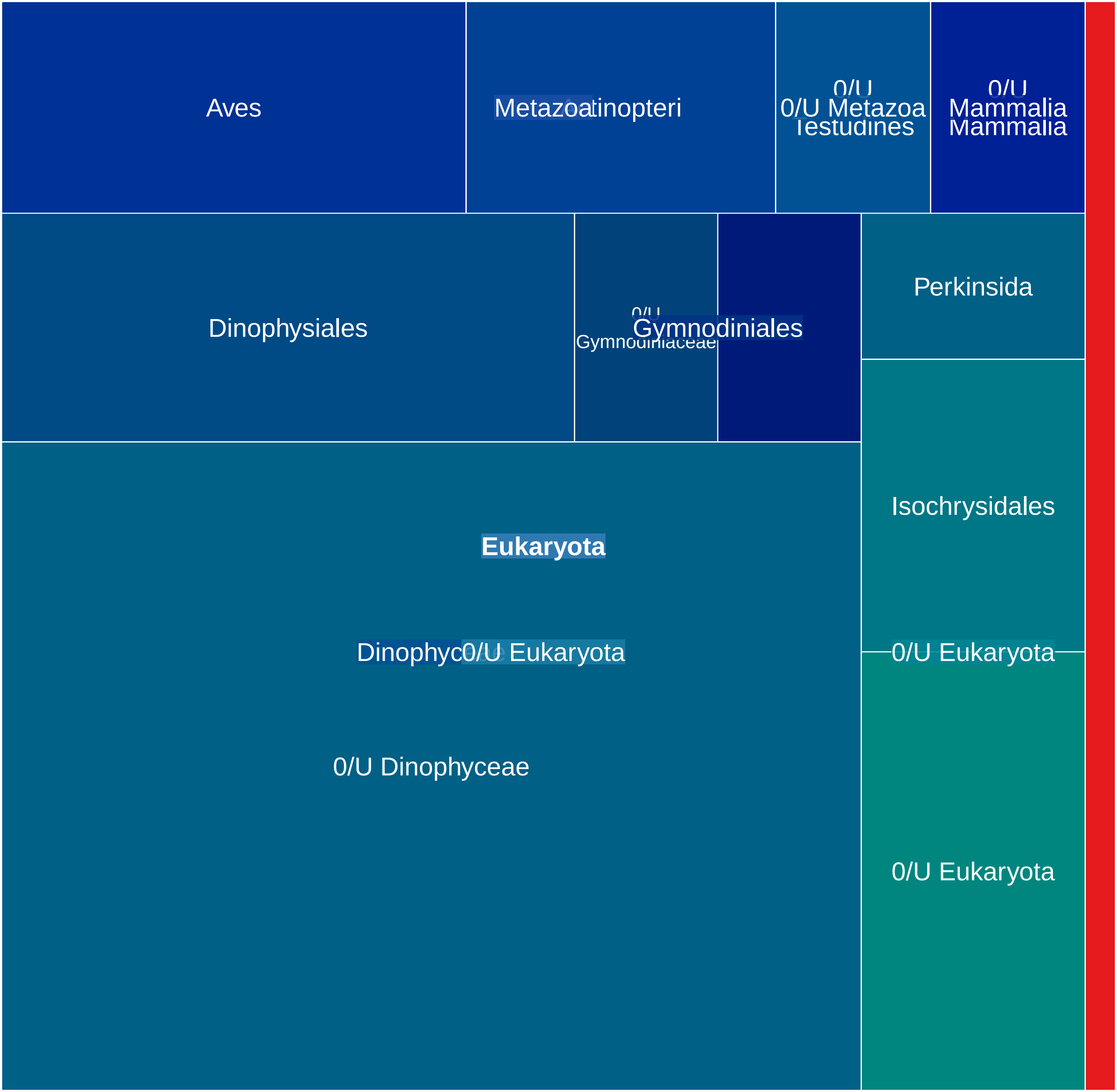

MGT−v1_872 : 550 unigenes, 390 (70.91%) taxonomically assigned

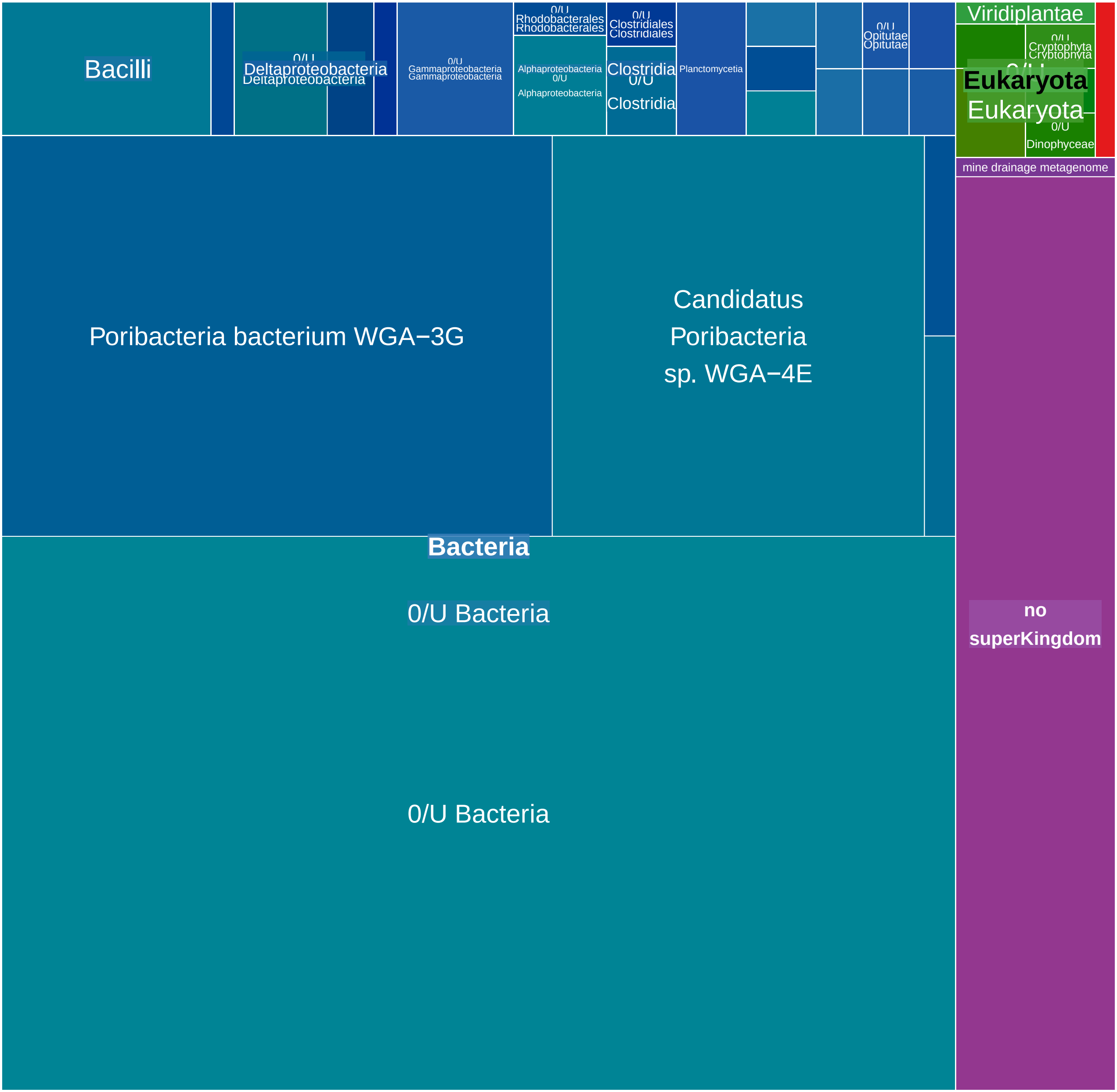

MGT−v1_873 : 544 unigenes, 55 (10.11%) taxonomically assigned

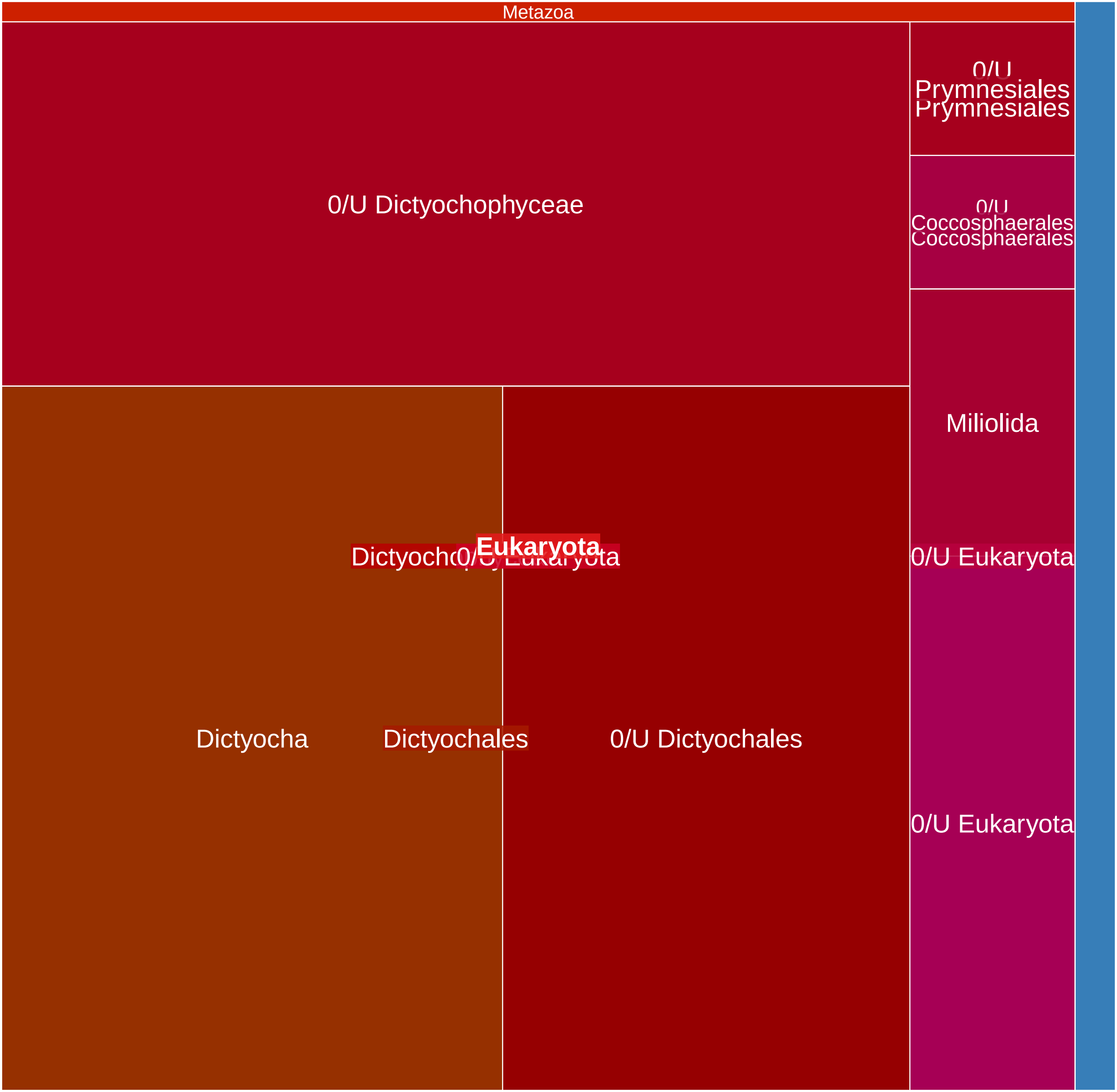

MGT−v1_874 : 542 unigenes, 22 (4.06%) taxonomically assigned

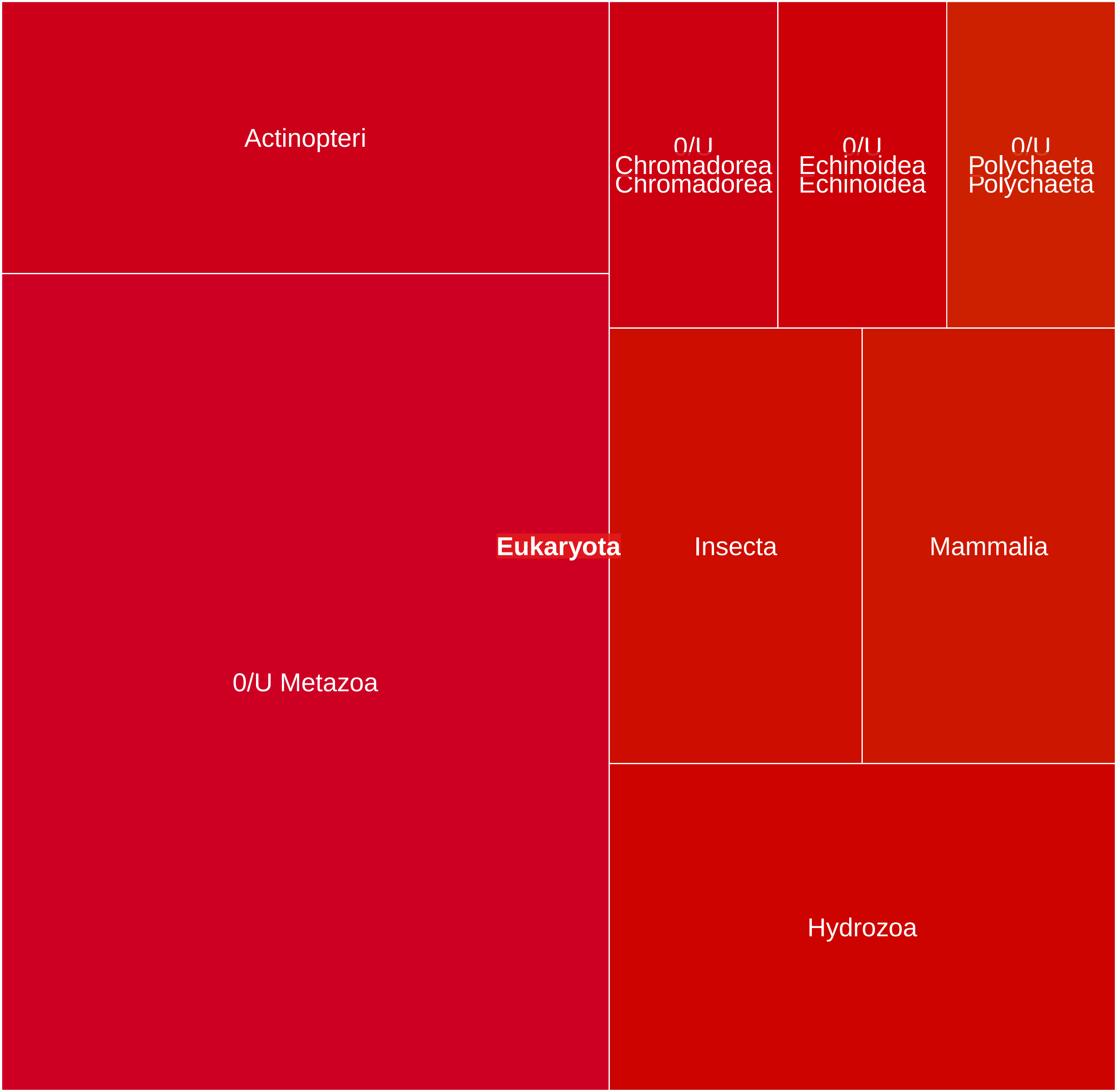

MGT−v1_875 : 542 unigenes, 56 (10.33%) taxonomically assigned

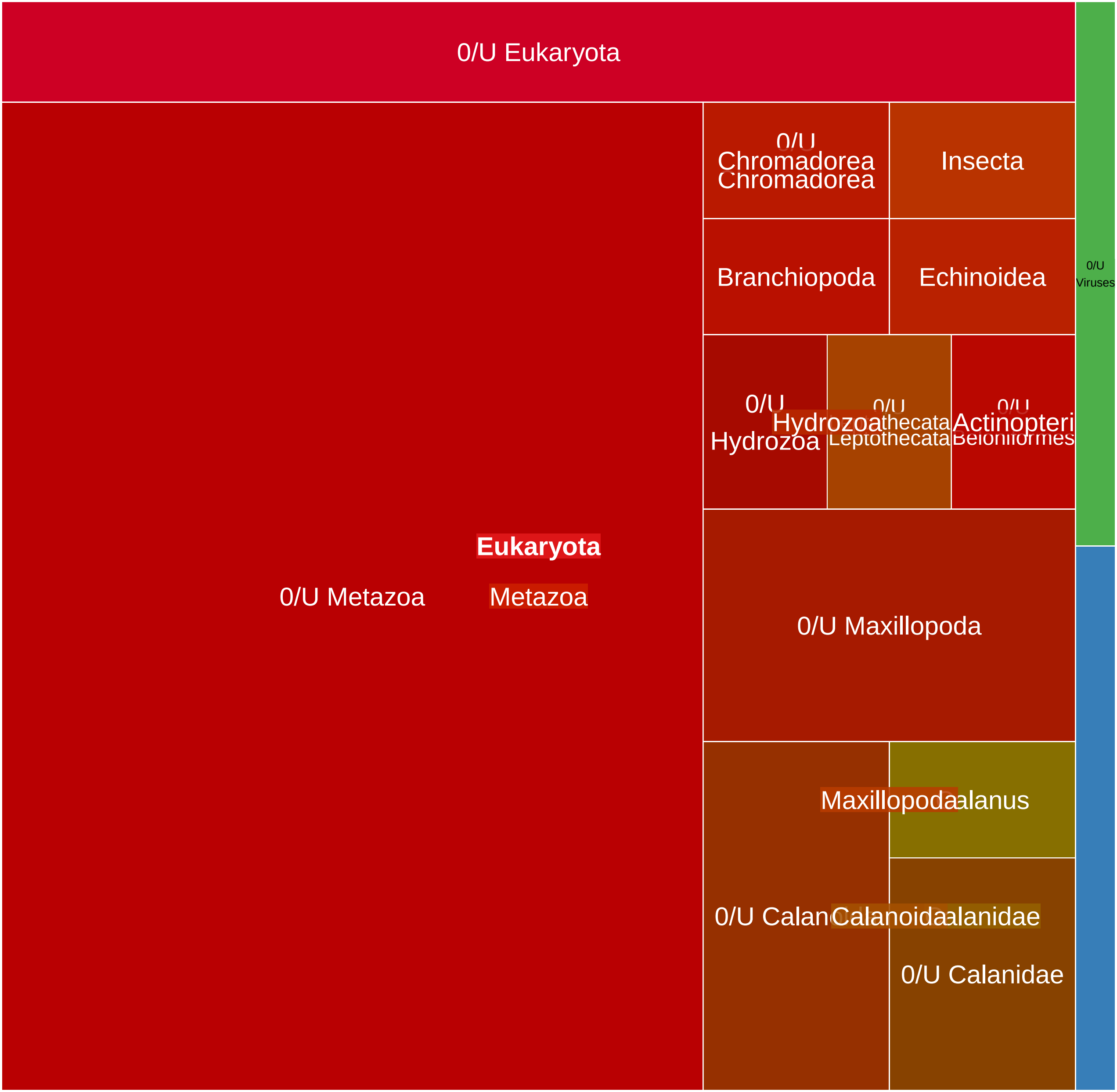

### MGT−v1_876 : 540 unigenes, 416 (77.04%) taxonomically assigned

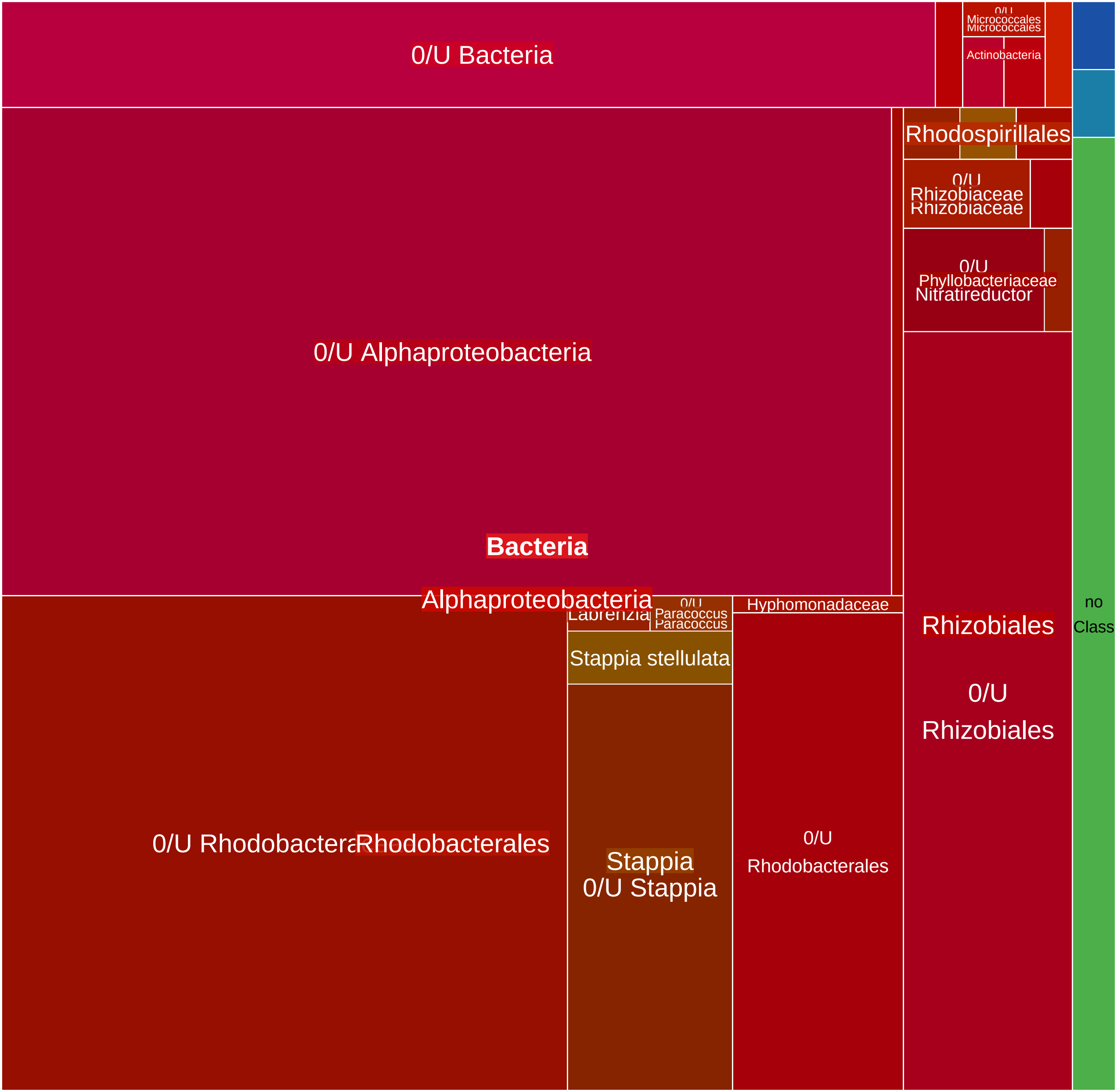

MGT−v1_877 : 539 unigenes, 463 (85.90%) taxonomically assigned

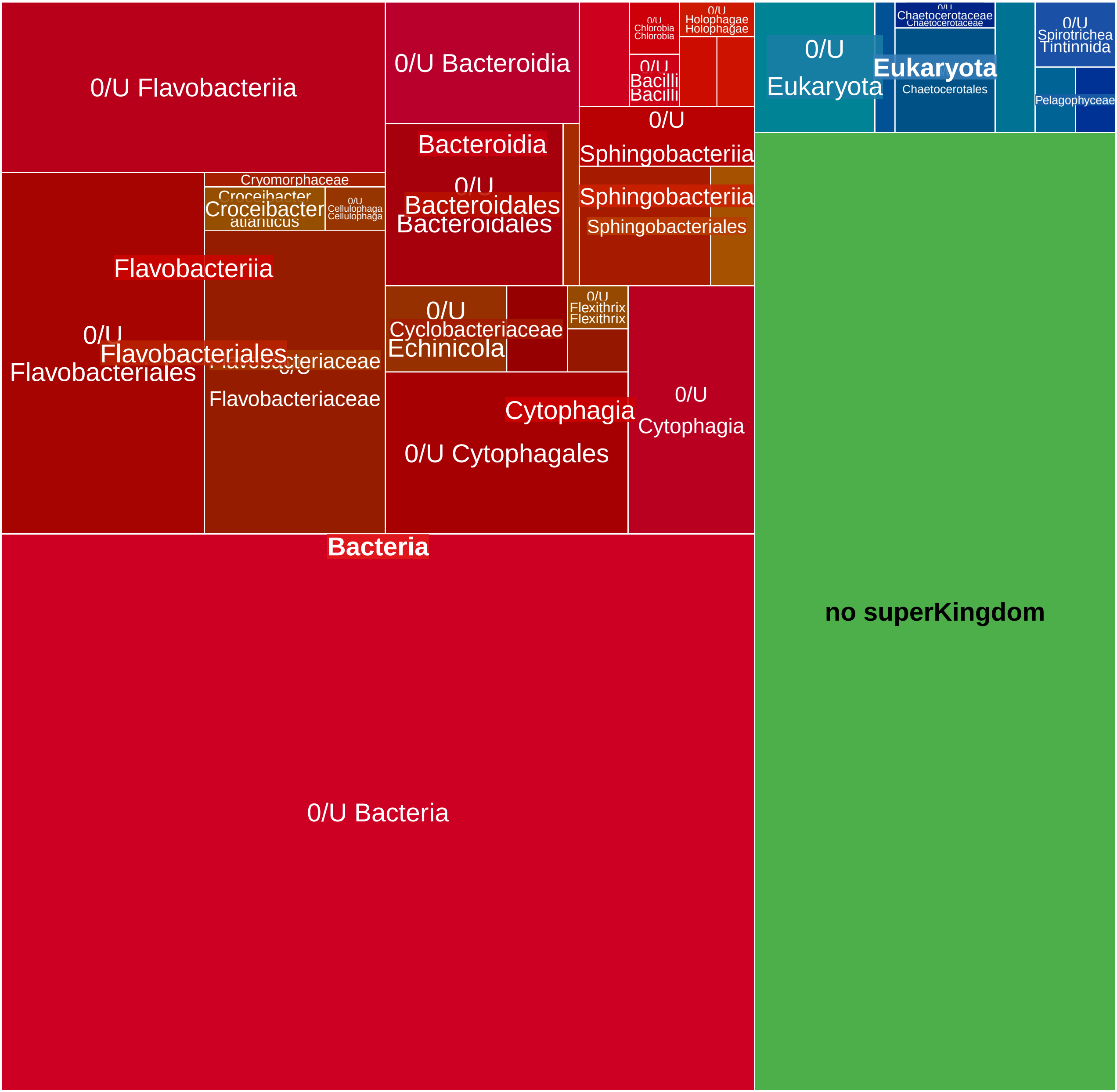

MGT−v1_878 : 537 unigenes, 309 (57.54%) taxonomically assigned

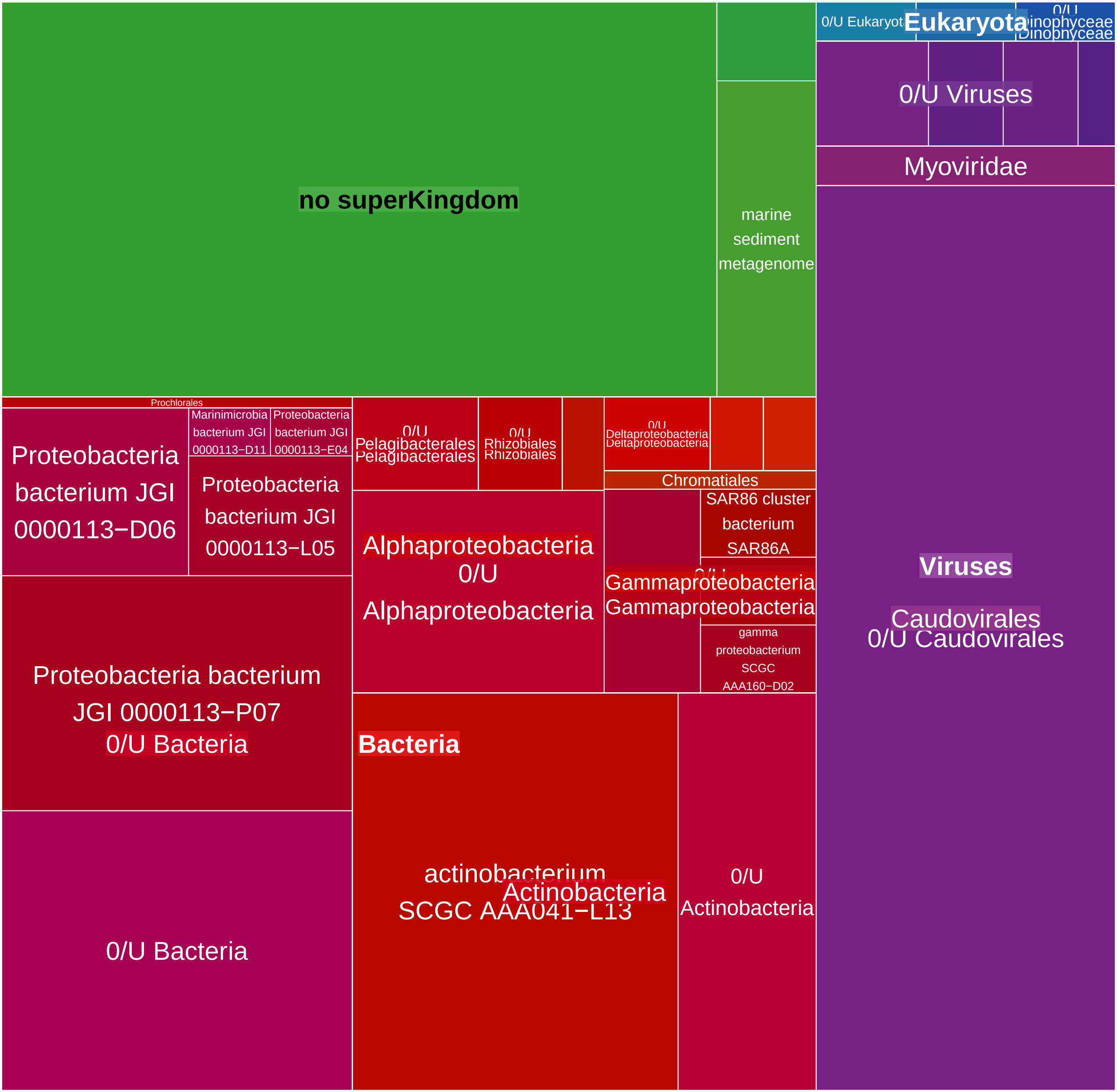

MGT−v1_879 : 536 unigenes, 43 (8.02%) taxonomically assigned

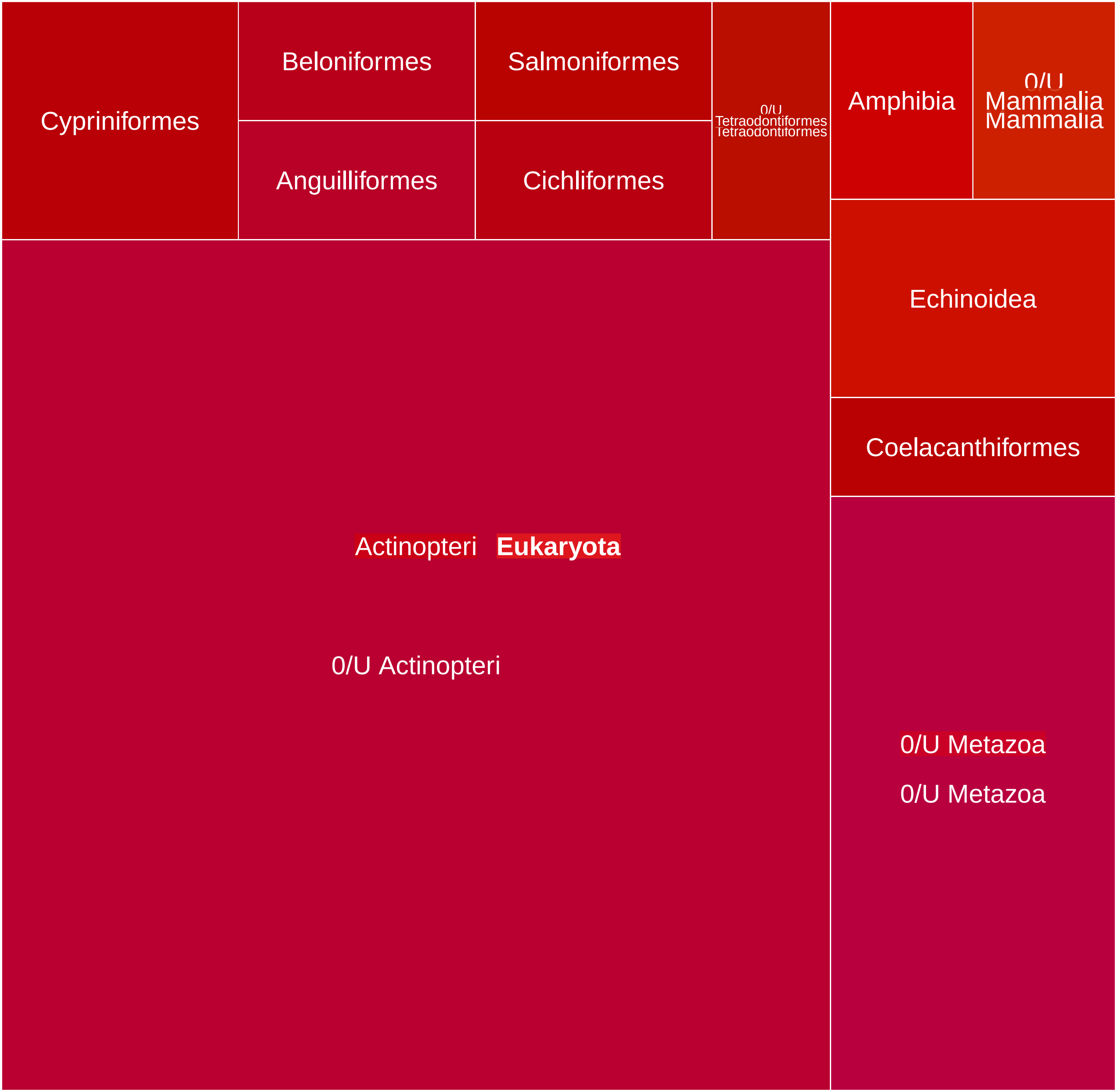

MGT−v1_880 : 532 unigenes, 71 (13.35%) taxonomically assigned

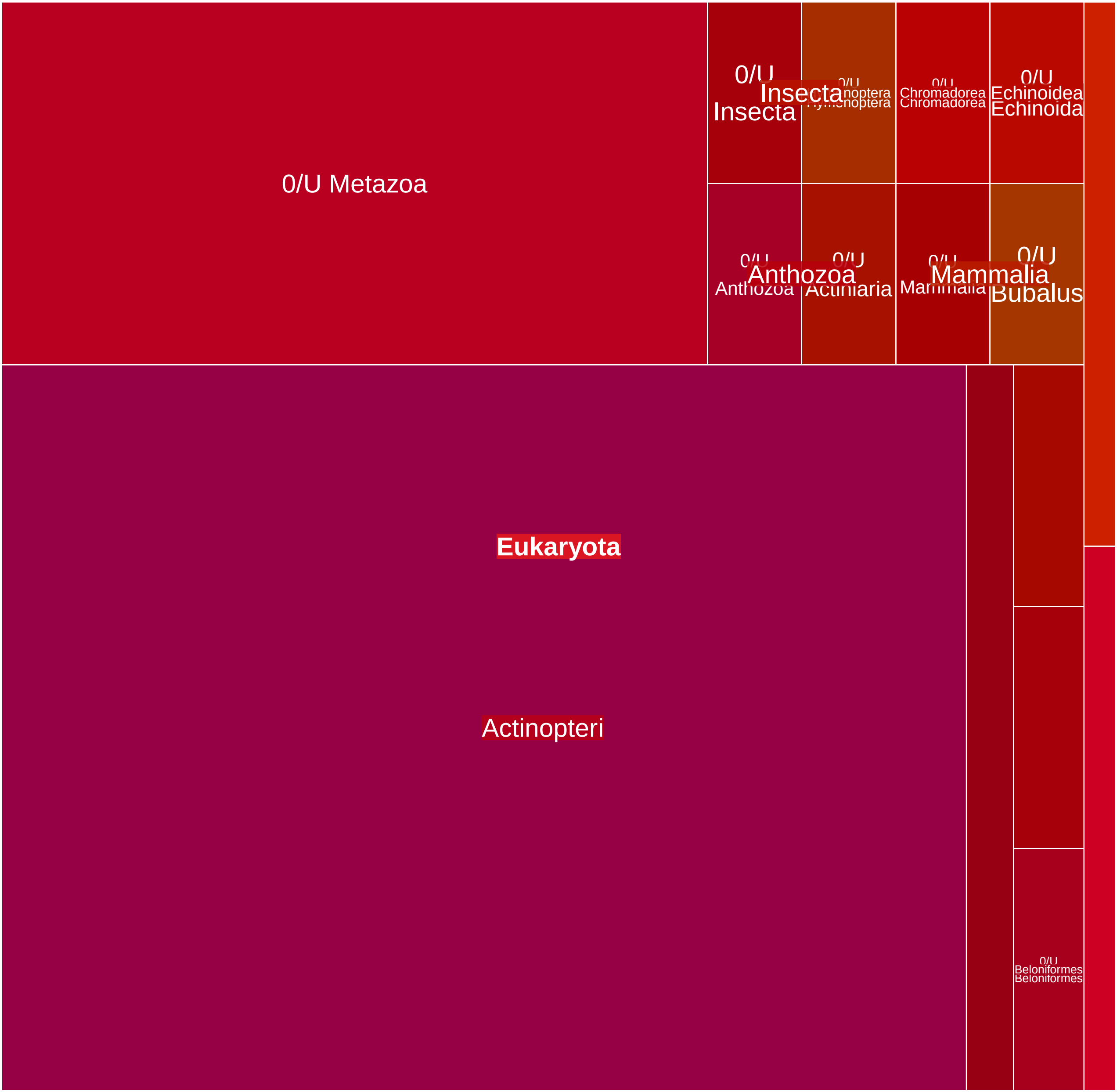

### MGT−v1_881 : 531 unigenes, 47 (8.85%) taxonomically assigned

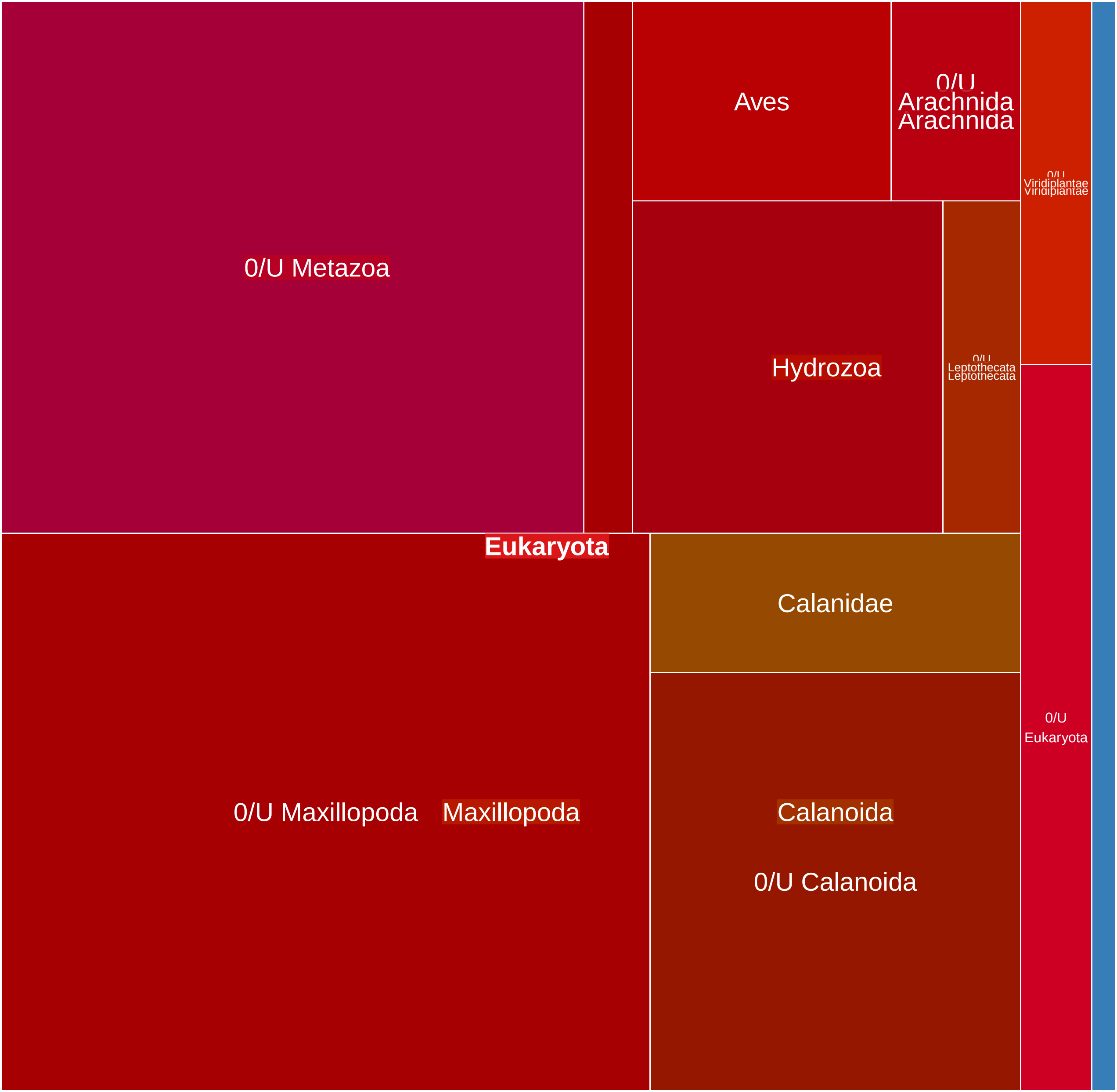

MGT−v1_882 : 530 unigenes, 87 (16.42%) taxonomically assigned

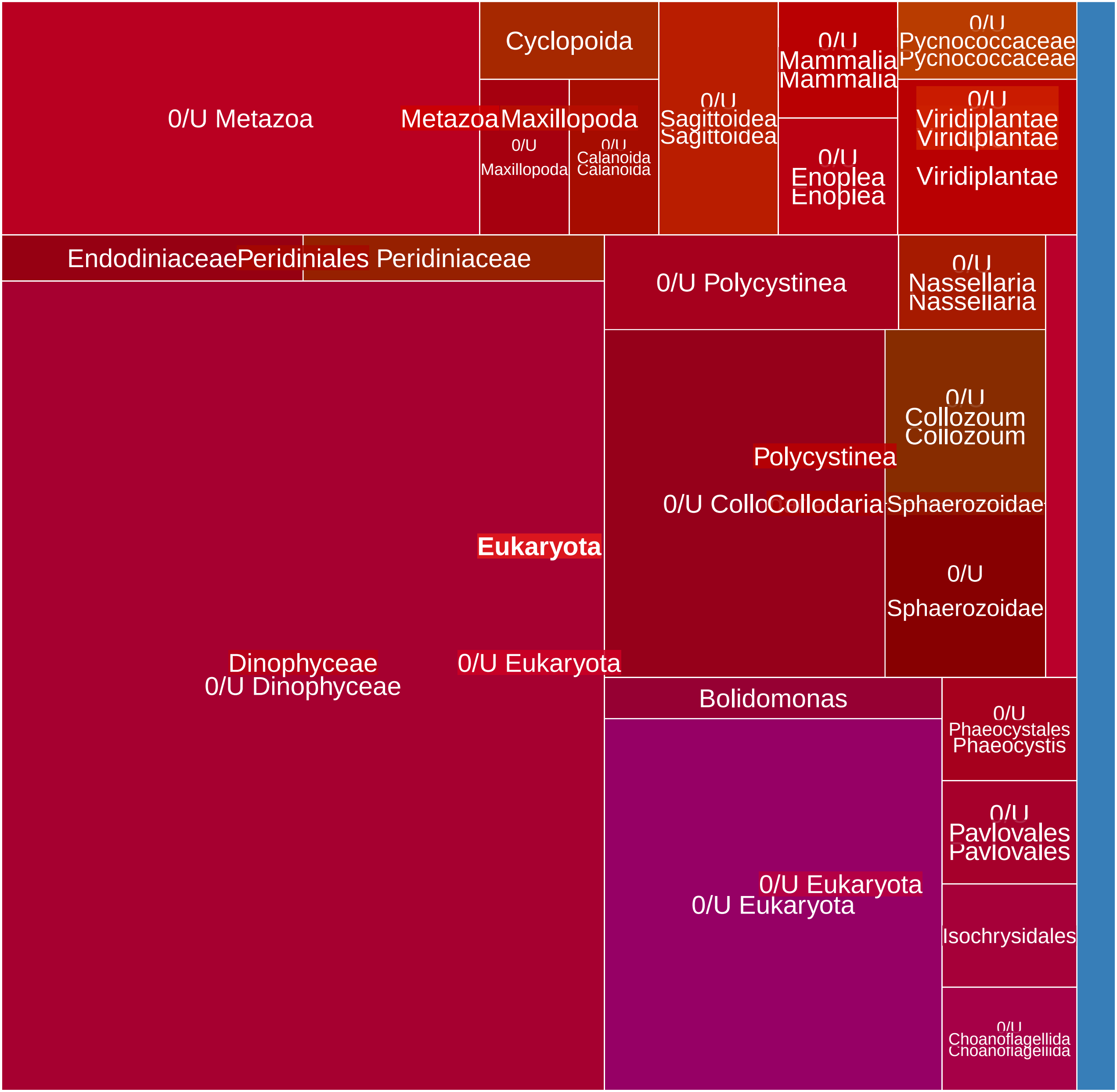

MGT−v1_883 : 529 unigenes, 62 (11.72%) taxonomically assigned

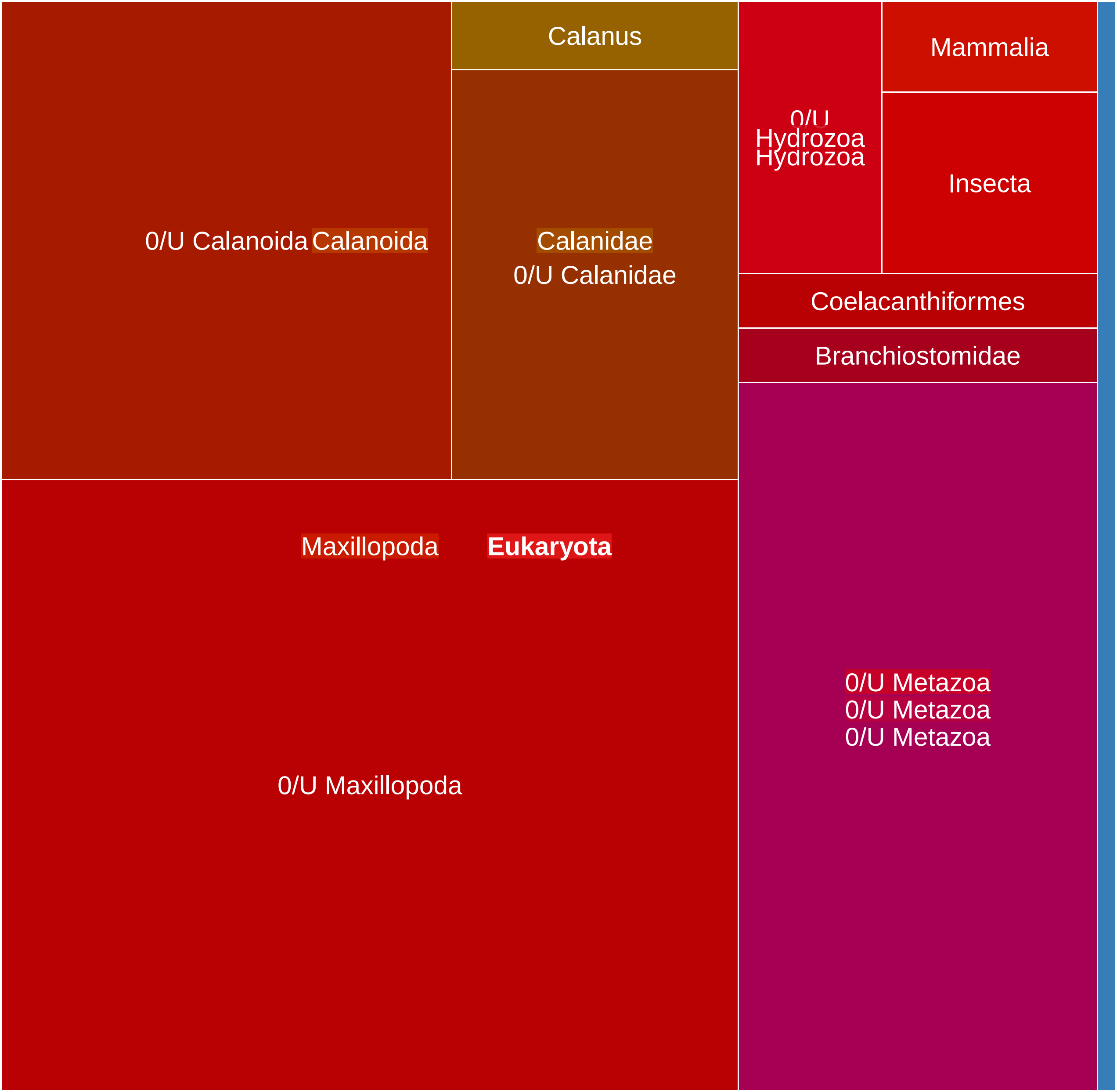

MGT−v1_884 : 529 unigenes, 15 (2.84%) taxonomically assigned

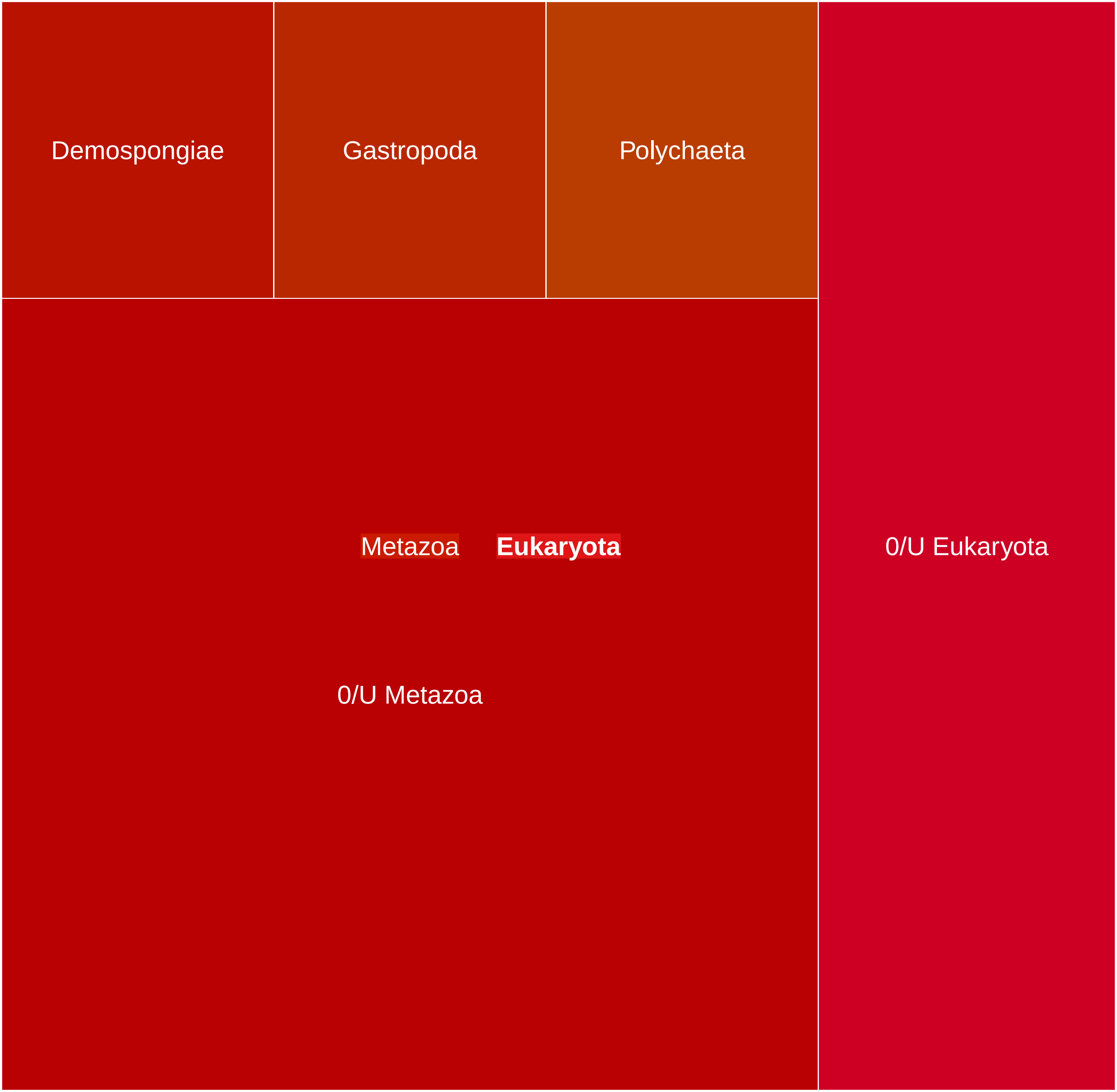

MGT−v1_885 : 528 unigenes, 16 (3.03%) taxonomically assigned

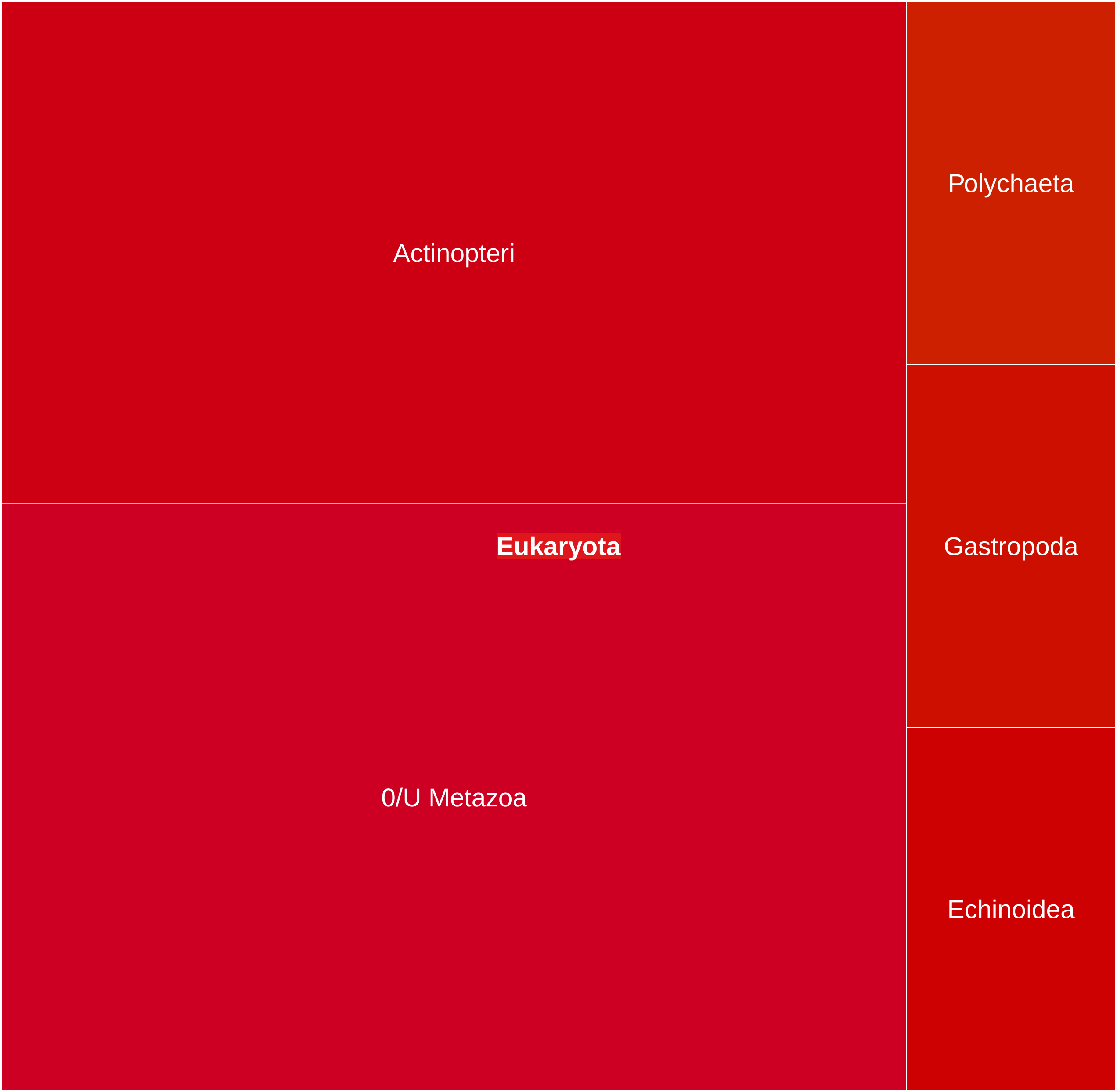

MGT−v1_886 : 528 unigenes, 80 (15.15%) taxonomically assigned

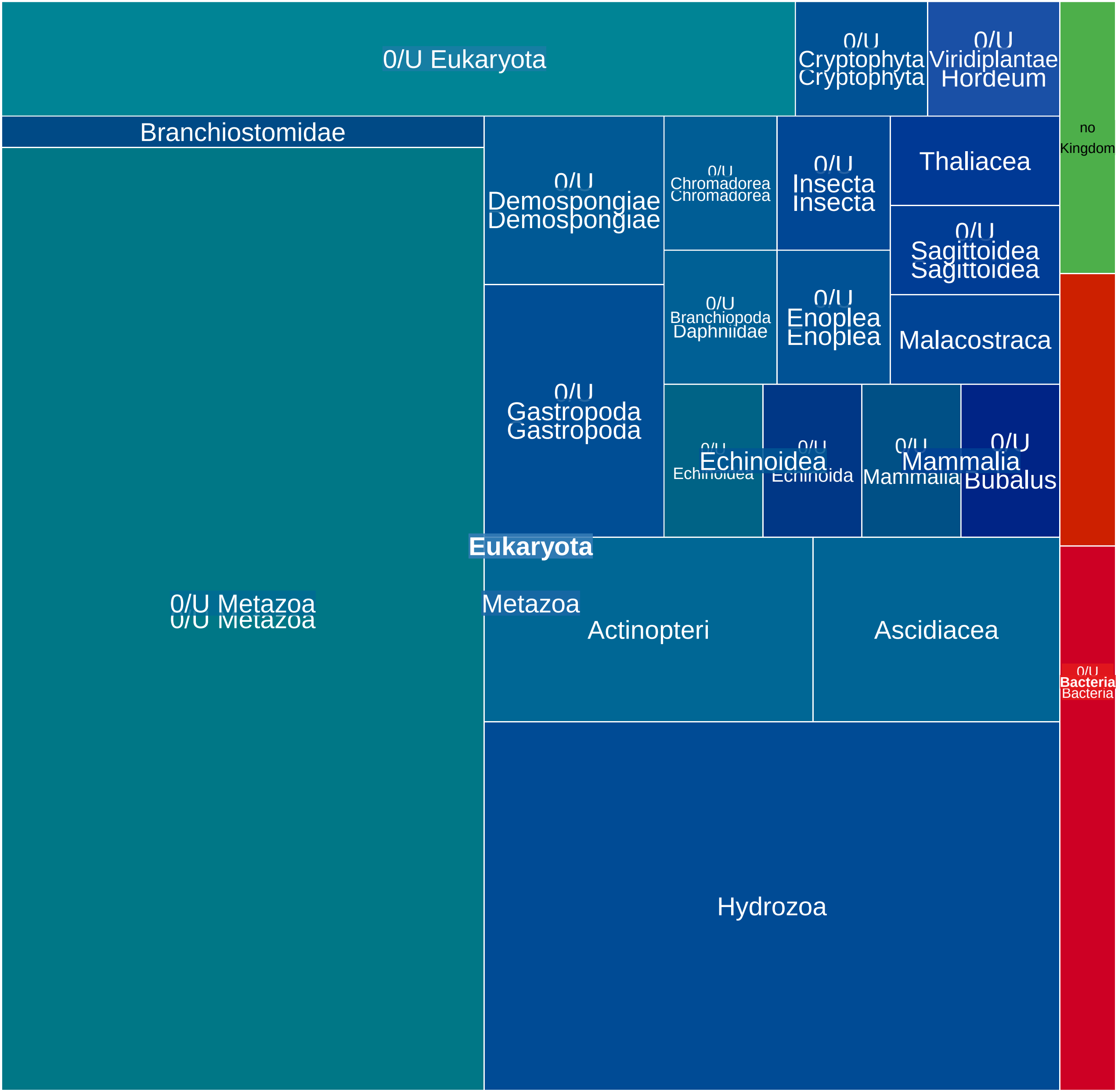

MGT−v1_887 : 526 unigenes, 323 (61.41%) taxonomically assigned

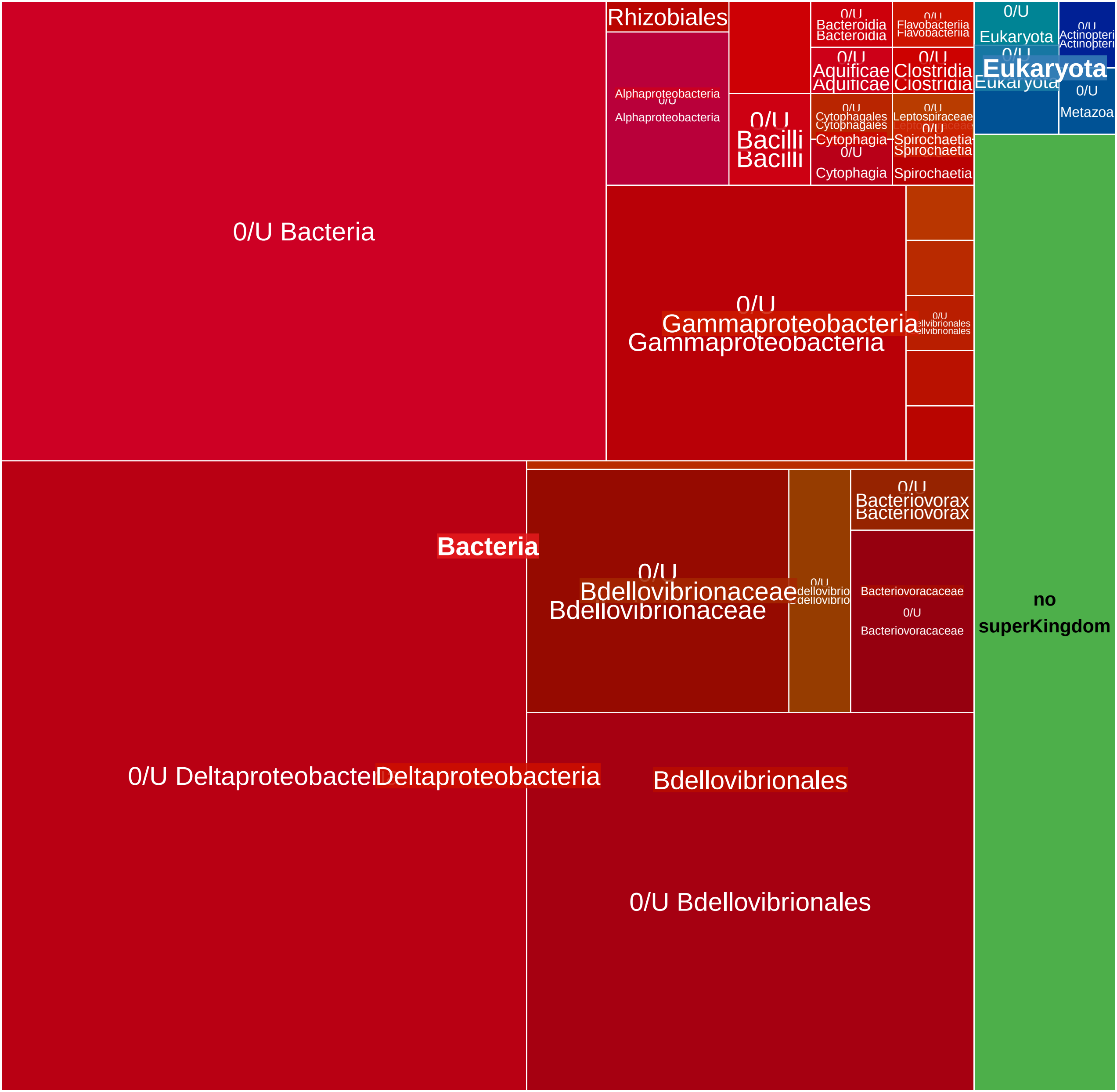

MGT−v1_888 : 526 unigenes, 225 (42.78%) taxonomically assigned

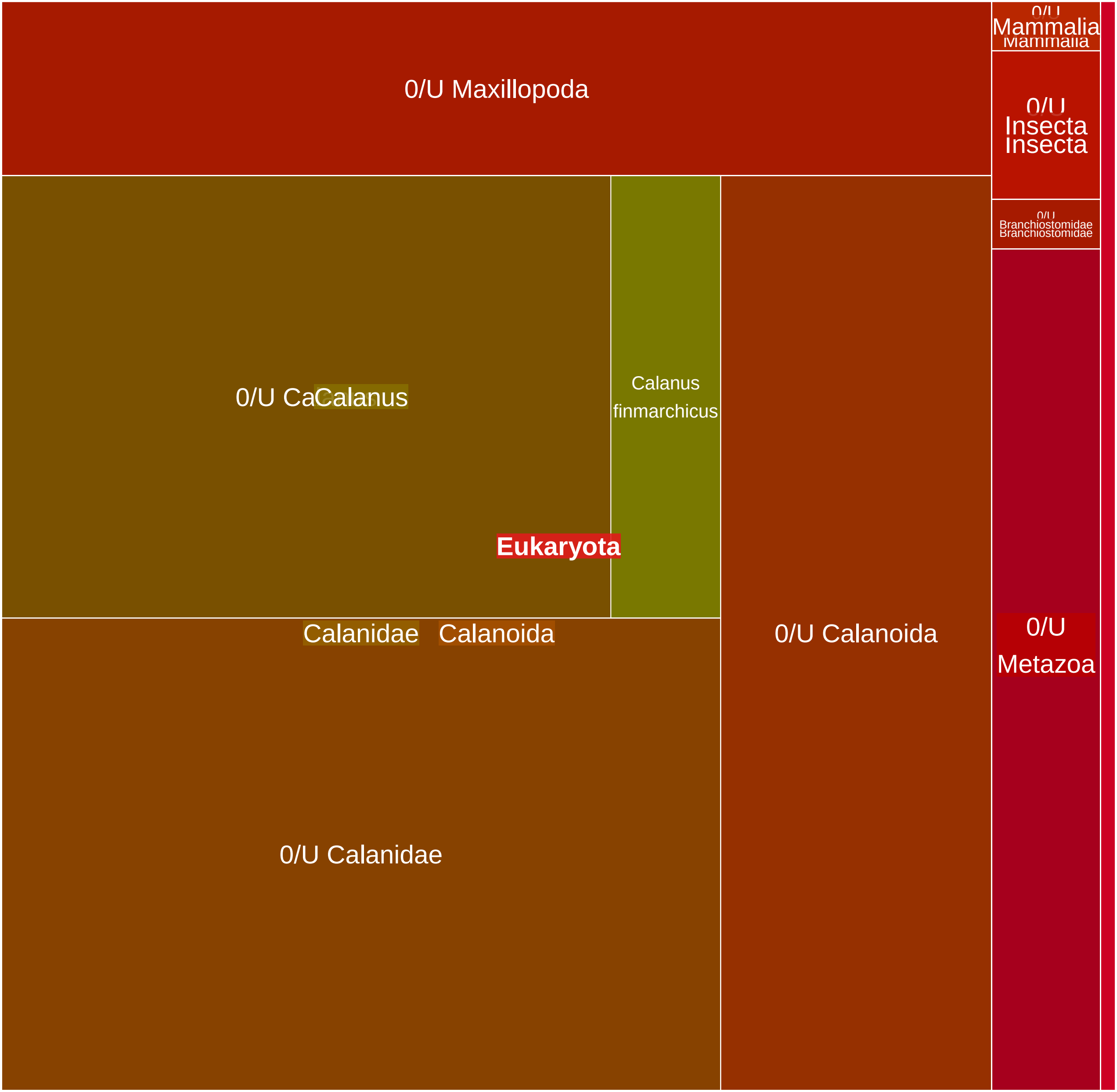

MGT−v1_889 : 521 unigenes, 185 (35.51%) taxonomically assigned

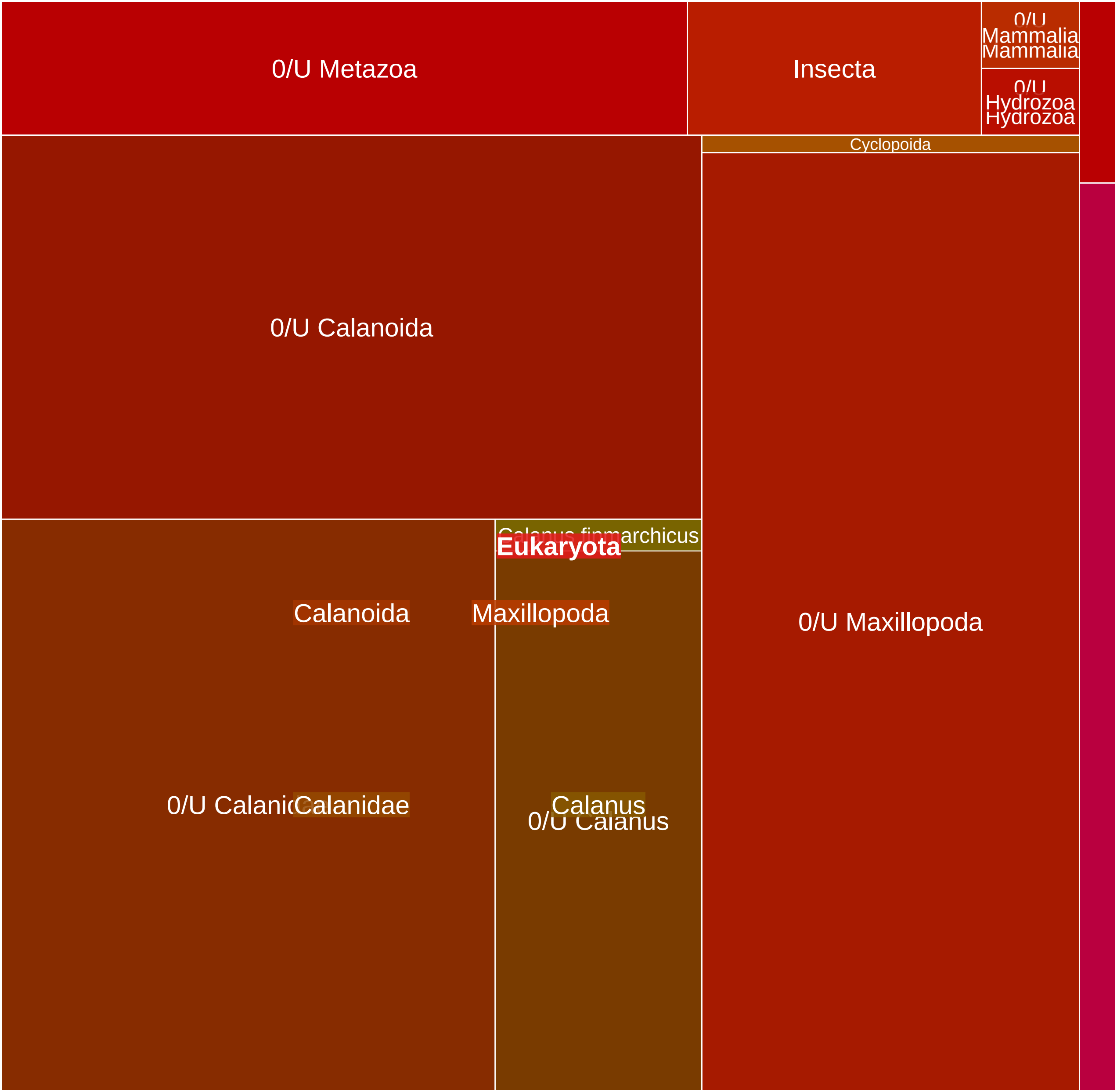

MGT−v1_890 : 521 unigenes, 24 (4.61%) taxonomically assigned

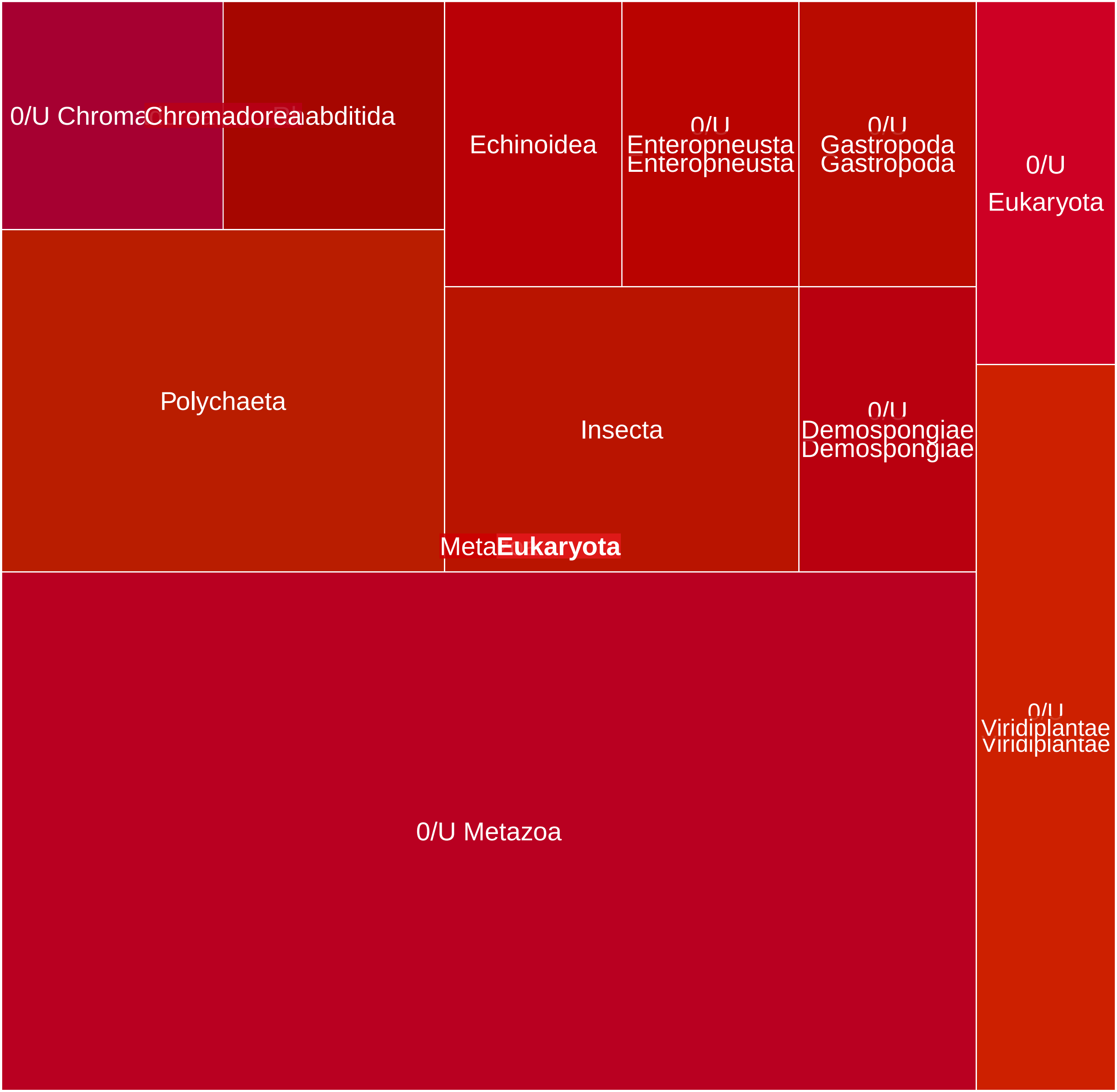

MGT−v1_891 : 521 unigenes, 59 (11.32%) taxonomically assigned

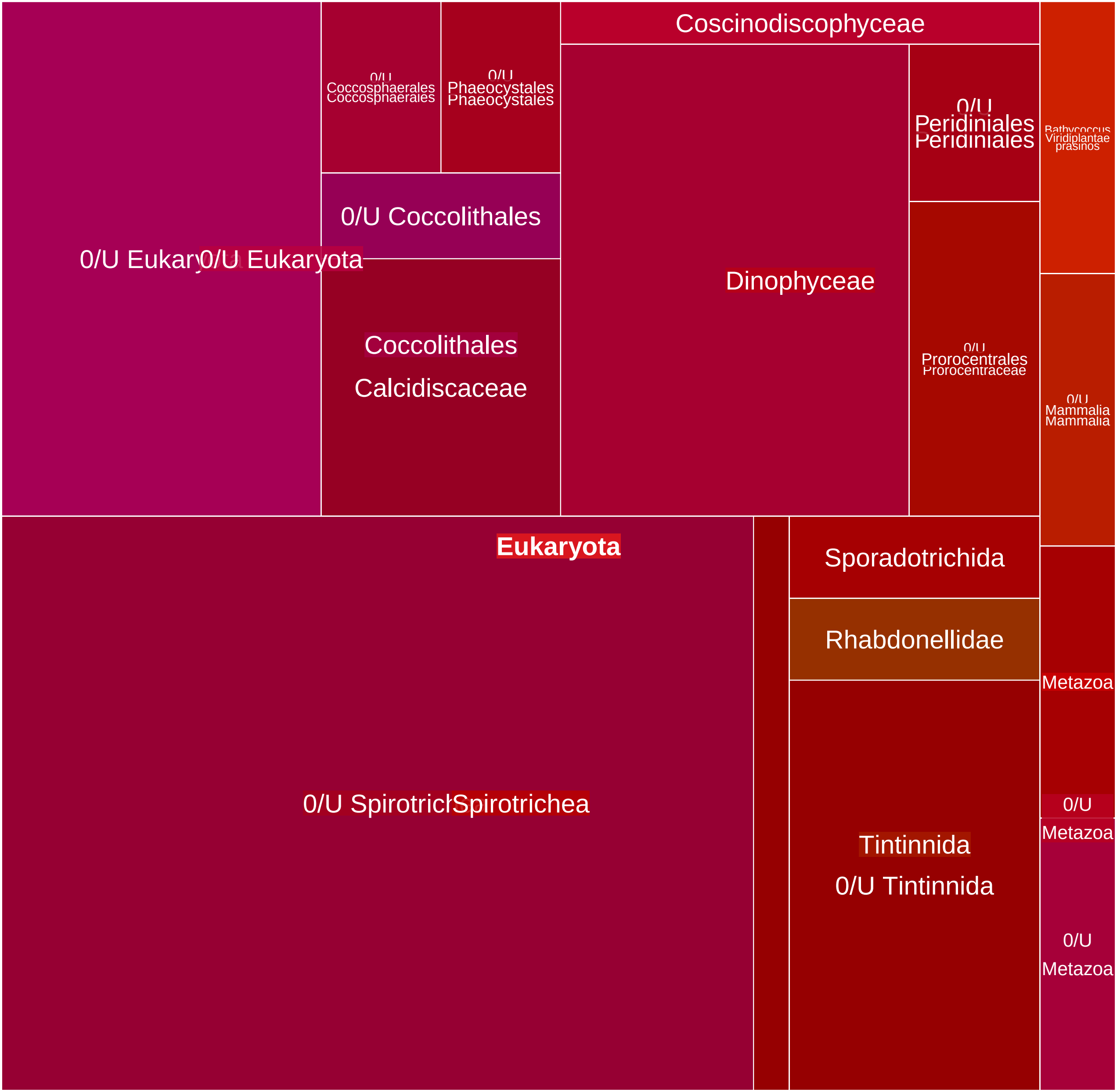

MGT−v1_892 : 521 unigenes, 24 (4.61%) taxonomically assigned

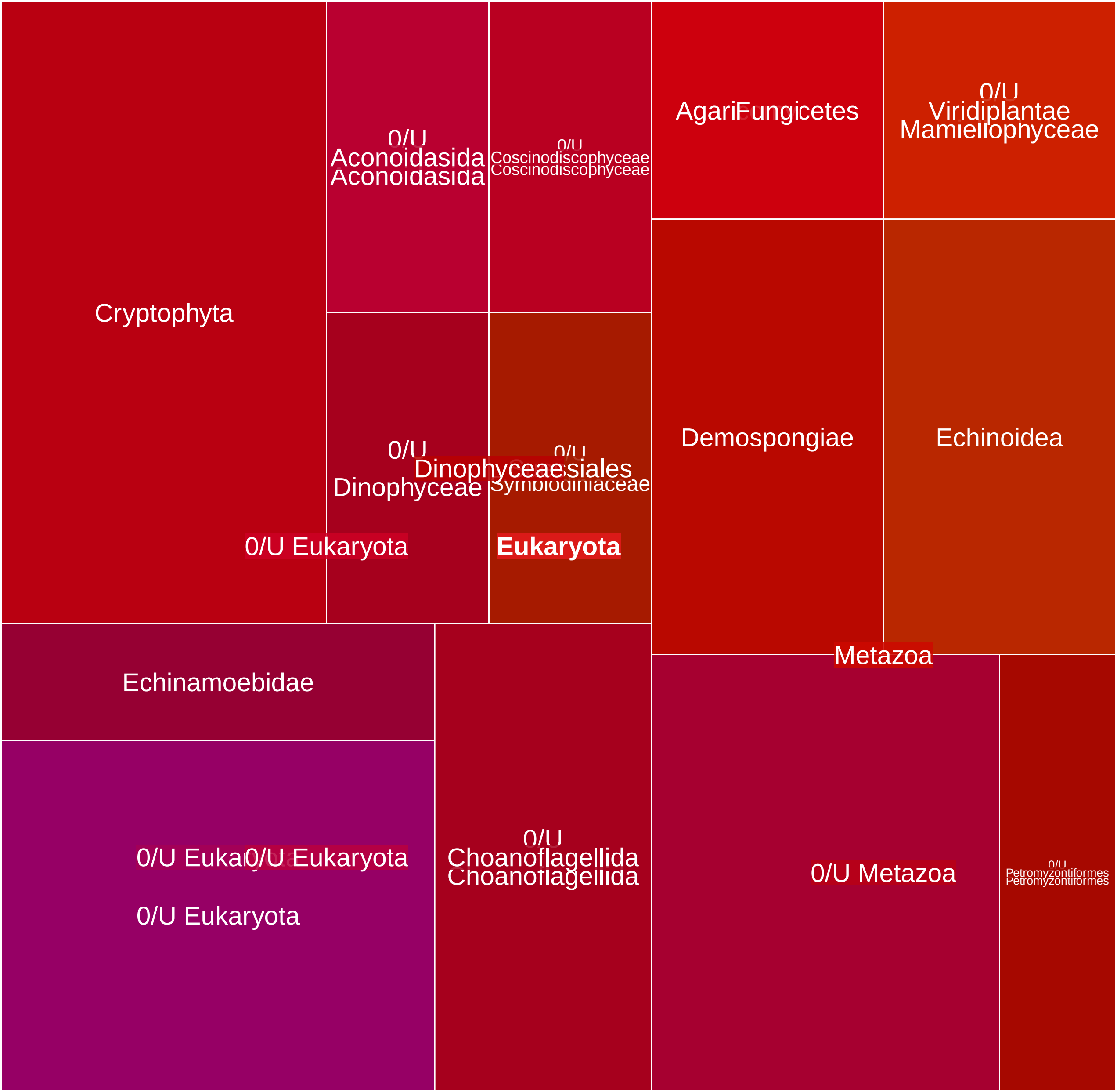

MGT−v1_893 : 520 unigenes, 489 (94.04%) taxonomically assigned

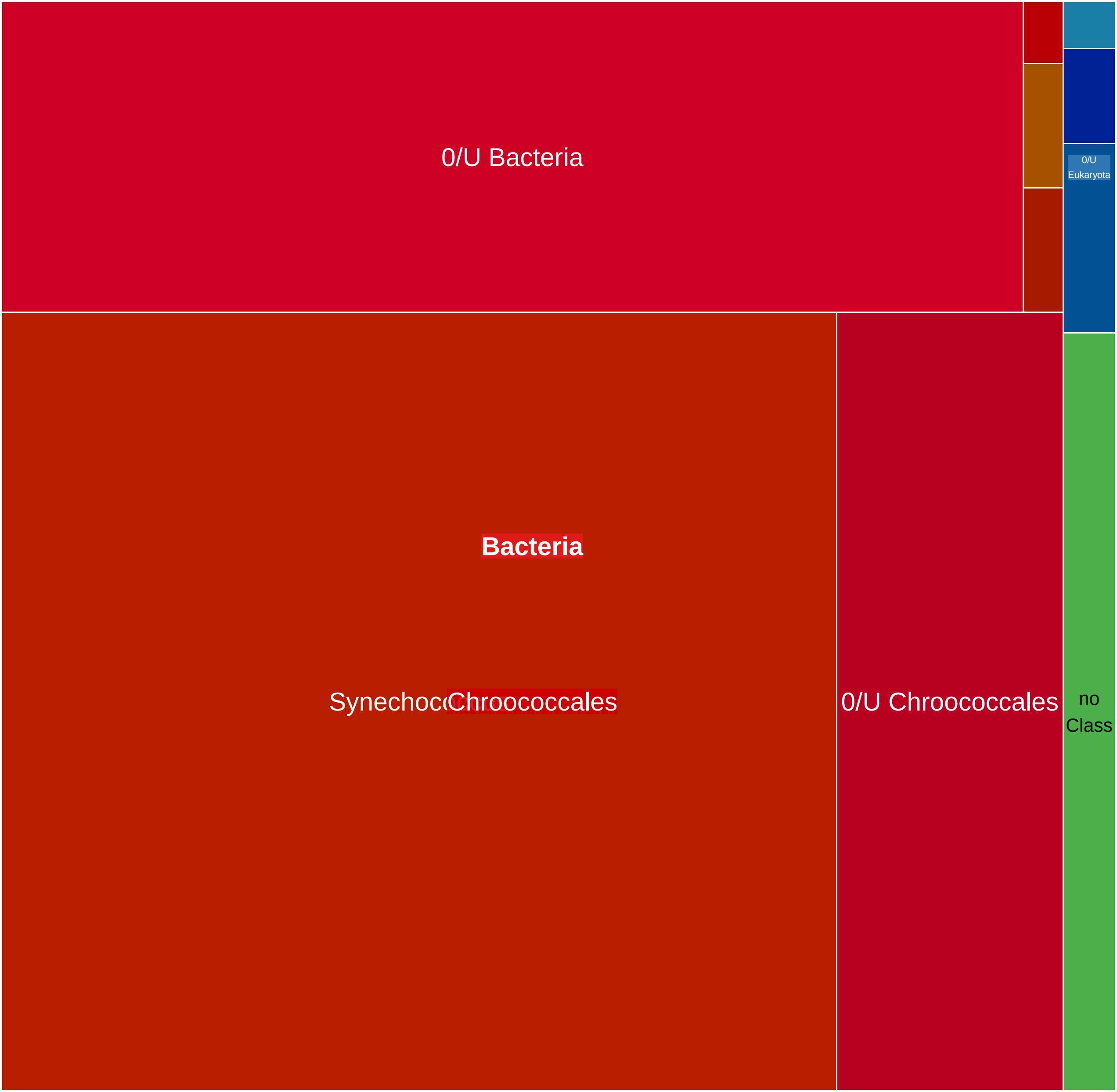

MGT−v1_894 : 519 unigenes, 32 (6.17%) taxonomically assigned

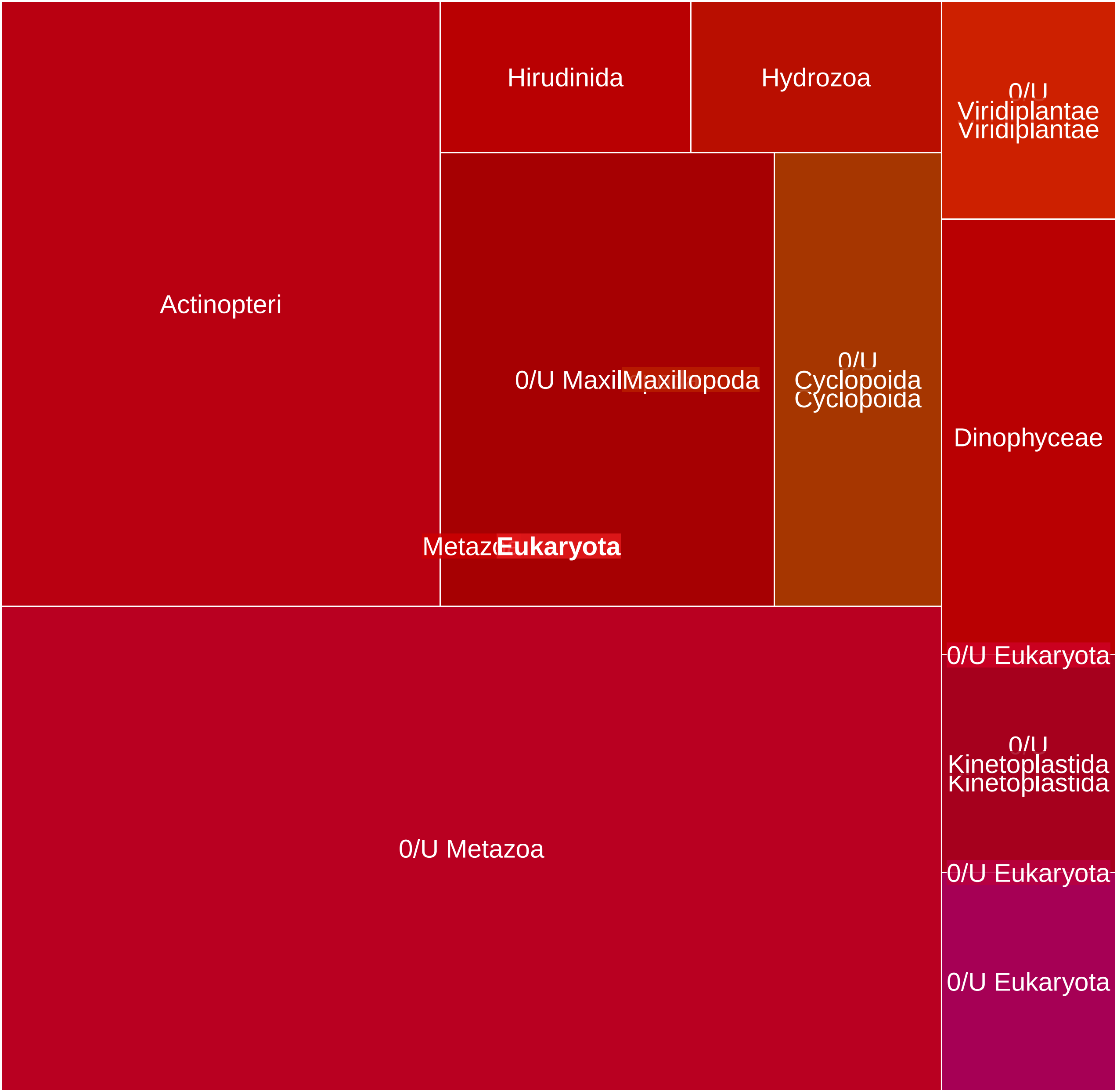

MGT−v1_895 : 518 unigenes, 149 (28.76%) taxonomically assigned

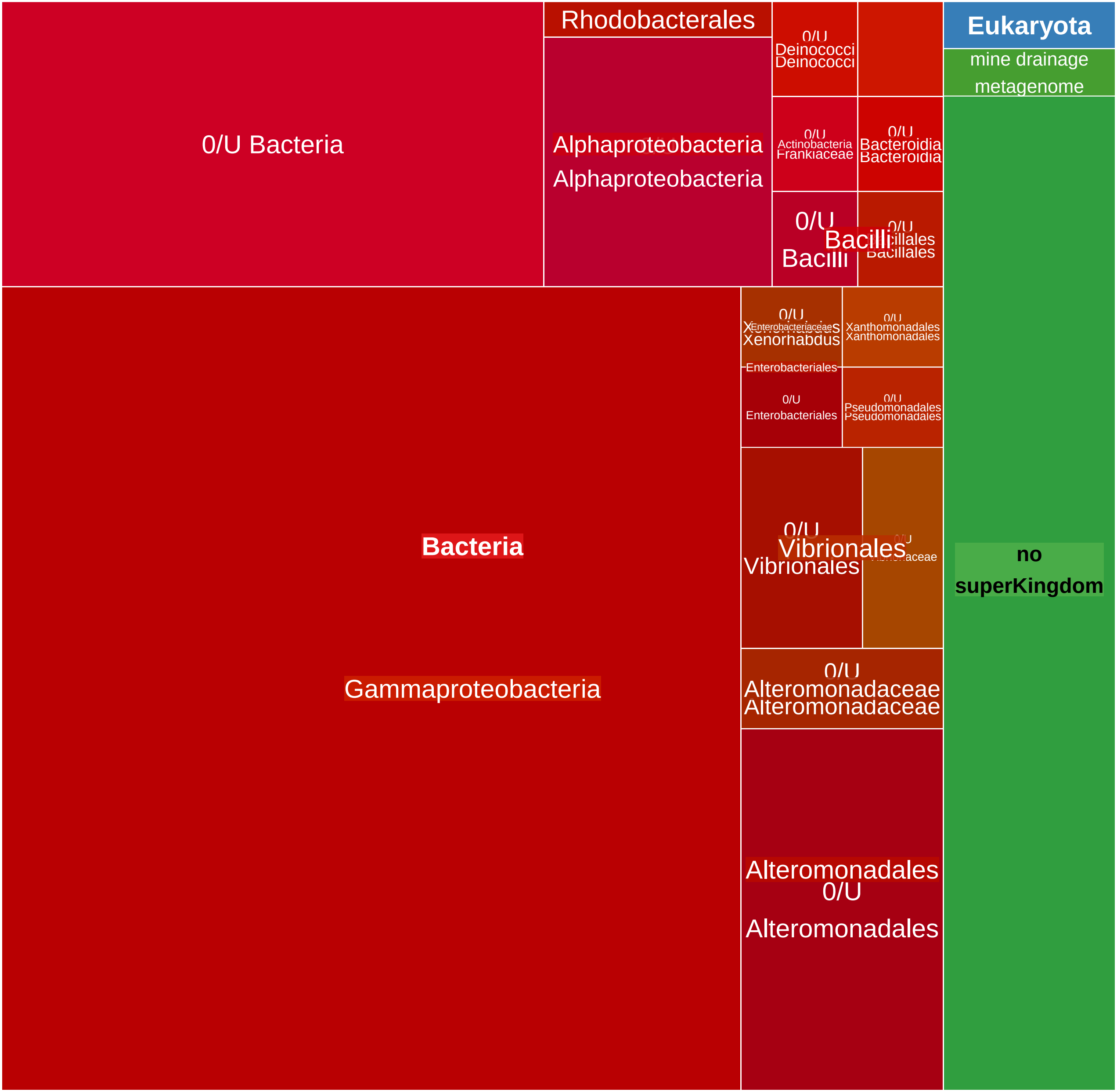

MGT−v1_896 : 517 unigenes, 342 (66.15%) taxonomically assigned

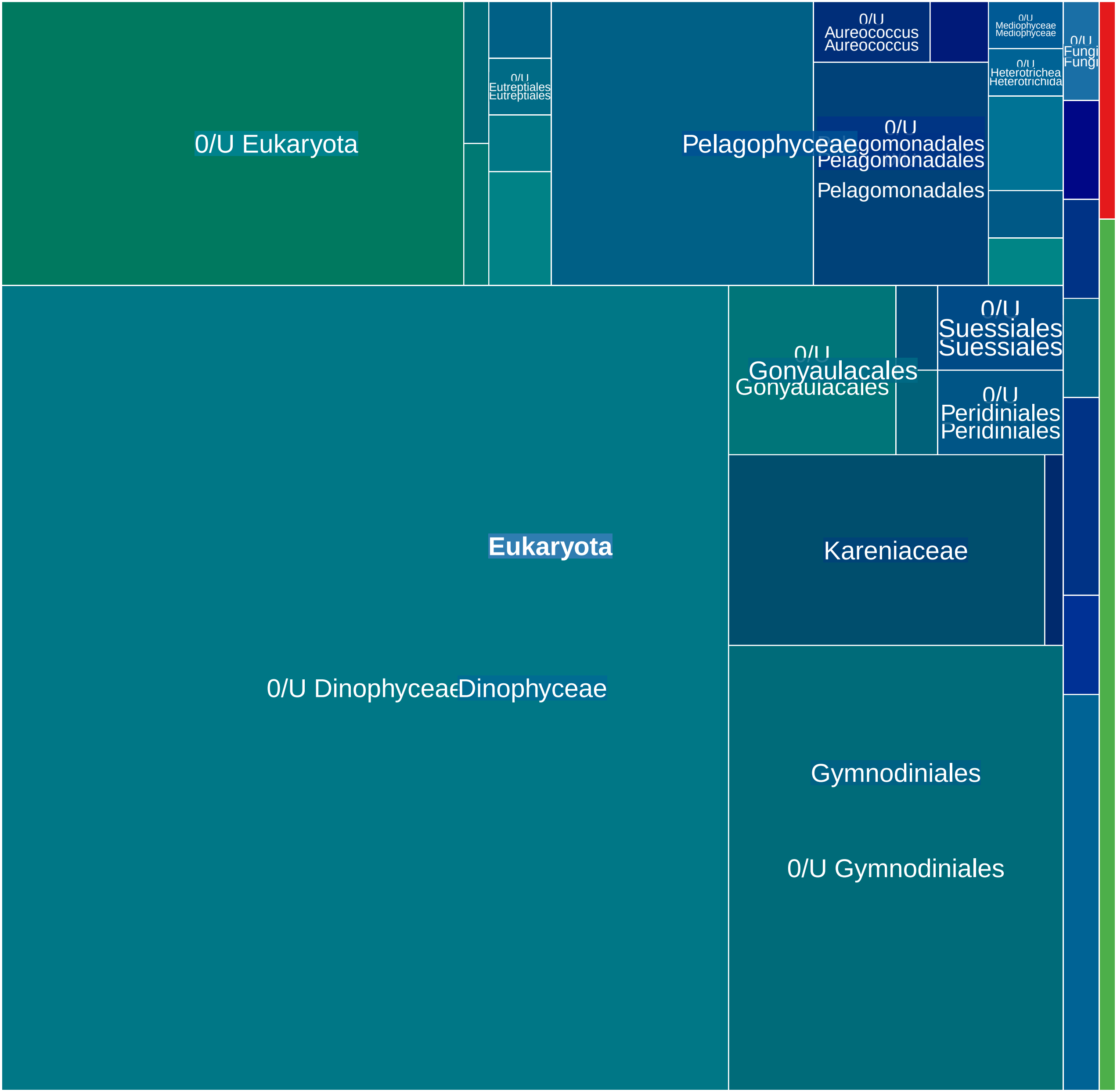

MGT−v1_897 : 517 unigenes, 61 (11.80%) taxonomically assigned

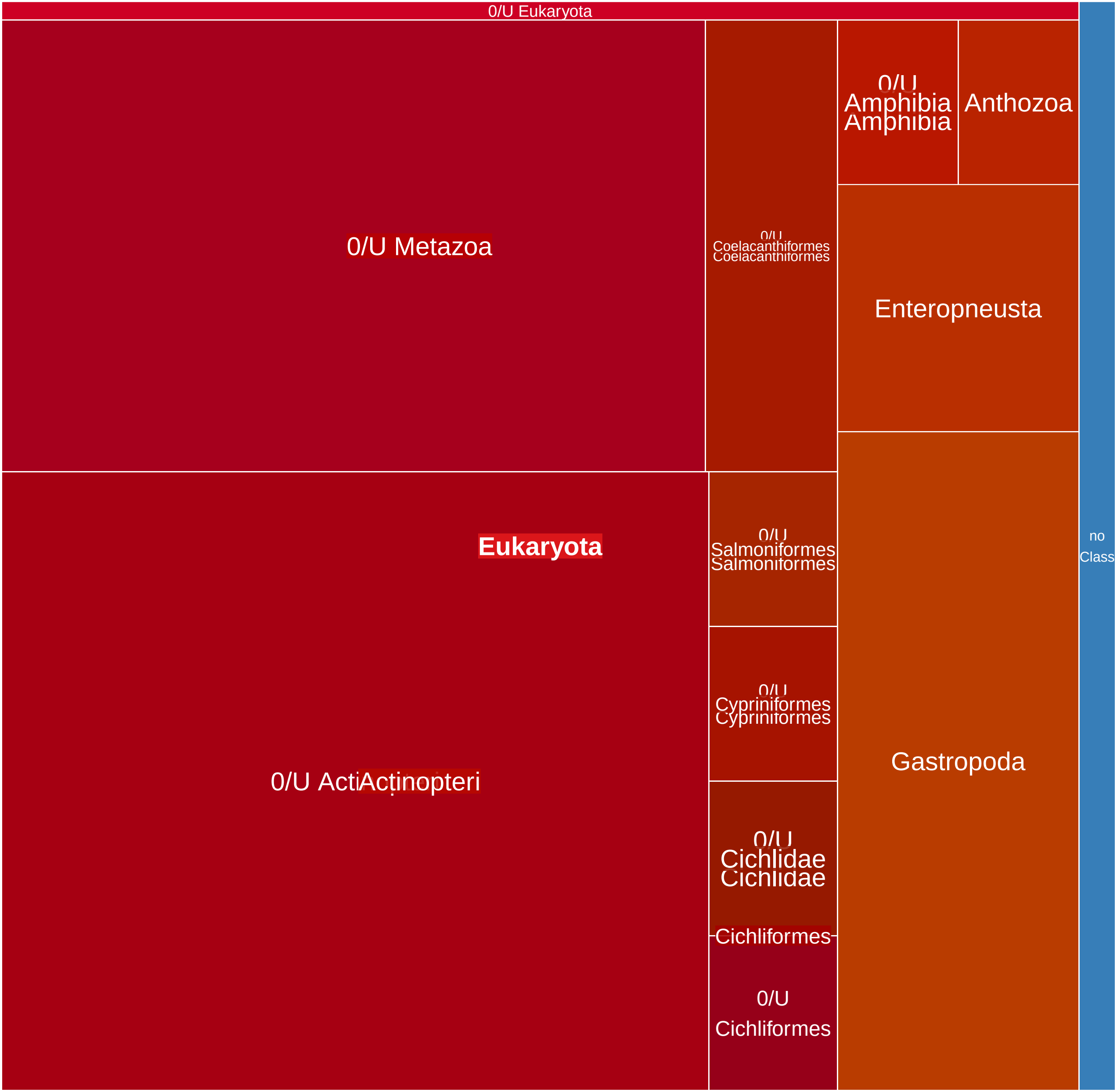

MGT−v1_898 : 517 unigenes, 58 (11.22%) taxonomically assigned

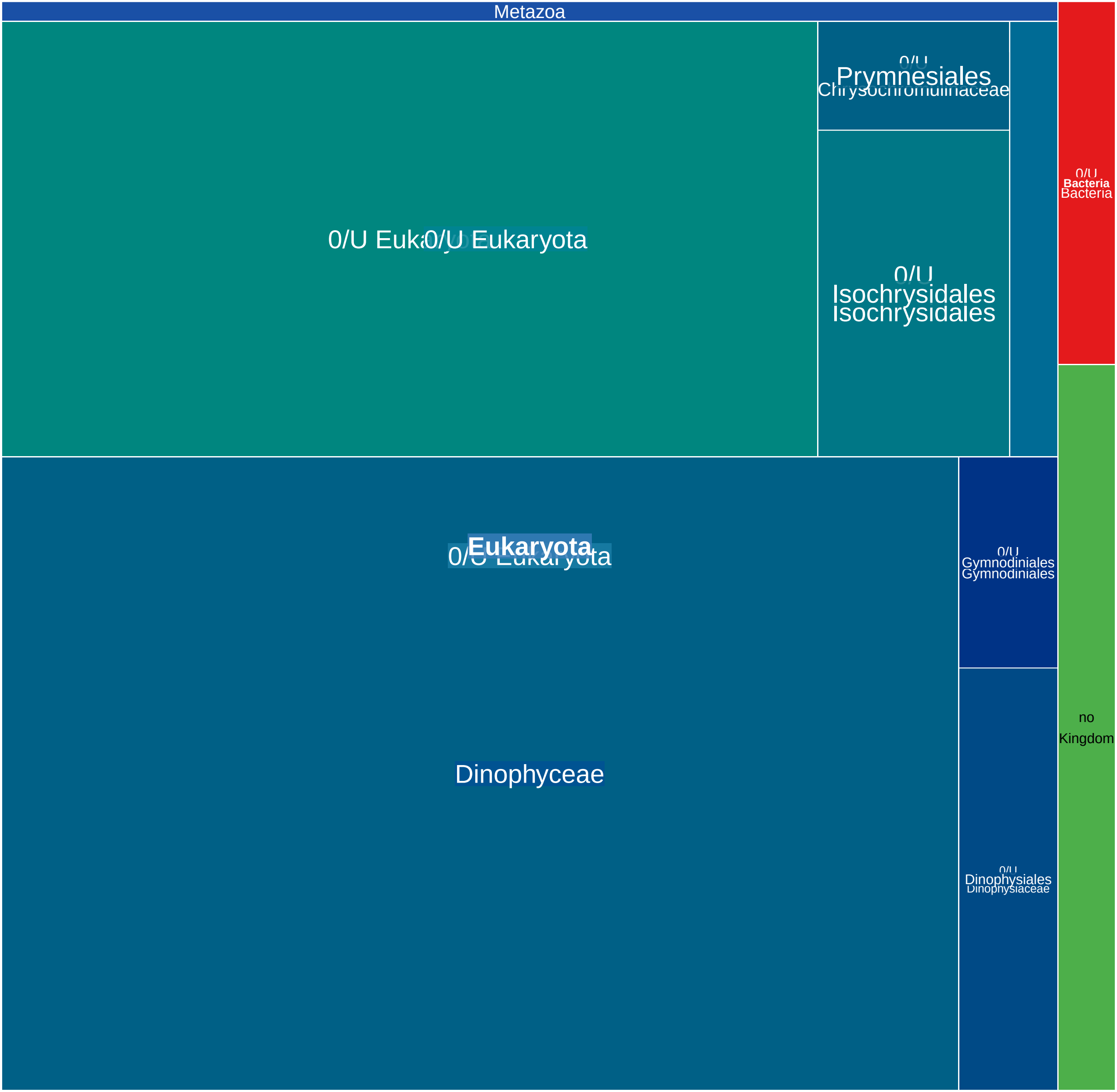

MGT−v1_899 : 516 unigenes, 153 (29.65%) taxonomically assigned

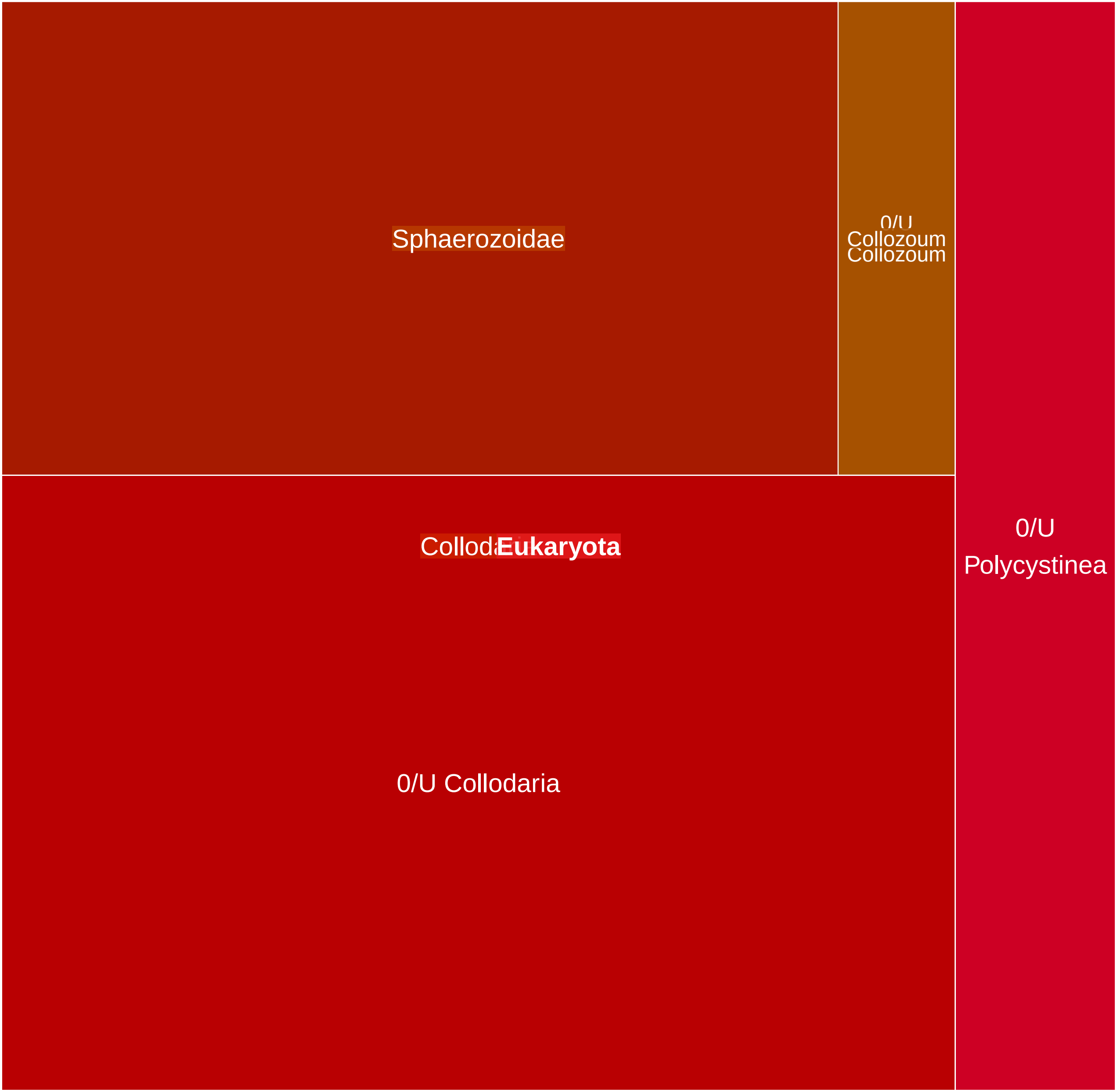

### MGT−v1_900 : 515 unigenes, 481 (93.40%) taxonomically assigned

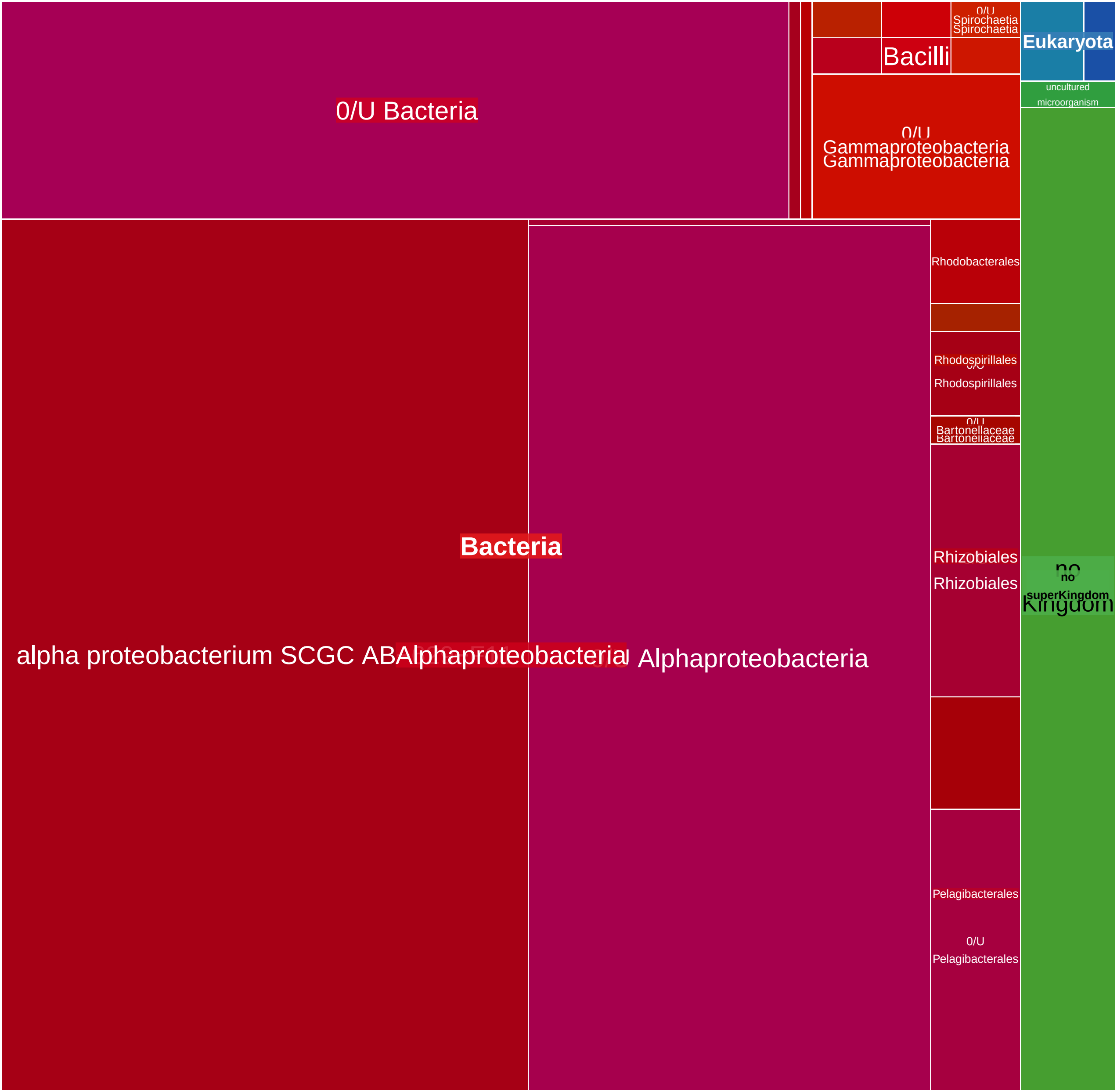

### MGT−v1_901 : 514 unigenes, 159 (30.93%) taxonomically assigned

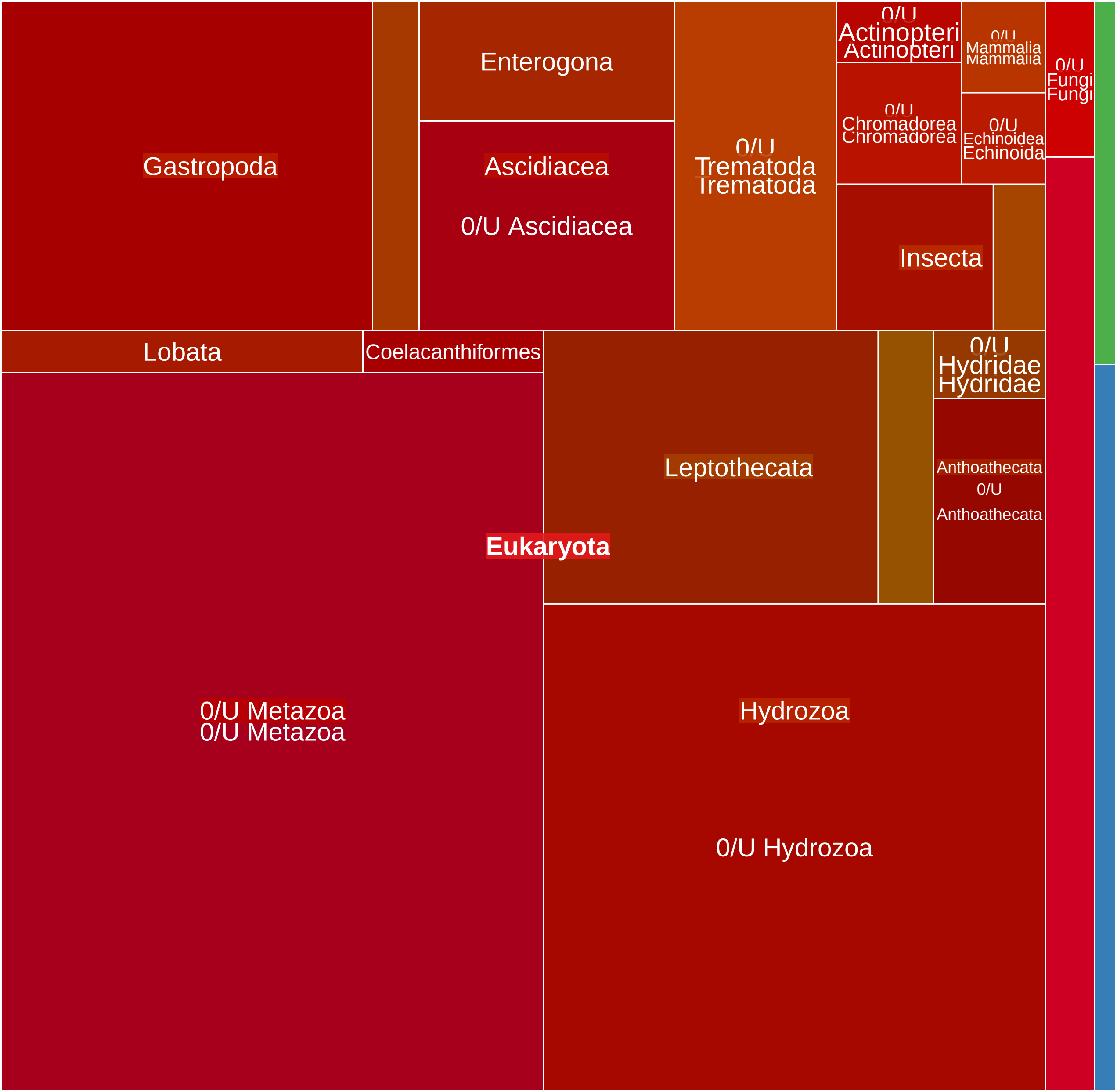

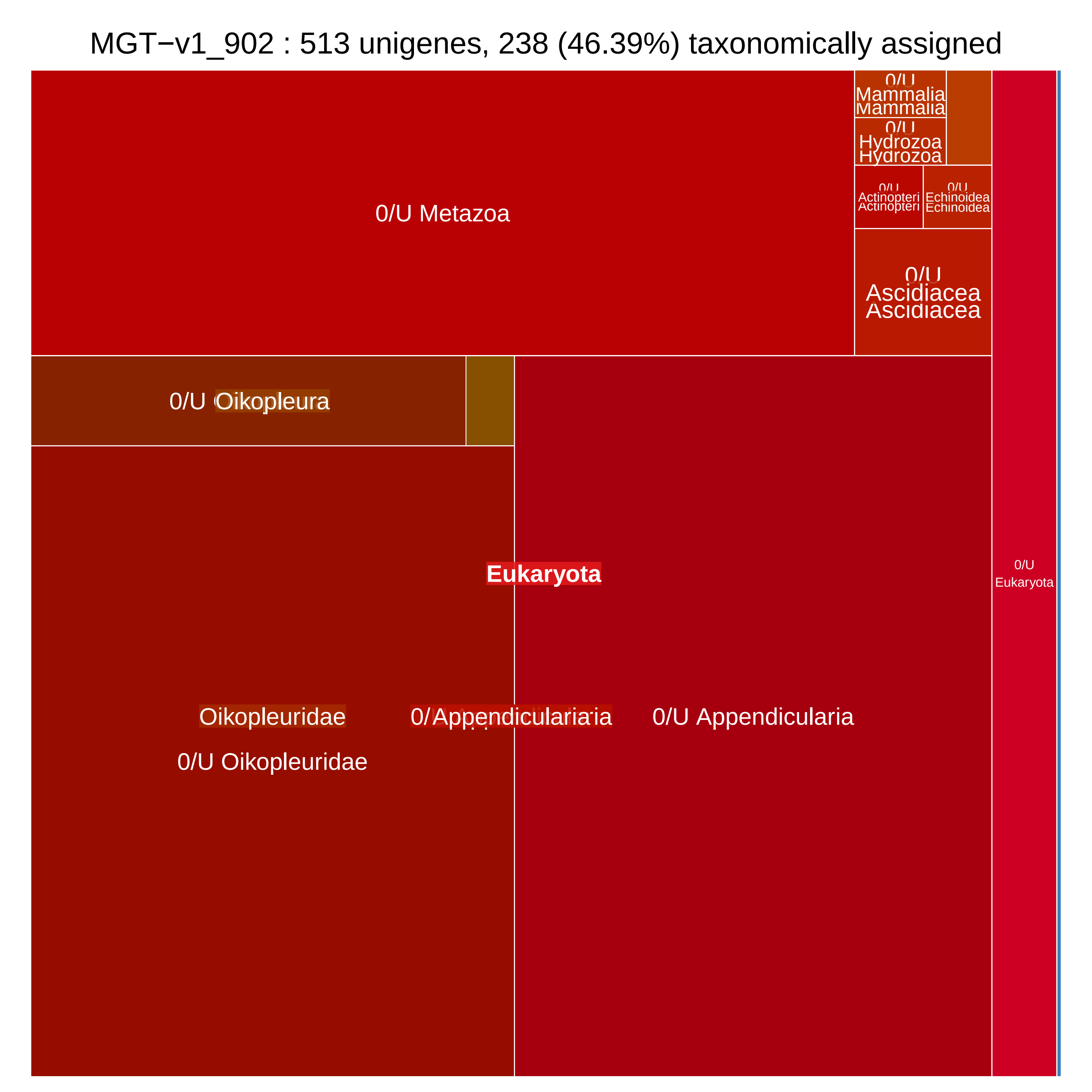
MGT−v1_902 : 513 unigenes, 238 (46.39%) taxonomically assigned
0/U Metazoa
0/U Mammalia
0/U Hydrozoa
0/U Actinopteri
0/U Echinoidea
0/U Ascidiacea
0/U Oikopleura
Eukaryota
Oikopleuridae
0/U Appendicularia
0/U Appendicularia
0/U Oikopleuridae
0/U Eukaryota

MGT−v1_903 : 512 unigenes, 235 (45.90%) taxonomically assigned

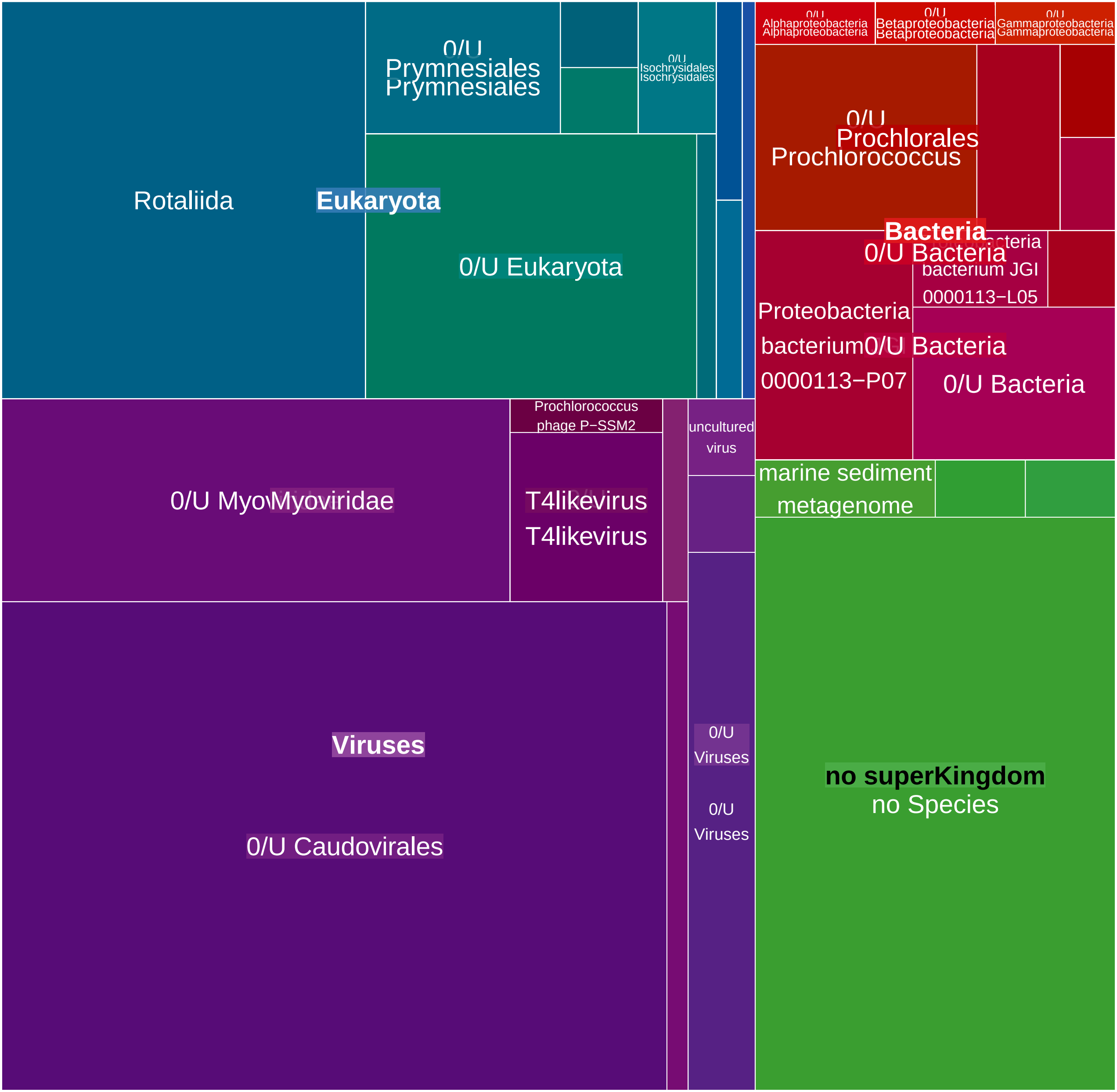

MGT−v1_904 : 512 unigenes, 485 (94.73%) taxonomically assigned

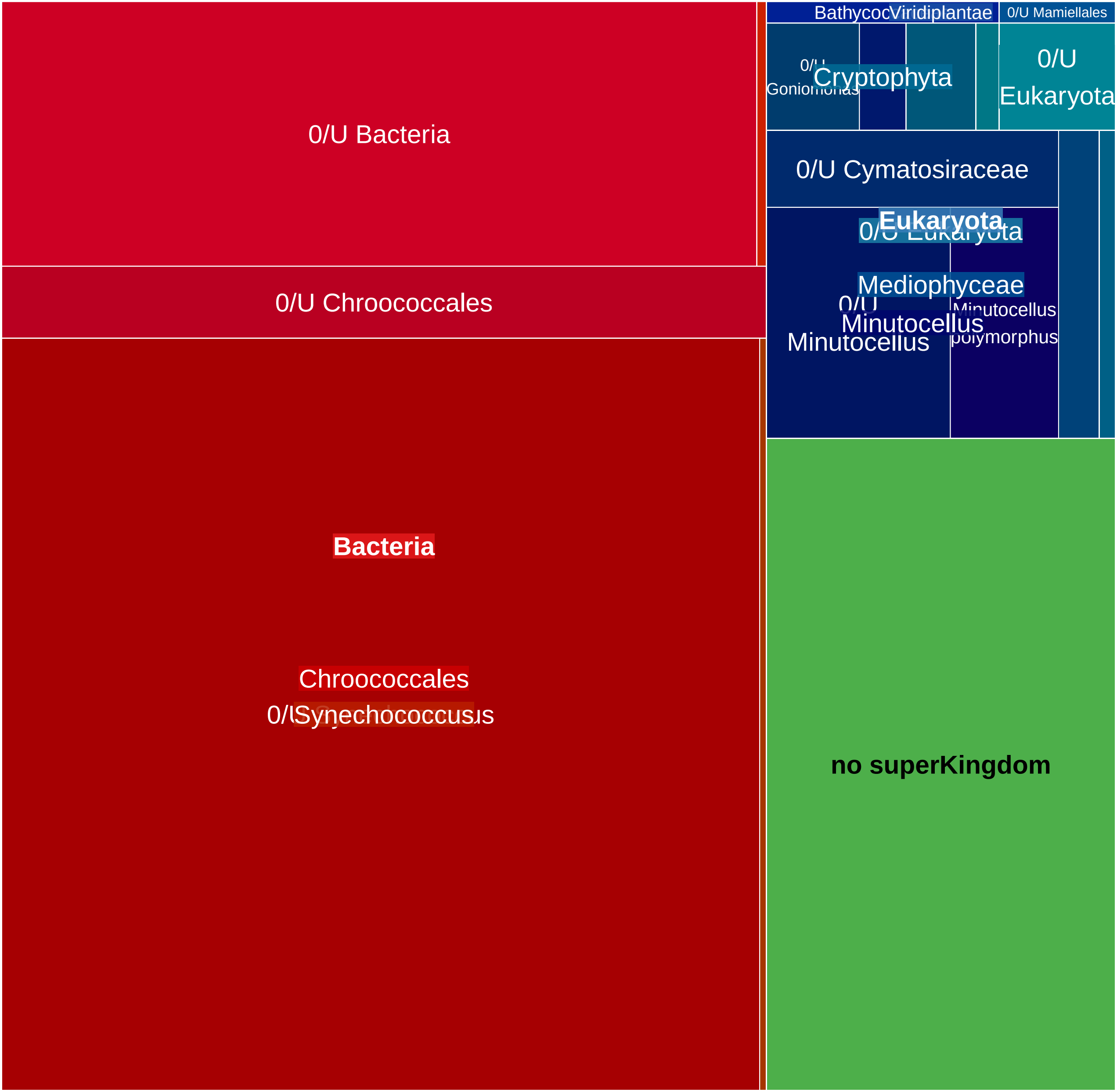

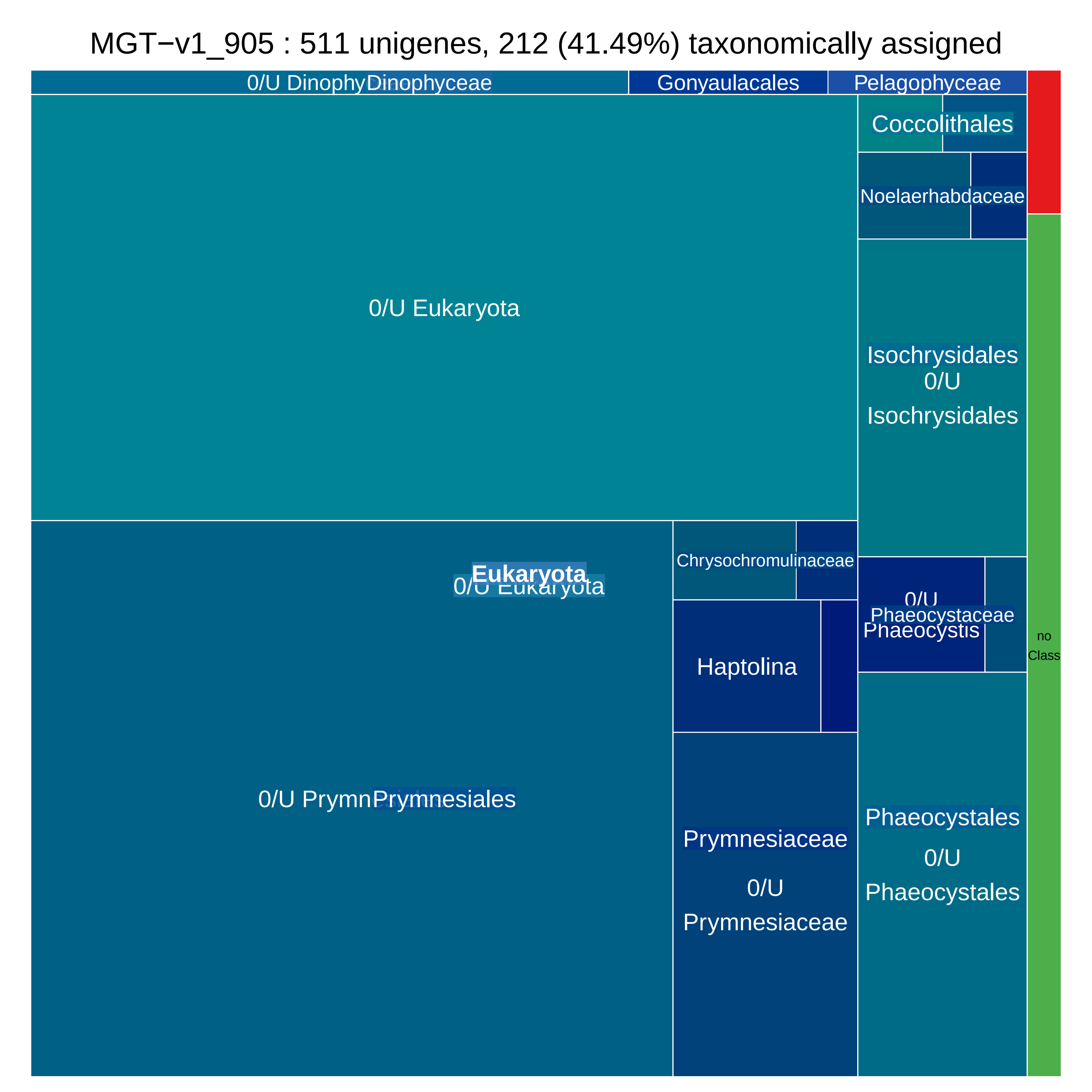
MGT−v1_905 : 511 unigenes, 212 (41.49%) taxonomically assigned
0/U Dinophy
Dinophyceae
Gonyaulacales
Pelagophyceae
0/U Eukaryota
Coccolithales
Noelaerhabdaceae
Isochrysidales
0/U
Isochrysidales
Eukaryota
0/U Eukaryota
Chrysochromulinaceae
0/U
Phaeocystaceae
Phaeocystis
Haptolina
no
Class
0/U Prymn
Prymnesiales
Prymnesiaceae
0/U
Prymnesiaceae
Phaeocystales
0/U
Phaeocystales

MGT−v1_906 : 510 unigenes, 260 (50.98%) taxonomically assigned

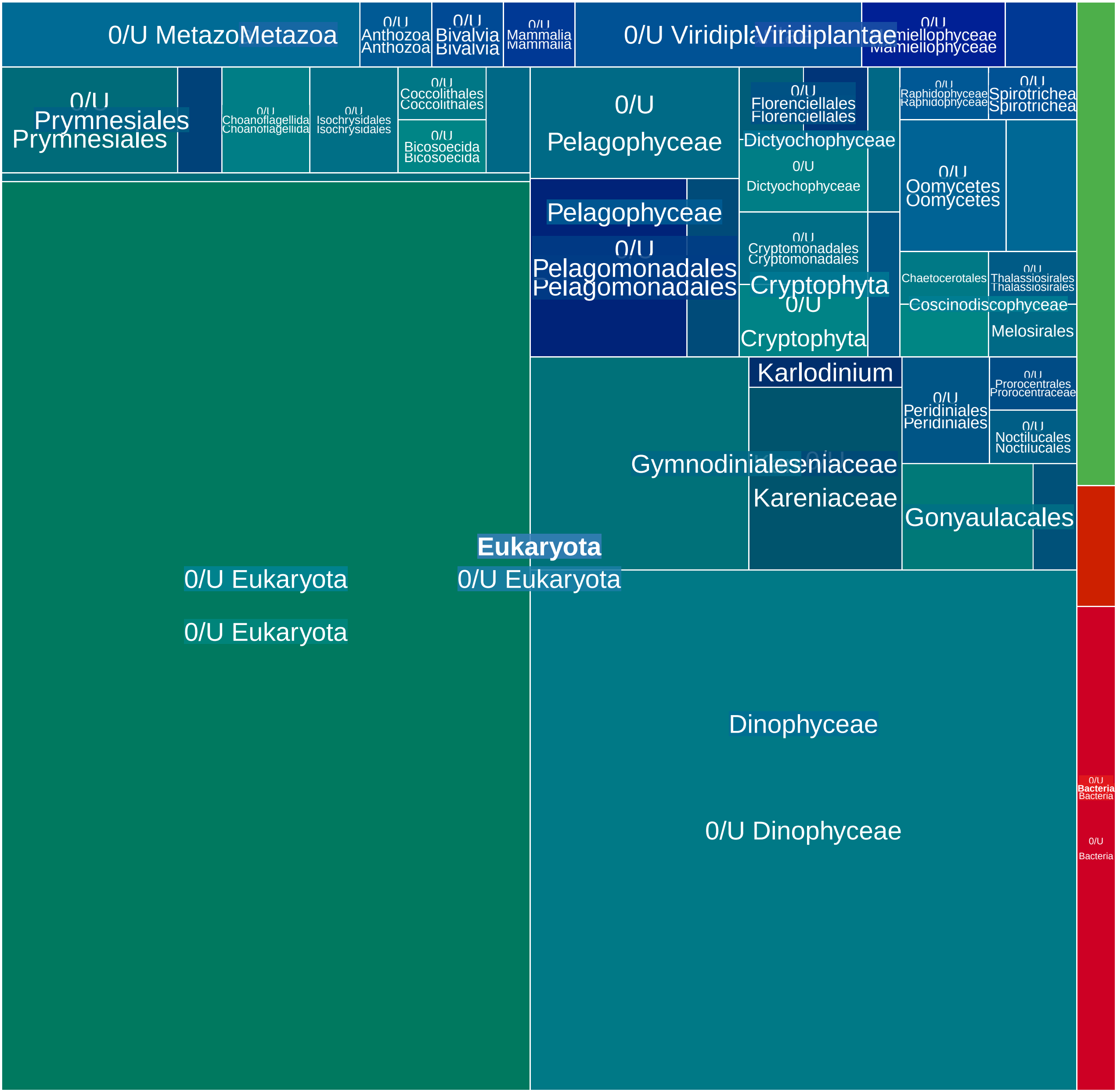

MGT−v1_907 : 509 unigenes, 13 (2.55%) taxonomically assigned

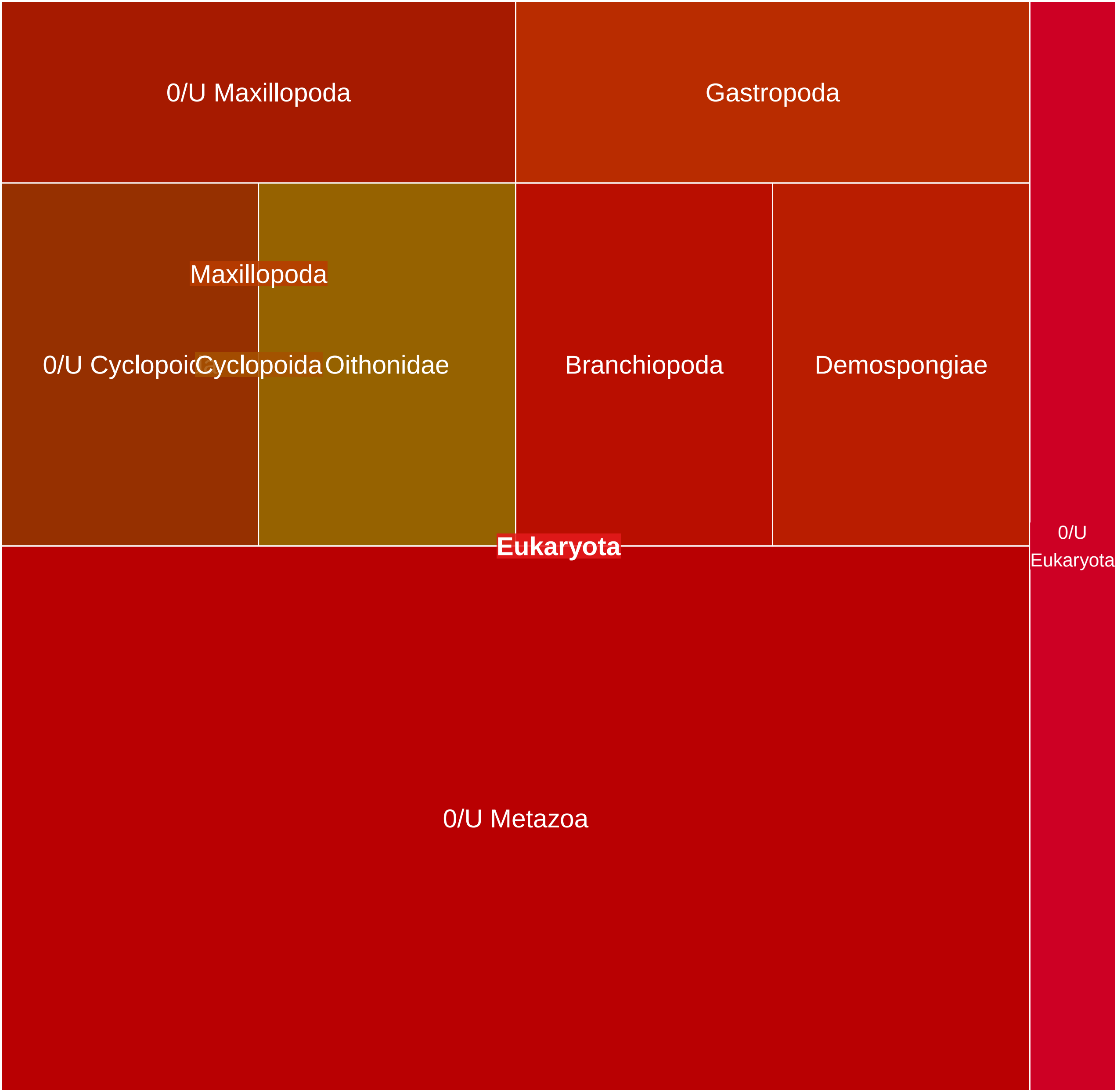

MGT−v1_908 : 508 unigenes, 5 (0.98%) taxonomically assigned

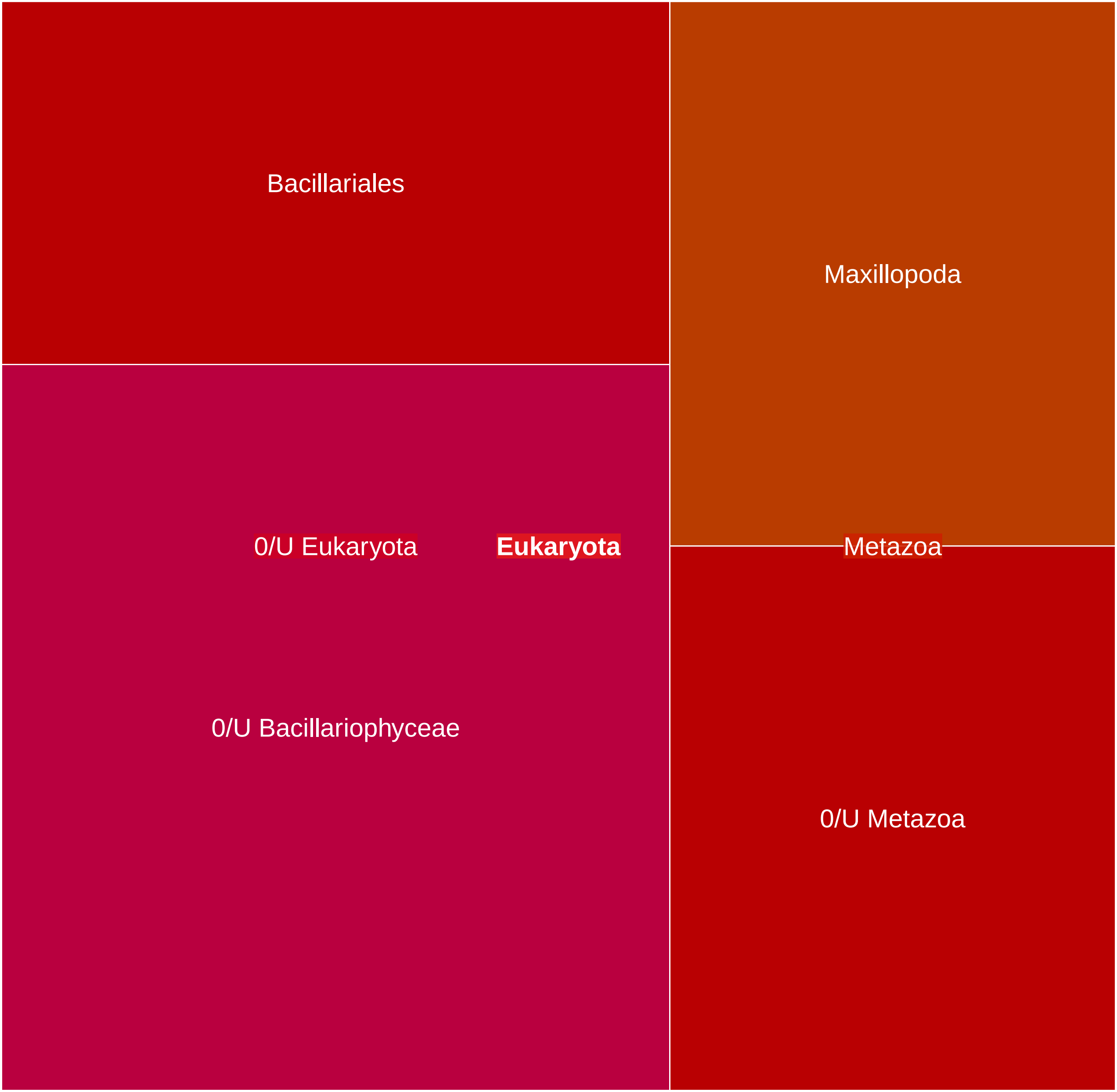

### MGT−v1_909 : 508 unigenes, 66 (12.99%) taxonomically assigned

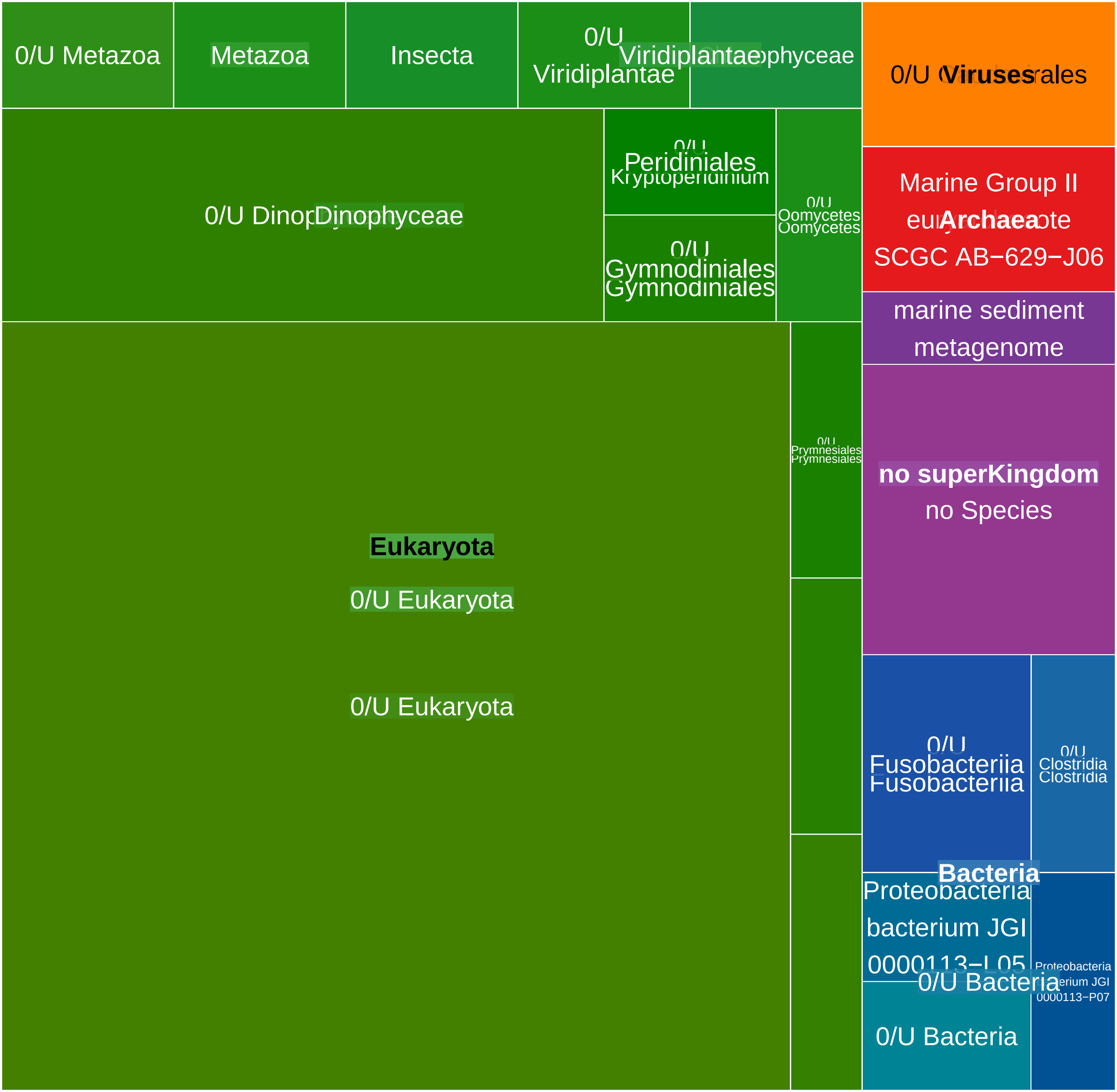

### MGT−v1_910 : 508 unigenes, 52 (10.24%) taxonomically assigned

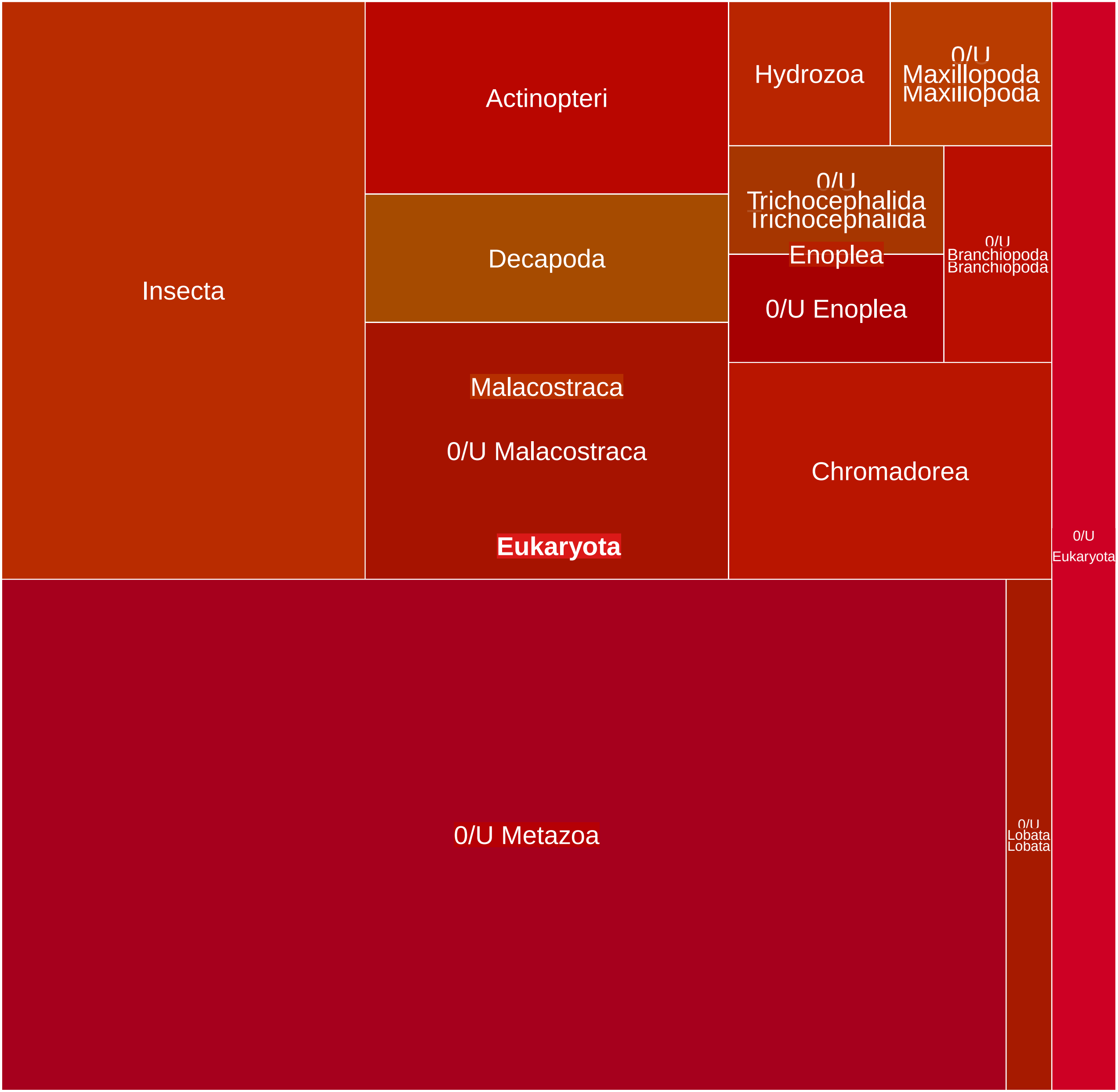

### MGT−v1_911 : 507 unigenes, 386 (76.13%) taxonomically assigned

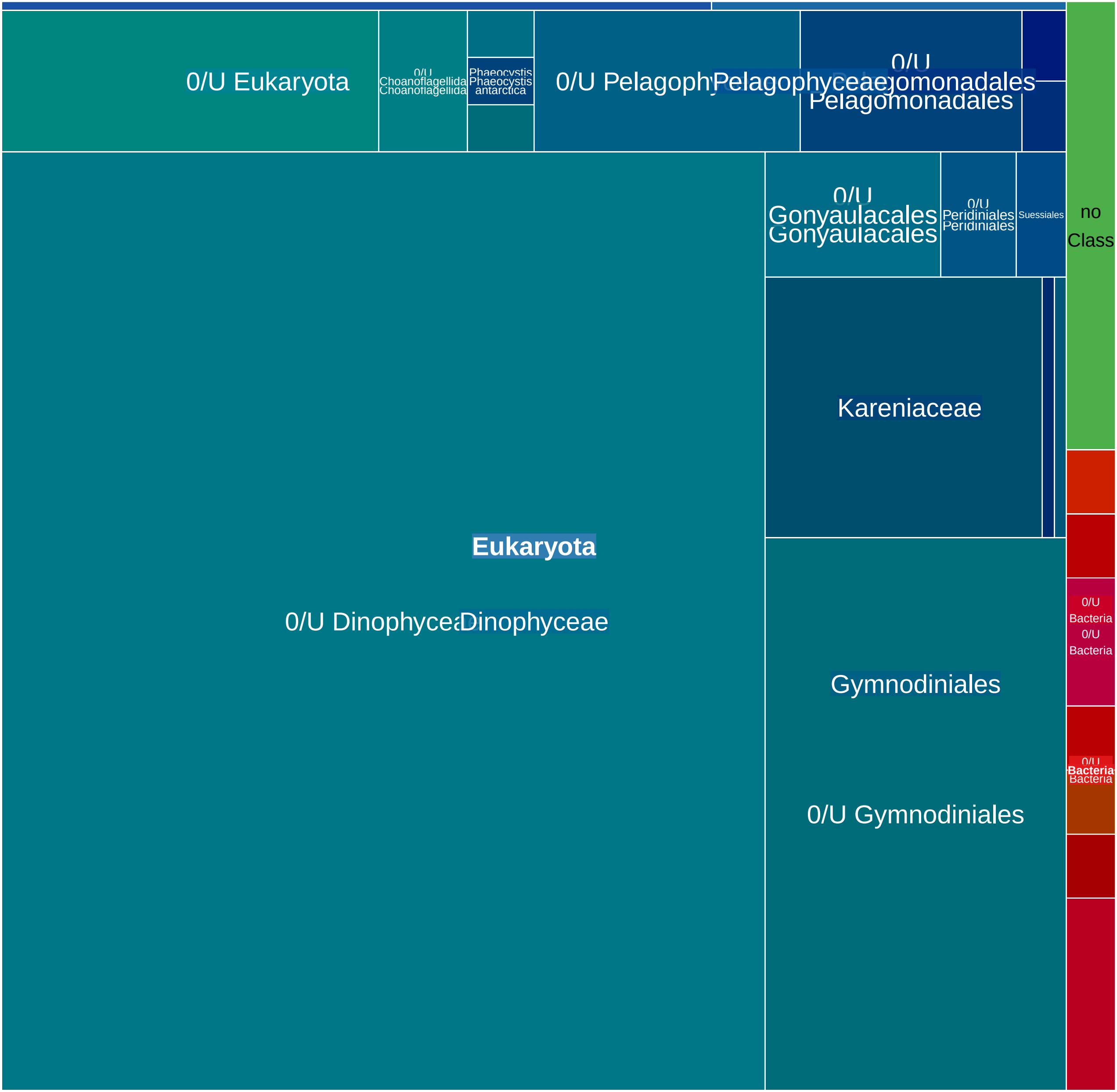

MGT−v1_912 : 506 unigenes, 198 (39.13%) taxonomically assigned

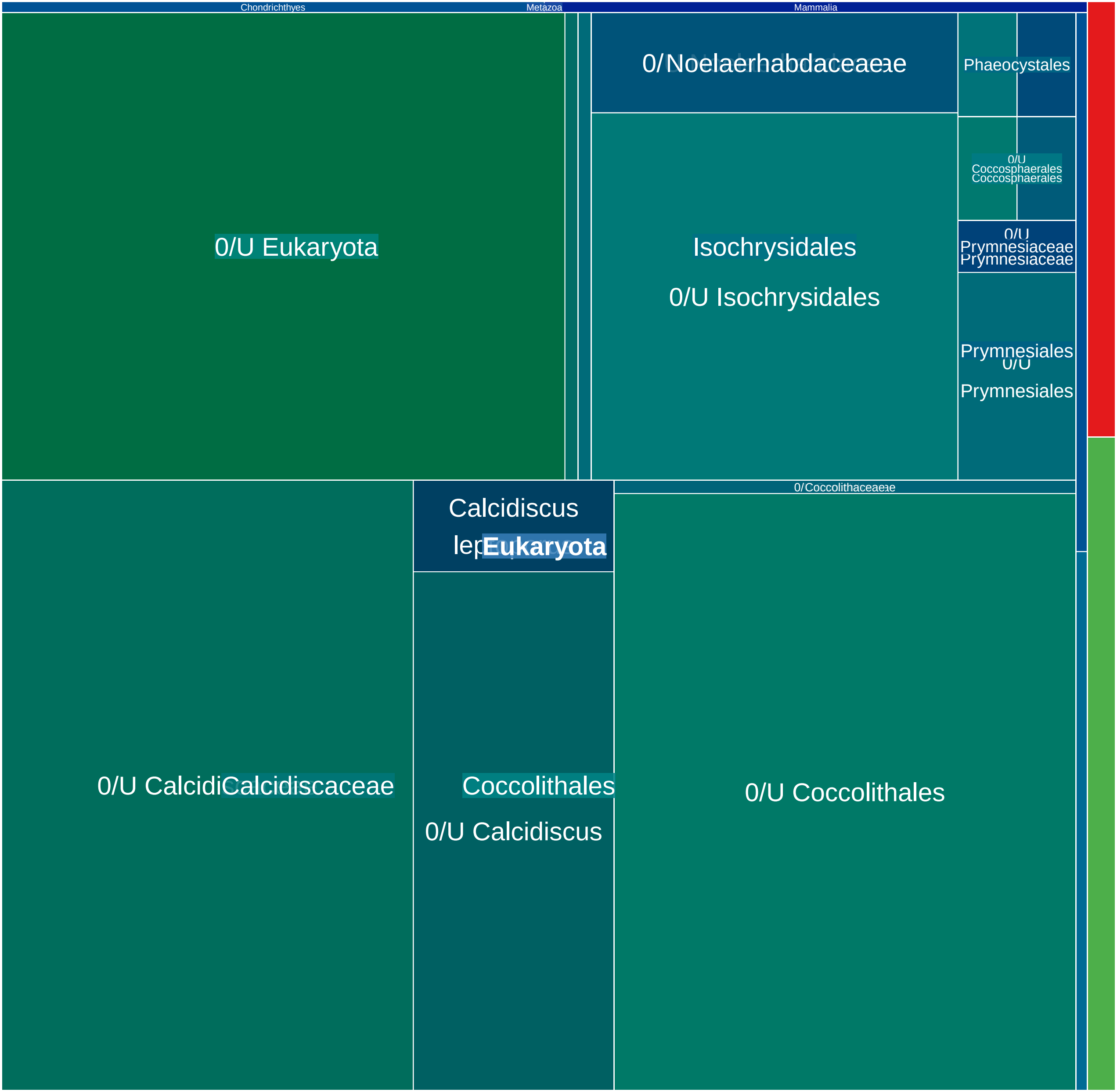

MGT−v1_913 : 505 unigenes, 489 (96.83%) taxonomically assigned

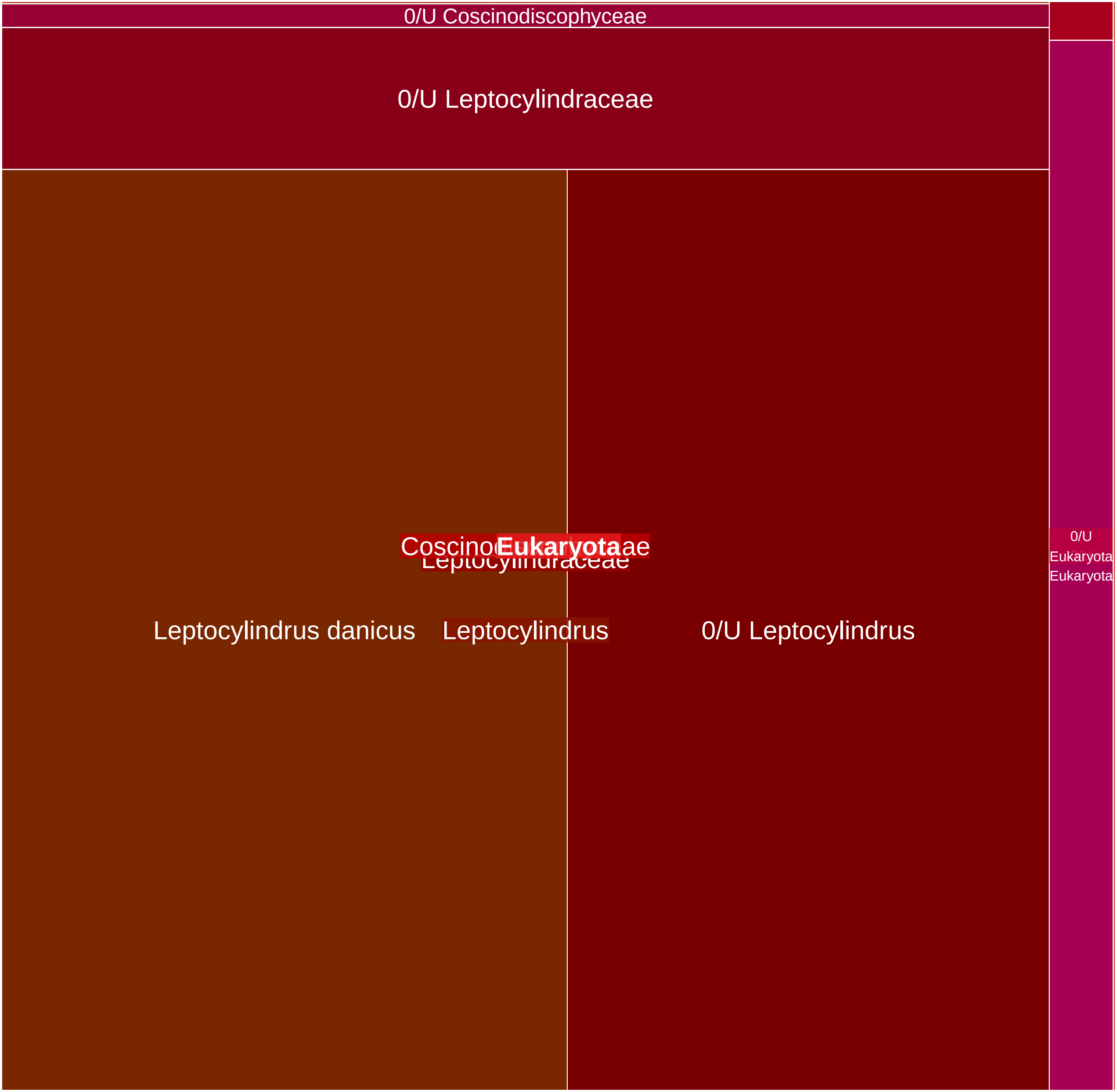

MGT−v1_914 : 505 unigenes, 370 (73.27%) taxonomically assigned

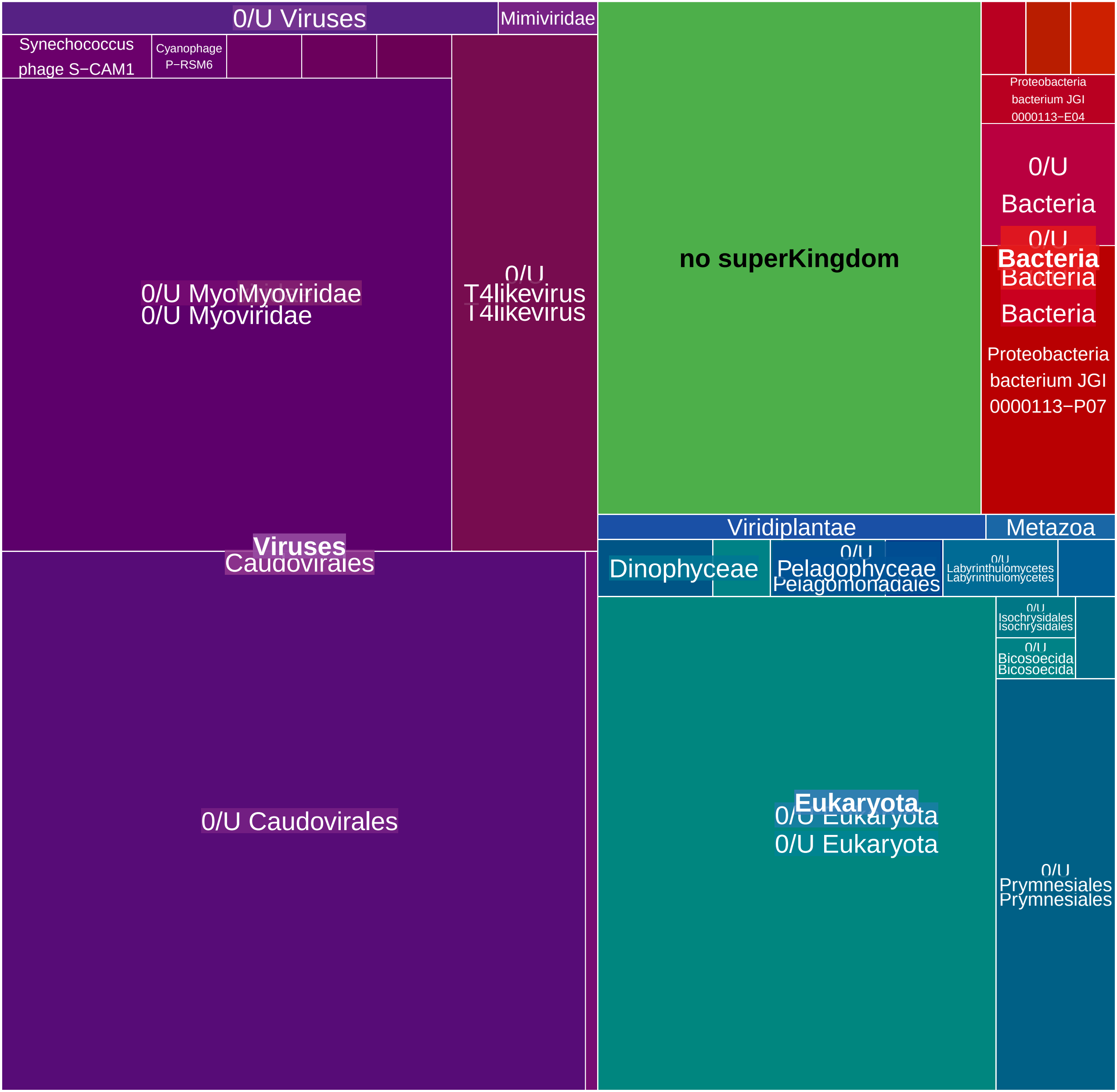

MGT−v1_915 : 505 unigenes, 294 (58.22%) taxonomically assigned

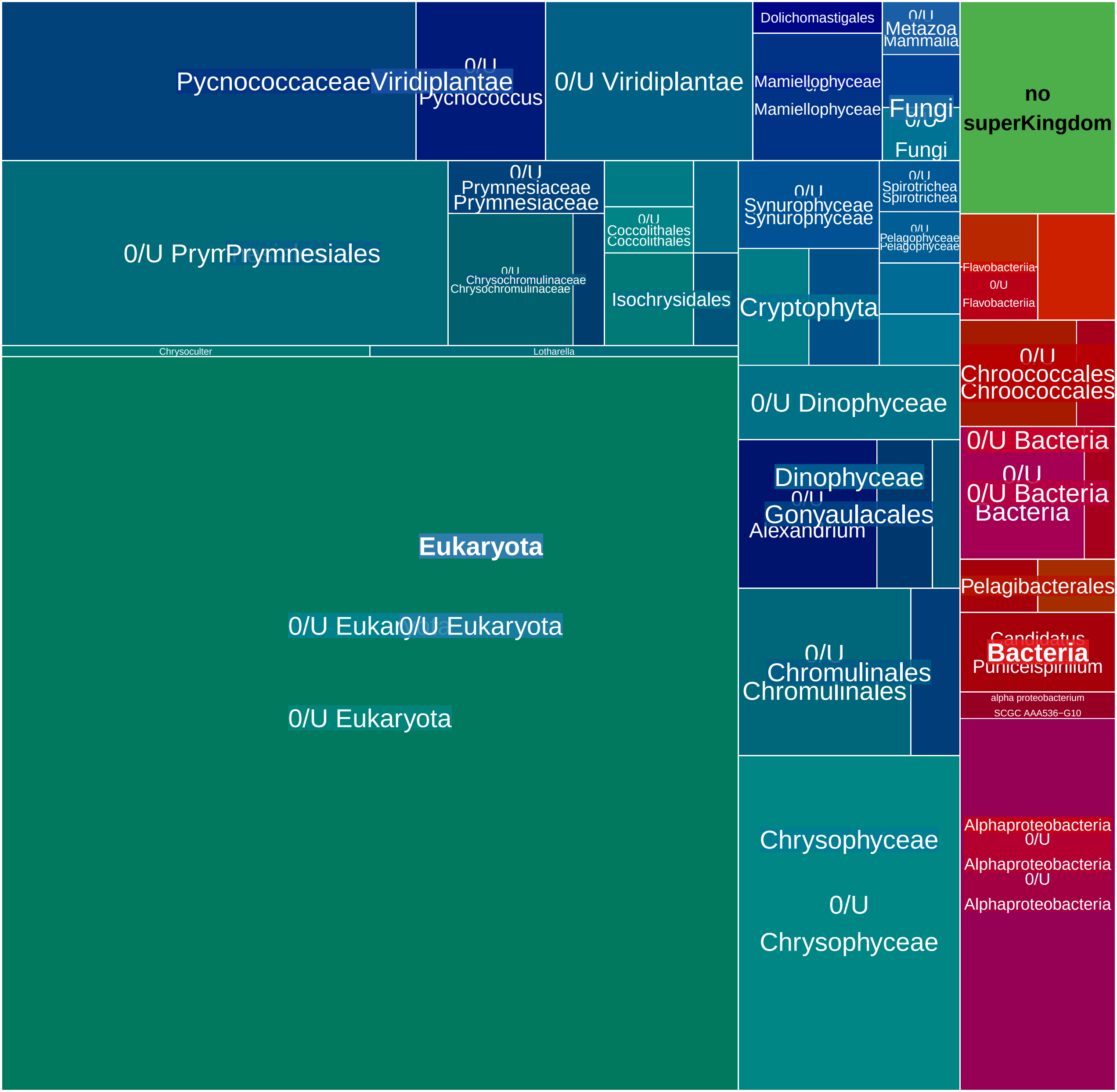

MGT−v1_916 : 504 unigenes, 16 (3.17%) taxonomically assigned

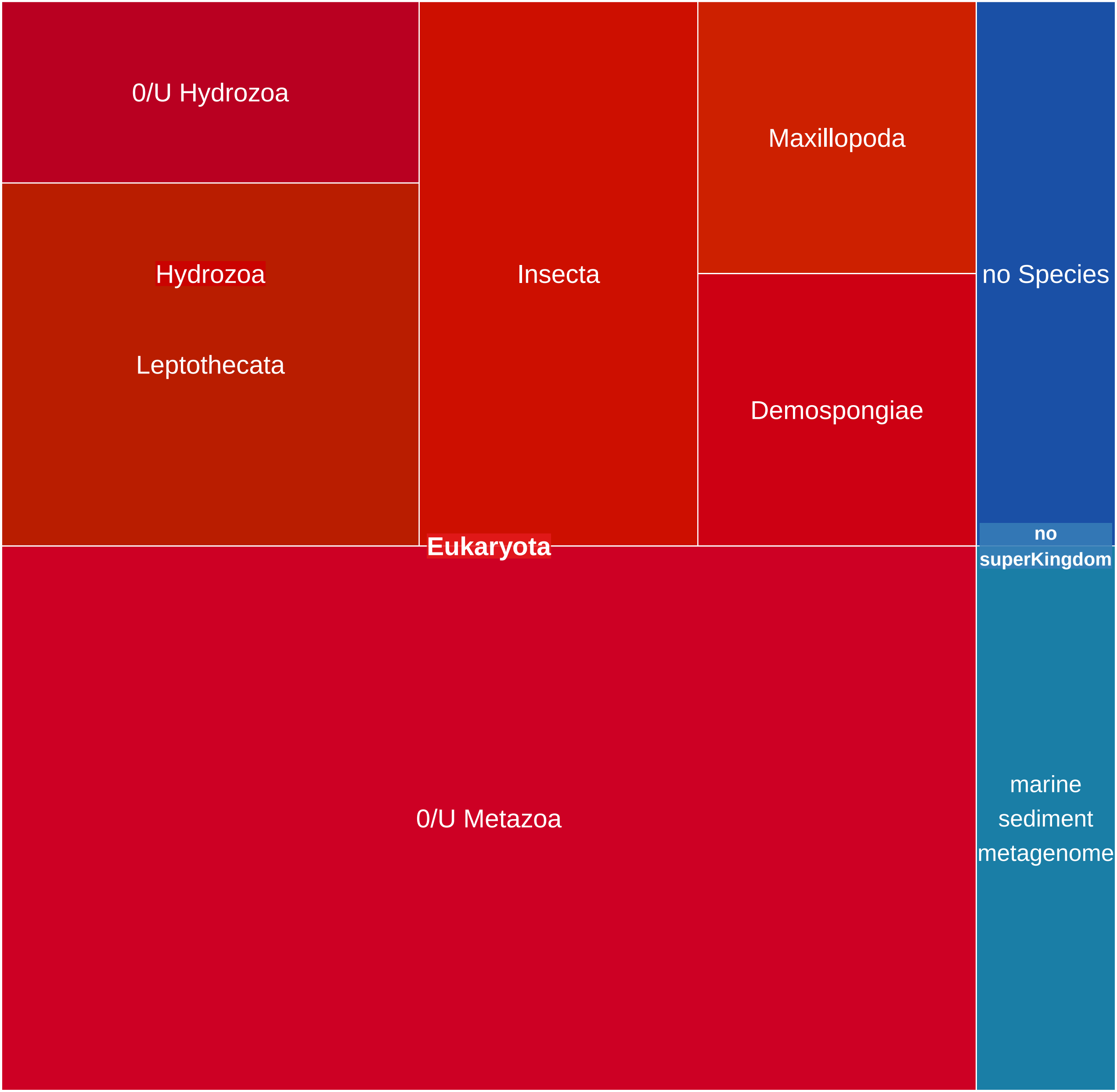

MGT−v1_917 : 504 unigenes, 438 (86.90%) taxonomically assigned

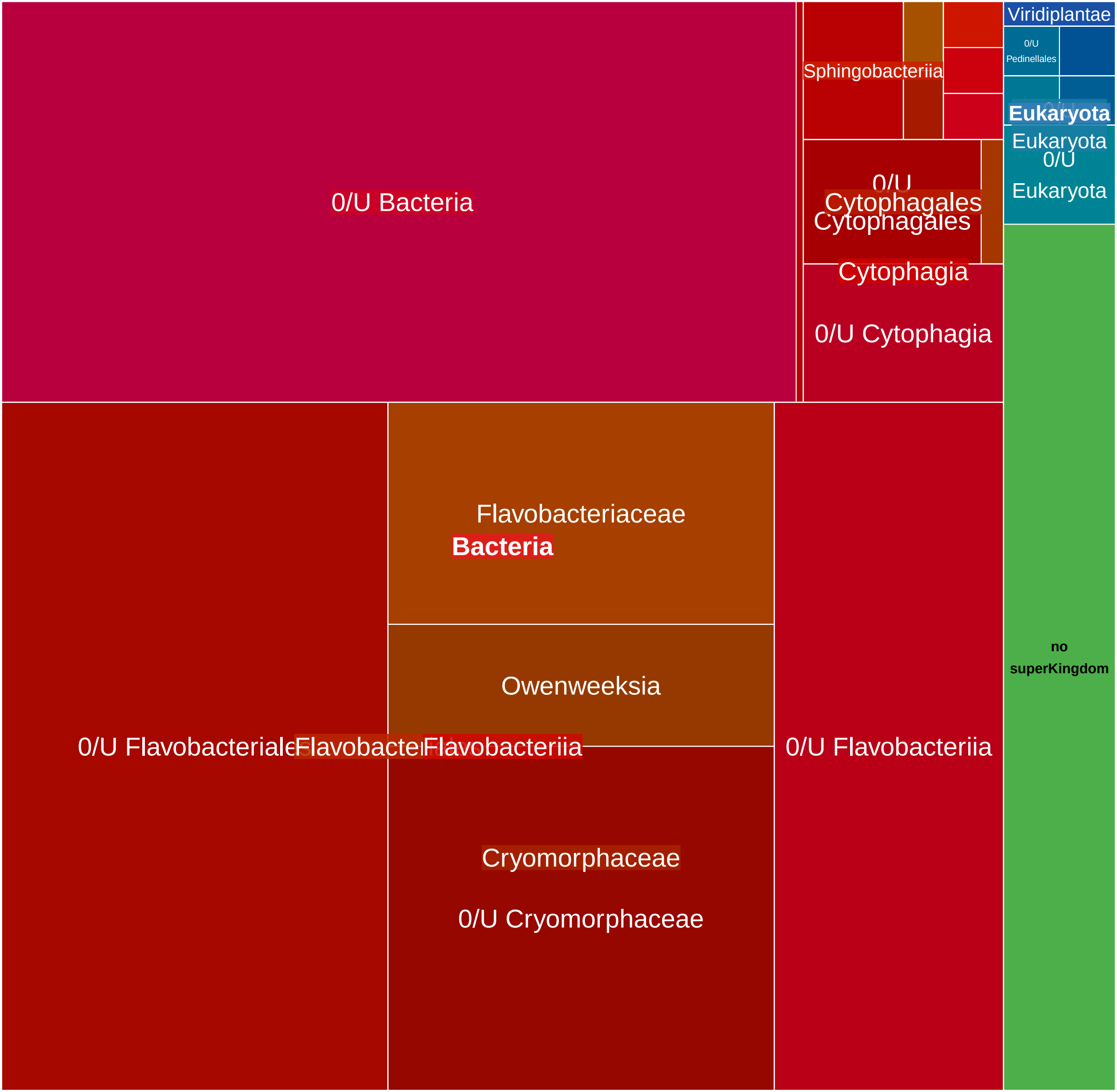

MGT−v1_918 : 503 unigenes, 366 (72.76%) taxonomically assigned

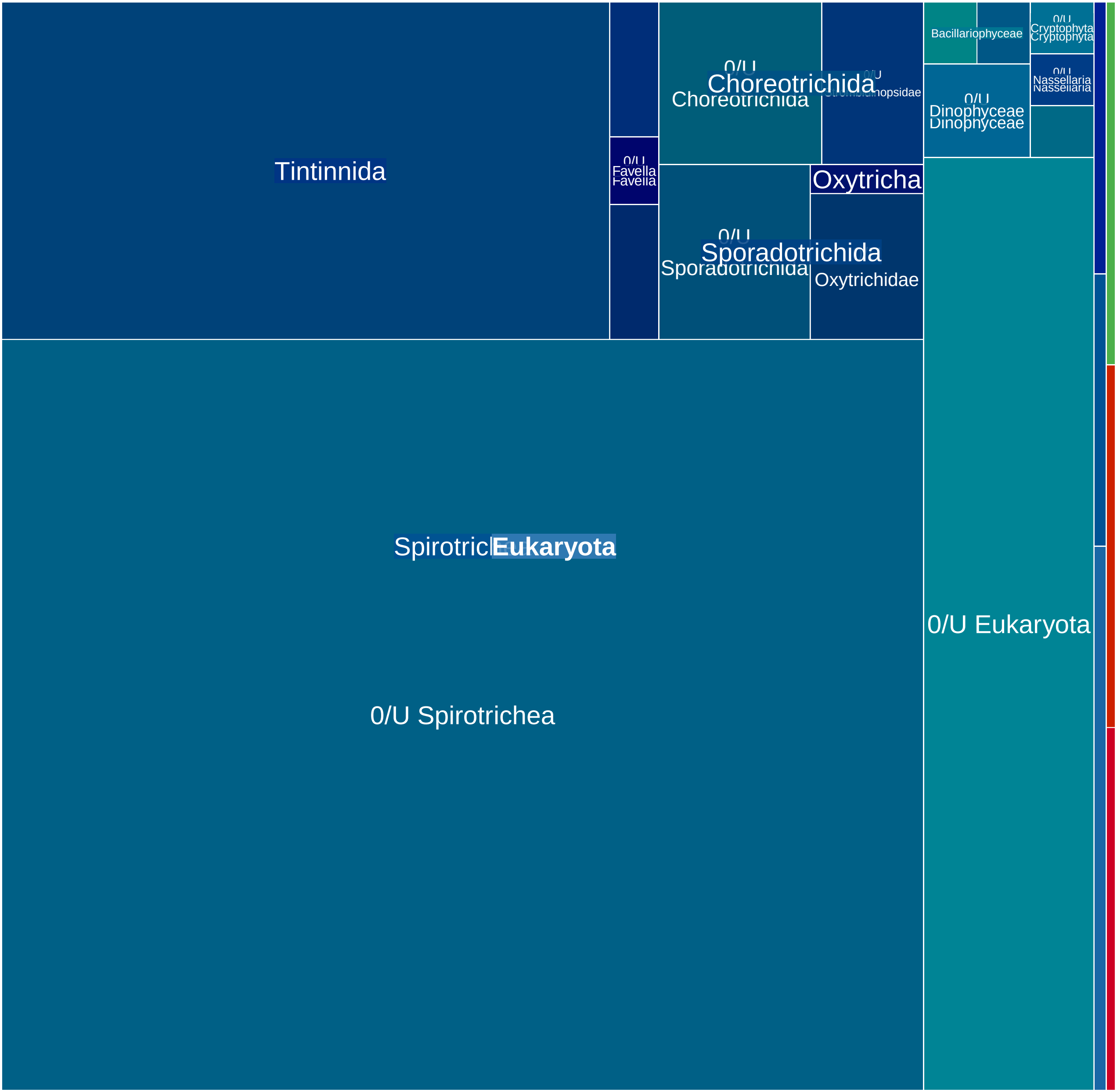

MGT−v1_919 : 503 unigenes, 49 (9.74%) taxonomically assigned

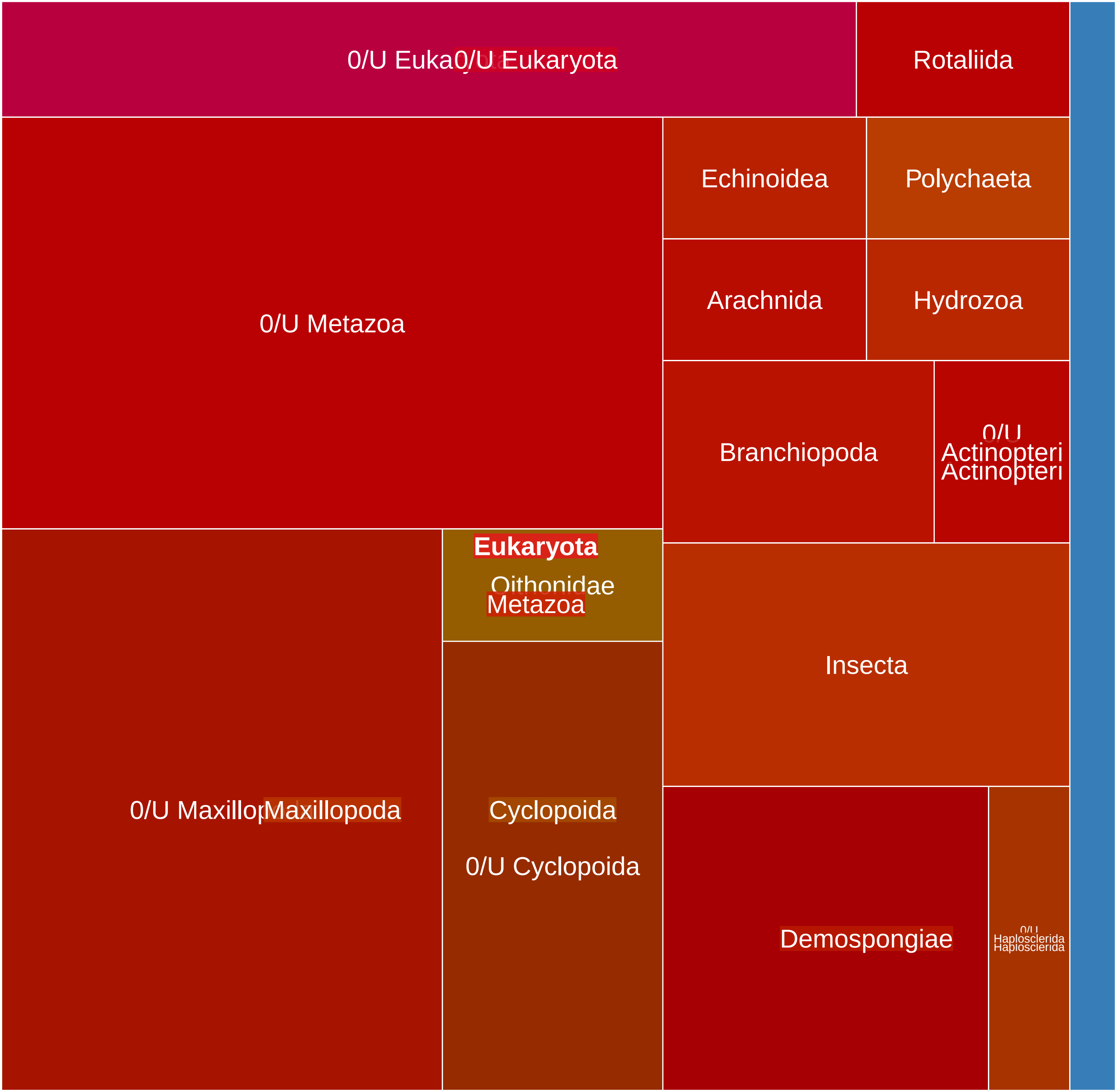

MGT−v1_920 : 503 unigenes, 29 (5.77%) taxonomically assigned

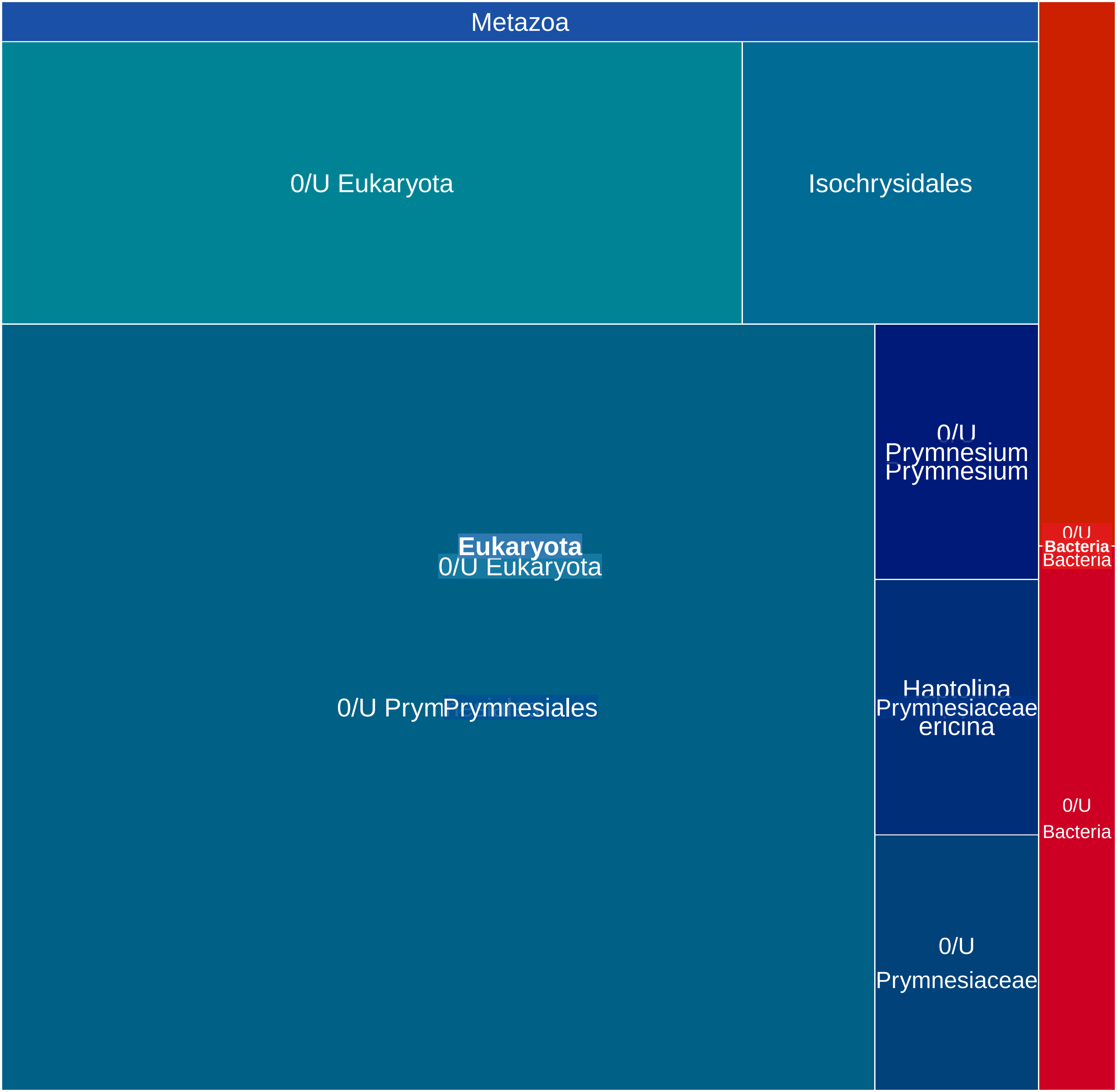

MGT−v1_921 : 502 unigenes, 235 (46.81%) taxonomically assigned

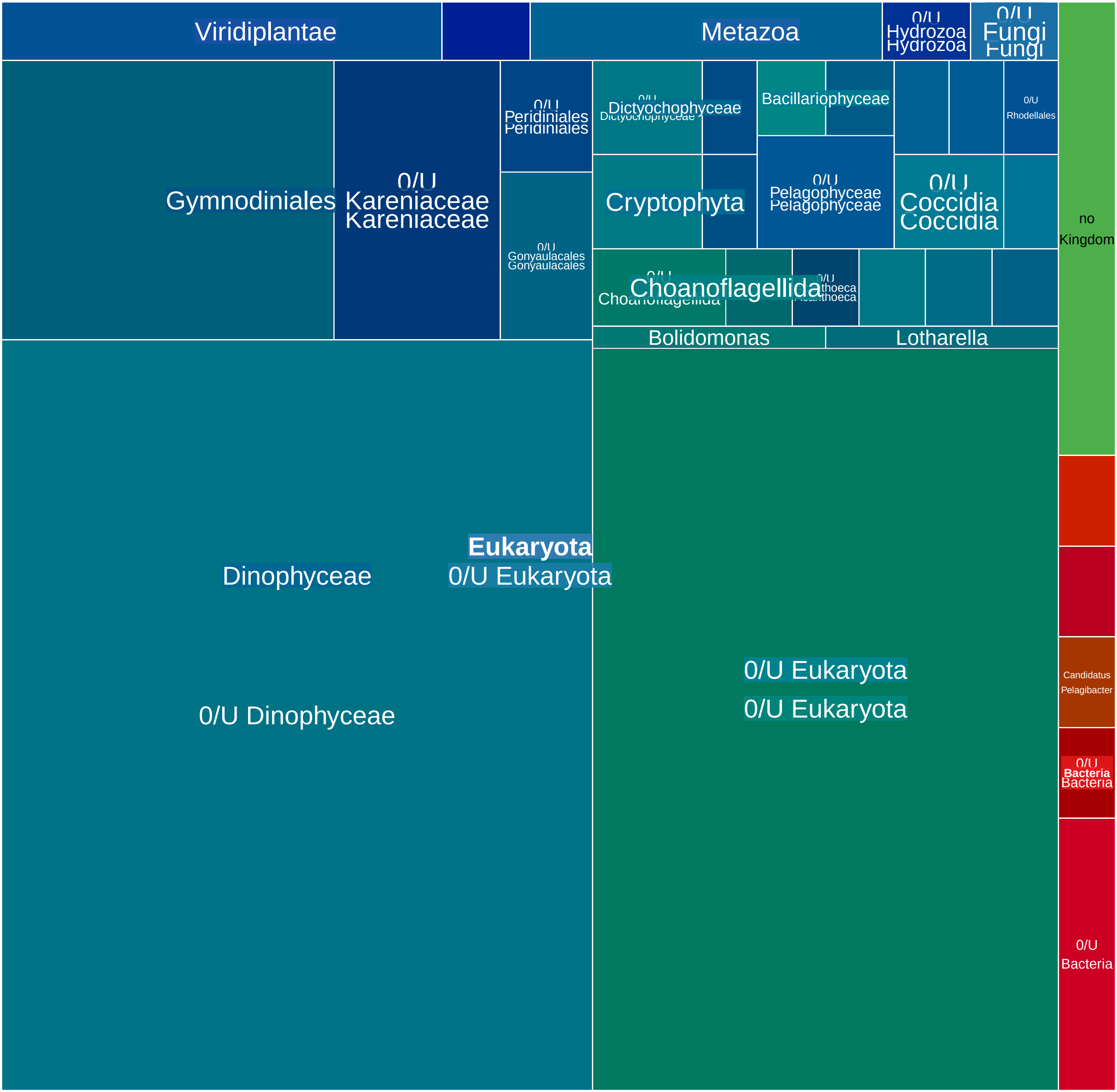

### MGT−v1_922 : 500 unigenes, 373 (74.60%) taxonomically assigned

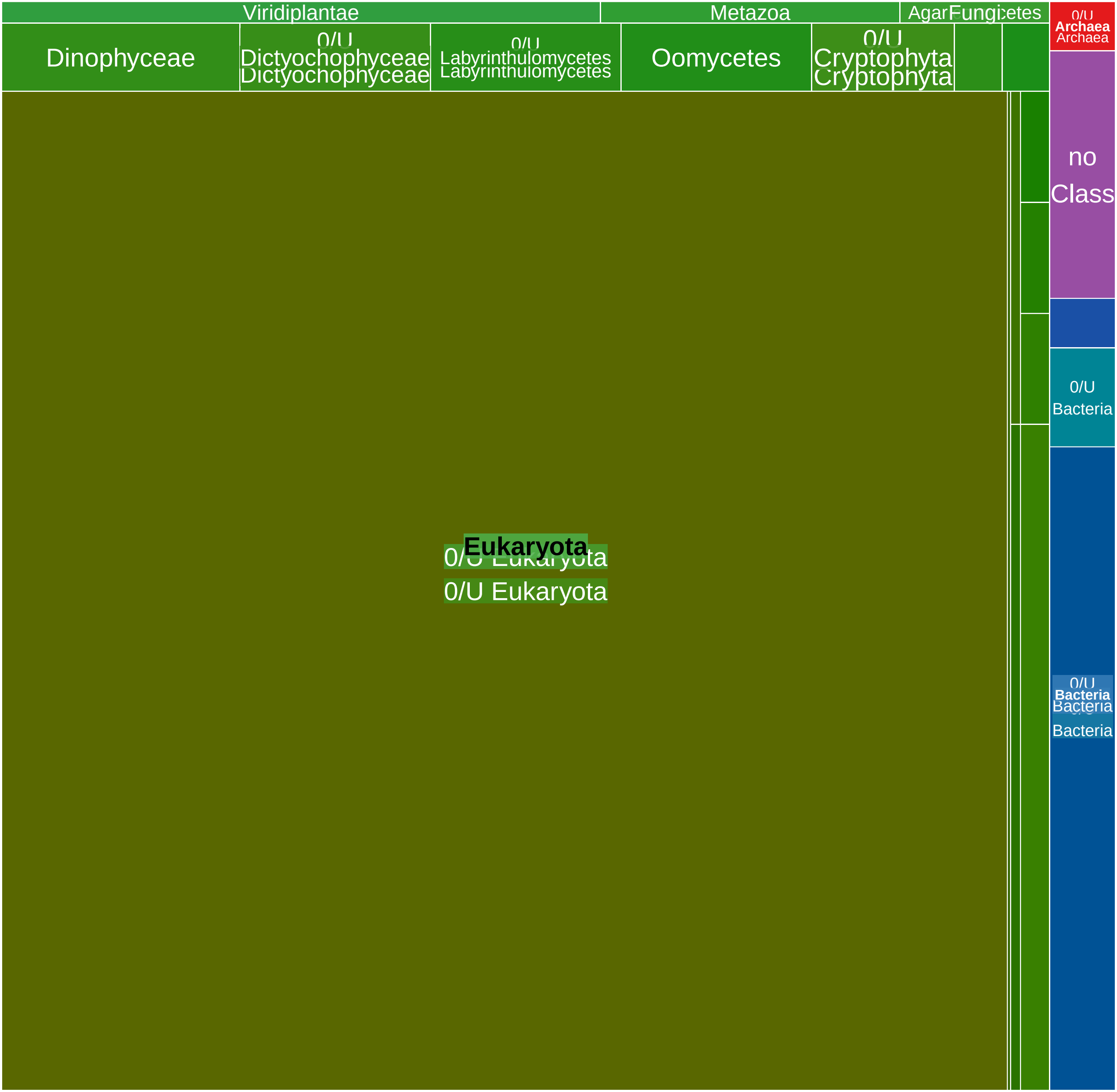

### MGT−v1_923 : 500 unigenes, 475 (95.00%) taxonomically assigned

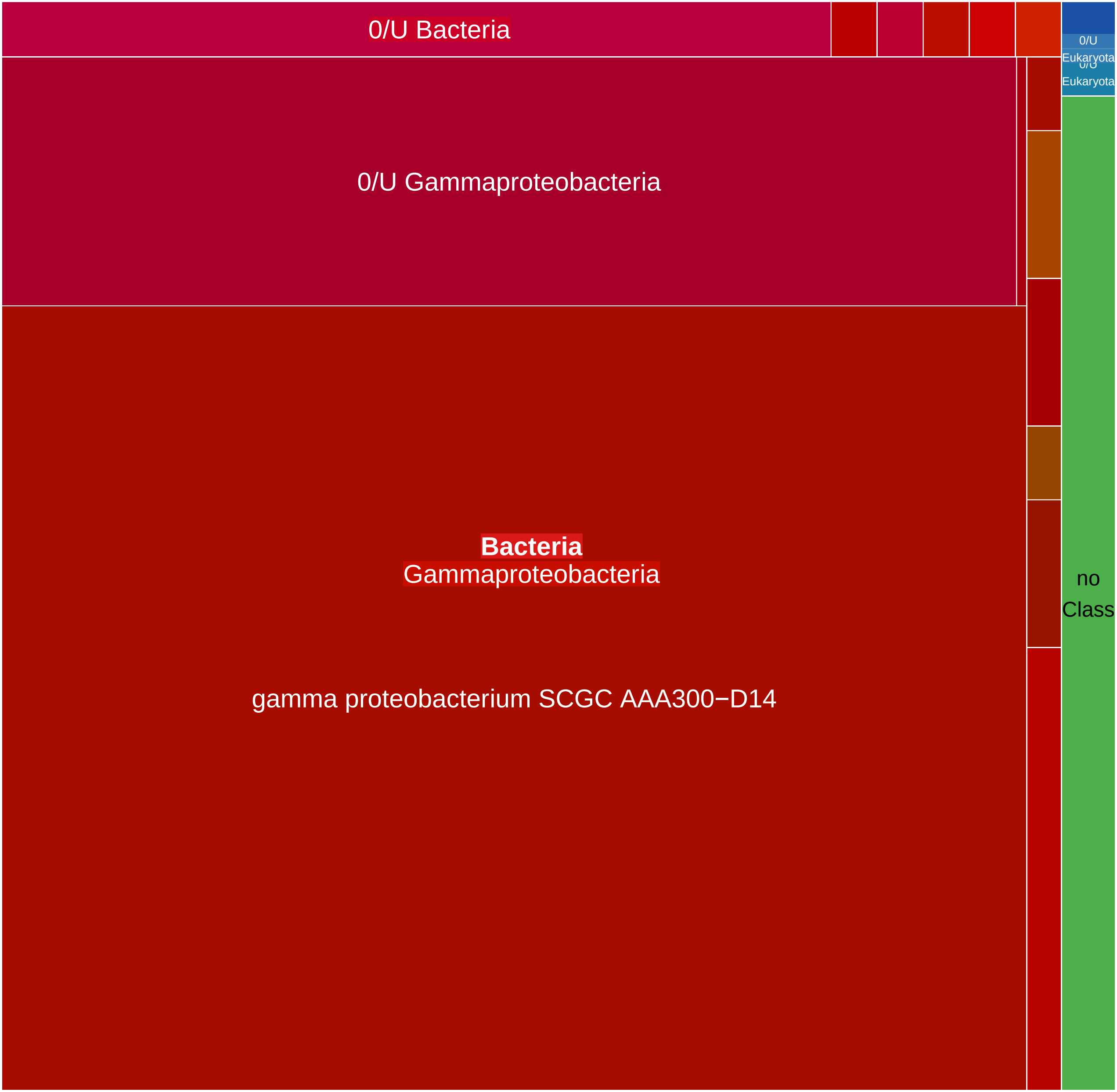

MGT−v1_924 : 500 unigenes, 11 (2.20%) taxonomically assigned

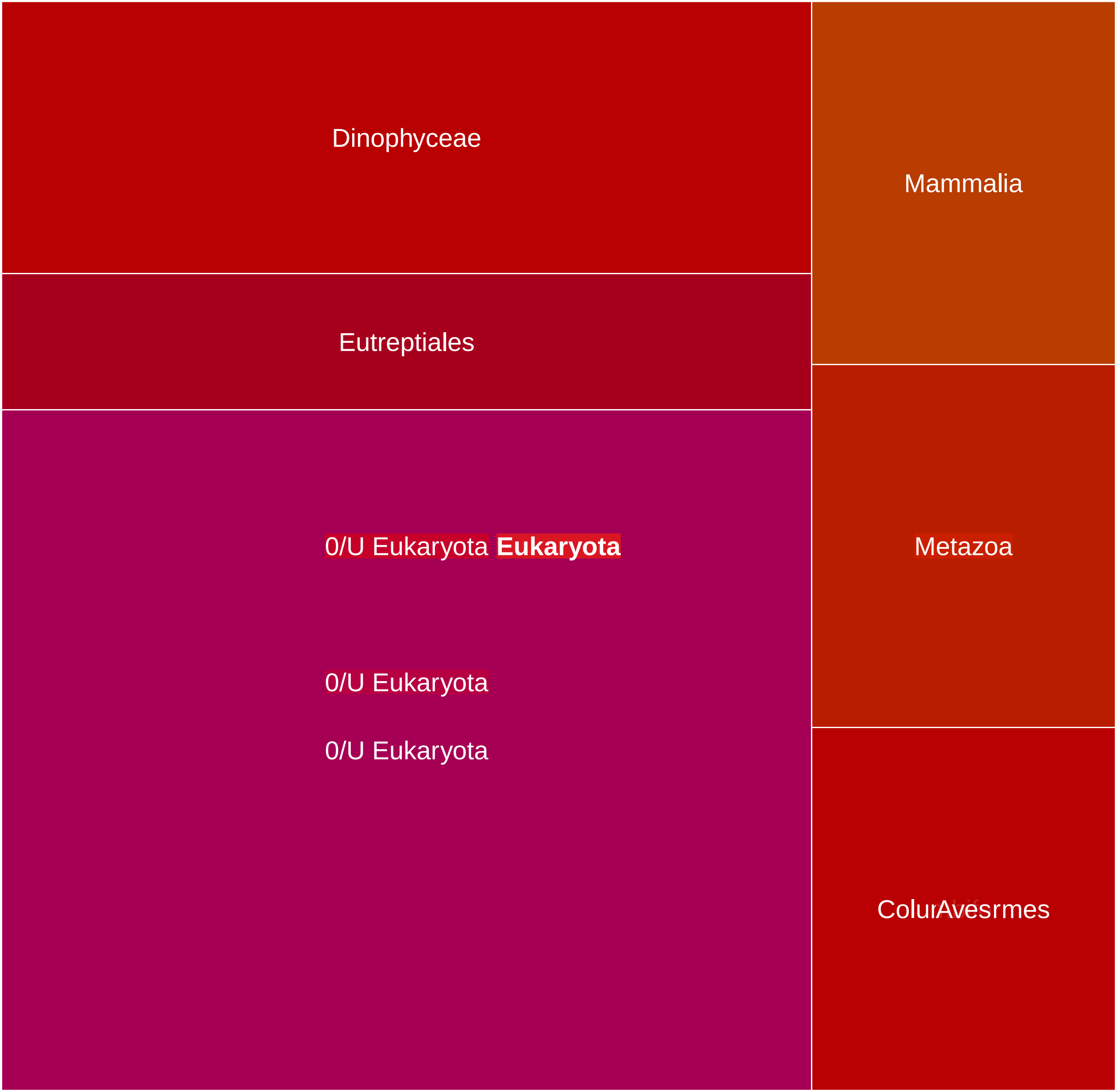
