## Additional supplemental Fig A2 for "Transcriptome reconstruction and functional analysis of eukaryotic marine plankton communities via high-throughput metagenomics and metatranscriptomics"

# Biogeolocalisation of MGT-v1 1   226807 unigenes, 71674 (31.6%) taxonomically assigned

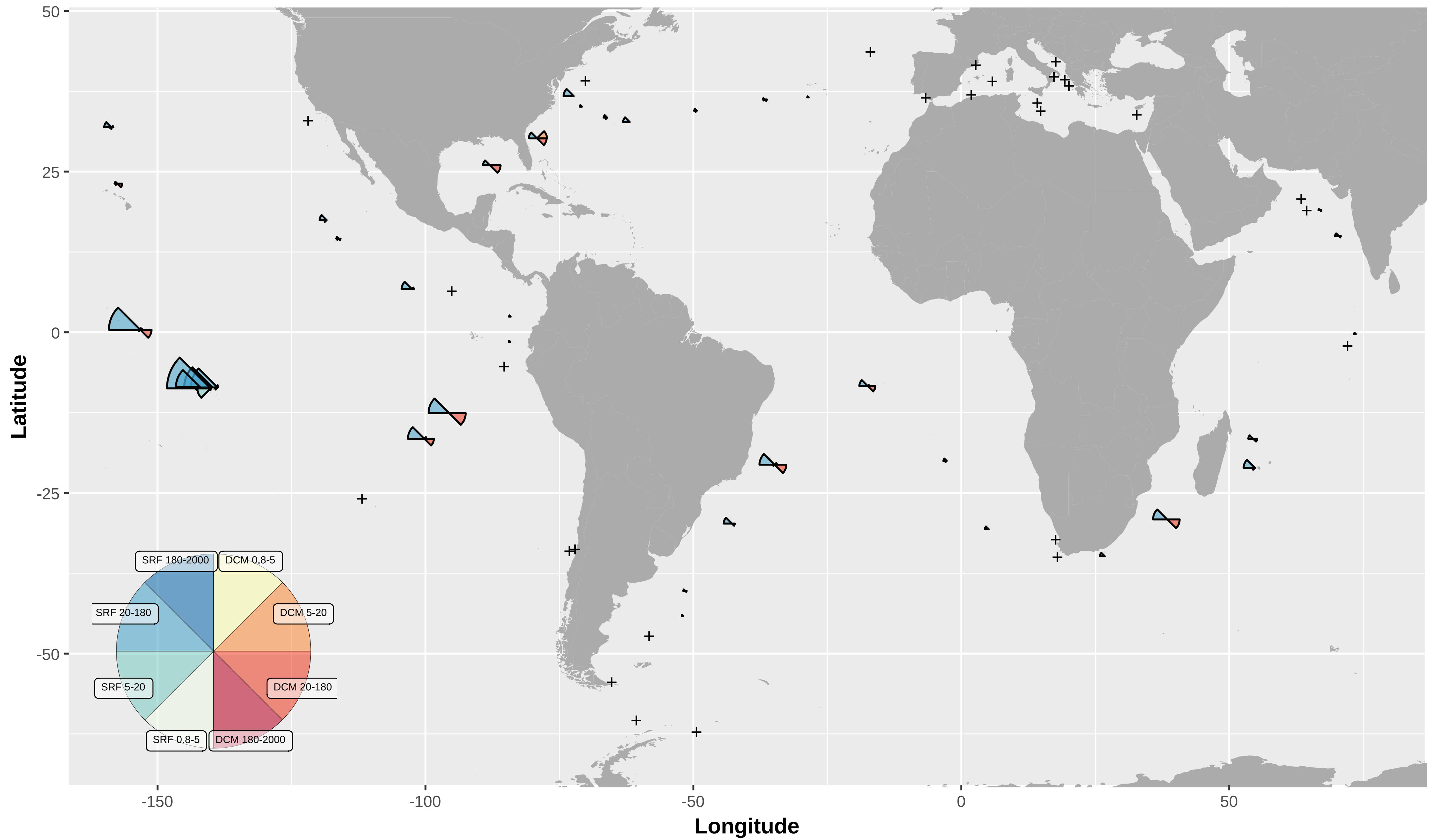


root;cellular organisms;Eukaryota;Opisthokonta;Metazoa;Eumetazoa;Bilateria;Protostomia;Ecdysozoa;Panarthropoda;Arthropoda;Mandibulata;Pancrustacea;Crustacea;Maxillopoda

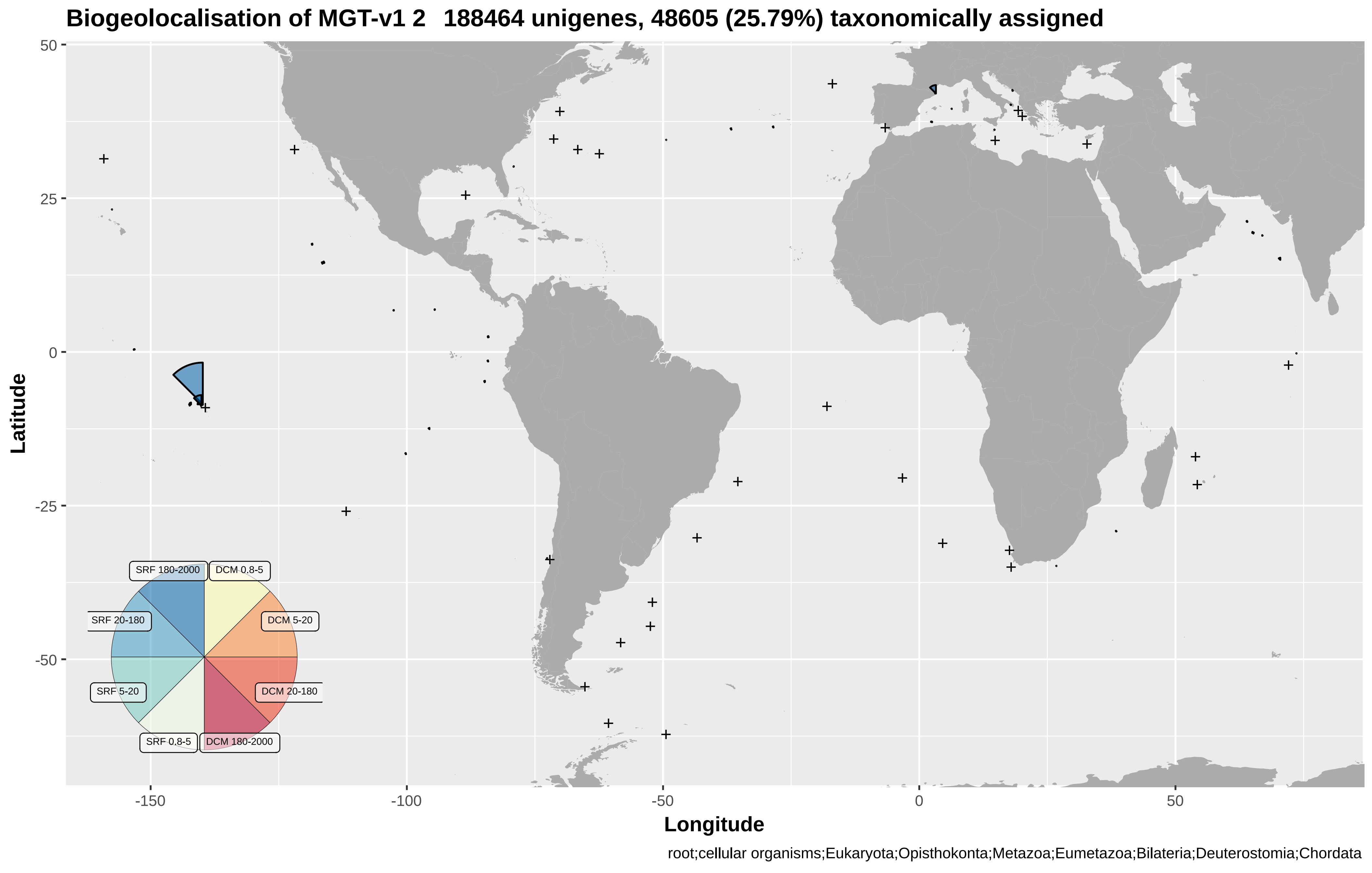

Biogeolocalisation of MGT-v1 2   188464 unigenes, 48605 (25.79%) taxonomically assigned
Latitude
50
25
0
-25
-50
-150
-100
-50
0
50
Longitude
SRF 180-2000
DCM 0.8-5
SRF 20-180
DCM 5-20
SRF 5-20
DCM 20-180
SRF 0.8-5
DCM 180-2000
root;cellular organisms;Eukaryota;Opisthokonta;Metazoa;Eumetazoa;Bilateria;Deuterostomia;Chordata

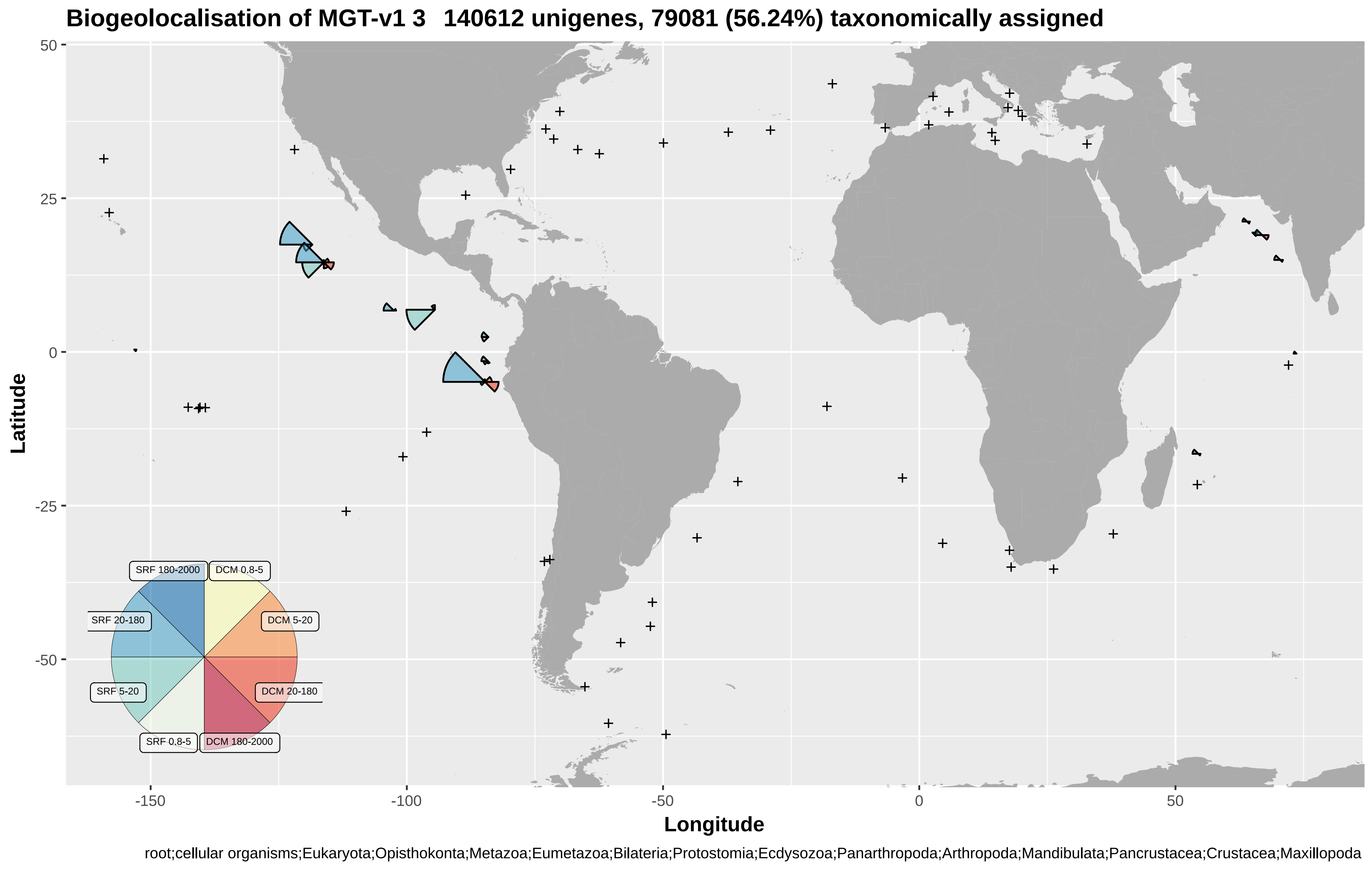
Biogeolocalisation of MGT-v1 3   140612 unigenes, 79081 (56.24%) taxonomically assigned
Latitude
Longitude
50
25
0
-25
-50
-150
-100
-50
0
50
SRF 180-2000
DCM 0.8-5
SRF 20-180
DCM 5-20
SRF 5-20
DCM 20-180
SRF 0.8-5
DCM 180-2000
root;cellular organisms;Eukaryota;Opisthokonta;Metazoa;Eumetazoa;Bilateria;Protostomia;Ecdysozoa;Panarthropoda;Arthropoda;Mandibulata;Pancrustacea;Crustacea;Maxillopoda

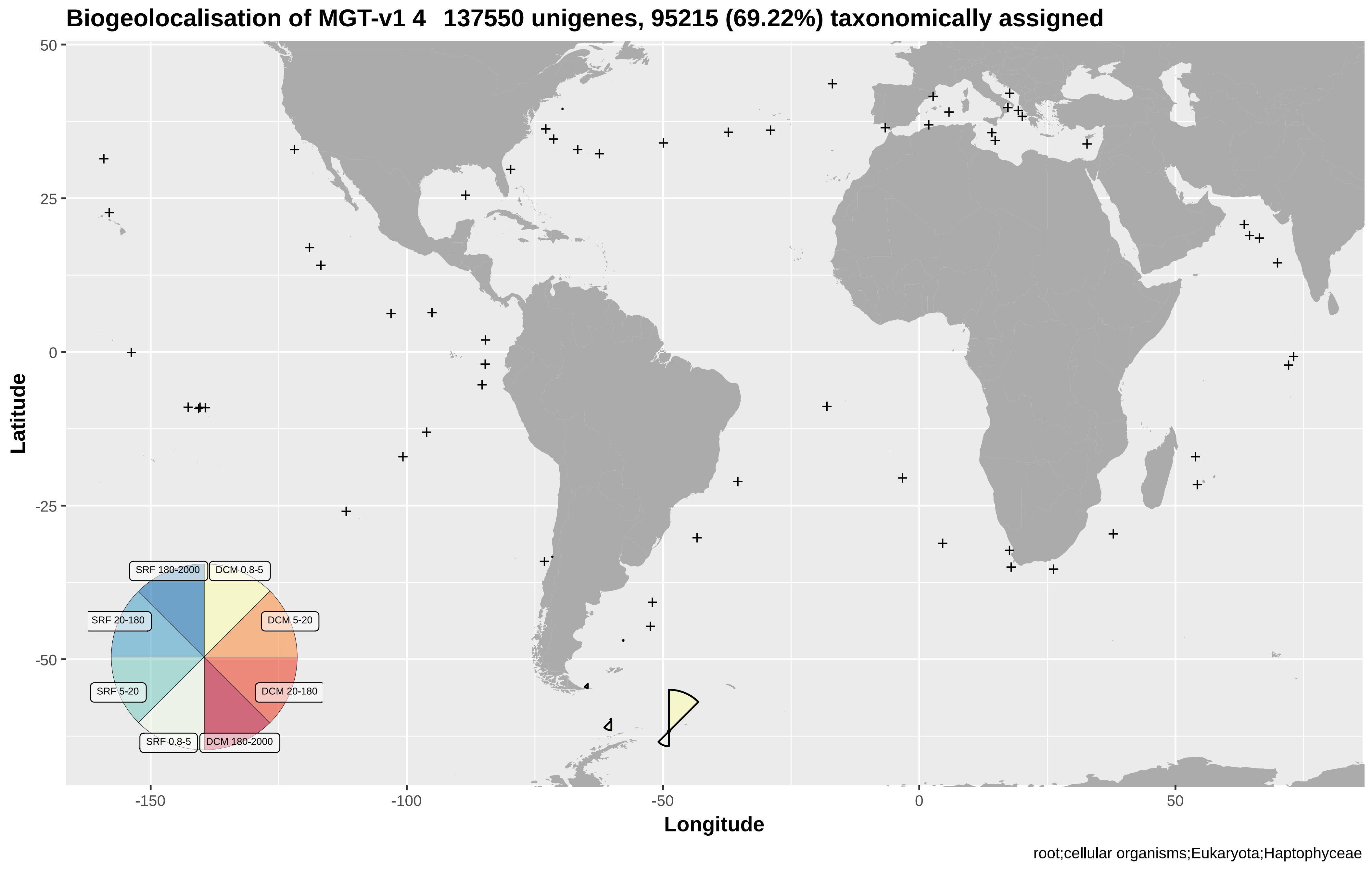
Biogeolocalisation of MGT-v1 4   137550 unigenes, 95215 (69.22%) taxonomically assigned
50
25
0
-25
-50
Latitude
-150
-100
-50
0
50
Longitude
SRF 180-2000
DCM 0.8-5
SRF 20-180
DCM 5-20
SRF 5-20
DCM 20-180
SRF 0.8-5
DCM 180-2000
root;cellular organisms;Eukaryota;Haptophyceae

# Biogeolocalisation of MGT-v1 5   136601 unigenes, 69325 (50.75%) taxonomically assigned

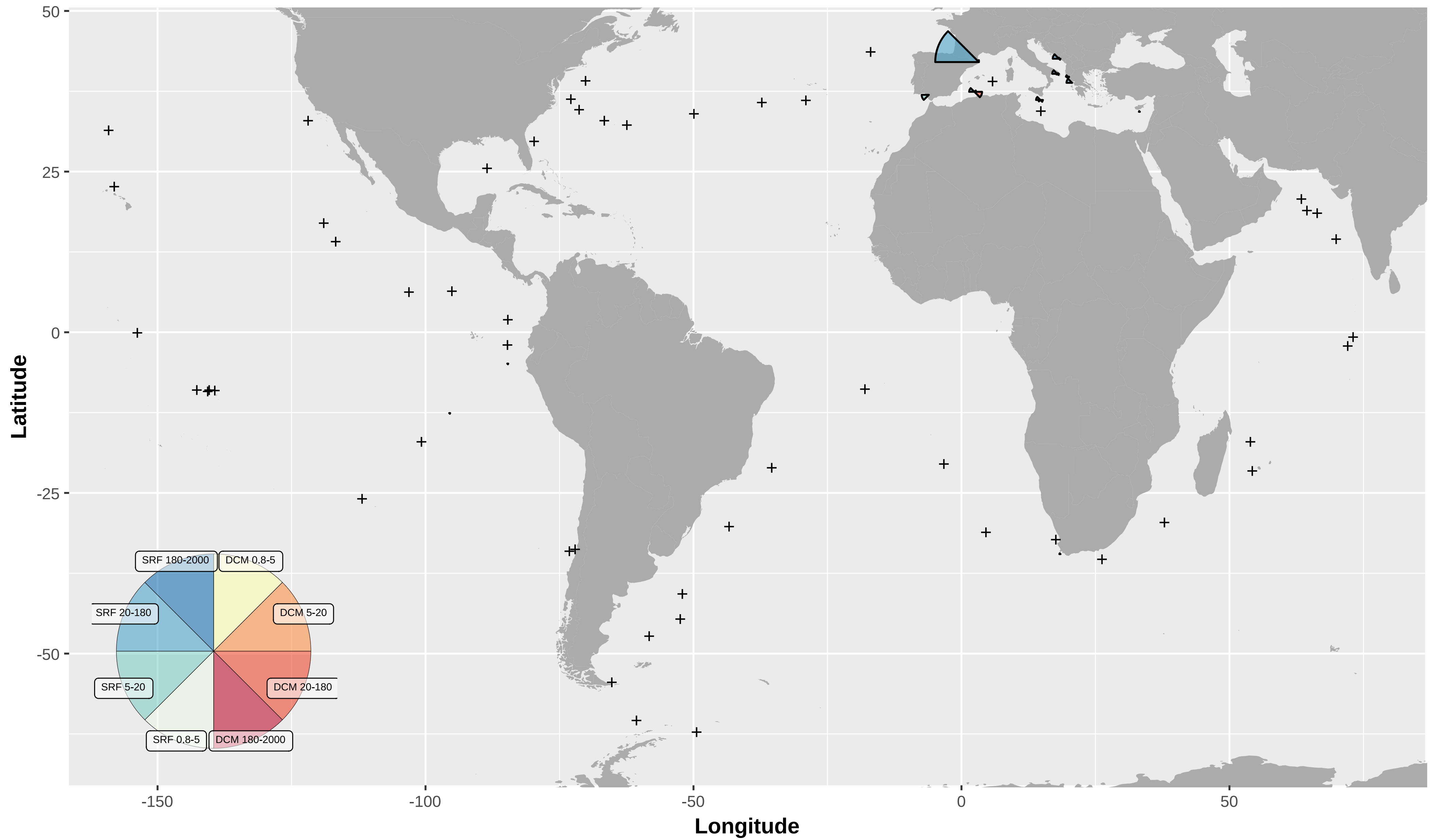


sthokonta;Metazoa;Eumetazoa;Bilateria;Protostomia;Ecdysozoa;Panarthropoda;Arthropoda;Mandibulata;Pancrustacea;Crustacea;Maxillopoda;Copepoda;Neocopepoda;Podoplea;Cyclopoida;Oithonidae

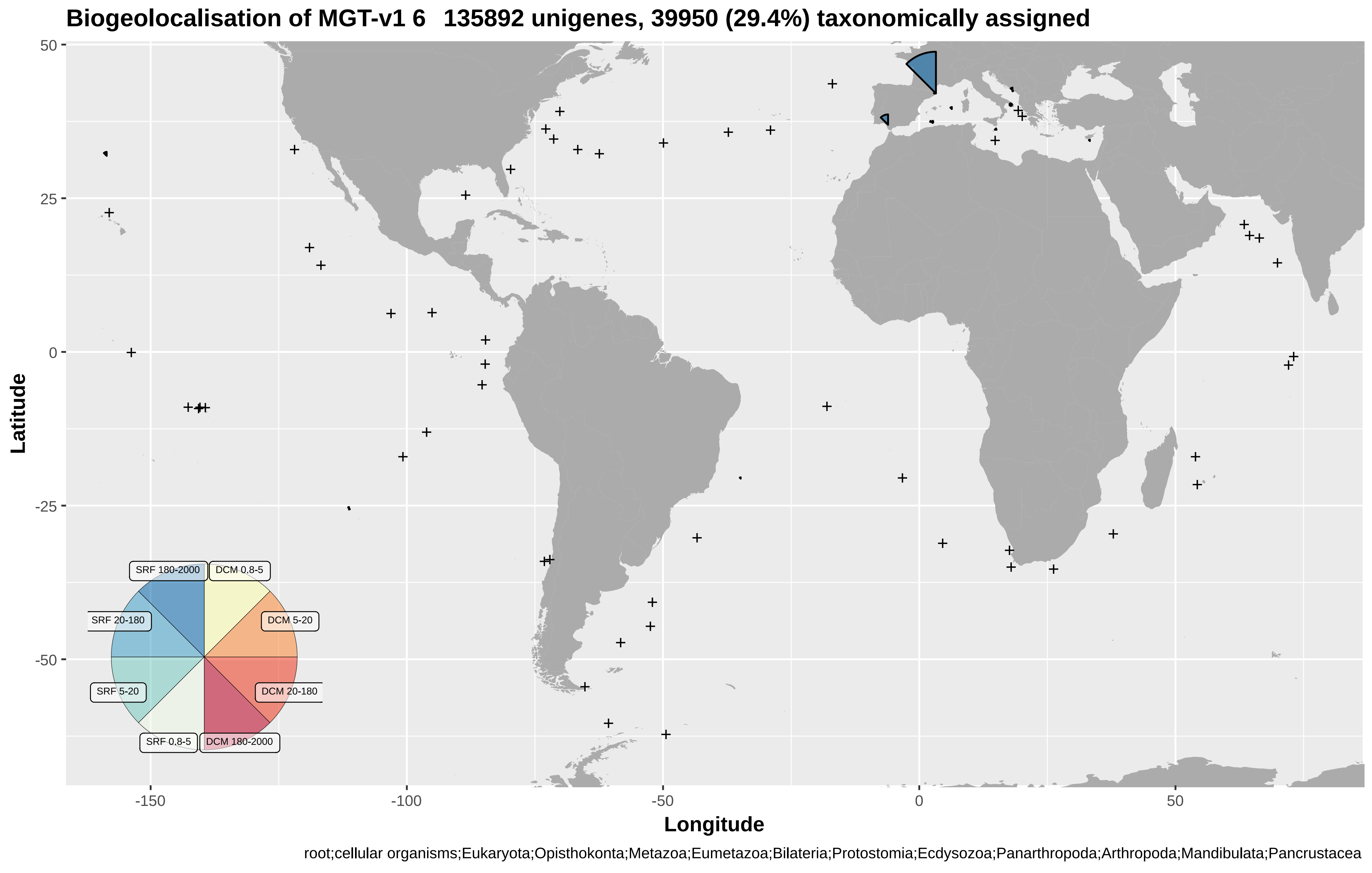
Biogeolocalisation of MGT-v1 6 135892 unigenes, 39950 (29.4%) taxonomically assigned
Latitude
50
25
0
-25
-50
Longitude
-150
-100
-50
0
50
SRF 180-2000
DCM 0.8-5
SRF 20-180
DCM 5-20
SRF 5-20
DCM 20-180
SRF 0.8-5
DCM 180-2000
root;cellular organisms;Eukaryota;Opisthokonta;Metazoa;Eumetazoa;Bilateria;Protostomia;Ecdysozoa;Panarthropoda;Arthropoda;Mandibulata;Pancrustacea

# Biogeolocalisation of MGT-v1 7   134359 unigenes, 28296 (21.06%) taxonomically assigned

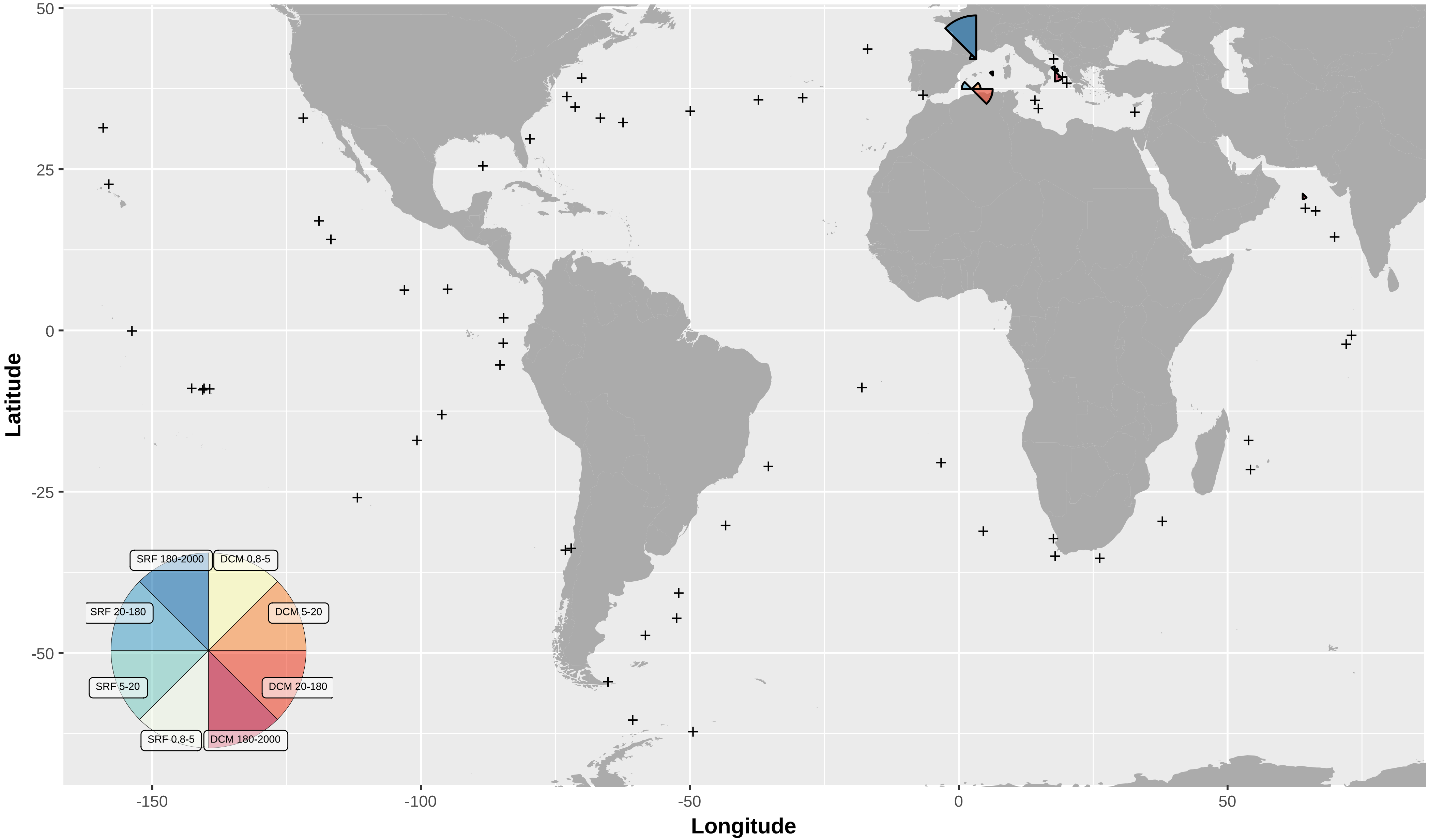


root;cellular organisms;Eukaryota;Opisthokonta;Metazoa;Eumetazoa;Bilateria;Protostomia;Ecdysozoa;Panarthropoda;Arthropoda;Mandibulata;Pancrustacea

## Biogeolocalisation of MGT-v1 8   123134 unigenes, 33117 (26.9%) taxonomically assigned

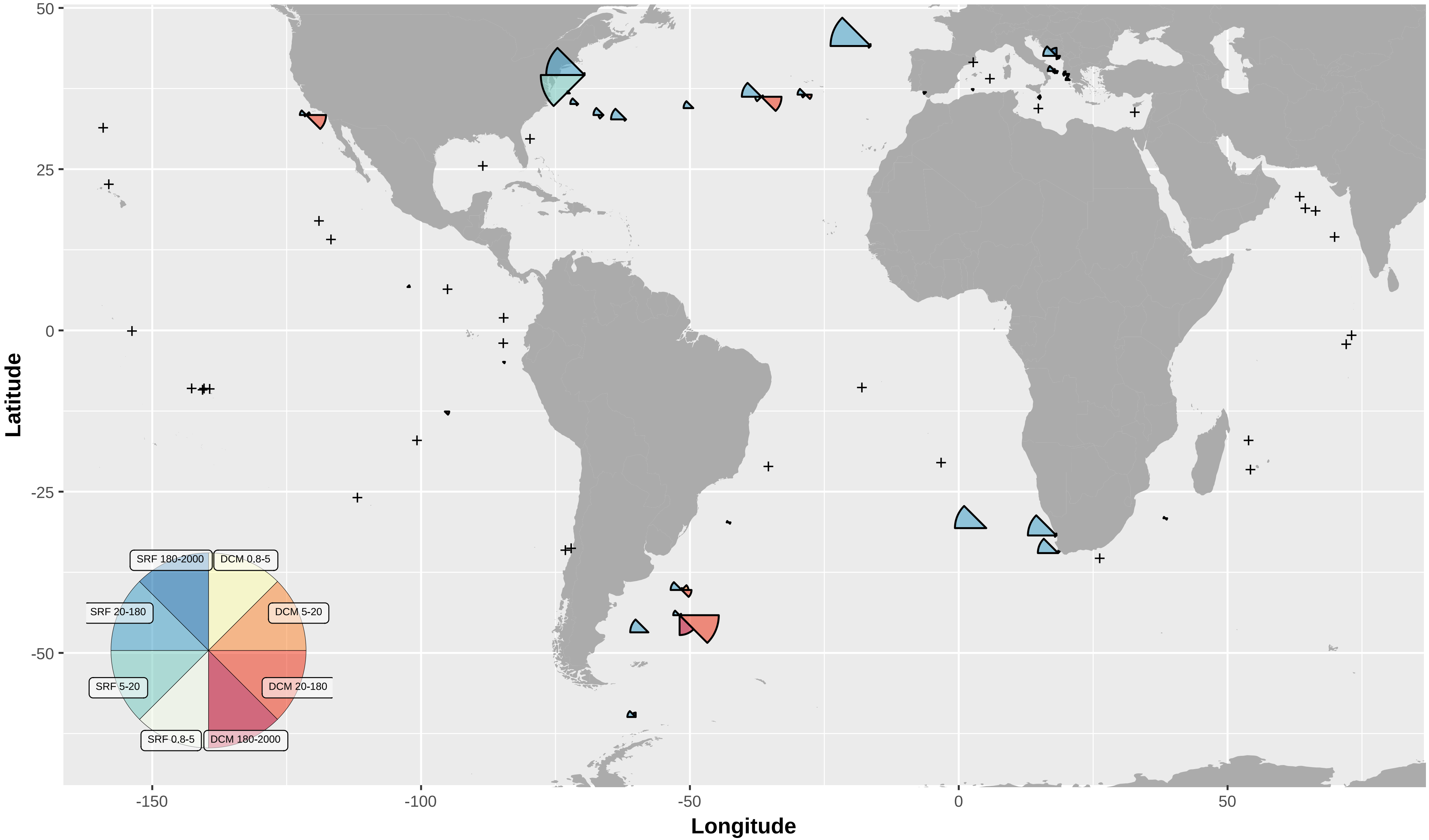


root;cellular organisms;Eukaryota;Opisthokonta;Metazoa;Eumetazoa;Bilateria;Protostomia;Ecdysozoa;Panarthropoda;Arthropoda;Mandibulata;Pancrustacea

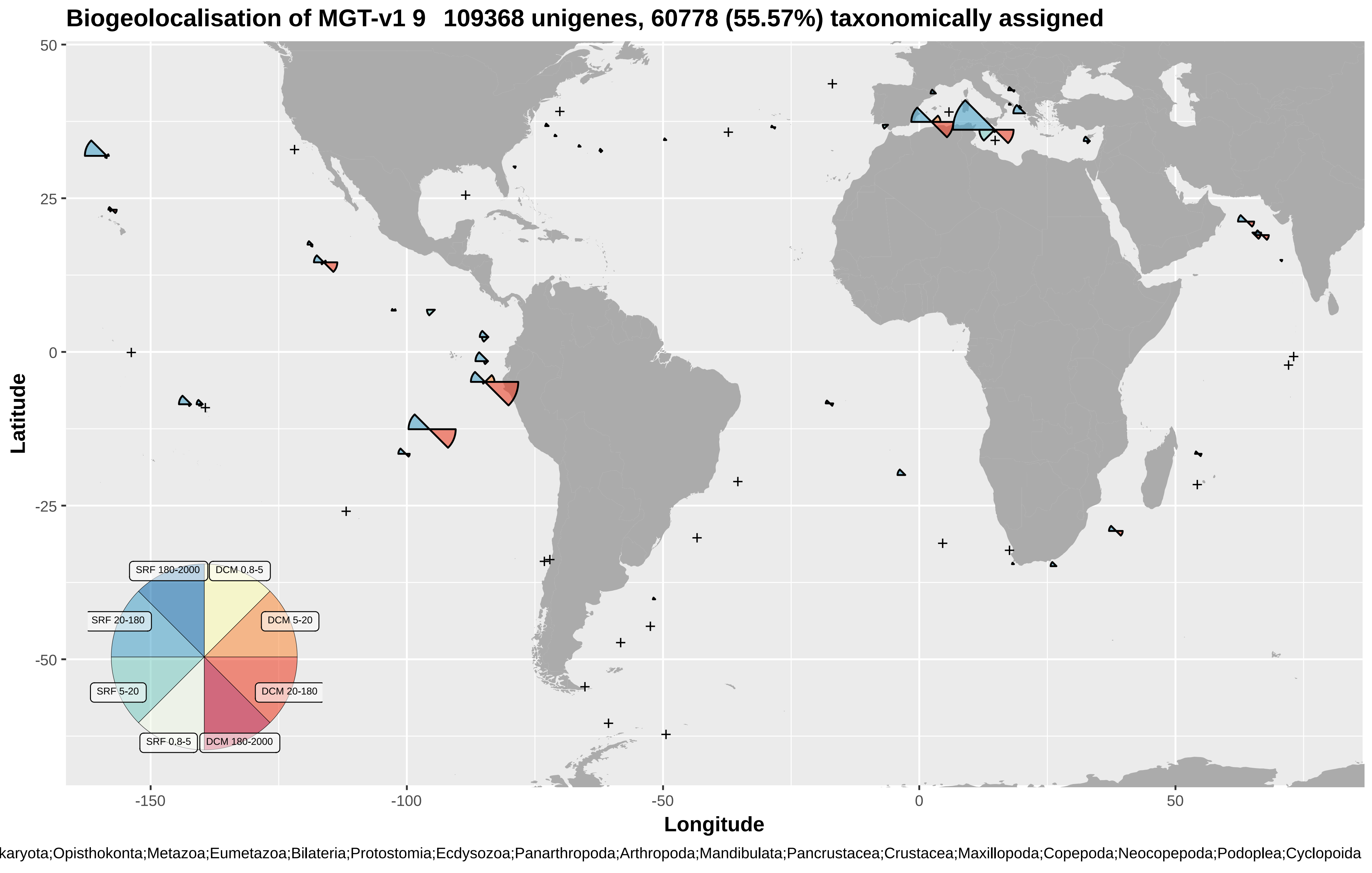

Biogeolocalisation of MGT-v1 9 109368 unigenes, 60778 (55.57%) taxonomically assigned
50
25
0
-25
-50
Latitude
-150
-100
-50
0
50
Longitude
SRF 180-2000
DCM 0.8-5
SRF 20-180
DCM 5-20
SRF 5-20
DCM 20-180
SRF 0.8-5
DCM 180-2000
karyota;Opisthokonta;Metazoa;Eumetazoa;Bilateria;Protostomia;Ecdysozoa;Panarthropoda;Arthropoda;Mandibulata;Pancrustacea;Crustacea;Maxillopoda;Copepoda;Neocopepoda;Podoplea;Cyclopoida

## Biogeolocalisation of MGT-v1 10        107664 unigenes, 36834 (34.21%) taxonomically assigned

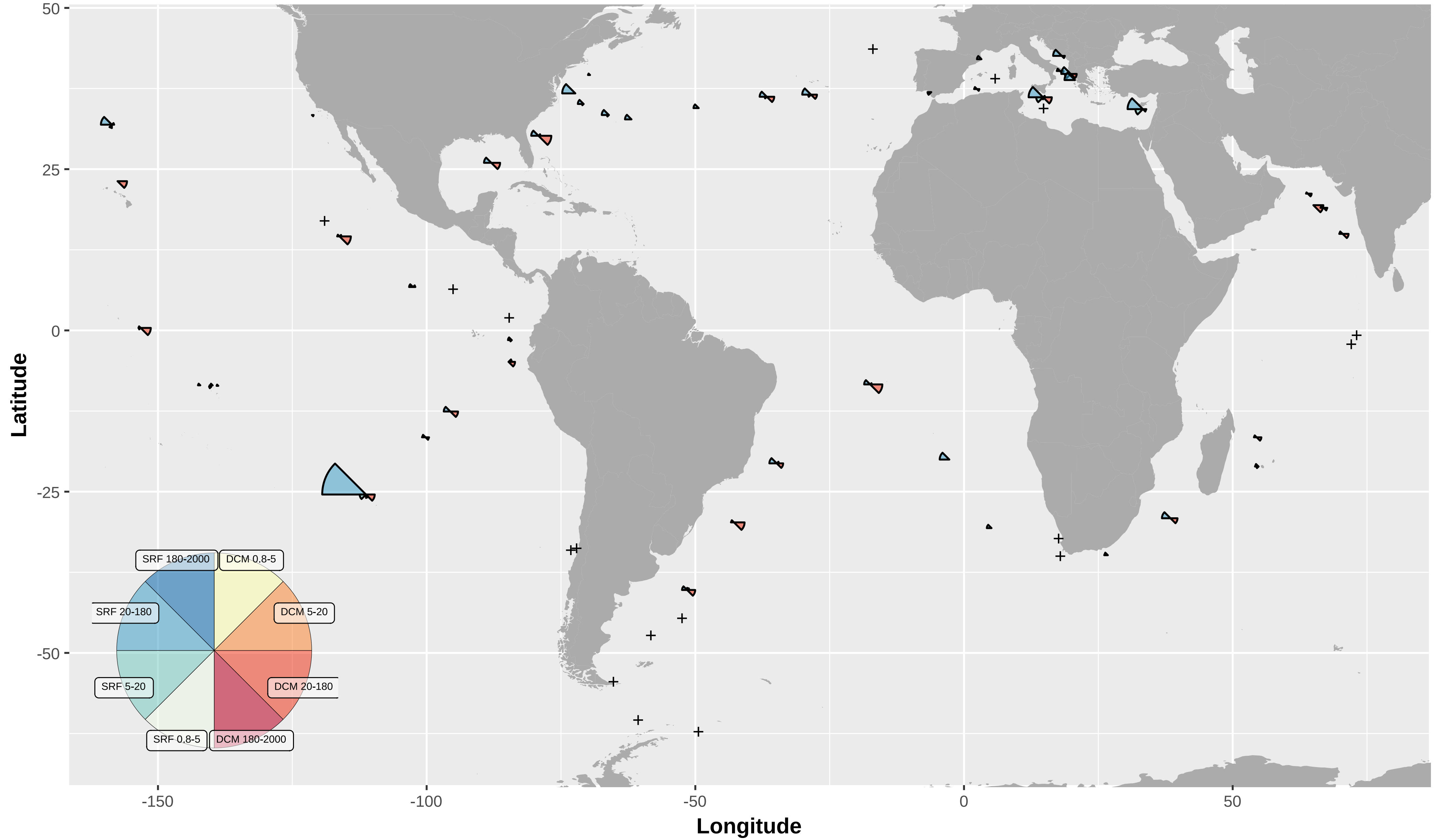


root;cellular organisms;Eukaryota;Opisthokonta;Metazoa;Eumetazoa;Bilateria;Protostomia;Ecdysozoa;Panarthropoda;Arthropoda;Mandibulata;Pancrustacea

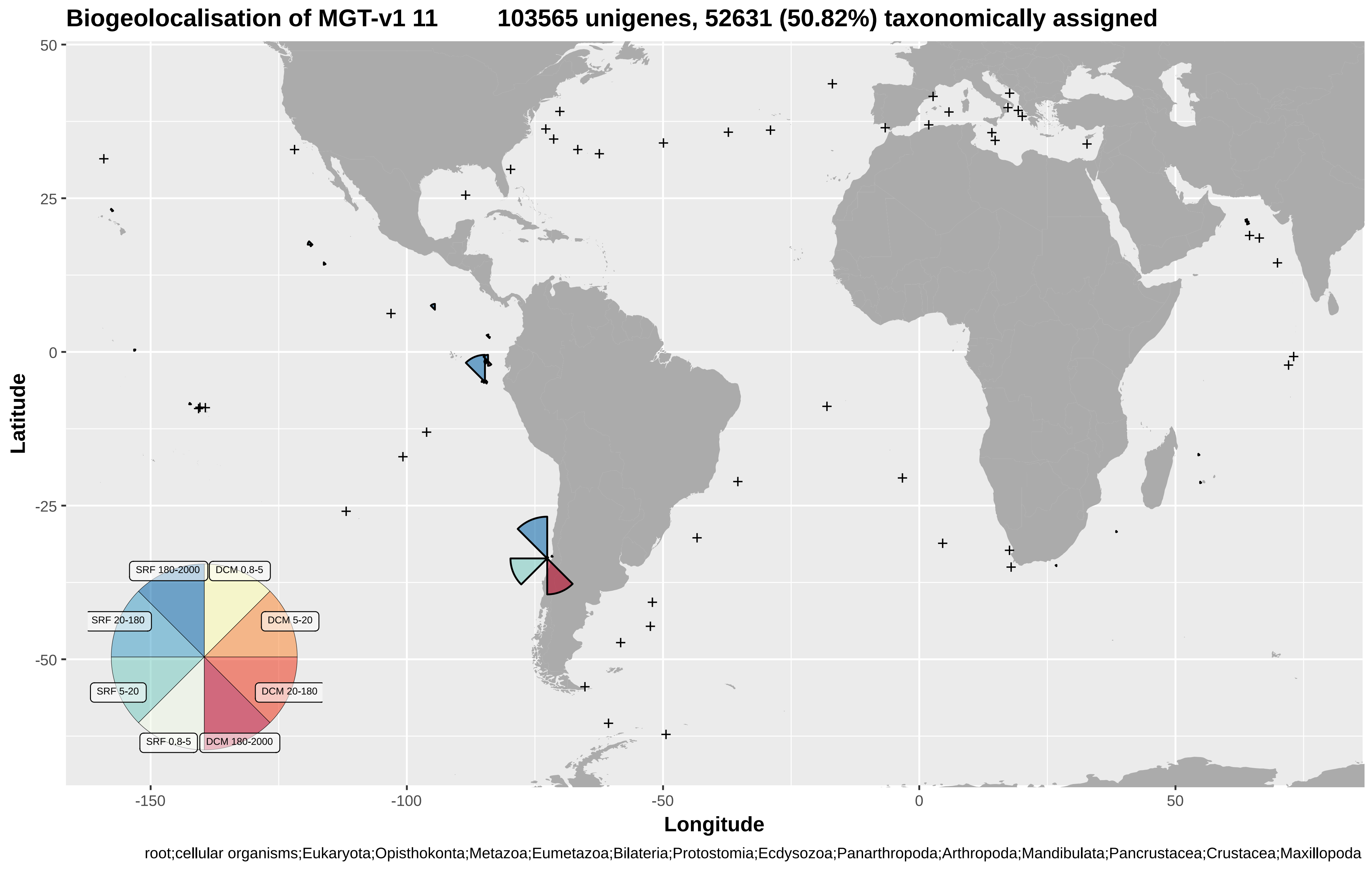
Biogeolocalisation of MGT-v1 11
103565 unigenes, 52631 (50.82%) taxonomically assigned
50
25
0
-25
-50
Latitude
-150
-100
-50
0
50
Longitude
SRF 180-2000
DCM 0.8-5
SRF 20-180
DCM 5-20
SRF 5-20
DCM 20-180
SRF 0.8-5
DCM 180-2000
root;cellular organisms;Eukaryota;Opisthokonta;Metazoa;Eumetazoa;Bilateria;Protostomia;Ecdysozoa;Panarthropoda;Arthropoda;Mandibulata;Pancrustacea;Crustacea;Maxillopoda

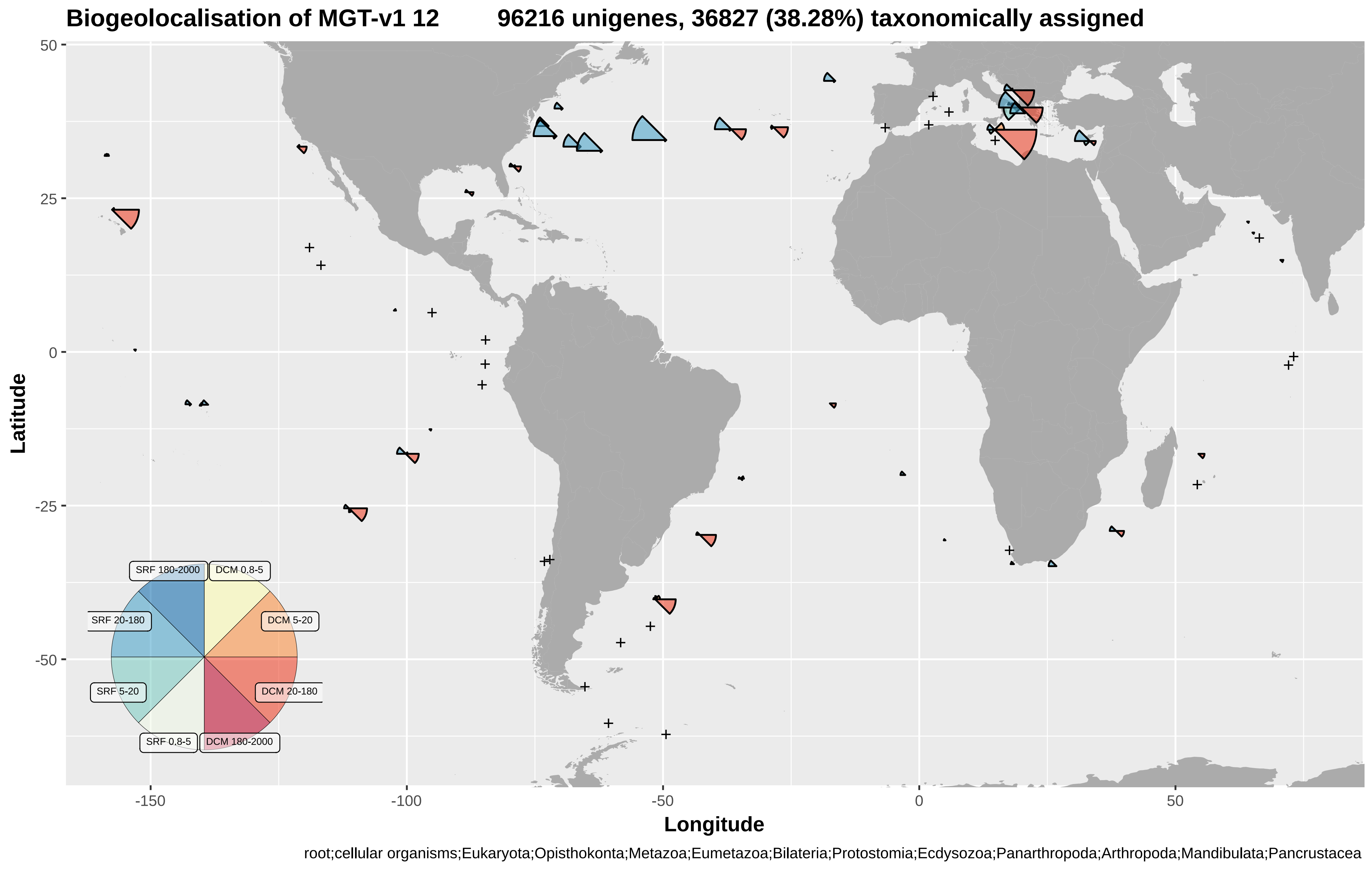
Biogeolocalisation of MGT-v1 12
96216 unigenes, 36827 (38.28%) taxonomically assigned
Latitude
50
25
0
-25
-50
Longitude
-150
-100
-50
0
50
SRF 180-2000
DCM 0.8-5
SRF 20-180
DCM 5-20
SRF 5-20
DCM 20-180
SRF 0.8-5
DCM 180-2000
root;cellular organisms;Eukaryota;Opisthokonta;Metazoa;Eumetazoa;Bilateria;Protostomia;Ecdysozoa;Panarthropoda;Arthropoda;Mandibulata;Pancrustacea

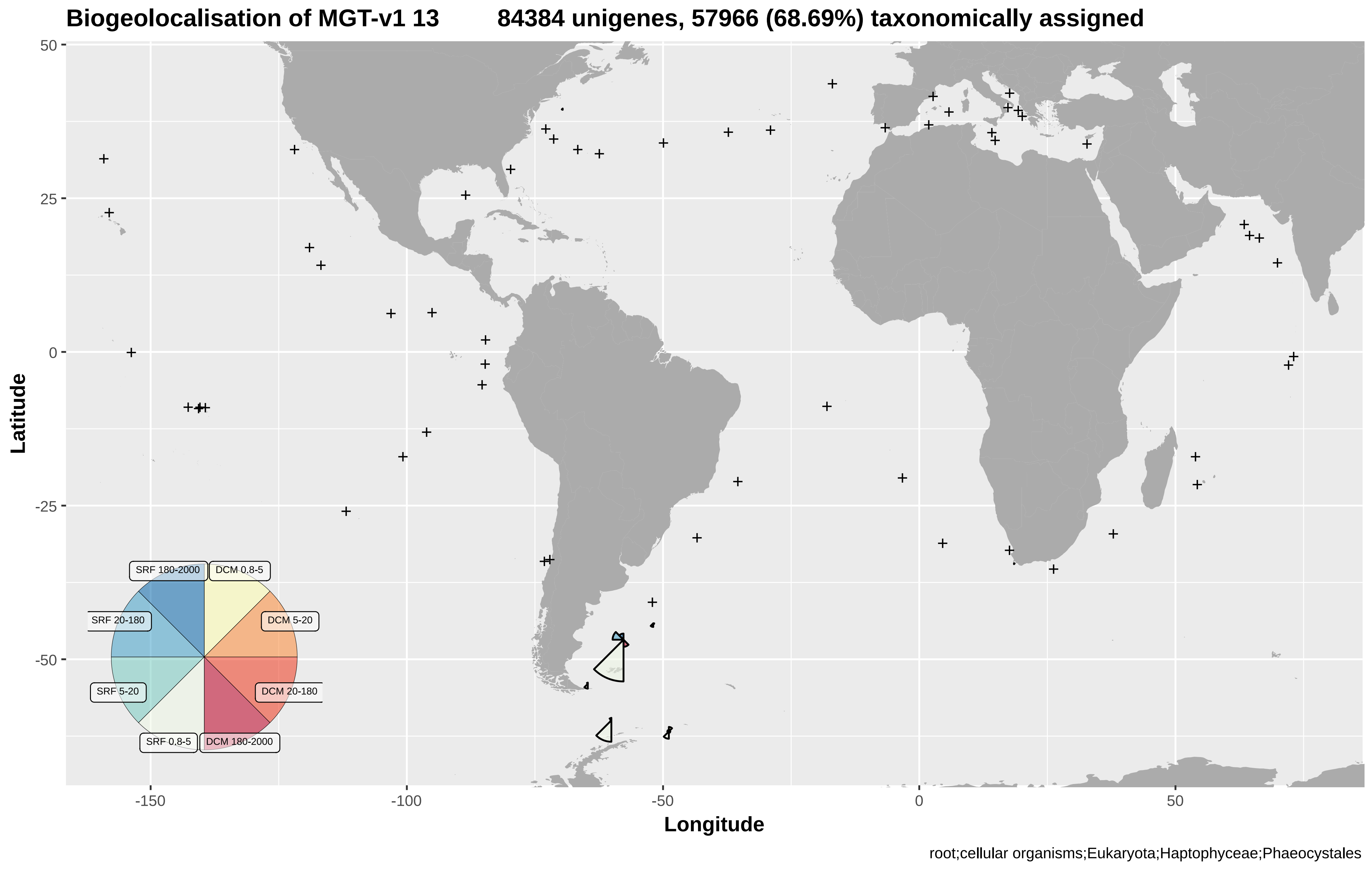

Biogeolocalisation of MGT-v1 13 84384 unigenes, 57966 (68.69%) taxonomically assigned
Latitude
Longitude
50
25
0
-25
-50
-150
-100
-50
0
50
SRF 180-2000
DCM 0.8-5
SRF 20-180
DCM 5-20
SRF 5-20
DCM 20-180
SRF 0.8-5
DCM 180-2000
root;cellular organisms;Eukaryota;Haptophyceae;Phaeocystales

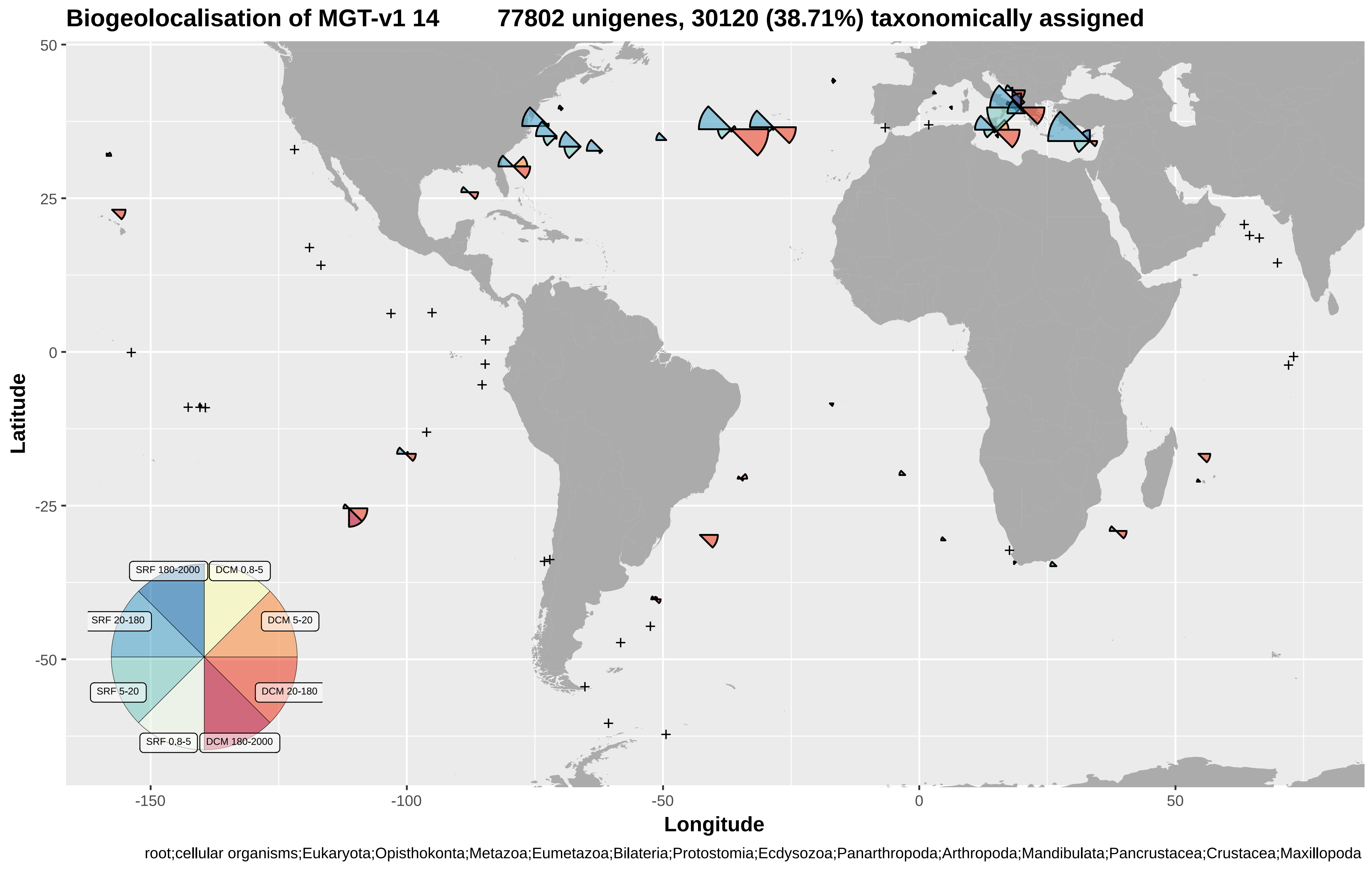

Biogeolocalisation of MGT-v1 14 77802 unigenes, 30120 (38.71%) taxonomically assigned
50
25
0
-25
-50
Latitude
-150
-100
-50
0
50
Longitude
SRF 180-2000
DCM 0.8-5
SRF 20-180
DCM 5-20
SRF 5-20
DCM 20-180
SRF 0.8-5
DCM 180-2000
root;cellular organisms;Eukaryota;Opisthokonta;Metazoa;Eumetazoa;Bilateria;Protostomia;Ecdysozoa;Panarthropoda;Arthropoda;Mandibulata;Pancrustacea;Crustacea;Maxillopoda

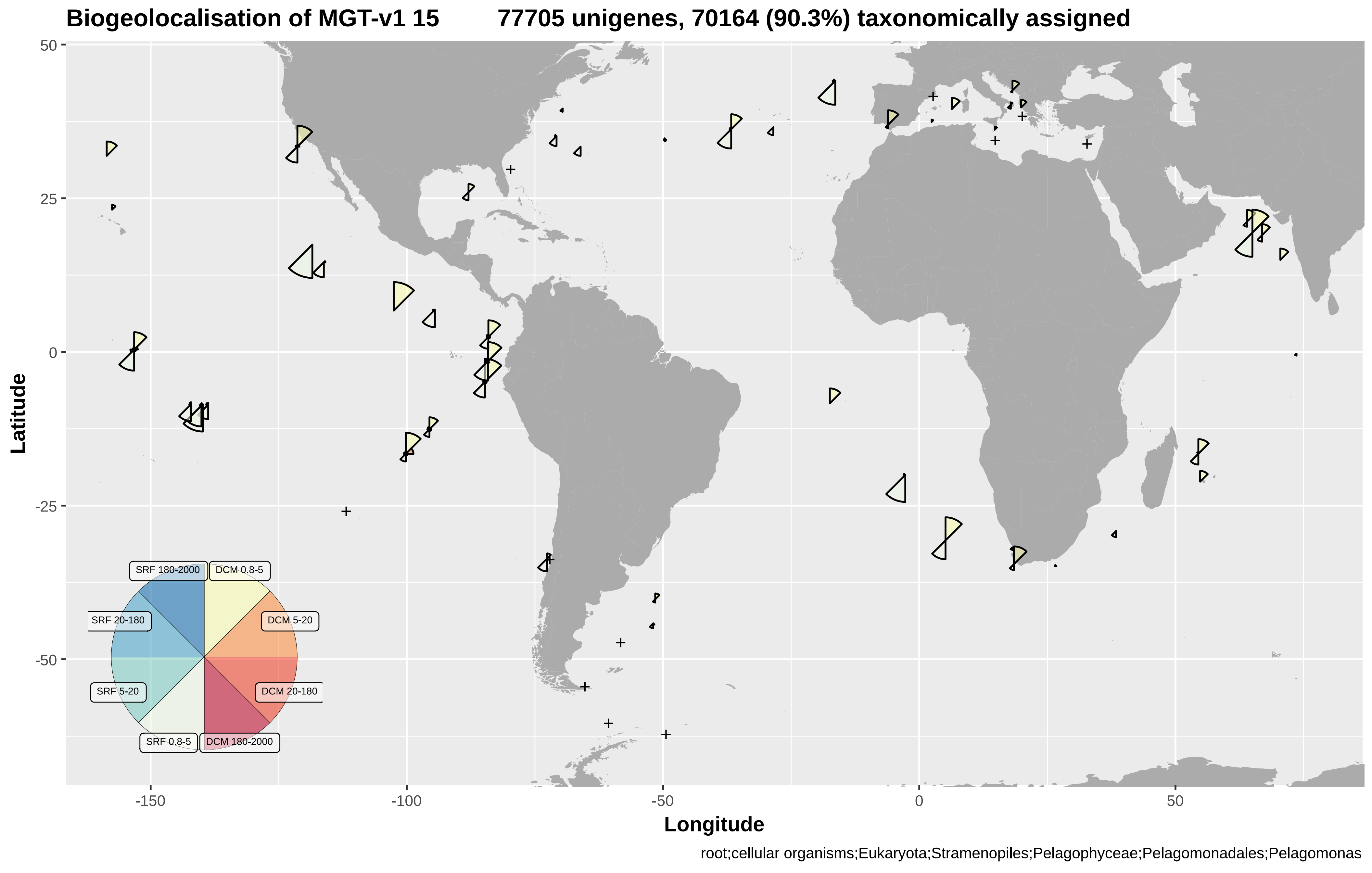

Biogeolocalisation of MGT-v1 15
77705 unigenes, 70164 (90.3%) taxonomically assigned
Latitude
Longitude
50
25
0
-25
-50
-150
-100
-50
0
50
SRF 180-2000
DCM 0.8-5
SRF 20-180
DCM 5-20
SRF 5-20
DCM 20-180
SRF 0.8-5
DCM 180-2000
root;cellular organisms;Eukaryota;Stramenopiles;Pelagophyceae;Pelagomonadales;Pelagomonas

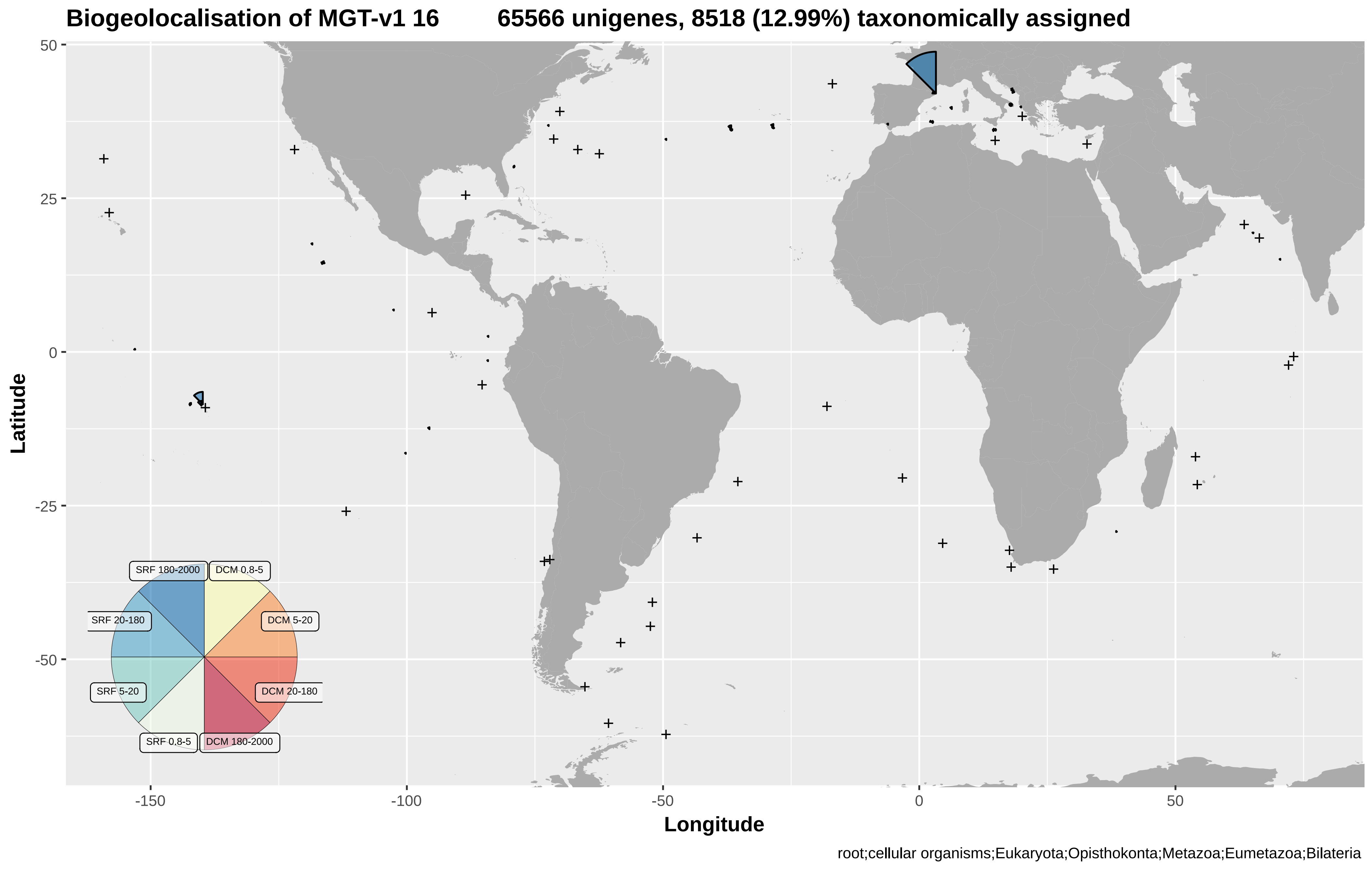

Biogeolocalisation of MGT-v1 16 65566 unigenes, 8518 (12.99%) taxonomically assigned
Latitude
50
25
0
-25
-50
Longitude
-150
-100
-50
0
50
SRF 180-2000
DCM 0.8-5
SRF 20-180
DCM 5-20
SRF 5-20
DCM 20-180
SRF 0.8-5
DCM 180-2000
root;cellular organisms;Eukaryota;Opisthokonta;Metazoa;Eumetazoa;Bilateria

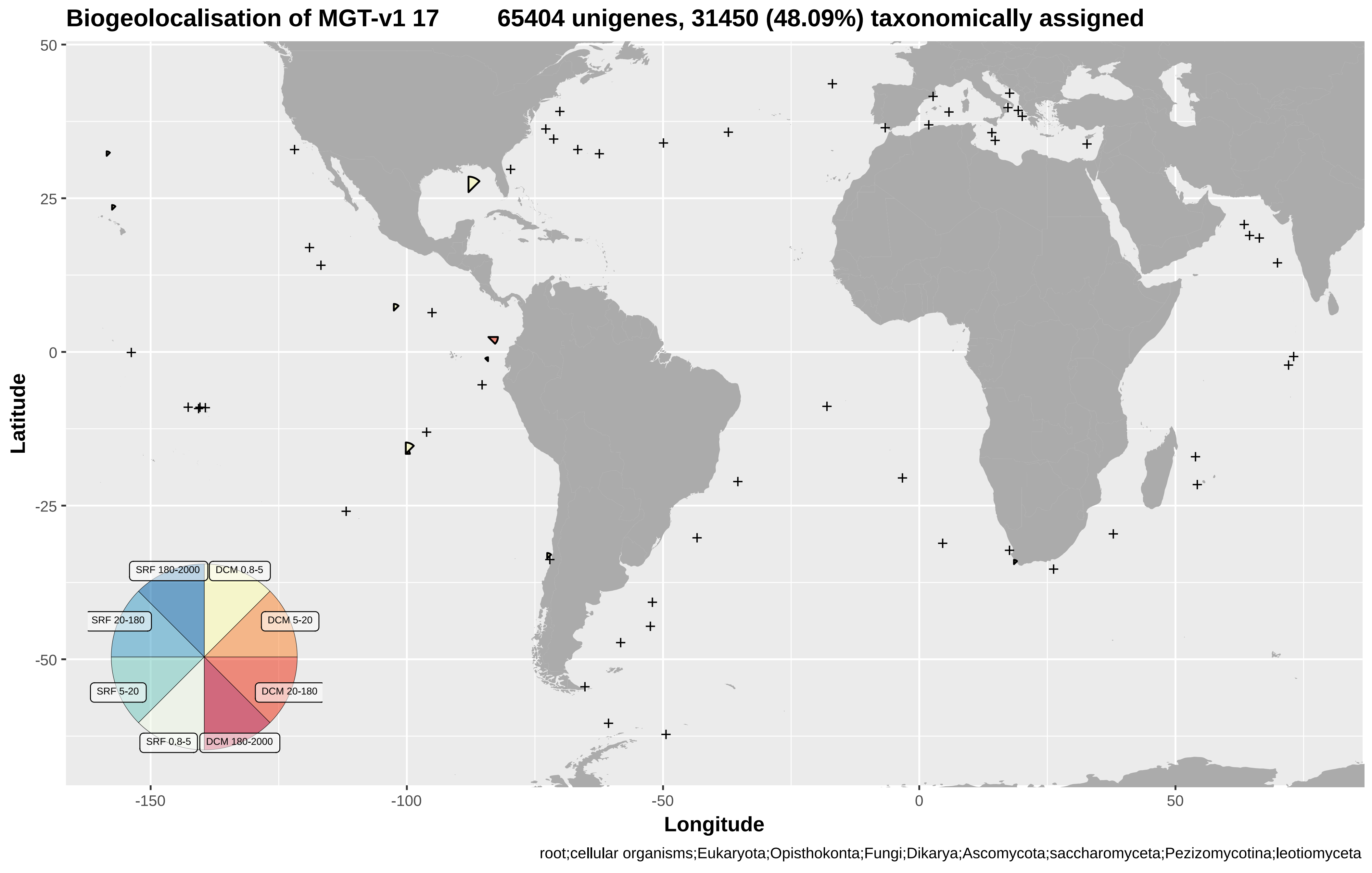
Biogeolocalisation of MGT-v1 17
65404 unigenes, 31450 (48.09%) taxonomically assigned
Latitude
Longitude
50
25
0
-25
-50
-150
-100
-50
0
50
SRF 180-2000
DCM 0.8-5
SRF 20-180
DCM 5-20
SRF 5-20
DCM 20-180
SRF 0.8-5
DCM 180-2000
root;cellular organisms;Eukaryota;Opisthokonta;Fungi;Dikarya;Ascomycota;saccharomyceta;Pezizomycotina;leotiomyceta

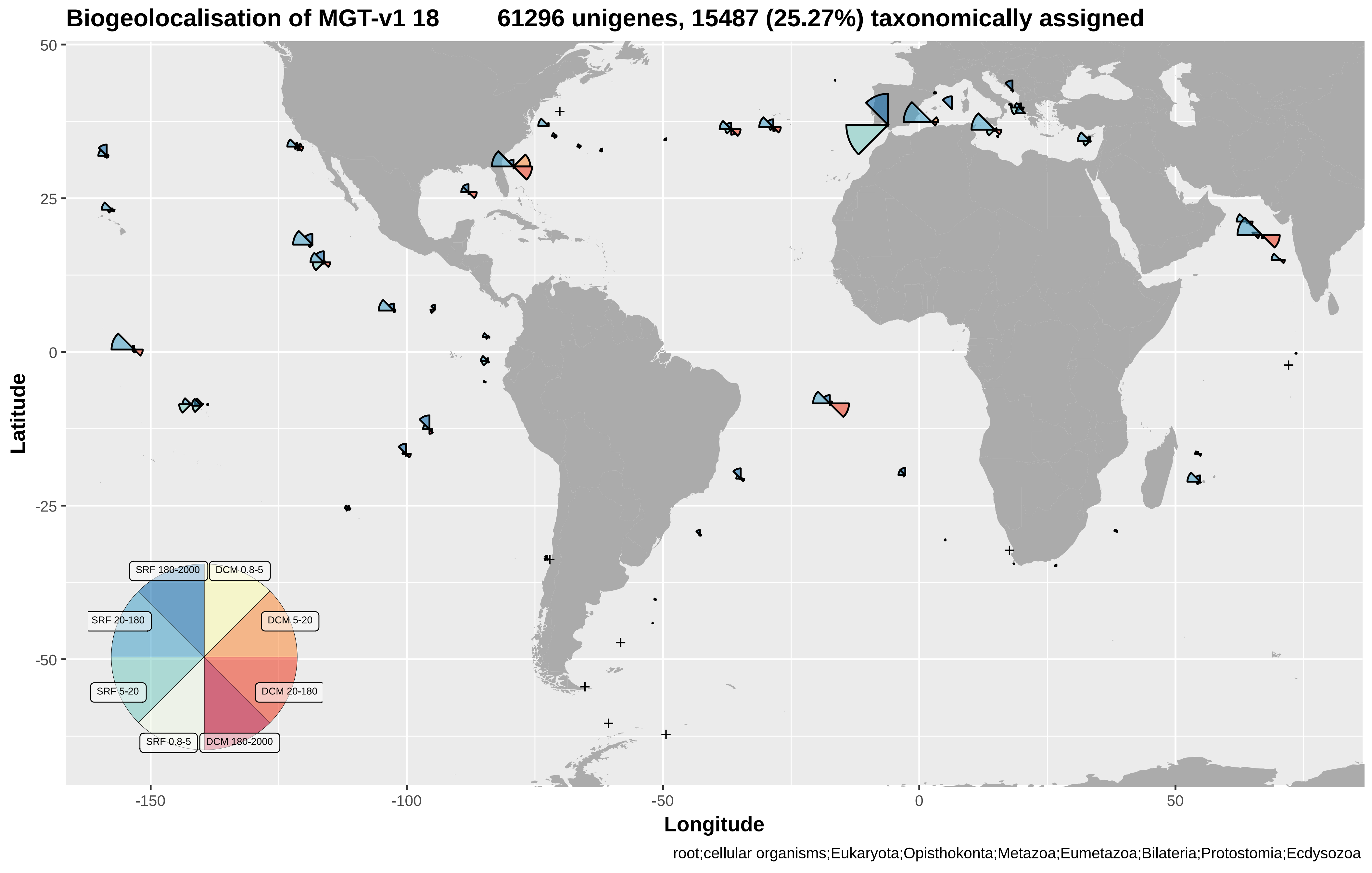

Biogeolocalisation of MGT-v1 18
61296 unigenes, 15487 (25.27%) taxonomically assigned
50
25
0
-25
-50
Latitude
-150
-100
-50
0
50
Longitude
SRF 180-2000
DCM 0.8-5
SRF 20-180
DCM 5-20
SRF 5-20
DCM 20-180
SRF 0.8-5
DCM 180-2000
root;cellular organisms;Eukaryota;Opisthokonta;Metazoa;Eumetazoa;Bilateria;Protostomia;Ecdysozoa

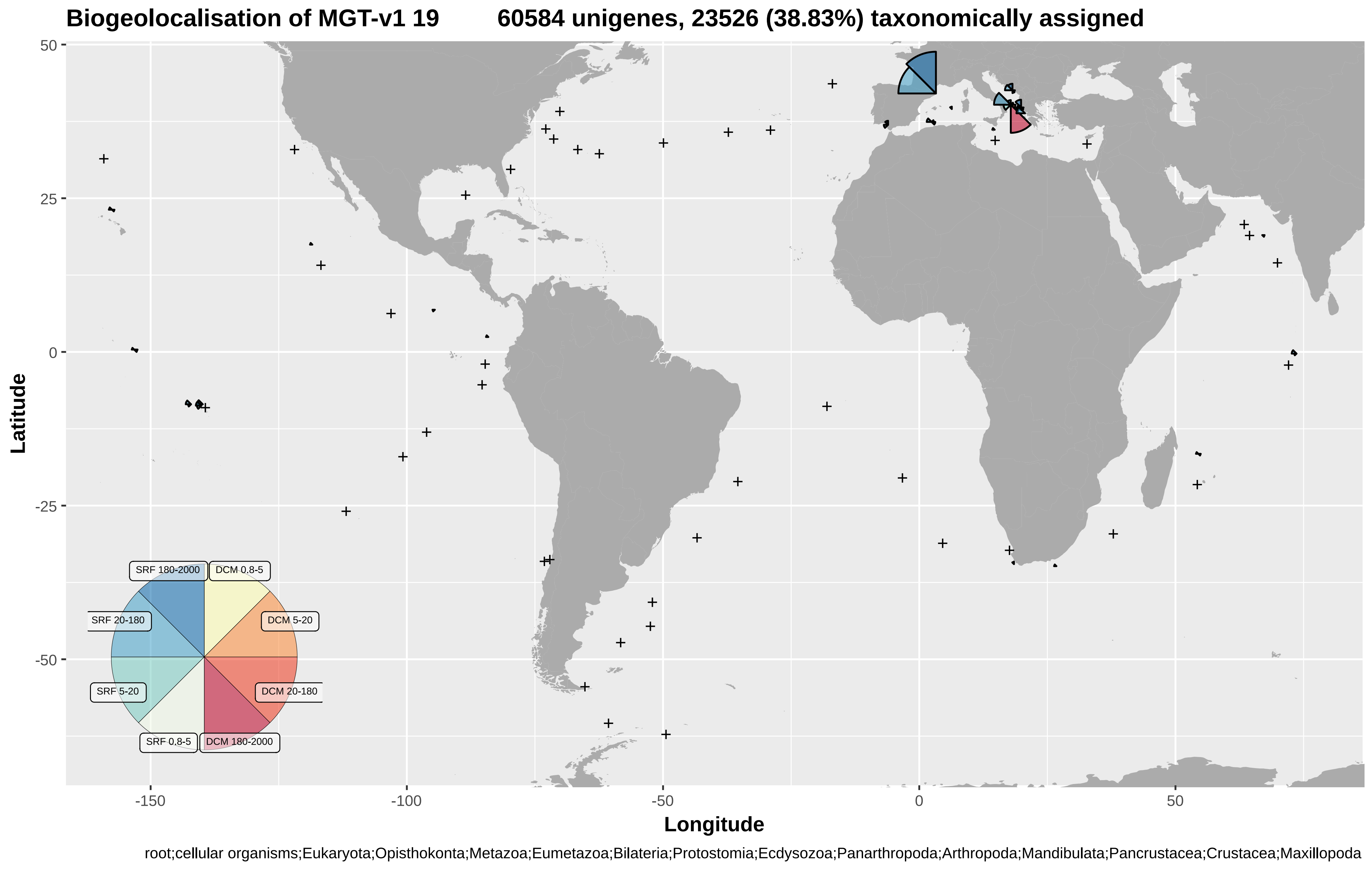
Biogeolocalisation of MGT-v1 19 60584 unigenes, 23526 (38.83%) taxonomically assigned
50
25
0
-25
-50
Latitude
-150
-100
-50
0
50
Longitude
SRF 180-2000
DCM 0.8-5
SRF 20-180
DCM 5-20
SRF 5-20
DCM 20-180
SRF 0.8-5
DCM 180-2000
root;cellular organisms;Eukaryota;Opisthokonta;Metazoa;Eumetazoa;Bilateria;Protostomia;Ecdysozoa;Panarthropoda;Arthropoda;Mandibulata;Pancrustacea;Crustacea;Maxillopoda

#### Biogeolocalisation of MGT-v1 20 58041 unigenes, 28047 (48.32%) taxonomically assigned

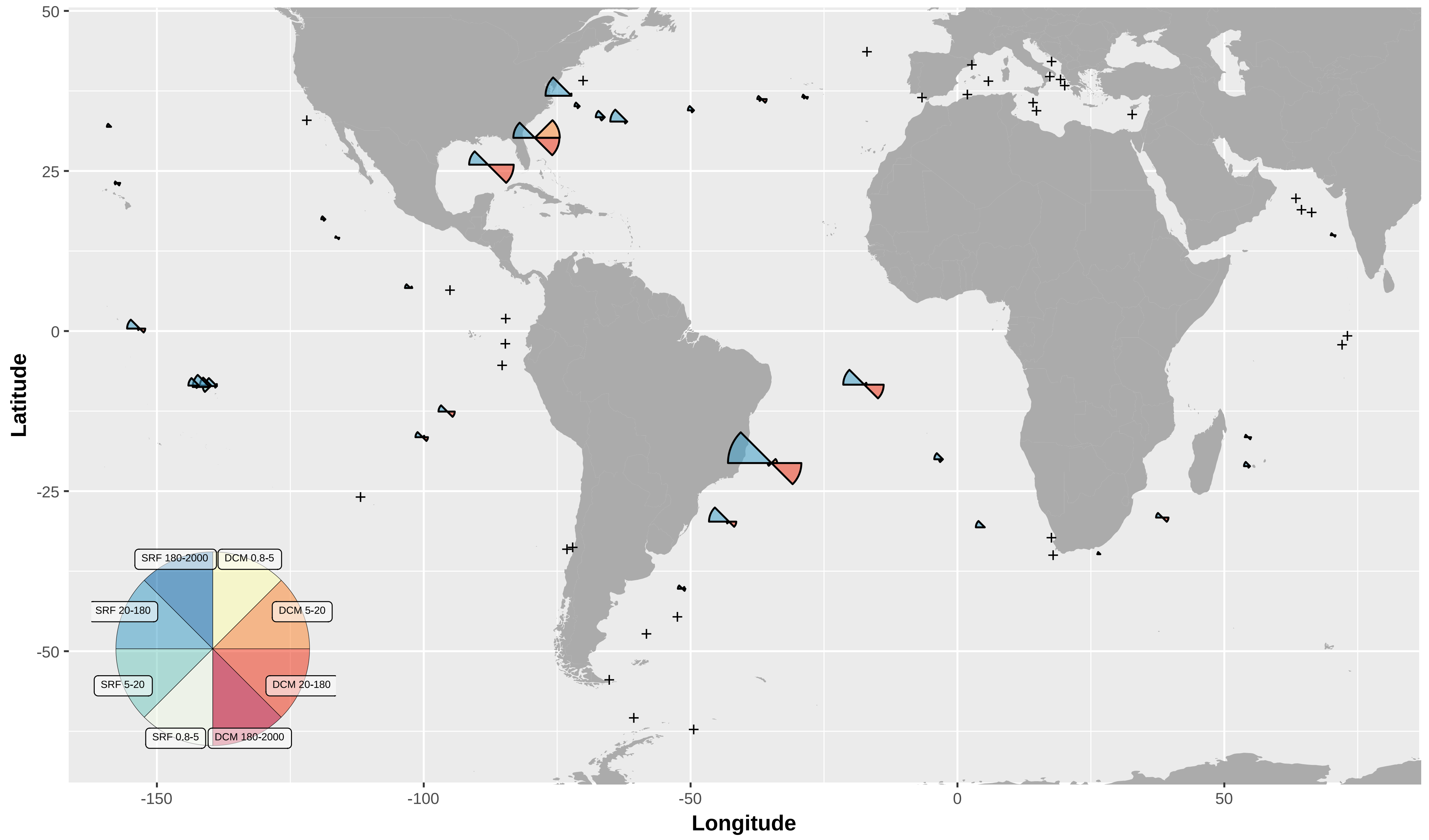


root;cellular organisms;Eukaryota;Opisthokonta;Metazoa;Eumetazoa;Bilateria;Protostomia;Ecdysozoa;Panarthropoda;Arthropoda;Mandibulata;Pancrustacea;Crustacea;Maxillopoda

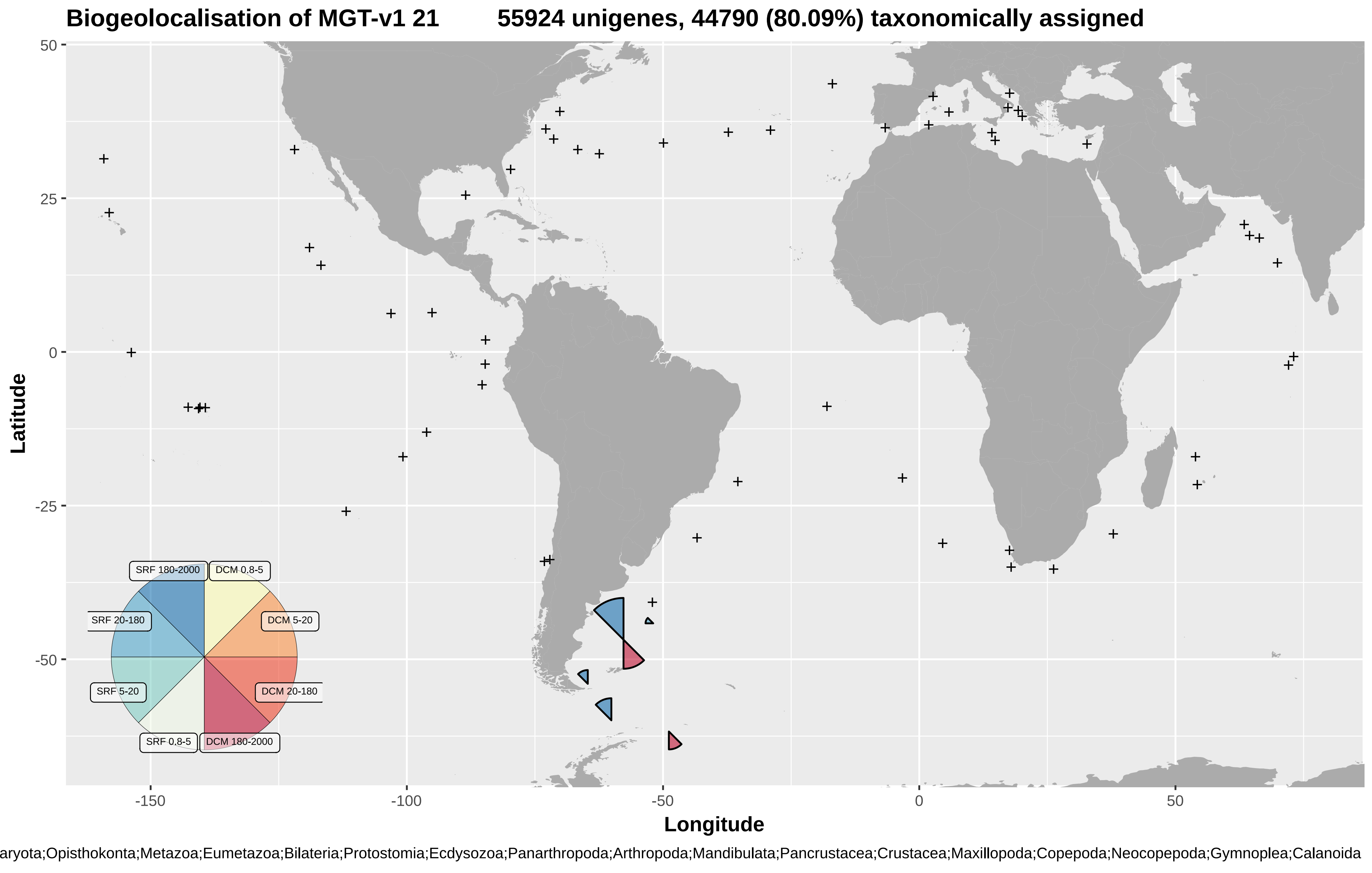
Biogeolocalisation of MGT-v1 21 55924 unigenes, 44790 (80.09%) taxonomically assigned
Latitude
Longitude
50
25
0
-25
-50
-150
-100
-50
0
50
SRF 180-2000
DCM 0.8-5
SRF 20-180
DCM 5-20
SRF 5-20
DCM 20-180
SRF 0.8-5
DCM 180-2000
aryota;Opisthokonta;Metazoa;Eumetazoa;Bilateria;Protostomia;Ecdysozoa;Panarthropoda;Arthropoda;Mandibulata;Pancrustacea;Crustacea;Maxillopoda;Copepoda;Neocopepoda;Gymnoplea;Calanoida

#### Biogeolocalisation of MGT-v1 22 55444 unigenes, 22551 (40.67%) taxonomically assigned

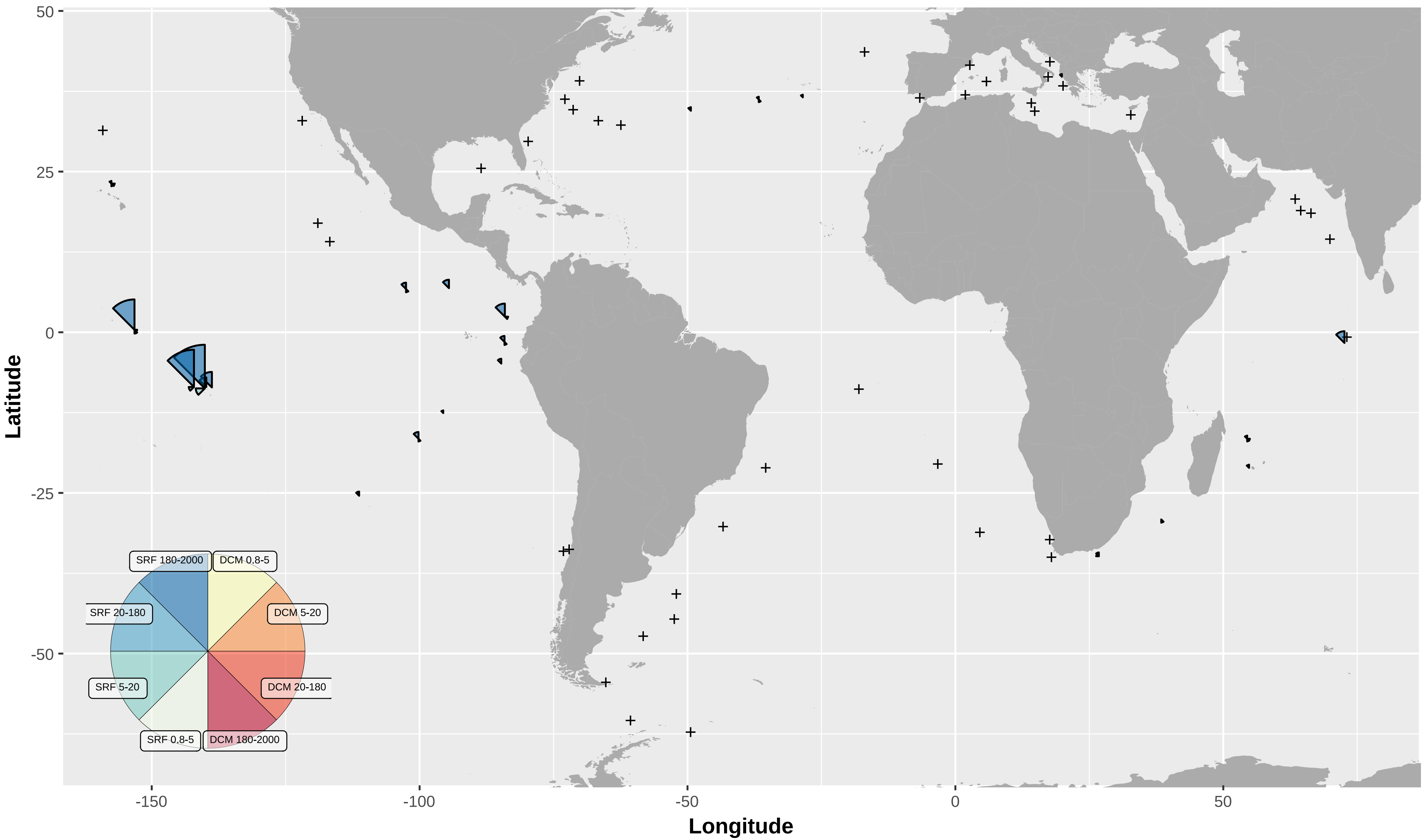


root;cellular organisms;Eukaryota;Opisthokonta;Metazoa;Eumetazoa;Bilateria;Protostomia;Ecdysozoa;Panarthropoda;Arthropoda;Mandibulata;Pancrustacea;Crustacea;Maxillopoda

#### Biogeolocalisation of MGT-v1 23 53910 unigenes, 28470 (52.81%) taxonomically assigned

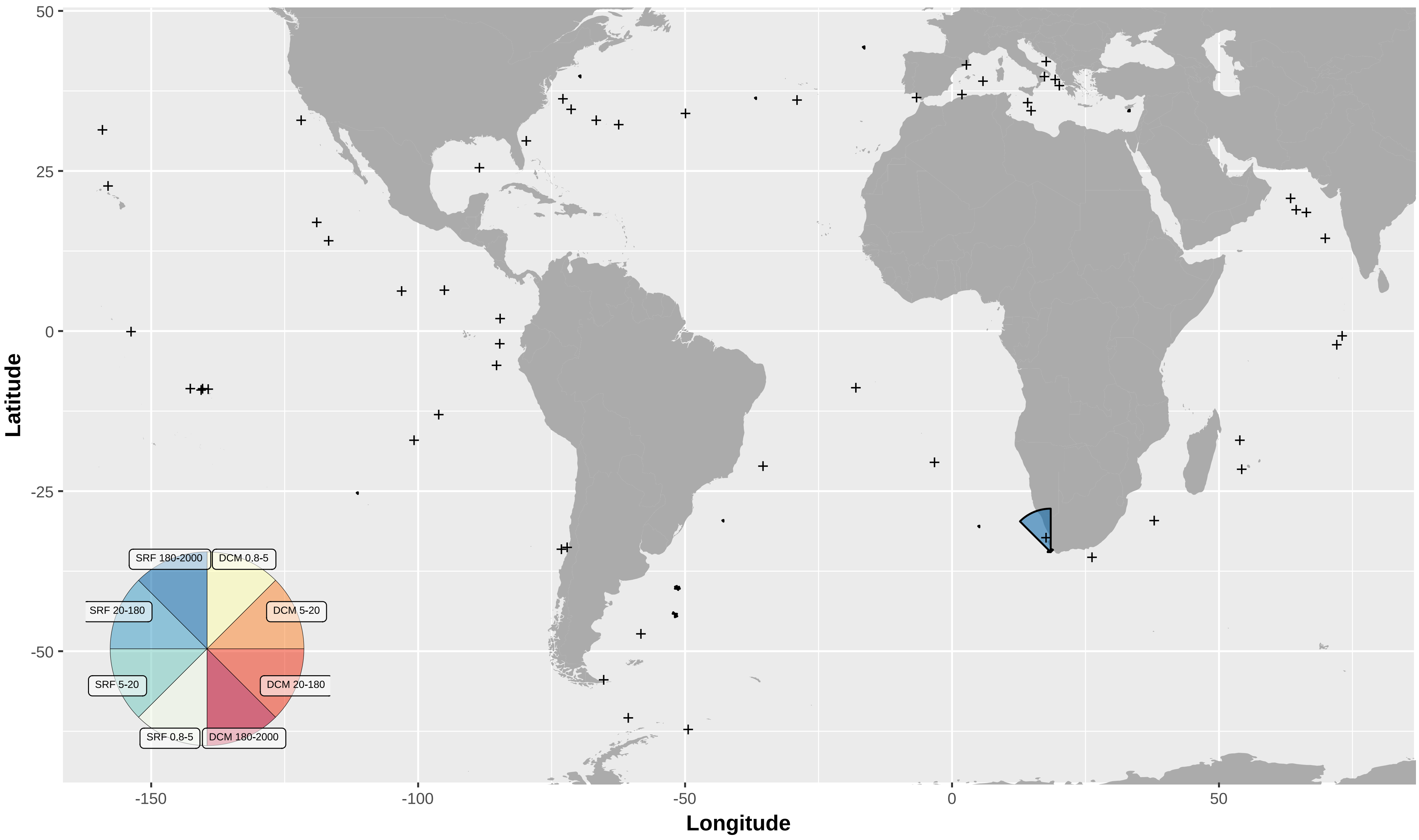


root;cellular organisms;Eukaryota;Opisthokonta;Metazoa;Eumetazoa;Bilateria;Protostomia;Ecdysozoa;Panarthropoda;Arthropoda;Mandibulata;Pancrustacea;Crustacea;Maxillopoda

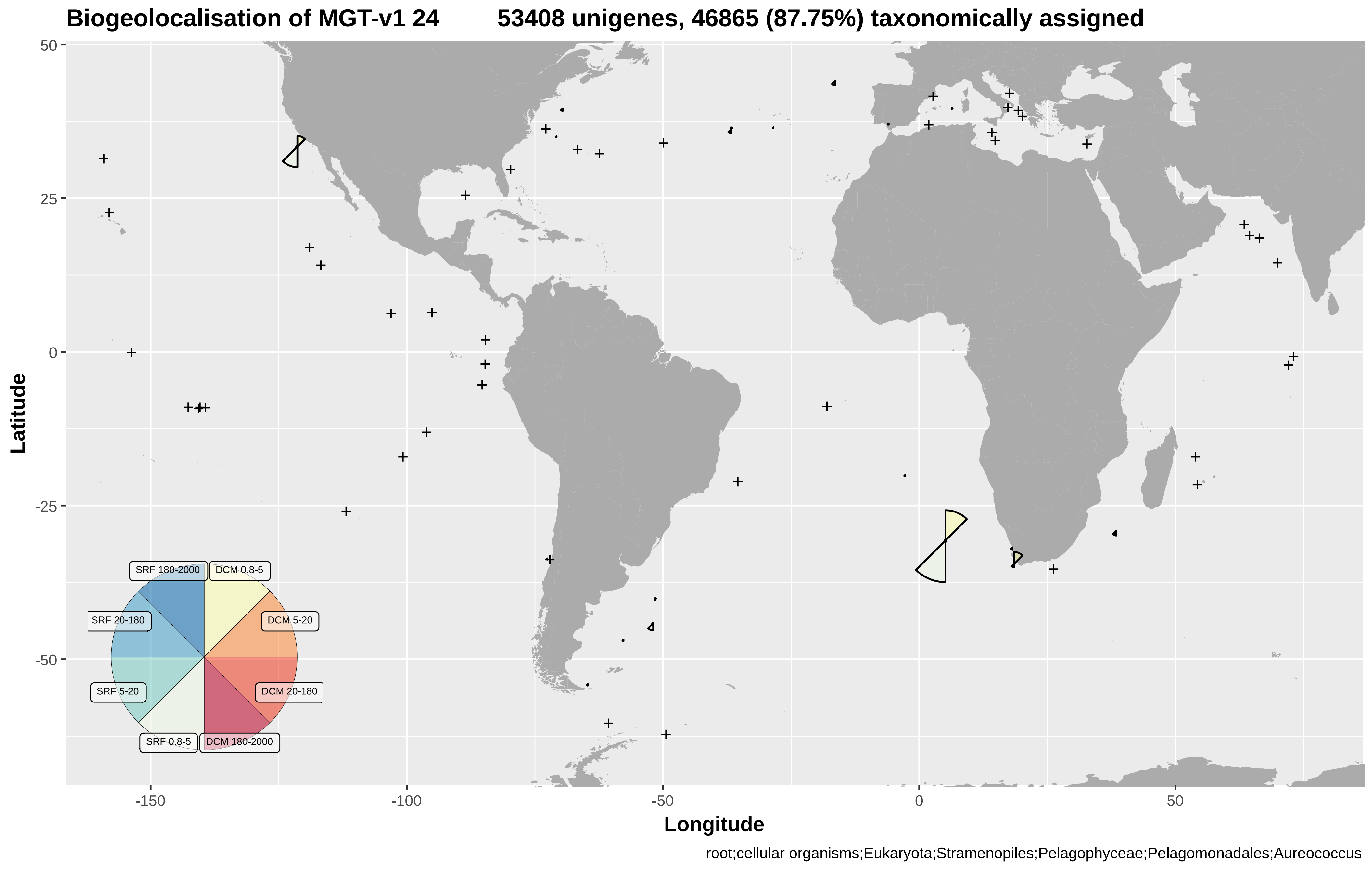

Biogeolocalisation of MGT-v1 24
53408 unigenes, 46865 (87.75%) taxonomically assigned
50
25
0
-25
-50
Latitude
-150
-100
-50
0
50
Longitude
SRF 180-2000
DCM 0.8-5
SRF 20-180
DCM 5-20
SRF 5-20
DCM 20-180
SRF 0.8-5
DCM 180-2000
root;cellular organisms;Eukaryota;Stramenopiles;Pelagophyceae;Pelagomonadales;Aureococcus

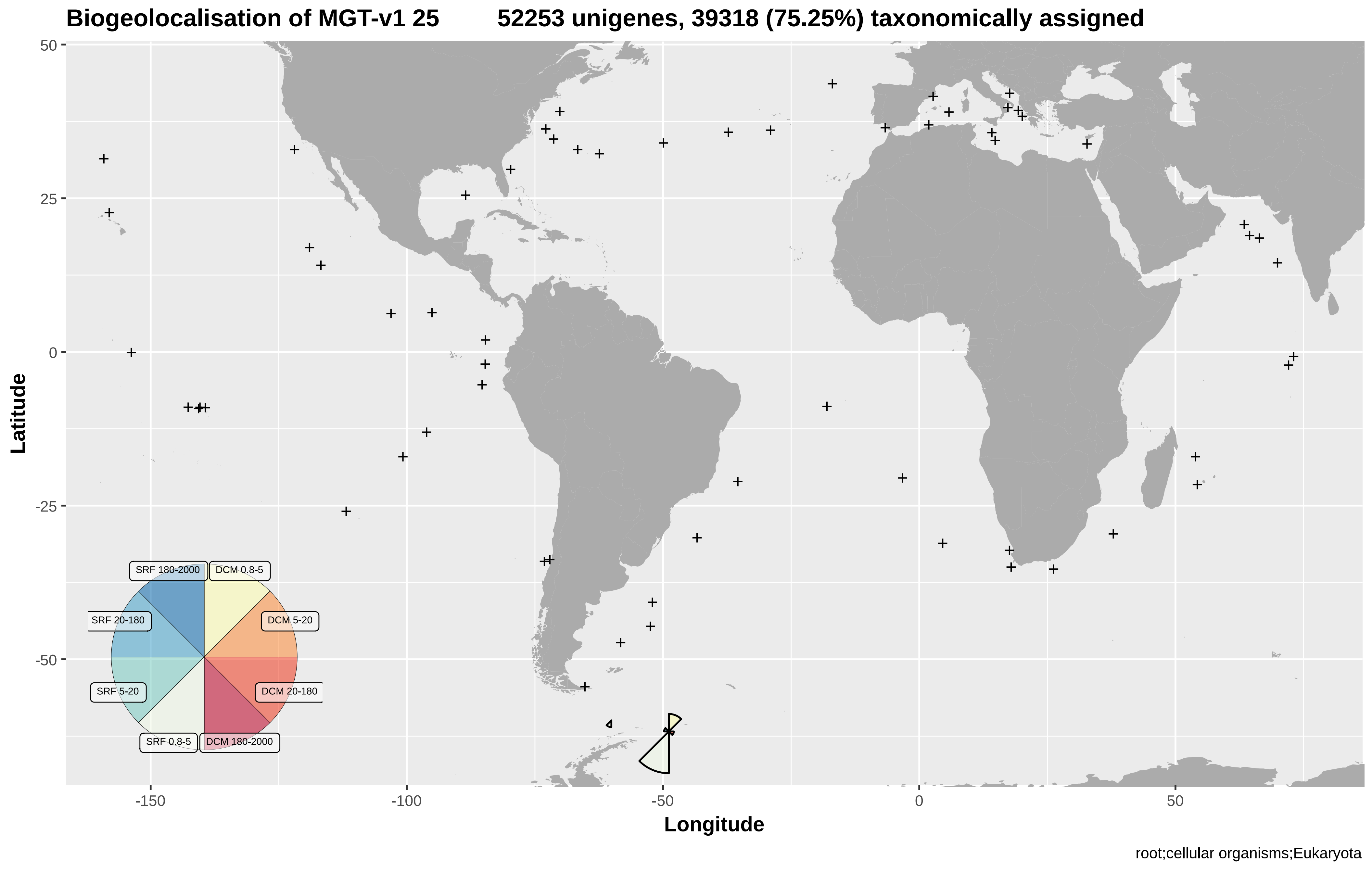

Biogeolocalisation of MGT-v1 25 52253 unigenes, 39318 (75.25%) taxonomically assigned
50
25
0
-25
-50
Latitude
-150
-100
-50
0
50
Longitude
SRF 180-2000
DCM 0.8-5
SRF 20-180
DCM 5-20
SRF 5-20
DCM 20-180
SRF 0.8-5
DCM 180-2000
root;cellular organisms;Eukaryota

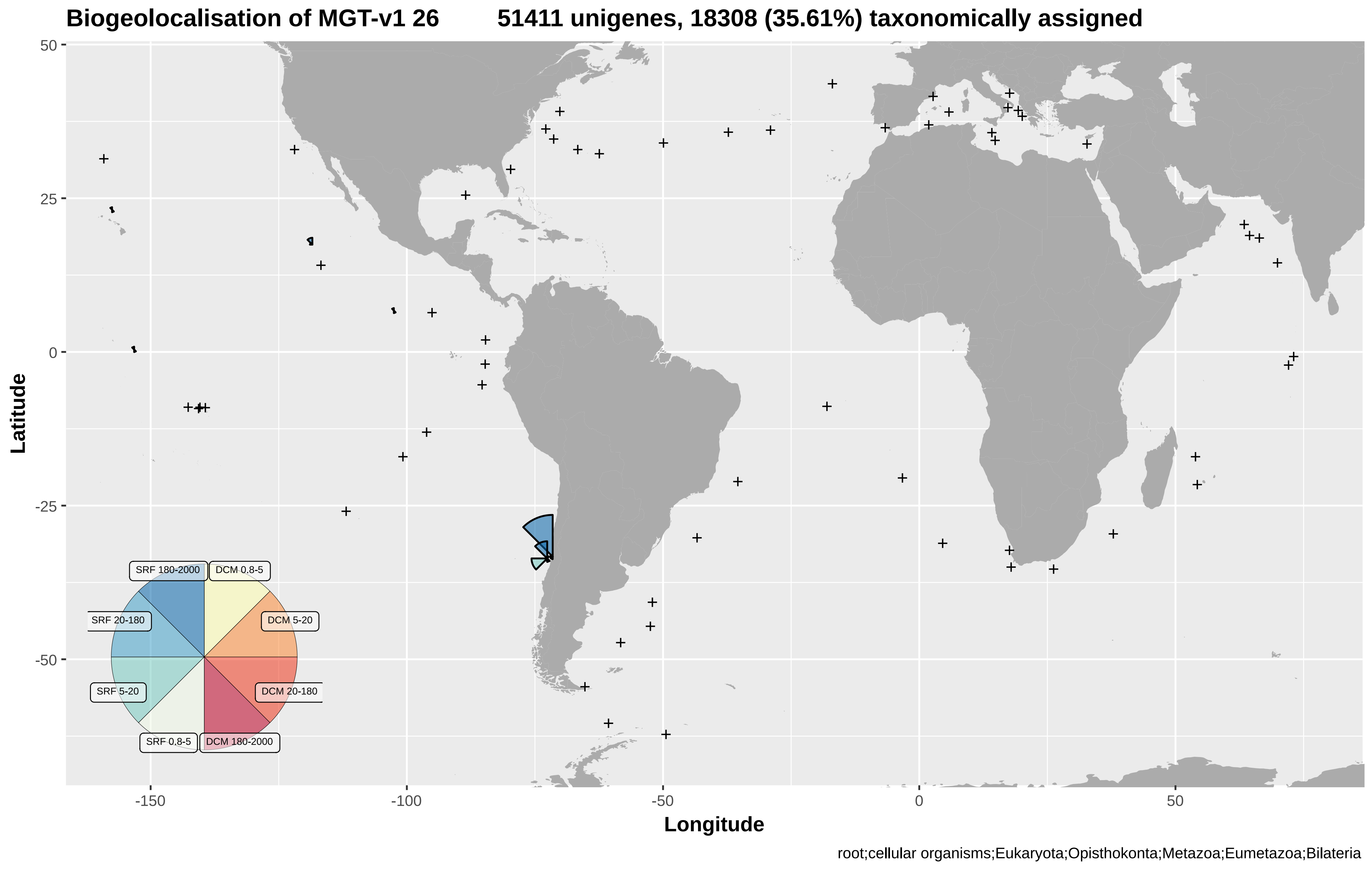
Biogeolocalisation of MGT-v1 26
51411 unigenes, 18308 (35.61%) taxonomically assigned
50
25
0
-25
-50
Latitude
-150
-100
-50
0
50
Longitude
SRF 180-2000
DCM 0.8-5
SRF 20-180
DCM 5-20
SRF 5-20
DCM 20-180
SRF 0.8-5
DCM 180-2000
root;cellular organisms;Eukaryota;Opisthokonta;Metazoa;Eumetazoa;Bilateria

#### Biogeolocalisation of MGT-v1 27 48515 unigenes, 17576 (36.23%) taxonomically assigned

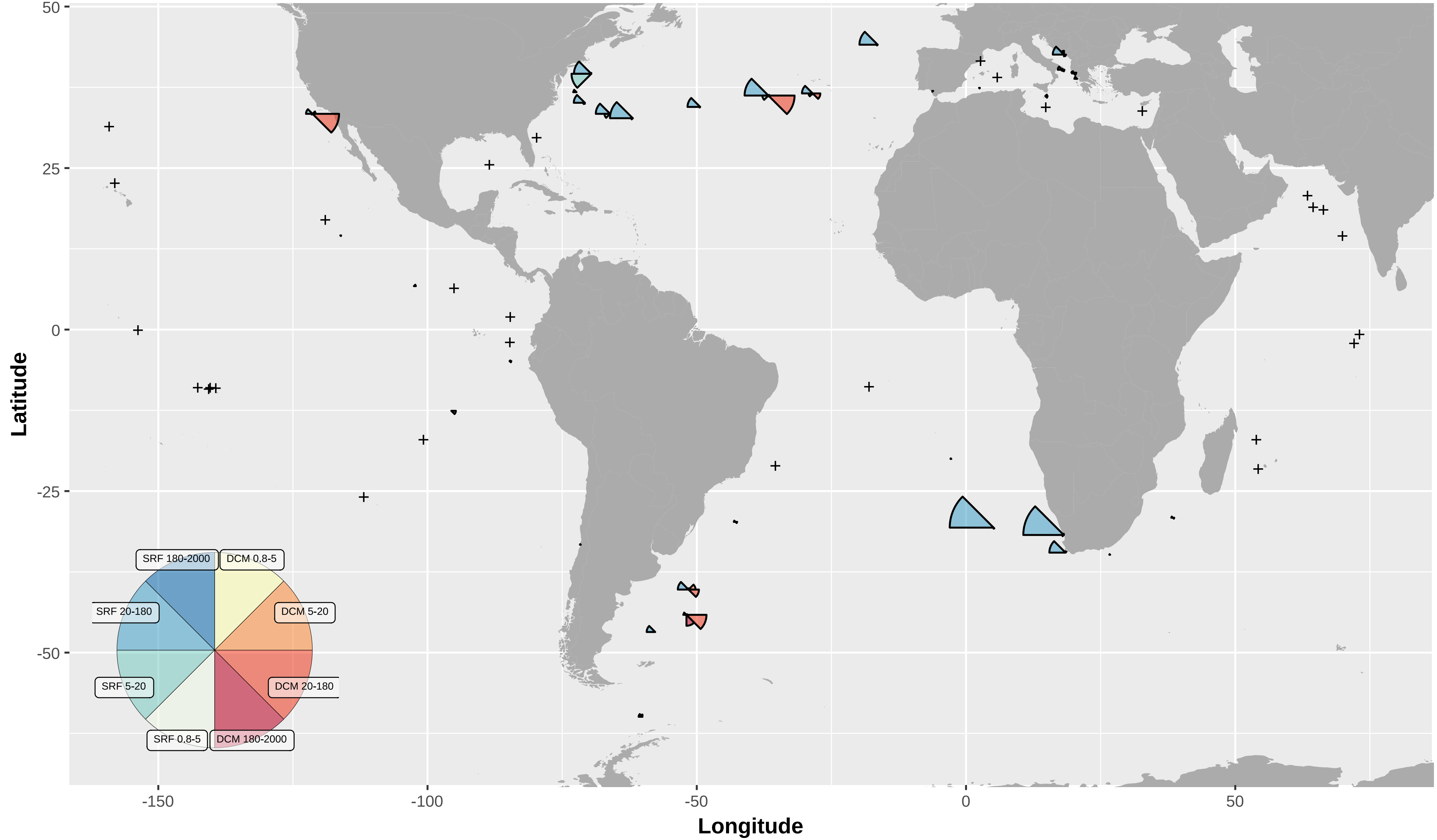


root;cellular organisms;Eukaryota;Opisthokonta;Metazoa;Eumetazoa;Bilateria;Protostomia;Ecdysozoa;Panarthropoda;Arthropoda;Mandibulata;Pancrustacea

Biogeolocalisation of MGT-v1 28 48509 unigenes, 3551 (7.32%) taxonomically assigned

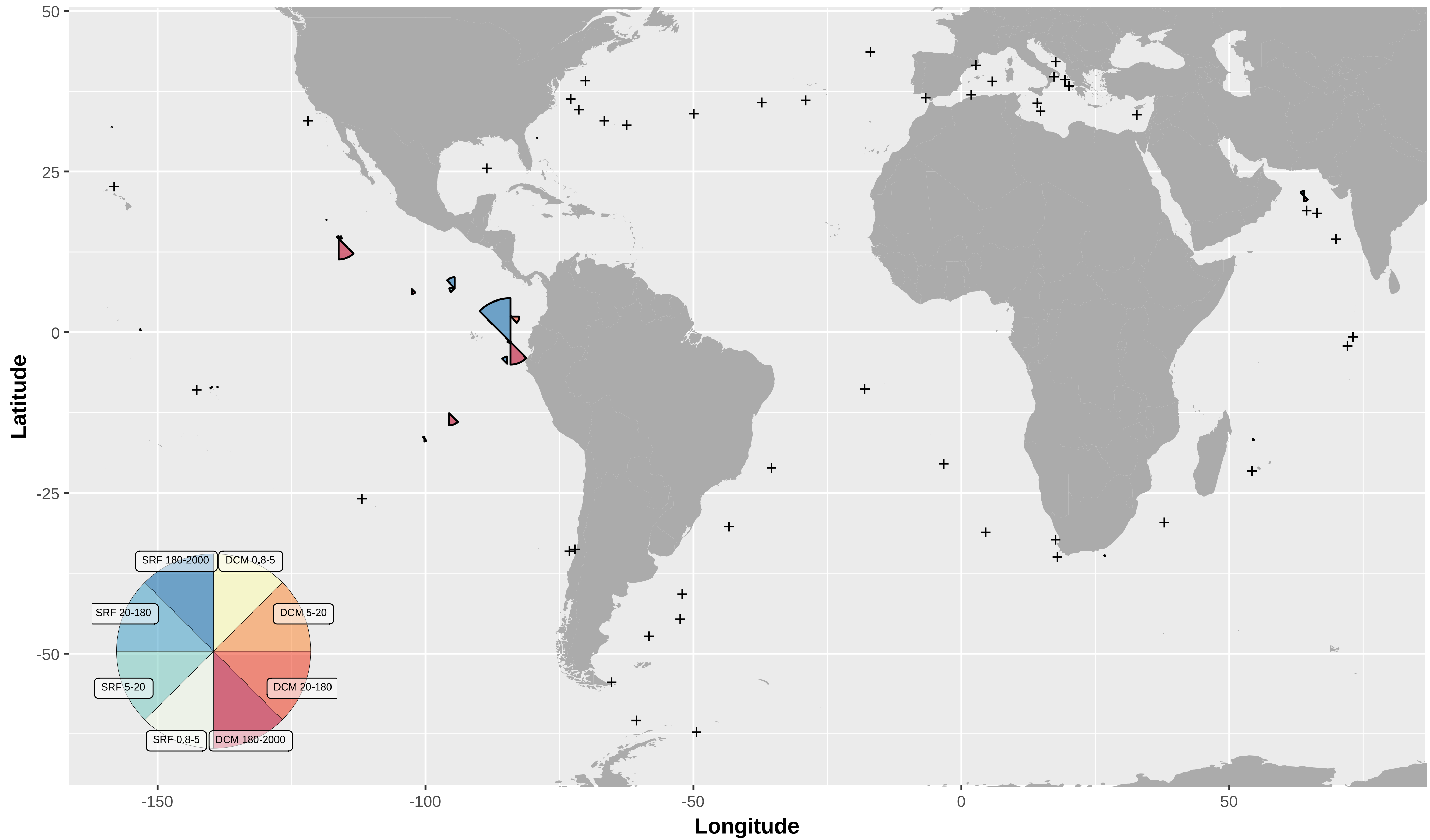


root;cellular organisms;Eukaryota;Opisthokonta;Metazoa;Eumetazoa;Bilateria

Biogeolocalisation of MGT-v1 29        48249 unigenes, 24942 (51.69%) taxonomically assigned

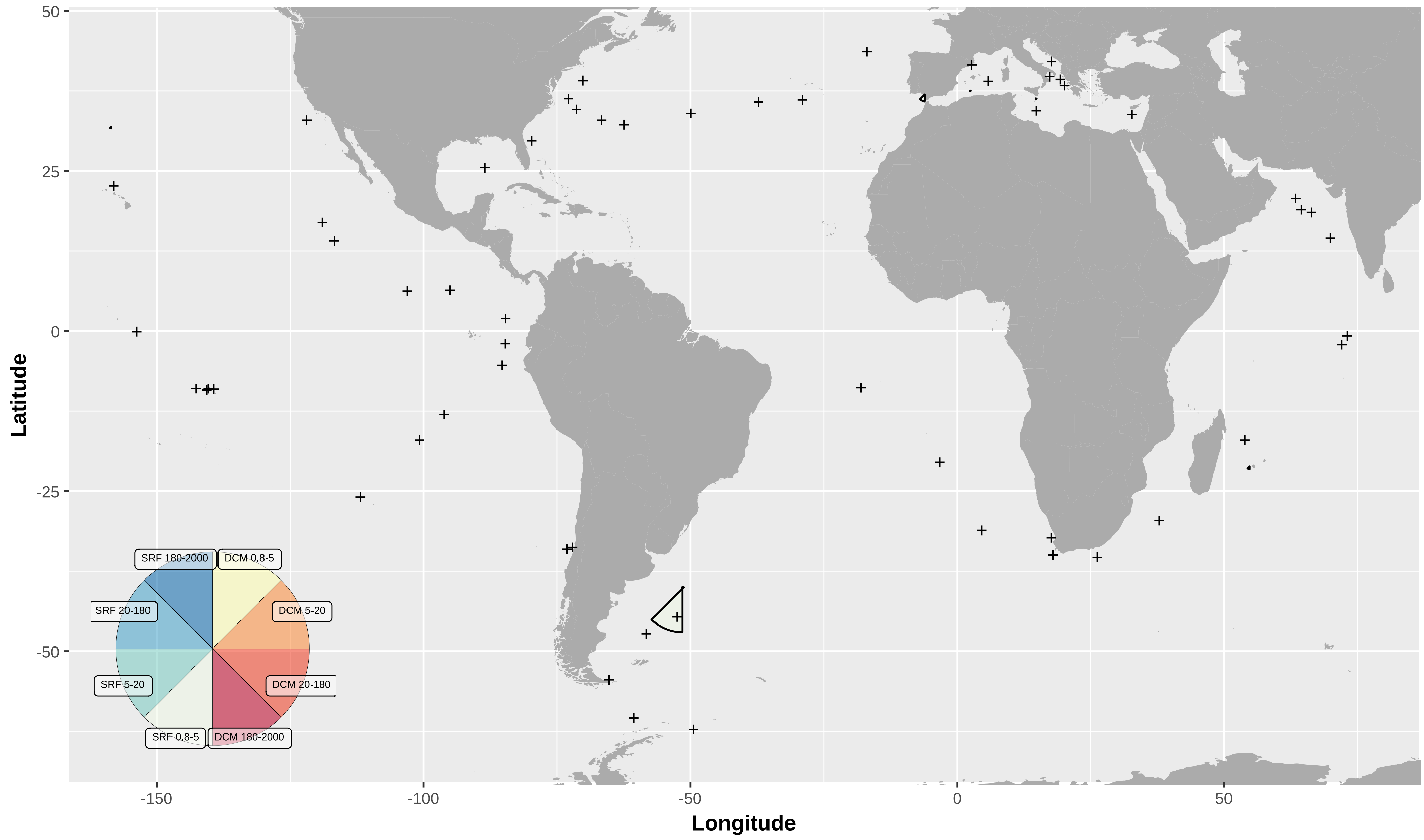


root;cellular organisms;Eukaryota;Haptophyceae

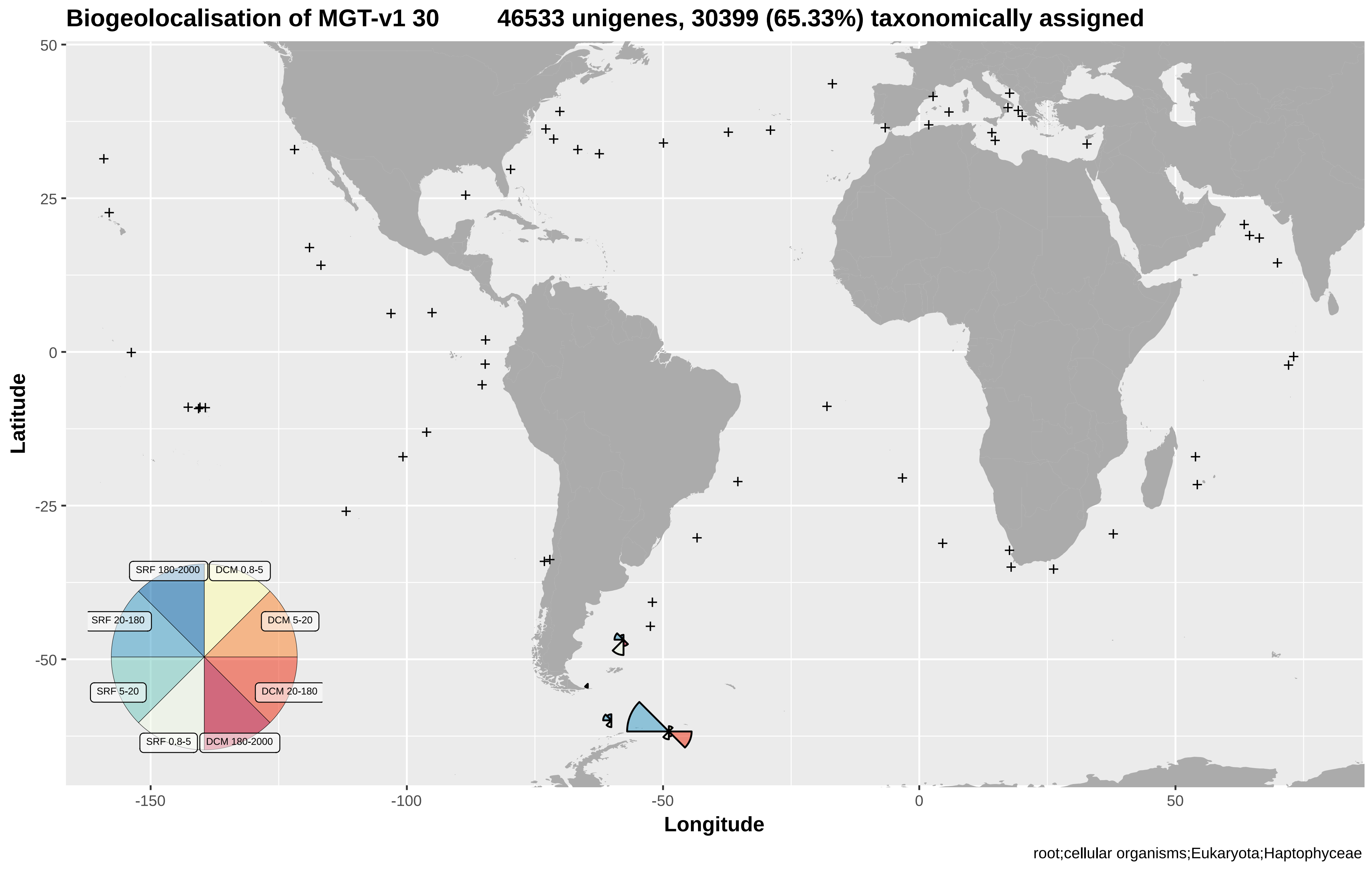

Biogeolocalisation of MGT-v1 30
46533 unigenes, 30399 (65.33%) taxonomically assigned
Latitude
Longitude
50
25
0
-25
-50
-150
-100
-50
0
50
SRF 180-2000
DCM 0.8-5
SRF 20-180
DCM 5-20
SRF 5-20
DCM 20-180
SRF 0.8-5
DCM 180-2000
root;cellular organisms;Eukaryota;Haptophyceae

# Biogeolocalisation of MGT-v1 31        45593 unigenes, 10587 (23.22%) taxonomically assigned

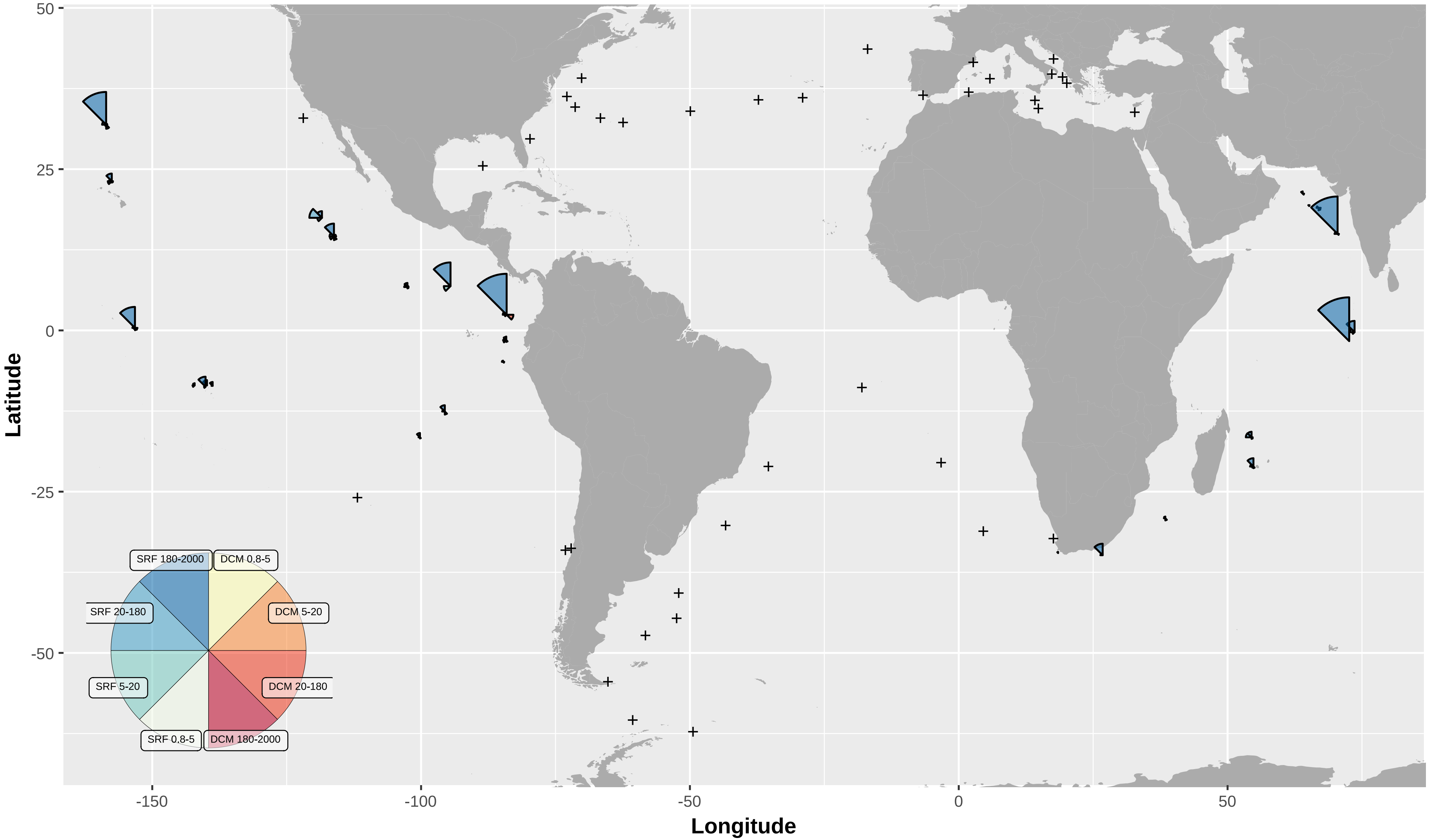


root;cellular organisms;Eukaryota;Opisthokonta;Metazoa;Eumetazoa;Bilateria;Protostomia;Ecdysozoa;Panarthropoda;Arthropoda;Mandibulata;Pancrustacea;Crustacea;Maxillopoda

# Biogeolocalisation of MGT-v1 32        43096 unigenes, 31508 (73.11%) taxonomically assigned

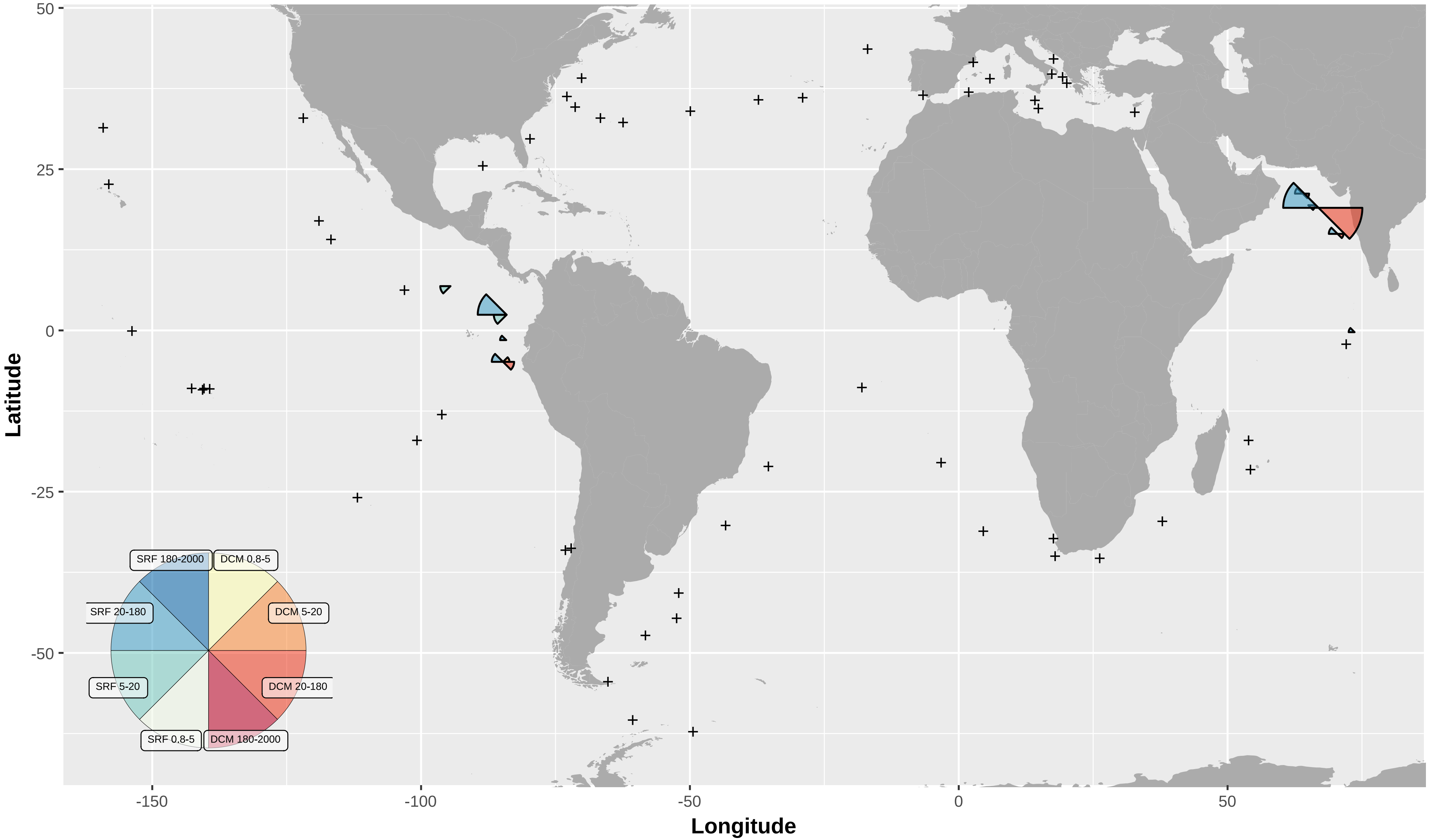


root;cellular organisms;Eukaryota;Opisthokonta;Metazoa;Eumetazoa;Bilateria;Protostomia;Ecdysozoa;Panarthropoda;Arthropoda;Mandibulata;Pancrustacea;Crustacea;Maxillopoda

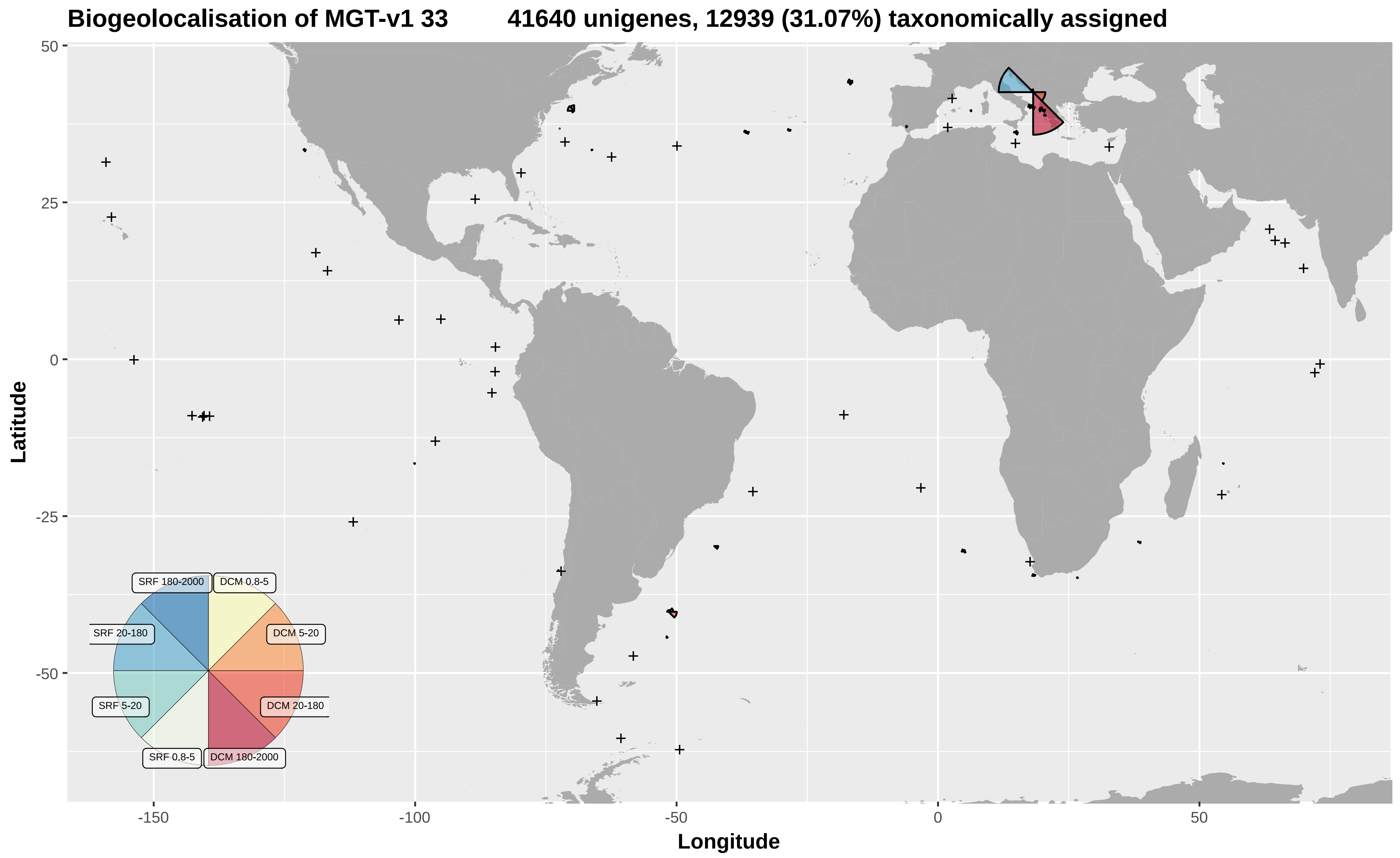

root;cellular organisms;Eukaryota;Opisthokonta;Metazoa;Eumetazoa;Bilateria;Protostomia;Ecdysozoa;Panarthropoda;Arthropoda;Mandibulata;Pancrustacea

#### Biogeolocalisation of MGT-v1 34 39353 unigenes, 22740 (57.78%) taxonomically assigned

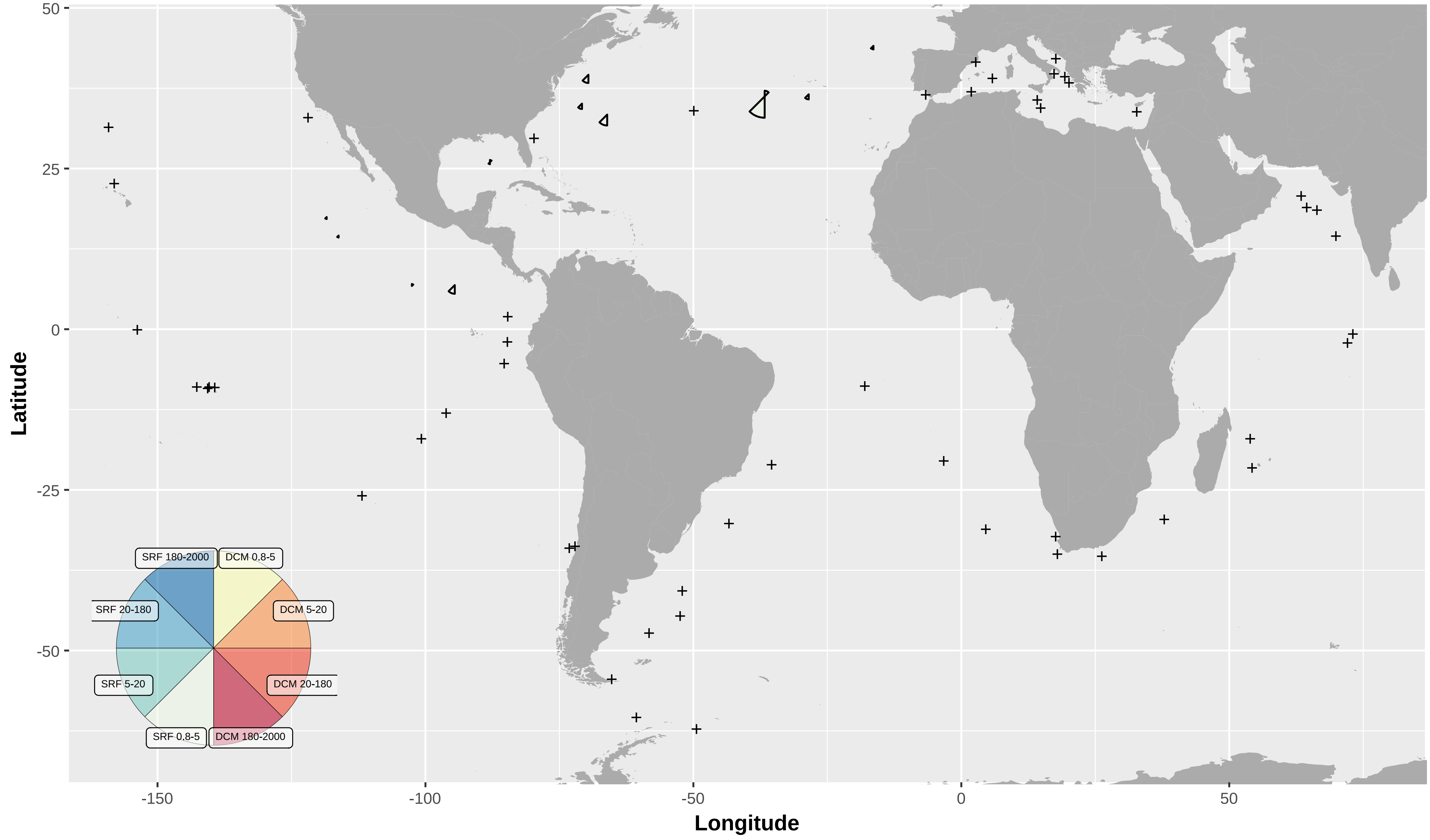


root;cellular organisms;Eukaryota

## Biogeolocalisation of MGT-v1 35        38940 unigenes, 19910 (51.13%) taxonomically assigned

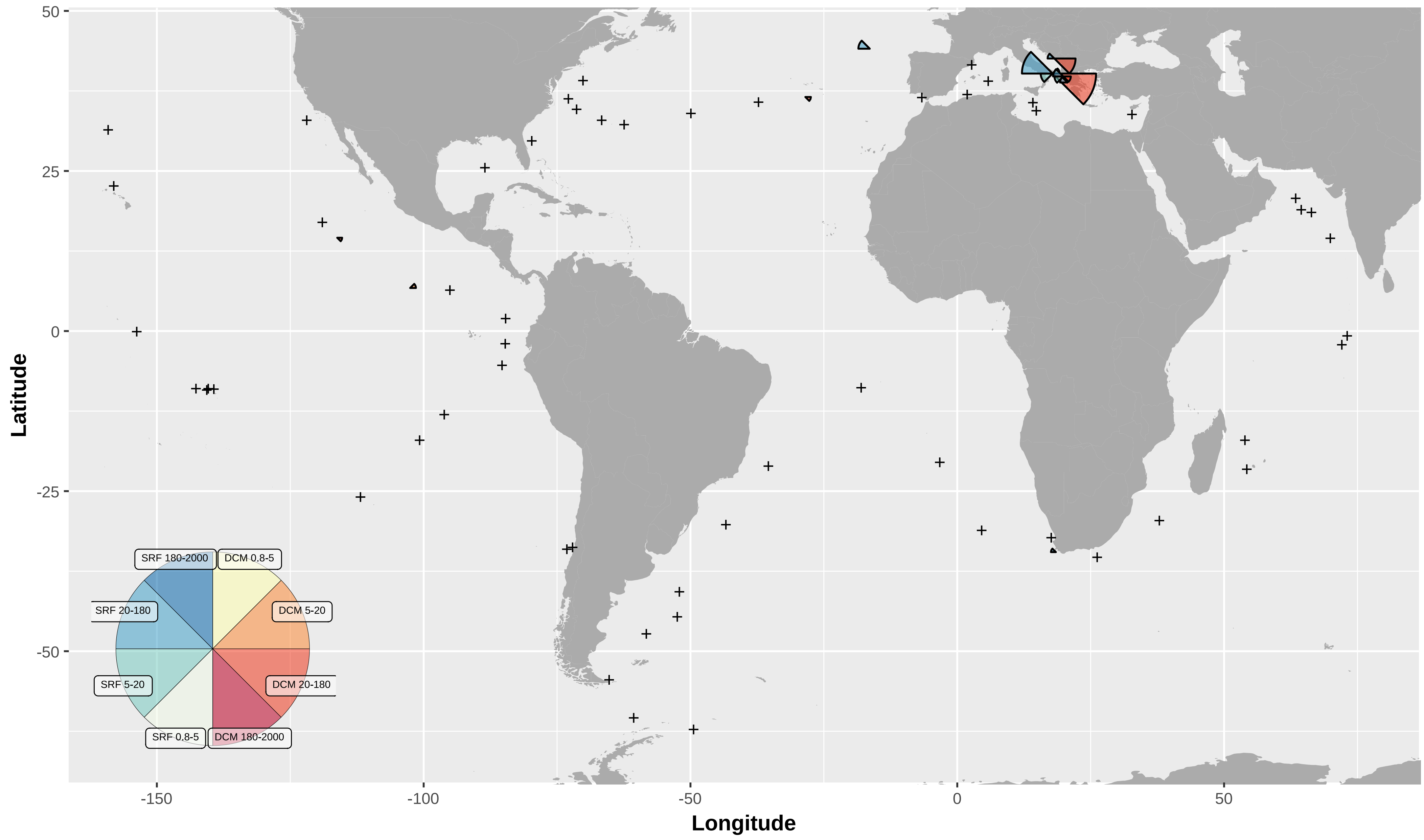


root;cellular organisms;Eukaryota;Opisthokonta;Metazoa;Eumetazoa;Bilateria;Protostomia;Ecdysozoa;Panarthropoda;Arthropoda

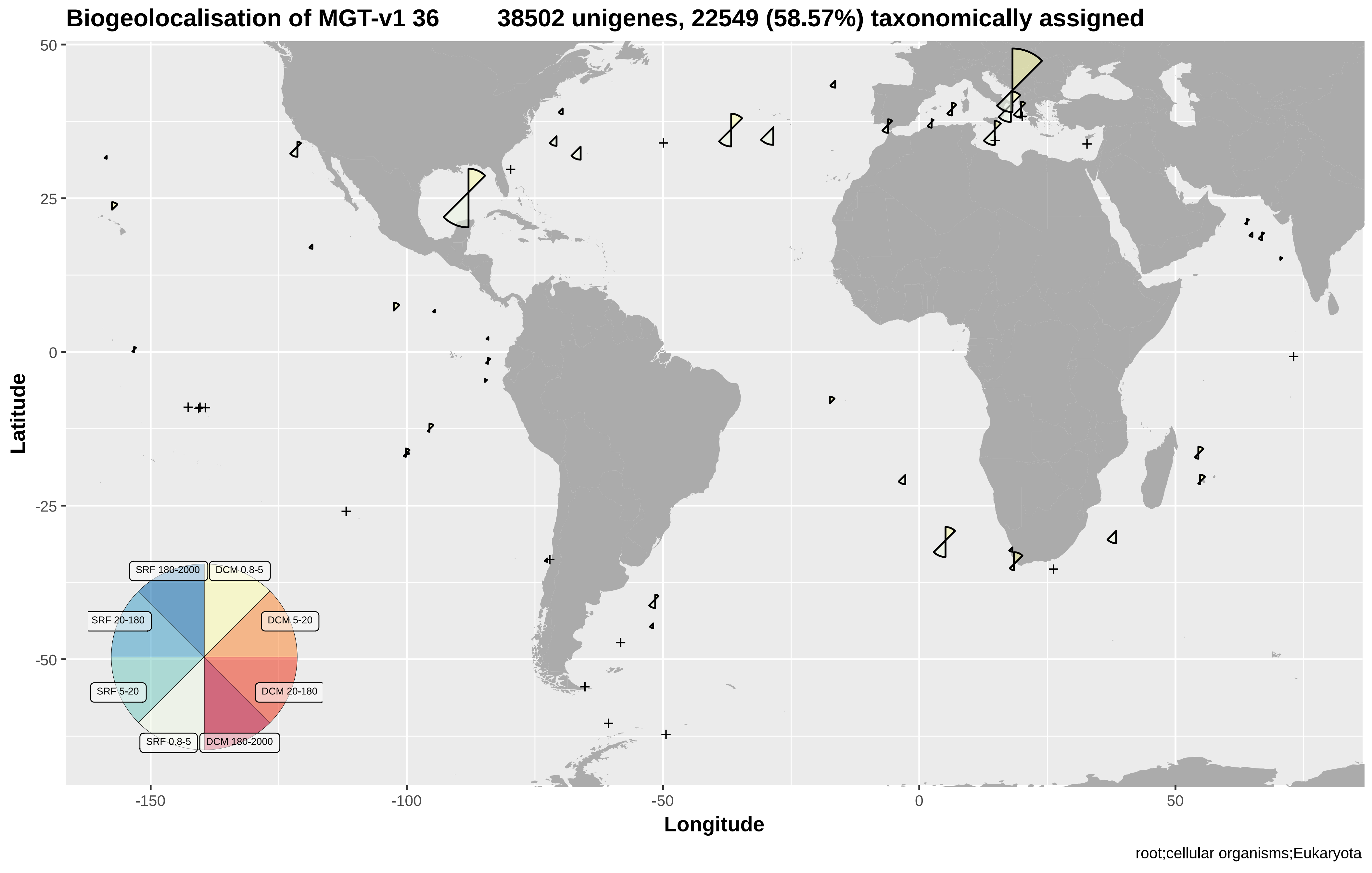

Biogeolocalisation of MGT-v1 36
38502 unigenes, 22549 (58.57%) taxonomically assigned
Latitude
50
25
0
-25
-50
Longitude
-150
-100
-50
0
50
SRF 180-2000
DCM 0.8-5
SRF 20-180
DCM 5-20
SRF 5-20
DCM 20-180
SRF 0.8-5
DCM 180-2000
root;cellular organisms;Eukaryota

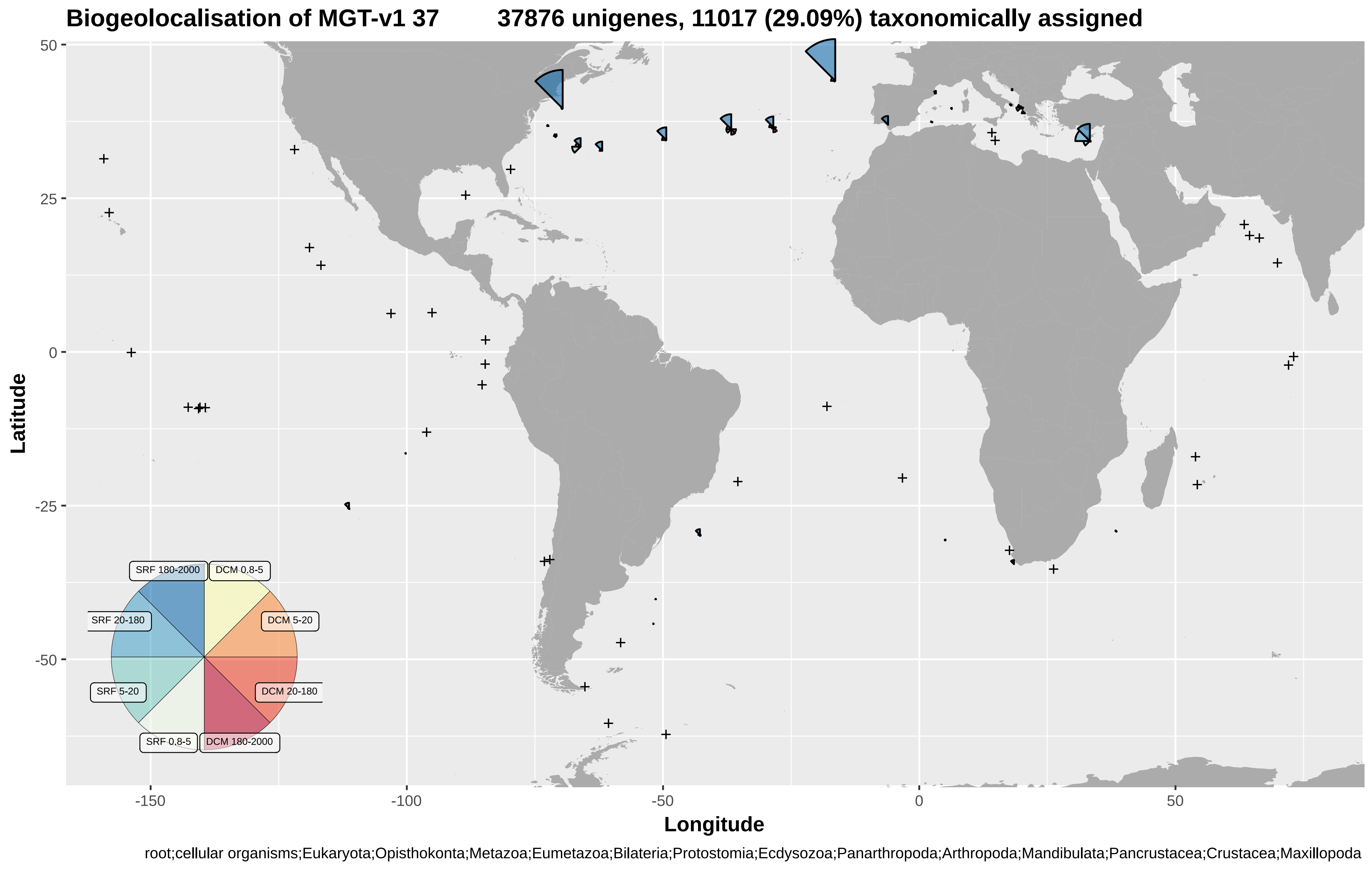
Biogeolocalisation of MGT-v1 37
37876 unigenes, 11017 (29.09%) taxonomically assigned
50
25
0
-25
-50
Latitude
-150
-100
-50
0
50
Longitude
SRF 180-2000
DCM 0.8-5
SRF 20-180
DCM 5-20
SRF 5-20
DCM 20-180
SRF 0.8-5
DCM 180-2000
root;cellular organisms;Eukaryota;Opisthokonta;Metazoa;Eumetazoa;Bilateria;Protostomia;Ecdysozoa;Panarthropoda;Arthropoda;Mandibulata;Pancrustacea;Crustacea;Maxillopoda

### Biogeolocalisation of MGT-v1 38 37557 unigenes, 13839 (36.85%) taxonomically assigned

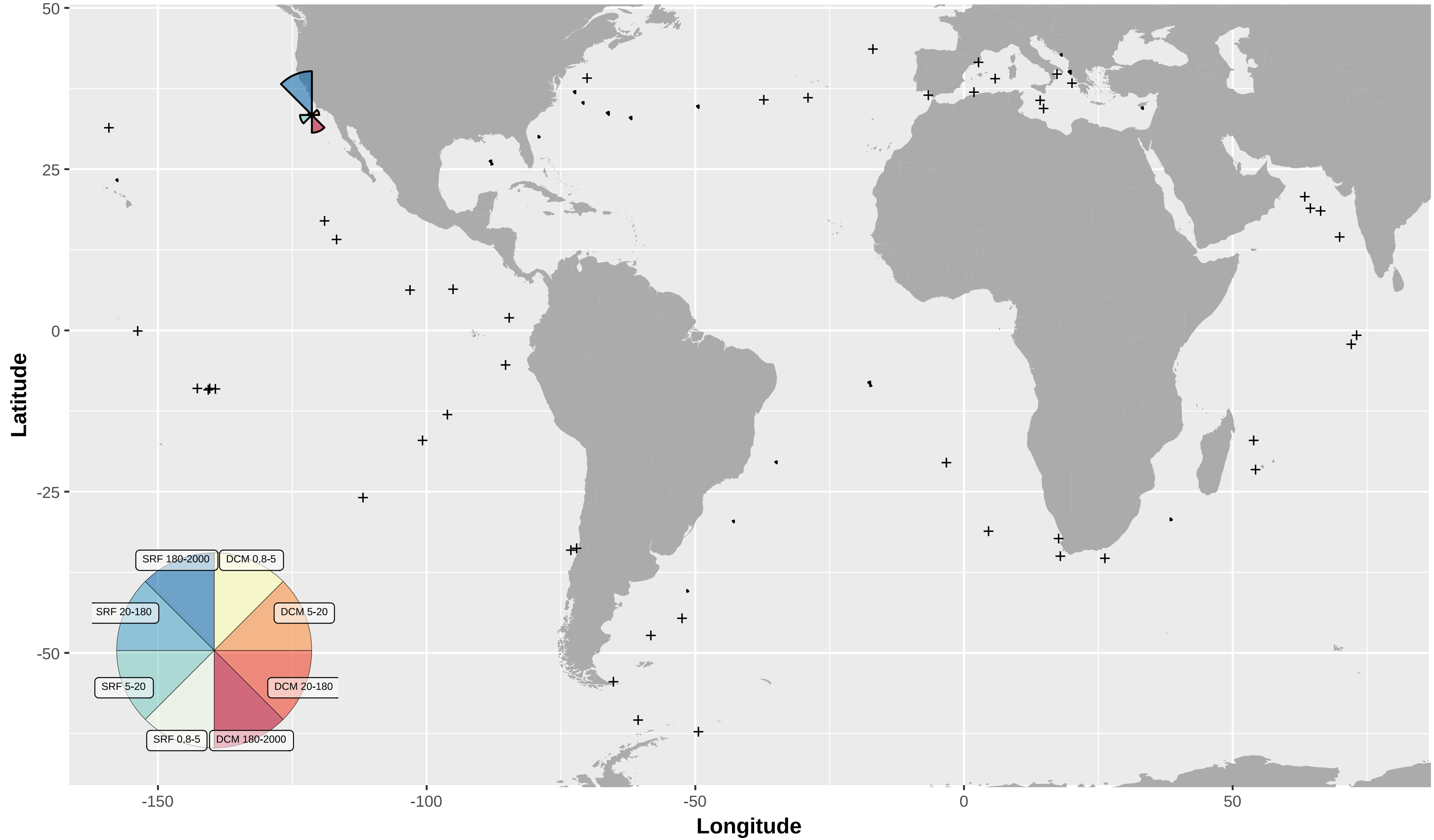


root;cellular organisms;Eukaryota;Opisthokonta;Metazoa;Eumetazoa

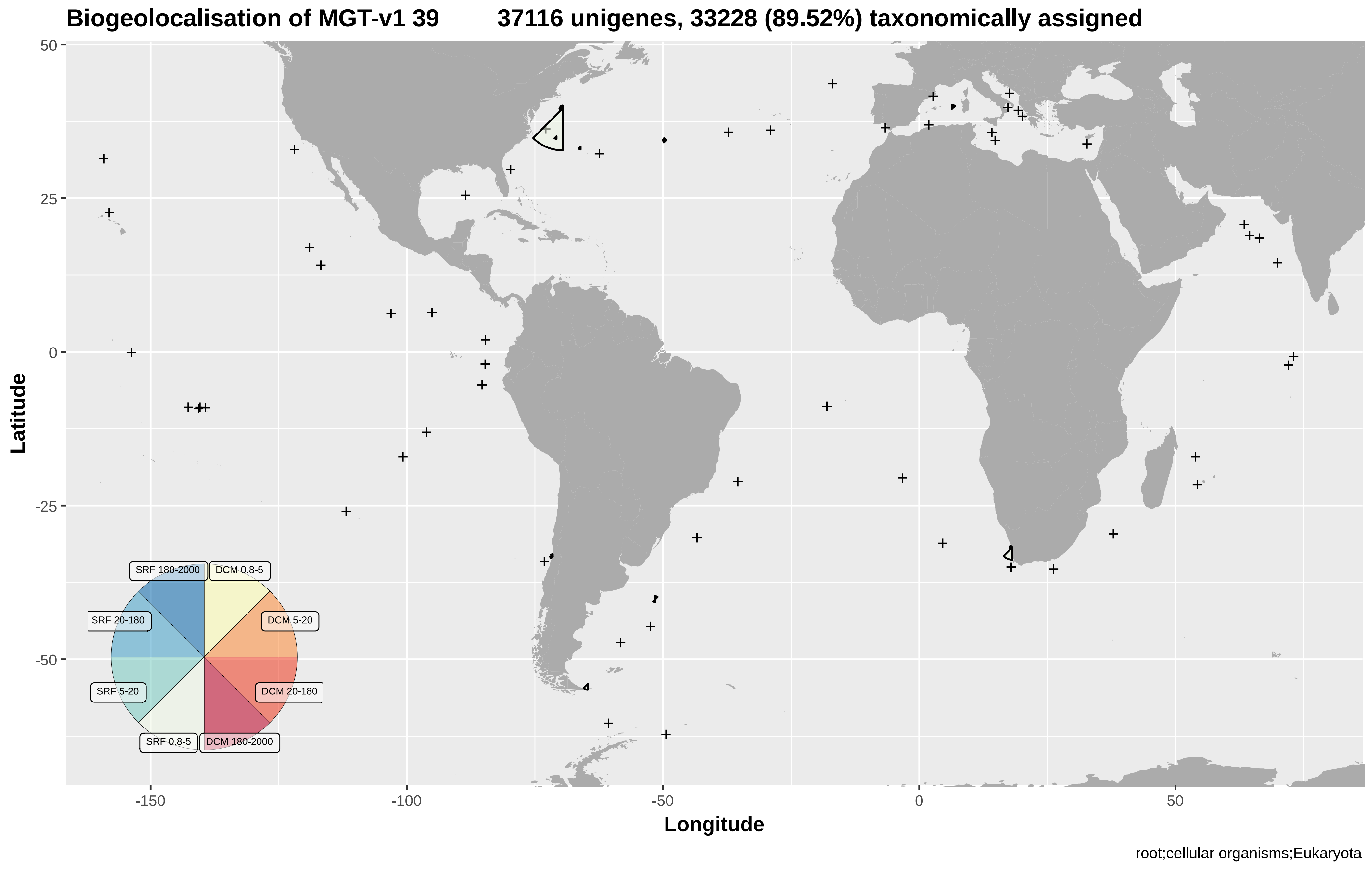
Biogeolocalisation of MGT-v1 39
37116 unigenes, 33228 (89.52%) taxonomically assigned
50
25
0
-25
-50
Latitude
-150
-100
-50
0
50
Longitude
SRF 180-2000
DCM 0.8-5
SRF 20-180
DCM 5-20
SRF 5-20
DCM 20-180
SRF 0.8-5
DCM 180-2000
root;cellular organisms;Eukaryota

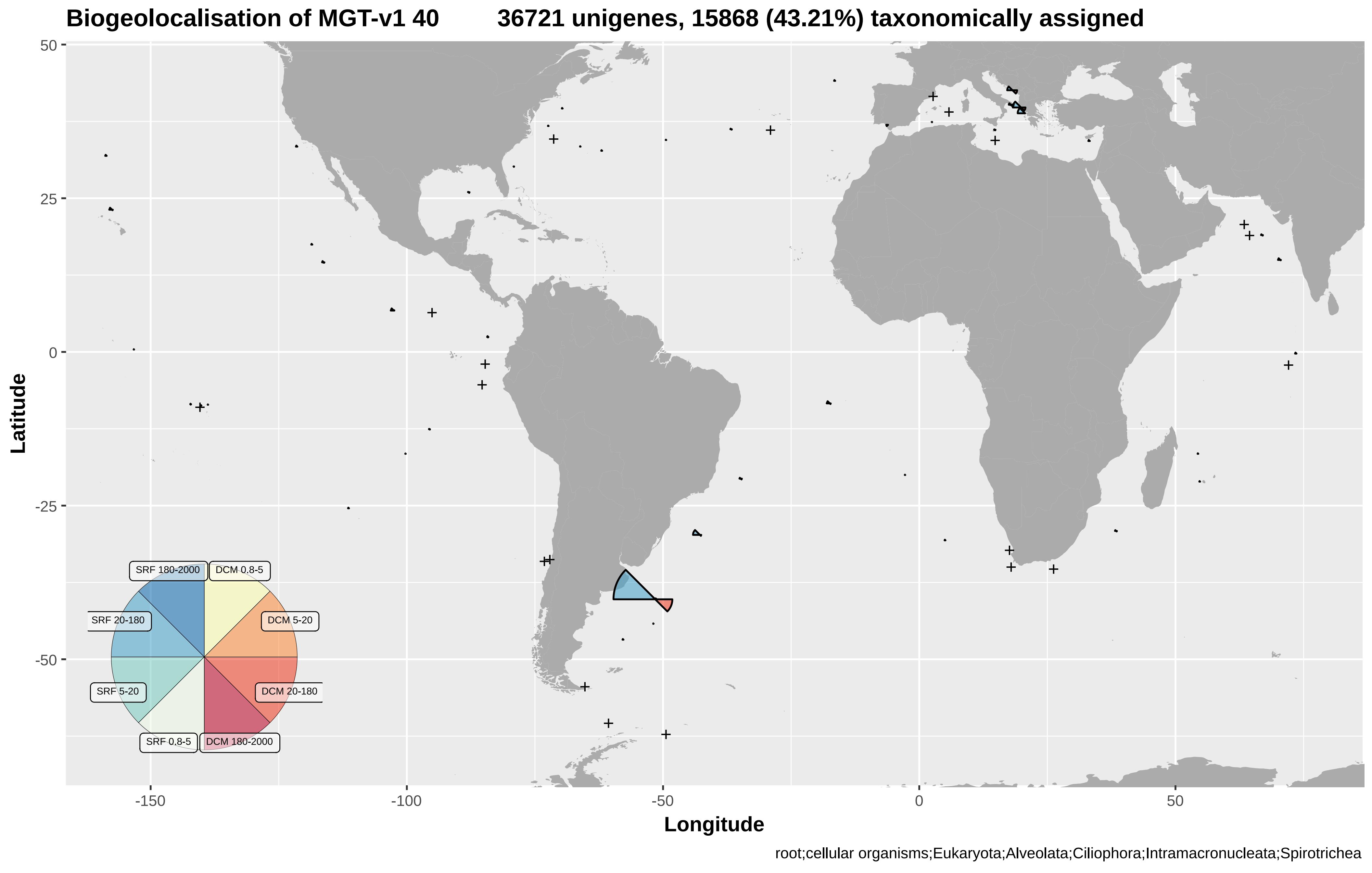

Biogeolocalisation of MGT-v1 40
36721 unigenes, 15868 (43.21%) taxonomically assigned
50
25
0
-25
-50
Latitude
-150
-100
-50
0
50
Longitude
SRF 180-2000
DCM 0.8-5
SRF 20-180
DCM 5-20
SRF 5-20
DCM 20-180
SRF 0.8-5
DCM 180-2000
root;cellular organisms;Eukaryota;Alveolata;Ciliophora;Intramacronucleata;Spirotrichea

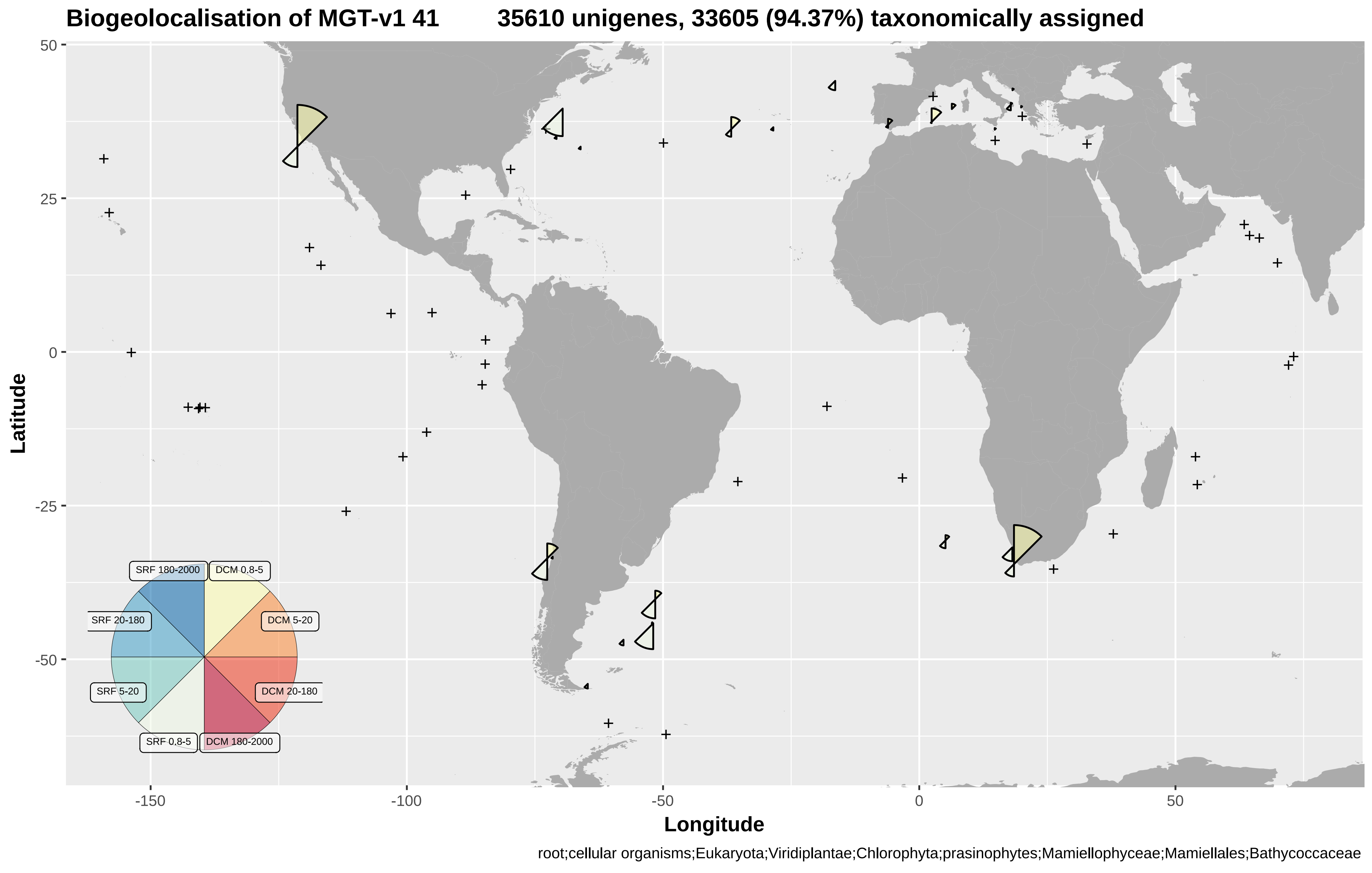
Biogeolocalisation of MGT-v1 41
35610 unigenes, 33605 (94.37%) taxonomically assigned
Latitude
50
25
0
-25
-50
Longitude
-150
-100
-50
0
50
SRF 180-2000
DCM 0.8-5
SRF 20-180
DCM 5-20
SRF 5-20
DCM 20-180
SRF 0.8-5
DCM 180-2000
root;cellular organisms;Eukaryota;Viridiplantae;Chlorophyta;prasinophytes;Mamiellophyceae;Mamiellales;Bathycoccaceae

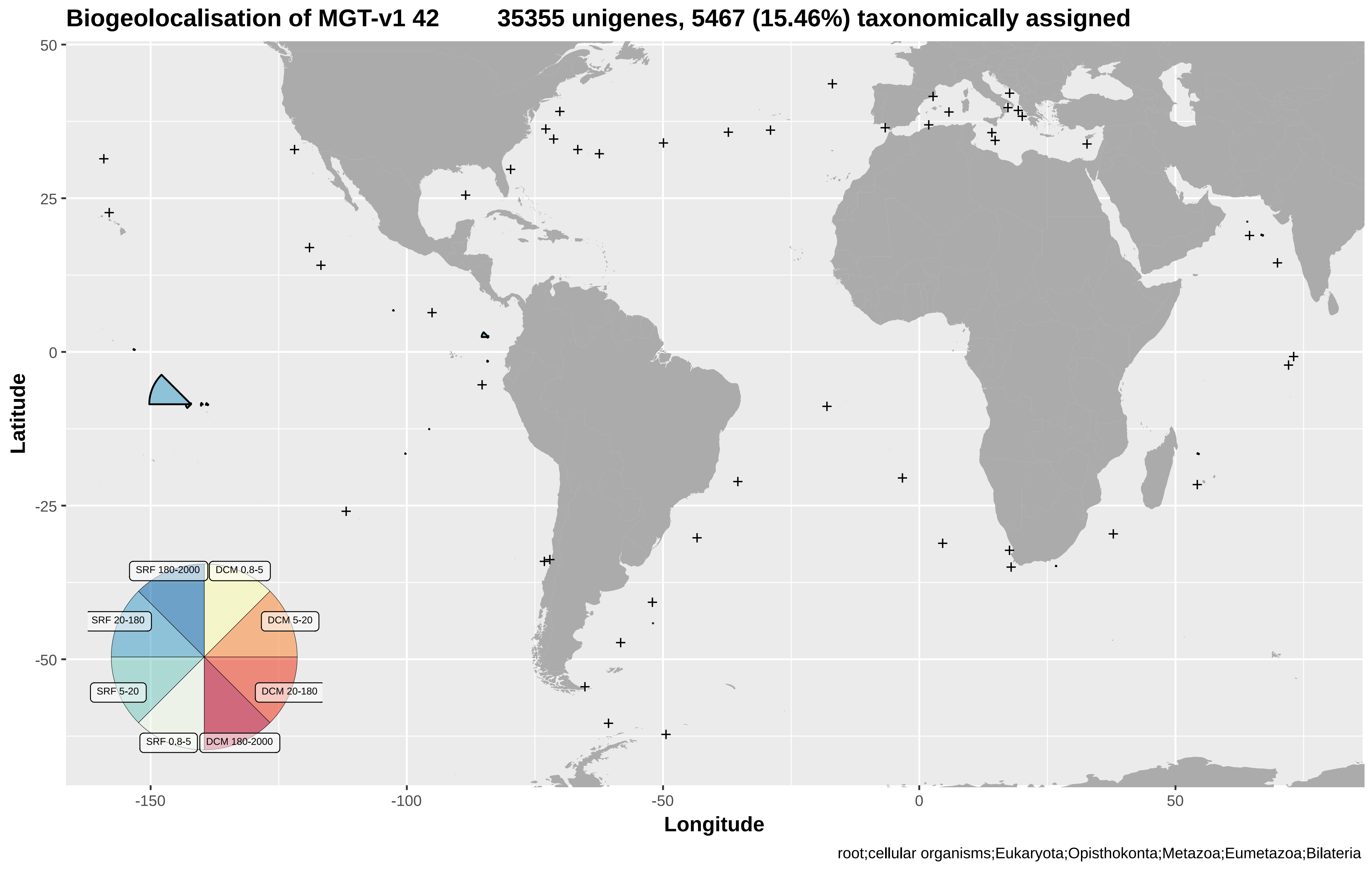
Biogeolocalisation of MGT-v1 42
35355 unigenes, 5467 (15.46%) taxonomically assigned
Latitude
Longitude
50
25
0
-25
-50
-150
-100
-50
0
50
SRF 180-2000
DCM 0.8-5
SRF 20-180
DCM 5-20
SRF 5-20
DCM 20-180
SRF 0.8-5
DCM 180-2000
root;cellular organisms;Eukaryota;Opisthokonta;Metazoa;Eumetazoa;Bilateria

## Biogeolocalisation of MGT-v1 43        35097 unigenes, 24909 (70.97%) taxonomically assigned

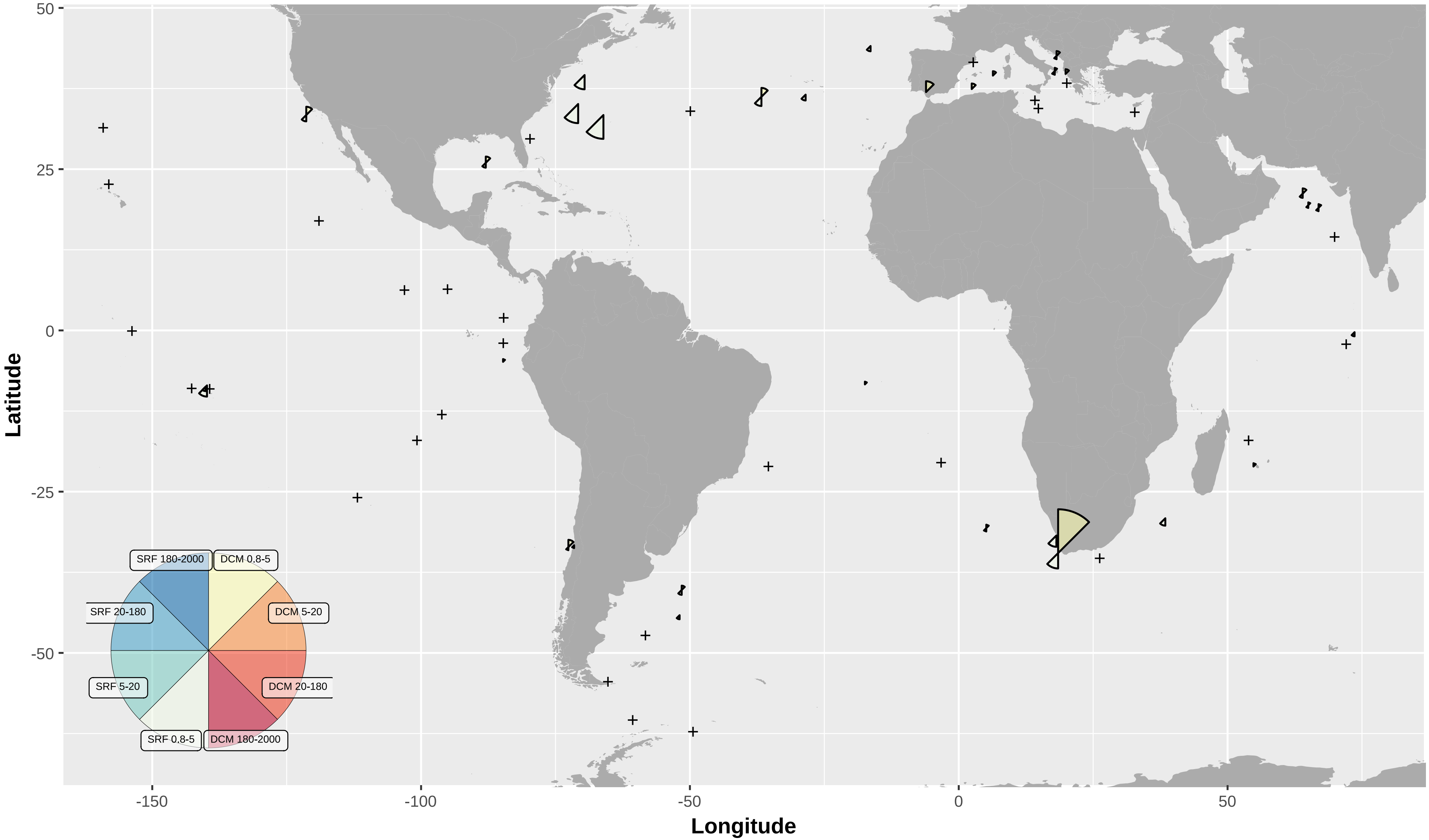


root;cellular organisms;Eukaryota

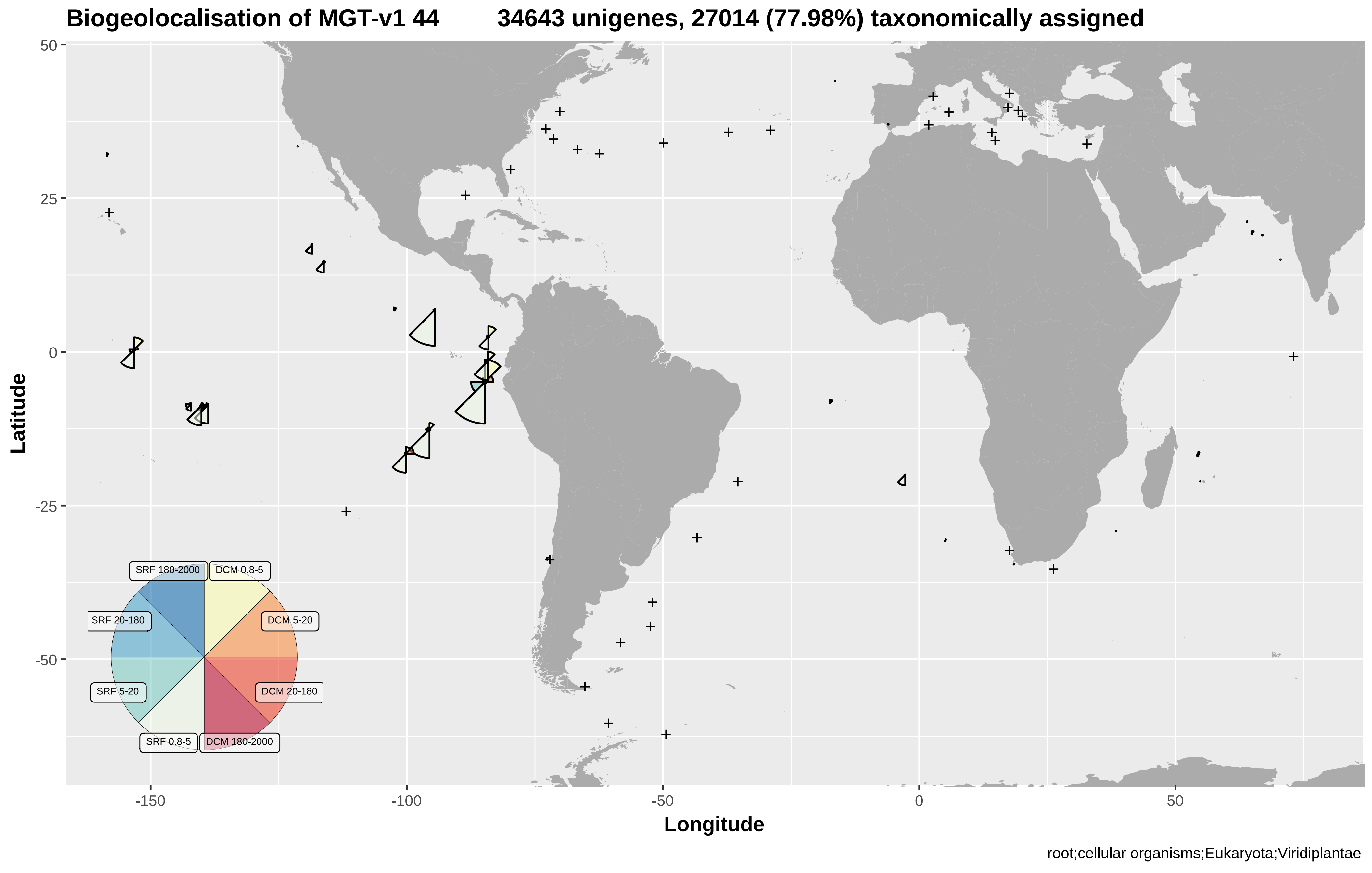
Biogeolocalisation of MGT-v1 44
34643 unigenes, 27014 (77.98%) taxonomically assigned
Latitude
Longitude
50
25
0
-25
-50
-150
-100
-50
0
50
SRF 180-2000
DCM 0.8-5
SRF 20-180
DCM 5-20
SRF 5-20
DCM 20-180
SRF 0.8-5
DCM 180-2000
root;cellular organisms;Eukaryota;Viridiplantae

## Biogeolocalisation of MGT-v1 45        33662 unigenes, 14831 (44.06%) taxonomically assigned

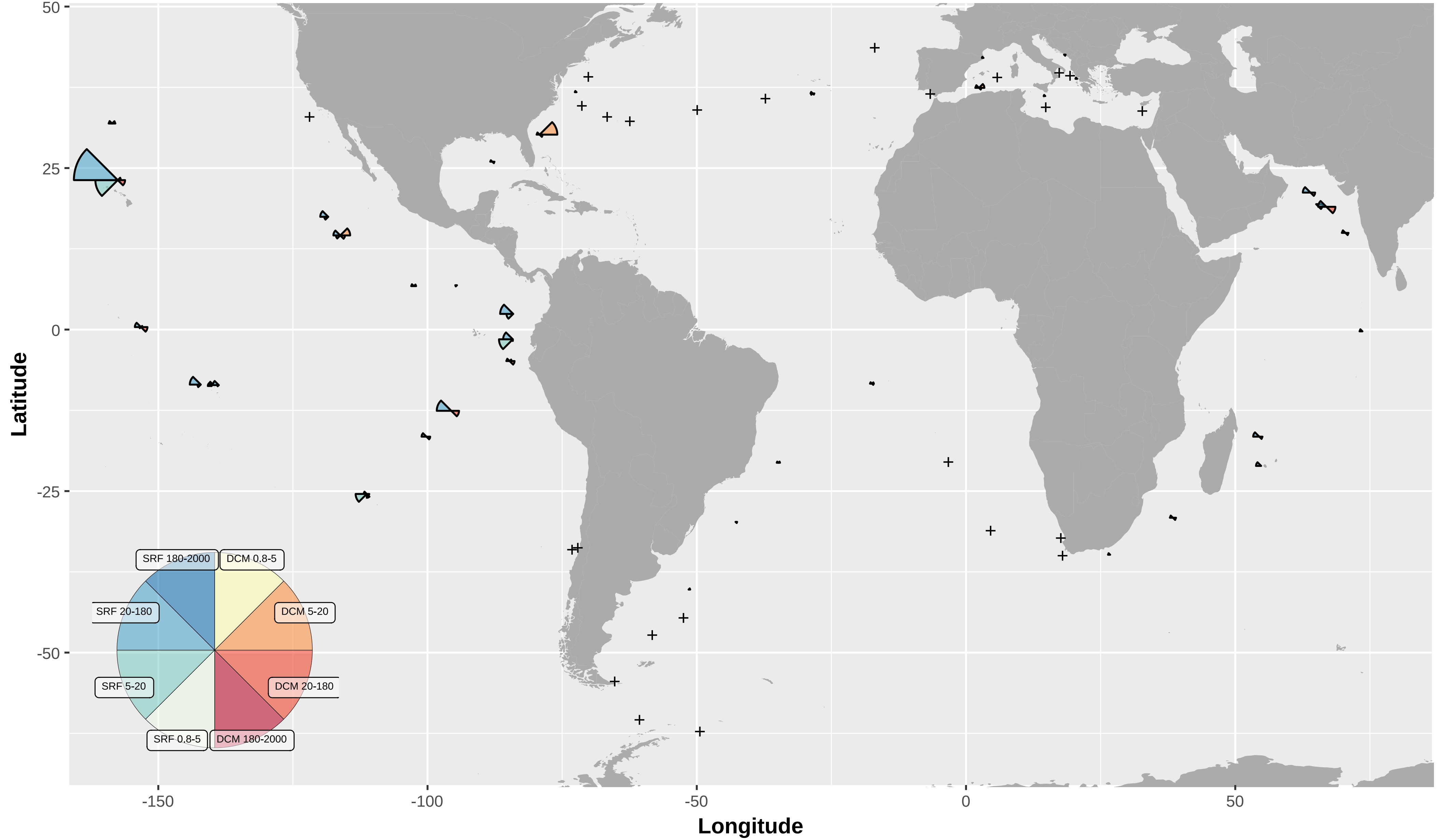


root;cellular organisms;Eukaryota;Opisthokonta;Metazoa;Eumetazoa;Bilateria;Protostomia

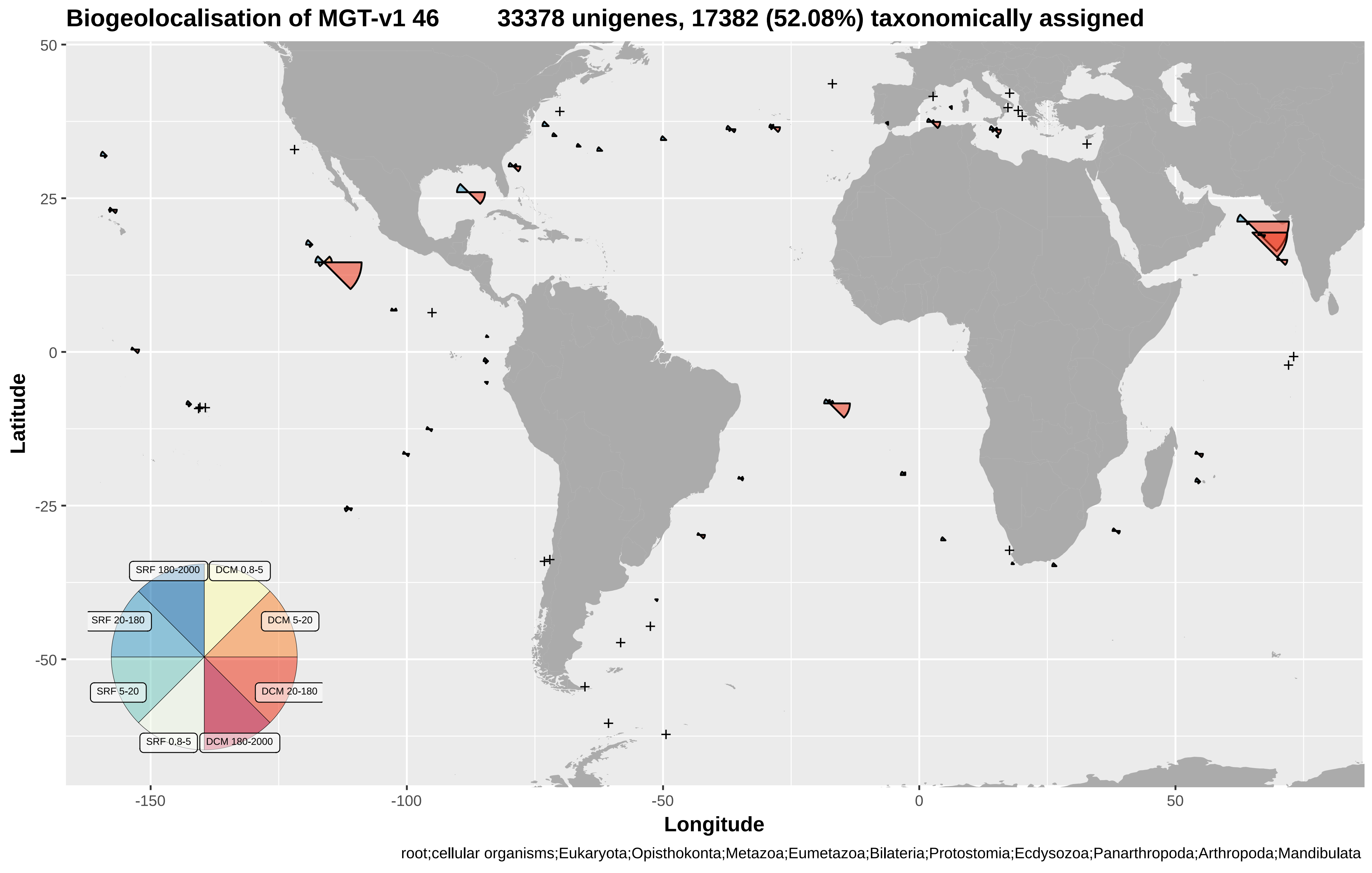
Biogeolocalisation of MGT-v1 46
33378 unigenes, 17382 (52.08%) taxonomically assigned
Latitude
Longitude
50
25
0
-25
-50
-150
-100
-50
0
50
SRF 180-2000
DCM 0.8-5
SRF 20-180
DCM 5-20
SRF 5-20
DCM 20-180
SRF 0.8-5
DCM 180-2000
root;cellular organisms;Eukaryota;Opisthokonta;Metazoa;Eumetazoa;Bilateria;Protostomia;Ecdysozoa;Panarthropoda;Arthropoda;Mandibulata

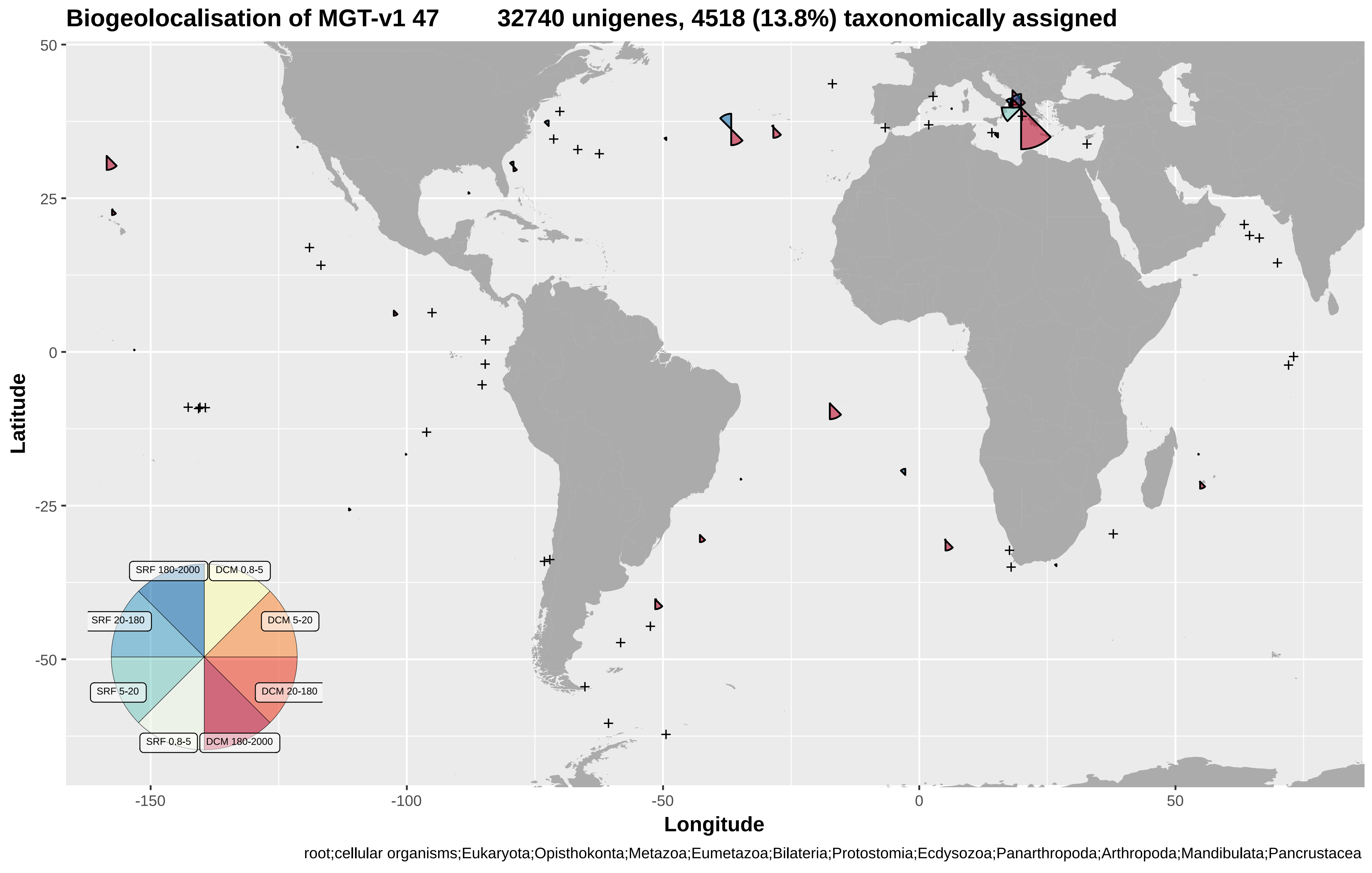

Biogeolocalisation of MGT-v1 47
32740 unigenes, 4518 (13.8%) taxonomically assigned
50
25
0
-25
-50
Latitude
-150
-100
-50
0
50
Longitude
SRF 180-2000
DCM 0.8-5
SRF 20-180
DCM 5-20
SRF 5-20
DCM 20-180
SRF 0.8-5
DCM 180-2000
root;cellular organisms;Eukaryota;Opisthokonta;Metazoa;Eumetazoa;Bilateria;Protostomia;Ecdysozoa;Panarthropoda;Arthropoda;Mandibulata;Pancrustacea

#### Biogeolocalisation of MGT-v1 48 31633 unigenes, 28011 (88.55%) taxonomically assigned

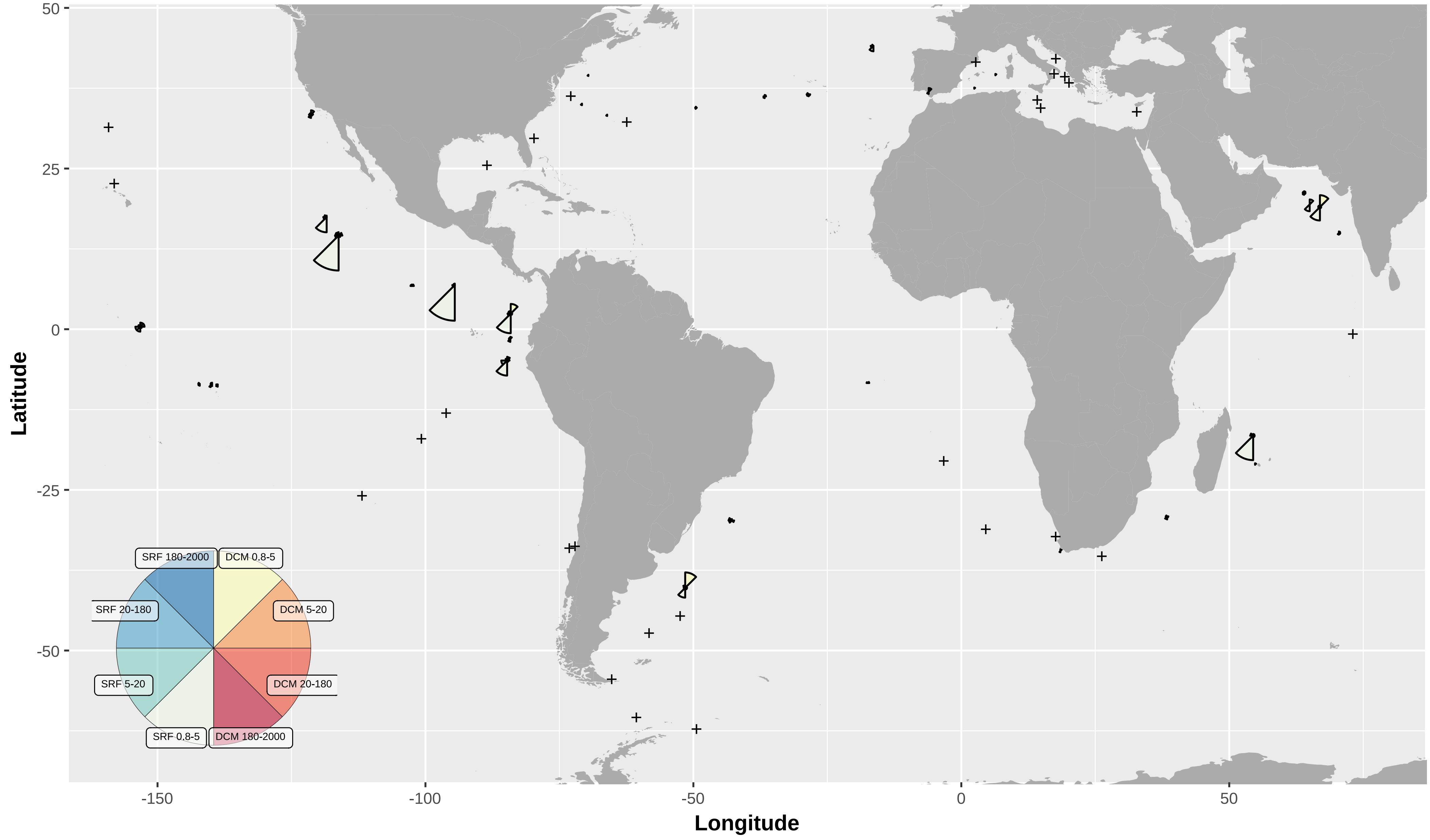


root;cellular organisms;Eukaryota;Viridiplantae;Chlorophyta;prasinophytes

#### Biogeolocalisation of MGT-v1 49 31155 unigenes, 11572 (37.14%) taxonomically assigned

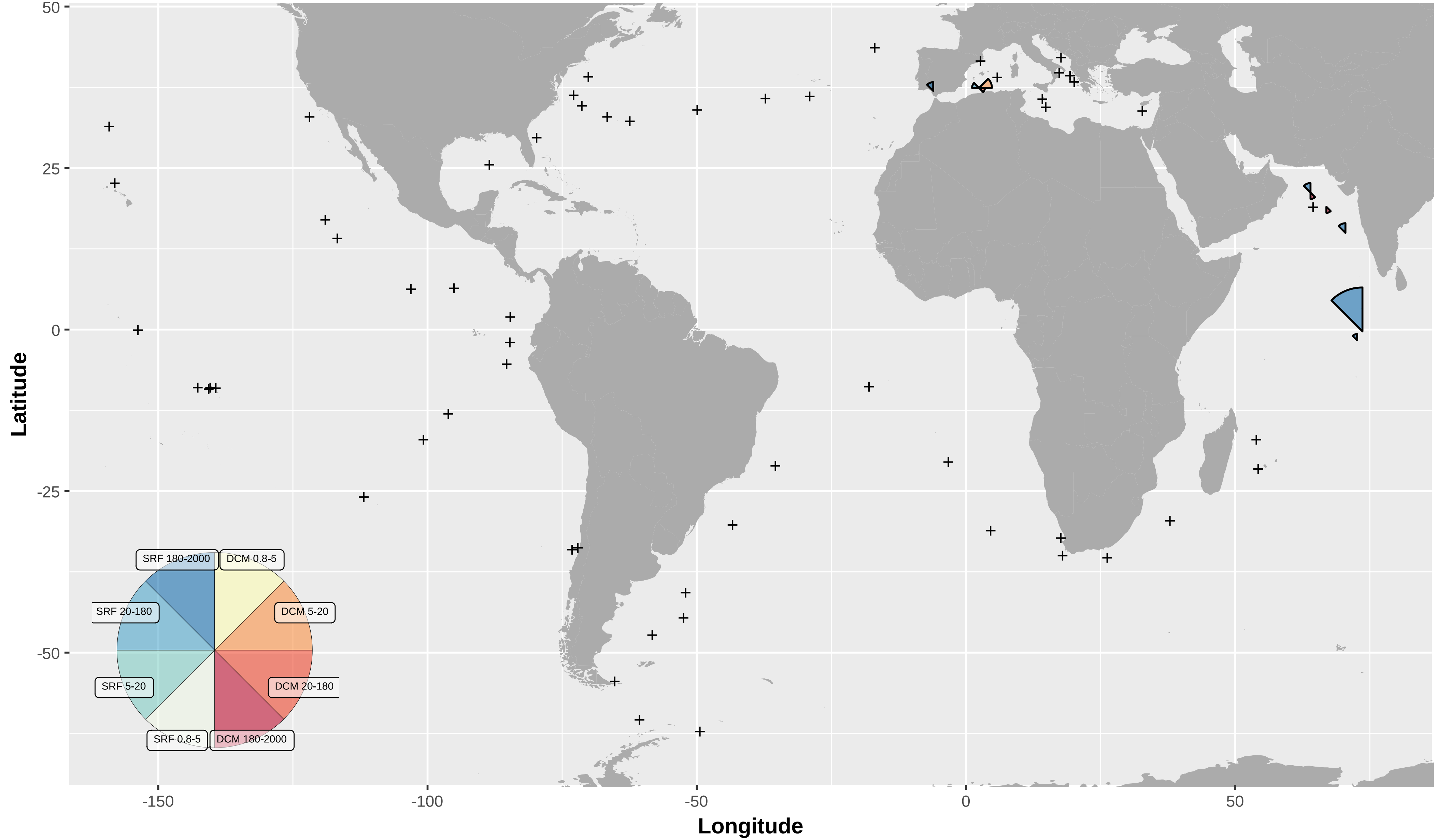


root;cellular organisms;Eukaryota;Opisthokonta;Metazoa;Eumetazoa;Bilateria;Protostomia;Ecdysozoa;Panarthropoda;Arthropoda;Mandibulata;Pancrustacea

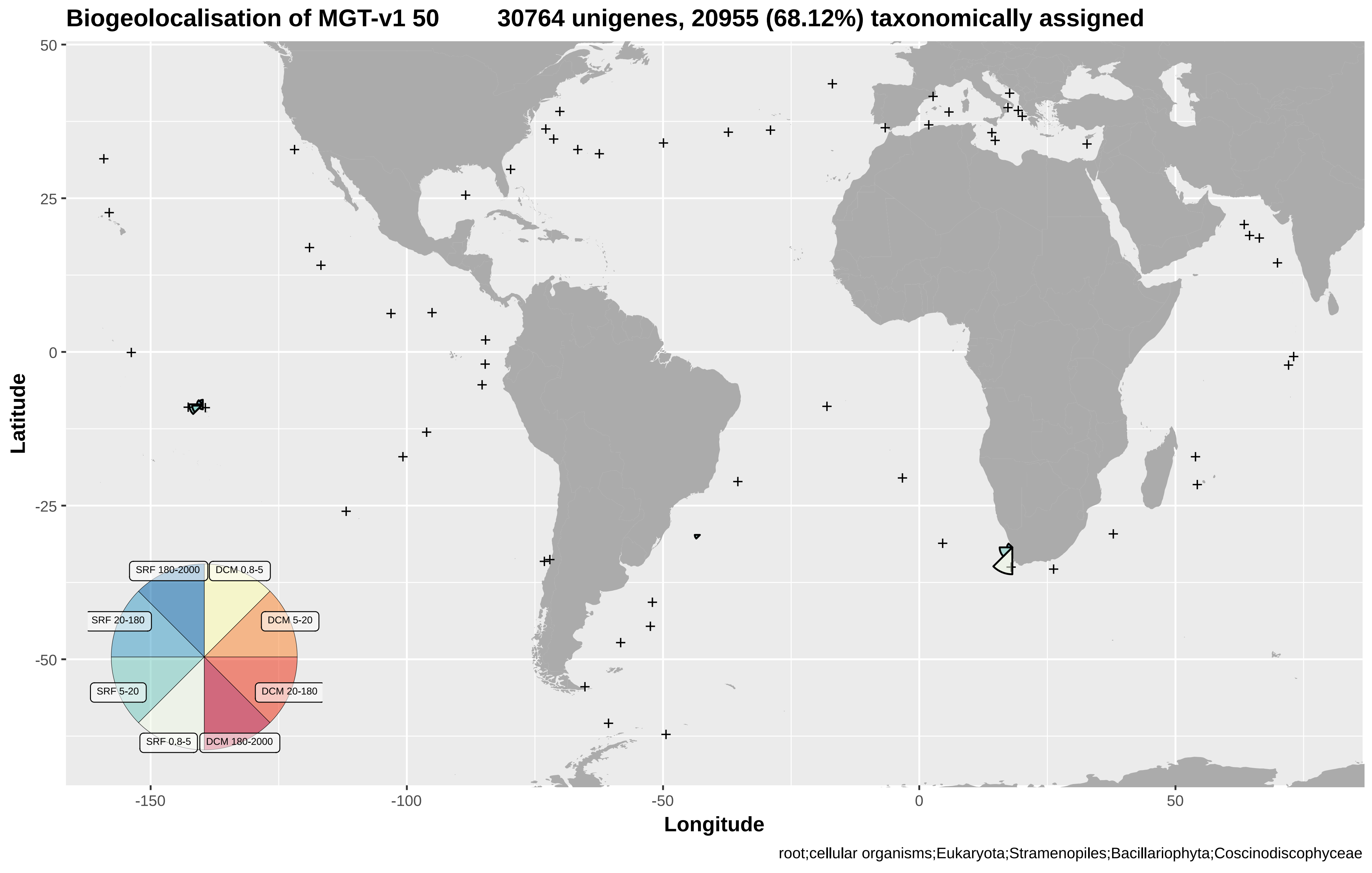
Biogeolocalisation of MGT-v1 50 30764 unigenes, 20955 (68.12%) taxonomically assigned
Latitude
Longitude
50
25
0
-25
-50
-150
-100
-50
0
50
SRF 180-2000
DCM 0.8-5
SRF 20-180
DCM 5-20
SRF 5-20
DCM 20-180
SRF 0.8-5
DCM 180-2000
root;cellular organisms;Eukaryota;Stramenopiles;Bacillariophyta;Coscinodiscophyceae

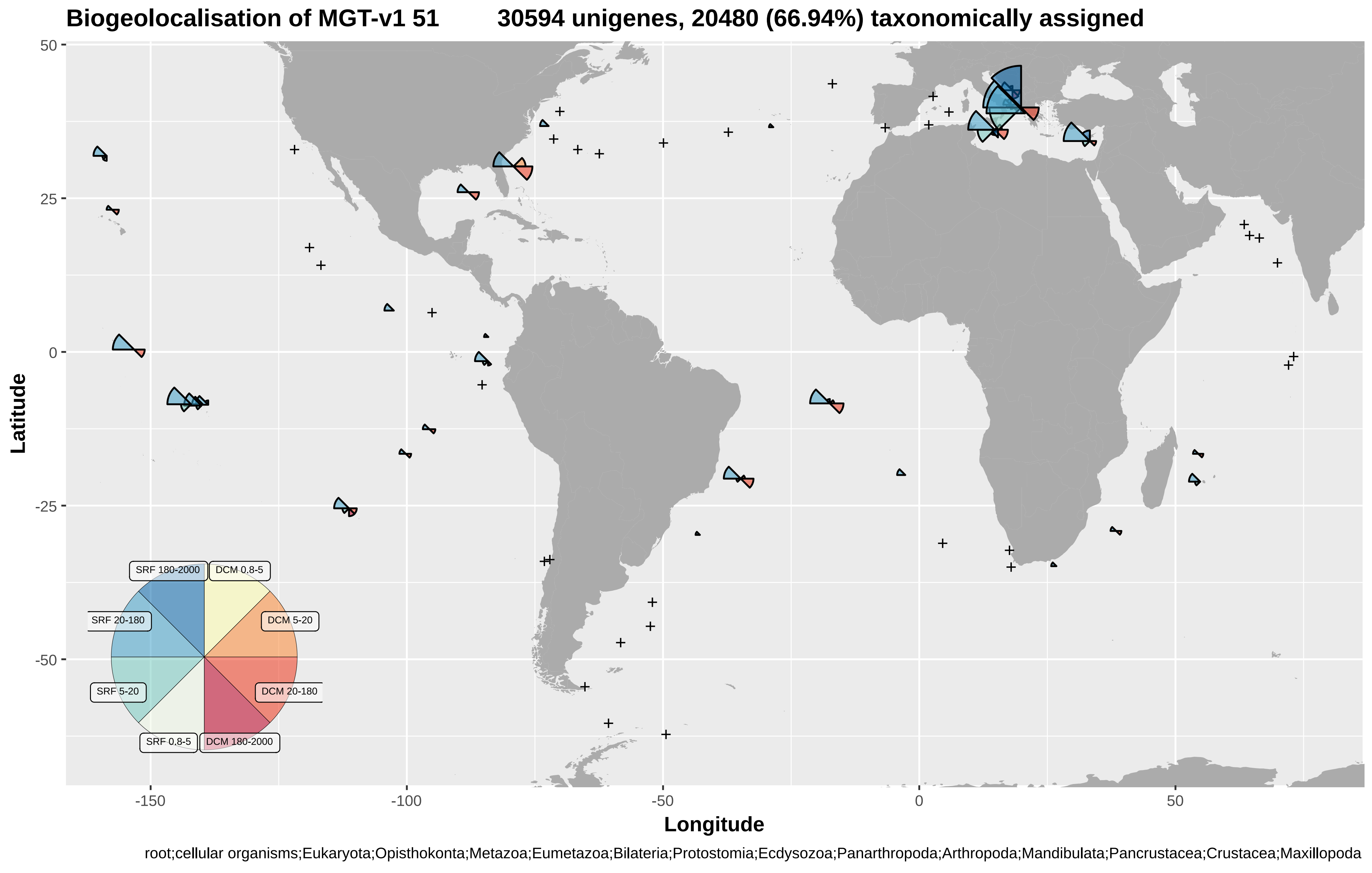
Biogeolocalisation of MGT-v1 51
30594 unigenes, 20480 (66.94%) taxonomically assigned
Latitude
Longitude
50
25
0
-25
-50
-150
-100
-50
0
50
SRF 180-2000
DCM 0.8-5
SRF 20-180
DCM 5-20
SRF 5-20
DCM 20-180
SRF 0.8-5
DCM 180-2000
root;cellular organisms;Eukaryota;Opisthokonta;Metazoa;Eumetazoa;Bilateria;Protostomia;Ecdysozoa;Panarthropoda;Arthropoda;Mandibulata;Pancrustacea;Crustacea;Maxillopoda

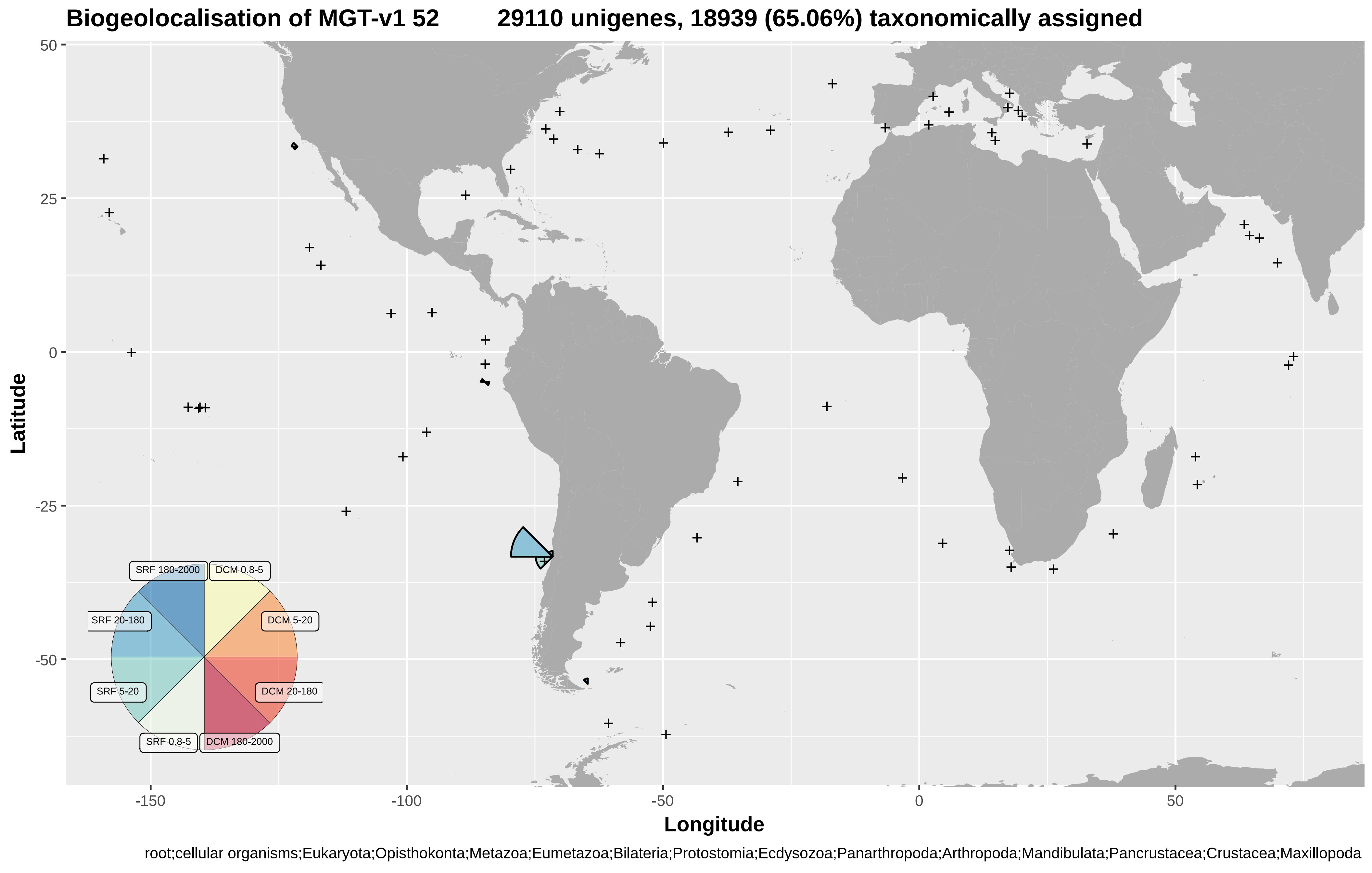
Biogeolocalisation of MGT-v1 52
29110 unigenes, 18939 (65.06%) taxonomically assigned
Latitude
Longitude
50
25
0
-25
-50
-150
-100
-50
0
50
SRF 180-2000
DCM 0.8-5
SRF 20-180
DCM 5-20
SRF 5-20
DCM 20-180
SRF 0.8-5
DCM 180-2000
root;cellular organisms;Eukaryota;Opisthokonta;Metazoa;Eumetazoa;Bilateria;Protostomia;Ecdysozoa;Panarthropoda;Arthropoda;Mandibulata;Pancrustacea;Crustacea;Maxillopoda

### Biogeolocalisation of MGT-v1 53 28858 unigenes, 18621 (64.53%) taxonomically assigned

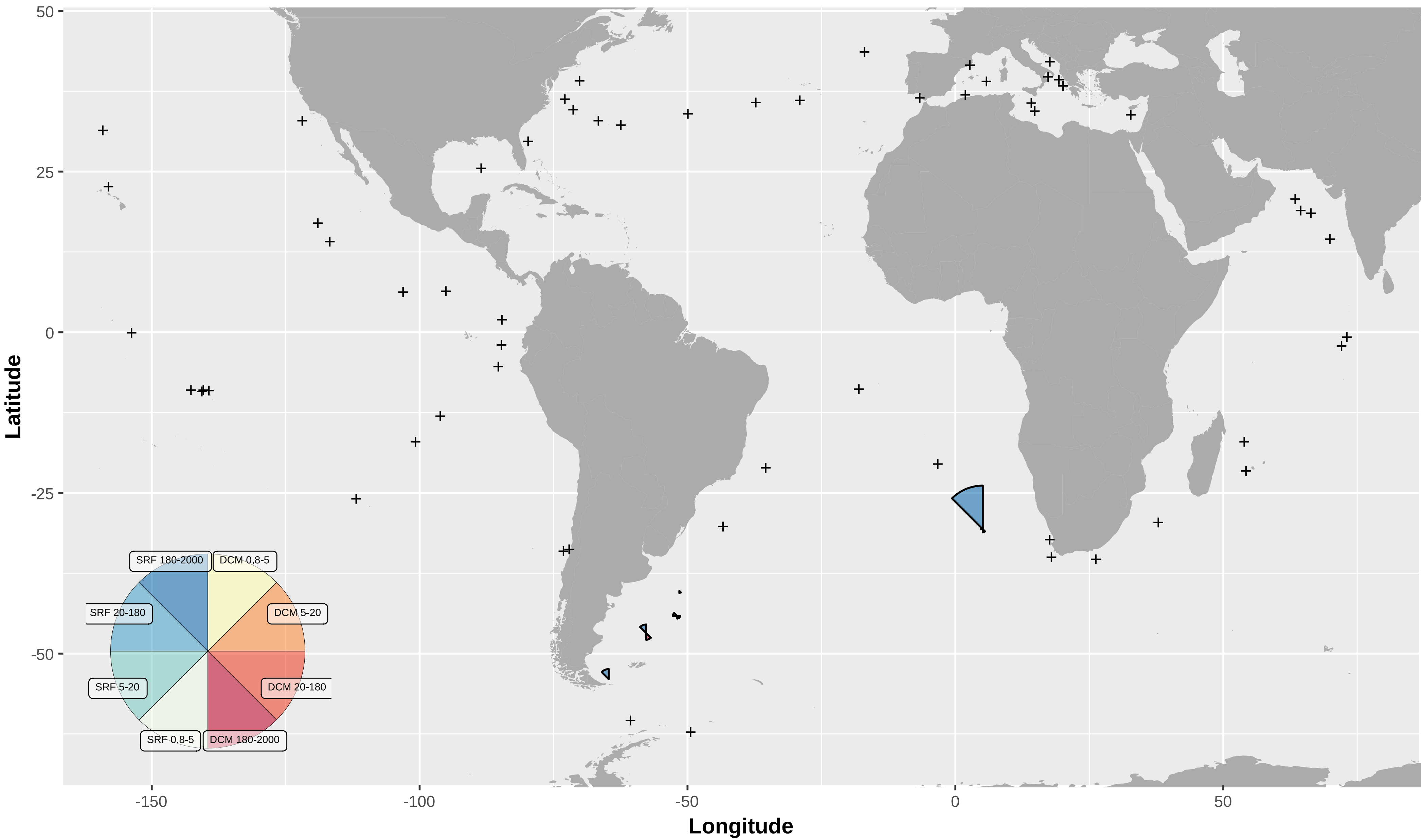


aryota;Opisthokonta;Metazoa;Eumetazoa;Bilateria;Protostomia;Ecdysozoa;Panarthropoda;Arthropoda;Mandibulata;Pancrustacea;Crustacea;Maxillopoda;Copepoda;Neocopepoda;Gymnoplea;Calanoida

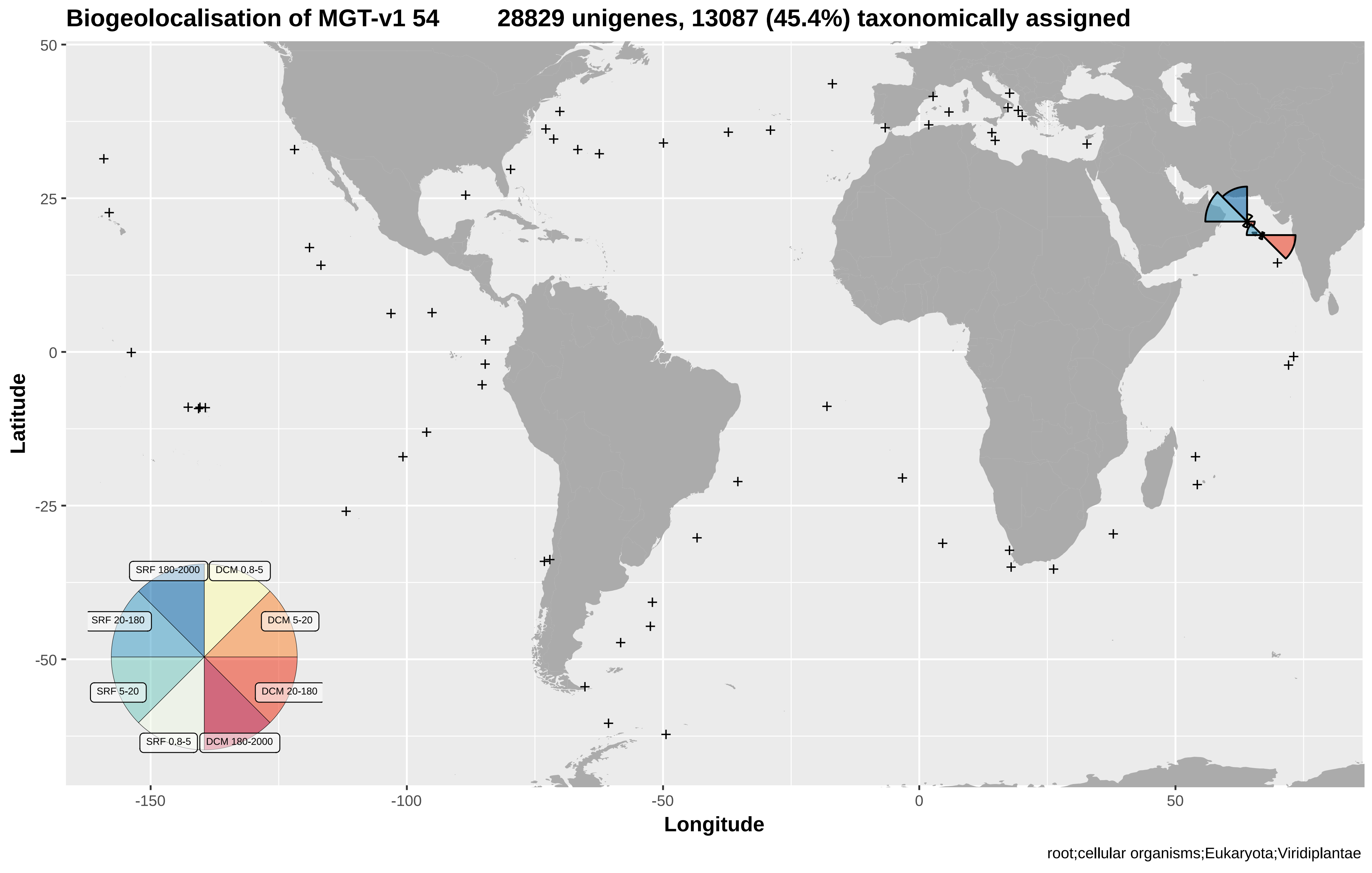
Biogeolocalisation of MGT-v1 54
28829 unigenes, 13087 (45.4%) taxonomically assigned
Latitude
Longitude
50
25
0
-25
-50
-150
-100
-50
0
50
SRF 180-2000
DCM 0.8-5
SRF 20-180
DCM 5-20
SRF 5-20
DCM 20-180
SRF 0.8-5
DCM 180-2000
root;cellular organisms;Eukaryota;Viridiplantae

#### Biogeolocalisation of MGT-v1 55 27304 unigenes, 2461 (9.01%) taxonomically assigned

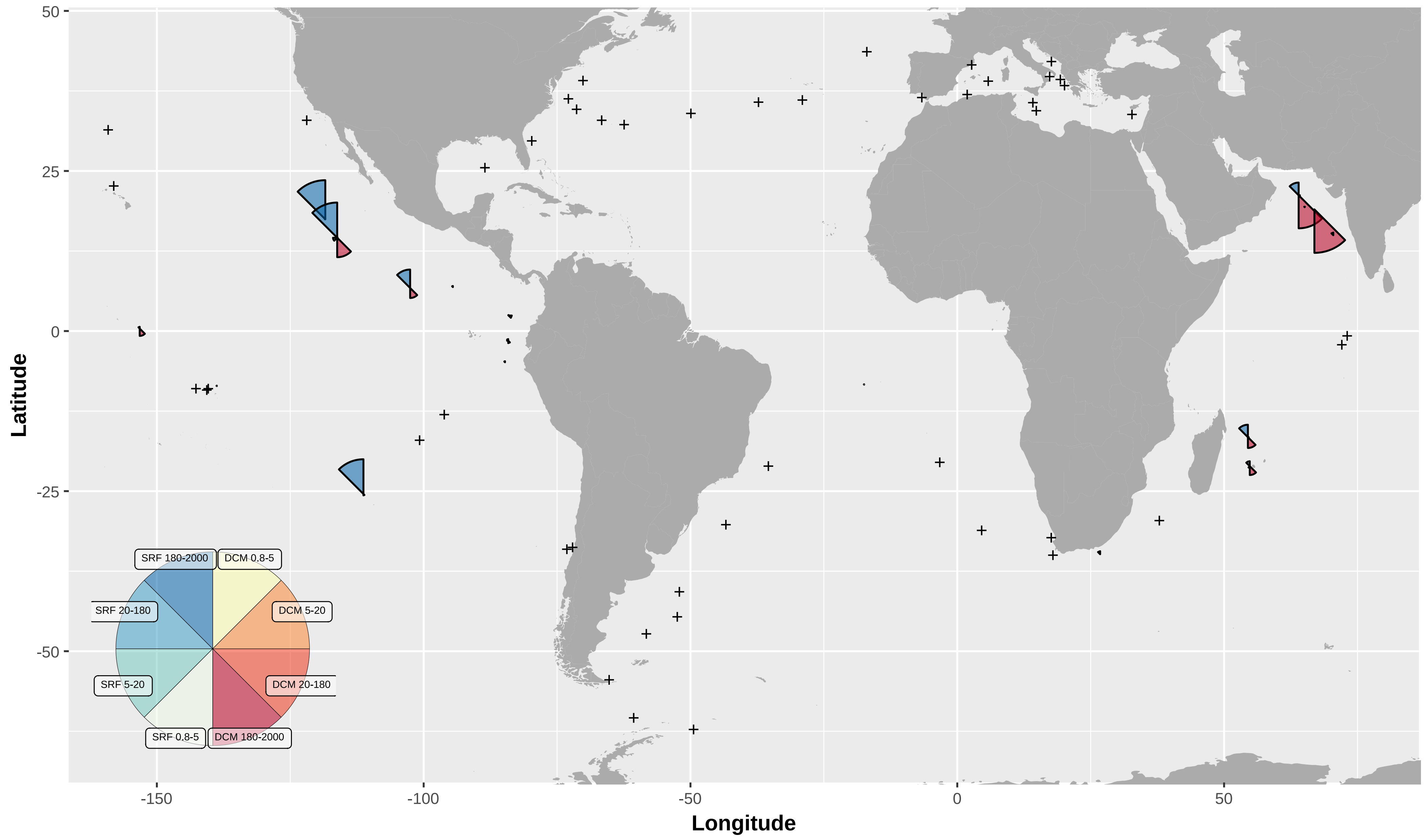


root;cellular organisms;Eukaryota;Opisthokonta;Metazoa;Eumetazoa;Bilateria

## Biogeolocalisation of MGT-v1 56        26118 unigenes, 10854 (41.56%) taxonomically assigned

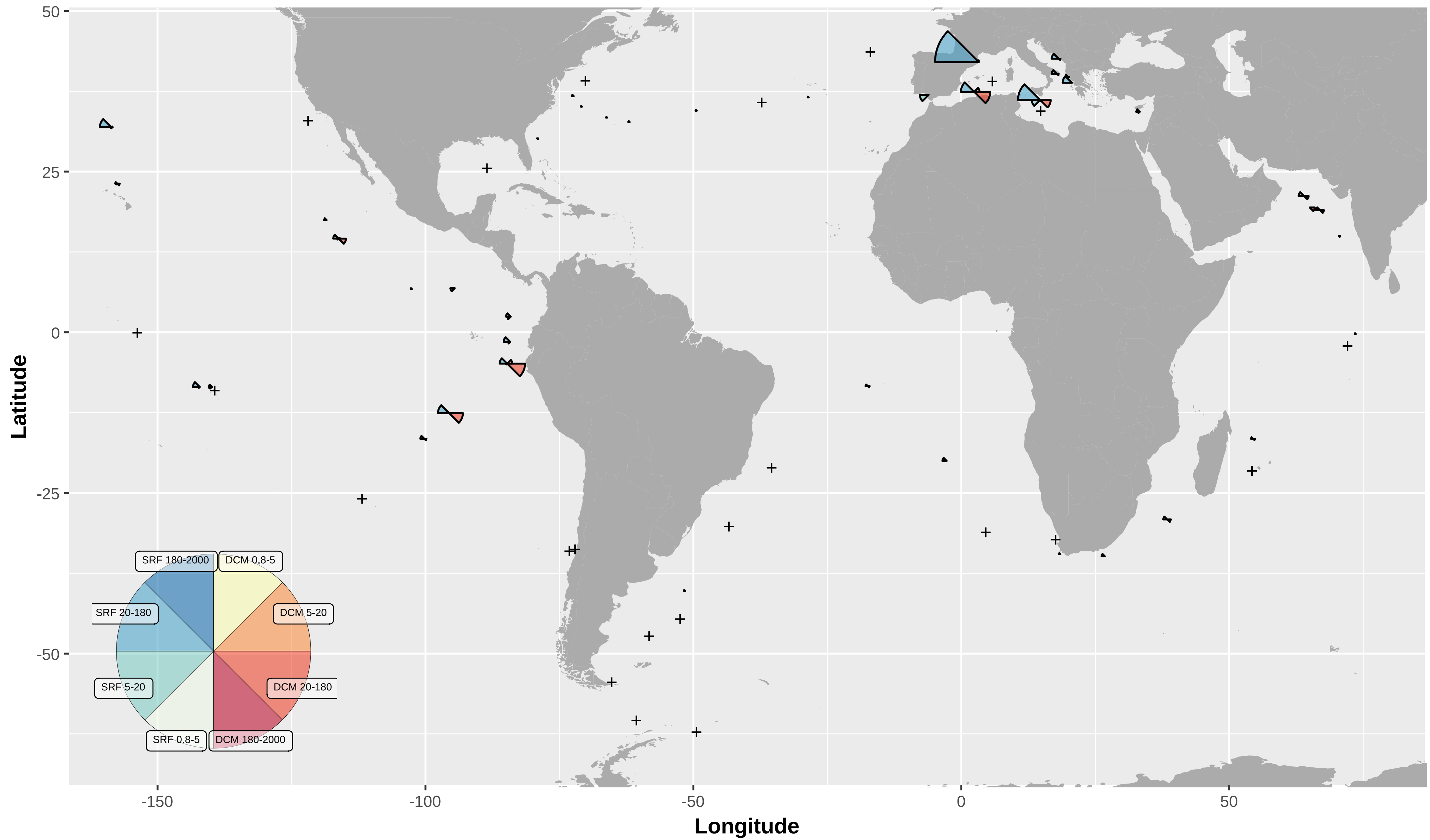


cellular organisms;Eukaryota;Opisthokonta;Metazoa;Eumetazoa;Bilateria;Protostomia;Ecdysozoa;Panarthropoda;Arthropoda;Mandibulata;Pancrustacea;Crustacea;Maxillopoda;Copepoda;Neocopepoda

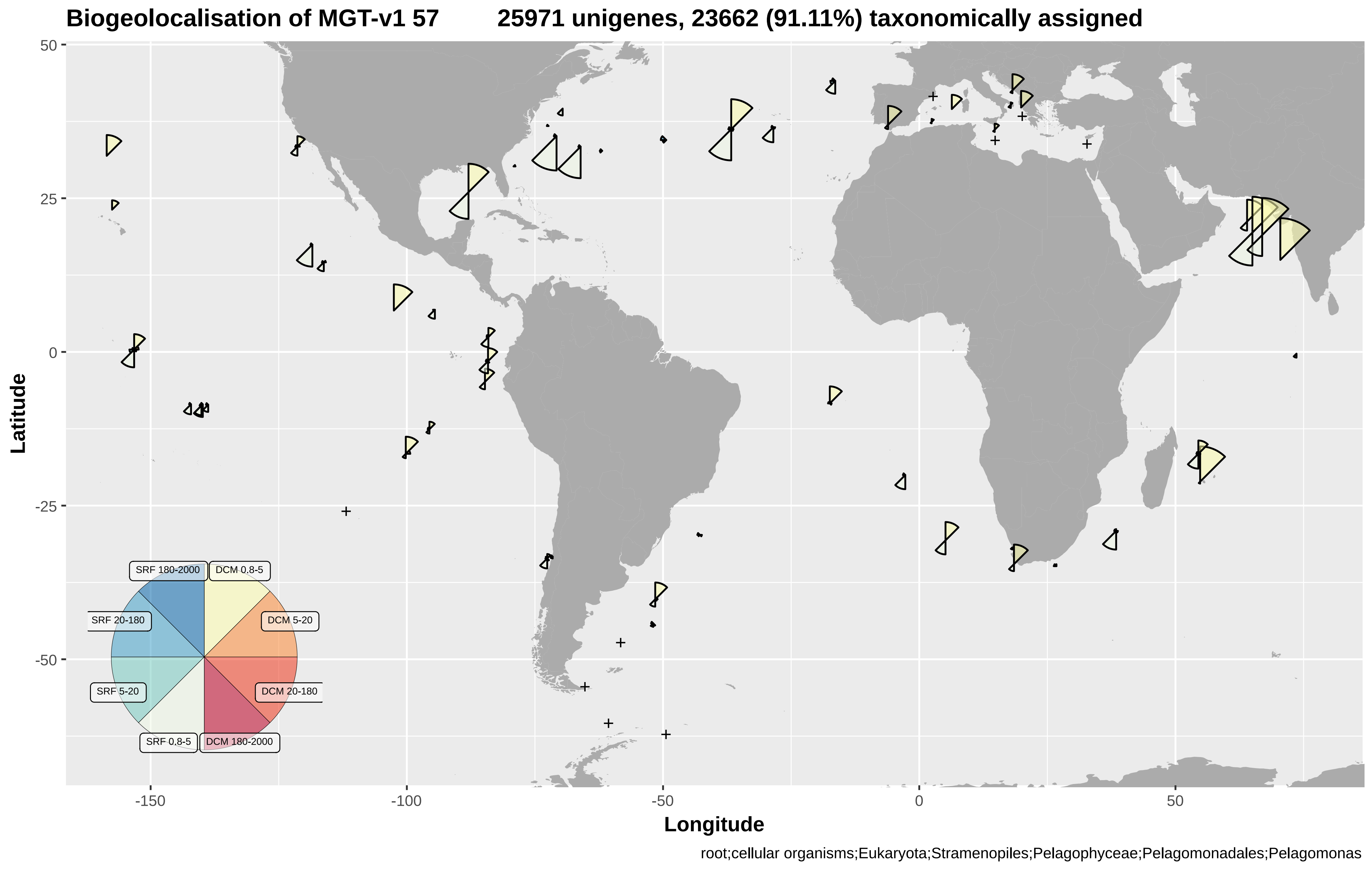
Biogeolocalisation of MGT-v1 57
25971 unigenes, 23662 (91.11%) taxonomically assigned
50
25
0
-25
-50
Latitude
-150
-100
-50
0
50
Longitude
SRF 180-2000
DCM 0.8-5
SRF 20-180
DCM 5-20
SRF 5-20
DCM 20-180
SRF 0.8-5
DCM 180-2000
root;cellular organisms;Eukaryota;Stramenopiles;Pelagophyceae;Pelagomonadales;Pelagomonas

#### Biogeolocalisation of MGT-v1 58 25964 unigenes, 22457 (86.49%) taxonomically assigned

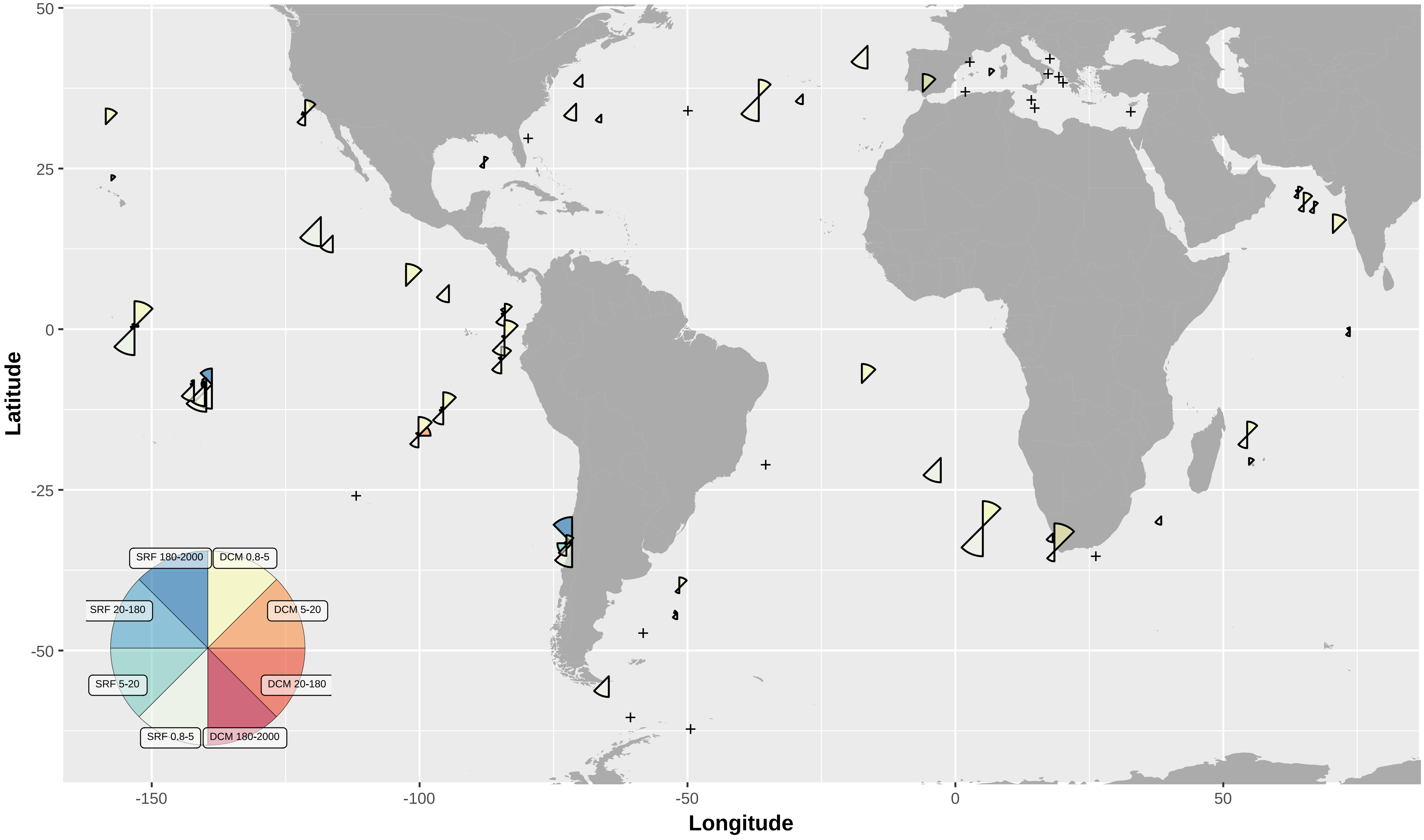


root;cellular organisms;Eukaryota;Haptophyceae;Phaeocystales

## Biogeolocalisation of MGT-v1 59        24885 unigenes, 16150 (64.9%) taxonomically assigned

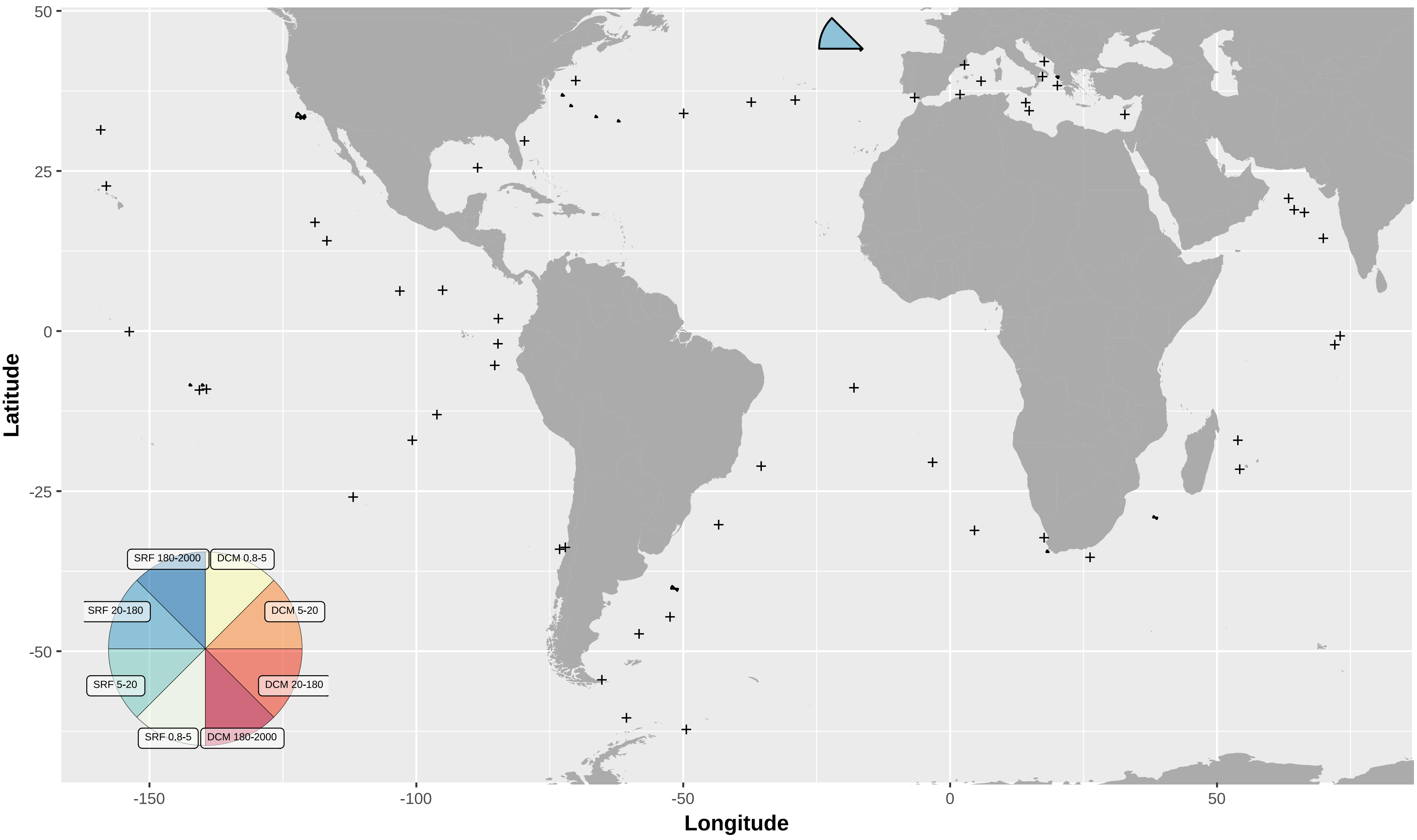


root;cellular organisms;Eukaryota

# Biogeolocalisation of MGT-v1 60        24590 unigenes, 20191 (82.11%) taxonomically assigned

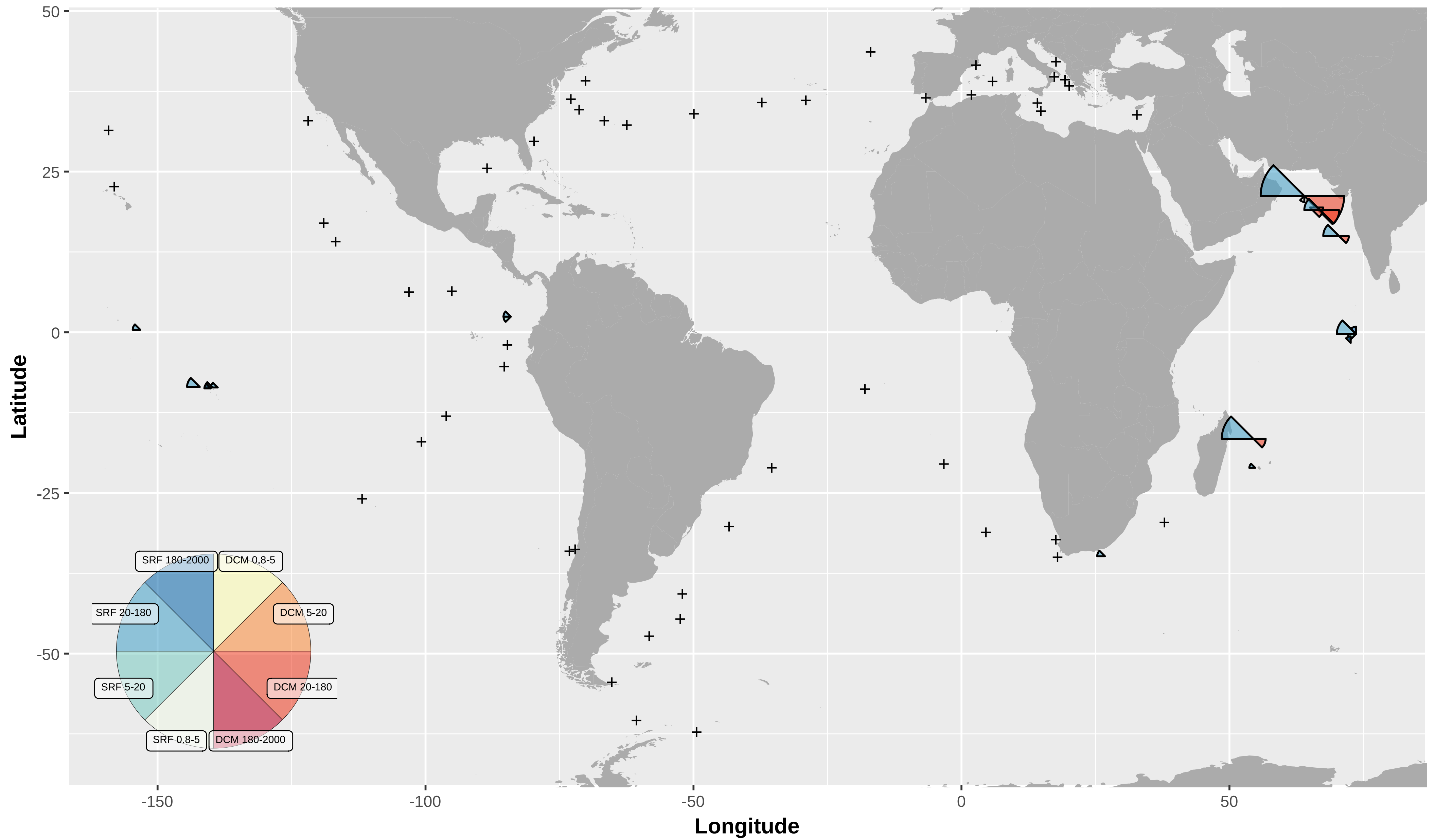


karyota;Opisthokonta;Metazoa;Eumetazoa;Bilateria;Protostomia;Ecdysozoa;Panarthropoda;Arthropoda;Mandibulata;Pancrustacea;Crustacea;Maxillopoda;Copepoda;Neocopepoda;Podoplea;Cyclopoida

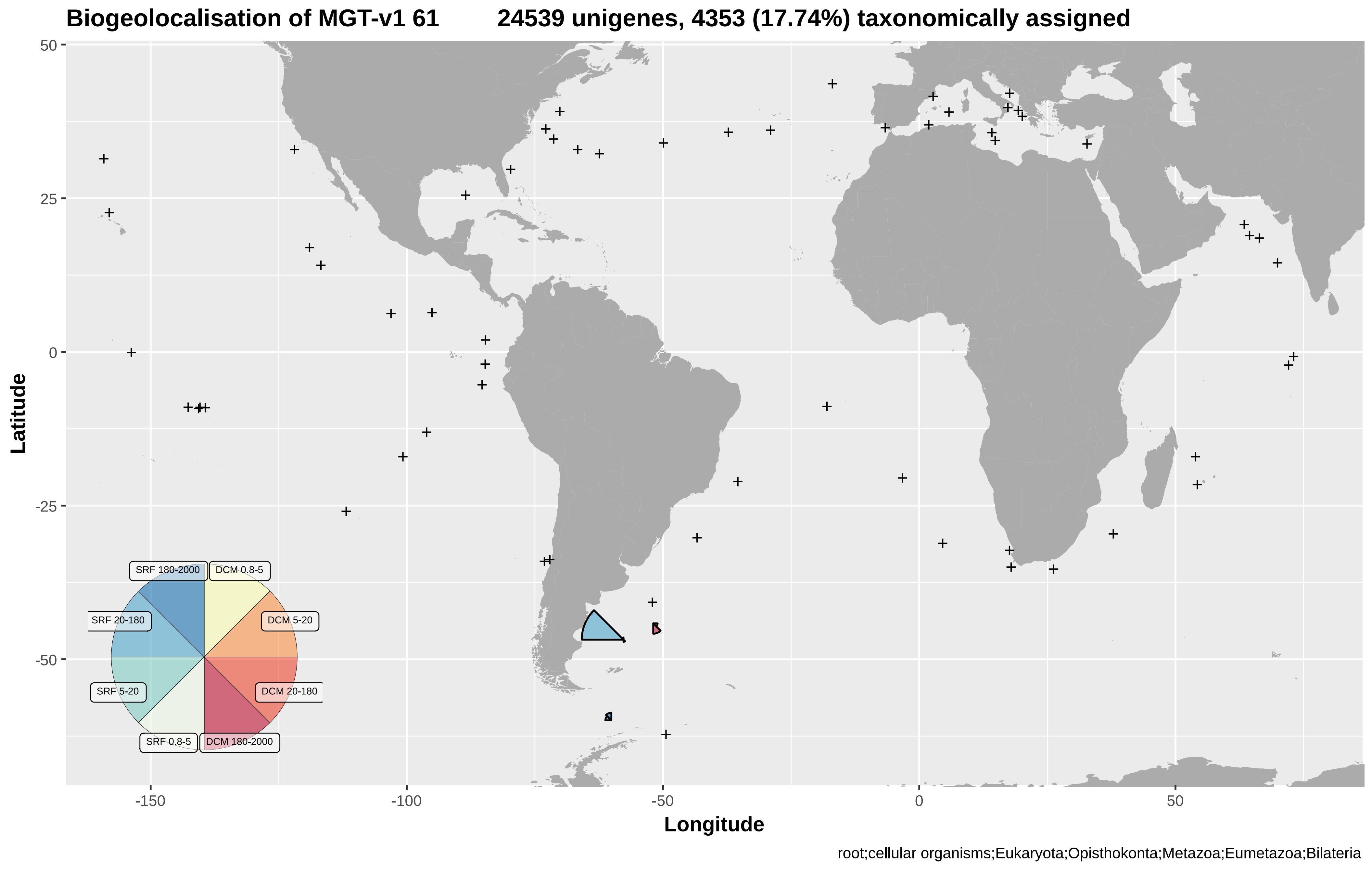

Biogeolocalisation of MGT-v1 61
24539 unigenes, 4353 (17.74%) taxonomically assigned
Latitude
Longitude
50
25
0
-25
-50
-150
-100
-50
0
50
SRF 180-2000
DCM 0.8-5
SRF 20-180
DCM 5-20
SRF 5-20
DCM 20-180
SRF 0.8-5
DCM 180-2000
root;cellular organisms;Eukaryota;Opisthokonta;Metazoa;Eumetazoa;Bilateria

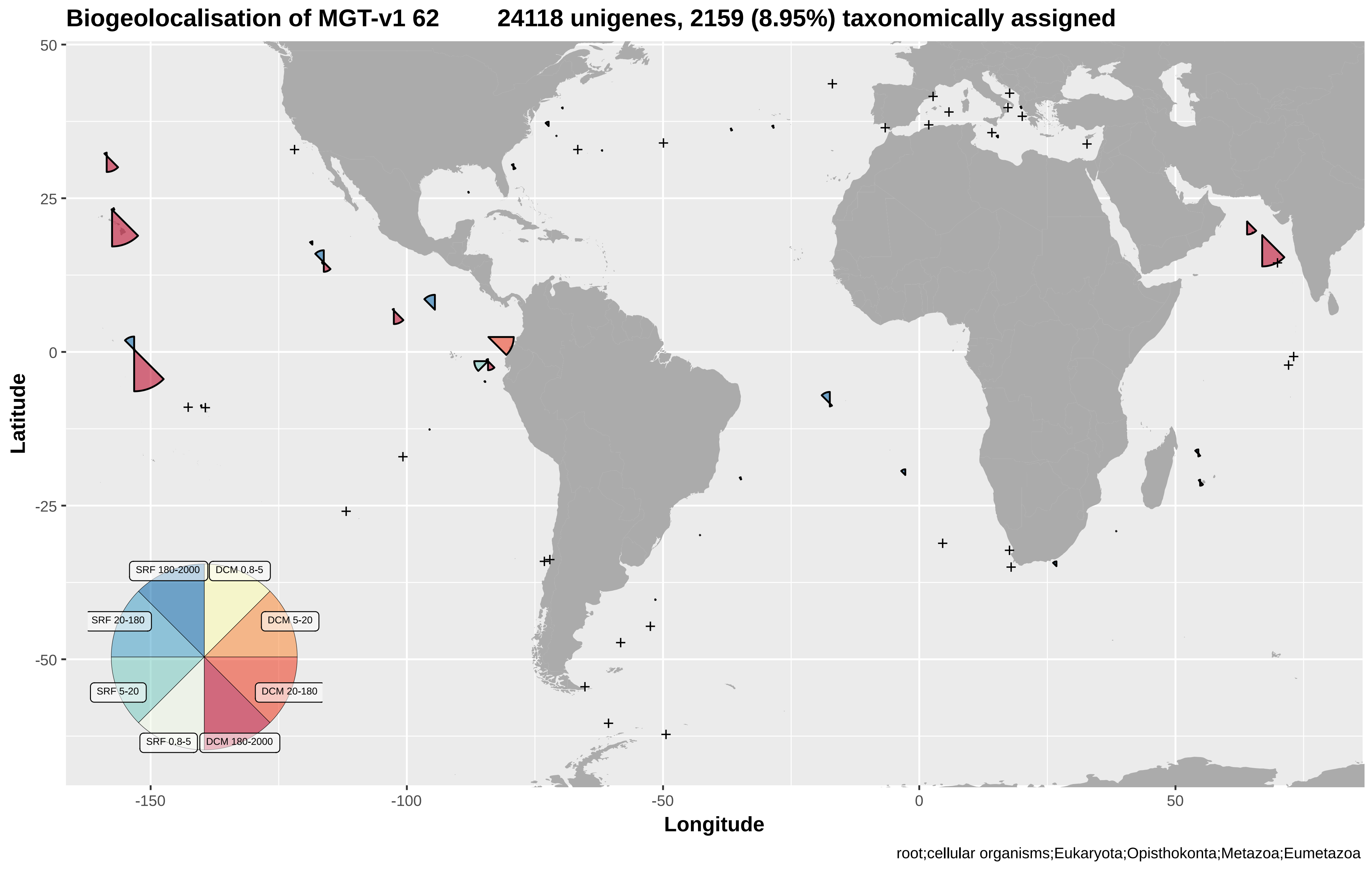
Biogeolocalisation of MGT-v1 62
24118 unigenes, 2159 (8.95%) taxonomically assigned
50
25
0
-25
-50
Latitude
-150
-100
-50
0
50
Longitude
SRF 180-2000
DCM 0.8-5
SRF 20-180
DCM 5-20
SRF 5-20
DCM 20-180
SRF 0.8-5
DCM 180-2000
root;cellular organisms;Eukaryota;Opisthokonta;Metazoa;Eumetazoa

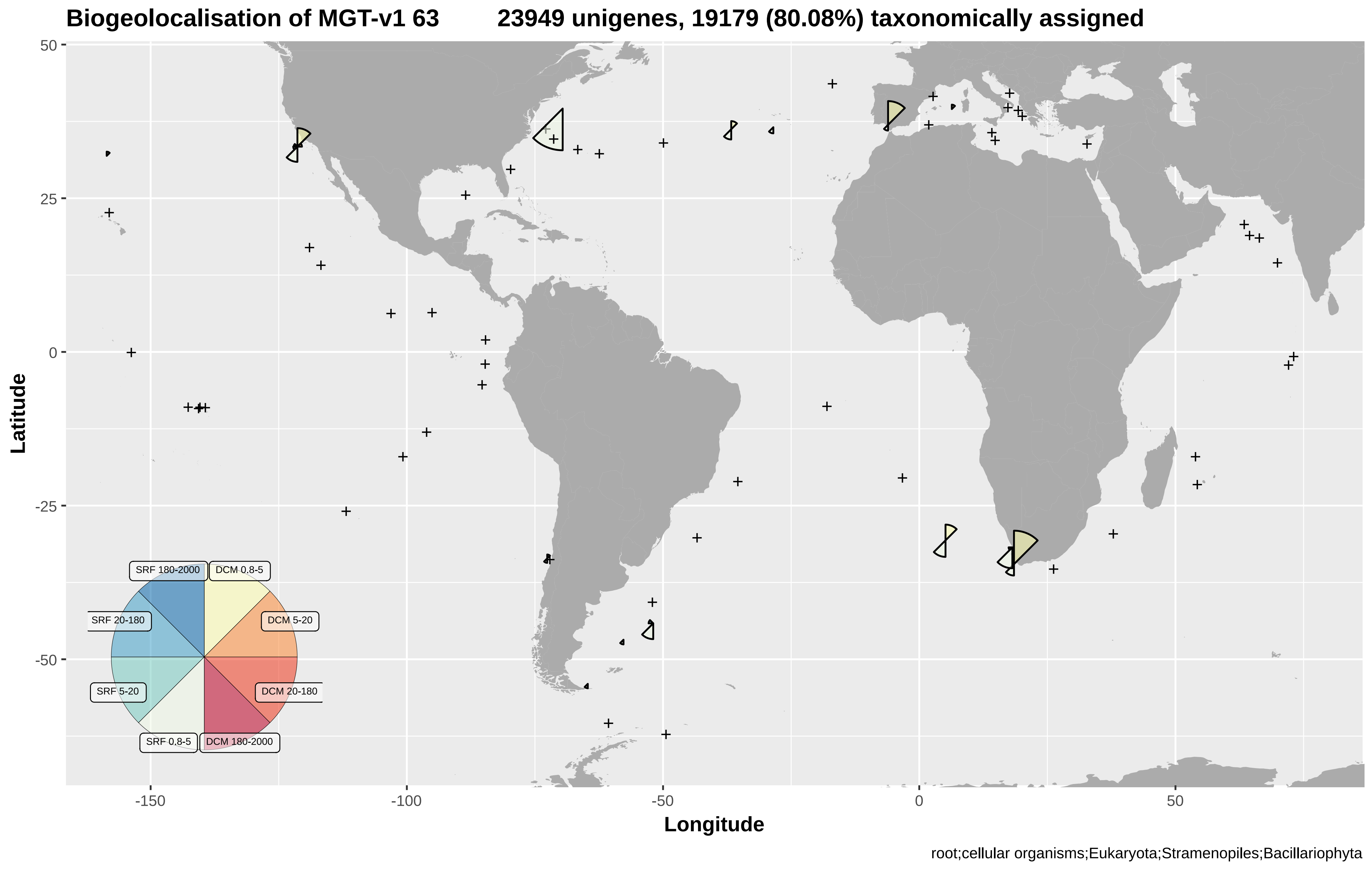
Biogeolocalisation of MGT-v1 63
23949 unigenes, 19179 (80.08%) taxonomically assigned
Latitude
Longitude
50
25
0
-25
-50
-150
-100
-50
0
50
SRF 180-2000
DCM 0.8-5
SRF 20-180
DCM 5-20
SRF 5-20
DCM 20-180
SRF 0.8-5
DCM 180-2000
root;cellular organisms;Eukaryota;Stramenopiles;Bacillariophyta

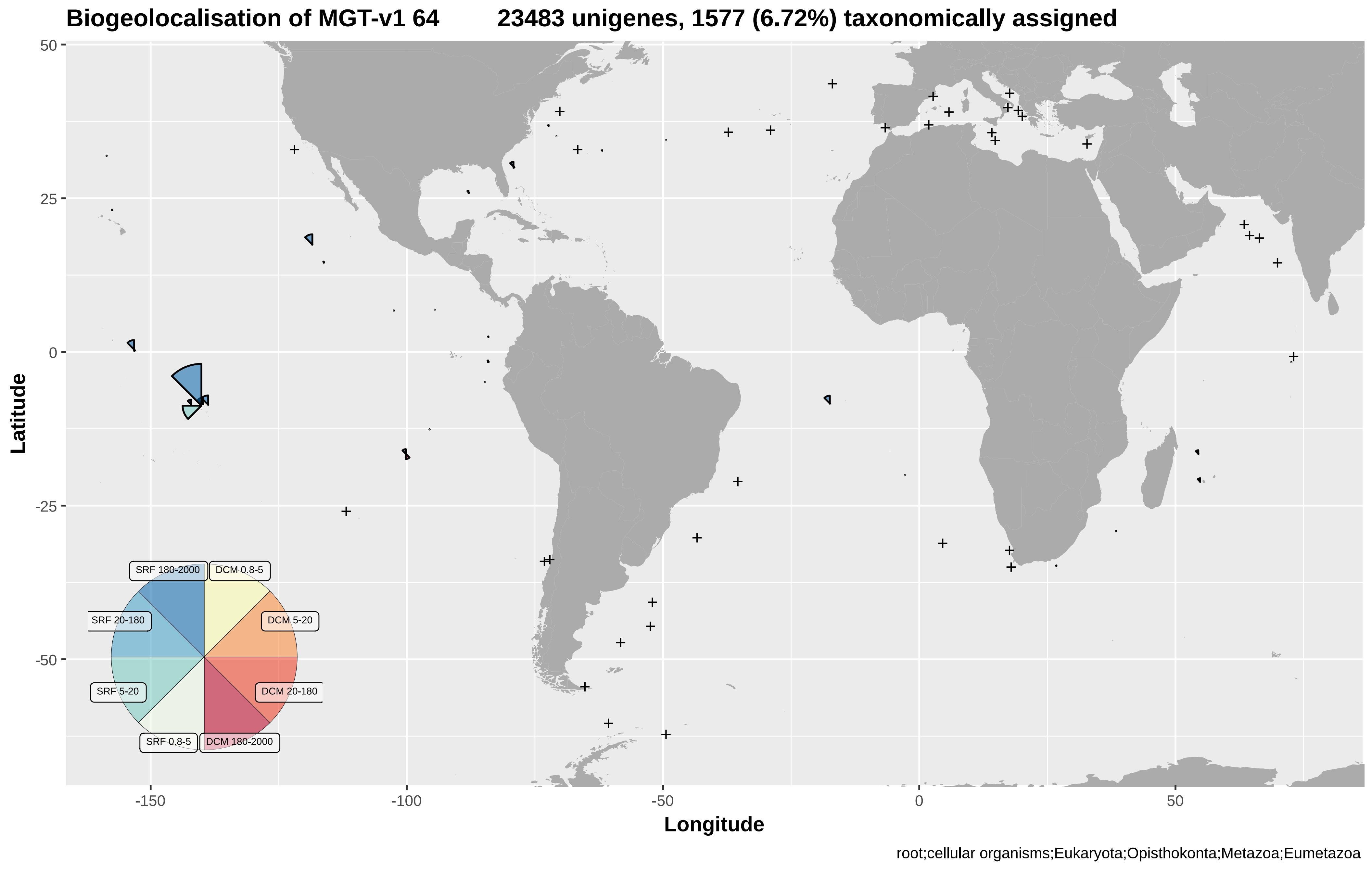
Biogeolocalisation of MGT-v1 64
23483 unigenes, 1577 (6.72%) taxonomically assigned
Latitude
Longitude
50
25
0
-25
-50
-150
-100
-50
0
50
SRF 180-2000
DCM 0.8-5
SRF 20-180
DCM 5-20
SRF 5-20
DCM 20-180
SRF 0.8-5
DCM 180-2000
root;cellular organisms;Eukaryota;Opisthokonta;Metazoa;Eumetazoa

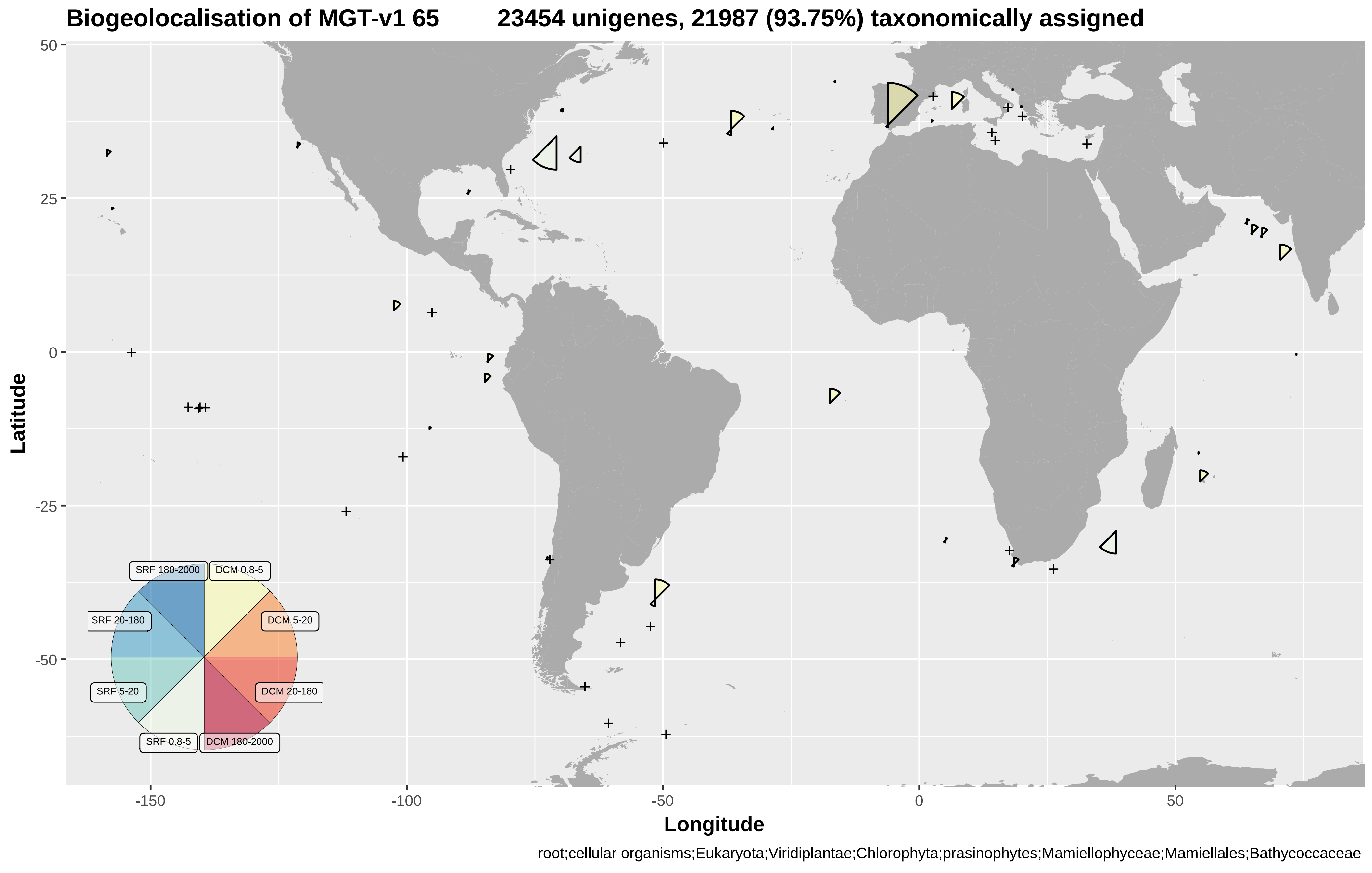
Biogeolocalisation of MGT-v1 65
23454 unigenes, 21987 (93.75%) taxonomically assigned
50
25
0
-25
-50
Latitude
-150
-100
-50
0
50
Longitude
SRF 180-2000
DCM 0.8-5
SRF 20-180
DCM 5-20
SRF 5-20
DCM 20-180
SRF 0.8-5
DCM 180-2000
root;cellular organisms;Eukaryota;Viridiplantae;Chlorophyta;prasinophytes;Mamiellophyceae;Mamiellales;Bathycoccaceae

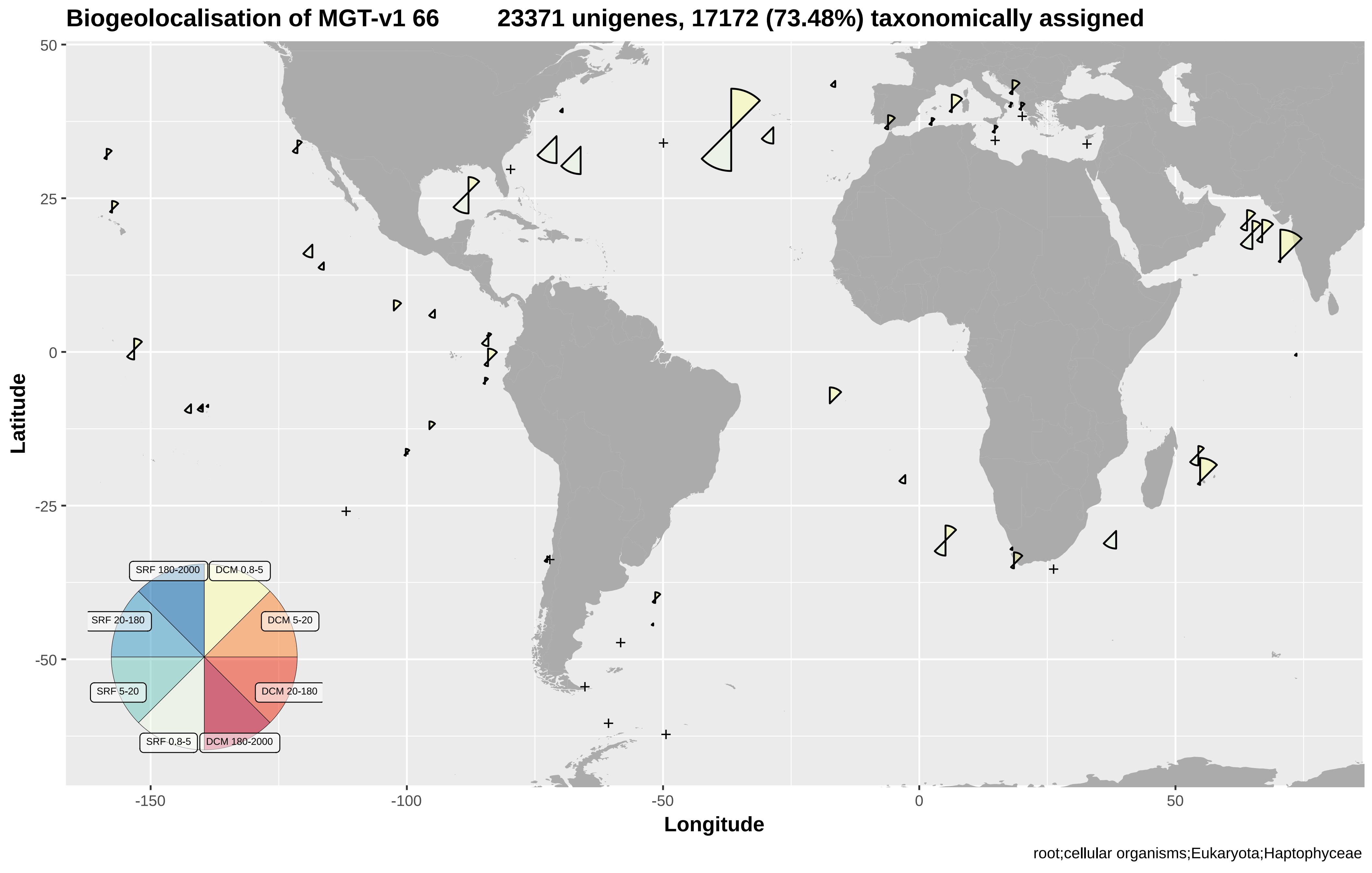

Biogeolocalisation of MGT-v1 66
23371 unigenes, 17172 (73.48%) taxonomically assigned
Latitude
Longitude
50
25
0
-25
-50
-150
-100
-50
0
50
SRF 180-2000
DCM 0.8-5
SRF 20-180
DCM 5-20
SRF 5-20
DCM 20-180
SRF 0.8-5
DCM 180-2000
root;cellular organisms;Eukaryota;Haptophyceae

**Biogeolocalisation of MGT-v1 67        23357 unigenes, 18717 (80.13%) taxonomically assigned**

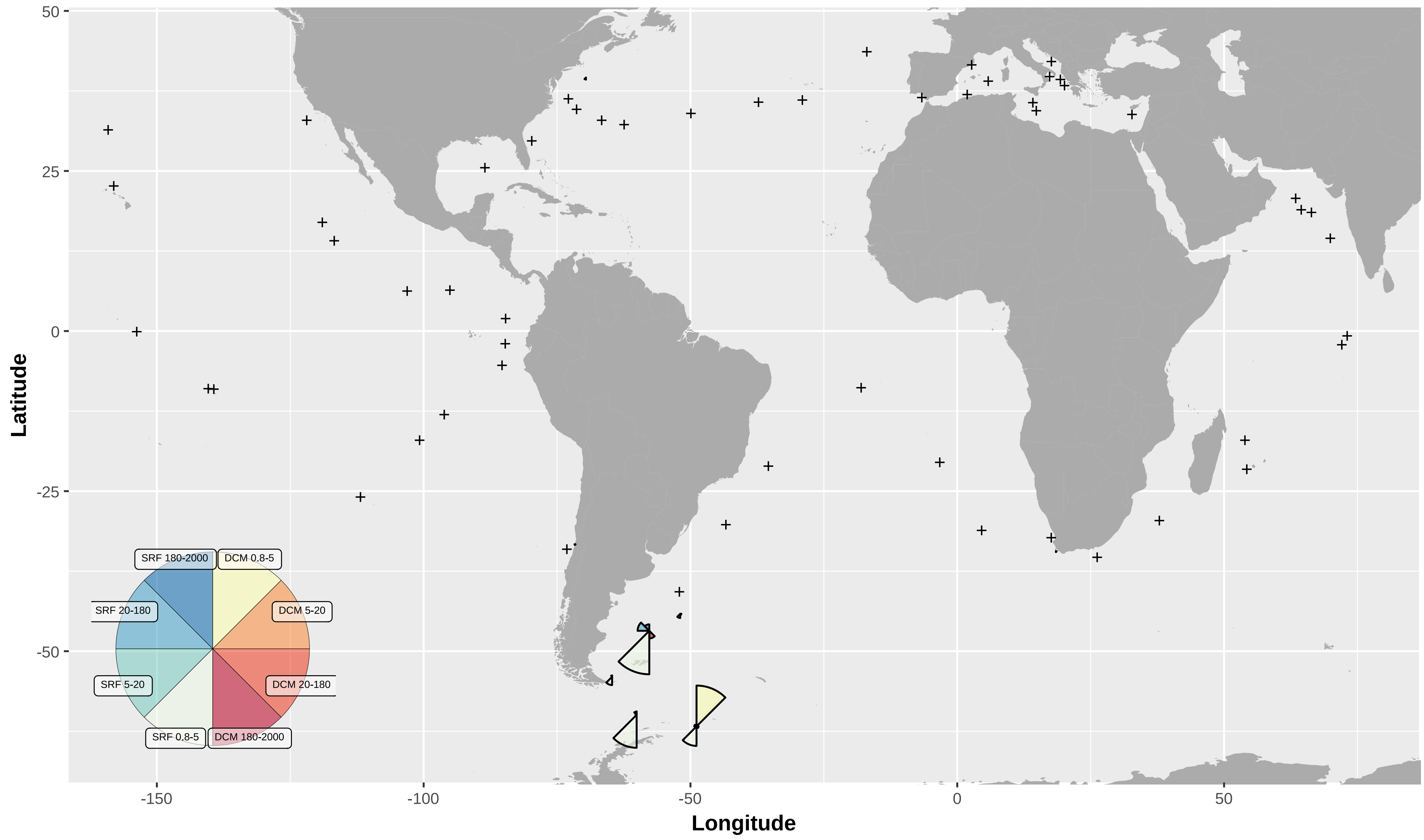


root;cellular organisms;Eukaryota;Haptophyceae

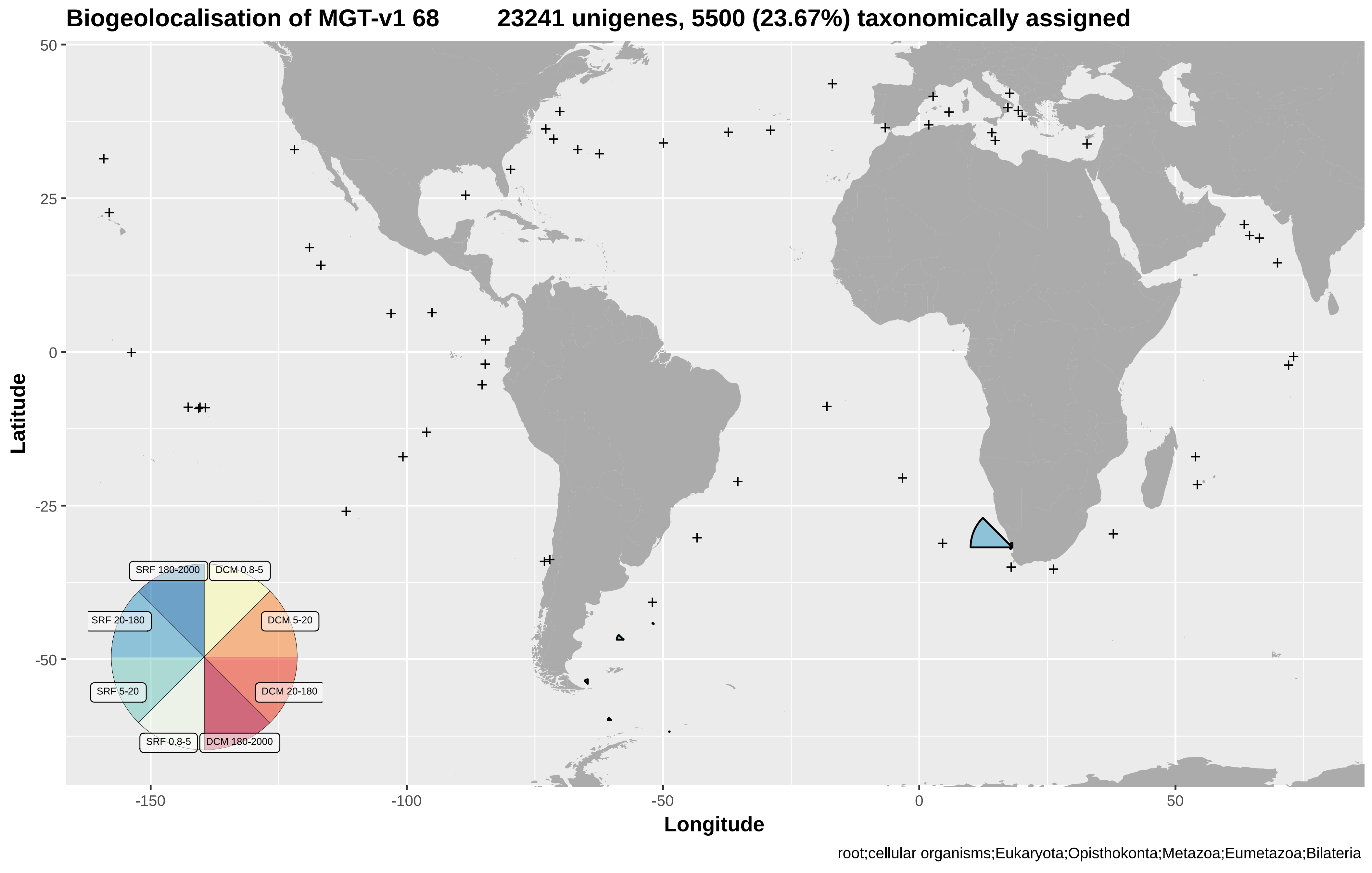

Biogeolocalisation of MGT-v1 68
23241 unigenes, 5500 (23.67%) taxonomically assigned
50
25
0
-25
-50
Latitude
-150
-100
-50
0
50
Longitude
SRF 180-2000
DCM 0.8-5
SRF 20-180
DCM 5-20
SRF 5-20
DCM 20-180
SRF 0.8-5
DCM 180-2000
root;cellular organisms;Eukaryota;Opisthokonta;Metazoa;Eumetazoa;Bilateria

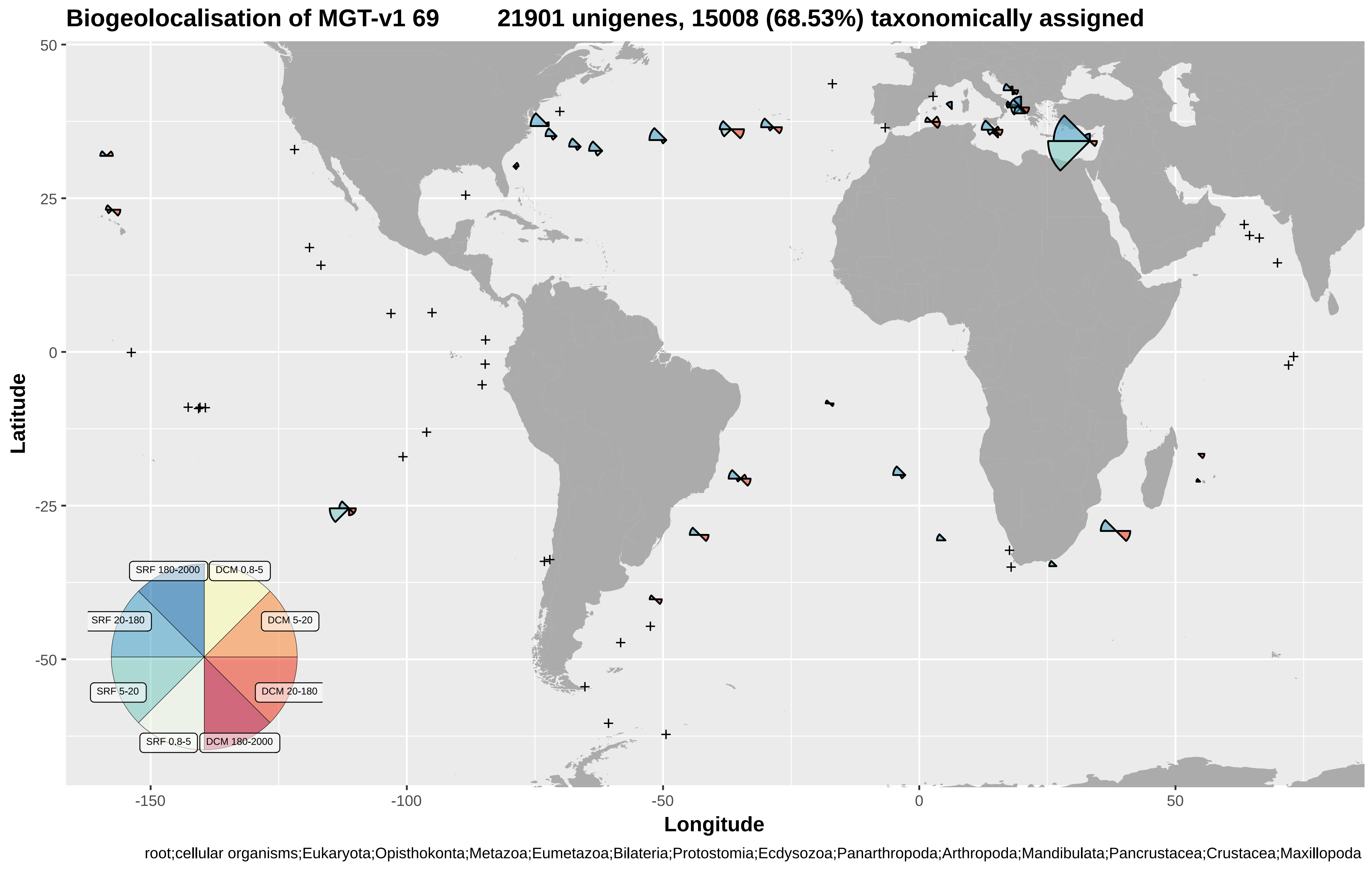

Biogeolocalisation of MGT-v1 69
21901 unigenes, 15008 (68.53%) taxonomically assigned
50
25
0
-25
-50
Latitude
-150
-100
-50
0
50
Longitude
SRF 180-2000
DCM 0.8-5
SRF 20-180
DCM 5-20
SRF 5-20
DCM 20-180
SRF 0.8-5
DCM 180-2000
root;cellular organisms;Eukaryota;Opisthokonta;Metazoa;Eumetazoa;Bilateria;Protostomia;Ecdysozoa;Panarthropoda;Arthropoda;Mandibulata;Pancrustacea;Crustacea;Maxillopoda

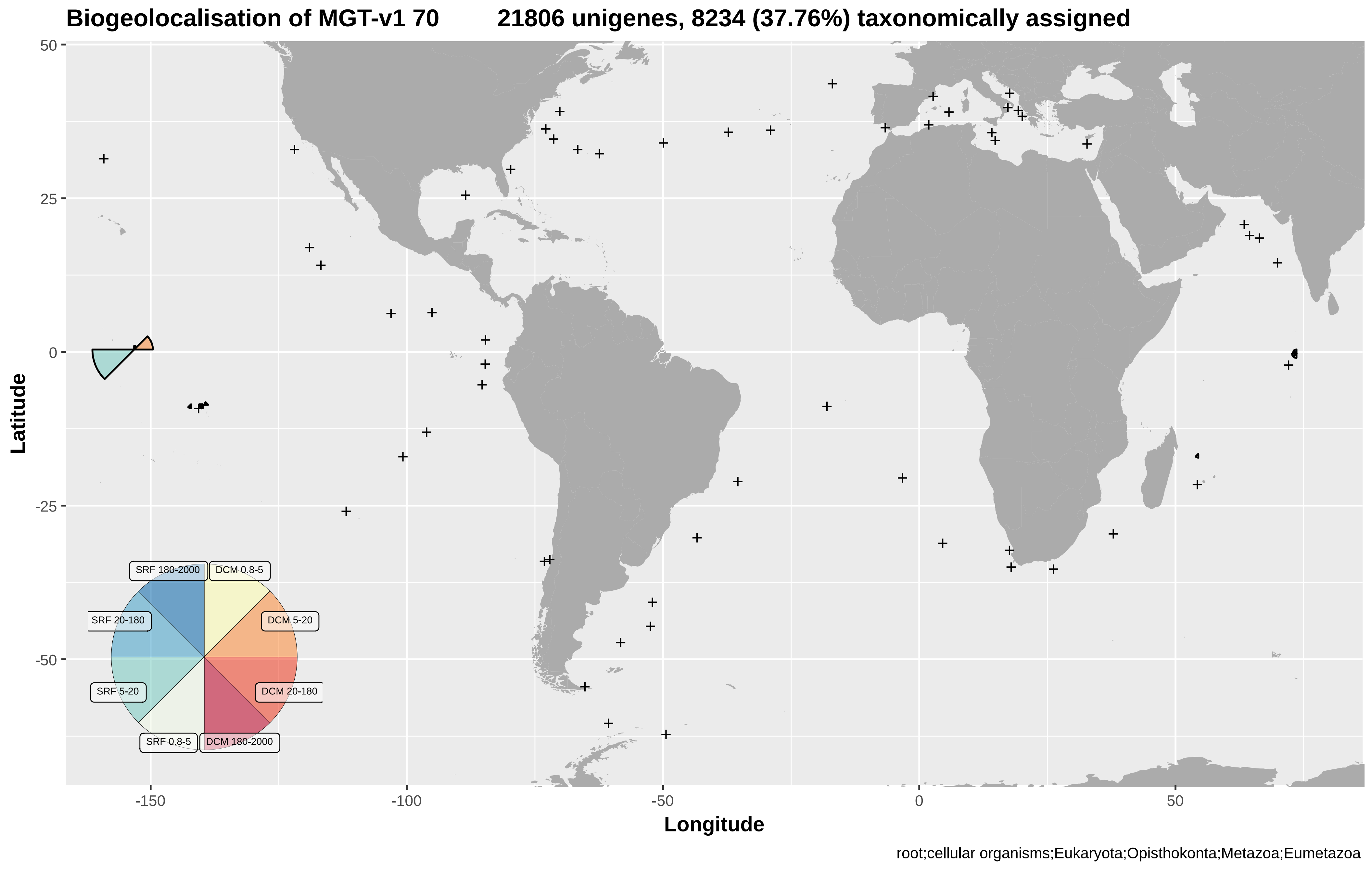

Biogeolocalisation of MGT-v1 70 21806 unigenes, 8234 (37.76%) taxonomically assigned
50
25
0
-25
-50
Latitude
-150
-100
-50
0
50
Longitude
SRF 180-2000
DCM 0.8-5
SRF 20-180
DCM 5-20
SRF 5-20
DCM 20-180
SRF 0.8-5
DCM 180-2000
root;cellular organisms;Eukaryota;Opisthokonta;Metazoa;Eumetazoa

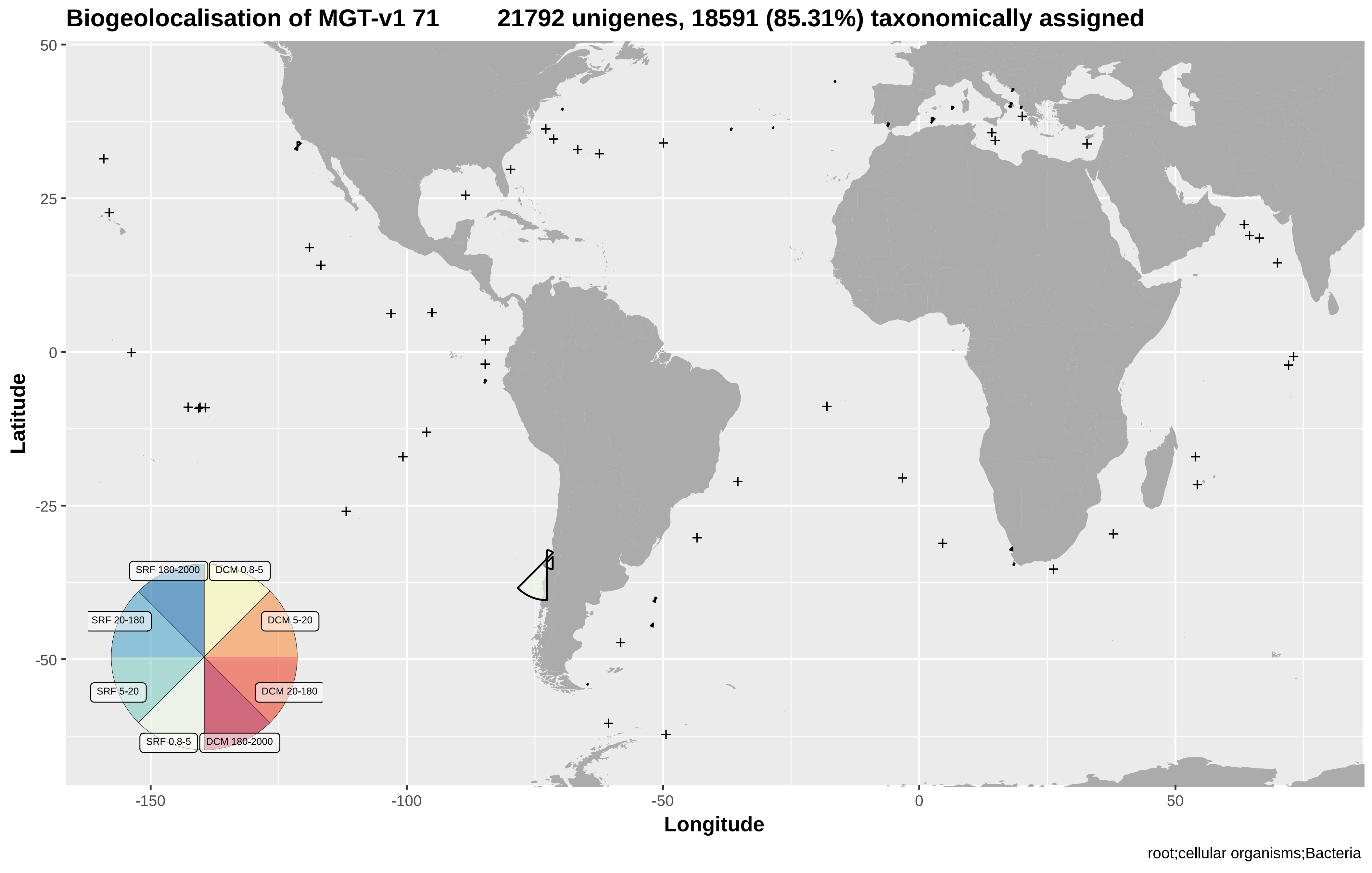
Biogeolocalisation of MGT-v1 71 21792 unigenes, 18591 (85.31%) taxonomically assigned
50
25
0
-25
-50
Latitude
-150
-100
-50
0
50
Longitude
SRF 180-2000
DCM 0.8-5
SRF 20-180
DCM 5-20
SRF 5-20
DCM 20-180
SRF 0.8-5
DCM 180-2000
root;cellular organisms;Bacteria

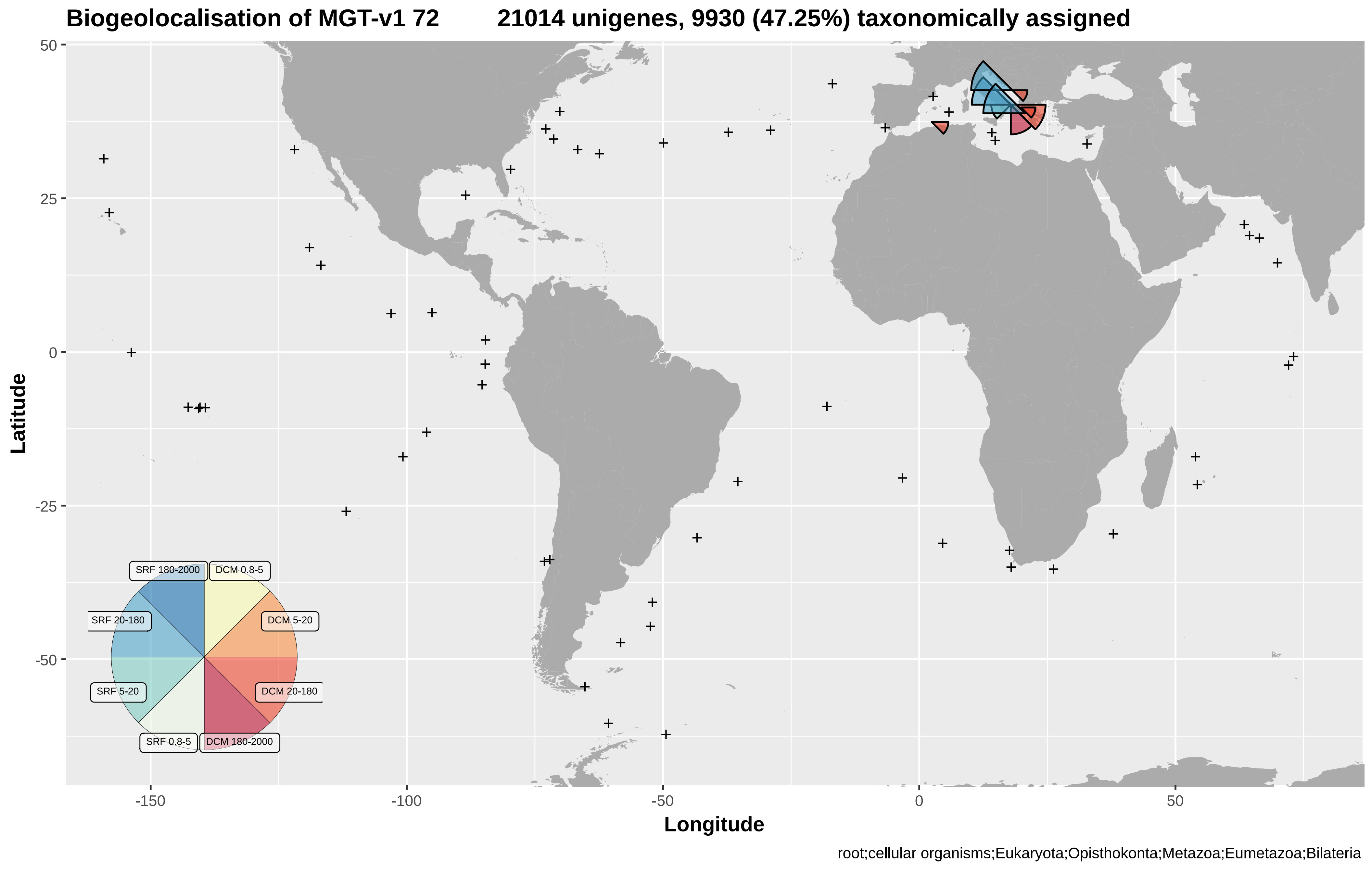
Biogeolocalisation of MGT-v1 72 21014 unigenes, 9930 (47.25%) taxonomically assigned
50
25
0
-25
-50
Latitude
-150
-100
-50
0
50
Longitude
SRF 180-2000
DCM 0.8-5
SRF 20-180
DCM 5-20
SRF 5-20
DCM 20-180
SRF 0.8-5
DCM 180-2000
root;cellular organisms;Eukaryota;Opisthokonta;Metazoa;Eumetazoa;Bilateria

#### Biogeolocalisation of MGT-v1 73 20936 unigenes, 1600 (7.64%) taxonomically assigned

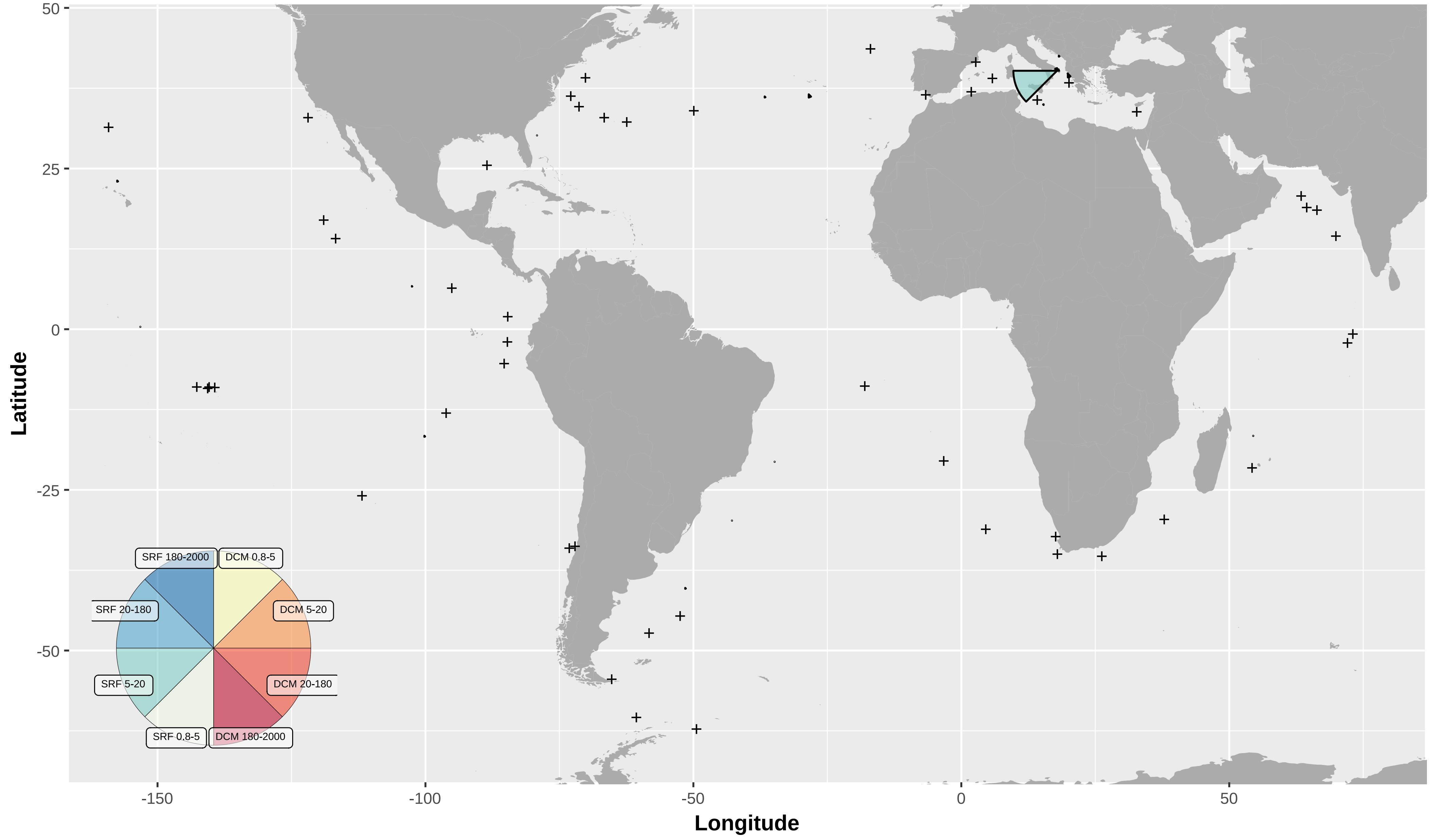


root;cellular organisms;Eukaryota;Opisthokonta;Metazoa;Eumetazoa

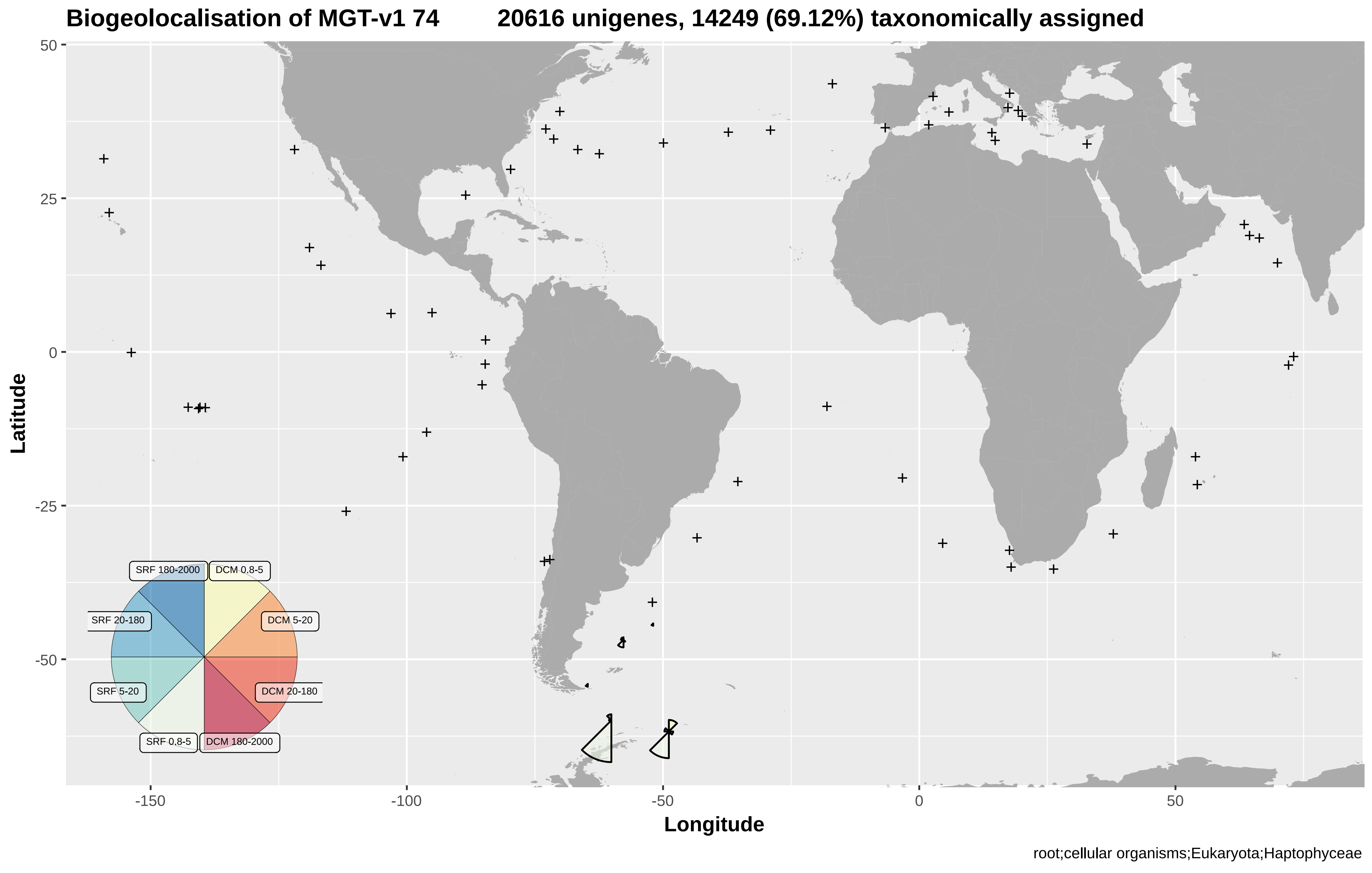

Biogeolocalisation of MGT-v1 74 20616 unigenes, 14249 (69.12%) taxonomically assigned
Latitude
Longitude
50
25
0
-25
-50
-150
-100
-50
0
50
SRF 180-2000
DCM 0.8-5
SRF 20-180
DCM 5-20
SRF 5-20
DCM 20-180
SRF 0.8-5
DCM 180-2000
root;cellular organisms;Eukaryota;Haptophyceae

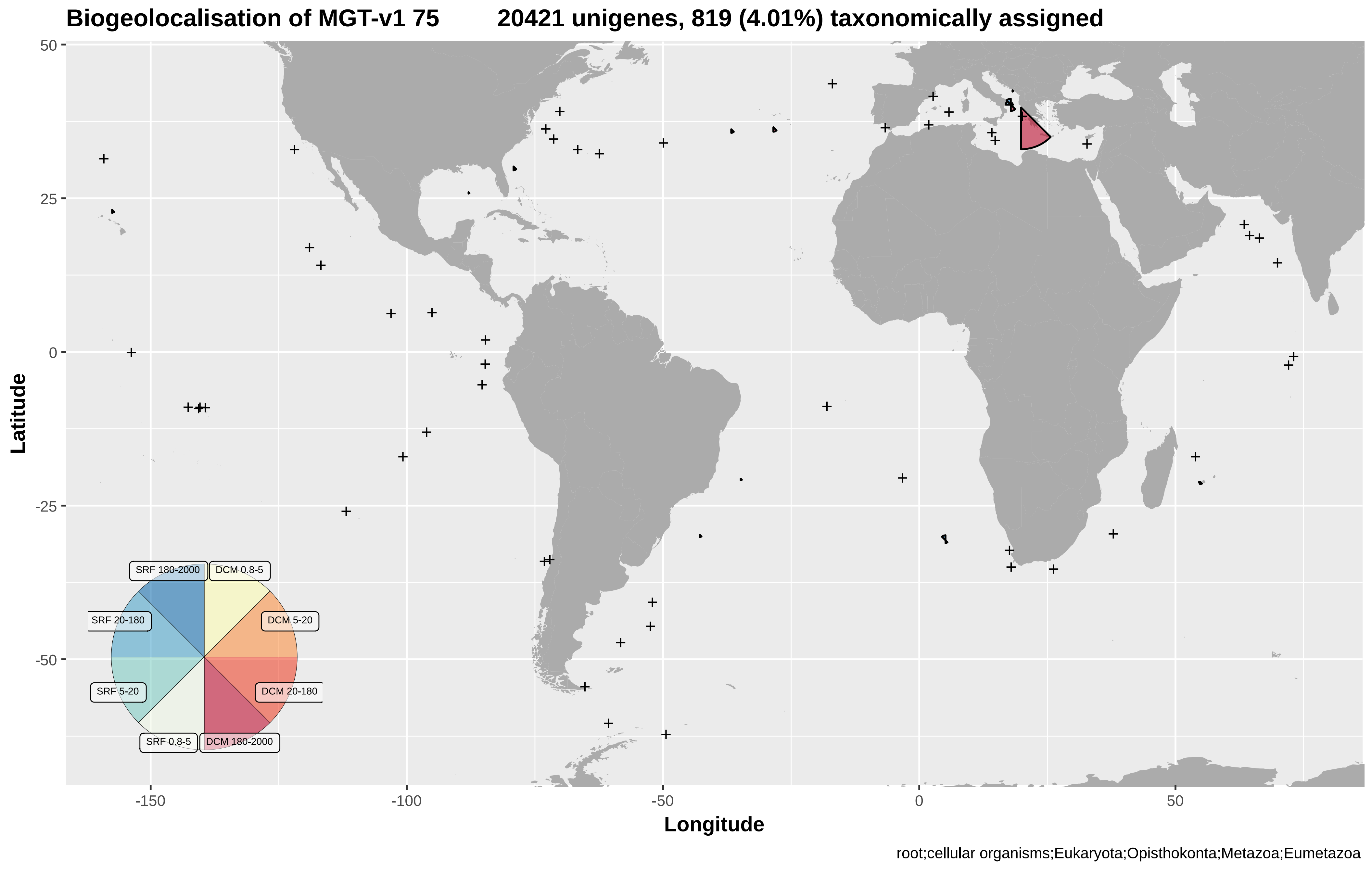

Biogeolocalisation of MGT-v1 75
20421 unigenes, 819 (4.01%) taxonomically assigned
Latitude
Longitude
50
25
0
-25
-50
-150
-100
-50
0
50
SRF 180-2000
DCM 0.8-5
SRF 20-180
DCM 5-20
SRF 5-20
DCM 20-180
SRF 0.8-5
DCM 180-2000
root;cellular organisms;Eukaryota;Opisthokonta;Metazoa;Eumetazoa

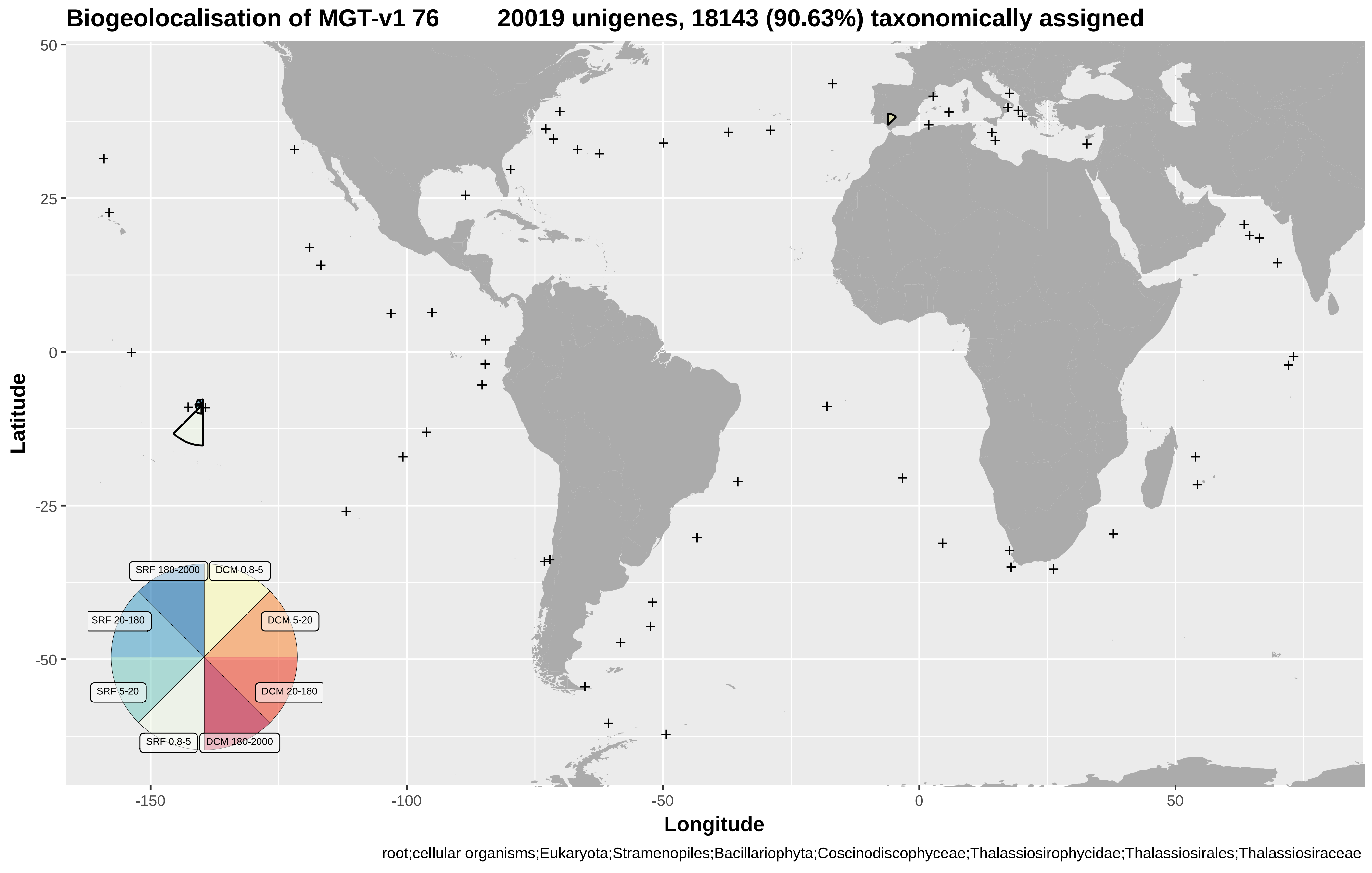

Biogeolocalisation of MGT-v1 76
20019 unigenes, 18143 (90.63%) taxonomically assigned
Latitude
Longitude
50
25
0
-25
-50
-150
-100
-50
0
50
SRF 180-2000
DCM 0.8-5
SRF 20-180
DCM 5-20
SRF 5-20
DCM 20-180
SRF 0.8-5
DCM 180-2000
root;cellular organisms;Eukaryota;Stramenopiles;Bacillariophyta;Coscinodiscophyceae;Thalassiosirophycidae;Thalassiosirales;Thalassiosiraceae

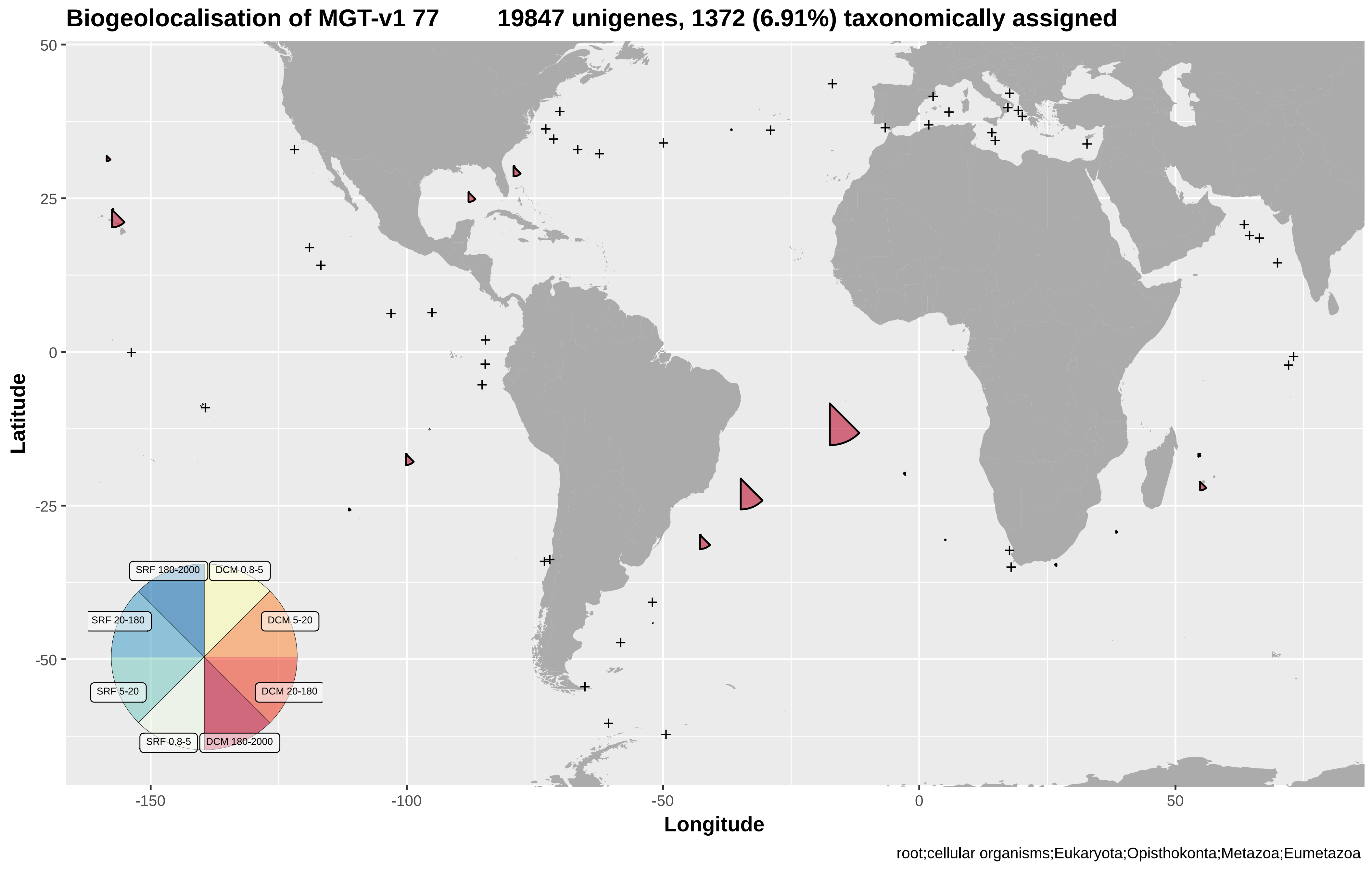

Biogeolocalisation of MGT-v1 77 19847 unigenes, 1372 (6.91%) taxonomically assigned
Latitude
Longitude
50
25
0
-25
-50
-150
-100
-50
0
50
SRF 180-2000
DCM 0.8-5
SRF 20-180
DCM 5-20
SRF 5-20
DCM 20-180
SRF 0.8-5
DCM 180-2000
root;cellular organisms;Eukaryota;Opisthokonta;Metazoa;Eumetazoa

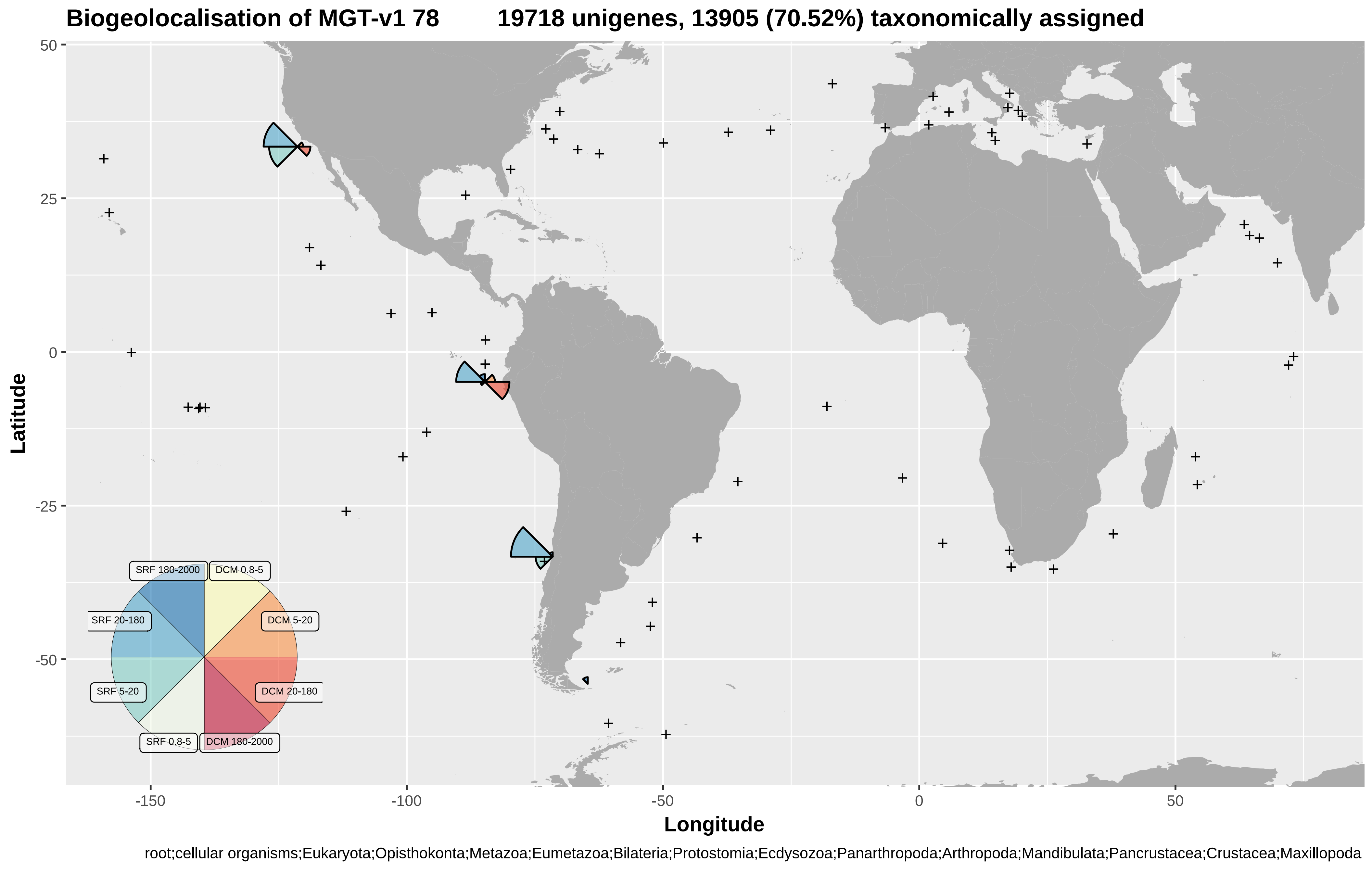
Biogeolocalisation of MGT-v1 78
19718 unigenes, 13905 (70.52%) taxonomically assigned
Latitude
Longitude
50
25
0
-25
-50
-150
-100
-50
0
50
SRF 180-2000
DCM 0.8-5
SRF 20-180
DCM 5-20
SRF 5-20
DCM 20-180
SRF 0.8-5
DCM 180-2000
root;cellular organisms;Eukaryota;Opisthokonta;Metazoa;Eumetazoa;Bilateria;Protostomia;Ecdysozoa;Panarthropoda;Arthropoda;Mandibulata;Pancrustacea;Crustacea;Maxillopoda

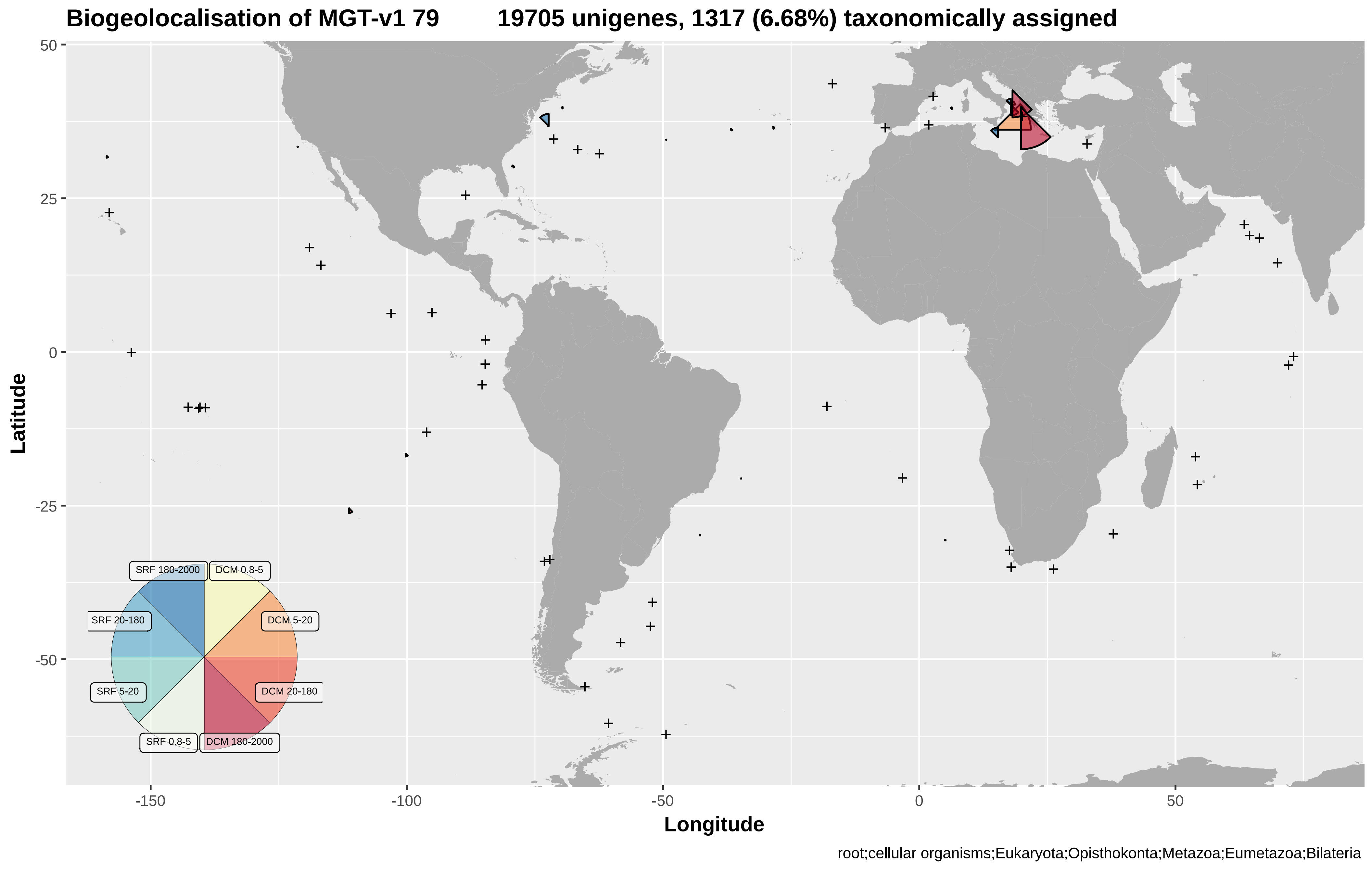
Biogeolocalisation of MGT-v1 79
19705 unigenes, 1317 (6.68%) taxonomically assigned
Latitude
Longitude
50
25
0
-25
-50
-150
-100
-50
0
50
SRF 180-2000
DCM 0.8-5
SRF 20-180
DCM 5-20
SRF 5-20
DCM 20-180
SRF 0.8-5
DCM 180-2000
root;cellular organisms;Eukaryota;Opisthokonta;Metazoa;Eumetazoa;Bilateria

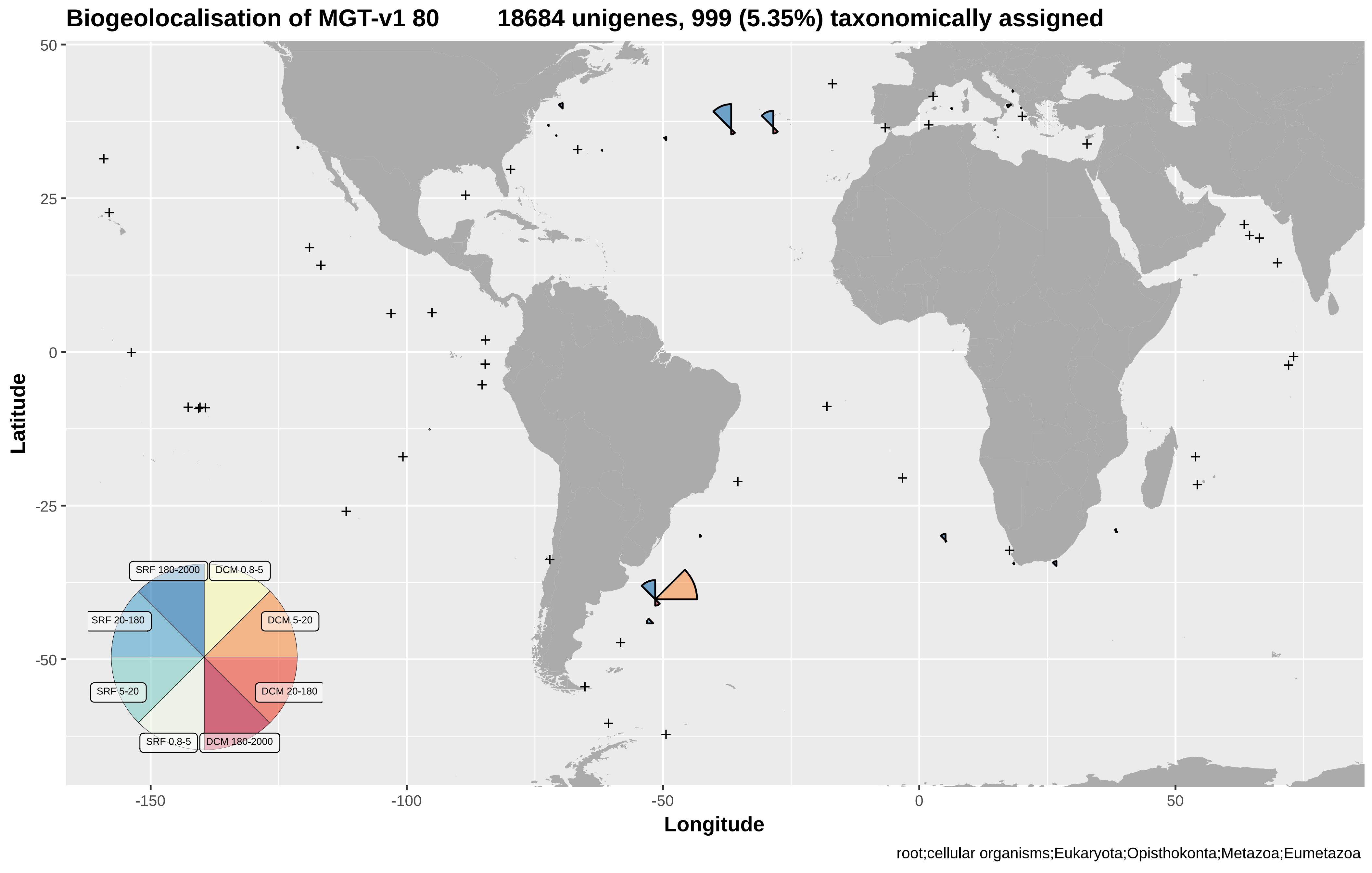
Biogeolocalisation of MGT-v1 80
18684 unigenes, 999 (5.35%) taxonomically assigned
Latitude
Longitude
50
25
0
-25
-50
-150
-100
-50
0
50
SRF 180-2000
DCM 0.8-5
SRF 20-180
DCM 5-20
SRF 5-20
DCM 20-180
SRF 0.8-5
DCM 180-2000
root;cellular organisms;Eukaryota;Opisthokonta;Metazoa;Eumetazoa

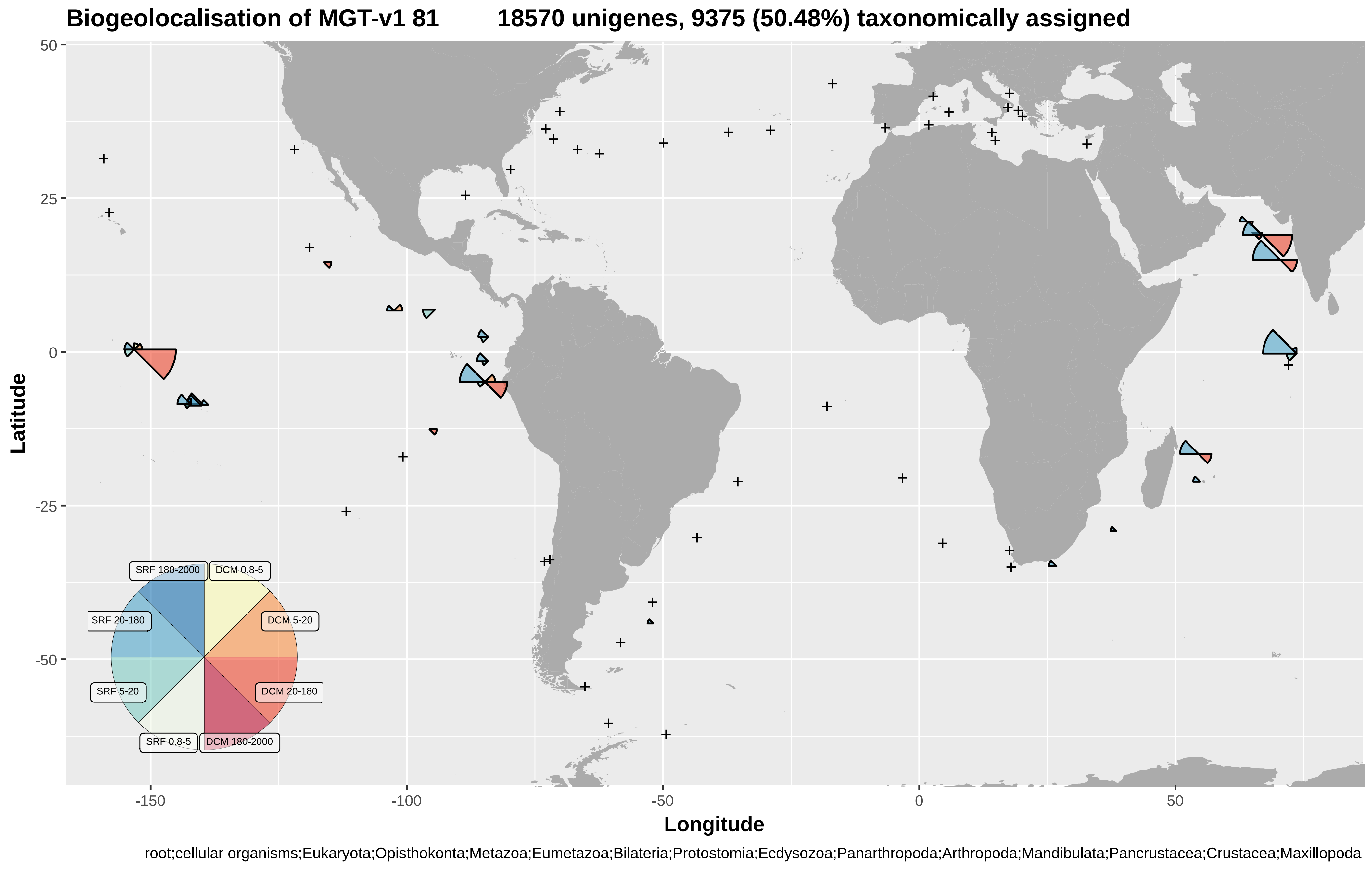

Biogeolocalisation of MGT-v1 81 18570 unigenes, 9375 (50.48%) taxonomically assigned
50
25
0
-25
-50
Latitude
-150
-100
-50
0
50
Longitude
SRF 180-2000
DCM 0.8-5
SRF 20-180
DCM 5-20
SRF 5-20
DCM 20-180
SRF 0.8-5
DCM 180-2000
root;cellular organisms;Eukaryota;Opisthokonta;Metazoa;Eumetazoa;Bilateria;Protostomia;Ecdysozoa;Panarthropoda;Arthropoda;Mandibulata;Pancrustacea;Crustacea;Maxillopoda

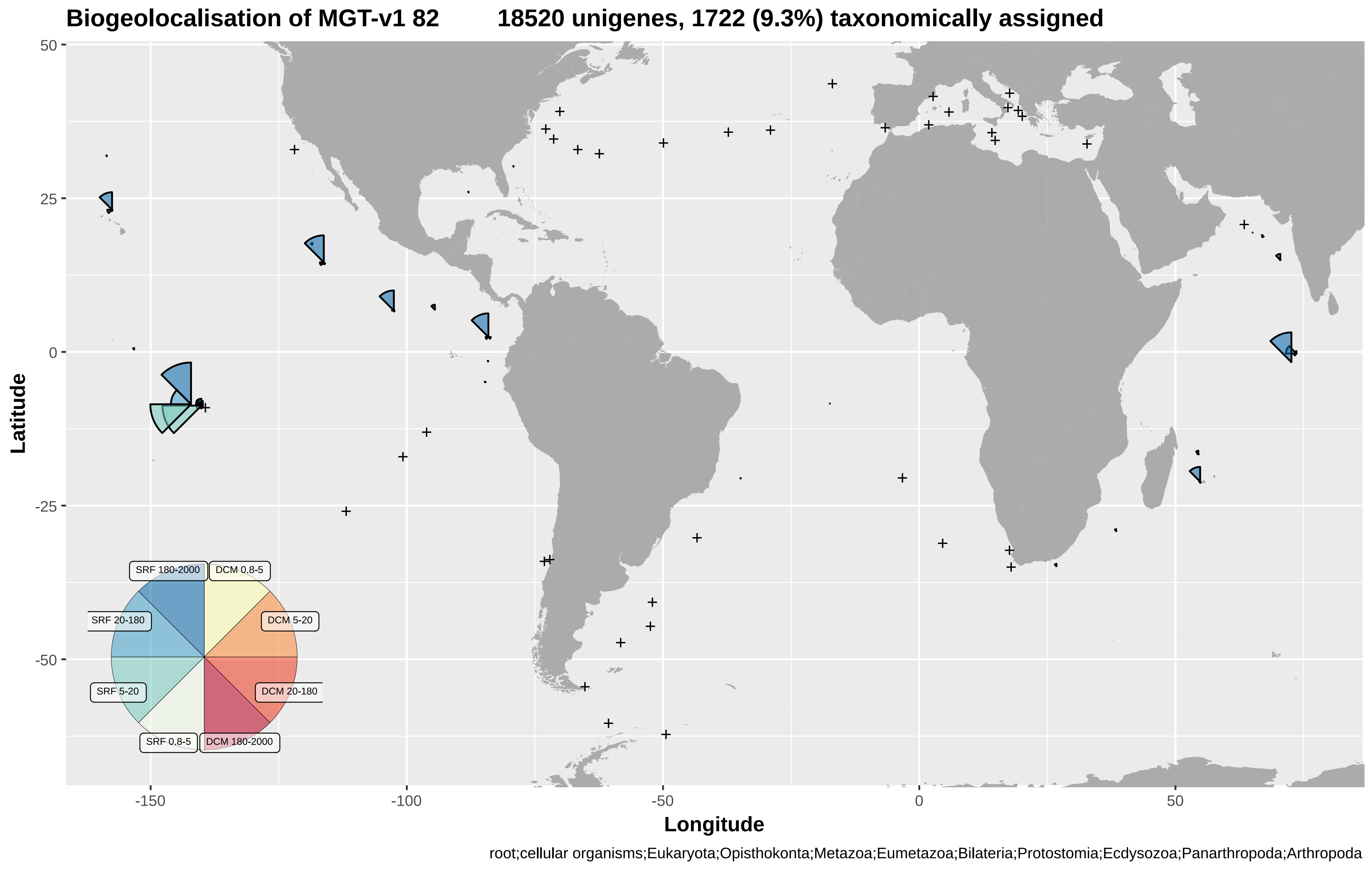
Biogeolocalisation of MGT-v1 82 18520 unigenes, 1722 (9.3%) taxonomically assigned
50
25
0
-25
-50
Latitude
-150
-100
-50
0
50
Longitude
SRF 180-2000
DCM 0.8-5
SRF 20-180
DCM 5-20
SRF 5-20
DCM 20-180
SRF 0.8-5
DCM 180-2000
root;cellular organisms;Eukaryota;Opisthokonta;Metazoa;Eumetazoa;Bilateria;Protostomia;Ecdysozoa;Panarthropoda;Arthropoda

# Biogeolocalisation of MGT-v1 83        18203 unigenes, 1900 (10.44%) taxonomically assigned

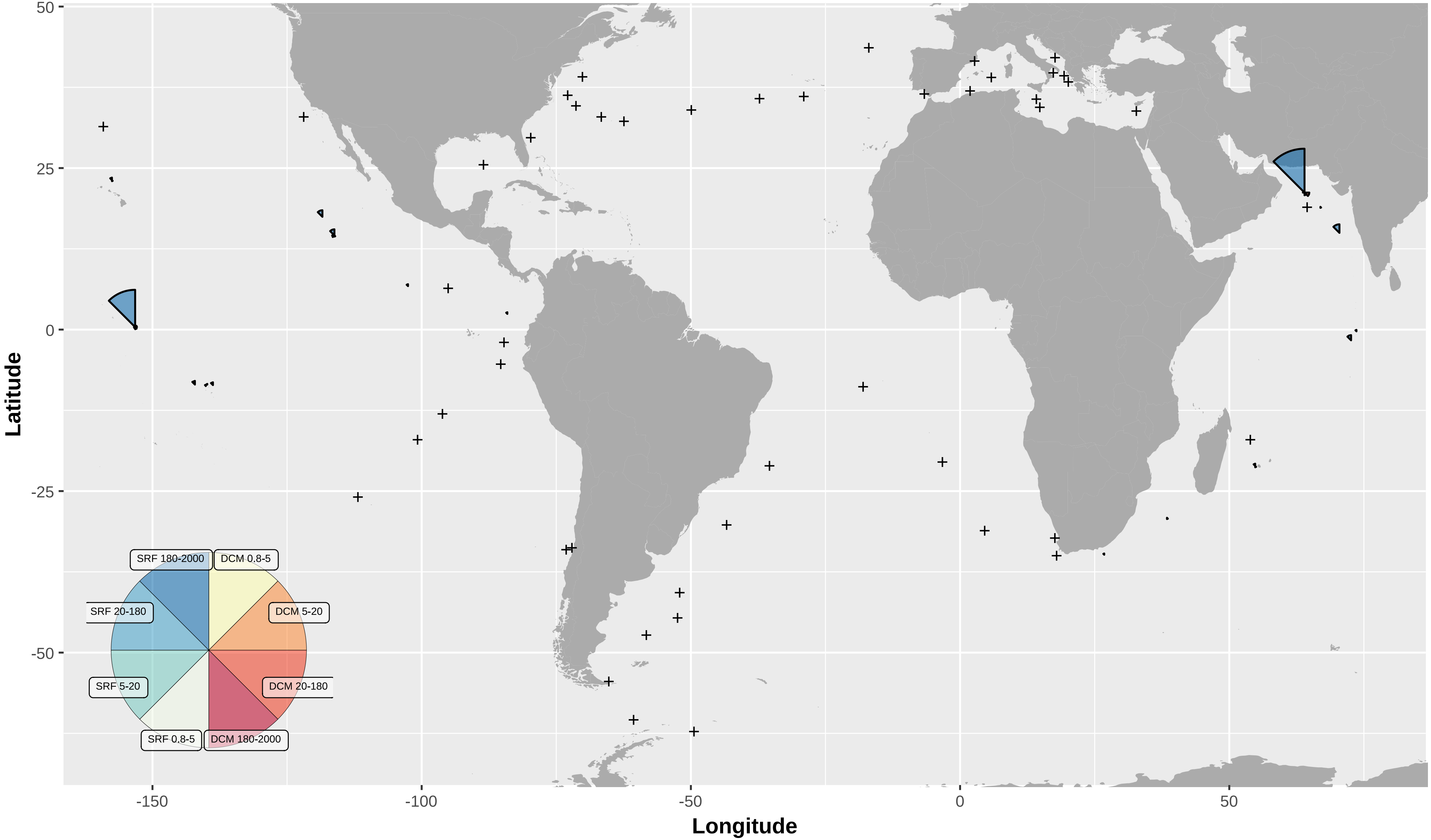


root;cellular organisms;Eukaryota;Opisthokonta;Metazoa;Eumetazoa;Bilateria

### Biogeolocalisation of MGT-v1 84 18092 unigenes, 17567 (97.1%) taxonomically assigned

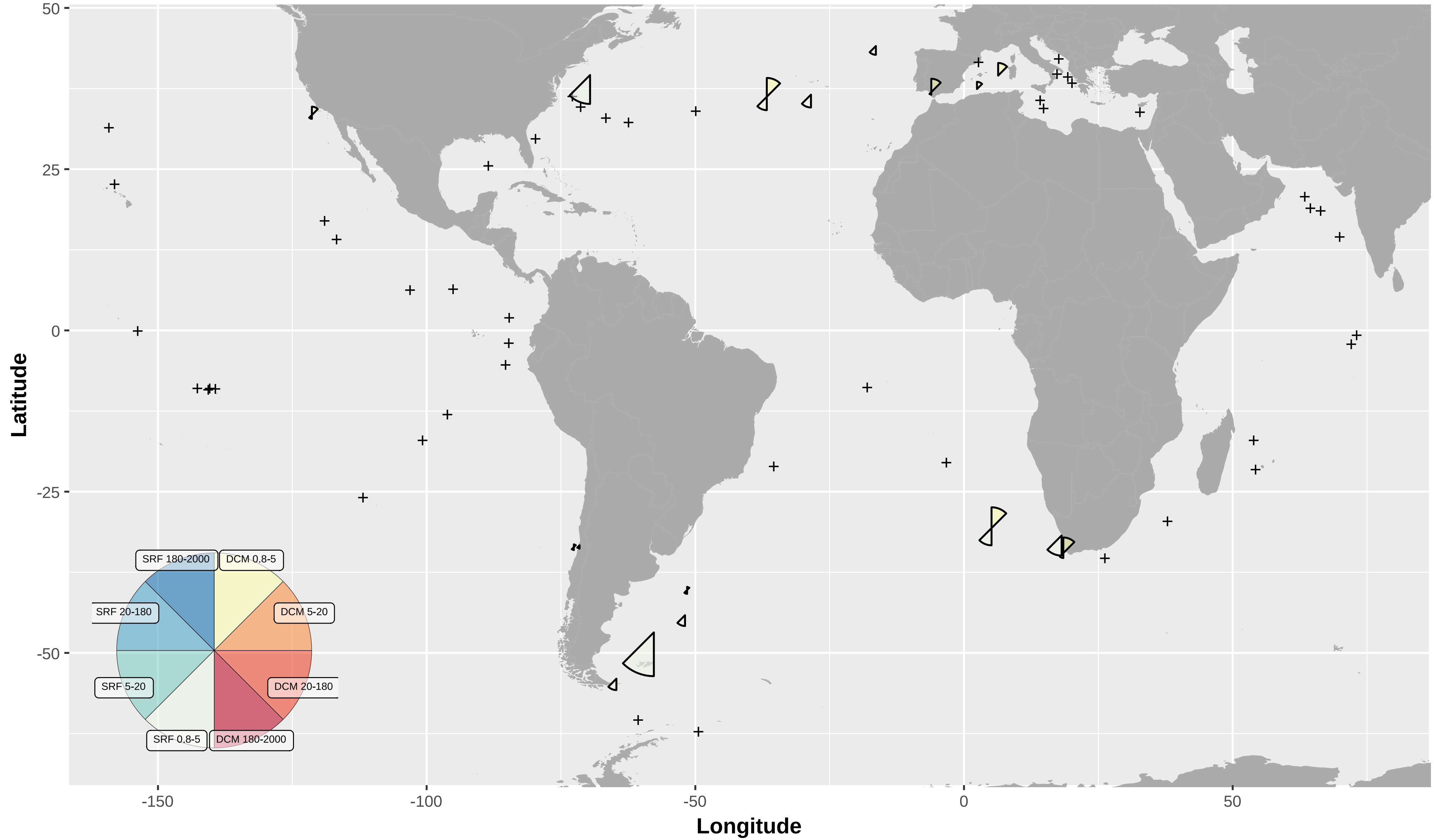


root;cellular organisms;Eukaryota;Viridiplantae;Chlorophyta;prasinophytes;Mamiellophyceae;Mamiellales;Mamiellaceae

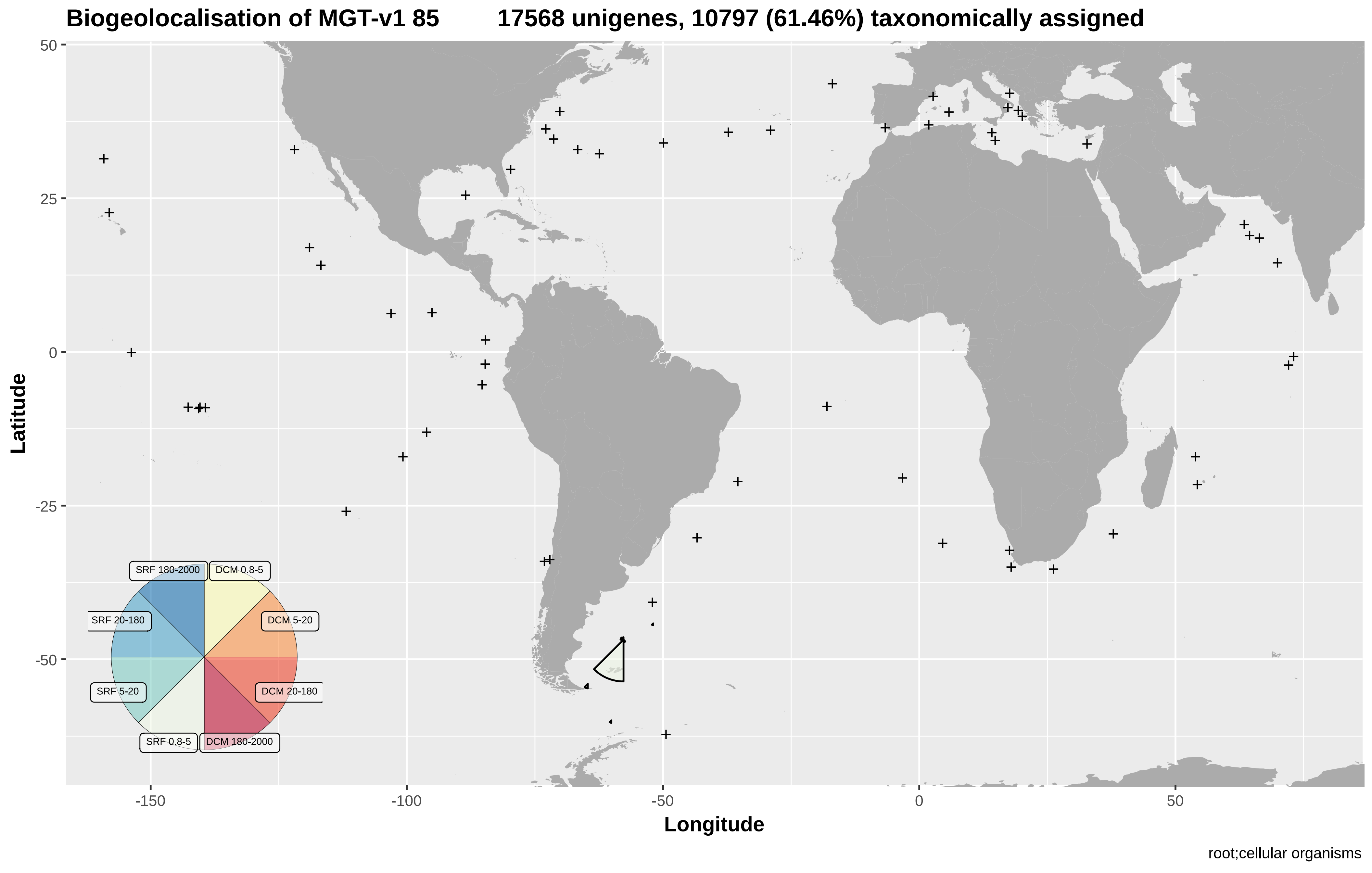

Biogeolocalisation of MGT-v1 85
17568 unigenes, 10797 (61.46%) taxonomically assigned
50
25
0
-25
-50
Latitude
-150
-100
-50
0
50
Longitude
SRF 180-2000
DCM 0.8-5
SRF 20-180
DCM 5-20
SRF 5-20
DCM 20-180
SRF 0.8-5
DCM 180-2000
root;cellular organisms

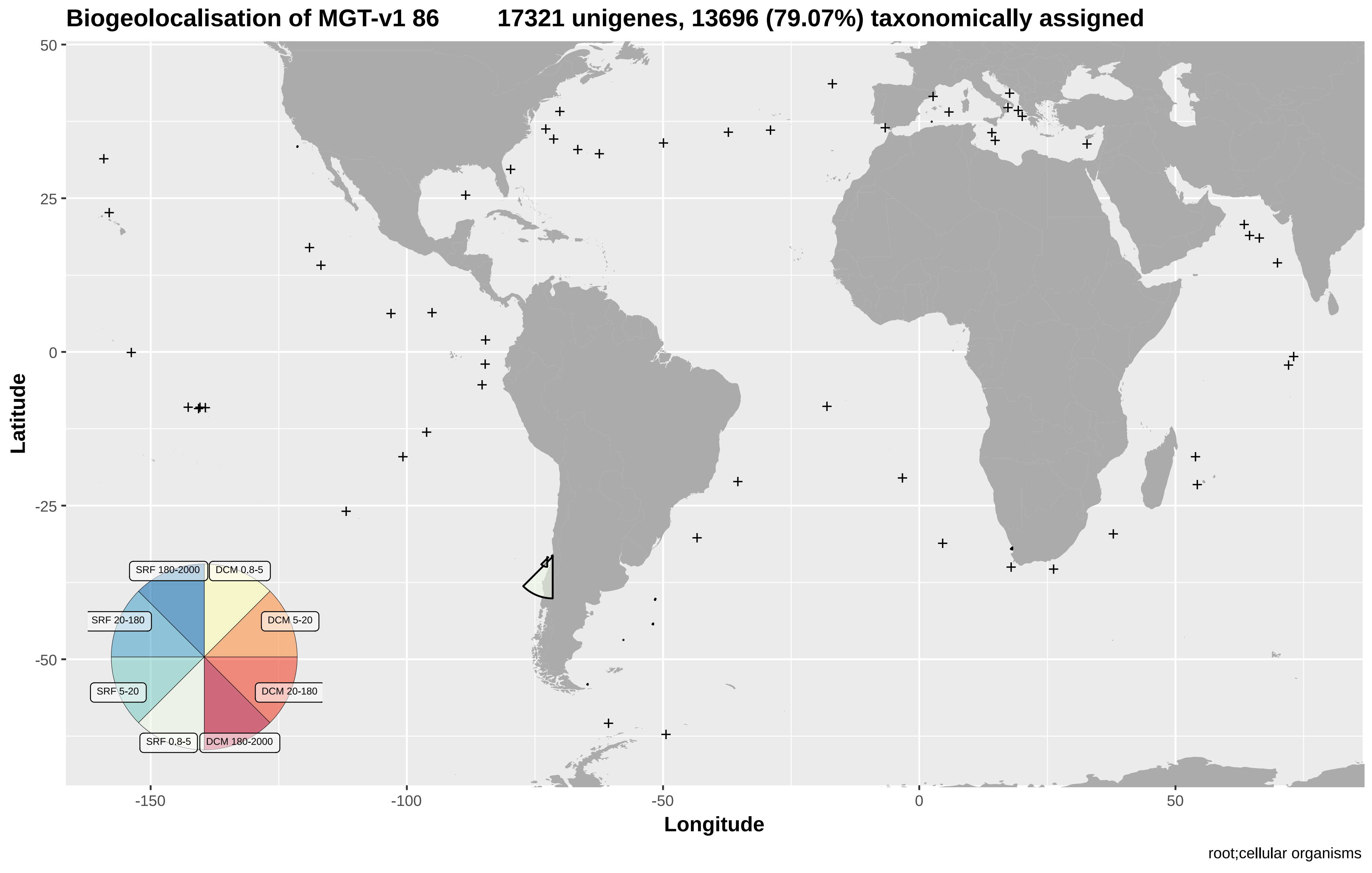

Biogeolocalisation of MGT-v1 86
17321 unigenes, 13696 (79.07%) taxonomically assigned
50
25
0
-25
-50
Latitude
-150
-100
-50
0
50
Longitude
SRF 180-2000
DCM 0.8-5
SRF 20-180
DCM 5-20
SRF 5-20
DCM 20-180
SRF 0.8-5
DCM 180-2000
root;cellular organisms

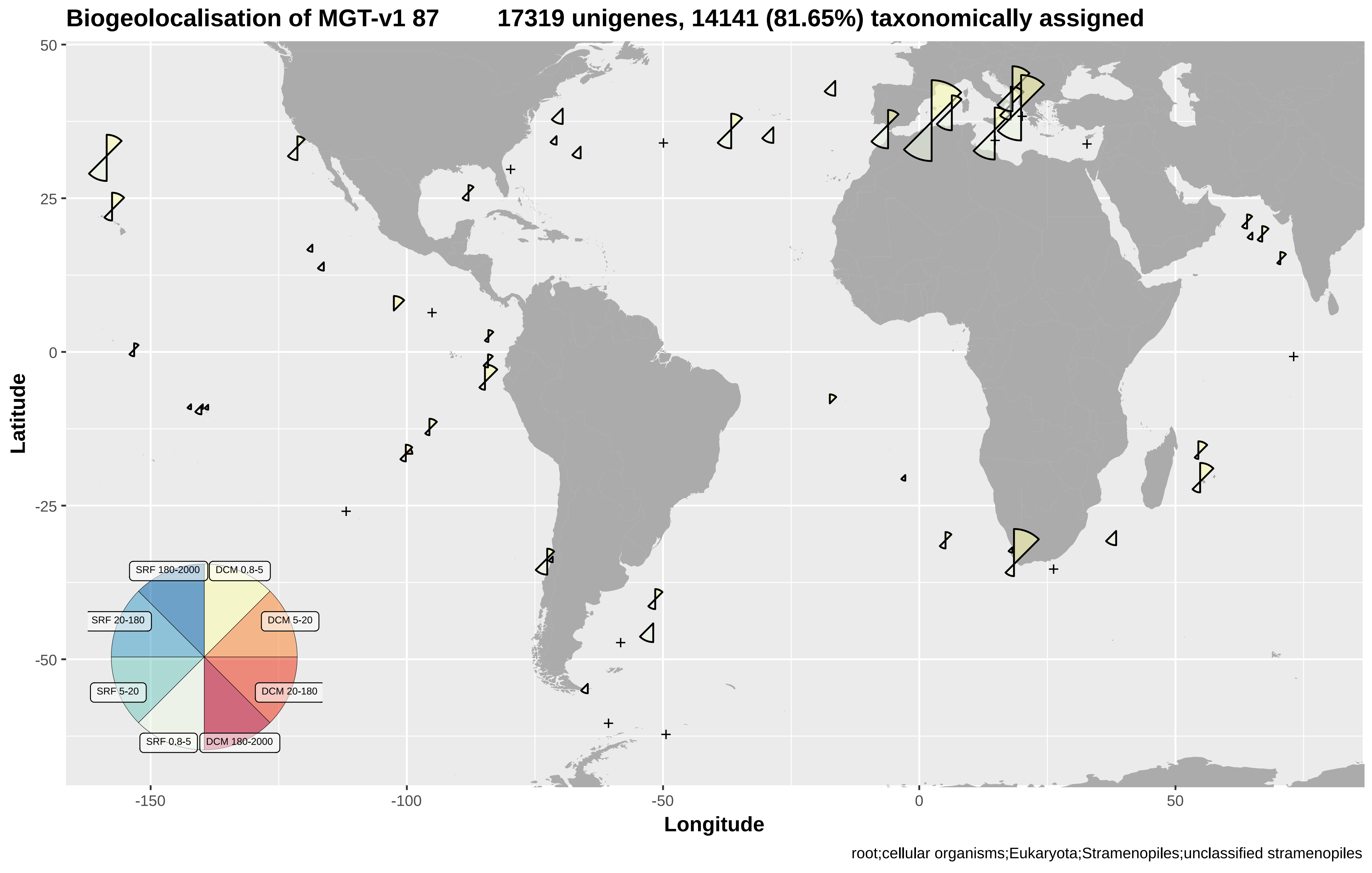
Biogeolocalisation of MGT-v1 87
17319 unigenes, 14141 (81.65%) taxonomically assigned
50
25
0
-25
-50
Latitude
-150
-100
-50
0
50
Longitude
SRF 180-2000
DCM 0.8-5
SRF 20-180
DCM 5-20
SRF 5-20
DCM 20-180
SRF 0.8-5
DCM 180-2000
root;cellular organisms;Eukaryota;Stramenopiles;unclassified stramenopiles

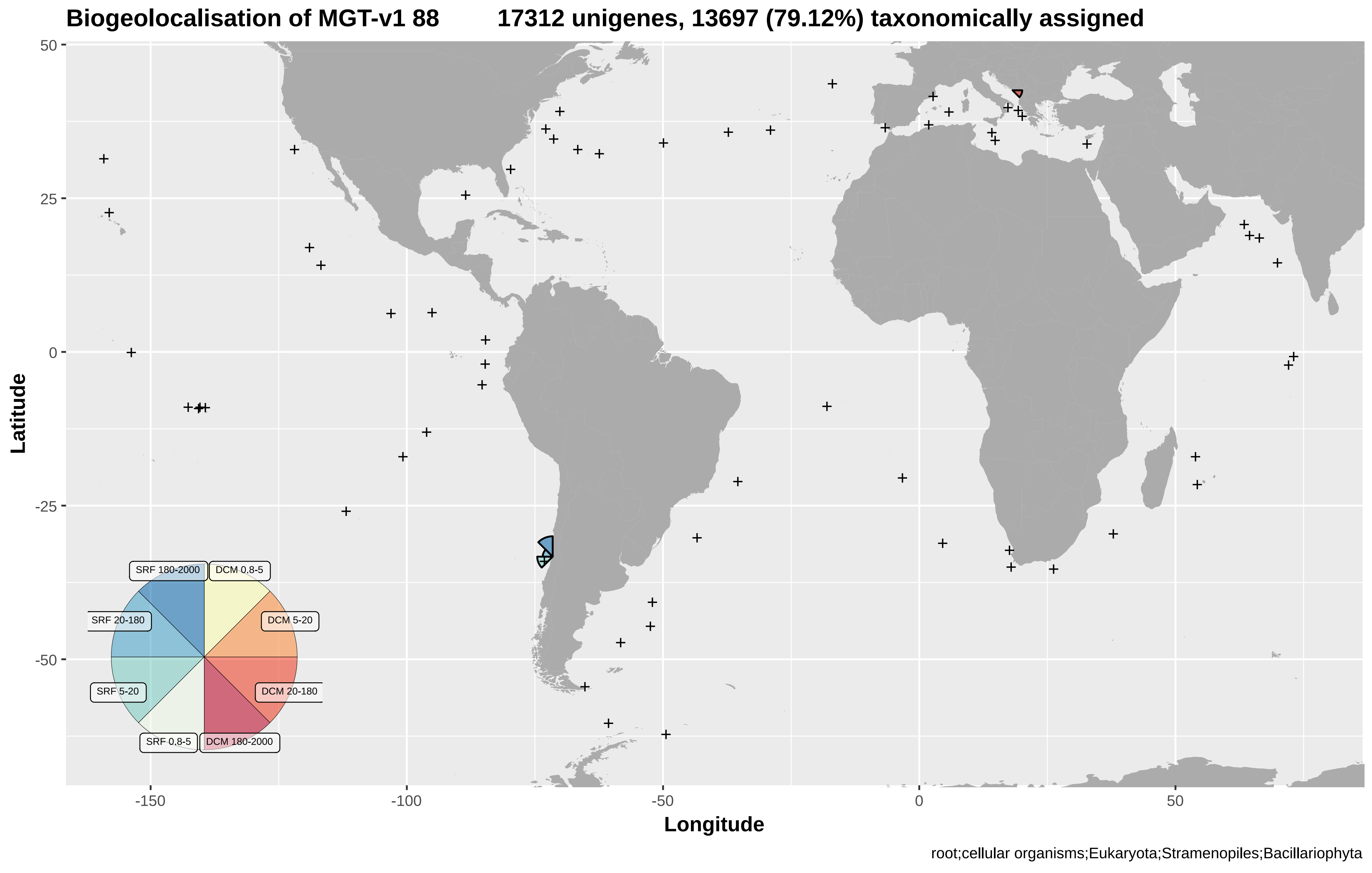

Biogeolocalisation of MGT-v1 88
17312 unigenes, 13697 (79.12%) taxonomically assigned
Latitude
Longitude
SRF 180-2000
DCM 0.8-5
SRF 20-180
DCM 5-20
SRF 5-20
DCM 20-180
SRF 0.8-5
DCM 180-2000
root;cellular organisms;Eukaryota;Stramenopiles;Bacillariophyta

# Biogeolocalisation of MGT-v1 89        17308 unigenes, 9914 (57.28%) taxonomically assigned

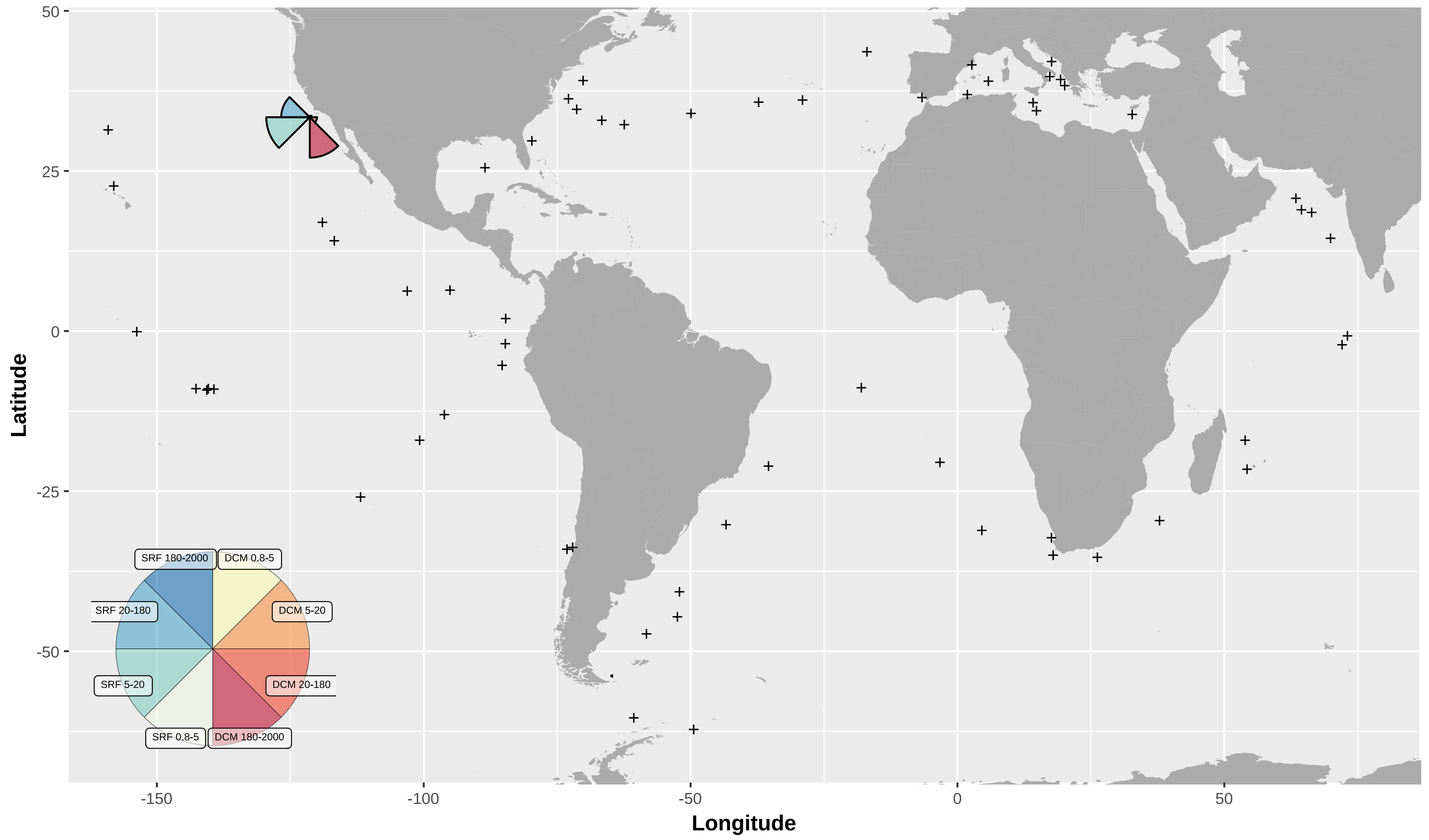


aryota;Opisthokonta;Metazoa;Eumetazoa;Bilateria;Protostomia;Ecdysozoa;Panarthropoda;Arthropoda;Mandibulata;Pancrustacea;Crustacea;Maxillopoda;Copepoda;Neocopepoda;Gymnoplea;Calanoida

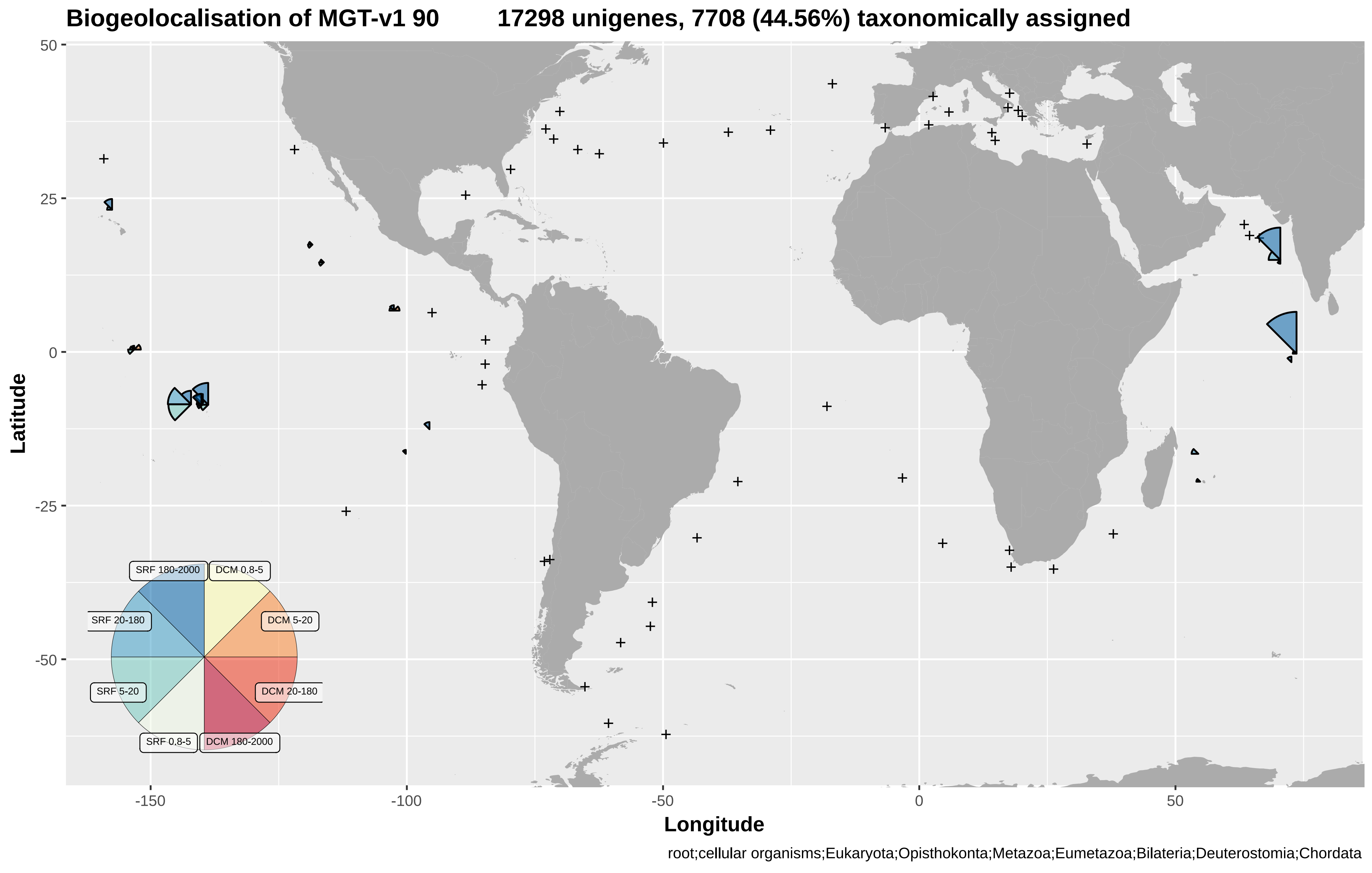
Biogeolocalisation of MGT-v1 90 17298 unigenes, 7708 (44.56%) taxonomically assigned
Latitude
Longitude
50
25
0
-25
-50
-150
-100
-50
0
50
SRF 180-2000
DCM 0.8-5
SRF 20-180
DCM 5-20
SRF 5-20
DCM 20-180
SRF 0.8-5
DCM 180-2000
root;cellular organisms;Eukaryota;Opisthokonta;Metazoa;Eumetazoa;Bilateria;Deuterostomia;Chordata

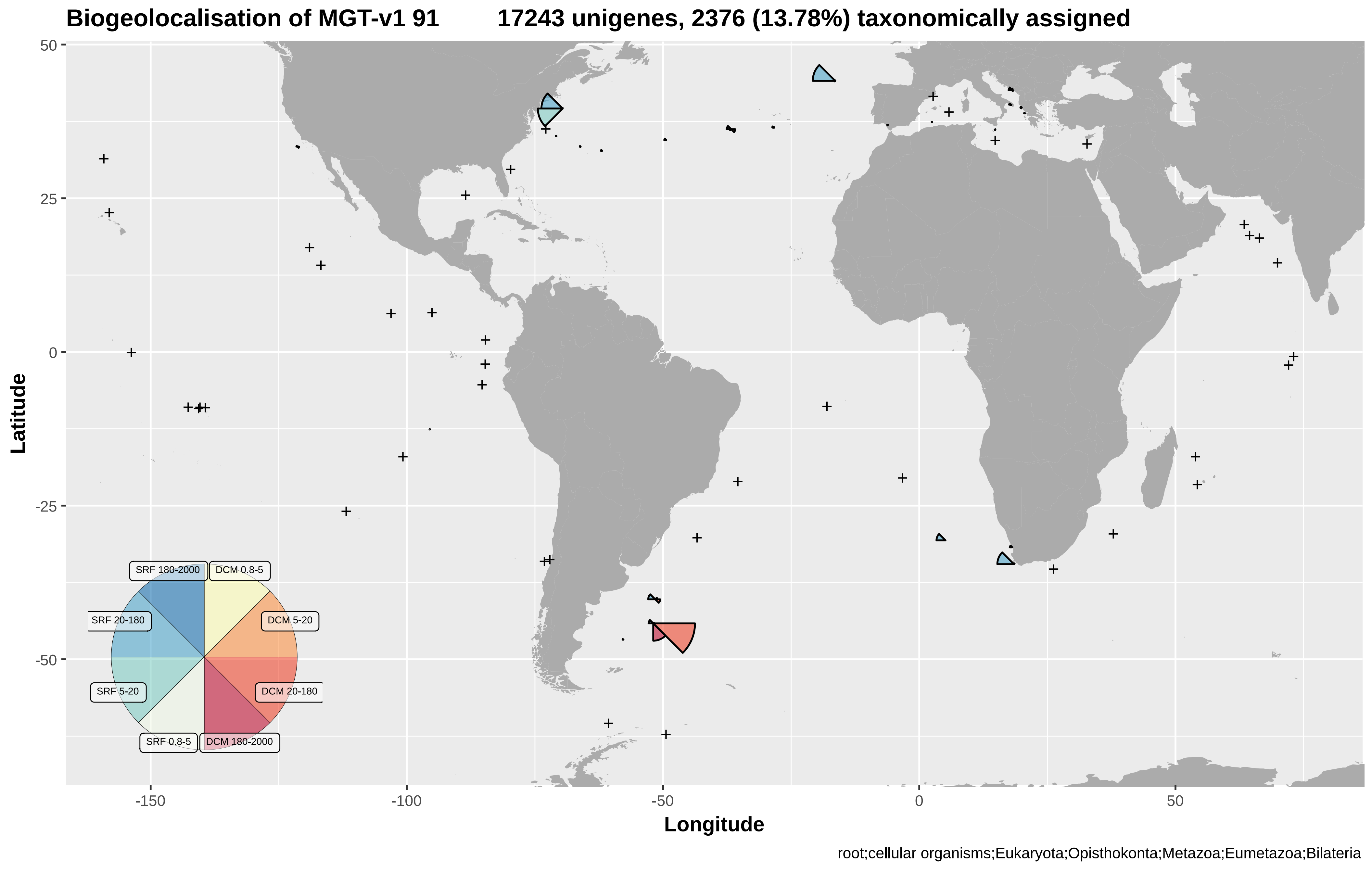

Biogeolocalisation of MGT-v1 91
17243 unigenes, 2376 (13.78%) taxonomically assigned
50
25
0
-25
-50
Latitude
-150
-100
-50
0
50
Longitude
SRF 180-2000
DCM 0.8-5
SRF 20-180
DCM 5-20
SRF 5-20
DCM 20-180
SRF 0.8-5
DCM 180-2000
root;cellular organisms;Eukaryota;Opisthokonta;Metazoa;Eumetazoa;Bilateria

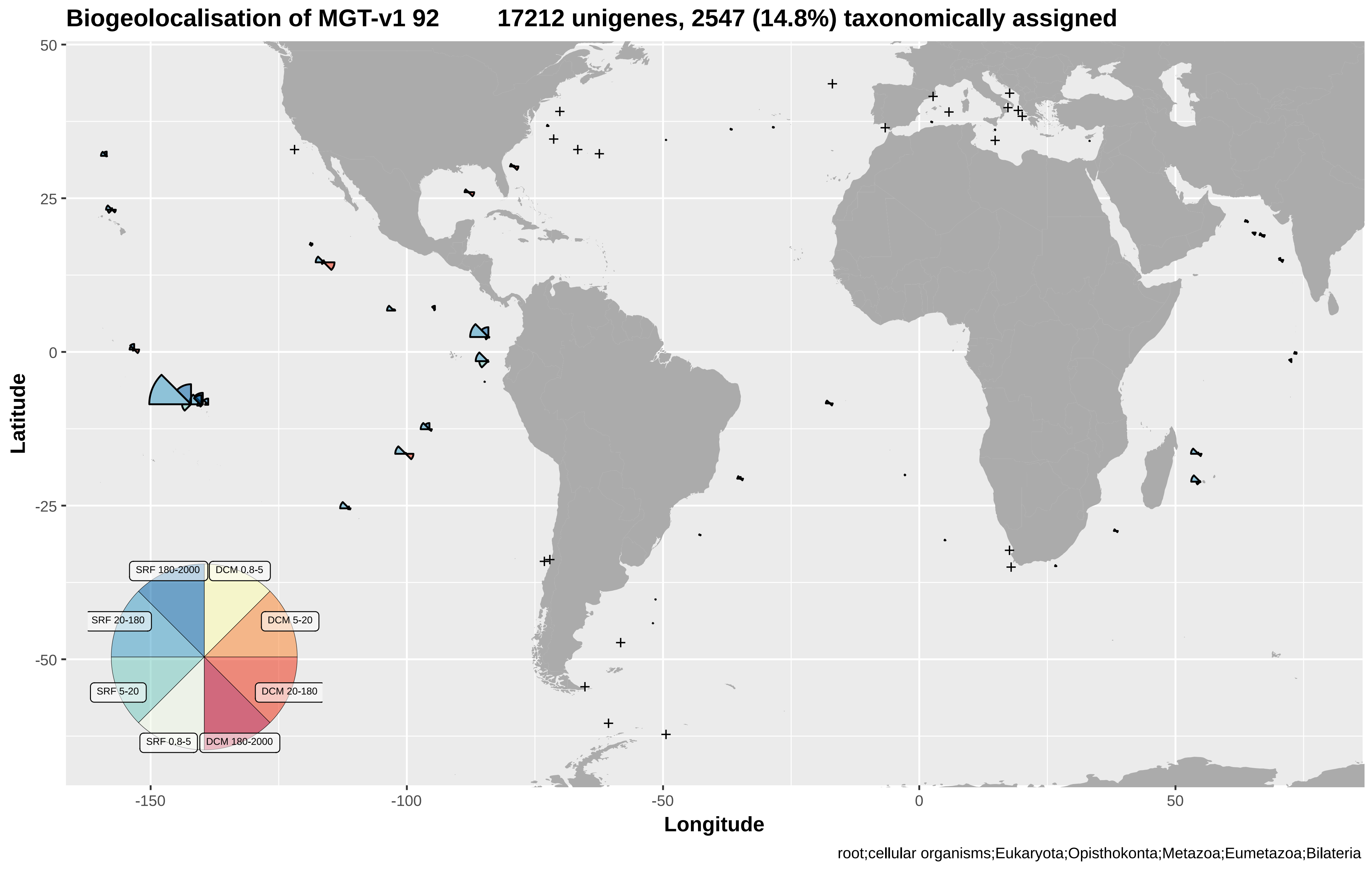
Biogeolocalisation of MGT-v1 92 17212 unigenes, 2547 (14.8%) taxonomically assigned
Latitude
Longitude
50
25
0
-25
-50
-150
-100
-50
0
50
SRF 180-2000
DCM 0.8-5
SRF 20-180
DCM 5-20
SRF 5-20
DCM 20-180
SRF 0.8-5
DCM 180-2000
root;cellular organisms;Eukaryota;Opisthokonta;Metazoa;Eumetazoa;Bilateria

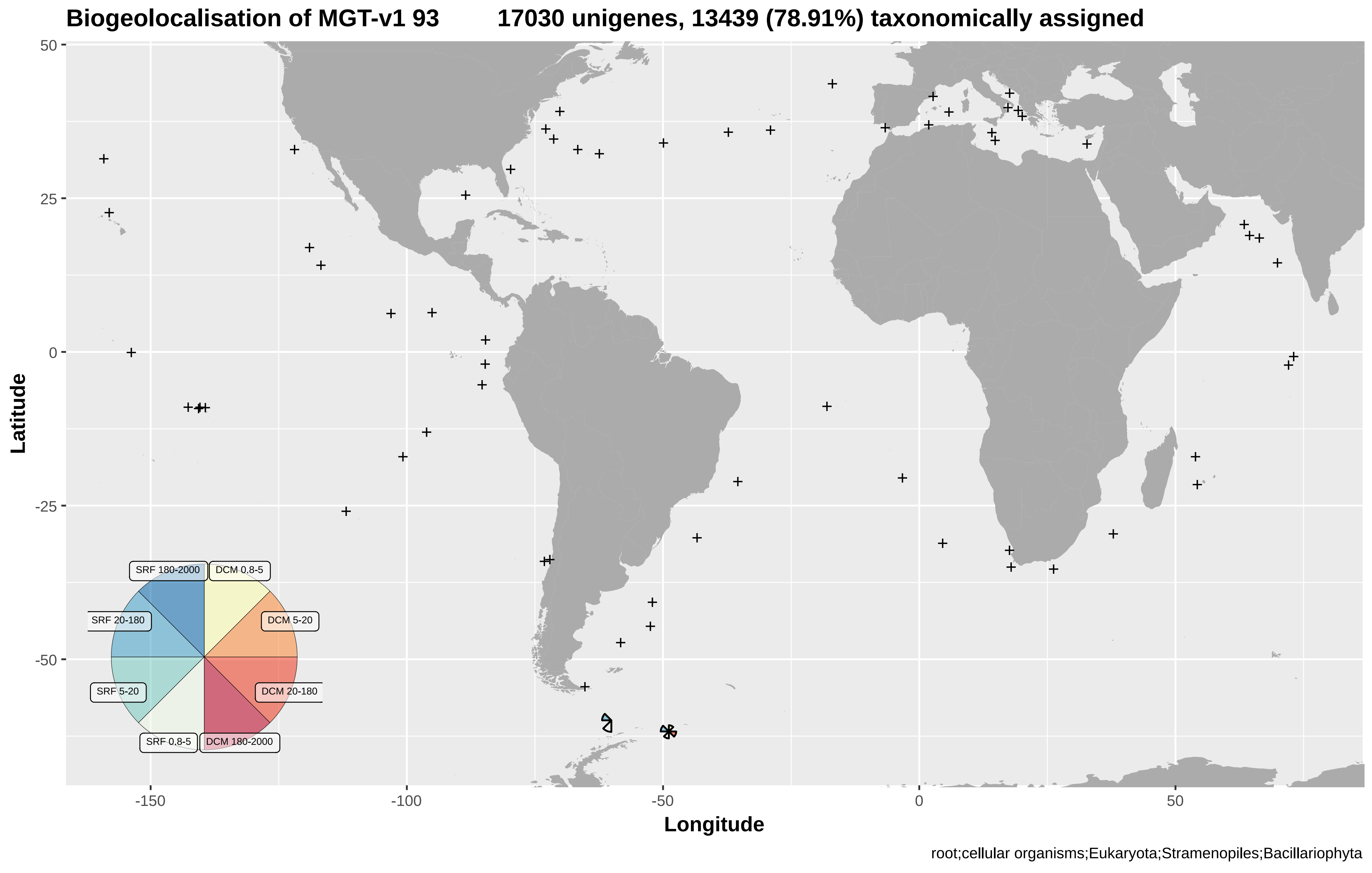
Biogeolocalisation of MGT-v1 93
17030 unigenes, 13439 (78.91%) taxonomically assigned
50
25
0
-25
-50
Latitude
-150
-100
-50
0
50
Longitude
SRF 180-2000
DCM 0.8-5
SRF 20-180
DCM 5-20
SRF 5-20
DCM 20-180
SRF 0.8-5
DCM 180-2000
root;cellular organisms;Eukaryota;Stramenopiles;Bacillariophyta

# Biogeolocalisation of MGT-v1 94        16921 unigenes, 10774 (63.67%) taxonomically assigned

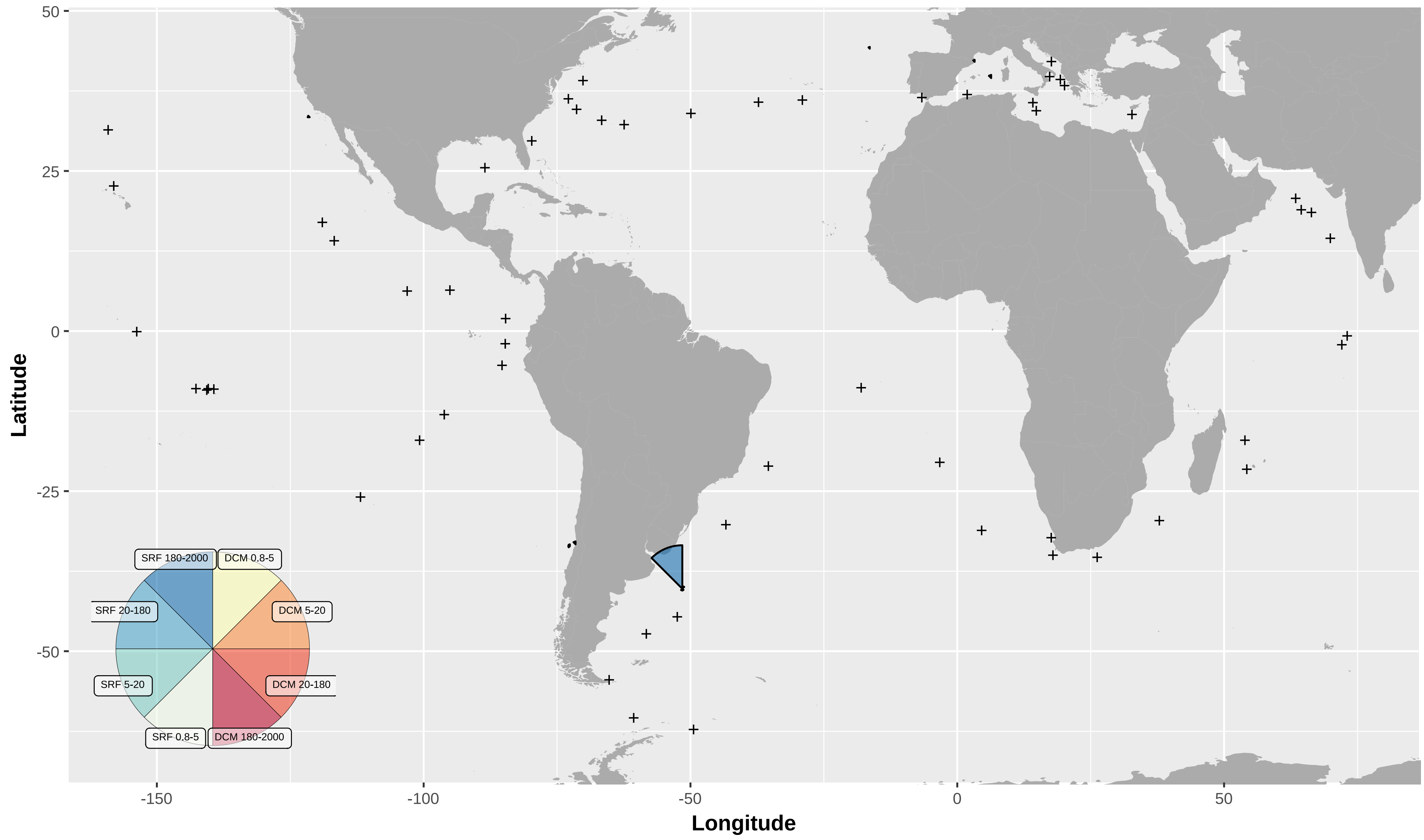


root;cellular organisms;Eukaryota;Opisthokonta;Metazoa;Eumetazoa;Bilateria

Biogeolocalisation of MGT-v1 95        16663 unigenes, 14010 (84.08%) taxonomically assigned

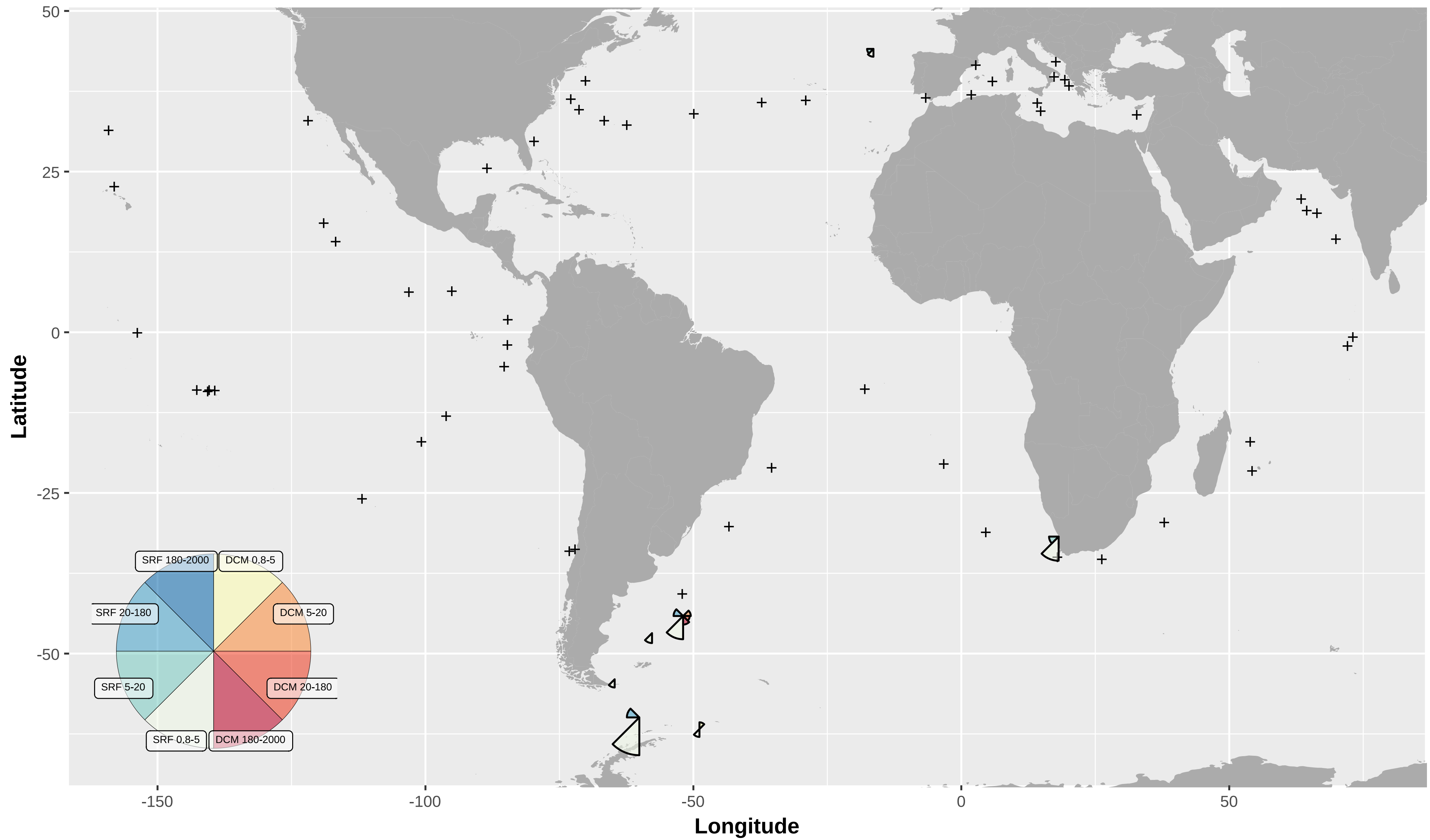


root;cellular organisms;Eukaryota;Stramenopiles;Bacillariophyta;Bacillariophyceae

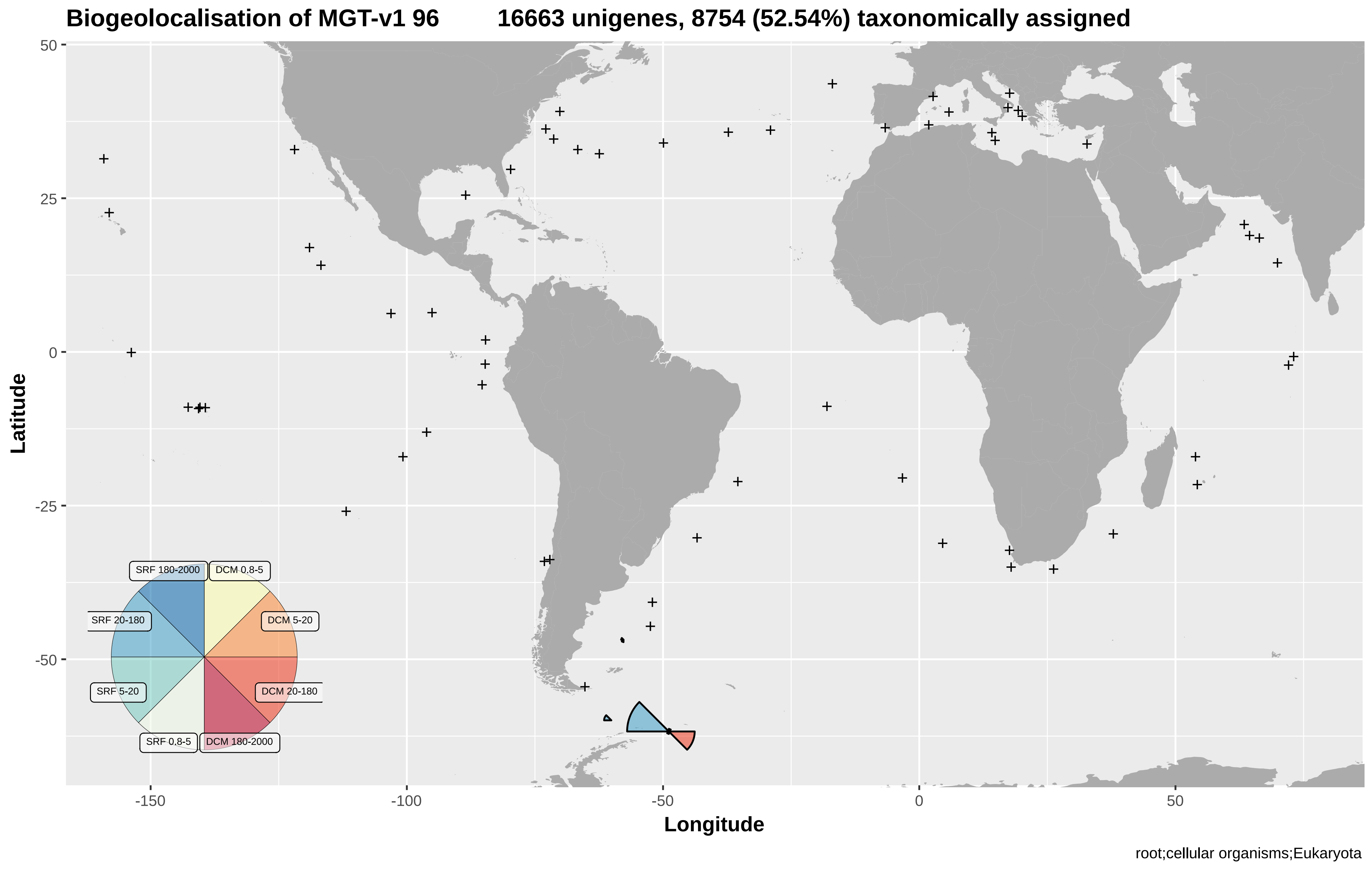
Biogeolocalisation of MGT-v1 96 16663 unigenes, 8754 (52.54%) taxonomically assigned
Latitude
Longitude
50
25
0
-25
-50
-150
-100
-50
0
50
SRF 180-2000
DCM 0.8-5
SRF 20-180
DCM 5-20
SRF 5-20
DCM 20-180
SRF 0.8-5
DCM 180-2000
root;cellular organisms;Eukaryota

## Biogeolocalisation of MGT-v1 97        16619 unigenes, 9608 (57.81%) taxonomically assigned

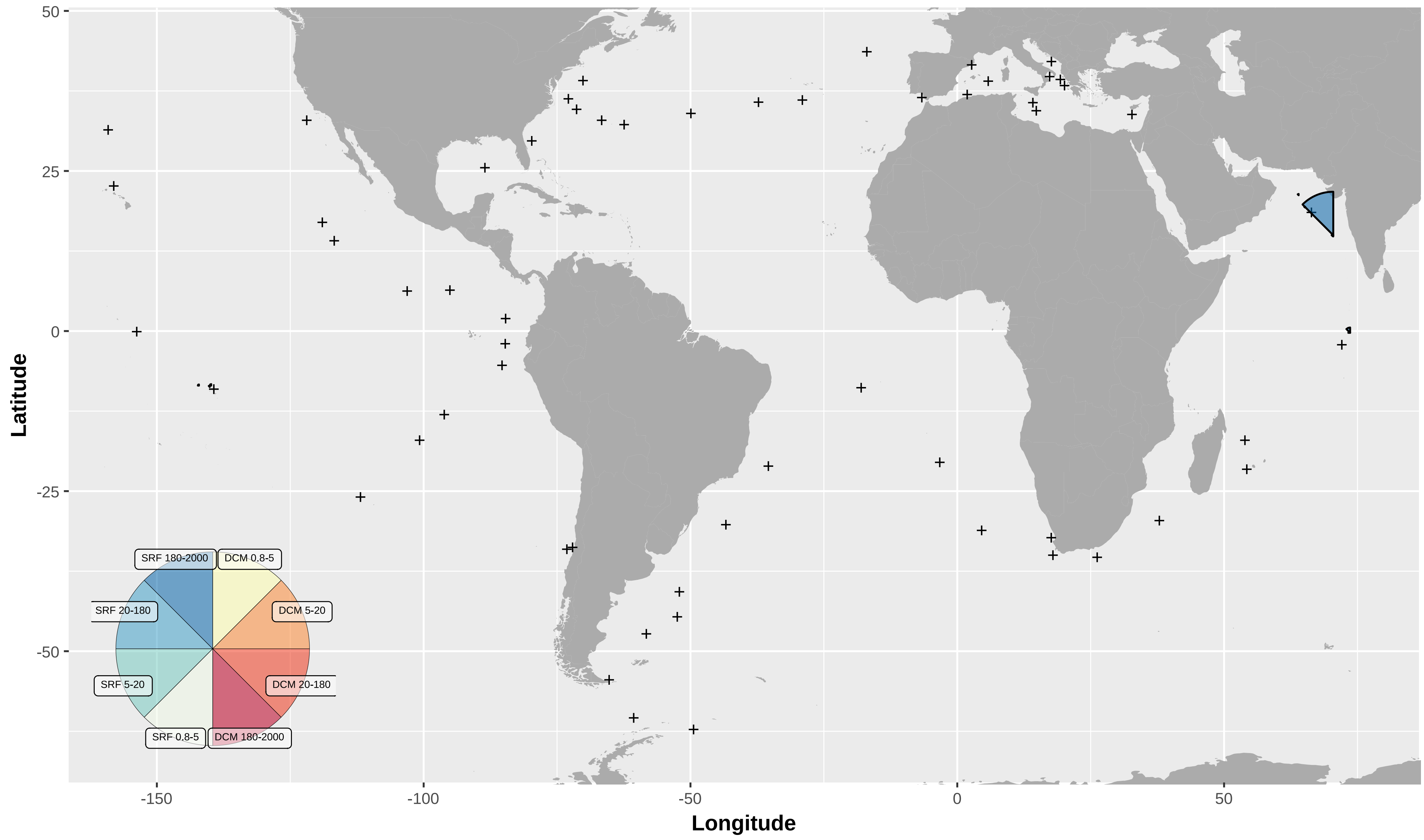


root;cellular organisms;Eukaryota;Opisthokonta;Metazoa;Eumetazoa;Bilateria;Deuterostomia;Chordata

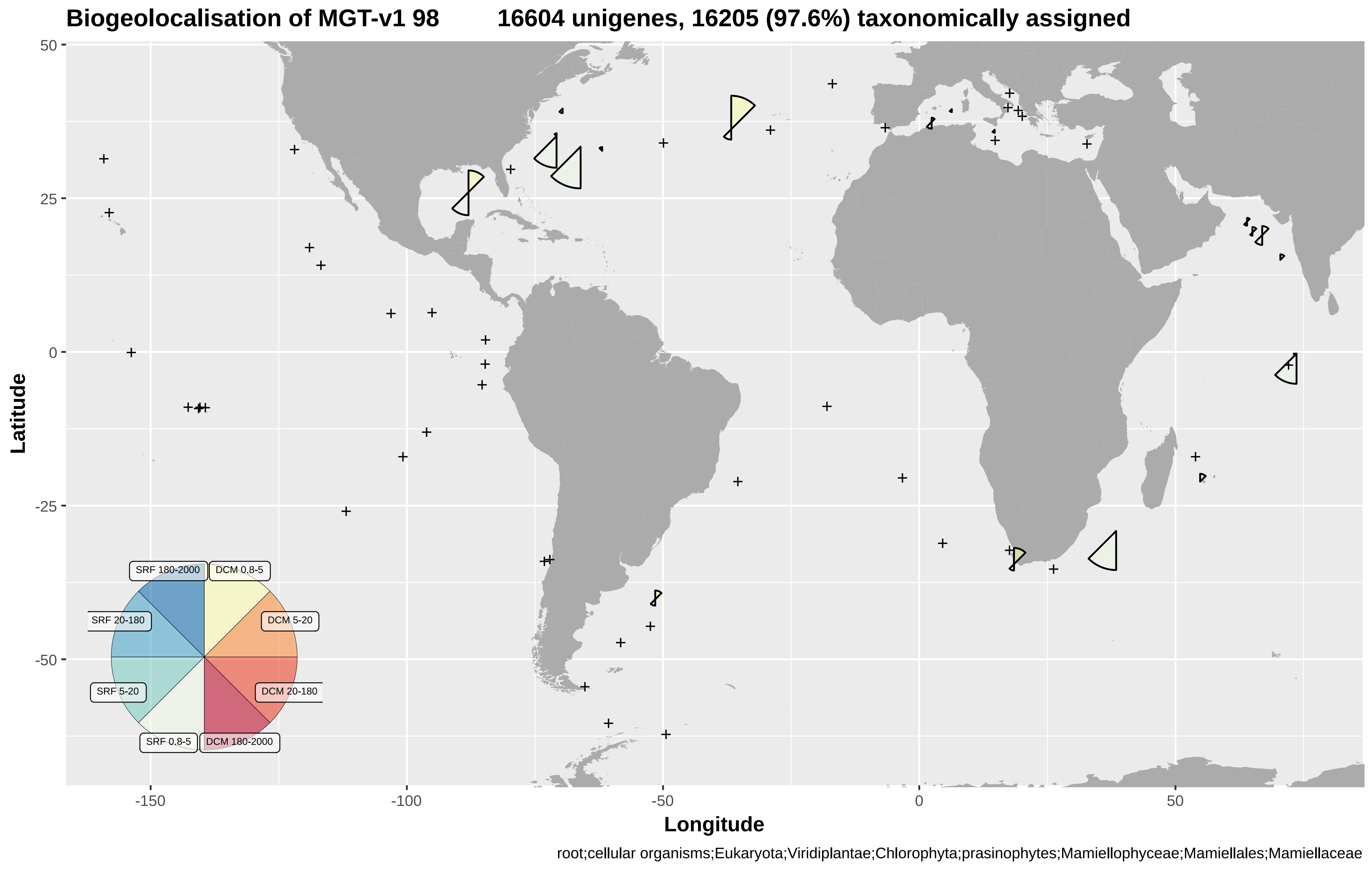
Biogeolocalisation of MGT-v1 98
16604 unigenes, 16205 (97.6%) taxonomically assigned
50
25
0
-25
-50
Latitude
-150
-100
-50
0
50
Longitude
SRF 180-2000
DCM 0.8-5
SRF 20-180
DCM 5-20
SRF 5-20
DCM 20-180
SRF 0.8-5
DCM 180-2000
root;cellular organisms;Eukaryota;Viridiplantae;Chlorophyta;prasinophytes;Mamiellophyceae;Mamiellales;Mamiellaceae

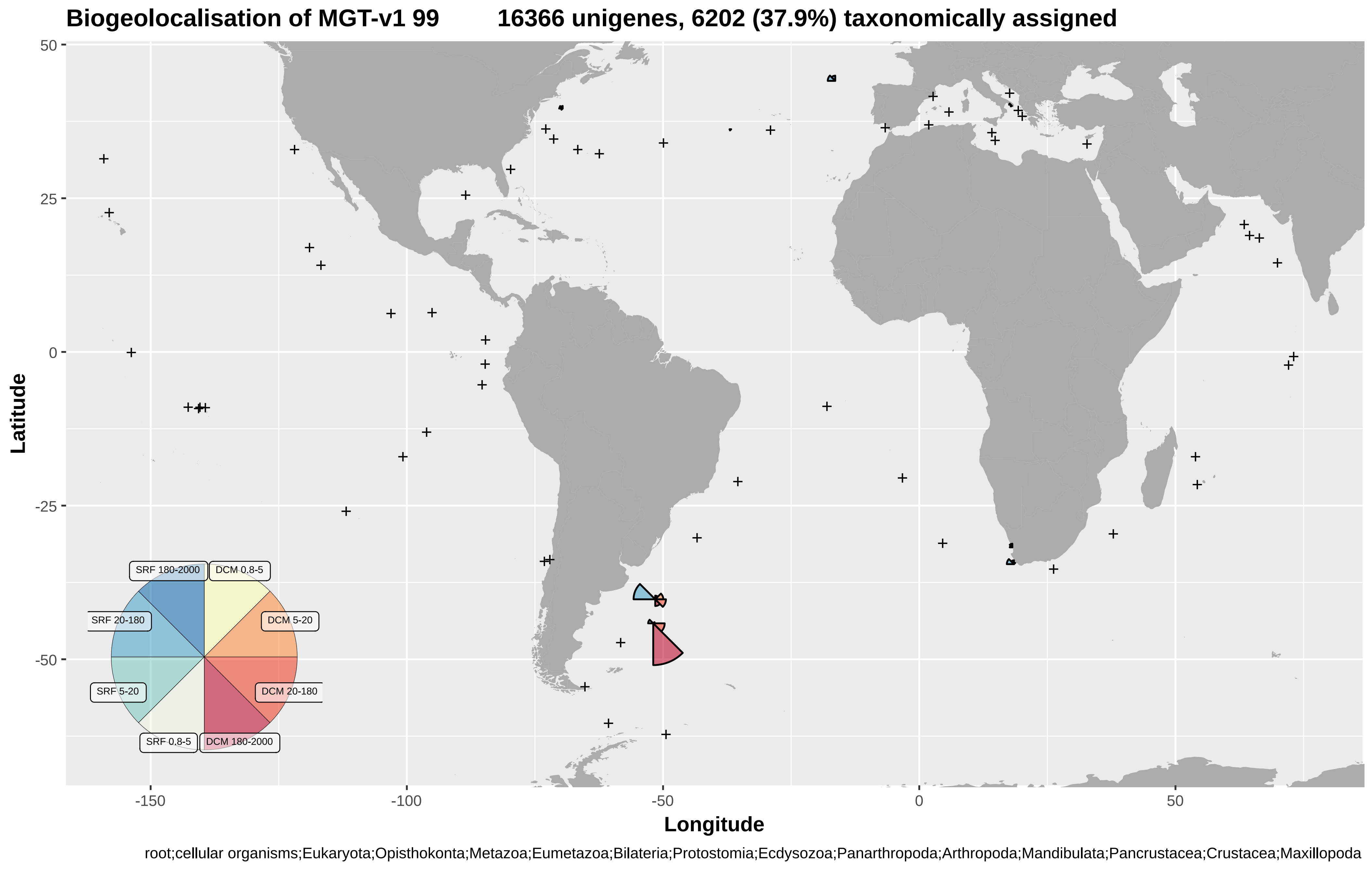

Biogeolocalisation of MGT-v1 99
16366 unigenes, 6202 (37.9%) taxonomically assigned
SRF 180-2000
DCM 0.8-5
SRF 20-180
DCM 5-20
SRF 5-20
DCM 20-180
SRF 0.8-5
DCM 180-2000
Latitude
Longitude
50
25
0
-25
-50
-150
-100
-50
0
50
root;cellular organisms;Eukaryota;Opisthokonta;Metazoa;Eumetazoa;Bilateria;Protostomia;Ecdysozoa;Panarthropoda;Arthropoda;Mandibulata;Pancrustacea;Crustacea;Maxillopoda

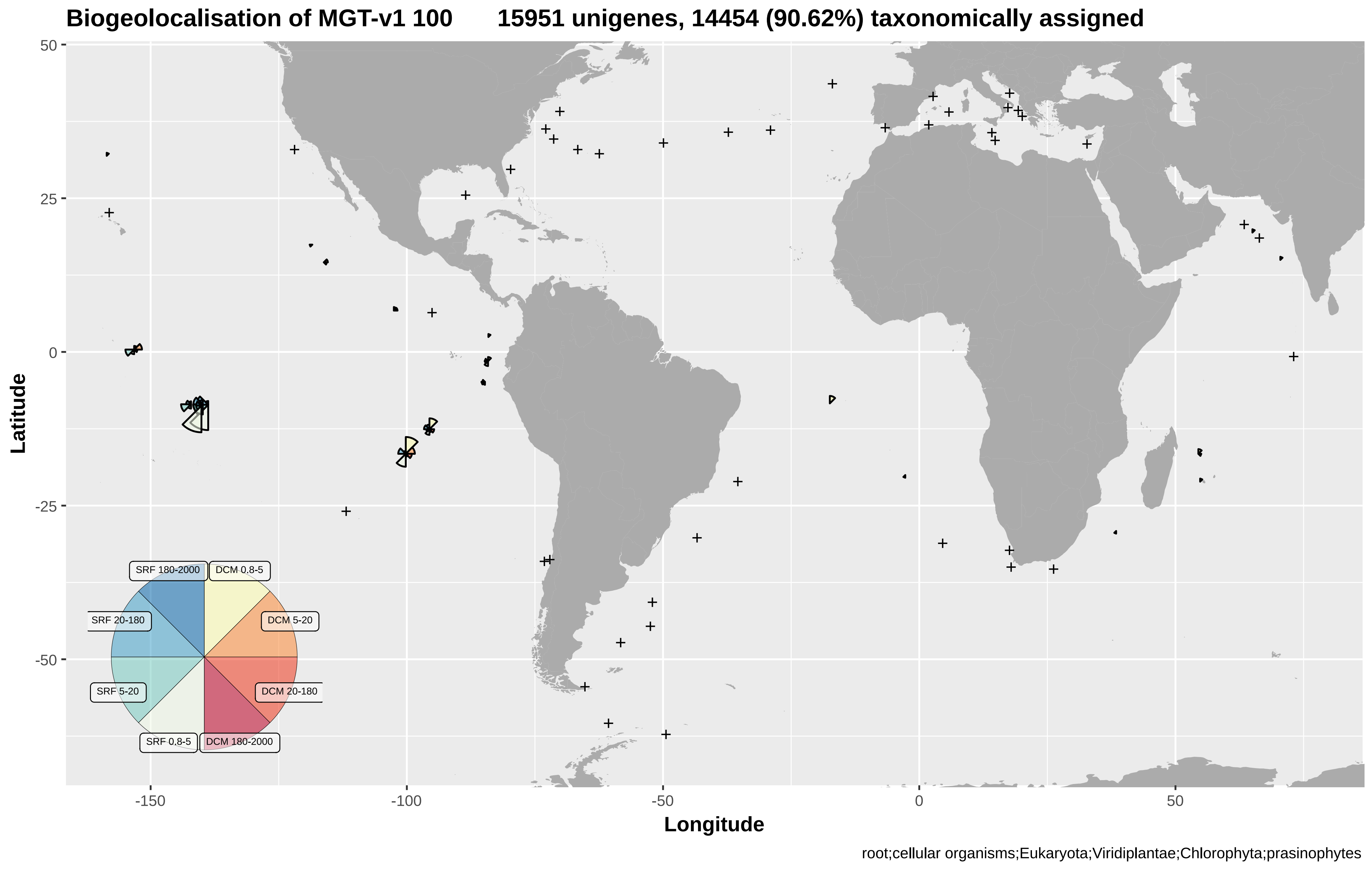
Biogeolocalisation of MGT-v1 100      15951 unigenes, 14454 (90.62%) taxonomically assigned
50
25
0
-25
-50
Latitude
-150
-100
-50
0
50
Longitude
SRF 180-2000
DCM 0.8-5
SRF 20-180
DCM 5-20
SRF 5-20
DCM 20-180
SRF 0.8-5
DCM 180-2000
root;cellular organisms;Eukaryota;Viridiplantae;Chlorophyta;prasinophytes

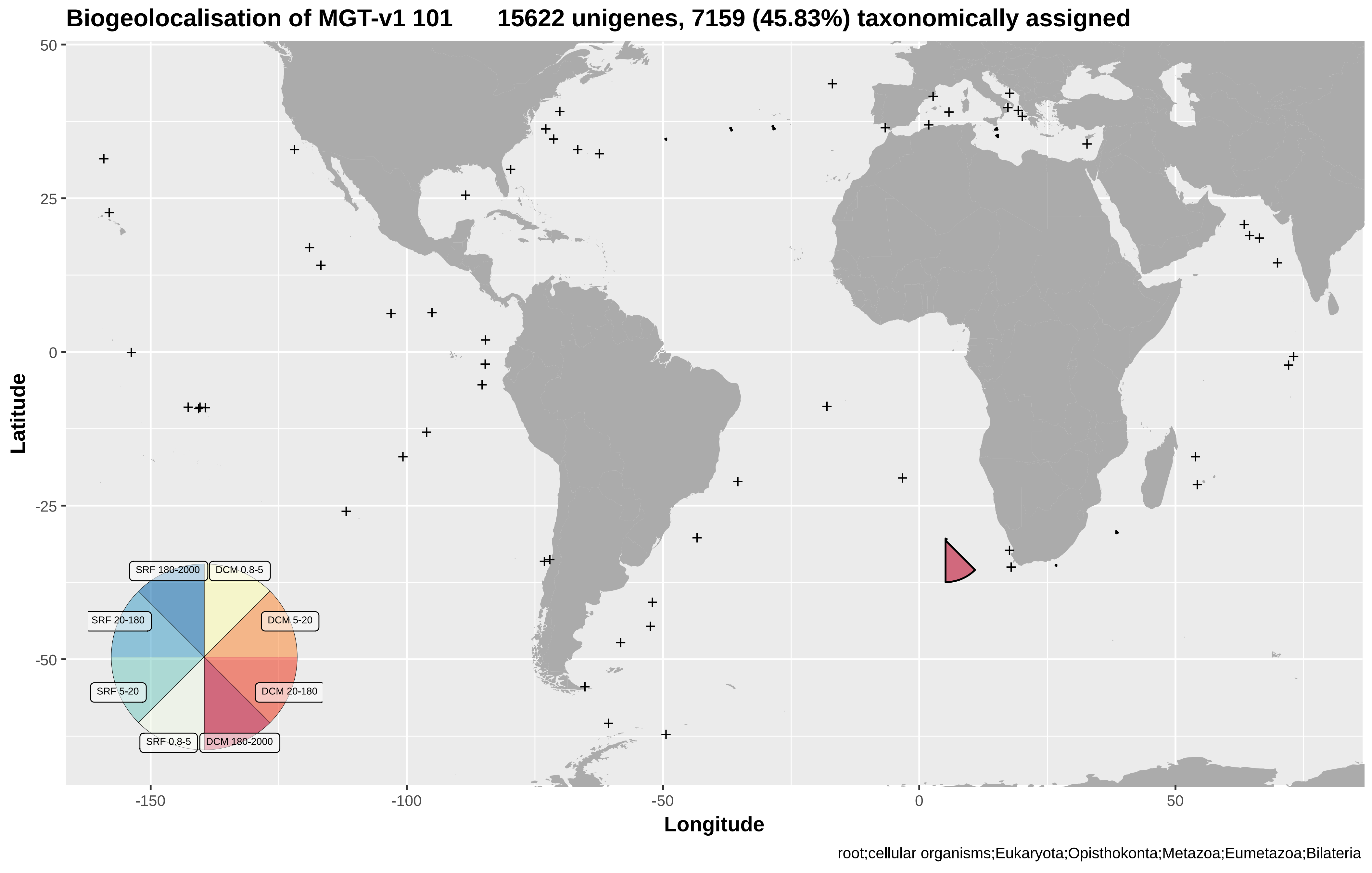
Biogeolocalisation of MGT-v1 101 15622 unigenes, 7159 (45.83%) taxonomically assigned
50
25
0
-25
-50
Latitude
-150
-100
-50
0
50
Longitude
SRF 180-2000
DCM 0.8-5
SRF 20-180
DCM 5-20
SRF 5-20
DCM 20-180
SRF 0.8-5
DCM 180-2000
root;cellular organisms;Eukaryota;Opisthokonta;Metazoa;Eumetazoa;Bilateria

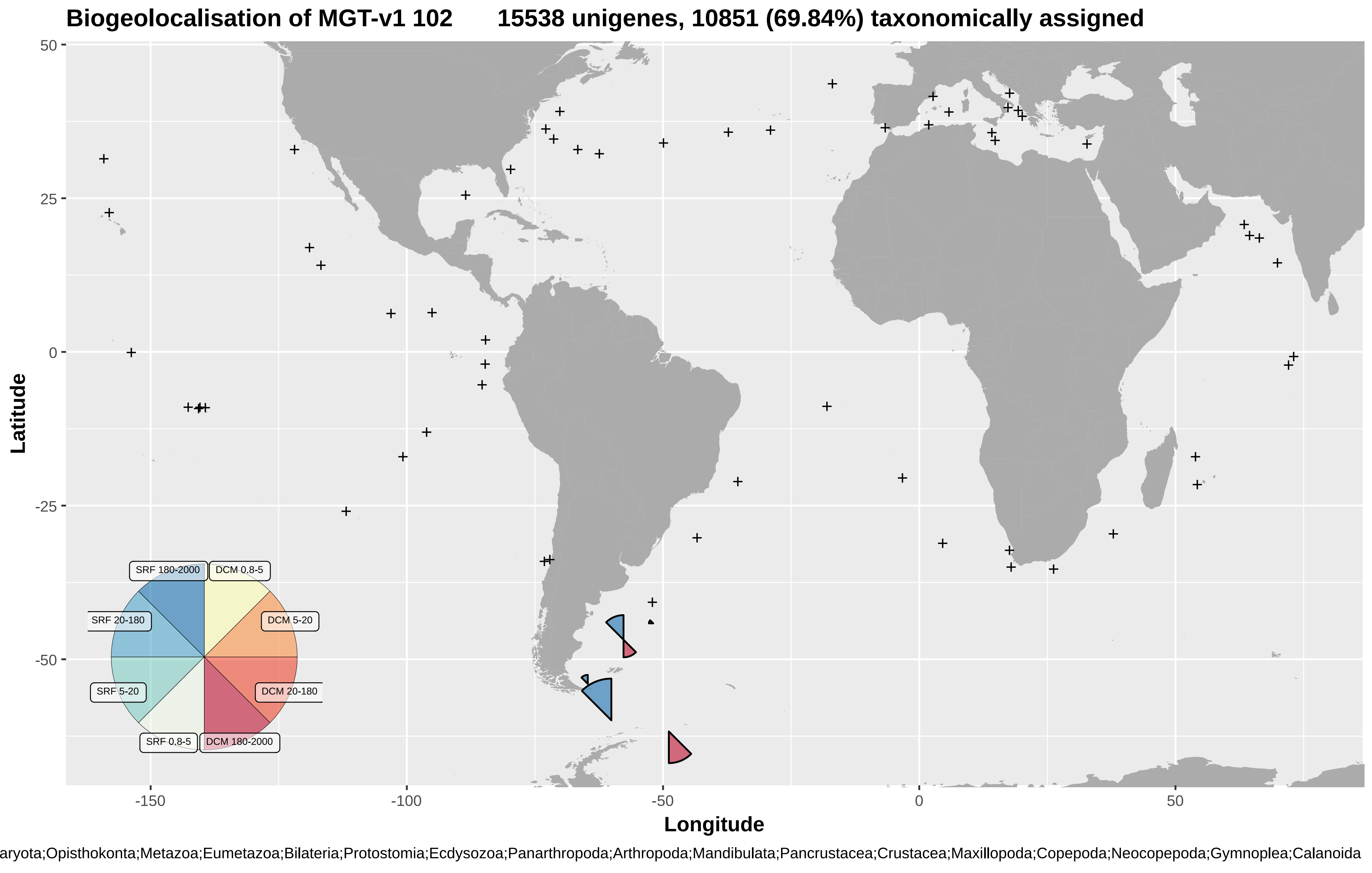
Biogeolocalisation of MGT-v1 102 15538 unigenes, 10851 (69.84%) taxonomically assigned
50
25
0
-25
-50
Latitude
-150
-100
-50
0
50
Longitude
SRF 180-2000
DCM 0.8-5
SRF 20-180
DCM 5-20
SRF 5-20
DCM 20-180
SRF 0.8-5
DCM 180-2000
aryota;Opisthokonta;Metazoa;Eumetazoa;Bilateria;Protostomia;Ecdysozoa;Panarthropoda;Arthropoda;Mandibulata;Pancrustacea;Crustacea;Maxillopoda;Copepoda;Neocopepoda;Gymnoplea;Calanoida

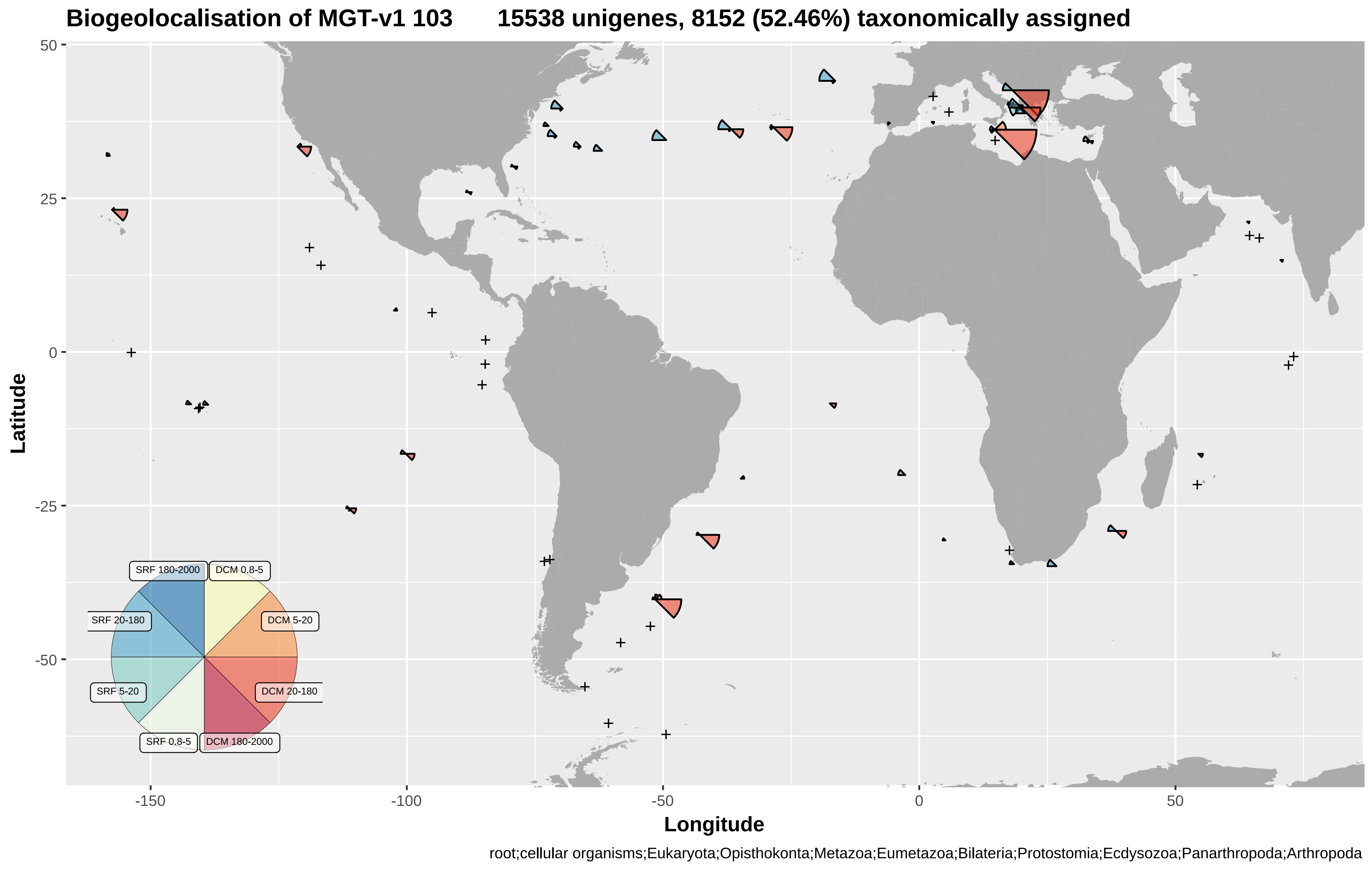
Biogeolocalisation of MGT-v1 103 15538 unigenes, 8152 (52.46%) taxonomically assigned
Latitude
Longitude
50
25
0
-25
-50
-150
-100
-50
0
50
SRF 180-2000
DCM 0.8-5
SRF 20-180
DCM 5-20
SRF 5-20
DCM 20-180
SRF 0.8-5
DCM 180-2000
root;cellular organisms;Eukaryota;Opisthokonta;Metazoa;Eumetazoa;Bilateria;Protostomia;Ecdysozoa;Panarthropoda;Arthropoda

Biogeolocalisation of MGT-v1 104      15010 unigenes, 13799 (91.93%) taxonomically assigned

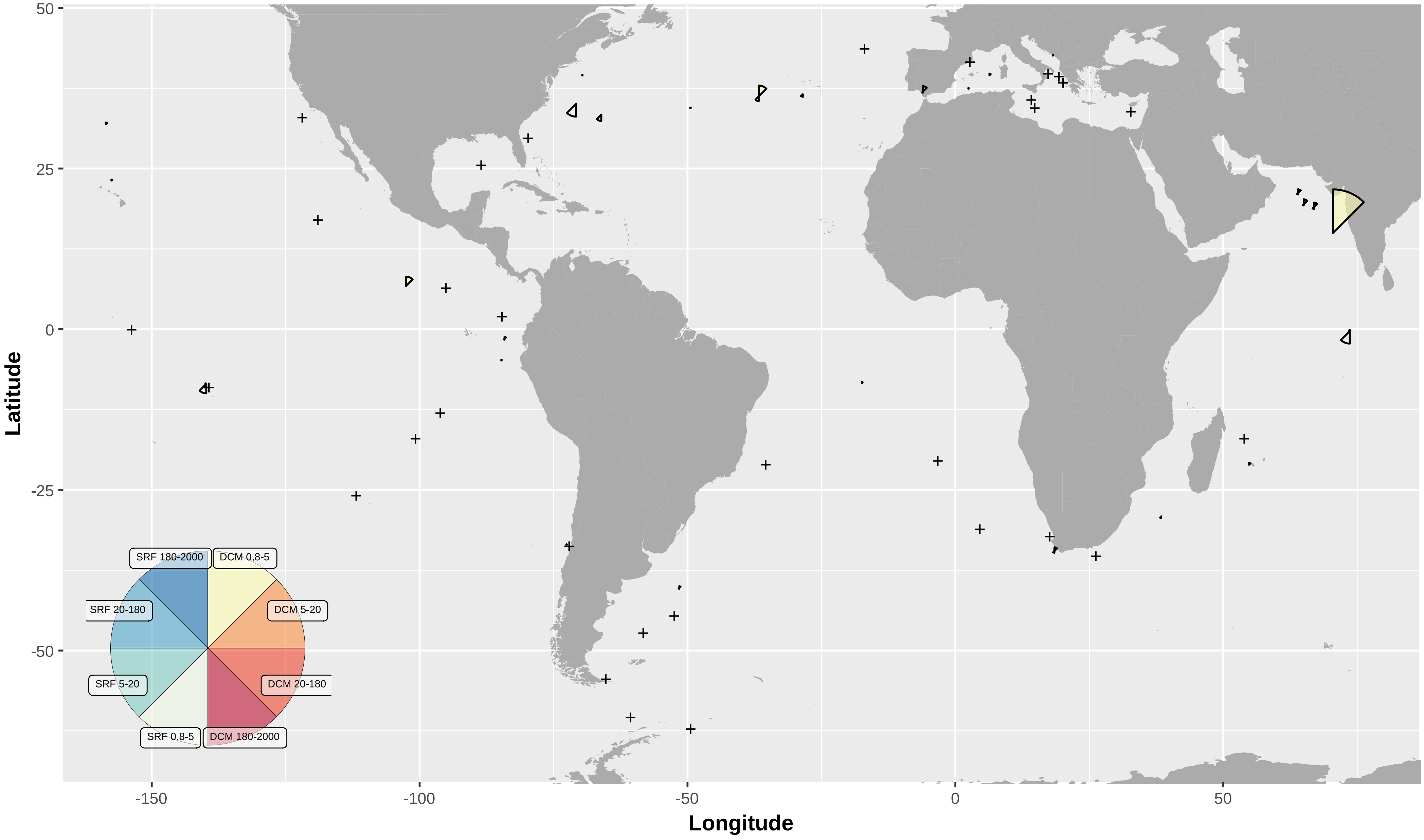


root;cellular organisms;Eukaryota;Viridiplantae;Chlorophyta;prasinophytes;Mamiellophyceae;Mamiellales

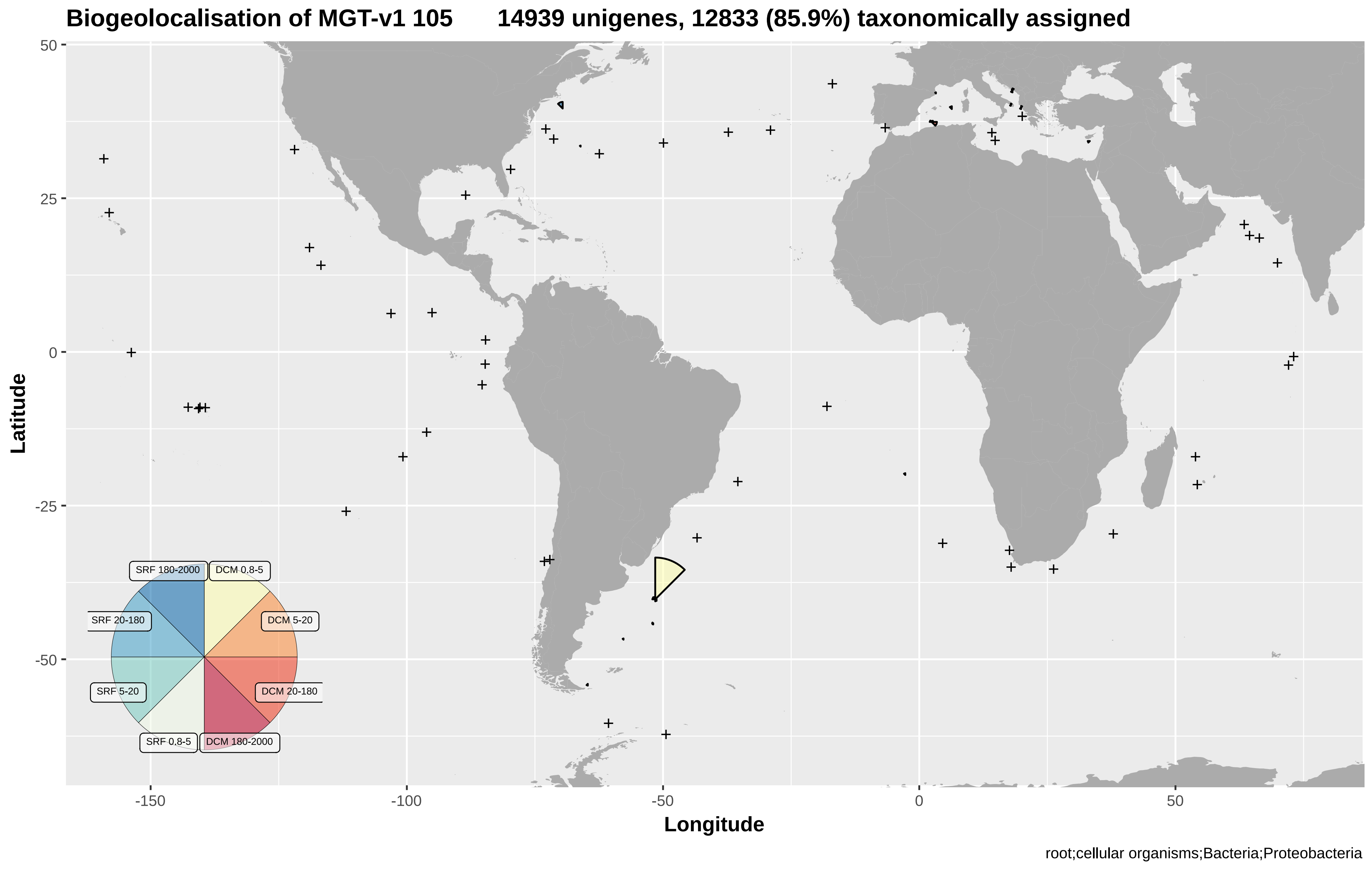
Biogeolocalisation of MGT-v1 105 14939 unigenes, 12833 (85.9%) taxonomically assigned
Latitude
Longitude
SRF 180-2000
DCM 0.8-5
SRF 20-180
DCM 5-20
SRF 5-20
DCM 20-180
SRF 0.8-5
DCM 180-2000
50
25
0
-25
-50
-150
-100
-50
0
50
root;cellular organisms;Bacteria;Proteobacteria

# Biogeolocalisation of MGT-v1 106      14931 unigenes, 8174 (54.75%) taxonomically assigned

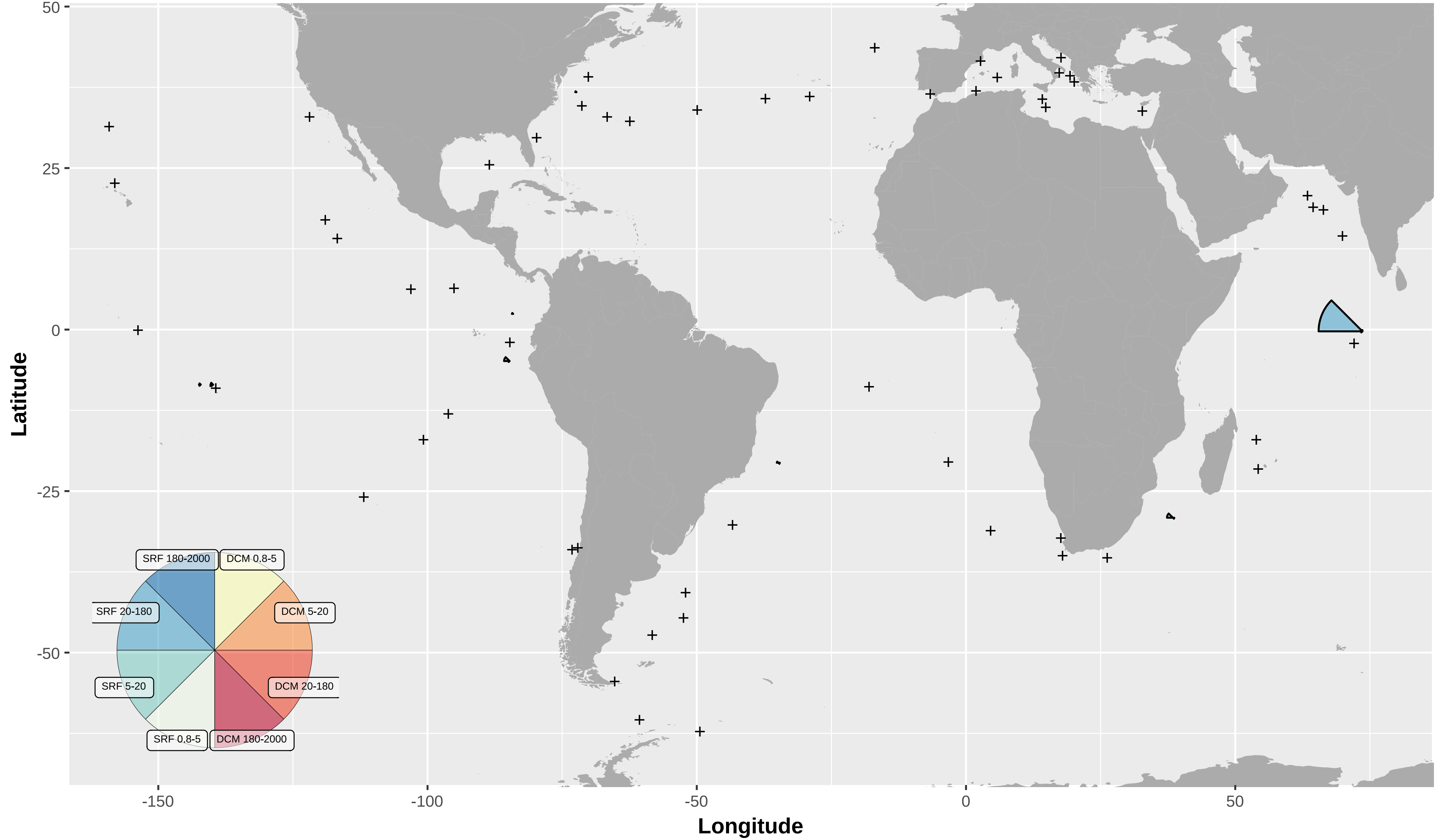


root;cellular organisms;Eukaryota;Alveolata;Ciliophora;Intramacronucleata;Spirotrichea

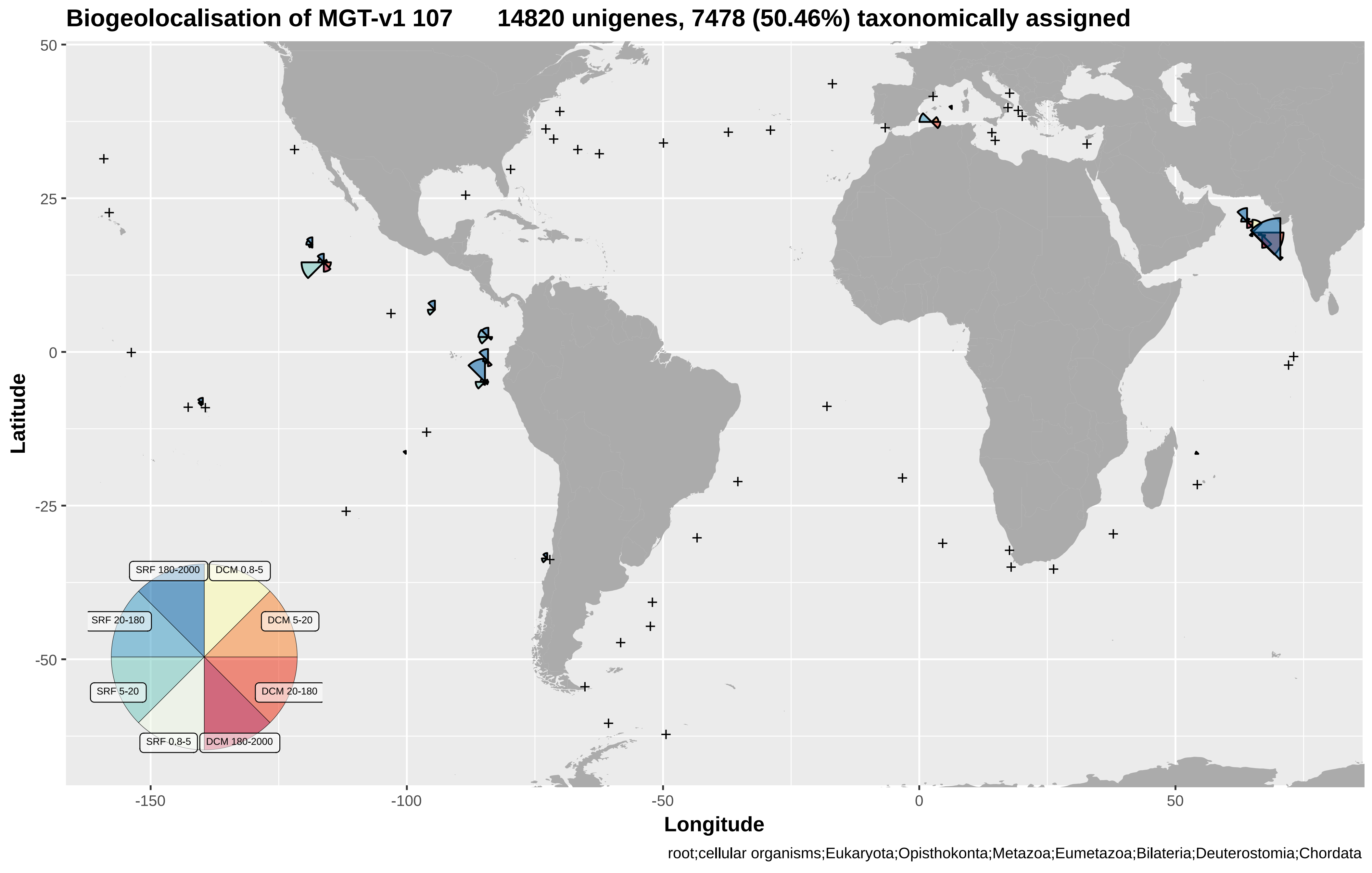
Biogeolocalisation of MGT-v1 107 14820 unigenes, 7478 (50.46%) taxonomically assigned
Latitude
50
25
0
-25
-50
Longitude
-150
-100
-50
0
50
SRF 180-2000
DCM 0.8-5
SRF 20-180
DCM 5-20
SRF 5-20
DCM 20-180
SRF 0.8-5
DCM 180-2000
root;cellular organisms;Eukaryota;Opisthokonta;Metazoa;Eumetazoa;Bilateria;Deuterostomia;Chordata

# Biogeolocalisation of MGT-v1 108      14792 unigenes, 10957 (74.07%) taxonomically assigned

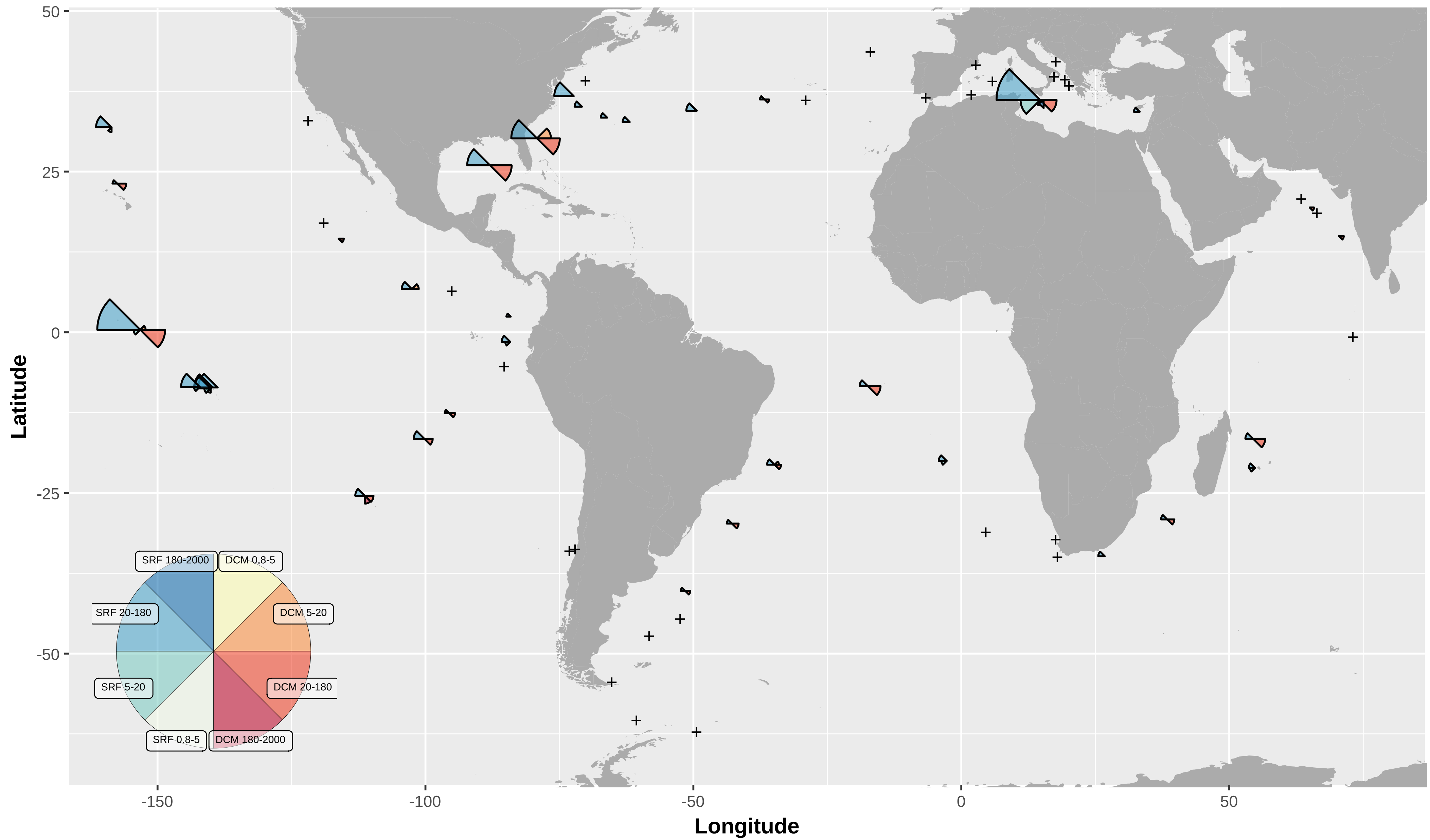


root;cellular organisms;Eukaryota;Opisthokonta;Metazoa;Eumetazoa;Bilateria;Protostomia;Ecdysozoa;Panarthropoda;Arthropoda;Mandibulata;Pancrustacea;Crustacea;Maxillopoda

### Biogeolocalisation of MGT-v1 109 14689 unigenes, 11897 (80.99%) taxonomically assigned

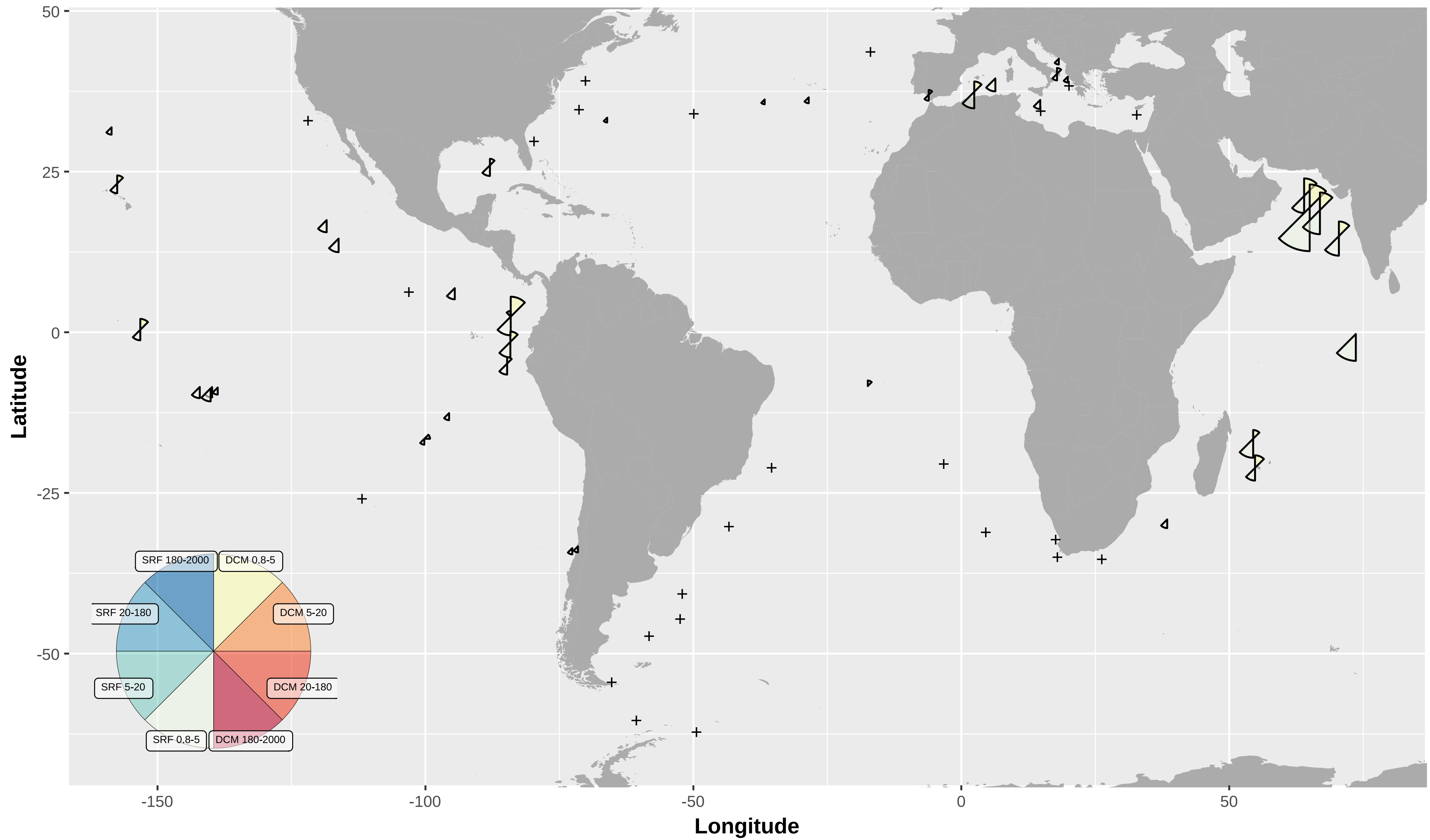


root;cellular organisms;Eukaryota;Stramenopiles

## Biogeolocalisation of MGT-v1 110      13810 unigenes, 2232 (16.16%) taxonomically assigned

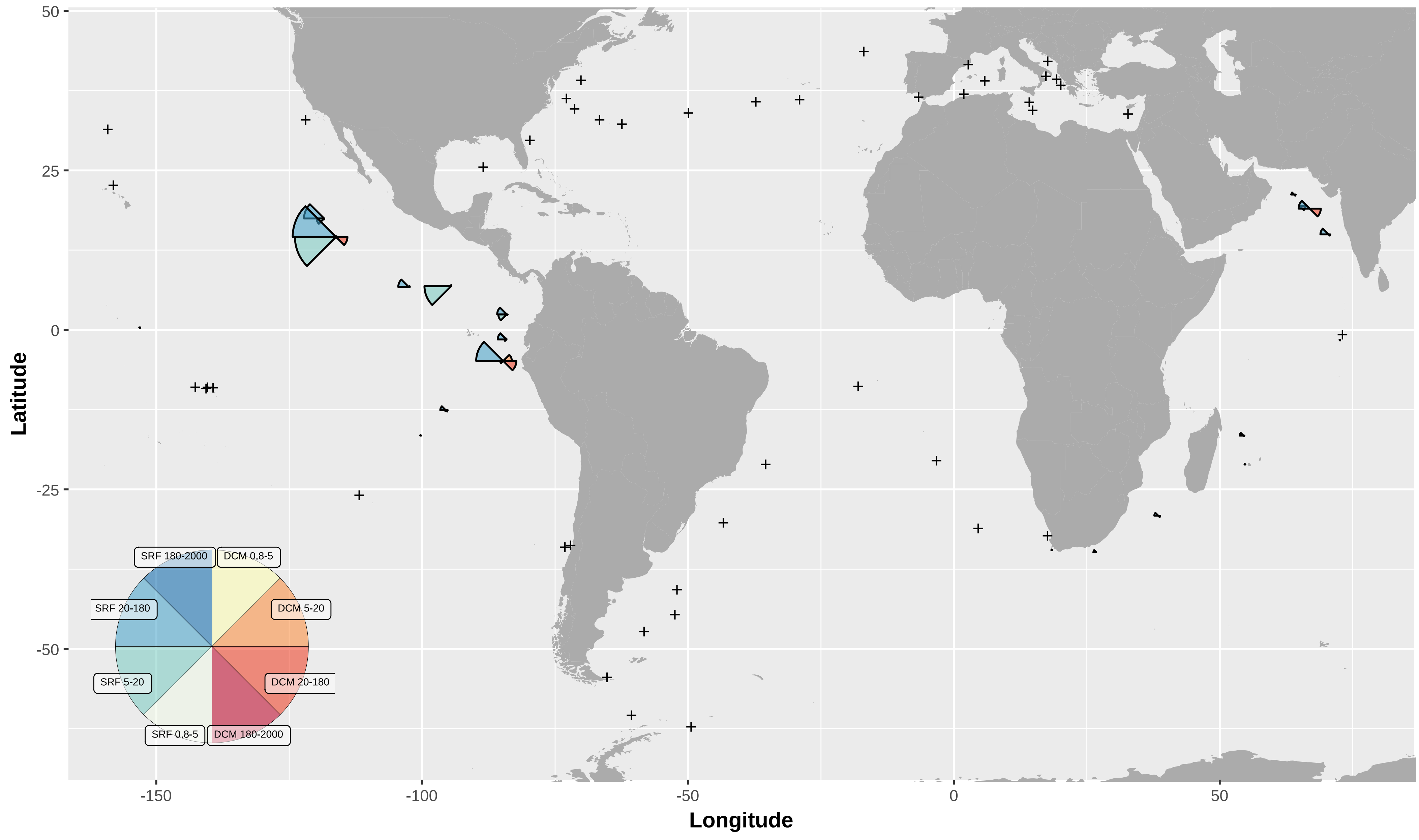


root;cellular organisms;Eukaryota;Opisthokonta;Metazoa;Eumetazoa;Bilateria;Protostomia;Ecdysozoa;Panarthropoda;Arthropoda;Mandibulata;Pancrustacea;Crustacea

## Biogeolocalisation of MGT-v1 111    13474 unigenes, 6676 (49.55%) taxonomically assigned

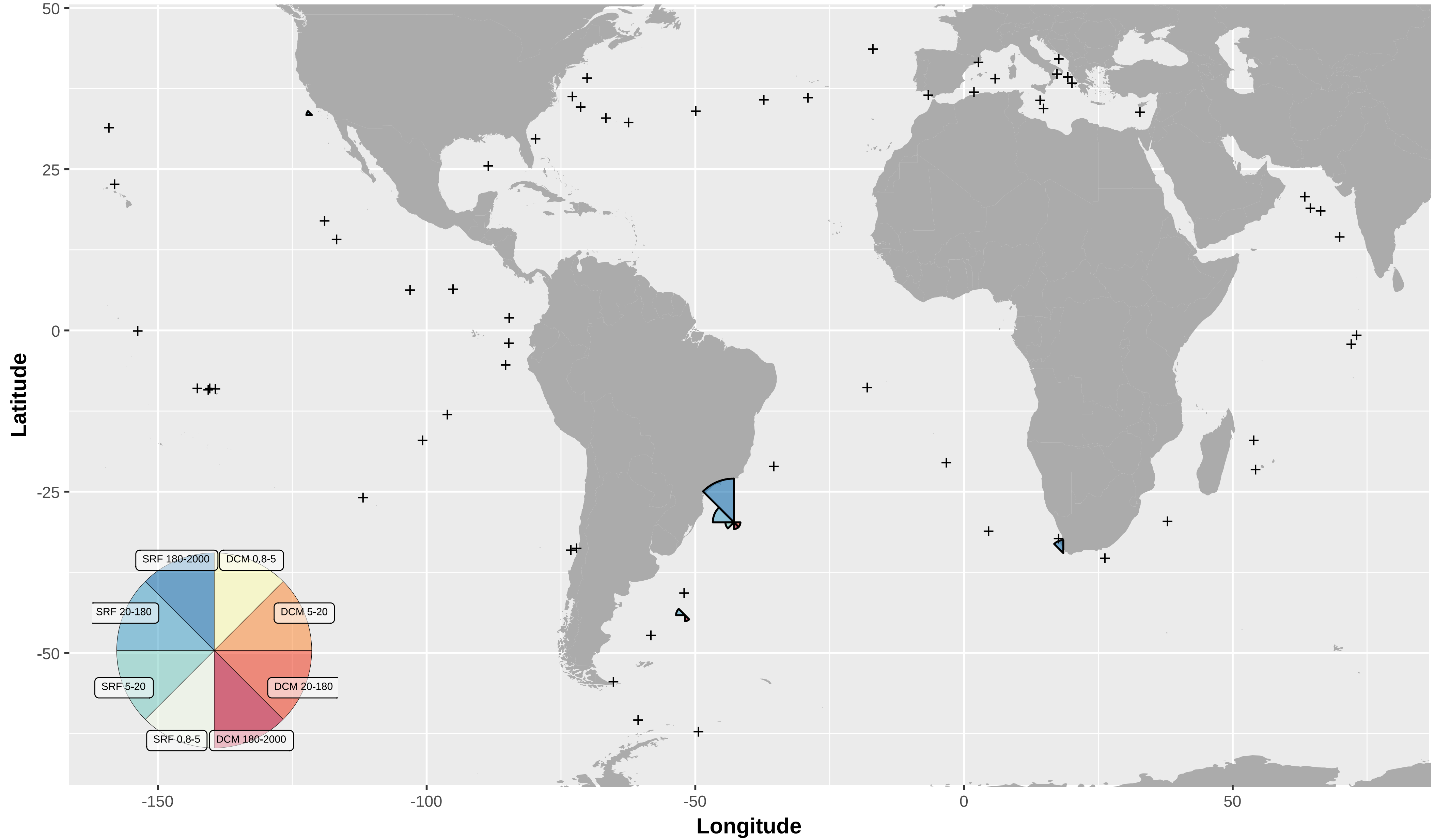


root;cellular organisms;Eukaryota;Opisthokonta;Metazoa;Eumetazoa;Bilateria;Deuterostomia;Chordata

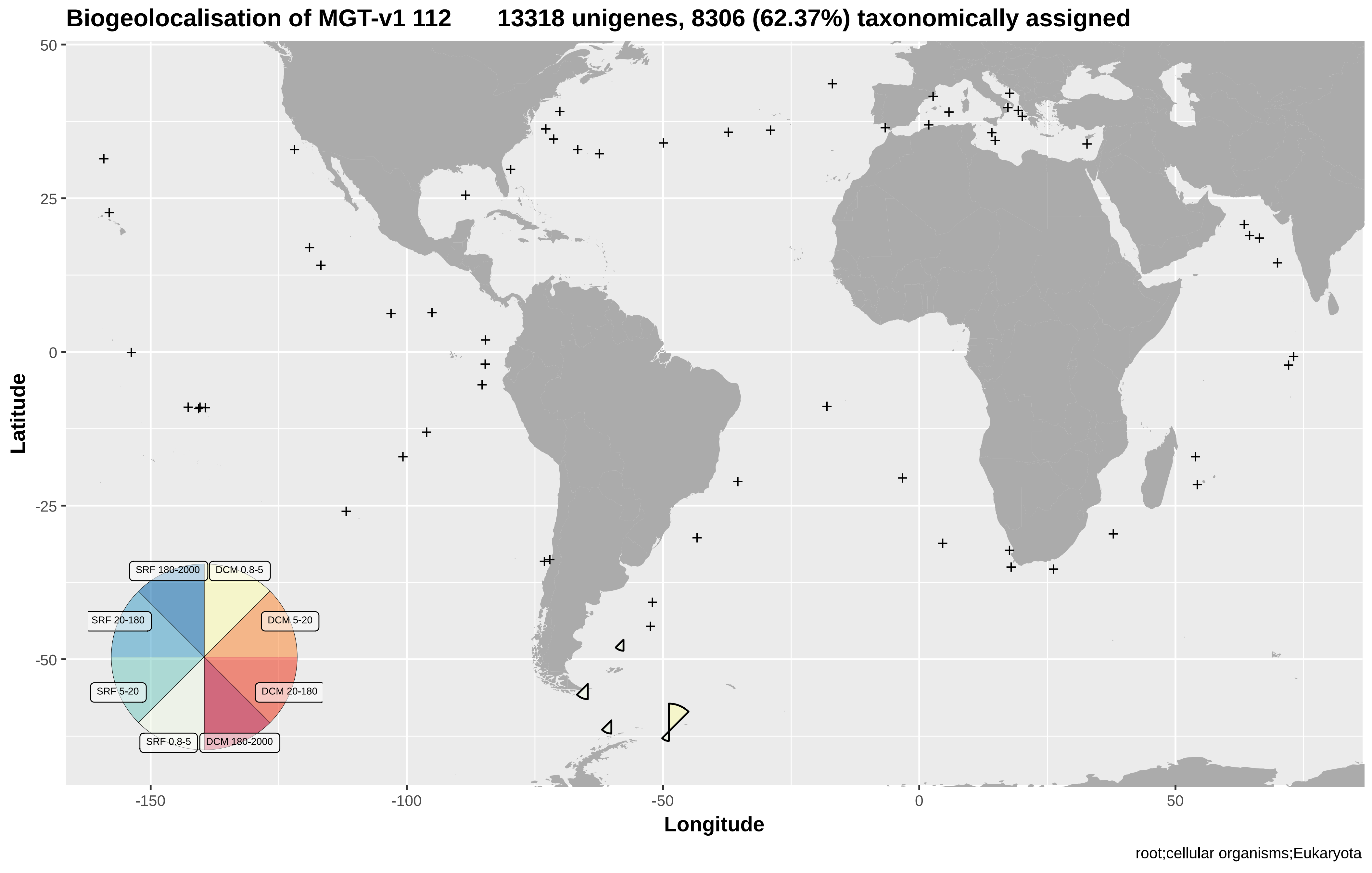
Biogeolocalisation of MGT-v1 112 13318 unigenes, 8306 (62.37%) taxonomically assigned
50
25
0
-25
-50
Latitude
-150
-100
-50
0
50
Longitude
SRF 180-2000
DCM 0.8-5
SRF 20-180
DCM 5-20
SRF 5-20
DCM 20-180
SRF 0.8-5
DCM 180-2000
root;cellular organisms;Eukaryota

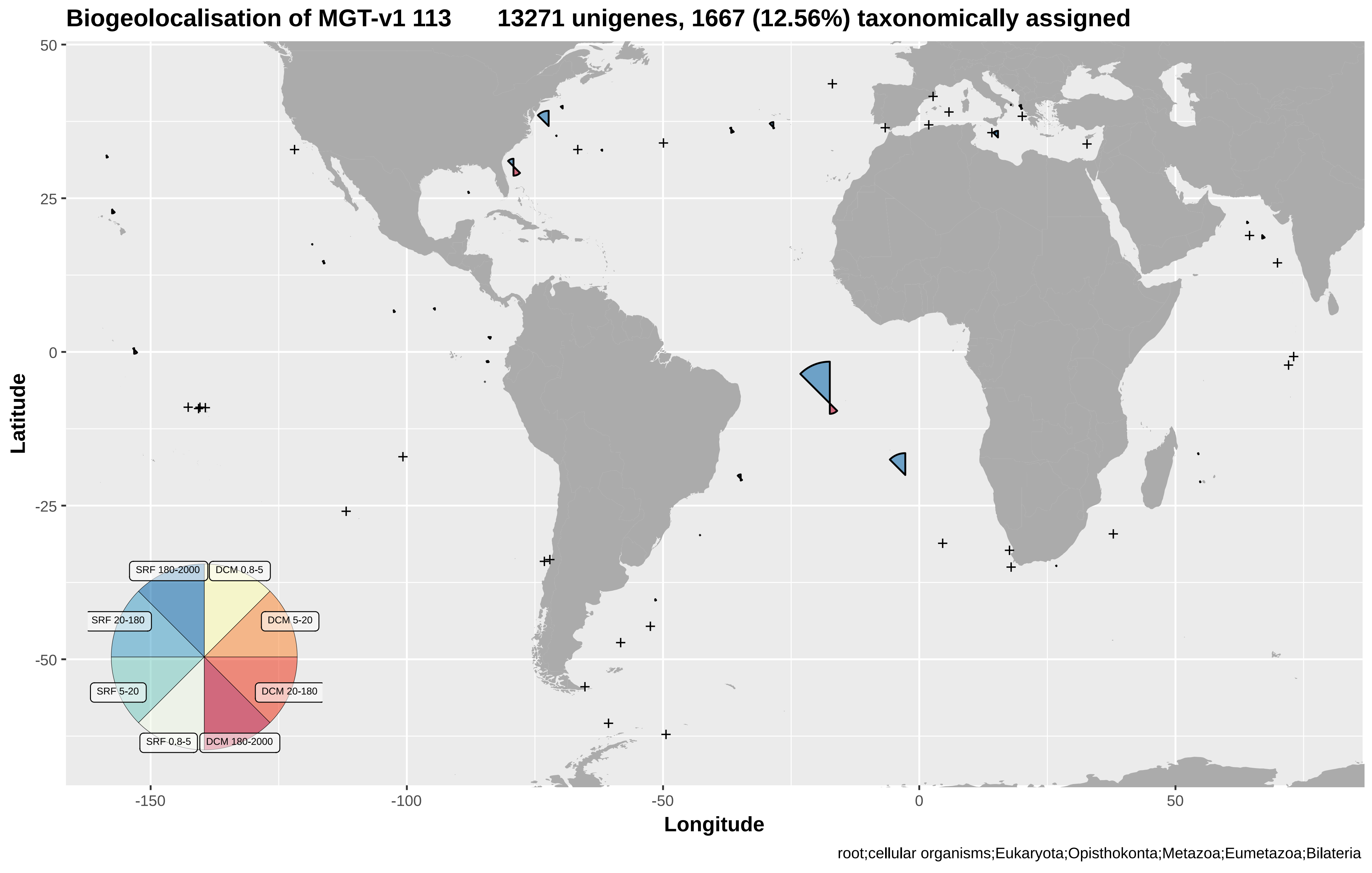

Biogeolocalisation of MGT-v1 113 13271 unigenes, 1667 (12.56%) taxonomically assigned
50
25
0
-25
-50
Latitude
-150
-100
-50
0
50
Longitude
SRF 180-2000
DCM 0.8-5
SRF 20-180
DCM 5-20
SRF 5-20
DCM 20-180
SRF 0.8-5
DCM 180-2000
root;cellular organisms;Eukaryota;Opisthokonta;Metazoa;Eumetazoa;Bilateria

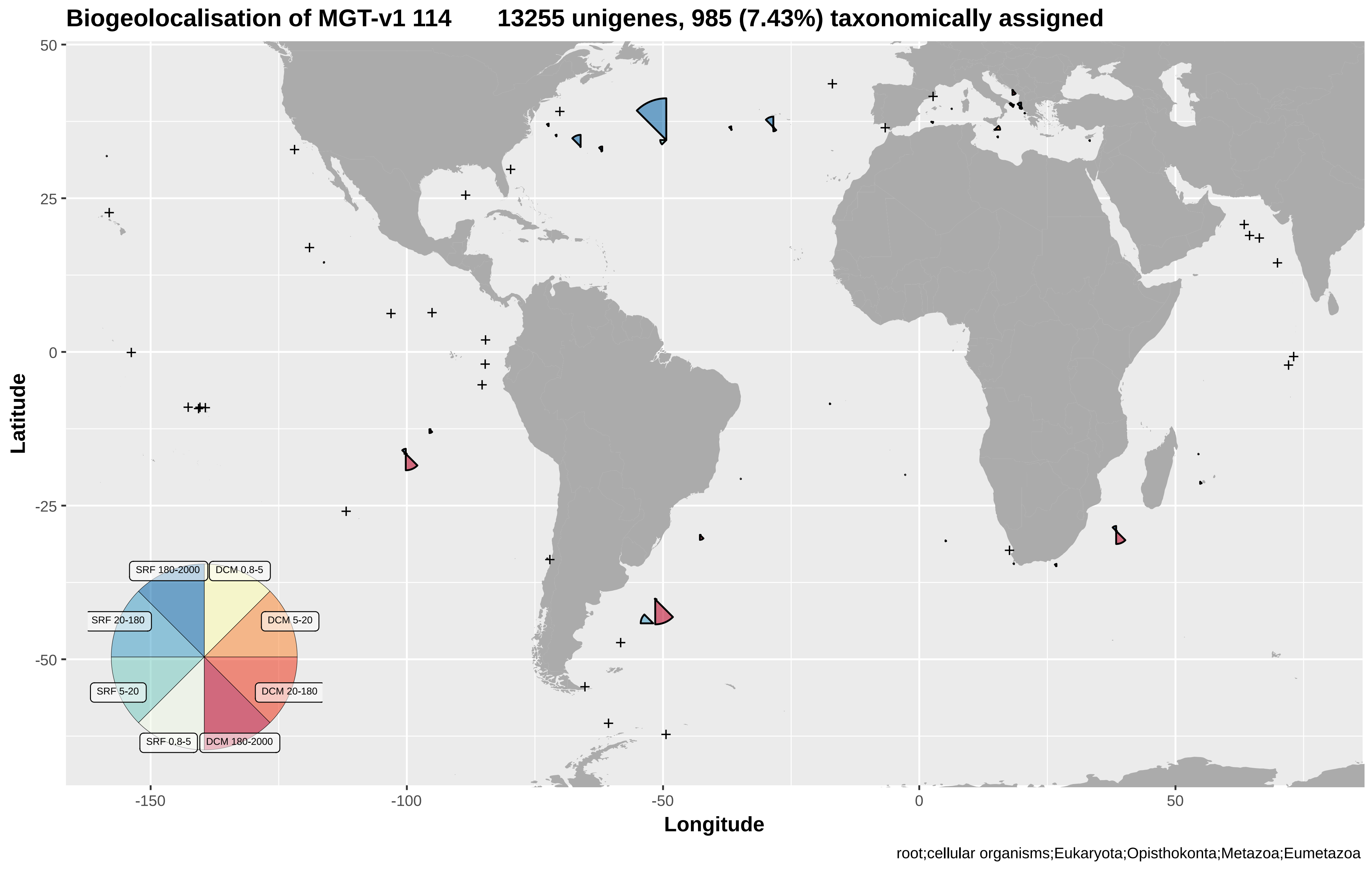

Biogeolocalisation of MGT-v1 114 13255 unigenes, 985 (7.43%) taxonomically assigned
50
25
0
-25
-50
Latitude
-150
-100
-50
0
50
Longitude
SRF 180-2000
DCM 0.8-5
SRF 20-180
DCM 5-20
SRF 5-20
DCM 20-180
SRF 0.8-5
DCM 180-2000
root;cellular organisms;Eukaryota;Opisthokonta;Metazoa;Eumetazoa

### Biogeolocalisation of MGT-v1 115 12923 unigenes, 11604 (89.79%) taxonomically assigned

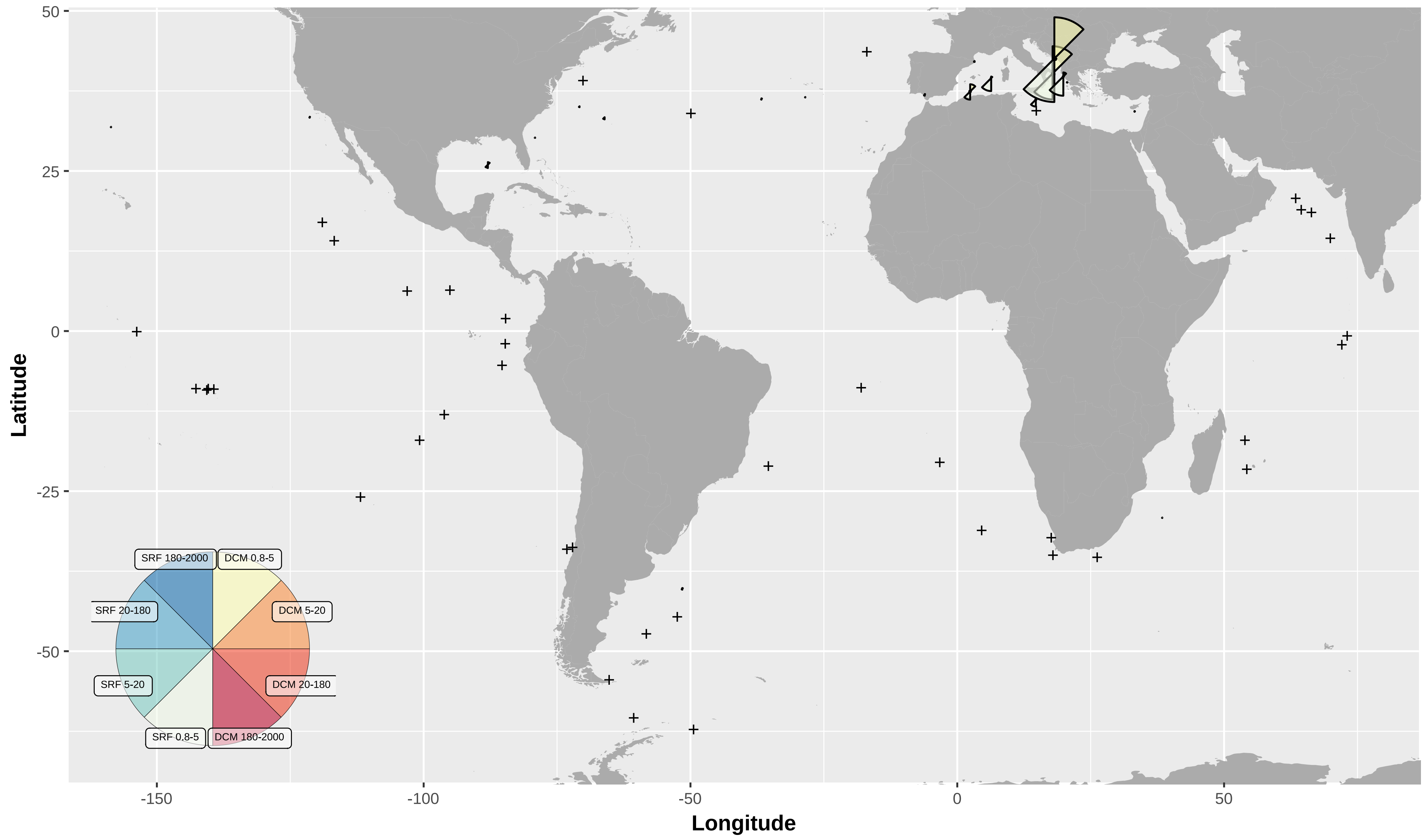


root;cellular organisms;Bacteria

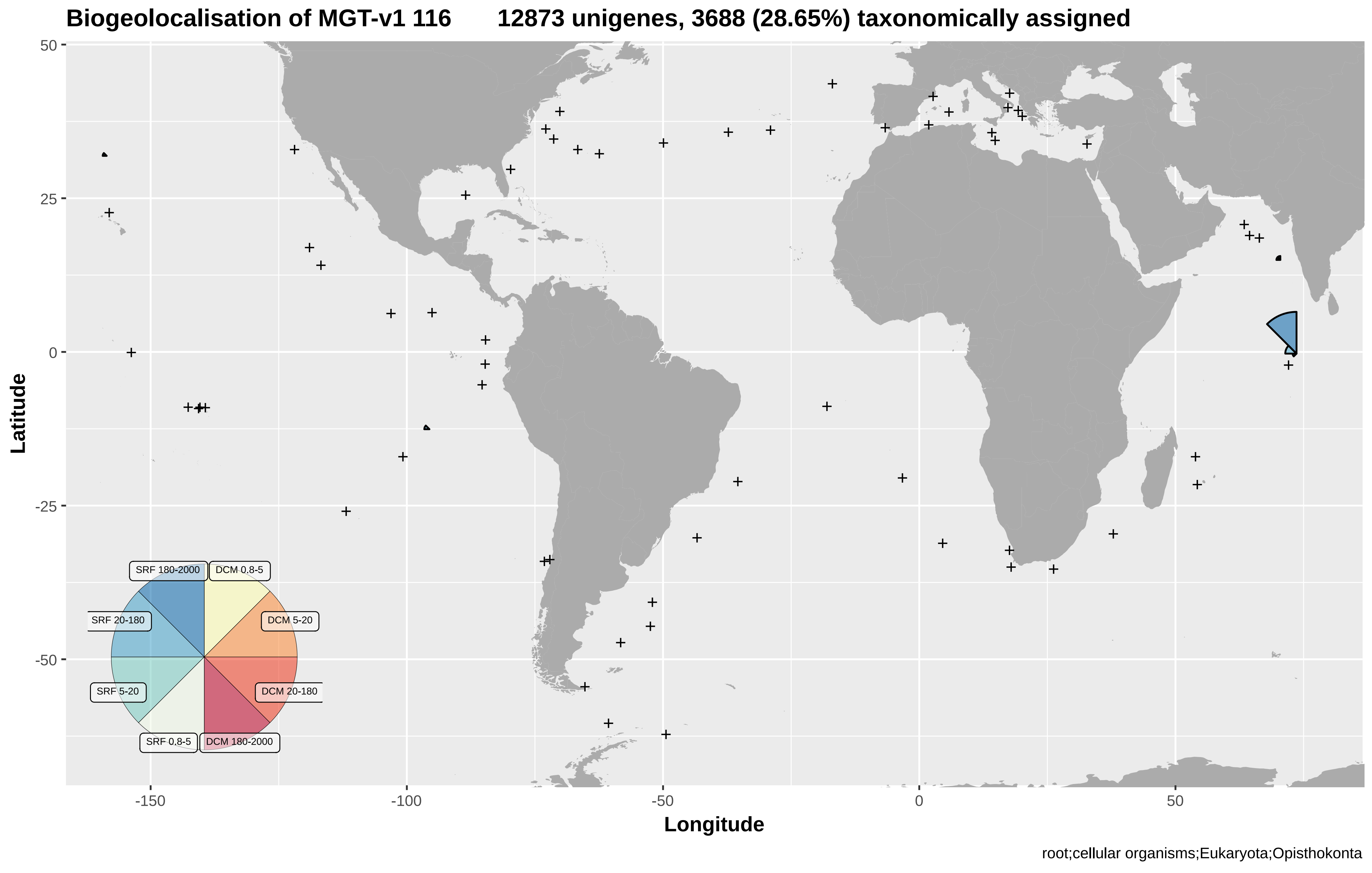

Biogeolocalisation of MGT-v1 116 12873 unigenes, 3688 (28.65%) taxonomically assigned
50
25
0
-25
-50
Latitude
-150
-100
-50
0
50
Longitude
SRF 180-2000
DCM 0.8-5
SRF 20-180
DCM 5-20
SRF 5-20
DCM 20-180
SRF 0.8-5
DCM 180-2000
root;cellular organisms;Eukaryota;Opisthokonta

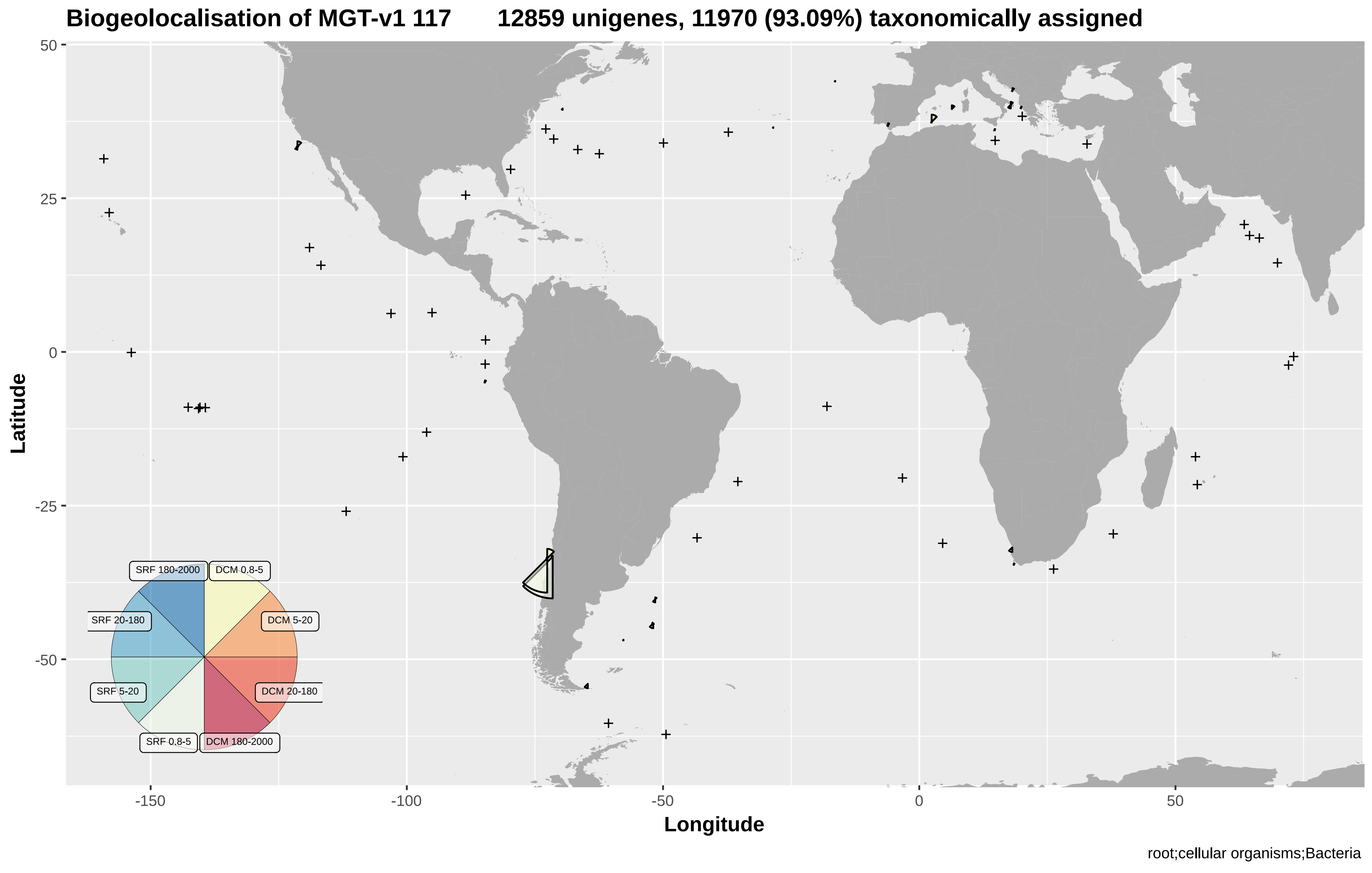

Biogeolocalisation of MGT-v1 117 12859 unigenes, 11970 (93.09%) taxonomically assigned
50
25
0
-25
-50
Latitude
-150
-100
-50
0
50
Longitude
SRF 180-2000
DCM 0.8-5
SRF 20-180
DCM 5-20
SRF 5-20
DCM 20-180
SRF 0.8-5
DCM 180-2000
root;cellular organisms;Bacteria

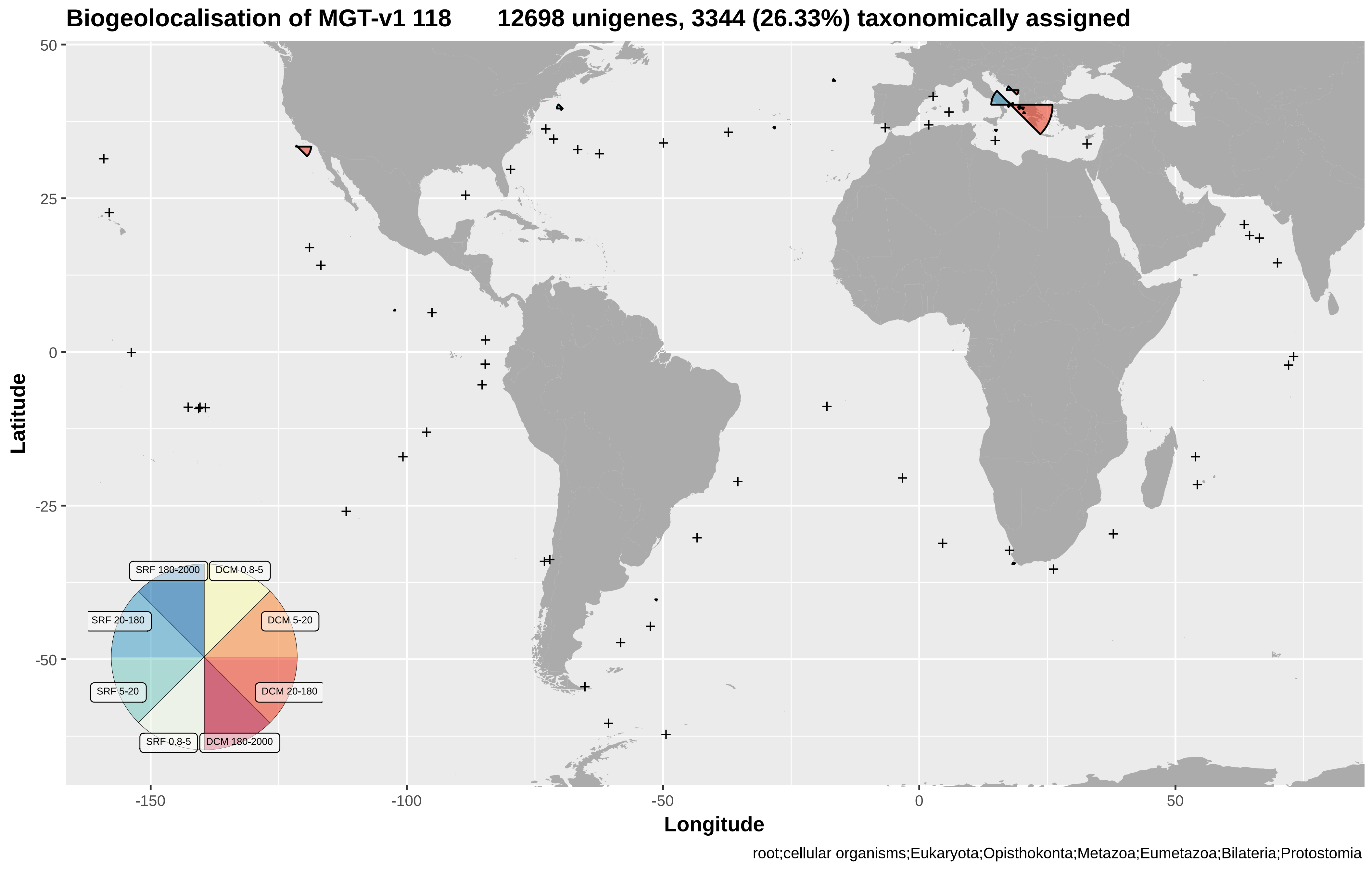
Biogeolocalisation of MGT-v1 118 12698 unigenes, 3344 (26.33%) taxonomically assigned
50
25
0
-25
-50
Latitude
-150
-100
-50
0
50
Longitude
SRF 180-2000
DCM 0.8-5
SRF 20-180
DCM 5-20
SRF 5-20
DCM 20-180
SRF 0.8-5
DCM 180-2000
root;cellular organisms;Eukaryota;Opisthokonta;Metazoa;Eumetazoa;Bilateria;Protostomia

## Biogeolocalisation of MGT-v1 119      12683 unigenes, 2354 (18.56%) taxonomically assigned

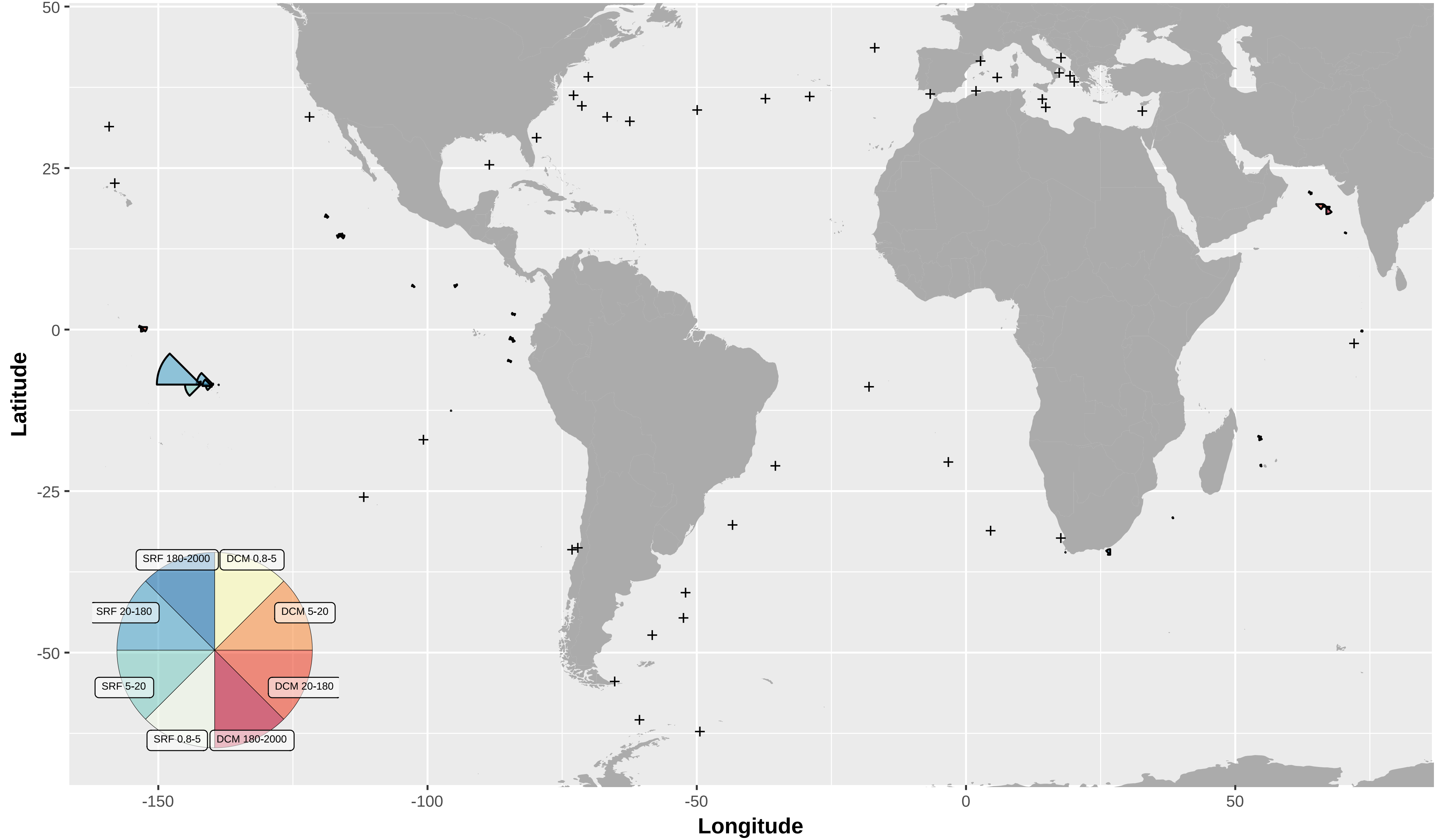


root;cellular organisms;Eukaryota;Opisthokonta;Metazoa;Eumetazoa;Bilateria;Protostomia;Ecdysozoa;Panarthropoda;Arthropoda;Mandibulata;Pancrustacea

# Biogeolocalisation of MGT-v1 120      12668 unigenes, 7381 (58.26%) taxonomically assigned

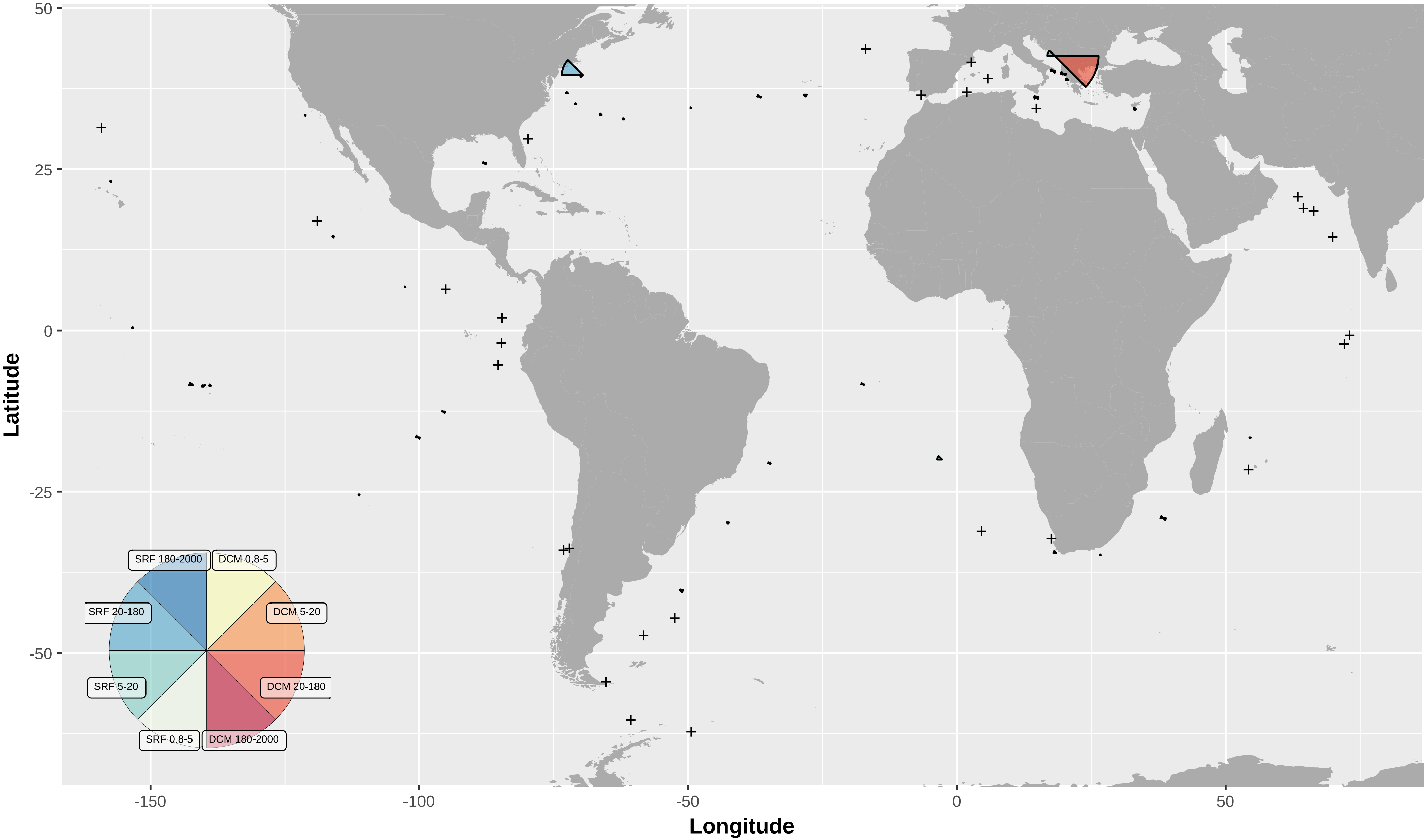


root;cellular organisms;Eukaryota;Alveolata;Ciliophora;Intramacronucleata;Spirotrichea

## Biogeolocalisation of MGT-v1 121      12643 unigenes, 2376 (18.79%) taxonomically assigned

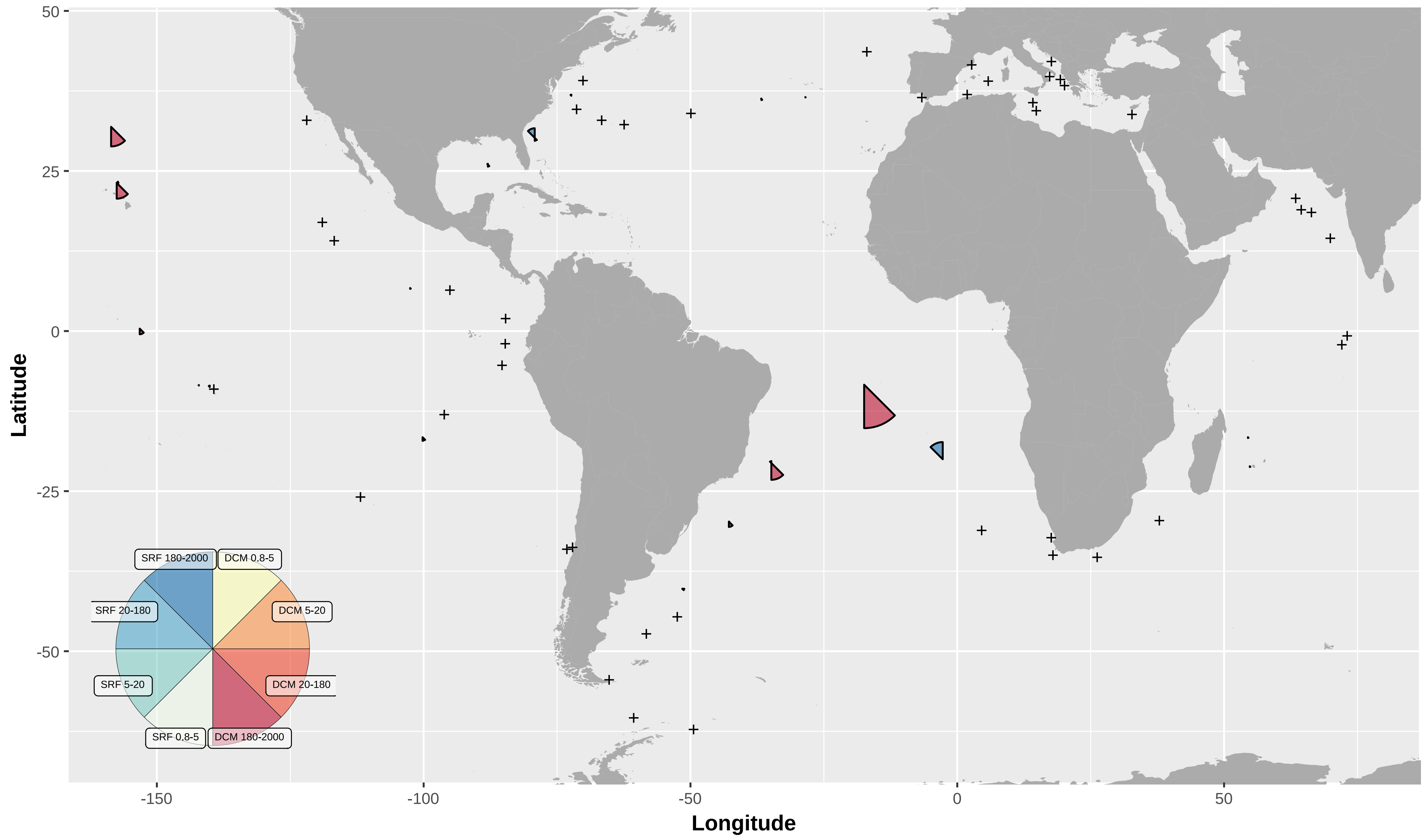


root;cellular organisms;Eukaryota;Opisthokonta;Metazoa;Eumetazoa;Bilateria;Protostomia

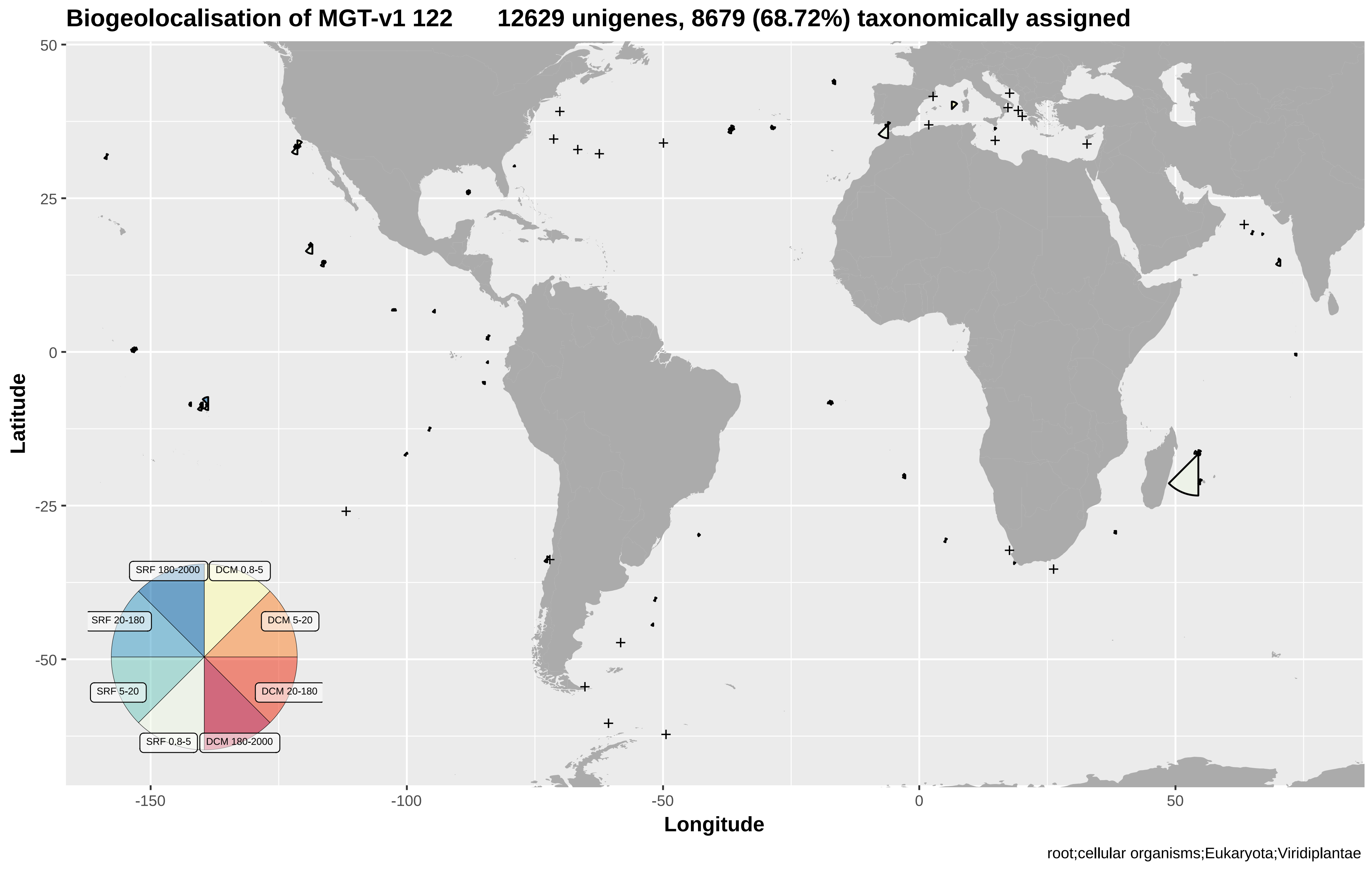
Biogeolocalisation of MGT-v1 122 12629 unigenes, 8679 (68.72%) taxonomically assigned
50
25
0
-25
-50
Latitude
-150
-100
-50
0
50
Longitude
SRF 180-2000
DCM 0.8-5
SRF 20-180
DCM 5-20
SRF 5-20
DCM 20-180
SRF 0.8-5
DCM 180-2000
root;cellular organisms;Eukaryota;Viridiplantae

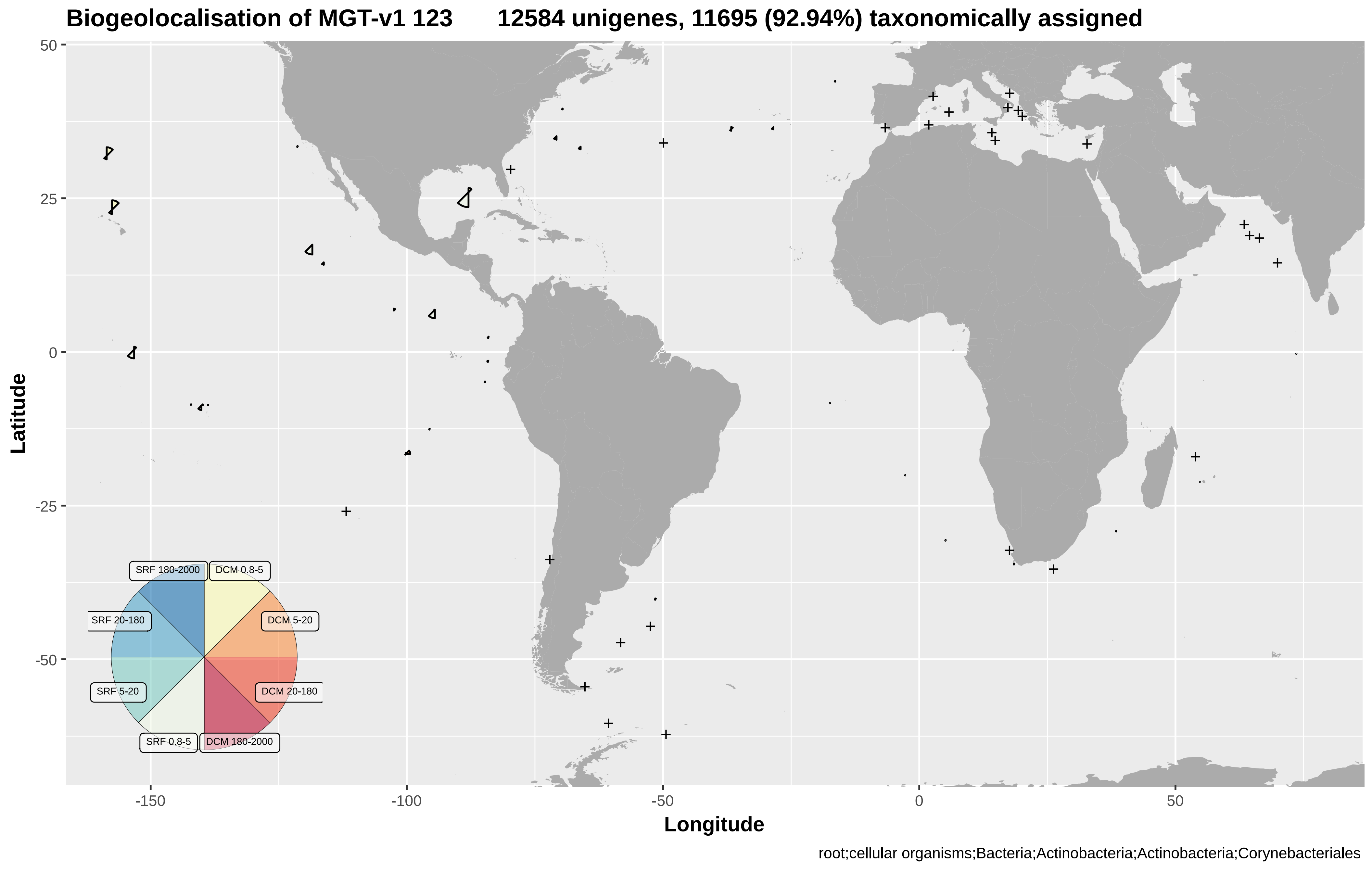

Biogeolocalisation of MGT-v1 123      12584 unigenes, 11695 (92.94%) taxonomically assigned
50
25
0
-25
-50
Latitude
-150
-100
-50
0
50
Longitude
SRF 180-2000
DCM 0.8-5
SRF 20-180
DCM 5-20
SRF 5-20
DCM 20-180
SRF 0.8-5
DCM 180-2000
root;cellular organisms;Bacteria;Actinobacteria;Actinobacteria;Corynebacteriales

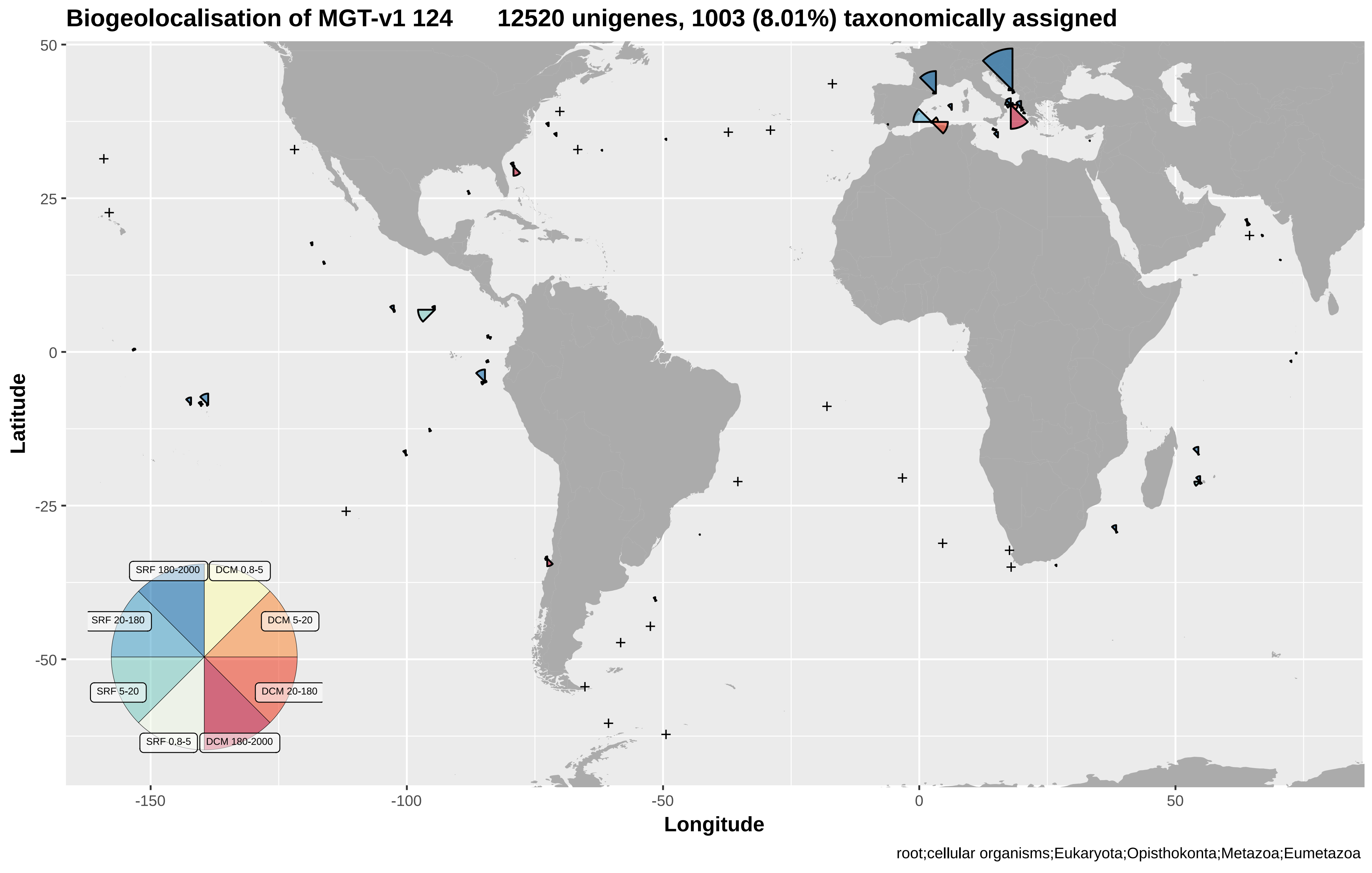
Biogeolocalisation of MGT-v1 124 12520 unigenes, 1003 (8.01%) taxonomically assigned
50
25
0
-25
-50
Latitude
-150
-100
-50
0
50
Longitude
SRF 180-2000
DCM 0.8-5
SRF 20-180
DCM 5-20
SRF 5-20
DCM 20-180
SRF 0.8-5
DCM 180-2000
root;cellular organisms;Eukaryota;Opisthokonta;Metazoa;Eumetazoa

# Biogeolocalisation of MGT-v1 125      12316 unigenes, 12026 (97.65%) taxonomically assigned

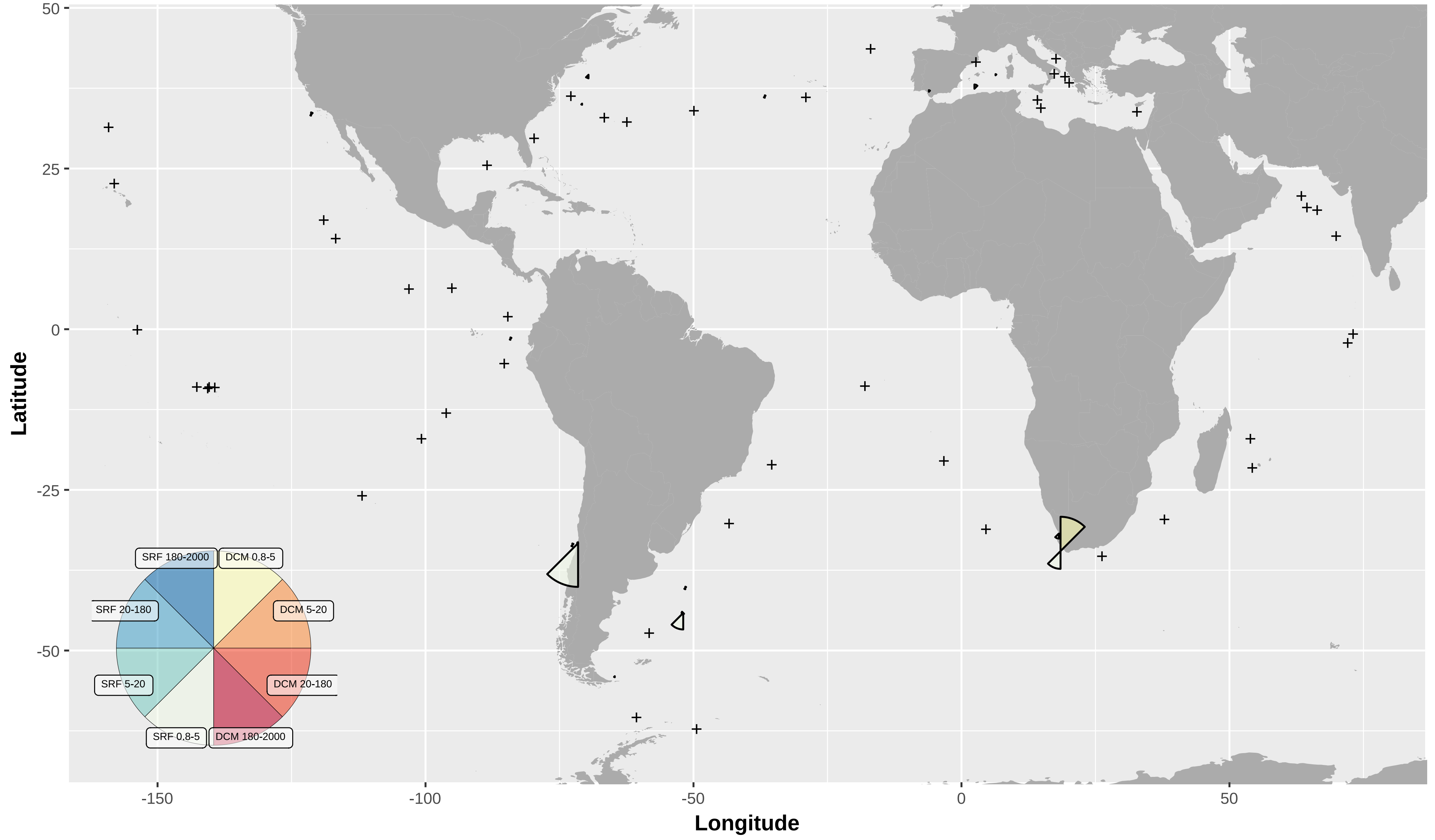


root;cellular organisms;Eukaryota;Viridiplantae;Chlorophyta;prasinophytes;Mamiellophyceae;Mamiellales;Bathycoccaceae;Ostreococcus

Biogeolocalisation of MGT-v1 126      12306 unigenes, 822 (6.68%) taxonomically assigned

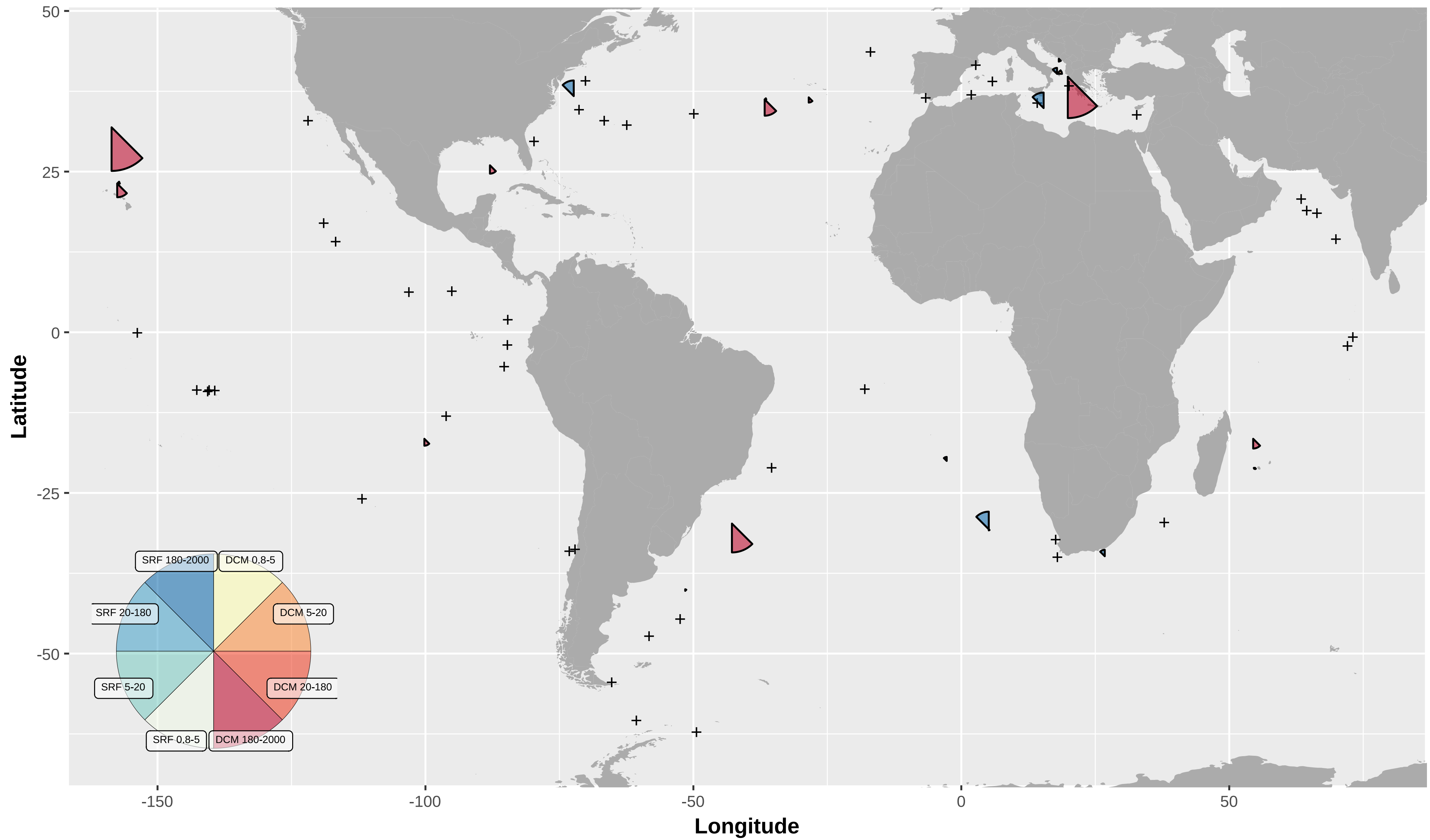


root;cellular organisms;Eukaryota;Opisthokonta;Metazoa;Eumetazoa;Bilateria

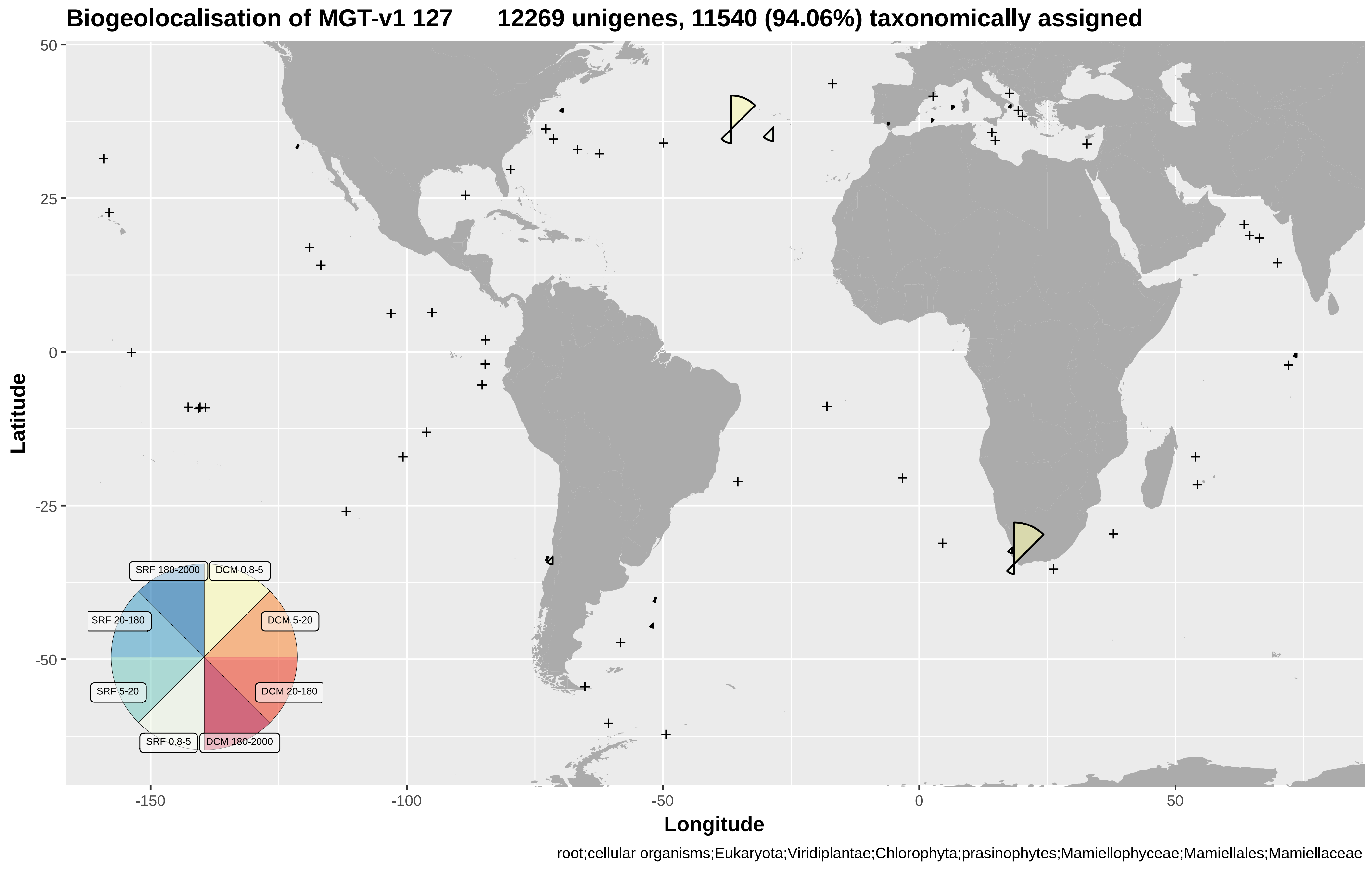
Biogeolocalisation of MGT-v1 127 12269 unigenes, 11540 (94.06%) taxonomically assigned
Latitude
50
25
0
-25
-50
Longitude
-150
-100
-50
0
50
SRF 180-2000
DCM 0.8-5
SRF 20-180
DCM 5-20
SRF 5-20
DCM 20-180
SRF 0.8-5
DCM 180-2000
root;cellular organisms;Eukaryota;Viridiplantae;Chlorophyta;prasinophytes;Mamiellophyceae;Mamiellales;Mamiellaceae

# Biogeolocalisation of MGT-v1 128      12153 unigenes, 6805 (55.99%) taxonomically assigned

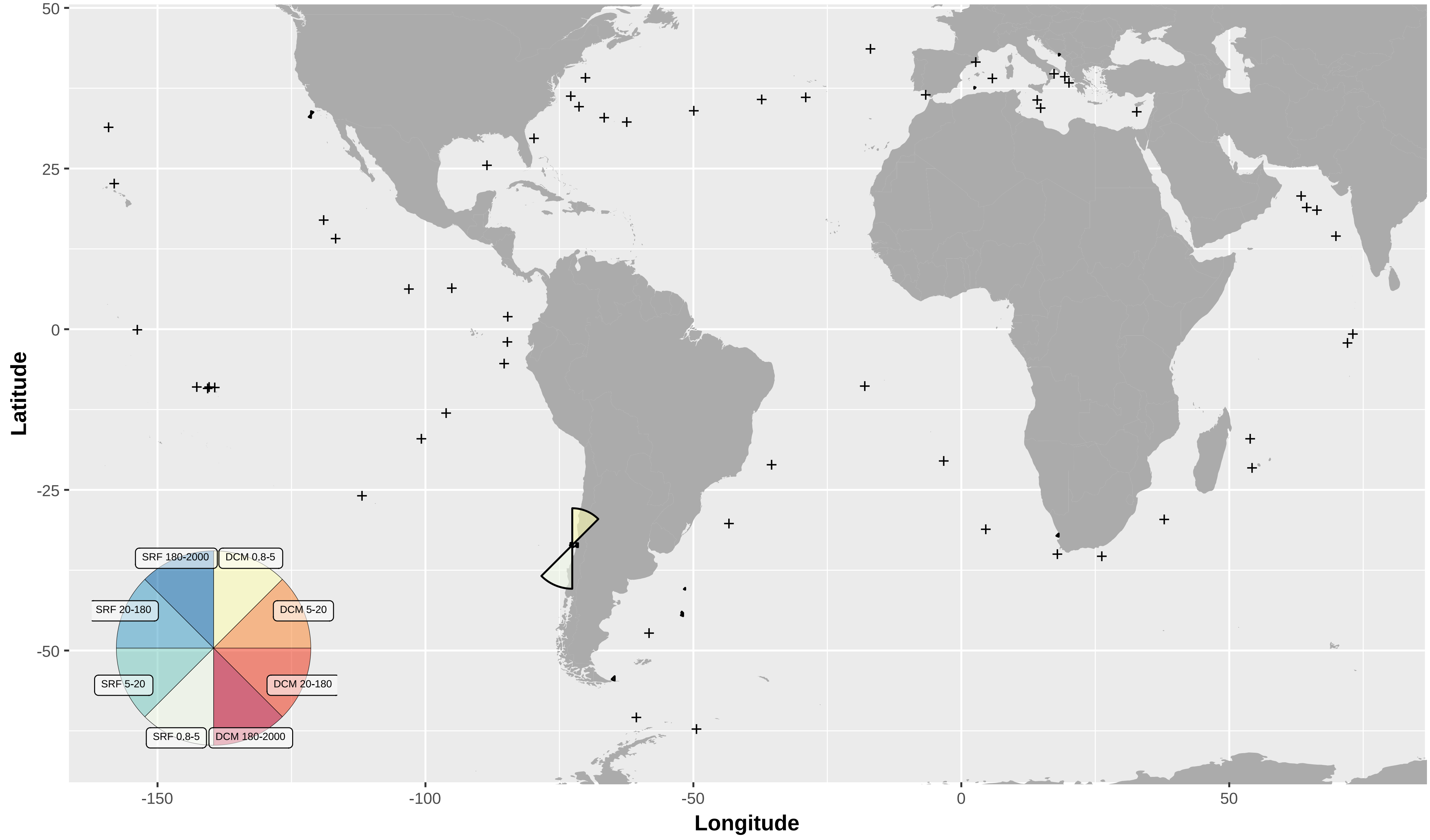


root;cellular organisms;Bacteria

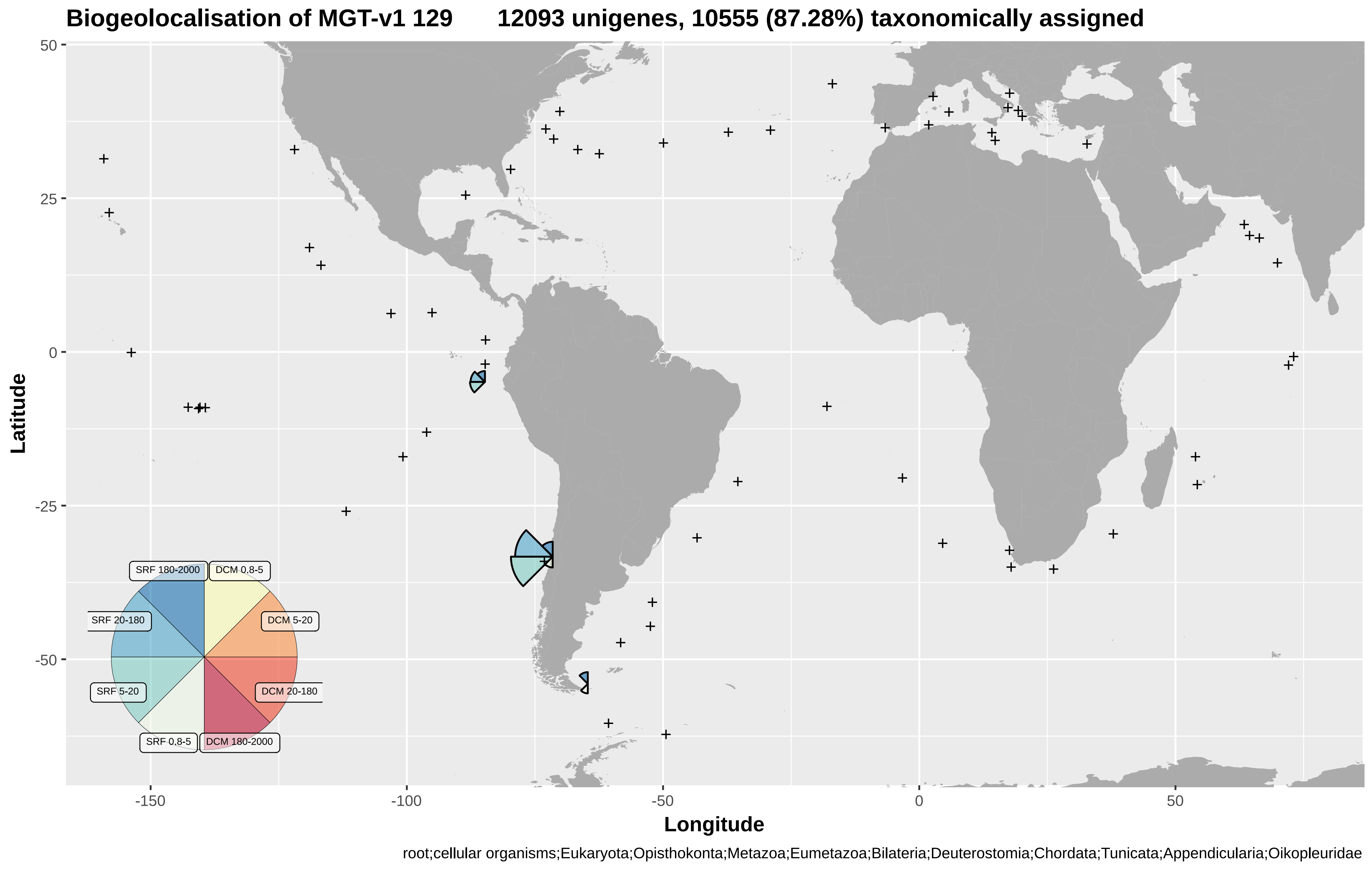

Biogeolocalisation of MGT-v1 129 12093 unigenes, 10555 (87.28%) taxonomically assigned
50
25
0
-25
-50
Latitude
-150
-100
-50
0
50
Longitude
SRF 180-2000
DCM 0.8-5
SRF 20-180
DCM 5-20
SRF 5-20
DCM 20-180
SRF 0.8-5
DCM 180-2000
root;cellular organisms;Eukaryota;Opisthokonta;Metazoa;Eumetazoa;Bilateria;Deuterostomia;Chordata;Tunicata;Appendicularia;Oikopleuridae

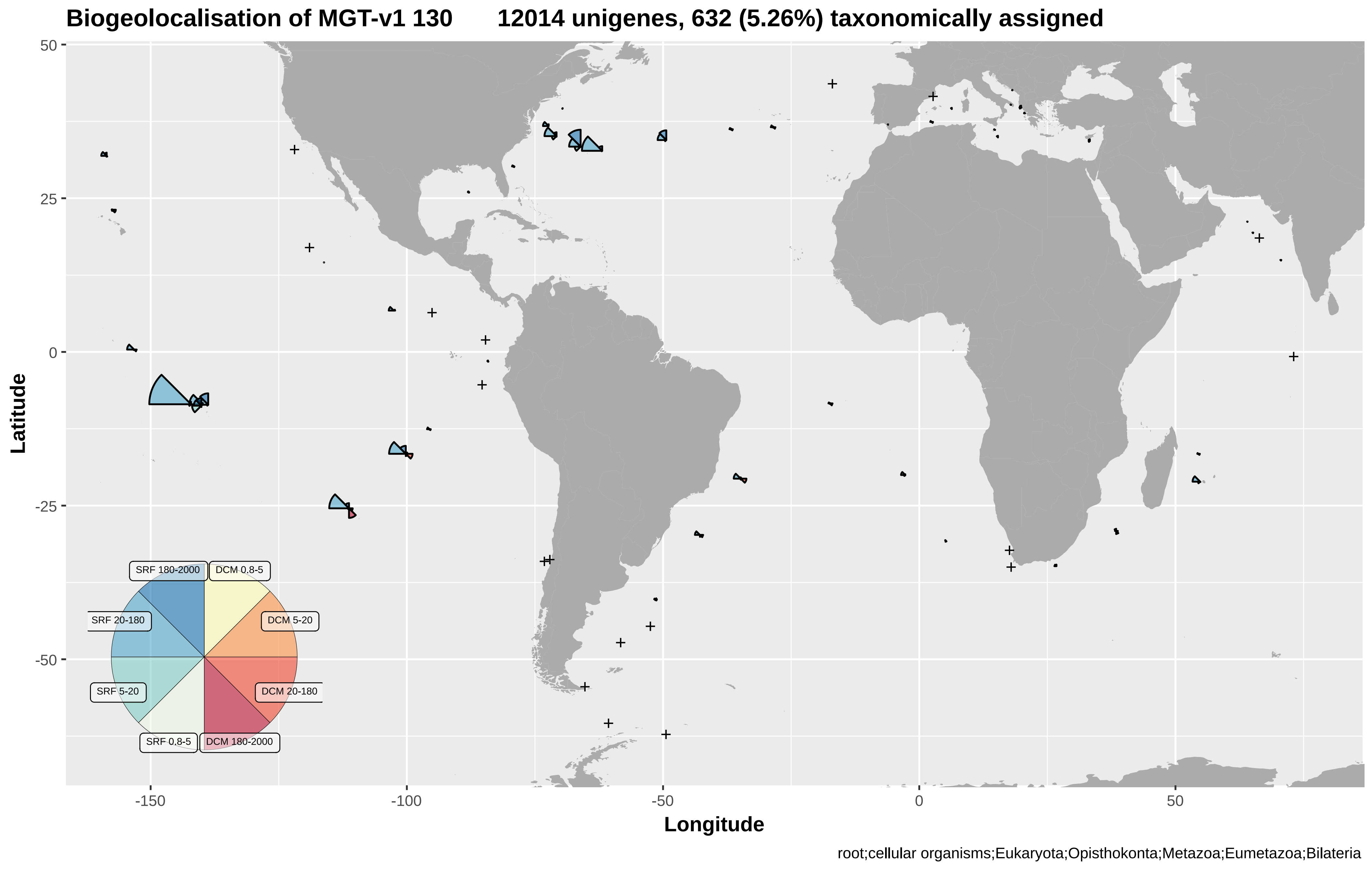

Biogeolocalisation of MGT-v1 130 12014 unigenes, 632 (5.26%) taxonomically assigned
Latitude
Longitude
50
25
0
-25
-50
-150
-100
-50
0
50
SRF 180-2000
DCM 0.8-5
SRF 20-180
DCM 5-20
SRF 5-20
DCM 20-180
SRF 0.8-5
DCM 180-2000
root;cellular organisms;Eukaryota;Opisthokonta;Metazoa;Eumetazoa;Bilateria

#### Biogeolocalisation of MGT-v1 131 11922 unigenes, 8692 (72.91%) taxonomically assigned

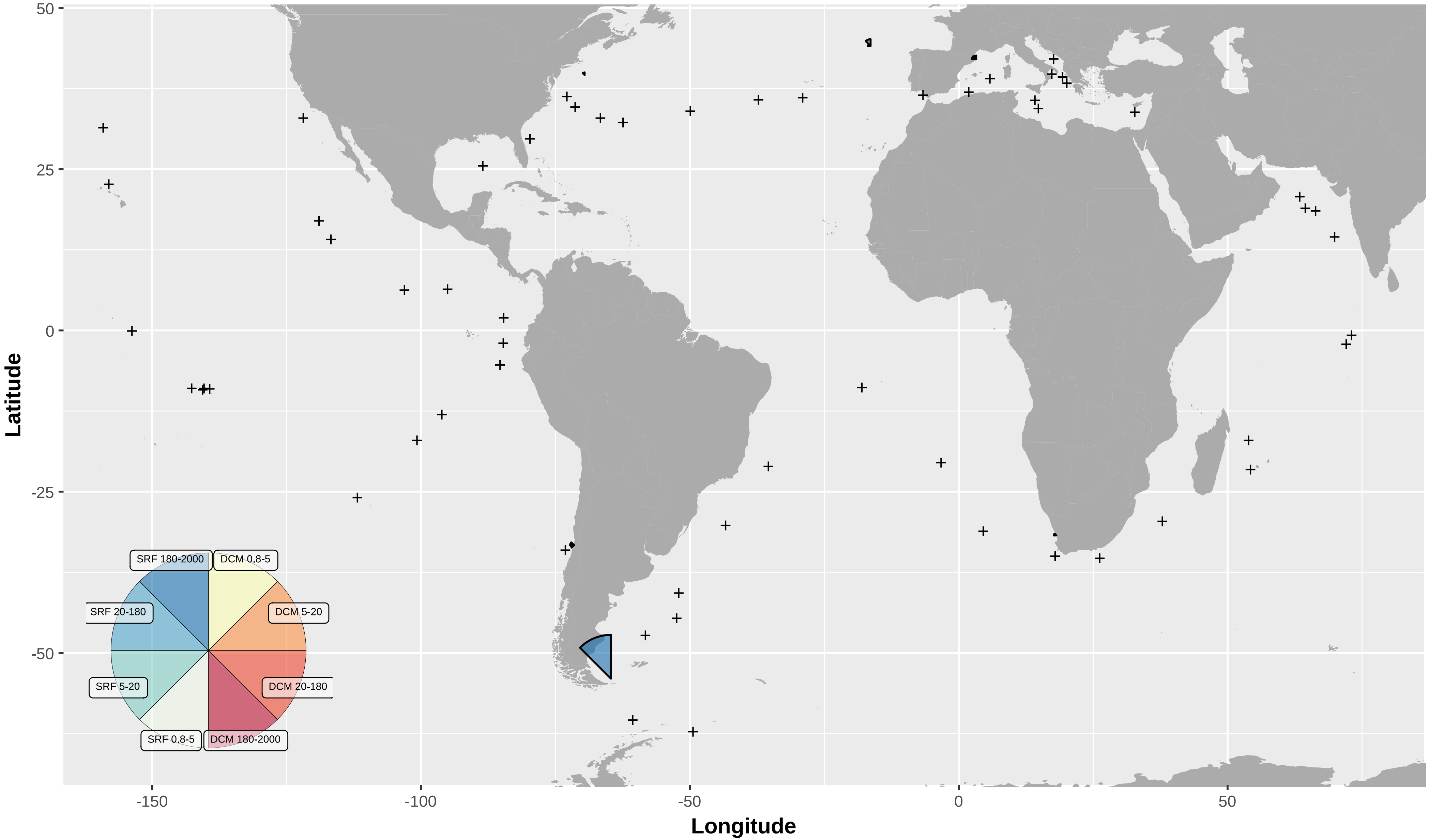


root;cellular organisms;Eukaryota;Opisthokonta;Metazoa;Eumetazoa;Bilateria;Protostomia;Ecdysozoa;Panarthropoda;Arthropoda;Mandibulata;Pancrustacea;Crustacea

## Biogeolocalisation of MGT-v1 132      11894 unigenes, 10226 (85.98%) taxonomically assigned

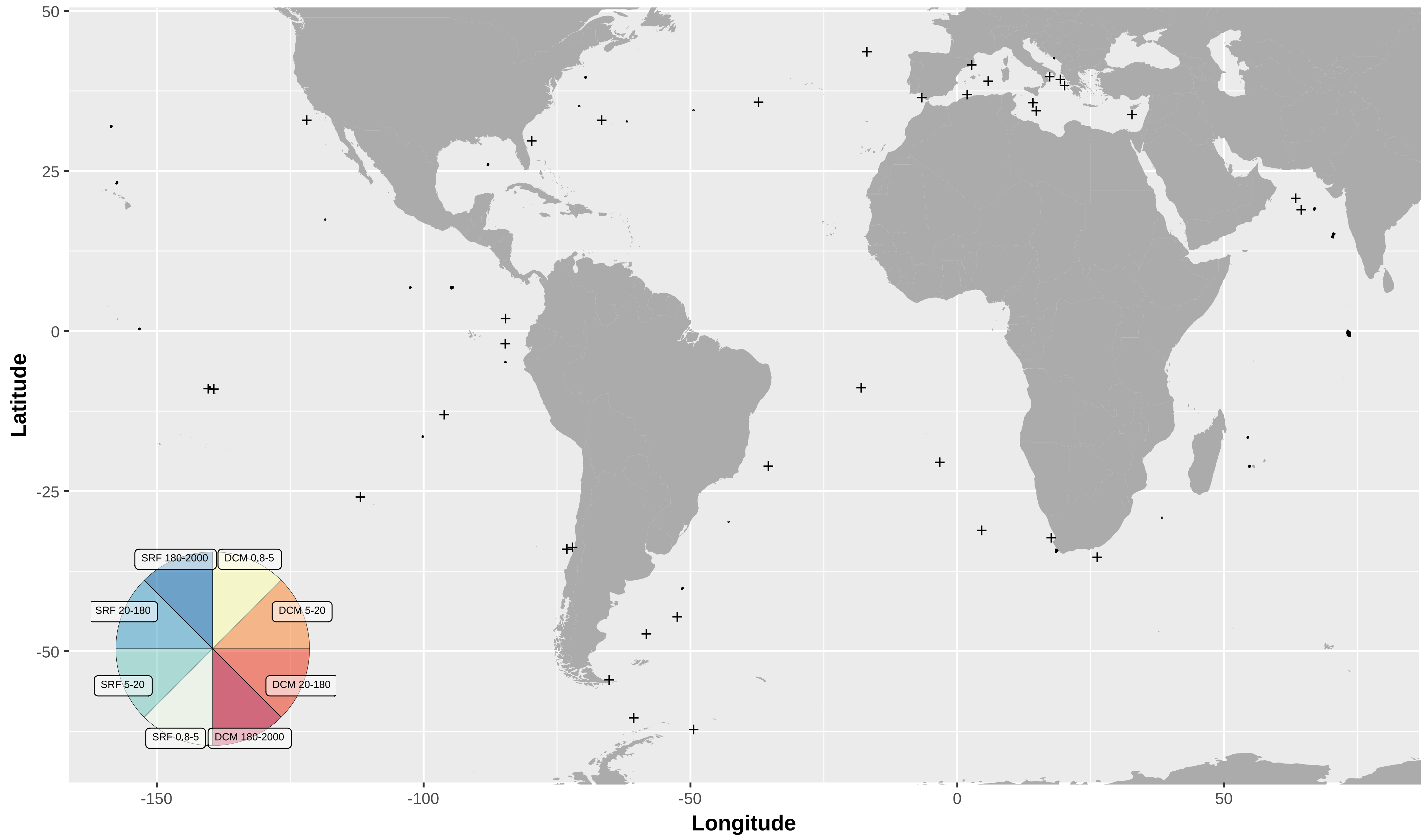


root;cellular organisms;Bacteria;Proteobacteria

## Biogeolocalisation of MGT-v1 133      11827 unigenes, 9413 (79.59%) taxonomically assigned

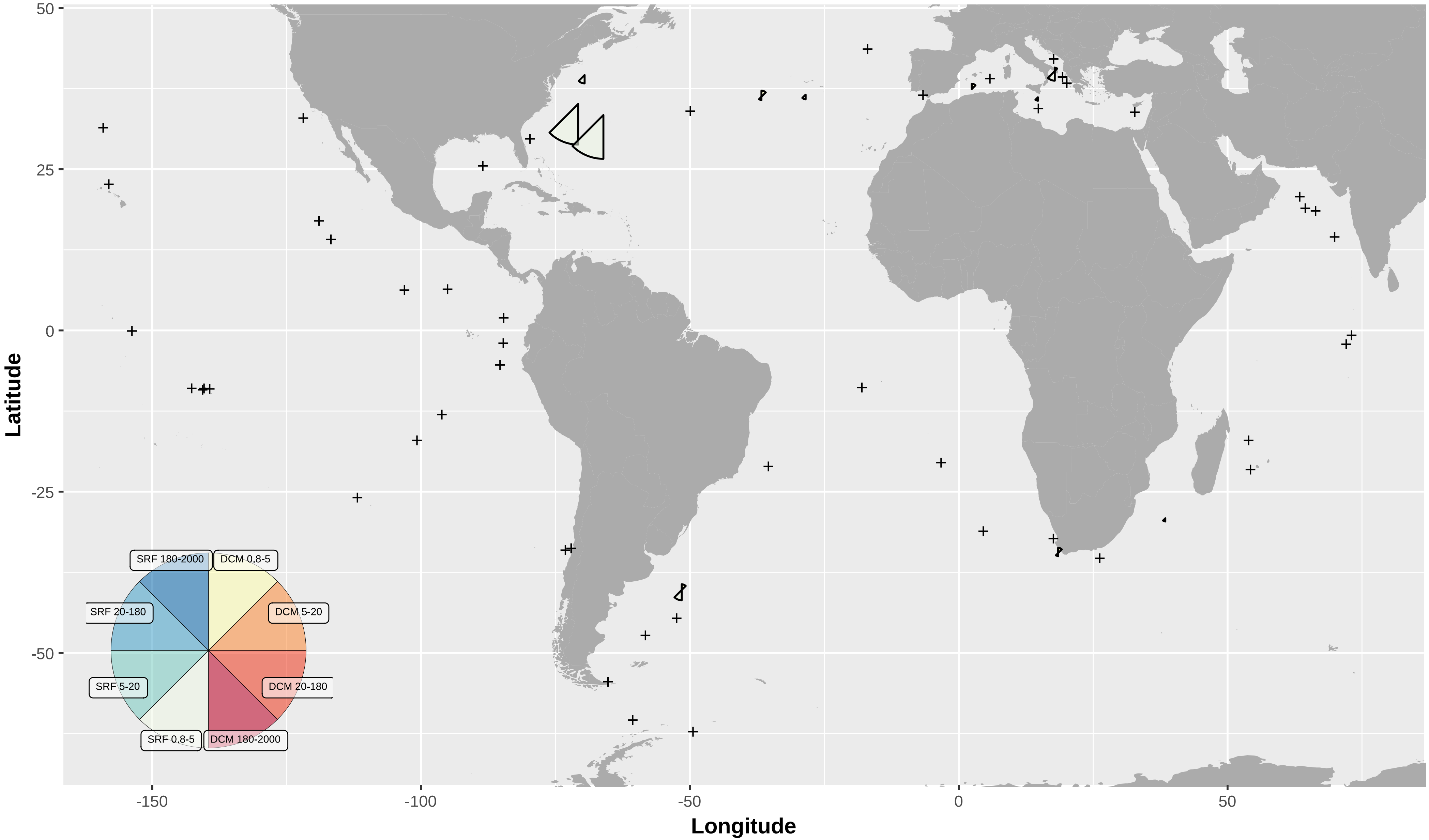


root;cellular organisms;Eukaryota;Viridiplantae;Chlorophyta;prasinophytes;Mamiellophyceae

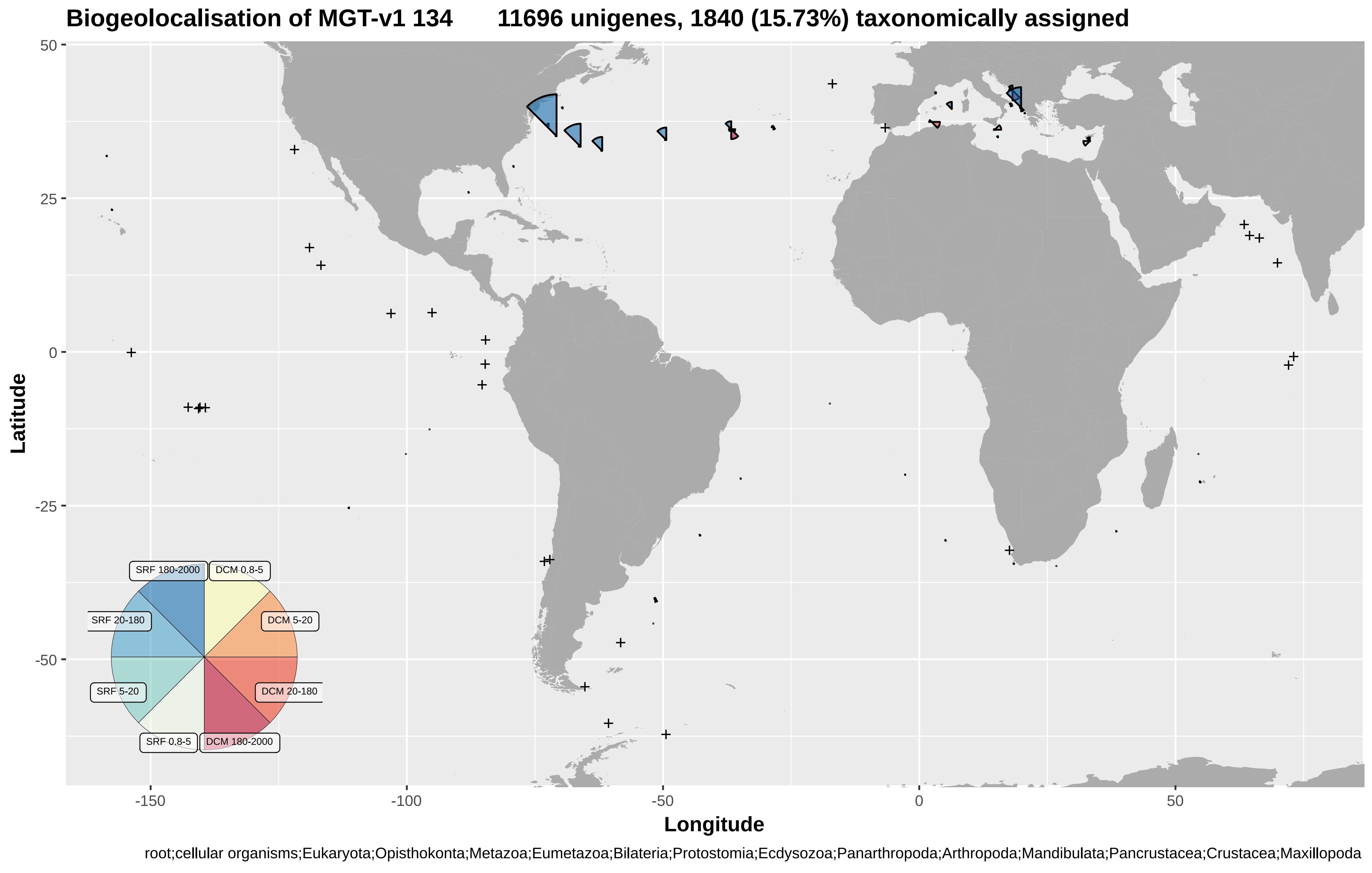
Biogeolocalisation of MGT-v1 134 11696 unigenes, 1840 (15.73%) taxonomically assigned
Latitude
50
25
0
-25
-50
Longitude
-150
-100
-50
0
50
SRF 180-2000
DCM 0.8-5
SRF 20-180
DCM 5-20
SRF 5-20
DCM 20-180
SRF 0.8-5
DCM 180-2000
root;cellular organisms;Eukaryota;Opisthokonta;Metazoa;Eumetazoa;Bilateria;Protostomia;Ecdysozoa;Panarthropoda;Arthropoda;Mandibulata;Pancrustacea;Crustacea;Maxillopoda

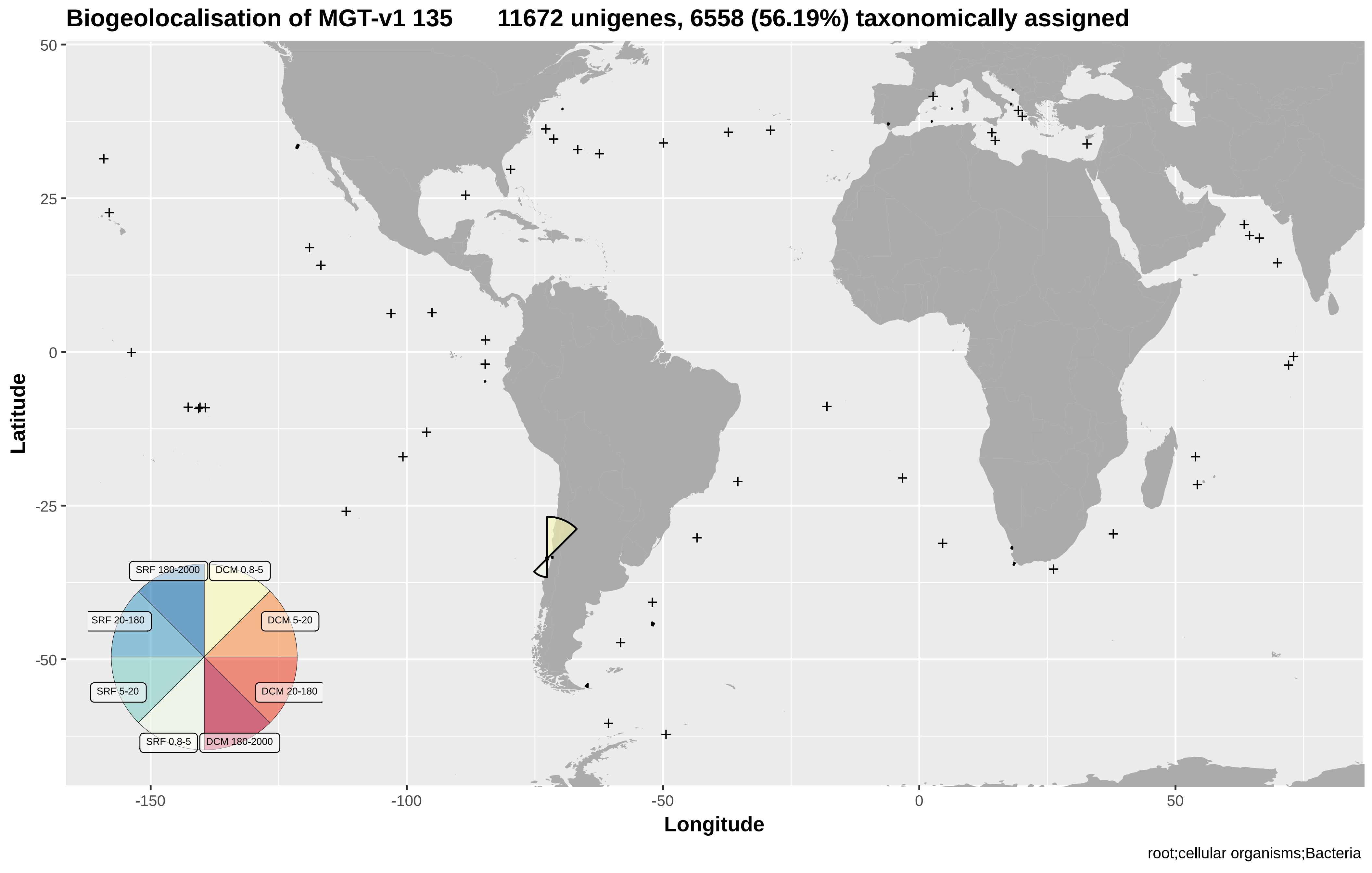

Biogeolocalisation of MGT-v1 135 11672 unigenes, 6558 (56.19%) taxonomically assigned
50
25
0
-25
-50
Latitude
-150
-100
-50
0
50
Longitude
SRF 180-2000
DCM 0.8-5
SRF 20-180
DCM 5-20
SRF 5-20
DCM 20-180
SRF 0.8-5
DCM 180-2000
root;cellular organisms;Bacteria

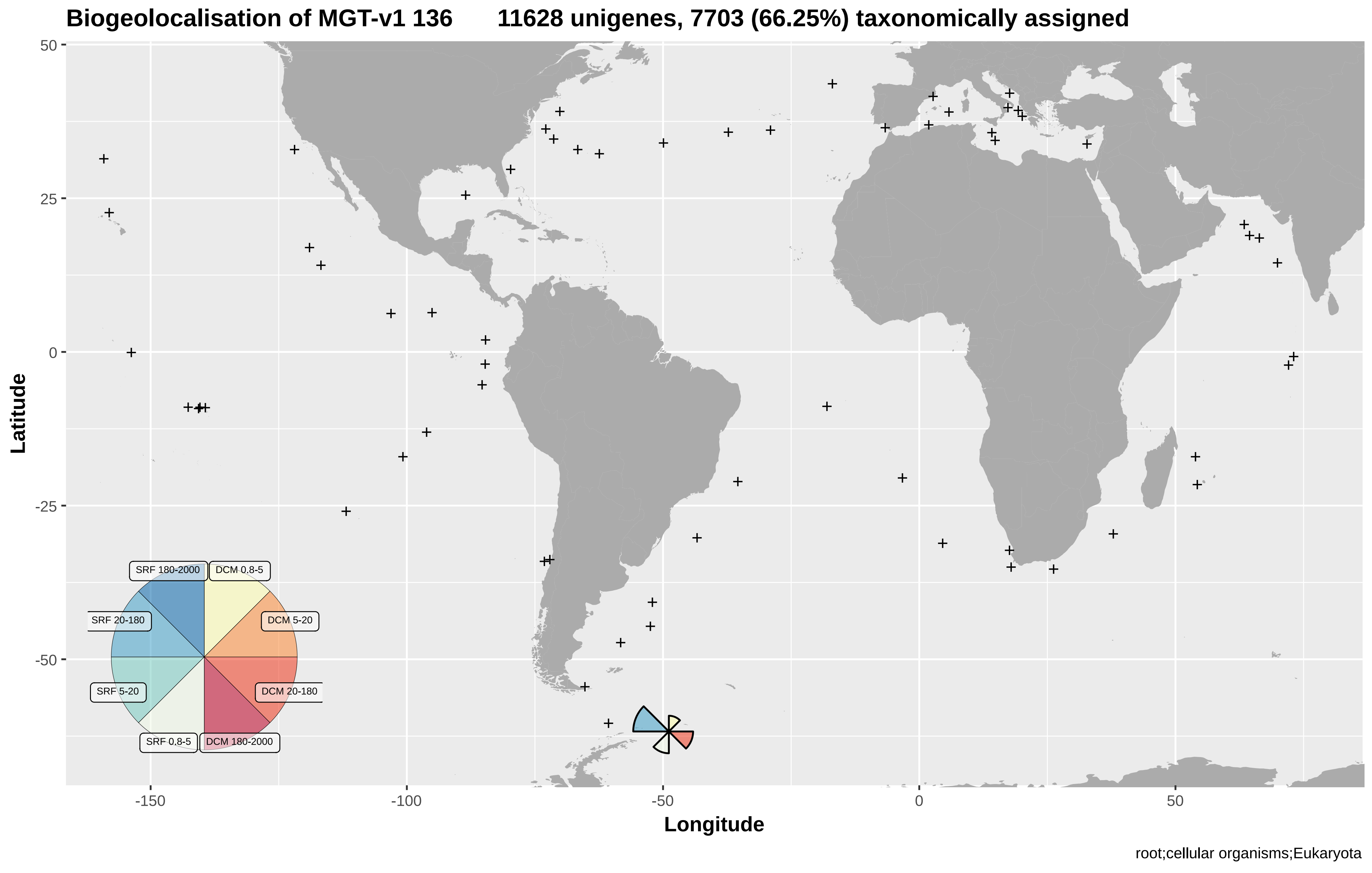
Biogeolocalisation of MGT-v1 136 11628 unigenes, 7703 (66.25%) taxonomically assigned
50
25
0
-25
-50
Latitude
-150
-100
-50
0
50
Longitude
SRF 180-2000
DCM 0.8-5
SRF 20-180
DCM 5-20
SRF 5-20
DCM 20-180
SRF 0.8-5
DCM 180-2000
root;cellular organisms;Eukaryota

## Biogeolocalisation of MGT-v1 137      11552 unigenes, 11225 (97.17%) taxonomically assigned

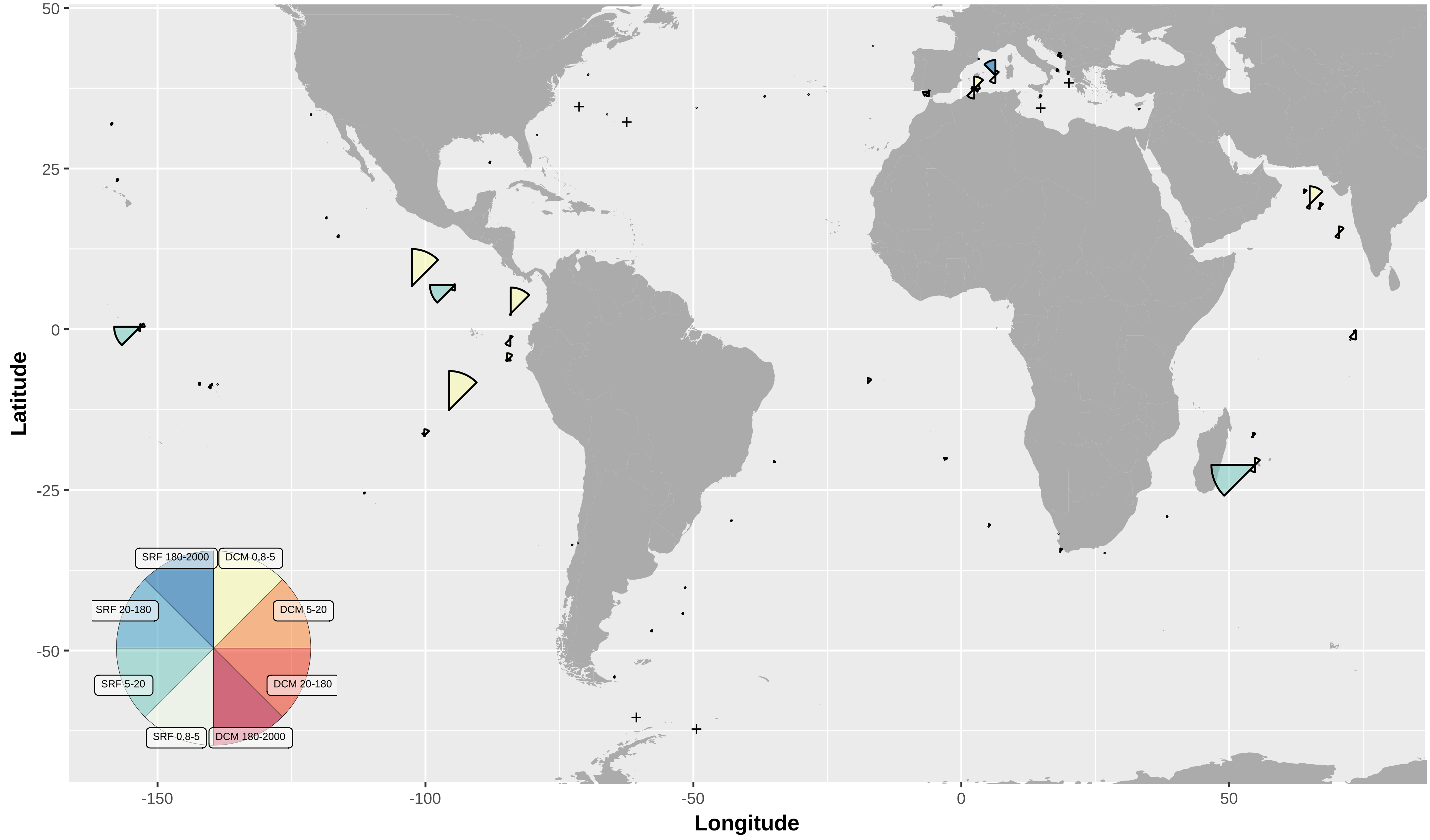


root;cellular organisms;Bacteria;Proteobacteria;Gammaproteobacteria;Alteromonadales;Alteromonadaceae

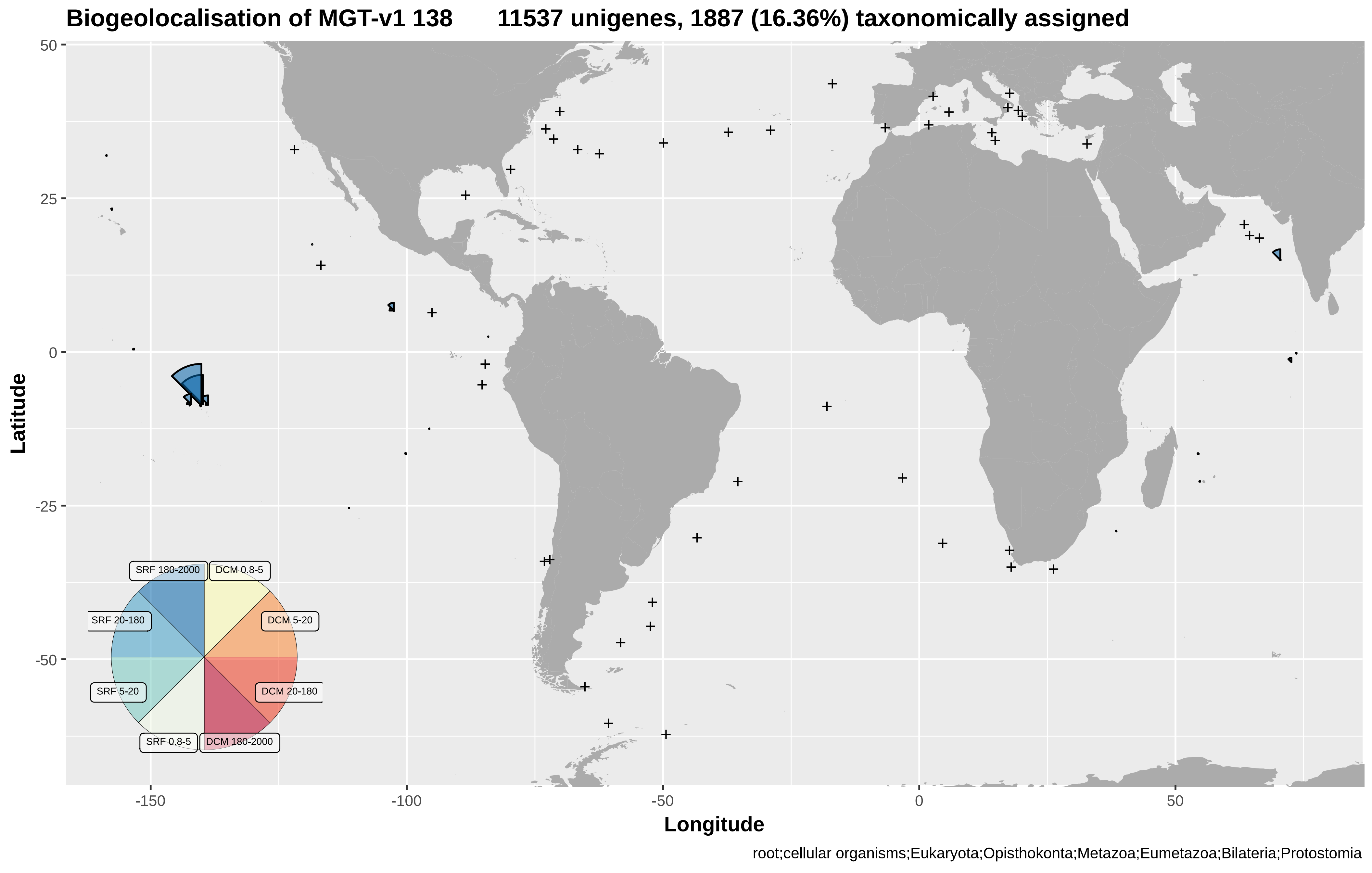
Biogeolocalisation of MGT-v1 138 11537 unigenes, 1887 (16.36%) taxonomically assigned
Latitude
Longitude
50
25
0
-25
-50
-150
-100
-50
0
50
SRF 180-2000
DCM 0.8-5
SRF 20-180
DCM 5-20
SRF 5-20
DCM 20-180
SRF 0.8-5
DCM 180-2000
root;cellular organisms;Eukaryota;Opisthokonta;Metazoa;Eumetazoa;Bilateria;Protostomia

## Biogeolocalisation of MGT-v1 139      11413 unigenes, 6319 (55.37%) taxonomically assigned

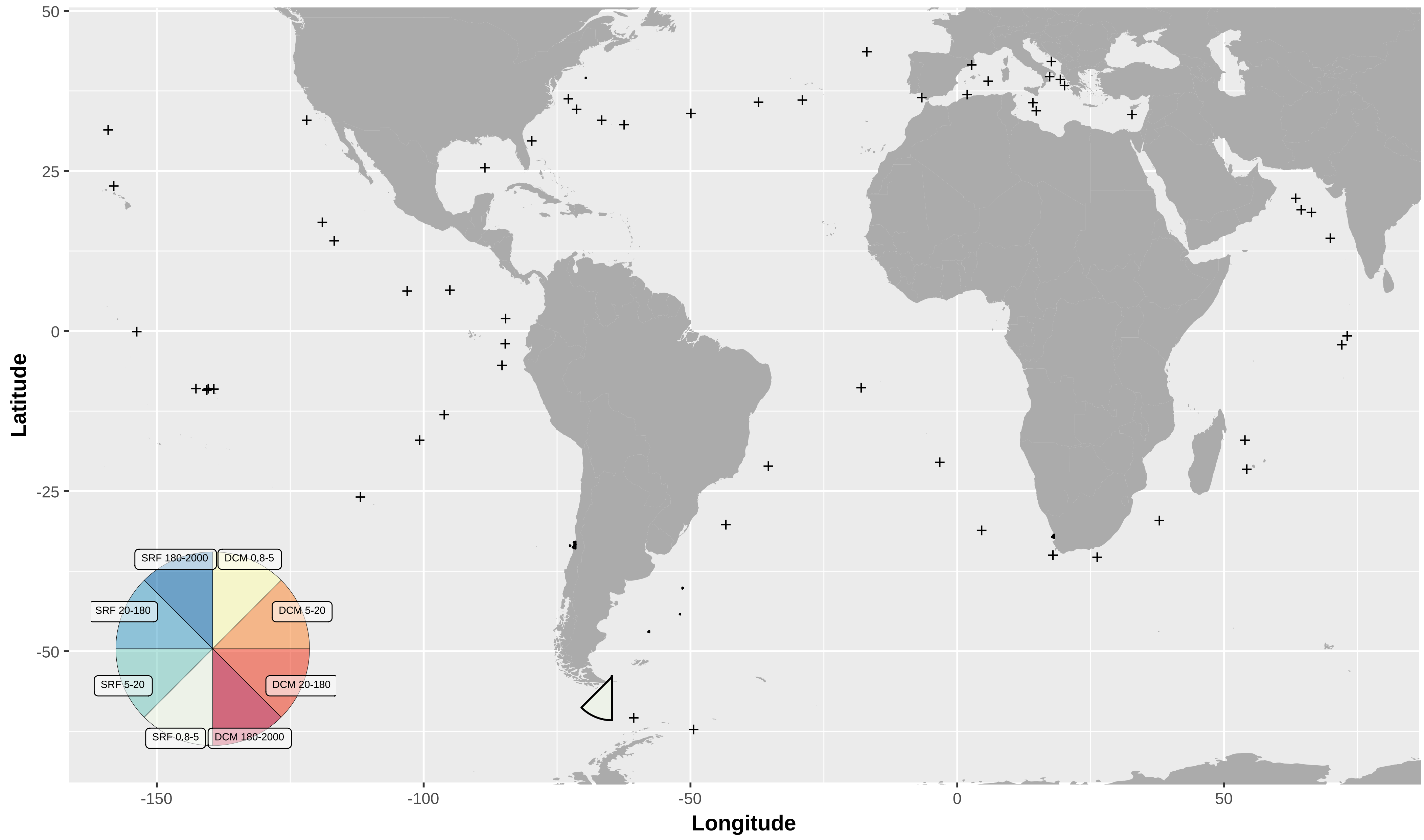


root;cellular organisms

#### Biogeolocalisation of MGT-v1 140 11221 unigenes, 8518 (75.91%) taxonomically assigned

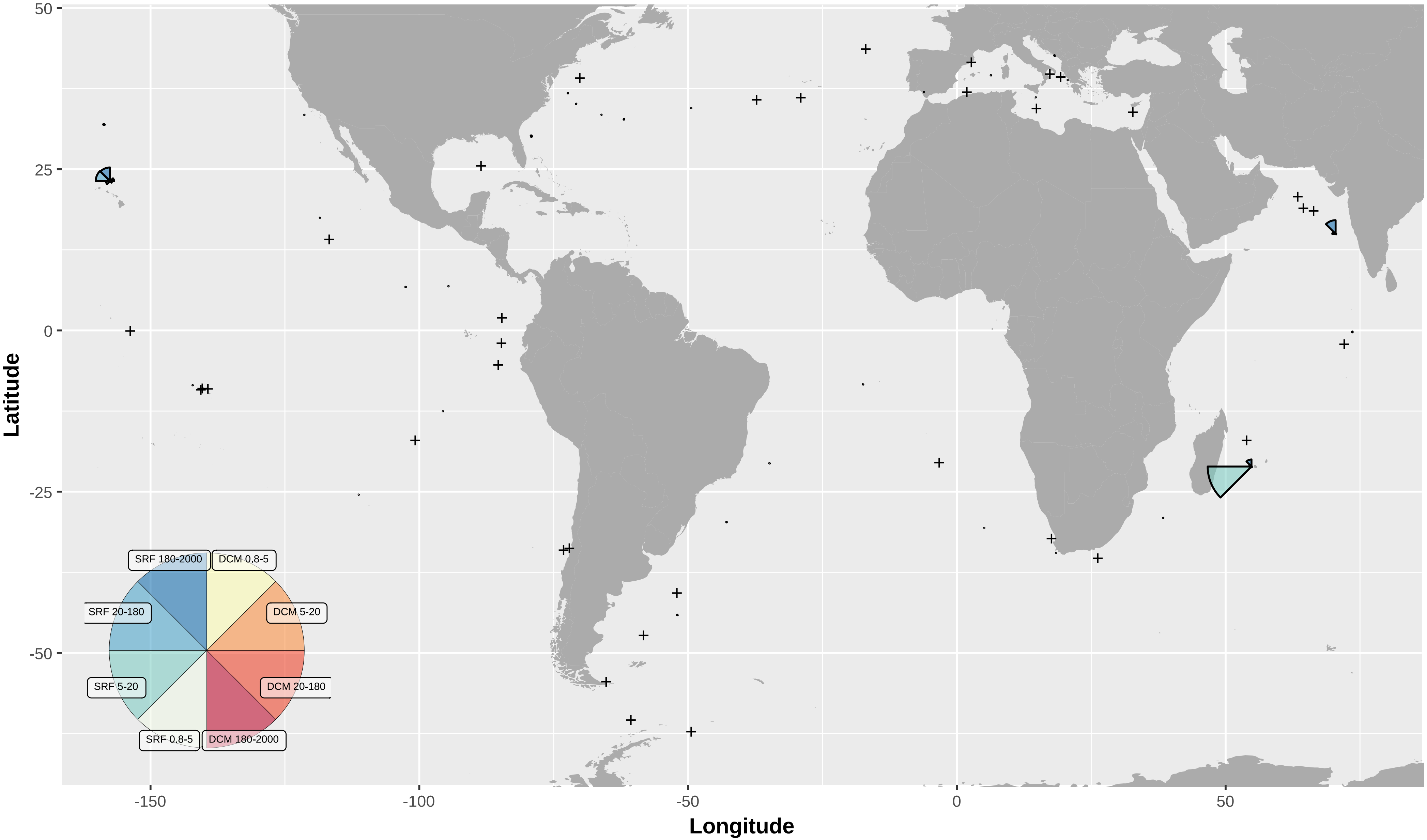


root;cellular organisms;Bacteria;Cyanobacteria;Oscillatoriophycideae

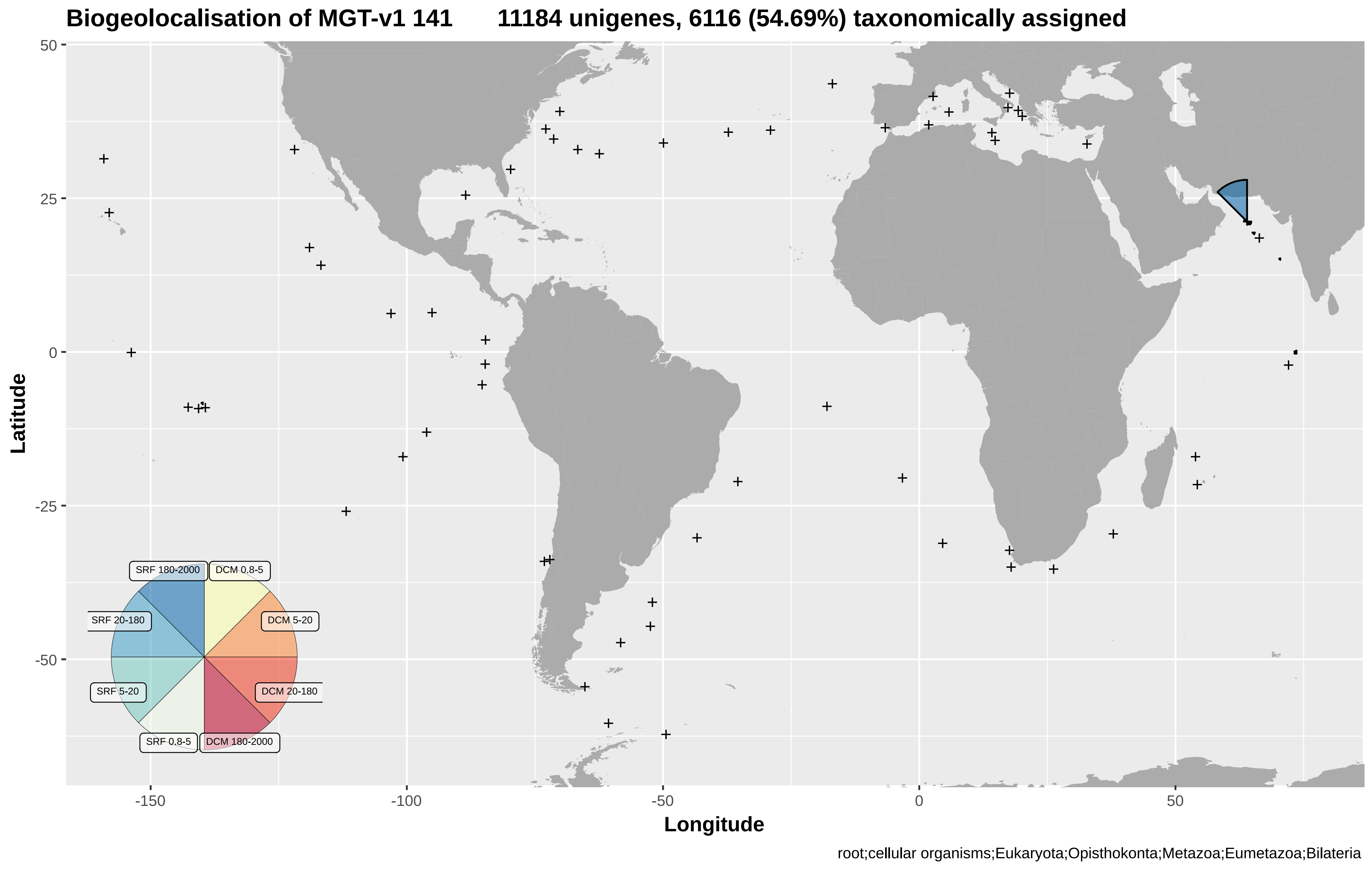
Biogeolocalisation of MGT-v1 141 11184 unigenes, 6116 (54.69%) taxonomically assigned
50
25
0
-25
-50
Latitude
-150
-100
-50
0
50
Longitude
SRF 180-2000
DCM 0.8-5
SRF 20-180
DCM 5-20
SRF 5-20
DCM 20-180
SRF 0.8-5
DCM 180-2000
root;cellular organisms;Eukaryota;Opisthokonta;Metazoa;Eumetazoa;Bilateria

#### Biogeolocalisation of MGT-v1 142 11167 unigenes, 1238 (11.09%) taxonomically assigned

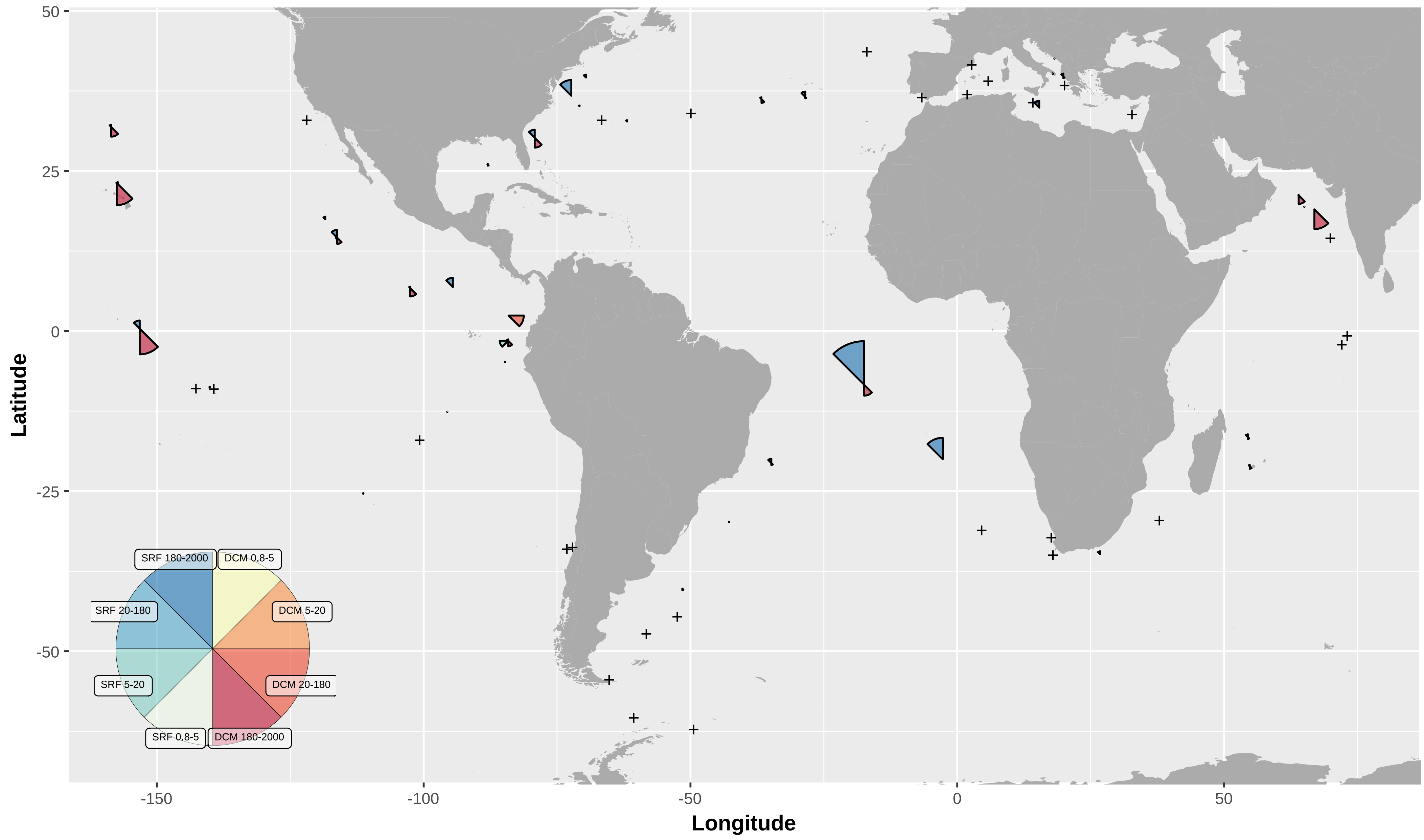


root;cellular organisms;Eukaryota;Opisthokonta;Metazoa;Eumetazoa;Bilateria

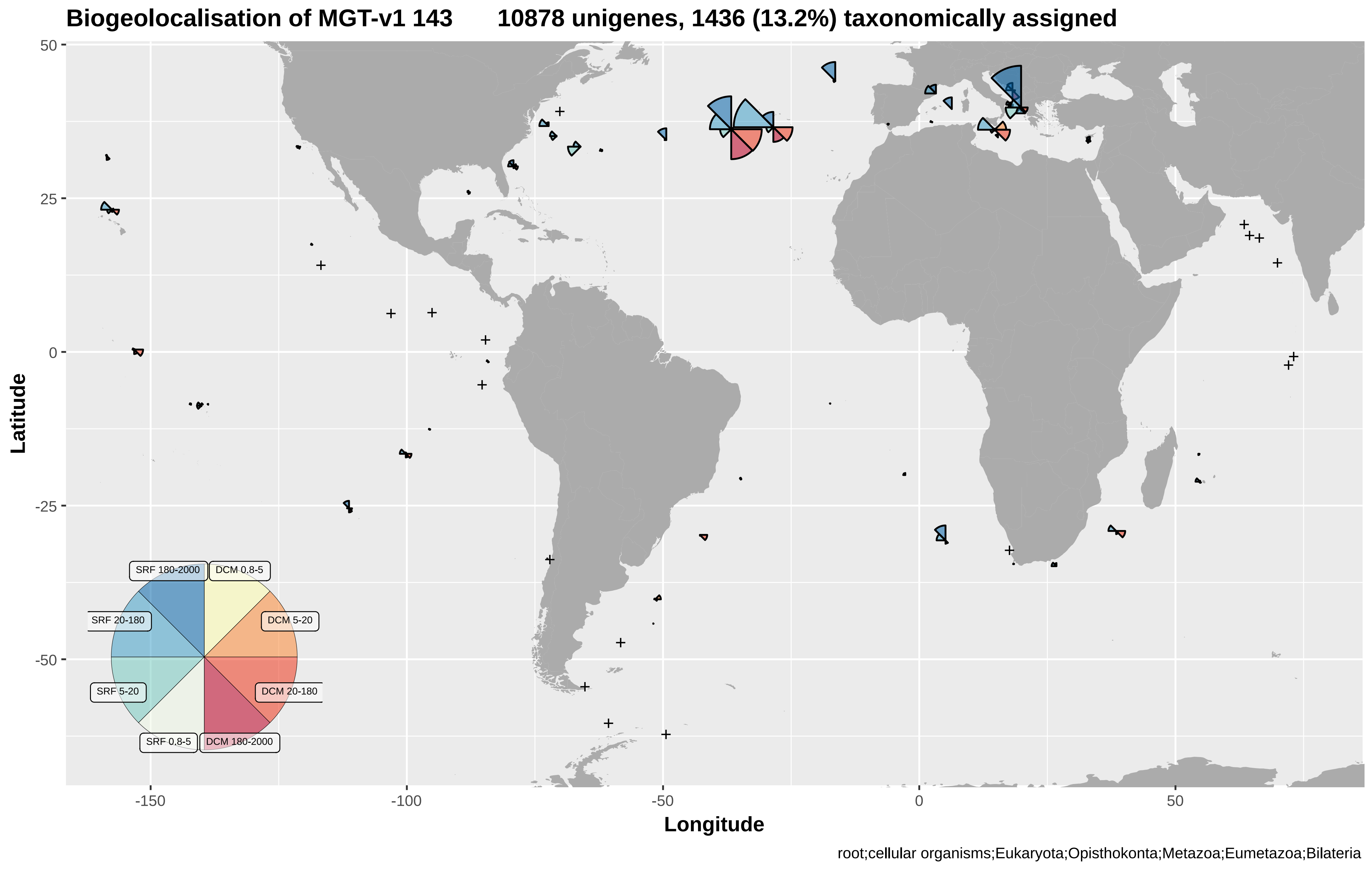

Biogeolocalisation of MGT-v1 143      10878 unigenes, 1436 (13.2%) taxonomically assigned
50
25
0
-25
-50
Latitude
-150
-100
-50
0
50
Longitude
SRF 180-2000
DCM 0.8-5
SRF 20-180
DCM 5-20
SRF 5-20
DCM 20-180
SRF 0.8-5
DCM 180-2000
root;cellular organisms;Eukaryota;Opisthokonta;Metazoa;Eumetazoa;Bilateria

#### Biogeolocalisation of MGT-v1 144 10820 unigenes, 7522 (69.52%) taxonomically assigned

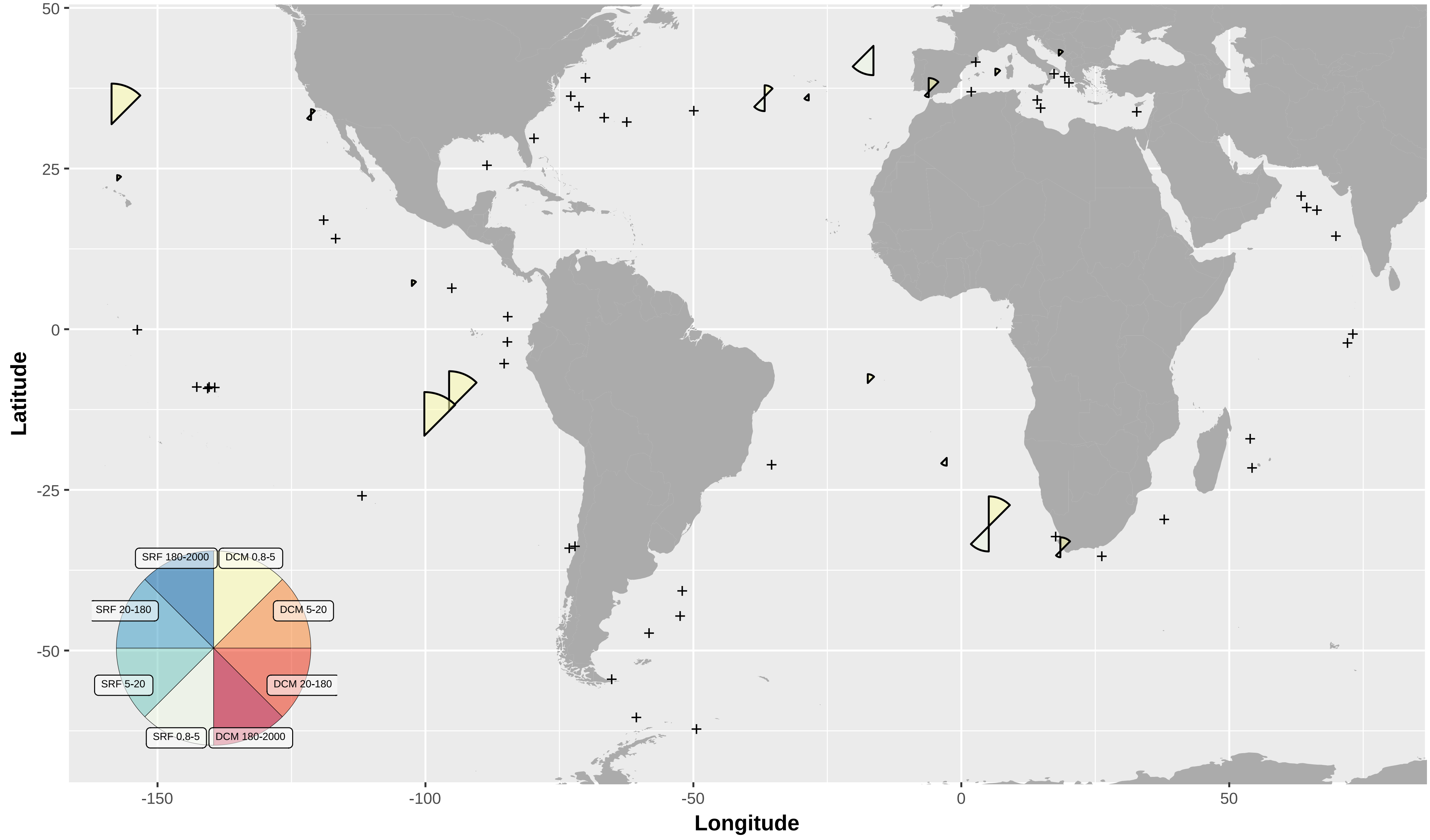


root;cellular organisms;Eukaryota;Haptophyceae

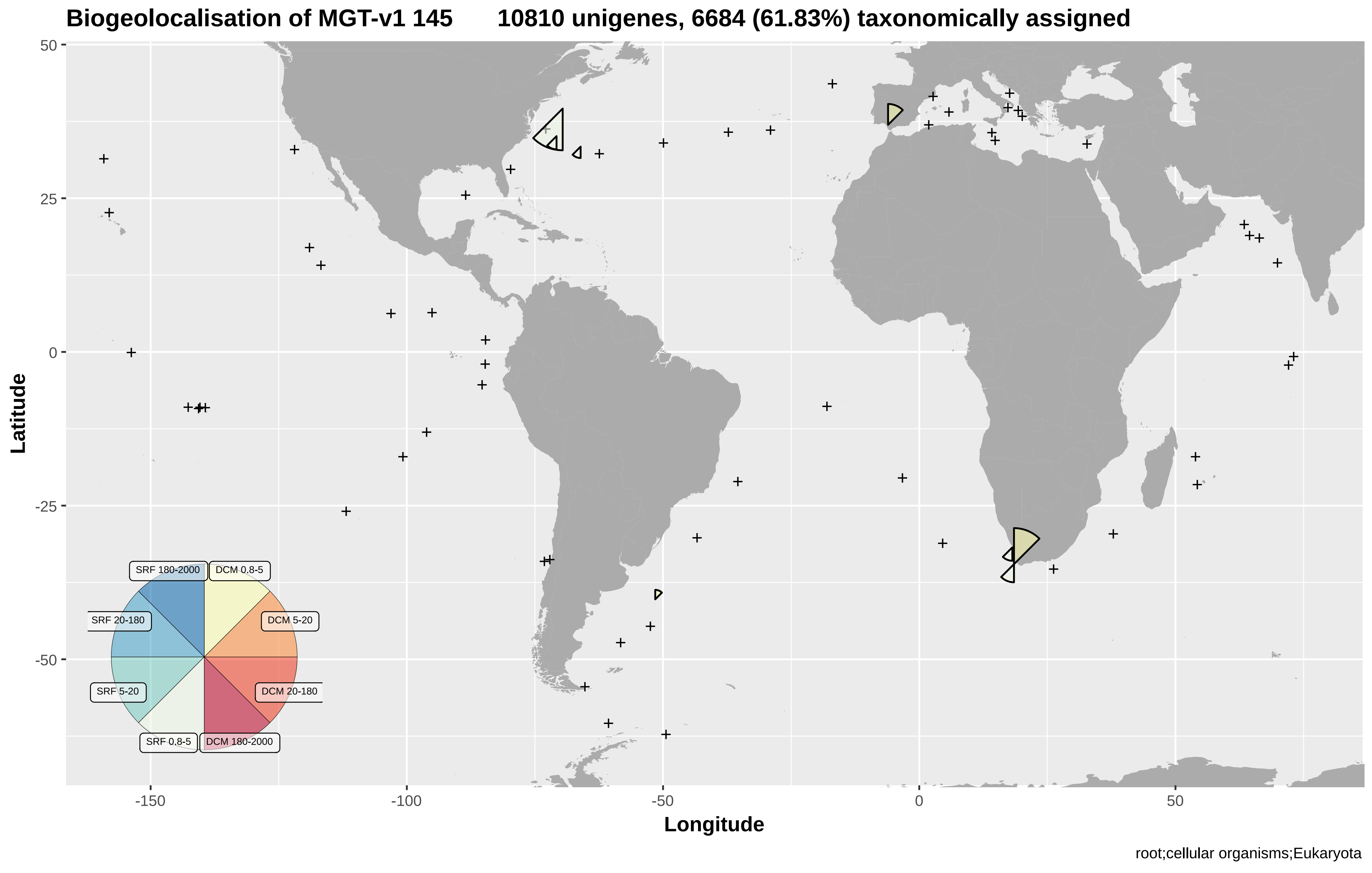
Biogeolocalisation of MGT-v1 145 10810 unigenes, 6684 (61.83%) taxonomically assigned
Latitude
Longitude
50
25
0
-25
-50
-150
-100
-50
0
50
SRF 180-2000
DCM 0.8-5
SRF 20-180
DCM 5-20
SRF 5-20
DCM 20-180
SRF 0.8-5
DCM 180-2000
root;cellular organisms;Eukaryota

### Biogeolocalisation of MGT-v1 146 10714 unigenes, 4454 (41.57%) taxonomically assigned

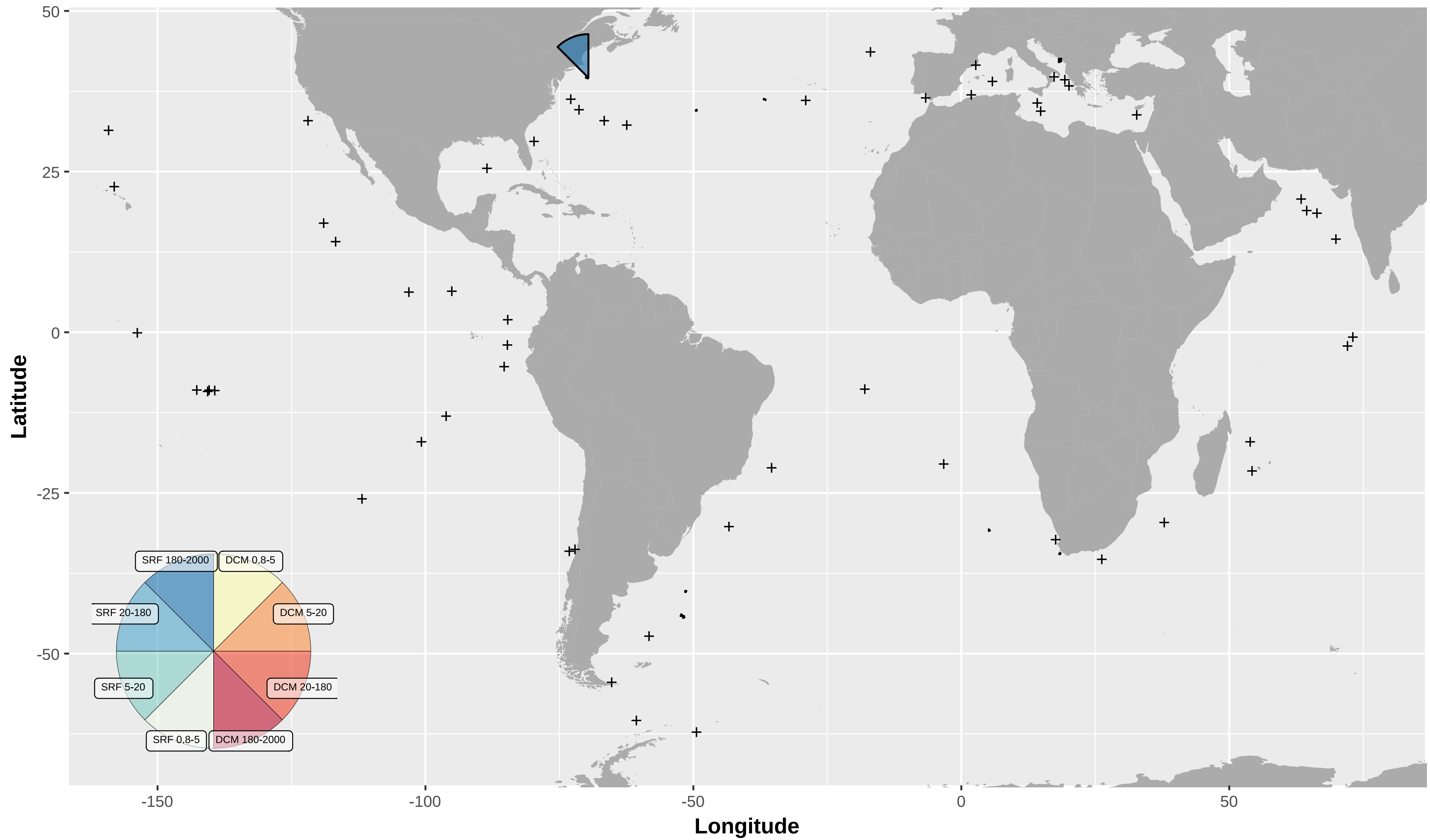


root;cellular organisms;Eukaryota;Opisthokonta;Metazoa;Eumetazoa;Bilateria;Protostomia;Ecdysozoa

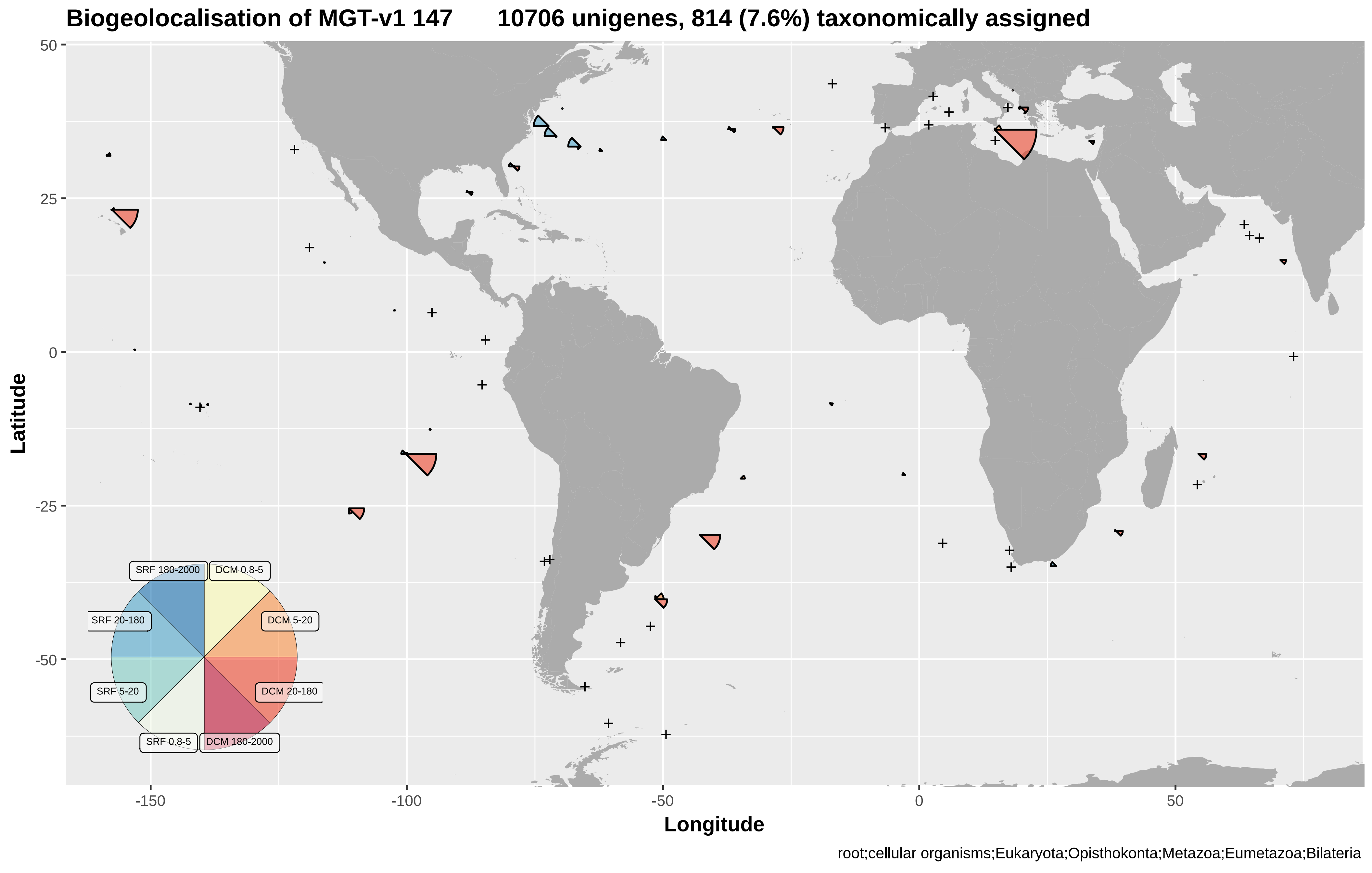

Biogeolocalisation of MGT-v1 147 10706 unigenes, 814 (7.6%) taxonomically assigned
50
25
0
-25
-50
Latitude
-150
-100
-50
0
50
Longitude
SRF 180-2000
DCM 0.8-5
SRF 20-180
DCM 5-20
SRF 5-20
DCM 20-180
SRF 0.8-5
DCM 180-2000
root;cellular organisms;Eukaryota;Opisthokonta;Metazoa;Eumetazoa;Bilateria

#### Biogeolocalisation of MGT-v1 148 10675 unigenes, 4052 (37.96%) taxonomically assigned

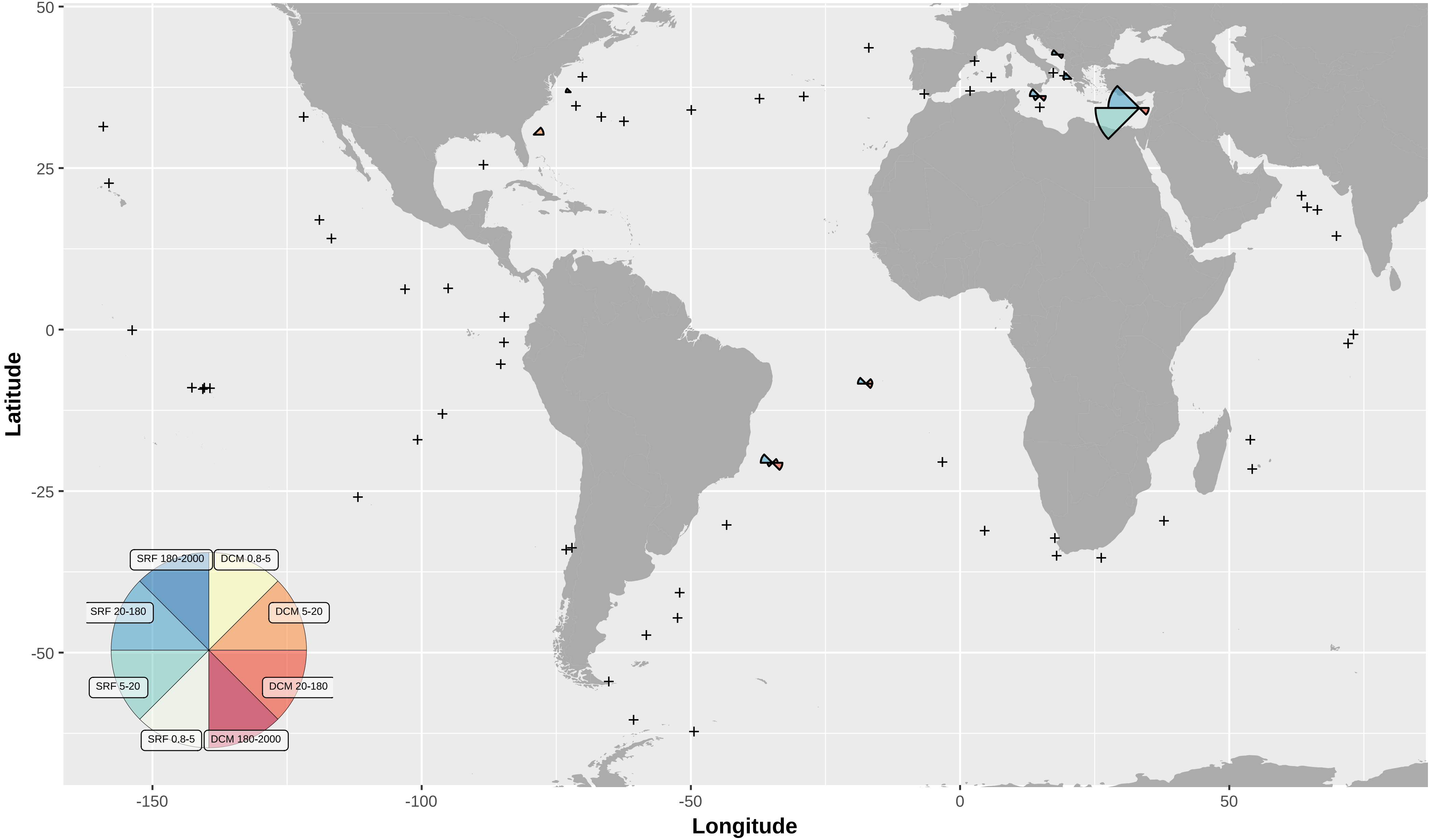


root;cellular organisms;Eukaryota;Opisthokonta;Metazoa;Eumetazoa;Bilateria;Protostomia

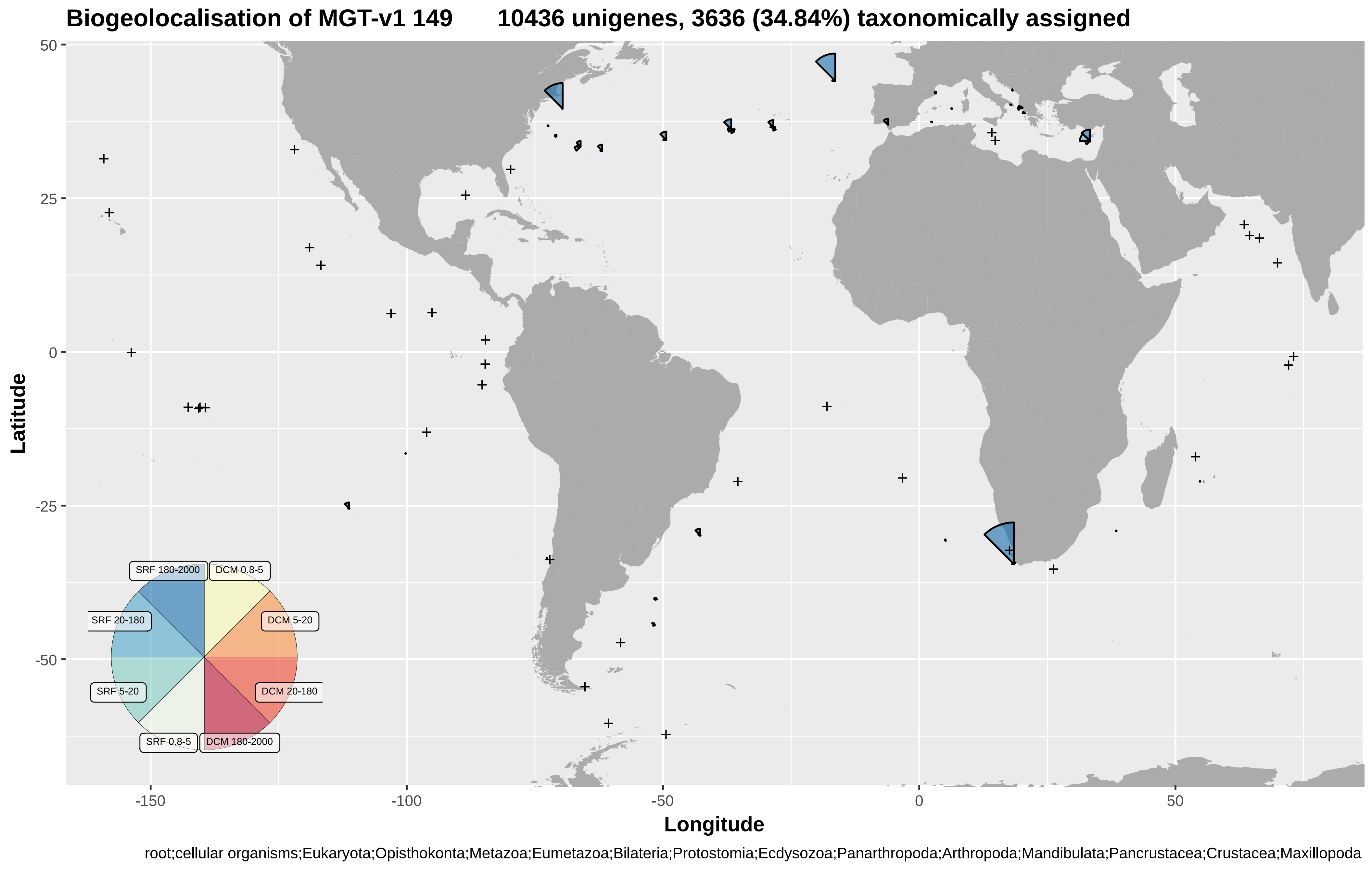
Biogeolocalisation of MGT-v1 149      10436 unigenes, 3636 (34.84%) taxonomically assigned
Latitude
Longitude
50
25
0
-25
-50
-150
-100
-50
0
50
SRF 180-2000
DCM 0.8-5
SRF 20-180
DCM 5-20
SRF 5-20
DCM 20-180
SRF 0.8-5
DCM 180-2000
root;cellular organisms;Eukaryota;Opisthokonta;Metazoa;Eumetazoa;Bilateria;Protostomia;Ecdysozoa;Panarthropoda;Arthropoda;Mandibulata;Pancrustacea;Crustacea;Maxillopoda

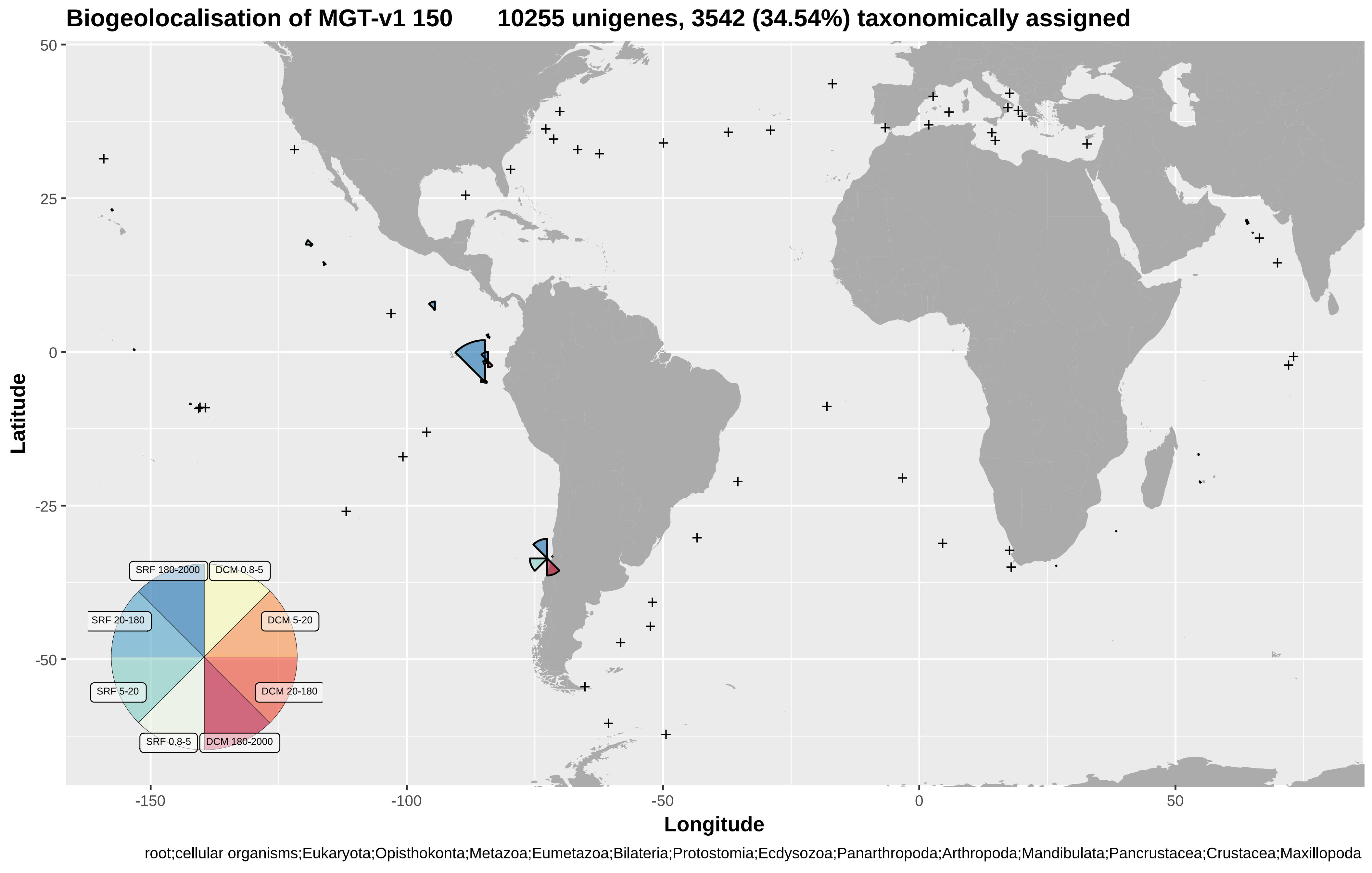
Biogeolocalisation of MGT-v1 150
10255 unigenes, 3542 (34.54%) taxonomically assigned
Latitude
Longitude
50
25
0
-25
-50
-150
-100
-50
0
50
SRF 180-2000
DCM 0.8-5
SRF 20-180
DCM 5-20
SRF 5-20
DCM 20-180
SRF 0.8-5
DCM 180-2000
root;cellular organisms;Eukaryota;Opisthokonta;Metazoa;Eumetazoa;Bilateria;Protostomia;Ecdysozoa;Panarthropoda;Arthropoda;Mandibulata;Pancrustacea;Crustacea;Maxillopoda

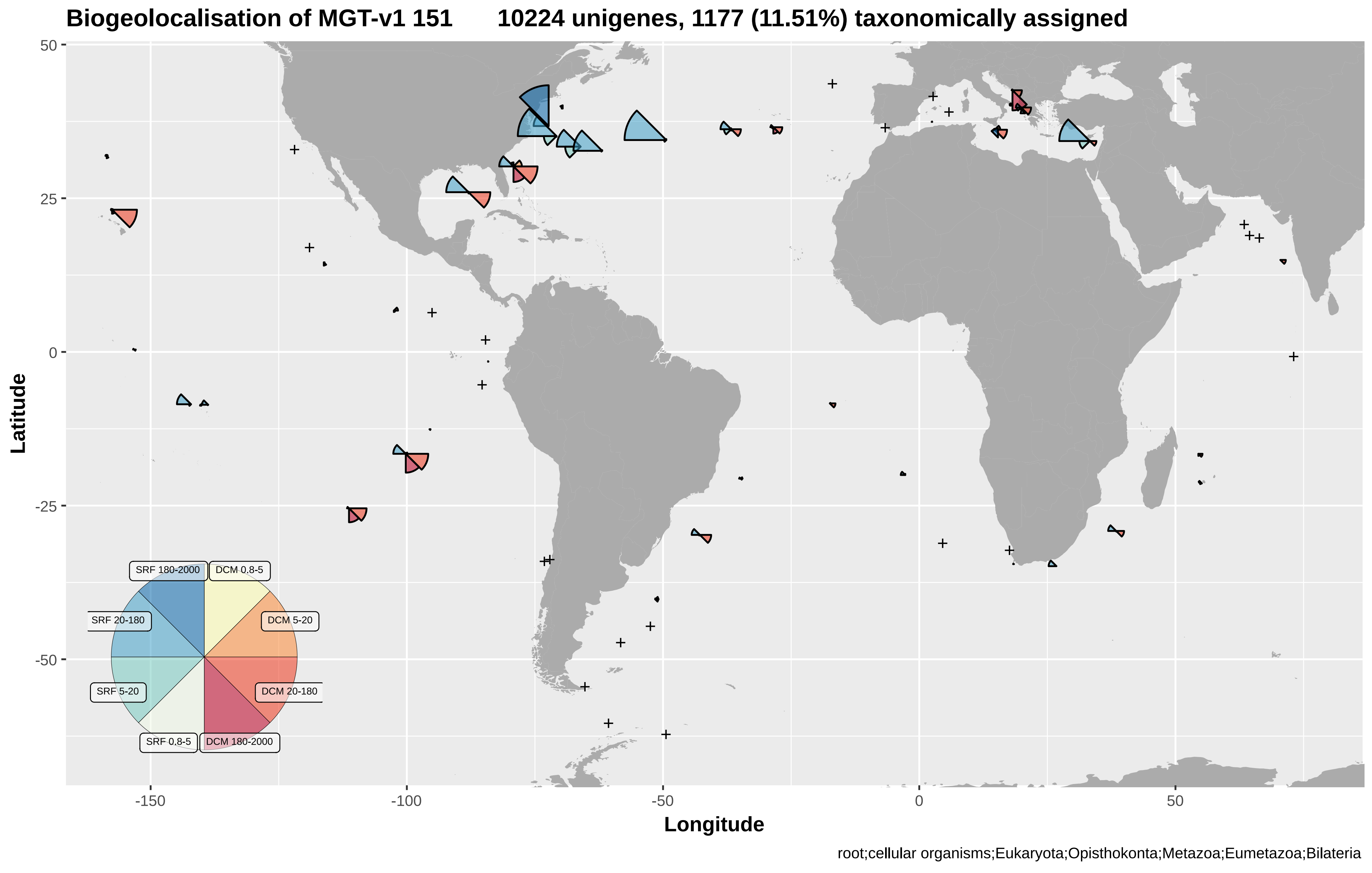

Biogeolocalisation of MGT-v1 151 10224 unigenes, 1177 (11.51%) taxonomically assigned
50
25
0
-25
-50
Latitude
-150
-100
-50
0
50
Longitude
SRF 180-2000
DCM 0.8-5
SRF 20-180
DCM 5-20
SRF 5-20
DCM 20-180
SRF 0.8-5
DCM 180-2000
root;cellular organisms;Eukaryota;Opisthokonta;Metazoa;Eumetazoa;Bilateria

## Biogeolocalisation of MGT-v1 152      10177 unigenes, 2530 (24.86%) taxonomically assigned

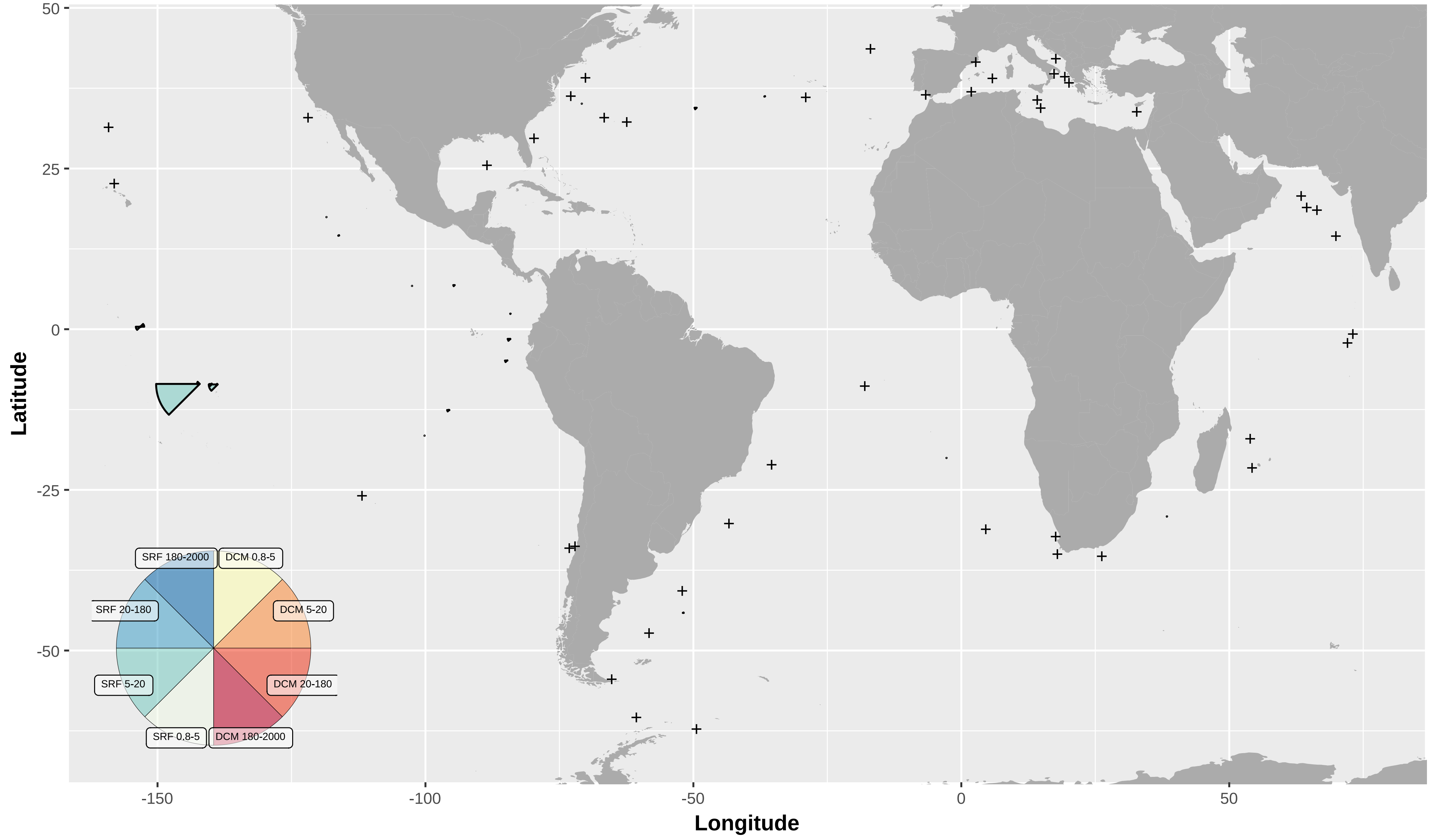


root;cellular organisms;Eukaryota;Stramenopiles;Bacillariophyta;Coscinodiscophyceae

## Biogeolocalisation of MGT-v1 153      9985 unigenes, 3643 (36.48%) taxonomically assigned

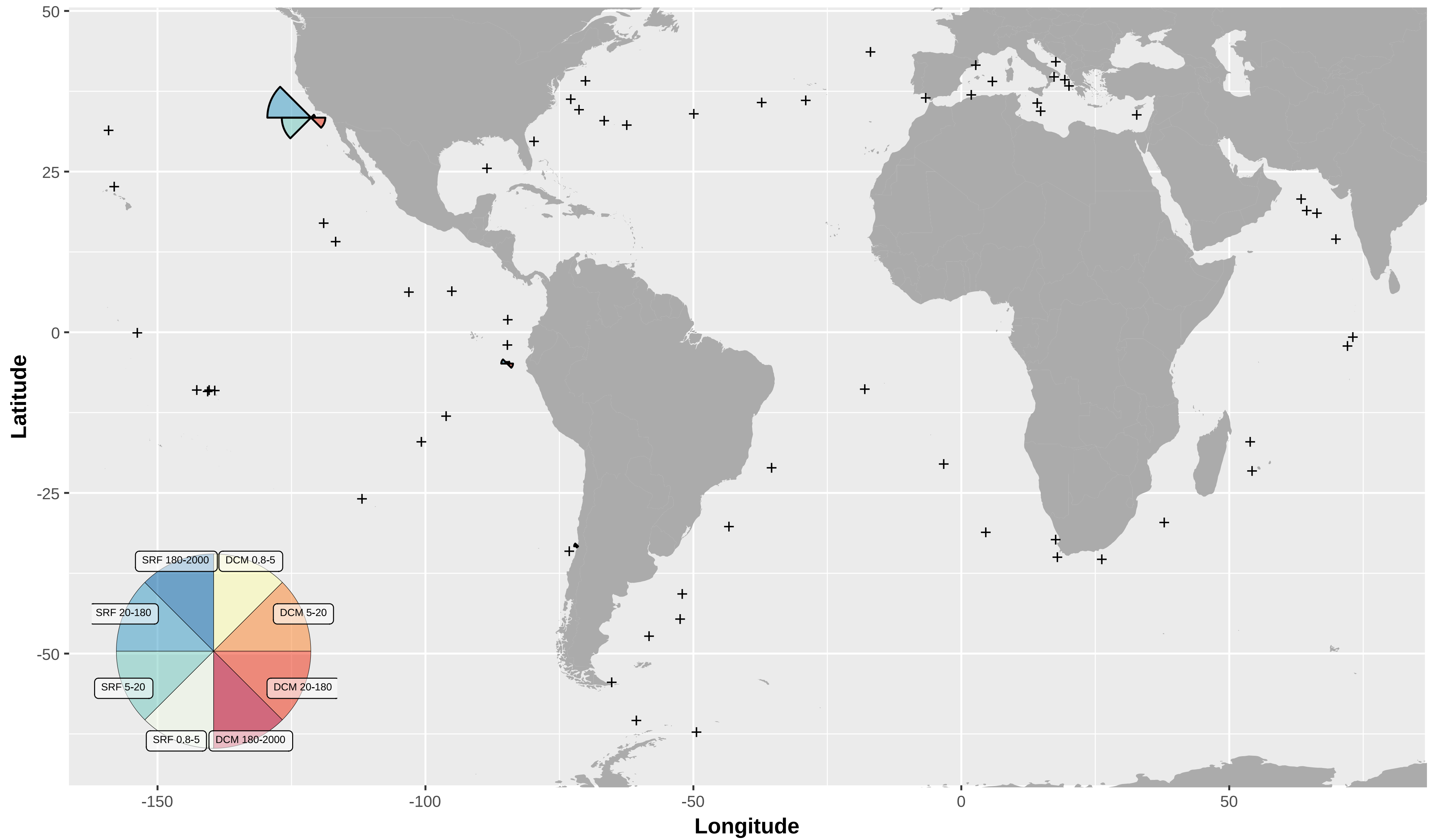


root;cellular organisms;Eukaryota;Opisthokonta;Metazoa;Eumetazoa;Bilateria;Protostomia;Ecdysozoa;Panarthropoda;Arthropoda;Mandibulata;Pancrustacea;Crustacea;Maxillopoda

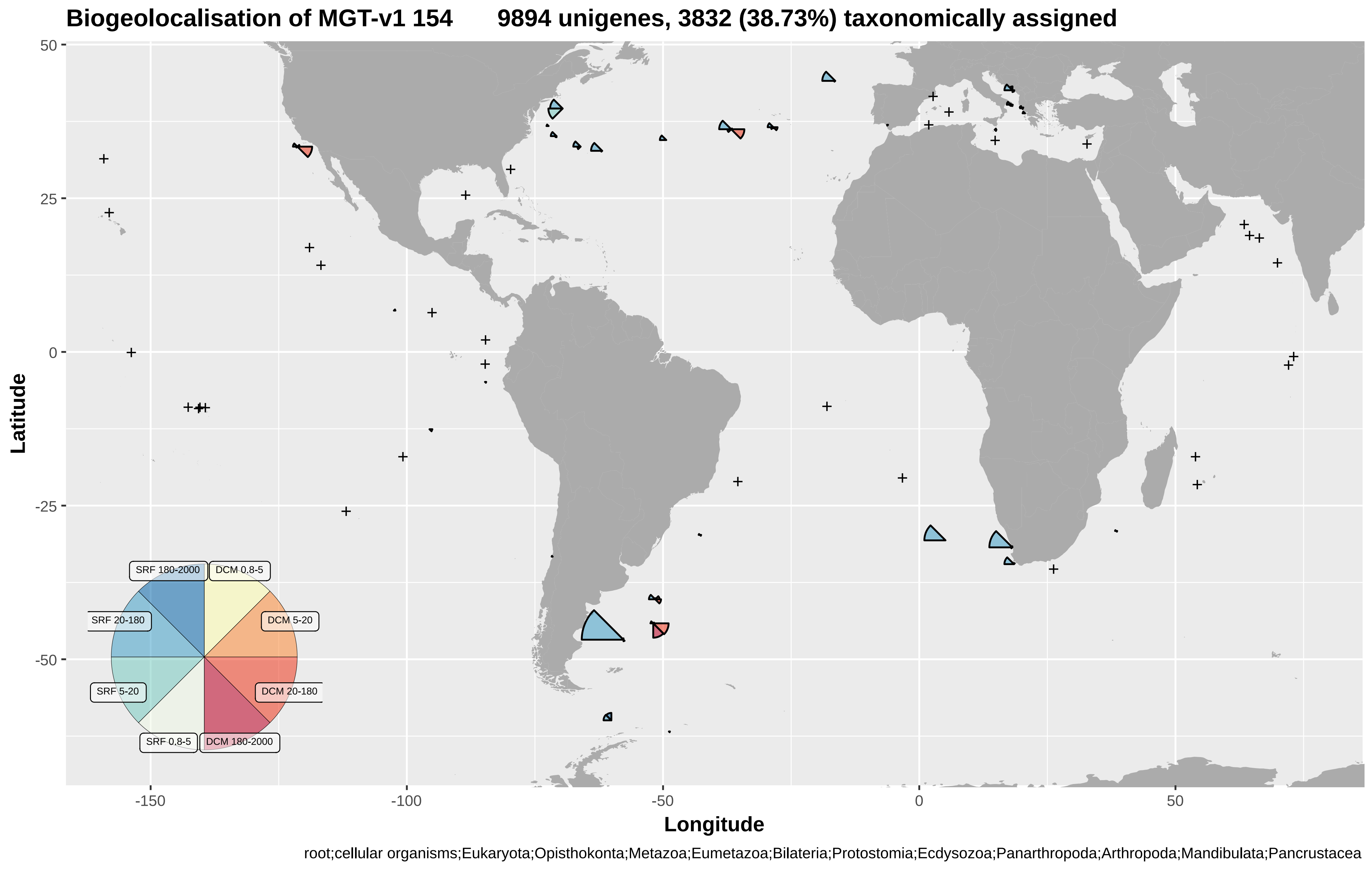
Biogeolocalisation of MGT-v1 154 9894 unigenes, 3832 (38.73%) taxonomically assigned
50
25
0
-25
-50
Latitude
-150
-100
-50
0
50
Longitude
SRF 180-2000
DCM 0.8-5
SRF 20-180
DCM 5-20
SRF 5-20
DCM 20-180
SRF 0.8-5
DCM 180-2000
root;cellular organisms;Eukaryota;Opisthokonta;Metazoa;Eumetazoa;Bilateria;Protostomia;Ecdysozoa;Panarthropoda;Arthropoda;Mandibulata;Pancrustacea

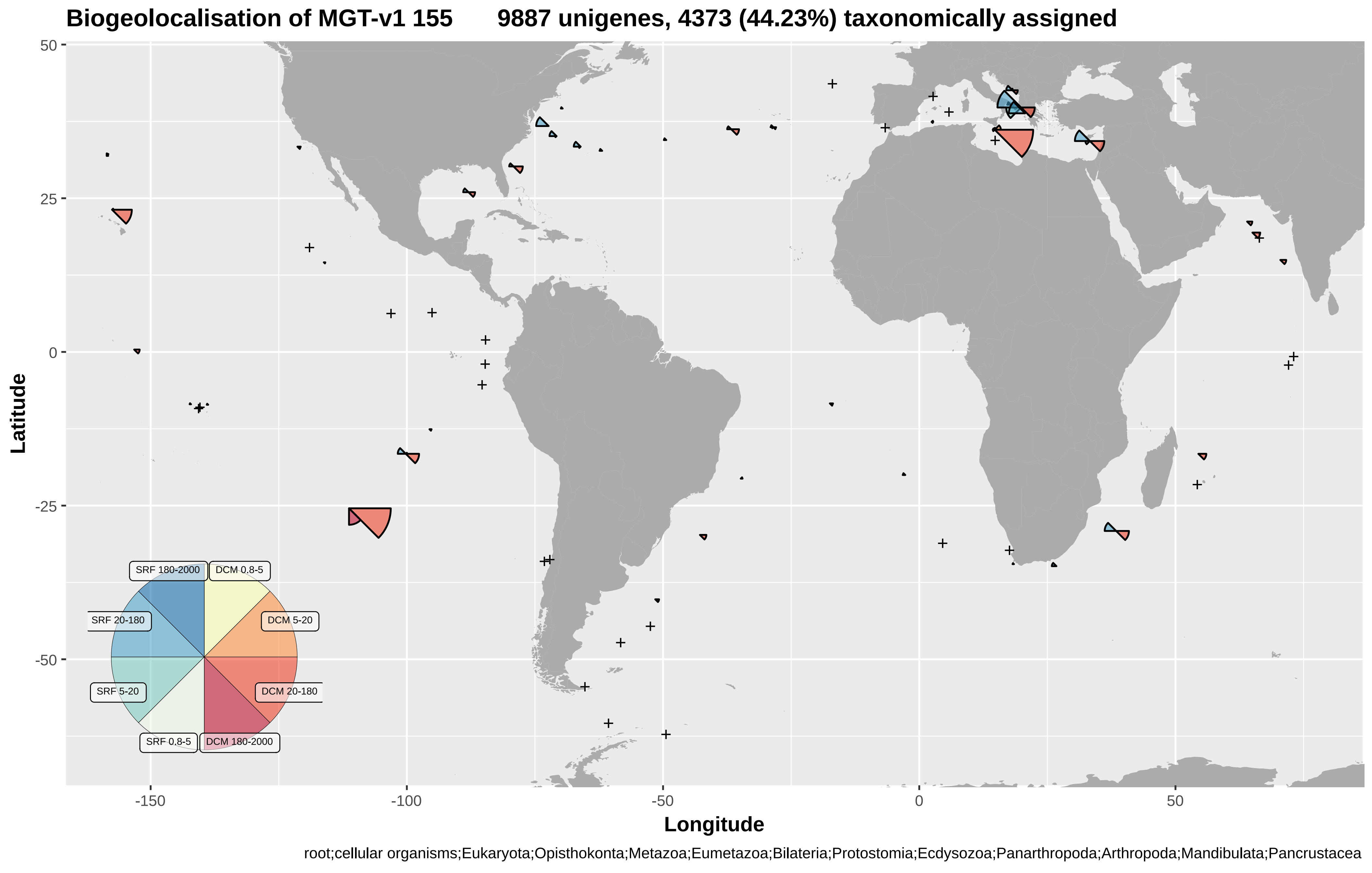
Biogeolocalisation of MGT-v1 155 9887 unigenes, 4373 (44.23%) taxonomically assigned
50
25
0
-25
-50
Latitude
-150
-100
-50
0
50
Longitude
SRF 180-2000
DCM 0.8-5
SRF 20-180
DCM 5-20
SRF 5-20
DCM 20-180
SRF 0.8-5
DCM 180-2000
root;cellular organisms;Eukaryota;Opisthokonta;Metazoa;Eumetazoa;Bilateria;Protostomia;Ecdysozoa;Panarthropoda;Arthropoda;Mandibulata;Pancrustacea

**Biogeolocalisation of MGT-v1 156      9787 unigenes, 6859 (70.08%) taxonomically assigned**

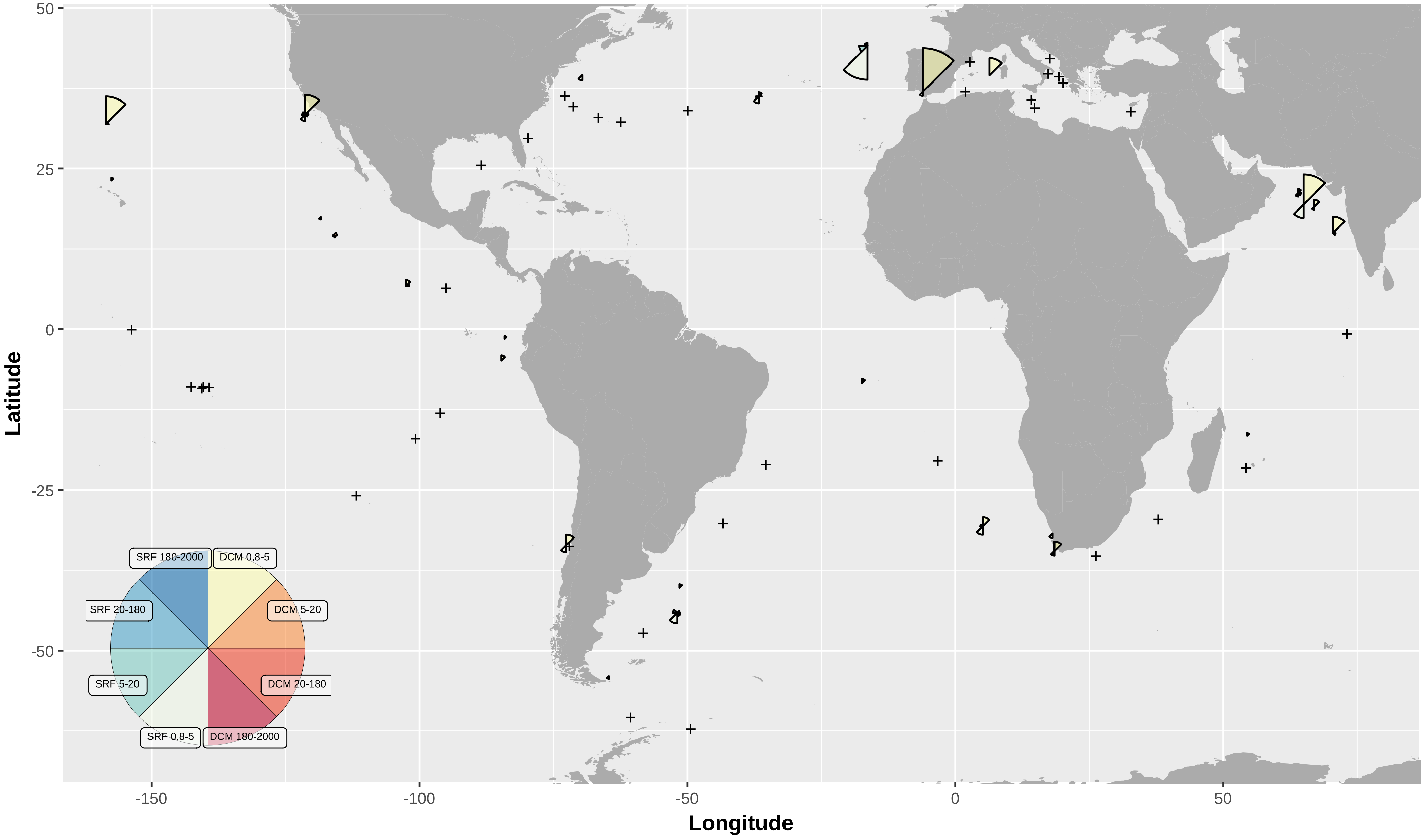


root;cellular organisms;Eukaryota;Viridiplantae

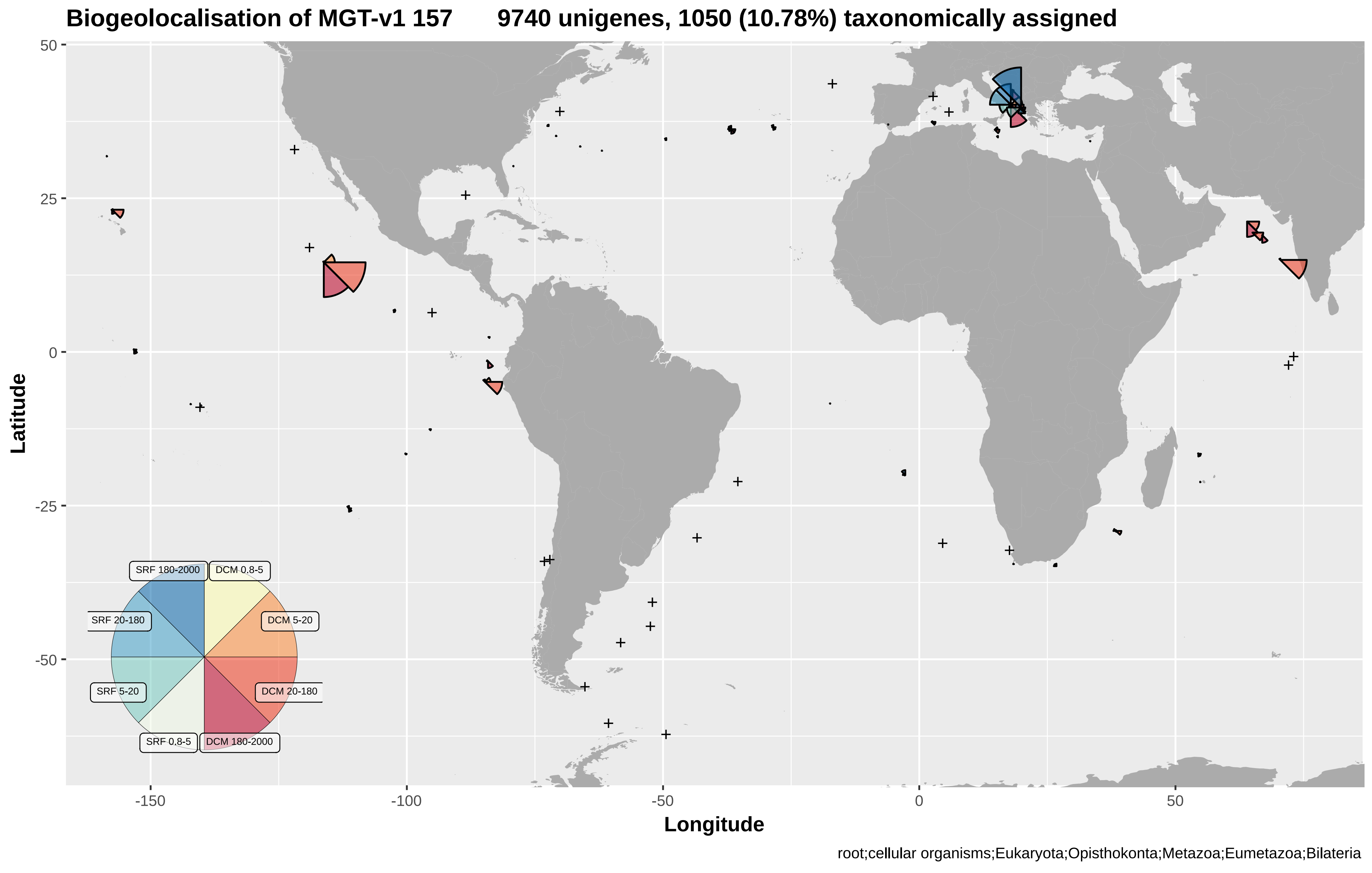
Biogeolocalisation of MGT-v1 157 9740 unigenes, 1050 (10.78%) taxonomically assigned
Latitude
Longitude
SRF 180-2000
DCM 0.8-5
SRF 20-180
DCM 5-20
SRF 5-20
DCM 20-180
SRF 0.8-5
DCM 180-2000
root;cellular organisms;Eukaryota;Opisthokonta;Metazoa;Eumetazoa;Bilateria

#### Biogeolocalisation of MGT-v1 158 9693 unigenes, 975 (10.06%) taxonomically assigned

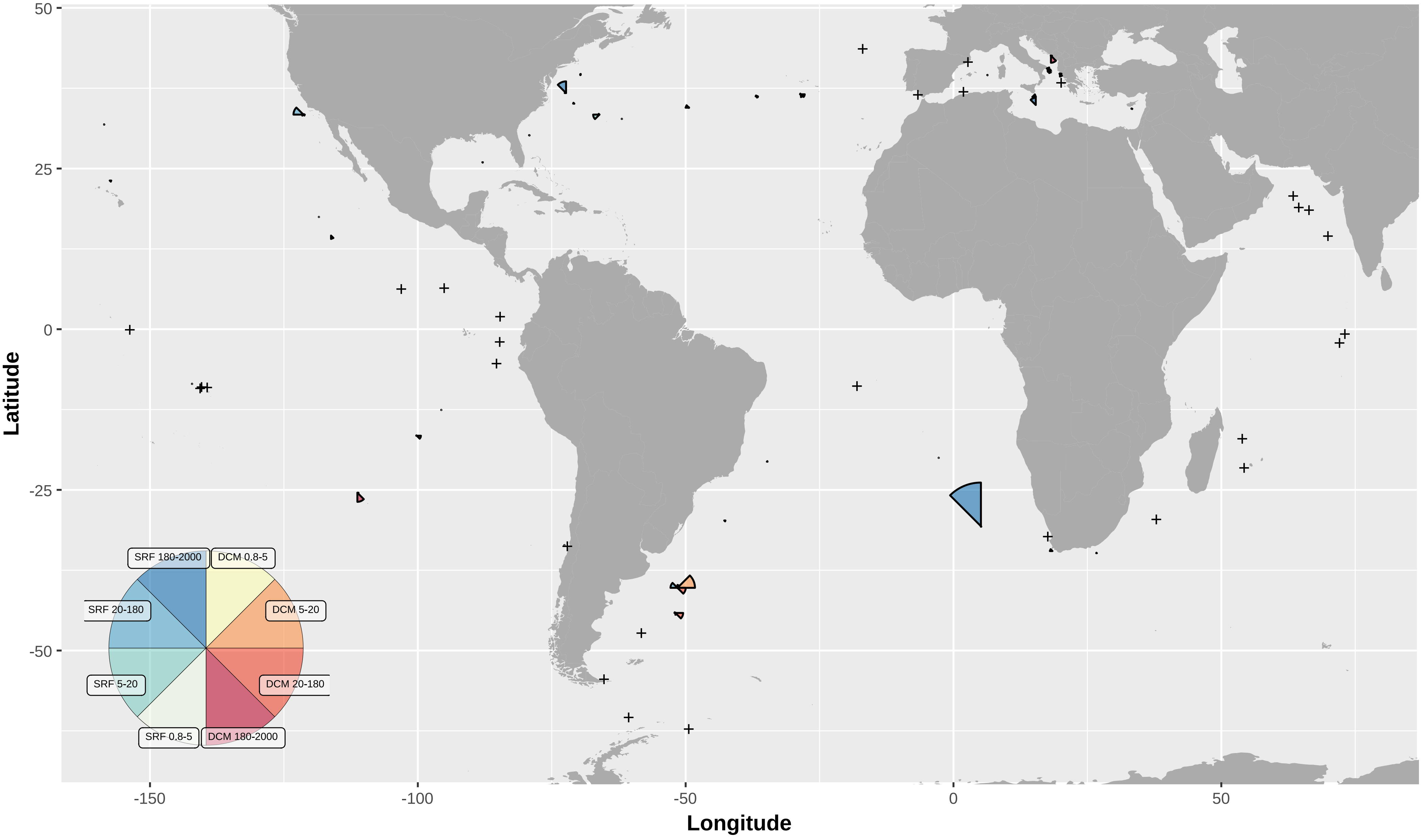


root;cellular organisms;Eukaryota;Opisthokonta;Metazoa;Eumetazoa;Bilateria

#### Biogeolocalisation of MGT-v1 159 9623 unigenes, 685 (7.12%) taxonomically assigned

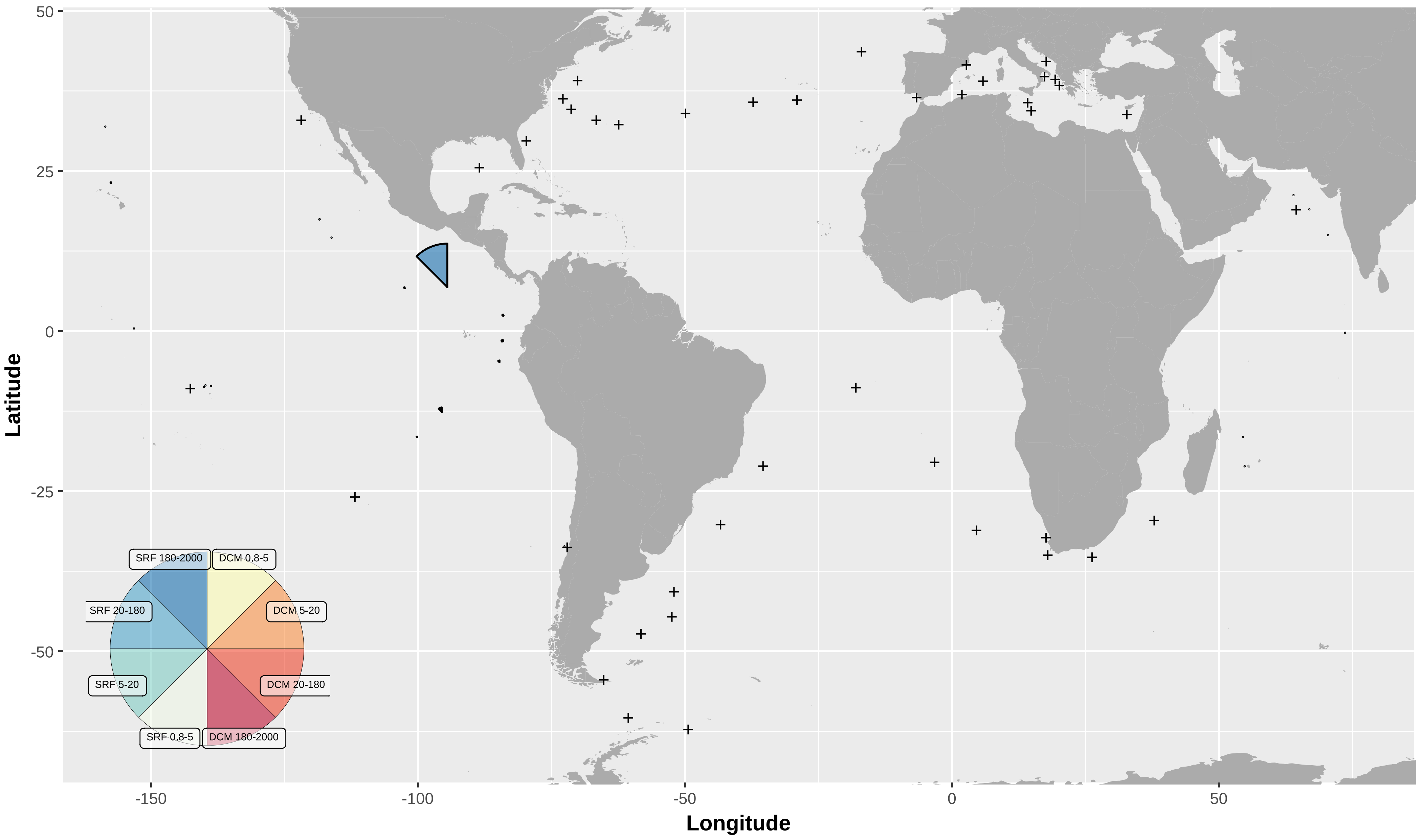


root;cellular organisms;Eukaryota;Opisthokonta;Metazoa;Eumetazoa;Bilateria

## Biogeolocalisation of MGT-v1 160      9494 unigenes, 3450 (36.34%) taxonomically assigned

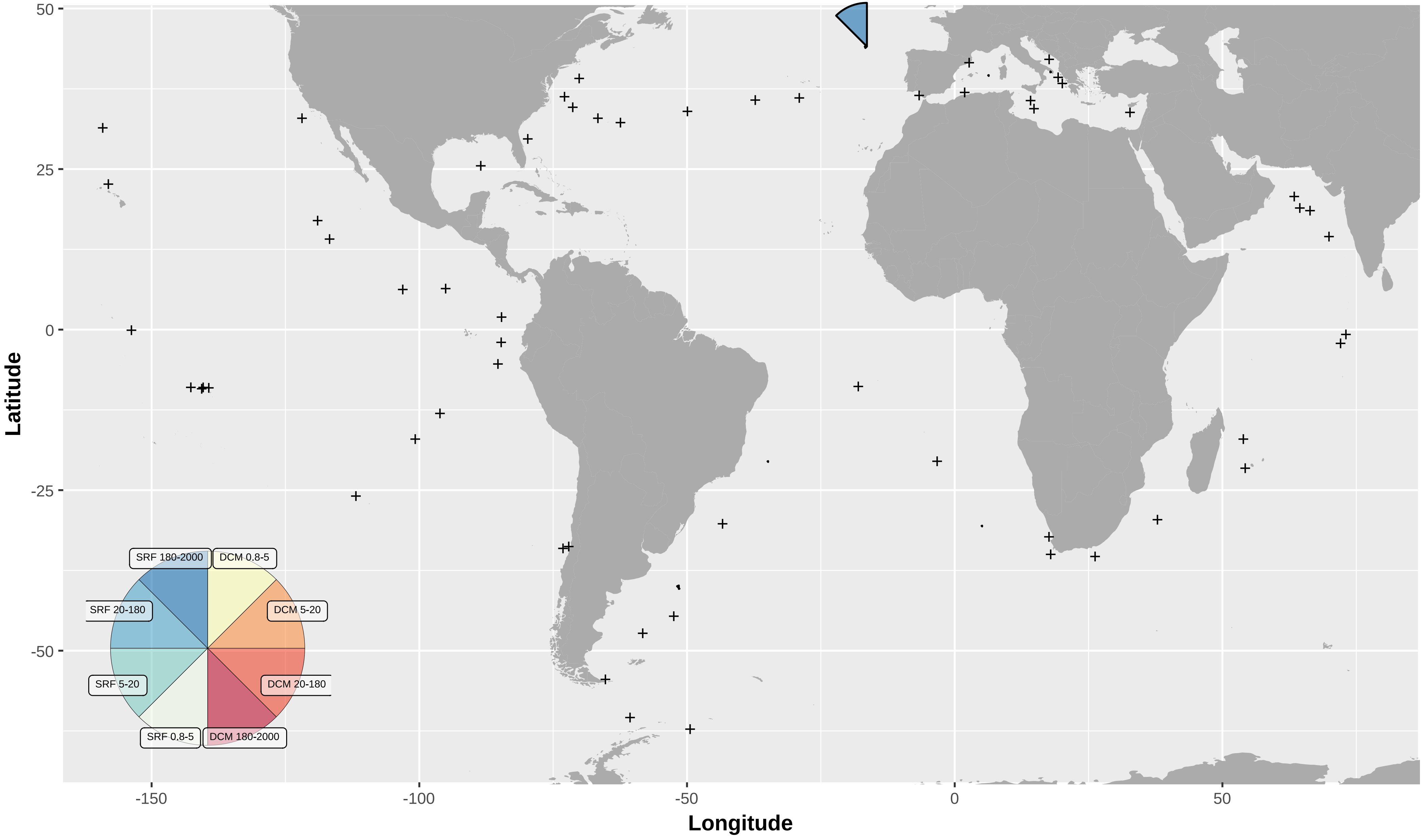


root;cellular organisms;Eukaryota;Opisthokonta;Metazoa;Eumetazoa;Bilateria

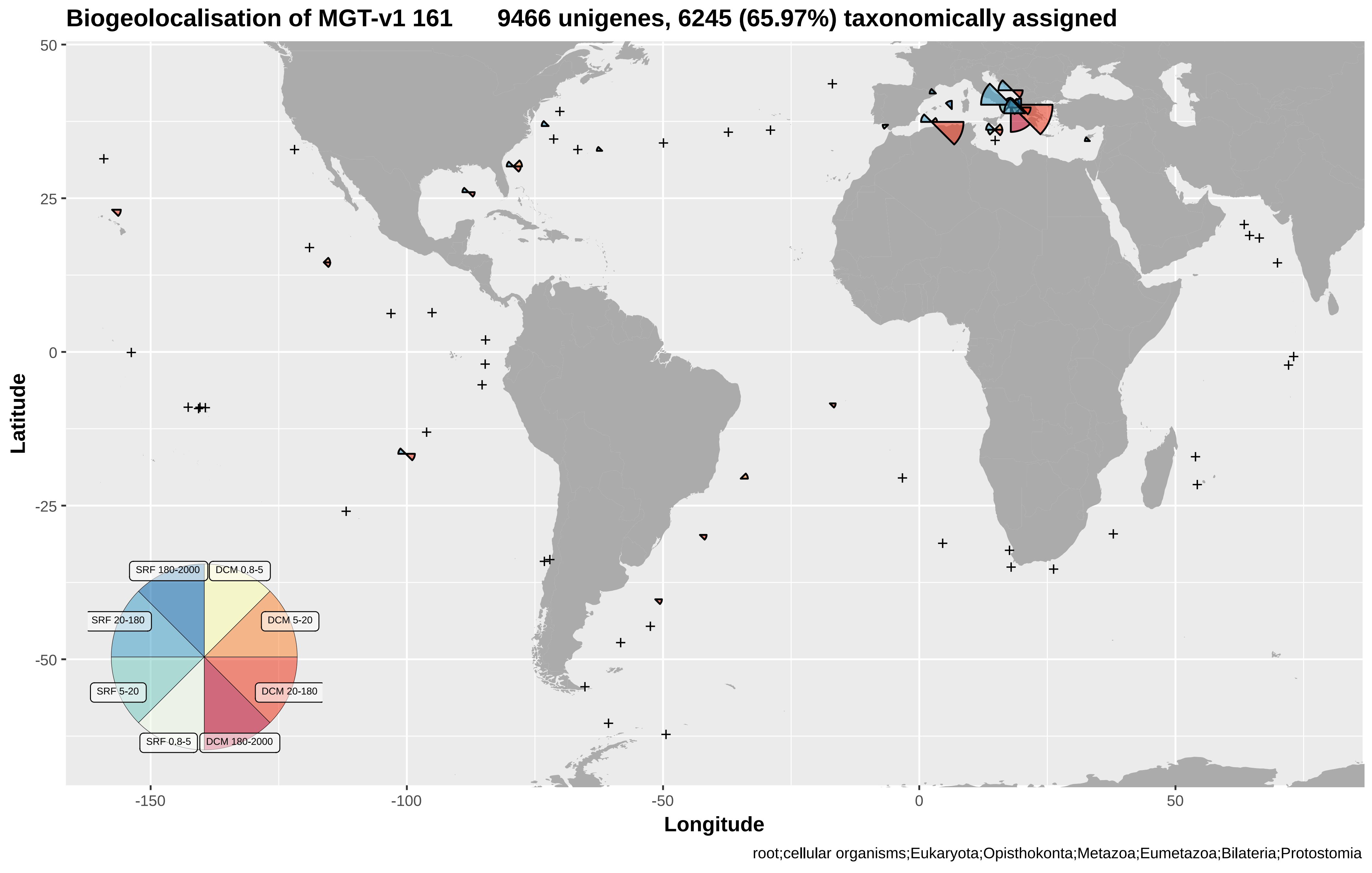
Biogeolocalisation of MGT-v1 161 9466 unigenes, 6245 (65.97%) taxonomically assigned
Latitude
50
25
0
-25
-50
Longitude
-150
-100
-50
0
50
SRF 180-2000
DCM 0.8-5
SRF 20-180
DCM 5-20
SRF 5-20
DCM 20-180
SRF 0.8-5
DCM 180-2000
root;cellular organisms;Eukaryota;Opisthokonta;Metazoa;Eumetazoa;Bilateria;Protostomia

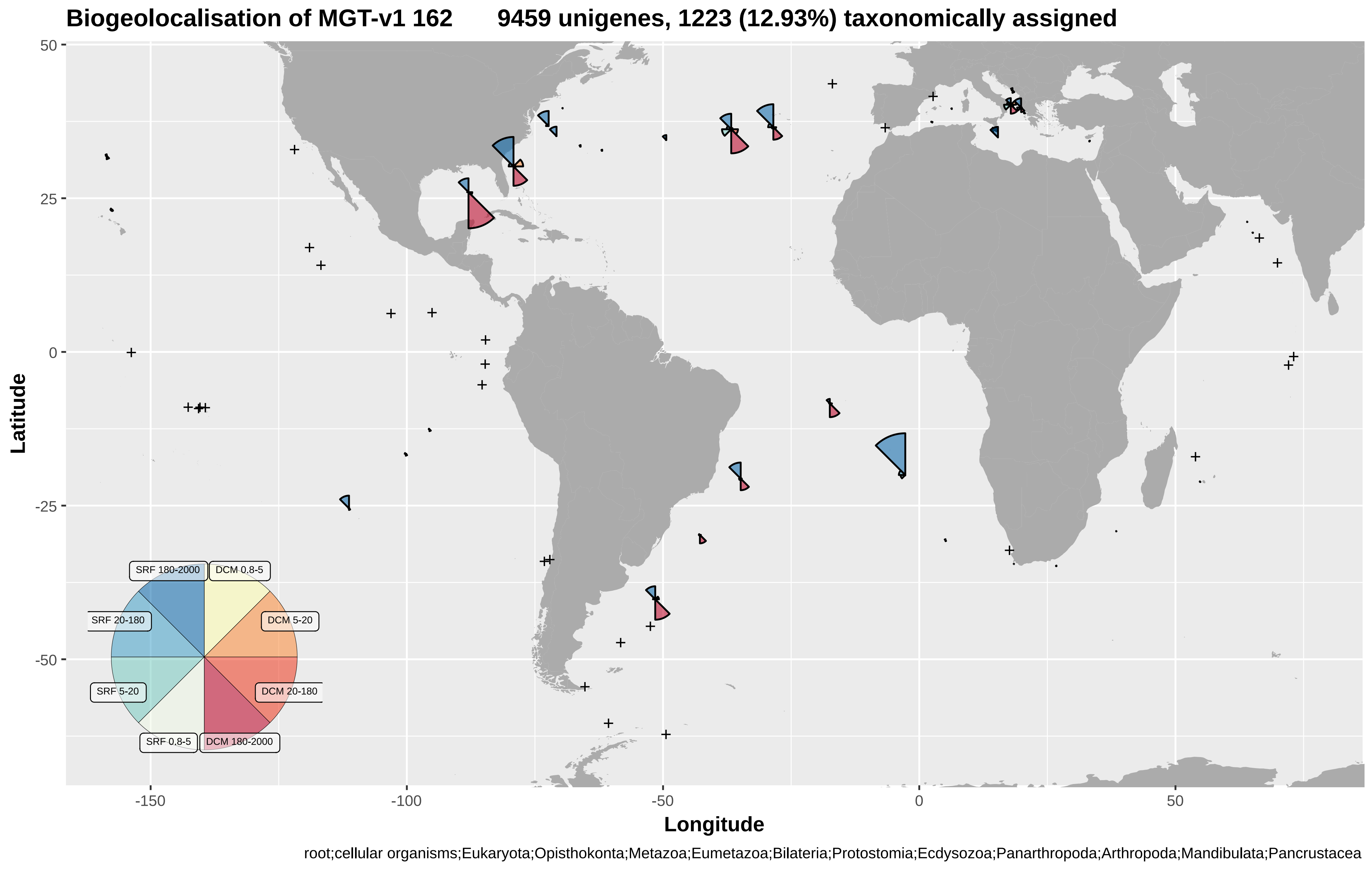

Biogeolocalisation of MGT-v1 162 9459 unigenes, 1223 (12.93%) taxonomically assigned
50
25
0
-25
-50
Latitude
-150
-100
-50
0
50
Longitude
SRF 180-2000
DCM 0.8-5
SRF 20-180
DCM 5-20
SRF 5-20
DCM 20-180
SRF 0.8-5
DCM 180-2000
root;cellular organisms;Eukaryota;Opisthokonta;Metazoa;Eumetazoa;Bilateria;Protostomia;Ecdysozoa;Panarthropoda;Arthropoda;Mandibulata;Pancrustacea

# Biogeolocalisation of MGT-v1 163      9400 unigenes, 1456 (15.49%) taxonomically assigned

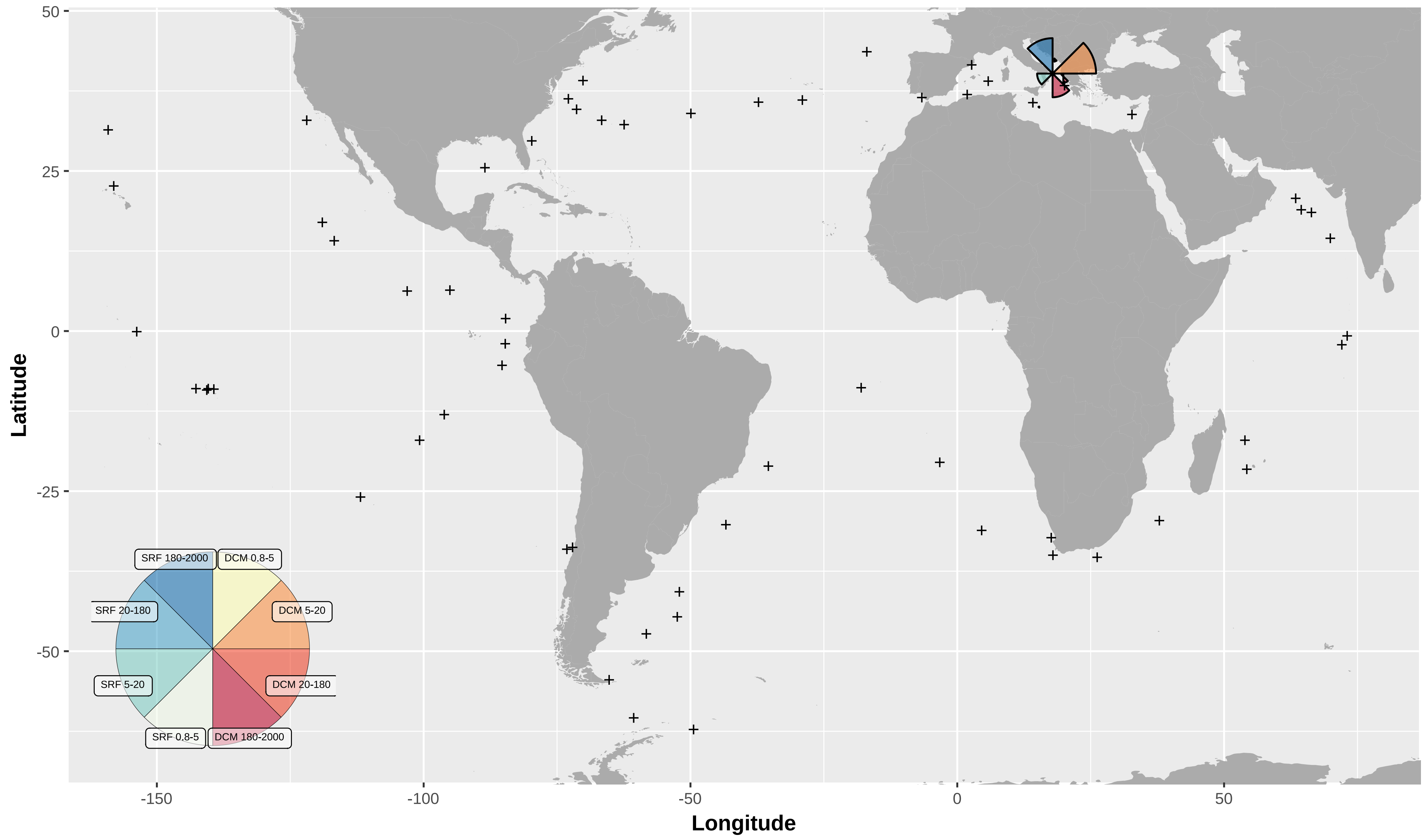


root;cellular organisms;Eukaryota;Opisthokonta;Metazoa;Eumetazoa;Bilateria

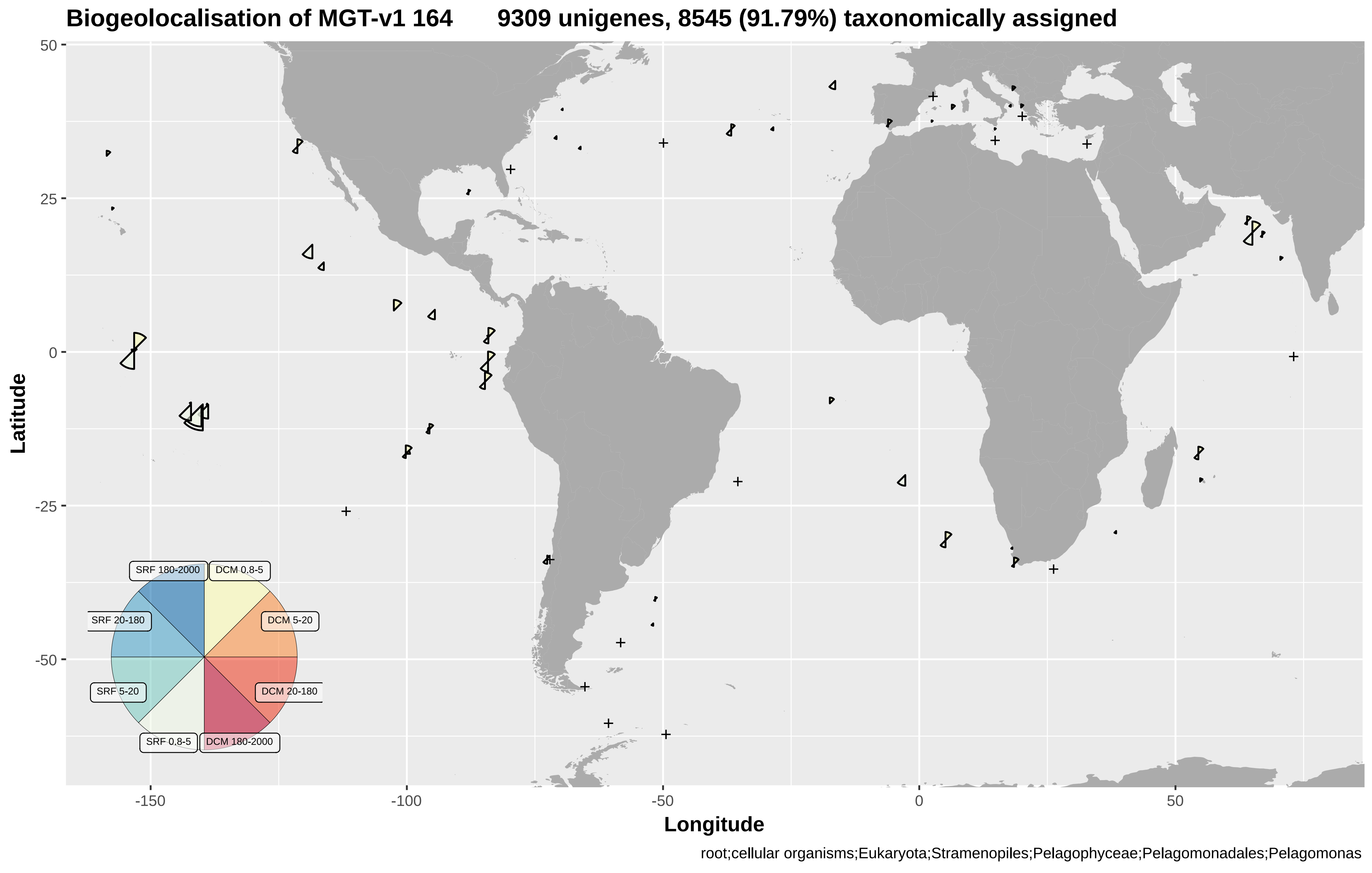

Biogeolocalisation of MGT-v1 164 9309 unigenes, 8545 (91.79%) taxonomically assigned
50
25
0
-25
-50
Latitude
-150
-100
-50
0
50
Longitude
SRF 180-2000
DCM 0.8-5
SRF 20-180
DCM 5-20
SRF 5-20
DCM 20-180
SRF 0.8-5
DCM 180-2000
root;cellular organisms;Eukaryota;Stramenopiles;Pelagophyceae;Pelagomonadales;Pelagomonas

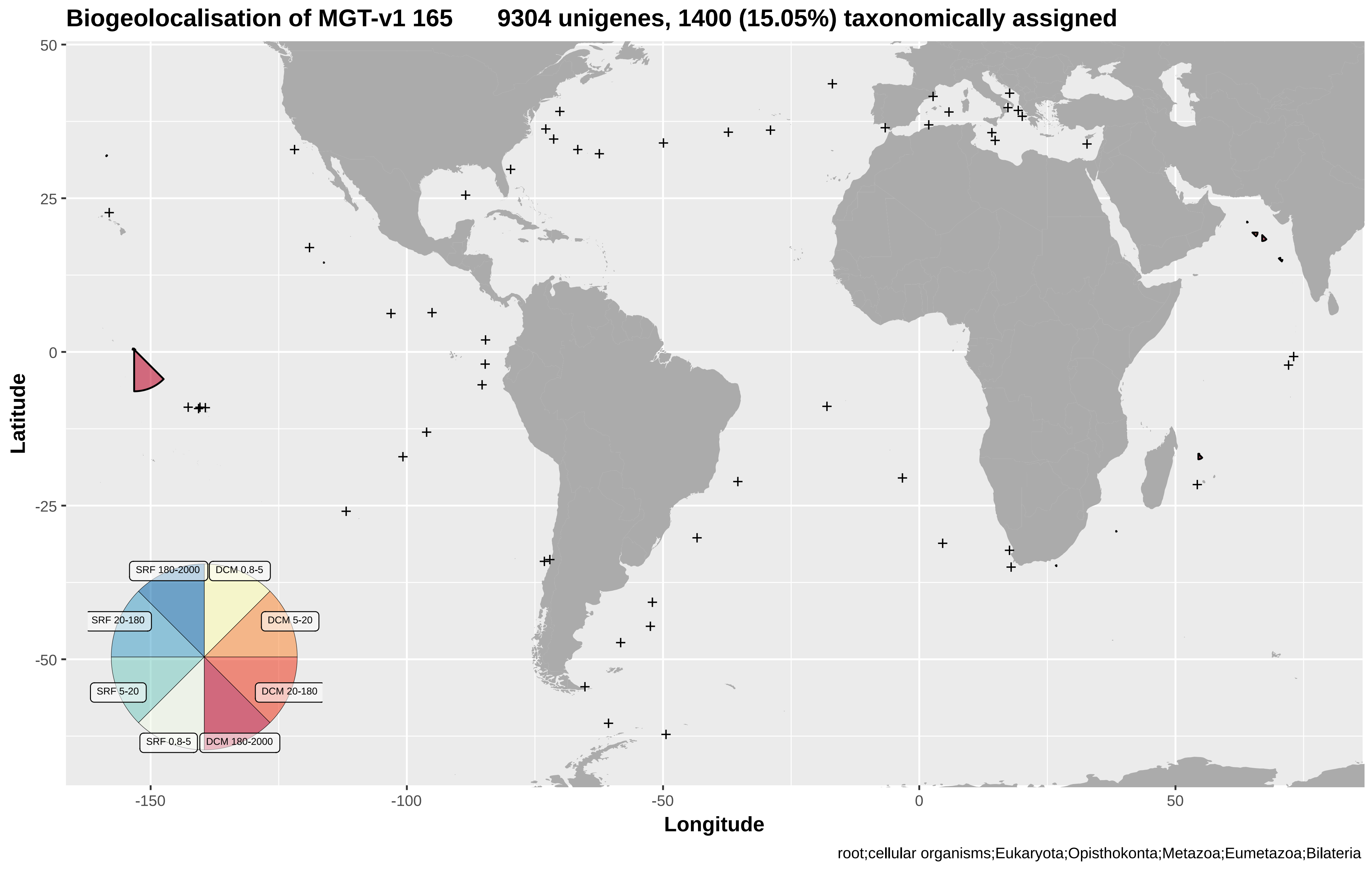

Biogeolocalisation of MGT-v1 165 9304 unigenes, 1400 (15.05%) taxonomically assigned
Latitude
Longitude
50
25
0
-25
-50
-150
-100
-50
0
50
SRF 180-2000
DCM 0.8-5
SRF 20-180
DCM 5-20
SRF 5-20
DCM 20-180
SRF 0.8-5
DCM 180-2000
root;cellular organisms;Eukaryota;Opisthokonta;Metazoa;Eumetazoa;Bilateria

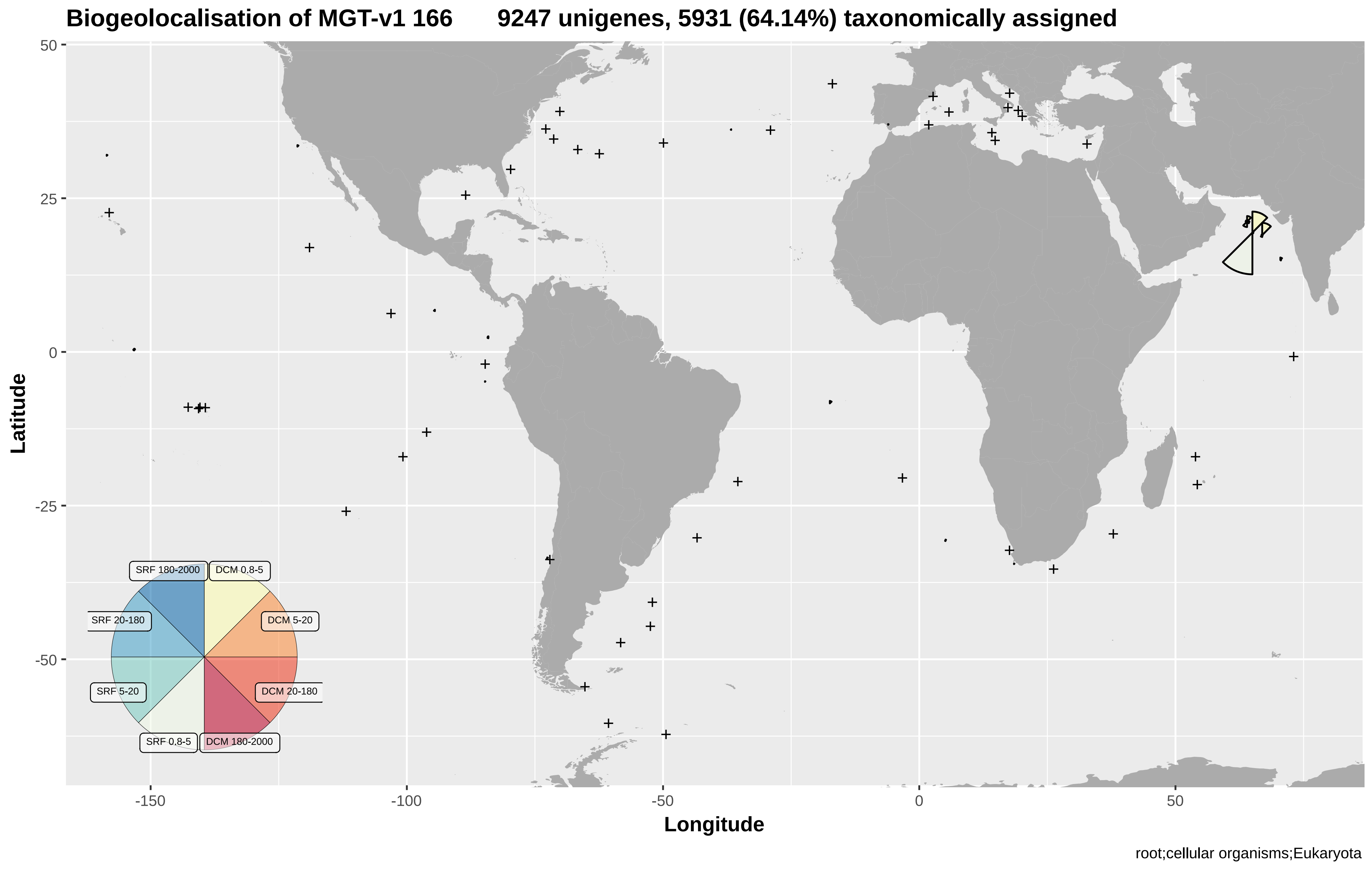
Biogeolocalisation of MGT-v1 166 9247 unigenes, 5931 (64.14%) taxonomically assigned
Latitude
Longitude
50
25
0
-25
-50
-150
-100
-50
0
50
SRF 180-2000
DCM 0.8-5
SRF 20-180
DCM 5-20
SRF 5-20
DCM 20-180
SRF 0.8-5
DCM 180-2000
root;cellular organisms;Eukaryota

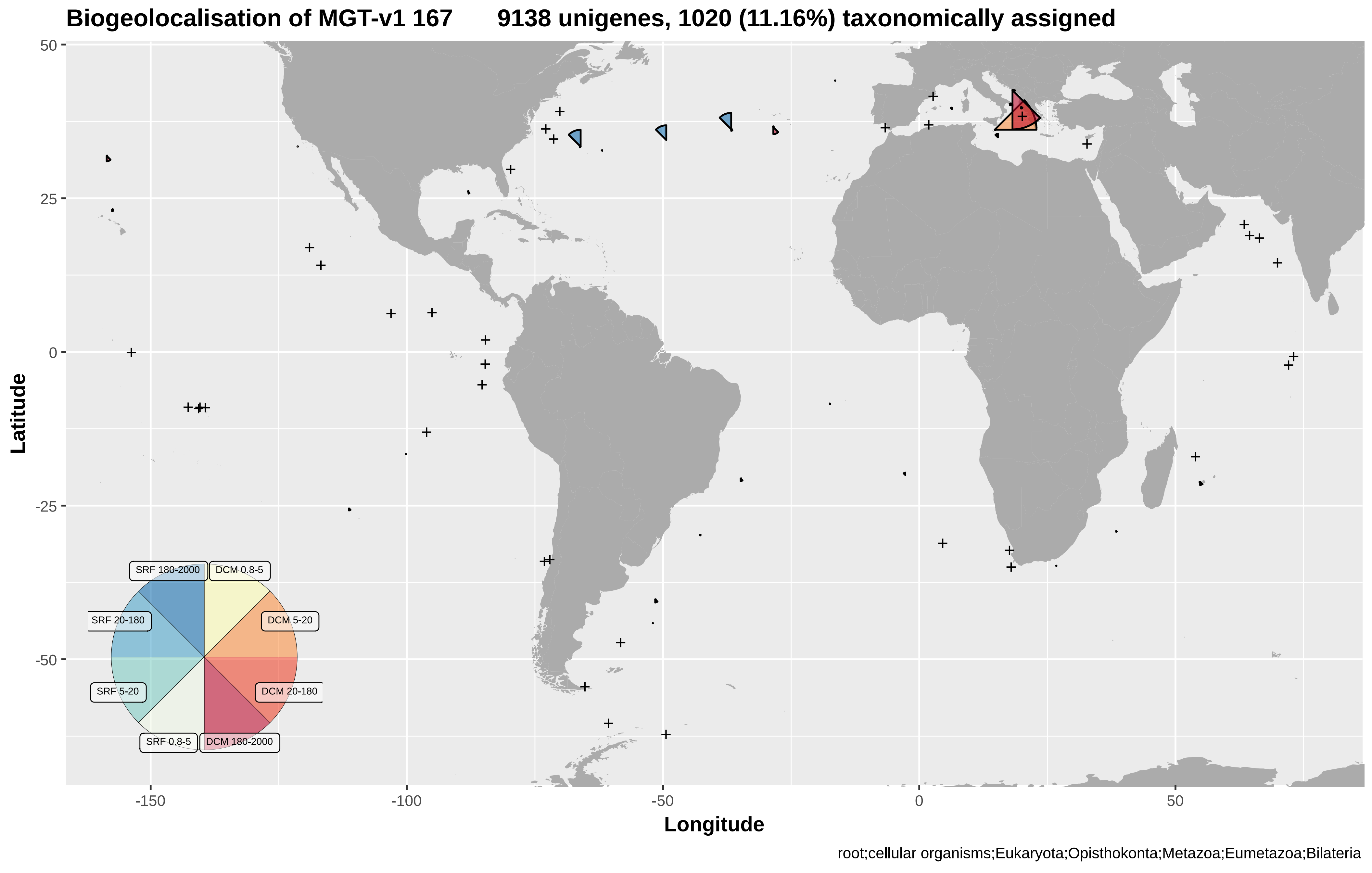

Biogeolocalisation of MGT-v1 167 9138 unigenes, 1020 (11.16%) taxonomically assigned
50
25
0
-25
-50
Latitude
-150
-100
-50
0
50
Longitude
SRF 180-2000
DCM 0.8-5
SRF 20-180
DCM 5-20
SRF 5-20
DCM 20-180
SRF 0.8-5
DCM 180-2000
root;cellular organisms;Eukaryota;Opisthokonta;Metazoa;Eumetazoa;Bilateria

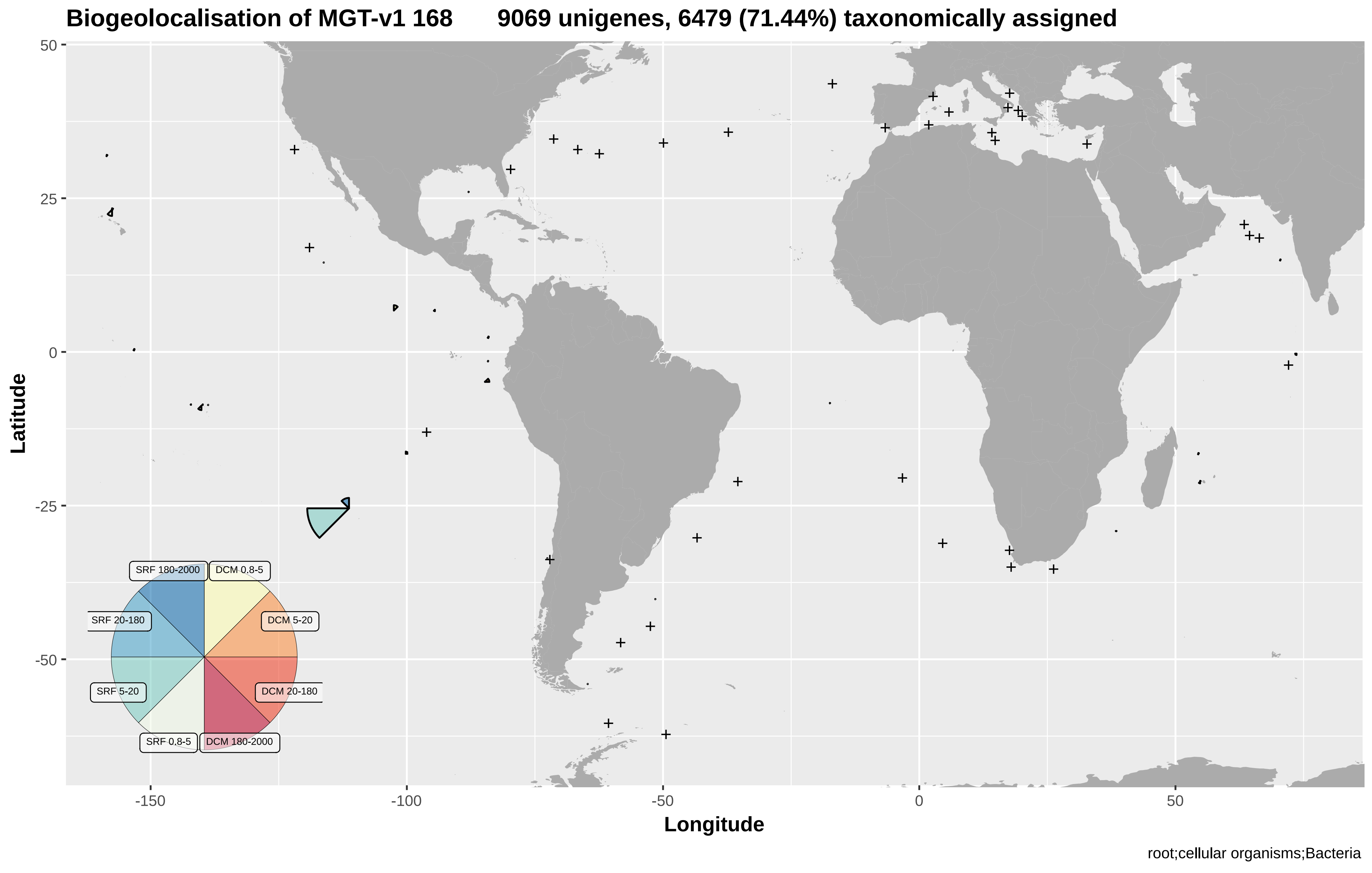
Biogeolocalisation of MGT-v1 168 9069 unigenes, 6479 (71.44%) taxonomically assigned
50
25
0
-25
-50
Latitude
-150
-100
-50
0
50
Longitude
SRF 180-2000
DCM 0.8-5
SRF 20-180
DCM 5-20
SRF 5-20
DCM 20-180
SRF 0.8-5
DCM 180-2000
root;cellular organisms;Bacteria

#### Biogeolocalisation of MGT-v1 169 9056 unigenes, 990 (10.93%) taxonomically assigned

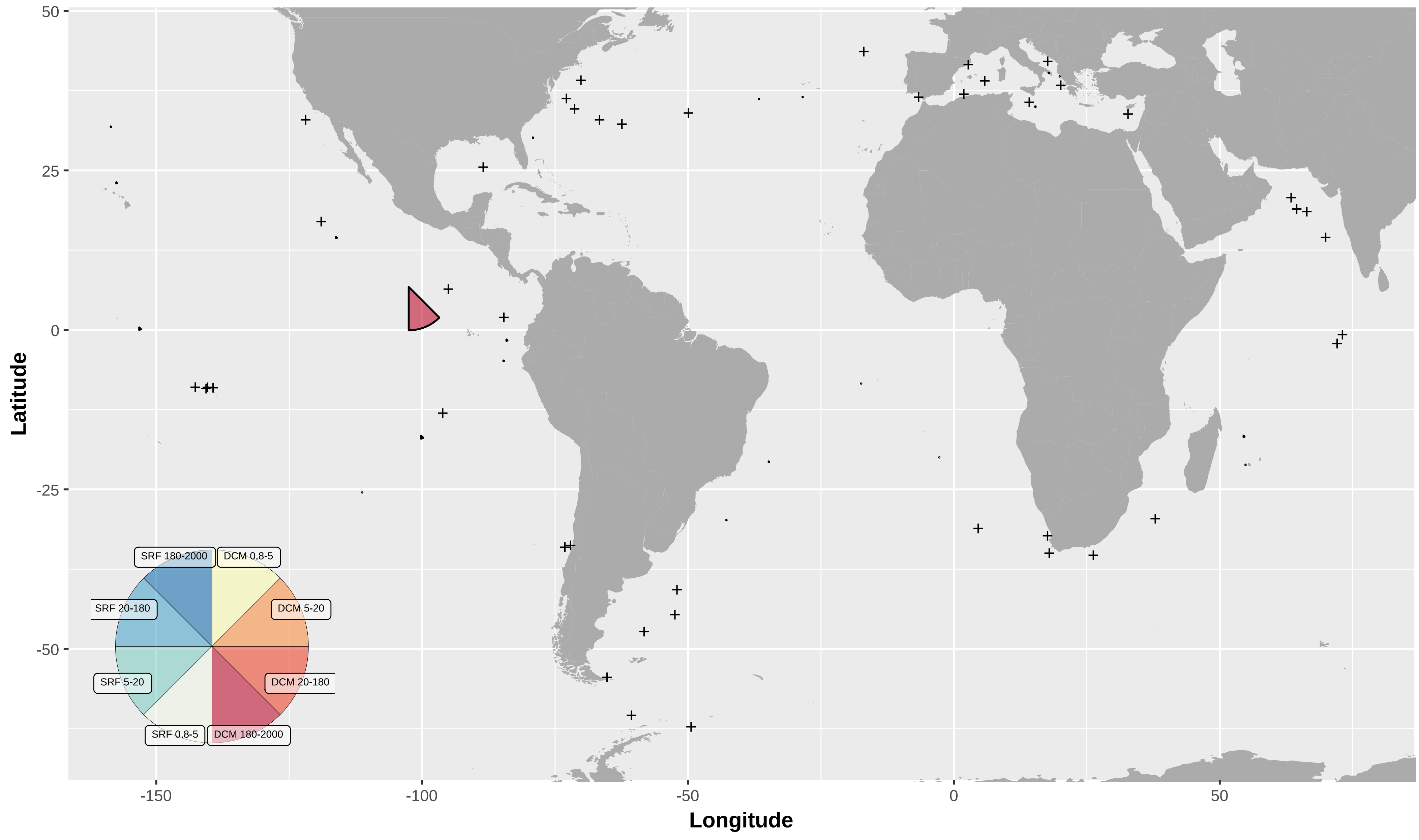


root;cellular organisms;Eukaryota;Opisthokonta;Metazoa;Eumetazoa

#### Biogeolocalisation of MGT-v1 170 9053 unigenes, 6712 (74.14%) taxonomically assigned

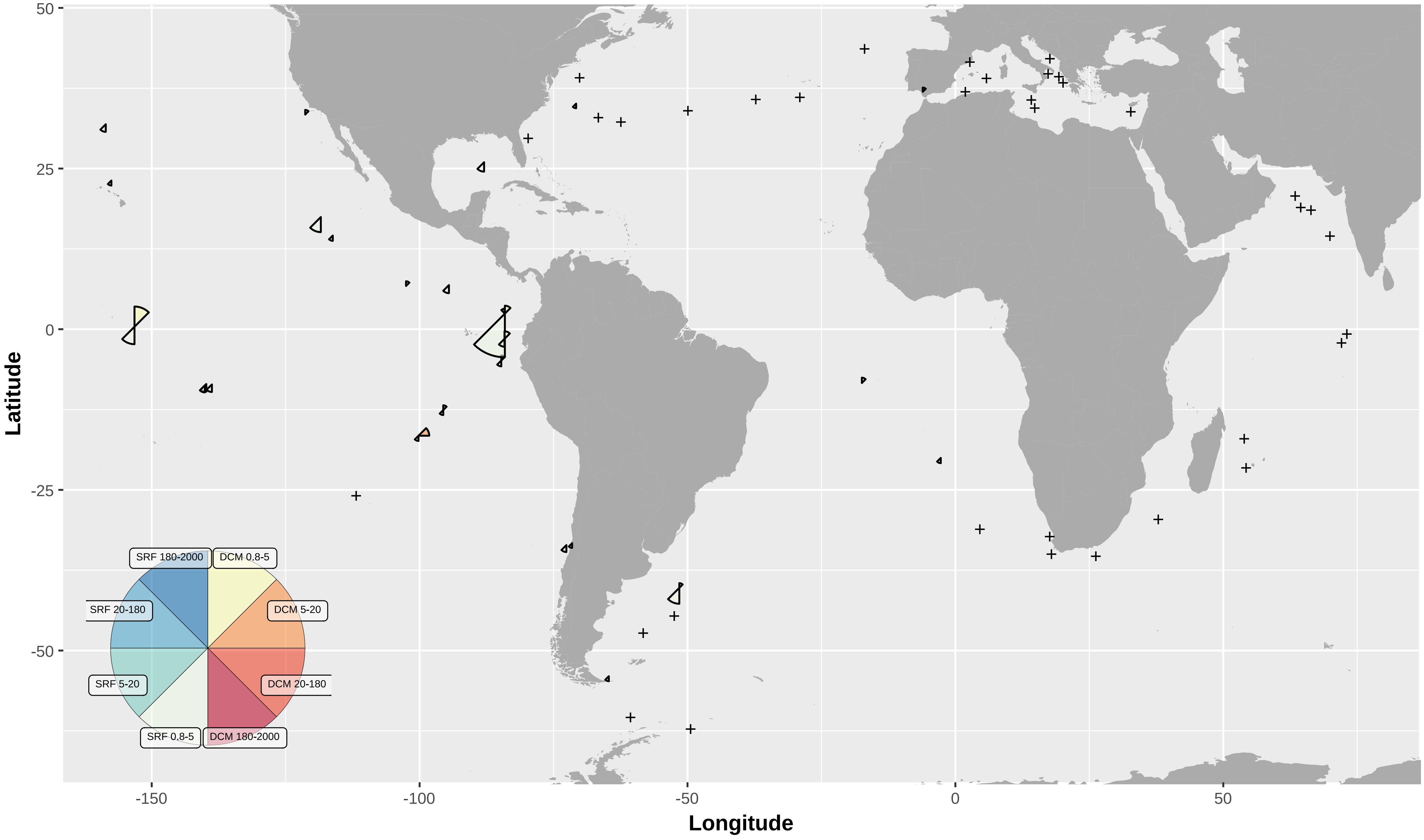


root;cellular organisms;Eukaryota;Opisthokonta;Choanoflagellida

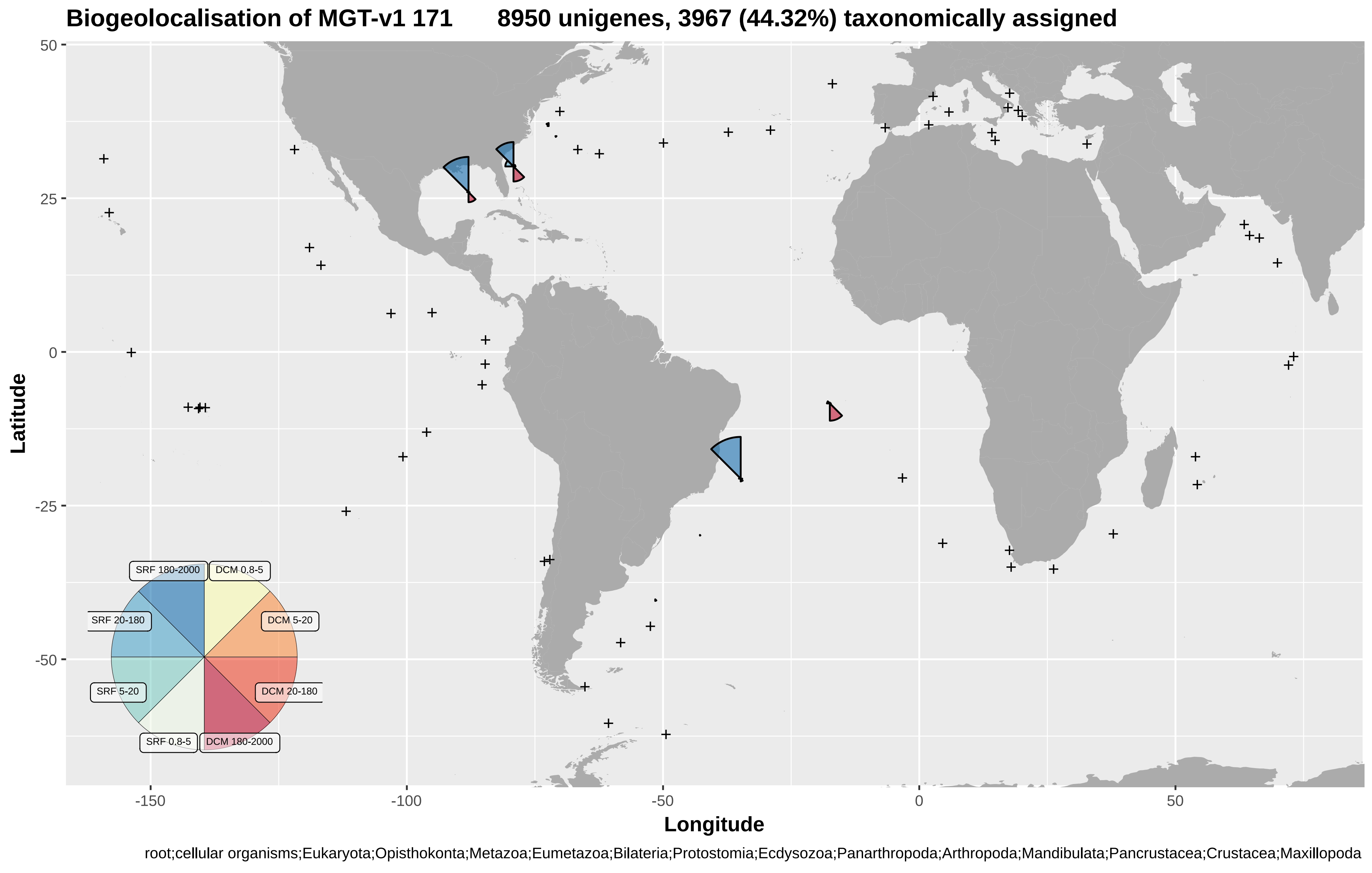
Biogeolocalisation of MGT-v1 171 8950 unigenes, 3967 (44.32%) taxonomically assigned
50
25
0
-25
-50
Latitude
-150
-100
-50
0
50
Longitude
SRF 180-2000
DCM 0.8-5
SRF 20-180
DCM 5-20
SRF 5-20
DCM 20-180
SRF 0.8-5
DCM 180-2000
root;cellular organisms;Eukaryota;Opisthokonta;Metazoa;Eumetazoa;Bilateria;Protostomia;Ecdysozoa;Panarthropoda;Arthropoda;Mandibulata;Pancrustacea;Crustacea;Maxillopoda

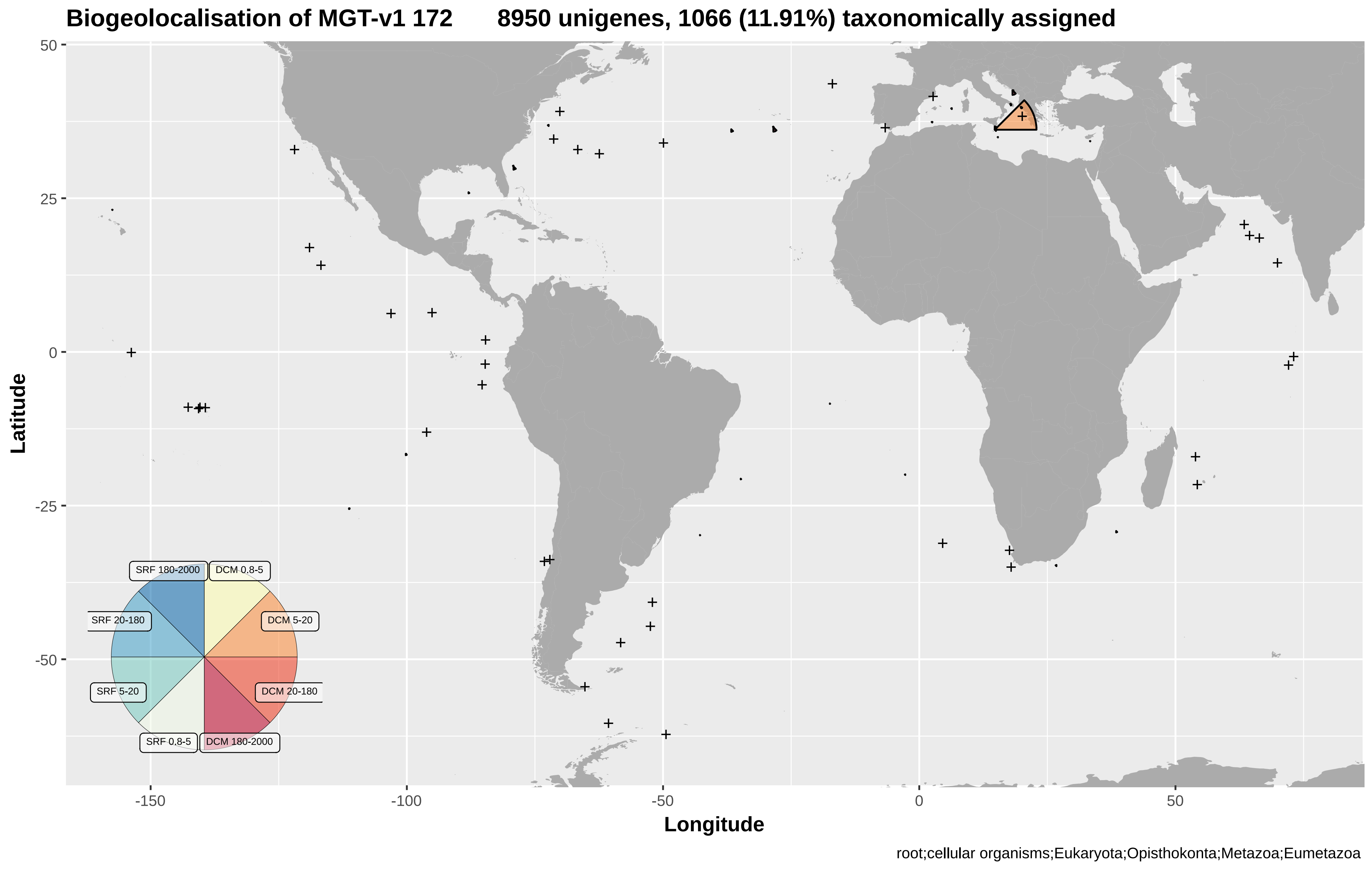

Biogeolocalisation of MGT-v1 172 8950 unigenes, 1066 (11.91%) taxonomically assigned
50
25
0
-25
-50
Latitude
-150
-100
-50
0
50
Longitude
SRF 180-2000
DCM 0.8-5
SRF 20-180
DCM 5-20
SRF 5-20
DCM 20-180
SRF 0.8-5
DCM 180-2000
root;cellular organisms;Eukaryota;Opisthokonta;Metazoa;Eumetazoa

## Biogeolocalisation of MGT-v1 173      8939 unigenes, 8315 (93.02%) taxonomically assigned

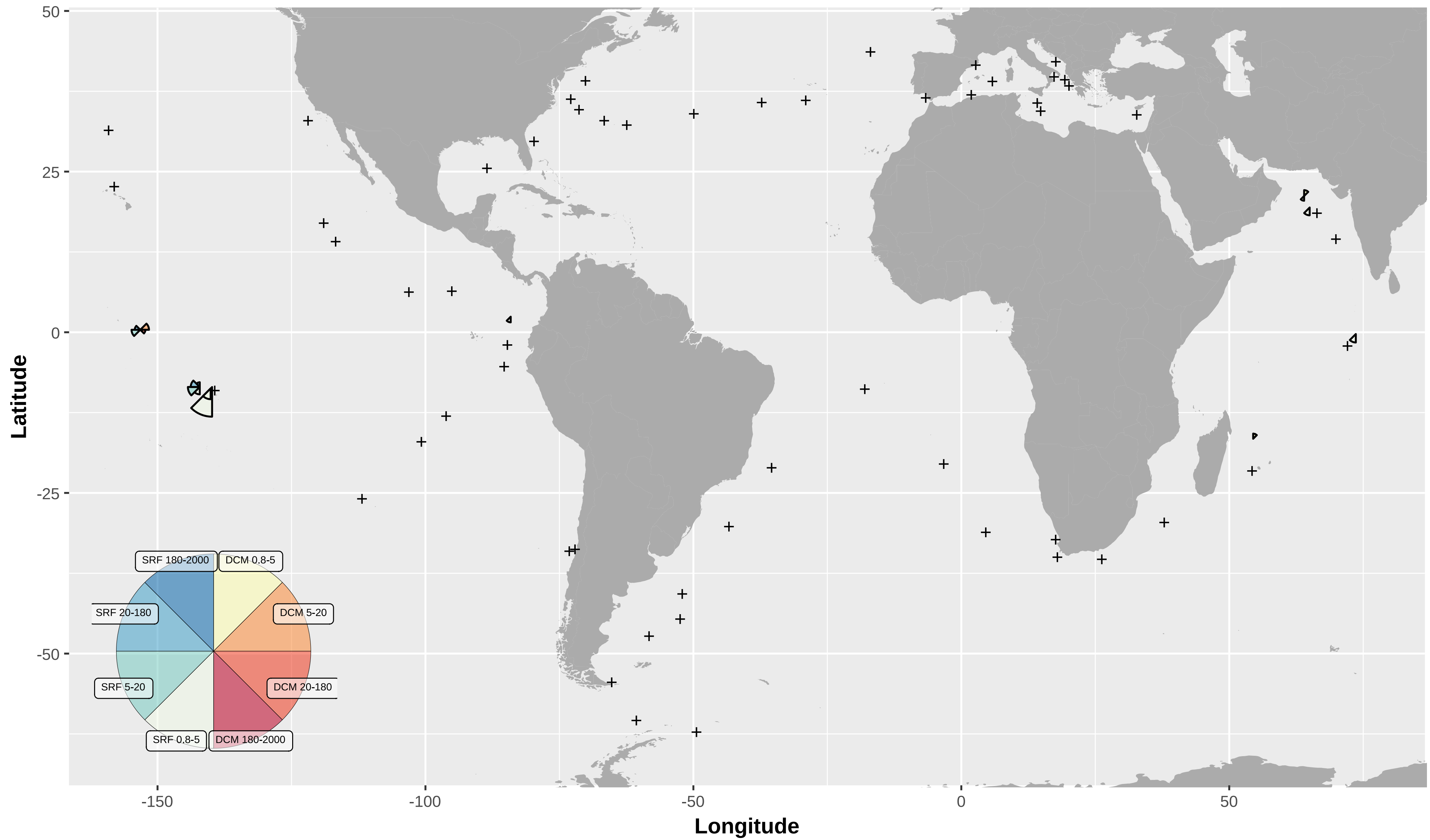


root;cellular organisms;Eukaryota;Viridiplantae;Chlorophyta;prasinophytes;Prasinococcales;Prasinoderma

## Biogeolocalisation of MGT-v1 174      8888 unigenes, 4186 (47.1%) taxonomically assigned

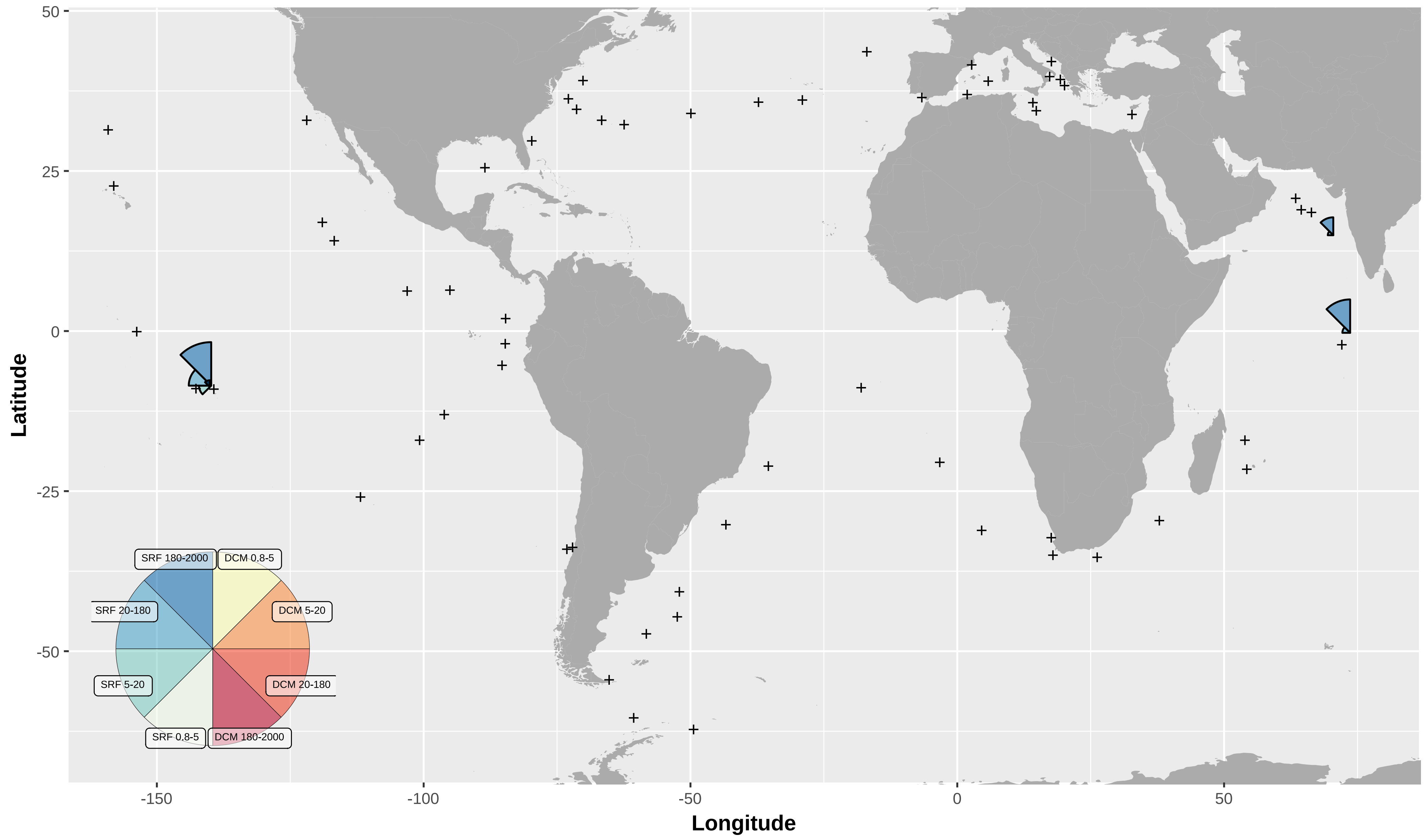


root;cellular organisms;Eukaryota;Opisthokonta;Metazoa;Eumetazoa;Bilateria;Deuterostomia;Chordata

#### Biogeolocalisation of MGT-v1 175 8797 unigenes, 7314 (83.14%) taxonomically assigned

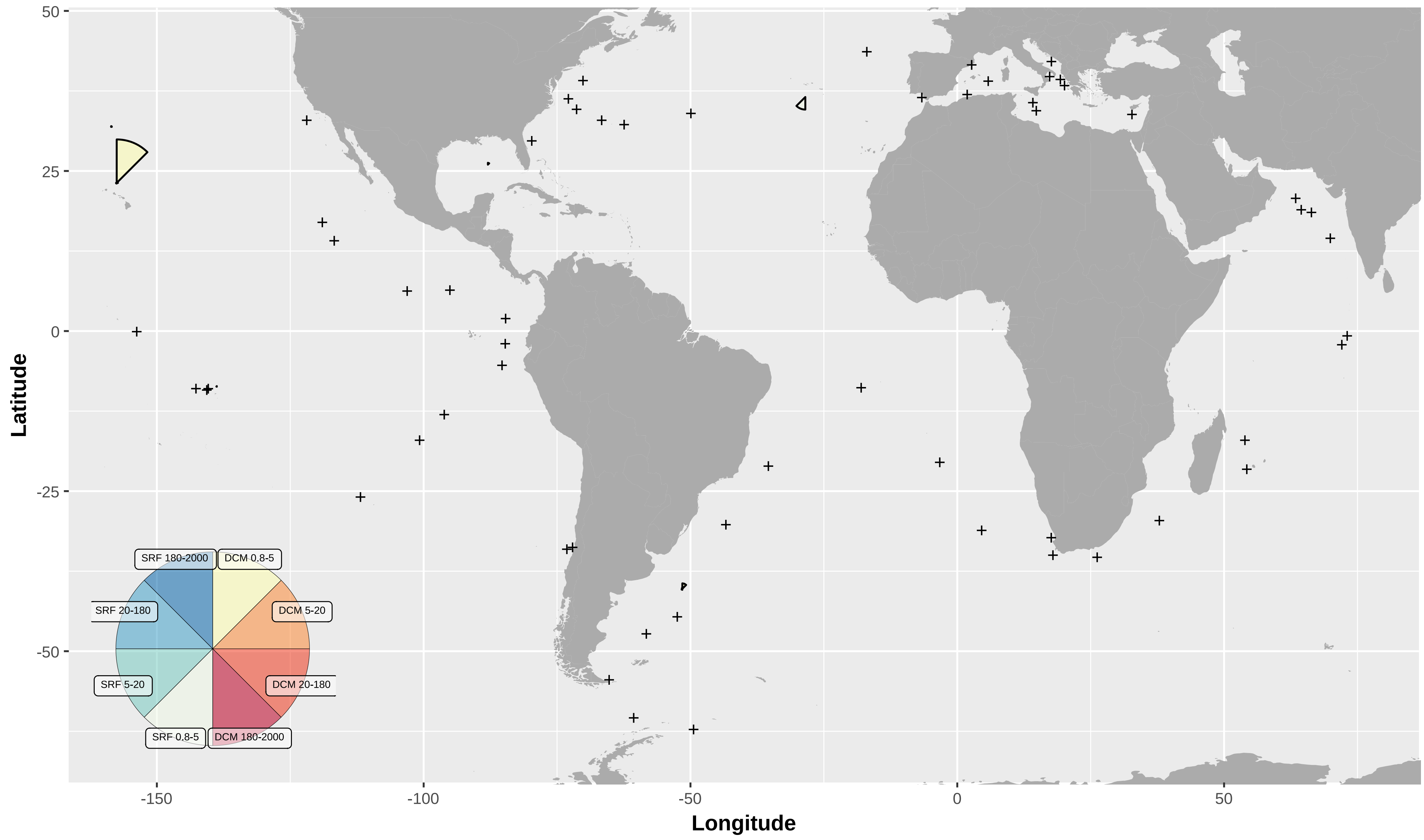


root;cellular organisms;Bacteria;Proteobacteria

## Biogeolocalisation of MGT-v1 176      8717 unigenes, 5410 (62.06%) taxonomically assigned

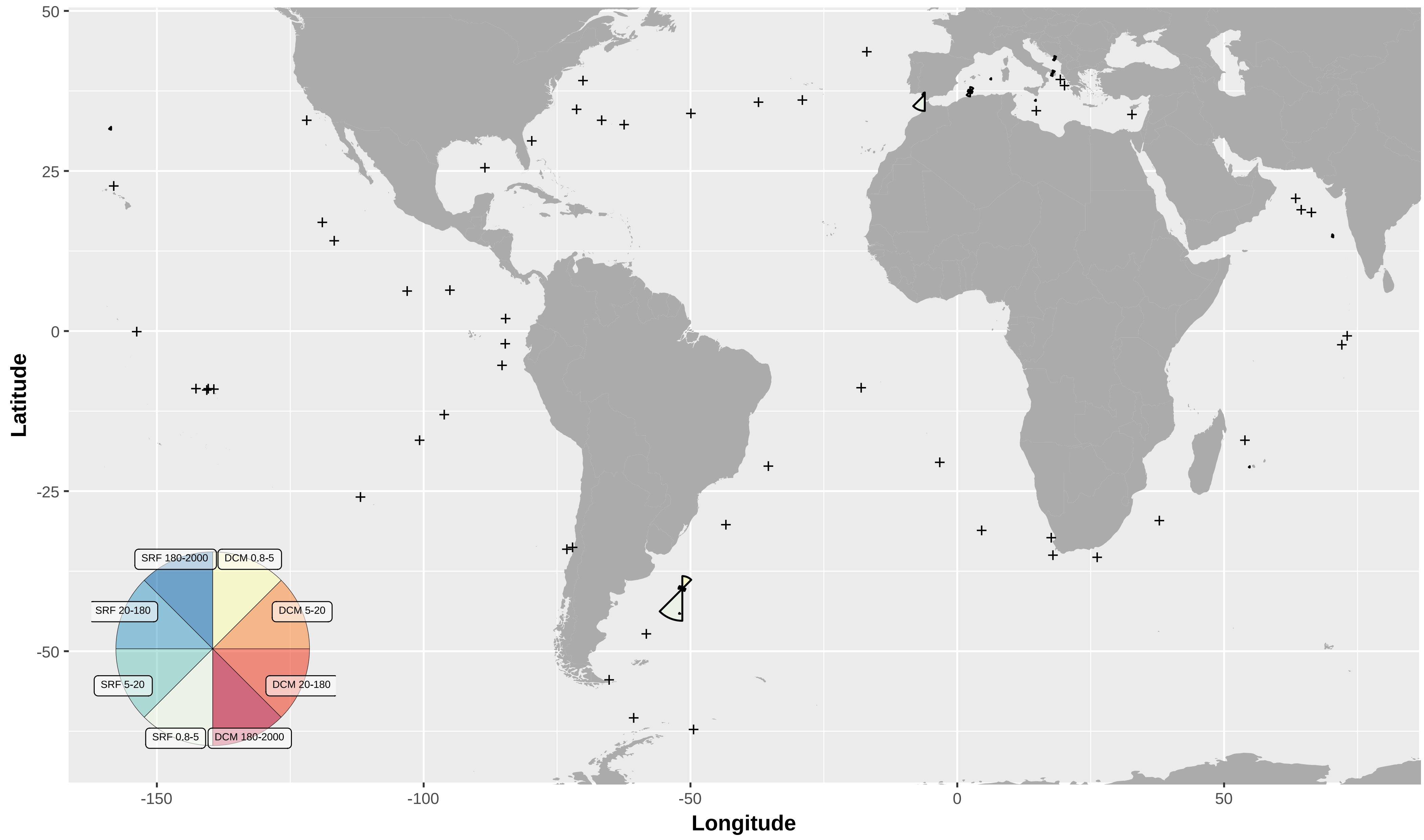


root;cellular organisms;Bacteria

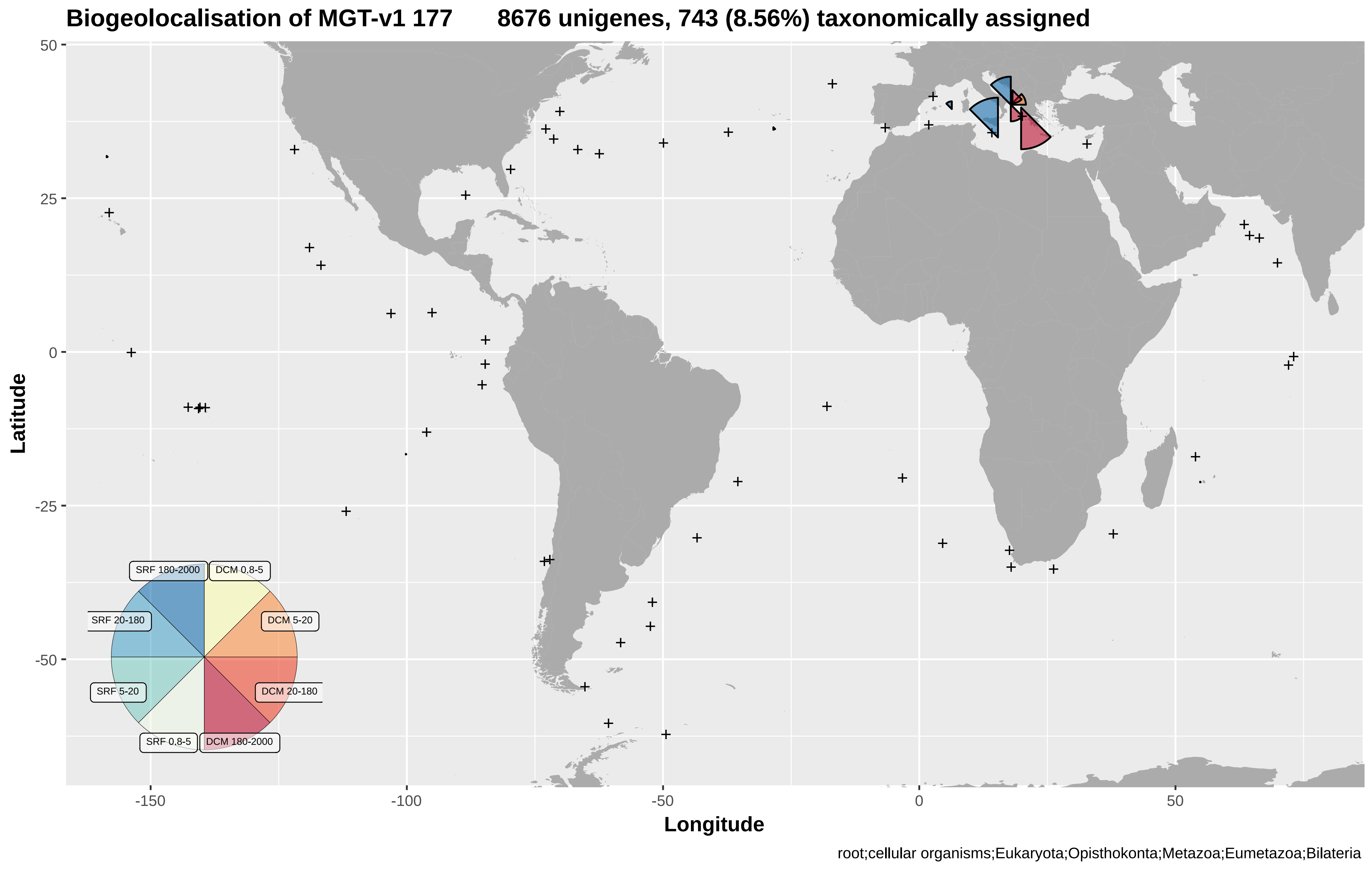

Biogeolocalisation of MGT-v1 177 8676 unigenes, 743 (8.56%) taxonomically assigned
Latitude
Longitude
50
25
0
-25
-50
-150
-100
-50
0
50
SRF 180-2000
DCM 0.8-5
SRF 20-180
DCM 5-20
SRF 5-20
DCM 20-180
SRF 0.8-5
DCM 180-2000
root;cellular organisms;Eukaryota;Opisthokonta;Metazoa;Eumetazoa;Bilateria

## Biogeolocalisation of MGT-v1 178      8653 unigenes, 7272 (84.04%) taxonomically assigned

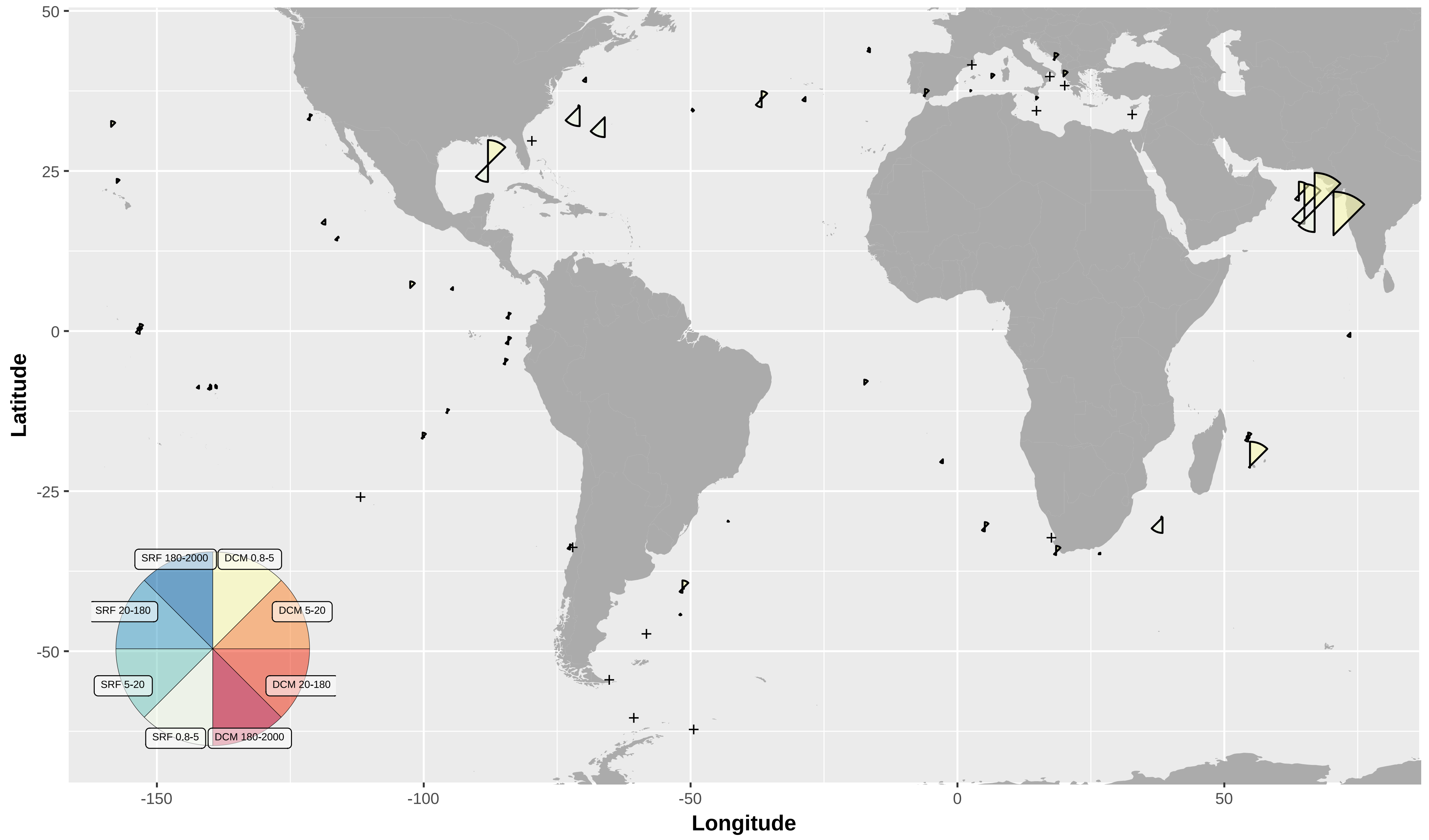


root;cellular organisms;Eukaryota;Stramenopiles;Pelagophyceae;Pelagomonadales;Pelagomonas

## Biogeolocalisation of MGT-v1 179      8603 unigenes, 7202 (83.71%) taxonomically assigned

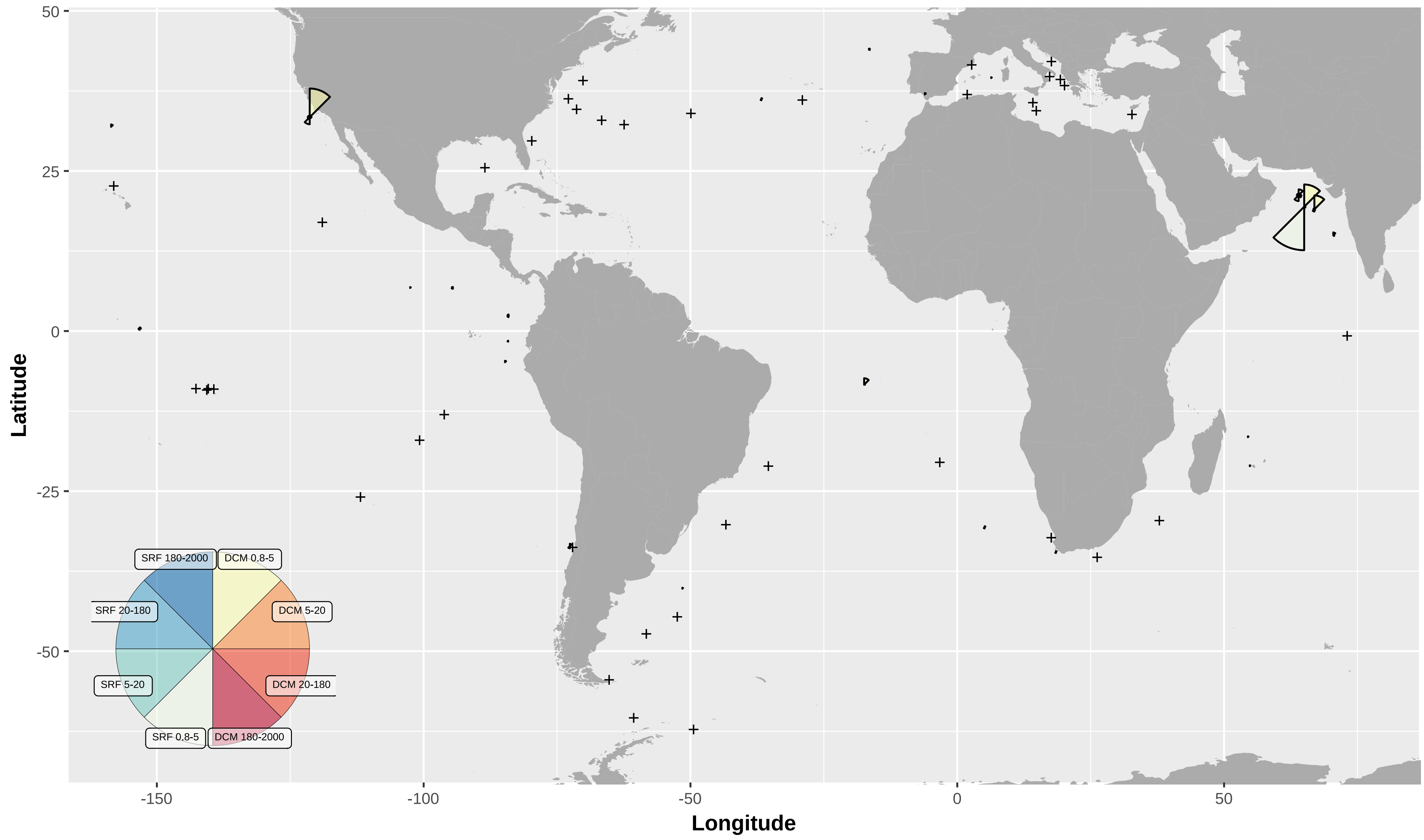


root;cellular organisms;Eukaryota;Viridiplantae

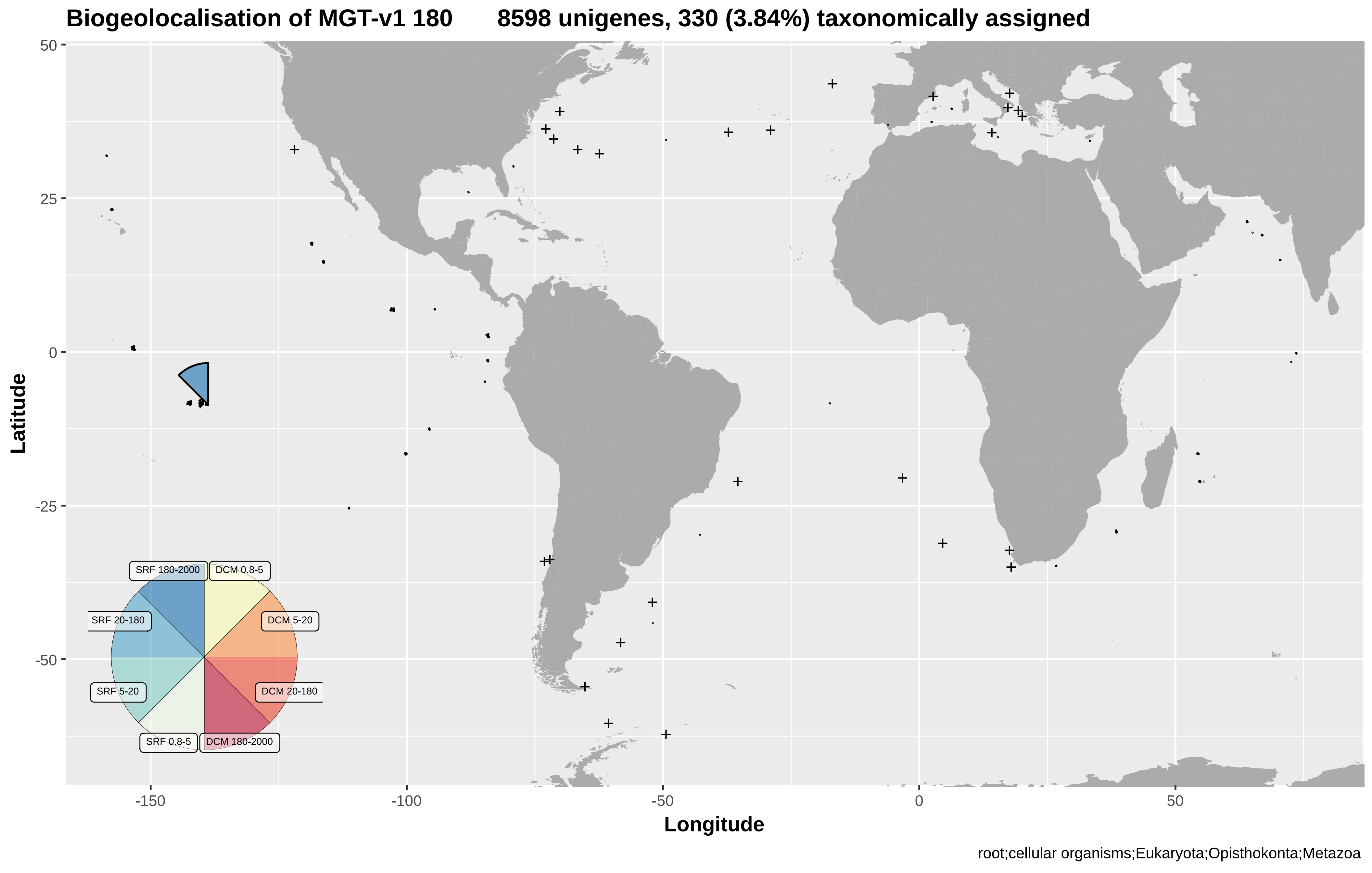

Biogeolocalisation of MGT-v1 180      8598 unigenes, 330 (3.84%) taxonomically assigned
50
25
0
-25
-50
Latitude
-150
-100
-50
0
50
Longitude
SRF 180-2000
DCM 0.8-5
SRF 20-180
DCM 5-20
SRF 5-20
DCM 20-180
SRF 0.8-5
DCM 180-2000
root;cellular organisms;Eukaryota;Opisthokonta;Metazoa

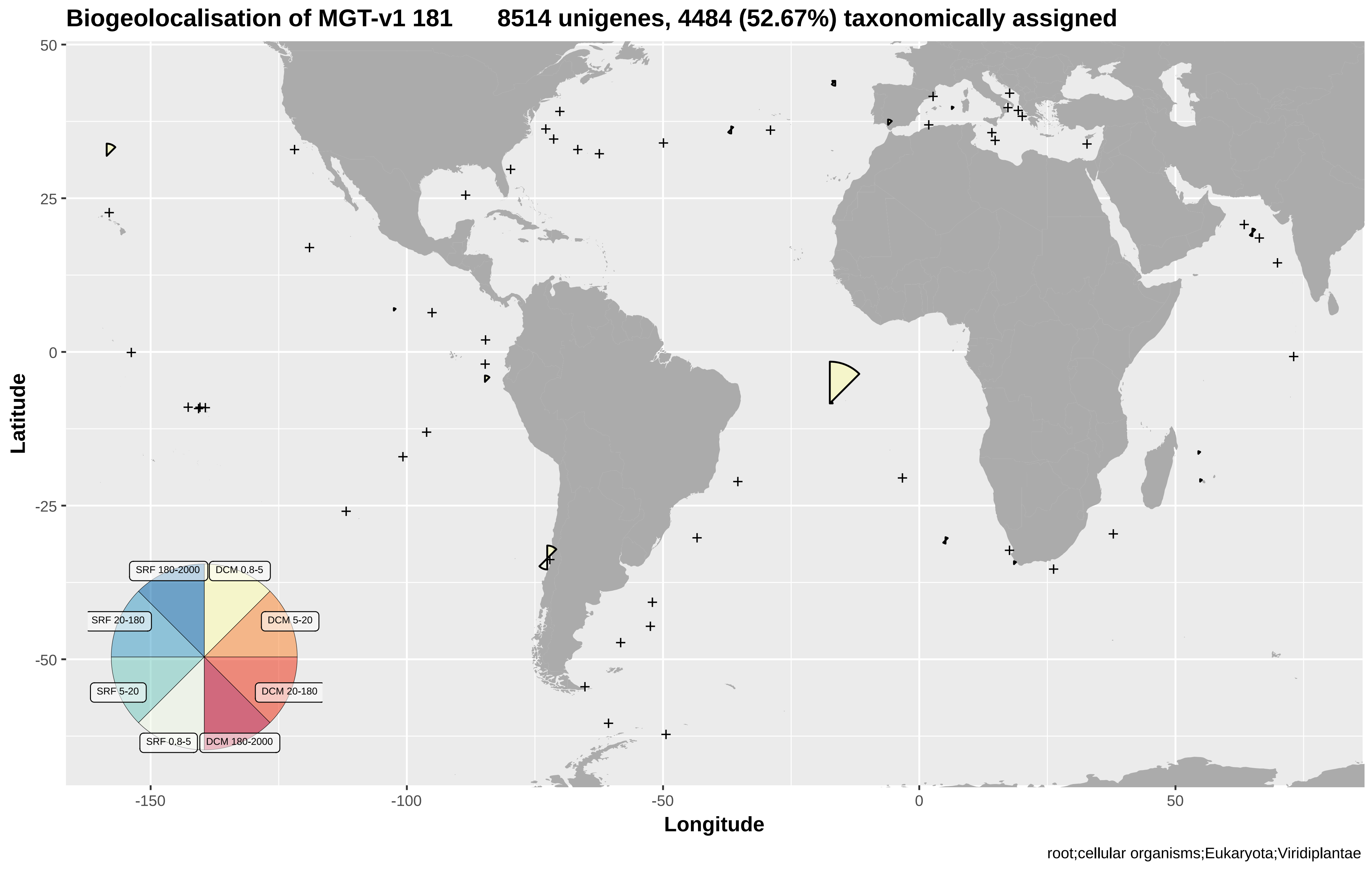
Biogeolocalisation of MGT-v1 181
8514 unigenes, 4484 (52.67%) taxonomically assigned
Latitude
Longitude
50
25
0
-25
-50
-150
-100
-50
0
50
SRF 180-2000
DCM 0.8-5
SRF 20-180
DCM 5-20
SRF 5-20
DCM 20-180
SRF 0.8-5
DCM 180-2000
root;cellular organisms;Eukaryota;Viridiplantae

# Biogeolocalisation of MGT-v1 182      8495 unigenes, 1454 (17.12%) taxonomically assigned

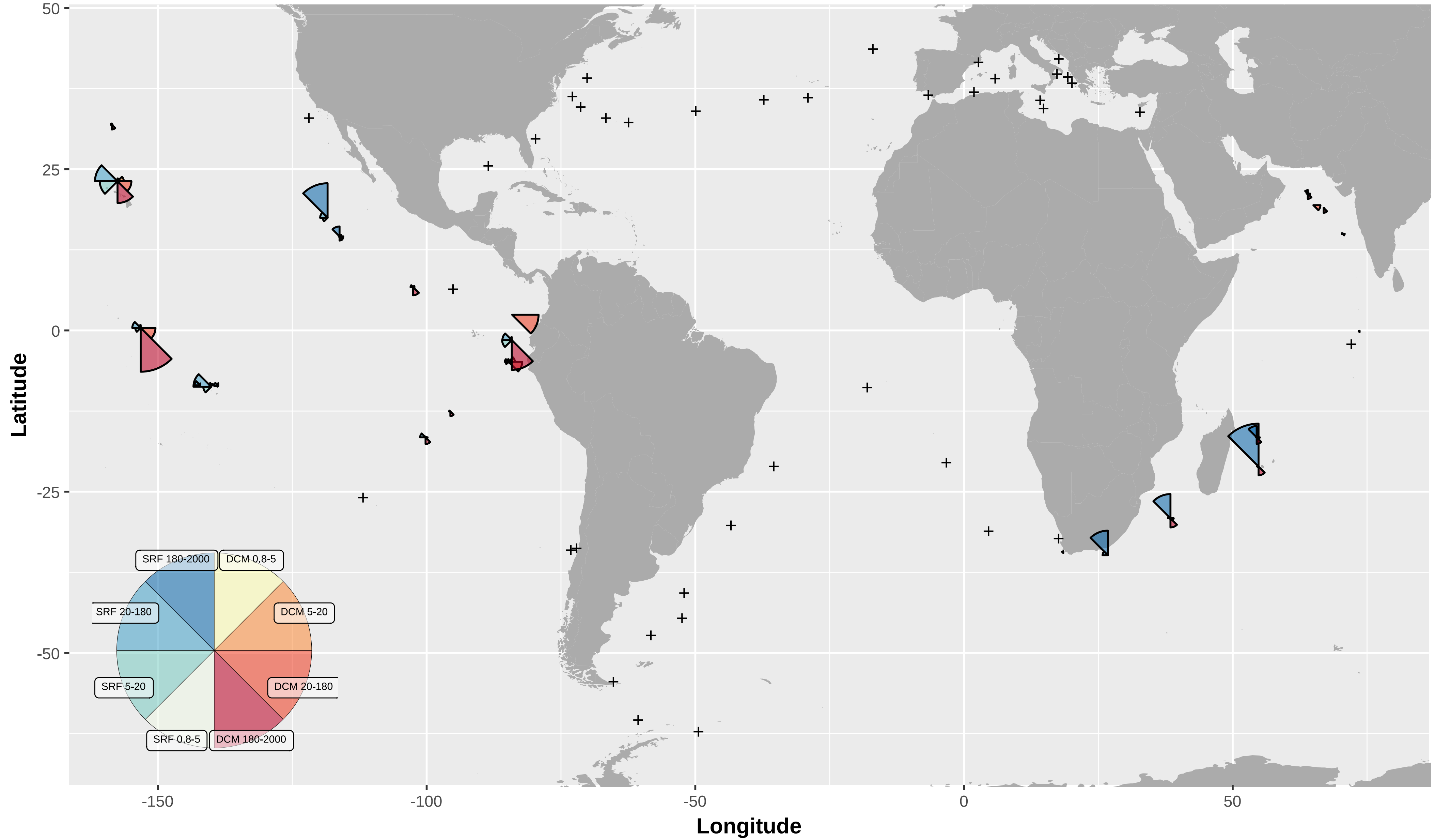


root;cellular organisms;Eukaryota;Opisthokonta;Metazoa;Eumetazoa;Bilateria;Protostomia;Ecdysozoa;Panarthropoda;Arthropoda;Mandibulata;Pancrustacea;Crustacea;Maxillopoda

Biogeolocalisation of MGT-v1 183 8433 unigenes, 3653 (43.32%) taxonomically assigned

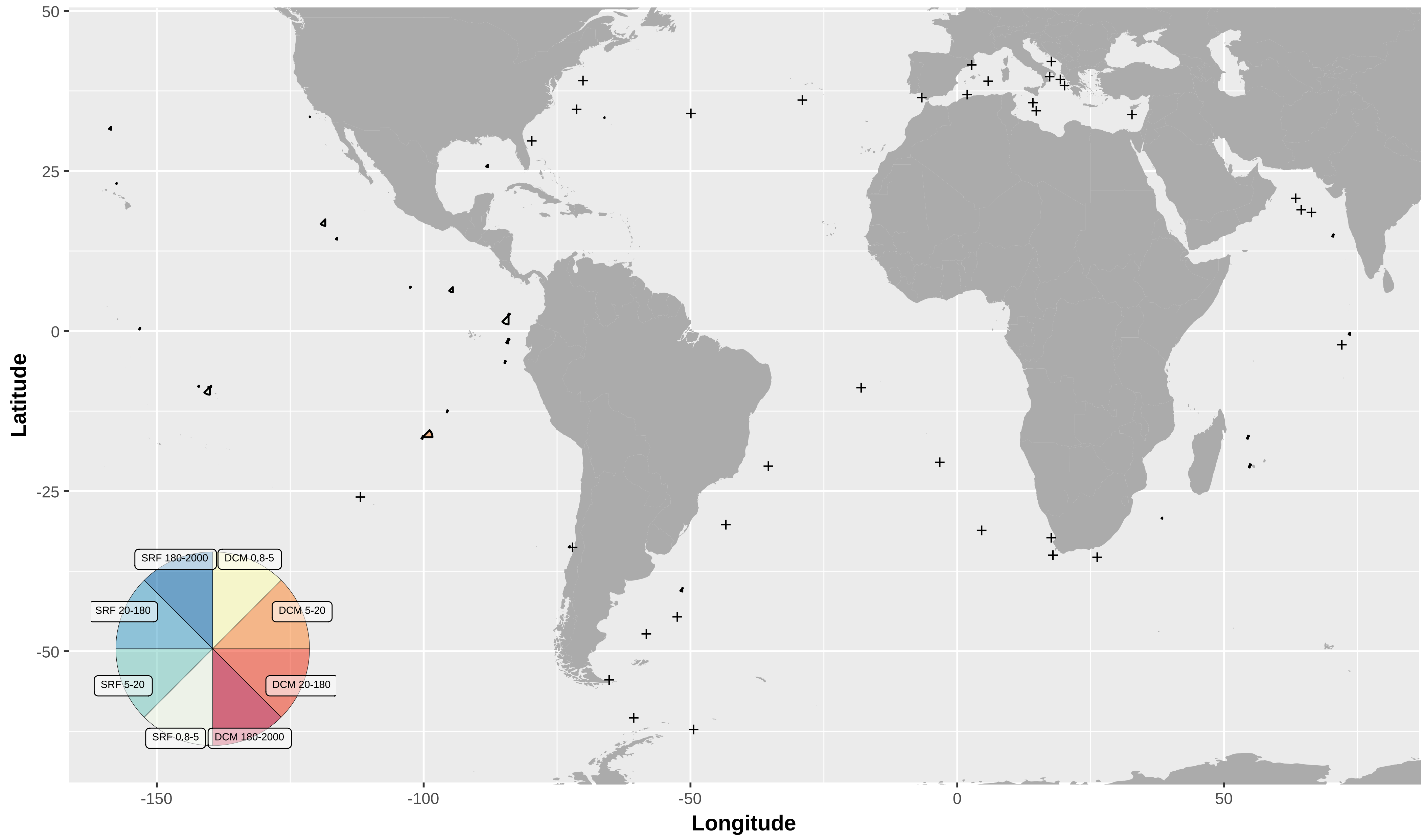


root;cellular organisms;Bacteria

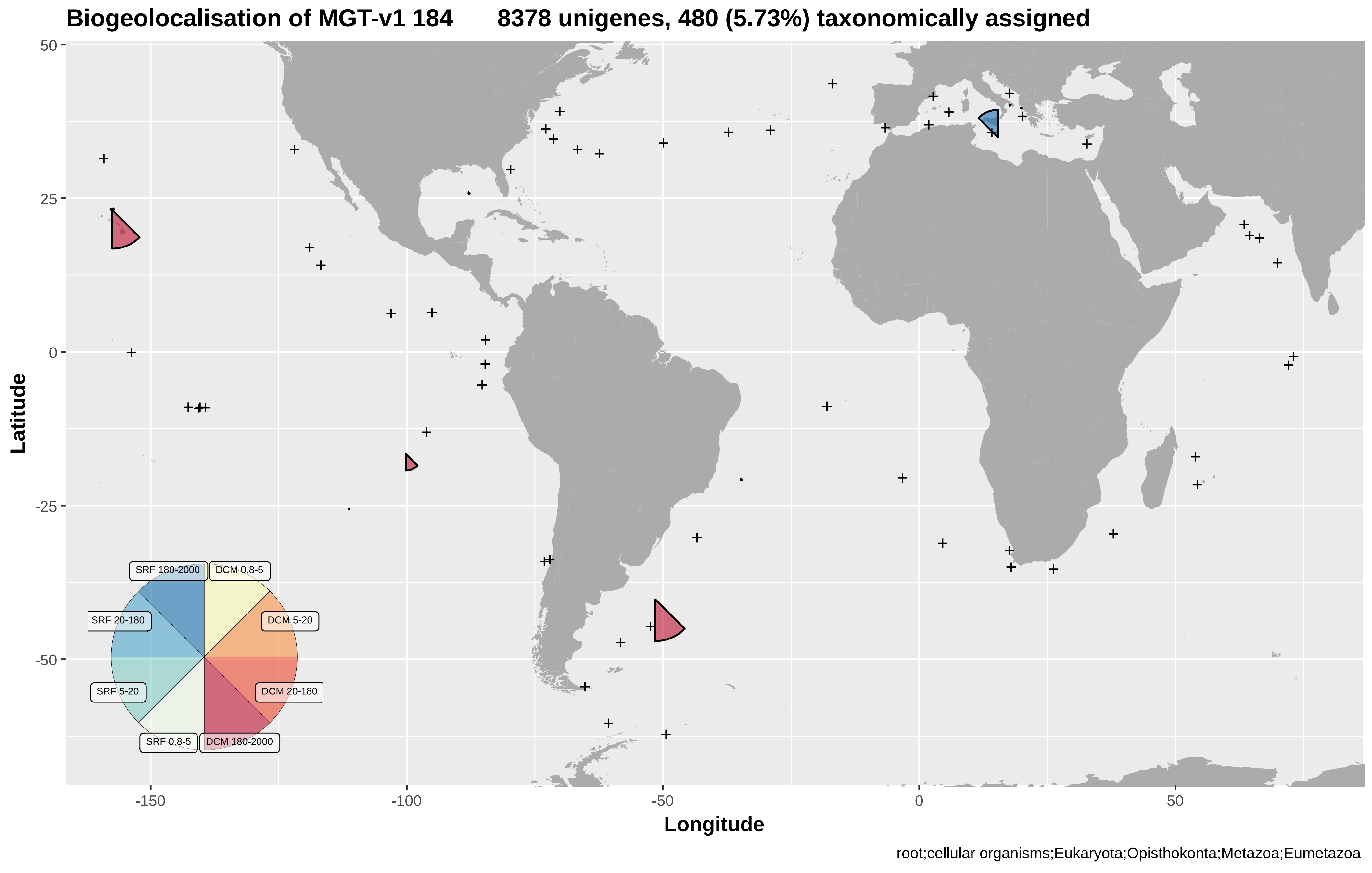

Biogeolocalisation of MGT-v1 184 8378 unigenes, 480 (5.73%) taxonomically assigned
50
25
0
-25
-50
Latitude
-150
-100
-50
0
50
Longitude
SRF 180-2000
DCM 0.8-5
SRF 20-180
DCM 5-20
SRF 5-20
DCM 20-180
SRF 0.8-5
DCM 180-2000
root;cellular organisms;Eukaryota;Opisthokonta;Metazoa;Eumetazoa

## Biogeolocalisation of MGT-v1 185      8243 unigenes, 627 (7.61%) taxonomically assigned

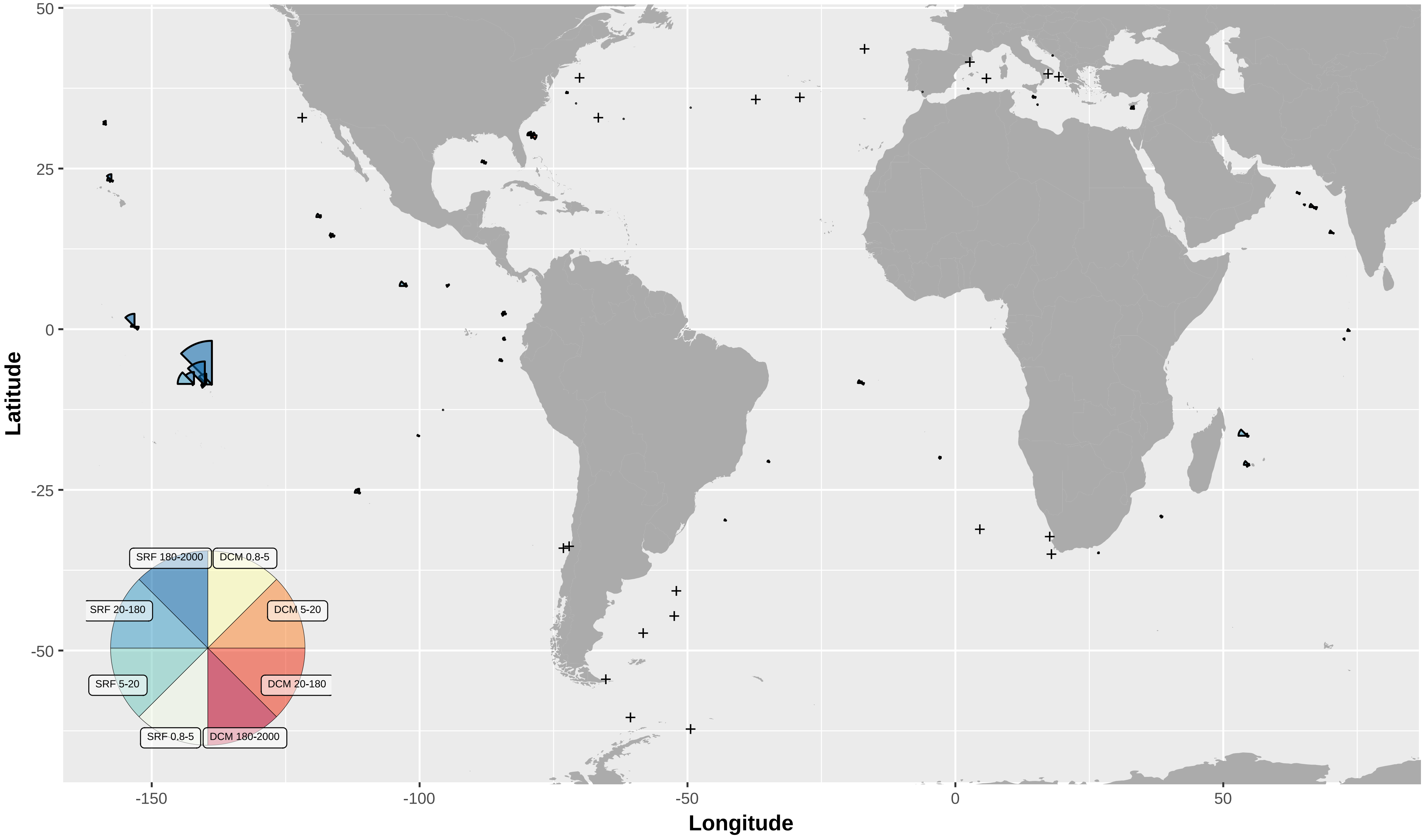


root;cellular organisms;Eukaryota;Opisthokonta;Metazoa;Eumetazoa;Bilateria

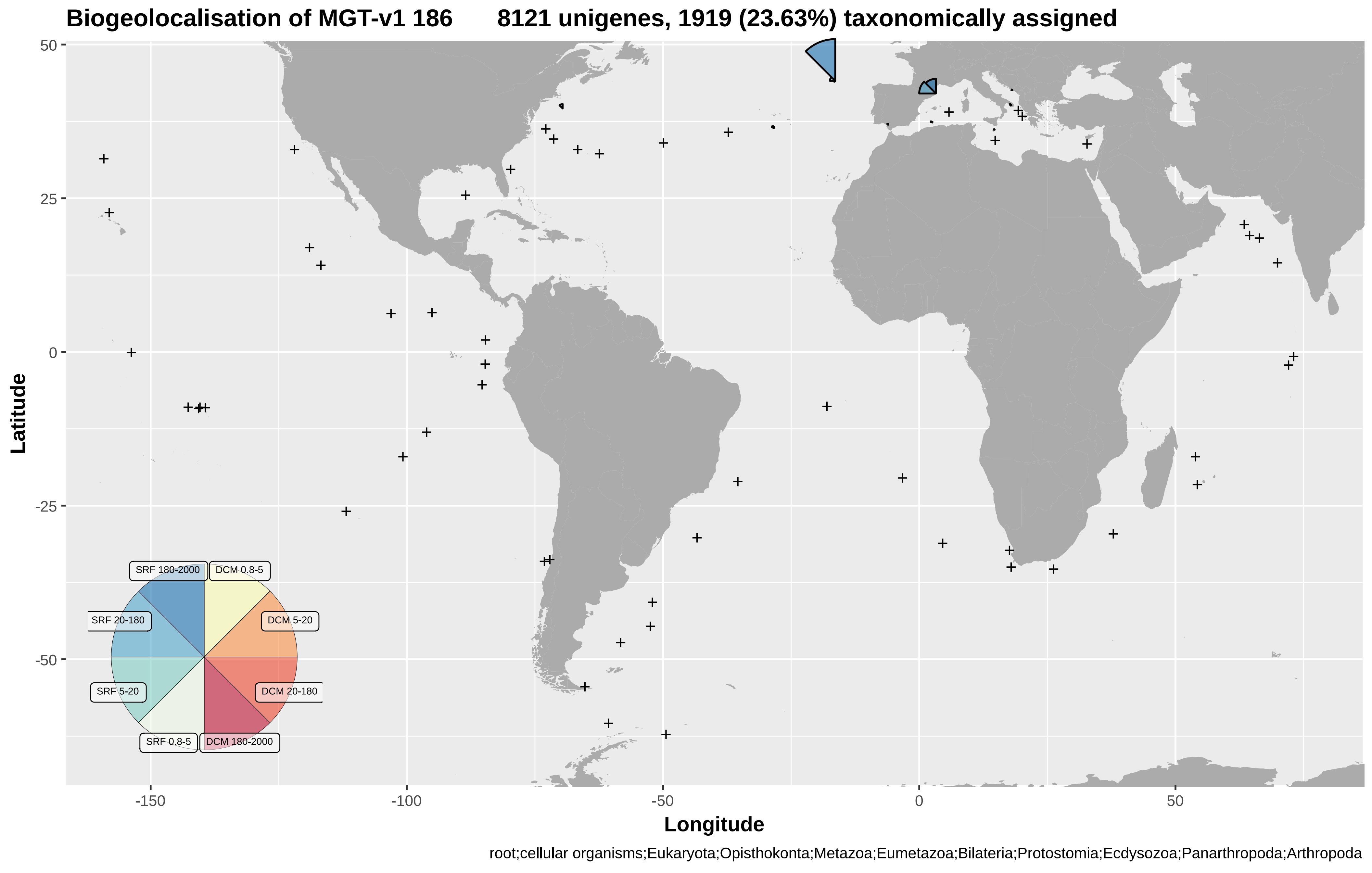

Biogeolocalisation of MGT-v1 186 8121 unigenes, 1919 (23.63%) taxonomically assigned
50
25
0
-25
-50
Latitude
-150
-100
-50
0
50
Longitude
SRF 180-2000
DCM 0.8-5
SRF 20-180
DCM 5-20
SRF 5-20
DCM 20-180
SRF 0.8-5
DCM 180-2000
root;cellular organisms;Eukaryota;Opisthokonta;Metazoa;Eumetazoa;Bilateria;Protostomia;Ecdysozoa;Panarthropoda;Arthropoda

**Biogeolocalisation of MGT-v1 187 8054 unigenes, 6974 (86.59%) taxonomically assigned**

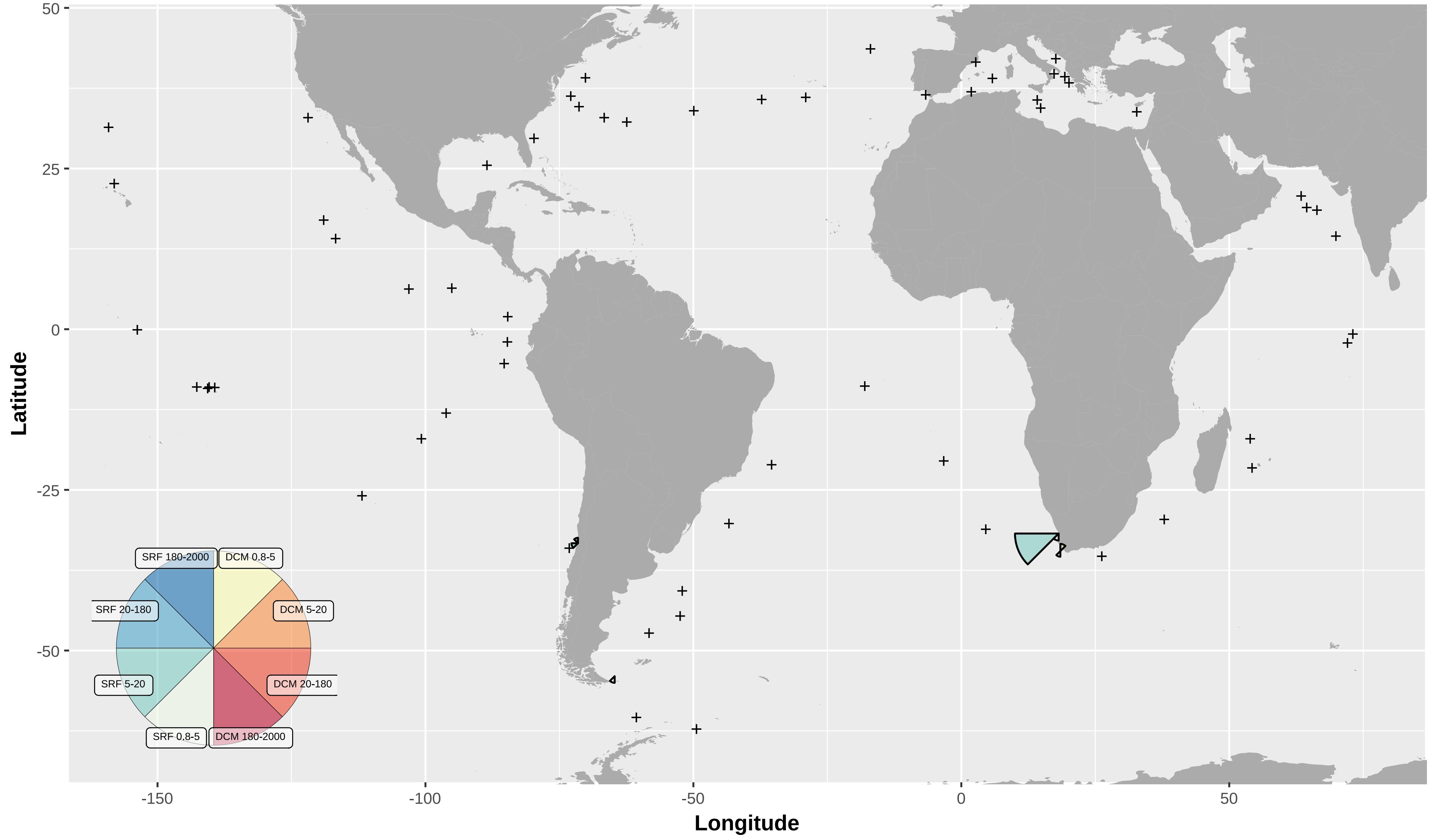


root;cellular organisms;Eukaryota;Stramenopiles;Bacillariophyta;Coscinodiscophyceae;Thalassiosirophycidae;Thalassiosirales;Thalassiosiraceae

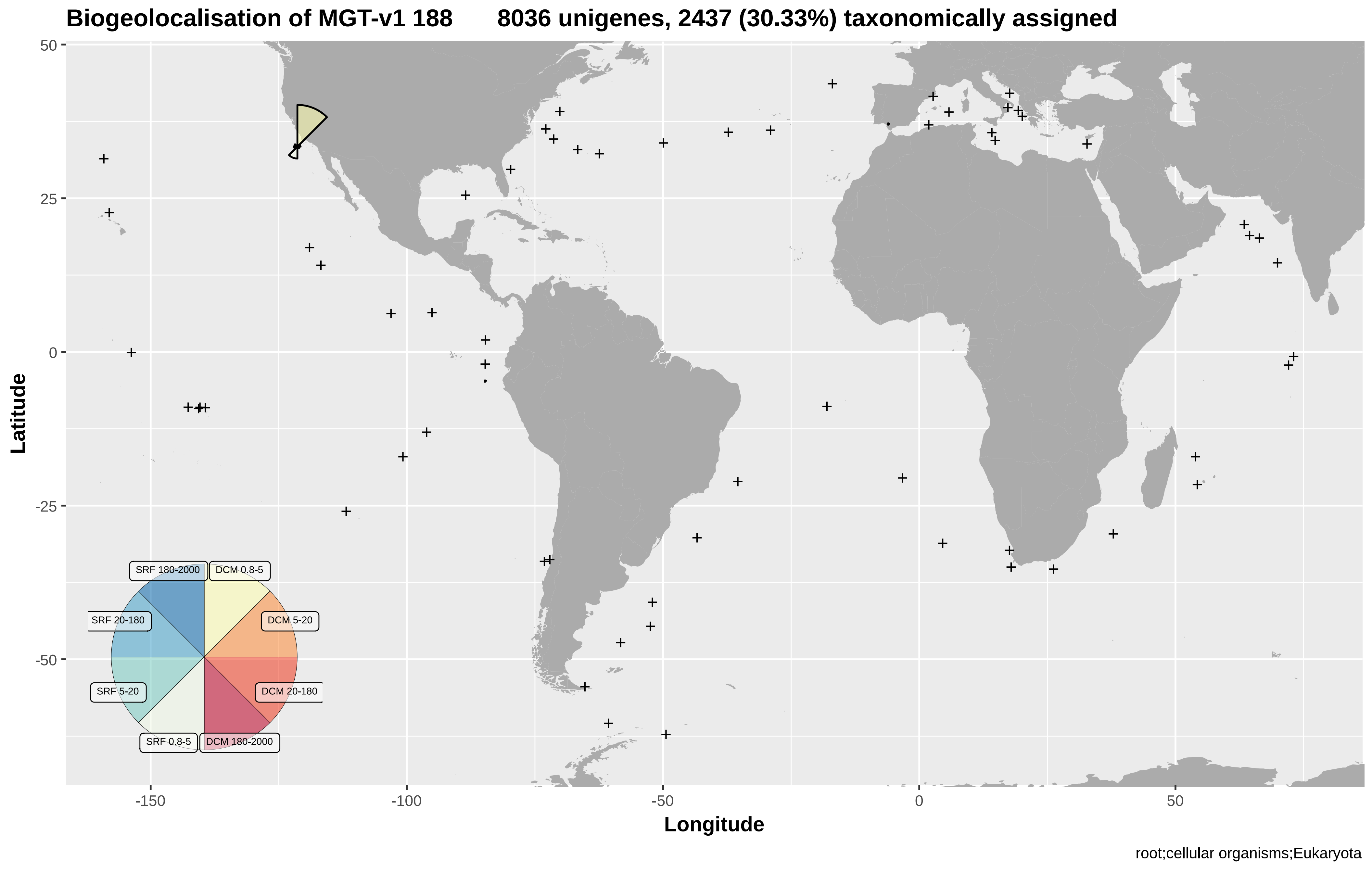

Biogeolocalisation of MGT-v1 188 8036 unigenes, 2437 (30.33%) taxonomically assigned
50
25
0
-25
-50
Latitude
-150
-100
-50
0
50
Longitude
SRF 180-2000
DCM 0.8-5
SRF 20-180
DCM 5-20
SRF 5-20
DCM 20-180
SRF 0.8-5
DCM 180-2000
root;cellular organisms;Eukaryota

Biogeolocalisation of MGT-v1 189      7758 unigenes, 605 (7.8%) taxonomically assigned

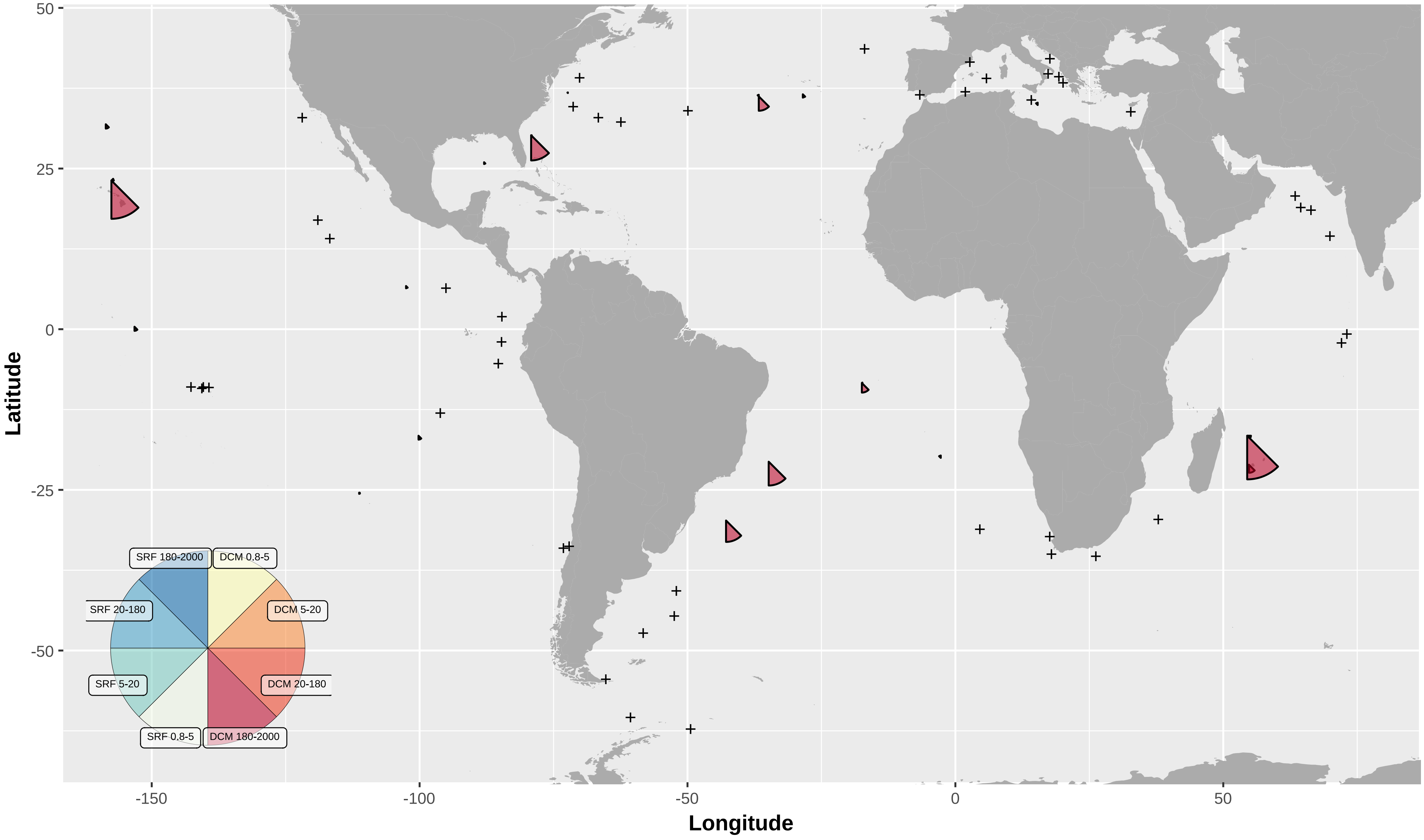


root;cellular organisms;Eukaryota;Opisthokonta;Metazoa;Eumetazoa

#### Biogeolocalisation of MGT-v1 190 7520 unigenes, 3679 (48.92%) taxonomically assigned

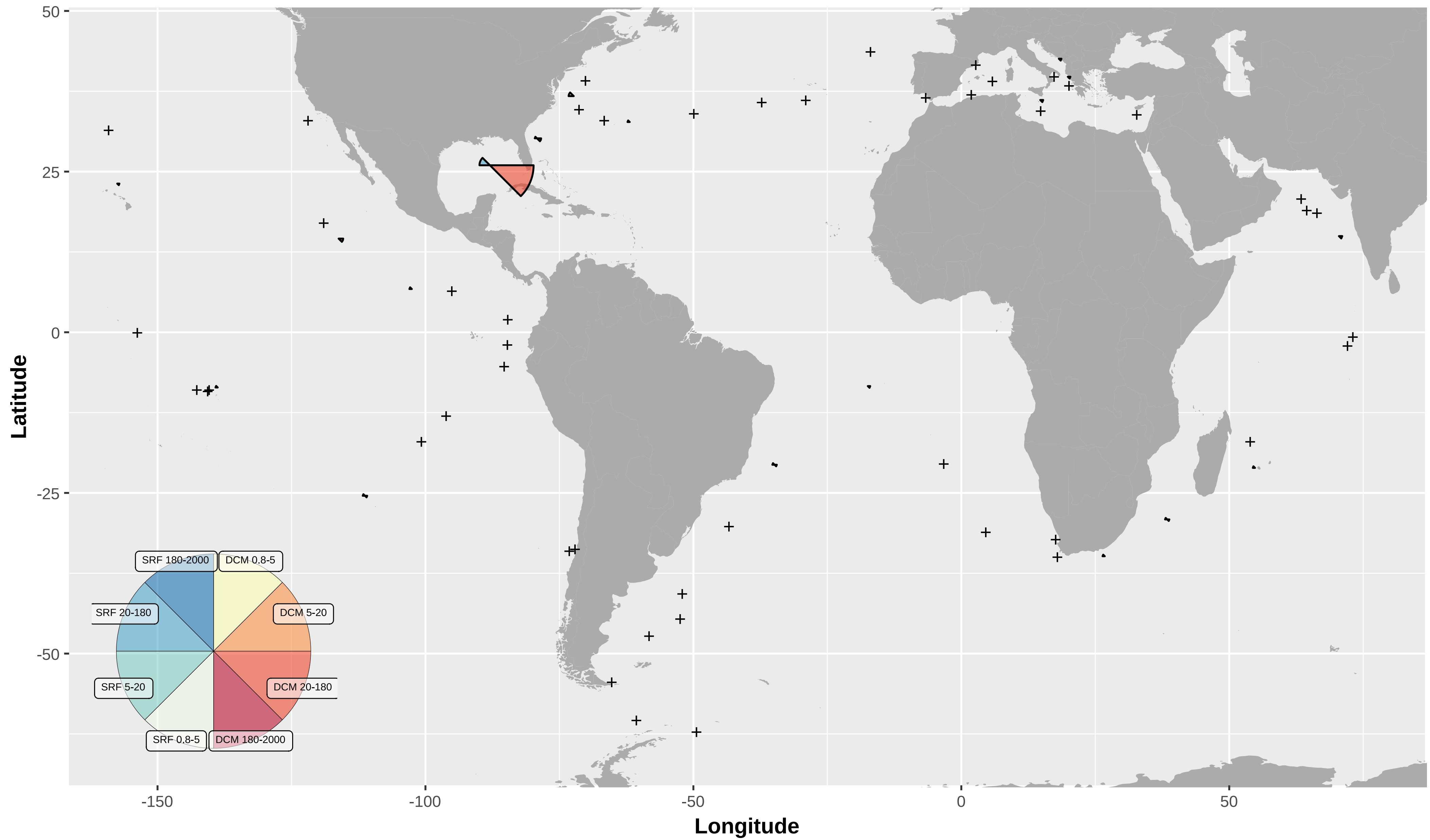


root;cellular organisms;Eukaryota;Alveolata;Ciliophora;Intramacronucleata;Spirotrichea

## Biogeolocalisation of MGT-v1 191      7517 unigenes, 5377 (71.53%) taxonomically assigned

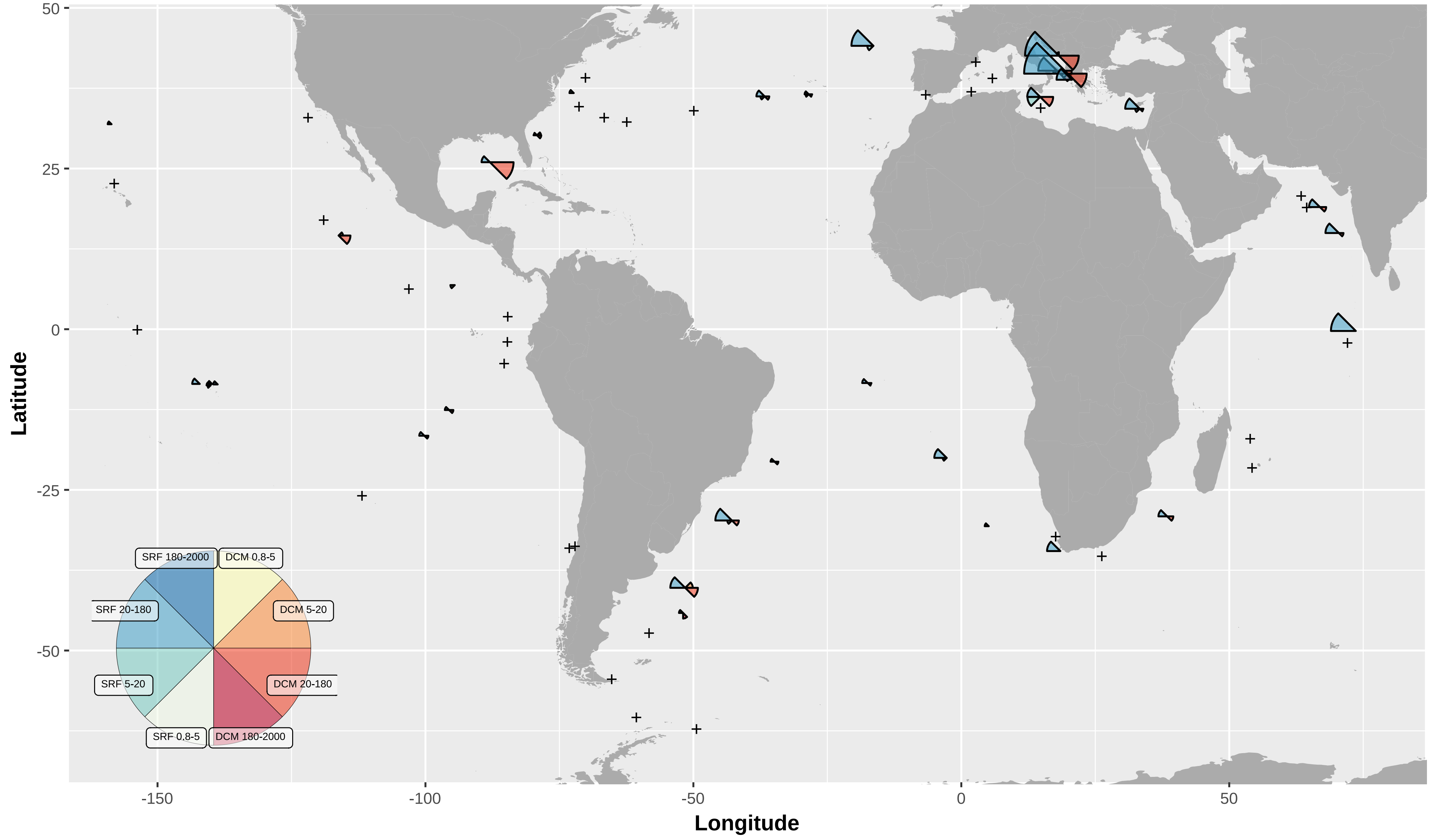


root;cellular organisms;Eukaryota;Alveolata;Ciliophora;Intramacronucleata;Spirotrichea

## Biogeolocalisation of MGT-v1 192      7455 unigenes, 758 (10.17%) taxonomically assigned

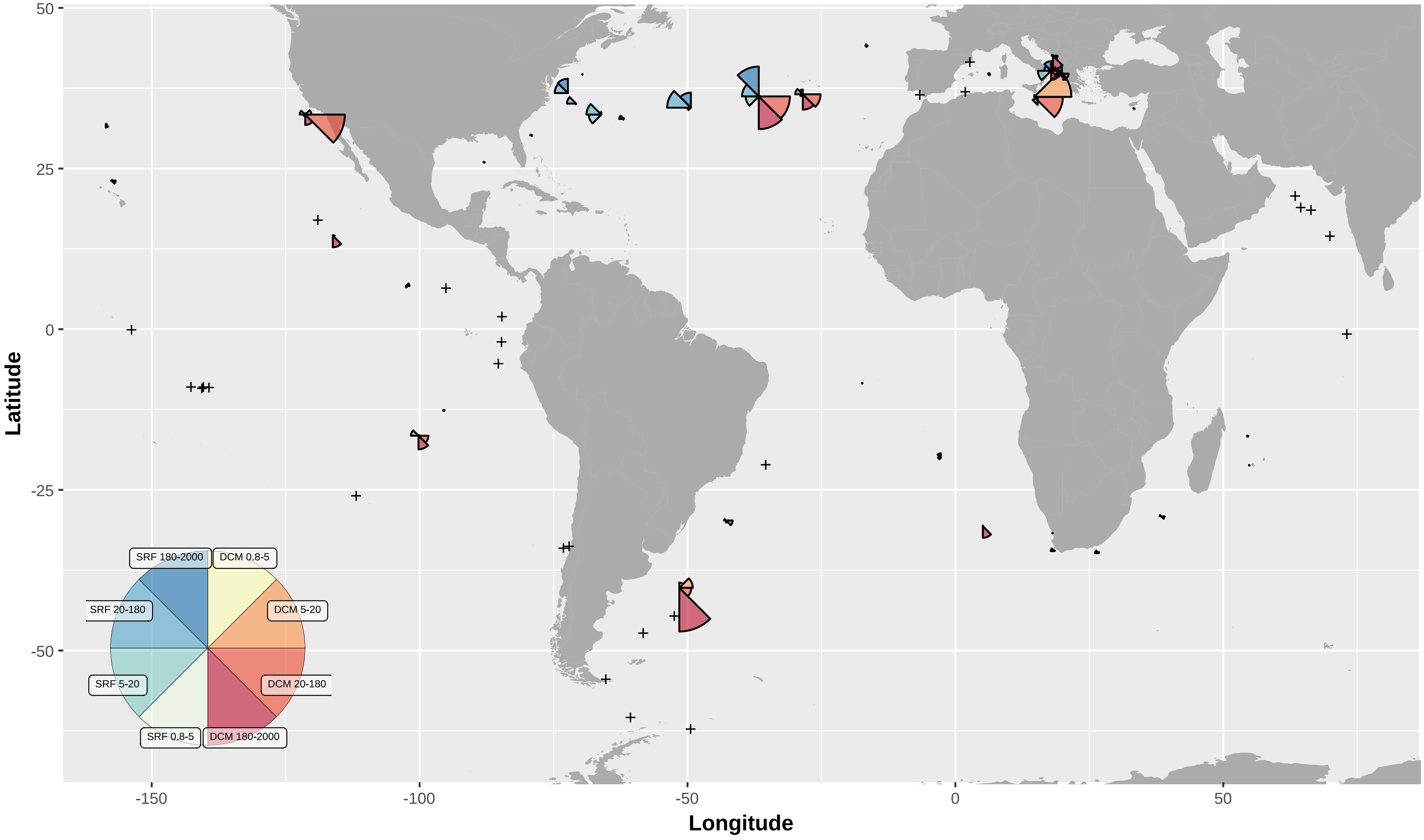


root;cellular organisms;Eukaryota;Opisthokonta;Metazoa;Eumetazoa;Bilateria

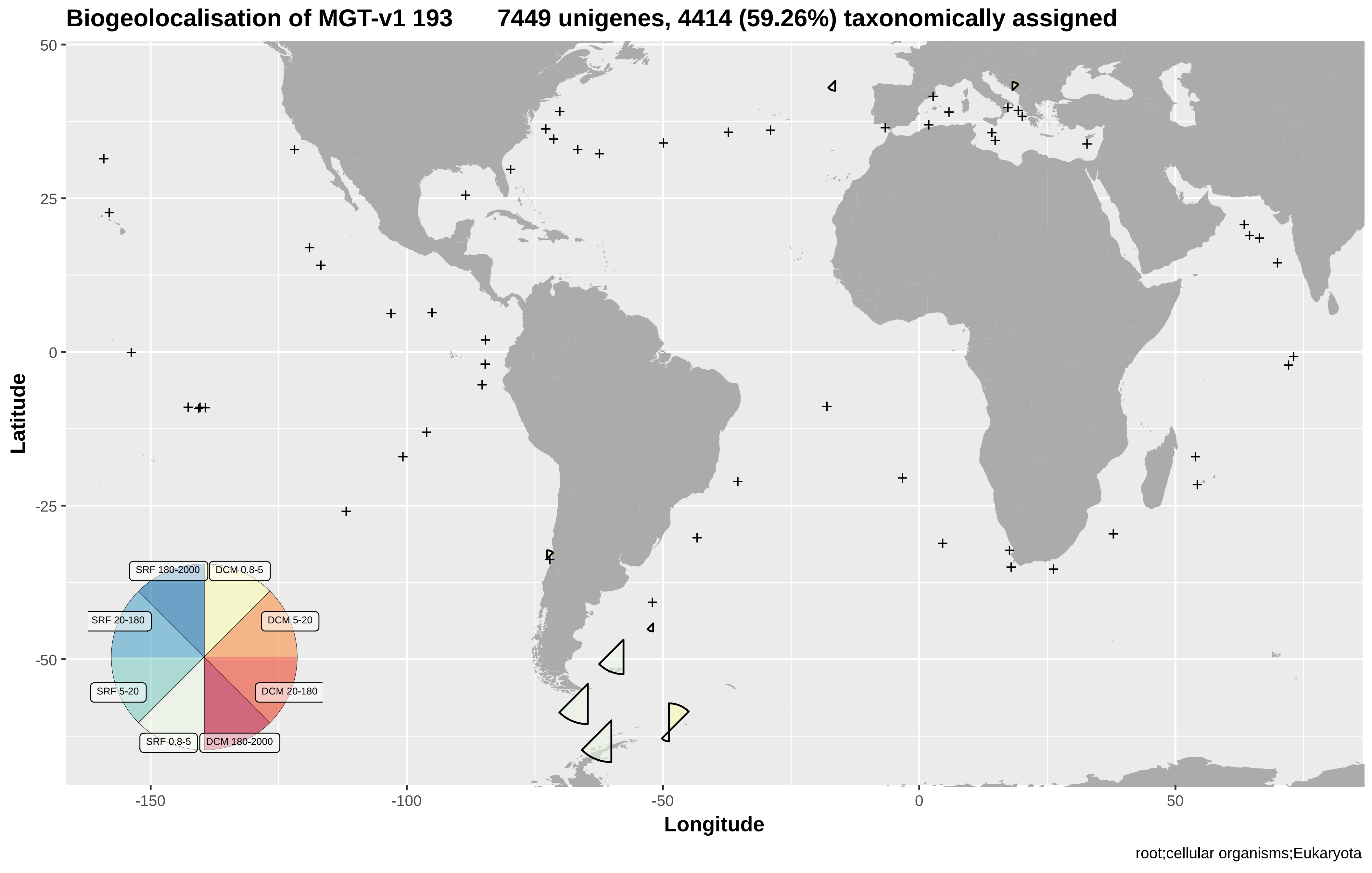

Biogeolocalisation of MGT-v1 193 7449 unigenes, 4414 (59.26%) taxonomically assigned
50
25
0
-25
-50
Latitude
-150
-100
-50
0
50
Longitude
SRF 180-2000
DCM 0.8-5
SRF 20-180
DCM 5-20
SRF 5-20
DCM 20-180
SRF 0.8-5
DCM 180-2000
root;cellular organisms;Eukaryota

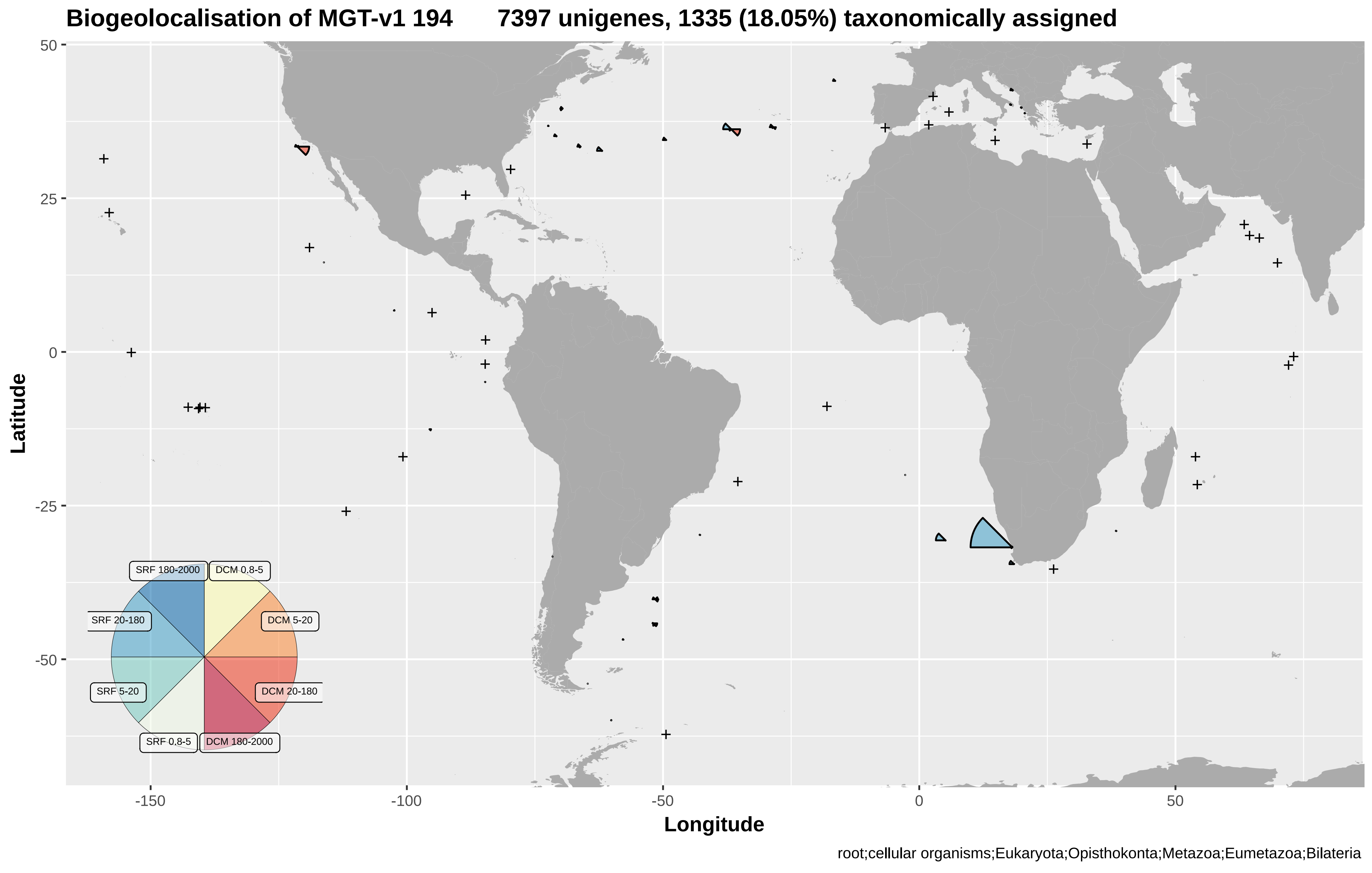

Biogeolocalisation of MGT-v1 194 7397 unigenes, 1335 (18.05%) taxonomically assigned
50
25
0
-25
-50
Latitude
-150
-100
-50
0
50
Longitude
SRF 180-2000
DCM 0.8-5
SRF 20-180
DCM 5-20
SRF 5-20
DCM 20-180
SRF 0.8-5
DCM 180-2000
root;cellular organisms;Eukaryota;Opisthokonta;Metazoa;Eumetazoa;Bilateria

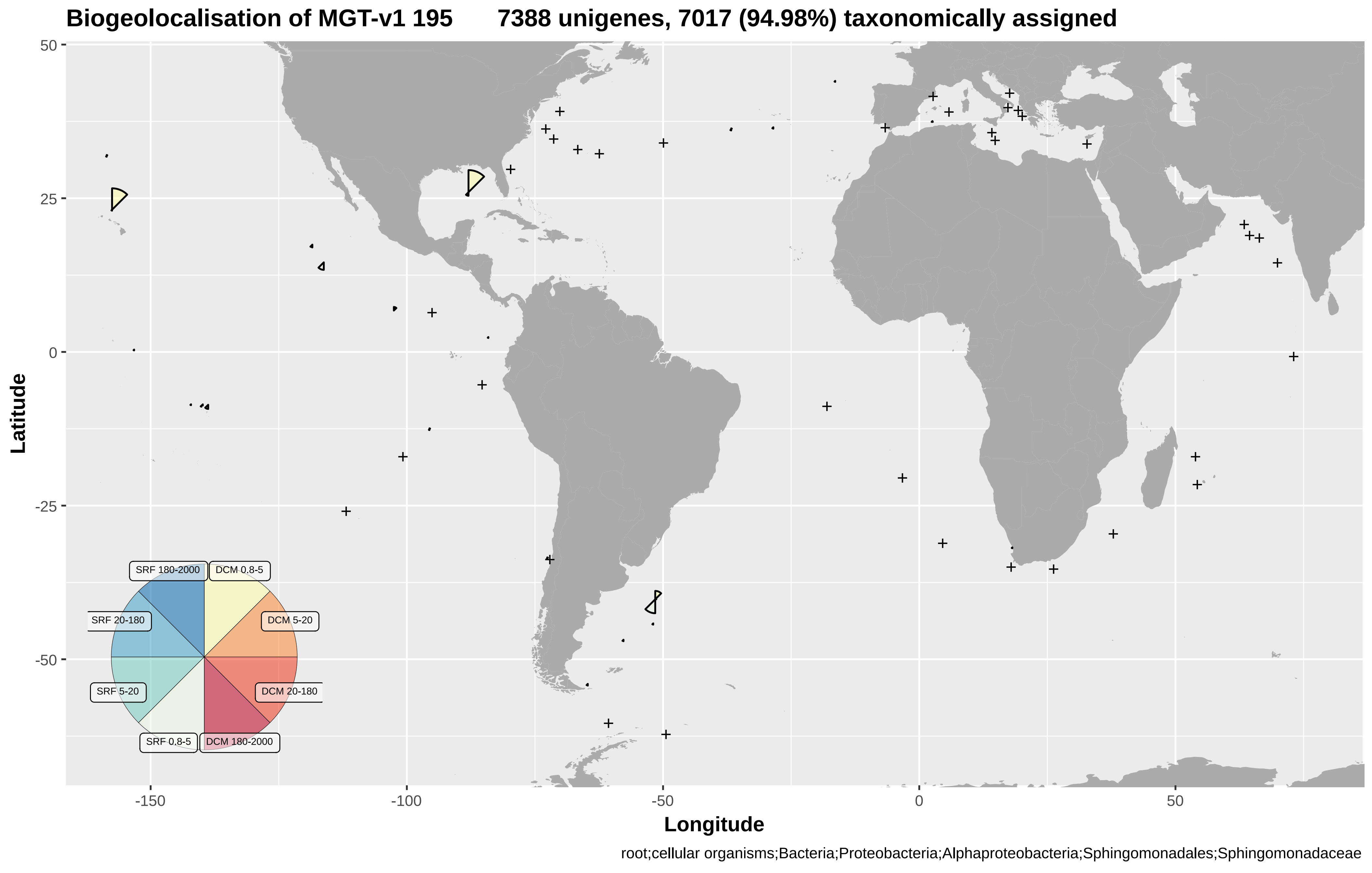

Biogeolocalisation of MGT-v1 195 7388 unigenes, 7017 (94.98%) taxonomically assigned
50
25
0
-25
-50
Latitude
-150
-100
-50
0
50
Longitude
SRF 180-2000
DCM 0.8-5
SRF 20-180
DCM 5-20
SRF 5-20
DCM 20-180
SRF 0.8-5
DCM 180-2000
root;cellular organisms;Bacteria;Proteobacteria;Alphaproteobacteria;Sphingomonadales;Sphingomonadaceae

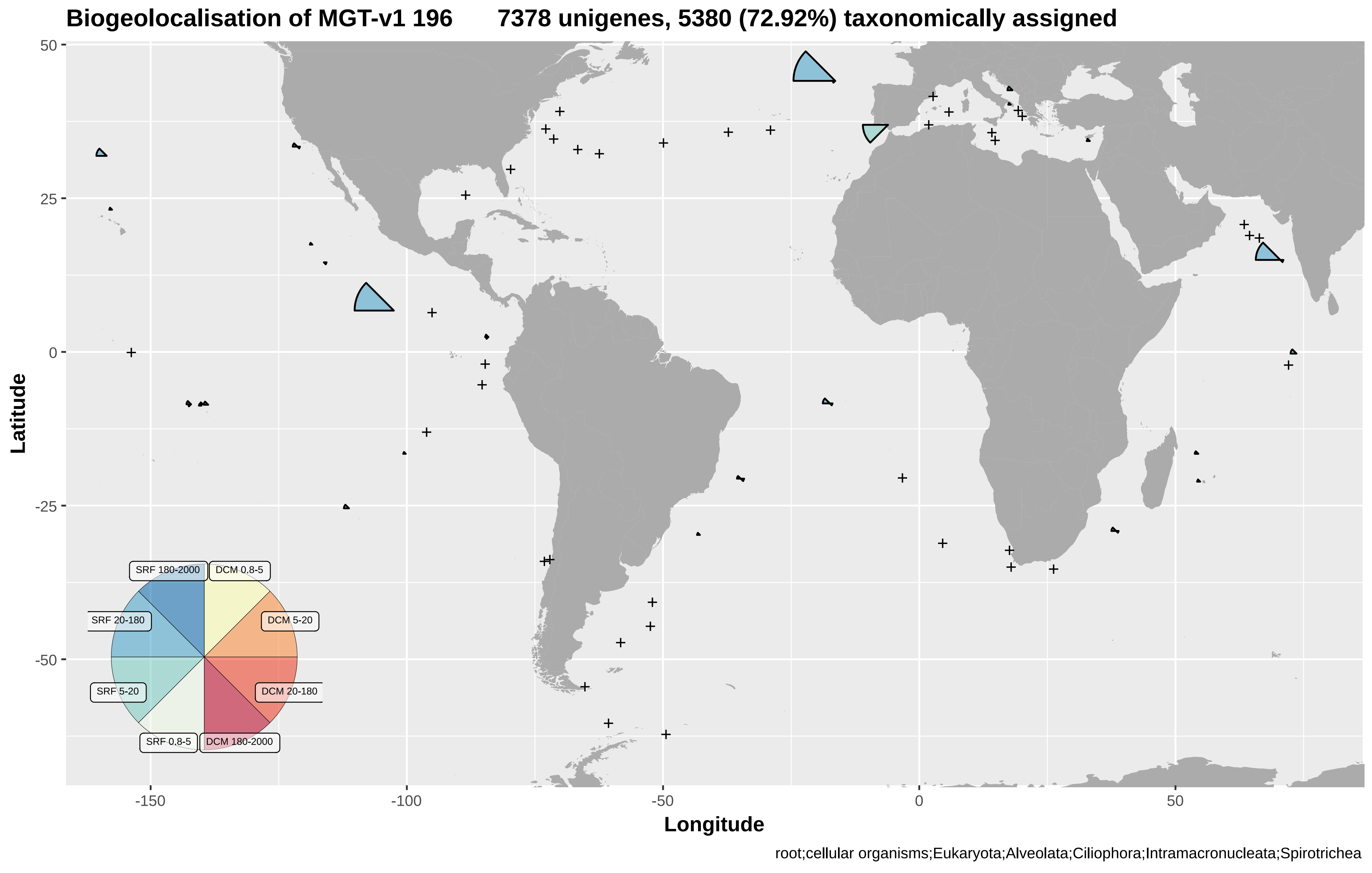

Biogeolocalisation of MGT-v1 196 7378 unigenes, 5380 (72.92%) taxonomically assigned
50
25
0
-25
-50
Latitude
-150
-100
-50
0
50
Longitude
SRF 180-2000
DCM 0.8-5
SRF 20-180
DCM 5-20
SRF 5-20
DCM 20-180
SRF 0.8-5
DCM 180-2000
root;cellular organisms;Eukaryota;Alveolata;Ciliophora;Intramacronucleata;Spirotrichea

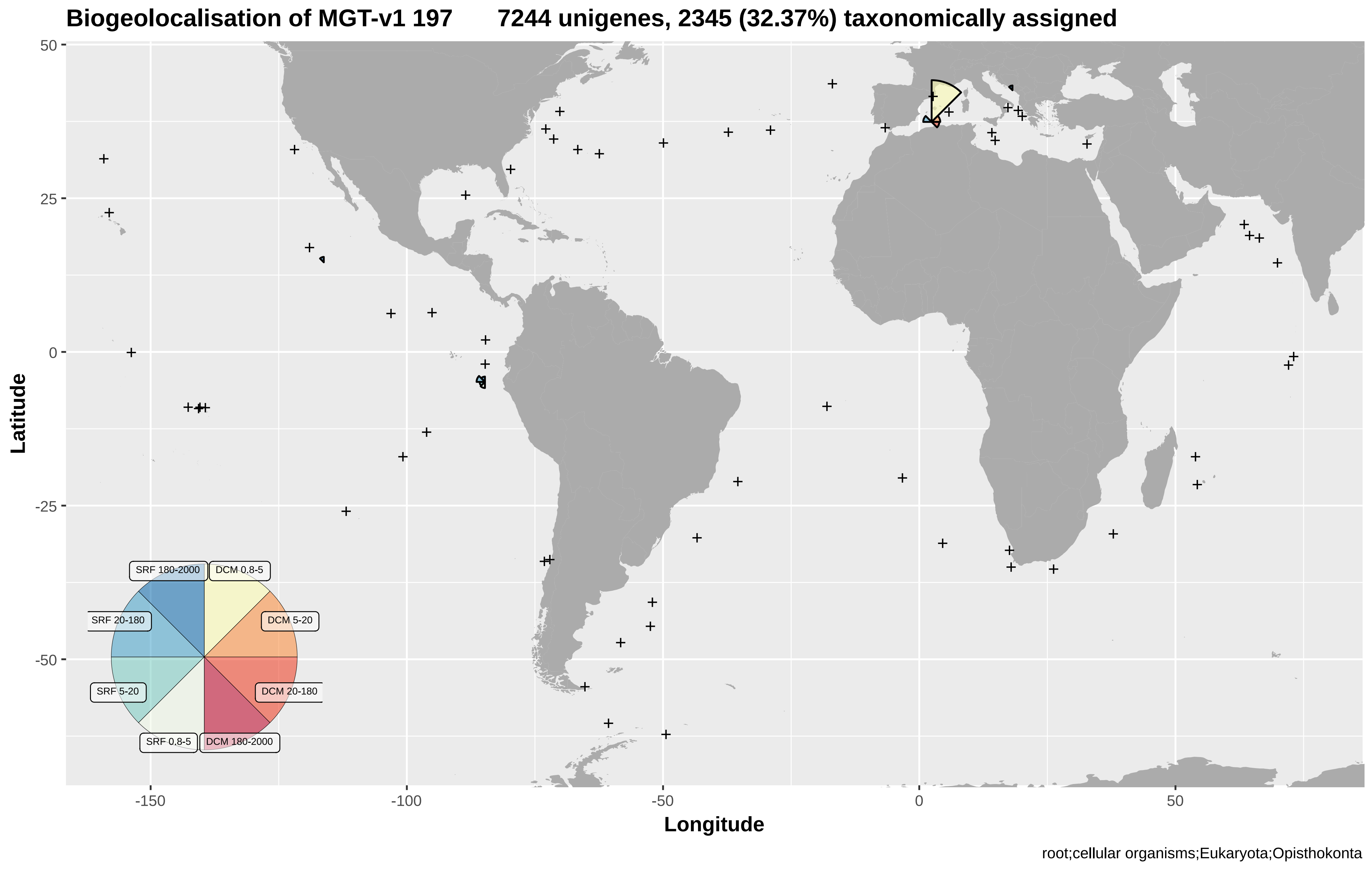
Biogeolocalisation of MGT-v1 197 7244 unigenes, 2345 (32.37%) taxonomically assigned
Latitude
Longitude
50
25
0
-25
-50
-150
-100
-50
0
50
SRF 180-2000
DCM 0.8-5
SRF 20-180
DCM 5-20
SRF 5-20
DCM 20-180
SRF 0.8-5
DCM 180-2000
root;cellular organisms;Eukaryota;Opisthokonta

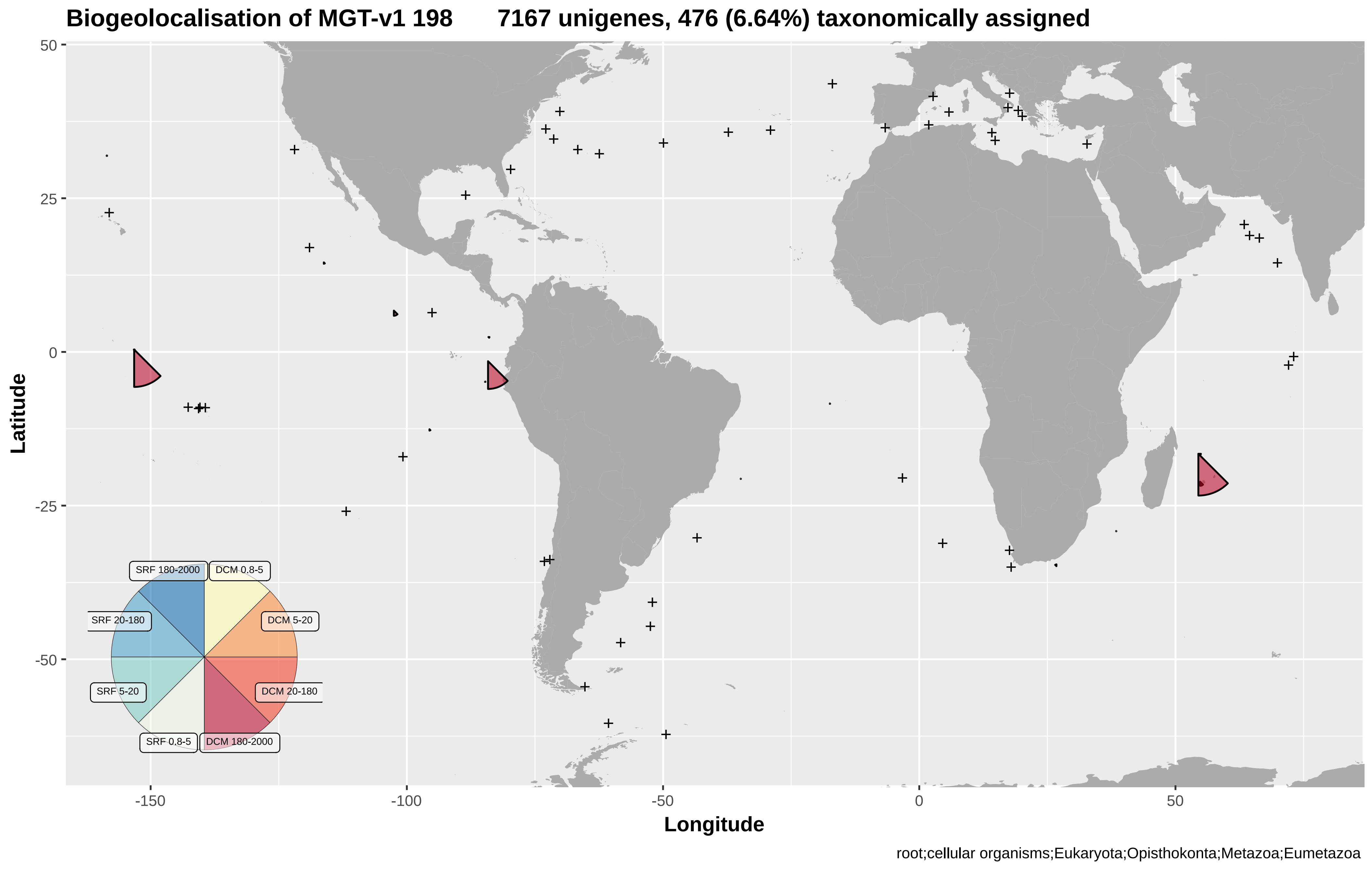
Biogeolocalisation of MGT-v1 198      7167 unigenes, 476 (6.64%) taxonomically assigned
50
25
0
-25
-50
Latitude
-150
-100
-50
0
50
Longitude
SRF 180-2000
DCM 0.8-5
SRF 20-180
DCM 5-20
SRF 5-20
DCM 20-180
SRF 0.8-5
DCM 180-2000
root;cellular organisms;Eukaryota;Opisthokonta;Metazoa;Eumetazoa

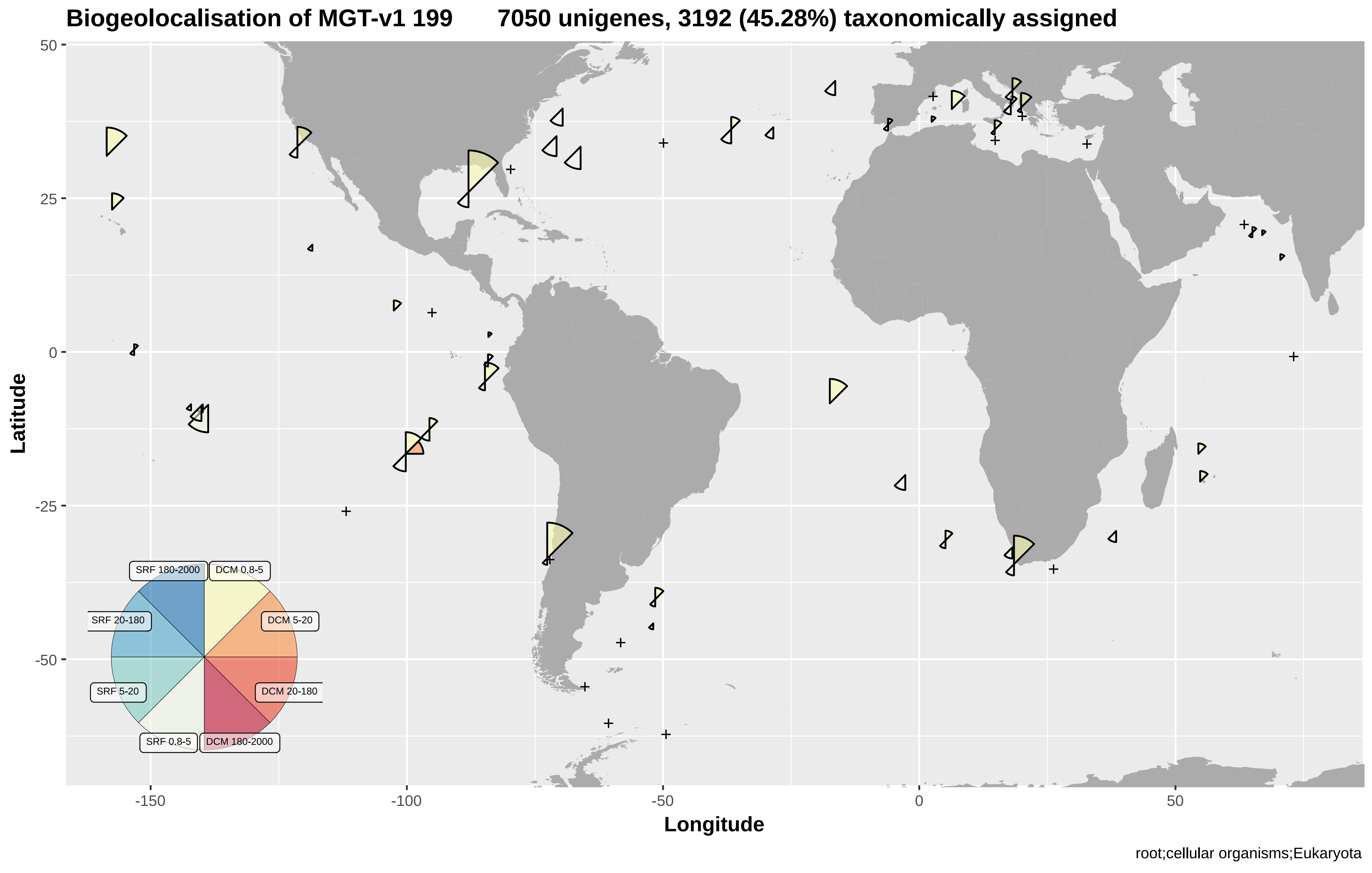

Biogeolocalisation of MGT-v1 199 7050 unigenes, 3192 (45.28%) taxonomically assigned
50
25
0
-25
-50
Latitude
-150
-100
-50
0
50
Longitude
SRF 180-2000
DCM 0.8-5
SRF 20-180
DCM 5-20
SRF 5-20
DCM 20-180
SRF 0.8-5
DCM 180-2000
root;cellular organisms;Eukaryota

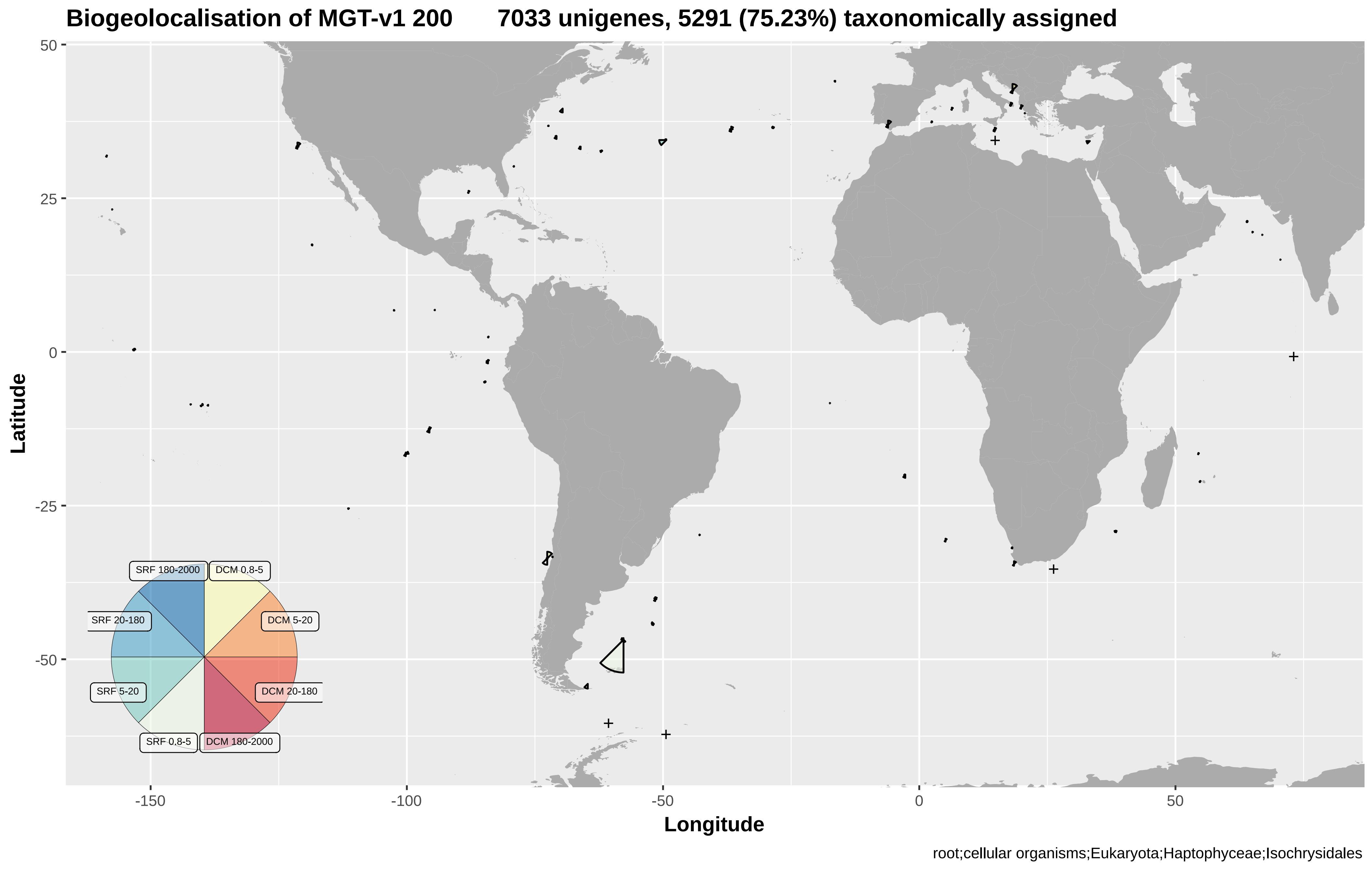

Biogeolocalisation of MGT-v1 200 7033 unigenes, 5291 (75.23%) taxonomically assigned
50
25
0
-25
-50
Latitude
-150
-100
-50
0
50
Longitude
SRF 180-2000
DCM 0.8-5
SRF 20-180
DCM 5-20
SRF 5-20
DCM 20-180
SRF 0.8-5
DCM 180-2000
root;cellular organisms;Eukaryota;Haptophyceae;Isochrysidales

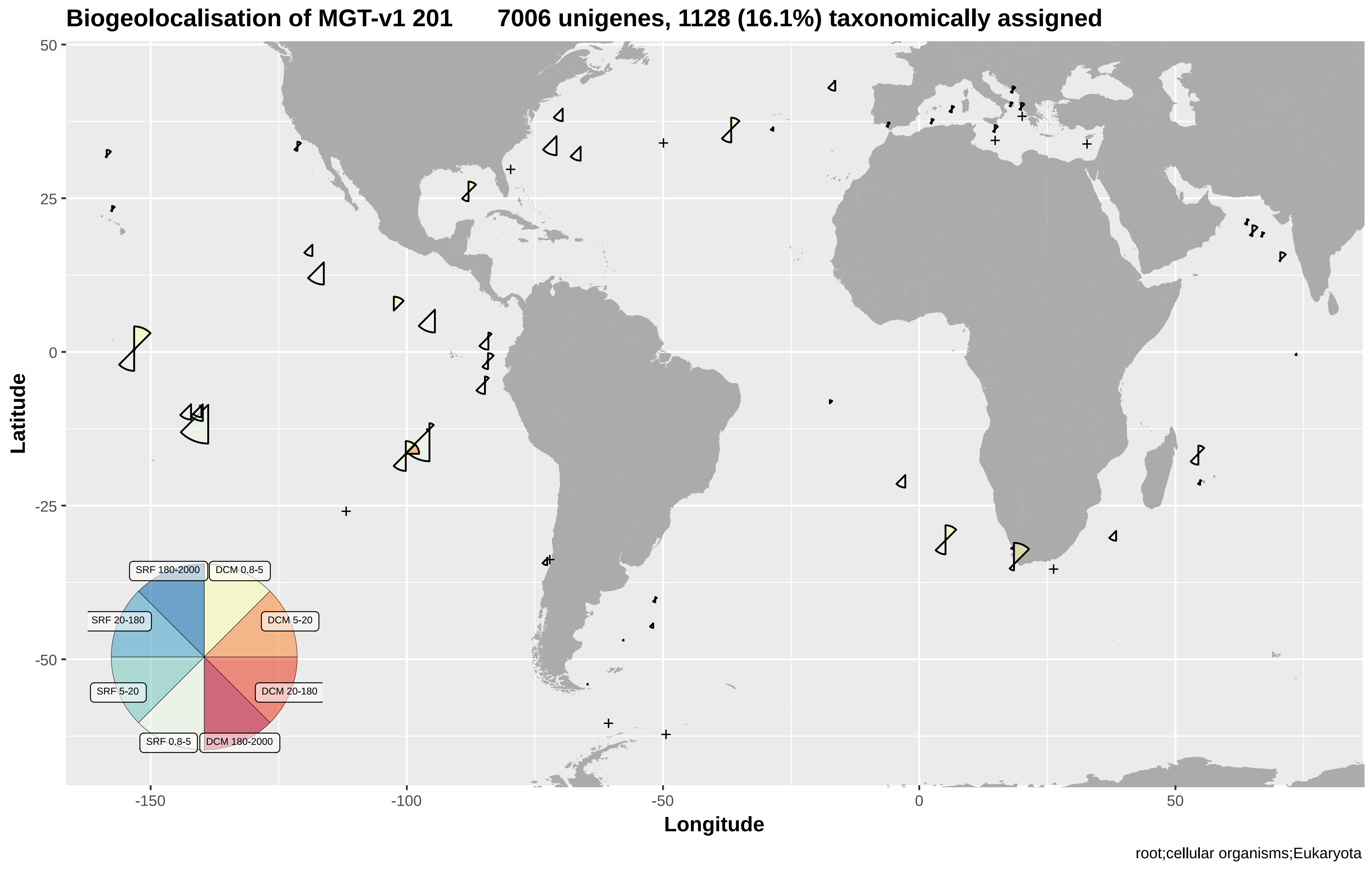

Biogeolocalisation of MGT-v1 201      7006 unigenes, 1128 (16.1%) taxonomically assigned
50
25
0
-25
-50
Latitude
-150
-100
-50
0
50
Longitude
SRF 180-2000
DCM 0.8-5
SRF 20-180
DCM 5-20
SRF 5-20
DCM 20-180
SRF 0.8-5
DCM 180-2000
root;cellular organisms;Eukaryota

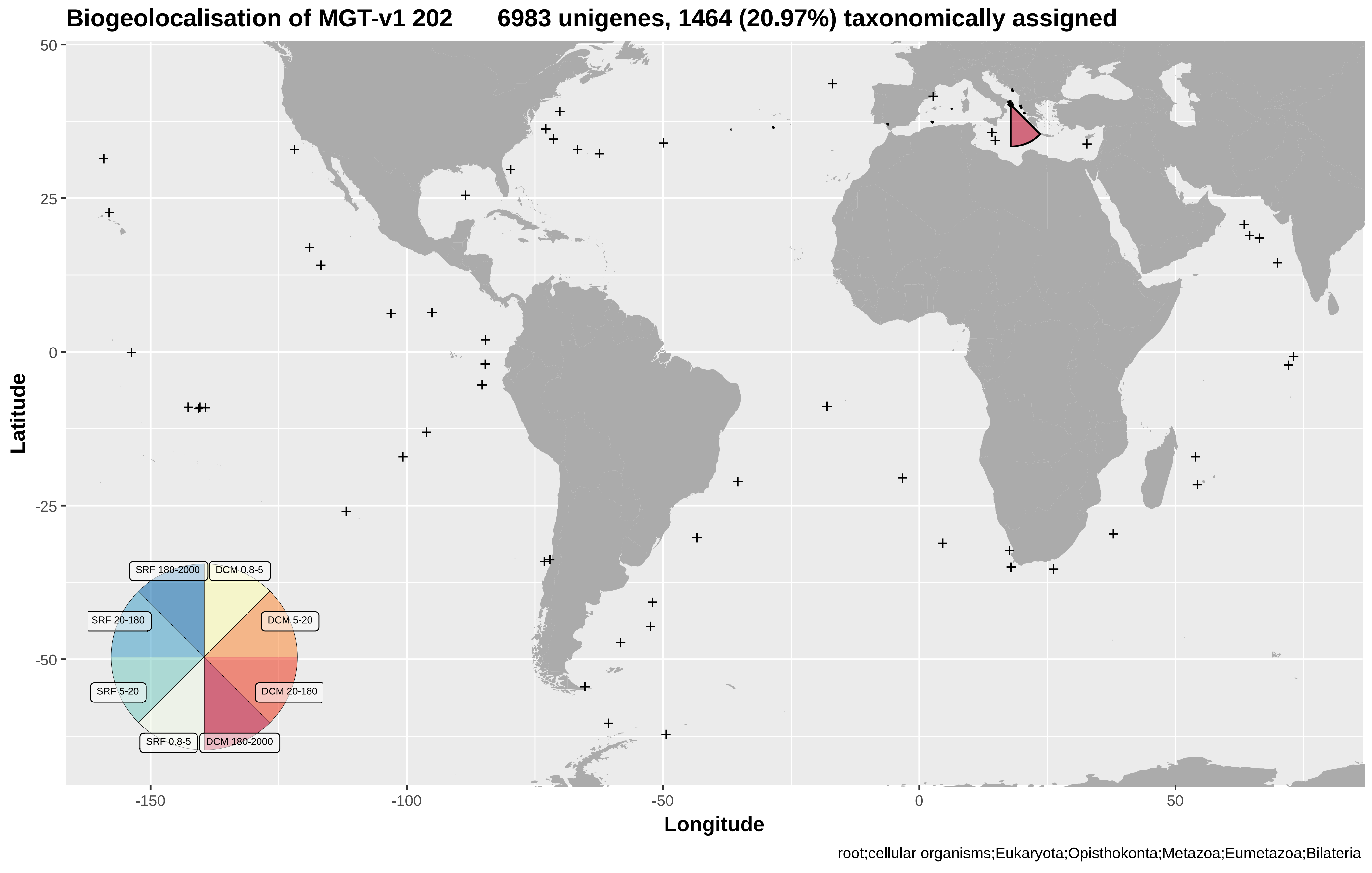

Biogeolocalisation of MGT-v1 202 6983 unigenes, 1464 (20.97%) taxonomically assigned
Latitude
Longitude
50
25
0
-25
-50
-150
-100
-50
0
50
SRF 180-2000
DCM 0.8-5
SRF 20-180
DCM 5-20
SRF 5-20
DCM 20-180
SRF 0.8-5
DCM 180-2000
root;cellular organisms;Eukaryota;Opisthokonta;Metazoa;Eumetazoa;Bilateria

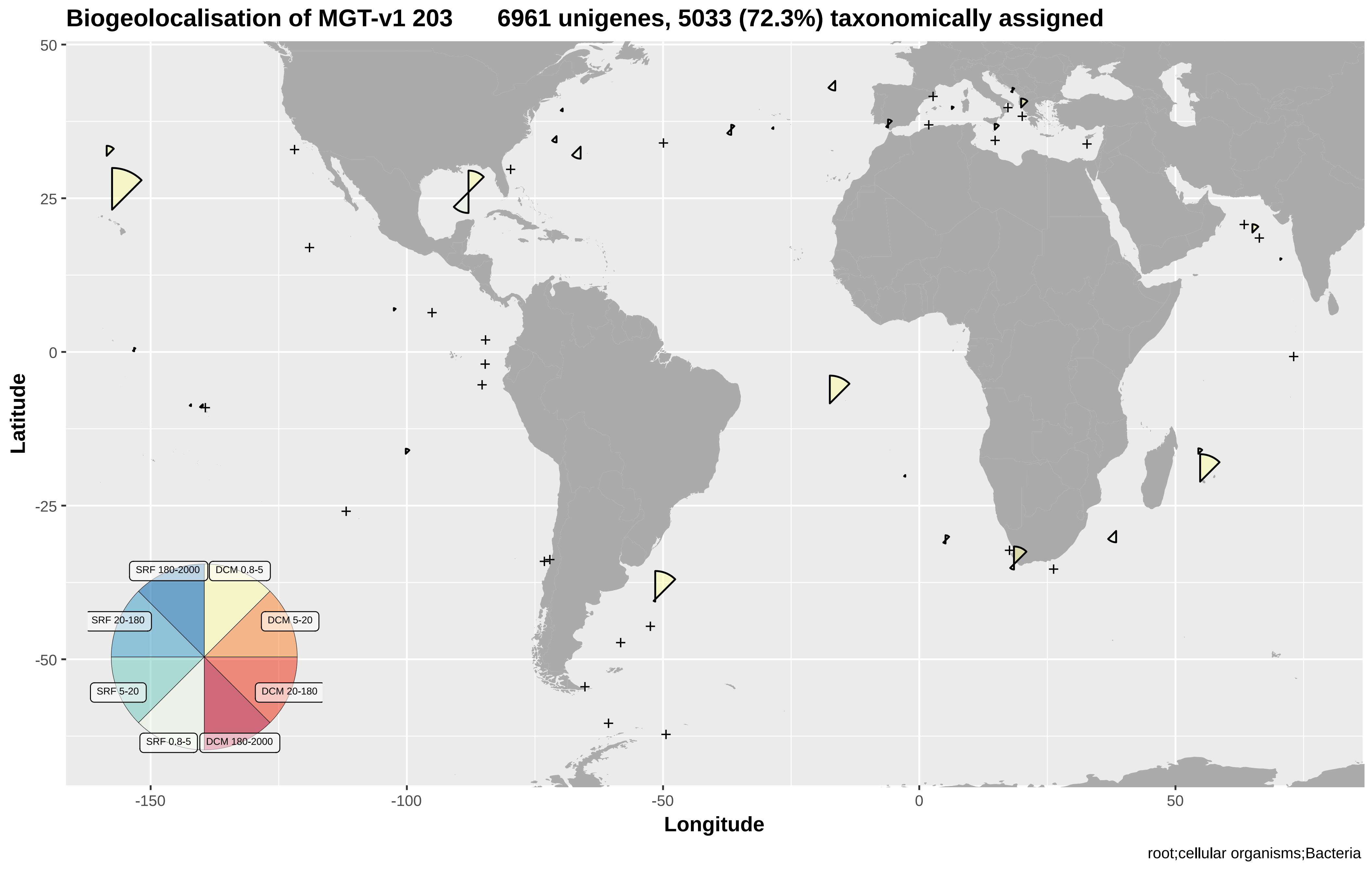
Biogeolocalisation of MGT-v1 203 6961 unigenes, 5033 (72.3%) taxonomically assigned
50
25
0
-25
-50
Latitude
-150
-100
-50
0
50
Longitude
SRF 180-2000
DCM 0.8-5
SRF 20-180
DCM 5-20
SRF 5-20
DCM 20-180
SRF 0.8-5
DCM 180-2000
root;cellular organisms;Bacteria

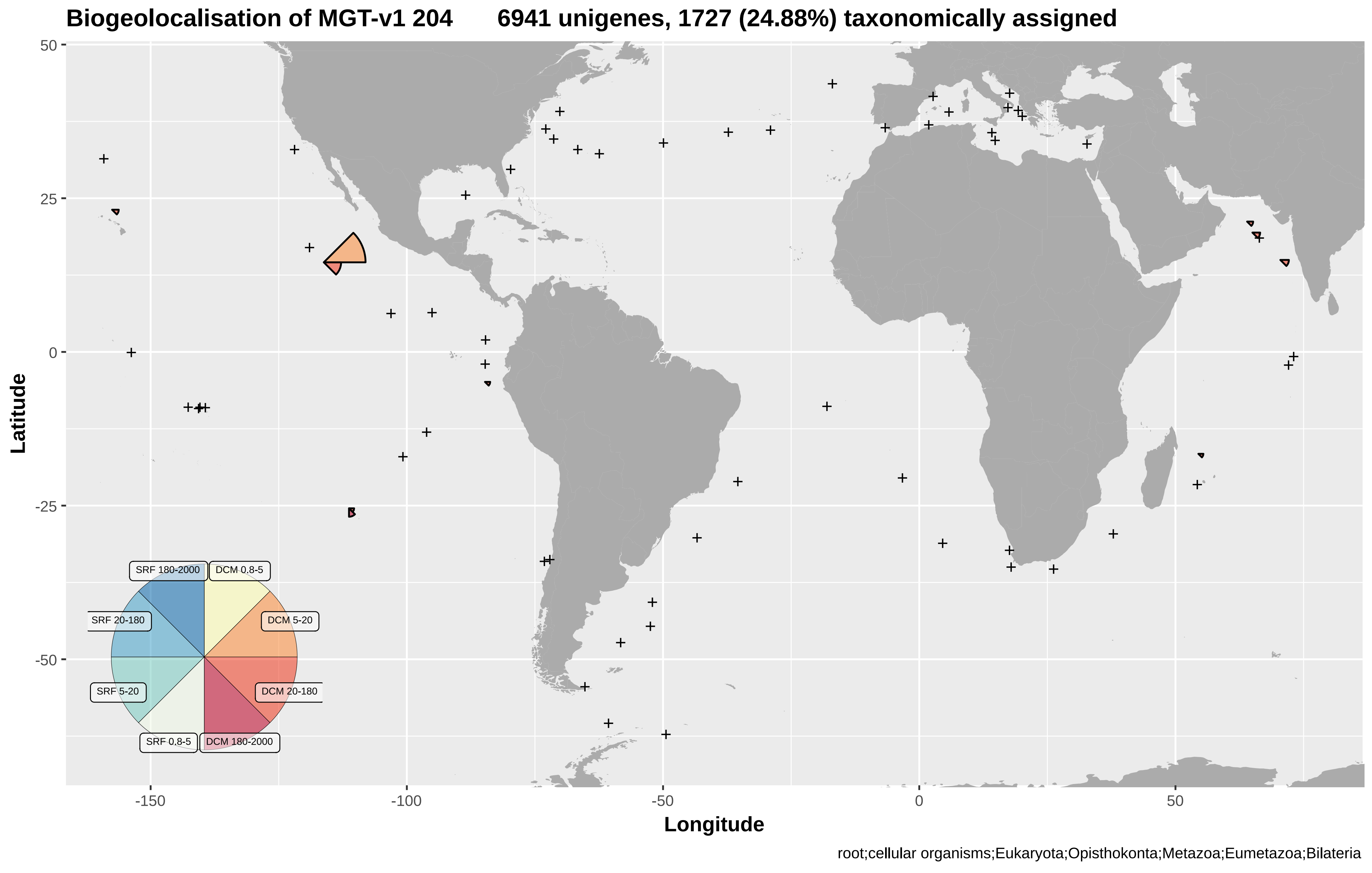

Biogeolocalisation of MGT-v1 204 6941 unigenes, 1727 (24.88%) taxonomically assigned
Latitude
Longitude
50
25
0
-25
-50
-150
-100
-50
0
50
SRF 180-2000
DCM 0.8-5
SRF 20-180
DCM 5-20
SRF 5-20
DCM 20-180
SRF 0.8-5
DCM 180-2000
root;cellular organisms;Eukaryota;Opisthokonta;Metazoa;Eumetazoa;Bilateria

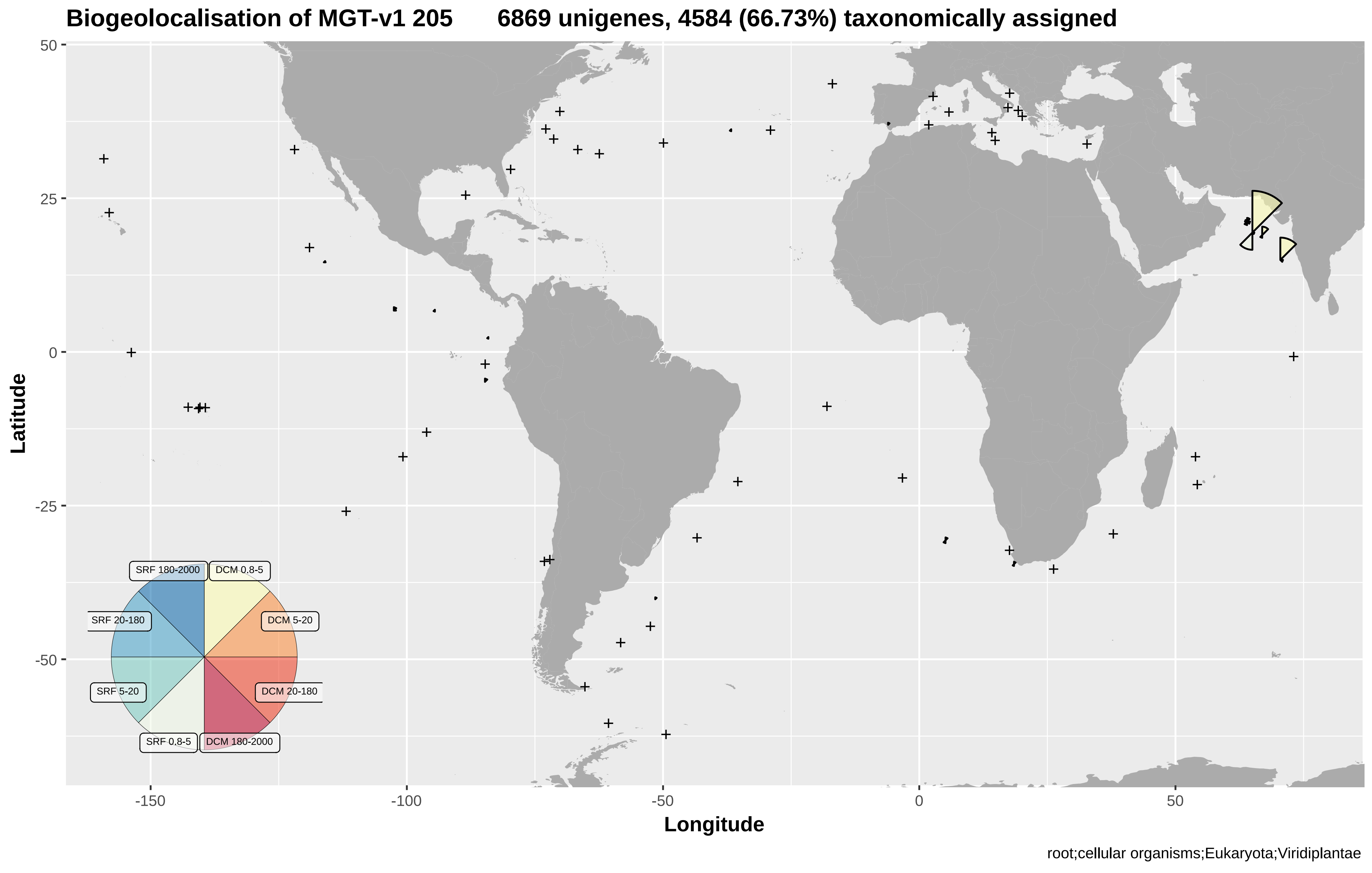

Biogeolocalisation of MGT-v1 205 6869 unigenes, 4584 (66.73%) taxonomically assigned
50
25
0
-25
-50
Latitude
-150
-100
-50
0
50
Longitude
SRF 180-2000
DCM 0.8-5
SRF 20-180
DCM 5-20
SRF 5-20
DCM 20-180
SRF 0.8-5
DCM 180-2000
root;cellular organisms;Eukaryota;Viridiplantae

### Biogeolocalisation of MGT-v1 206 6804 unigenes, 5341 (78.5%) taxonomically assigned

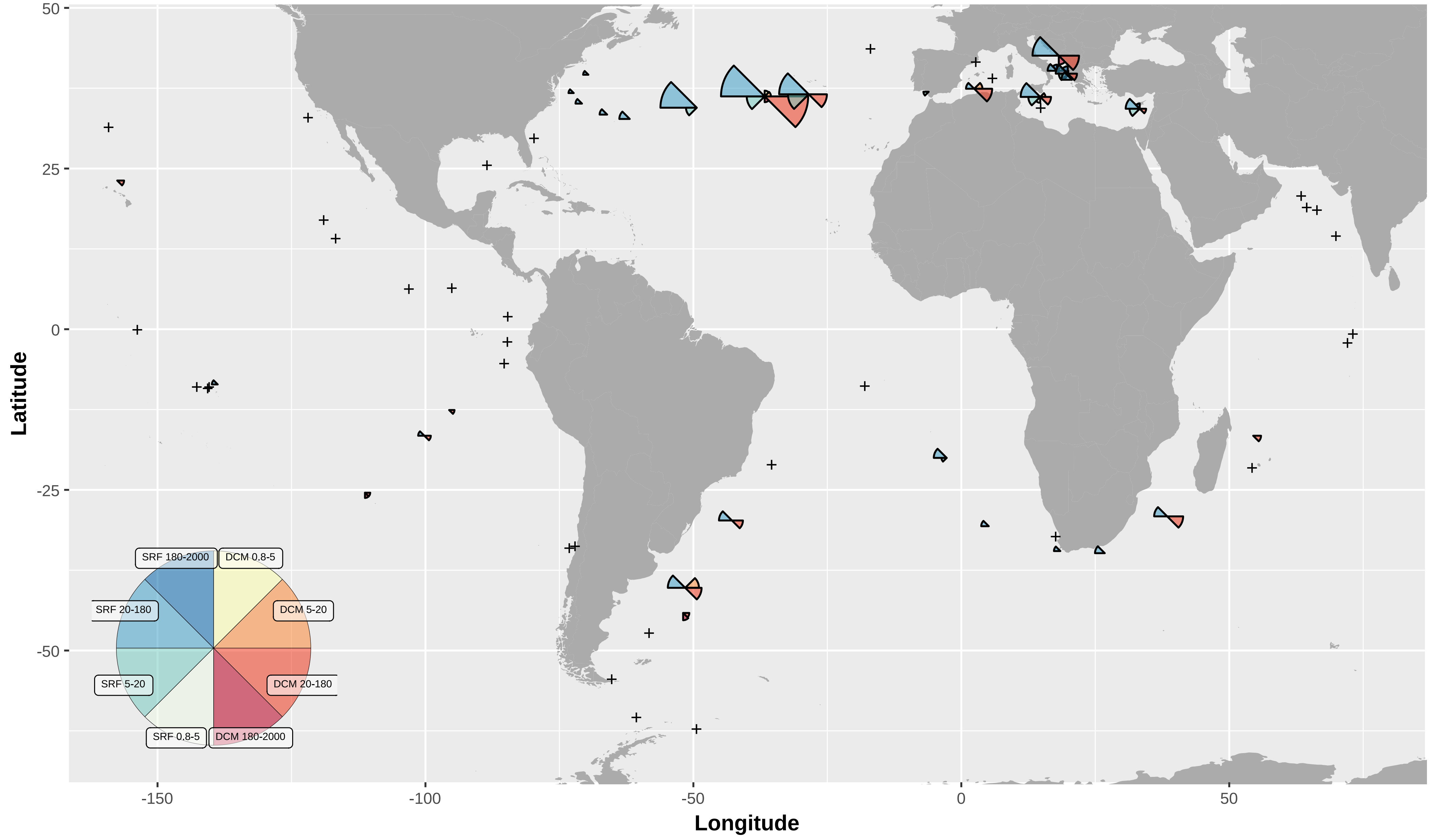


root;cellular organisms;Eukaryota;Opisthokonta;Metazoa;Eumetazoa;Bilateria;Protostomia;Ecdysozoa;Panarthropoda;Arthropoda;Mandibulata;Pancrustacea;Crustacea;Maxillopoda

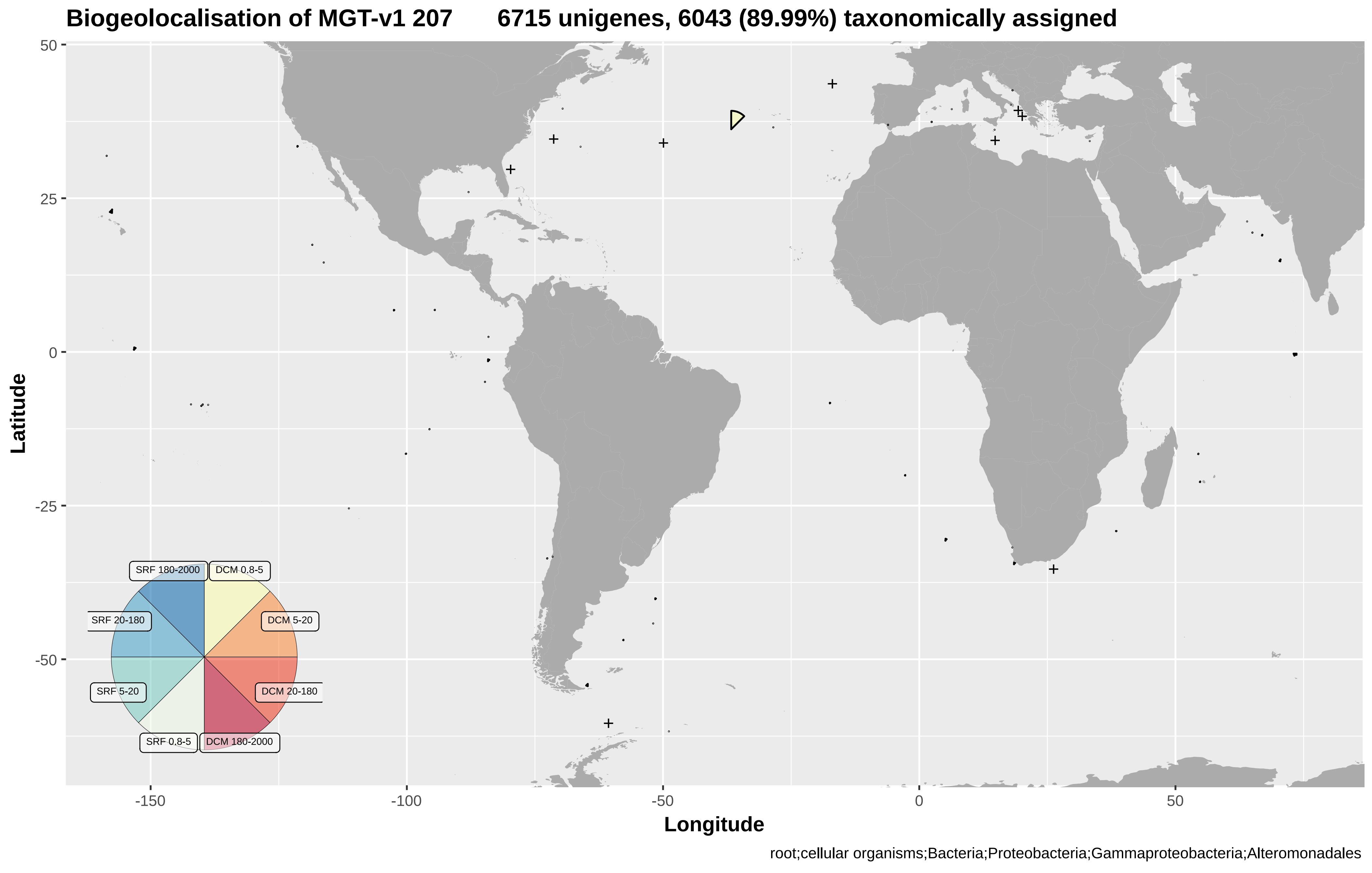
Biogeolocalisation of MGT-v1 207 6715 unigenes, 6043 (89.99%) taxonomically assigned
Latitude
Longitude
50
25
0
-25
-50
-150
-100
-50
0
50
SRF 180-2000
DCM 0.8-5
SRF 20-180
DCM 5-20
SRF 5-20
DCM 20-180
SRF 0.8-5
DCM 180-2000
root;cellular organisms;Bacteria;Proteobacteria;Gammaproteobacteria;Alteromonadales

Biogeolocalisation of MGT-v1 208 6656 unigenes, 1776 (26.68%) taxonomically assigned

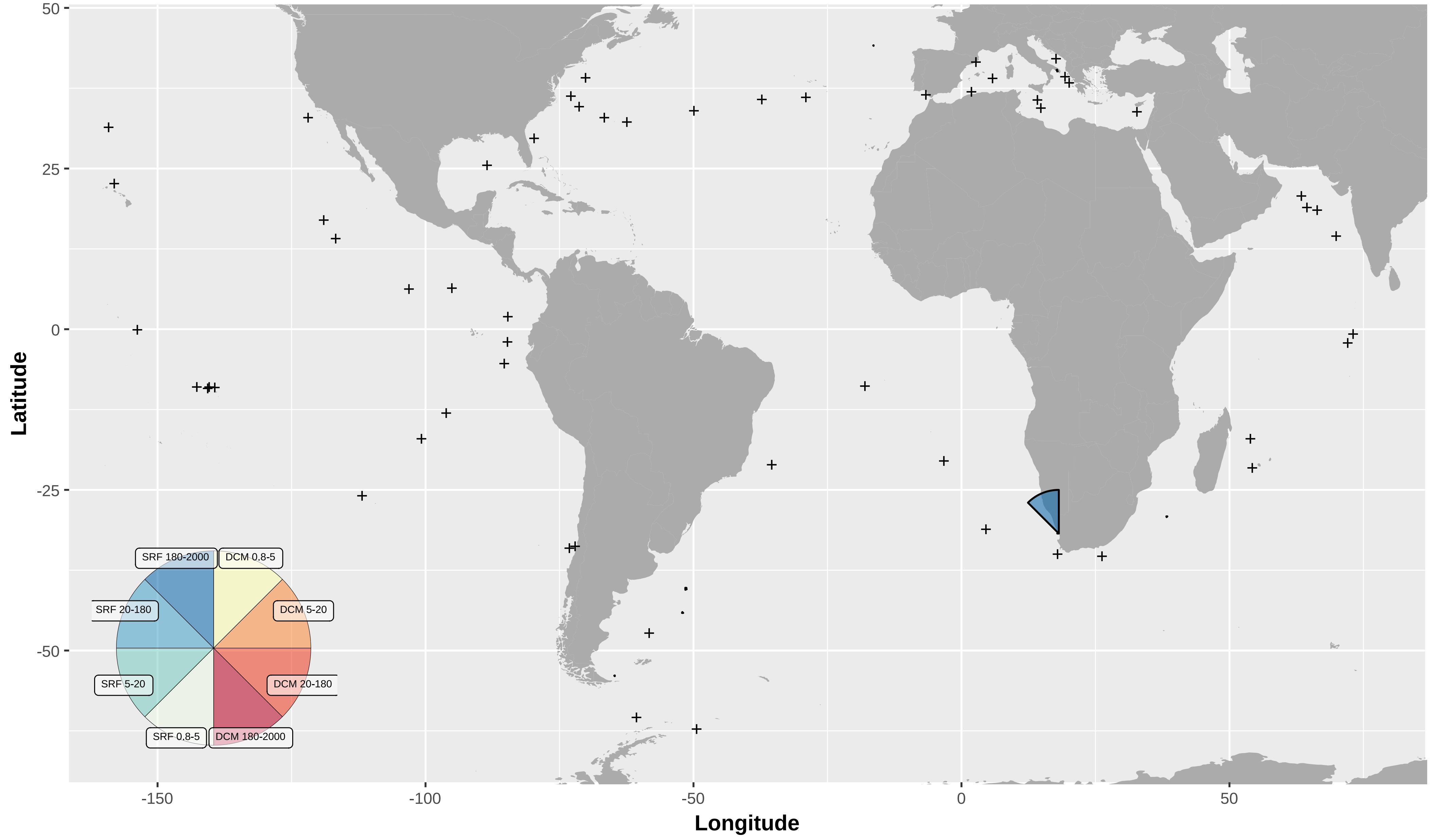


root;cellular organisms;Eukaryota;Opisthokonta;Metazoa;Eumetazoa;Bilateria

### Biogeolocalisation of MGT-v1 209 6345 unigenes, 830 (13.08%) taxonomically assigned

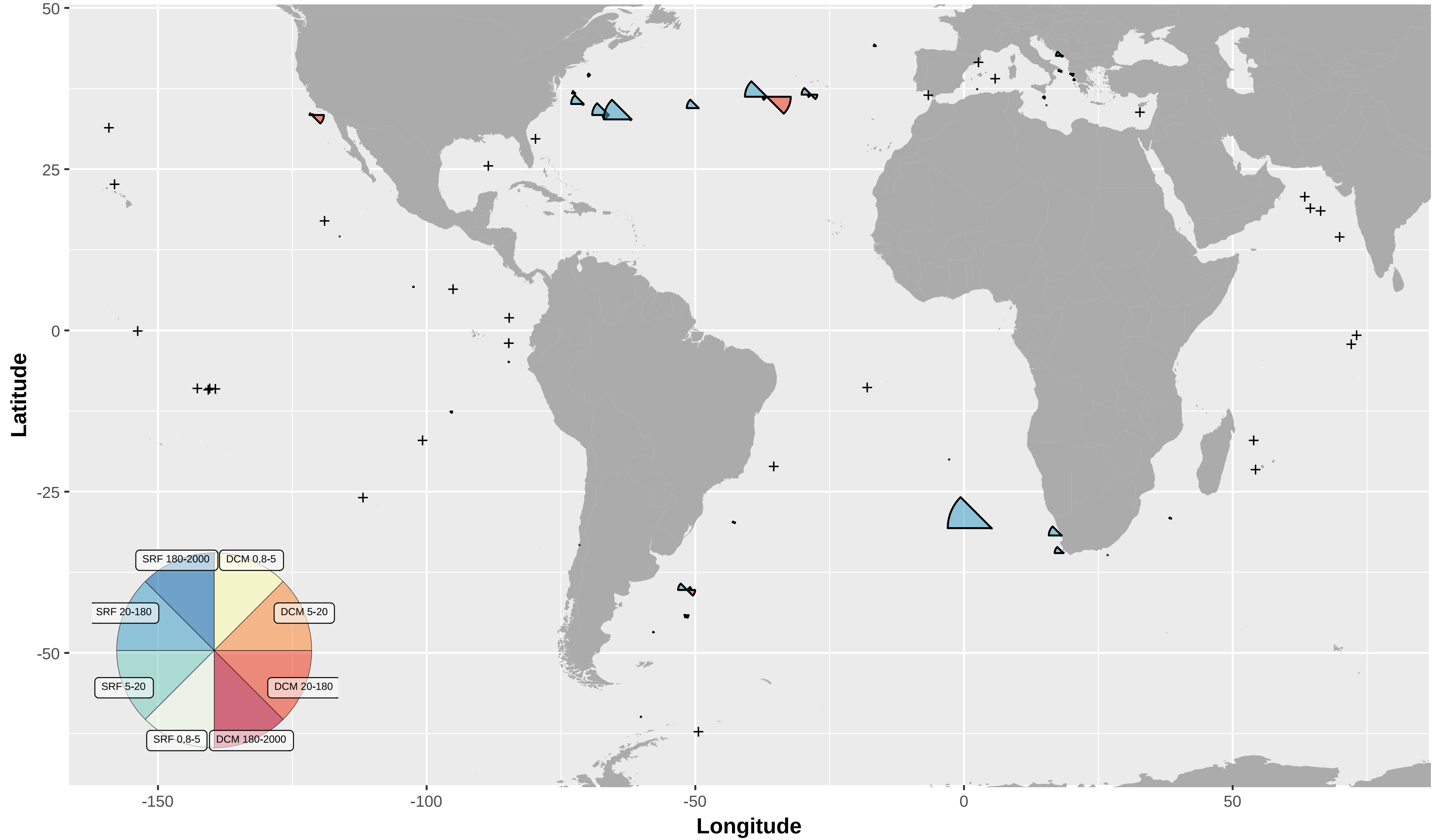


root;cellular organisms;Eukaryota;Opisthokonta;Metazoa;Eumetazoa;Bilateria

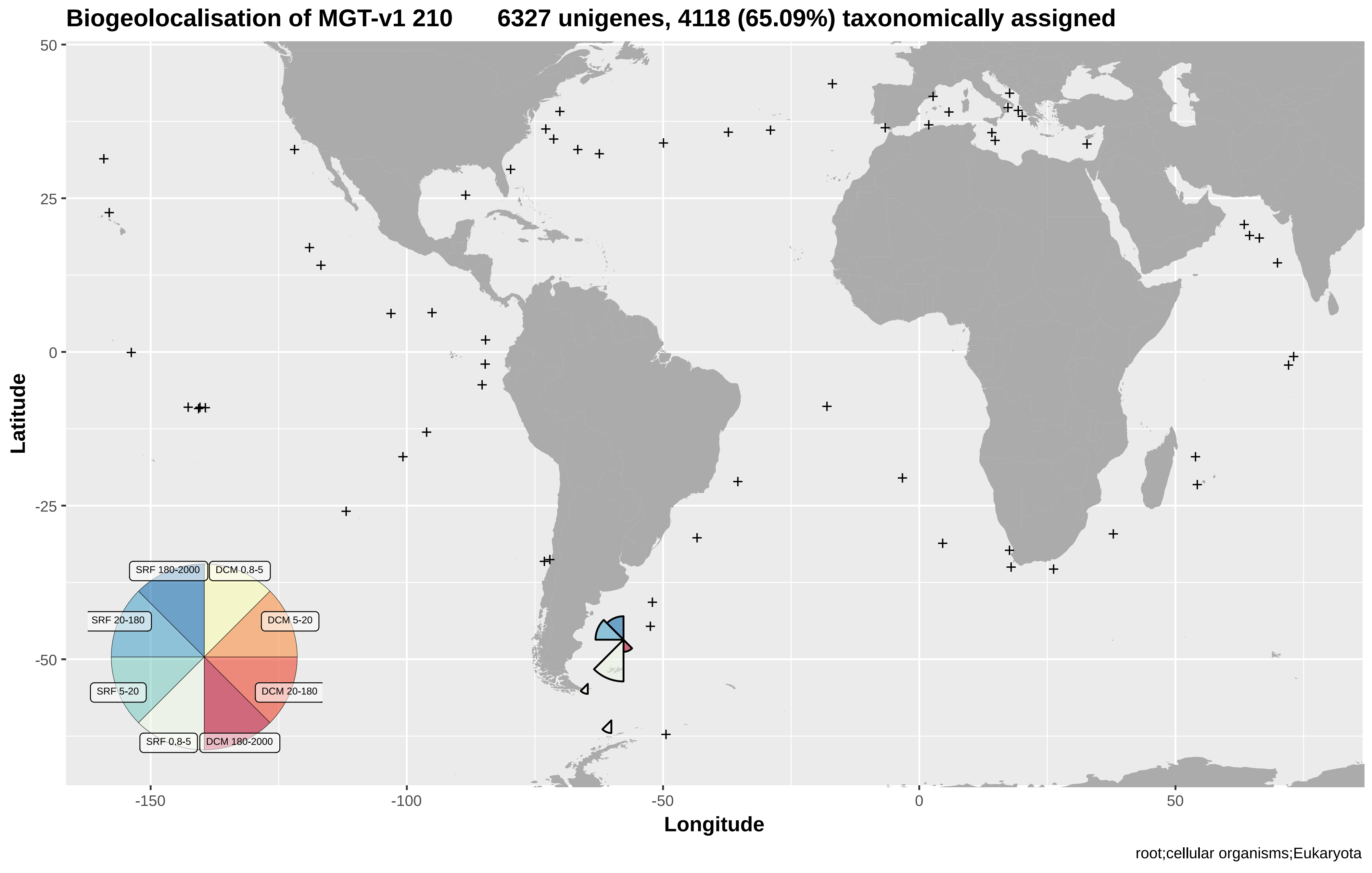
Biogeolocalisation of MGT-v1 210
6327 unigenes, 4118 (65.09%) taxonomically assigned
Latitude
Longitude
50
25
0
-25
-50
-150
-100
-50
0
50
SRF 180-2000
DCM 0.8-5
SRF 20-180
DCM 5-20
SRF 5-20
DCM 20-180
SRF 0.8-5
DCM 180-2000
root;cellular organisms;Eukaryota

**Biogeolocalisation of MGT-v1 211      6304 unigenes, 4492 (71.26%) taxonomically assigned**

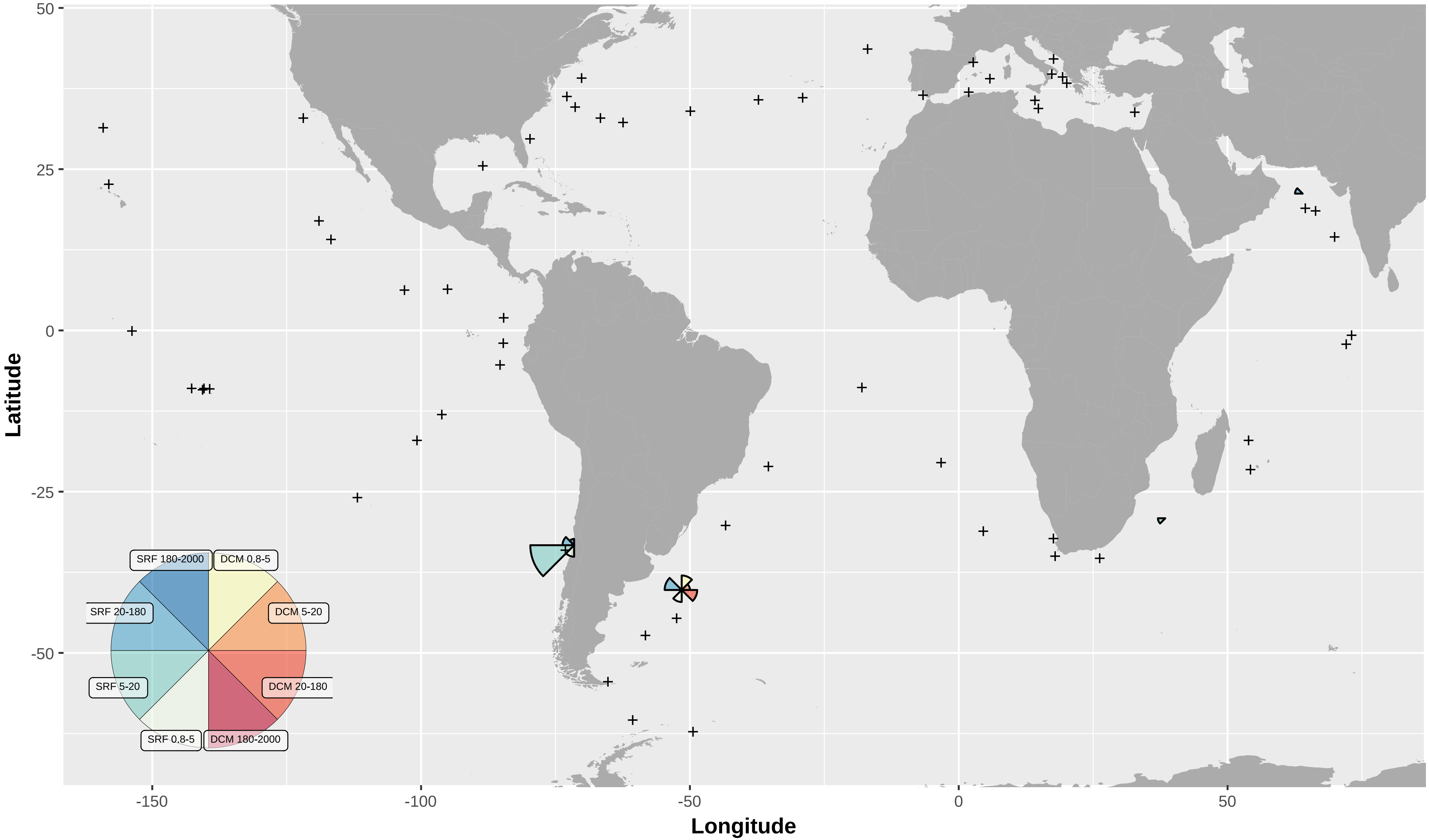


root;cellular organisms;Eukaryota;Stramenopiles;Bacillariophyta

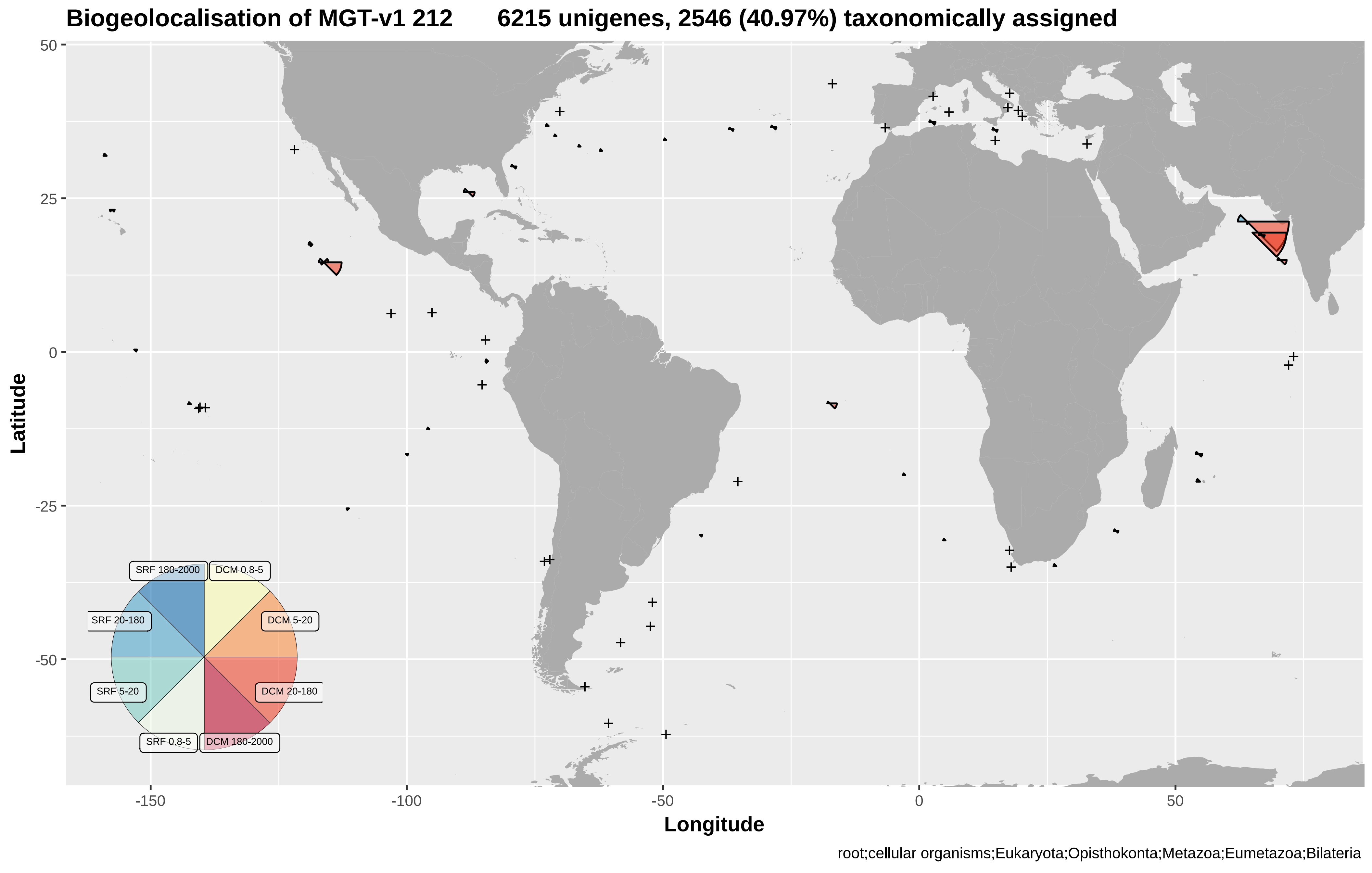

Biogeolocalisation of MGT-v1 212 6215 unigenes, 2546 (40.97%) taxonomically assigned
50
25
0
-25
-50
Latitude
-150
-100
-50
0
50
Longitude
SRF 180-2000
DCM 0.8-5
SRF 20-180
DCM 5-20
SRF 5-20
DCM 20-180
SRF 0.8-5
DCM 180-2000
root;cellular organisms;Eukaryota;Opisthokonta;Metazoa;Eumetazoa;Bilateria

#### Biogeolocalisation of MGT-v1 213 6203 unigenes, 2406 (38.79%) taxonomically assigned

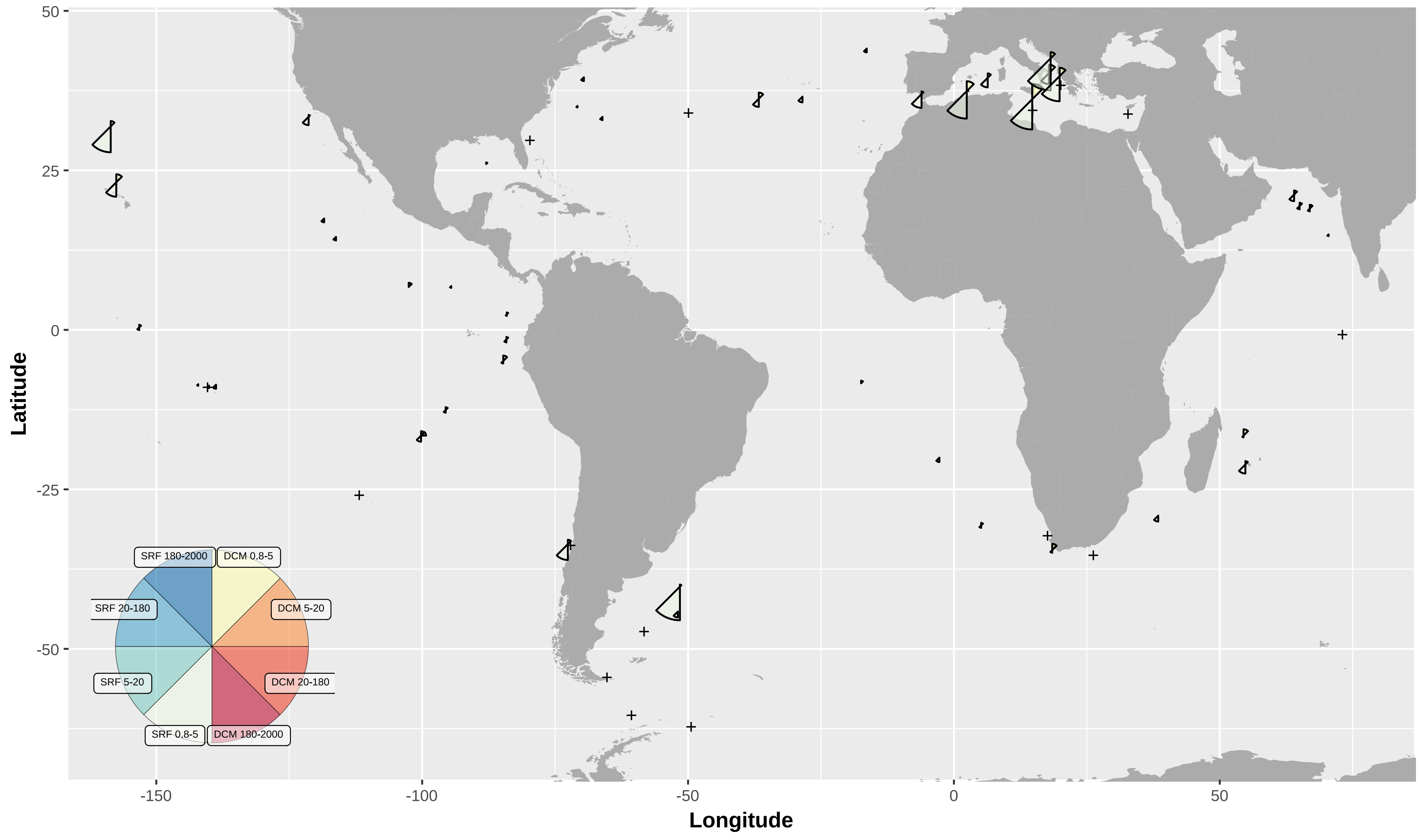


root;cellular organisms;Eukaryota;Stramenopiles

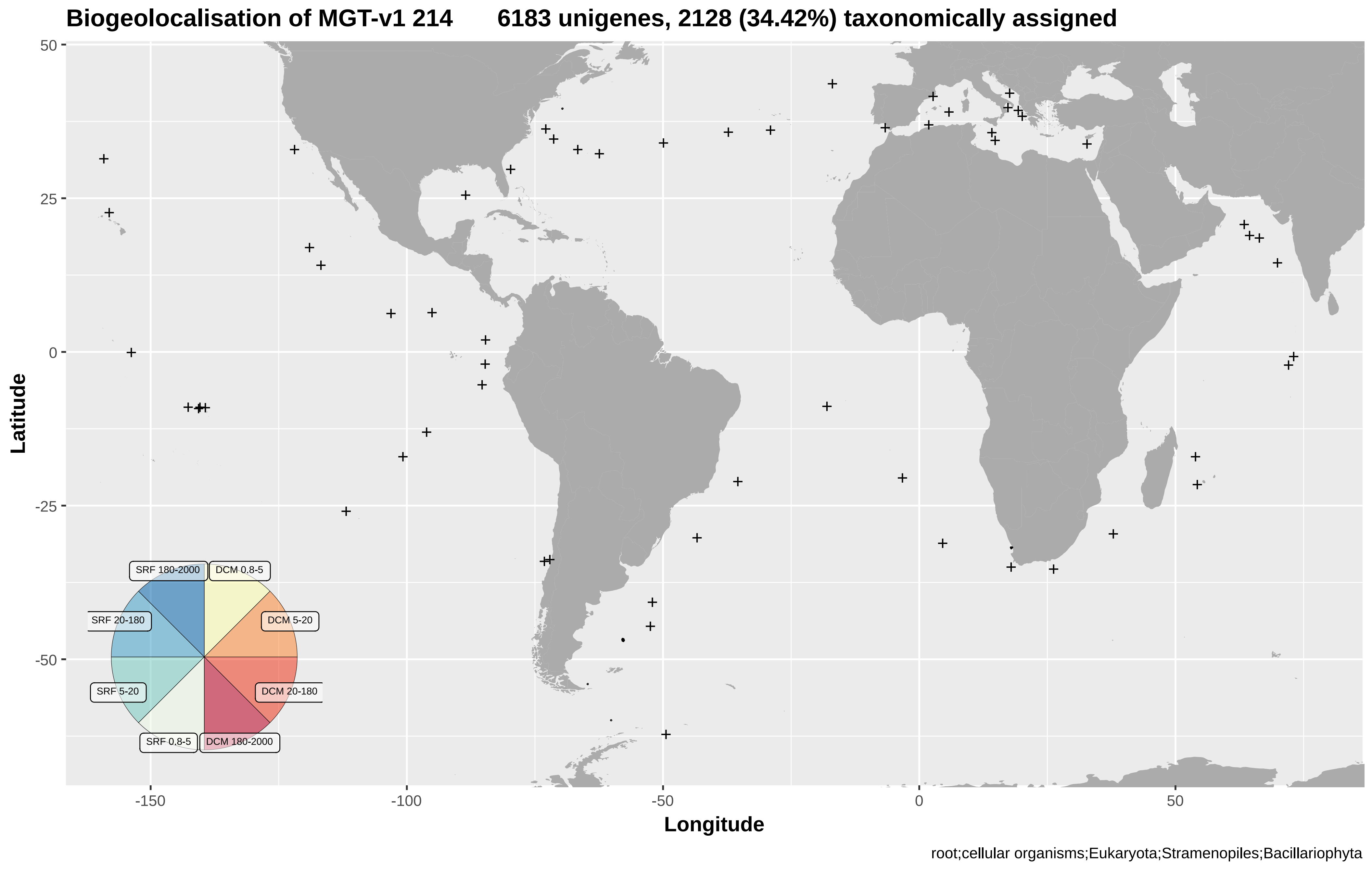
Biogeolocalisation of MGT-v1 214 6183 unigenes, 2128 (34.42%) taxonomically assigned
50
25
0
-25
-50
Latitude
-150
-100
-50
0
50
Longitude
SRF 180-2000
DCM 0.8-5
SRF 20-180
DCM 5-20
SRF 5-20
DCM 20-180
SRF 0.8-5
DCM 180-2000
root;cellular organisms;Eukaryota;Stramenopiles;Bacillariophyta

#### Biogeolocalisation of MGT-v1 215 6182 unigenes, 4780 (77.32%) taxonomically assigned

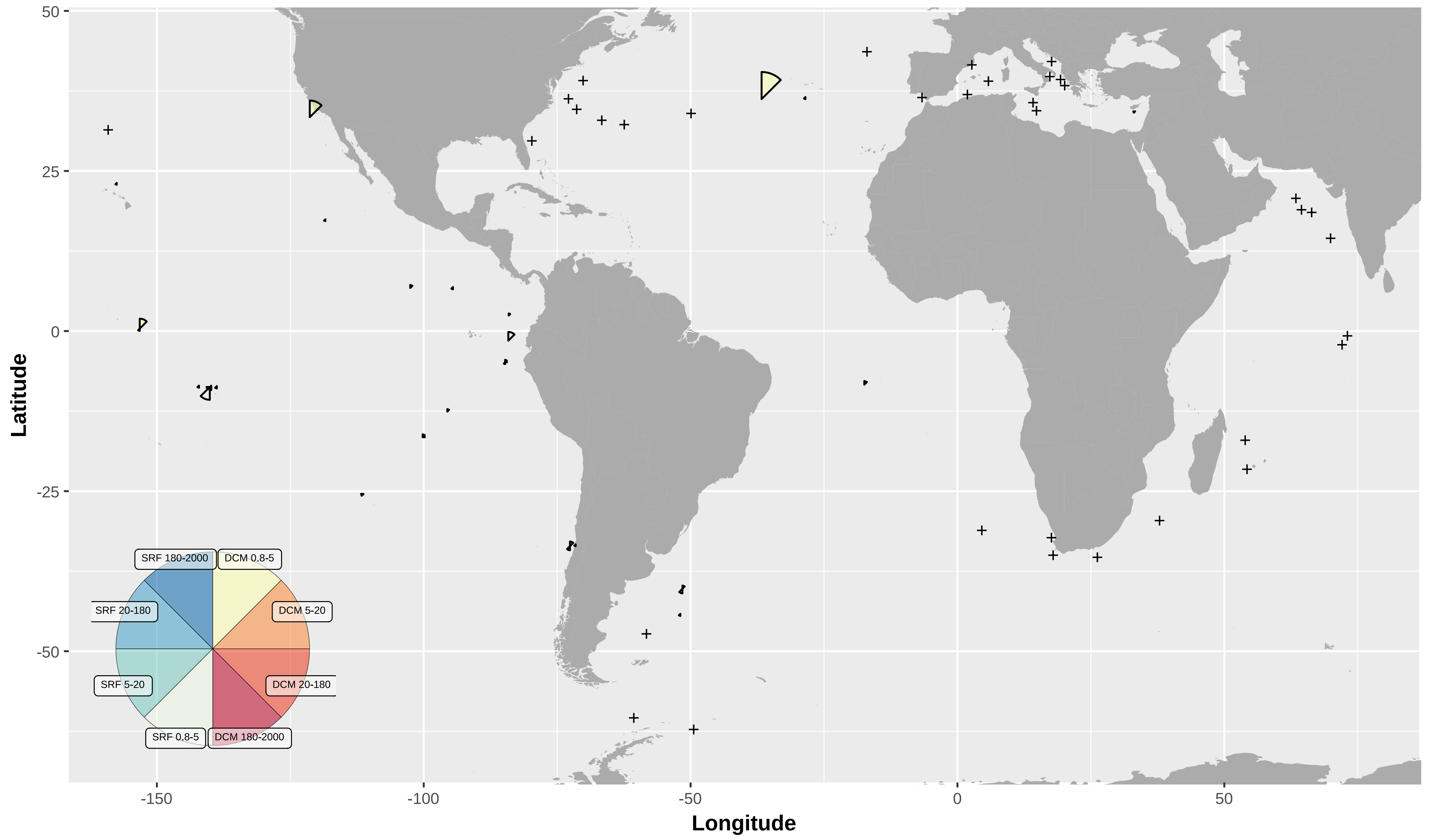


root;cellular organisms;Bacteria;Proteobacteria;Alphaproteobacteria

**Biogeolocalisation of MGT-v1 216      6139 unigenes, 1723 (28.07%) taxonomically assigned**

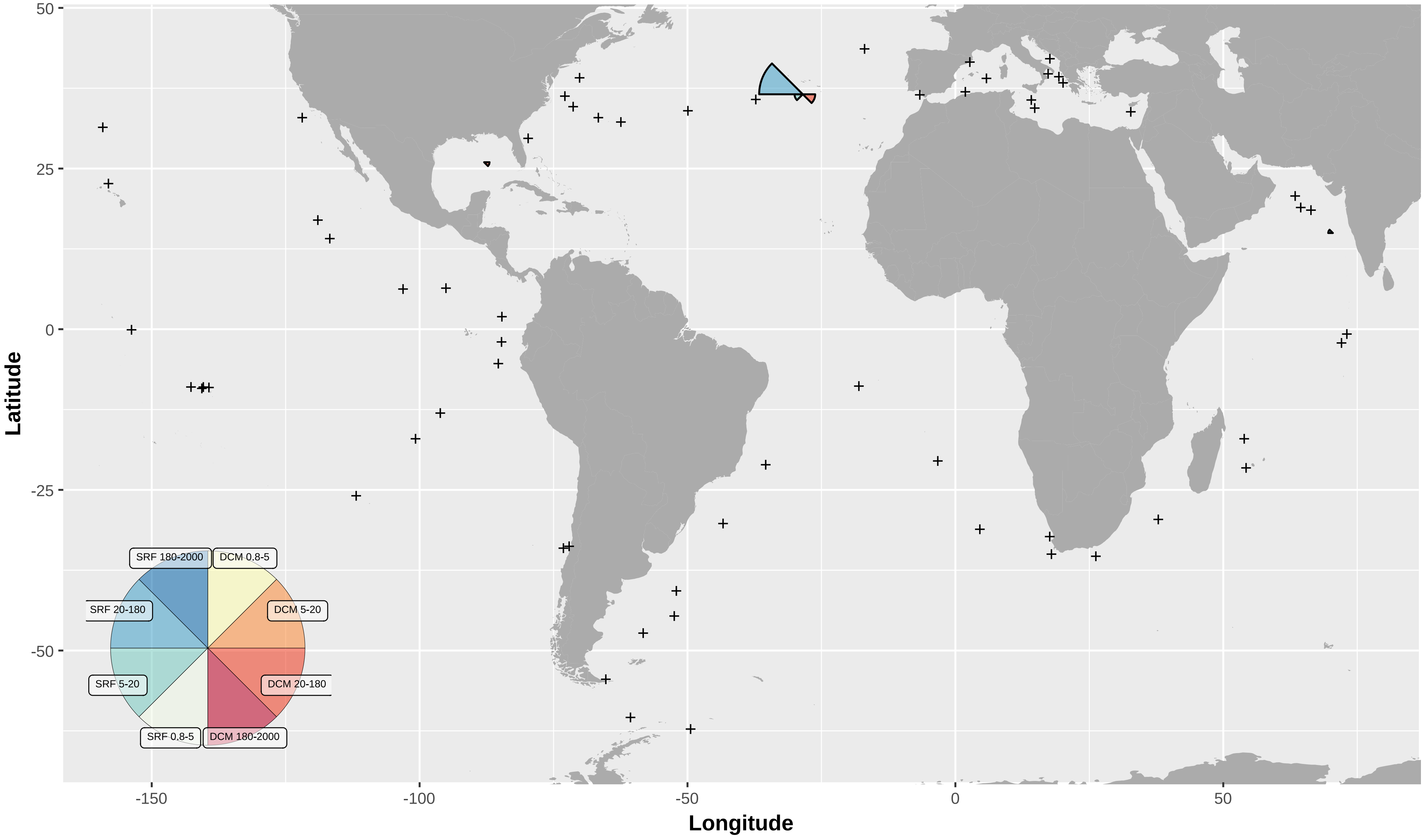


root;cellular organisms;Eukaryota;Opisthokonta

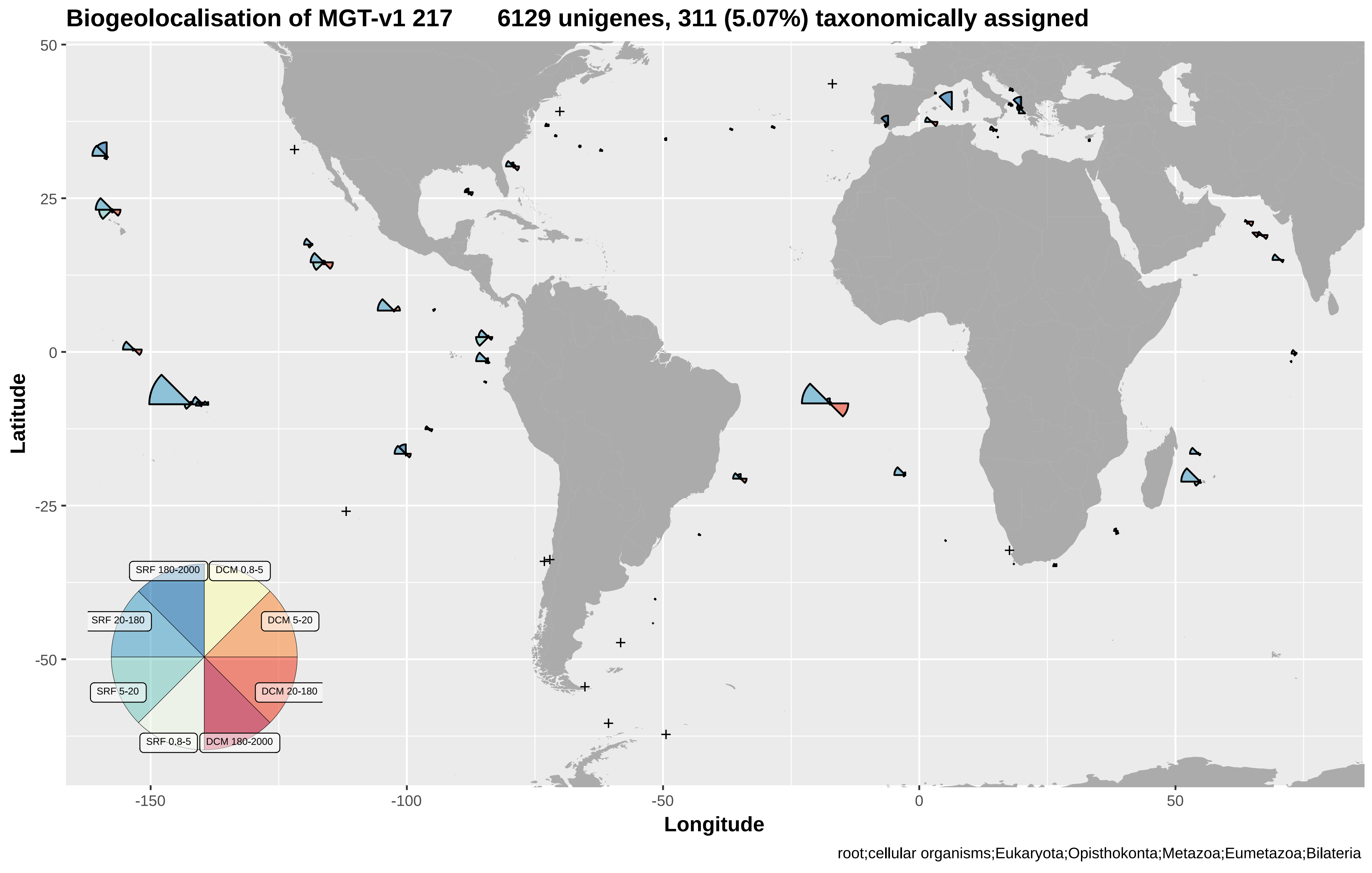
Biogeolocalisation of MGT-v1 217 6129 unigenes, 311 (5.07%) taxonomically assigned
50
25
0
-25
-50
Latitude
-150
-100
-50
0
50
Longitude
SRF 180-2000
DCM 0.8-5
SRF 20-180
DCM 5-20
SRF 5-20
DCM 20-180
SRF 0.8-5
DCM 180-2000
root;cellular organisms;Eukaryota;Opisthokonta;Metazoa;Eumetazoa;Bilateria

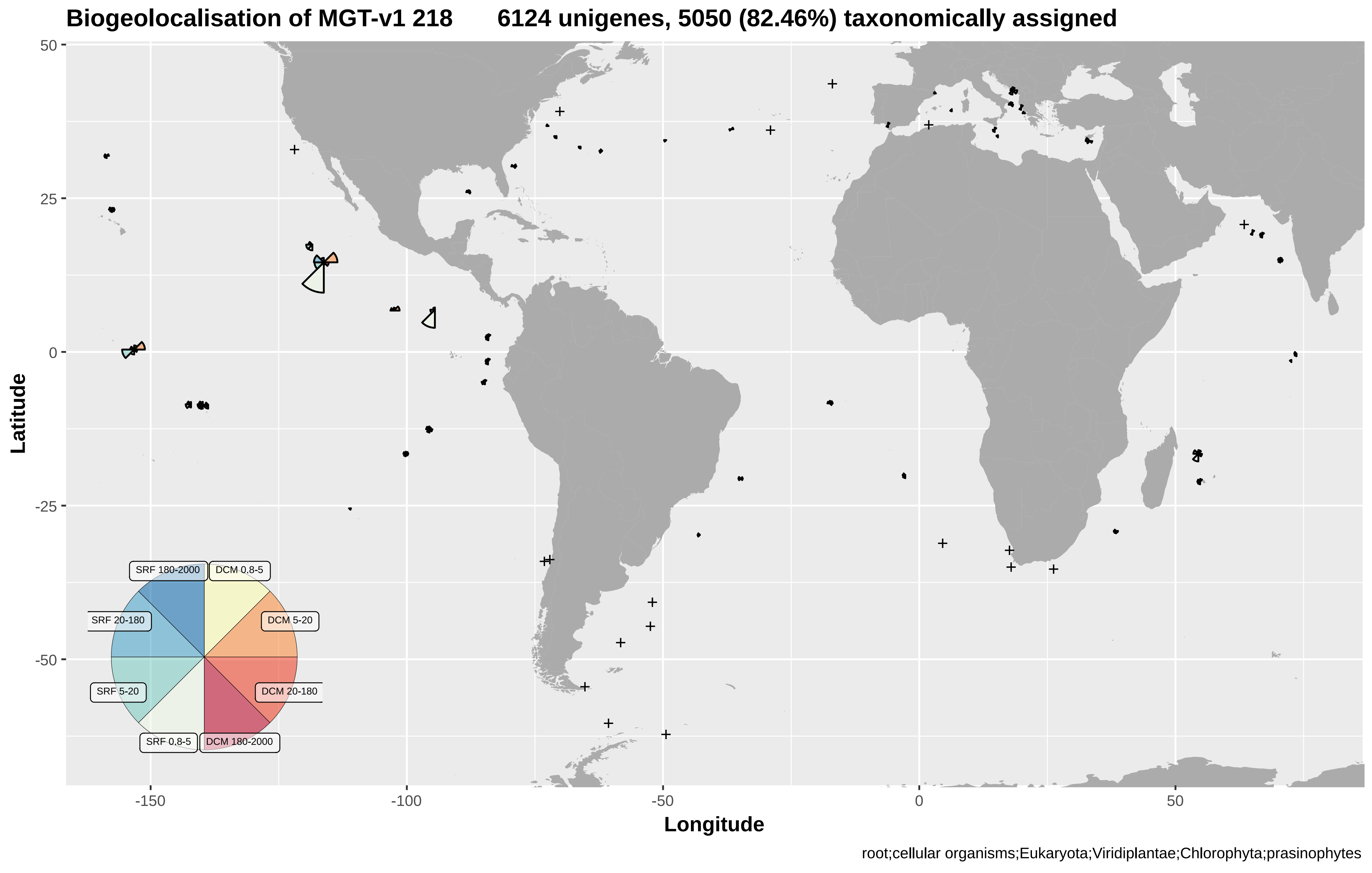
Biogeolocalisation of MGT-v1 218 6124 unigenes, 5050 (82.46%) taxonomically assigned
Latitude
Longitude
50
25
0
-25
-50
-150
-100
-50
0
50
SRF 180-2000
DCM 0.8-5
SRF 20-180
DCM 5-20
SRF 5-20
DCM 20-180
SRF 0.8-5
DCM 180-2000
root;cellular organisms;Eukaryota;Viridiplantae;Chlorophyta;prasinophytes

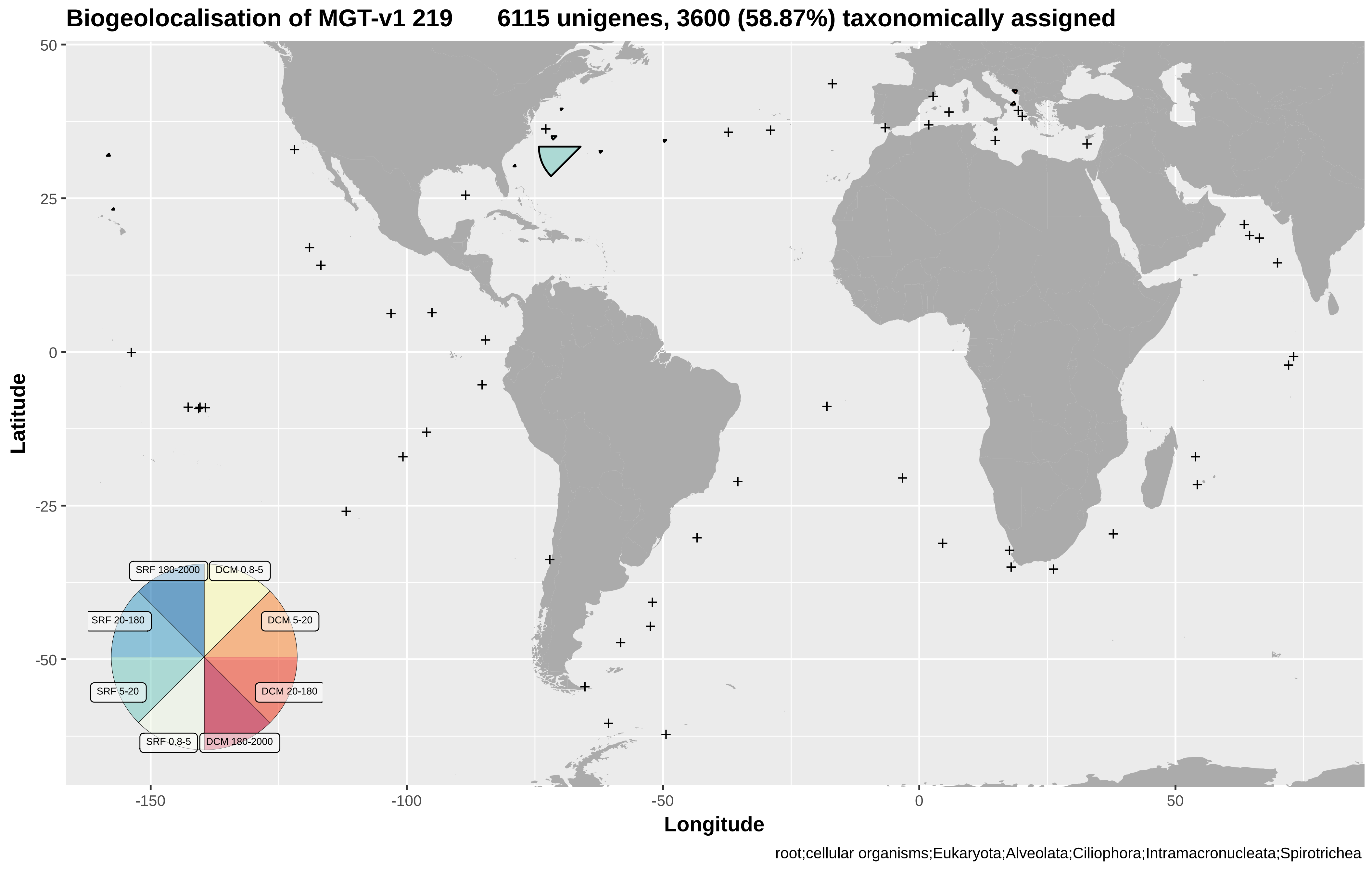

Biogeolocalisation of MGT-v1 219 6115 unigenes, 3600 (58.87%) taxonomically assigned
Latitude
Longitude
50
25
0
-25
-50
-150
-100
-50
0
50
SRF 180-2000
DCM 0.8-5
SRF 20-180
DCM 5-20
SRF 5-20
DCM 20-180
SRF 0.8-5
DCM 180-2000
root;cellular organisms;Eukaryota;Alveolata;Ciliophora;Intramacronucleata;Spirotrichea

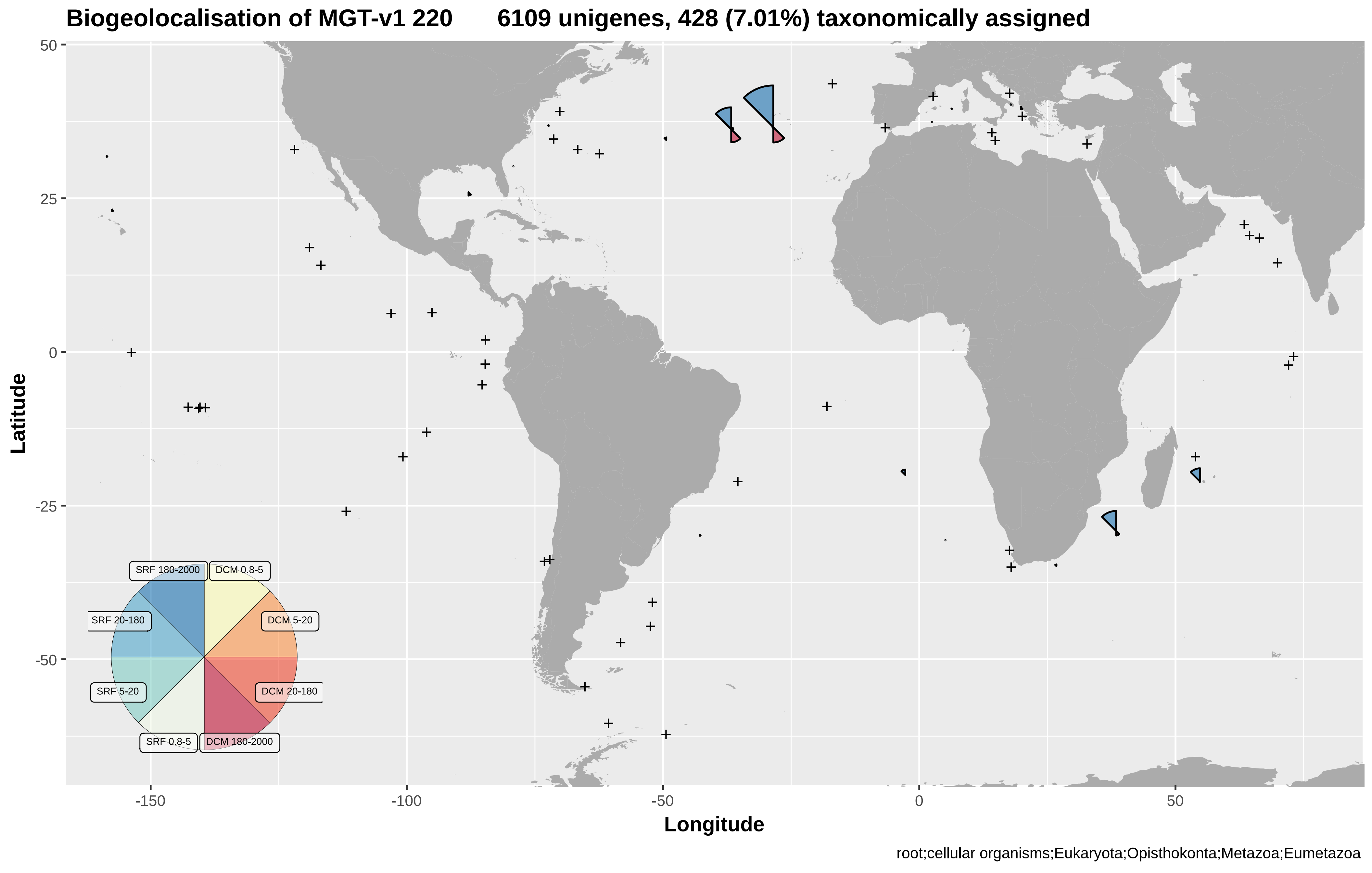

Biogeolocalisation of MGT-v1 220 6109 unigenes, 428 (7.01%) taxonomically assigned
Latitude
Longitude
50
25
0
-25
-50
-150
-100
-50
0
50
SRF 180-2000
DCM 0.8-5
SRF 20-180
DCM 5-20
SRF 5-20
DCM 20-180
SRF 0.8-5
DCM 180-2000
root;cellular organisms;Eukaryota;Opisthokonta;Metazoa;Eumetazoa

# Biogeolocalisation of MGT-v1 221      6033 unigenes, 753 (12.48%) taxonomically assigned

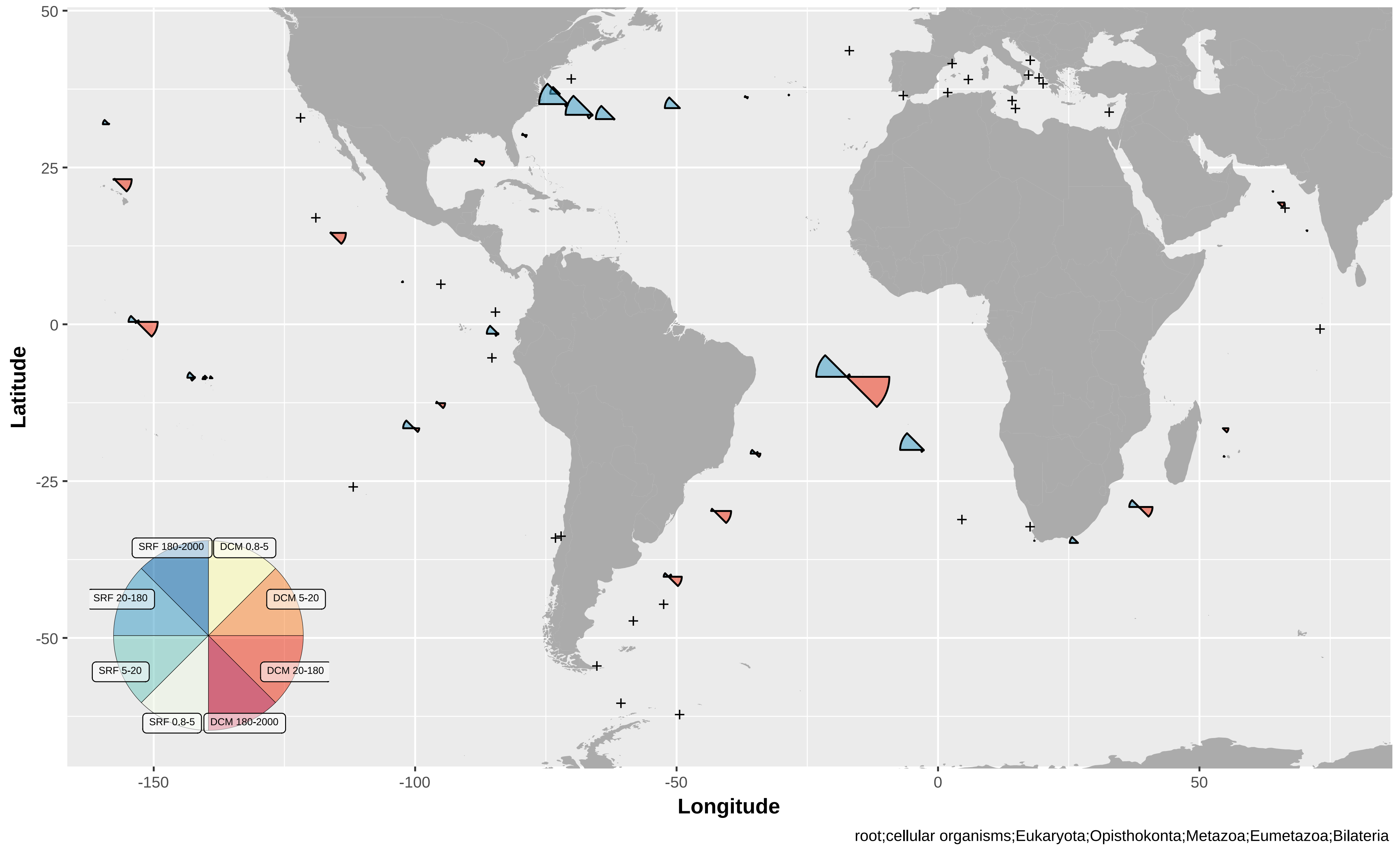

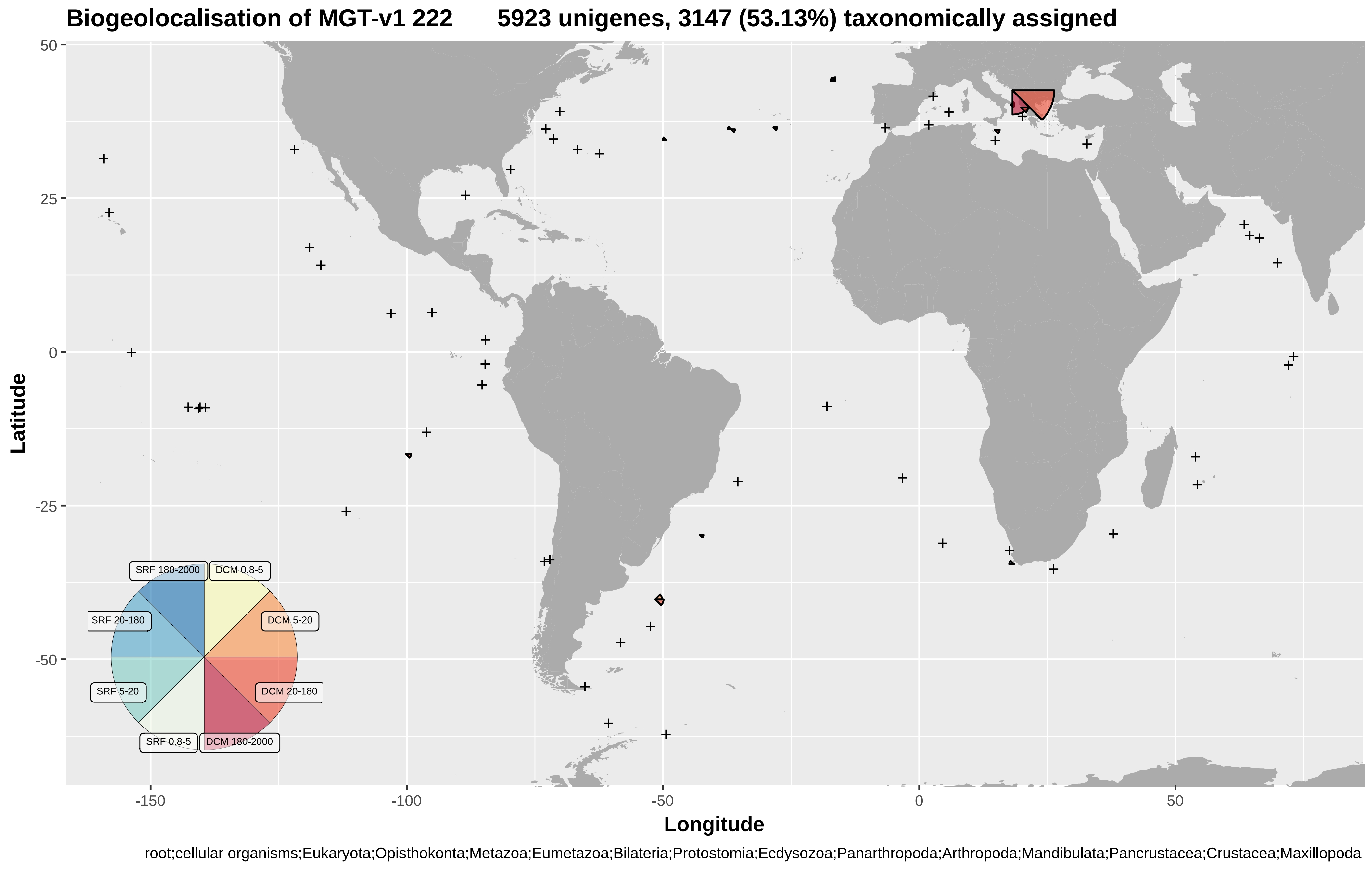
Biogeolocalisation of MGT-v1 222      5923 unigenes, 3147 (53.13%) taxonomically assigned
Latitude
Longitude
50
25
0
-25
-50
-150
-100
-50
0
50
SRF 180-2000
DCM 0.8-5
SRF 20-180
DCM 5-20
SRF 5-20
DCM 20-180
SRF 0.8-5
DCM 180-2000
root;cellular organisms;Eukaryota;Opisthokonta;Metazoa;Eumetazoa;Bilateria;Protostomia;Ecdysozoa;Panarthropoda;Arthropoda;Mandibulata;Pancrustacea;Crustacea;Maxillopoda

# Biogeolocalisation of MGT-v1 223      5916 unigenes, 5207 (88.02%) taxonomically assigned

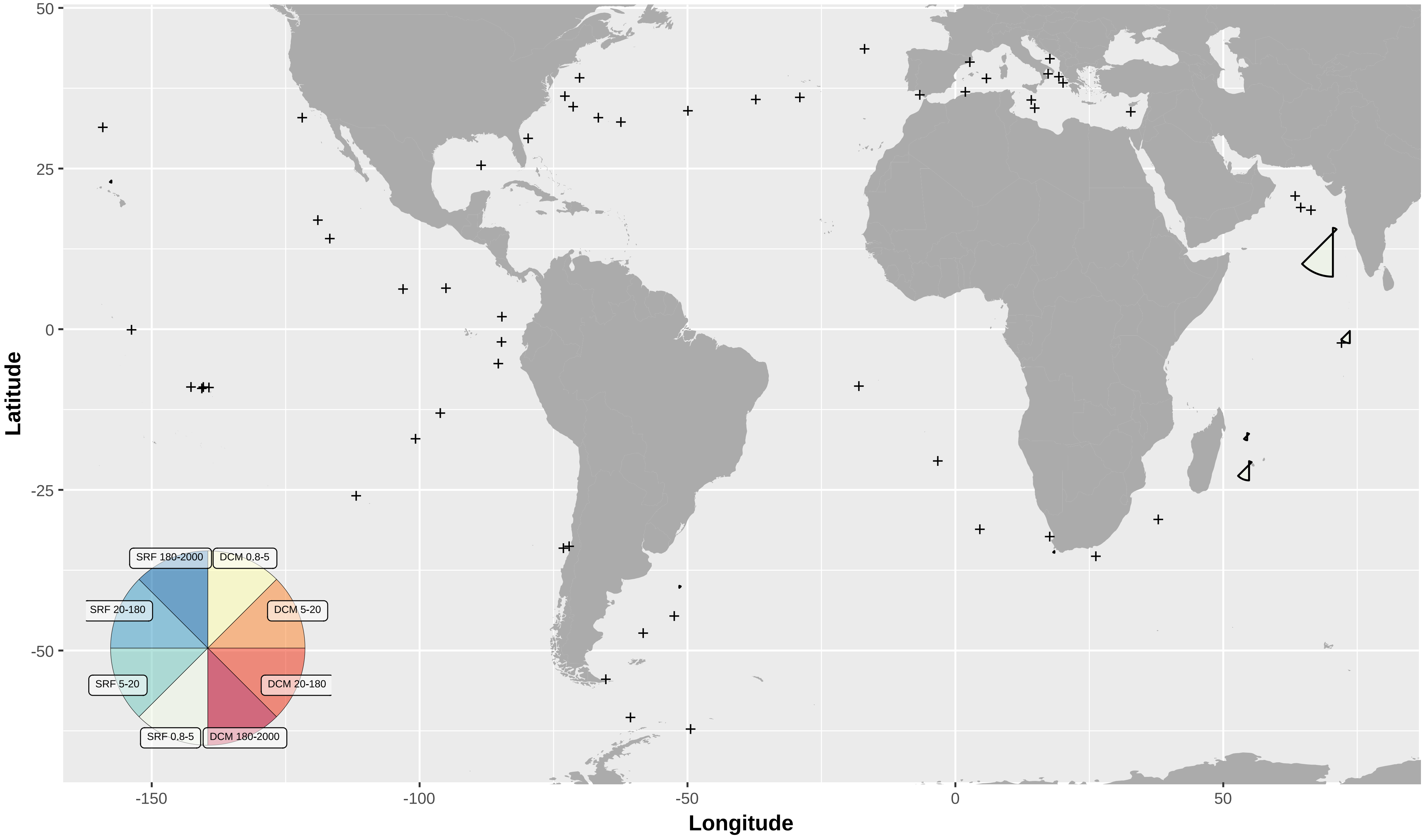


root;cellular organisms;Bacteria;Proteobacteria;Gammaproteobacteria

# Biogeolocalisation of MGT-v1 224      5900 unigenes, 3454 (58.54%) taxonomically assigned

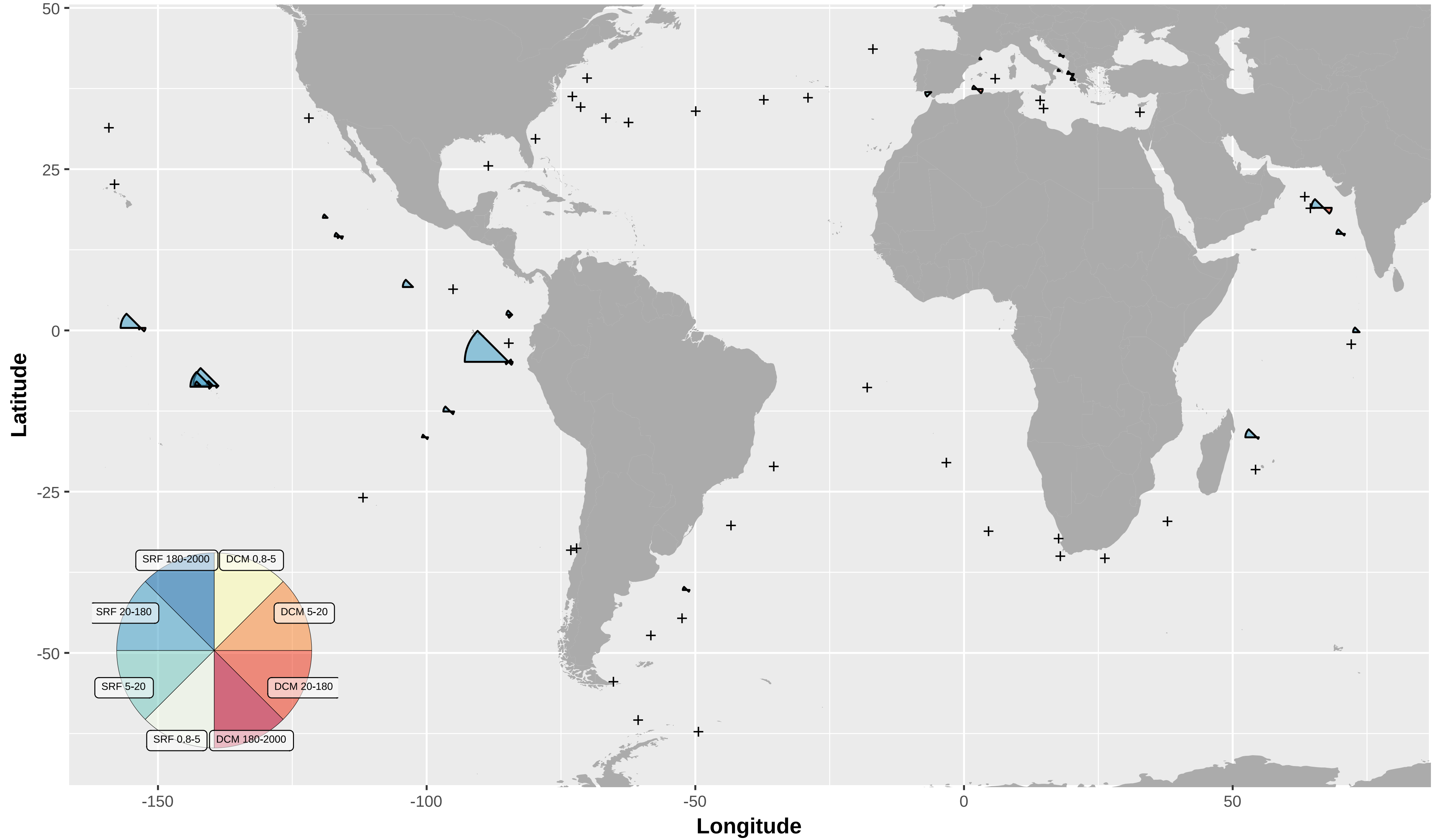


root;cellular organisms;Eukaryota;Alveolata;Ciliophora;Intramacronucleata;Spirotrichea

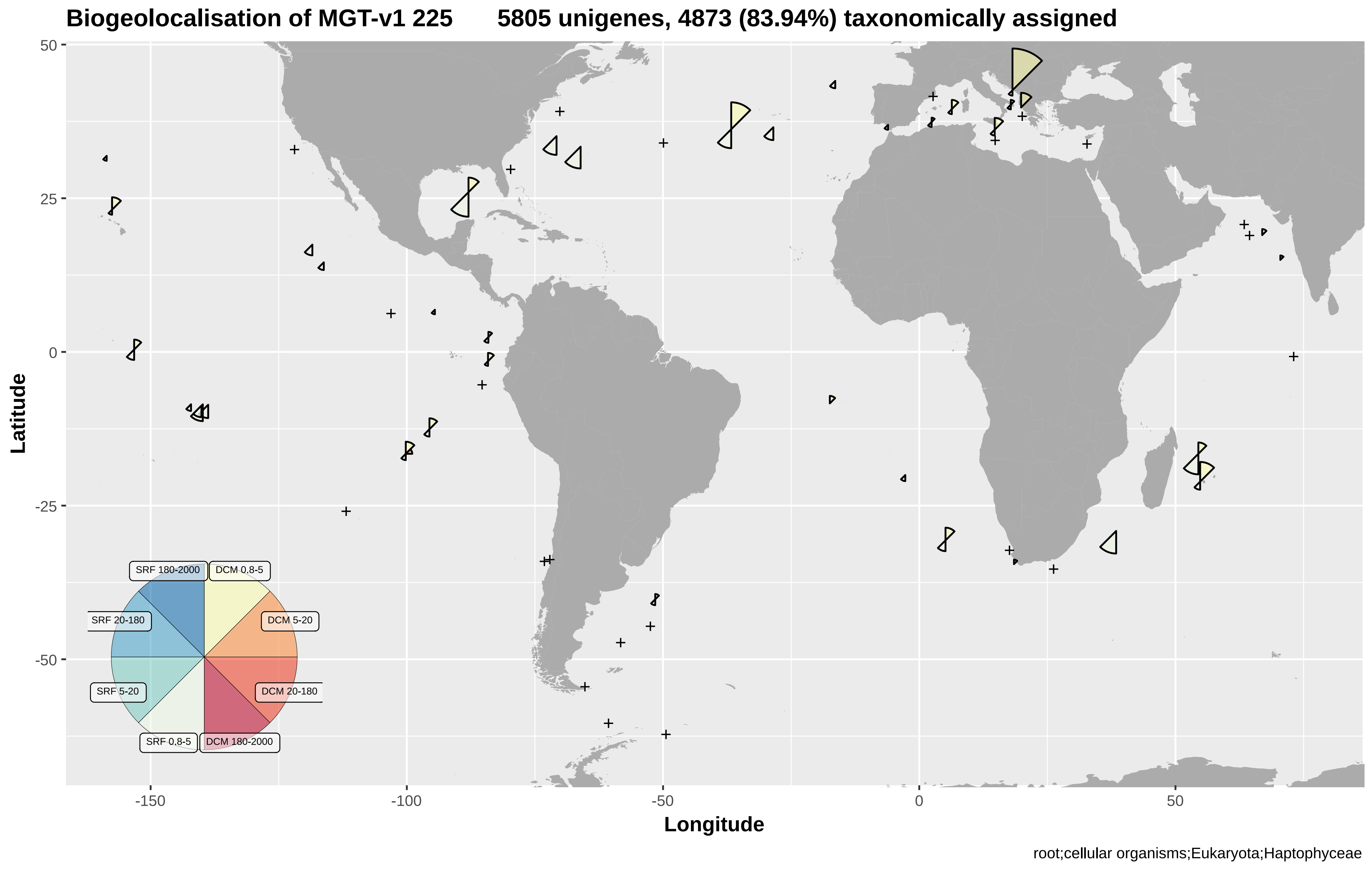
Biogeolocalisation of MGT-v1 225      5805 unigenes, 4873 (83.94%) taxonomically assigned
50
25
0
-25
-50
Latitude
-150
-100
-50
0
50
Longitude
SRF 180-2000
DCM 0.8-5
SRF 20-180
DCM 5-20
SRF 5-20
DCM 20-180
SRF 0.8-5
DCM 180-2000
root;cellular organisms;Eukaryota;Haptophyceae

### Biogeolocalisation of MGT-v1 226 5748 unigenes, 1706 (29.68%) taxonomically assigned

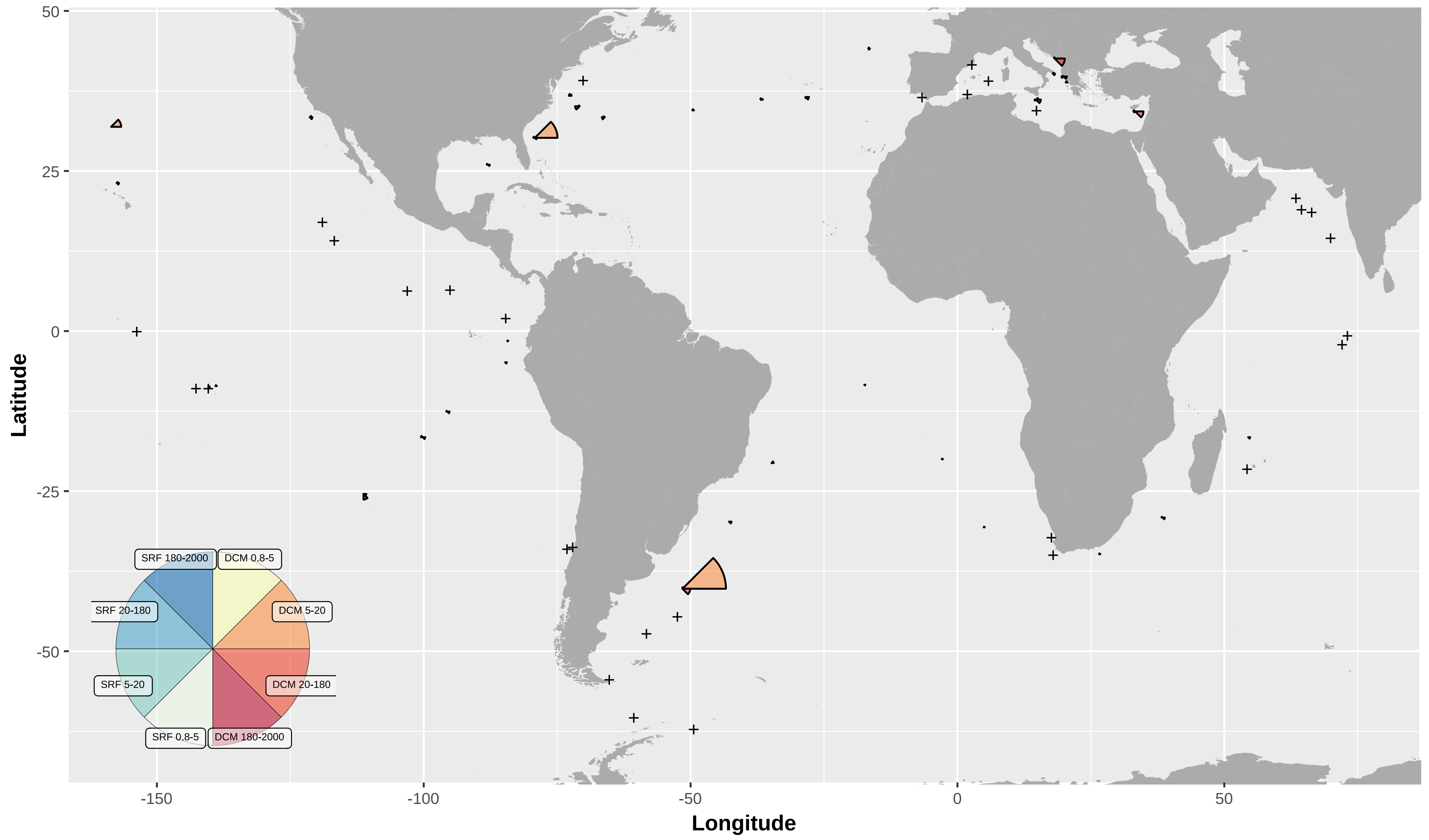


root;cellular organisms;Eukaryota;Opisthokonta;Metazoa;Eumetazoa;Bilateria

## Biogeolocalisation of MGT-v1 227      5738 unigenes, 5641 (98.31%) taxonomically assigned

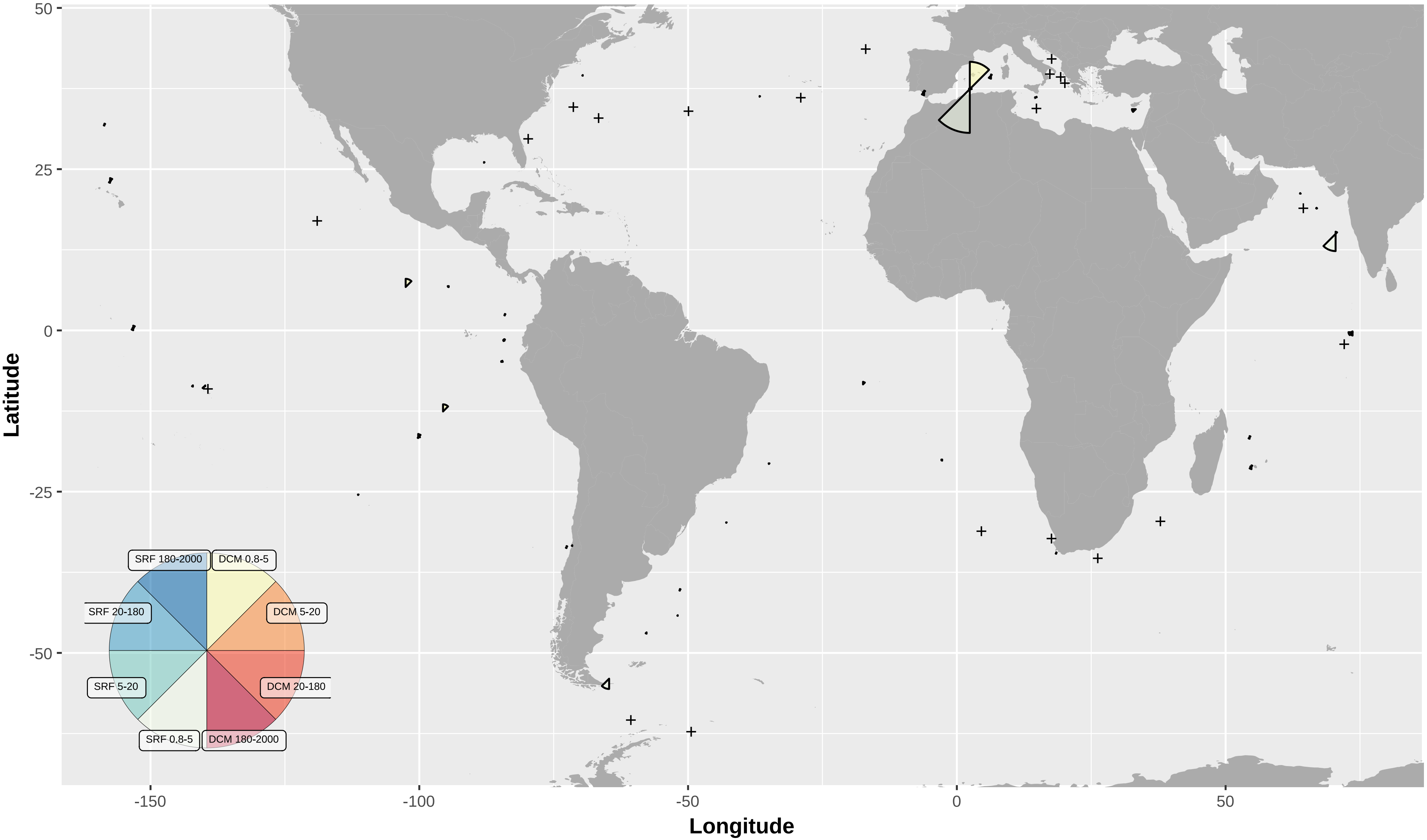


root;cellular organisms;Bacteria;Proteobacteria;Gammaproteobacteria;Alteromonadales;Alteromonadaceae

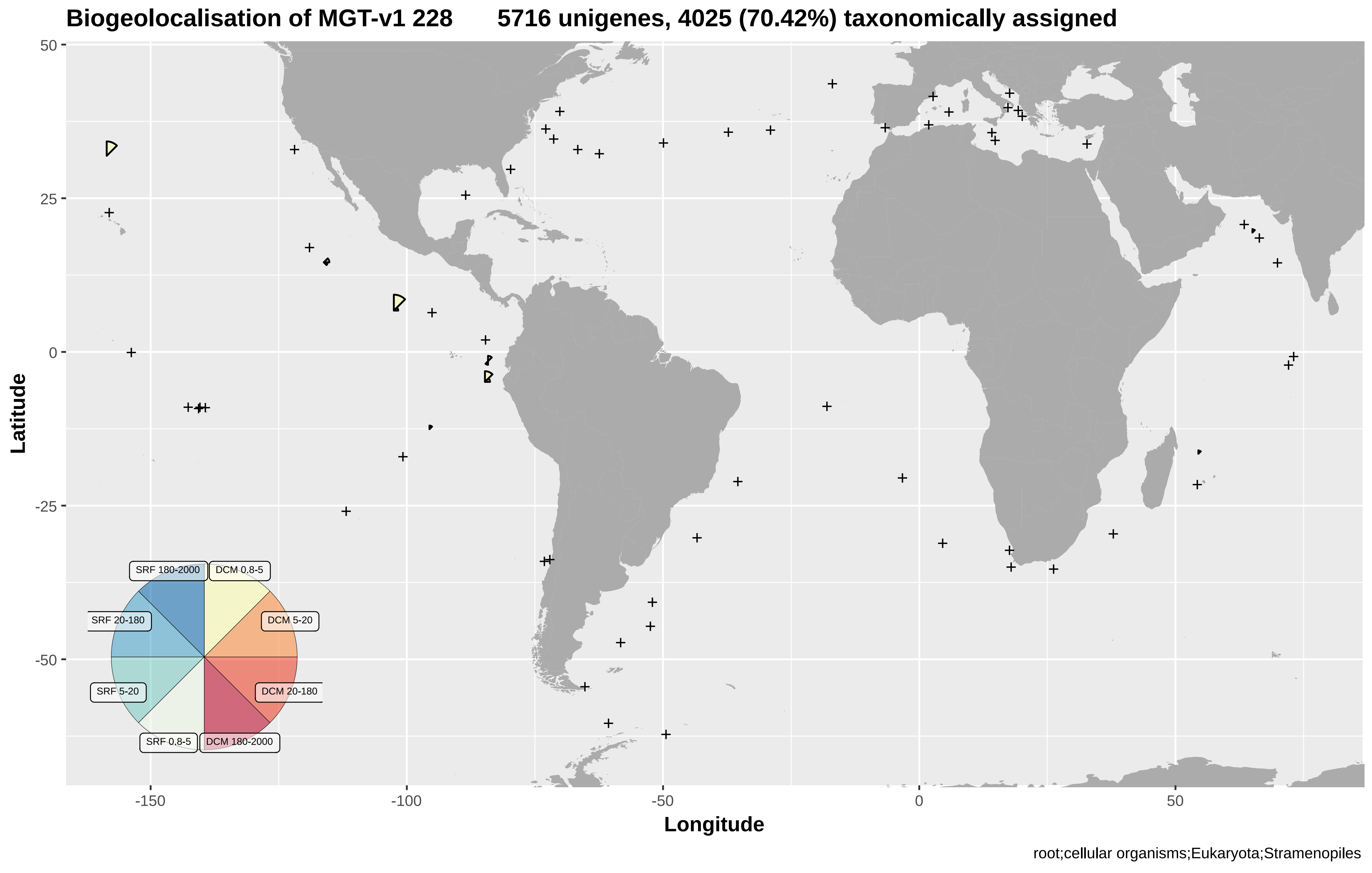

Biogeolocalisation of MGT-v1 228 5716 unigenes, 4025 (70.42%) taxonomically assigned
50
25
0
-25
-50
Latitude
-150
-100
-50
0
50
Longitude
SRF 180-2000
DCM 0.8-5
SRF 20-180
DCM 5-20
SRF 5-20
DCM 20-180
SRF 0.8-5
DCM 180-2000
root;cellular organisms;Eukaryota;Stramenopiles

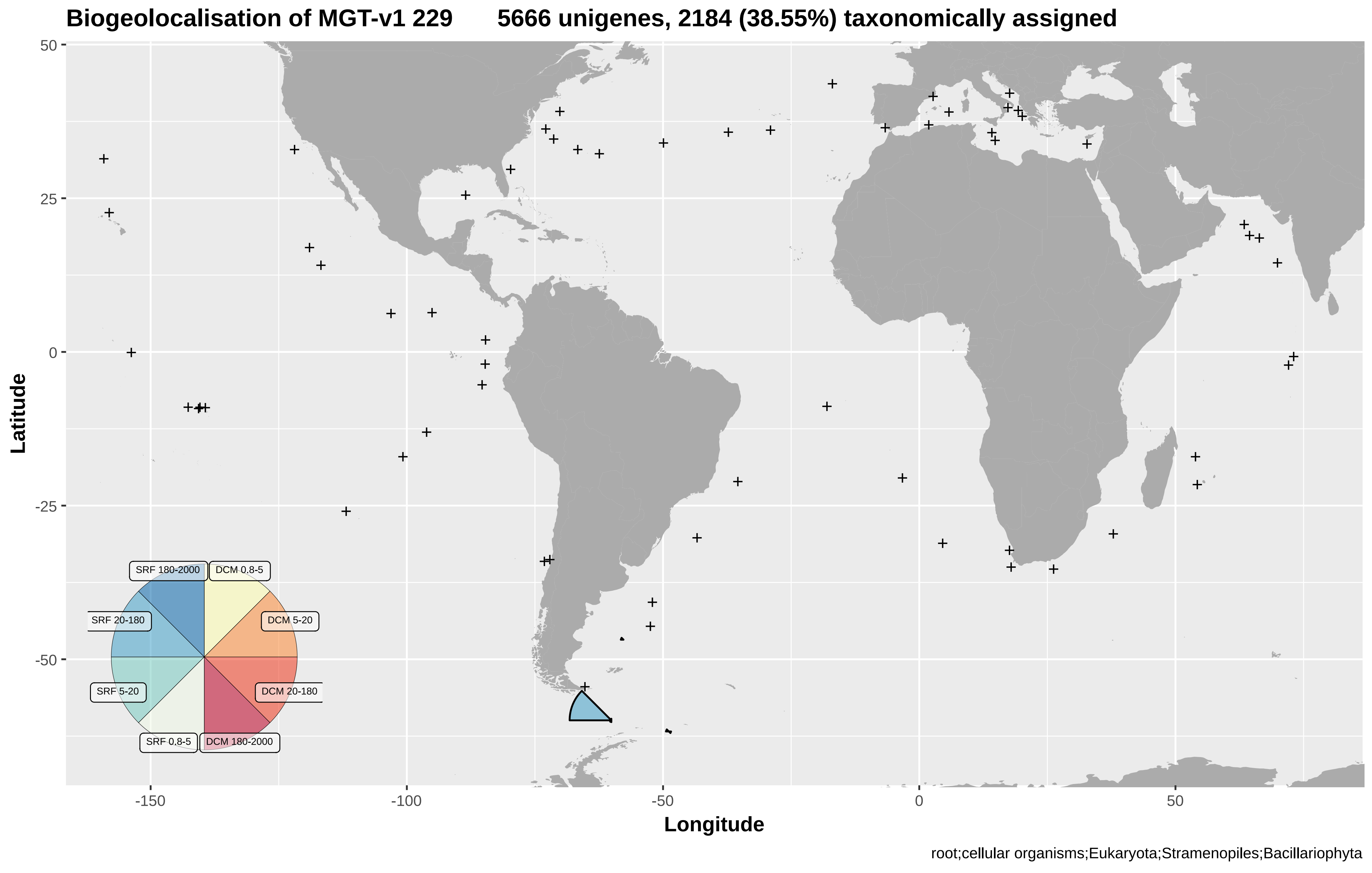
Biogeolocalisation of MGT-v1 229 5666 unigenes, 2184 (38.55%) taxonomically assigned
50
25
0
-25
-50
Latitude
-150
-100
-50
0
50
Longitude
SRF 180-2000
DCM 0.8-5
SRF 20-180
DCM 5-20
SRF 5-20
DCM 20-180
SRF 0.8-5
DCM 180-2000
root;cellular organisms;Eukaryota;Stramenopiles;Bacillariophyta

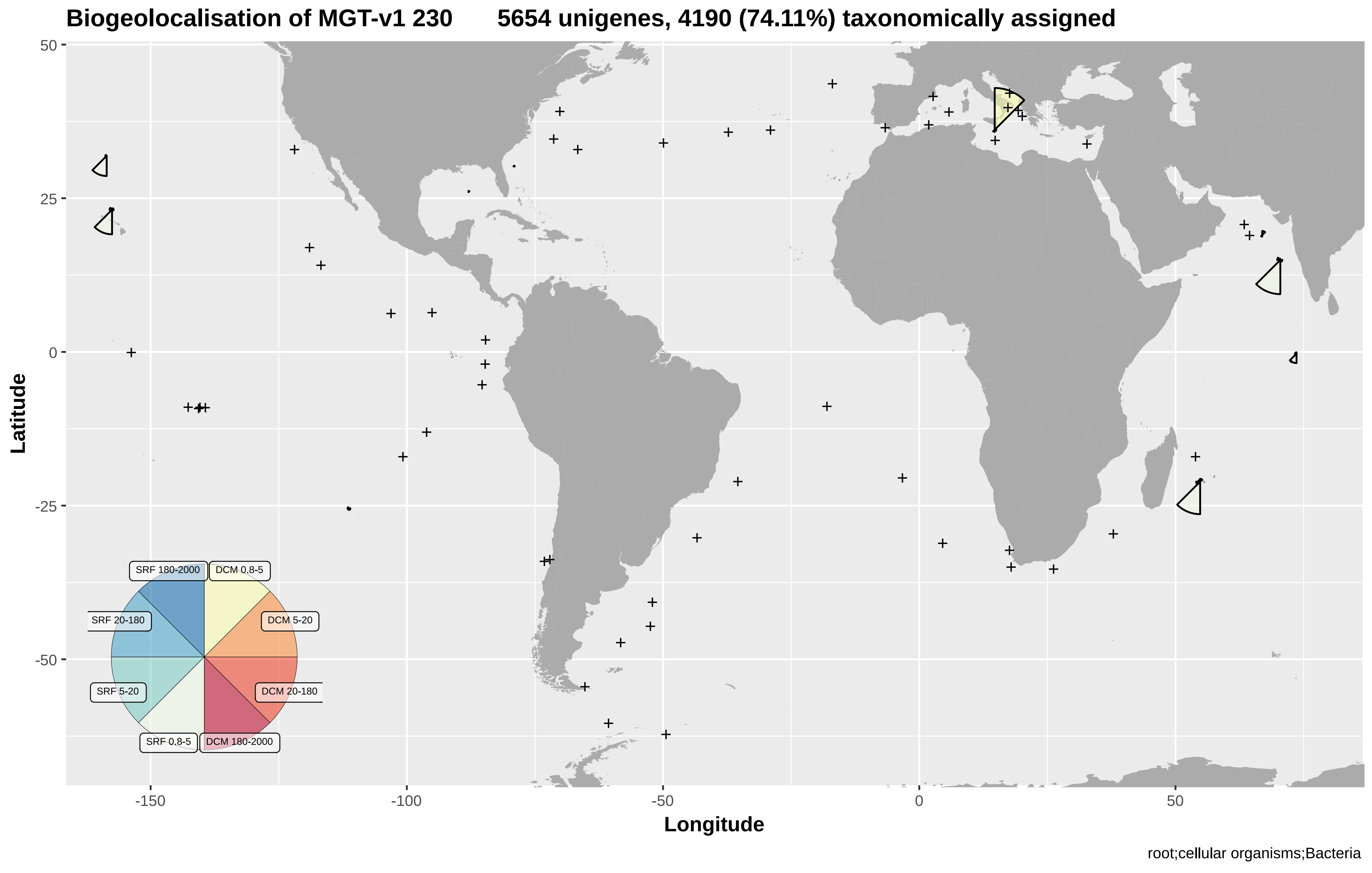
Biogeolocalisation of MGT-v1 230 5654 unigenes, 4190 (74.11%) taxonomically assigned
Latitude
Longitude
50
25
0
-25
-50
-150
-100
-50
0
50
SRF 180-2000
DCM 0.8-5
SRF 20-180
DCM 5-20
SRF 5-20
DCM 20-180
SRF 0.8-5
DCM 180-2000
root;cellular organisms;Bacteria

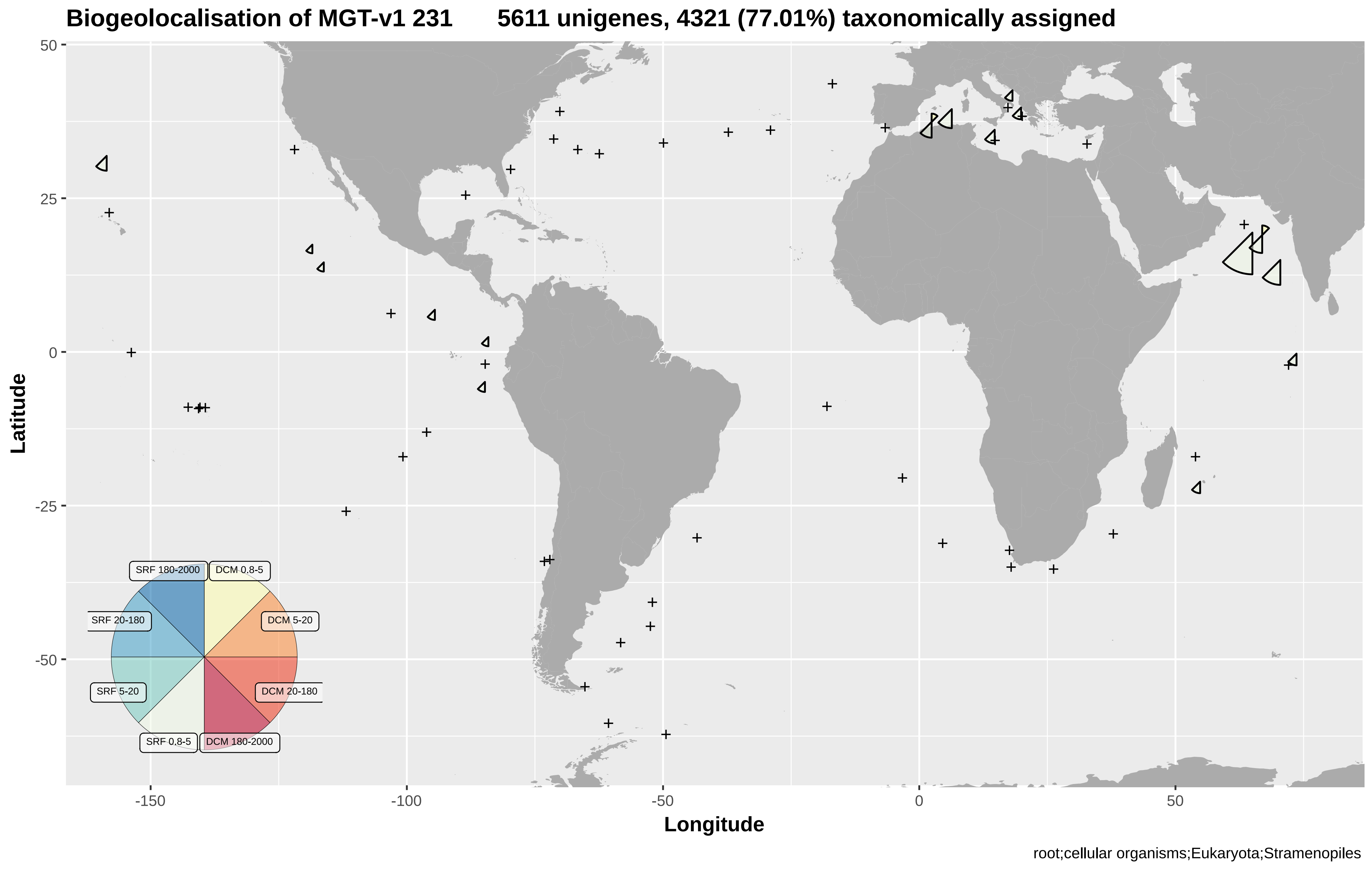
Biogeolocalisation of MGT-v1 231 5611 unigenes, 4321 (77.01%) taxonomically assigned
Latitude
50
25
0
-25
-50
Longitude
-150
-100
-50
0
50
SRF 180-2000
DCM 0.8-5
SRF 20-180
DCM 5-20
SRF 5-20
DCM 20-180
SRF 0.8-5
DCM 180-2000
root;cellular organisms;Eukaryota;Stramenopiles

#### Biogeolocalisation of MGT-v1 232 5587 unigenes, 536 (9.59%) taxonomically assigned

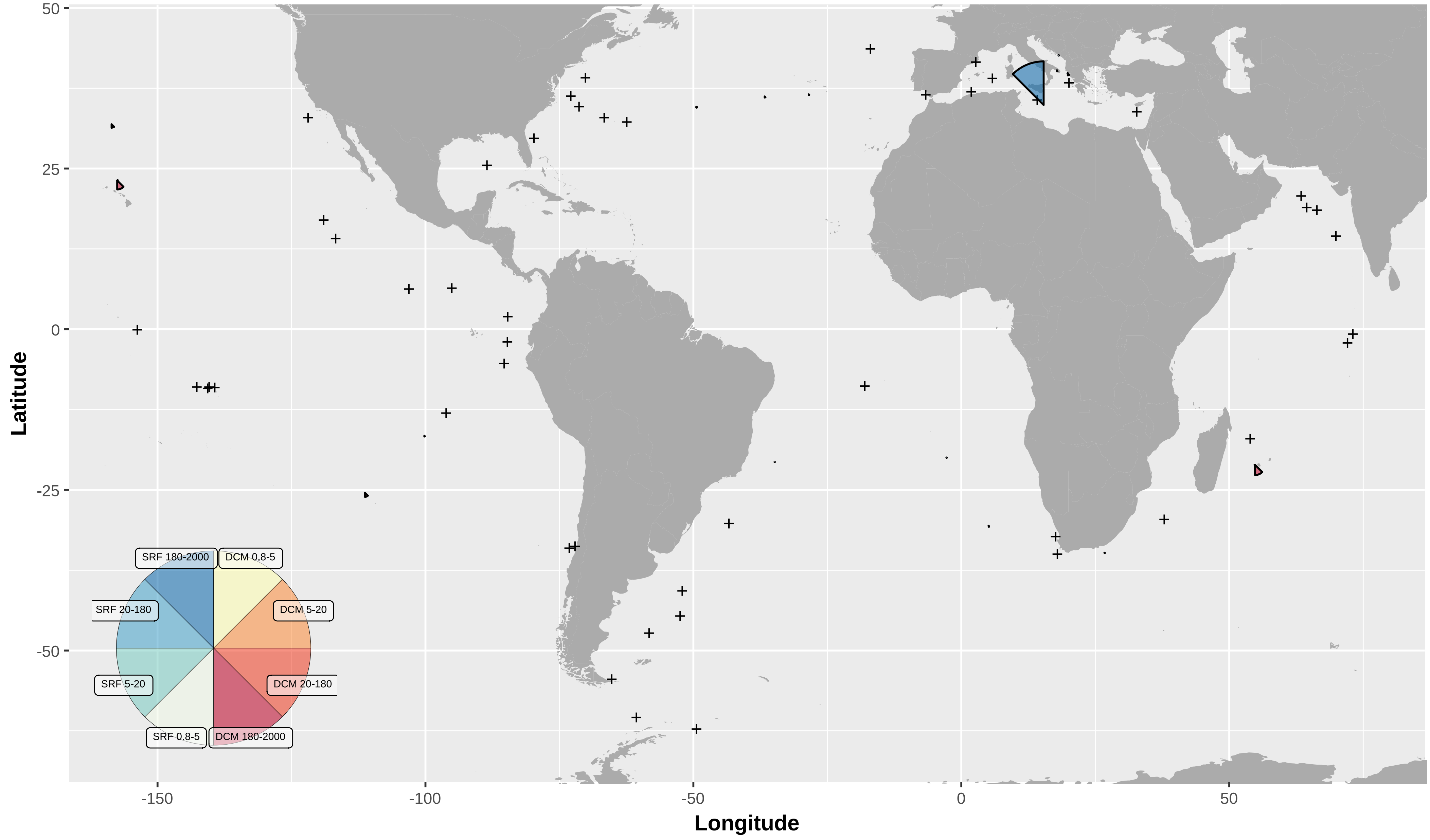


root;cellular organisms;Eukaryota;Opisthokonta;Metazoa;Eumetazoa;Bilateria

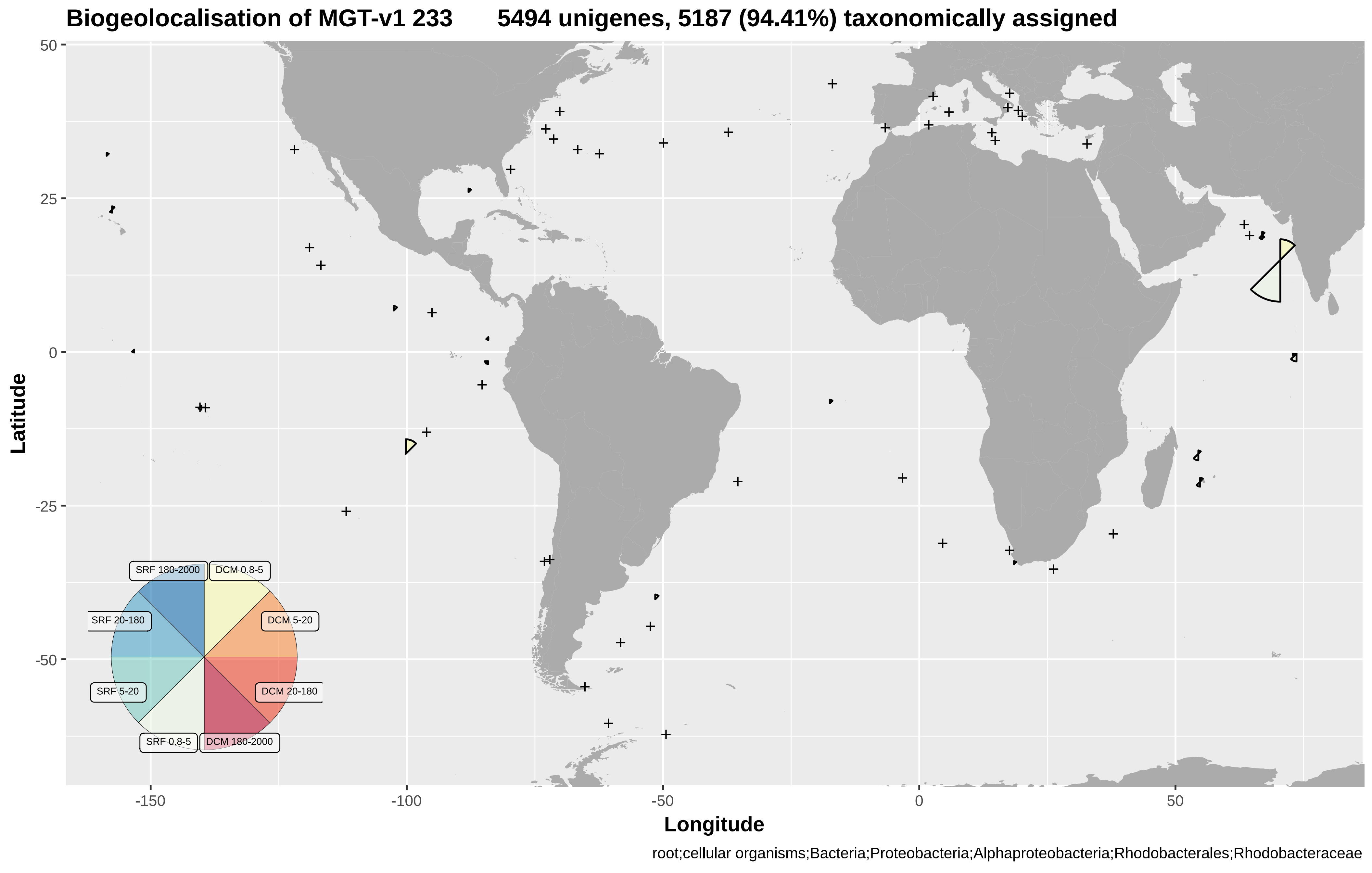

Biogeolocalisation of MGT-v1 233 5494 unigenes, 5187 (94.41%) taxonomically assigned
Latitude
Longitude
50
25
0
-25
-50
-150
-100
-50
0
50
SRF 180-2000
DCM 0.8-5
SRF 20-180
DCM 5-20
SRF 5-20
DCM 20-180
SRF 0.8-5
DCM 180-2000
root;cellular organisms;Bacteria;Proteobacteria;Alphaproteobacteria;Rhodobacterales;Rhodobacteraceae

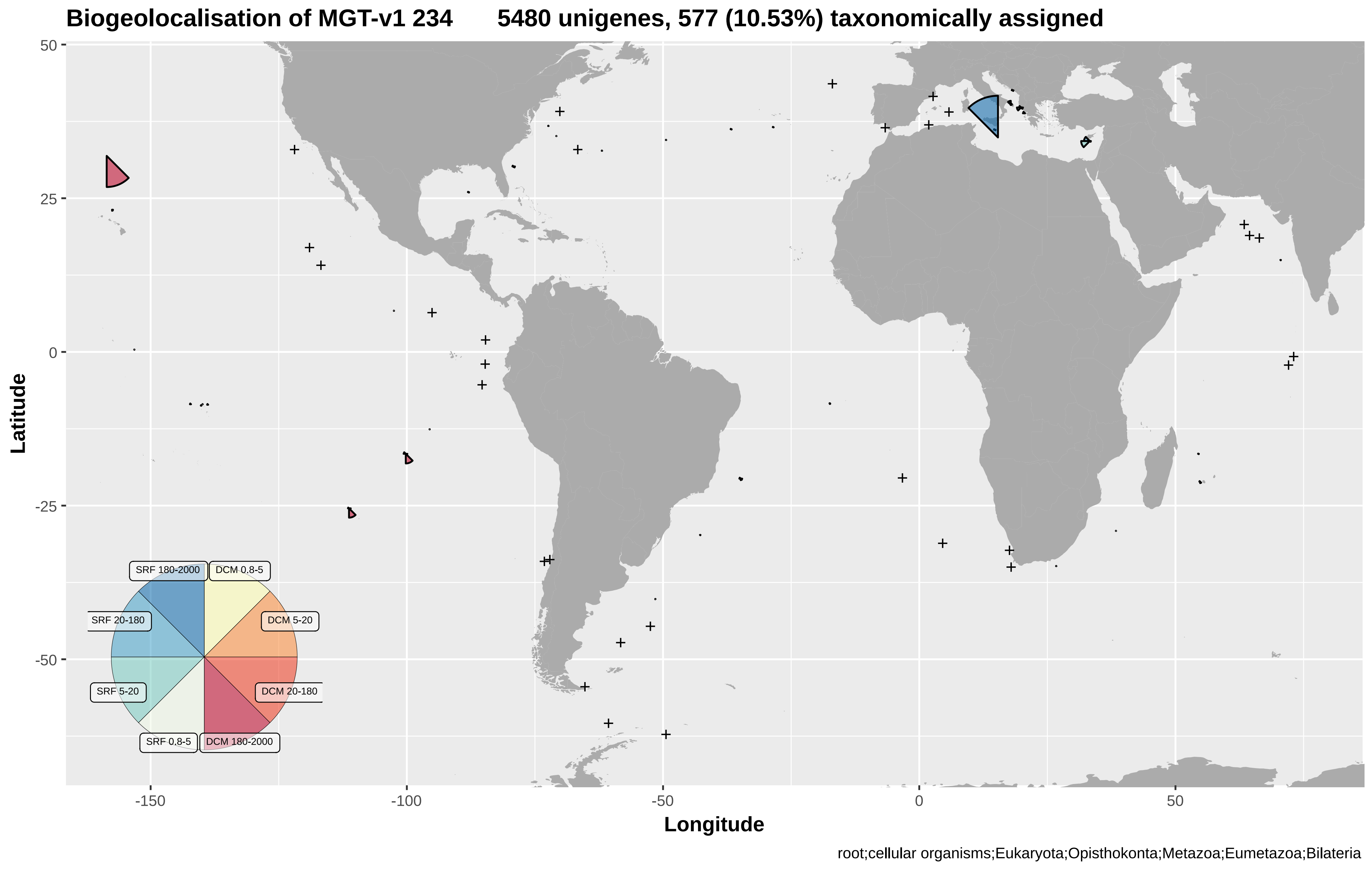
Biogeolocalisation of MGT-v1 234 5480 unigenes, 577 (10.53%) taxonomically assigned
50
25
0
-25
-50
Latitude
-150
-100
-50
0
50
Longitude
SRF 180-2000
DCM 0.8-5
SRF 20-180
DCM 5-20
SRF 5-20
DCM 20-180
SRF 0.8-5
DCM 180-2000
root;cellular organisms;Eukaryota;Opisthokonta;Metazoa;Eumetazoa;Bilateria

# Biogeolocalisation of MGT-v1 235      5473 unigenes, 4740 (86.61%) taxonomically assigned

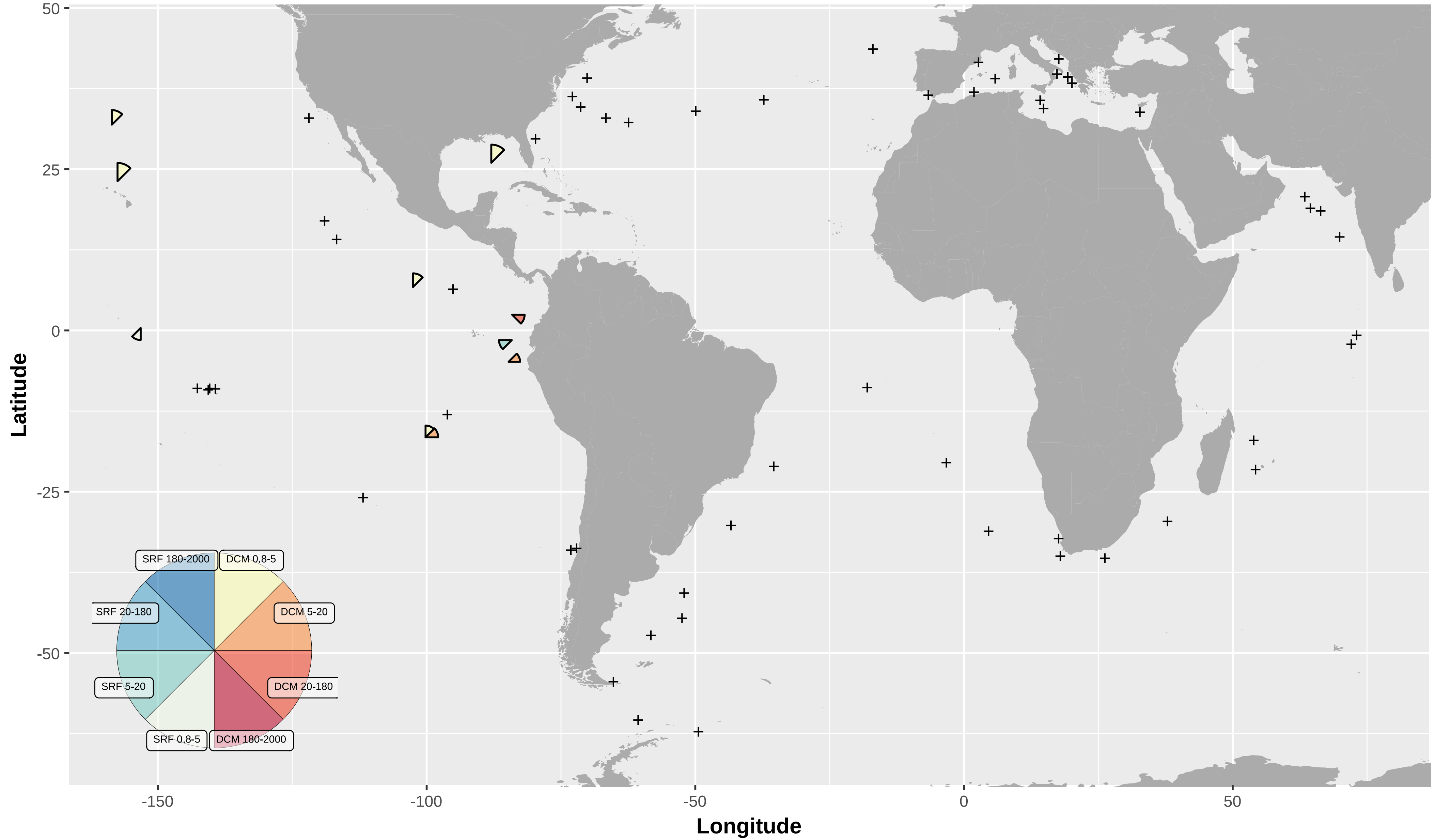


root;cellular organisms;Eukaryota;Opisthokonta;Fungi;Dikarya;Ascomycota;saccharomyceta;Pezizomycotina;leotiomyceta

## Biogeolocalisation of MGT-v1 236      5456 unigenes, 356 (6.52%) taxonomically assigned

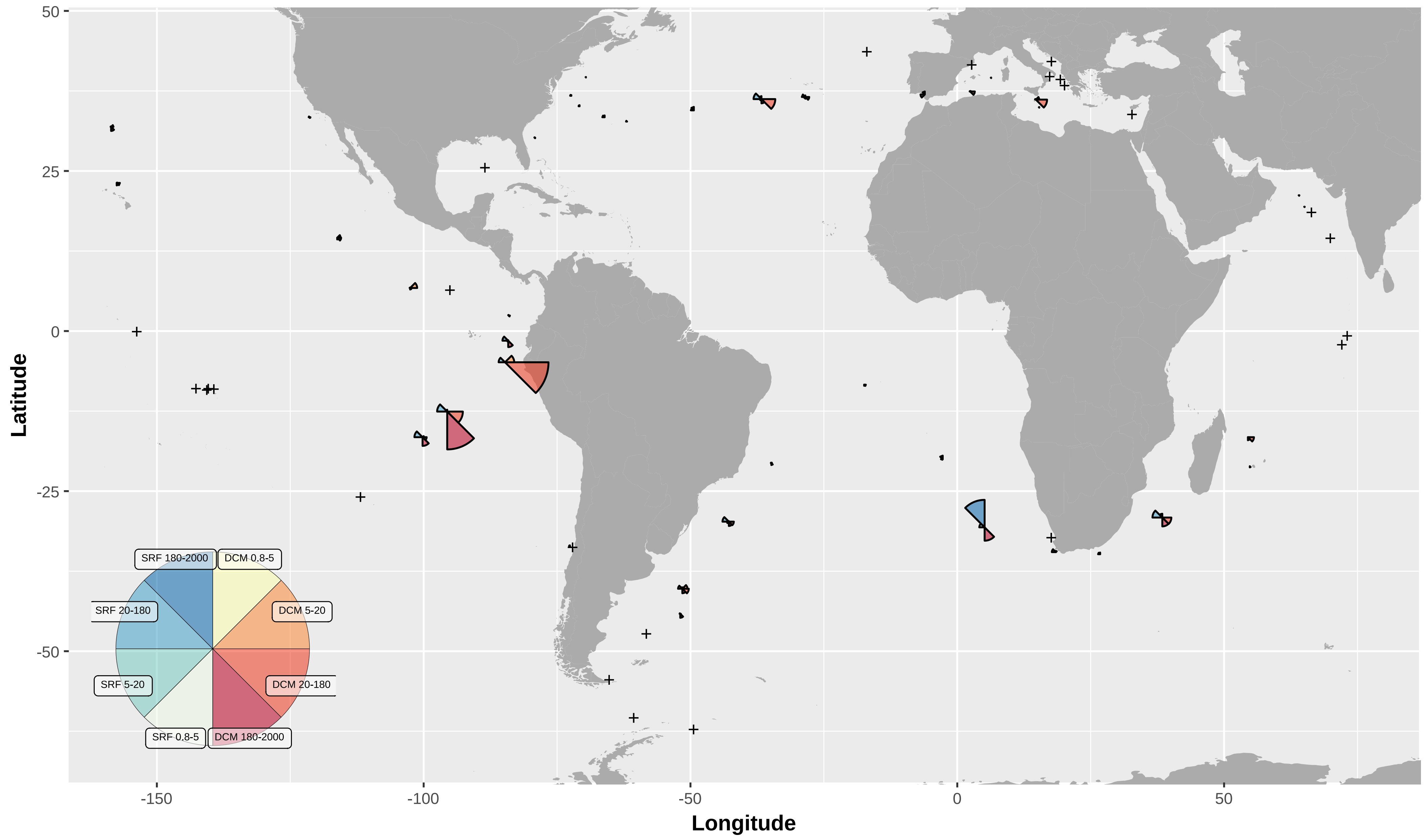


root;cellular organisms;Eukaryota;Opisthokonta;Metazoa;Eumetazoa;Bilateria

## Biogeolocalisation of MGT-v1 237      5449 unigenes, 373 (6.85%) taxonomically assigned

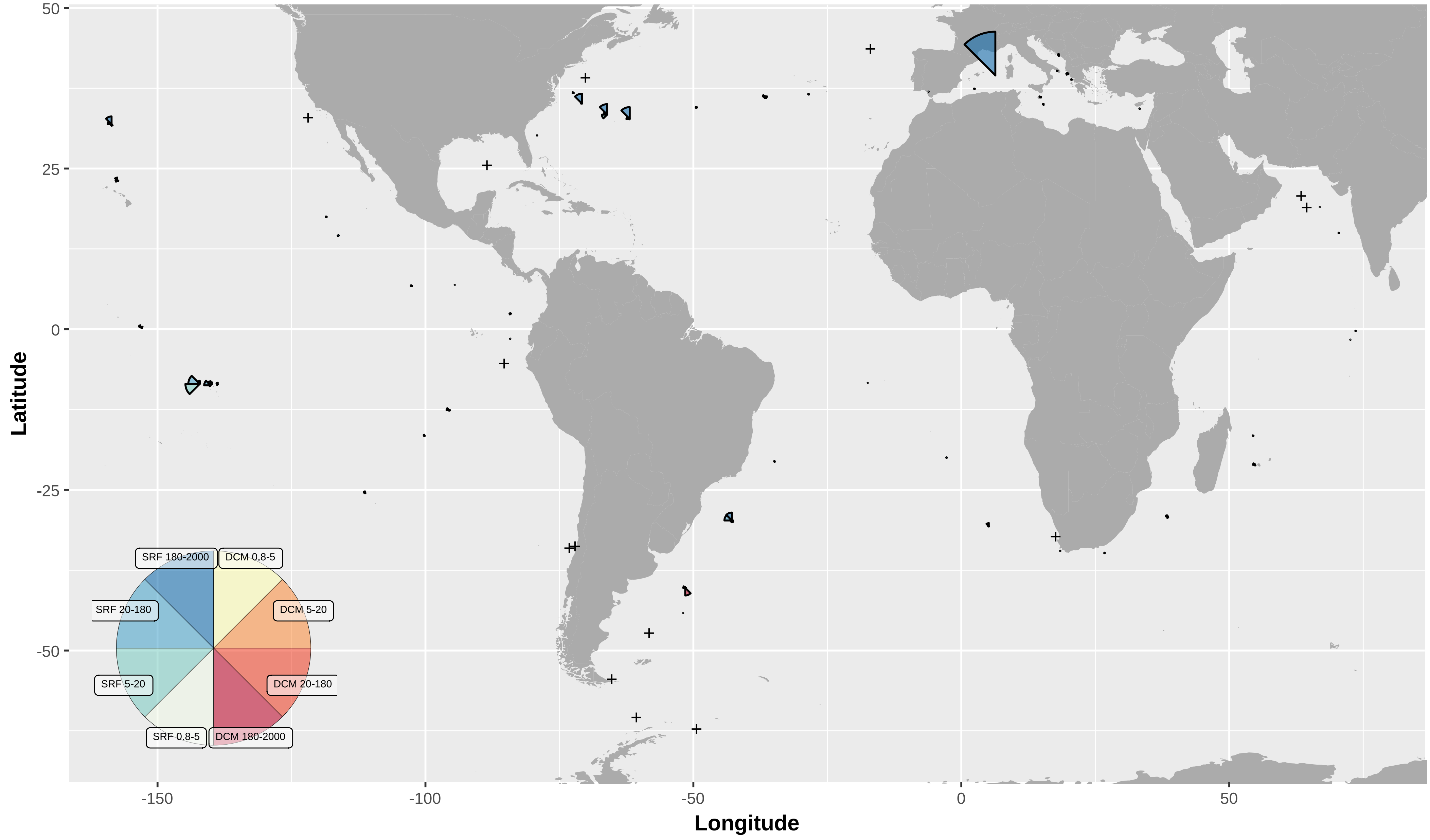


root;cellular organisms;Eukaryota;Opisthokonta;Metazoa;Eumetazoa;Bilateria

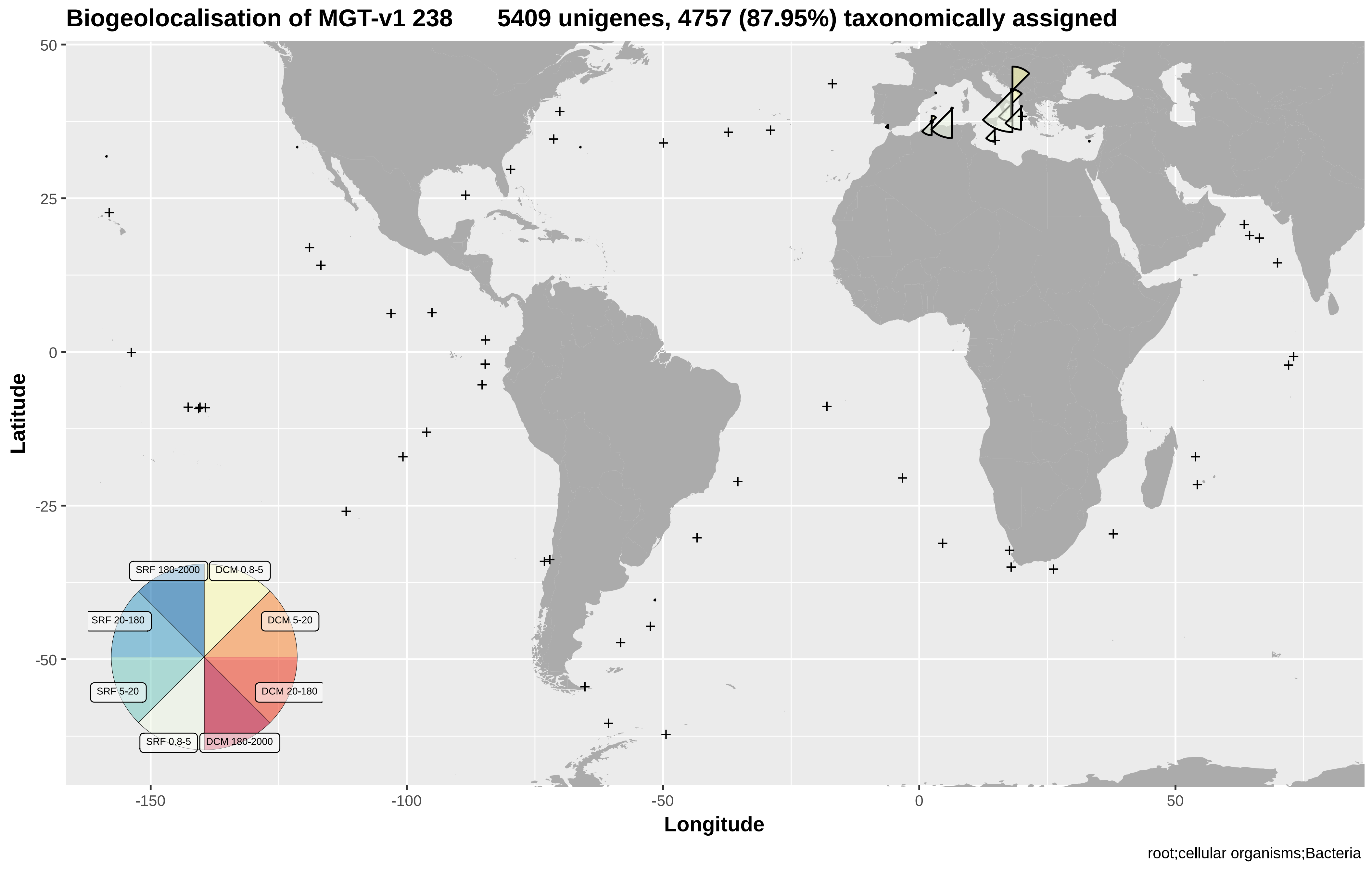
Biogeolocalisation of MGT-v1 238 5409 unigenes, 4757 (87.95%) taxonomically assigned
Latitude
Longitude
50
25
0
-25
-50
-150
-100
-50
0
50
SRF 180-2000
DCM 0.8-5
SRF 20-180
DCM 5-20
SRF 5-20
DCM 20-180
SRF 0.8-5
DCM 180-2000
root;cellular organisms;Bacteria

## Biogeolocalisation of MGT-v1 239      5408 unigenes, 4931 (91.18%) taxonomically assigned

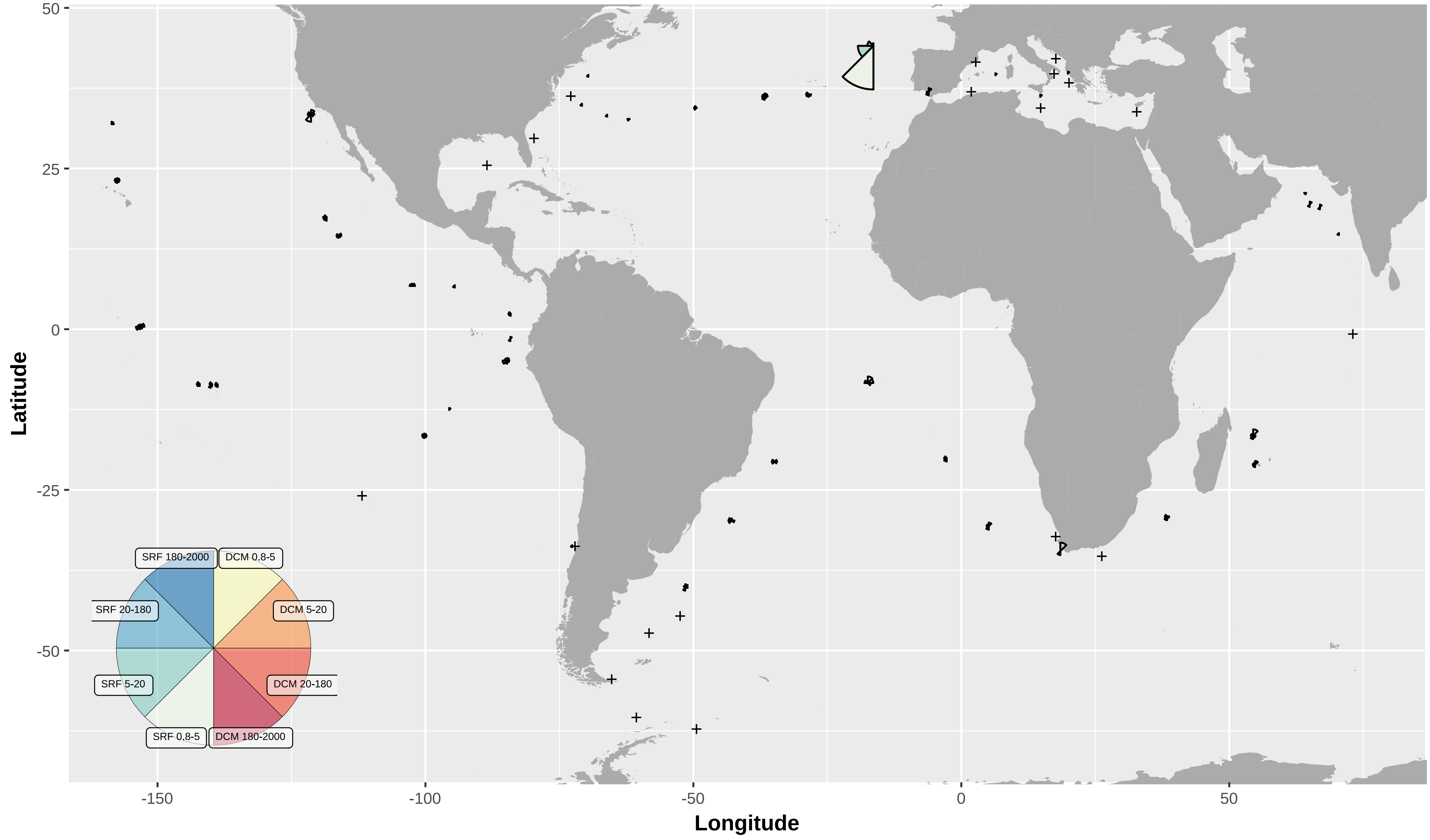


root;cellular organisms;Eukaryota;Viridiplantae;Chlorophyta;prasinophytes

## Biogeolocalisation of MGT-v1 240      5292 unigenes, 4781 (90.34%) taxonomically assigned

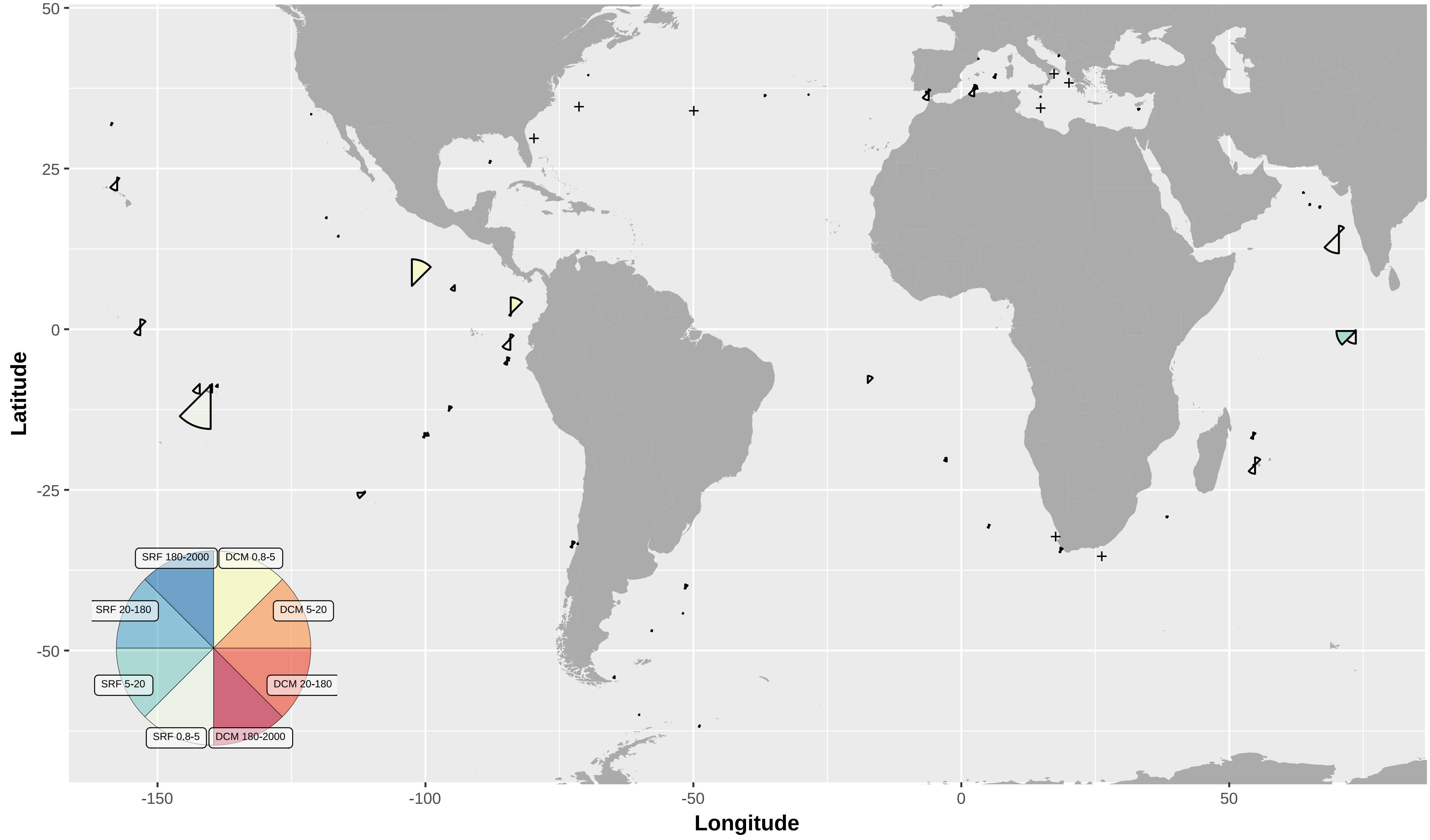


root;cellular organisms;Bacteria;Proteobacteria;Gammaproteobacteria

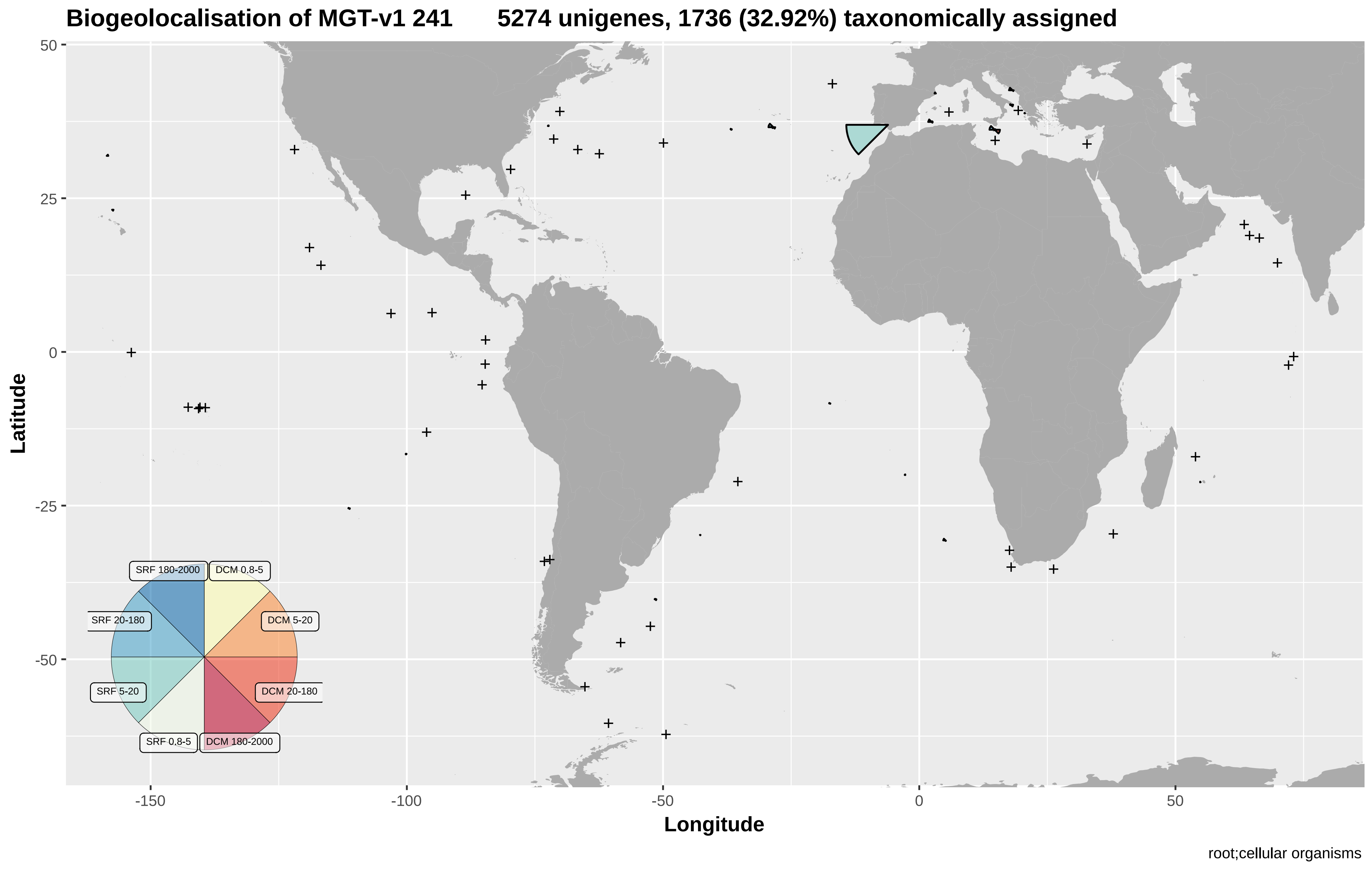
Biogeolocalisation of MGT-v1 241 5274 unigenes, 1736 (32.92%) taxonomically assigned
Latitude
Longitude
50
25
0
-25
-50
-150
-100
-50
0
50
SRF 180-2000
DCM 0.8-5
SRF 20-180
DCM 5-20
SRF 5-20
DCM 20-180
SRF 0.8-5
DCM 180-2000
root;cellular organisms

### Biogeolocalisation of MGT-v1 242 5257 unigenes, 1361 (25.89%) taxonomically assigned

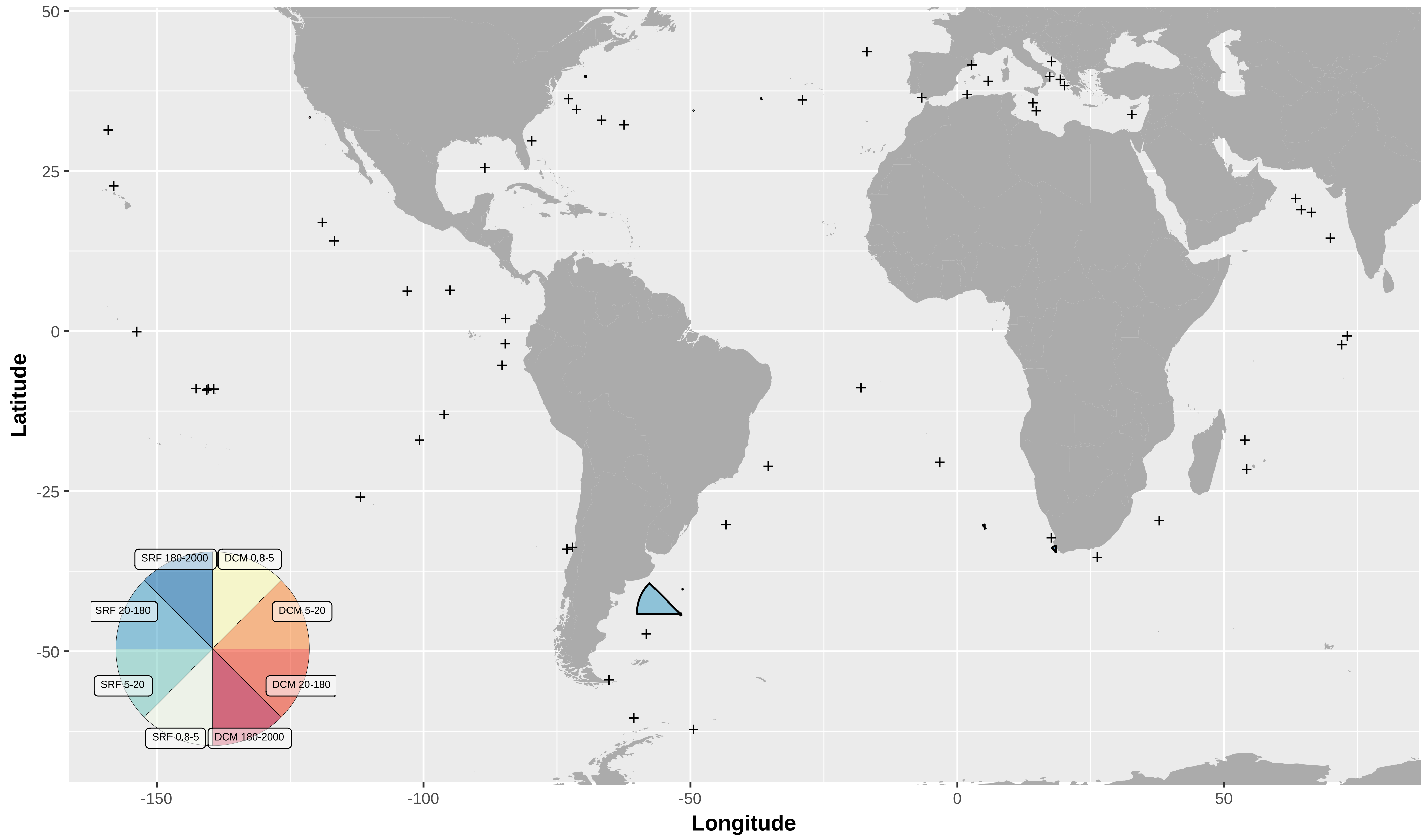


root;cellular organisms

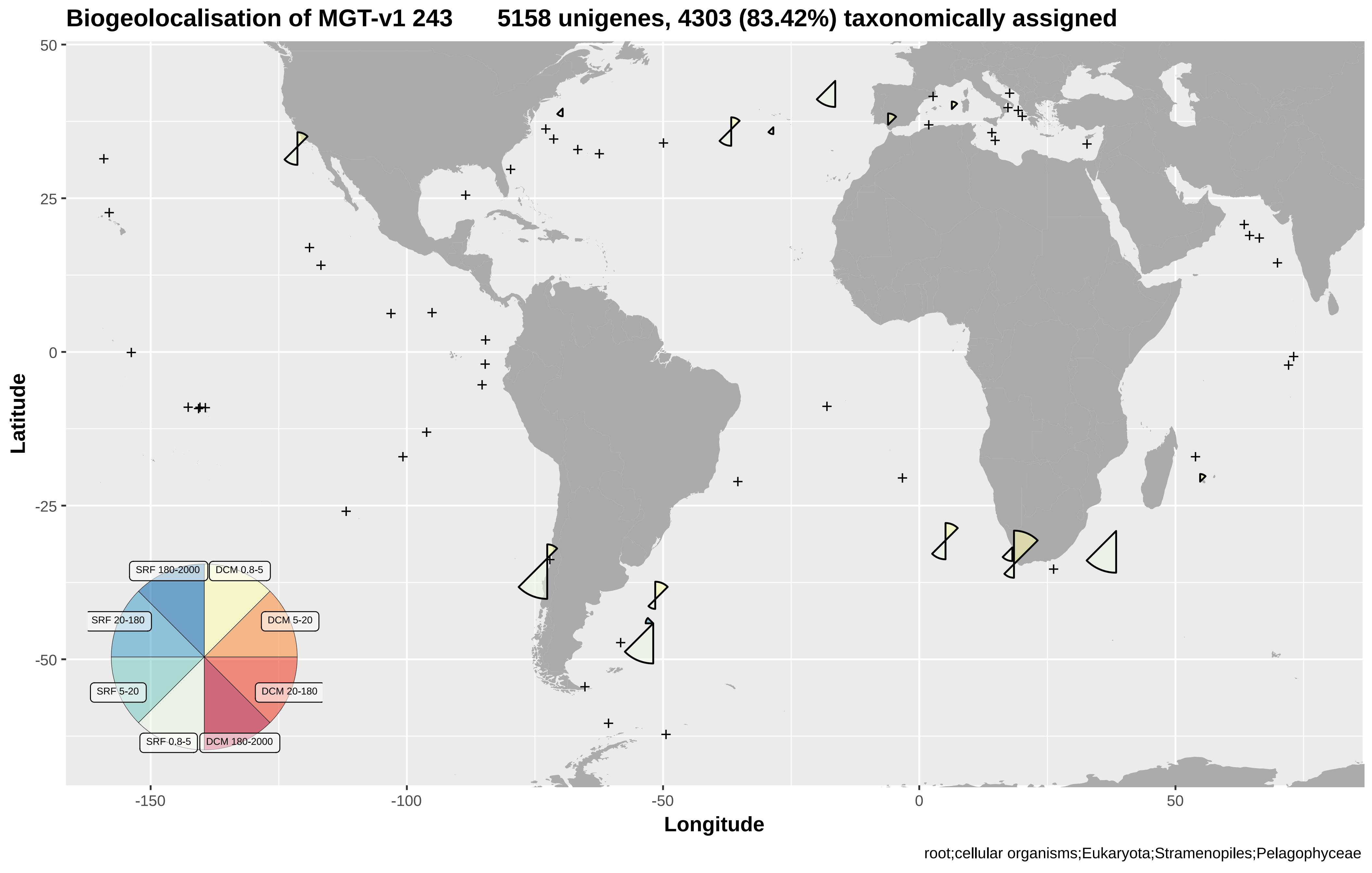

Biogeolocalisation of MGT-v1 243 5158 unigenes, 4303 (83.42%) taxonomically assigned
50
25
0
-25
-50
Latitude
-150
-100
-50
0
50
Longitude
SRF 180-2000
DCM 0.8-5
SRF 20-180
DCM 5-20
SRF 5-20
DCM 20-180
SRF 0.8-5
DCM 180-2000
root;cellular organisms;Eukaryota;Stramenopiles;Pelagophyceae

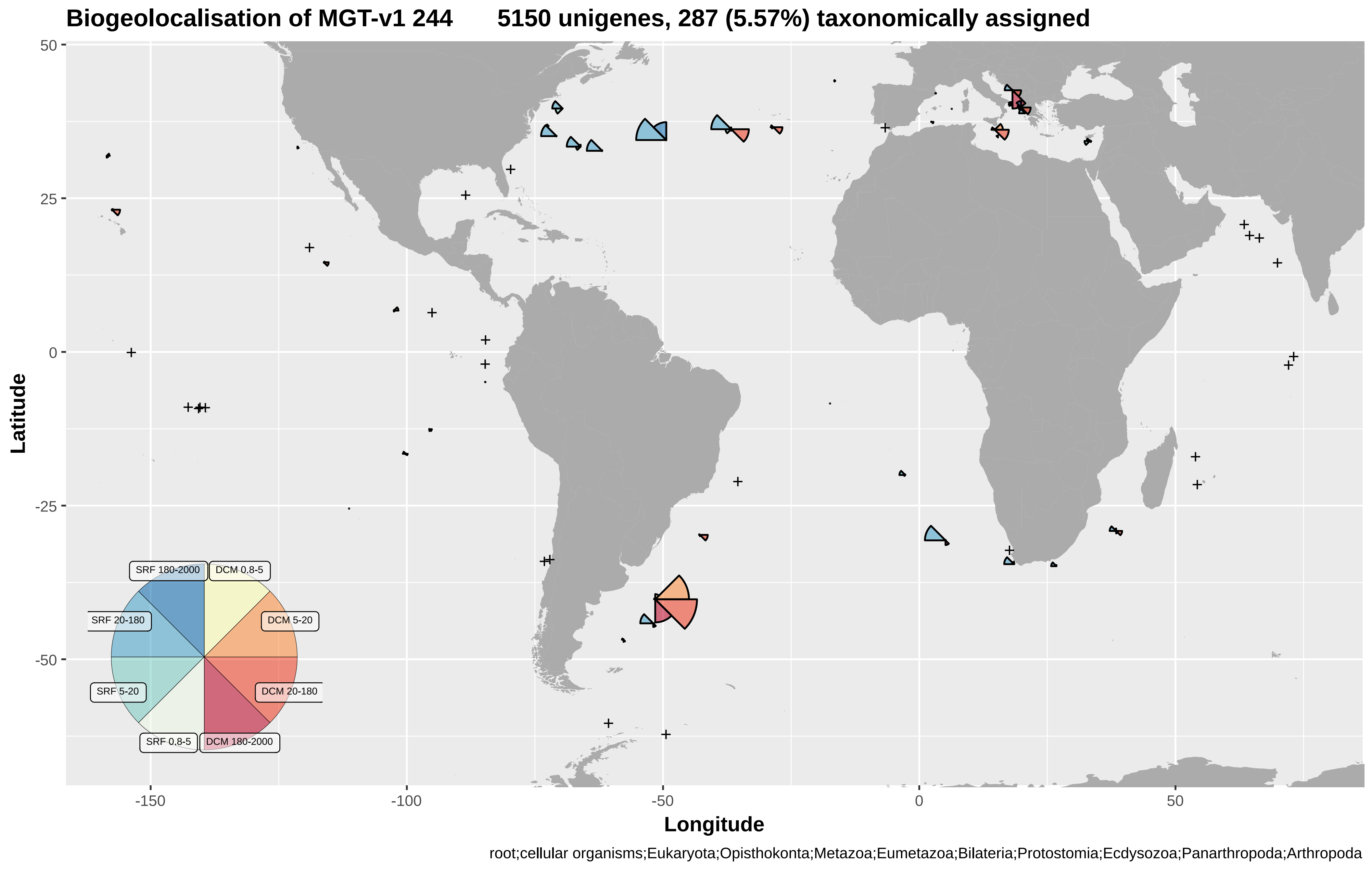

Biogeolocalisation of MGT-v1 244 5150 unigenes, 287 (5.57%) taxonomically assigned
50
25
0
-25
-50
Latitude
-150
-100
-50
0
50
Longitude
SRF 180-2000
DCM 0.8-5
SRF 20-180
DCM 5-20
SRF 5-20
DCM 20-180
SRF 0.8-5
DCM 180-2000
root;cellular organisms;Eukaryota;Opisthokonta;Metazoa;Eumetazoa;Bilateria;Protostomia;Ecdysozoa;Panarthropoda;Arthropoda

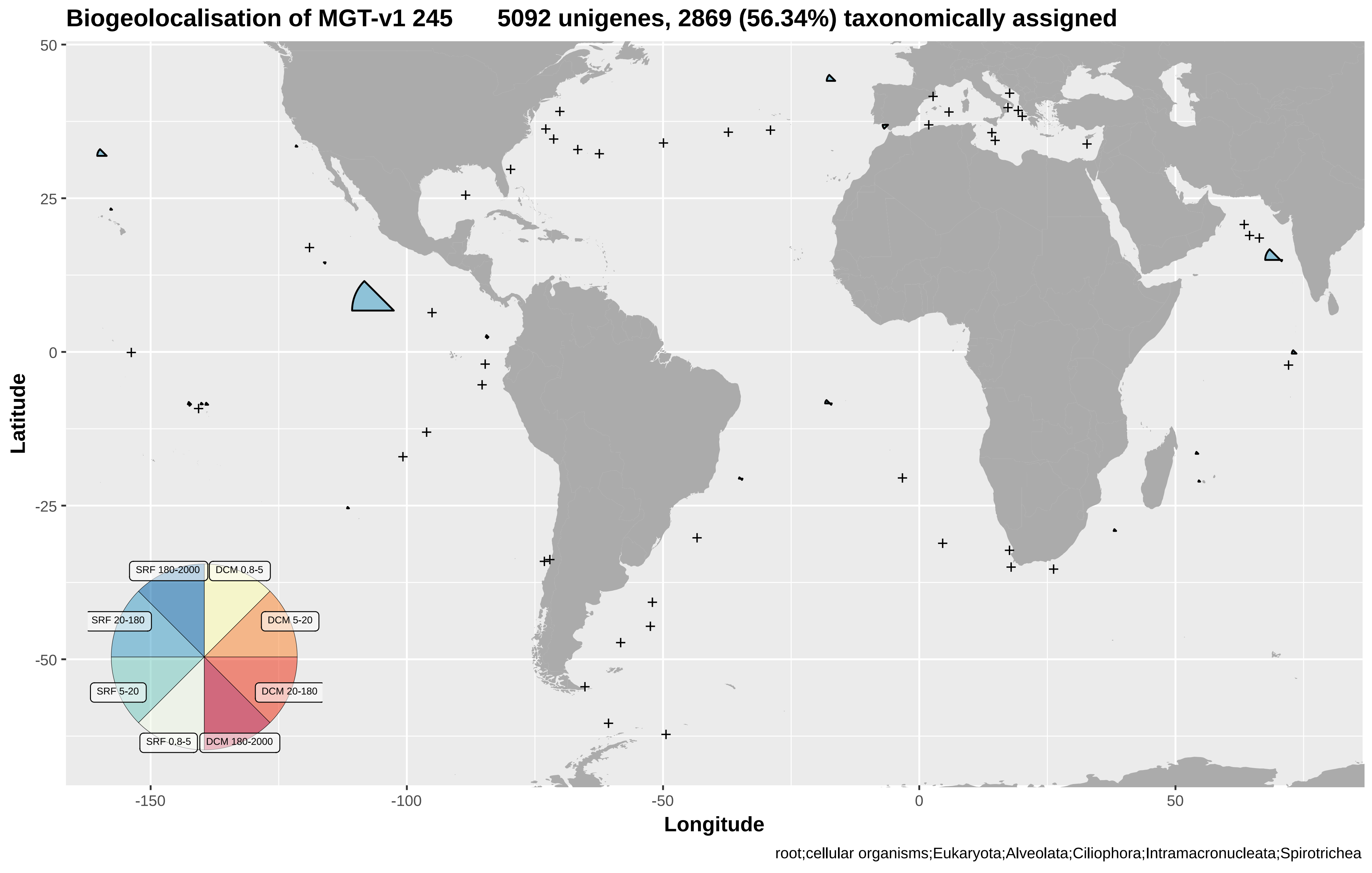

Biogeolocalisation of MGT-v1 245 5092 unigenes, 2869 (56.34%) taxonomically assigned
50
25
0
-25
-50
Latitude
-150
-100
-50
0
50
Longitude
SRF 180-2000
DCM 0.8-5
SRF 20-180
DCM 5-20
SRF 5-20
DCM 20-180
SRF 0.8-5
DCM 180-2000
root;cellular organisms;Eukaryota;Alveolata;Ciliophora;Intramacronucleata;Spirotrichea

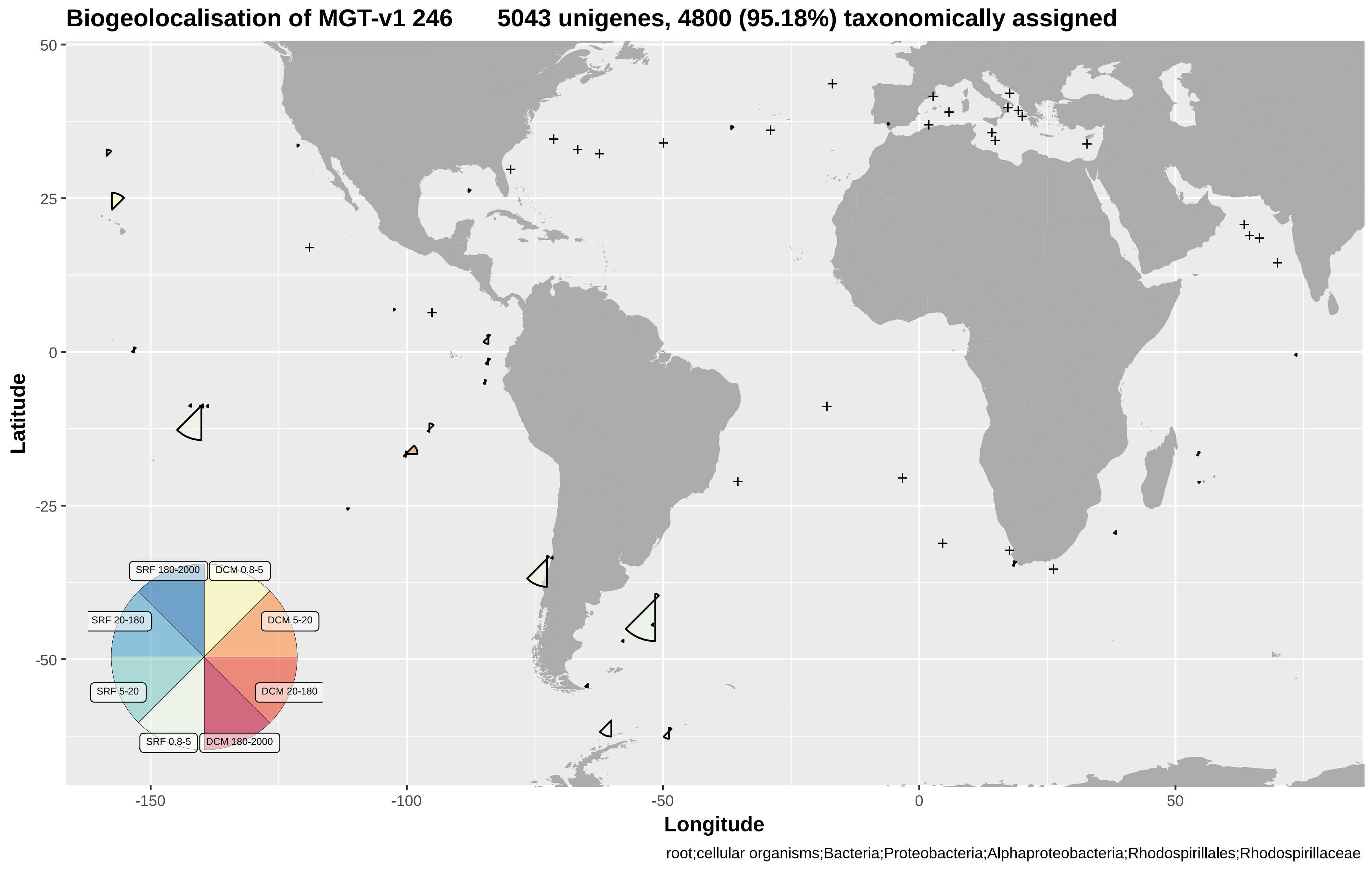

Biogeolocalisation of MGT-v1 246 5043 unigenes, 4800 (95.18%) taxonomically assigned
50
25
0
-25
-50
Latitude
-150
-100
-50
0
50
Longitude
SRF 180-2000
DCM 0.8-5
SRF 20-180
DCM 5-20
SRF 5-20
DCM 20-180
SRF 0.8-5
DCM 180-2000
root;cellular organisms;Bacteria;Proteobacteria;Alphaproteobacteria;Rhodospirillales;Rhodospirillaceae

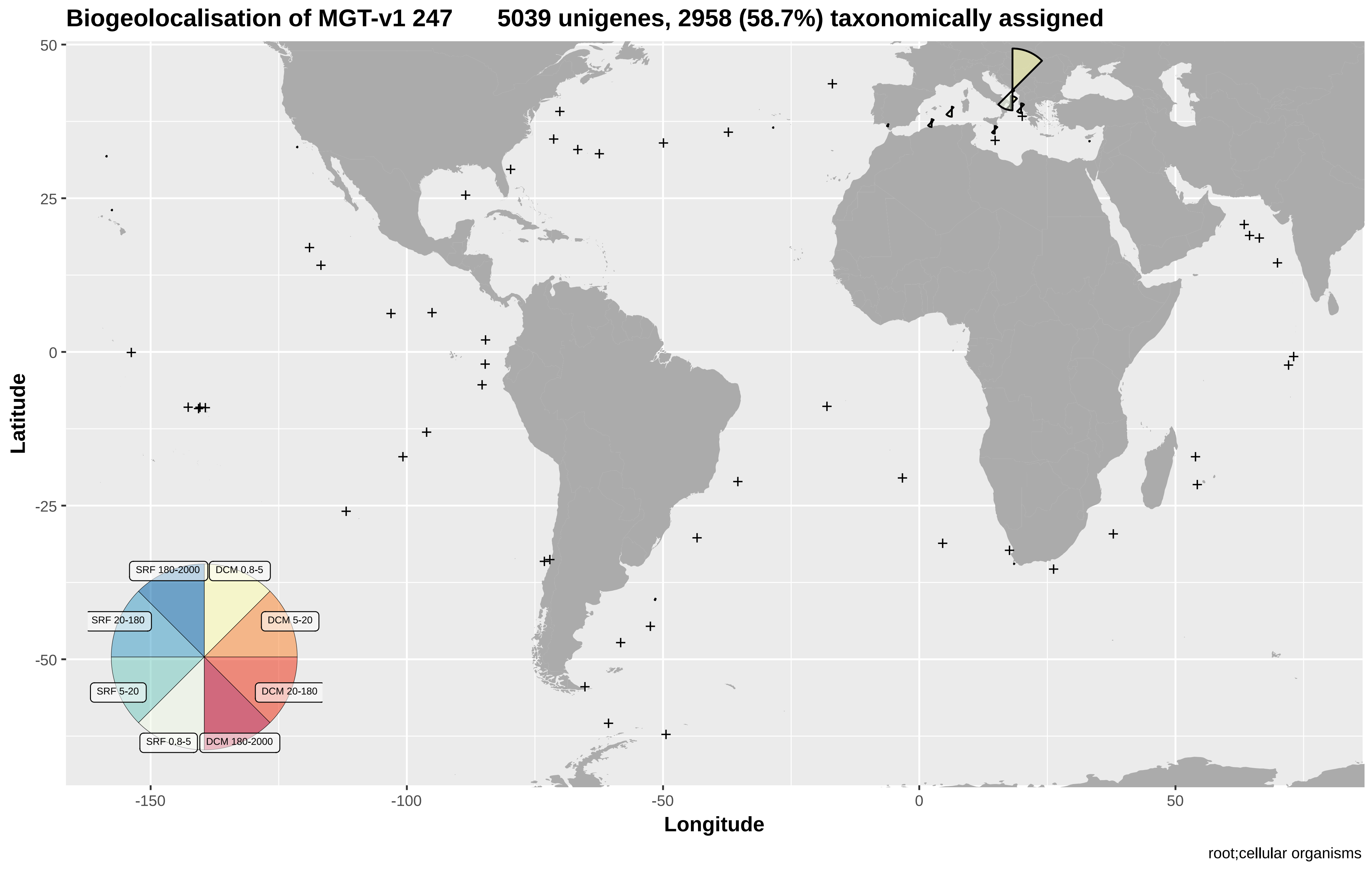
Biogeolocalisation of MGT-v1 247 5039 unigenes, 2958 (58.7%) taxonomically assigned
50
25
0
-25
-50
Latitude
-150
-100
-50
0
50
Longitude
SRF 180-2000
DCM 0.8-5
SRF 20-180
DCM 5-20
SRF 5-20
DCM 20-180
SRF 0.8-5
DCM 180-2000
root;cellular organisms

#### Biogeolocalisation of MGT-v1 248 4981 unigenes, 488 (9.8%) taxonomically assigned

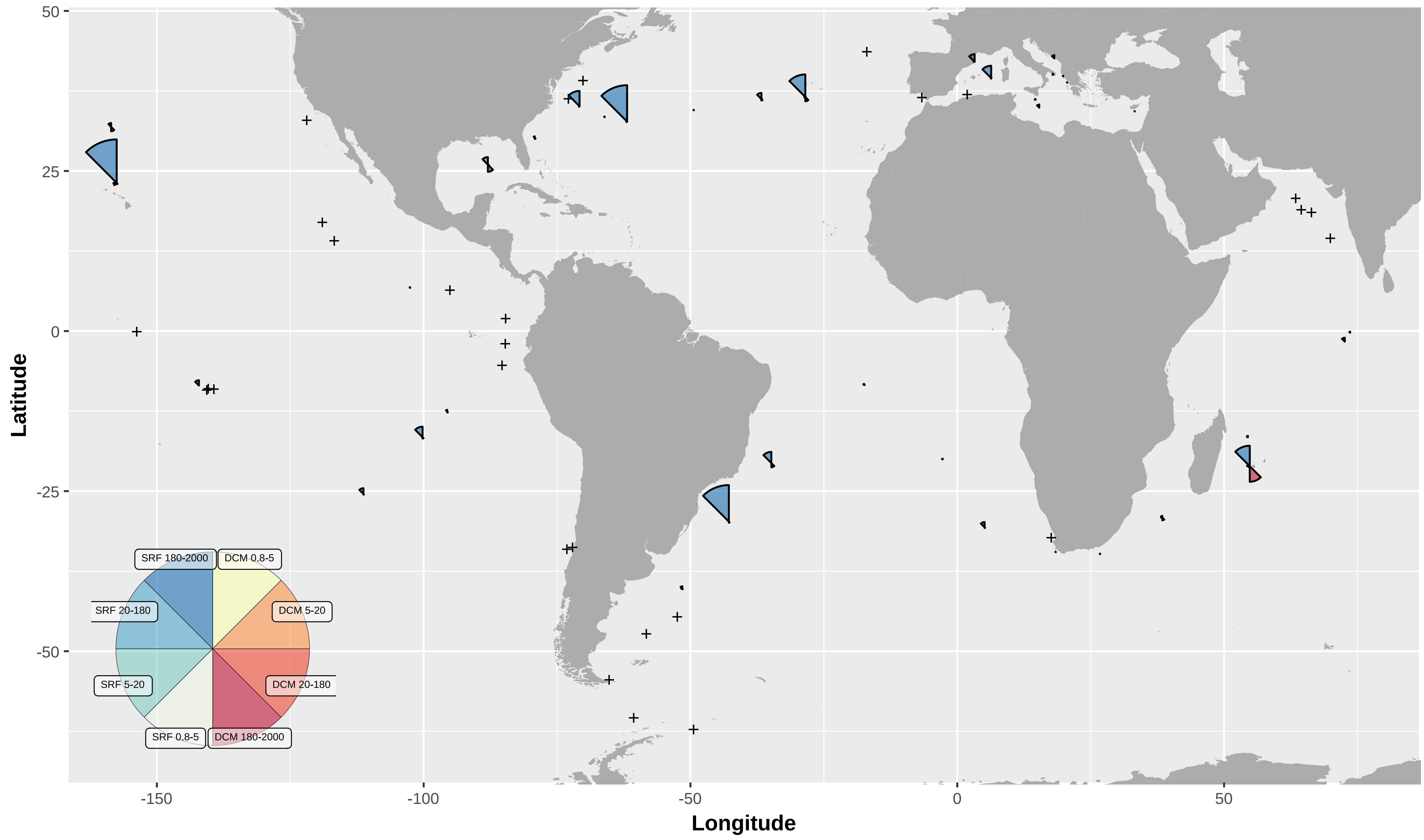


root;cellular organisms;Eukaryota;Opisthokonta

# Biogeolocalisation of MGT-v1 249      4980 unigenes, 3882 (77.95%) taxonomically assigned

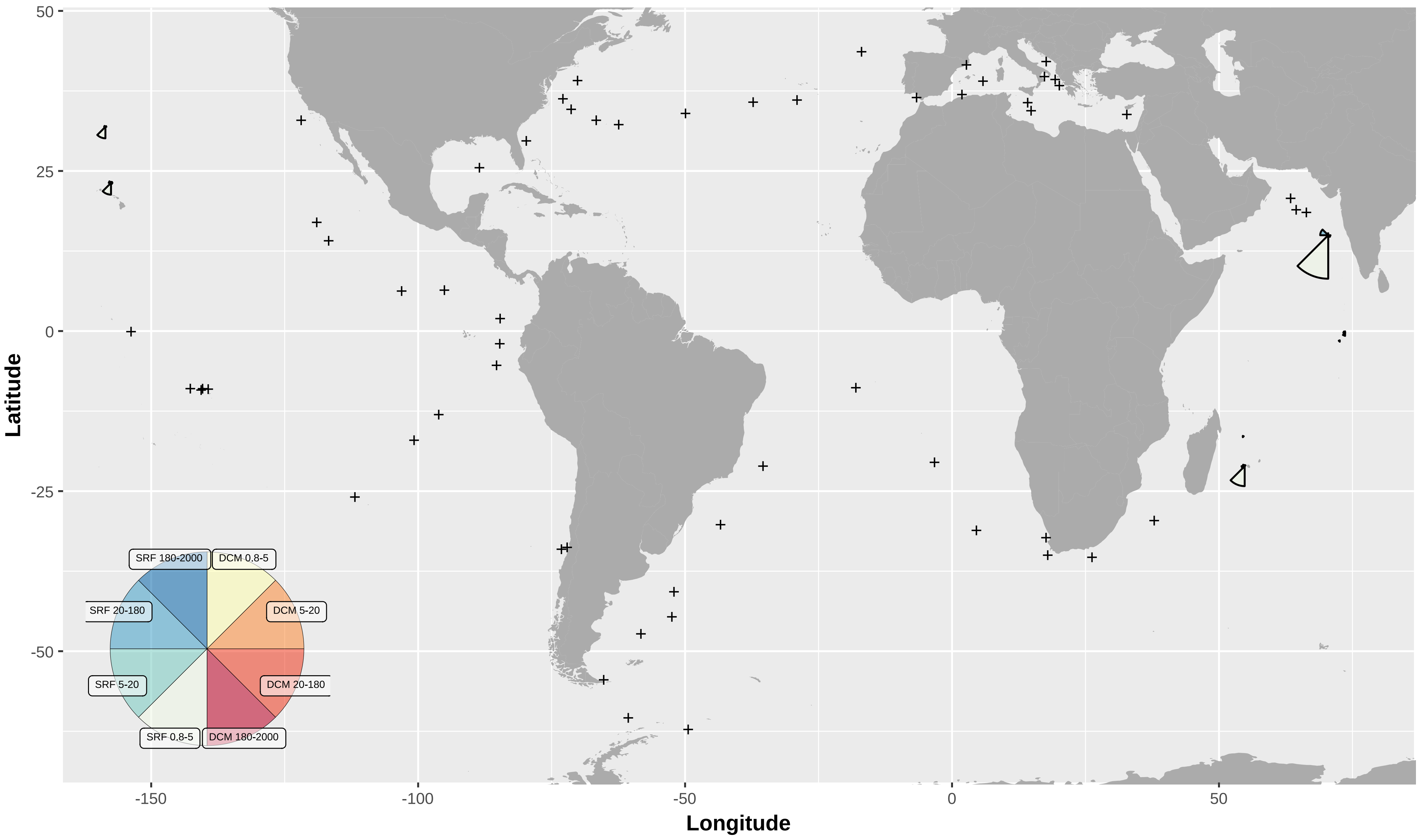


root;cellular organisms;Bacteria

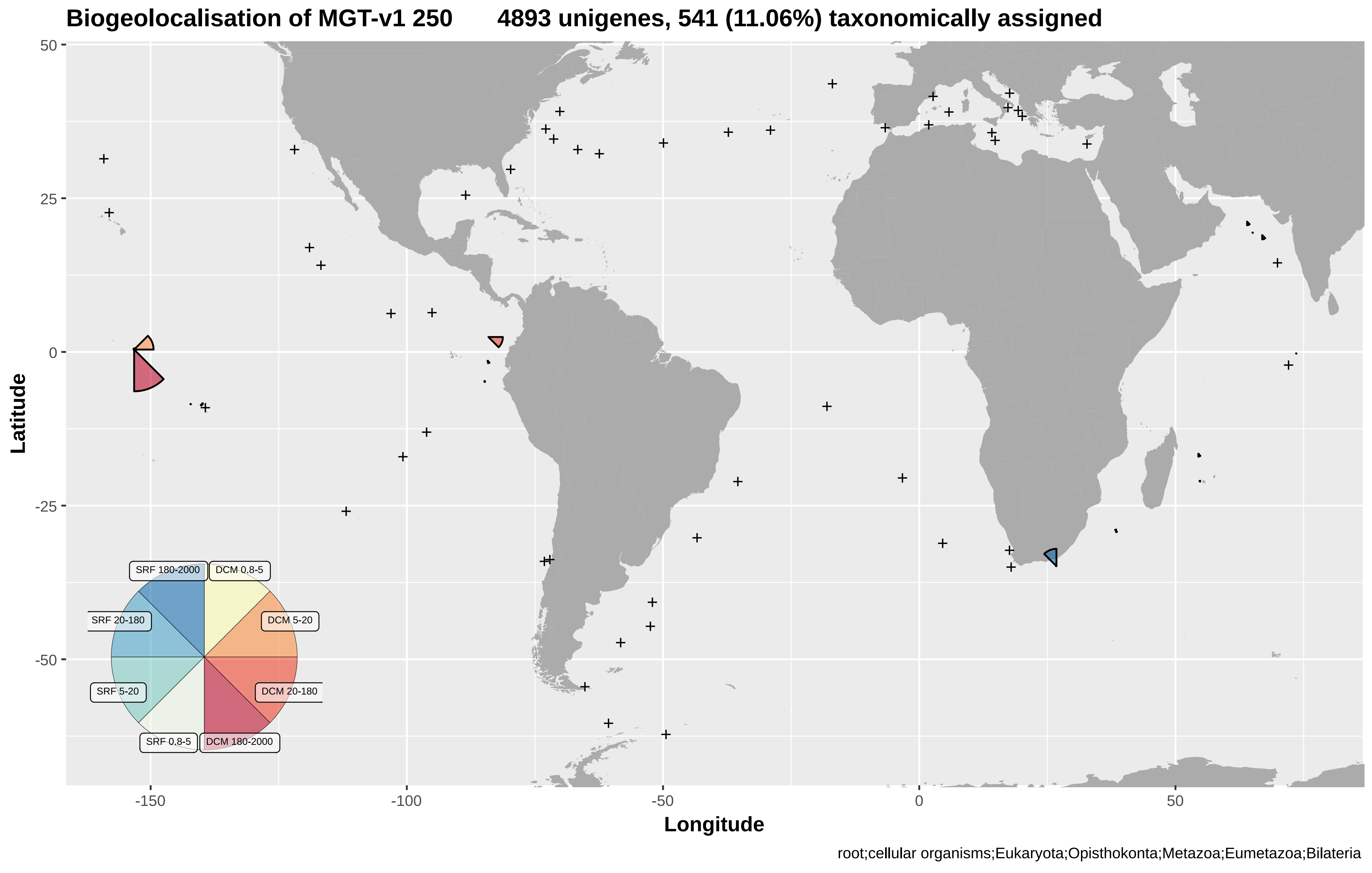
Biogeolocalisation of MGT-v1 250 4893 unigenes, 541 (11.06%) taxonomically assigned
50
25
0
-25
-50
Latitude
-150
-100
-50
0
50
Longitude
SRF 180-2000
DCM 0.8-5
SRF 20-180
DCM 5-20
SRF 5-20
DCM 20-180
SRF 0.8-5
DCM 180-2000
root;cellular organisms;Eukaryota;Opisthokonta;Metazoa;Eumetazoa;Bilateria

### Biogeolocalisation of MGT-v1 251 4889 unigenes, 866 (17.71%) taxonomically assigned

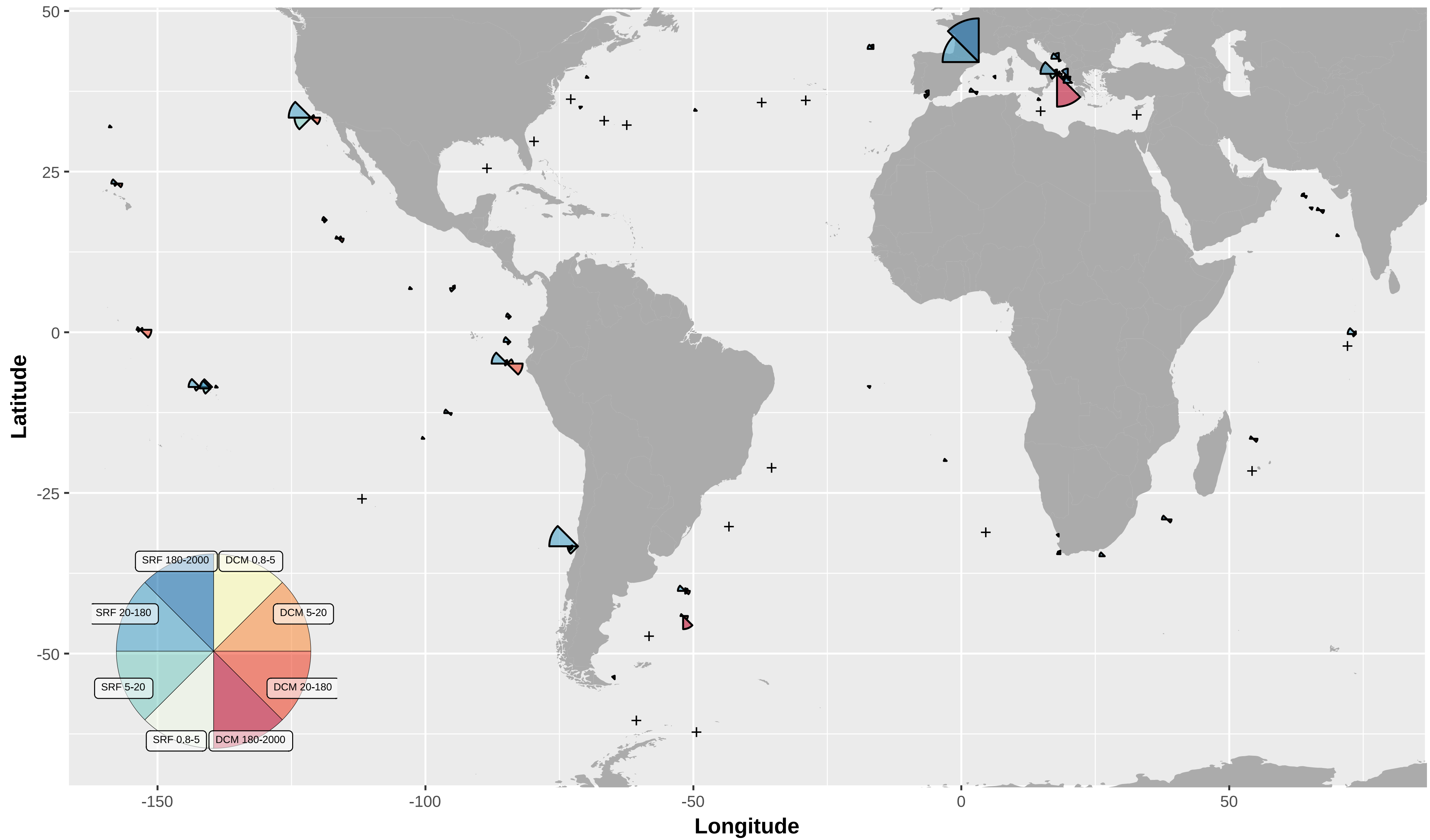


root;cellular organisms;Eukaryota;Opisthokonta;Metazoa;Eumetazoa;Bilateria;Protostomia;Ecdysozoa;Panarthropoda;Arthropoda;Mandibulata;Pancrustacea;Crustacea;Maxillopoda

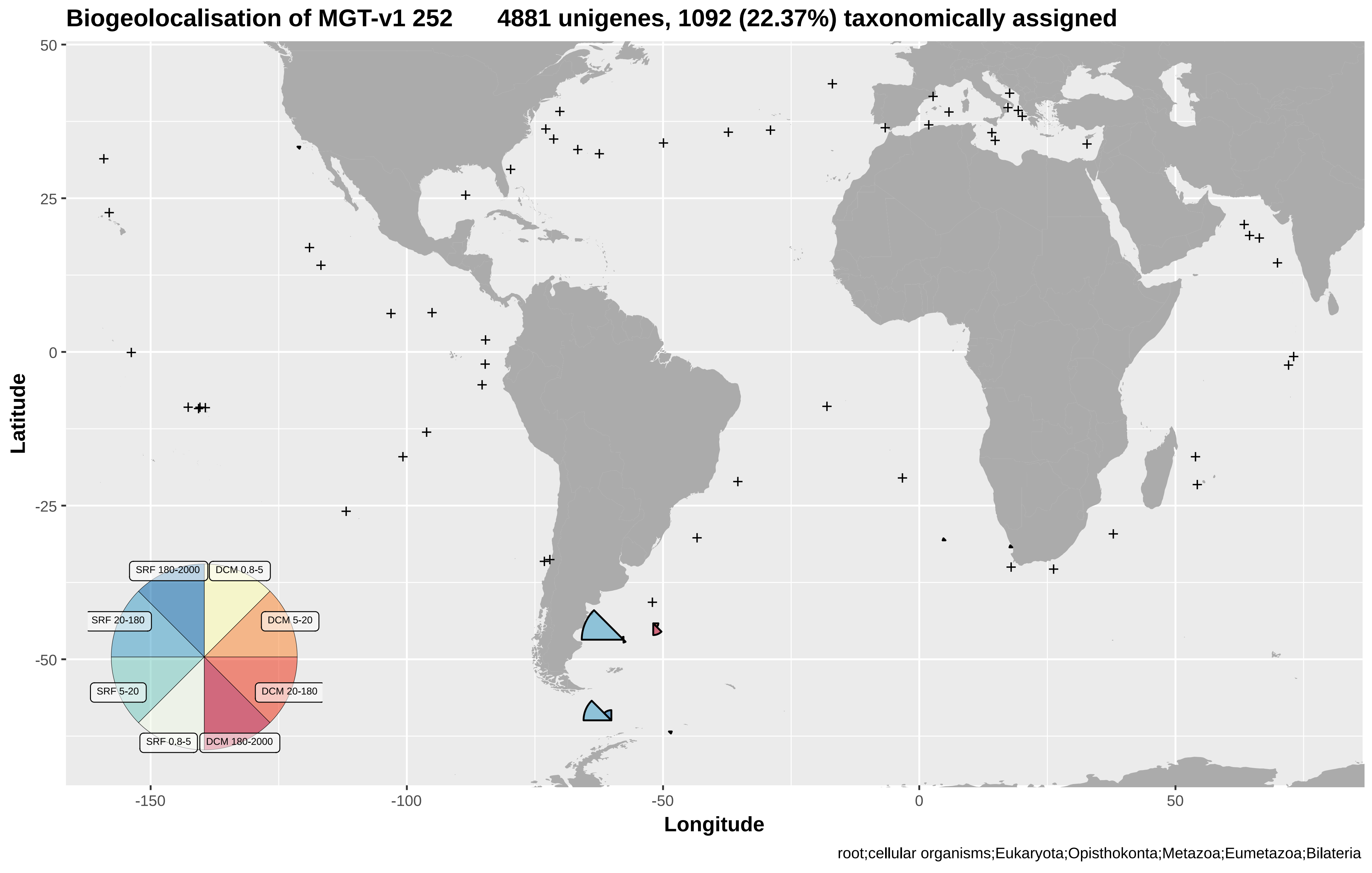

Biogeolocalisation of MGT-v1 252 4881 unigenes, 1092 (22.37%) taxonomically assigned
50
25
0
-25
-50
Latitude
-150
-100
-50
0
50
Longitude
SRF 180-2000
DCM 0.8-5
SRF 20-180
DCM 5-20
SRF 5-20
DCM 20-180
SRF 0.8-5
DCM 180-2000
root;cellular organisms;Eukaryota;Opisthokonta;Metazoa;Eumetazoa;Bilateria

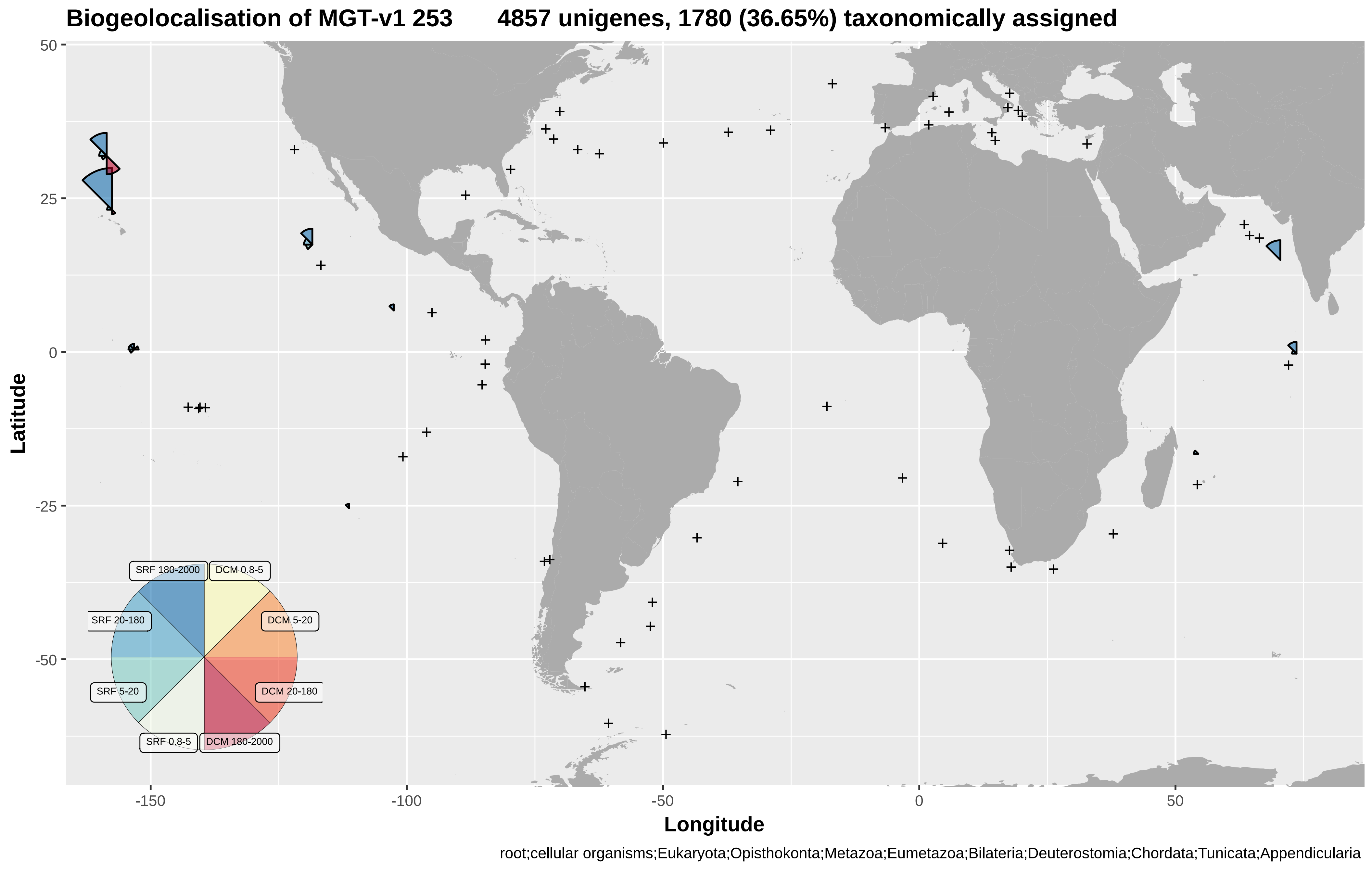

Biogeolocalisation of MGT-v1 253 4857 unigenes, 1780 (36.65%) taxonomically assigned
50
25
0
-25
-50
Latitude
-150
-100
-50
0
50
Longitude
SRF 180-2000
DCM 0.8-5
SRF 20-180
DCM 5-20
SRF 5-20
DCM 20-180
SRF 0.8-5
DCM 180-2000
root;cellular organisms;Eukaryota;Opisthokonta;Metazoa;Eumetazoa;Bilateria;Deuterostomia;Chordata;Tunicata;Appendicularia

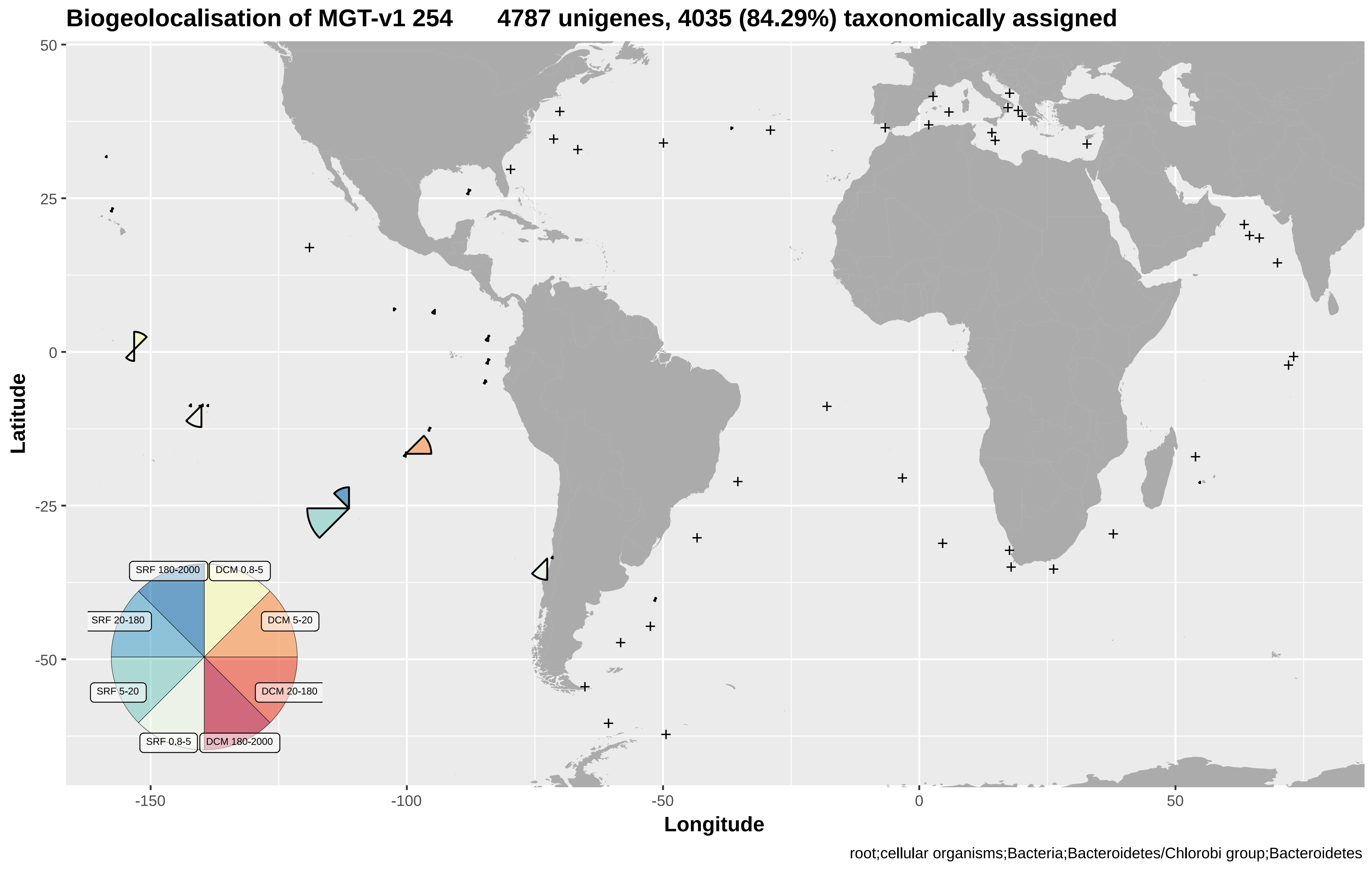

Biogeolocalisation of MGT-v1 254 4787 unigenes, 4035 (84.29%) taxonomically assigned
50
25
0
-25
-50
Latitude
-150
-100
-50
0
50
Longitude
SRF 180-2000
DCM 0.8-5
SRF 20-180
DCM 5-20
SRF 5-20
DCM 20-180
SRF 0.8-5
DCM 180-2000
root;cellular organisms;Bacteria;Bacteroidetes/Chlorobi group;Bacteroidetes

Biogeolocalisation of MGT-v1 255 4743 unigenes, 962 (20.28%) taxonomically assigned

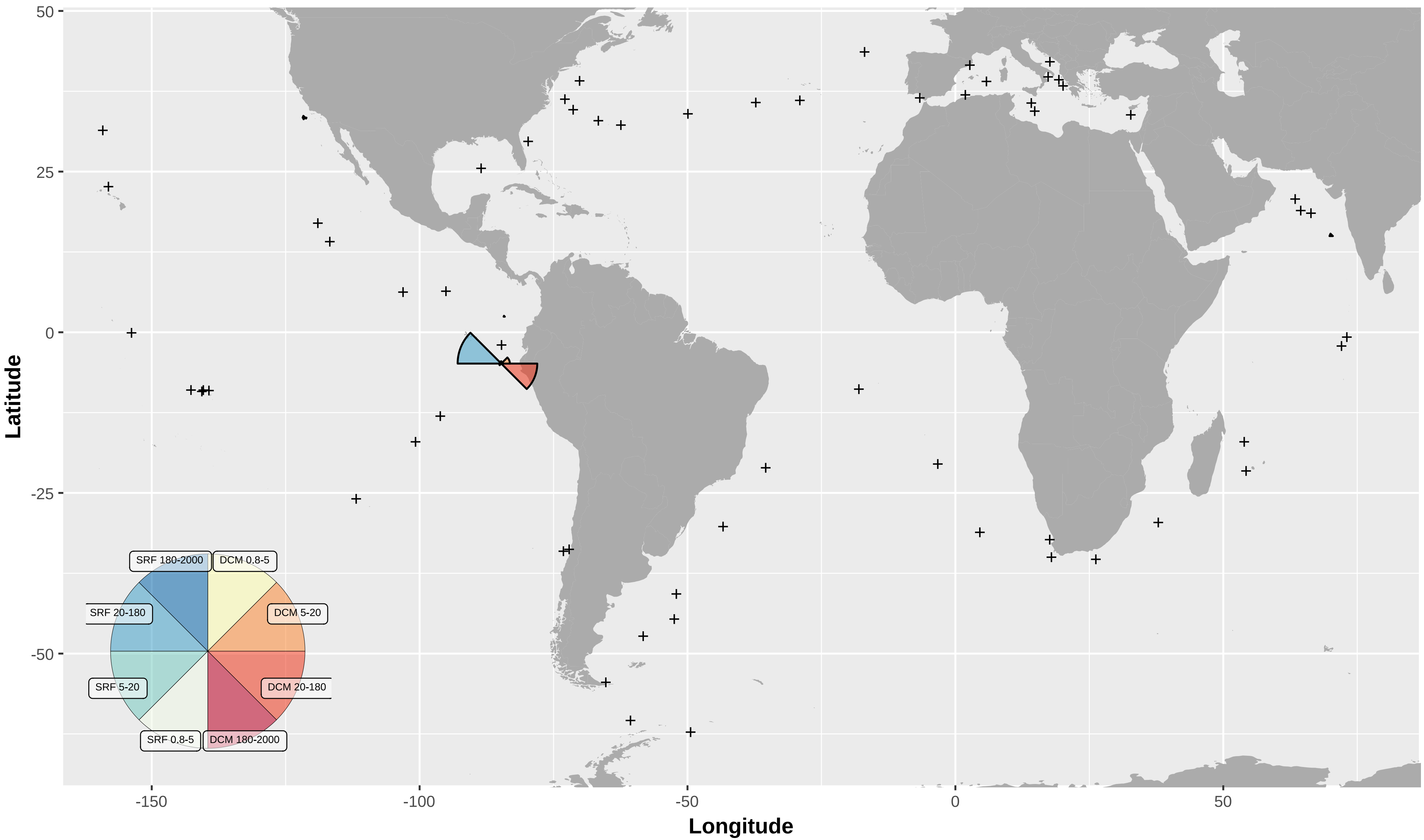


root;cellular organisms;Eukaryota;Opisthokonta;Metazoa;Eumetazoa;Bilateria;Protostomia;Ecdysozoa;Panarthropoda;Arthropoda;Mandibulata;Pancrustacea;Crustacea;Maxillopoda

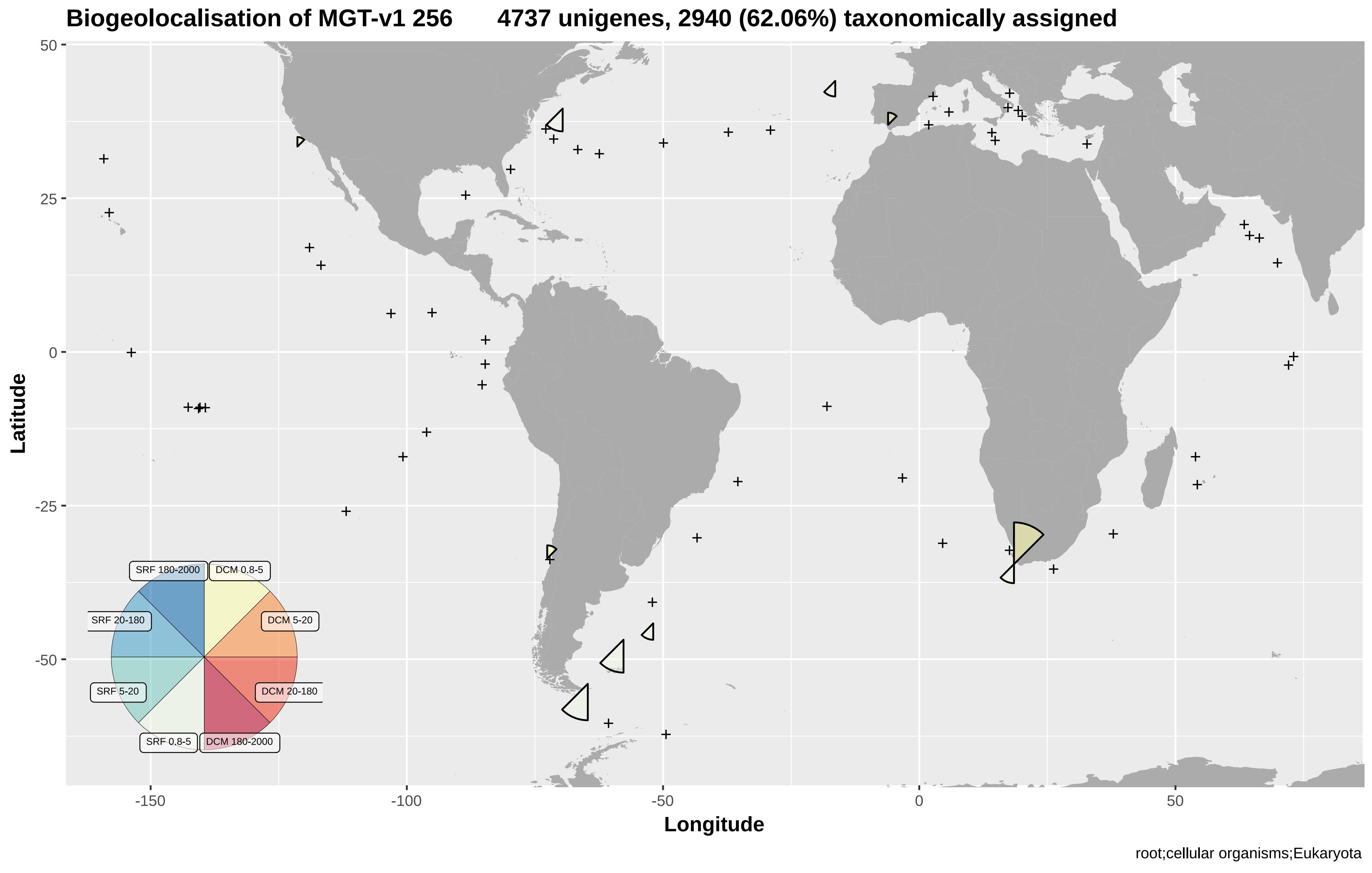

Biogeolocalisation of MGT-v1 256 4737 unigenes, 2940 (62.06%) taxonomically assigned
50
25
0
-25
-50
Latitude
-150
-100
-50
0
50
Longitude
SRF 180-2000
DCM 0.8-5
SRF 20-180
DCM 5-20
SRF 5-20
DCM 20-180
SRF 0.8-5
DCM 180-2000
root;cellular organisms;Eukaryota

## Biogeolocalisation of MGT-v1 257      4729 unigenes, 2839 (60.03%) taxonomically assigned

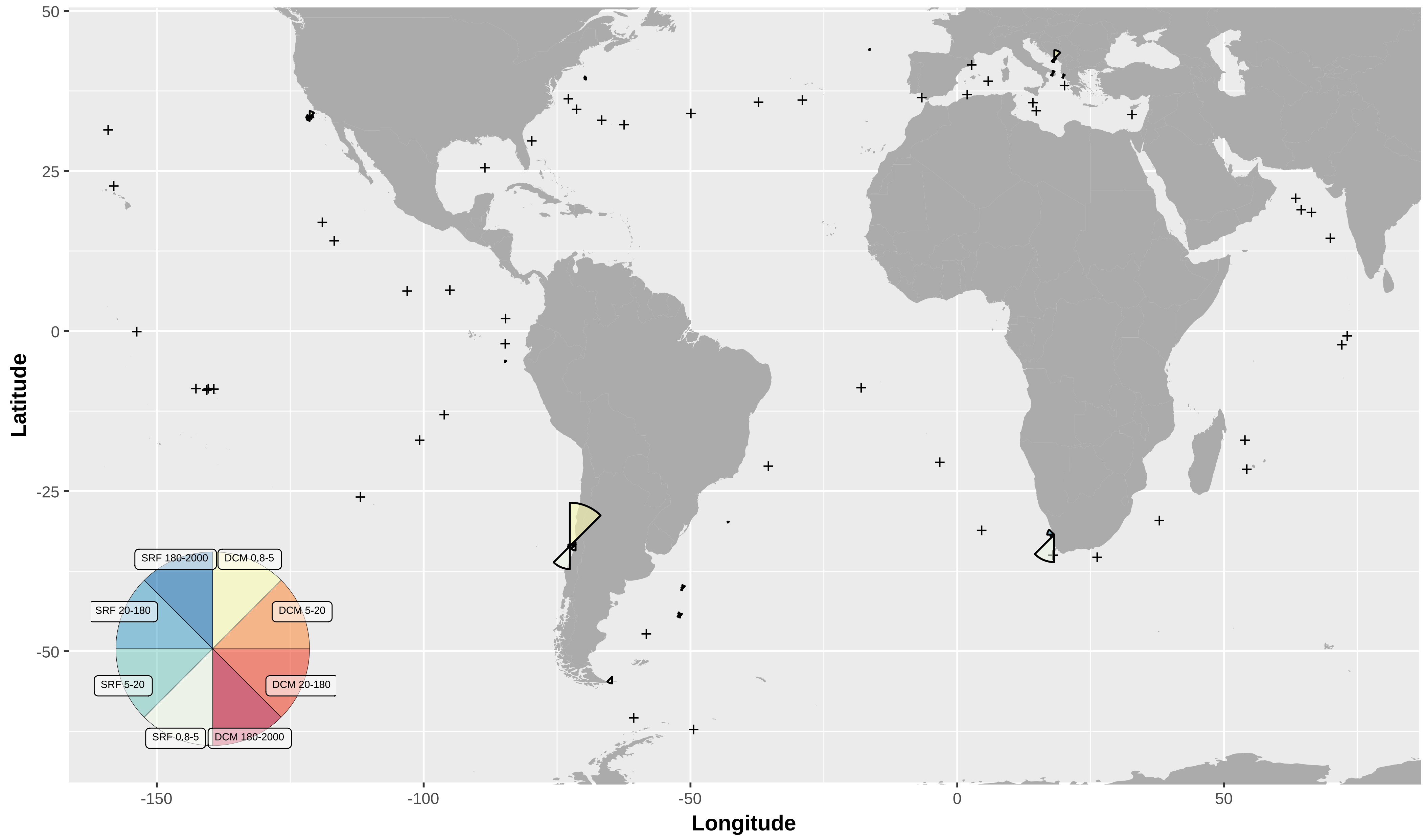


root;cellular organisms

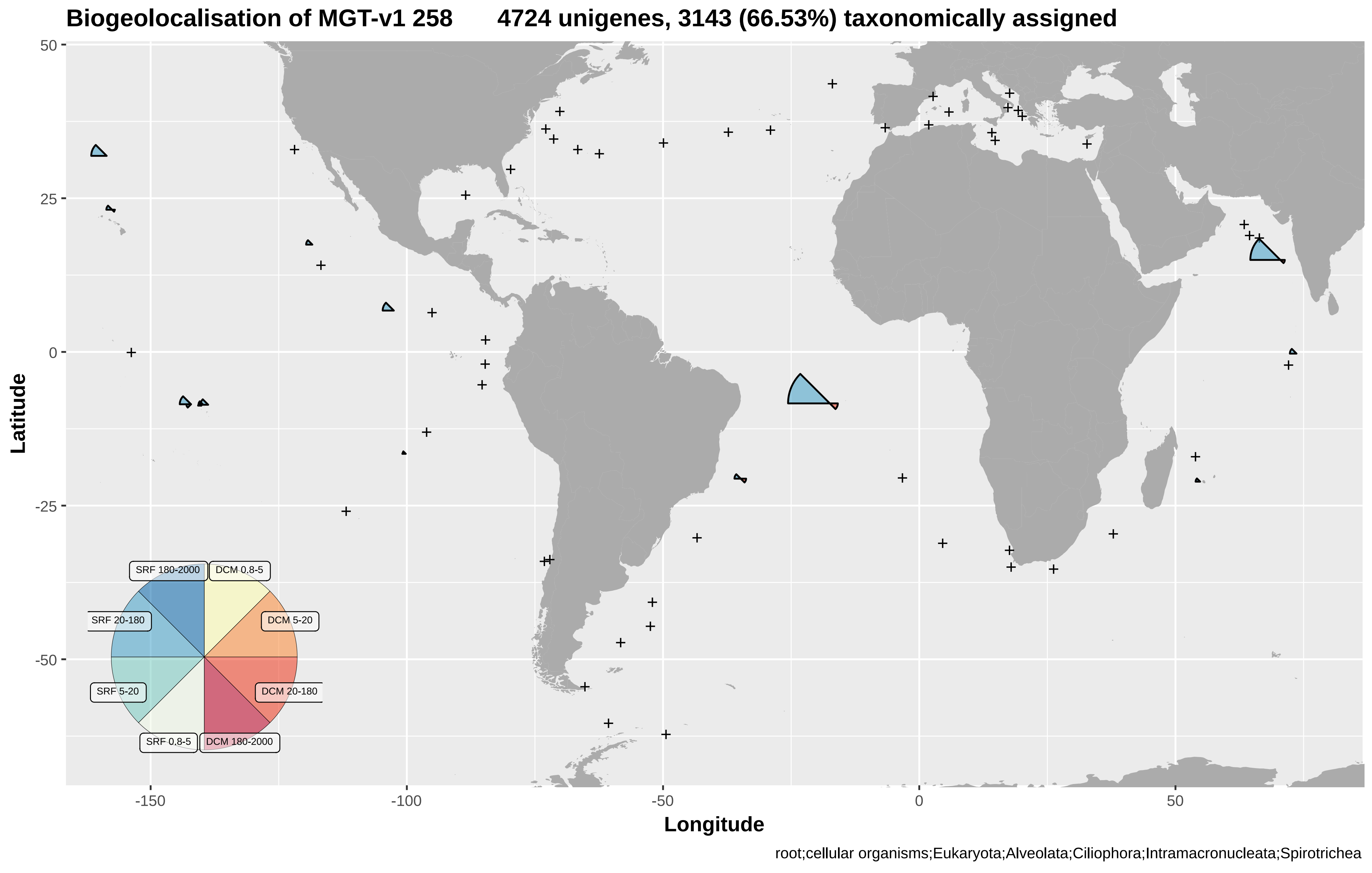

Biogeolocalisation of MGT-v1 258 4724 unigenes, 3143 (66.53%) taxonomically assigned
Latitude
Longitude
50
25
0
-25
-50
-150
-100
-50
0
50
SRF 180-2000
DCM 0.8-5
SRF 20-180
DCM 5-20
SRF 5-20
DCM 20-180
SRF 0.8-5
DCM 180-2000
root;cellular organisms;Eukaryota;Alveolata;Ciliophora;Intramacronucleata;Spirotrichea

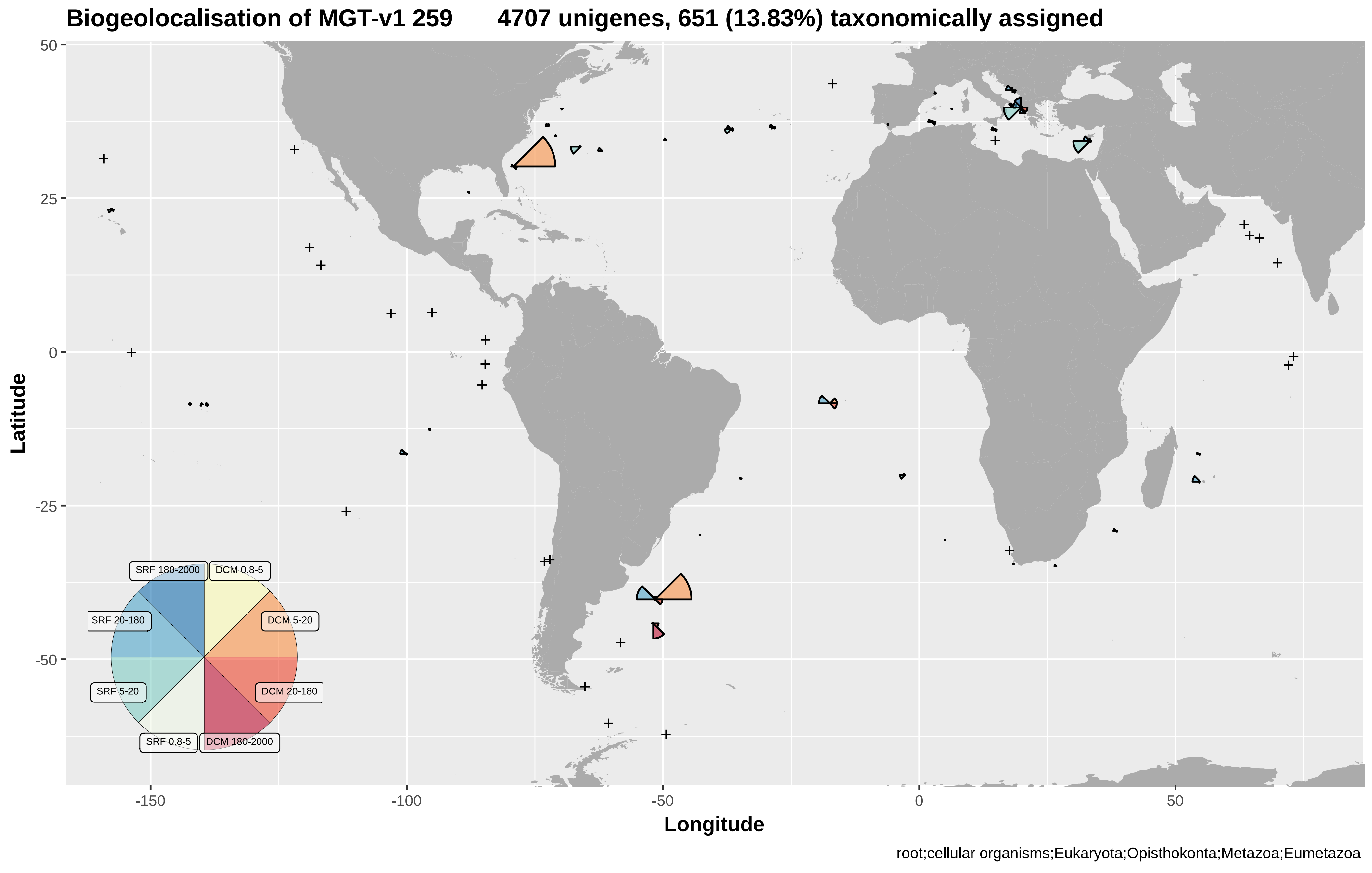
Biogeolocalisation of MGT-v1 259
4707 unigenes, 651 (13.83%) taxonomically assigned
Latitude
Longitude
50
25
0
-25
-50
-150
-100
-50
0
50
SRF 180-2000
DCM 0.8-5
SRF 20-180
DCM 5-20
SRF 5-20
DCM 20-180
SRF 0.8-5
DCM 180-2000
root;cellular organisms;Eukaryota;Opisthokonta;Metazoa;Eumetazoa

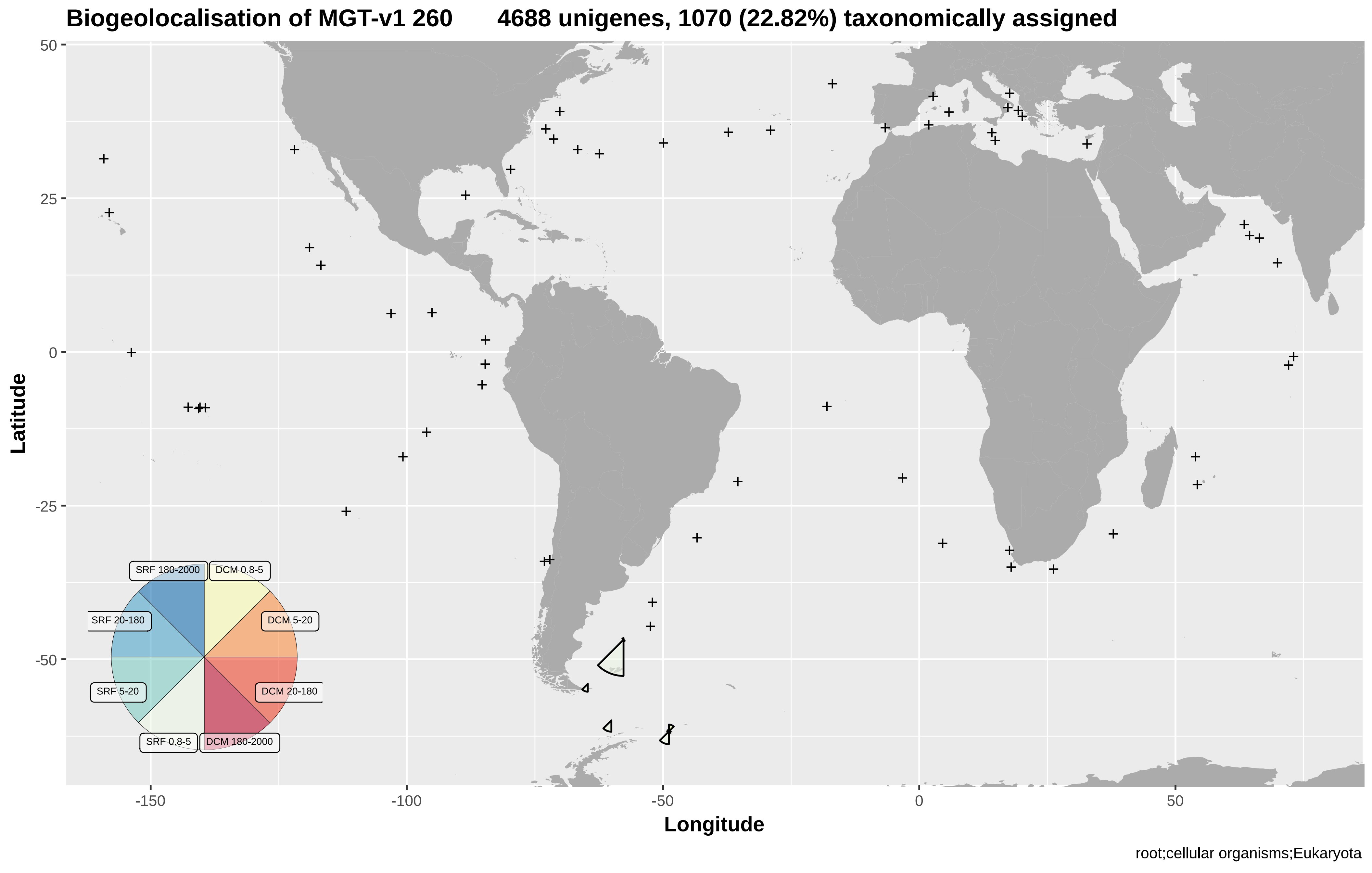
Biogeolocalisation of MGT-v1 260 4688 unigenes, 1070 (22.82%) taxonomically assigned
Latitude
Longitude
50
25
0
-25
-50
-150
-100
-50
0
50
SRF 180-2000
DCM 0.8-5
SRF 20-180
DCM 5-20
SRF 5-20
DCM 20-180
SRF 0.8-5
DCM 180-2000
root;cellular organisms;Eukaryota

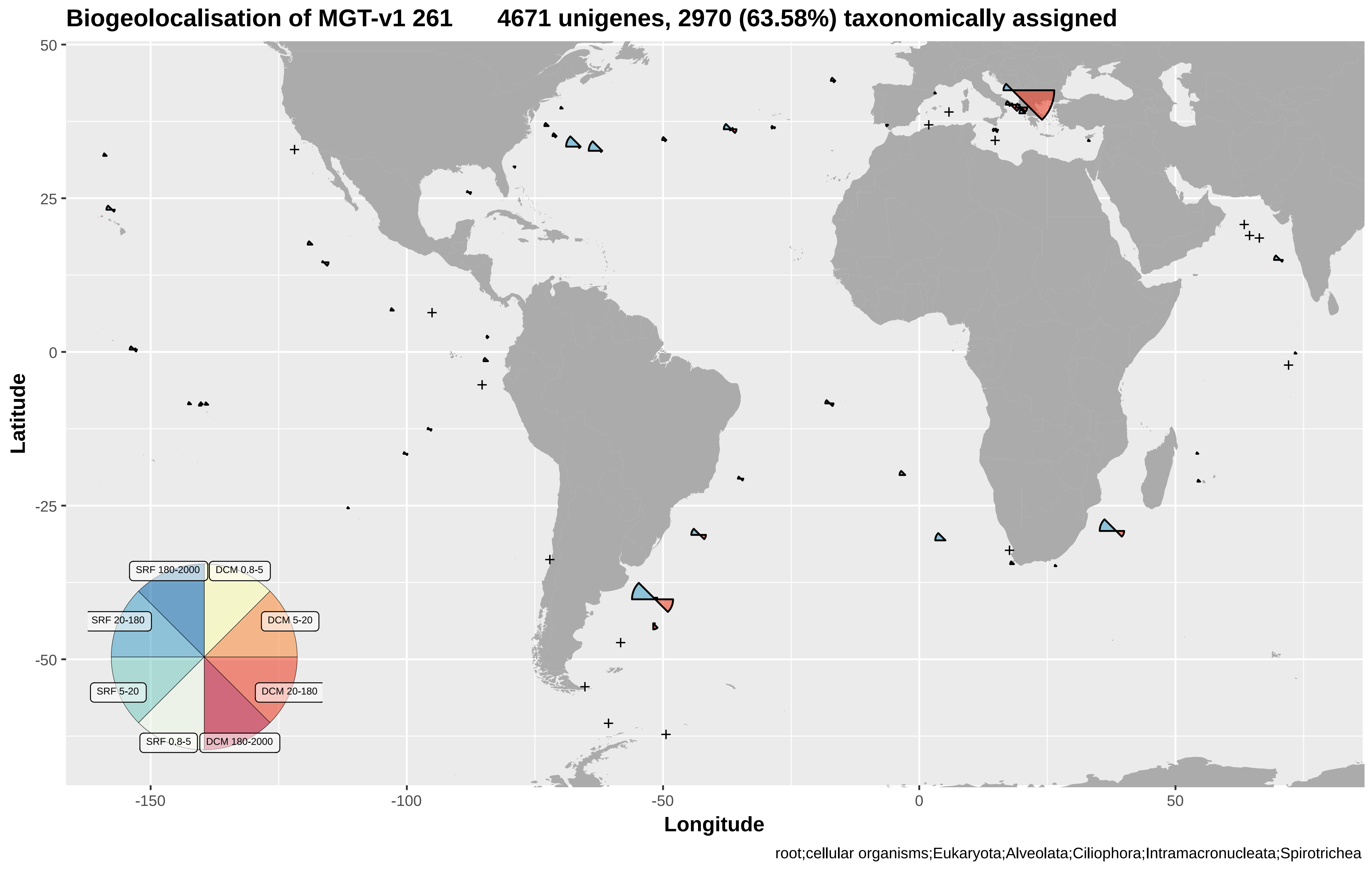

Biogeolocalisation of MGT-v1 261
4671 unigenes, 2970 (63.58%) taxonomically assigned
50
25
0
-25
-50
Latitude
-150
-100
-50
0
50
Longitude
SRF 180-2000
DCM 0.8-5
SRF 20-180
DCM 5-20
SRF 5-20
DCM 20-180
SRF 0.8-5
DCM 180-2000
root;cellular organisms;Eukaryota;Alveolata;Ciliophora;Intramacronucleata;Spirotrichea

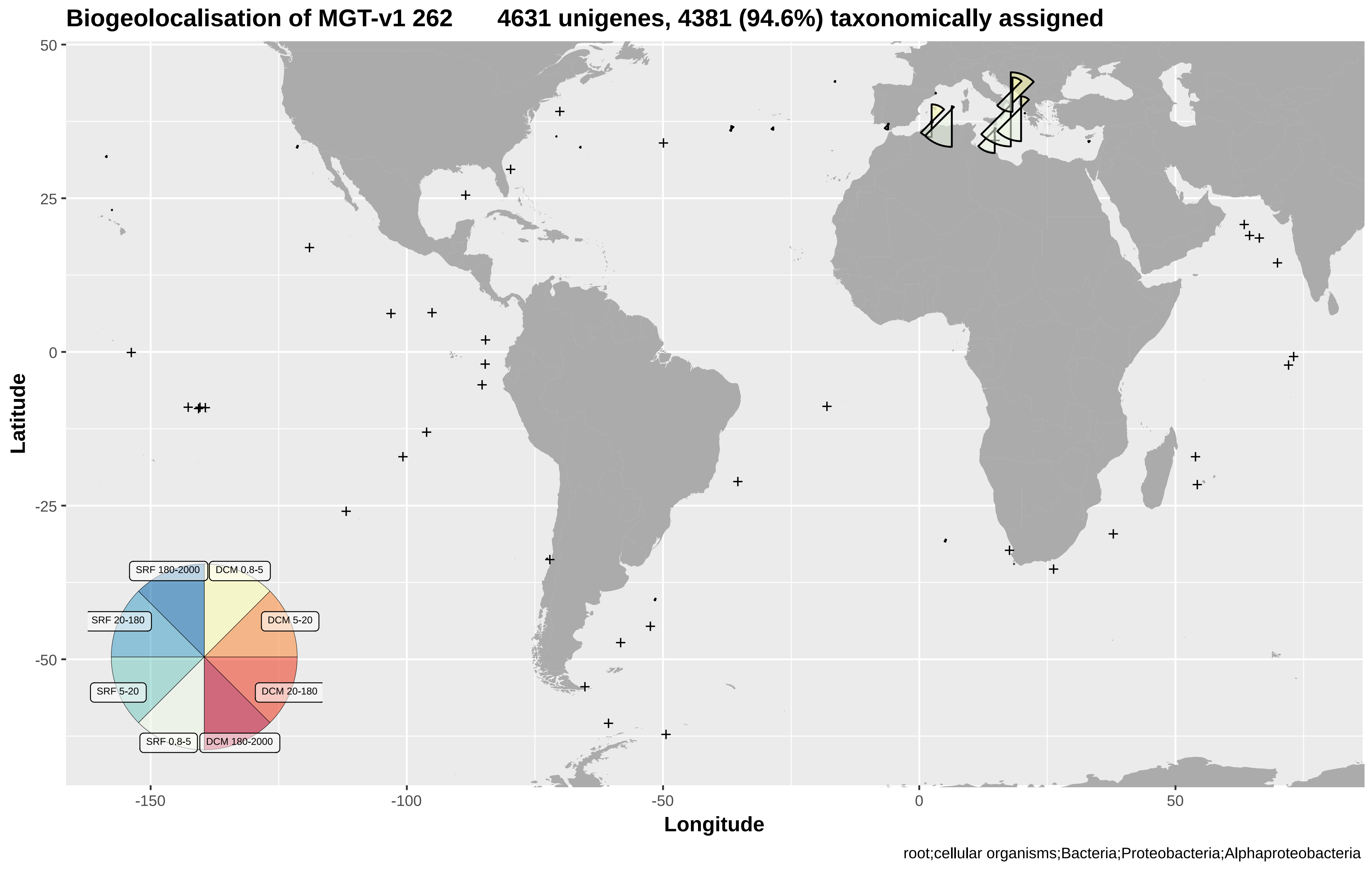

Biogeolocalisation of MGT-v1 262 4631 unigenes, 4381 (94.6%) taxonomically assigned
50
25
0
-25
-50
Latitude
-150
-100
-50
0
50
Longitude
SRF 180-2000
DCM 0.8-5
SRF 20-180
DCM 5-20
SRF 5-20
DCM 20-180
SRF 0.8-5
DCM 180-2000
root;cellular organisms;Bacteria;Proteobacteria;Alphaproteobacteria

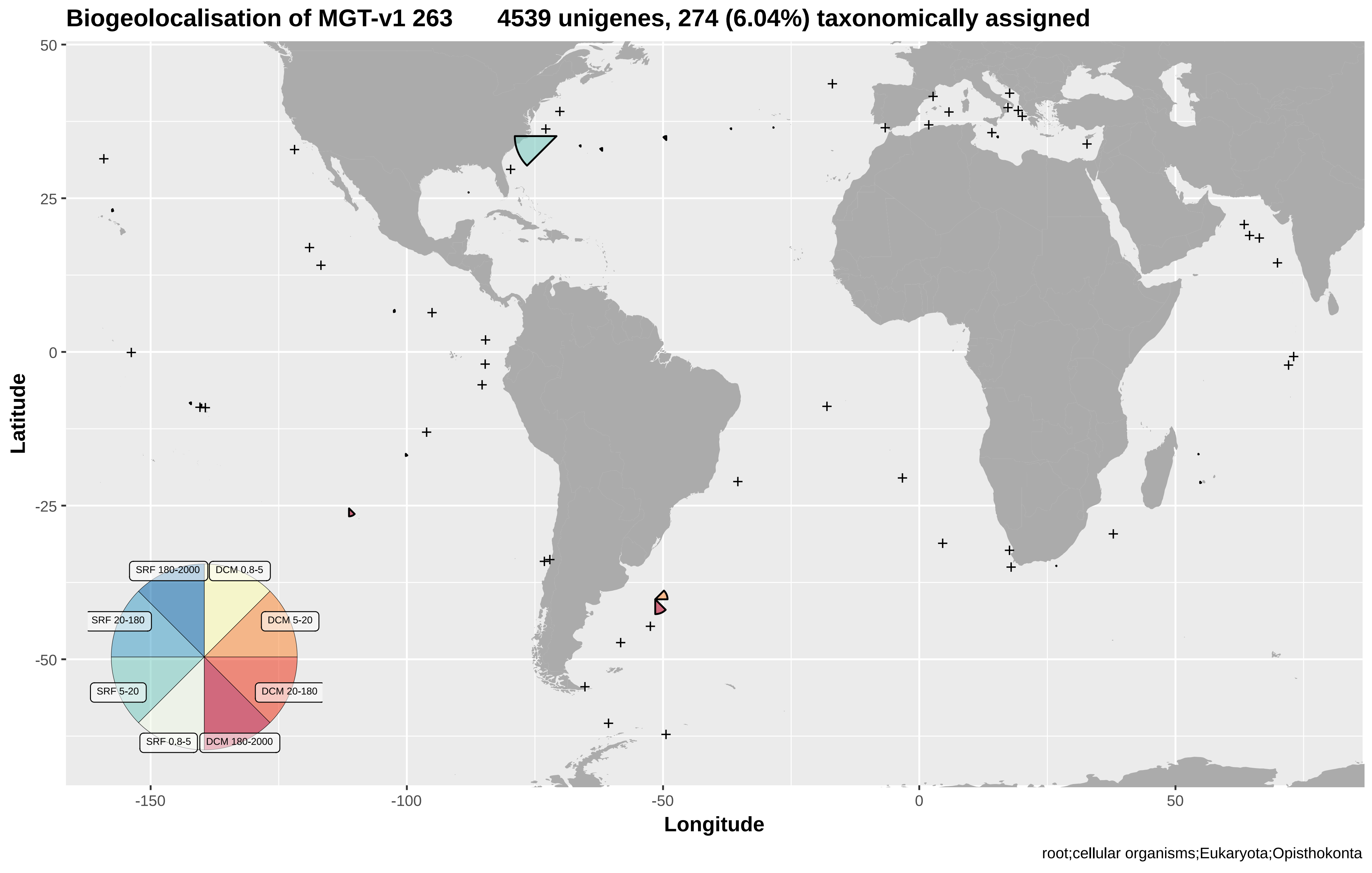
Biogeolocalisation of MGT-v1 263
4539 unigenes, 274 (6.04%) taxonomically assigned
Latitude
Longitude
50
25
0
-25
-50
-150
-100
-50
0
50
SRF 180-2000
DCM 0.8-5
SRF 20-180
DCM 5-20
SRF 5-20
DCM 20-180
SRF 0.8-5
DCM 180-2000
root;cellular organisms;Eukaryota;Opisthokonta

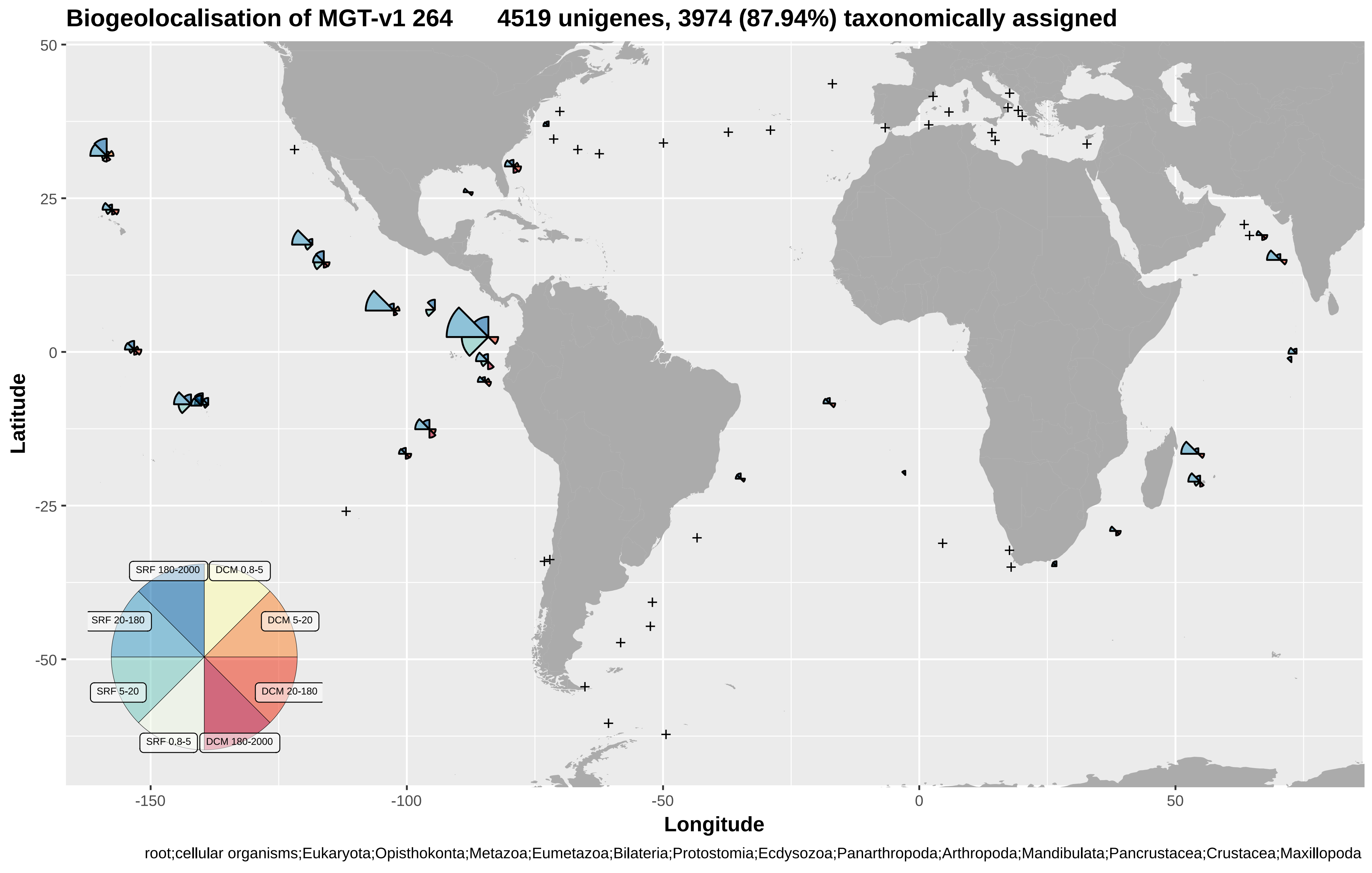

Biogeolocalisation of MGT-v1 264 4519 unigenes, 3974 (87.94%) taxonomically assigned
50
25
0
-25
-50
Latitude
-150
-100
-50
0
50
Longitude
SRF 180-2000
DCM 0.8-5
SRF 20-180
DCM 5-20
SRF 5-20
DCM 20-180
SRF 0.8-5
DCM 180-2000
root;cellular organisms;Eukaryota;Opisthokonta;Metazoa;Eumetazoa;Bilateria;Protostomia;Ecdysozoa;Panarthropoda;Arthropoda;Mandibulata;Pancrustacea;Crustacea;Maxillopoda

## Biogeolocalisation of MGT-v1 265      4491 unigenes, 620 (13.81%) taxonomically assigned

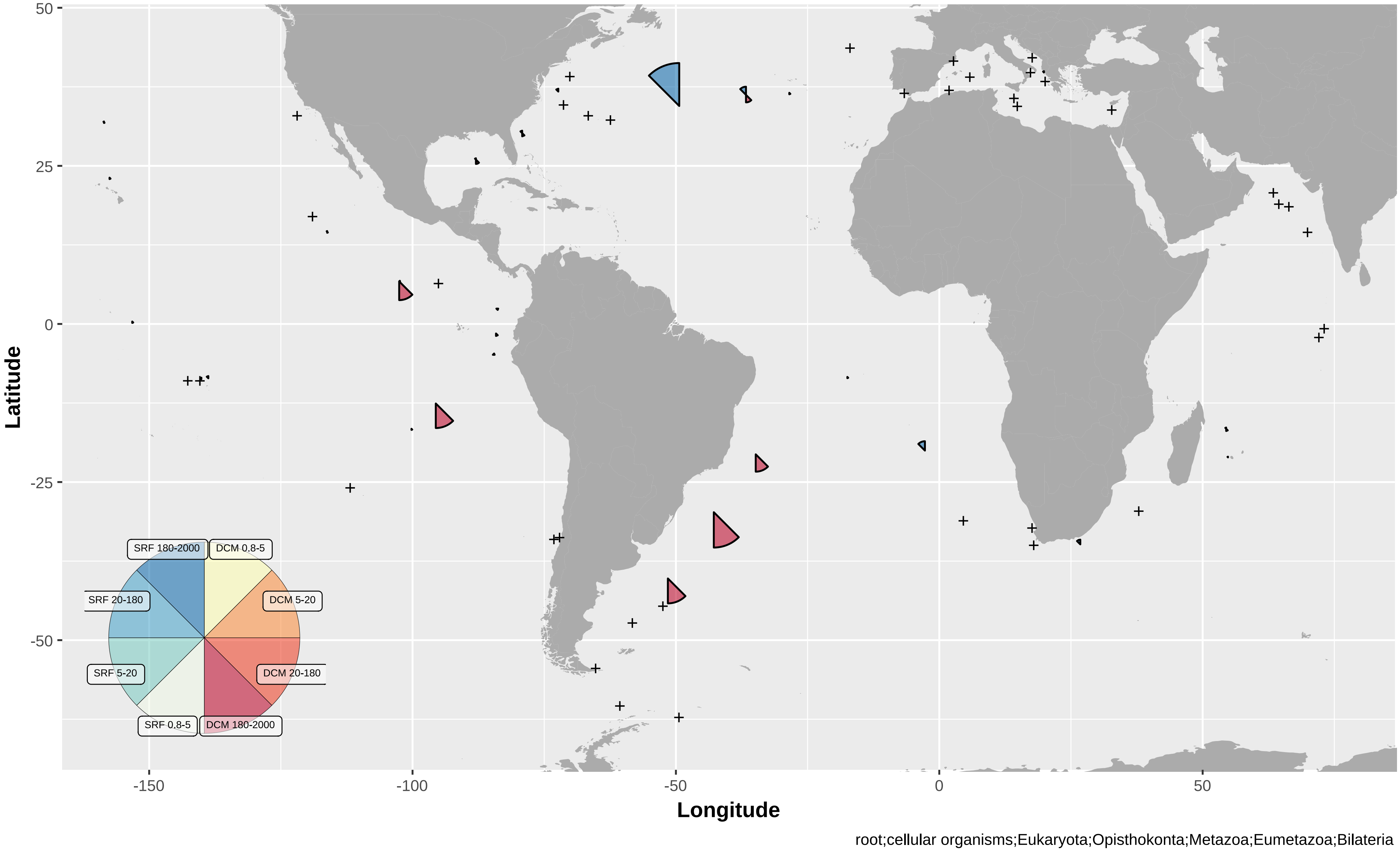

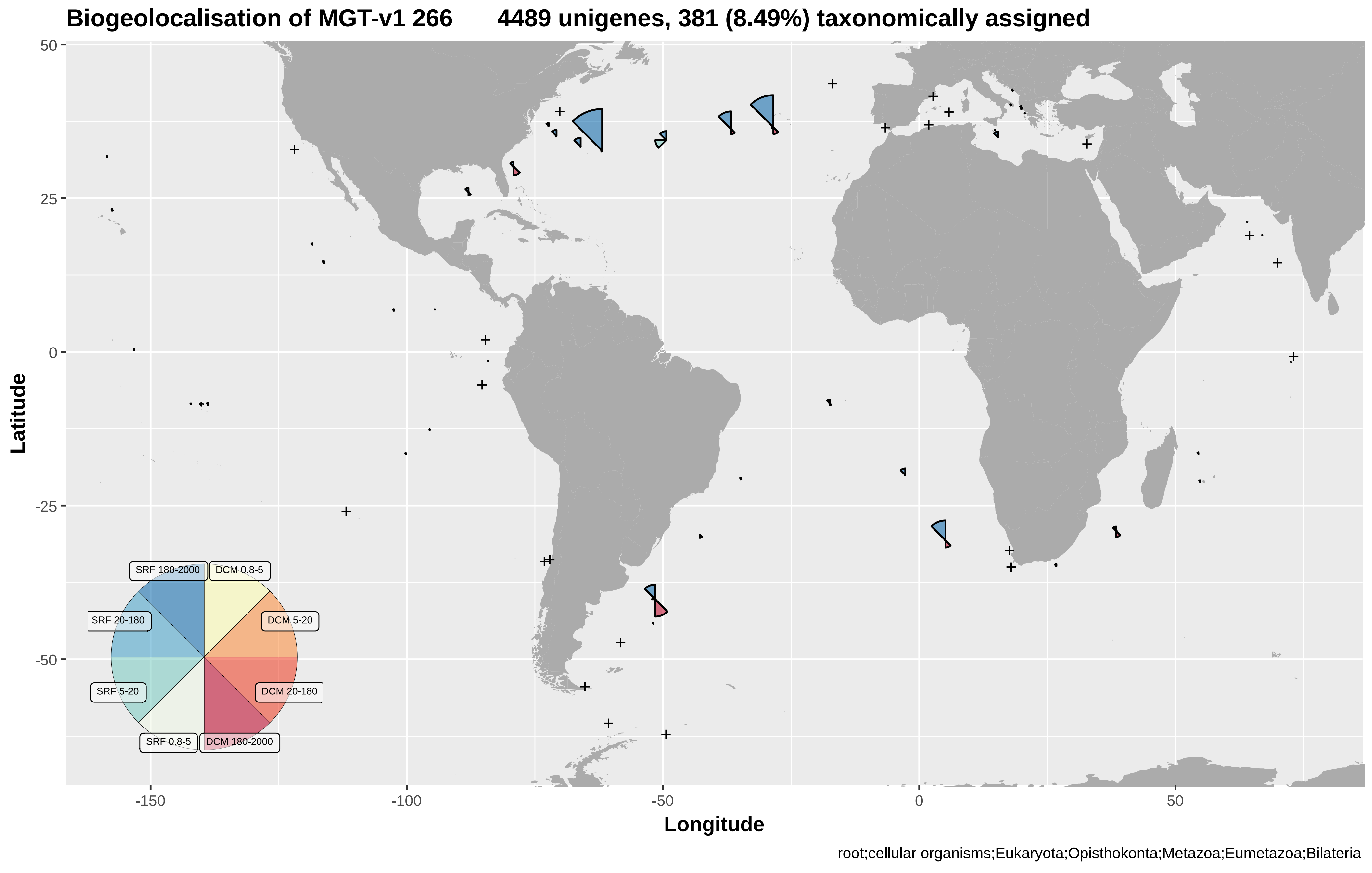
Biogeolocalisation of MGT-v1 266      4489 unigenes, 381 (8.49%) taxonomically assigned
50
25
0
-25
-50
Latitude
-150
-100
-50
0
50
Longitude
SRF 180-2000
DCM 0.8-5
SRF 20-180
DCM 5-20
SRF 5-20
DCM 20-180
SRF 0.8-5
DCM 180-2000
root;cellular organisms;Eukaryota;Opisthokonta;Metazoa;Eumetazoa;Bilateria

# Biogeolocalisation of MGT-v1 267      4470 unigenes, 921 (20.6%) taxonomically assigned

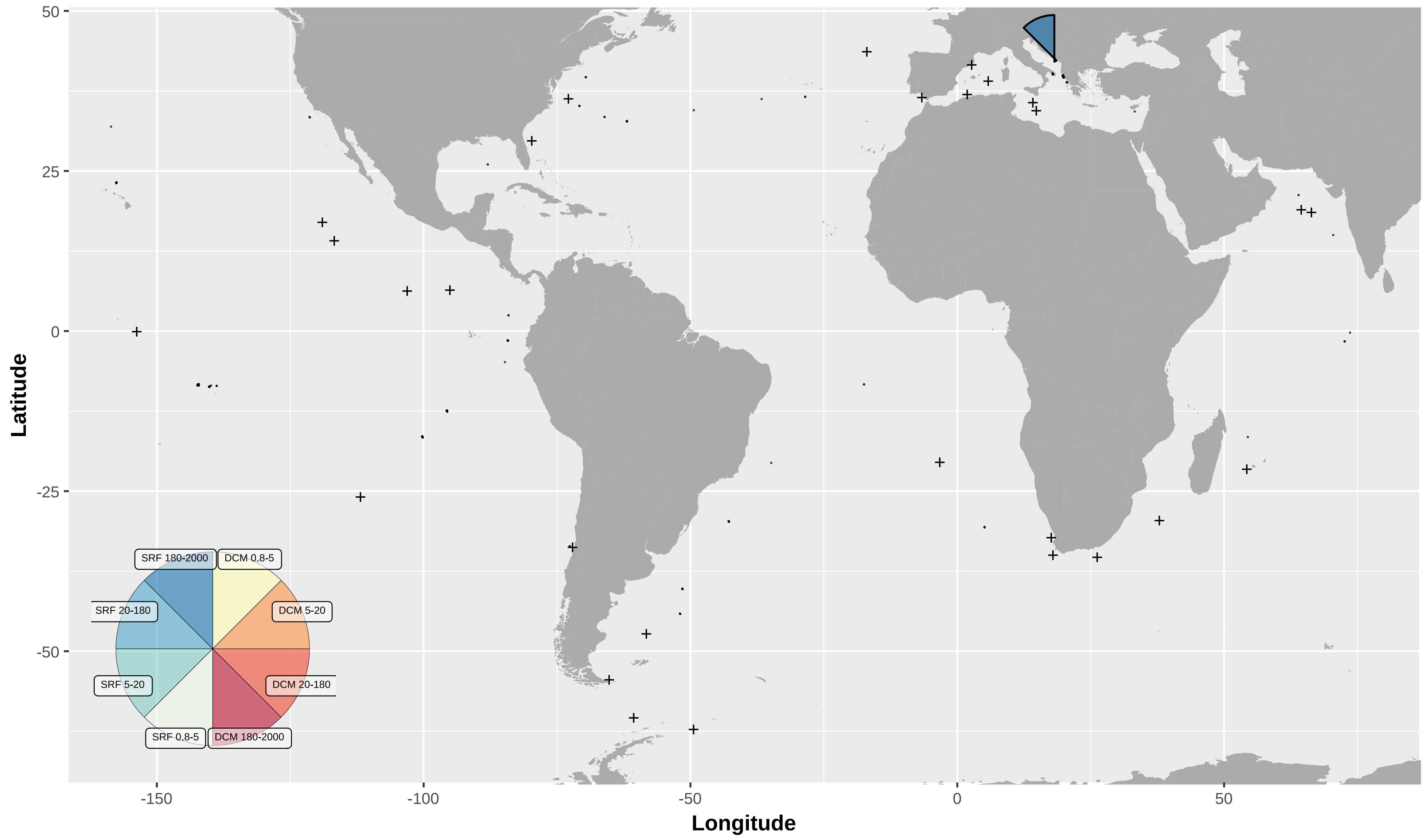


root;cellular organisms;Eukaryota

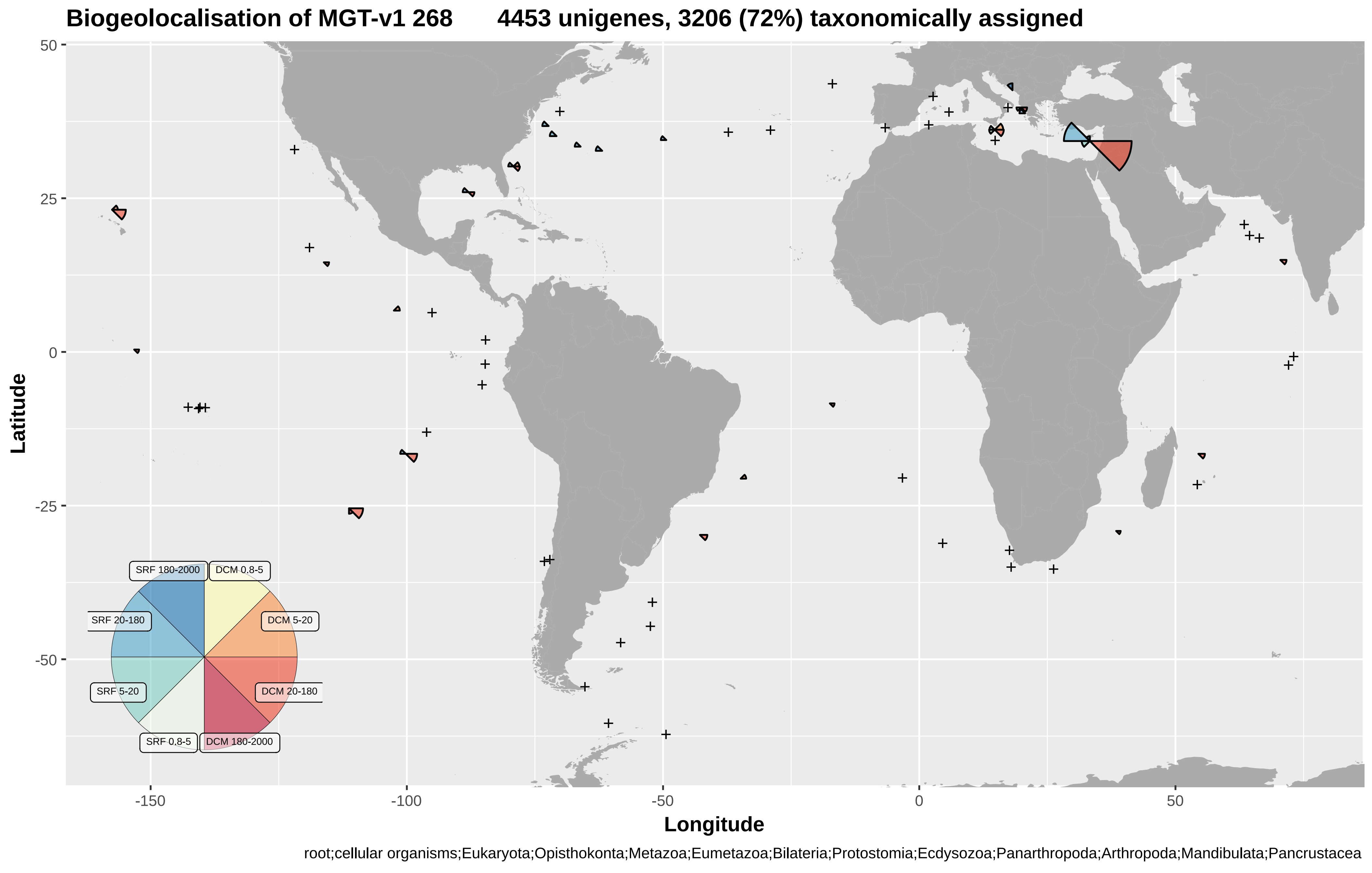

Biogeolocalisation of MGT-v1 268 4453 unigenes, 3206 (72%) taxonomically assigned
50
25
0
-25
-50
Latitude
-150
-100
-50
0
50
Longitude
SRF 180-2000
DCM 0.8-5
SRF 20-180
DCM 5-20
SRF 5-20
DCM 20-180
SRF 0.8-5
DCM 180-2000
root;cellular organisms;Eukaryota;Opisthokonta;Metazoa;Eumetazoa;Bilateria;Protostomia;Ecdysozoa;Panarthropoda;Arthropoda;Mandibulata;Pancrustacea

**Biogeolocalisation of MGT-v1 269      4436 unigenes, 1249 (28.16%) taxonomically assigned**

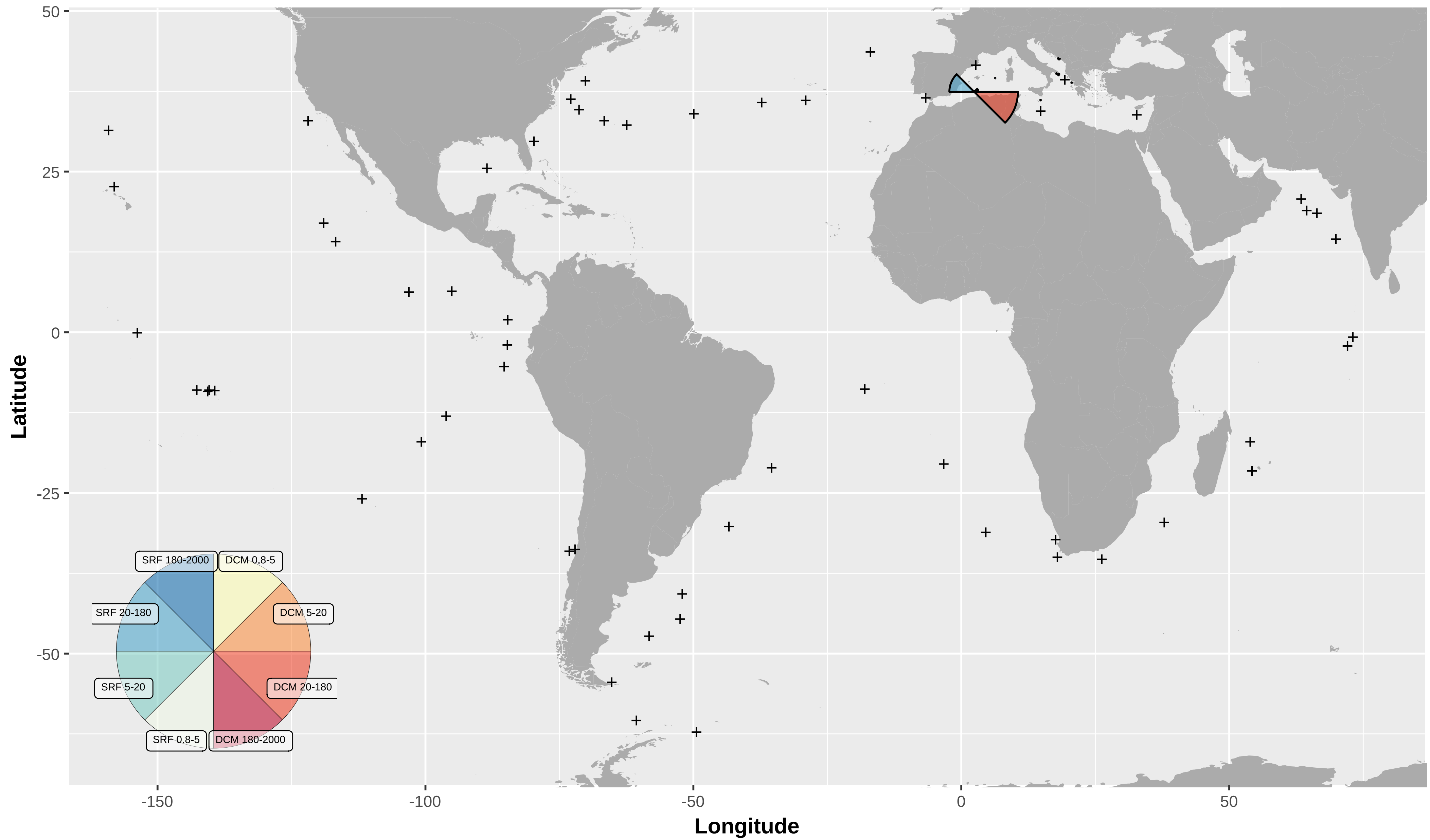


root;cellular organisms;Eukaryota;Opisthokonta

## Biogeolocalisation of MGT-v1 270    4415 unigenes, 4283 (97.01%) taxonomically assigned

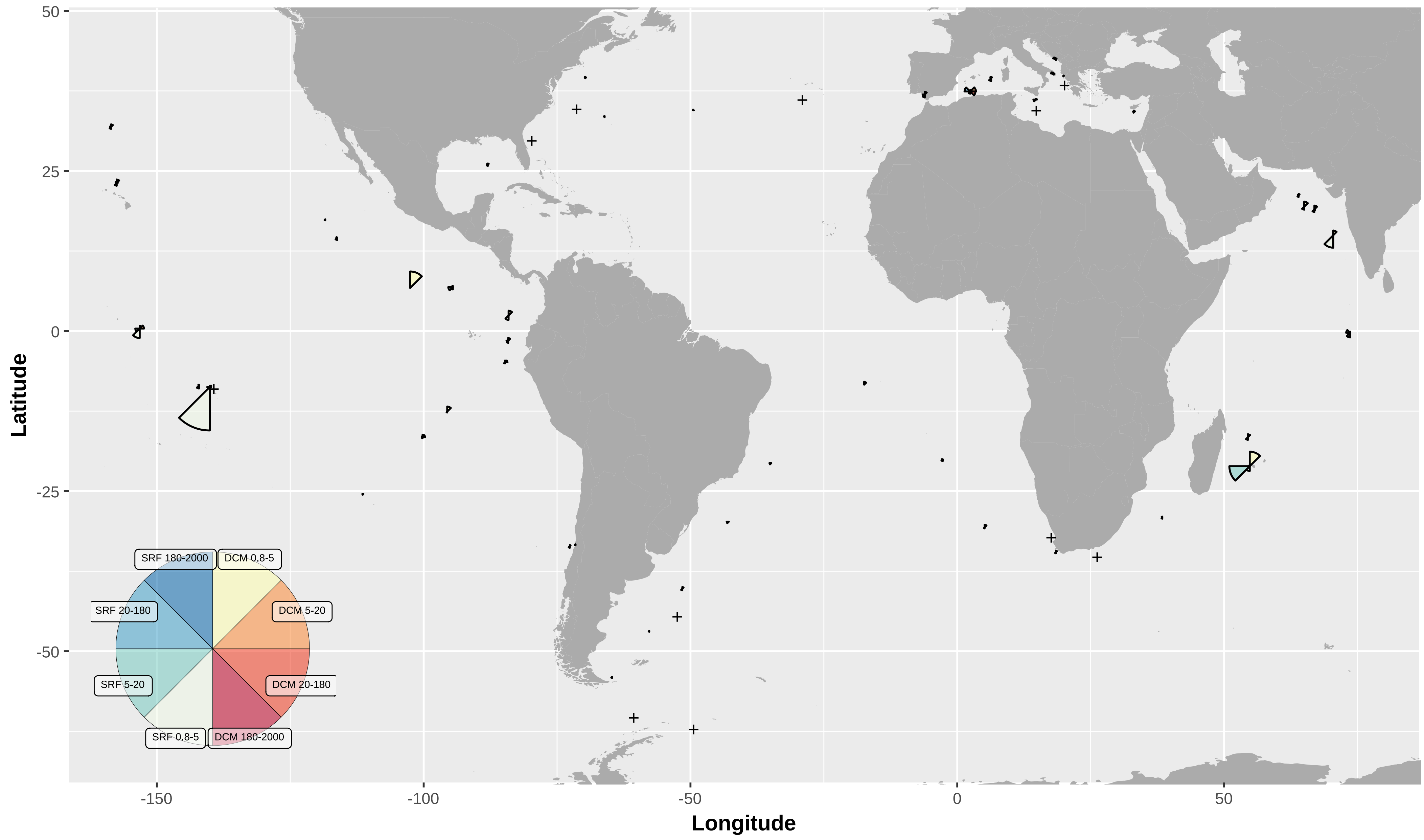


root;cellular organisms;Bacteria;Proteobacteria;Gammaproteobacteria;Alteromonadales;Alteromonadaceae;Marinobacter

## Biogeolocalisation of MGT-v1 271      4385 unigenes, 3259 (74.32%) taxonomically assigned

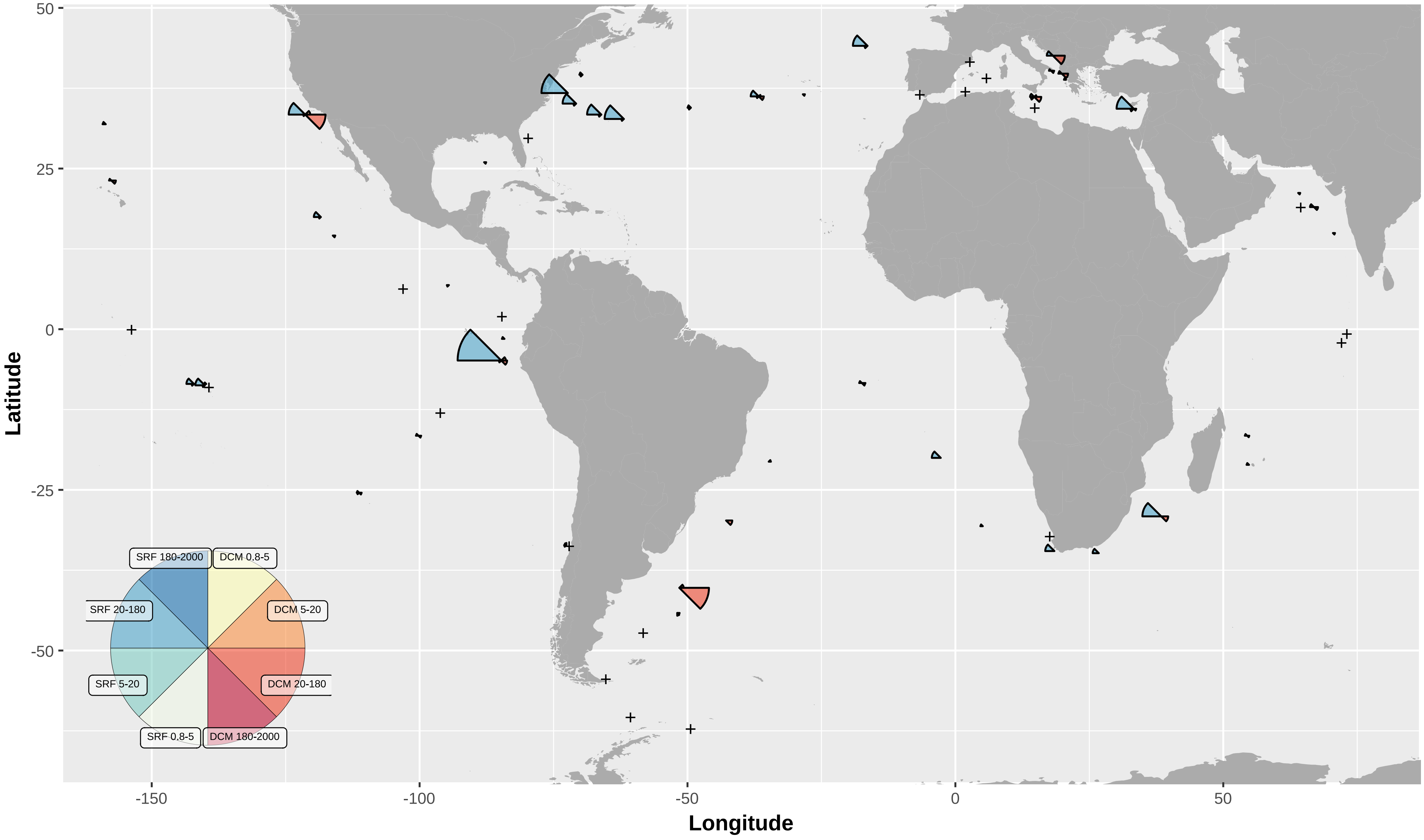


root;cellular organisms;Eukaryota;Alveolata;Ciliophora;Intramacronucleata;Spirotrichea

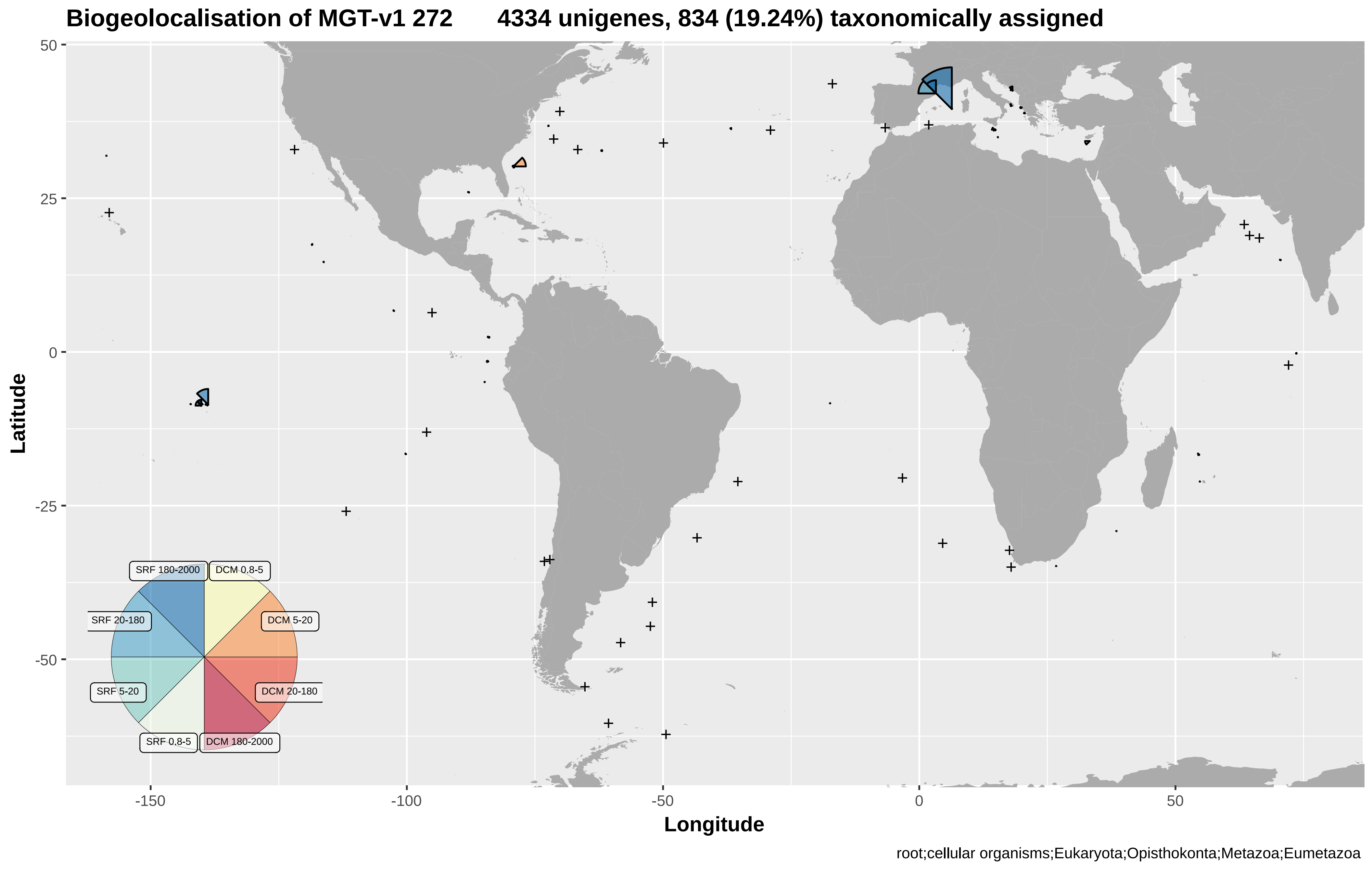
Biogeolocalisation of MGT-v1 272 4334 unigenes, 834 (19.24%) taxonomically assigned
Latitude
Longitude
50
25
0
-25
-50
-150
-100
-50
0
50
SRF 180-2000
DCM 0.8-5
SRF 20-180
DCM 5-20
SRF 5-20
DCM 20-180
SRF 0.8-5
DCM 180-2000
root;cellular organisms;Eukaryota;Opisthokonta;Metazoa;Eumetazoa

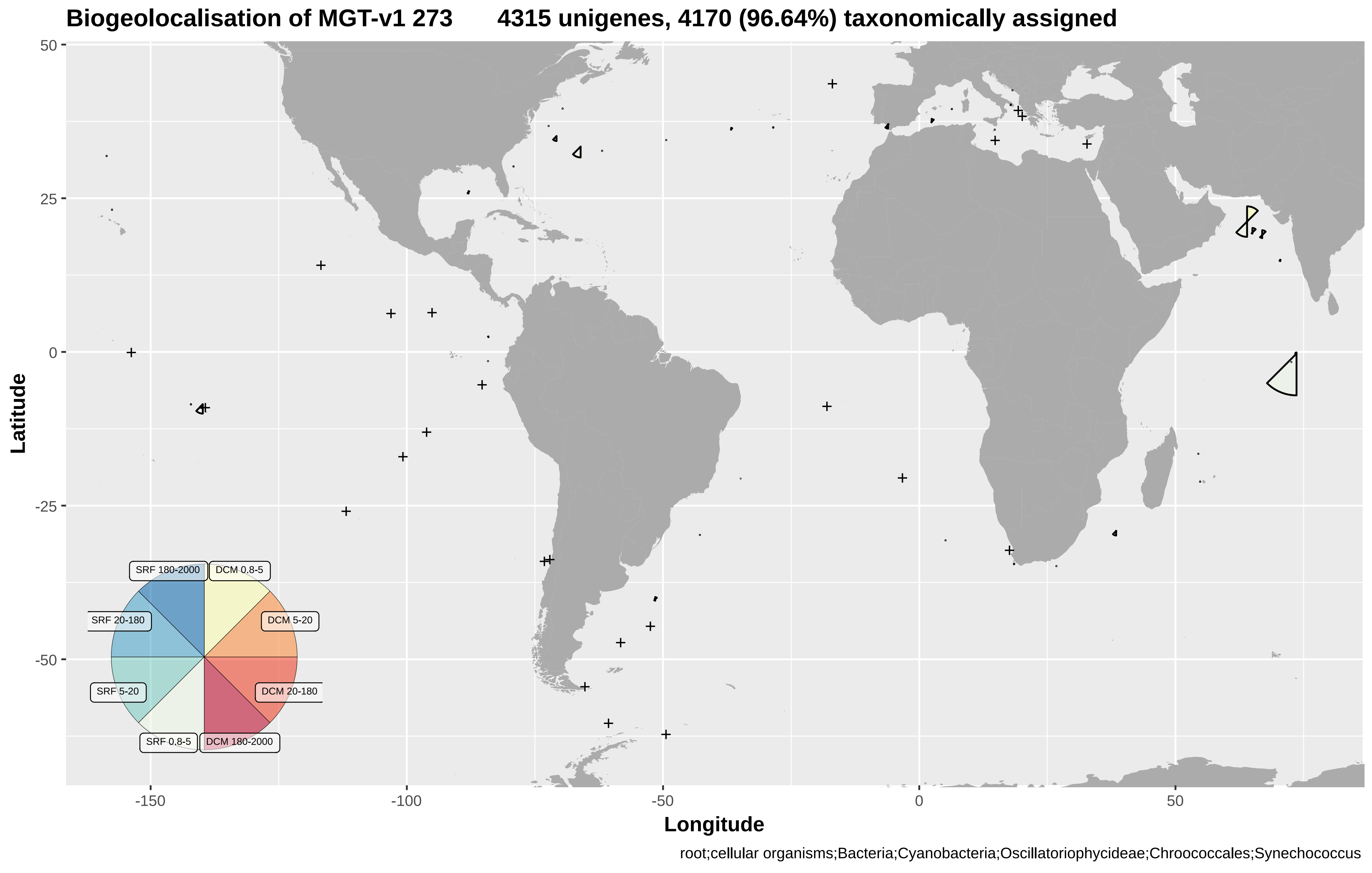
Biogeolocalisation of MGT-v1 273
4315 unigenes, 4170 (96.64%) taxonomically assigned
Latitude
Longitude
50
25
0
-25
-50
-150
-100
-50
0
50
SRF 180-2000
DCM 0.8-5
SRF 20-180
DCM 5-20
SRF 5-20
DCM 20-180
SRF 0.8-5
DCM 180-2000
root;cellular organisms;Bacteria;Cyanobacteria;Oscillatoriophycideae;Chroococcales;Synechococcus

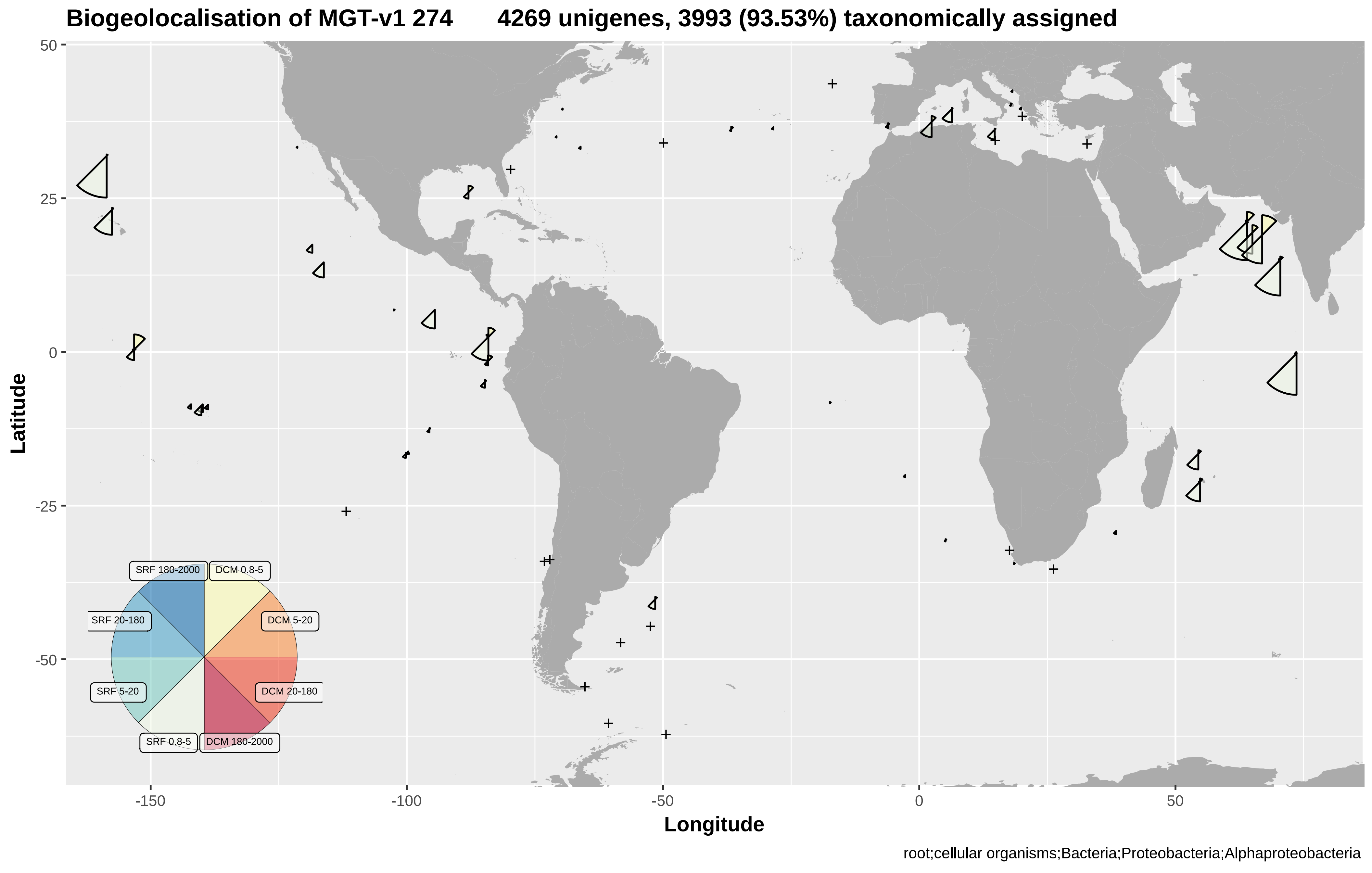

Biogeolocalisation of MGT-v1 274 4269 unigenes, 3993 (93.53%) taxonomically assigned
Latitude
Longitude
50
25
0
-25
-50
-150
-100
-50
0
50
SRF 180-2000
DCM 0.8-5
SRF 20-180
DCM 5-20
SRF 5-20
DCM 20-180
SRF 0.8-5
DCM 180-2000
root;cellular organisms;Bacteria;Proteobacteria;Alphaproteobacteria

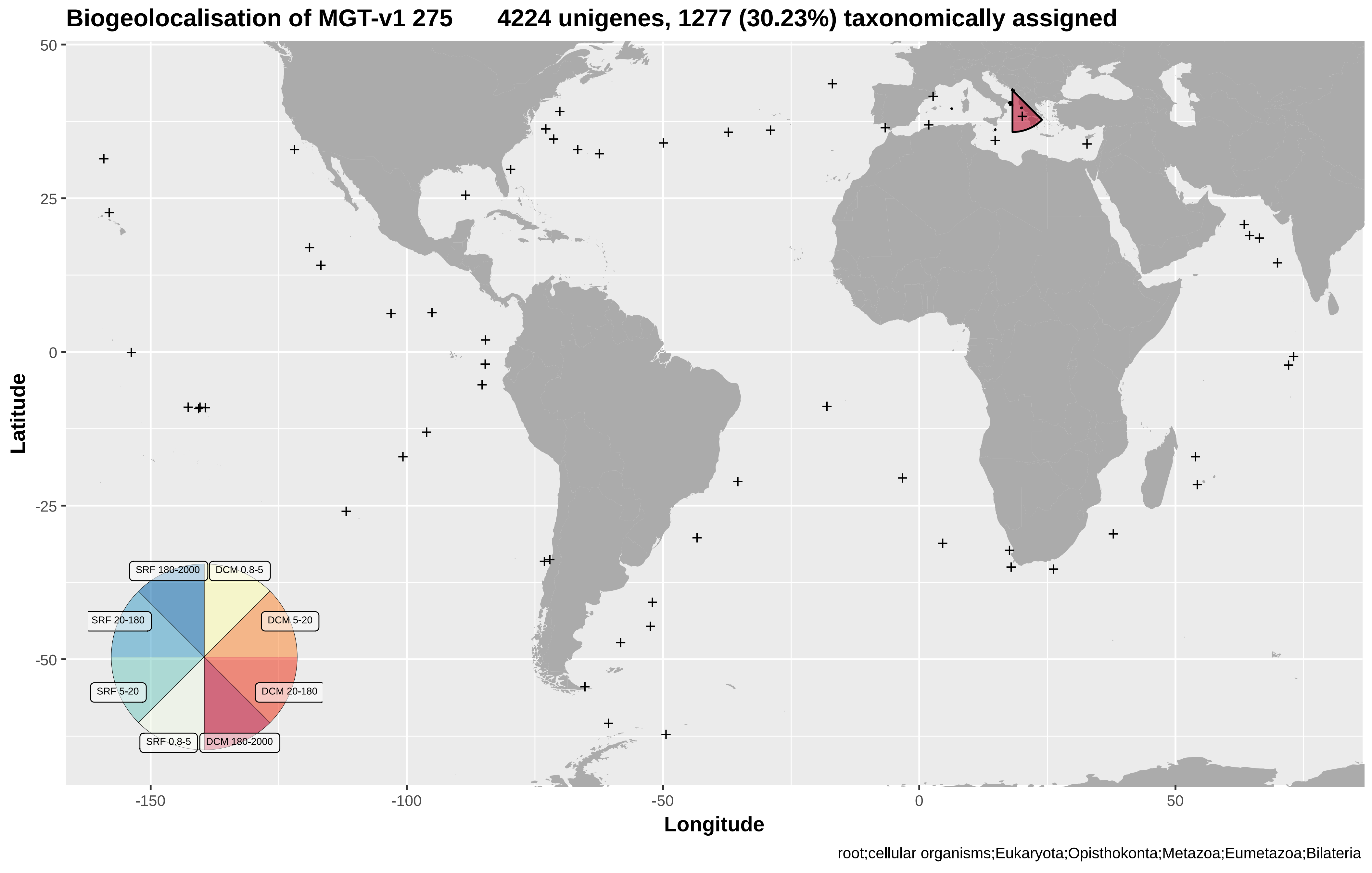

Biogeolocalisation of MGT-v1 275 4224 unigenes, 1277 (30.23%) taxonomically assigned
50
25
0
-25
-50
Latitude
-150
-100
-50
0
50
Longitude
SRF 180-2000
DCM 0.8-5
SRF 20-180
DCM 5-20
SRF 5-20
DCM 20-180
SRF 0.8-5
DCM 180-2000
root;cellular organisms;Eukaryota;Opisthokonta;Metazoa;Eumetazoa;Bilateria

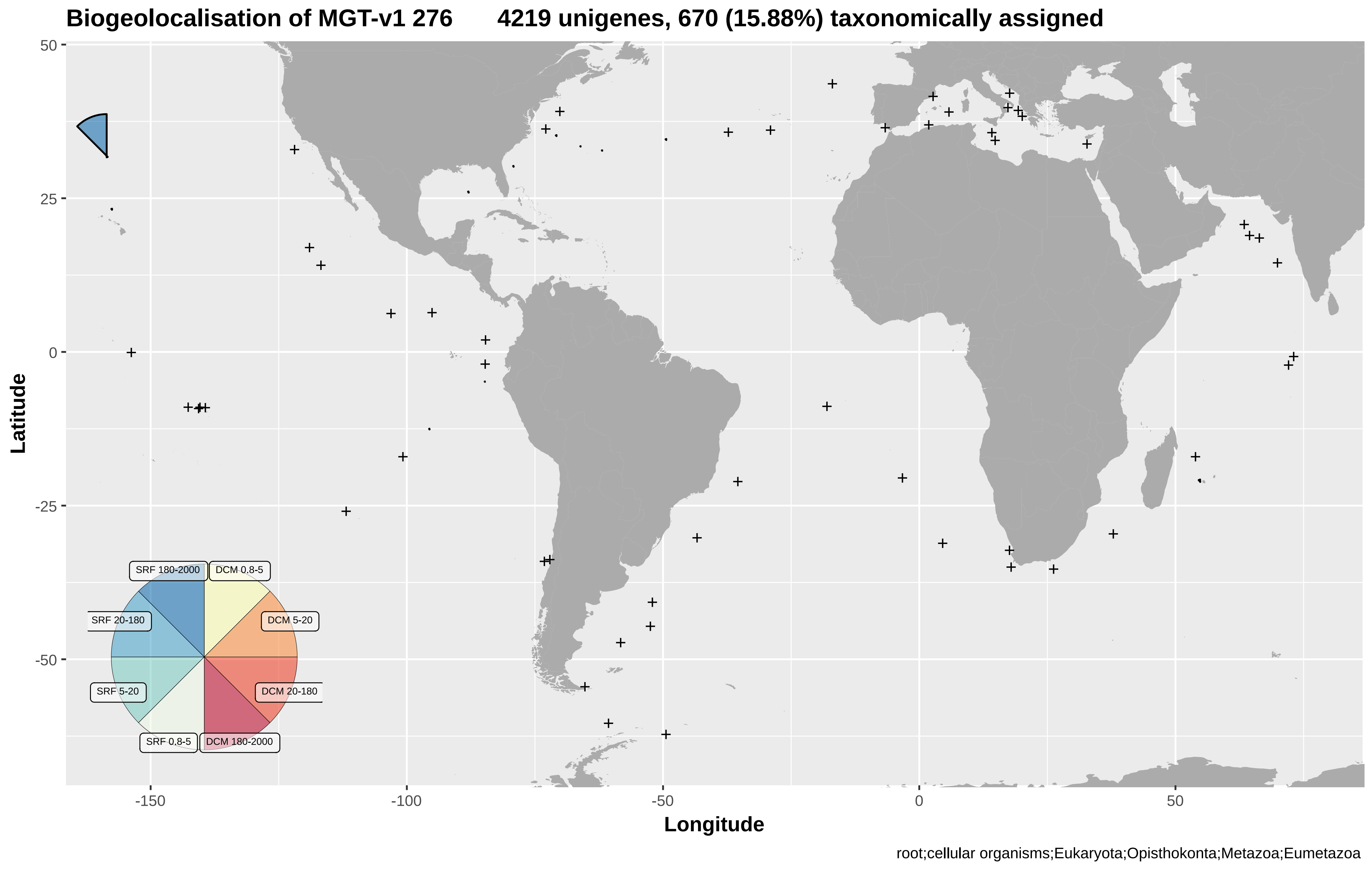
Biogeolocalisation of MGT-v1 276
4219 unigenes, 670 (15.88%) taxonomically assigned
50
25
0
-25
-50
Latitude
-150
-100
-50
0
50
Longitude
SRF 180-2000
DCM 0.8-5
SRF 20-180
DCM 5-20
SRF 5-20
DCM 20-180
SRF 0.8-5
DCM 180-2000
root;cellular organisms;Eukaryota;Opisthokonta;Metazoa;Eumetazoa

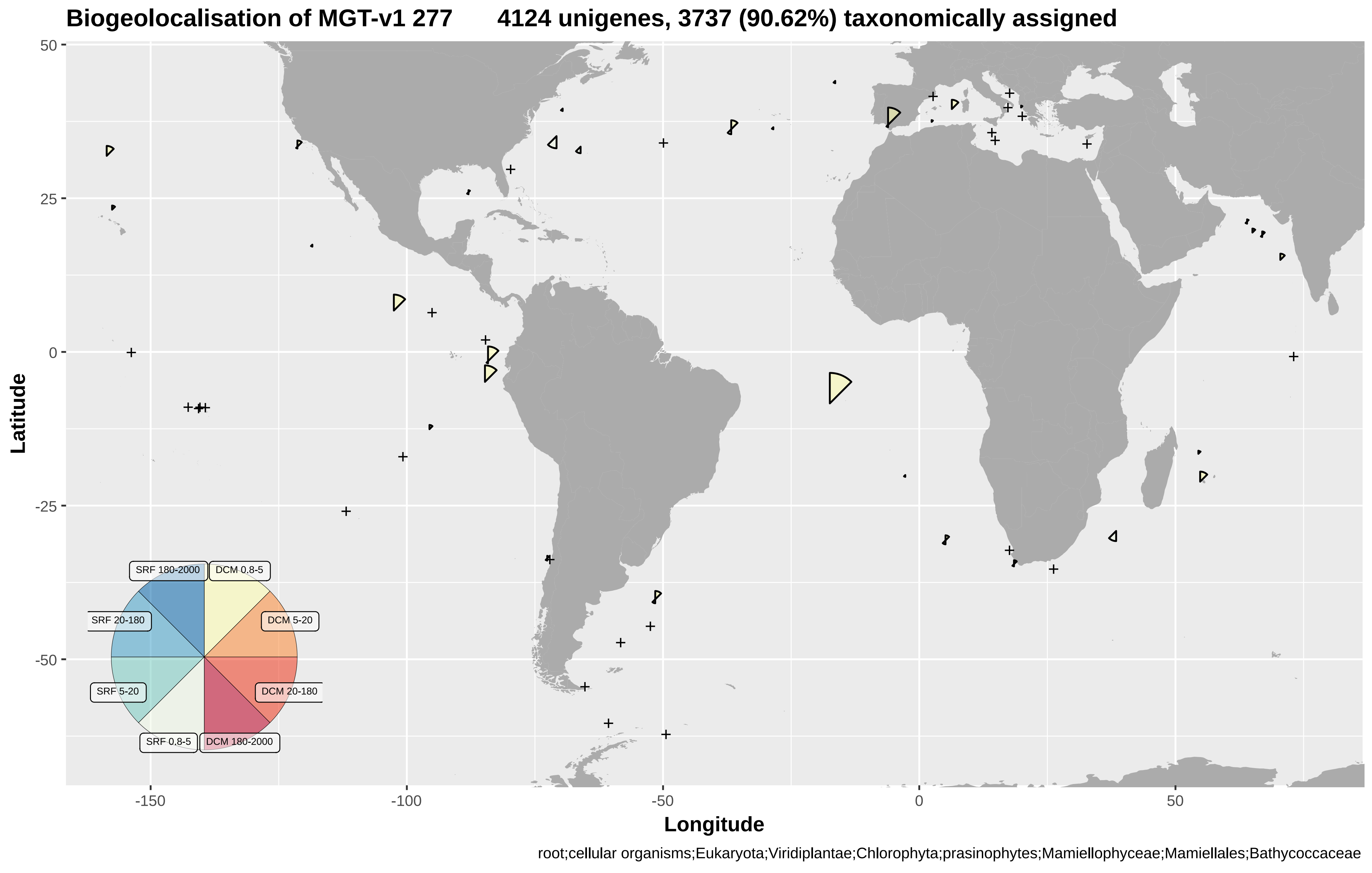

Biogeolocalisation of MGT-v1 277 4124 unigenes, 3737 (90.62%) taxonomically assigned
50
25
0
-25
-50
Latitude
-150
-100
-50
0
50
Longitude
SRF 180-2000
DCM 0.8-5
SRF 20-180
DCM 5-20
SRF 5-20
DCM 20-180
SRF 0.8-5
DCM 180-2000
root;cellular organisms;Eukaryota;Viridiplantae;Chlorophyta;prasinophytes;Mamiellophyceae;Mamiellales;Bathycoccaceae

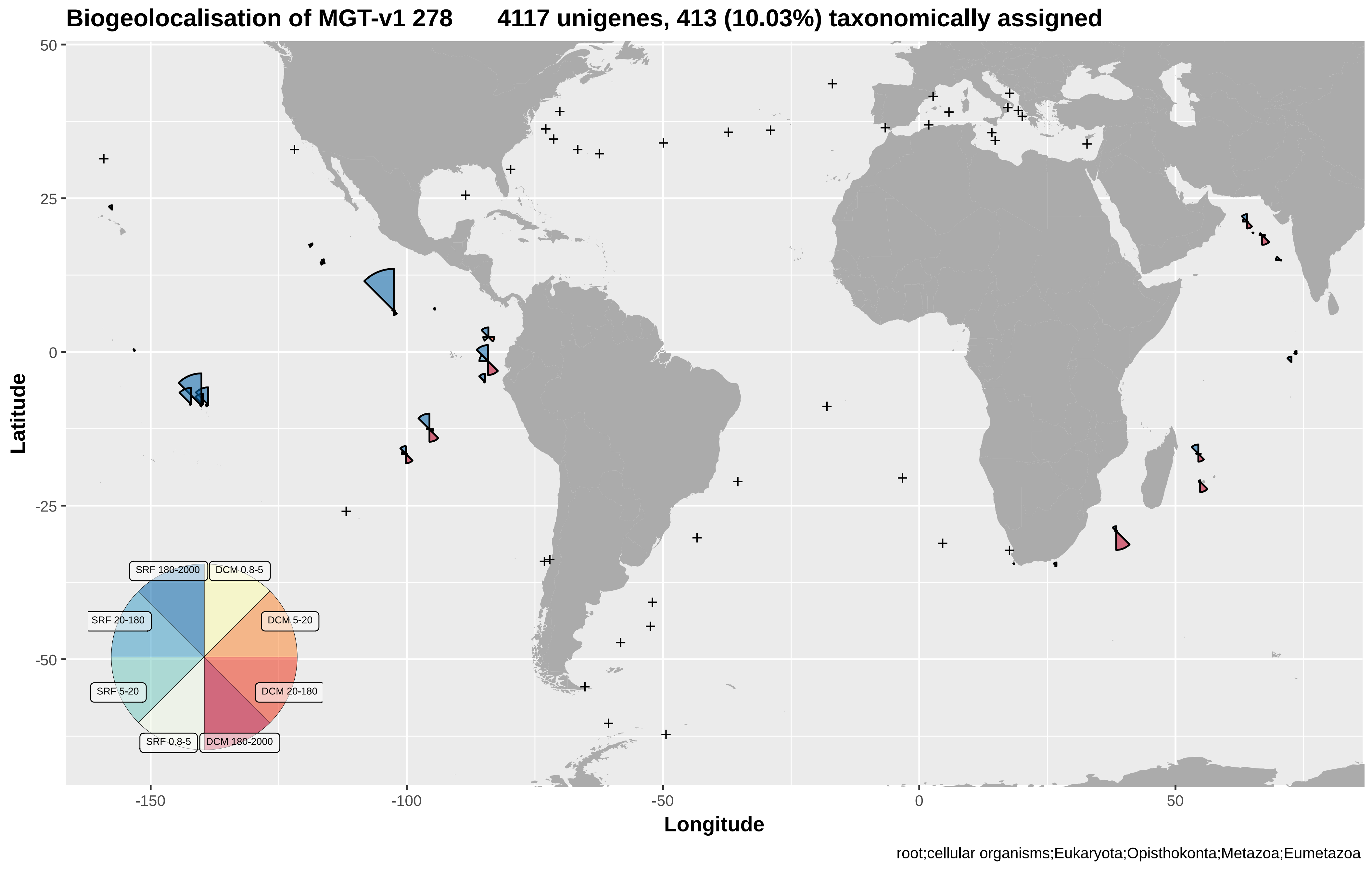

Biogeolocalisation of MGT-v1 278
4117 unigenes, 413 (10.03%) taxonomically assigned
Latitude
Longitude
50
25
0
-25
-50
-150
-100
-50
0
50
SRF 180-2000
DCM 0.8-5
SRF 20-180
DCM 5-20
SRF 5-20
DCM 20-180
SRF 0.8-5
DCM 180-2000
root;cellular organisms;Eukaryota;Opisthokonta;Metazoa;Eumetazoa

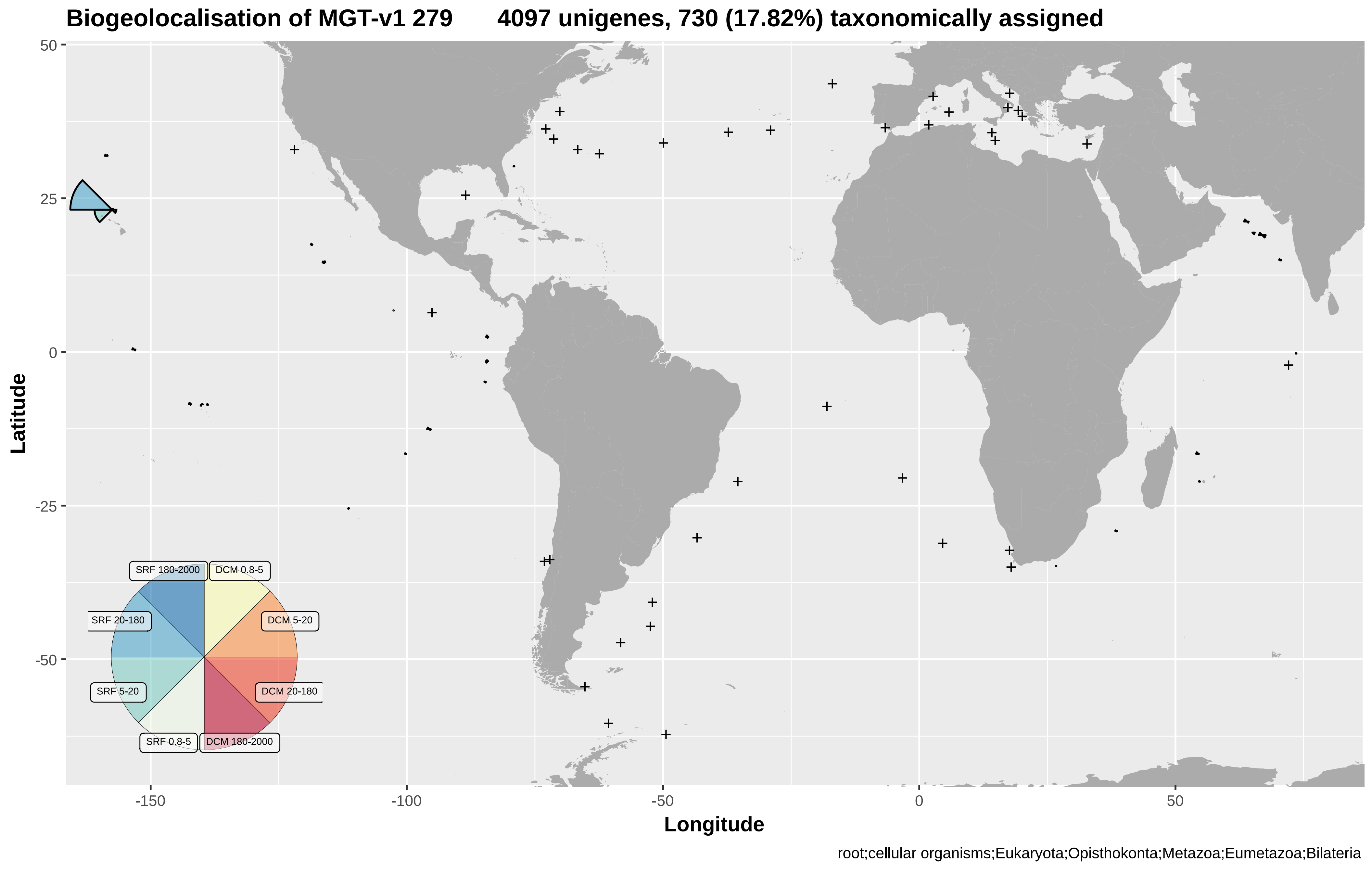
Biogeolocalisation of MGT-v1 279 4097 unigenes, 730 (17.82%) taxonomically assigned
50
25
0
-25
-50
Latitude
-150
-100
-50
0
50
Longitude
SRF 180-2000
DCM 0.8-5
SRF 20-180
DCM 5-20
SRF 5-20
DCM 20-180
SRF 0.8-5
DCM 180-2000
root;cellular organisms;Eukaryota;Opisthokonta;Metazoa;Eumetazoa;Bilateria

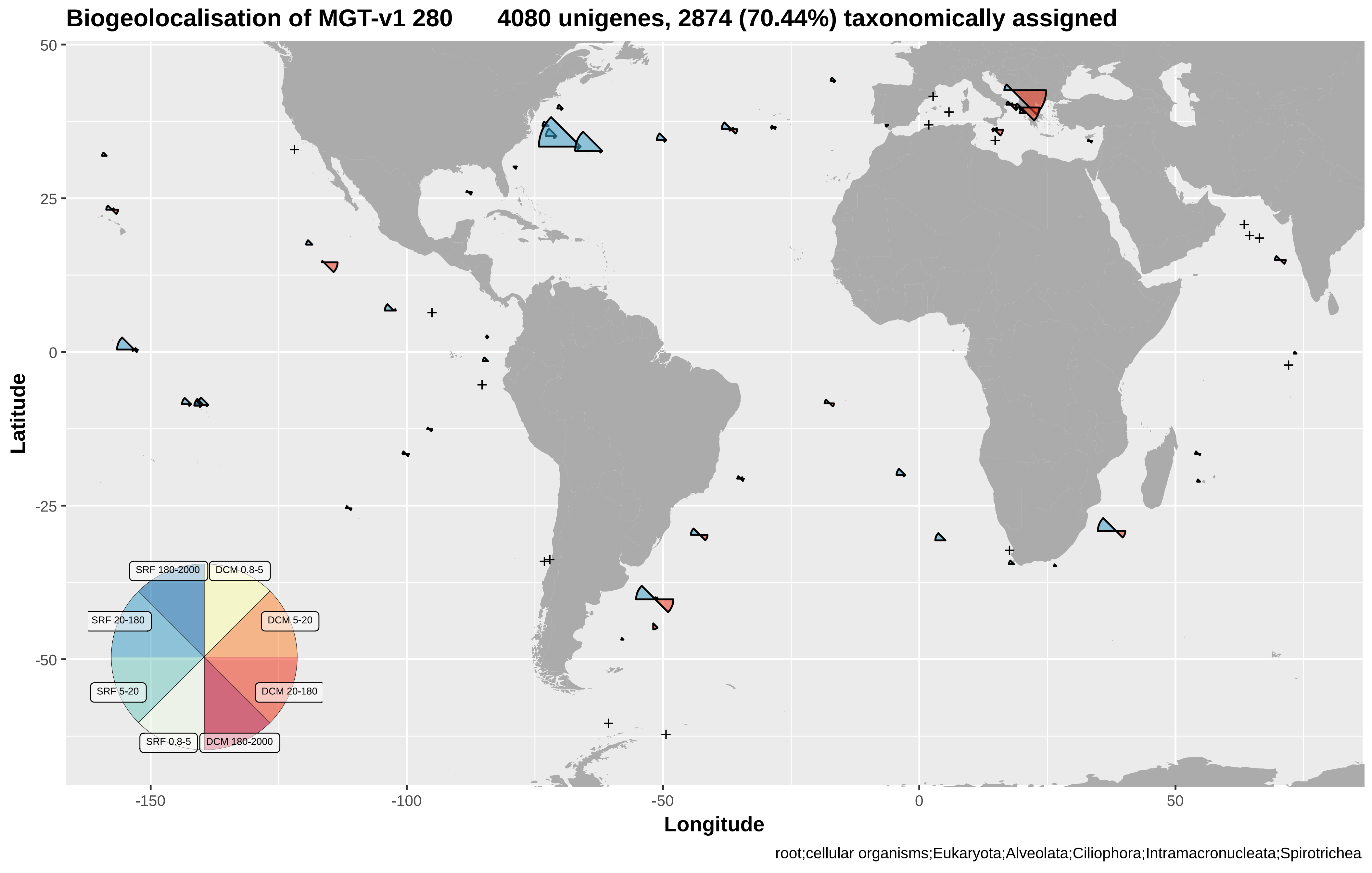

Biogeolocalisation of MGT-v1 280
4080 unigenes, 2874 (70.44%) taxonomically assigned
50
25
0
-25
-50
Latitude
-150
-100
-50
0
50
Longitude
SRF 180-2000
DCM 0.8-5
SRF 20-180
DCM 5-20
SRF 5-20
DCM 20-180
SRF 0.8-5
DCM 180-2000
root;cellular organisms;Eukaryota;Alveolata;Ciliophora;Intramacronucleata;Spirotrichea

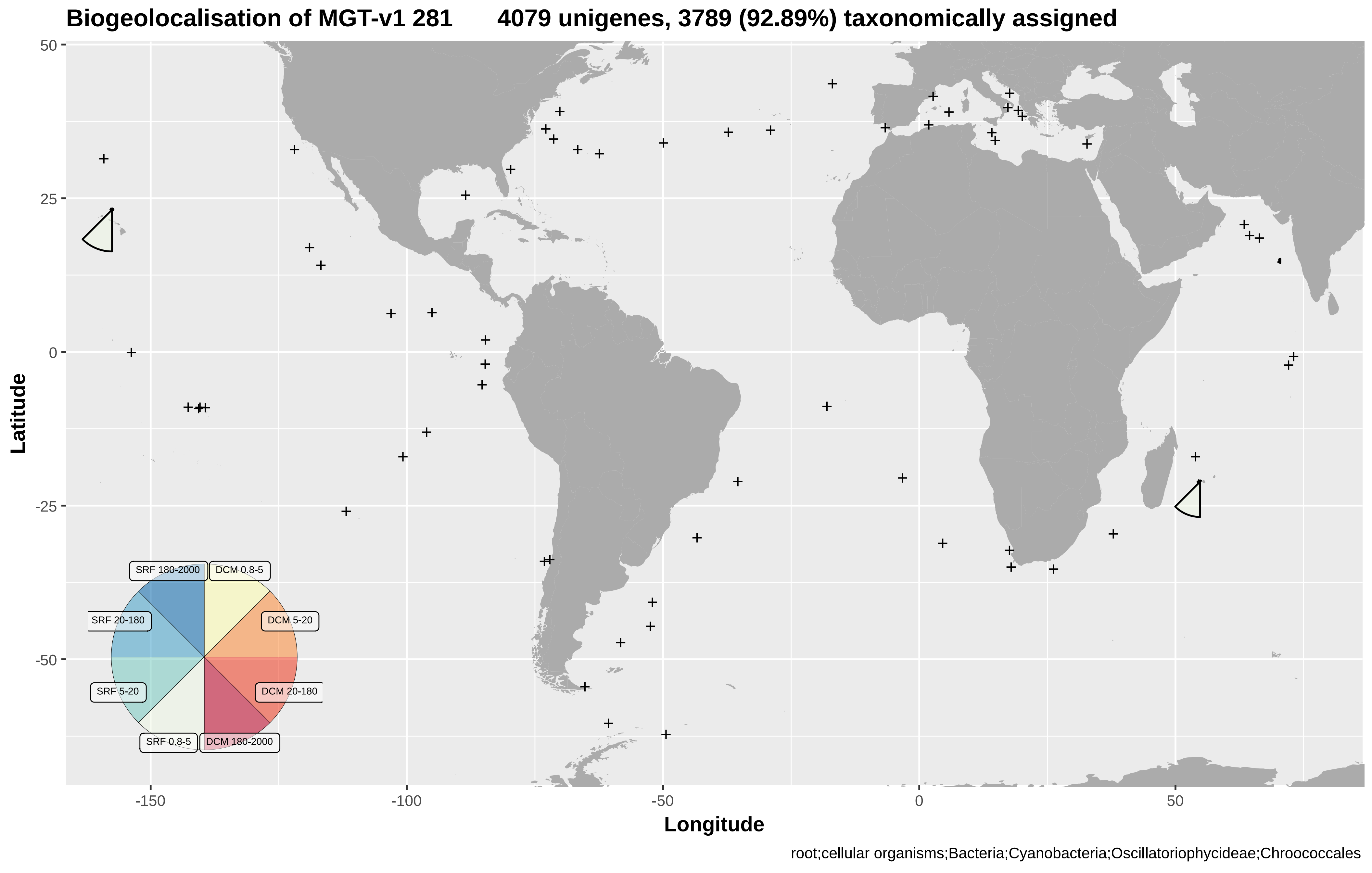

Biogeolocalisation of MGT-v1 281 4079 unigenes, 3789 (92.89%) taxonomically assigned
Latitude
50
25
0
-25
-50
-150
-100
-50
0
50
Longitude
SRF 180-2000
DCM 0.8-5
SRF 20-180
DCM 5-20
SRF 5-20
DCM 20-180
SRF 0.8-5
DCM 180-2000
root;cellular organisms;Bacteria;Cyanobacteria;Oscillatoriophycideae;Chroococcales

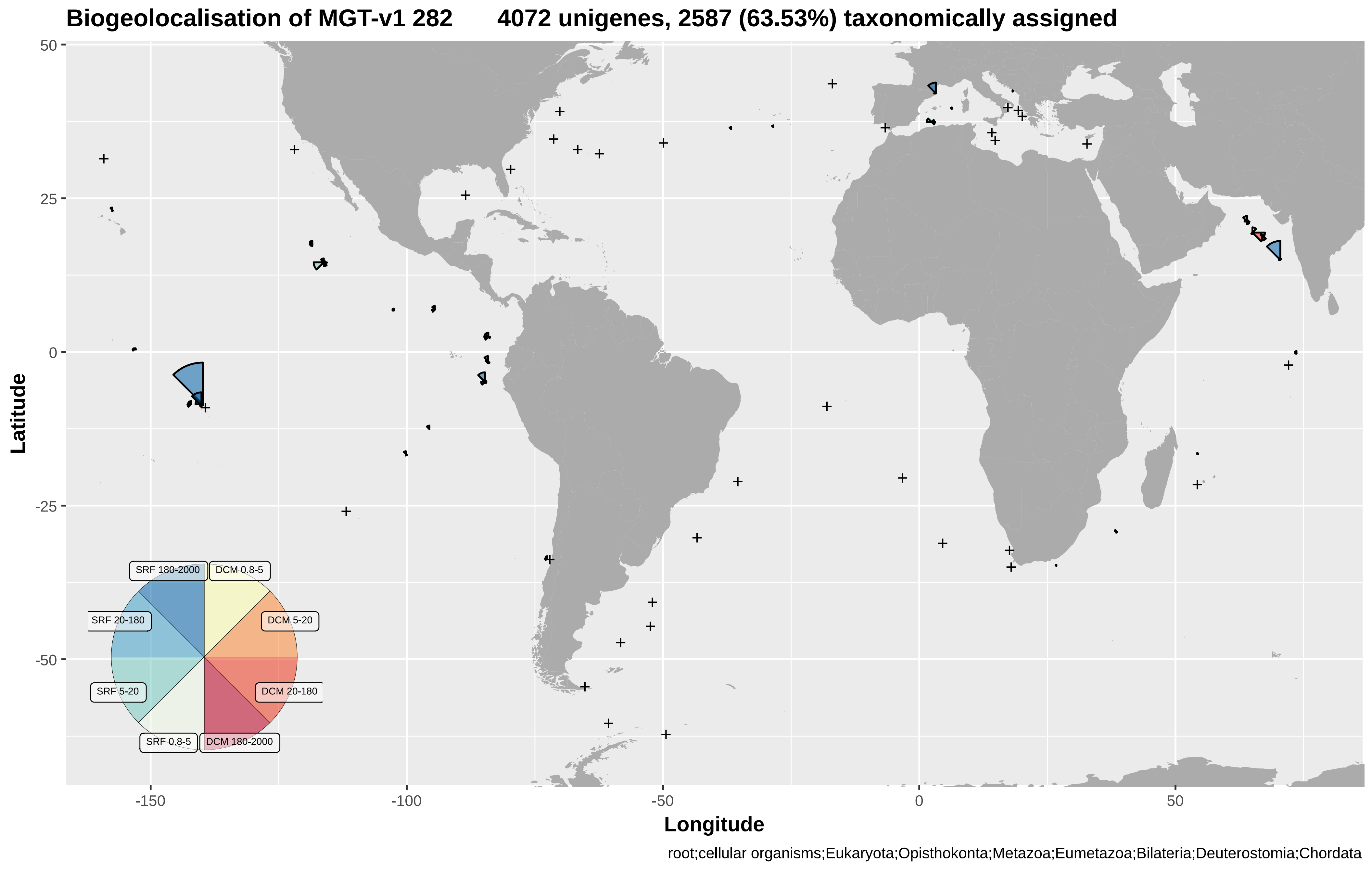

Biogeolocalisation of MGT-v1 282 4072 unigenes, 2587 (63.53%) taxonomically assigned
50
25
0
-25
-50
Latitude
-150
-100
-50
0
50
Longitude
SRF 180-2000
DCM 0.8-5
SRF 20-180
DCM 5-20
SRF 5-20
DCM 20-180
SRF 0.8-5
DCM 180-2000
root;cellular organisms;Eukaryota;Opisthokonta;Metazoa;Eumetazoa;Bilateria;Deuterostomia;Chordata

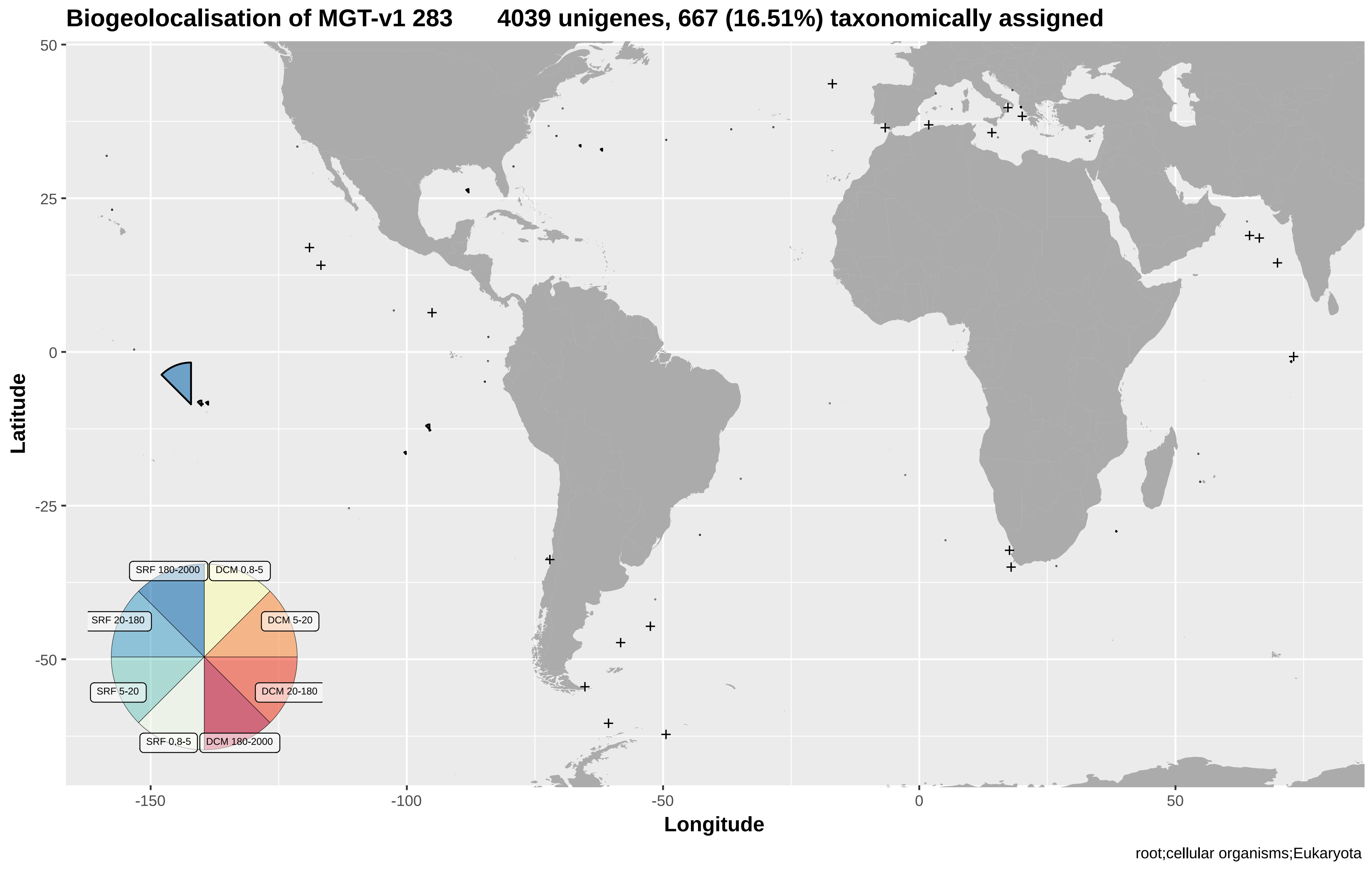

Biogeolocalisation of MGT-v1 283
4039 unigenes, 667 (16.51%) taxonomically assigned
50
25
0
-25
-50
Latitude
-150
-100
-50
0
50
Longitude
SRF 180-2000
DCM 0.8-5
SRF 20-180
DCM 5-20
SRF 5-20
DCM 20-180
SRF 0.8-5
DCM 180-2000
root;cellular organisms;Eukaryota

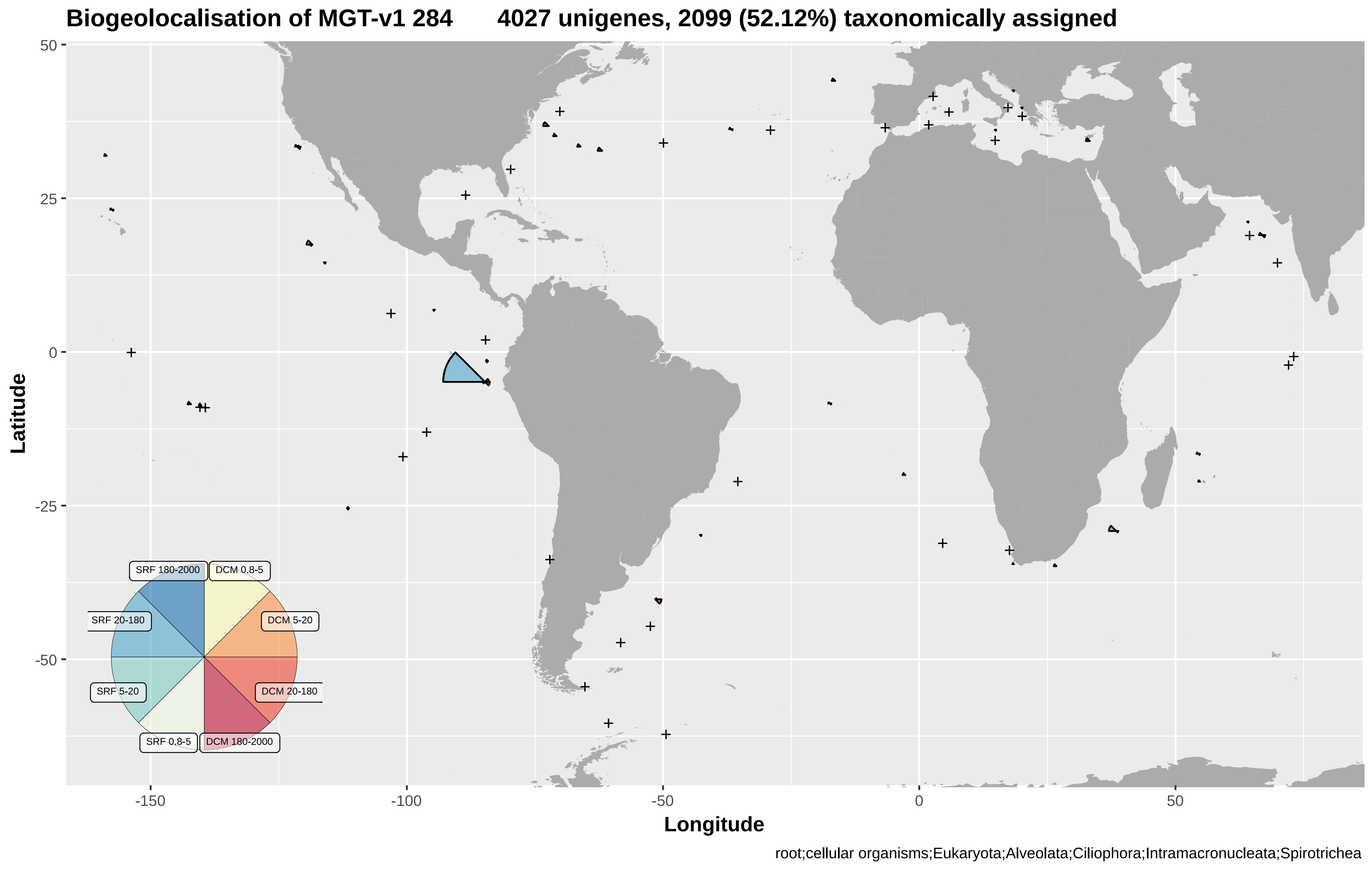

Biogeolocalisation of MGT-v1 284 4027 unigenes, 2099 (52.12%) taxonomically assigned
Latitude
Longitude
50
25
0
-25
-50
-150
-100
-50
0
50
SRF 180-2000
DCM 0.8-5
SRF 20-180
DCM 5-20
SRF 5-20
DCM 20-180
SRF 0.8-5
DCM 180-2000
root;cellular organisms;Eukaryota;Alveolata;Ciliophora;Intramacronucleata;Spirotrichea

## Biogeolocalisation of MGT-v1 285      3949 unigenes, 460 (11.65%) taxonomically assigned

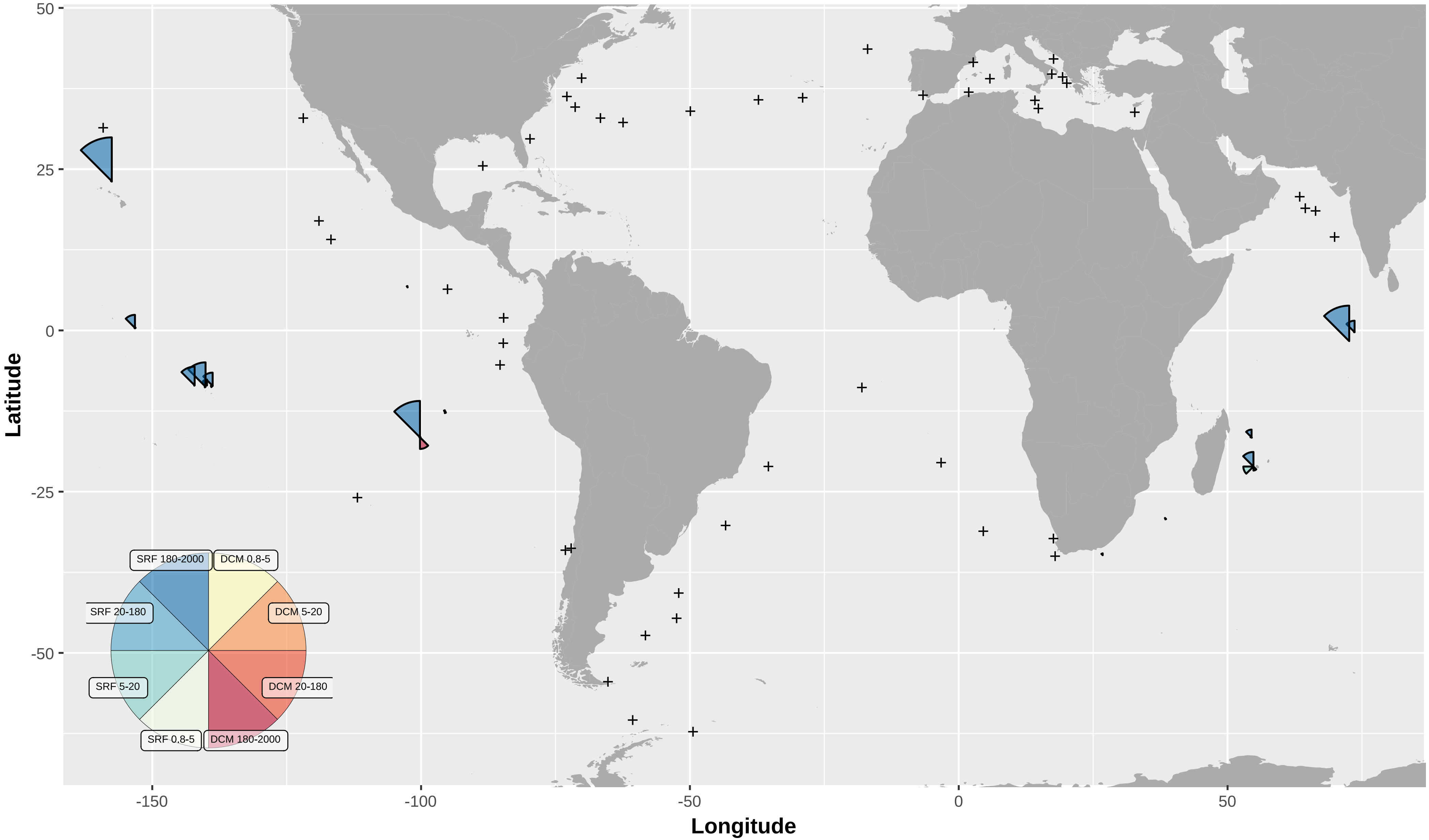


root;cellular organisms;Eukaryota;Opisthokonta;Metazoa;Eumetazoa;Bilateria

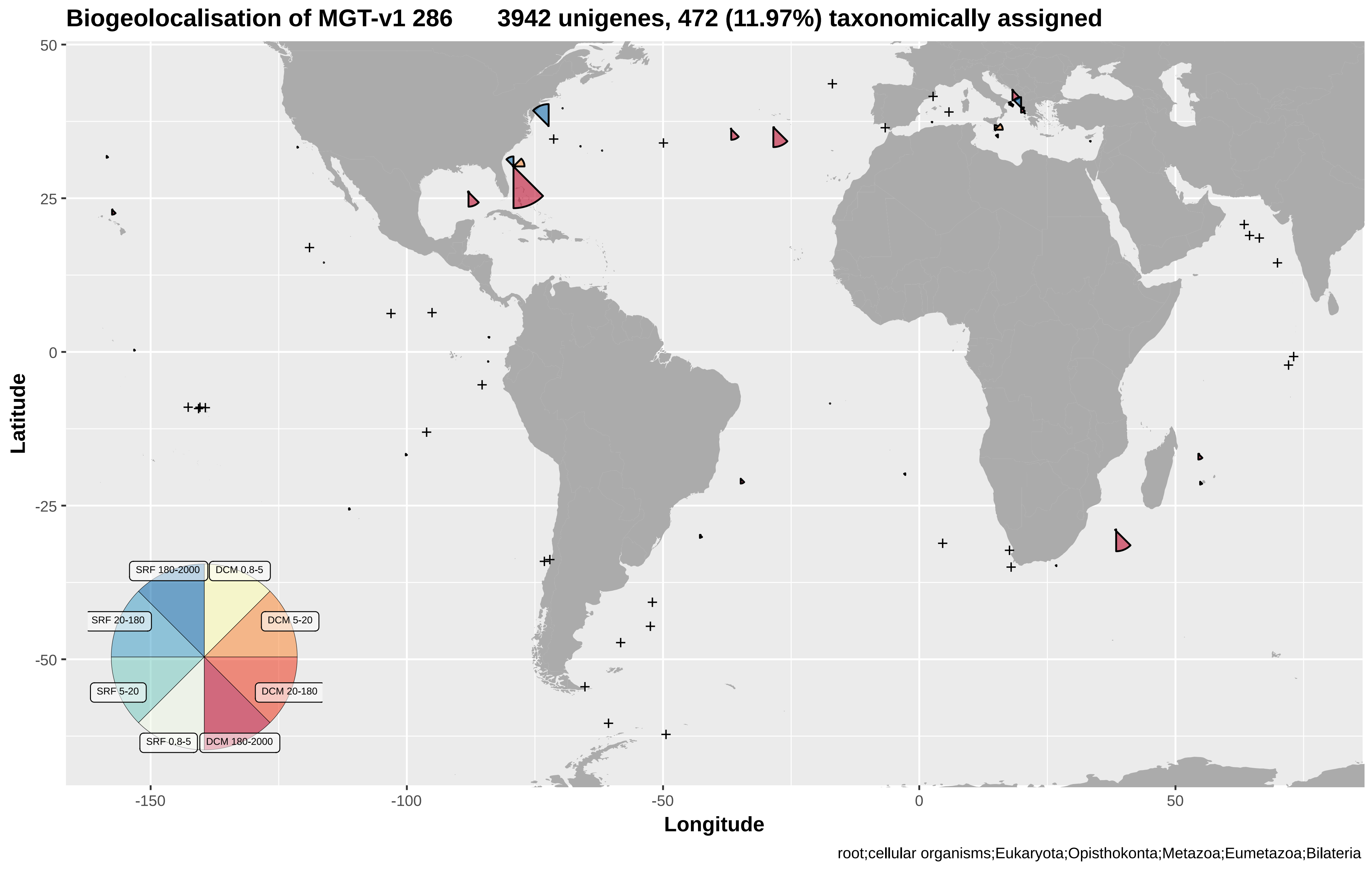

Biogeolocalisation of MGT-v1 286      3942 unigenes, 472 (11.97%) taxonomically assigned
50
25
0
-25
-50
Latitude
-150
-100
-50
0
50
Longitude
SRF 180-2000
DCM 0.8-5
SRF 20-180
DCM 5-20
SRF 5-20
DCM 20-180
SRF 0.8-5
DCM 180-2000
root;cellular organisms;Eukaryota;Opisthokonta;Metazoa;Eumetazoa;Bilateria

### Biogeolocalisation of MGT-v1 287 3910 unigenes, 2377 (60.79%) taxonomically assigned

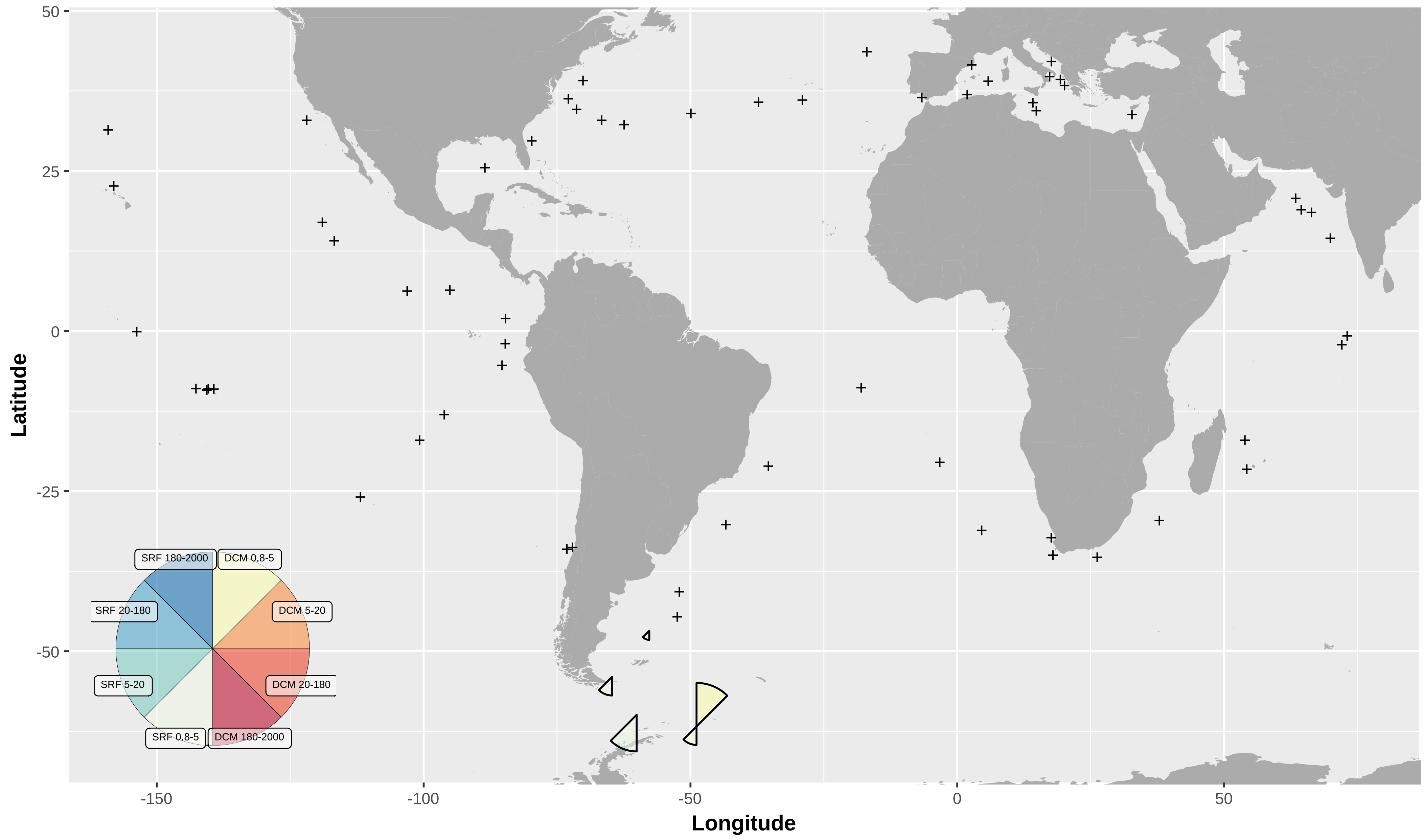


root;cellular organisms;Eukaryota

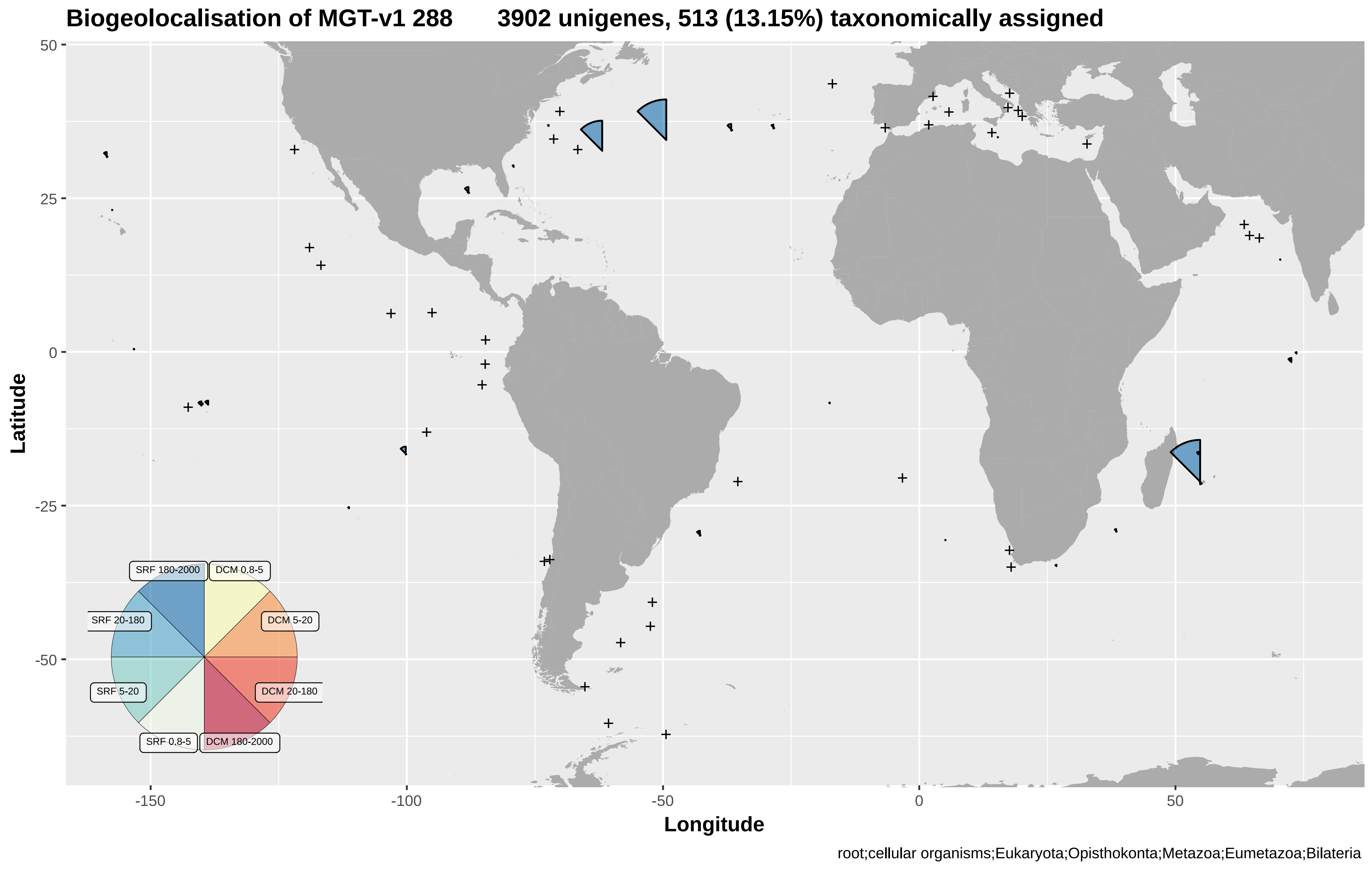
Biogeolocalisation of MGT-v1 288 3902 unigenes, 513 (13.15%) taxonomically assigned
Latitude
Longitude
50
25
0
-25
-50
-150
-100
-50
0
50
SRF 180-2000
DCM 0.8-5
SRF 20-180
DCM 5-20
SRF 5-20
DCM 20-180
SRF 0.8-5
DCM 180-2000
root;cellular organisms;Eukaryota;Opisthokonta;Metazoa;Eumetazoa;Bilateria

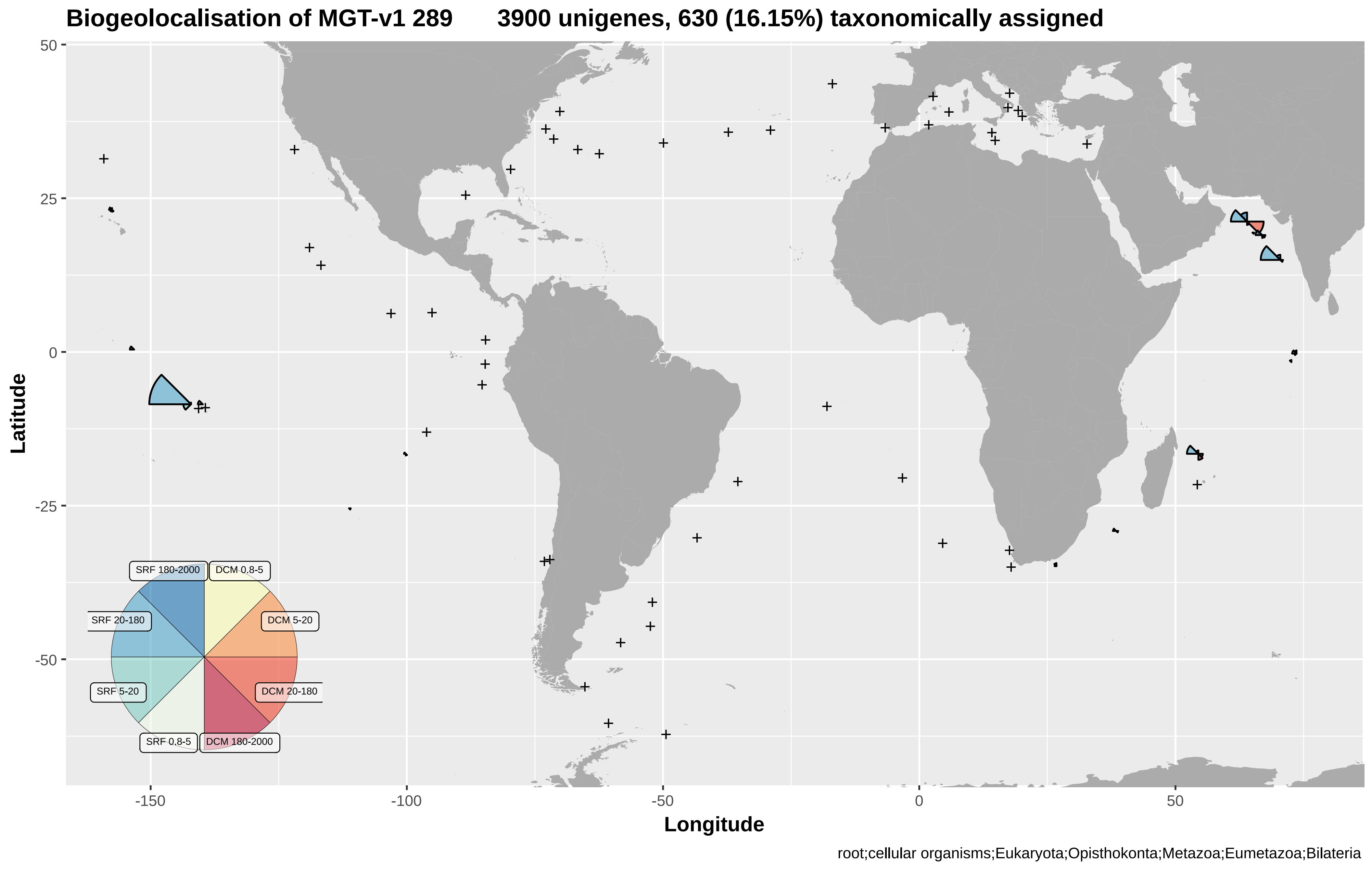
Biogeolocalisation of MGT-v1 289      3900 unigenes, 630 (16.15%) taxonomically assigned
50
25
0
-25
-50
Latitude
-150
-100
-50
0
50
Longitude
SRF 180-2000
DCM 0.8-5
SRF 20-180
DCM 5-20
SRF 5-20
DCM 20-180
SRF 0.8-5
DCM 180-2000
root;cellular organisms;Eukaryota;Opisthokonta;Metazoa;Eumetazoa;Bilateria

#### Biogeolocalisation of MGT-v1 290 3899 unigenes, 3199 (82.05%) taxonomically assigned

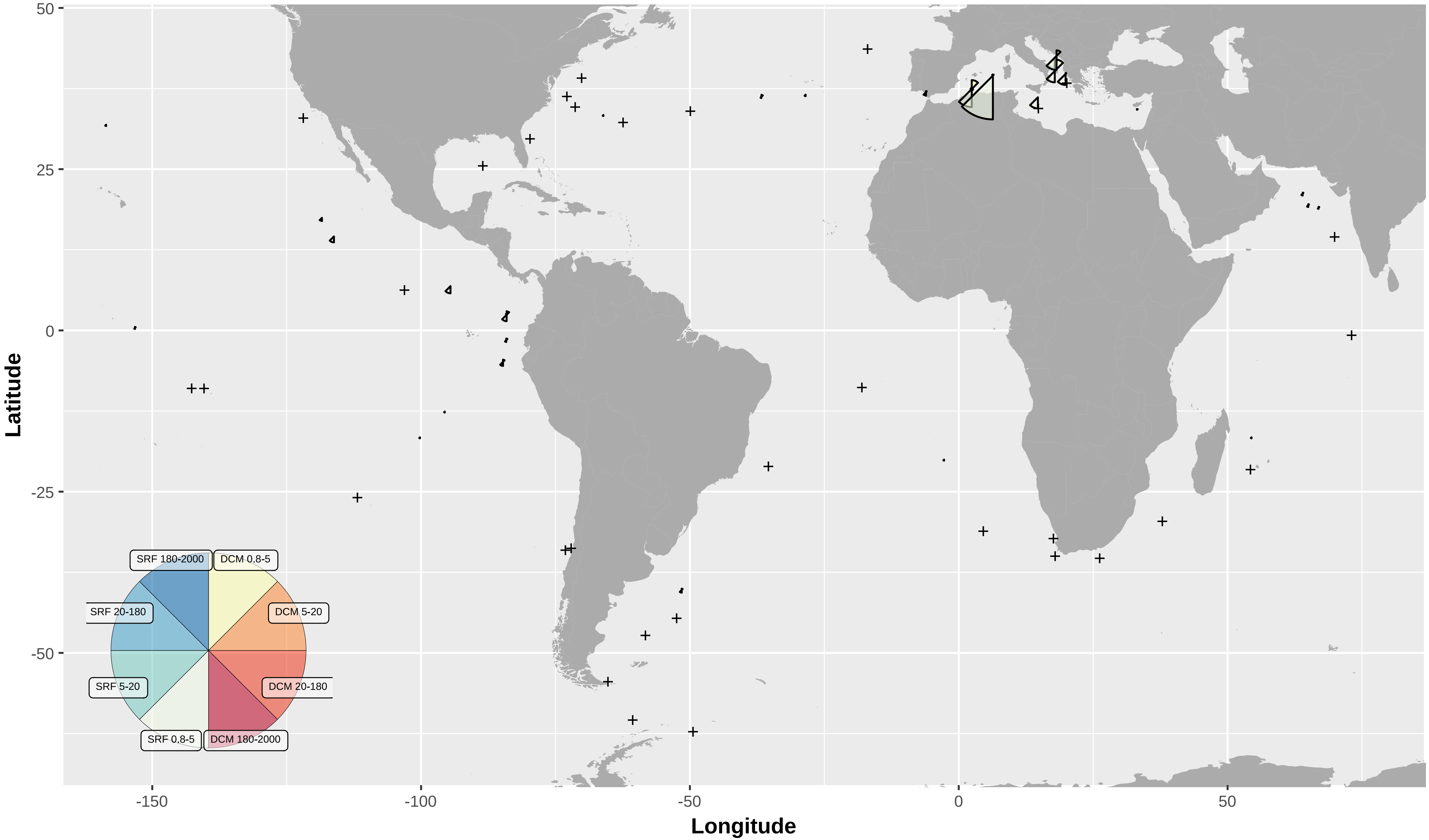


root;cellular organisms;Bacteria;Proteobacteria

#### Biogeolocalisation of MGT-v1 291 3887 unigenes, 410 (10.55%) taxonomically assigned

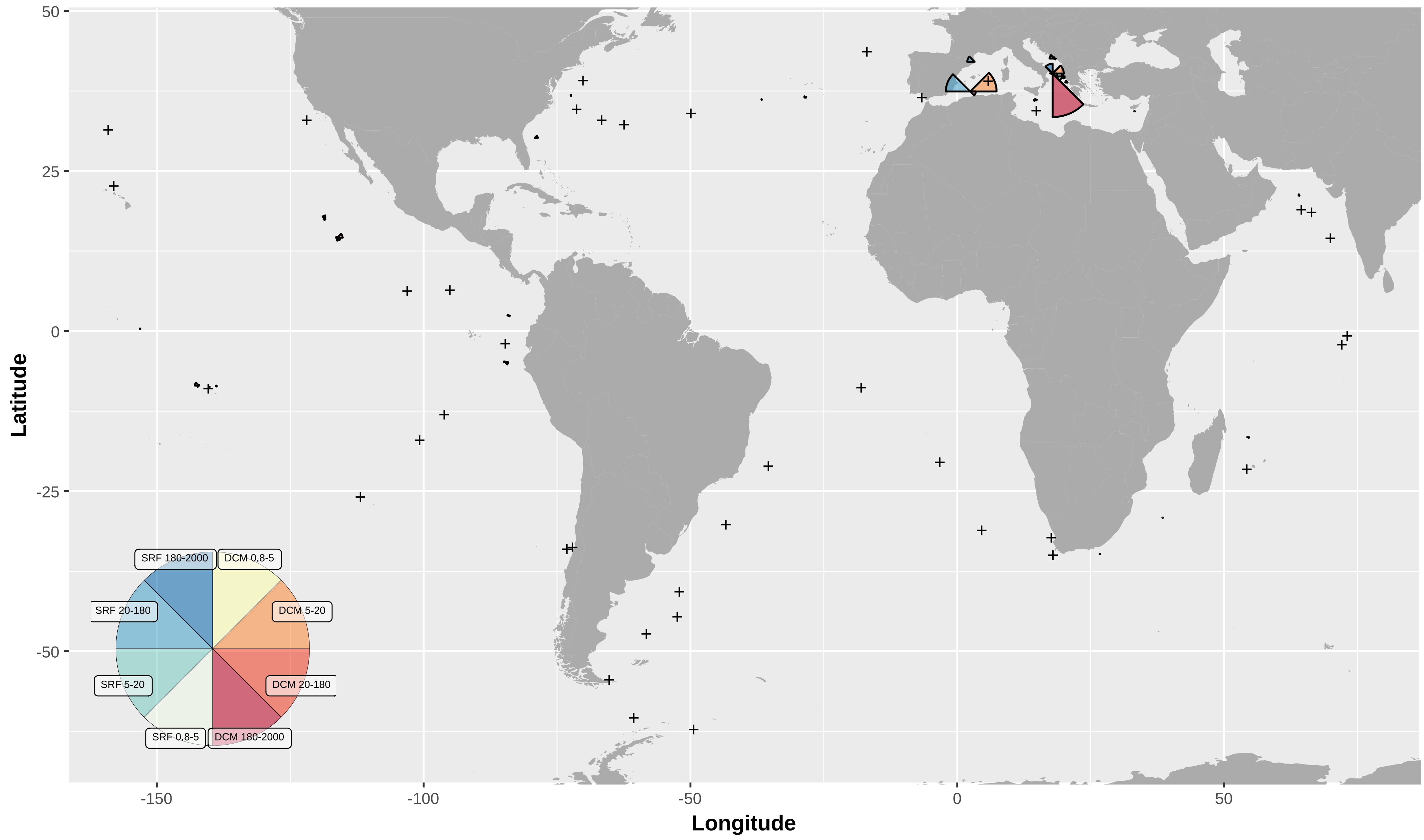


root;cellular organisms;Eukaryota;Opisthokonta;Metazoa;Eumetazoa;Bilateria

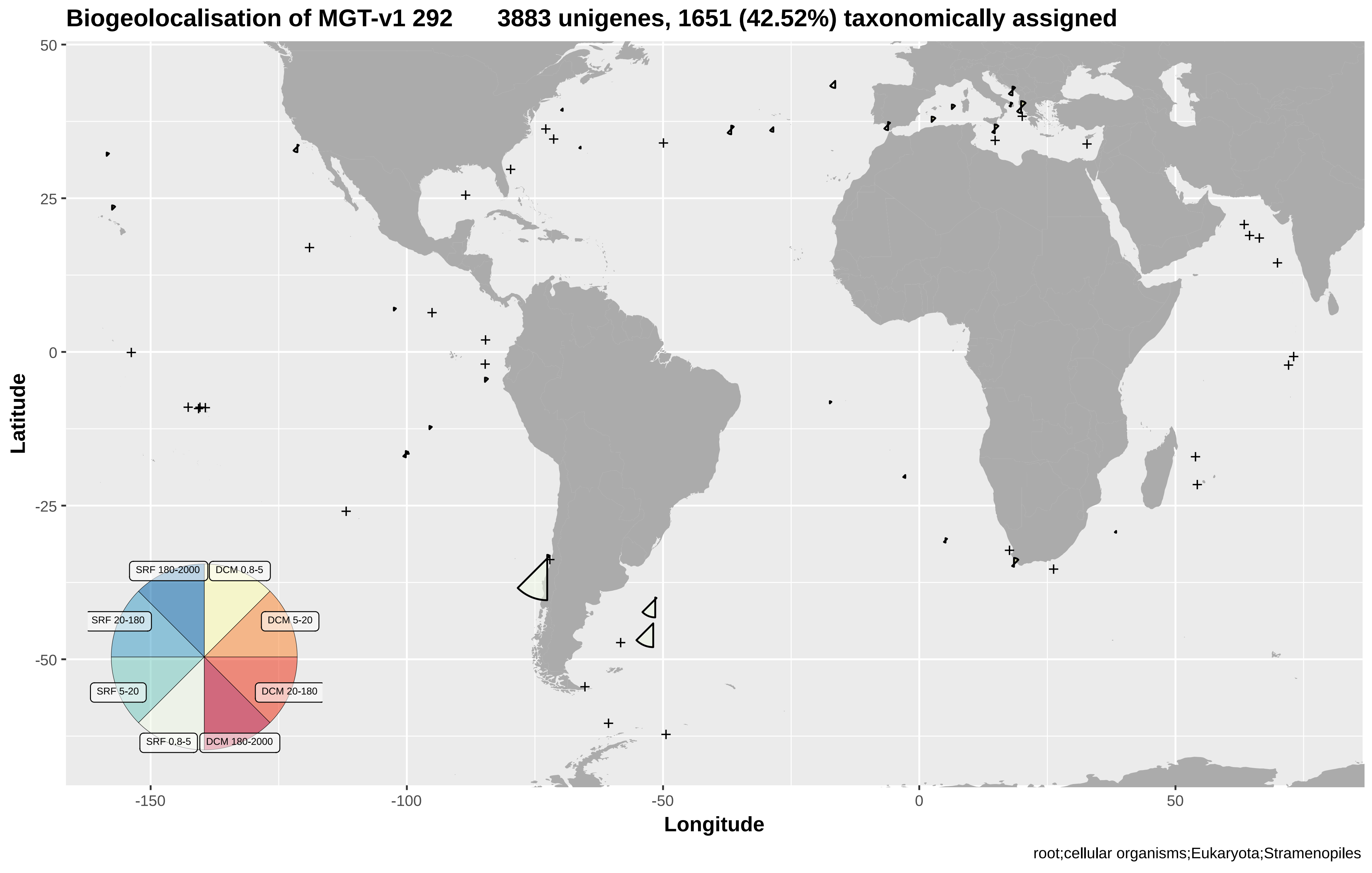
Biogeolocalisation of MGT-v1 292 3883 unigenes, 1651 (42.52%) taxonomically assigned
Latitude
50
25
0
-25
-50
Longitude
-150
-100
-50
0
50
SRF 180-2000
DCM 0.8-5
SRF 20-180
DCM 5-20
SRF 5-20
DCM 20-180
SRF 0.8-5
DCM 180-2000
root;cellular organisms;Eukaryota;Stramenopiles

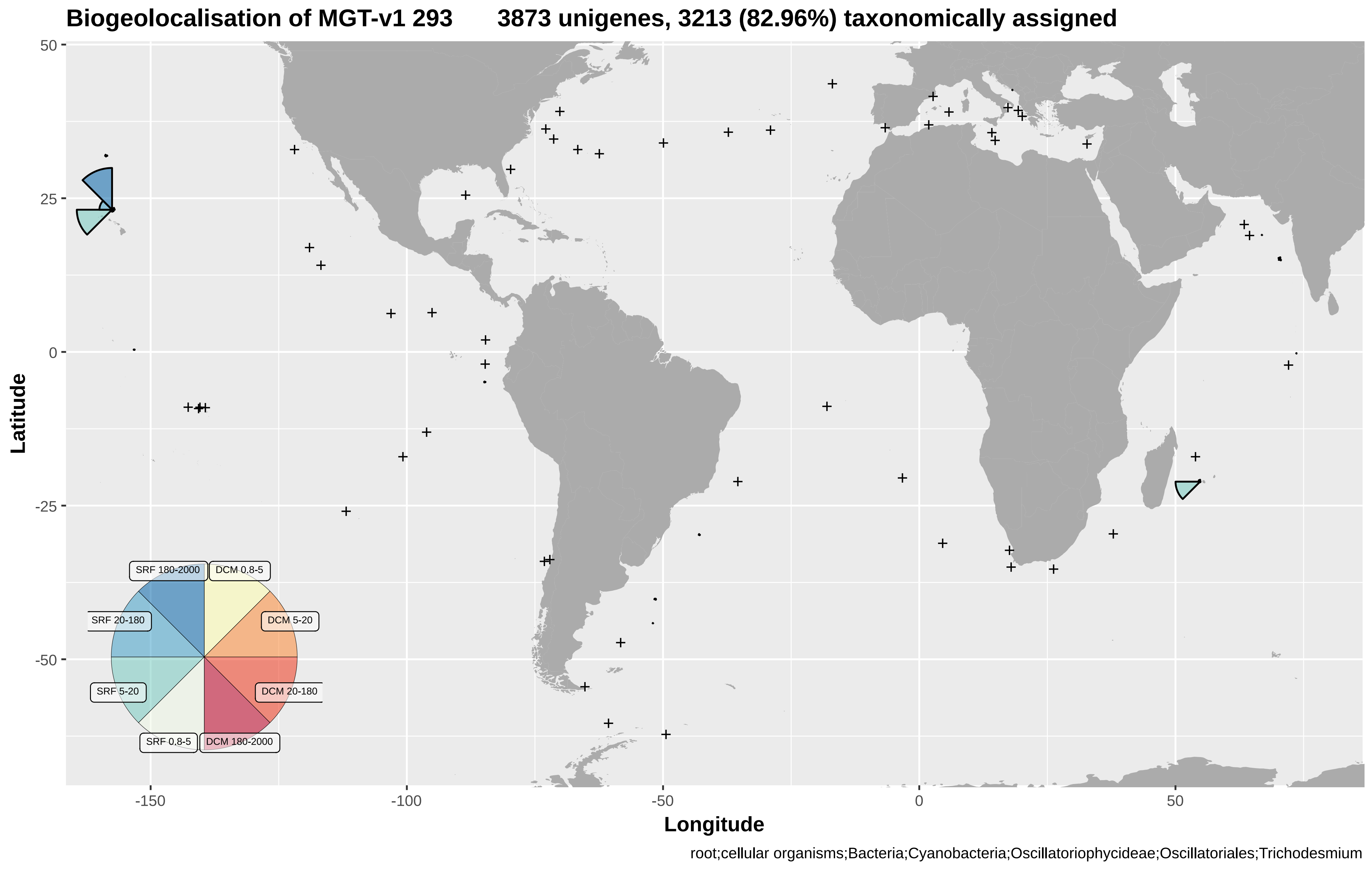

Biogeolocalisation of MGT-v1 293 3873 unigenes, 3213 (82.96%) taxonomically assigned
Latitude
Longitude
50
25
0
-25
-50
-150
-100
-50
0
50
SRF 180-2000
DCM 0.8-5
SRF 20-180
DCM 5-20
SRF 5-20
DCM 20-180
SRF 0.8-5
DCM 180-2000
root;cellular organisms;Bacteria;Cyanobacteria;Oscillatoriophycideae;Oscillatoriales;Trichodesmium

## Biogeolocalisation of MGT-v1 294      3865 unigenes, 862 (22.3%) taxonomically assigned

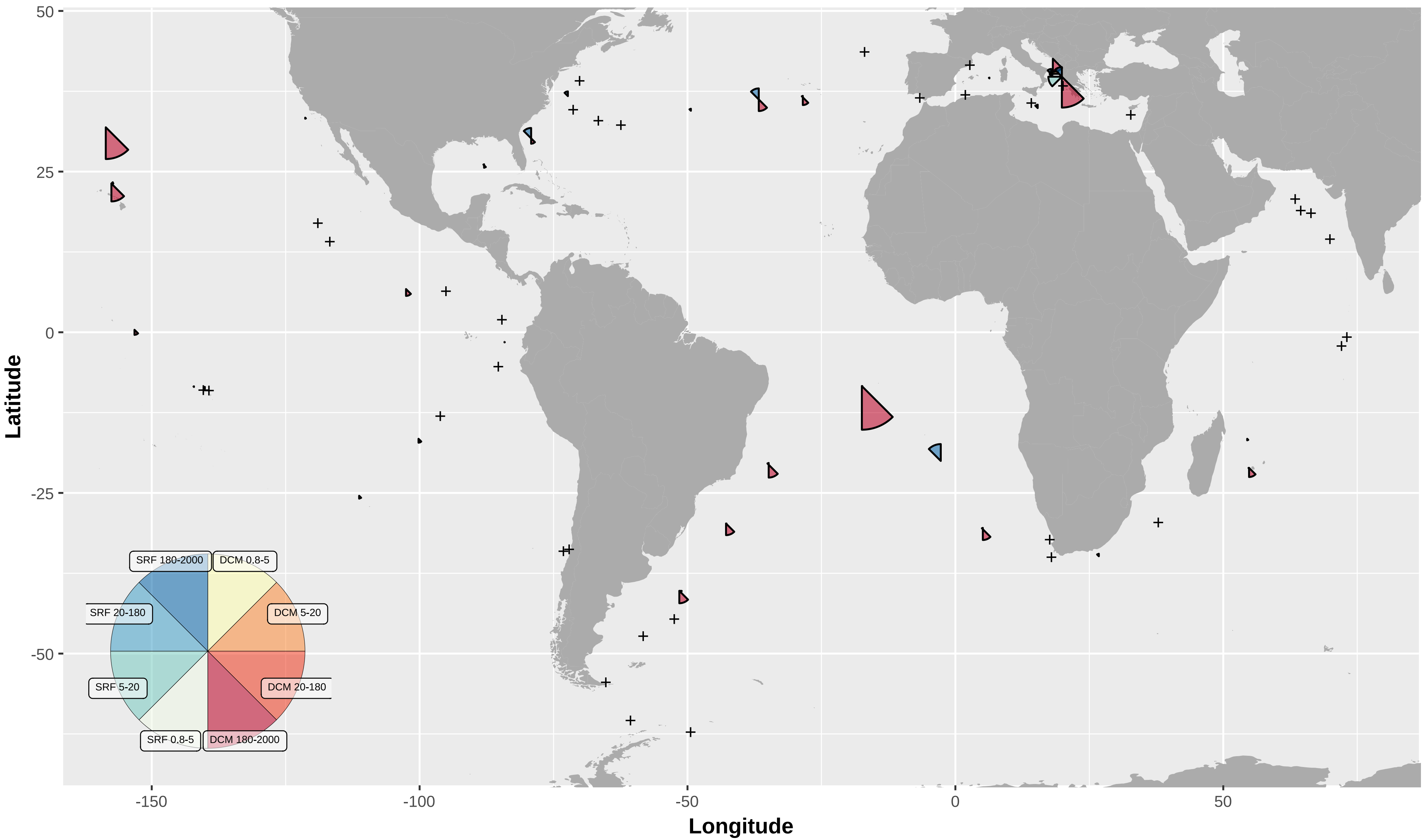


root;cellular organisms;Eukaryota;Opisthokonta;Metazoa;Eumetazoa;Bilateria

### Biogeolocalisation of MGT-v1 295 3819 unigenes, 387 (10.13%) taxonomically assigned

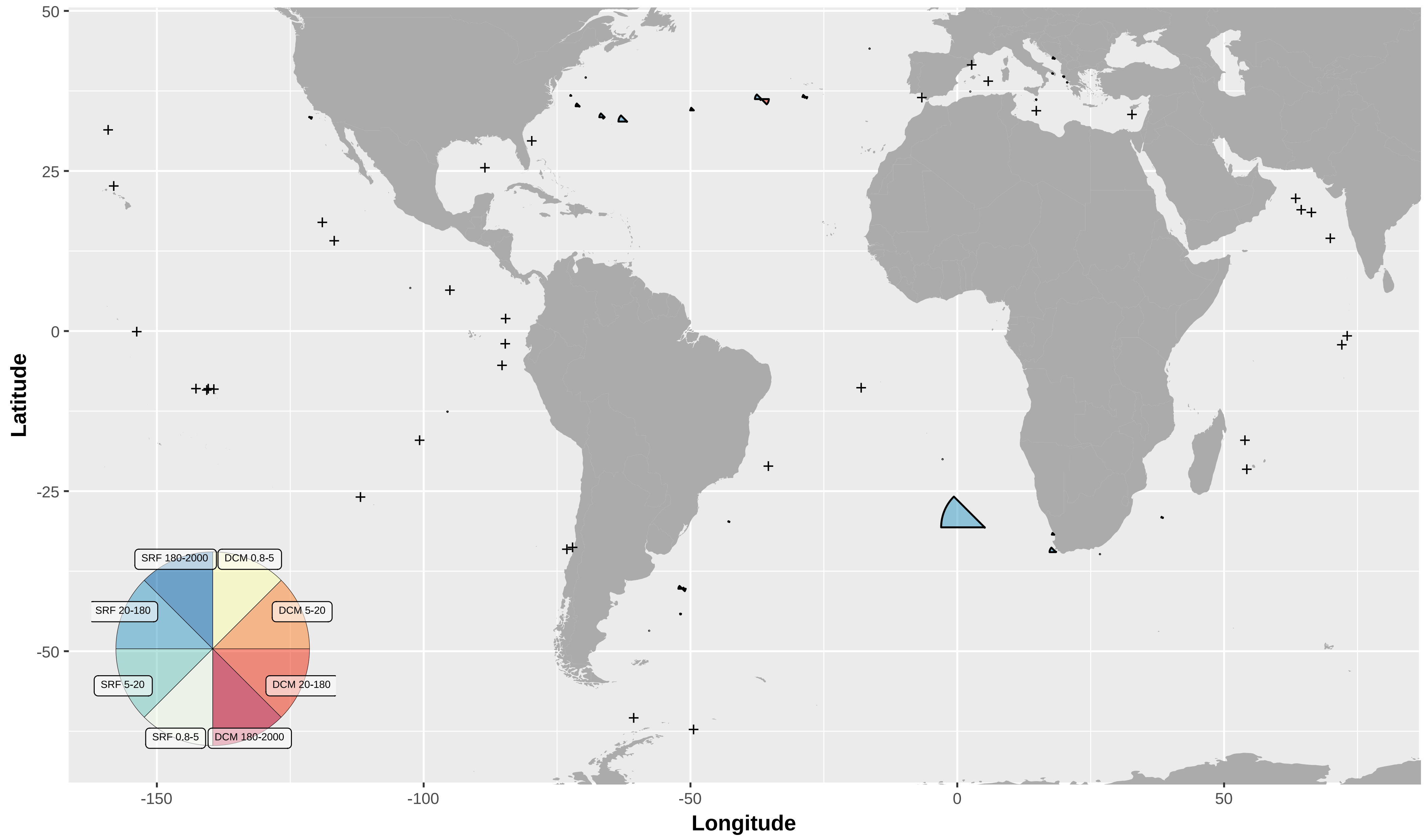


root;cellular organisms;Eukaryota;Opisthokonta;Metazoa;Eumetazoa;Bilateria

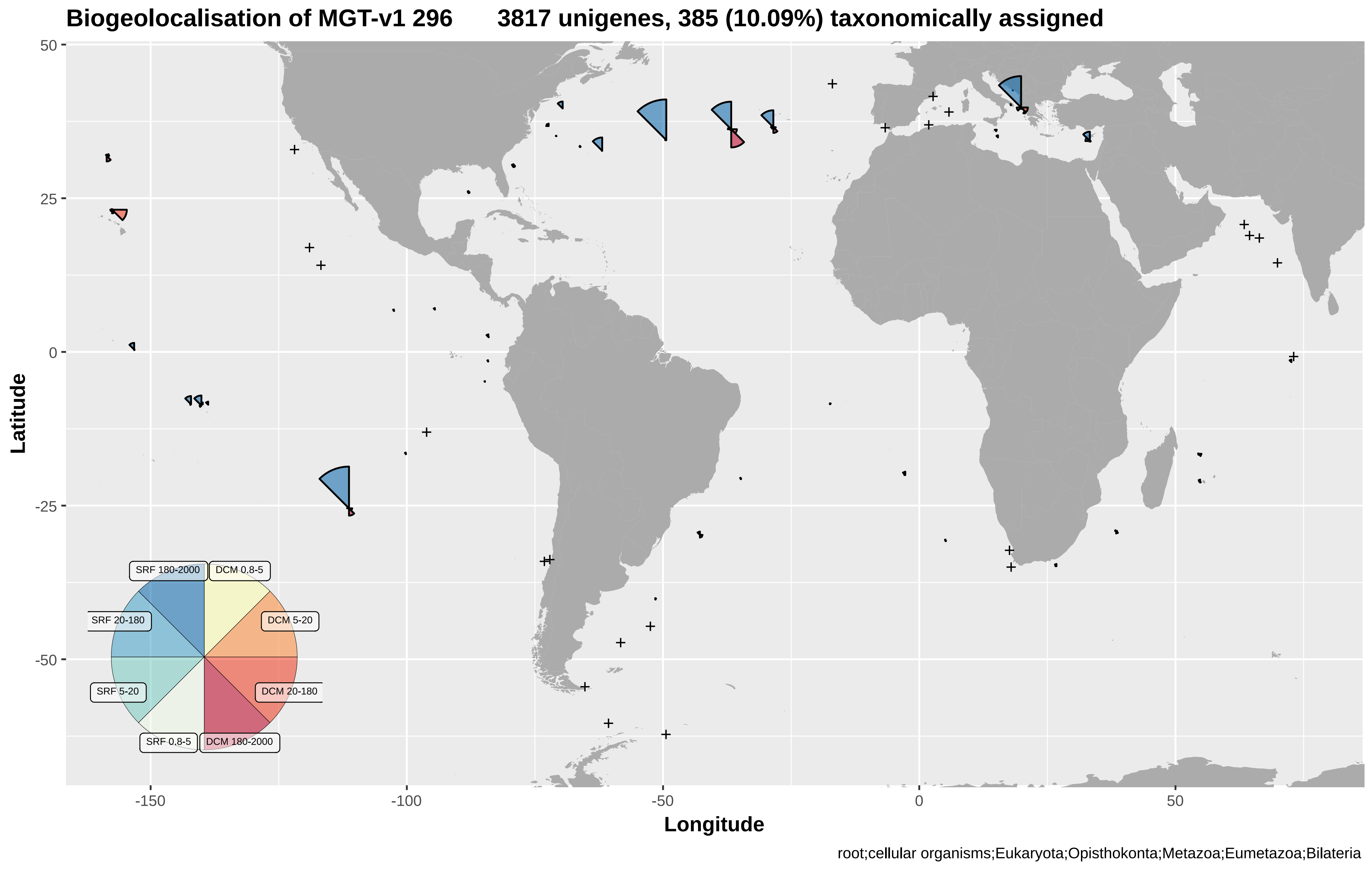
Biogeolocalisation of MGT-v1 296 3817 unigenes, 385 (10.09%) taxonomically assigned
Latitude
Longitude
50
25
0
-25
-50
-150
-100
-50
0
50
SRF 180-2000
DCM 0.8-5
SRF 20-180
DCM 5-20
SRF 5-20
DCM 20-180
SRF 0.8-5
DCM 180-2000
root;cellular organisms;Eukaryota;Opisthokonta;Metazoa;Eumetazoa;Bilateria

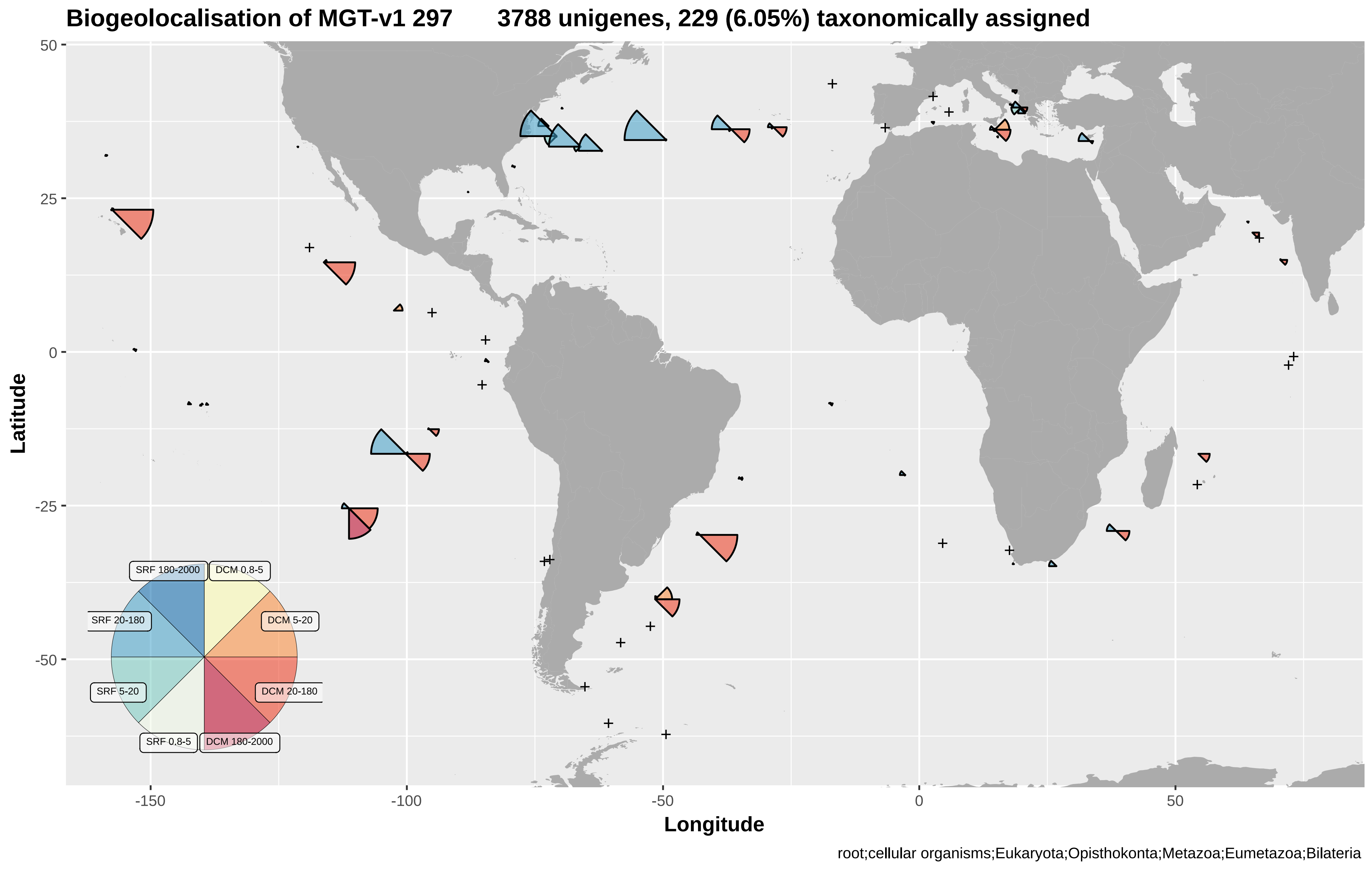
Biogeolocalisation of MGT-v1 297 3788 unigenes, 229 (6.05%) taxonomically assigned
50
25
0
-25
-50
Latitude
-150
-100
-50
0
50
Longitude
SRF 180-2000
DCM 0.8-5
SRF 20-180
DCM 5-20
SRF 5-20
DCM 20-180
SRF 0.8-5
DCM 180-2000
root;cellular organisms;Eukaryota;Opisthokonta;Metazoa;Eumetazoa;Bilateria

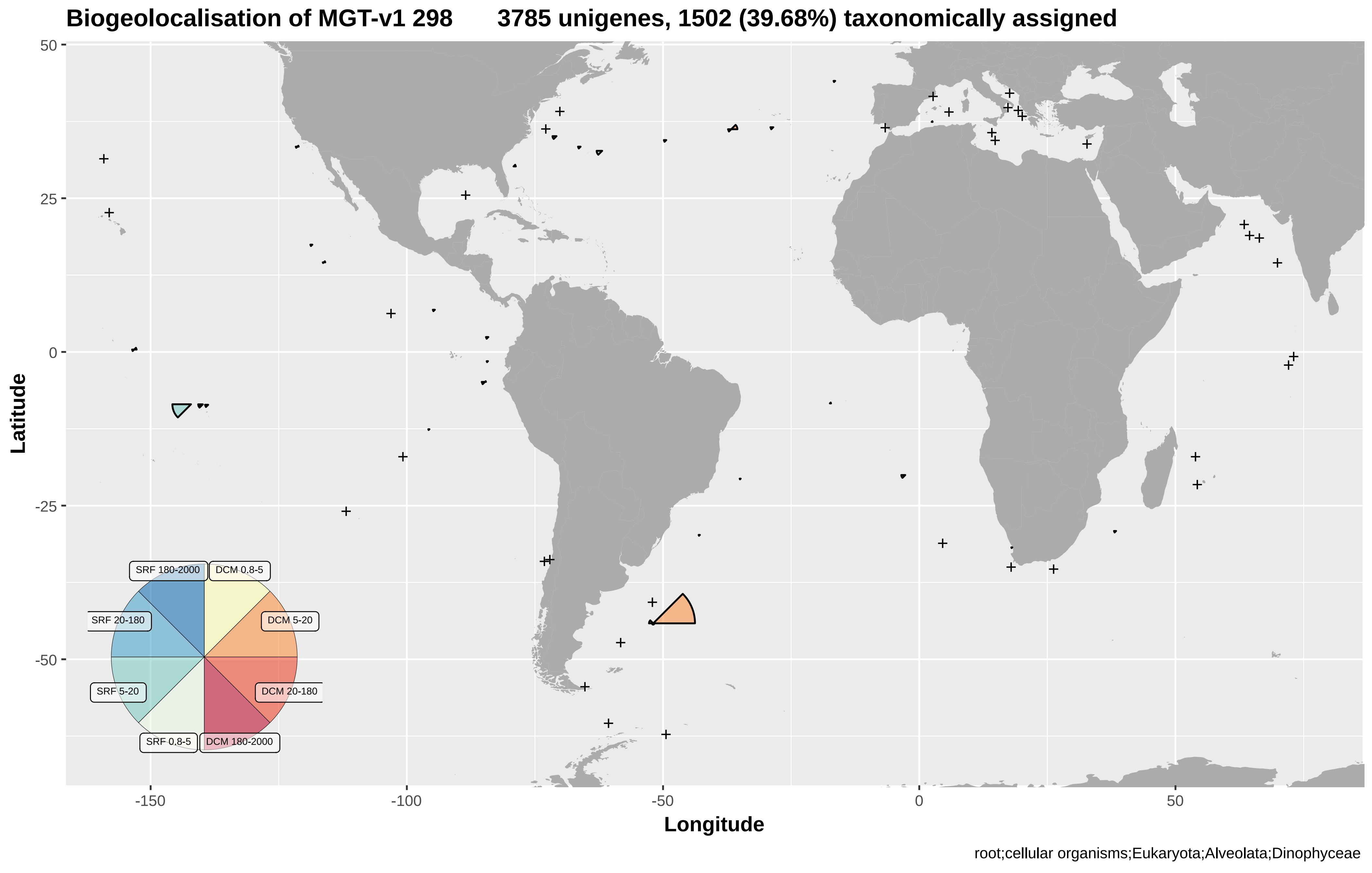
Biogeolocalisation of MGT-v1 298 3785 unigenes, 1502 (39.68%) taxonomically assigned
Latitude
Longitude
50
25
0
-25
-50
-150
-100
-50
0
50
SRF 180-2000
DCM 0.8-5
SRF 20-180
DCM 5-20
SRF 5-20
DCM 20-180
SRF 0.8-5
DCM 180-2000
root;cellular organisms;Eukaryota;Alveolata;Dinophyceae

## Biogeolocalisation of MGT-v1 299      3784 unigenes, 433 (11.44%) taxonomically assigned

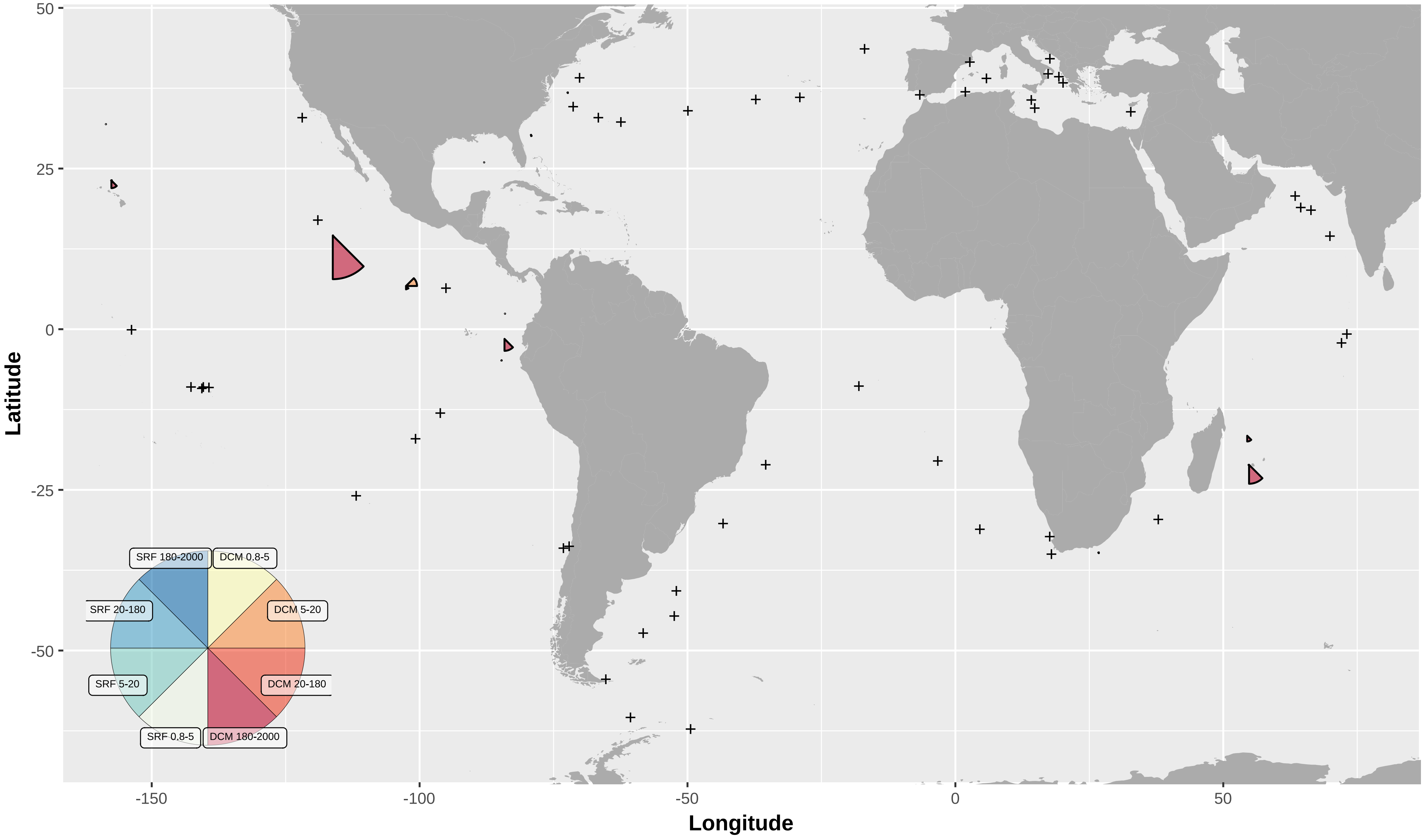


root;cellular organisms;Eukaryota;Opisthokonta;Metazoa;Eumetazoa;Bilateria

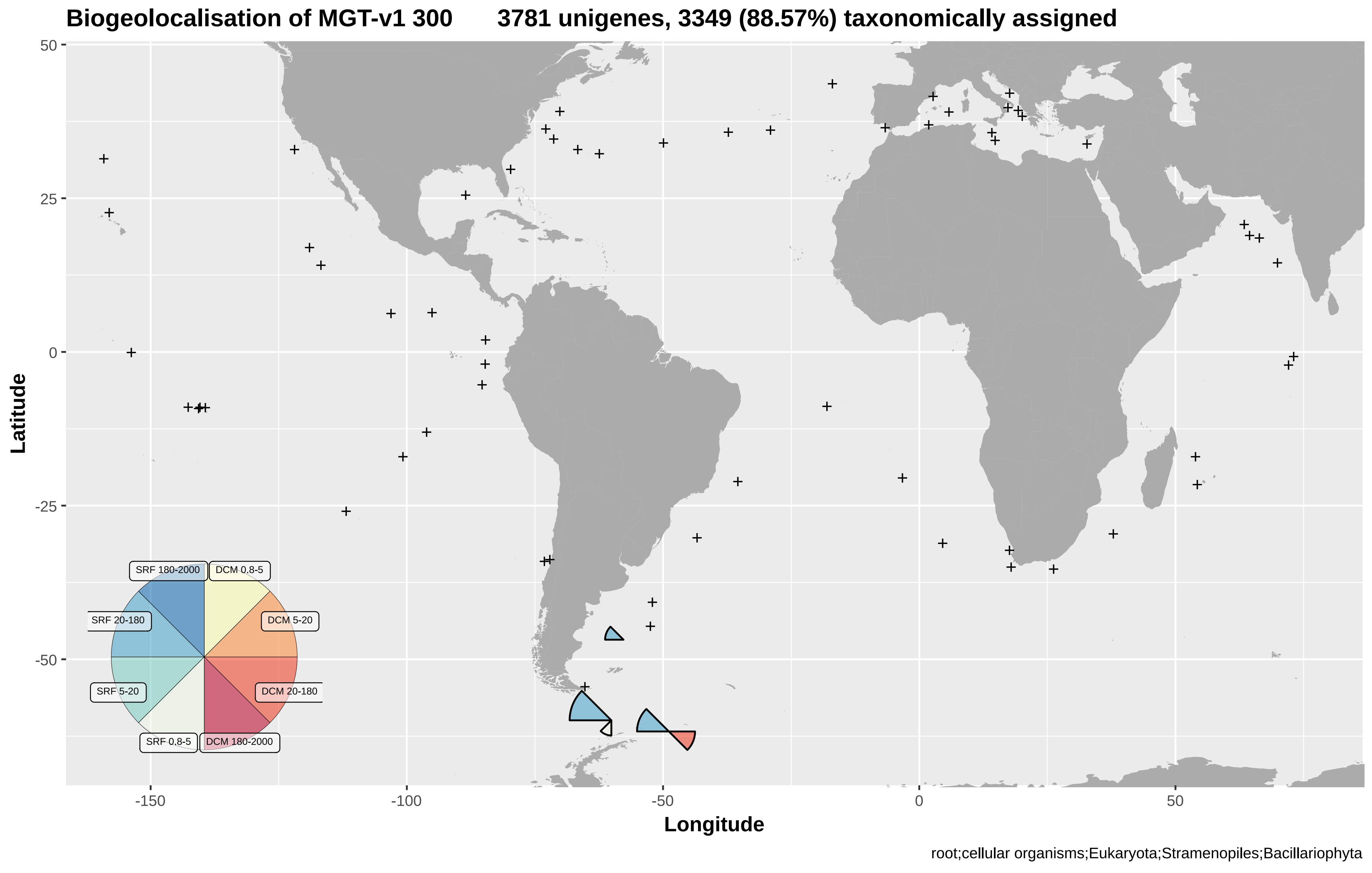

Biogeolocalisation of MGT-v1 300 3781 unigenes, 3349 (88.57%) taxonomically assigned
50
25
0
-25
-50
Latitude
-150
-100
-50
0
50
Longitude
SRF 180-2000
DCM 0.8-5
SRF 20-180
DCM 5-20
SRF 5-20
DCM 20-180
SRF 0.8-5
DCM 180-2000
root;cellular organisms;Eukaryota;Stramenopiles;Bacillariophyta

#### Biogeolocalisation of MGT-v1 301 3735 unigenes, 2201 (58.93%) taxonomically assigned

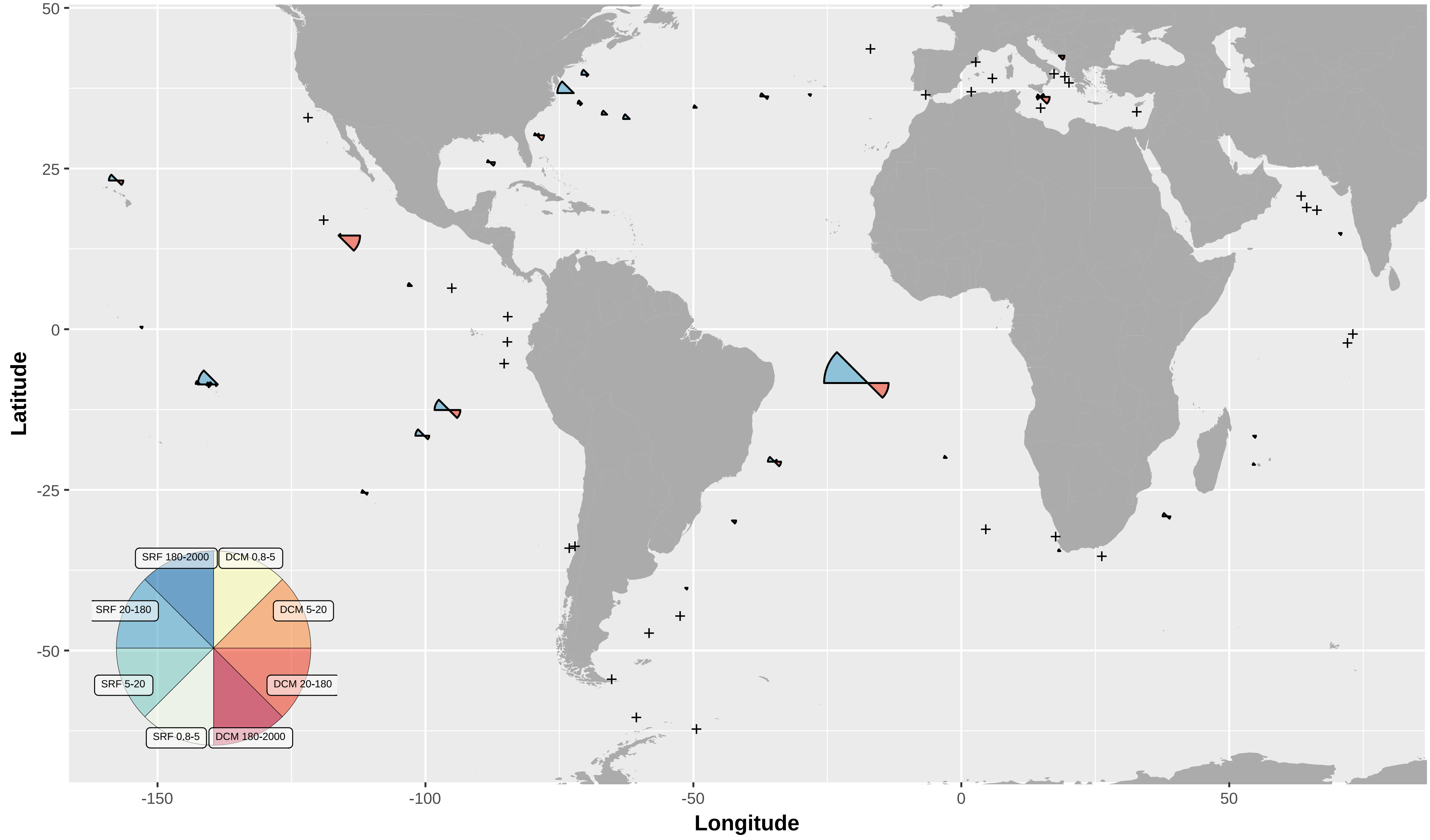


root;cellular organisms;Eukaryota;Alveolata;Ciliophora;Intramacronucleata;Spirotrichea

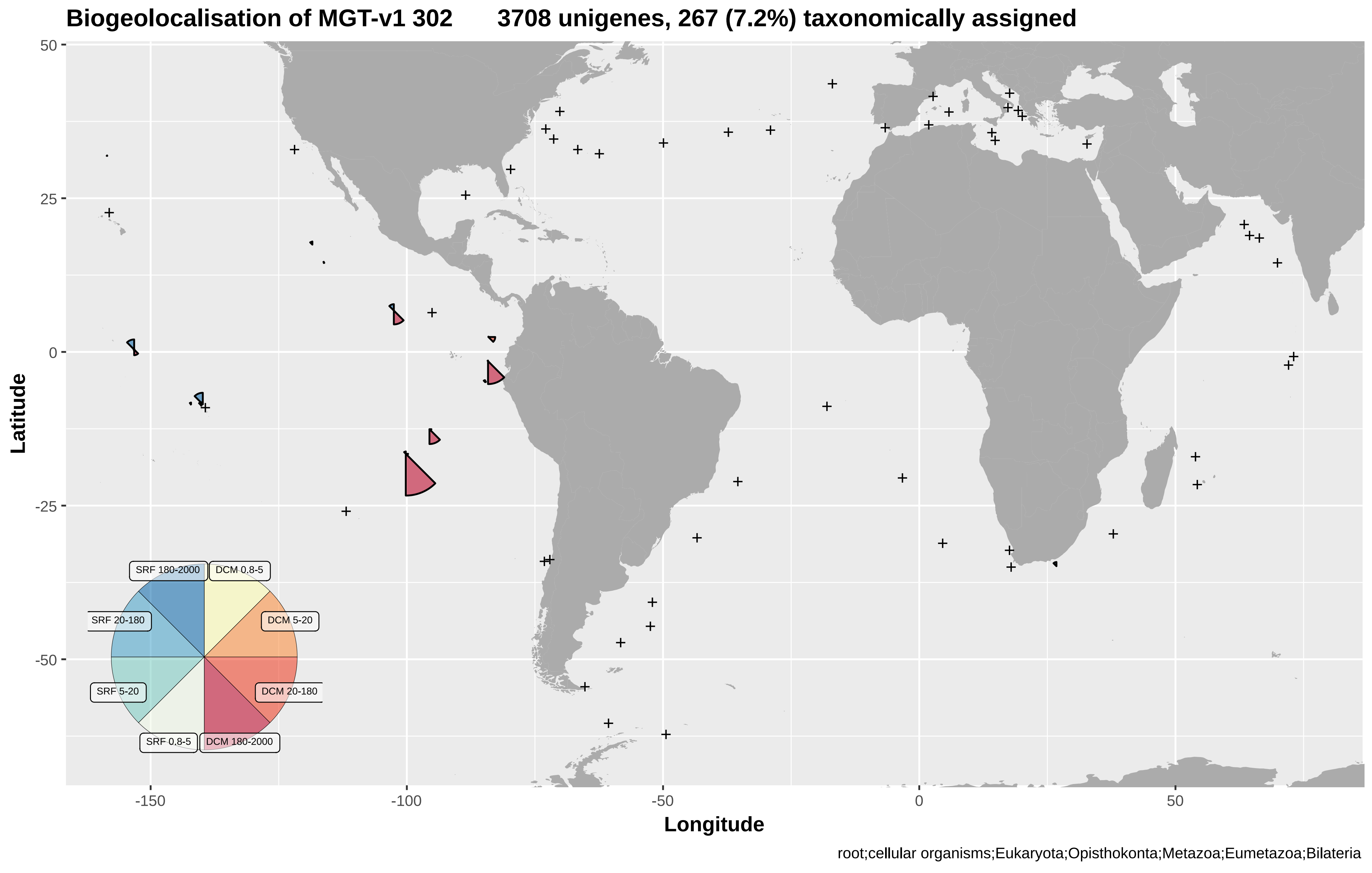
Biogeolocalisation of MGT-v1 302 3708 unigenes, 267 (7.2%) taxonomically assigned
Latitude
Longitude
50
25
0
-25
-50
-150
-100
-50
0
50
SRF 180-2000
DCM 0.8-5
SRF 20-180
DCM 5-20
SRF 5-20
DCM 20-180
SRF 0.8-5
DCM 180-2000
root;cellular organisms;Eukaryota;Opisthokonta;Metazoa;Eumetazoa;Bilateria

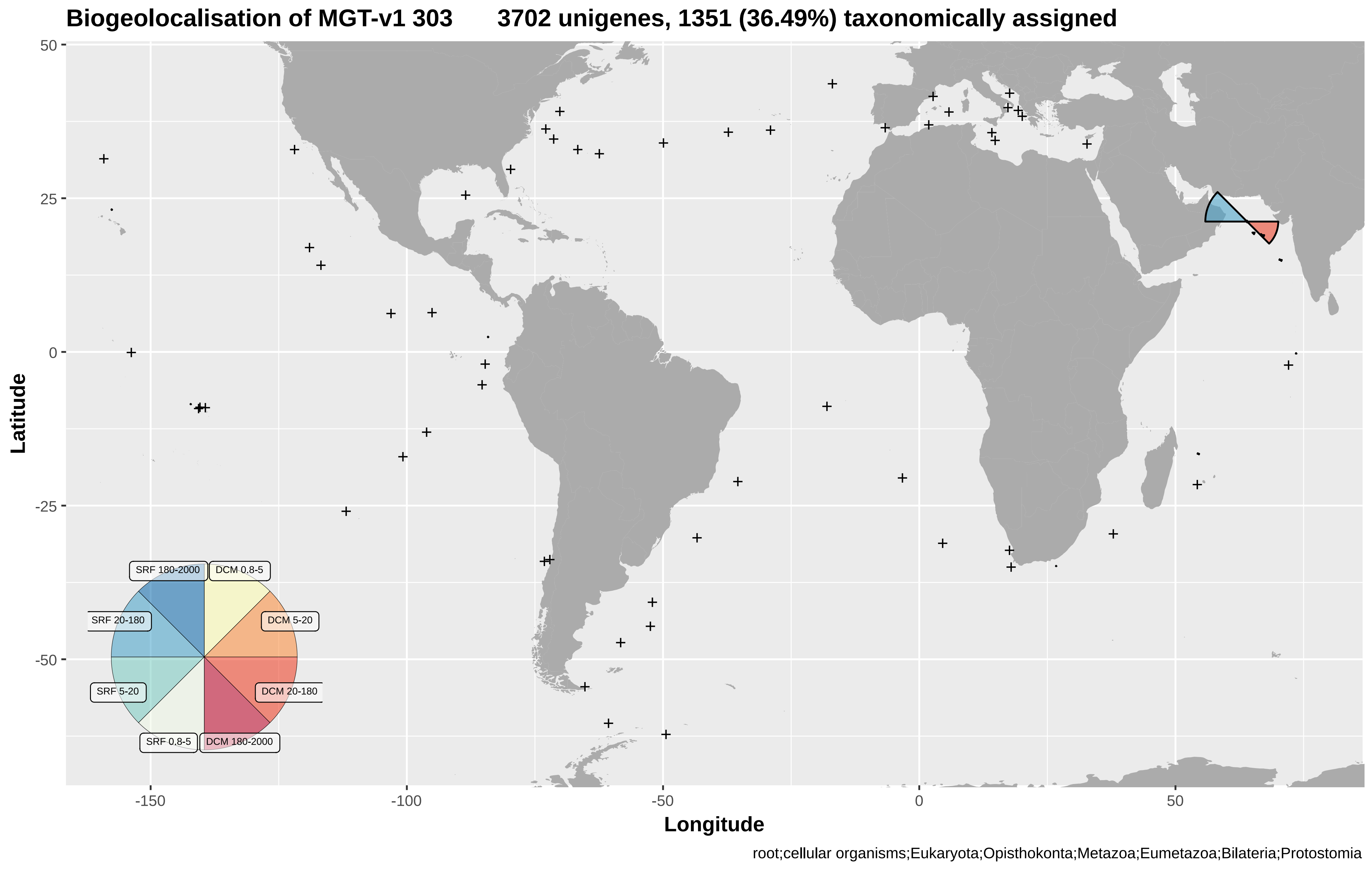
Biogeolocalisation of MGT-v1 303 3702 unigenes, 1351 (36.49%) taxonomically assigned
Latitude
Longitude
50
25
0
-25
-50
-150
-100
-50
0
50
SRF 180-2000
DCM 0.8-5
SRF 20-180
DCM 5-20
SRF 5-20
DCM 20-180
SRF 0.8-5
DCM 180-2000
root;cellular organisms;Eukaryota;Opisthokonta;Metazoa;Eumetazoa;Bilateria;Protostomia

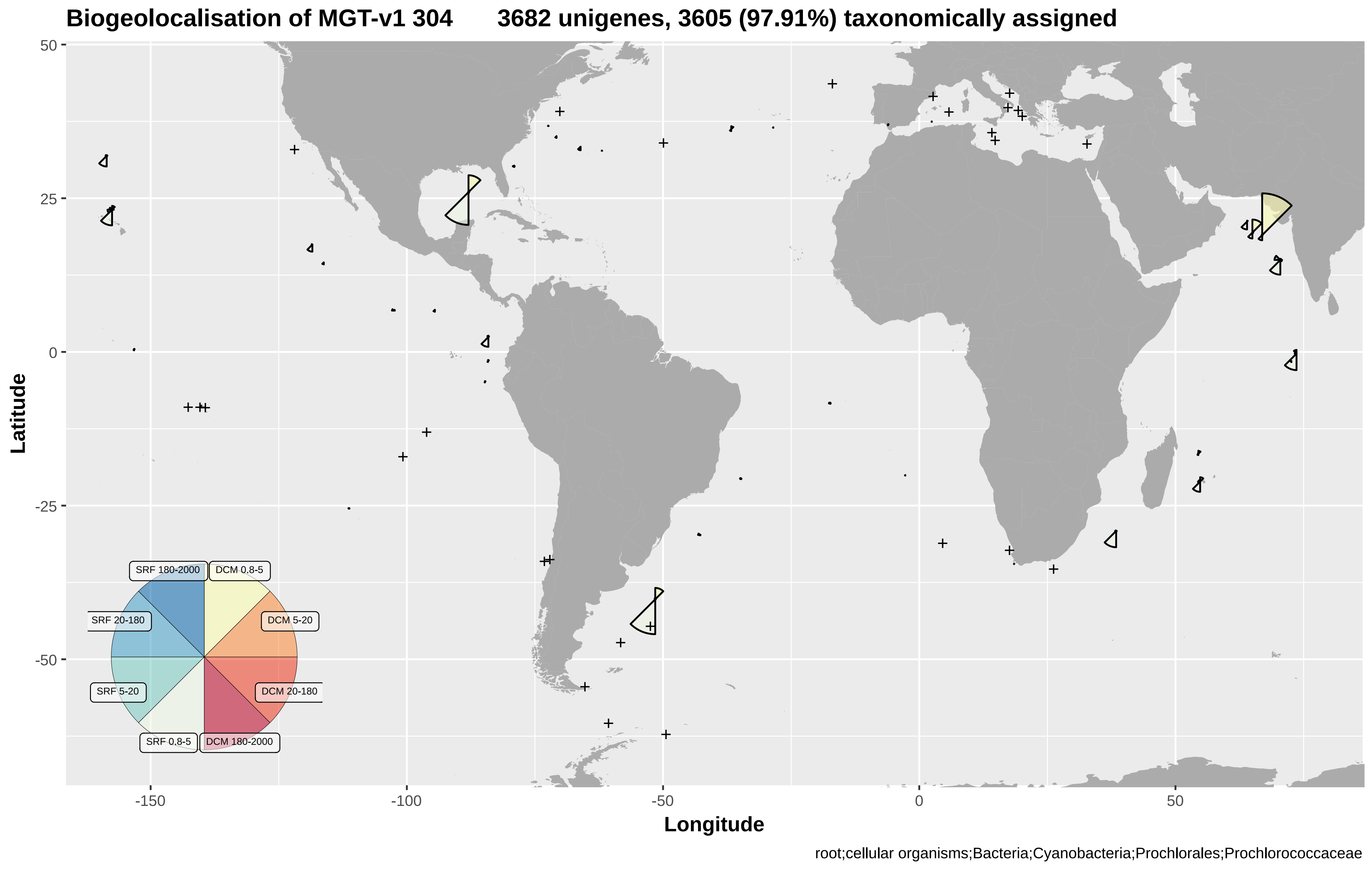
Biogeolocalisation of MGT-v1 304 3682 unigenes, 3605 (97.91%) taxonomically assigned
50
25
0
-25
-50
Latitude
-150
-100
-50
0
50
Longitude
SRF 180-2000
DCM 0.8-5
SRF 20-180
DCM 5-20
SRF 5-20
DCM 20-180
SRF 0.8-5
DCM 180-2000
root;cellular organisms;Bacteria;Cyanobacteria;Prochlorales;Prochlorococcaceae

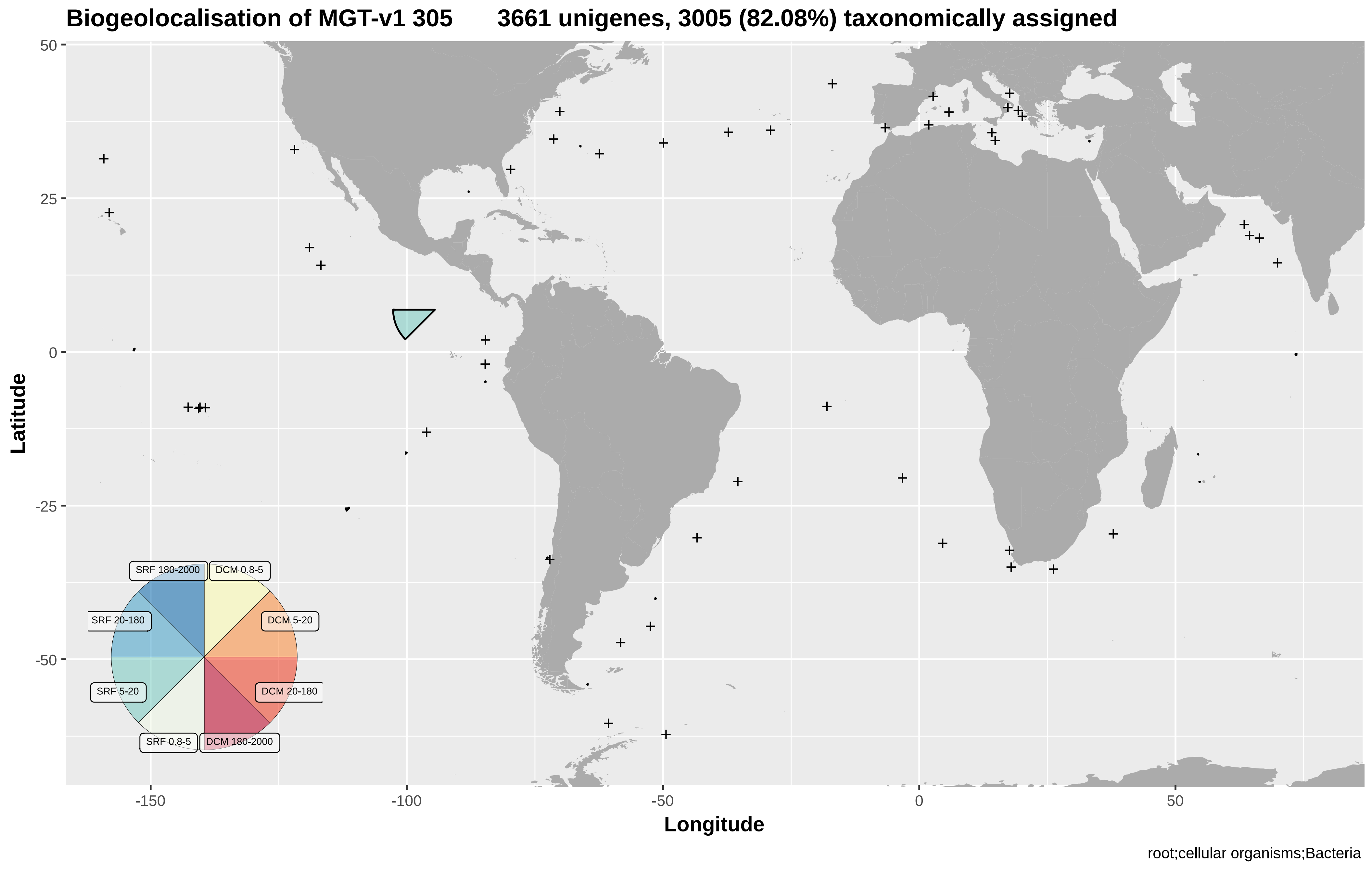

Biogeolocalisation of MGT-v1 305 3661 unigenes, 3005 (82.08%) taxonomically assigned
Latitude
Longitude
50
25
0
-25
-50
-150
-100
-50
0
50
SRF 180-2000
DCM 0.8-5
SRF 20-180
DCM 5-20
SRF 5-20
DCM 20-180
SRF 0.8-5
DCM 180-2000
root;cellular organisms;Bacteria

### Biogeolocalisation of MGT-v1 306 3659 unigenes, 495 (13.53%) taxonomically assigned

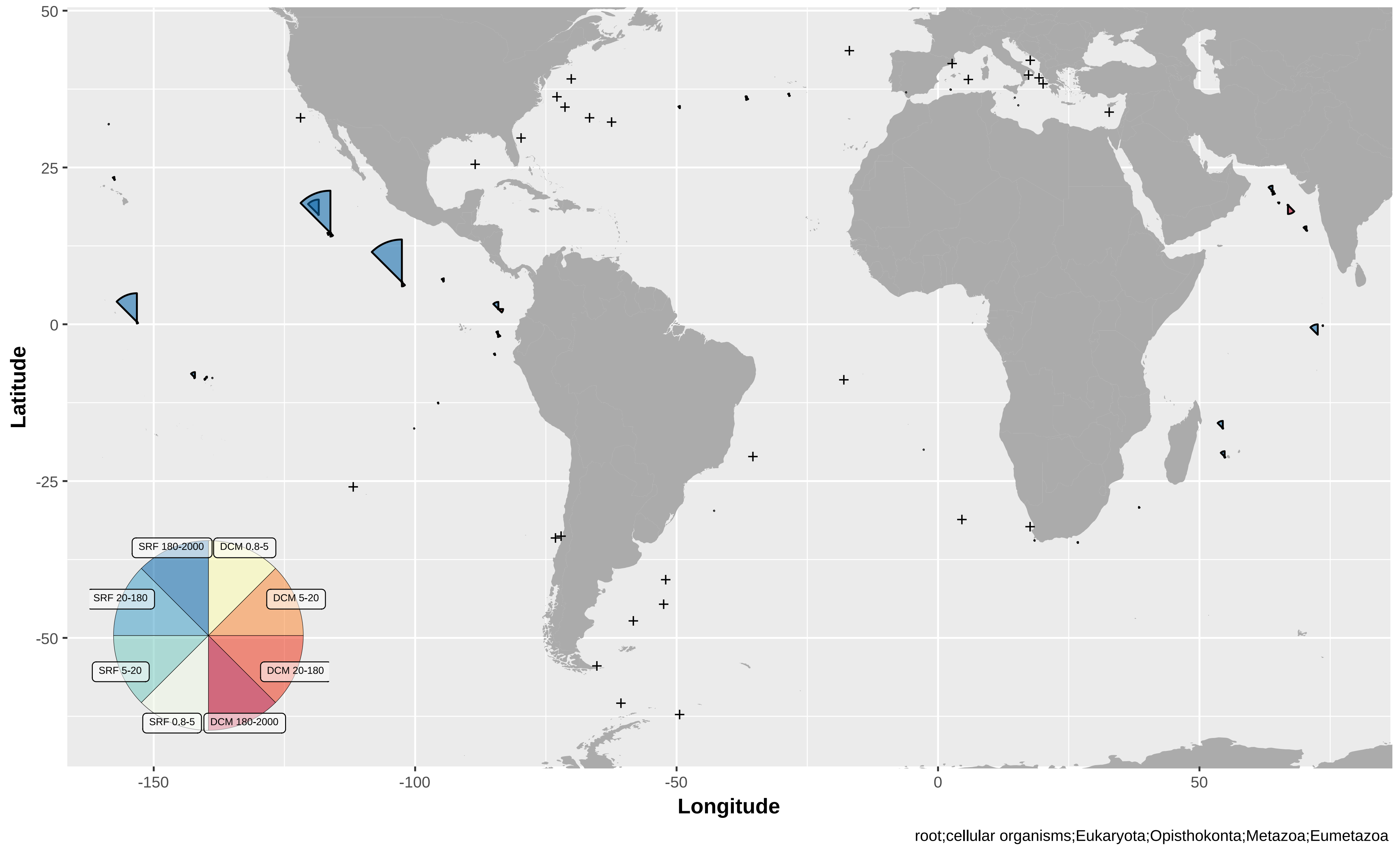

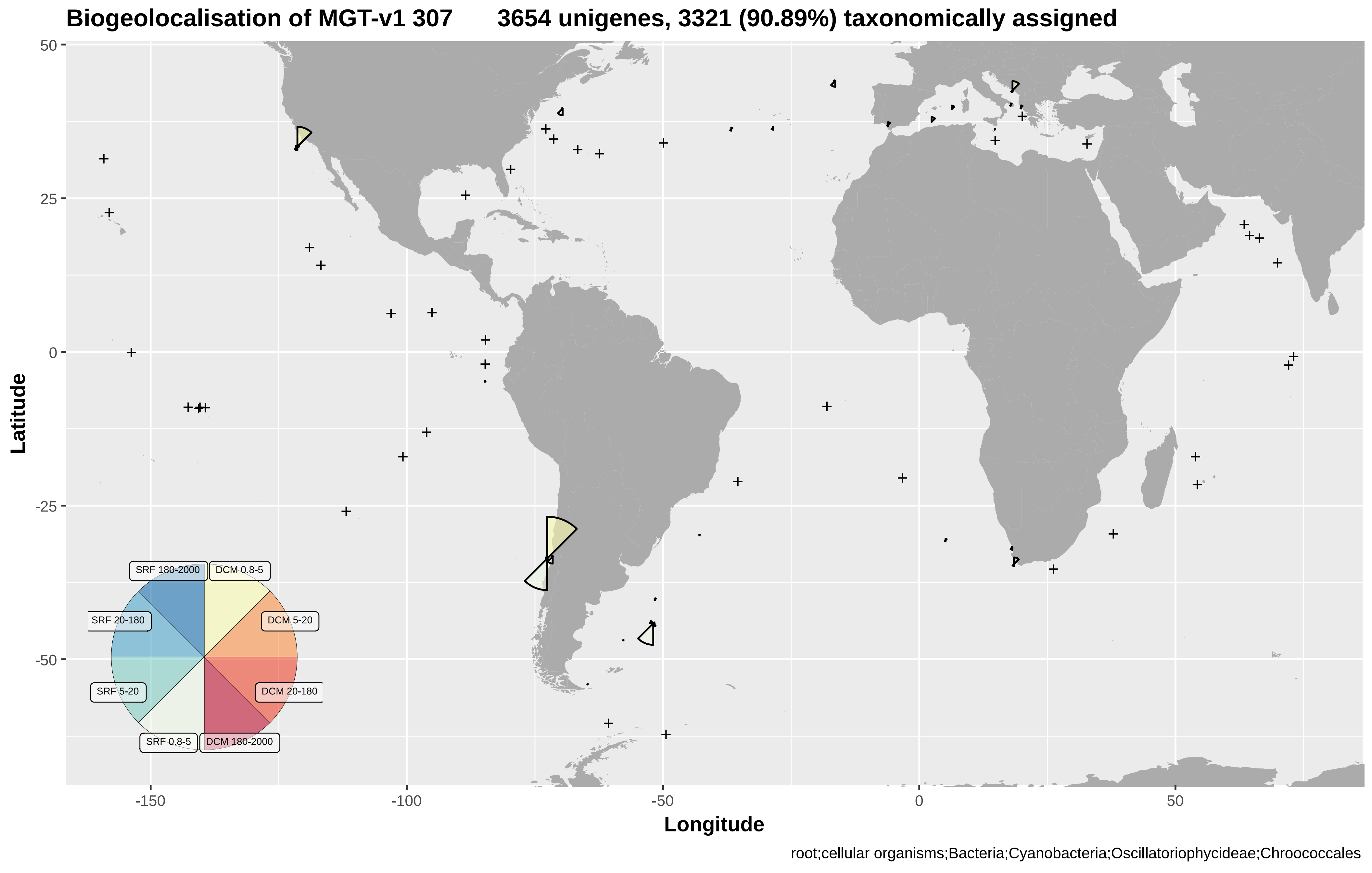
Biogeolocalisation of MGT-v1 307 3654 unigenes, 3321 (90.89%) taxonomically assigned
Latitude
Longitude
50
25
0
-25
-50
-150
-100
-50
0
50
SRF 180-2000
DCM 0.8-5
SRF 20-180
DCM 5-20
SRF 5-20
DCM 20-180
SRF 0.8-5
DCM 180-2000
root;cellular organisms;Bacteria;Cyanobacteria;Oscillatoriophycideae;Chroococcales

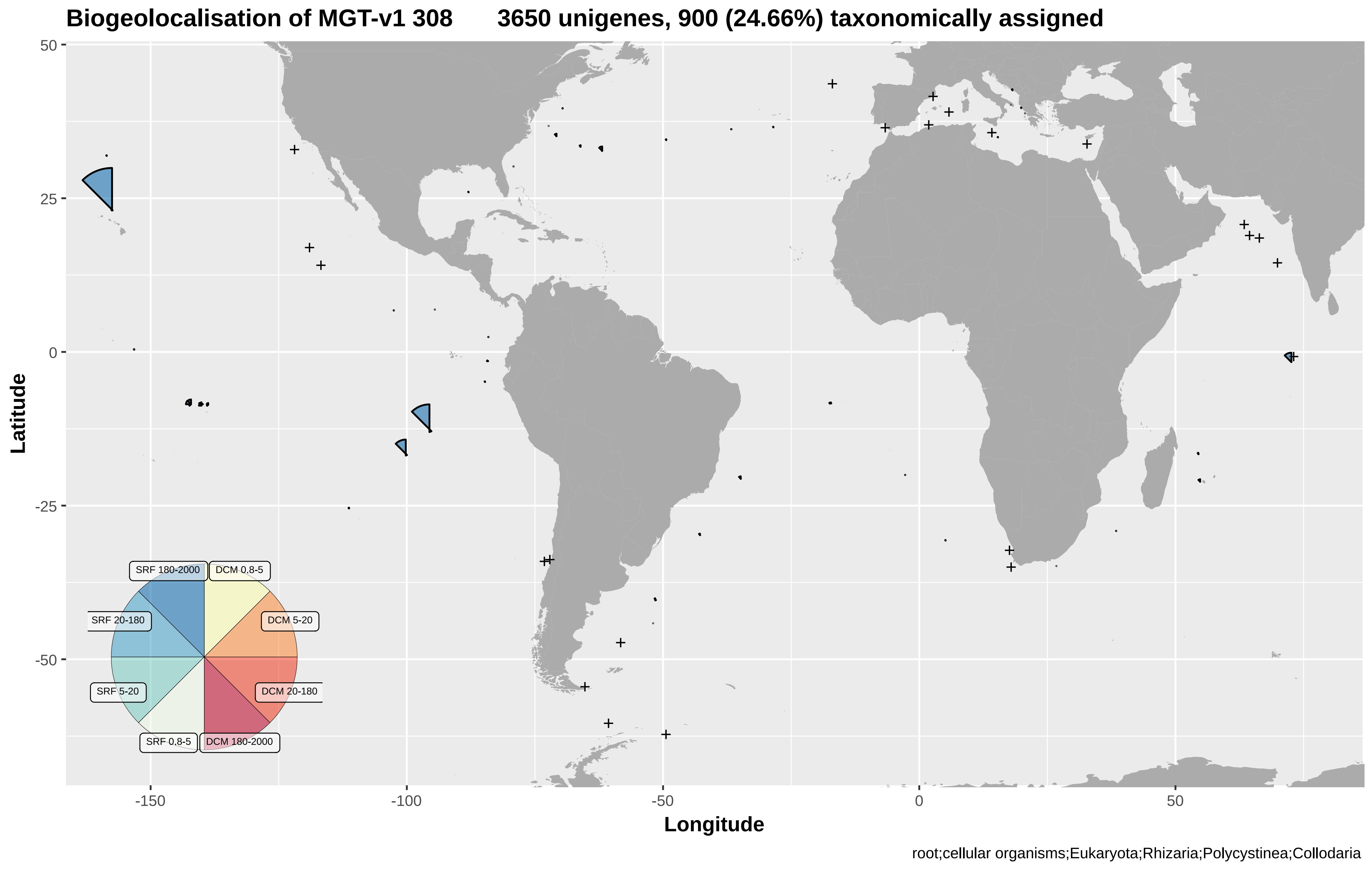

Biogeolocalisation of MGT-v1 308 3650 unigenes, 900 (24.66%) taxonomically assigned
50
25
0
-25
-50
Latitude
-150
-100
-50
0
50
Longitude
SRF 180-2000
DCM 0.8-5
SRF 20-180
DCM 5-20
SRF 5-20
DCM 20-180
SRF 0.8-5
DCM 180-2000
root;cellular organisms;Eukaryota;Rhizaria;Polycystinea;Collodaria

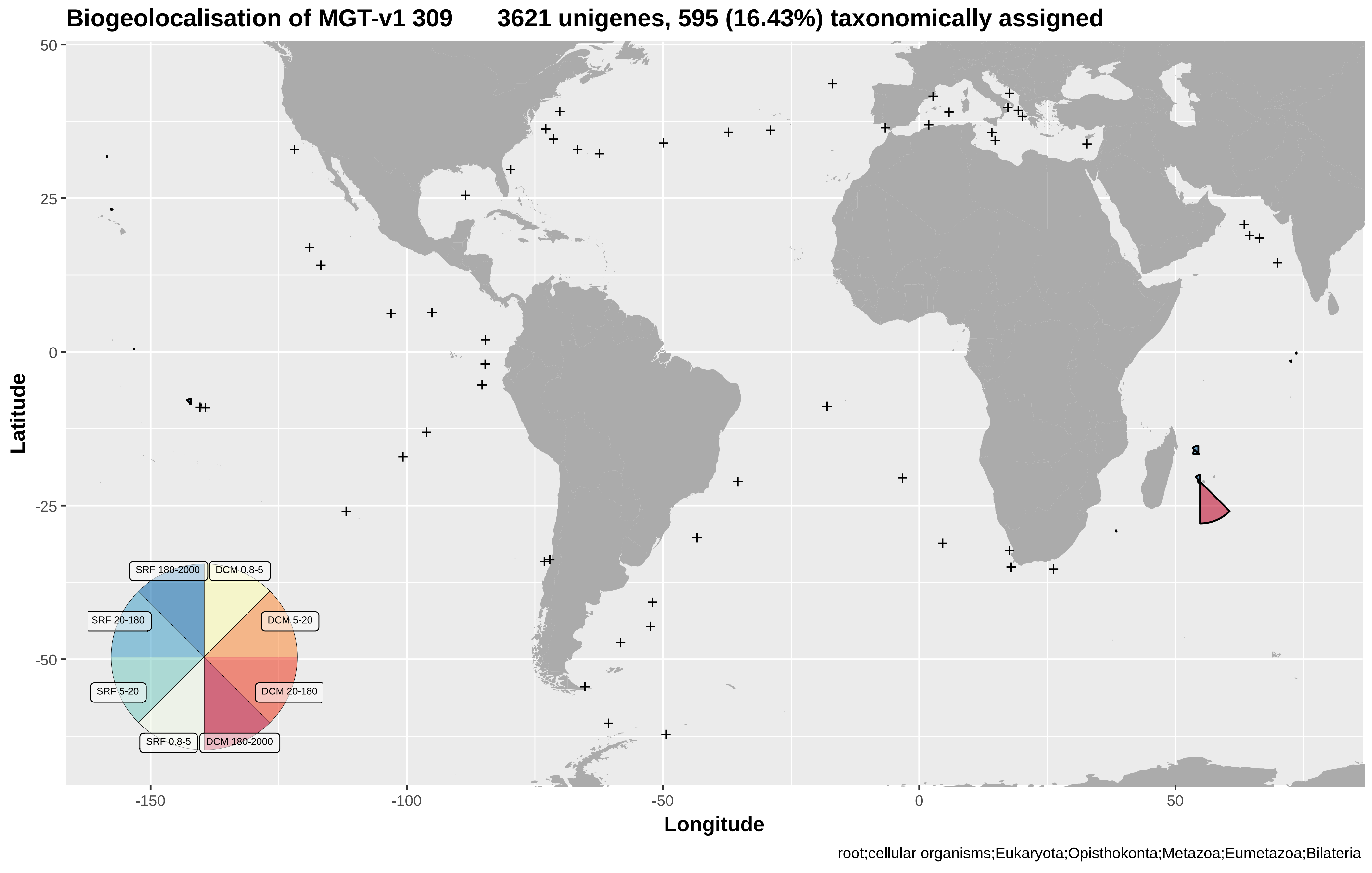

Biogeolocalisation of MGT-v1 309 3621 unigenes, 595 (16.43%) taxonomically assigned
Latitude
Longitude
50
25
0
-25
-50
-150
-100
-50
0
50
SRF 180-2000
DCM 0.8-5
SRF 20-180
DCM 5-20
SRF 5-20
DCM 20-180
SRF 0.8-5
DCM 180-2000
root;cellular organisms;Eukaryota;Opisthokonta;Metazoa;Eumetazoa;Bilateria

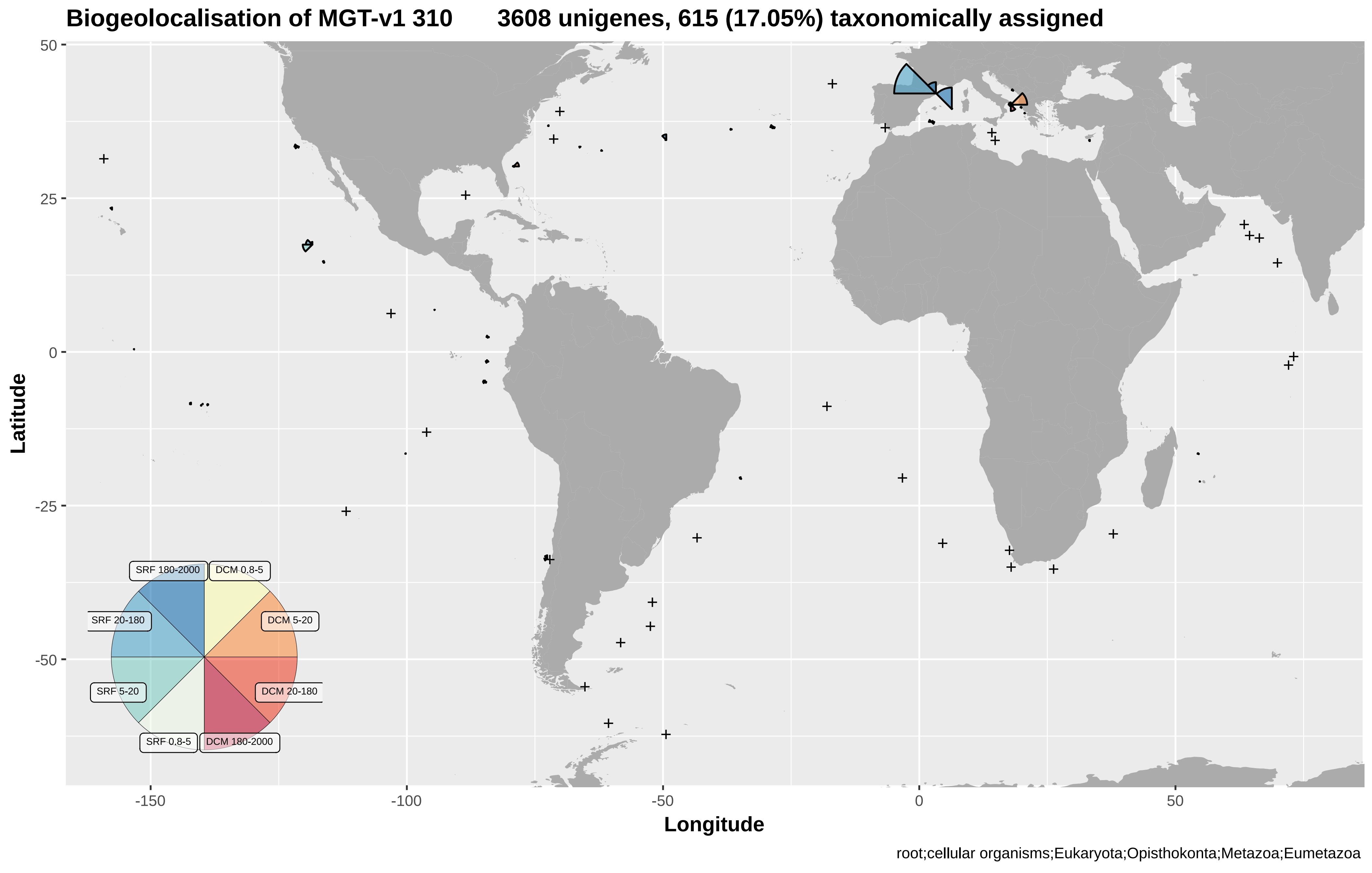

Biogeolocalisation of MGT-v1 310 3608 unigenes, 615 (17.05%) taxonomically assigned
50
25
0
-25
-50
Latitude
-150
-100
-50
0
50
Longitude
SRF 180-2000
DCM 0.8-5
SRF 20-180
DCM 5-20
SRF 5-20
DCM 20-180
SRF 0.8-5
DCM 180-2000
root;cellular organisms;Eukaryota;Opisthokonta;Metazoa;Eumetazoa

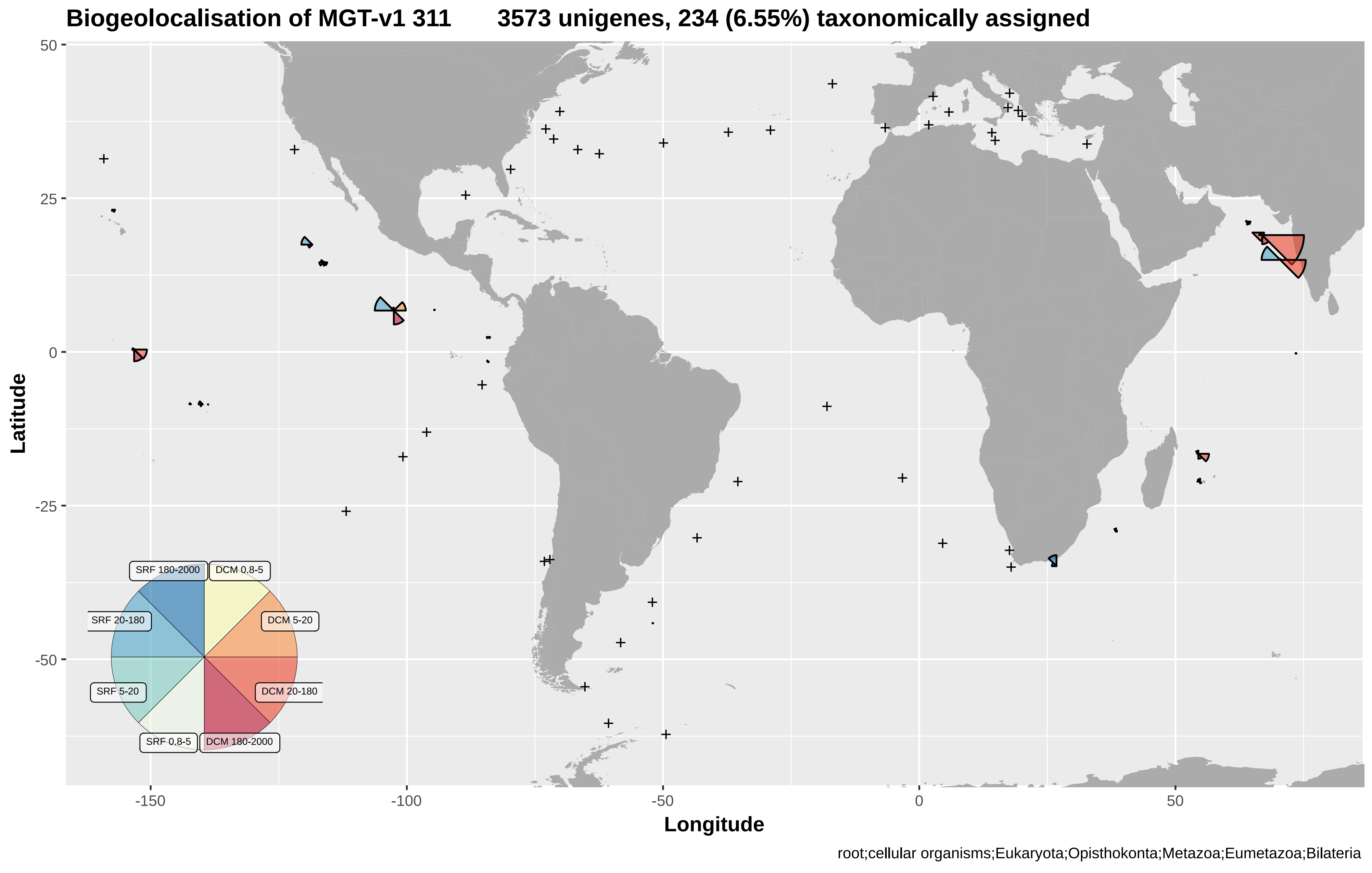

Biogeolocalisation of MGT-v1 311      3573 unigenes, 234 (6.55%) taxonomically assigned
50
25
0
-25
-50
Latitude
-150
-100
-50
0
50
Longitude
SRF 180-2000
DCM 0.8-5
SRF 20-180
DCM 5-20
SRF 5-20
DCM 20-180
SRF 0.8-5
DCM 180-2000
root;cellular organisms;Eukaryota;Opisthokonta;Metazoa;Eumetazoa;Bilateria

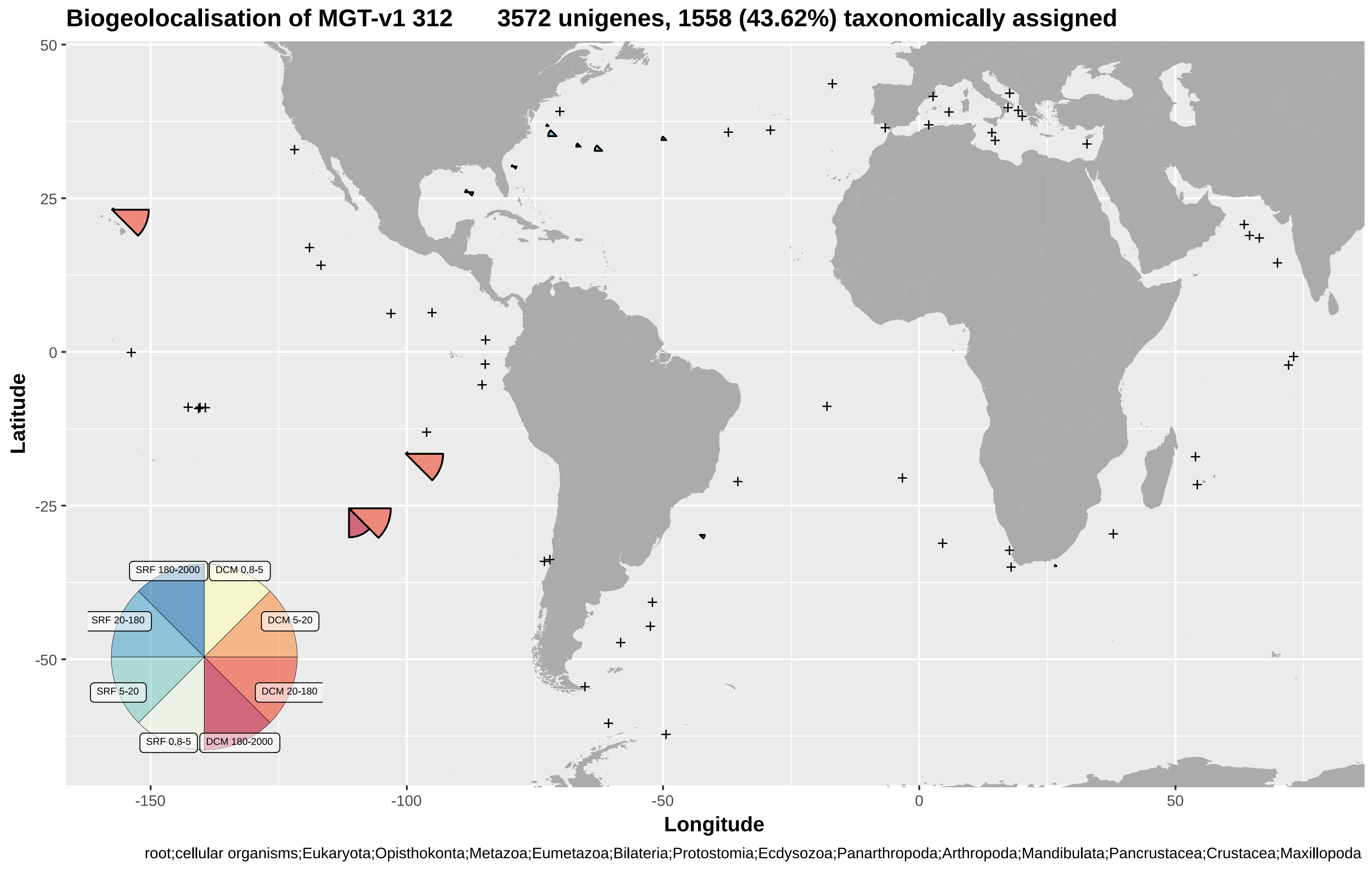
Biogeolocalisation of MGT-v1 312 3572 unigenes, 1558 (43.62%) taxonomically assigned
Latitude
Longitude
50
25
0
-25
-50
-150
-100
-50
0
50
SRF 180-2000
DCM 0.8-5
SRF 20-180
DCM 5-20
SRF 5-20
DCM 20-180
SRF 0.8-5
DCM 180-2000
root;cellular organisms;Eukaryota;Opisthokonta;Metazoa;Eumetazoa;Bilateria;Protostomia;Ecdysozoa;Panarthropoda;Arthropoda;Mandibulata;Pancrustacea;Crustacea;Maxillopoda

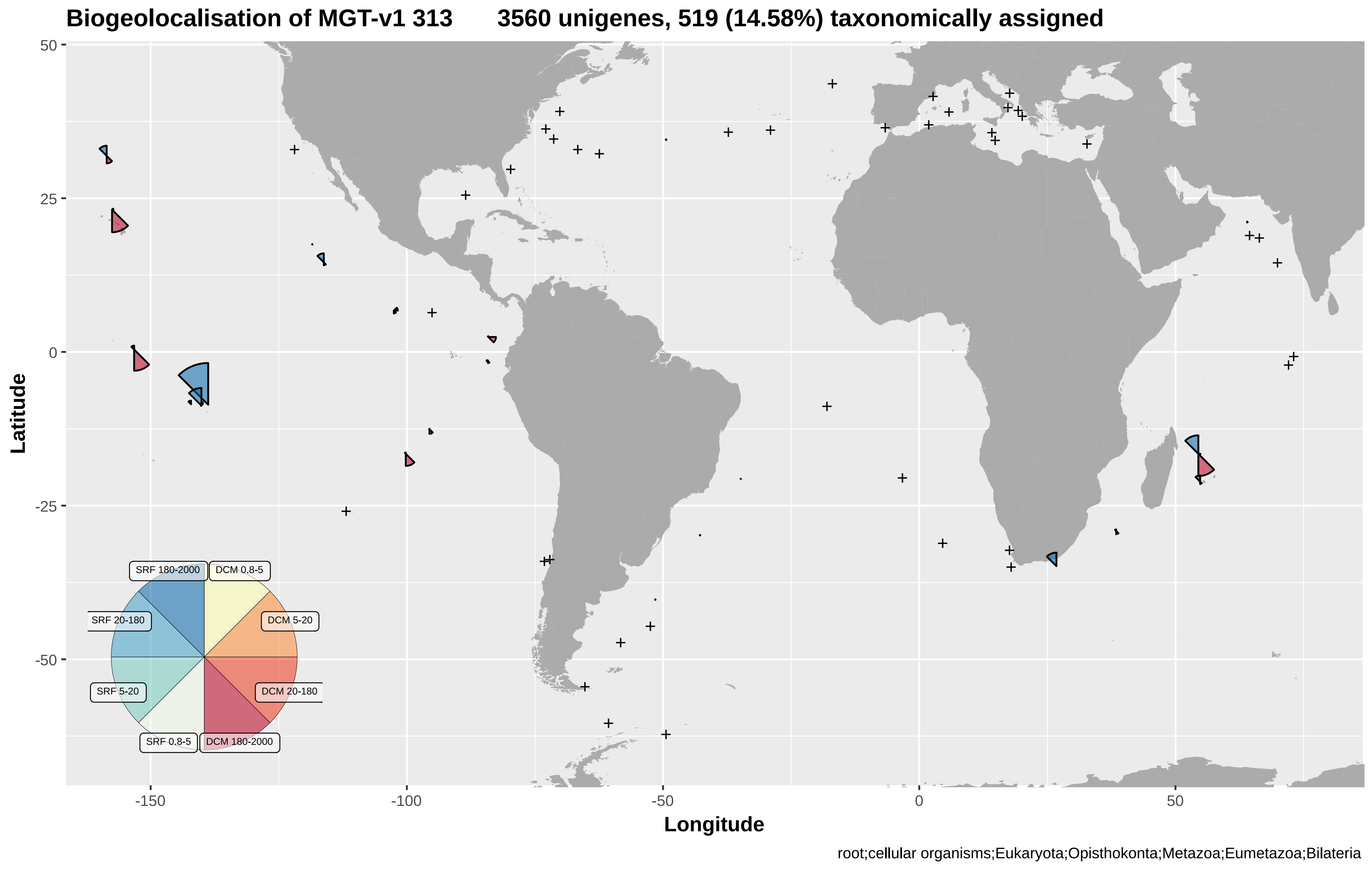

Biogeolocalisation of MGT-v1 313
3560 unigenes, 519 (14.58%) taxonomically assigned
50
25
0
-25
-50
Latitude
-150
-100
-50
0
50
Longitude
SRF 180-2000
DCM 0.8-5
SRF 20-180
DCM 5-20
SRF 5-20
DCM 20-180
SRF 0.8-5
DCM 180-2000
root;cellular organisms;Eukaryota;Opisthokonta;Metazoa;Eumetazoa;Bilateria

#### Biogeolocalisation of MGT-v1 314 3555 unigenes, 455 (12.8%) taxonomically assigned

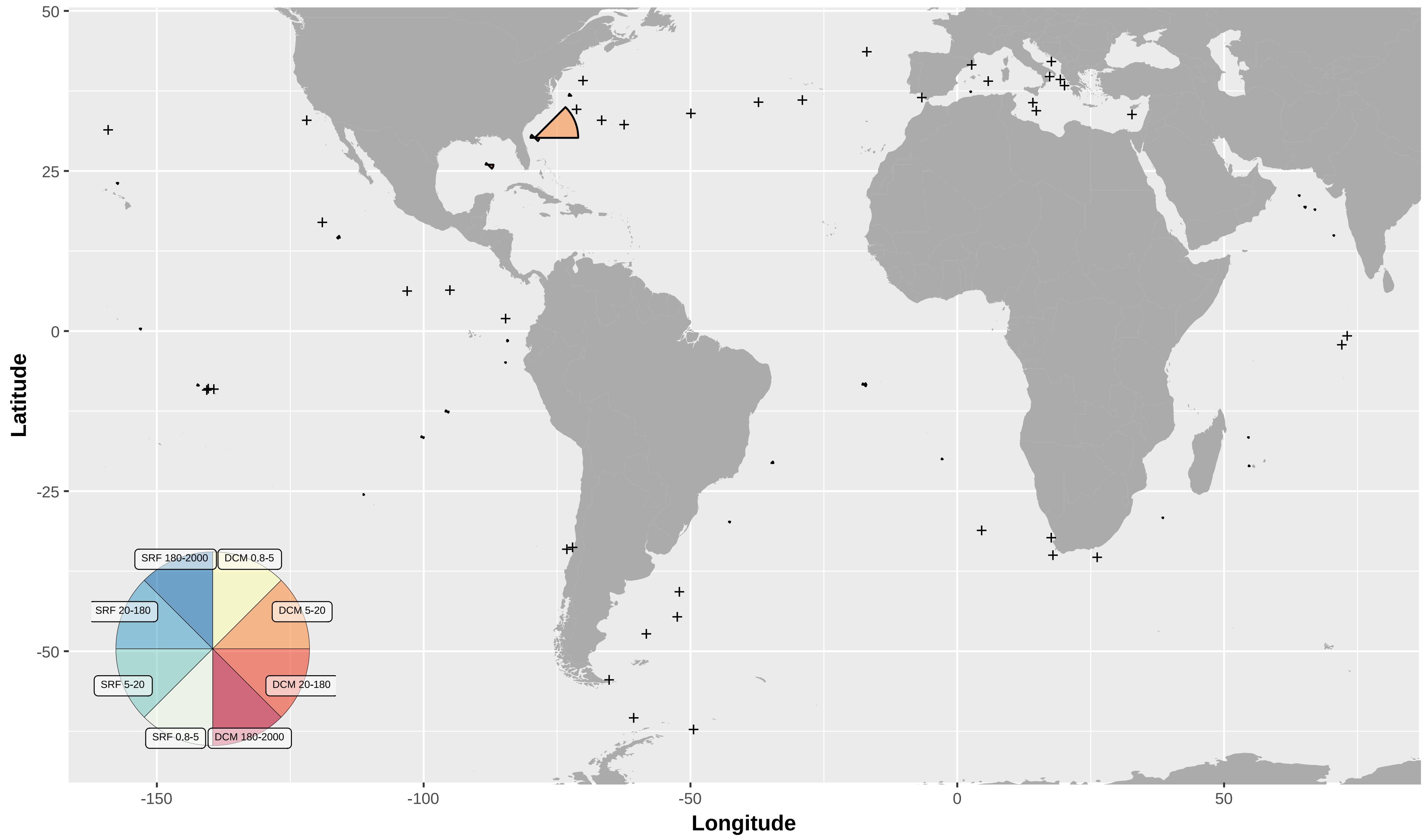


root;cellular organisms;Eukaryota;Opisthokonta;Metazoa;Eumetazoa;Bilateria

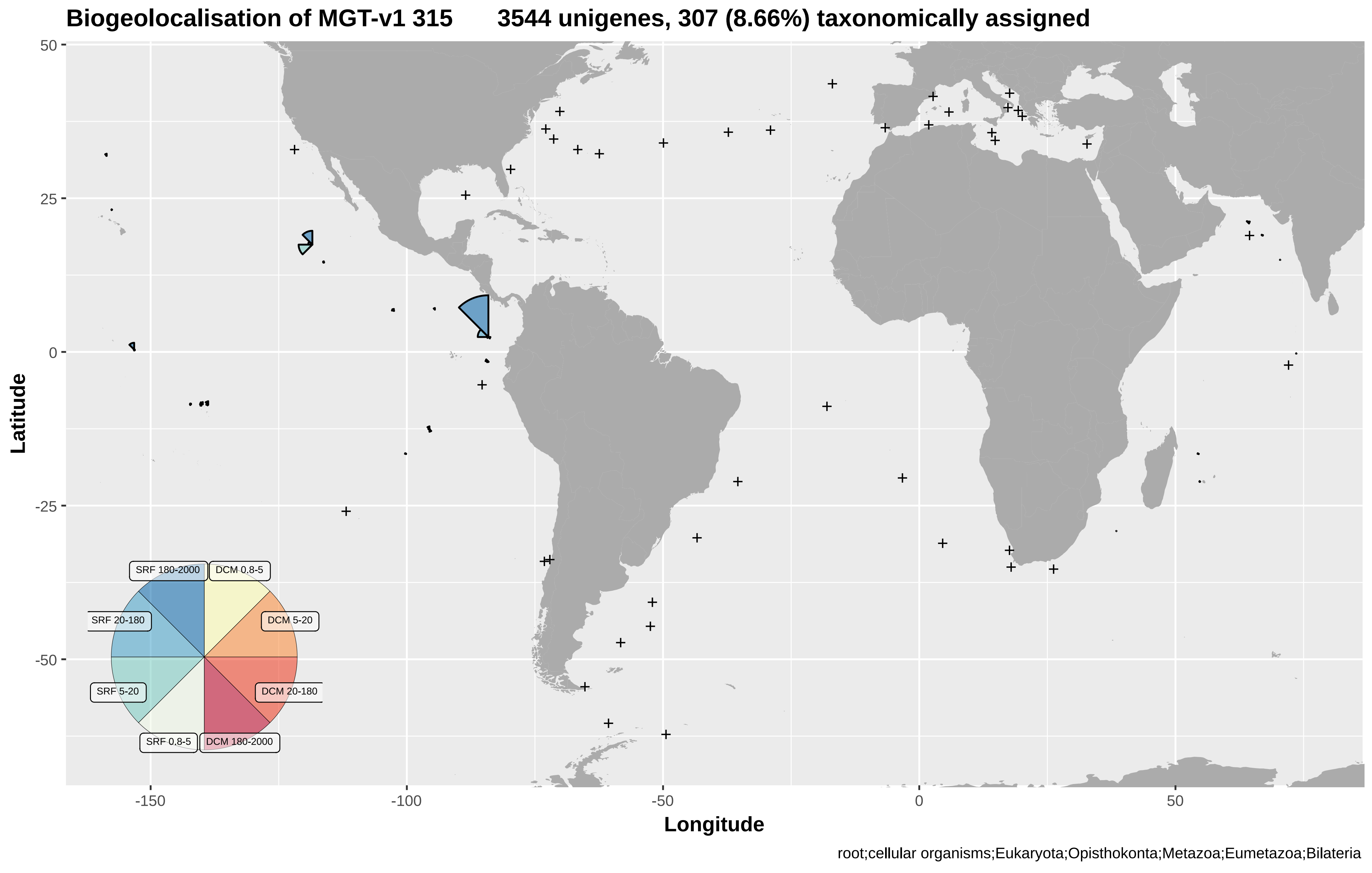

Biogeolocalisation of MGT-v1 315 3544 unigenes, 307 (8.66%) taxonomically assigned
50
25
0
-25
-50
Latitude
-150
-100
-50
0
50
Longitude
SRF 180-2000
DCM 0.8-5
SRF 20-180
DCM 5-20
SRF 5-20
DCM 20-180
SRF 0.8-5
DCM 180-2000
root;cellular organisms;Eukaryota;Opisthokonta;Metazoa;Eumetazoa;Bilateria

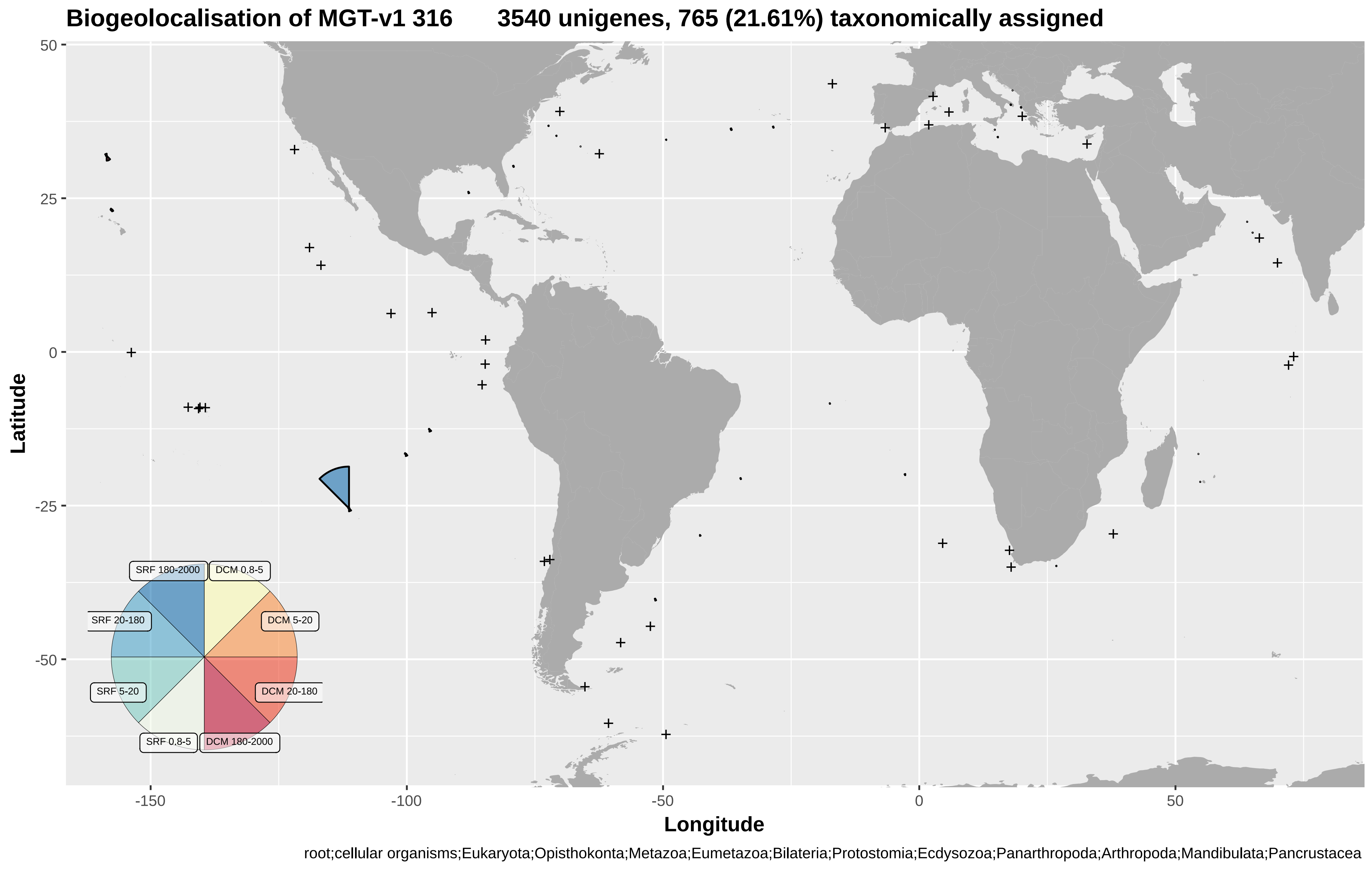
Biogeolocalisation of MGT-v1 316 3540 unigenes, 765 (21.61%) taxonomically assigned
50
25
0
-25
-50
Latitude
-150
-100
-50
0
50
Longitude
SRF 180-2000
DCM 0.8-5
SRF 20-180
DCM 5-20
SRF 5-20
DCM 20-180
SRF 0.8-5
DCM 180-2000
root;cellular organisms;Eukaryota;Opisthokonta;Metazoa;Eumetazoa;Bilateria;Protostomia;Ecdysozoa;Panarthropoda;Arthropoda;Mandibulata;Pancrustacea

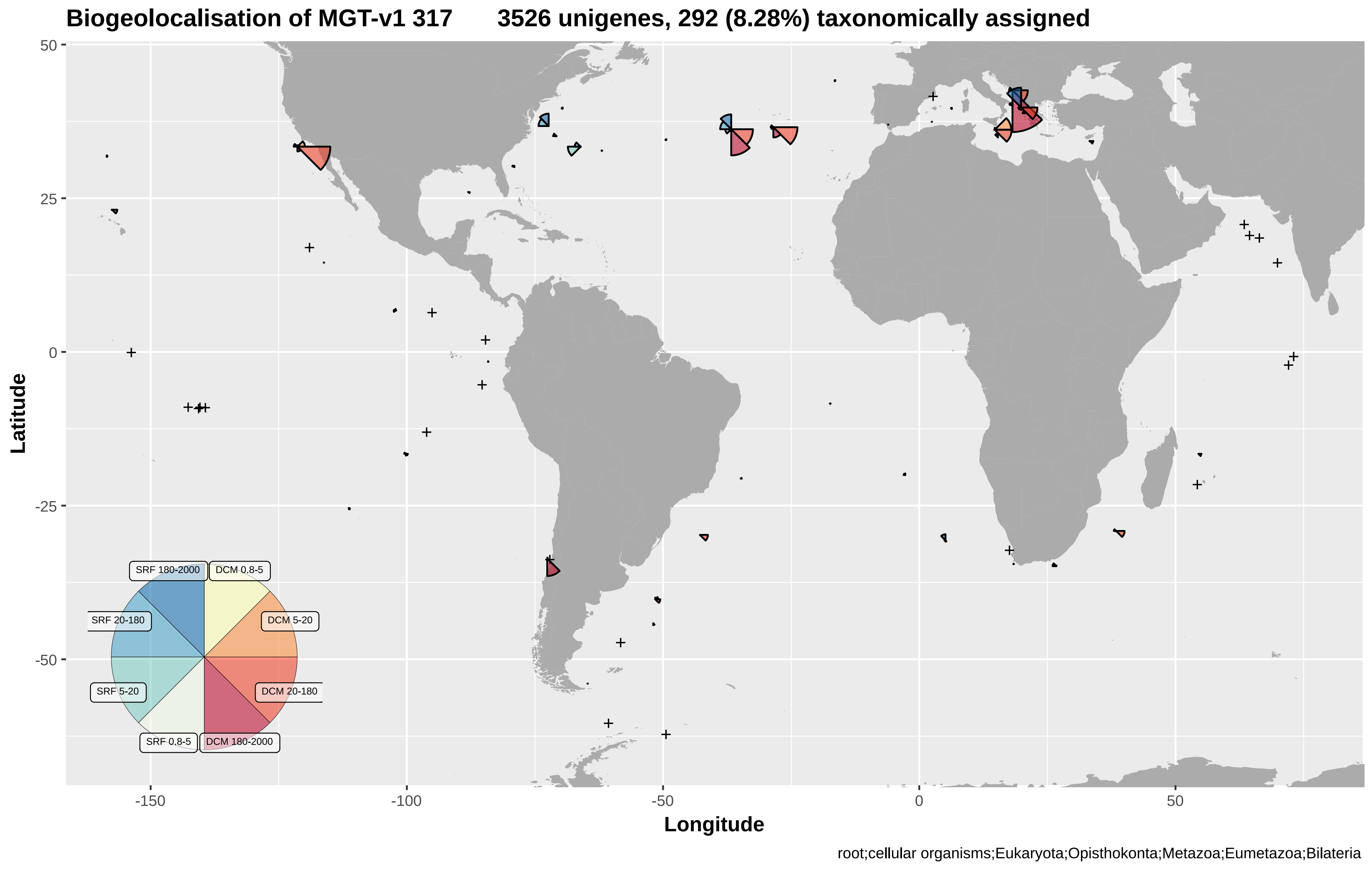
Biogeolocalisation of MGT-v1 317 3526 unigenes, 292 (8.28%) taxonomically assigned
Latitude
Longitude
50
25
0
-25
-50
-150
-100
-50
0
50
SRF 180-2000
DCM 0.8-5
SRF 20-180
DCM 5-20
SRF 5-20
DCM 20-180
SRF 0.8-5
DCM 180-2000
root;cellular organisms;Eukaryota;Opisthokonta;Metazoa;Eumetazoa;Bilateria

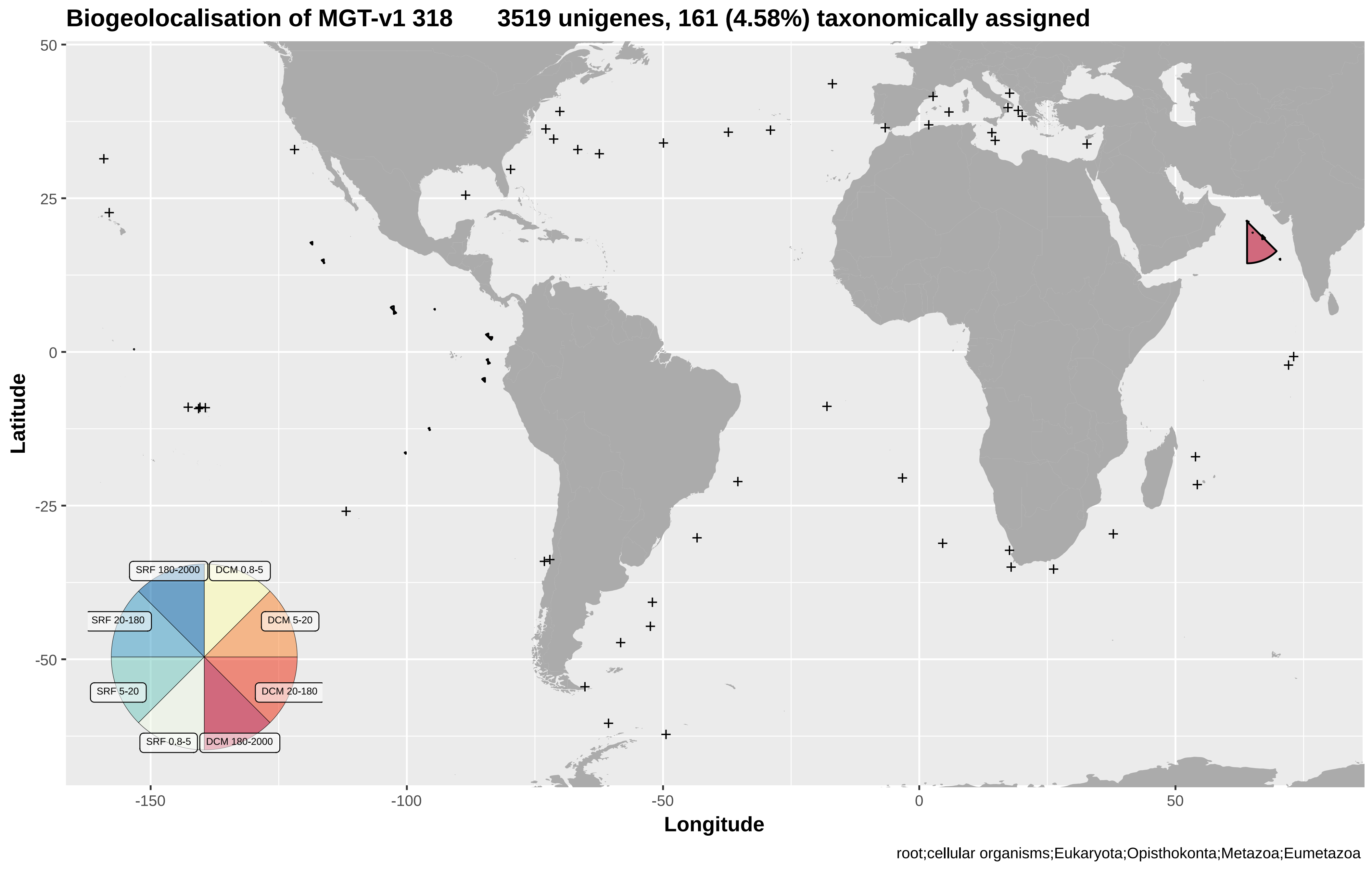
Biogeolocalisation of MGT-v1 318 3519 unigenes, 161 (4.58%) taxonomically assigned
50
25
0
-25
-50
Latitude
-150
-100
-50
0
50
Longitude
SRF 180-2000
DCM 0.8-5
SRF 20-180
DCM 5-20
SRF 5-20
DCM 20-180
SRF 0.8-5
DCM 180-2000
root;cellular organisms;Eukaryota;Opisthokonta;Metazoa;Eumetazoa

## Biogeolocalisation of MGT-v1 319      3513 unigenes, 3301 (93.97%) taxonomically assigned

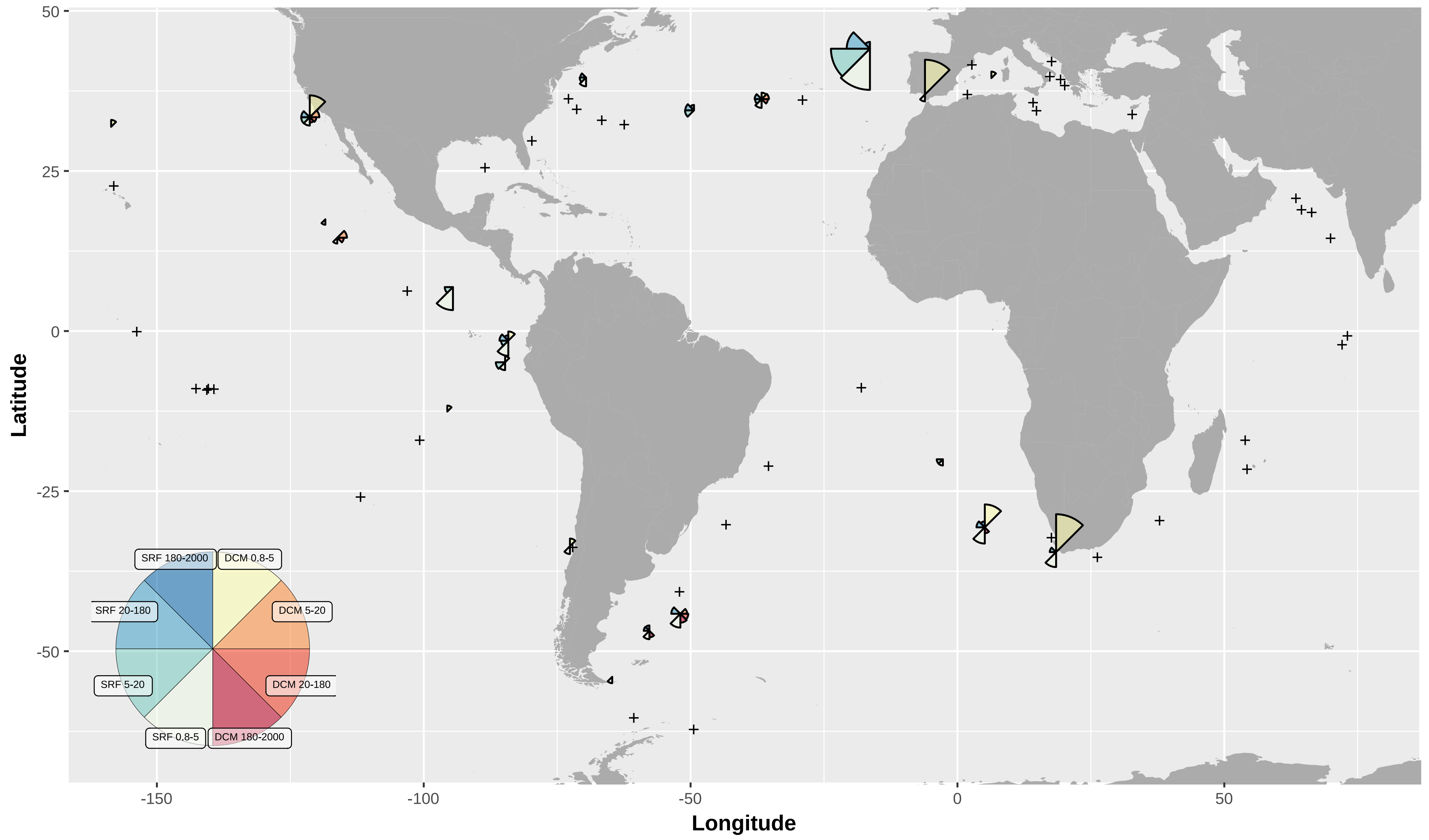


root;cellular organisms;Eukaryota;Viridiplantae;Chlorophyta;prasinophytes;Prasinococcales;Prasinoderma

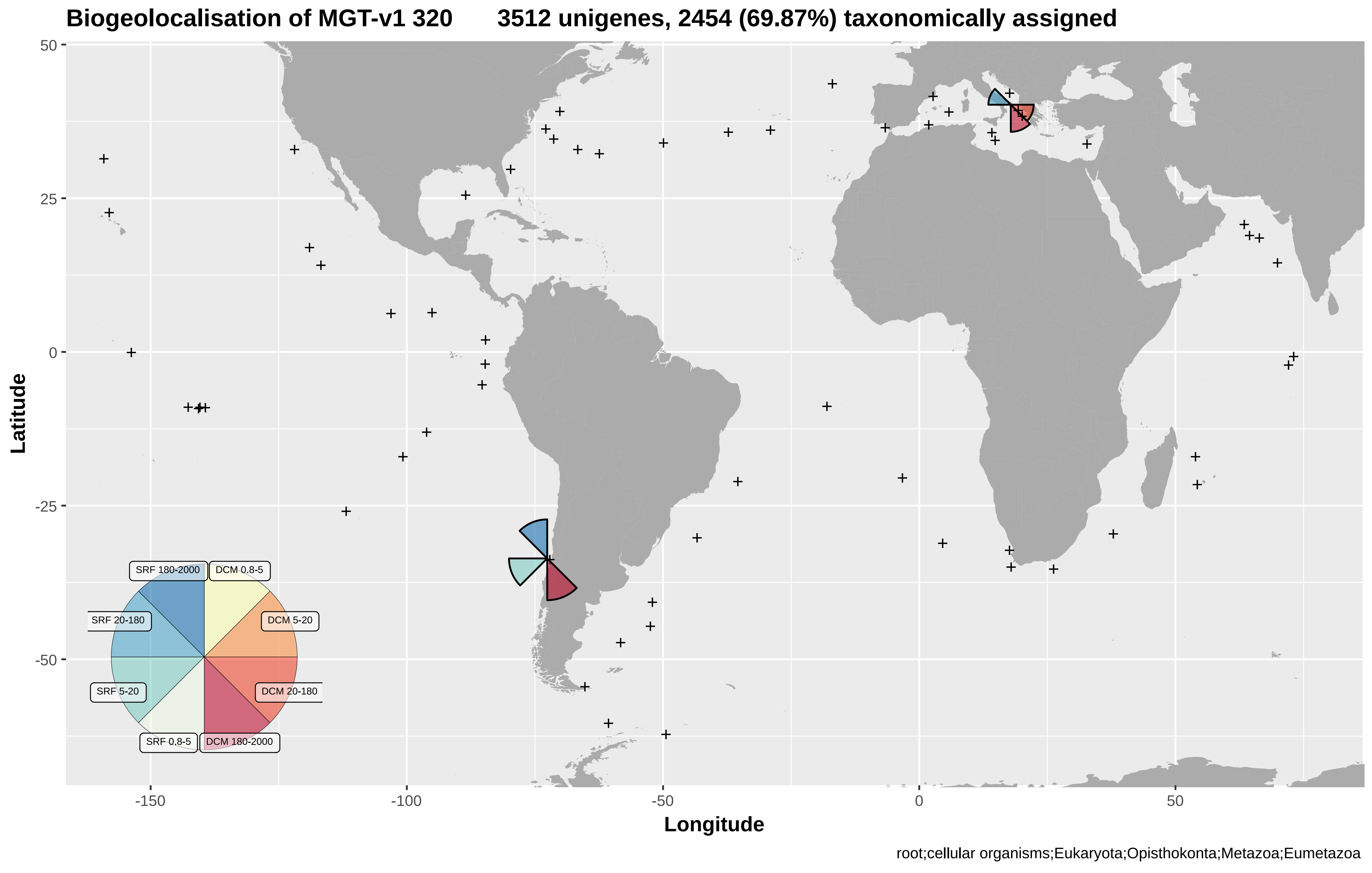
Biogeolocalisation of MGT-v1 320
3512 unigenes, 2454 (69.87%) taxonomically assigned
Latitude
Longitude
50
25
0
-25
-50
-150
-100
-50
0
50
SRF 180-2000
DCM 0.8-5
SRF 20-180
DCM 5-20
SRF 5-20
DCM 20-180
SRF 0.8-5
DCM 180-2000
root;cellular organisms;Eukaryota;Opisthokonta;Metazoa;Eumetazoa

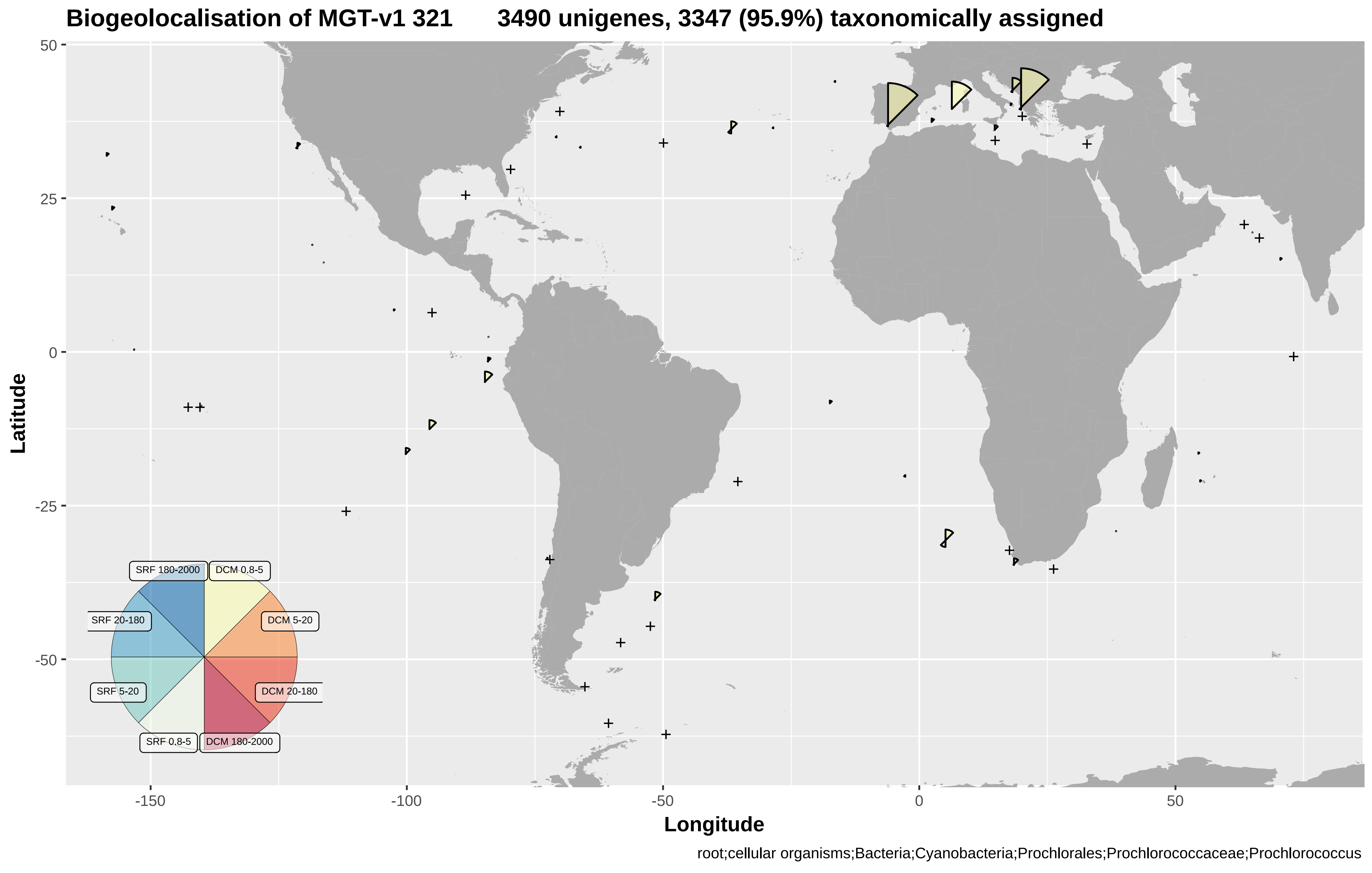
Biogeolocalisation of MGT-v1 321 3490 unigenes, 3347 (95.9%) taxonomically assigned
50
25
0
-25
-50
Latitude
-150
-100
-50
0
50
Longitude
SRF 180-2000
DCM 0.8-5
SRF 20-180
DCM 5-20
SRF 5-20
DCM 20-180
SRF 0.8-5
DCM 180-2000
root;cellular organisms;Bacteria;Cyanobacteria;Prochlorales;Prochlorococcaceae;Prochlorococcus

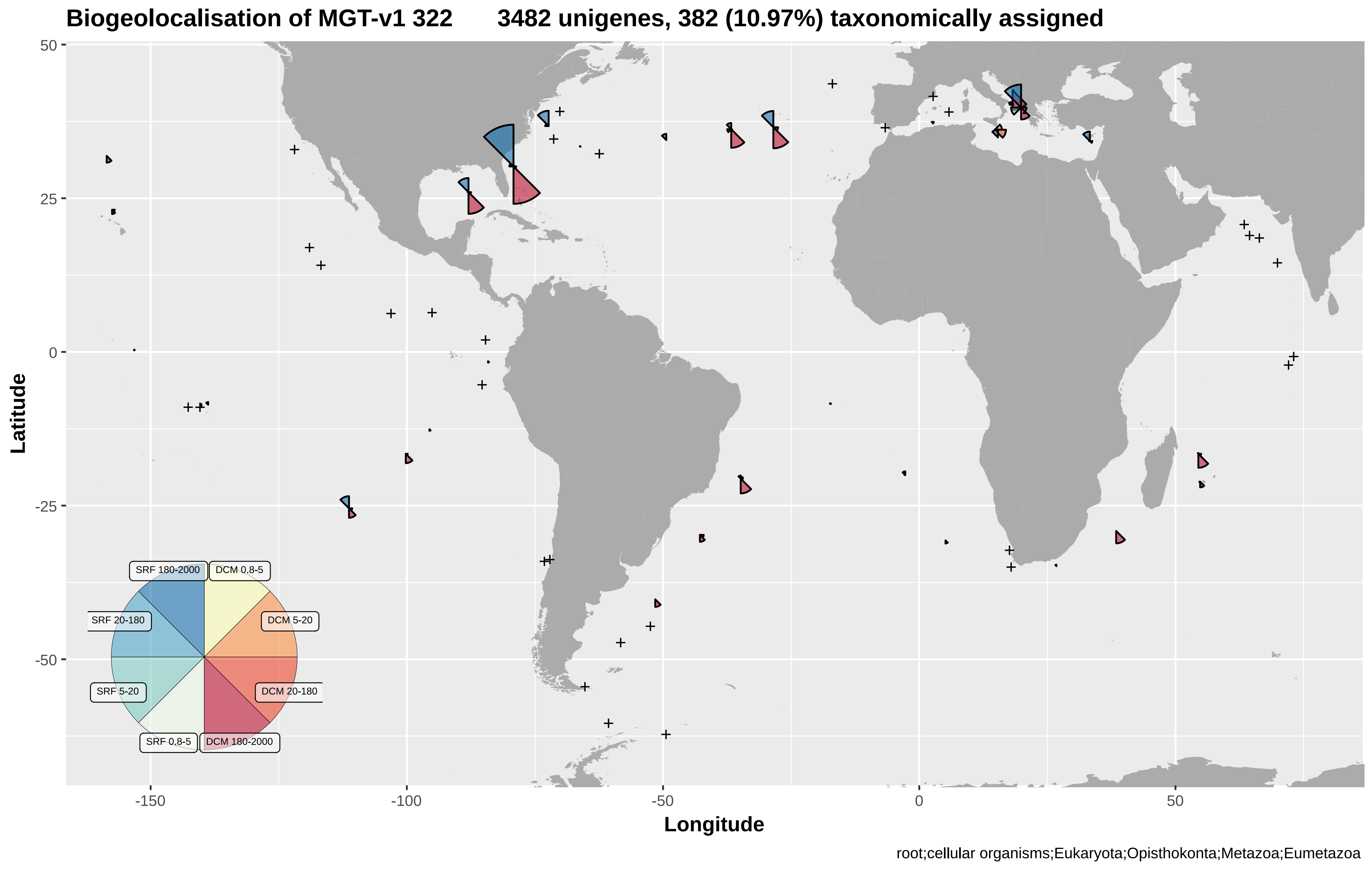

Biogeolocalisation of MGT-v1 322
3482 unigenes, 382 (10.97%) taxonomically assigned
50
25
0
-25
-50
Latitude
-150
-100
-50
0
50
Longitude
SRF 180-2000
DCM 0.8-5
SRF 20-180
DCM 5-20
SRF 5-20
DCM 20-180
SRF 0.8-5
DCM 180-2000
root;cellular organisms;Eukaryota;Opisthokonta;Metazoa;Eumetazoa

## Biogeolocalisation of MGT-v1 323      3472 unigenes, 1961 (56.48%) taxonomically assigned

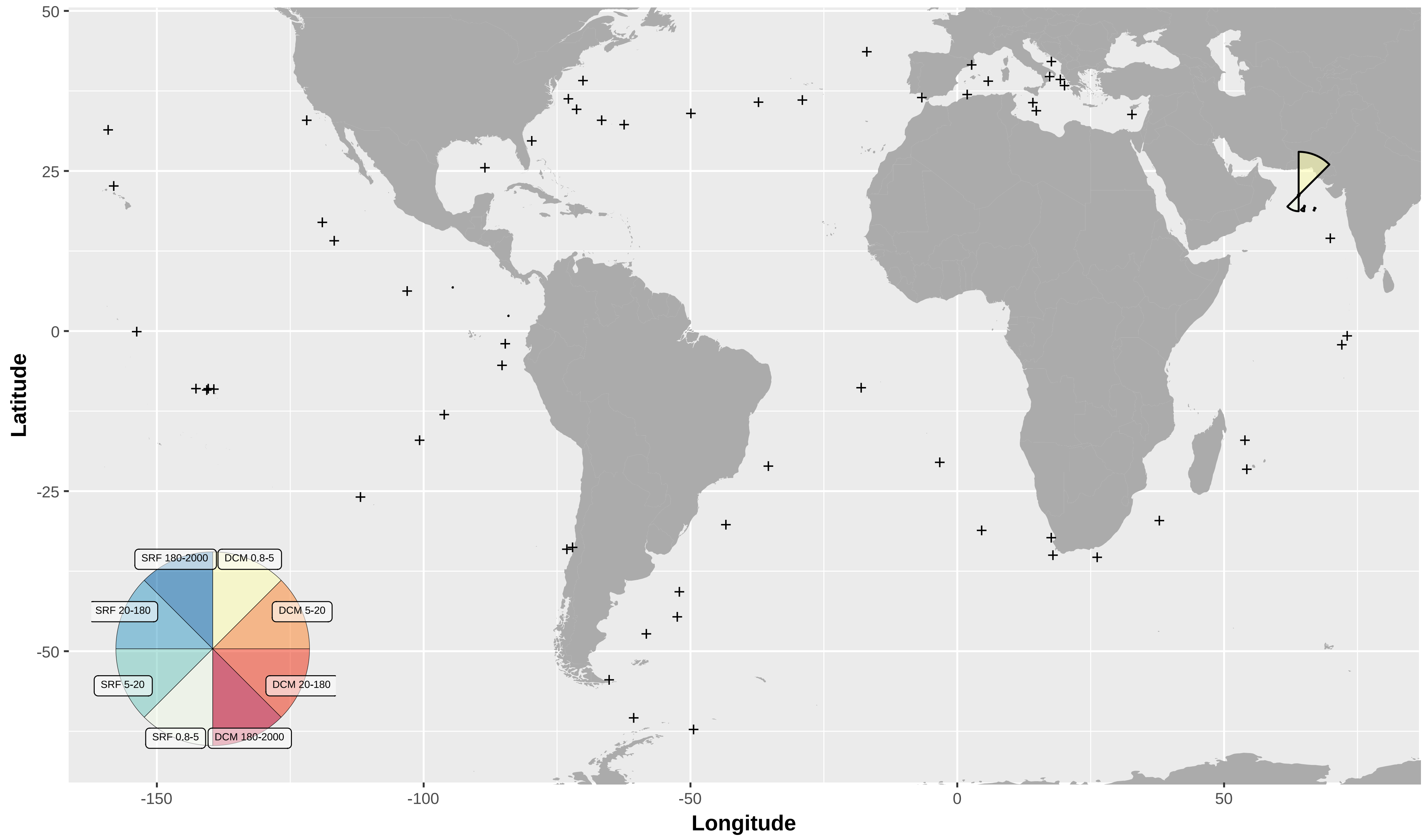


root

## Biogeolocalisation of MGT-v1 324      3458 unigenes, 2216 (64.08%) taxonomically assigned

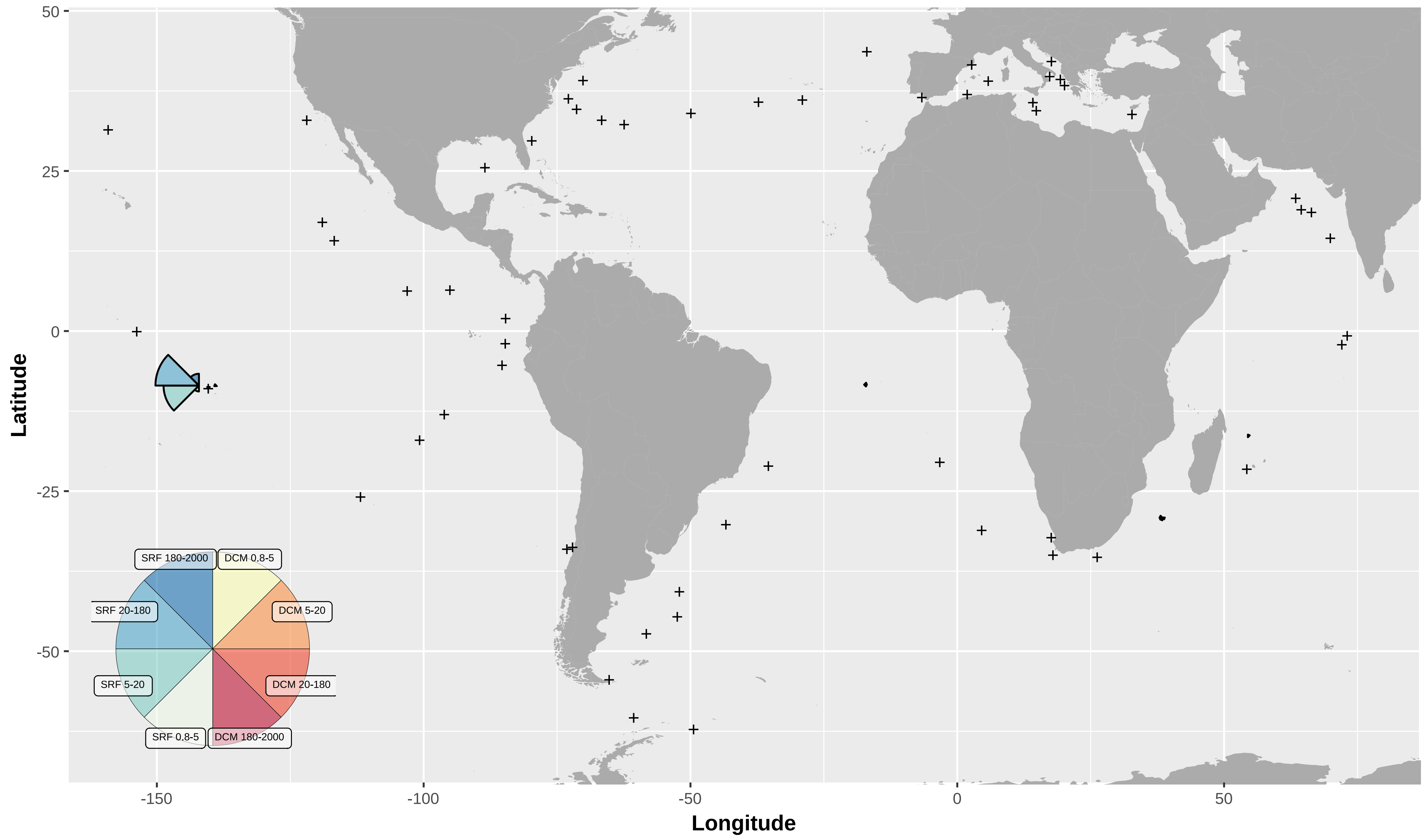


root;cellular organisms

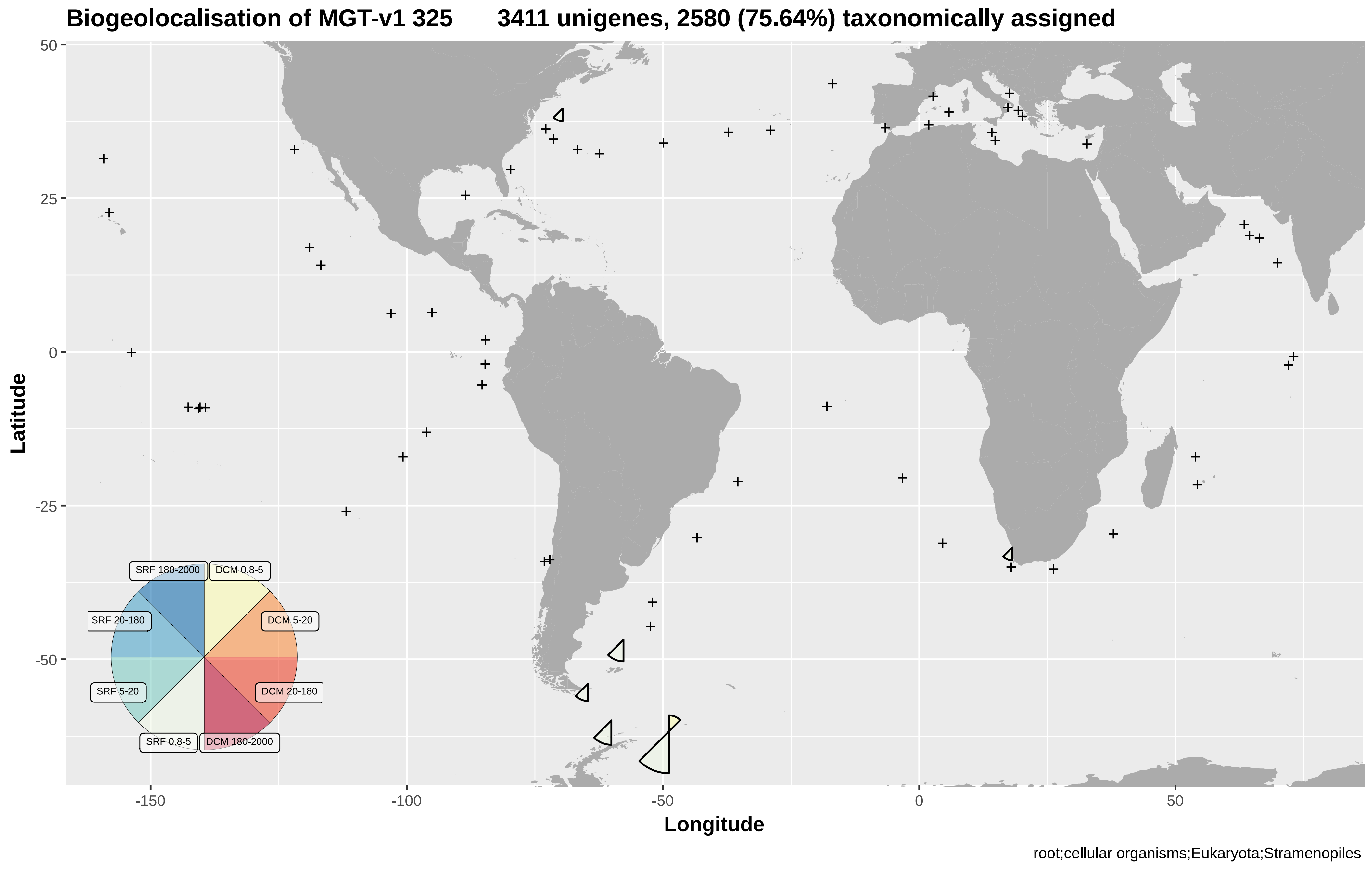

Biogeolocalisation of MGT-v1 325 3411 unigenes, 2580 (75.64%) taxonomically assigned
50
25
0
-25
-50
Latitude
-150
-100
-50
0
50
Longitude
SRF 180-2000
DCM 0.8-5
SRF 20-180
DCM 5-20
SRF 5-20
DCM 20-180
SRF 0.8-5
DCM 180-2000
root;cellular organisms;Eukaryota;Stramenopiles

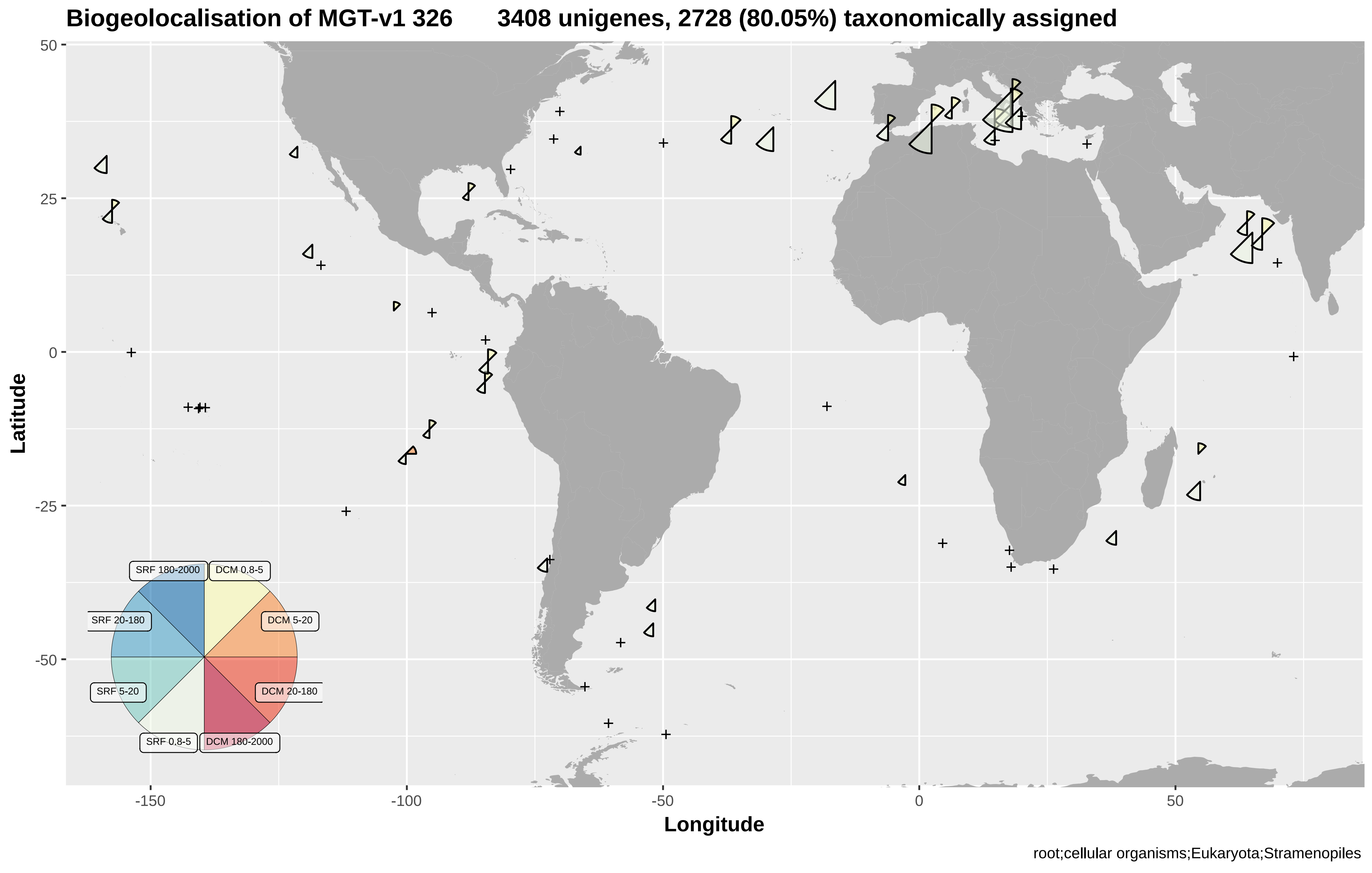
Biogeolocalisation of MGT-v1 326 3408 unigenes, 2728 (80.05%) taxonomically assigned
Latitude
50
25
0
-25
-50
-150
-100
-50
0
50
Longitude
SRF 180-2000
DCM 0.8-5
SRF 20-180
DCM 5-20
SRF 5-20
DCM 20-180
SRF 0.8-5
DCM 180-2000
root;cellular organisms;Eukaryota;Stramenopiles

## Biogeolocalisation of MGT-v1 327      3406 unigenes, 1894 (55.61%) taxonomically assigned

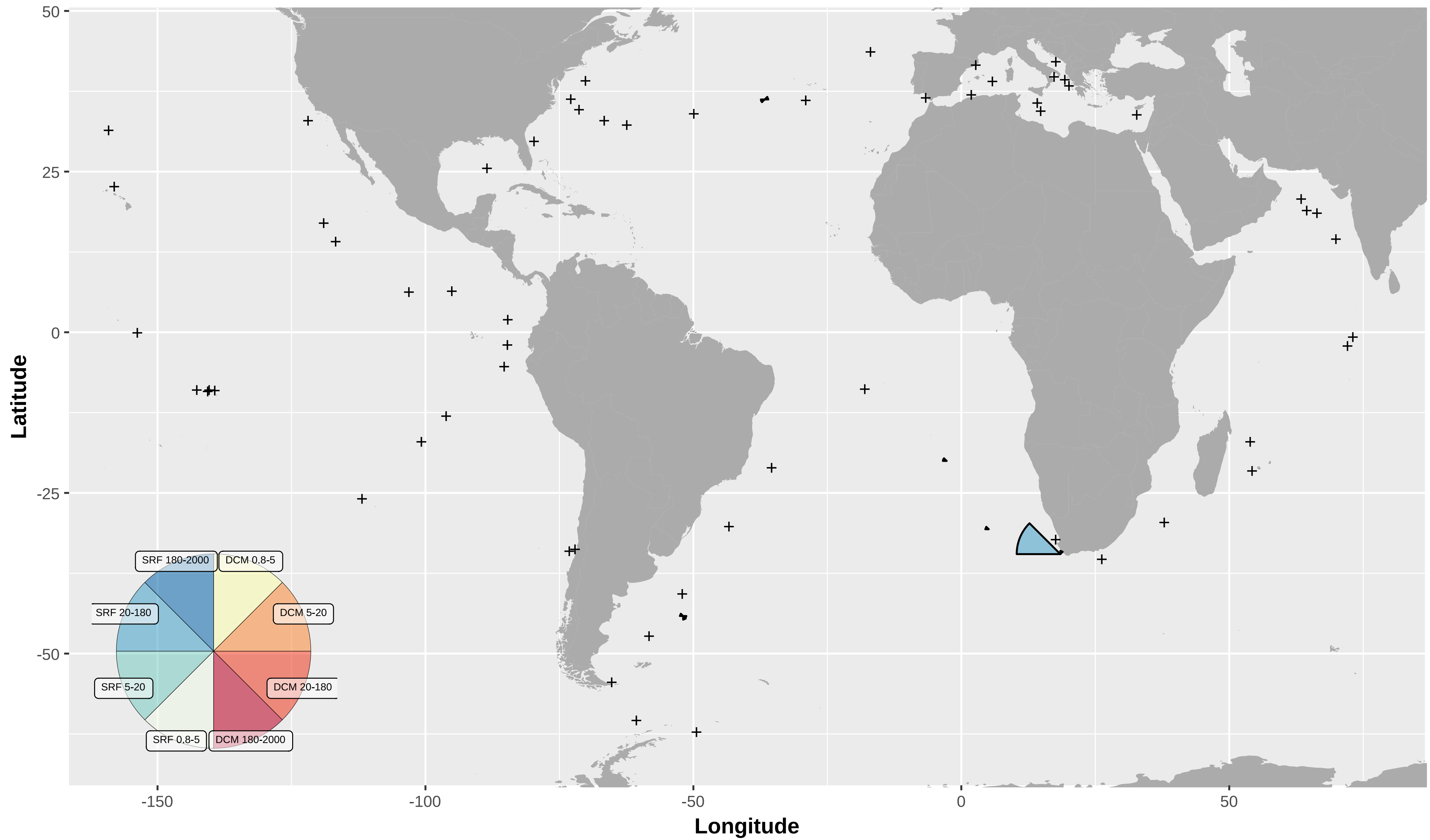


root;cellular organisms;Eukaryota;Alveolata;Ciliophora;Intramacronucleata;Spirotrichea

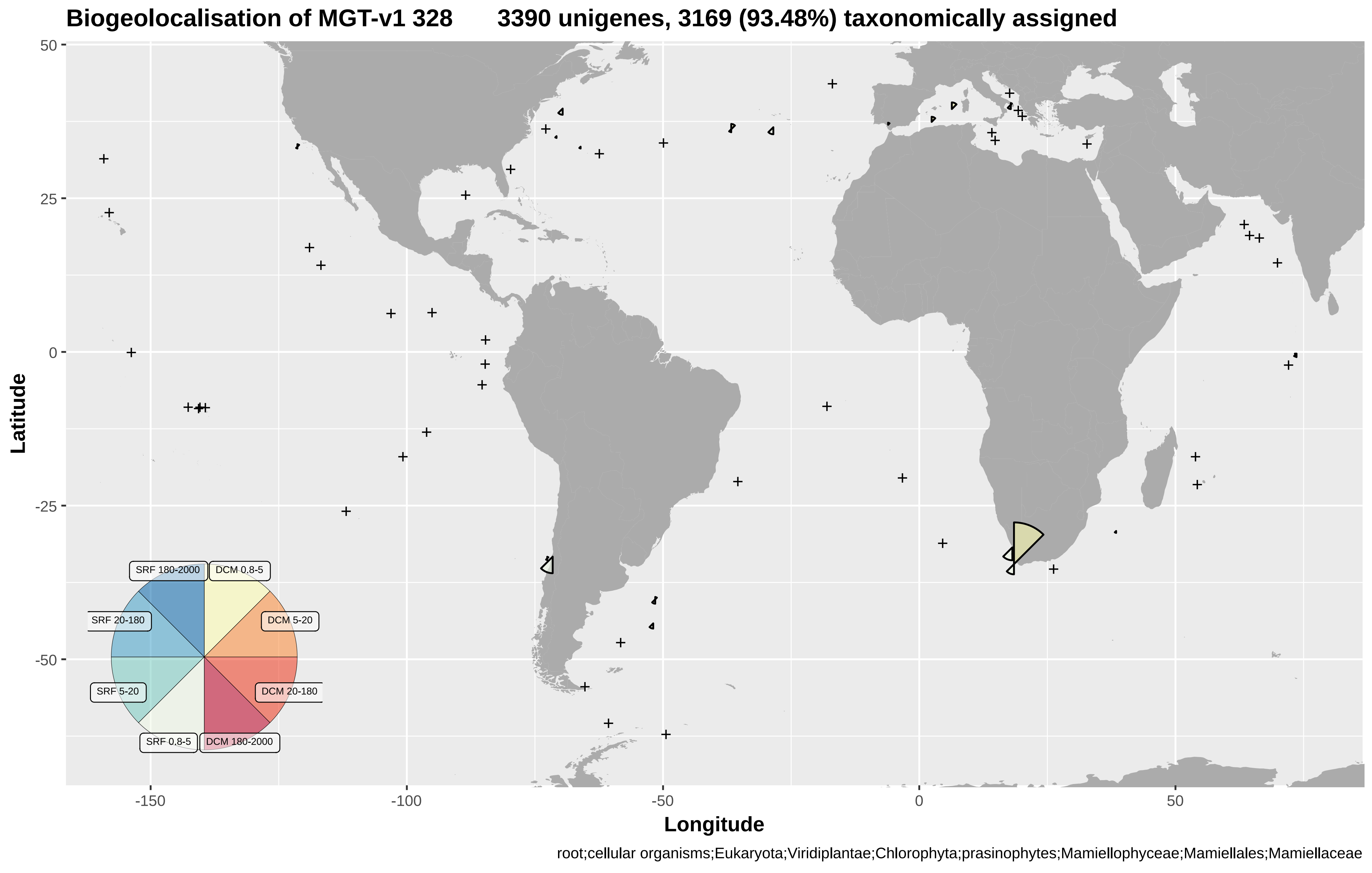
Biogeolocalisation of MGT-v1 328 3390 unigenes, 3169 (93.48%) taxonomically assigned
Latitude
Longitude
50
25
0
-25
-50
-150
-100
-50
0
50
SRF 180-2000
DCM 0.8-5
SRF 20-180
DCM 5-20
SRF 5-20
DCM 20-180
SRF 0.8-5
DCM 180-2000
root;cellular organisms;Eukaryota;Viridiplantae;Chlorophyta;prasinophytes;Mamiellophyceae;Mamiellales;Mamiellaceae

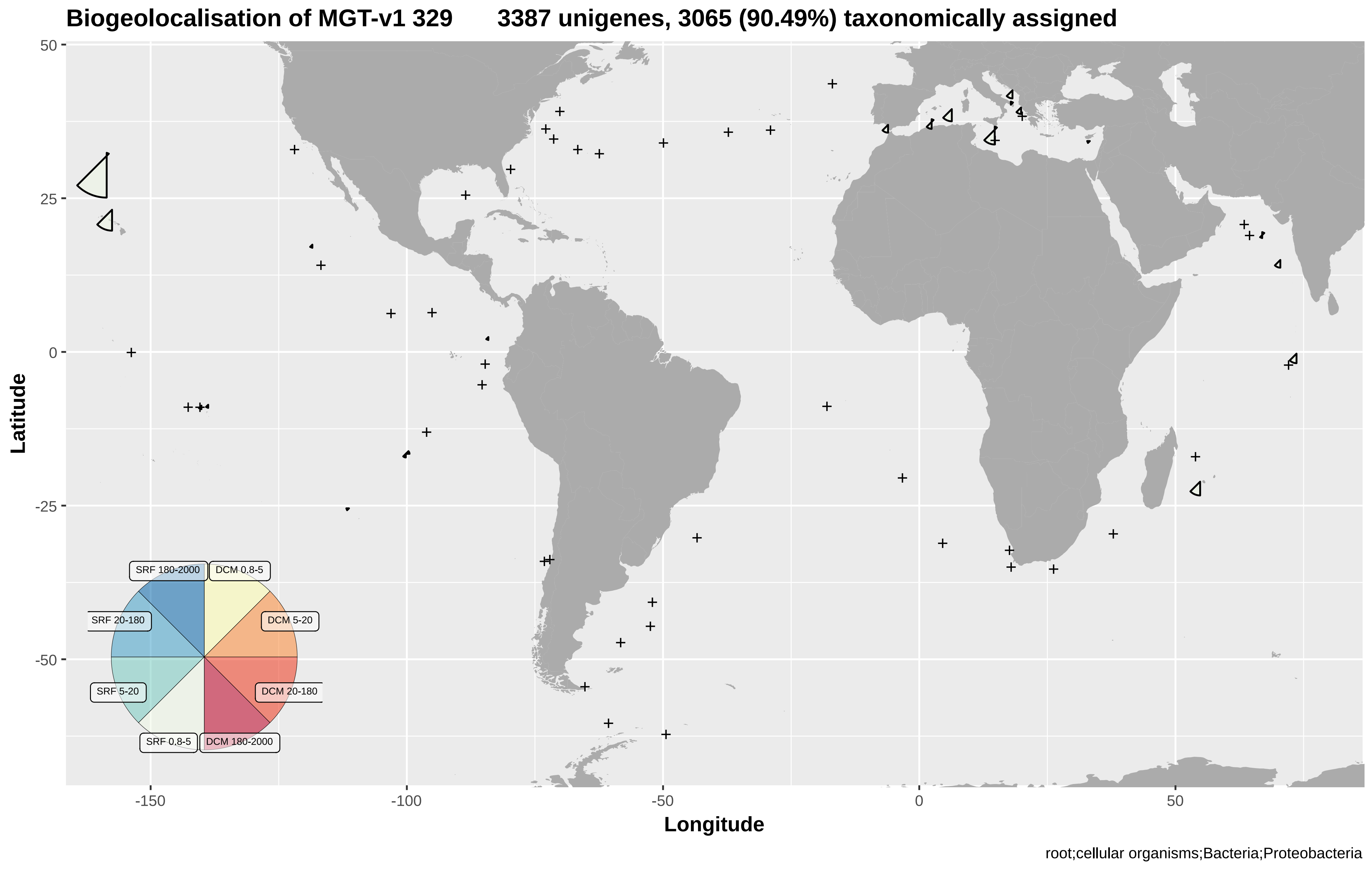

Biogeolocalisation of MGT-v1 329 3387 unigenes, 3065 (90.49%) taxonomically assigned
50
25
0
-25
-50
Latitude
-150
-100
-50
0
50
Longitude
SRF 180-2000
DCM 0.8-5
SRF 20-180
DCM 5-20
SRF 5-20
DCM 20-180
SRF 0.8-5
DCM 180-2000
root;cellular organisms;Bacteria;Proteobacteria

# Biogeolocalisation of MGT-v1 330      3376 unigenes, 1691 (50.09%) taxonomically assigned

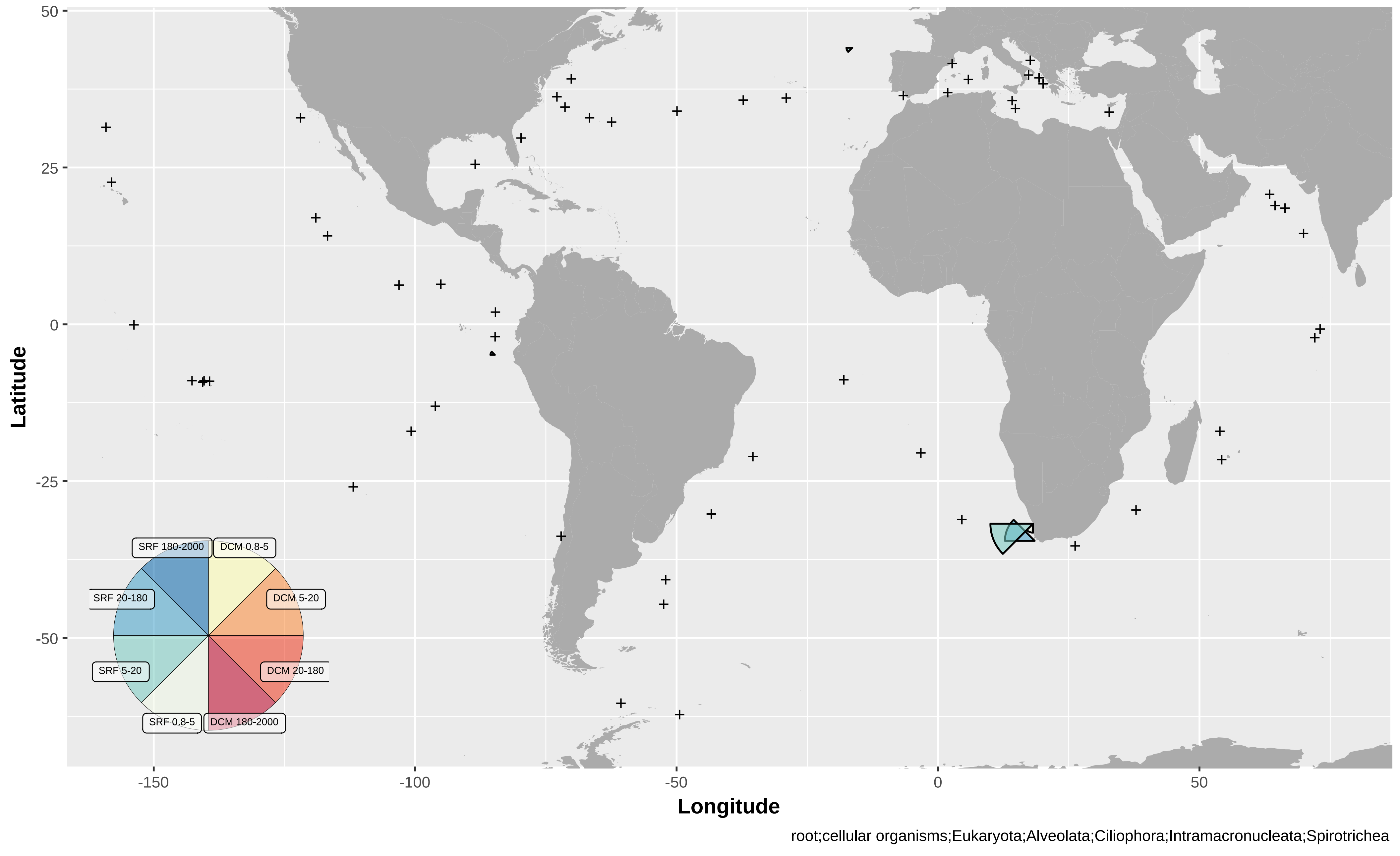

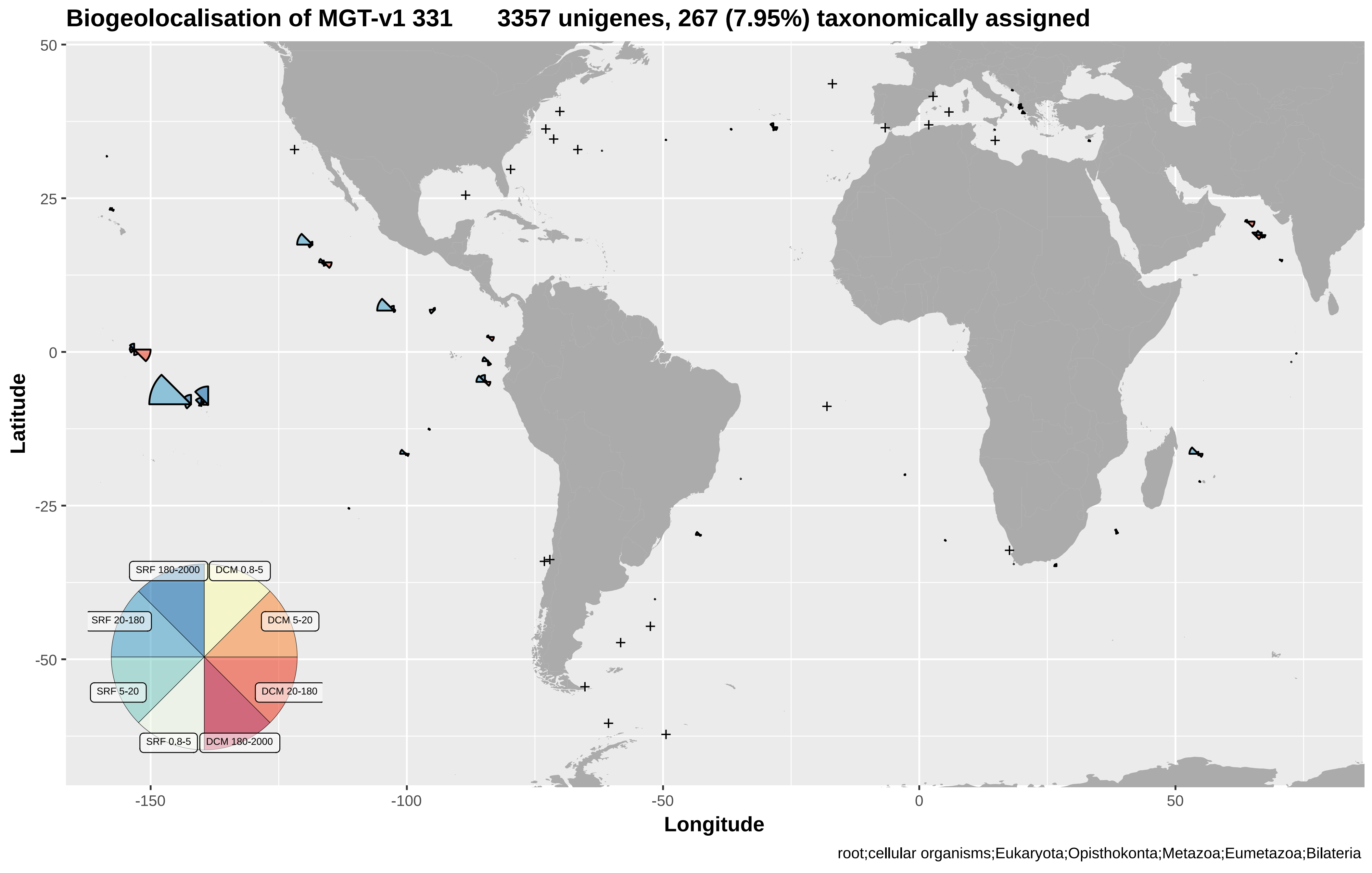
Biogeolocalisation of MGT-v1 331 3357 unigenes, 267 (7.95%) taxonomically assigned
Latitude
Longitude
50
25
0
-25
-50
-150
-100
-50
0
50
SRF 180-2000
DCM 0.8-5
SRF 20-180
DCM 5-20
SRF 5-20
DCM 20-180
SRF 0.8-5
DCM 180-2000
root;cellular organisms;Eukaryota;Opisthokonta;Metazoa;Eumetazoa;Bilateria

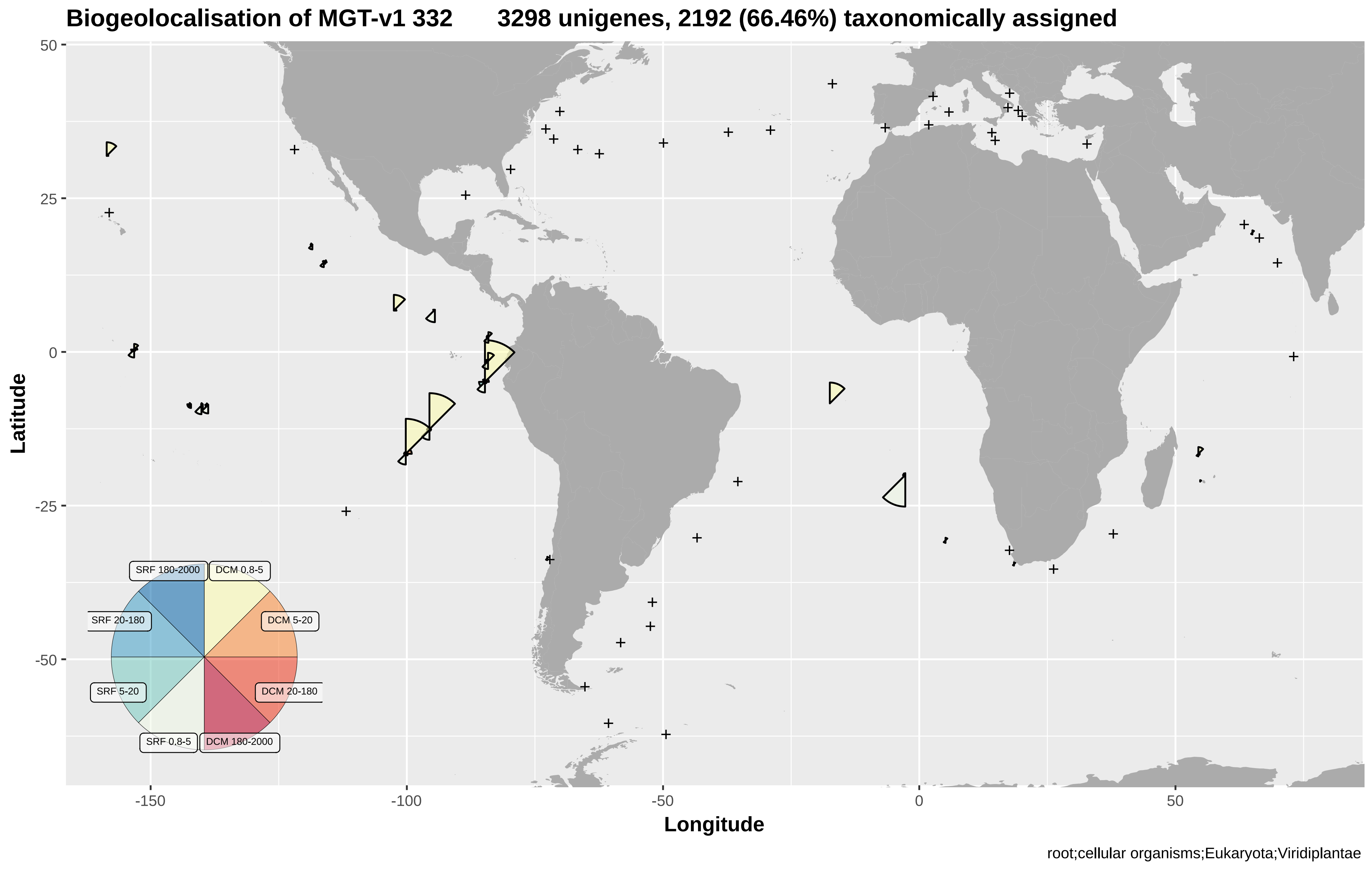
Biogeolocalisation of MGT-v1 332 3298 unigenes, 2192 (66.46%) taxonomically assigned
50
25
0
-25
-50
Latitude
-150
-100
-50
0
50
Longitude
SRF 180-2000
DCM 0.8-5
SRF 20-180
DCM 5-20
SRF 5-20
DCM 20-180
SRF 0.8-5
DCM 180-2000
root;cellular organisms;Eukaryota;Viridiplantae

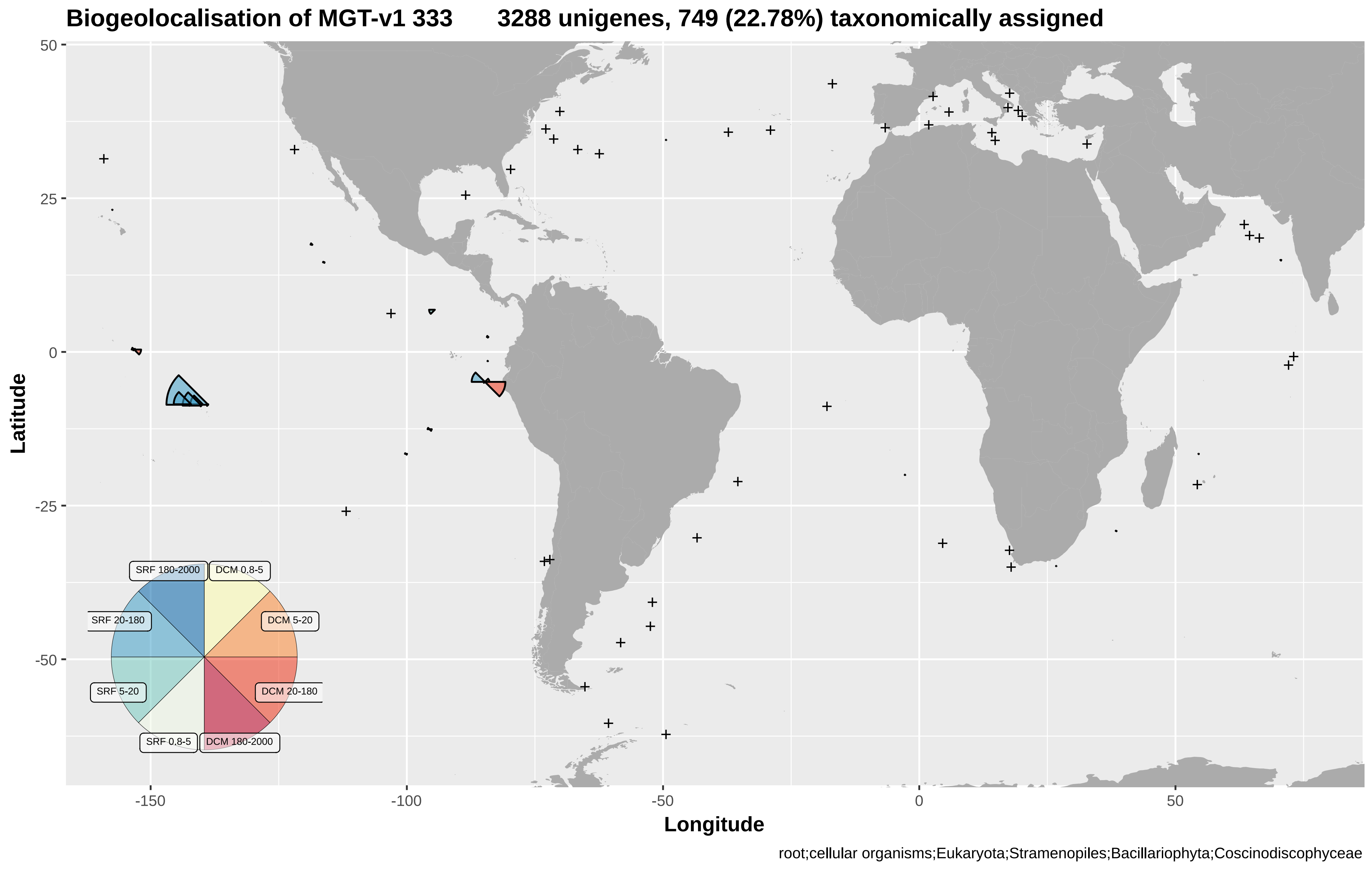
Biogeolocalisation of MGT-v1 333
3288 unigenes, 749 (22.78%) taxonomically assigned
Latitude
Longitude
50
25
0
-25
-50
-150
-100
-50
0
50
SRF 180-2000
DCM 0.8-5
SRF 20-180
DCM 5-20
SRF 5-20
DCM 20-180
SRF 0.8-5
DCM 180-2000
root;cellular organisms;Eukaryota;Stramenopiles;Bacillariophyta;Coscinodiscophyceae

#### Biogeolocalisation of MGT-v1 334 3273 unigenes, 1692 (51.7%) taxonomically assigned

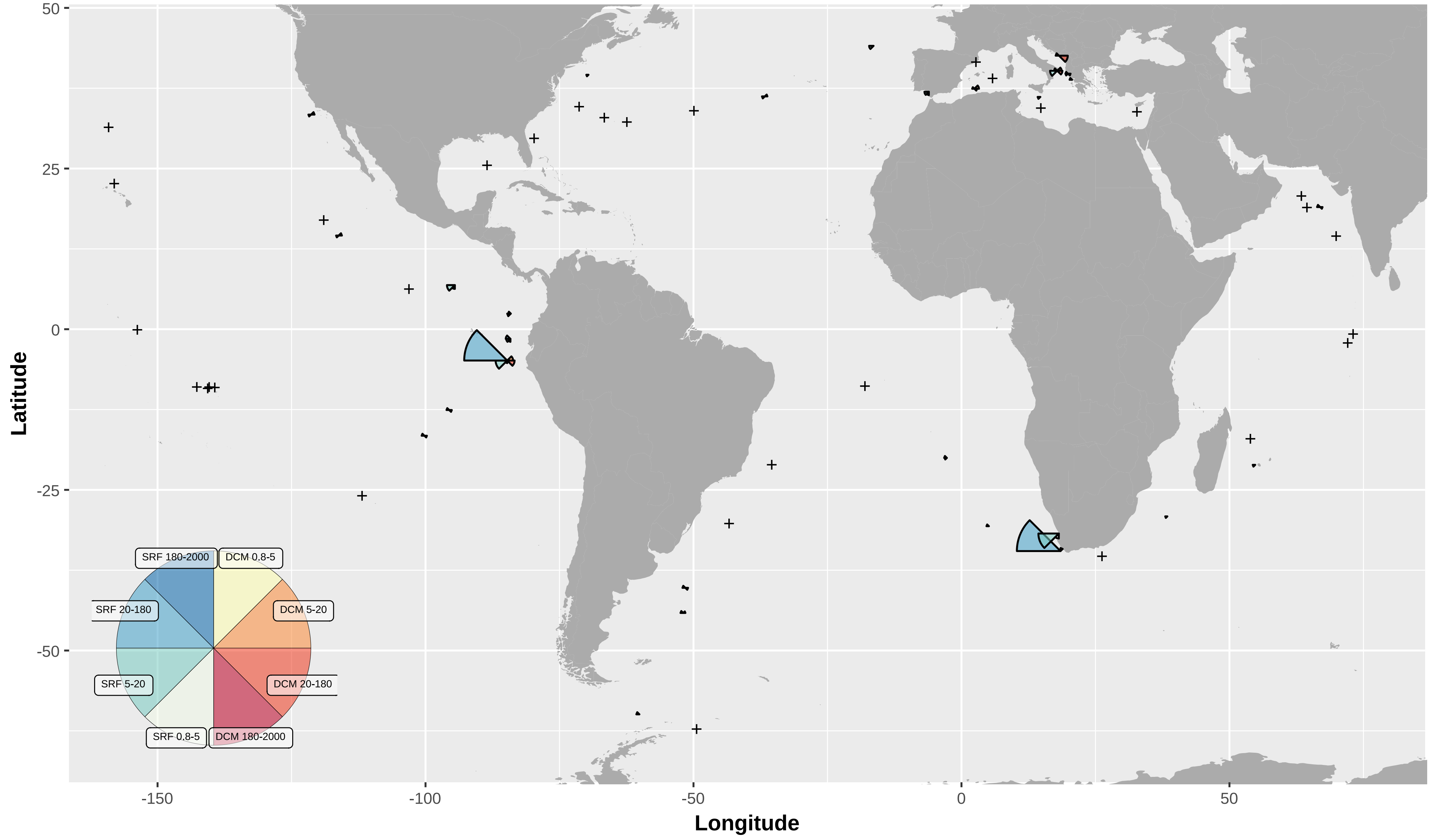


root;cellular organisms;Eukaryota;Alveolata;Ciliophora;Intramacronucleata;Spirotrichea

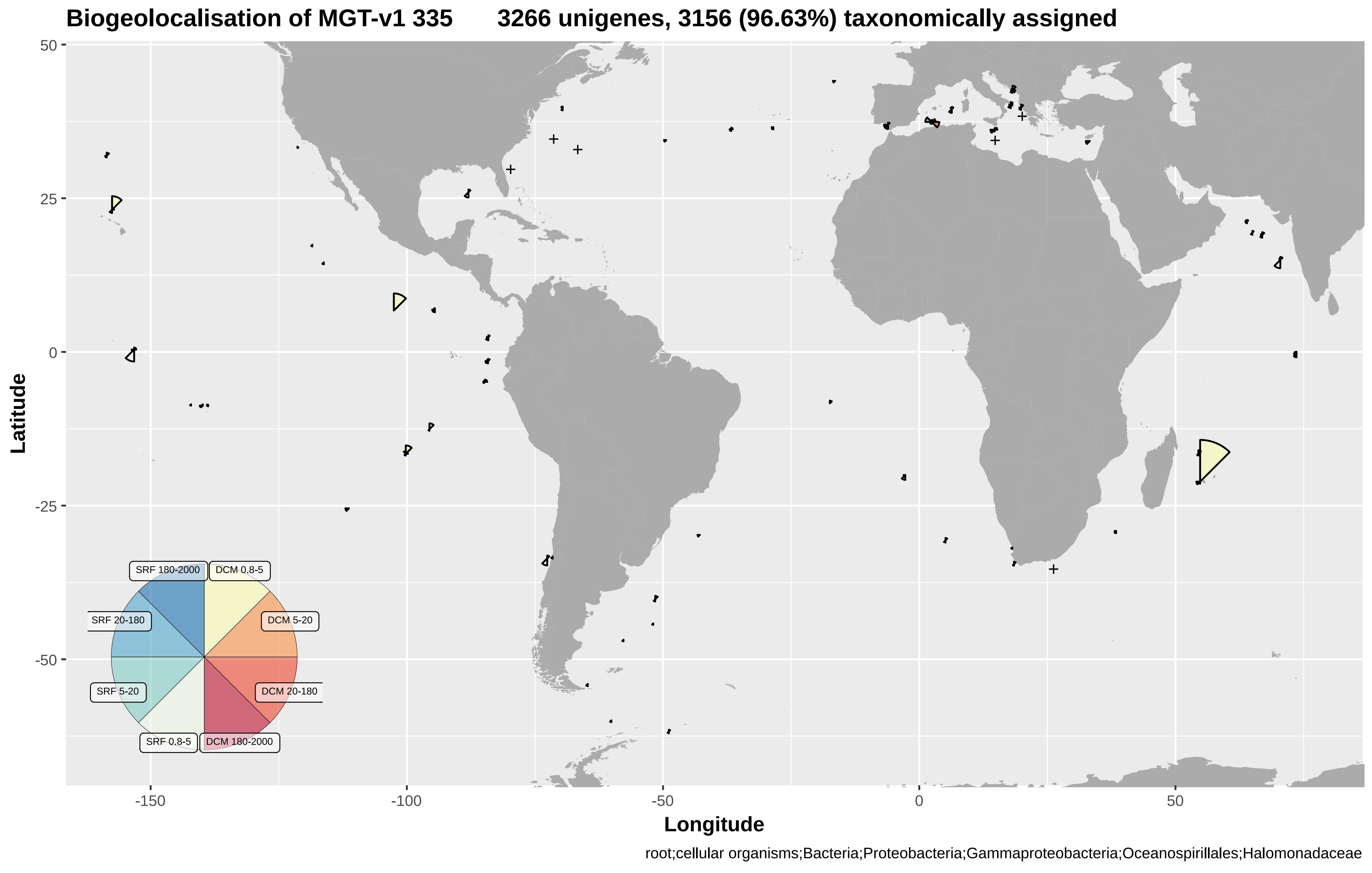

Biogeolocalisation of MGT-v1 335 3266 unigenes, 3156 (96.63%) taxonomically assigned
50
25
0
-25
-50
Latitude
-150
-100
-50
0
50
Longitude
SRF 180-2000
DCM 0.8-5
SRF 20-180
DCM 5-20
SRF 5-20
DCM 20-180
SRF 0.8-5
DCM 180-2000
root;cellular organisms;Bacteria;Proteobacteria;Gammaproteobacteria;Oceanospirillales;Halomonadaceae

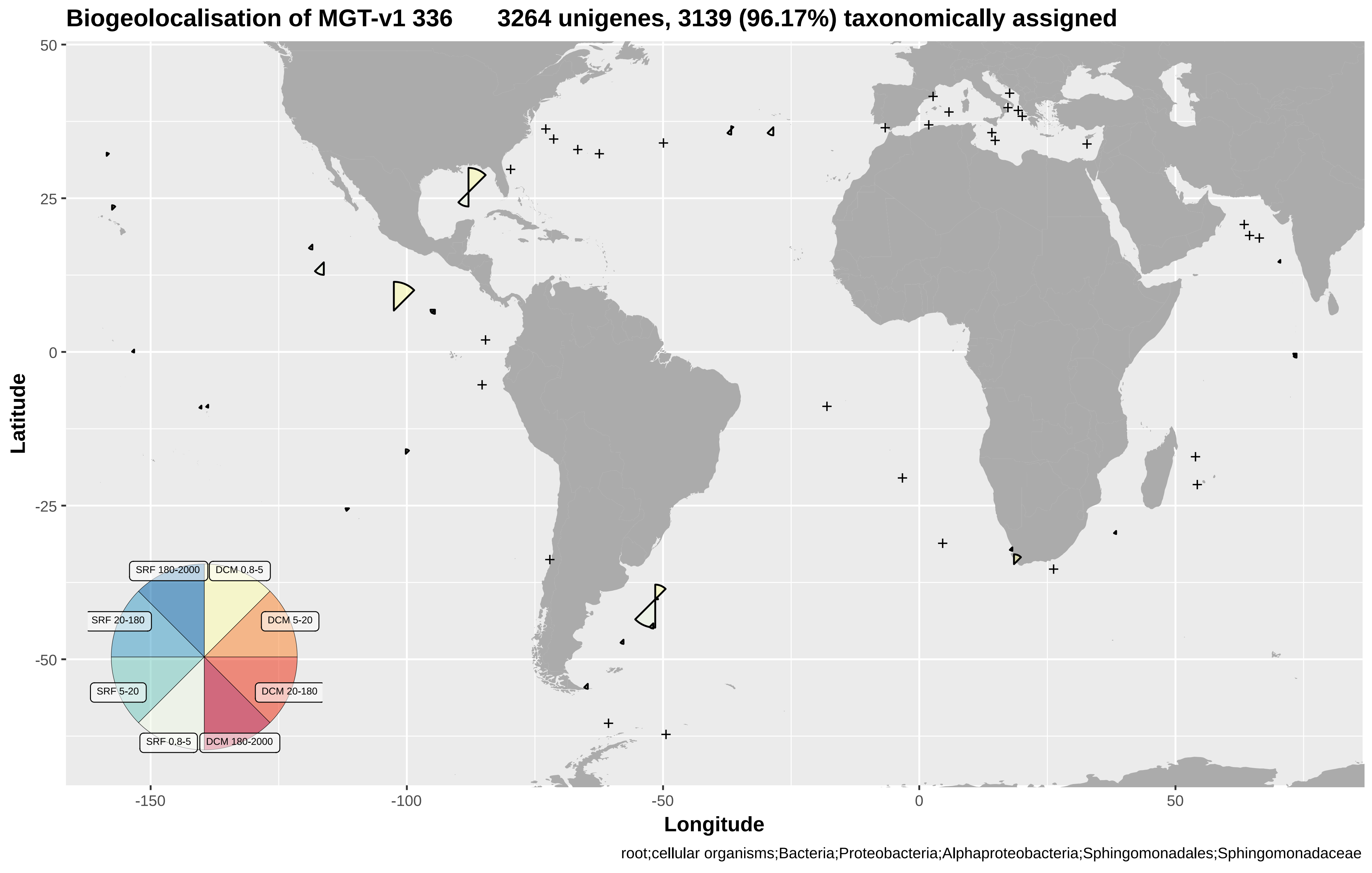
Biogeolocalisation of MGT-v1 336 3264 unigenes, 3139 (96.17%) taxonomically assigned
50
25
0
-25
-50
Latitude
-150
-100
-50
0
50
Longitude
SRF 180-2000
DCM 0.8-5
SRF 20-180
DCM 5-20
SRF 5-20
DCM 20-180
SRF 0.8-5
DCM 180-2000
root;cellular organisms;Bacteria;Proteobacteria;Alphaproteobacteria;Sphingomonadales;Sphingomonadaceae

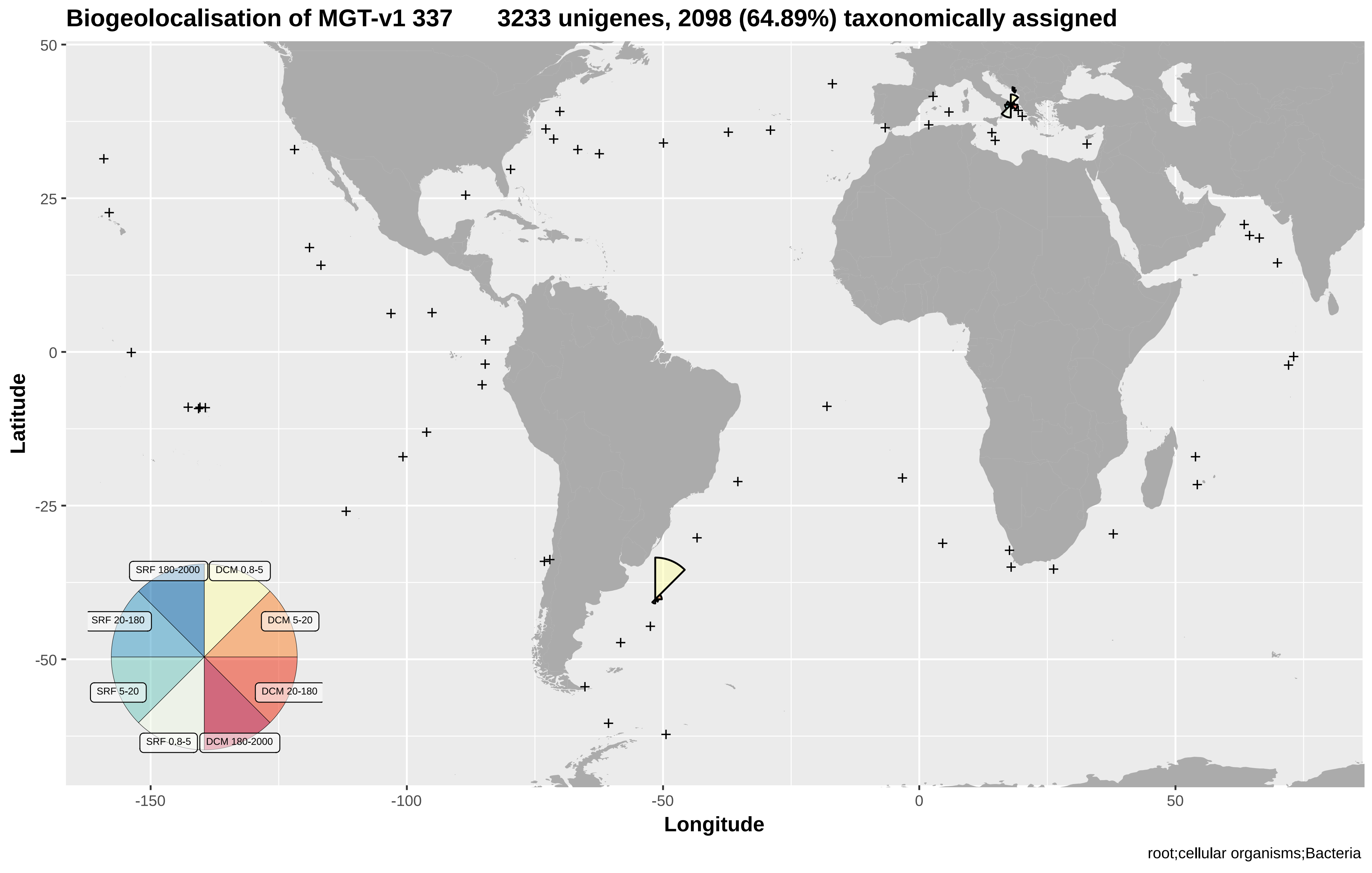

Biogeolocalisation of MGT-v1 337 3233 unigenes, 2098 (64.89%) taxonomically assigned
50
25
0
-25
-50
Latitude
-150
-100
-50
0
50
Longitude
SRF 180-2000
DCM 0.8-5
SRF 20-180
DCM 5-20
SRF 5-20
DCM 20-180
SRF 0.8-5
DCM 180-2000
root;cellular organisms;Bacteria

### Biogeolocalisation of MGT-v1 338 3222 unigenes, 3146 (97.64%) taxonomically assigned

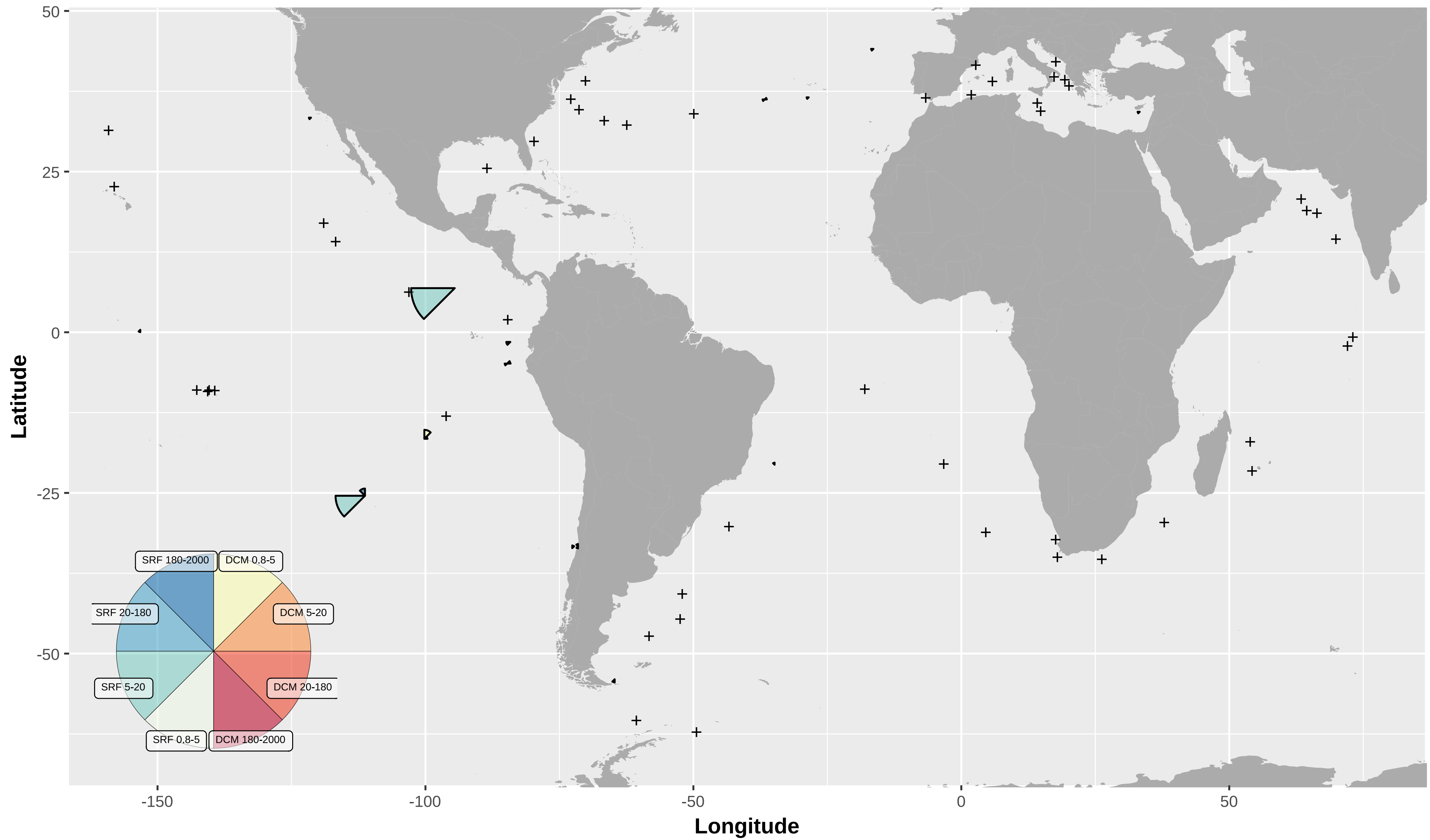


root;cellular organisms;Bacteria;Bacteroidetes/Chlorobi group;Bacteroidetes;Flavobacteriia;Flavobacteriales;Flavobacteriaceae;Croceibacter

# Biogeolocalisation of MGT-v1 339      3196 unigenes, 2642 (82.67%) taxonomically assigned

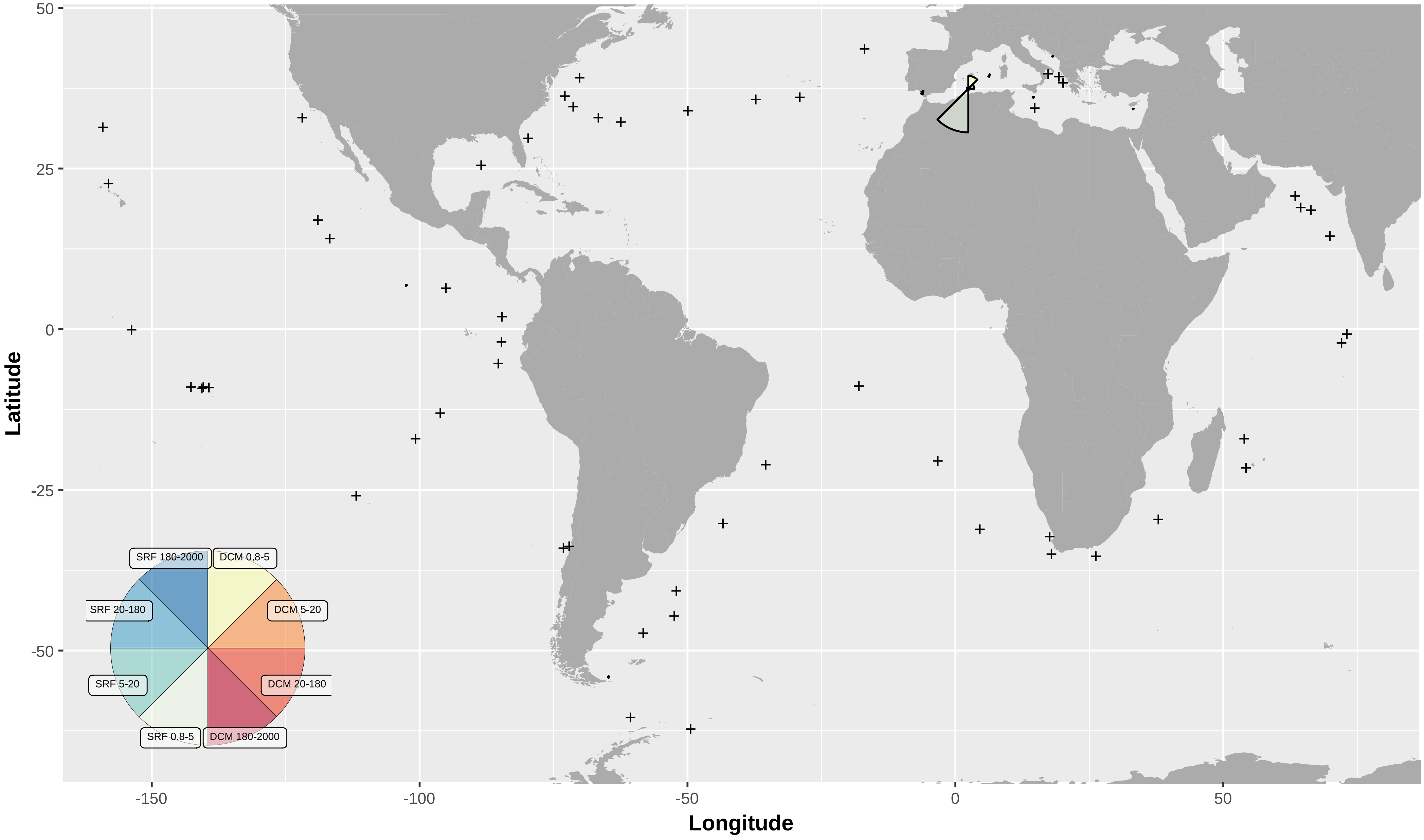


root;cellular organisms;Bacteria;Proteobacteria

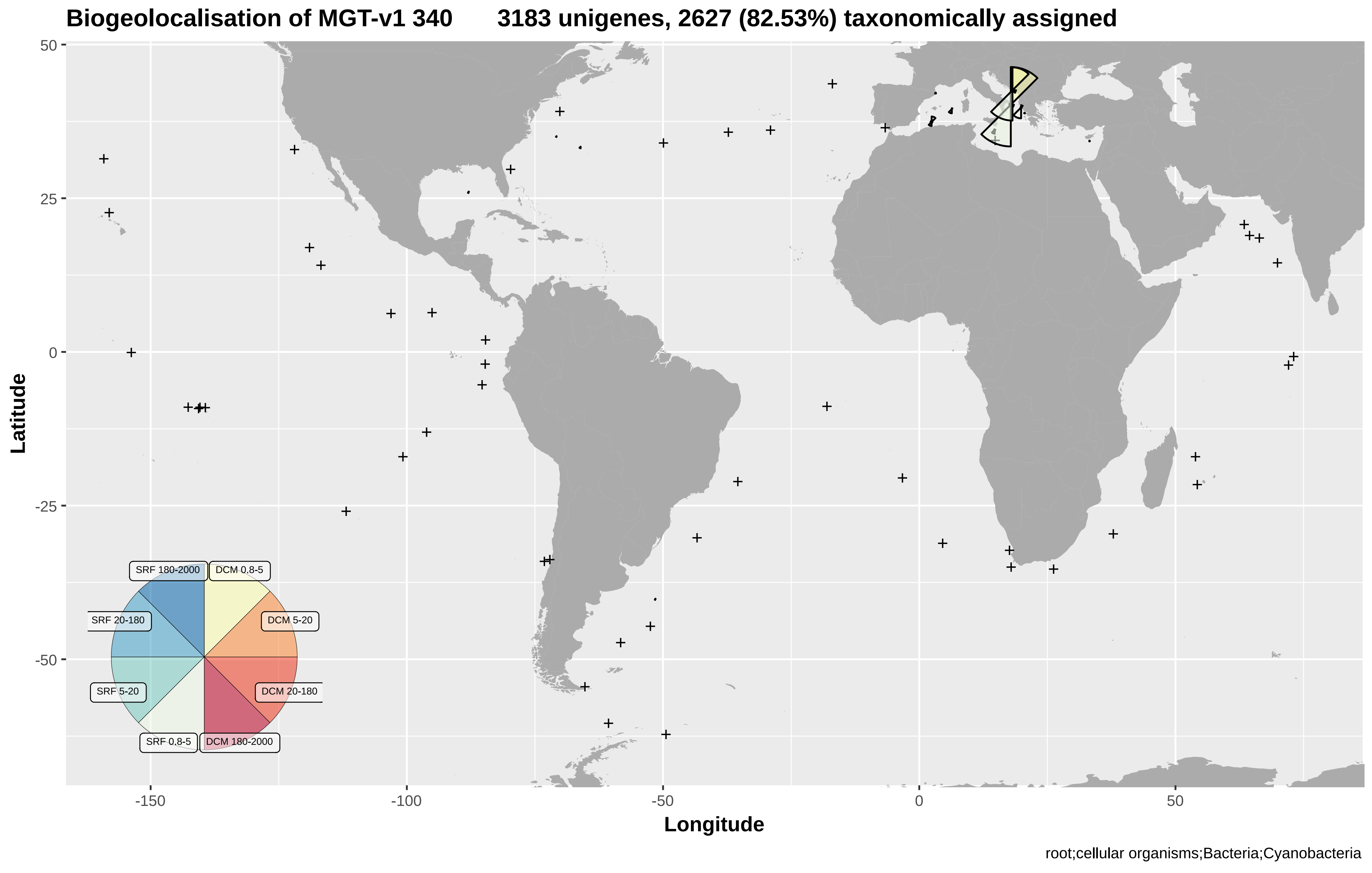
Biogeolocalisation of MGT-v1 340 3183 unigenes, 2627 (82.53%) taxonomically assigned
Latitude
Longitude
50
25
0
-25
-50
-150
-100
-50
0
50
SRF 180-2000
DCM 0.8-5
SRF 20-180
DCM 5-20
SRF 5-20
DCM 20-180
SRF 0.8-5
DCM 180-2000
root;cellular organisms;Bacteria;Cyanobacteria

# Biogeolocalisation of MGT-v1 341      3176 unigenes, 2913 (91.72%) taxonomically assigned

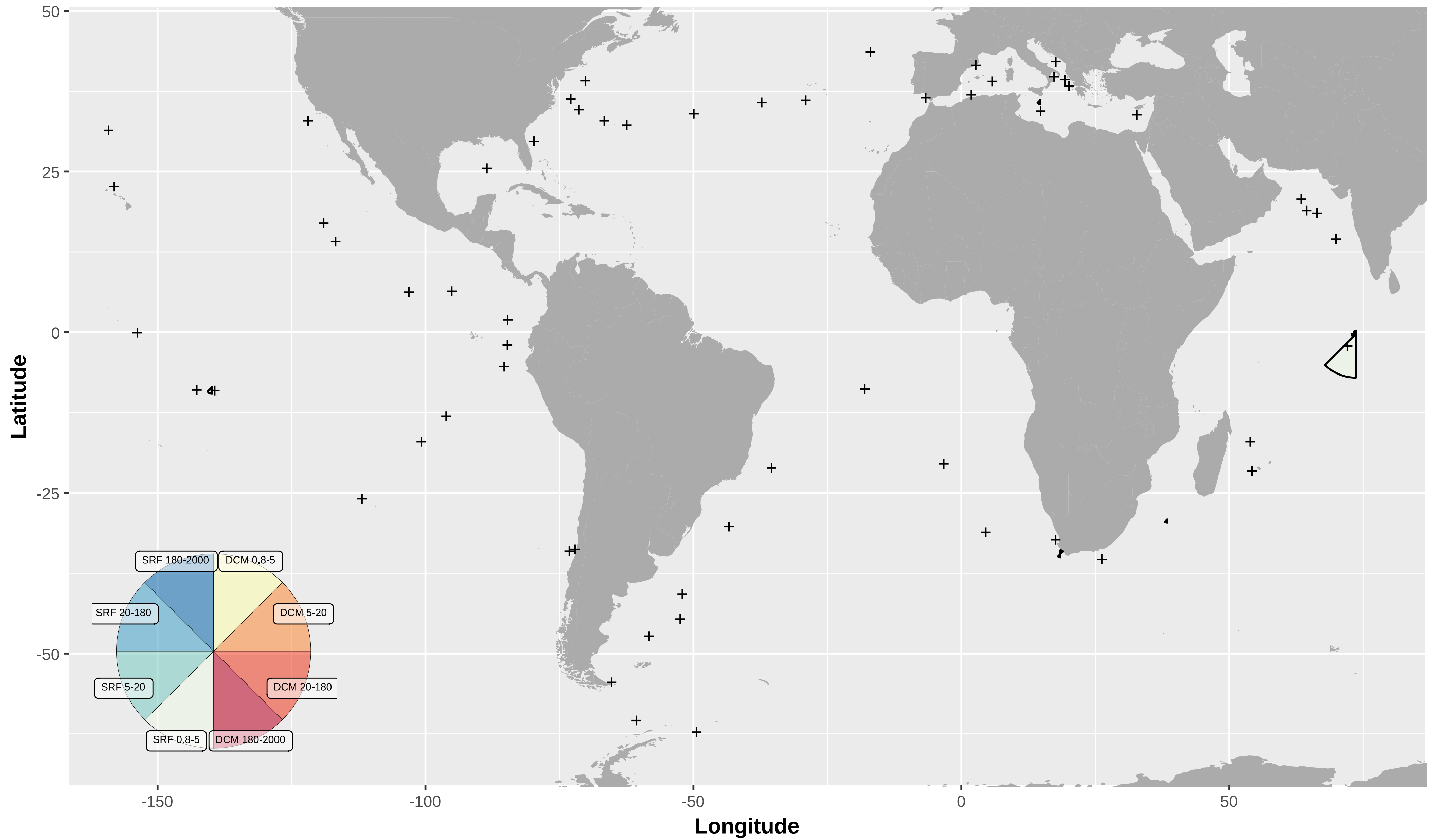


root;cellular organisms;Eukaryota;Viridiplantae;Chlorophyta;prasinophytes;Mamiellophyceae

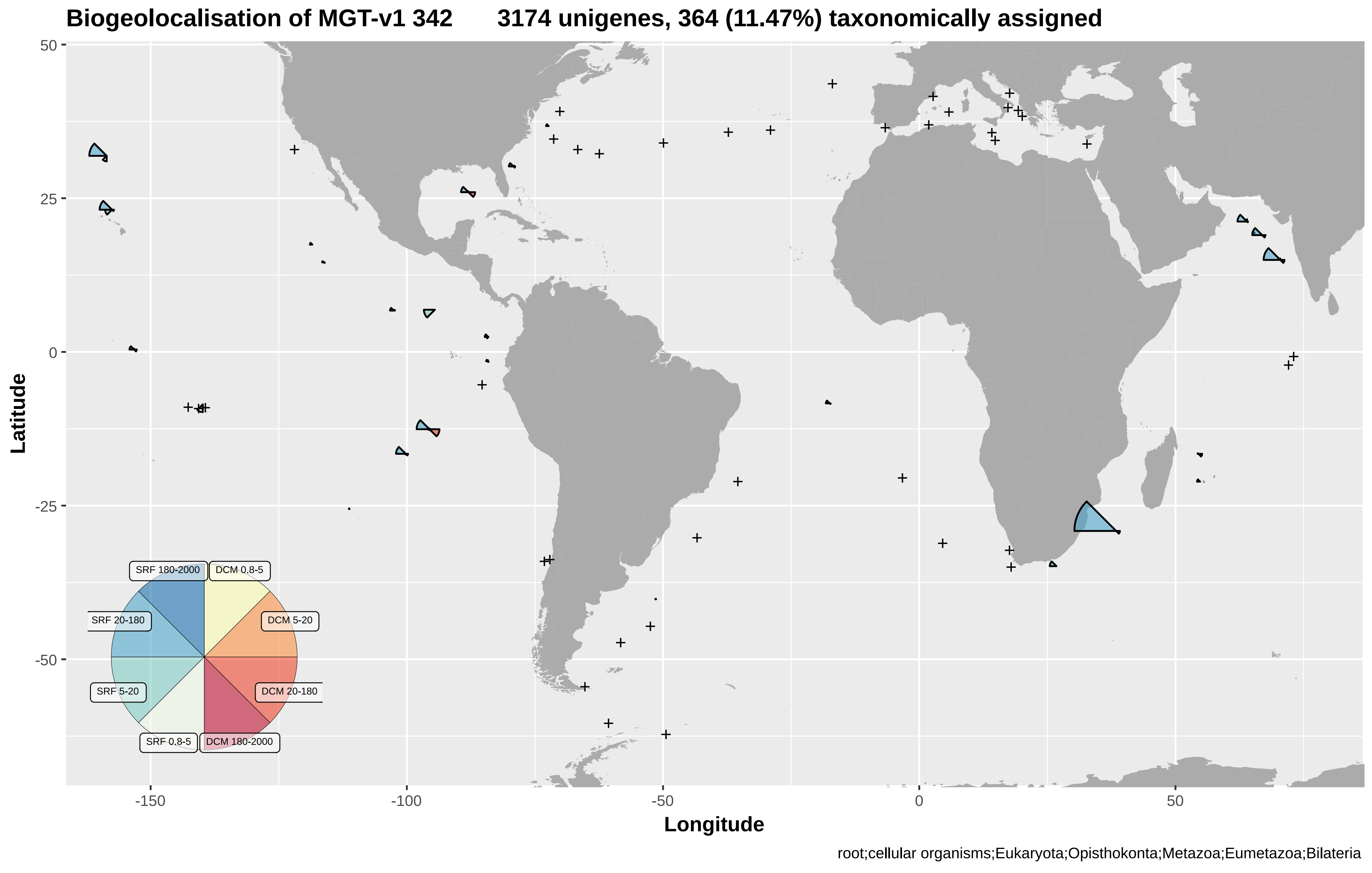
Biogeolocalisation of MGT-v1 342 3174 unigenes, 364 (11.47%) taxonomically assigned
Latitude
Longitude
50
25
0
-25
-50
-150
-100
-50
0
50
SRF 180-2000
DCM 0.8-5
SRF 20-180
DCM 5-20
SRF 5-20
DCM 20-180
SRF 0.8-5
DCM 180-2000
root;cellular organisms;Eukaryota;Opisthokonta;Metazoa;Eumetazoa;Bilateria

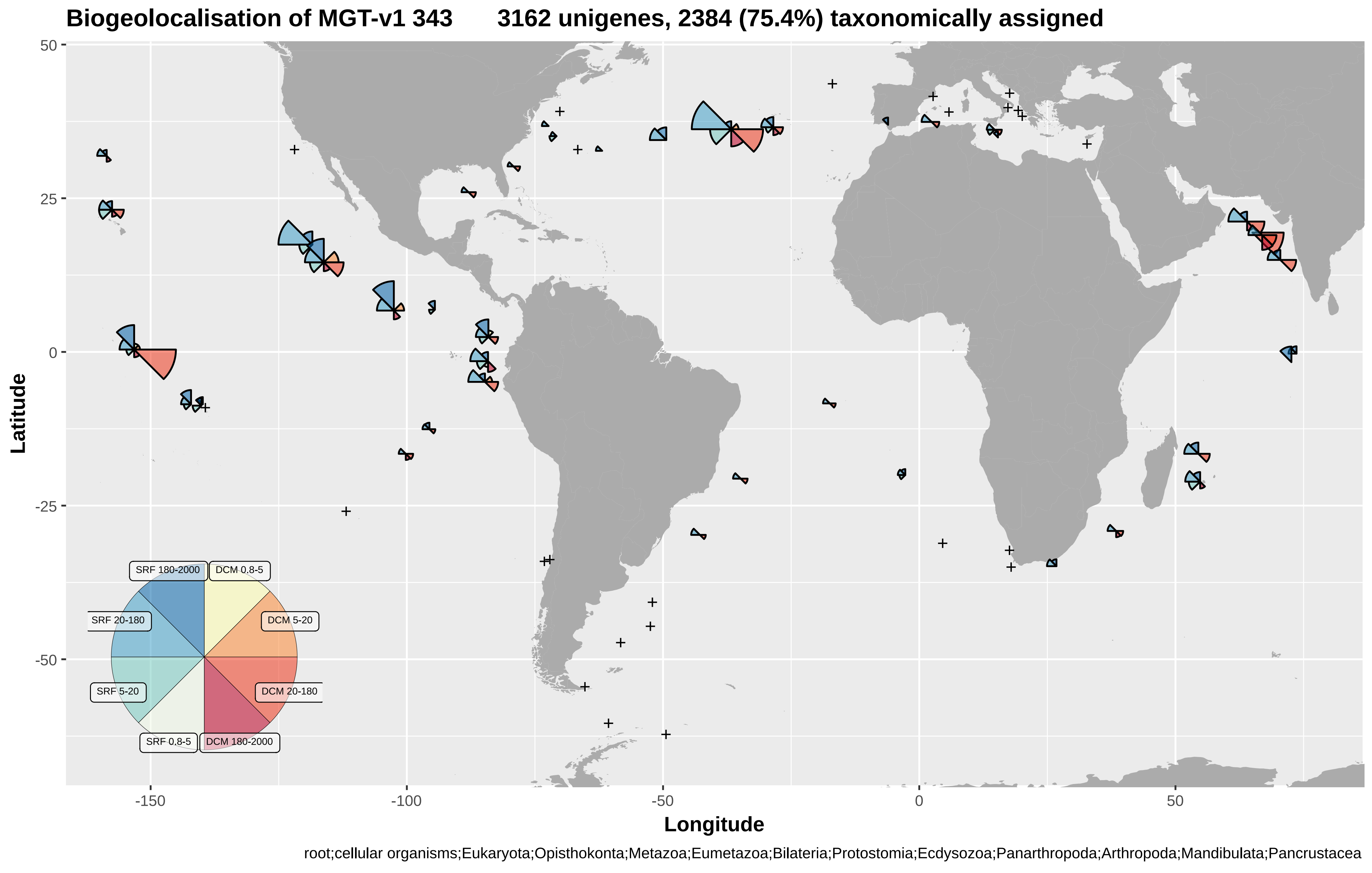
Biogeolocalisation of MGT-v1 343 3162 unigenes, 2384 (75.4%) taxonomically assigned
Latitude
Longitude
50
25
0
-25
-50
-150
-100
-50
0
50
SRF 180-2000
DCM 0.8-5
SRF 20-180
DCM 5-20
SRF 5-20
DCM 20-180
SRF 0.8-5
DCM 180-2000
root;cellular organisms;Eukaryota;Opisthokonta;Metazoa;Eumetazoa;Bilateria;Protostomia;Ecdysozoa;Panarthropoda;Arthropoda;Mandibulata;Pancrustacea

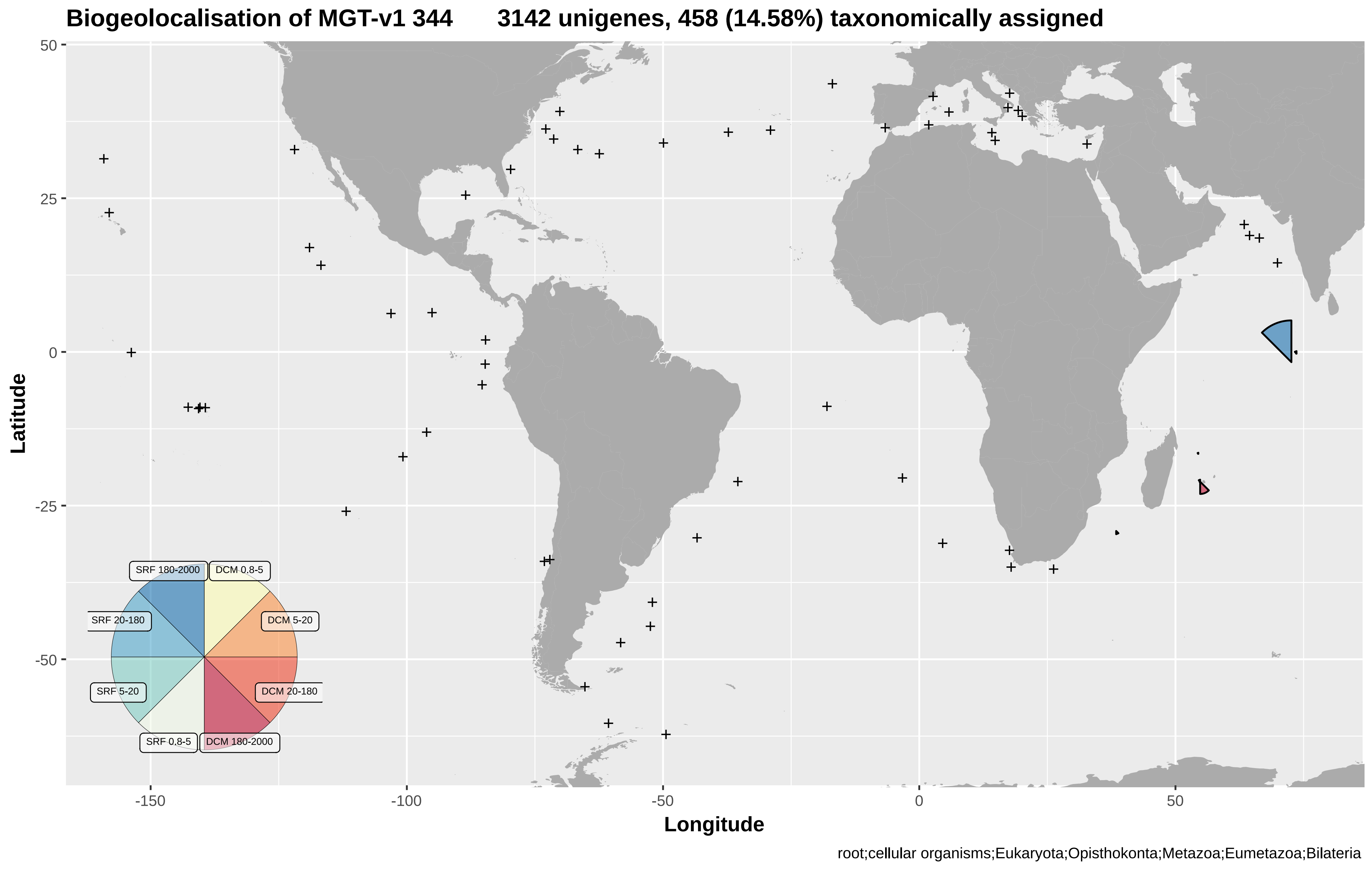
Biogeolocalisation of MGT-v1 344
3142 unigenes, 458 (14.58%) taxonomically assigned
50
25
0
-25
-50
Latitude
-150
-100
-50
0
50
Longitude
SRF 180-2000
DCM 0.8-5
SRF 20-180
DCM 5-20
SRF 5-20
DCM 20-180
SRF 0.8-5
DCM 180-2000
root;cellular organisms;Eukaryota;Opisthokonta;Metazoa;Eumetazoa;Bilateria

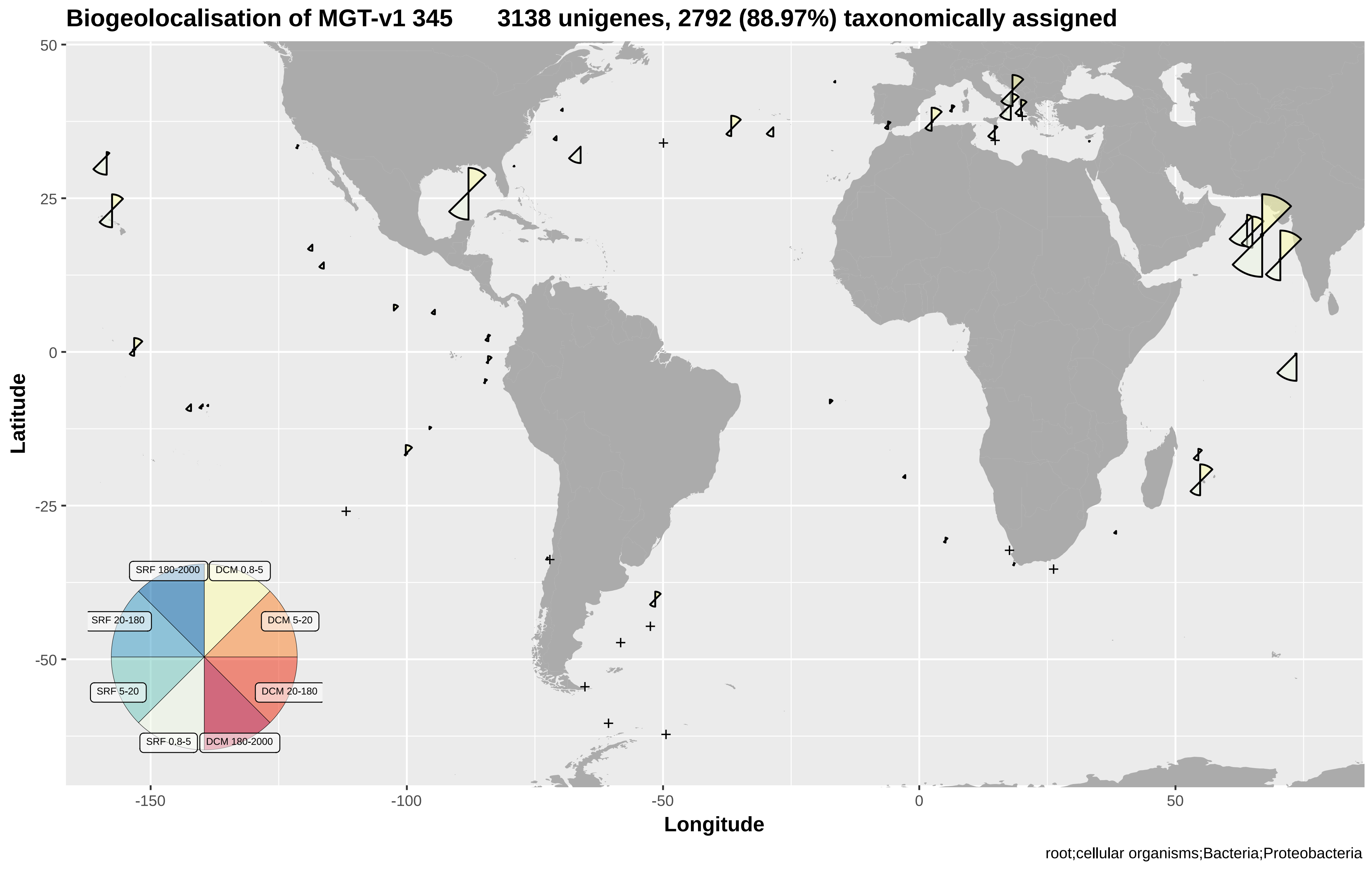

Biogeolocalisation of MGT-v1 345 3138 unigenes, 2792 (88.97%) taxonomically assigned
50
25
0
-25
-50
Latitude
-150
-100
-50
0
50
Longitude
SRF 180-2000
DCM 0.8-5
SRF 20-180
DCM 5-20
SRF 5-20
DCM 20-180
SRF 0.8-5
DCM 180-2000
root;cellular organisms;Bacteria;Proteobacteria

#### Biogeolocalisation of MGT-v1 346 3088 unigenes, 2955 (95.69%) taxonomically assigned

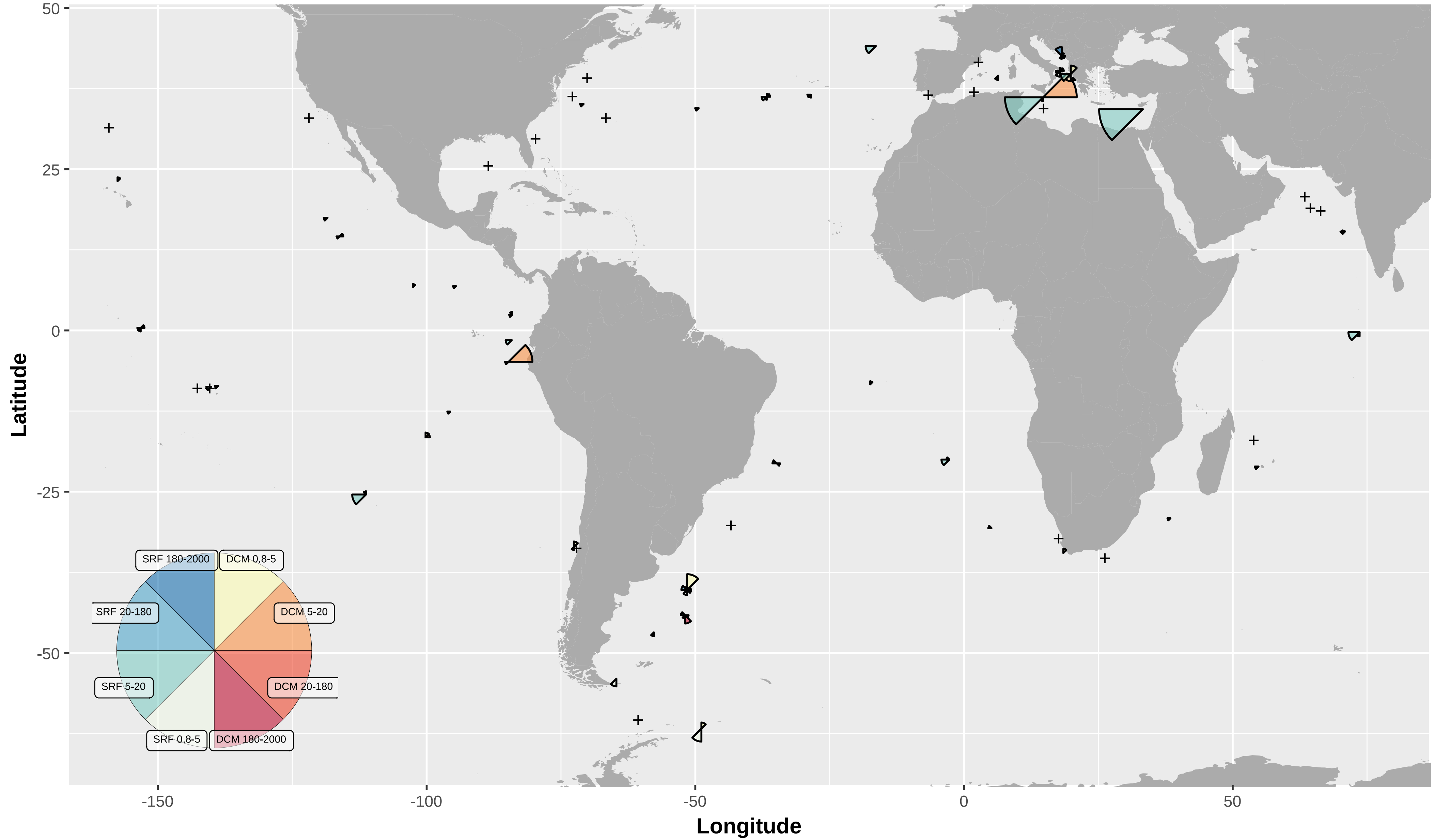


root;cellular organisms;Bacteria;Proteobacteria;Gammaproteobacteria;Pseudomonadales;Pseudomonadaceae

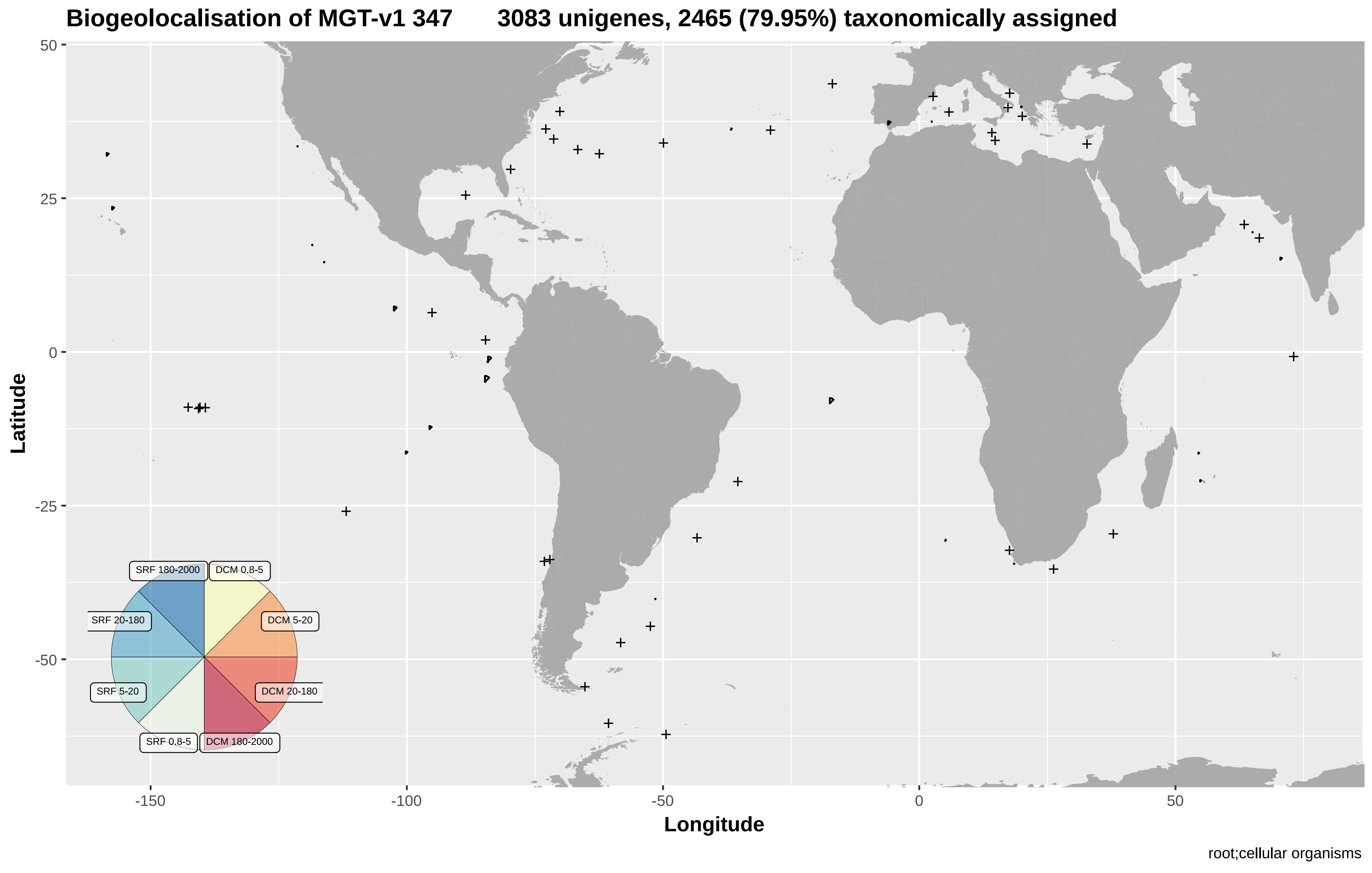
Biogeolocalisation of MGT-v1 347 3083 unigenes, 2465 (79.95%) taxonomically assigned
50
25
0
-25
-50
Latitude
-150
-100
-50
0
50
Longitude
SRF 180-2000
DCM 0.8-5
SRF 20-180
DCM 5-20
SRF 5-20
DCM 20-180
SRF 0.8-5
DCM 180-2000
root;cellular organisms

# Biogeolocalisation of MGT-v1 348      3042 unigenes, 452 (14.86%) taxonomically assigned

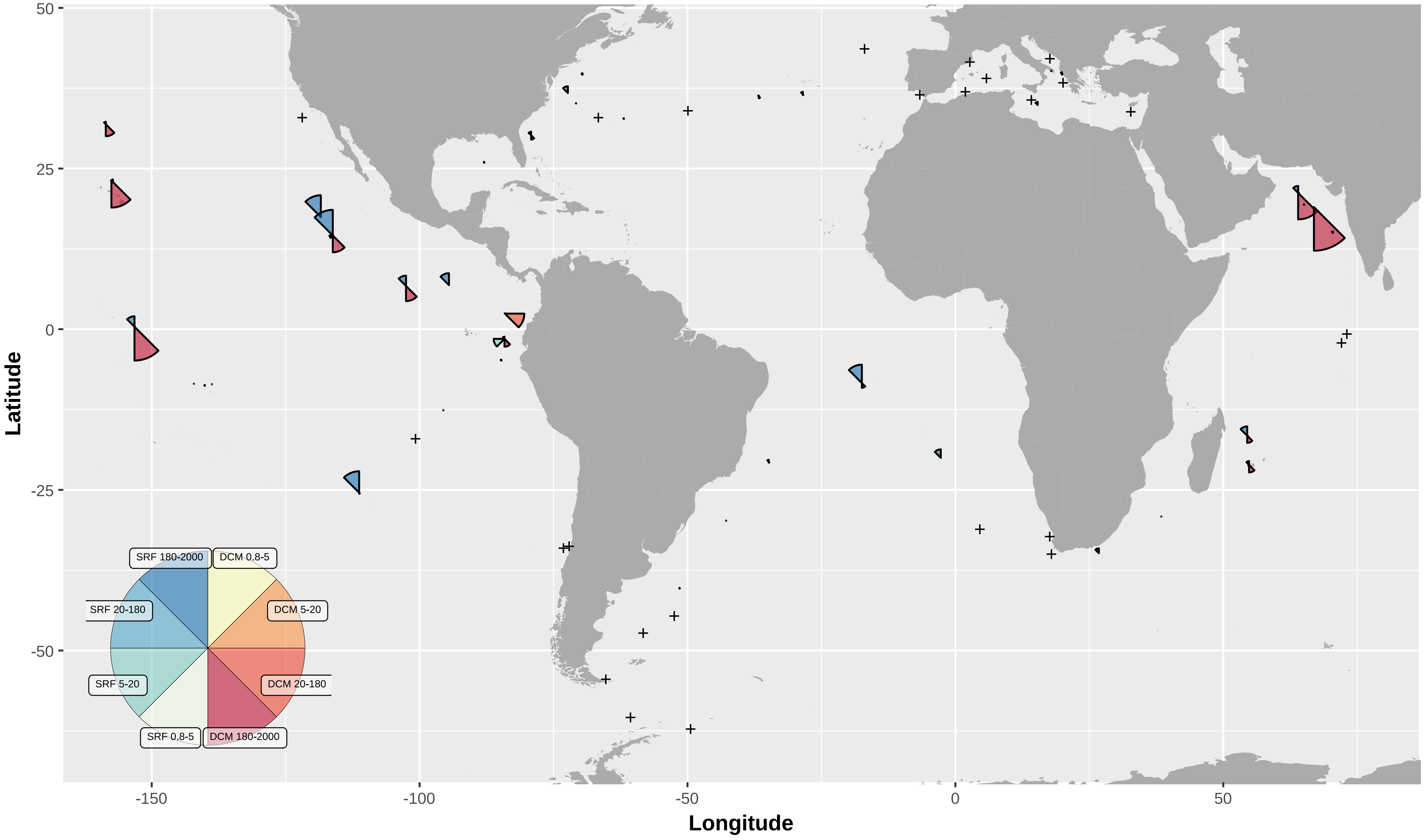


root;cellular organisms;Eukaryota;Opisthokonta;Metazoa;Eumetazoa;Bilateria

# Biogeolocalisation of MGT-v1 349      3032 unigenes, 193 (6.37%) taxonomically assigned

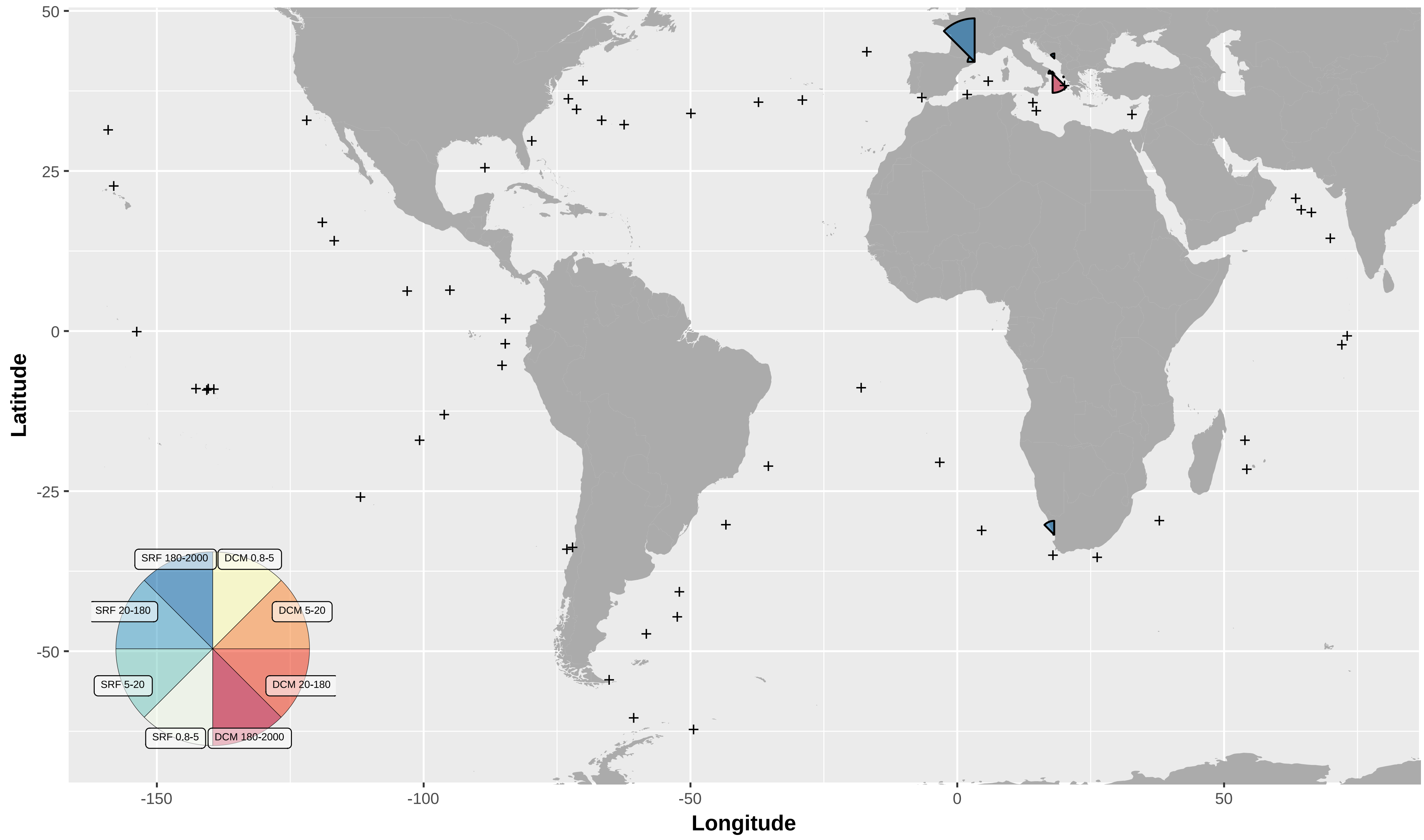


root;cellular organisms;Eukaryota;Opisthokonta;Metazoa;Eumetazoa

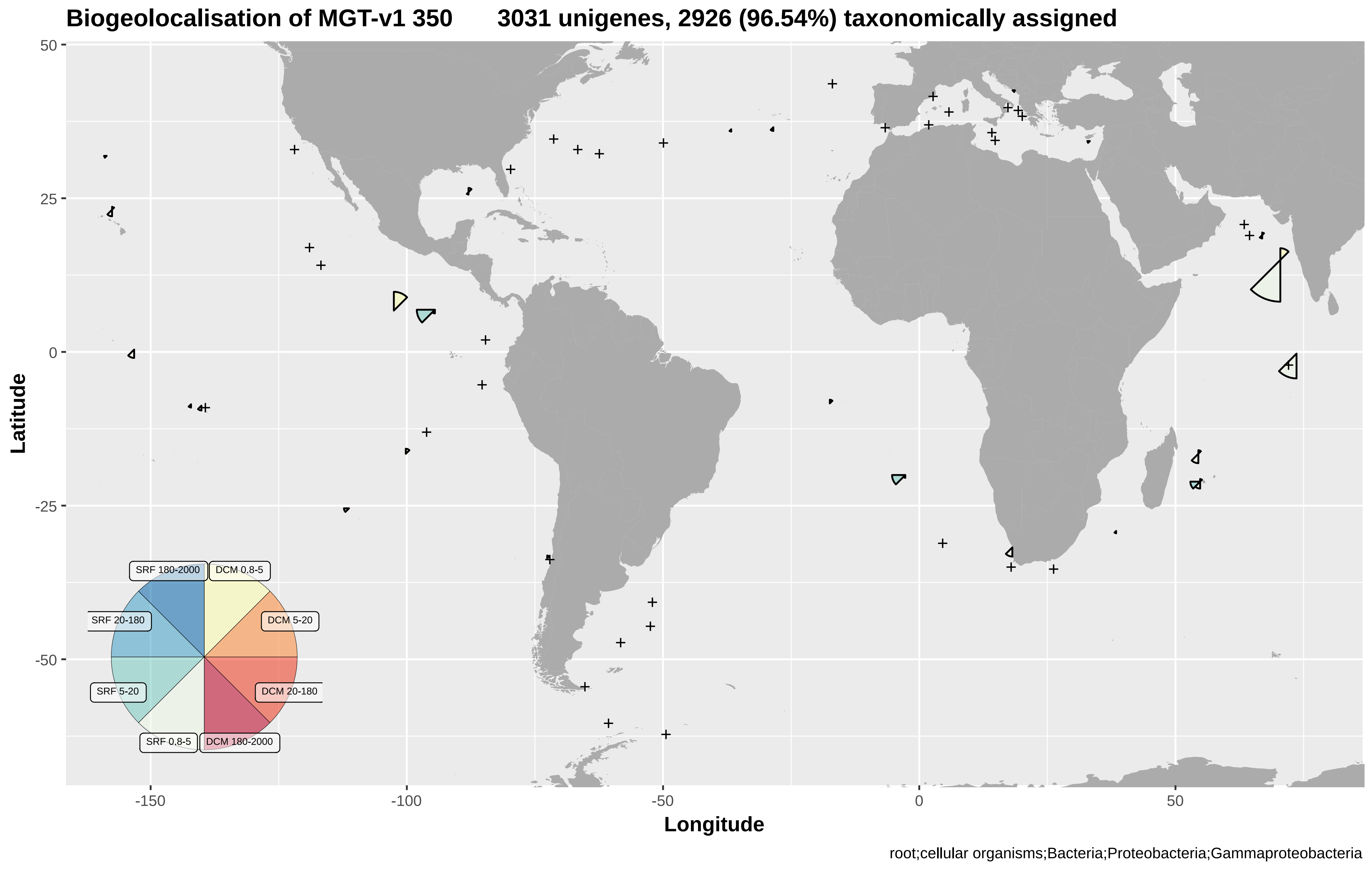

Biogeolocalisation of MGT-v1 350 3031 unigenes, 2926 (96.54%) taxonomically assigned
Latitude
Longitude
50
25
0
-25
-50
-150
-100
-50
0
50
SRF 180-2000
DCM 0.8-5
SRF 20-180
DCM 5-20
SRF 5-20
DCM 20-180
SRF 0.8-5
DCM 180-2000
root;cellular organisms;Bacteria;Proteobacteria;Gammaproteobacteria

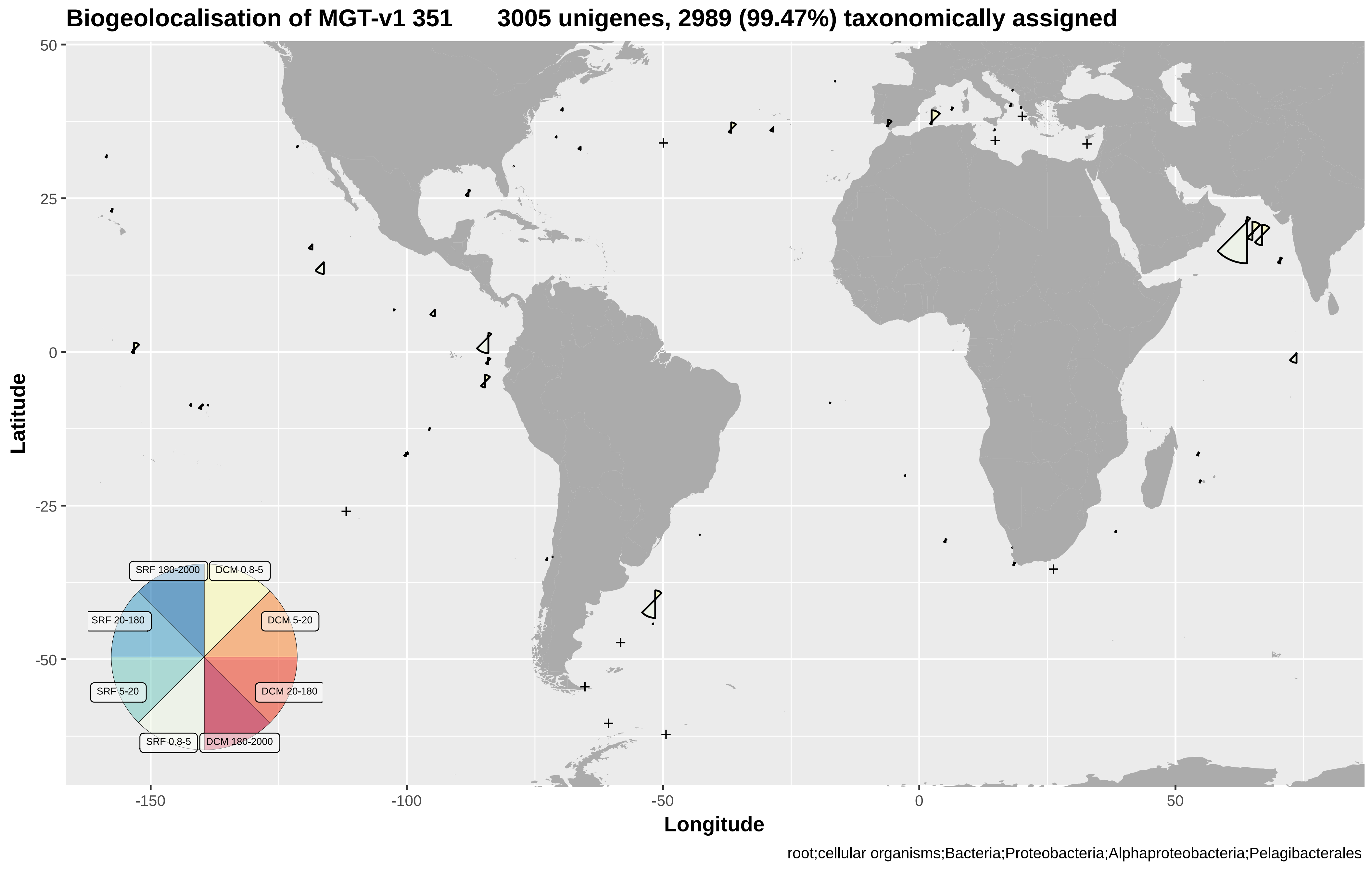

Biogeolocalisation of MGT-v1 351 3005 unigenes, 2989 (99.47%) taxonomically assigned
Latitude
Longitude
50
25
0
-25
-50
-150
-100
-50
0
50
SRF 180-2000
DCM 0.8-5
SRF 20-180
DCM 5-20
SRF 5-20
DCM 20-180
SRF 0.8-5
DCM 180-2000
root;cellular organisms;Bacteria;Proteobacteria;Alphaproteobacteria;Pelagibacterales

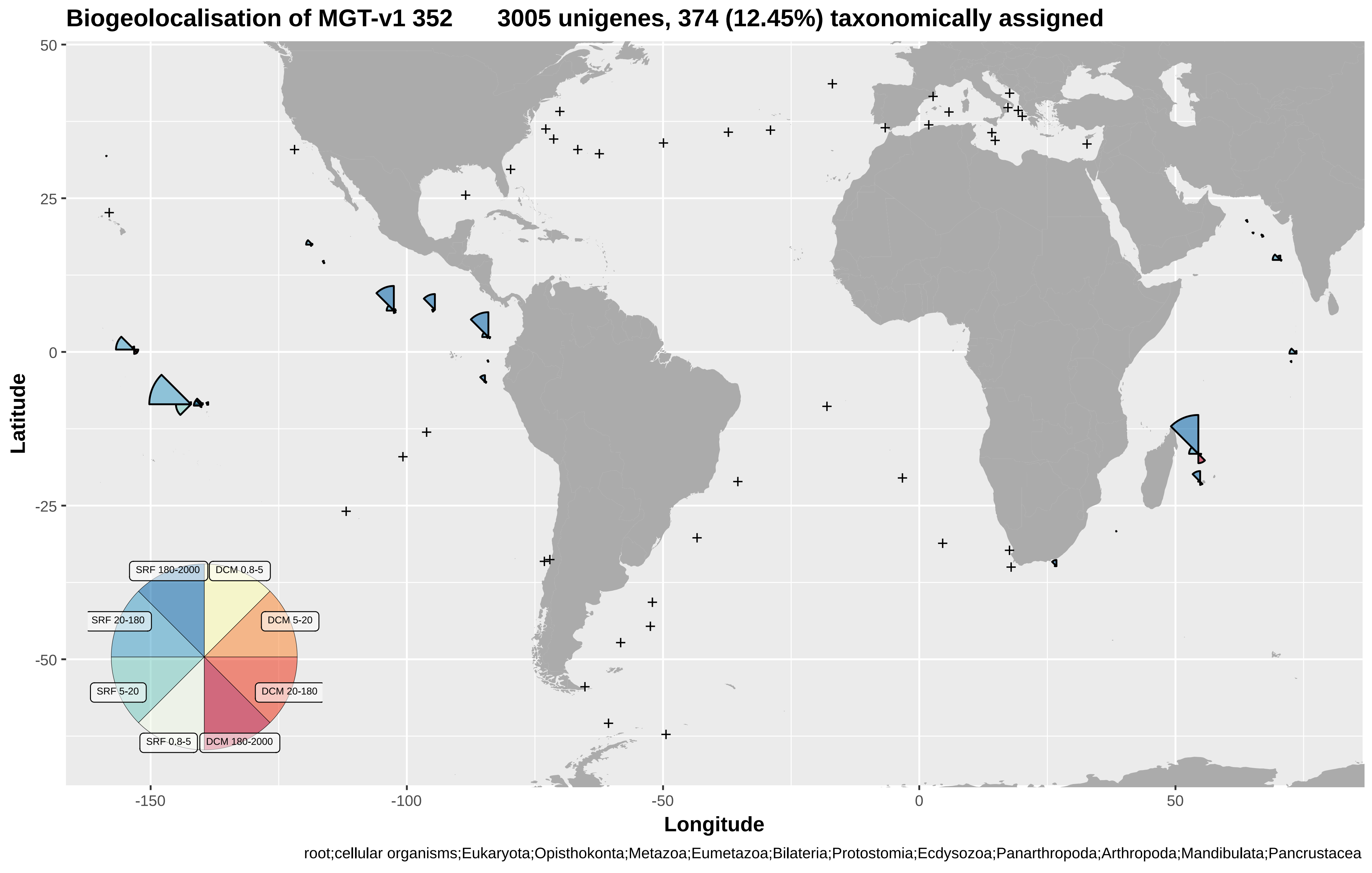
Biogeolocalisation of MGT-v1 352
3005 unigenes, 374 (12.45%) taxonomically assigned
50
25
0
-25
-50
Latitude
-150
-100
-50
0
50
Longitude
SRF 180-2000
DCM 0.8-5
SRF 20-180
DCM 5-20
SRF 5-20
DCM 20-180
SRF 0.8-5
DCM 180-2000
root;cellular organisms;Eukaryota;Opisthokonta;Metazoa;Eumetazoa;Bilateria;Protostomia;Ecdysozoa;Panarthropoda;Arthropoda;Mandibulata;Pancrustacea

**Biogeolocalisation of MGT-v1 353      2995 unigenes, 2789 (93.12%) taxonomically assigned**

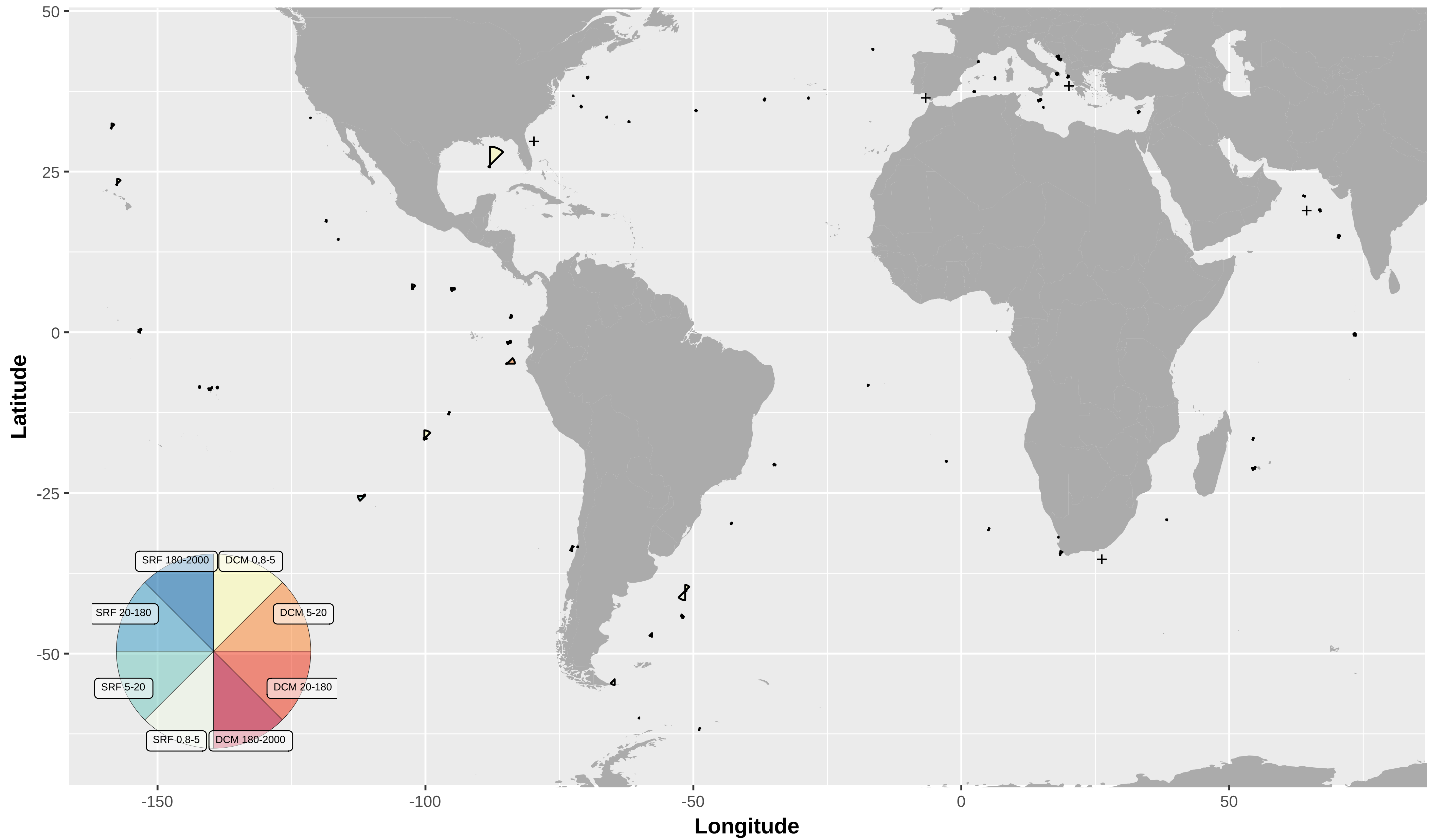


root;cellular organisms;Bacteria;Proteobacteria;Alphaproteobacteria;Sphingomonadales

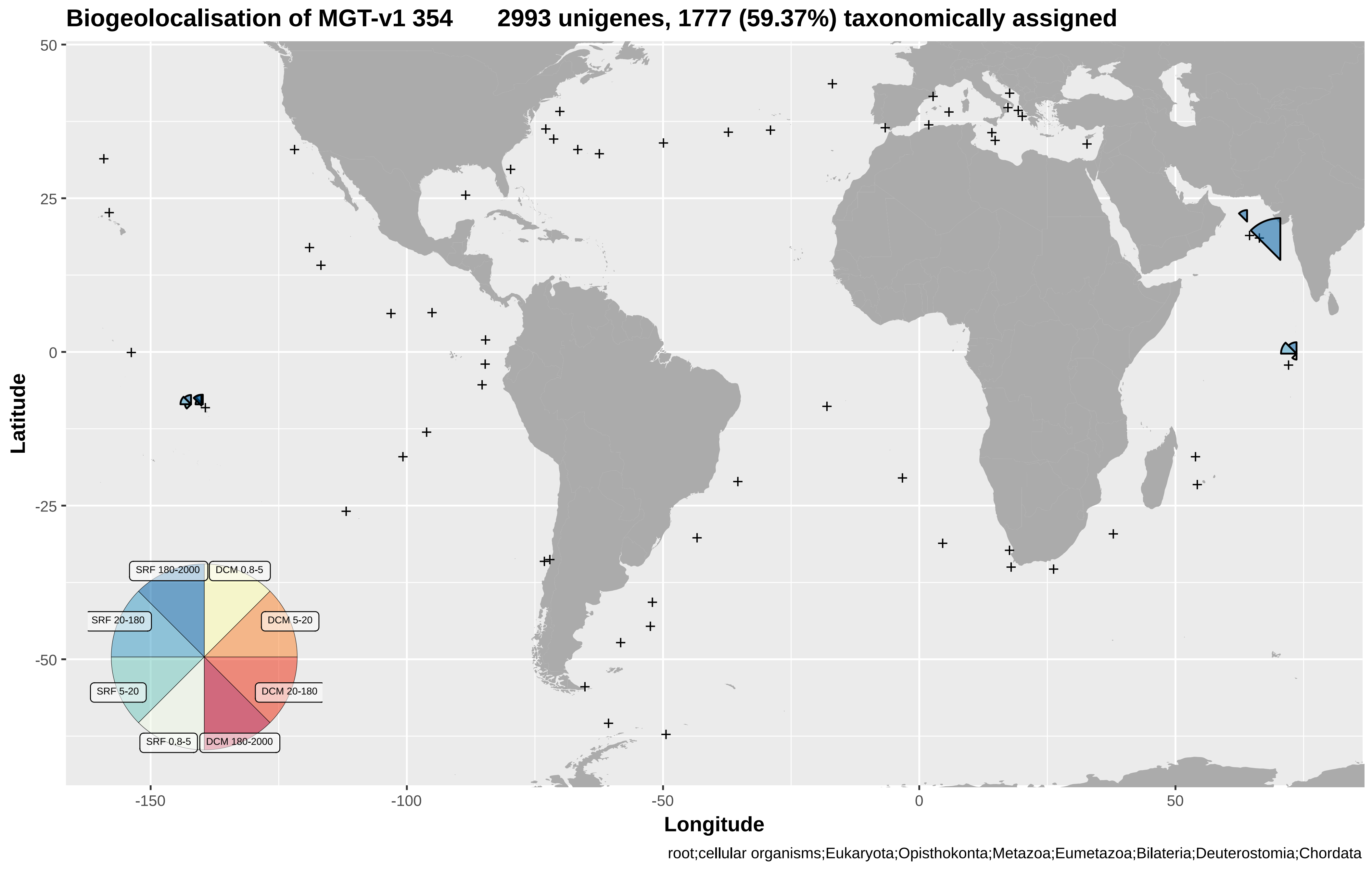
Biogeolocalisation of MGT-v1 354 2993 unigenes, 1777 (59.37%) taxonomically assigned
50
25
0
-25
-50
Latitude
-150
-100
-50
0
50
Longitude
SRF 180-2000
DCM 0.8-5
SRF 20-180
DCM 5-20
SRF 5-20
DCM 20-180
SRF 0.8-5
DCM 180-2000
root;cellular organisms;Eukaryota;Opisthokonta;Metazoa;Eumetazoa;Bilateria;Deuterostomia;Chordata

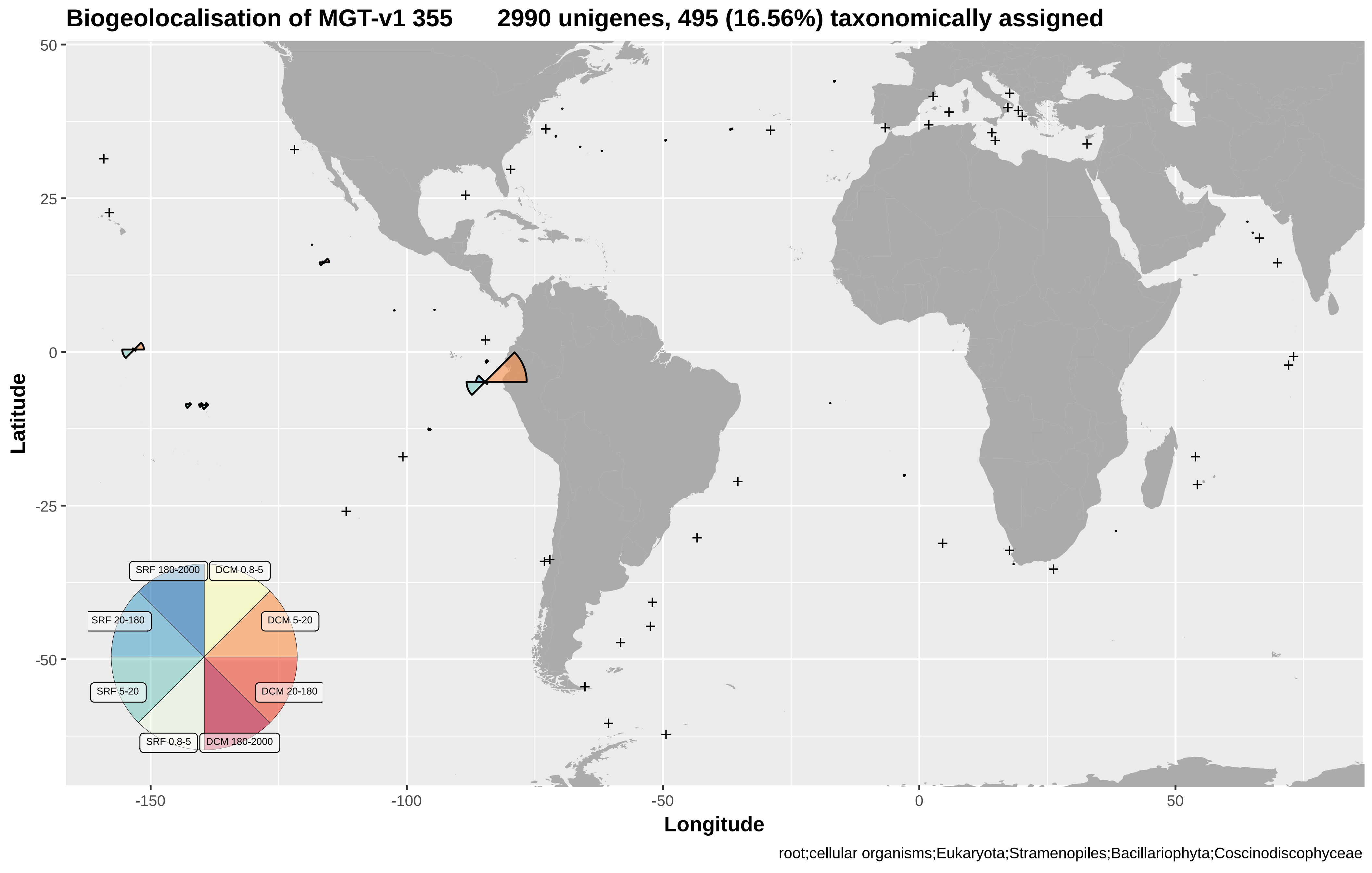

Biogeolocalisation of MGT-v1 355 2990 unigenes, 495 (16.56%) taxonomically assigned
Latitude
Longitude
50
25
0
-25
-50
-150
-100
-50
0
50
SRF 180-2000
DCM 0.8-5
SRF 20-180
DCM 5-20
SRF 5-20
DCM 20-180
SRF 0.8-5
DCM 180-2000
root;cellular organisms;Eukaryota;Stramenopiles;Bacillariophyta;Coscinodiscophyceae

#### Biogeolocalisation of MGT-v1 356 2968 unigenes, 584 (19.68%) taxonomically assigned

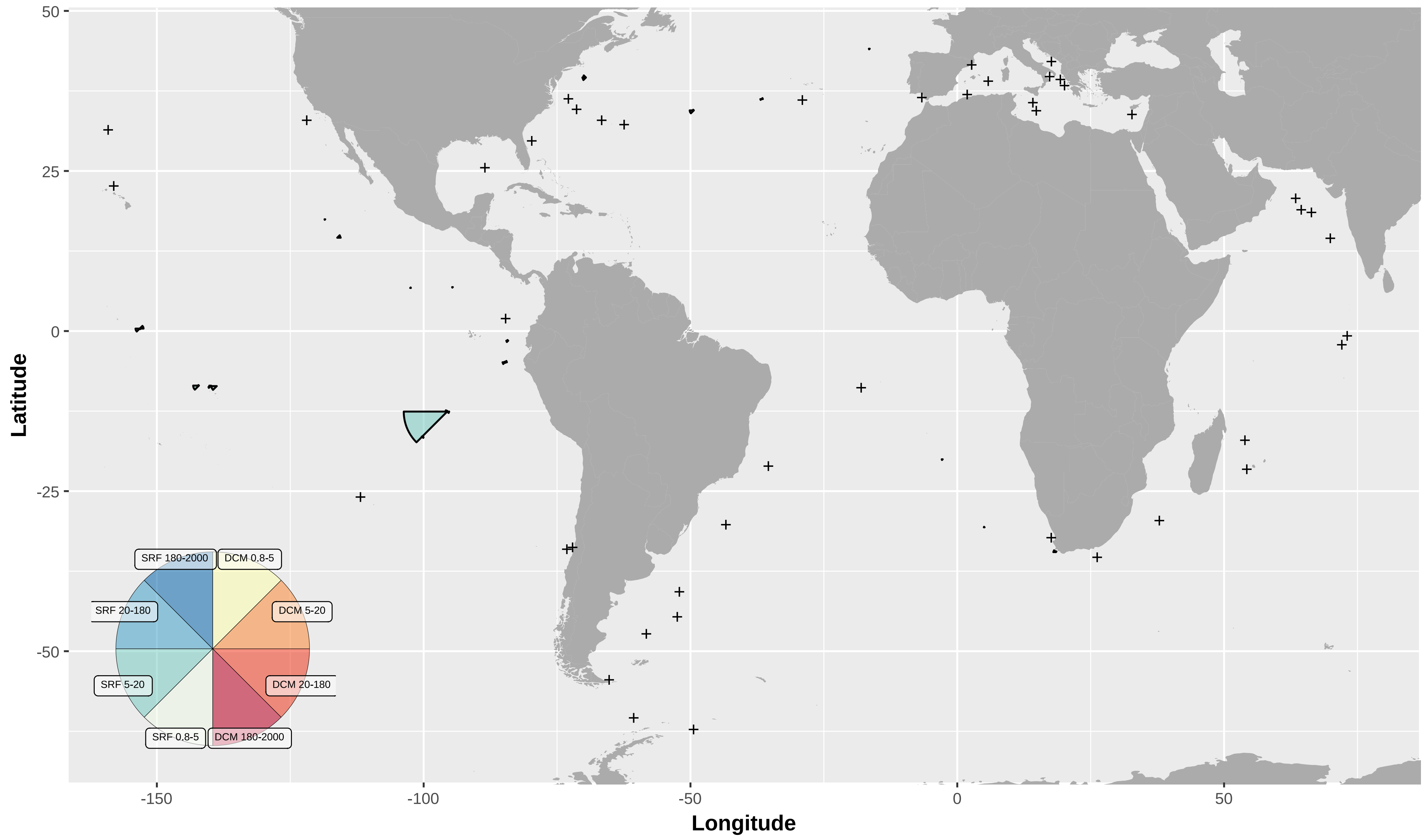


root;cellular organisms;Eukaryota;Stramenopiles;Bacillariophyta

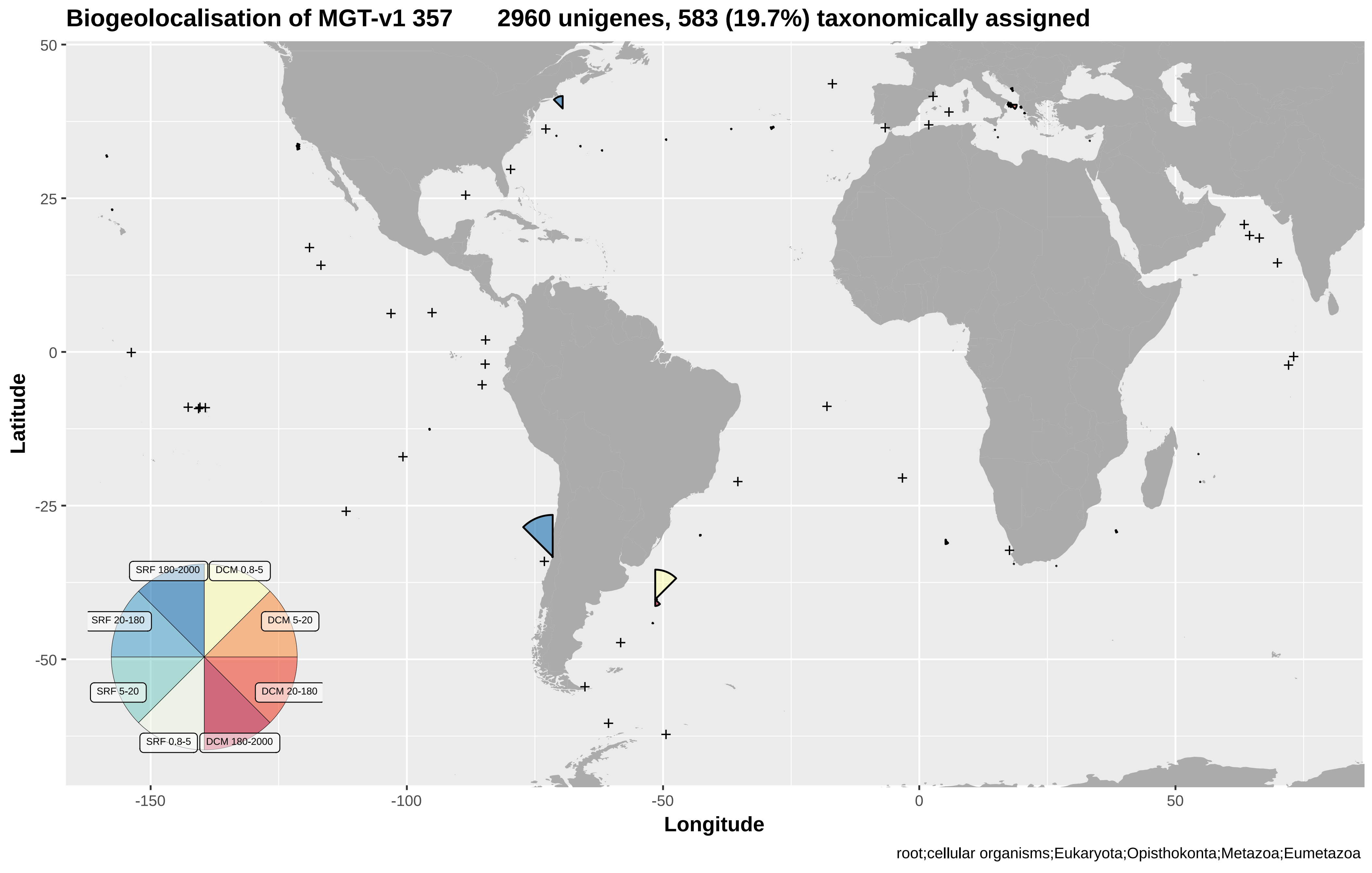
Biogeolocalisation of MGT-v1 357 2960 unigenes, 583 (19.7%) taxonomically assigned
50
25
0
-25
-50
Latitude
-150
-100
-50
0
50
Longitude
SRF 180-2000
DCM 0.8-5
SRF 20-180
DCM 5-20
SRF 5-20
DCM 20-180
SRF 0.8-5
DCM 180-2000
root;cellular organisms;Eukaryota;Opisthokonta;Metazoa;Eumetazoa

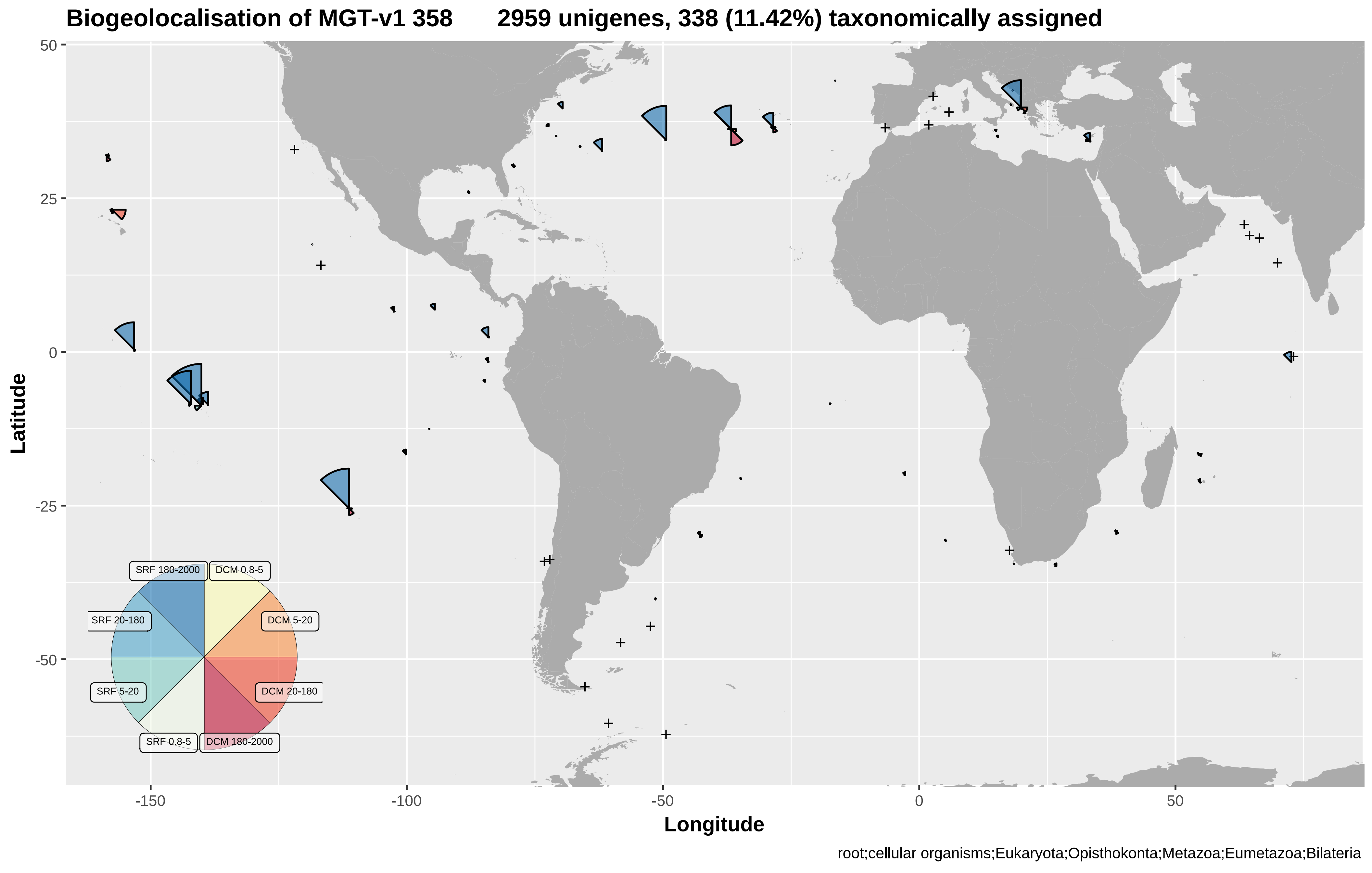
Biogeolocalisation of MGT-v1 358 2959 unigenes, 338 (11.42%) taxonomically assigned
50
25
0
-25
-50
Latitude
-150
-100
-50
0
50
Longitude
SRF 180-2000
DCM 0.8-5
SRF 20-180
DCM 5-20
SRF 5-20
DCM 20-180
SRF 0.8-5
DCM 180-2000
root;cellular organisms;Eukaryota;Opisthokonta;Metazoa;Eumetazoa;Bilateria

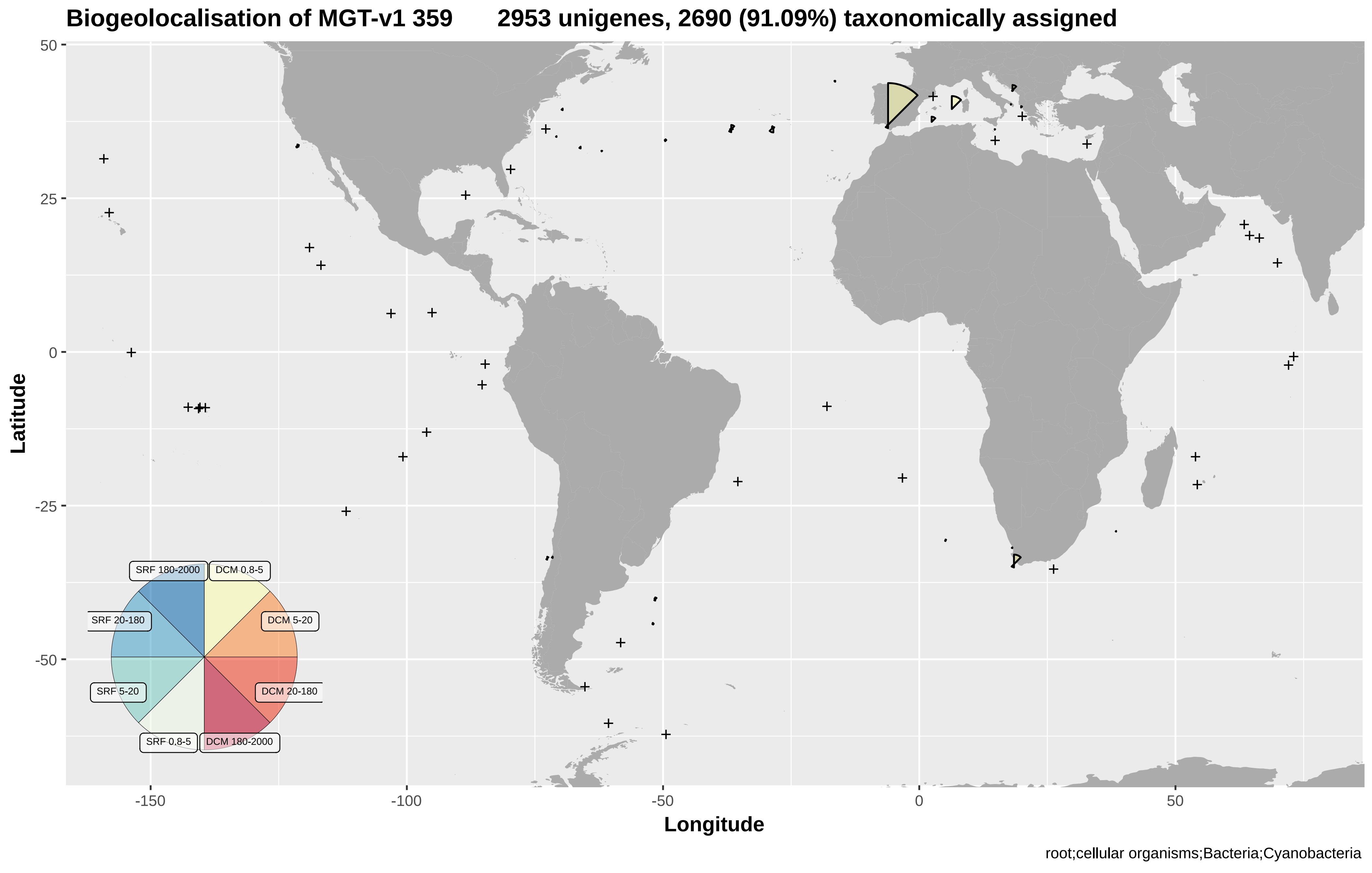
Biogeolocalisation of MGT-v1 359 2953 unigenes, 2690 (91.09%) taxonomically assigned
50
25
0
-25
-50
Latitude
-150
-100
-50
0
50
Longitude
SRF 180-2000
DCM 0.8-5
SRF 20-180
DCM 5-20
SRF 5-20
DCM 20-180
SRF 0.8-5
DCM 180-2000
root;cellular organisms;Bacteria;Cyanobacteria

## Biogeolocalisation of MGT-v1 360      2940 unigenes, 561 (19.08%) taxonomically assigned

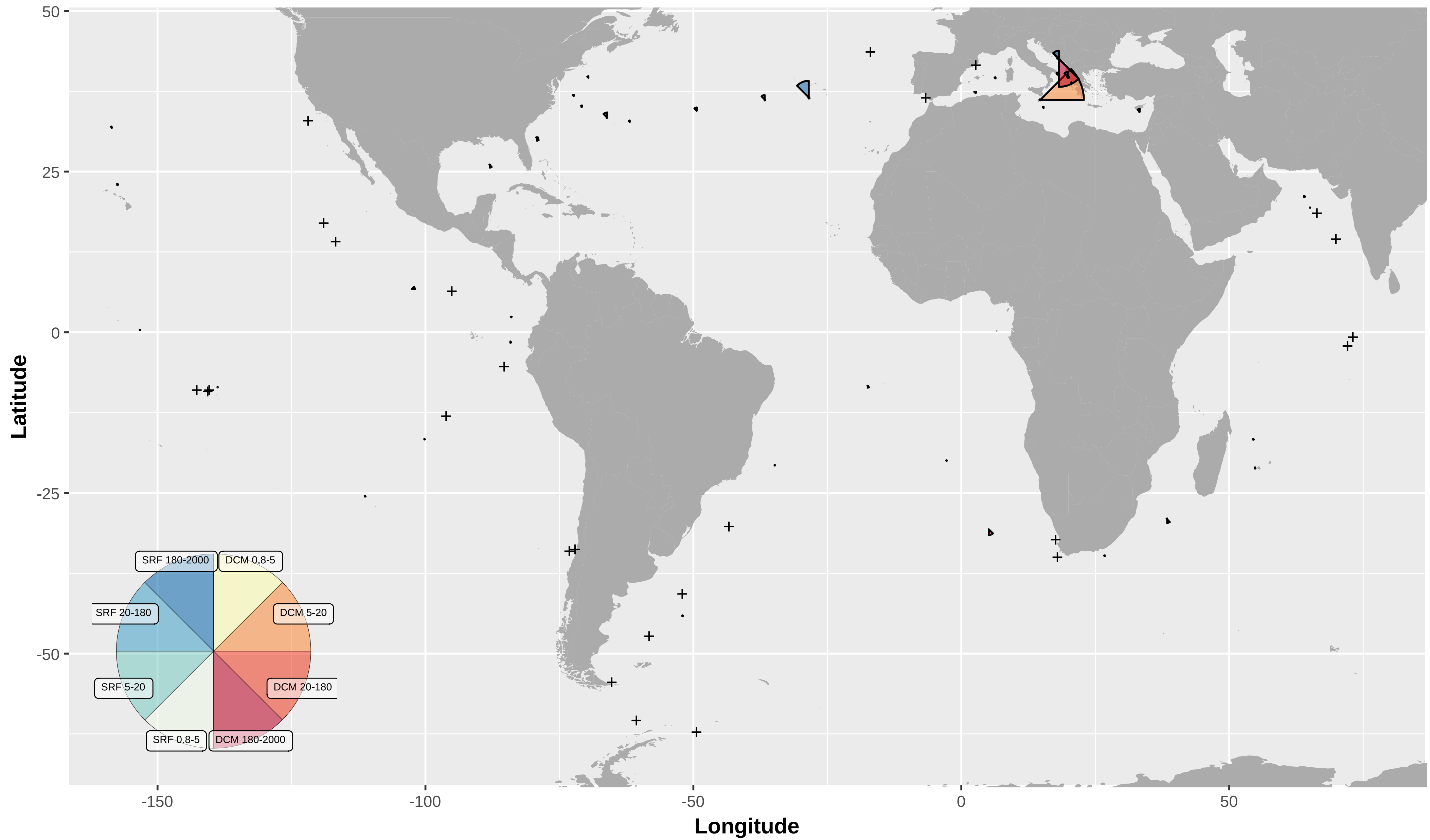


root;cellular organisms;Eukaryota;Opisthokonta;Metazoa;Eumetazoa

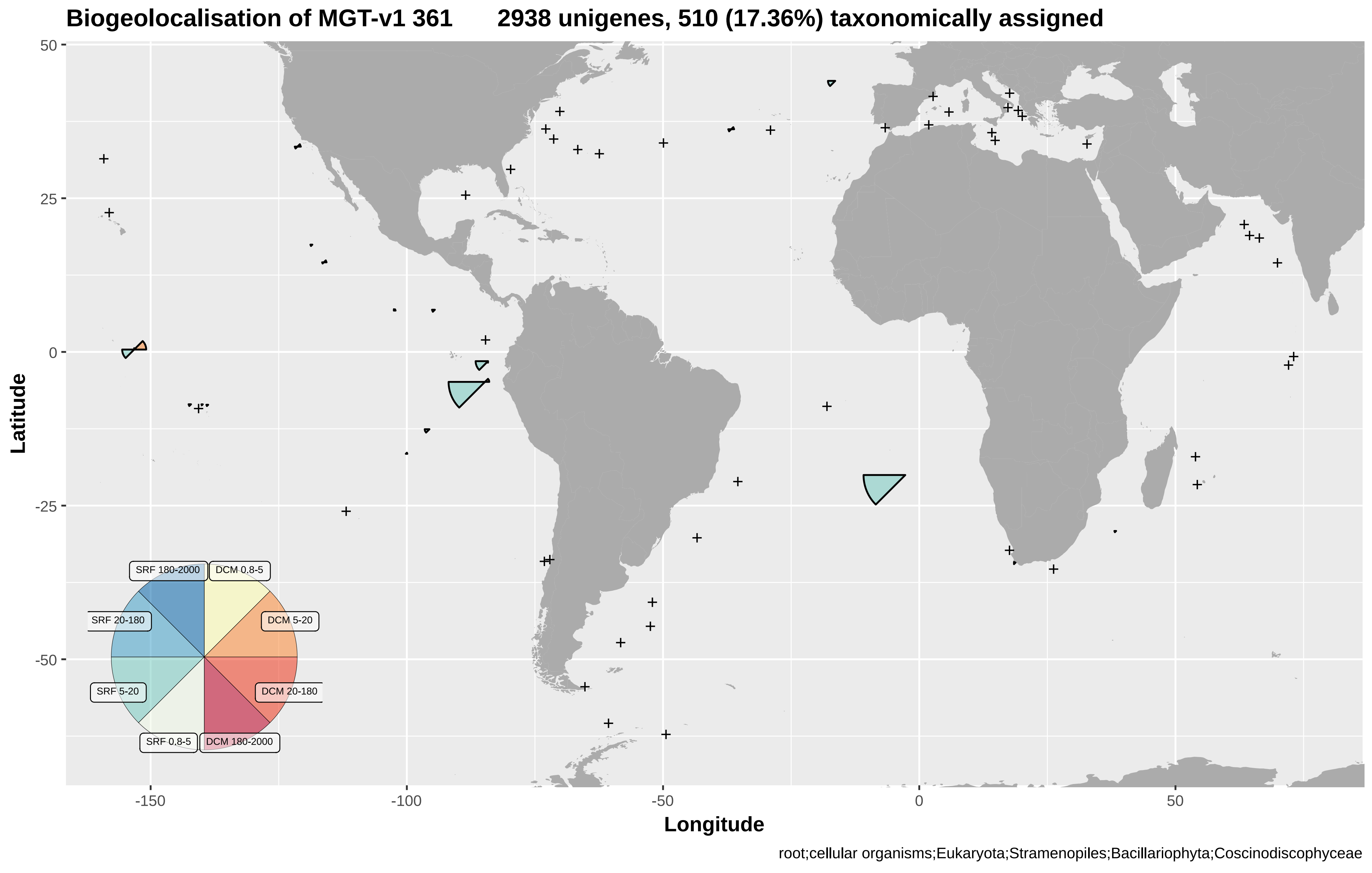

Biogeolocalisation of MGT-v1 361 2938 unigenes, 510 (17.36%) taxonomically assigned
50
25
0
-25
-50
Latitude
-150
-100
-50
0
50
Longitude
SRF 180-2000
DCM 0.8-5
SRF 20-180
DCM 5-20
SRF 5-20
DCM 20-180
SRF 0.8-5
DCM 180-2000
root;cellular organisms;Eukaryota;Stramenopiles;Bacillariophyta;Coscinodiscophyceae

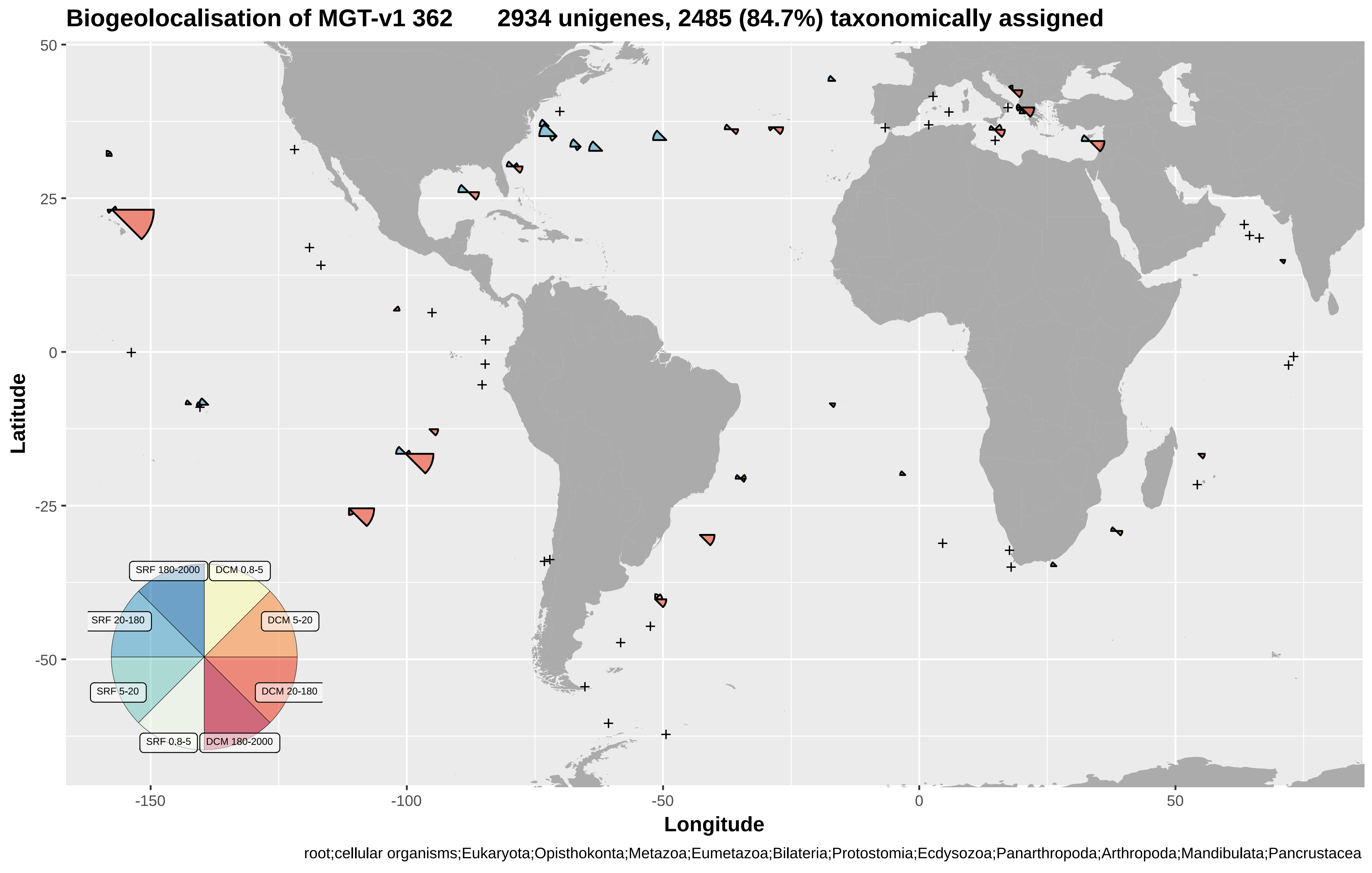
Biogeolocalisation of MGT-v1 362 2934 unigenes, 2485 (84.7%) taxonomically assigned
Latitude
50
25
0
-25
-50
Longitude
-150
-100
-50
0
50
SRF 180-2000
DCM 0.8-5
SRF 20-180
DCM 5-20
SRF 5-20
DCM 20-180
SRF 0.8-5
DCM 180-2000
root;cellular organisms;Eukaryota;Opisthokonta;Metazoa;Eumetazoa;Bilateria;Protostomia;Ecdysozoa;Panarthropoda;Arthropoda;Mandibulata;Pancrustacea

#### Biogeolocalisation of MGT-v1 363 2932 unigenes, 573 (19.54%) taxonomically assigned

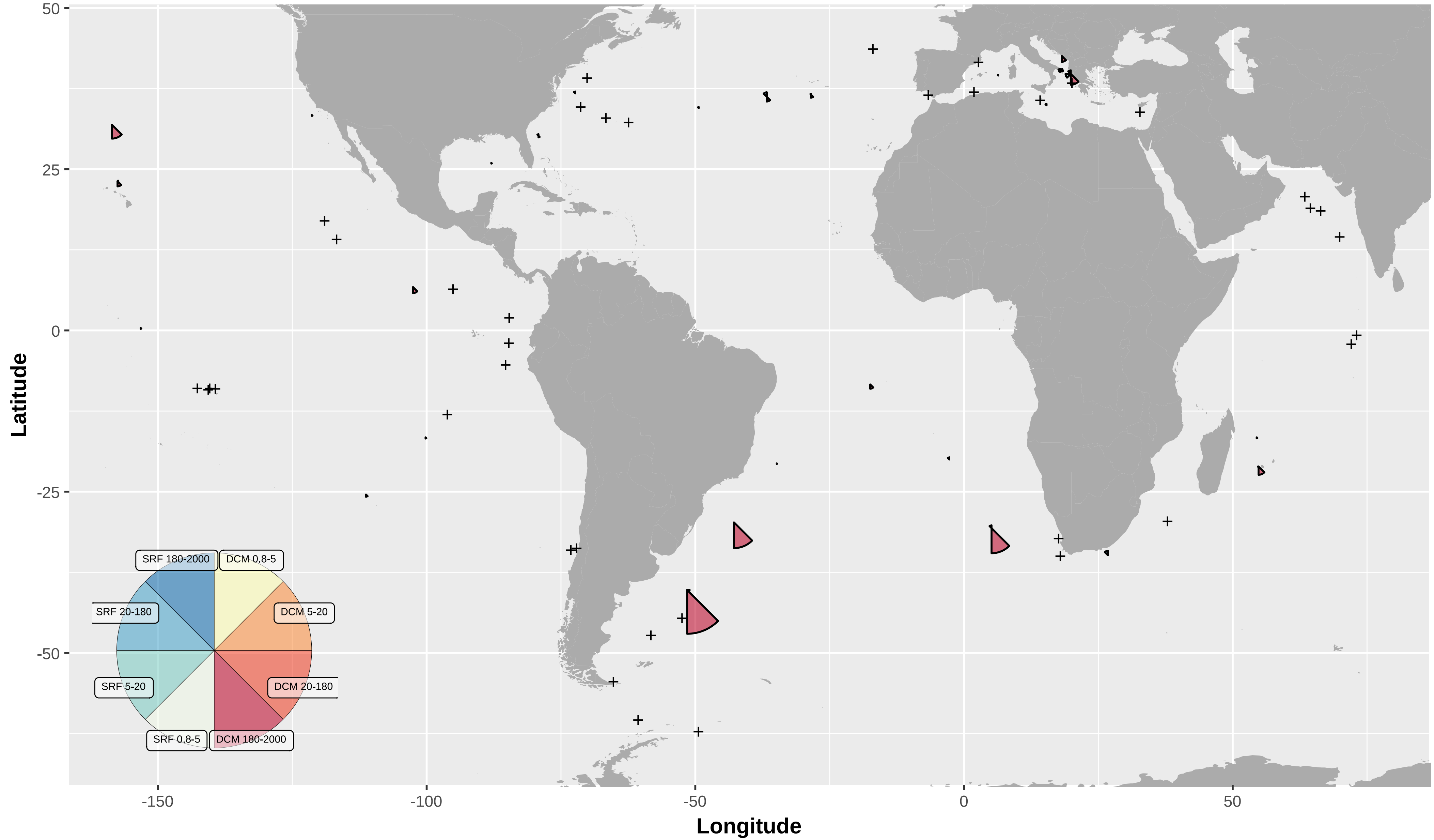


root;cellular organisms;Eukaryota;Opisthokonta;Metazoa;Eumetazoa;Bilateria;Protostomia;Ecdysozoa;Panarthropoda;Arthropoda;Mandibulata;Pancrustacea;Crustacea;Maxillopoda

**Biogeolocalisation of MGT-v1 364      2923 unigenes, 2117 (72.43%) taxonomically assigned**

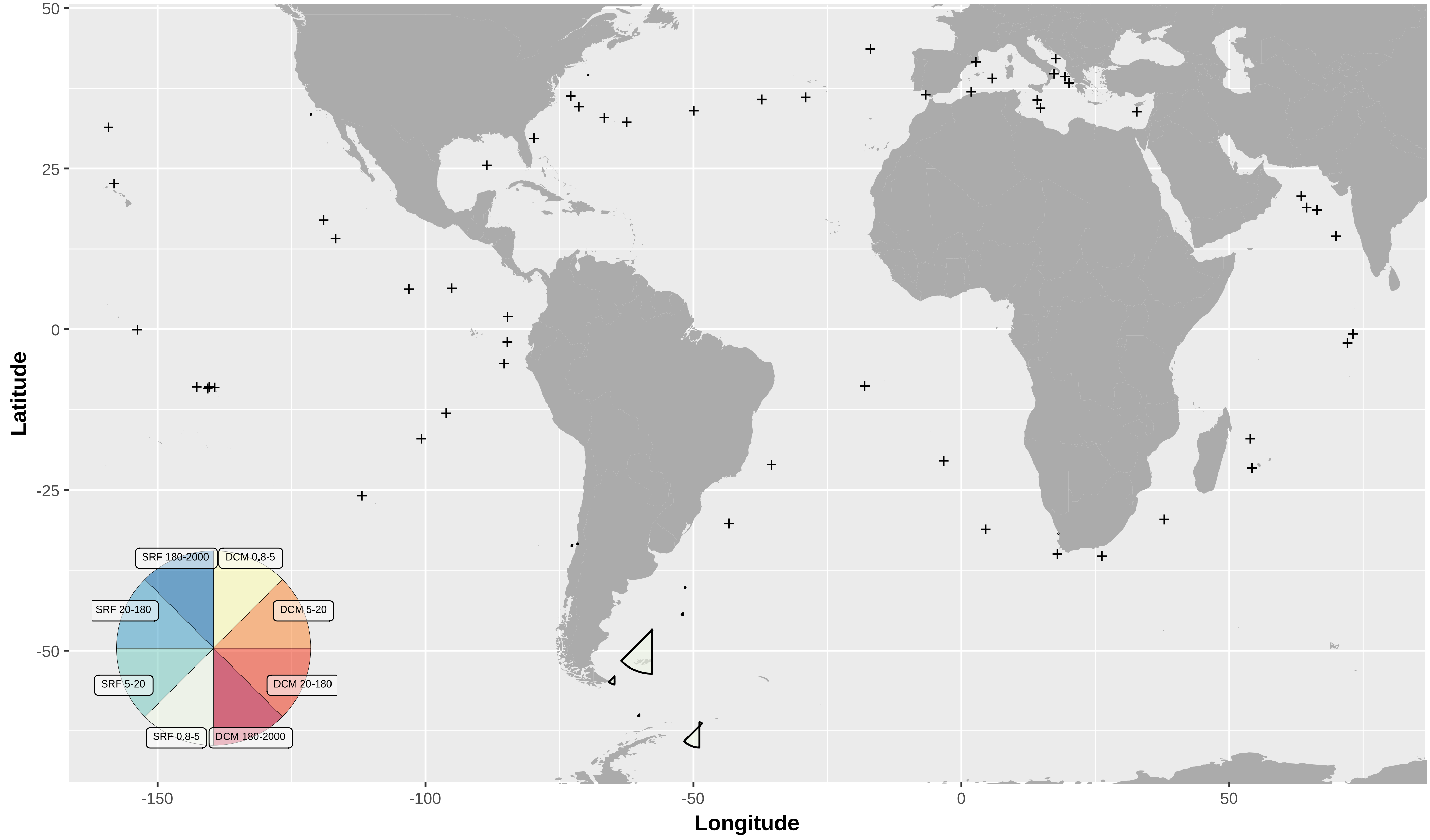


root;cellular organisms;Bacteria

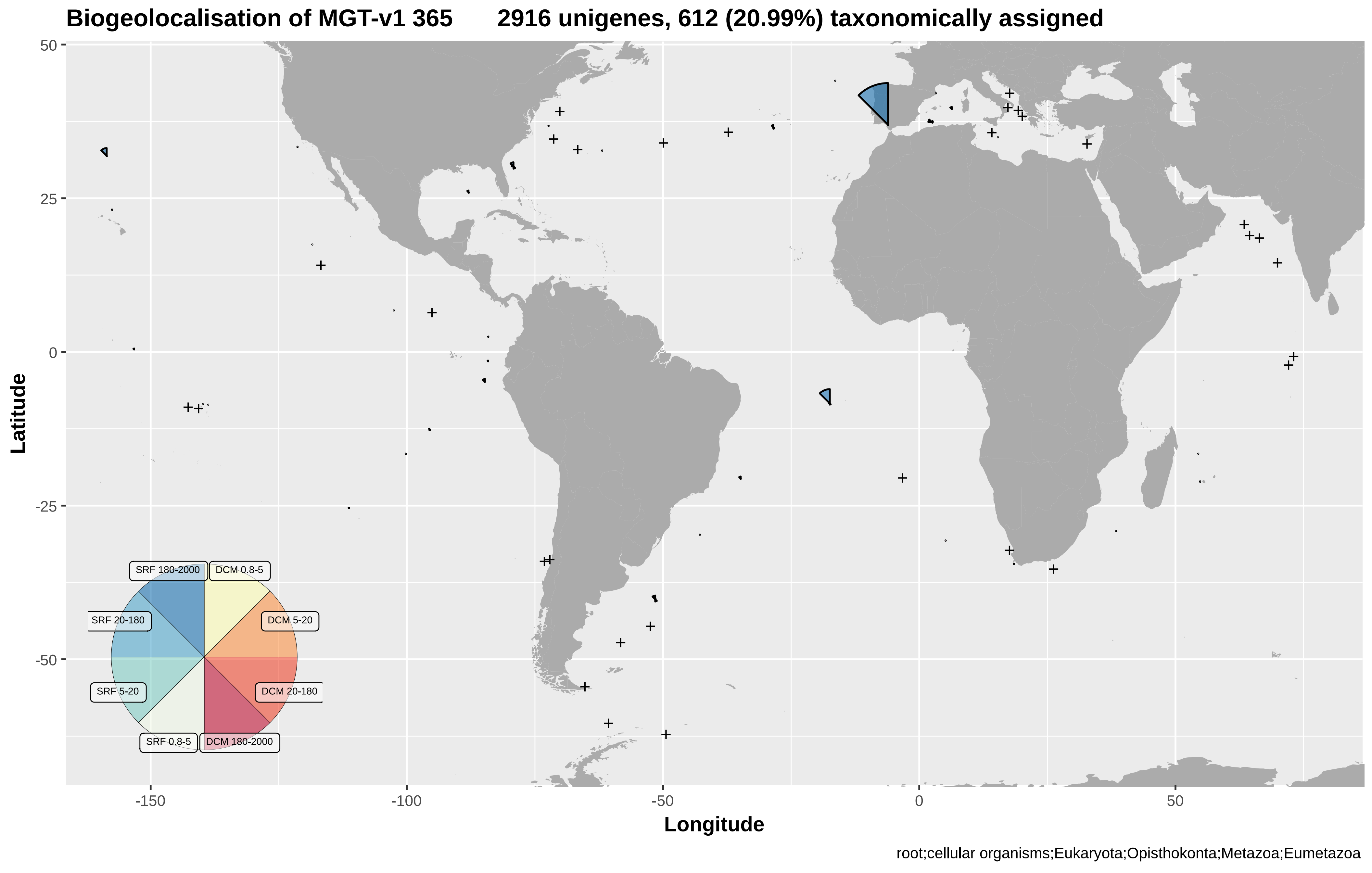

Biogeolocalisation of MGT-v1 365 2916 unigenes, 612 (20.99%) taxonomically assigned
50
25
0
-25
-50
Latitude
-150
-100
-50
0
50
Longitude
SRF 180-2000
DCM 0.8-5
SRF 20-180
DCM 5-20
SRF 5-20
DCM 20-180
SRF 0.8-5
DCM 180-2000
root;cellular organisms;Eukaryota;Opisthokonta;Metazoa;Eumetazoa

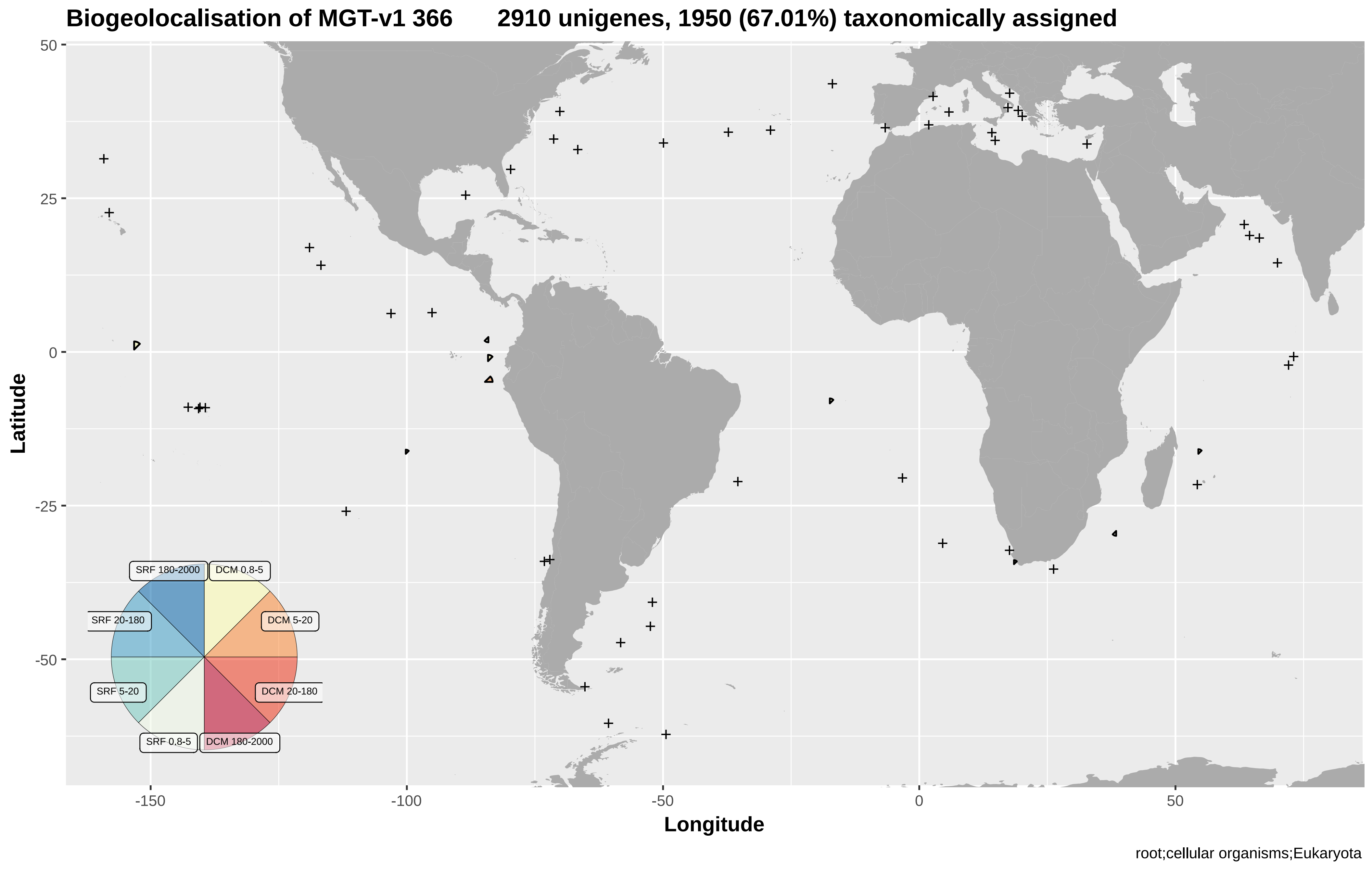

Biogeolocalisation of MGT-v1 366 2910 unigenes, 1950 (67.01%) taxonomically assigned
50
25
0
-25
-50
Latitude
-150
-100
-50
0
50
Longitude
SRF 180-2000
DCM 0.8-5
SRF 20-180
DCM 5-20
SRF 5-20
DCM 20-180
SRF 0.8-5
DCM 180-2000
root;cellular organisms;Eukaryota

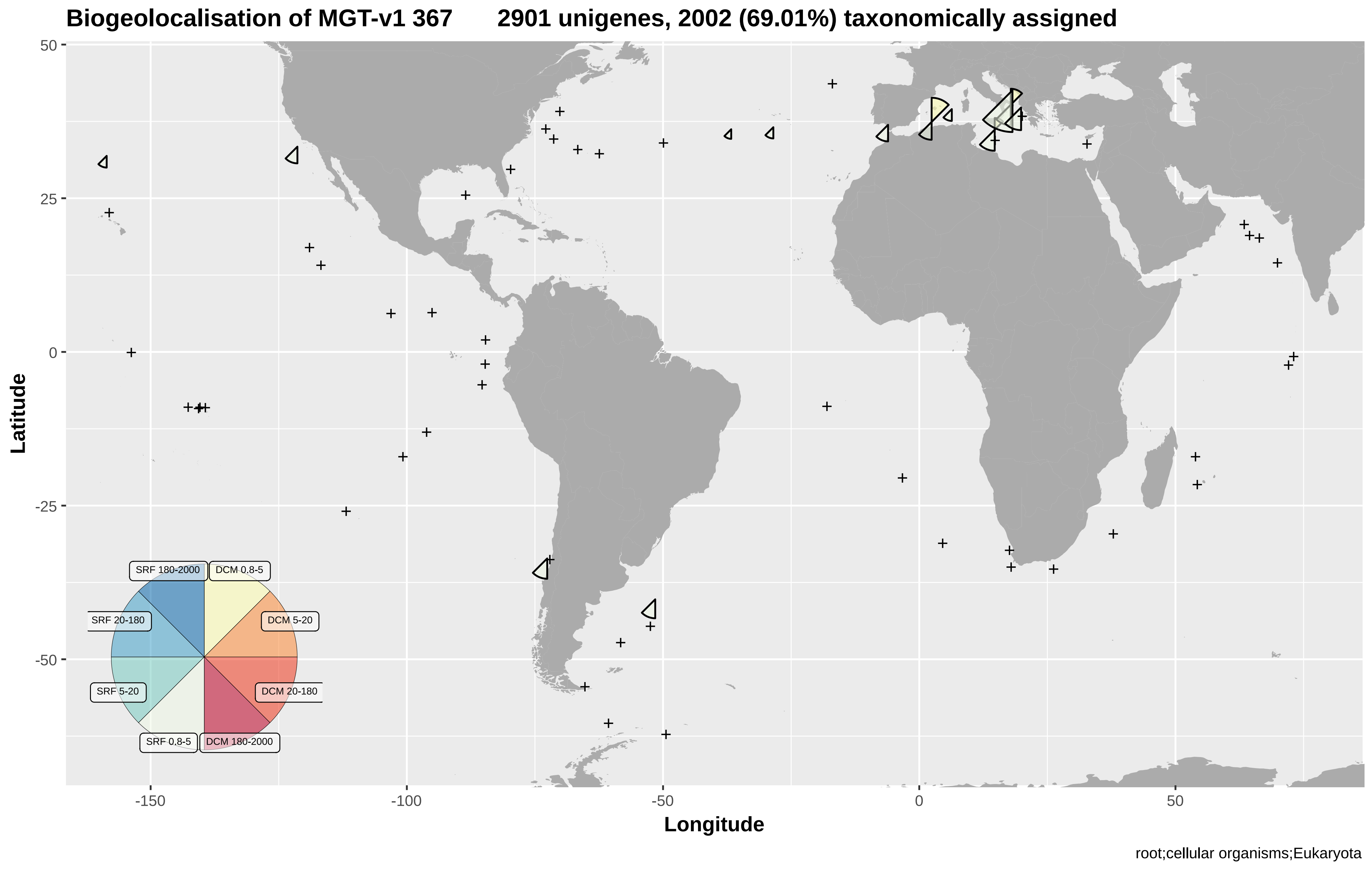

Biogeolocalisation of MGT-v1 367 2901 unigenes, 2002 (69.01%) taxonomically assigned
Latitude
Longitude
50
25
0
-25
-50
-150
-100
-50
0
50
SRF 180-2000
DCM 0.8-5
SRF 20-180
DCM 5-20
SRF 5-20
DCM 20-180
SRF 0.8-5
DCM 180-2000
root;cellular organisms;Eukaryota

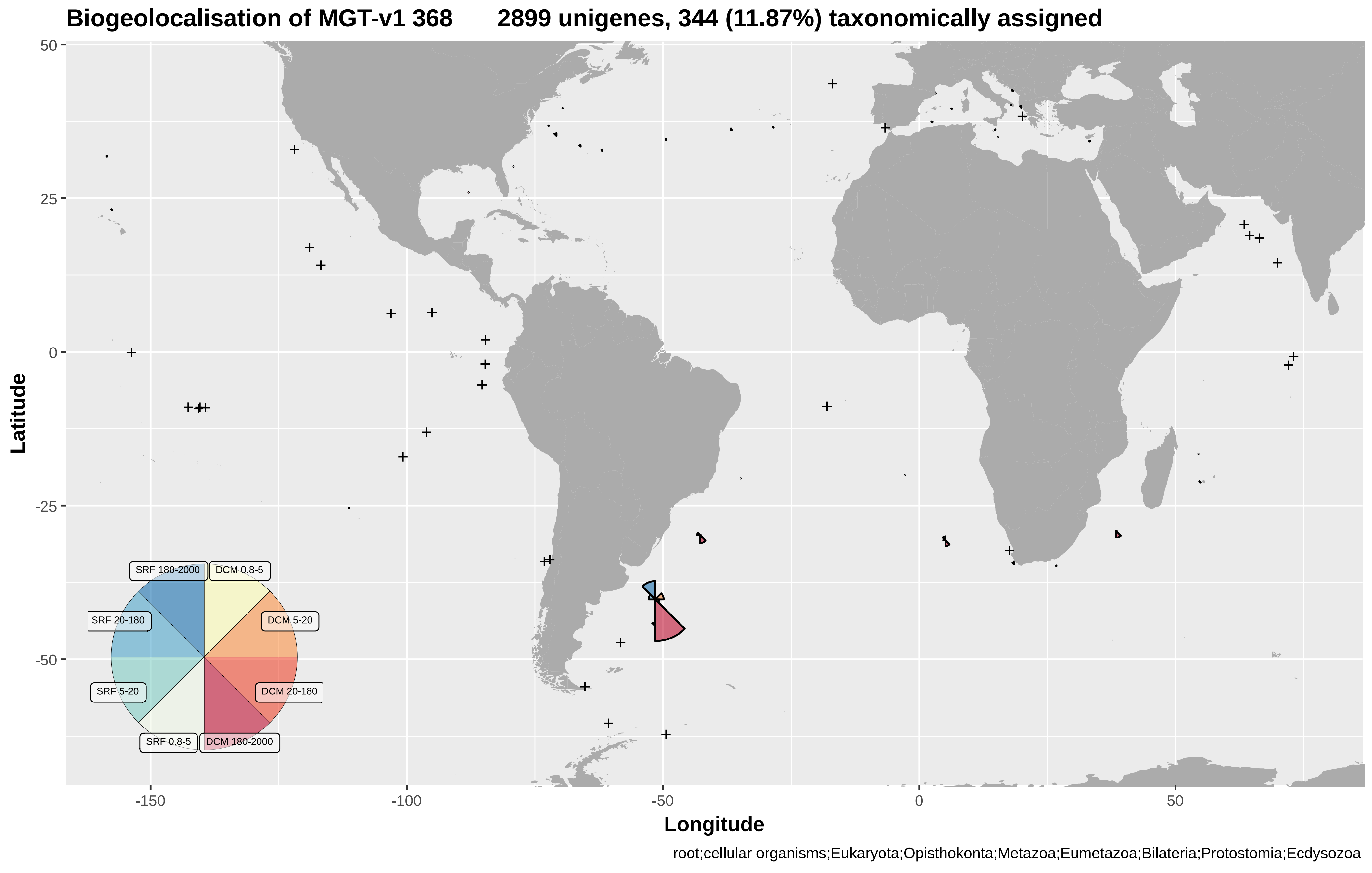

Biogeolocalisation of MGT-v1 368 2899 unigenes, 344 (11.87%) taxonomically assigned
50
25
0
-25
-50
Latitude
-150
-100
-50
0
50
Longitude
SRF 180-2000
DCM 0.8-5
SRF 20-180
DCM 5-20
SRF 5-20
DCM 20-180
SRF 0.8-5
DCM 180-2000
root;cellular organisms;Eukaryota;Opisthokonta;Metazoa;Eumetazoa;Bilateria;Protostomia;Ecdysozoa

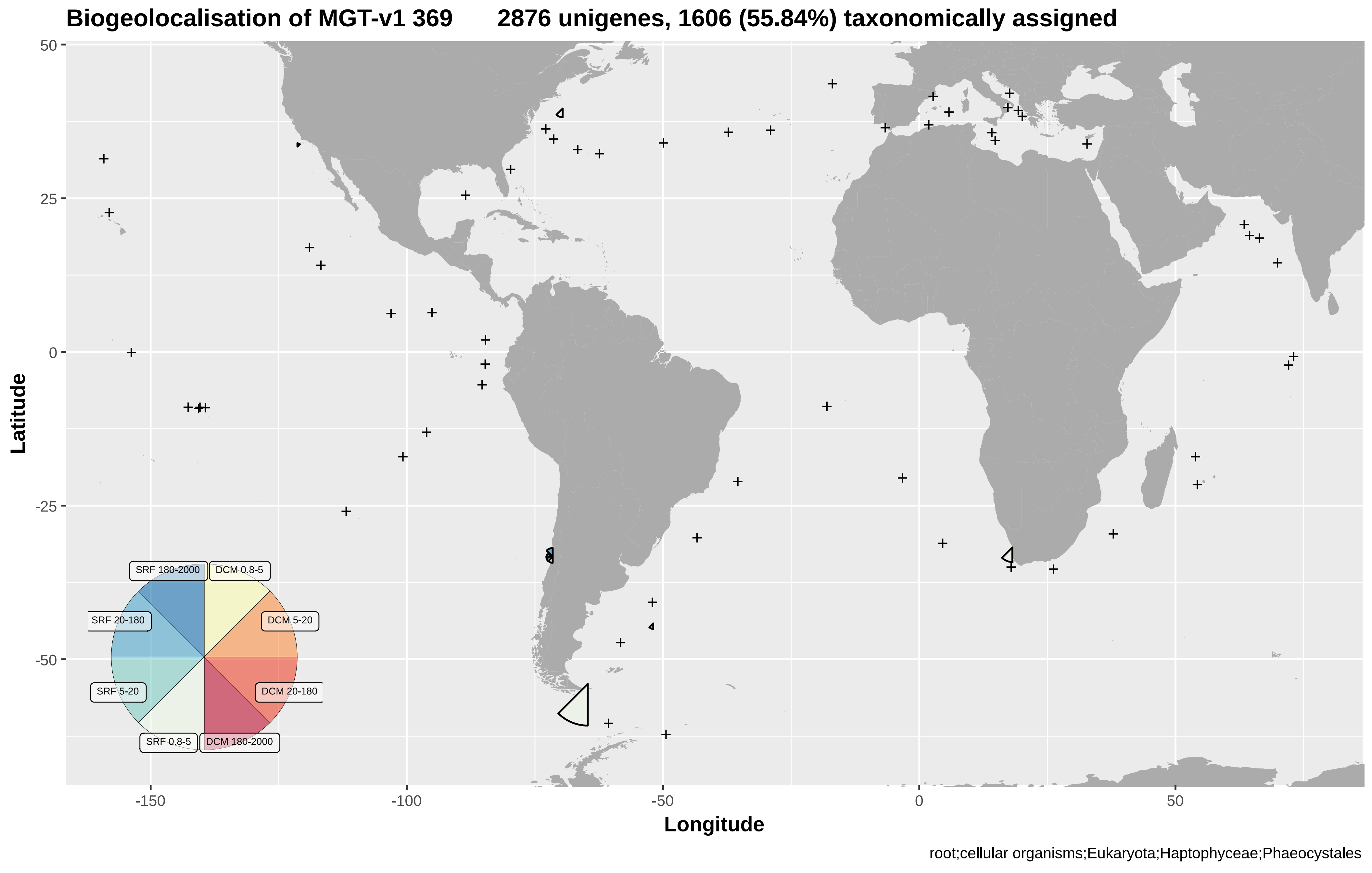
Biogeolocalisation of MGT-v1 369 2876 unigenes, 1606 (55.84%) taxonomically assigned
50
25
0
-25
-50
Latitude
-150
-100
-50
0
50
Longitude
SRF 180-2000
DCM 0.8-5
SRF 20-180
DCM 5-20
SRF 5-20
DCM 20-180
SRF 0.8-5
DCM 180-2000
root;cellular organisms;Eukaryota;Haptophyceae;Phaeocystales

#### Biogeolocalisation of MGT-v1 370  2875 unigenes, 2639 (91.79%) taxonomically assigned

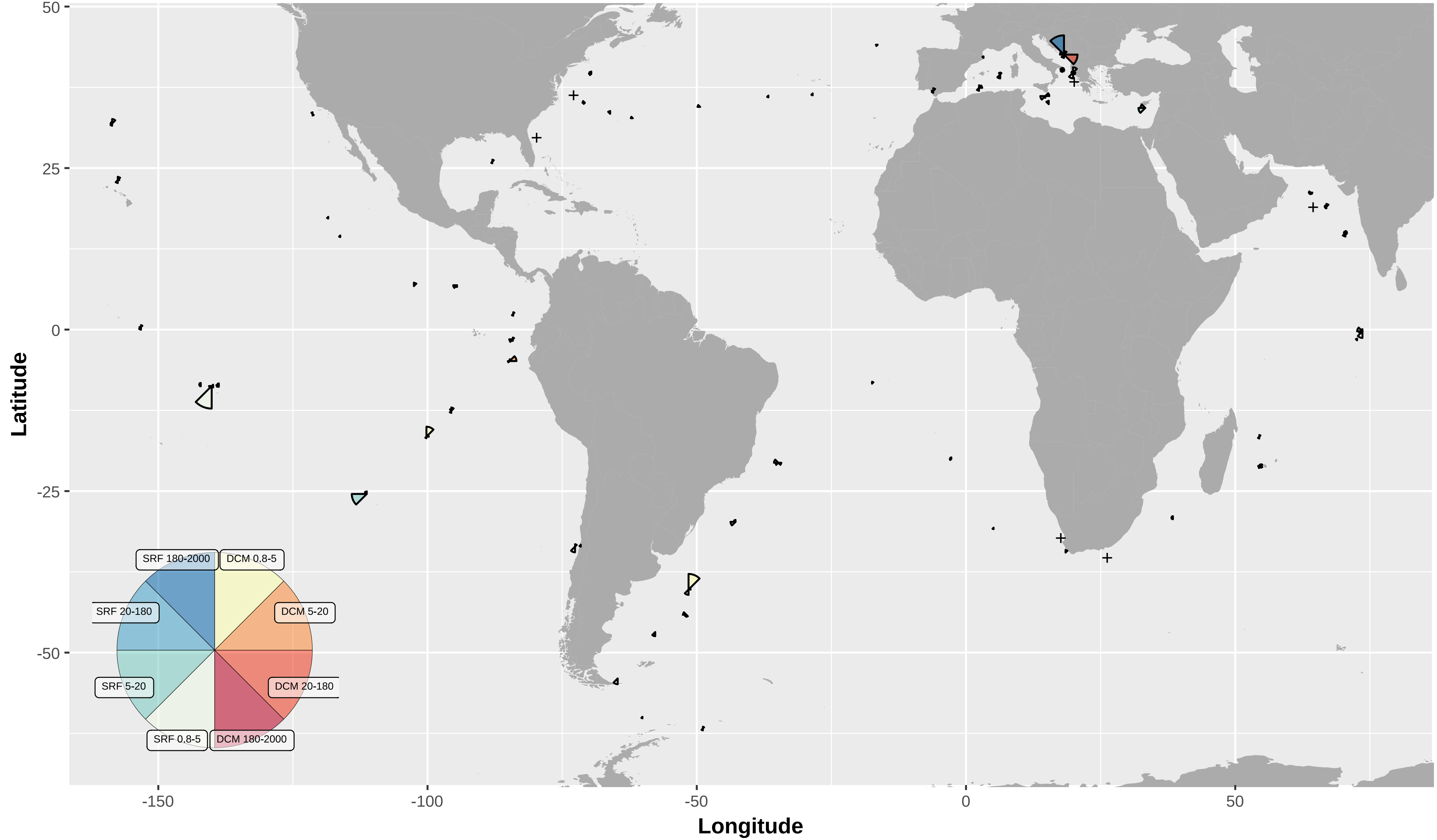


root;cellular organisms;Bacteria;Proteobacteria;Alphaproteobacteria;Sphingomonadales

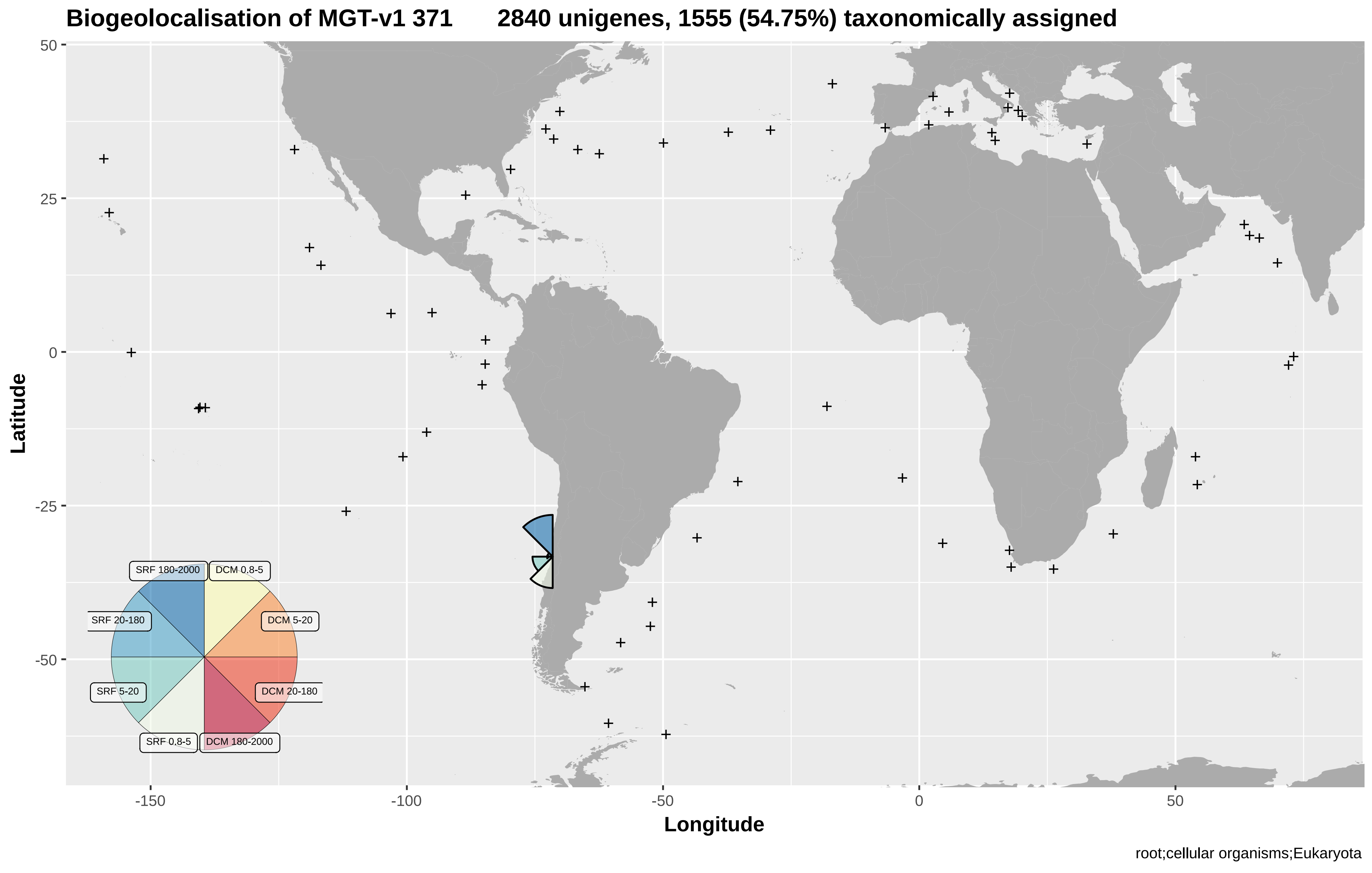

Biogeolocalisation of MGT-v1 371 2840 unigenes, 1555 (54.75%) taxonomically assigned
50
25
0
-25
-50
Latitude
-150
-100
-50
0
50
Longitude
SRF 180-2000
DCM 0.8-5
SRF 20-180
DCM 5-20
SRF 5-20
DCM 20-180
SRF 0.8-5
DCM 180-2000
root;cellular organisms;Eukaryota

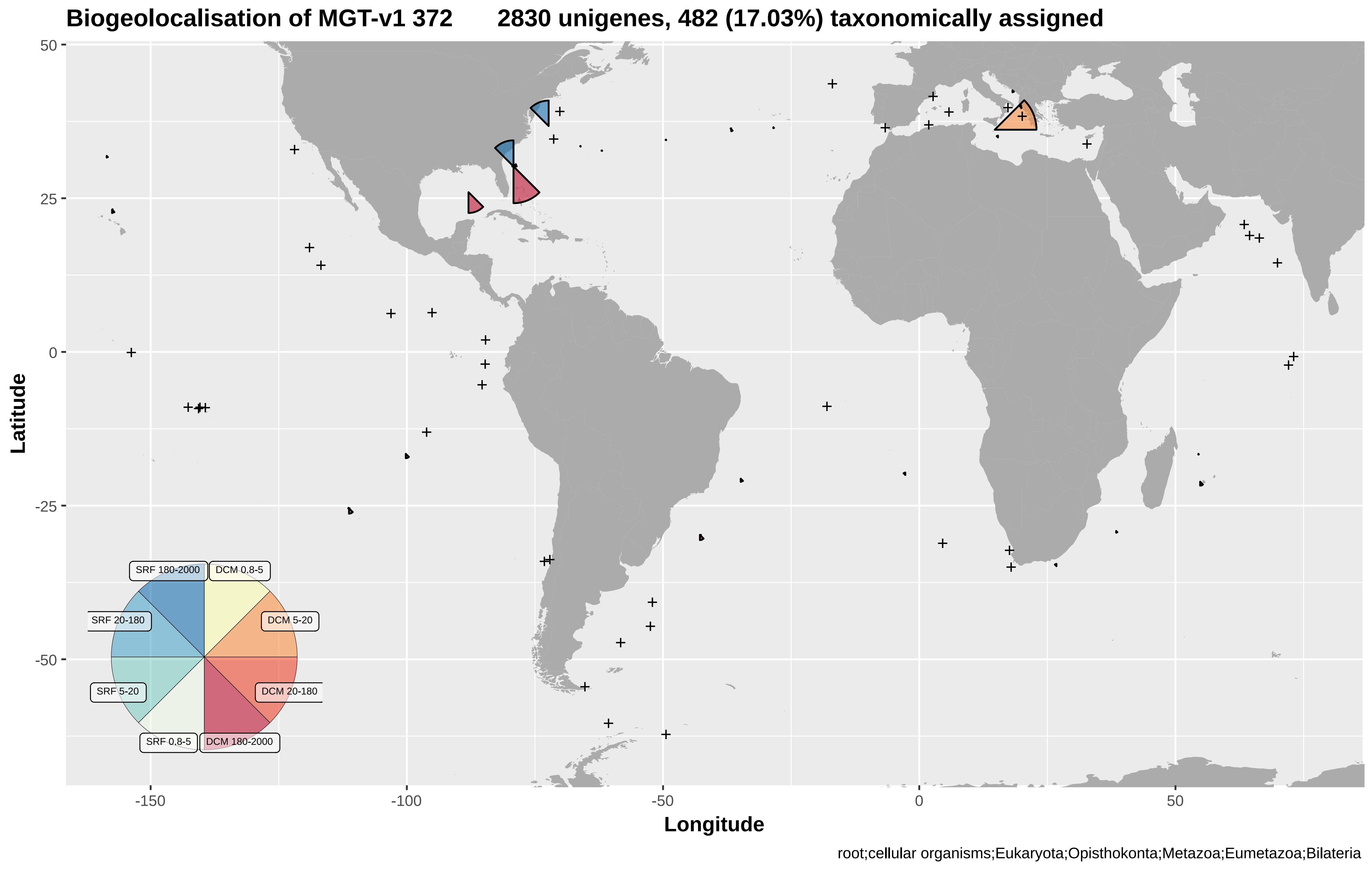

Biogeolocalisation of MGT-v1 372 2830 unigenes, 482 (17.03%) taxonomically assigned
50
25
0
-25
-50
Latitude
-150
-100
-50
0
50
Longitude
SRF 180-2000
DCM 0.8-5
SRF 20-180
DCM 5-20
SRF 5-20
DCM 20-180
SRF 0.8-5
DCM 180-2000
root;cellular organisms;Eukaryota;Opisthokonta;Metazoa;Eumetazoa;Bilateria

# Biogeolocalisation of MGT-v1 373      2830 unigenes, 2601 (91.91%) taxonomically assigned

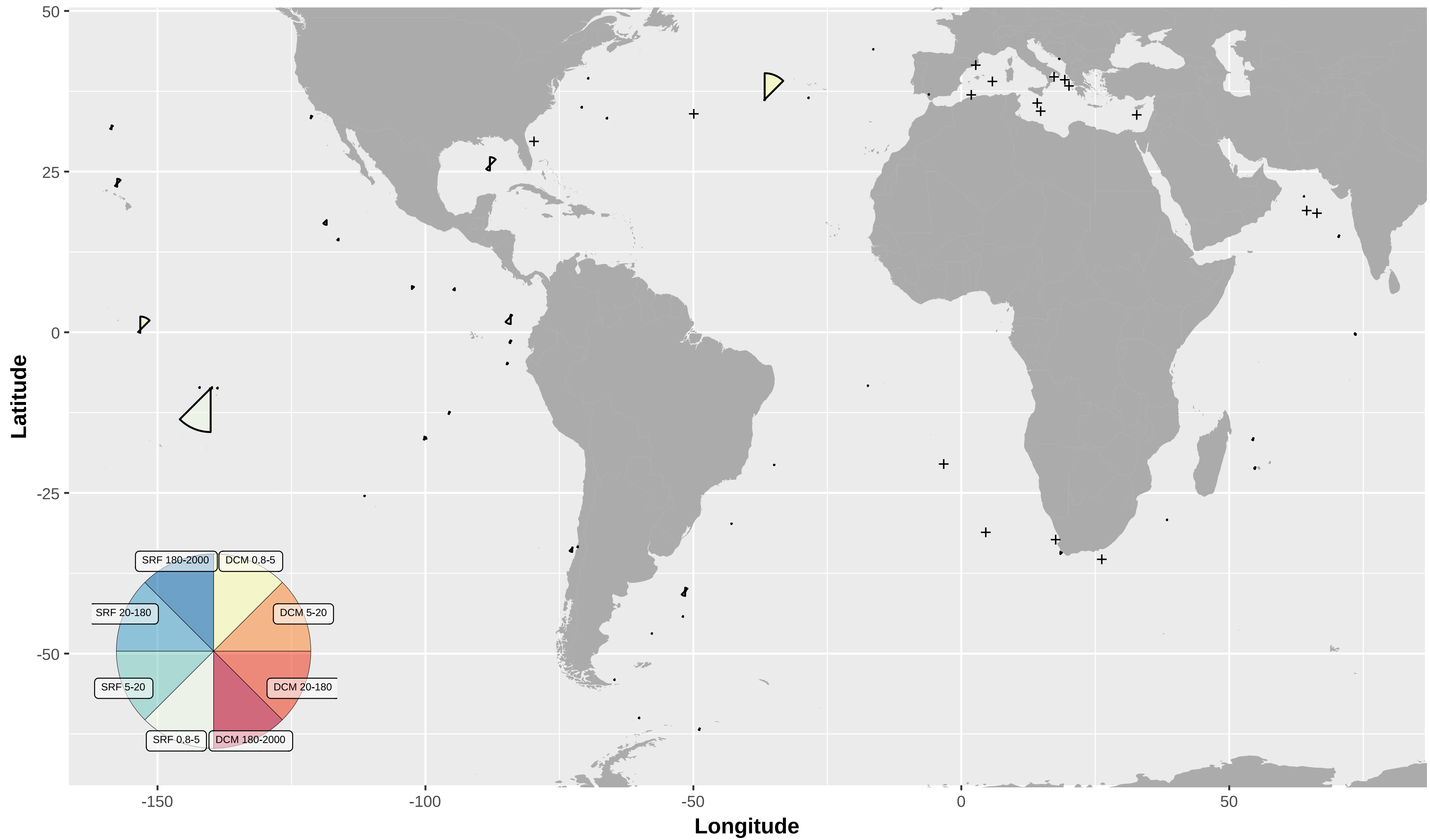


root;cellular organisms;Bacteria;Actinobacteria;Actinobacteria;Micrococcales

#### Biogeolocalisation of MGT-v1 374 2830 unigenes, 2355 (83.22%) taxonomically assigned

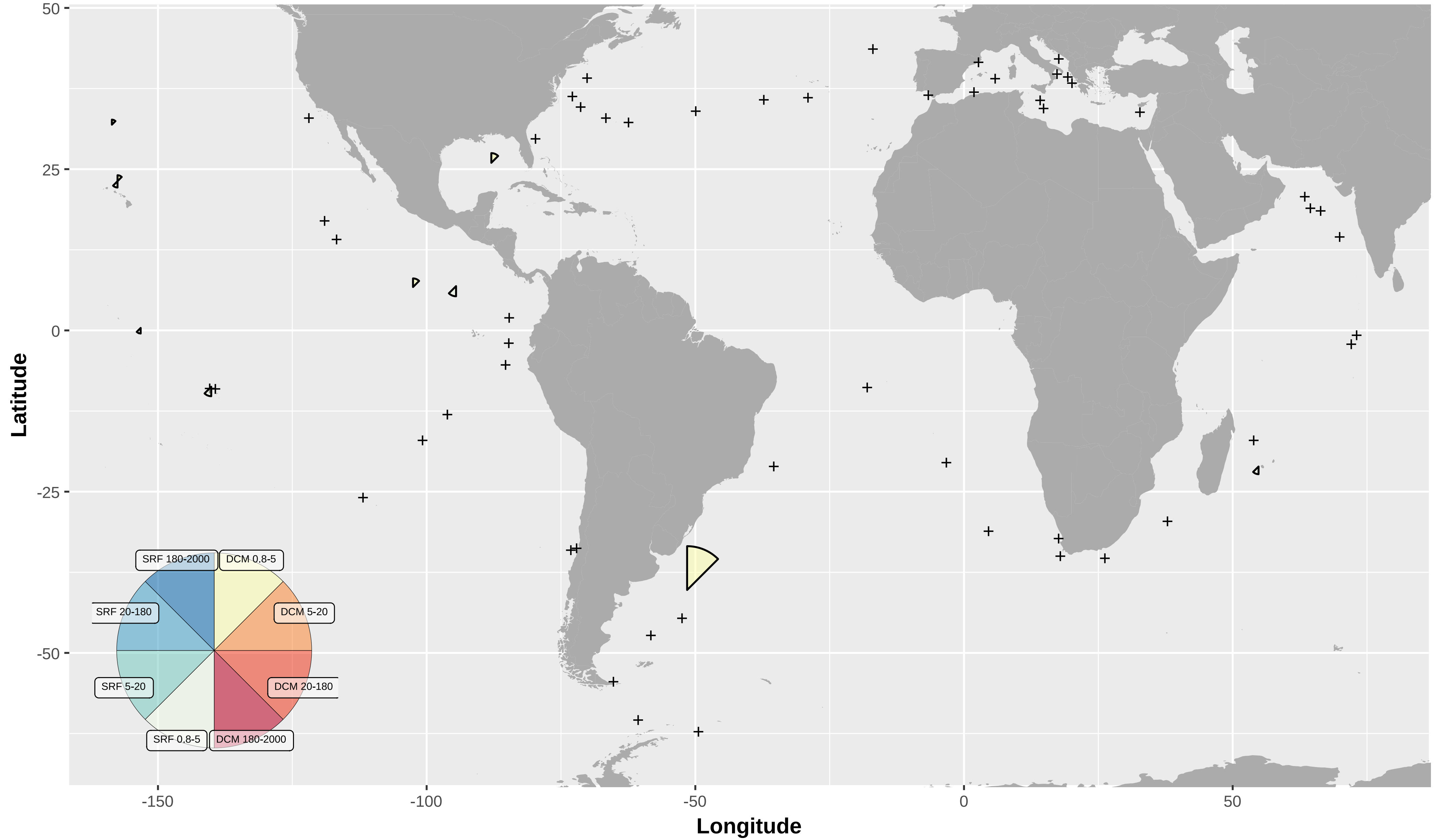


root;cellular organisms;Bacteria;Proteobacteria;Alphaproteobacteria

# Biogeolocalisation of MGT-v1 375      2826 unigenes, 188 (6.65%) taxonomically assigned

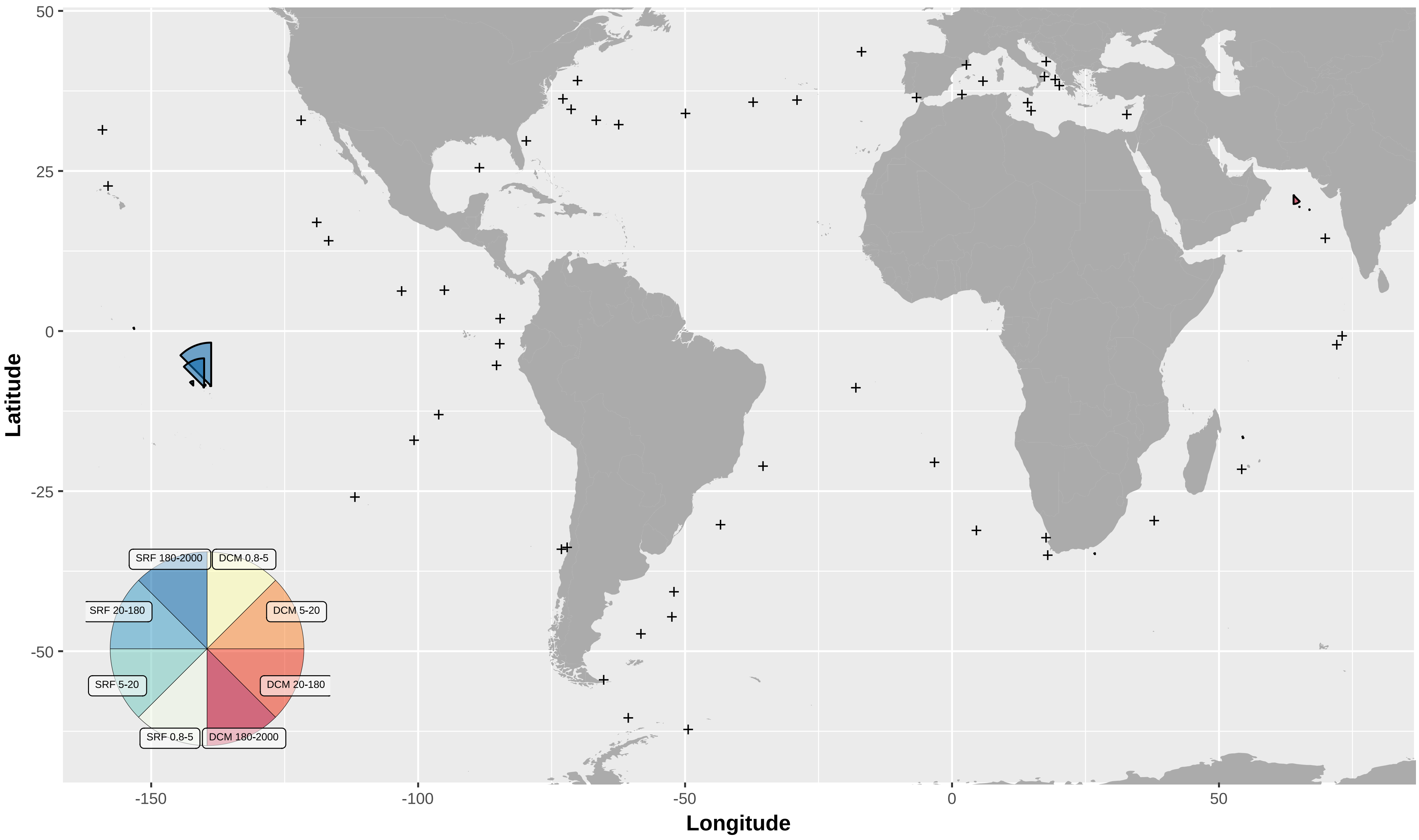


root;cellular organisms;Eukaryota;Opisthokonta;Metazoa;Eumetazoa;Bilateria

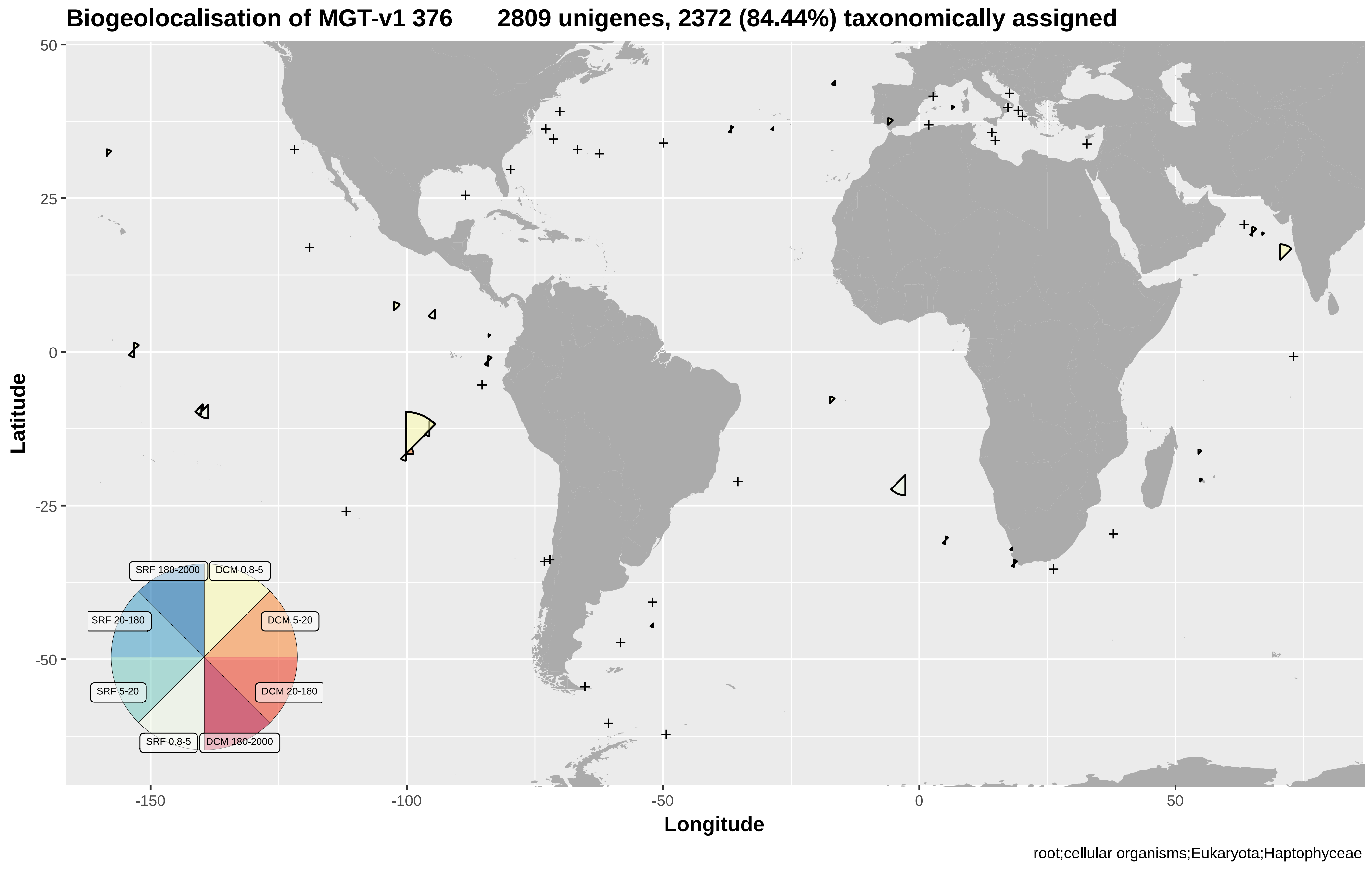

Biogeolocalisation of MGT-v1 376 2809 unigenes, 2372 (84.44%) taxonomically assigned
50
25
0
-25
-50
Latitude
-150
-100
-50
0
50
Longitude
SRF 180-2000
DCM 0.8-5
SRF 20-180
DCM 5-20
SRF 5-20
DCM 20-180
SRF 0.8-5
DCM 180-2000
root;cellular organisms;Eukaryota;Haptophyceae

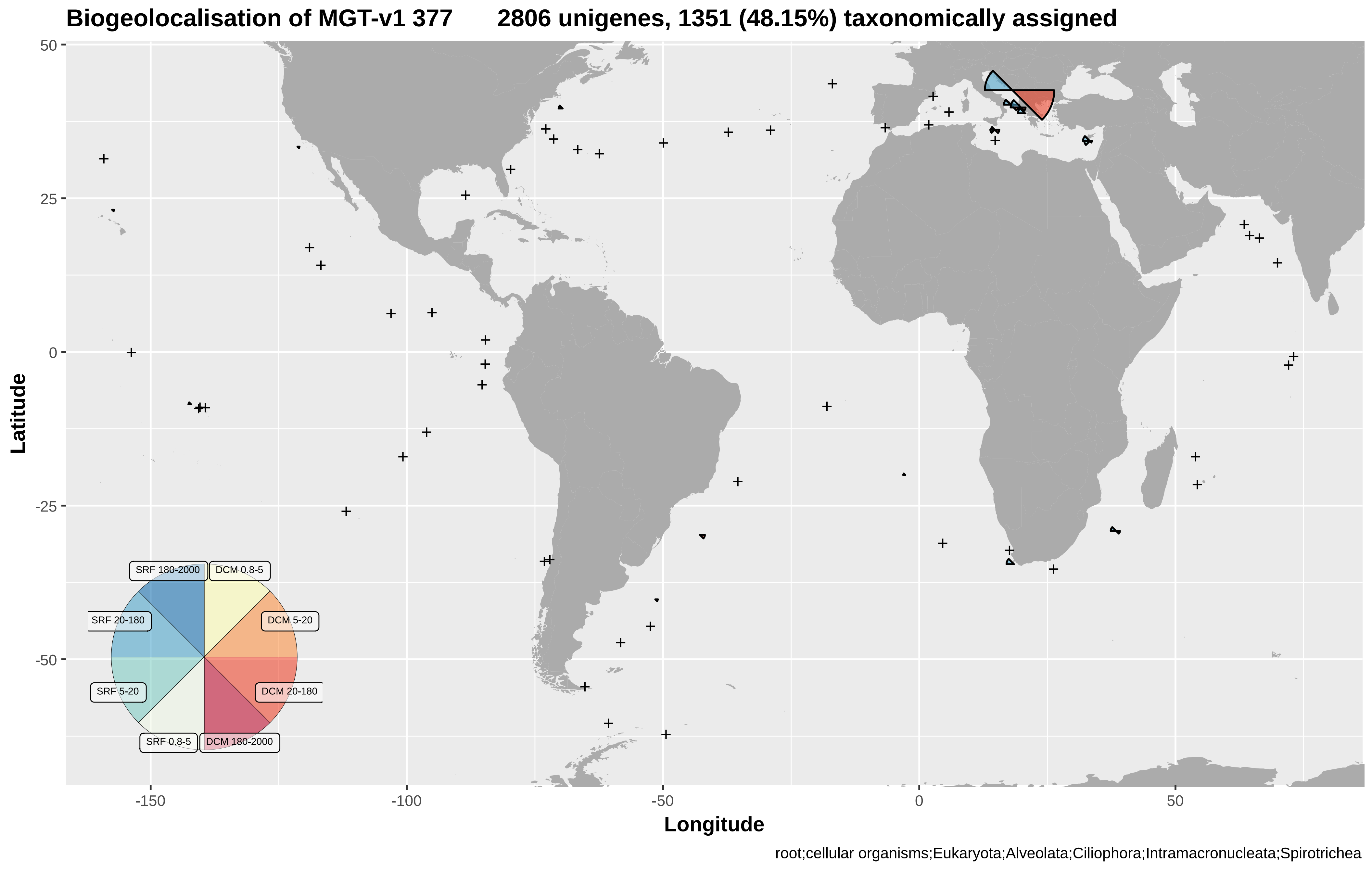

Biogeolocalisation of MGT-v1 377 2806 unigenes, 1351 (48.15%) taxonomically assigned
50
25
0
-25
-50
Latitude
-150
-100
-50
0
50
Longitude
SRF 180-2000
DCM 0.8-5
SRF 20-180
DCM 5-20
SRF 5-20
DCM 20-180
SRF 0.8-5
DCM 180-2000
root;cellular organisms;Eukaryota;Alveolata;Ciliophora;Intramacronucleata;Spirotrichea

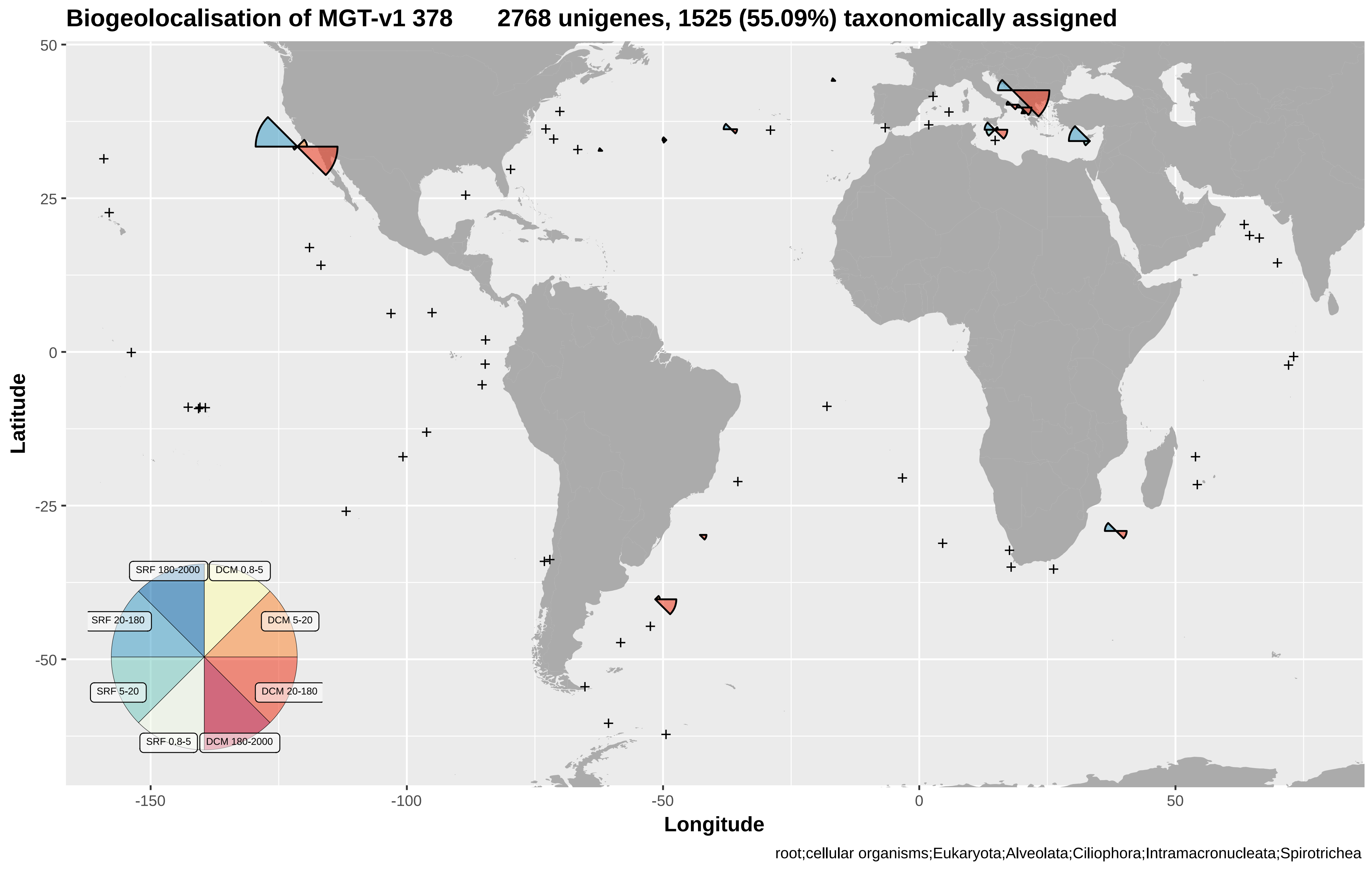
Biogeolocalisation of MGT-v1 378      2768 unigenes, 1525 (55.09%) taxonomically assigned
50
25
0
-25
-50
Latitude
-150
-100
-50
0
50
Longitude
SRF 180-2000
DCM 0.8-5
SRF 20-180
DCM 5-20
SRF 5-20
DCM 20-180
SRF 0.8-5
DCM 180-2000
root;cellular organisms;Eukaryota;Alveolata;Ciliophora;Intramacronucleata;Spirotrichea

**Biogeolocalisation of MGT-v1 379      2765 unigenes, 2067 (74.76%) taxonomically assigned**

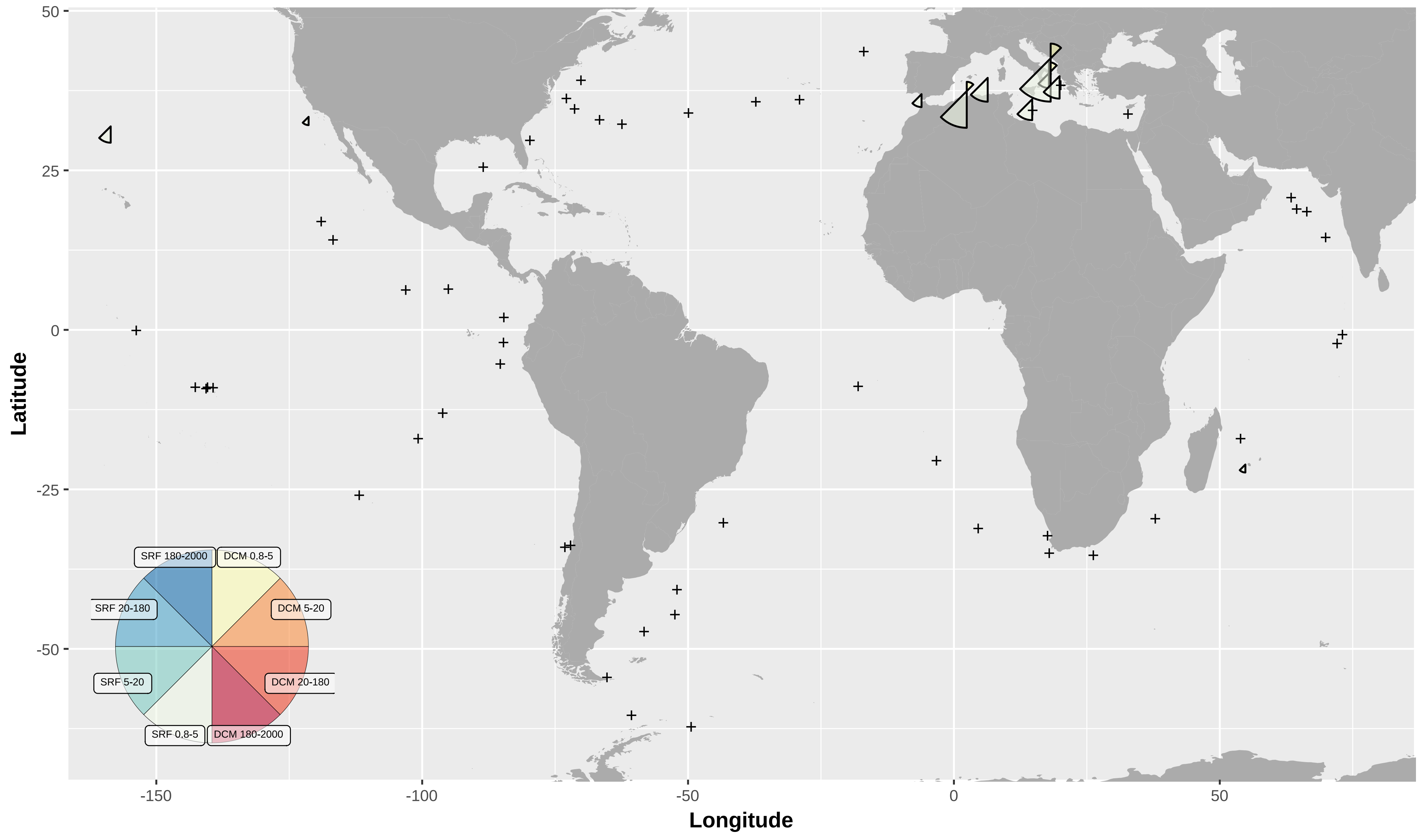


root;cellular organisms;Eukaryota

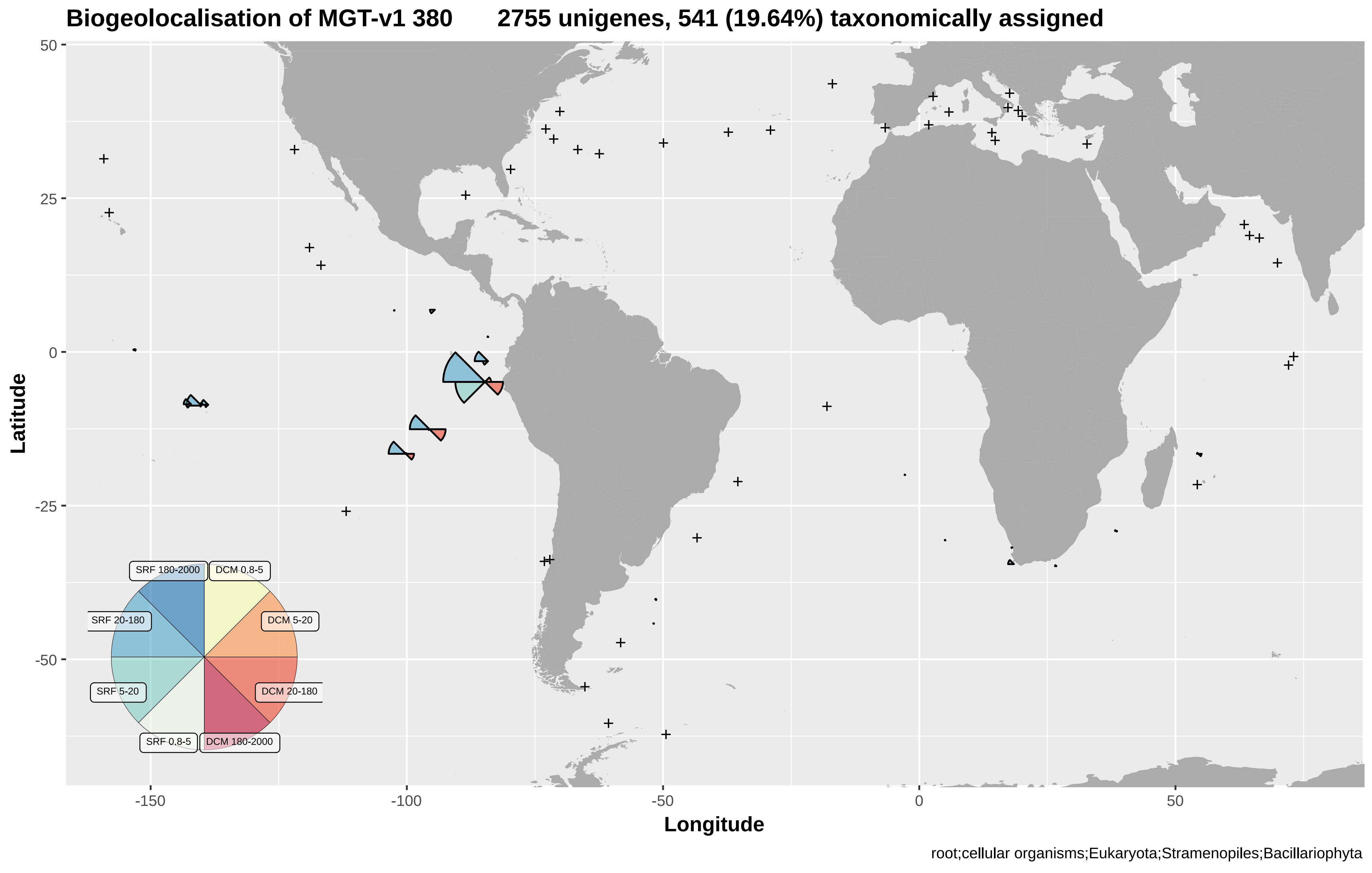

Biogeolocalisation of MGT-v1 380 2755 unigenes, 541 (19.64%) taxonomically assigned
Latitude
50
25
0
-25
-50
Longitude
-150
-100
-50
0
50
SRF 180-2000
DCM 0.8-5
SRF 20-180
DCM 5-20
SRF 5-20
DCM 20-180
SRF 0.8-5
DCM 180-2000
root;cellular organisms;Eukaryota;Stramenopiles;Bacillariophyta

# Biogeolocalisation of MGT-v1 381      2745 unigenes, 288 (10.49%) taxonomically assigned

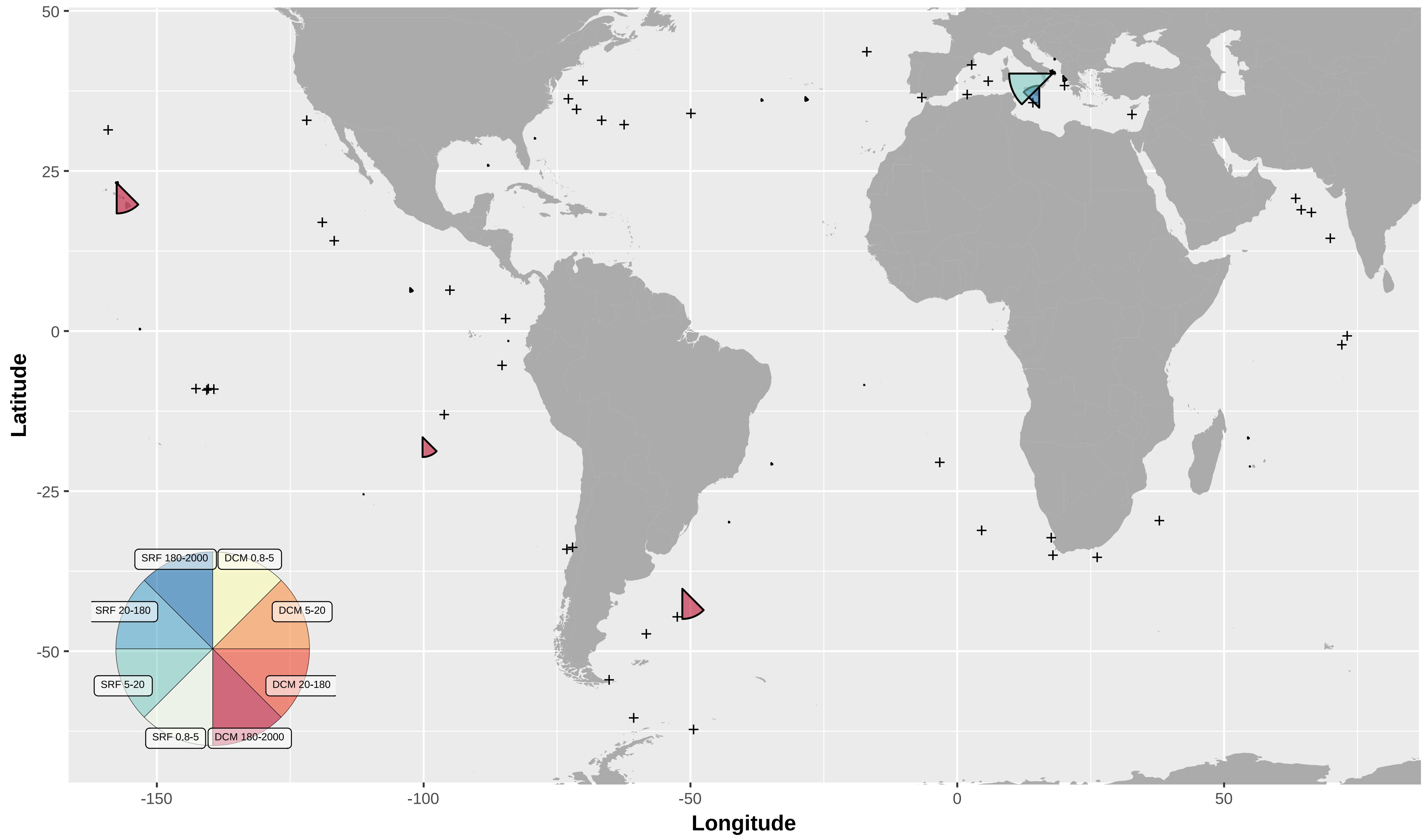


root;cellular organisms;Eukaryota;Opisthokonta;Metazoa;Eumetazoa

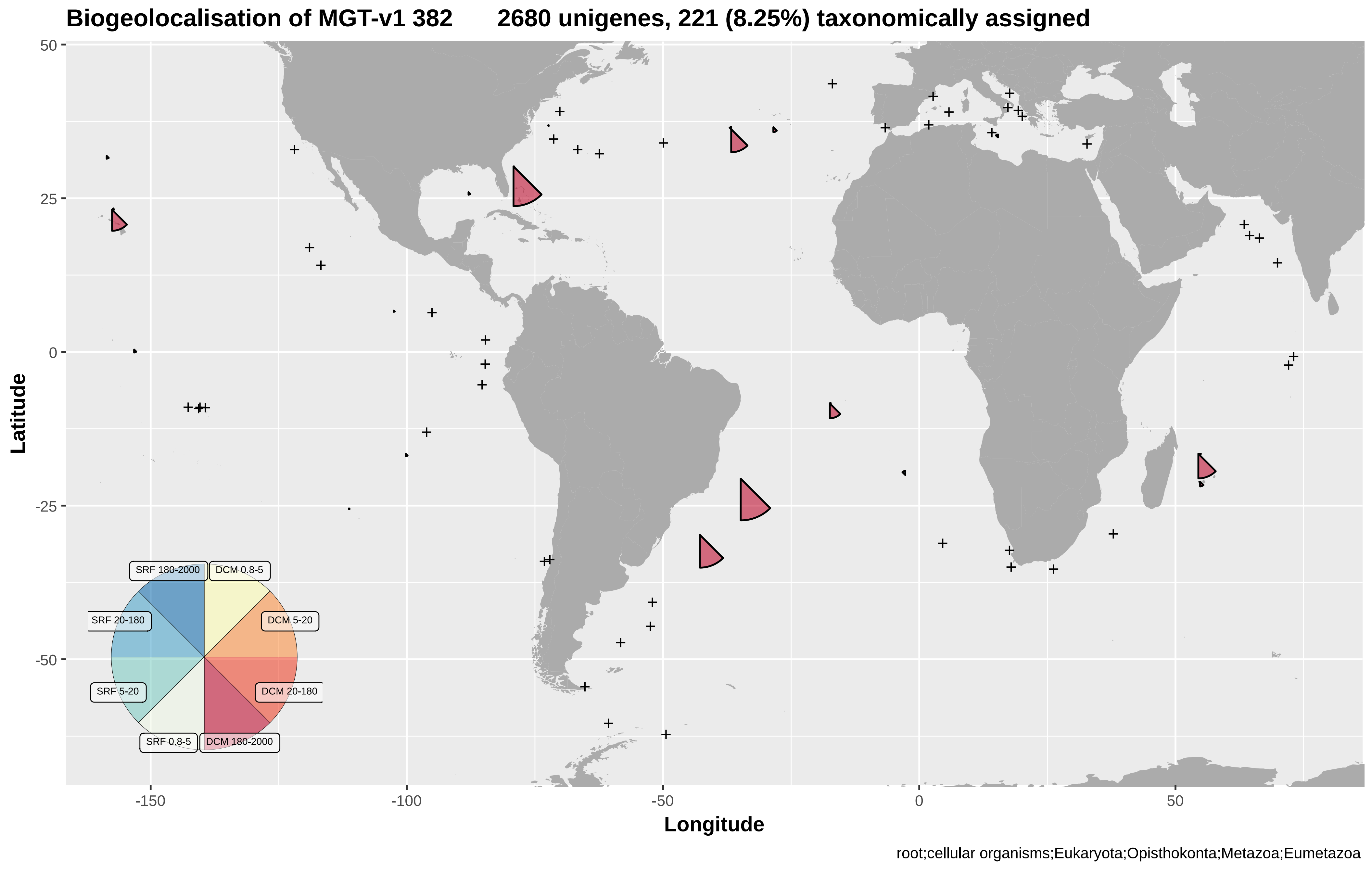
Biogeolocalisation of MGT-v1 382 2680 unigenes, 221 (8.25%) taxonomically assigned
Latitude
Longitude
50
25
0
-25
-50
-150
-100
-50
0
50
SRF 180-2000
DCM 0.8-5
SRF 20-180
DCM 5-20
SRF 5-20
DCM 20-180
SRF 0.8-5
DCM 180-2000
root;cellular organisms;Eukaryota;Opisthokonta;Metazoa;Eumetazoa

## Biogeolocalisation of MGT-v1 383      2670 unigenes, 2475 (92.7%) taxonomically assigned

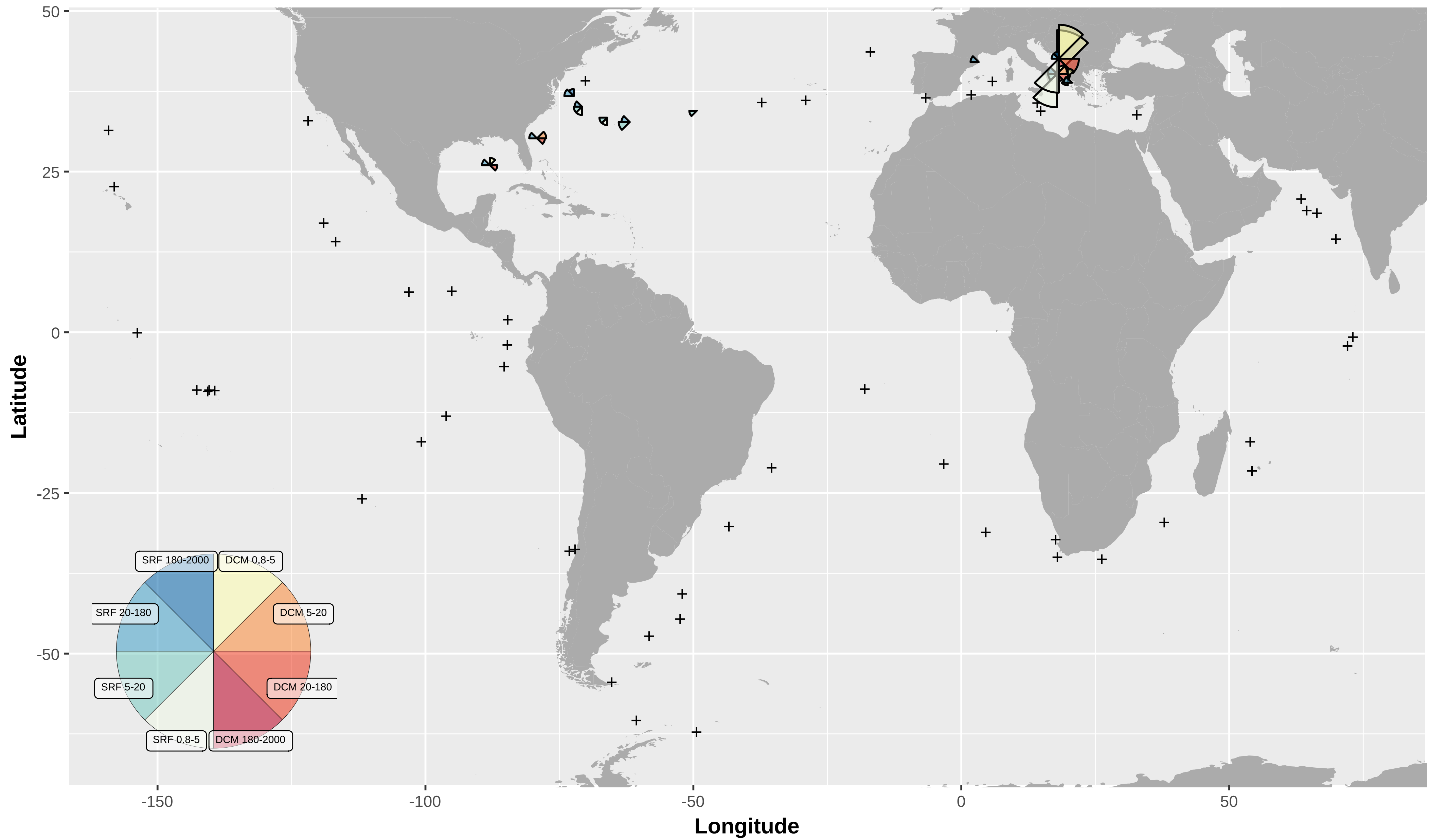


root;cellular organisms;Eukaryota;Viridiplantae;Chlorophyta;prasinophytes;Pycnococcaceae

Biogeolocalisation of MGT-v1 384      2665 unigenes, 2549 (95.65%) taxonomically assigned

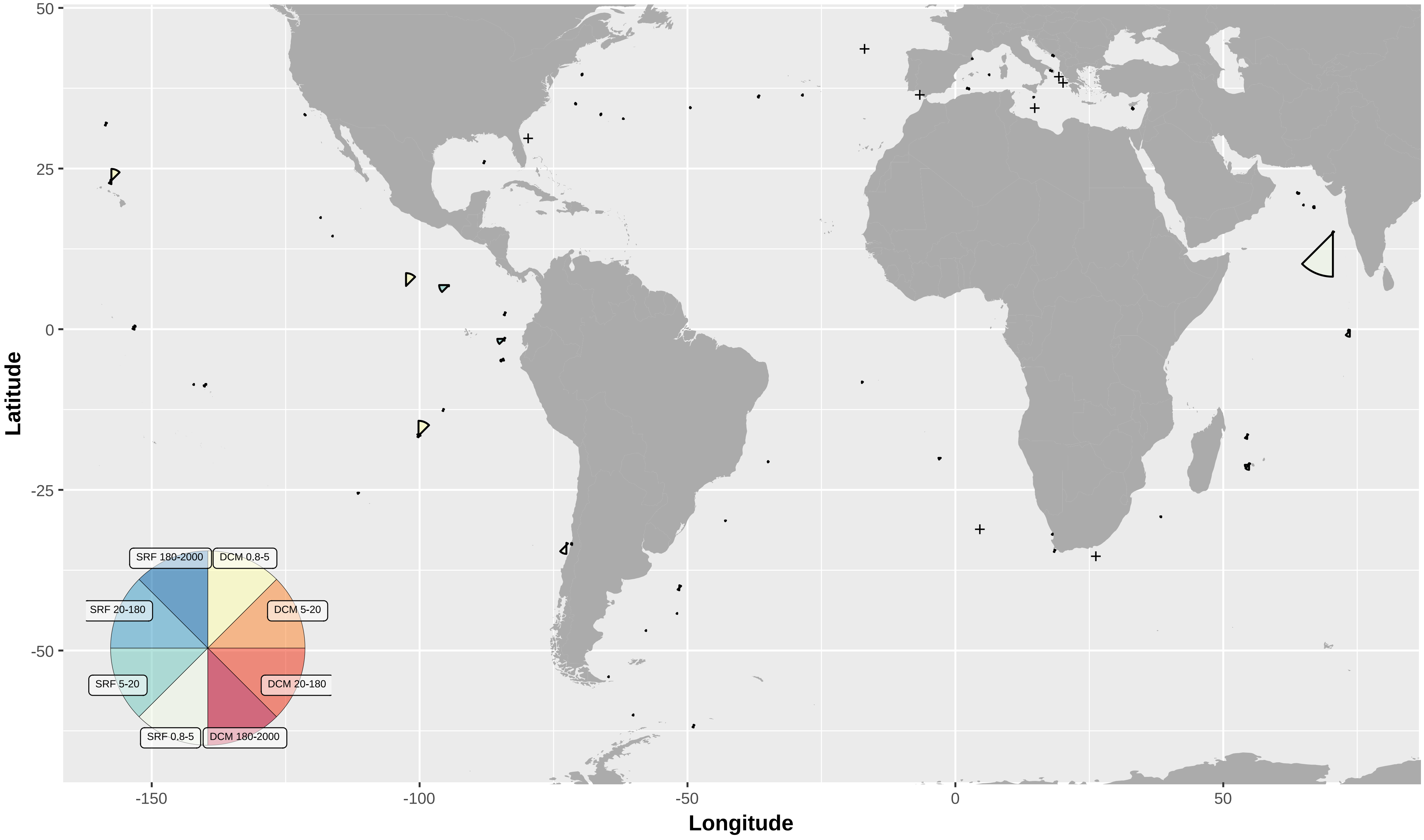


root;cellular organisms;Bacteria;Proteobacteria;Gammaproteobacteria

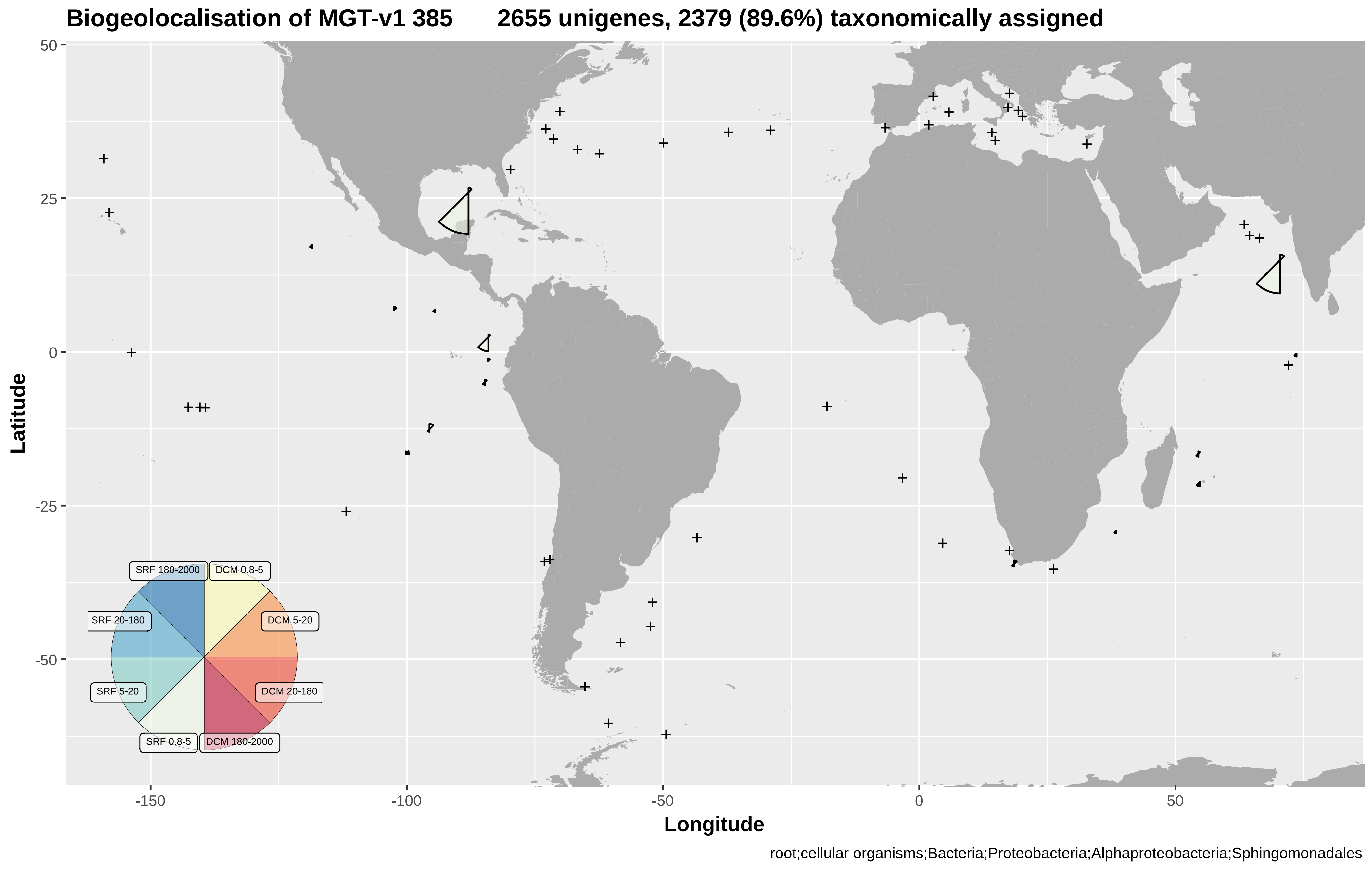

Biogeolocalisation of MGT-v1 385 2655 unigenes, 2379 (89.6%) taxonomically assigned
50
25
0
-25
-50
Latitude
-150
-100
-50
0
50
Longitude
SRF 180-2000
DCM 0.8-5
SRF 20-180
DCM 5-20
SRF 5-20
DCM 20-180
SRF 0.8-5
DCM 180-2000
root;cellular organisms;Bacteria;Proteobacteria;Alphaproteobacteria;Sphingomonadales

#### Biogeolocalisation of MGT-v1 386 2652 unigenes, 113 (4.26%) taxonomically assigned

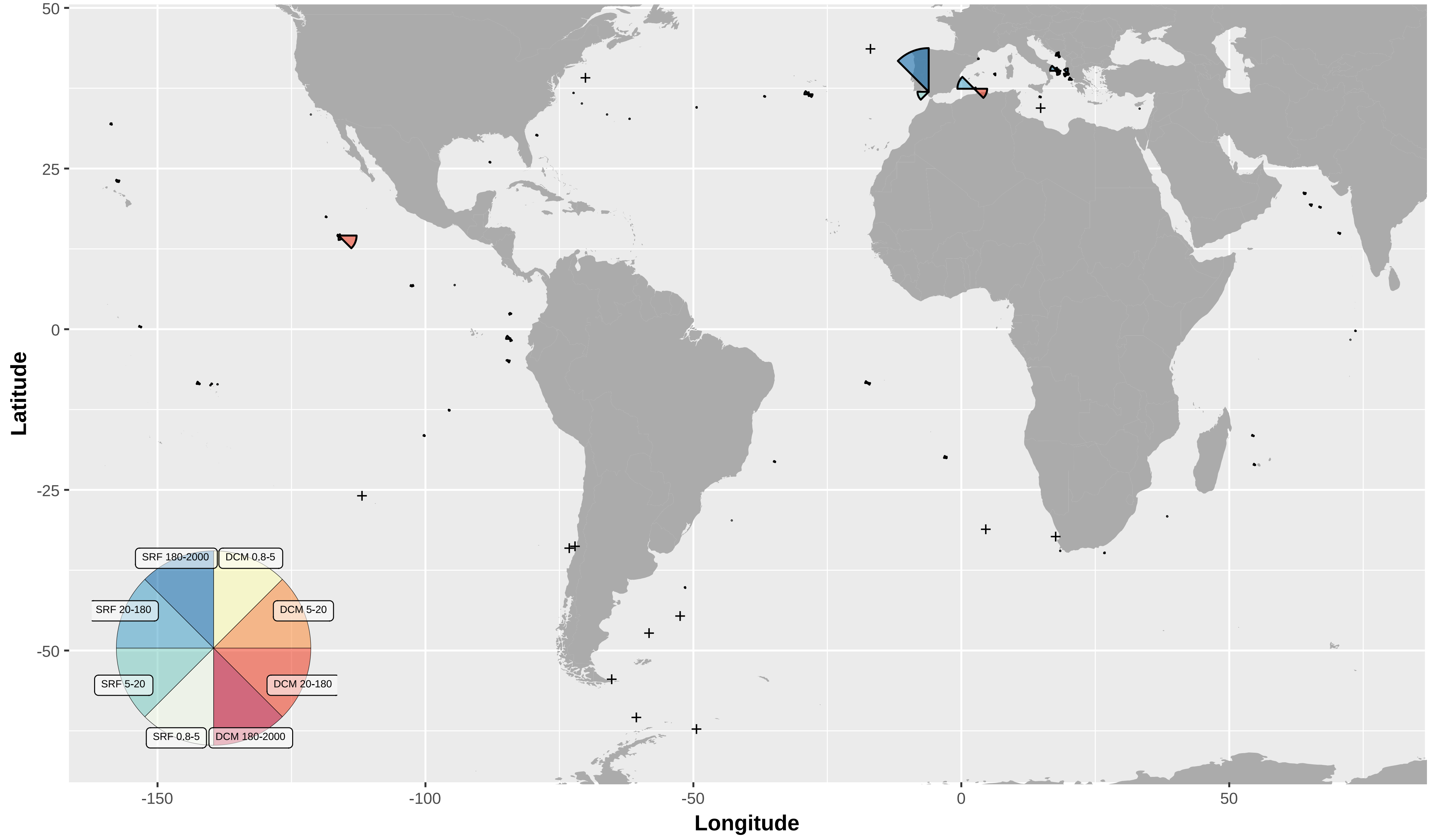


root;cellular organisms;Eukaryota;Opisthokonta;Metazoa;Eumetazoa

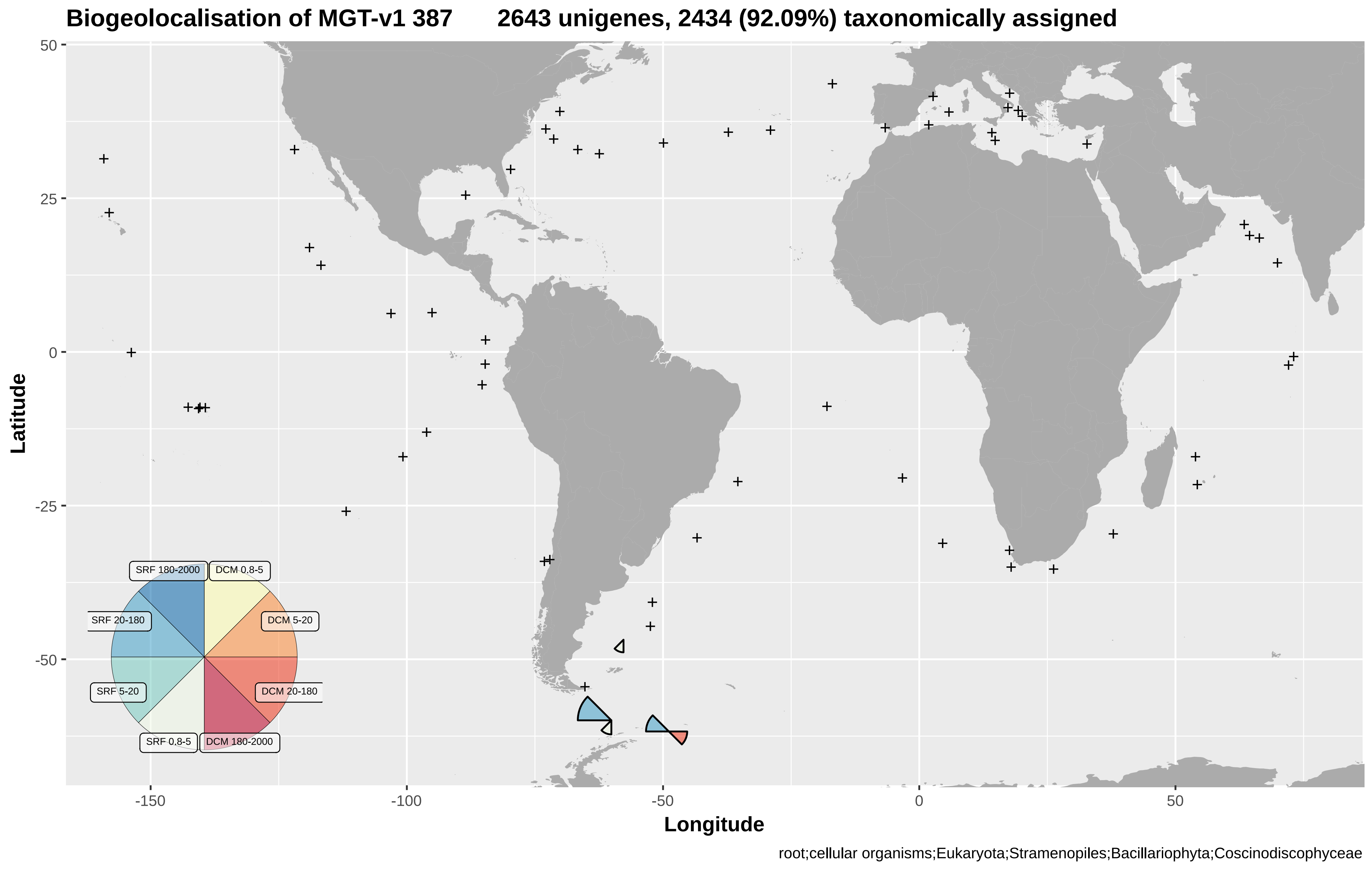

Biogeolocalisation of MGT-v1 387 2643 unigenes, 2434 (92.09%) taxonomically assigned
50
25
0
-25
-50
Latitude
-150
-100
-50
0
50
Longitude
SRF 180-2000
DCM 0.8-5
SRF 20-180
DCM 5-20
SRF 5-20
DCM 20-180
SRF 0.8-5
DCM 180-2000
root;cellular organisms;Eukaryota;Stramenopiles;Bacillariophyta;Coscinodiscophyceae

## Biogeolocalisation of MGT-v1 388      2642 unigenes, 975 (36.9%) taxonomically assigned

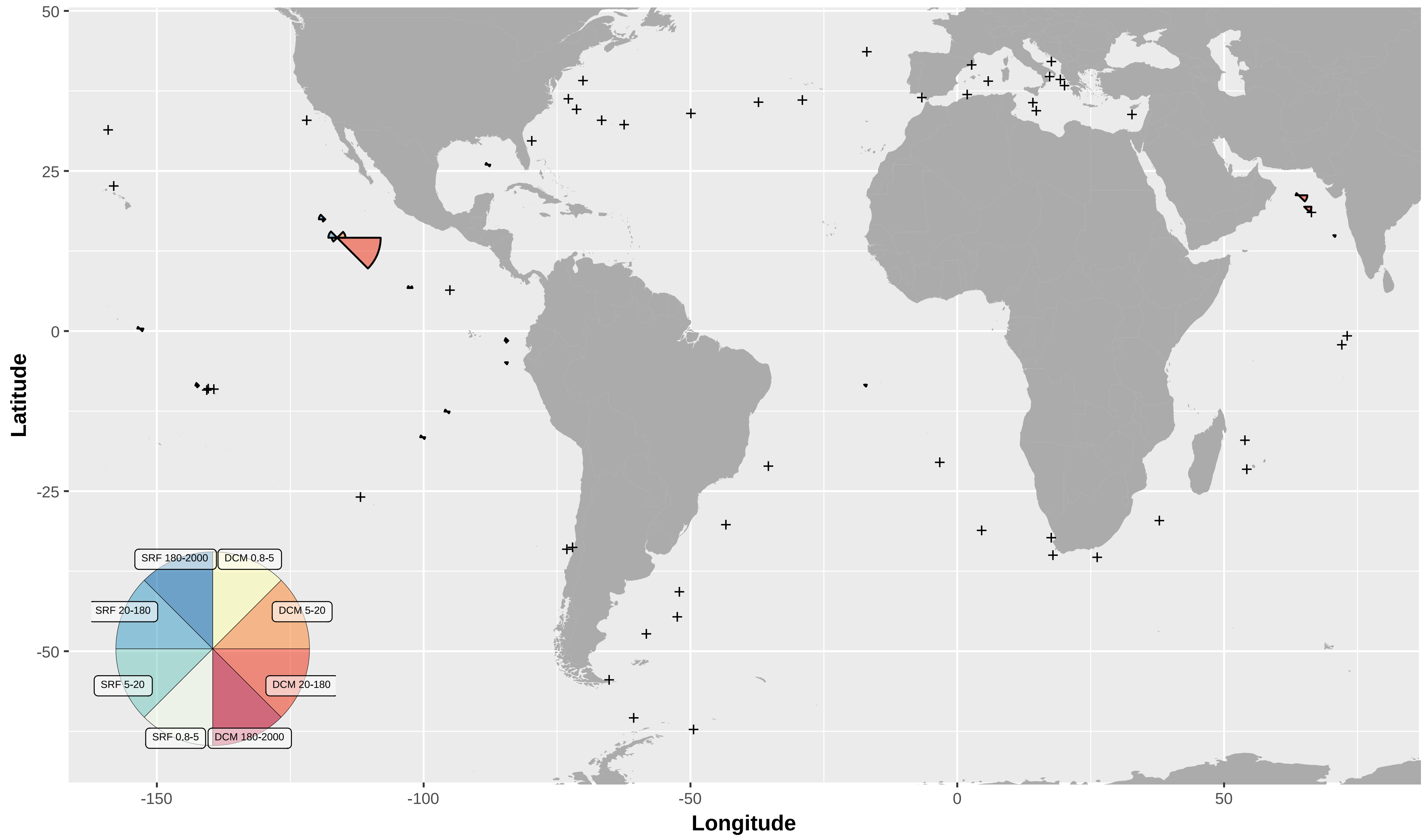


root;cellular organisms;Eukaryota;Opisthokonta;Metazoa;Eumetazoa;Bilateria

Biogeolocalisation of MGT-v1 389 2632 unigenes, 352 (13.37%) taxonomically assigned

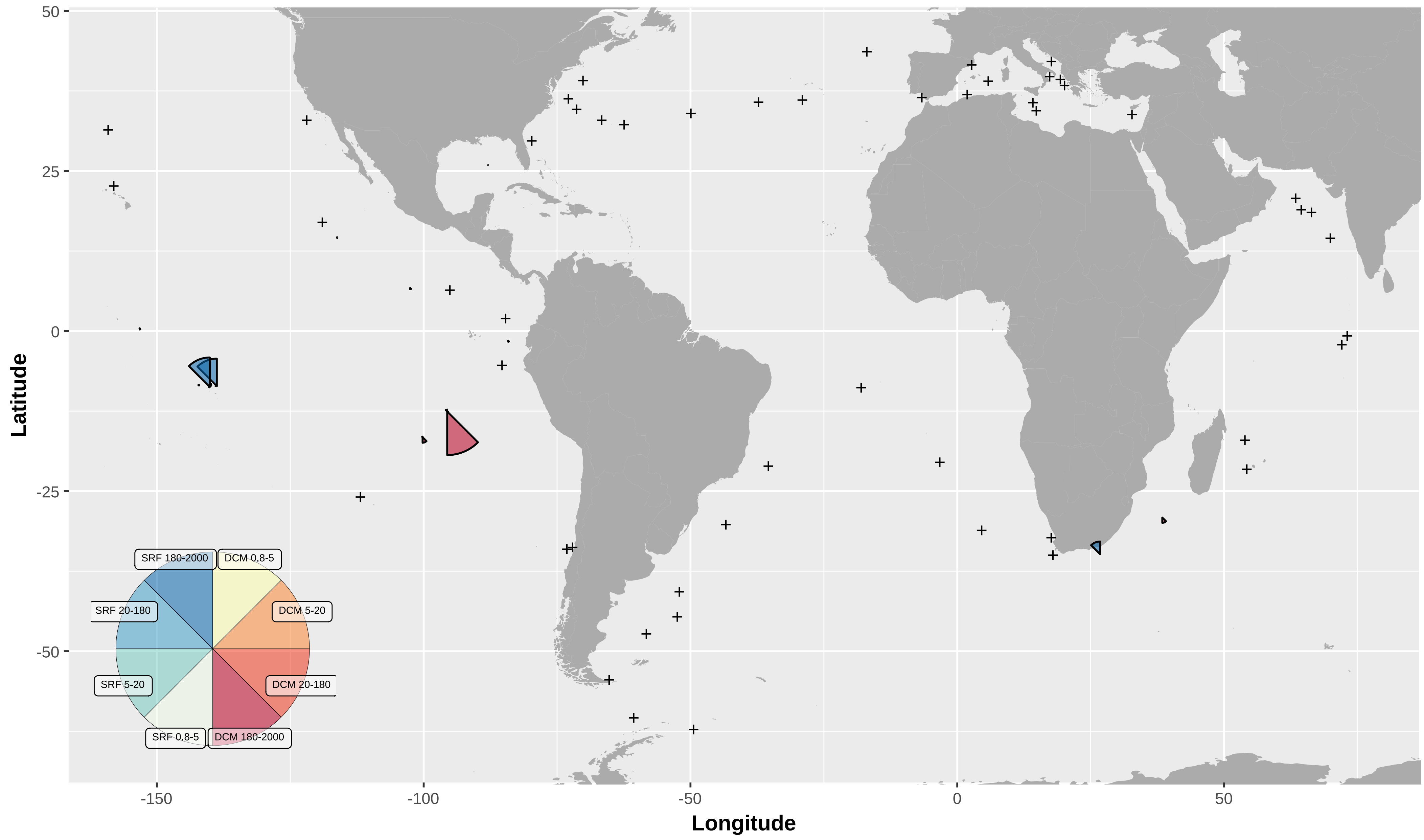


root;cellular organisms;Eukaryota;Opisthokonta;Metazoa;Eumetazoa;Bilateria

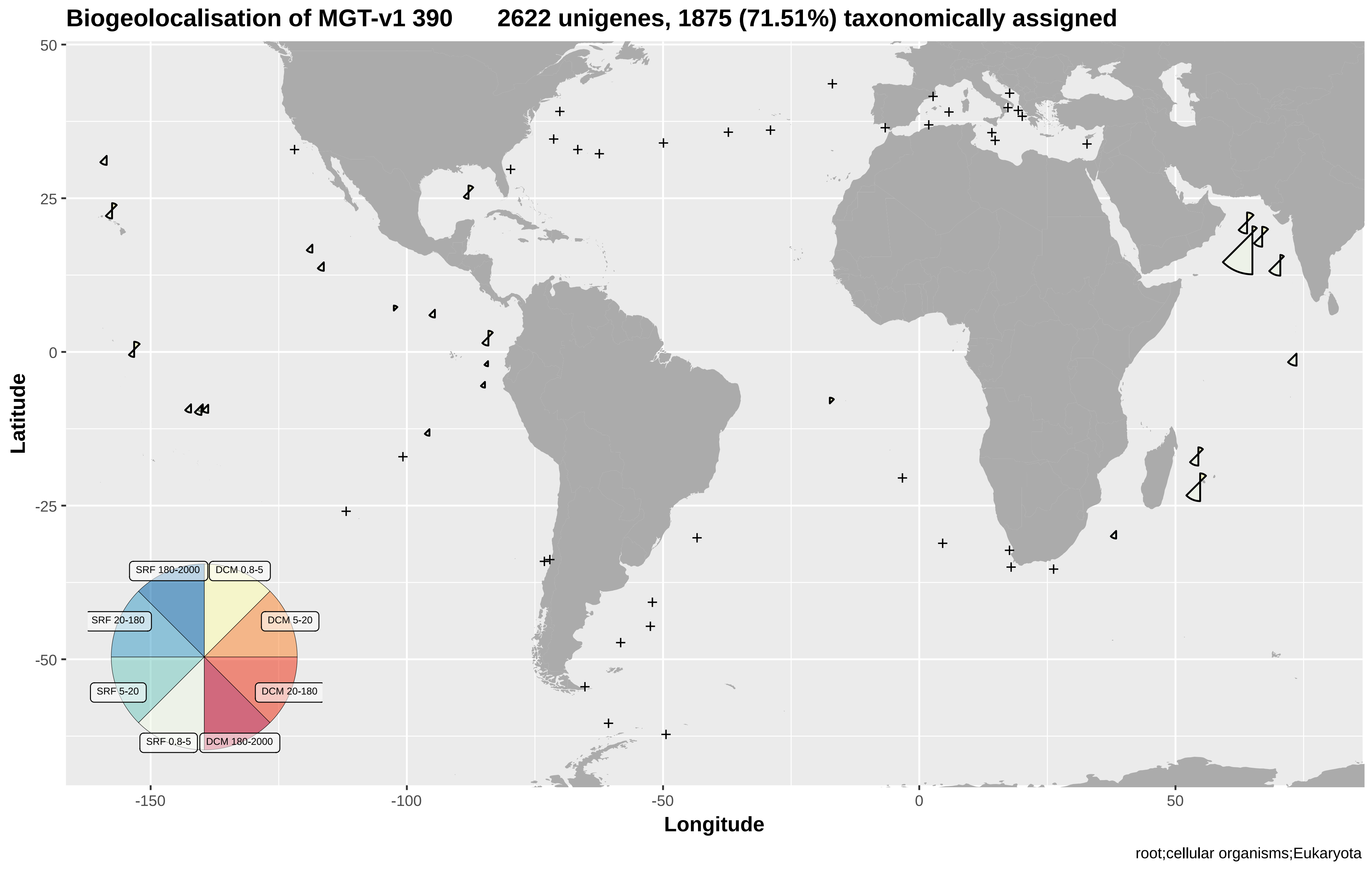

Biogeolocalisation of MGT-v1 390 2622 unigenes, 1875 (71.51%) taxonomically assigned
50
25
0
-25
-50
Latitude
-150
-100
-50
0
50
Longitude
SRF 180-2000
DCM 0.8-5
SRF 20-180
DCM 5-20
SRF 5-20
DCM 20-180
SRF 0.8-5
DCM 180-2000
root;cellular organisms;Eukaryota

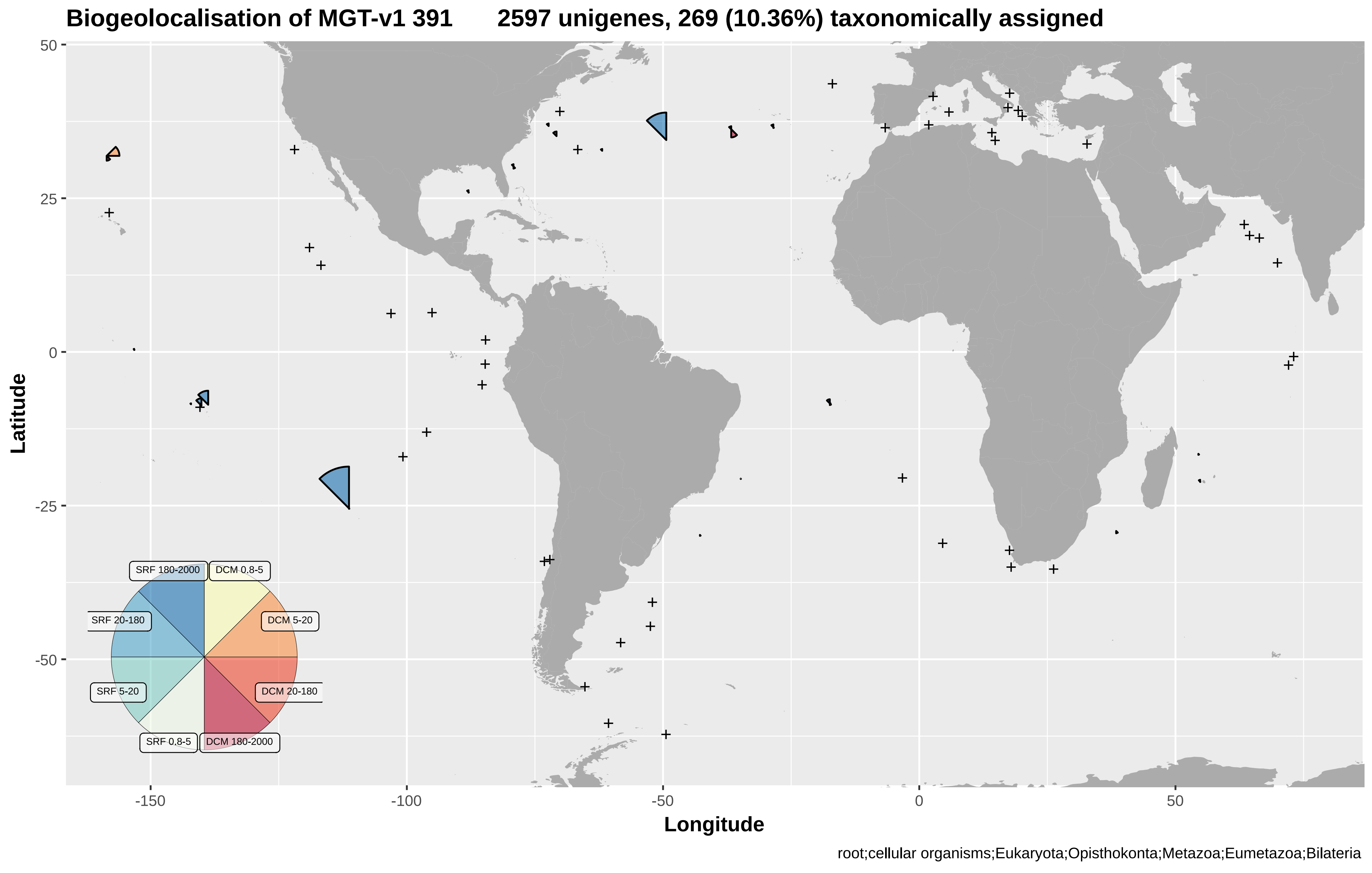

Biogeolocalisation of MGT-v1 391 2597 unigenes, 269 (10.36%) taxonomically assigned
50
25
0
-25
-50
Latitude
-150
-100
-50
0
50
Longitude
SRF 180-2000
DCM 0.8-5
SRF 20-180
DCM 5-20
SRF 5-20
DCM 20-180
SRF 0.8-5
DCM 180-2000
root;cellular organisms;Eukaryota;Opisthokonta;Metazoa;Eumetazoa;Bilateria

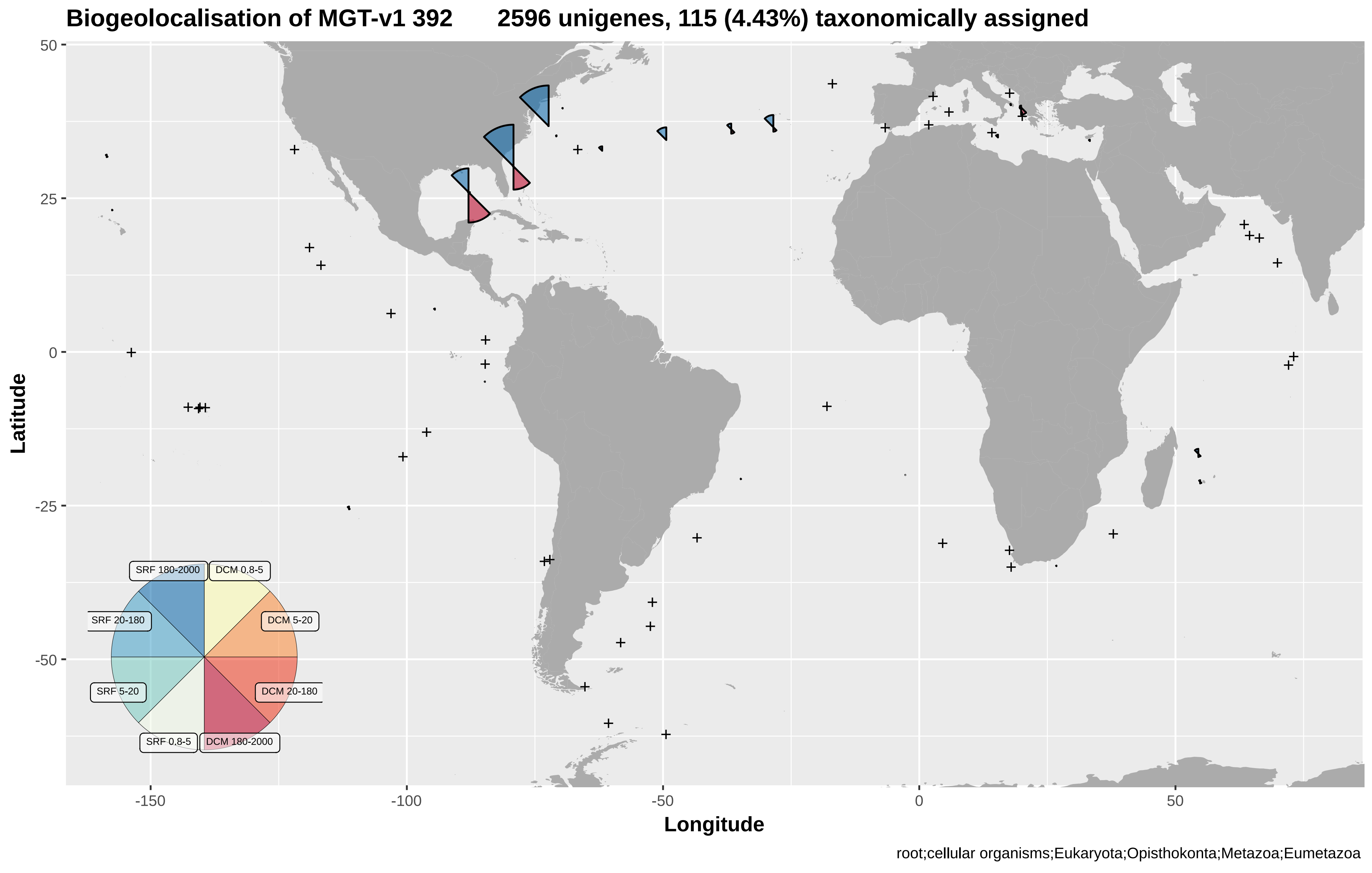
Biogeolocalisation of MGT-v1 392
2596 unigenes, 115 (4.43%) taxonomically assigned
Latitude
Longitude
50
25
0
-25
-50
-150
-100
-50
0
50
SRF 180-2000
DCM 0.8-5
SRF 20-180
DCM 5-20
SRF 5-20
DCM 20-180
SRF 0.8-5
DCM 180-2000
root;cellular organisms;Eukaryota;Opisthokonta;Metazoa;Eumetazoa

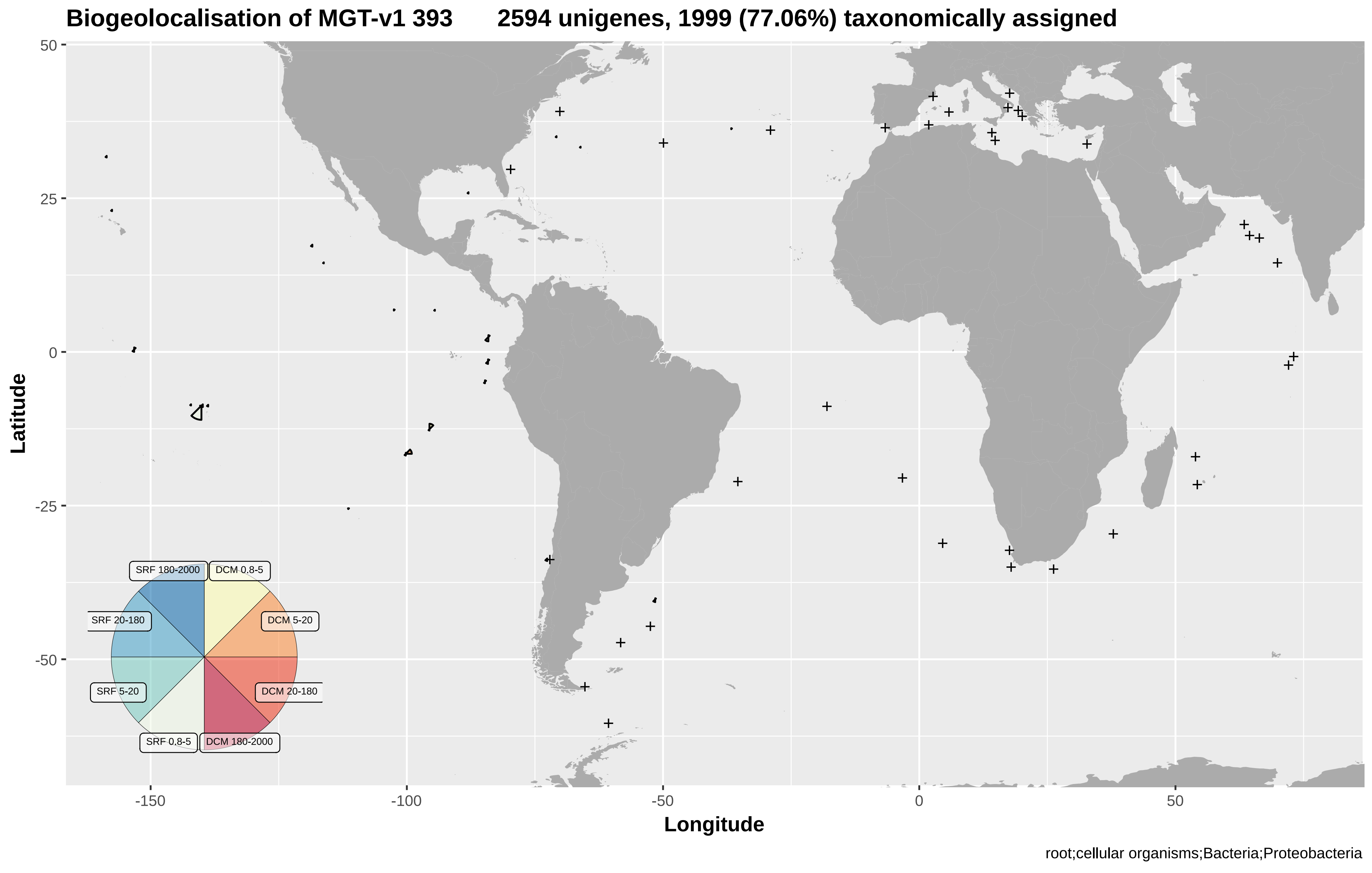
Biogeolocalisation of MGT-v1 393 2594 unigenes, 1999 (77.06%) taxonomically assigned
50
25
0
-25
-50
Latitude
-150
-100
-50
0
50
Longitude
SRF 180-2000
DCM 0.8-5
SRF 20-180
DCM 5-20
SRF 5-20
DCM 20-180
SRF 0.8-5
DCM 180-2000
root;cellular organisms;Bacteria;Proteobacteria

#### Biogeolocalisation of MGT-v1 394 2583 unigenes, 328 (12.7%) taxonomically assigned

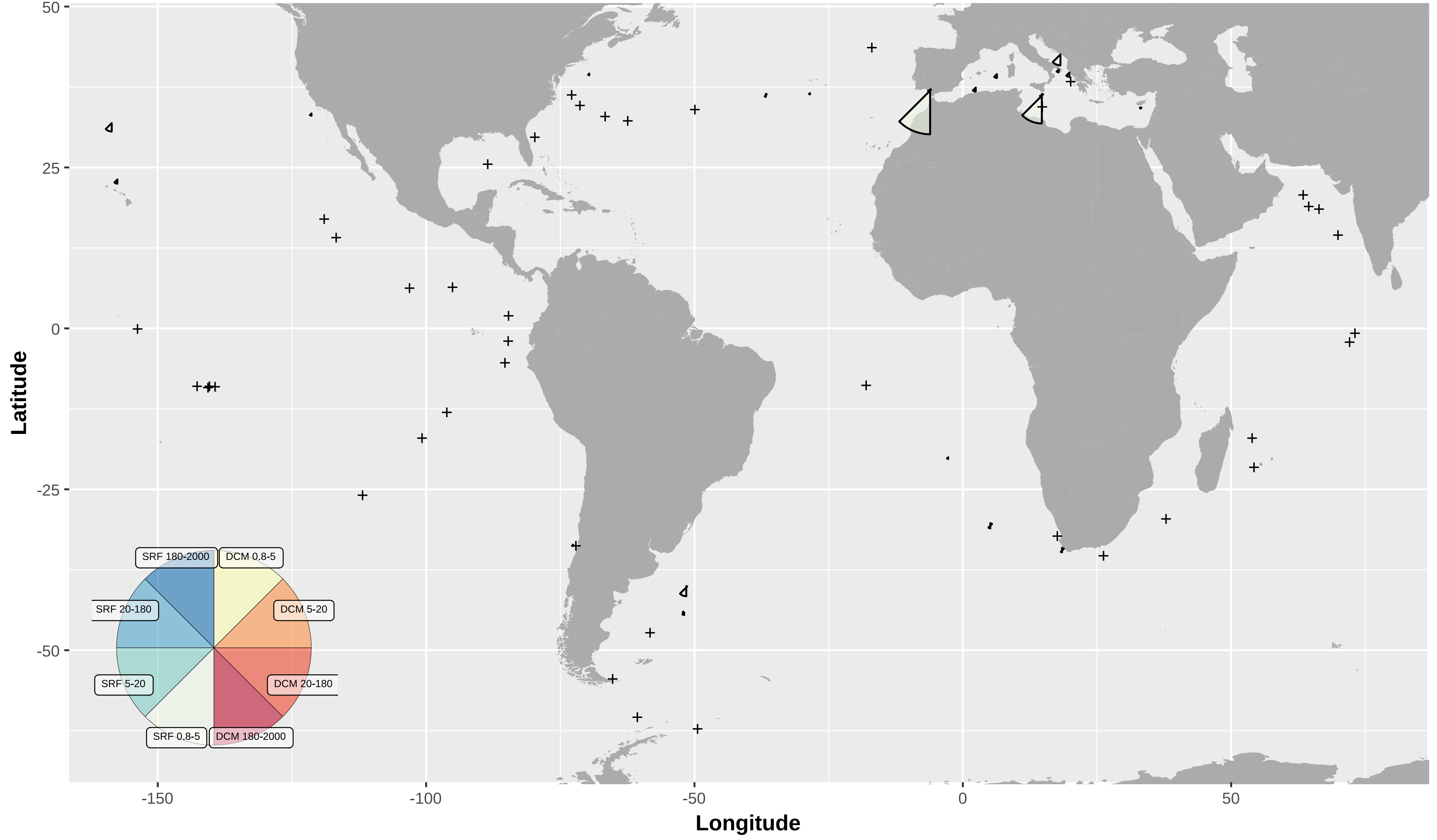


root;cellular organisms;Eukaryota

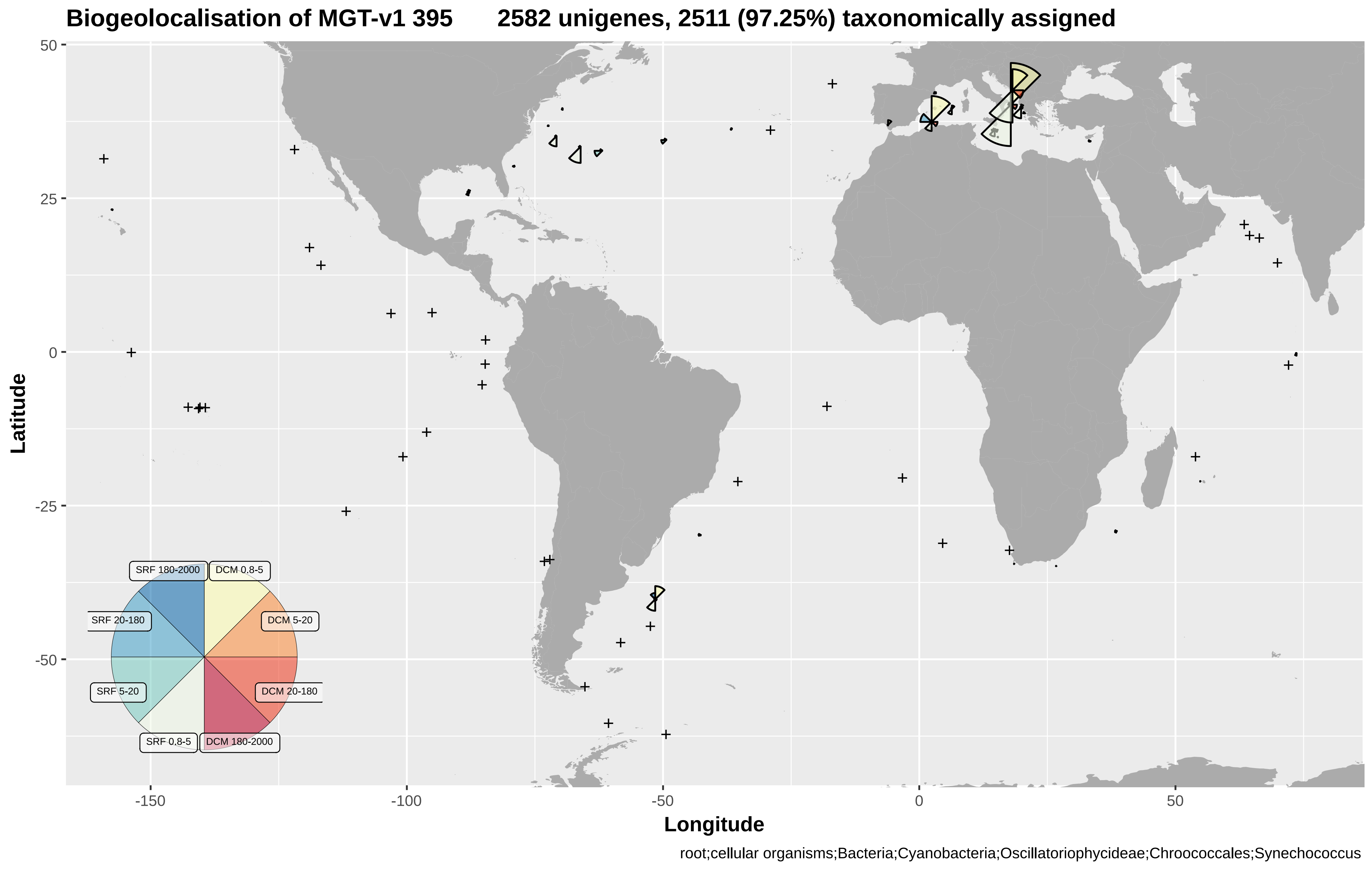

Biogeolocalisation of MGT-v1 395 2582 unigenes, 2511 (97.25%) taxonomically assigned
Latitude
Longitude
50
25
0
-25
-50
-150
-100
-50
0
50
SRF 180-2000
DCM 0.8-5
SRF 20-180
DCM 5-20
SRF 5-20
DCM 20-180
SRF 0.8-5
DCM 180-2000
root;cellular organisms;Bacteria;Cyanobacteria;Oscillatoriophycideae;Chroococcales;Synechococcus

### Biogeolocalisation of MGT-v1 396 2579 unigenes, 324 (12.56%) taxonomically assigned

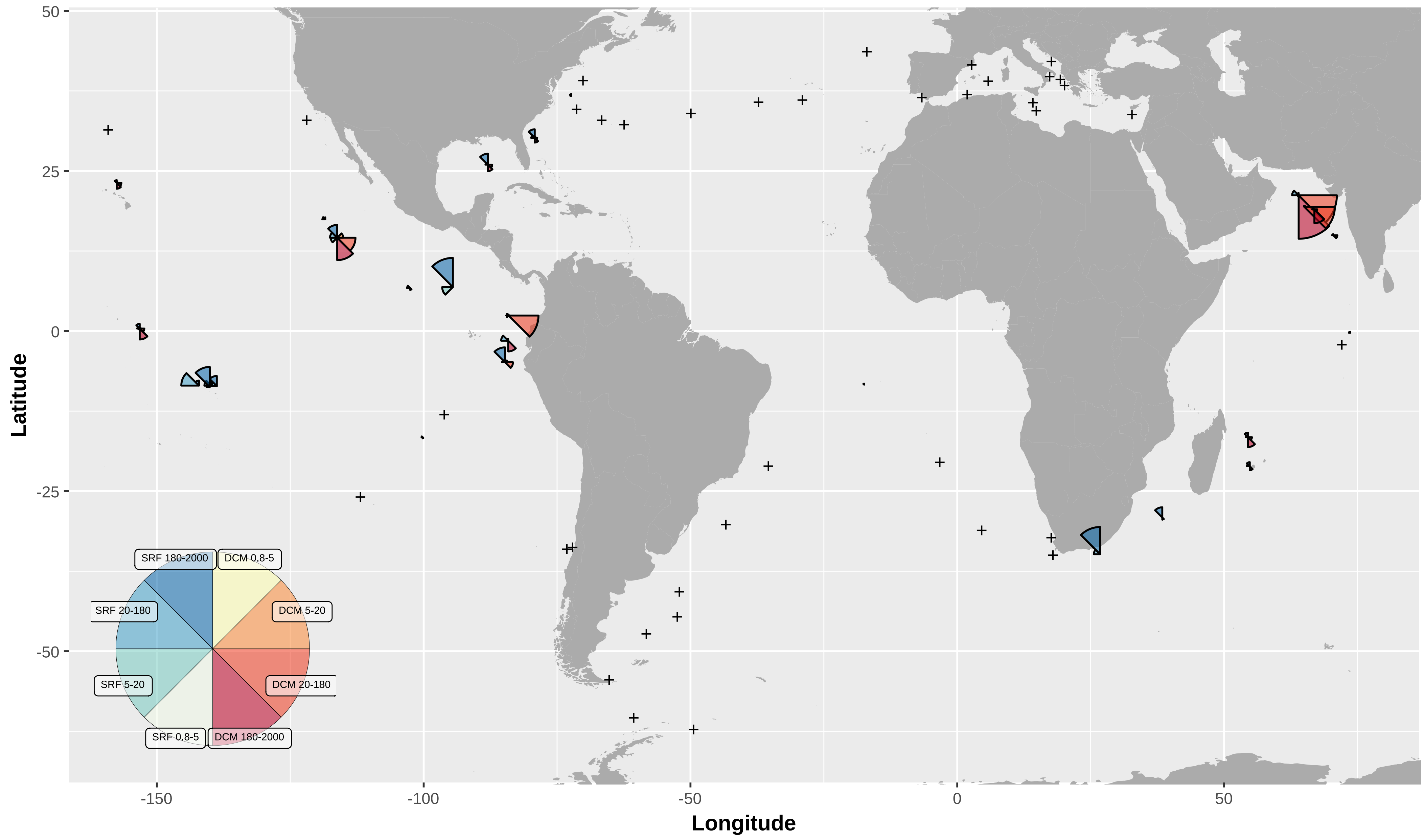


root;cellular organisms;Eukaryota;Opisthokonta;Metazoa;Eumetazoa;Bilateria;Protostomia;Ecdysozoa;Panarthropoda;Arthropoda;Mandibulata;Pancrustacea;Crustacea;Maxillopoda

# Biogeolocalisation of MGT-v1 397      2578 unigenes, 260 (10.09%) taxonomically assigned

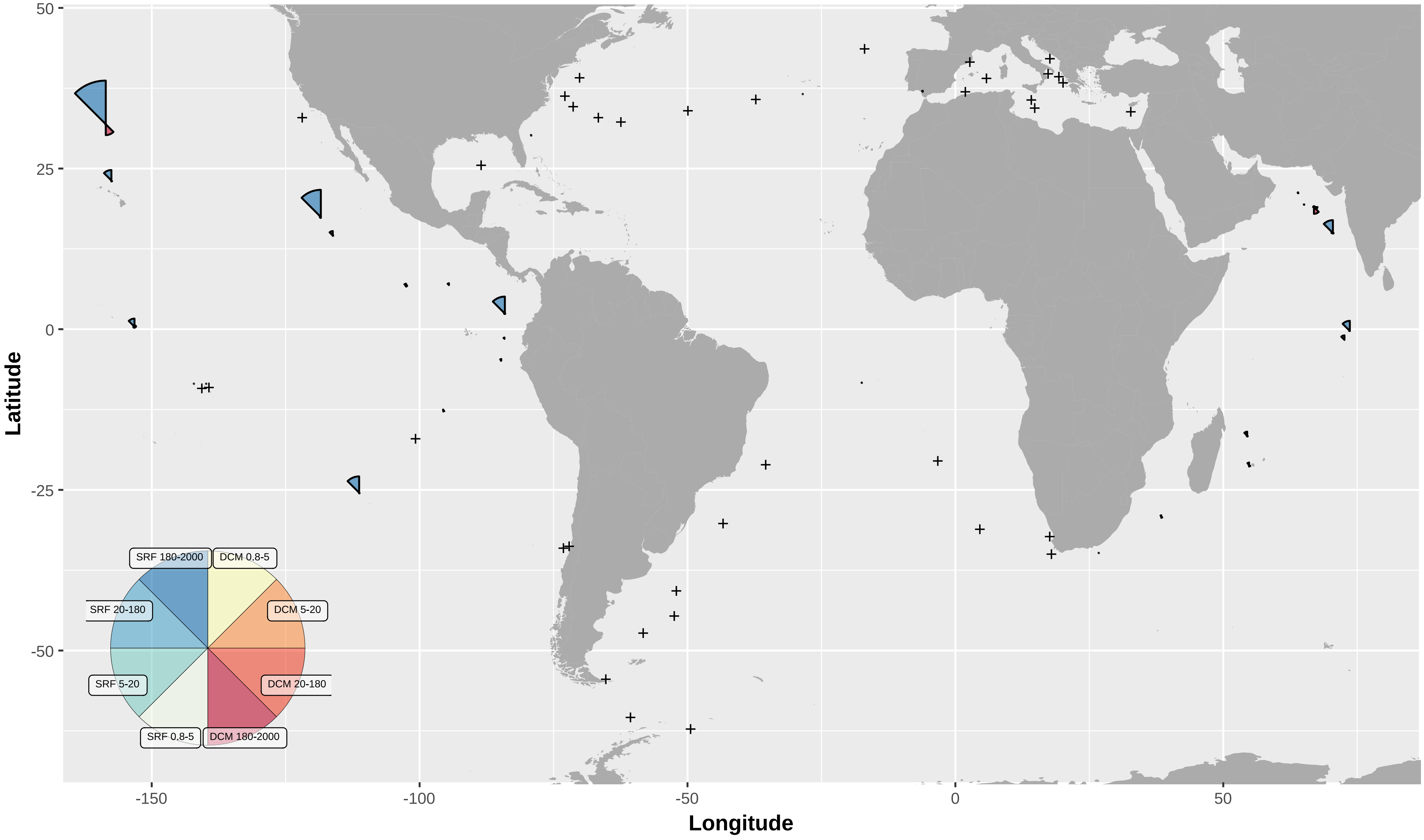


root;cellular organisms;Eukaryota;Opisthokonta;Metazoa;Eumetazoa

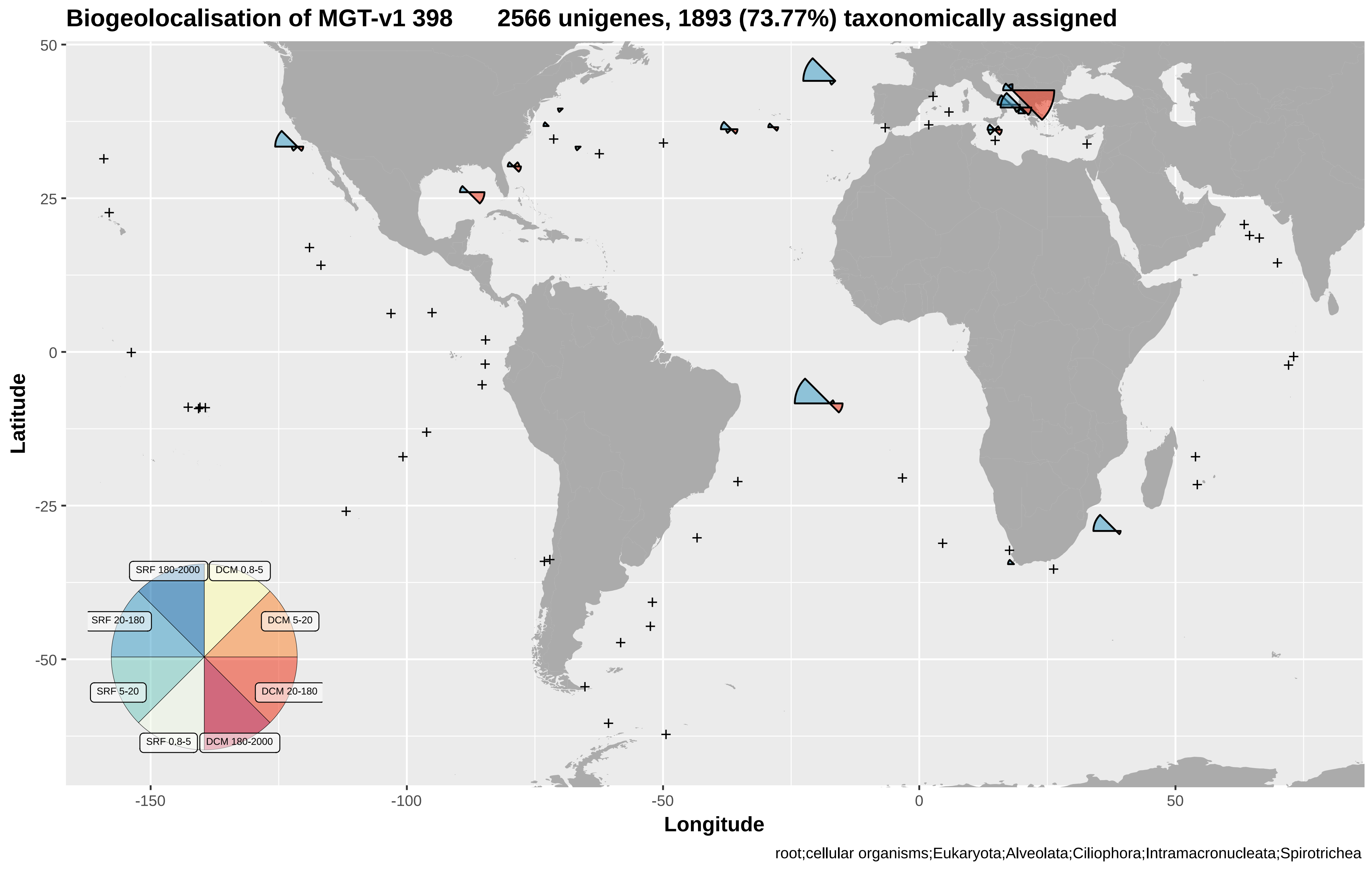
Biogeolocalisation of MGT-v1 398
2566 unigenes, 1893 (73.77%) taxonomically assigned
50
25
0
-25
-50
Latitude
-150
-100
-50
0
50
Longitude
SRF 180-2000
DCM 0.8-5
SRF 20-180
DCM 5-20
SRF 5-20
DCM 20-180
SRF 0.8-5
DCM 180-2000
root;cellular organisms;Eukaryota;Alveolata;Ciliophora;Intramacronucleata;Spirotrichea

Biogeolocalisation of MGT-v1 399      2550 unigenes, 253 (9.92%) taxonomically assigned

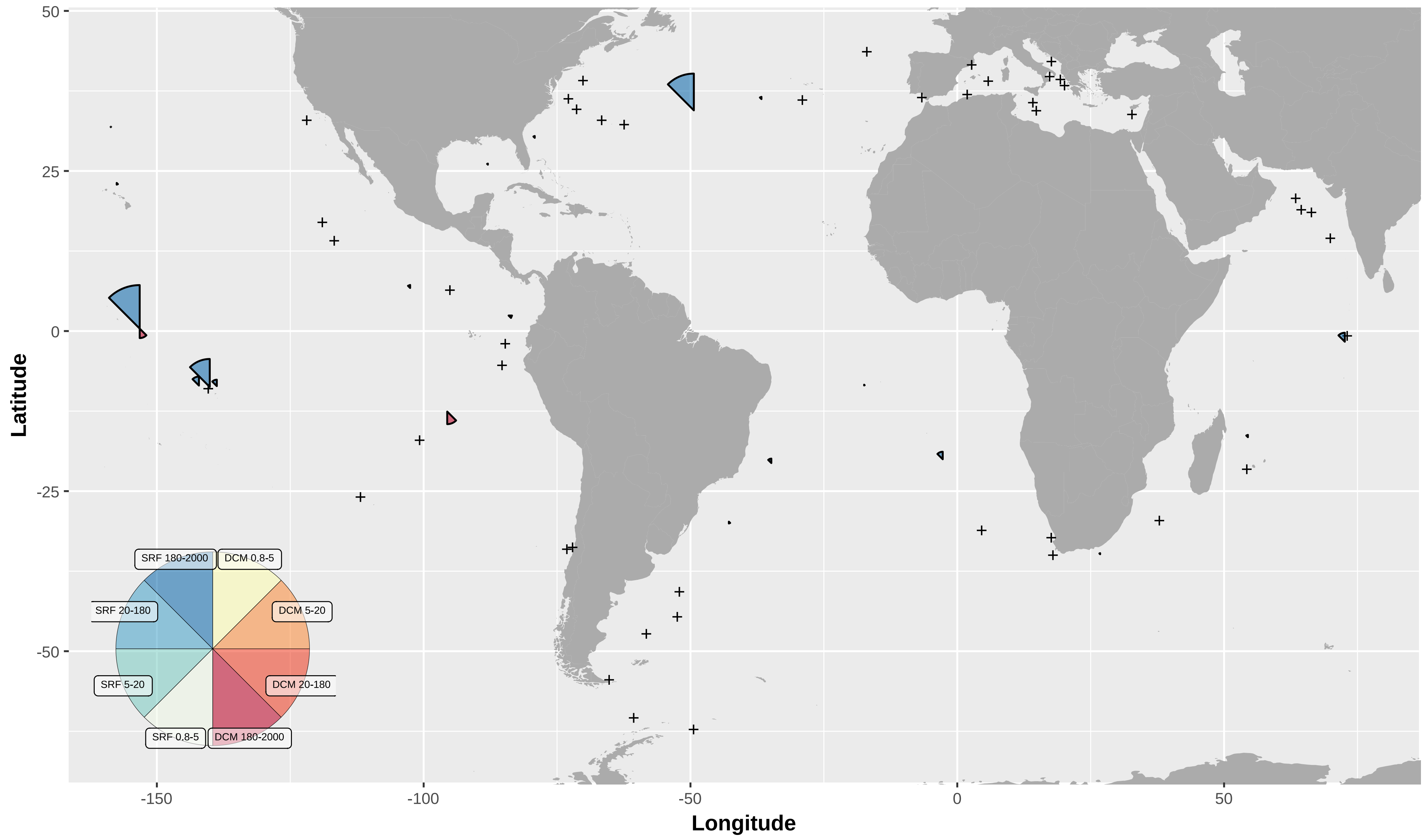


root;cellular organisms;Eukaryota;Opisthokonta;Metazoa;Eumetazoa

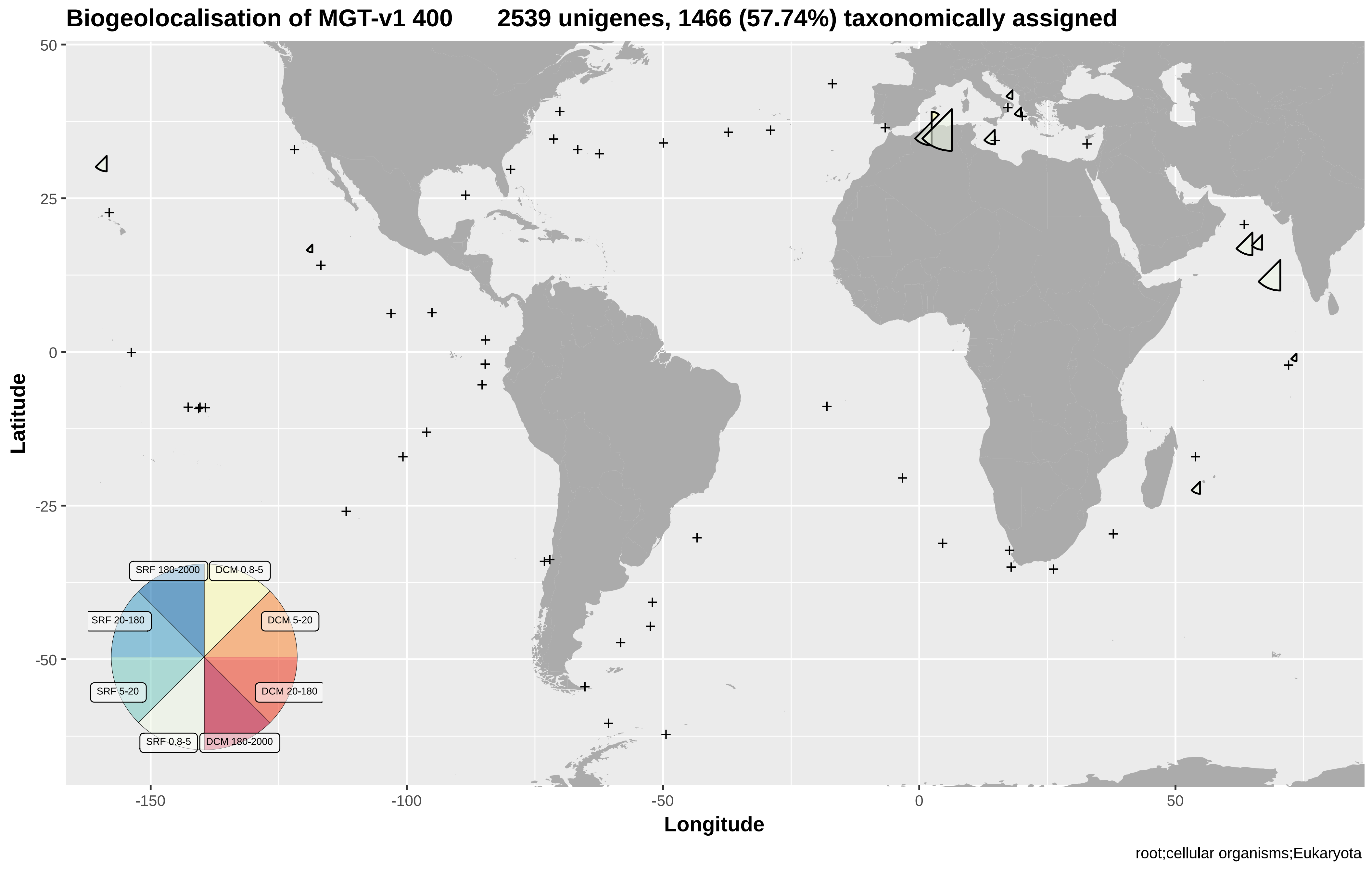

Biogeolocalisation of MGT-v1 400 2539 unigenes, 1466 (57.74%) taxonomically assigned
Latitude
Longitude
50
25
0
-25
-50
-150
-100
-50
0
50
SRF 180-2000
DCM 0.8-5
SRF 20-180
DCM 5-20
SRF 5-20
DCM 20-180
SRF 0.8-5
DCM 180-2000
root;cellular organisms;Eukaryota

# Biogeolocalisation of MGT-v1 401      2536 unigenes, 198 (7.81%) taxonomically assigned

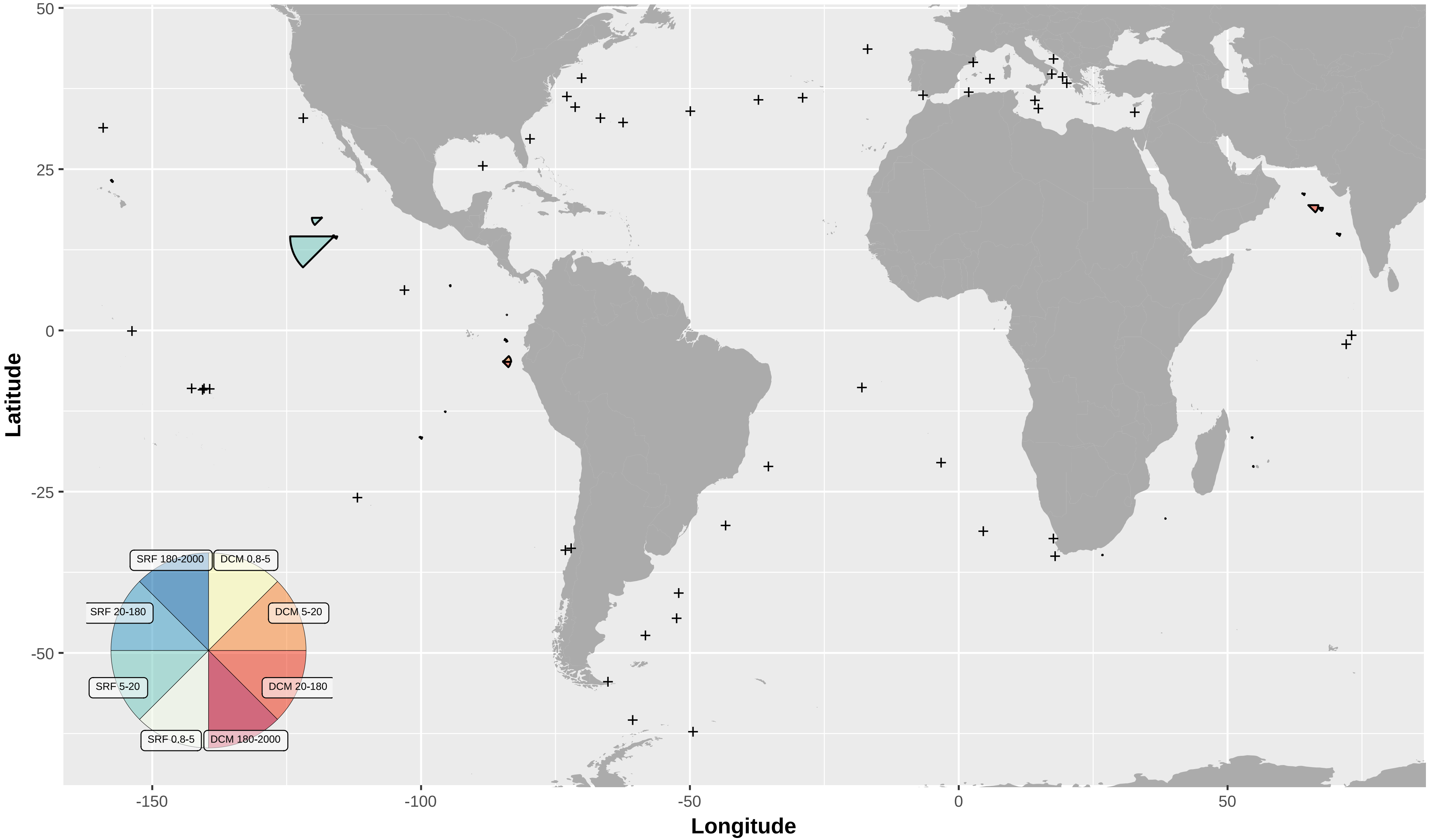


root;cellular organisms;Eukaryota;Opisthokonta;Metazoa;Eumetazoa

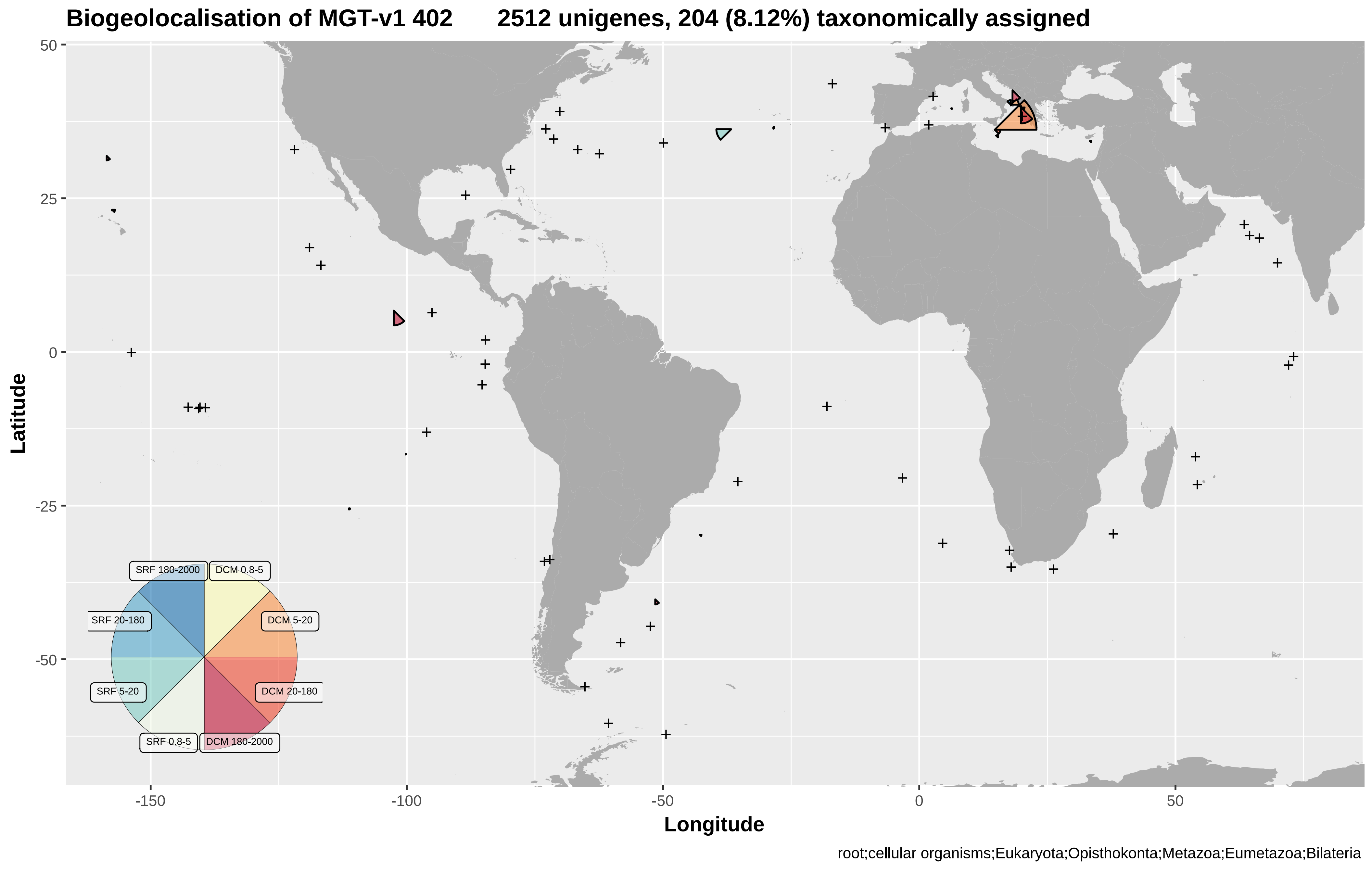
Biogeolocalisation of MGT-v1 402
2512 unigenes, 204 (8.12%) taxonomically assigned
Latitude
Longitude
50
25
0
-25
-50
-150
-100
-50
0
50
SRF 180-2000
DCM 0.8-5
SRF 20-180
DCM 5-20
SRF 5-20
DCM 20-180
SRF 0.8-5
DCM 180-2000
root;cellular organisms;Eukaryota;Opisthokonta;Metazoa;Eumetazoa;Bilateria

#### Biogeolocalisation of MGT-v1 403 2498 unigenes, 1514 (60.61%) taxonomically assigned

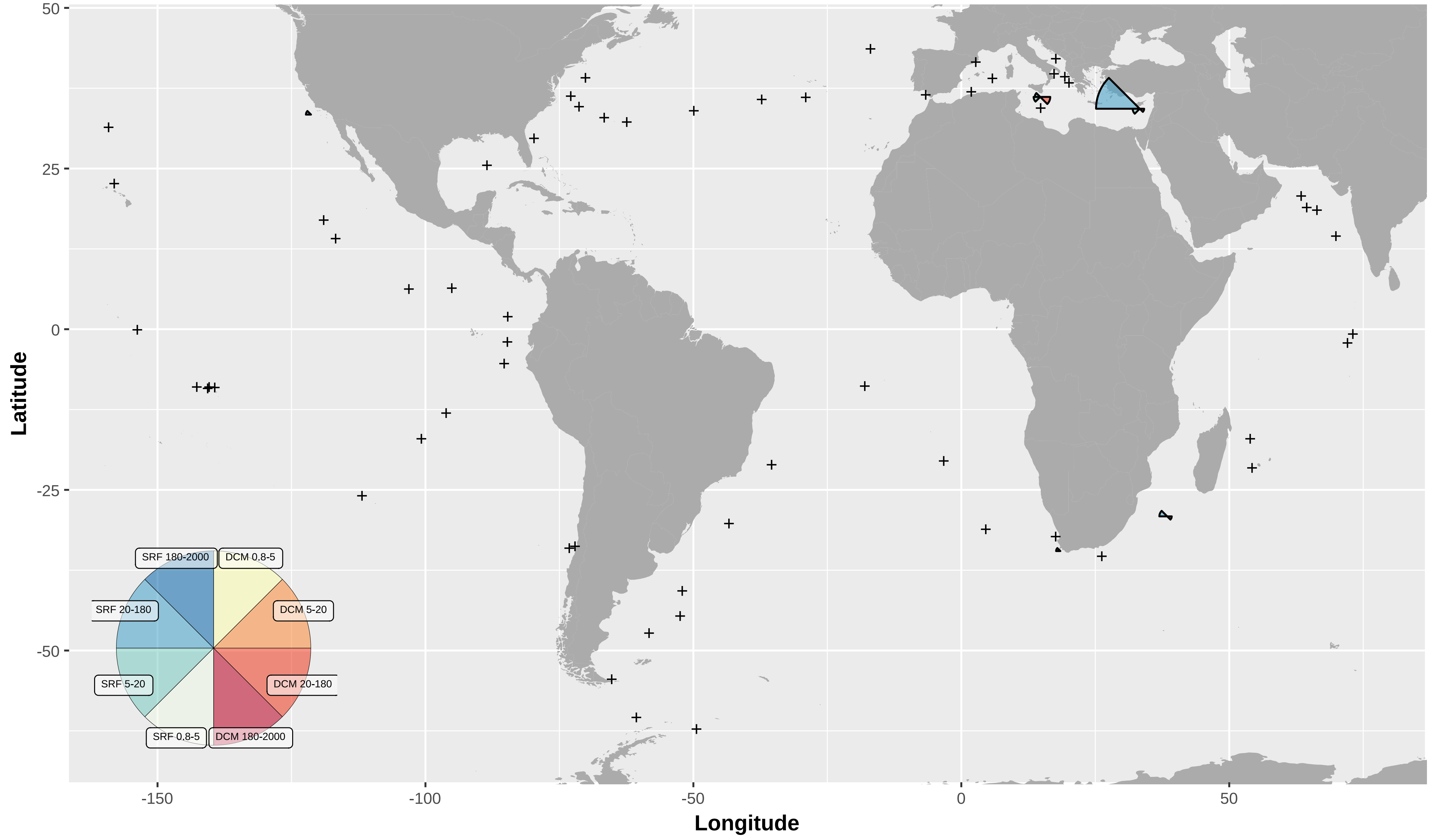


root;cellular organisms;Eukaryota;Alveolata;Ciliophora;Intramacronucleata;Spirotrichea

### Biogeolocalisation of MGT-v1 404 2488 unigenes, 1327 (53.34%) taxonomically assigned

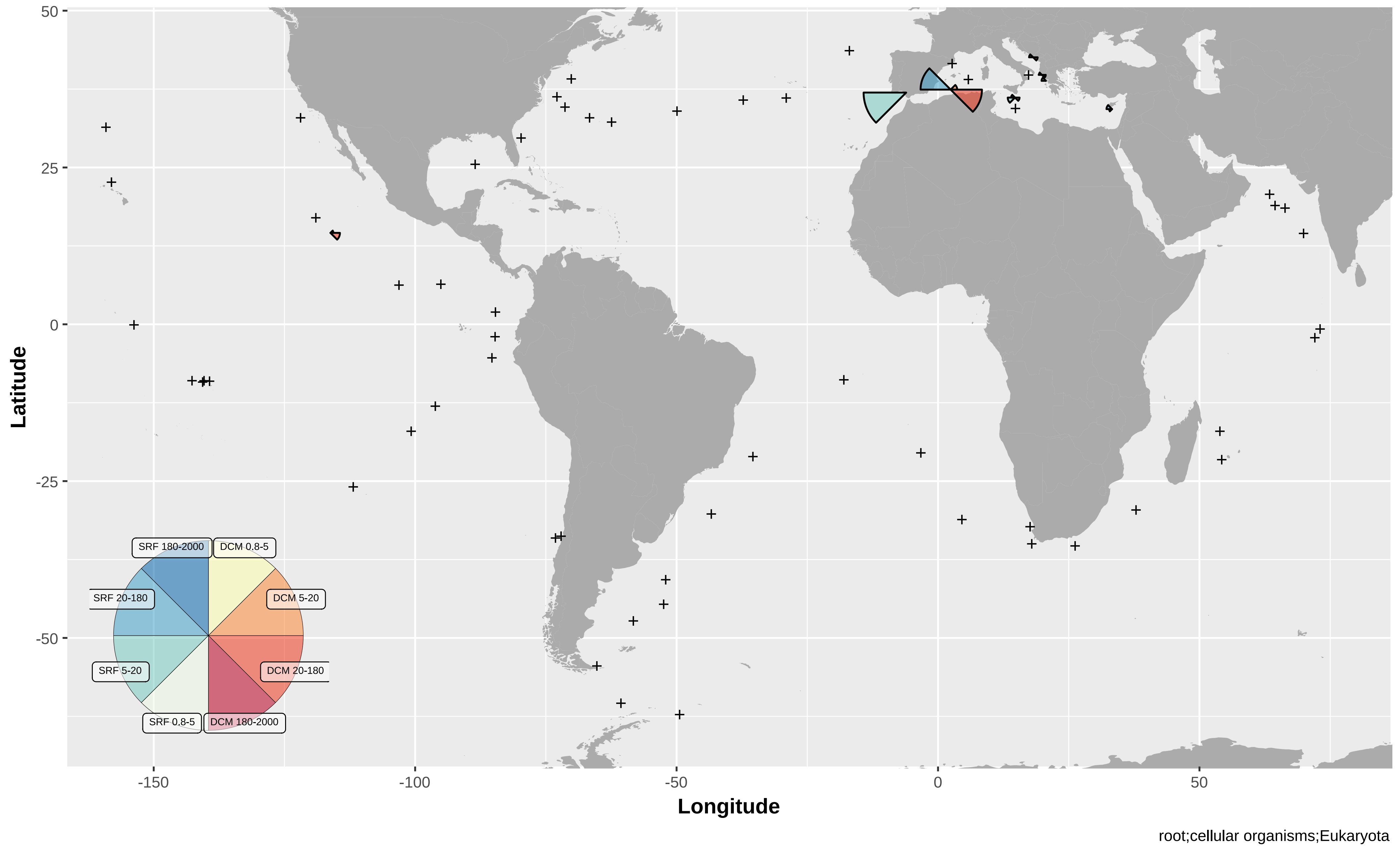

#### Biogeolocalisation of MGT-v1 405 2483 unigenes, 371 (14.94%) taxonomically assigned

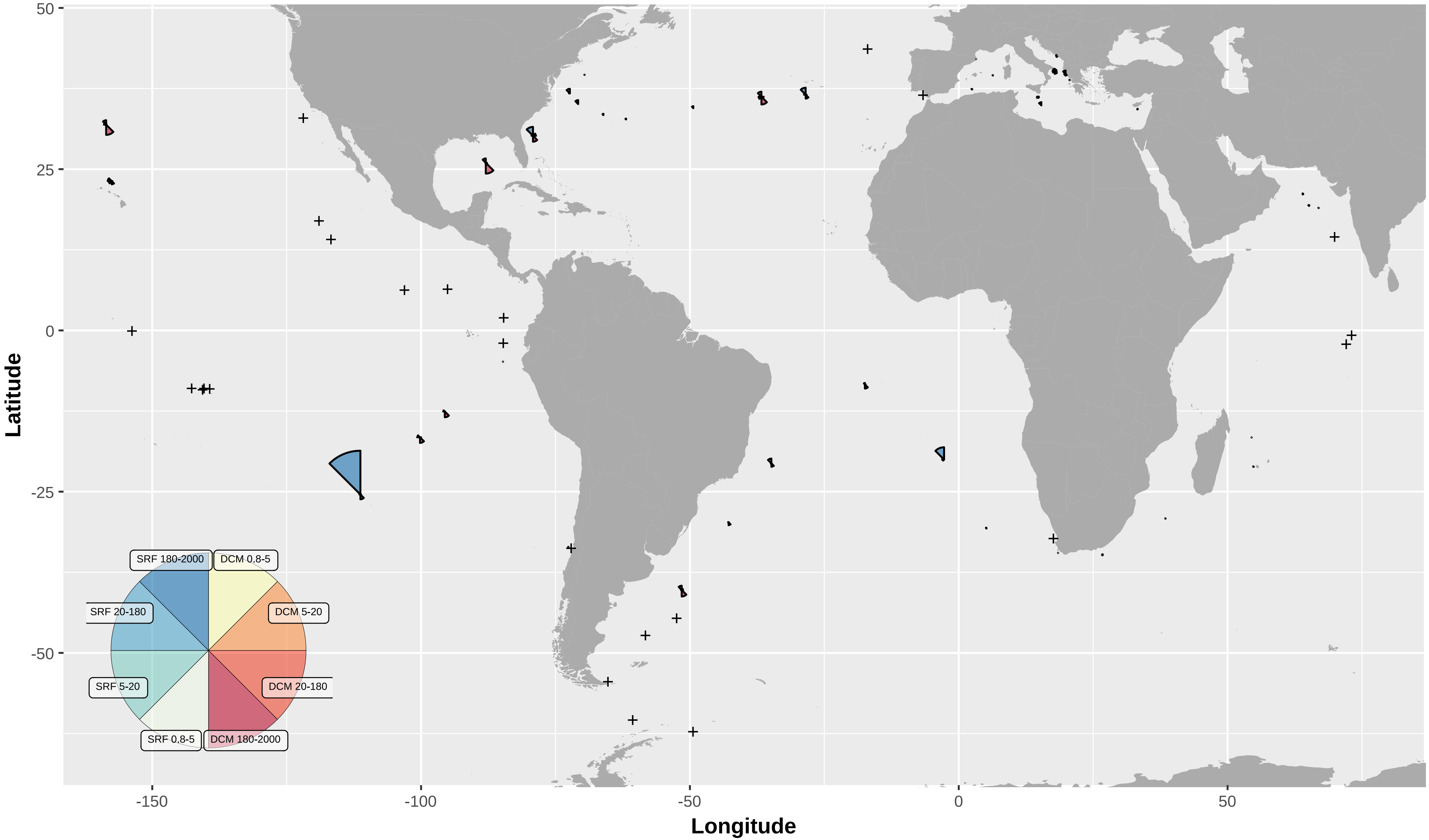


root;cellular organisms;Eukaryota;Opisthokonta;Metazoa;Eumetazoa;Bilateria;Protostomia;Ecdysozoa;Panarthropoda;Arthropoda;Mandibulata;Pancrustacea;Crustacea;Maxillopoda

# Biogeolocalisation of MGT-v1 406      2469 unigenes, 639 (25.88%) taxonomically assigned

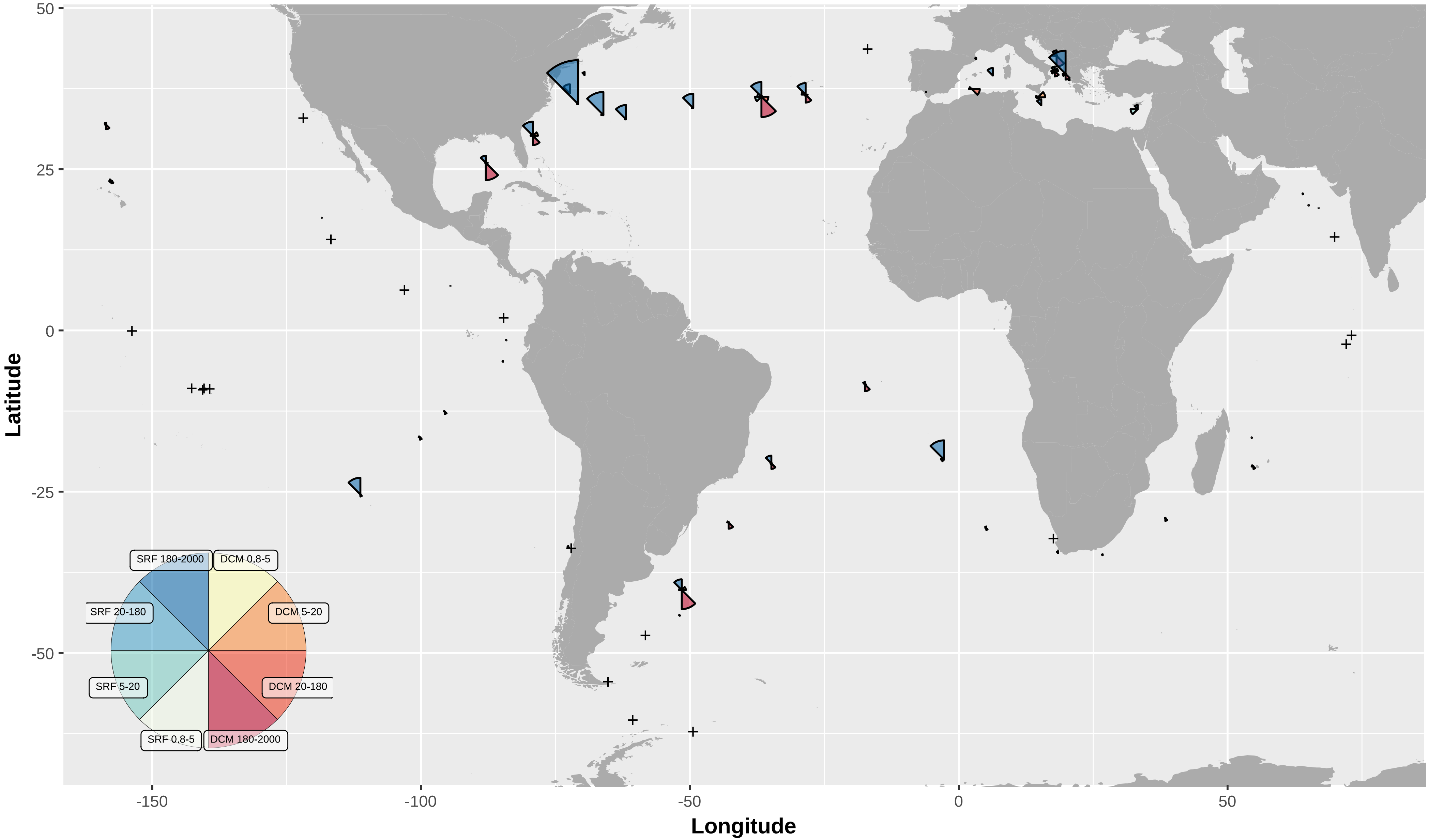


root;cellular organisms;Eukaryota;Opisthokonta;Metazoa;Eumetazoa;Bilateria;Protostomia;Ecdysozoa;Panarthropoda;Arthropoda;Mandibulata;Pancrustacea

## Biogeolocalisation of MGT-v1 407      2443 unigenes, 244 (9.99%) taxonomically assigned

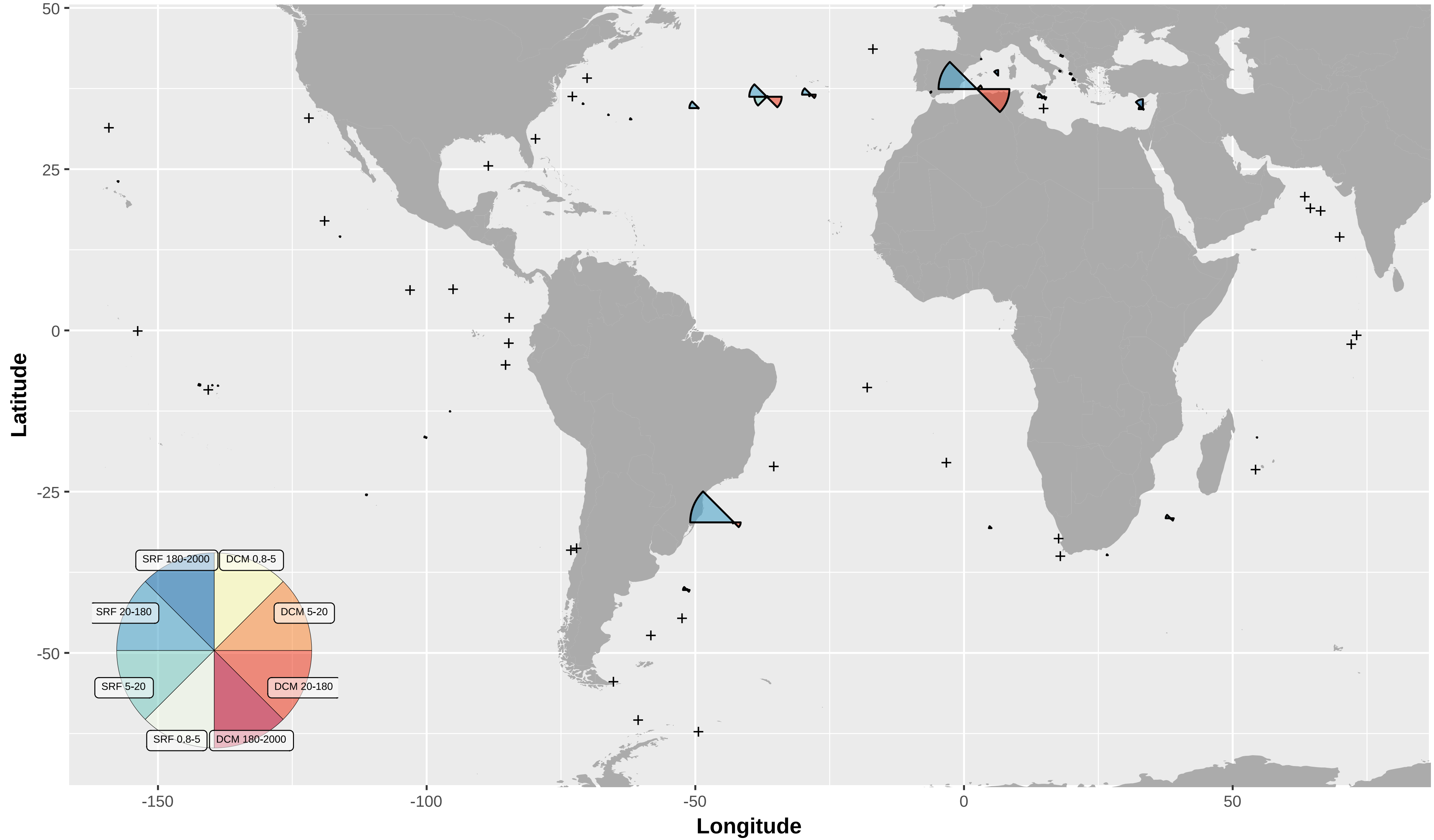


root;cellular organisms;Eukaryota;Opisthokonta;Metazoa;Eumetazoa;Bilateria

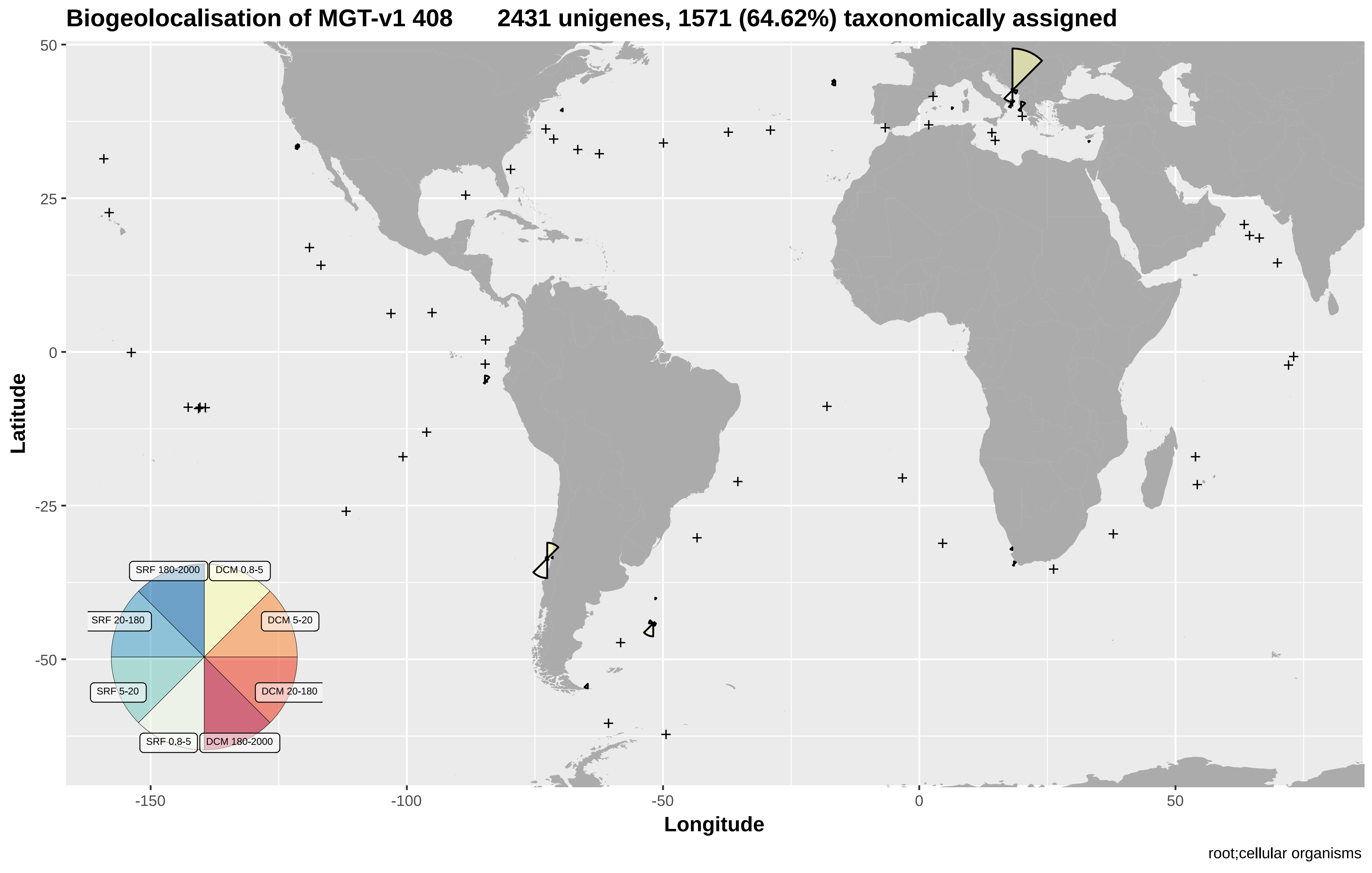
Biogeolocalisation of MGT-v1 408 2431 unigenes, 1571 (64.62%) taxonomically assigned
50
25
0
-25
-50
Latitude
-150
-100
-50
0
50
Longitude
SRF 180-2000
DCM 0.8-5
SRF 20-180
DCM 5-20
SRF 5-20
DCM 20-180
SRF 0.8-5
DCM 180-2000
root;cellular organisms

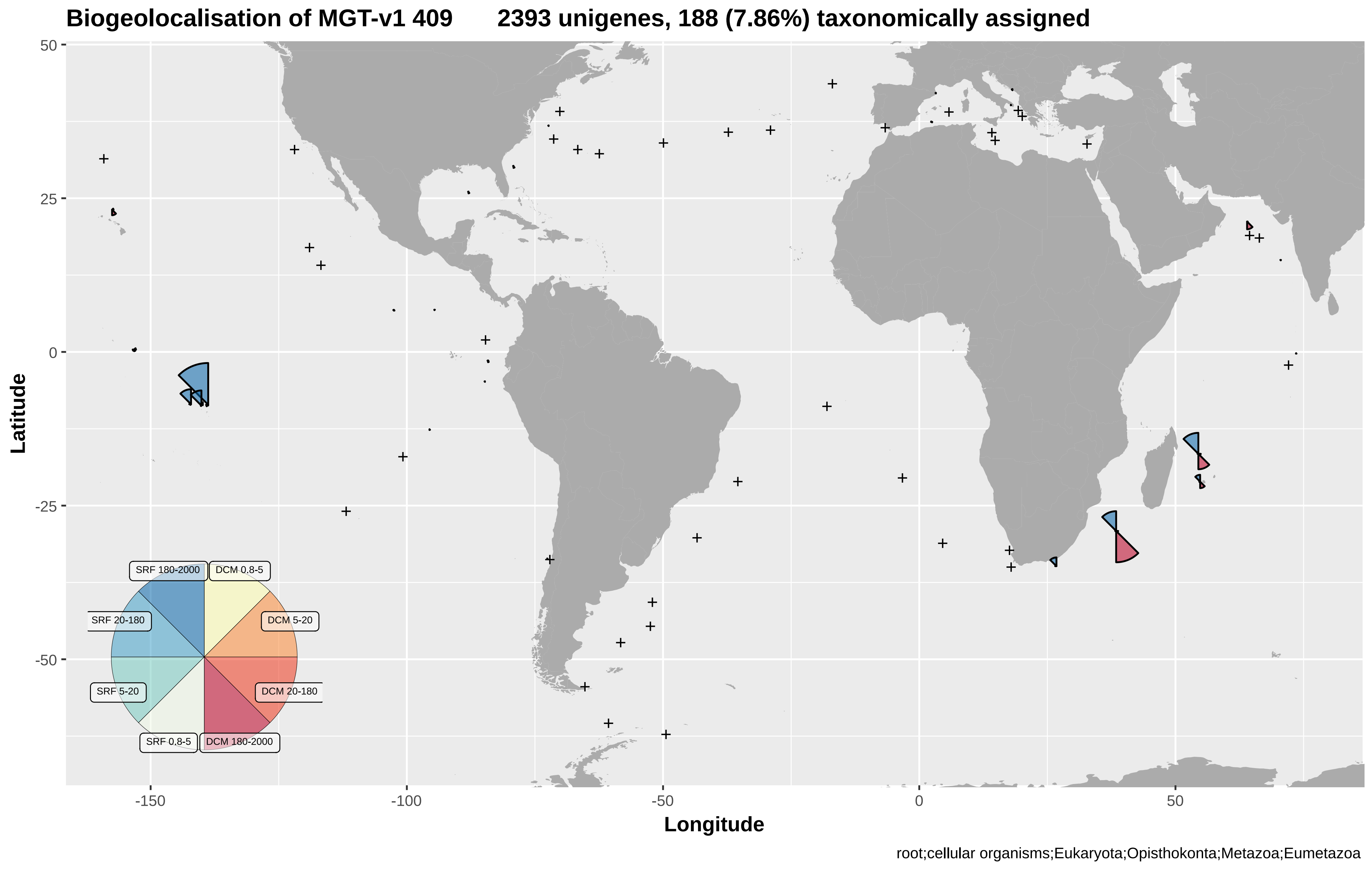

Biogeolocalisation of MGT-v1 409 2393 unigenes, 188 (7.86%) taxonomically assigned
50
25
0
-25
-50
Latitude
-150
-100
-50
0
50
Longitude
SRF 180-2000
DCM 0.8-5
SRF 20-180
DCM 5-20
SRF 5-20
DCM 20-180
SRF 0.8-5
DCM 180-2000
root;cellular organisms;Eukaryota;Opisthokonta;Metazoa;Eumetazoa

## Biogeolocalisation of MGT-v1 410      2391 unigenes, 1638 (68.51%) taxonomically assigned

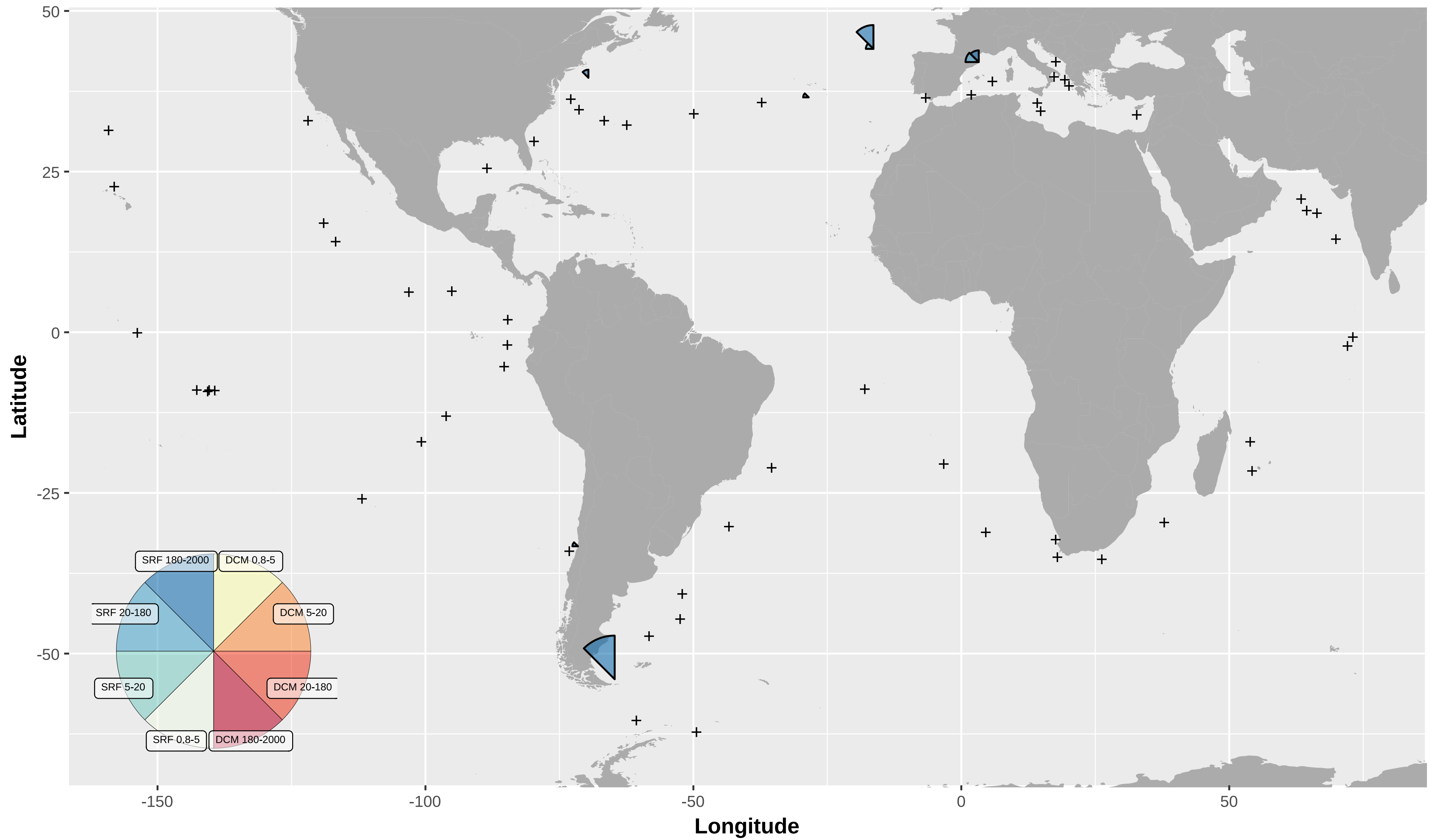


root;cellular organisms;Eukaryota;Opisthokonta;Metazoa;Eumetazoa;Bilateria;Protostomia;Ecdysozoa;Panarthropoda;Arthropoda;Mandibulata;Pancrustacea;Crustacea;Maxillopoda

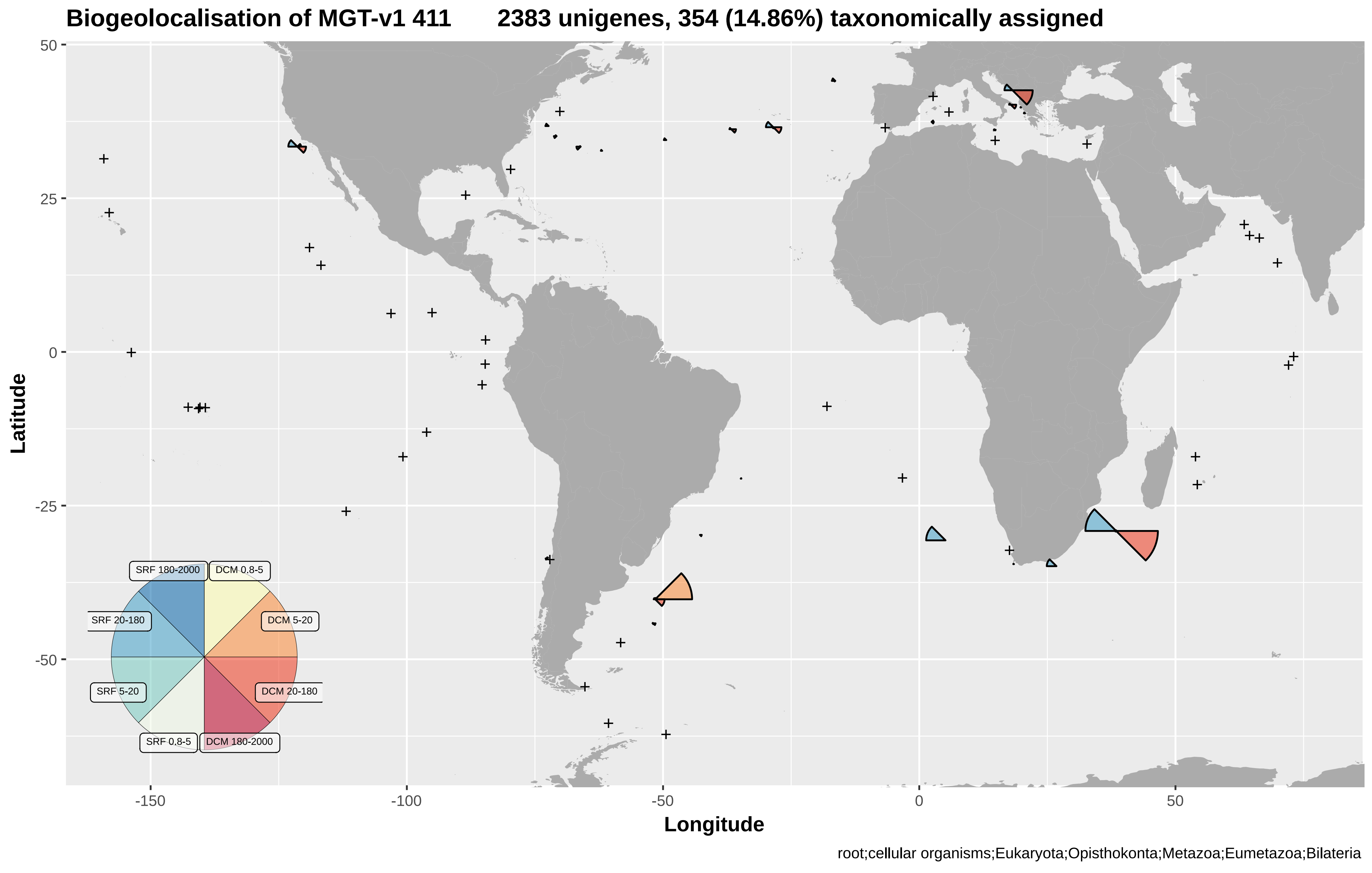

Biogeolocalisation of MGT-v1 411
2383 unigenes, 354 (14.86%) taxonomically assigned
Latitude
50
25
0
-25
-50
Longitude
-150
-100
-50
0
50
SRF 180-2000
DCM 0.8-5
SRF 20-180
DCM 5-20
SRF 5-20
DCM 20-180
SRF 0.8-5
DCM 180-2000
root;cellular organisms;Eukaryota;Opisthokonta;Metazoa;Eumetazoa;Bilateria

### Biogeolocalisation of MGT-v1 412 2361 unigenes, 2287 (96.87%) taxonomically assigned

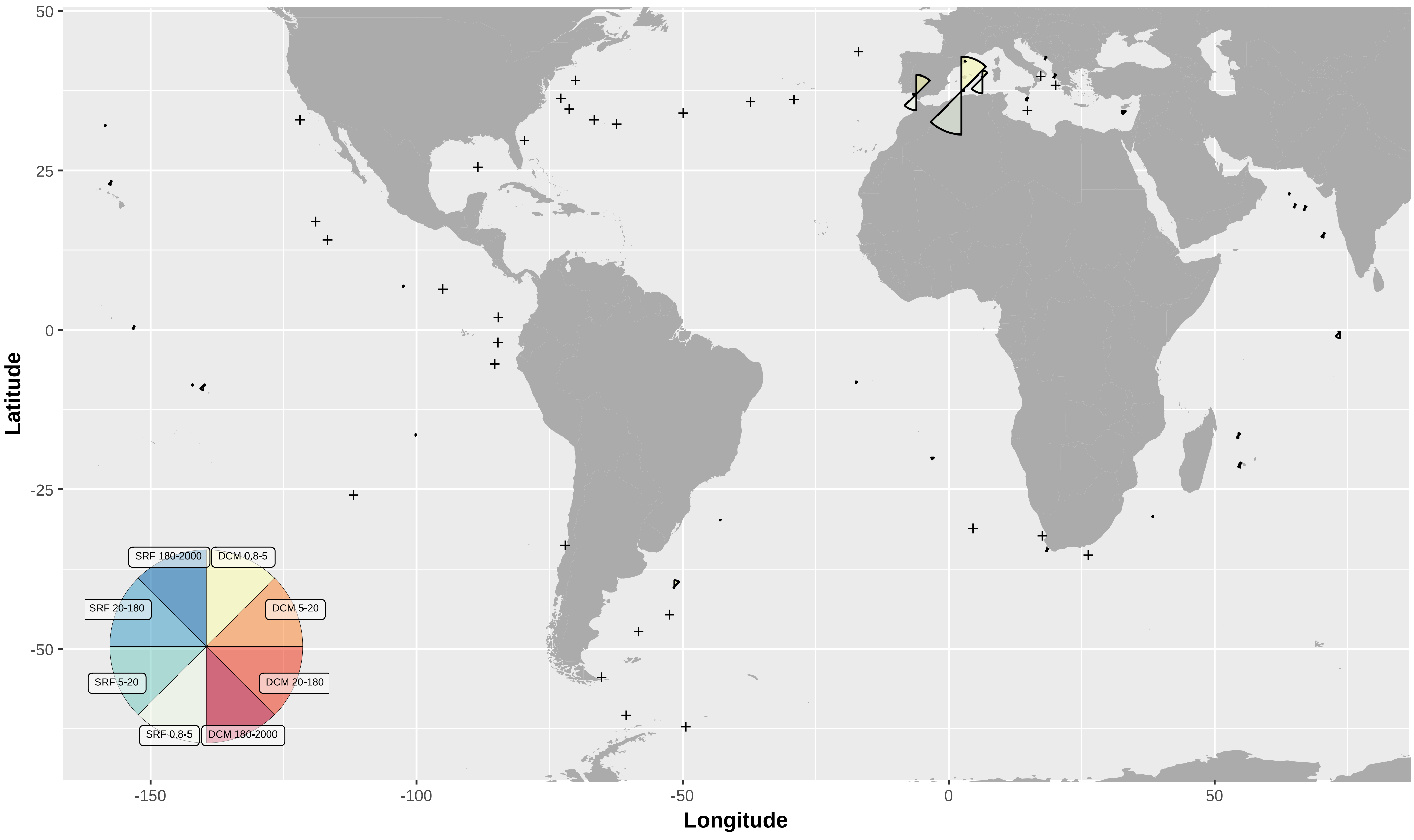


root;cellular organisms;Bacteria

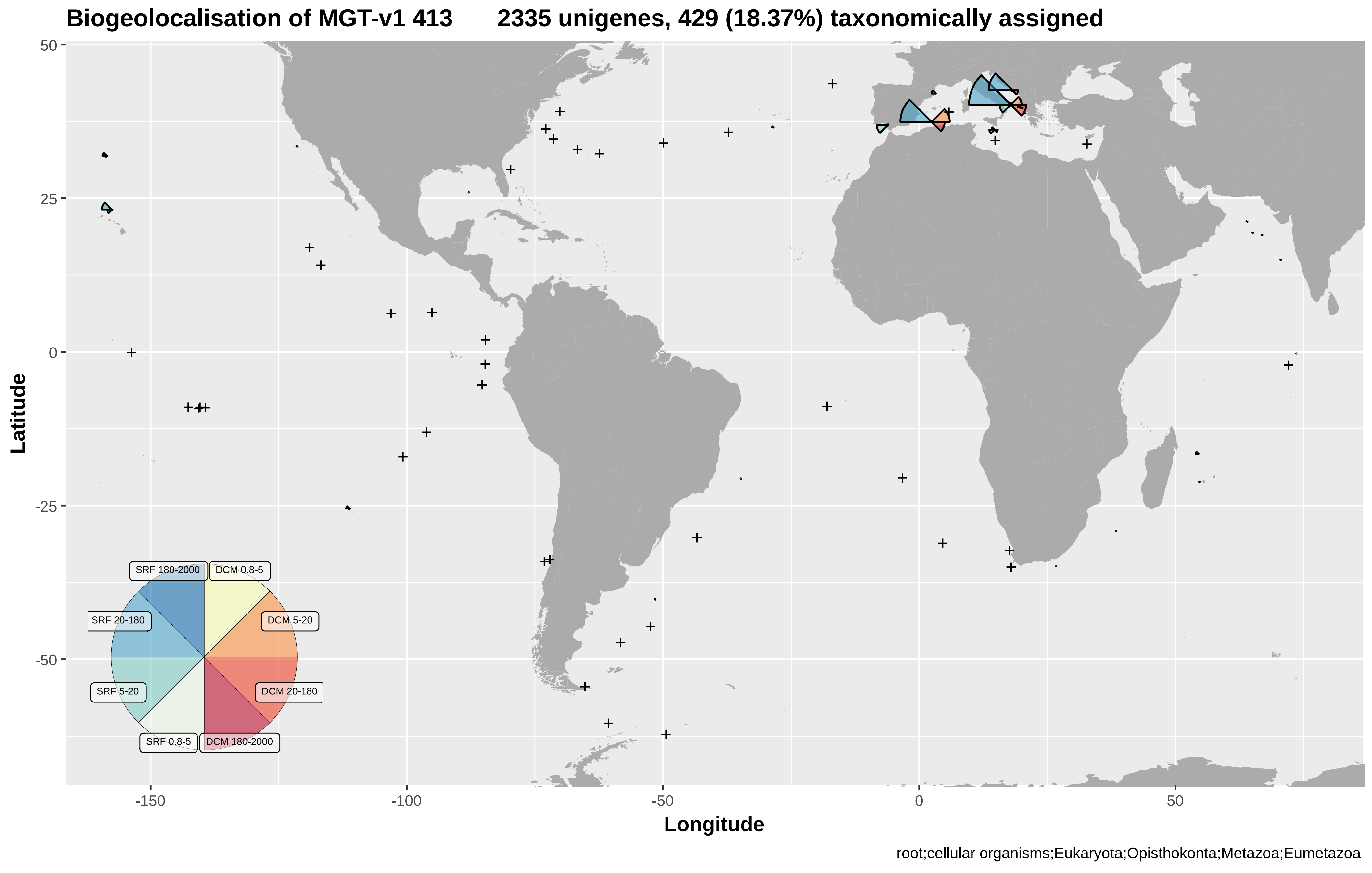
Biogeolocalisation of MGT-v1 413
2335 unigenes, 429 (18.37%) taxonomically assigned
50
25
0
-25
-50
Latitude
-150
-100
-50
0
50
Longitude
SRF 180-2000
DCM 0.8-5
SRF 20-180
DCM 5-20
SRF 5-20
DCM 20-180
SRF 0.8-5
DCM 180-2000
root;cellular organisms;Eukaryota;Opisthokonta;Metazoa;Eumetazoa

#### Biogeolocalisation of MGT-v1 414 2326 unigenes, 1689 (72.61%) taxonomically assigned

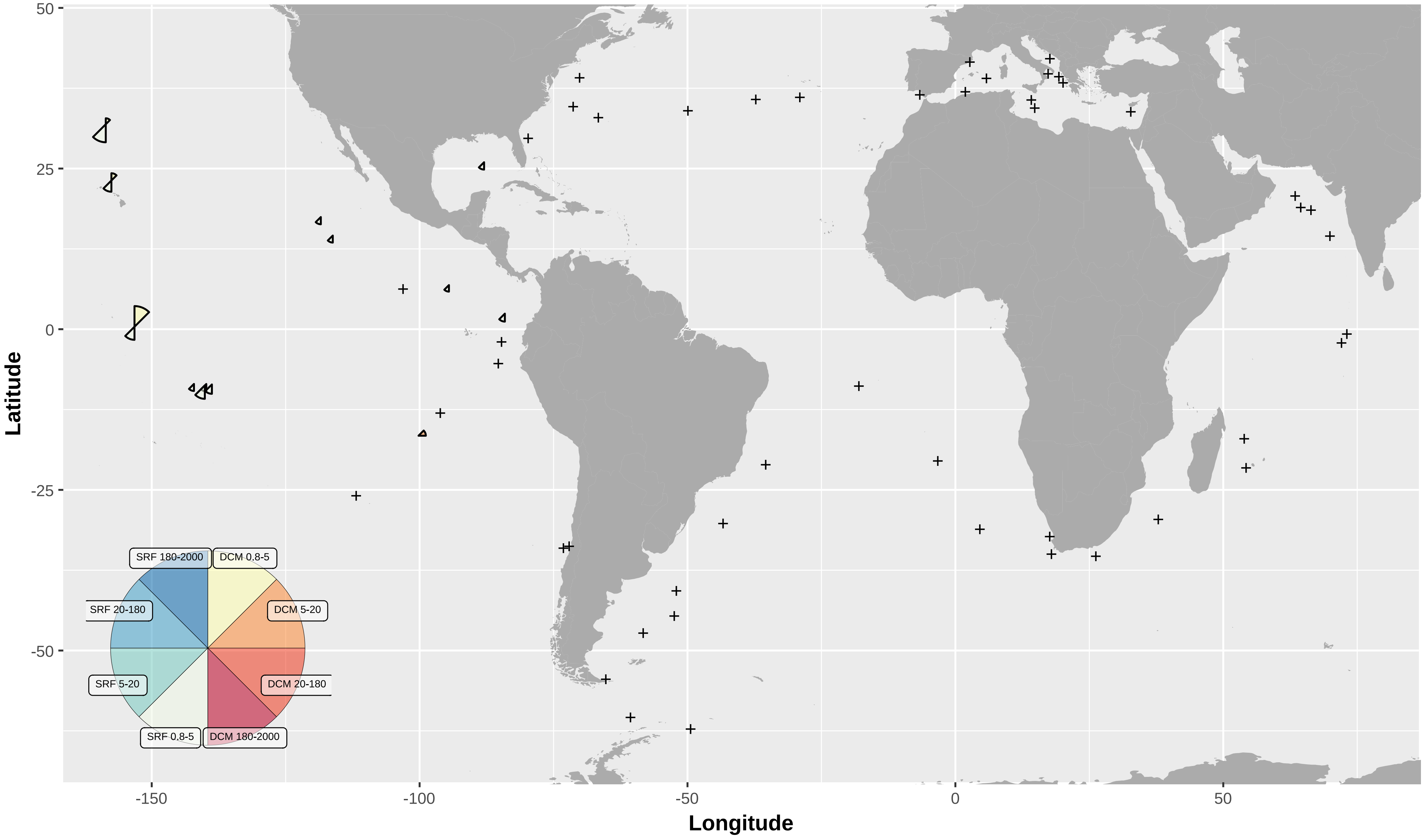


root;cellular organisms;Eukaryota

# Biogeolocalisation of MGT-v1 415      2316 unigenes, 583 (25.17%) taxonomically assigned

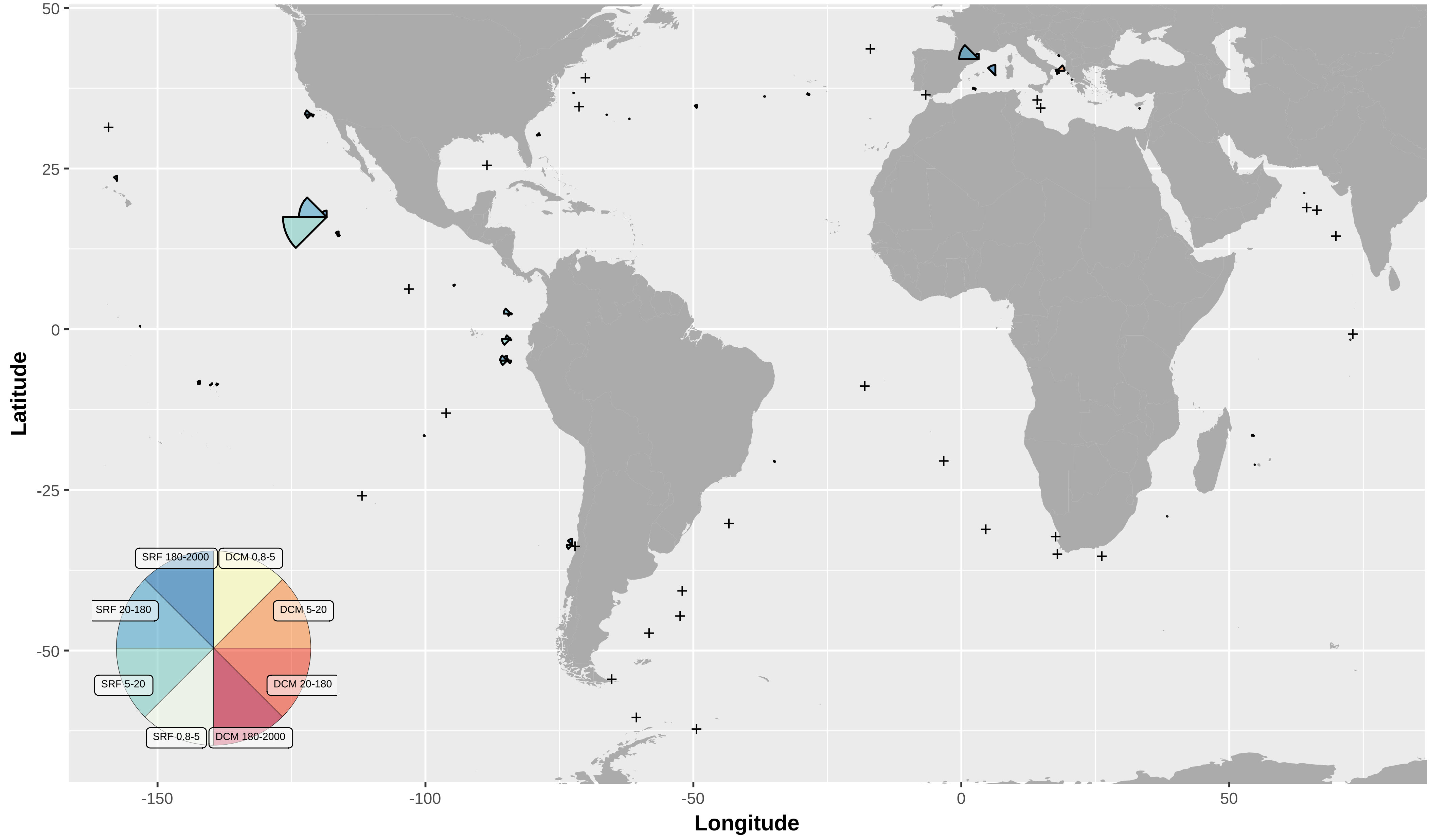


root;cellular organisms;Eukaryota;Opisthokonta;Metazoa;Eumetazoa;Bilateria

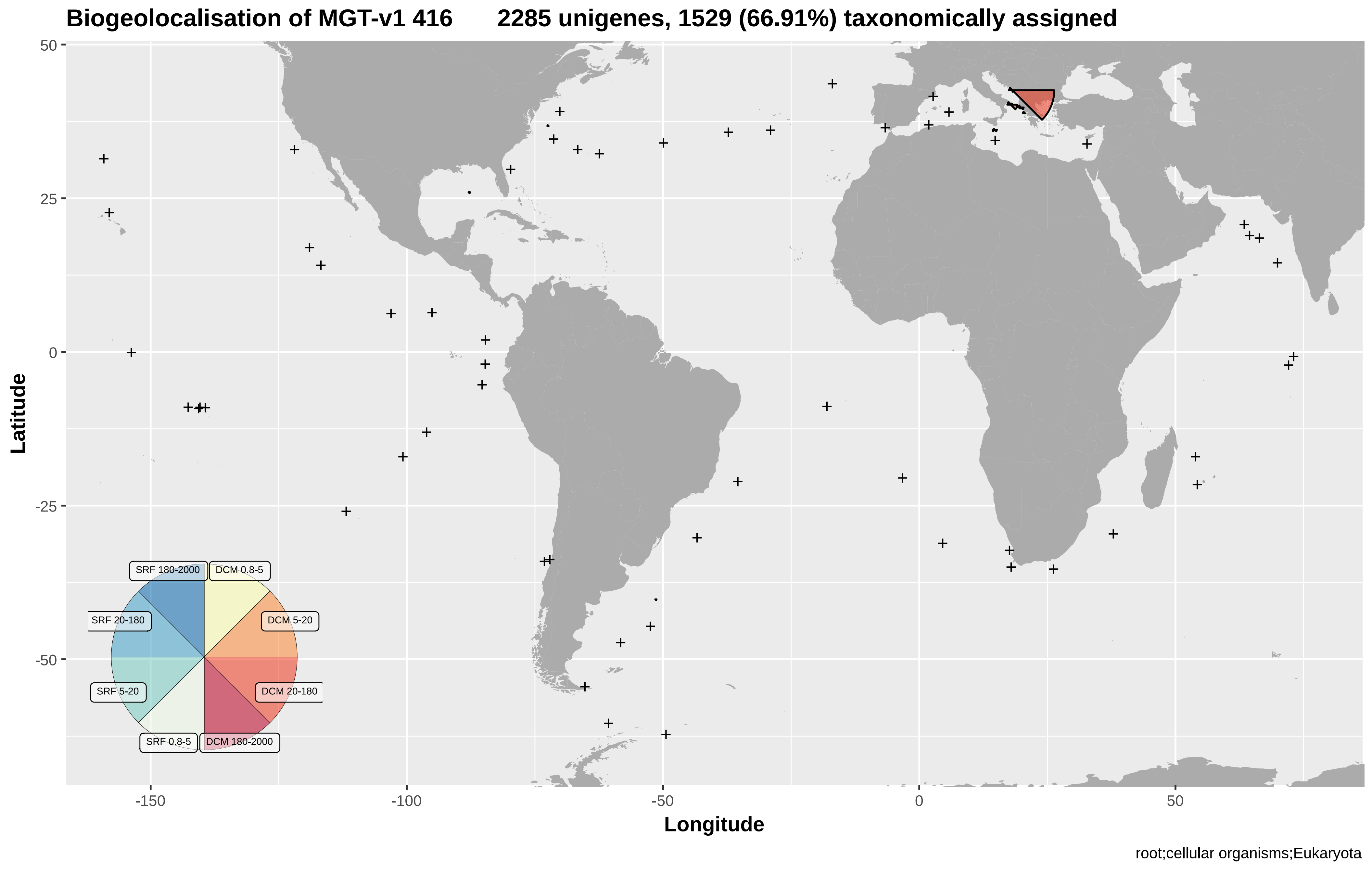

Biogeolocalisation of MGT-v1 416 2285 unigenes, 1529 (66.91%) taxonomically assigned
Latitude
Longitude
50
25
0
-25
-50
-150
-100
-50
0
50
SRF 180-2000
DCM 0.8-5
SRF 20-180
DCM 5-20
SRF 5-20
DCM 20-180
SRF 0.8-5
DCM 180-2000
root;cellular organisms;Eukaryota

# Biogeolocalisation of MGT-v1 417      2271 unigenes, 629 (27.7%) taxonomically assigned

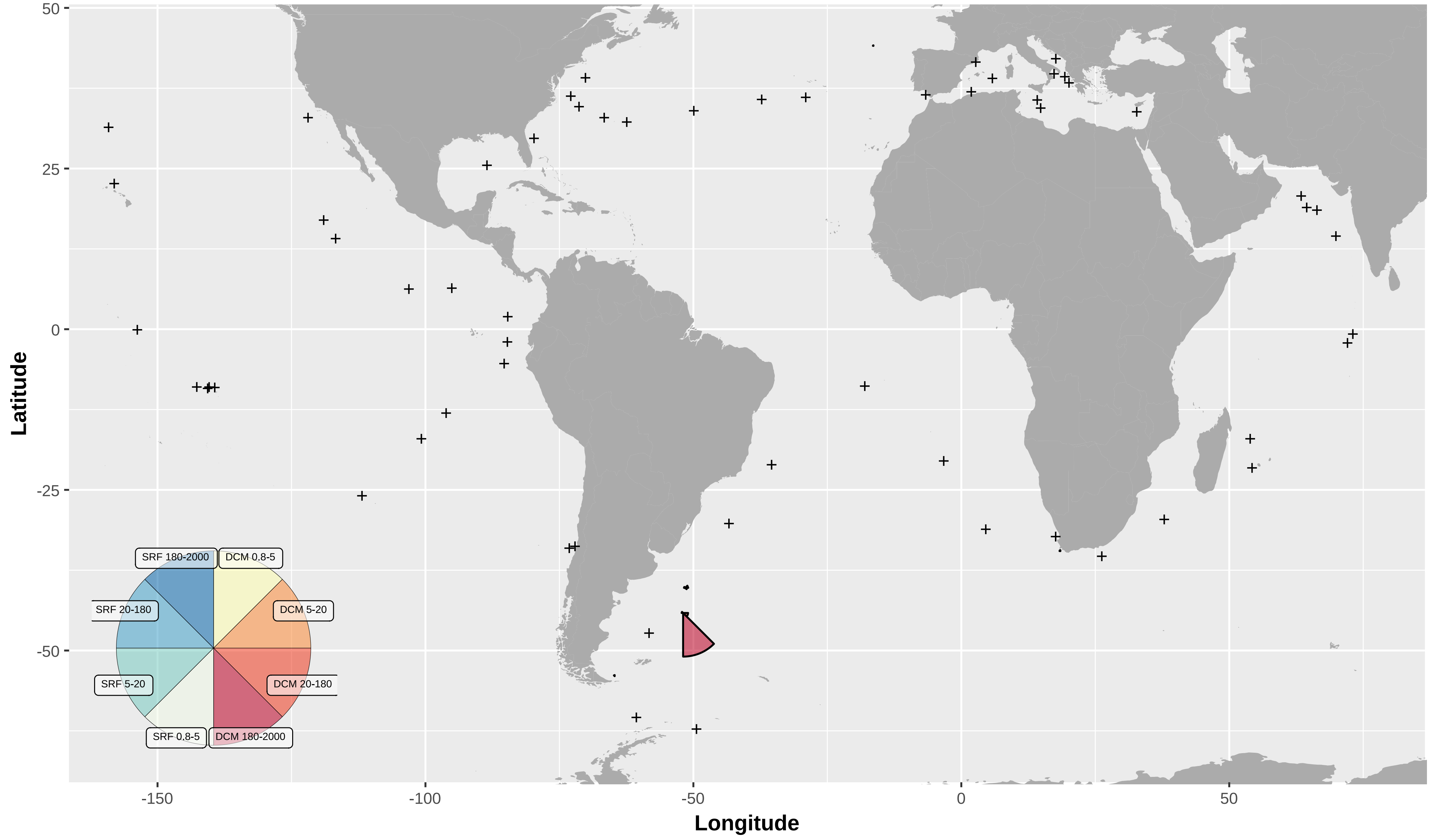


root;cellular organisms;Eukaryota;Opisthokonta;Metazoa;Eumetazoa;Bilateria;Protostomia;Ecdysozoa;Panarthropoda;Arthropoda;Mandibulata;Pancrustacea

## Biogeolocalisation of MGT-v1 418      2261 unigenes, 267 (11.81%) taxonomically assigned

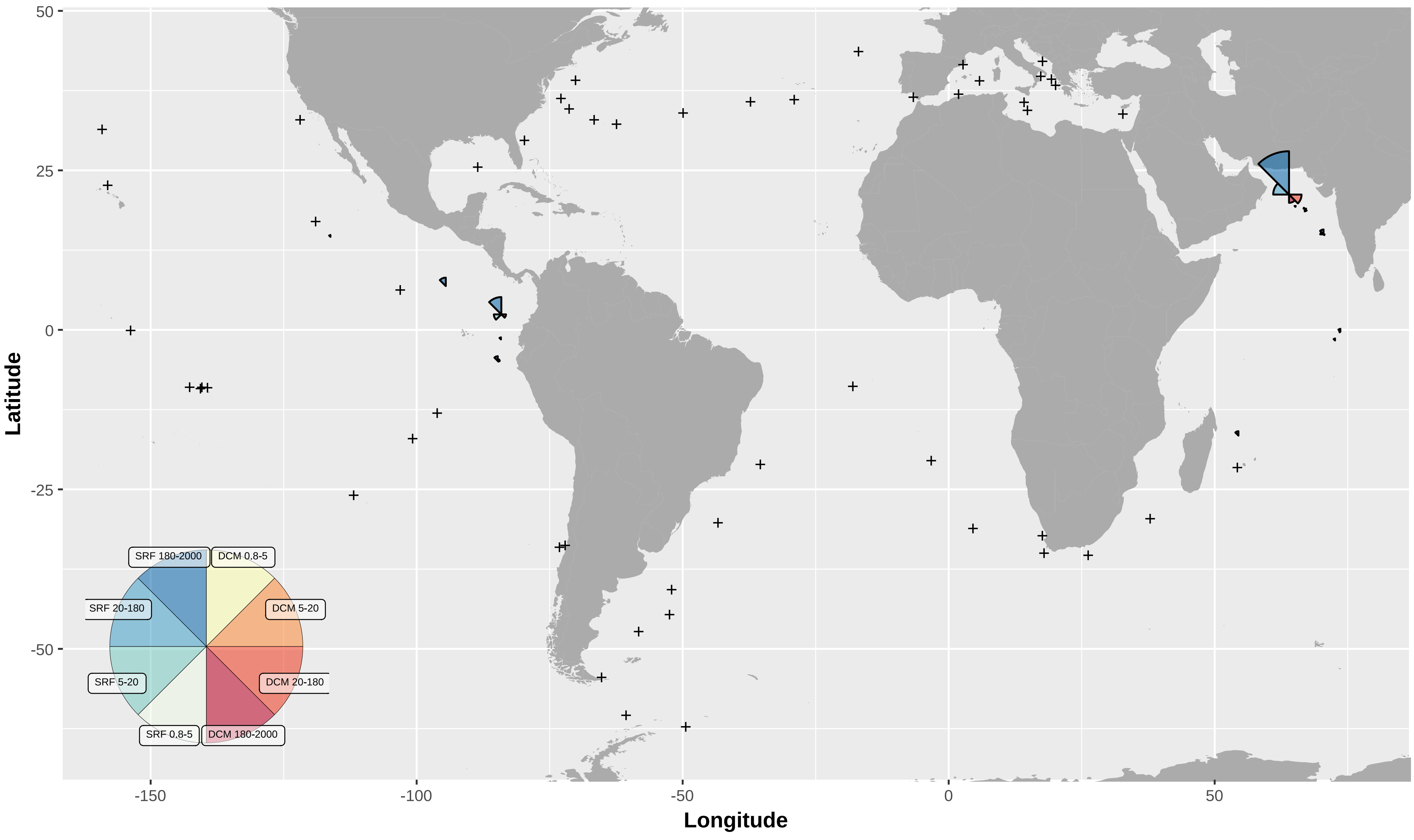


root;cellular organisms;Eukaryota;Opisthokonta;Metazoa;Eumetazoa;Bilateria

# Biogeolocalisation of MGT-v1 419      2252 unigenes, 480 (21.31%) taxonomically assigned

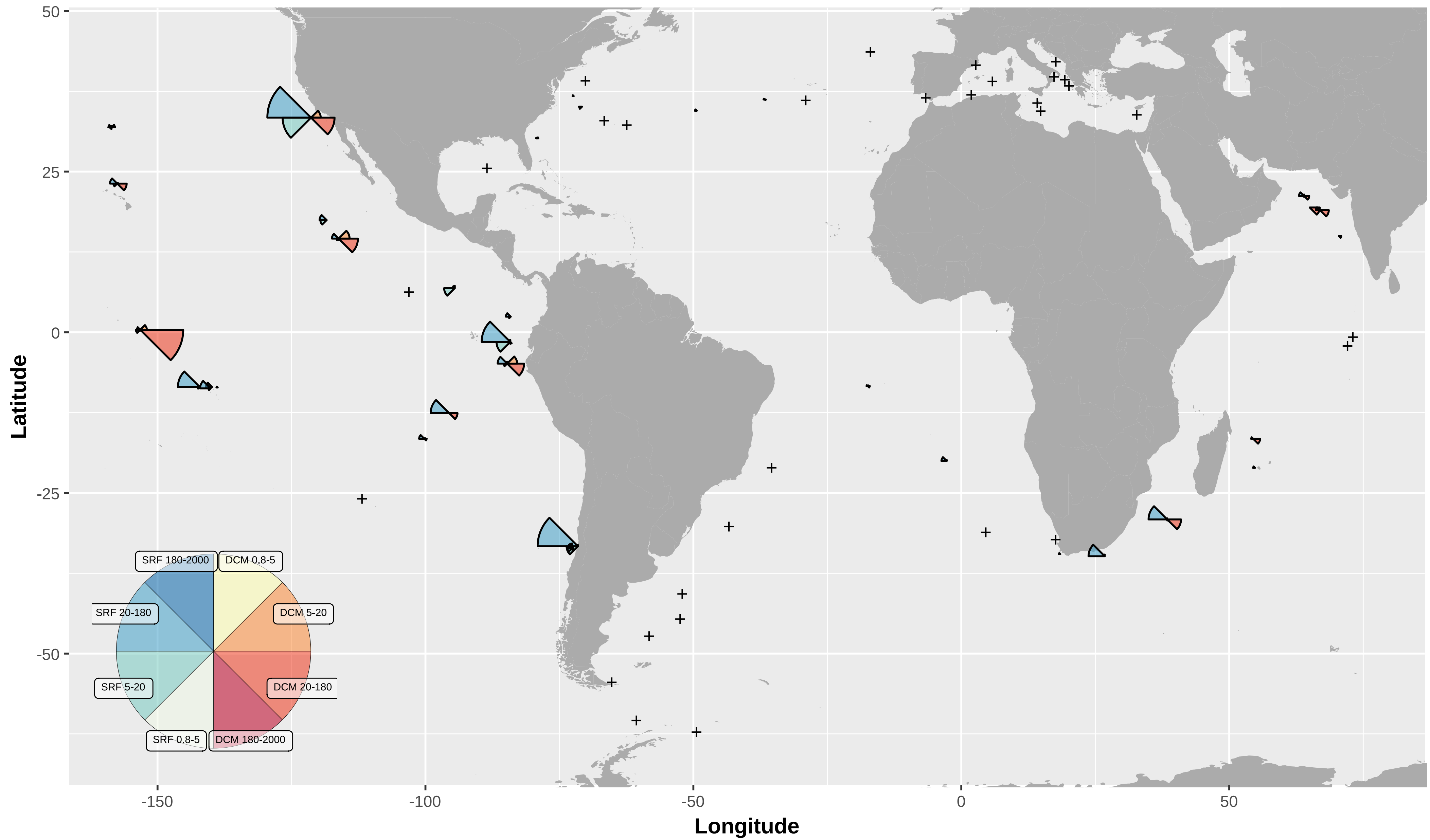


root;cellular organisms;Eukaryota;Opisthokonta;Metazoa;Eumetazoa;Bilateria;Protostomia;Ecdysozoa;Panarthropoda;Arthropoda;Mandibulata;Pancrustacea;Crustacea;Maxillopoda

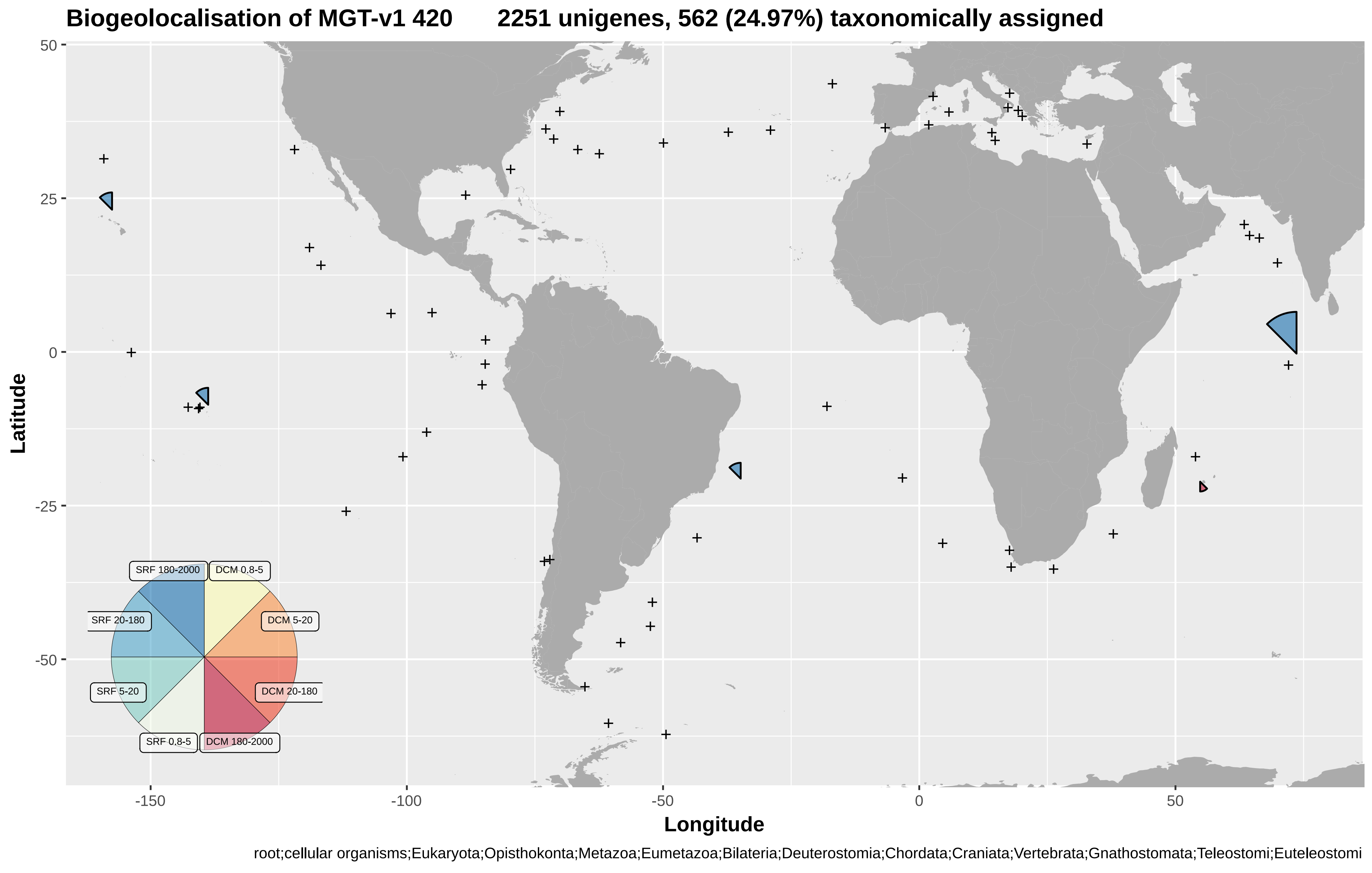
Biogeolocalisation of MGT-v1 420 2251 unigenes, 562 (24.97%) taxonomically assigned
50
25
0
-25
-50
Latitude
-150
-100
-50
0
50
Longitude
SRF 180-2000
DCM 0.8-5
SRF 20-180
DCM 5-20
SRF 5-20
DCM 20-180
SRF 0.8-5
DCM 180-2000
root;cellular organisms;Eukaryota;Opisthokonta;Metazoa;Eumetazoa;Bilateria;Deuterostomia;Chordata;Craniata;Vertebrata;Gnathostomata;Teleostomi;Euteleostomi

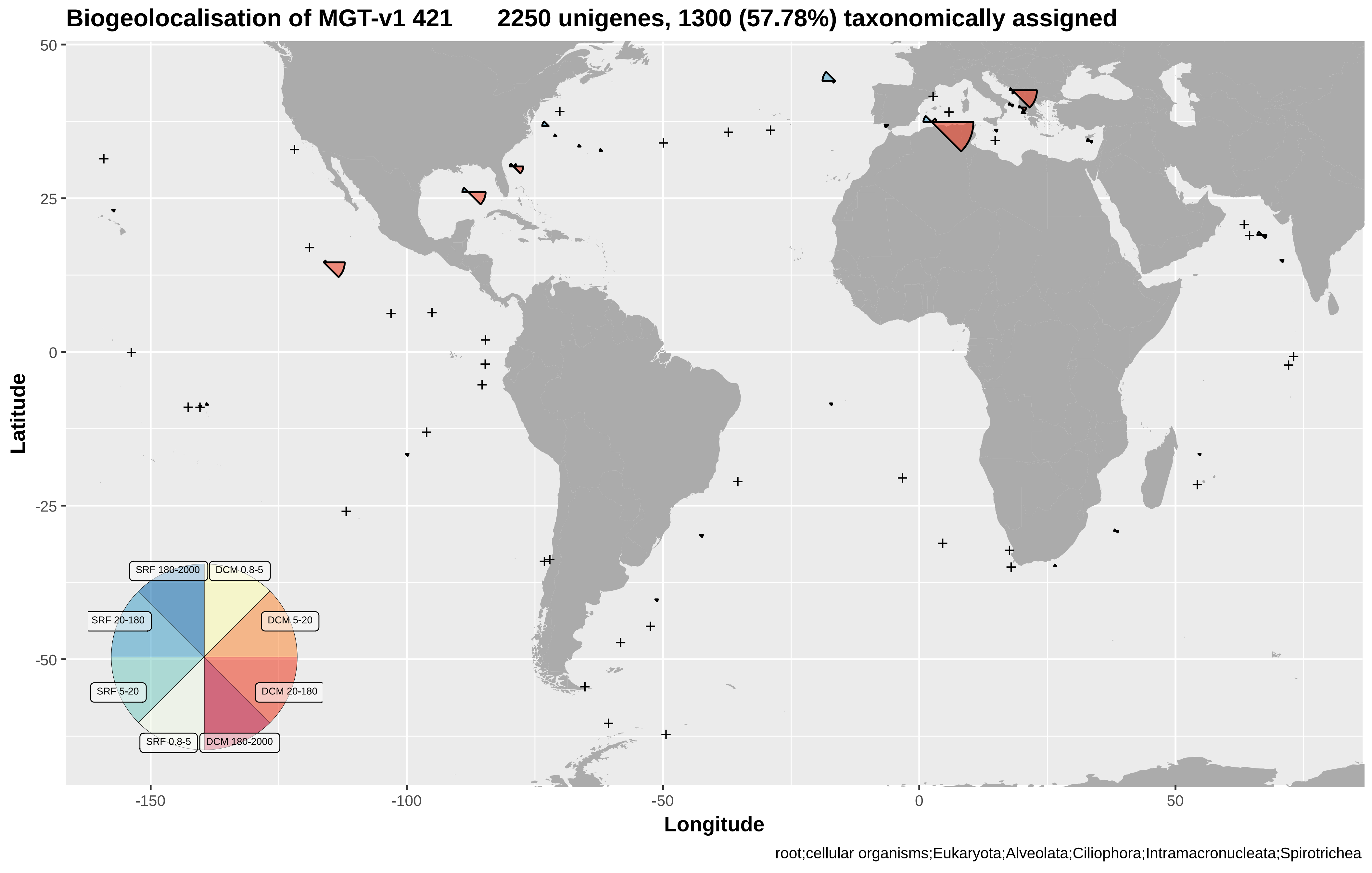
Biogeolocalisation of MGT-v1 421
2250 unigenes, 1300 (57.78%) taxonomically assigned
Latitude
Longitude
50
25
0
-25
-50
-150
-100
-50
0
50
SRF 180-2000
DCM 0.8-5
SRF 20-180
DCM 5-20
SRF 5-20
DCM 20-180
SRF 0.8-5
DCM 180-2000
root;cellular organisms;Eukaryota;Alveolata;Ciliophora;Intramacronucleata;Spirotrichea

#### Biogeolocalisation of MGT-v1 422 2247 unigenes, 144 (6.41%) taxonomically assigned

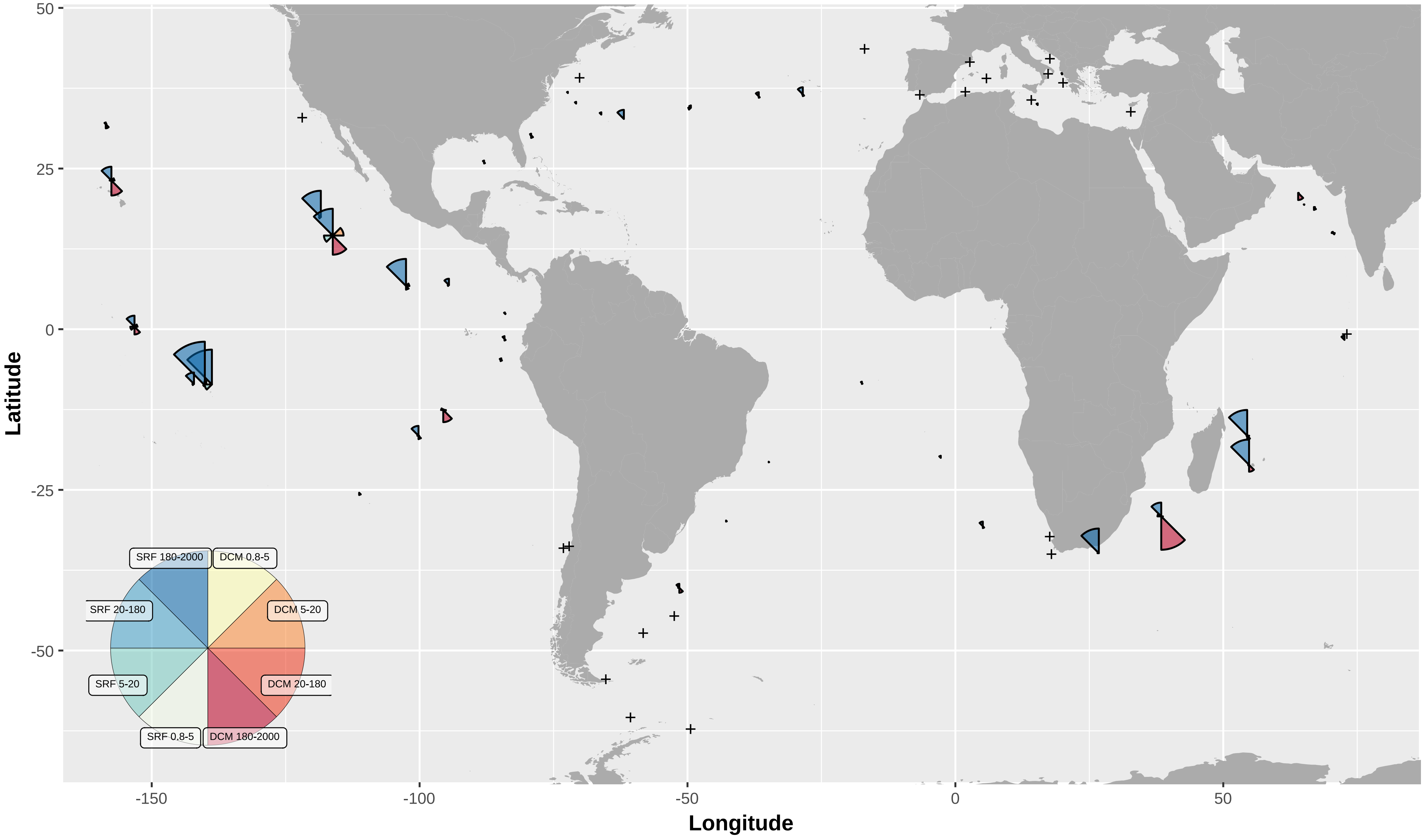


root;cellular organisms;Eukaryota;Opisthokonta;Metazoa;Eumetazoa;Bilateria

Biogeolocalisation of MGT-v1 423      2239 unigenes, 110 (4.91%) taxonomically assigned

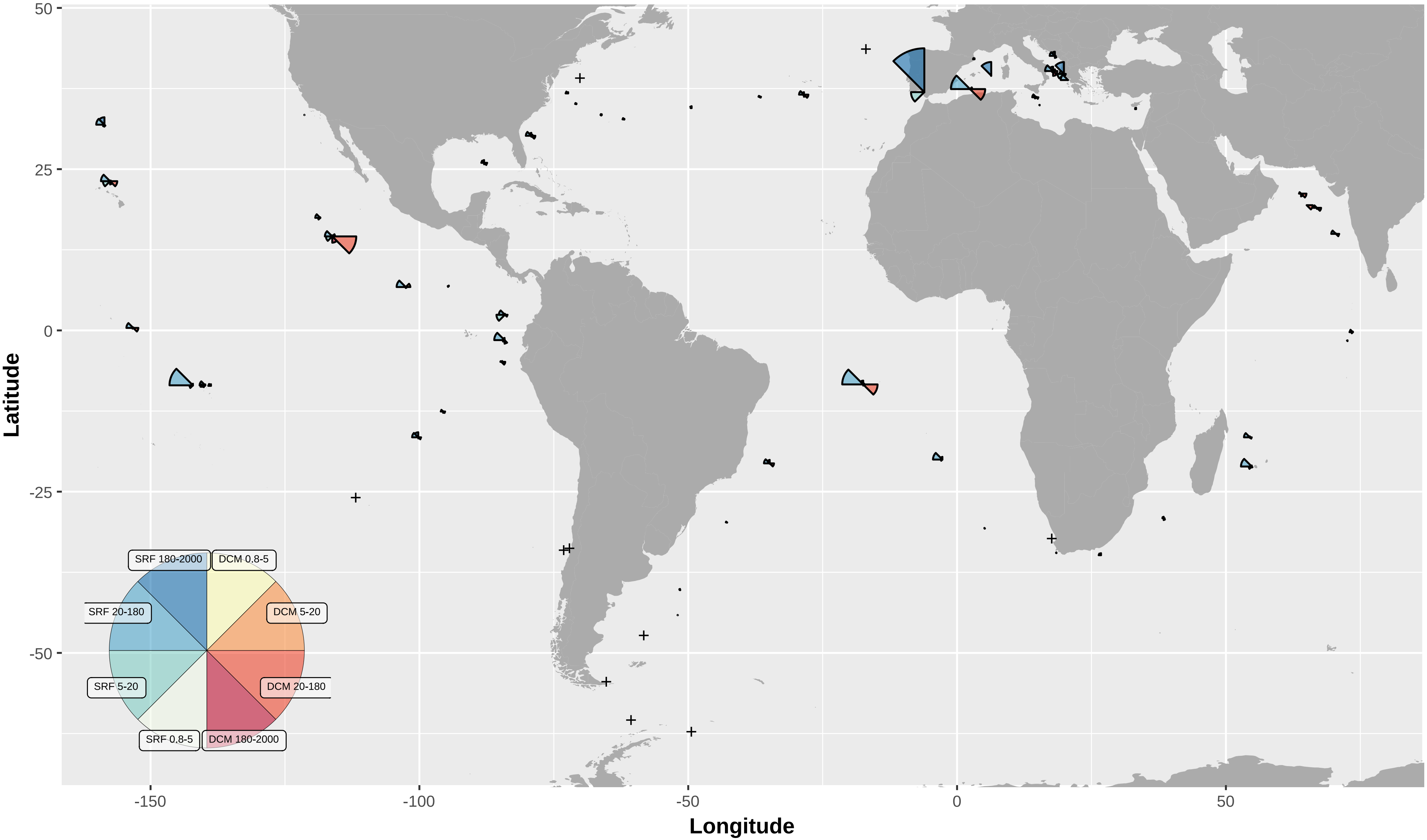


root;cellular organisms;Eukaryota;Opisthokonta;Metazoa;Eumetazoa

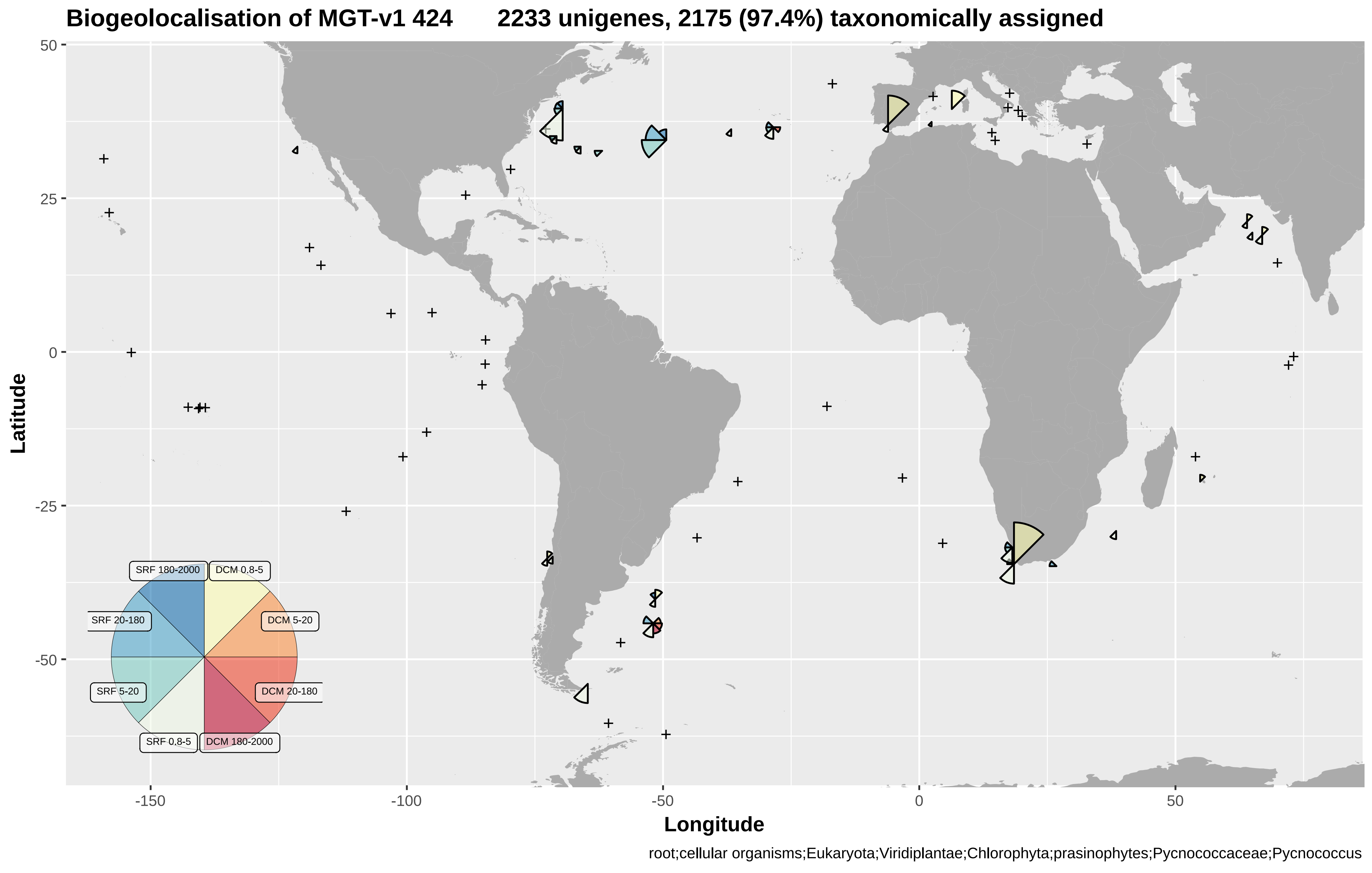

Biogeolocalisation of MGT-v1 424 2233 unigenes, 2175 (97.4%) taxonomically assigned
50
25
0
-25
-50
Latitude
-150
-100
-50
0
50
Longitude
SRF 180-2000
DCM 0.8-5
SRF 20-180
DCM 5-20
SRF 5-20
DCM 20-180
SRF 0.8-5
DCM 180-2000
root;cellular organisms;Eukaryota;Viridiplantae;Chlorophyta;prasinophytes;Pycnococcaceae;Pycnococcus

#### Biogeolocalisation of MGT-v1 425 2226 unigenes, 2146 (96.41%) taxonomically assigned

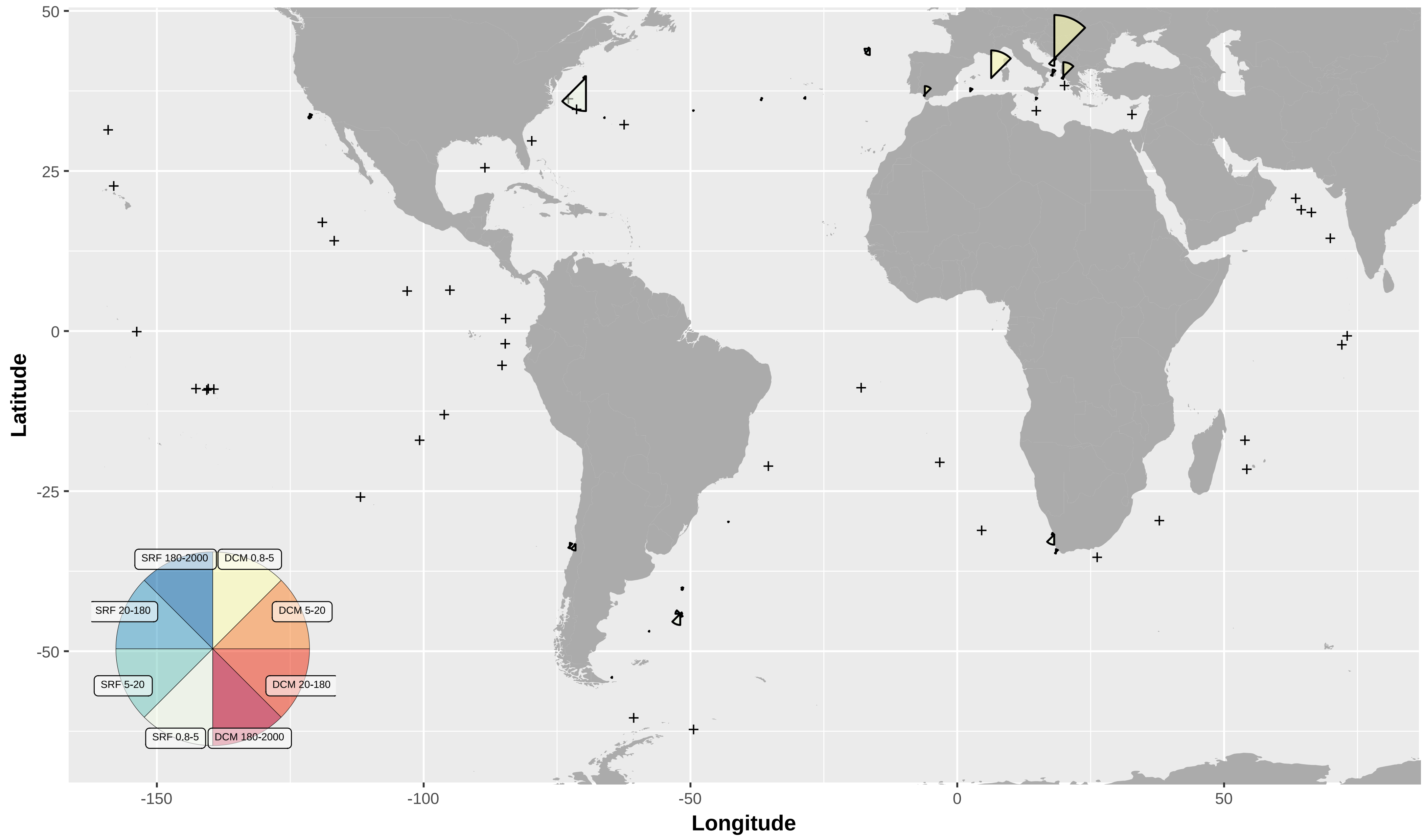


root;cellular organisms;Bacteria;Cyanobacteria;Oscillatoriophycideae;Chroococcales;Synechococcus

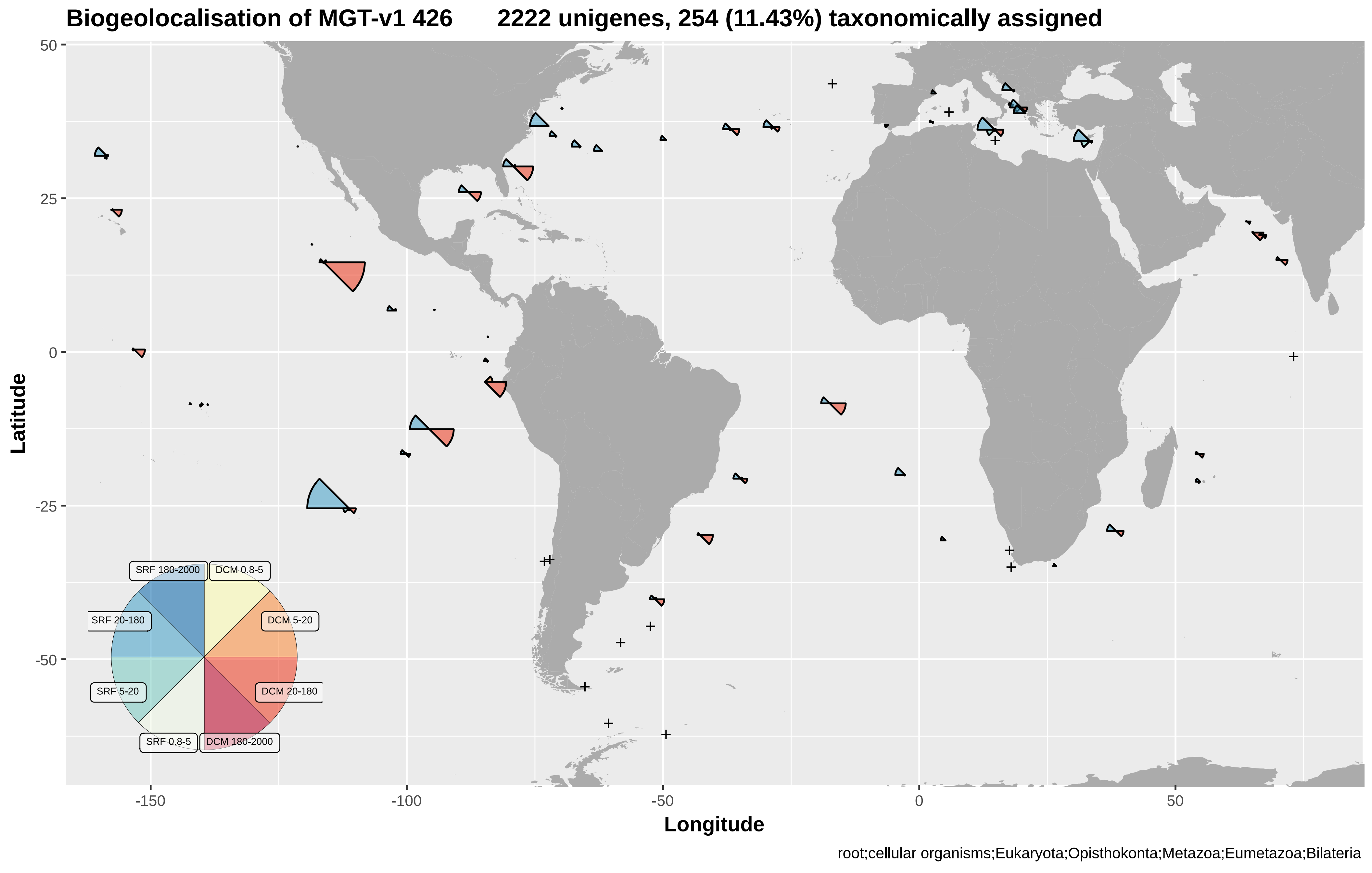

Biogeolocalisation of MGT-v1 426 2222 unigenes, 254 (11.43%) taxonomically assigned
Latitude
50
25
0
-25
-50
-150
-100
-50
0
50
Longitude
SRF 180-2000
DCM 0.8-5
SRF 20-180
DCM 5-20
SRF 5-20
DCM 20-180
SRF 0.8-5
DCM 180-2000
root;cellular organisms;Eukaryota;Opisthokonta;Metazoa;Eumetazoa;Bilateria

Biogeolocalisation of MGT-v1 427      2222 unigenes, 530 (23.85%) taxonomically assigned

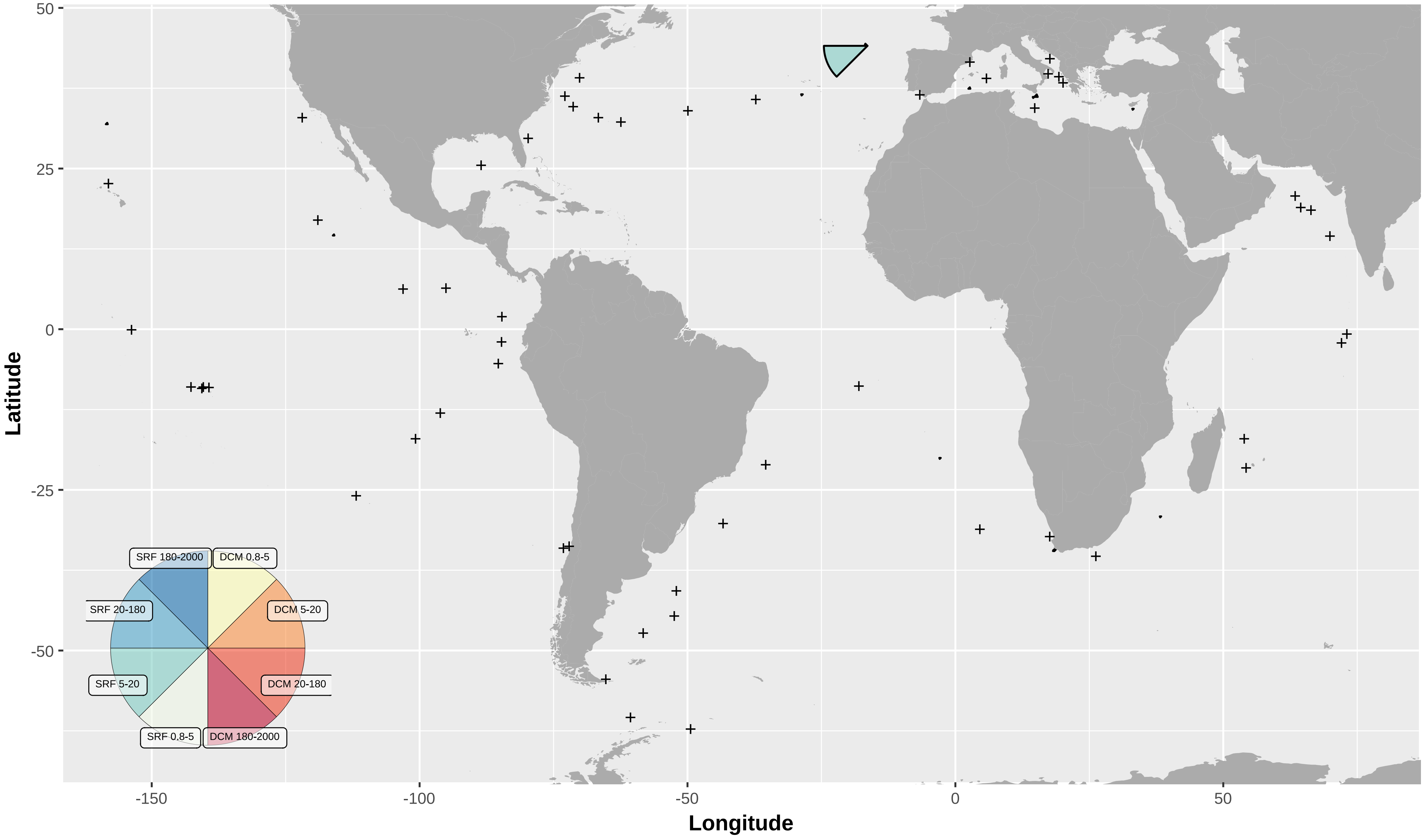


root;cellular organisms;Eukaryota

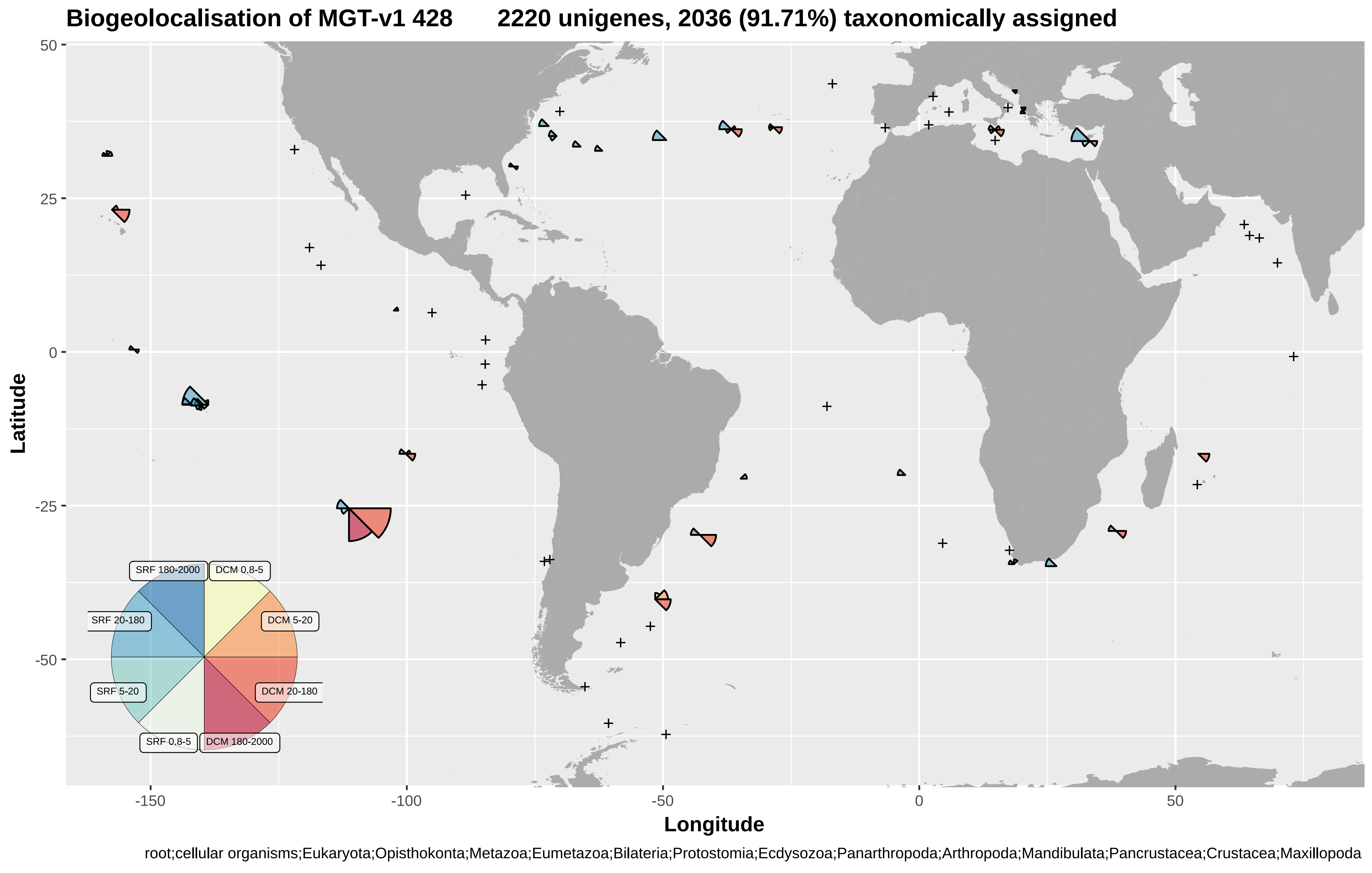

Biogeolocalisation of MGT-v1 428
2220 unigenes, 2036 (91.71%) taxonomically assigned
Latitude
50
25
0
-25
-50
Longitude
-150
-100
-50
0
50
SRF 180-2000
DCM 0.8-5
SRF 20-180
DCM 5-20
SRF 5-20
DCM 20-180
SRF 0.8-5
DCM 180-2000
root;cellular organisms;Eukaryota;Opisthokonta;Metazoa;Eumetazoa;Bilateria;Protostomia;Ecdysozoa;Panarthropoda;Arthropoda;Mandibulata;Pancrustacea;Crustacea;Maxillopoda

# Biogeolocalisation of MGT-v1 429      2208 unigenes, 255 (11.55%) taxonomically assigned

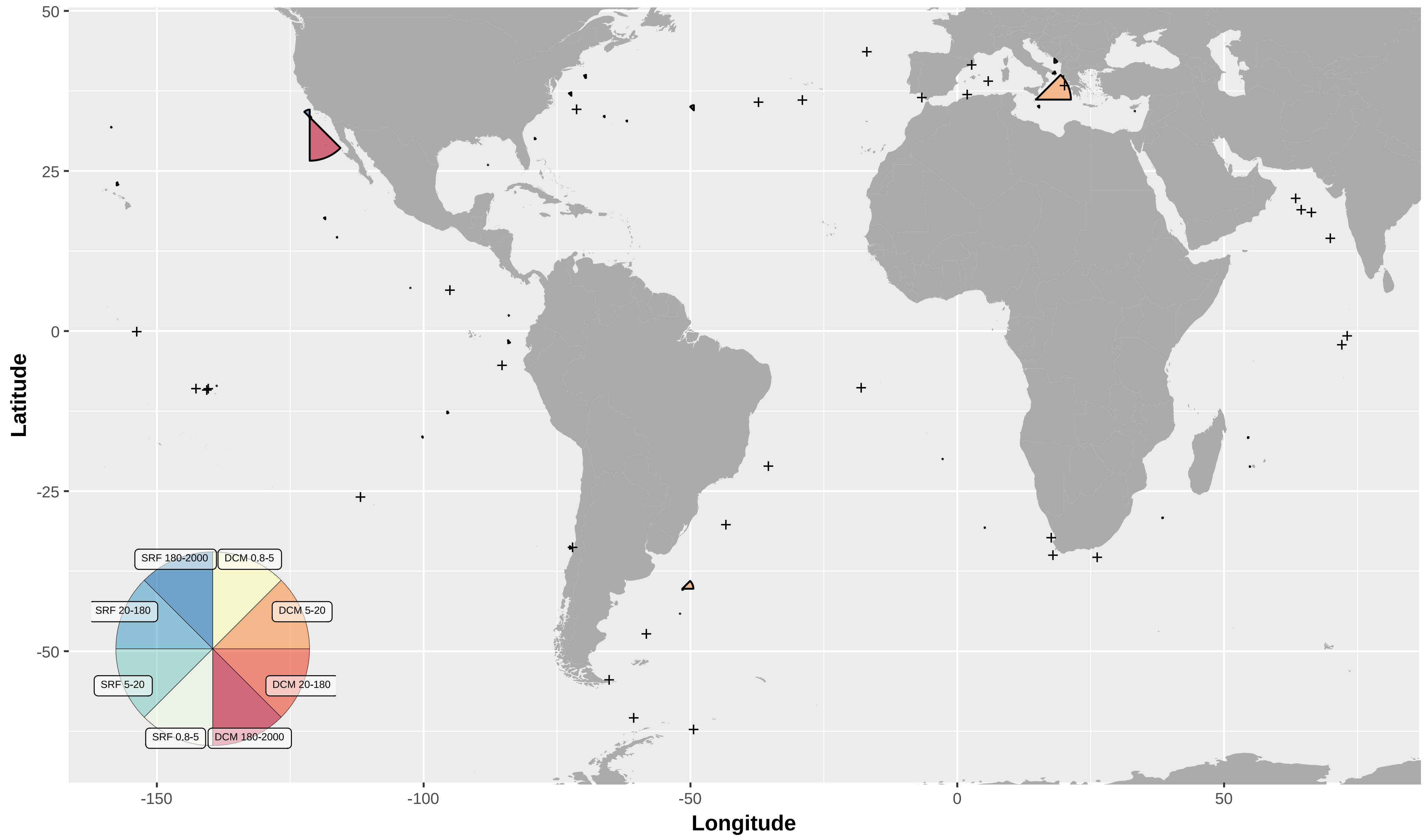


root;cellular organisms;Eukaryota;Opisthokonta;Metazoa;Eumetazoa

## Biogeolocalisation of MGT-v1 430      2203 unigenes, 1841 (83.57%) taxonomically assigned

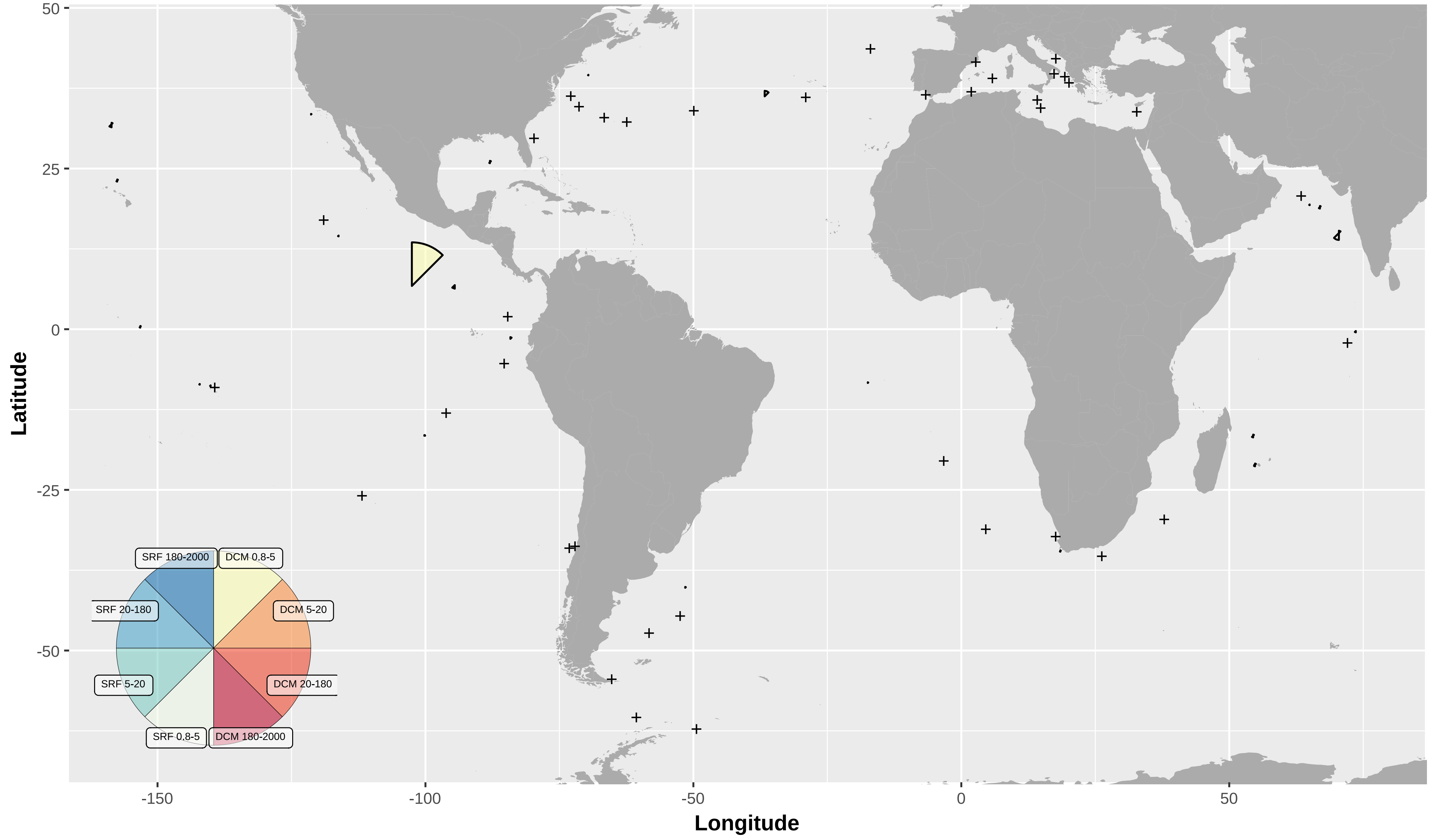


root;cellular organisms;Bacteria;Proteobacteria;Gammaproteobacteria

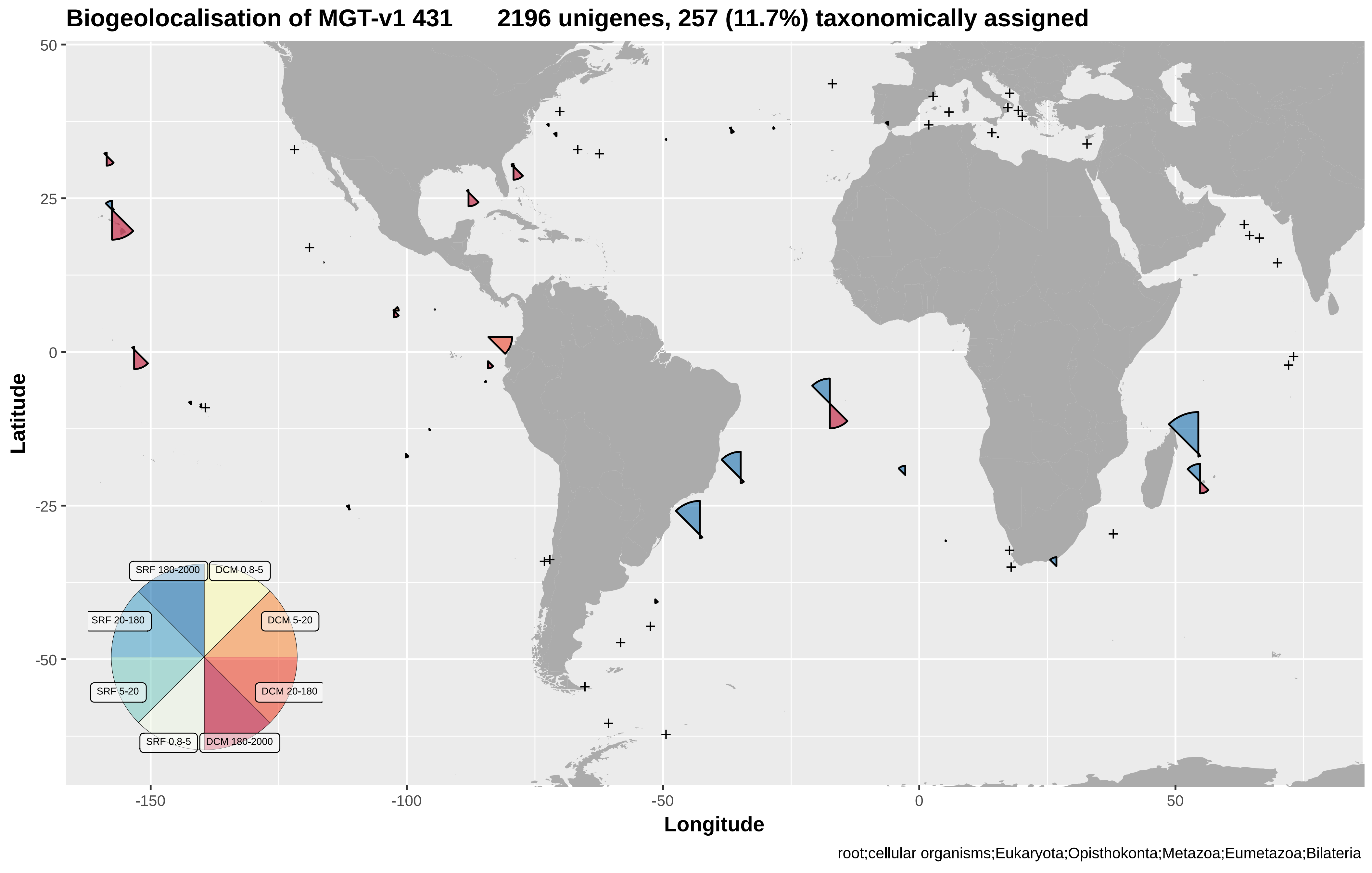

Biogeolocalisation of MGT-v1 431 2196 unigenes, 257 (11.7%) taxonomically assigned
50
25
0
-25
-50
Latitude
-150
-100
-50
0
50
Longitude
SRF 180-2000
DCM 0.8-5
SRF 20-180
DCM 5-20
SRF 5-20
DCM 20-180
SRF 0.8-5
DCM 180-2000
root;cellular organisms;Eukaryota;Opisthokonta;Metazoa;Eumetazoa;Bilateria

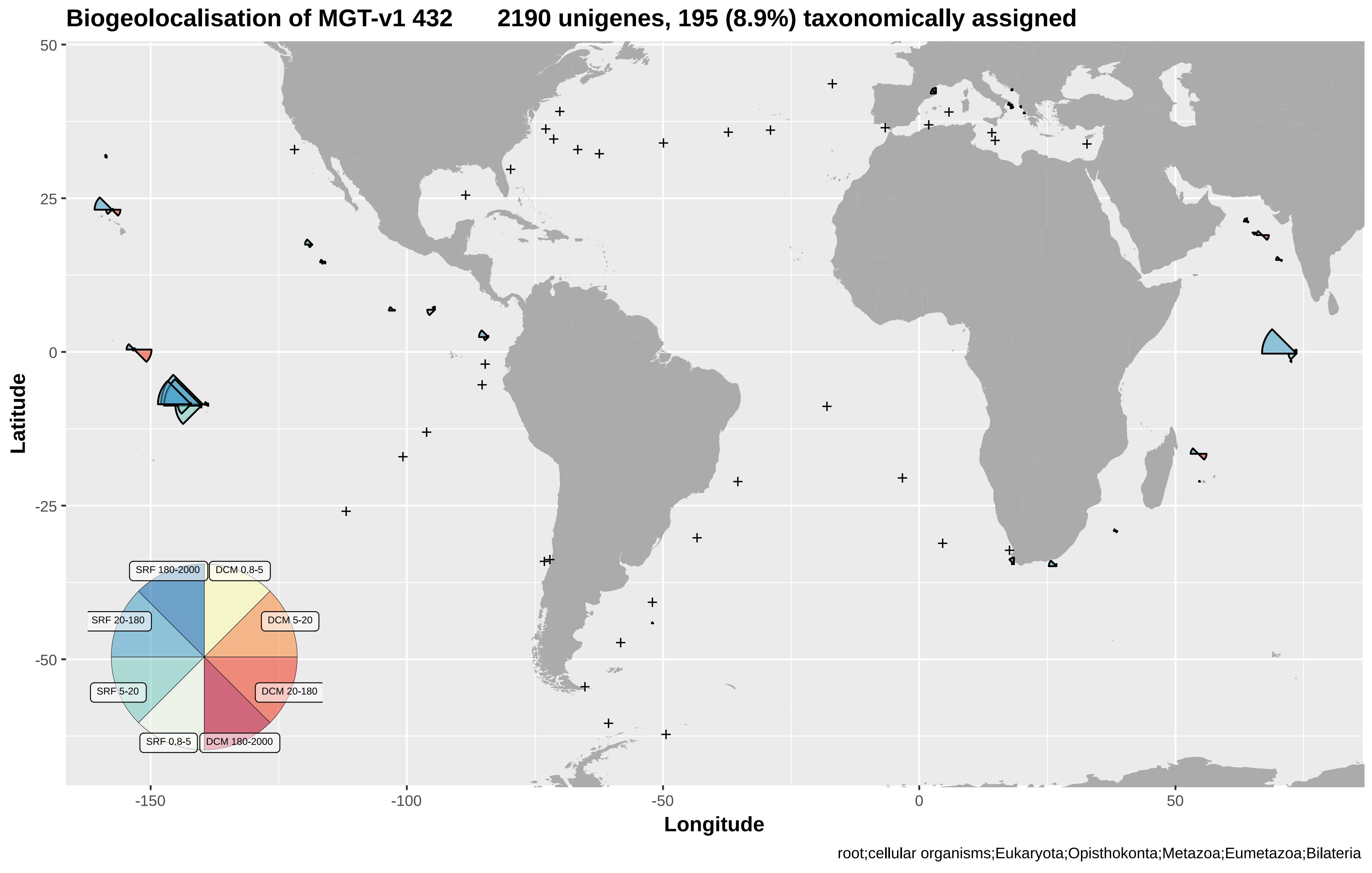
Biogeolocalisation of MGT-v1 432 2190 unigenes, 195 (8.9%) taxonomically assigned
50
25
0
-25
-50
Latitude
-150
-100
-50
0
50
Longitude
SRF 180-2000
DCM 0.8-5
SRF 20-180
DCM 5-20
SRF 5-20
DCM 20-180
SRF 0.8-5
DCM 180-2000
root;cellular organisms;Eukaryota;Opisthokonta;Metazoa;Eumetazoa;Bilateria

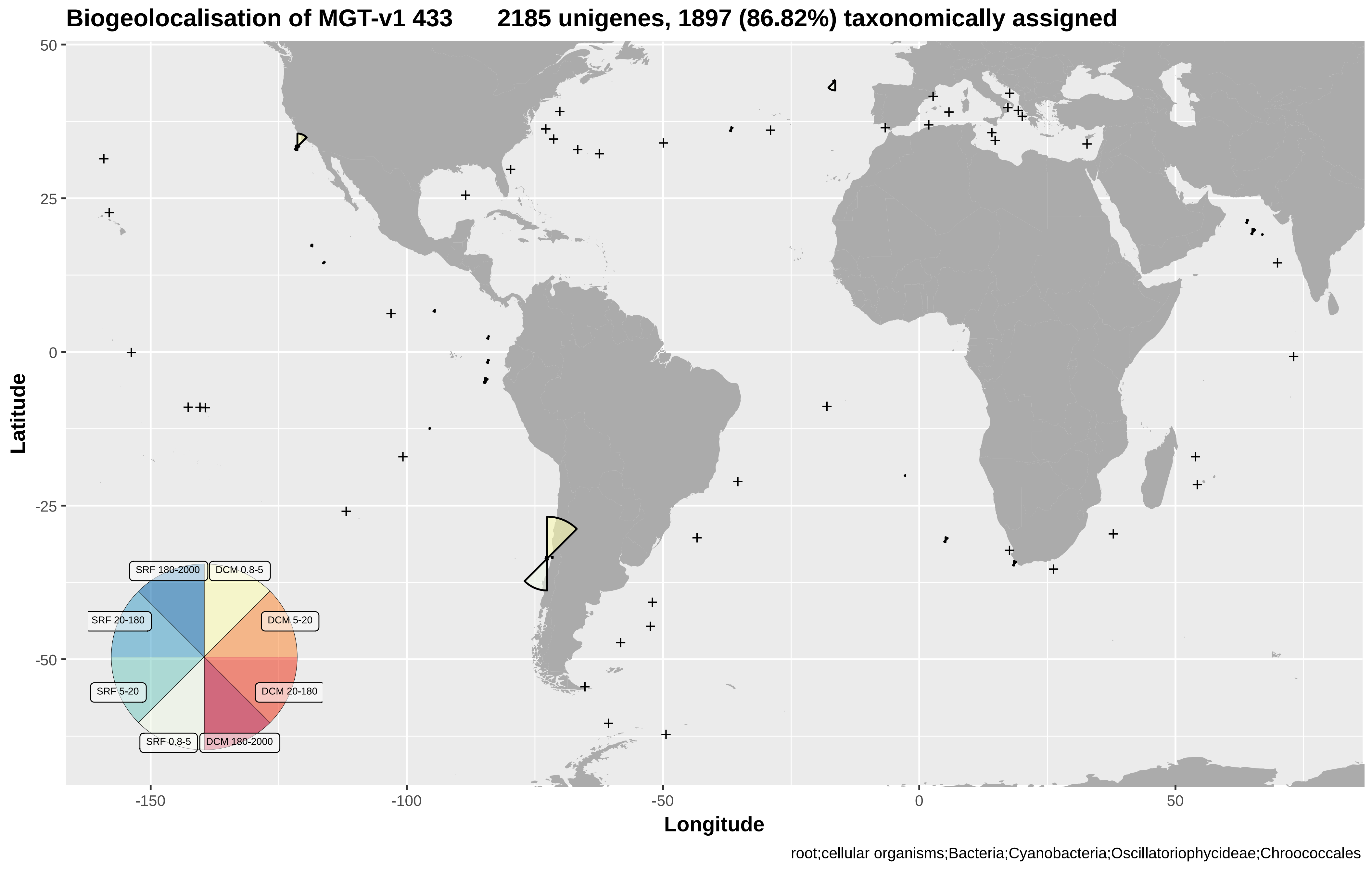

Biogeolocalisation of MGT-v1 433 2185 unigenes, 1897 (86.82%) taxonomically assigned
Latitude
Longitude
50
25
0
-25
-50
-150
-100
-50
0
50
SRF 180-2000
DCM 0.8-5
SRF 20-180
DCM 5-20
SRF 5-20
DCM 20-180
SRF 0.8-5
DCM 180-2000
root;cellular organisms;Bacteria;Cyanobacteria;Oscillatoriophycideae;Chroococcales

## Biogeolocalisation of MGT-v1 434      2149 unigenes, 241 (11.21%) taxonomically assigned

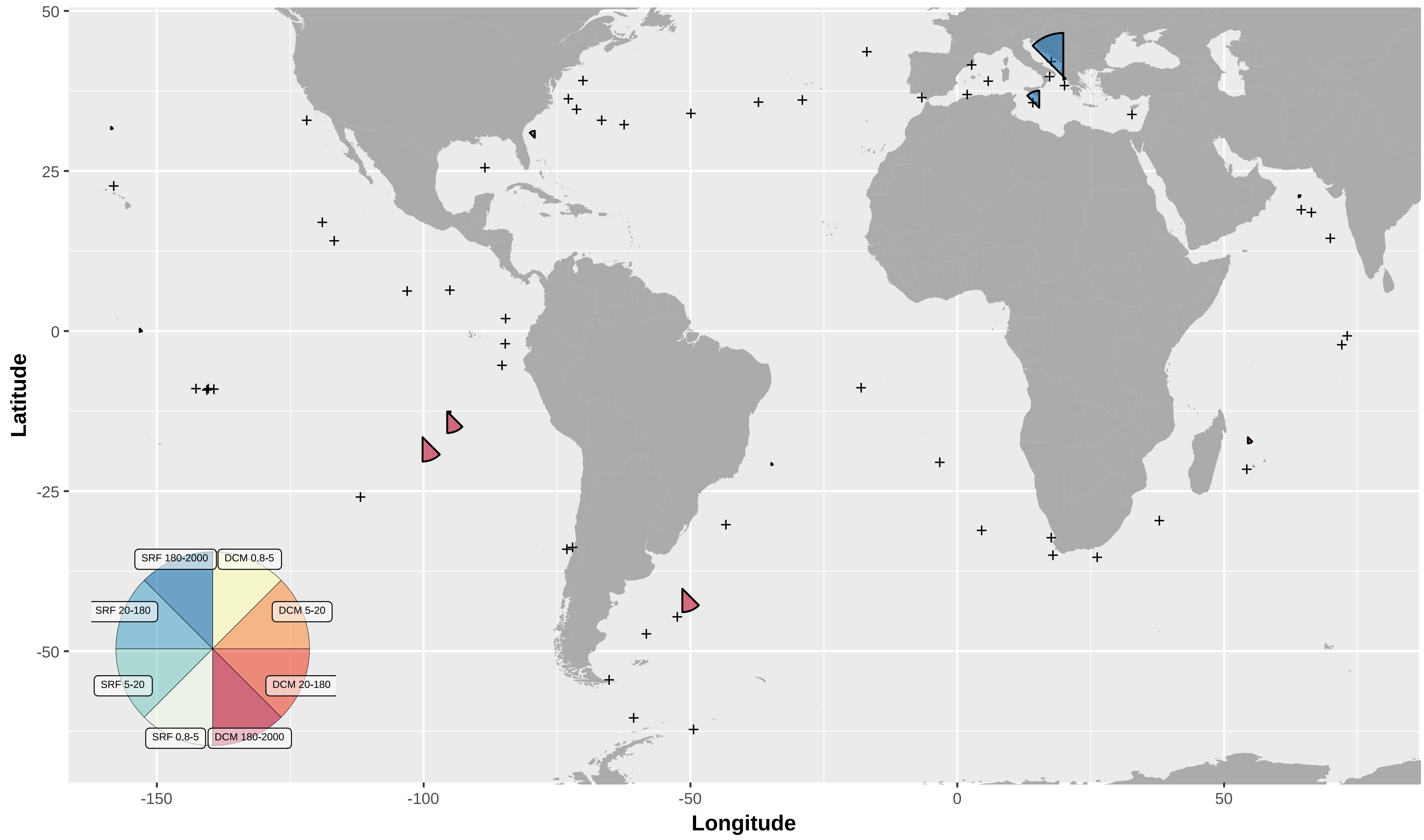


root;cellular organisms;Eukaryota;Opisthokonta;Metazoa;Eumetazoa

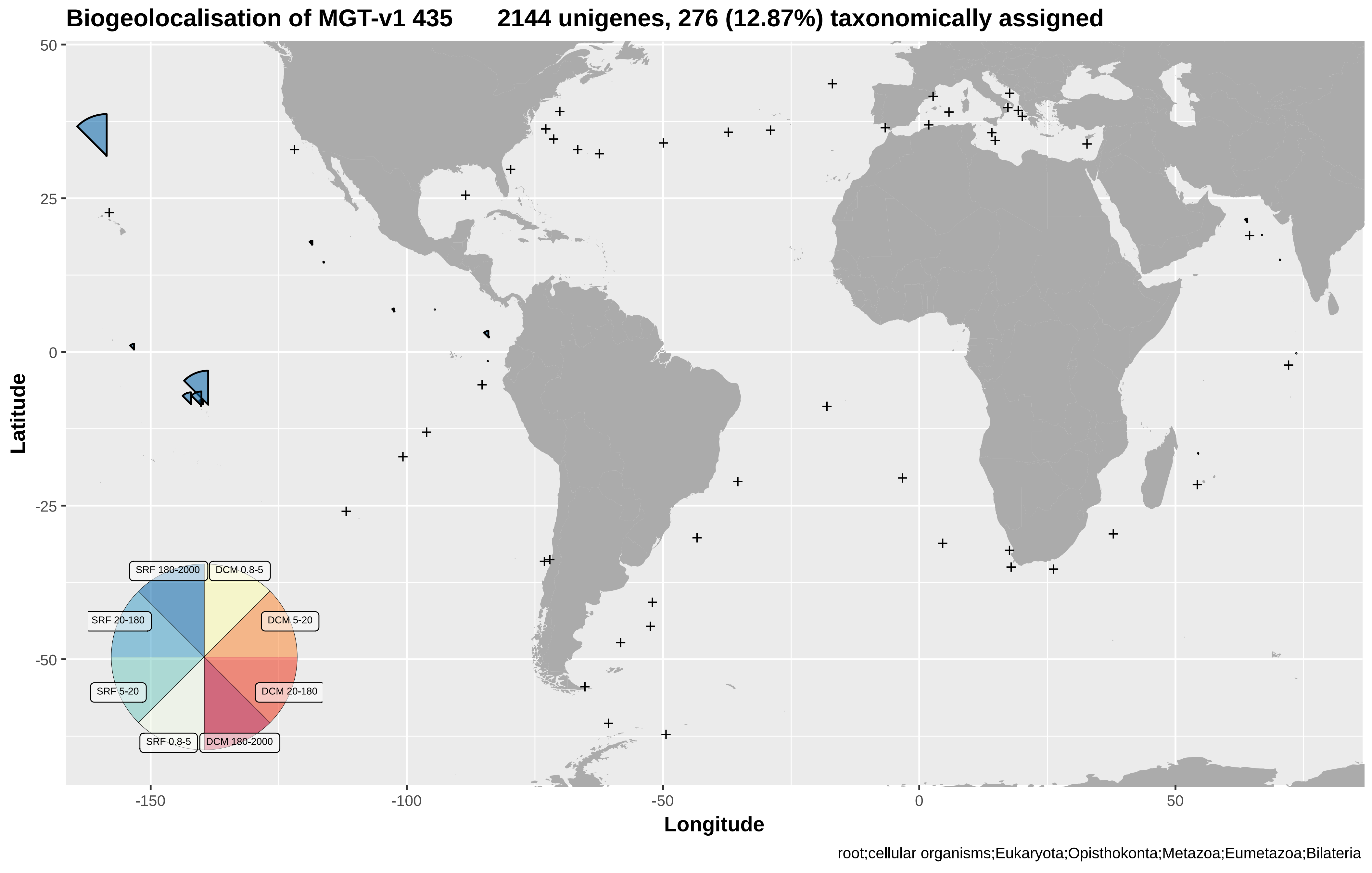

Biogeolocalisation of MGT-v1 435 2144 unigenes, 276 (12.87%) taxonomically assigned
50
25
0
-25
-50
Latitude
-150
-100
-50
0
50
Longitude
SRF 180-2000
DCM 0.8-5
SRF 20-180
DCM 5-20
SRF 5-20
DCM 20-180
SRF 0.8-5
DCM 180-2000
root;cellular organisms;Eukaryota;Opisthokonta;Metazoa;Eumetazoa;Bilateria

#### Biogeolocalisation of MGT-v1 436 2140 unigenes, 591 (27.62%) taxonomically assigned

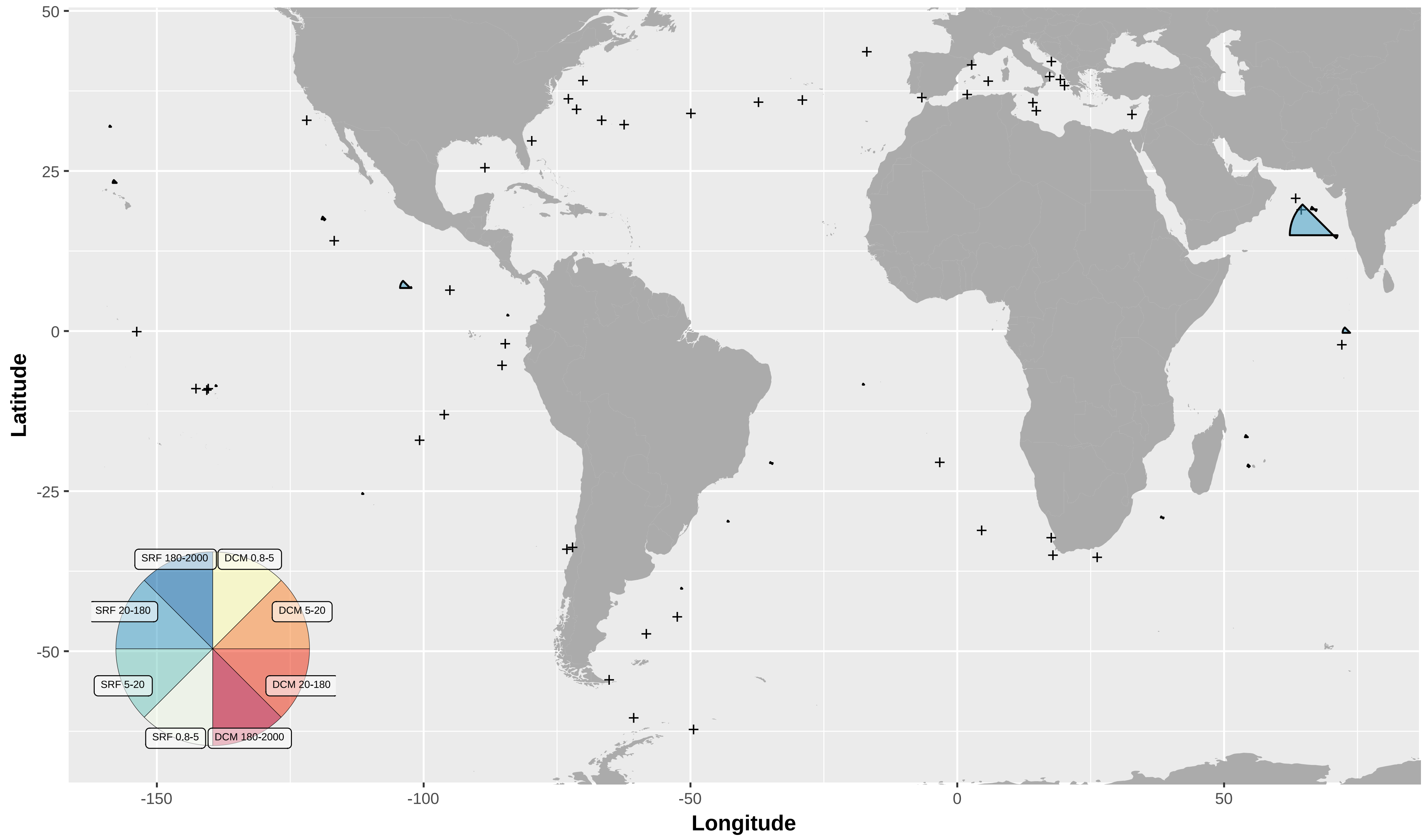


root;cellular organisms;Eukaryota;Alveolata;Ciliophora

# Biogeolocalisation of MGT-v1 437      2136 unigenes, 468 (21.91%) taxonomically assigned

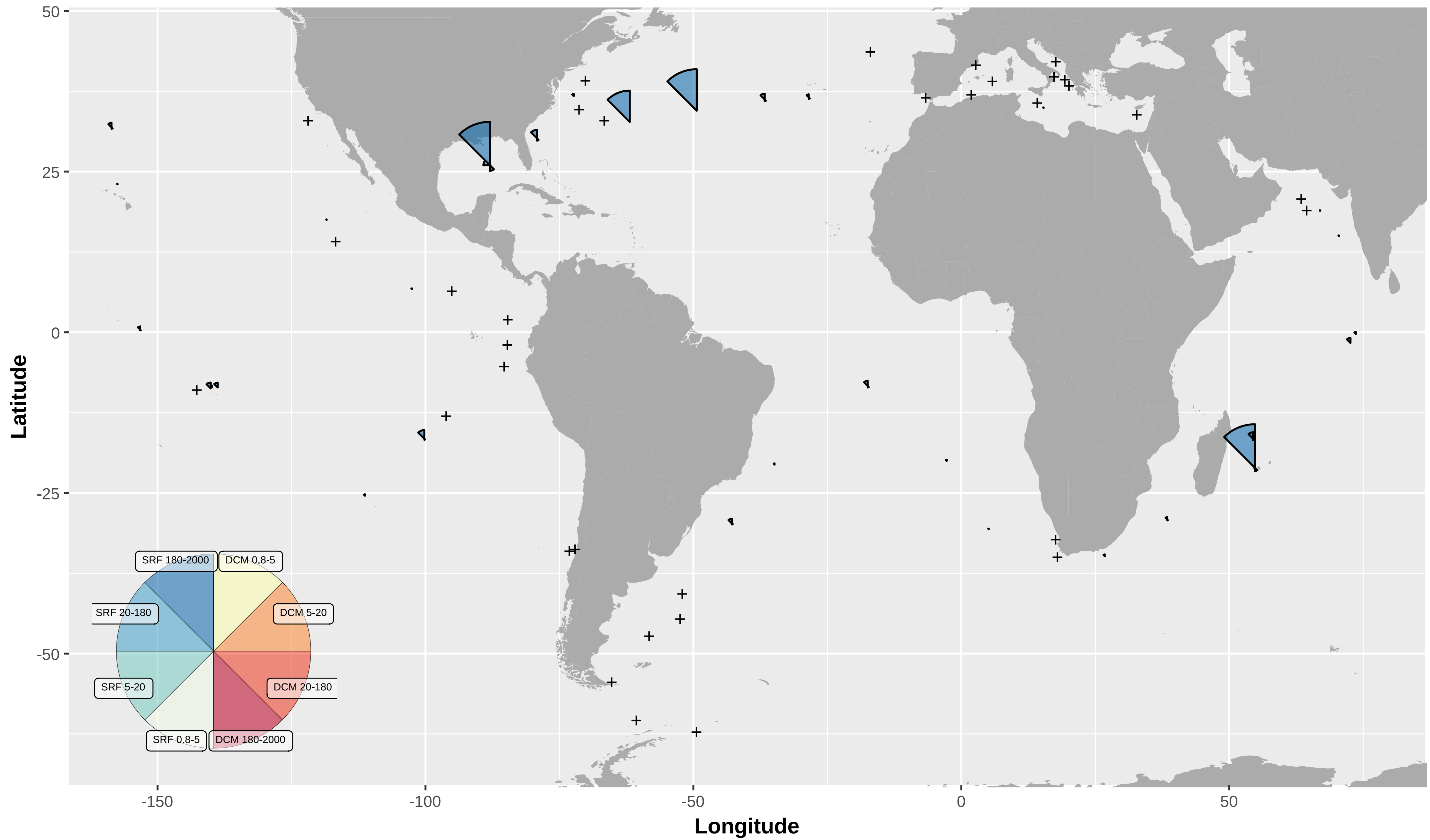


root;cellular organisms;Eukaryota;Opisthokonta;Metazoa;Eumetazoa;Bilateria;Protostomia;Ecdysozoa;Panarthropoda;Arthropoda

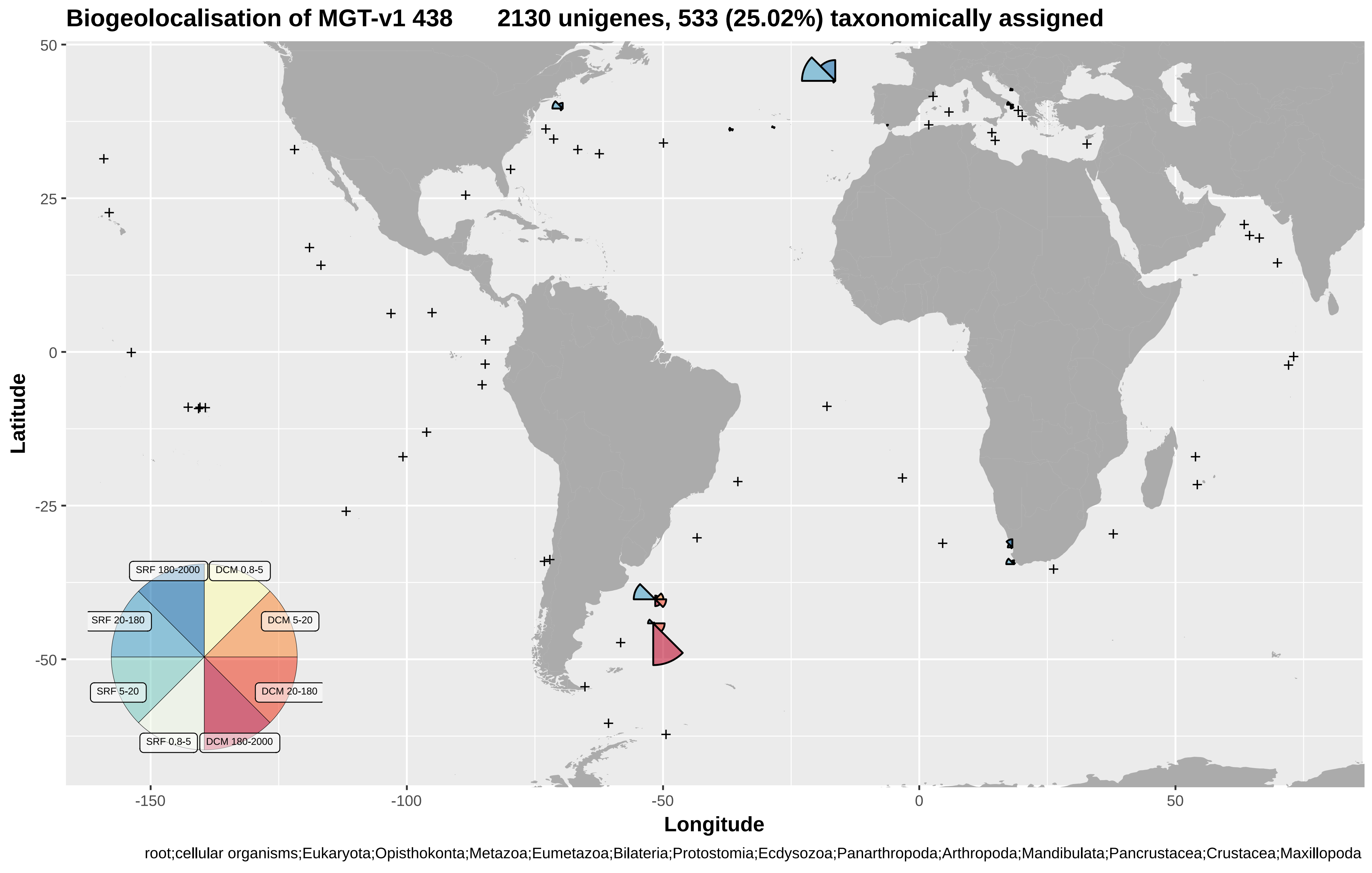

Biogeolocalisation of MGT-v1 438 2130 unigenes, 533 (25.02%) taxonomically assigned
50
25
0
-25
-50
Latitude
-150
-100
-50
0
50
Longitude
SRF 180-2000
DCM 0.8-5
SRF 20-180
DCM 5-20
SRF 5-20
DCM 20-180
SRF 0.8-5
DCM 180-2000
root;cellular organisms;Eukaryota;Opisthokonta;Metazoa;Eumetazoa;Bilateria;Protostomia;Ecdysozoa;Panarthropoda;Arthropoda;Mandibulata;Pancrustacea;Crustacea;Maxillopoda

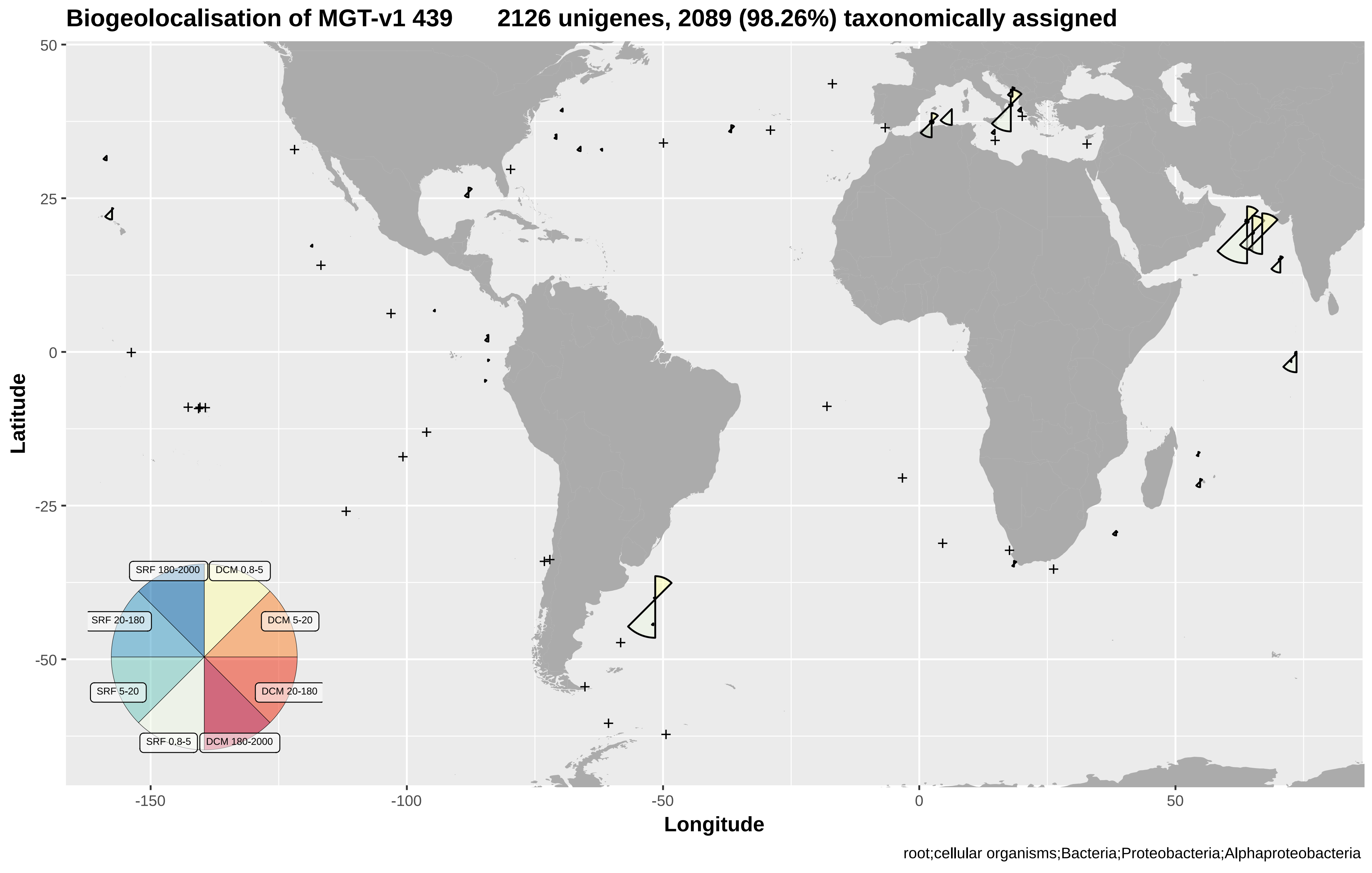
Biogeolocalisation of MGT-v1 439 2126 unigenes, 2089 (98.26%) taxonomically assigned
Latitude
Longitude
50
25
0
-25
-50
-150
-100
-50
0
50
SRF 180-2000
DCM 0.8-5
SRF 20-180
DCM 5-20
SRF 5-20
DCM 20-180
SRF 0.8-5
DCM 180-2000
root;cellular organisms;Bacteria;Proteobacteria;Alphaproteobacteria

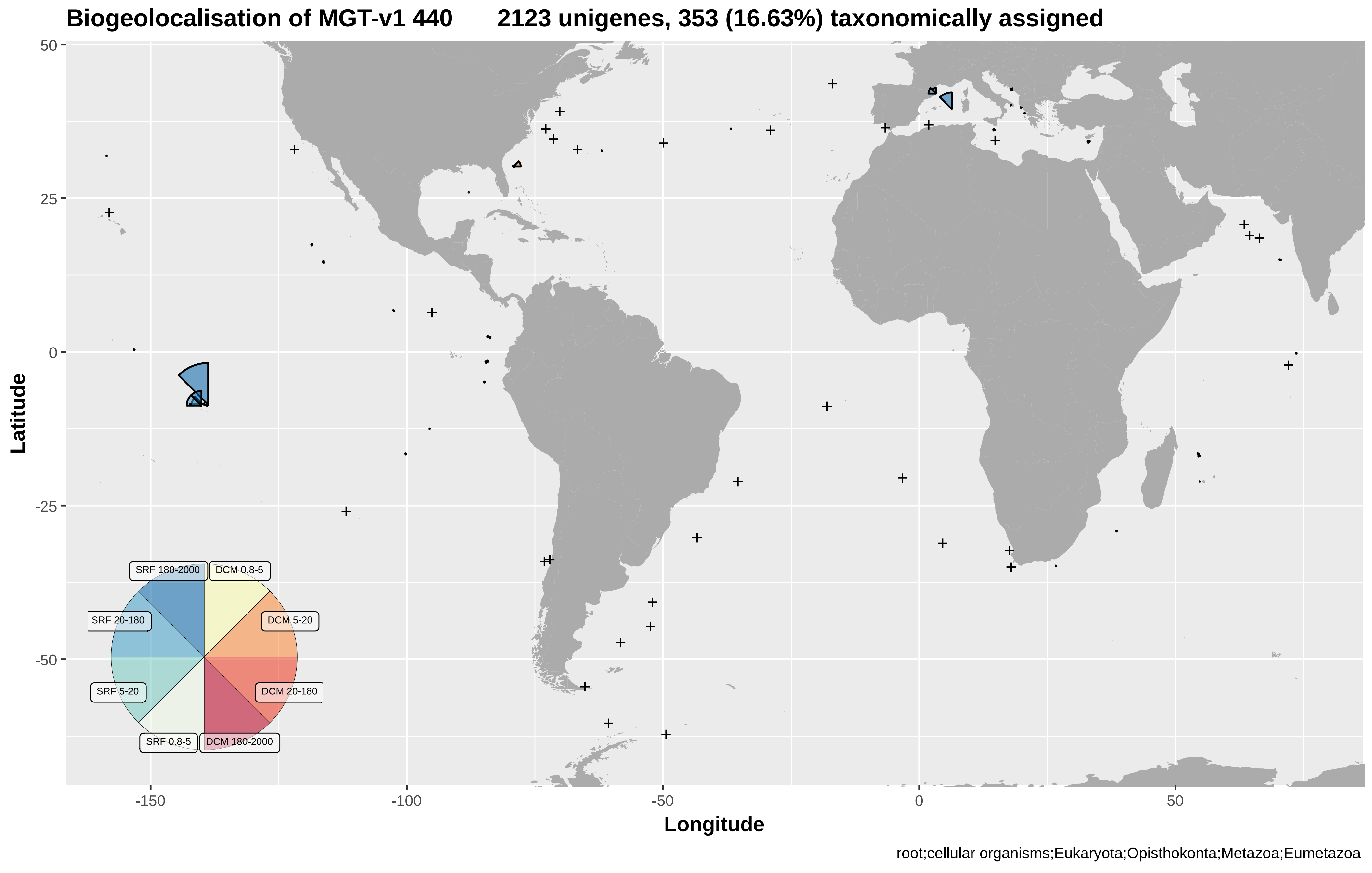
Biogeolocalisation of MGT-v1 440
2123 unigenes, 353 (16.63%) taxonomically assigned
Latitude
Longitude
50
25
0
-25
-50
-150
-100
-50
0
50
SRF 180-2000
DCM 0.8-5
SRF 20-180
DCM 5-20
SRF 5-20
DCM 20-180
SRF 0.8-5
DCM 180-2000
root;cellular organisms;Eukaryota;Opisthokonta;Metazoa;Eumetazoa

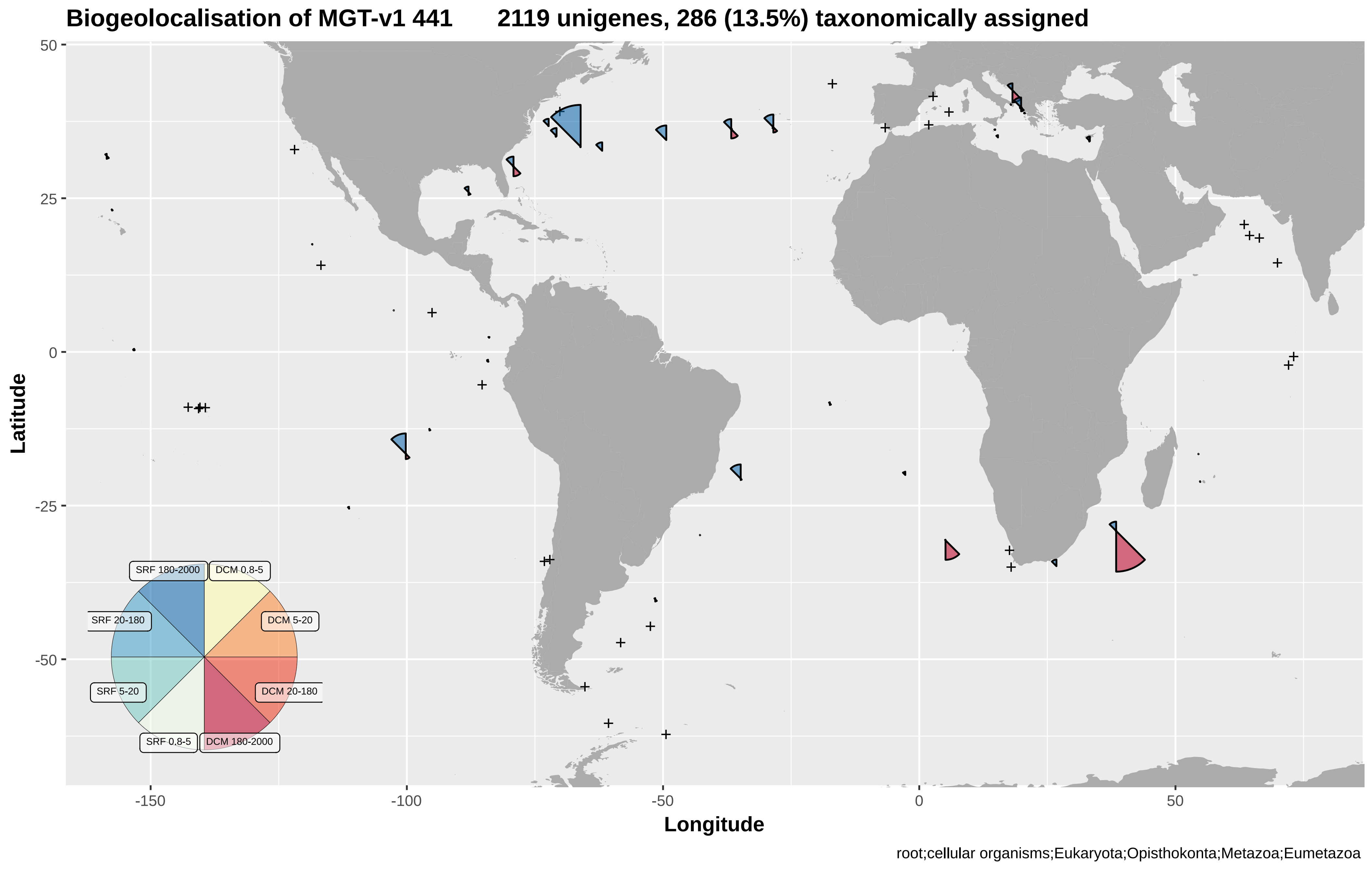
Biogeolocalisation of MGT-v1 441 2119 unigenes, 286 (13.5%) taxonomically assigned
Latitude
Longitude
50
25
0
-25
-50
-150
-100
-50
0
50
SRF 180-2000
DCM 0.8-5
SRF 20-180
DCM 5-20
SRF 5-20
DCM 20-180
SRF 0.8-5
DCM 180-2000
root;cellular organisms;Eukaryota;Opisthokonta;Metazoa;Eumetazoa

#### Biogeolocalisation of MGT-v1 442 2098 unigenes, 1833 (87.37%) taxonomically assigned

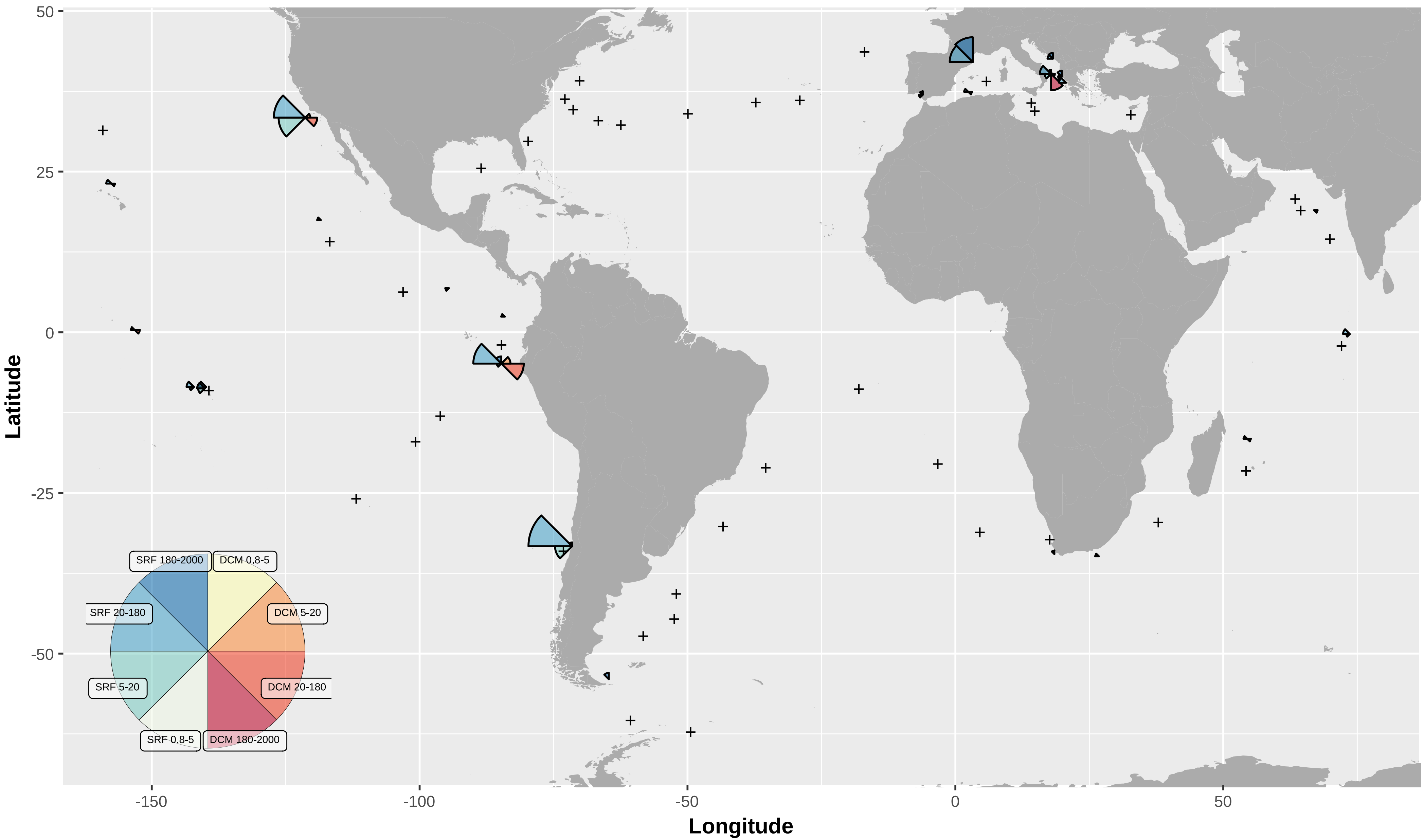


root;cellular organisms;Eukaryota;Opisthokonta;Metazoa;Eumetazoa;Bilateria;Protostomia;Ecdysozoa;Panarthropoda;Arthropoda;Mandibulata;Pancrustacea;Crustacea;Maxillopoda

## Biogeolocalisation of MGT-v1 443      2097 unigenes, 374 (17.84%) taxonomically assigned

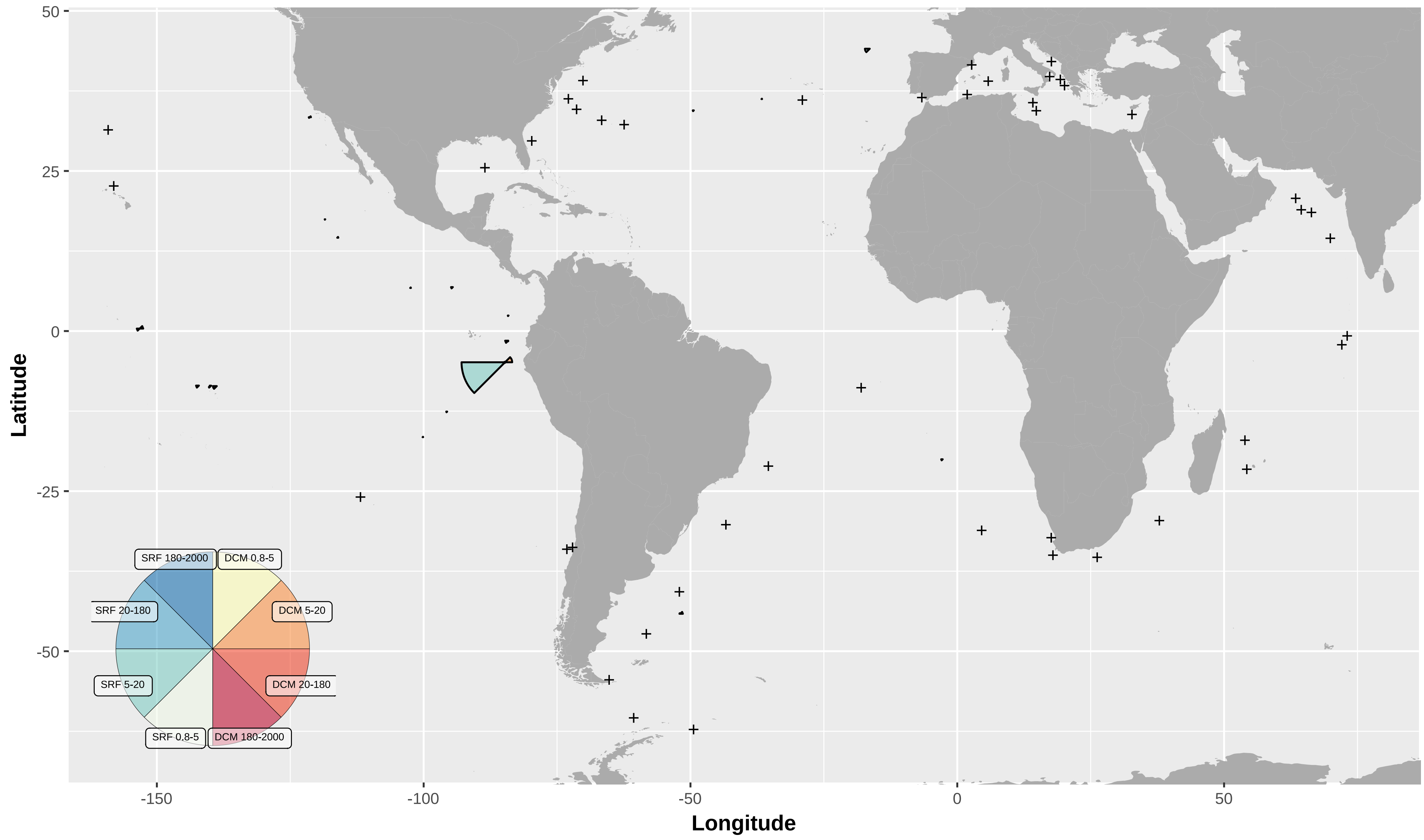


root;cellular organisms;Eukaryota;Stramenopiles;Bacillariophyta

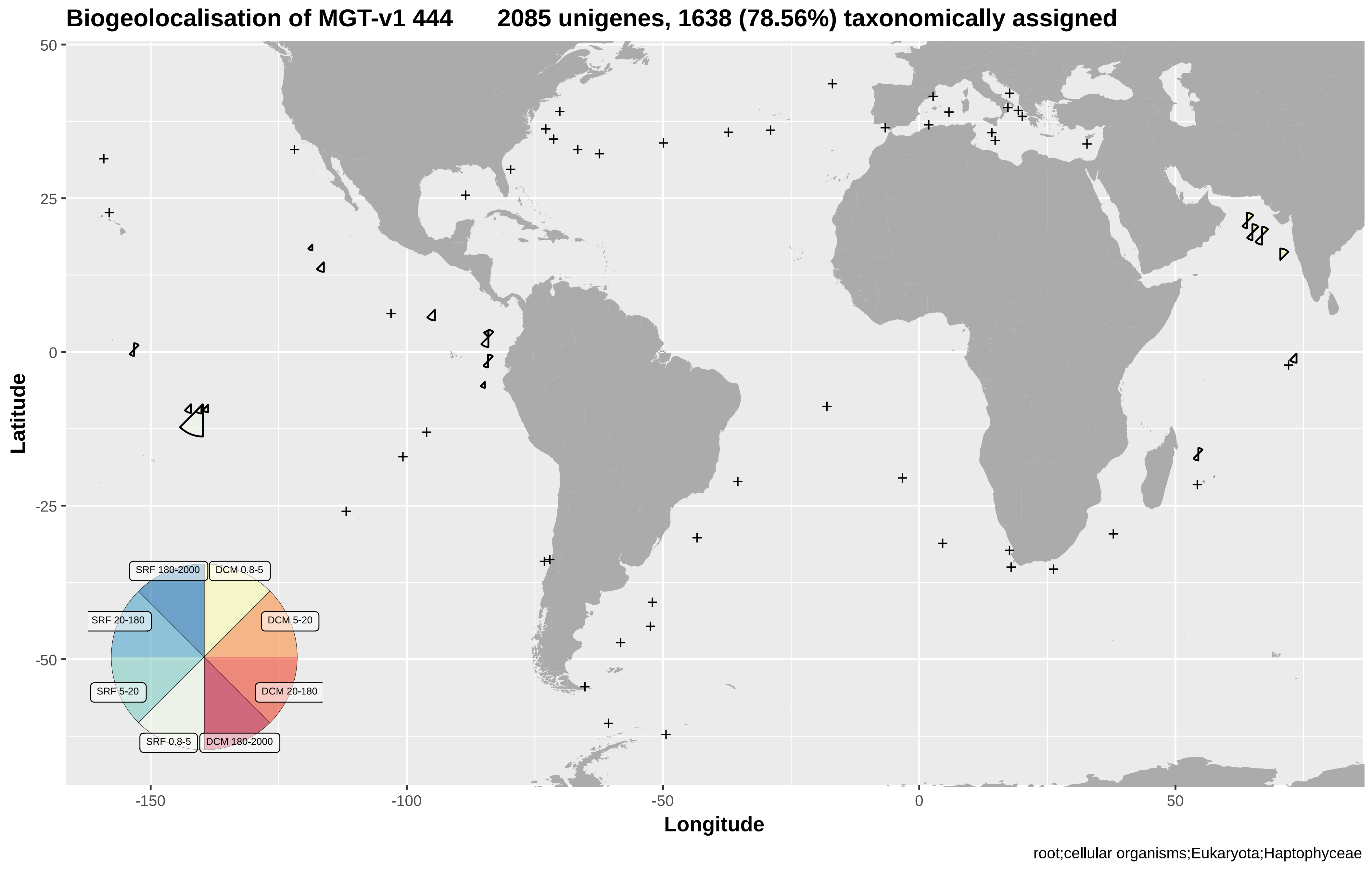

Biogeolocalisation of MGT-v1 444      2085 unigenes, 1638 (78.56%) taxonomically assigned
50
25
0
-25
-50
Latitude
-150
-100
-50
0
50
Longitude
SRF 180-2000
DCM 0.8-5
SRF 20-180
DCM 5-20
SRF 5-20
DCM 20-180
SRF 0.8-5
DCM 180-2000
root;cellular organisms;Eukaryota;Haptophyceae

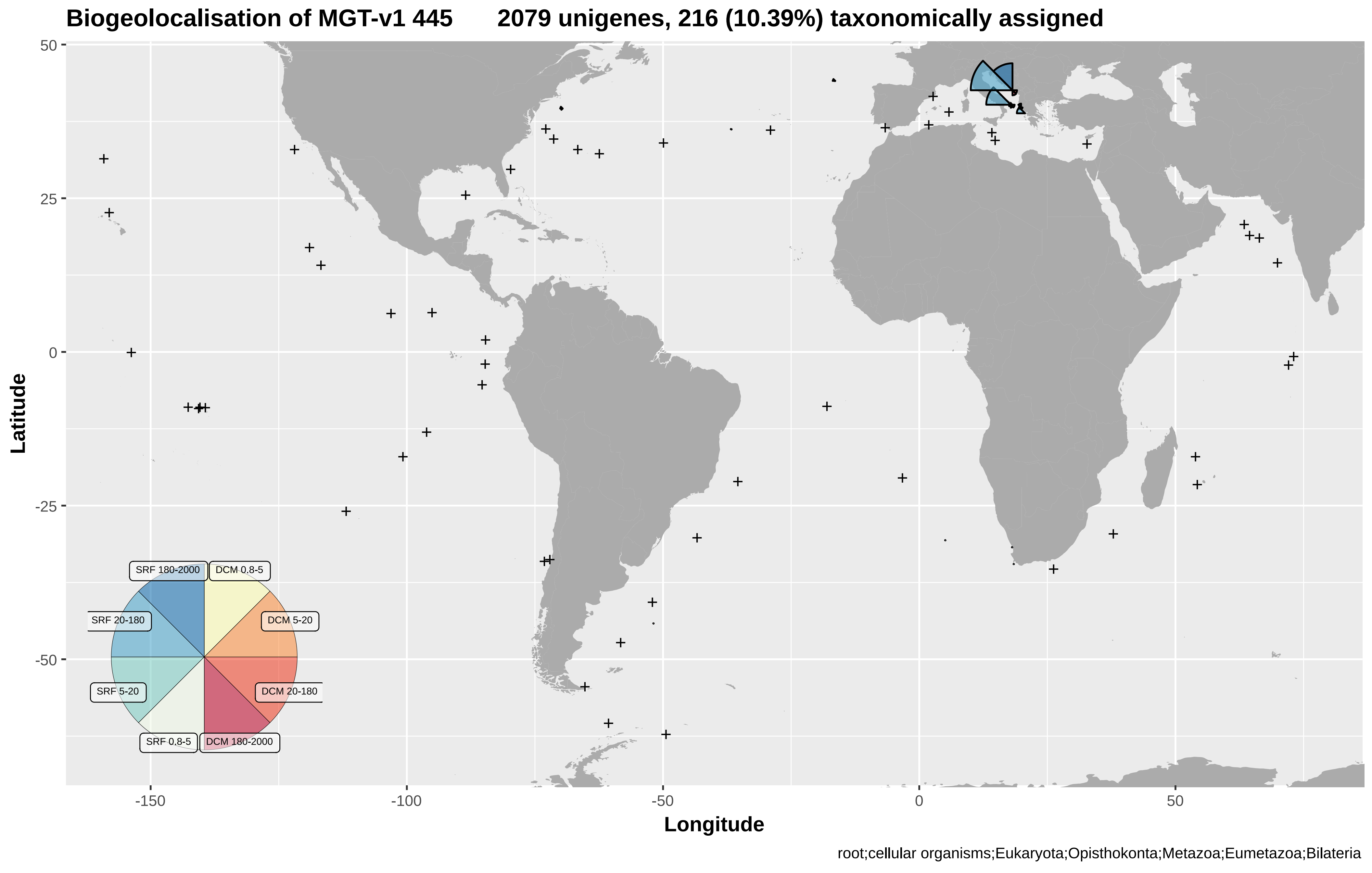

Biogeolocalisation of MGT-v1 445 2079 unigenes, 216 (10.39%) taxonomically assigned
Latitude
Longitude
50
25
0
-25
-50
-150
-100
-50
0
50
SRF 180-2000
DCM 0.8-5
SRF 20-180
DCM 5-20
SRF 5-20
DCM 20-180
SRF 0.8-5
DCM 180-2000
root;cellular organisms;Eukaryota;Opisthokonta;Metazoa;Eumetazoa;Bilateria

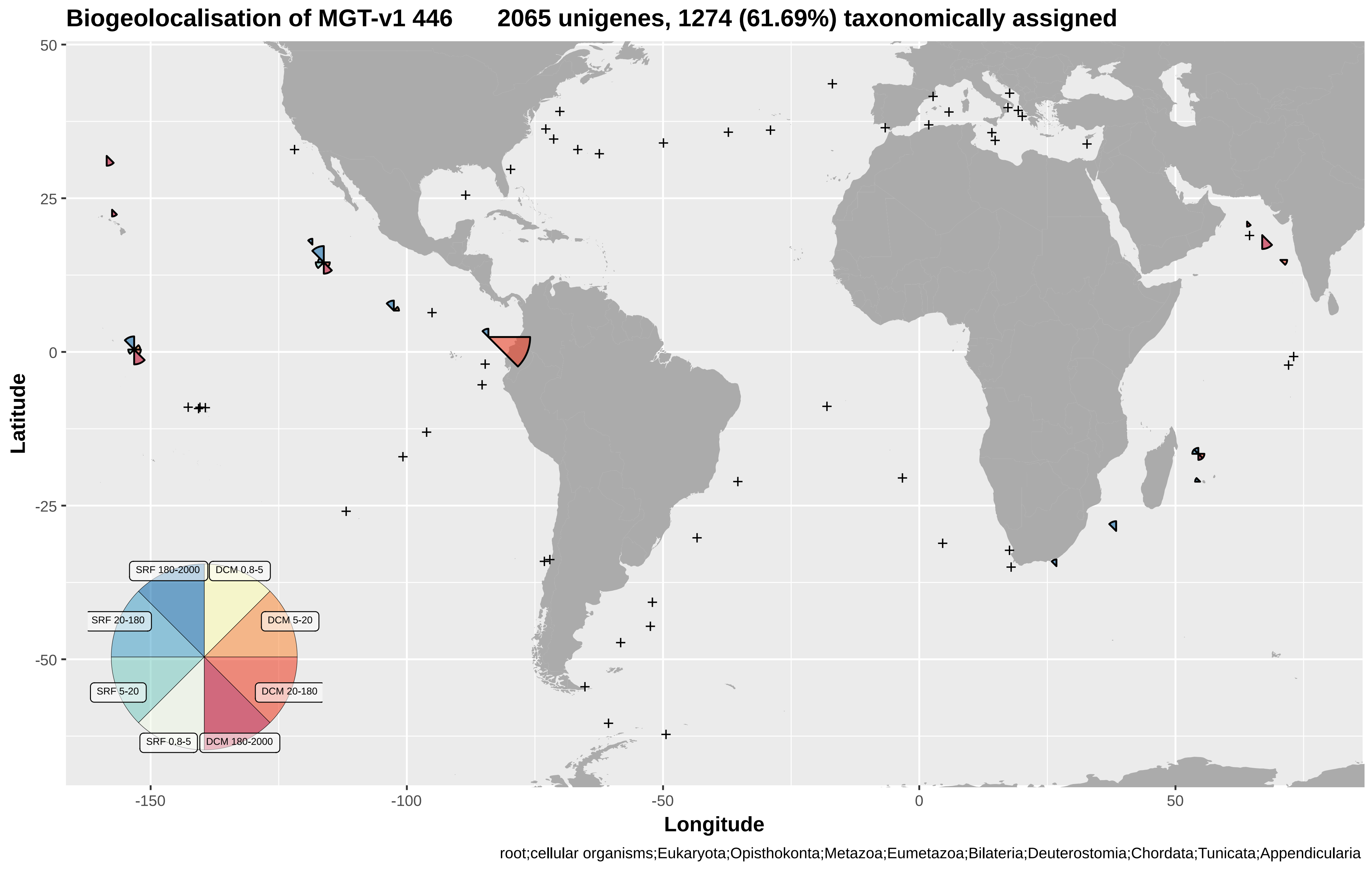

Biogeolocalisation of MGT-v1 446 2065 unigenes, 1274 (61.69%) taxonomically assigned
50
25
0
-25
-50
Latitude
-150
-100
-50
0
50
Longitude
SRF 180-2000
DCM 0.8-5
SRF 20-180
DCM 5-20
SRF 5-20
DCM 20-180
SRF 0.8-5
DCM 180-2000
root;cellular organisms;Eukaryota;Opisthokonta;Metazoa;Eumetazoa;Bilateria;Deuterostomia;Chordata;Tunicata;Appendicularia

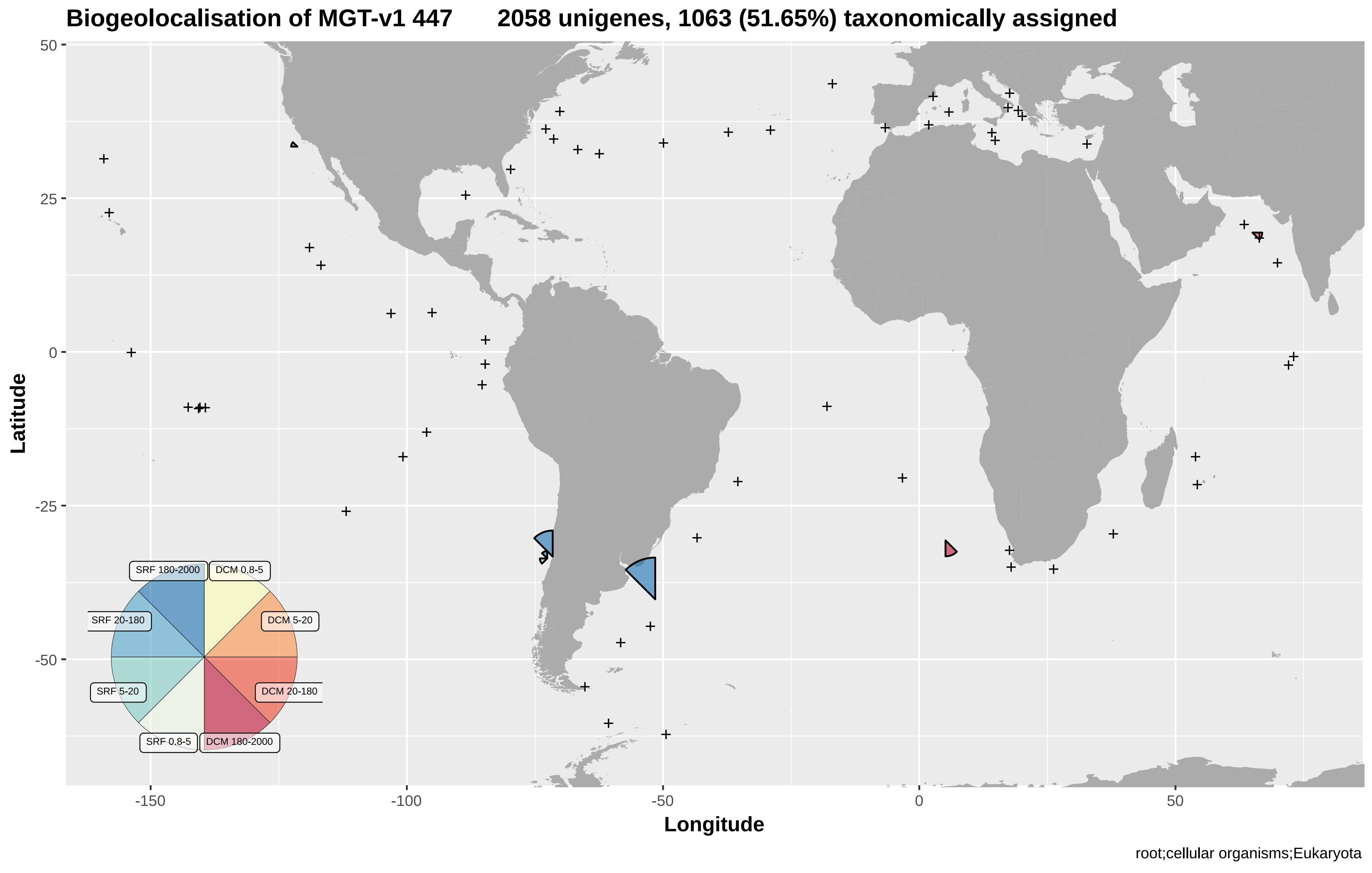

Biogeolocalisation of MGT-v1 447 2058 unigenes, 1063 (51.65%) taxonomically assigned
50
25
0
-25
-50
Latitude
-150
-100
-50
0
50
Longitude
SRF 180-2000
DCM 0.8-5
SRF 20-180
DCM 5-20
SRF 5-20
DCM 20-180
SRF 0.8-5
DCM 180-2000
root;cellular organisms;Eukaryota

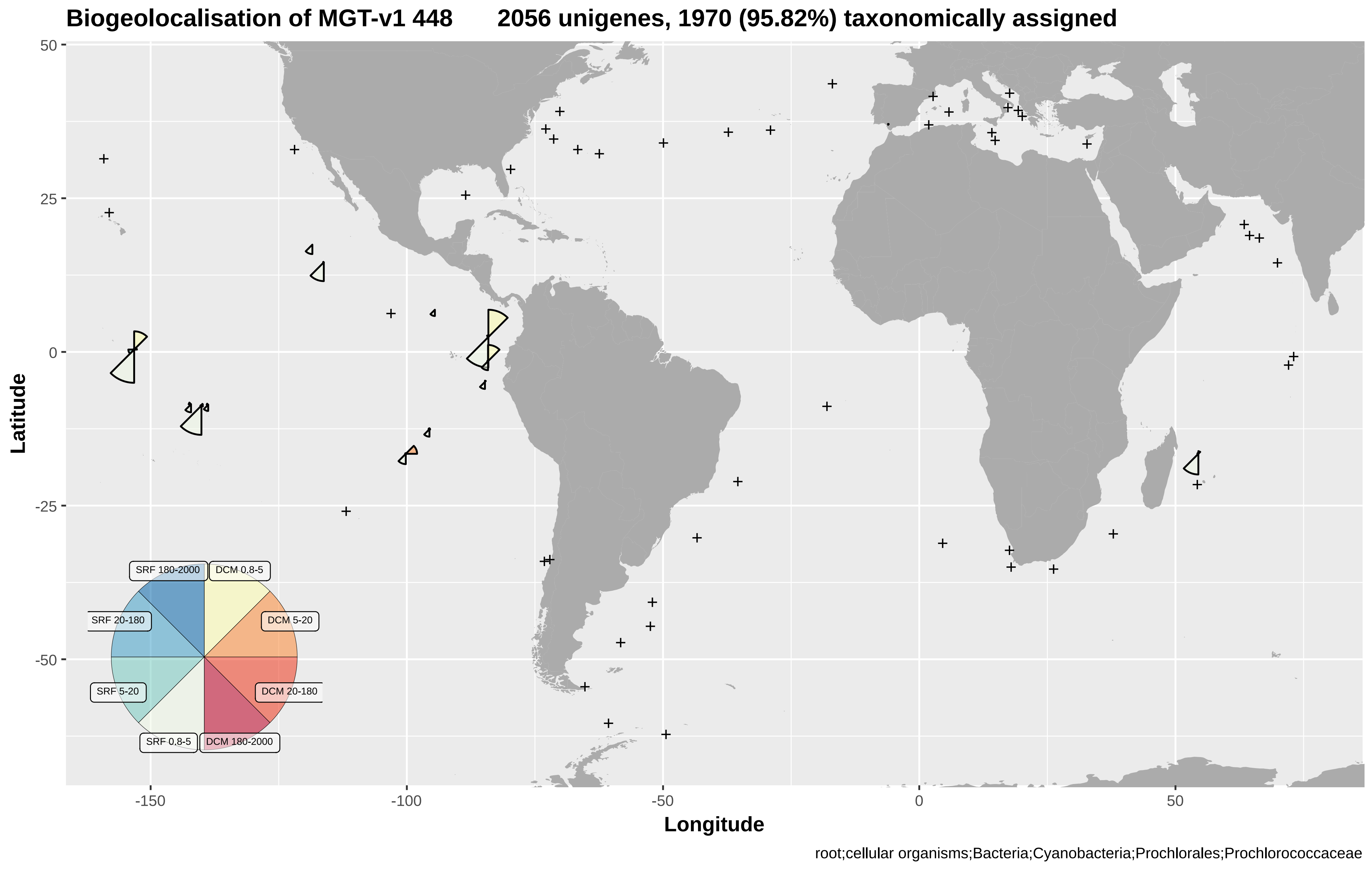

Biogeolocalisation of MGT-v1 448 2056 unigenes, 1970 (95.82%) taxonomically assigned
50
25
0
-25
-50
Latitude
-150
-100
-50
0
50
Longitude
SRF 180-2000
DCM 0.8-5
SRF 20-180
DCM 5-20
SRF 5-20
DCM 20-180
SRF 0.8-5
DCM 180-2000
root;cellular organisms;Bacteria;Cyanobacteria;Prochlorales;Prochlorococcaceae

# Biogeolocalisation of MGT-v1 449      2040 unigenes, 1912 (93.73%) taxonomically assigned

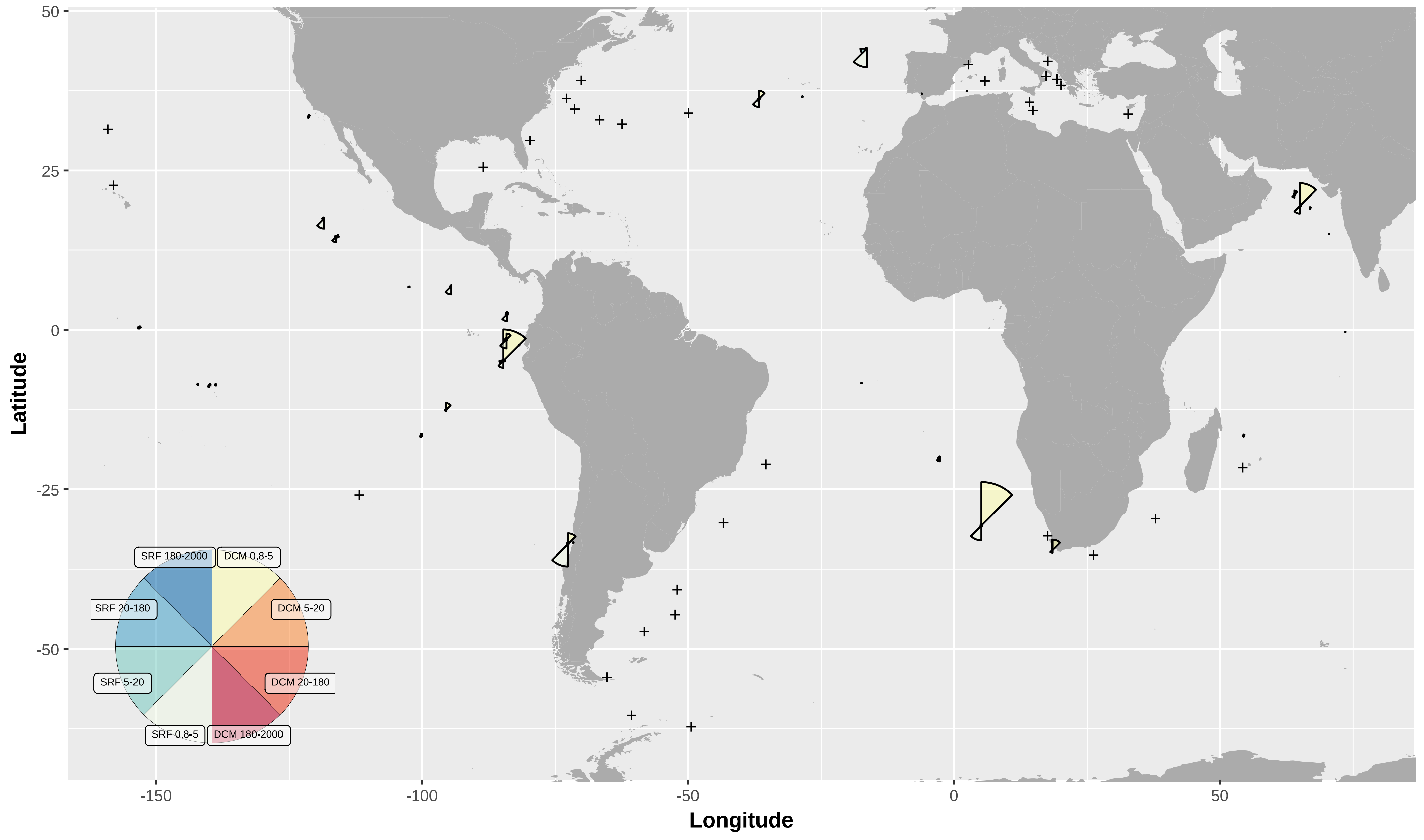


root;cellular organisms;Bacteria;Cyanobacteria;Oscillatoriophycideae;Chroococcales

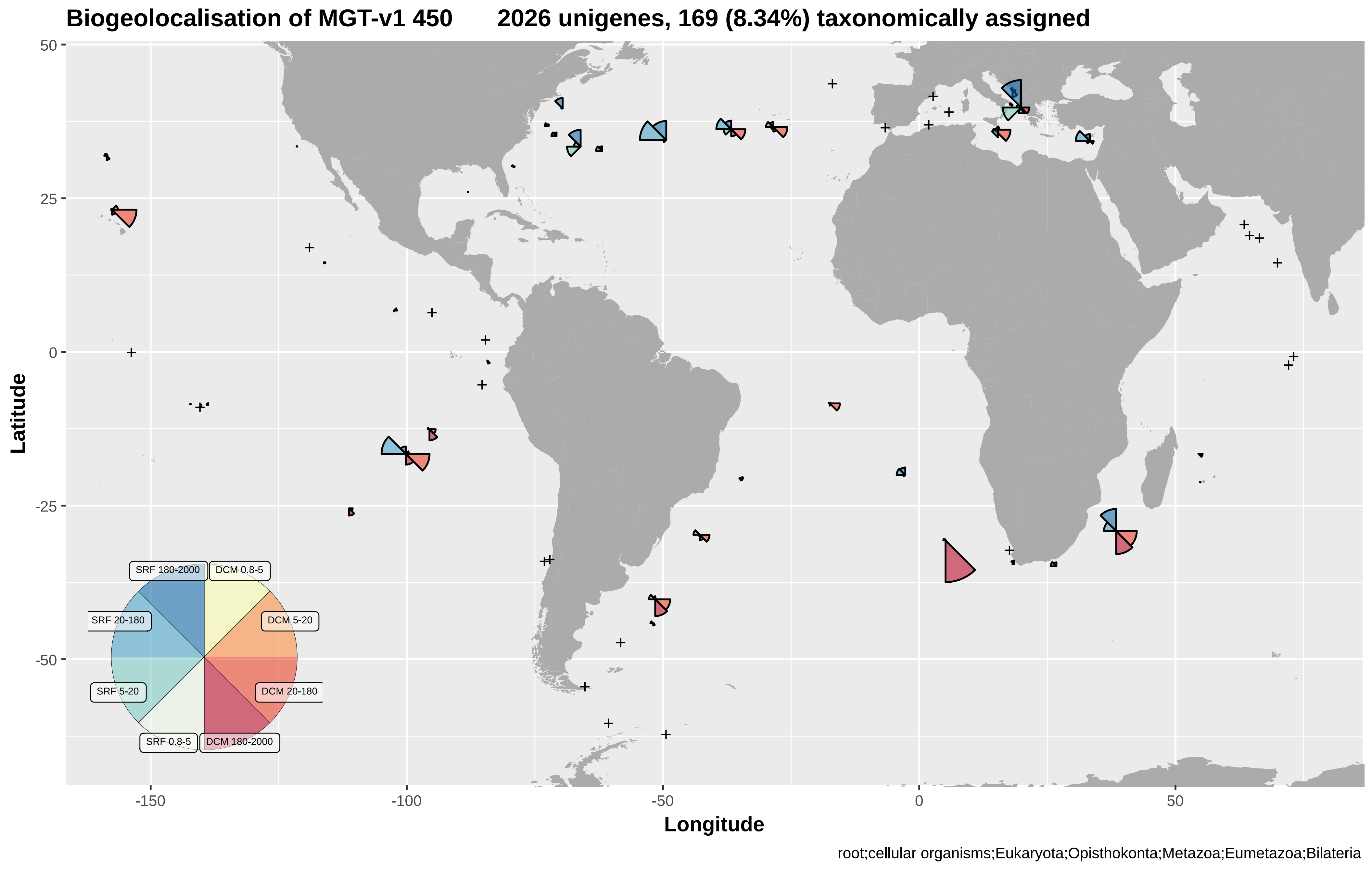

Biogeolocalisation of MGT-v1 450
2026 unigenes, 169 (8.34%) taxonomically assigned
50
25
0
-25
-50
Latitude
-150
-100
-50
0
50
Longitude
SRF 180-2000
DCM 0.8-5
SRF 20-180
DCM 5-20
SRF 5-20
DCM 20-180
SRF 0.8-5
DCM 180-2000
root;cellular organisms;Eukaryota;Opisthokonta;Metazoa;Eumetazoa;Bilateria

## Biogeolocalisation of MGT-v1 451      2021 unigenes, 1595 (78.92%) taxonomically assigned

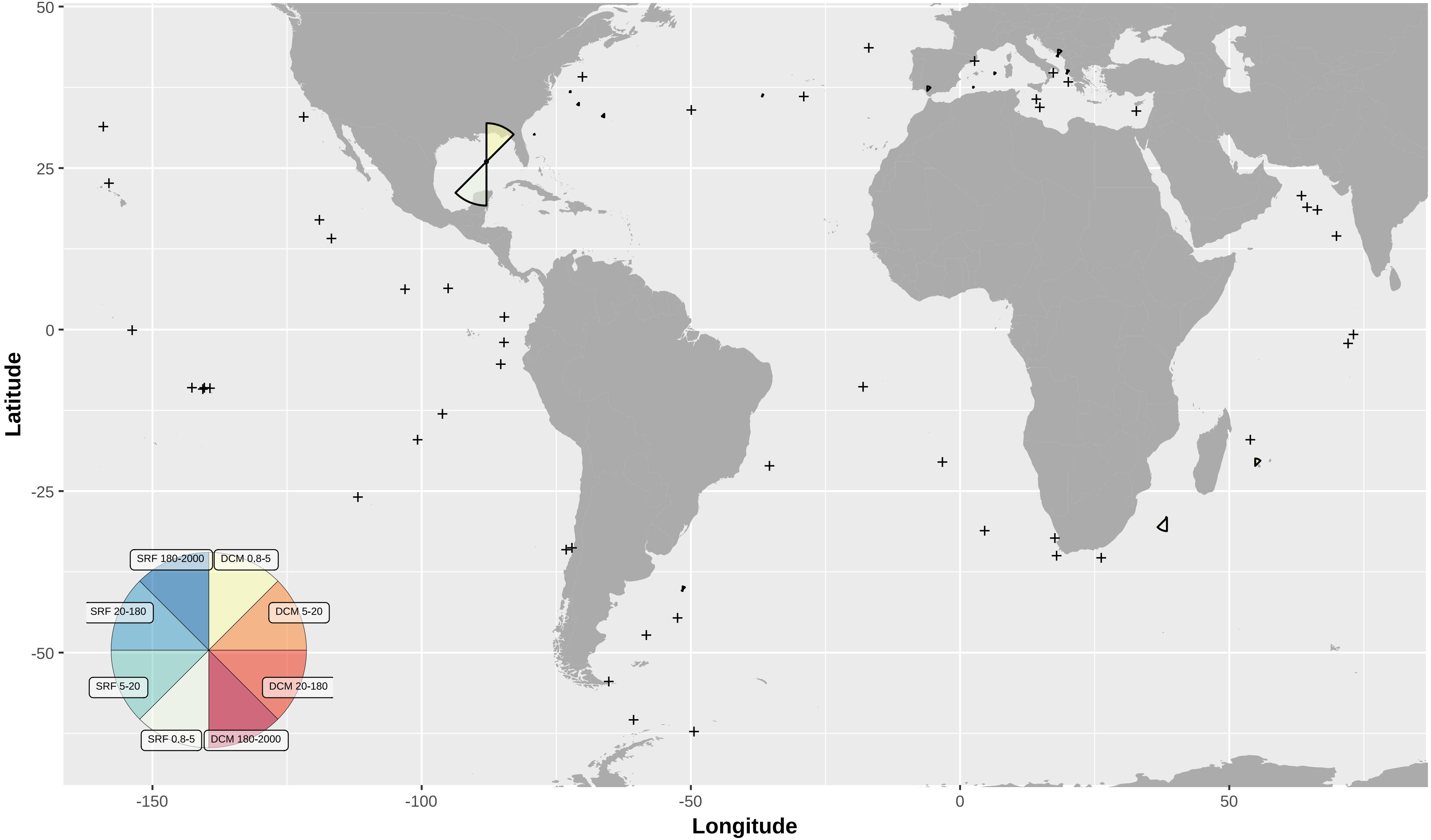


root;cellular organisms;Bacteria;Cyanobacteria;Oscillatoriophycideae

#### Biogeolocalisation of MGT-v1 452 2013 unigenes, 195 (9.69%) taxonomically assigned

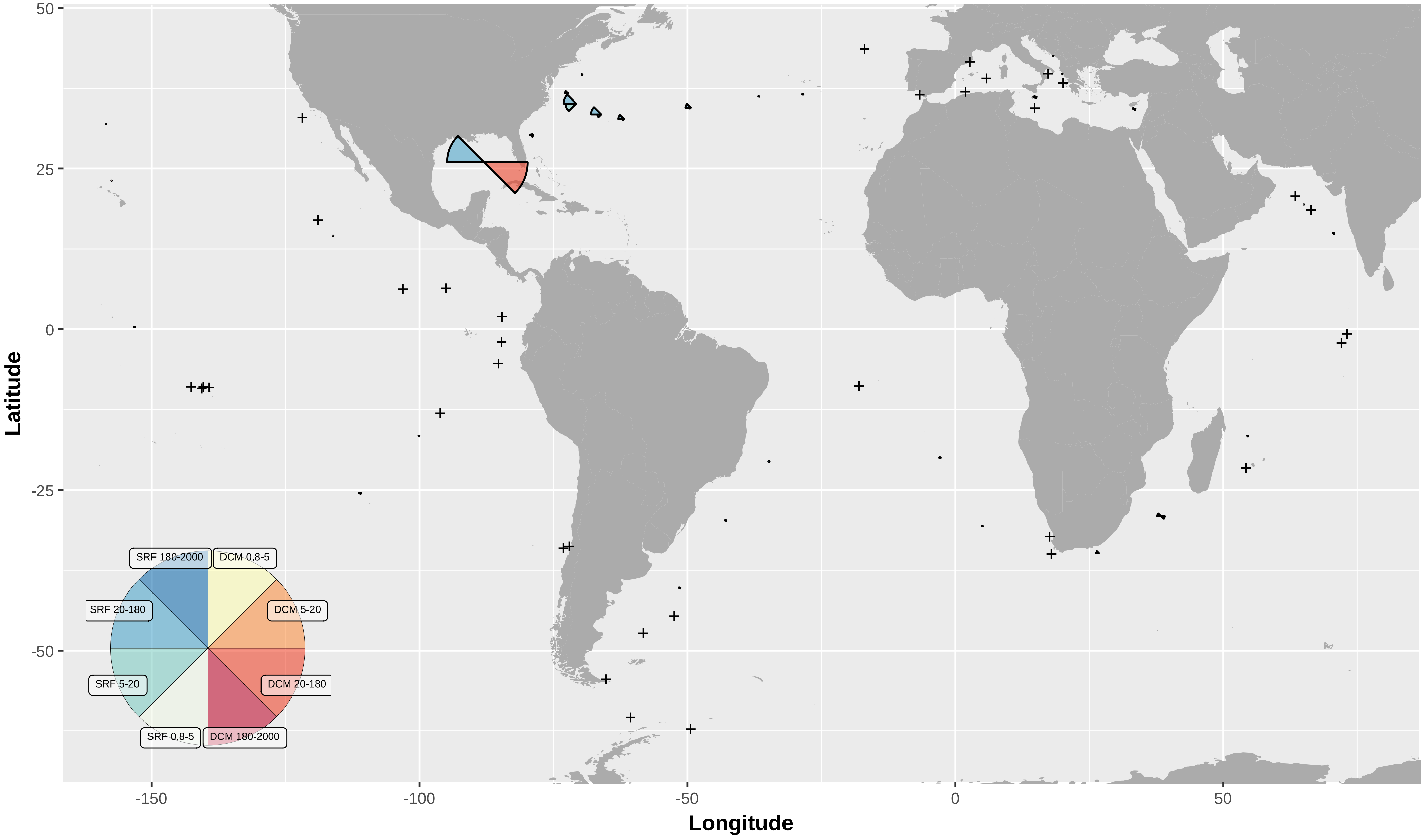


root;cellular organisms;Eukaryota

## Biogeolocalisation of MGT-v1 453      1991 unigenes, 211 (10.6%) taxonomically assigned

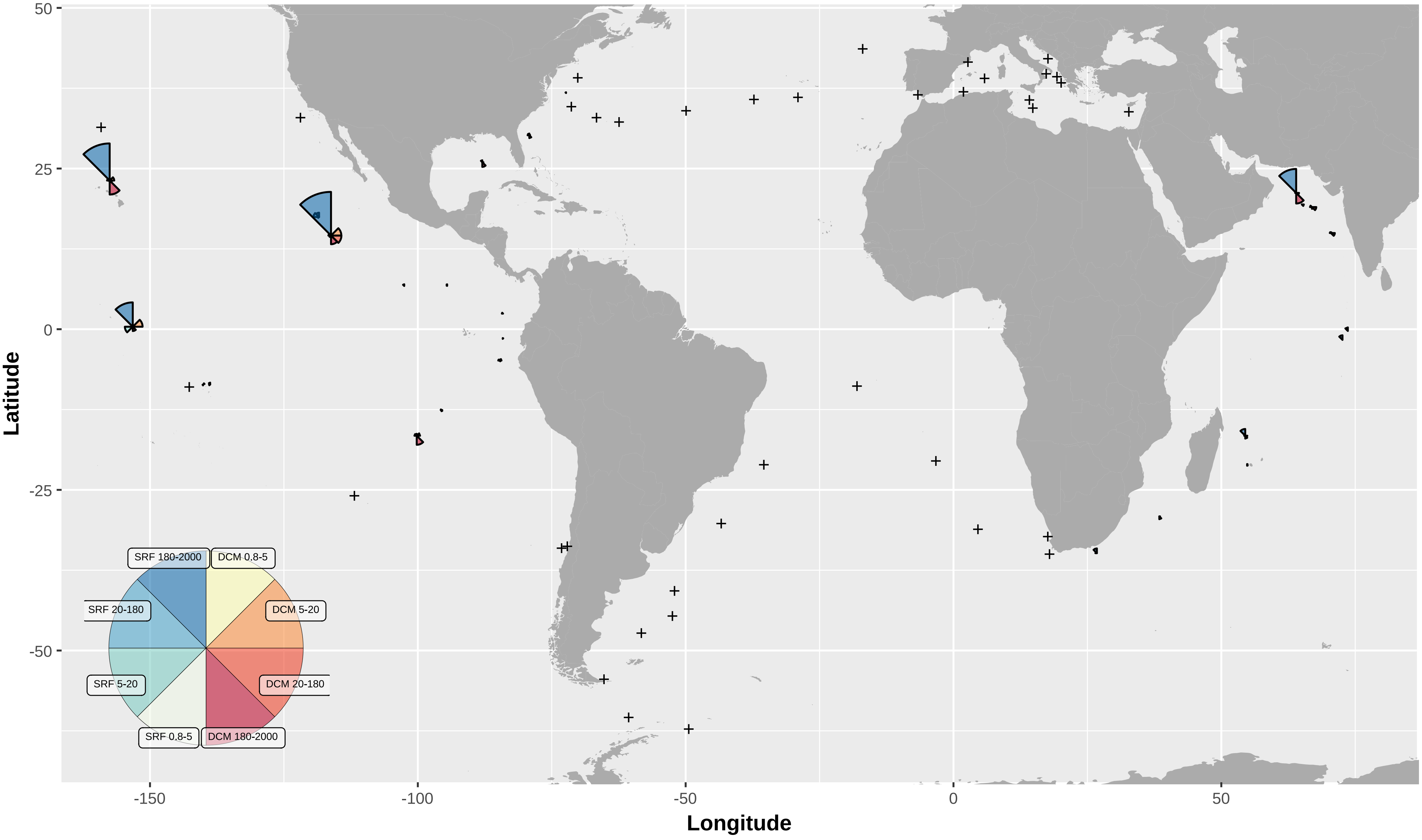


root;cellular organisms;Eukaryota;Opisthokonta;Metazoa;Eumetazoa

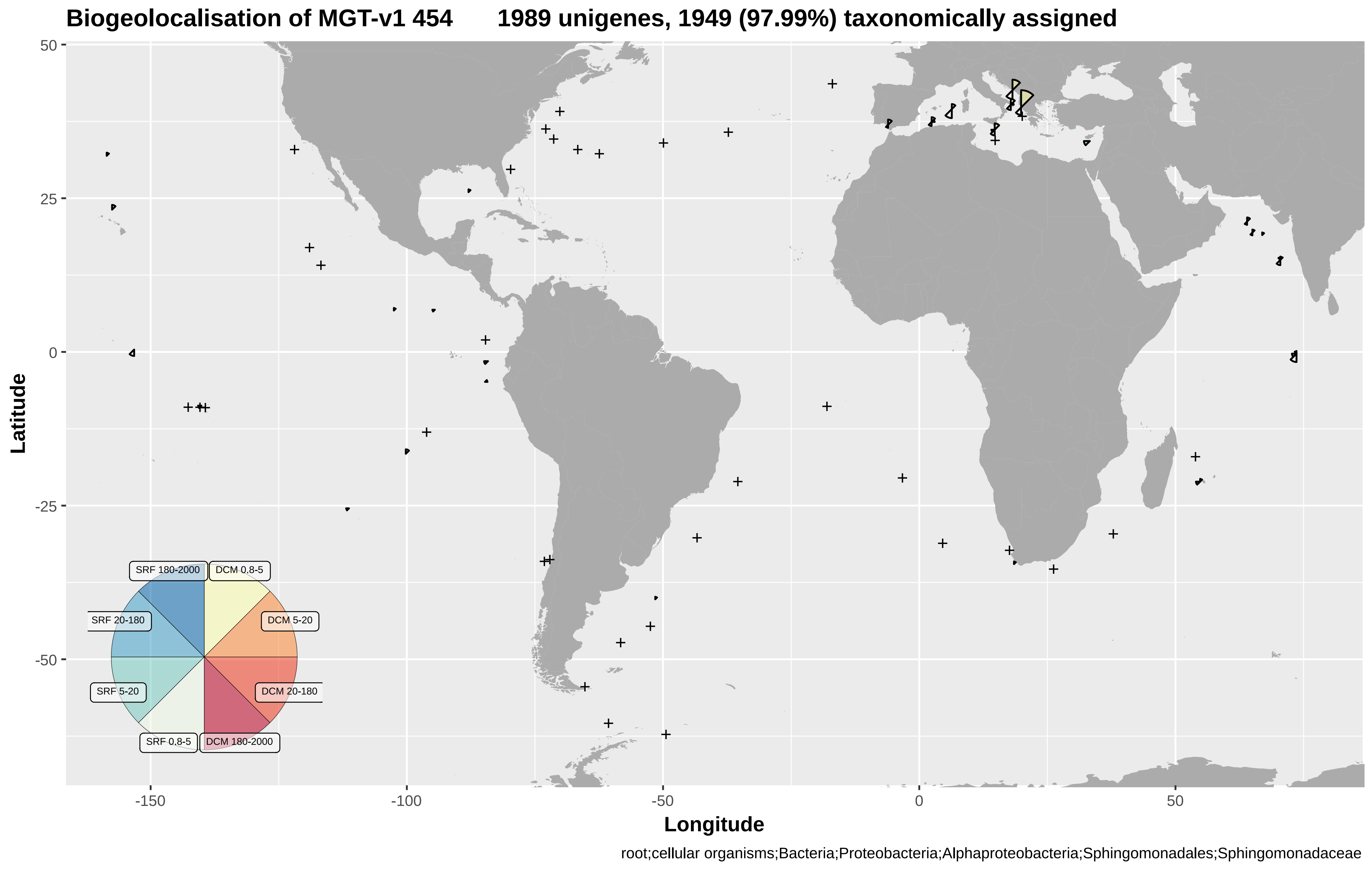

Biogeolocalisation of MGT-v1 454 1989 unigenes, 1949 (97.99%) taxonomically assigned
Latitude
Longitude
50
25
0
-25
-50
-150
-100
-50
0
50
SRF 180-2000
DCM 0.8-5
SRF 20-180
DCM 5-20
SRF 5-20
DCM 20-180
SRF 0.8-5
DCM 180-2000
root;cellular organisms;Bacteria;Proteobacteria;Alphaproteobacteria;Sphingomonadales;Sphingomonadaceae

#### Biogeolocalisation of MGT-v1 455 1986 unigenes, 296 (14.9%) taxonomically assigned

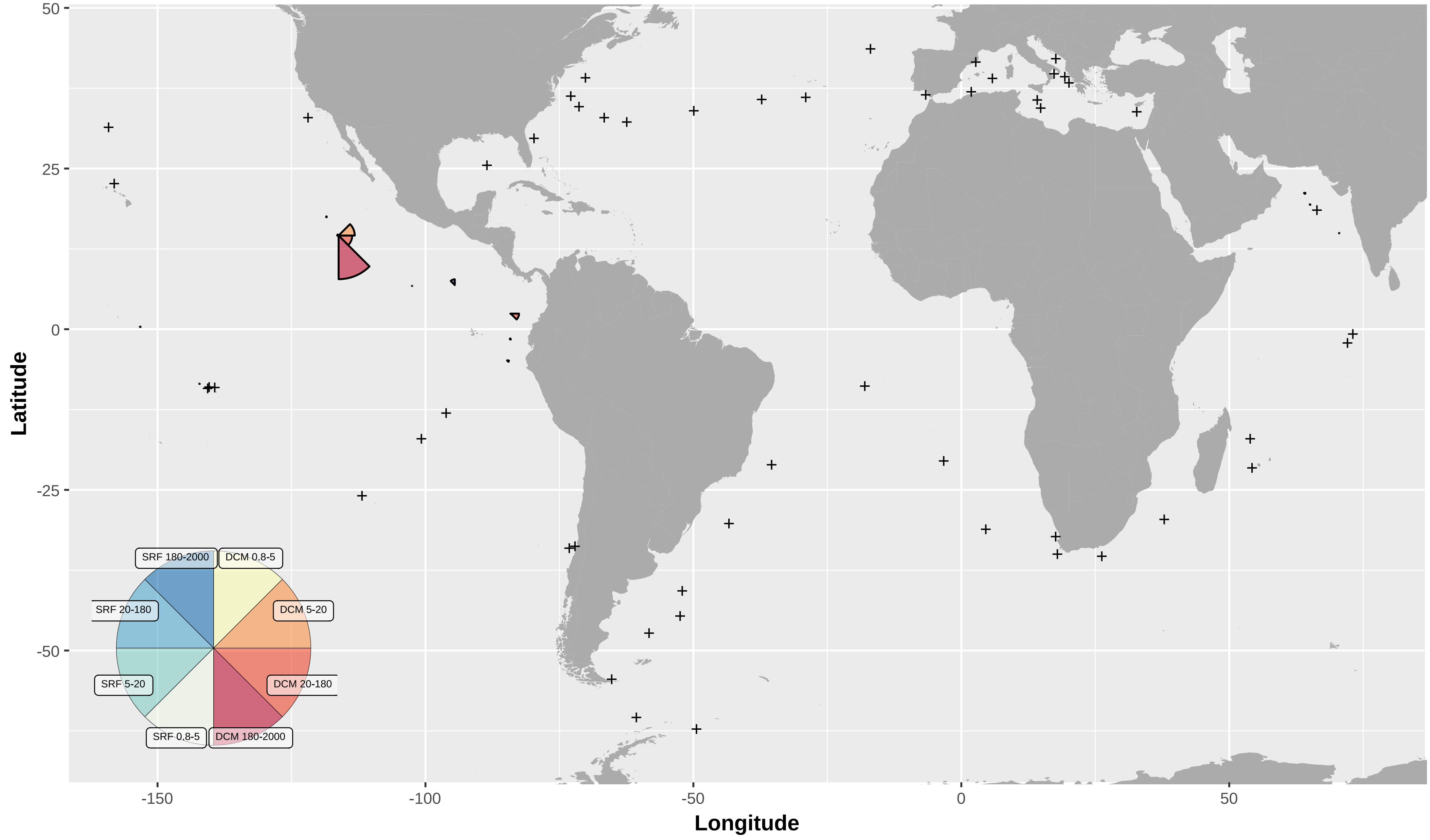


root;cellular organisms;Eukaryota;Opisthokonta;Metazoa;Eumetazoa;Bilateria

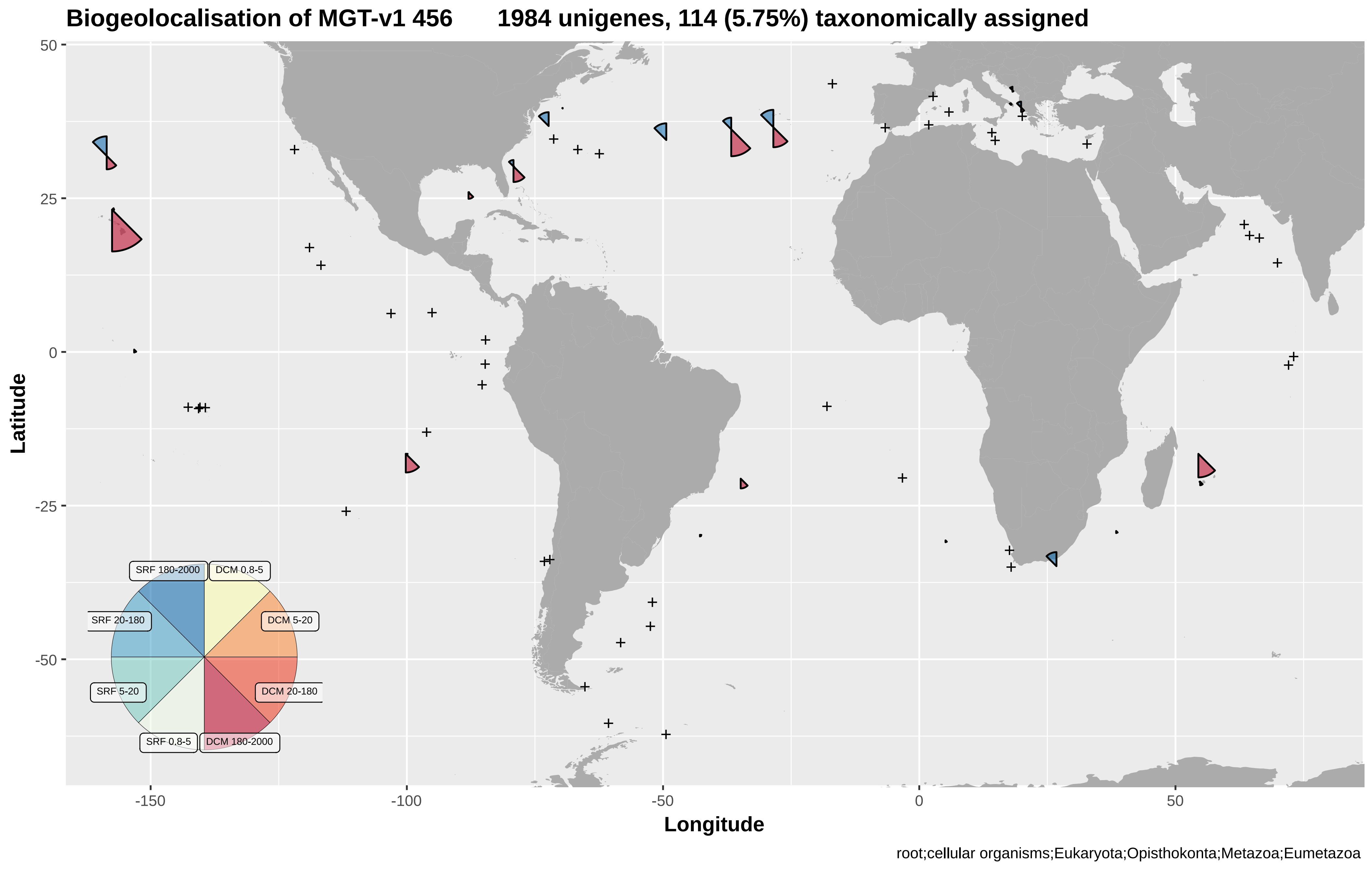

Biogeolocalisation of MGT-v1 456 1984 unigenes, 114 (5.75%) taxonomically assigned
50
25
0
-25
-50
Latitude
-150
-100
-50
0
50
Longitude
SRF 180-2000
DCM 0.8-5
SRF 20-180
DCM 5-20
SRF 5-20
DCM 20-180
SRF 0.8-5
DCM 180-2000
root;cellular organisms;Eukaryota;Opisthokonta;Metazoa;Eumetazoa

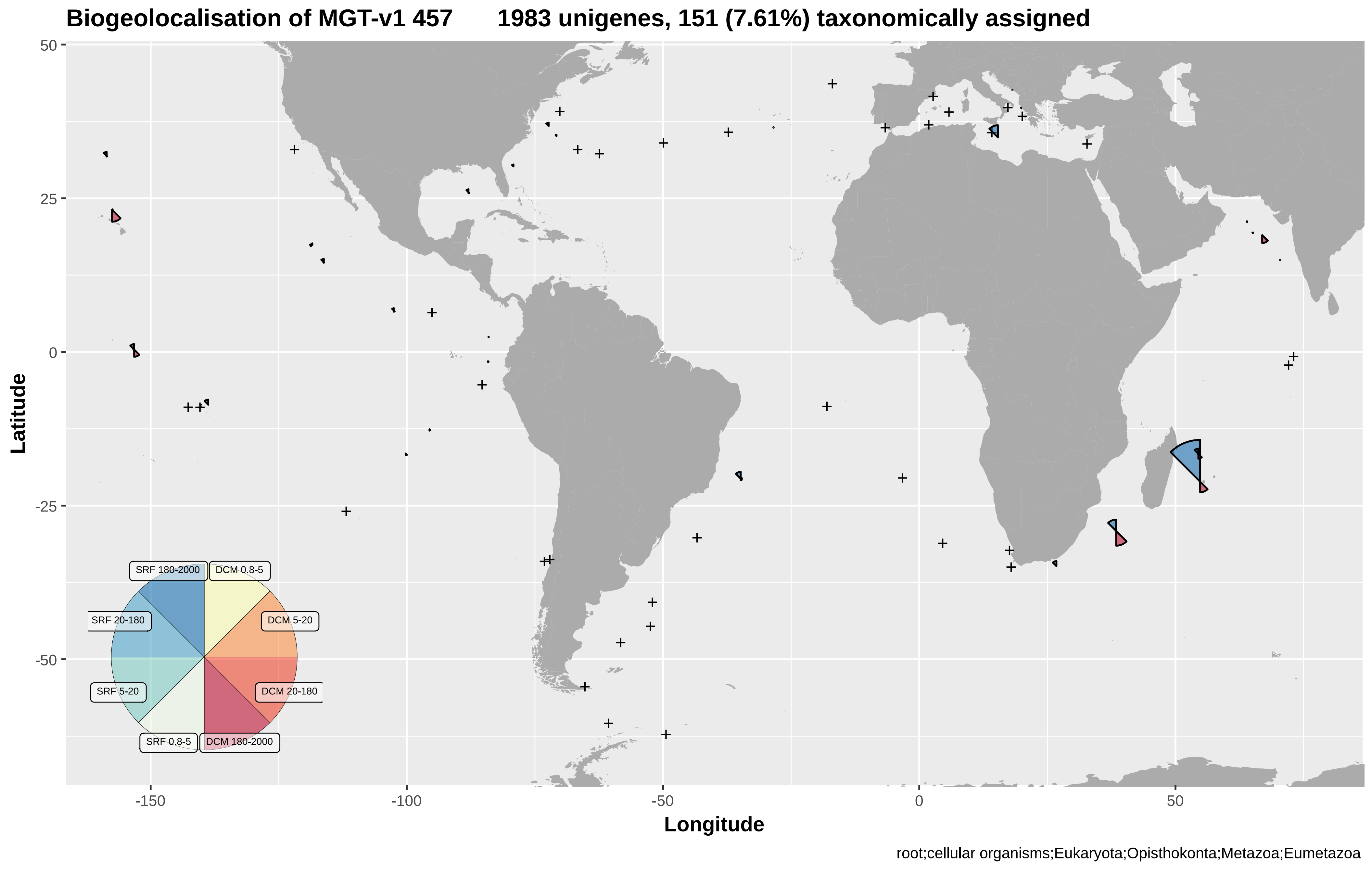

Biogeolocalisation of MGT-v1 457 1983 unigenes, 151 (7.61%) taxonomically assigned
Latitude
50
25
0
-25
-50
Longitude
-150
-100
-50
0
50
SRF 180-2000
DCM 0.8-5
SRF 20-180
DCM 5-20
SRF 5-20
DCM 20-180
SRF 0.8-5
DCM 180-2000
root;cellular organisms;Eukaryota;Opisthokonta;Metazoa;Eumetazoa

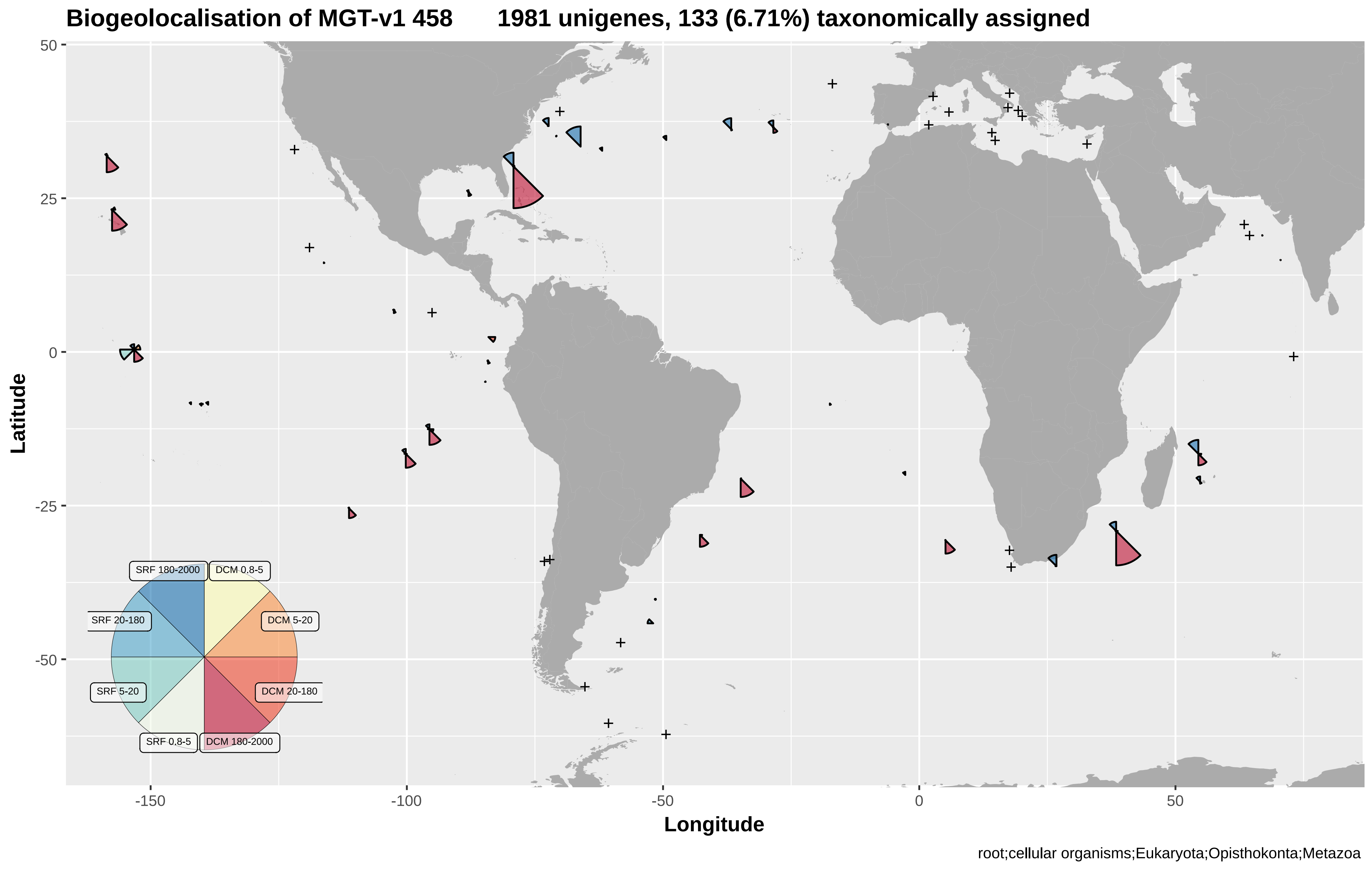

Biogeolocalisation of MGT-v1 458 1981 unigenes, 133 (6.71%) taxonomically assigned
50
25
0
-25
-50
Latitude
-150
-100
-50
0
50
Longitude
SRF 180-2000
DCM 0.8-5
SRF 20-180
DCM 5-20
SRF 5-20
DCM 20-180
SRF 0.8-5
DCM 180-2000
root;cellular organisms;Eukaryota;Opisthokonta;Metazoa

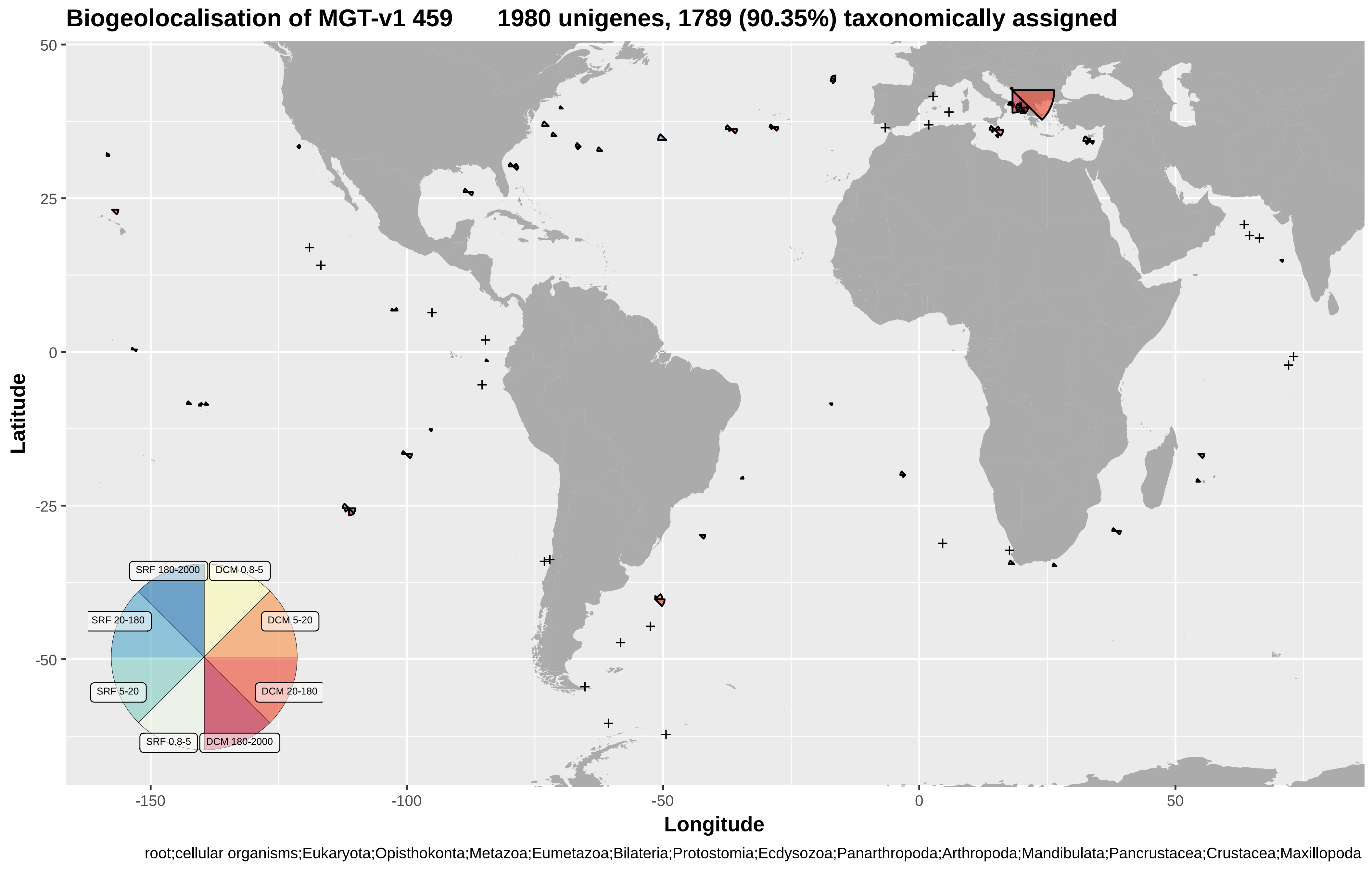
Biogeolocalisation of MGT-v1 459 1980 unigenes, 1789 (90.35%) taxonomically assigned
50
25
0
-25
-50
Latitude
-150
-100
-50
0
50
Longitude
SRF 180-2000
DCM 0.8-5
SRF 20-180
DCM 5-20
SRF 5-20
DCM 20-180
SRF 0.8-5
DCM 180-2000
root;cellular organisms;Eukaryota;Opisthokonta;Metazoa;Eumetazoa;Bilateria;Protostomia;Ecdysozoa;Panarthropoda;Arthropoda;Mandibulata;Pancrustacea;Crustacea;Maxillopoda

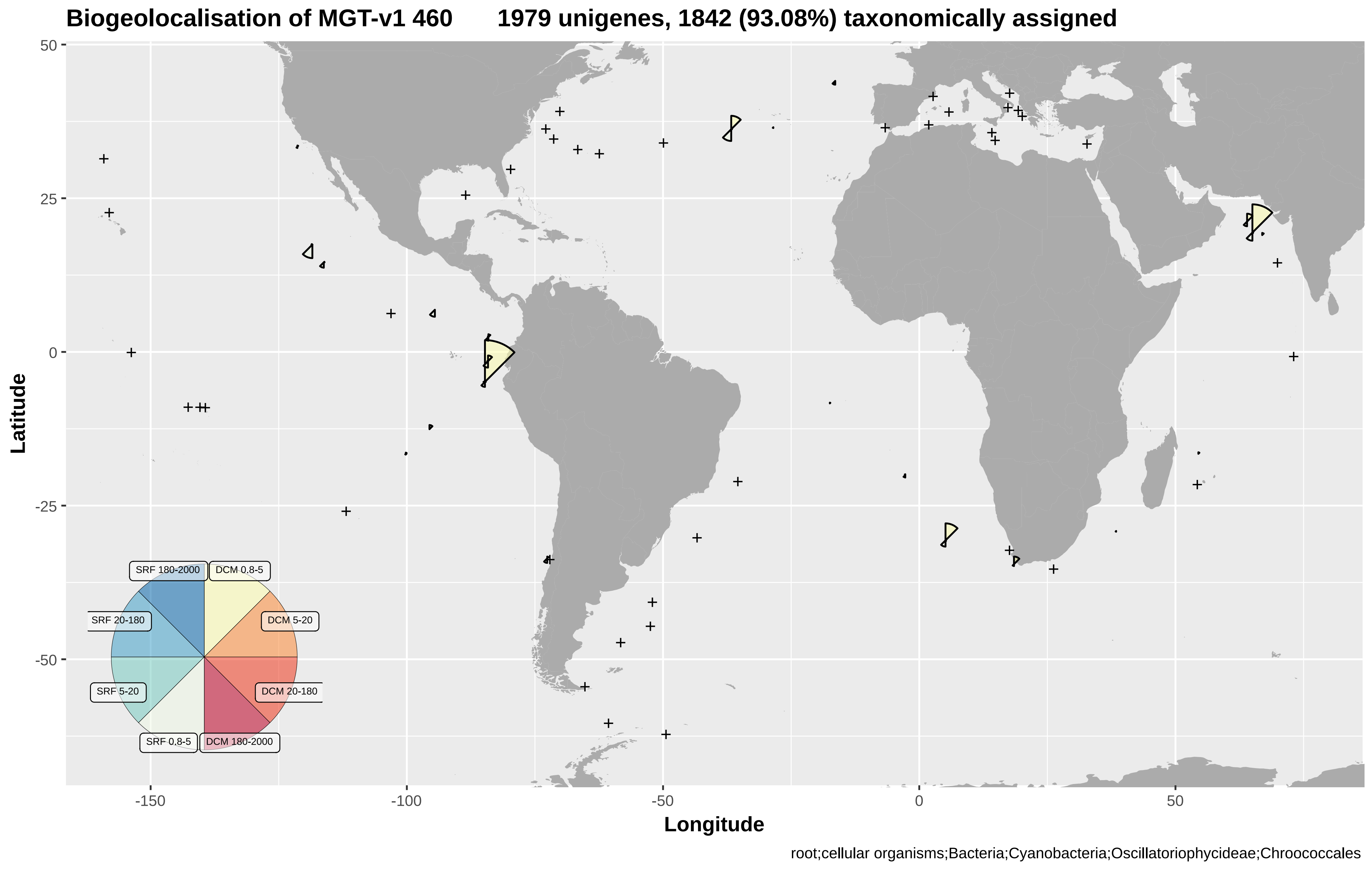
Biogeolocalisation of MGT-v1 460 1979 unigenes, 1842 (93.08%) taxonomically assigned
Latitude
Longitude
50
25
0
-25
-50
-150
-100
-50
0
50
SRF 180-2000
DCM 0.8-5
SRF 20-180
DCM 5-20
SRF 5-20
DCM 20-180
SRF 0.8-5
DCM 180-2000
root;cellular organisms;Bacteria;Cyanobacteria;Oscillatoriophycideae;Chroococcales

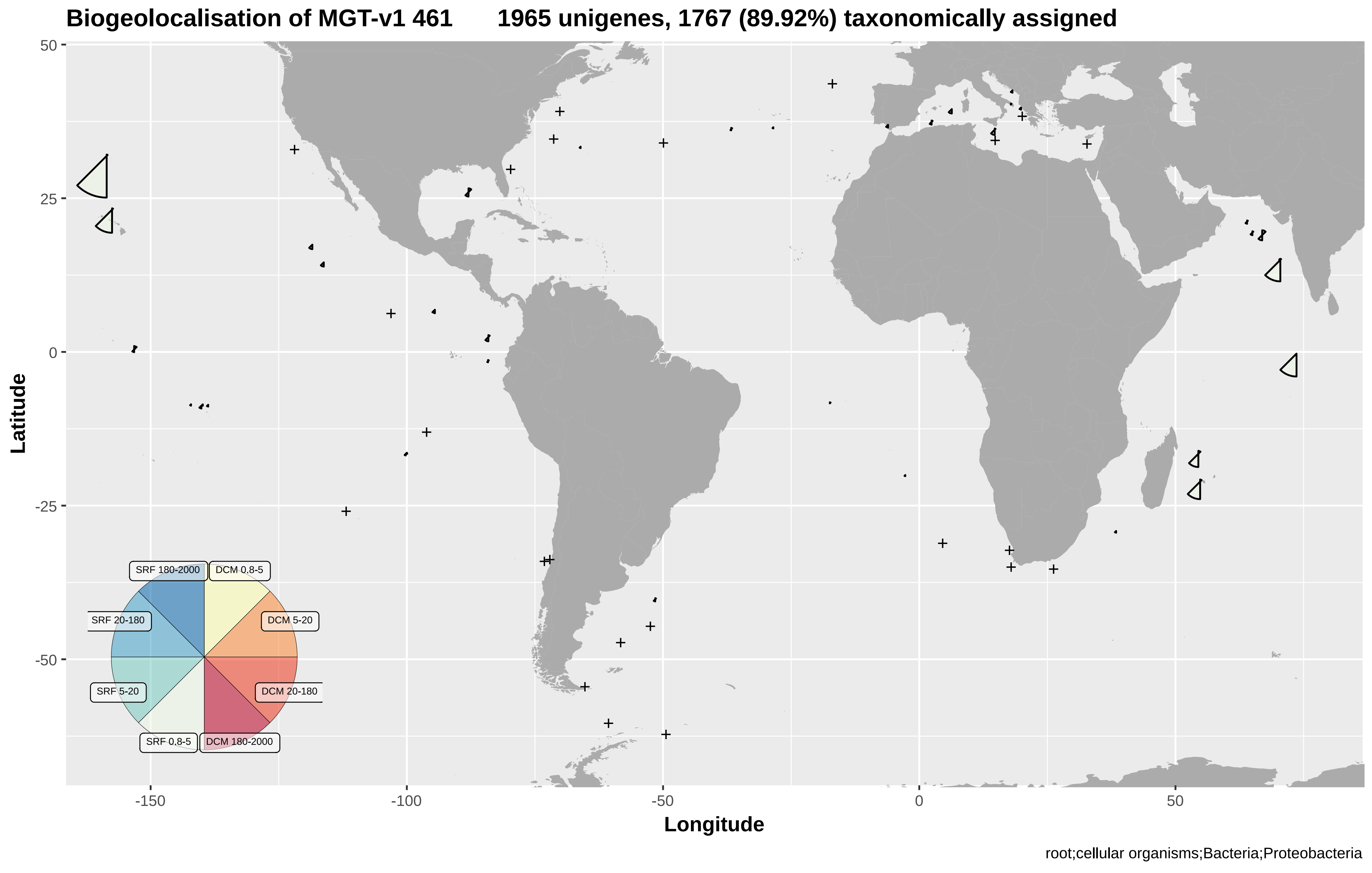

Biogeolocalisation of MGT-v1 461
1965 unigenes, 1767 (89.92%) taxonomically assigned
Latitude
Longitude
50
25
0
-25
-50
-150
-100
-50
0
50
SRF 180-2000
DCM 0.8-5
SRF 20-180
DCM 5-20
SRF 5-20
DCM 20-180
SRF 0.8-5
DCM 180-2000
root;cellular organisms;Bacteria;Proteobacteria

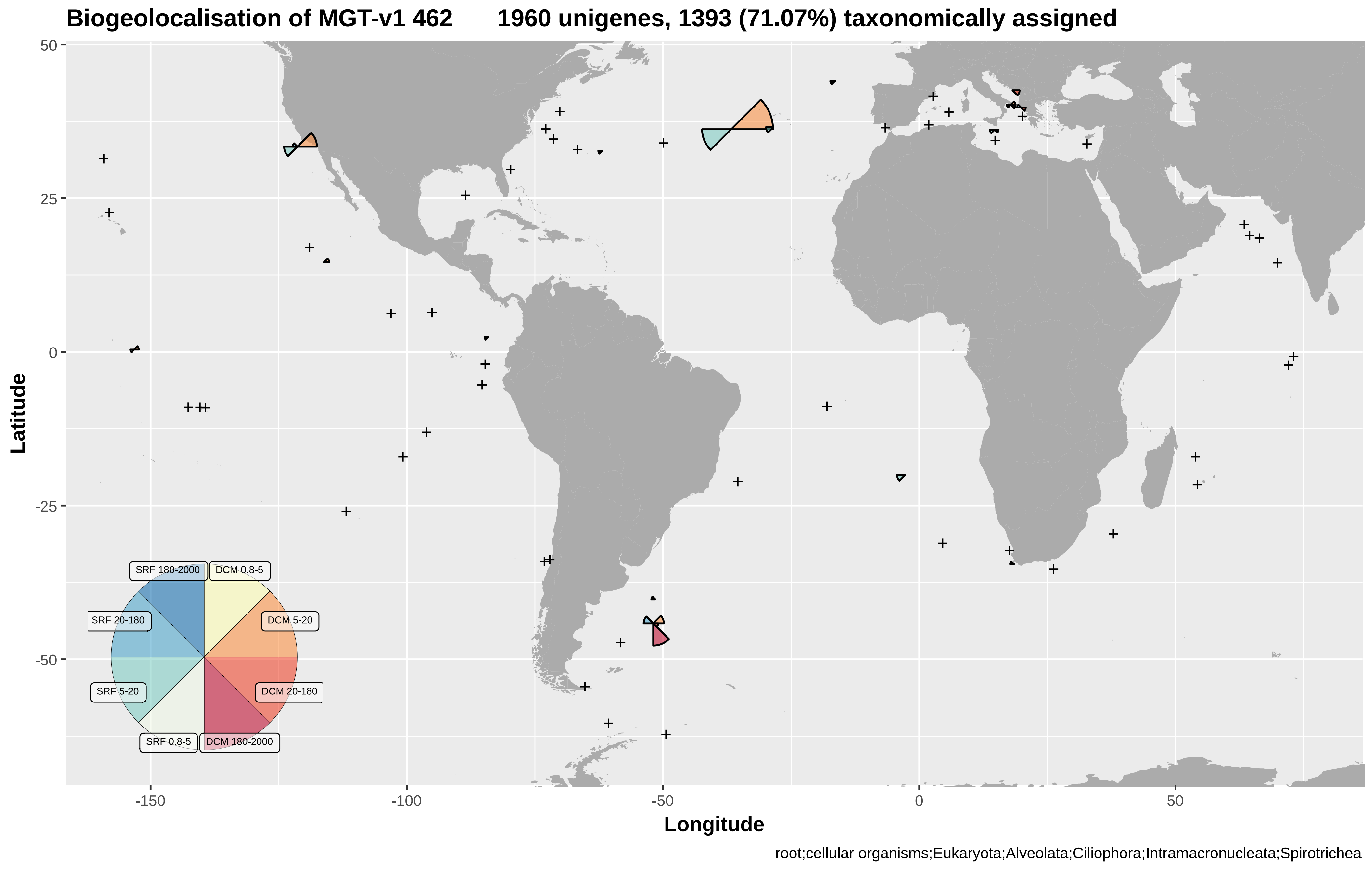
Biogeolocalisation of MGT-v1 462
1960 unigenes, 1393 (71.07%) taxonomically assigned
Latitude
Longitude
50
25
0
-25
-50
-150
-100
-50
0
50
SRF 180-2000
DCM 0.8-5
SRF 20-180
DCM 5-20
SRF 5-20
DCM 20-180
SRF 0.8-5
DCM 180-2000
root;cellular organisms;Eukaryota;Alveolata;Ciliophora;Intramacronucleata;Spirotrichea

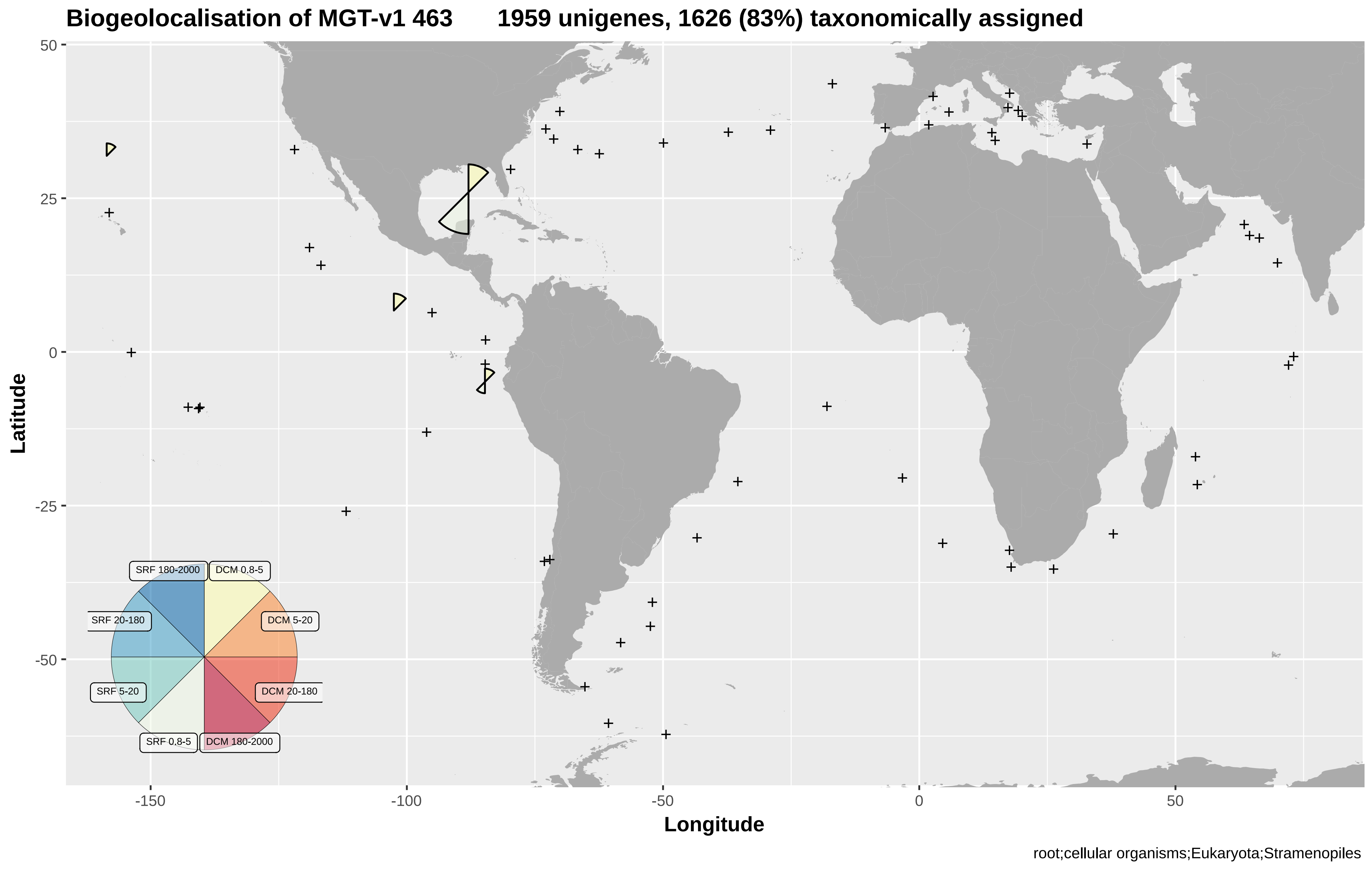

Biogeolocalisation of MGT-v1 463
1959 unigenes, 1626 (83%) taxonomically assigned
Latitude
Longitude
50
25
0
-25
-50
-150
-100
-50
0
50
SRF 180-2000
DCM 0.8-5
SRF 20-180
DCM 5-20
SRF 5-20
DCM 20-180
SRF 0.8-5
DCM 180-2000
root;cellular organisms;Eukaryota;Stramenopiles

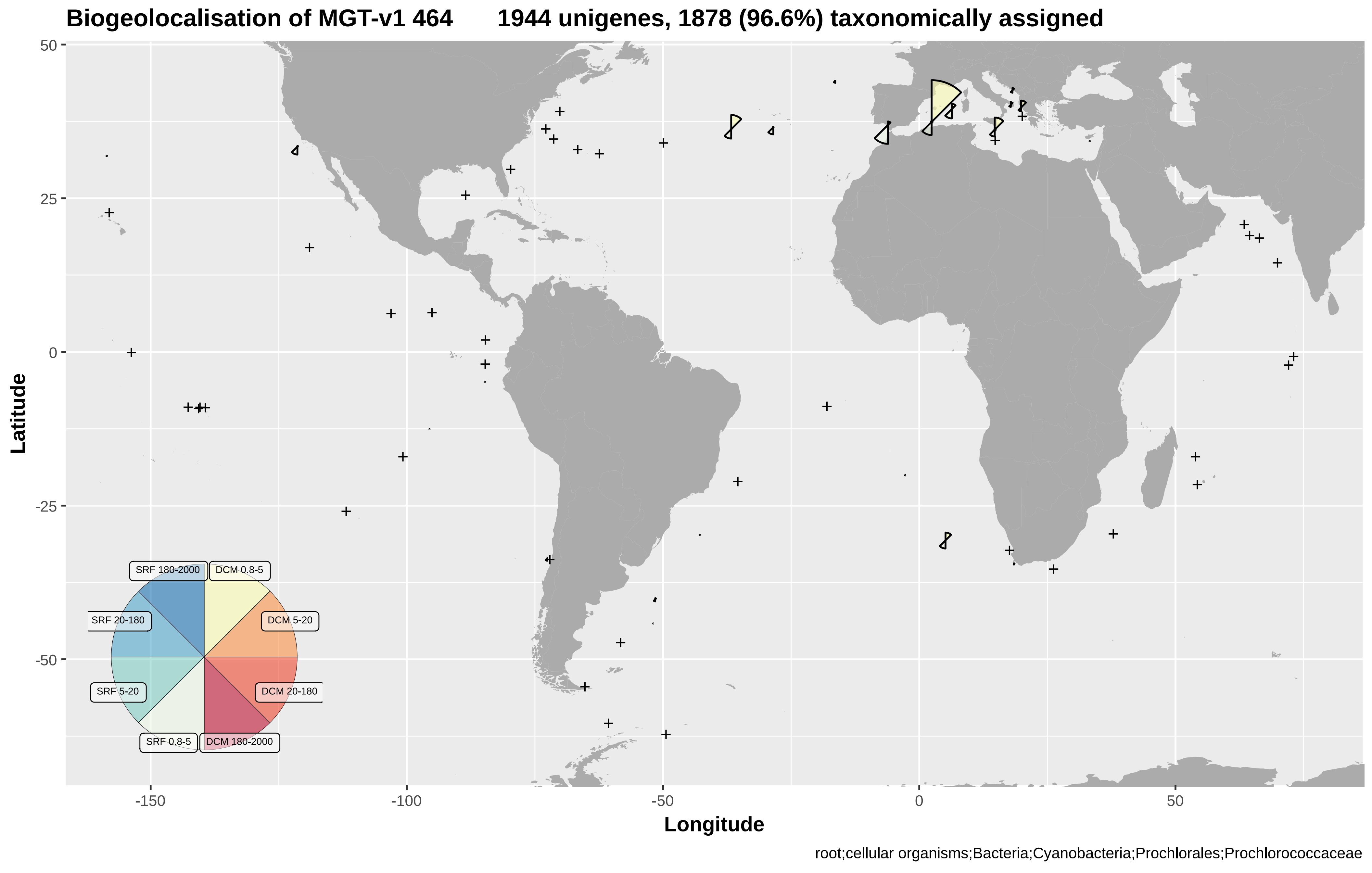

Biogeolocalisation of MGT-v1 464 1944 unigenes, 1878 (96.6%) taxonomically assigned
50
25
0
-25
-50
Latitude
-150
-100
-50
0
50
Longitude
SRF 180-2000
DCM 0.8-5
SRF 20-180
DCM 5-20
SRF 5-20
DCM 20-180
SRF 0.8-5
DCM 180-2000
root;cellular organisms;Bacteria;Cyanobacteria;Prochlorales;Prochlorococcaceae

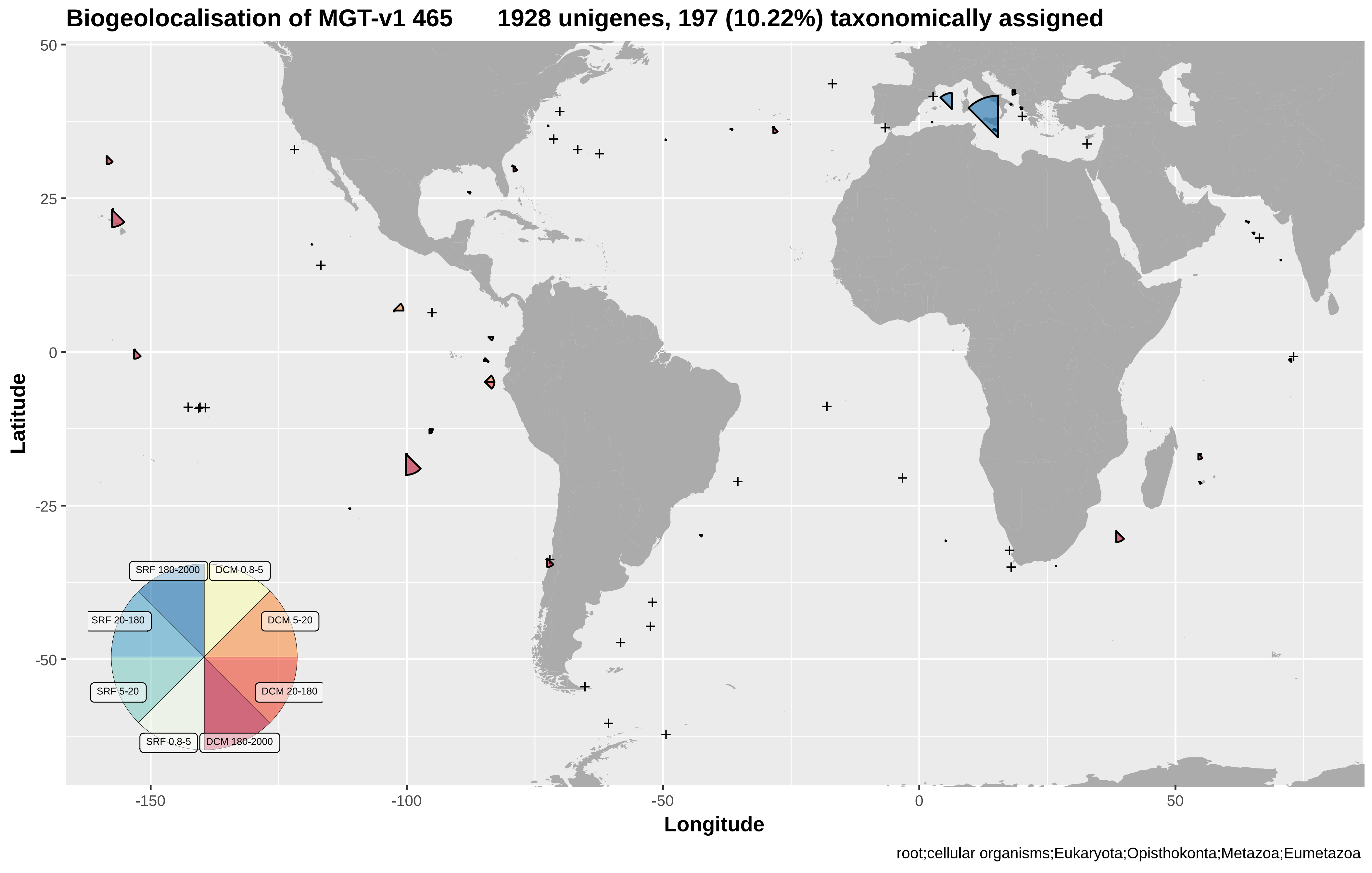

Biogeolocalisation of MGT-v1 465
1928 unigenes, 197 (10.22%) taxonomically assigned
50
25
0
-25
-50
Latitude
-150
-100
-50
0
50
Longitude
SRF 180-2000
DCM 0.8-5
SRF 20-180
DCM 5-20
SRF 5-20
DCM 20-180
SRF 0.8-5
DCM 180-2000
root;cellular organisms;Eukaryota;Opisthokonta;Metazoa;Eumetazoa

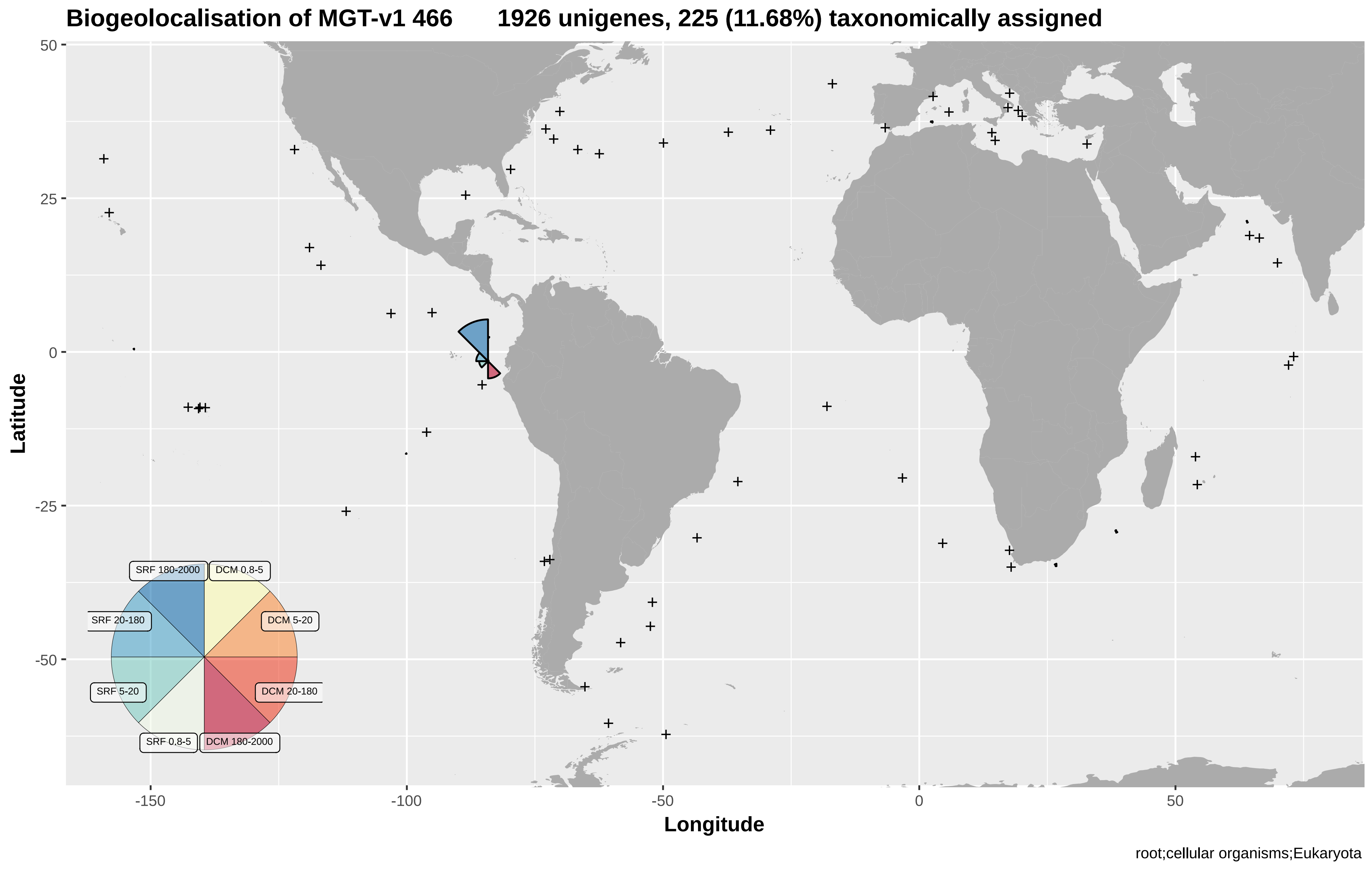
Biogeolocalisation of MGT-v1 466
1926 unigenes, 225 (11.68%) taxonomically assigned
Latitude
Longitude
50
25
0
-25
-50
-150
-100
-50
0
50
SRF 180-2000
DCM 0.8-5
SRF 20-180
DCM 5-20
SRF 5-20
DCM 20-180
SRF 0.8-5
DCM 180-2000
root;cellular organisms;Eukaryota

Biogeolocalisation of MGT-v1 467      1917 unigenes, 1734 (90.45%) taxonomically assigned

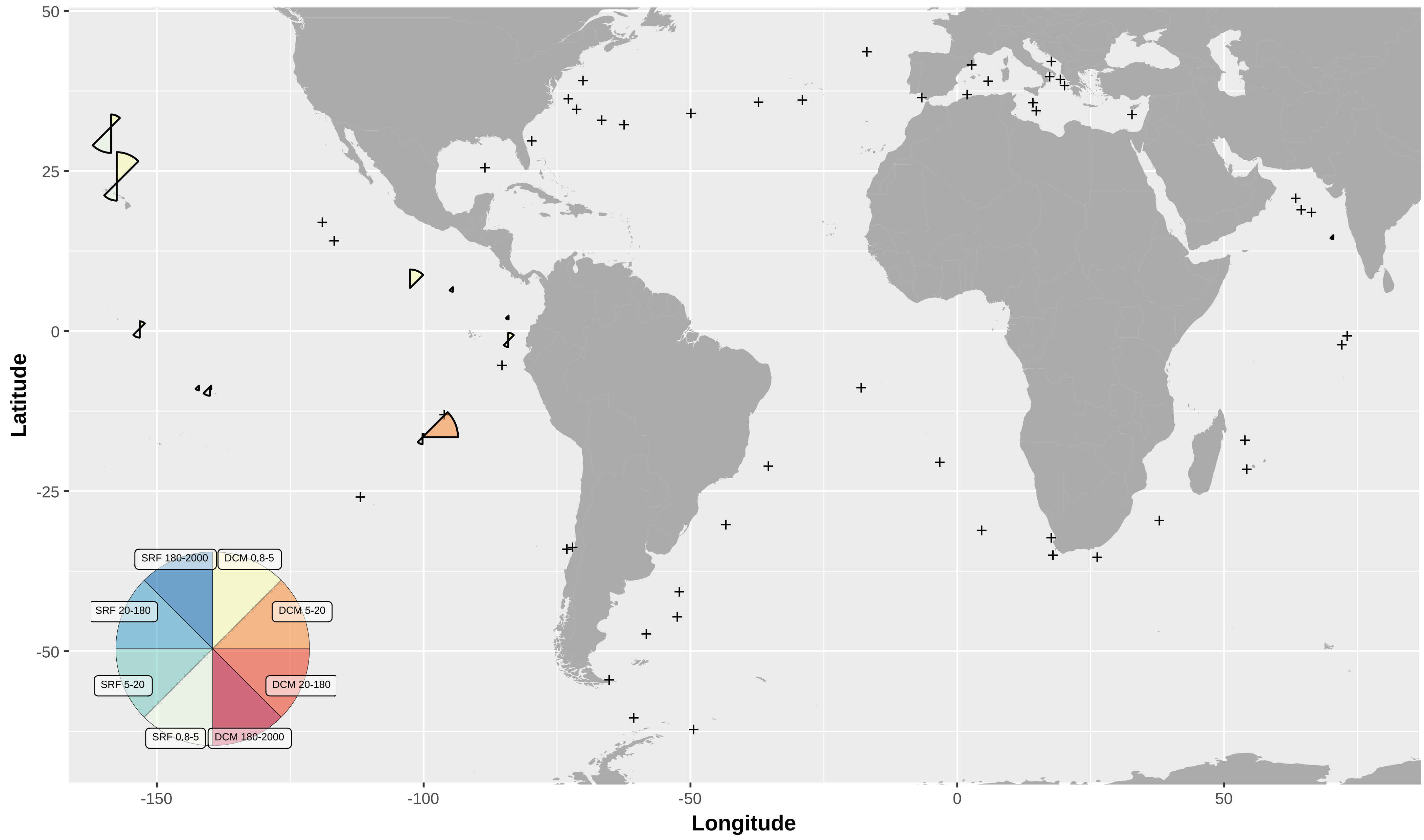


root;cellular organisms;Bacteria;Proteobacteria;Gammaproteobacteria

#### Biogeolocalisation of MGT-v1 468 1908 unigenes, 277 (14.52%) taxonomically assigned

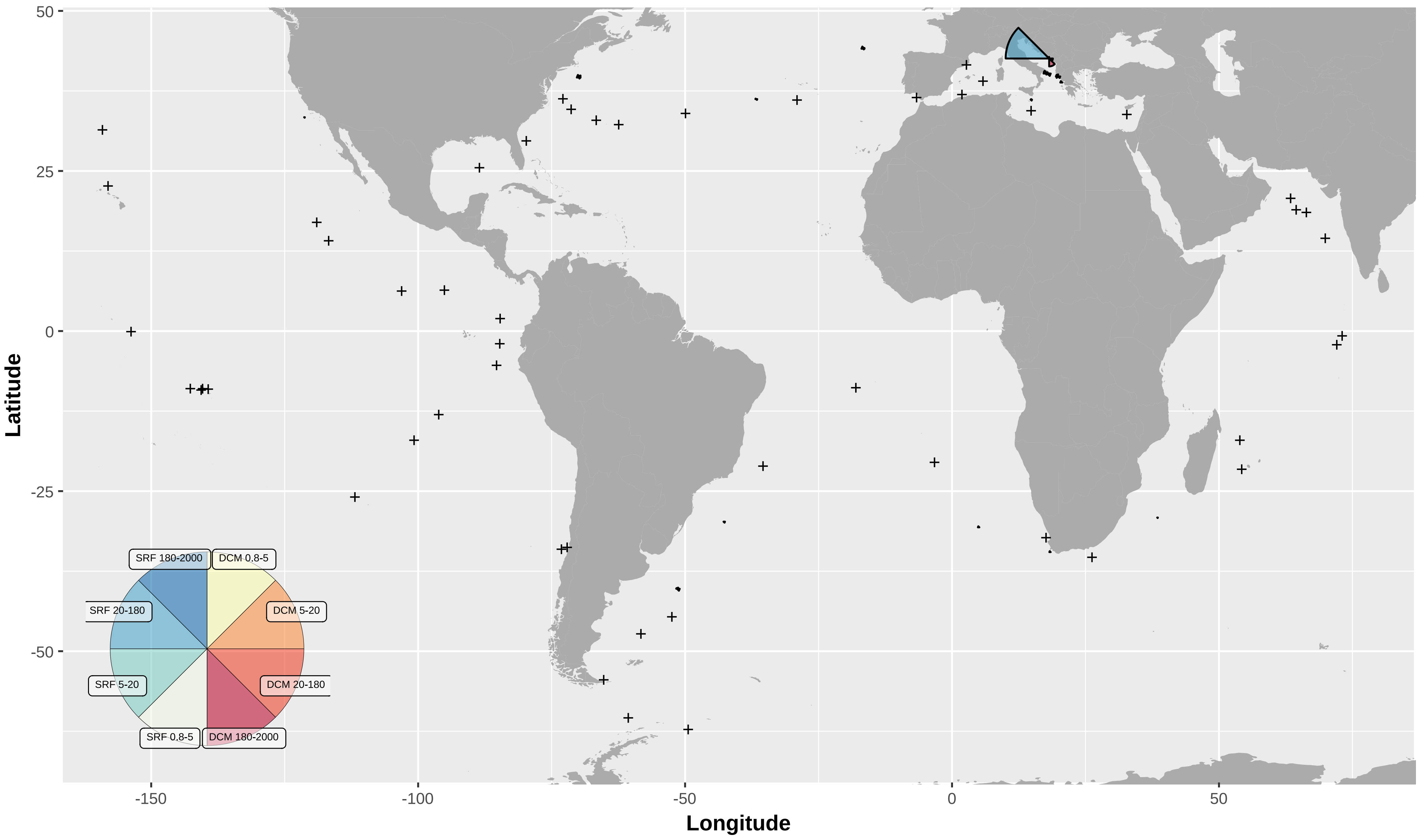


root;cellular organisms;Eukaryota;Opisthokonta;Metazoa;Eumetazoa;Bilateria

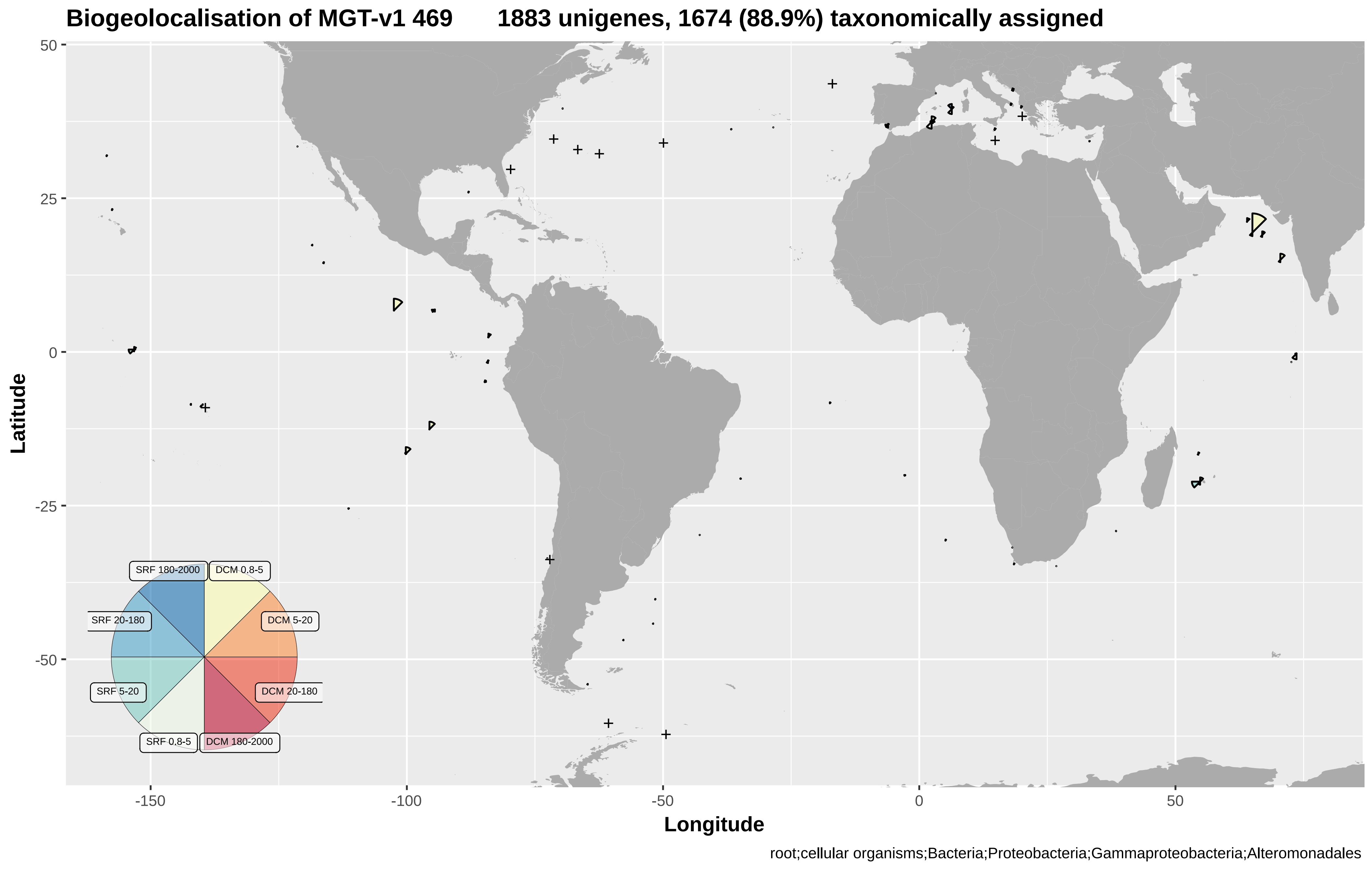

Biogeolocalisation of MGT-v1 469 1883 unigenes, 1674 (88.9%) taxonomically assigned
Latitude
Longitude
50
25
0
-25
-50
-150
-100
-50
0
50
SRF 180-2000
DCM 0.8-5
SRF 20-180
DCM 5-20
SRF 5-20
DCM 20-180
SRF 0.8-5
DCM 180-2000
root;cellular organisms;Bacteria;Proteobacteria;Gammaproteobacteria;Alteromonadales

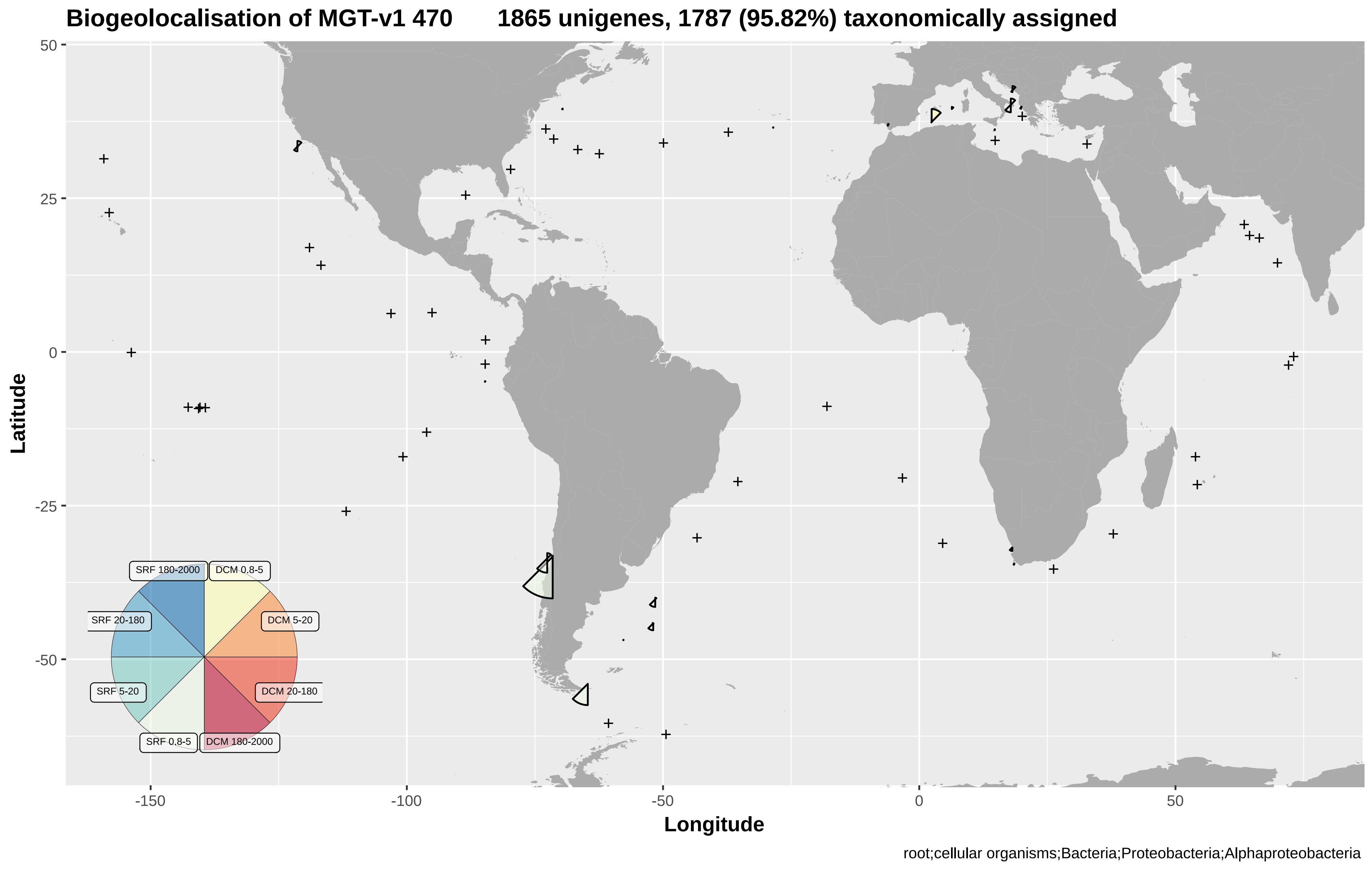

Biogeolocalisation of MGT-v1 470
1865 unigenes, 1787 (95.82%) taxonomically assigned
Latitude
Longitude
50
25
0
-25
-50
-150
-100
-50
0
50
SRF 180-2000
DCM 0.8-5
SRF 20-180
DCM 5-20
SRF 5-20
DCM 20-180
SRF 0.8-5
DCM 180-2000
root;cellular organisms;Bacteria;Proteobacteria;Alphaproteobacteria

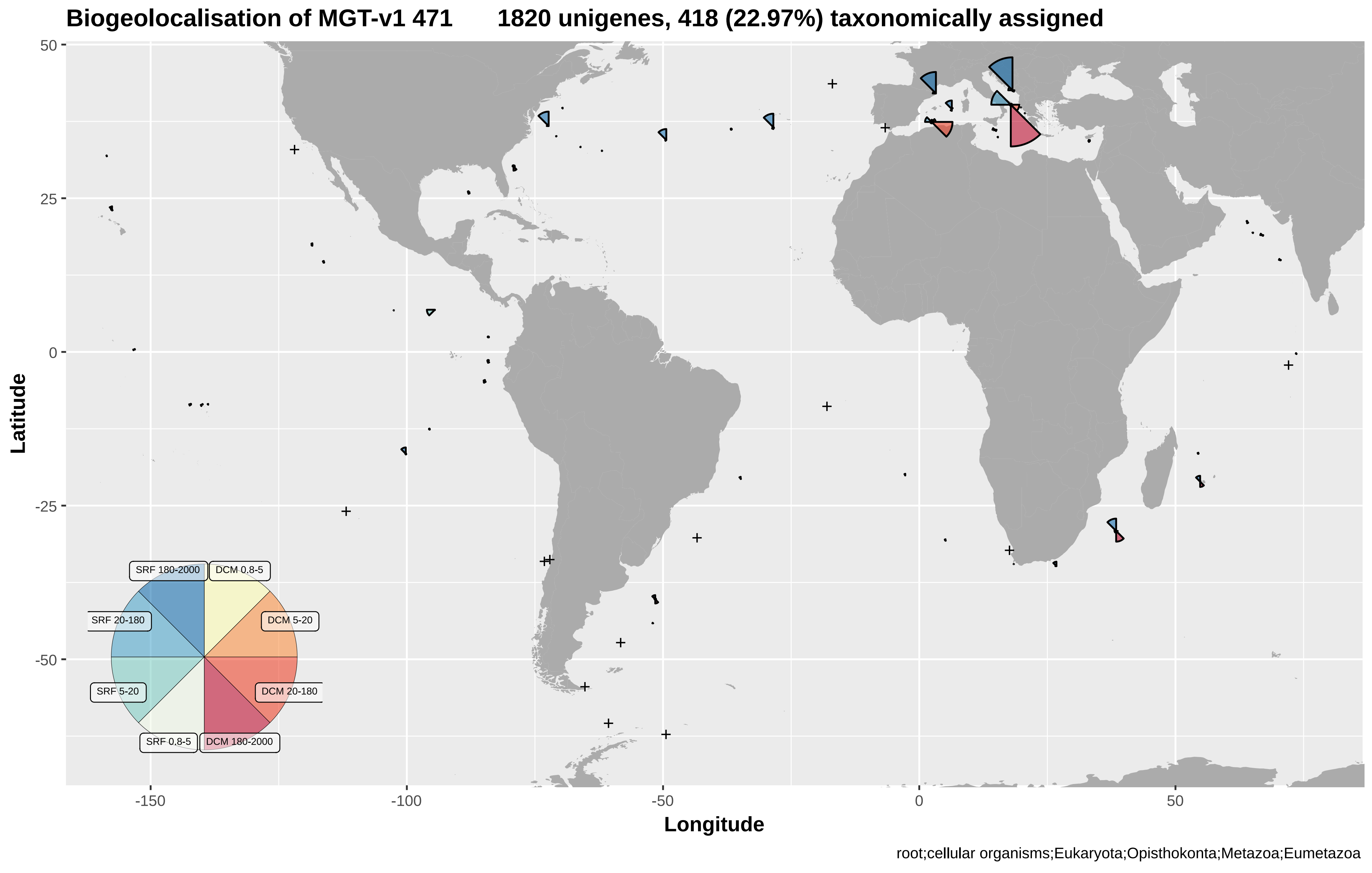

Biogeolocalisation of MGT-v1 471 1820 unigenes, 418 (22.97%) taxonomically assigned
50
25
0
-25
-50
Latitude
-150
-100
-50
0
50
Longitude
SRF 180-2000
DCM 0.8-5
SRF 20-180
DCM 5-20
SRF 5-20
DCM 20-180
SRF 0.8-5
DCM 180-2000
root;cellular organisms;Eukaryota;Opisthokonta;Metazoa;Eumetazoa

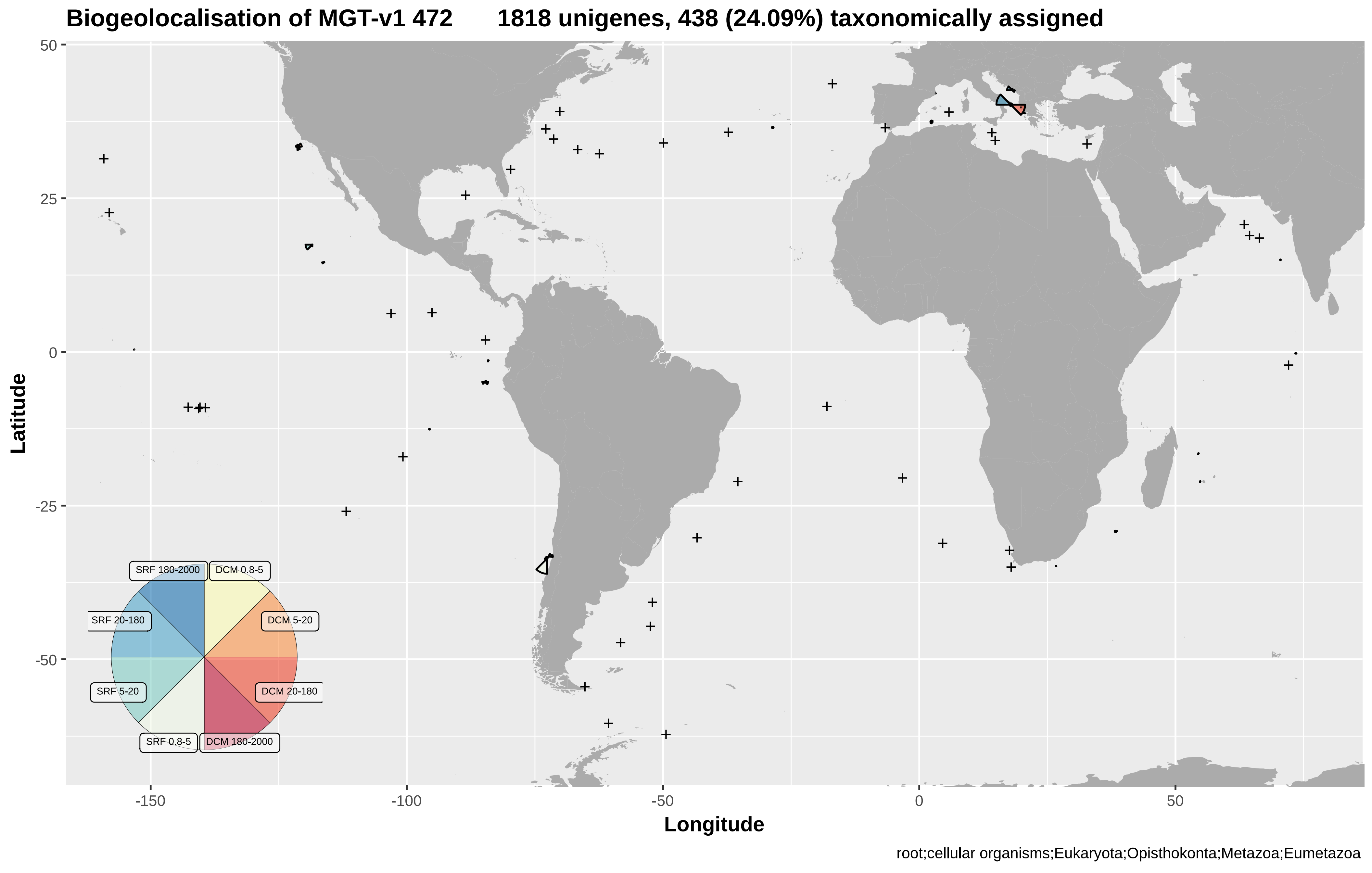

Biogeolocalisation of MGT-v1 472 1818 unigenes, 438 (24.09%) taxonomically assigned
50
25
0
-25
-50
Latitude
-150
-100
-50
0
50
Longitude
SRF 180-2000
DCM 0.8-5
SRF 20-180
DCM 5-20
SRF 5-20
DCM 20-180
SRF 0.8-5
DCM 180-2000
root;cellular organisms;Eukaryota;Opisthokonta;Metazoa;Eumetazoa

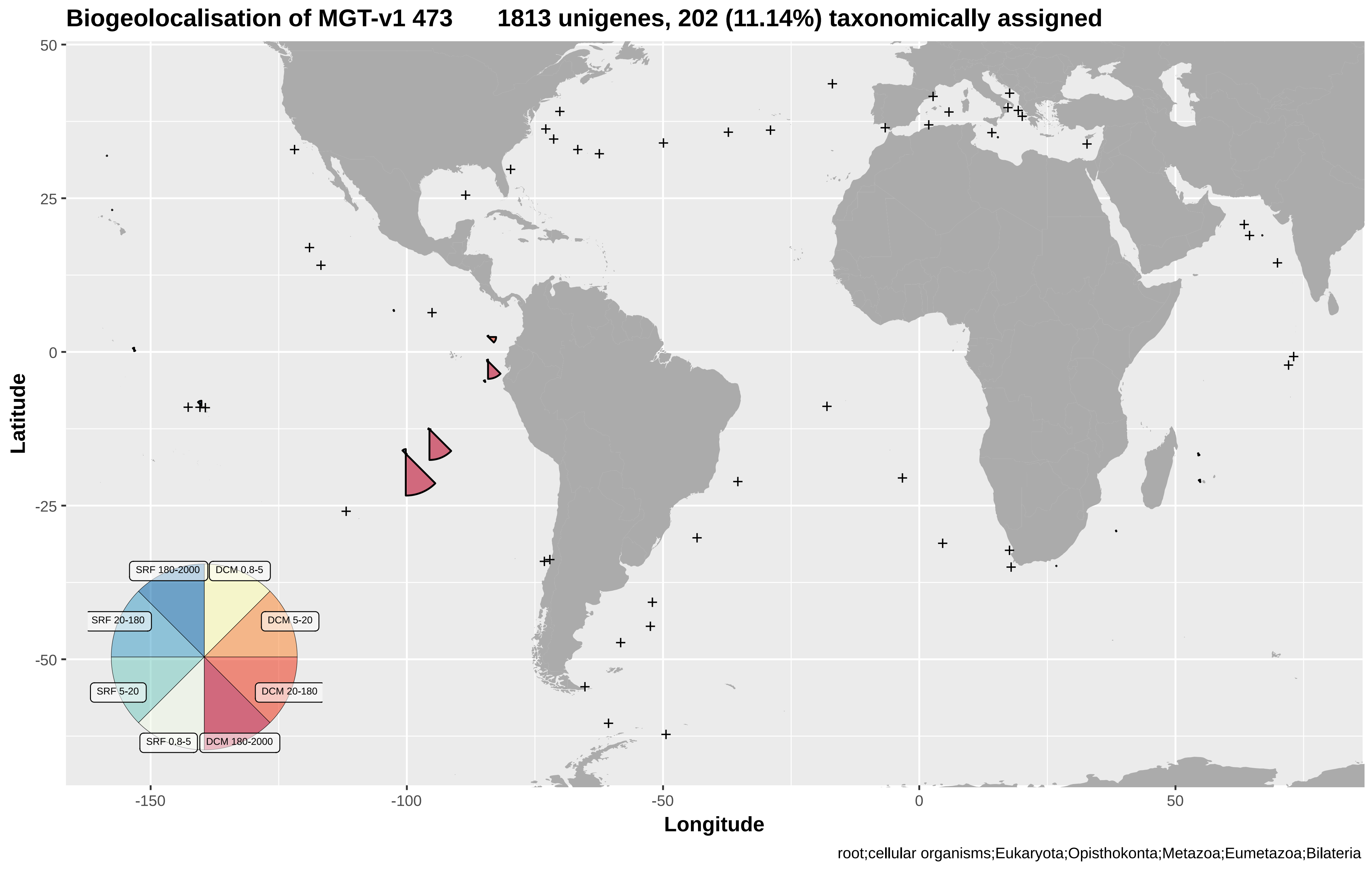
Biogeolocalisation of MGT-v1 473 1813 unigenes, 202 (11.14%) taxonomically assigned
50
25
0
-25
-50
Latitude
-150
-100
-50
0
50
Longitude
SRF 180-2000
DCM 0.8-5
SRF 20-180
DCM 5-20
SRF 5-20
DCM 20-180
SRF 0.8-5
DCM 180-2000
root;cellular organisms;Eukaryota;Opisthokonta;Metazoa;Eumetazoa;Bilateria

# Biogeolocalisation of MGT-v1 474      1812 unigenes, 727 (40.12%) taxonomically assigned

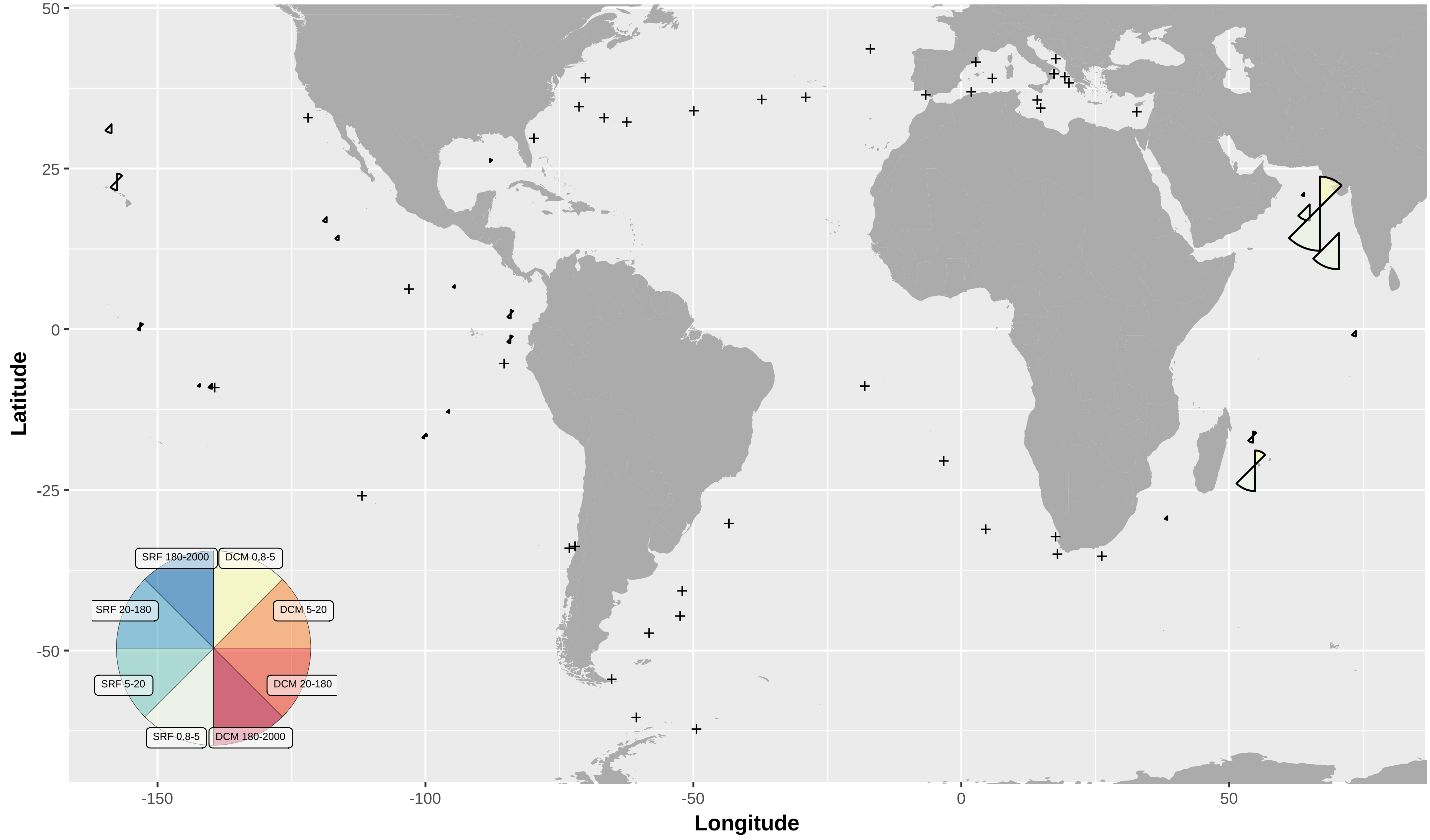


root;cellular organisms;Eukaryota;Haptophyceae

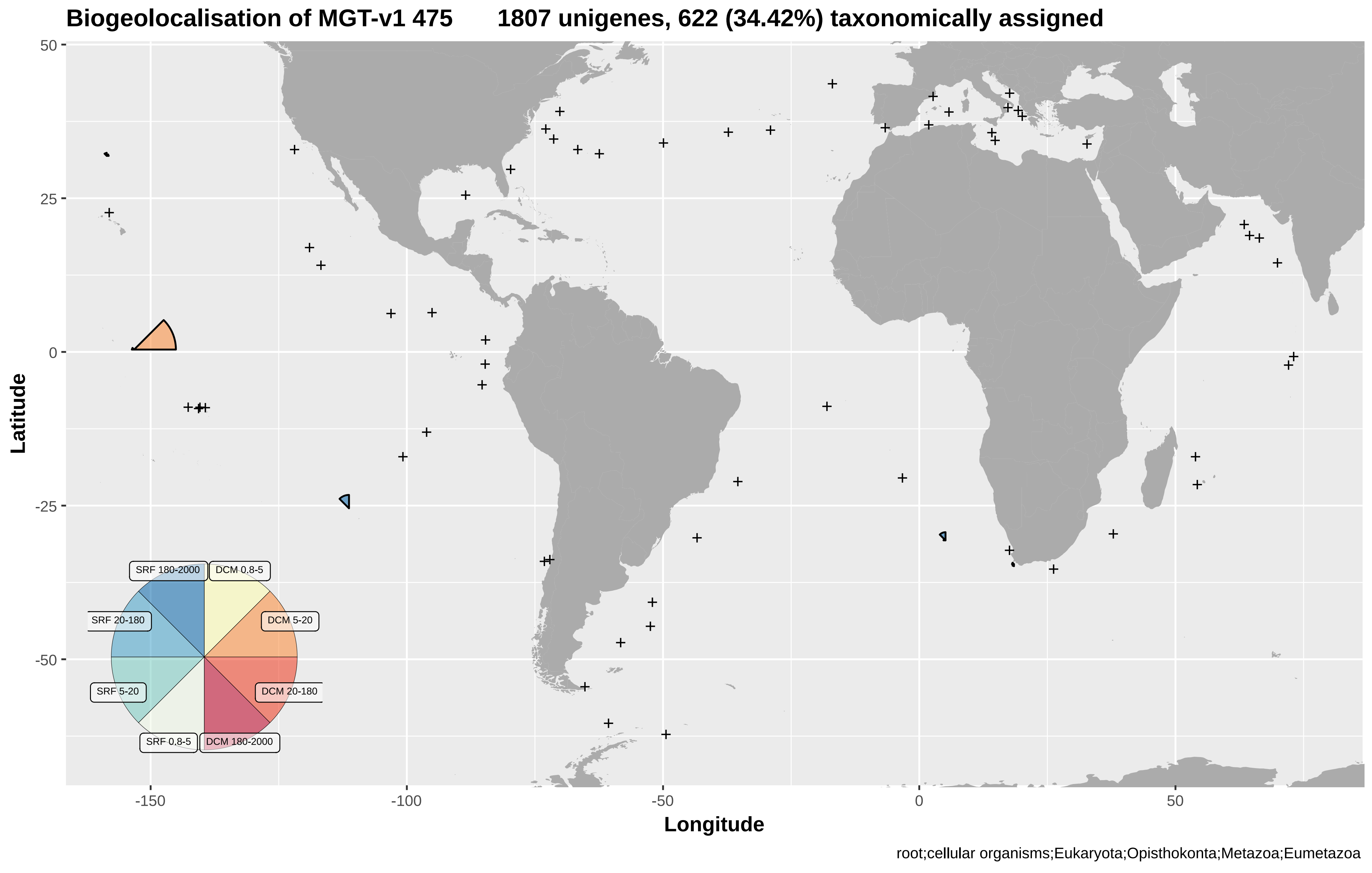
Biogeolocalisation of MGT-v1 475 1807 unigenes, 622 (34.42%) taxonomically assigned
50
25
0
-25
-50
Latitude
-150
-100
-50
0
50
Longitude
SRF 180-2000
DCM 0.8-5
SRF 20-180
DCM 5-20
SRF 5-20
DCM 20-180
SRF 0.8-5
DCM 180-2000
root;cellular organisms;Eukaryota;Opisthokonta;Metazoa;Eumetazoa

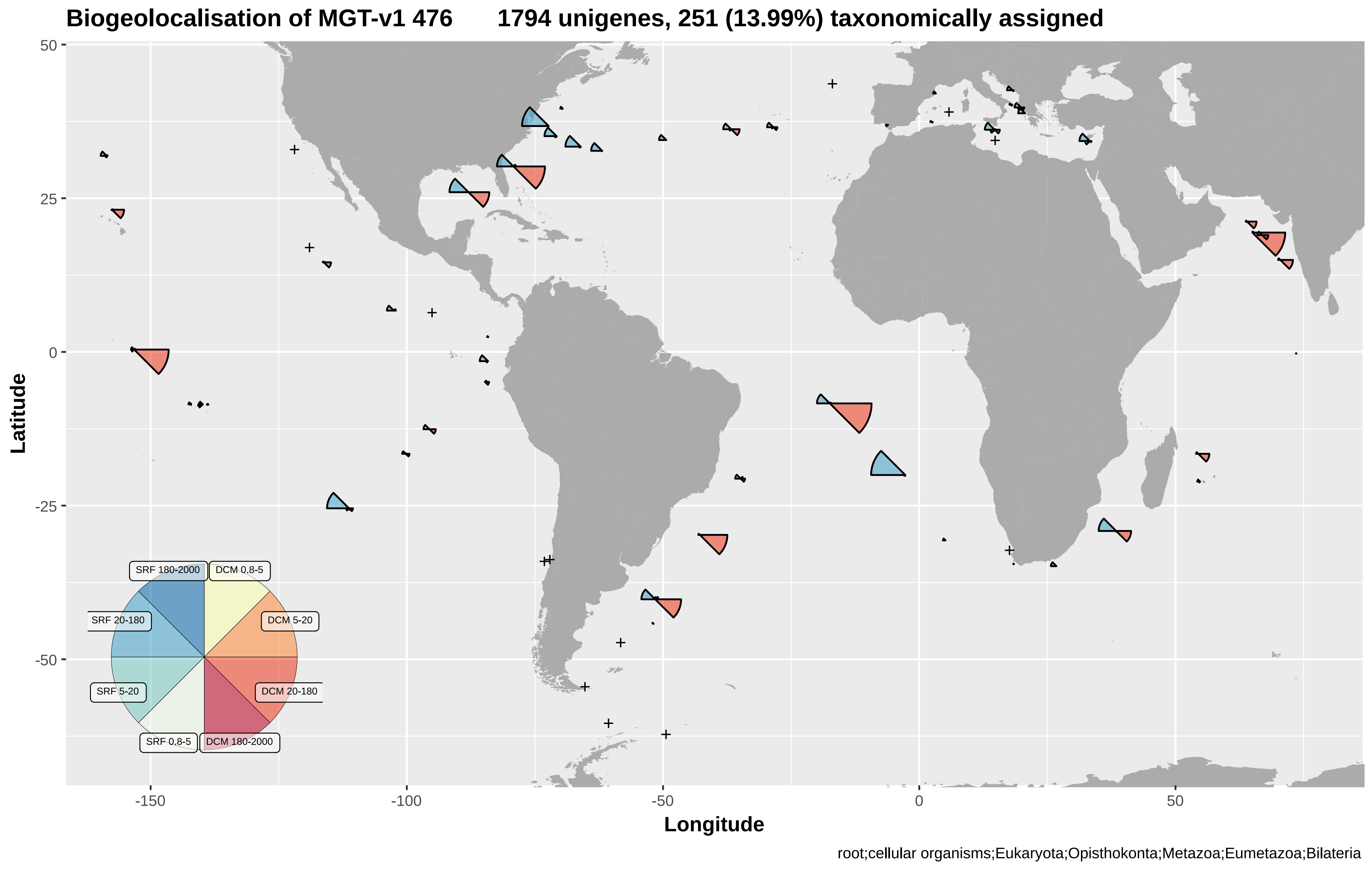
Biogeolocalisation of MGT-v1 476
1794 unigenes, 251 (13.99%) taxonomically assigned
50
25
0
-25
-50
Latitude
-150
-100
-50
0
50
Longitude
SRF 180-2000
DCM 0.8-5
SRF 20-180
DCM 5-20
SRF 5-20
DCM 20-180
SRF 0.8-5
DCM 180-2000
root;cellular organisms;Eukaryota;Opisthokonta;Metazoa;Eumetazoa;Bilateria

Biogeolocalisation of MGT-v1 477    1794 unigenes, 191 (10.65%) taxonomically assigned

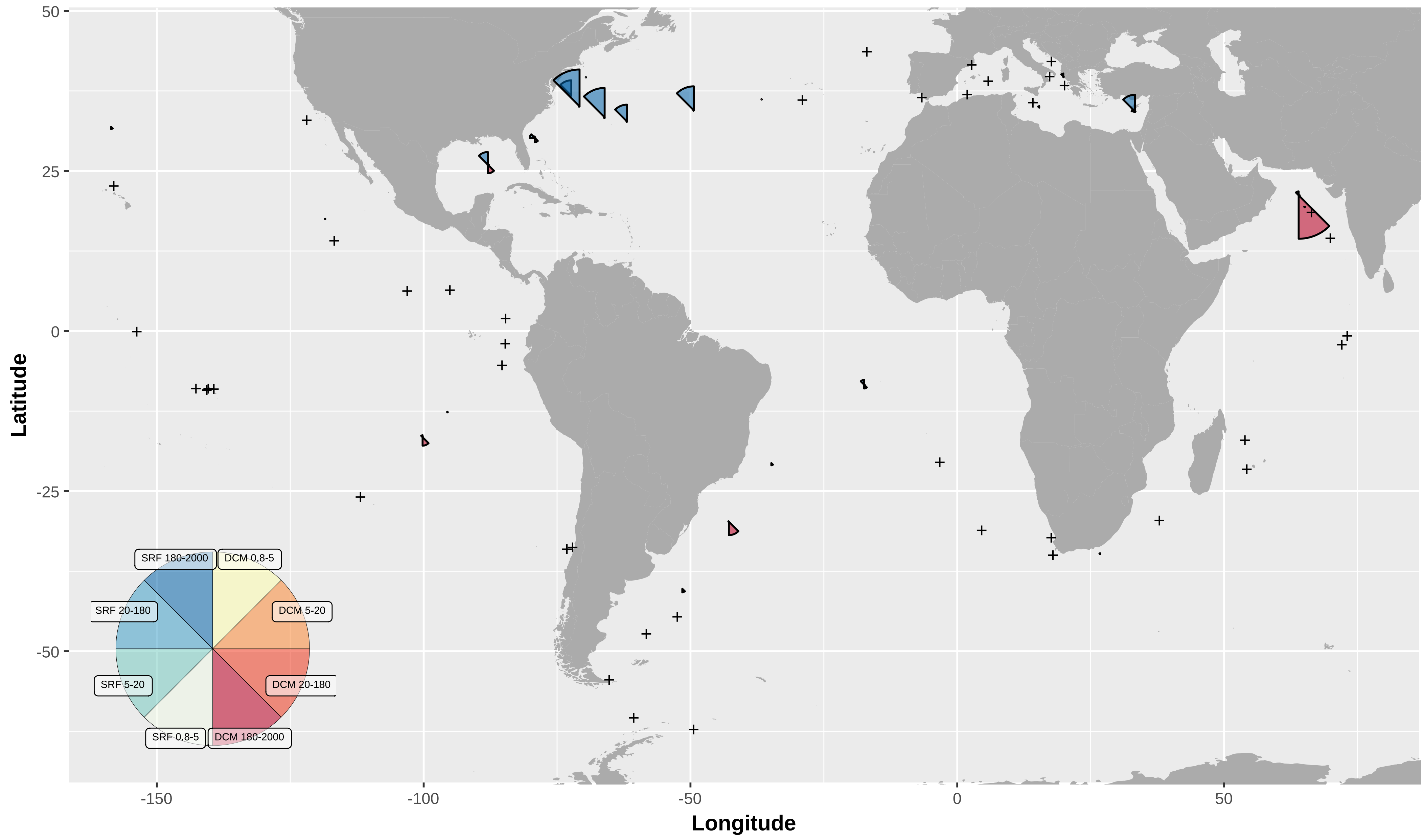


root;cellular organisms;Eukaryota;Opisthokonta;Metazoa;Eumetazoa;Bilateria

## Biogeolocalisation of MGT-v1 478      1792 unigenes, 119 (6.64%) taxonomically assigned

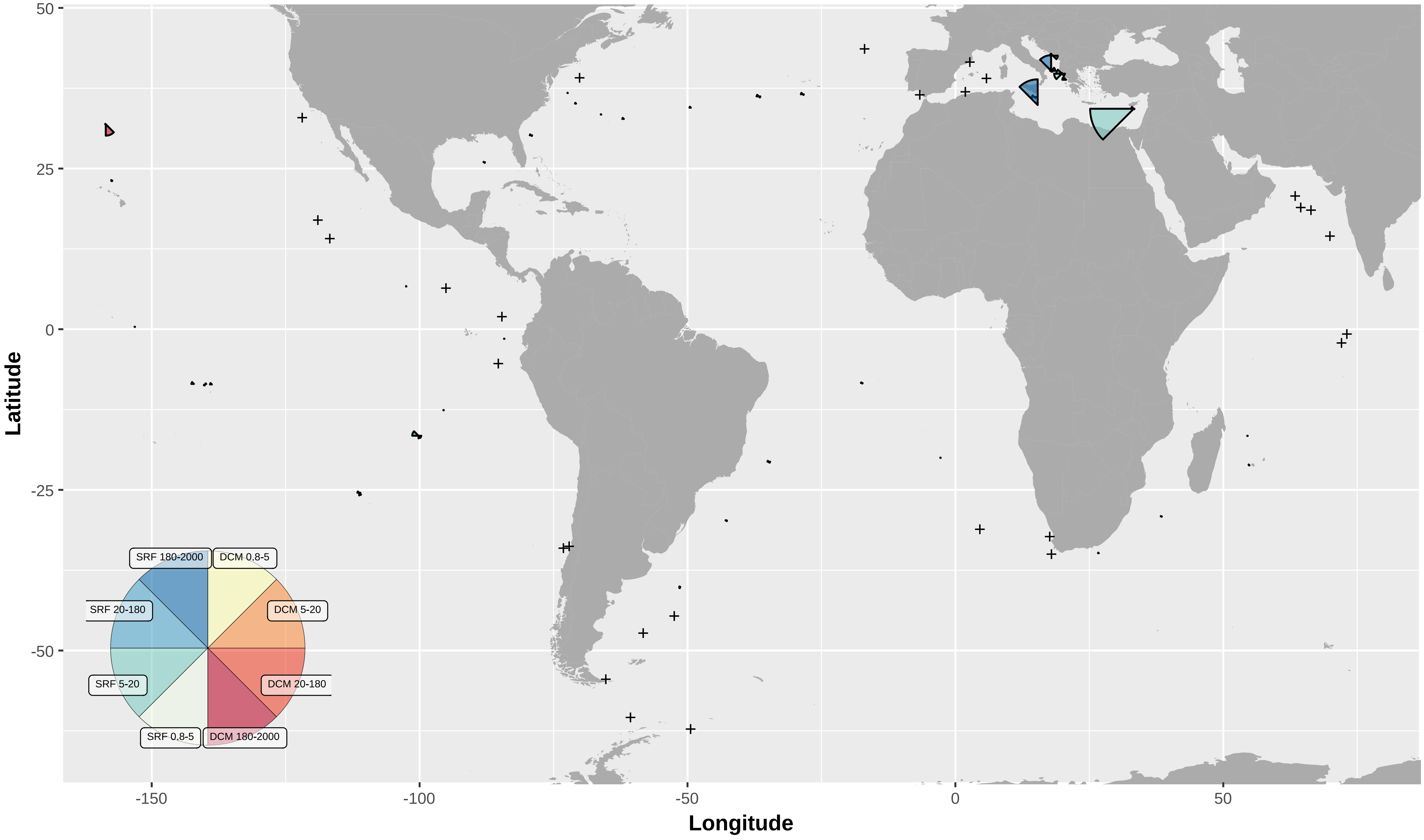


root;cellular organisms;Eukaryota;Opisthokonta;Metazoa;Eumetazoa;Bilateria

#### Biogeolocalisation of MGT-v1 479 1780 unigenes, 1594 (89.55%) taxonomically assigned

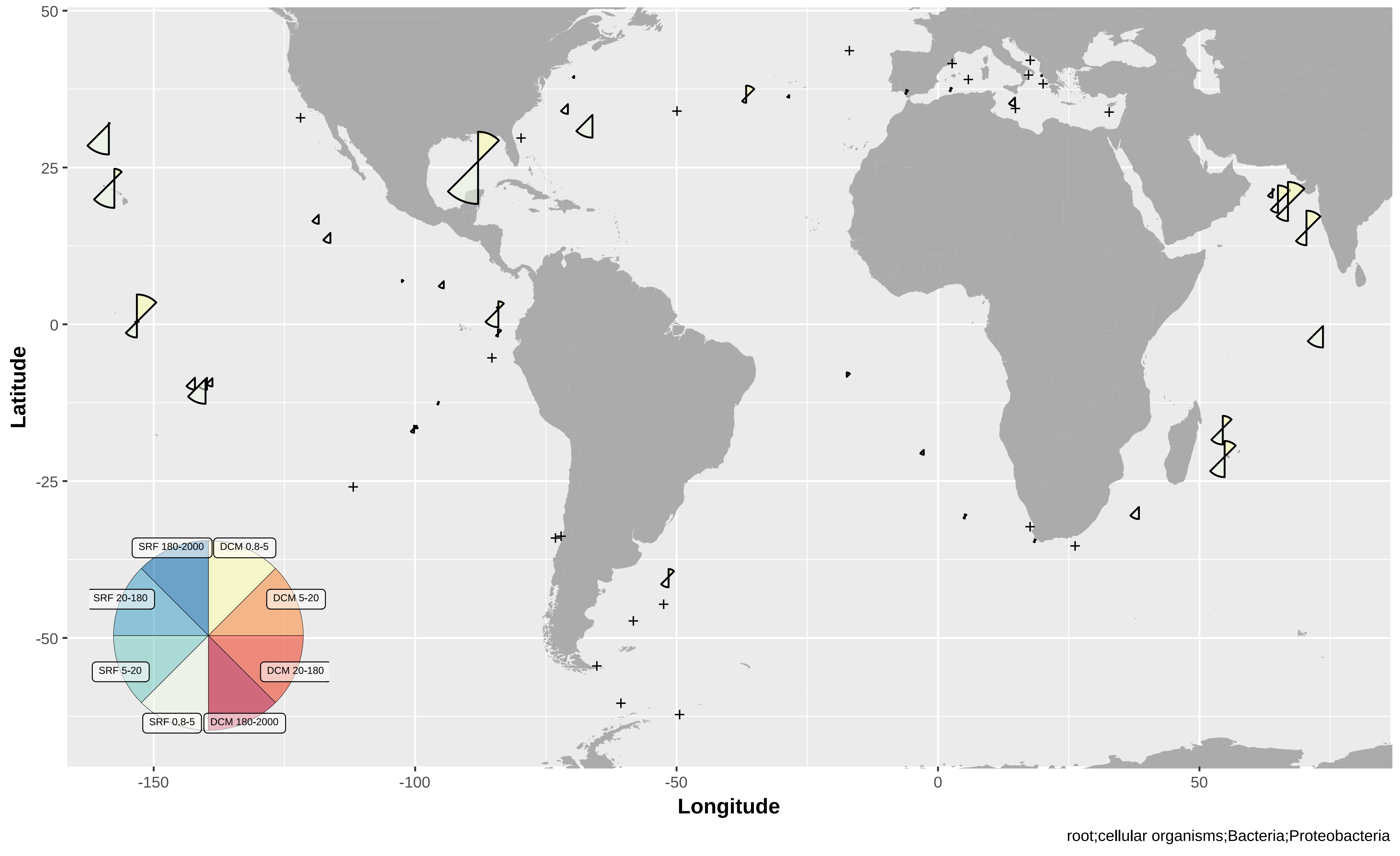

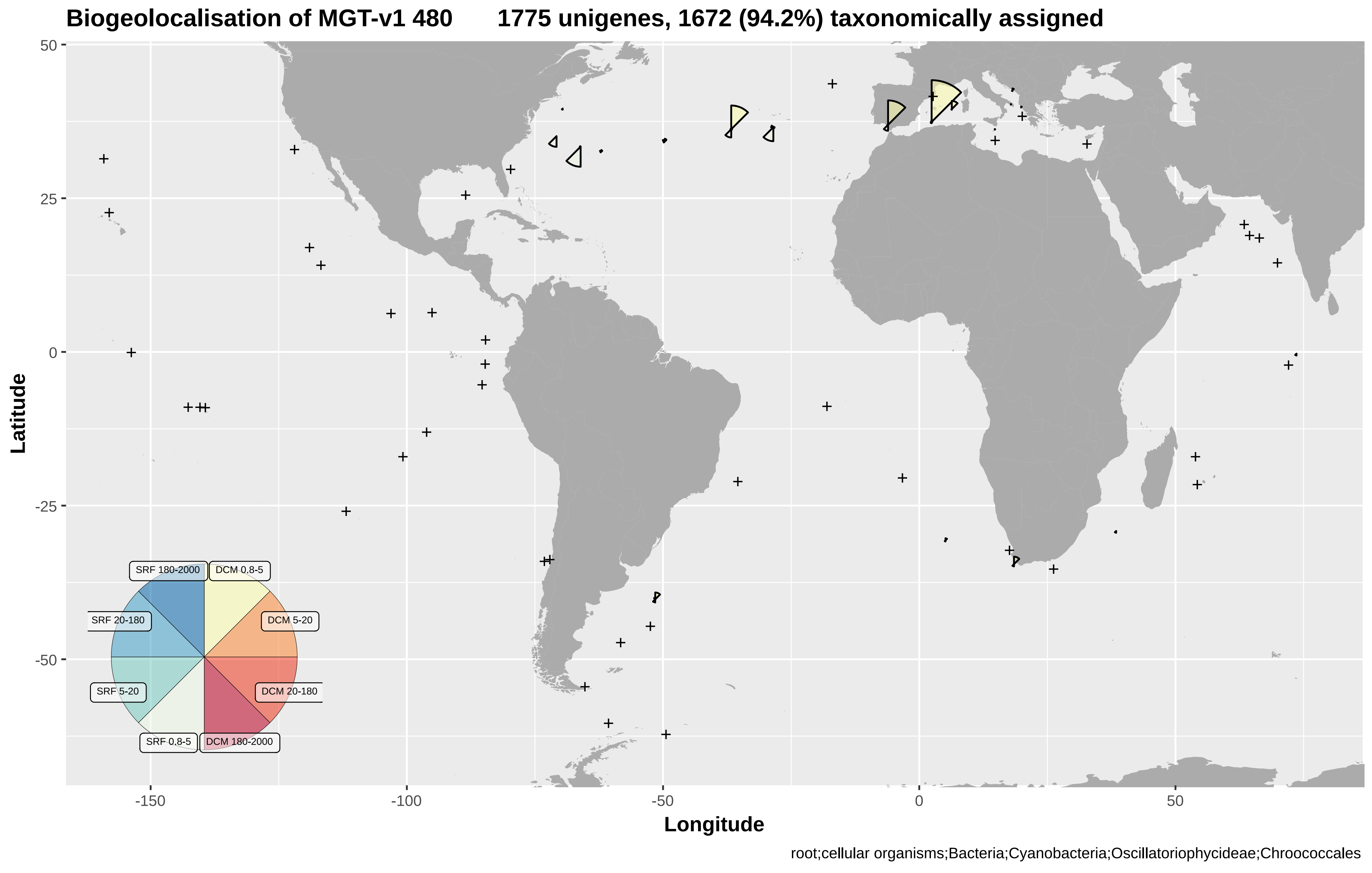

Biogeolocalisation of MGT-v1 480 1775 unigenes, 1672 (94.2%) taxonomically assigned
50
25
0
-25
-50
Latitude
-150
-100
-50
0
50
Longitude
SRF 180-2000
DCM 0.8-5
SRF 20-180
DCM 5-20
SRF 5-20
DCM 20-180
SRF 0.8-5
DCM 180-2000
root;cellular organisms;Bacteria;Cyanobacteria;Oscillatoriophycideae;Chroococcales

# Biogeolocalisation of MGT-v1 481      1767 unigenes, 137 (7.75%) taxonomically assigned

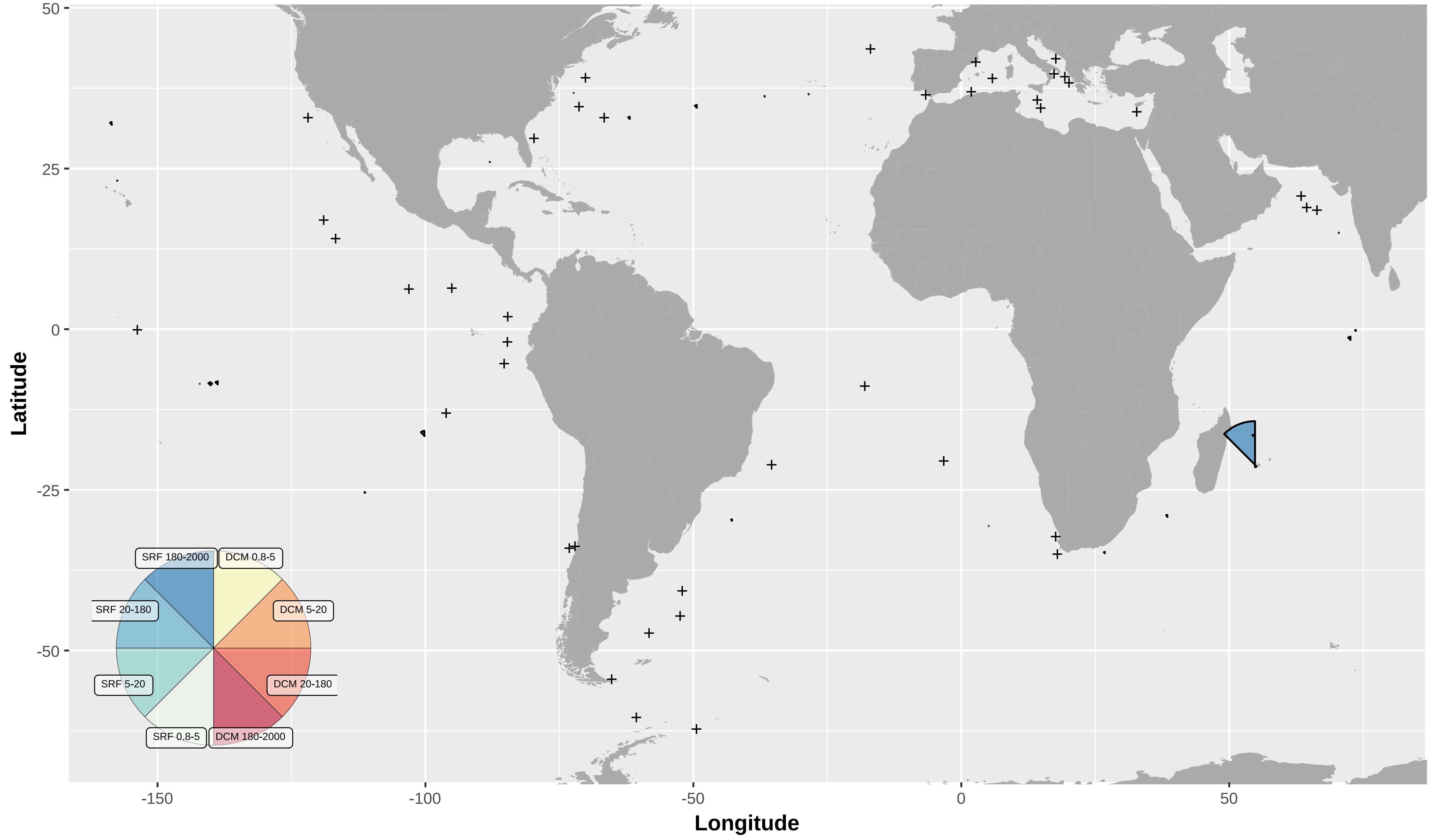


root;cellular organisms;Eukaryota;Opisthokonta;Metazoa;Eumetazoa

**Biogeolocalisation of MGT-v1 482 1767 unigenes, 1538 (87.04%) taxonomically assigned**

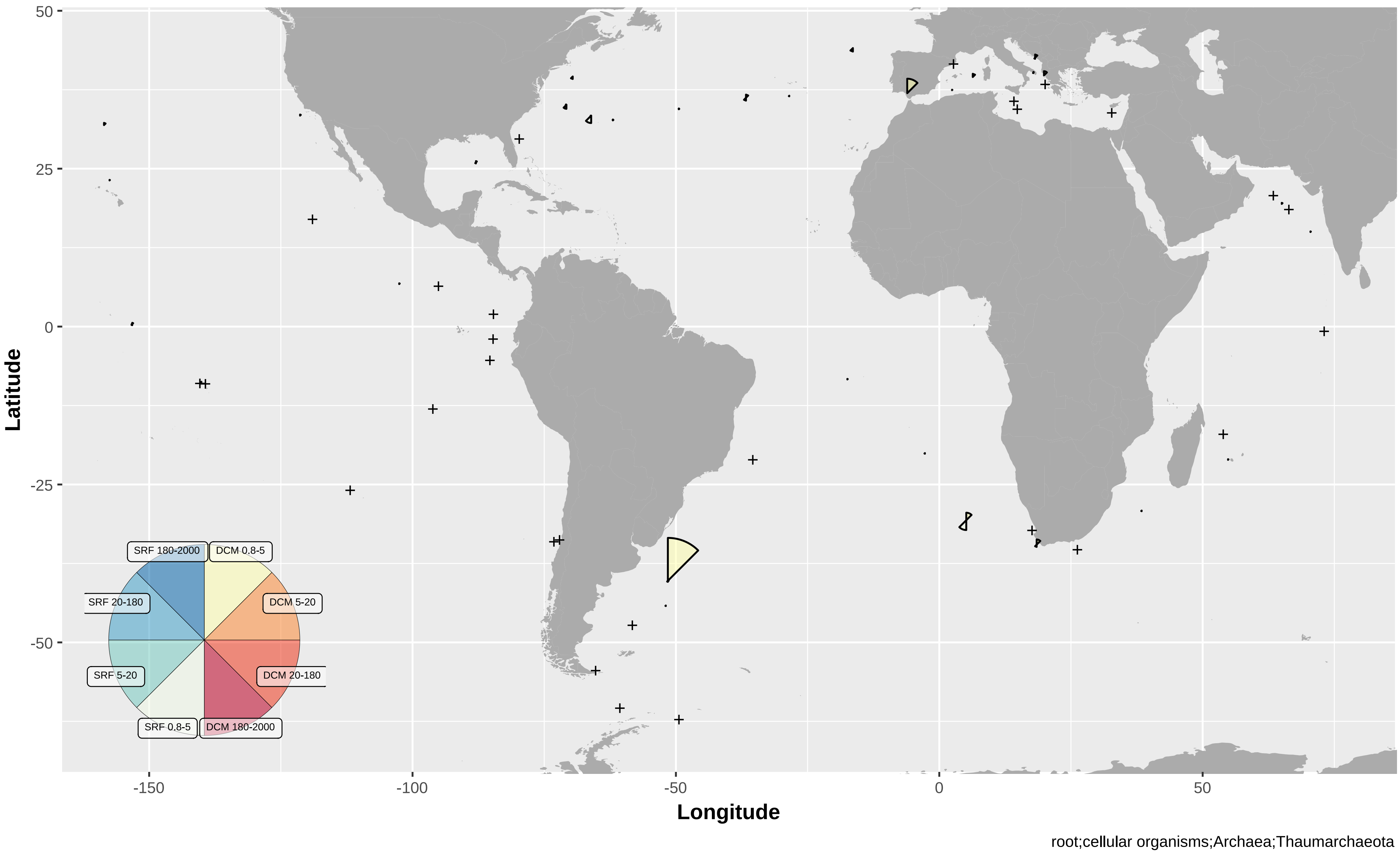

#### Biogeolocalisation of MGT-v1 483 1756 unigenes, 98 (5.58%) taxonomically assigned

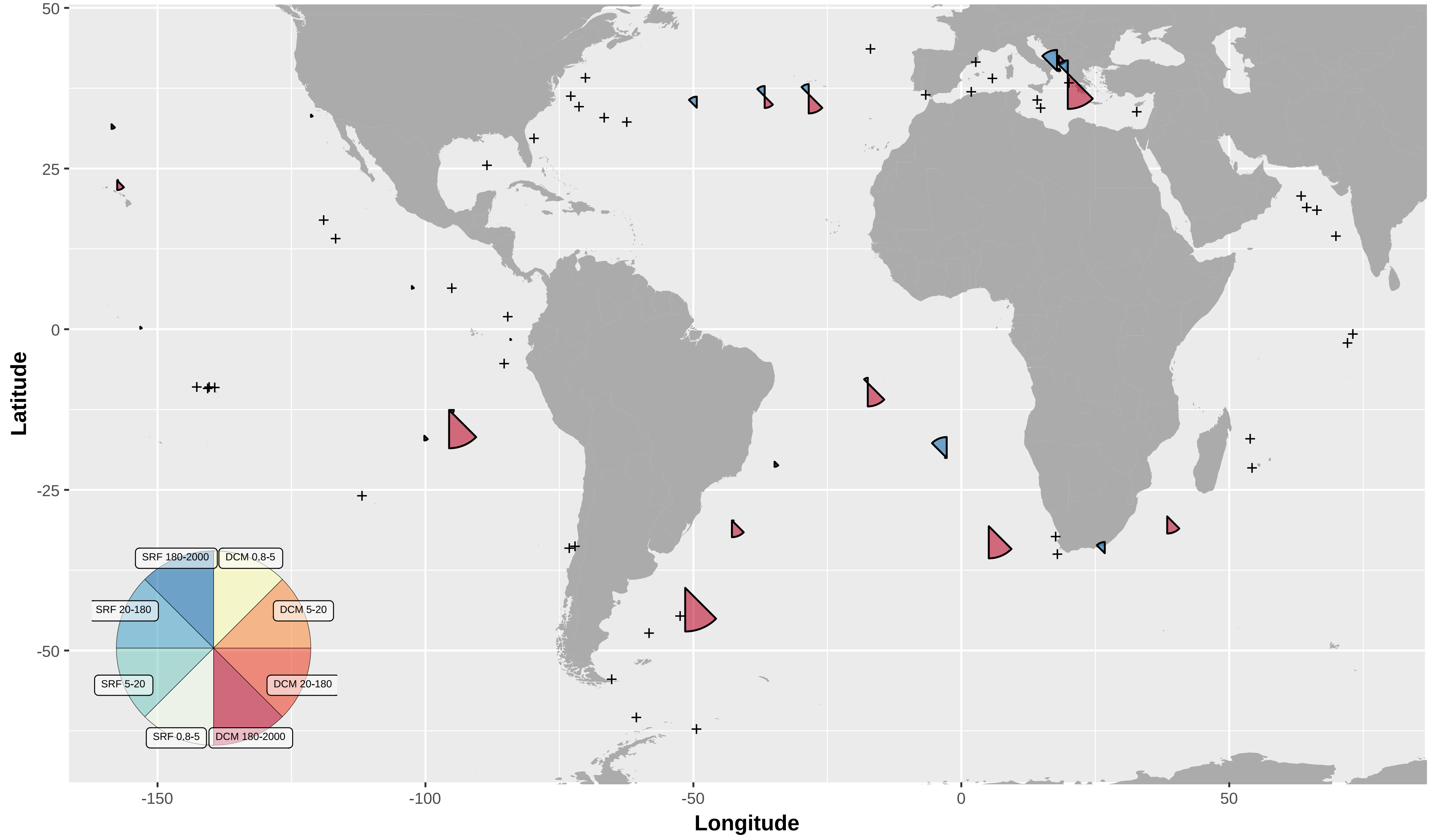


root;cellular organisms;Eukaryota;Opisthokonta;Metazoa;Eumetazoa

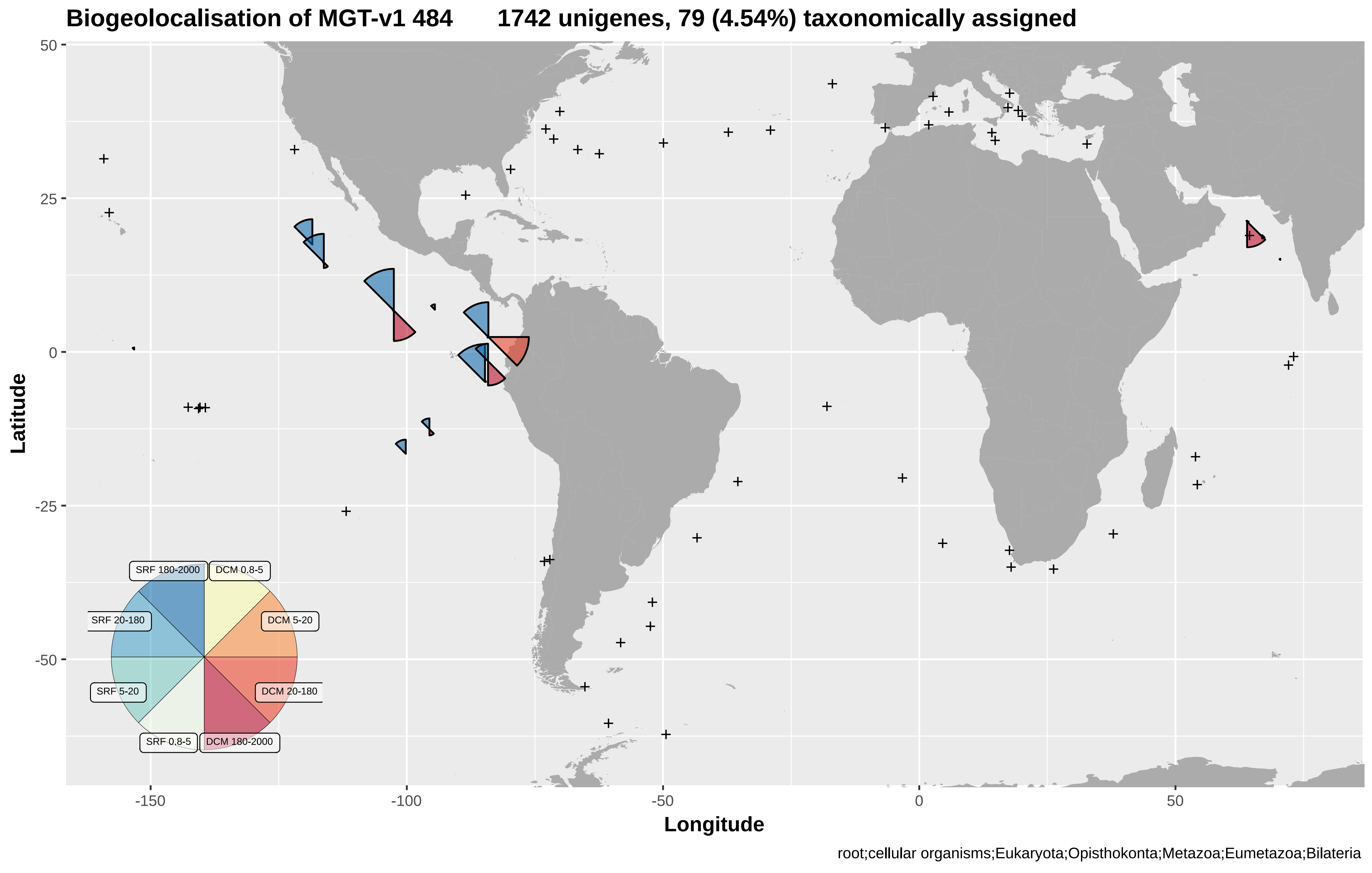
Biogeolocalisation of MGT-v1 484 1742 unigenes, 79 (4.54%) taxonomically assigned
50
25
0
-25
-50
Latitude
-150
-100
-50
0
50
Longitude
SRF 180-2000
DCM 0.8-5
SRF 20-180
DCM 5-20
SRF 5-20
DCM 20-180
SRF 0.8-5
DCM 180-2000
root;cellular organisms;Eukaryota;Opisthokonta;Metazoa;Eumetazoa;Bilateria

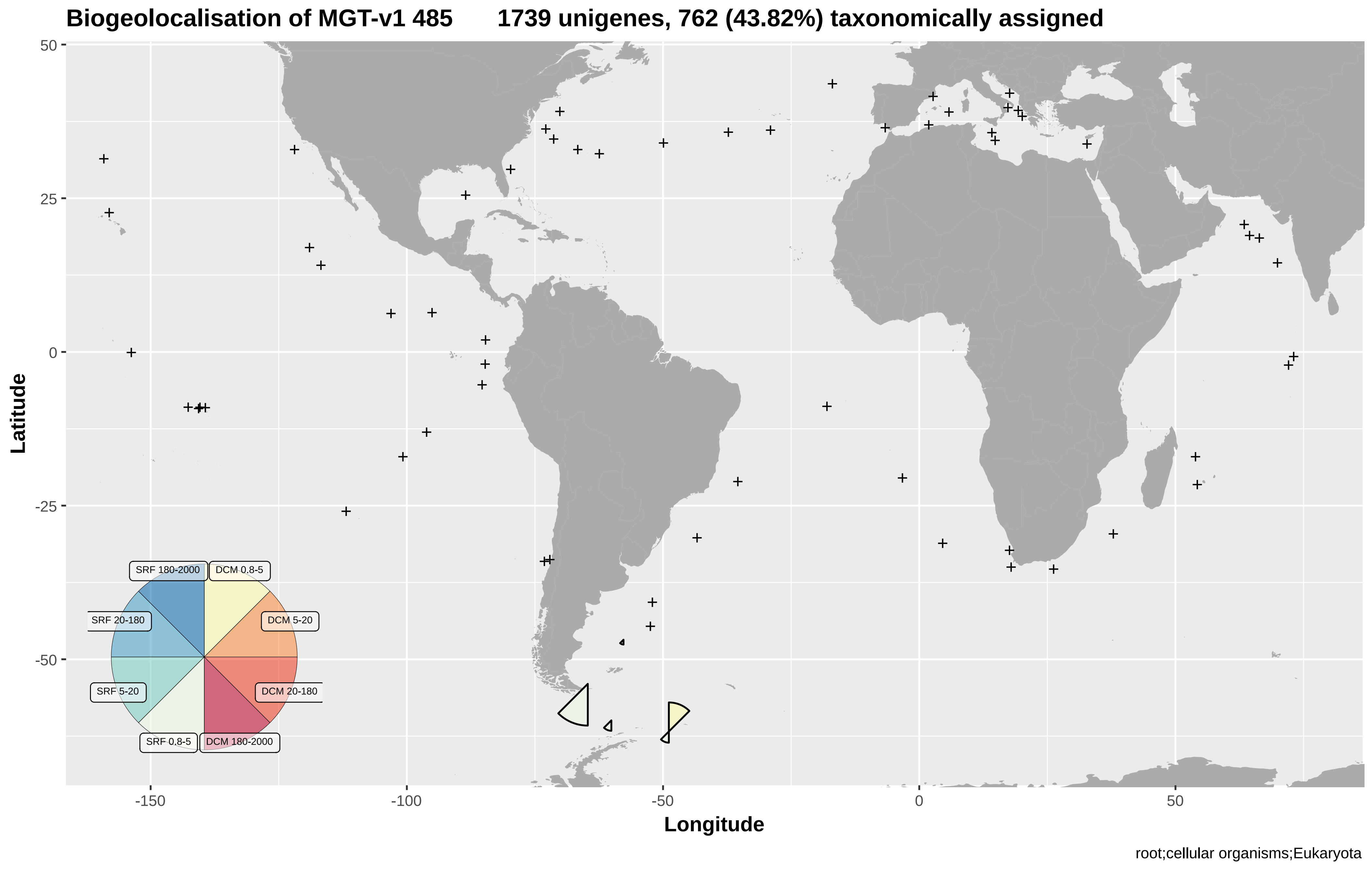
Biogeolocalisation of MGT-v1 485 1739 unigenes, 762 (43.82%) taxonomically assigned
50
25
0
-25
-50
Latitude
-150
-100
-50
0
50
Longitude
SRF 180-2000
DCM 0.8-5
SRF 20-180
DCM 5-20
SRF 5-20
DCM 20-180
SRF 0.8-5
DCM 180-2000
root;cellular organisms;Eukaryota

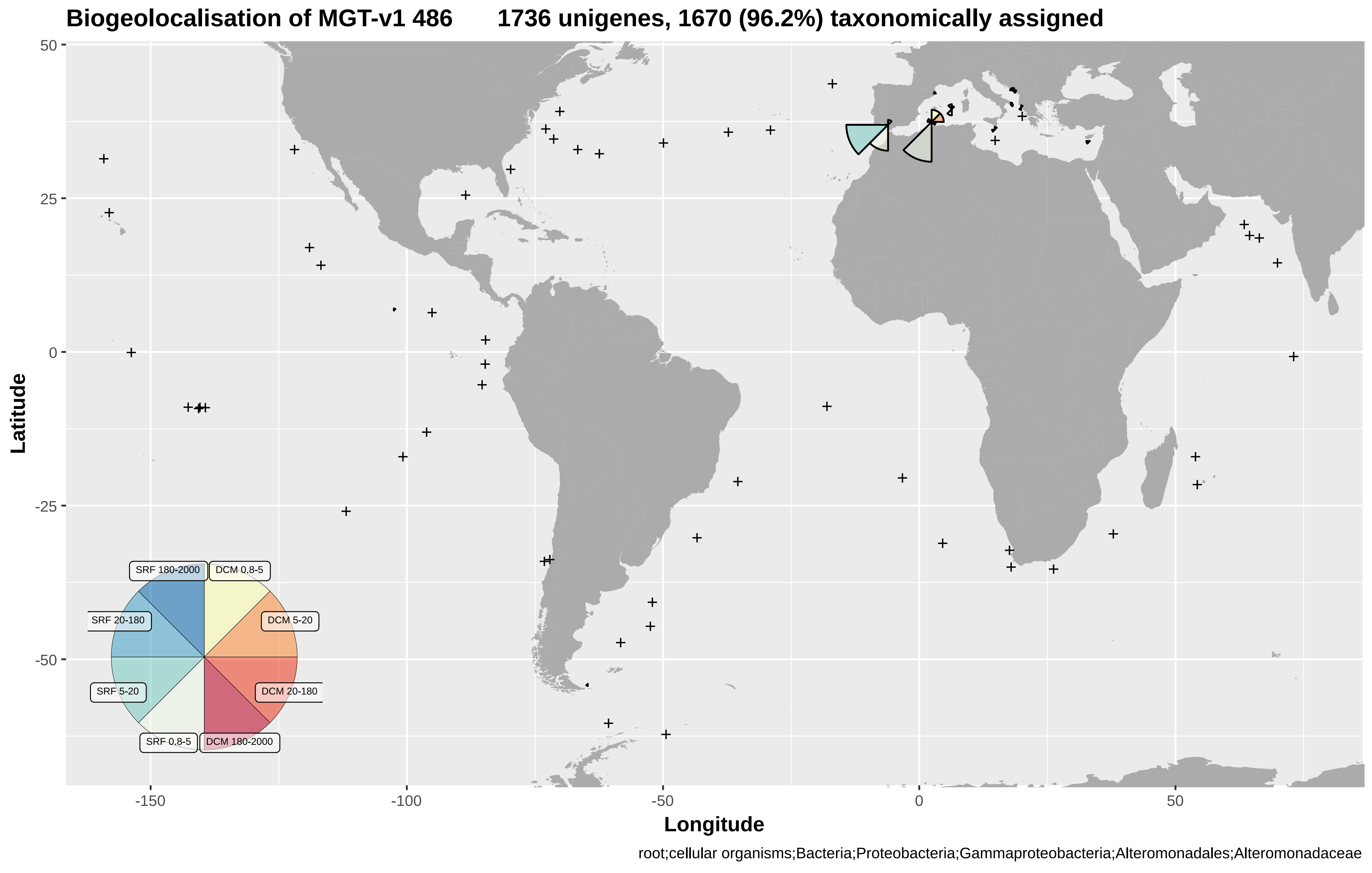

Biogeolocalisation of MGT-v1 486 1736 unigenes, 1670 (96.2%) taxonomically assigned
50
25
0
-25
-50
Latitude
-150
-100
-50
0
50
Longitude
SRF 180-2000
DCM 0.8-5
SRF 20-180
DCM 5-20
SRF 5-20
DCM 20-180
SRF 0.8-5
DCM 180-2000
root;cellular organisms;Bacteria;Proteobacteria;Gammaproteobacteria;Alteromonadales;Alteromonadaceae

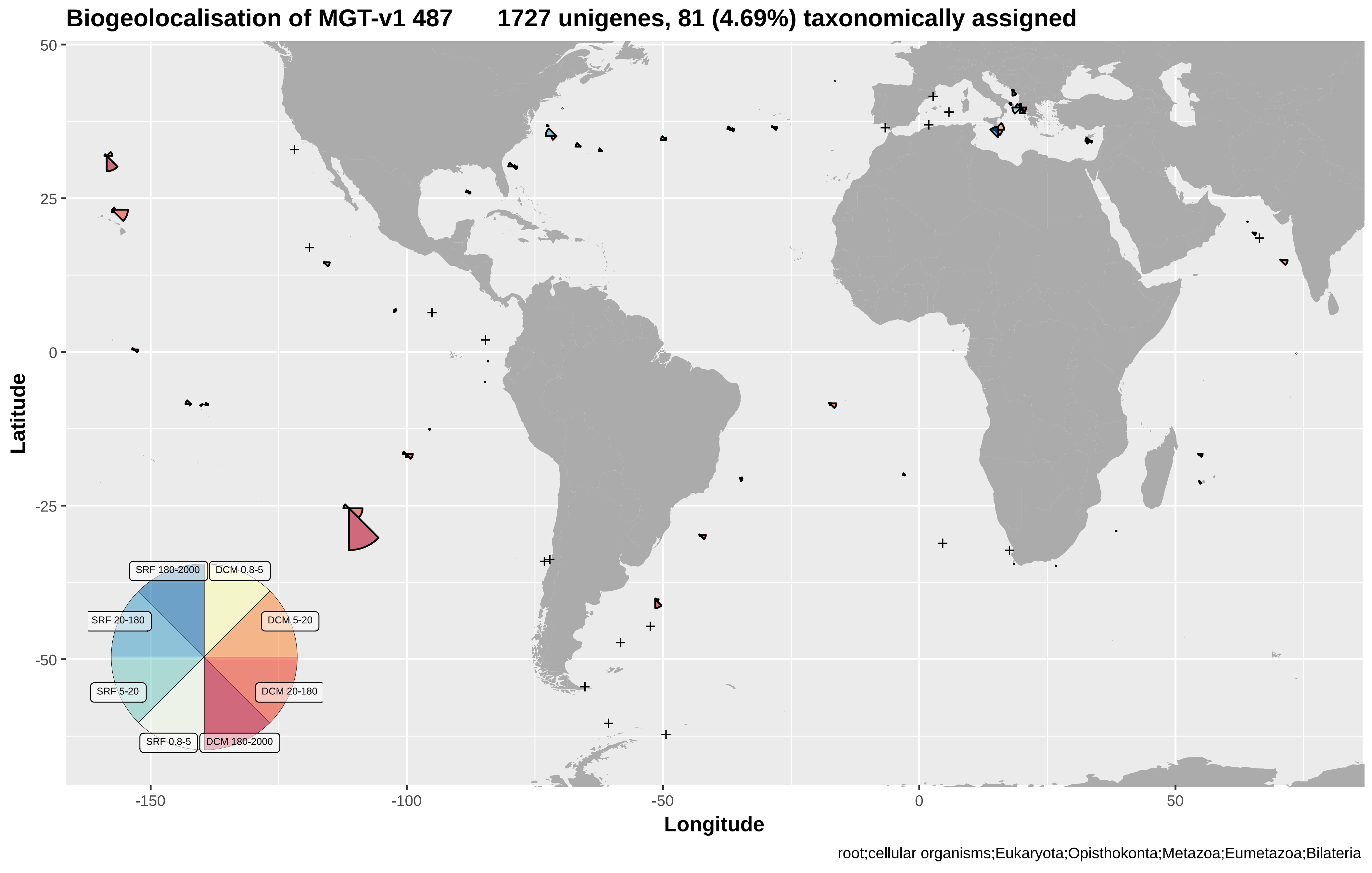

Biogeolocalisation of MGT-v1 487 1727 unigenes, 81 (4.69%) taxonomically assigned
Latitude
Longitude
50
25
0
-25
-50
-150
-100
-50
0
50
SRF 180-2000
DCM 0.8-5
SRF 20-180
DCM 5-20
SRF 5-20
DCM 20-180
SRF 0.8-5
DCM 180-2000
root;cellular organisms;Eukaryota;Opisthokonta;Metazoa;Eumetazoa;Bilateria

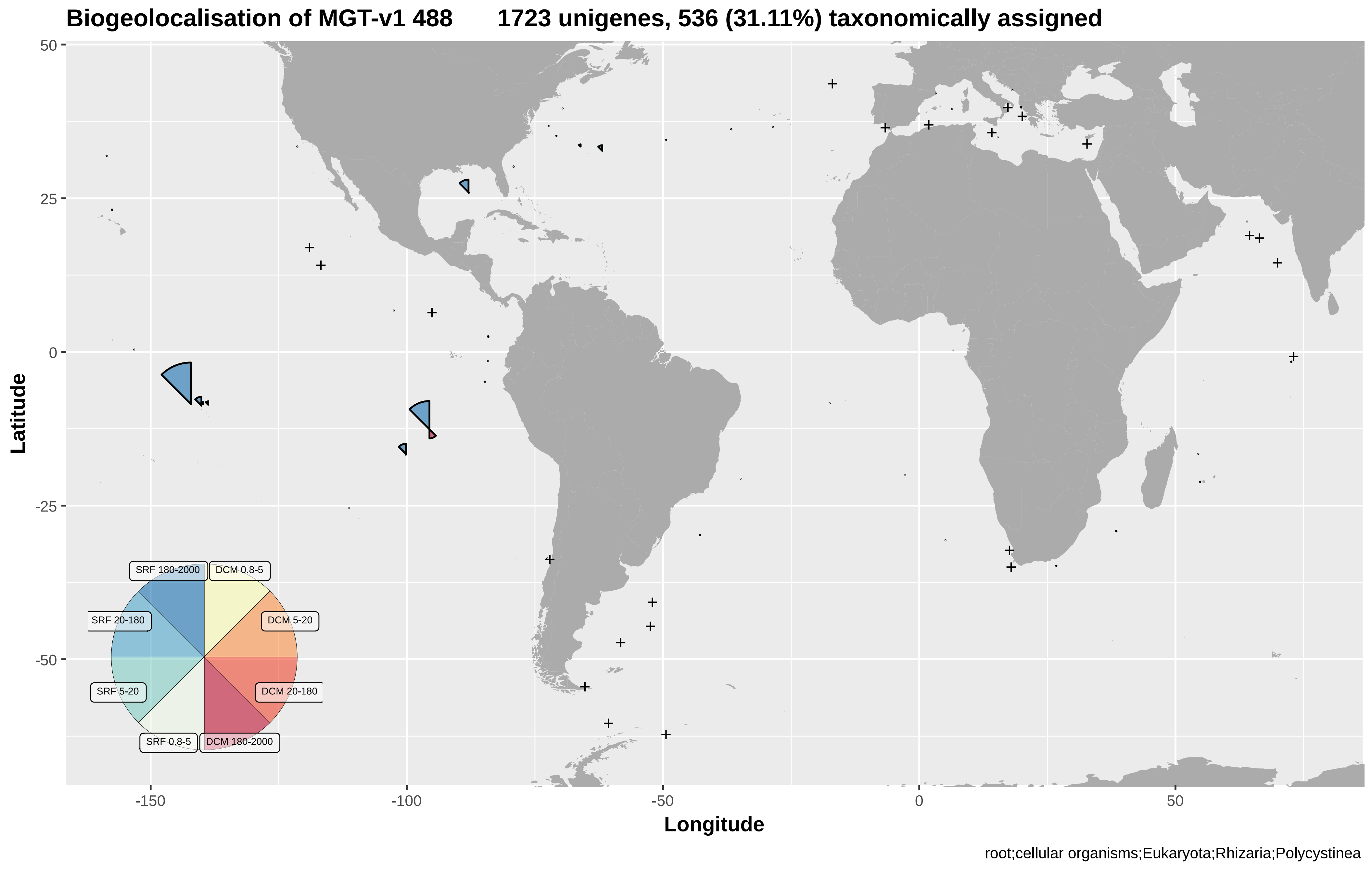

Biogeolocalisation of MGT-v1 488 1723 unigenes, 536 (31.11%) taxonomically assigned
Latitude
Longitude
50
25
0
-25
-50
-150
-100
-50
0
50
SRF 180-2000
DCM 0.8-5
SRF 20-180
DCM 5-20
SRF 5-20
DCM 20-180
SRF 0.8-5
DCM 180-2000
root;cellular organisms;Eukaryota;Rhizaria;Polycystinea

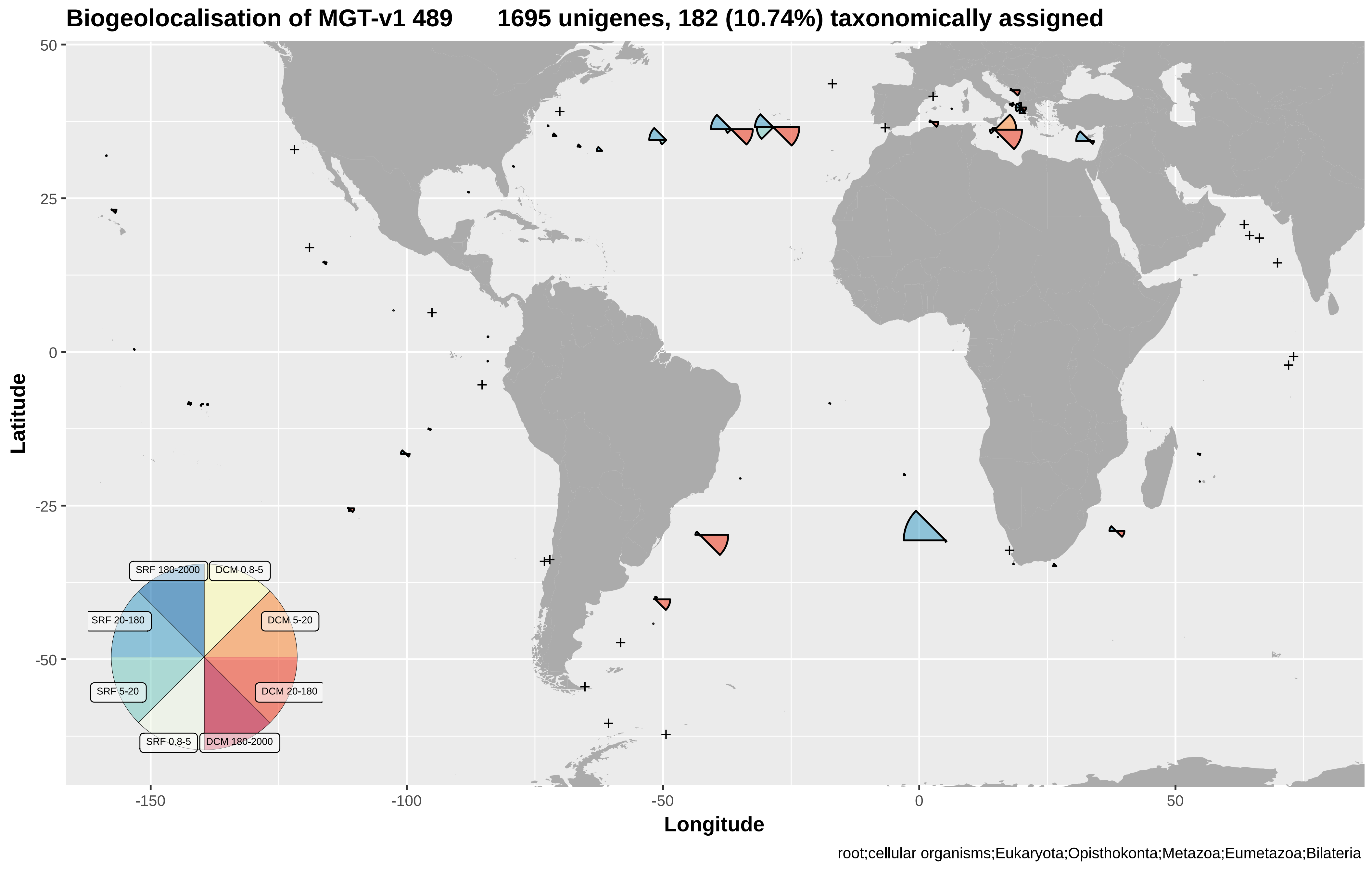
Biogeolocalisation of MGT-v1 489      1695 unigenes, 182 (10.74%) taxonomically assigned
Latitude
Longitude
50
25
0
-25
-50
-150
-100
-50
0
50
SRF 180-2000
DCM 0.8-5
SRF 20-180
DCM 5-20
SRF 5-20
DCM 20-180
SRF 0.8-5
DCM 180-2000
root;cellular organisms;Eukaryota;Opisthokonta;Metazoa;Eumetazoa;Bilateria

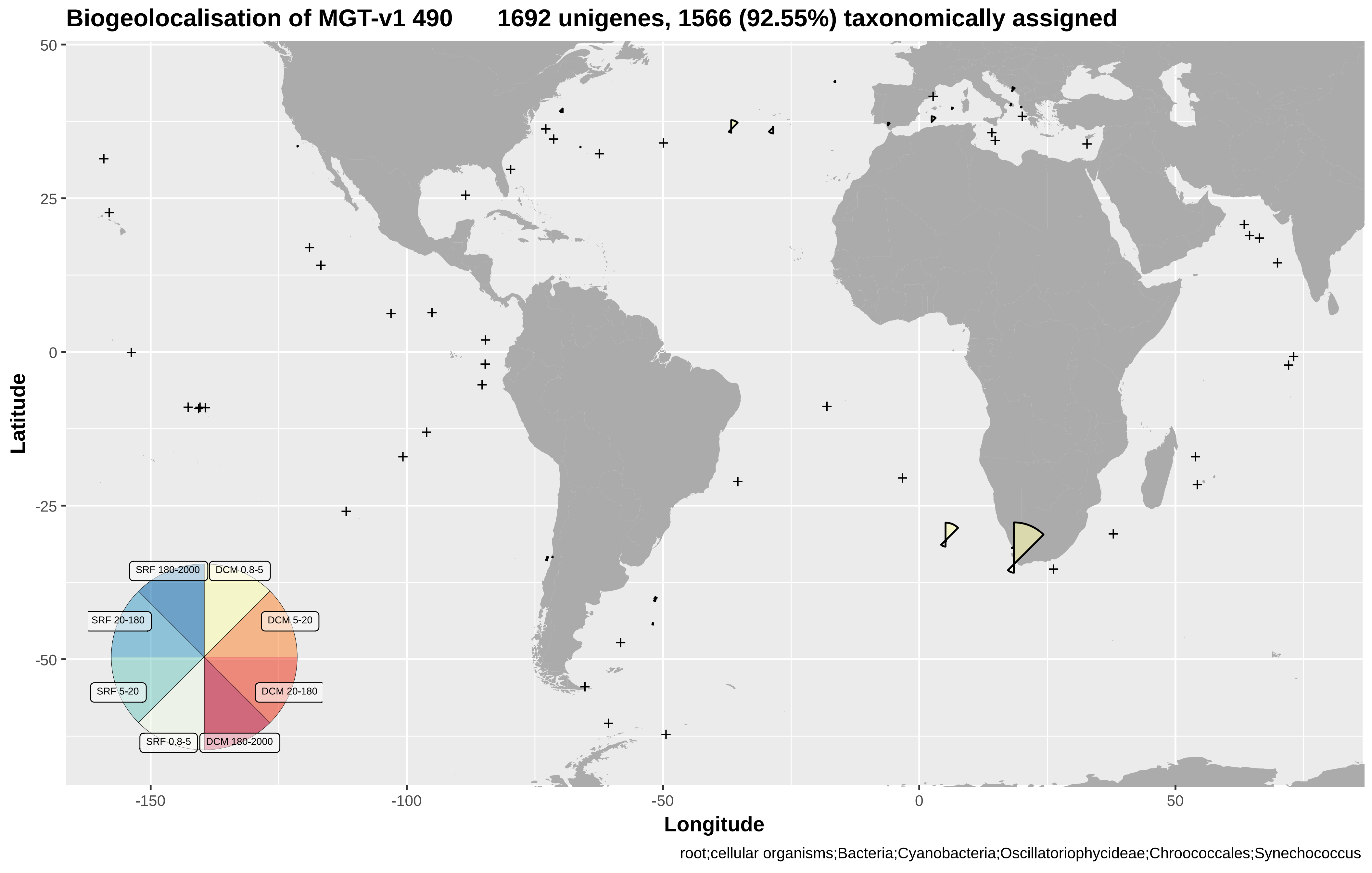
Biogeolocalisation of MGT-v1 490 1692 unigenes, 1566 (92.55%) taxonomically assigned
50
25
0
-25
-50
Latitude
-150
-100
-50
0
50
Longitude
SRF 180-2000
DCM 0.8-5
SRF 20-180
DCM 5-20
SRF 5-20
DCM 20-180
SRF 0.8-5
DCM 180-2000
root;cellular organisms;Bacteria;Cyanobacteria;Oscillatoriophycideae;Chroococcales;Synechococcus

# Biogeolocalisation of MGT-v1 491      1691 unigenes, 1594 (94.26%) taxonomically assigned

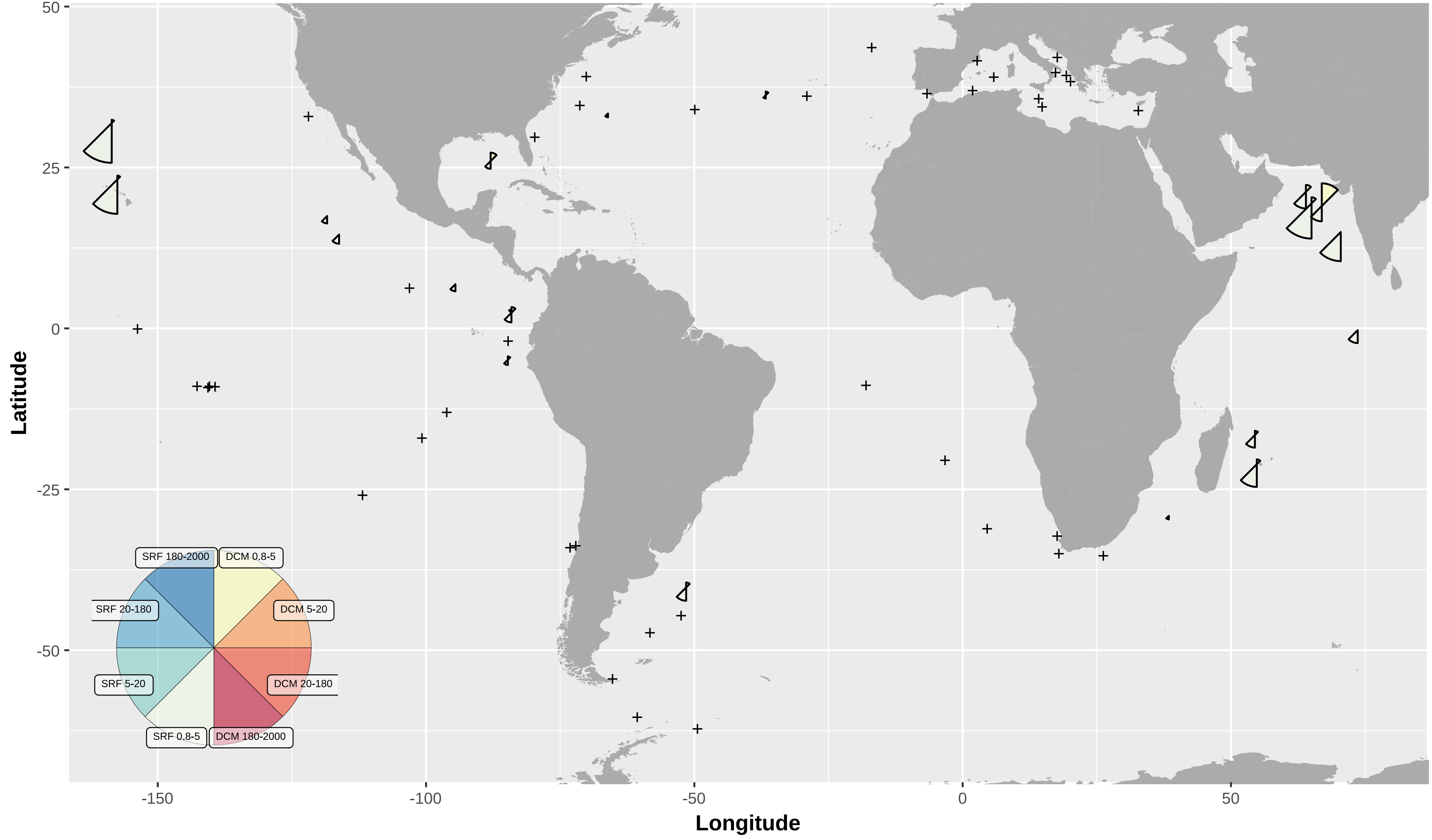


root;cellular organisms;Bacteria;Proteobacteria;Alphaproteobacteria

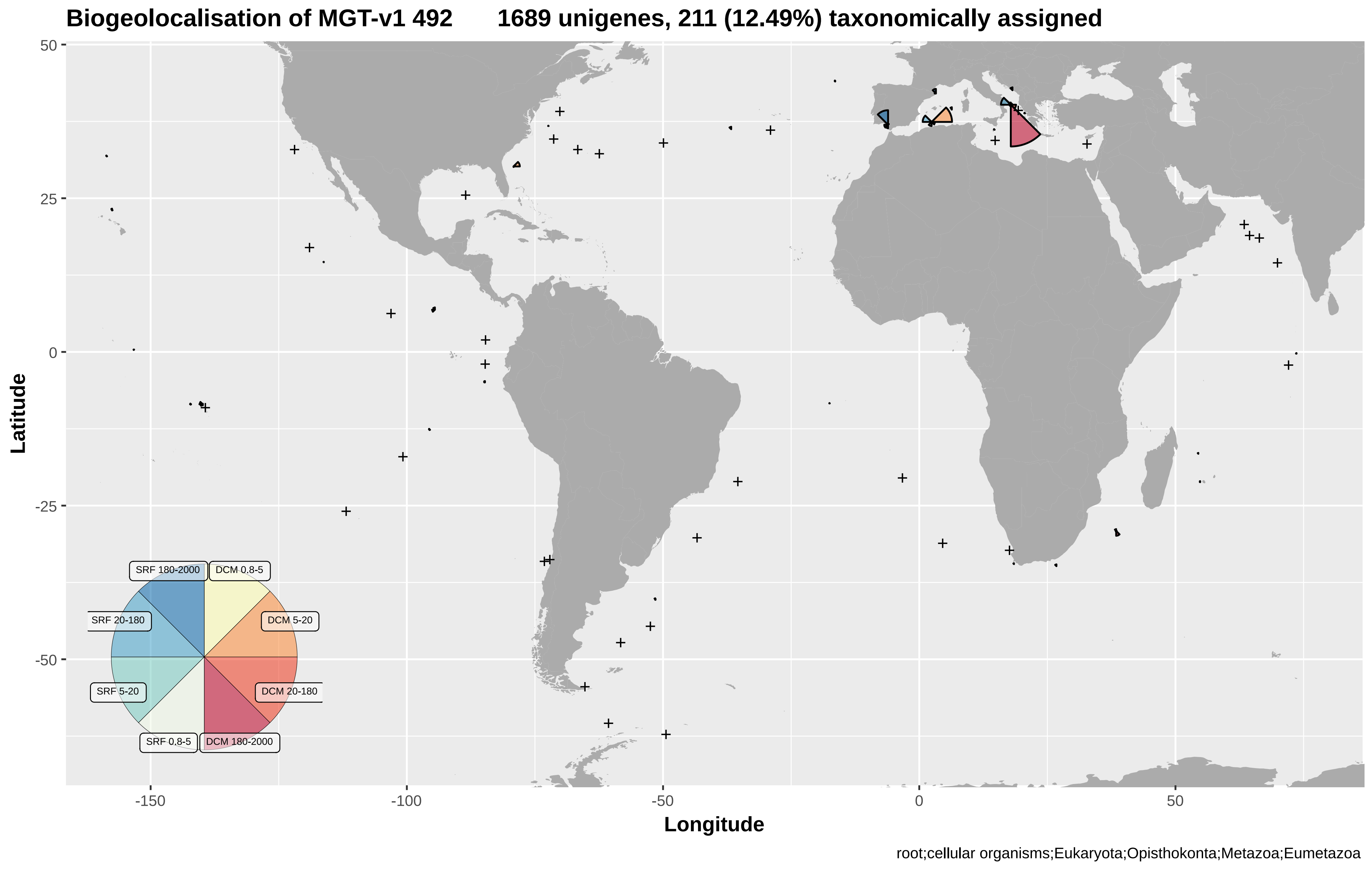

Biogeolocalisation of MGT-v1 492 1689 unigenes, 211 (12.49%) taxonomically assigned
Latitude
Longitude
50
25
0
-25
-50
-150
-100
-50
0
50
SRF 180-2000
DCM 0.8-5
SRF 20-180
DCM 5-20
SRF 5-20
DCM 20-180
SRF 0.8-5
DCM 180-2000
root;cellular organisms;Eukaryota;Opisthokonta;Metazoa;Eumetazoa

# Biogeolocalisation of MGT-v1 493      1681 unigenes, 1227 (72.99%) taxonomically assigned

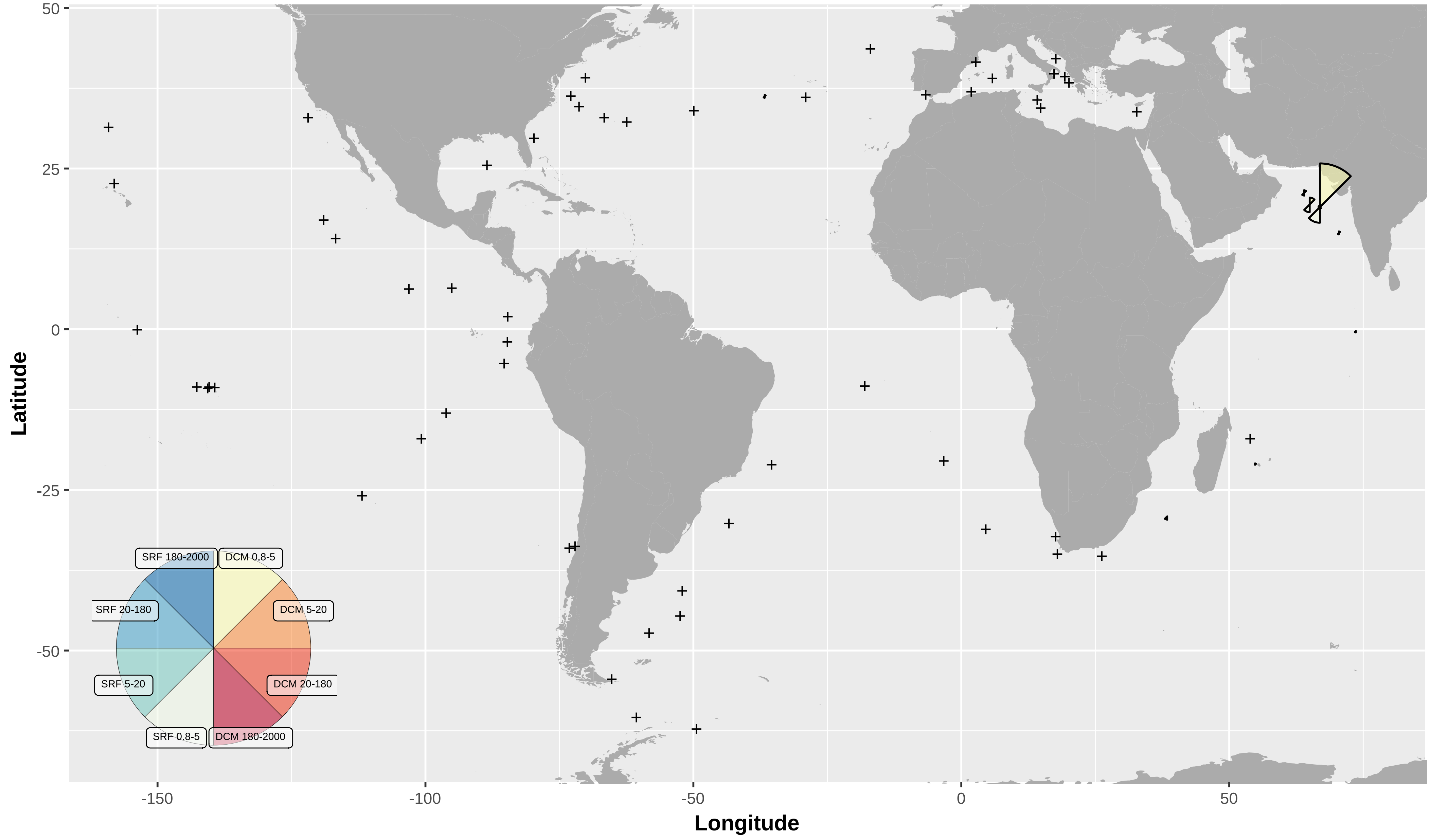


root;cellular organisms;Bacteria

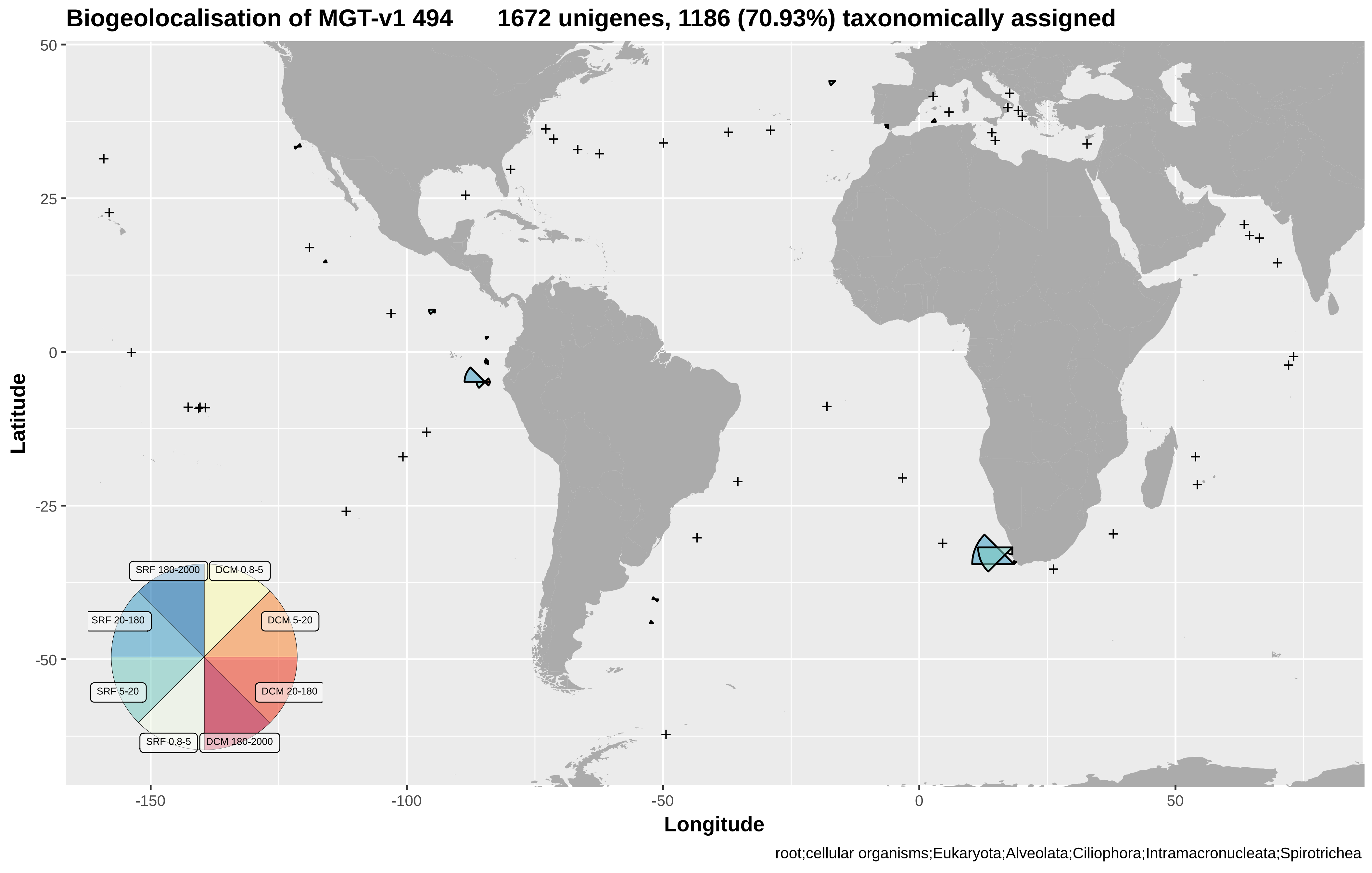

Biogeolocalisation of MGT-v1 494 1672 unigenes, 1186 (70.93%) taxonomically assigned
Latitude
50
25
0
-25
-50
Longitude
-150
-100
-50
0
50
SRF 180-2000
DCM 0.8-5
SRF 20-180
DCM 5-20
SRF 5-20
DCM 20-180
SRF 0.8-5
DCM 180-2000
root;cellular organisms;Eukaryota;Alveolata;Ciliophora;Intramacronucleata;Spirotrichea

## Biogeolocalisation of MGT-v1 495      1664 unigenes, 592 (35.58%) taxonomically assigned

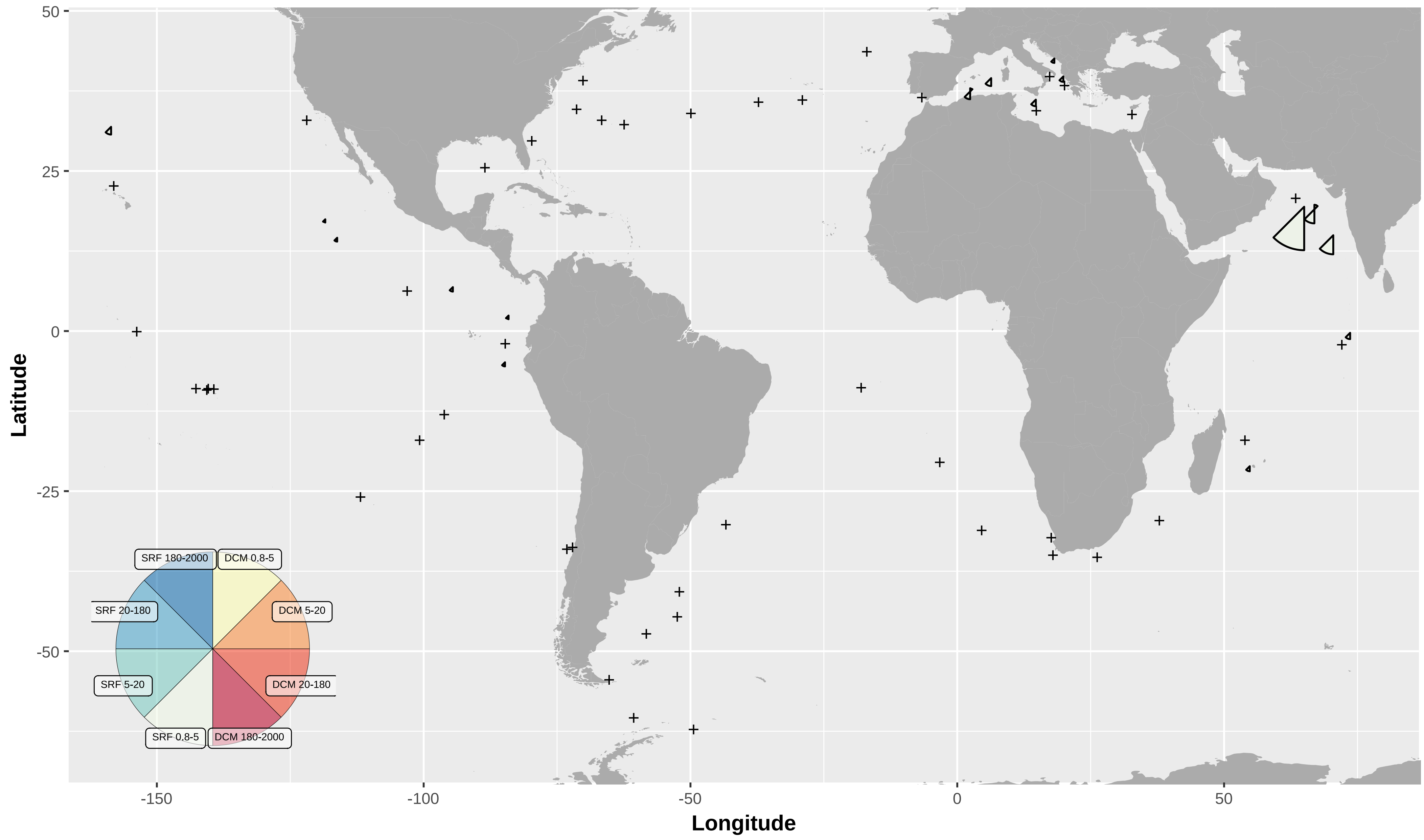


root;cellular organisms;Eukaryota

# Biogeolocalisation of MGT-v1 496      1653 unigenes, 214 (12.95%) taxonomically assigned

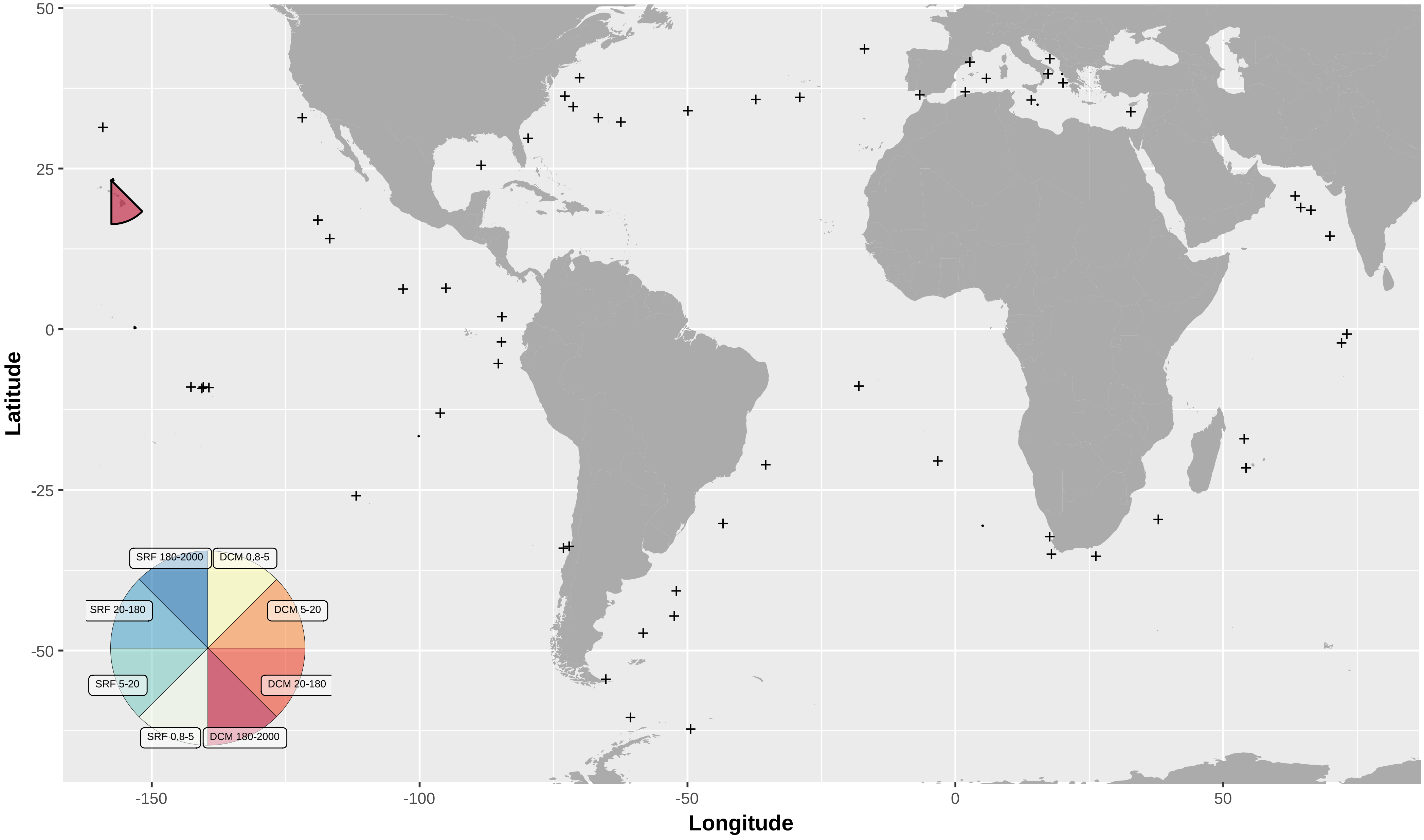


root;cellular organisms;Eukaryota;Opisthokonta;Metazoa;Eumetazoa;Bilateria

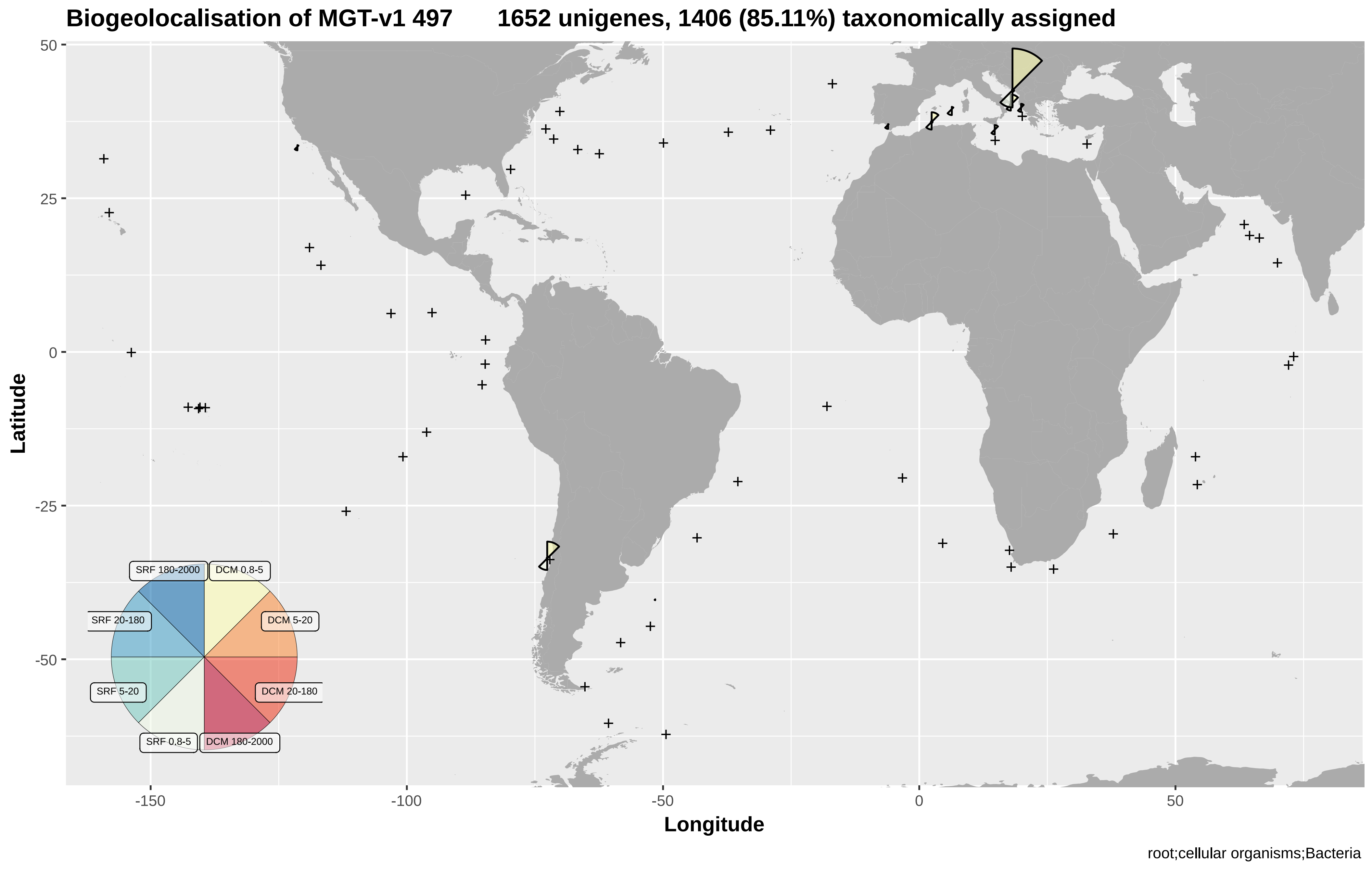

Biogeolocalisation of MGT-v1 497 1652 unigenes, 1406 (85.11%) taxonomically assigned
SRF 180-2000
DCM 0.8-5
SRF 20-180
DCM 5-20
SRF 5-20
DCM 20-180
SRF 0.8-5
DCM 180-2000
Latitude
Longitude
50
25
0
-25
-50
-150
-100
-50
0
50
root;cellular organisms;Bacteria

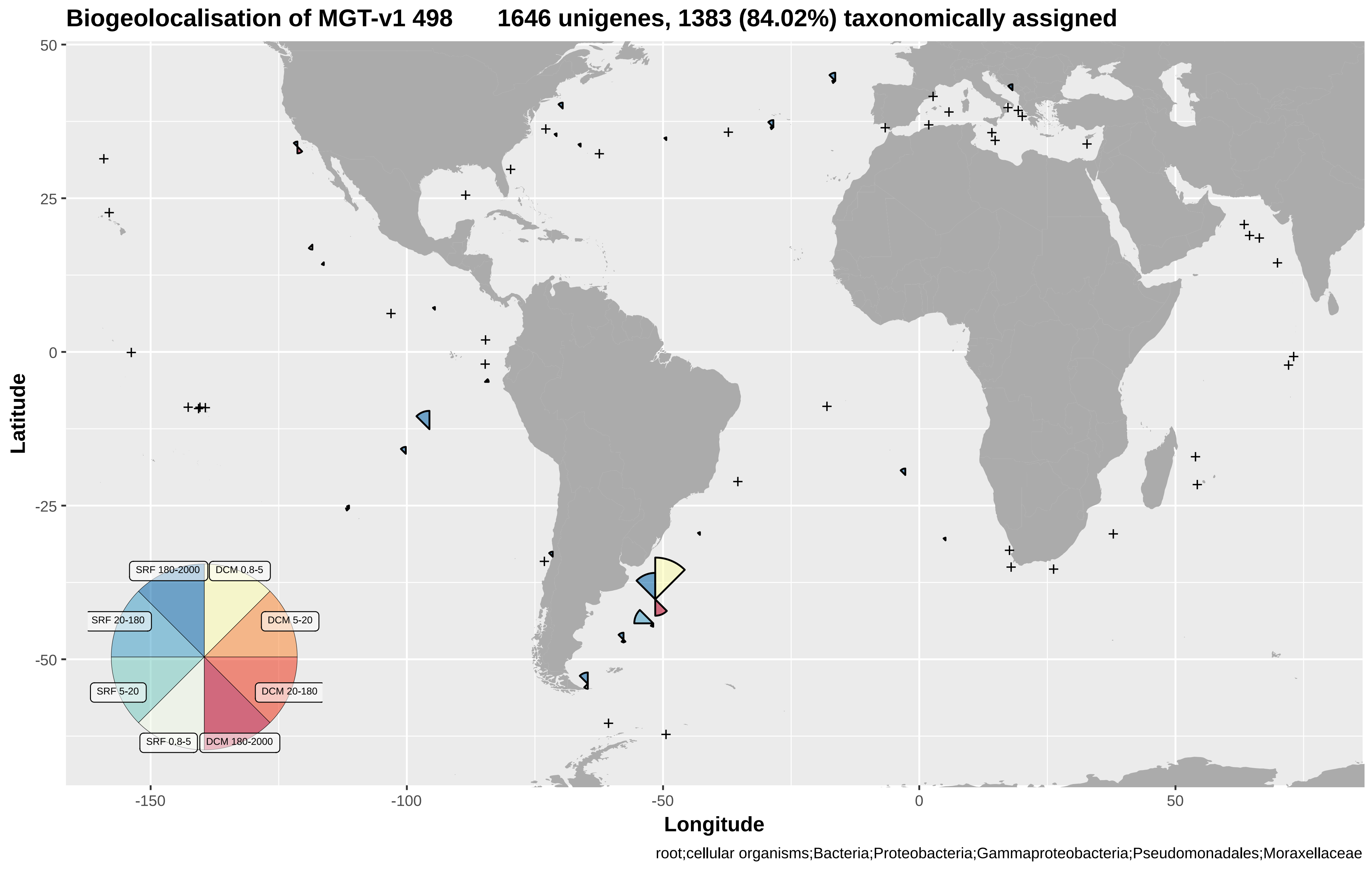
Biogeolocalisation of MGT-v1 498 1646 unigenes, 1383 (84.02%) taxonomically assigned
50
25
0
-25
-50
Latitude
-150
-100
-50
0
50
Longitude
SRF 180-2000
DCM 0.8-5
SRF 20-180
DCM 5-20
SRF 5-20
DCM 20-180
SRF 0.8-5
DCM 180-2000
root;cellular organisms;Bacteria;Proteobacteria;Gammaproteobacteria;Pseudomonadales;Moraxellaceae

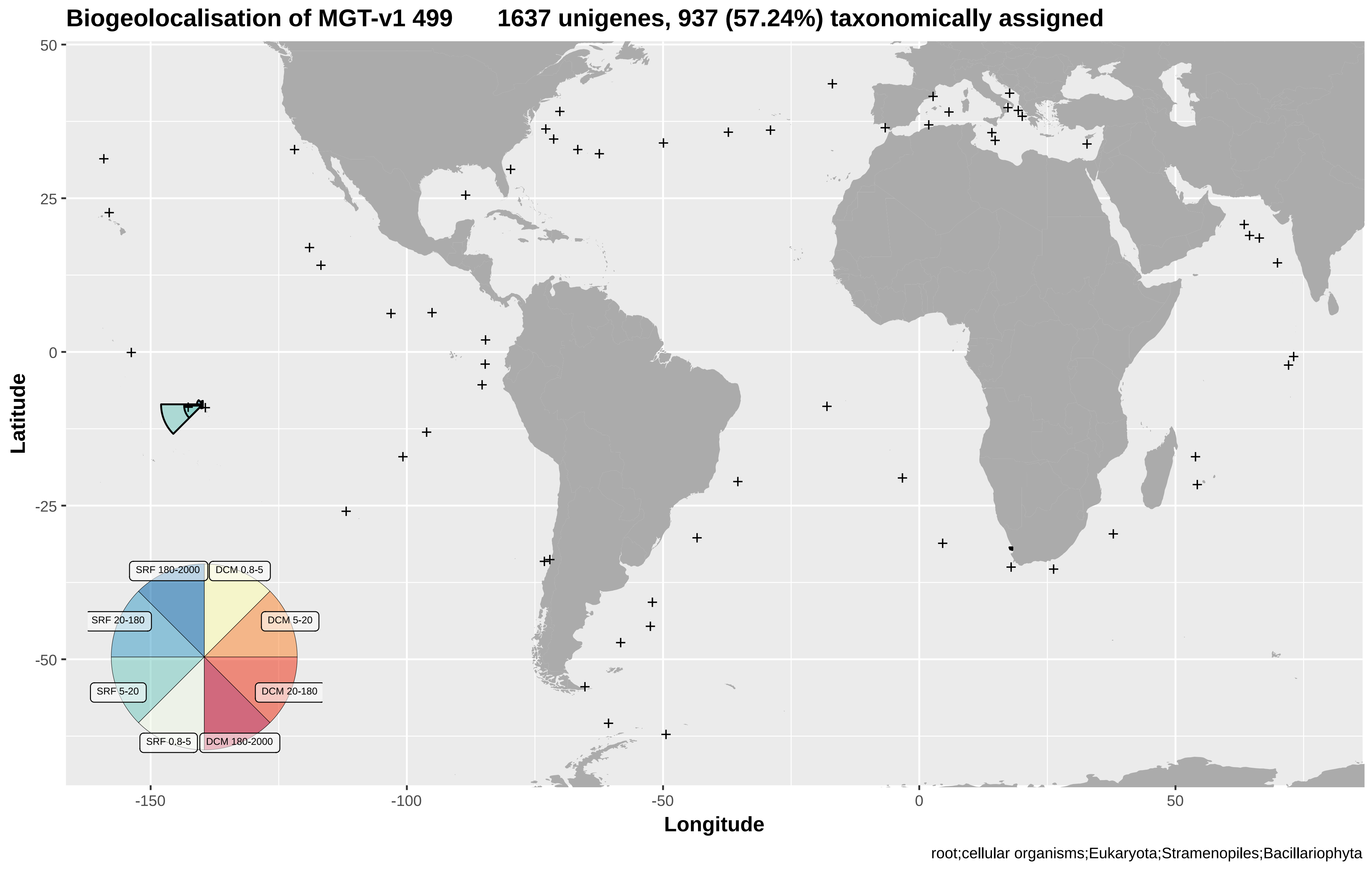
Biogeolocalisation of MGT-v1 499 1637 unigenes, 937 (57.24%) taxonomically assigned
Latitude
50
25
0
-25
-50
Longitude
-150
-100
-50
0
50
SRF 180-2000
DCM 0.8-5
SRF 20-180
DCM 5-20
SRF 5-20
DCM 20-180
SRF 0.8-5
DCM 180-2000
root;cellular organisms;Eukaryota;Stramenopiles;Bacillariophyta

## Biogeolocalisation of MGT-v1 500      1637 unigenes, 119 (7.27%) taxonomically assigned

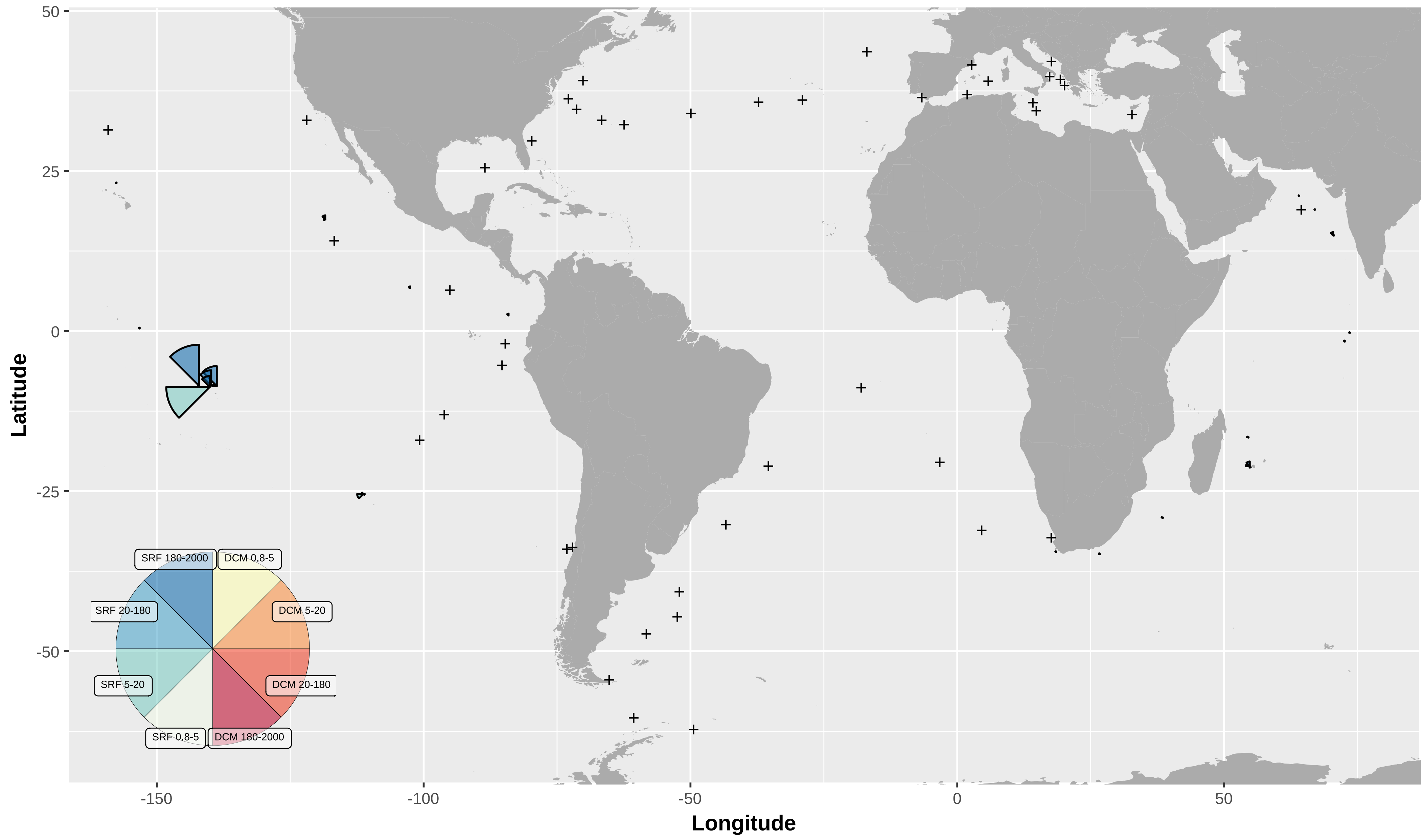


root;cellular organisms;Eukaryota;Opisthokonta;Metazoa;Eumetazoa

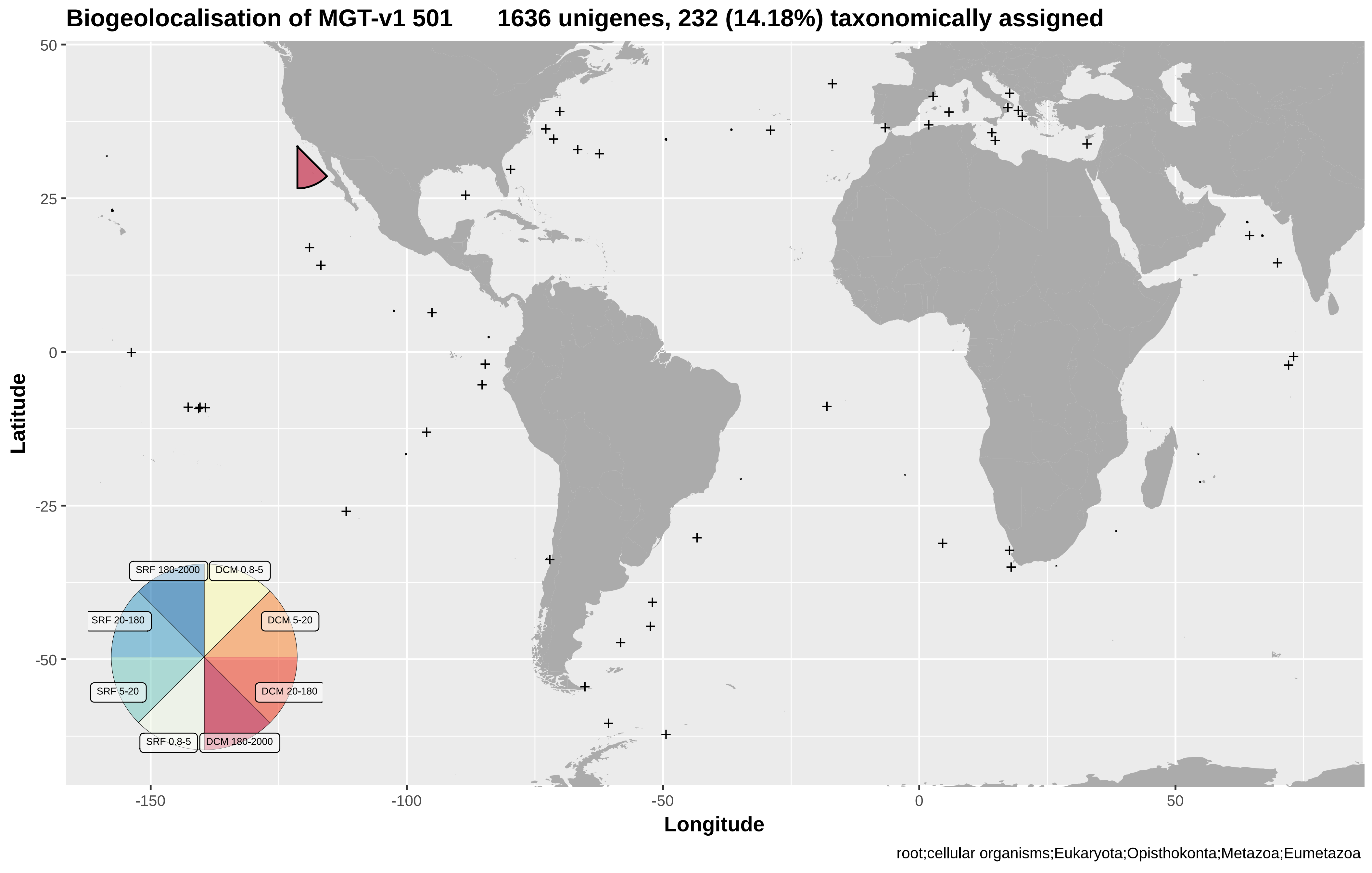

Biogeolocalisation of MGT-v1 501
1636 unigenes, 232 (14.18%) taxonomically assigned
50
25
0
-25
-50
Latitude
-150
-100
-50
0
50
Longitude
SRF 180-2000
DCM 0.8-5
SRF 20-180
DCM 5-20
SRF 5-20
DCM 20-180
SRF 0.8-5
DCM 180-2000
root;cellular organisms;Eukaryota;Opisthokonta;Metazoa;Eumetazoa

# Biogeolocalisation of MGT-v1 502      1633 unigenes, 93 (5.7%) taxonomically assigned

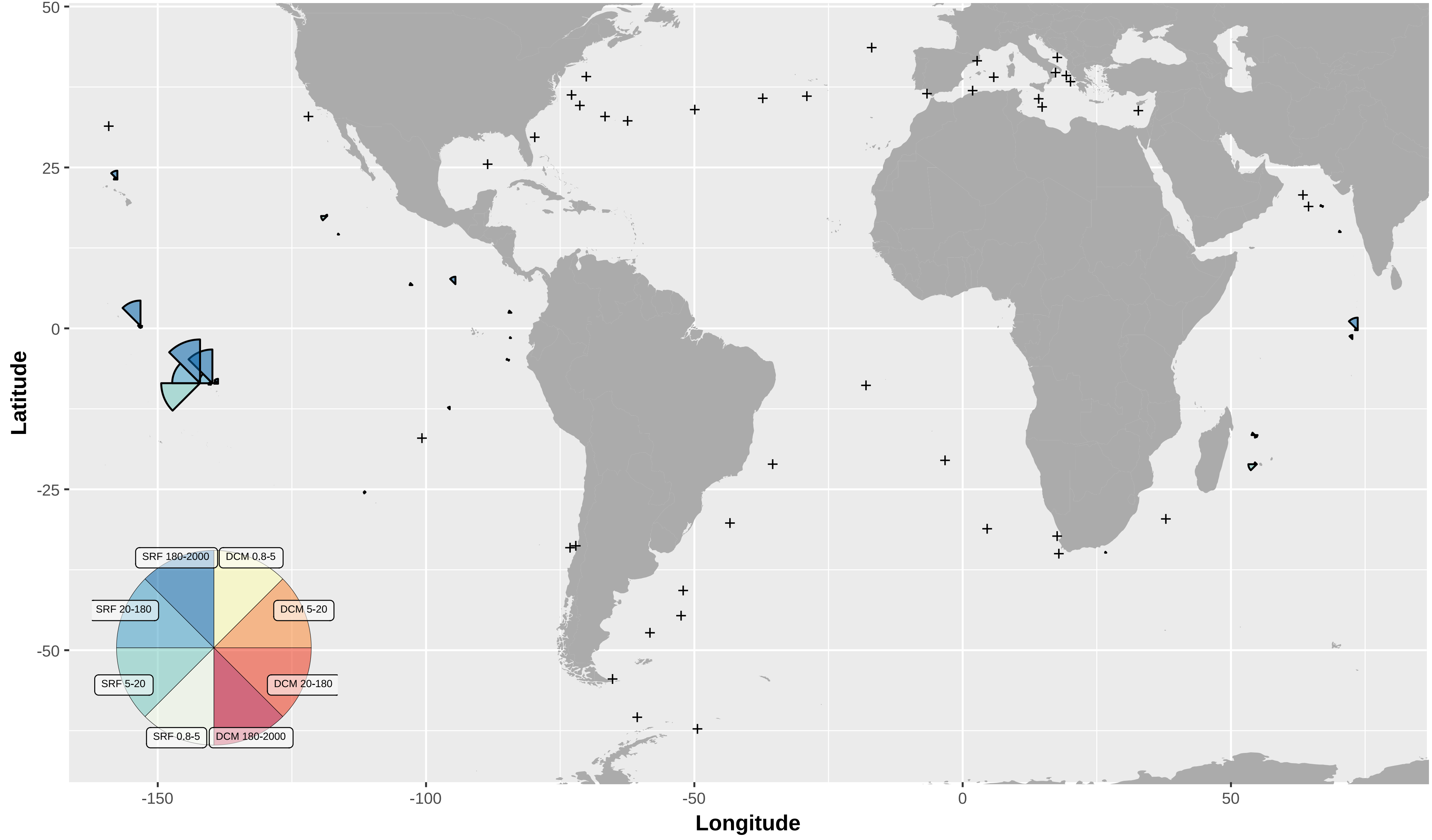


root;cellular organisms;Eukaryota;Opisthokonta;Metazoa;Eumetazoa;Bilateria

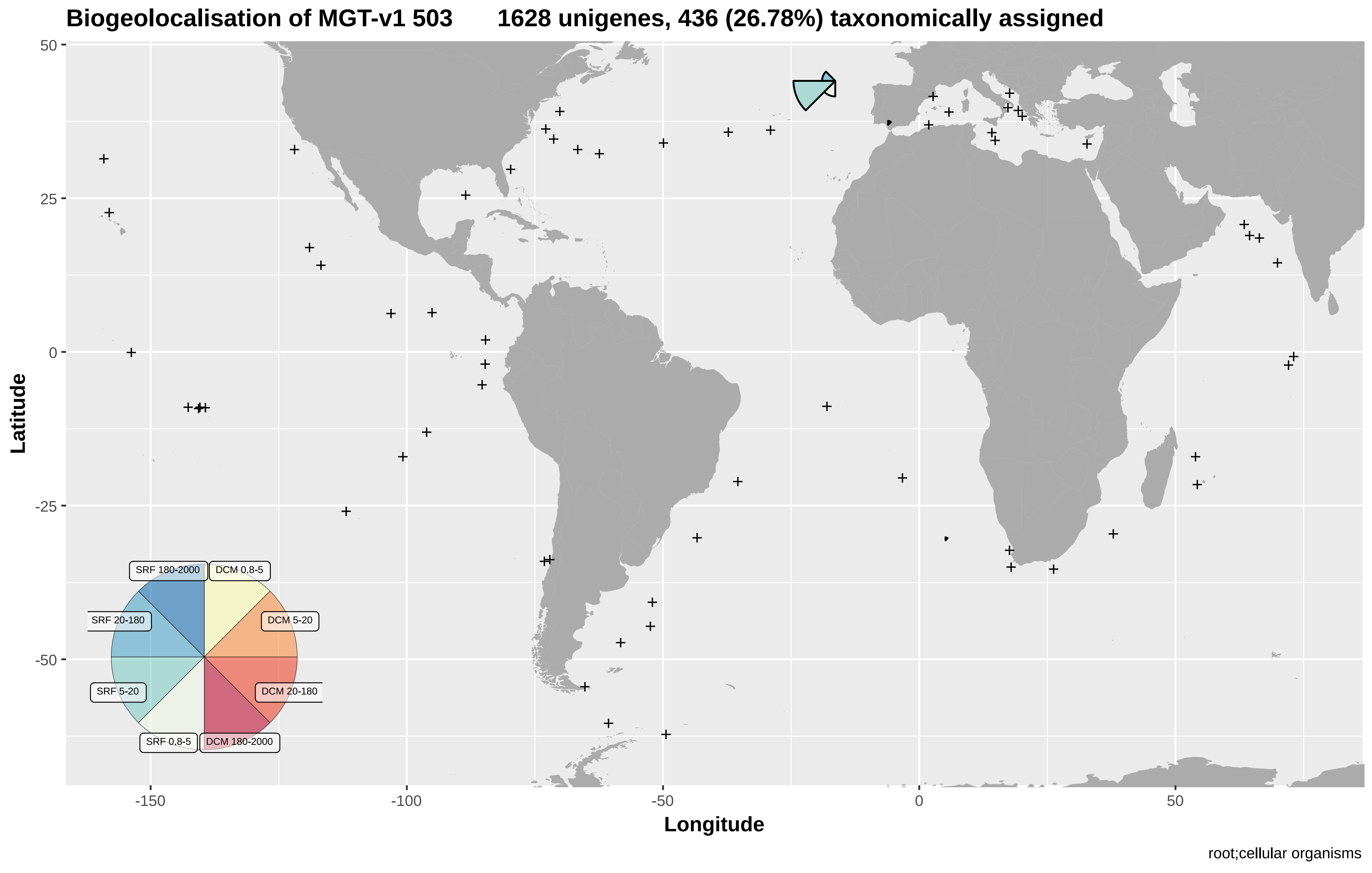

Biogeolocalisation of MGT-v1 503 1628 unigenes, 436 (26.78%) taxonomically assigned
Latitude
Longitude
50
25
0
-25
-50
-150
-100
-50
0
50
SRF 180-2000
DCM 0.8-5
SRF 20-180
DCM 5-20
SRF 5-20
DCM 20-180
SRF 0.8-5
DCM 180-2000
root;cellular organisms

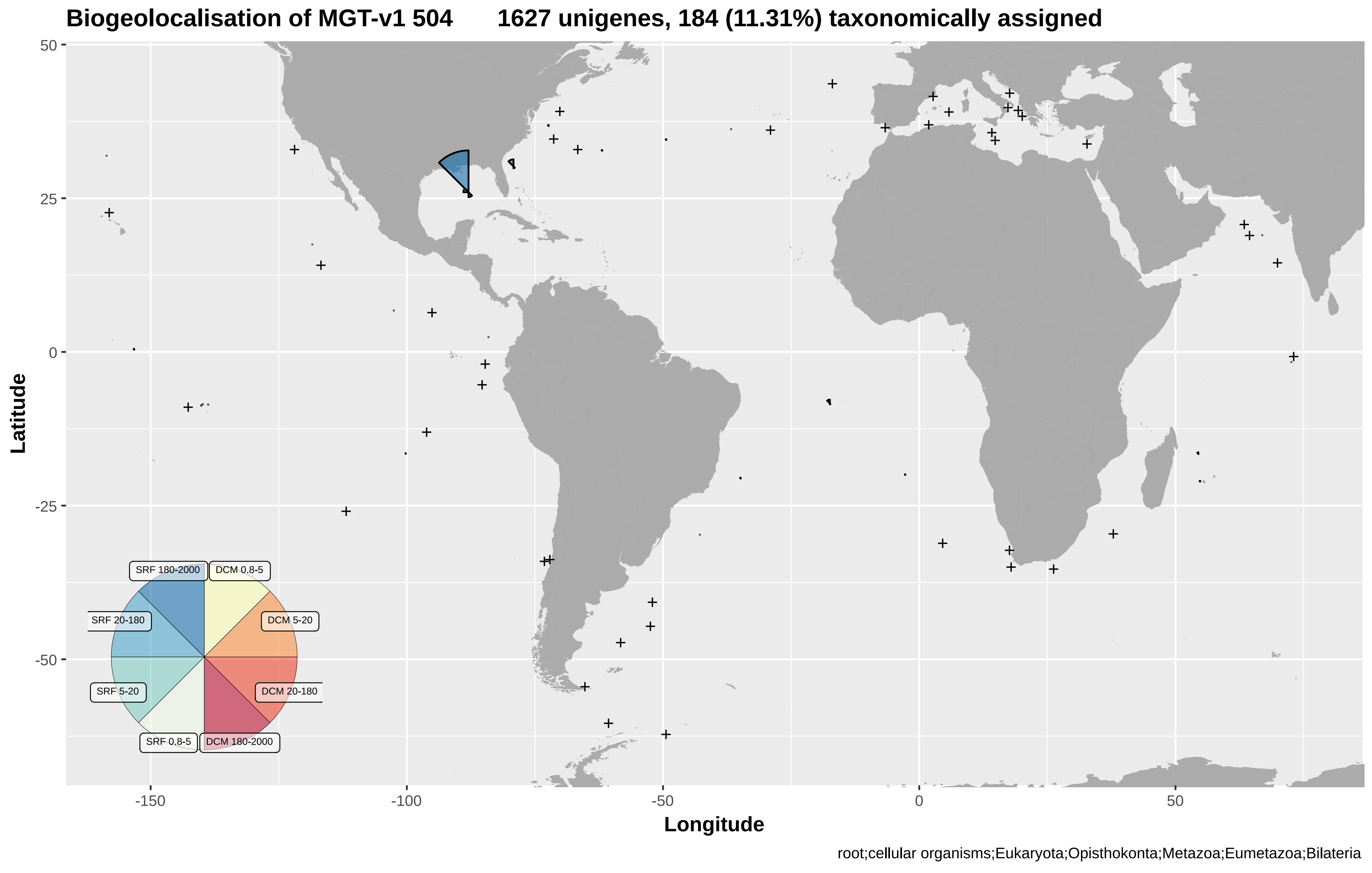

Biogeolocalisation of MGT-v1 504 1627 unigenes, 184 (11.31%) taxonomically assigned
Latitude
Longitude
SRF 180-2000
DCM 0.8-5
SRF 20-180
DCM 5-20
SRF 5-20
DCM 20-180
SRF 0.8-5
DCM 180-2000
root;cellular organisms;Eukaryota;Opisthokonta;Metazoa;Eumetazoa;Bilateria

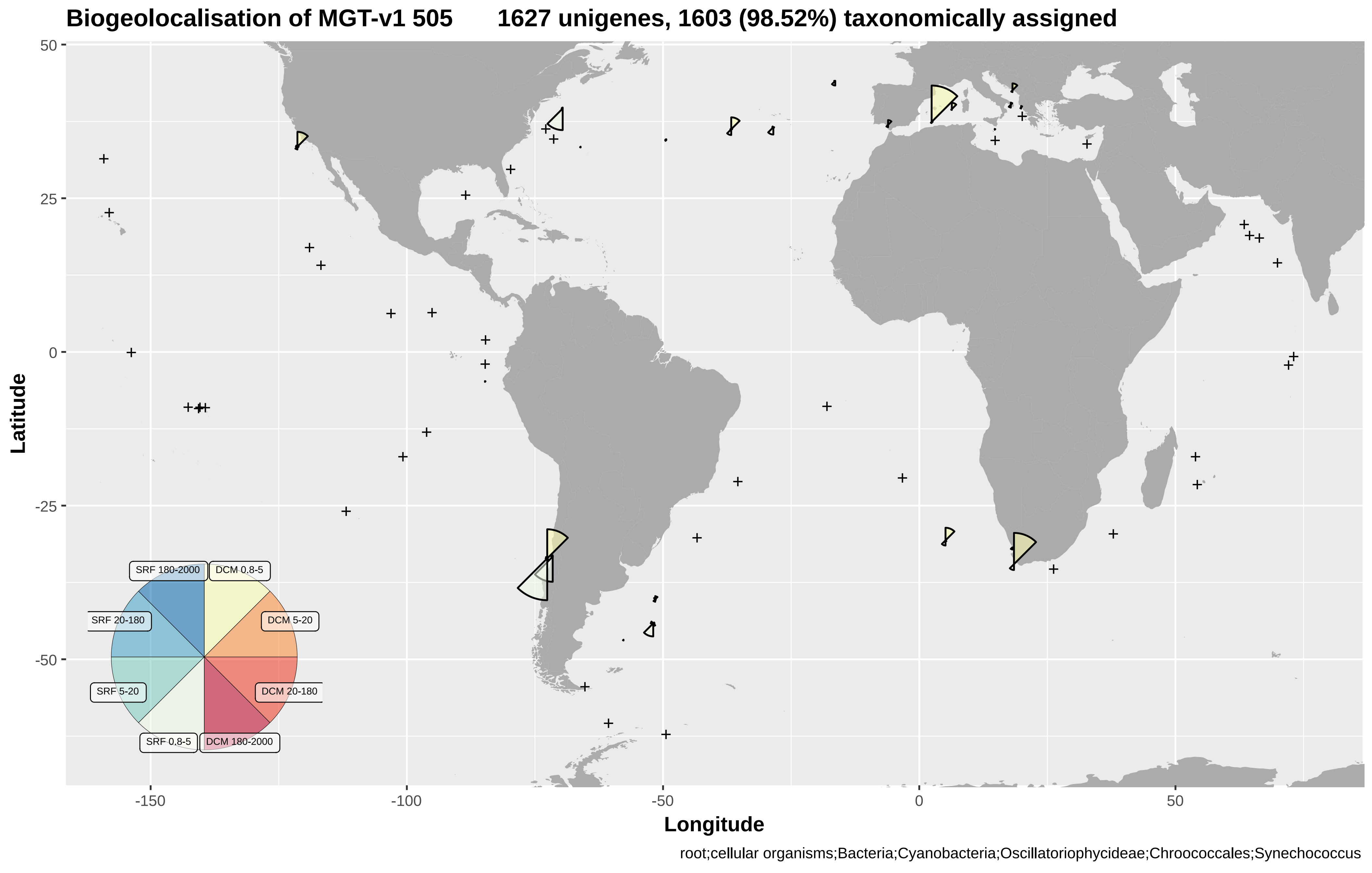

Biogeolocalisation of MGT-v1 505      1627 unigenes, 1603 (98.52%) taxonomically assigned
Latitude
Longitude
50
25
0
-25
-50
-150
-100
-50
0
50
SRF 180-2000
DCM 0.8-5
SRF 20-180
DCM 5-20
SRF 5-20
DCM 20-180
SRF 0.8-5
DCM 180-2000
root;cellular organisms;Bacteria;Cyanobacteria;Oscillatoriophycideae;Chroococcales;Synechococcus

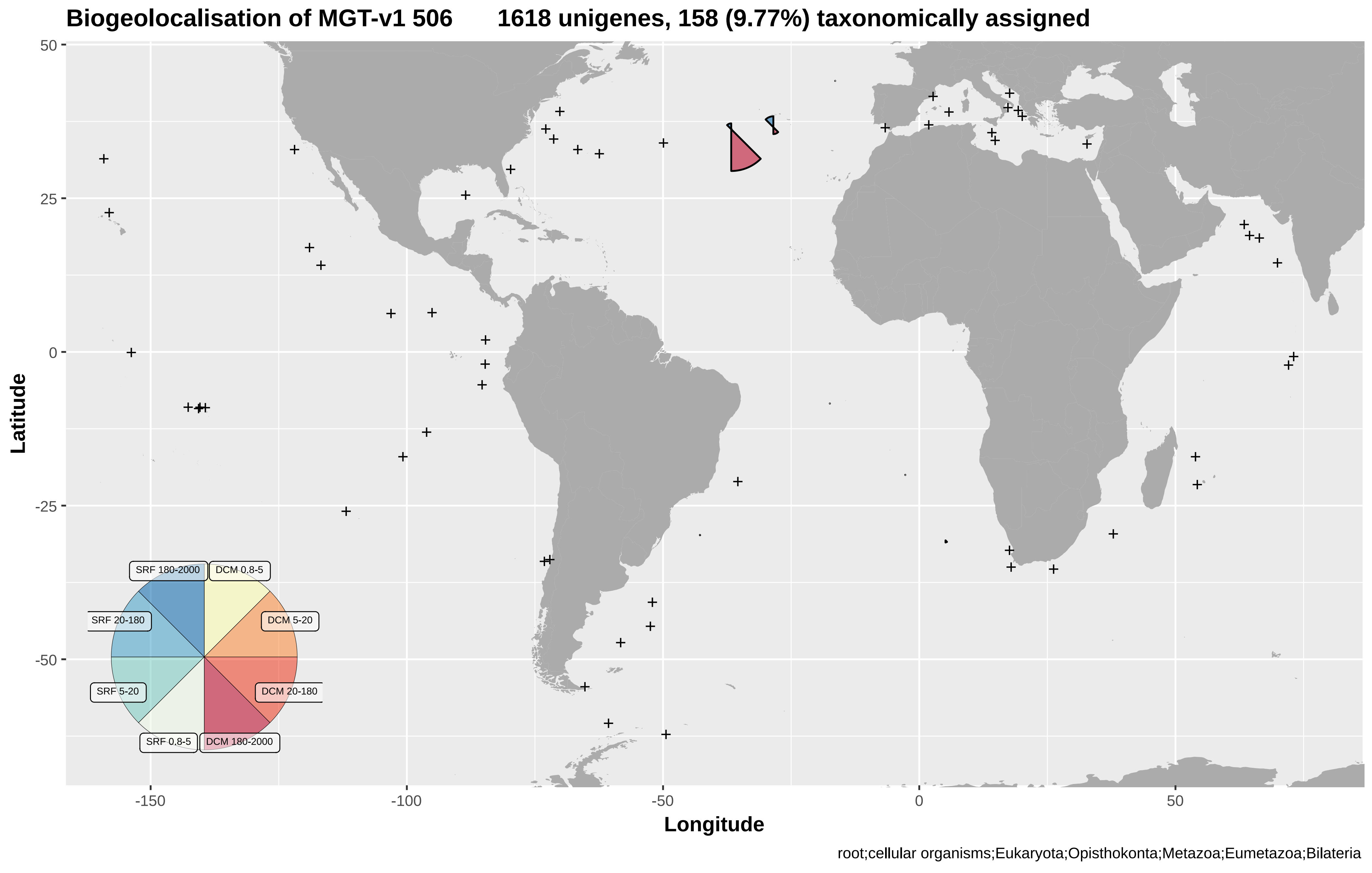
Biogeolocalisation of MGT-v1 506
1618 unigenes, 158 (9.77%) taxonomically assigned
Latitude
Longitude
50
25
0
-25
-50
-150
-100
-50
0
50
SRF 180-2000
DCM 0.8-5
SRF 20-180
DCM 5-20
SRF 5-20
DCM 20-180
SRF 0.8-5
DCM 180-2000
root;cellular organisms;Eukaryota;Opisthokonta;Metazoa;Eumetazoa;Bilateria

## Biogeolocalisation of MGT-v1 507    1601 unigenes, 202 (12.62%) taxonomically assigned

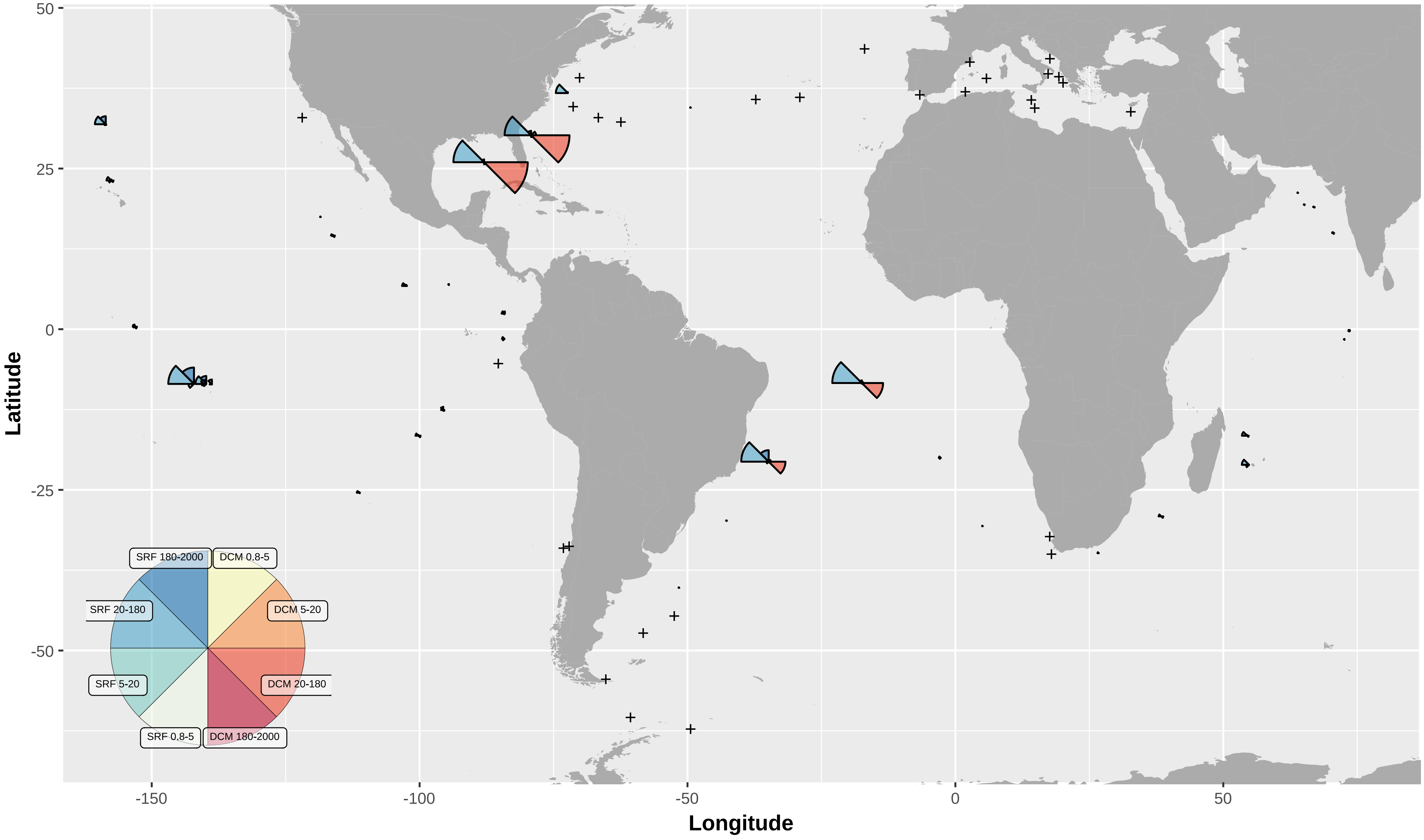


root;cellular organisms;Eukaryota;Opisthokonta;Metazoa;Eumetazoa;Bilateria

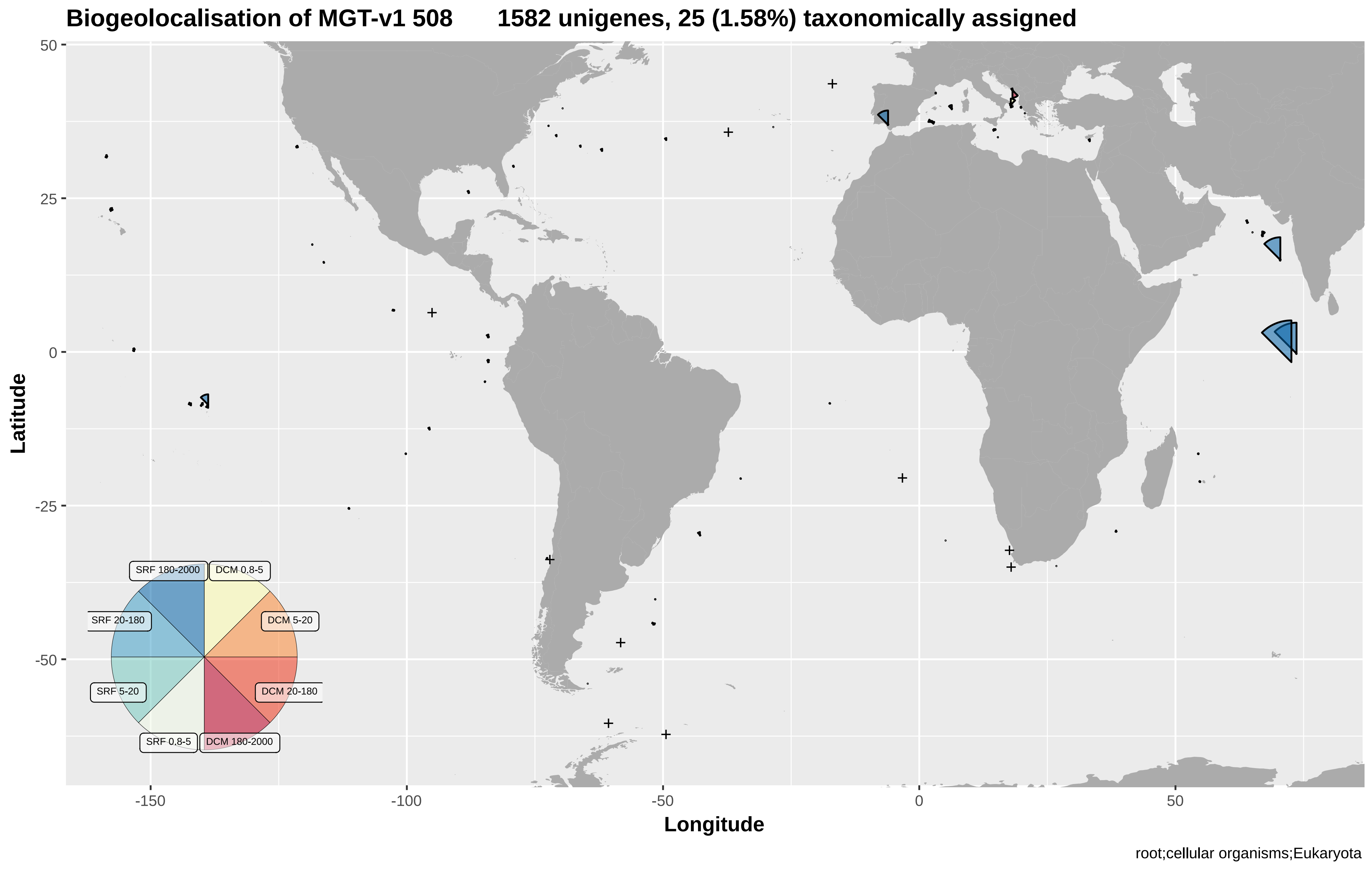

Biogeolocalisation of MGT-v1 508 1582 unigenes, 25 (1.58%) taxonomically assigned
Latitude
Longitude
50
25
0
-25
-50
-150
-100
-50
0
50
SRF 180-2000
DCM 0.8-5
SRF 20-180
DCM 5-20
SRF 5-20
DCM 20-180
SRF 0.8-5
DCM 180-2000
root;cellular organisms;Eukaryota

# Biogeolocalisation of MGT-v1 509      1578 unigenes, 974 (61.72%) taxonomically assigned

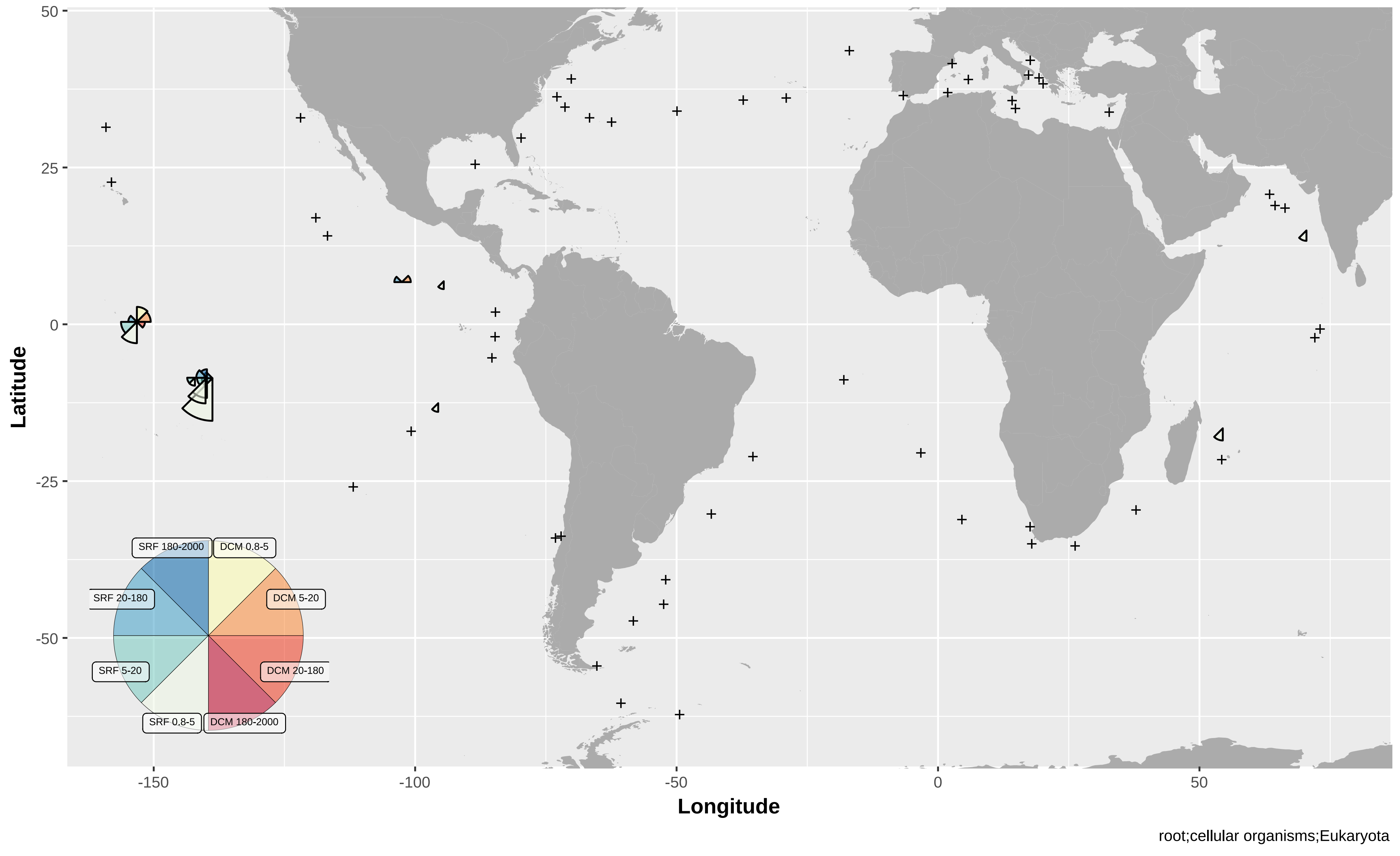

# Biogeolocalisation of MGT-v1 510      1575 unigenes, 728 (46.22%) taxonomically assigned

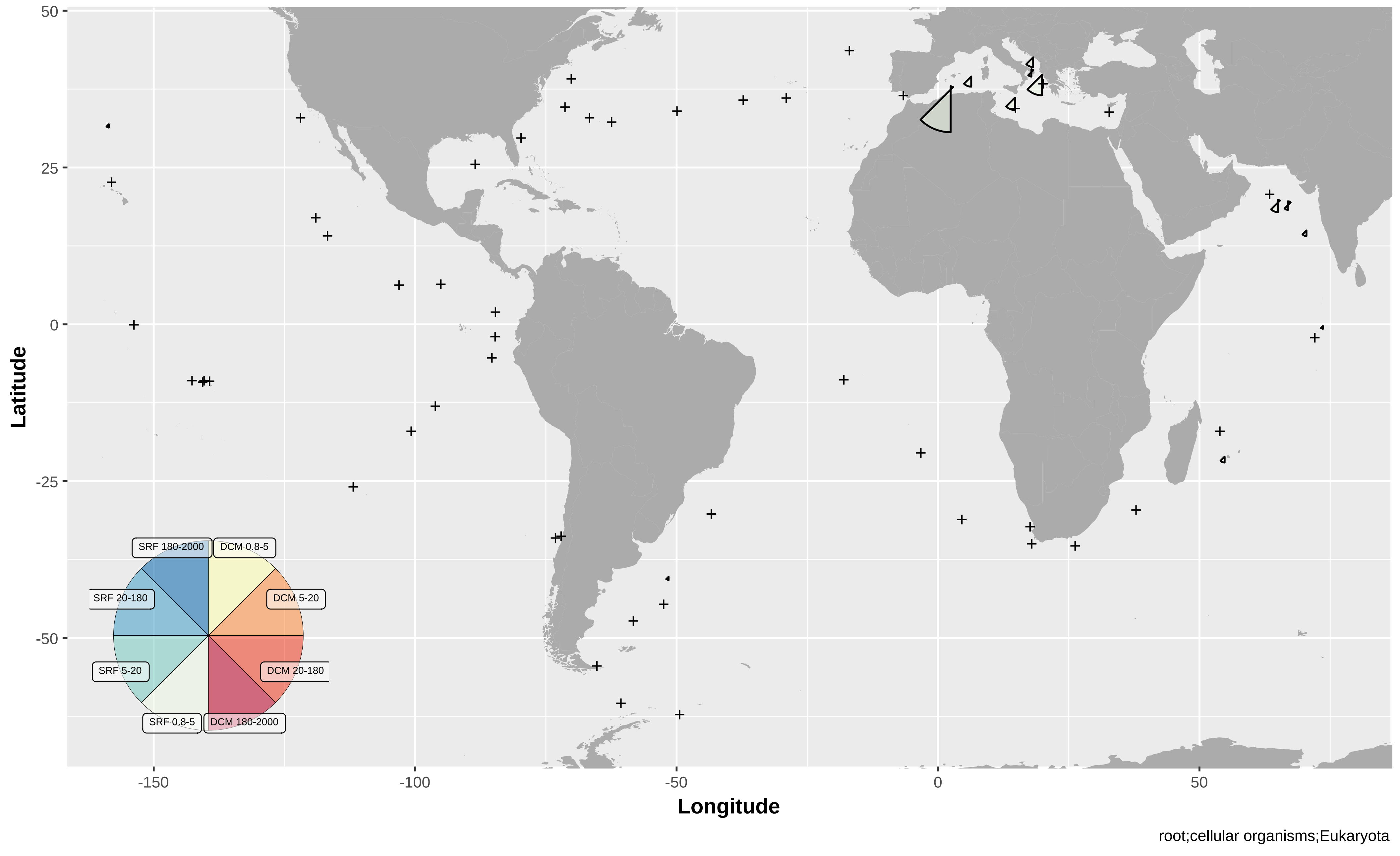

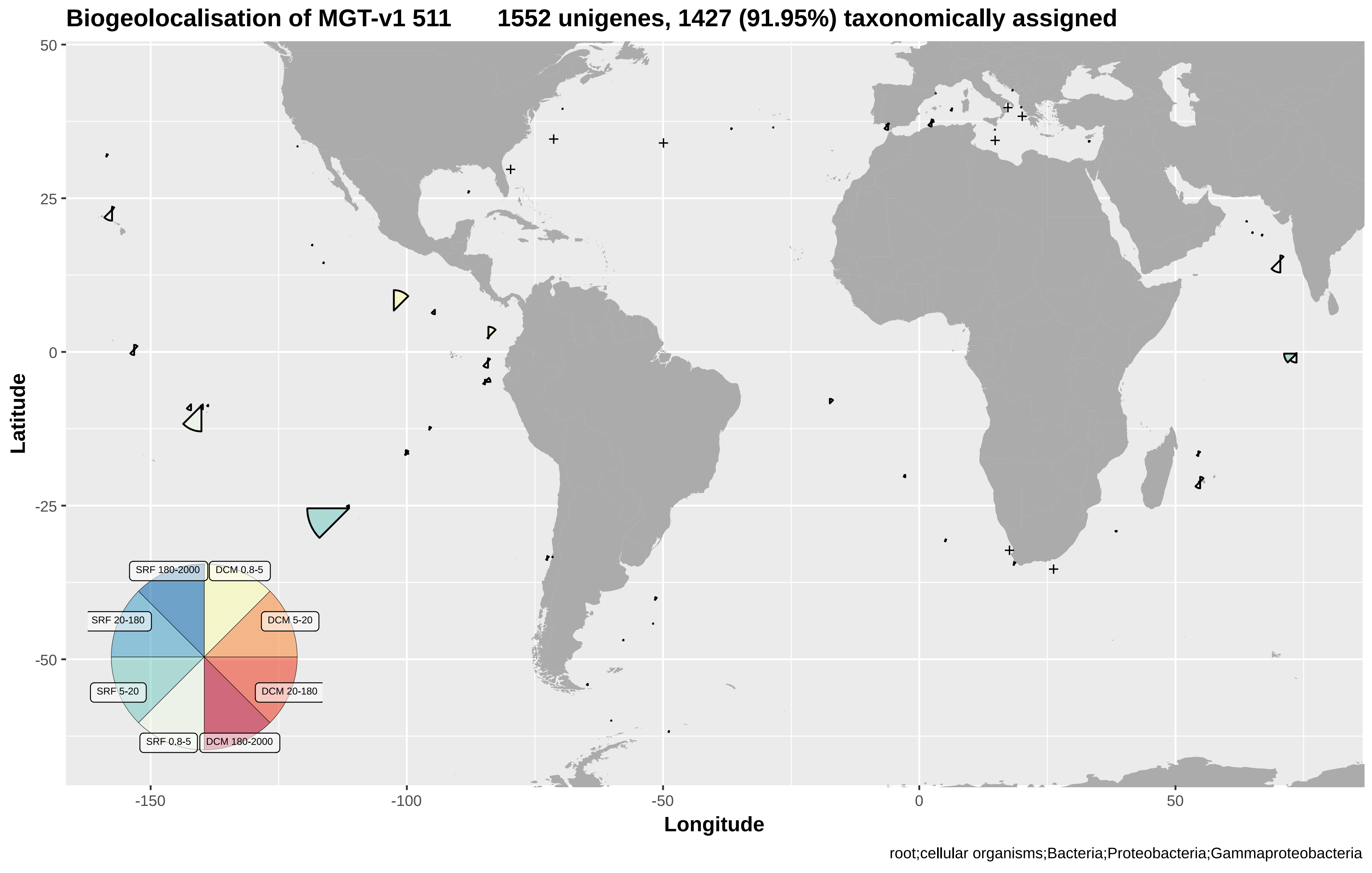
Biogeolocalisation of MGT-v1 511
1552 unigenes, 1427 (91.95%) taxonomically assigned
Latitude
Longitude
50
25
0
-25
-50
-150
-100
-50
0
50
SRF 180-2000
DCM 0.8-5
SRF 20-180
DCM 5-20
SRF 5-20
DCM 20-180
SRF 0.8-5
DCM 180-2000
root;cellular organisms;Bacteria;Proteobacteria;Gammaproteobacteria

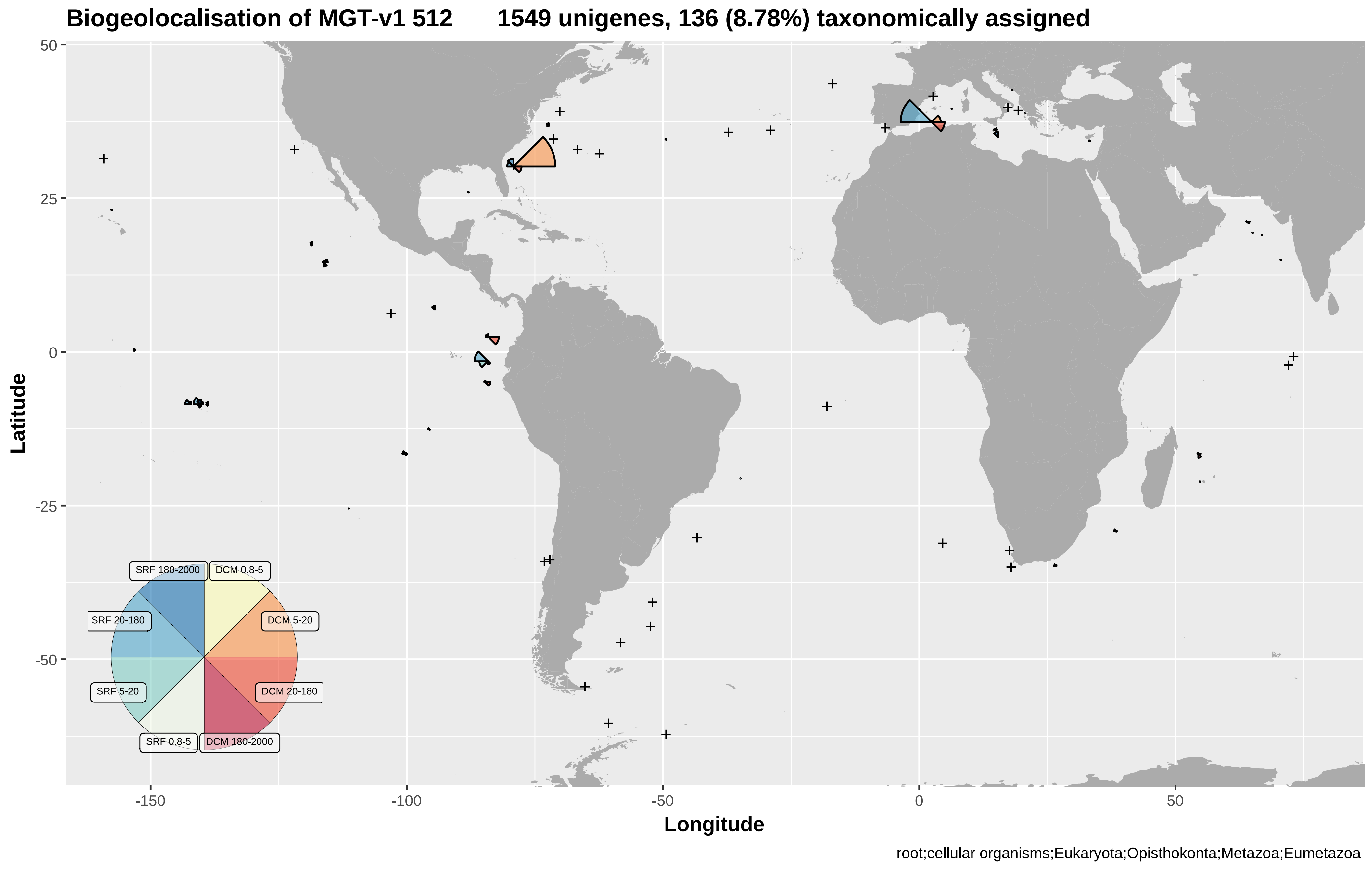
Biogeolocalisation of MGT-v1 512
1549 unigenes, 136 (8.78%) taxonomically assigned
50
25
0
-25
-50
Latitude
-150
-100
-50
0
50
Longitude
SRF 180-2000
DCM 0.8-5
SRF 20-180
DCM 5-20
SRF 5-20
DCM 20-180
SRF 0.8-5
DCM 180-2000
root;cellular organisms;Eukaryota;Opisthokonta;Metazoa;Eumetazoa

### Biogeolocalisation of MGT-v1 513 1546 unigenes, 161 (10.41%) taxonomically assigned

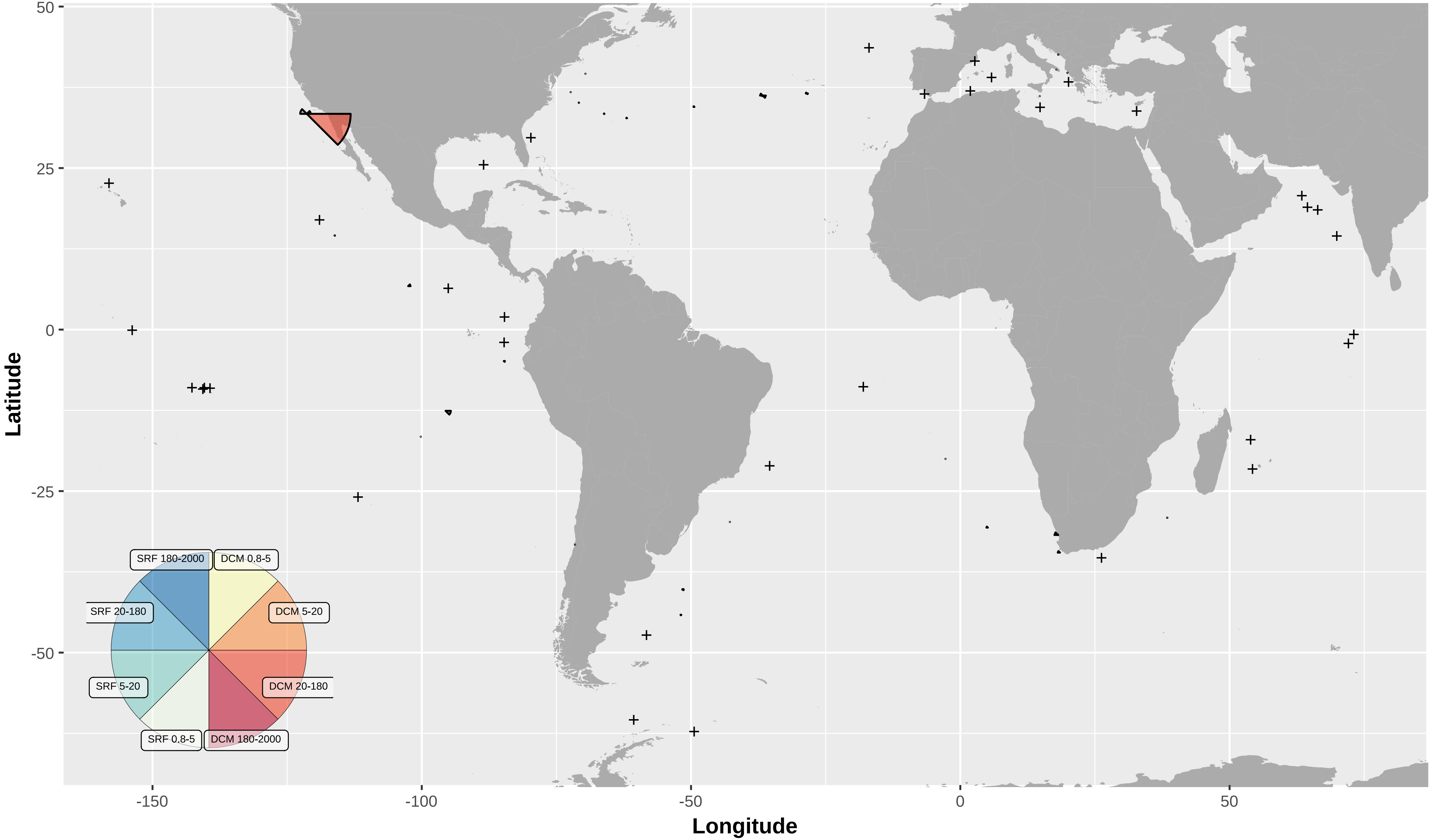


root;cellular organisms;Eukaryota

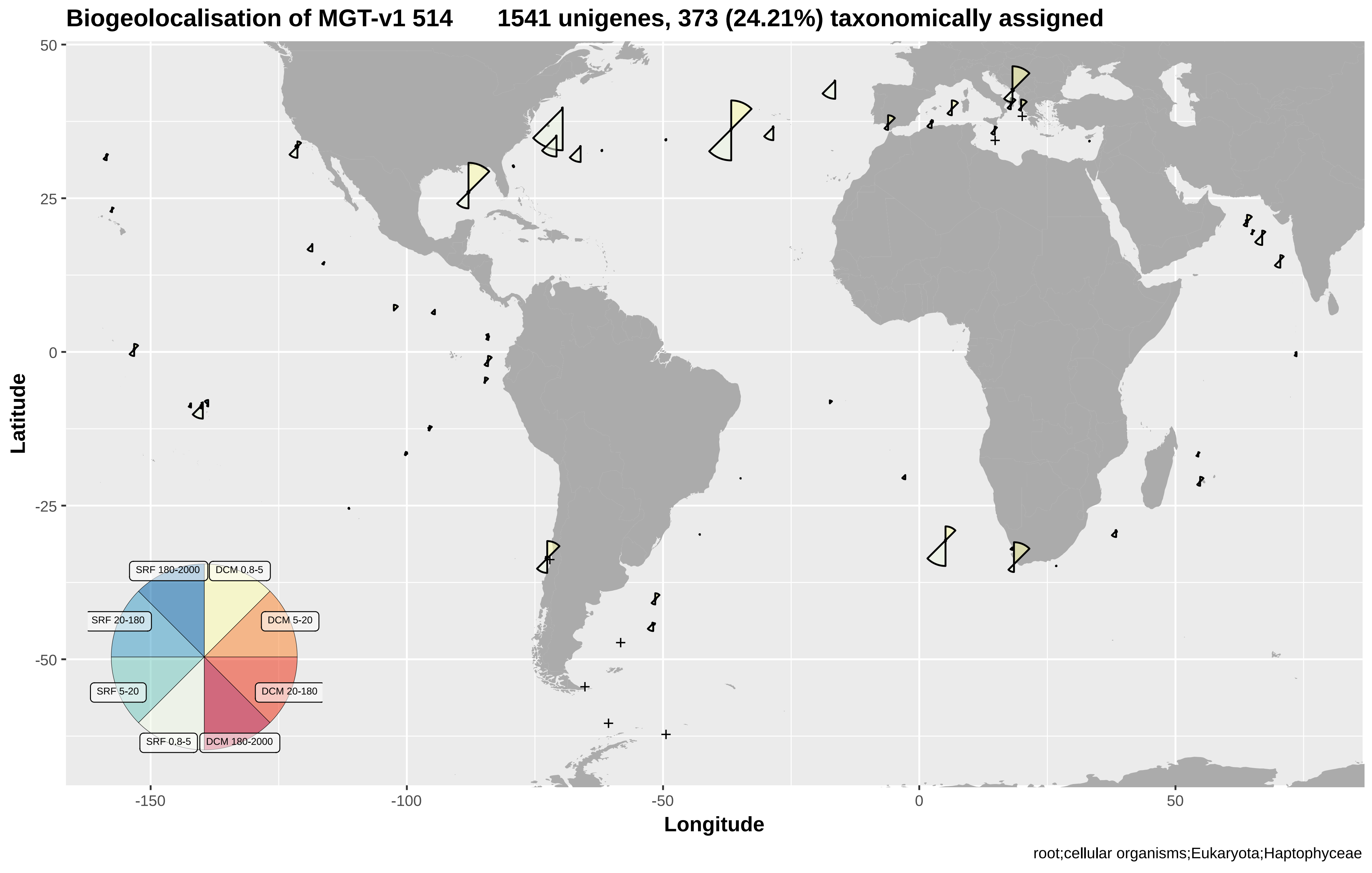
Biogeolocalisation of MGT-v1 514 1541 unigenes, 373 (24.21%) taxonomically assigned
50
25
0
-25
-50
Latitude
-150
-100
-50
0
50
Longitude
SRF 180-2000
DCM 0.8-5
SRF 20-180
DCM 5-20
SRF 5-20
DCM 20-180
SRF 0.8-5
DCM 180-2000
root;cellular organisms;Eukaryota;Haptophyceae

## Biogeolocalisation of MGT-v1 515      1534 unigenes, 205 (13.36%) taxonomically assigned

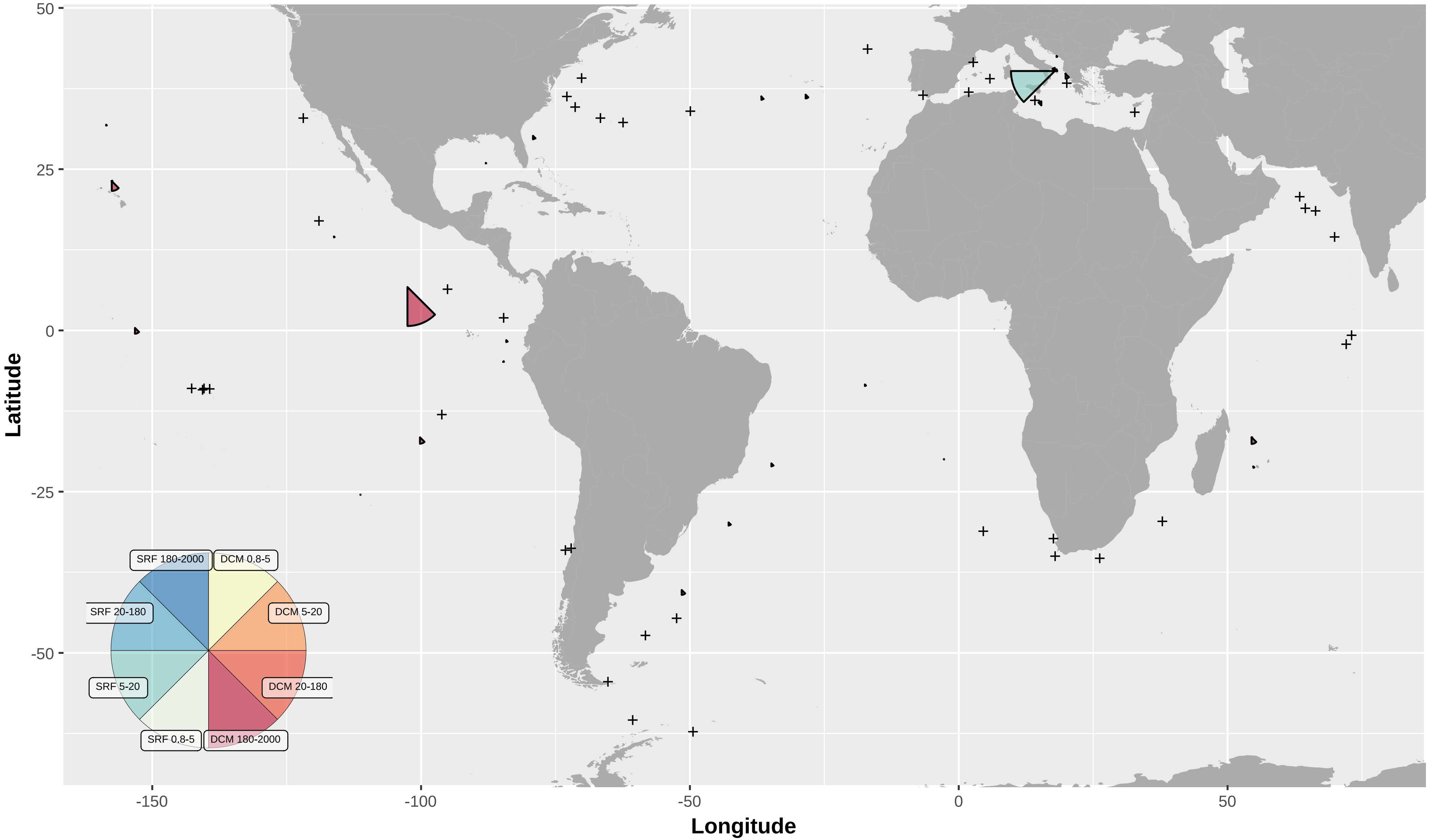


root;cellular organisms;Eukaryota;Opisthokonta;Metazoa;Eumetazoa;Bilateria

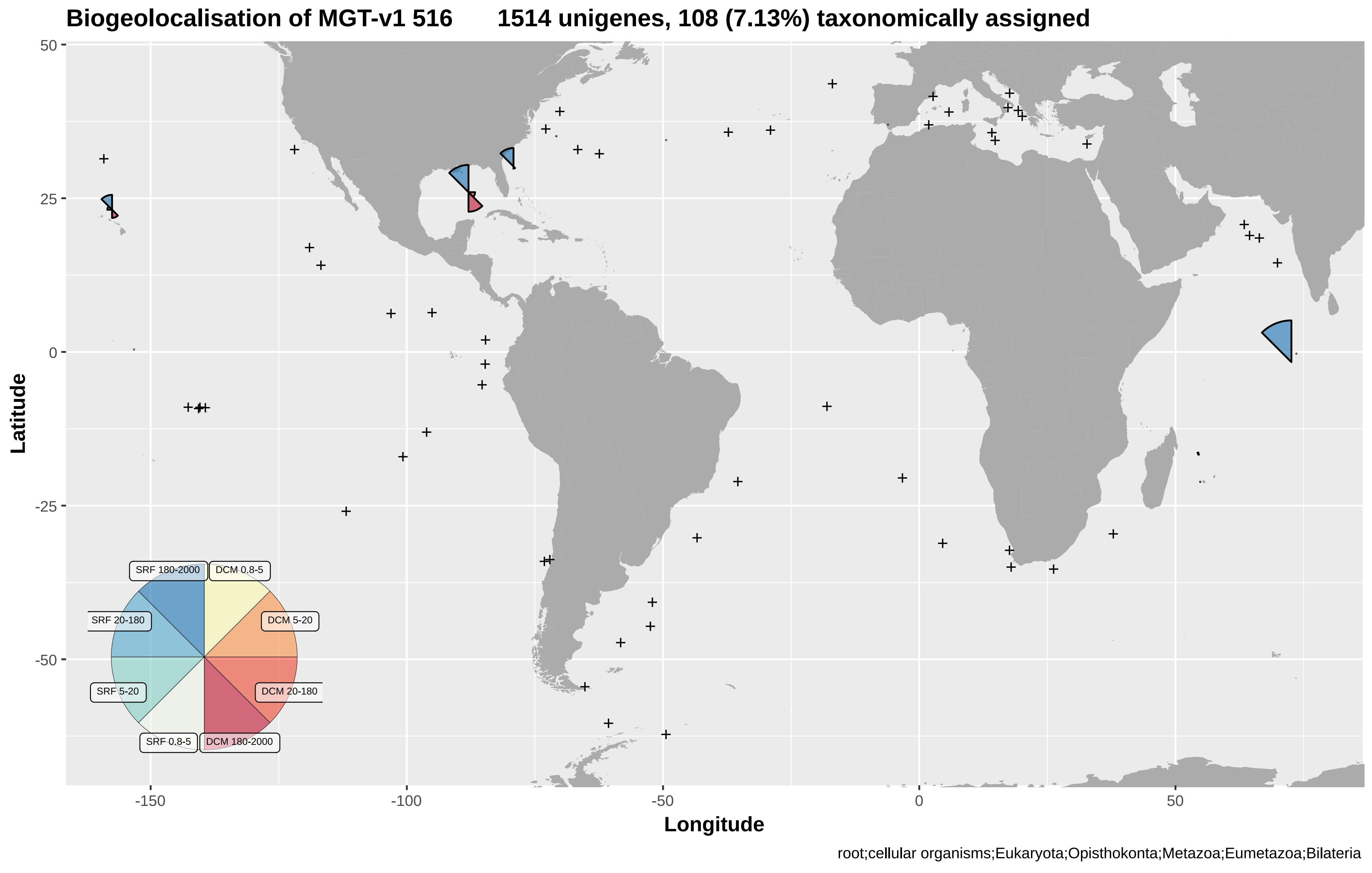
Biogeolocalisation of MGT-v1 516
1514 unigenes, 108 (7.13%) taxonomically assigned
50
25
0
-25
-50
Latitude
-150
-100
-50
0
50
Longitude
SRF 180-2000
DCM 0.8-5
SRF 20-180
DCM 5-20
SRF 5-20
DCM 20-180
SRF 0.8-5
DCM 180-2000
root;cellular organisms;Eukaryota;Opisthokonta;Metazoa;Eumetazoa;Bilateria

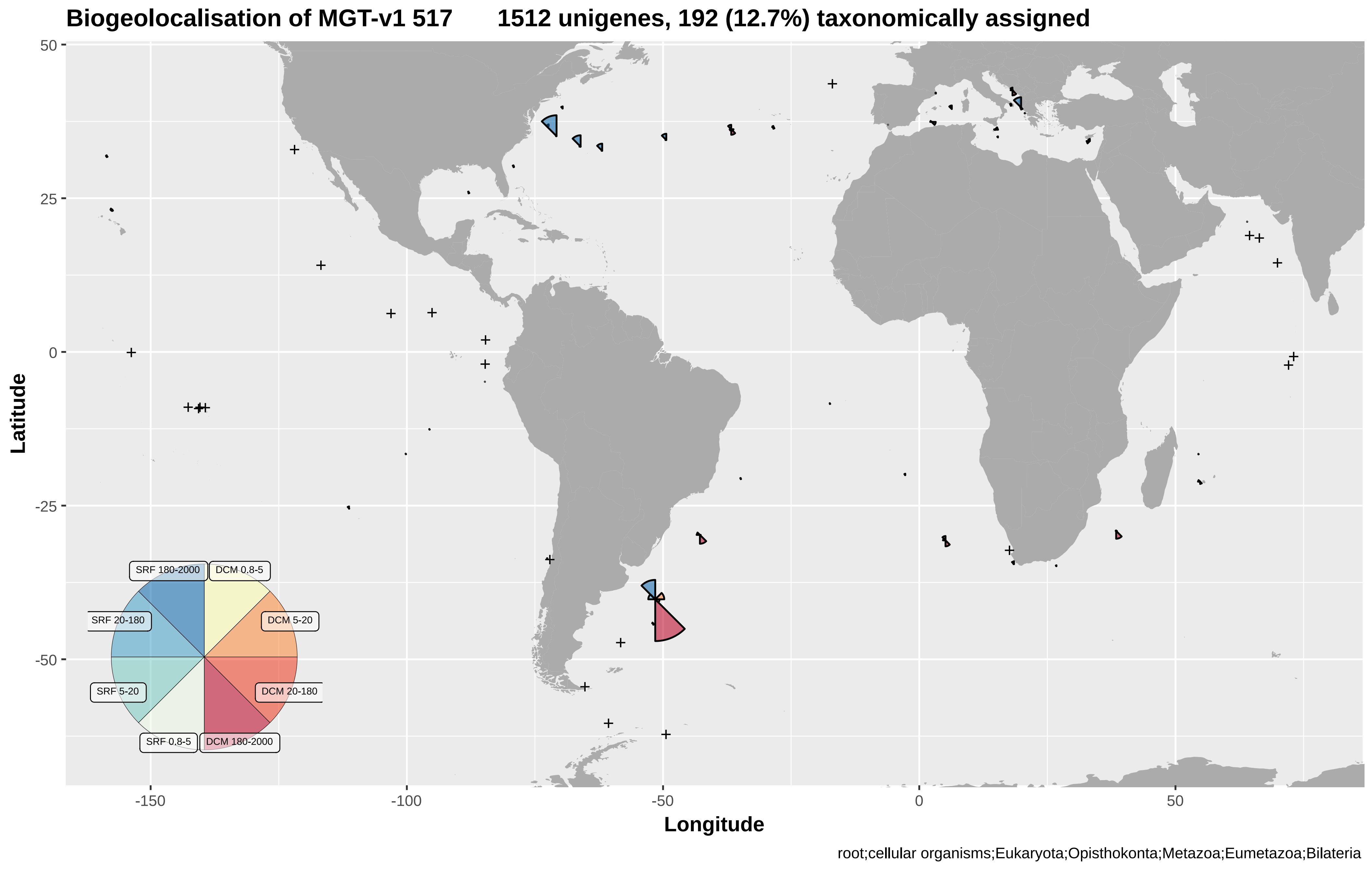
Biogeolocalisation of MGT-v1 517 1512 unigenes, 192 (12.7%) taxonomically assigned
Latitude
50
25
0
-25
-50
Longitude
-150
-100
-50
0
50
SRF 180-2000
DCM 0.8-5
SRF 20-180
DCM 5-20
SRF 5-20
DCM 20-180
SRF 0.8-5
DCM 180-2000
root;cellular organisms;Eukaryota;Opisthokonta;Metazoa;Eumetazoa;Bilateria

## Biogeolocalisation of MGT-v1 518      1506 unigenes, 294 (19.52%) taxonomically assigned

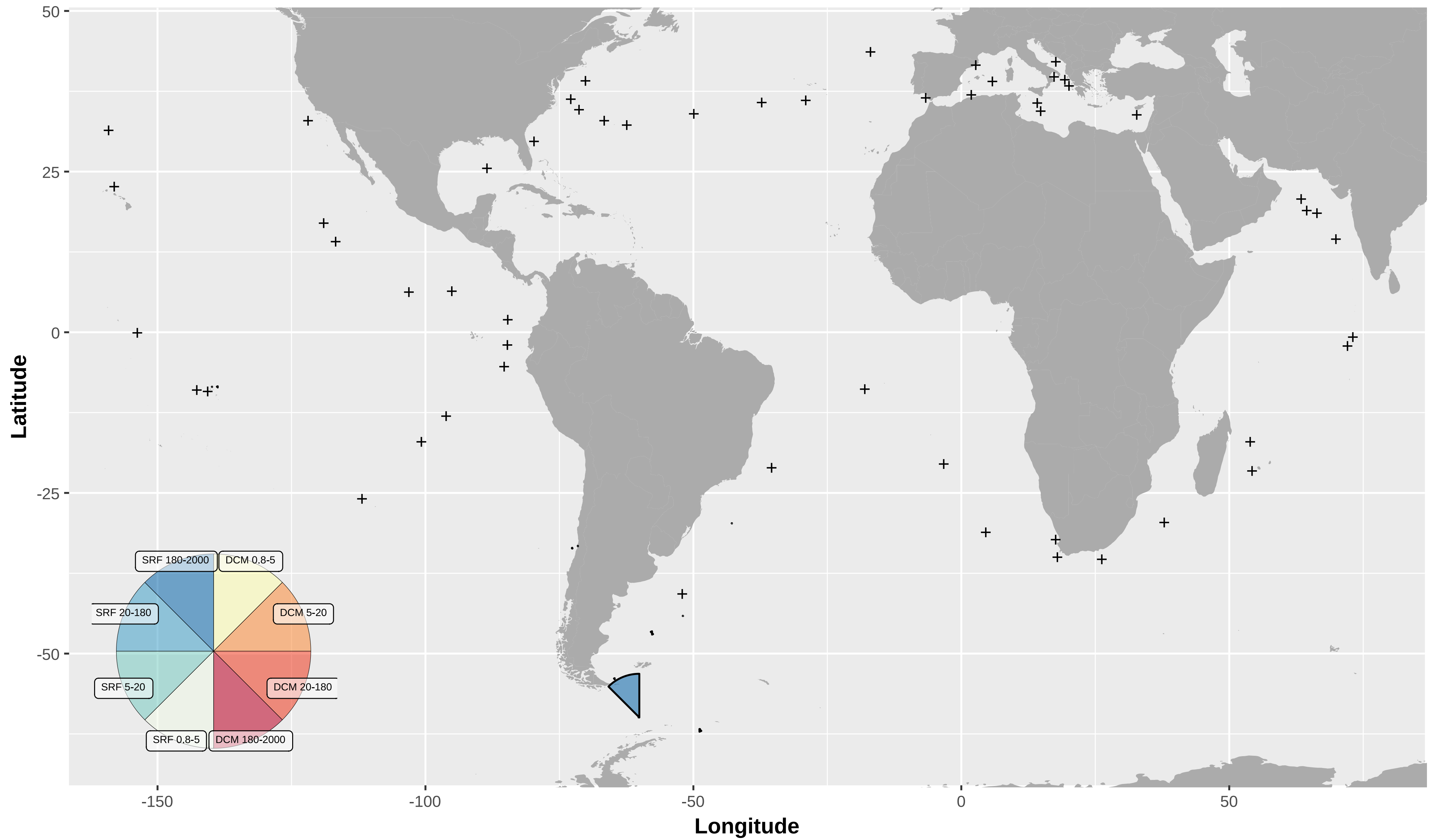


root;cellular organisms;Eukaryota;Opisthokonta;Metazoa;Eumetazoa;Bilateria

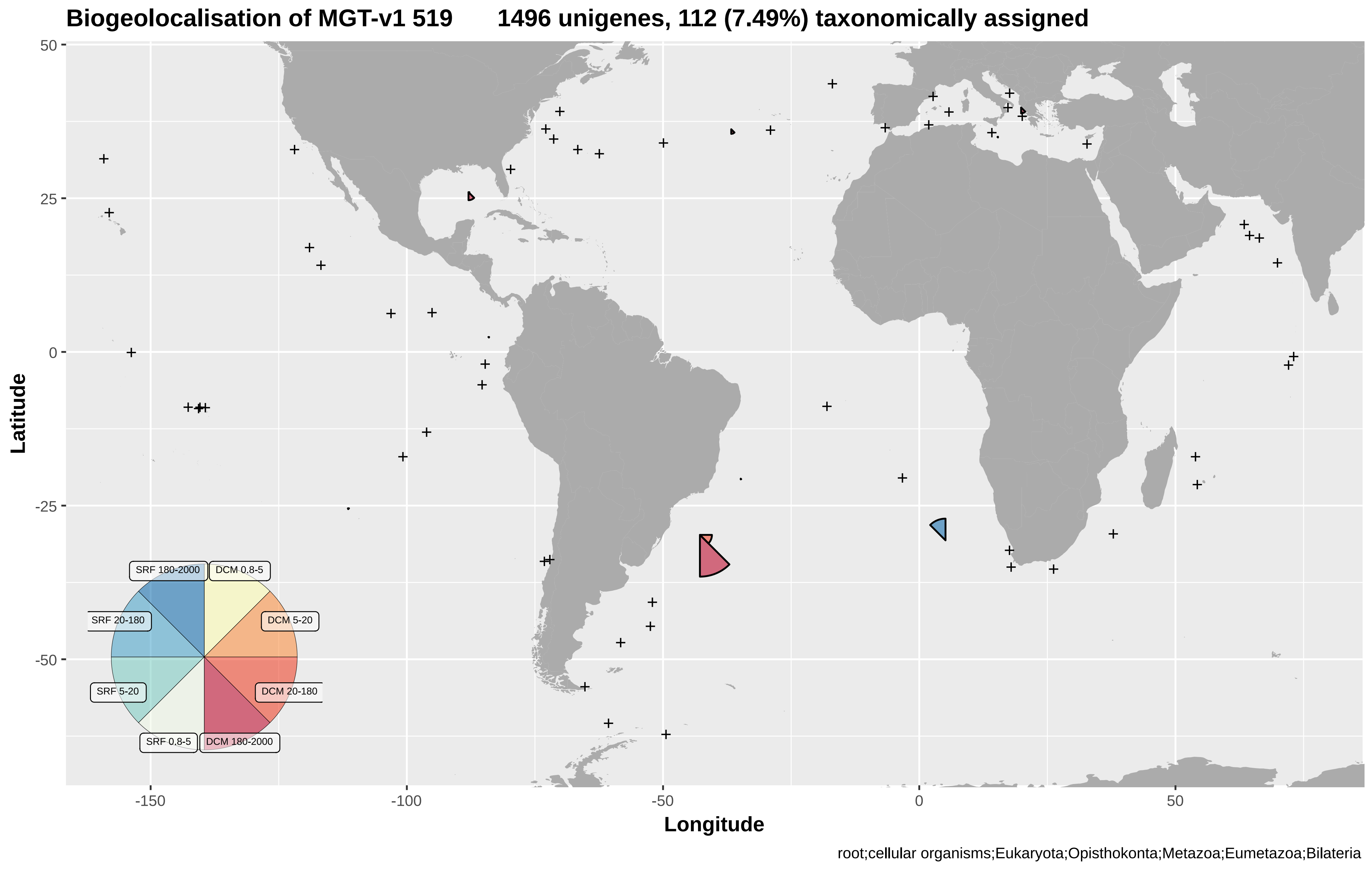

Biogeolocalisation of MGT-v1 519 1496 unigenes, 112 (7.49%) taxonomically assigned
Latitude
Longitude
50
25
0
-25
-50
-150
-100
-50
0
50
SRF 180-2000
DCM 0.8-5
SRF 20-180
DCM 5-20
SRF 5-20
DCM 20-180
SRF 0.8-5
DCM 180-2000
root;cellular organisms;Eukaryota;Opisthokonta;Metazoa;Eumetazoa;Bilateria

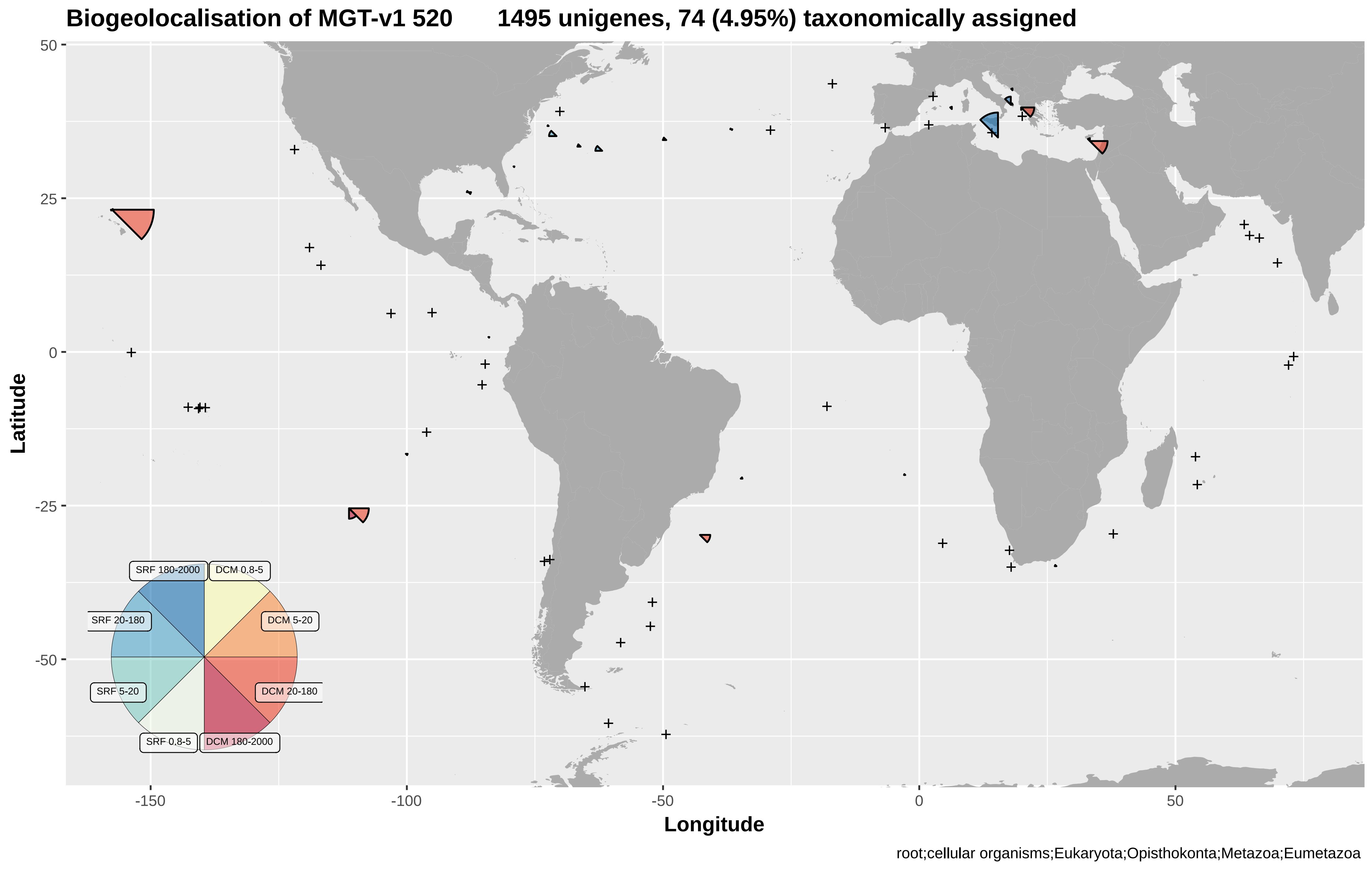
Biogeolocalisation of MGT-v1 520 1495 unigenes, 74 (4.95%) taxonomically assigned
Latitude
Longitude
50
25
0
-25
-50
-150
-100
-50
0
50
SRF 180-2000
DCM 0.8-5
SRF 20-180
DCM 5-20
SRF 5-20
DCM 20-180
SRF 0.8-5
DCM 180-2000
root;cellular organisms;Eukaryota;Opisthokonta;Metazoa;Eumetazoa

#### Biogeolocalisation of MGT-v1 521 1493 unigenes, 184 (12.32%) taxonomically assigned

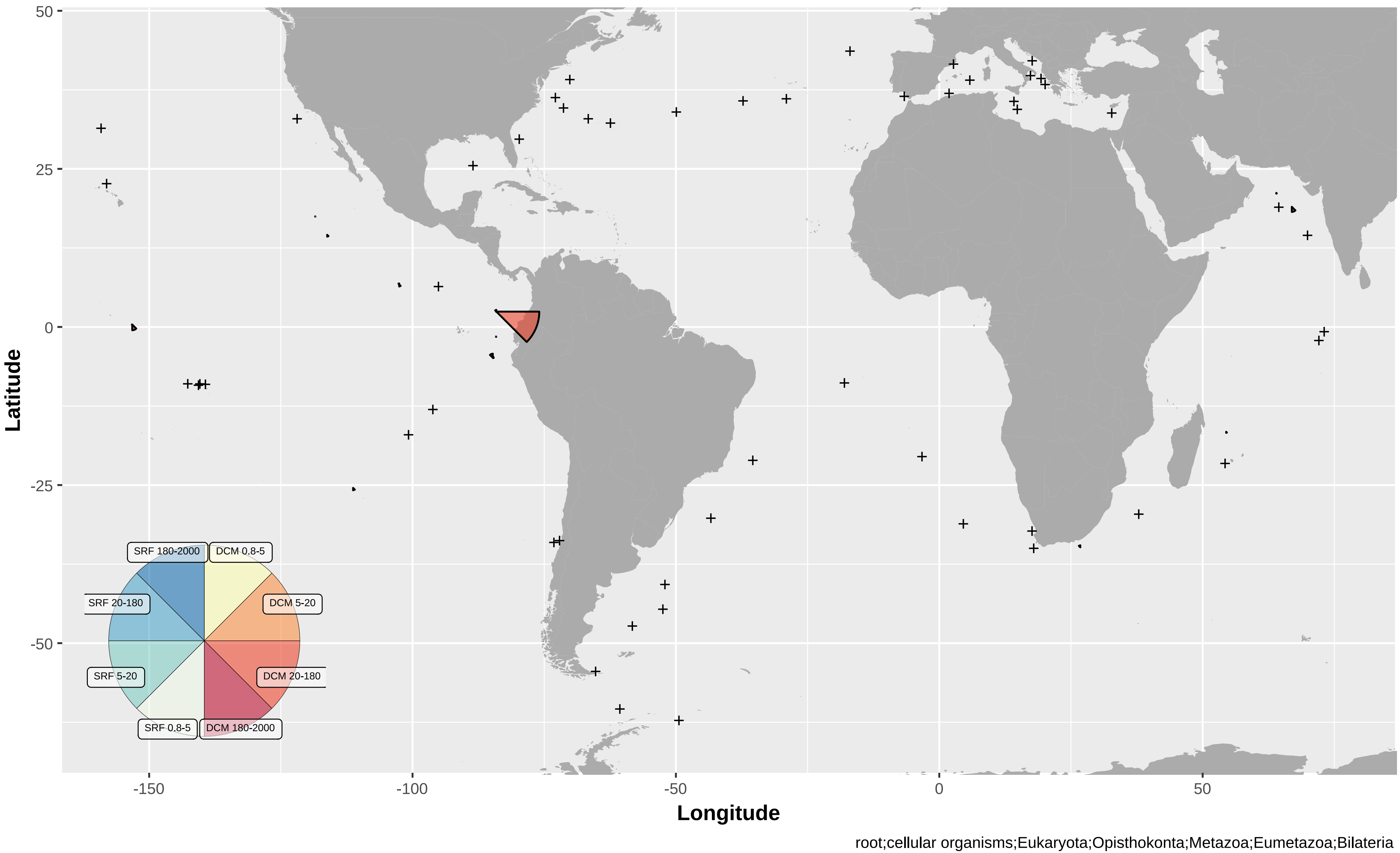

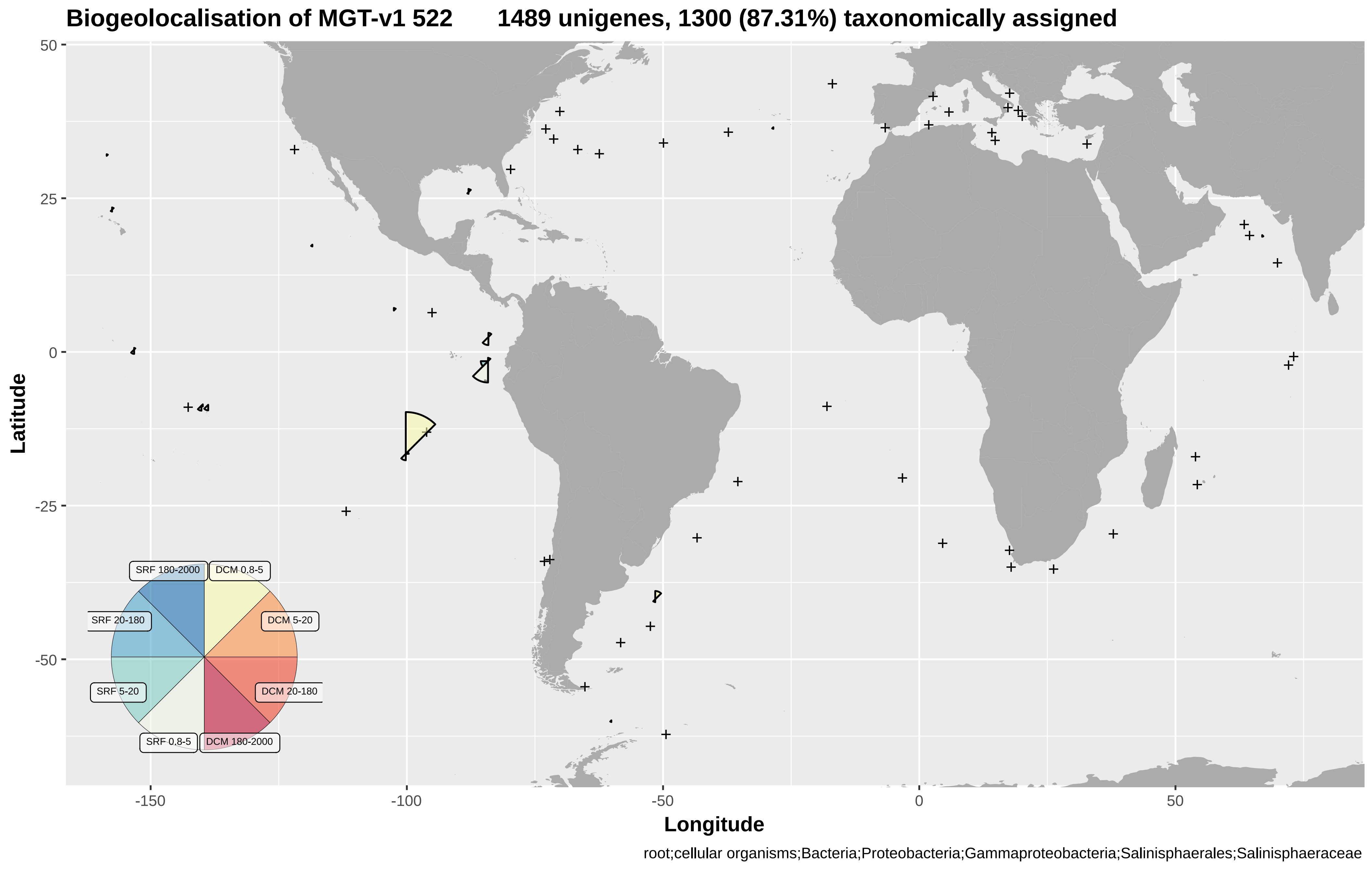

Biogeolocalisation of MGT-v1 522 1489 unigenes, 1300 (87.31%) taxonomically assigned
50
25
0
-25
-50
Latitude
-150
-100
-50
0
50
Longitude
SRF 180-2000
DCM 0.8-5
SRF 20-180
DCM 5-20
SRF 5-20
DCM 20-180
SRF 0.8-5
DCM 180-2000
root;cellular organisms;Bacteria;Proteobacteria;Gammaproteobacteria;Salinisphaerales;Salinisphaeraceae

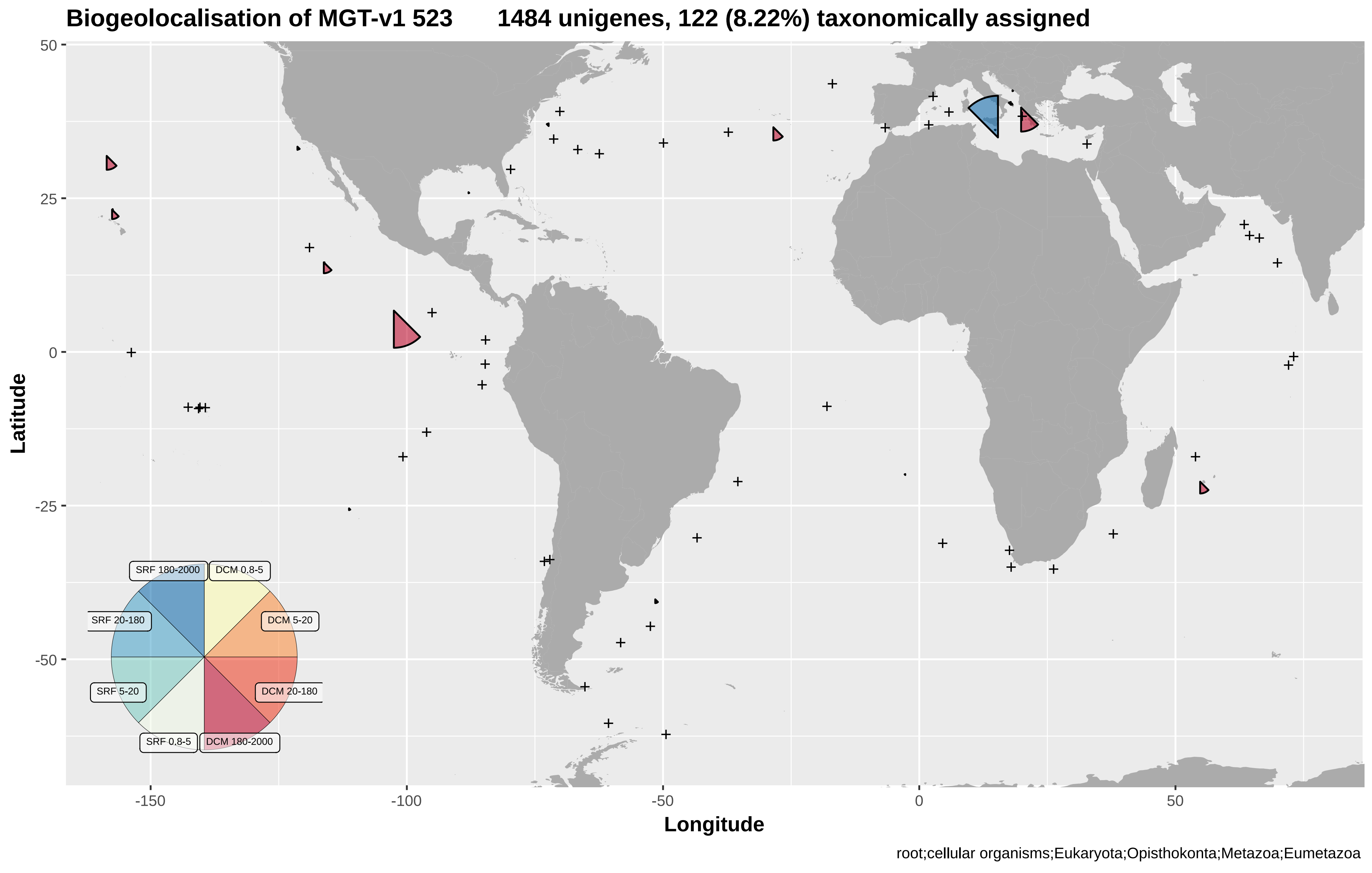
Biogeolocalisation of MGT-v1 523 1484 unigenes, 122 (8.22%) taxonomically assigned
Latitude
Longitude
50
25
0
-25
-50
-150
-100
-50
0
50
SRF 180-2000
DCM 0.8-5
SRF 20-180
DCM 5-20
SRF 5-20
DCM 20-180
SRF 0.8-5
DCM 180-2000
root;cellular organisms;Eukaryota;Opisthokonta;Metazoa;Eumetazoa

#### Biogeolocalisation of MGT-v1 524 1482 unigenes, 1357 (91.57%) taxonomically assigned

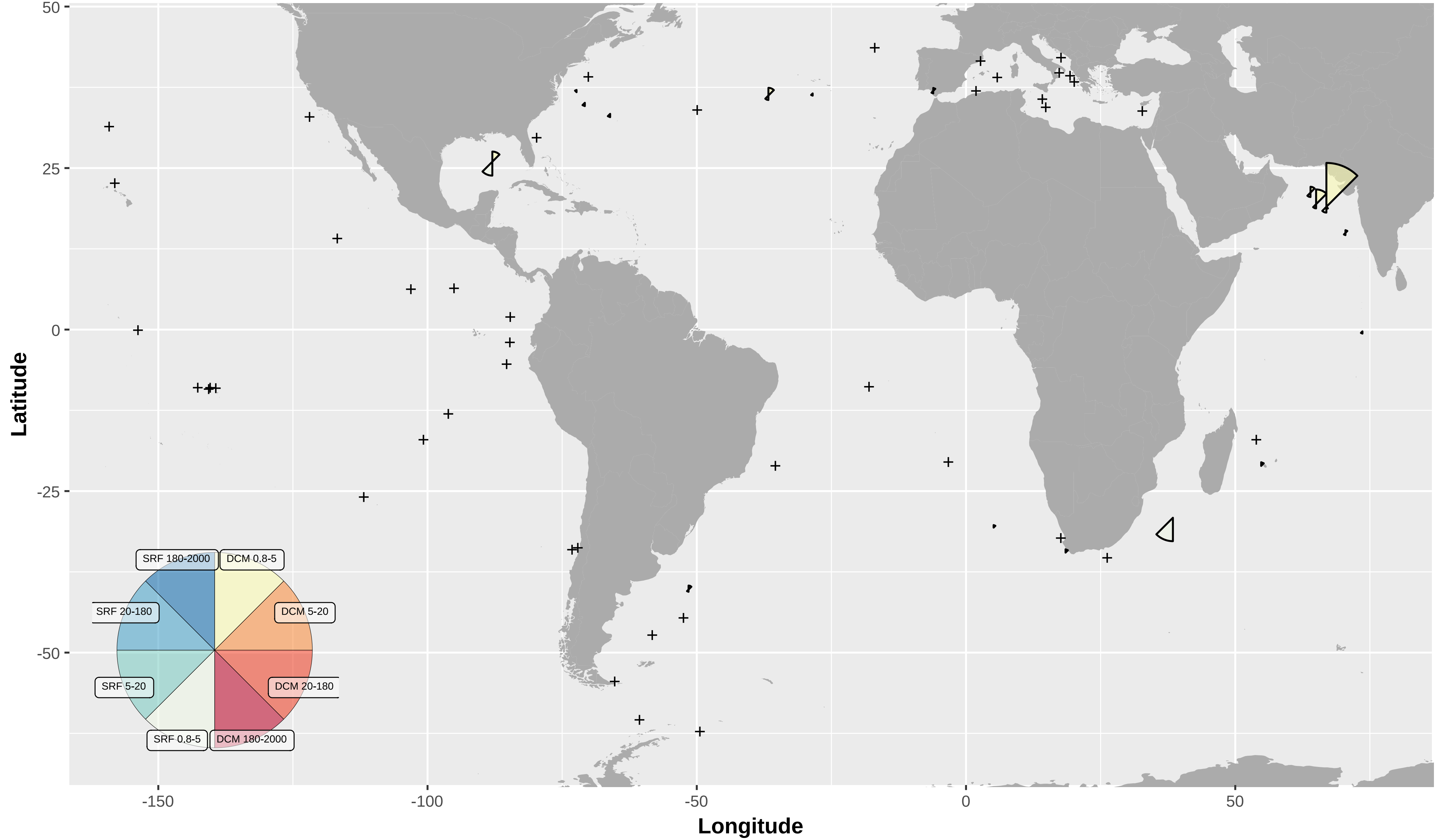


root;cellular organisms;Bacteria;Cyanobacteria;Oscillatoriophycideae;Chroococcales

# Biogeolocalisation of MGT-v1 525      1449 unigenes, 92 (6.35%) taxonomically assigned

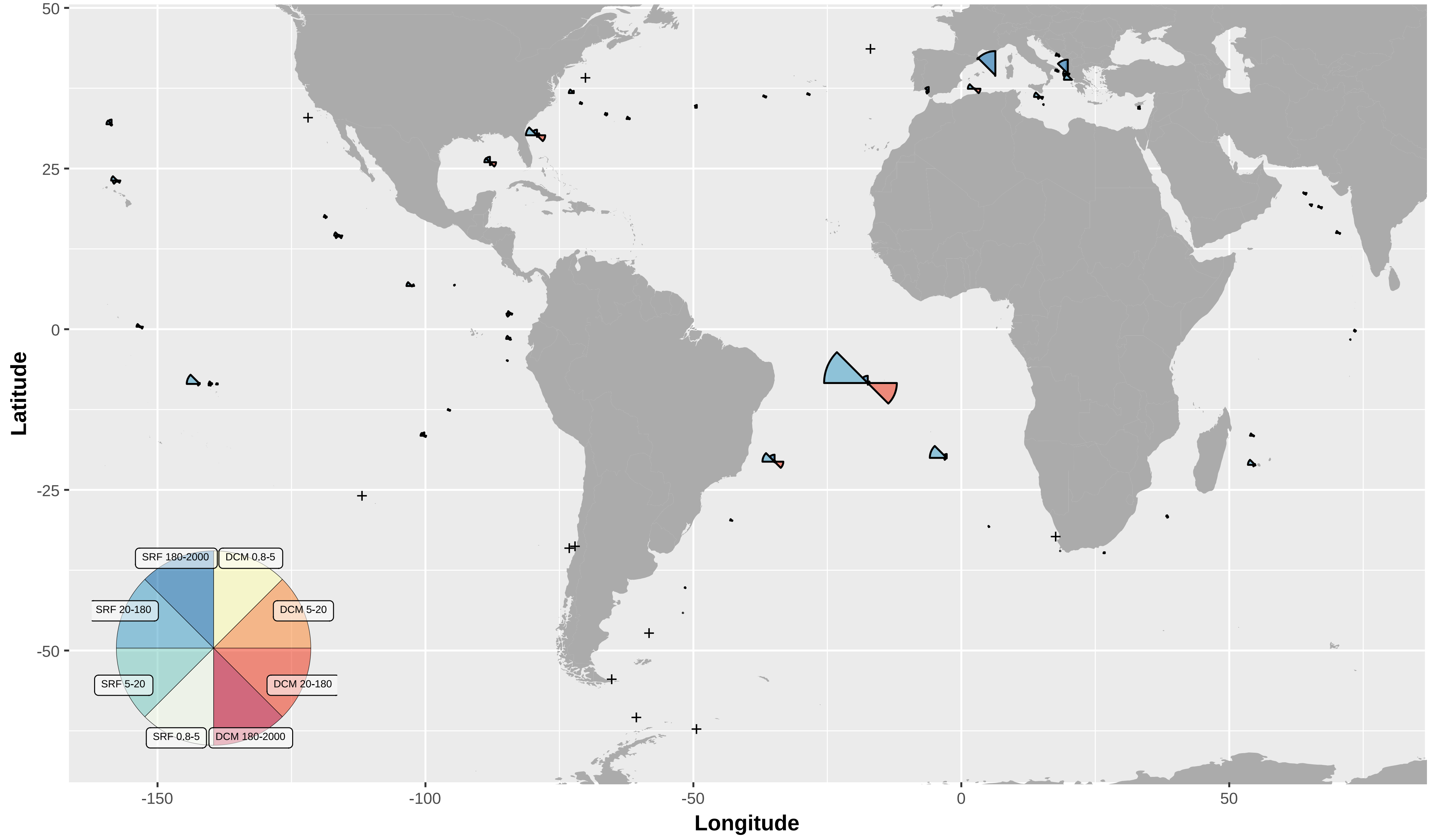


root;cellular organisms;Eukaryota;Opisthokonta

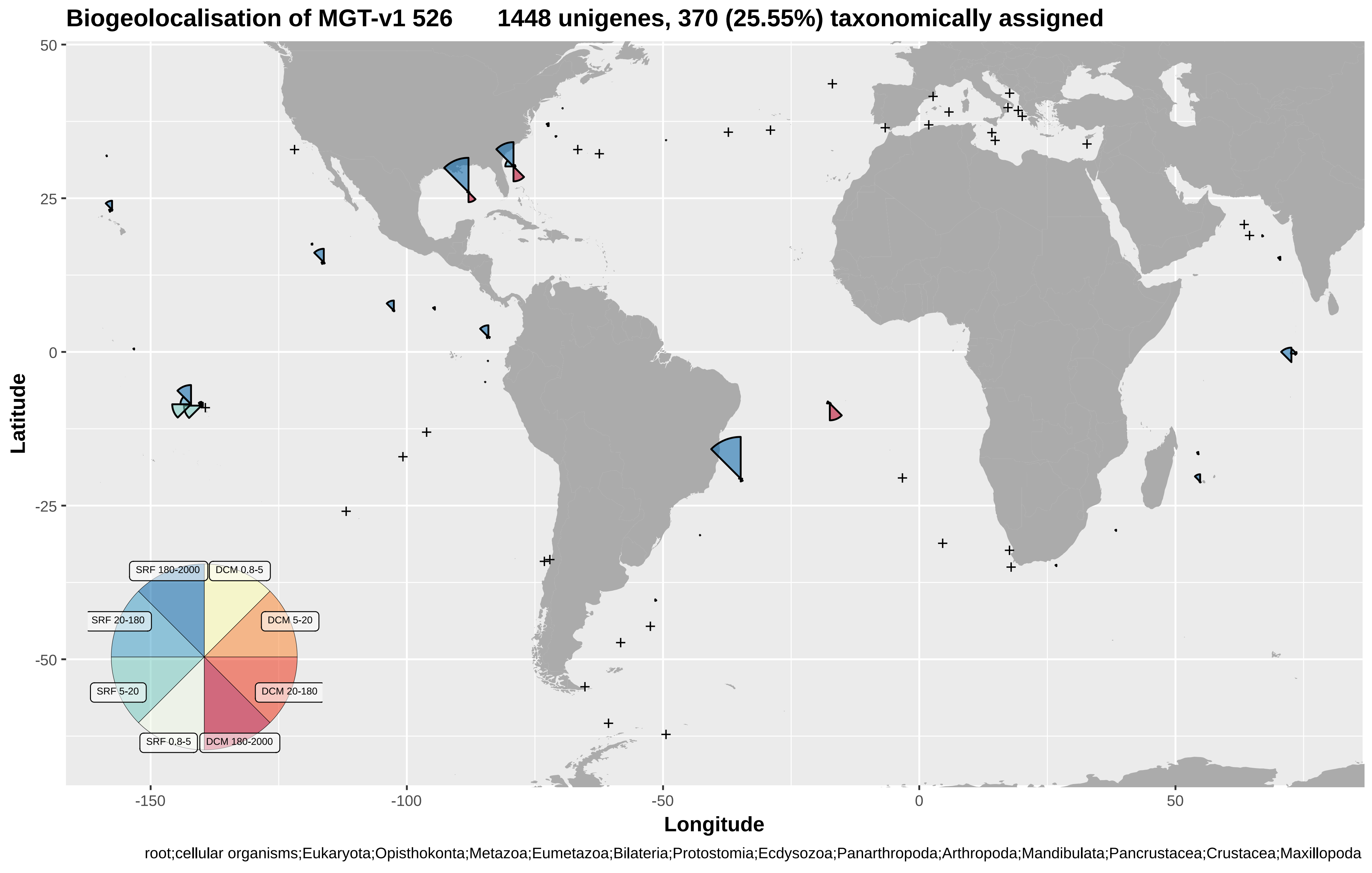
Biogeolocalisation of MGT-v1 526      1448 unigenes, 370 (25.55%) taxonomically assigned
50
25
0
-25
-50
Latitude
-150
-100
-50
0
50
Longitude
SRF 180-2000
DCM 0.8-5
SRF 20-180
DCM 5-20
SRF 5-20
DCM 20-180
SRF 0.8-5
DCM 180-2000
root;cellular organisms;Eukaryota;Opisthokonta;Metazoa;Eumetazoa;Bilateria;Protostomia;Ecdysozoa;Panarthropoda;Arthropoda;Mandibulata;Pancrustacea;Crustacea;Maxillopoda

### Biogeolocalisation of MGT-v1 527 1431 unigenes, 1391 (97.2%) taxonomically assigned

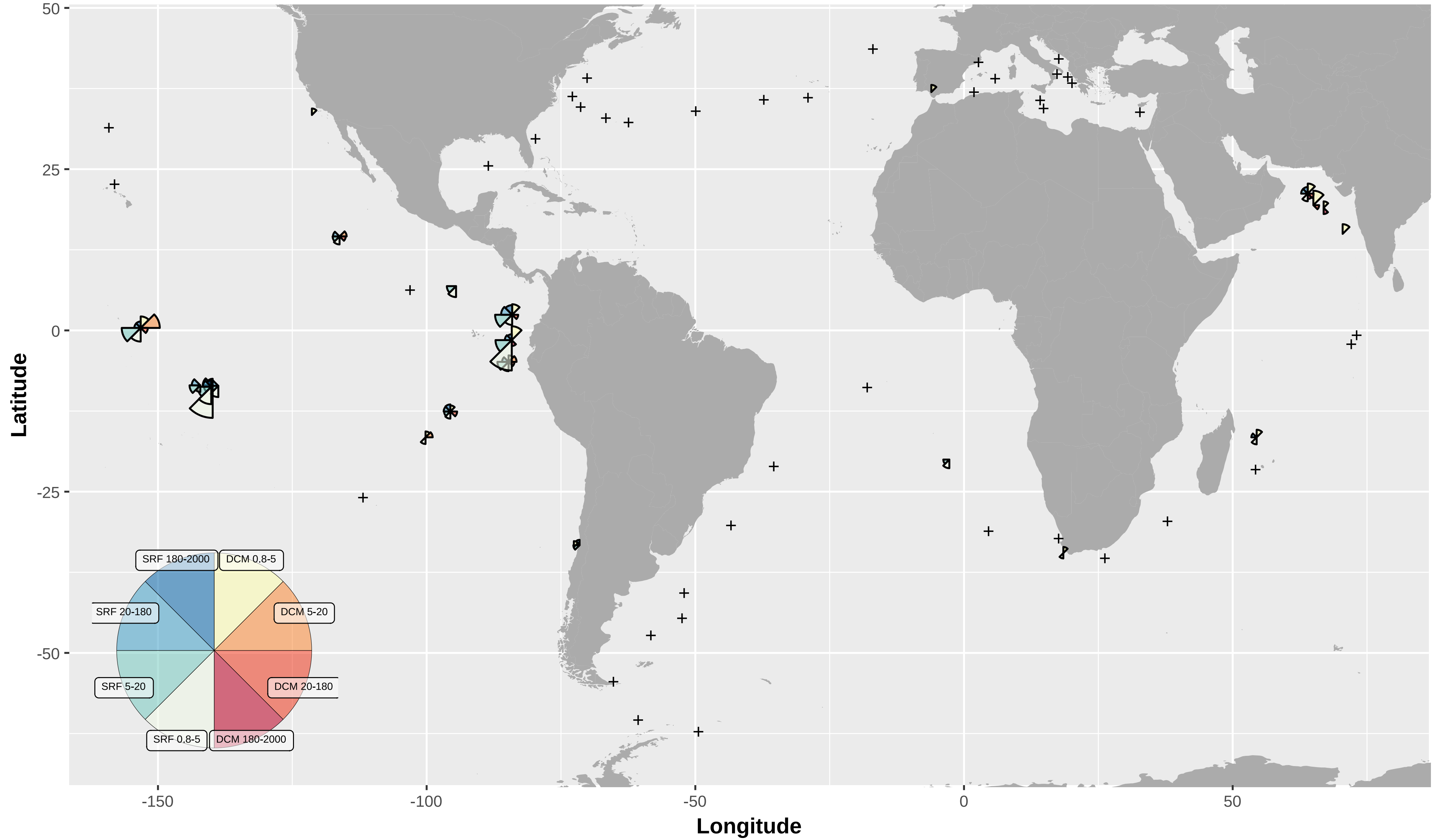


root;cellular organisms;Eukaryota;Stramenopiles;Bacillariophyta;Bacillariophyceae;Bacillariophycidae;Bacillariales

## Biogeolocalisation of MGT-v1 528      1430 unigenes, 90 (6.29%) taxonomically assigned

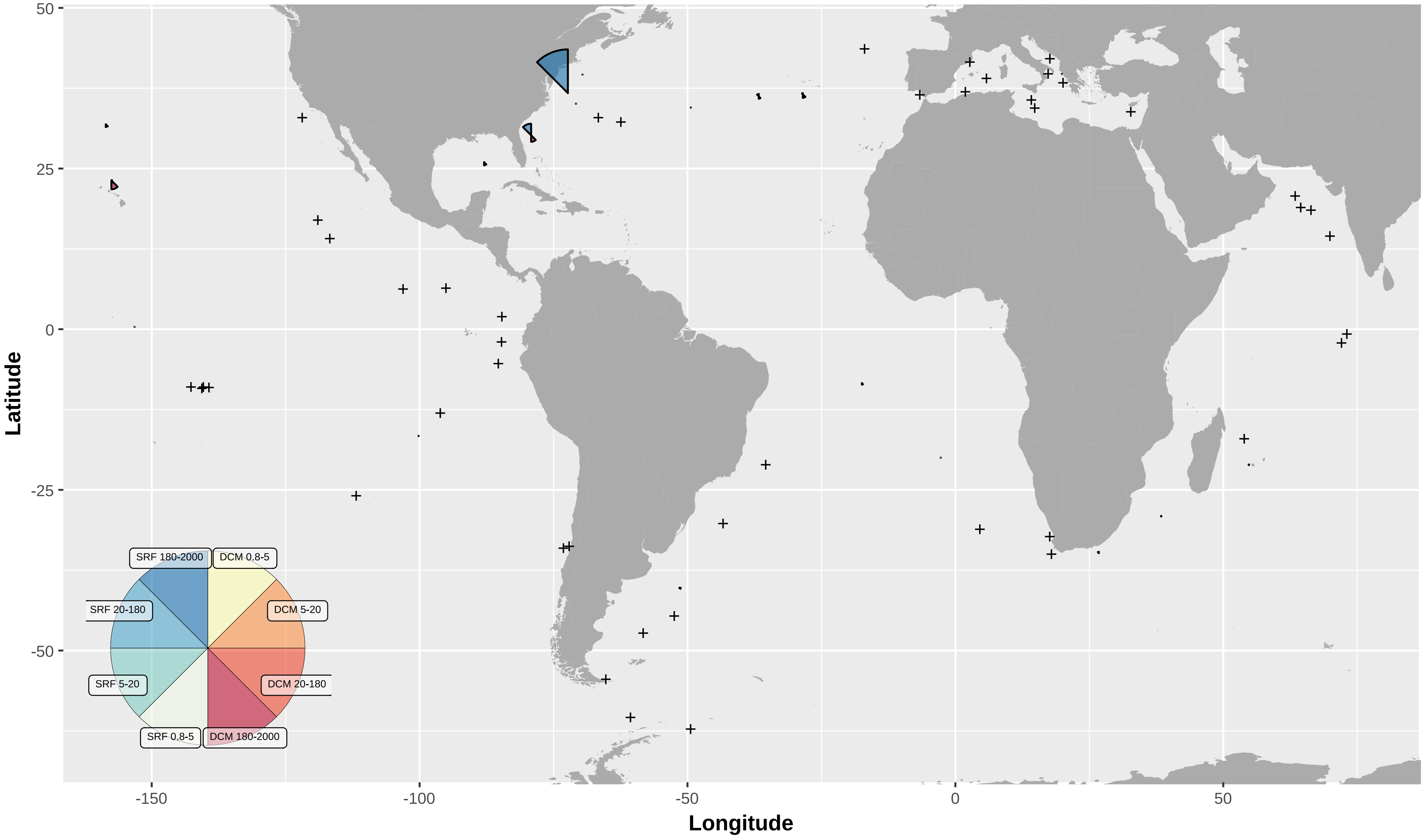


root;cellular organisms;Eukaryota;Opisthokonta;Metazoa;Eumetazoa

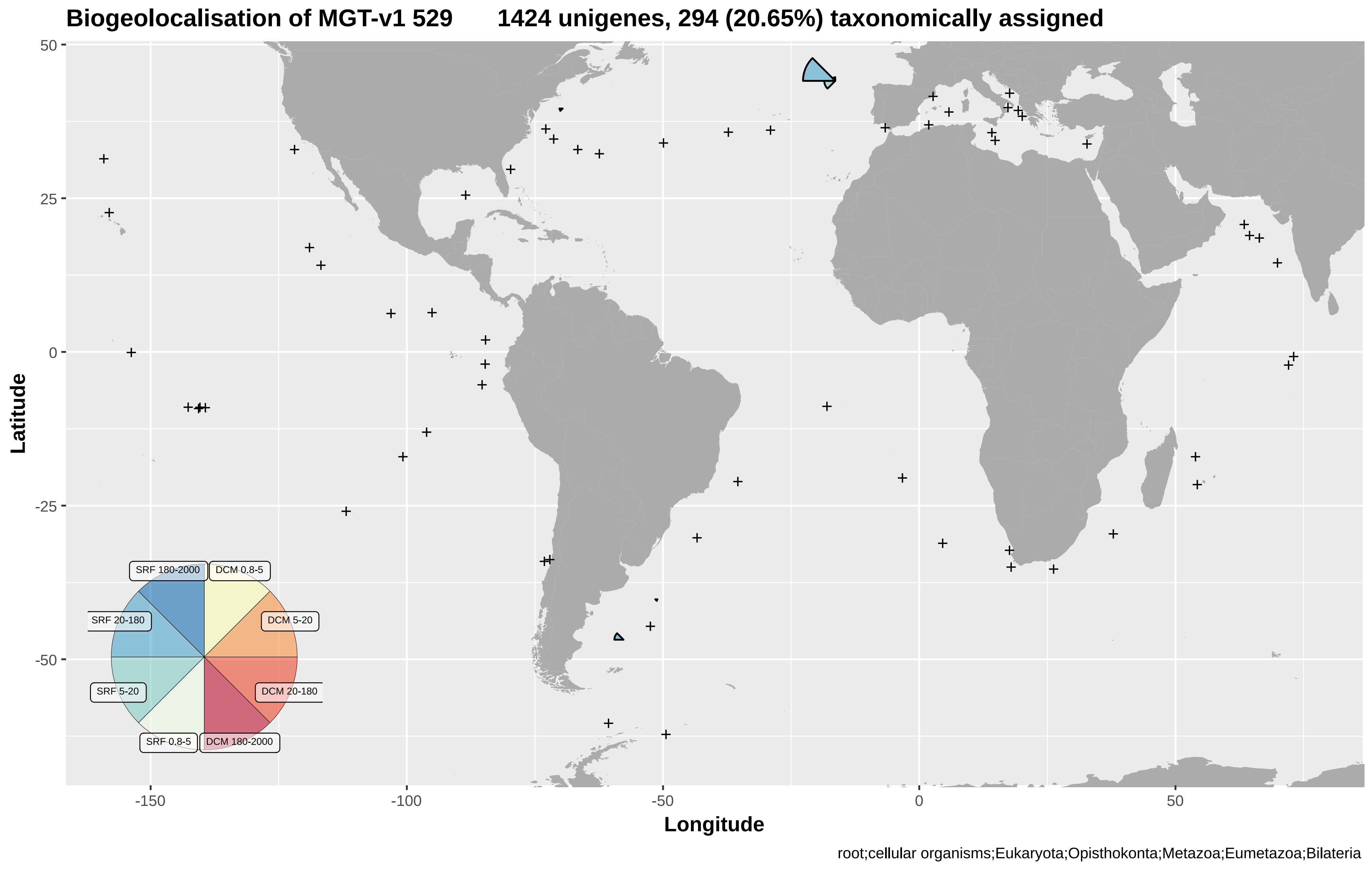
Biogeolocalisation of MGT-v1 529 1424 unigenes, 294 (20.65%) taxonomically assigned
50
25
0
-25
-50
Latitude
-150
-100
-50
0
50
Longitude
SRF 180-2000
DCM 0.8-5
SRF 20-180
DCM 5-20
SRF 5-20
DCM 20-180
SRF 0.8-5
DCM 180-2000
root;cellular organisms;Eukaryota;Opisthokonta;Metazoa;Eumetazoa;Bilateria

#### Biogeolocalisation of MGT-v1 530 1423 unigenes, 686 (48.21%) taxonomically assigned

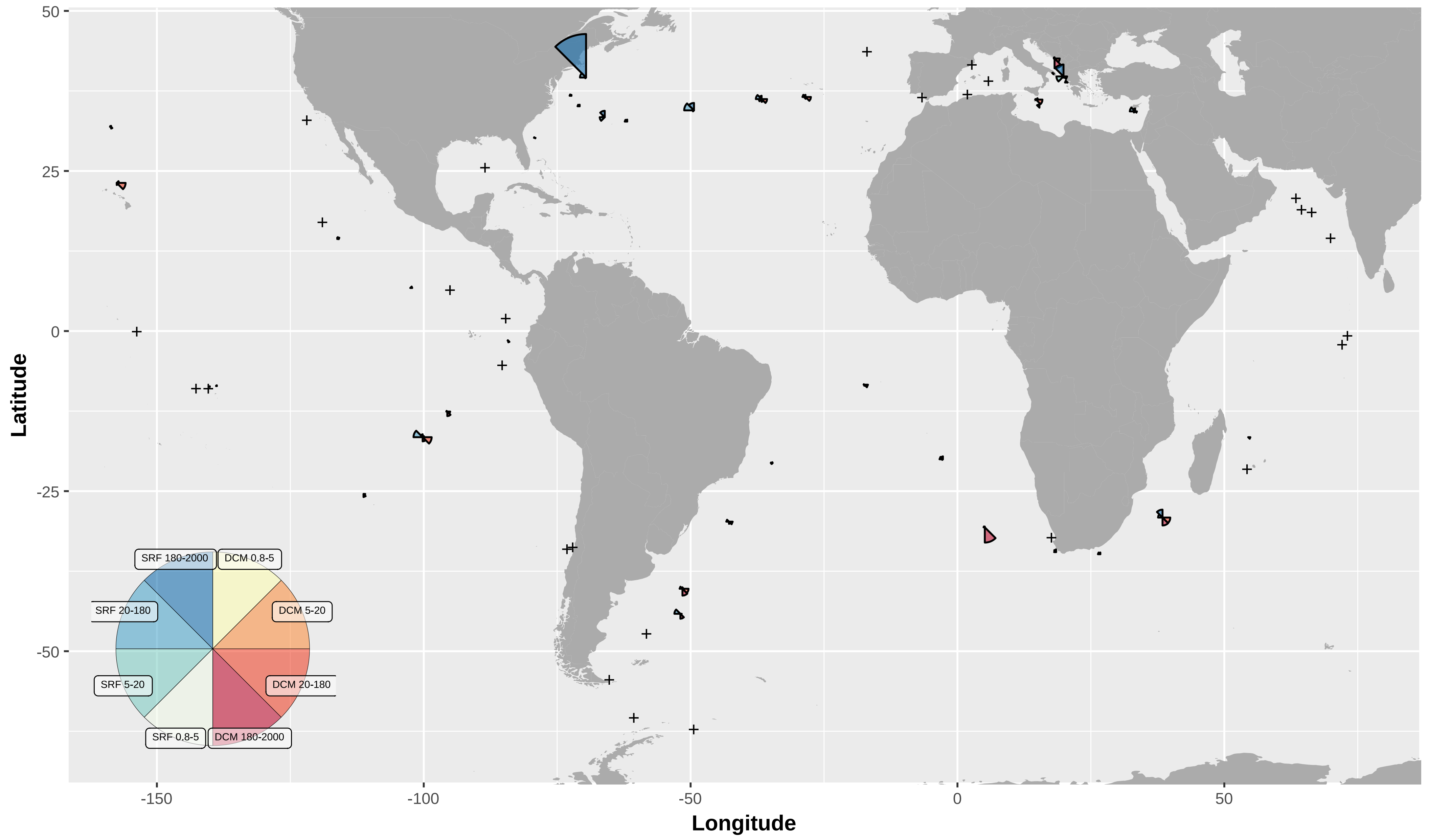


root;cellular organisms;Eukaryota;Opisthokonta;Metazoa;Eumetazoa;Bilateria;Protostomia;Ecdysozoa;Panarthropoda;Arthropoda;Mandibulata;Pancrustacea

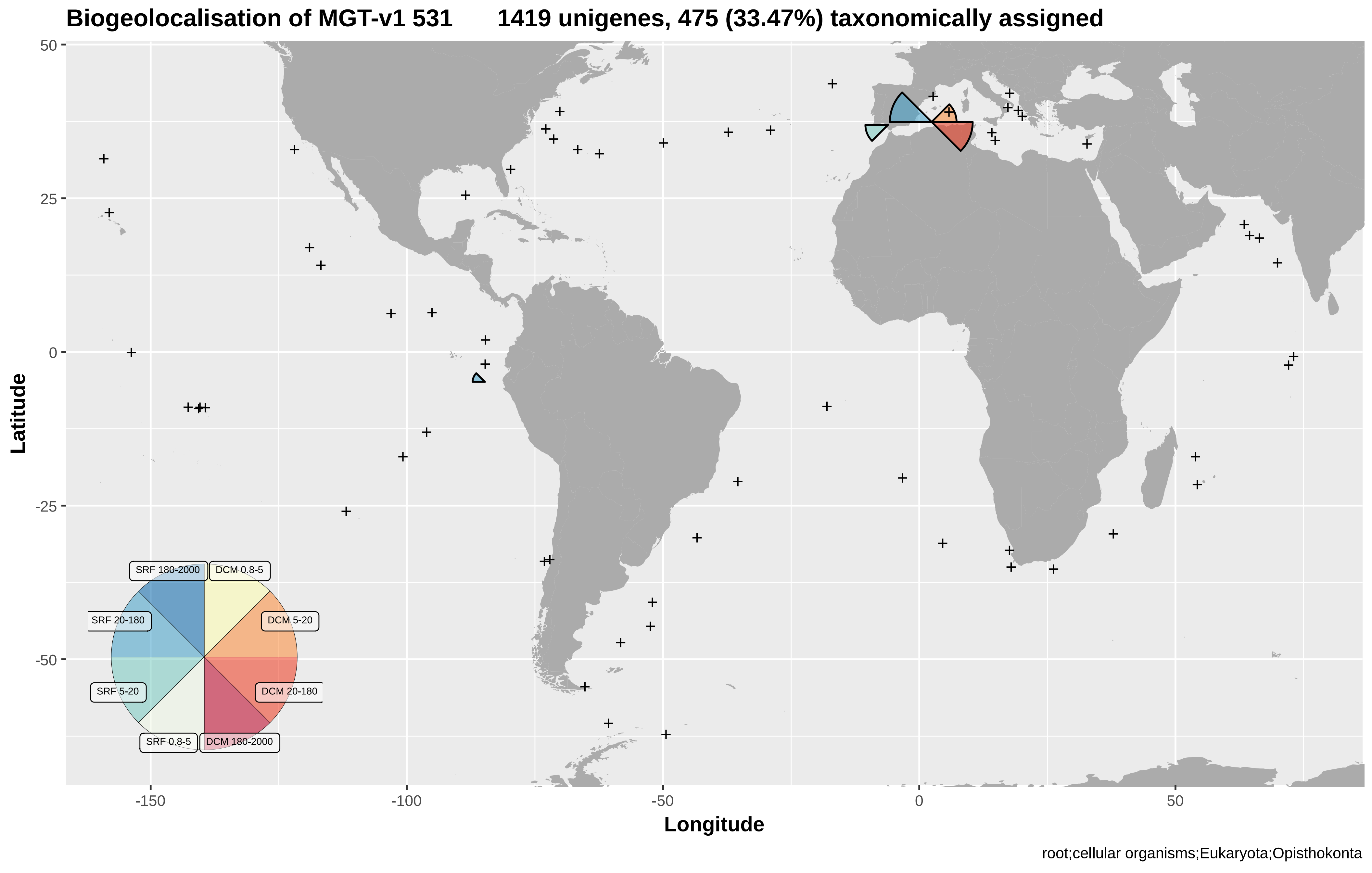

Biogeolocalisation of MGT-v1 531
1419 unigenes, 475 (33.47%) taxonomically assigned
Latitude
Longitude
50
25
0
-25
-50
-150
-100
-50
0
50
SRF 180-2000
DCM 0.8-5
SRF 20-180
DCM 5-20
SRF 5-20
DCM 20-180
SRF 0.8-5
DCM 180-2000
root;cellular organisms;Eukaryota;Opisthokonta

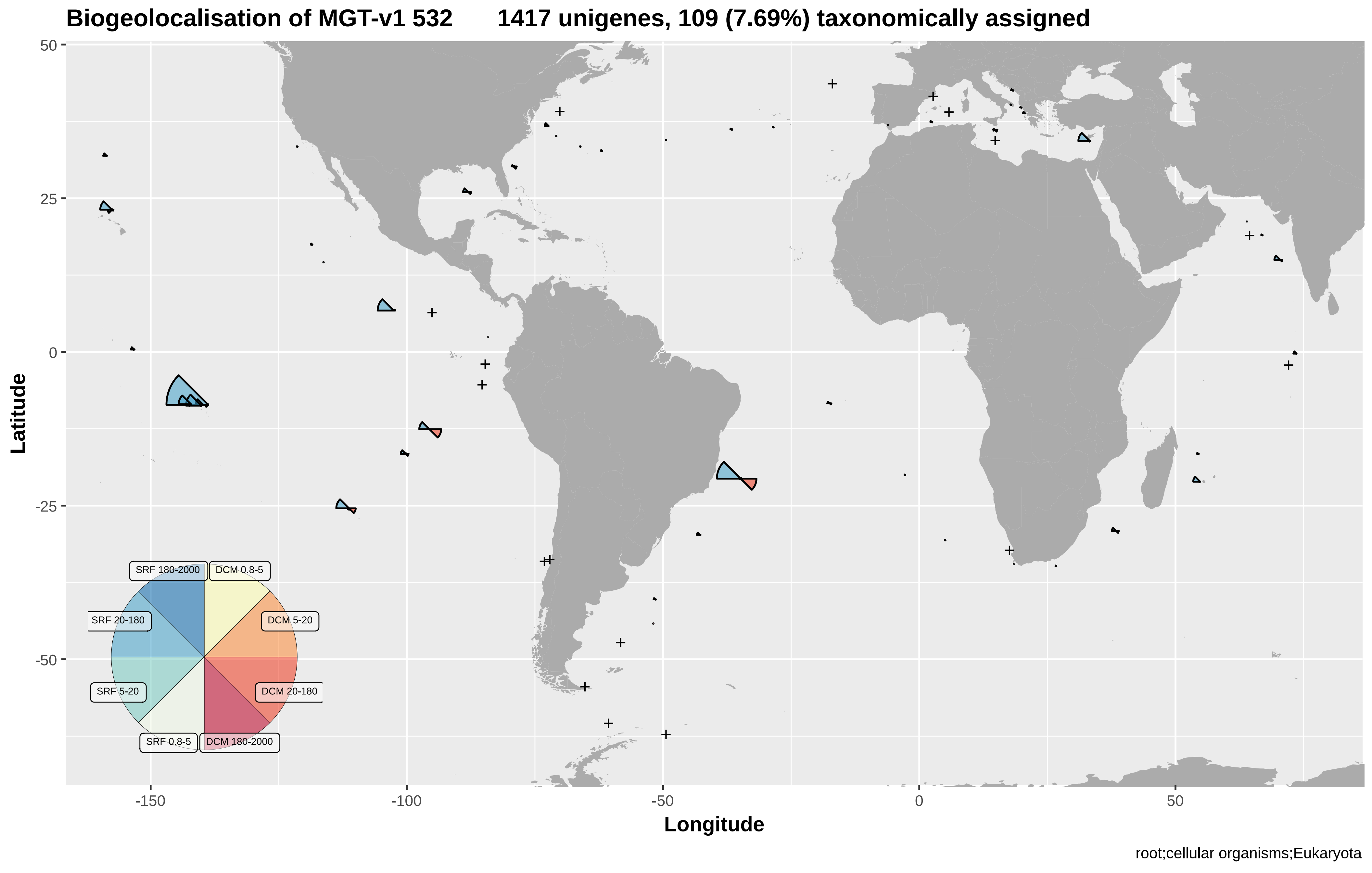

Biogeolocalisation of MGT-v1 532
1417 unigenes, 109 (7.69%) taxonomically assigned
Latitude
Longitude
50
25
0
-25
-50
-150
-100
-50
0
50
SRF 180-2000
DCM 0.8-5
SRF 20-180
DCM 5-20
SRF 5-20
DCM 20-180
SRF 0.8-5
DCM 180-2000
root;cellular organisms;Eukaryota

### Biogeolocalisation of MGT-v1 533 1410 unigenes, 198 (14.04%) taxonomically assigned

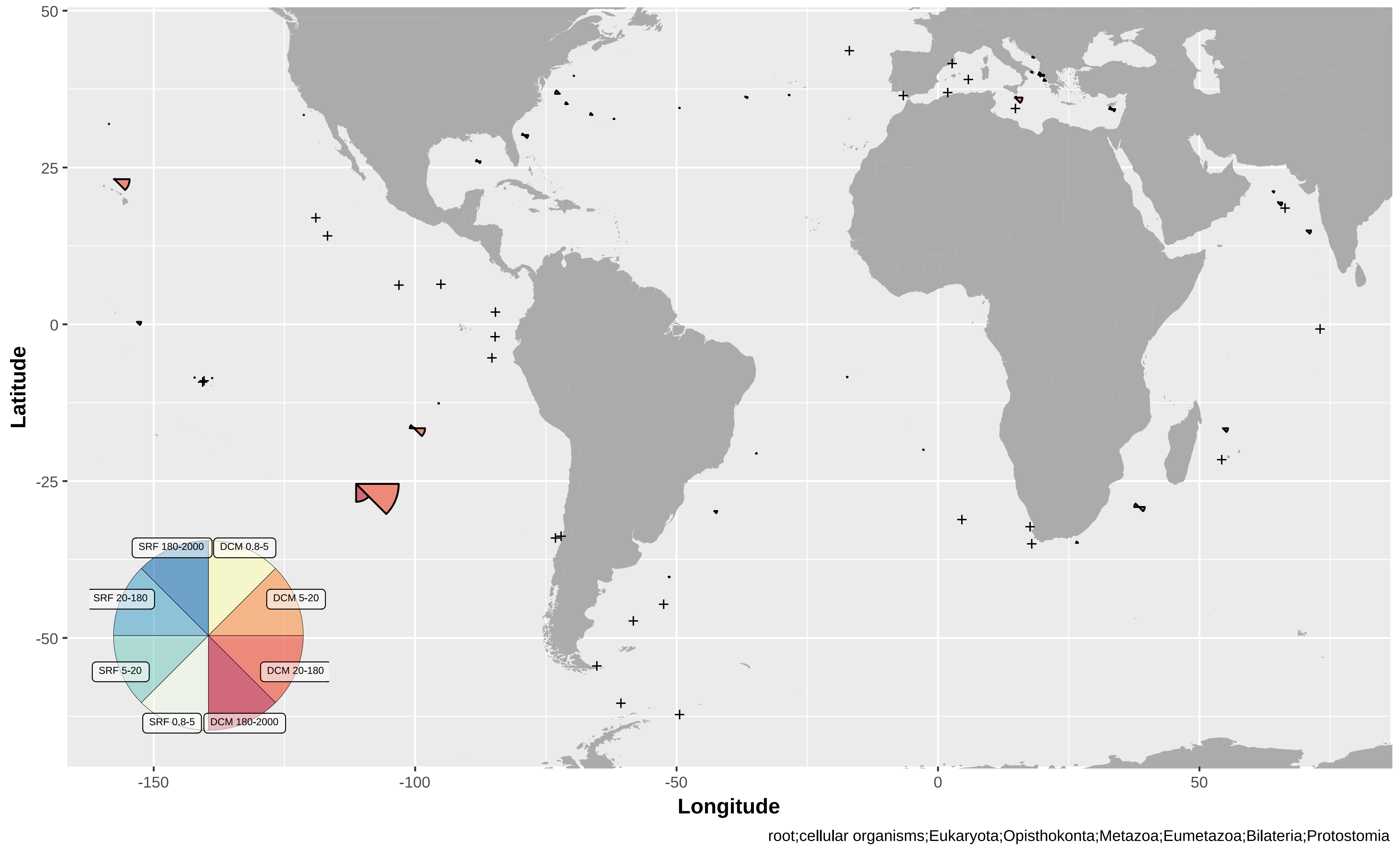

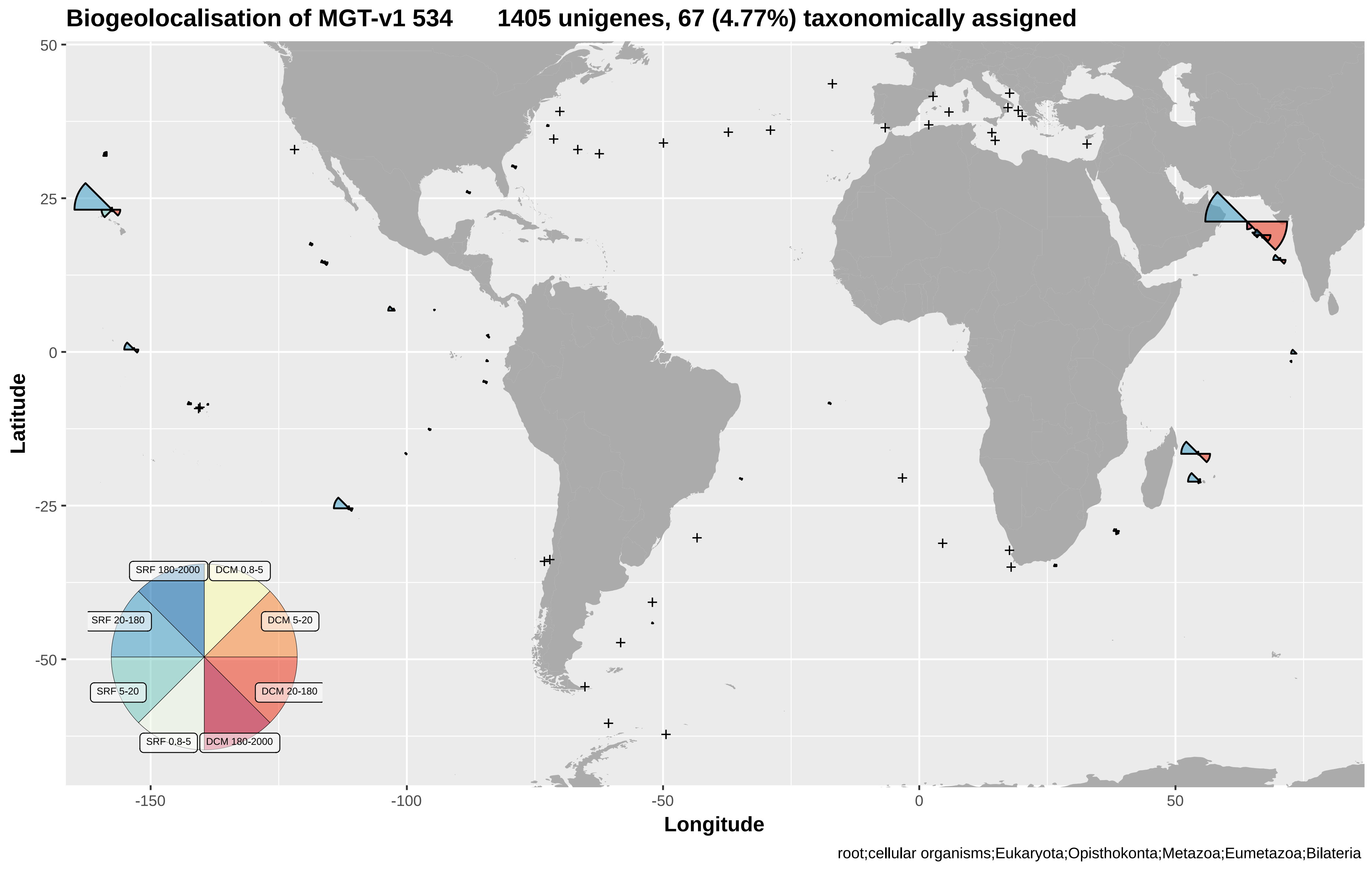
Biogeolocalisation of MGT-v1 534 1405 unigenes, 67 (4.77%) taxonomically assigned
Latitude
50
25
0
-25
-50
Longitude
-150
-100
-50
0
50
SRF 180-2000
DCM 0.8-5
SRF 20-180
DCM 5-20
SRF 5-20
DCM 20-180
SRF 0.8-5
DCM 180-2000
root;cellular organisms;Eukaryota;Opisthokonta;Metazoa;Eumetazoa;Bilateria

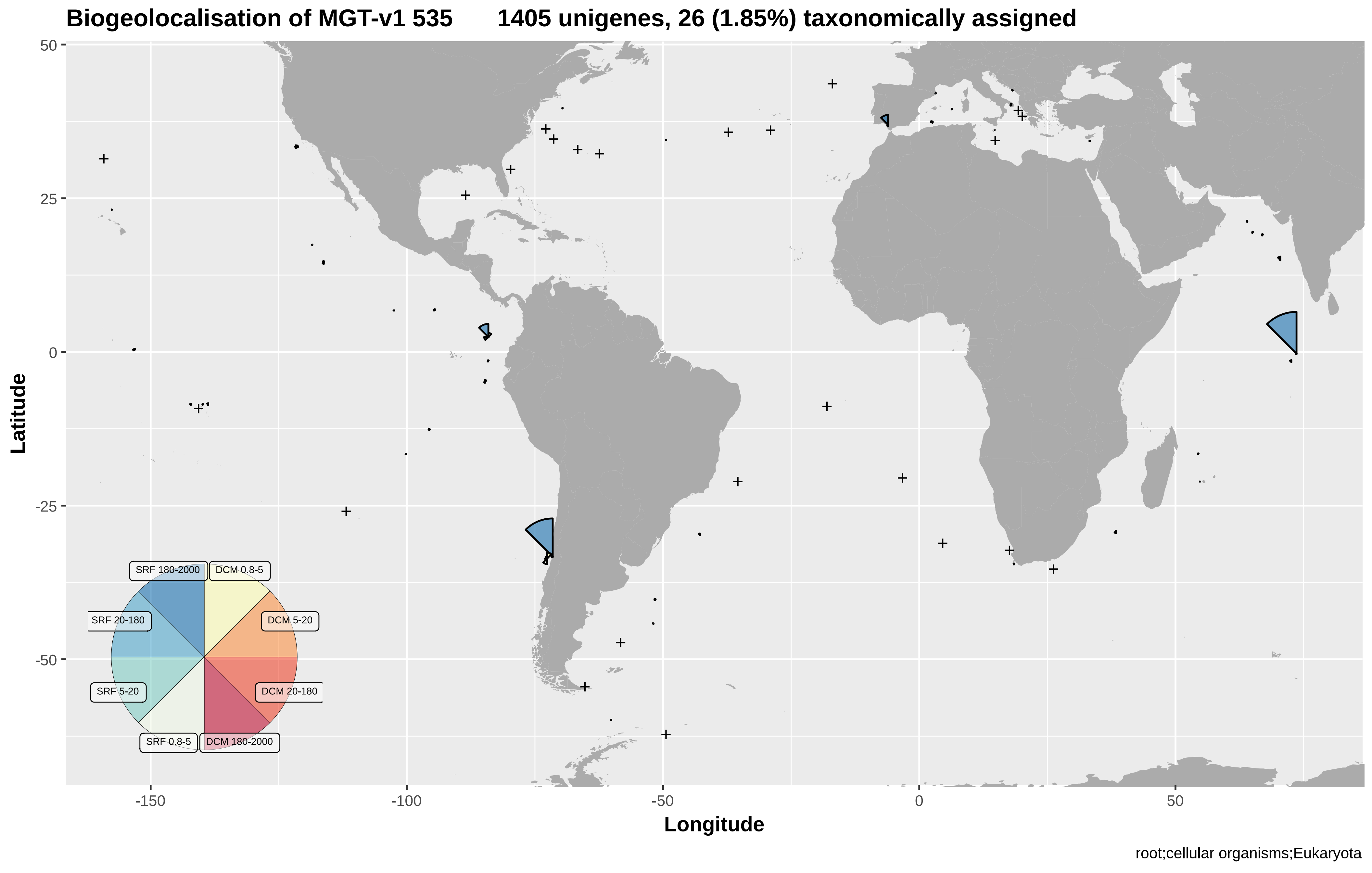
Biogeolocalisation of MGT-v1 535 1405 unigenes, 26 (1.85%) taxonomically assigned
Latitude
Longitude
50
25
0
-25
-50
-150
-100
-50
0
50
SRF 180-2000
DCM 0.8-5
SRF 20-180
DCM 5-20
SRF 5-20
DCM 20-180
SRF 0.8-5
DCM 180-2000
root;cellular organisms;Eukaryota

Biogeolocalisation of MGT-v1 536 1404 unigenes, 1218 (86.75%) taxonomically assigned

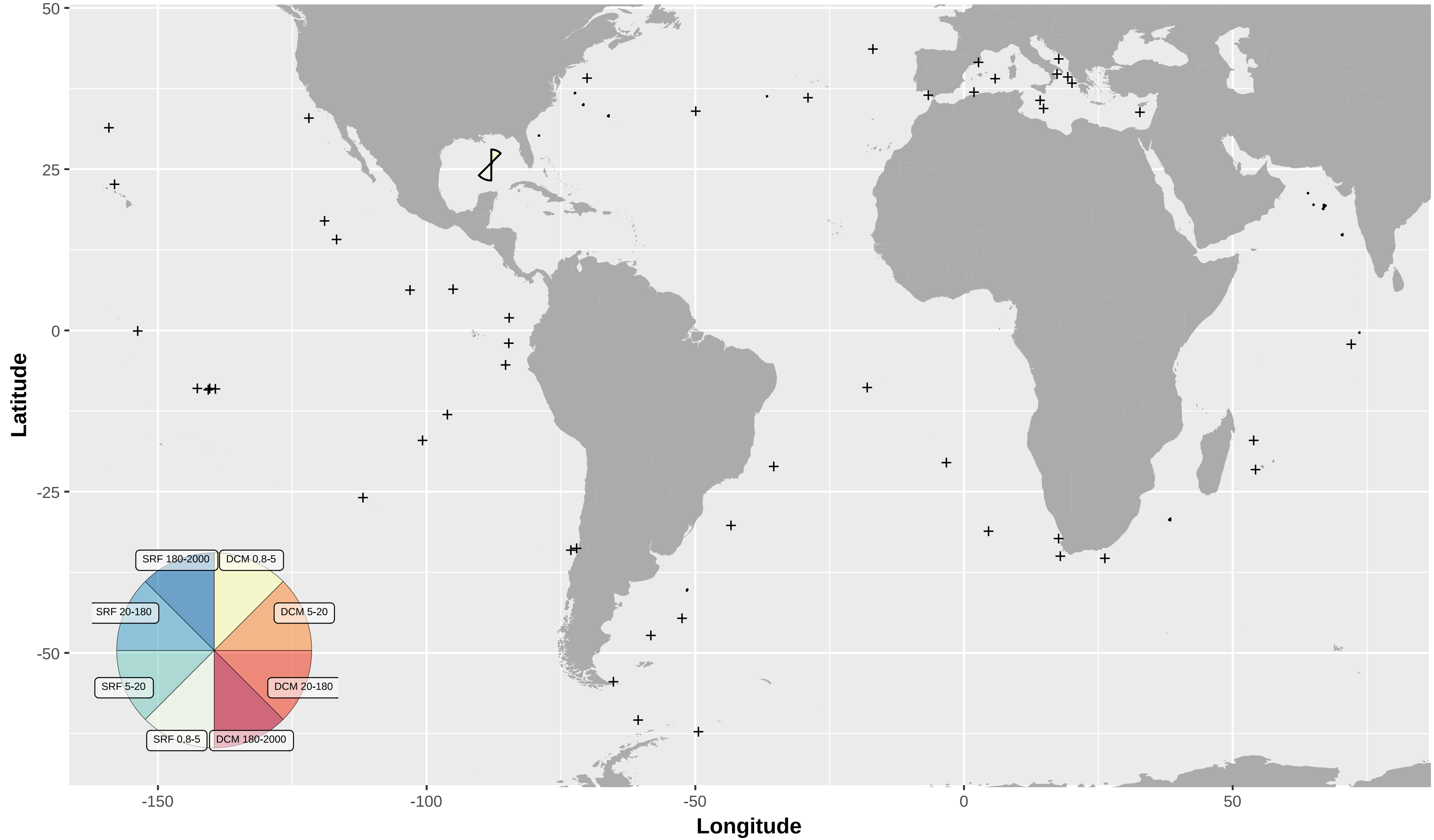


root;cellular organisms;Bacteria;Cyanobacteria;Oscillatoriophycideae

#### Biogeolocalisation of MGT-v1 537 1404 unigenes, 791 (56.34%) taxonomically assigned

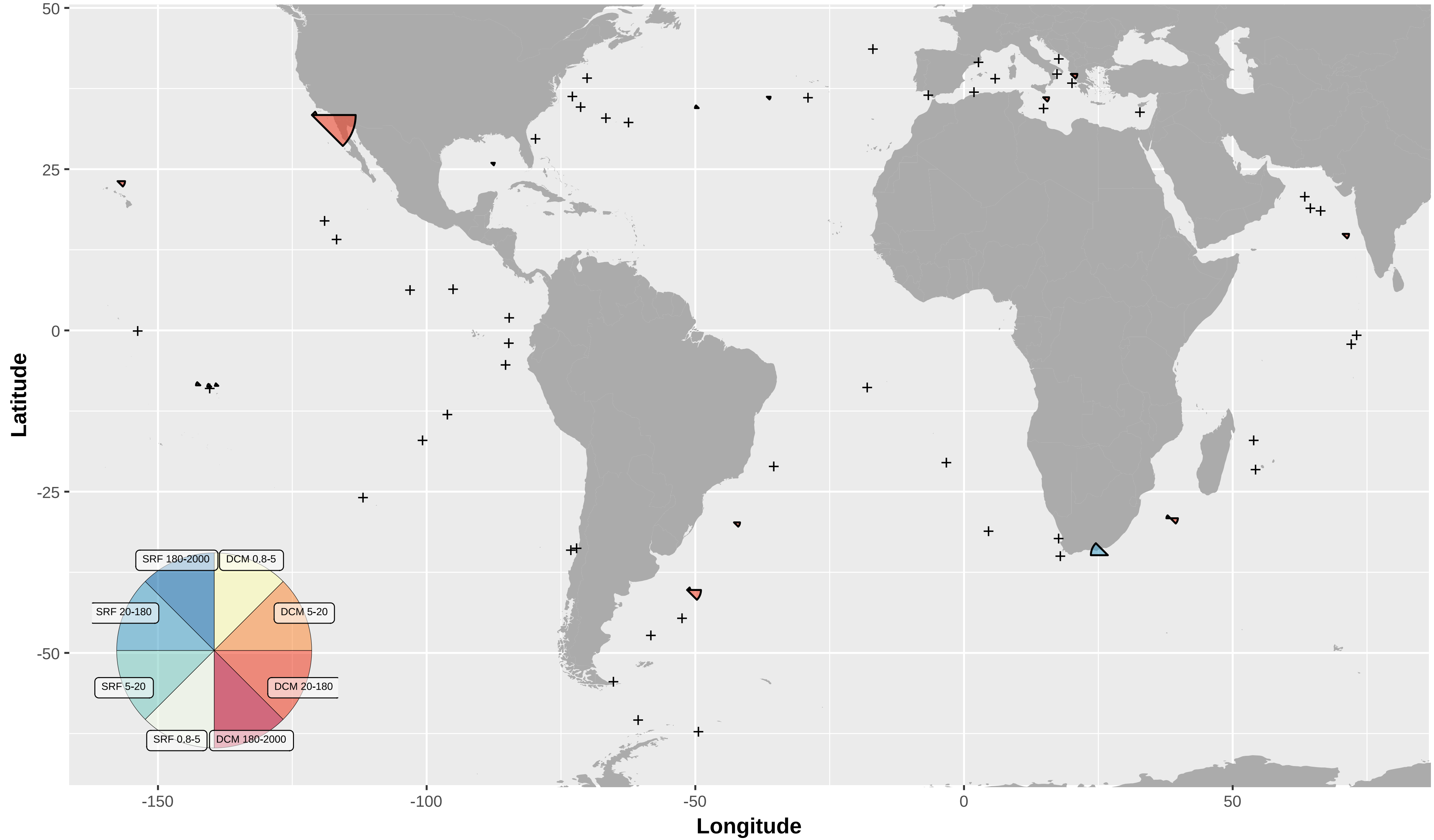


root;cellular organisms;Eukaryota;Alveolata;Ciliophora;Intramacronucleata;Spirotrichea

#### Biogeolocalisation of MGT-v1 538 1403 unigenes, 1178 (83.96%) taxonomically assigned

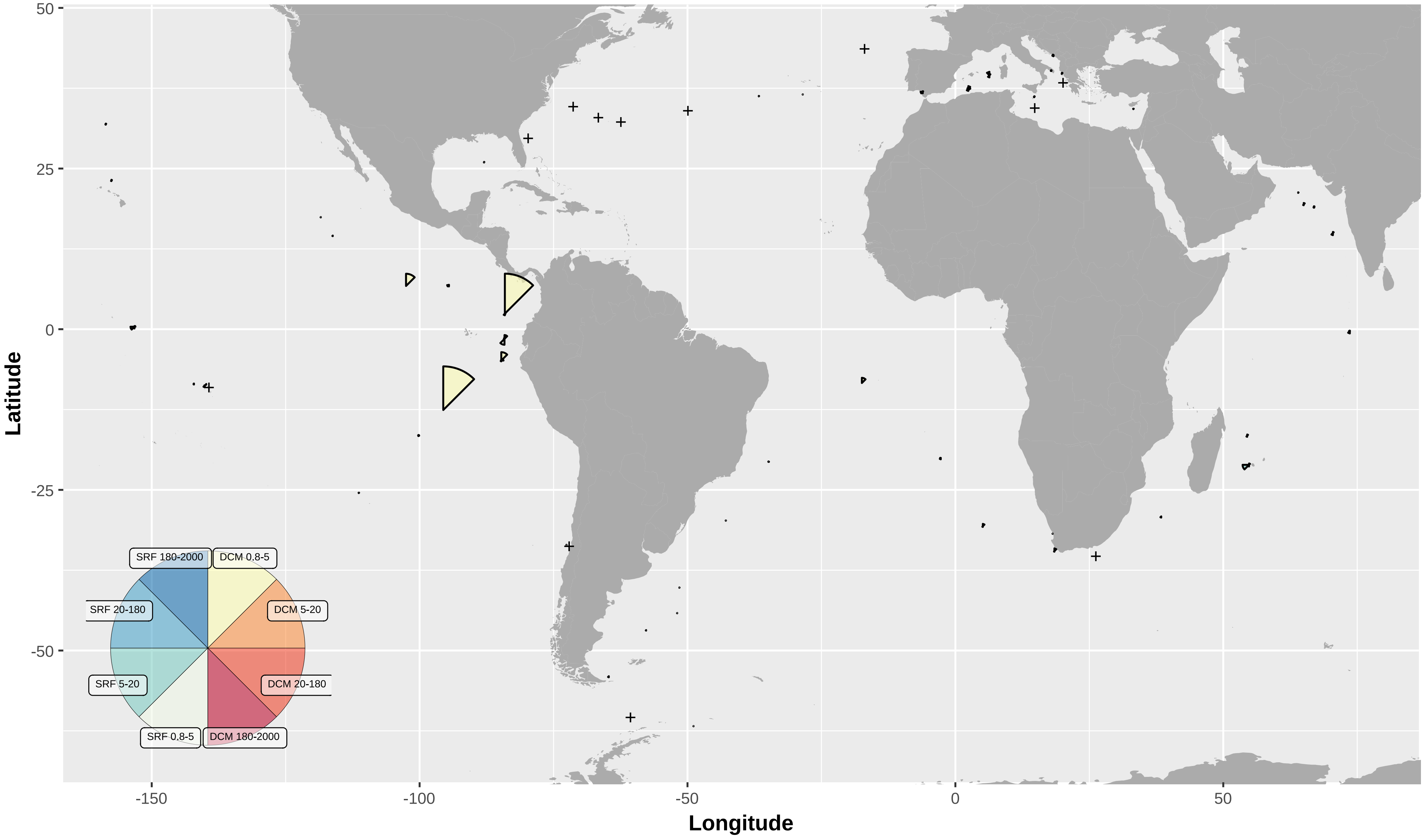


root;cellular organisms;Bacteria;Proteobacteria;Gammaproteobacteria

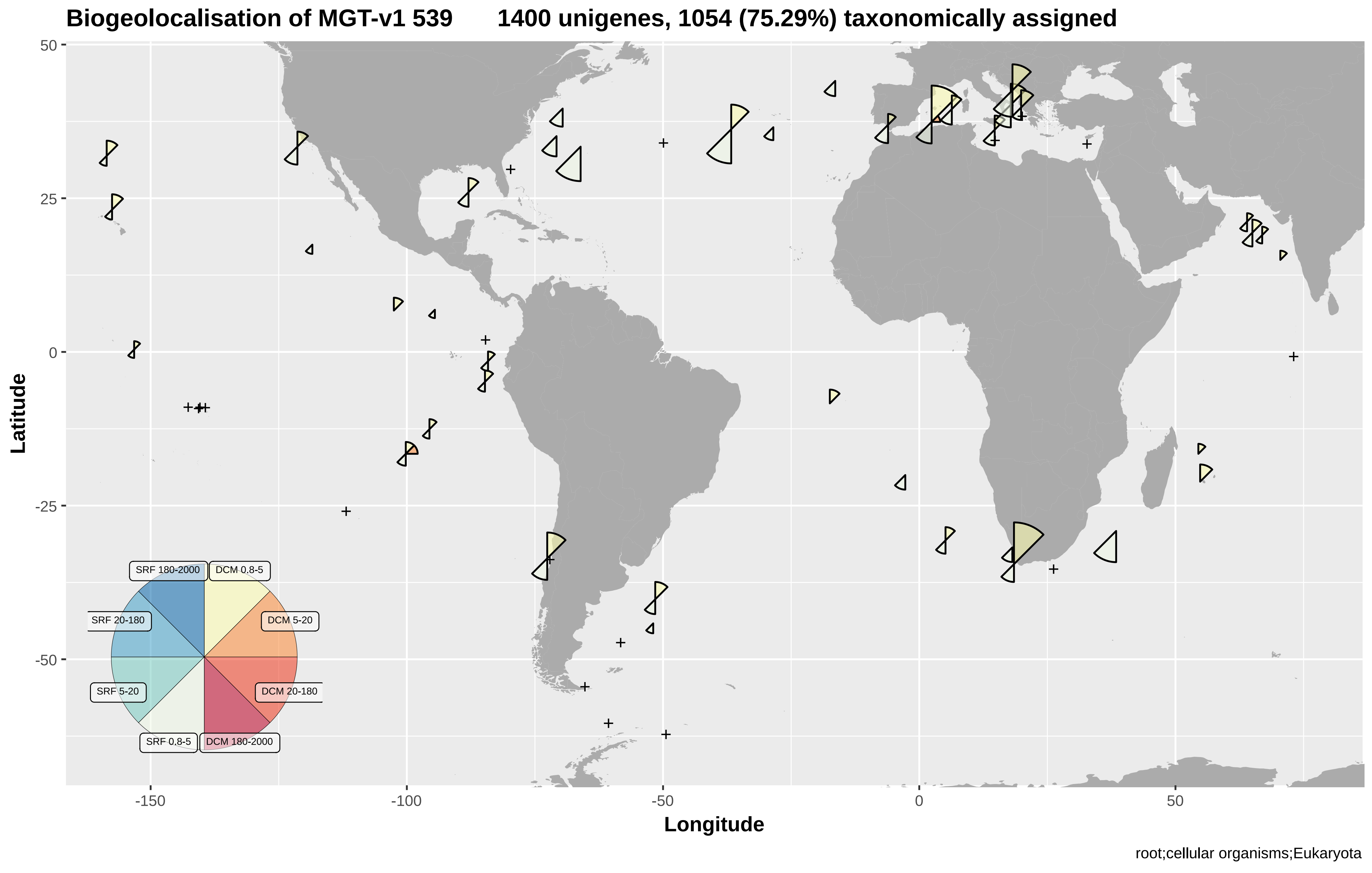

Biogeolocalisation of MGT-v1 539 1400 unigenes, 1054 (75.29%) taxonomically assigned
50
25
0
-25
-50
Latitude
-150
-100
-50
0
50
Longitude
SRF 180-2000
DCM 0.8-5
SRF 20-180
DCM 5-20
SRF 5-20
DCM 20-180
SRF 0.8-5
DCM 180-2000
root;cellular organisms;Eukaryota

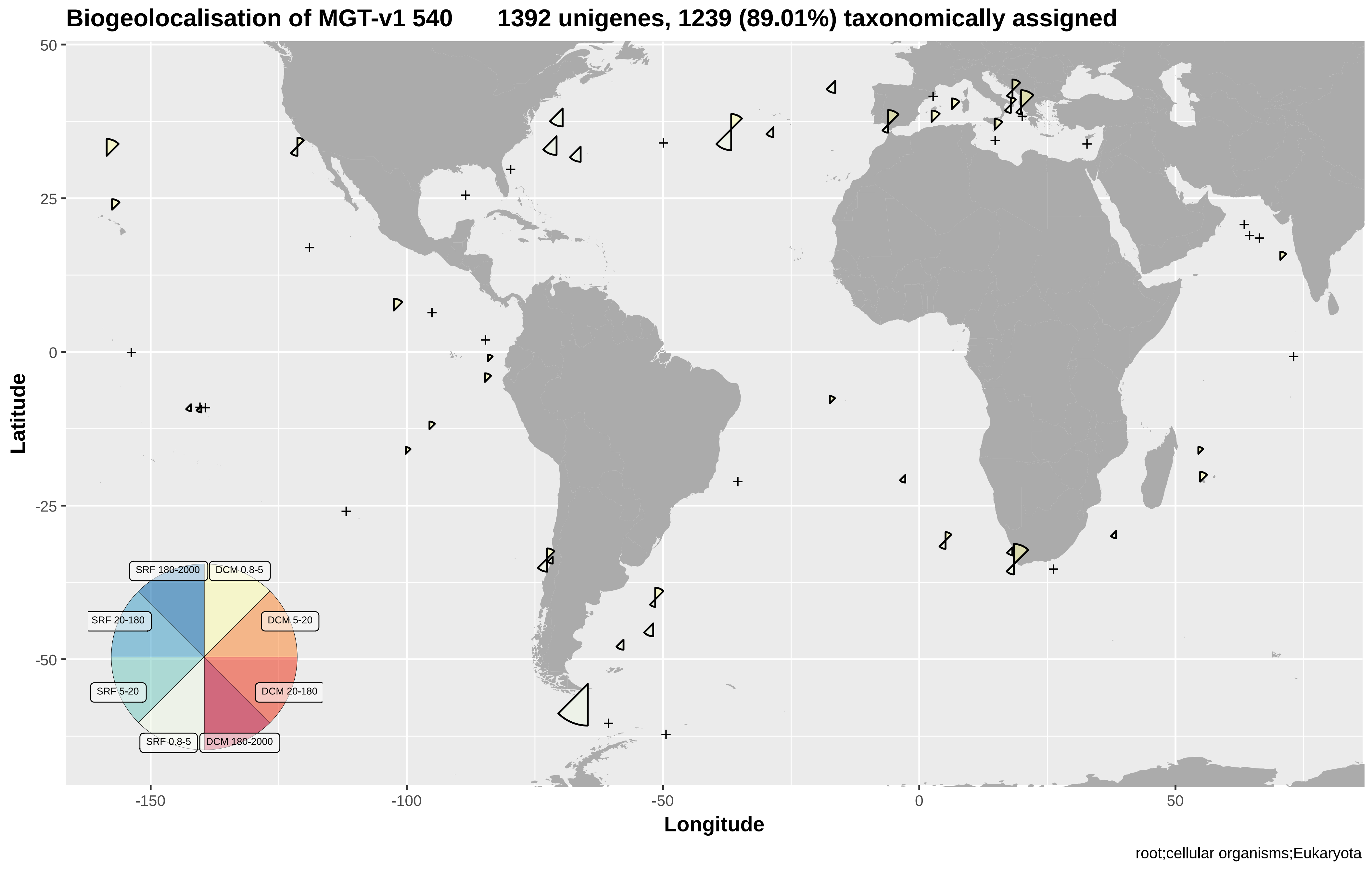

Biogeolocalisation of MGT-v1 540 1392 unigenes, 1239 (89.01%) taxonomically assigned
50
25
0
-25
-50
Latitude
-150
-100
-50
0
50
Longitude
SRF 180-2000
DCM 0.8-5
SRF 20-180
DCM 5-20
SRF 5-20
DCM 20-180
SRF 0.8-5
DCM 180-2000
root;cellular organisms;Eukaryota

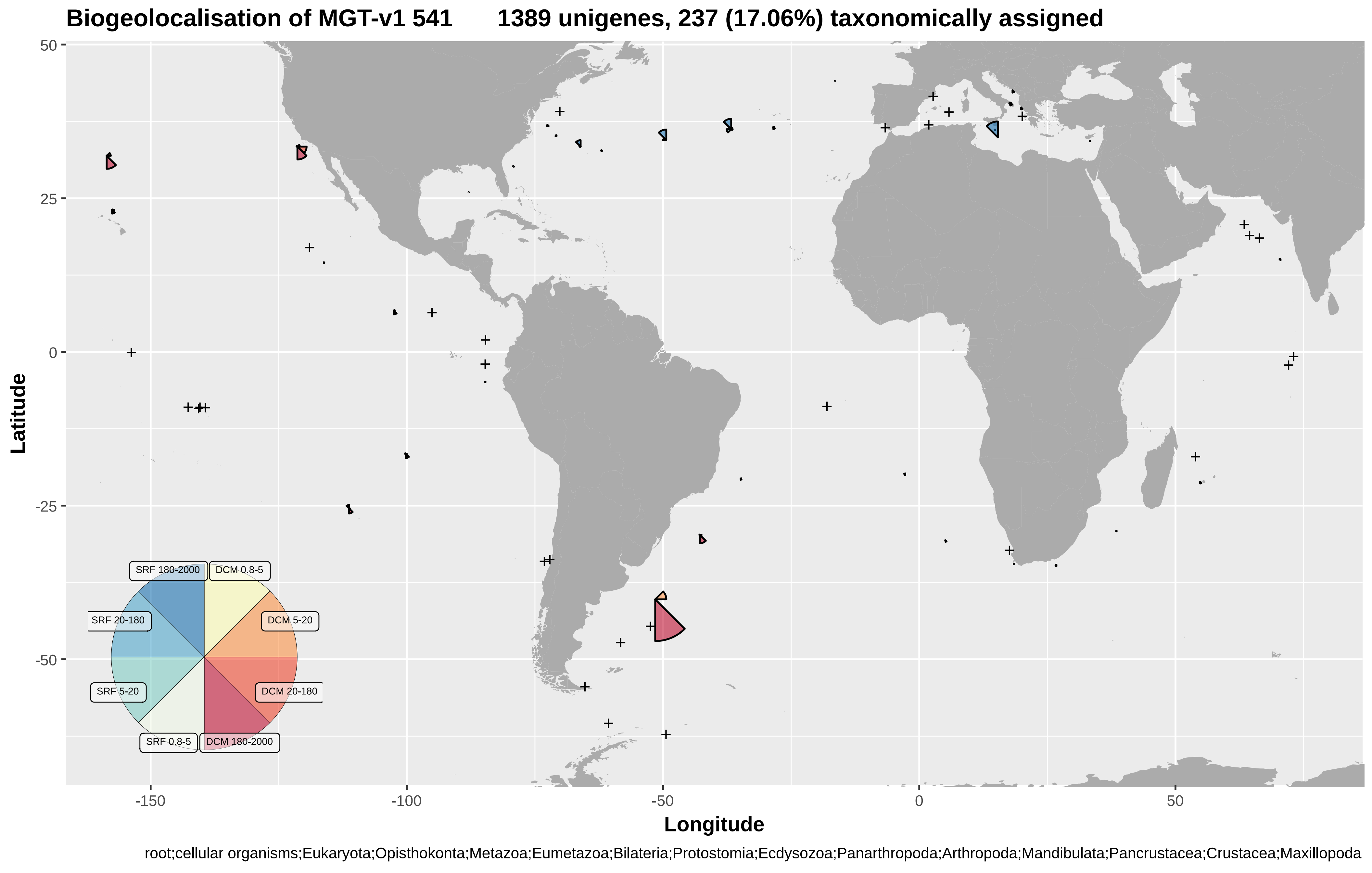
Biogeolocalisation of MGT-v1 541 1389 unigenes, 237 (17.06%) taxonomically assigned
Latitude
Longitude
50
25
0
-25
-50
-150
-100
-50
0
50
SRF 180-2000
DCM 0.8-5
SRF 20-180
DCM 5-20
SRF 5-20
DCM 20-180
SRF 0.8-5
DCM 180-2000
root;cellular organisms;Eukaryota;Opisthokonta;Metazoa;Eumetazoa;Bilateria;Protostomia;Ecdysozoa;Panarthropoda;Arthropoda;Mandibulata;Pancrustacea;Crustacea;Maxillopoda

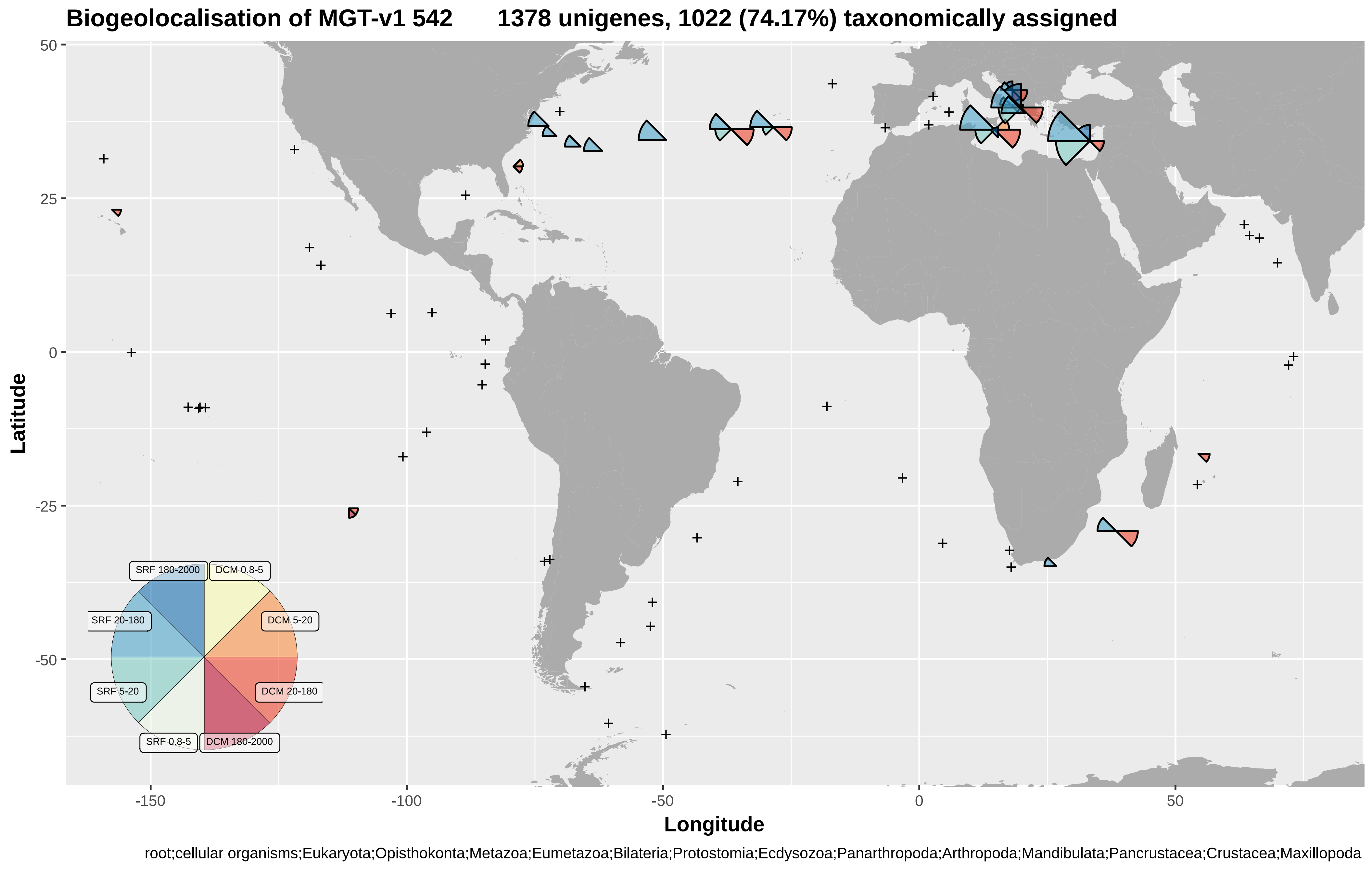

Biogeolocalisation of MGT-v1 542 1378 unigenes, 1022 (74.17%) taxonomically assigned
50
25
0
-25
-50
Latitude
-150
-100
-50
0
50
Longitude
SRF 180-2000
DCM 0.8-5
SRF 20-180
DCM 5-20
SRF 5-20
DCM 20-180
SRF 0.8-5
DCM 180-2000
root;cellular organisms;Eukaryota;Opisthokonta;Metazoa;Eumetazoa;Bilateria;Protostomia;Ecdysozoa;Panarthropoda;Arthropoda;Mandibulata;Pancrustacea;Crustacea;Maxillopoda

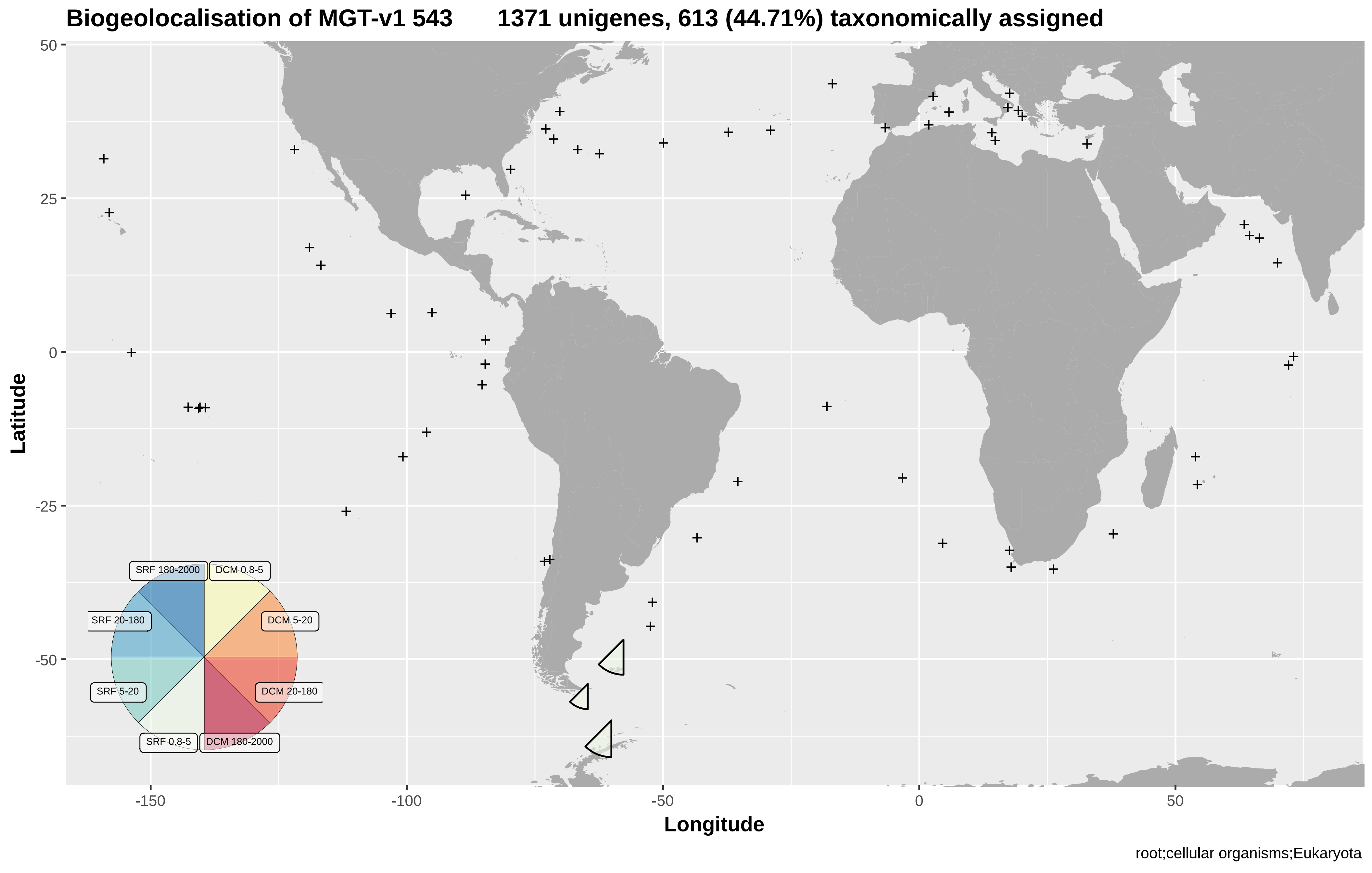

Biogeolocalisation of MGT-v1 543 1371 unigenes, 613 (44.71%) taxonomically assigned
Latitude
Longitude
50
25
0
-25
-50
-150
-100
-50
0
50
SRF 180-2000
DCM 0.8-5
SRF 20-180
DCM 5-20
SRF 5-20
DCM 20-180
SRF 0.8-5
DCM 180-2000
root;cellular organisms;Eukaryota

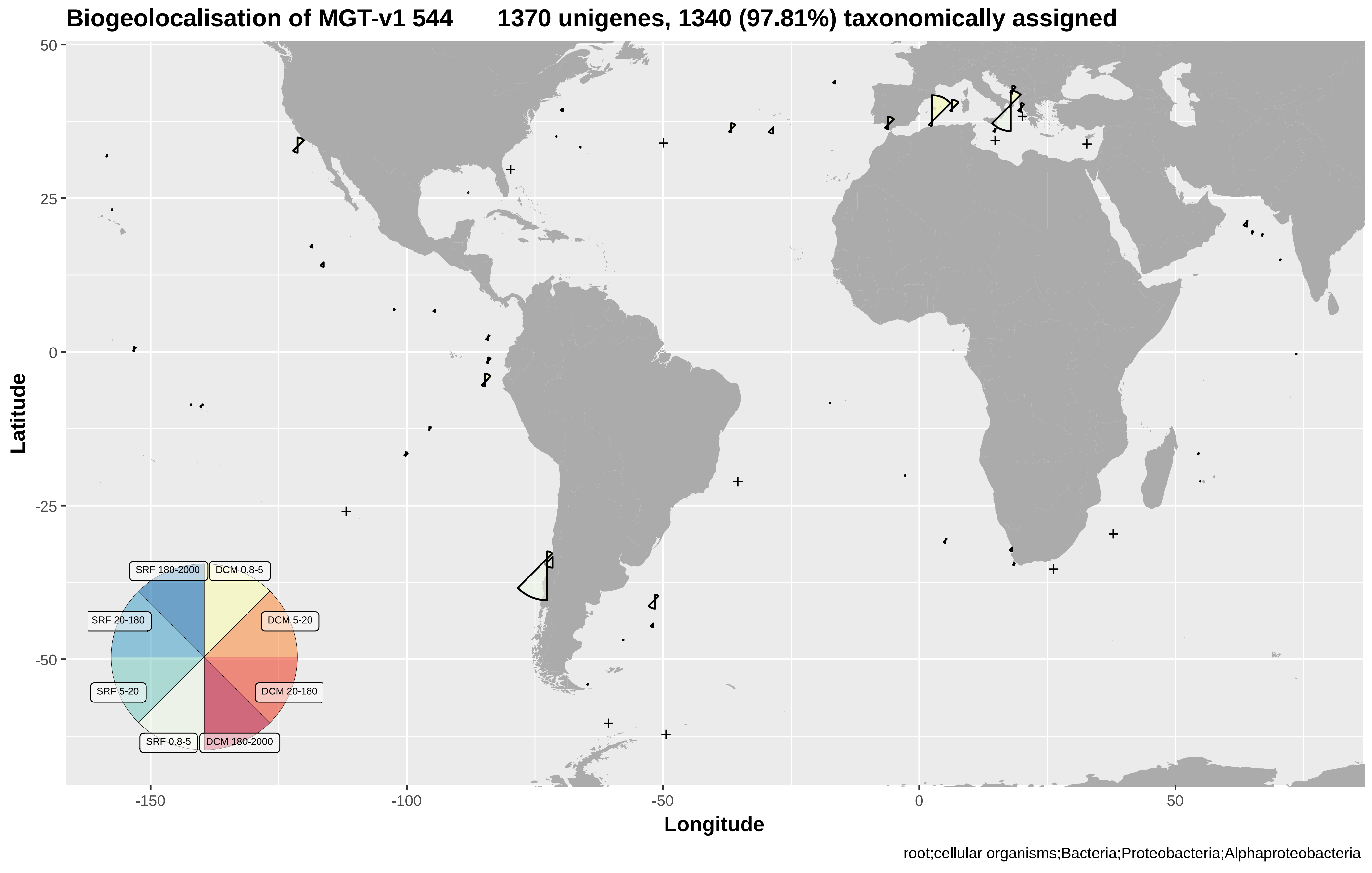
Biogeolocalisation of MGT-v1 544 1370 unigenes, 1340 (97.81%) taxonomically assigned
Latitude
Longitude
50
25
0
-25
-50
-150
-100
-50
0
50
SRF 180-2000
DCM 0.8-5
SRF 20-180
DCM 5-20
SRF 5-20
DCM 20-180
SRF 0.8-5
DCM 180-2000
root;cellular organisms;Bacteria;Proteobacteria;Alphaproteobacteria

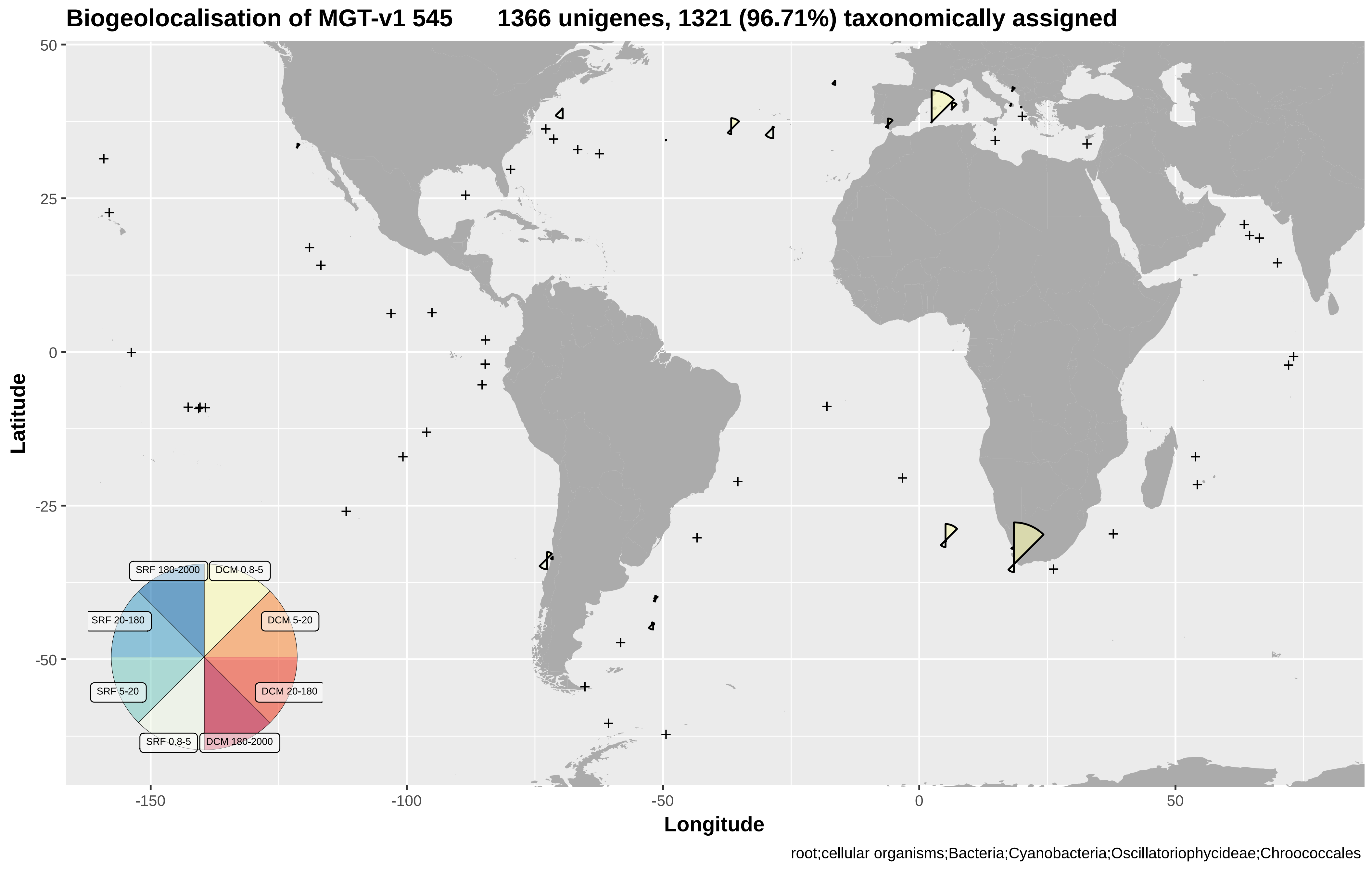
Biogeolocalisation of MGT-v1 545
1366 unigenes, 1321 (96.71%) taxonomically assigned
Latitude
Longitude
50
25
0
-25
-50
-150
-100
-50
0
50
SRF 180-2000
DCM 0.8-5
SRF 20-180
DCM 5-20
SRF 5-20
DCM 20-180
SRF 0.8-5
DCM 180-2000
root;cellular organisms;Bacteria;Cyanobacteria;Oscillatoriophycideae;Chroococcales

#### Biogeolocalisation of MGT-v1 546 1359 unigenes, 194 (14.28%) taxonomically assigned

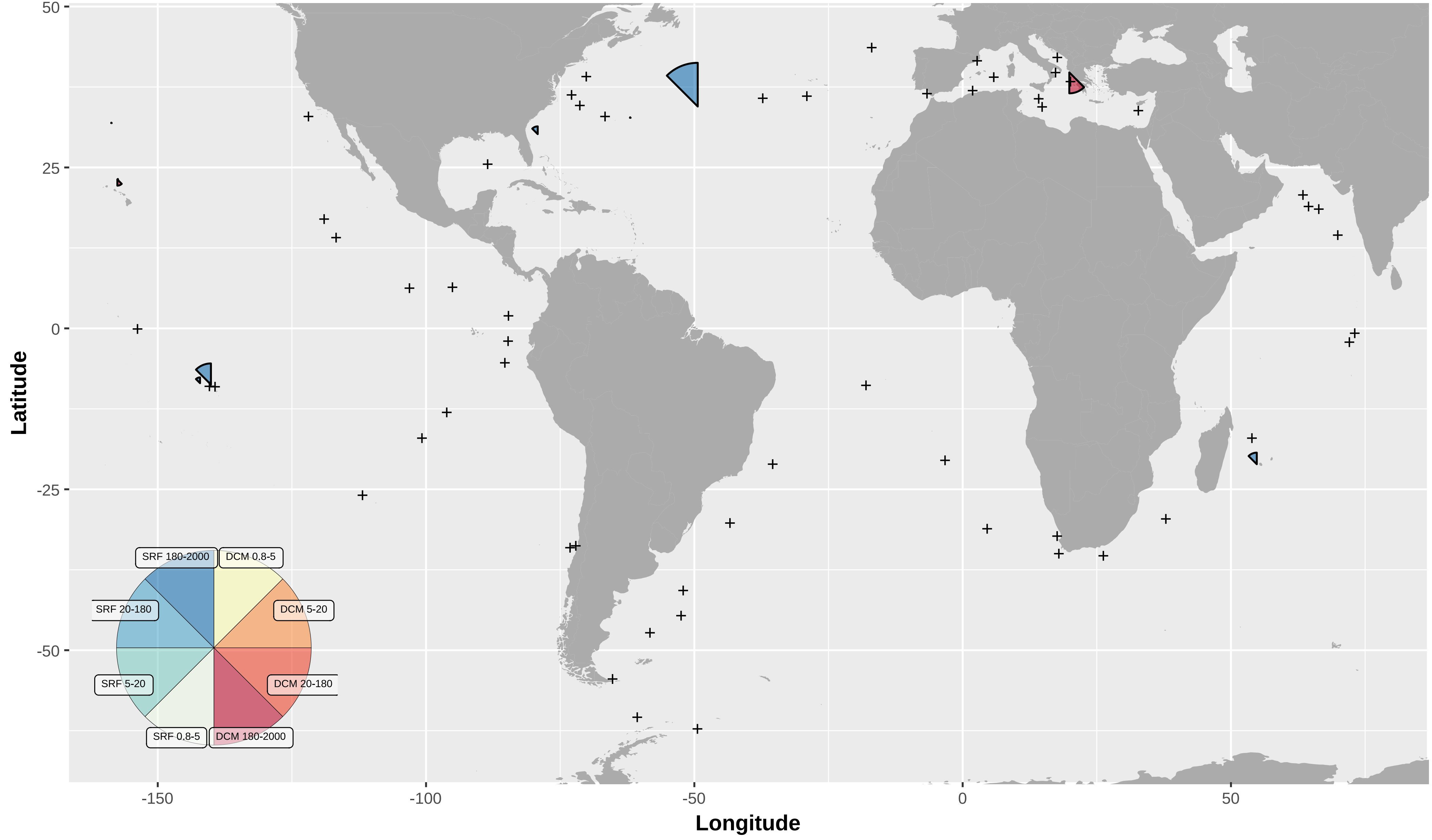


root;cellular organisms;Eukaryota;Opisthokonta;Metazoa;Eumetazoa

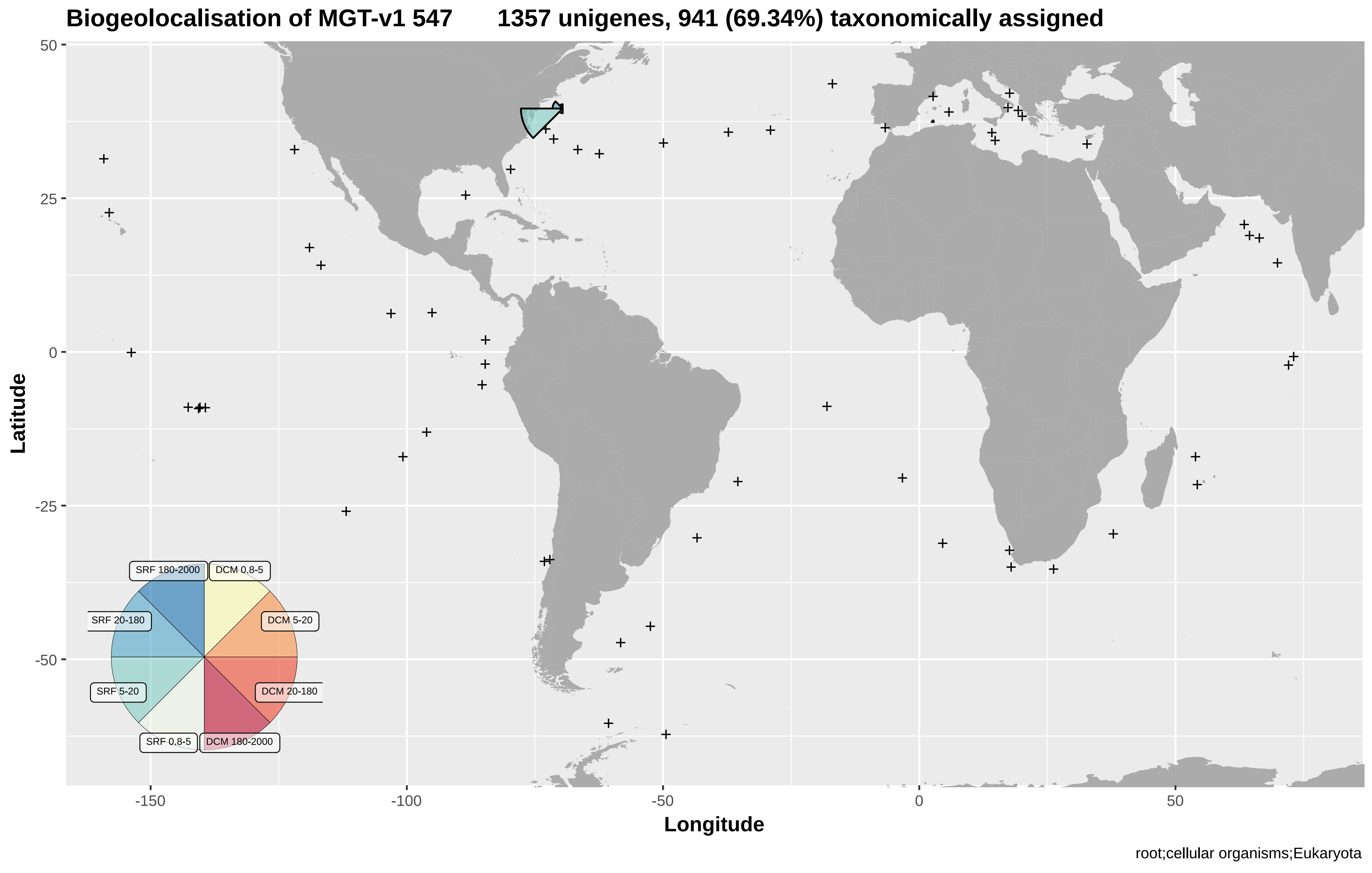

Biogeolocalisation of MGT-v1 547 1357 unigenes, 941 (69.34%) taxonomically assigned
Latitude
Longitude
50
25
0
-25
-50
-150
-100
-50
0
50
SRF 180-2000
DCM 0.8-5
SRF 20-180
DCM 5-20
SRF 5-20
DCM 20-180
SRF 0.8-5
DCM 180-2000
root;cellular organisms;Eukaryota

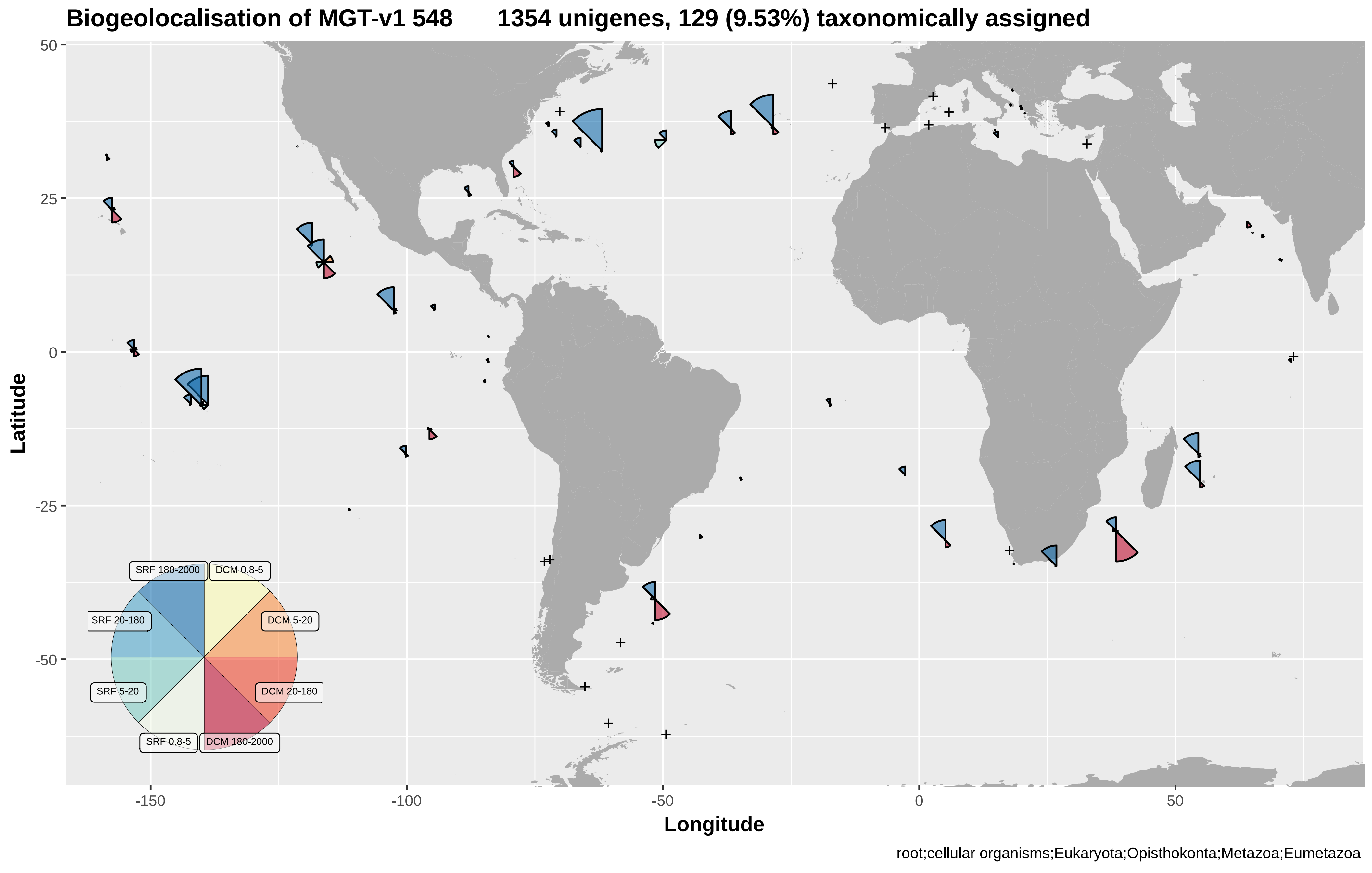

Biogeolocalisation of MGT-v1 548
1354 unigenes, 129 (9.53%) taxonomically assigned
50
25
0
-25
-50
Latitude
-150
-100
-50
0
50
Longitude
SRF 180-2000
DCM 0.8-5
SRF 20-180
DCM 5-20
SRF 5-20
DCM 20-180
SRF 0.8-5
DCM 180-2000
root;cellular organisms;Eukaryota;Opisthokonta;Metazoa;Eumetazoa

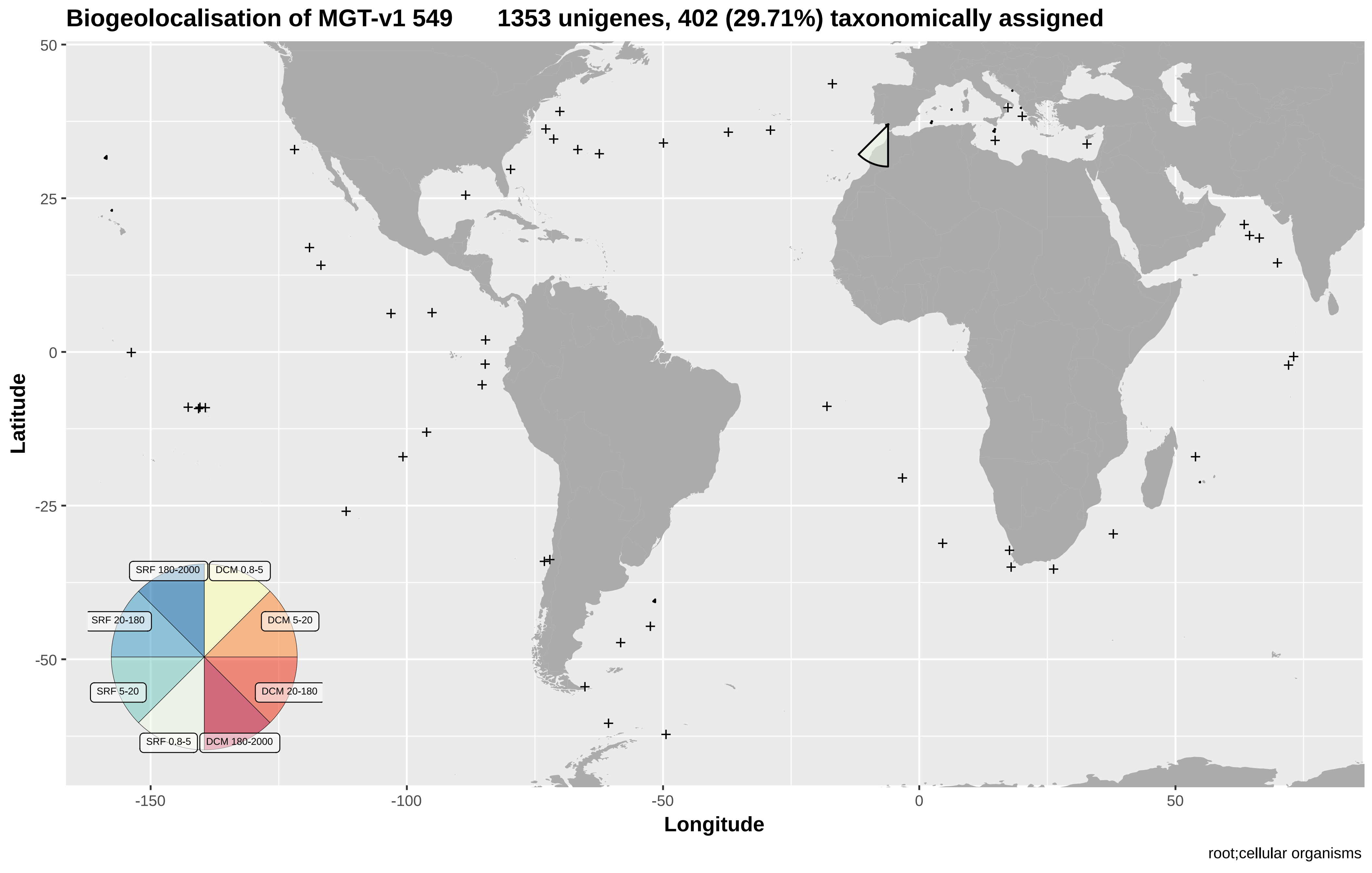
Biogeolocalisation of MGT-v1 549 1353 unigenes, 402 (29.71%) taxonomically assigned
Latitude
Longitude
50
25
0
-25
-50
-150
-100
-50
0
50
SRF 180-2000
DCM 0.8-5
SRF 20-180
DCM 5-20
SRF 5-20
DCM 20-180
SRF 0.8-5
DCM 180-2000
root;cellular organisms

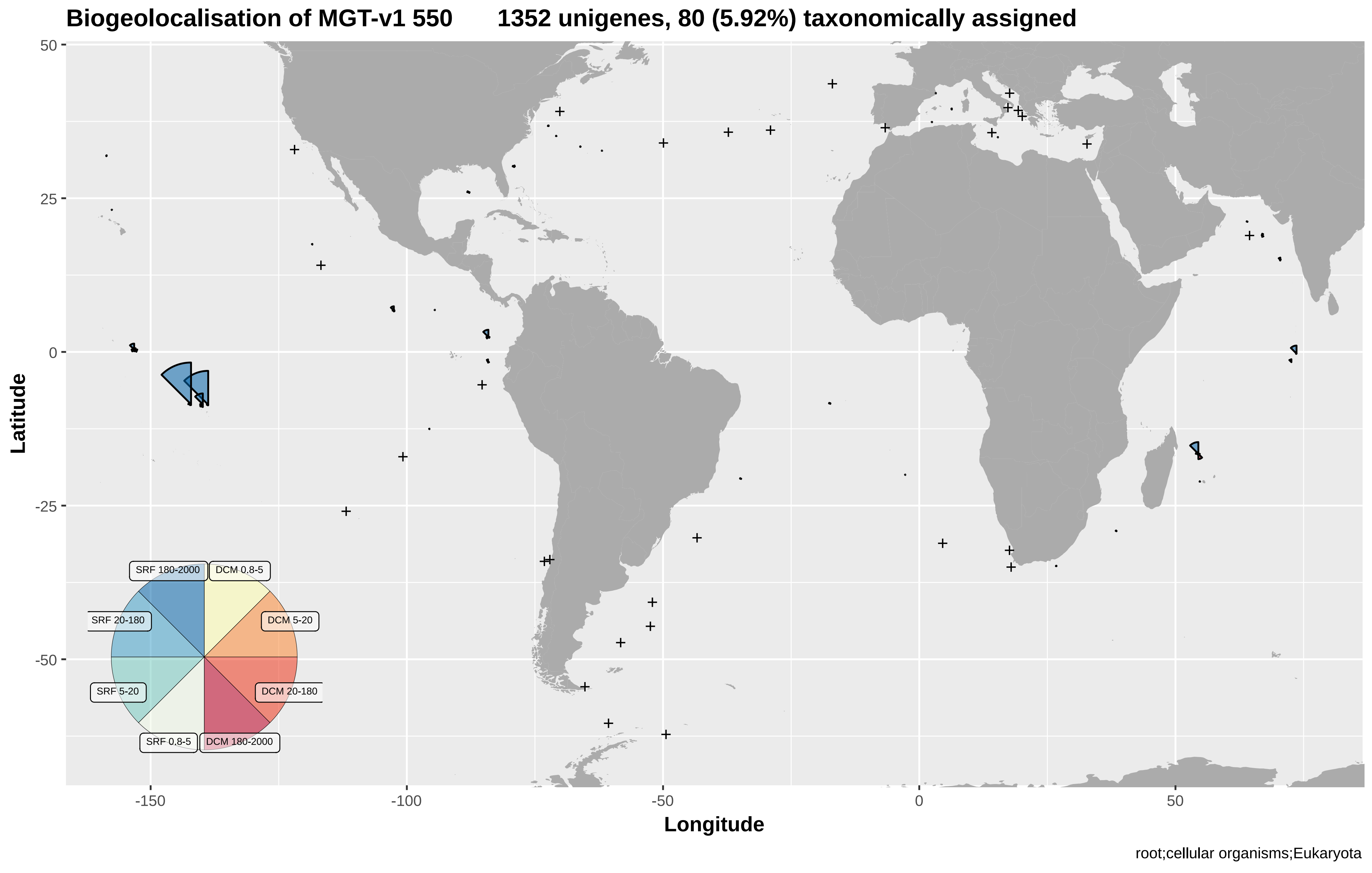

Biogeolocalisation of MGT-v1 550 1352 unigenes, 80 (5.92%) taxonomically assigned
50
25
0
-25
-50
Latitude
-150
-100
-50
0
50
Longitude
SRF 180-2000
DCM 0.8-5
SRF 20-180
DCM 5-20
SRF 5-20
DCM 20-180
SRF 0.8-5
DCM 180-2000
root;cellular organisms;Eukaryota

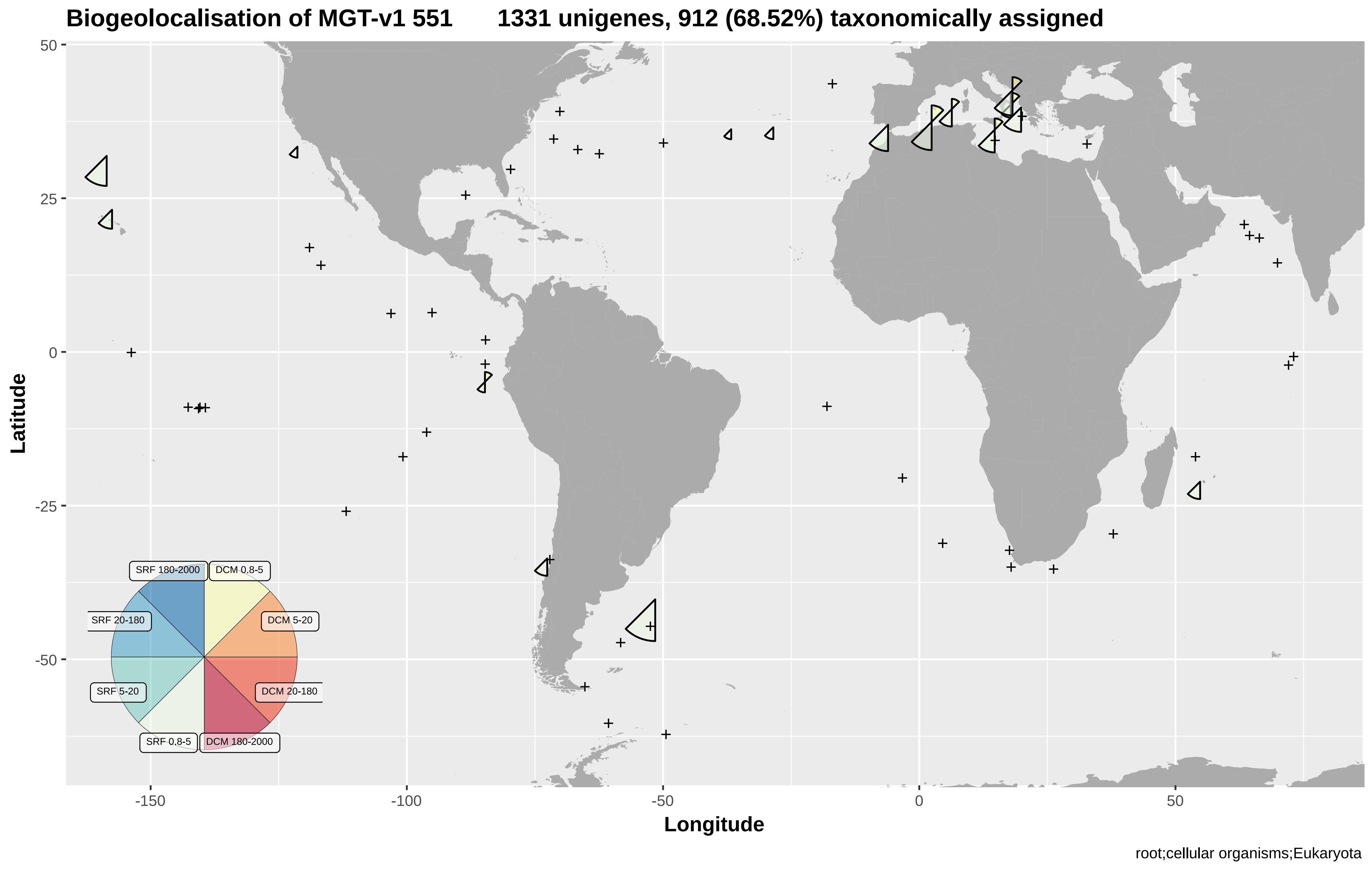

Biogeolocalisation of MGT-v1 551
1331 unigenes, 912 (68.52%) taxonomically assigned
50
25
0
-25
-50
Latitude
-150
-100
-50
0
50
Longitude
SRF 180-2000
DCM 0.8-5
SRF 20-180
DCM 5-20
SRF 5-20
DCM 20-180
SRF 0.8-5
DCM 180-2000
root;cellular organisms;Eukaryota

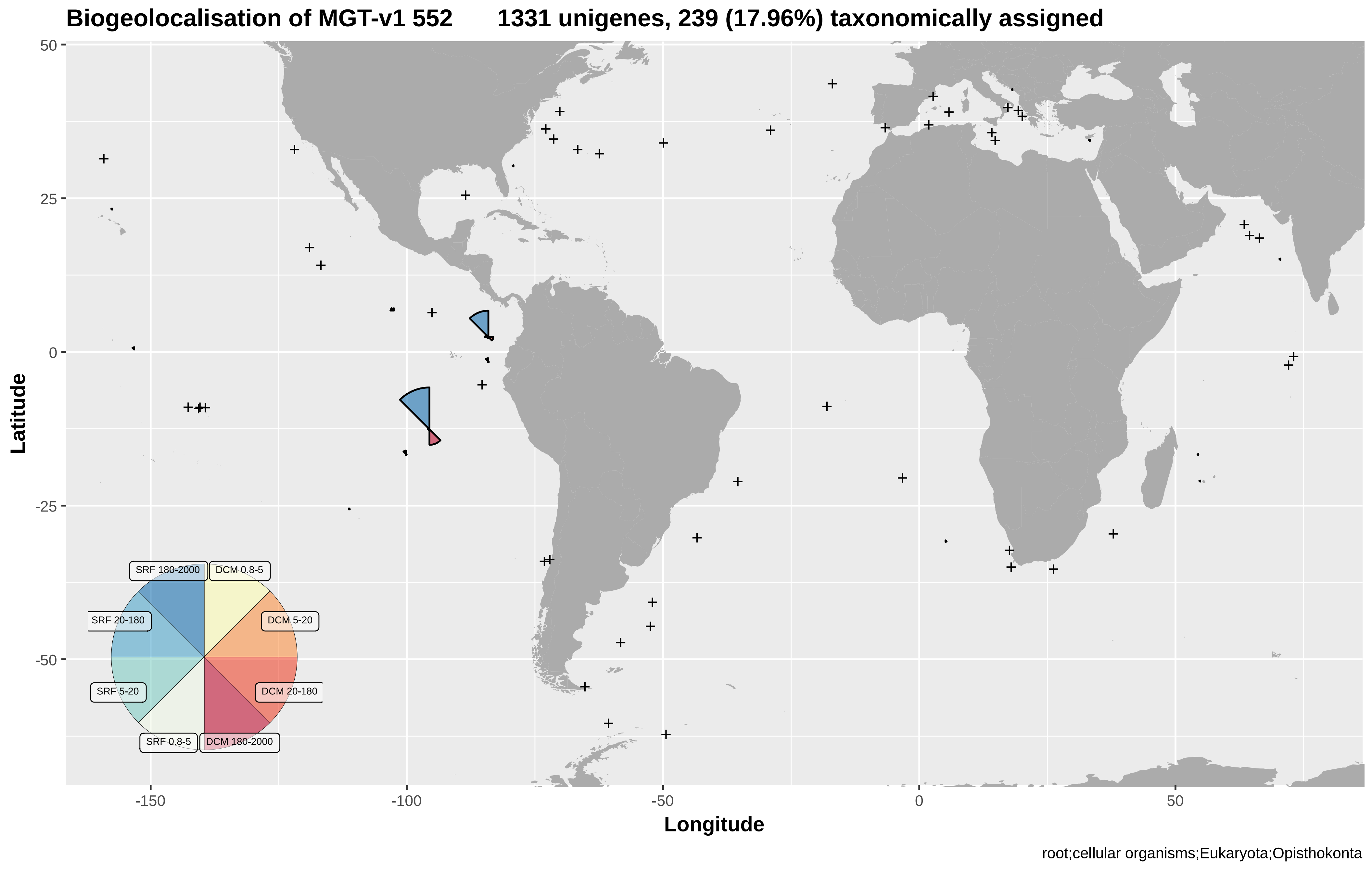

Biogeolocalisation of MGT-v1 552 1331 unigenes, 239 (17.96%) taxonomically assigned
50
25
0
-25
-50
Latitude
-150
-100
-50
0
50
Longitude
SRF 180-2000
DCM 0.8-5
SRF 20-180
DCM 5-20
SRF 5-20
DCM 20-180
SRF 0.8-5
DCM 180-2000
root;cellular organisms;Eukaryota;Opisthokonta

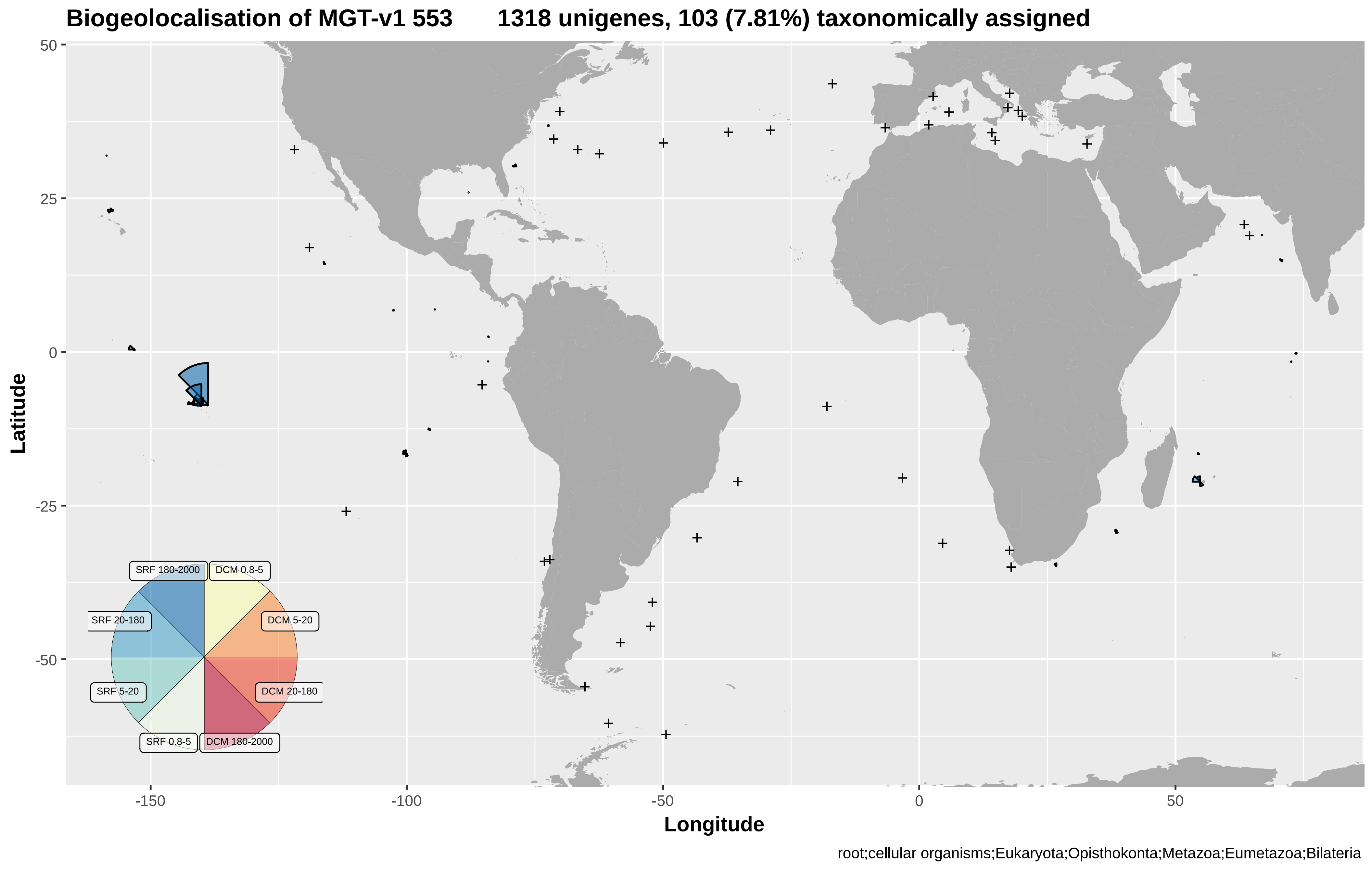
Biogeolocalisation of MGT-v1 553 1318 unigenes, 103 (7.81%) taxonomically assigned
50
25
0
-25
-50
Latitude
-150
-100
-50
0
50
Longitude
SRF 180-2000
DCM 0.8-5
SRF 20-180
DCM 5-20
SRF 5-20
DCM 20-180
SRF 0.8-5
DCM 180-2000
root;cellular organisms;Eukaryota;Opisthokonta;Metazoa;Eumetazoa;Bilateria

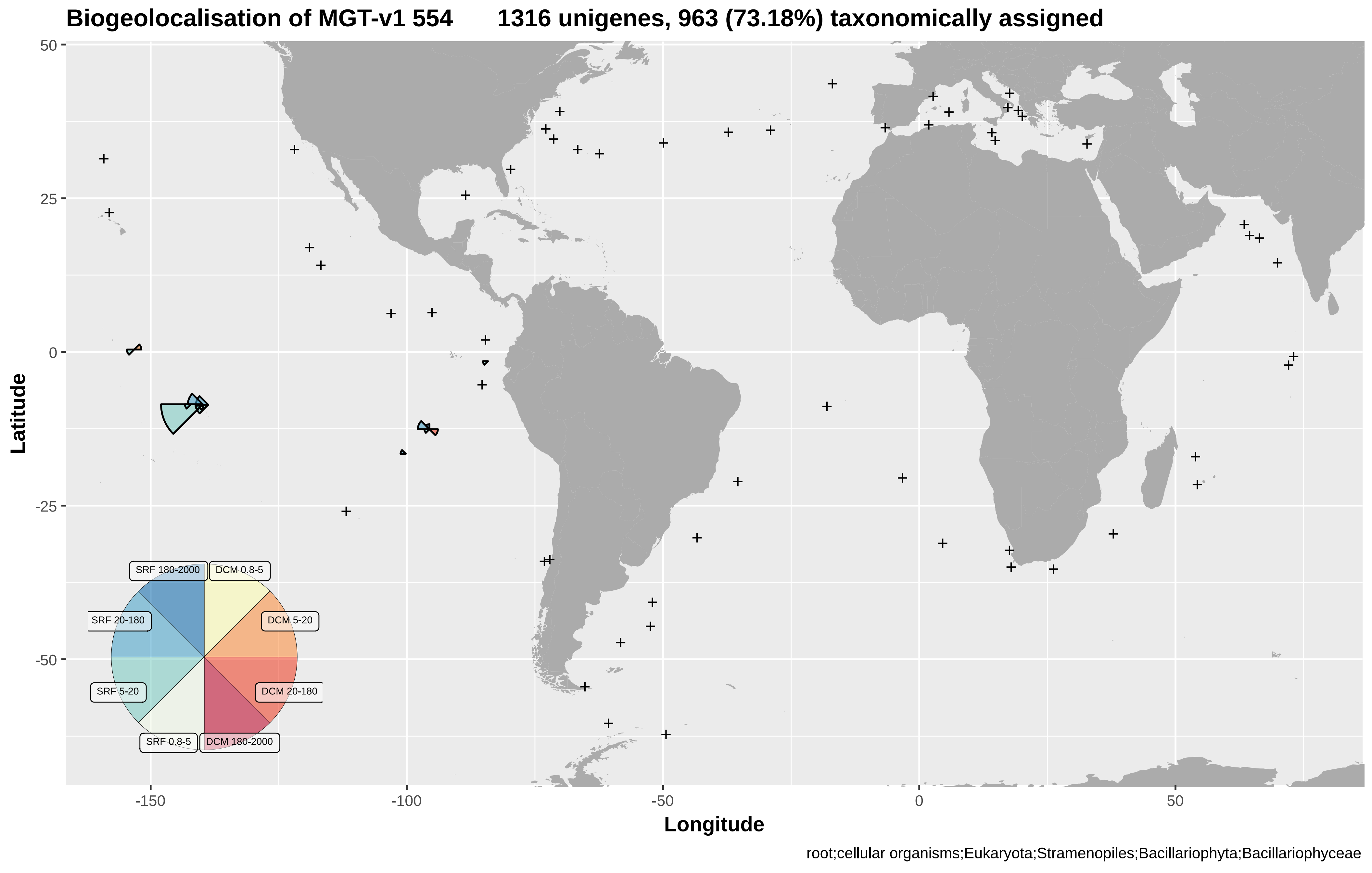

Biogeolocalisation of MGT-v1 554 1316 unigenes, 963 (73.18%) taxonomically assigned
Latitude
50
25
0
-25
-50
Longitude
-150
-100
-50
0
50
SRF 180-2000
DCM 0.8-5
SRF 20-180
DCM 5-20
SRF 5-20
DCM 20-180
SRF 0.8-5
DCM 180-2000
root;cellular organisms;Eukaryota;Stramenopiles;Bacillariophyta;Bacillariophyceae

Biogeolocalisation of MGT-v1 555      1311 unigenes, 292 (22.27%) taxonomically assigned

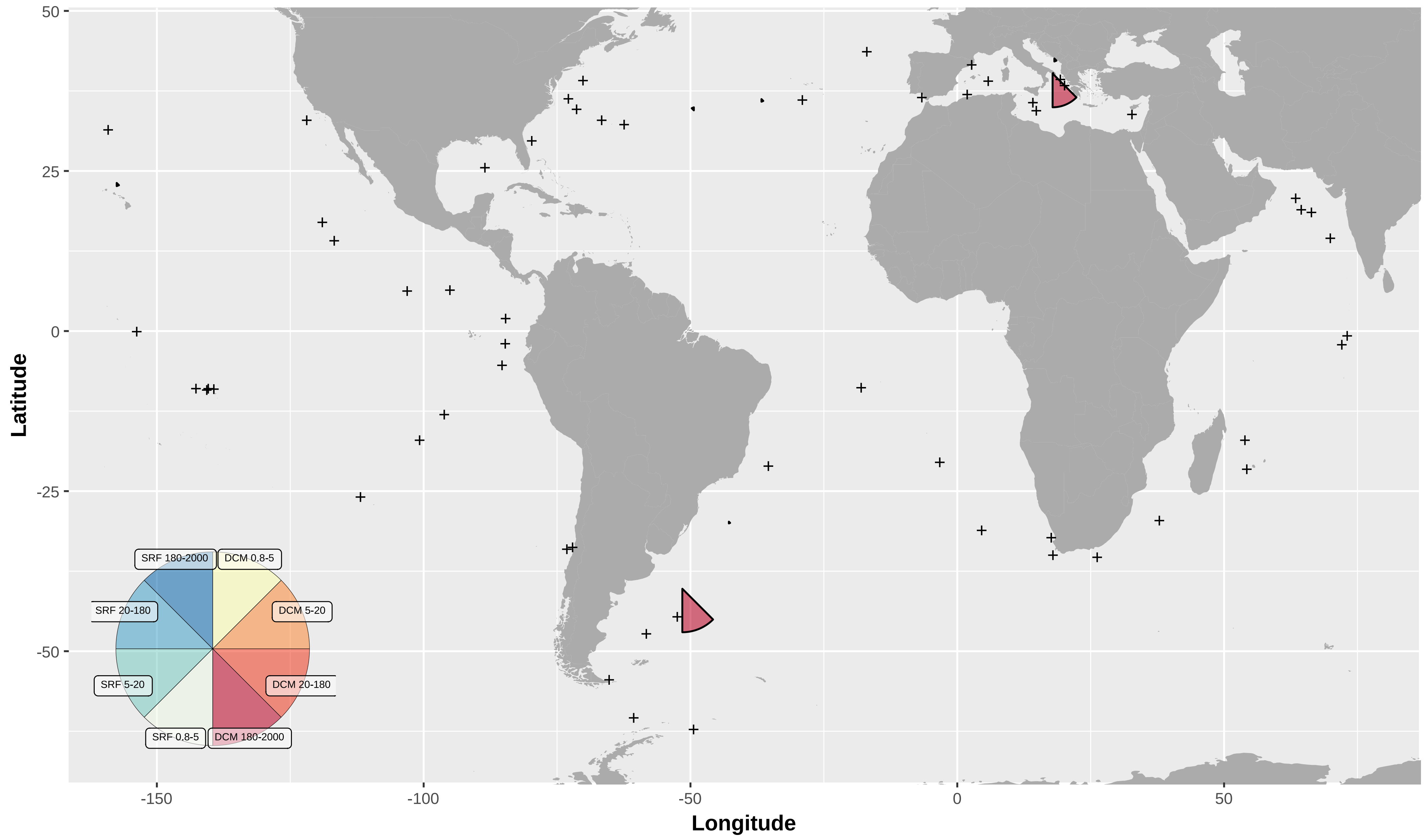


root;cellular organisms;Eukaryota;Opisthokonta;Metazoa;Eumetazoa;Bilateria

## Biogeolocalisation of MGT-v1 556      1311 unigenes, 114 (8.7%) taxonomically assigned

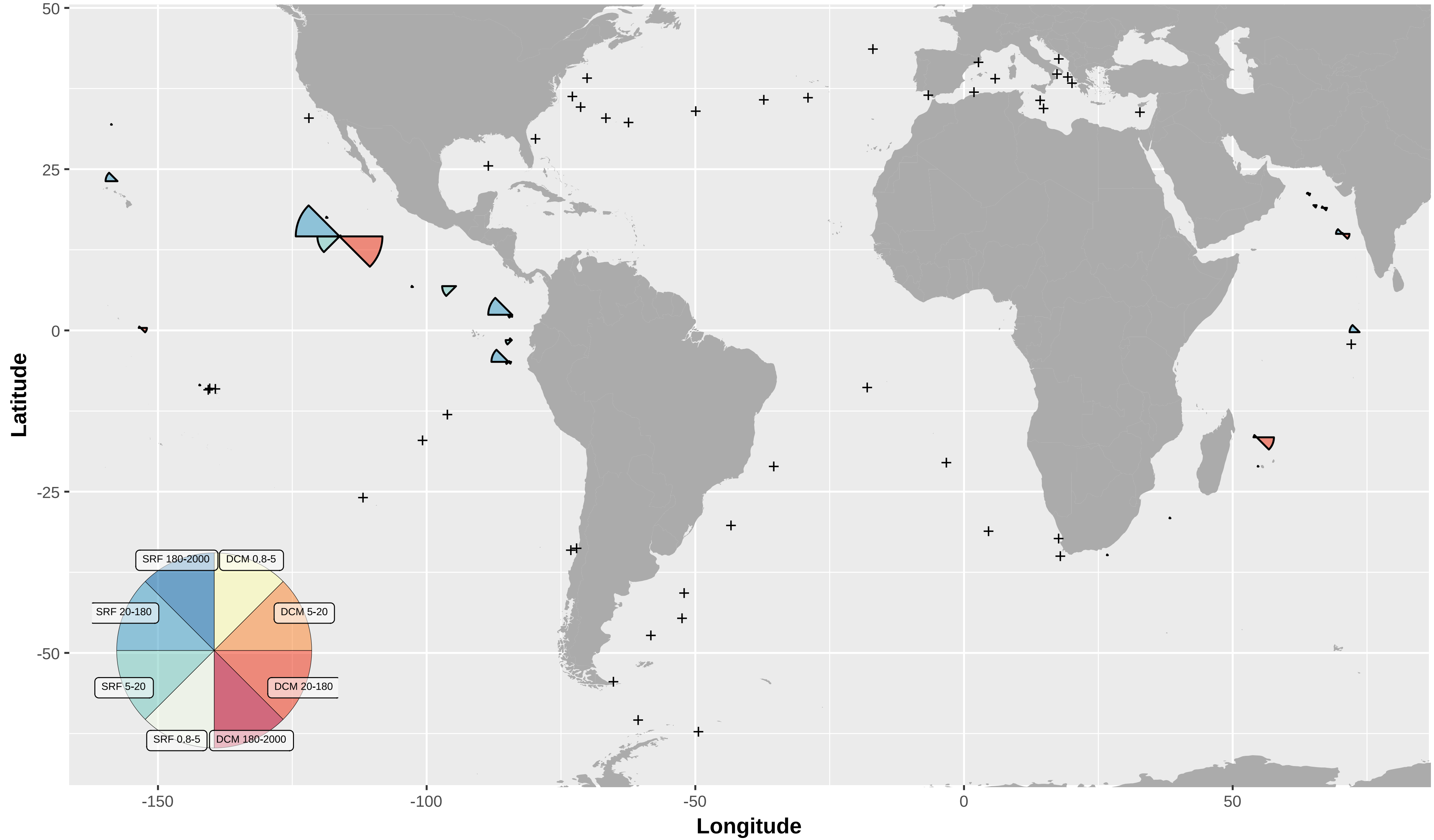


root;cellular organisms;Eukaryota;Opisthokonta;Metazoa;Eumetazoa;Bilateria

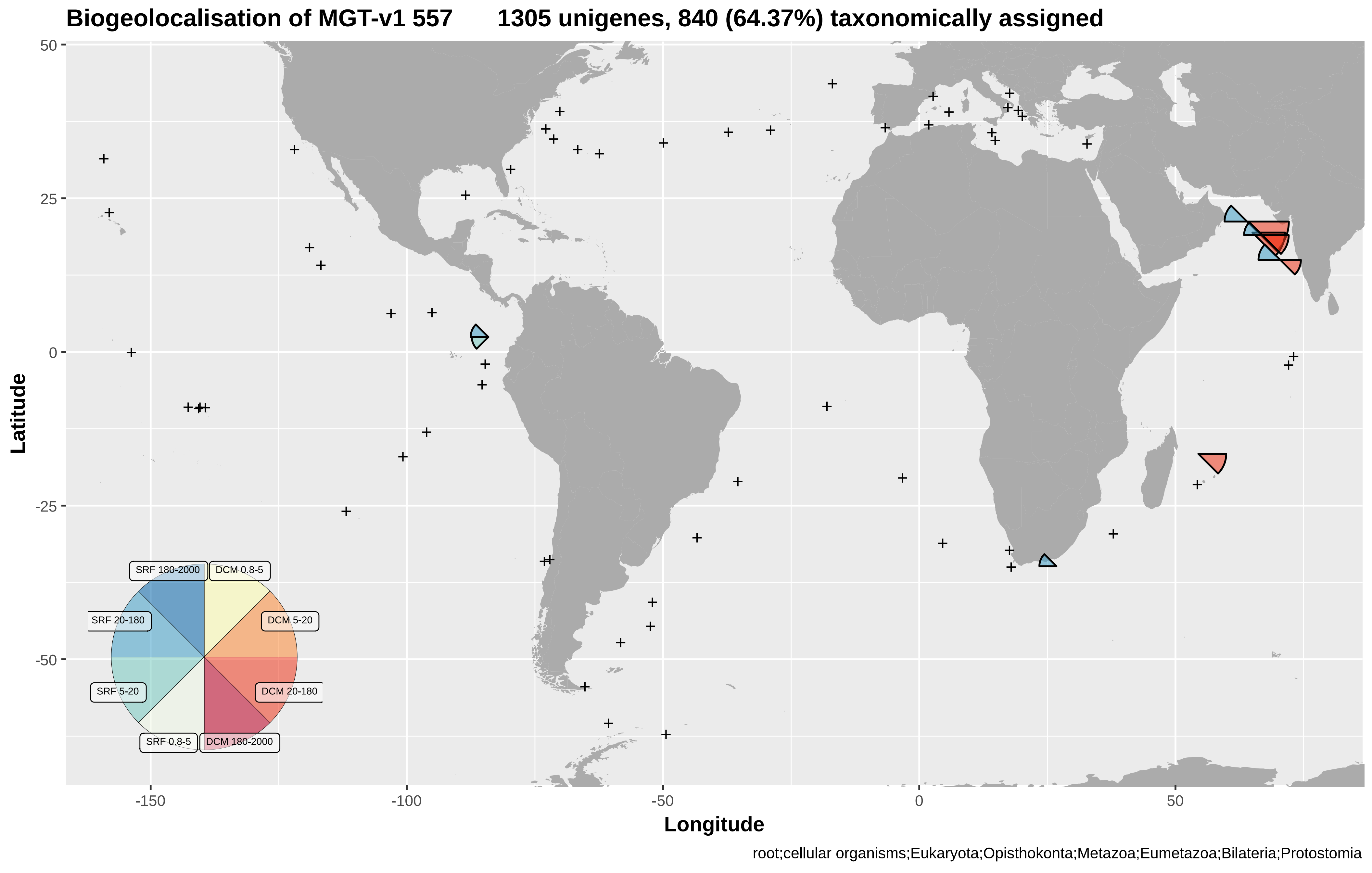

Biogeolocalisation of MGT-v1 557 1305 unigenes, 840 (64.37%) taxonomically assigned
50
25
0
-25
-50
Latitude
-150
-100
-50
0
50
Longitude
SRF 180-2000
DCM 0.8-5
SRF 20-180
DCM 5-20
SRF 5-20
DCM 20-180
SRF 0.8-5
DCM 180-2000
root;cellular organisms;Eukaryota;Opisthokonta;Metazoa;Eumetazoa;Bilateria;Protostomia

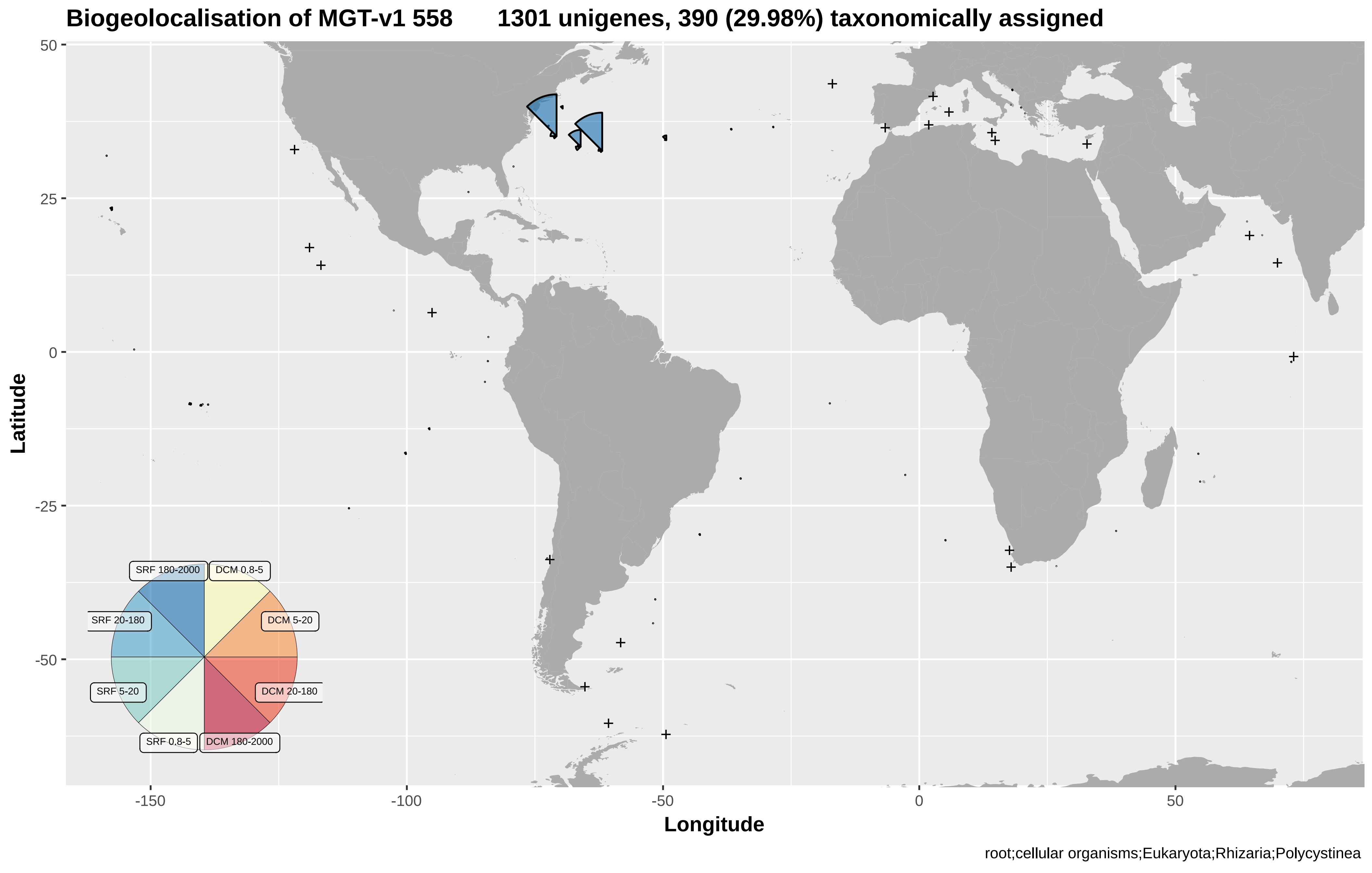
Biogeolocalisation of MGT-v1 558 1301 unigenes, 390 (29.98%) taxonomically assigned
Latitude
50
25
0
-25
-50
Longitude
-150
-100
-50
0
50
SRF 180-2000
DCM 0.8-5
SRF 20-180
DCM 5-20
SRF 5-20
DCM 20-180
SRF 0.8-5
DCM 180-2000
root;cellular organisms;Eukaryota;Rhizaria;Polycystinea

# Biogeolocalisation of MGT-v1 559      1289 unigenes, 1123 (87.12%) taxonomically assigned

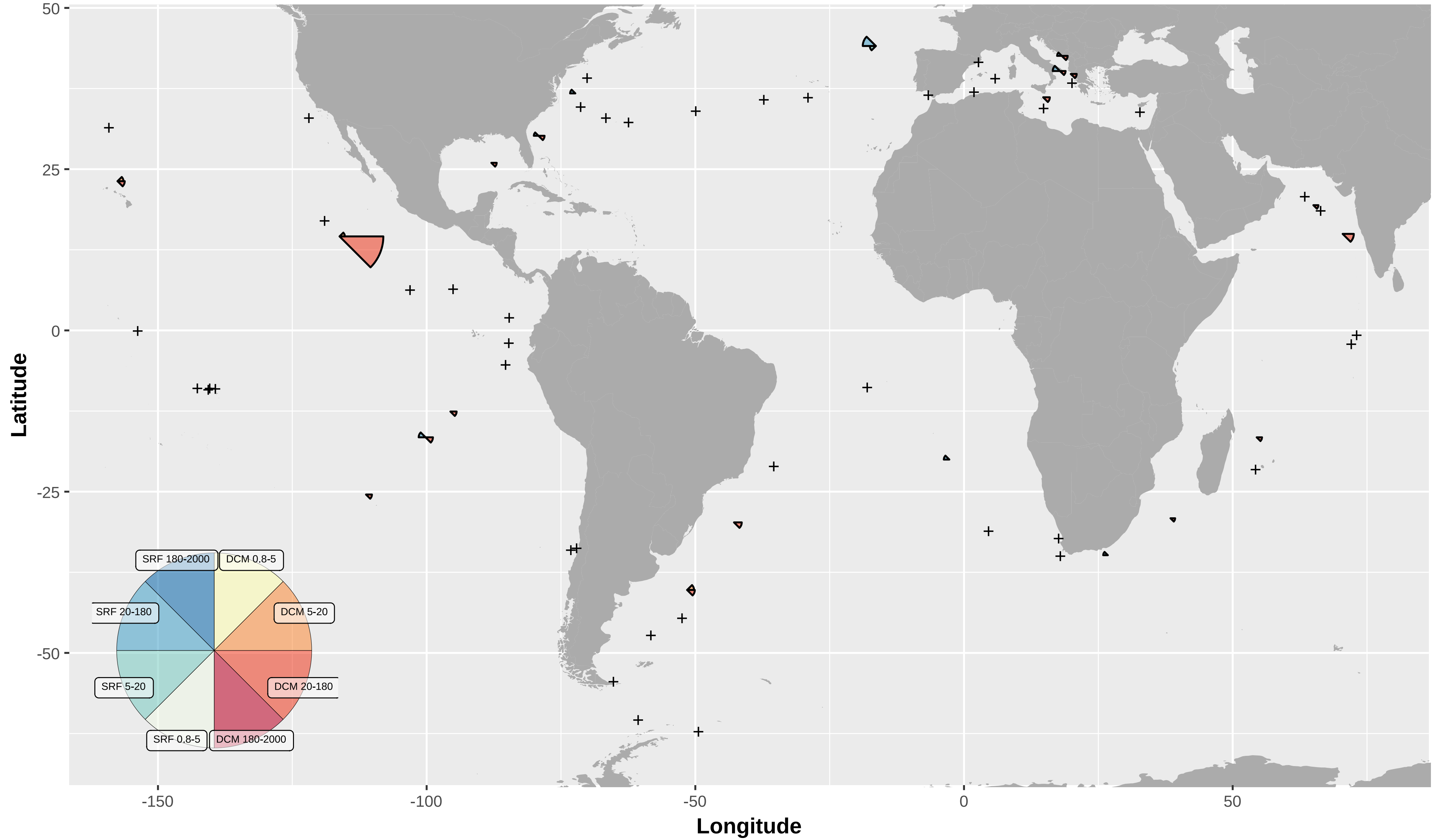


root;cellular organisms;Eukaryota;Alveolata;Ciliophora;Intramacronucleata;Spirotrichea

# Biogeolocalisation of MGT-v1 560      1285 unigenes, 151 (11.75%) taxonomically assigned

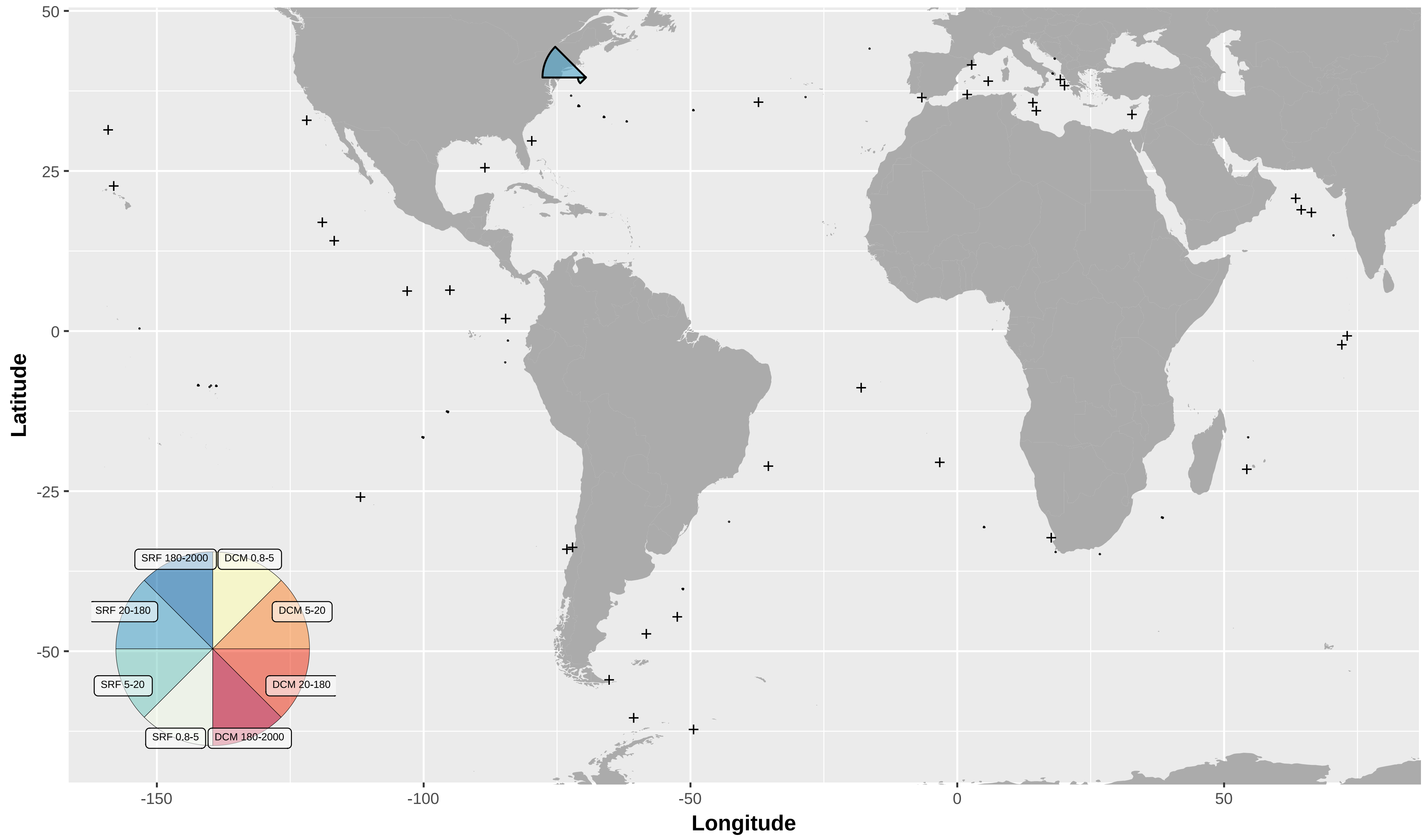


root;cellular organisms

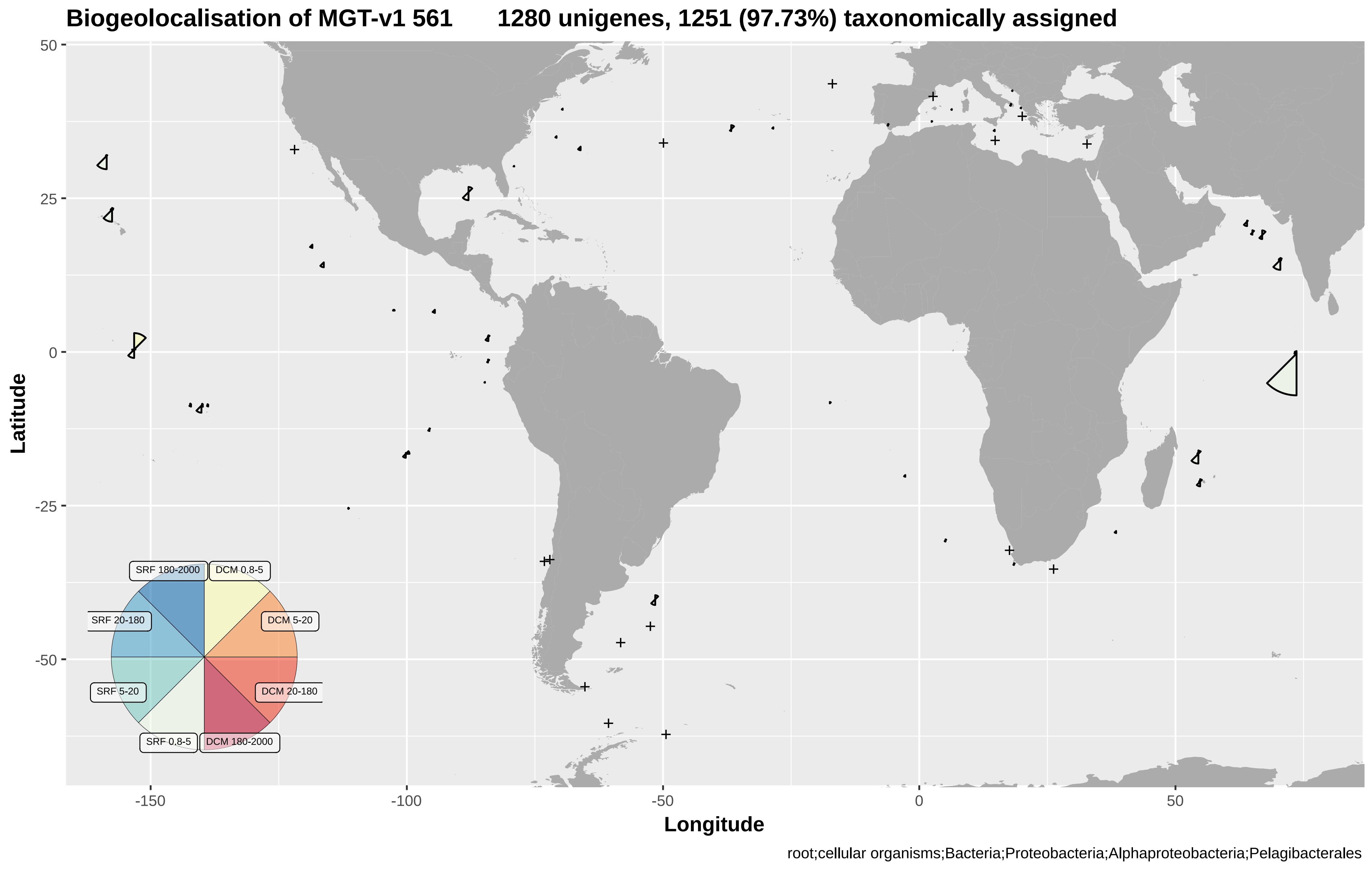

Biogeolocalisation of MGT-v1 561 1280 unigenes, 1251 (97.73%) taxonomically assigned
50
25
0
-25
-50
Latitude
-150
-100
-50
0
50
Longitude
SRF 180-2000
DCM 0.8-5
SRF 20-180
DCM 5-20
SRF 5-20
DCM 20-180
SRF 0.8-5
DCM 180-2000
root;cellular organisms;Bacteria;Proteobacteria;Alphaproteobacteria;Pelagibacterales

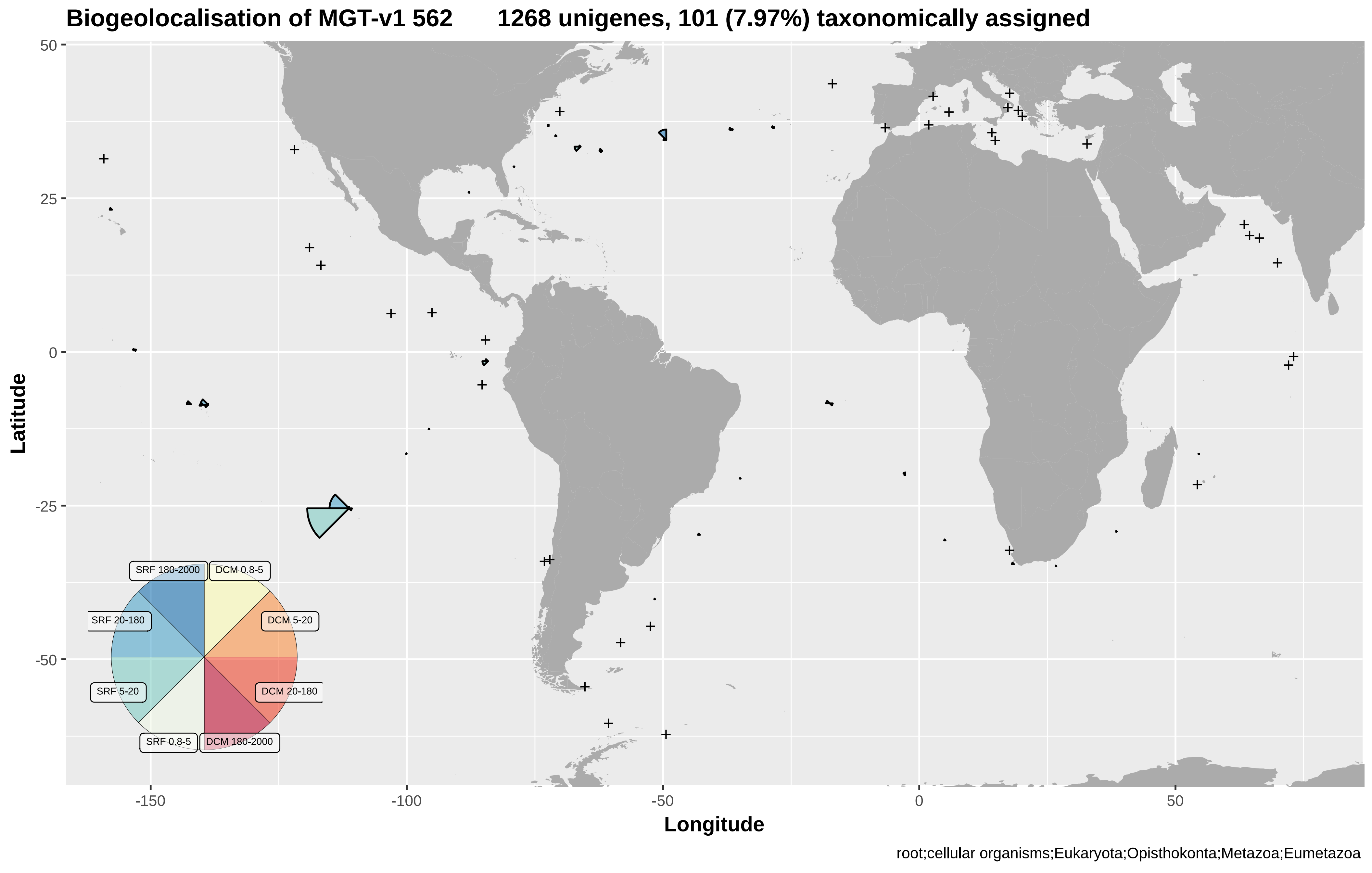

Biogeolocalisation of MGT-v1 562 1268 unigenes, 101 (7.97%) taxonomically assigned
Latitude
Longitude
50
25
0
-25
-50
-150
-100
-50
0
50
SRF 180-2000
DCM 0.8-5
SRF 20-180
DCM 5-20
SRF 5-20
DCM 20-180
SRF 0.8-5
DCM 180-2000
root;cellular organisms;Eukaryota;Opisthokonta;Metazoa;Eumetazoa

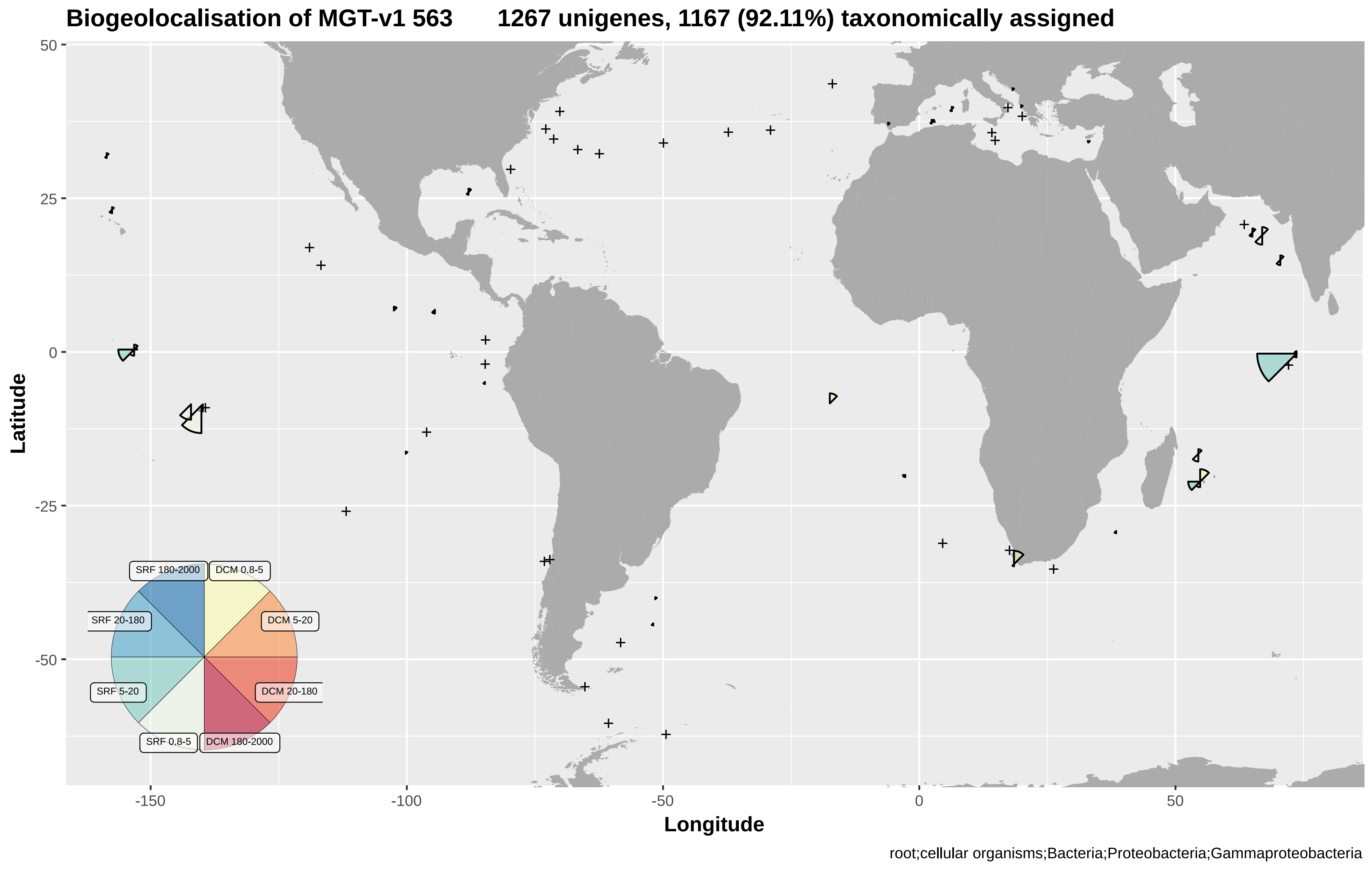

Biogeolocalisation of MGT-v1 563 1267 unigenes, 1167 (92.11%) taxonomically assigned
50
25
0
-25
-50
Latitude
-150
-100
-50
0
50
Longitude
SRF 180-2000
DCM 0.8-5
SRF 20-180
DCM 5-20
SRF 5-20
DCM 20-180
SRF 0.8-5
DCM 180-2000
root;cellular organisms;Bacteria;Proteobacteria;Gammaproteobacteria

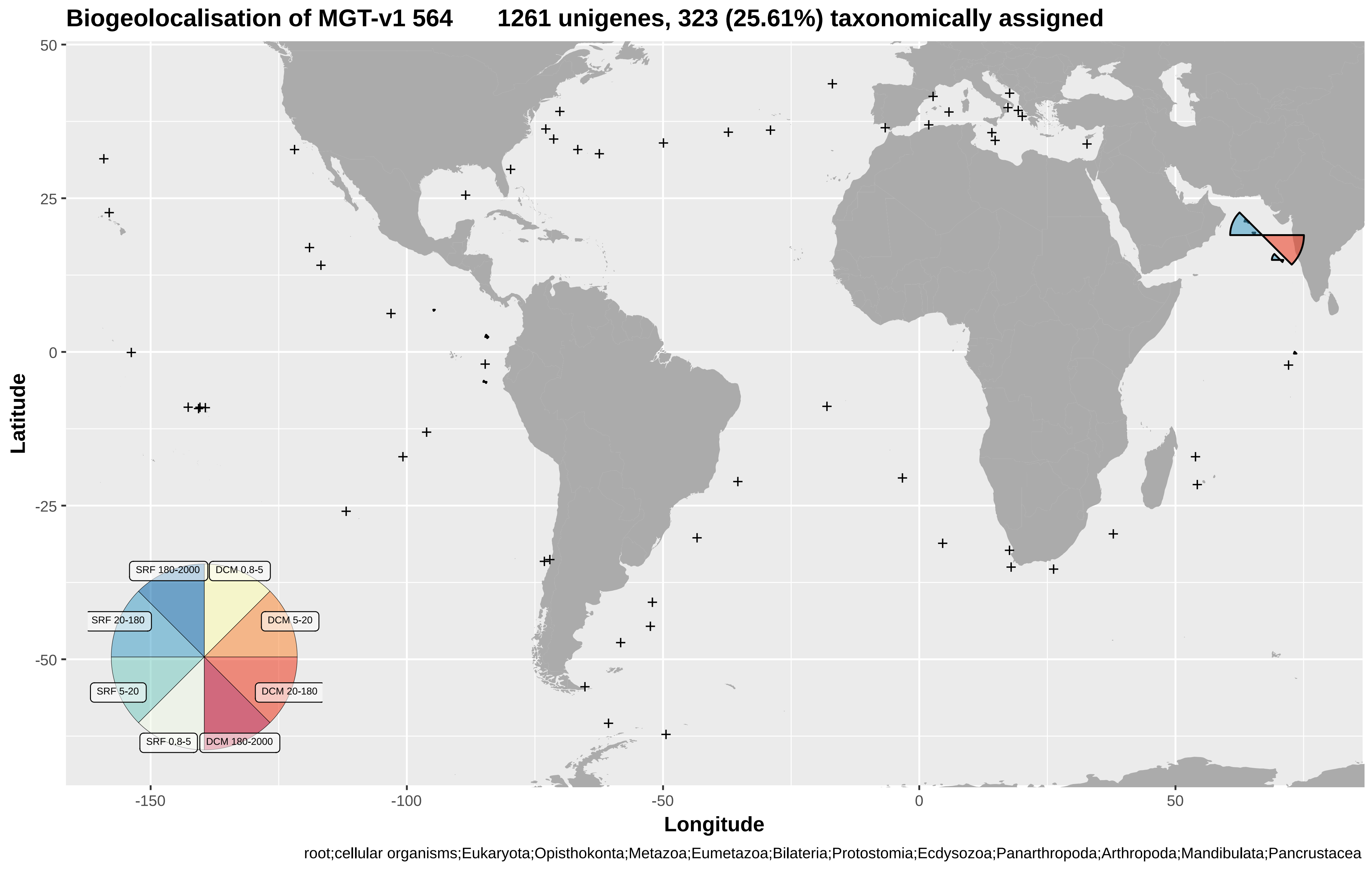

Biogeolocalisation of MGT-v1 564 1261 unigenes, 323 (25.61%) taxonomically assigned
Latitude
Longitude
50
25
0
-25
-50
-150
-100
-50
0
50
SRF 180-2000
DCM 0.8-5
SRF 20-180
DCM 5-20
SRF 5-20
DCM 20-180
SRF 0.8-5
DCM 180-2000
root;cellular organisms;Eukaryota;Opisthokonta;Metazoa;Eumetazoa;Bilateria;Protostomia;Ecdysozoa;Panarthropoda;Arthropoda;Mandibulata;Pancrustacea

## Biogeolocalisation of MGT-v1 565      1259 unigenes, 393 (31.22%) taxonomically assigned

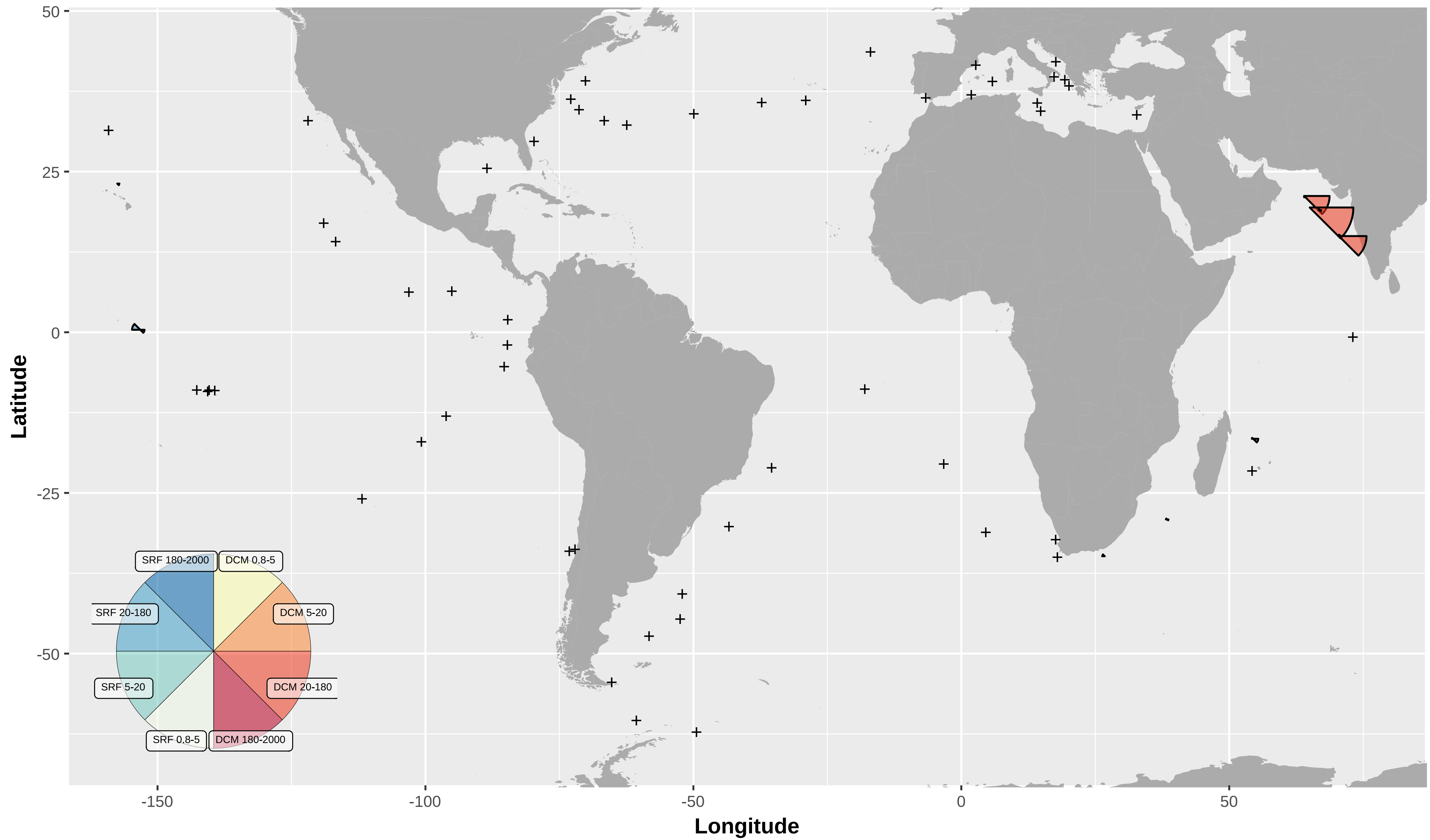


root;cellular organisms;Eukaryota;Opisthokonta;Metazoa;Eumetazoa;Bilateria;Protostomia;Ecdysozoa;Panarthropoda;Arthropoda;Mandibulata;Pancrustacea;Crustacea;Maxillopoda

## Biogeolocalisation of MGT-v1 566      1258 unigenes, 212 (16.85%) taxonomically assigned

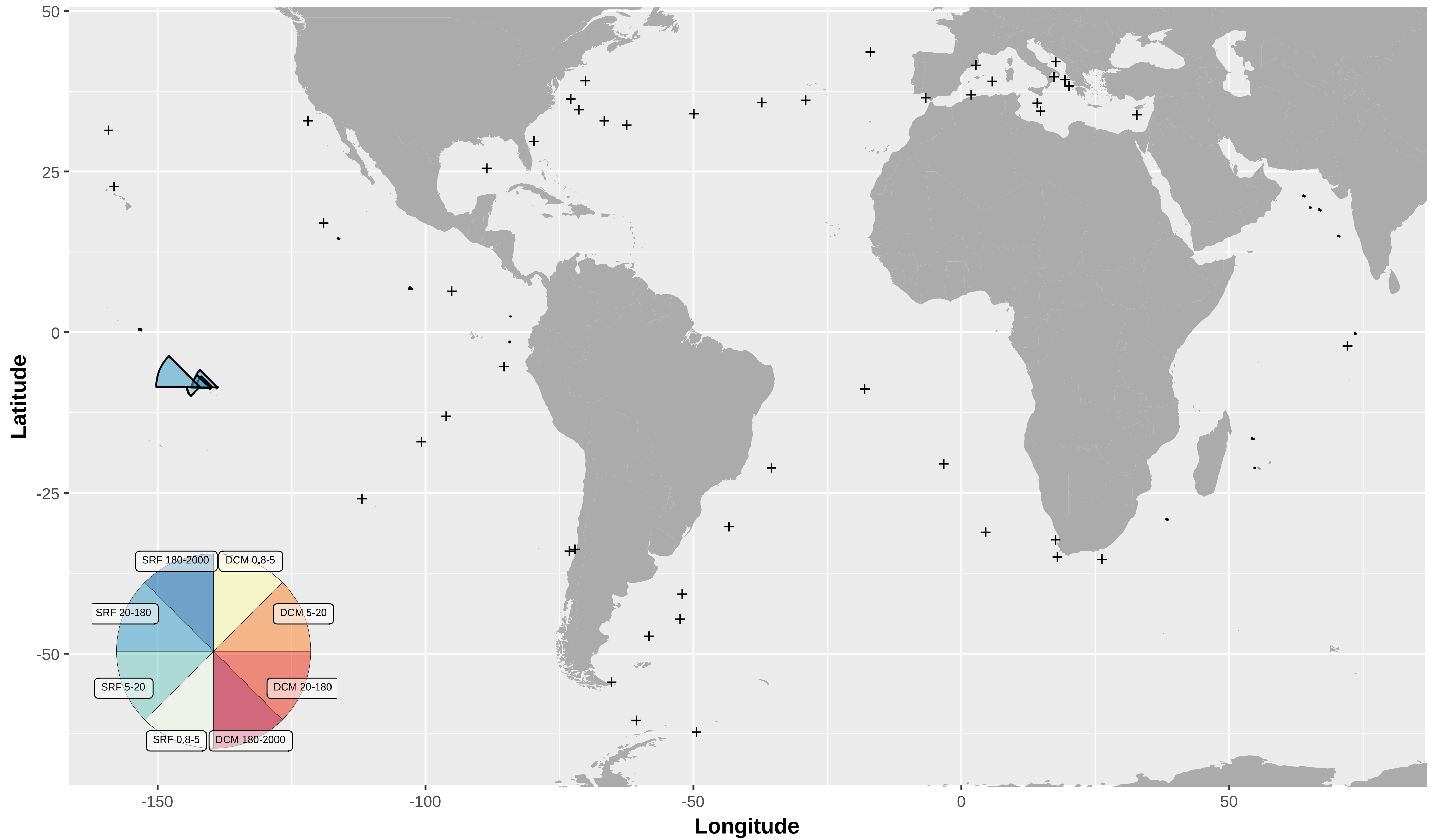


root;cellular organisms;Eukaryota

## Biogeolocalisation of MGT-v1 567      1258 unigenes, 202 (16.06%) taxonomically assigned

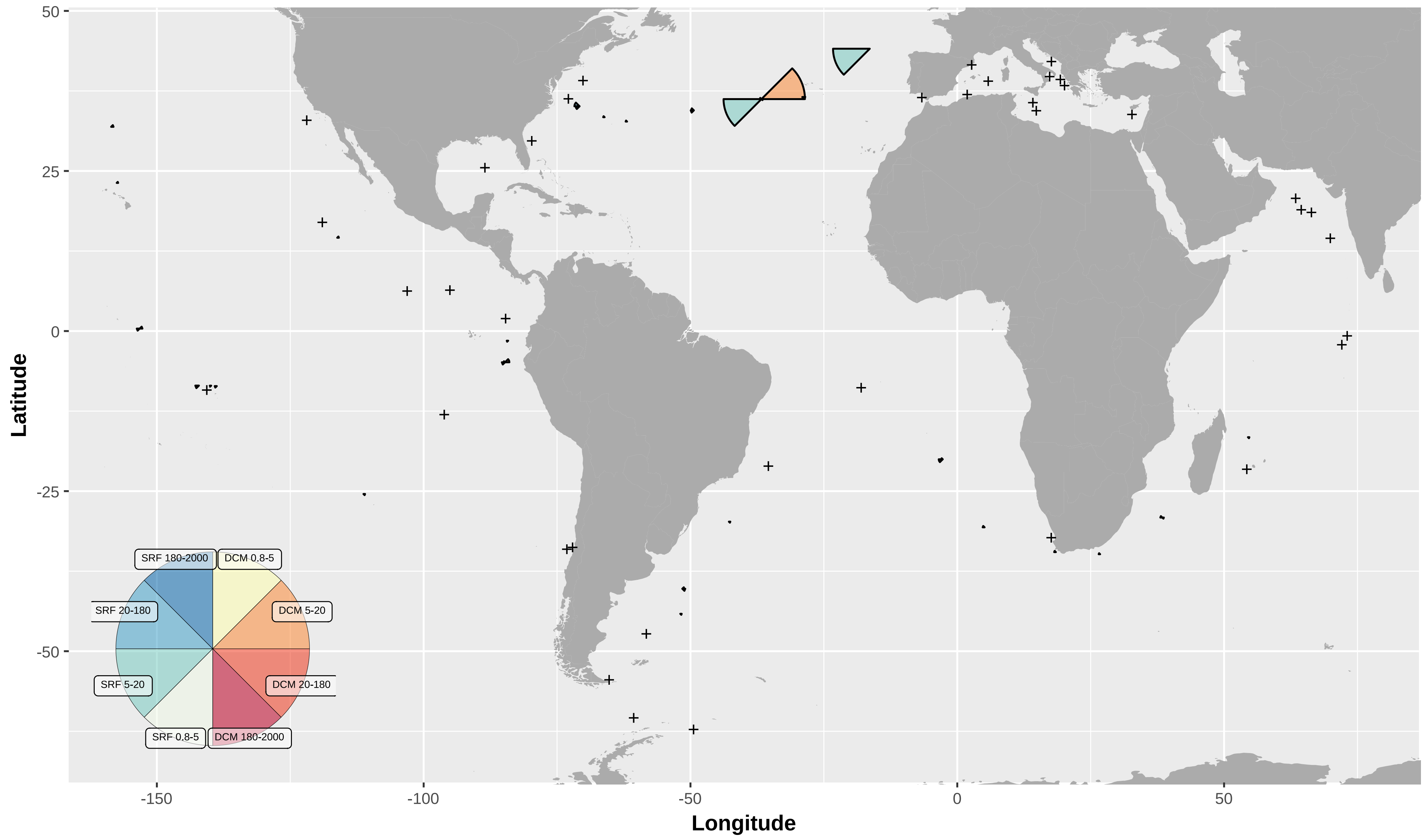


root;cellular organisms;Eukaryota;Stramenopiles;Bacillariophyta

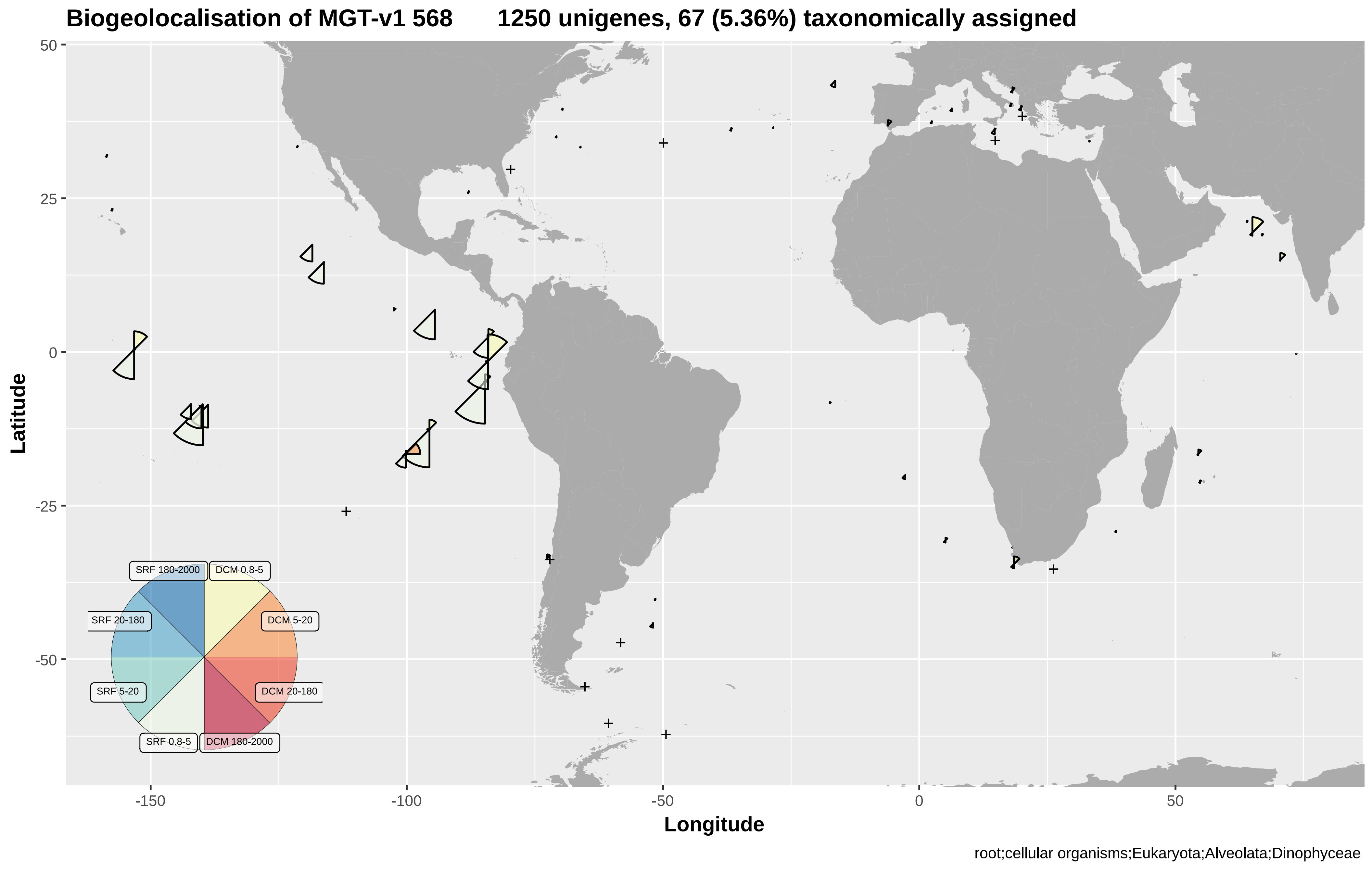

Biogeolocalisation of MGT-v1 568 1250 unigenes, 67 (5.36%) taxonomically assigned
50
25
0
-25
-50
Latitude
-150
-100
-50
0
50
Longitude
SRF 180-2000
DCM 0.8-5
SRF 20-180
DCM 5-20
SRF 5-20
DCM 20-180
SRF 0.8-5
DCM 180-2000
root;cellular organisms;Eukaryota;Alveolata;Dinophyceae

## Biogeolocalisation of MGT-v1 569      1237 unigenes, 116 (9.38%) taxonomically assigned

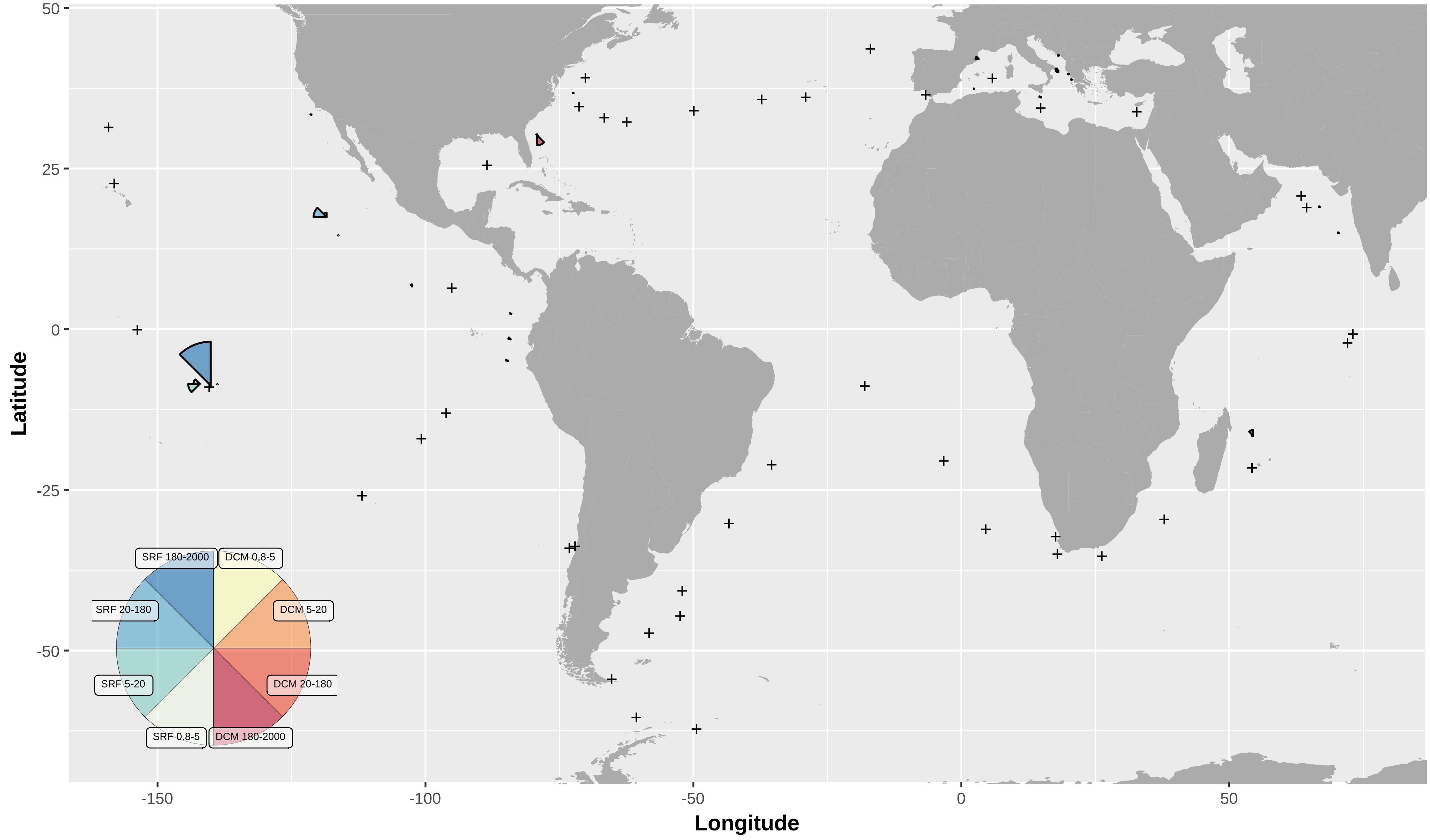


root;cellular organisms;Eukaryota;Opisthokonta;Metazoa;Eumetazoa;Bilateria

#### Biogeolocalisation of MGT-v1 570 1232 unigenes, 130 (10.55%) taxonomically assigned

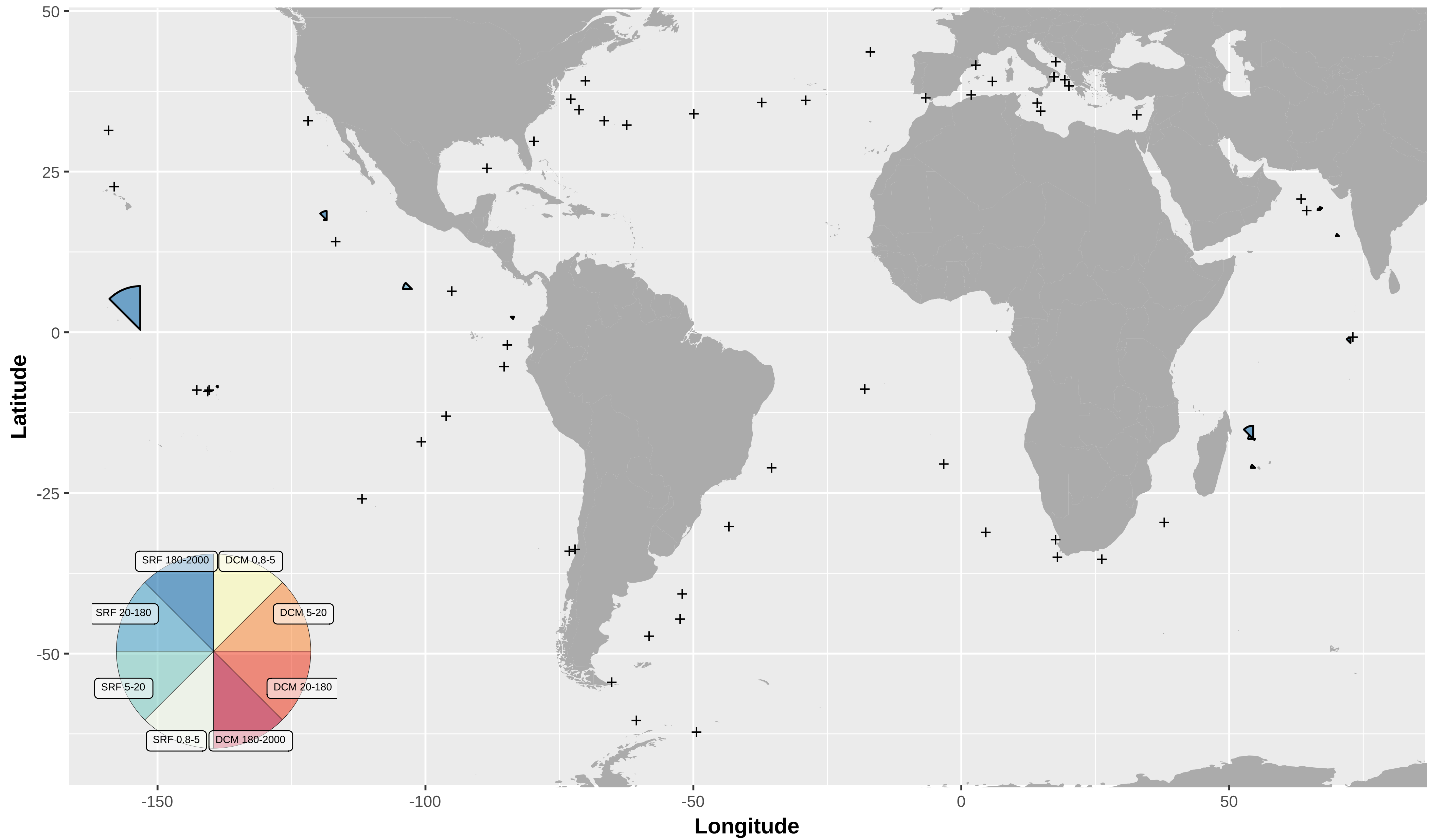


root;cellular organisms;Eukaryota;Opisthokonta;Metazoa;Eumetazoa;Bilateria

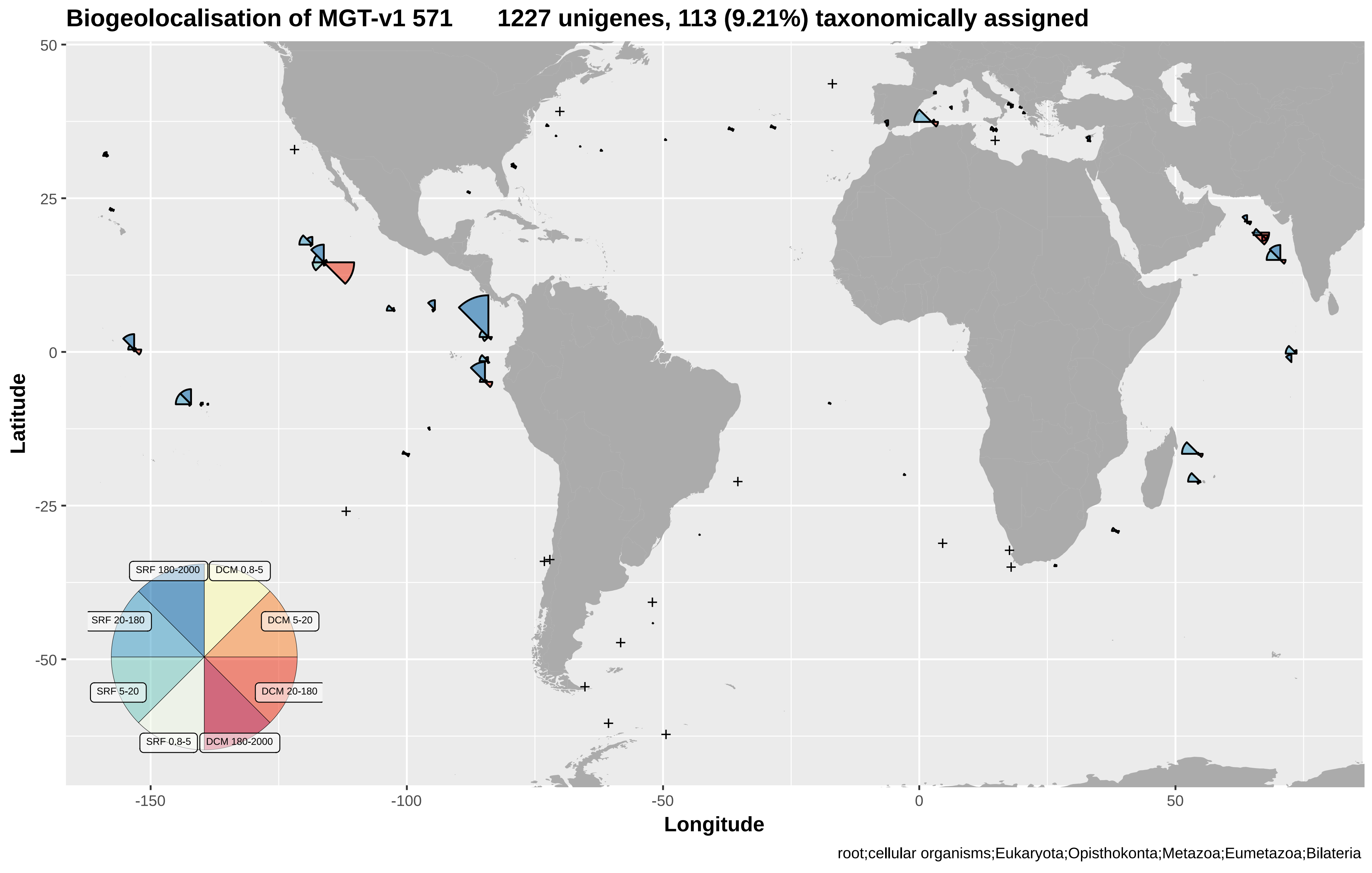

Biogeolocalisation of MGT-v1 571 1227 unigenes, 113 (9.21%) taxonomically assigned
50
25
0
-25
-50
Latitude
-150
-100
-50
0
50
Longitude
SRF 180-2000
DCM 0.8-5
SRF 20-180
DCM 5-20
SRF 5-20
DCM 20-180
SRF 0.8-5
DCM 180-2000
root;cellular organisms;Eukaryota;Opisthokonta;Metazoa;Eumetazoa;Bilateria

## Biogeolocalisation of MGT-v1 572      1223 unigenes, 922 (75.39%) taxonomically assigned

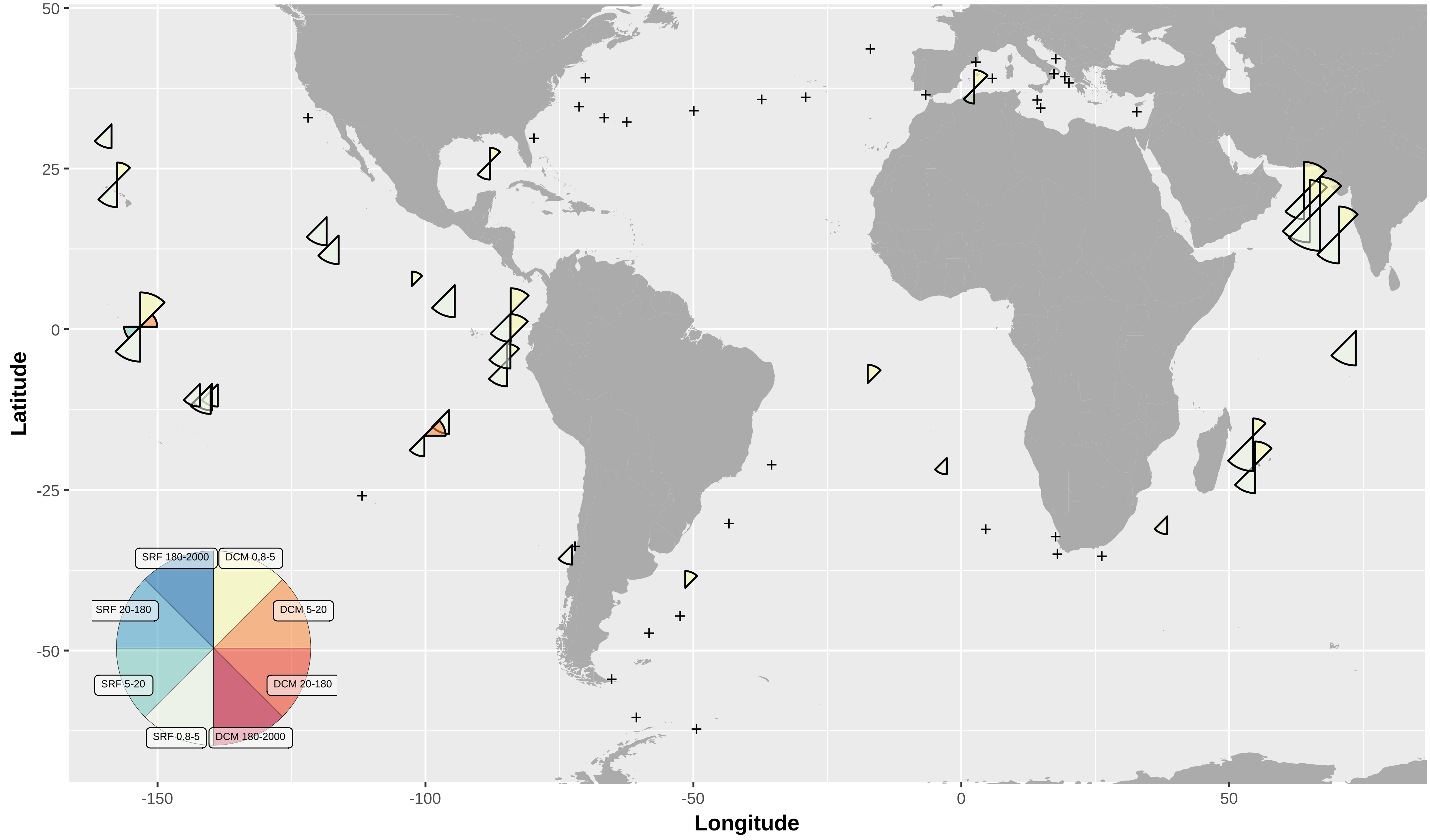


root;cellular organisms;Eukaryota

## Biogeolocalisation of MGT-v1 573      1223 unigenes, 248 (20.28%) taxonomically assigned

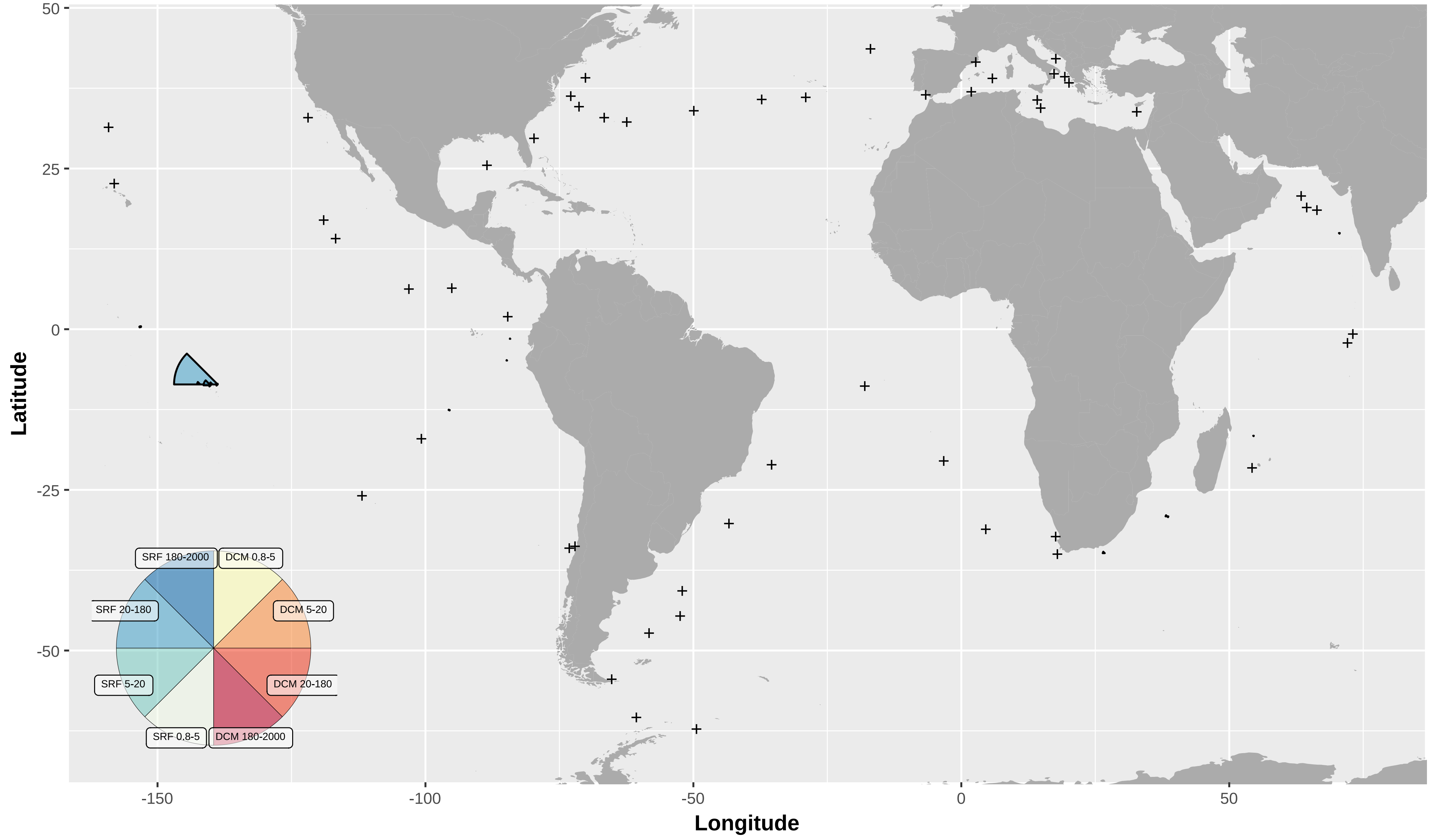


root;cellular organisms;Eukaryota

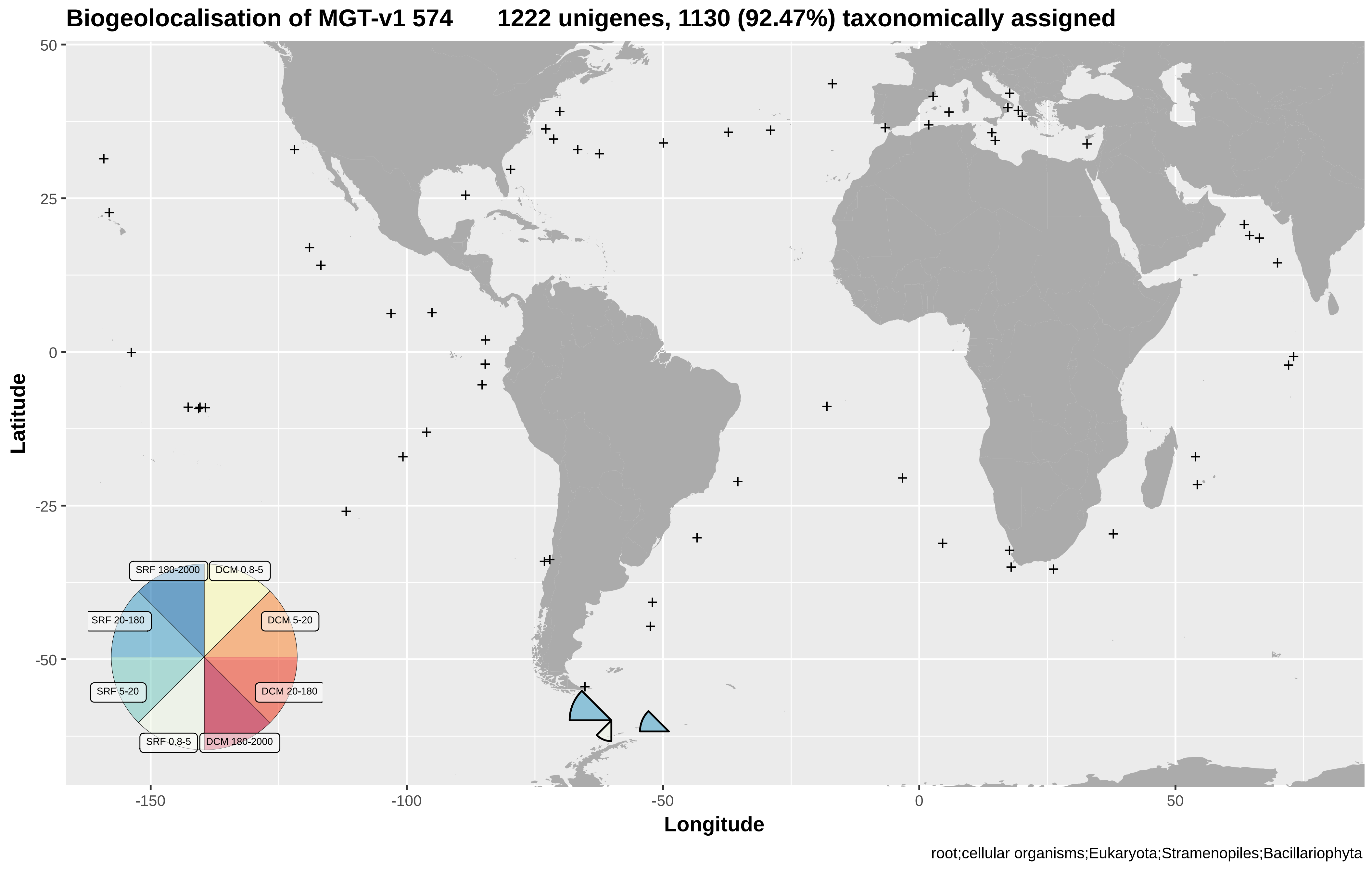
Biogeolocalisation of MGT-v1 574 1222 unigenes, 1130 (92.47%) taxonomically assigned
Latitude
Longitude
50
25
0
-25
-50
-150
-100
-50
0
50
SRF 180-2000
DCM 0.8-5
SRF 20-180
DCM 5-20
SRF 5-20
DCM 20-180
SRF 0.8-5
DCM 180-2000
root;cellular organisms;Eukaryota;Stramenopiles;Bacillariophyta

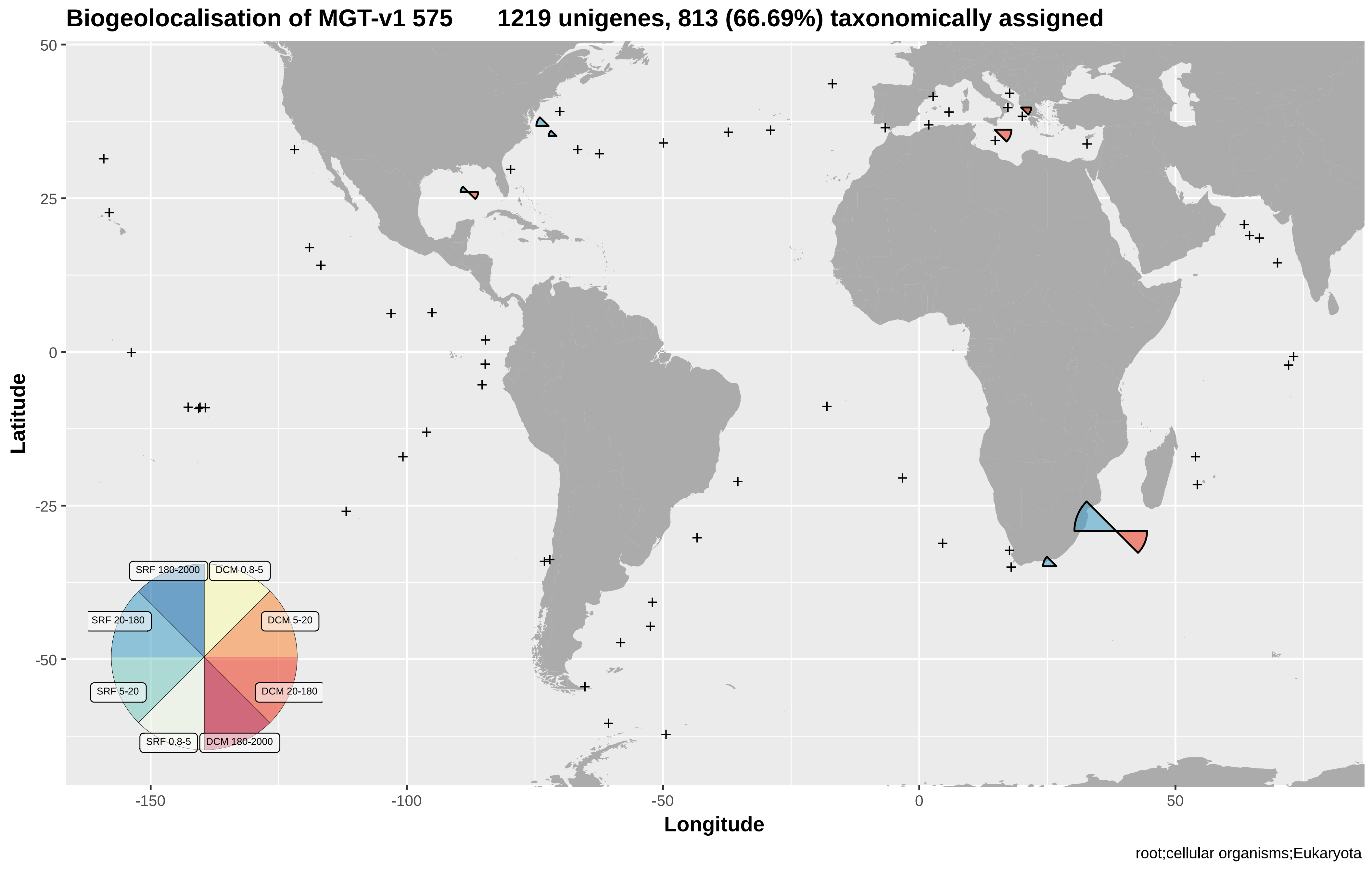
Biogeolocalisation of MGT-v1 575 1219 unigenes, 813 (66.69%) taxonomically assigned
50
25
0
-25
-50
Latitude
-150
-100
-50
0
50
Longitude
SRF 180-2000
DCM 0.8-5
SRF 20-180
DCM 5-20
SRF 5-20
DCM 20-180
SRF 0.8-5
DCM 180-2000
root;cellular organisms;Eukaryota

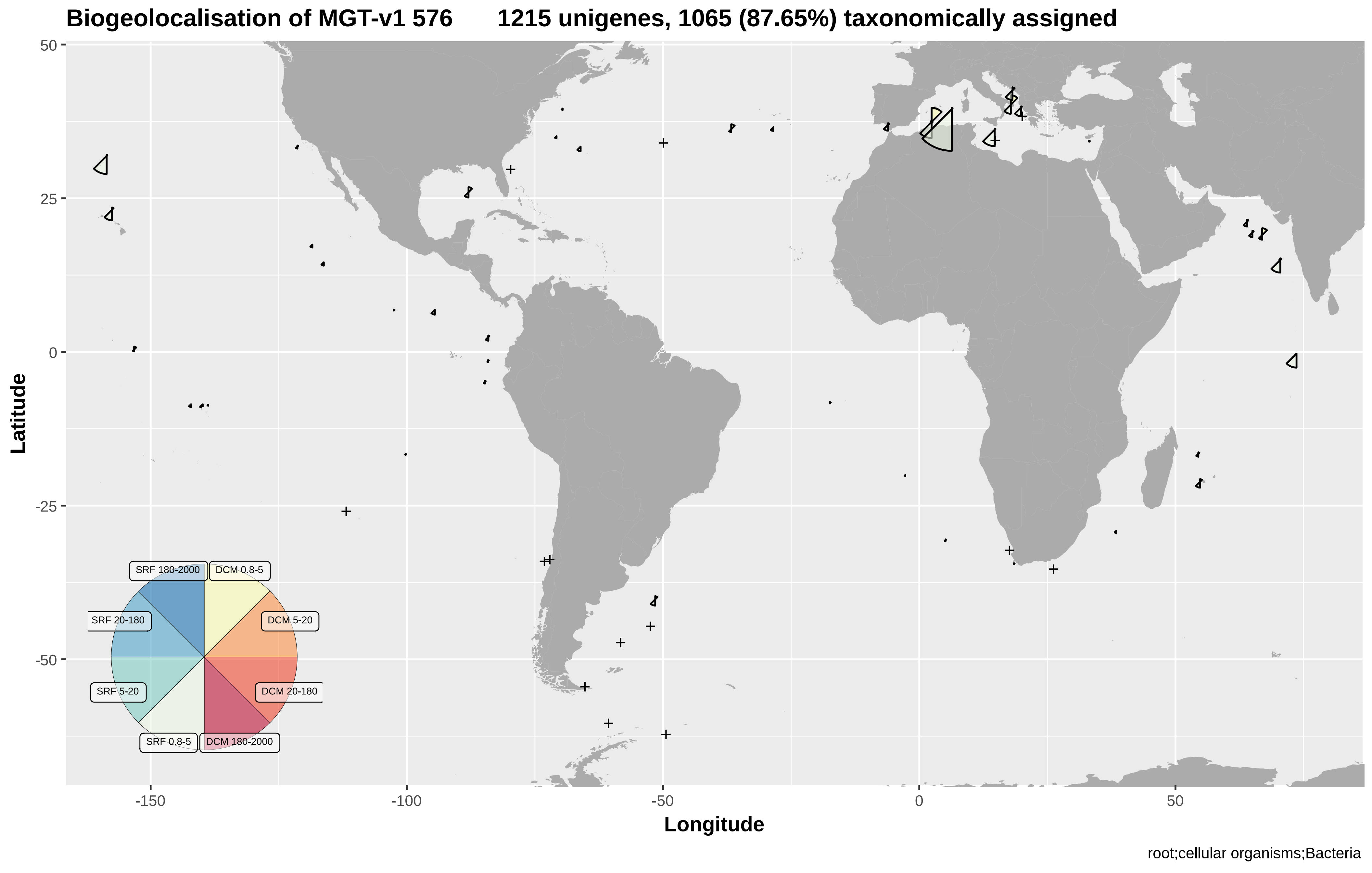

Biogeolocalisation of MGT-v1 576 1215 unigenes, 1065 (87.65%) taxonomically assigned
50
25
0
-25
-50
Latitude
-150
-100
-50
0
50
Longitude
SRF 180-2000
DCM 0.8-5
SRF 20-180
DCM 5-20
SRF 5-20
DCM 20-180
SRF 0.8-5
DCM 180-2000
root;cellular organisms;Bacteria

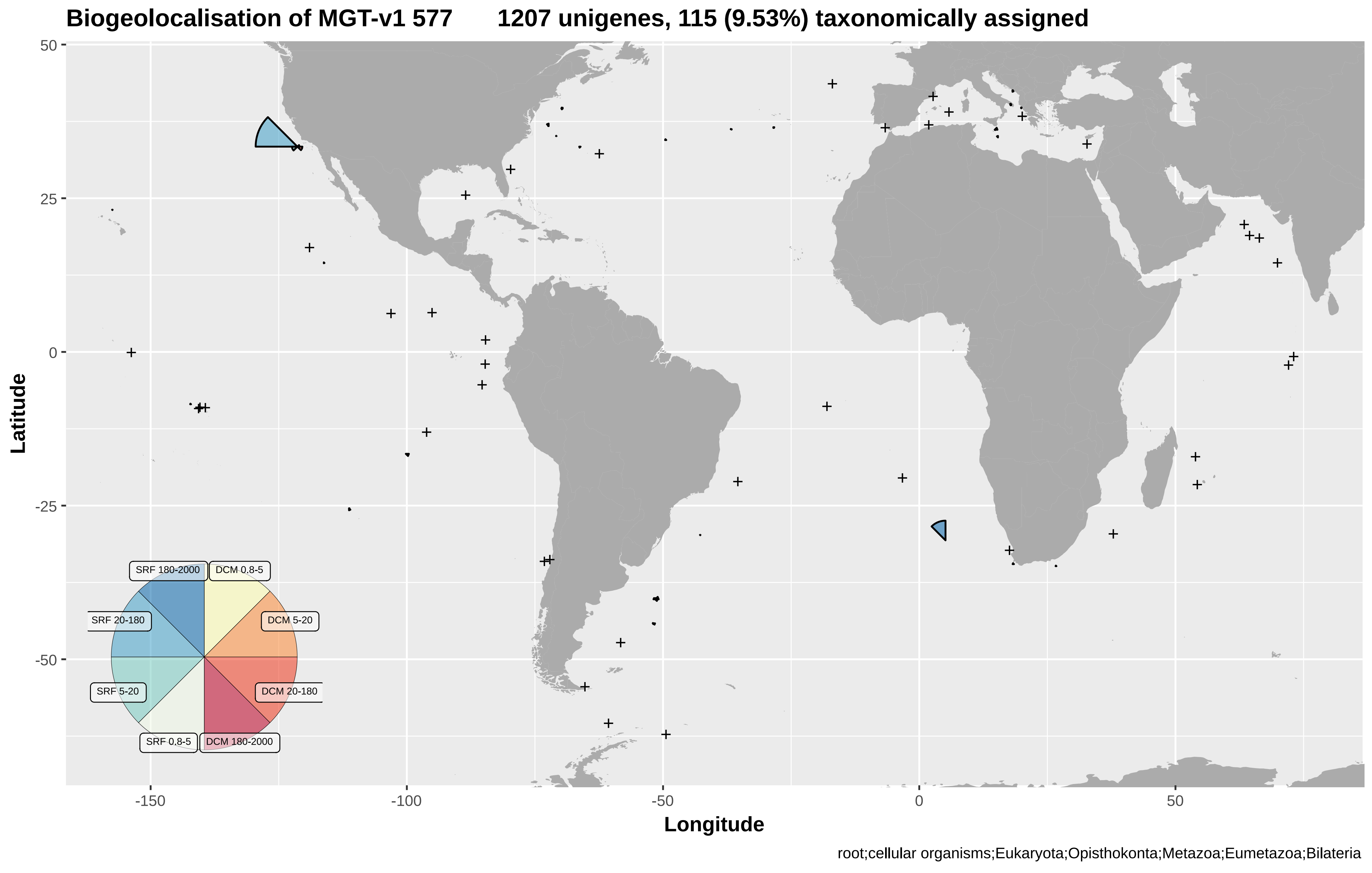

Biogeolocalisation of MGT-v1 577 1207 unigenes, 115 (9.53%) taxonomically assigned
Latitude
Longitude
50
25
0
-25
-50
-150
-100
-50
0
50
SRF 180-2000
DCM 0.8-5
SRF 20-180
DCM 5-20
SRF 5-20
DCM 20-180
SRF 0.8-5
DCM 180-2000
root;cellular organisms;Eukaryota;Opisthokonta;Metazoa;Eumetazoa;Bilateria

Biogeolocalisation of MGT-v1 578 1197 unigenes, 275 (22.97%) taxonomically assigned

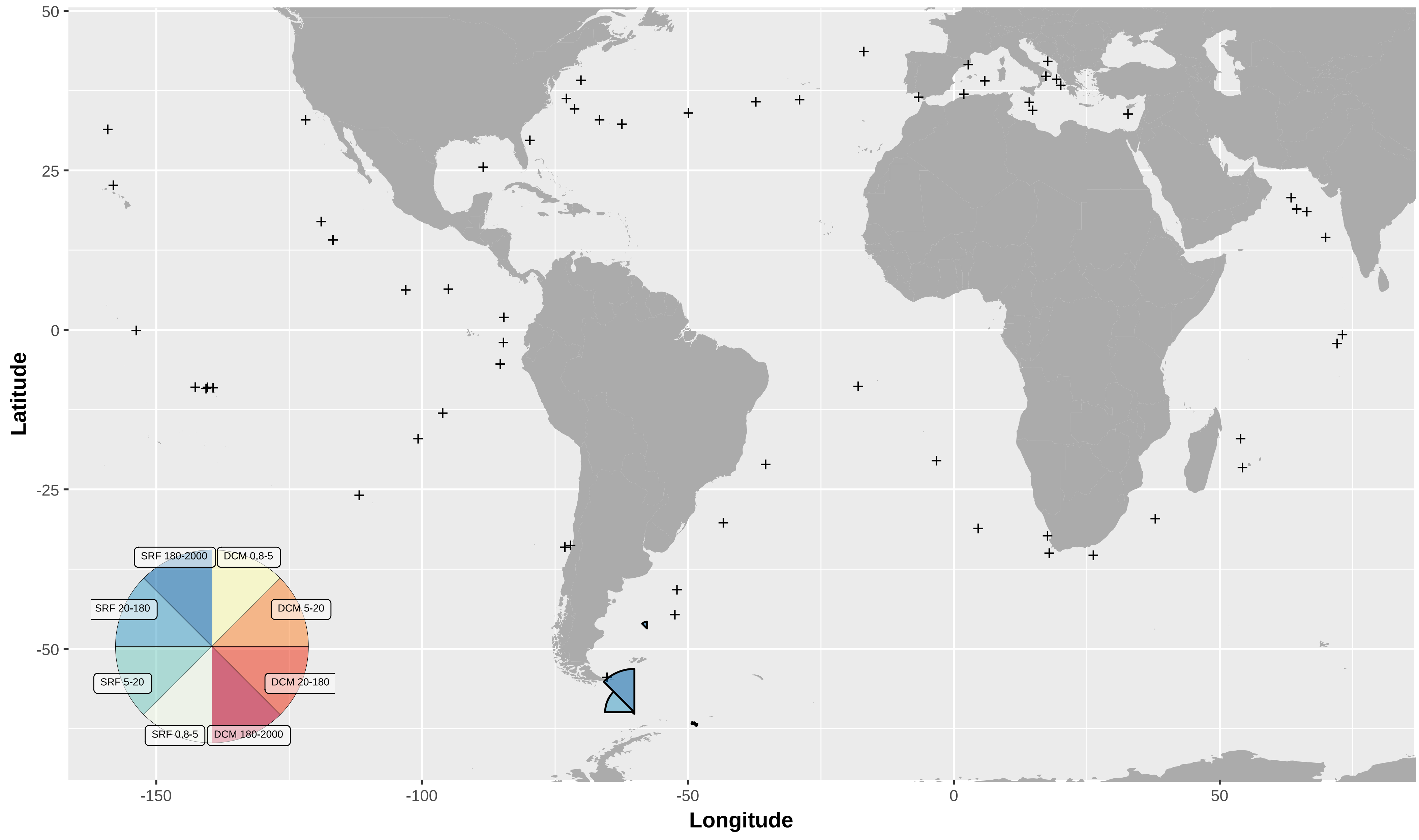


root;cellular organisms;Eukaryota;Opisthokonta;Metazoa;Eumetazoa;Bilateria

**Biogeolocalisation of MGT-v1 579      1194 unigenes, 1025 (85.85%) taxonomically assigned**

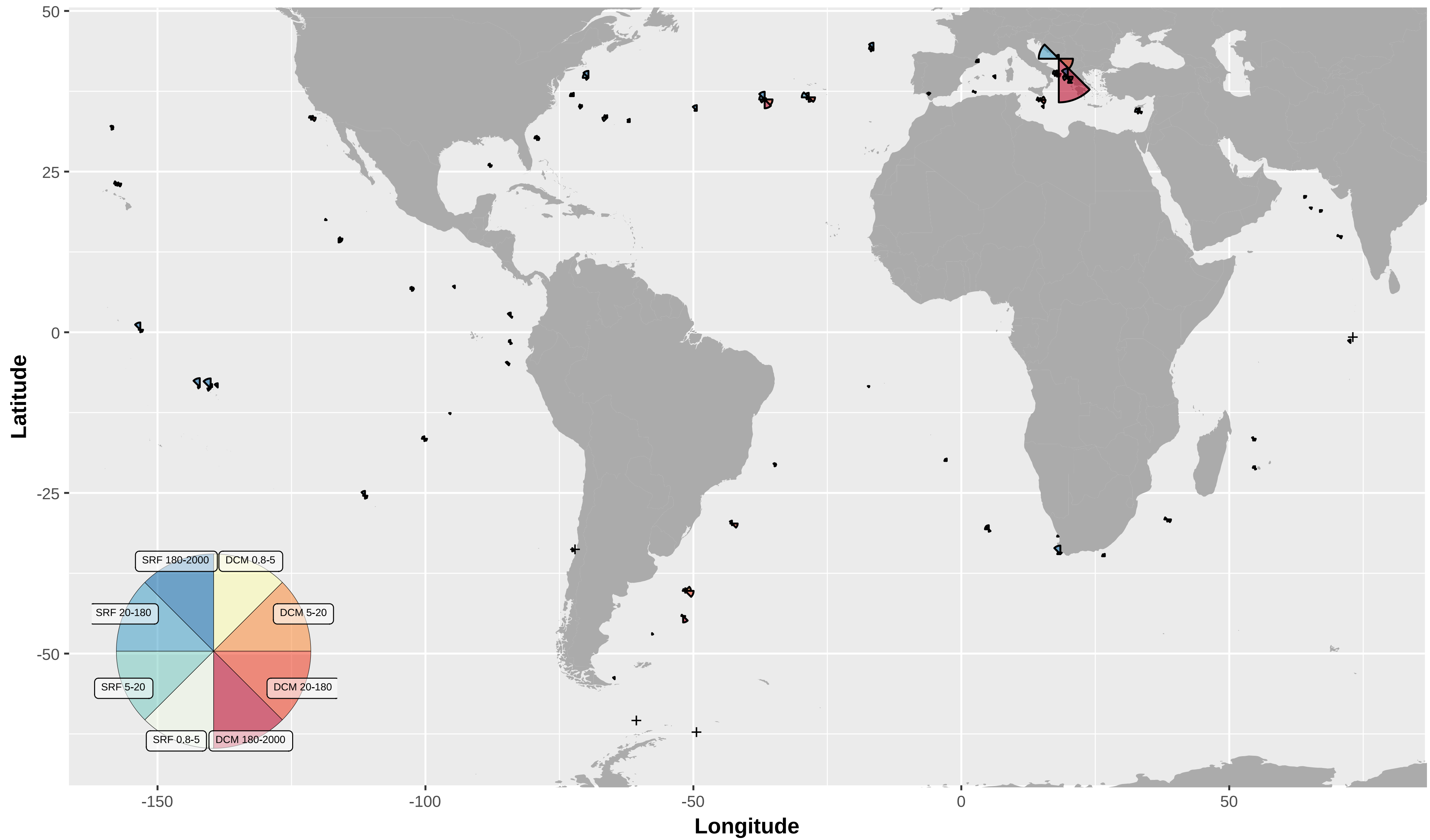


root;cellular organisms;Eukaryota;Opisthokonta;Metazoa;Eumetazoa;Bilateria;Protostomia;Ecdysozoa;Panarthropoda;Arthropoda;Mandibulata;Pancrustacea;Crustacea;Maxillopoda

# Biogeolocalisation of MGT-v1 580      1189 unigenes, 1117 (93.94%) taxonomically assigned

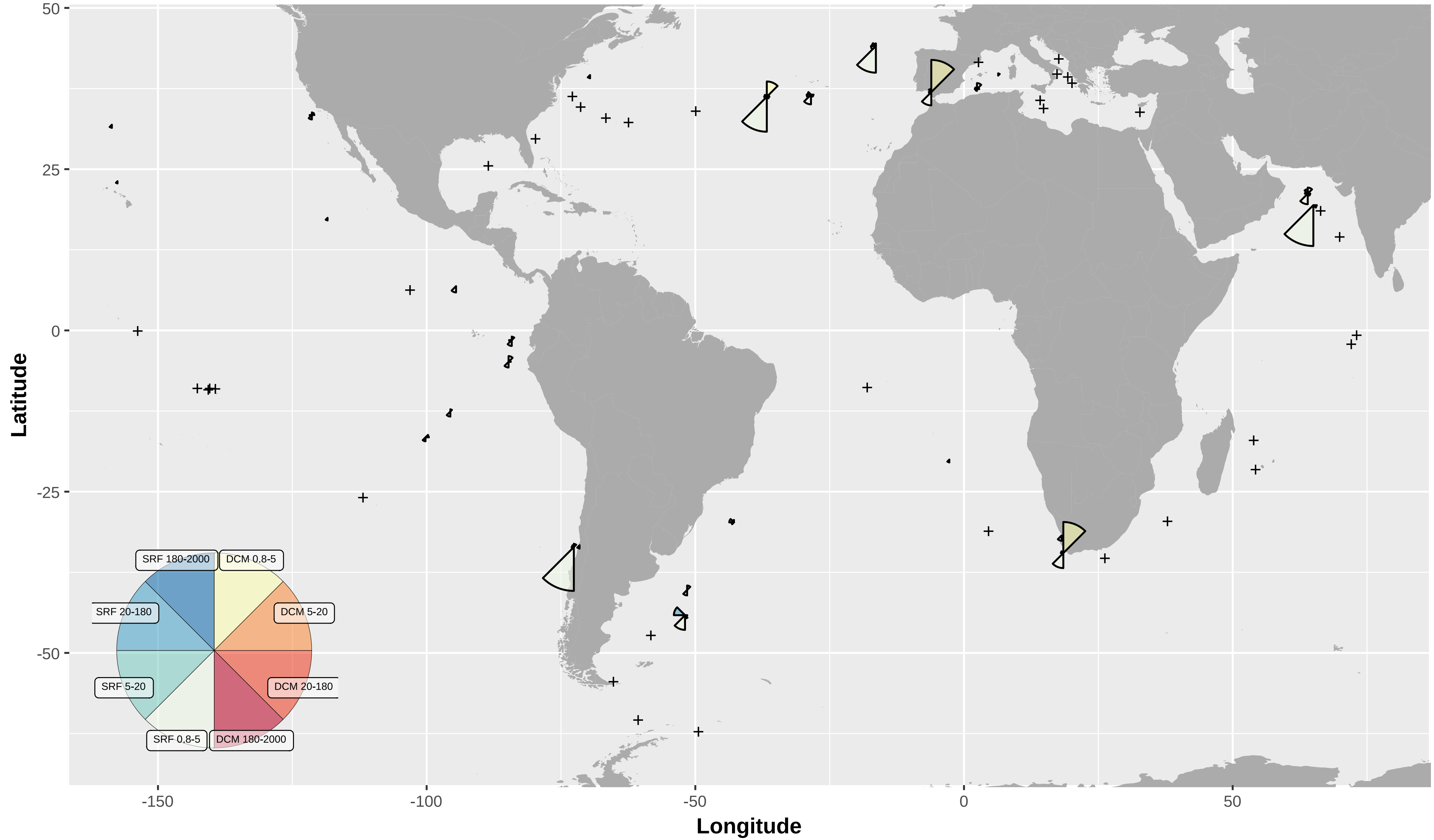


root;cellular organisms;Bacteria;Bacteroidetes/Chlorobi group;Bacteroidetes;Flavobacteriia;Flavobacteriales

## Biogeolocalisation of MGT-v1 581      1188 unigenes, 7 (0.59%) taxonomically assigned

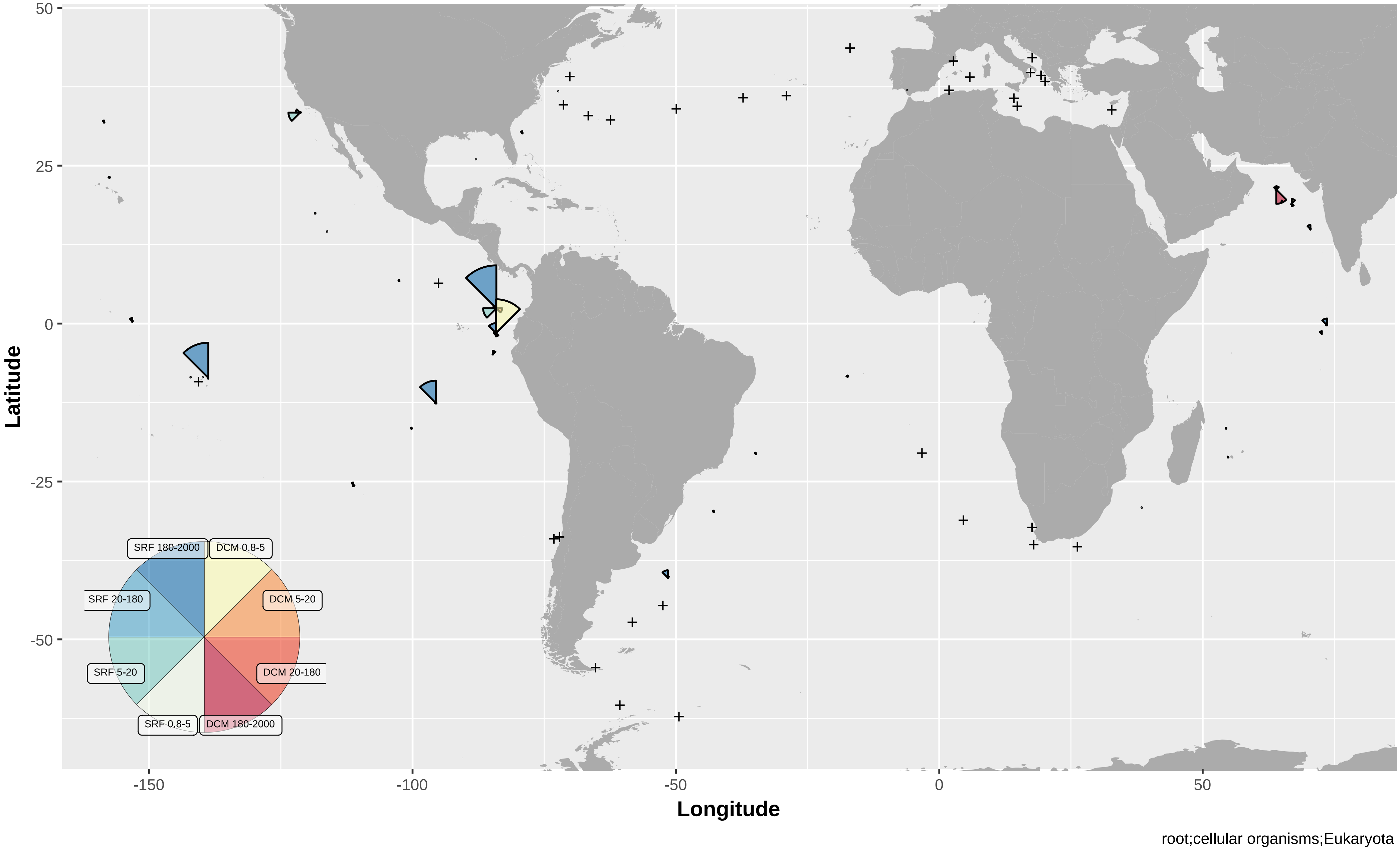

### Biogeolocalisation of MGT-v1 582 1187 unigenes, 921 (77.59%) taxonomically assigned

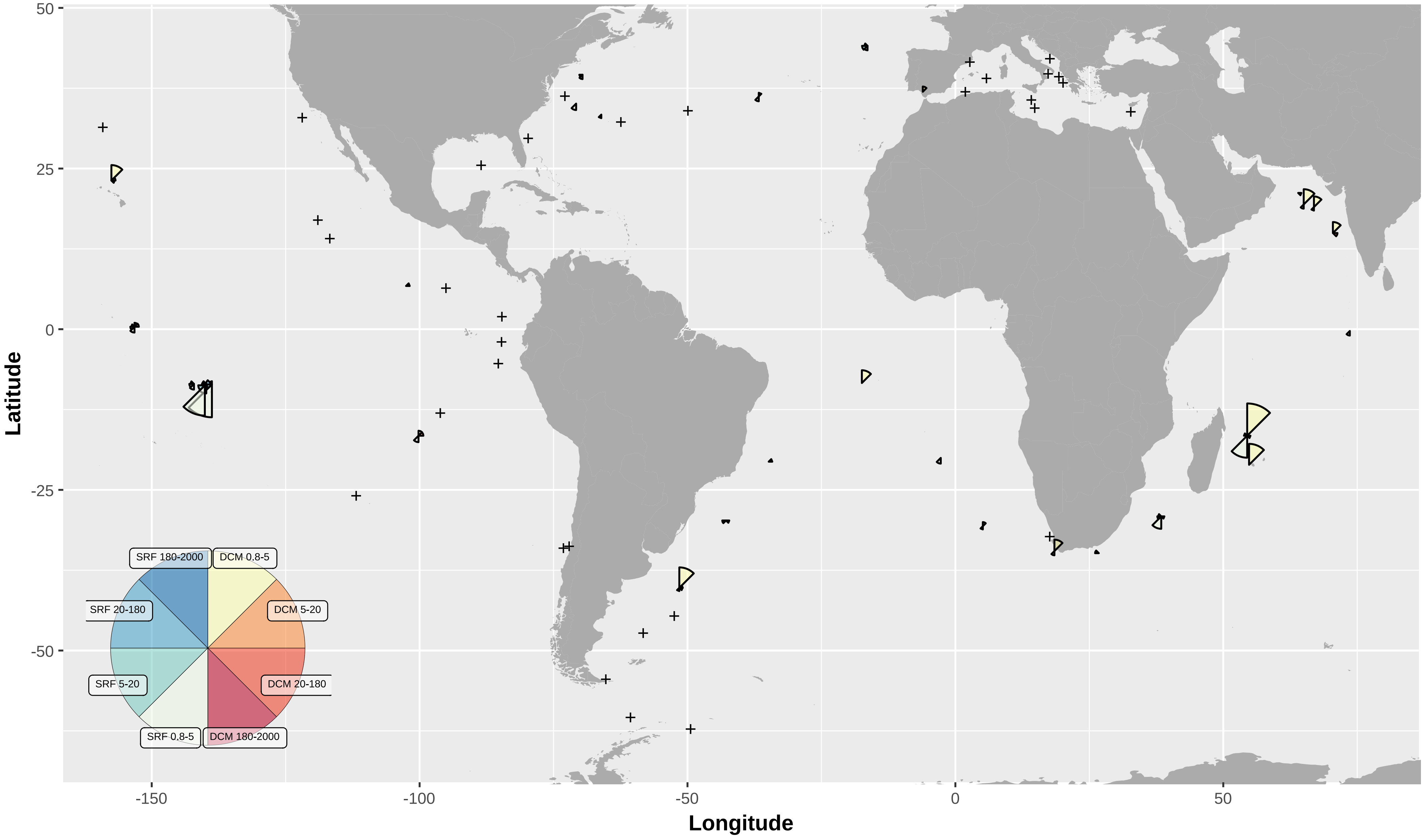


root;cellular organisms;Bacteria

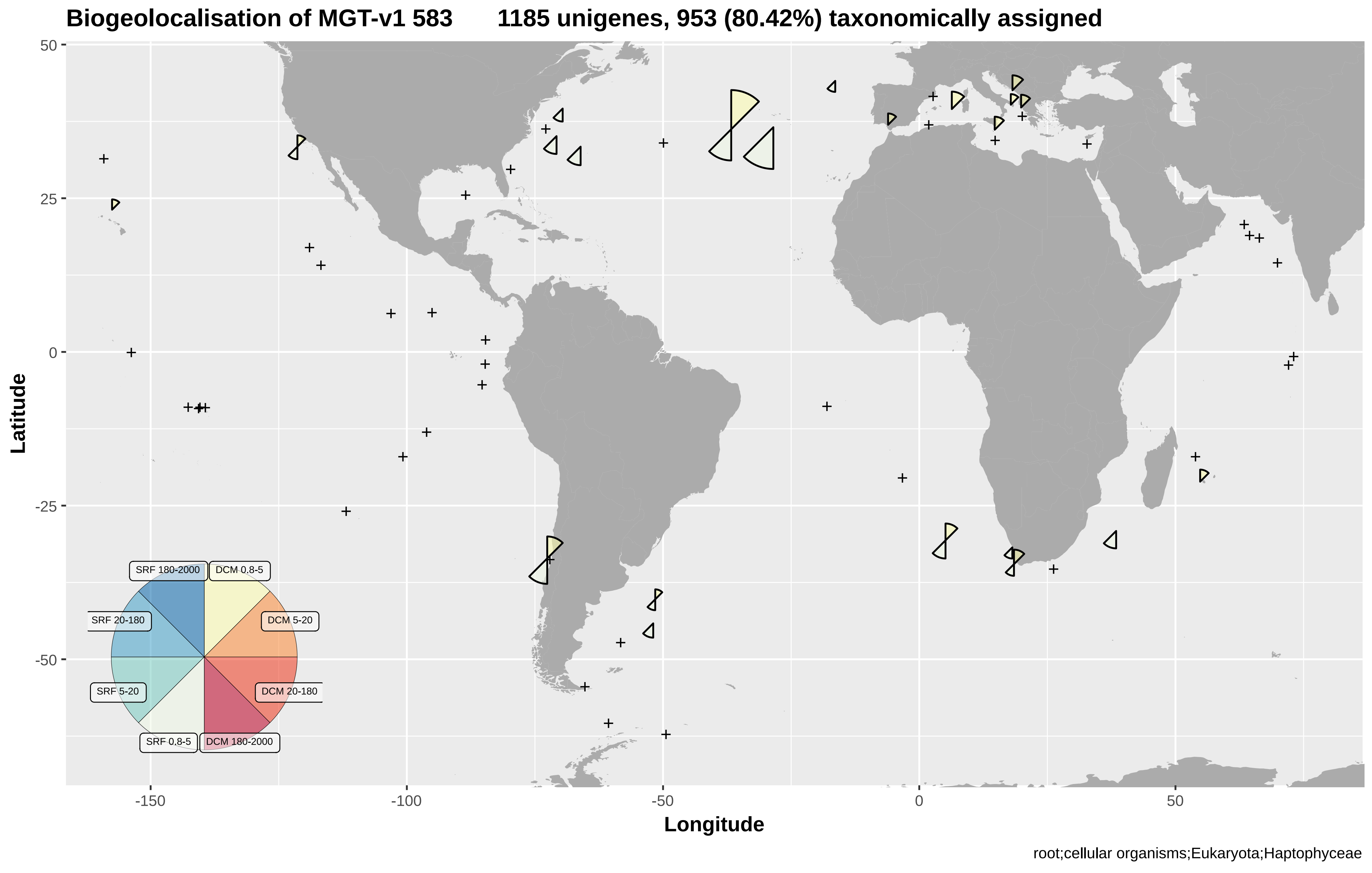
Biogeolocalisation of MGT-v1 583 1185 unigenes, 953 (80.42%) taxonomically assigned
Latitude
Longitude
50
25
0
-25
-50
-150
-100
-50
0
50
SRF 180-2000
DCM 0.8-5
SRF 20-180
DCM 5-20
SRF 5-20
DCM 20-180
SRF 0.8-5
DCM 180-2000
root;cellular organisms;Eukaryota;Haptophyceae

# Biogeolocalisation of MGT-v1 584      1175 unigenes, 150 (12.77%) taxonomically assigned

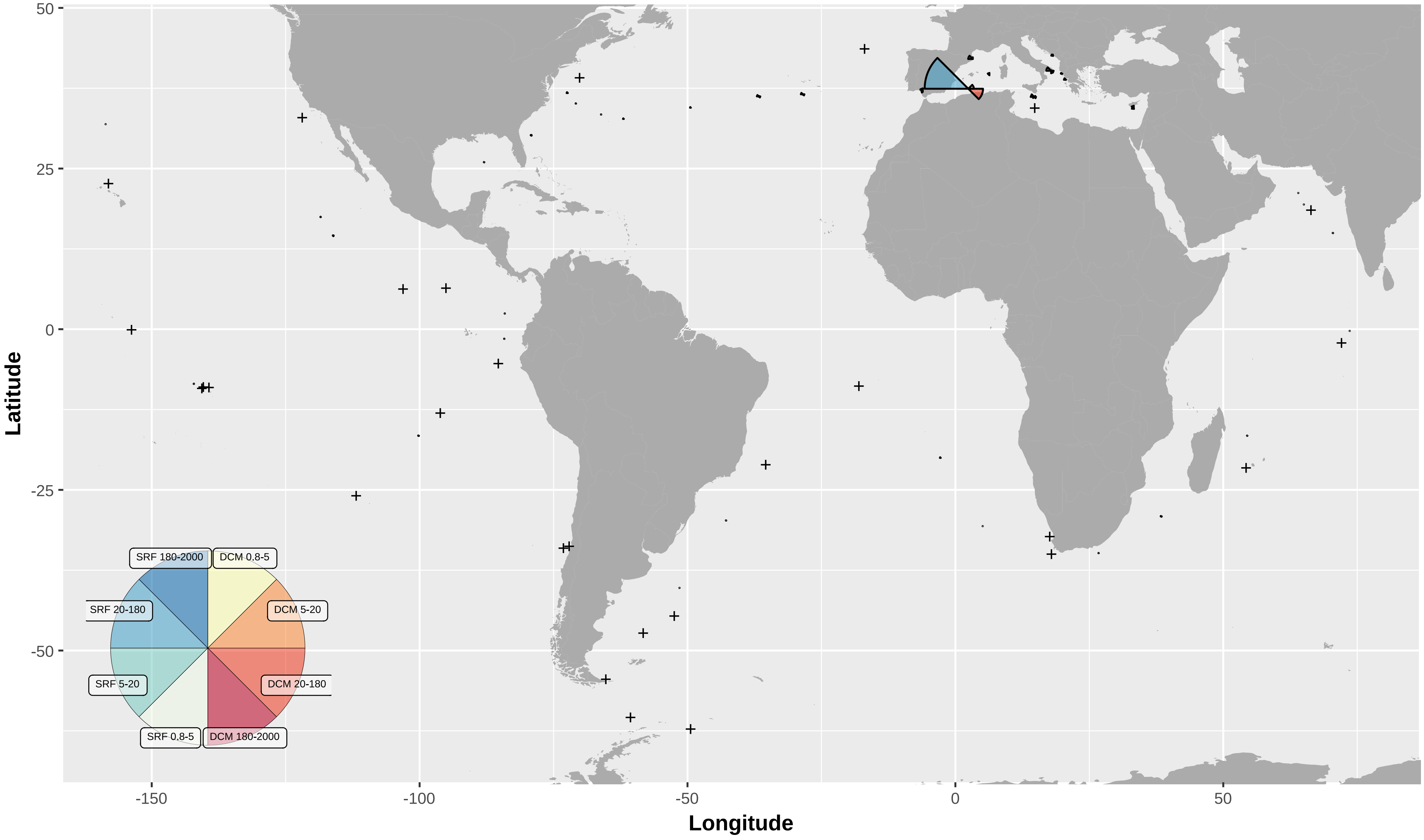


root;cellular organisms;Eukaryota

#### Biogeolocalisation of MGT-v1 585 1175 unigenes, 157 (13.36%) taxonomically assigned

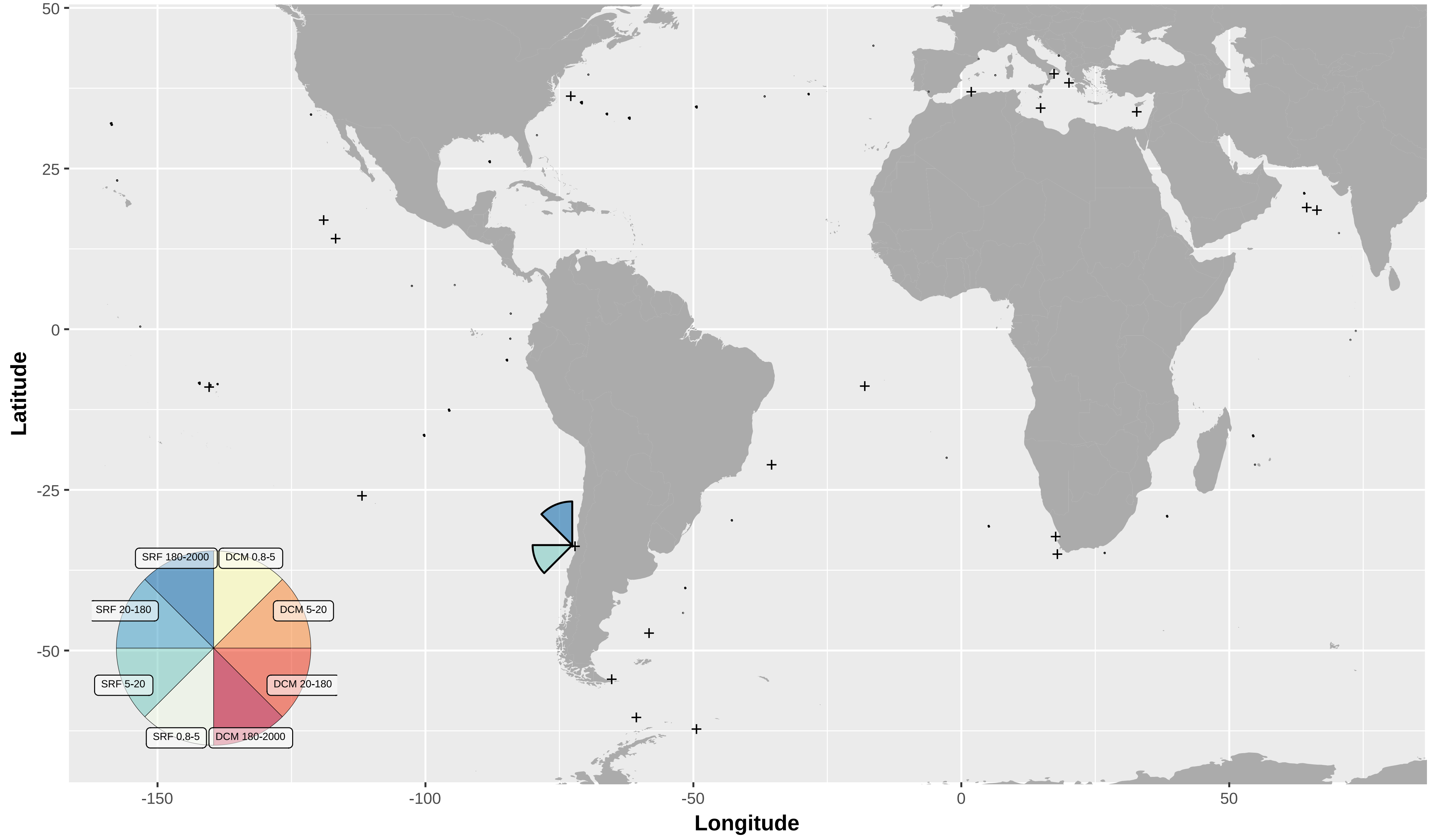


root;cellular organisms;Eukaryota

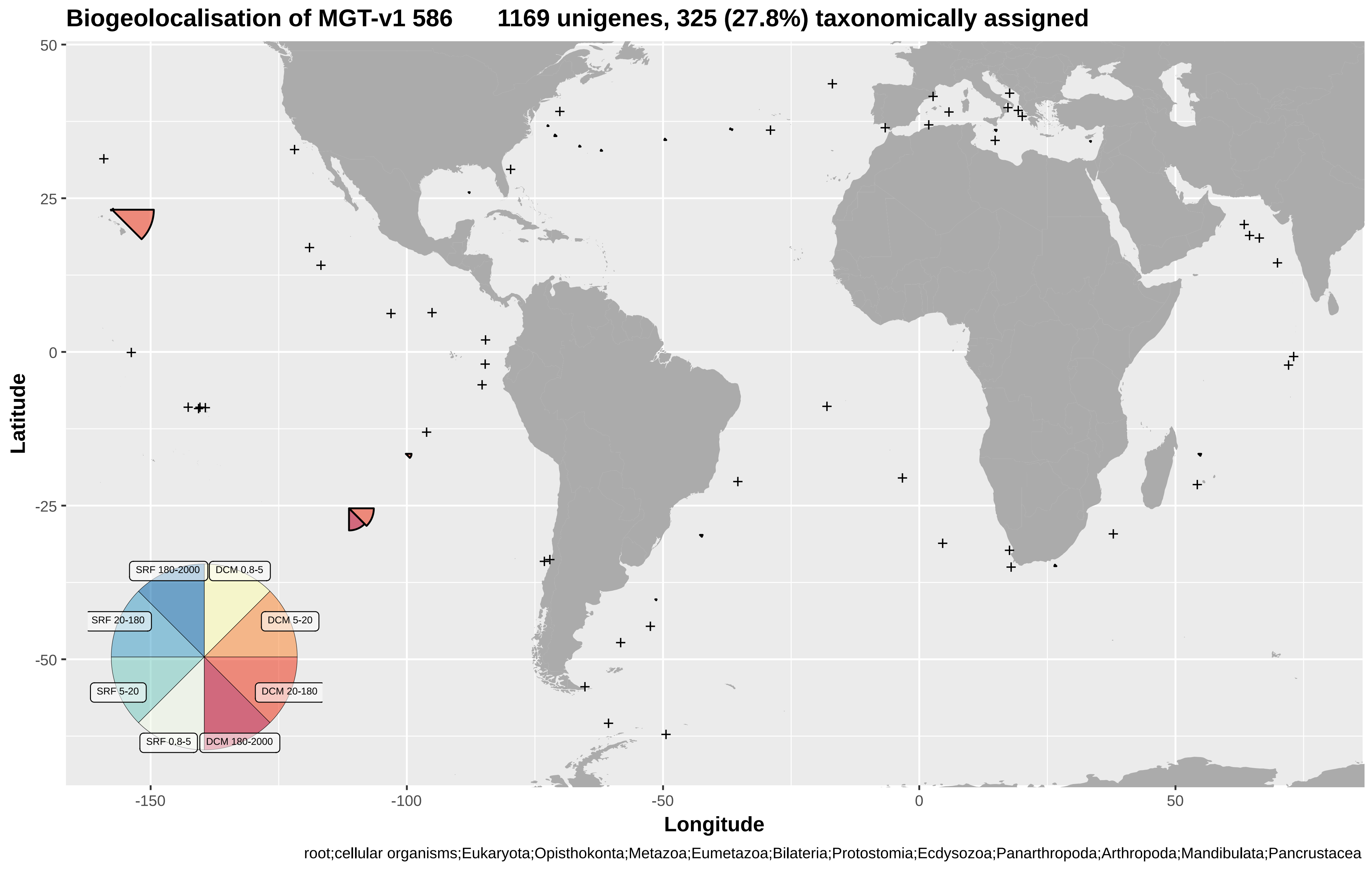

Biogeolocalisation of MGT-v1 586
1169 unigenes, 325 (27.8%) taxonomically assigned
Latitude
Longitude
50
25
0
-25
-50
-150
-100
-50
0
50
SRF 180-2000
DCM 0.8-5
SRF 20-180
DCM 5-20
SRF 5-20
DCM 20-180
SRF 0.8-5
DCM 180-2000
root;cellular organisms;Eukaryota;Opisthokonta;Metazoa;Eumetazoa;Bilateria;Protostomia;Ecdysozoa;Panarthropoda;Arthropoda;Mandibulata;Pancrustacea

#### Biogeolocalisation of MGT-v1 587 1168 unigenes, 1120 (95.89%) taxonomically assigned

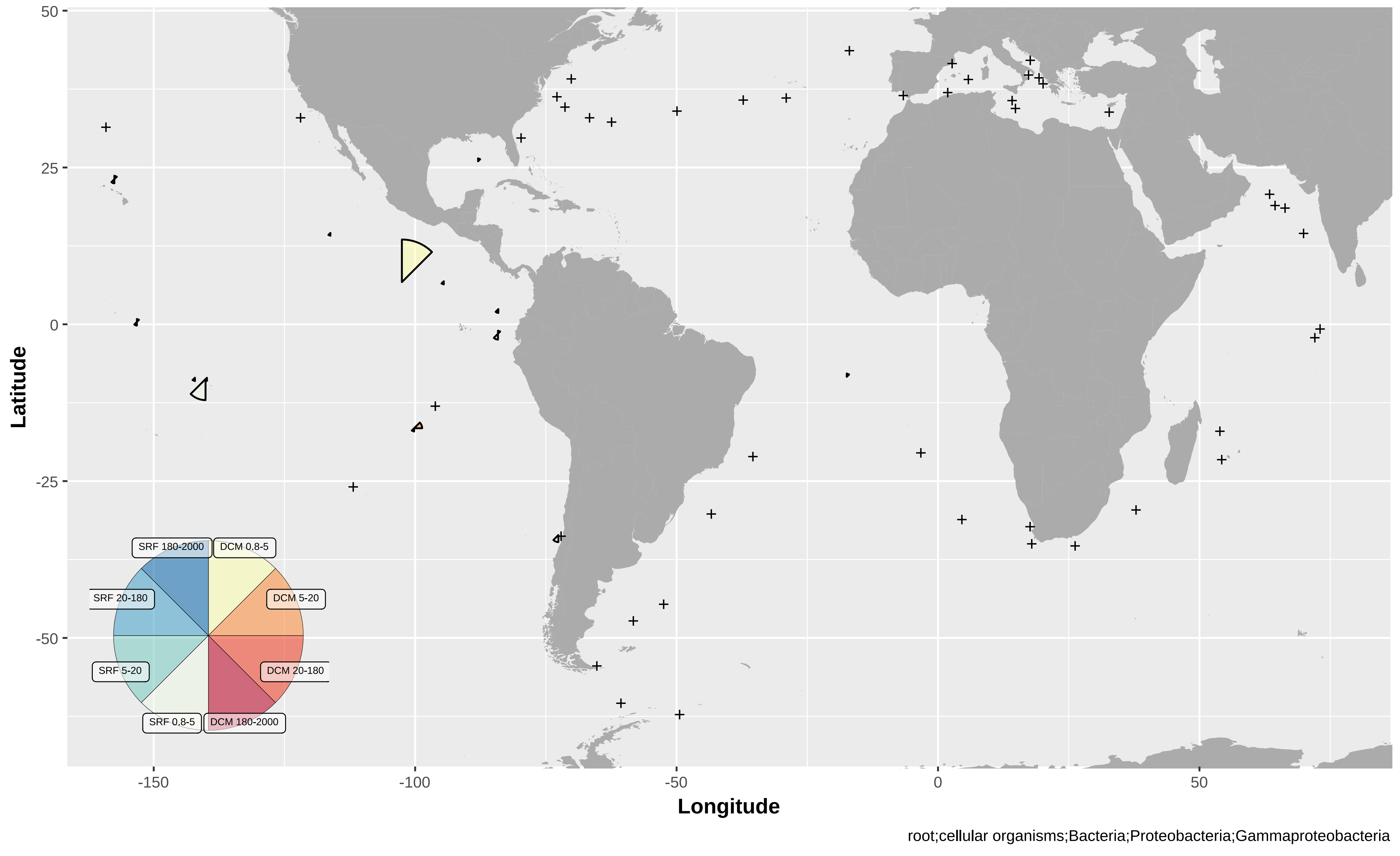

Biogeolocalisation of MGT-v1 588 1161 unigenes, 69 (5.94%) taxonomically assigned

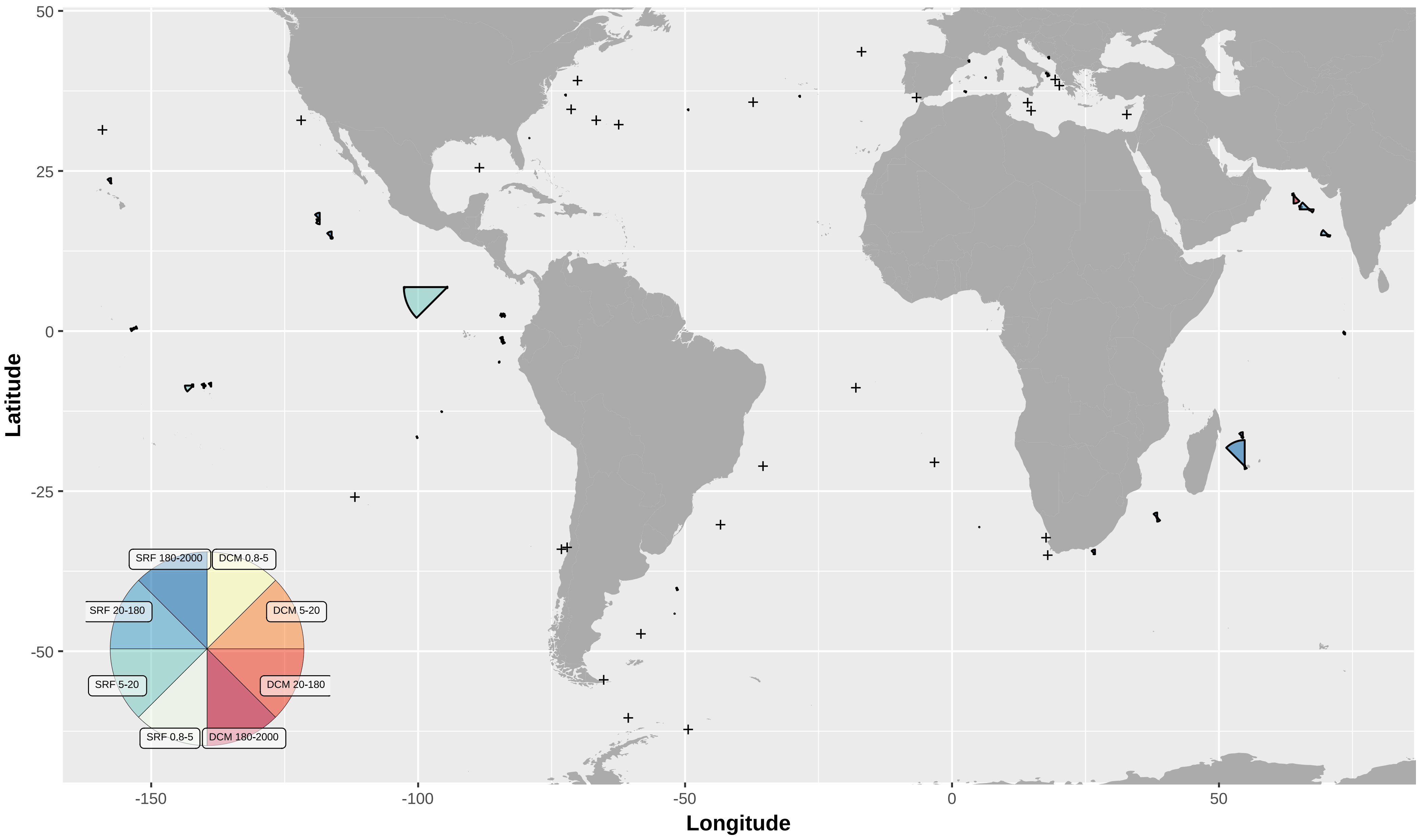


root;cellular organisms;Eukaryota;Opisthokonta;Metazoa;Eumetazoa

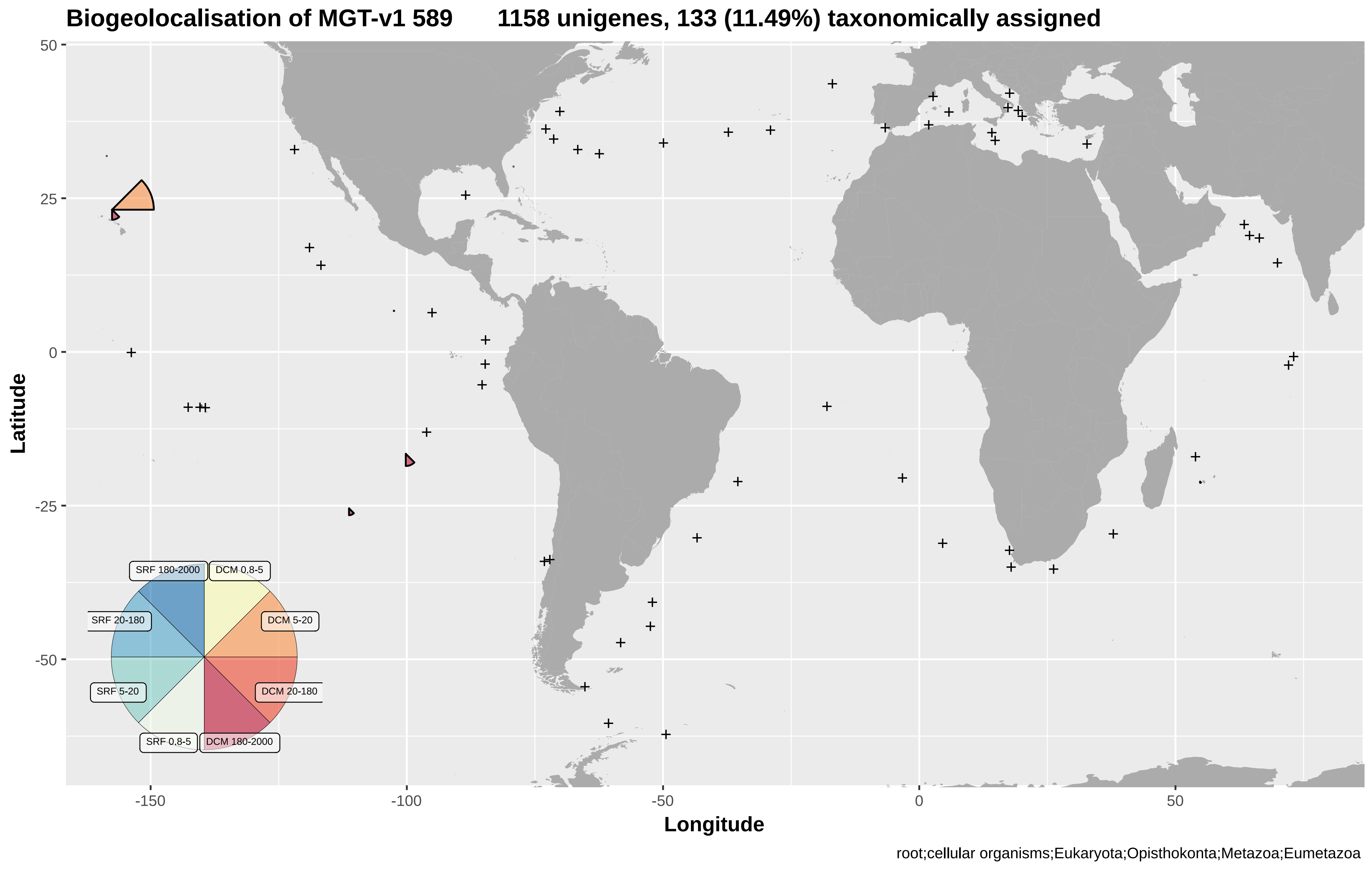

Biogeolocalisation of MGT-v1 589 1158 unigenes, 133 (11.49%) taxonomically assigned
Latitude
Longitude
50
25
0
-25
-50
-150
-100
-50
0
50
SRF 180-2000
DCM 0.8-5
SRF 20-180
DCM 5-20
SRF 5-20
DCM 20-180
SRF 0.8-5
DCM 180-2000
root;cellular organisms;Eukaryota;Opisthokonta;Metazoa;Eumetazoa

Biogeolocalisation of MGT-v1 590      1157 unigenes, 111 (9.59%) taxonomically assigned

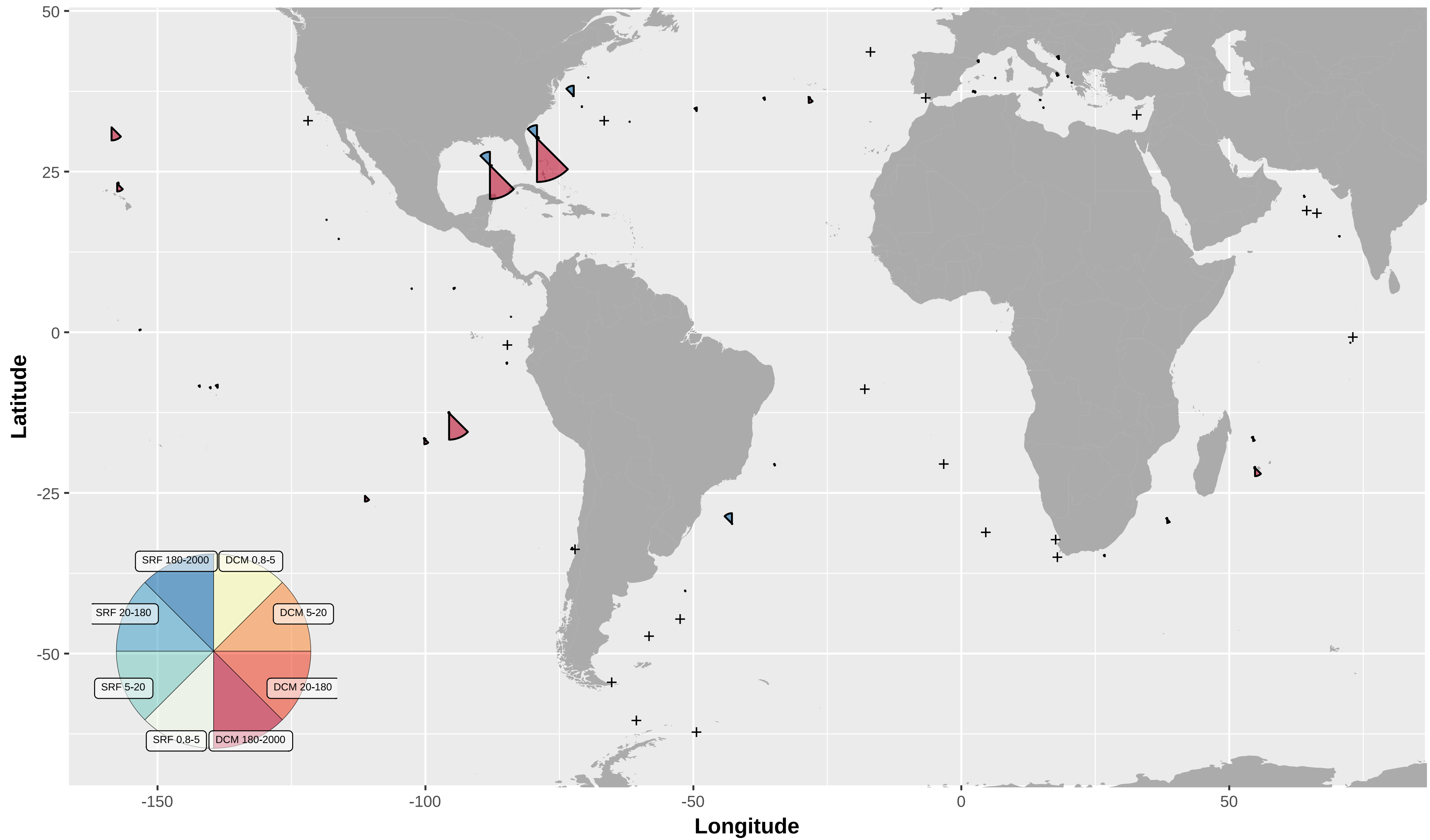


root;cellular organisms;Eukaryota;Opisthokonta;Metazoa;Eumetazoa;Bilateria

# Biogeolocalisation of MGT-v1 591      1154 unigenes, 76 (6.59%) taxonomically assigned

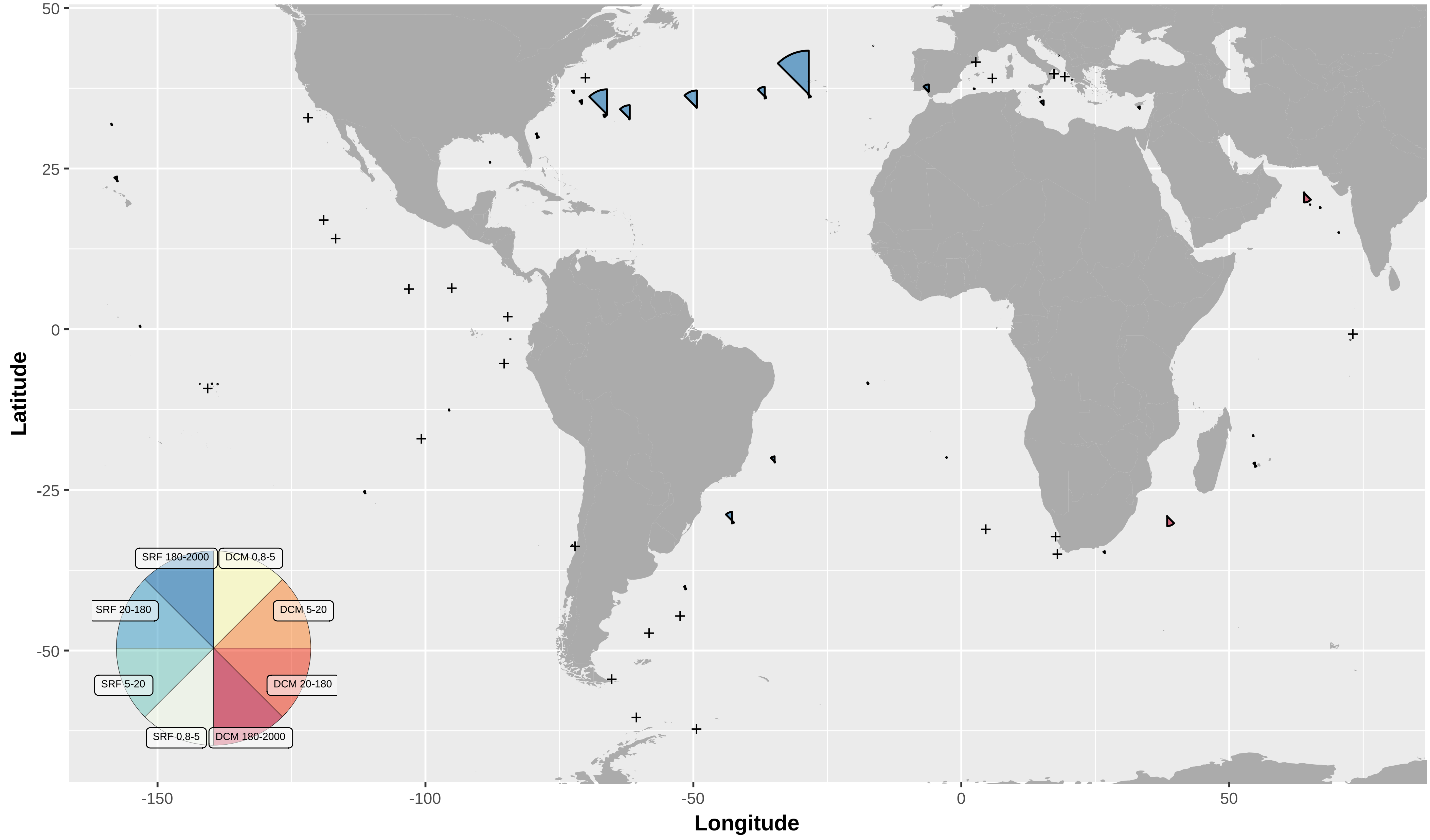


root;cellular organisms;Eukaryota;Opisthokonta;Metazoa;Eumetazoa

Biogeolocalisation of MGT-v1 592 1153 unigenes, 132 (11.45%) taxonomically assigned

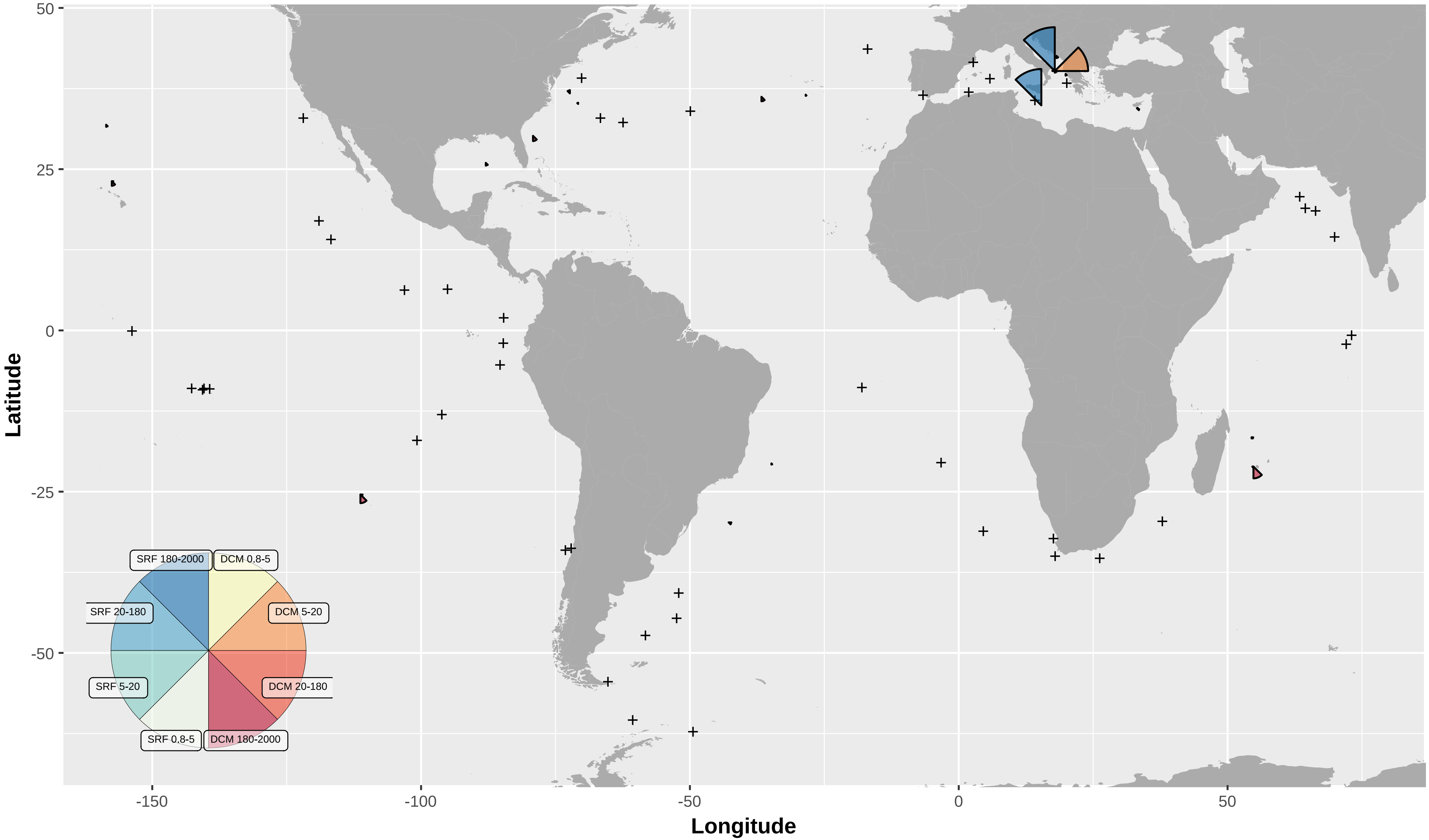


root;cellular organisms;Eukaryota;Opisthokonta;Metazoa;Eumetazoa;Bilateria

## Biogeolocalisation of MGT-v1 593      1147 unigenes, 1040 (90.67%) taxonomically assigned

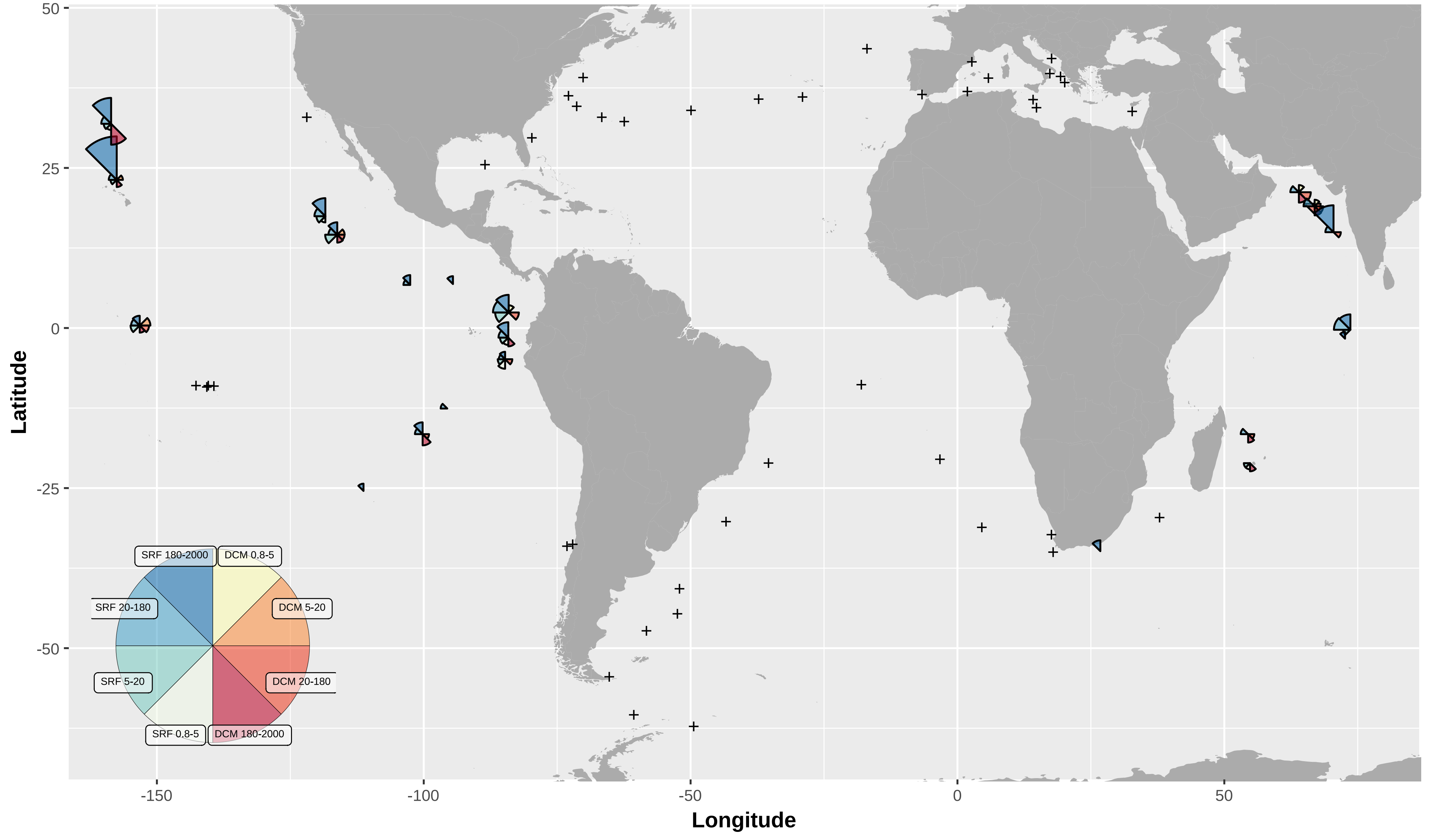


root;cellular organisms;Eukaryota;Opisthokonta;Metazoa;Eumetazoa;Bilateria;Deuterostomia;Chordata;Tunicata;Appendicularia

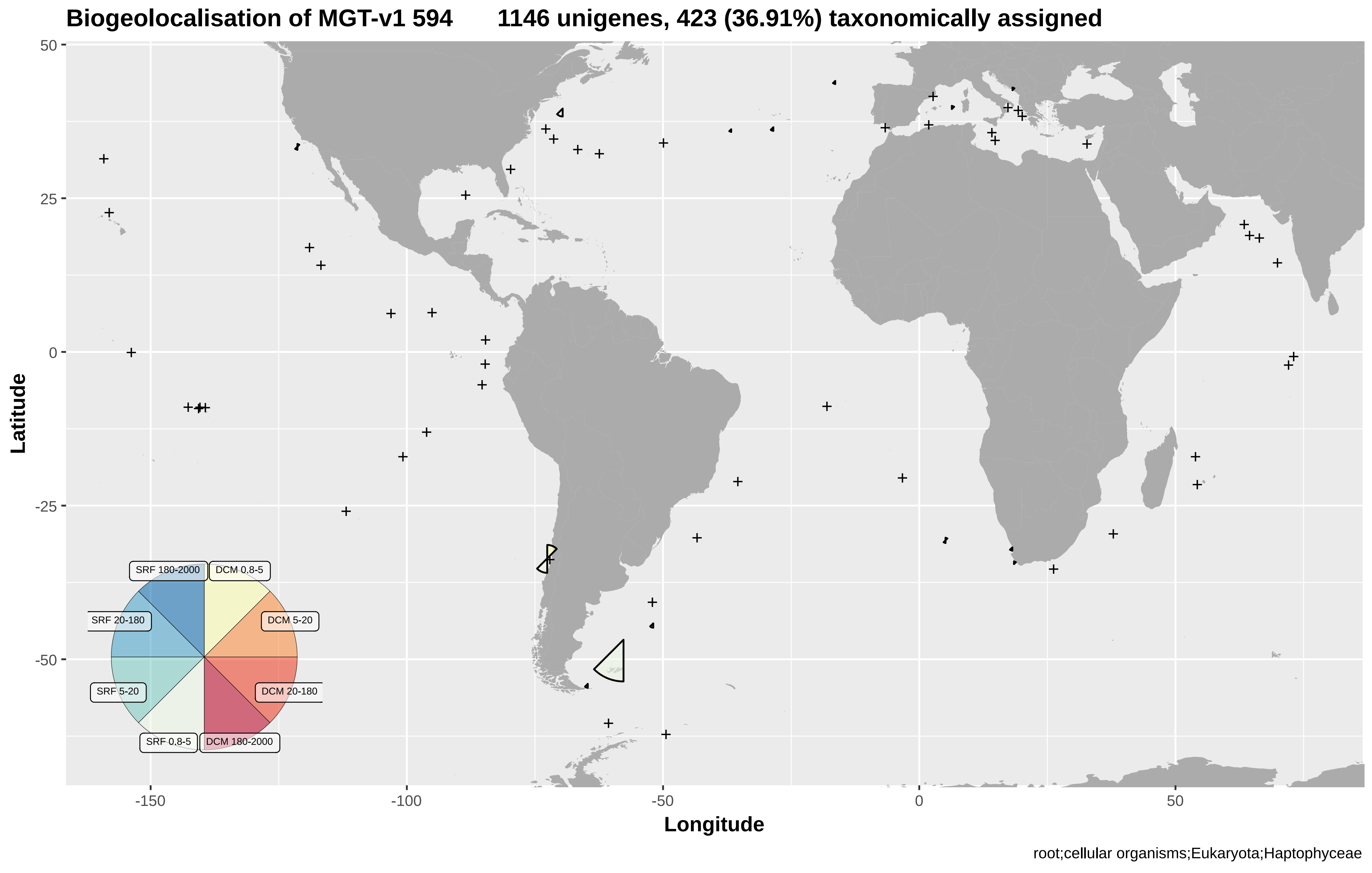
Biogeolocalisation of MGT-v1 594 1146 unigenes, 423 (36.91%) taxonomically assigned
Latitude
Longitude
50
25
0
-25
-50
-150
-100
-50
0
50
SRF 180-2000
DCM 0.8-5
SRF 20-180
DCM 5-20
SRF 5-20
DCM 20-180
SRF 0.8-5
DCM 180-2000
root;cellular organisms;Eukaryota;Haptophyceae

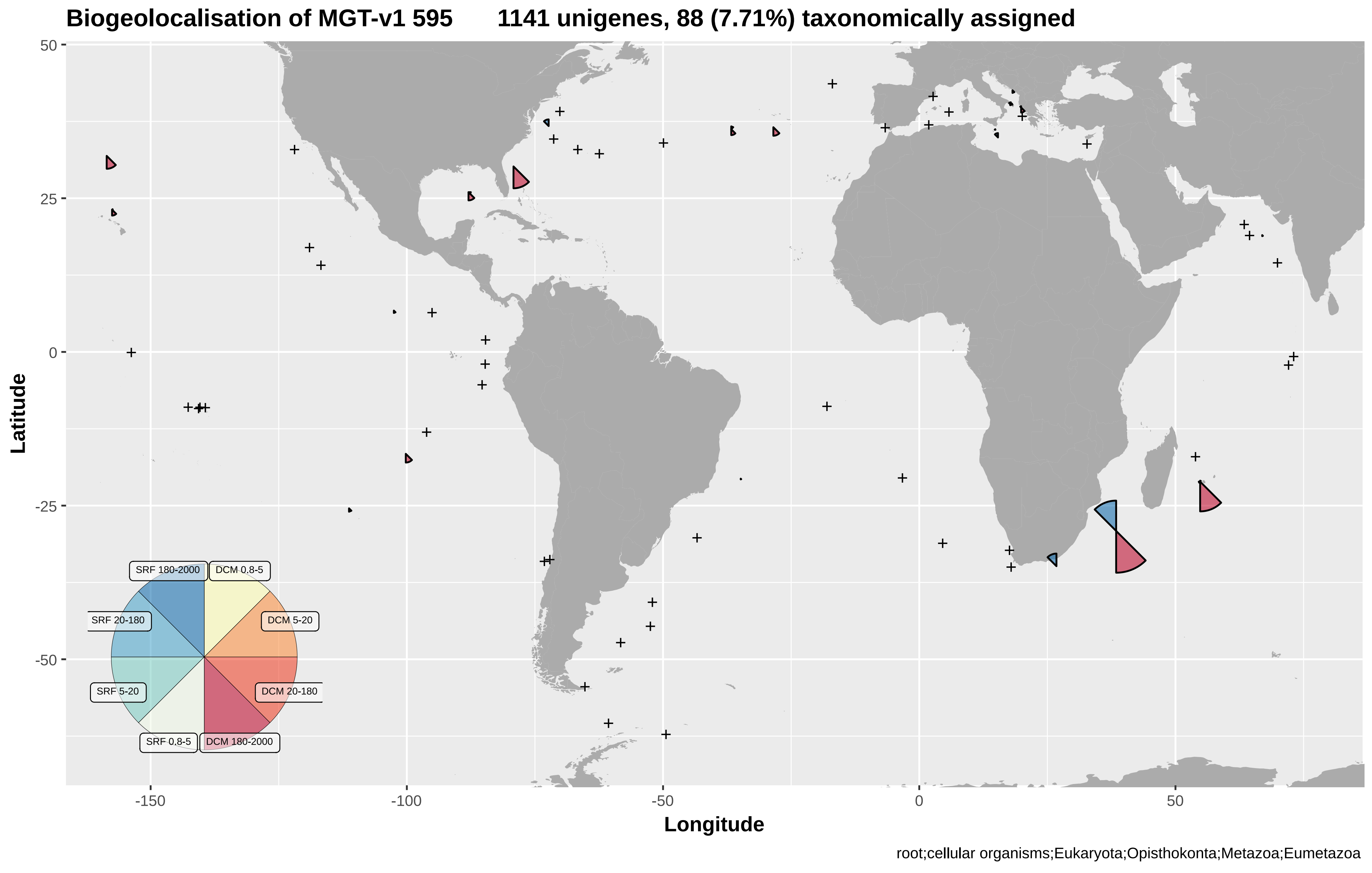
Biogeolocalisation of MGT-v1 595 1141 unigenes, 88 (7.71%) taxonomically assigned
50
25
0
-25
-50
Latitude
-150
-100
-50
0
50
Longitude
SRF 180-2000
DCM 0.8-5
SRF 20-180
DCM 5-20
SRF 5-20
DCM 20-180
SRF 0.8-5
DCM 180-2000
root;cellular organisms;Eukaryota;Opisthokonta;Metazoa;Eumetazoa

#### Biogeolocalisation of MGT-v1 596 1137 unigenes, 533 (46.88%) taxonomically assigned

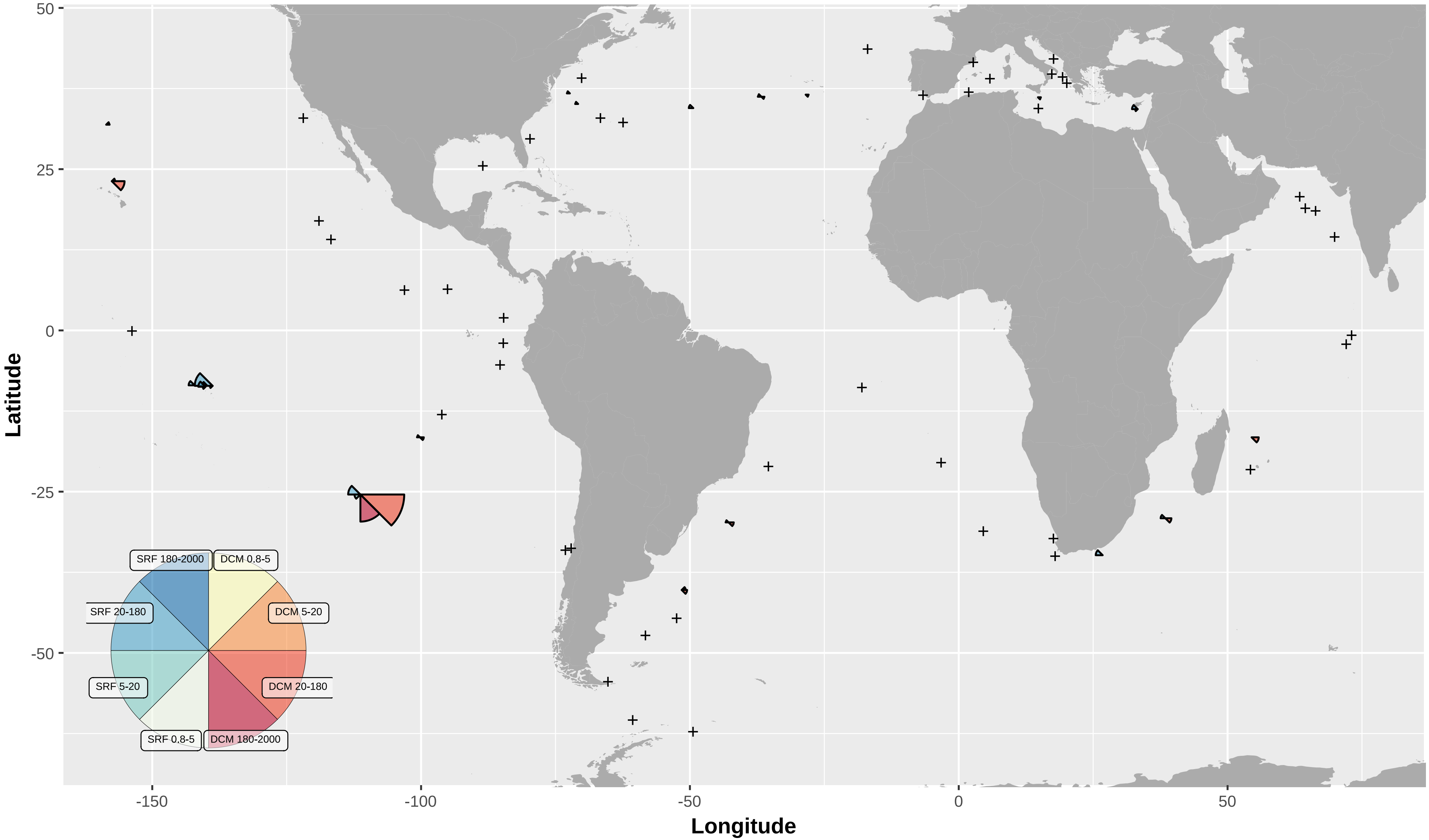


root;cellular organisms;Eukaryota;Opisthokonta;Metazoa;Eumetazoa;Bilateria;Protostomia;Ecdysozoa;Panarthropoda;Arthropoda;Mandibulata;Pancrustacea;Crustacea;Maxillopoda

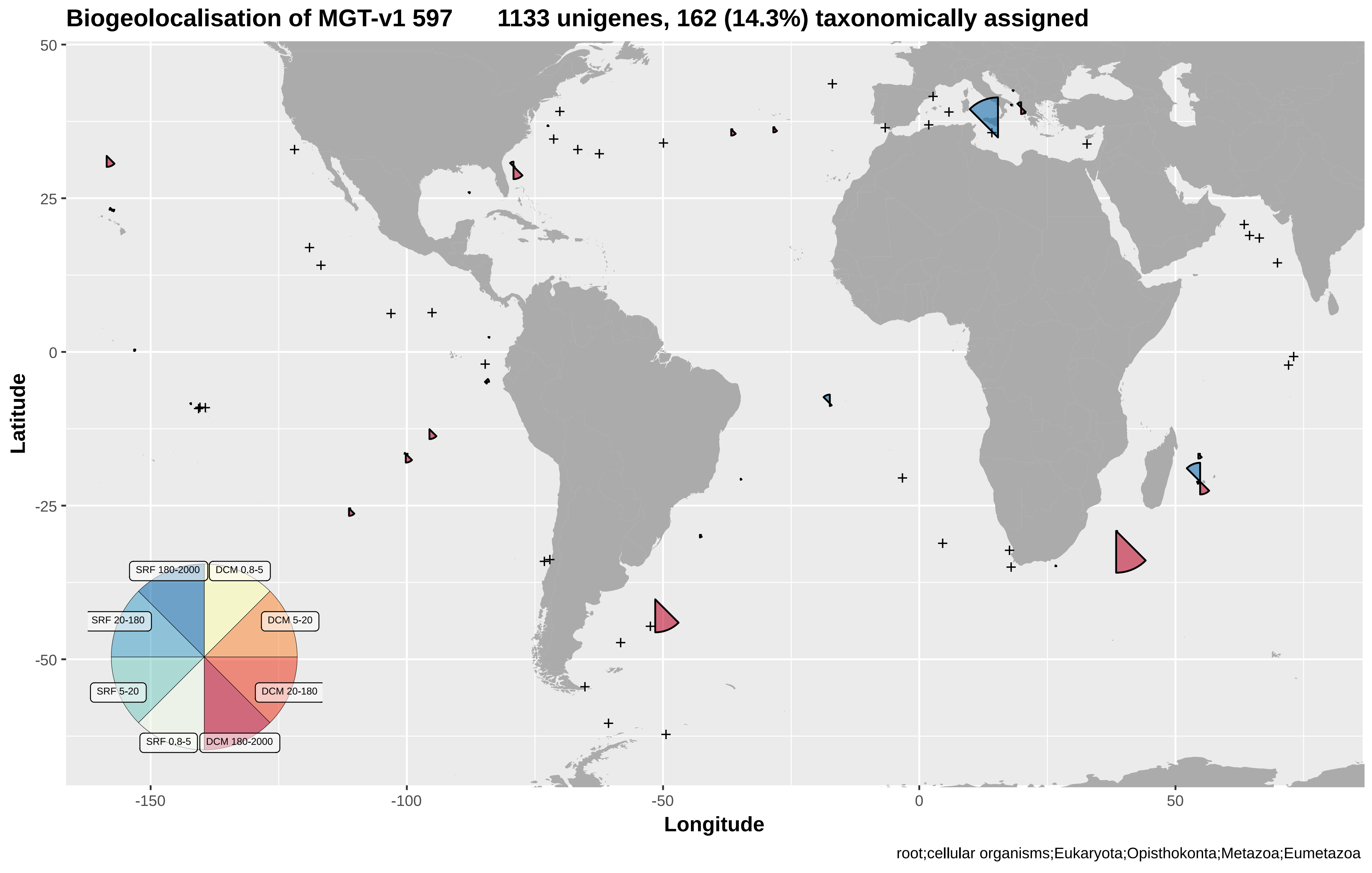
Biogeolocalisation of MGT-v1 597
1133 unigenes, 162 (14.3%) taxonomically assigned
Latitude
Longitude
50
25
0
-25
-50
-150
-100
-50
0
50
SRF 180-2000
DCM 0.8-5
SRF 20-180
DCM 5-20
SRF 5-20
DCM 20-180
SRF 0.8-5
DCM 180-2000
root;cellular organisms;Eukaryota;Opisthokonta;Metazoa;Eumetazoa

# Biogeolocalisation of MGT-v1 598      1132 unigenes, 324 (28.62%) taxonomically assigned

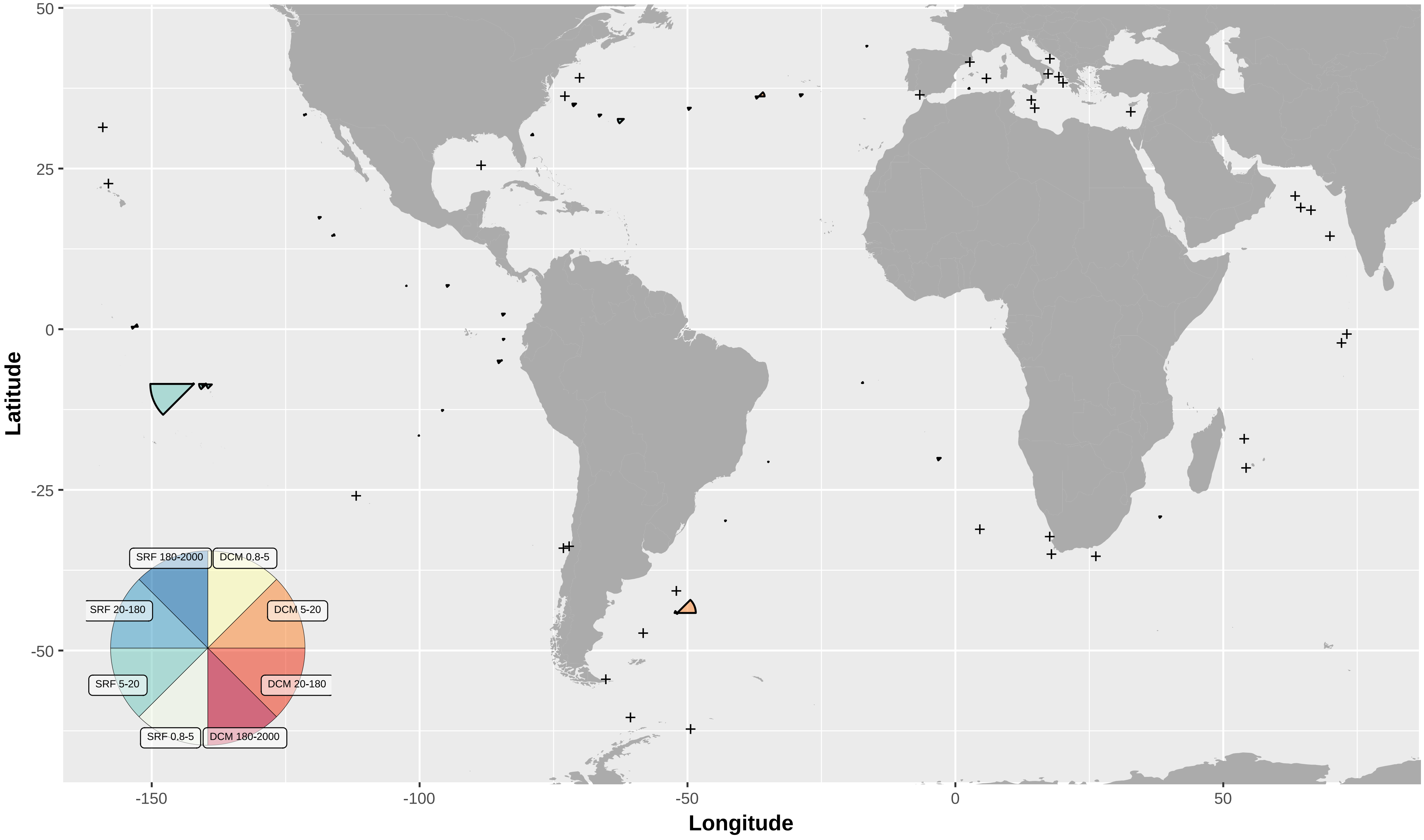


root;cellular organisms;Eukaryota;Alveolata;Dinophyceae

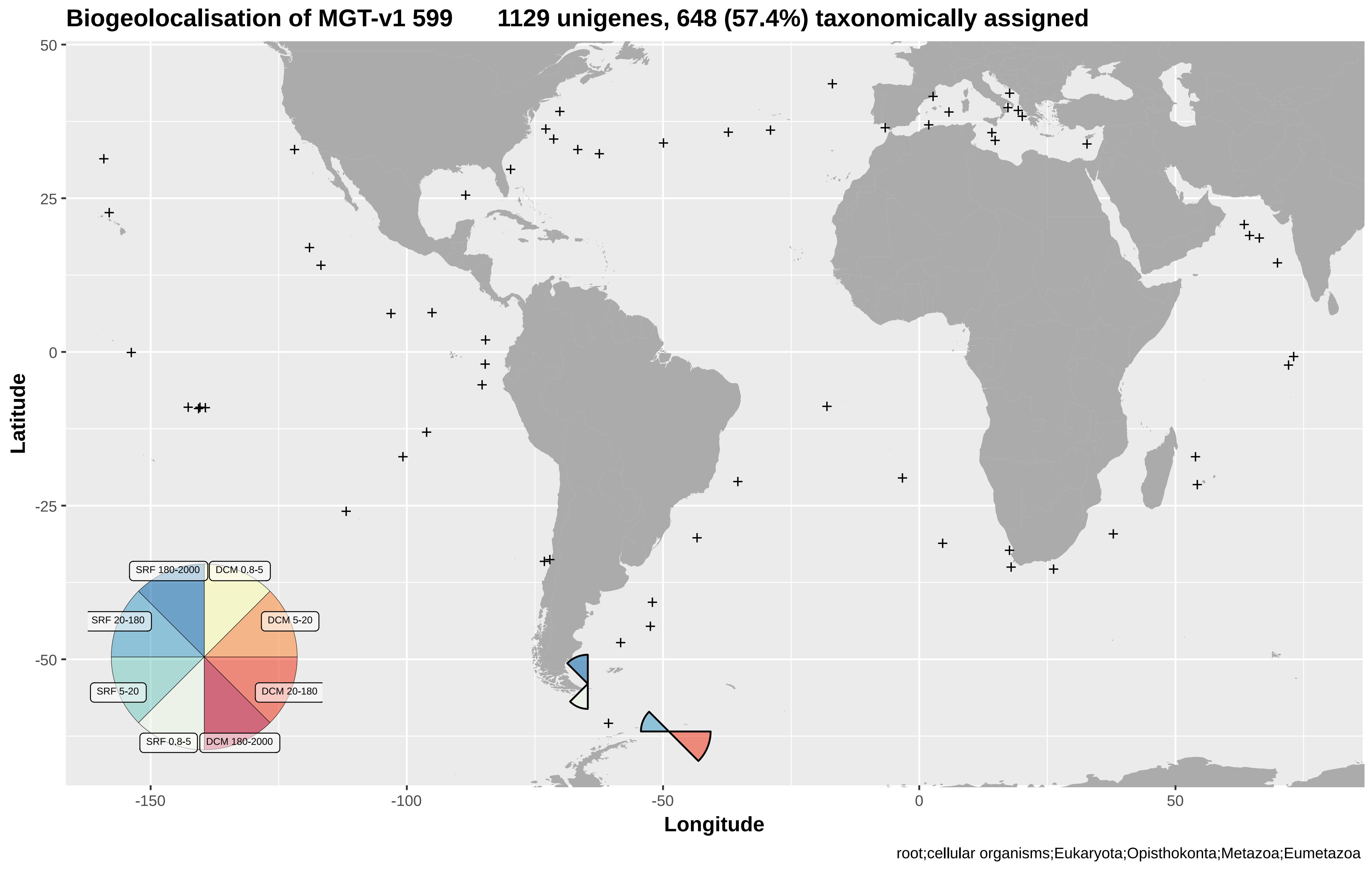
Biogeolocalisation of MGT-v1 599
1129 unigenes, 648 (57.4%) taxonomically assigned
Latitude
Longitude
50
25
0
-25
-50
-150
-100
-50
0
50
SRF 180-2000
DCM 0.8-5
SRF 20-180
DCM 5-20
SRF 5-20
DCM 20-180
SRF 0.8-5
DCM 180-2000
root;cellular organisms;Eukaryota;Opisthokonta;Metazoa;Eumetazoa

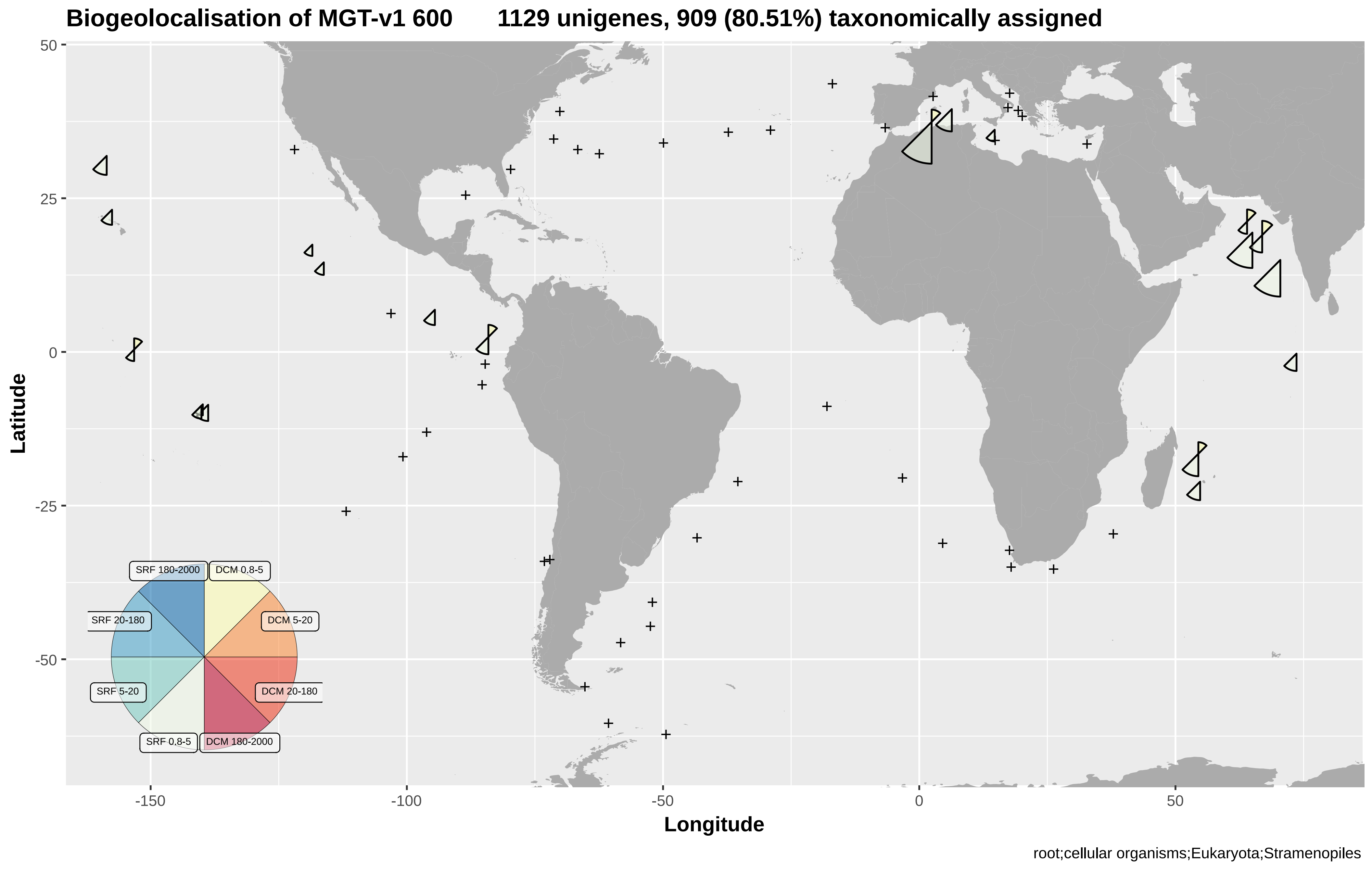

Biogeolocalisation of MGT-v1 600 1129 unigenes, 909 (80.51%) taxonomically assigned
50
25
0
-25
-50
Latitude
-150
-100
-50
0
50
Longitude
SRF 180-2000
DCM 0.8-5
SRF 20-180
DCM 5-20
SRF 5-20
DCM 20-180
SRF 0.8-5
DCM 180-2000
root;cellular organisms;Eukaryota;Stramenopiles

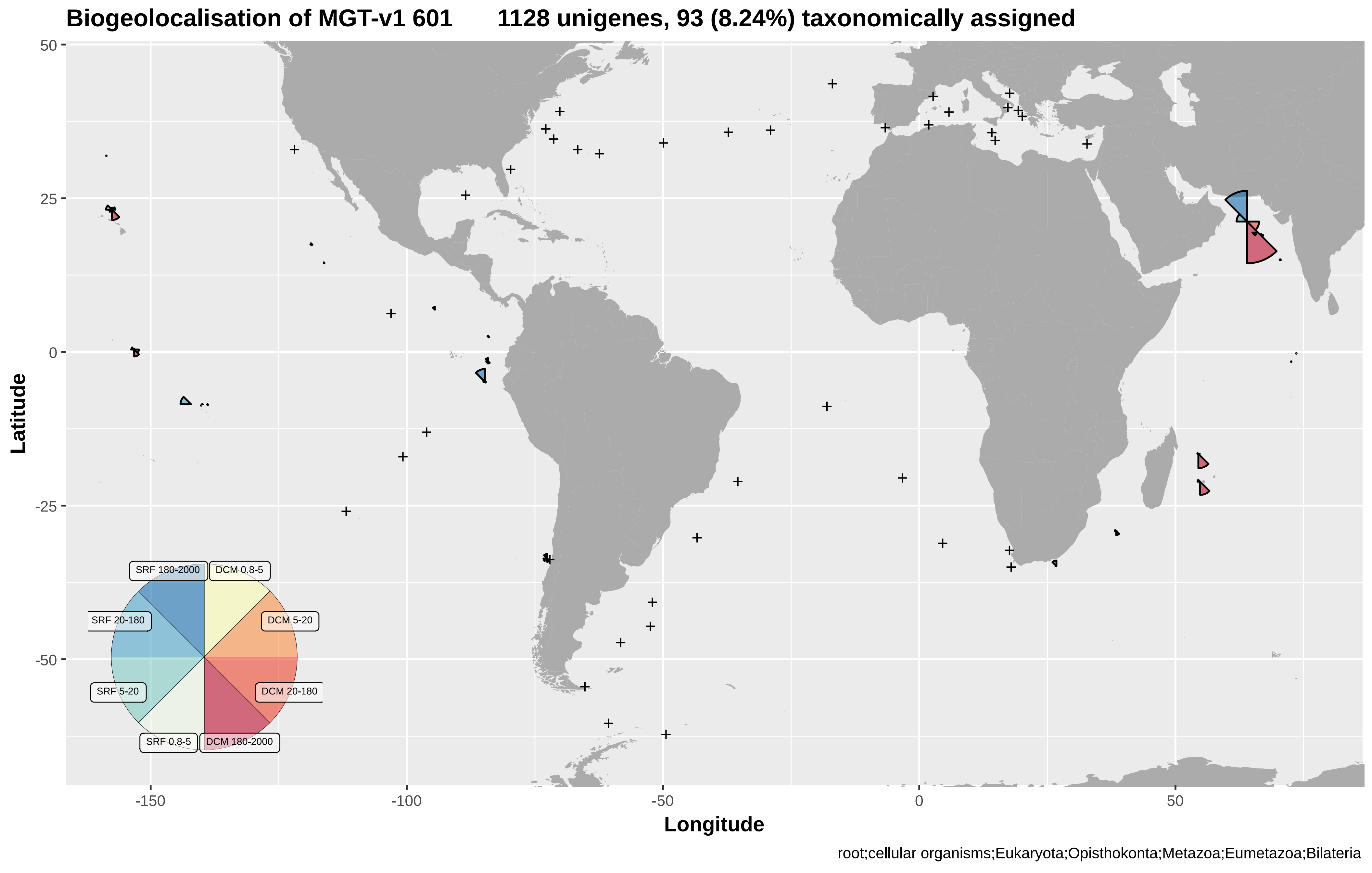
Biogeolocalisation of MGT-v1 601 1128 unigenes, 93 (8.24%) taxonomically assigned
Latitude
Longitude
50
25
0
-25
-50
-150
-100
-50
0
50
SRF 180-2000
DCM 0.8-5
SRF 20-180
DCM 5-20
SRF 5-20
DCM 20-180
SRF 0.8-5
DCM 180-2000
root;cellular organisms;Eukaryota;Opisthokonta;Metazoa;Eumetazoa;Bilateria

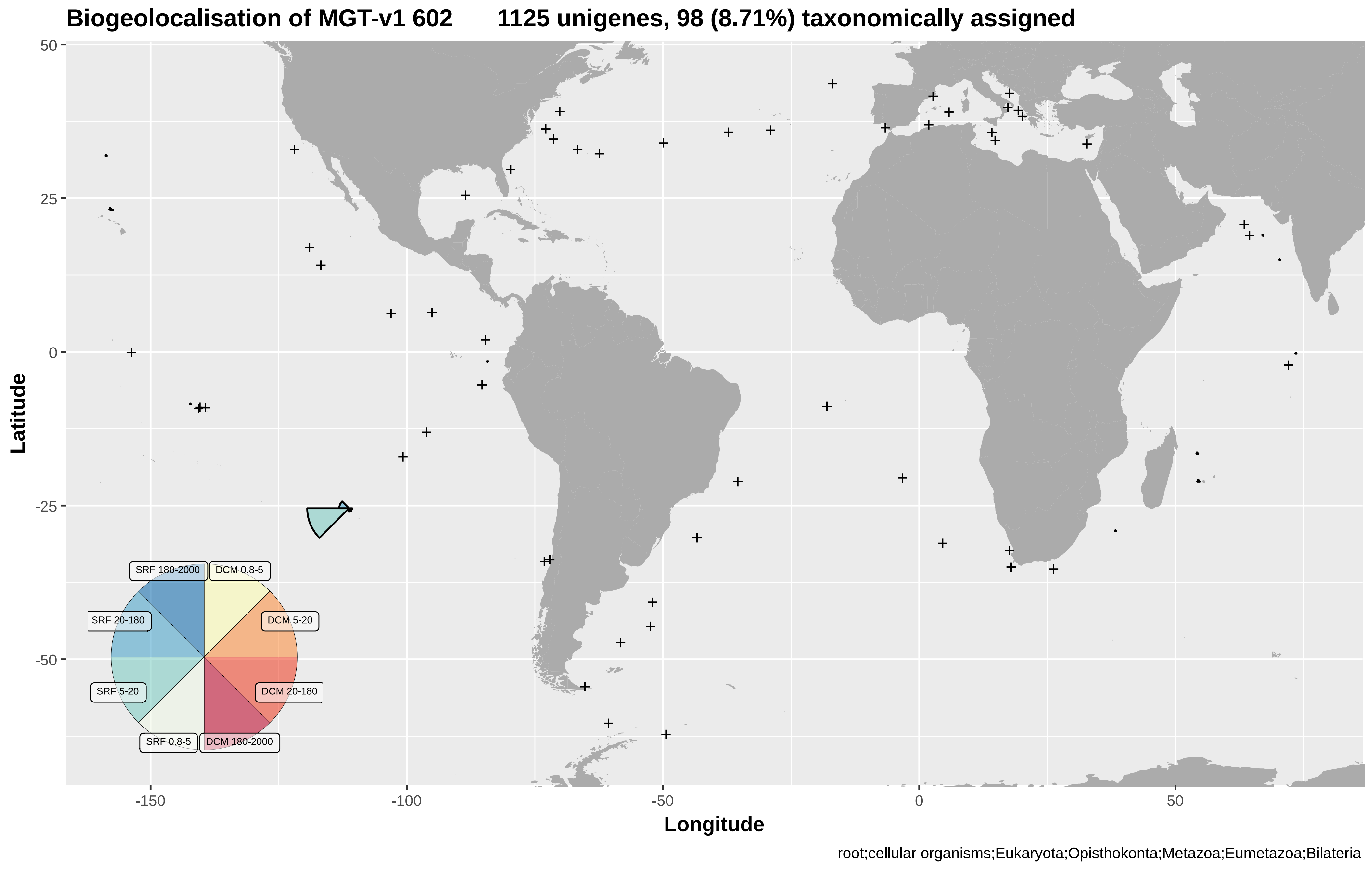

Biogeolocalisation of MGT-v1 602 1125 unigenes, 98 (8.71%) taxonomically assigned
50
25
0
-25
-50
Latitude
-150
-100
-50
0
50
Longitude
SRF 180-2000
DCM 0.8-5
SRF 20-180
DCM 5-20
SRF 5-20
DCM 20-180
SRF 0.8-5
DCM 180-2000
root;cellular organisms;Eukaryota;Opisthokonta;Metazoa;Eumetazoa;Bilateria

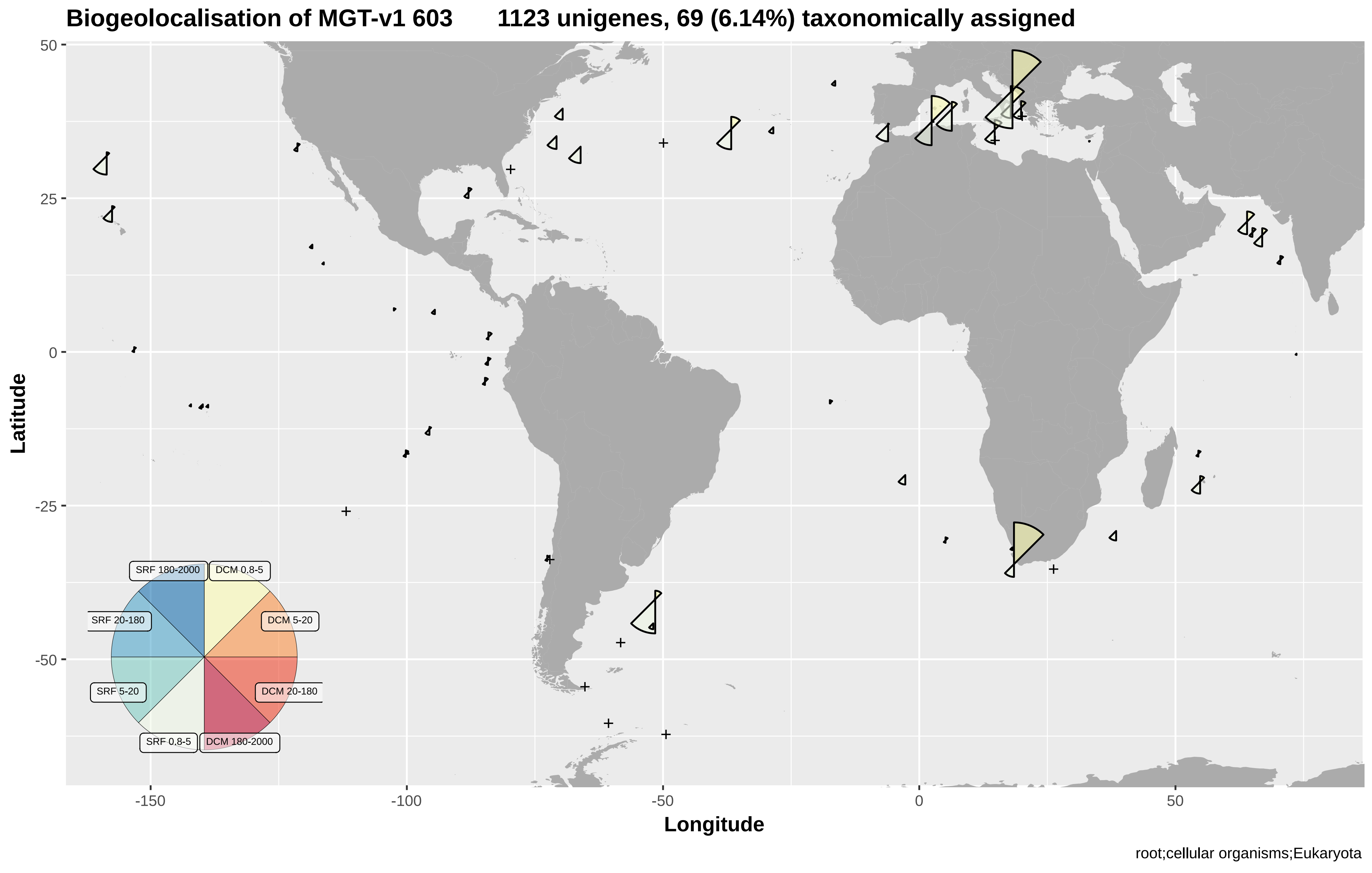

Biogeolocalisation of MGT-v1 603 1123 unigenes, 69 (6.14%) taxonomically assigned
Latitude
Longitude
50
25
0
-25
-50
-150
-100
-50
0
50
SRF 180-2000
DCM 0.8-5
SRF 20-180
DCM 5-20
SRF 5-20
DCM 20-180
SRF 0.8-5
DCM 180-2000
root;cellular organisms;Eukaryota

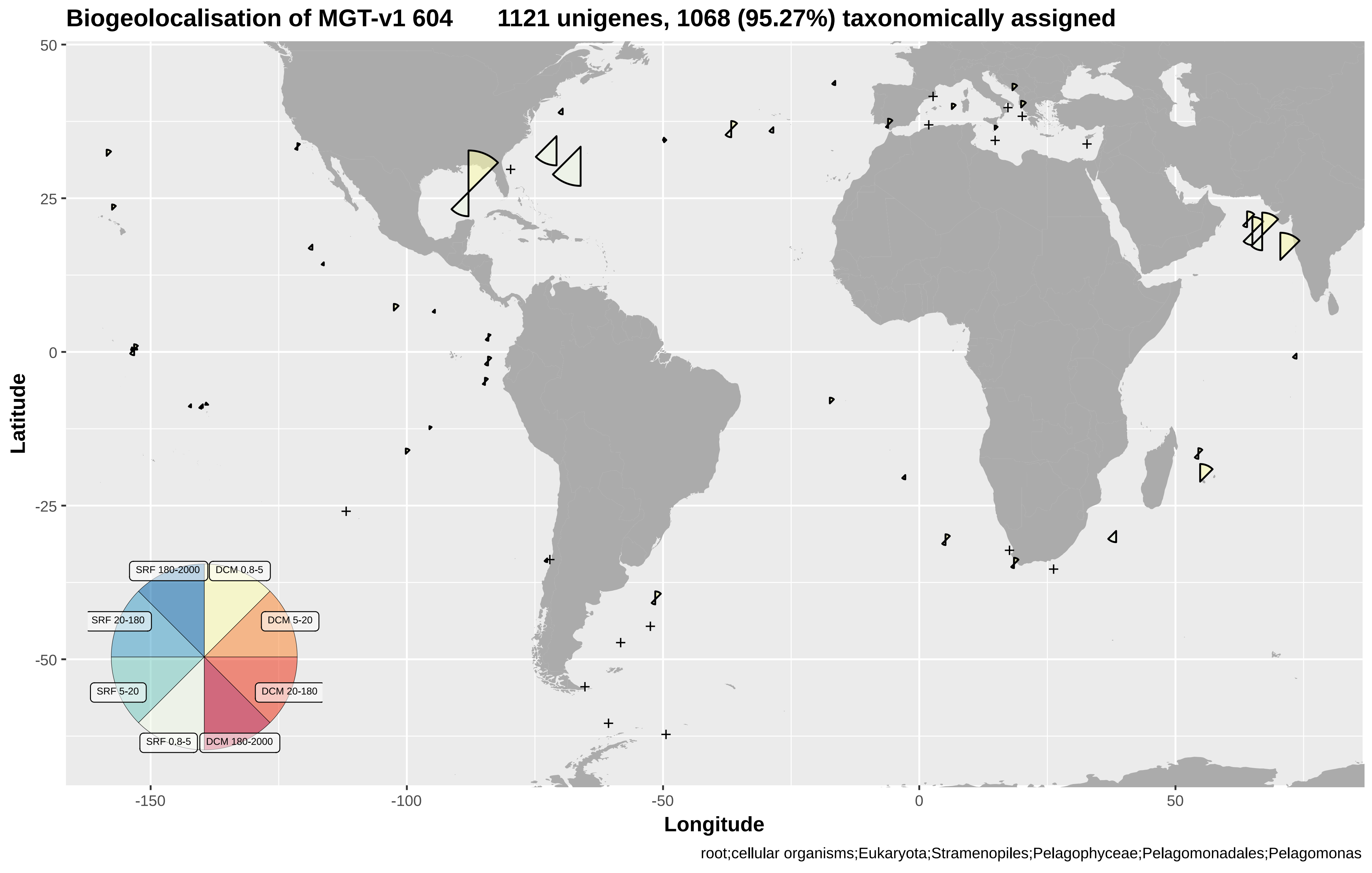

Biogeolocalisation of MGT-v1 604 1121 unigenes, 1068 (95.27%) taxonomically assigned
Latitude
50
25
0
-25
-50
Longitude
-150
-100
-50
0
50
SRF 180-2000
DCM 0.8-5
SRF 20-180
DCM 5-20
SRF 5-20
DCM 20-180
SRF 0.8-5
DCM 180-2000
root;cellular organisms;Eukaryota;Stramenopiles;Pelagophyceae;Pelagomonadales;Pelagomonas

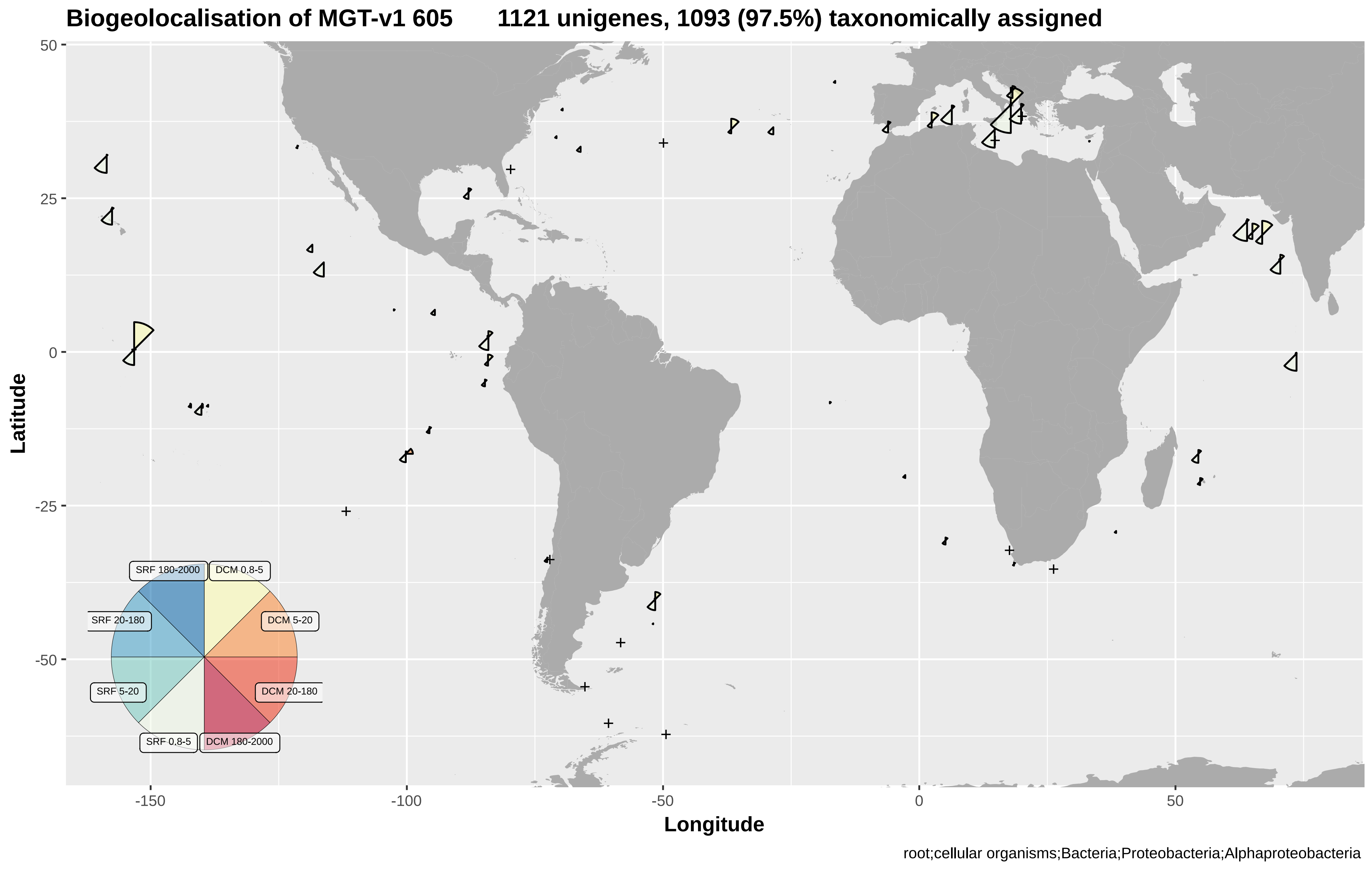

Biogeolocalisation of MGT-v1 605 1121 unigenes, 1093 (97.5%) taxonomically assigned
Latitude
50
25
0
-25
-50
Longitude
-150
-100
-50
0
50
SRF 180-2000
DCM 0.8-5
SRF 20-180
DCM 5-20
SRF 5-20
DCM 20-180
SRF 0.8-5
DCM 180-2000
root;cellular organisms;Bacteria;Proteobacteria;Alphaproteobacteria

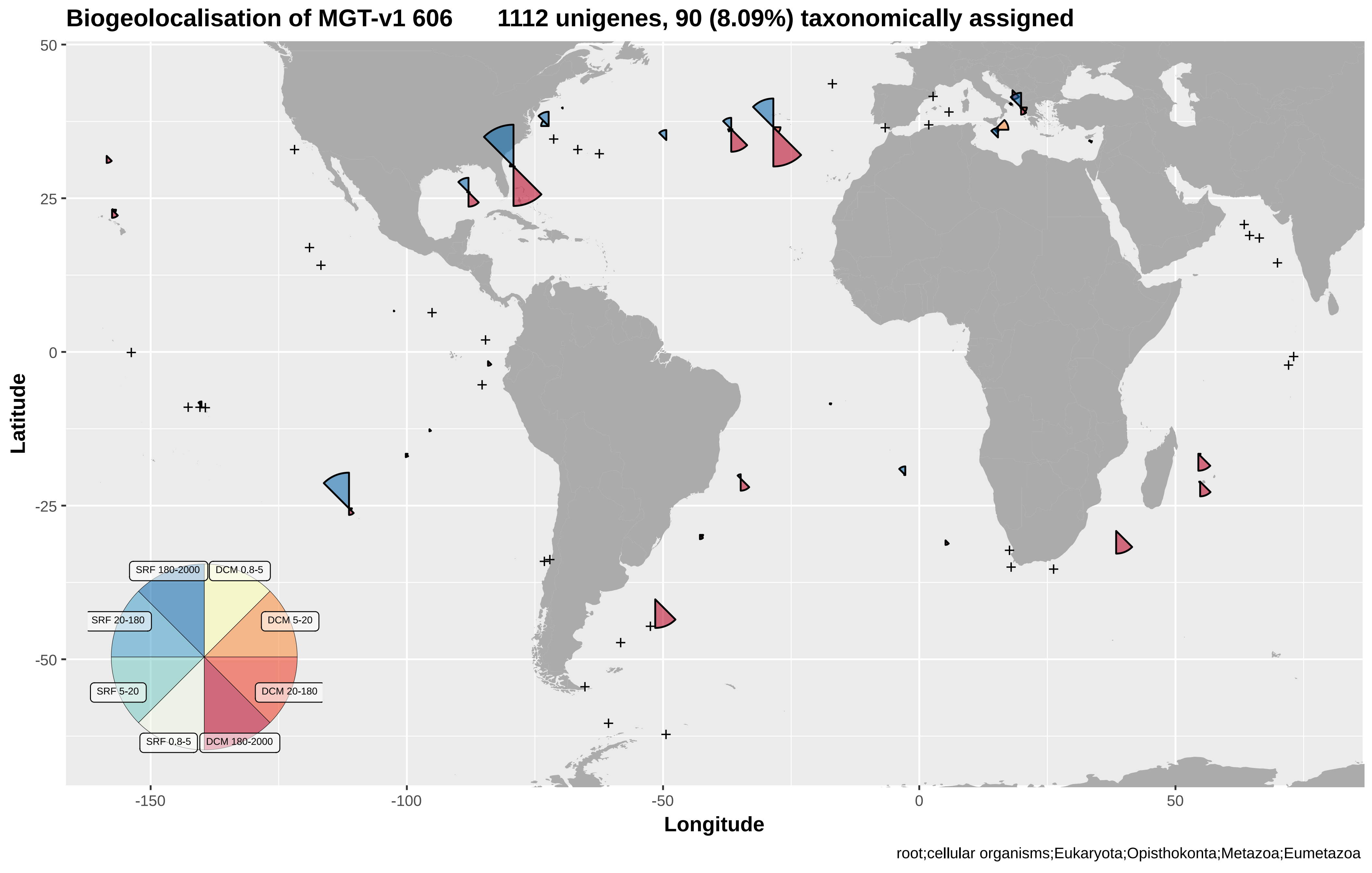

Biogeolocalisation of MGT-v1 606 1112 unigenes, 90 (8.09%) taxonomically assigned
Latitude
Longitude
50
25
0
-25
-50
-150
-100
-50
0
50
SRF 180-2000
DCM 0.8-5
SRF 20-180
DCM 5-20
SRF 5-20
DCM 20-180
SRF 0.8-5
DCM 180-2000
root;cellular organisms;Eukaryota;Opisthokonta;Metazoa;Eumetazoa

## Biogeolocalisation of MGT-v1 607      1109 unigenes, 88 (7.94%) taxonomically assigned

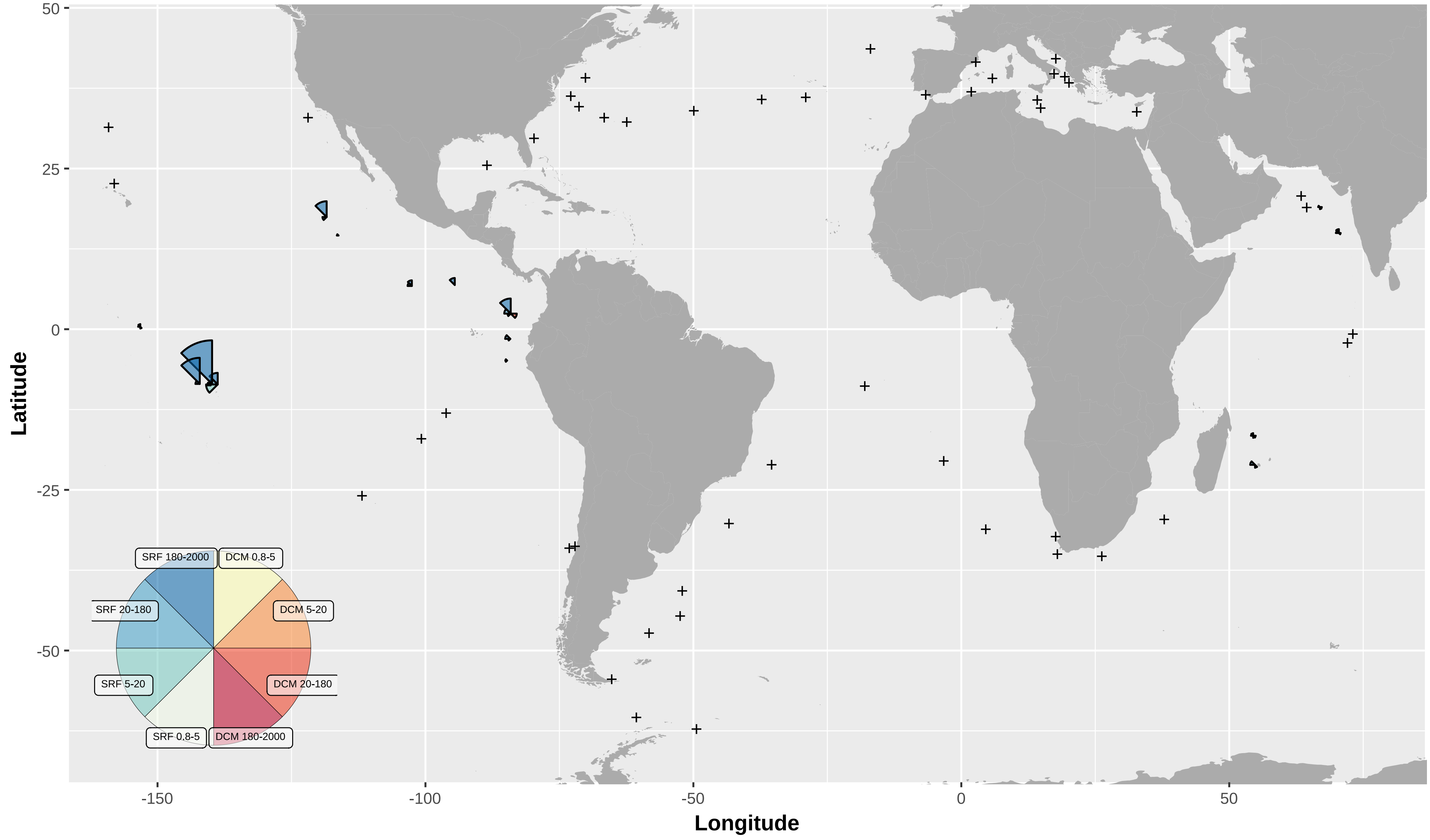


root;cellular organisms;Eukaryota;Opisthokonta;Metazoa;Eumetazoa;Bilateria

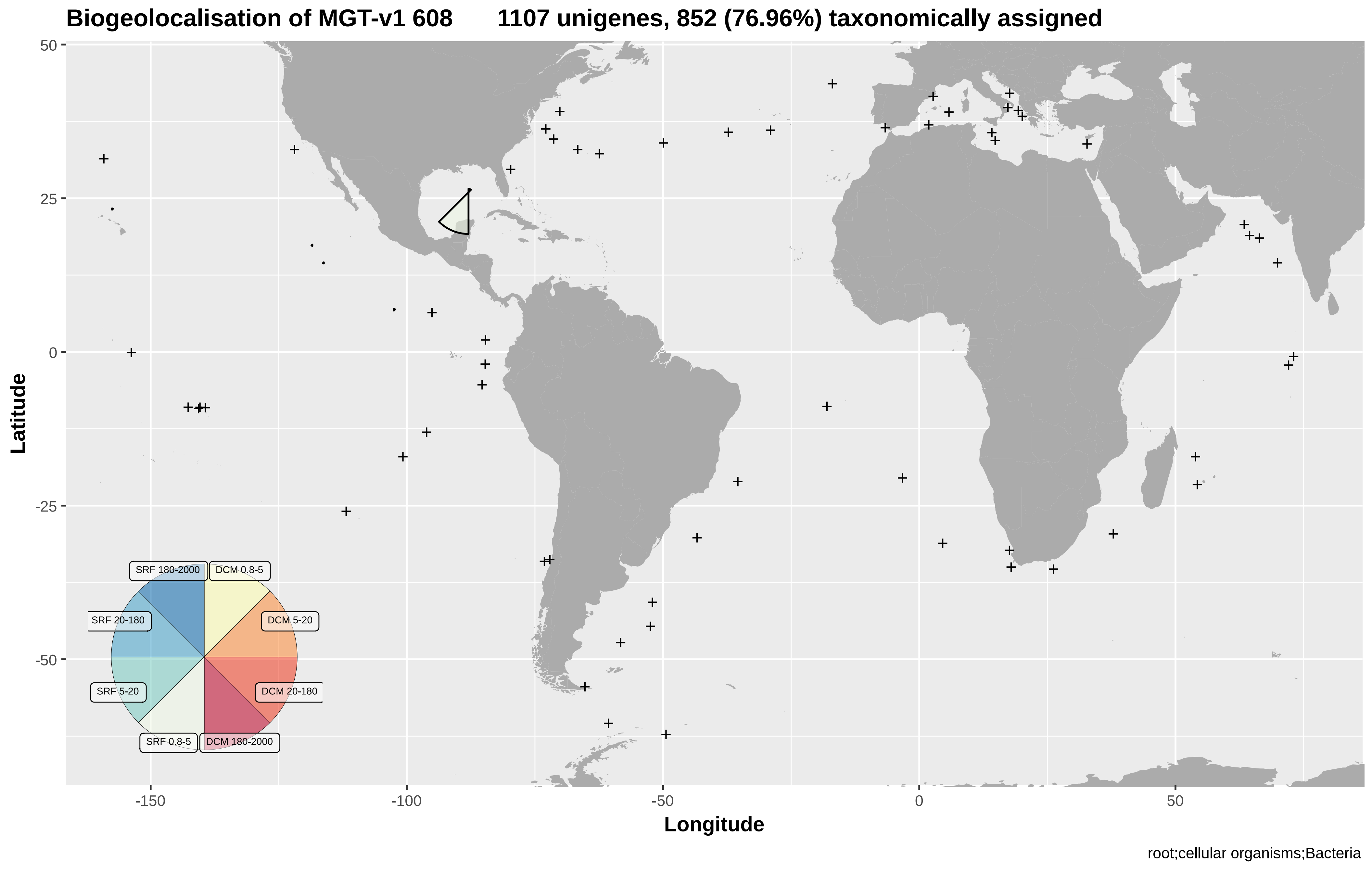
Biogeolocalisation of MGT-v1 608
1107 unigenes, 852 (76.96%) taxonomically assigned
Latitude
Longitude
50
25
0
-25
-50
-150
-100
-50
0
50
SRF 180-2000
DCM 0.8-5
SRF 20-180
DCM 5-20
SRF 5-20
DCM 20-180
SRF 0.8-5
DCM 180-2000
root;cellular organisms;Bacteria

### Biogeolocalisation of MGT-v1 609 1106 unigenes, 799 (72.24%) taxonomically assigned

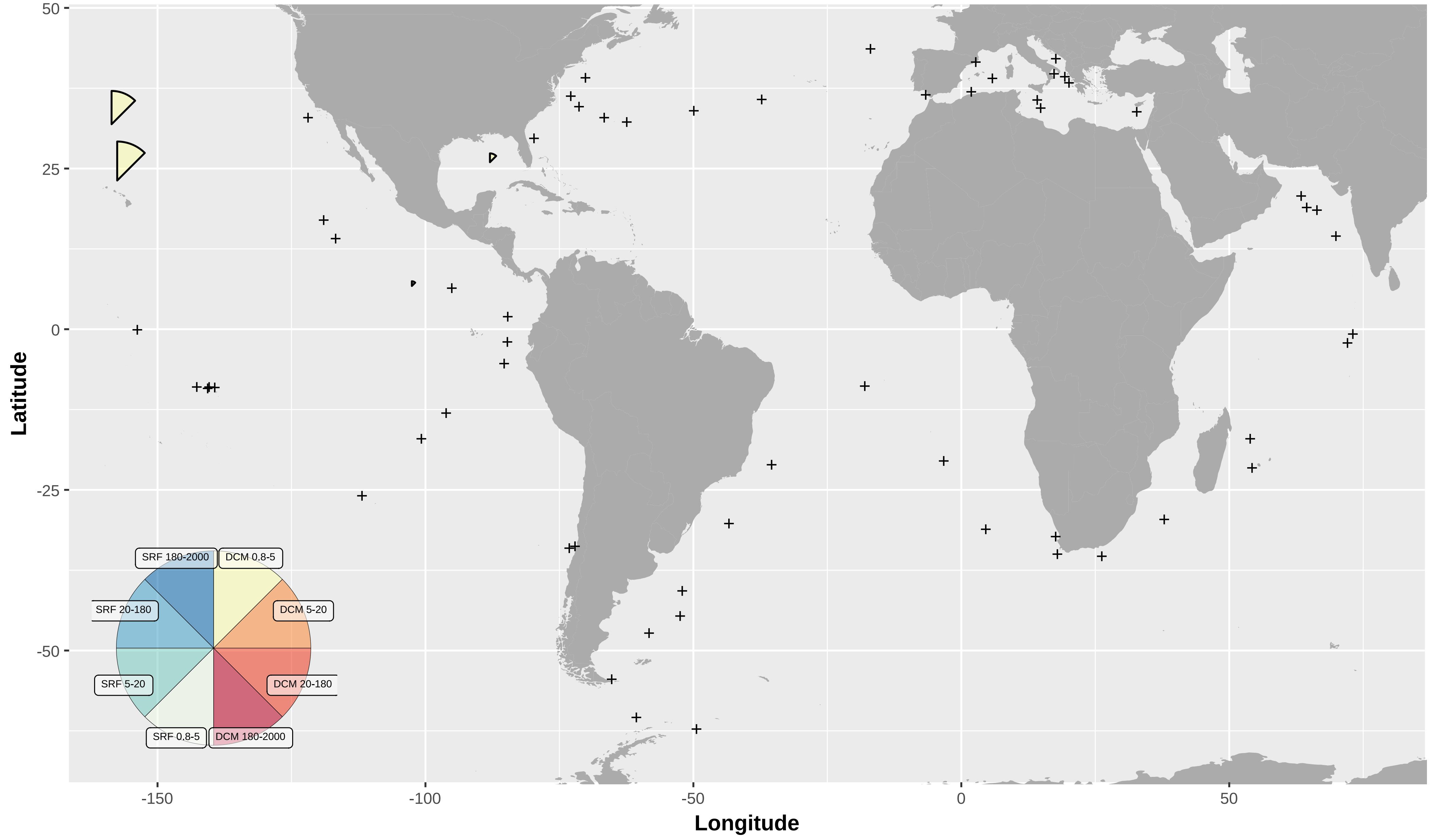


root;cellular organisms;Bacteria

## Biogeolocalisation of MGT-v1 610      1104 unigenes, 94 (8.51%) taxonomically assigned

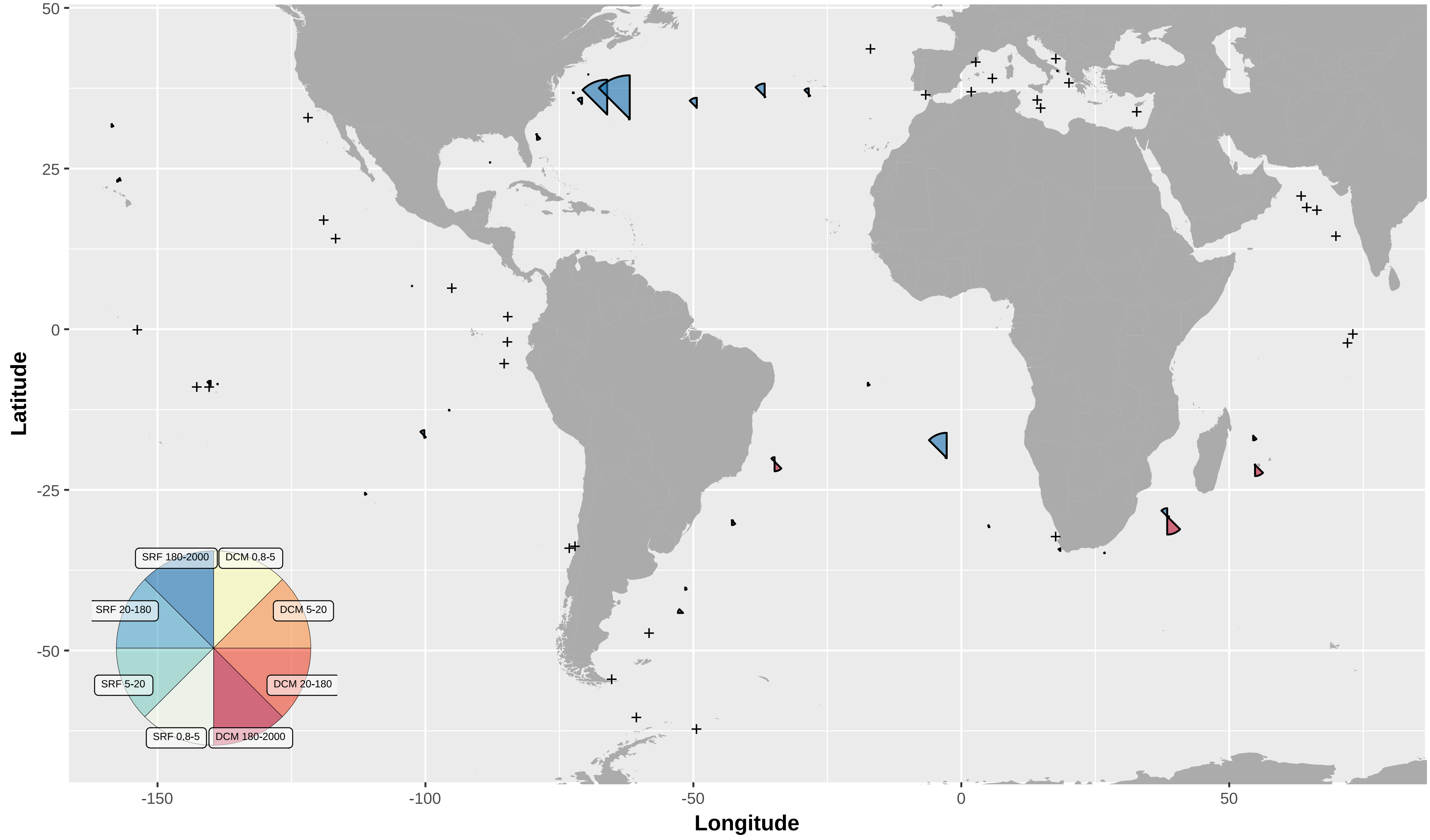


root;cellular organisms;Eukaryota

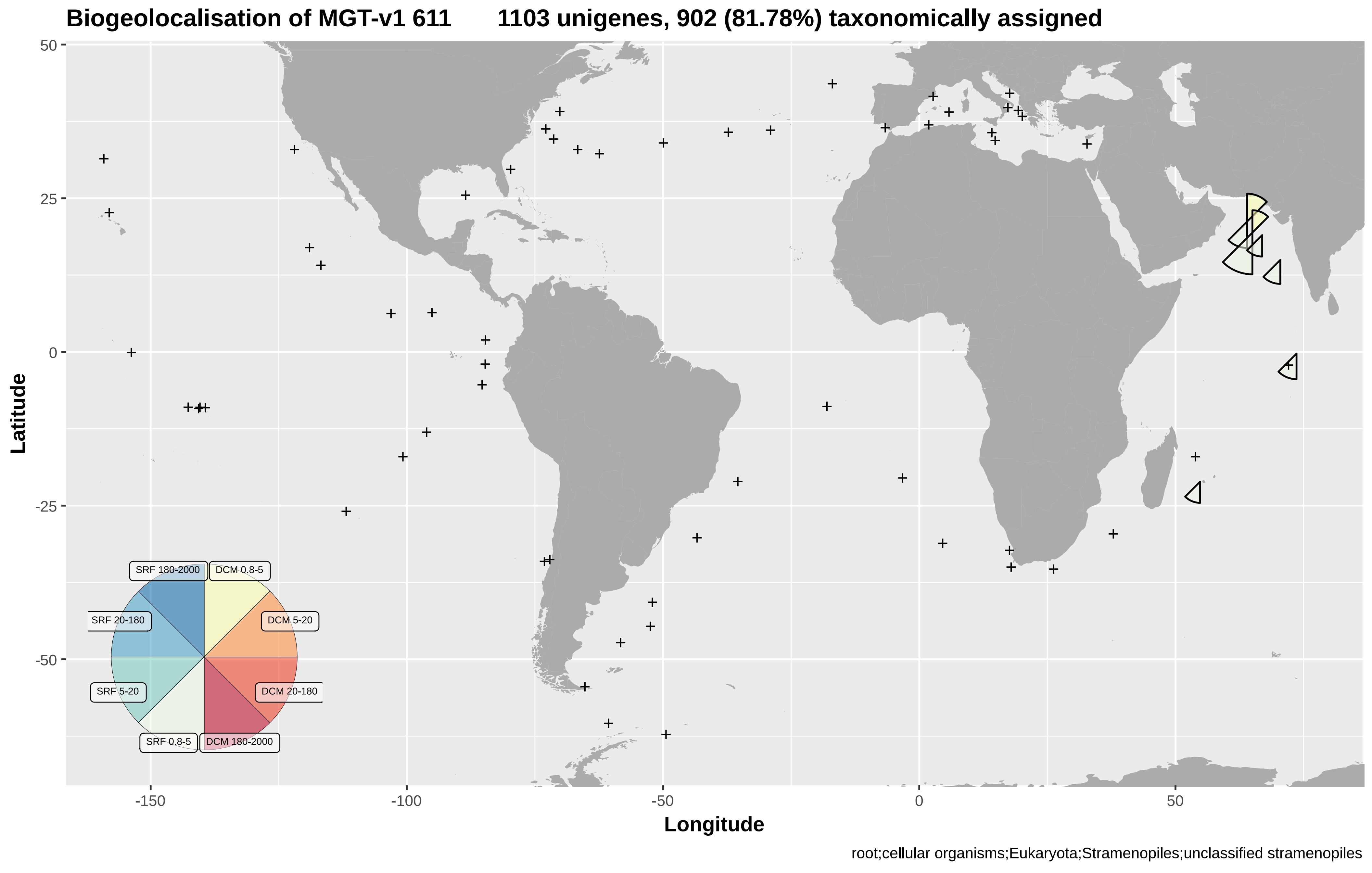

Biogeolocalisation of MGT-v1 611 1103 unigenes, 902 (81.78%) taxonomically assigned
50
25
0
-25
-50
Latitude
-150
-100
-50
0
50
Longitude
SRF 180-2000
DCM 0.8-5
SRF 20-180
DCM 5-20
SRF 5-20
DCM 20-180
SRF 0.8-5
DCM 180-2000
root;cellular organisms;Eukaryota;Stramenopiles;unclassified stramenopiles

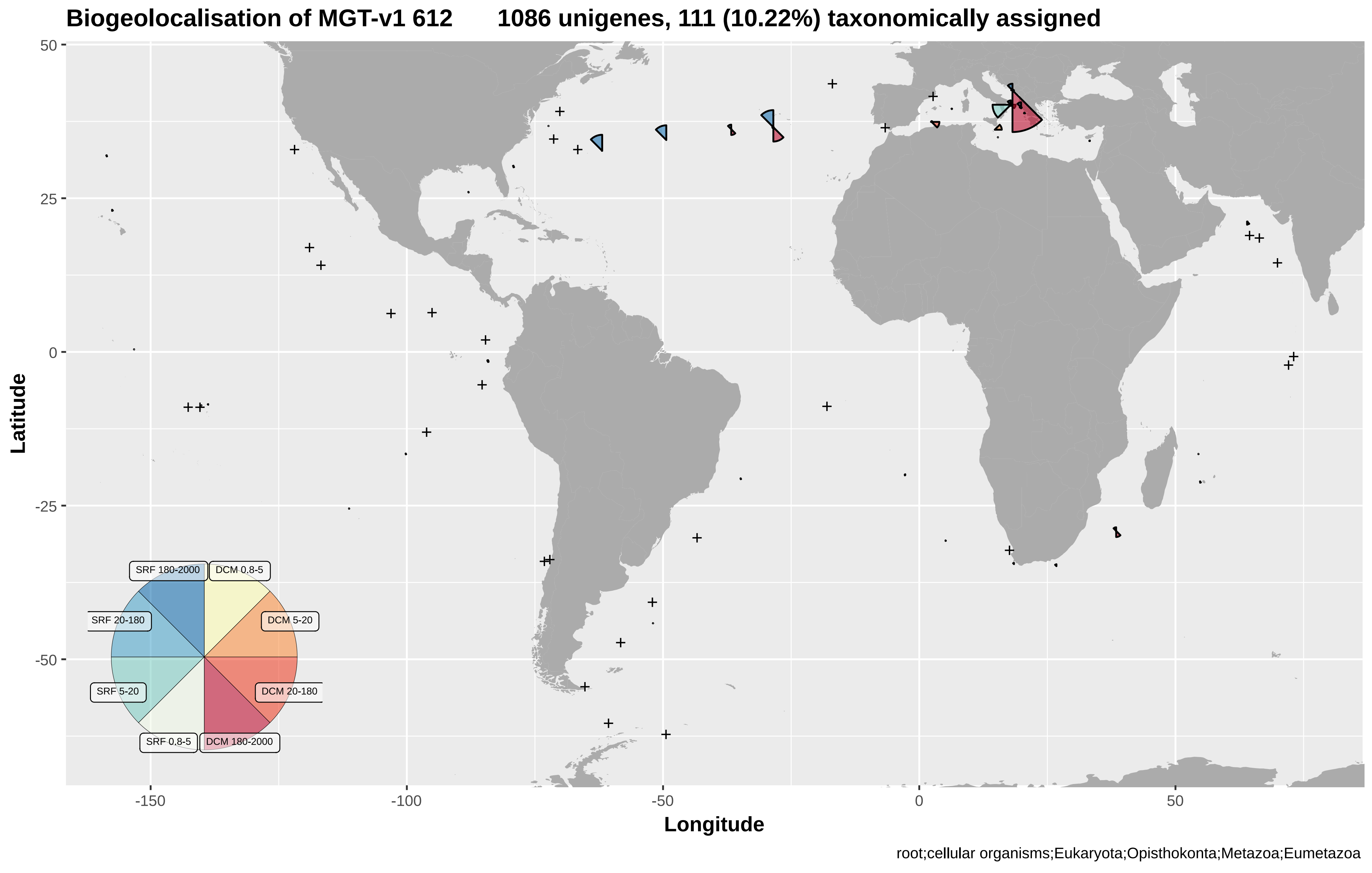
Biogeolocalisation of MGT-v1 612 1086 unigenes, 111 (10.22%) taxonomically assigned
Latitude
50
25
0
-25
-50
Longitude
-150
-100
-50
0
50
SRF 180-2000
DCM 0.8-5
SRF 20-180
DCM 5-20
SRF 5-20
DCM 20-180
SRF 0.8-5
DCM 180-2000
root;cellular organisms;Eukaryota;Opisthokonta;Metazoa;Eumetazoa

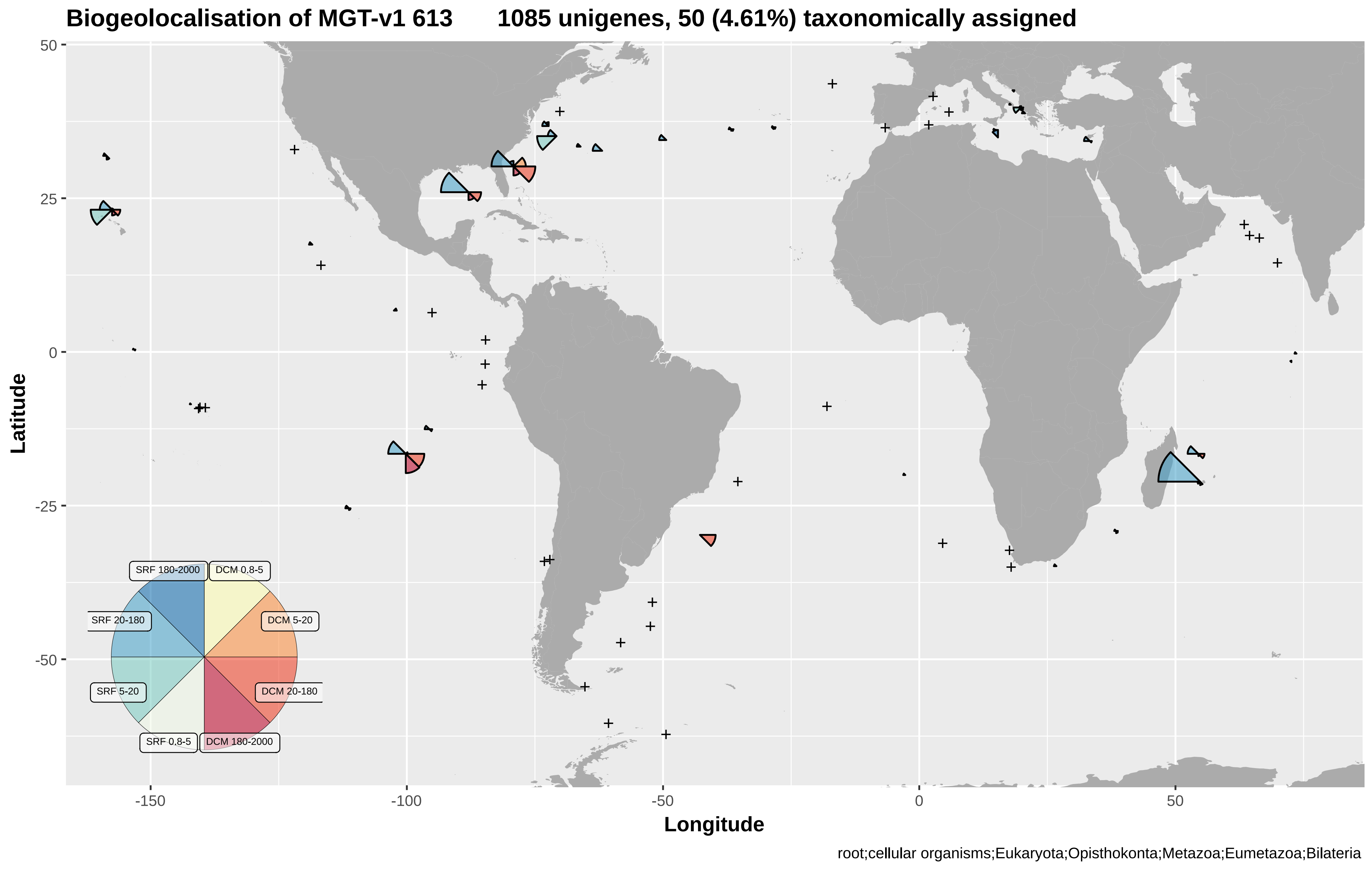

Biogeolocalisation of MGT-v1 613 1085 unigenes, 50 (4.61%) taxonomically assigned
Latitude
Longitude
50
25
0
-25
-50
-150
-100
-50
0
50
SRF 180-2000
DCM 0.8-5
SRF 20-180
DCM 5-20
SRF 5-20
DCM 20-180
SRF 0.8-5
DCM 180-2000
root;cellular organisms;Eukaryota;Opisthokonta;Metazoa;Eumetazoa;Bilateria

#### Biogeolocalisation of MGT-v1 614 1082 unigenes, 128 (11.83%) taxonomically assigned

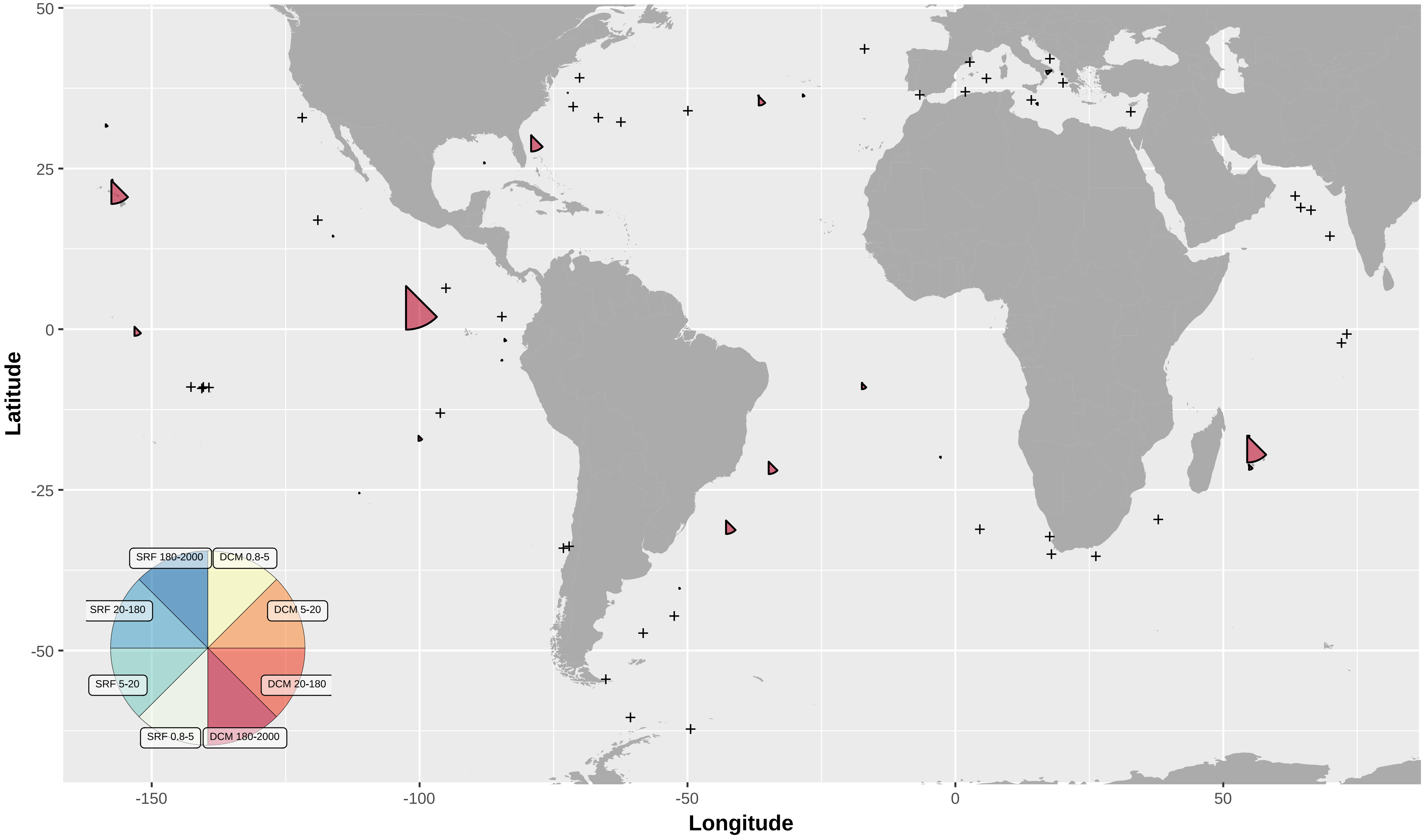


root;cellular organisms;Eukaryota;Opisthokonta;Metazoa;Eumetazoa

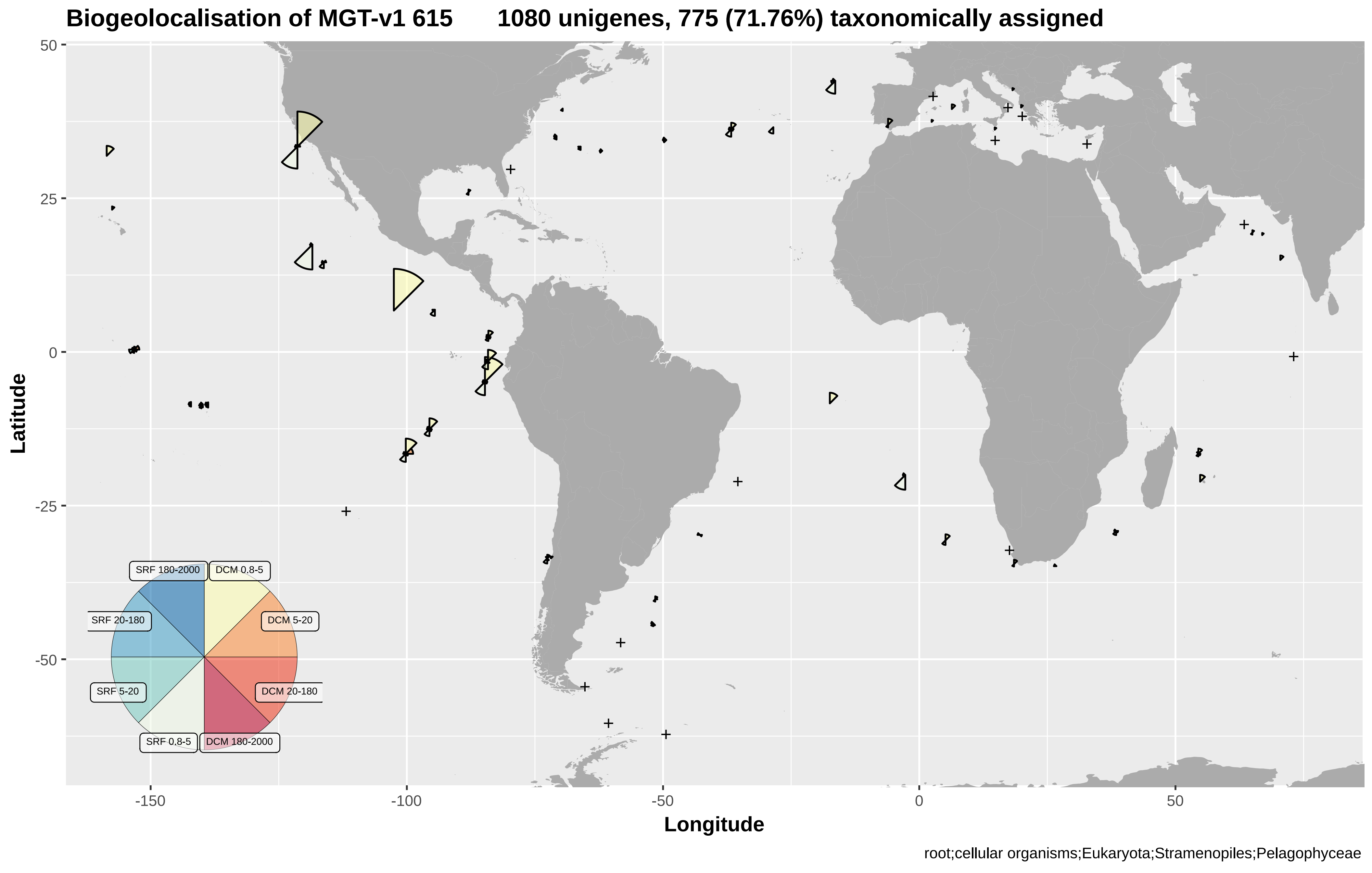
Biogeolocalisation of MGT-v1 615 1080 unigenes, 775 (71.76%) taxonomically assigned
50
25
0
-25
-50
Latitude
-150
-100
-50
0
50
Longitude
SRF 180-2000
DCM 0.8-5
SRF 20-180
DCM 5-20
SRF 5-20
DCM 20-180
SRF 0.8-5
DCM 180-2000
root;cellular organisms;Eukaryota;Stramenopiles;Pelagophyceae

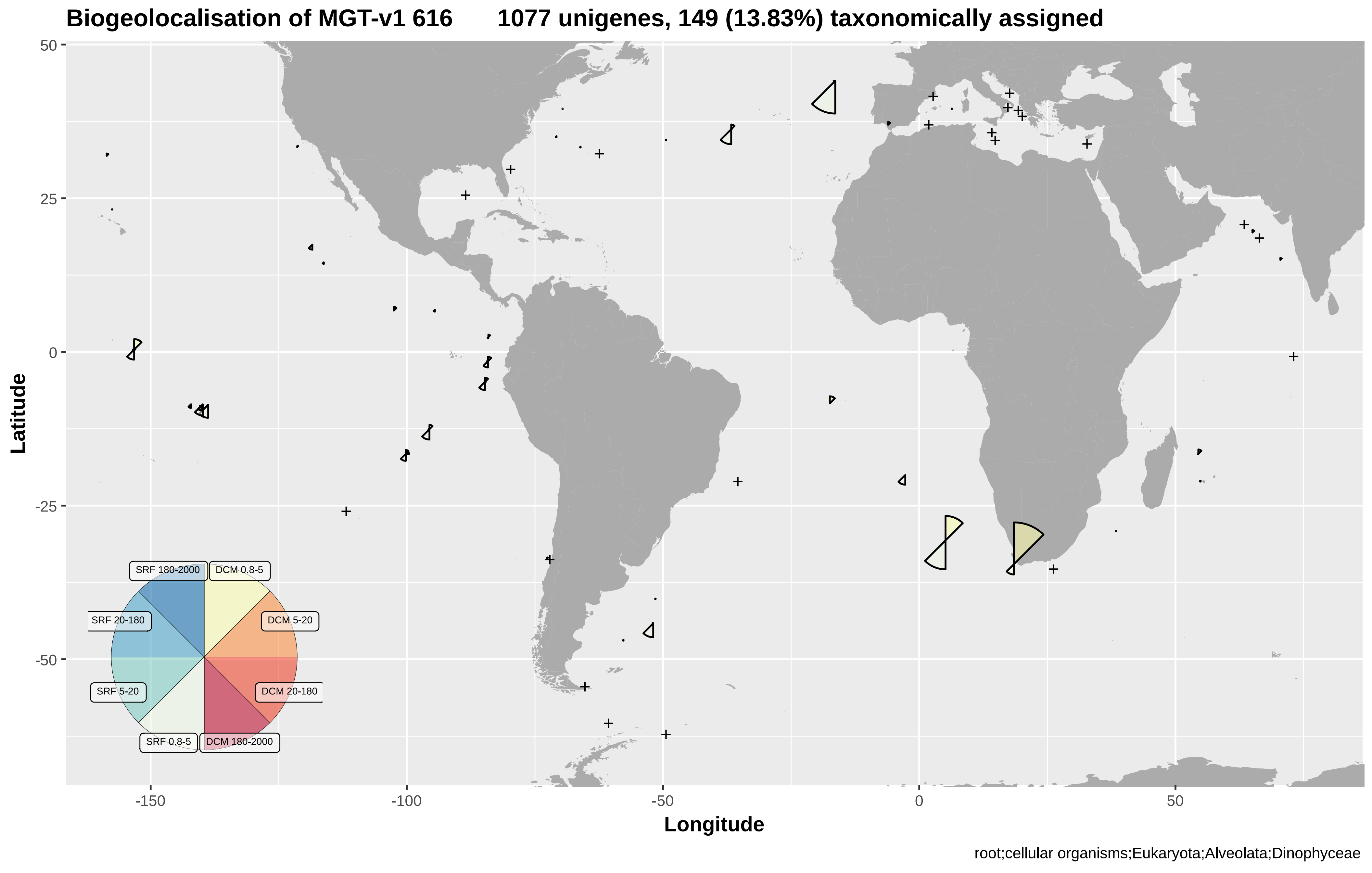
Biogeolocalisation of MGT-v1 616 1077 unigenes, 149 (13.83%) taxonomically assigned
Latitude
Longitude
50
25
0
-25
-50
-150
-100
-50
0
50
SRF 180-2000
DCM 0.8-5
SRF 20-180
DCM 5-20
SRF 5-20
DCM 20-180
SRF 0.8-5
DCM 180-2000
root;cellular organisms;Eukaryota;Alveolata;Dinophyceae

# Biogeolocalisation of MGT-v1 617      1077 unigenes, 658 (61.1%) taxonomically assigned

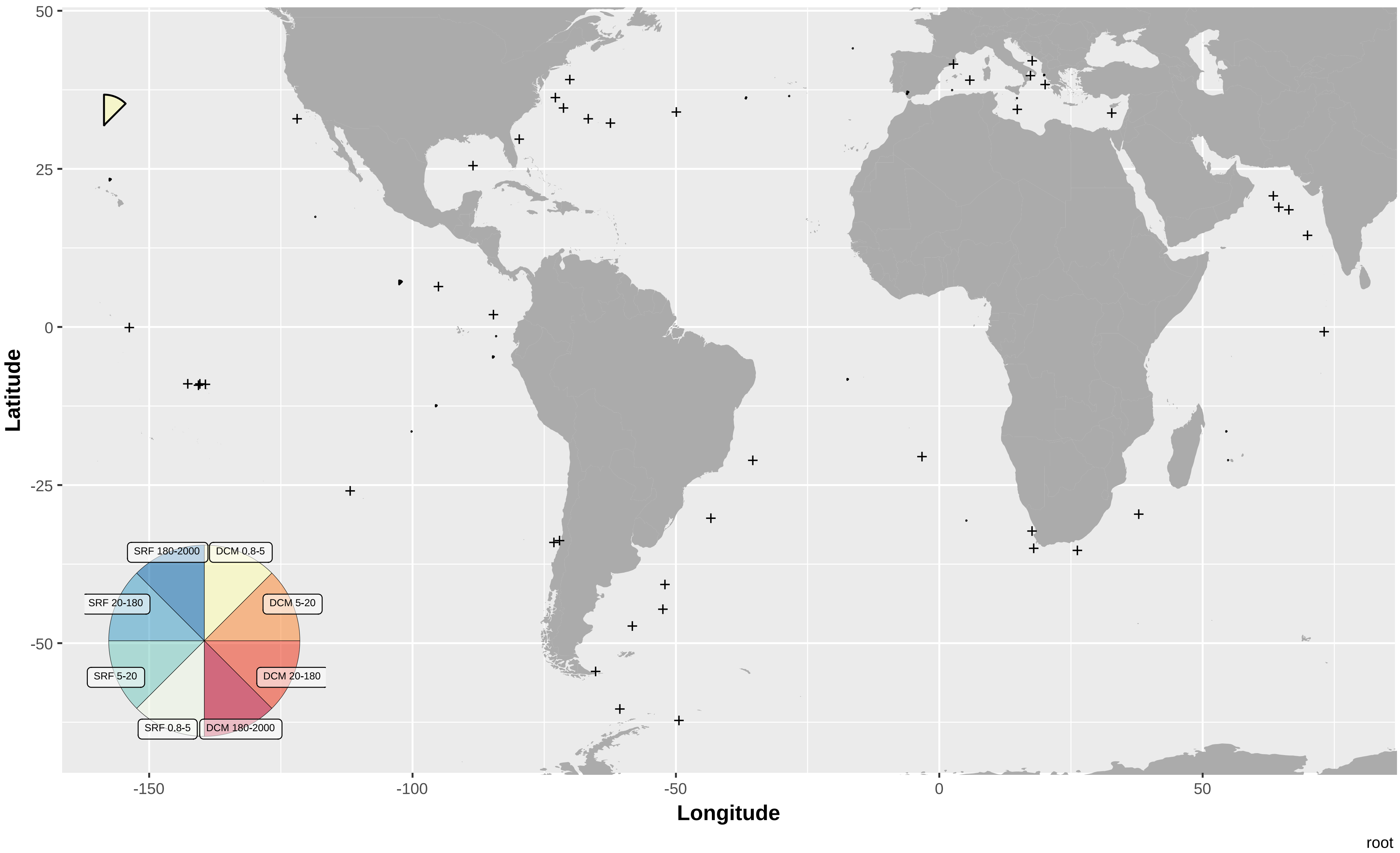

#### Biogeolocalisation of MGT-v1 618 1077 unigenes, 971 (90.16%) taxonomically assigned

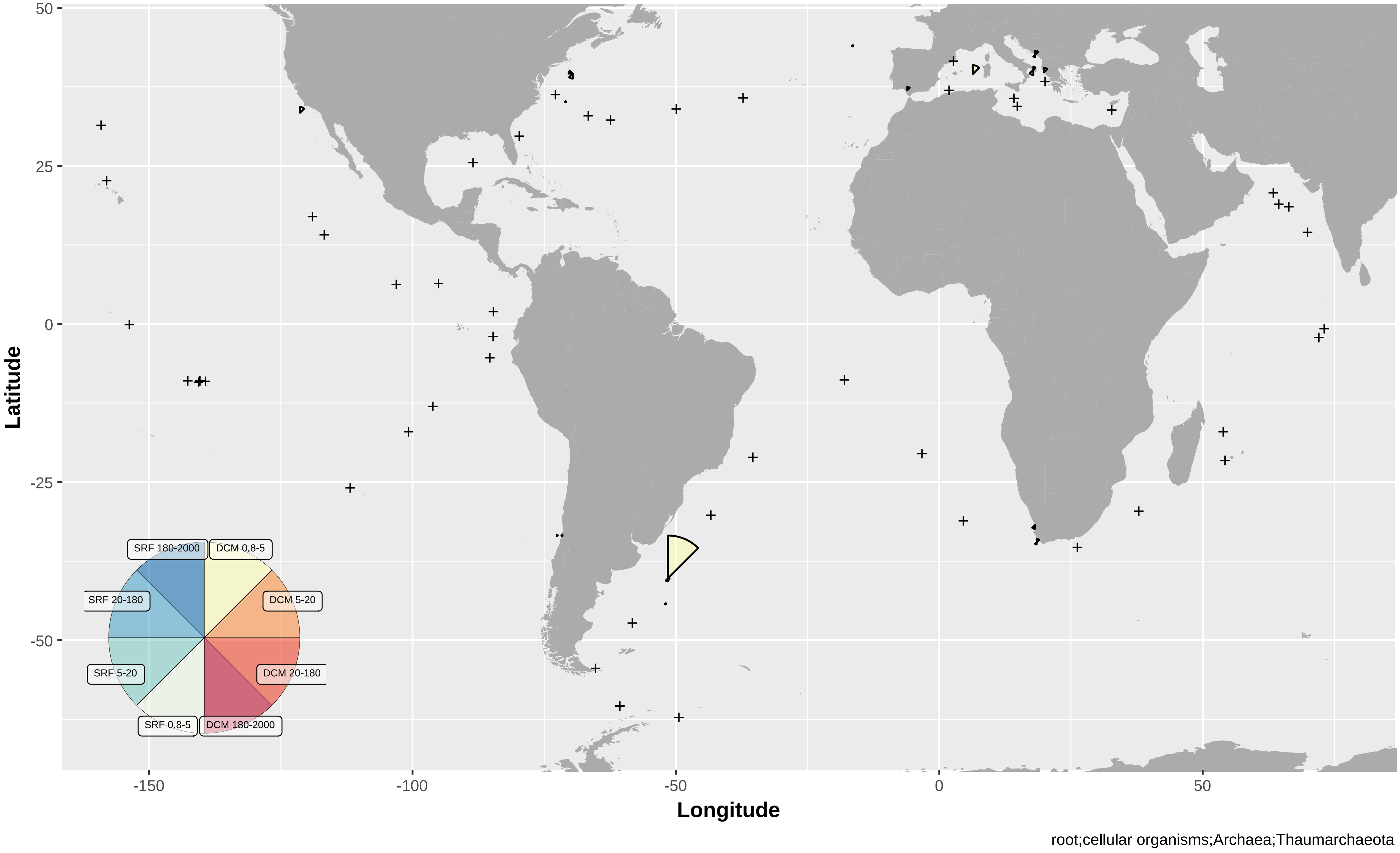

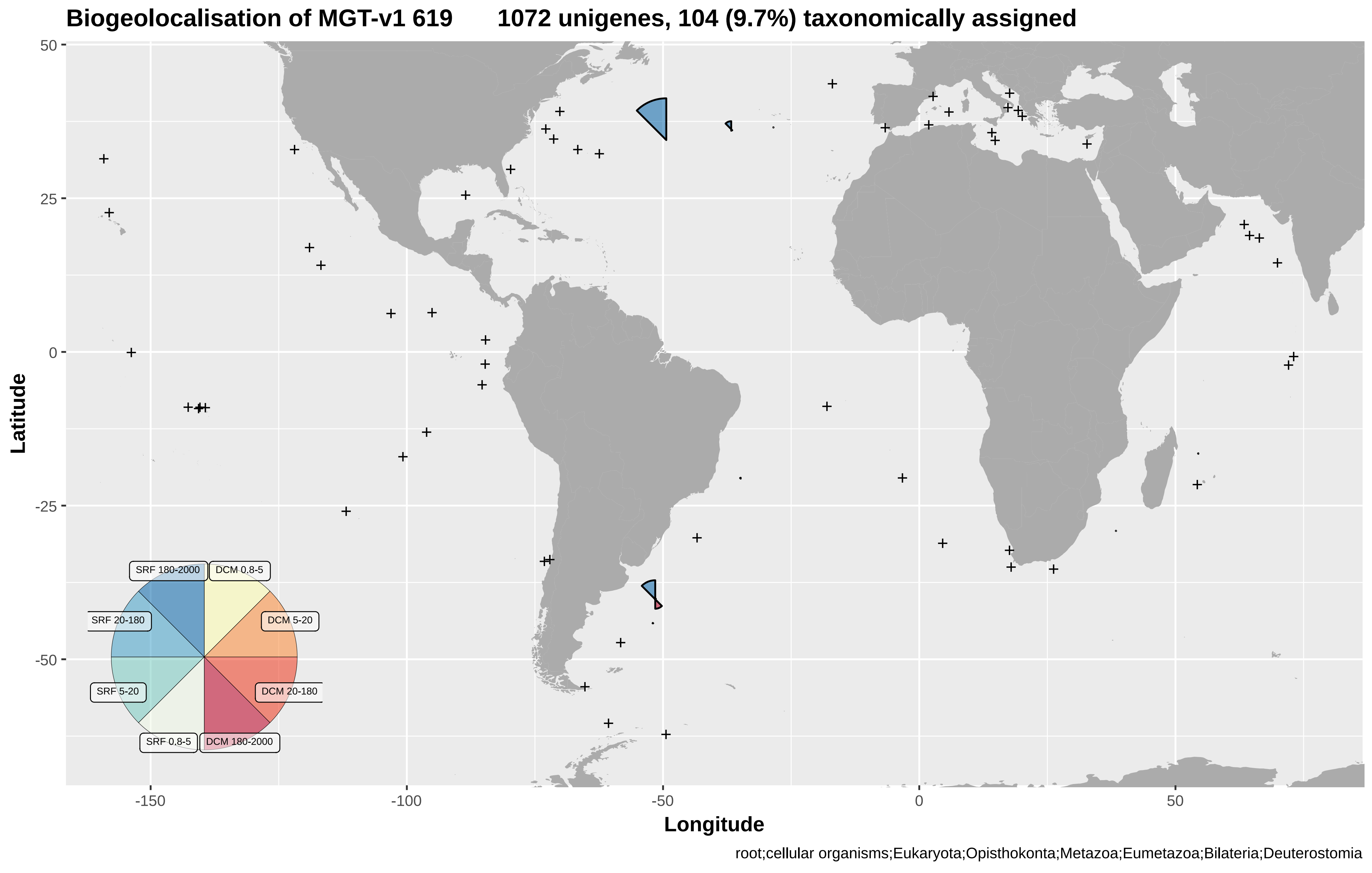
Biogeolocalisation of MGT-v1 619 1072 unigenes, 104 (9.7%) taxonomically assigned
Latitude
Longitude
50
25
0
-25
-50
-150
-100
-50
0
50
SRF 180-2000
DCM 0.8-5
SRF 20-180
DCM 5-20
SRF 5-20
DCM 20-180
SRF 0.8-5
DCM 180-2000
root;cellular organisms;Eukaryota;Opisthokonta;Metazoa;Eumetazoa;Bilateria;Deuterostomia

# Biogeolocalisation of MGT-v1 620      1071 unigenes, 510 (47.62%) taxonomically assigned

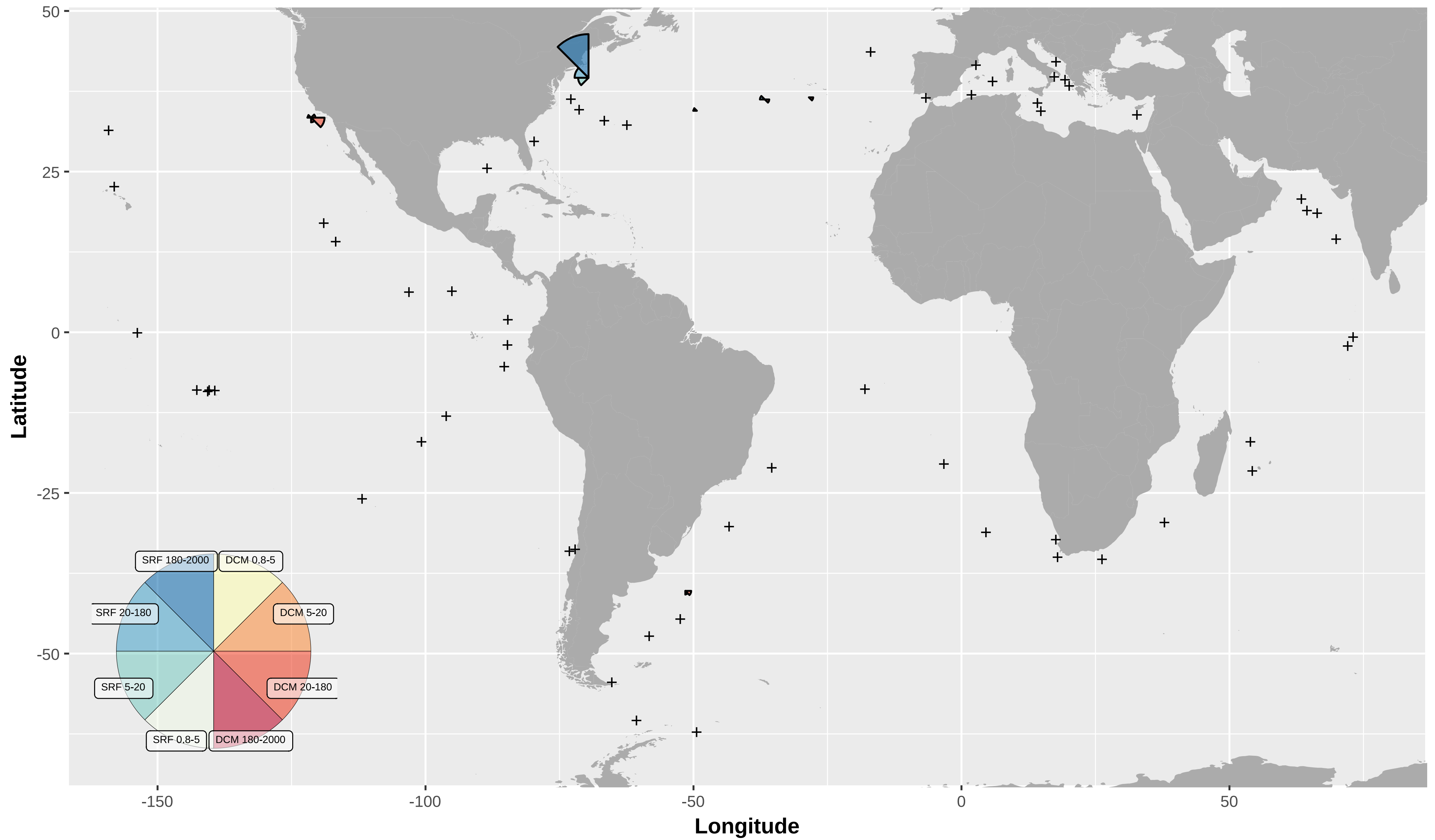


root;cellular organisms;Eukaryota;Opisthokonta;Metazoa;Eumetazoa;Bilateria;Protostomia;Ecdysozoa;Panarthropoda;Arthropoda;Mandibulata;Pancrustacea;Crustacea;Maxillopoda

## Biogeolocalisation of MGT-v1 621      1071 unigenes, 678 (63.31%) taxonomically assigned

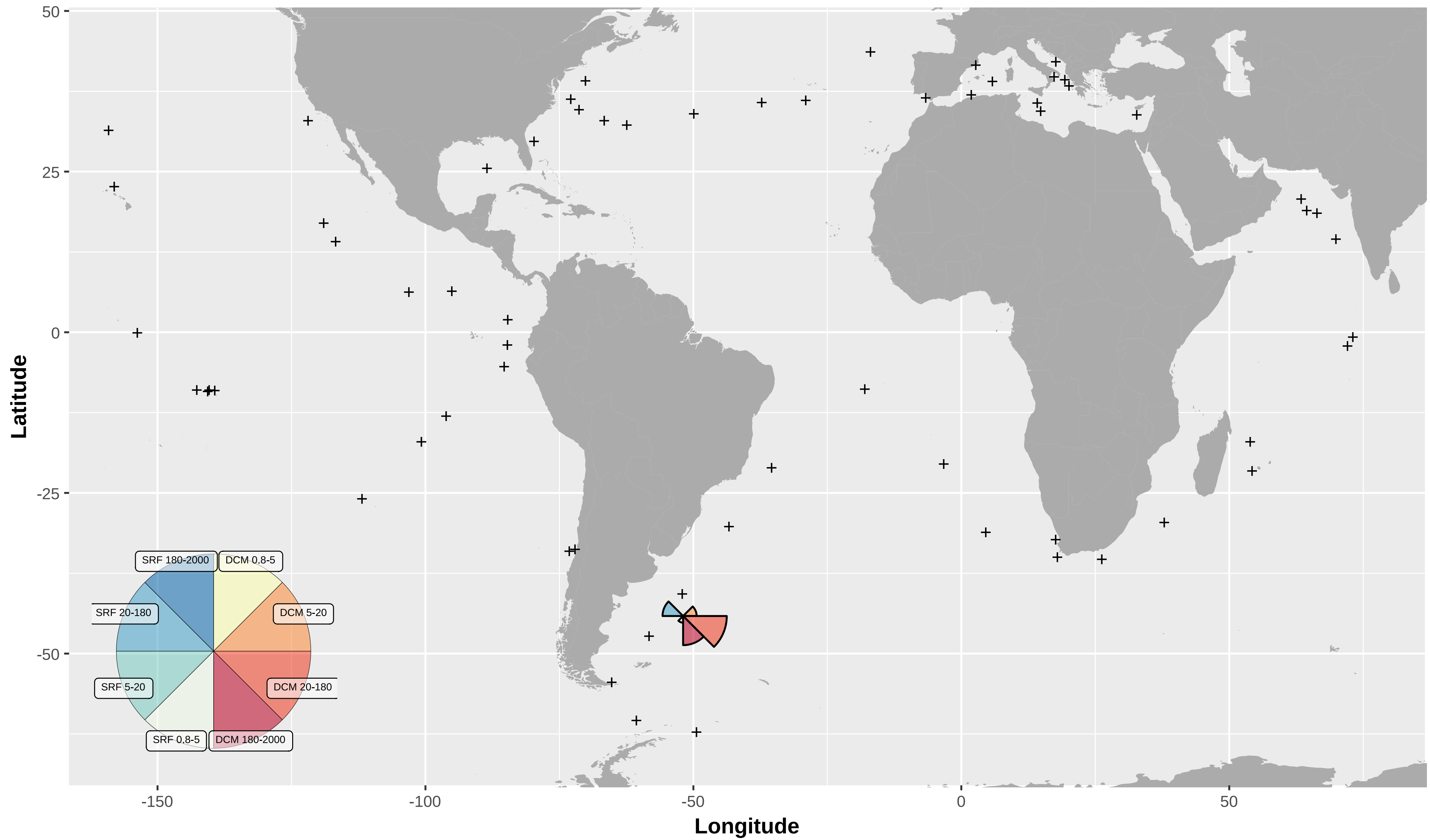


root;cellular organisms;Eukaryota;Alveolata;Ciliophora;Intramacronucleata;Spirotrichea

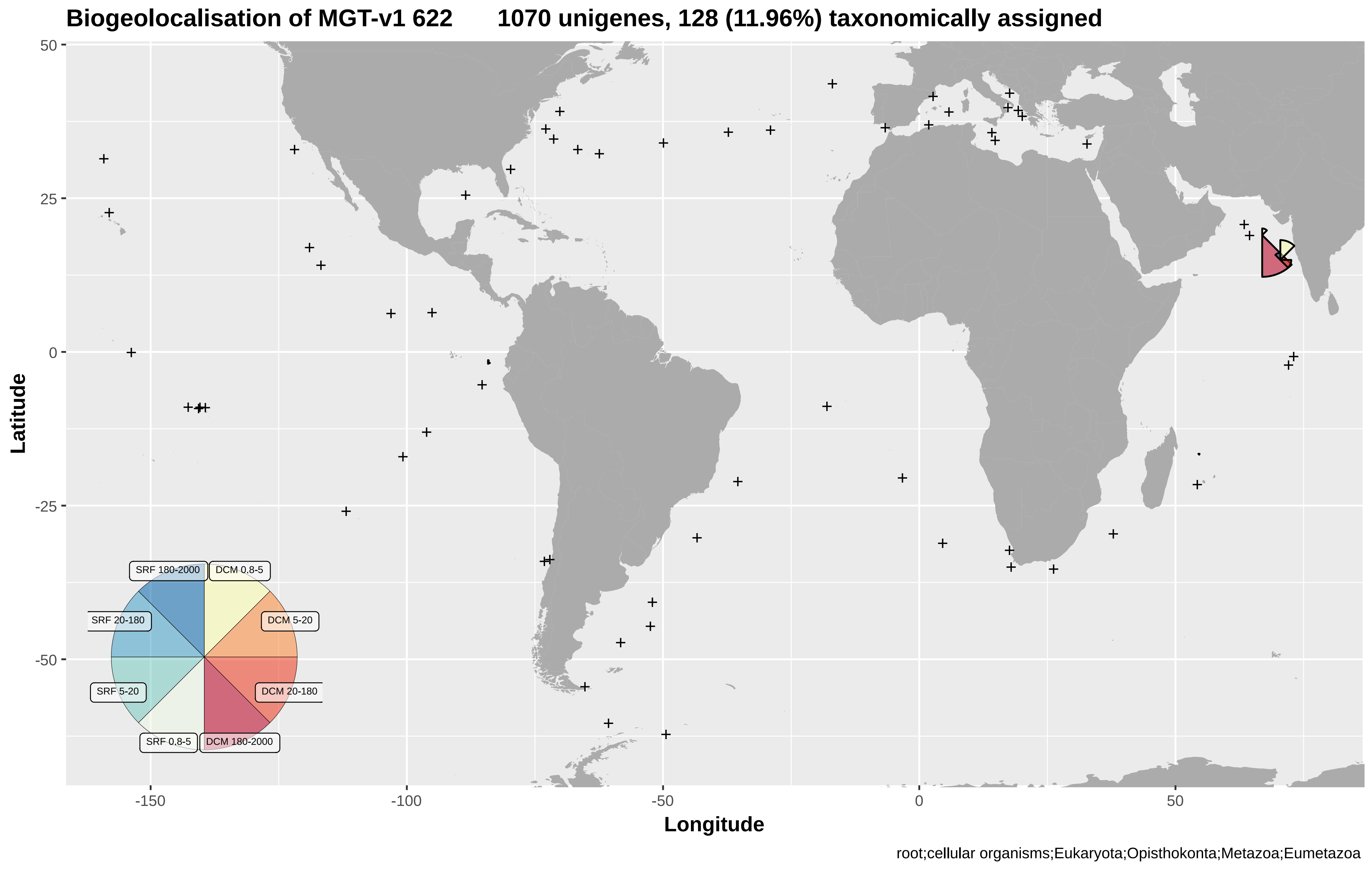

Biogeolocalisation of MGT-v1 622 1070 unigenes, 128 (11.96%) taxonomically assigned
Latitude
Longitude
50
25
0
-25
-50
-150
-100
-50
0
50
SRF 180-2000
DCM 0.8-5
SRF 20-180
DCM 5-20
SRF 5-20
DCM 20-180
SRF 0.8-5
DCM 180-2000
root;cellular organisms;Eukaryota;Opisthokonta;Metazoa;Eumetazoa

#### Biogeolocalisation of MGT-v1 623 1066 unigenes, 580 (54.41%) taxonomically assigned

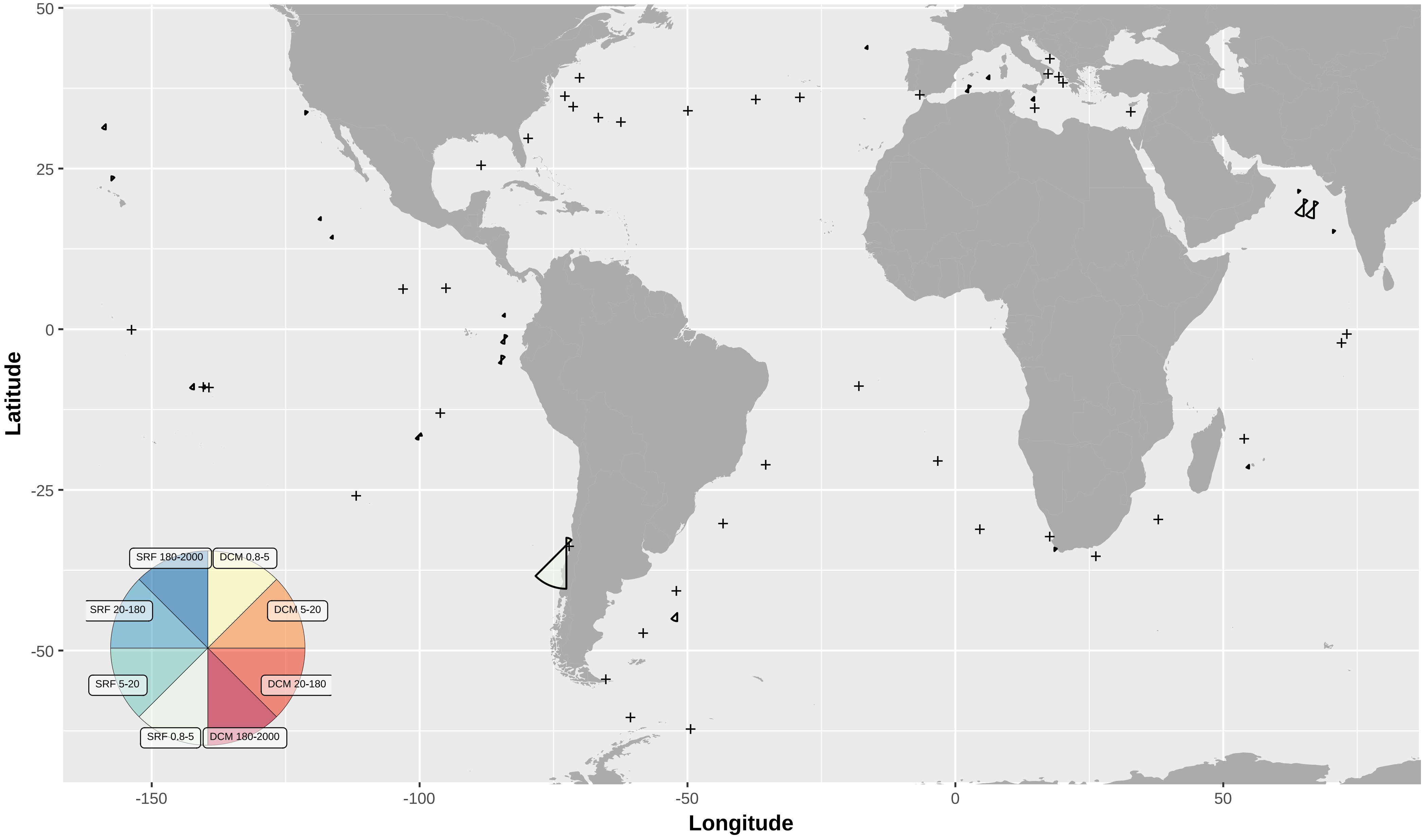


root;cellular organisms;Eukaryota

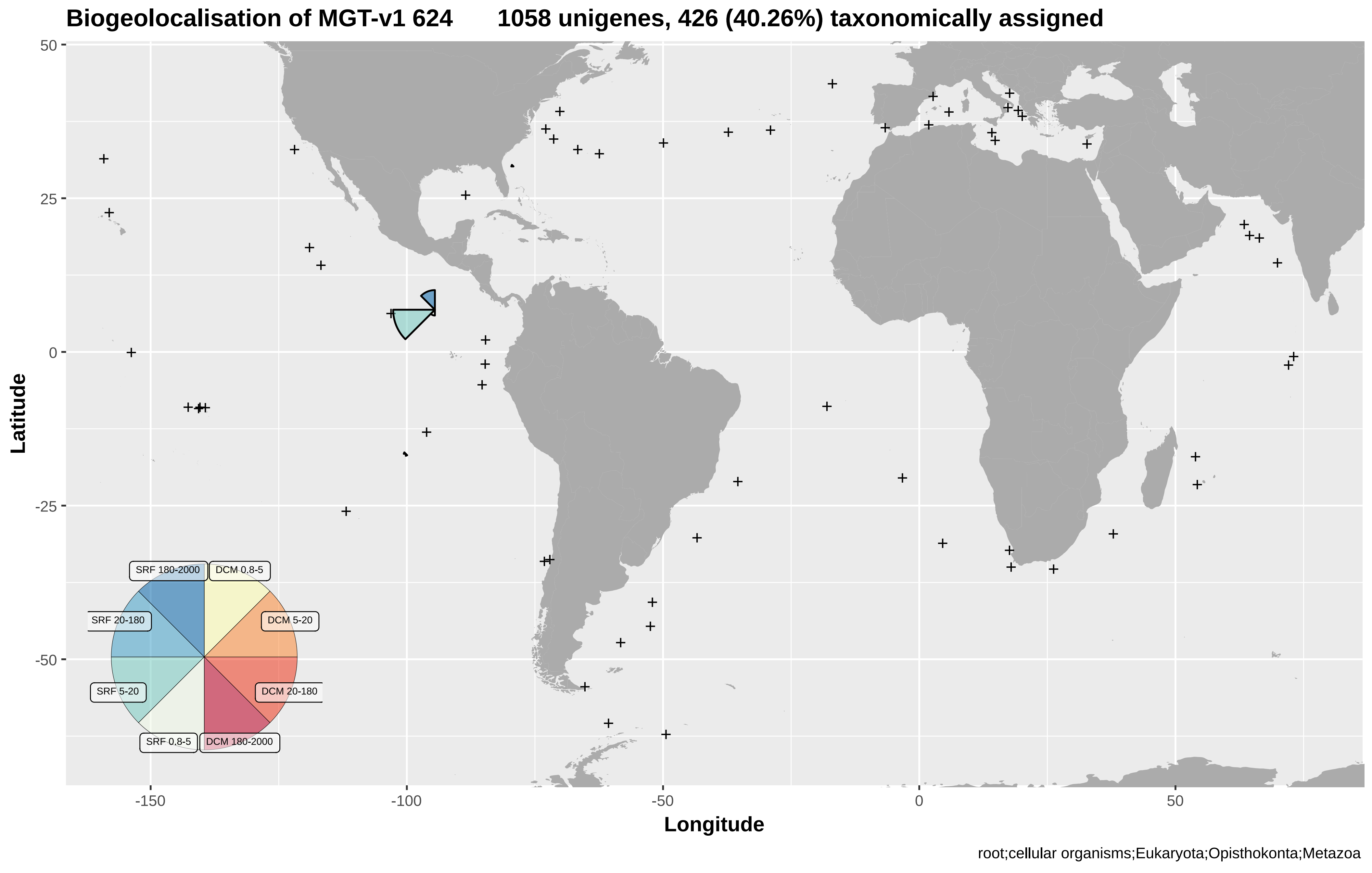

Biogeolocalisation of MGT-v1 624 1058 unigenes, 426 (40.26%) taxonomically assigned
50
25
0
-25
-50
Latitude
-150
-100
-50
0
50
Longitude
SRF 180-2000
DCM 0.8-5
SRF 20-180
DCM 5-20
SRF 5-20
DCM 20-180
SRF 0.8-5
DCM 180-2000
root;cellular organisms;Eukaryota;Opisthokonta;Metazoa

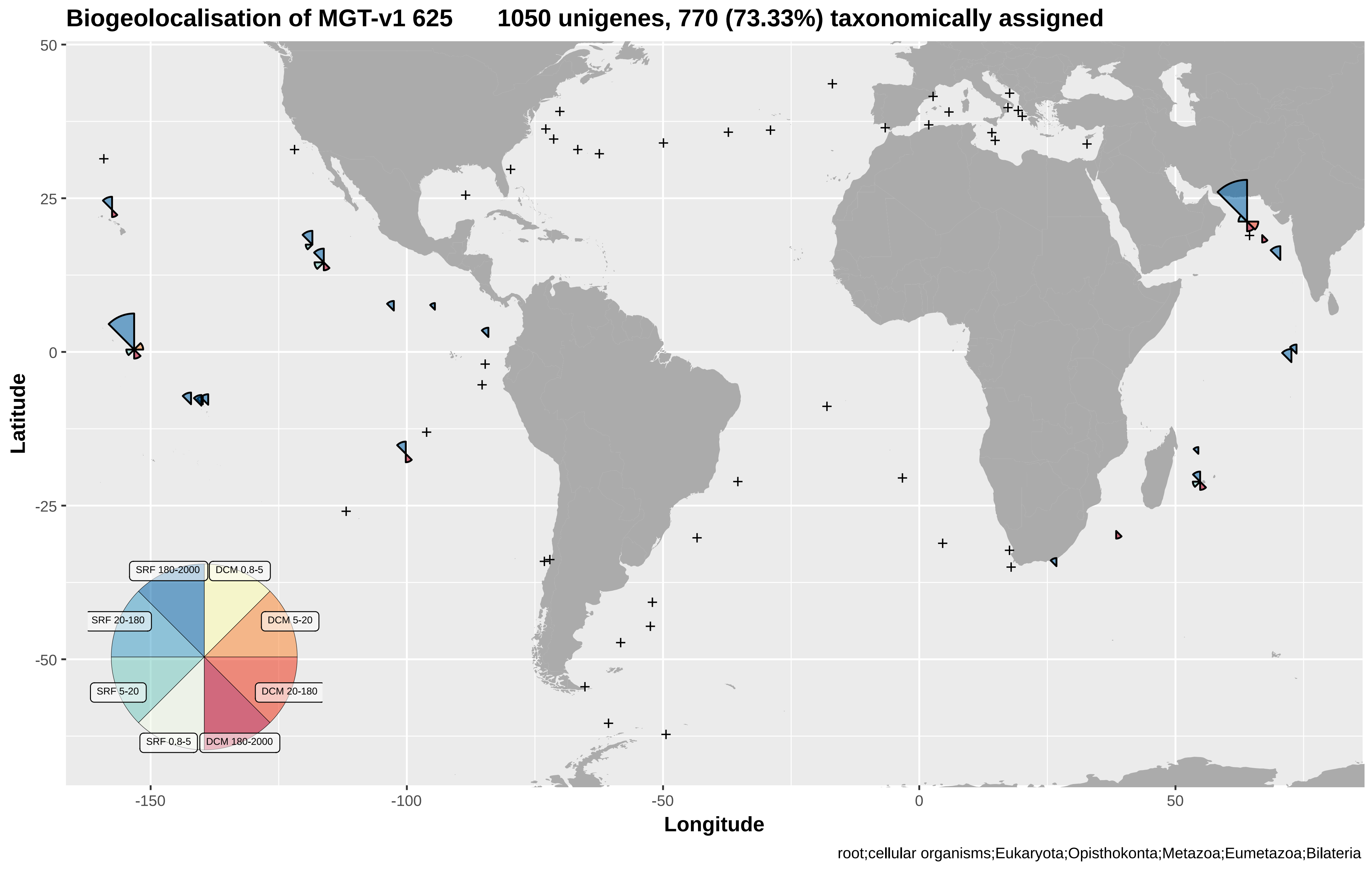
Biogeolocalisation of MGT-v1 625 1050 unigenes, 770 (73.33%) taxonomically assigned
Latitude
Longitude
50
25
0
-25
-50
-150
-100
-50
0
50
SRF 180-2000
DCM 0.8-5
SRF 20-180
DCM 5-20
SRF 5-20
DCM 20-180
SRF 0.8-5
DCM 180-2000
root;cellular organisms;Eukaryota;Opisthokonta;Metazoa;Eumetazoa;Bilateria

#### Biogeolocalisation of MGT-v1 626 1045 unigenes, 85 (8.13%) taxonomically assigned

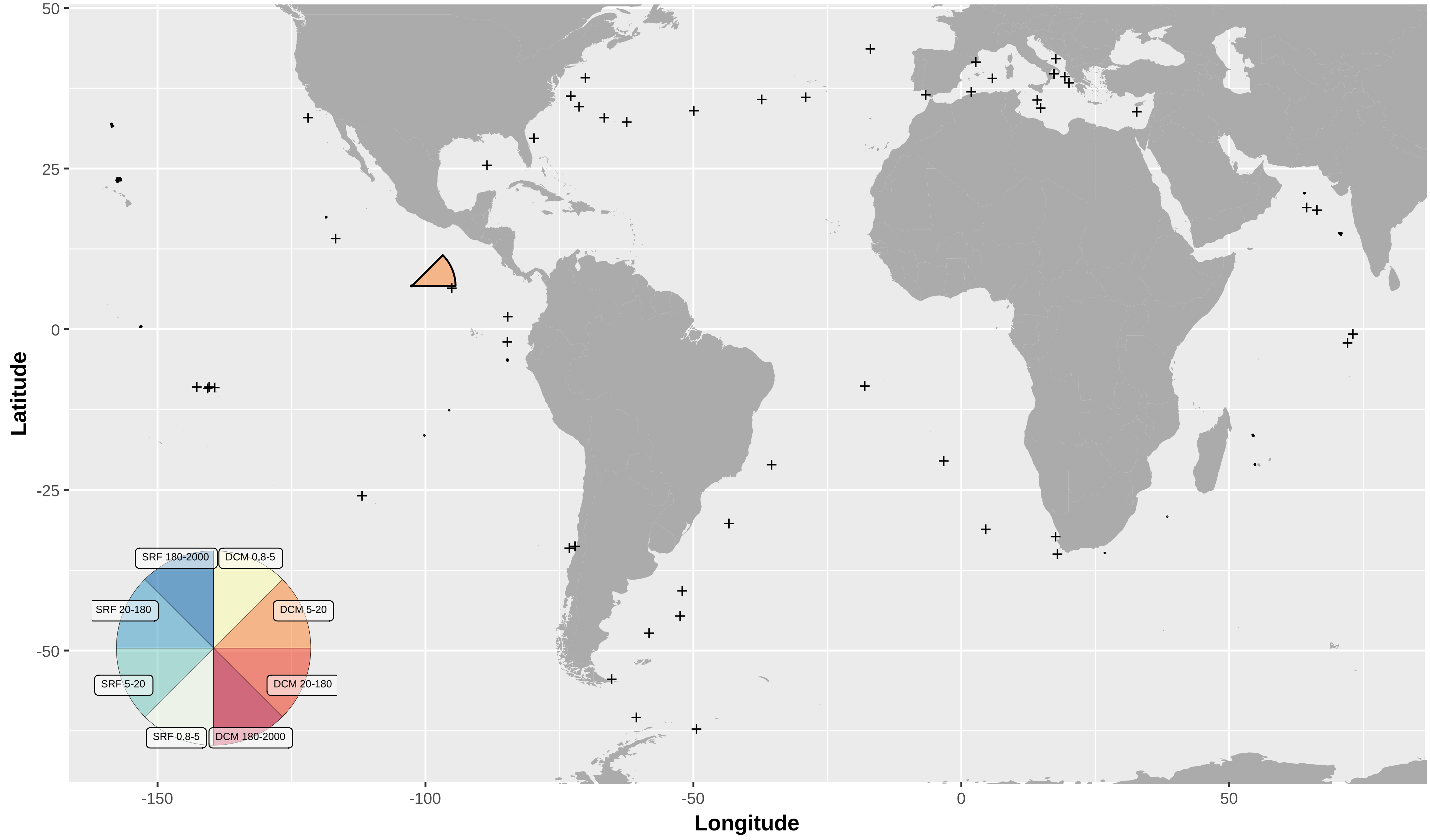


root;cellular organisms;Eukaryota;Opisthokonta;Metazoa;Eumetazoa

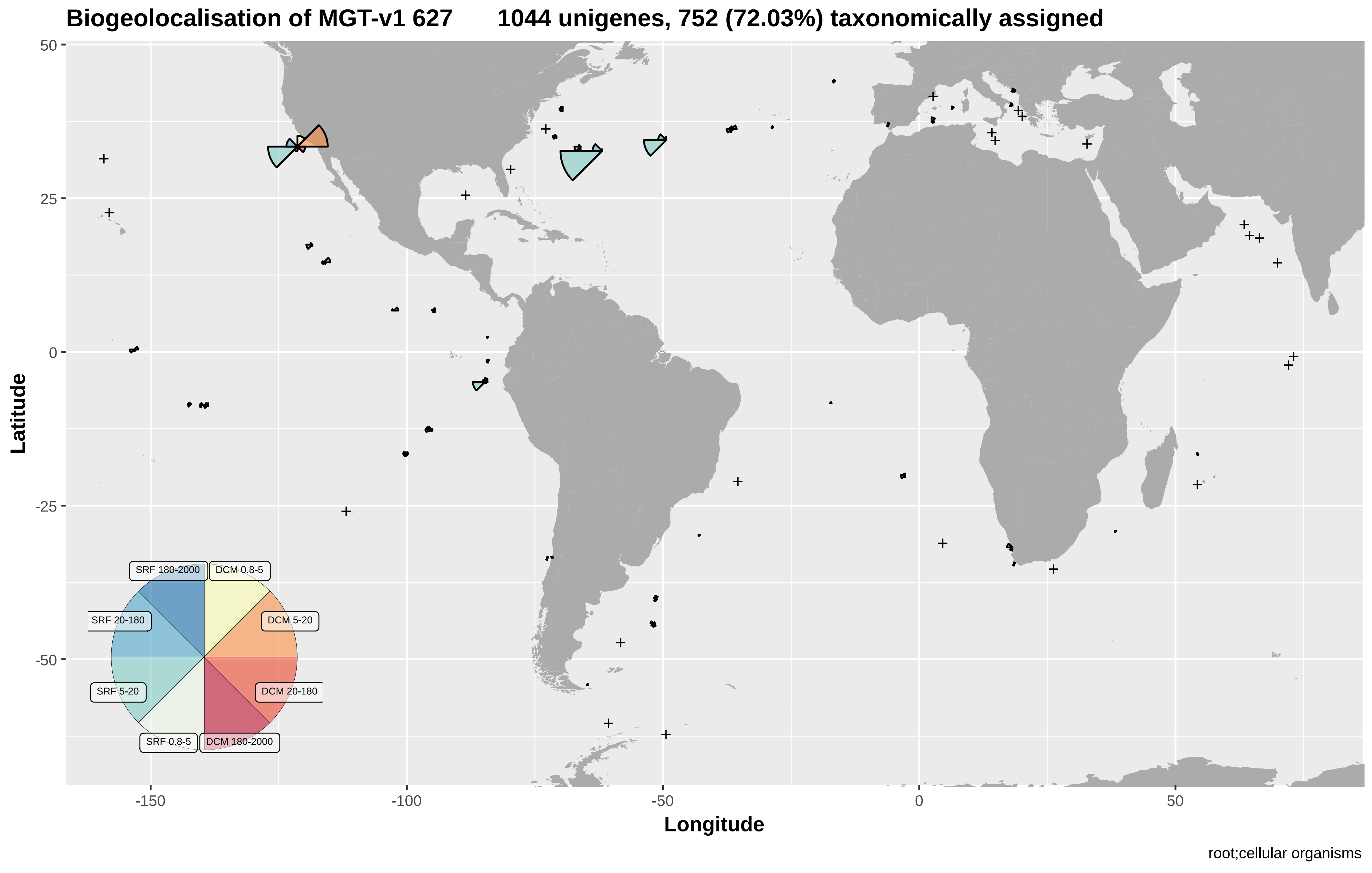
Biogeolocalisation of MGT-v1 627 1044 unigenes, 752 (72.03%) taxonomically assigned
50
25
0
-25
-50
Latitude
-150
-100
-50
0
50
Longitude
SRF 180-2000
DCM 0.8-5
SRF 20-180
DCM 5-20
SRF 5-20
DCM 20-180
SRF 0.8-5
DCM 180-2000
root;cellular organisms

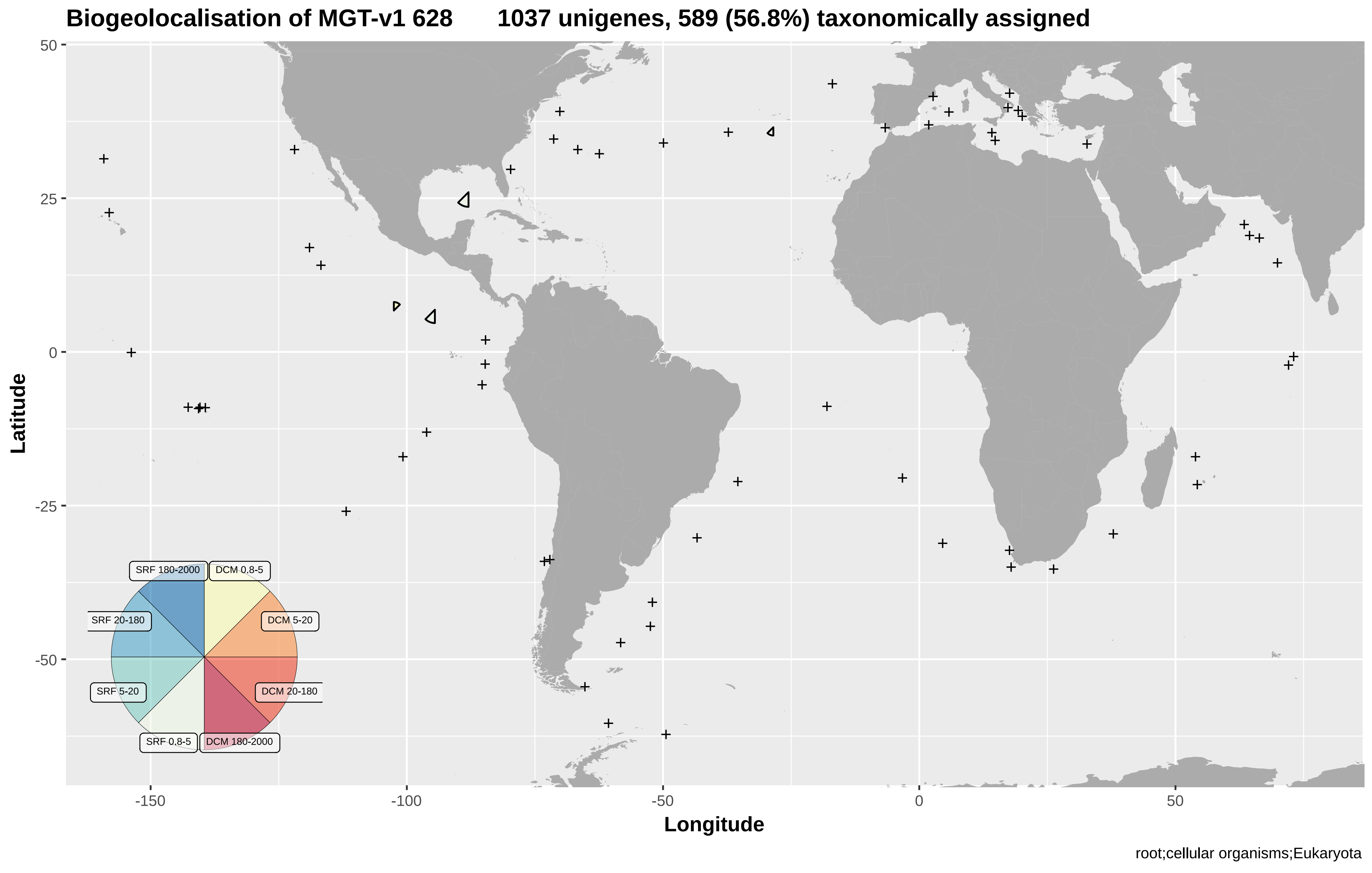
Biogeolocalisation of MGT-v1 628 1037 unigenes, 589 (56.8%) taxonomically assigned
Latitude
Longitude
50
25
0
-25
-50
-150
-100
-50
0
50
SRF 180-2000
DCM 0.8-5
SRF 20-180
DCM 5-20
SRF 5-20
DCM 20-180
SRF 0.8-5
DCM 180-2000
root;cellular organisms;Eukaryota

Biogeolocalisation of MGT-v1 629 1027 unigenes, 984 (95.81%) taxonomically assigned

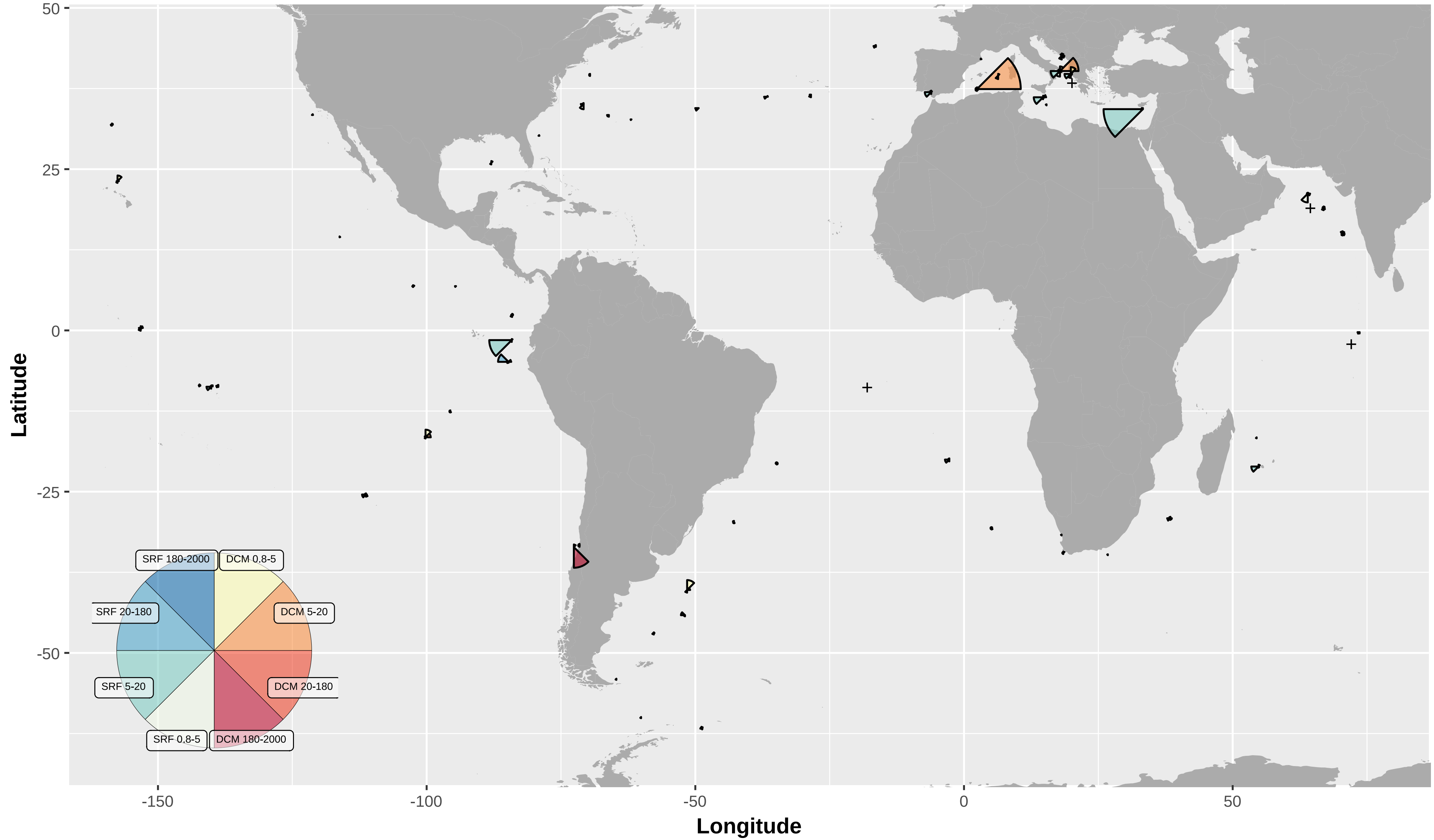


root;cellular organisms;Eukaryota

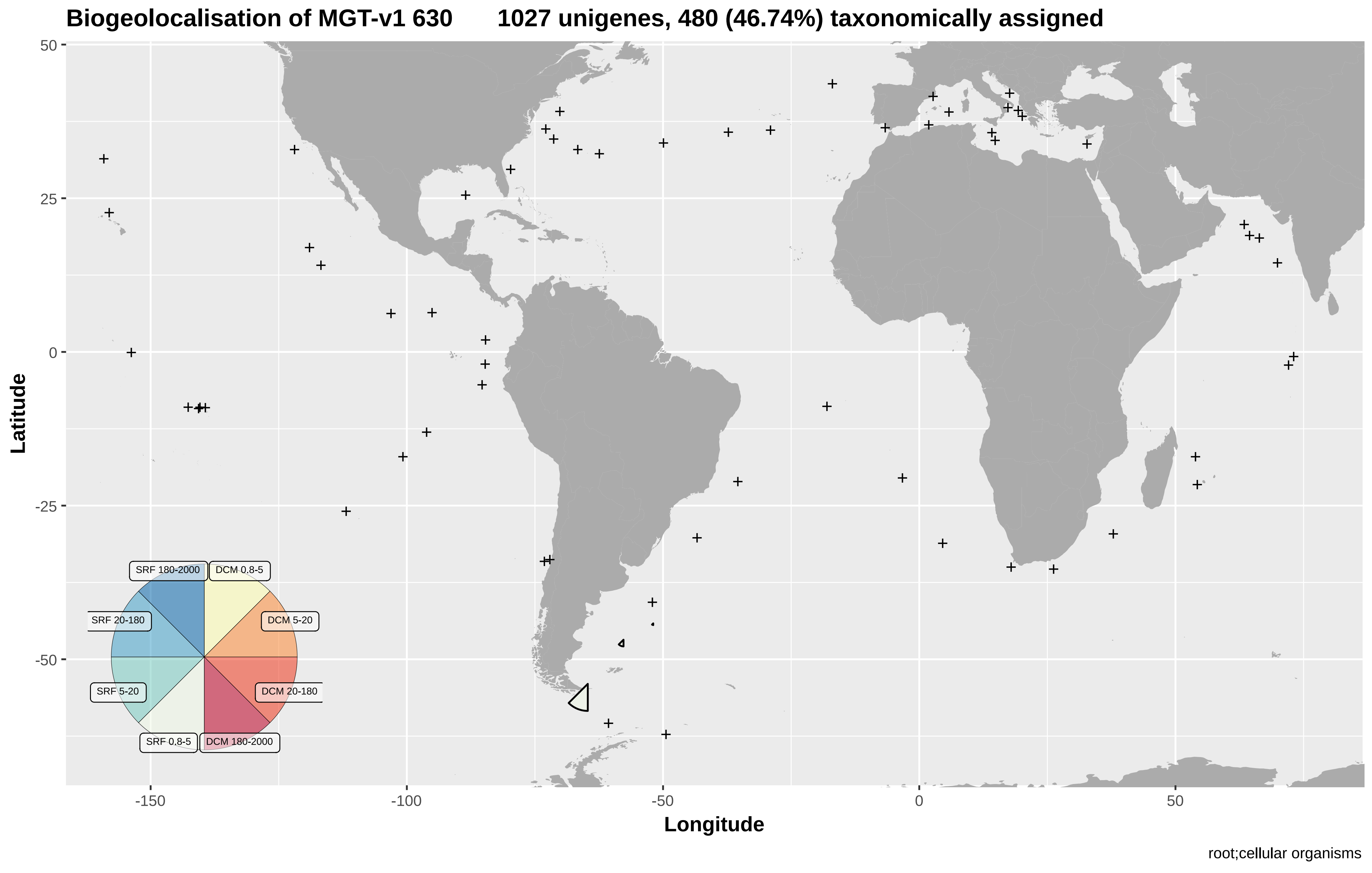

Biogeolocalisation of MGT-v1 630 1027 unigenes, 480 (46.74%) taxonomically assigned
50
25
0
-25
-50
Latitude
-150
-100
-50
0
50
Longitude
SRF 180-2000
DCM 0.8-5
SRF 20-180
DCM 5-20
SRF 5-20
DCM 20-180
SRF 0.8-5
DCM 180-2000
root;cellular organisms

## Biogeolocalisation of MGT-v1 631      1023 unigenes, 743 (72.63%) taxonomically assigned

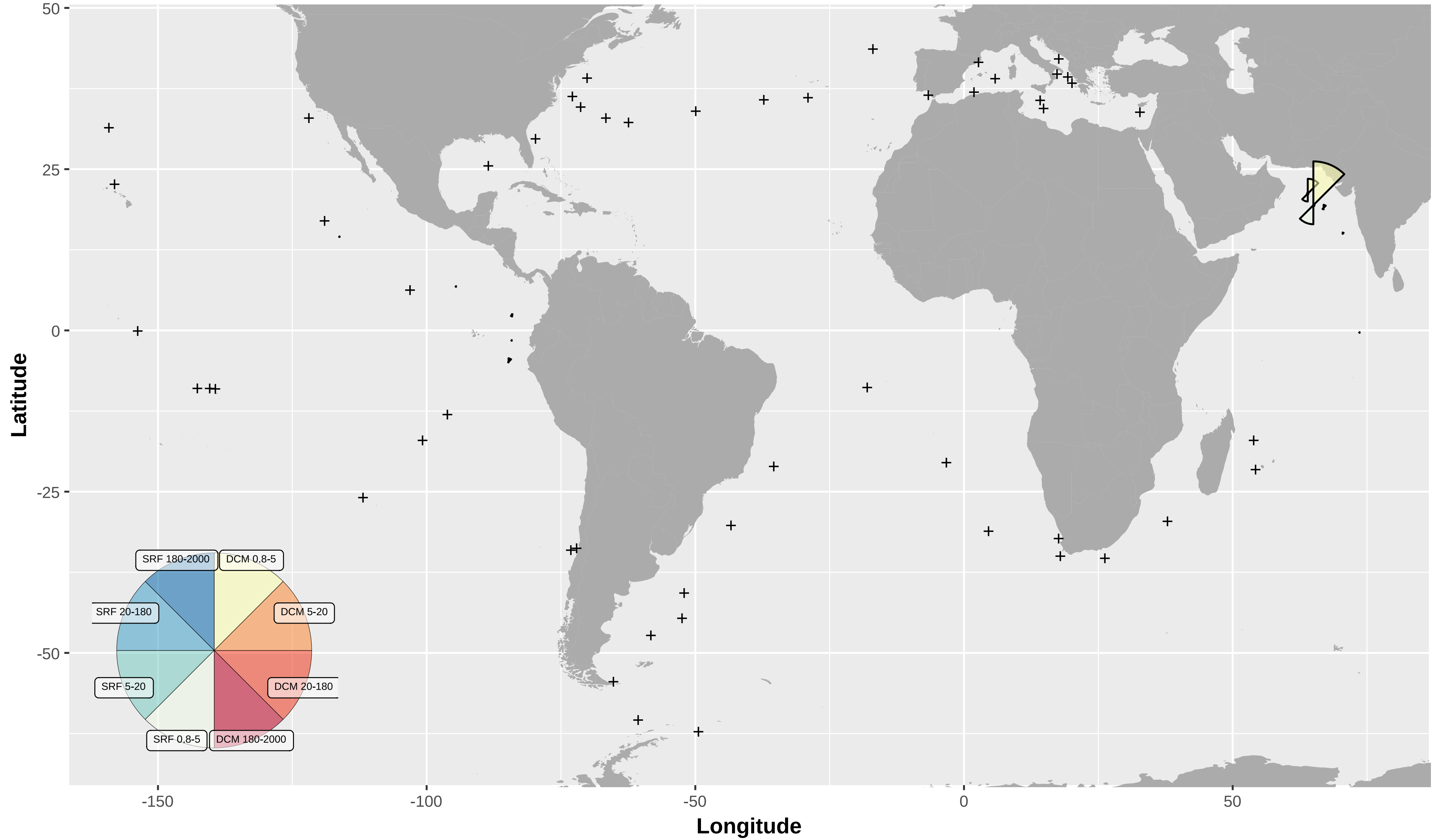


root

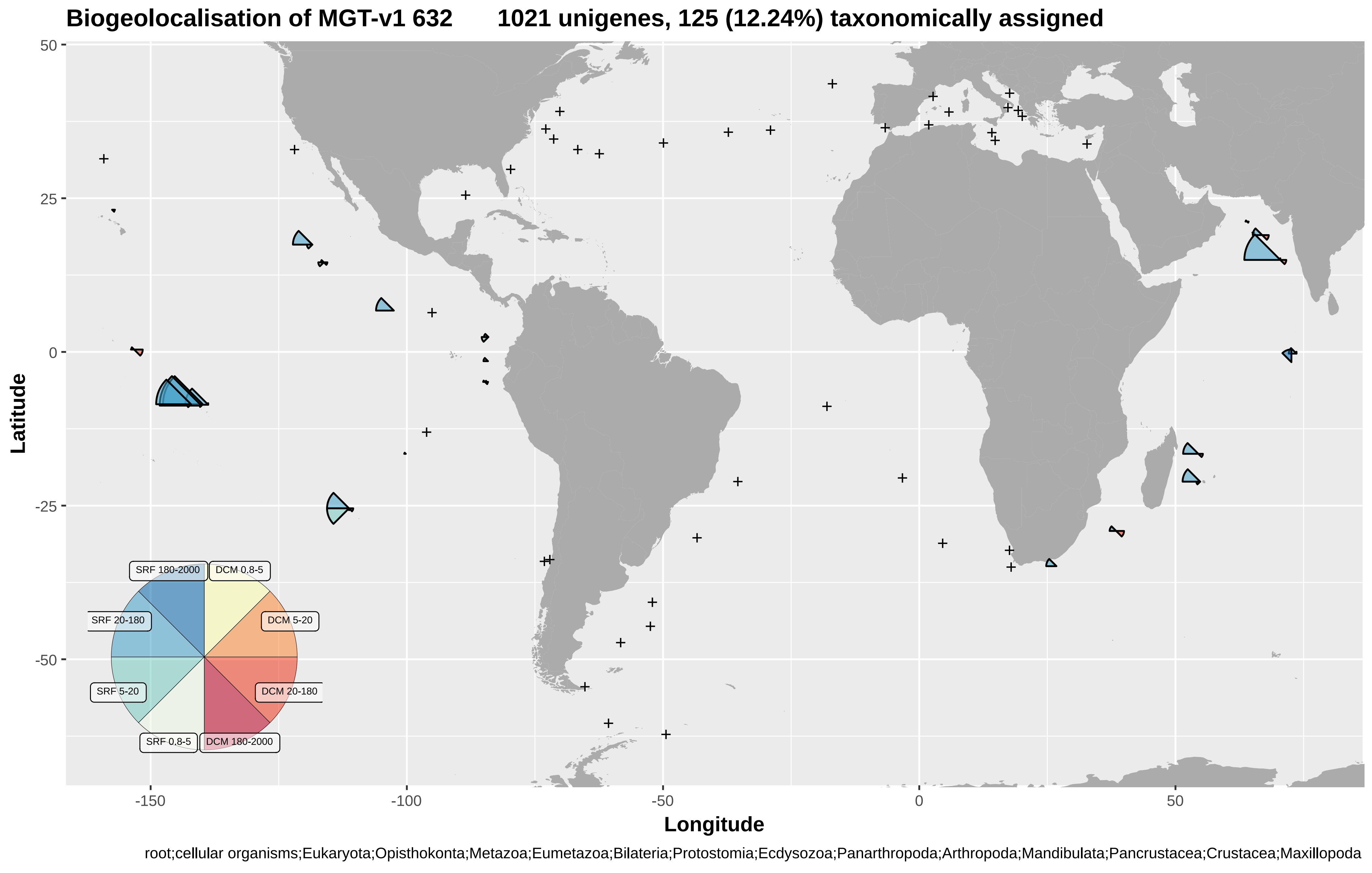

Biogeolocalisation of MGT-v1 632 1021 unigenes, 125 (12.24%) taxonomically assigned
Latitude
Longitude
50
25
0
-25
-50
-150
-100
-50
0
50
SRF 180-2000
DCM 0.8-5
SRF 20-180
DCM 5-20
SRF 5-20
DCM 20-180
SRF 0.8-5
DCM 180-2000
root;cellular organisms;Eukaryota;Opisthokonta;Metazoa;Eumetazoa;Bilateria;Protostomia;Ecdysozoa;Panarthropoda;Arthropoda;Mandibulata;Pancrustacea;Crustacea;Maxillopoda

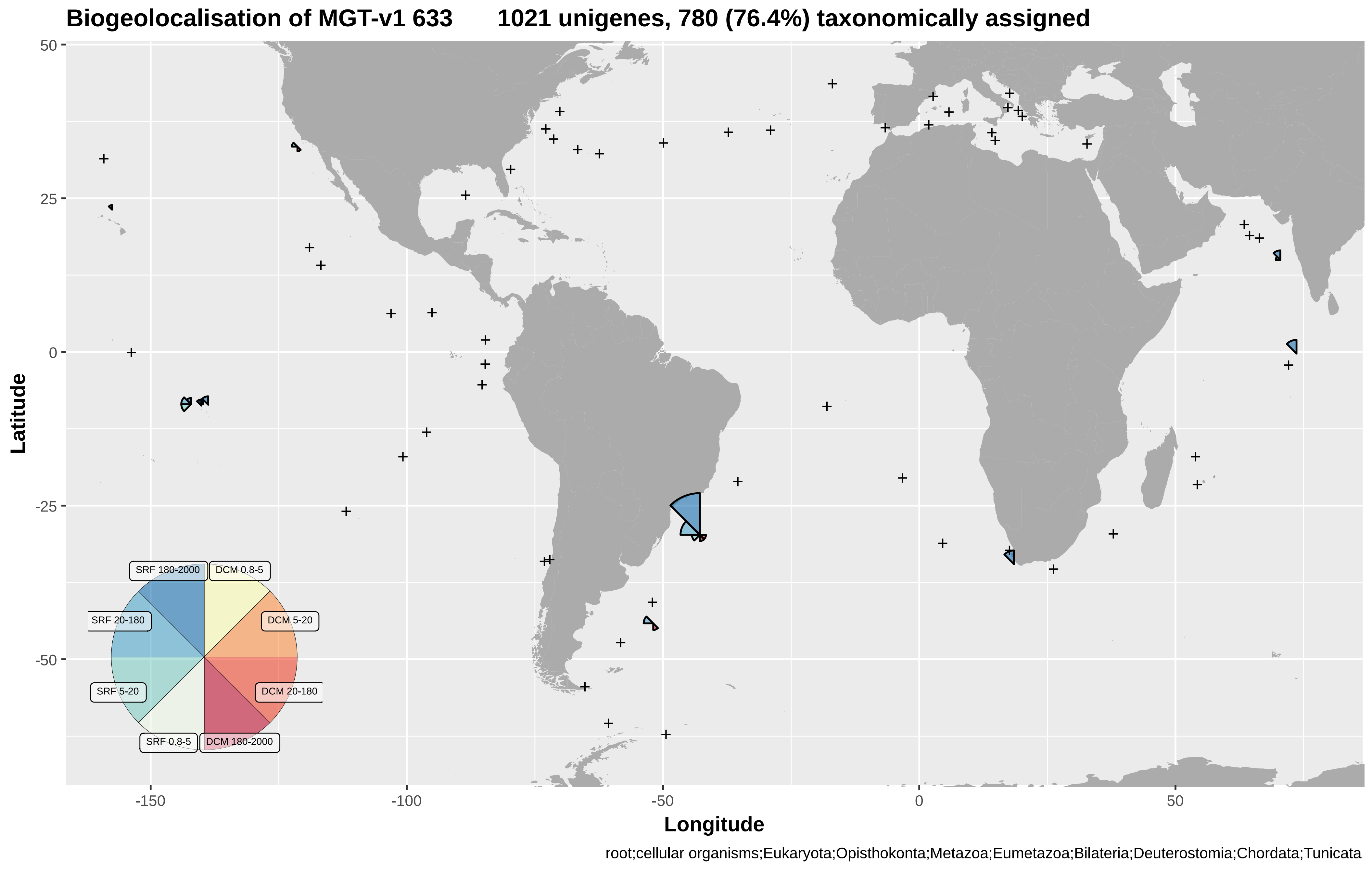
Biogeolocalisation of MGT-v1 633 1021 unigenes, 780 (76.4%) taxonomically assigned
Latitude
Longitude
50
25
0
-25
-50
-150
-100
-50
0
50
SRF 180-2000
DCM 0.8-5
SRF 20-180
DCM 5-20
SRF 5-20
DCM 20-180
SRF 0.8-5
DCM 180-2000
root;cellular organisms;Eukaryota;Opisthokonta;Metazoa;Eumetazoa;Bilateria;Deuterostomia;Chordata;Tunicata

## Biogeolocalisation of MGT-v1 634    1018 unigenes, 59 (5.8%) taxonomically assigned

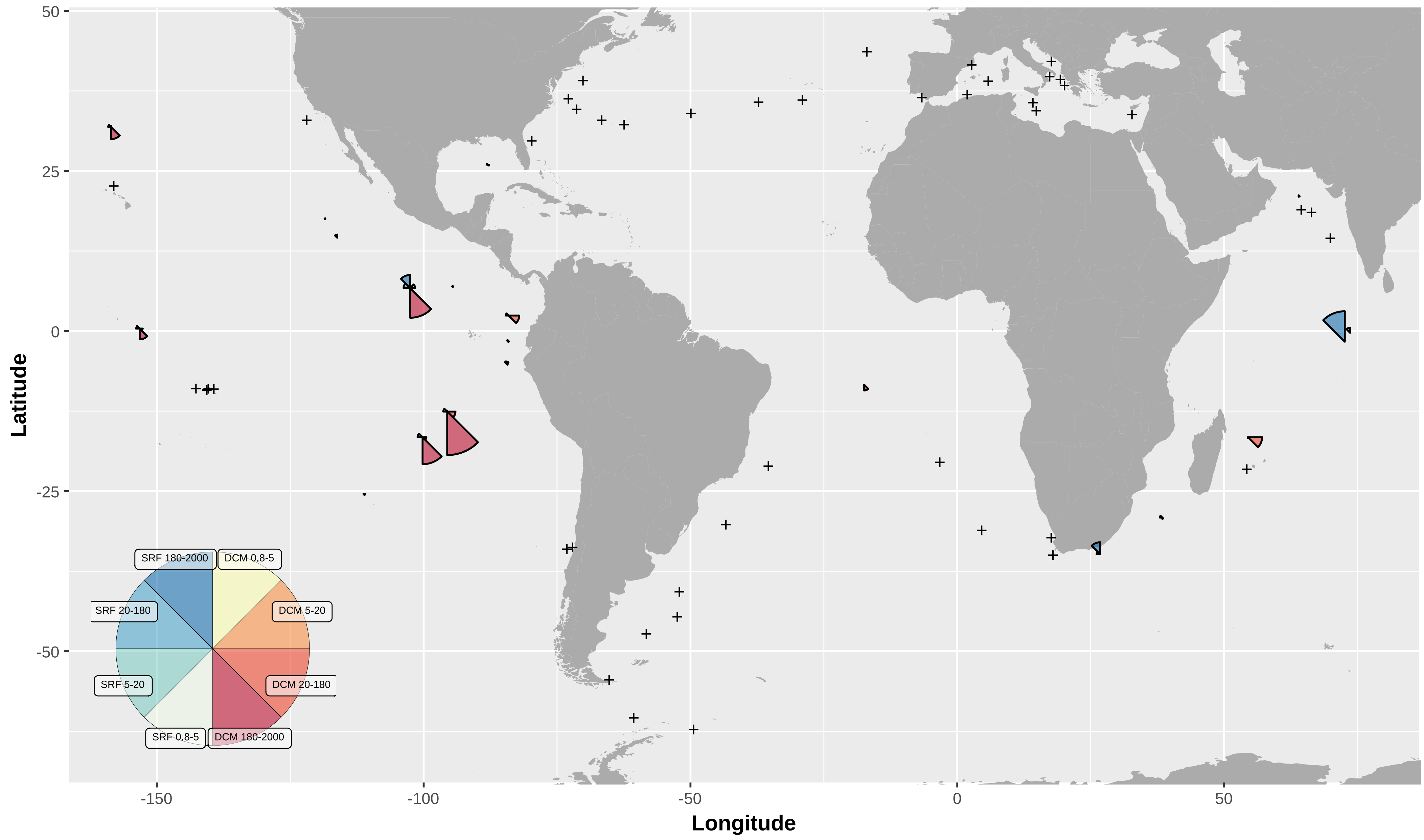


root;cellular organisms;Eukaryota;Opisthokonta;Metazoa;Eumetazoa

#### Biogeolocalisation of MGT-v1 635 1015 unigenes, 71 (7%) taxonomically assigned

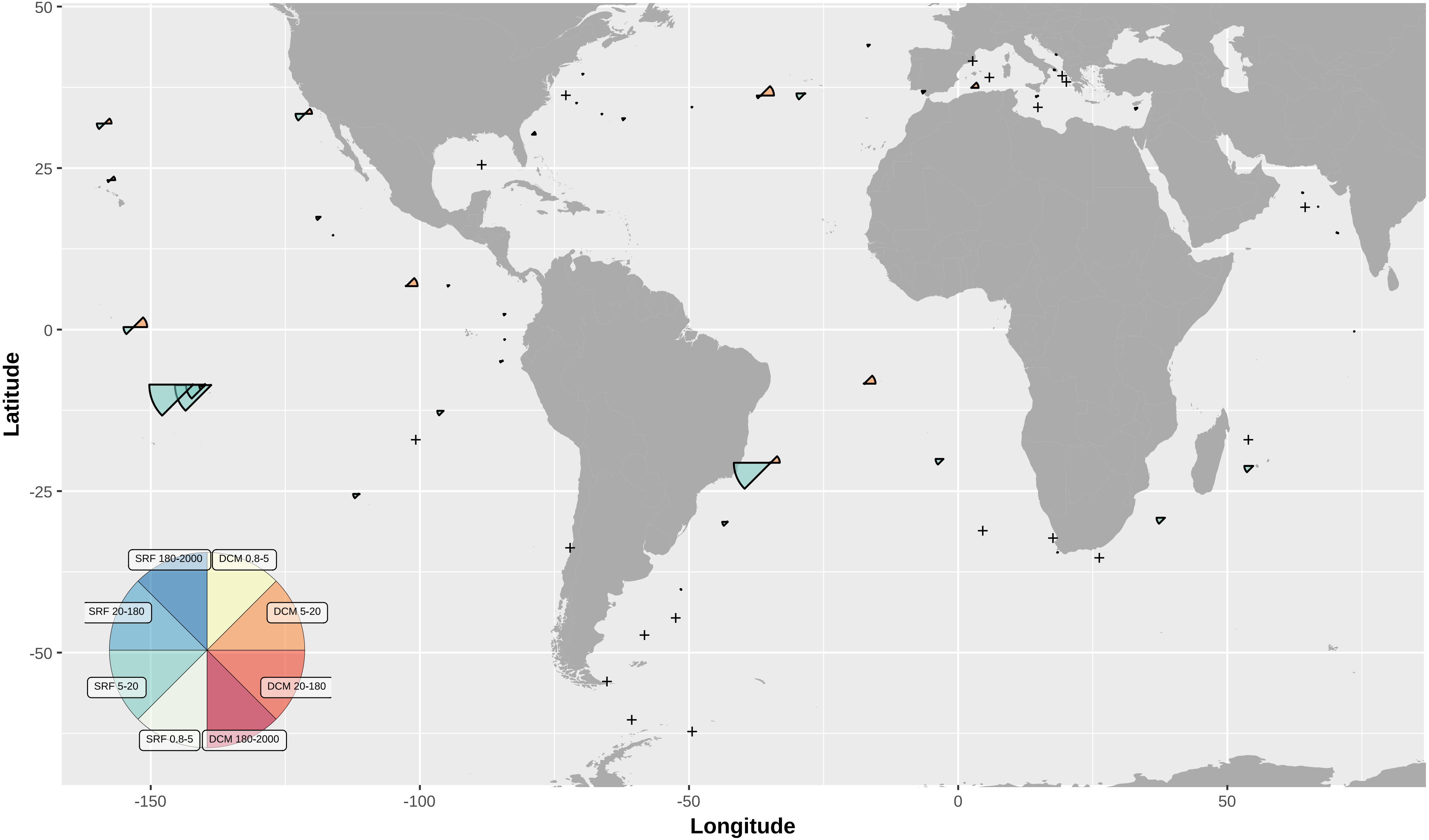


root;cellular organisms;Eukaryota

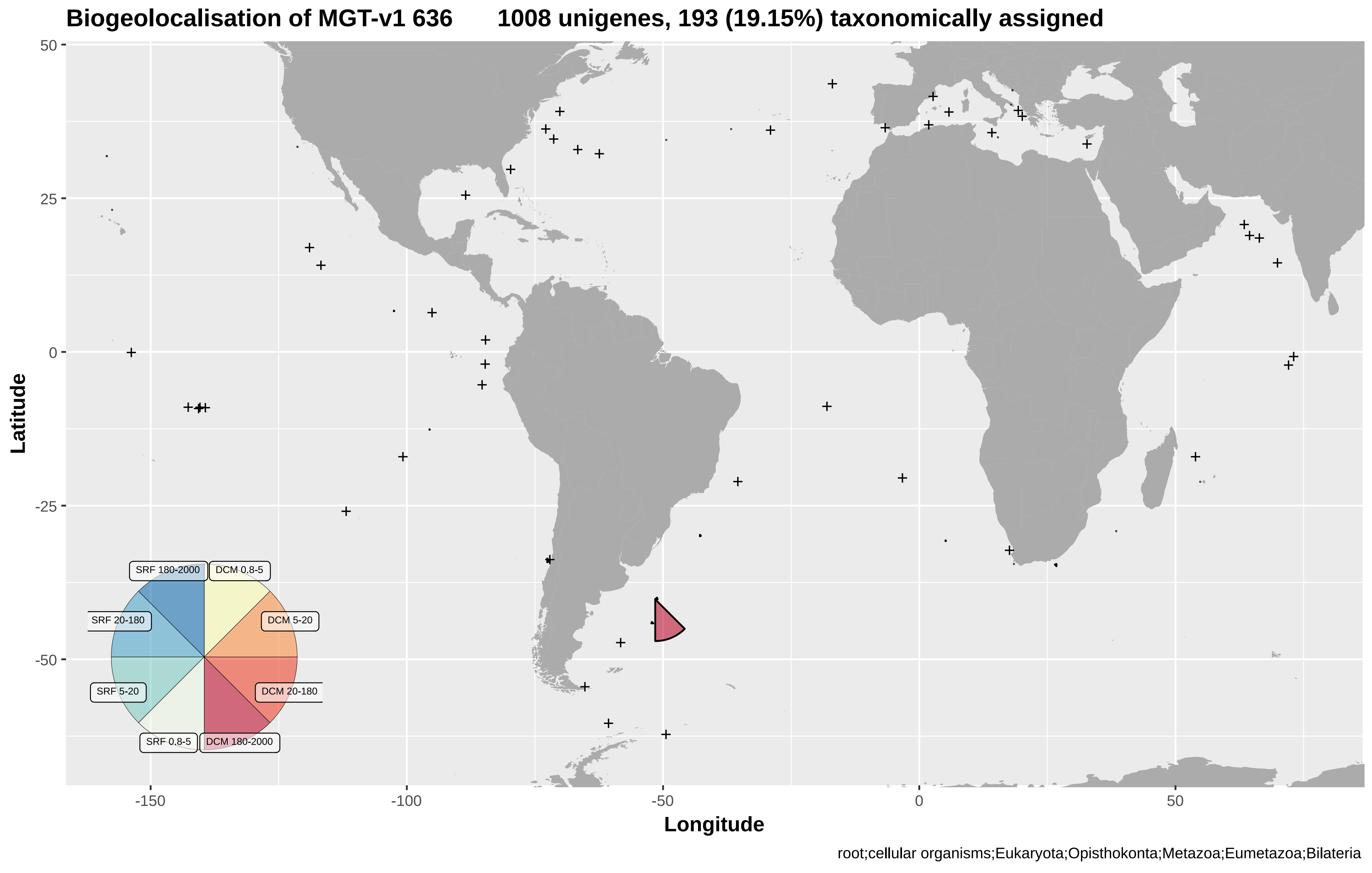
Biogeolocalisation of MGT-v1 636
1008 unigenes, 193 (19.15%) taxonomically assigned
Latitude
Longitude
50
25
0
-25
-50
-150
-100
-50
0
50
SRF 180-2000
DCM 0.8-5
SRF 20-180
DCM 5-20
SRF 5-20
DCM 20-180
SRF 0.8-5
DCM 180-2000
root;cellular organisms;Eukaryota;Opisthokonta;Metazoa;Eumetazoa;Bilateria

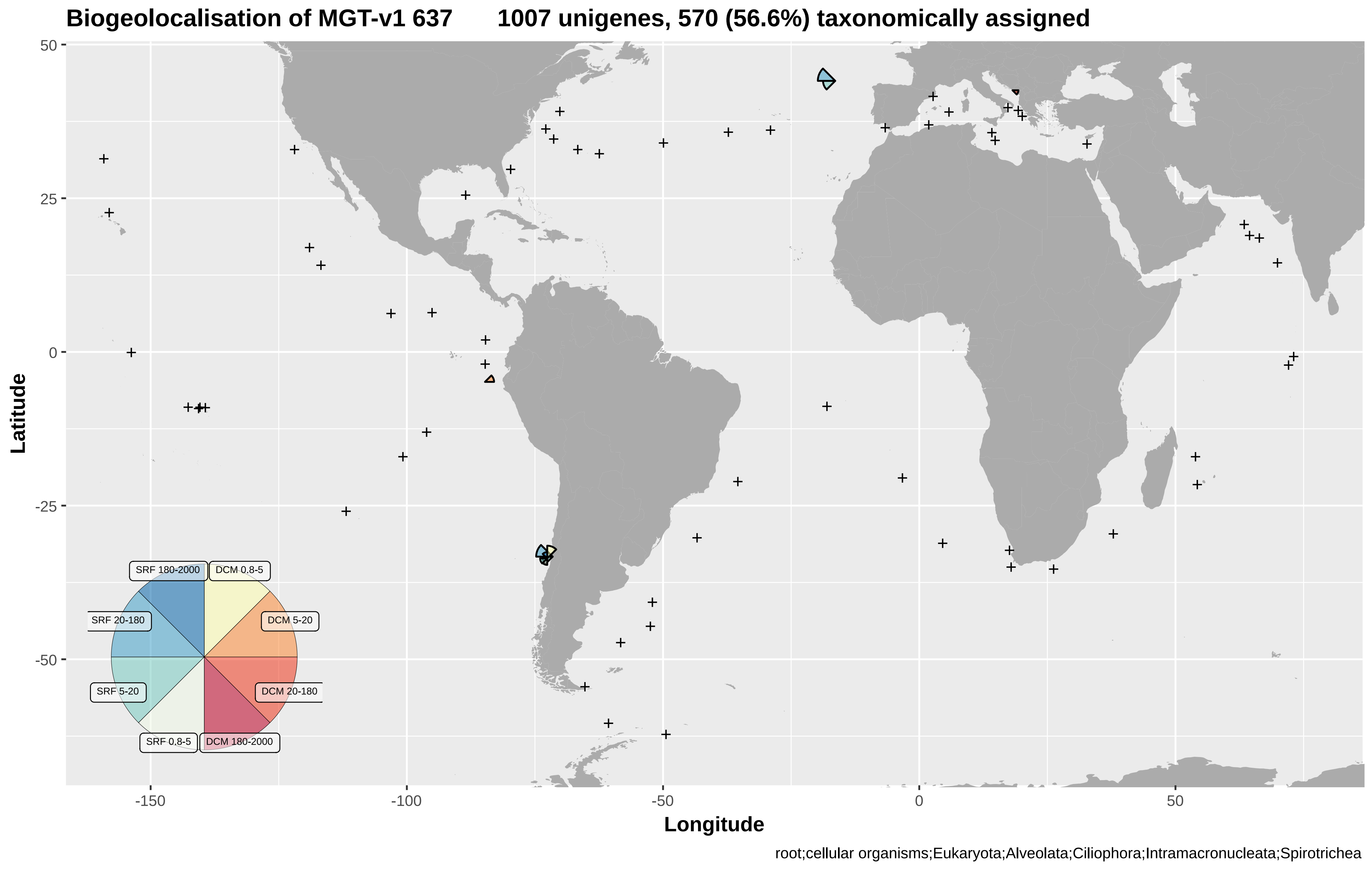
Biogeolocalisation of MGT-v1 637 1007 unigenes, 570 (56.6%) taxonomically assigned
50
25
0
-25
-50
Latitude
-150
-100
-50
0
50
Longitude
SRF 180-2000
DCM 0.8-5
SRF 20-180
DCM 5-20
SRF 5-20
DCM 20-180
SRF 0.8-5
DCM 180-2000
root;cellular organisms;Eukaryota;Alveolata;Ciliophora;Intramacronucleata;Spirotrichea

#### Biogeolocalisation of MGT-v1 638 995 unigenes, 108 (10.85%) taxonomically assigned

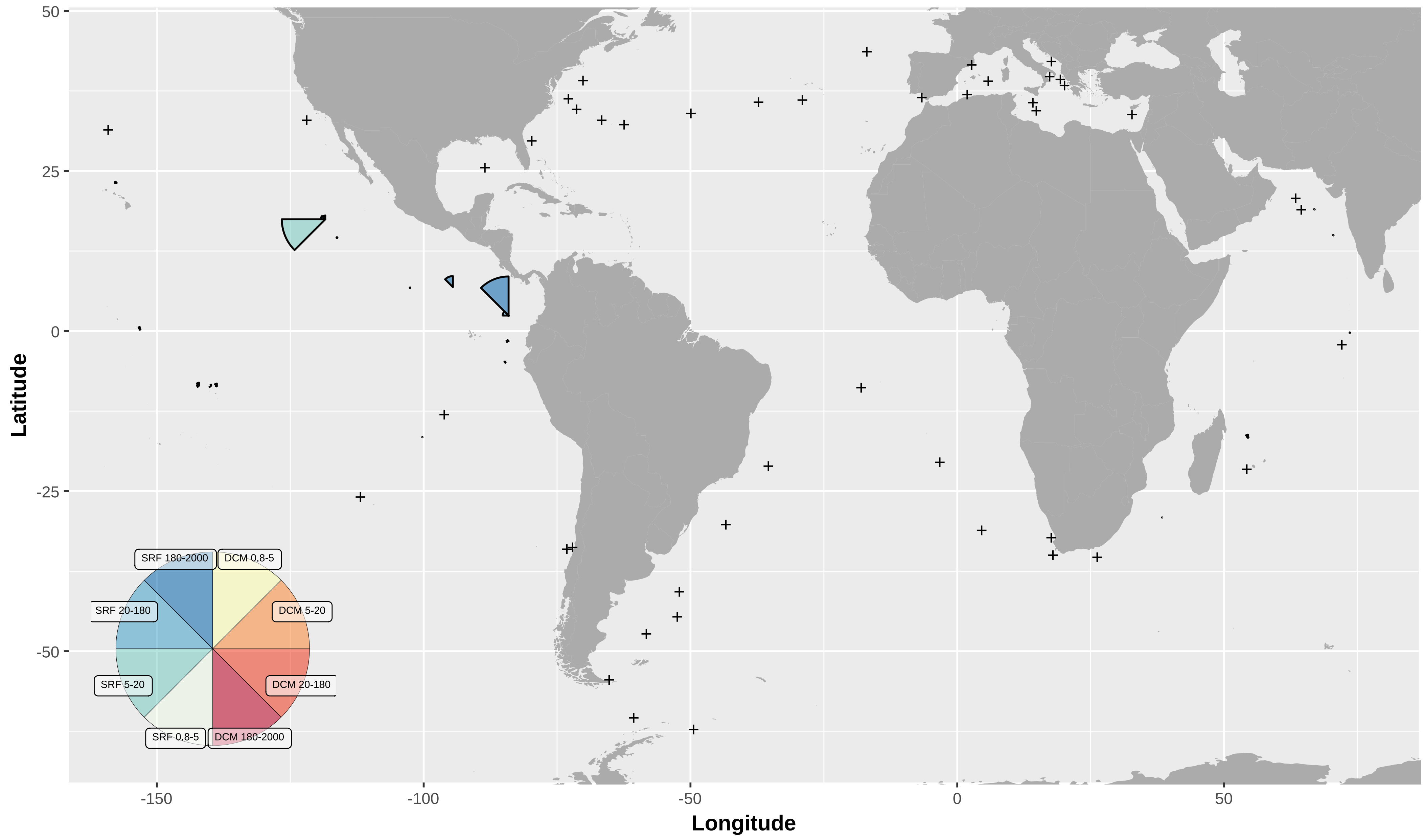


root;cellular organisms;Eukaryota;Opisthokonta;Metazoa;Eumetazoa;Bilateria

## Biogeolocalisation of MGT-v1 639      993 unigenes, 141 (14.2%) taxonomically assigned

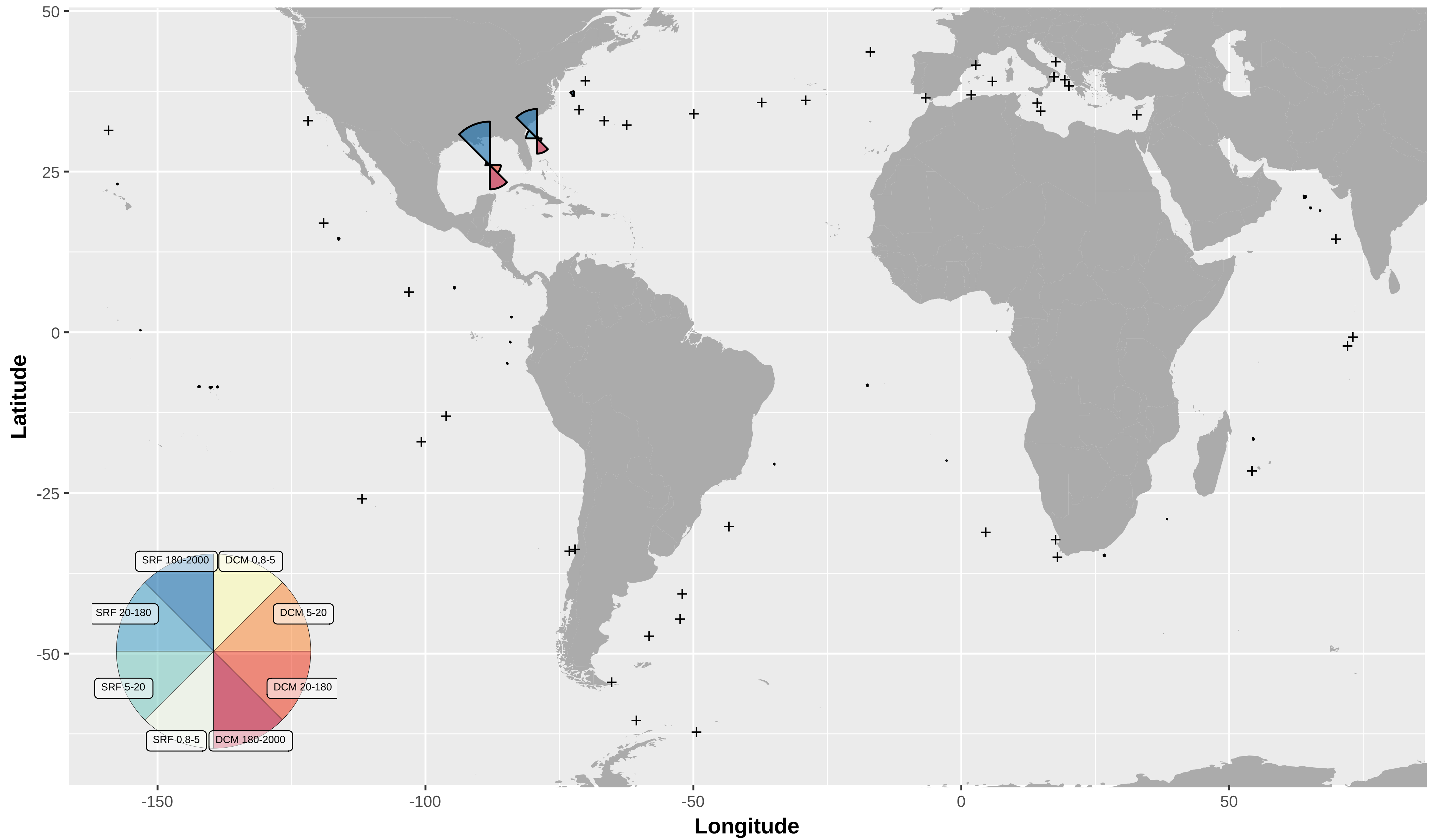


root;cellular organisms;Eukaryota;Opisthokonta;Metazoa;Eumetazoa;Bilateria

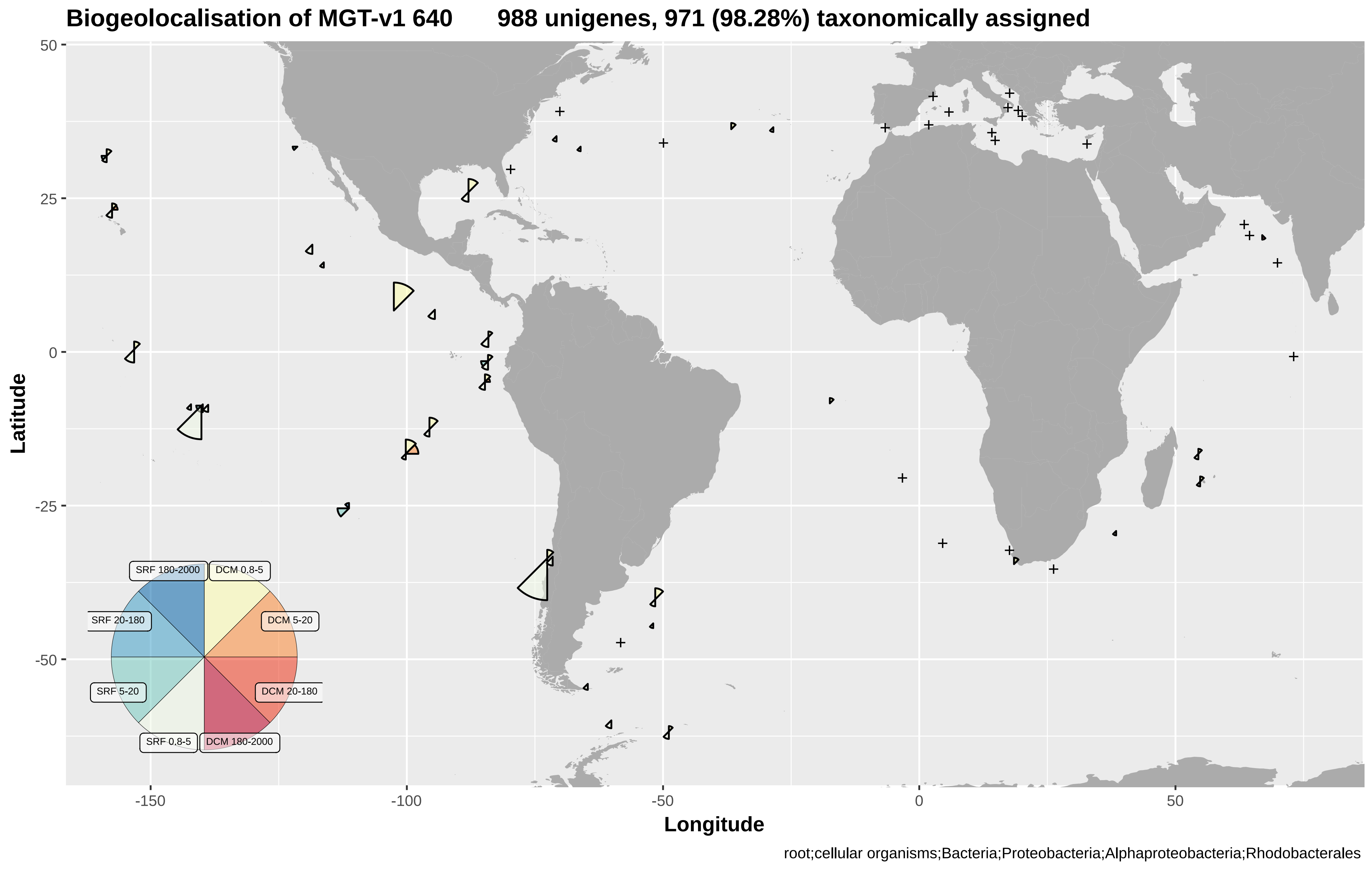
Biogeolocalisation of MGT-v1 640 988 unigenes, 971 (98.28%) taxonomically assigned
Latitude
Longitude
50
25
0
-25
-50
-150
-100
-50
0
50
SRF 180-2000
DCM 0.8-5
SRF 20-180
DCM 5-20
SRF 5-20
DCM 20-180
SRF 0.8-5
DCM 180-2000
root;cellular organisms;Bacteria;Proteobacteria;Alphaproteobacteria;Rhodobacterales

Biogeolocalisation of MGT-v1 641      980 unigenes, 64 (6.53%) taxonomically assigned

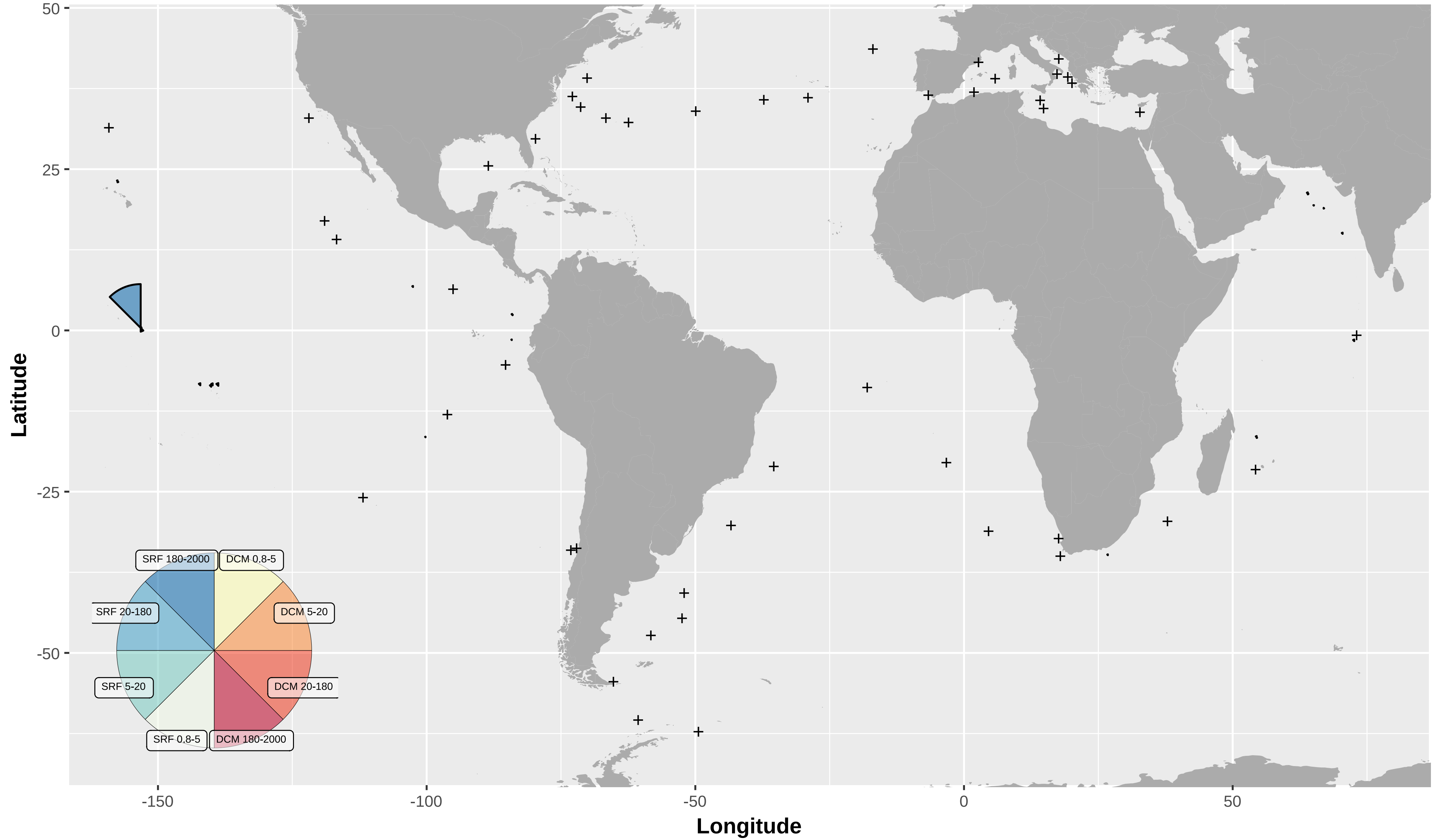


root;cellular organisms;Eukaryota;Opisthokonta;Metazoa;Eumetazoa

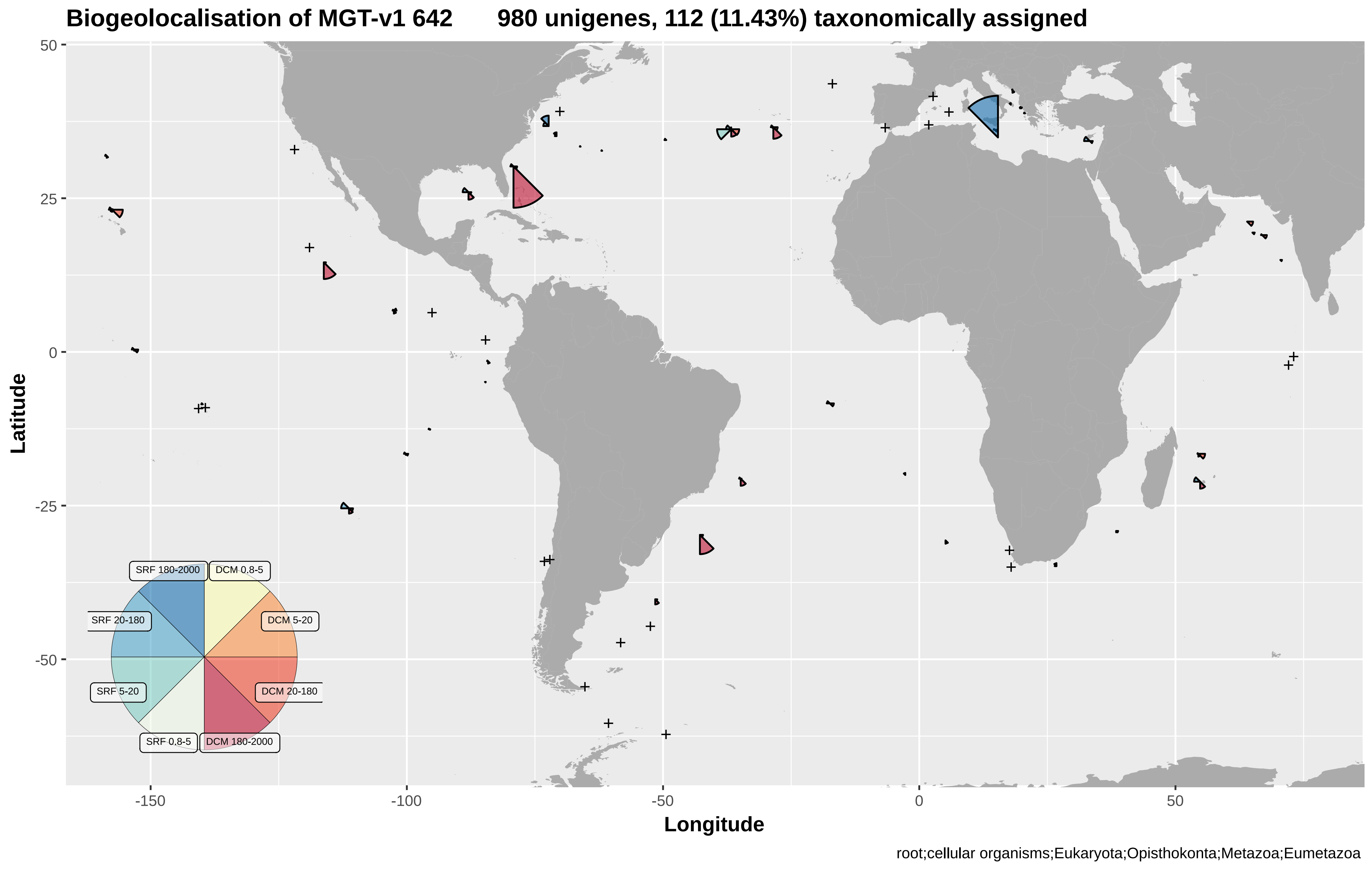

Biogeolocalisation of MGT-v1 642 980 unigenes, 112 (11.43%) taxonomically assigned
Latitude
Longitude
50
25
0
-25
-50
-150
-100
-50
0
50
SRF 180-2000
DCM 0.8-5
SRF 20-180
DCM 5-20
SRF 5-20
DCM 20-180
SRF 0.8-5
DCM 180-2000
root;cellular organisms;Eukaryota;Opisthokonta;Metazoa;Eumetazoa

Biogeolocalisation of MGT-v1 643      978 unigenes, 907 (92.74%) taxonomically assigned

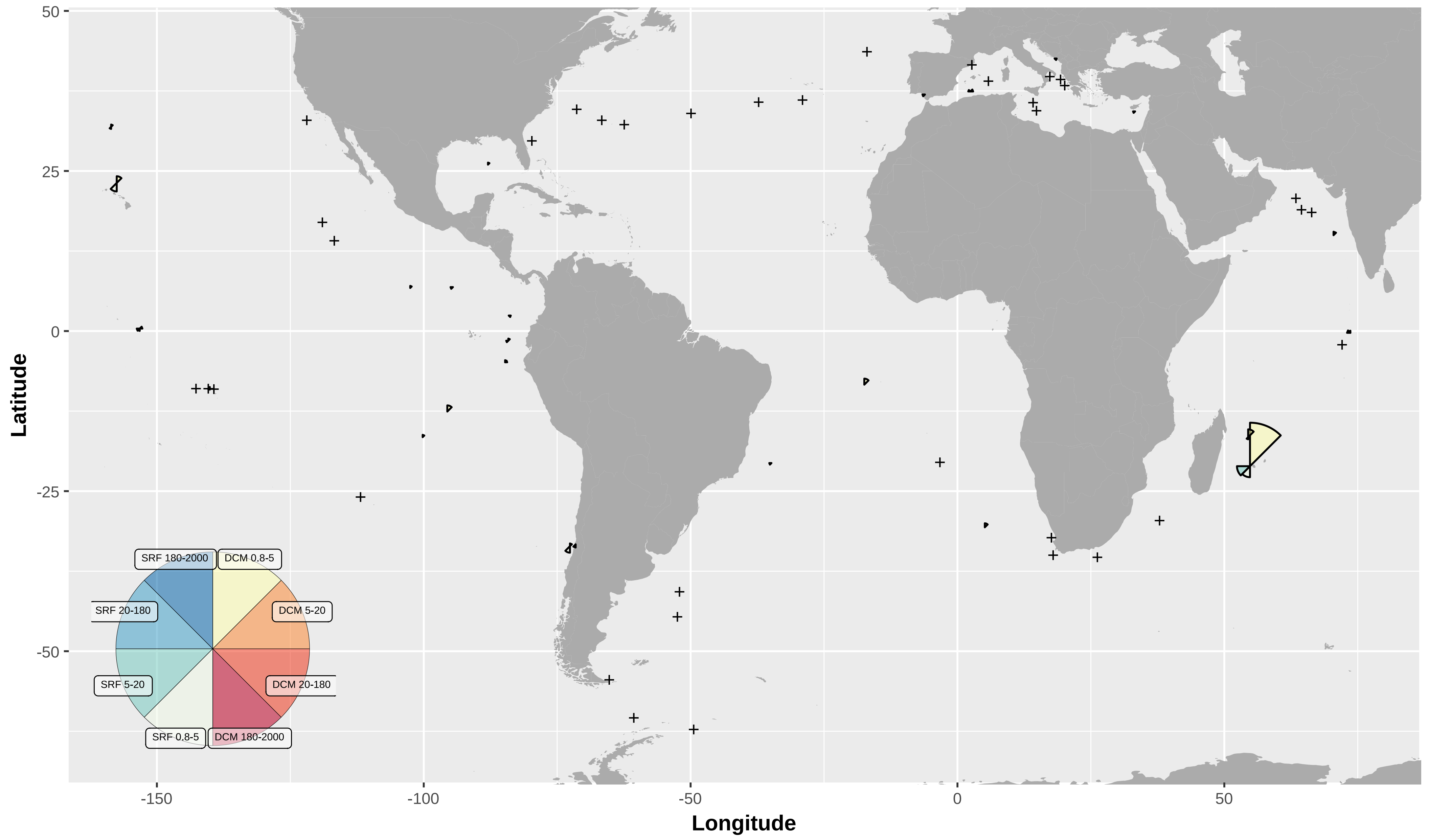


root;cellular organisms;Bacteria;Proteobacteria;Gammaproteobacteria;Alteromonadales;Alteromonadaceae

## Biogeolocalisation of MGT-v1 644      978 unigenes, 679 (69.43%) taxonomically assigned

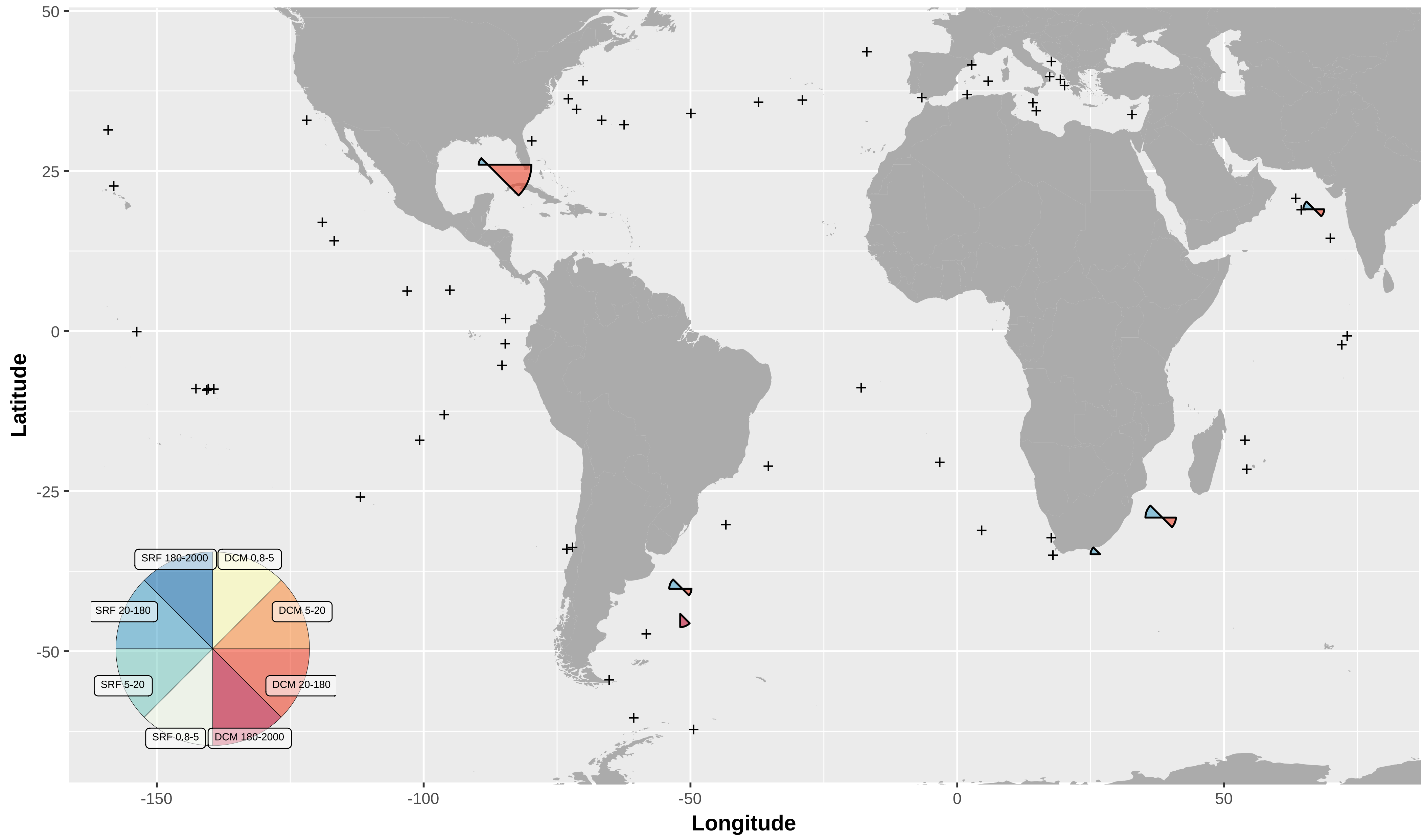


root;cellular organisms;Eukaryota;Stramenopiles;Bacillariophyta

# Biogeolocalisation of MGT-v1 645      978 unigenes, 297 (30.37%) taxonomically assigned

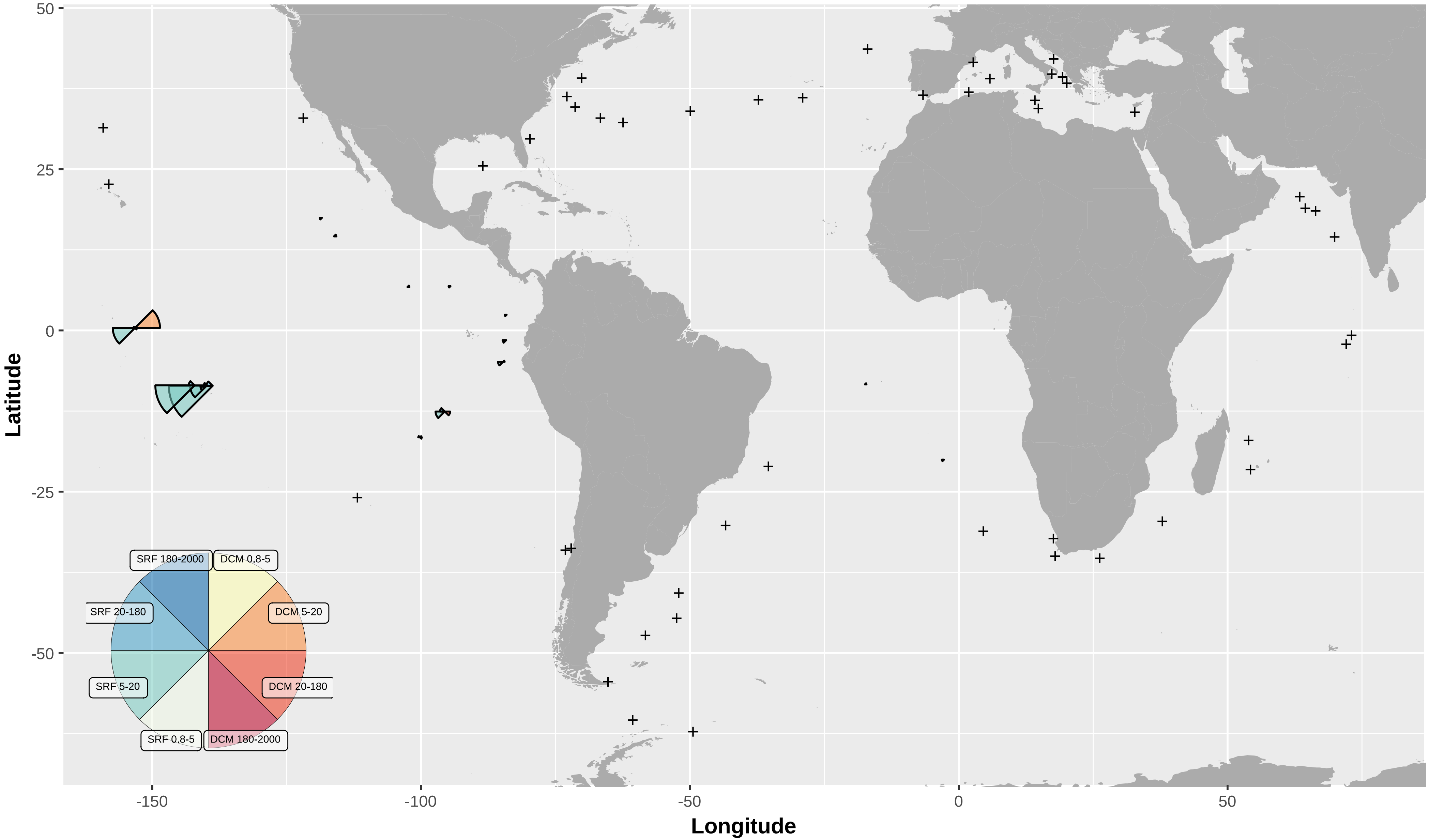


root;cellular organisms;Eukaryota;Stramenopiles;Bacillariophyta

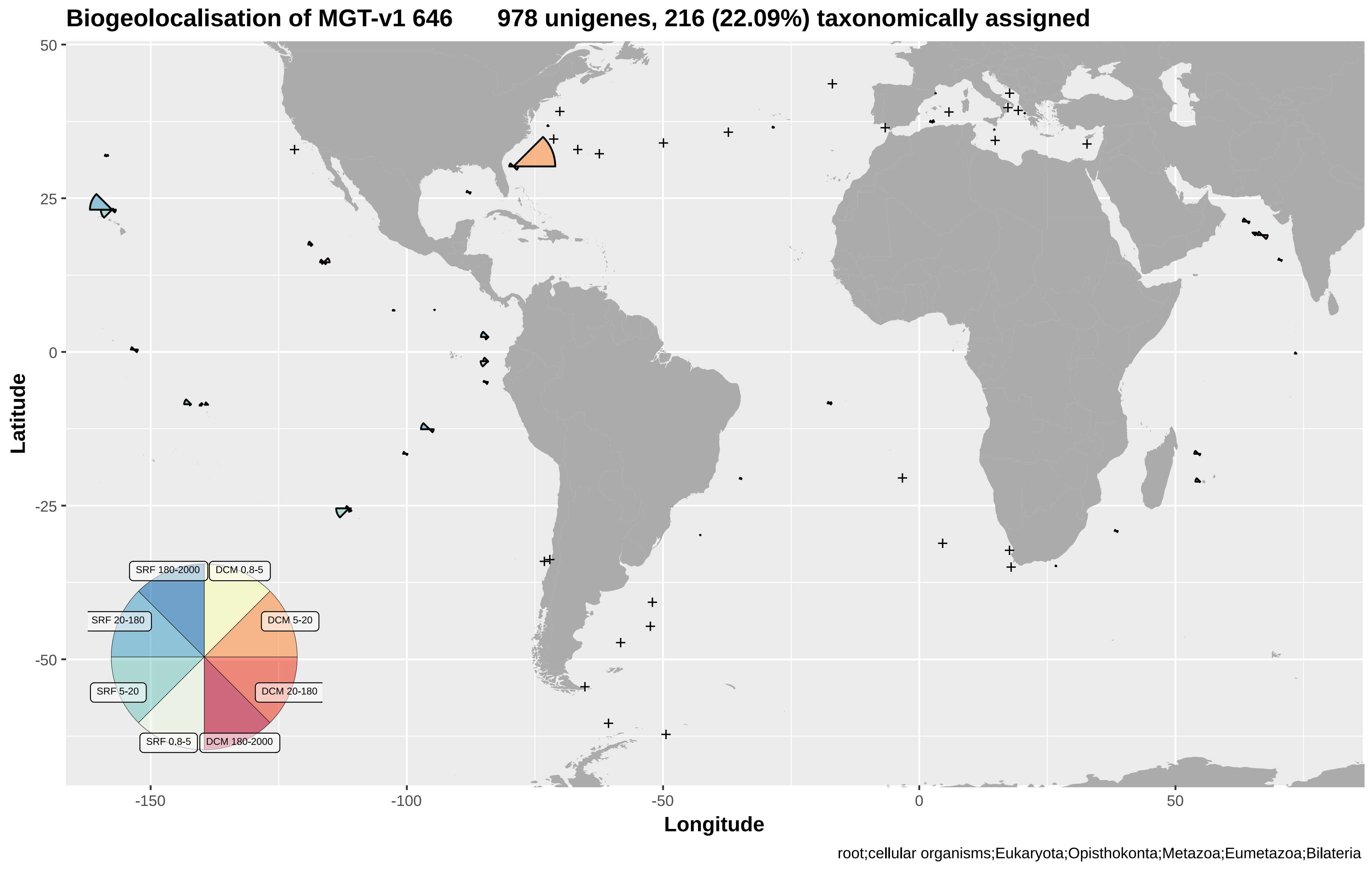
Biogeolocalisation of MGT-v1 646 978 unigenes, 216 (22.09%) taxonomically assigned
Latitude
Longitude
50
25
0
-25
-50
-150
-100
-50
0
50
SRF 180-2000
DCM 0.8-5
SRF 20-180
DCM 5-20
SRF 5-20
DCM 20-180
SRF 0.8-5
DCM 180-2000
root;cellular organisms;Eukaryota;Opisthokonta;Metazoa;Eumetazoa;Bilateria

## Biogeolocalisation of MGT-v1 647      977 unigenes, 962 (98.46%) taxonomically assigned

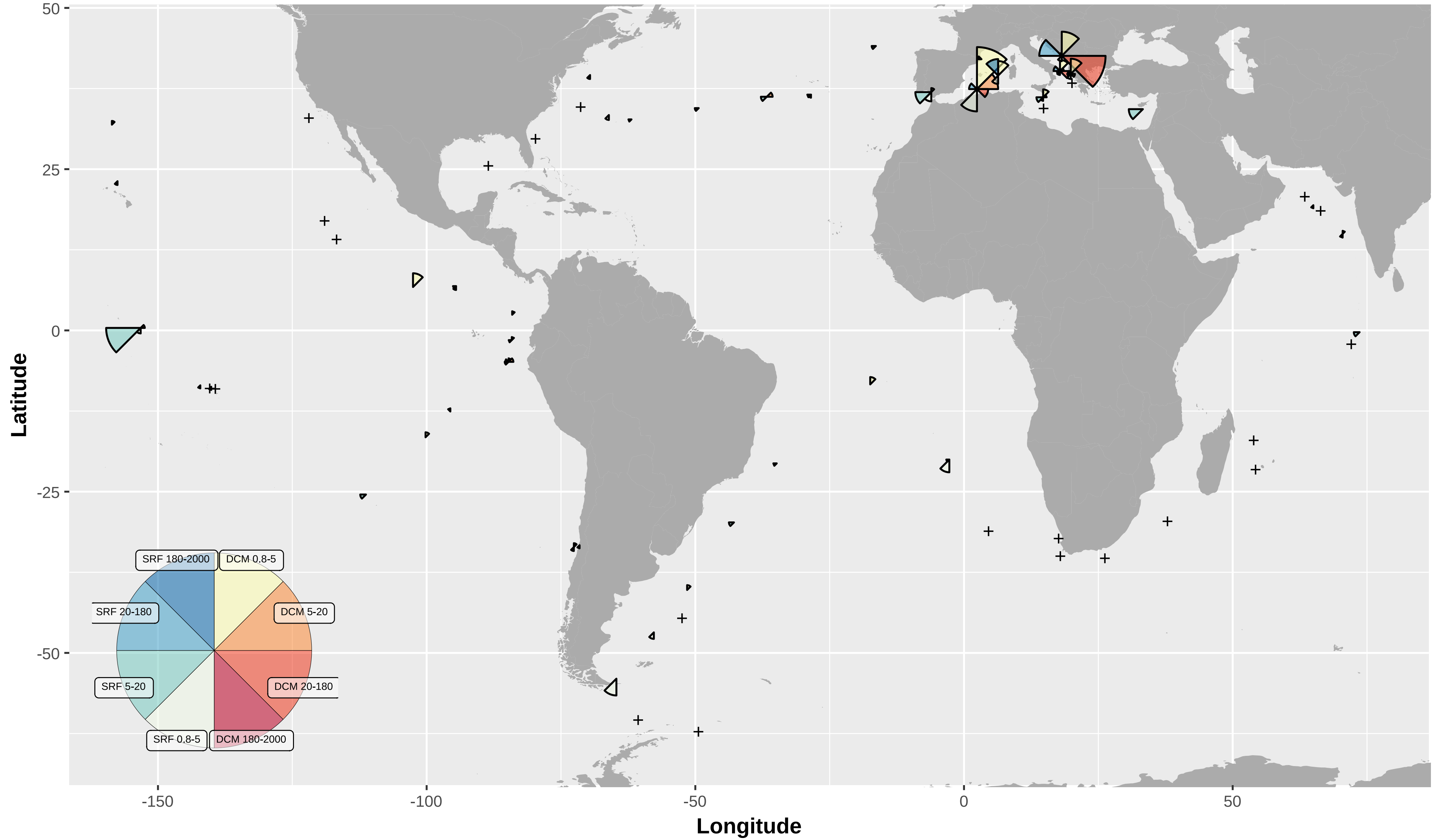


root;cellular organisms;Bacteria;Proteobacteria;Gammaproteobacteria;Alteromonadales;Alteromonadaceae

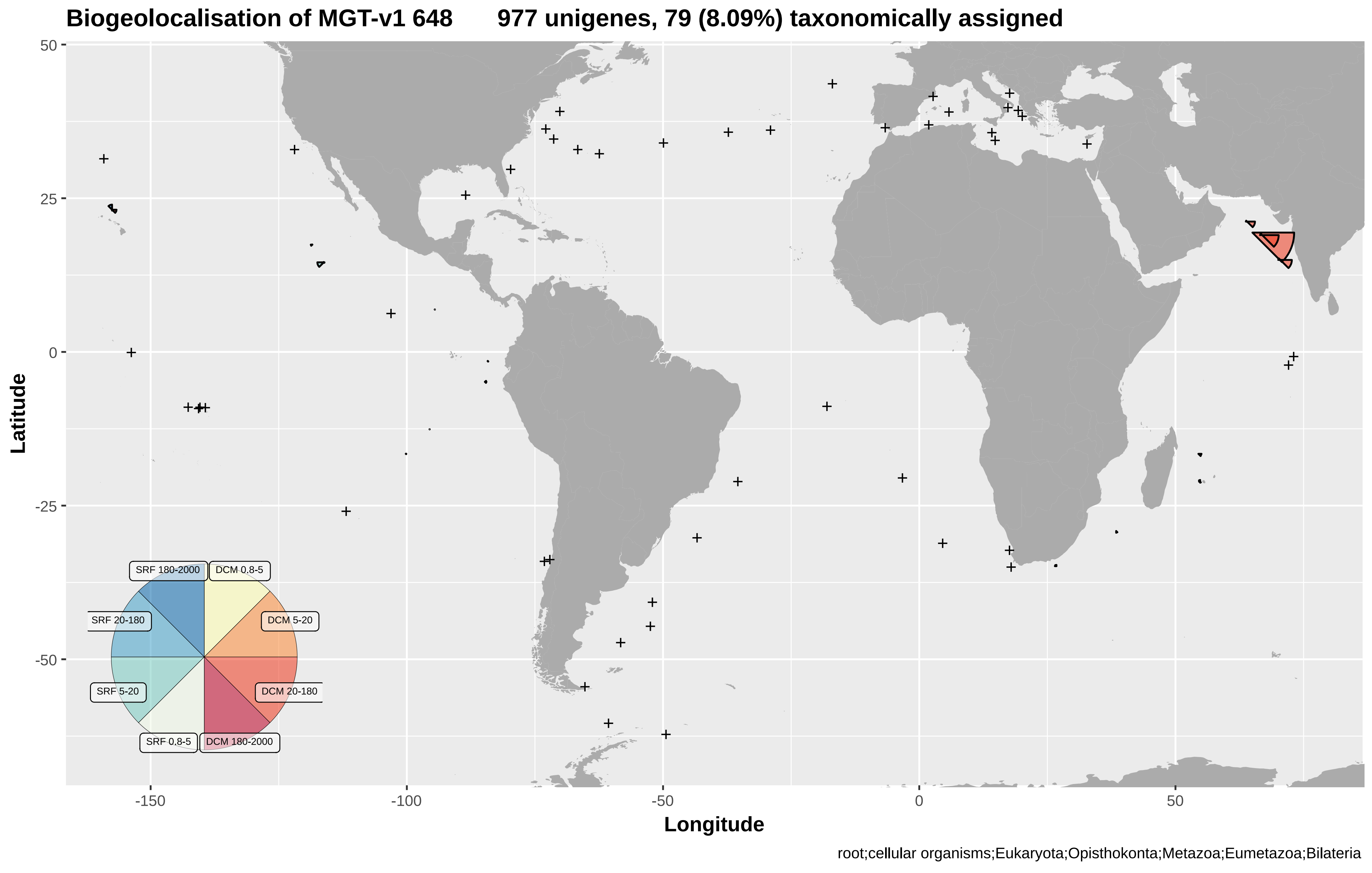
Biogeolocalisation of MGT-v1 648 977 unigenes, 79 (8.09%) taxonomically assigned
50
25
0
-25
-50
Latitude
-150
-100
-50
0
50
Longitude
SRF 180-2000
DCM 0.8-5
SRF 20-180
DCM 5-20
SRF 5-20
DCM 20-180
SRF 0.8-5
DCM 180-2000
root;cellular organisms;Eukaryota;Opisthokonta;Metazoa;Eumetazoa;Bilateria

**Biogeolocalisation of MGT-v1 649      976 unigenes, 944 (96.72%) taxonomically assigned**

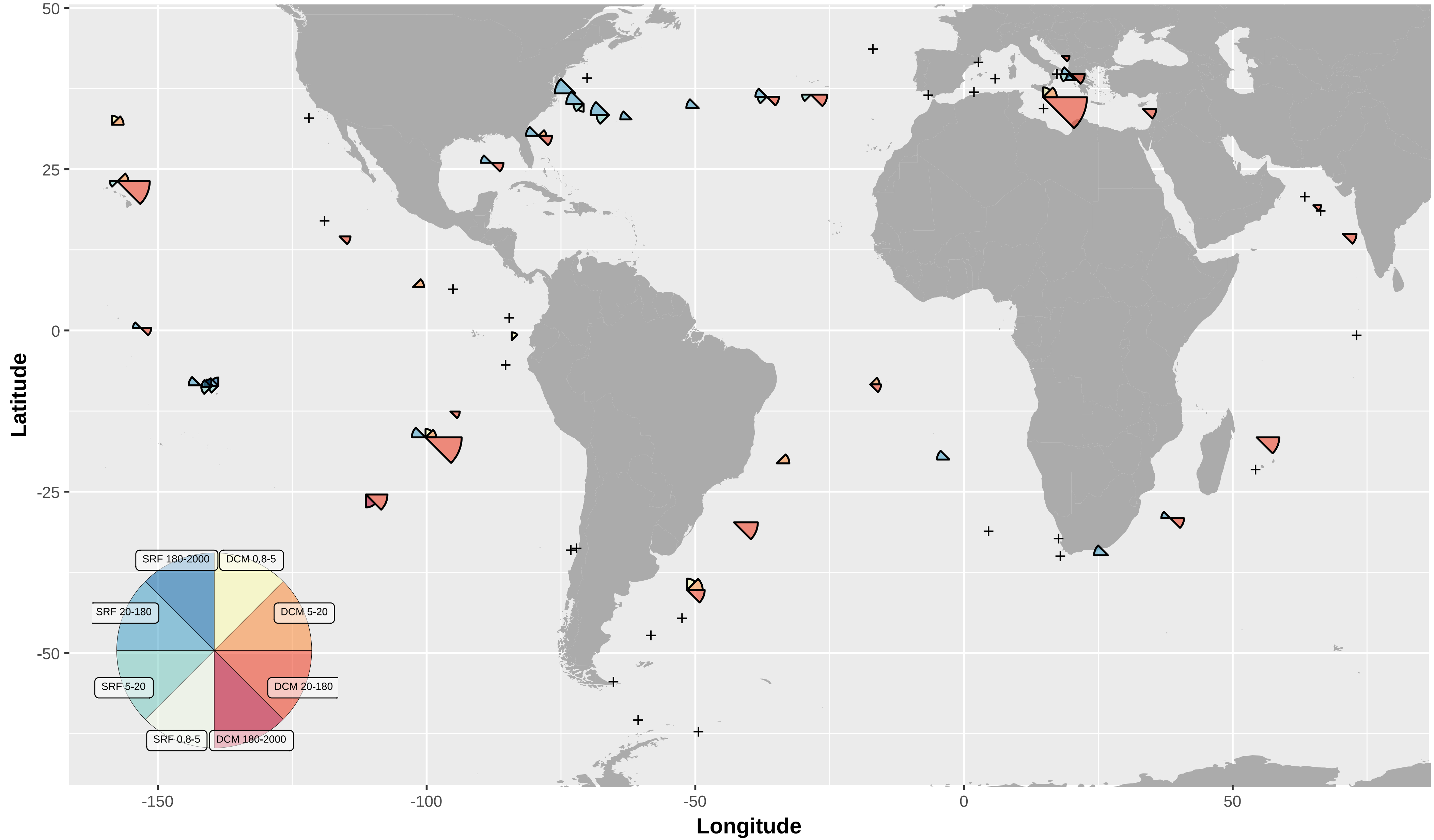


root;cellular organisms;Eukaryota;Opisthokonta;Metazoa;Eumetazoa;Bilateria;Protostomia;Ecdysozoa;Panarthropoda;Arthropoda;Mandibulata;Pancrustacea;Crustacea;Maxillopoda

# Biogeolocalisation of MGT-v1 650      976 unigenes, 300 (30.74%) taxonomically assigned

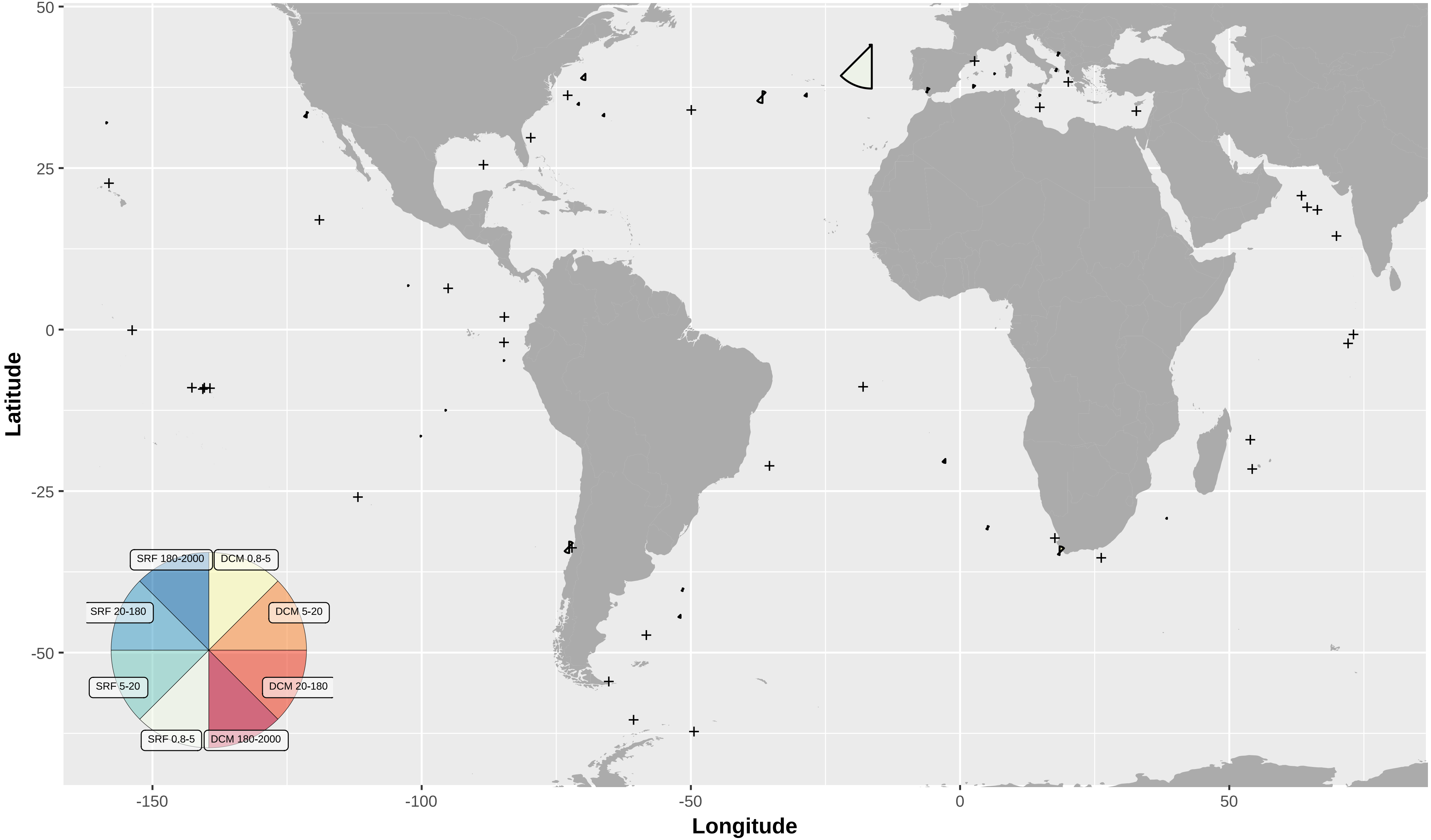


root;cellular organisms;Eukaryota

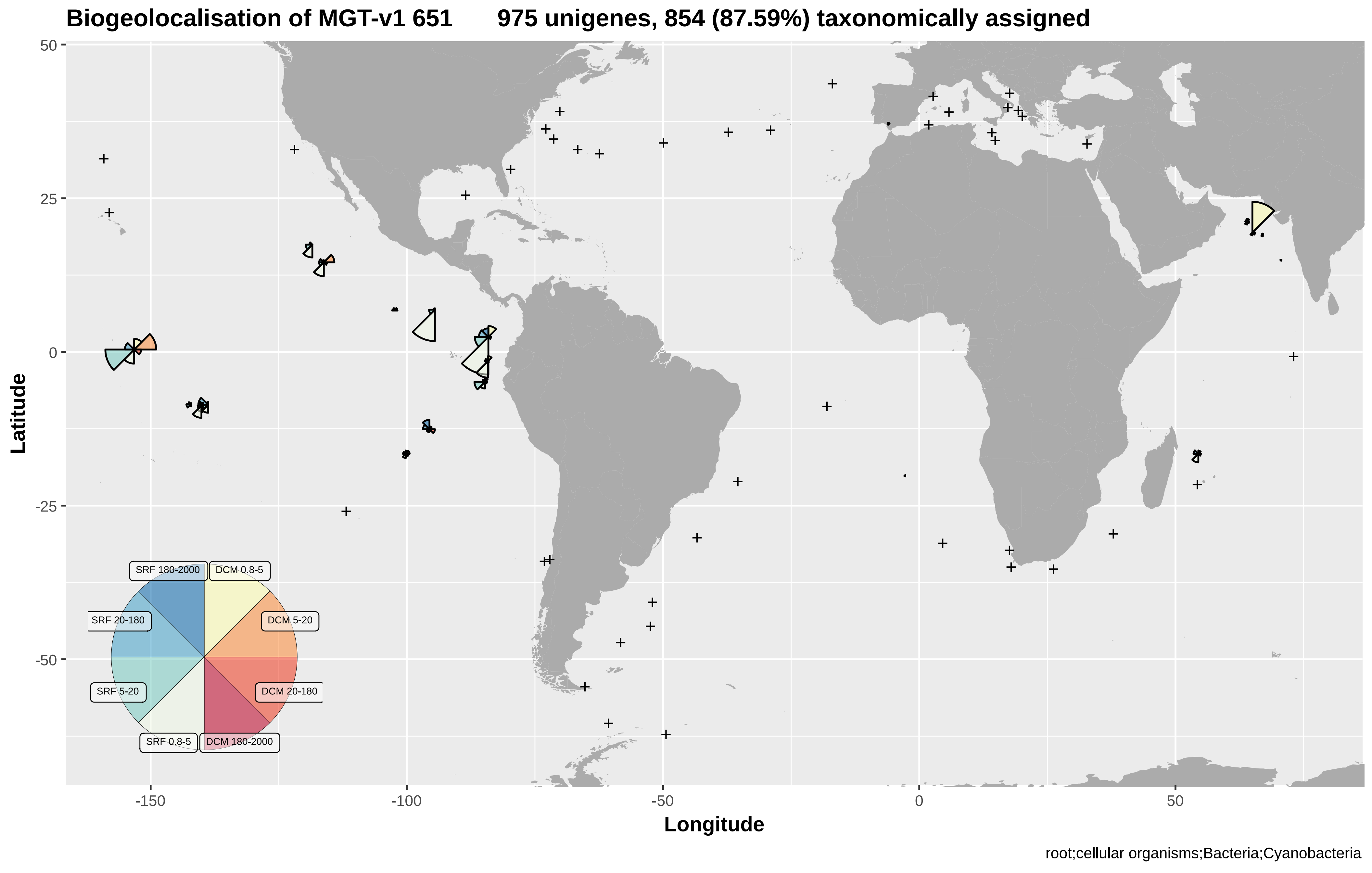
Biogeolocalisation of MGT-v1 651 975 unigenes, 854 (87.59%) taxonomically assigned
50
25
0
-25
-50
Latitude
-150
-100
-50
0
50
Longitude
SRF 180-2000
DCM 0.8-5
SRF 20-180
DCM 5-20
SRF 5-20
DCM 20-180
SRF 0.8-5
DCM 180-2000
root;cellular organisms;Bacteria;Cyanobacteria

## Biogeolocalisation of MGT-v1 652      974 unigenes, 782 (80.29%) taxonomically assigned

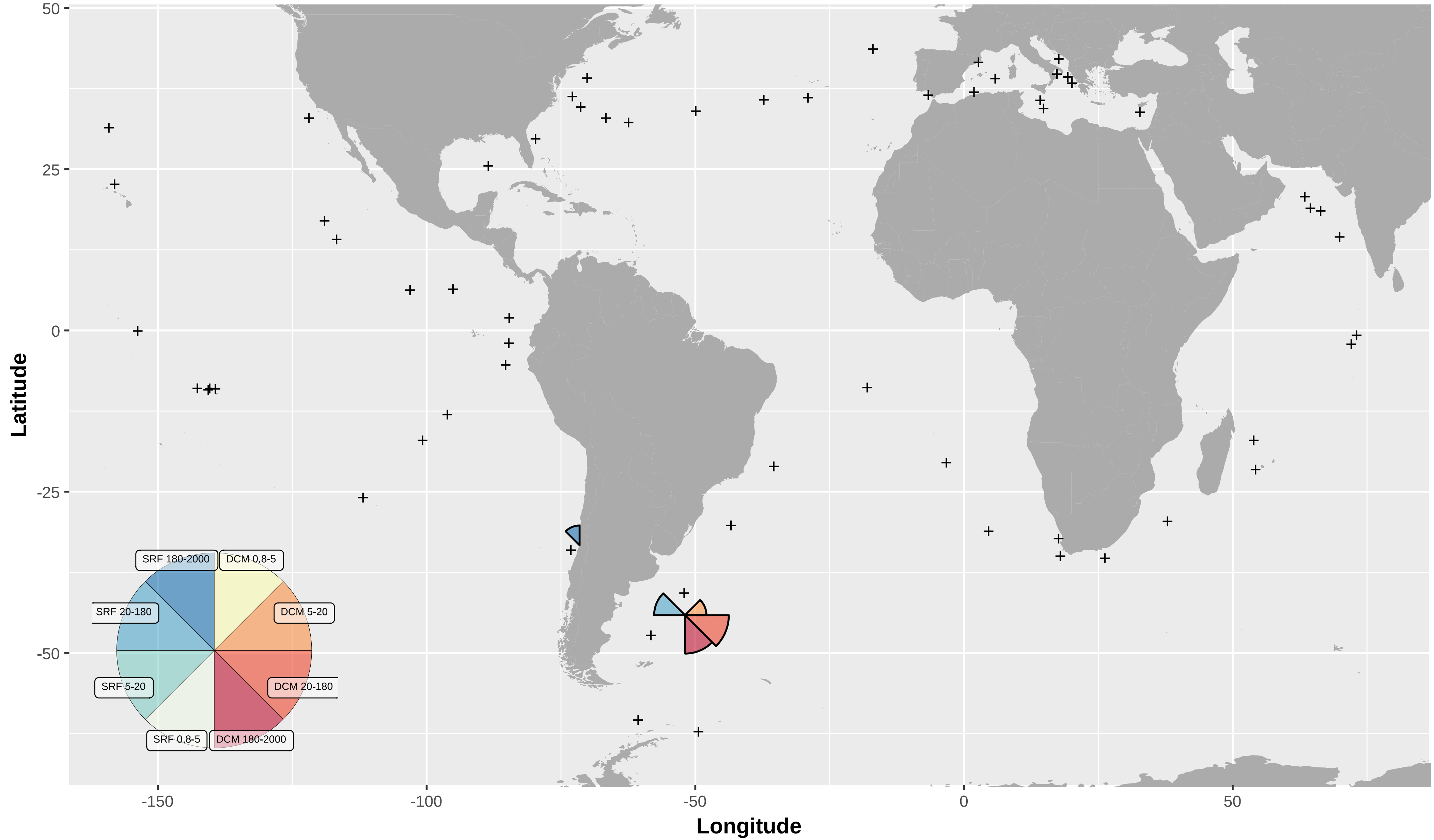


root;cellular organisms;Eukaryota;Alveolata;Ciliophora;Intramacronucleata;Spirotrichea

# Biogeolocalisation of MGT-v1 653      973 unigenes, 232 (23.84%) taxonomically assigned

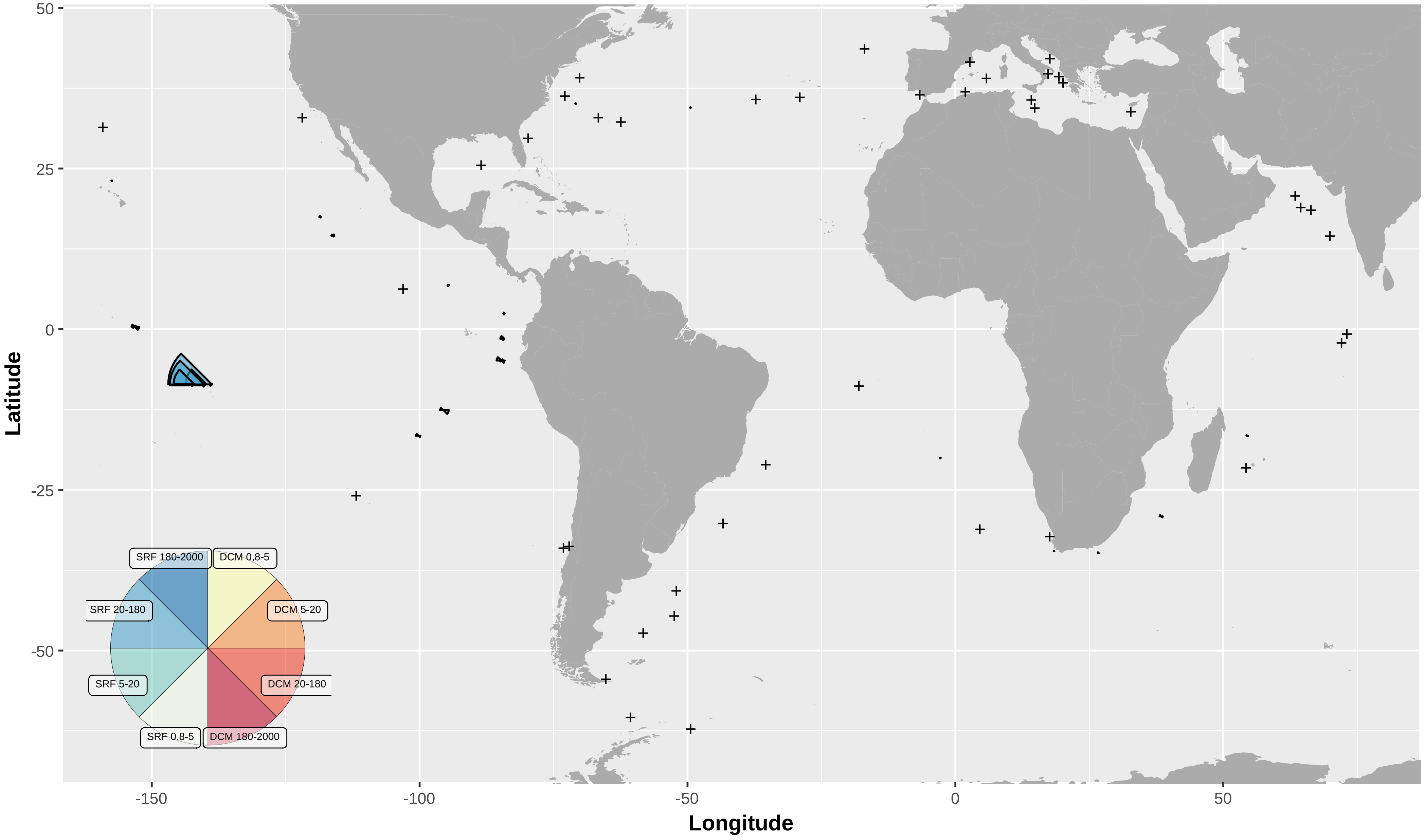


root;cellular organisms;Eukaryota

## Biogeolocalisation of MGT-v1 654      970 unigenes, 134 (13.81%) taxonomically assigned

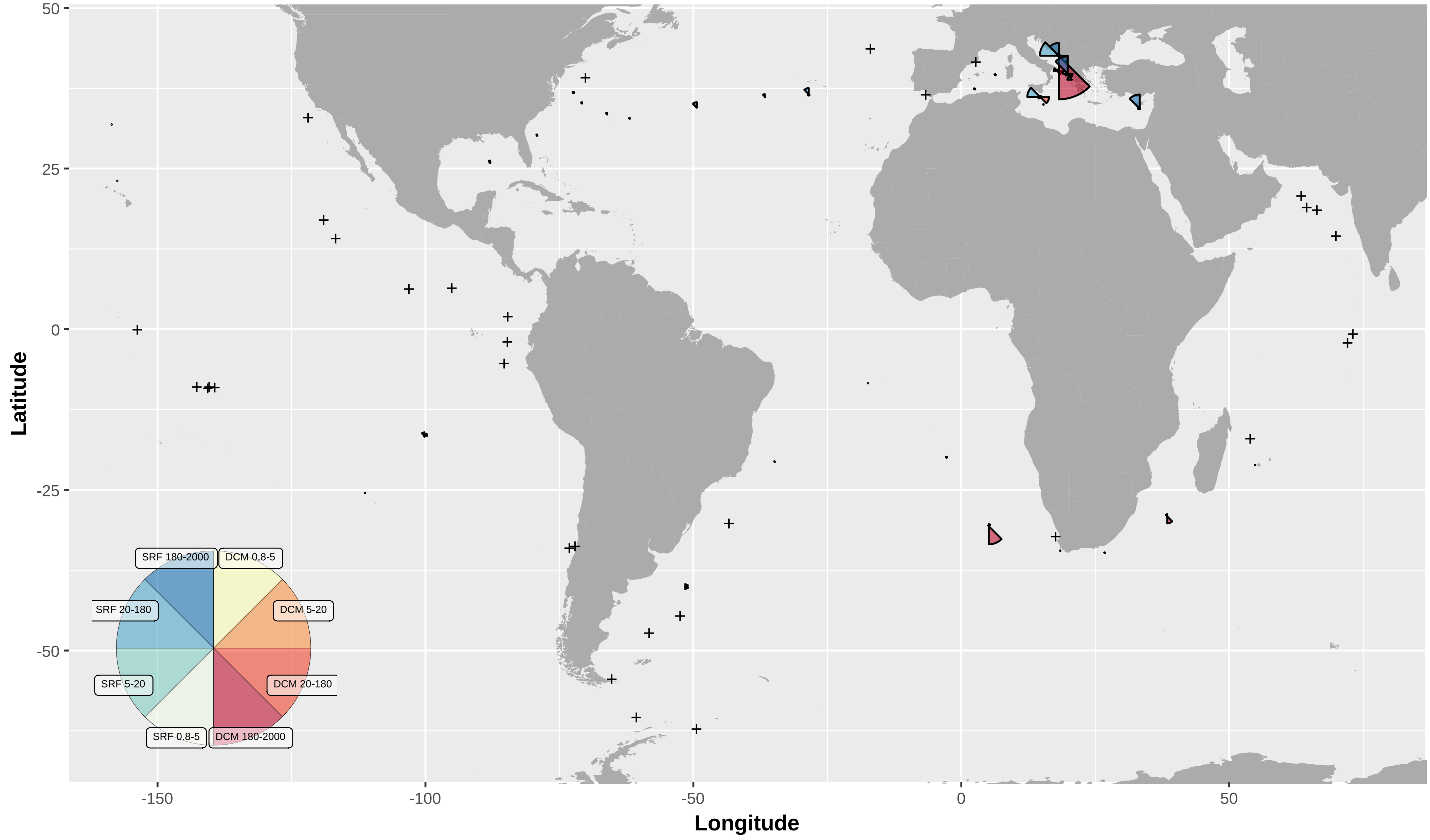


root;cellular organisms;Eukaryota;Opisthokonta;Metazoa;Eumetazoa

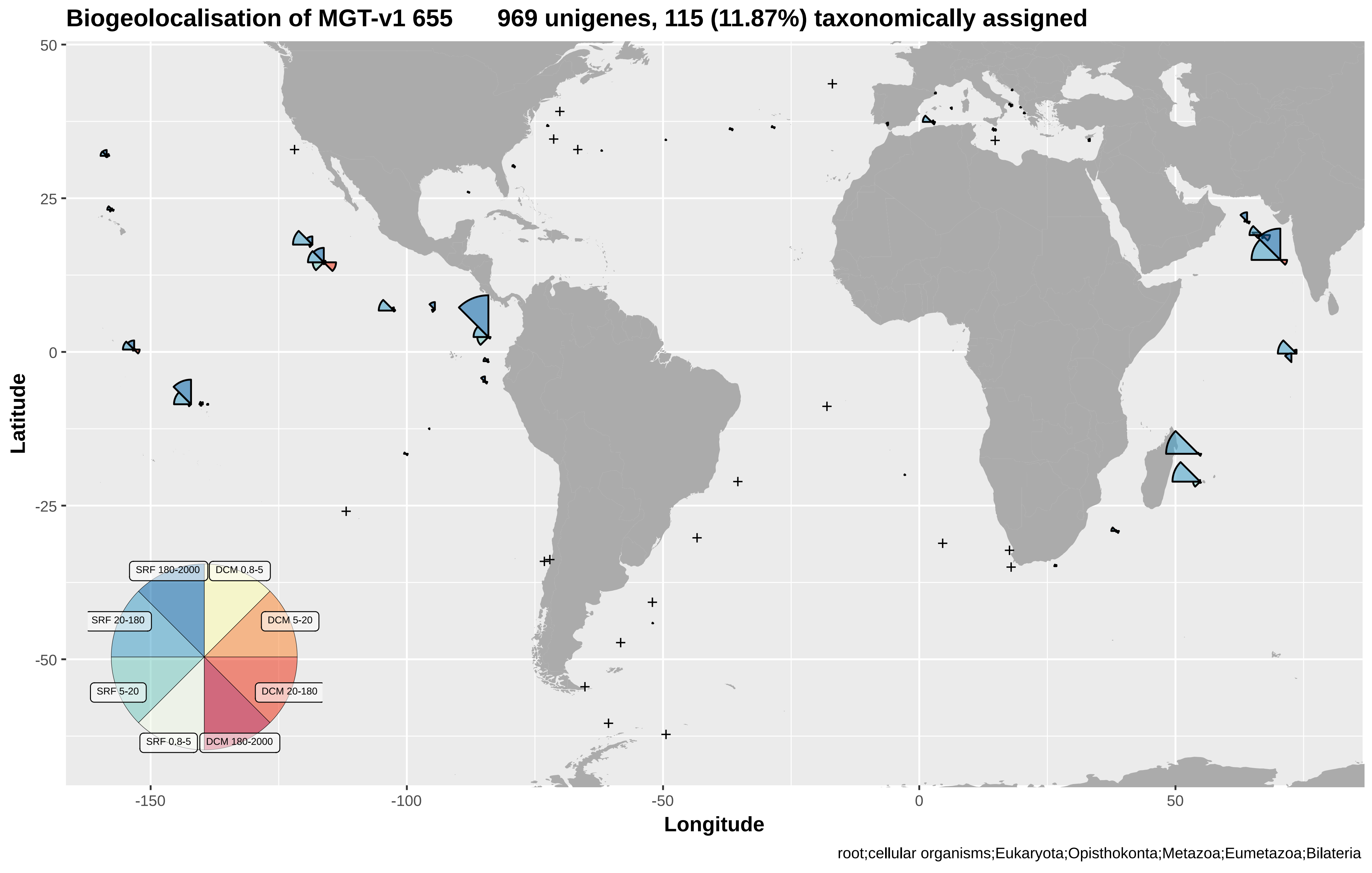

Biogeolocalisation of MGT-v1 655 969 unigenes, 115 (11.87%) taxonomically assigned
Latitude
Longitude
50
25
0
-25
-50
-150
-100
-50
0
50
SRF 180-2000
DCM 0.8-5
SRF 20-180
DCM 5-20
SRF 5-20
DCM 20-180
SRF 0.8-5
DCM 180-2000
root;cellular organisms;Eukaryota;Opisthokonta;Metazoa;Eumetazoa;Bilateria

#### Biogeolocalisation of MGT-v1 656 968 unigenes, 714 (73.76%) taxonomically assigned

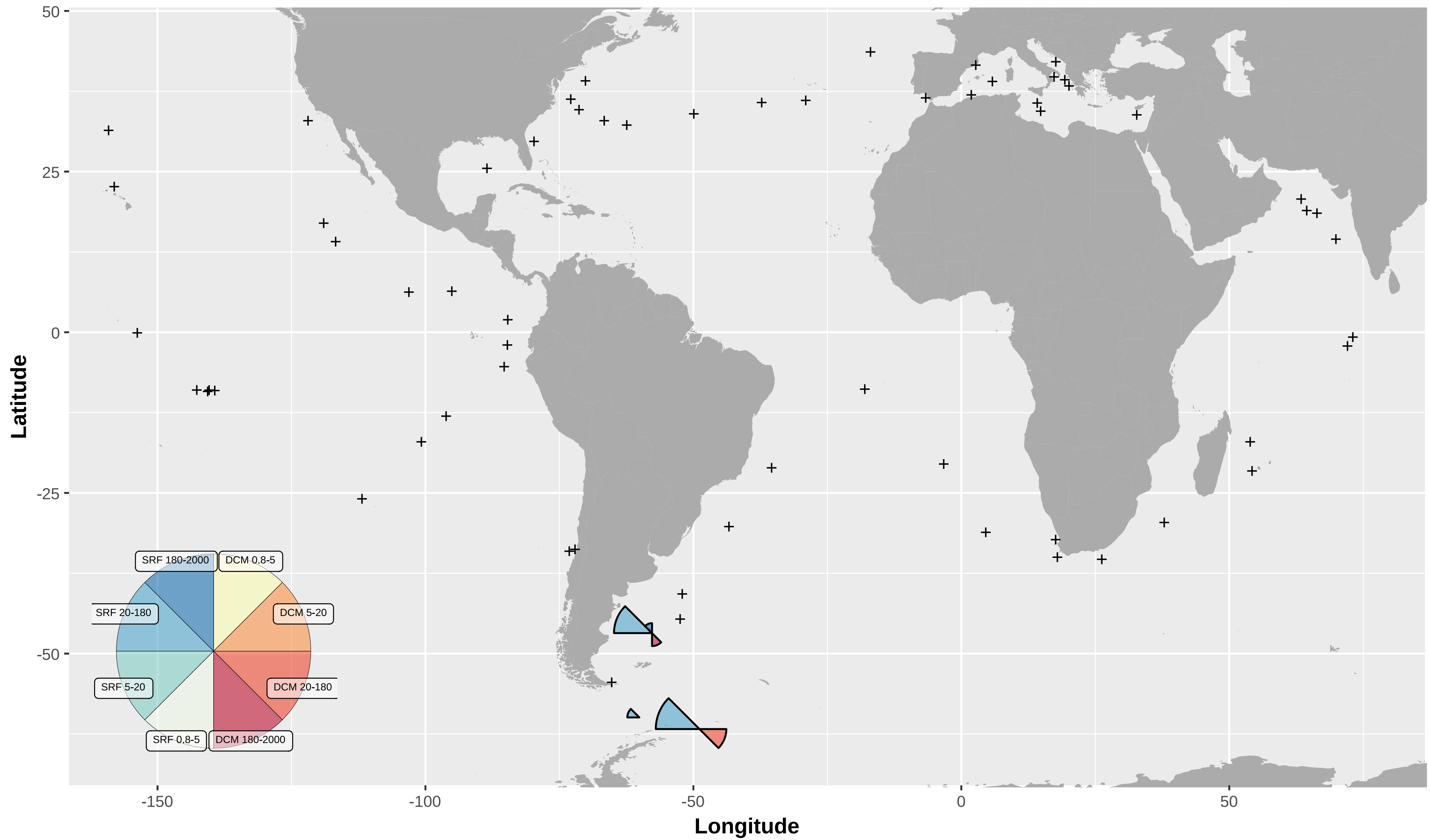


root;cellular organisms;Eukaryota;Stramenopiles;Bacillariophyta

### Biogeolocalisation of MGT-v1 657 967 unigenes, 603 (62.36%) taxonomically assigned

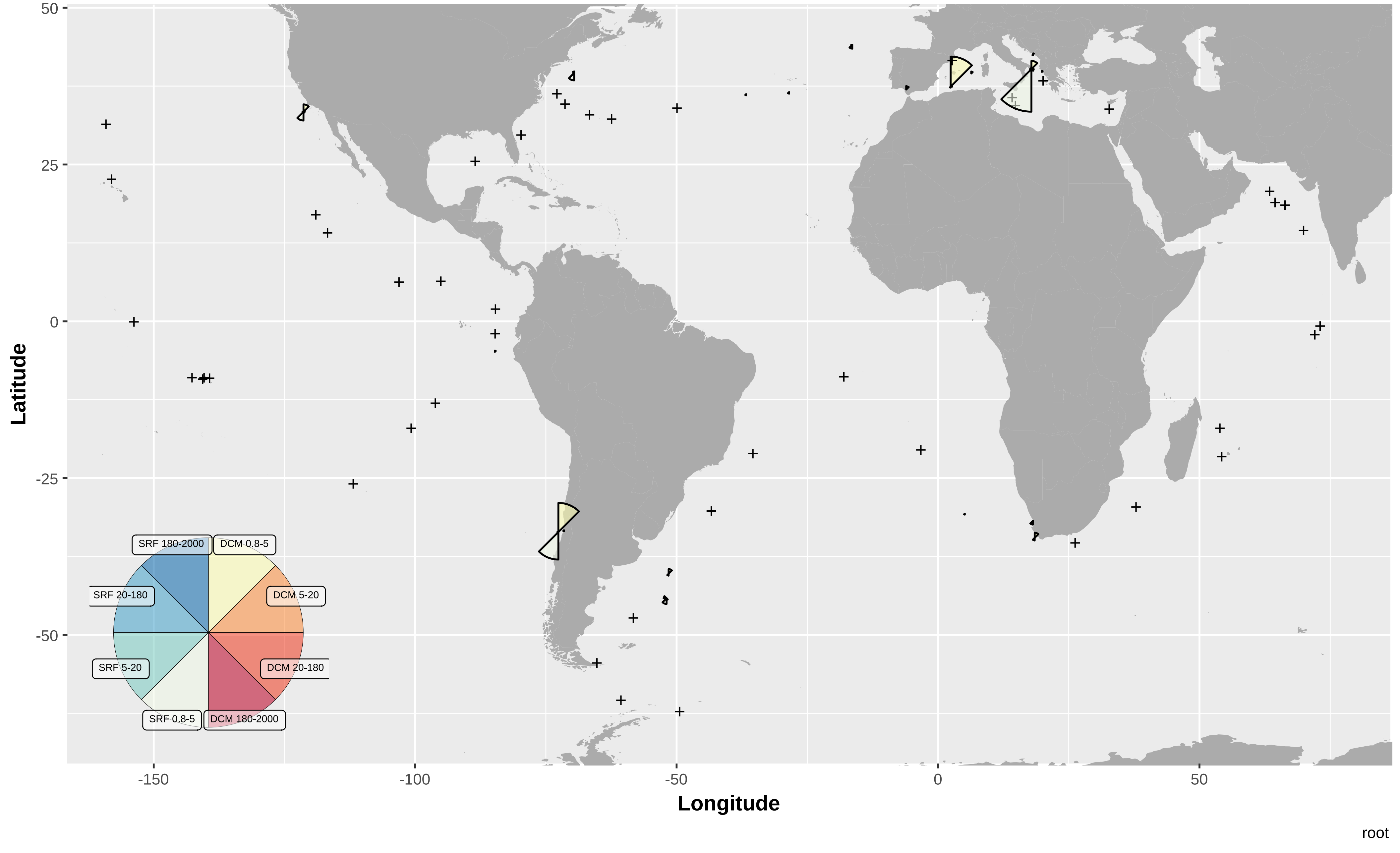

# Biogeolocalisation of MGT-v1 658      966 unigenes, 692 (71.64%) taxonomically assigned

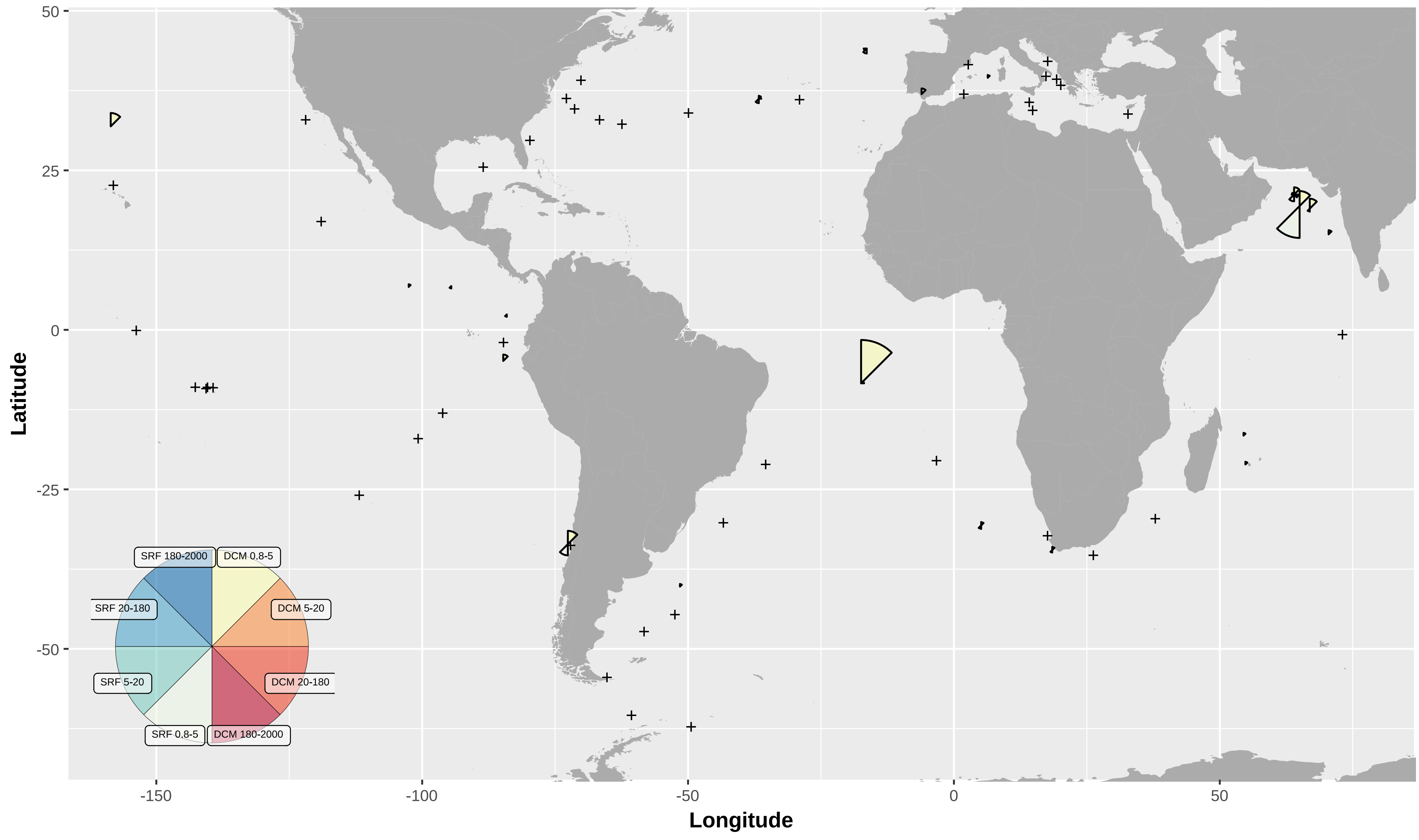


root;cellular organisms;Eukaryota;Viridiplantae

## Biogeolocalisation of MGT-v1 659      964 unigenes, 878 (91.08%) taxonomically assigned

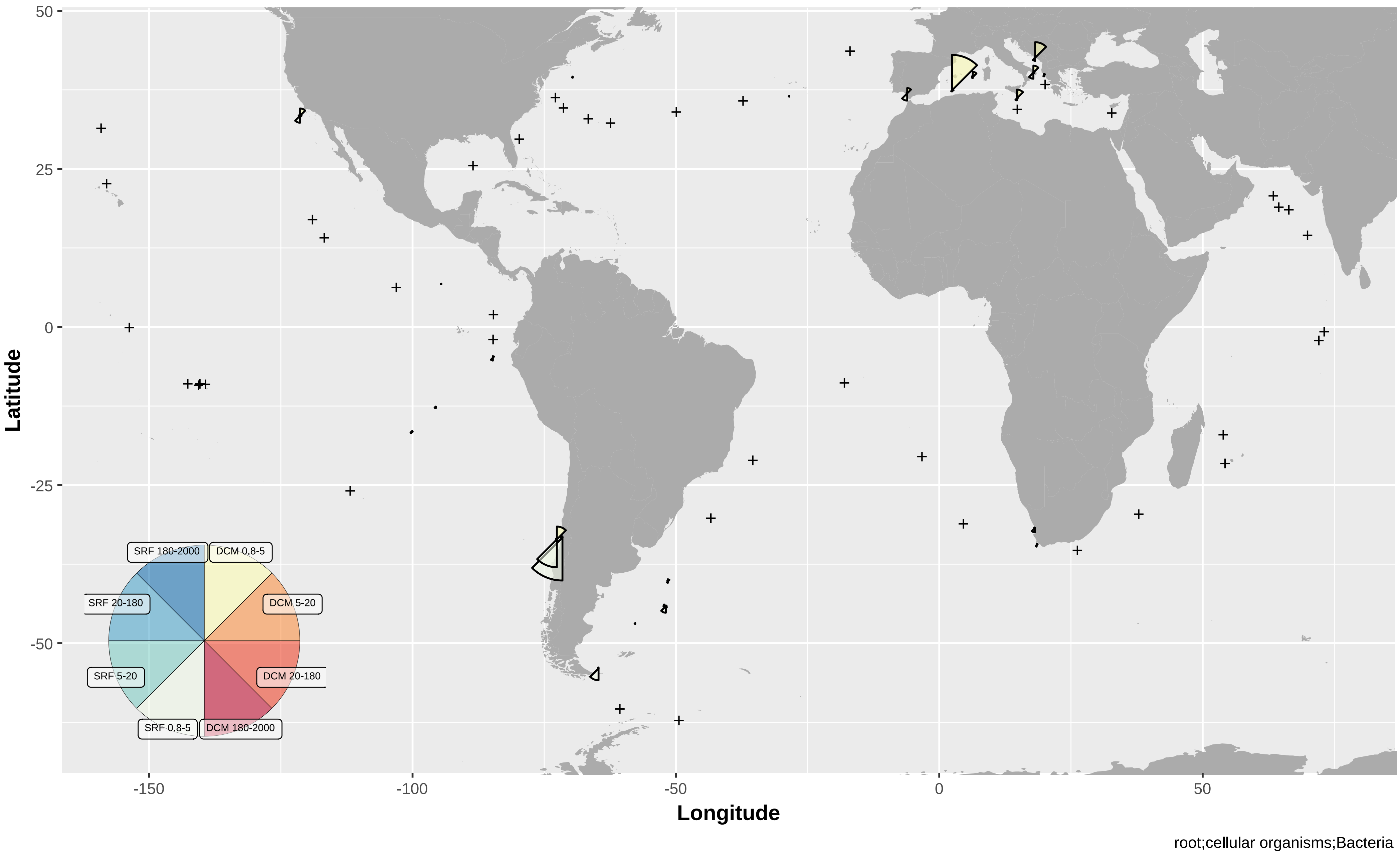

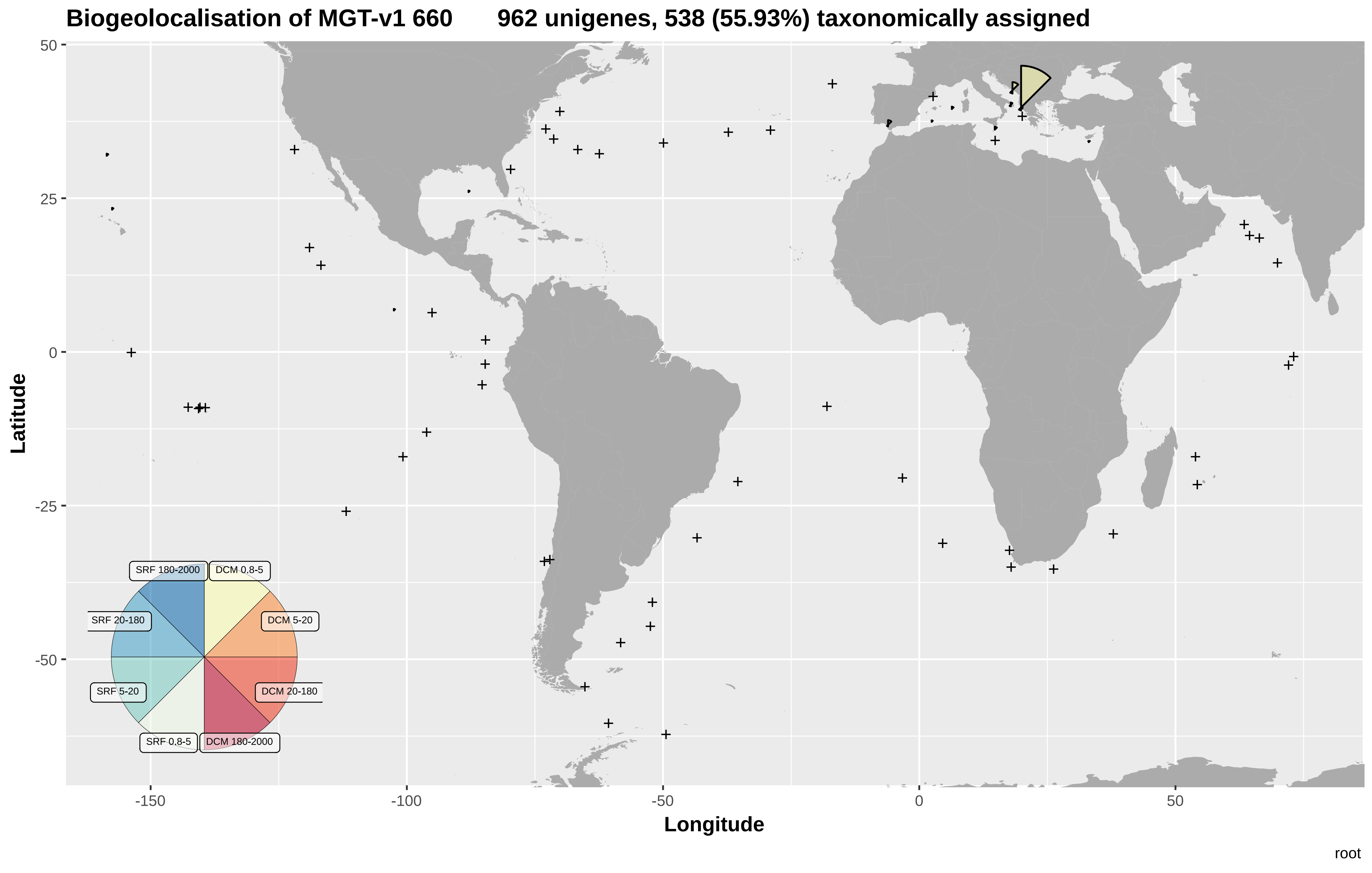
Biogeolocalisation of MGT-v1 660 962 unigenes, 538 (55.93%) taxonomically assigned
Latitude
Longitude
50
25
0
-25
-50
-150
-100
-50
0
50
SRF 180-2000
DCM 0.8-5
SRF 20-180
DCM 5-20
SRF 5-20
DCM 20-180
SRF 0.8-5
DCM 180-2000
root

Biogeolocalisation of MGT-v1 661      959 unigenes, 63 (6.57%) taxonomically assigned

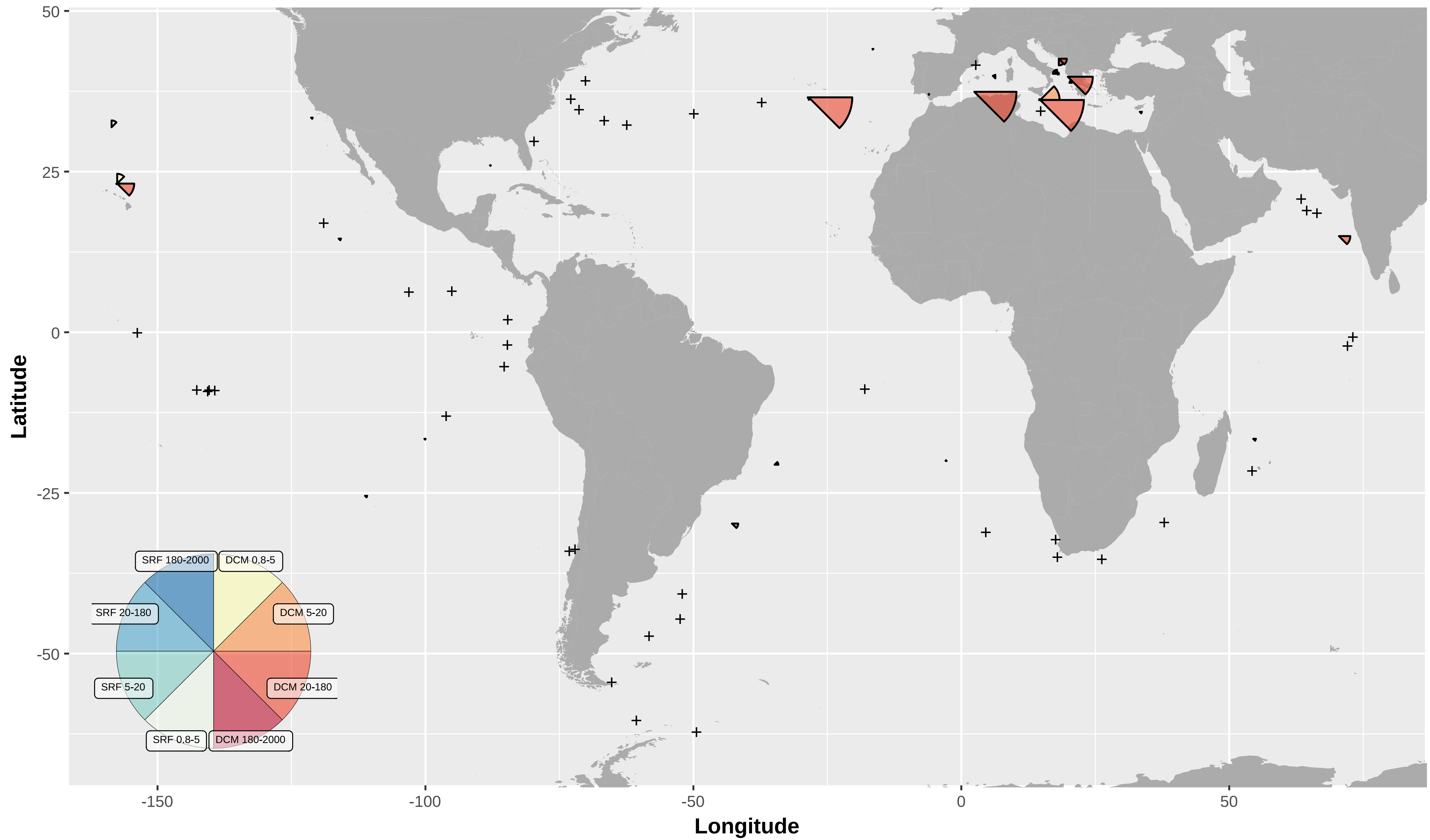


root;cellular organisms;Eukaryota;Opisthokonta;Metazoa;Eumetazoa

#### Biogeolocalisation of MGT-v1 662 958 unigenes, 104 (10.86%) taxonomically assigned

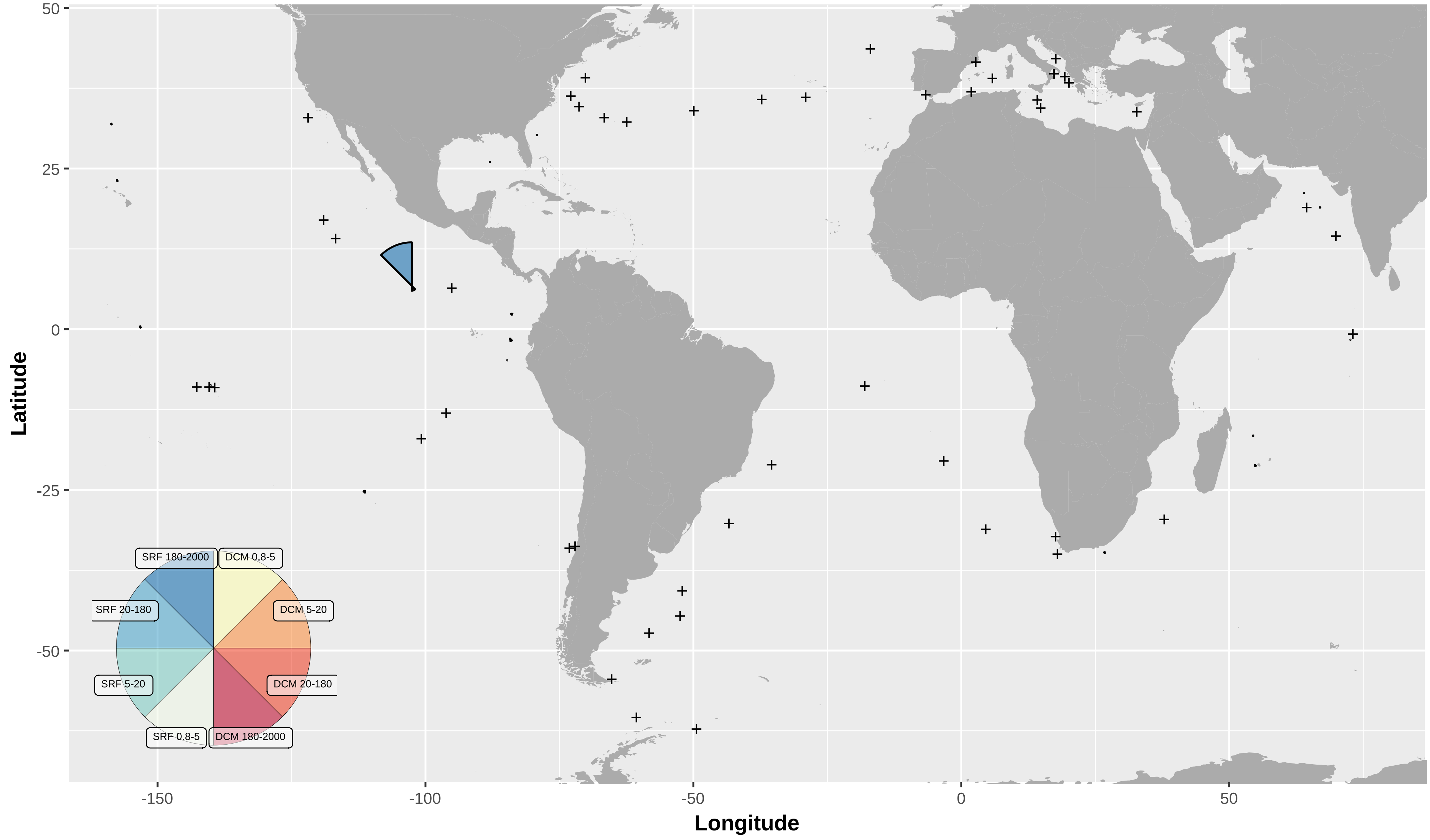


root;cellular organisms;Eukaryota;Opisthokonta;Metazoa;Eumetazoa

Biogeolocalisation of MGT-v1 663 953 unigenes, 910 (95.49%) taxonomically assigned

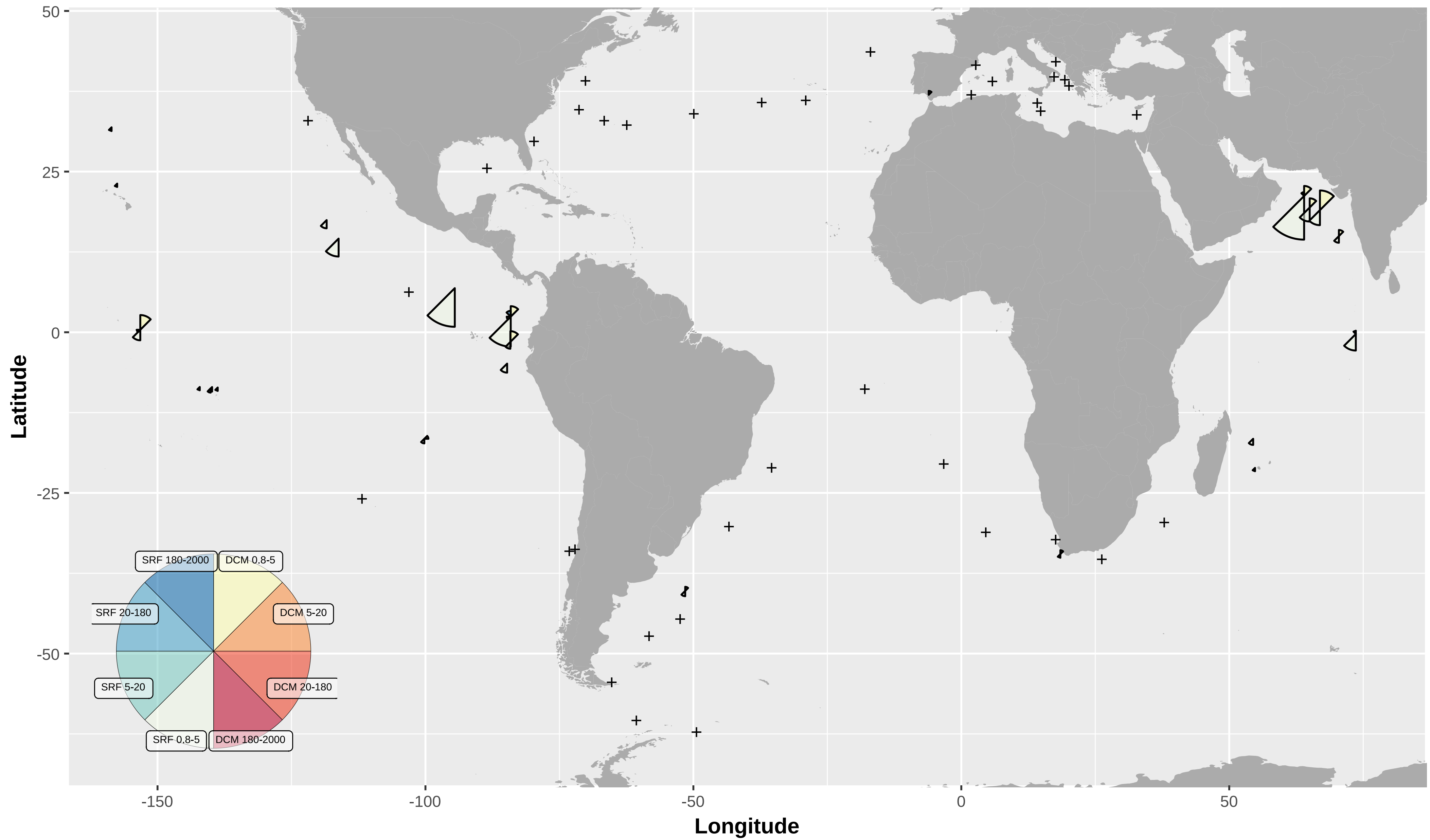


root;cellular organisms;Bacteria;Bacteroidetes/Chlorobi group;Bacteroidetes;Flavobacteriia

**Biogeolocalisation of MGT-v1 664      950 unigenes, 436 (45.89%) taxonomically assigned**

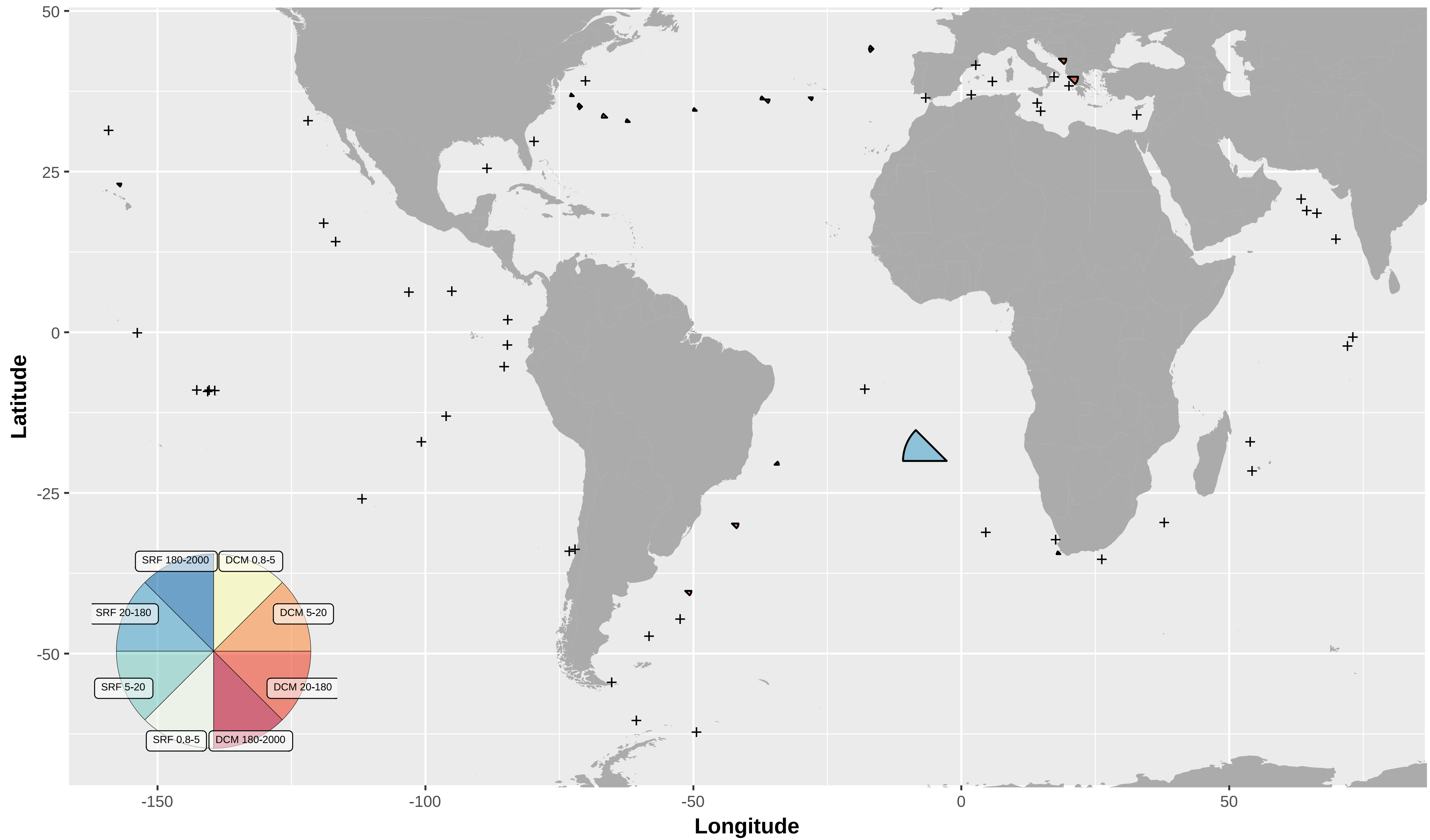


root;cellular organisms;Eukaryota;Alveolata;Ciliophora;Intramacronucleata;Spirotrichea

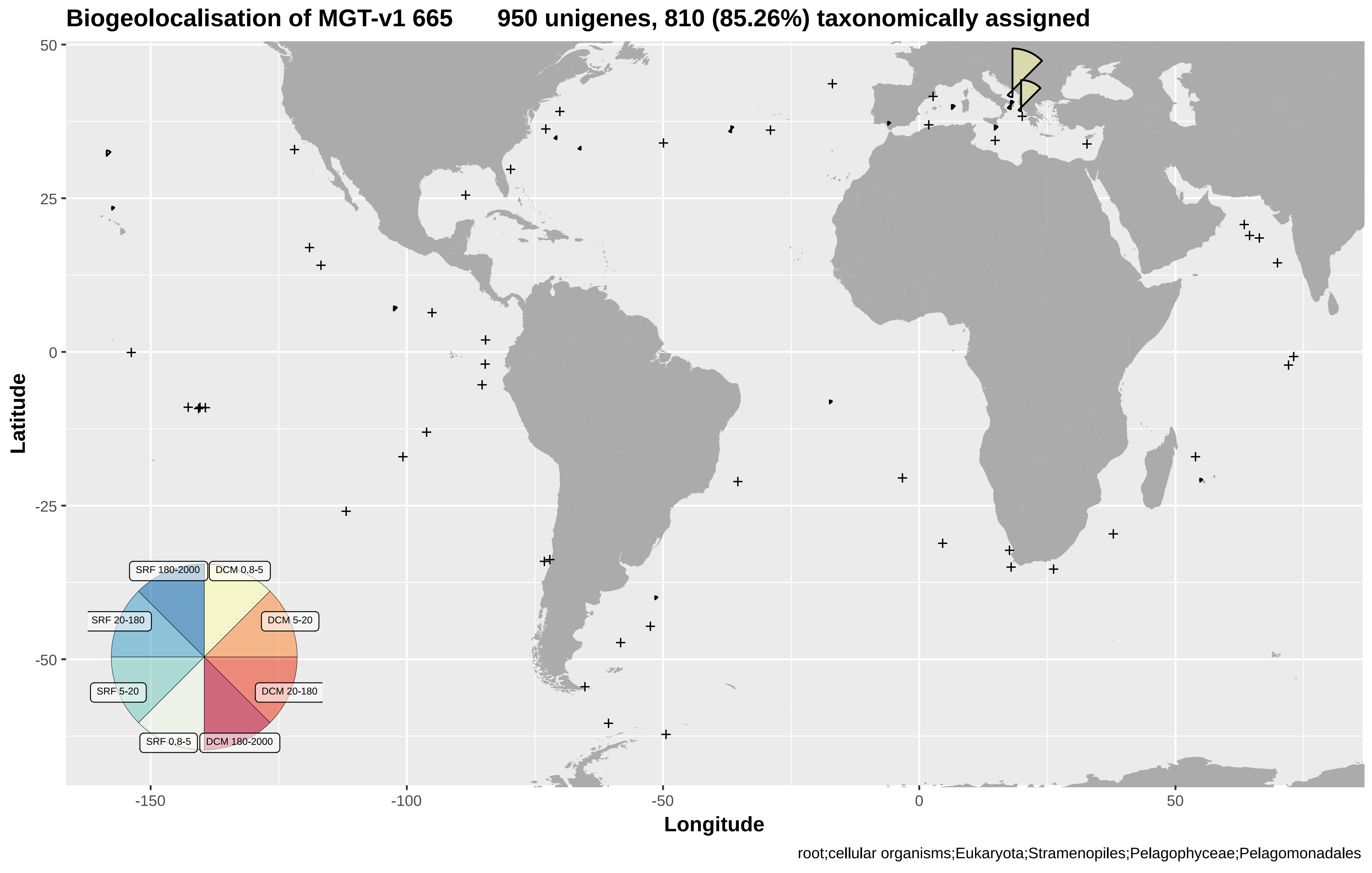
Biogeolocalisation of MGT-v1 665
950 unigenes, 810 (85.26%) taxonomically assigned
Latitude
Longitude
50
25
0
-25
-50
-150
-100
-50
0
50
SRF 180-2000
DCM 0.8-5
SRF 20-180
DCM 5-20
SRF 5-20
DCM 20-180
SRF 0.8-5
DCM 180-2000
root;cellular organisms;Eukaryota;Stramenopiles;Pelagophyceae;Pelagomonadales

# Biogeolocalisation of MGT-v1 666      946 unigenes, 894 (94.5%) taxonomically assigned

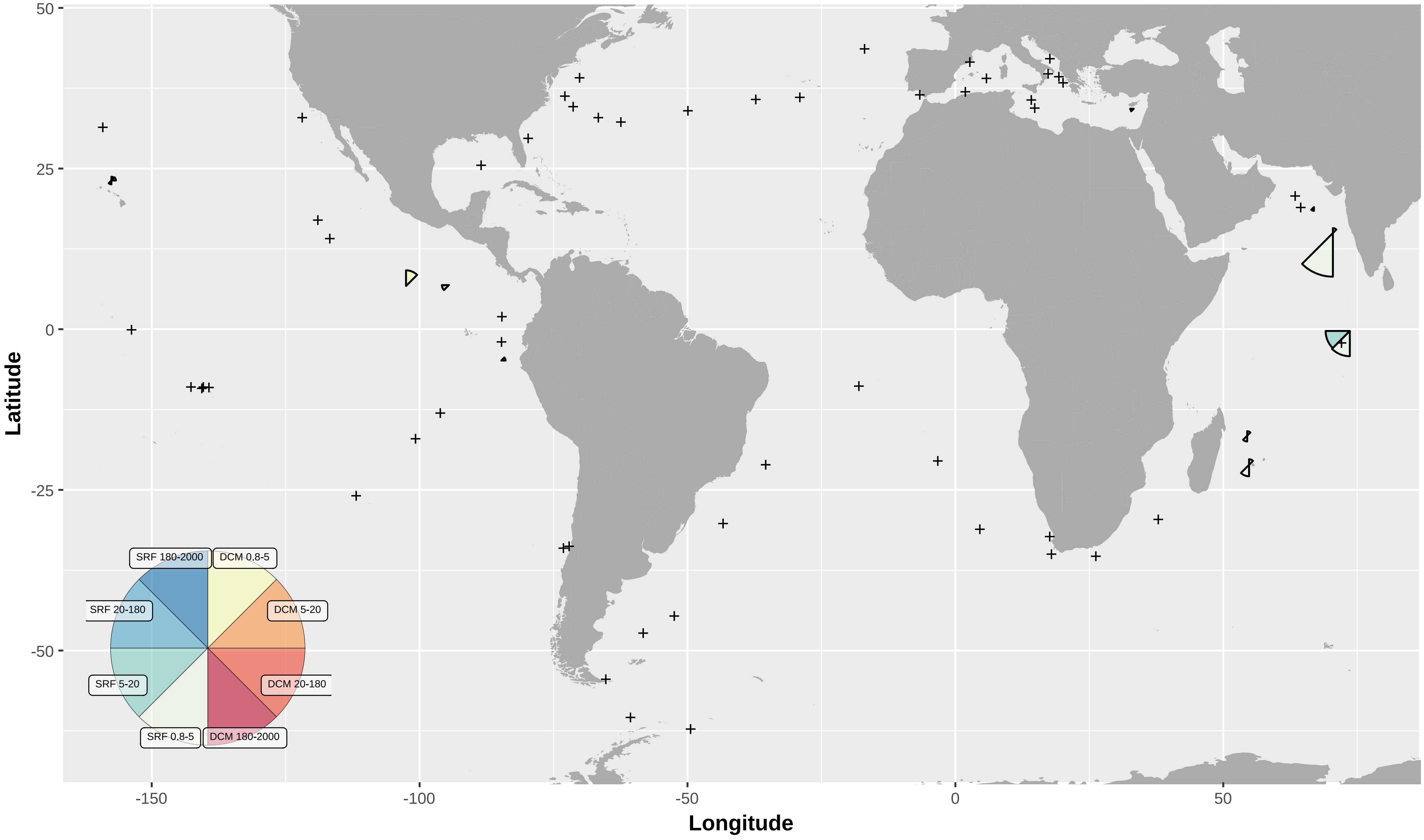


root;cellular organisms;Bacteria;Bacteroidetes/Chlorobi group;Bacteroidetes;Flavobacteriia;Flavobacteriales;Flavobacteriaceae

#### Biogeolocalisation of MGT-v1 667 945 unigenes, 906 (95.87%) taxonomically assigned

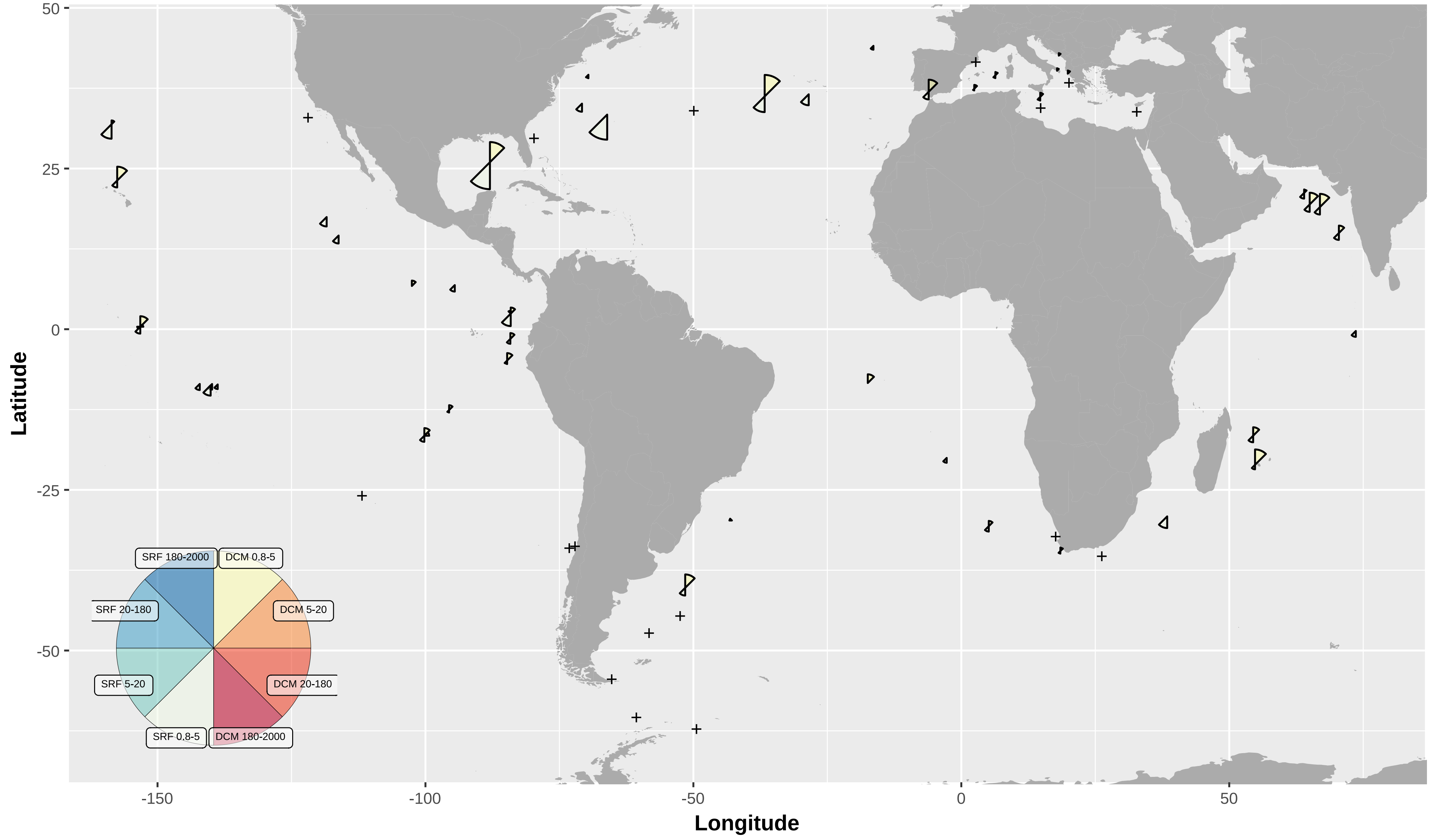


root;cellular organisms;Bacteria;Proteobacteria;Alphaproteobacteria

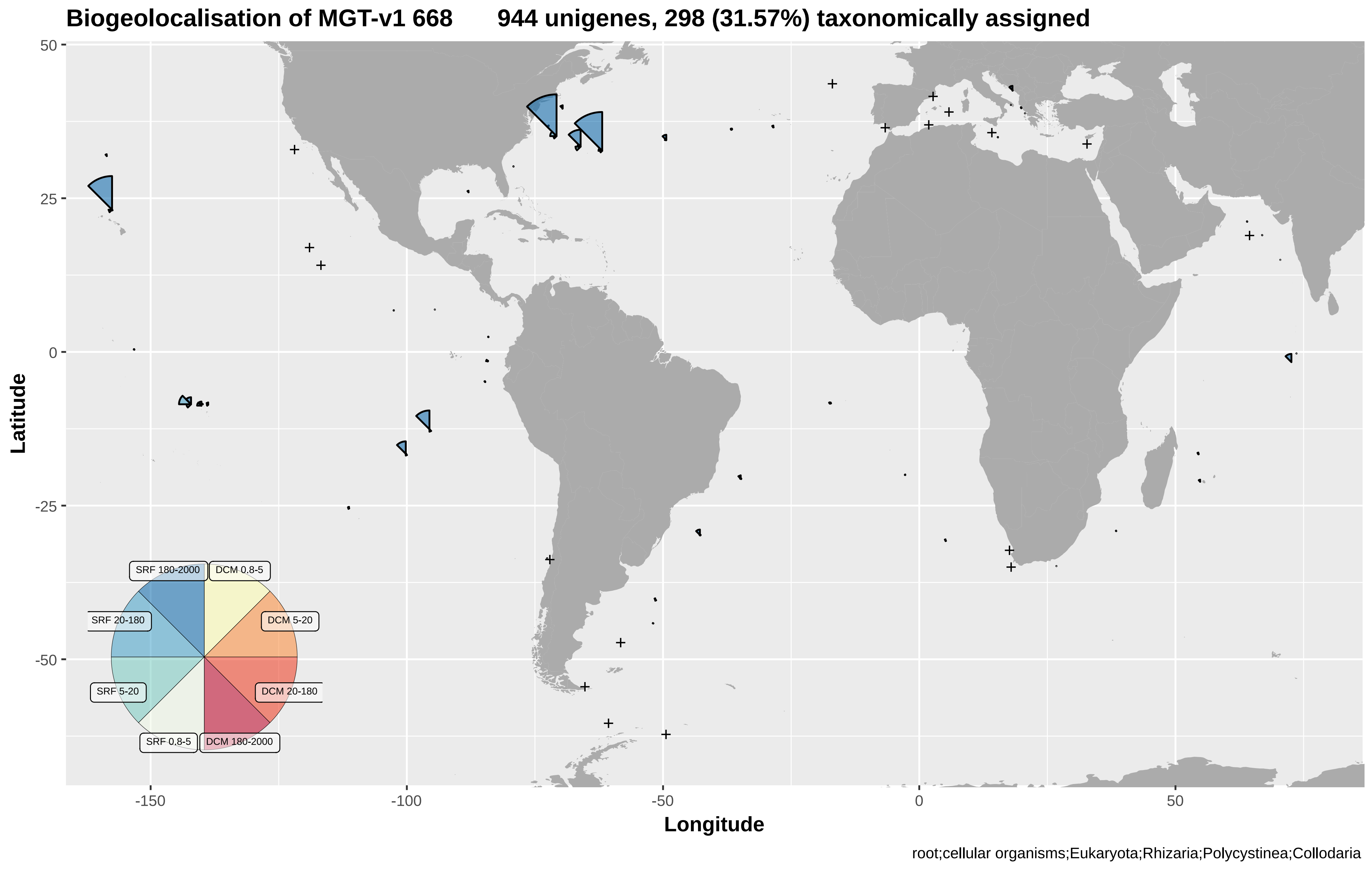
Biogeolocalisation of MGT-v1 668      944 unigenes, 298 (31.57%) taxonomically assigned
Latitude
Longitude
50
25
0
-25
-50
-150
-100
-50
0
50
SRF 180-2000
DCM 0.8-5
SRF 20-180
DCM 5-20
SRF 5-20
DCM 20-180
SRF 0.8-5
DCM 180-2000
root;cellular organisms;Eukaryota;Rhizaria;Polycystinea;Collodaria

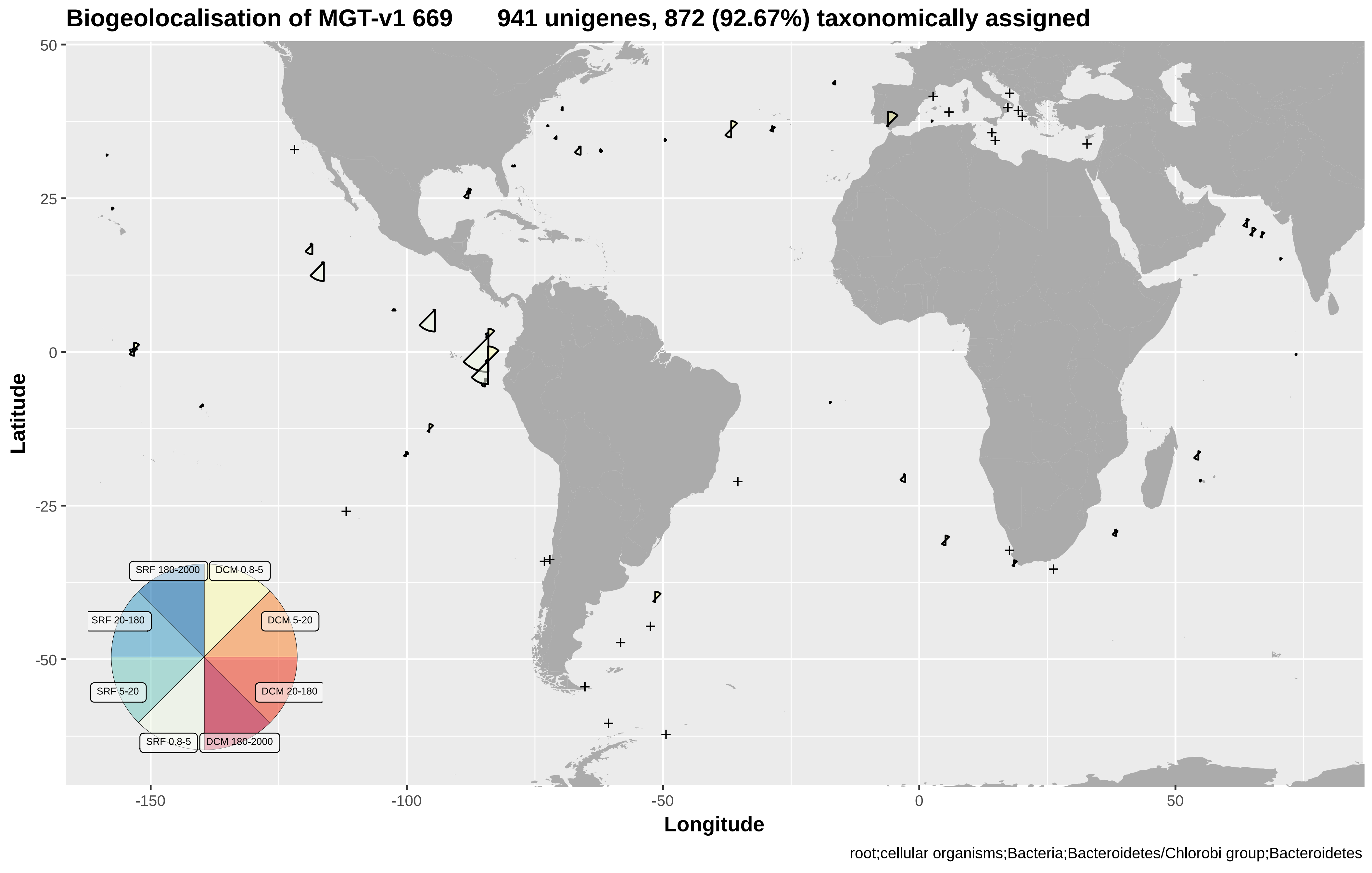

Biogeolocalisation of MGT-v1 669      941 unigenes, 872 (92.67%) taxonomically assigned
50
25
0
-25
-50
Latitude
-150
-100
-50
0
50
Longitude
SRF 180-2000
DCM 0.8-5
SRF 20-180
DCM 5-20
SRF 5-20
DCM 20-180
SRF 0.8-5
DCM 180-2000
root;cellular organisms;Bacteria;Bacteroidetes/Chlorobi group;Bacteroidetes

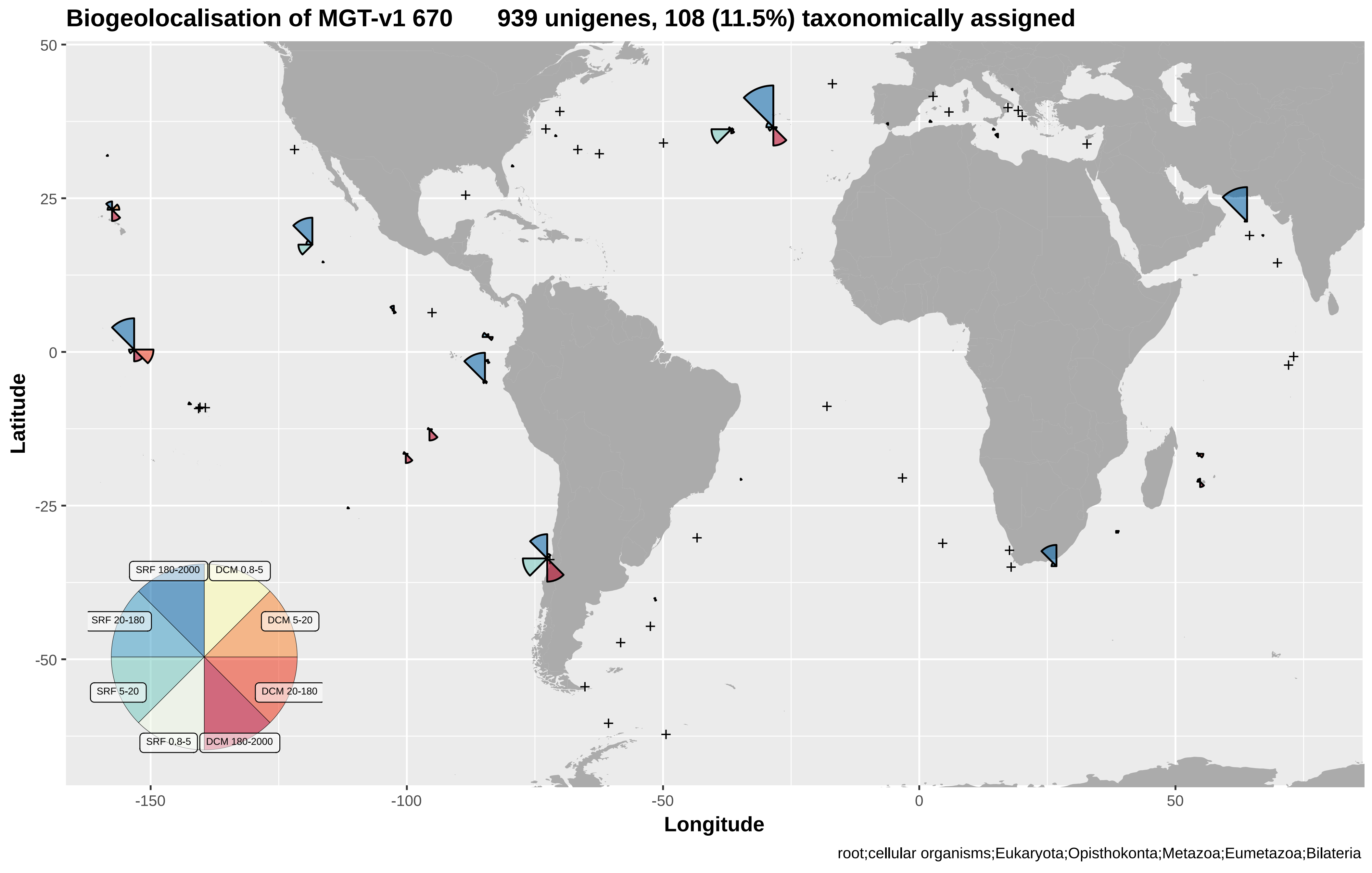

Biogeolocalisation of MGT-v1 670 939 unigenes, 108 (11.5%) taxonomically assigned
50
25
0
-25
-50
Latitude
-150
-100
-50
0
50
Longitude
SRF 180-2000
DCM 0.8-5
SRF 20-180
DCM 5-20
SRF 5-20
DCM 20-180
SRF 0.8-5
DCM 180-2000
root;cellular organisms;Eukaryota;Opisthokonta;Metazoa;Eumetazoa;Bilateria

### Biogeolocalisation of MGT-v1 671 937 unigenes, 80 (8.54%) taxonomically assigned

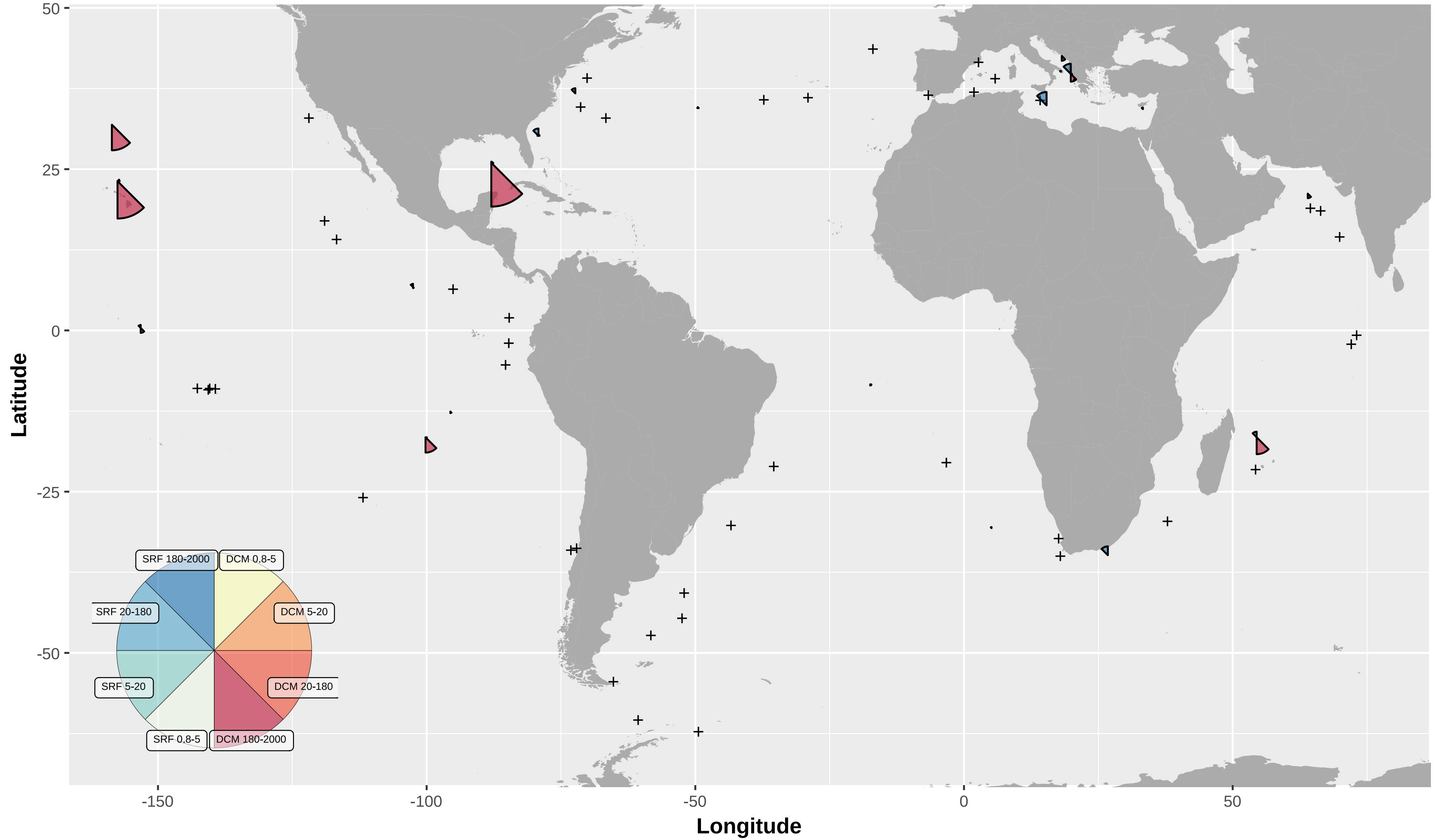


root;cellular organisms;Eukaryota;Opisthokonta;Metazoa;Eumetazoa

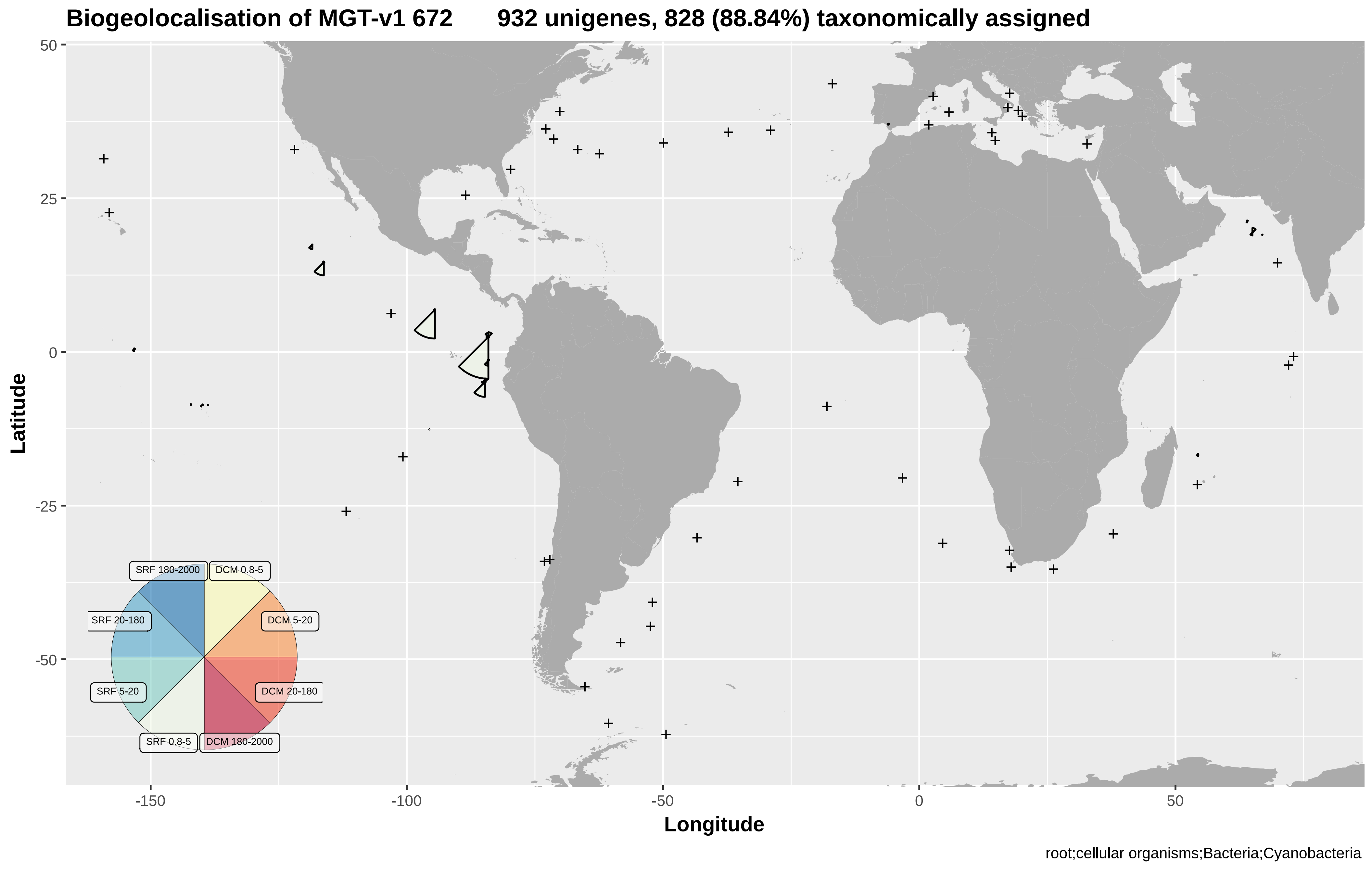

Biogeolocalisation of MGT-v1 672 932 unigenes, 828 (88.84%) taxonomically assigned
Latitude
Longitude
50
25
0
-25
-50
-150
-100
-50
0
50
SRF 180-2000
DCM 0.8-5
SRF 20-180
DCM 5-20
SRF 5-20
DCM 20-180
SRF 0.8-5
DCM 180-2000
root;cellular organisms;Bacteria;Cyanobacteria

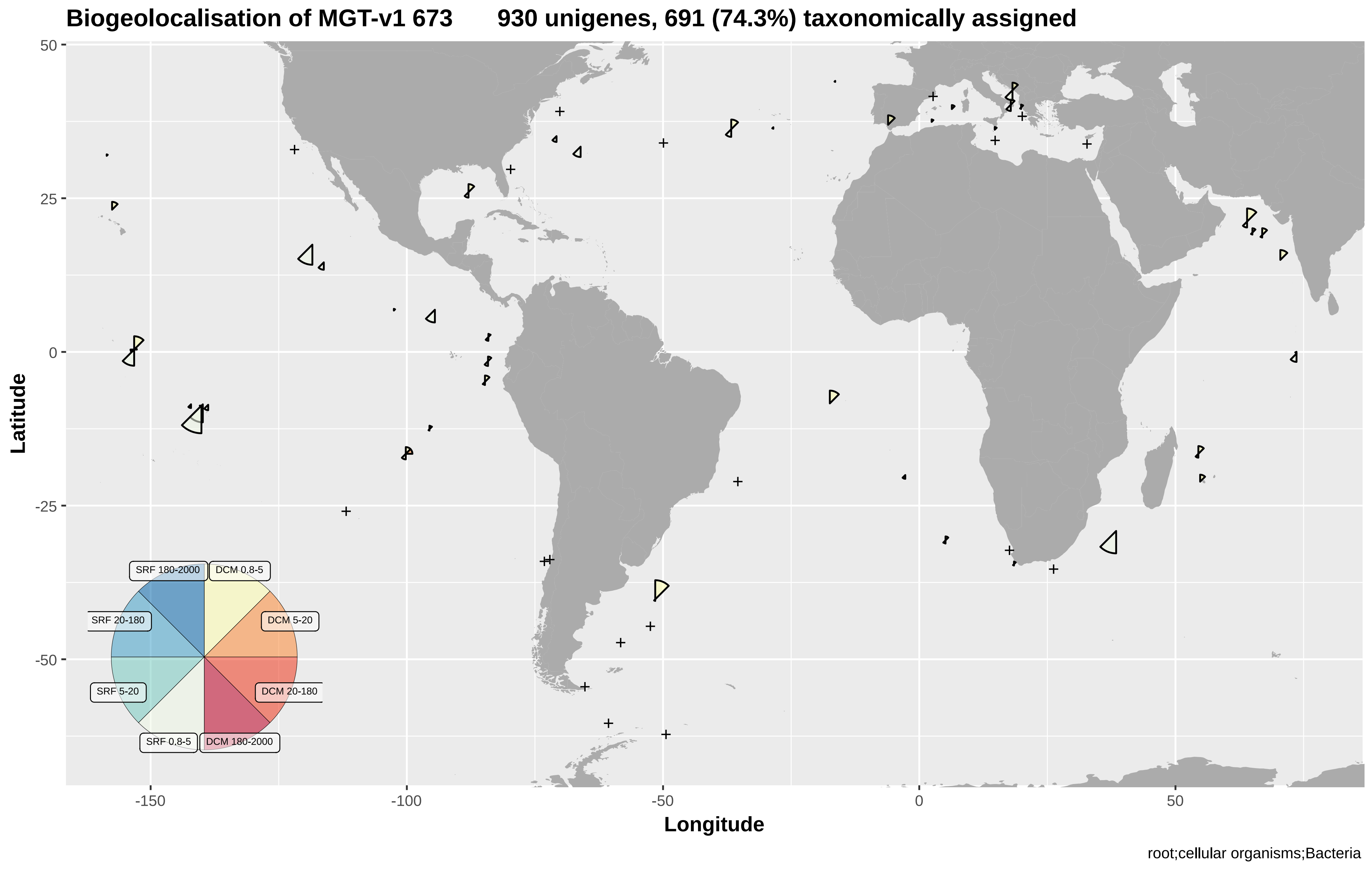
Biogeolocalisation of MGT-v1 673 930 unigenes, 691 (74.3%) taxonomically assigned
50
25
0
-25
-50
Latitude
-150
-100
-50
0
50
Longitude
SRF 180-2000
DCM 0.8-5
SRF 20-180
DCM 5-20
SRF 5-20
DCM 20-180
SRF 0.8-5
DCM 180-2000
root;cellular organisms;Bacteria

## Biogeolocalisation of MGT-v1 674      928 unigenes, 920 (99.14%) taxonomically assigned

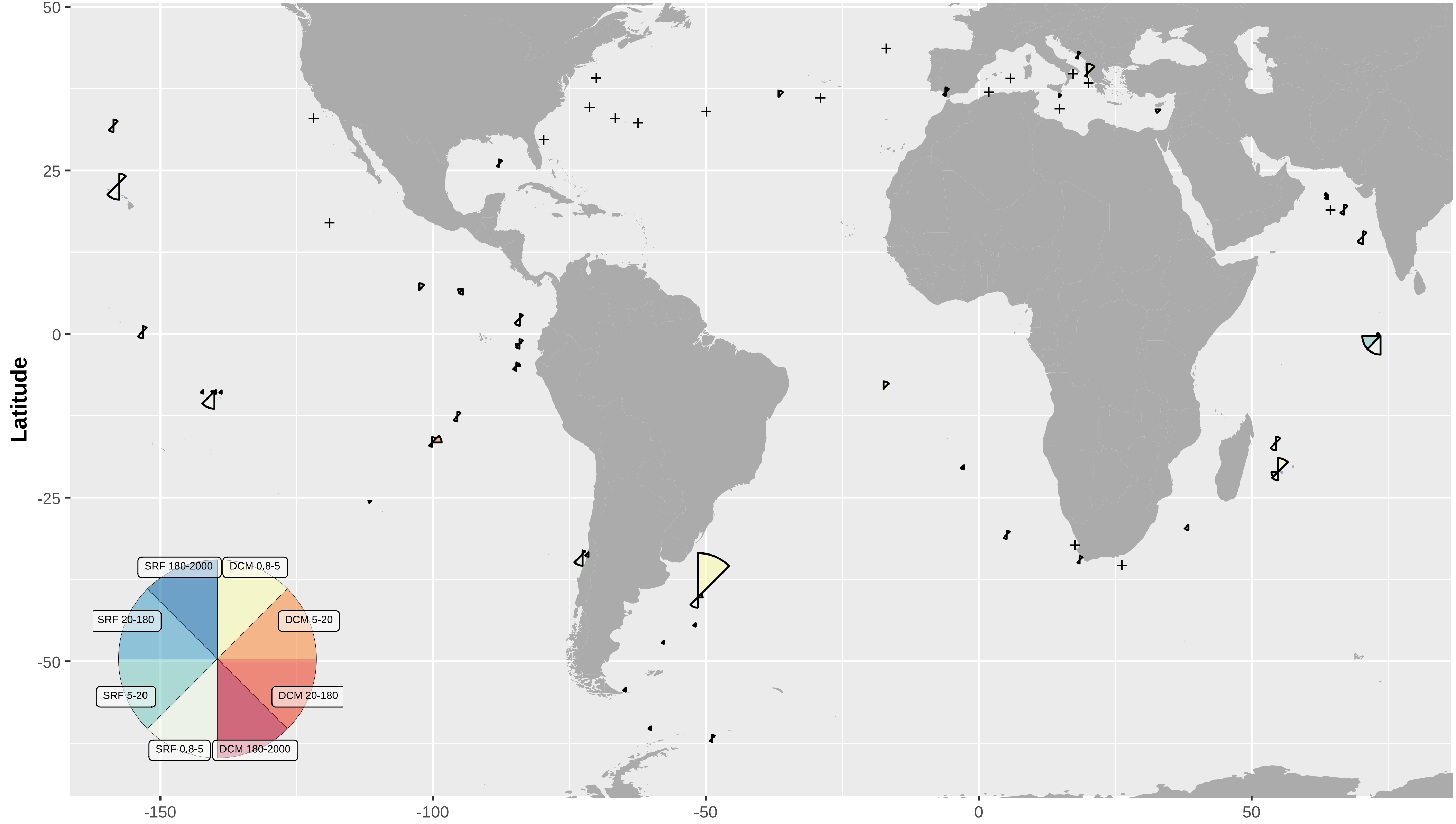


root;cellular organisms;Bacteria;Proteobacteria;Alphaproteobacteria;Rhodobacterales;Hyphomonadaceae;Oceanicaulis

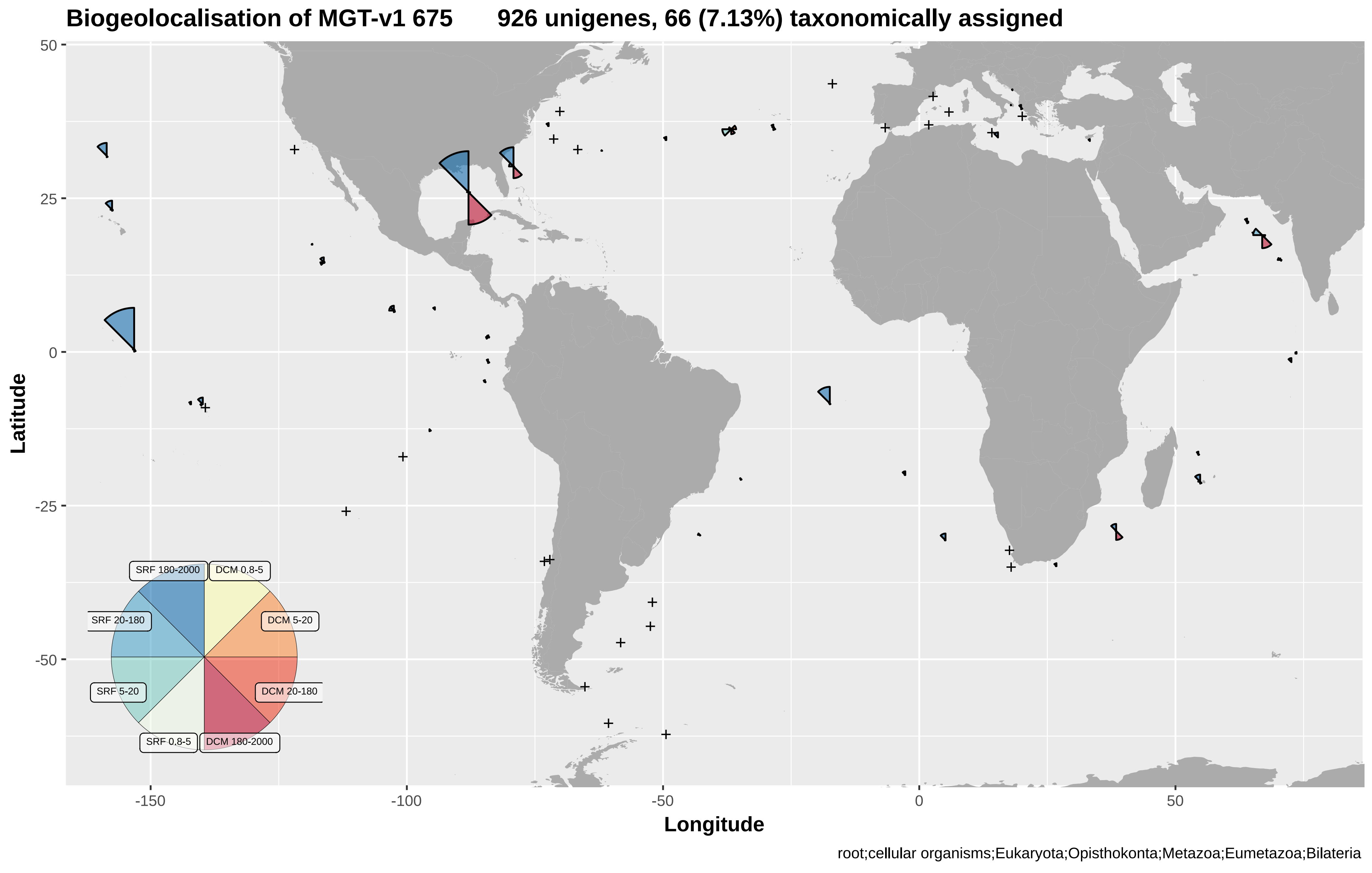

Biogeolocalisation of MGT-v1 675 926 unigenes, 66 (7.13%) taxonomically assigned
50
25
0
-25
-50
Latitude
-150
-100
-50
0
50
Longitude
SRF 180-2000
DCM 0.8-5
SRF 20-180
DCM 5-20
SRF 5-20
DCM 20-180
SRF 0.8-5
DCM 180-2000
root;cellular organisms;Eukaryota;Opisthokonta;Metazoa;Eumetazoa;Bilateria

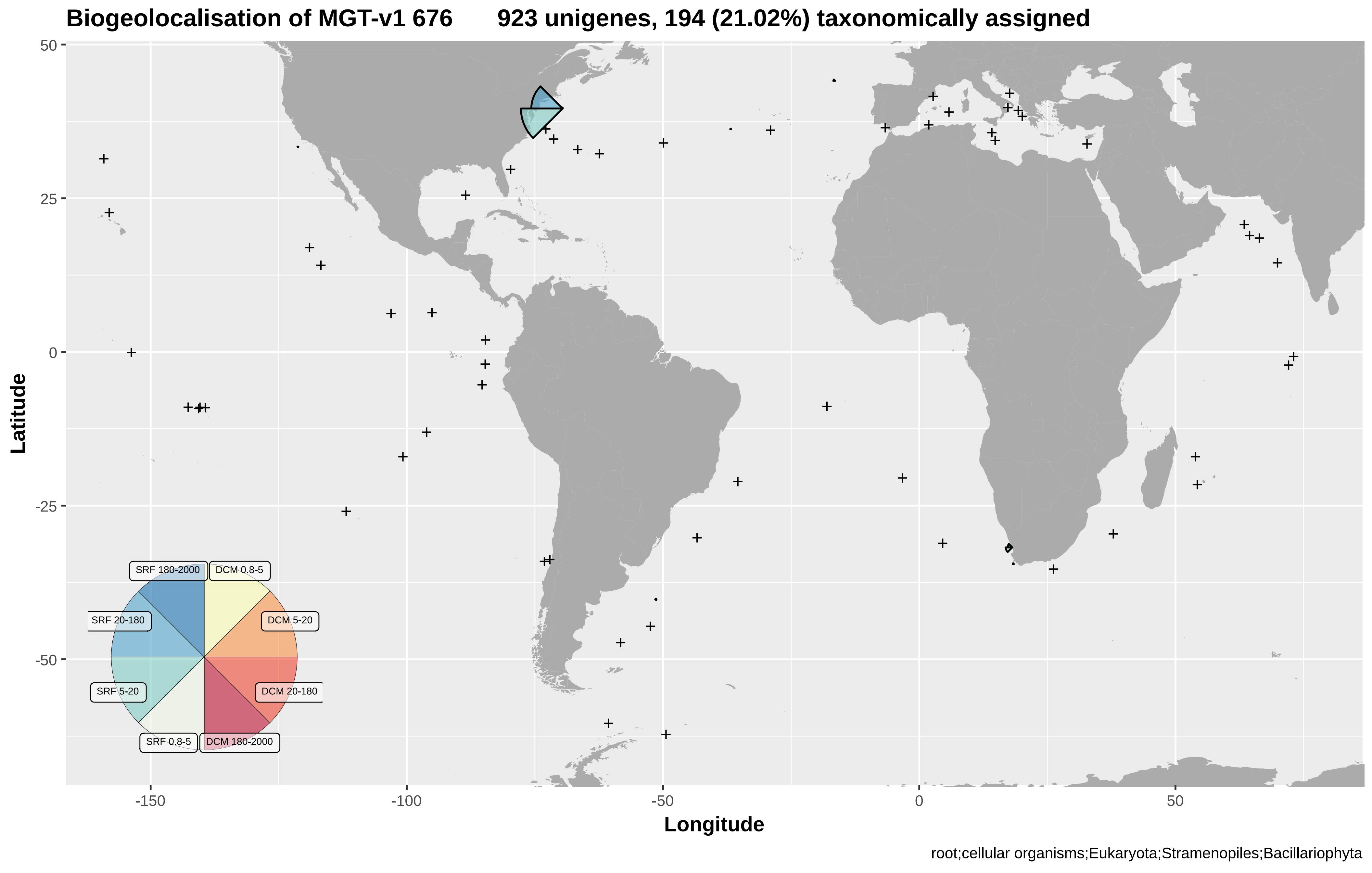

Biogeolocalisation of MGT-v1 676 923 unigenes, 194 (21.02%) taxonomically assigned
Latitude
50
25
0
-25
-50
Longitude
-150
-100
-50
0
50
SRF 180-2000
DCM 0.8-5
SRF 20-180
DCM 5-20
SRF 5-20
DCM 20-180
SRF 0.8-5
DCM 180-2000
root;cellular organisms;Eukaryota;Stramenopiles;Bacillariophyta

## Biogeolocalisation of MGT-v1 677      922 unigenes, 863 (93.6%) taxonomically assigned

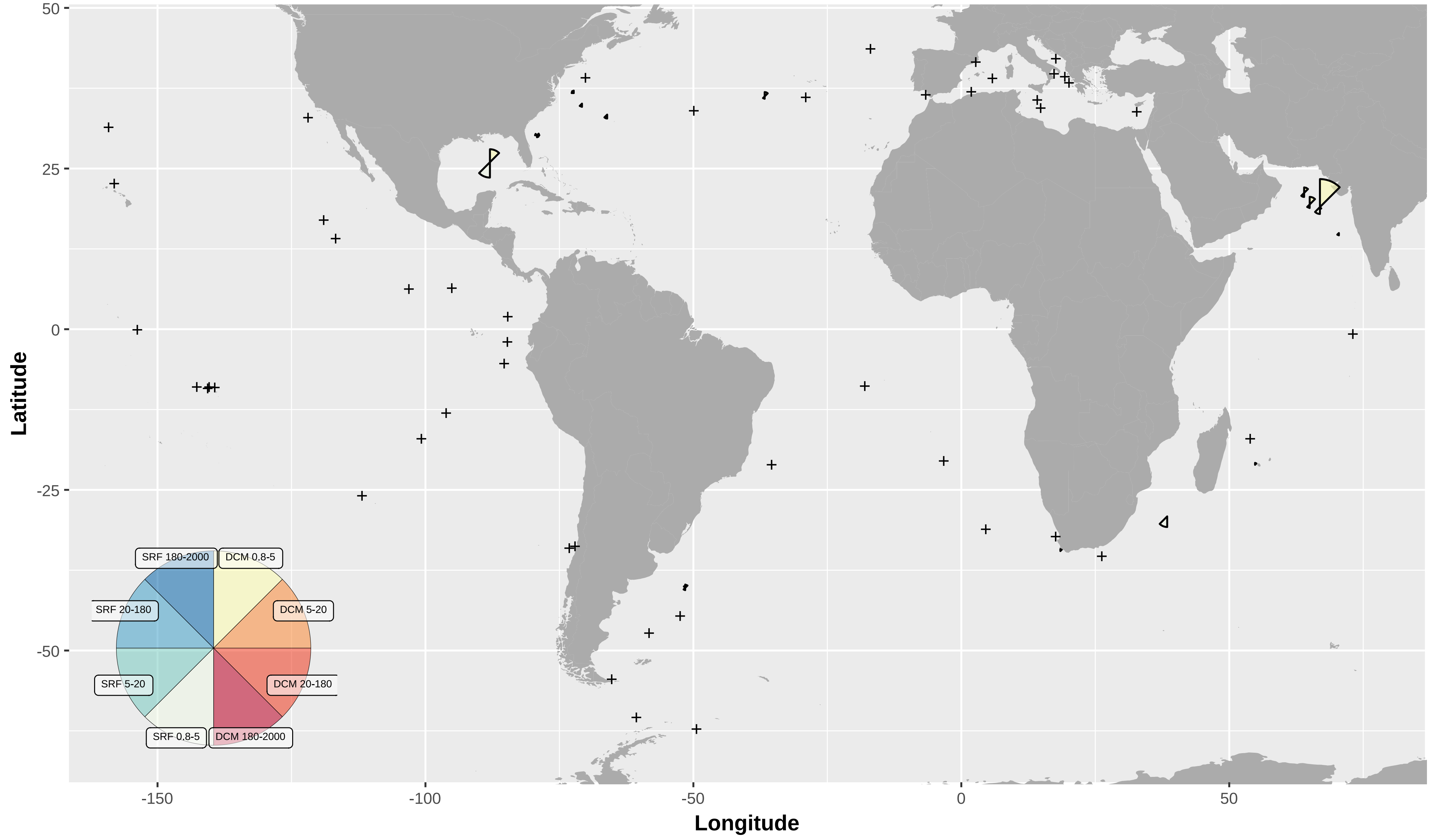


root;cellular organisms;Bacteria;Cyanobacteria;Oscillatoriophycideae;Chroococcales

## Biogeolocalisation of MGT-v1 678      922 unigenes, 390 (42.3%) taxonomically assigned

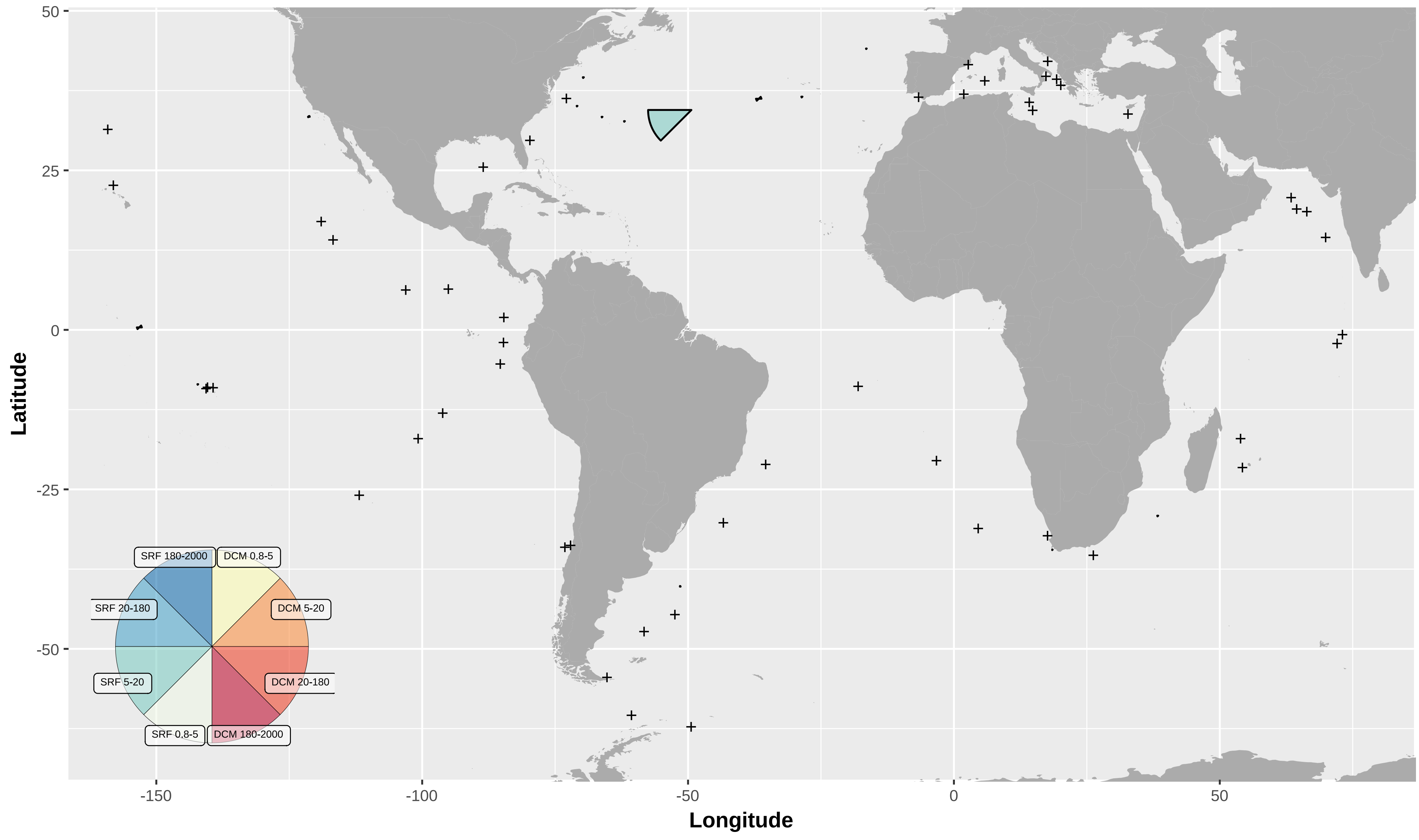


root;cellular organisms;Eukaryota;Haptophyceae

# Biogeolocalisation of MGT-v1 679      914 unigenes, 535 (58.53%) taxonomically assigned

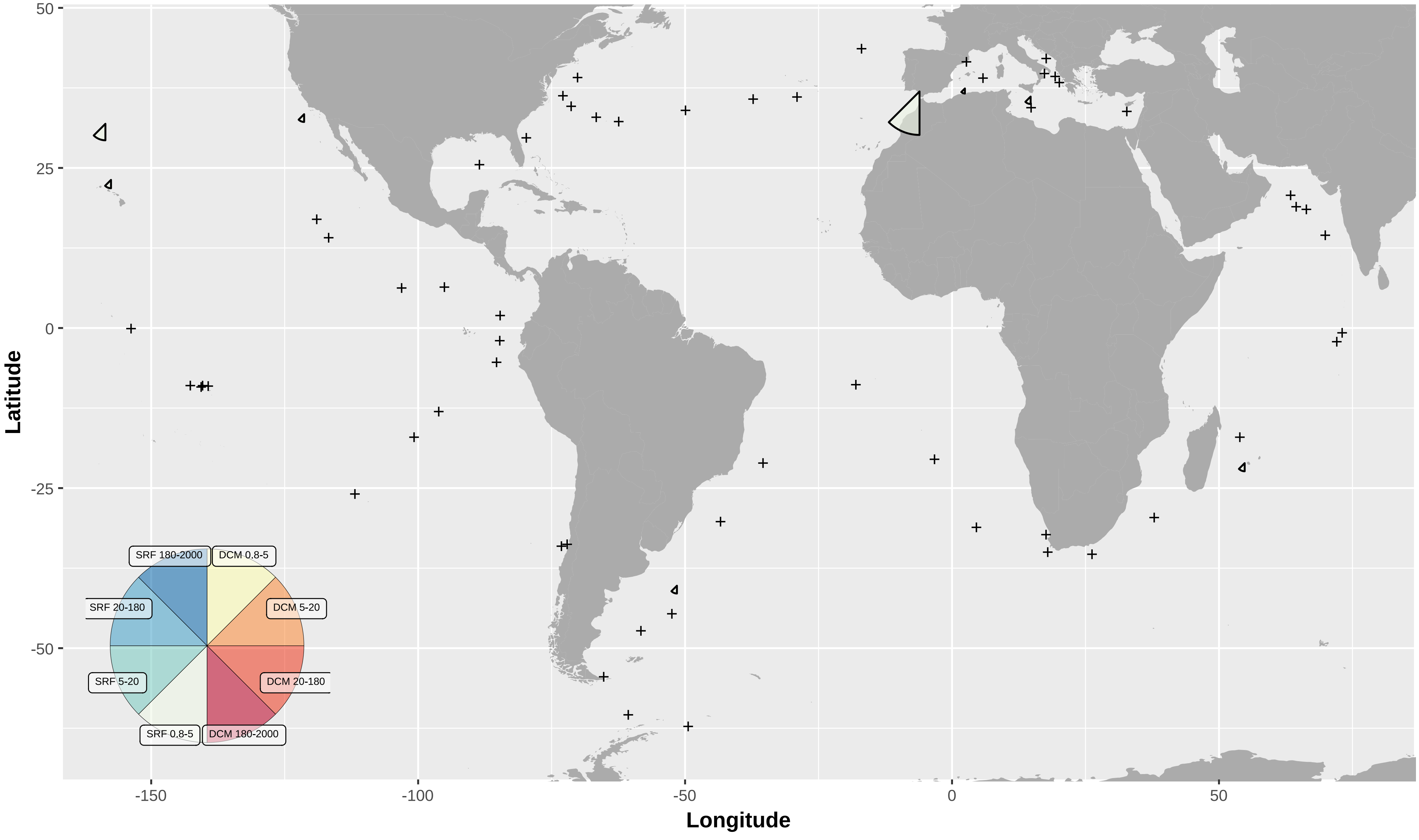


root;cellular organisms;Eukaryota

#### Biogeolocalisation of MGT-v1 680 913 unigenes, 58 (6.35%) taxonomically assigned

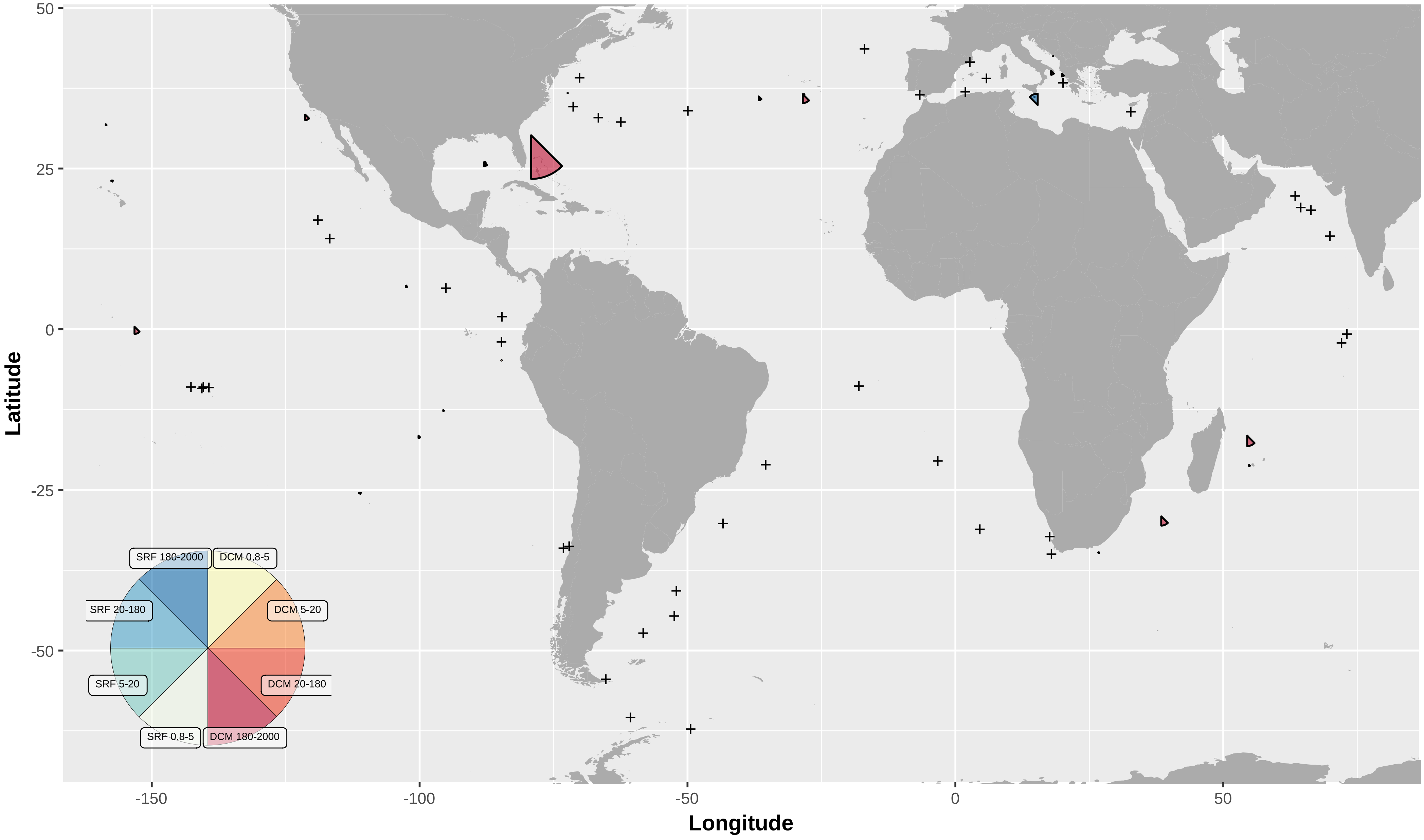


root;cellular organisms;Eukaryota;Opisthokonta;Metazoa;Eumetazoa;Bilateria

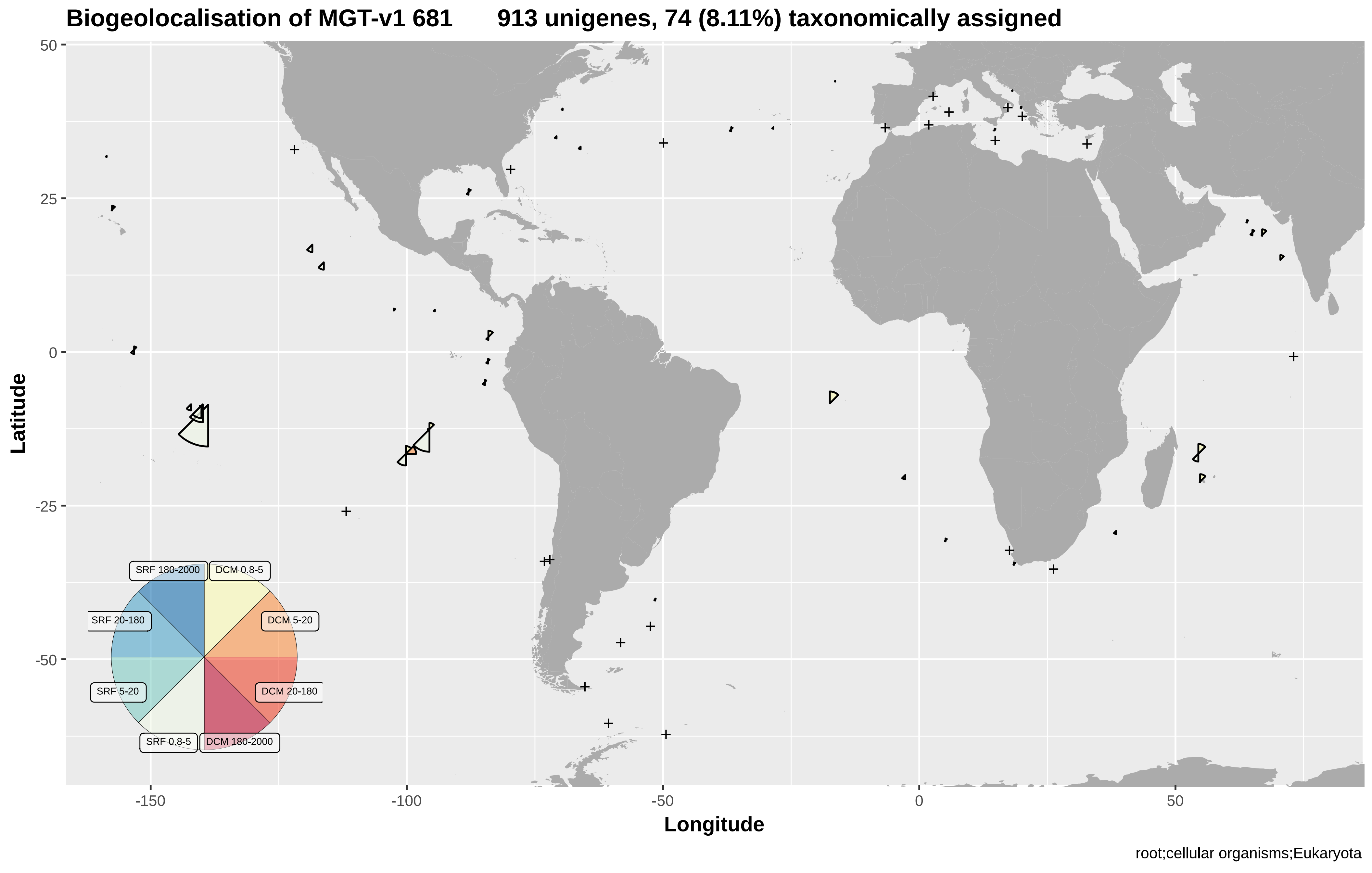

Biogeolocalisation of MGT-v1 681 913 unigenes, 74 (8.11%) taxonomically assigned
Latitude
Longitude
50
25
0
-25
-50
-150
-100
-50
0
50
SRF 180-2000
DCM 0.8-5
SRF 20-180
DCM 5-20
SRF 5-20
DCM 20-180
SRF 0.8-5
DCM 180-2000
root;cellular organisms;Eukaryota

# Biogeolocalisation of MGT-v1 682      912 unigenes, 256 (28.07%) taxonomically assigned

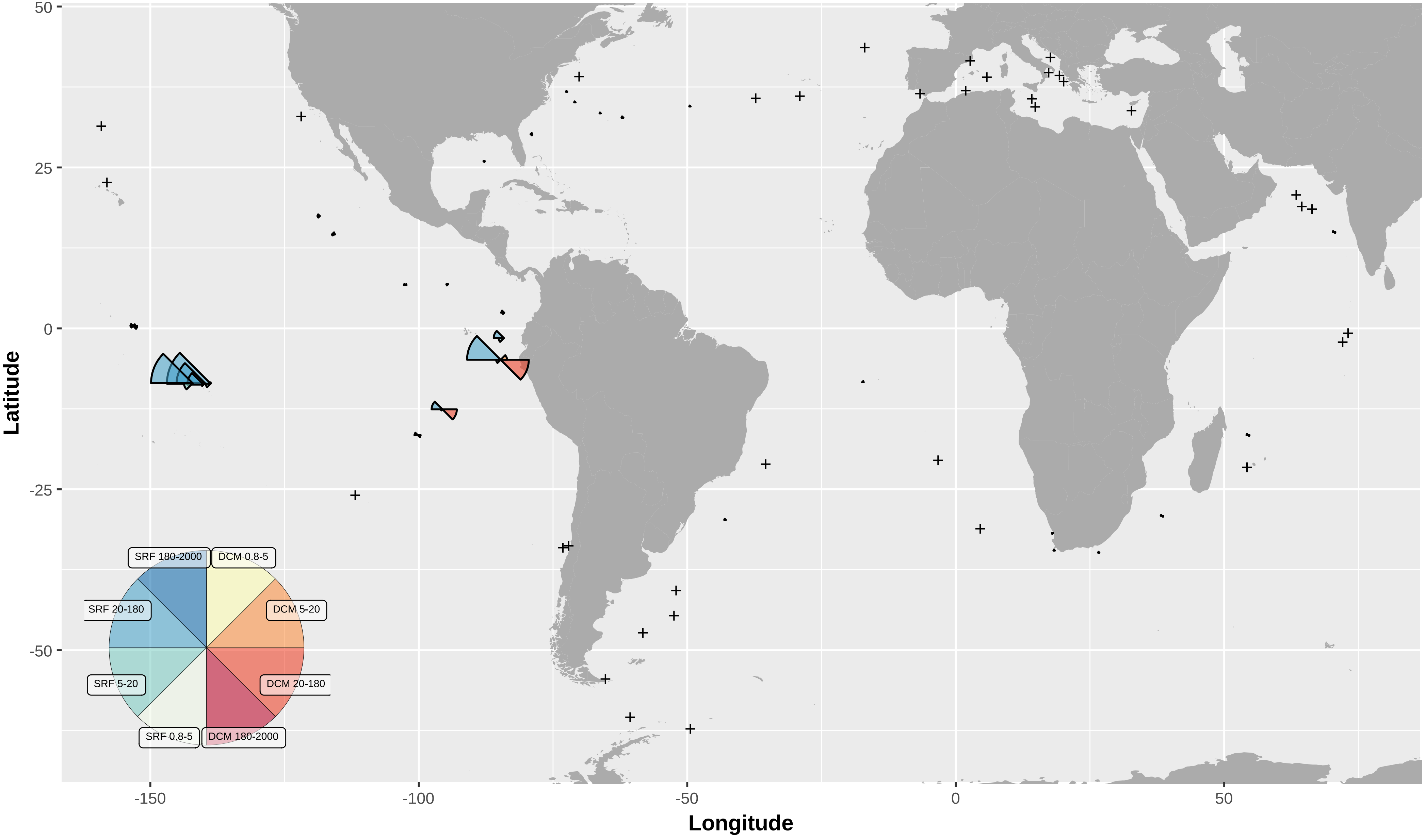


root;cellular organisms;Eukaryota;Stramenopiles;Bacillariophyta

## Biogeolocalisation of MGT-v1 683      910 unigenes, 75 (8.24%) taxonomically assigned

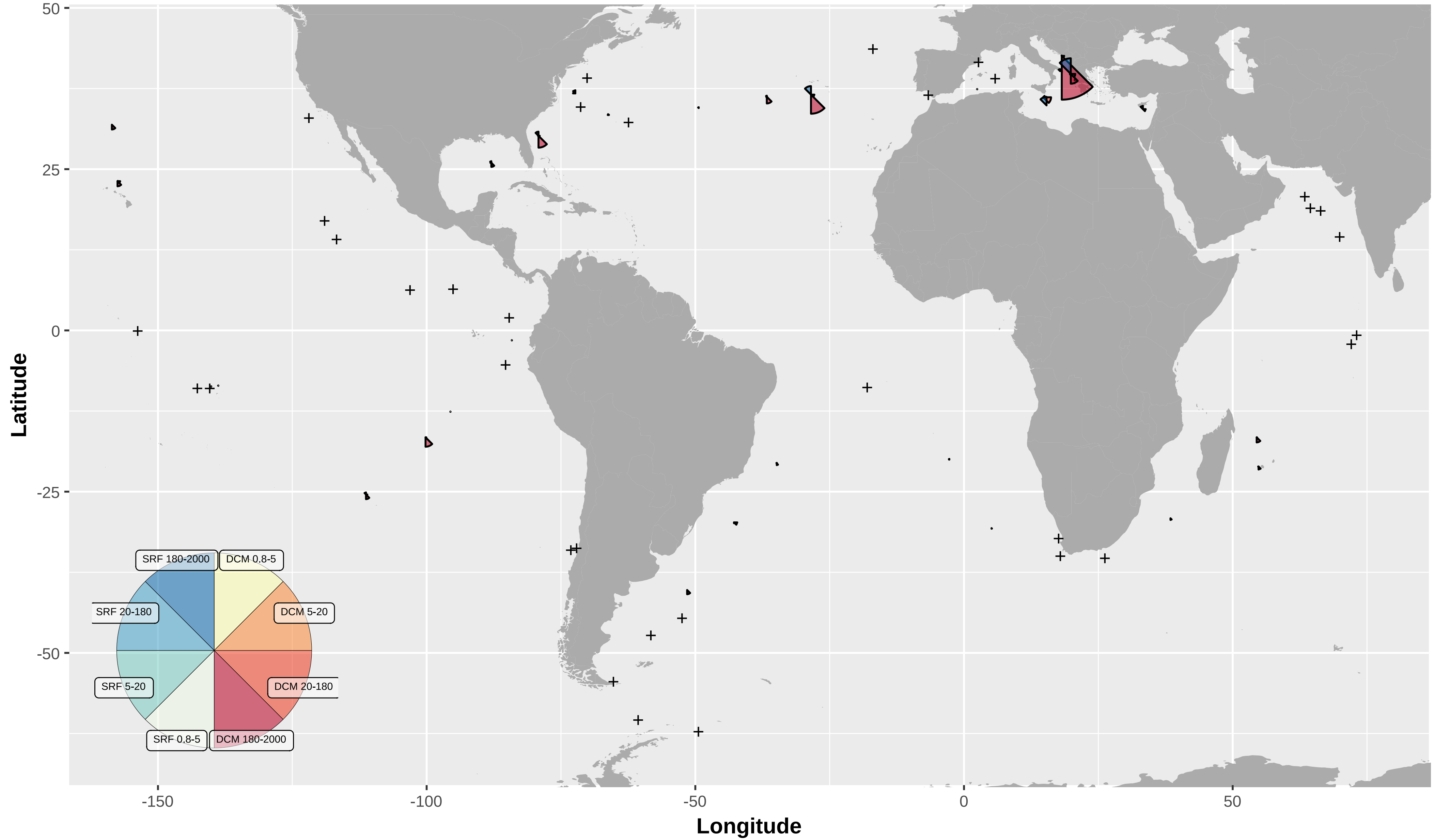


root;cellular organisms;Eukaryota;Opisthokonta;Metazoa;Eumetazoa

### Biogeolocalisation of MGT-v1 684 909 unigenes, 50 (5.5%) taxonomically assigned

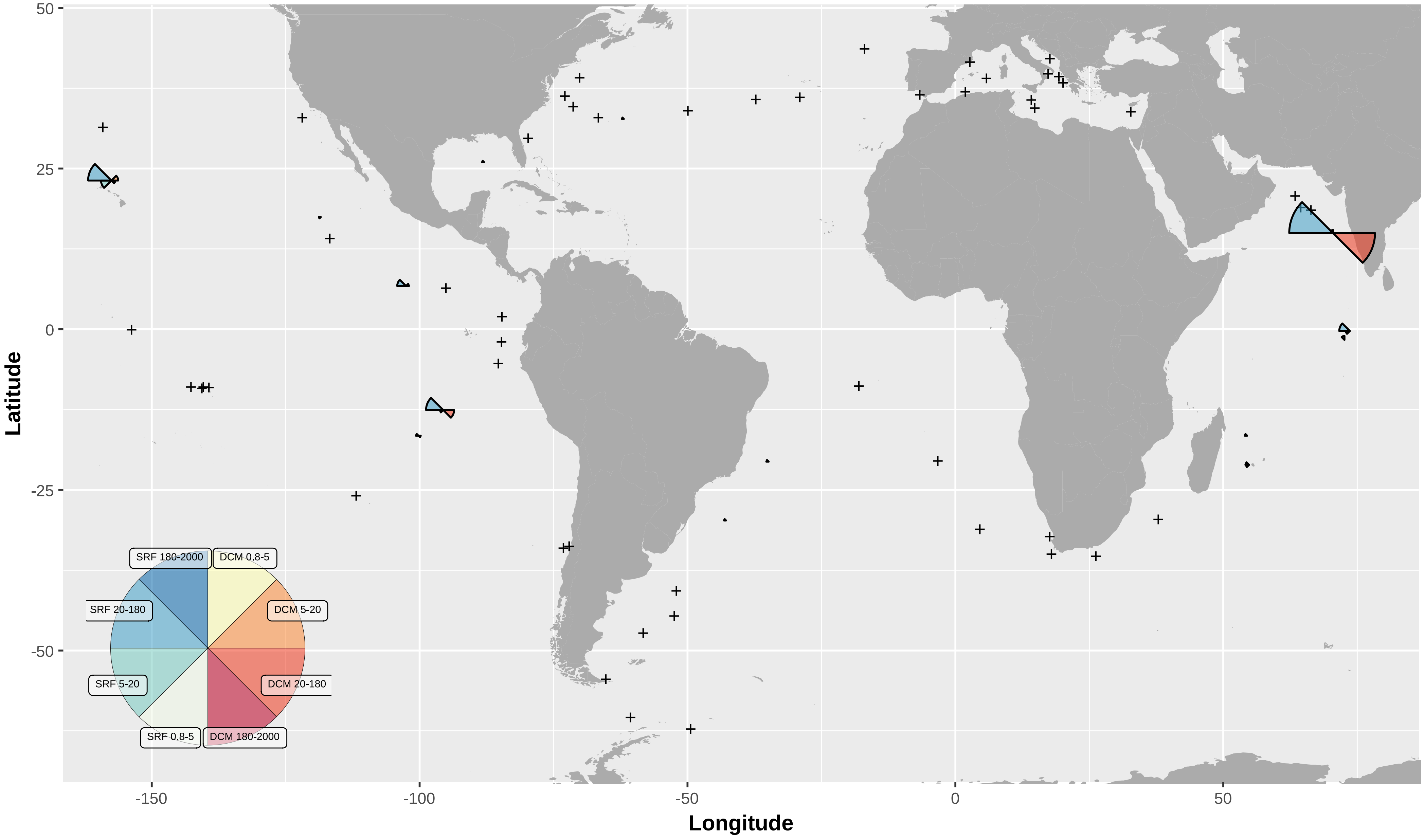


root;cellular organisms;Eukaryota

## Biogeolocalisation of MGT-v1 685      907 unigenes, 140 (15.44%) taxonomically assigned

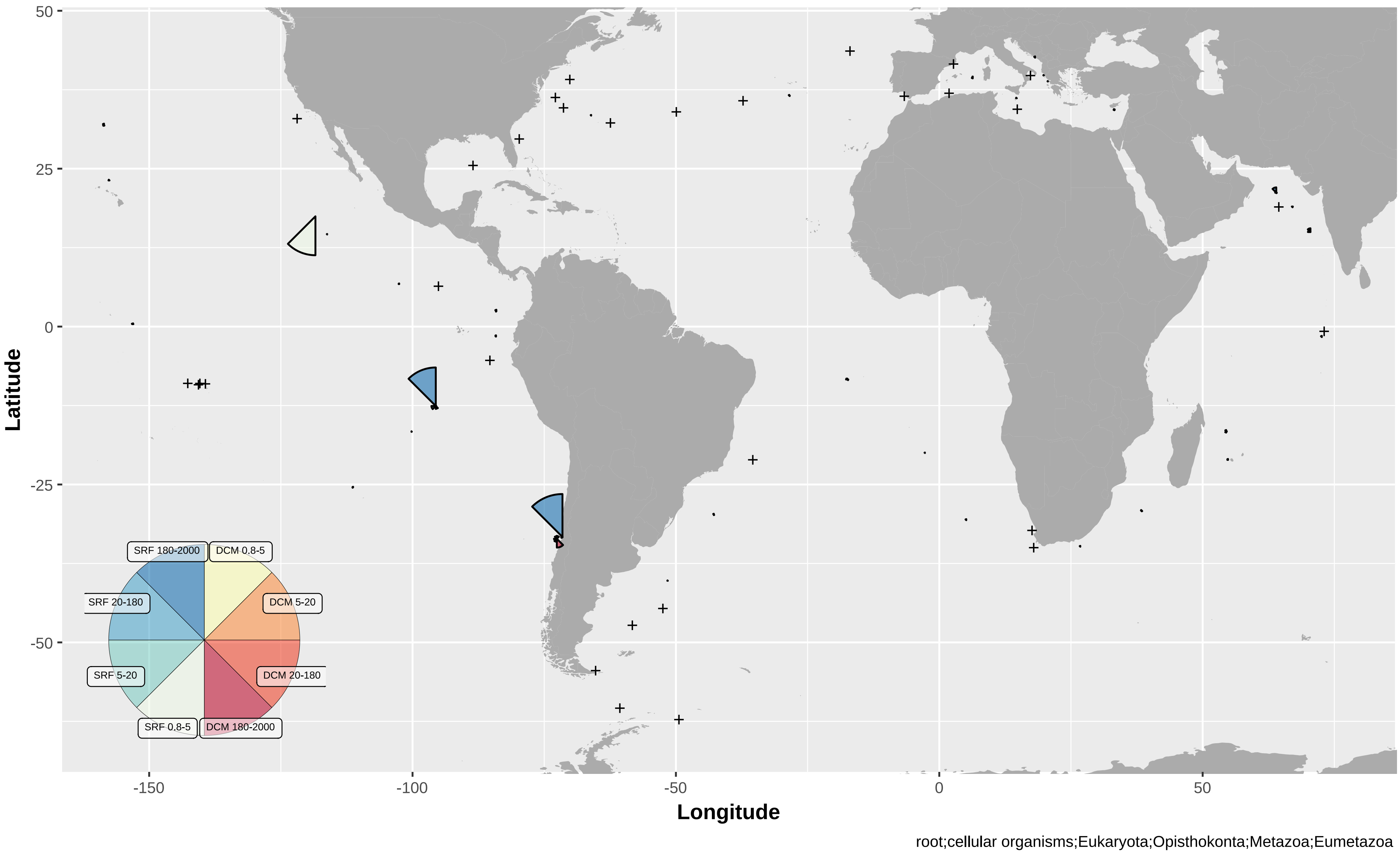

## Biogeolocalisation of MGT-v1 686      901 unigenes, 47 (5.22%) taxonomically assigned

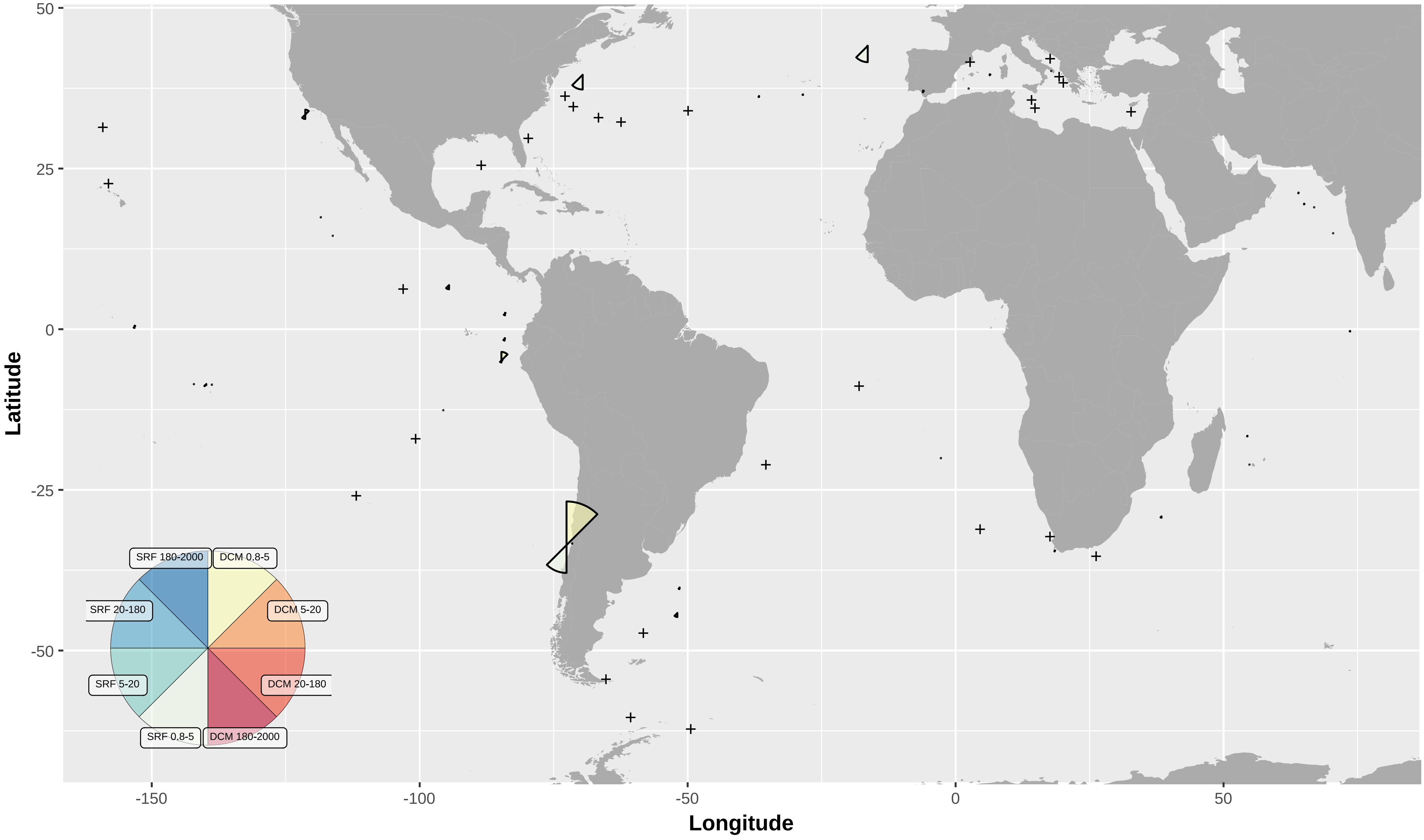


root;cellular organisms

## Biogeolocalisation of MGT-v1 687      900 unigenes, 72 (8%) taxonomically assigned

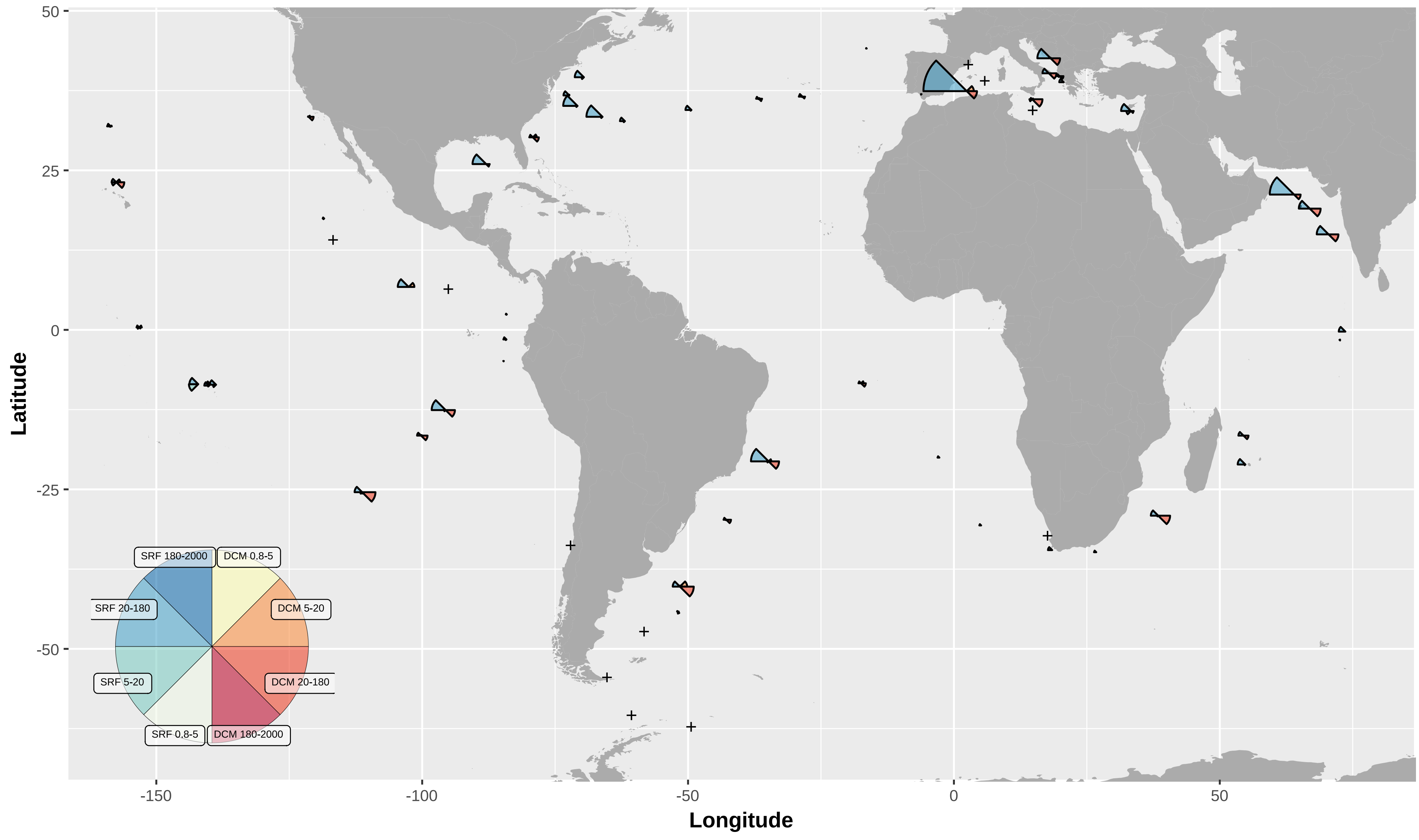


root;cellular organisms;Eukaryota

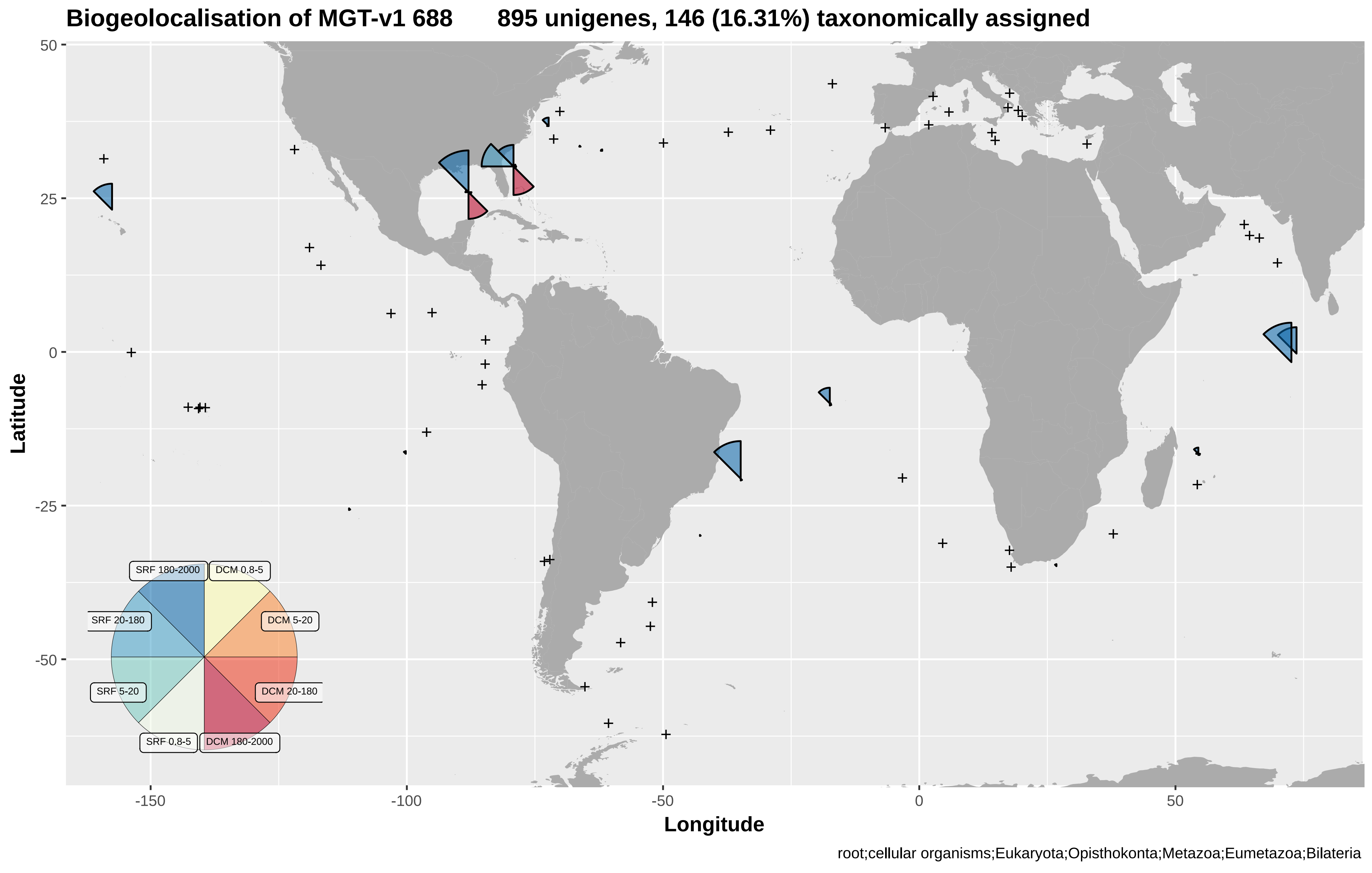
Biogeolocalisation of MGT-v1 688 895 unigenes, 146 (16.31%) taxonomically assigned
50
25
0
-25
-50
Latitude
-150
-100
-50
0
50
Longitude
SRF 180-2000
DCM 0.8-5
SRF 20-180
DCM 5-20
SRF 5-20
DCM 20-180
SRF 0.8-5
DCM 180-2000
root;cellular organisms;Eukaryota;Opisthokonta;Metazoa;Eumetazoa;Bilateria

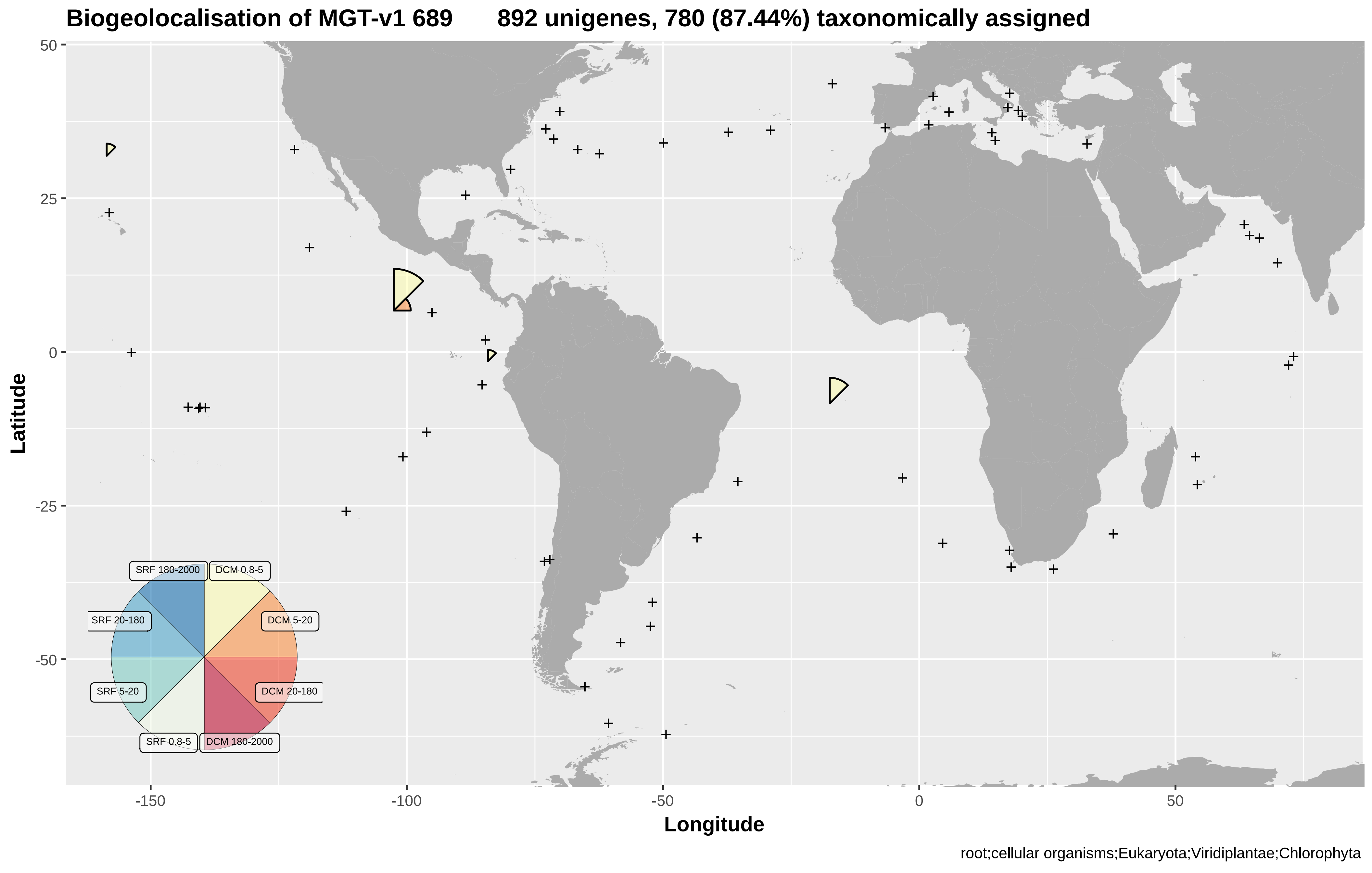
Biogeolocalisation of MGT-v1 689 892 unigenes, 780 (87.44%) taxonomically assigned
50
25
0
-25
-50
Latitude
-150
-100
-50
0
50
Longitude
SRF 180-2000
DCM 0.8-5
SRF 20-180
DCM 5-20
SRF 5-20
DCM 20-180
SRF 0.8-5
DCM 180-2000
root;cellular organisms;Eukaryota;Viridiplantae;Chlorophyta

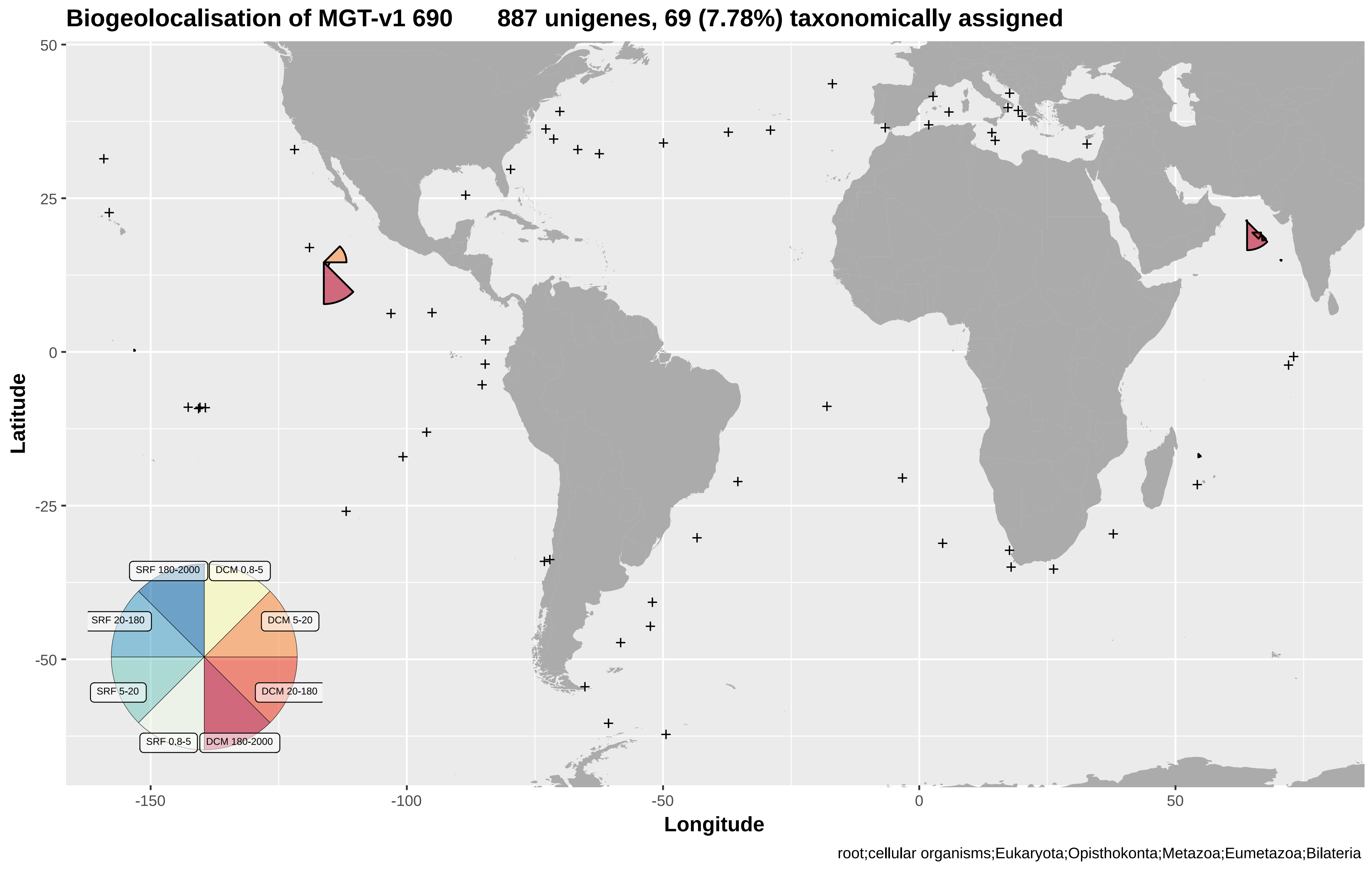
Biogeolocalisation of MGT-v1 690
887 unigenes, 69 (7.78%) taxonomically assigned
50
25
0
-25
-50
Latitude
-150
-100
-50
0
50
Longitude
SRF 180-2000
DCM 0.8-5
SRF 20-180
DCM 5-20
SRF 5-20
DCM 20-180
SRF 0.8-5
DCM 180-2000
root;cellular organisms;Eukaryota;Opisthokonta;Metazoa;Eumetazoa;Bilateria

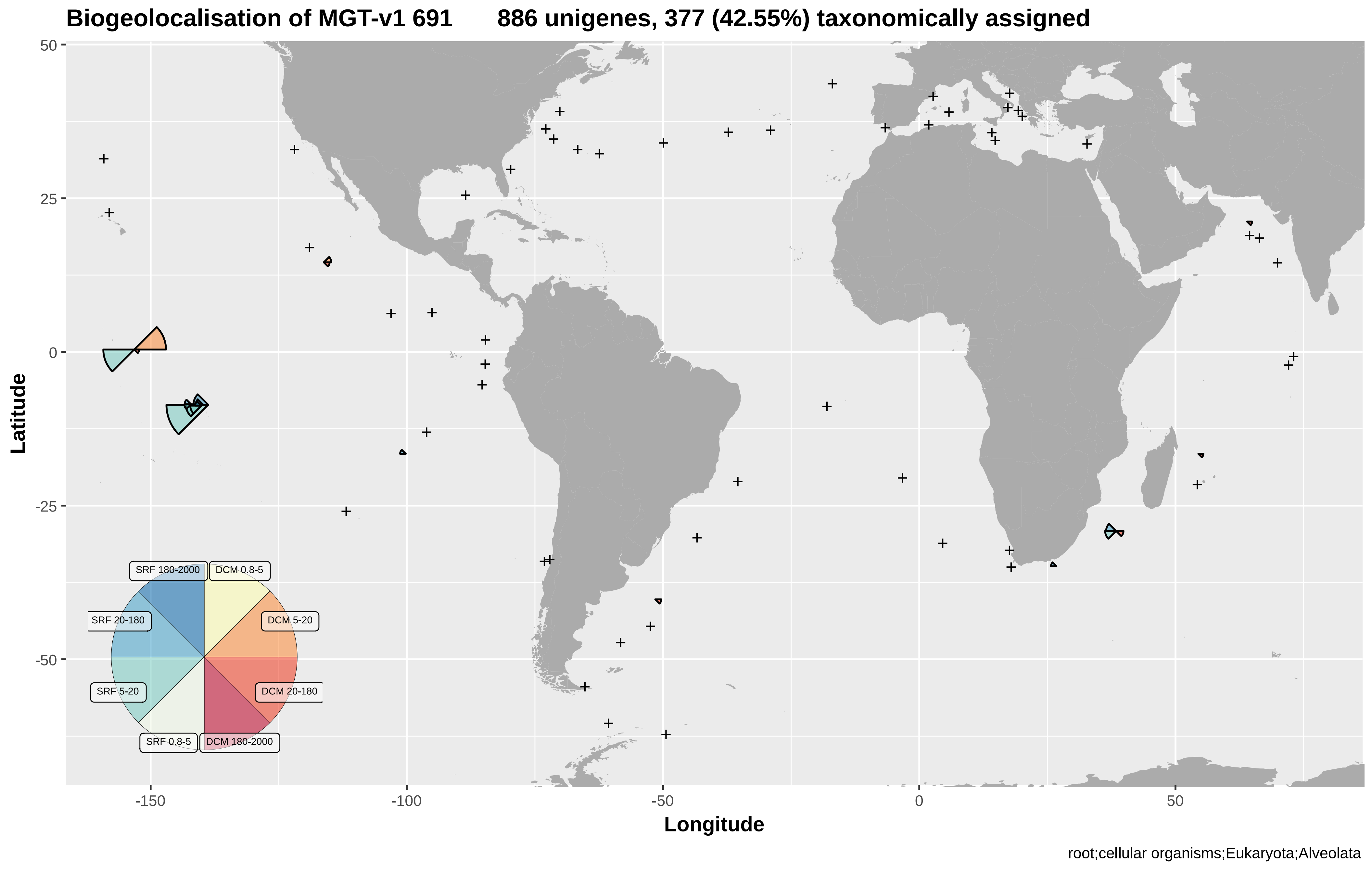
Biogeolocalisation of MGT-v1 691 886 unigenes, 377 (42.55%) taxonomically assigned
50
25
0
-25
-50
Latitude
-150
-100
-50
0
50
Longitude
SRF 180-2000
DCM 0.8-5
SRF 20-180
DCM 5-20
SRF 5-20
DCM 20-180
SRF 0.8-5
DCM 180-2000
root;cellular organisms;Eukaryota;Alveolata

### Biogeolocalisation of MGT-v1 692 885 unigenes, 830 (93.79%) taxonomically assigned

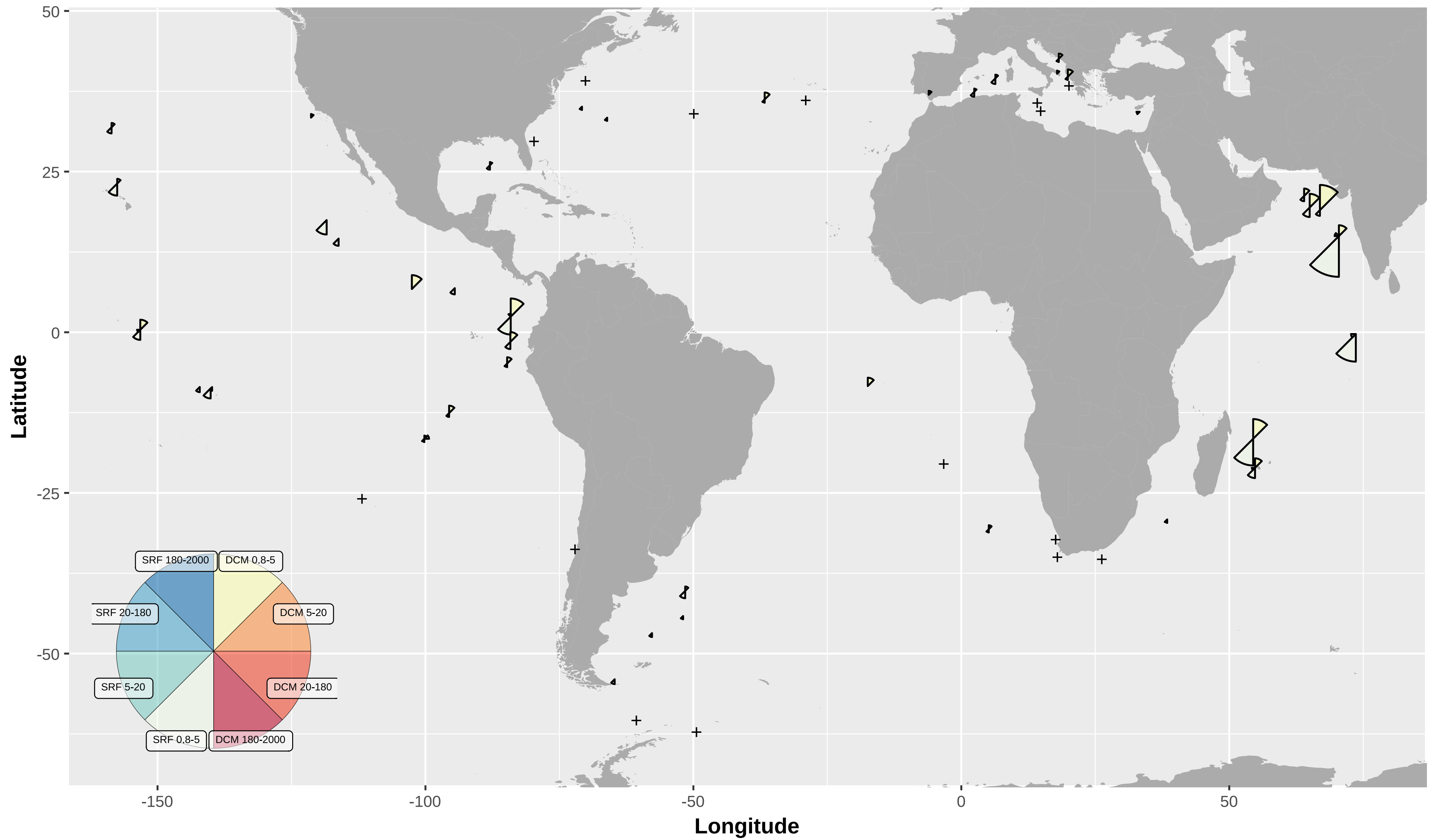


root;cellular organisms;Bacteria;Proteobacteria;Alphaproteobacteria;Sphingomonadales

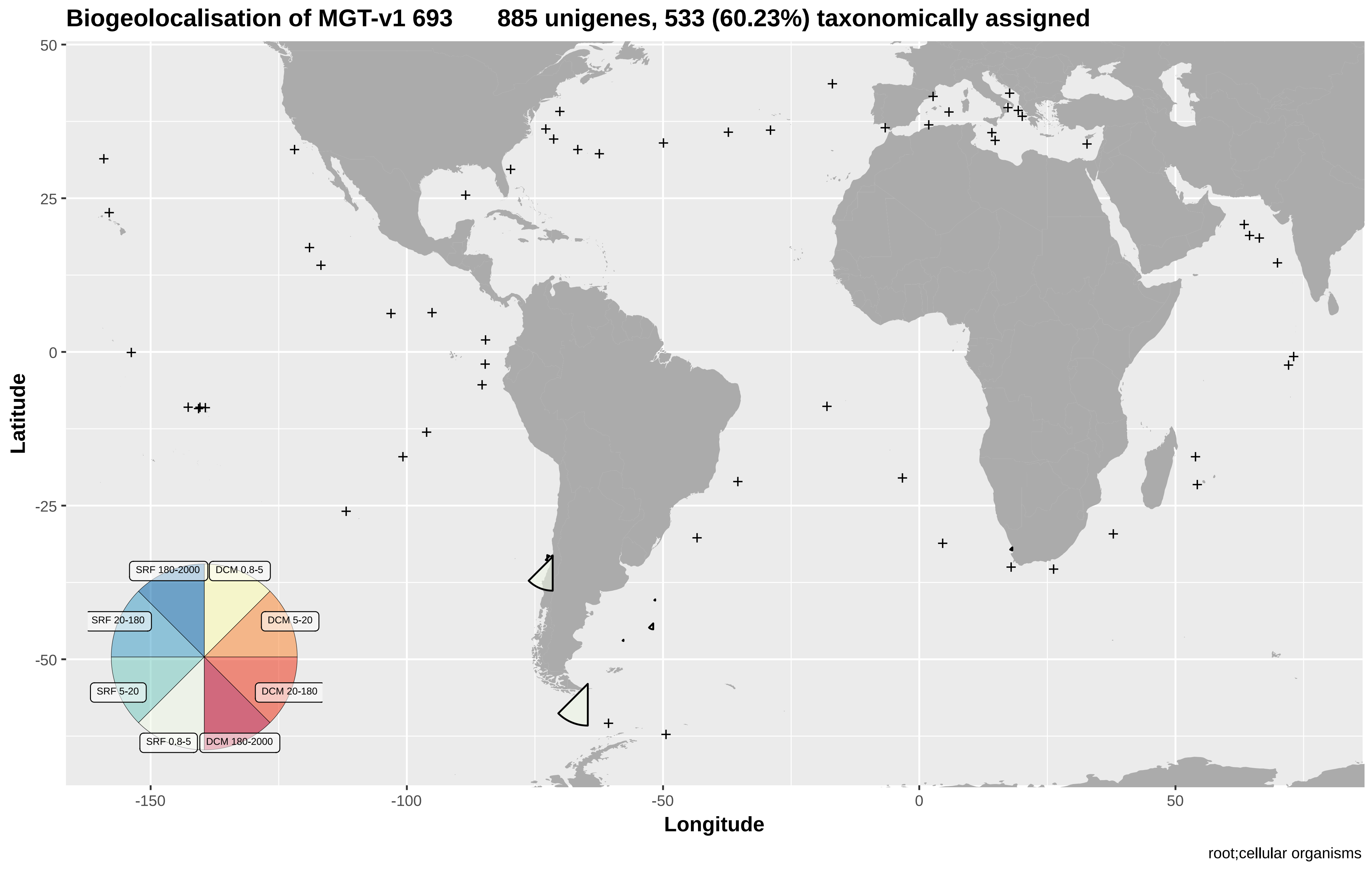

Biogeolocalisation of MGT-v1 693 885 unigenes, 533 (60.23%) taxonomically assigned
50
25
0
-25
-50
Latitude
-150
-100
-50
0
50
Longitude
SRF 180-2000
DCM 0.8-5
SRF 20-180
DCM 5-20
SRF 5-20
DCM 20-180
SRF 0.8-5
DCM 180-2000
root;cellular organisms

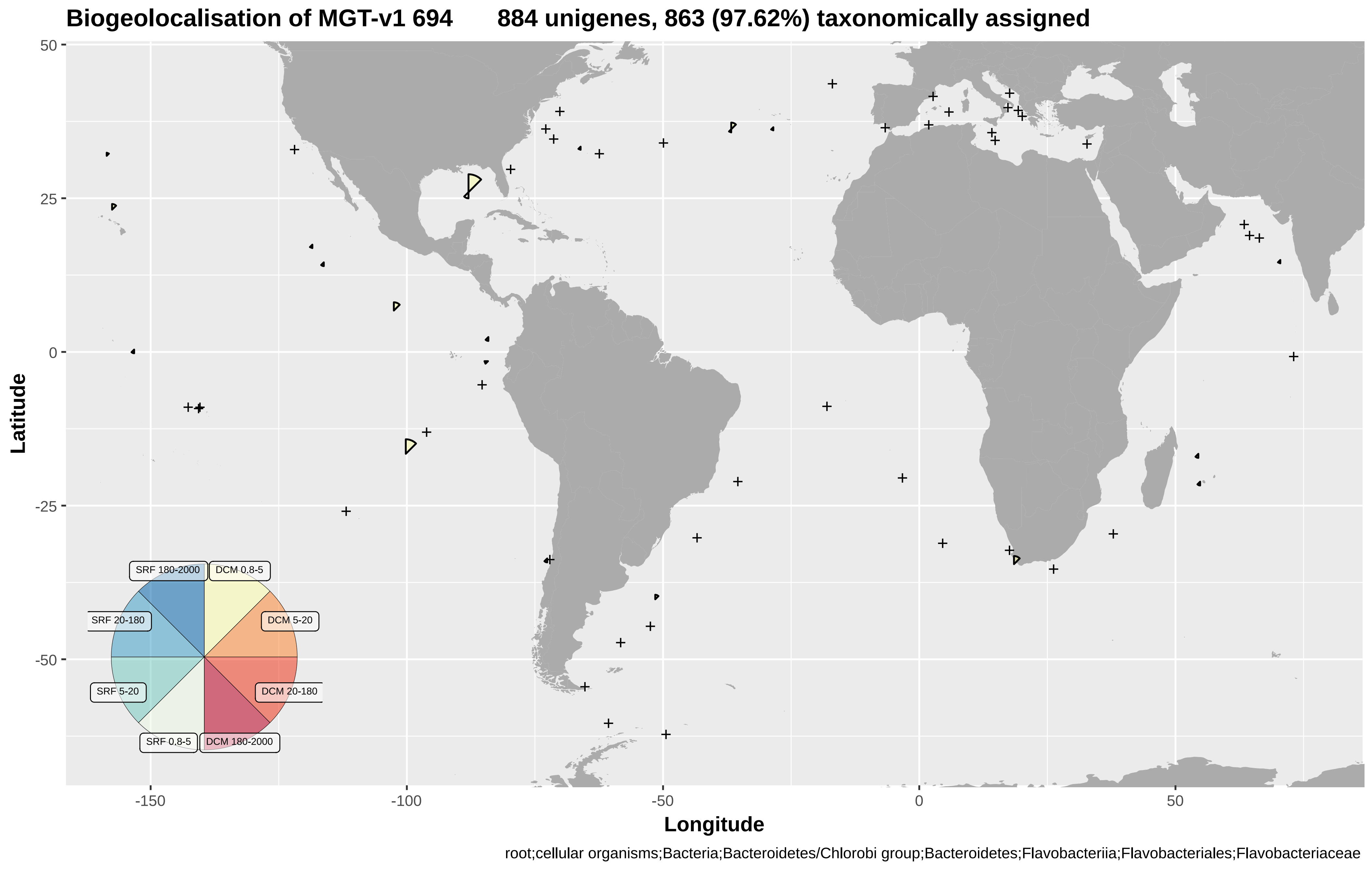

Biogeolocalisation of MGT-v1 694
884 unigenes, 863 (97.62%) taxonomically assigned
50
25
0
-25
-50
Latitude
-150
-100
-50
0
50
Longitude
SRF 180-2000
DCM 0.8-5
SRF 20-180
DCM 5-20
SRF 5-20
DCM 20-180
SRF 0.8-5
DCM 180-2000
root;cellular organisms;Bacteria;Bacteroidetes/Chlorobi group;Bacteroidetes;Flavobacteriia;Flavobacteriales;Flavobacteriaceae

# Biogeolocalisation of MGT-v1 695      883 unigenes, 63 (7.13%) taxonomically assigned

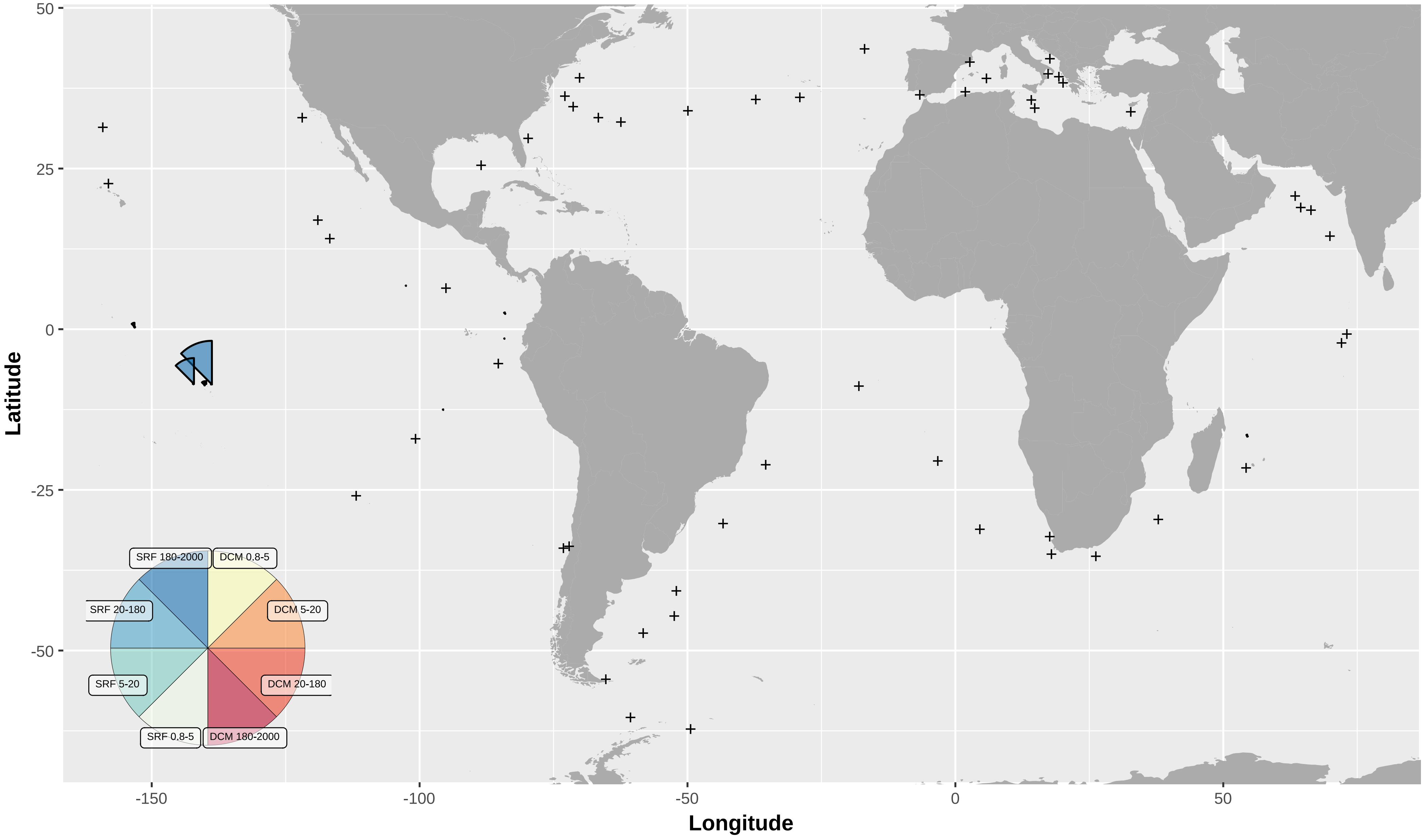


root;cellular organisms;Eukaryota

## Biogeolocalisation of MGT-v1 696      883 unigenes, 652 (73.84%) taxonomically assigned

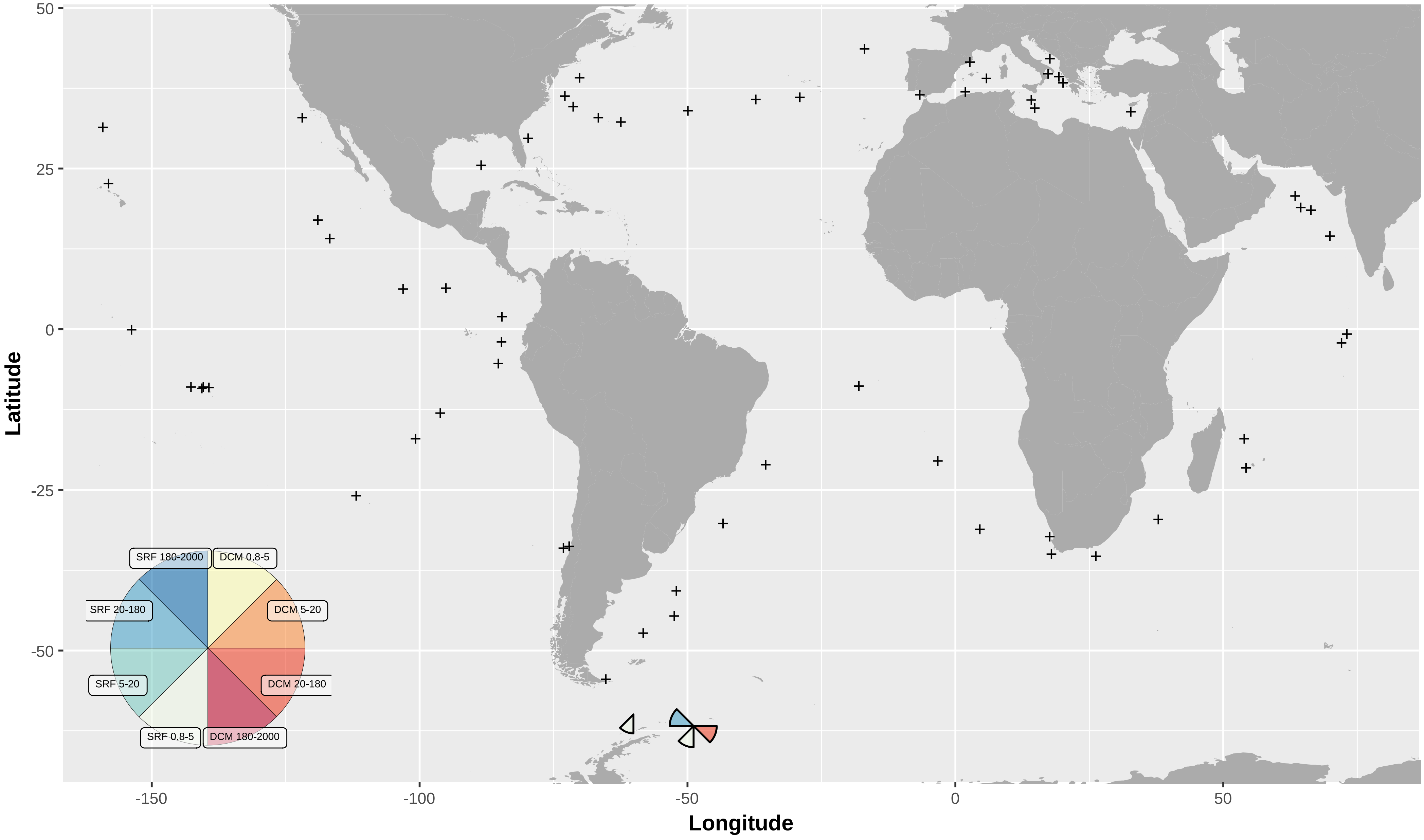


root;cellular organisms;Eukaryota

#### Biogeolocalisation of MGT-v1 697 877 unigenes, 688 (78.45%) taxonomically assigned

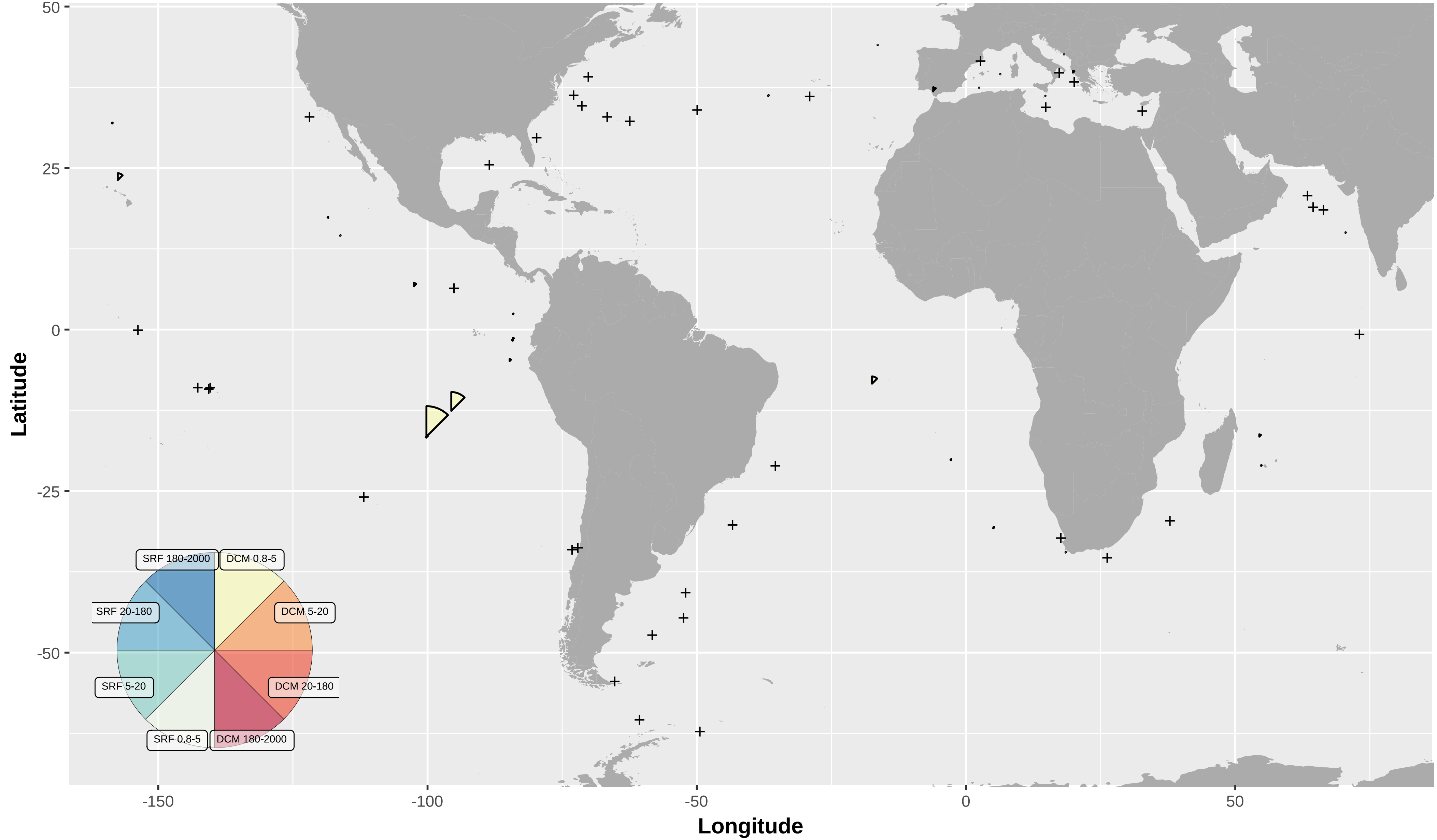


root;Viruses;dsDNA viruses, no RNA stage;Caudovirales

## Biogeolocalisation of MGT-v1 698      873 unigenes, 792 (90.72%) taxonomically assigned

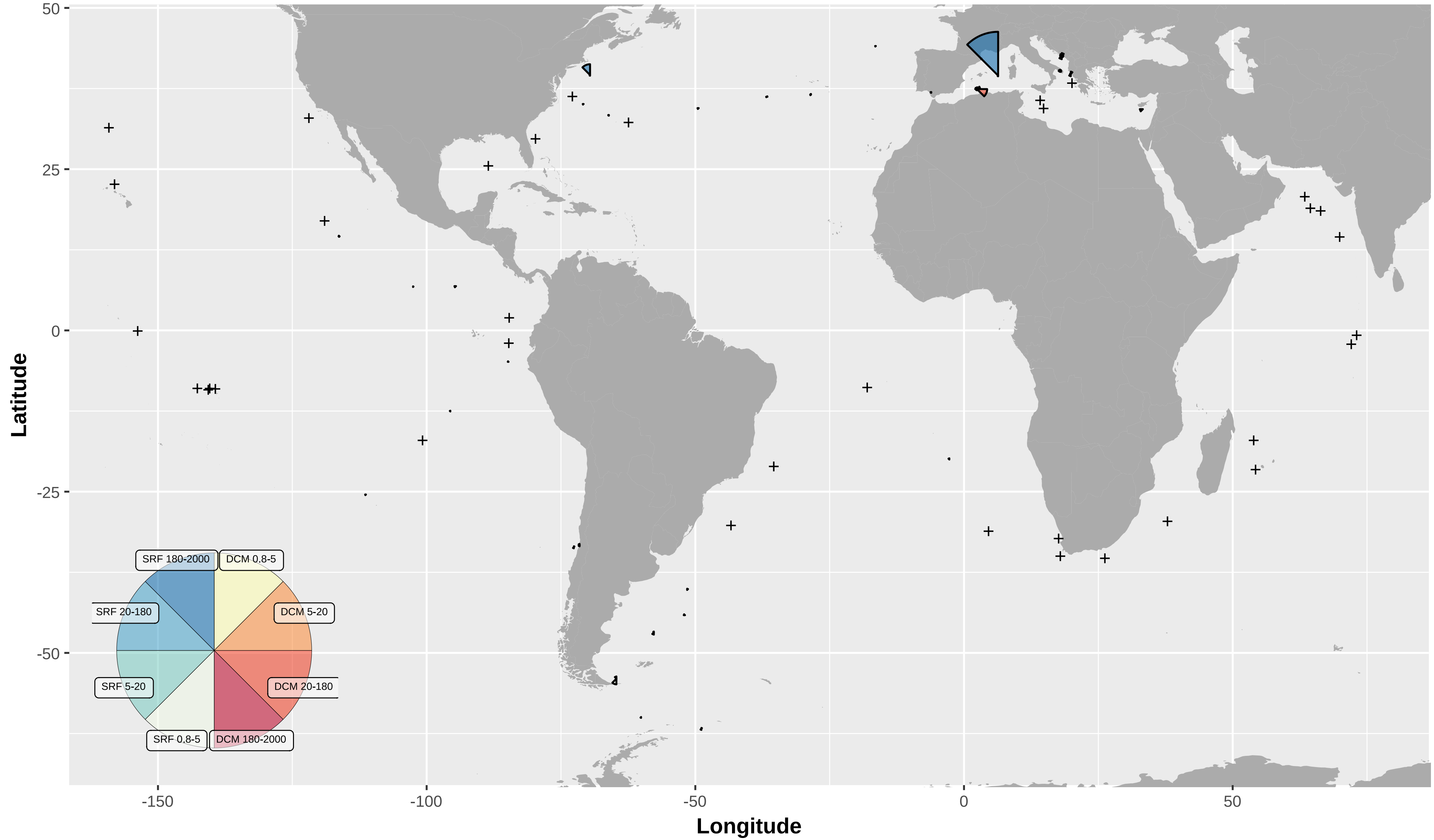


root;cellular organisms;Bacteria;Proteobacteria;Gammaproteobacteria;Alteromonadales

Biogeolocalisation of MGT-v1 699      873 unigenes, 79 (9.05%) taxonomically assigned

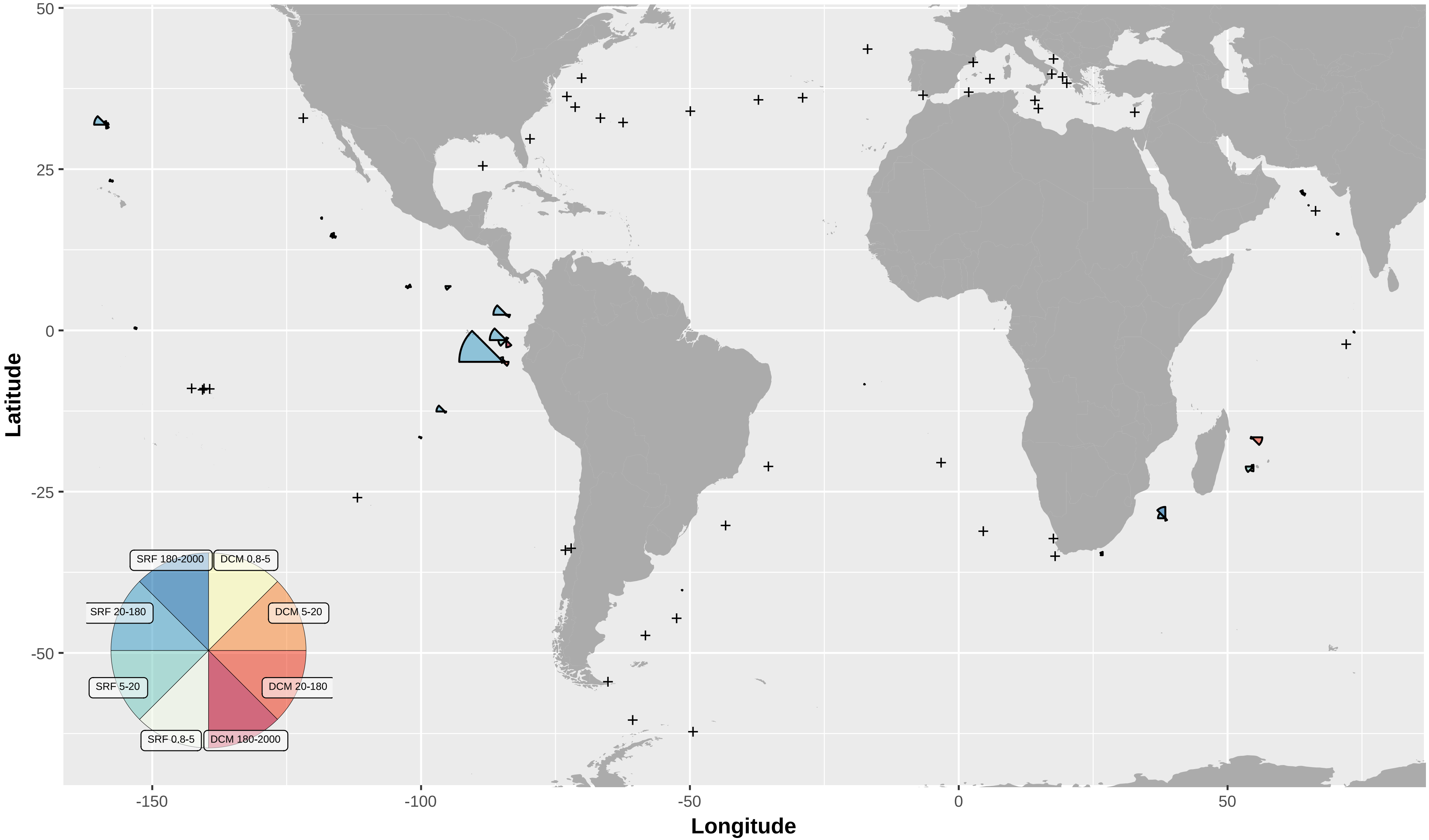


root;cellular organisms;Eukaryota;Opisthokonta;Metazoa;Eumetazoa;Bilateria

## Biogeolocalisation of MGT-v1 700      871 unigenes, 606 (69.58%) taxonomically assigned

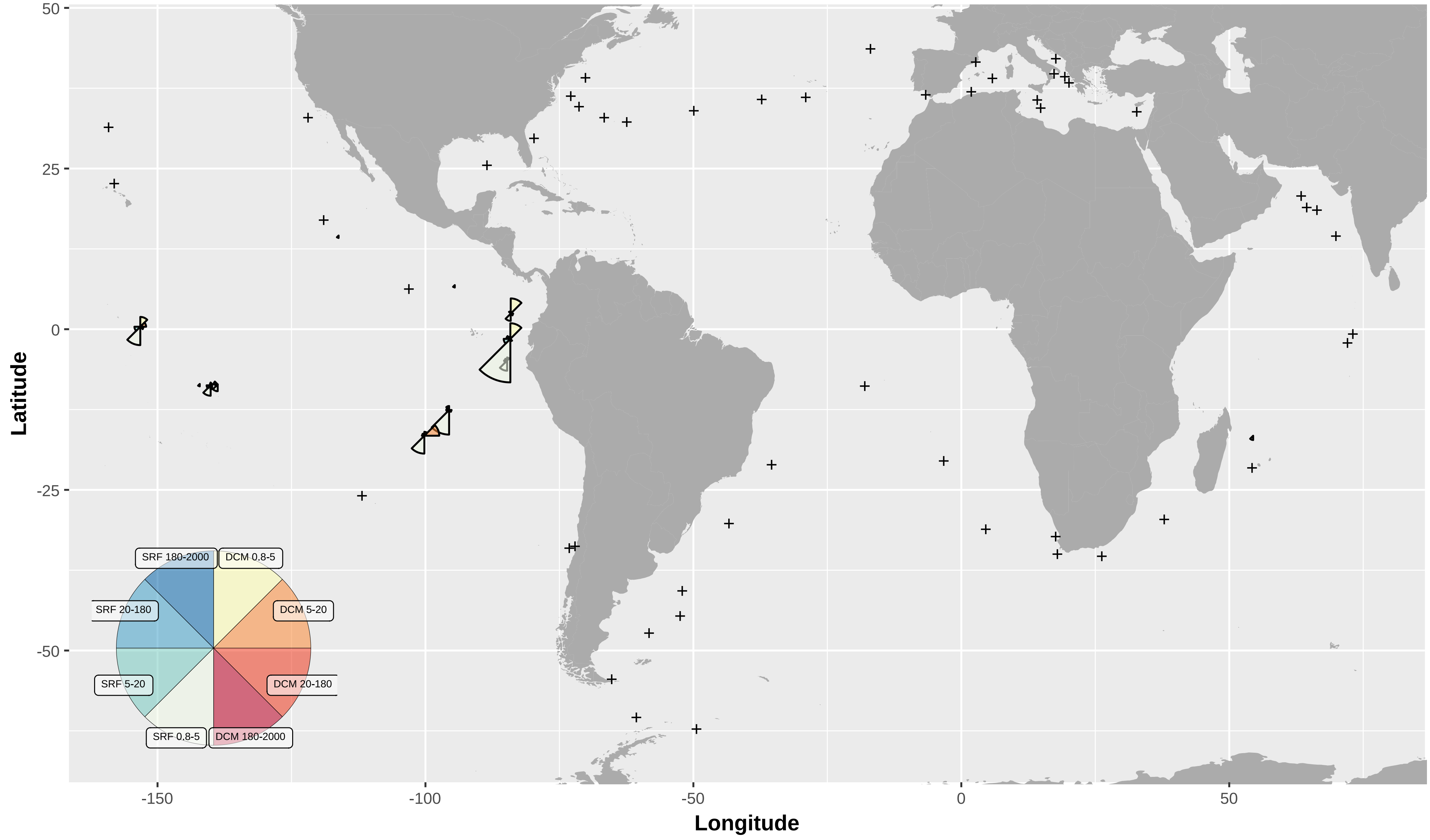


root;cellular organisms;Bacteria

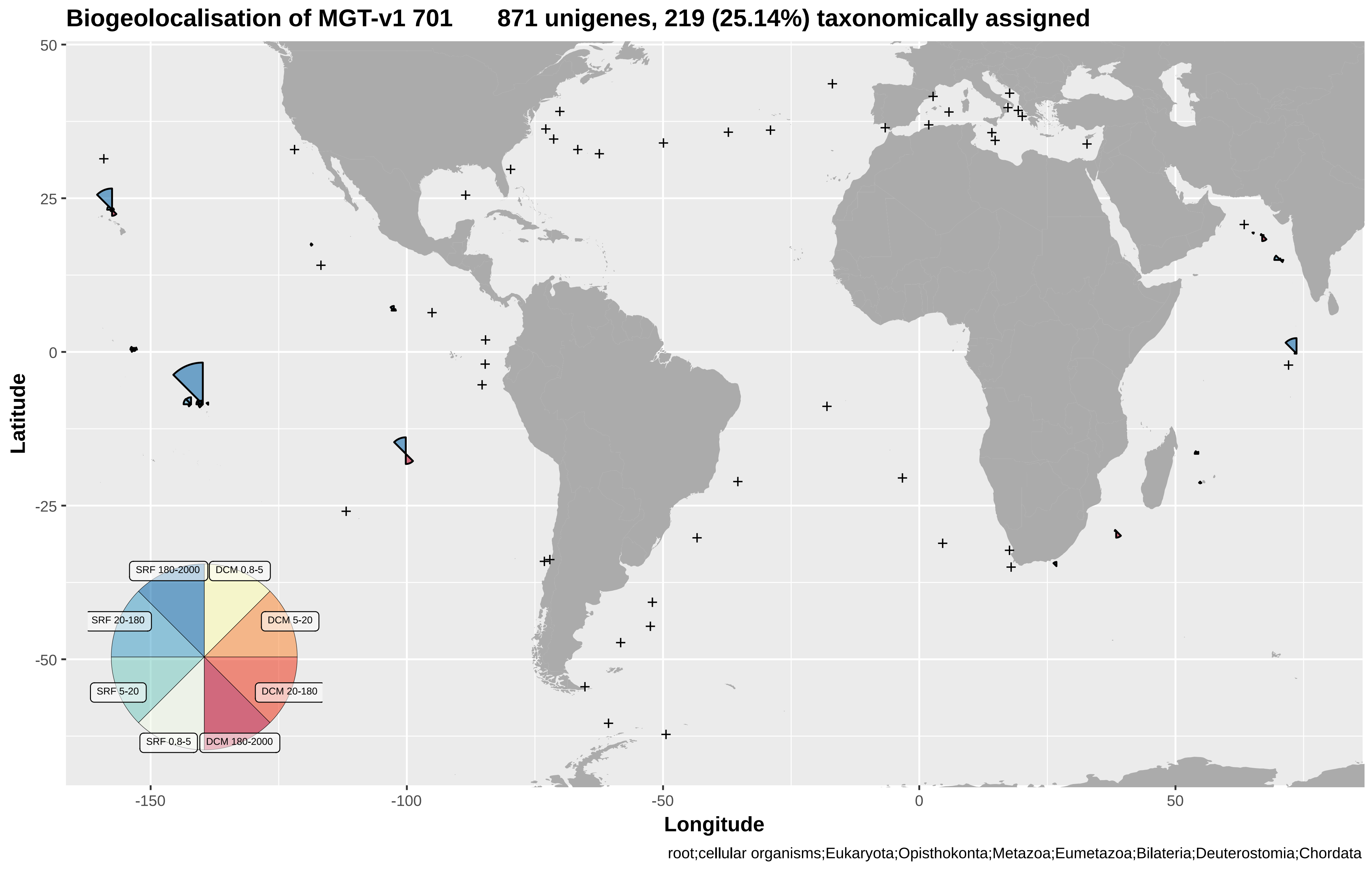
Biogeolocalisation of MGT-v1 701 871 unigenes, 219 (25.14%) taxonomically assigned
Latitude
Longitude
50
25
0
-25
-50
-150
-100
-50
0
50
SRF 180-2000
DCM 0.8-5
SRF 20-180
DCM 5-20
SRF 5-20
DCM 20-180
SRF 0.8-5
DCM 180-2000
root;cellular organisms;Eukaryota;Opisthokonta;Metazoa;Eumetazoa;Bilateria;Deuterostomia;Chordata

# Biogeolocalisation of MGT-v1 702      870 unigenes, 754 (86.67%) taxonomically assigned

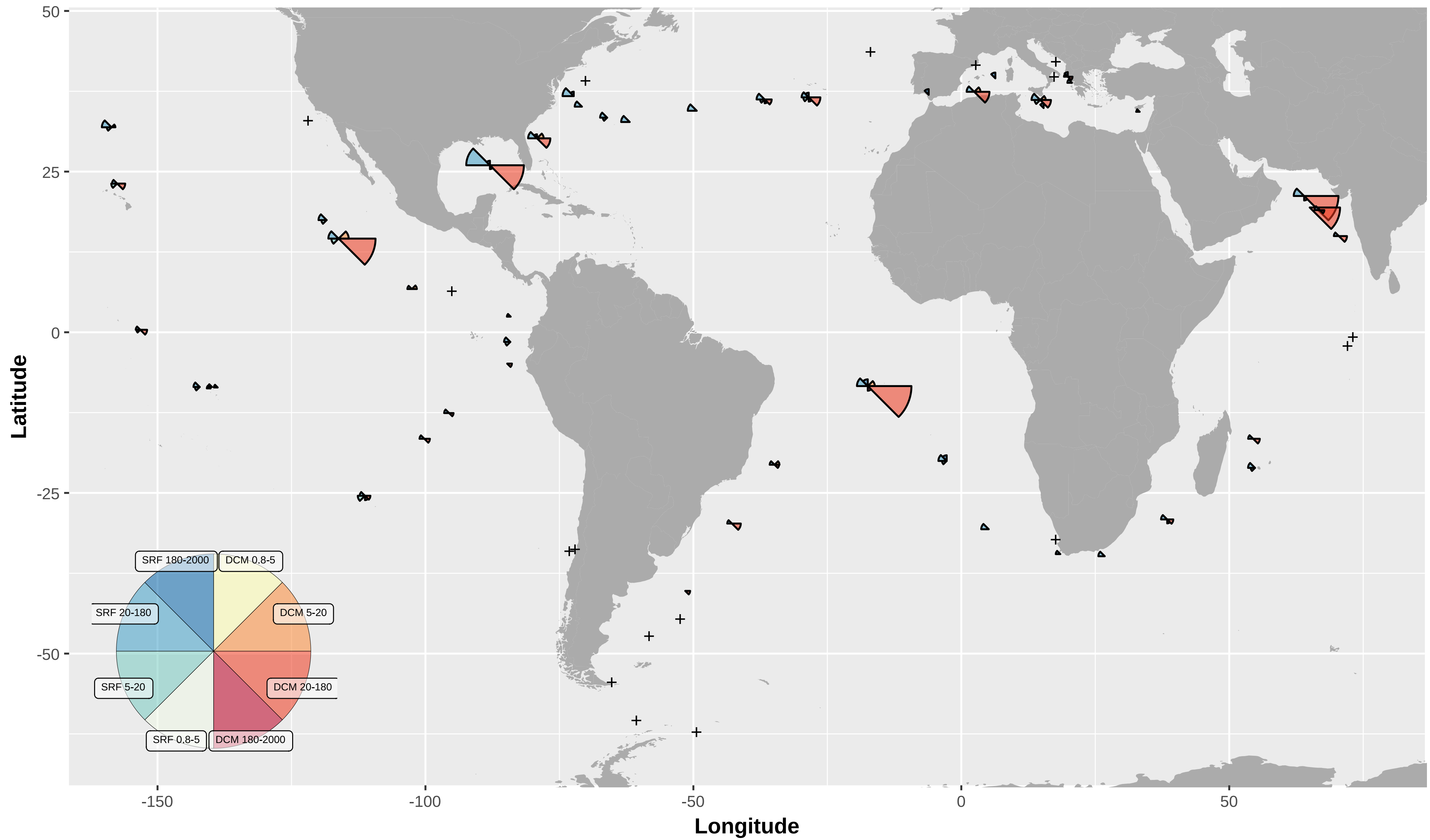


root;cellular organisms;Eukaryota;Opisthokonta;Metazoa;Eumetazoa;Bilateria;Protostomia;Ecdysozoa;Panarthropoda;Arthropoda;Mandibulata;Pancrustacea

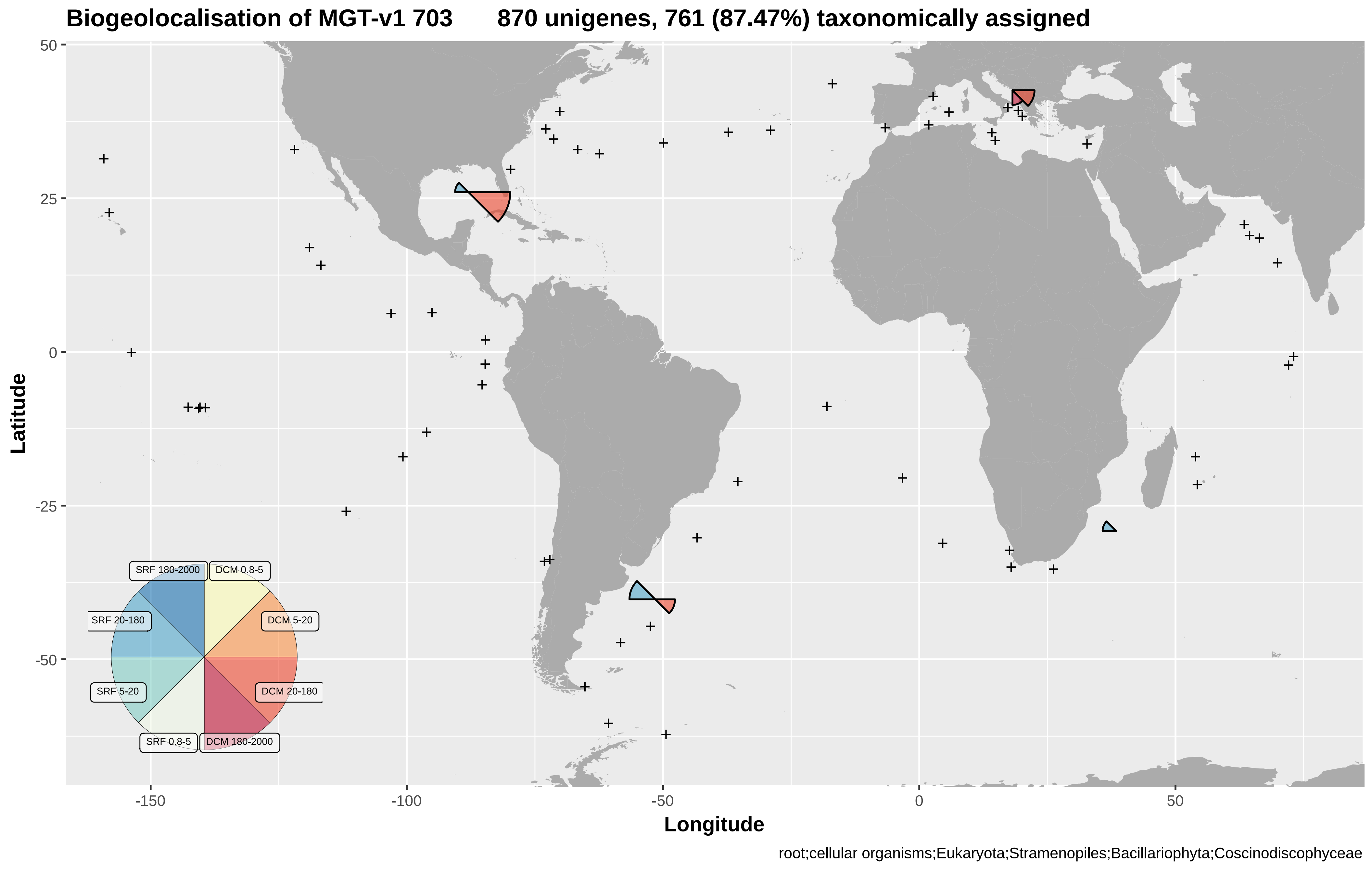
Biogeolocalisation of MGT-v1 703
870 unigenes, 761 (87.47%) taxonomically assigned
Latitude
Longitude
50
25
0
-25
-50
-150
-100
-50
0
50
SRF 180-2000
DCM 0.8-5
SRF 20-180
DCM 5-20
SRF 5-20
DCM 20-180
SRF 0.8-5
DCM 180-2000
root;cellular organisms;Eukaryota;Stramenopiles;Bacillariophyta;Coscinodiscophyceae

# Biogeolocalisation of MGT-v1 704      868 unigenes, 857 (98.73%) taxonomically assigned

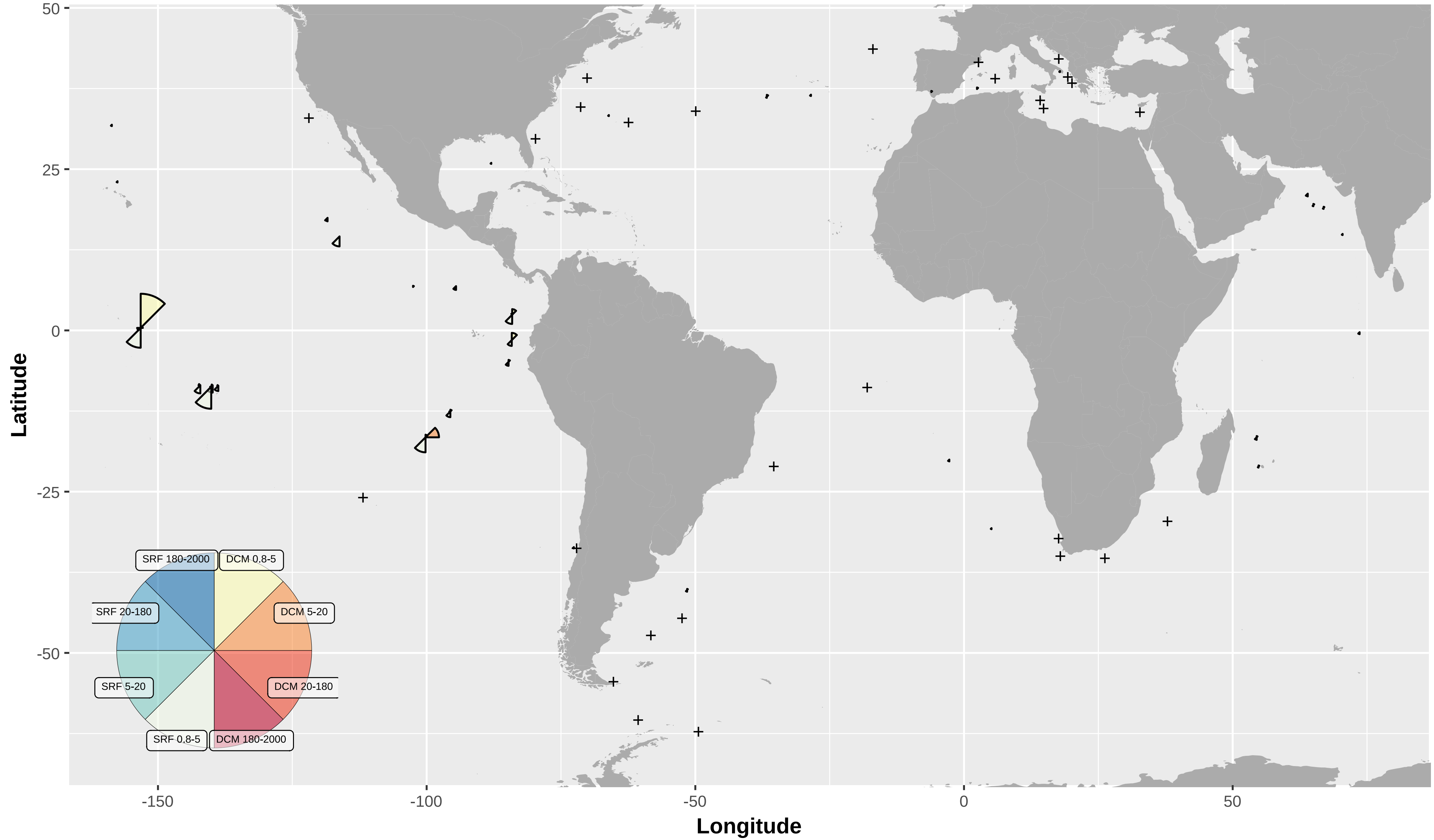


root;cellular organisms;Bacteria;Proteobacteria;Alphaproteobacteria

## Biogeolocalisation of MGT-v1 705      867 unigenes, 107 (12.34%) taxonomically assigned

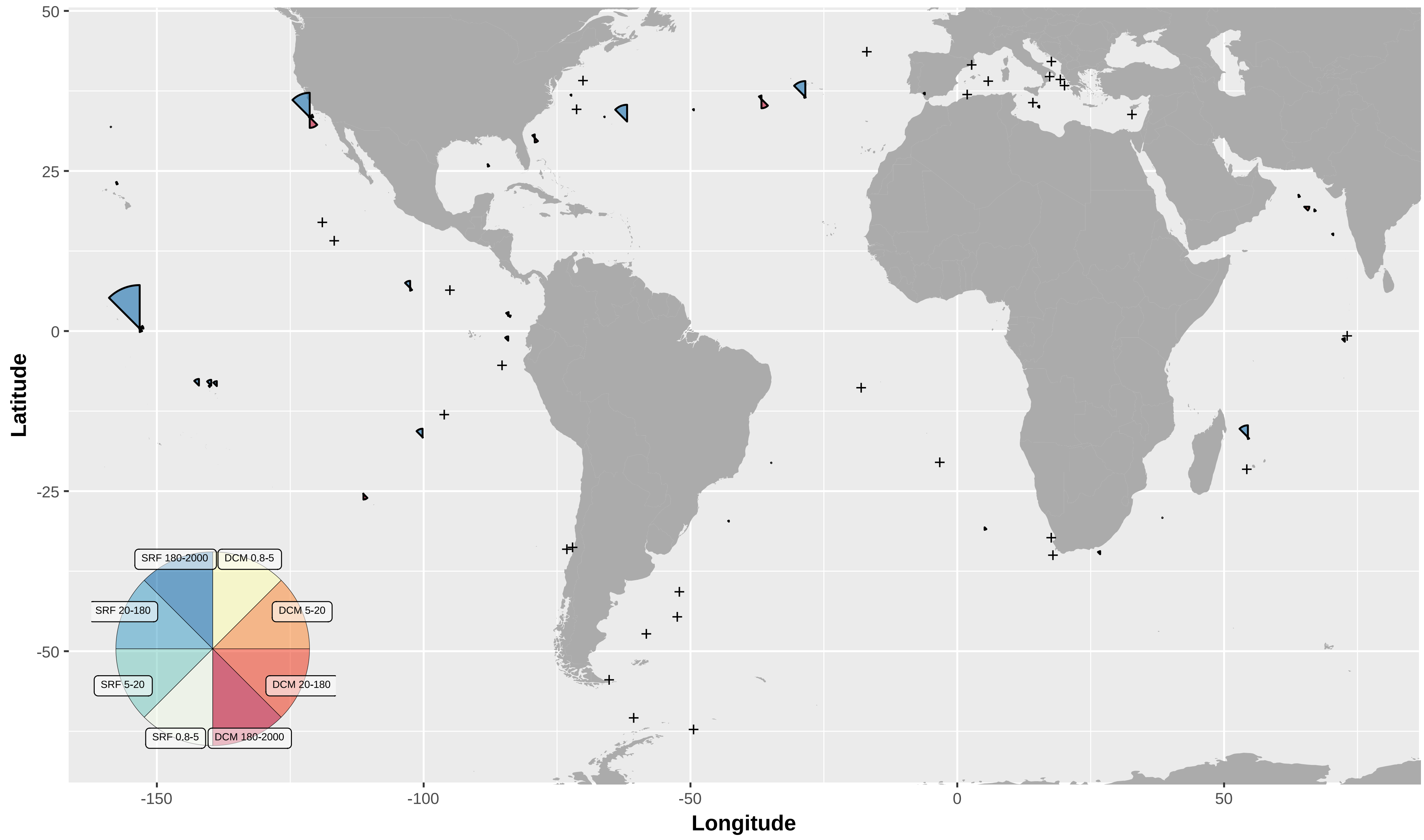


root;cellular organisms;Eukaryota;Opisthokonta;Metazoa

# Biogeolocalisation of MGT-v1 706      853 unigenes, 230 (26.96%) taxonomically assigned

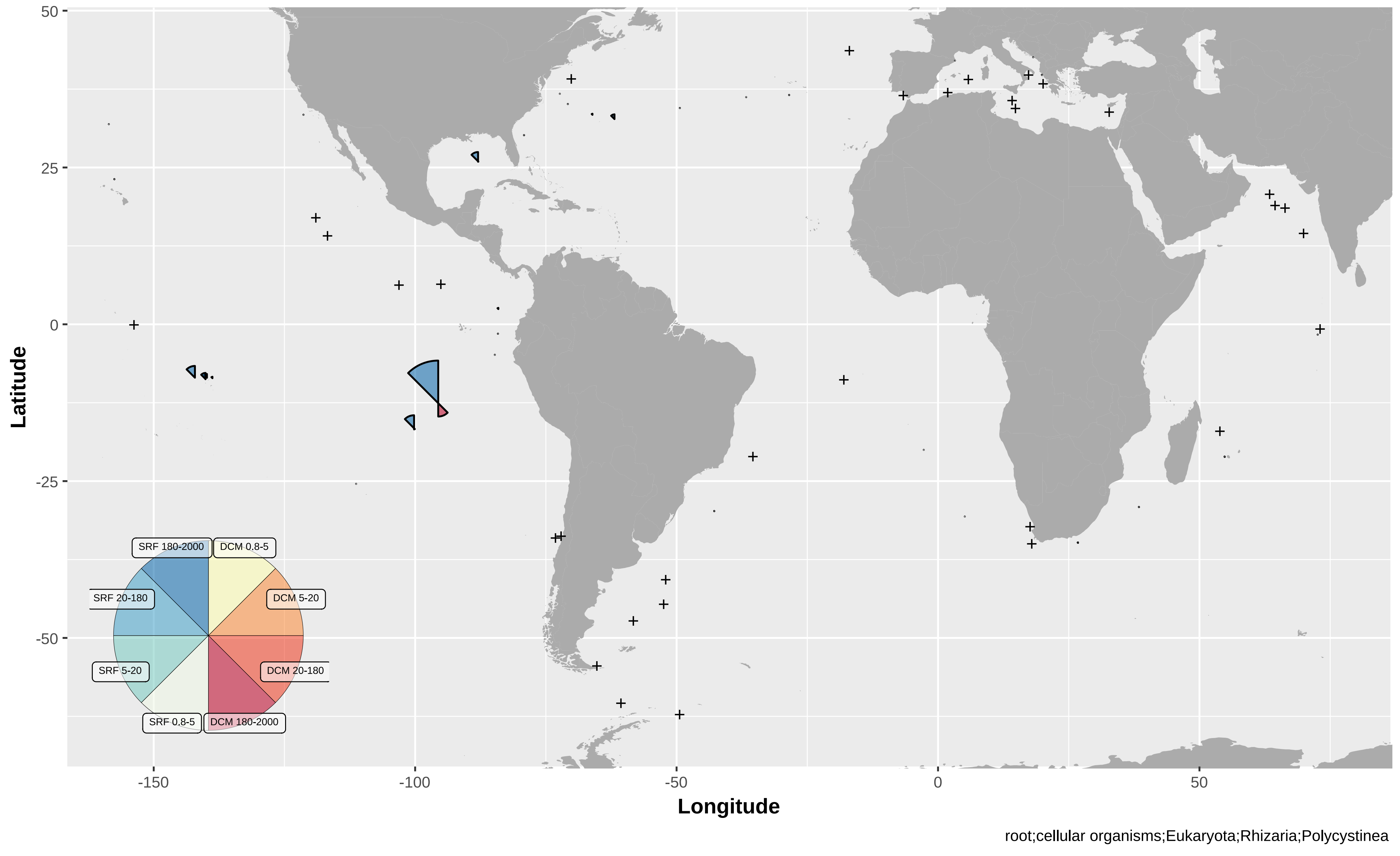

# Biogeolocalisation of MGT-v1 707      849 unigenes, 760 (89.52%) taxonomically assigned

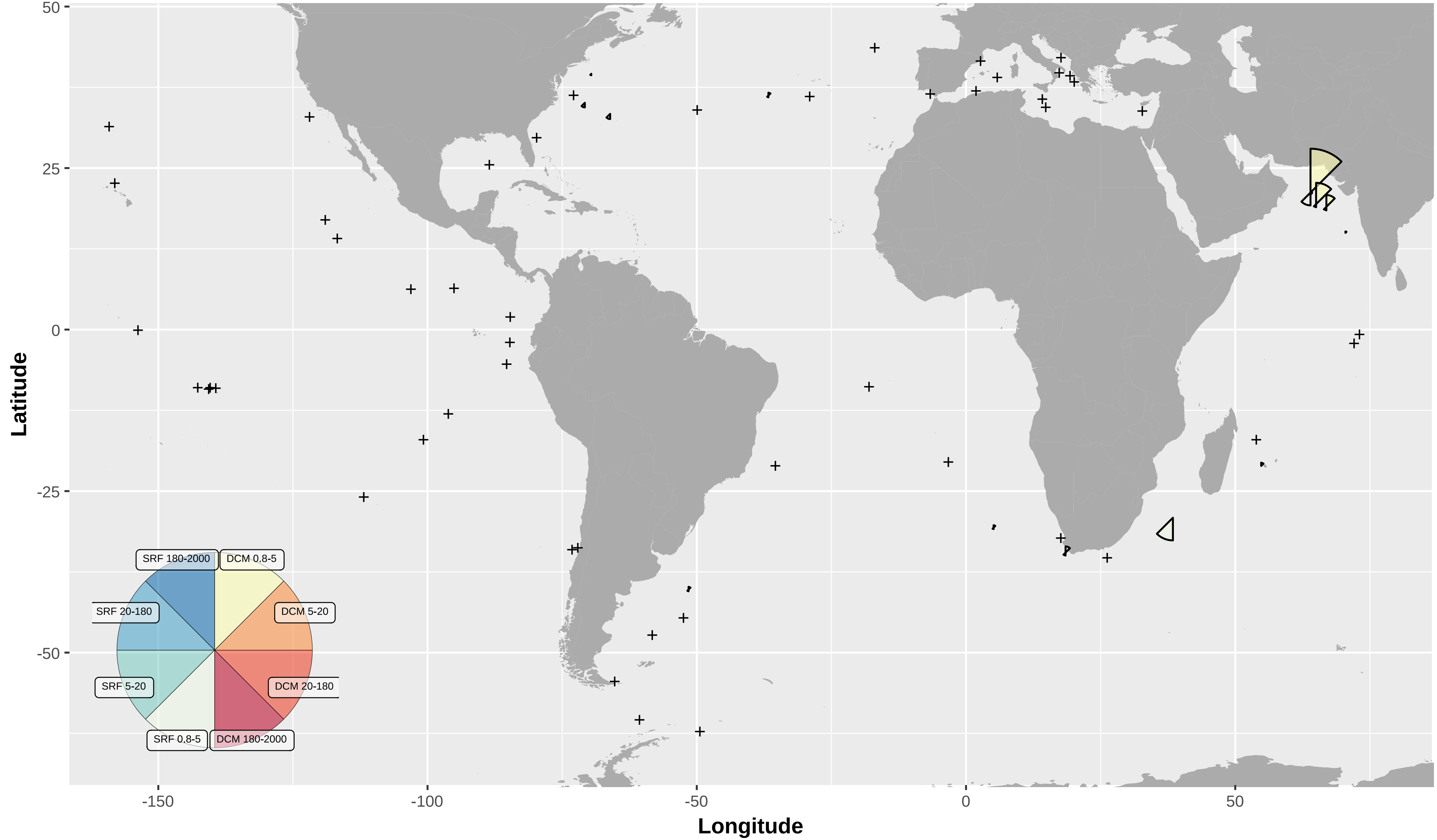


root;cellular organisms;Bacteria;Cyanobacteria;Oscillatoriophycideae

## Biogeolocalisation of MGT-v1 708      849 unigenes, 87 (10.25%) taxonomically assigned

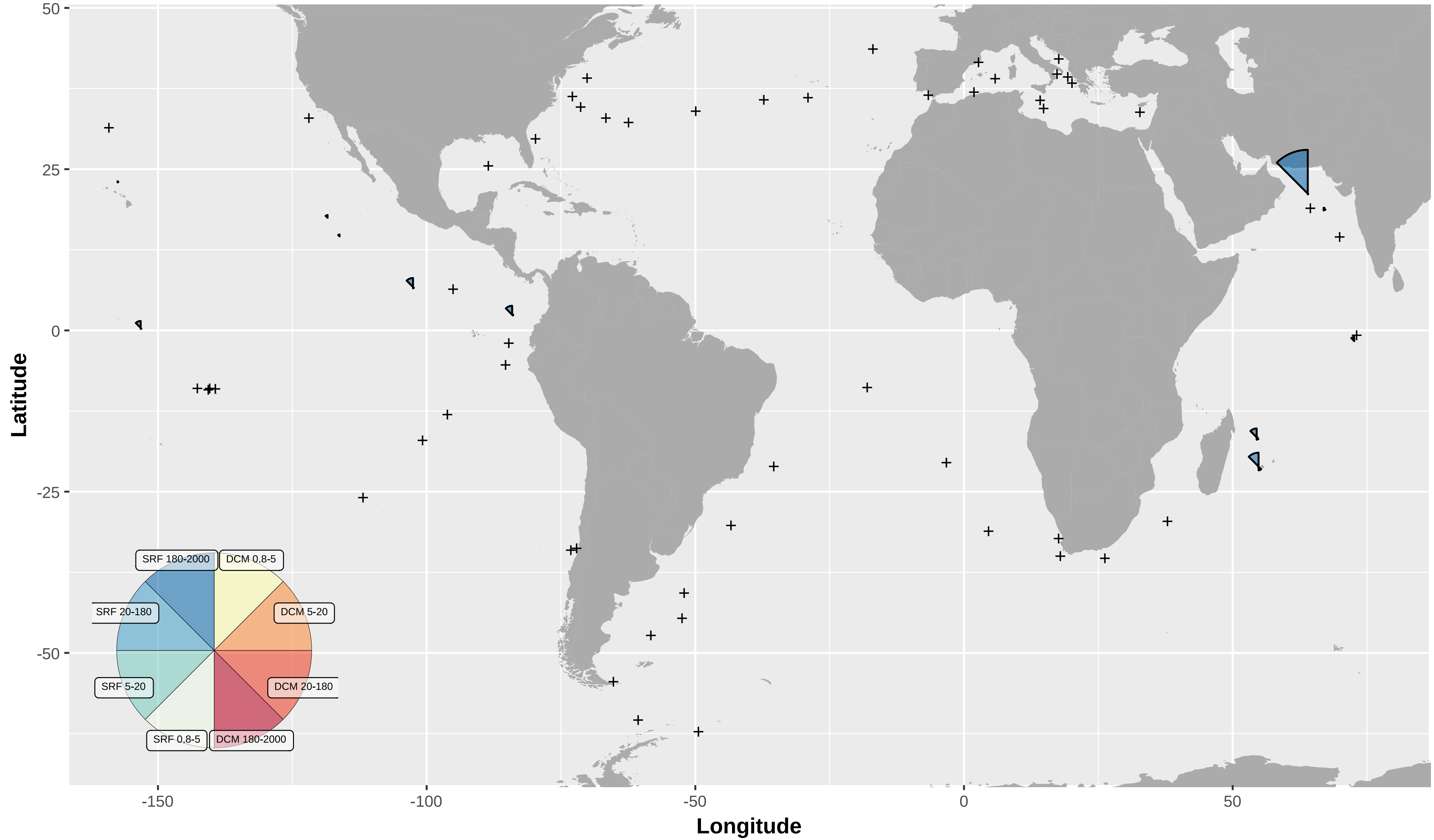


root;cellular organisms;Eukaryota;Opisthokonta;Metazoa;Eumetazoa;Bilateria

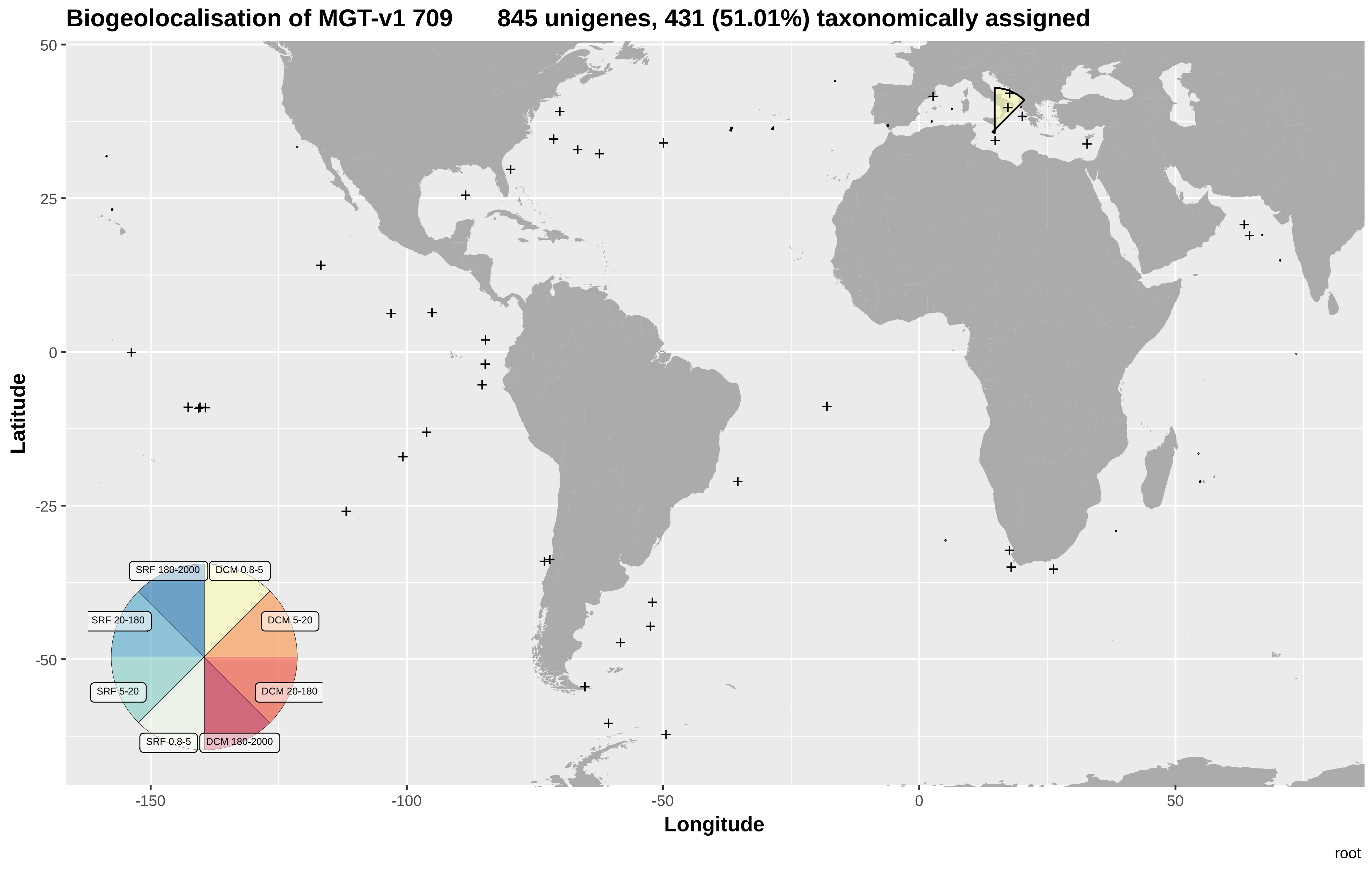
Biogeolocalisation of MGT-v1 709 845 unigenes, 431 (51.01%) taxonomically assigned
Latitude
Longitude
50
25
0
-25
-50
-150
-100
-50
0
50
SRF 180-2000
DCM 0.8-5
SRF 20-180
DCM 5-20
SRF 5-20
DCM 20-180
SRF 0.8-5
DCM 180-2000
root

# Biogeolocalisation of MGT-v1 710      844 unigenes, 600 (71.09%) taxonomically assigned

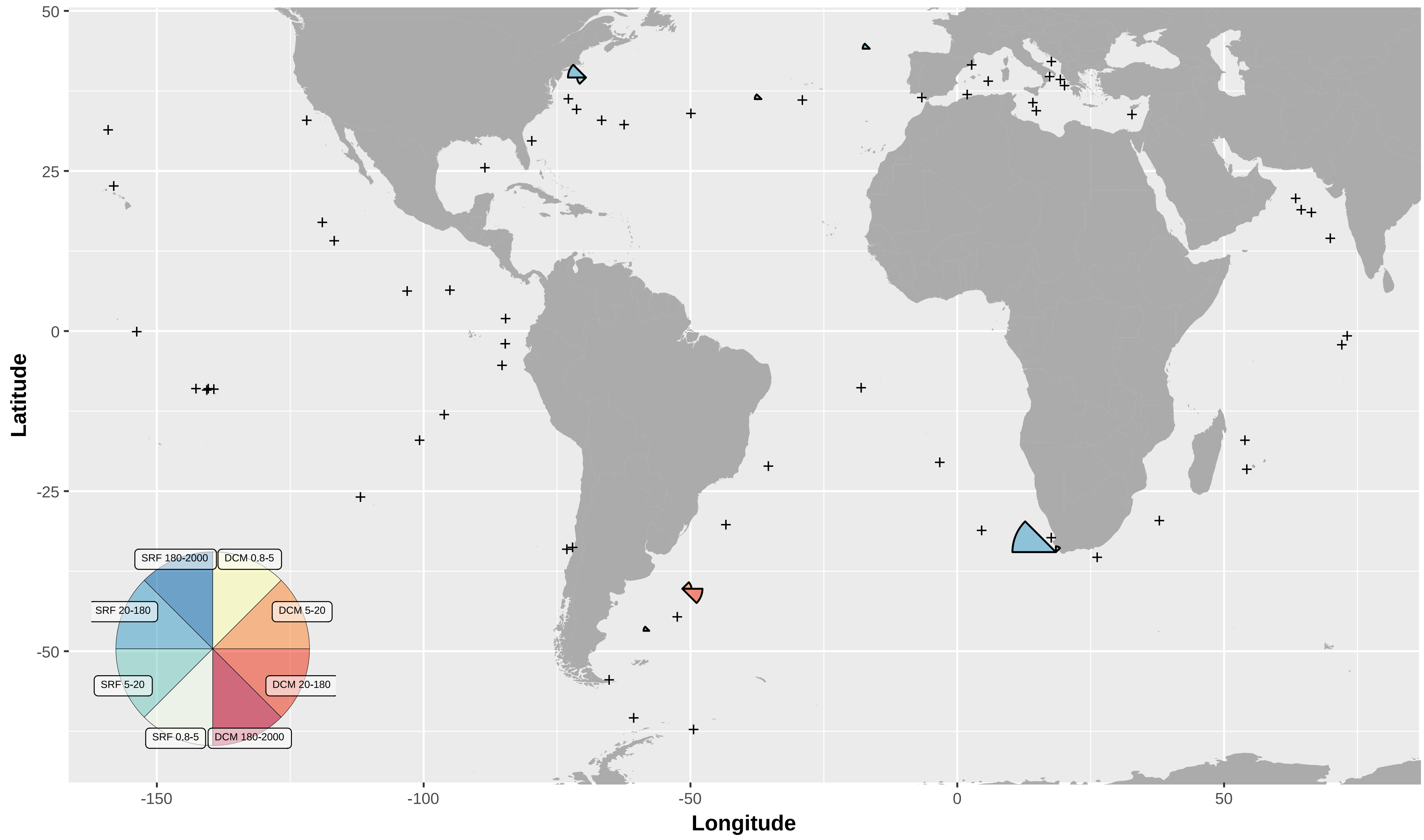


root;cellular organisms;Eukaryota;Alveolata;Ciliophora;Intramacronucleata;Spirotrichea

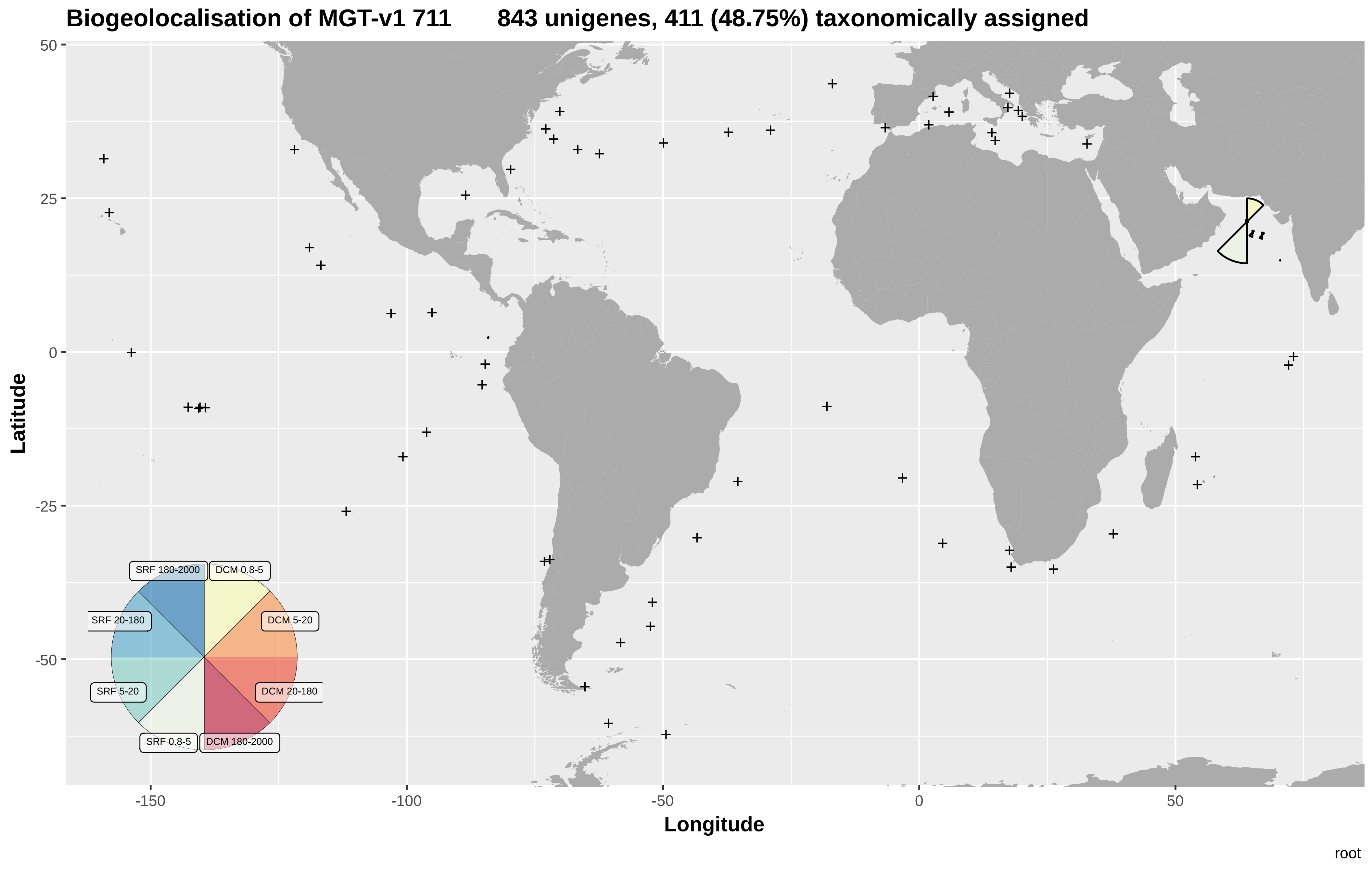

Biogeolocalisation of MGT-v1 711 843 unigenes, 411 (48.75%) taxonomically assigned
Latitude
Longitude
50
25
0
-25
-50
-150
-100
-50
0
50
SRF 180-2000
DCM 0.8-5
SRF 20-180
DCM 5-20
SRF 5-20
DCM 20-180
SRF 0.8-5
DCM 180-2000
root

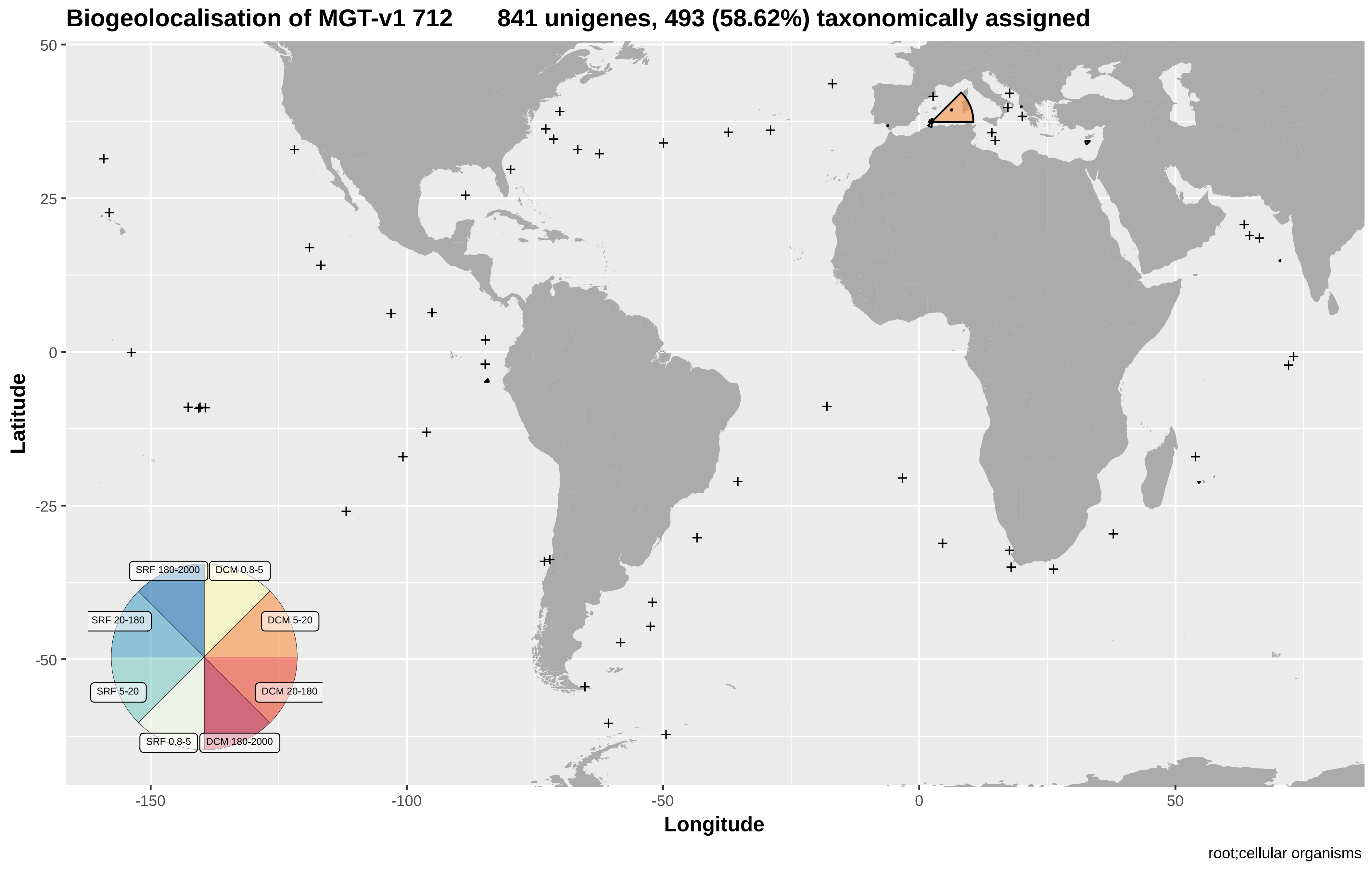

Biogeolocalisation of MGT-v1 712 841 unigenes, 493 (58.62%) taxonomically assigned
50
25
0
-25
-50
Latitude
-150
-100
-50
0
50
Longitude
SRF 180-2000
DCM 0.8-5
SRF 20-180
DCM 5-20
SRF 5-20
DCM 20-180
SRF 0.8-5
DCM 180-2000
root;cellular organisms

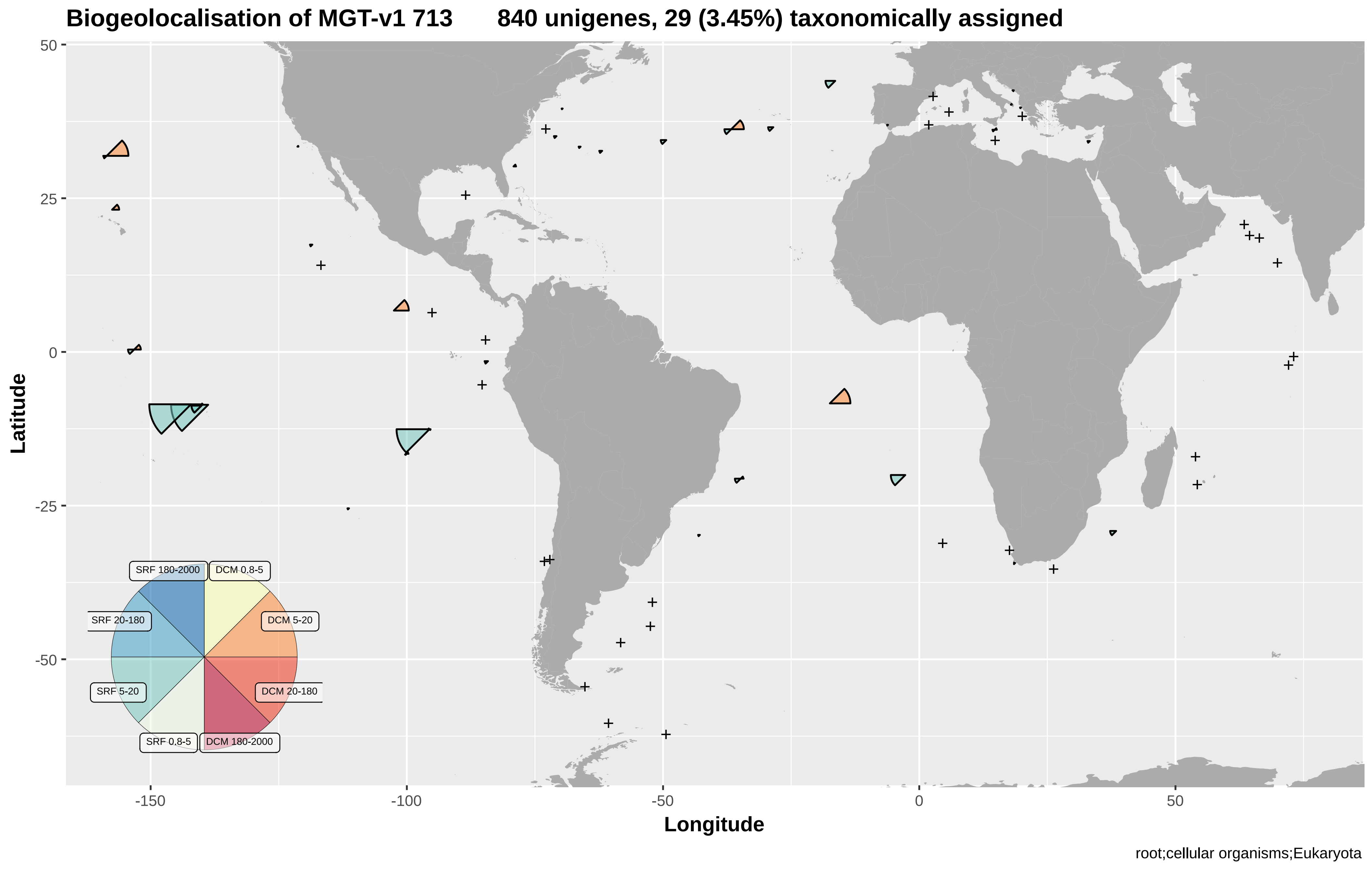

Biogeolocalisation of MGT-v1 713 840 unigenes, 29 (3.45%) taxonomically assigned
Latitude
Longitude
50
25
0
-25
-50
-150
-100
-50
0
50
SRF 180-2000
DCM 0.8-5
SRF 20-180
DCM 5-20
SRF 5-20
DCM 20-180
SRF 0.8-5
DCM 180-2000
root;cellular organisms;Eukaryota

#### Biogeolocalisation of MGT-v1 714 836 unigenes, 603 (72.13%) taxonomically assigned

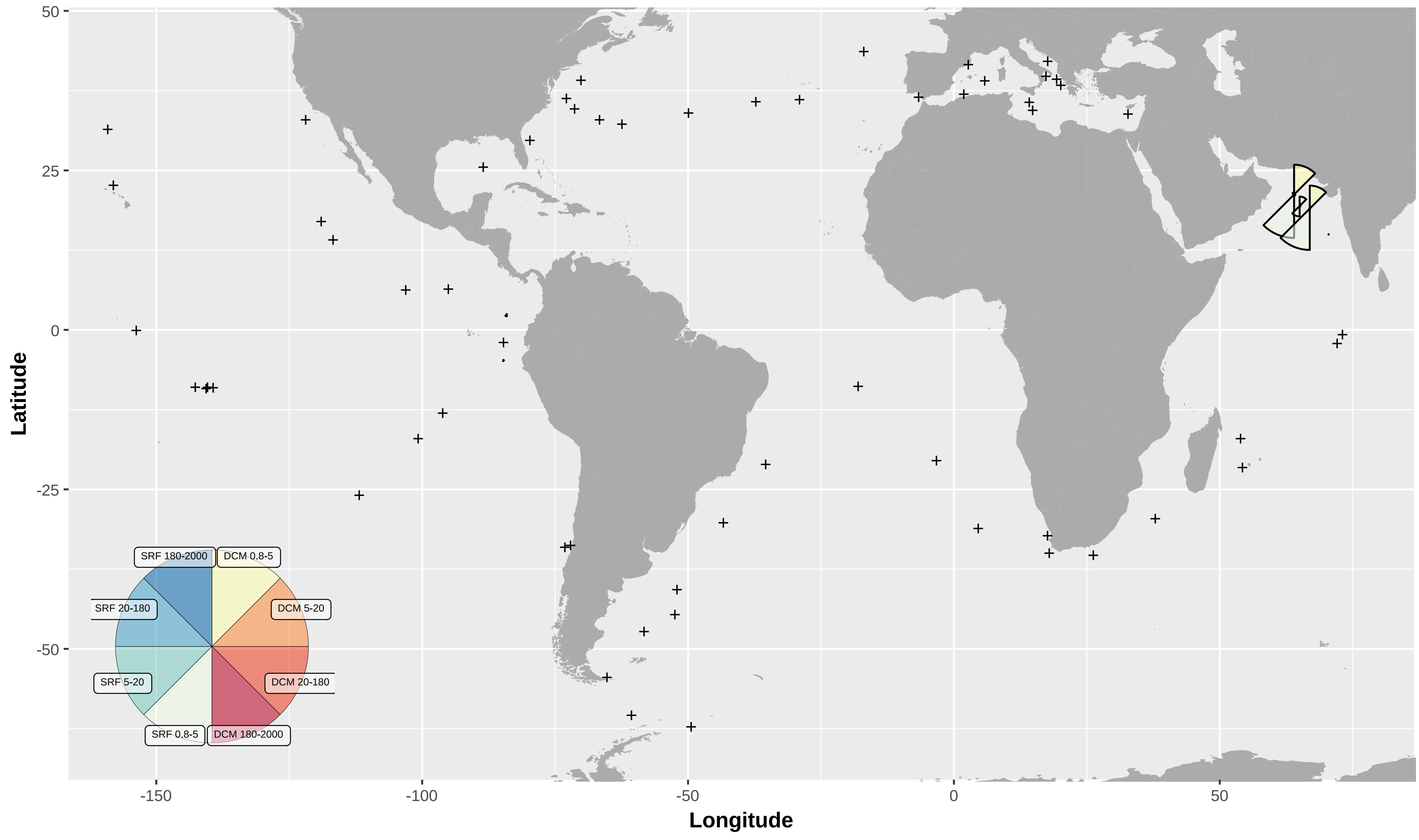


root

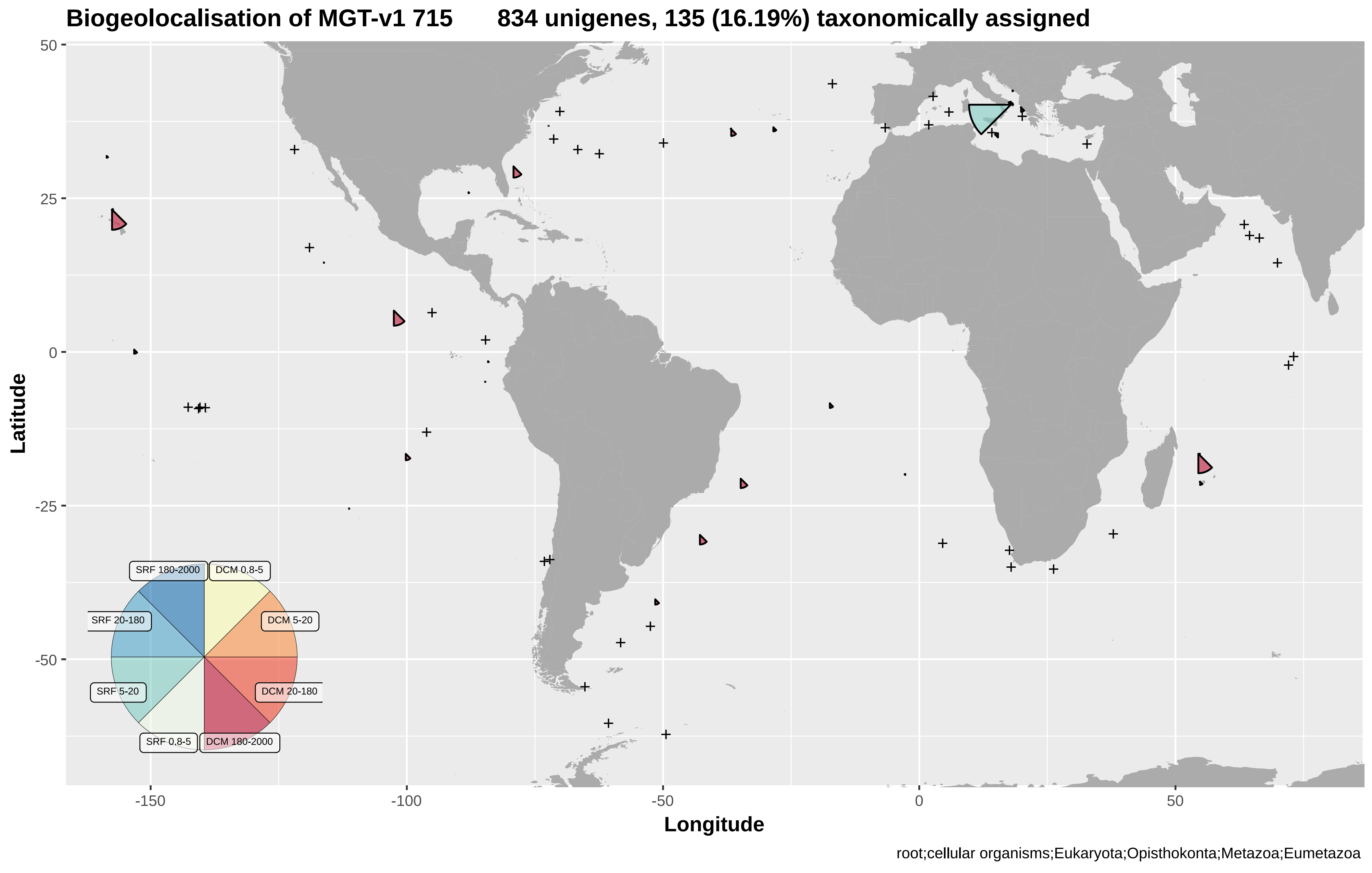

Biogeolocalisation of MGT-v1 715 834 unigenes, 135 (16.19%) taxonomically assigned
Latitude
Longitude
50
25
0
-25
-50
-150
-100
-50
0
50
SRF 180-2000
DCM 0.8-5
SRF 20-180
DCM 5-20
SRF 5-20
DCM 20-180
SRF 0.8-5
DCM 180-2000
root;cellular organisms;Eukaryota;Opisthokonta;Metazoa;Eumetazoa

# Biogeolocalisation of MGT-v1 716      831 unigenes, 601 (72.32%) taxonomically assigned

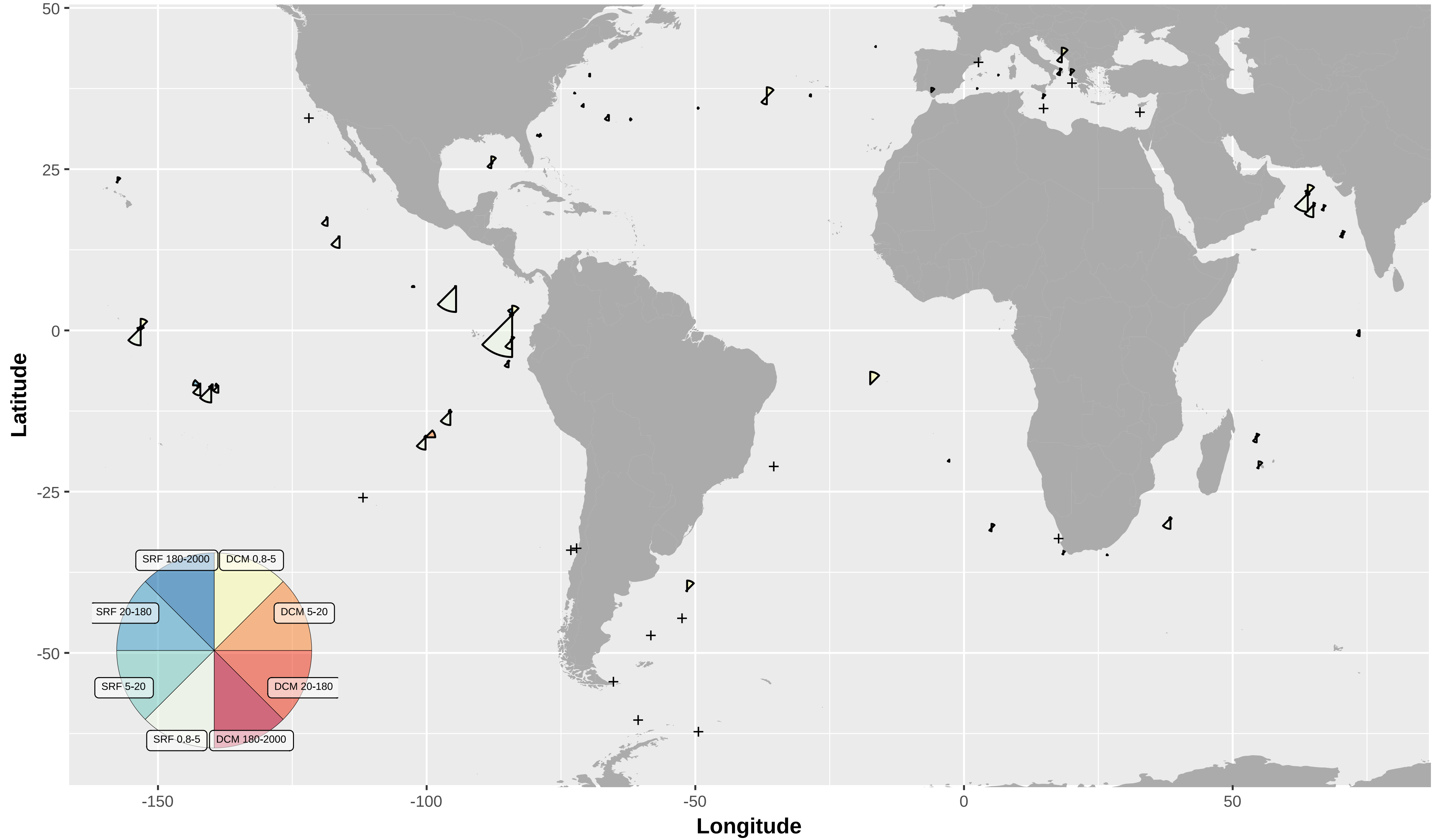


root;cellular organisms;Bacteria

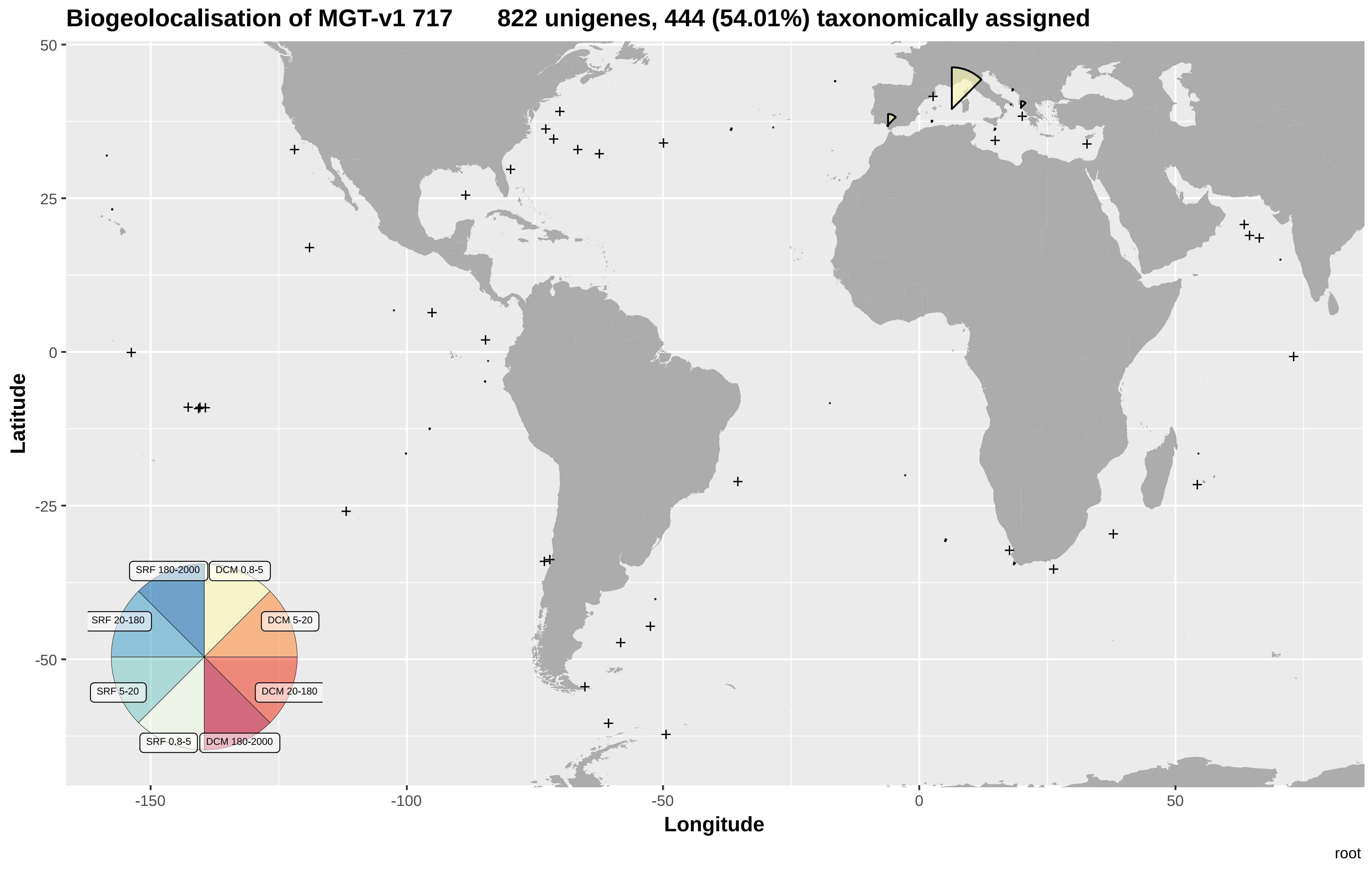
Biogeolocalisation of MGT-v1 717
822 unigenes, 444 (54.01%) taxonomically assigned
50
25
0
-25
-50
Latitude
-150
-100
-50
0
50
Longitude
SRF 180-2000
DCM 0.8-5
SRF 20-180
DCM 5-20
SRF 5-20
DCM 20-180
SRF 0.8-5
DCM 180-2000
root

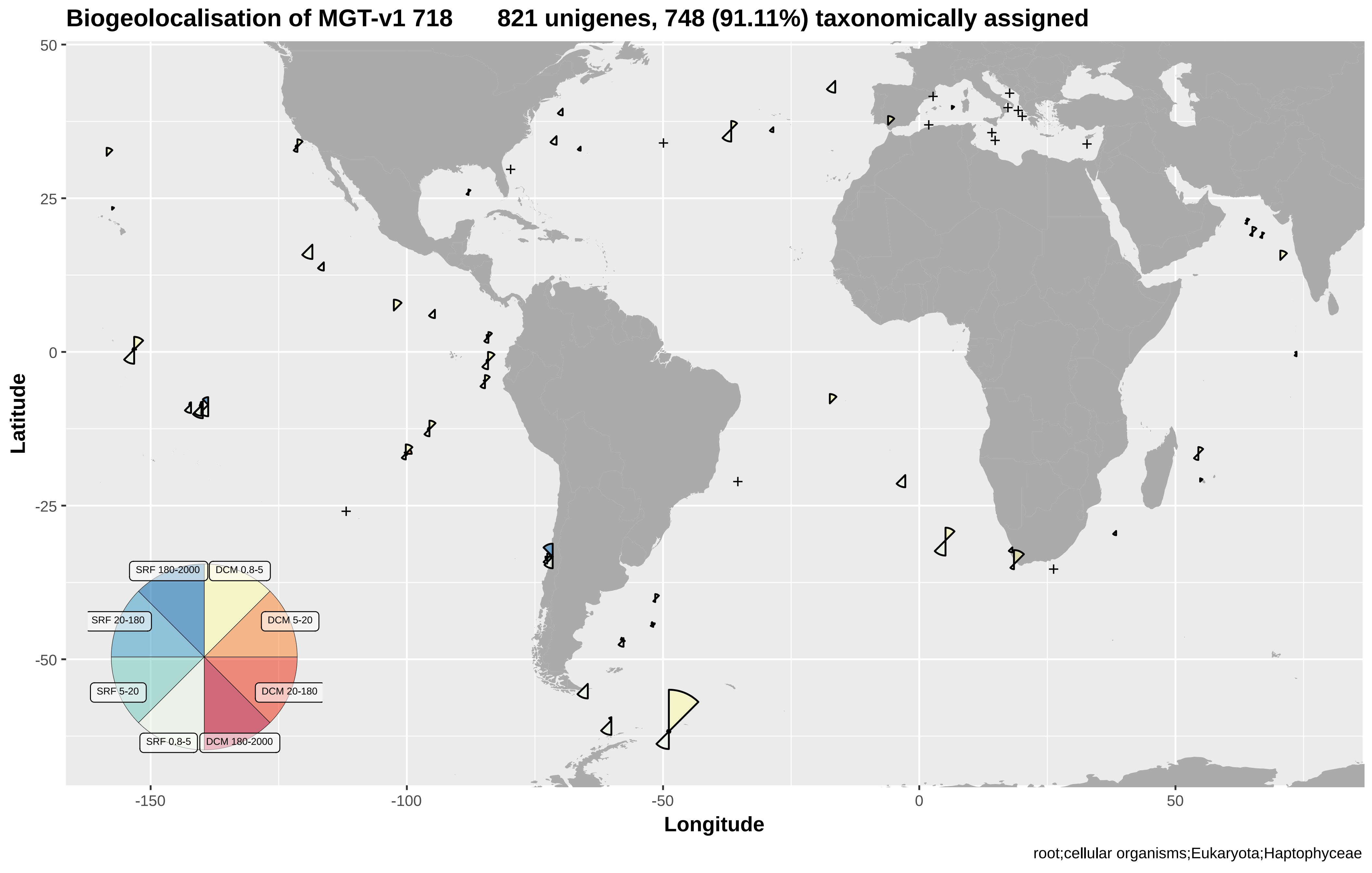

Biogeolocalisation of MGT-v1 718 821 unigenes, 748 (91.11%) taxonomically assigned
50
25
0
-25
-50
Latitude
-150
-100
-50
0
50
Longitude
SRF 180-2000
DCM 0.8-5
SRF 20-180
DCM 5-20
SRF 5-20
DCM 20-180
SRF 0.8-5
DCM 180-2000
root;cellular organisms;Eukaryota;Haptophyceae

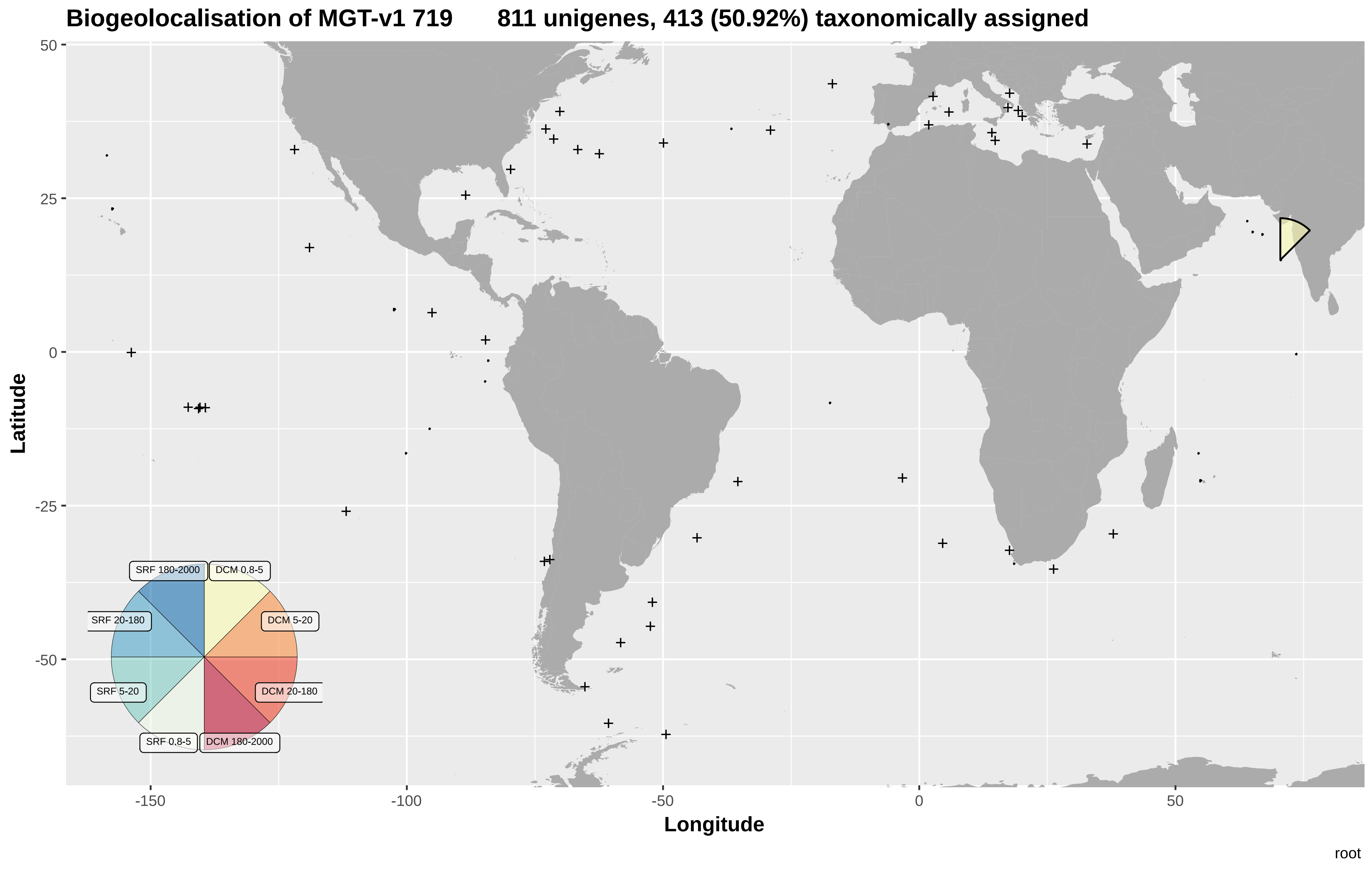

Biogeolocalisation of MGT-v1 719 811 unigenes, 413 (50.92%) taxonomically assigned
50
25
0
-25
-50
Latitude
-150
-100
-50
0
50
Longitude
SRF 180-2000
DCM 0.8-5
SRF 20-180
DCM 5-20
SRF 5-20
DCM 20-180
SRF 0.8-5
DCM 180-2000
root

# Biogeolocalisation of MGT-v1 720      806 unigenes, 32 (3.97%) taxonomically assigned

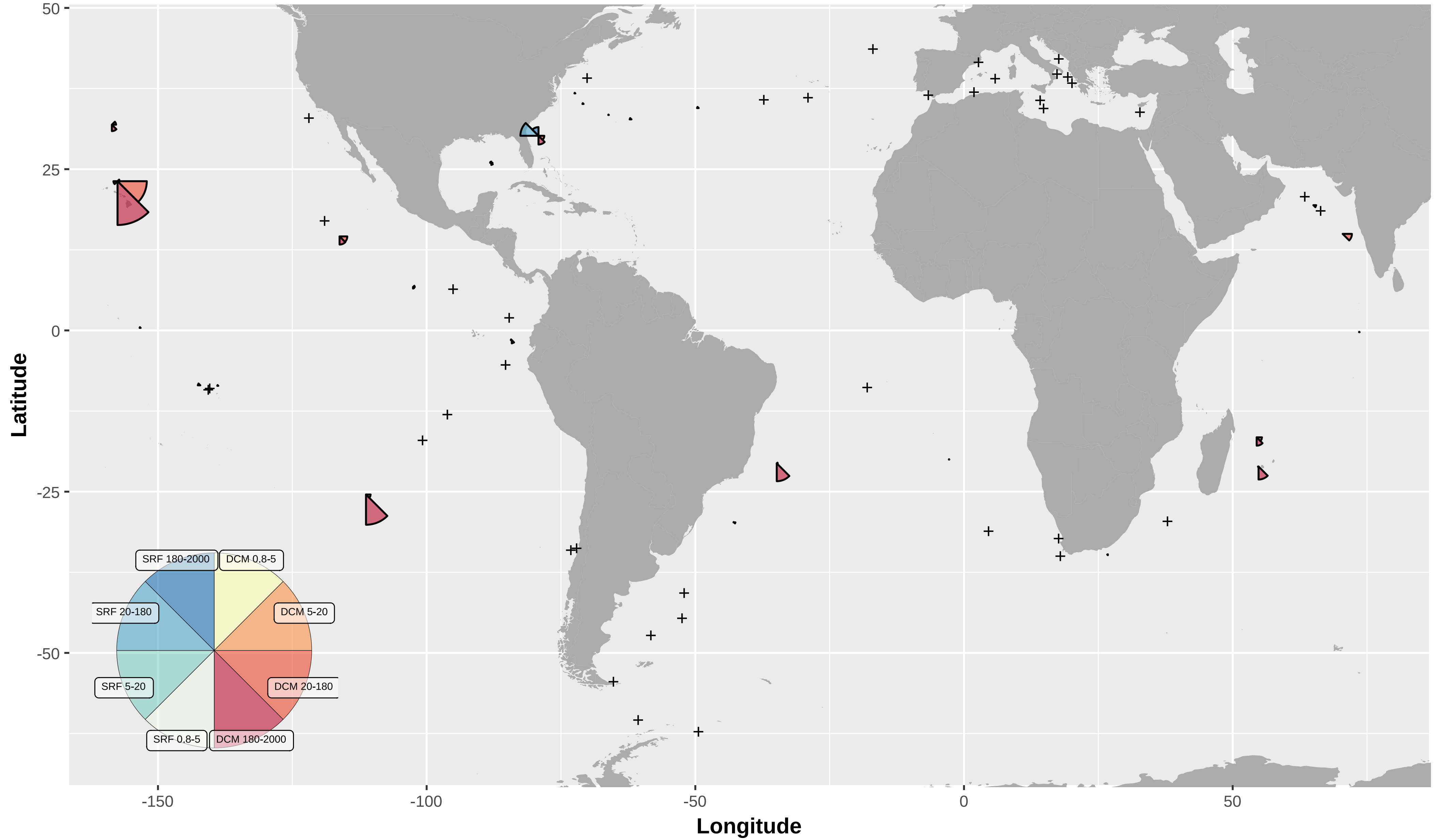


root;cellular organisms;Eukaryota;Opisthokonta;Metazoa;Eumetazoa;Bilateria

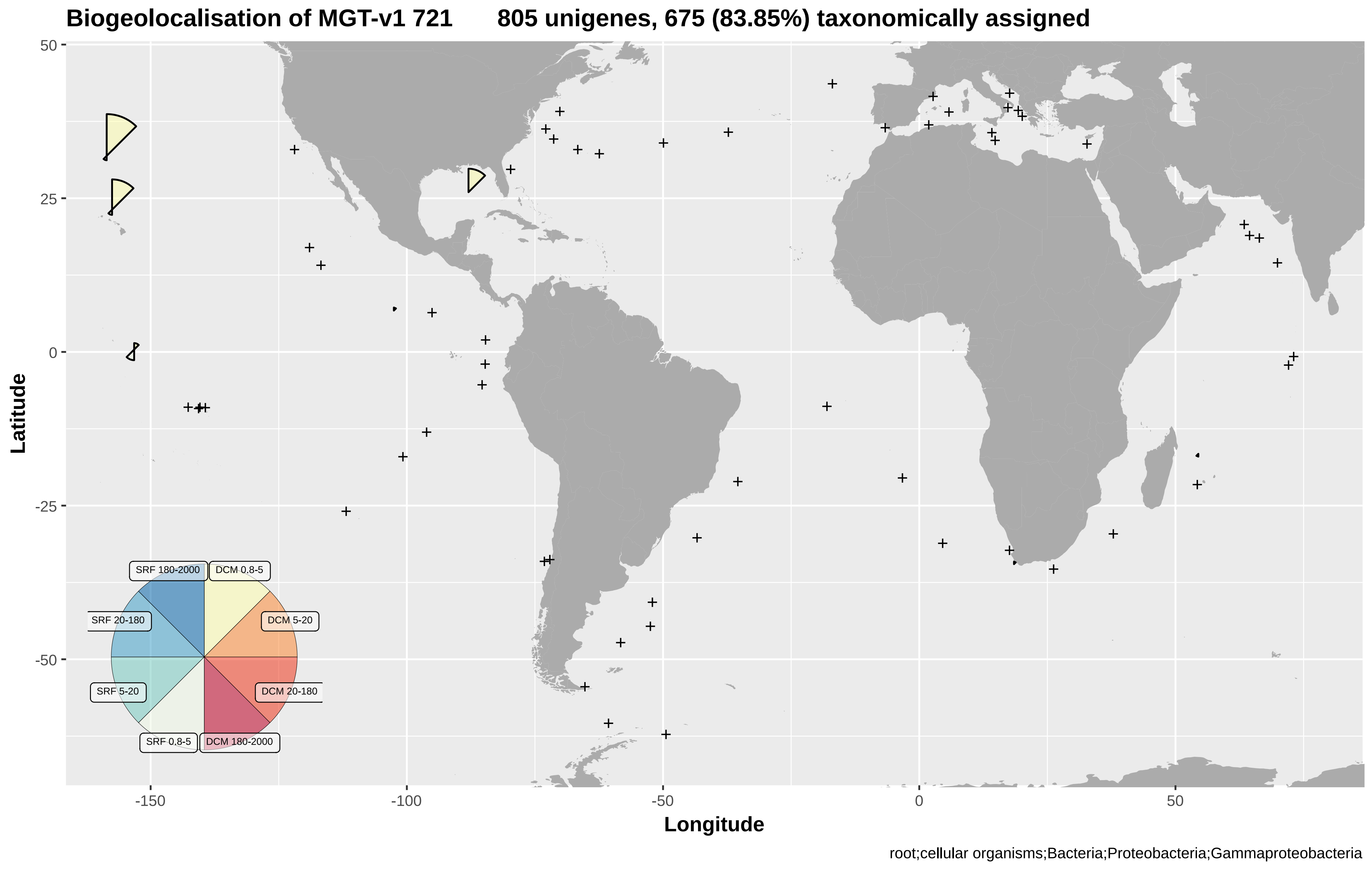
Biogeolocalisation of MGT-v1 721      805 unigenes, 675 (83.85%) taxonomically assigned
50
25
0
-25
-50
Latitude
-150
-100
-50
0
50
Longitude
SRF 180-2000
DCM 0.8-5
SRF 20-180
DCM 5-20
SRF 5-20
DCM 20-180
SRF 0.8-5
DCM 180-2000
root;cellular organisms;Bacteria;Proteobacteria;Gammaproteobacteria

### Biogeolocalisation of MGT-v1 722 798 unigenes, 171 (21.43%) taxonomically assigned

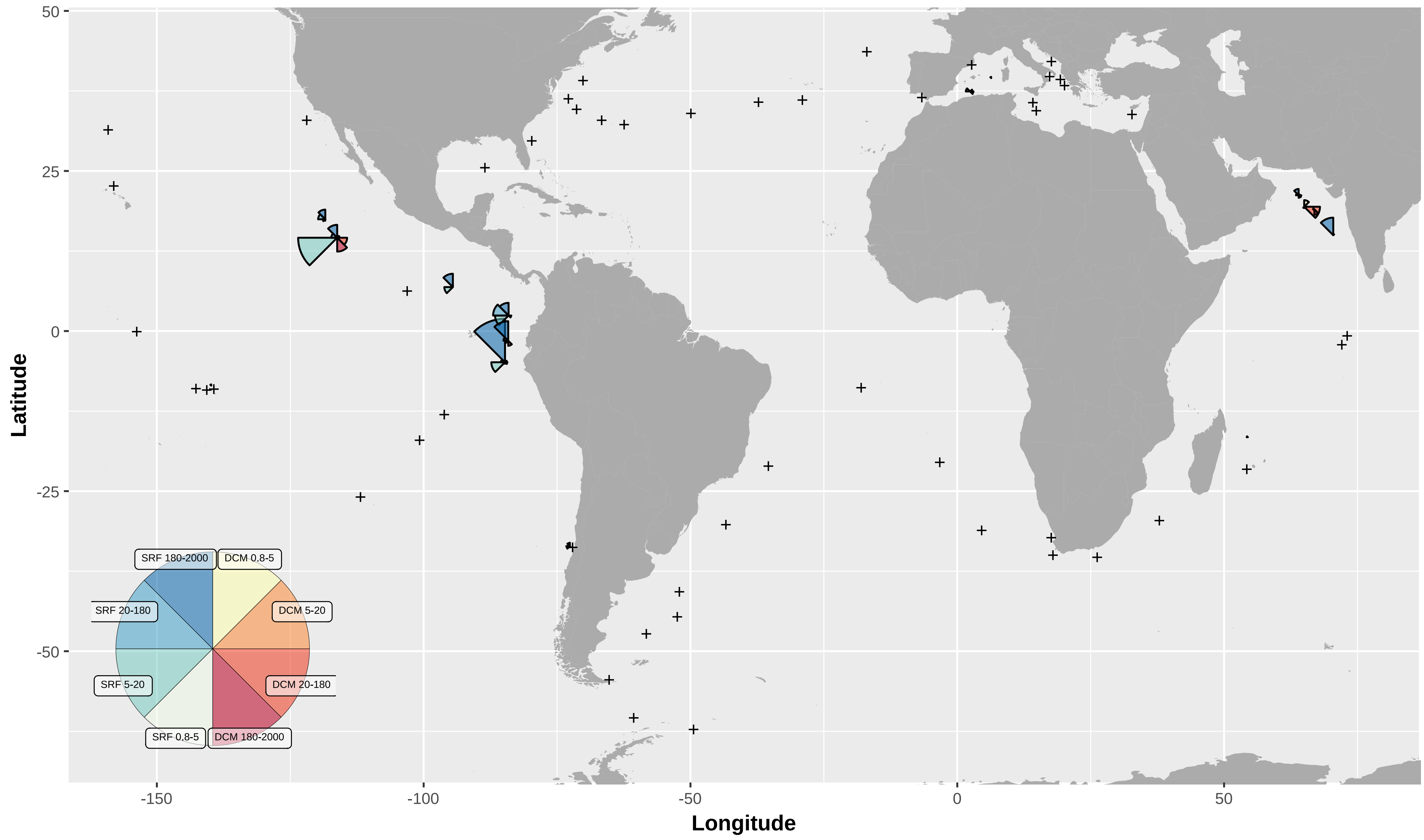


root;cellular organisms;Eukaryota;Opisthokonta;Metazoa;Eumetazoa;Bilateria;Deuterostomia;Chordata

## Biogeolocalisation of MGT-v1 723      797 unigenes, 757 (94.98%) taxonomically assigned

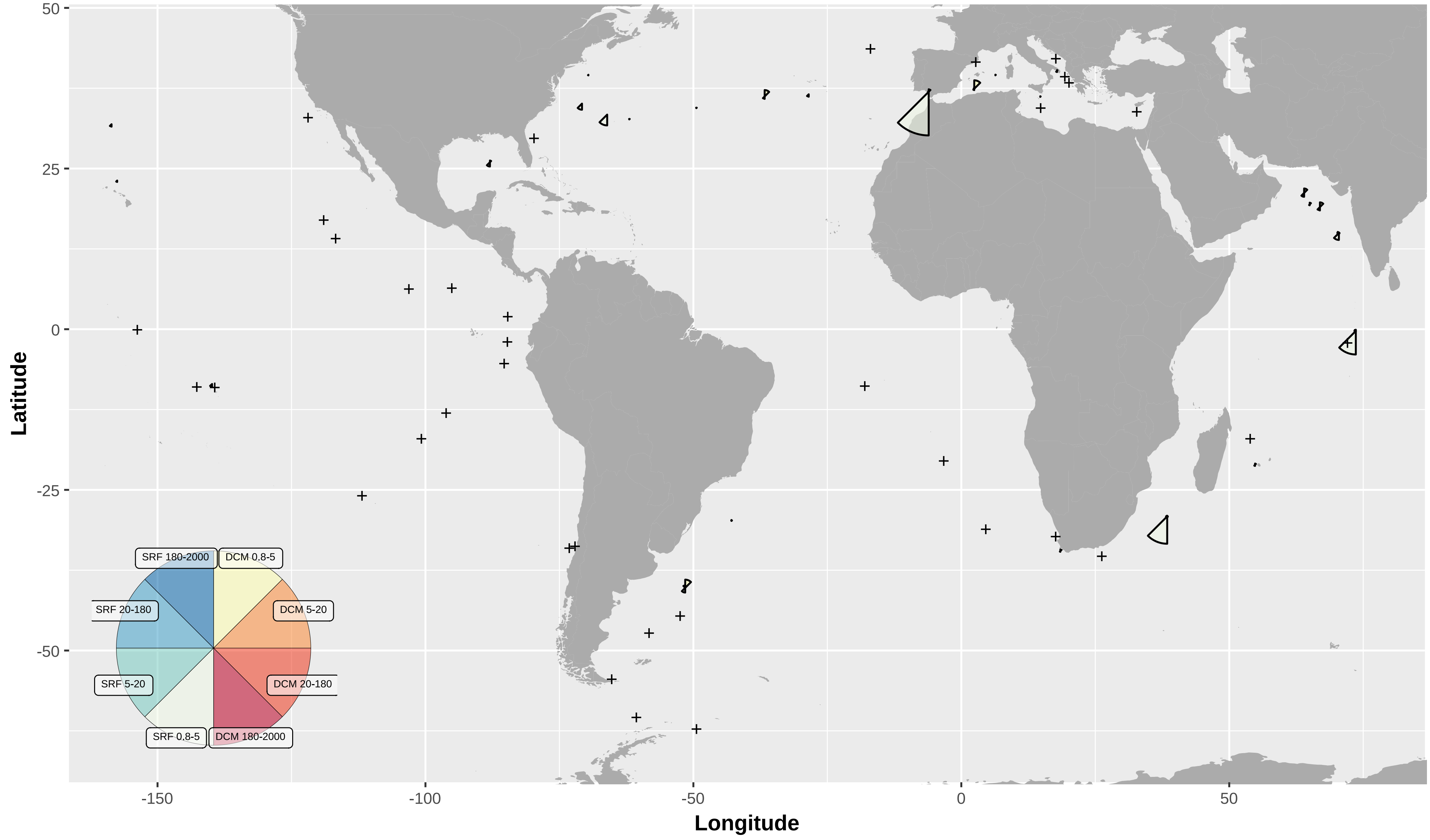


root;cellular organisms;Bacteria;Cyanobacteria;Oscillatoriophycideae;Chroococcales;Synechococcus

## Biogeolocalisation of MGT-v1 724      797 unigenes, 511 (64.12%) taxonomically assigned

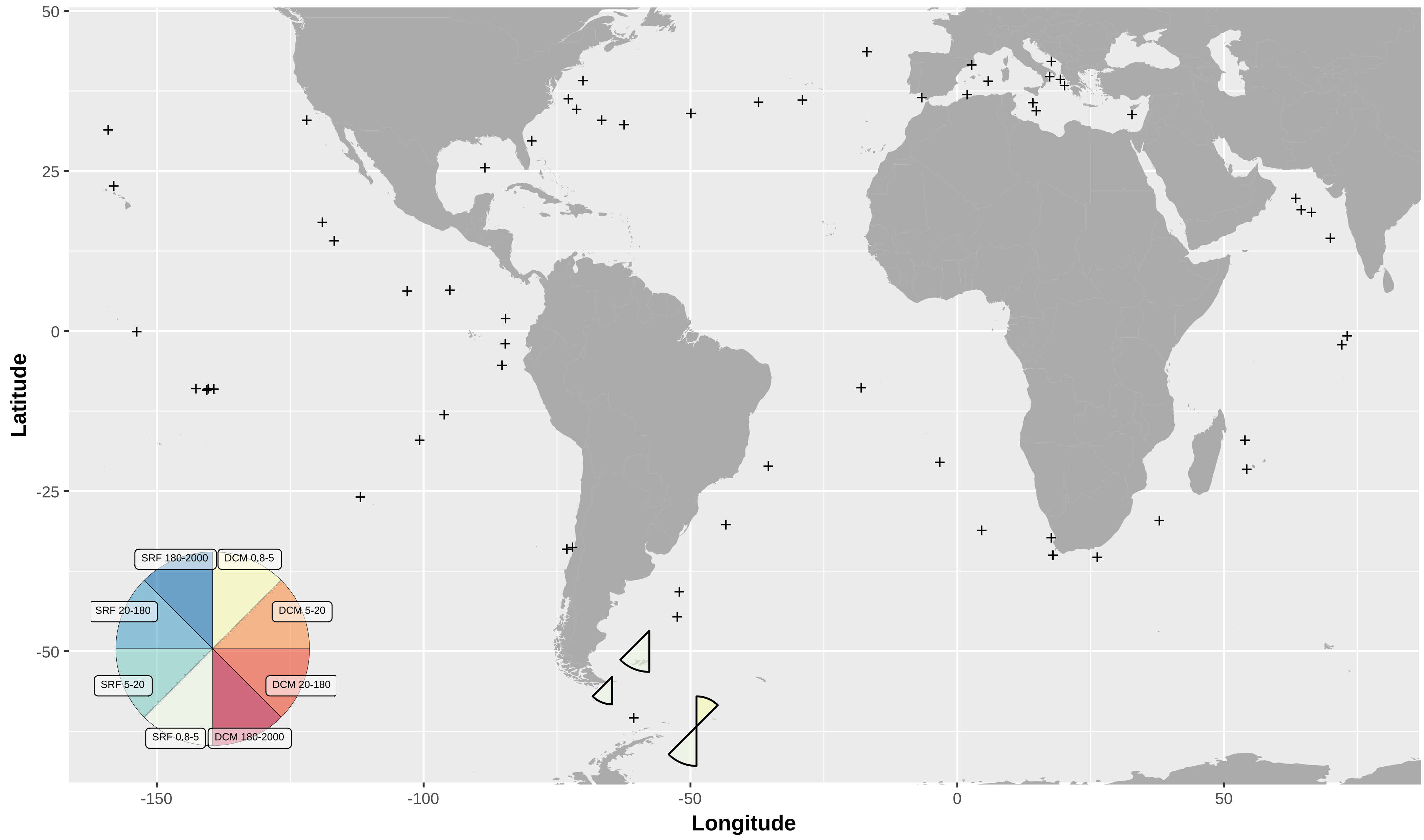


root;cellular organisms;Eukaryota

#### Biogeolocalisation of MGT-v1 725 797 unigenes, 404 (50.69%) taxonomically assigned

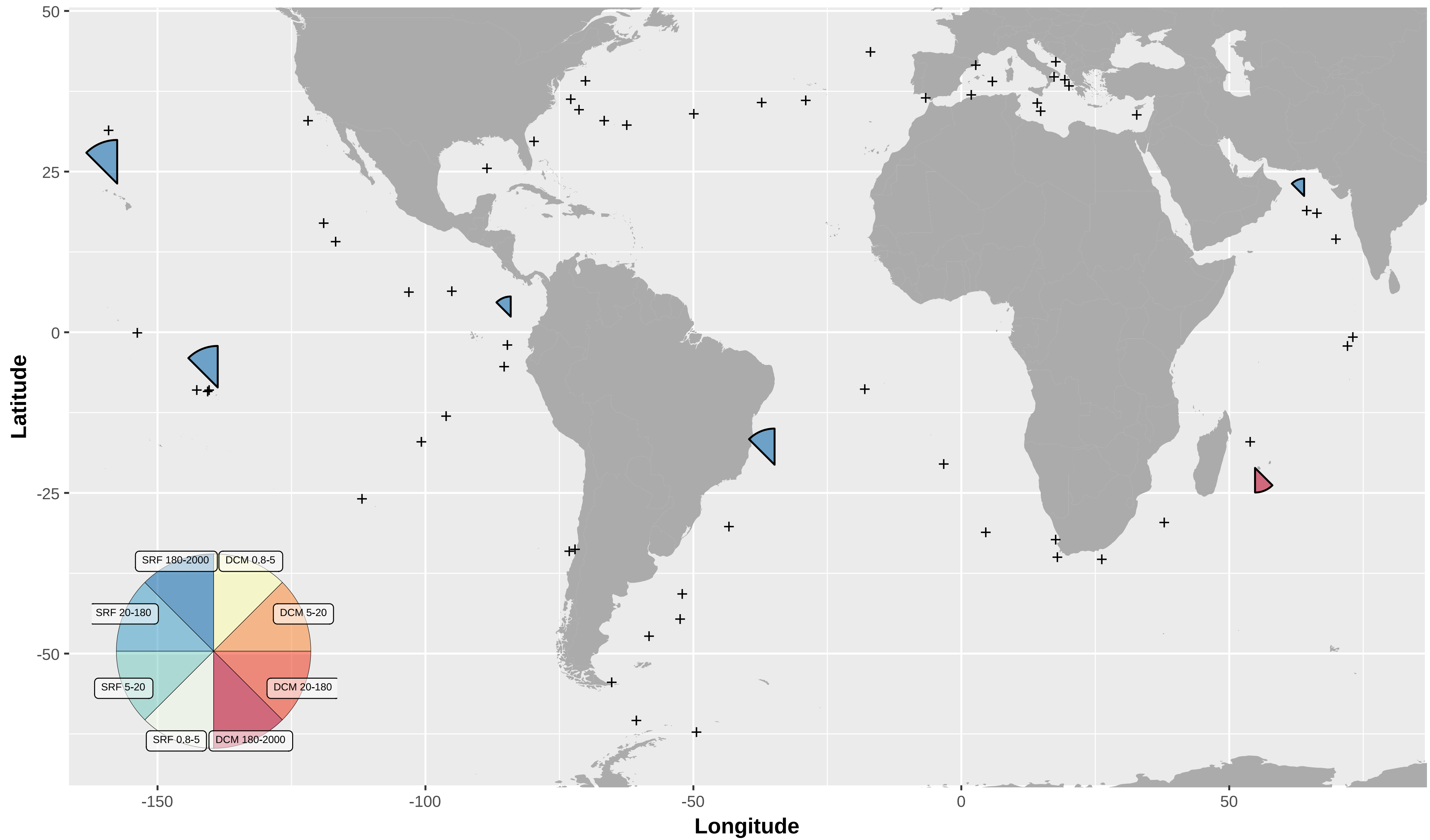


oa;Bilateria;Deuterostomia;Chordata;Craniata;Vertebrata;Gnathostomata;Teleostomi;Euteleostomi;Actinopterygii;Actinopteri;Neopterygii;Teleostei;Osteoglossocephalai;Clupeocephala;Euteleosteomorpha

# Biogeolocalisation of MGT-v1 726      796 unigenes, 465 (58.42%) taxonomically assigned

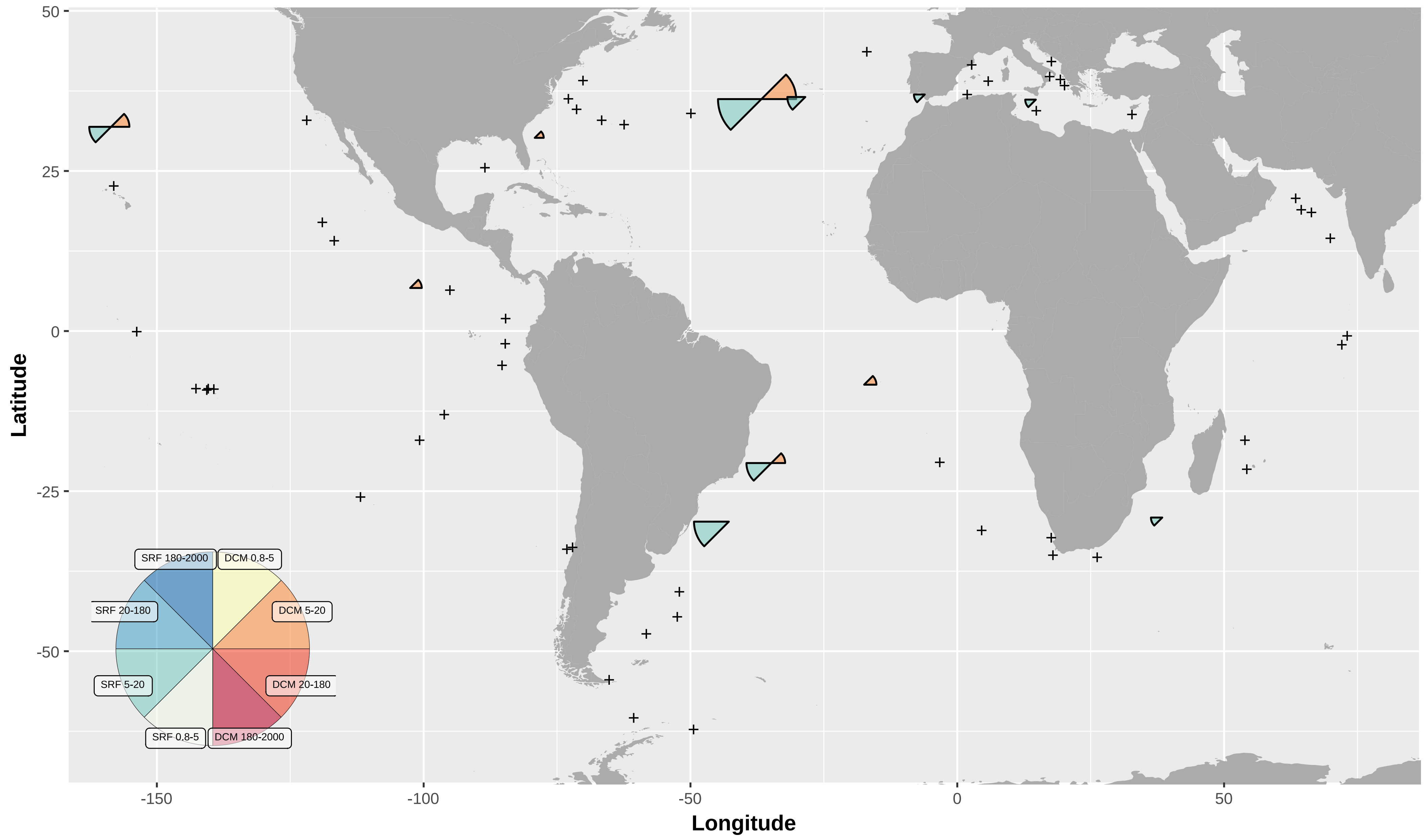


root;cellular organisms;Eukaryota;Haptophyceae

# Biogeolocalisation of MGT-v1 727      793 unigenes, 183 (23.08%) taxonomically assigned

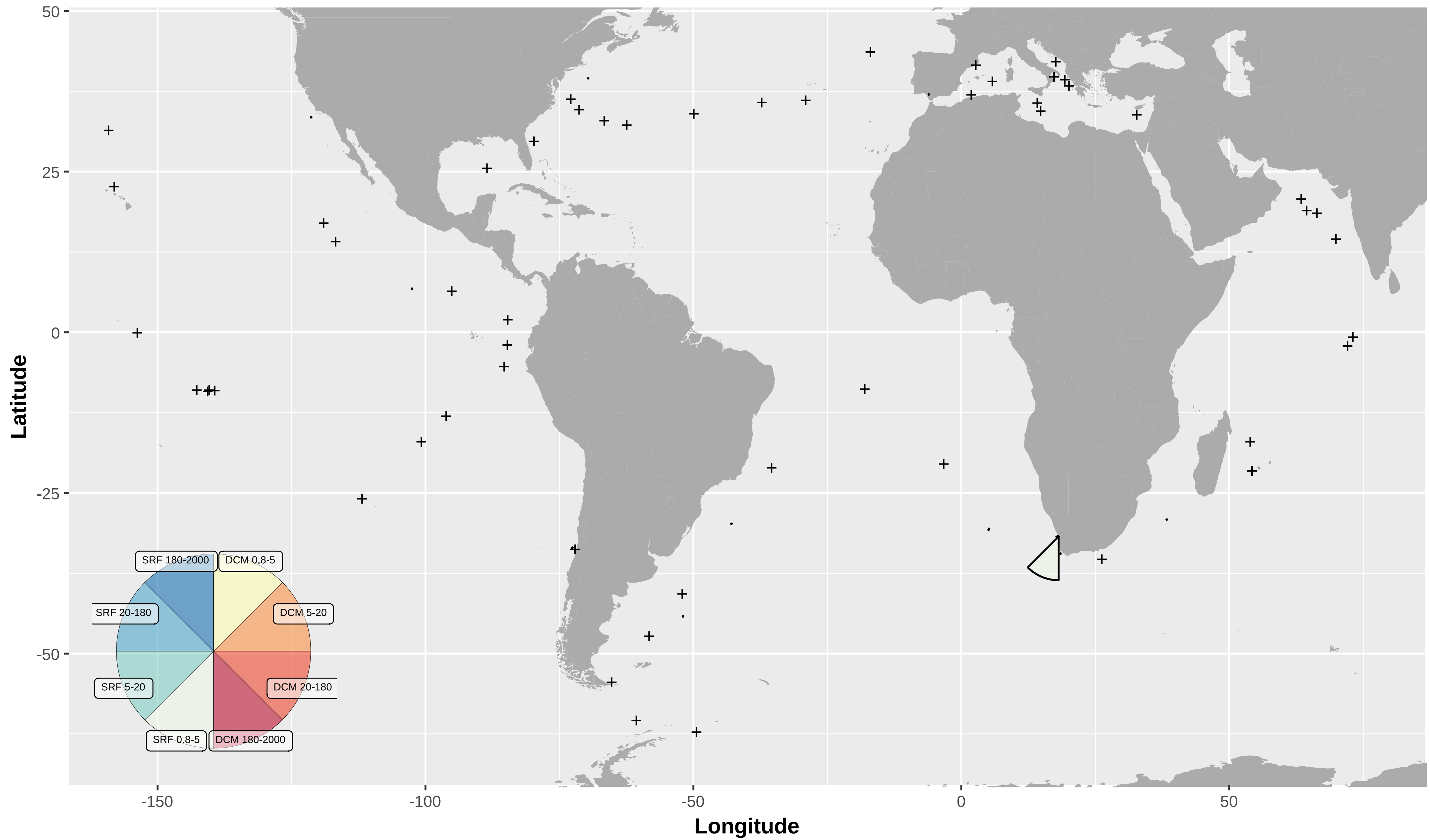


root;cellular organisms

## Biogeolocalisation of MGT-v1 728      789 unigenes, 563 (71.36%) taxonomically assigned

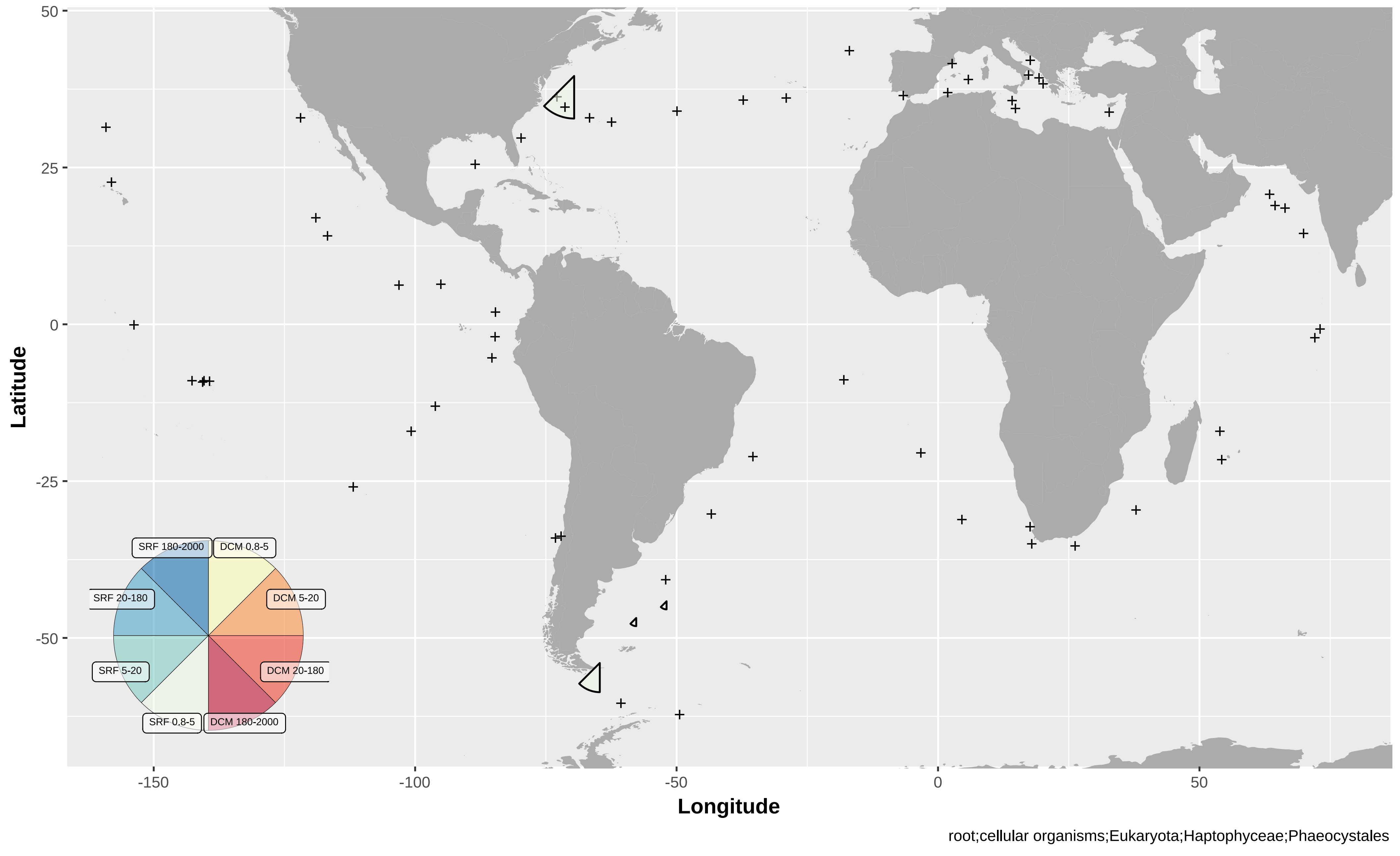

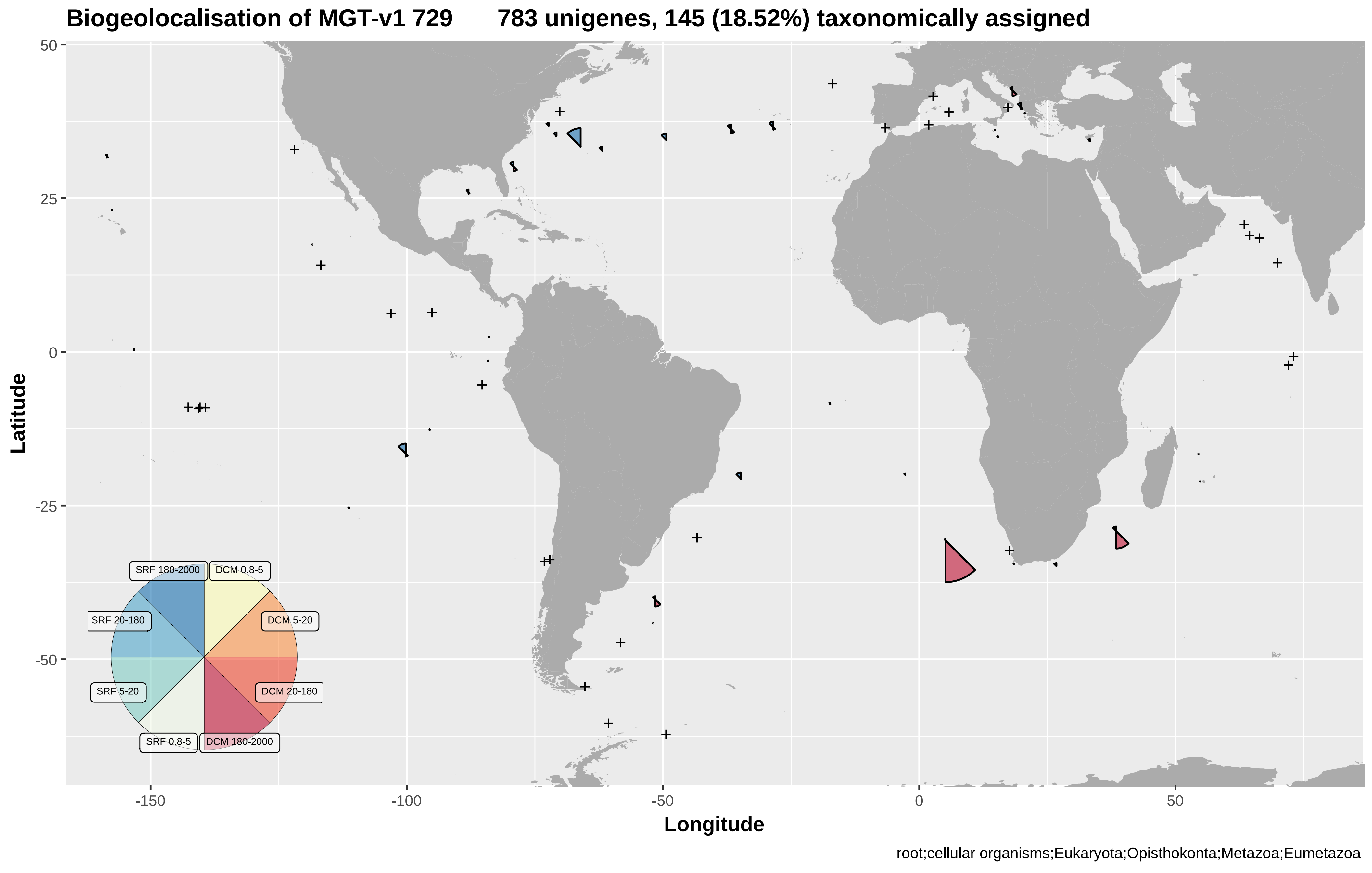

Biogeolocalisation of MGT-v1 729 783 unigenes, 145 (18.52%) taxonomically assigned
Latitude
50
25
0
-25
-50
Longitude
-150
-100
-50
0
50
SRF 180-2000
DCM 0.8-5
SRF 20-180
DCM 5-20
SRF 5-20
DCM 20-180
SRF 0.8-5
DCM 180-2000
root;cellular organisms;Eukaryota;Opisthokonta;Metazoa;Eumetazoa

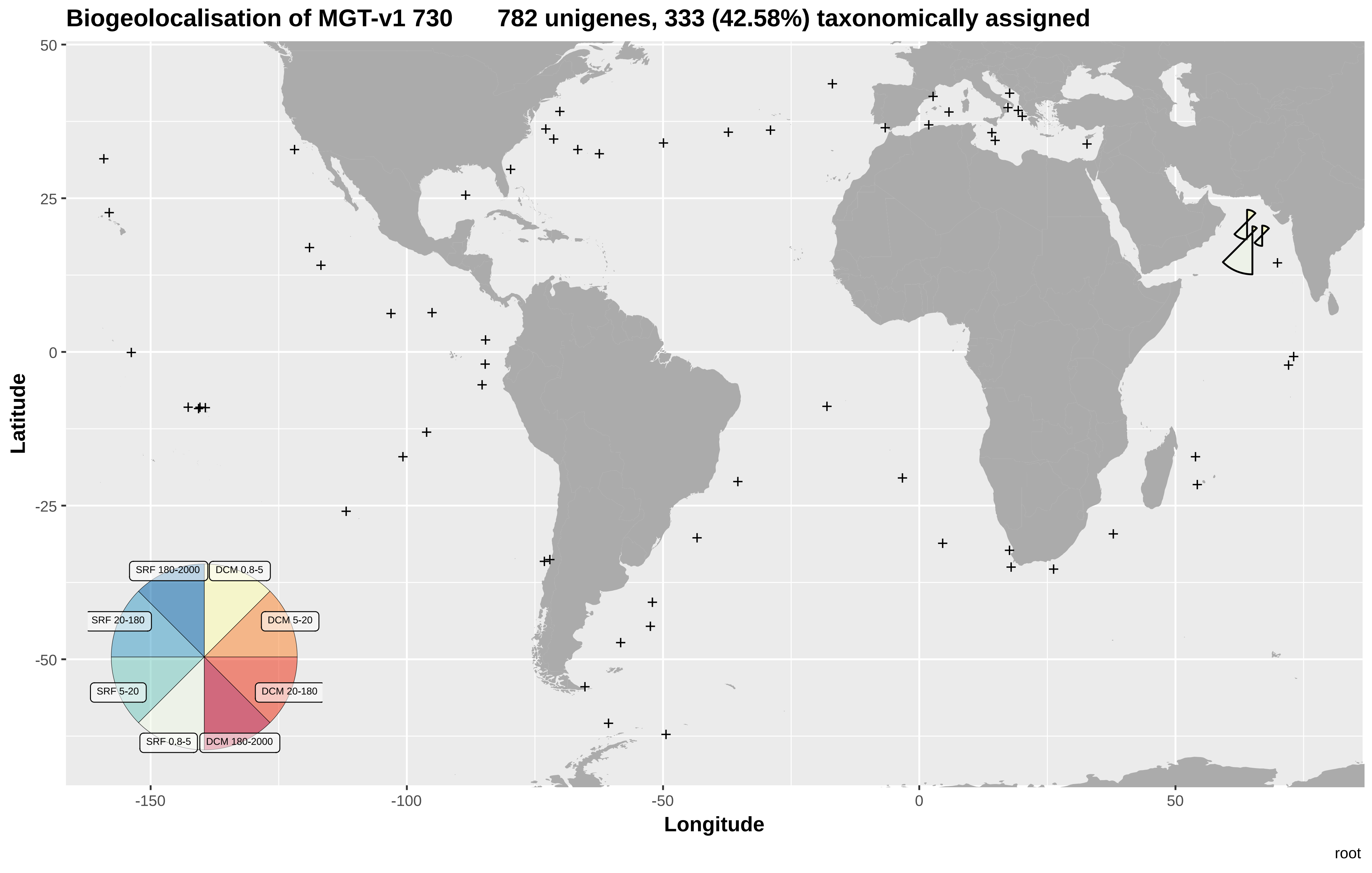
Biogeolocalisation of MGT-v1 730 782 unigenes, 333 (42.58%) taxonomically assigned
50
25
0
-25
-50
Latitude
-150
-100
-50
0
50
Longitude
SRF 180-2000
DCM 0.8-5
SRF 20-180
DCM 5-20
SRF 5-20
DCM 20-180
SRF 0.8-5
DCM 180-2000
root

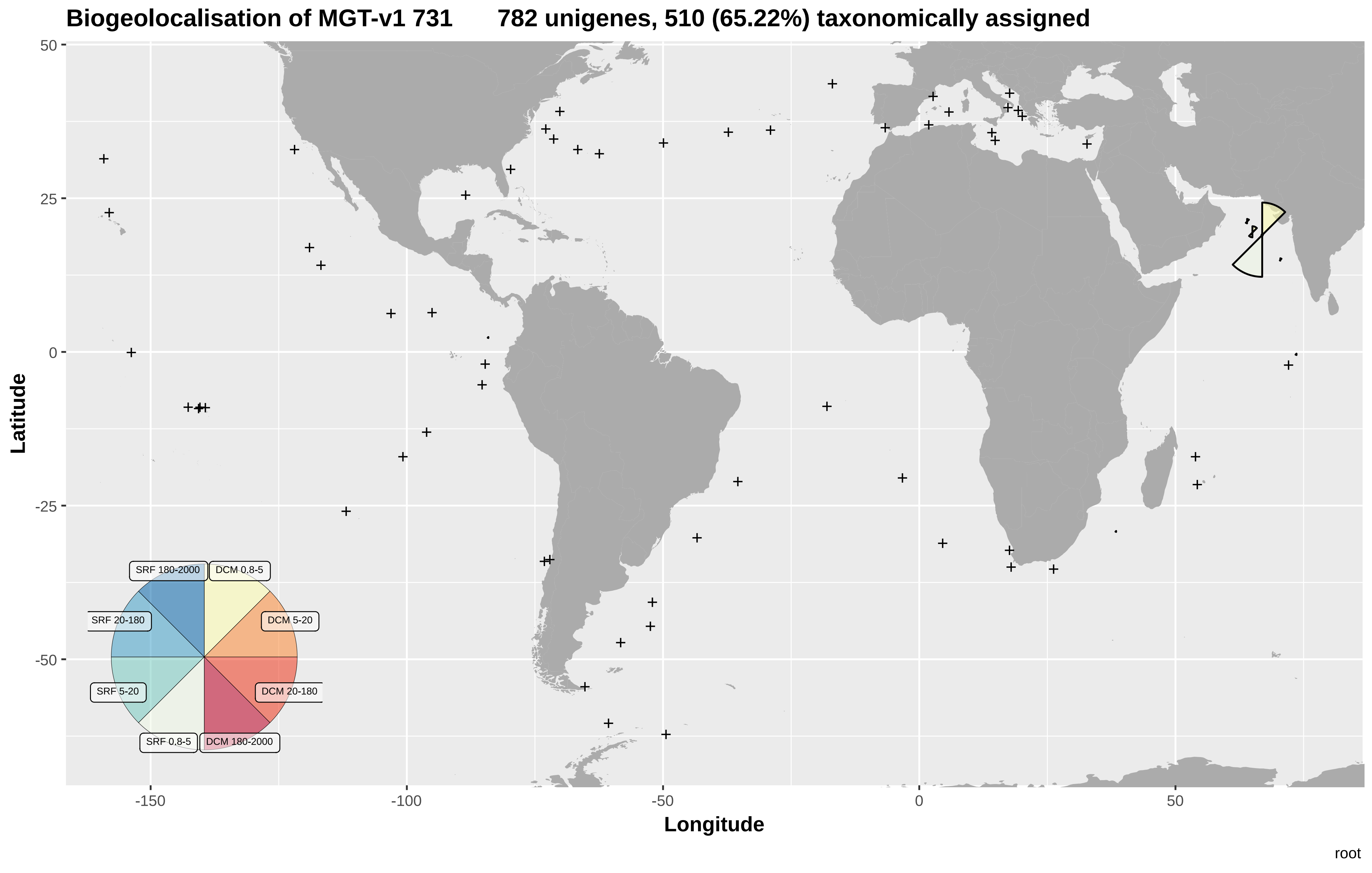
Biogeolocalisation of MGT-v1 731      782 unigenes, 510 (65.22%) taxonomically assigned
50
25
0
-25
-50
Latitude
-150
-100
-50
0
50
Longitude
SRF 180-2000
DCM 0.8-5
SRF 20-180
DCM 5-20
SRF 5-20
DCM 20-180
SRF 0.8-5
DCM 180-2000
root

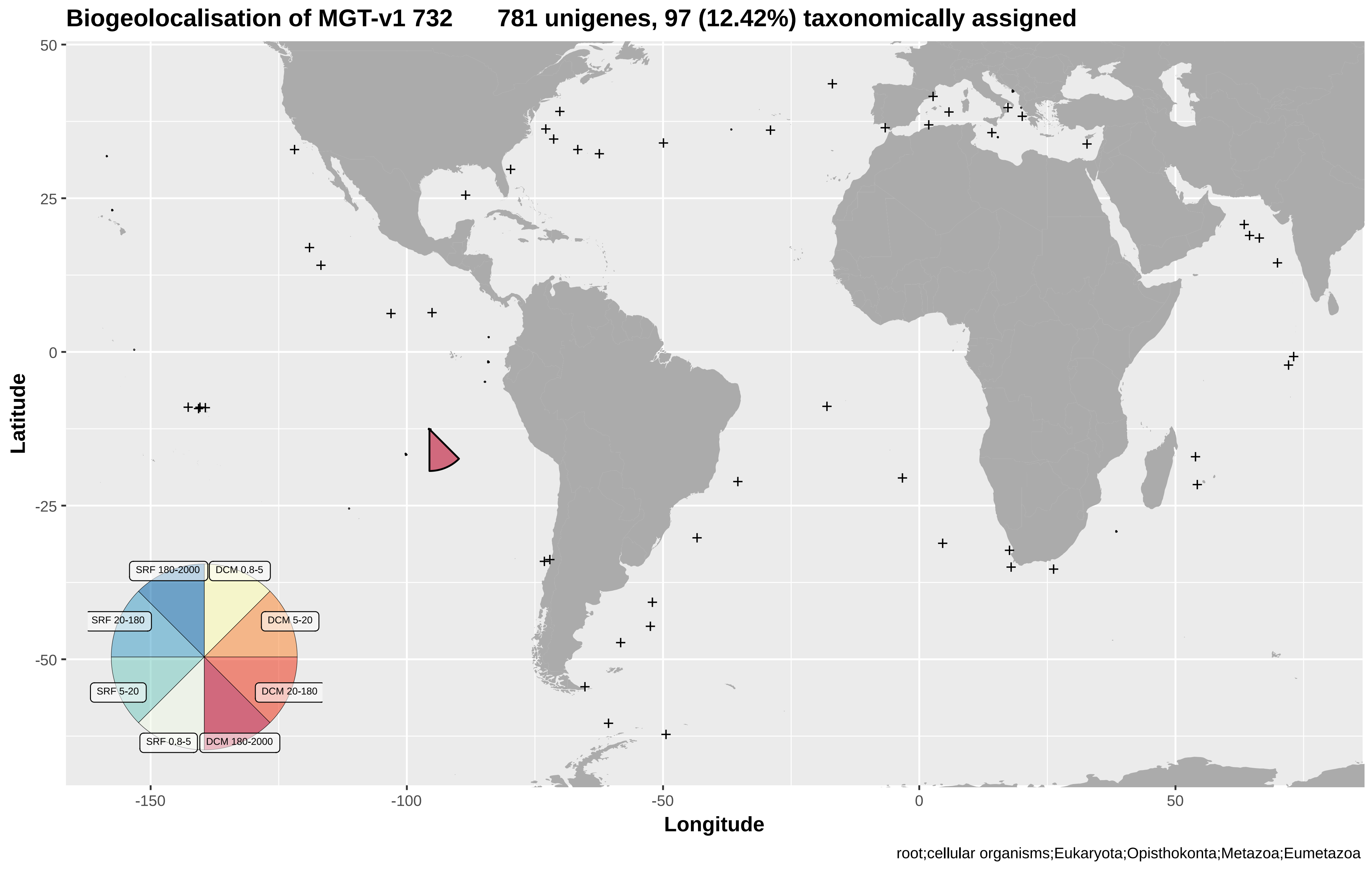

Biogeolocalisation of MGT-v1 732 781 unigenes, 97 (12.42%) taxonomically assigned
50
25
0
-25
-50
Latitude
-150
-100
-50
0
50
Longitude
SRF 180-2000
DCM 0.8-5
SRF 20-180
DCM 5-20
SRF 5-20
DCM 20-180
SRF 0.8-5
DCM 180-2000
root;cellular organisms;Eukaryota;Opisthokonta;Metazoa;Eumetazoa

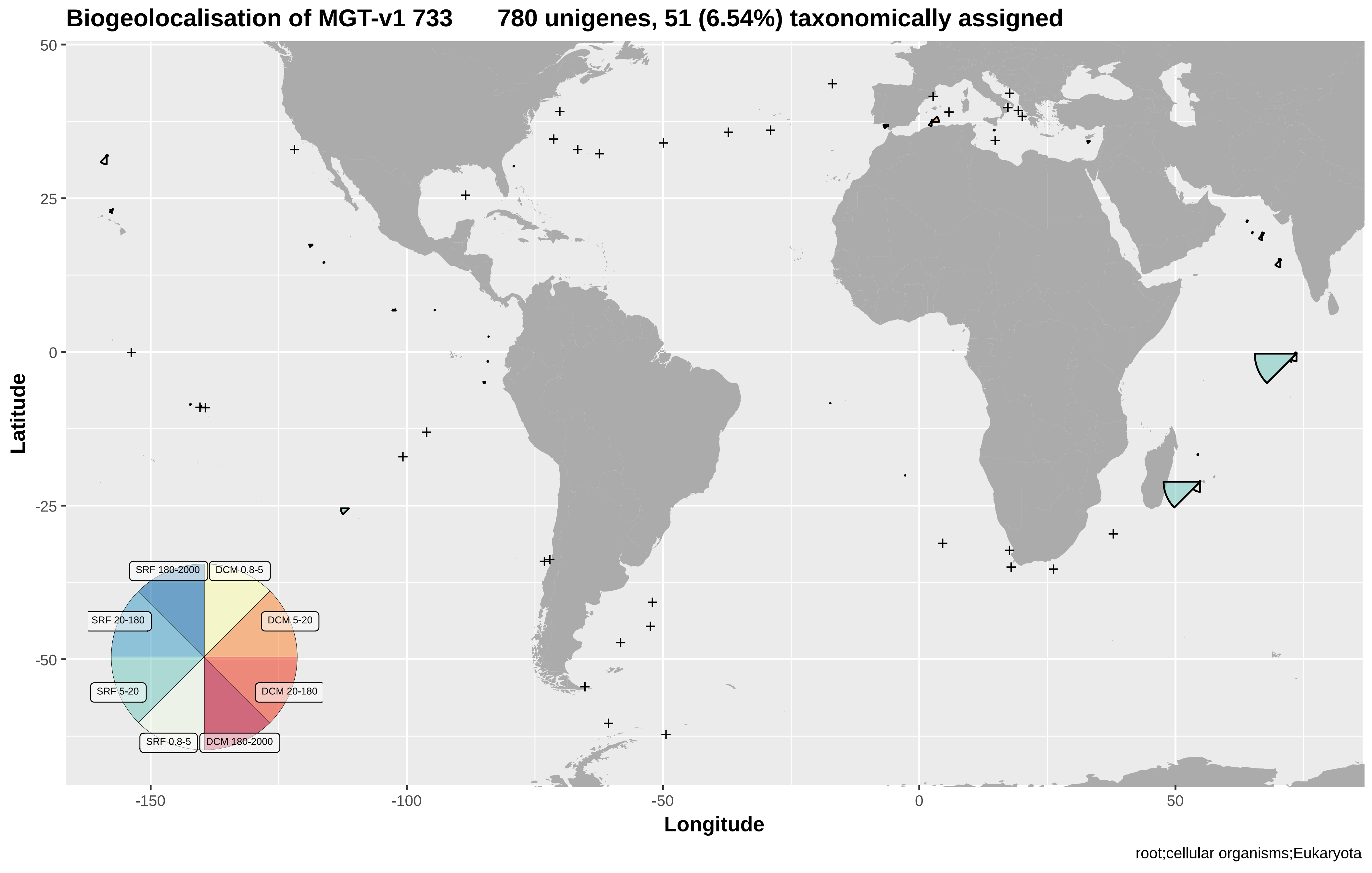
Biogeolocalisation of MGT-v1 733 780 unigenes, 51 (6.54%) taxonomically assigned
Latitude
Longitude
50
25
0
-25
-50
-150
-100
-50
0
50
SRF 180-2000
DCM 0.8-5
SRF 20-180
DCM 5-20
SRF 5-20
DCM 20-180
SRF 0.8-5
DCM 180-2000
root;cellular organisms;Eukaryota

# Biogeolocalisation of MGT-v1 734      780 unigenes, 142 (18.21%) taxonomically assigned

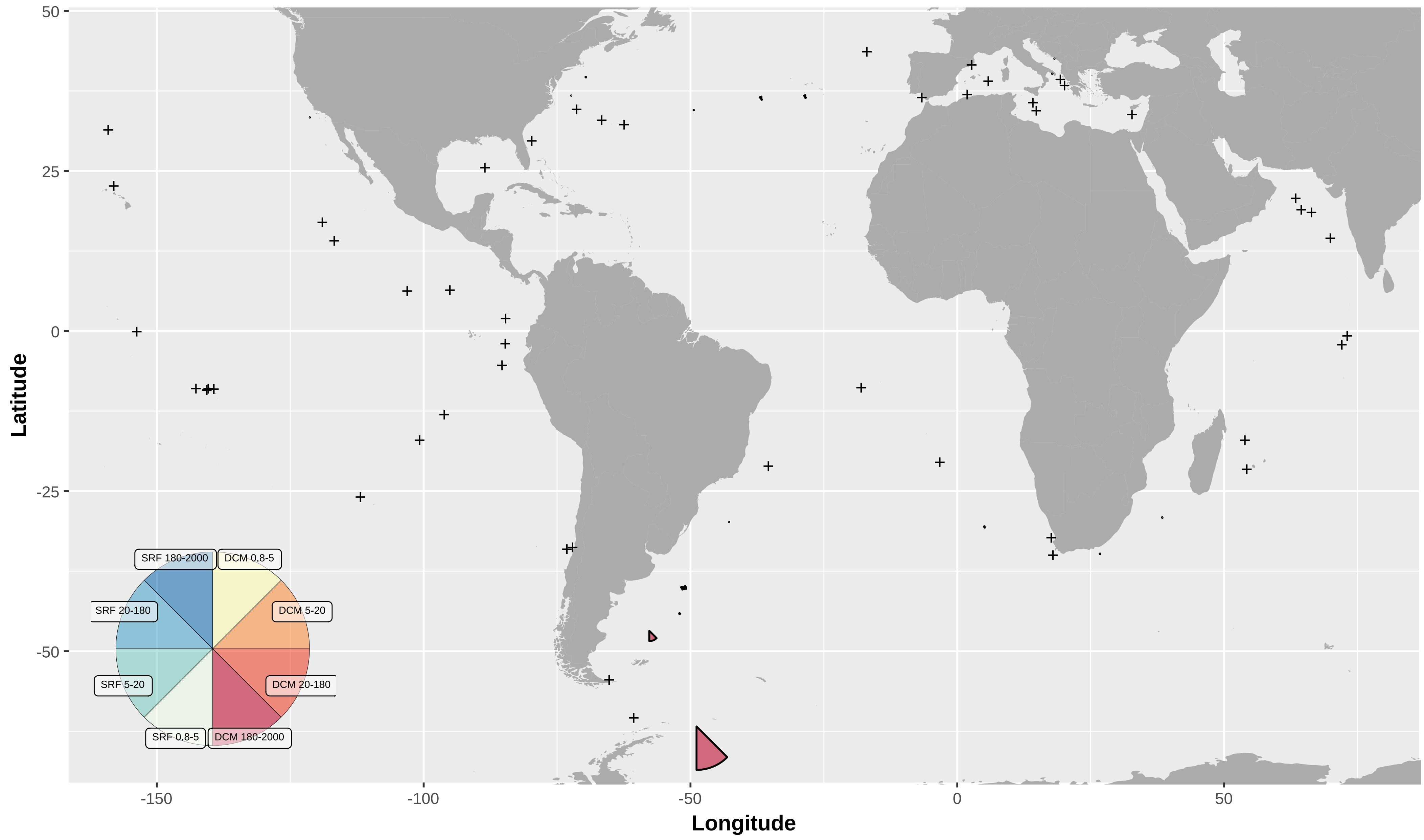


root;cellular organisms;Eukaryota;Opisthokonta;Metazoa;Eumetazoa

## Biogeolocalisation of MGT-v1 735      777 unigenes, 97 (12.48%) taxonomically assigned

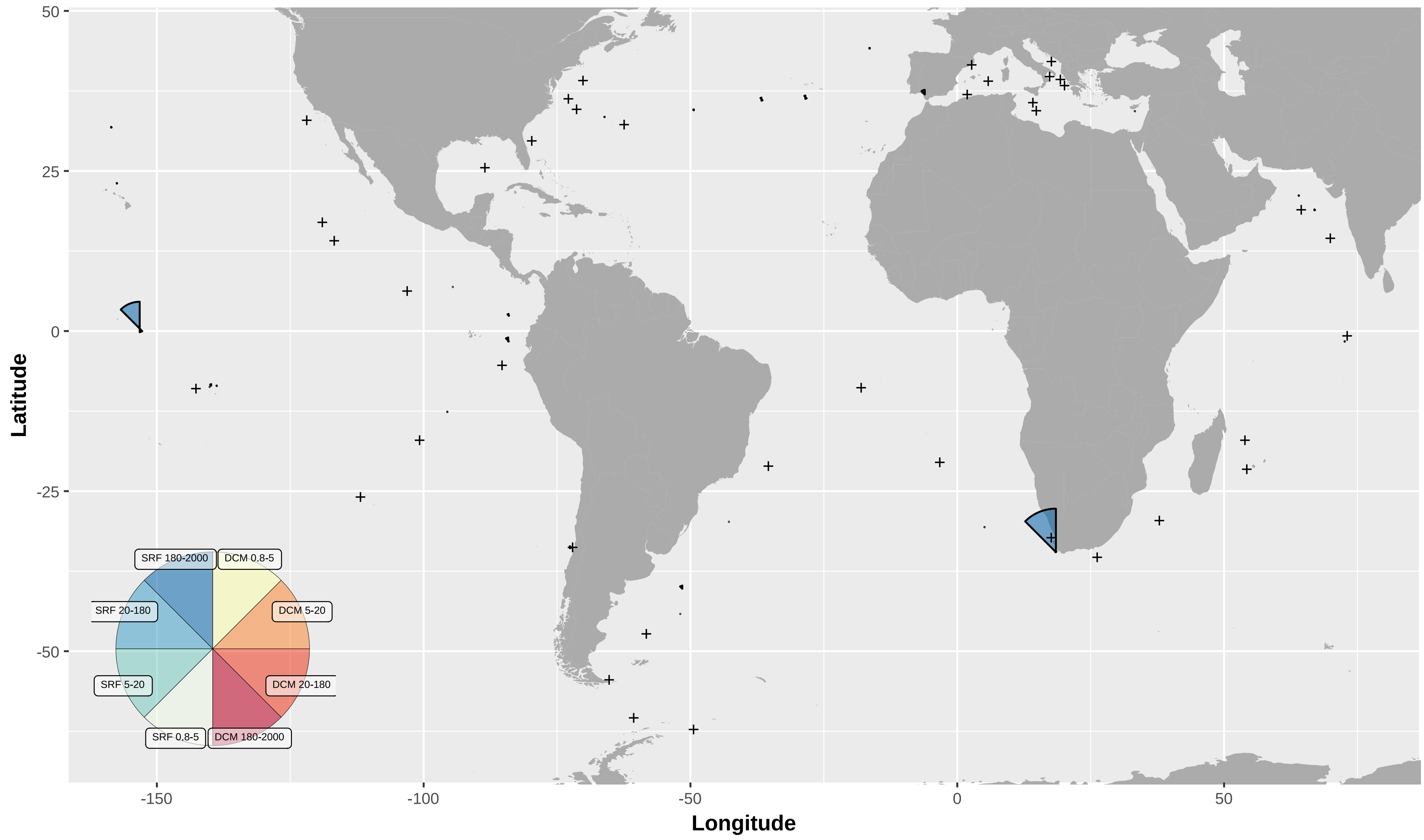


root;cellular organisms;Eukaryota

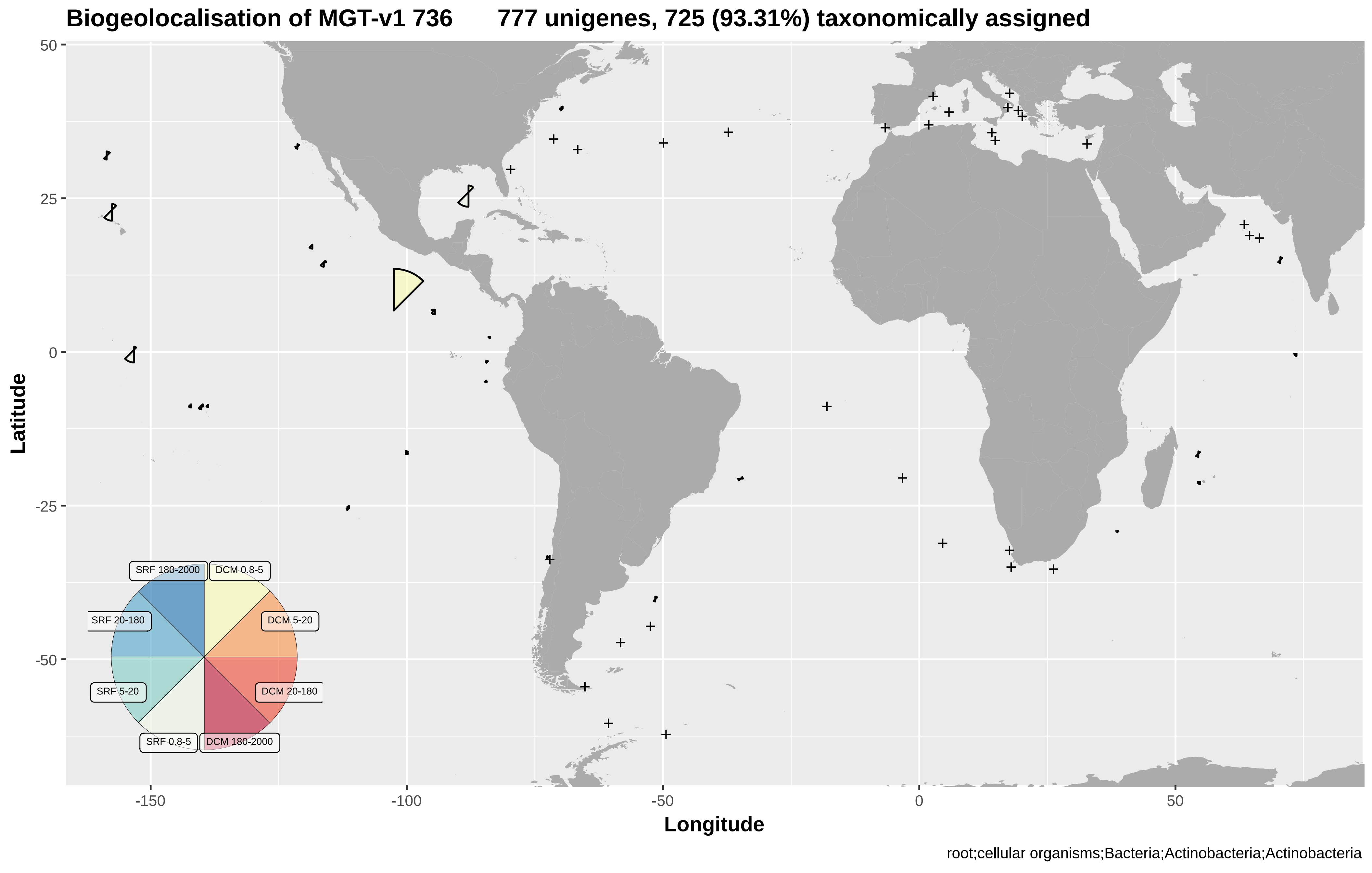

Biogeolocalisation of MGT-v1 736 777 unigenes, 725 (93.31%) taxonomically assigned
Latitude
50
25
0
-25
-50
Longitude
-150
-100
-50
0
50
SRF 180-2000
DCM 0.8-5
SRF 20-180
DCM 5-20
SRF 5-20
DCM 20-180
SRF 0.8-5
DCM 180-2000
root;cellular organisms;Bacteria;Actinobacteria;Actinobacteria

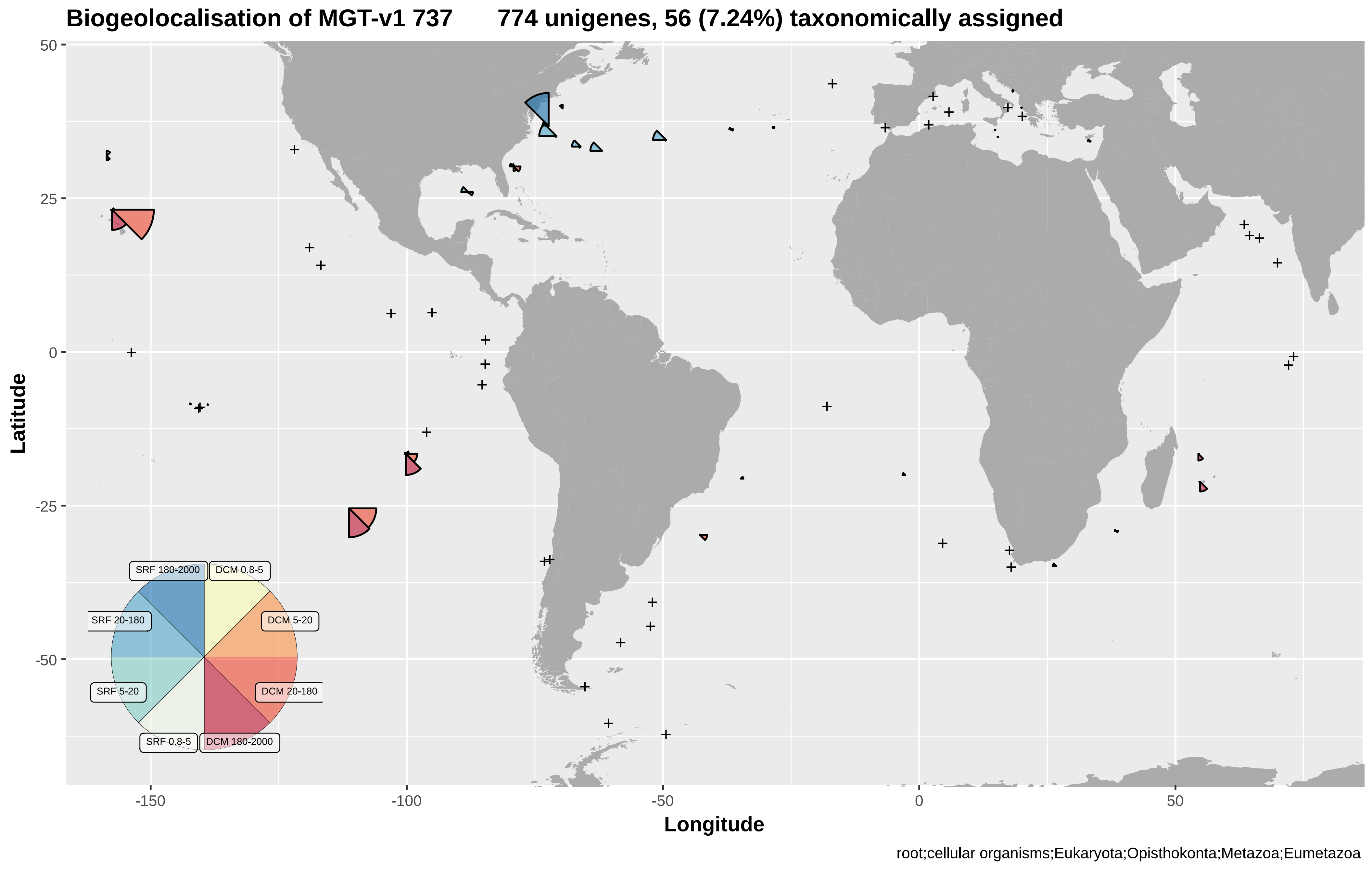
Biogeolocalisation of MGT-v1 737 774 unigenes, 56 (7.24%) taxonomically assigned
Latitude
Longitude
50
25
0
-25
-50
-150
-100
-50
0
50
SRF 180-2000
DCM 0.8-5
SRF 20-180
DCM 5-20
SRF 5-20
DCM 20-180
SRF 0.8-5
DCM 180-2000
root;cellular organisms;Eukaryota;Opisthokonta;Metazoa;Eumetazoa

## Biogeolocalisation of MGT-v1 738      770 unigenes, 423 (54.94%) taxonomically assigned

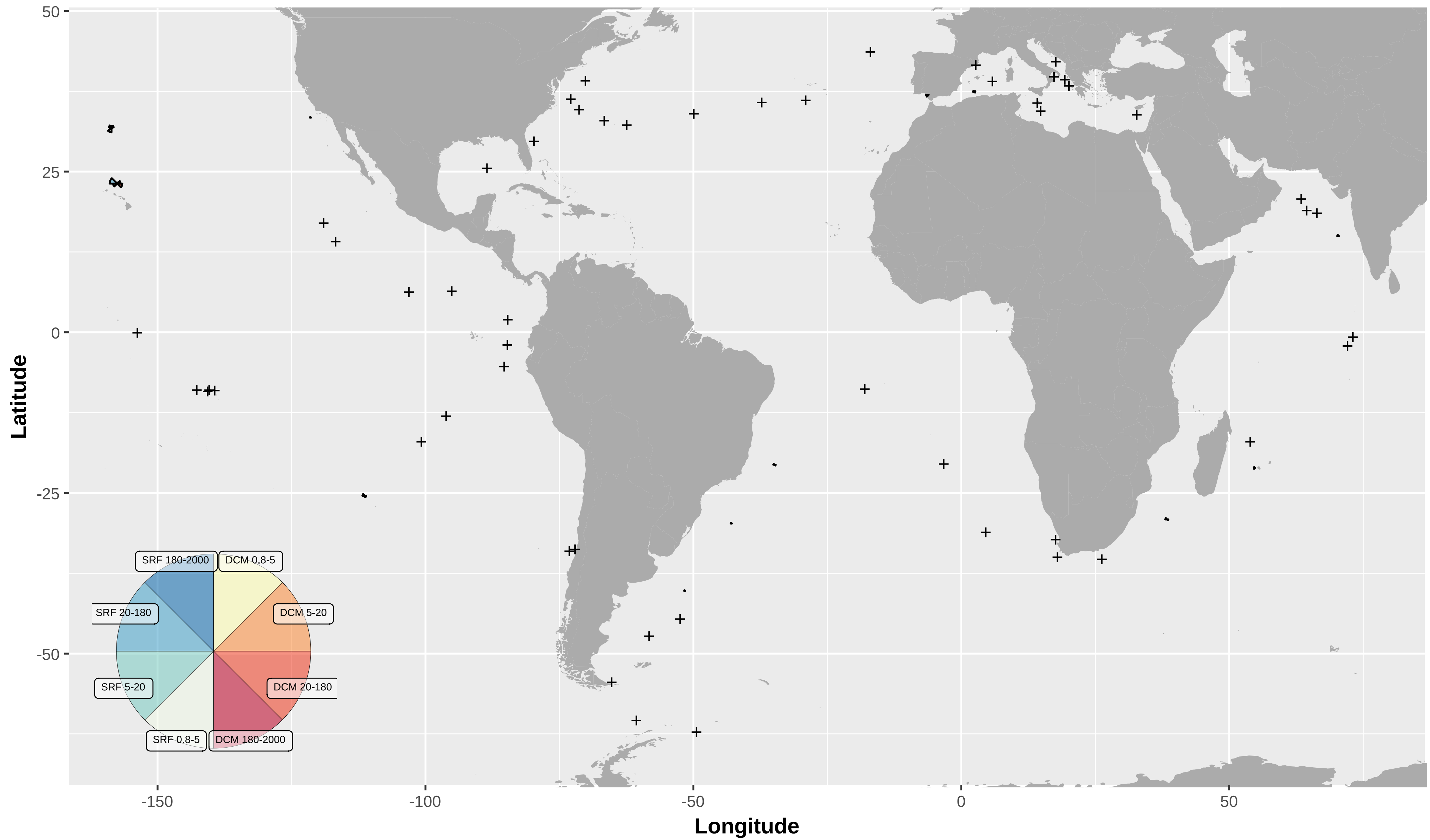


root;cellular organisms

# Biogeolocalisation of MGT-v1 739      769 unigenes, 128 (16.64%) taxonomically assigned

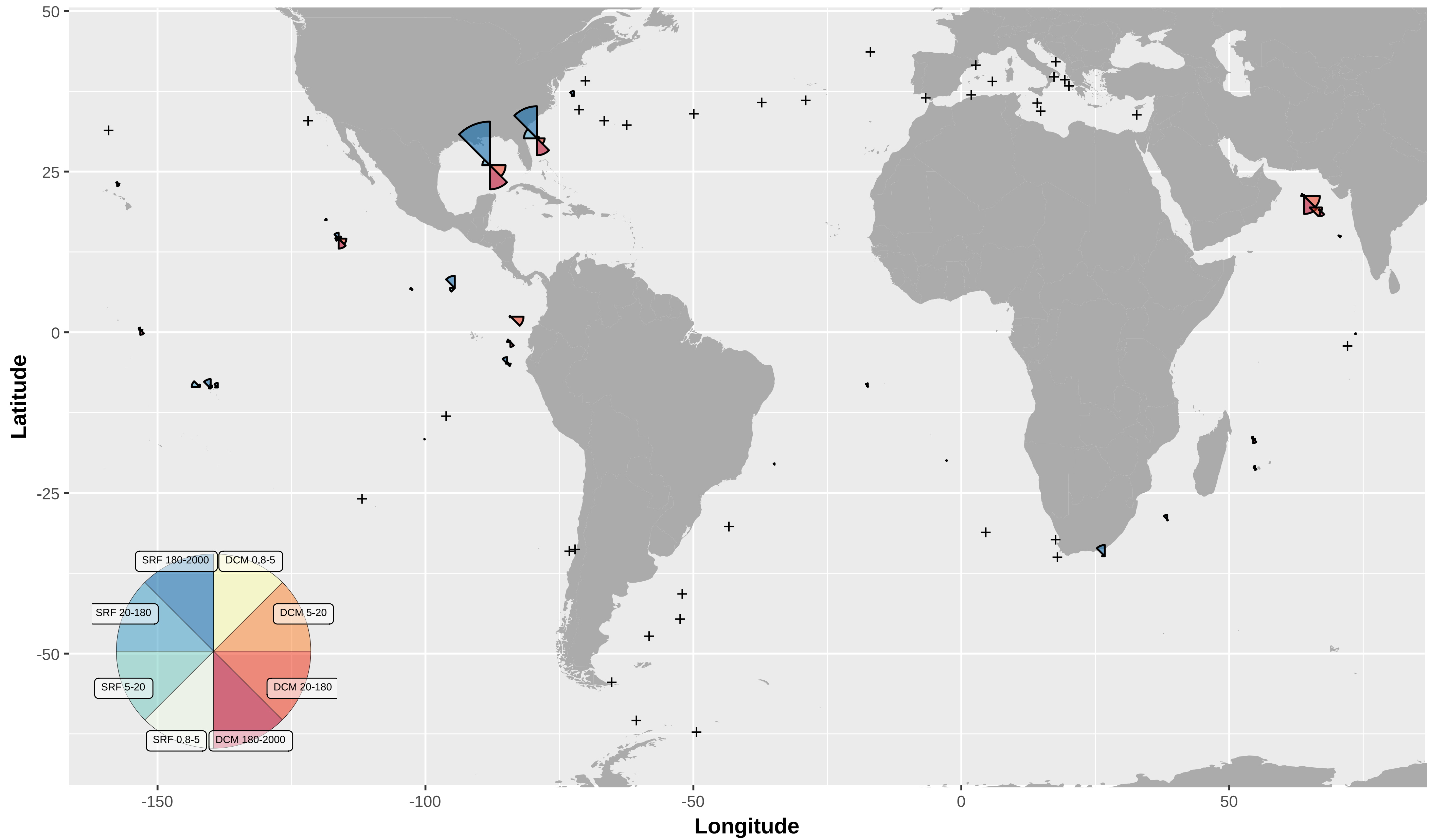


root;cellular organisms;Eukaryota;Opisthokonta;Metazoa;Eumetazoa;Bilateria;Protostomia;Ecdysozoa;Panarthropoda;Arthropoda;Mandibulata;Pancrustacea;Crustacea;Maxillopoda

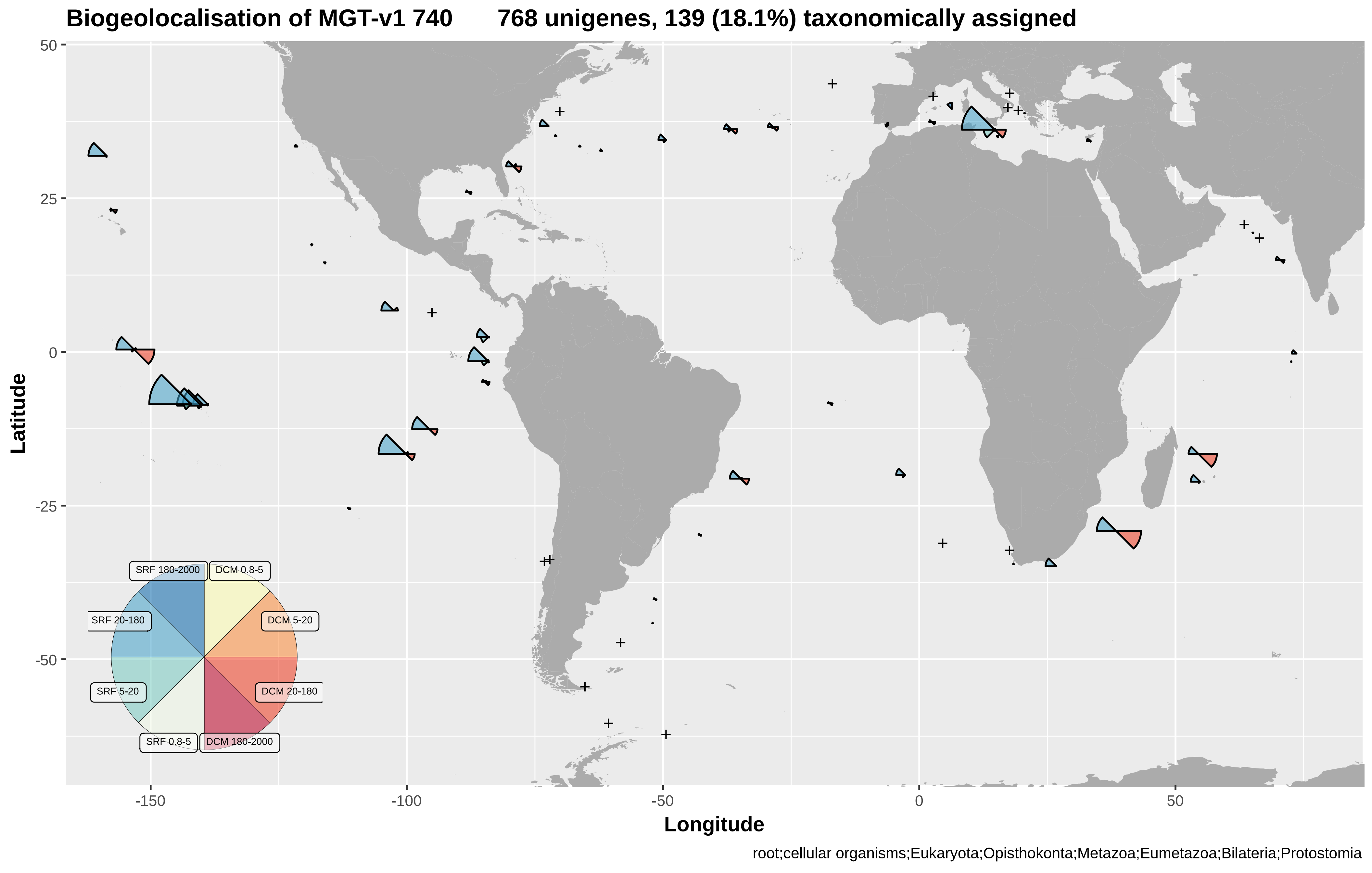
Biogeolocalisation of MGT-v1 740 768 unigenes, 139 (18.1%) taxonomically assigned
50
25
0
-25
-50
Latitude
-150
-100
-50
0
50
Longitude
SRF 180-2000
DCM 0.8-5
SRF 20-180
DCM 5-20
SRF 5-20
DCM 20-180
SRF 0.8-5
DCM 180-2000
root;cellular organisms;Eukaryota;Opisthokonta;Metazoa;Eumetazoa;Bilateria;Protostomia

### Biogeolocalisation of MGT-v1 741 766 unigenes, 120 (15.67%) taxonomically assigned

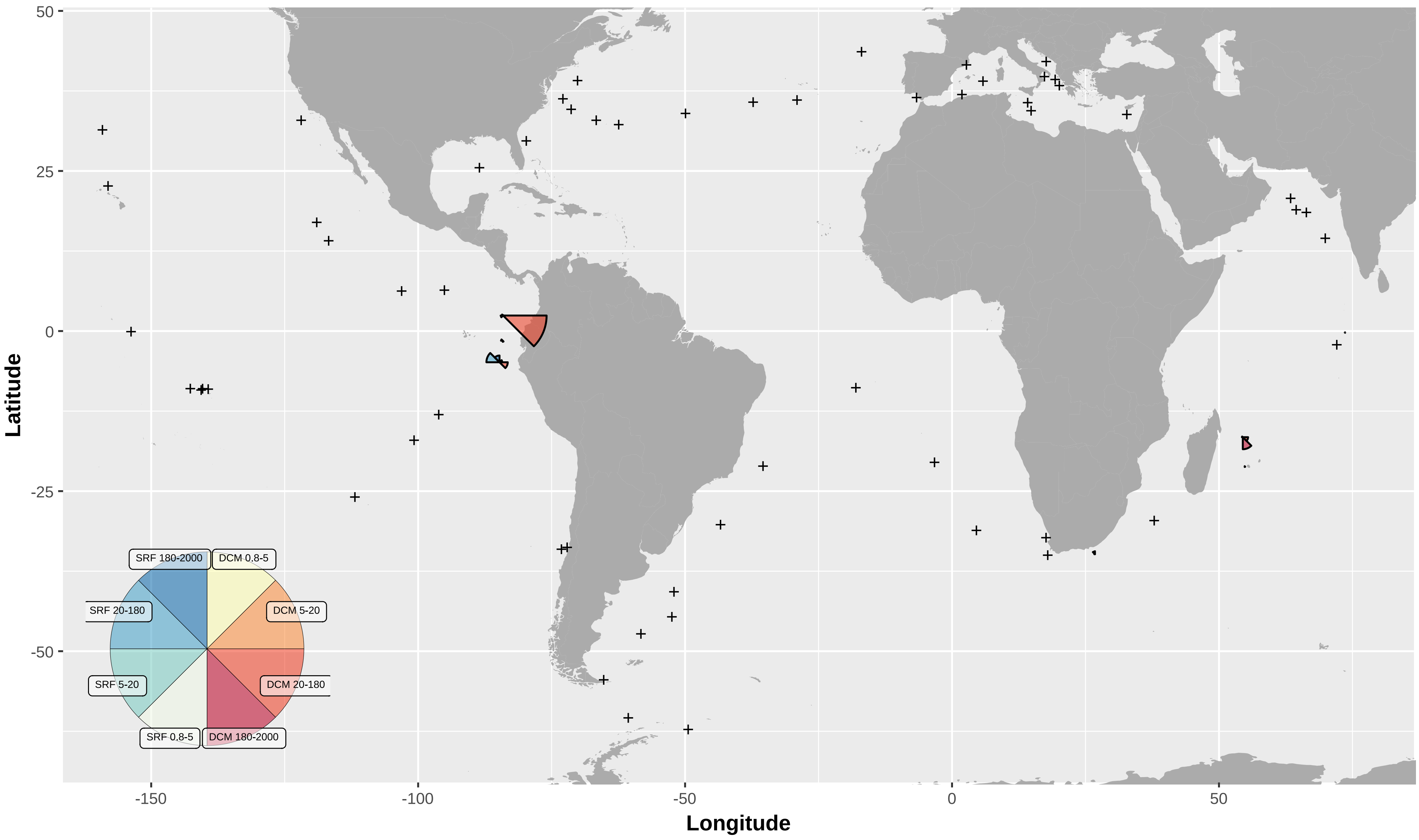


root;cellular organisms;Eukaryota;Opisthokonta;Metazoa;Eumetazoa;Bilateria

### Biogeolocalisation of MGT-v1 742 765 unigenes, 49 (6.41%) taxonomically assigned

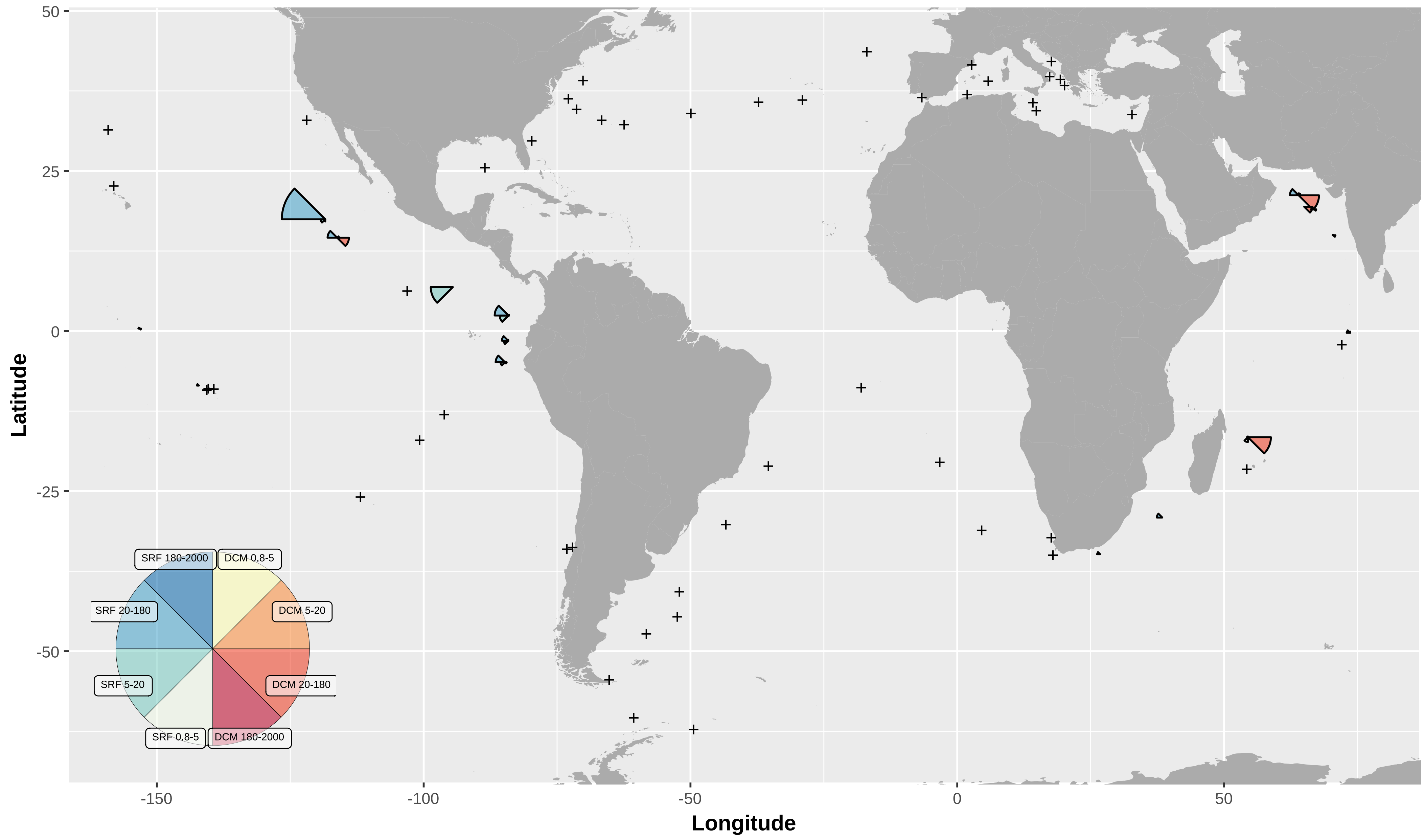


root;cellular organisms;Eukaryota;Opisthokonta;Metazoa;Eumetazoa;Bilateria

### Biogeolocalisation of MGT-v1 743 764 unigenes, 512 (67.02%) taxonomically assigned

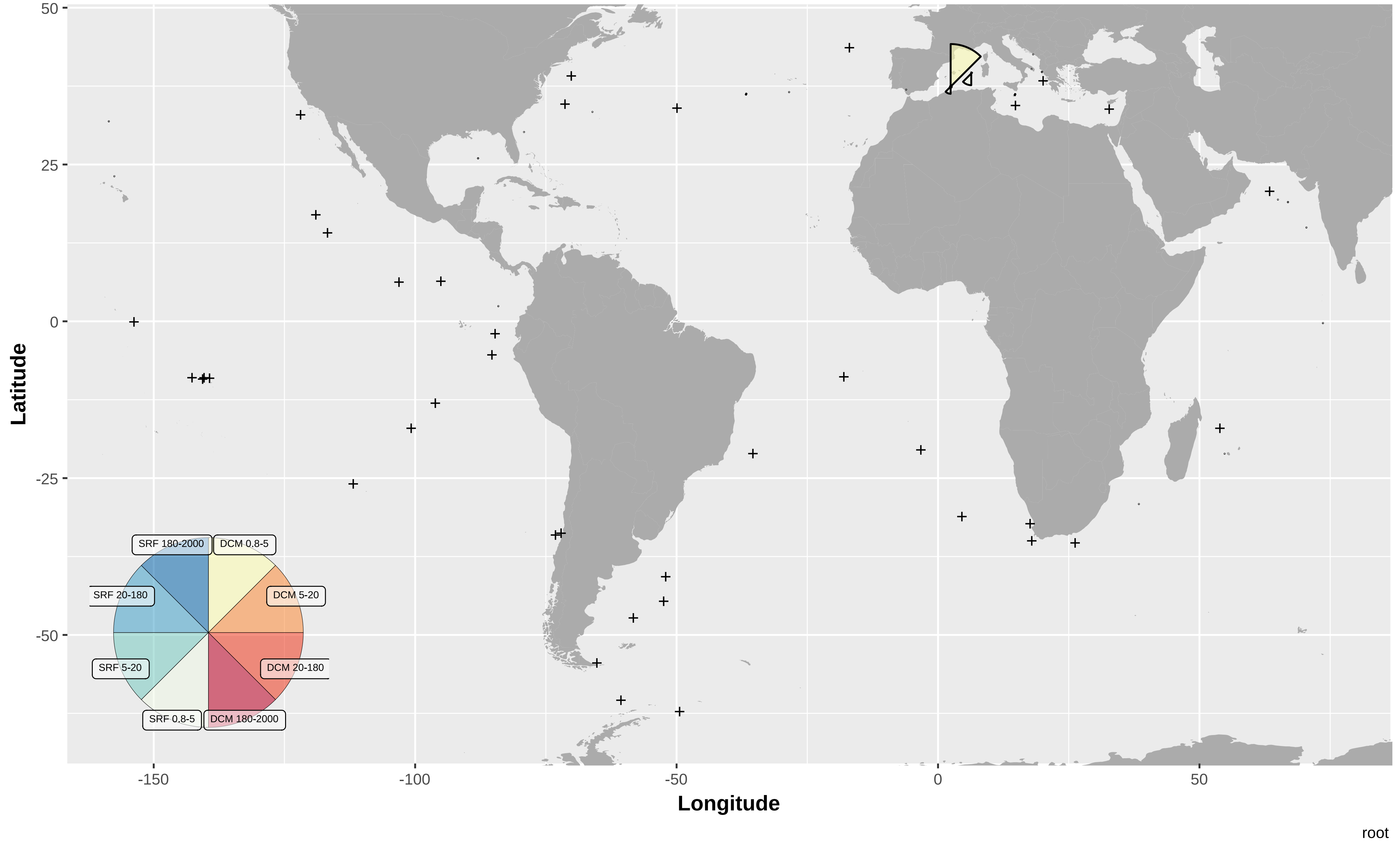

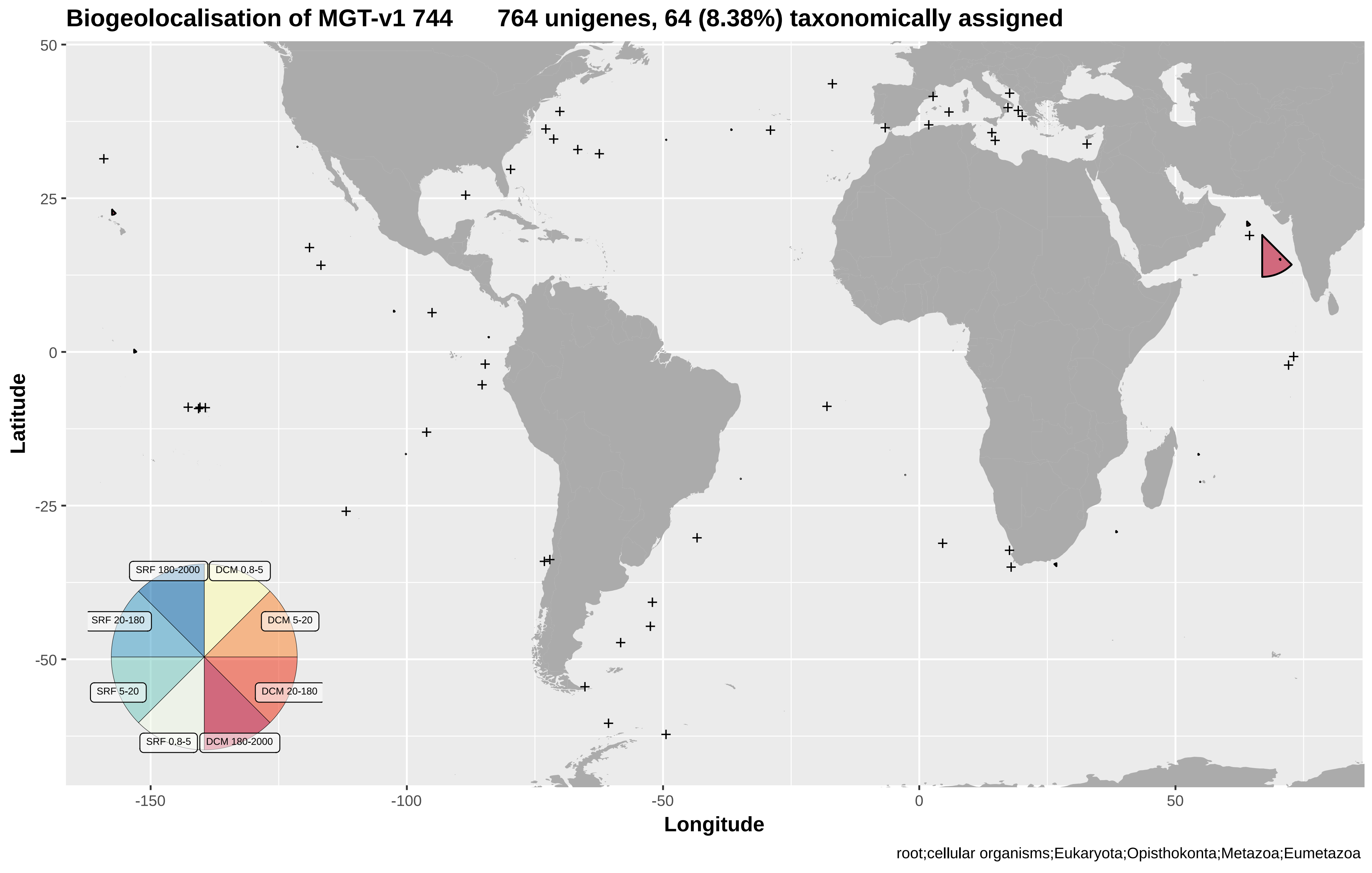
Biogeolocalisation of MGT-v1 744
764 unigenes, 64 (8.38%) taxonomically assigned
50
25
0
-25
-50
Latitude
-150
-100
-50
0
50
Longitude
SRF 180-2000
DCM 0.8-5
SRF 20-180
DCM 5-20
SRF 5-20
DCM 20-180
SRF 0.8-5
DCM 180-2000
root;cellular organisms;Eukaryota;Opisthokonta;Metazoa;Eumetazoa

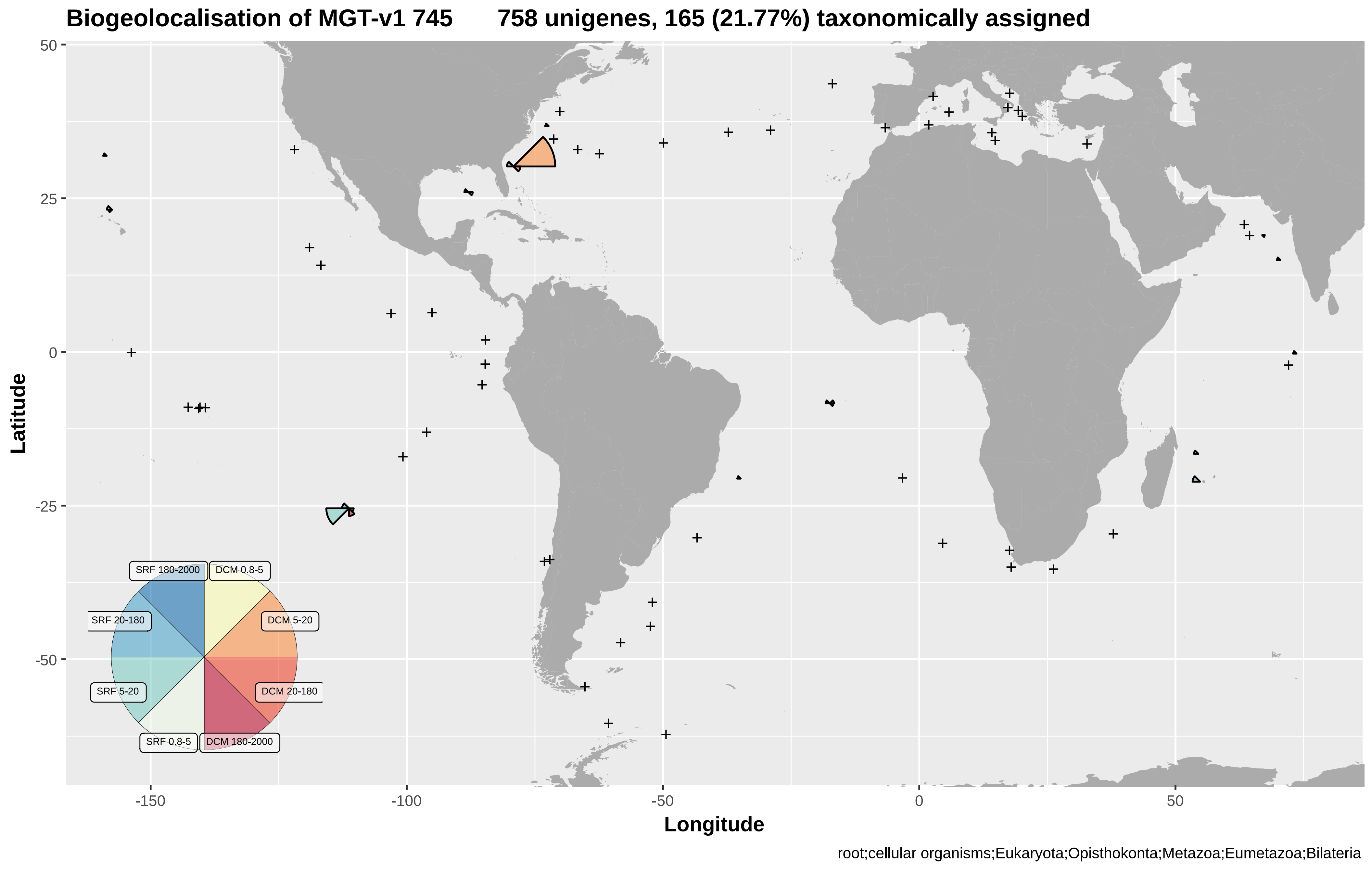

Biogeolocalisation of MGT-v1 745 758 unigenes, 165 (21.77%) taxonomically assigned
50
25
0
-25
-50
Latitude
-150
-100
-50
0
50
Longitude
SRF 180-2000
DCM 0.8-5
SRF 20-180
DCM 5-20
SRF 5-20
DCM 20-180
SRF 0.8-5
DCM 180-2000
root;cellular organisms;Eukaryota;Opisthokonta;Metazoa;Eumetazoa;Bilateria

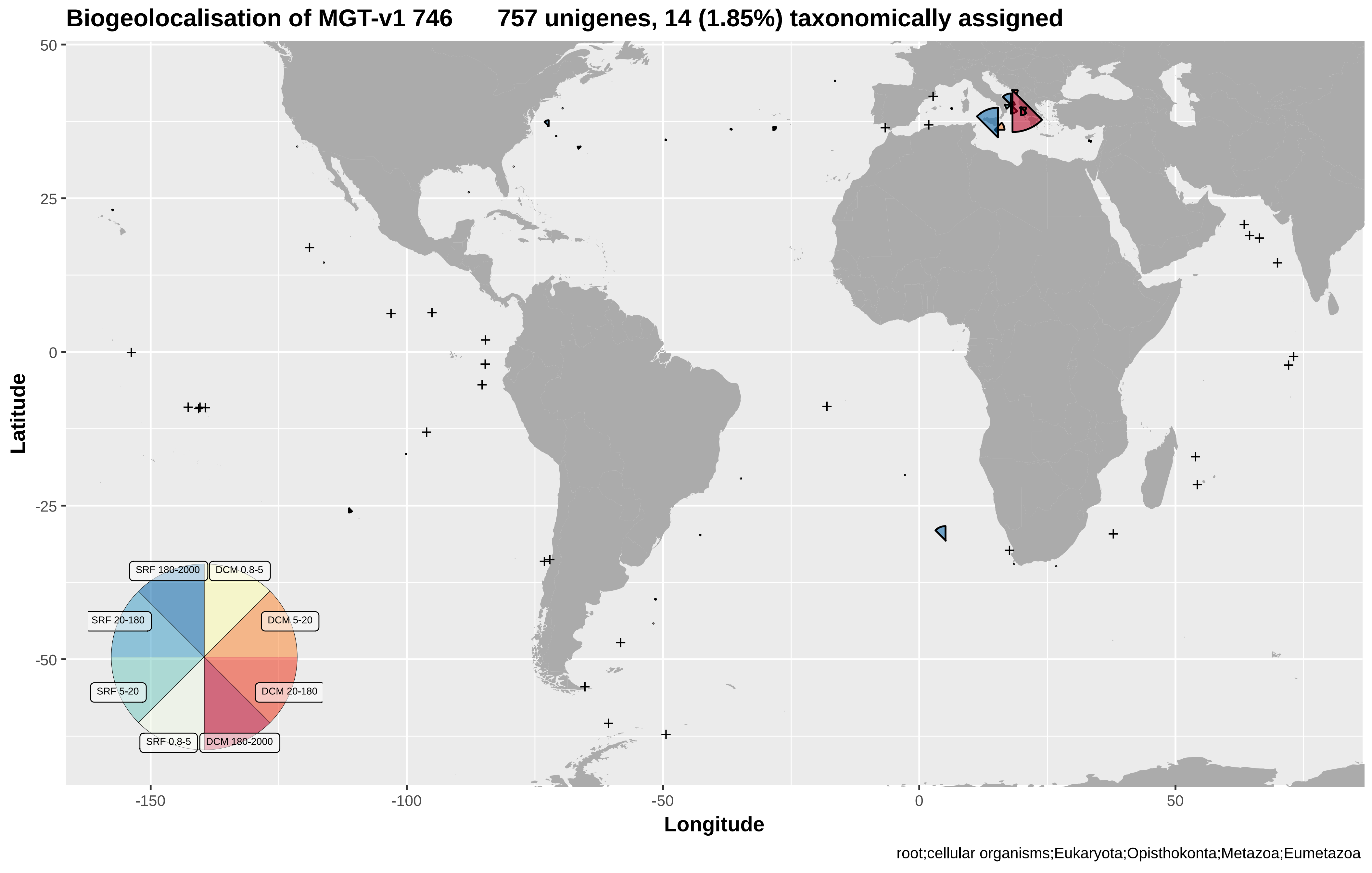
Biogeolocalisation of MGT-v1 746 757 unigenes, 14 (1.85%) taxonomically assigned
50
25
0
-25
-50
Latitude
-150
-100
-50
0
50
Longitude
SRF 180-2000
DCM 0.8-5
SRF 20-180
DCM 5-20
SRF 5-20
DCM 20-180
SRF 0.8-5
DCM 180-2000
root;cellular organisms;Eukaryota;Opisthokonta;Metazoa;Eumetazoa

## Biogeolocalisation of MGT-v1 747      756 unigenes, 540 (71.43%) taxonomically assigned

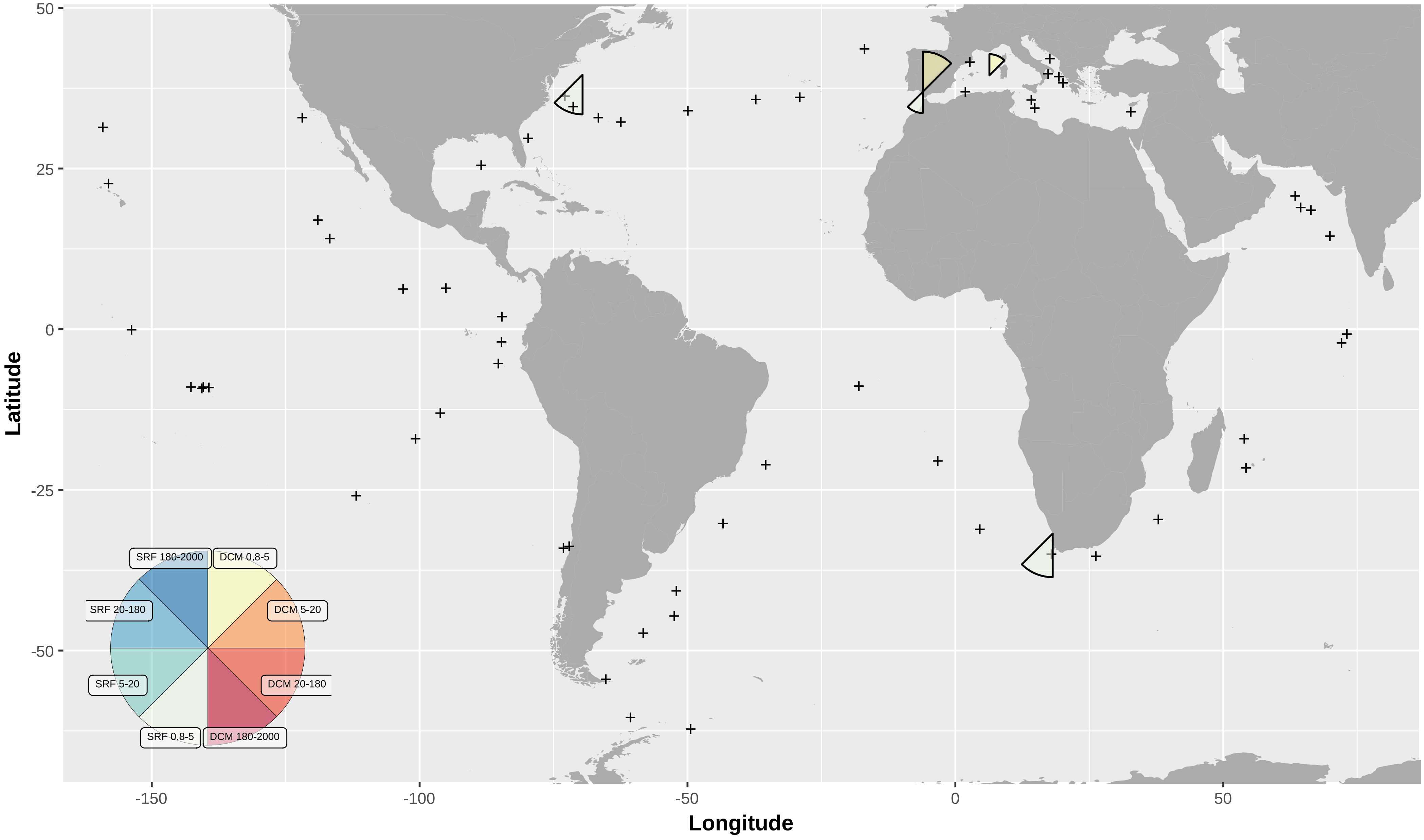


root;cellular organisms;Eukaryota;Haptophyceae

## Biogeolocalisation of MGT-v1 748      750 unigenes, 150 (20%) taxonomically assigned

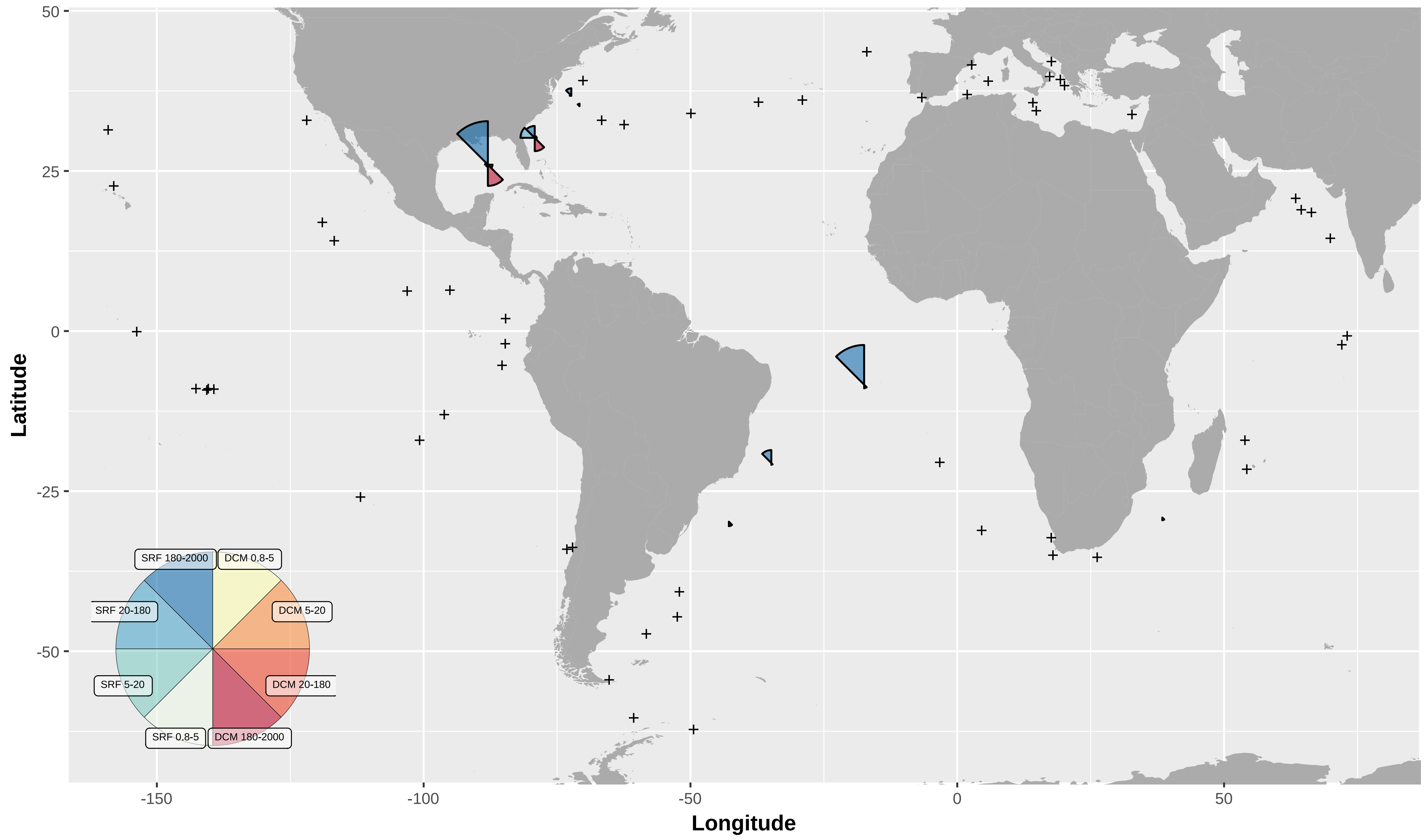


root;cellular organisms;Eukaryota;Opisthokonta;Metazoa;Eumetazoa;Bilateria

# Biogeolocalisation of MGT-v1 749      746 unigenes, 167 (22.39%) taxonomically assigned

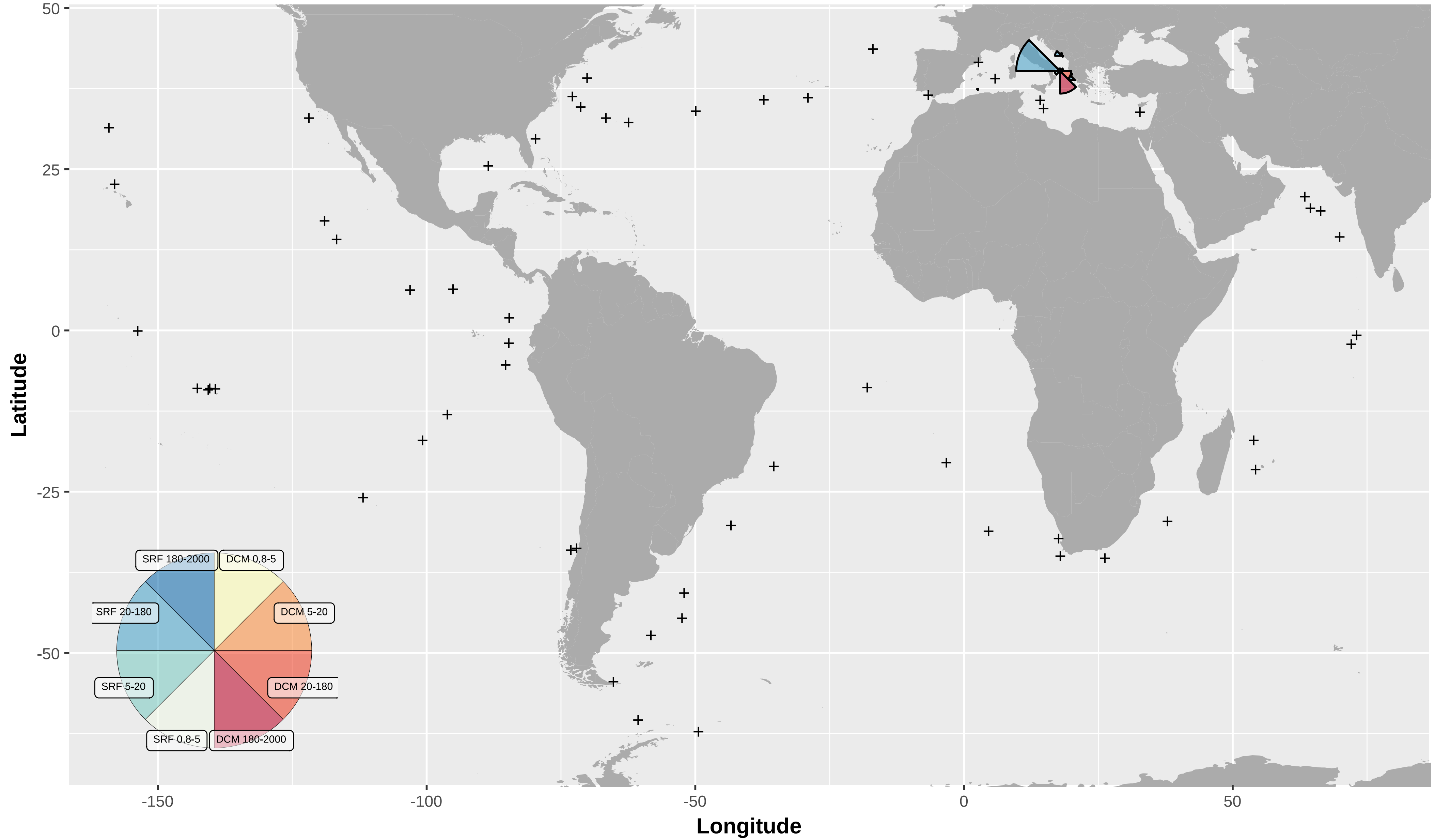


root;cellular organisms;Eukaryota;Opisthokonta;Metazoa;Eumetazoa;Bilateria

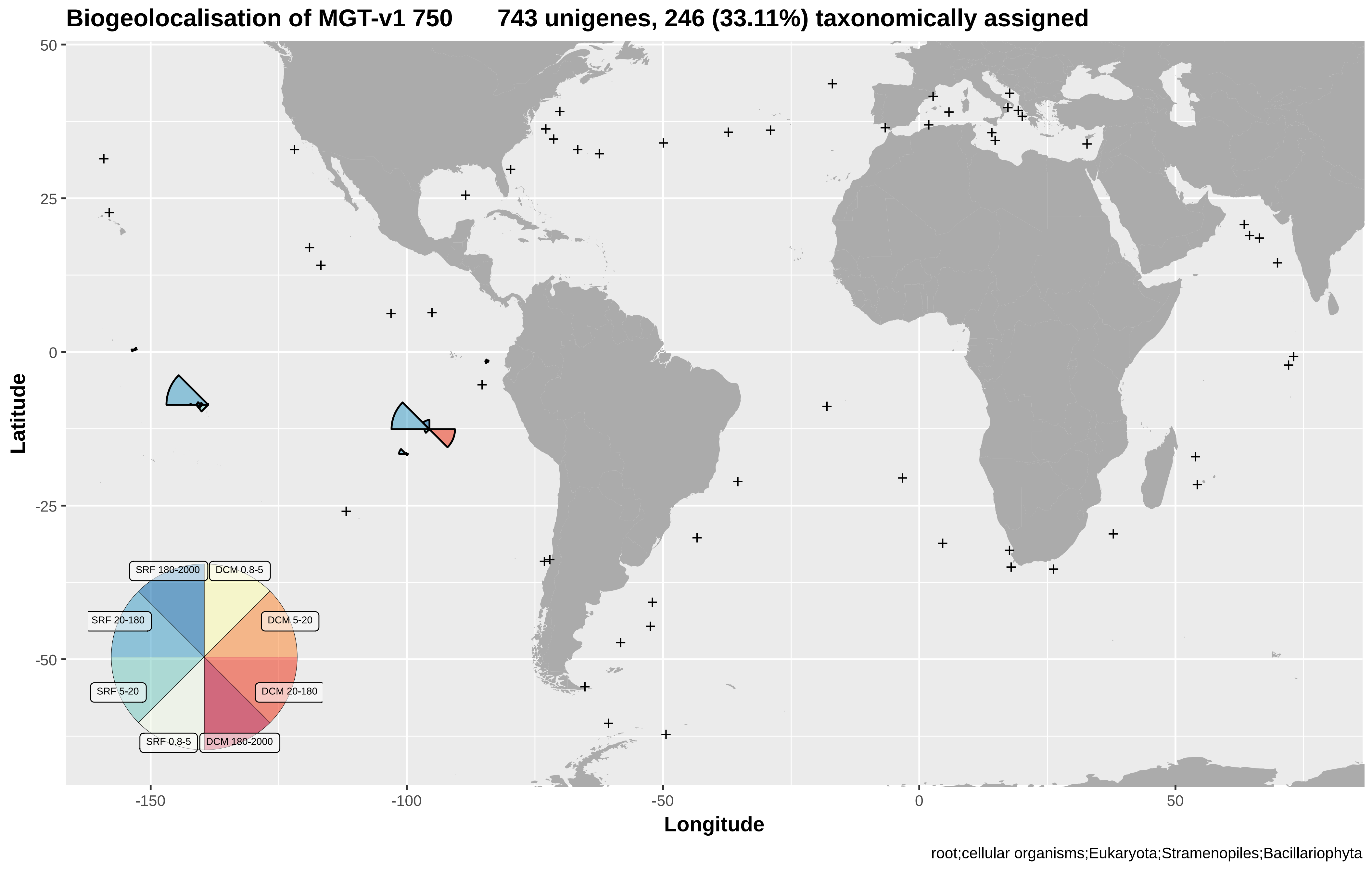

Biogeolocalisation of MGT-v1 750 743 unigenes, 246 (33.11%) taxonomically assigned
50
25
0
-25
-50
Latitude
-150
-100
-50
0
50
Longitude
SRF 180-2000
DCM 0.8-5
SRF 20-180
DCM 5-20
SRF 5-20
DCM 20-180
SRF 0.8-5
DCM 180-2000
root;cellular organisms;Eukaryota;Stramenopiles;Bacillariophyta

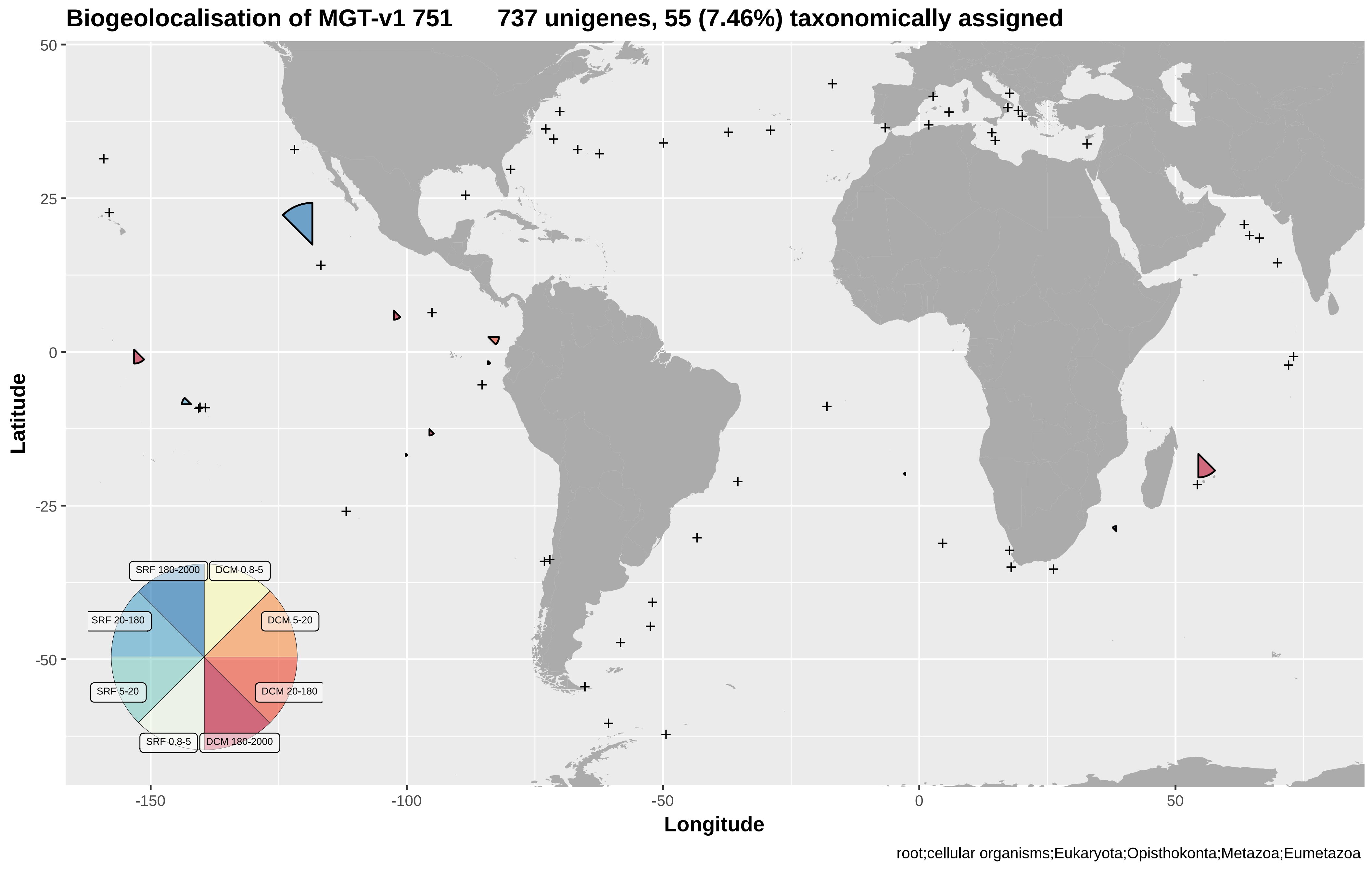
Biogeolocalisation of MGT-v1 751 737 unigenes, 55 (7.46%) taxonomically assigned
Latitude
Longitude
50
25
0
-25
-50
-150
-100
-50
0
50
SRF 180-2000
DCM 0.8-5
SRF 20-180
DCM 5-20
SRF 5-20
DCM 20-180
SRF 0.8-5
DCM 180-2000
root;cellular organisms;Eukaryota;Opisthokonta;Metazoa;Eumetazoa

## Biogeolocalisation of MGT-v1 752      734 unigenes, 33 (4.5%) taxonomically assigned

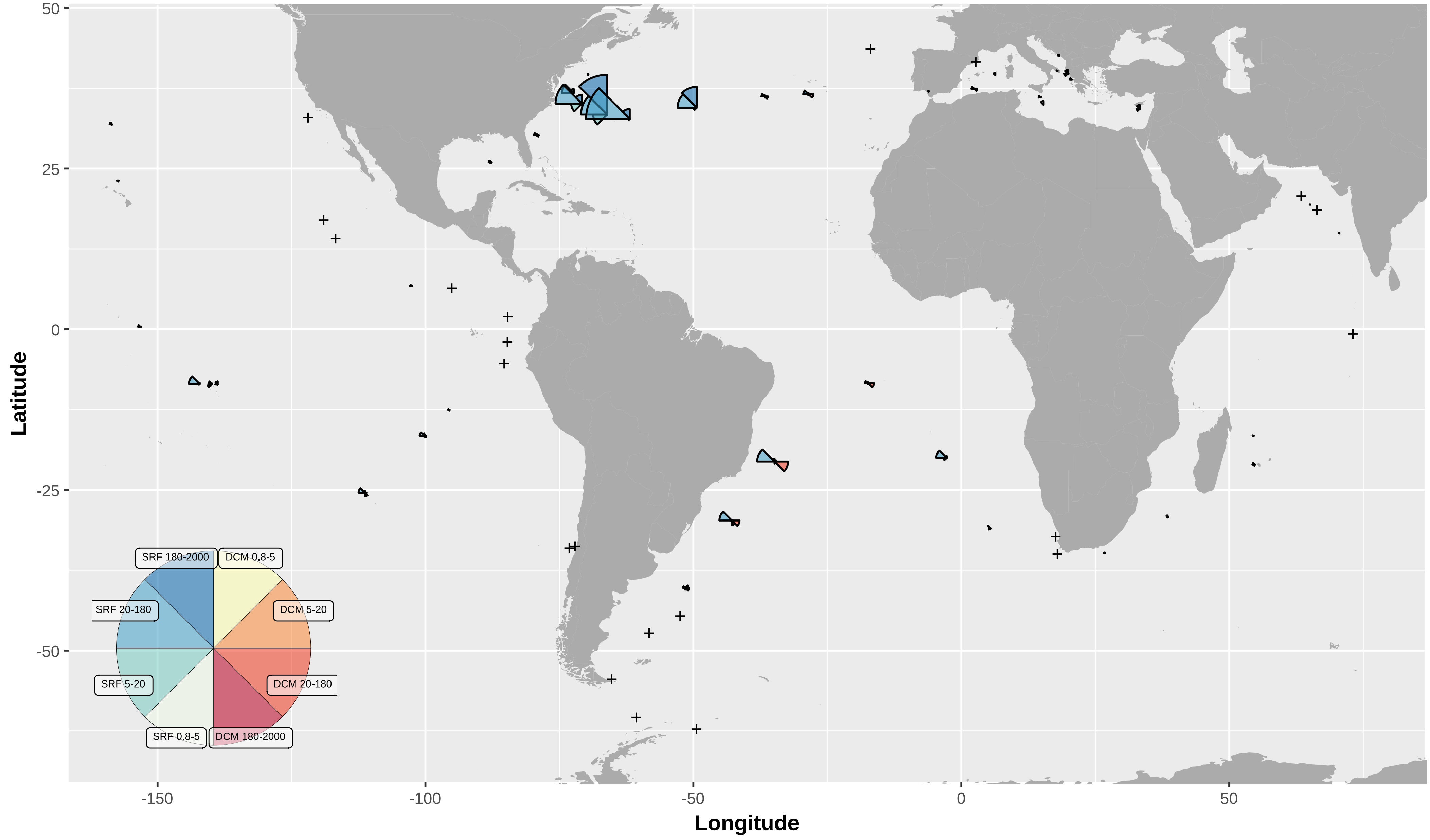


root;cellular organisms;Eukaryota;Opisthokonta;Metazoa;Eumetazoa;Bilateria

# Biogeolocalisation of MGT-v1 753      734 unigenes, 715 (97.41%) taxonomically assigned

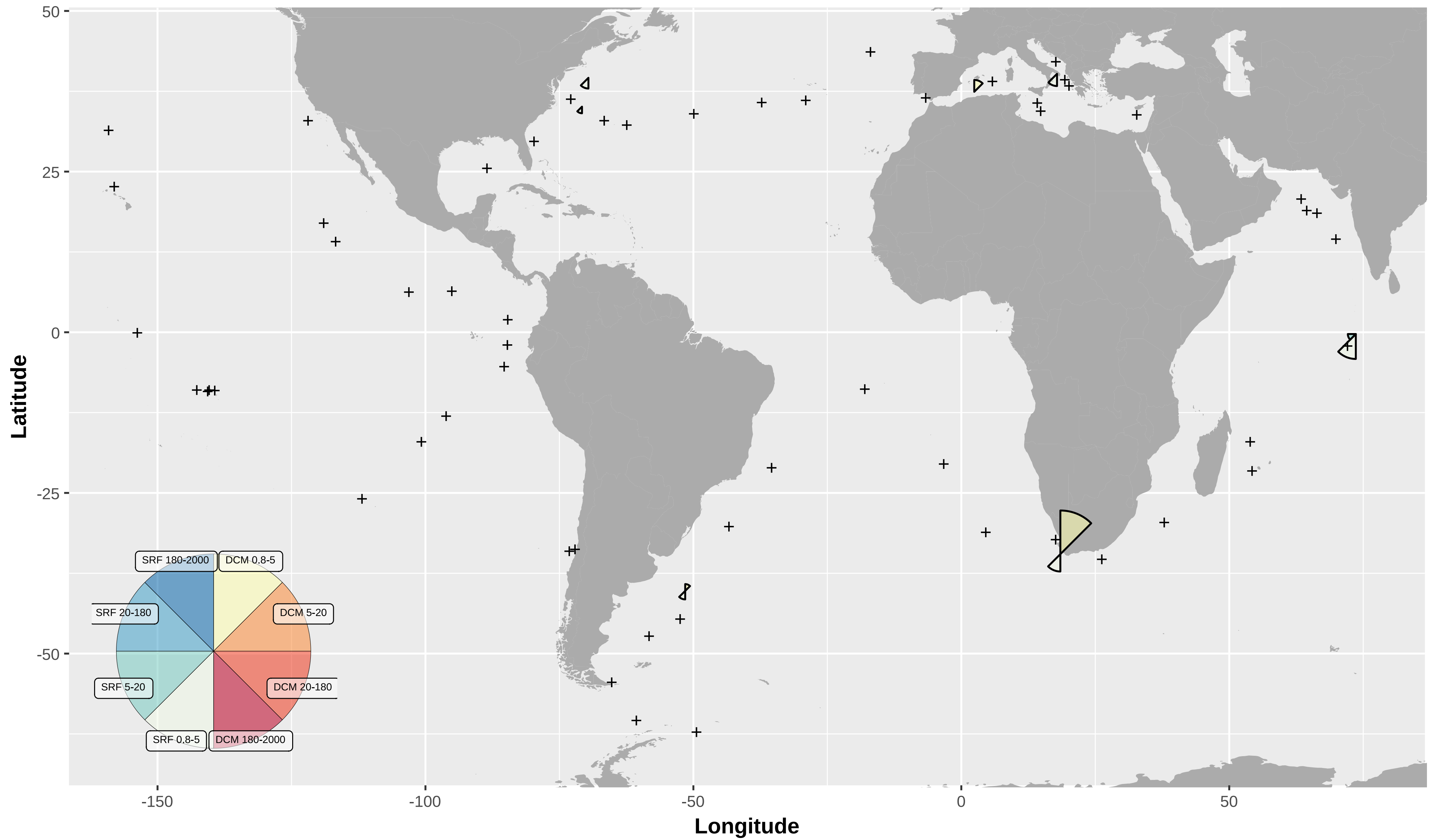


root;cellular organisms;Eukaryota;Viridiplantae;Chlorophyta;prasinophytes;Mamiellophyceae;Mamiellales;Mamiellaceae

## Biogeolocalisation of MGT-v1 754      733 unigenes, 724 (98.77%) taxonomically assigned

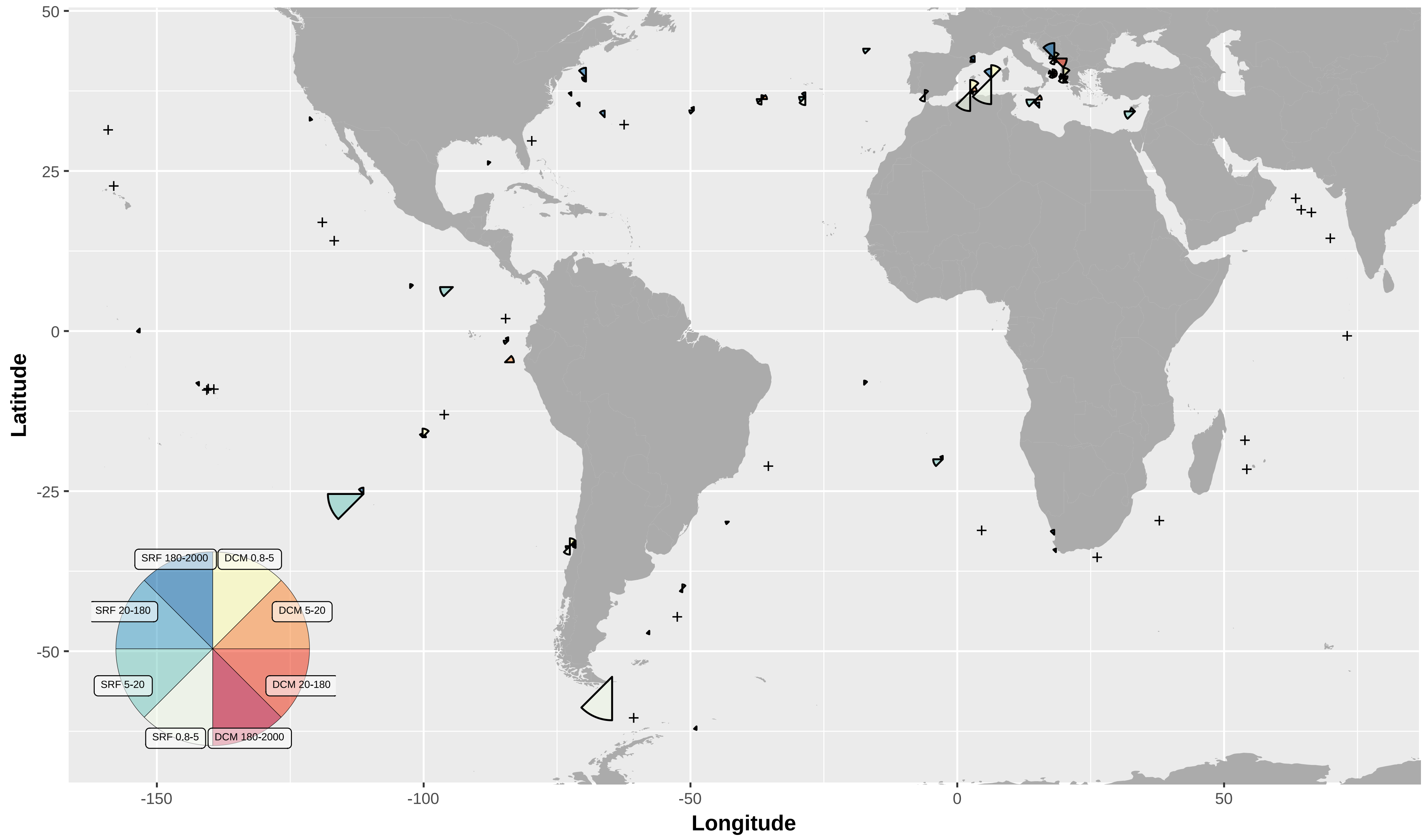


root;cellular organisms;Bacteria;Proteobacteria;Alphaproteobacteria;Rhodobacterales;Rhodobacteraceae

Biogeolocalisation of MGT-v1 755      726 unigenes, 674 (92.84%) taxonomically assigned

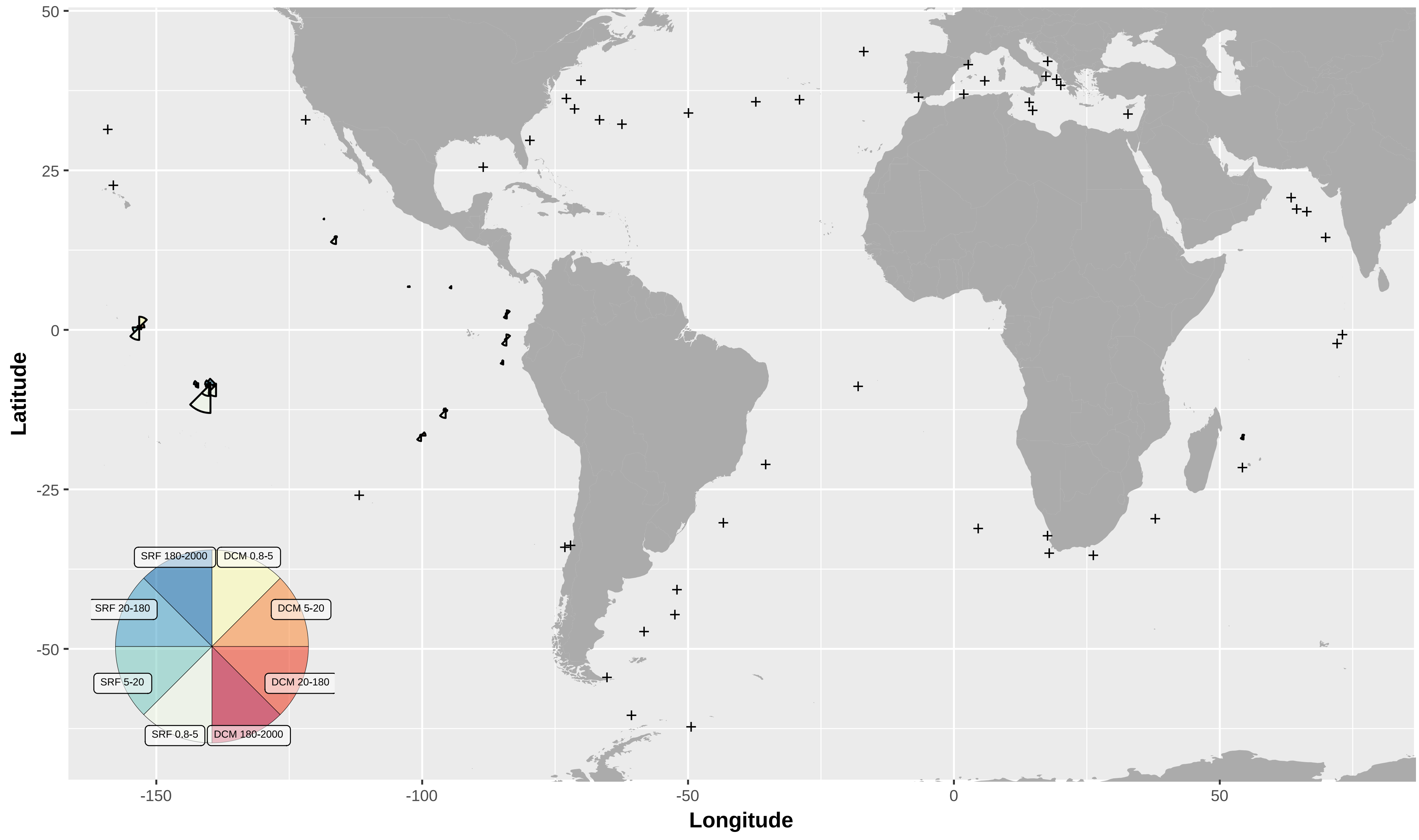


root;cellular organisms;Bacteria

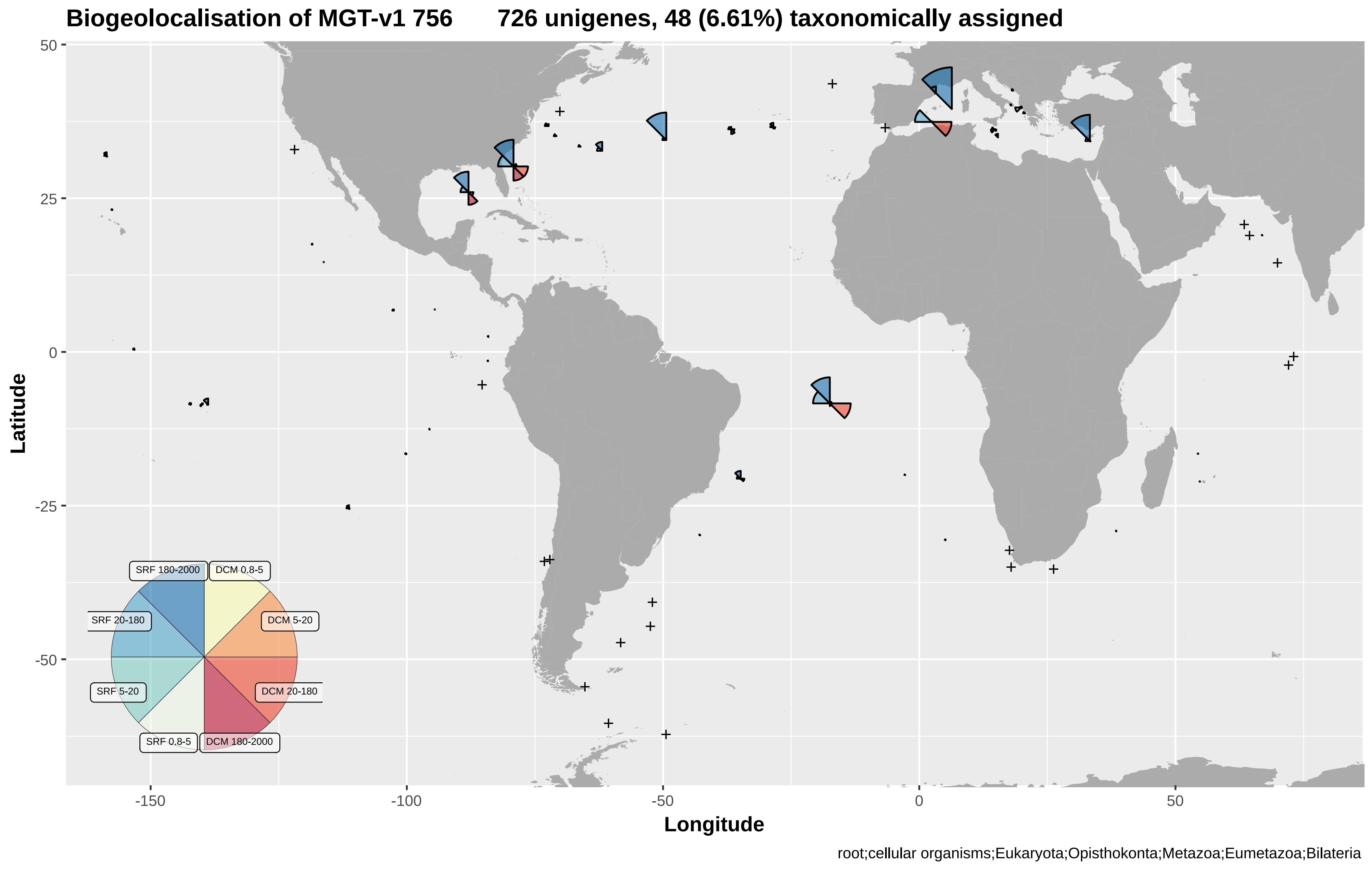
Biogeolocalisation of MGT-v1 756 726 unigenes, 48 (6.61%) taxonomically assigned
50
25
0
-25
-50
Latitude
-150
-100
-50
0
50
Longitude
SRF 180-2000
DCM 0.8-5
SRF 20-180
DCM 5-20
SRF 5-20
DCM 20-180
SRF 0.8-5
DCM 180-2000
root;cellular organisms;Eukaryota;Opisthokonta;Metazoa;Eumetazoa;Bilateria

Biogeolocalisation of MGT-v1 757 725 unigenes, 67 (9.24%) taxonomically assigned

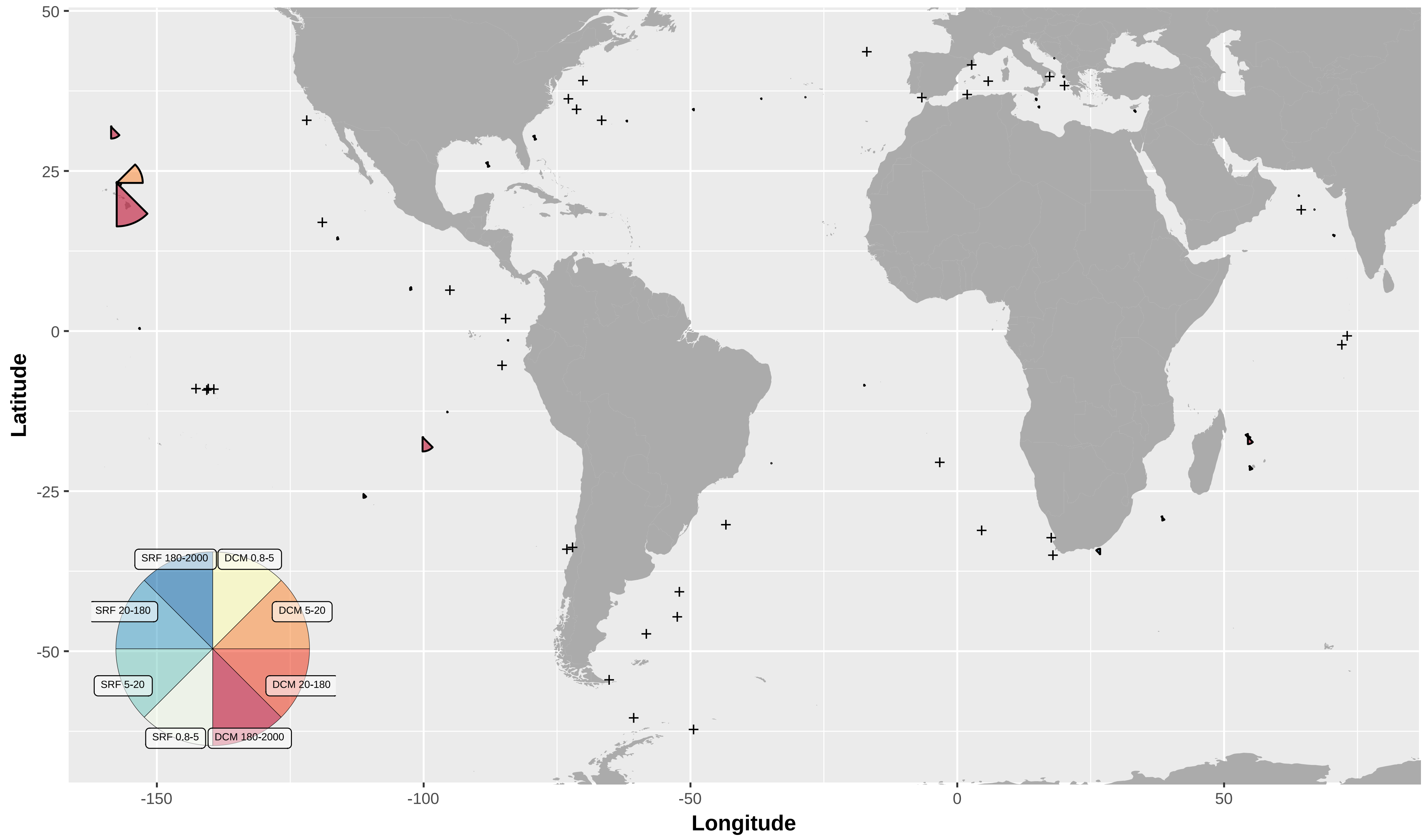


root;cellular organisms;Eukaryota;Opisthokonta;Metazoa;Eumetazoa

## Biogeolocalisation of MGT-v1 758      724 unigenes, 123 (16.99%) taxonomically assigned

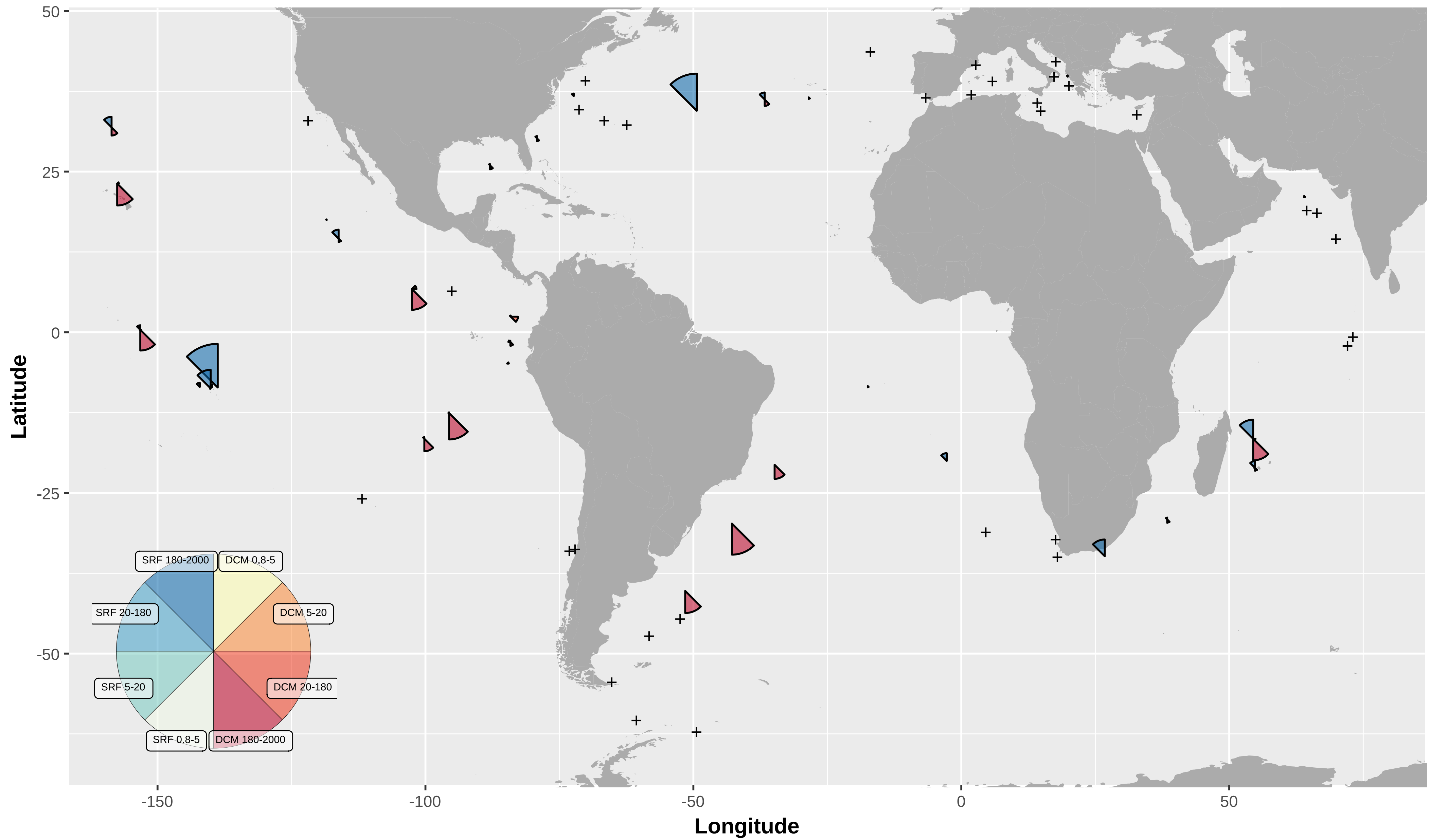


root;cellular organisms;Eukaryota;Opisthokonta;Metazoa;Eumetazoa;Bilateria

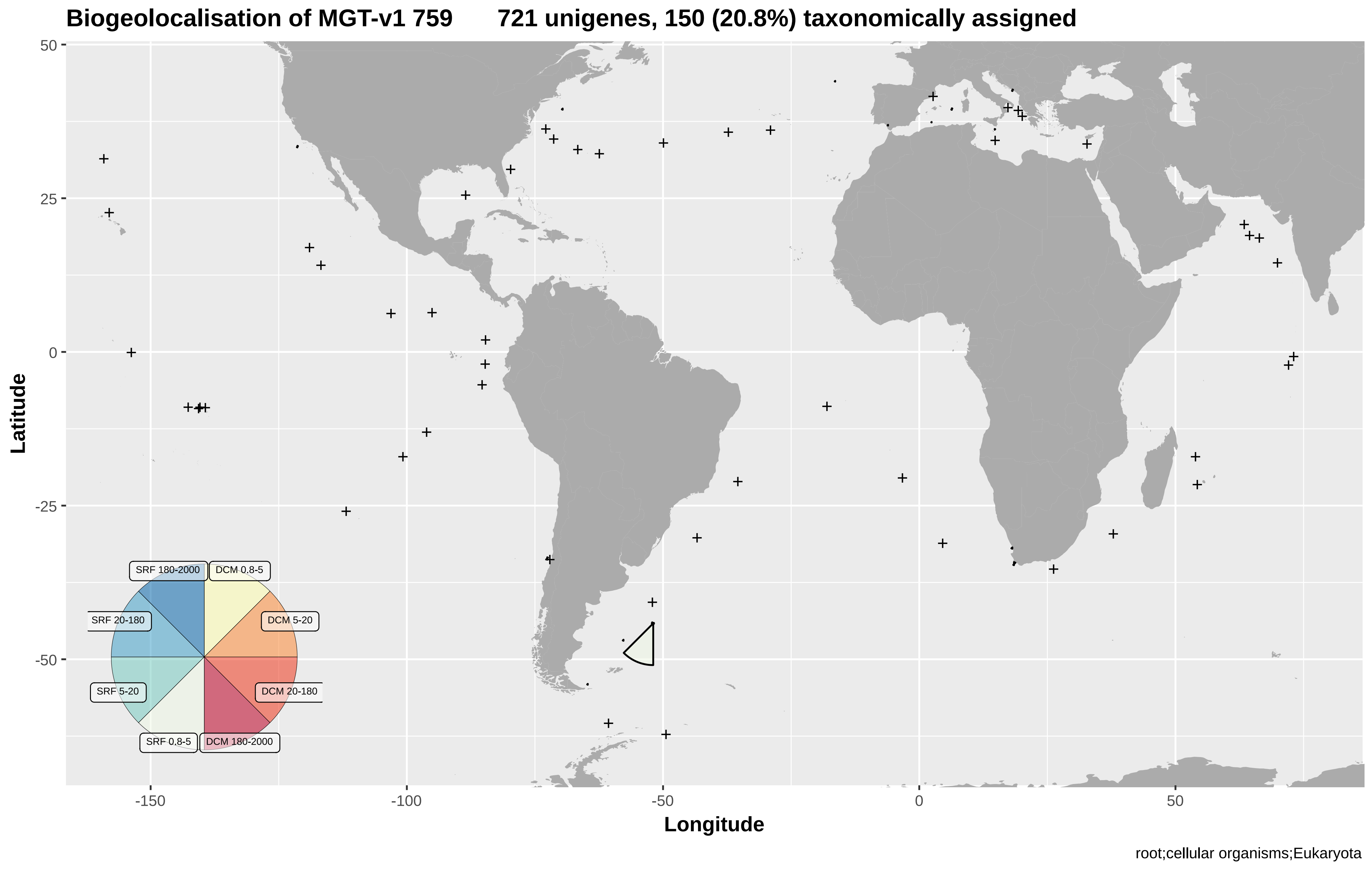
Biogeolocalisation of MGT-v1 759 721 unigenes, 150 (20.8%) taxonomically assigned
50
25
0
-25
-50
Latitude
-150
-100
-50
0
50
Longitude
SRF 180-2000
DCM 0.8-5
SRF 20-180
DCM 5-20
SRF 5-20
DCM 20-180
SRF 0.8-5
DCM 180-2000
root;cellular organisms;Eukaryota

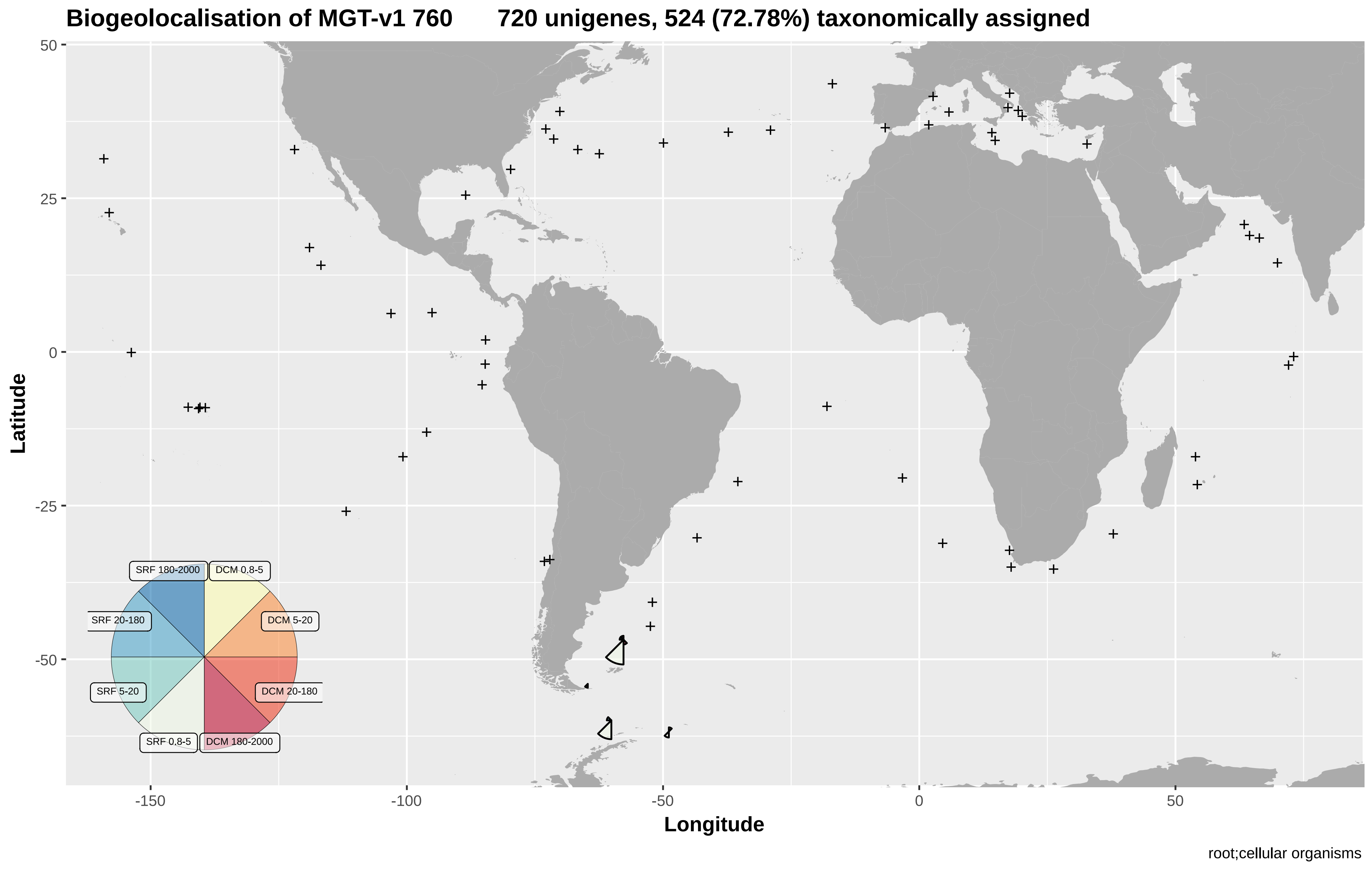

Biogeolocalisation of MGT-v1 760 720 unigenes, 524 (72.78%) taxonomically assigned
Latitude
50
25
0
-25
-50
Longitude
-150
-100
-50
0
50
SRF 180-2000
DCM 0.8-5
SRF 20-180
DCM 5-20
SRF 5-20
DCM 20-180
SRF 0.8-5
DCM 180-2000
root;cellular organisms

#### Biogeolocalisation of MGT-v1 761 719 unigenes, 632 (87.9%) taxonomically assigned

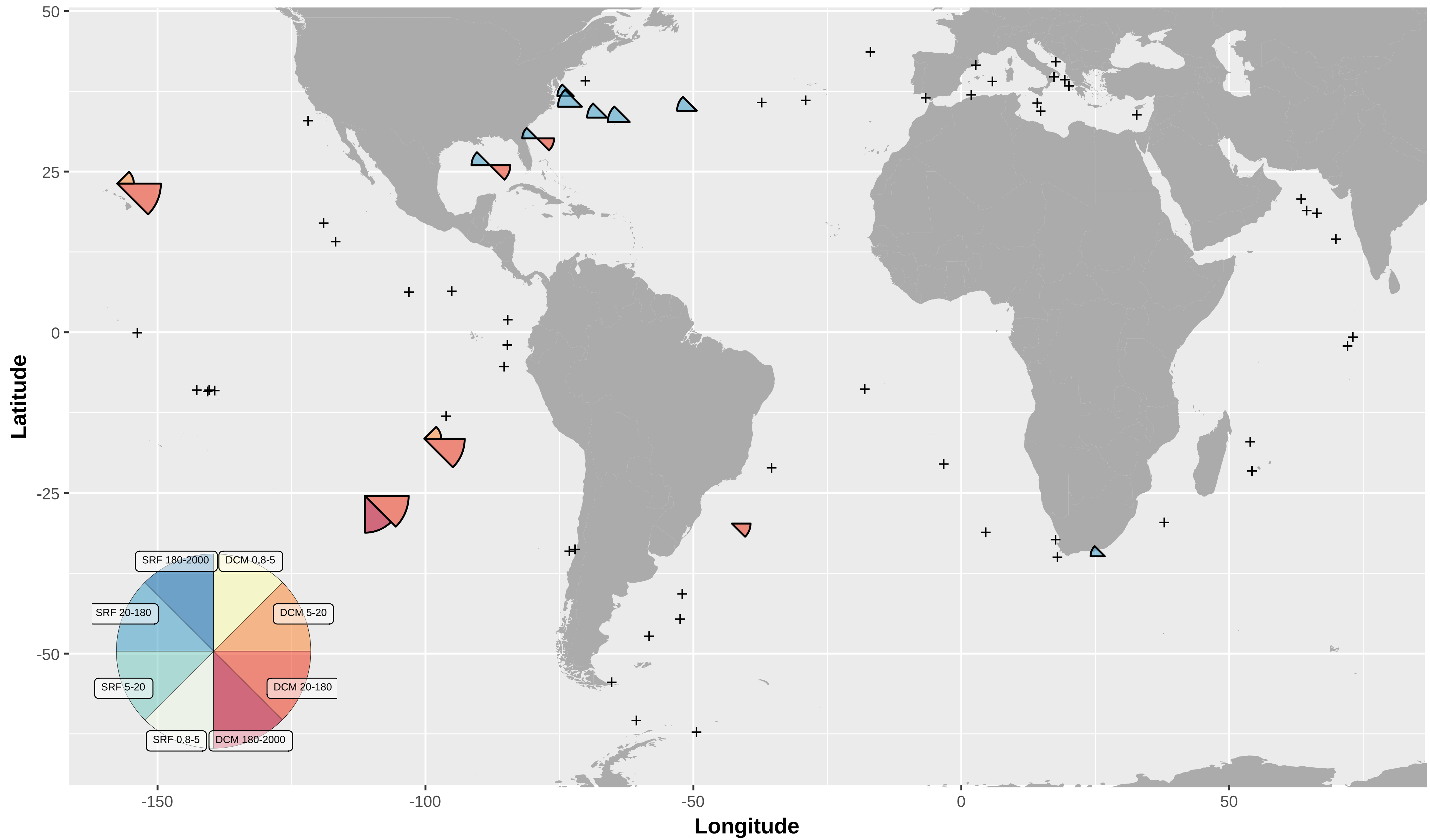


root;cellular organisms;Eukaryota;Opisthokonta;Metazoa;Eumetazoa;Bilateria;Protostomia;Ecdysozoa;Panarthropoda;Arthropoda;Mandibulata;Pancrustacea;Crustacea;Maxillopoda

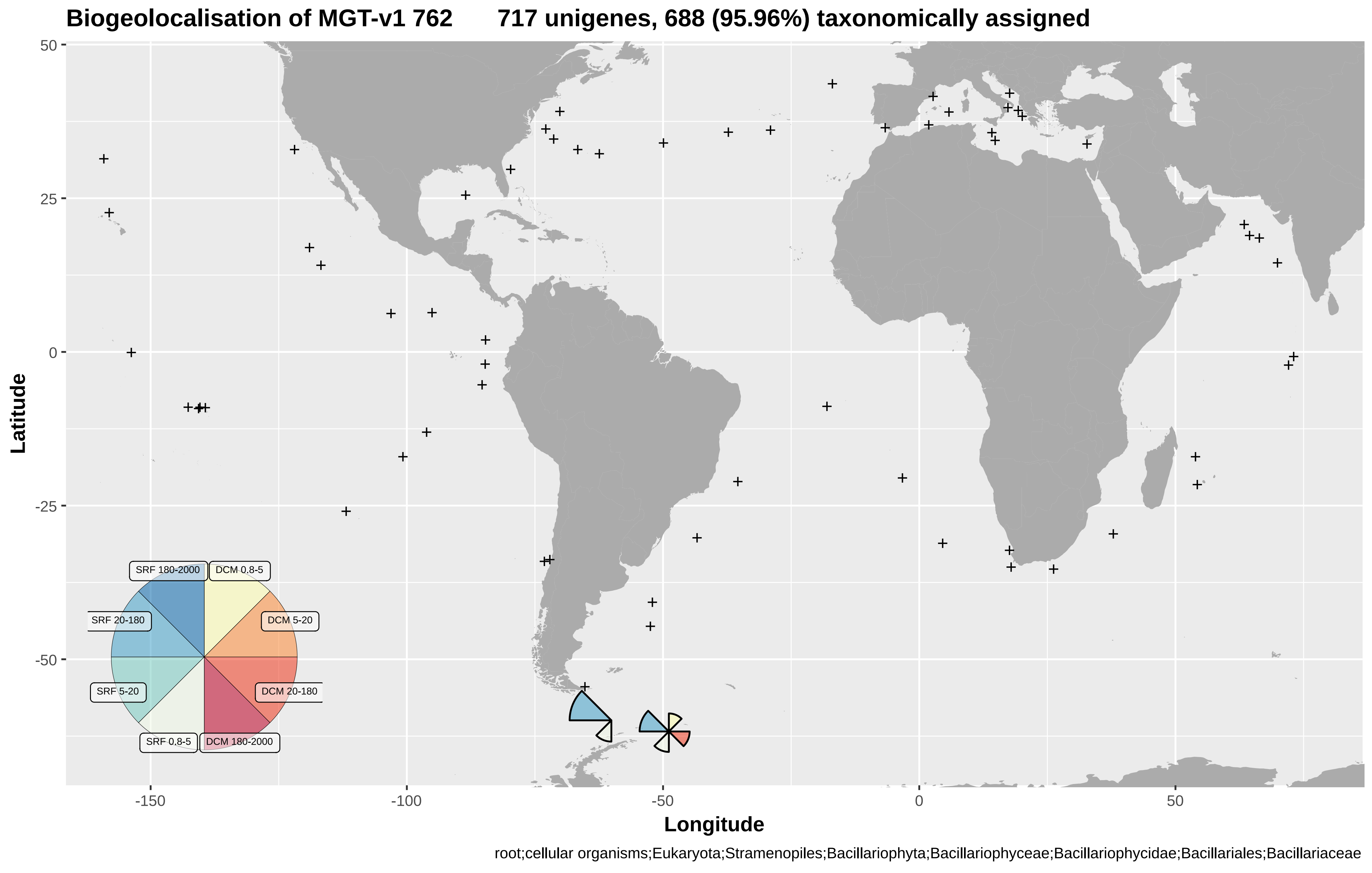

Biogeolocalisation of MGT-v1 762 717 unigenes, 688 (95.96%) taxonomically assigned
50
25
0
-25
-50
Latitude
-150
-100
-50
0
50
Longitude
SRF 180-2000
DCM 0.8-5
SRF 20-180
DCM 5-20
SRF 5-20
DCM 20-180
SRF 0.8-5
DCM 180-2000
root;cellular organisms;Eukaryota;Stramenopiles;Bacillariophyta;Bacillariophyceae;Bacillariophycidae;Bacillariales;Bacillariaceae

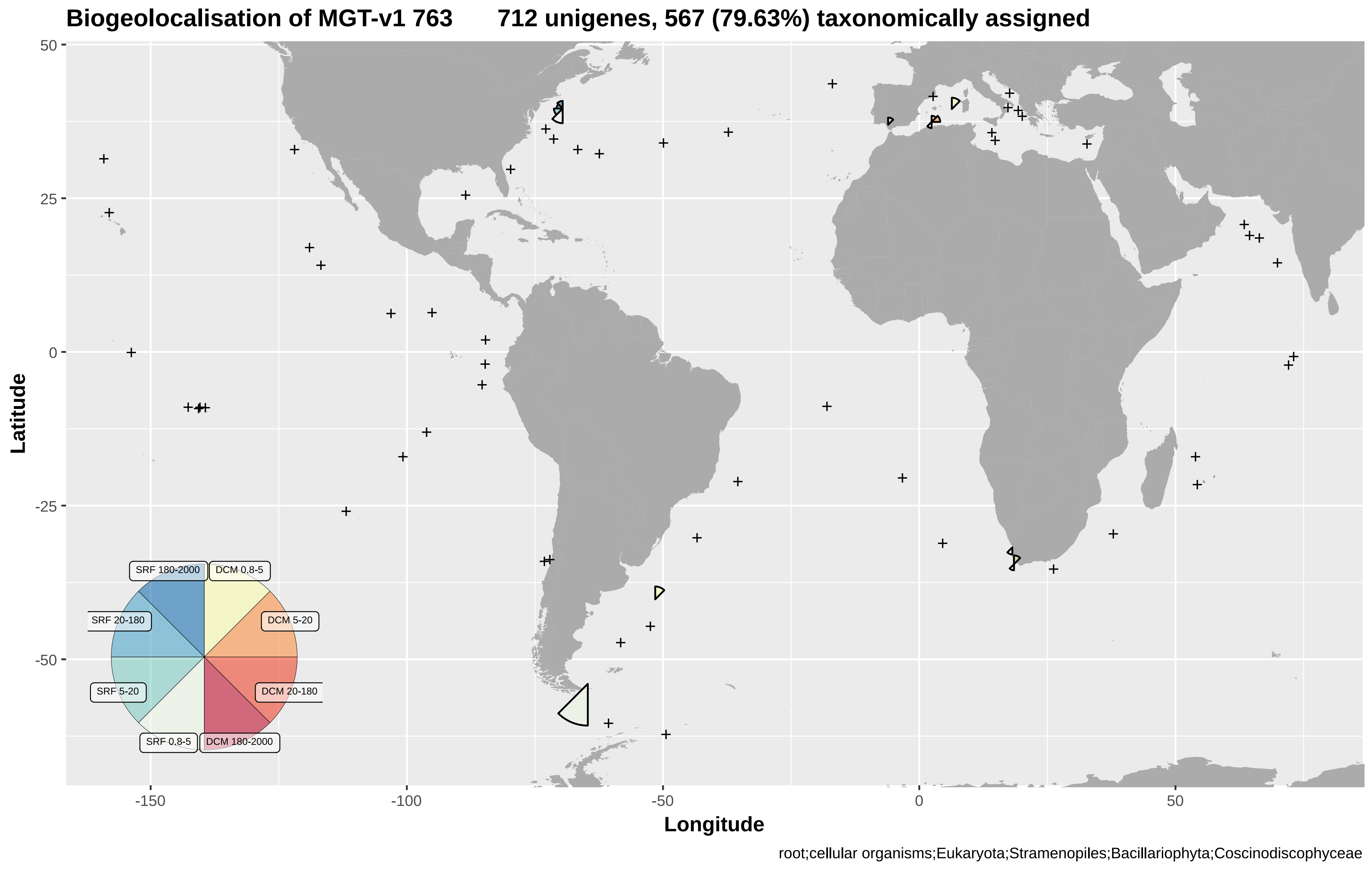

Biogeolocalisation of MGT-v1 763 712 unigenes, 567 (79.63%) taxonomically assigned
Latitude
50
25
0
-25
-50
-150
-100
-50
0
50
Longitude
SRF 180-2000
DCM 0.8-5
SRF 20-180
DCM 5-20
SRF 5-20
DCM 20-180
SRF 0.8-5
DCM 180-2000
root;cellular organisms;Eukaryota;Stramenopiles;Bacillariophyta;Coscinodiscophyceae

**Biogeolocalisation of MGT-v1 764      711 unigenes, 54 (7.59%) taxonomically assigned**

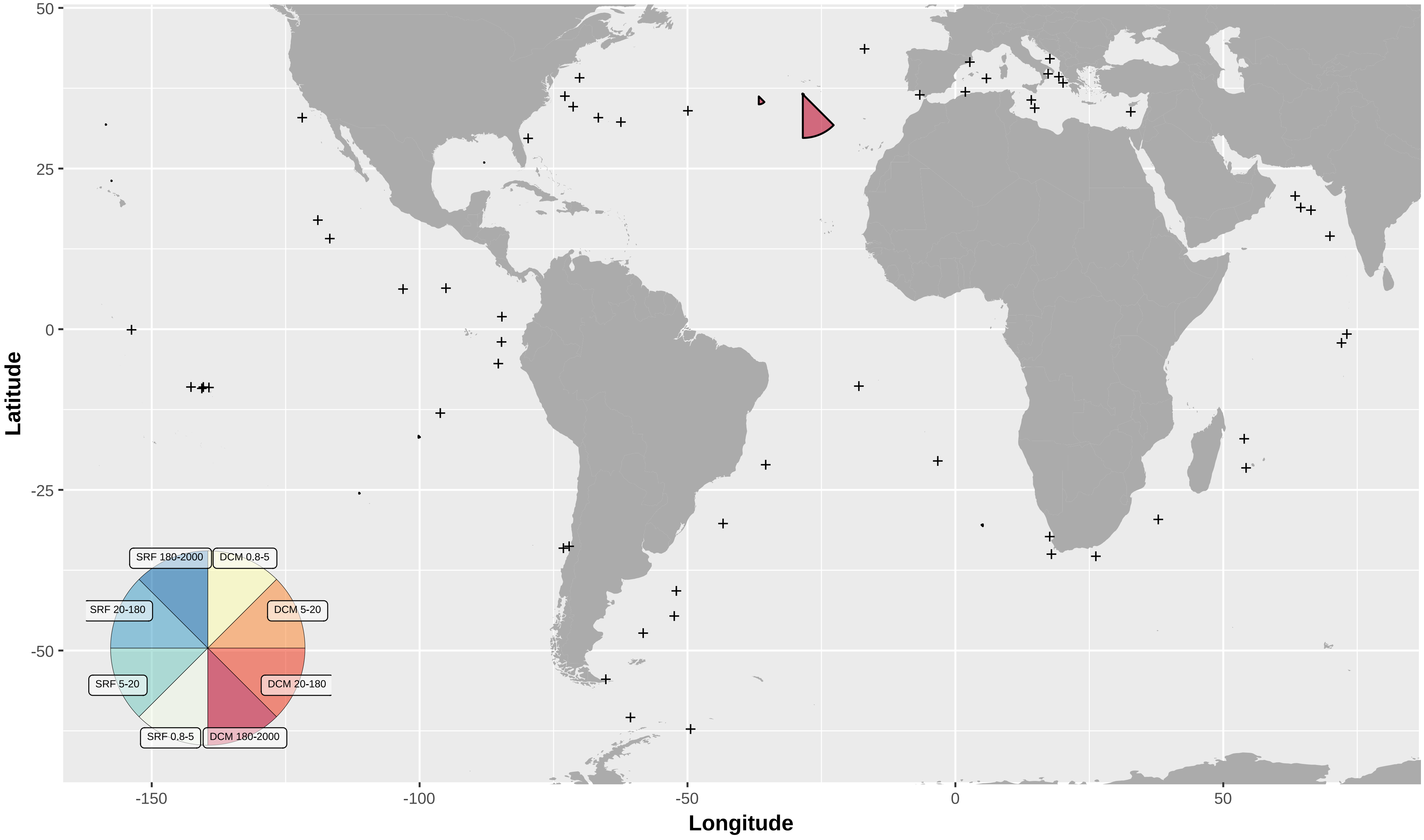


root;cellular organisms;Eukaryota;Opisthokonta;Metazoa;Eumetazoa

## Biogeolocalisation of MGT-v1 765      711 unigenes, 40 (5.63%) taxonomically assigned

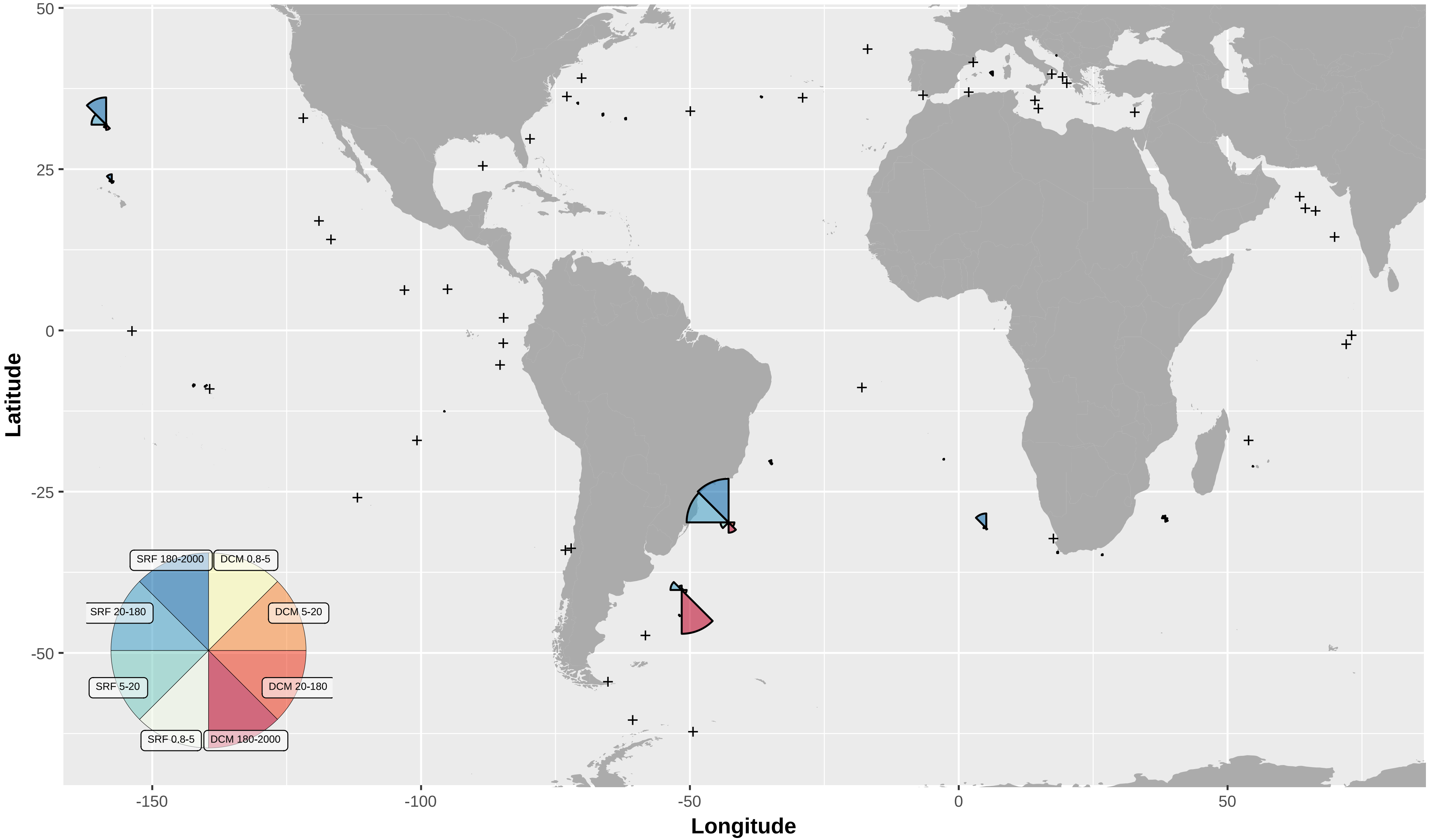


root;cellular organisms;Eukaryota;Opisthokonta;Metazoa;Eumetazoa;Bilateria

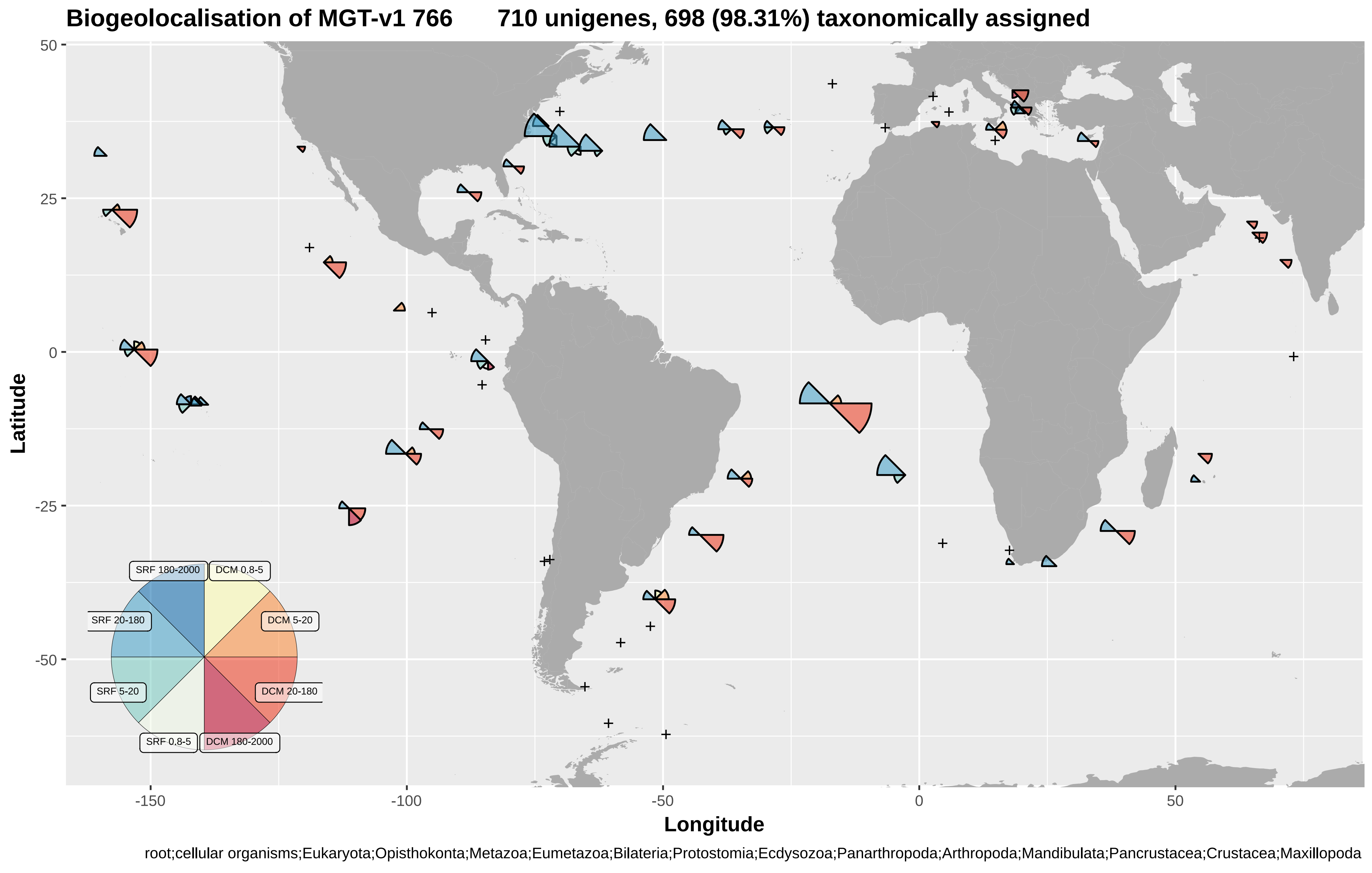
Biogeolocalisation of MGT-v1 766
710 unigenes, 698 (98.31%) taxonomically assigned
Latitude
Longitude
50
25
0
-25
-50
-150
-100
-50
0
50
SRF 180-2000
DCM 0.8-5
SRF 20-180
DCM 5-20
SRF 5-20
DCM 20-180
SRF 0.8-5
DCM 180-2000
root;cellular organisms;Eukaryota;Opisthokonta;Metazoa;Eumetazoa;Bilateria;Protostomia;Ecdysozoa;Panarthropoda;Arthropoda;Mandibulata;Pancrustacea;Crustacea;Maxillopoda

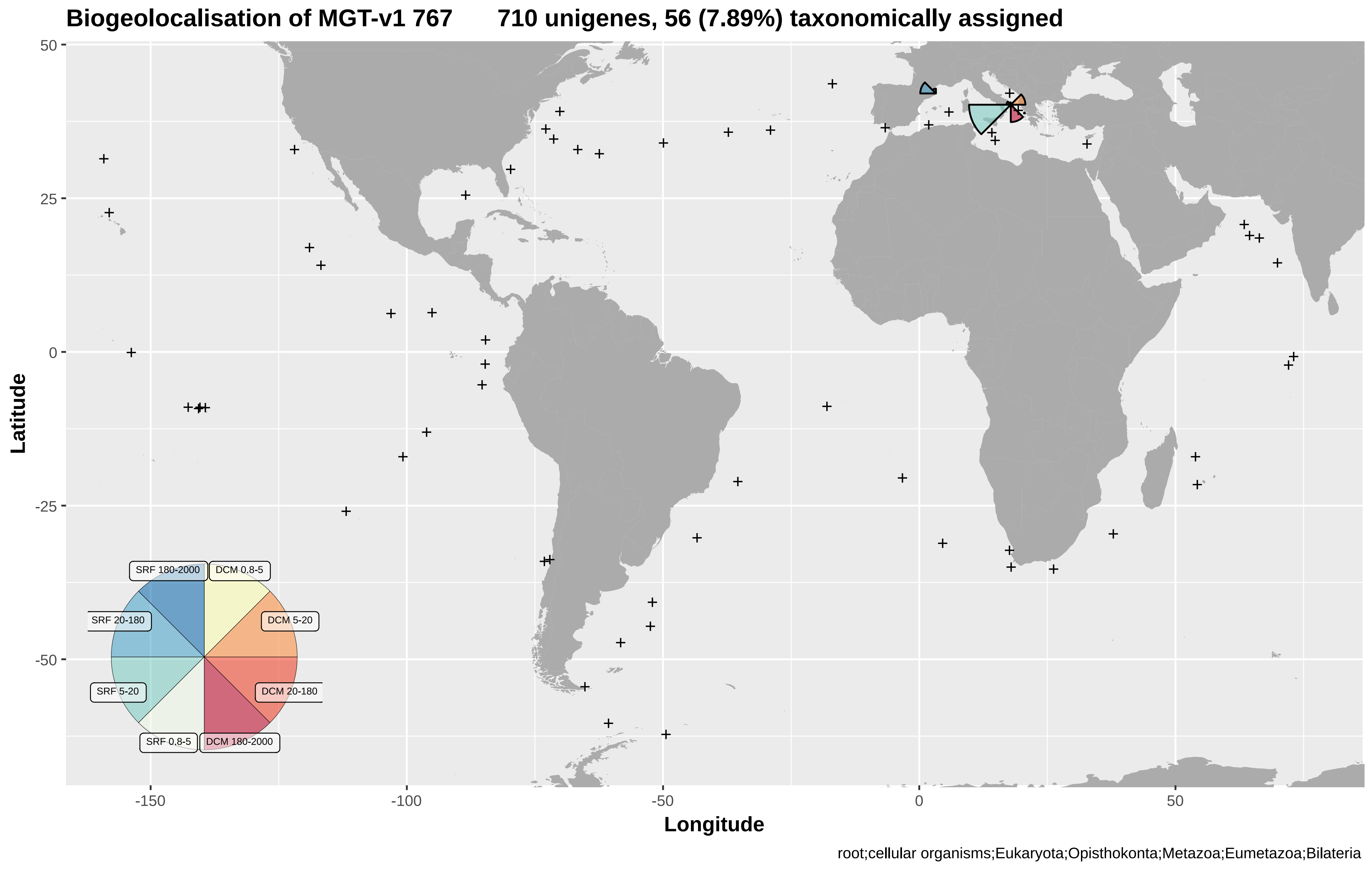

Biogeolocalisation of MGT-v1 767      710 unigenes, 56 (7.89%) taxonomically assigned
50
25
0
-25
-50
Latitude
-150
-100
-50
0
50
Longitude
SRF 180-2000
DCM 0.8-5
SRF 20-180
DCM 5-20
SRF 5-20
DCM 20-180
SRF 0.8-5
DCM 180-2000
root;cellular organisms;Eukaryota;Opisthokonta;Metazoa;Eumetazoa;Bilateria

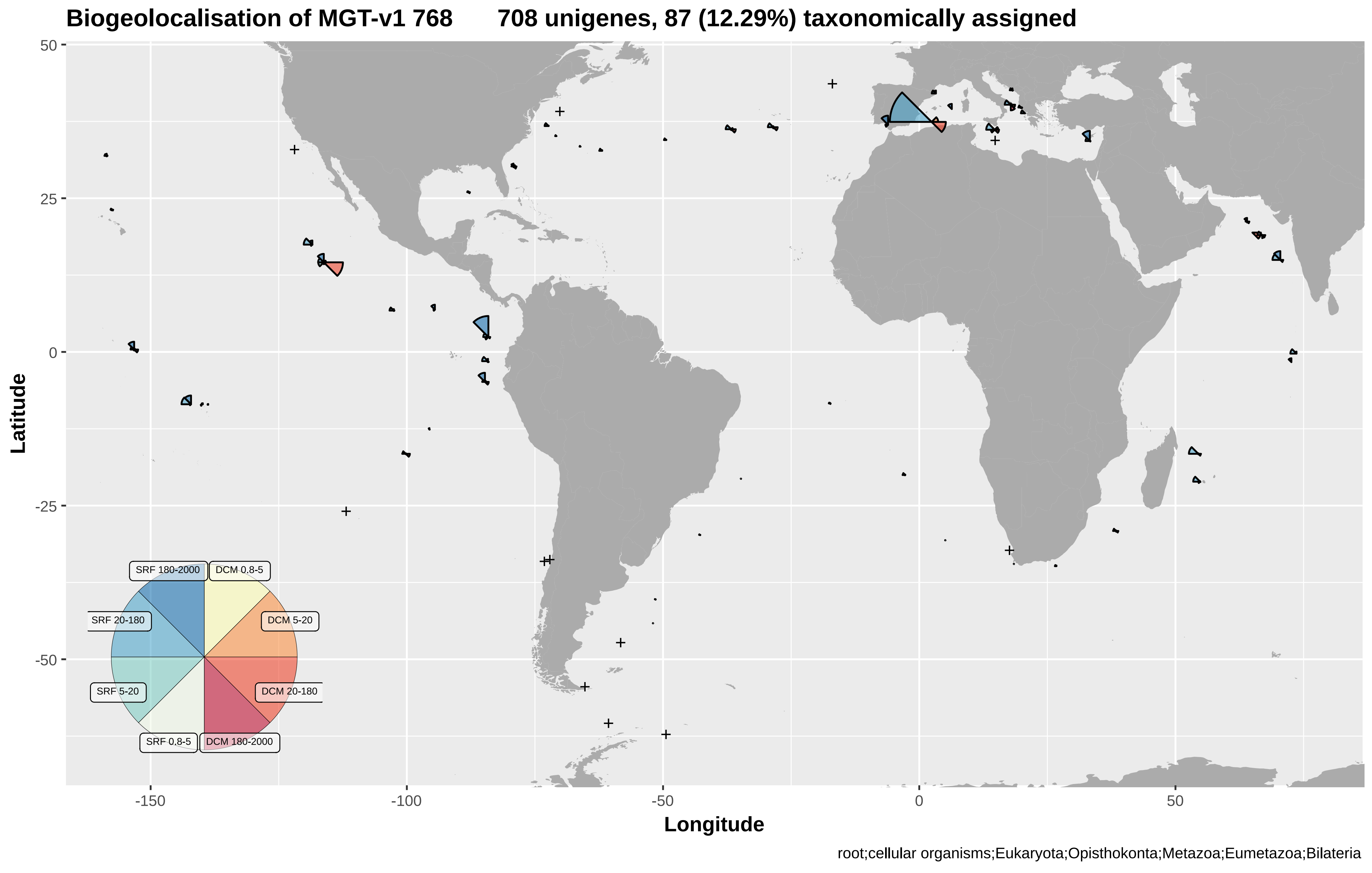

Biogeolocalisation of MGT-v1 768 708 unigenes, 87 (12.29%) taxonomically assigned
50
25
0
-25
-50
Latitude
-150
-100
-50
0
50
Longitude
SRF 180-2000
DCM 0.8-5
SRF 20-180
DCM 5-20
SRF 5-20
DCM 20-180
SRF 0.8-5
DCM 180-2000
root;cellular organisms;Eukaryota;Opisthokonta;Metazoa;Eumetazoa;Bilateria

## Biogeolocalisation of MGT-v1 769      705 unigenes, 142 (20.14%) taxonomically assigned

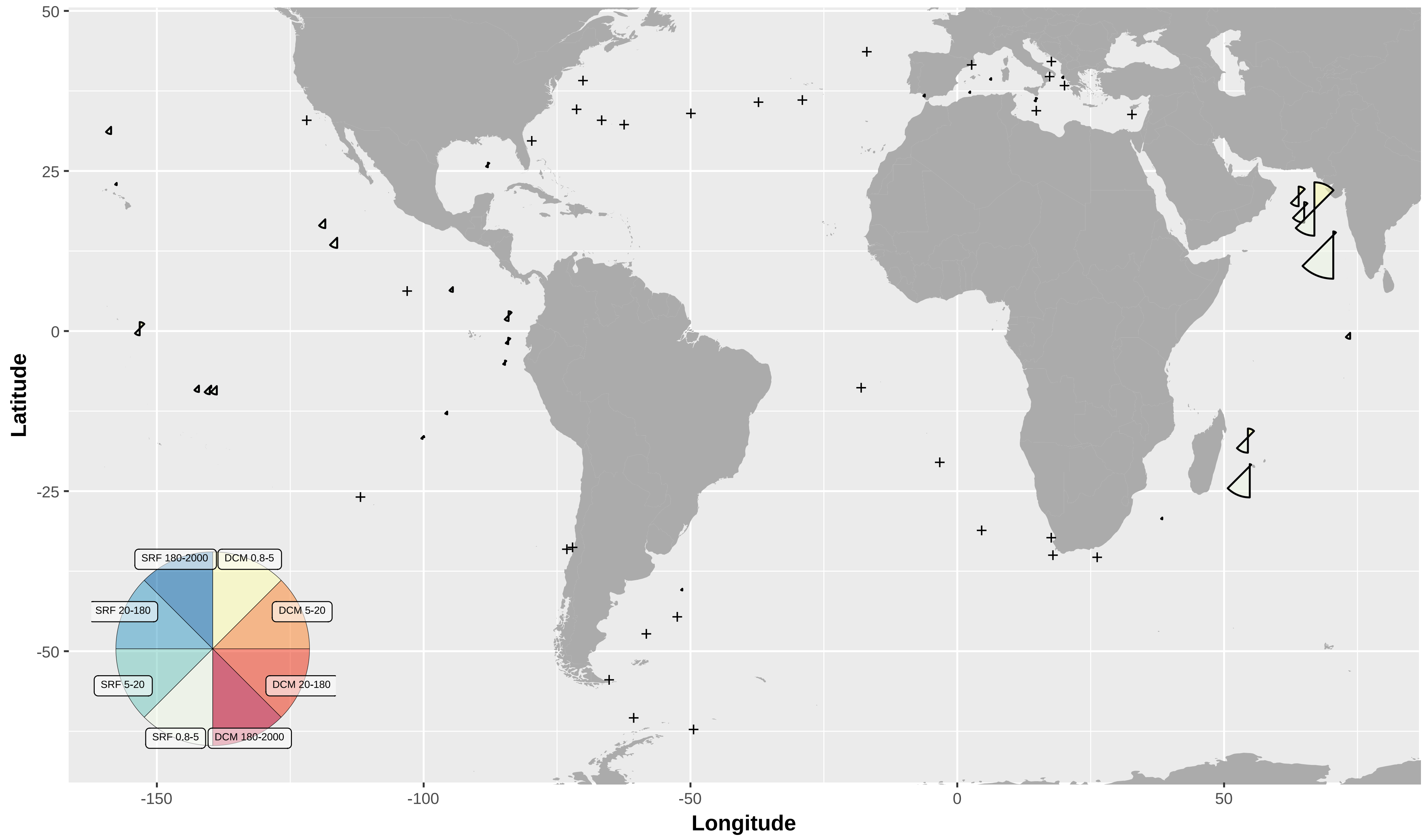


root;cellular organisms;Eukaryota;Stramenopiles

## Biogeolocalisation of MGT-v1 770      705 unigenes, 365 (51.77%) taxonomically assigned

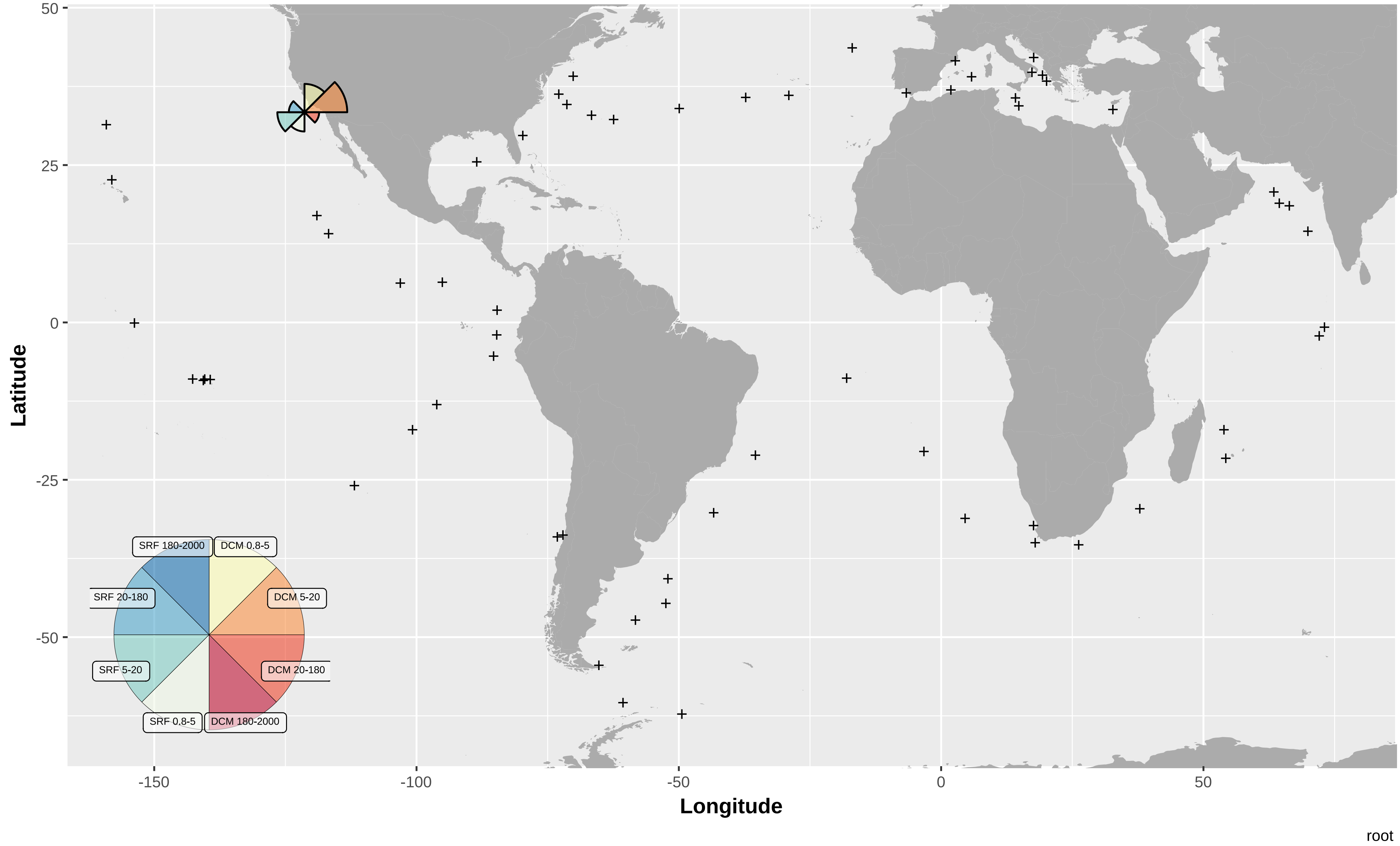

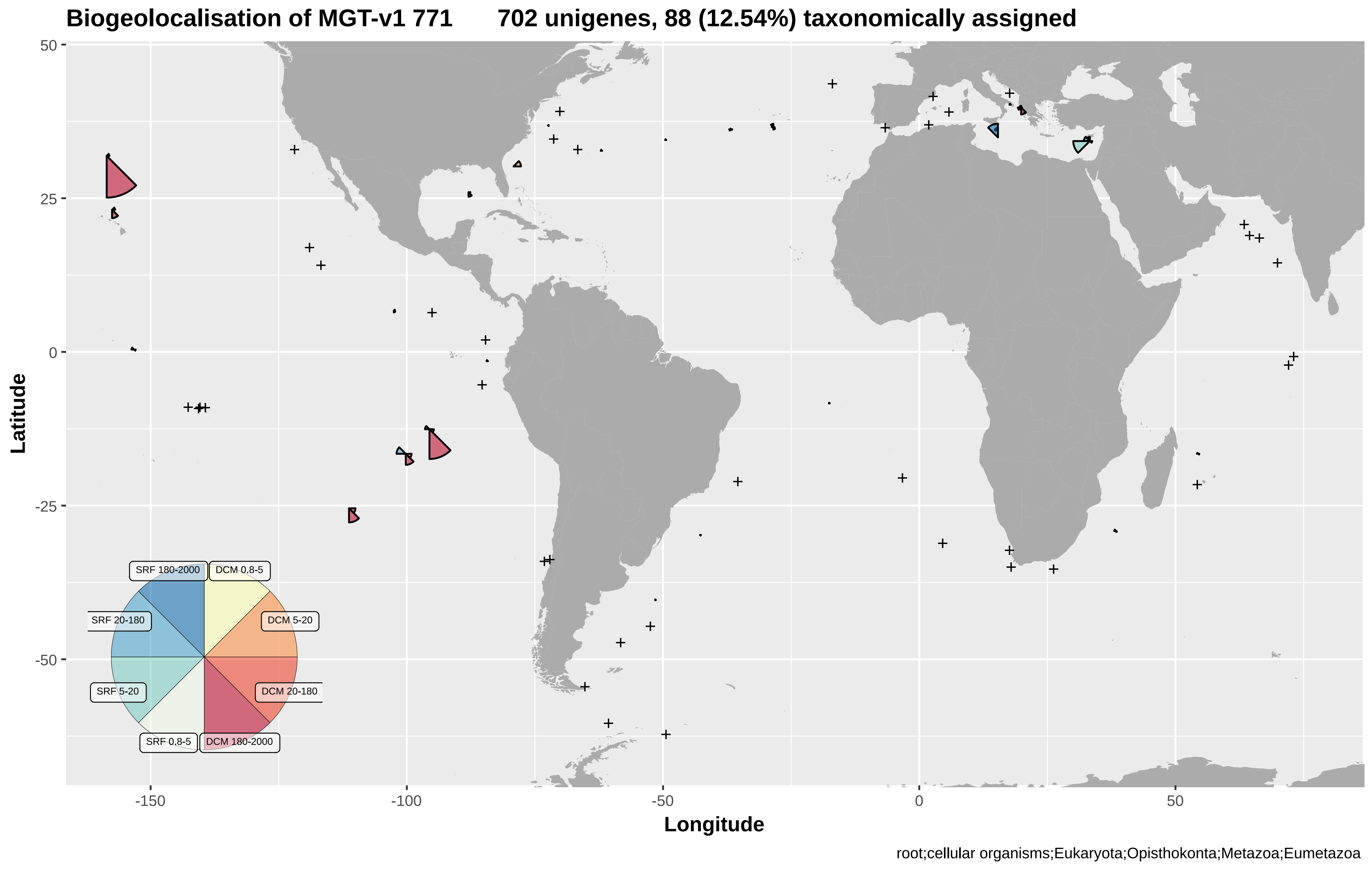
Biogeolocalisation of MGT-v1 771 702 unigenes, 88 (12.54%) taxonomically assigned
50
25
0
-25
-50
Latitude
-150
-100
-50
0
50
Longitude
SRF 180-2000
DCM 0.8-5
SRF 20-180
DCM 5-20
SRF 5-20
DCM 20-180
SRF 0.8-5
DCM 180-2000
root;cellular organisms;Eukaryota;Opisthokonta;Metazoa;Eumetazoa

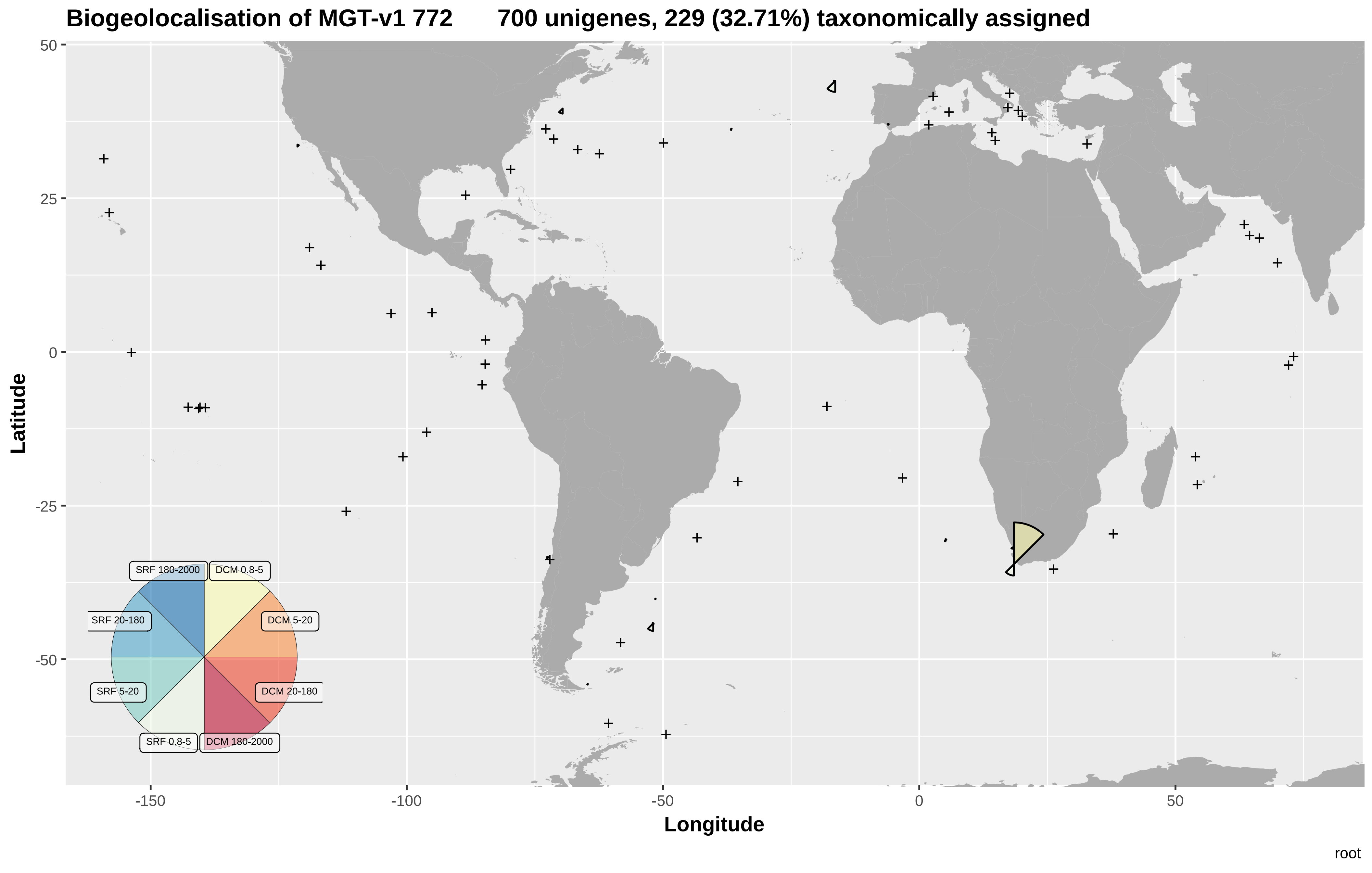
Biogeolocalisation of MGT-v1 772 700 unigenes, 229 (32.71%) taxonomically assigned
50
25
0
-25
-50
Latitude
-150
-100
-50
0
50
Longitude
SRF 180-2000
DCM 0.8-5
SRF 20-180
DCM 5-20
SRF 5-20
DCM 20-180
SRF 0.8-5
DCM 180-2000
root

#### Biogeolocalisation of MGT-v1 773 699 unigenes, 312 (44.64%) taxonomically assigned

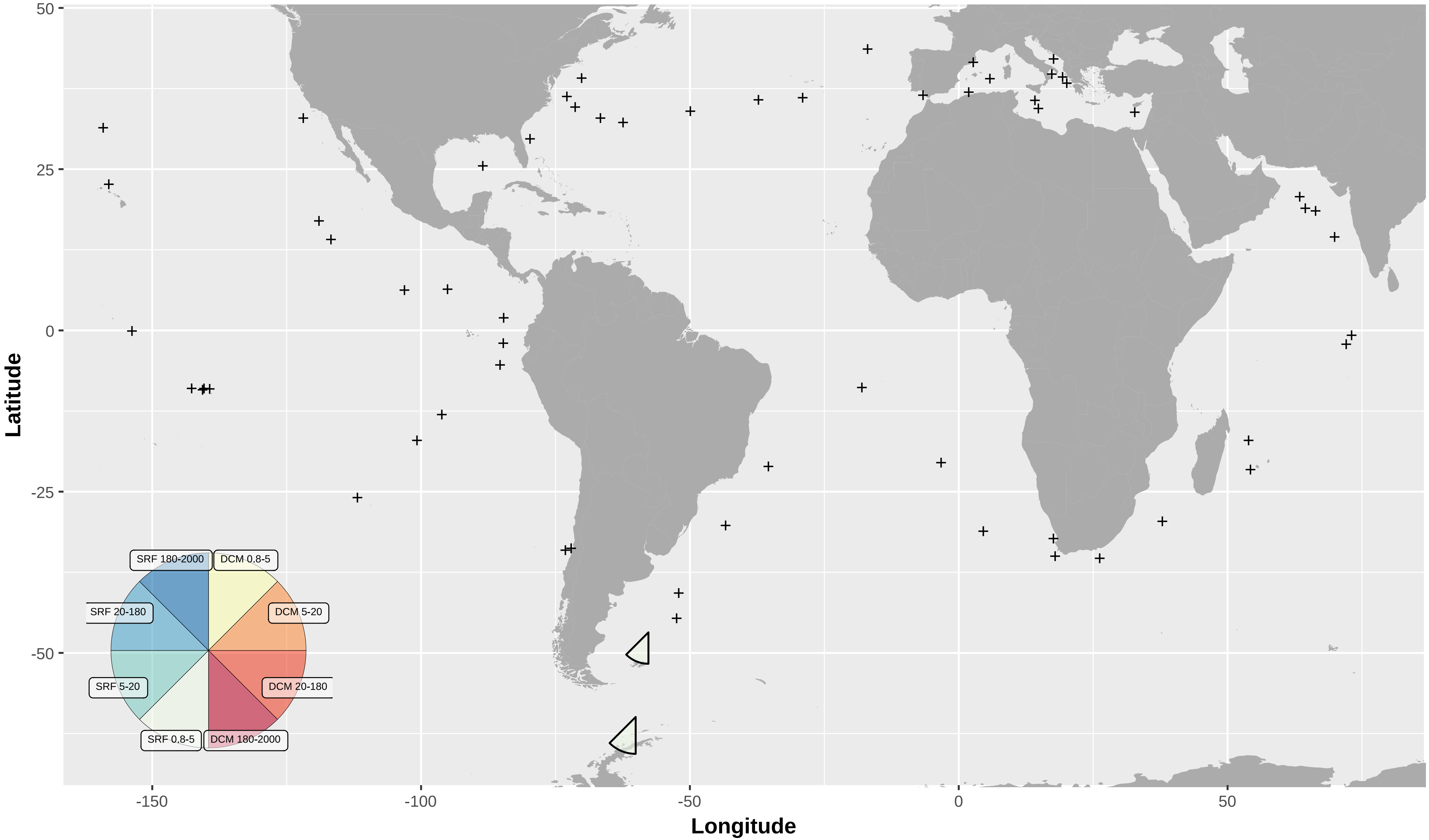


root;cellular organisms;Eukaryota

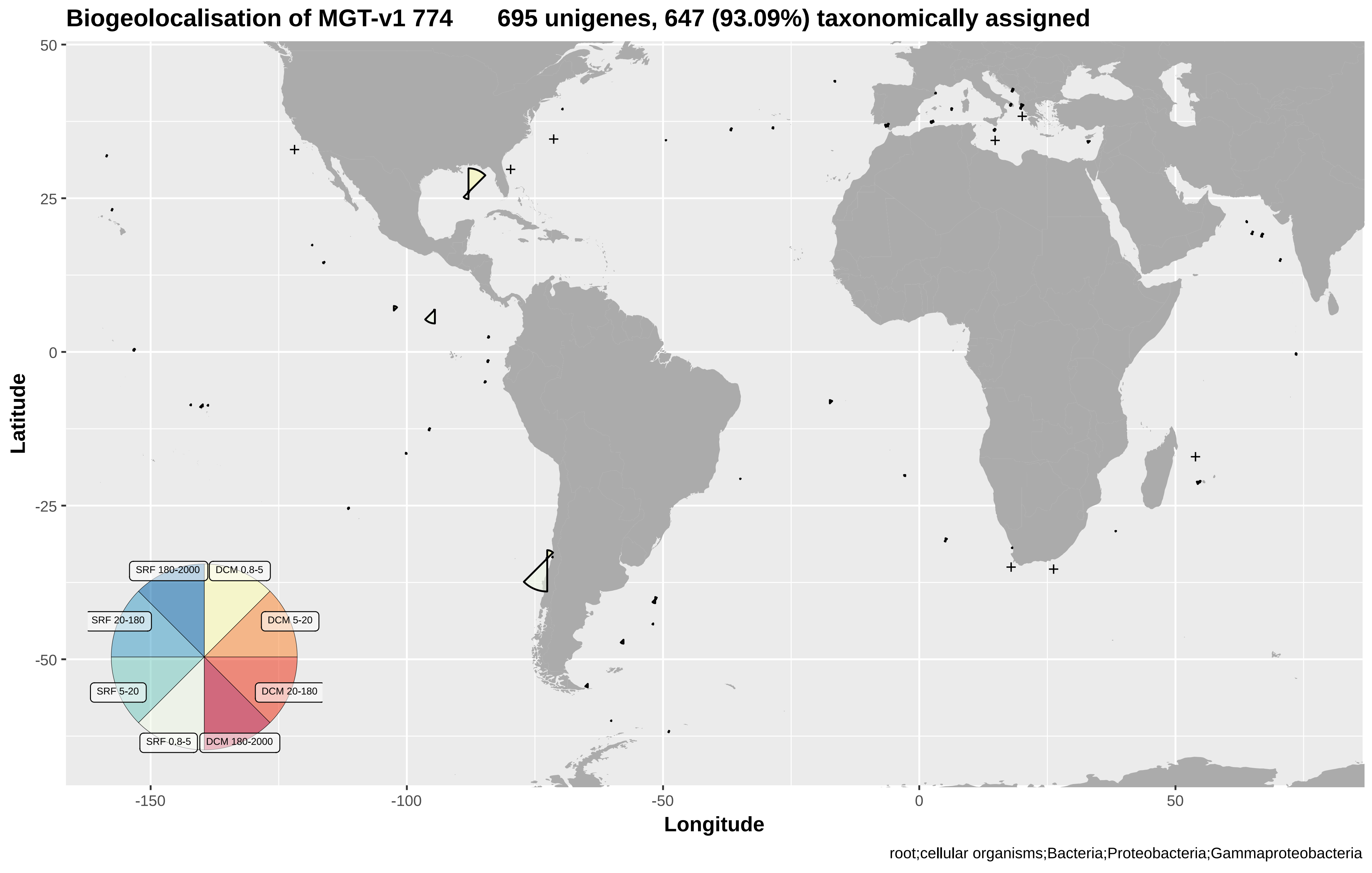
Biogeolocalisation of MGT-v1 774 695 unigenes, 647 (93.09%) taxonomically assigned
50
25
0
-25
-50
Latitude
-150
-100
-50
0
50
Longitude
SRF 180-2000
DCM 0.8-5
SRF 20-180
DCM 5-20
SRF 5-20
DCM 20-180
SRF 0.8-5
DCM 180-2000
root;cellular organisms;Bacteria;Proteobacteria;Gammaproteobacteria

## Biogeolocalisation of MGT-v1 775      694 unigenes, 51 (7.35%) taxonomically assigned

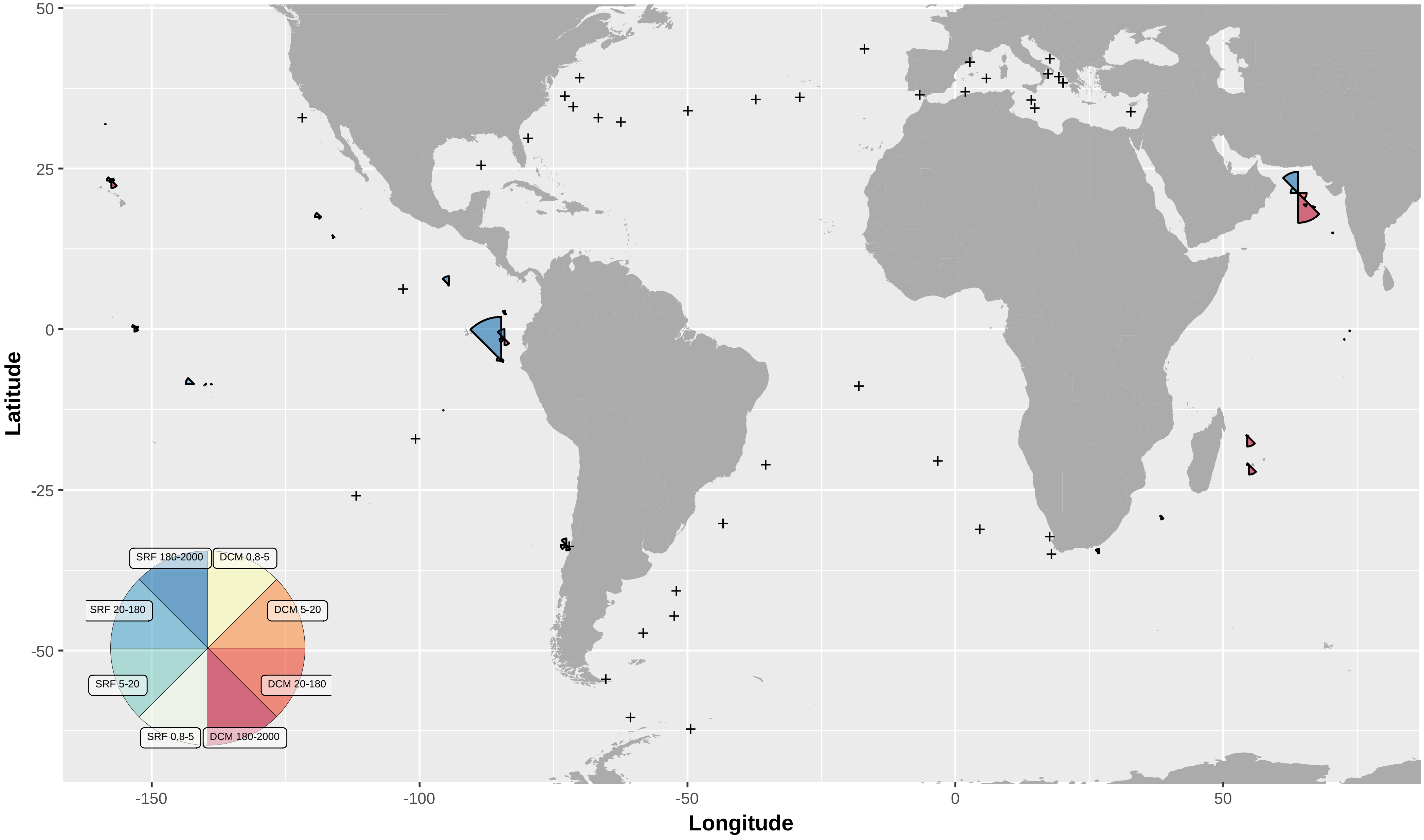


root;cellular organisms;Eukaryota;Opisthokonta;Metazoa;Eumetazoa;Bilateria

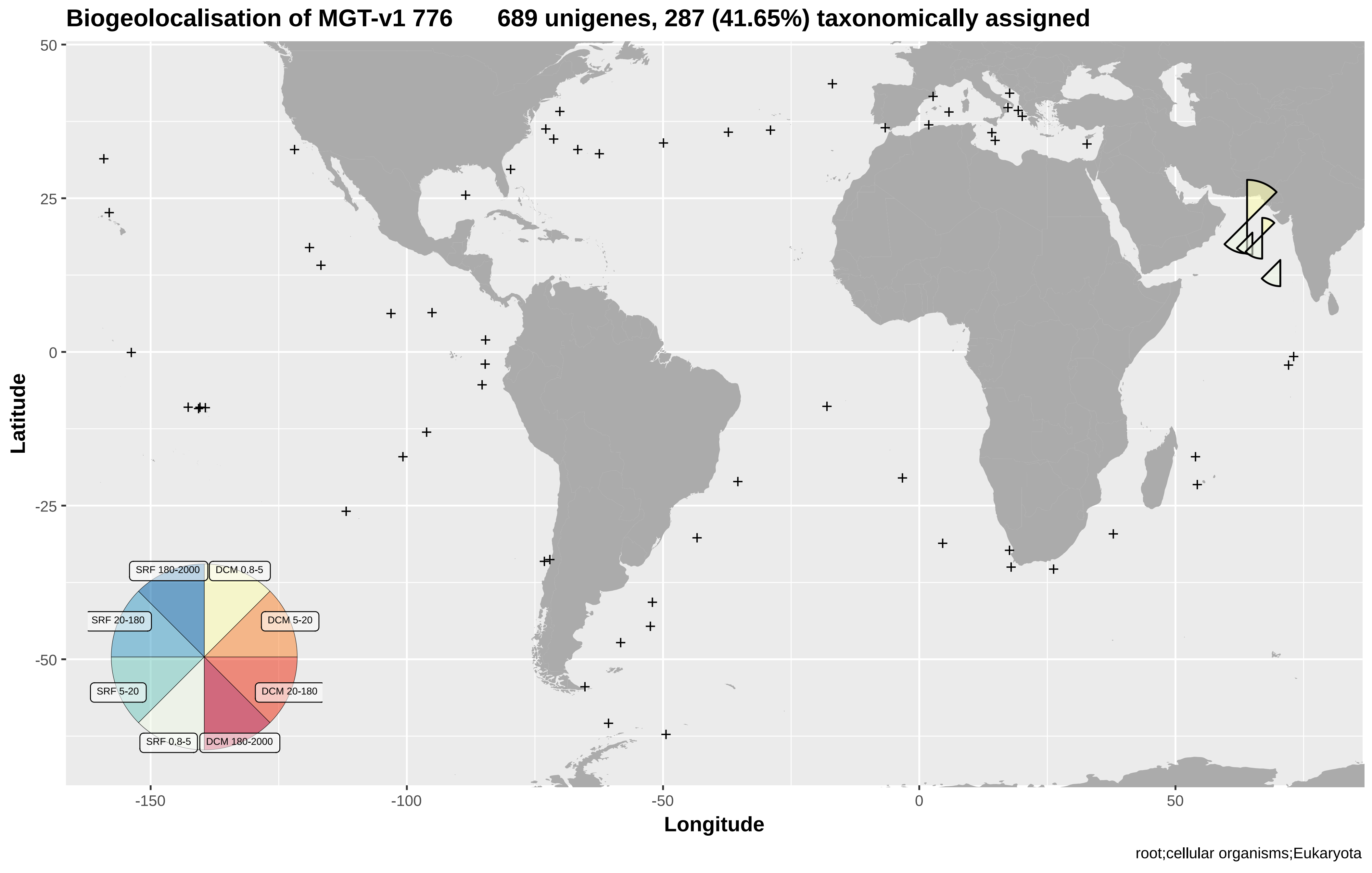

Biogeolocalisation of MGT-v1 776 689 unigenes, 287 (41.65%) taxonomically assigned
Latitude
Longitude
50
25
0
-25
-50
-150
-100
-50
0
50
SRF 180-2000
DCM 0.8-5
SRF 20-180
DCM 5-20
SRF 5-20
DCM 20-180
SRF 0.8-5
DCM 180-2000
root;cellular organisms;Eukaryota

#### Biogeolocalisation of MGT-v1 777 689 unigenes, 668 (96.95%) taxonomically assigned

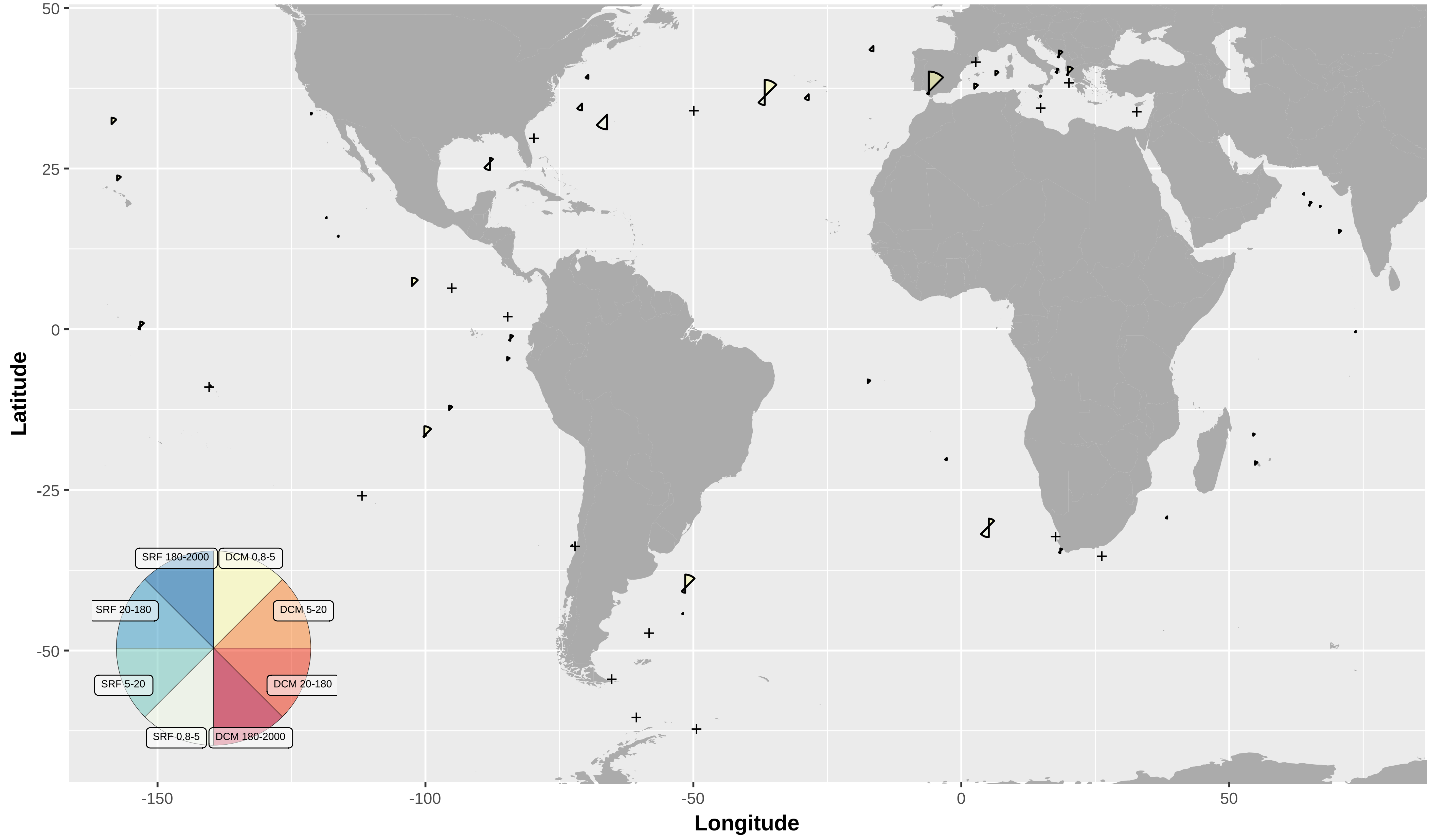


root;cellular organisms;Bacteria;Proteobacteria;Alphaproteobacteria

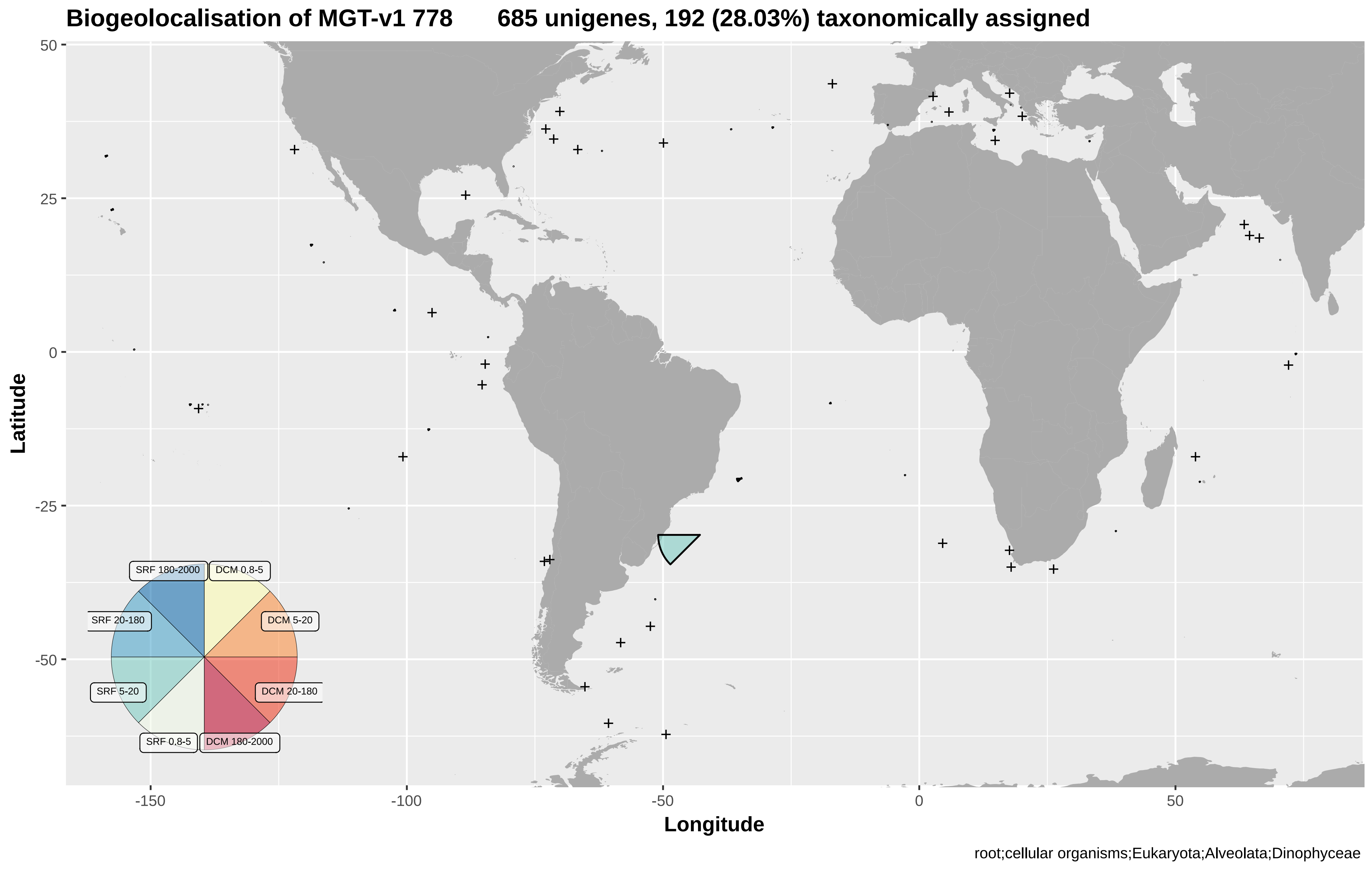
Biogeolocalisation of MGT-v1 778 685 unigenes, 192 (28.03%) taxonomically assigned
Latitude
Longitude
50
25
0
-25
-50
-150
-100
-50
0
50
SRF 180-2000
DCM 0.8-5
SRF 20-180
DCM 5-20
SRF 5-20
DCM 20-180
SRF 0.8-5
DCM 180-2000
root;cellular organisms;Eukaryota;Alveolata;Dinophyceae

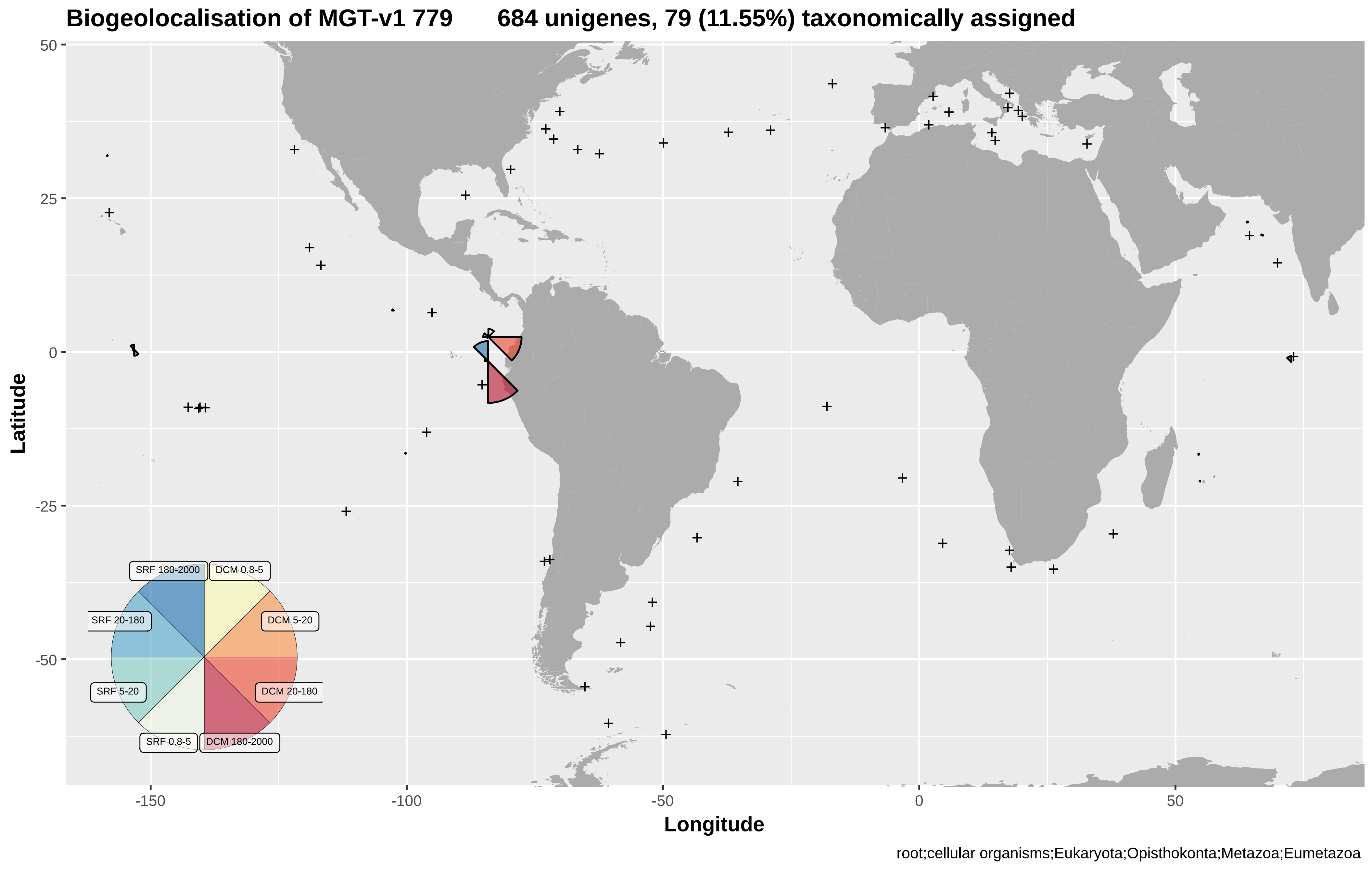

Biogeolocalisation of MGT-v1 779 684 unigenes, 79 (11.55%) taxonomically assigned
50
25
0
-25
-50
Latitude
-150
-100
-50
0
50
Longitude
SRF 180-2000
DCM 0.8-5
SRF 20-180
DCM 5-20
SRF 5-20
DCM 20-180
SRF 0.8-5
DCM 180-2000
root;cellular organisms;Eukaryota;Opisthokonta;Metazoa;Eumetazoa

## Biogeolocalisation of MGT-v1 780      683 unigenes, 75 (10.98%) taxonomically assigned

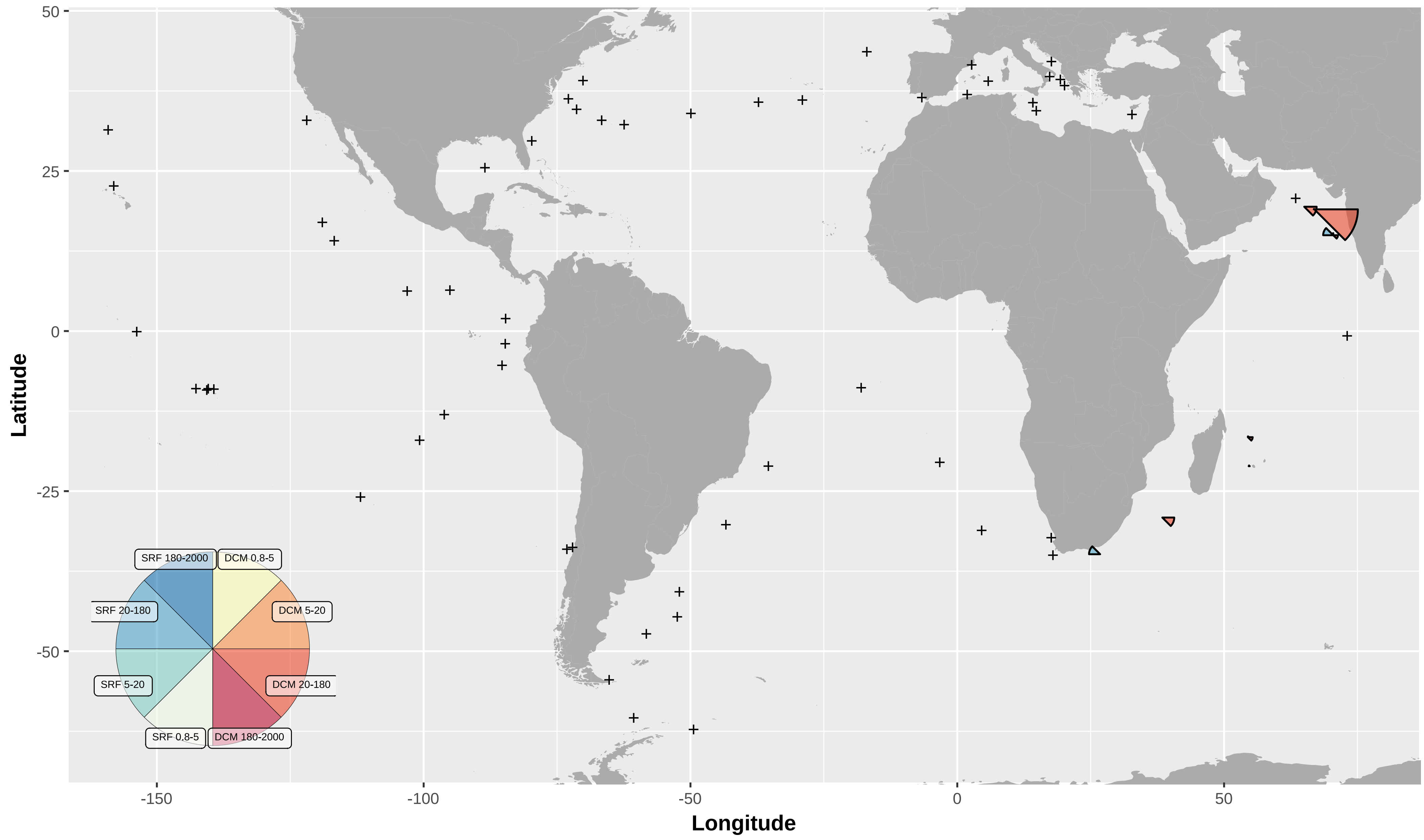


root;cellular organisms;Eukaryota;Opisthokonta;Metazoa;Eumetazoa

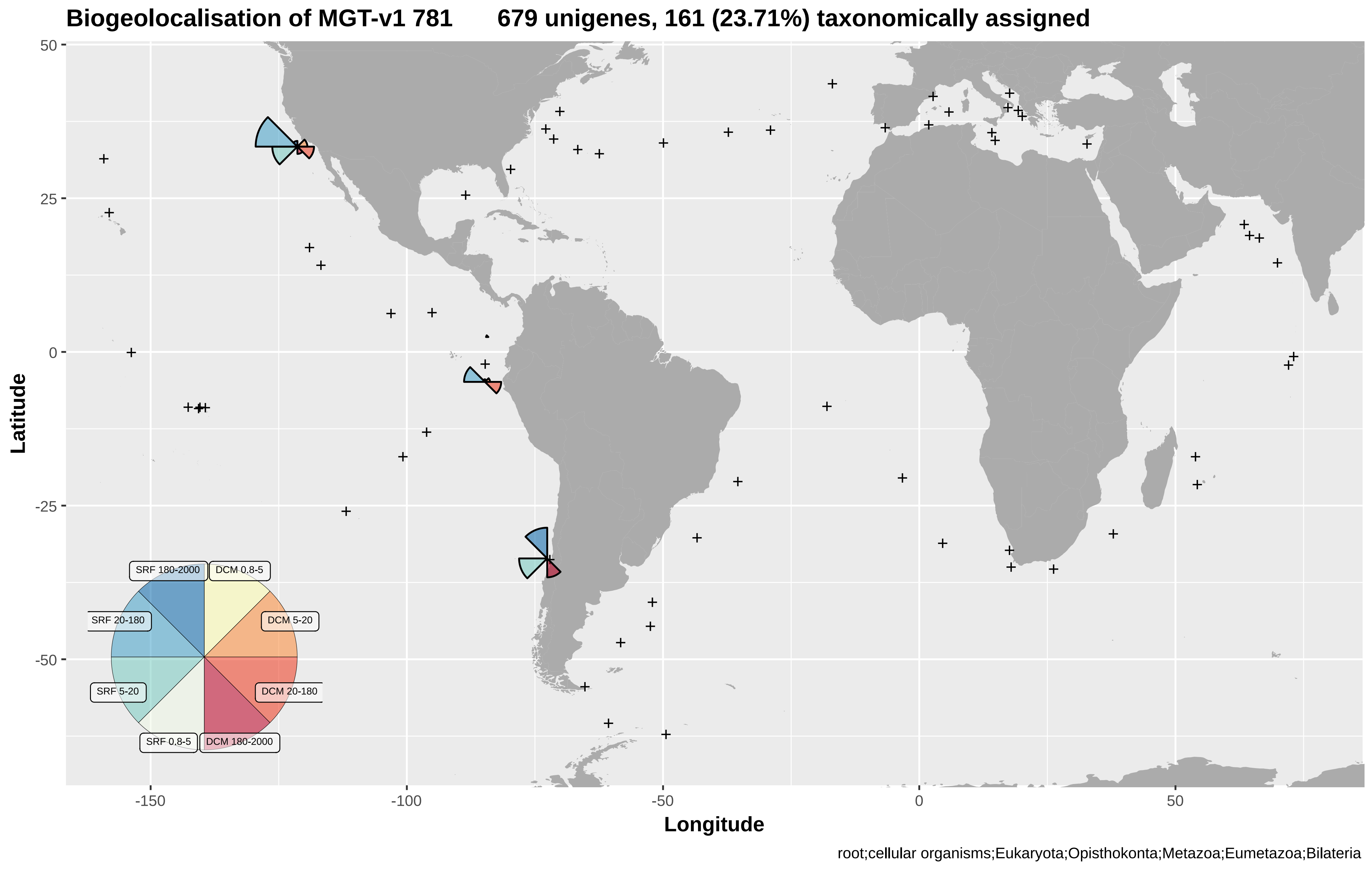
Biogeolocalisation of MGT-v1 781 679 unigenes, 161 (23.71%) taxonomically assigned
50
25
0
-25
-50
Latitude
-150
-100
-50
0
50
Longitude
SRF 180-2000
DCM 0.8-5
SRF 20-180
DCM 5-20
SRF 5-20
DCM 20-180
SRF 0.8-5
DCM 180-2000
root;cellular organisms;Eukaryota;Opisthokonta;Metazoa;Eumetazoa;Bilateria

# Biogeolocalisation of MGT-v1 782      675 unigenes, 665 (98.52%) taxonomically assigned

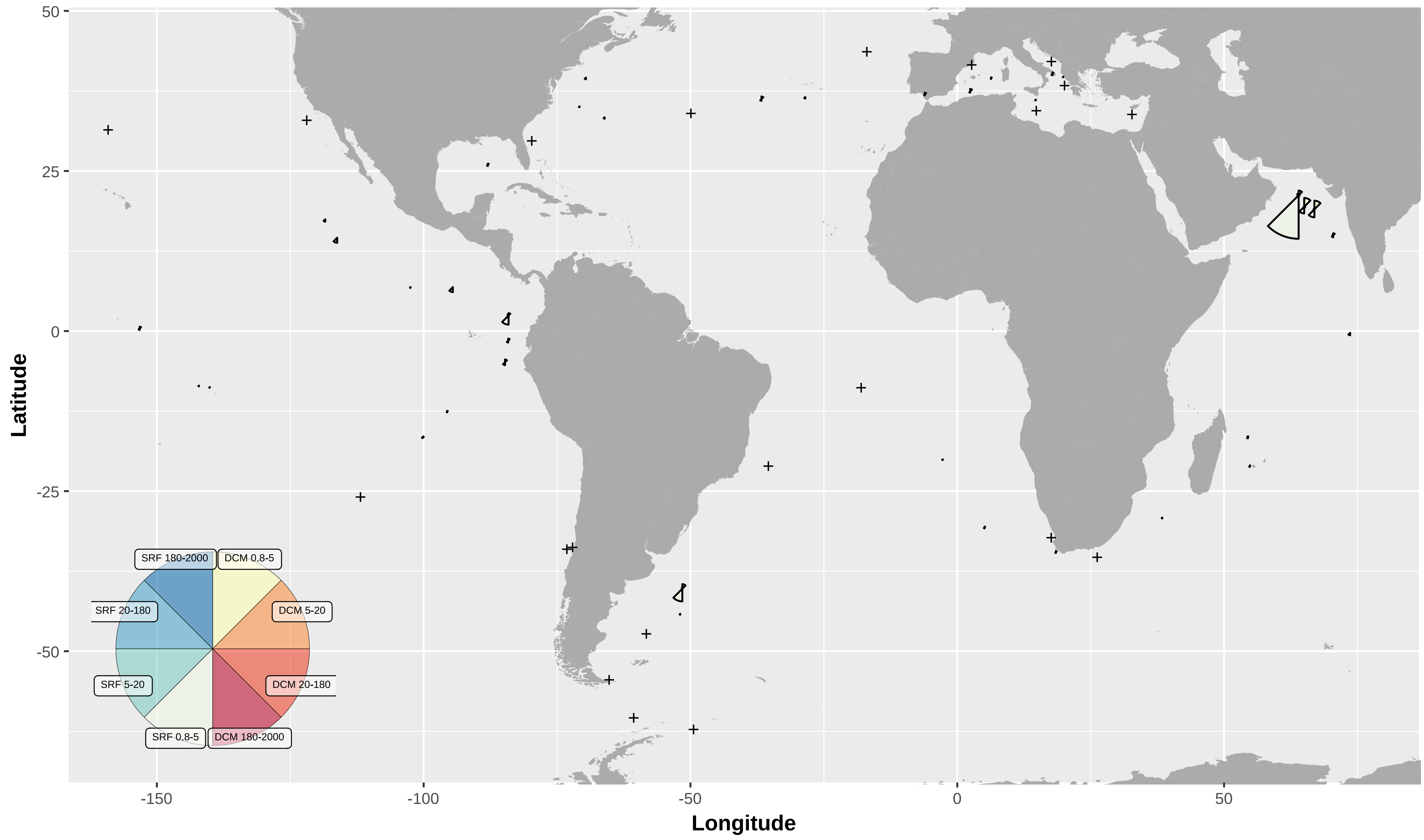


root;cellular organisms;Bacteria;Proteobacteria;Alphaproteobacteria;Pelagibacterales

## Biogeolocalisation of MGT-v1 783      675 unigenes, 112 (16.59%) taxonomically assigned

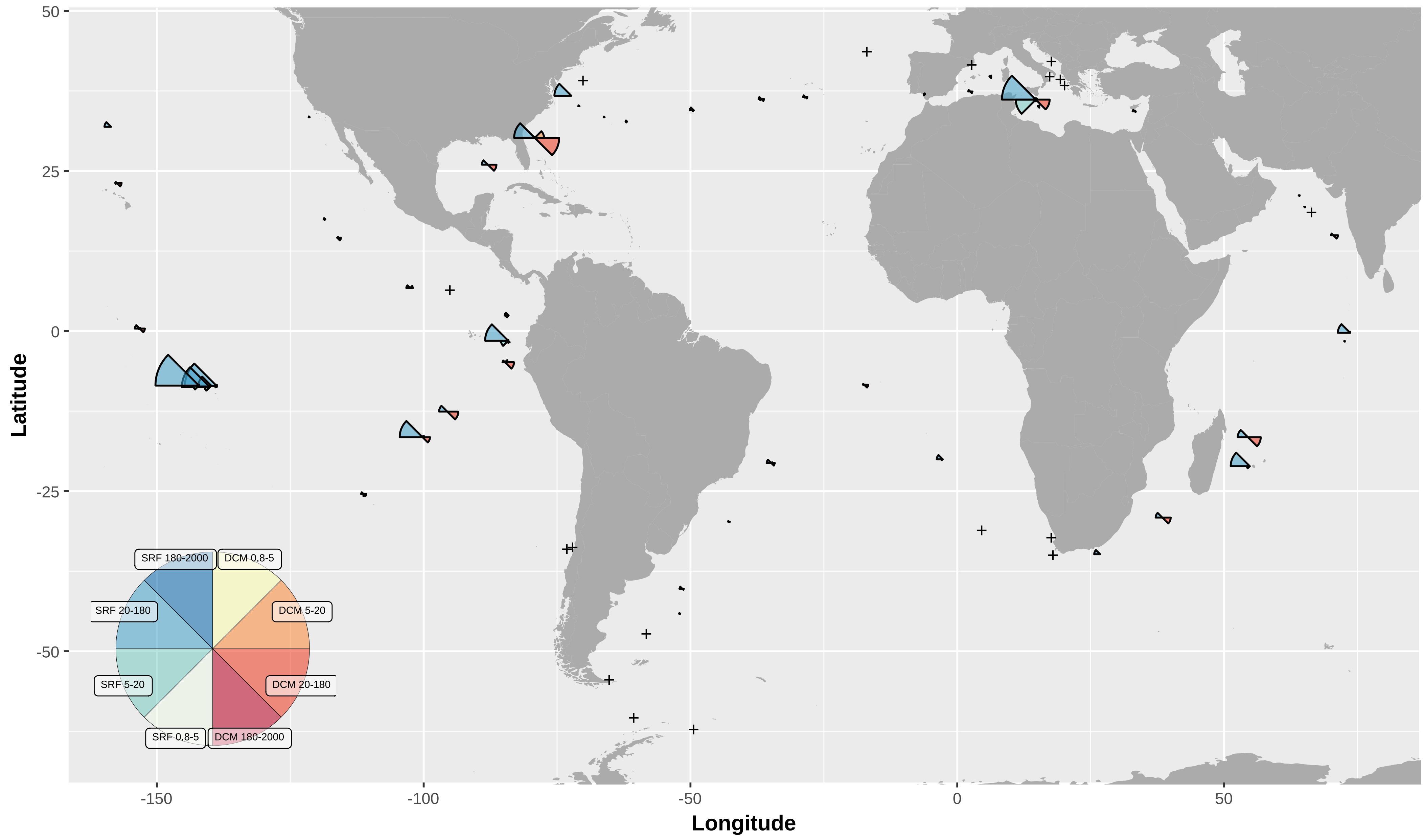


root;cellular organisms;Eukaryota;Opisthokonta;Metazoa;Eumetazoa;Bilateria;Protostomia;Ecdysozoa

# Biogeolocalisation of MGT-v1 784      675 unigenes, 59 (8.74%) taxonomically assigned

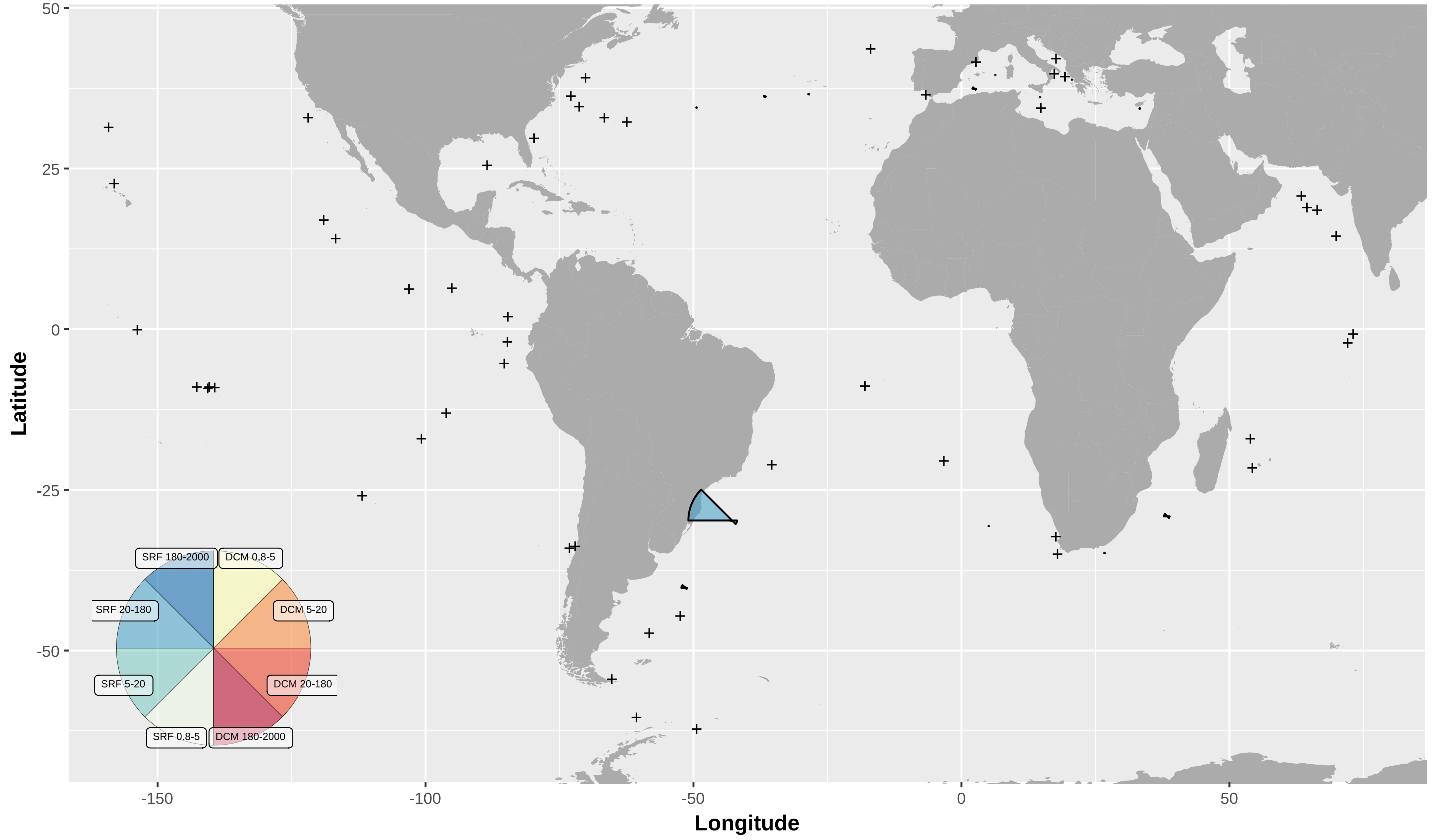


root;cellular organisms;Eukaryota;Opisthokonta;Metazoa

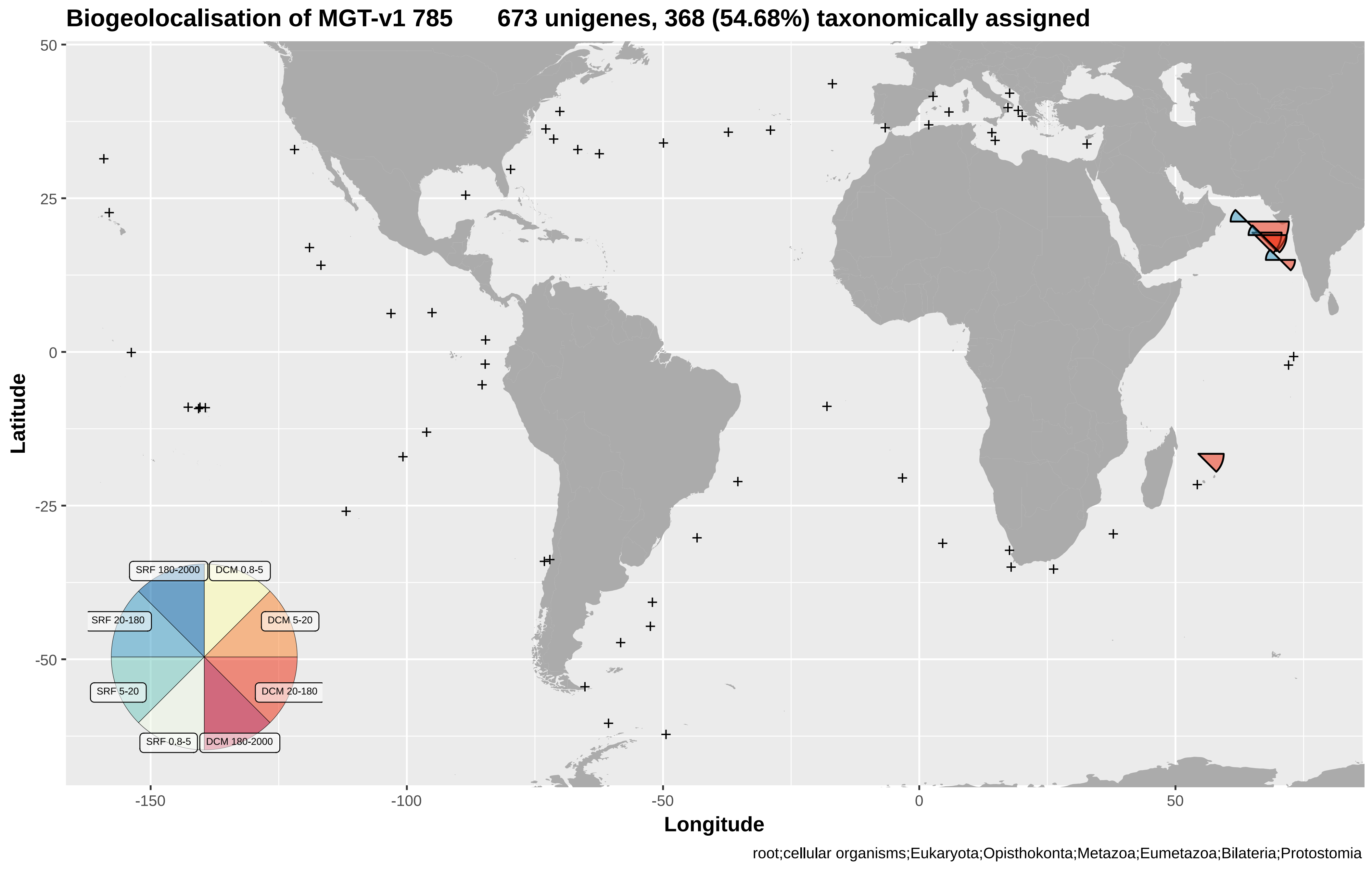

Biogeolocalisation of MGT-v1 785 673 unigenes, 368 (54.68%) taxonomically assigned
50
25
0
-25
-50
Latitude
-150
-100
-50
0
50
Longitude
SRF 180-2000
DCM 0.8-5
SRF 20-180
DCM 5-20
SRF 5-20
DCM 20-180
SRF 0.8-5
DCM 180-2000
root;cellular organisms;Eukaryota;Opisthokonta;Metazoa;Eumetazoa;Bilateria;Protostomia

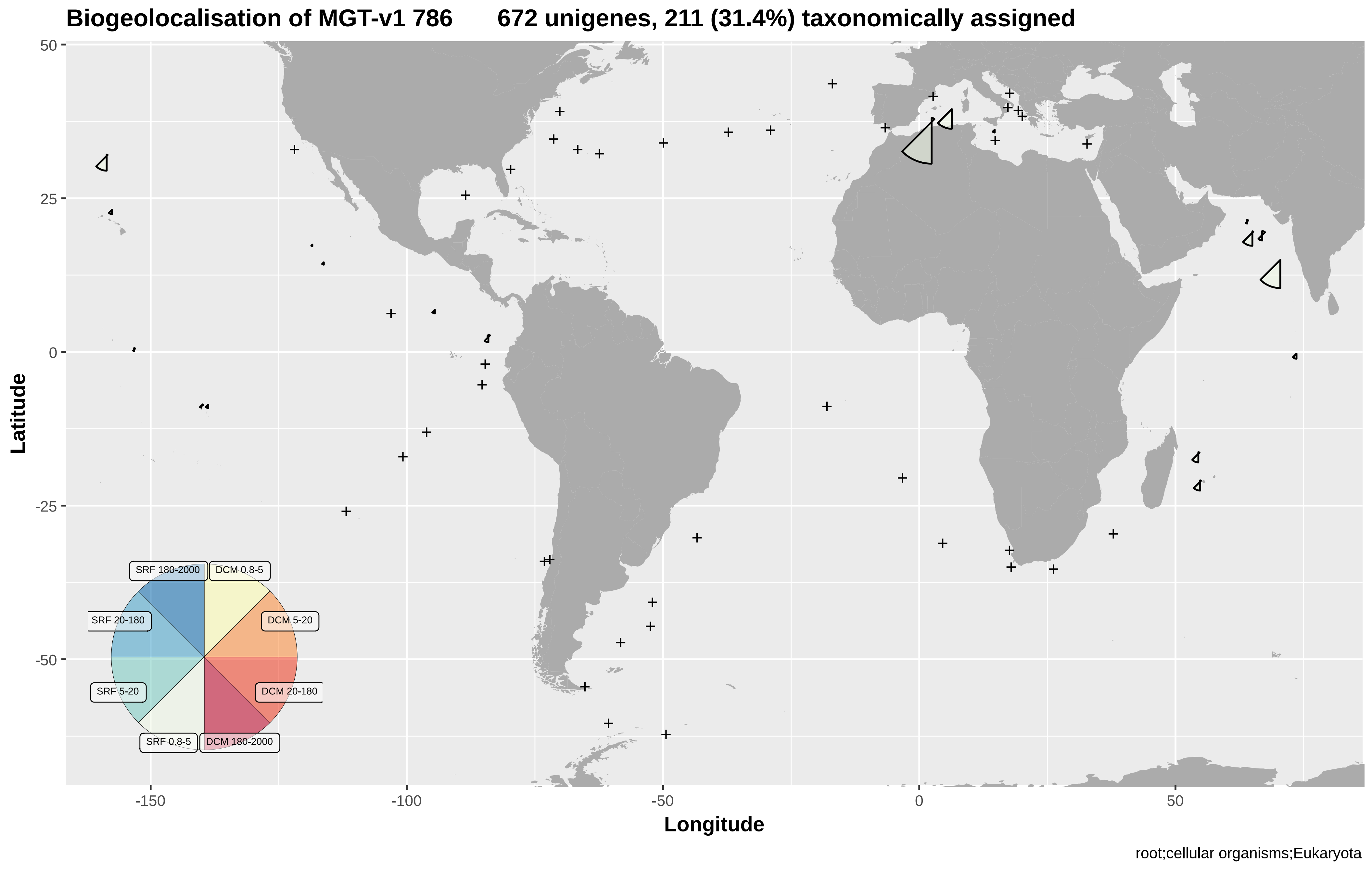
Biogeolocalisation of MGT-v1 786 672 unigenes, 211 (31.4%) taxonomically assigned
50
25
0
-25
-50
Latitude
-150
-100
-50
0
50
Longitude
SRF 180-2000
DCM 0.8-5
SRF 20-180
DCM 5-20
SRF 5-20
DCM 20-180
SRF 0.8-5
DCM 180-2000
root;cellular organisms;Eukaryota

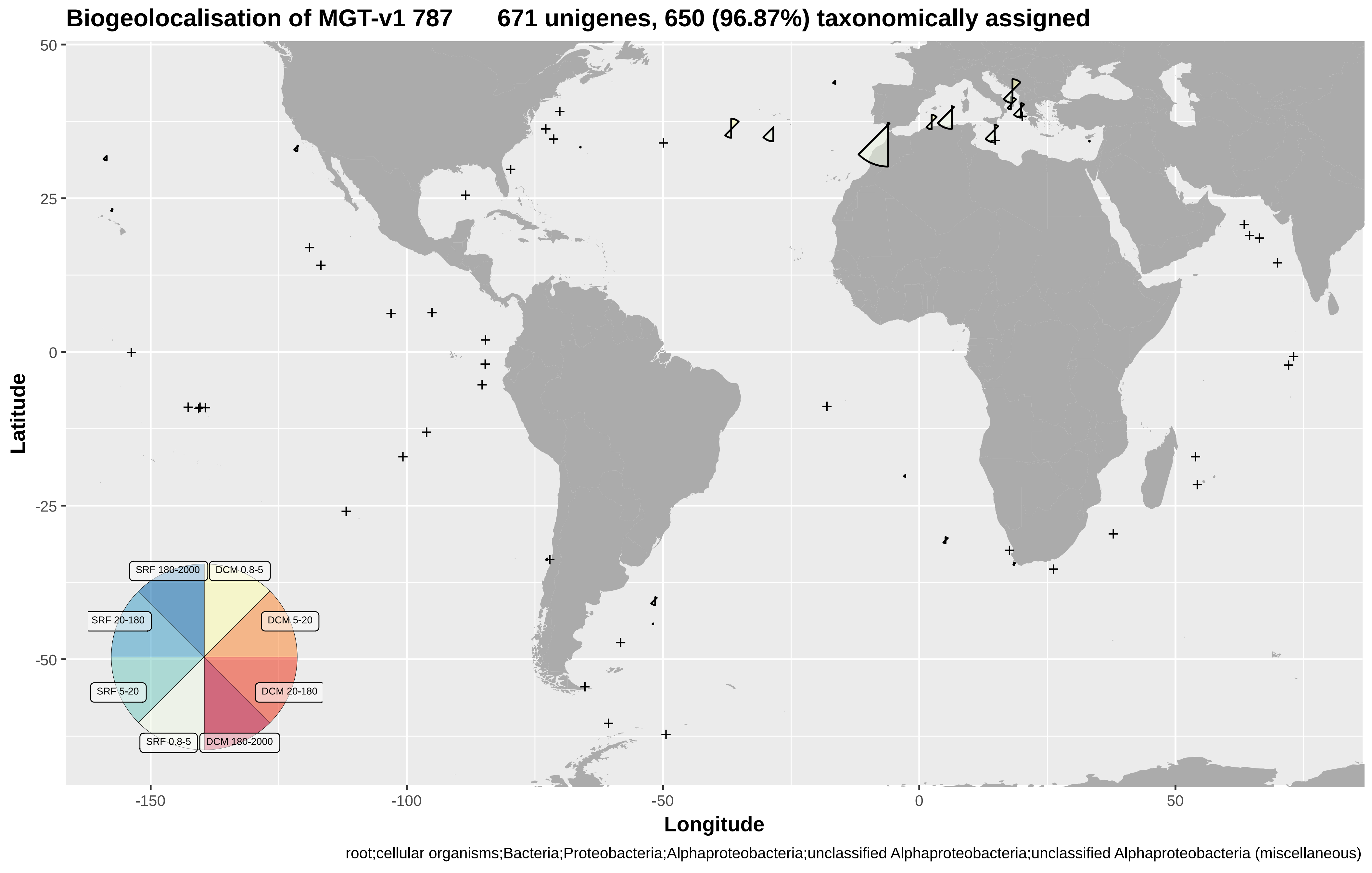

Biogeolocalisation of MGT-v1 787      671 unigenes, 650 (96.87%) taxonomically assigned
Latitude
Longitude
50
25
0
-25
-50
-150
-100
-50
0
50
SRF 180-2000
DCM 0.8-5
SRF 20-180
DCM 5-20
SRF 5-20
DCM 20-180
SRF 0.8-5
DCM 180-2000
root;cellular organisms;Bacteria;Proteobacteria;Alphaproteobacteria;unclassified Alphaproteobacteria;unclassified Alphaproteobacteria (miscellaneous)

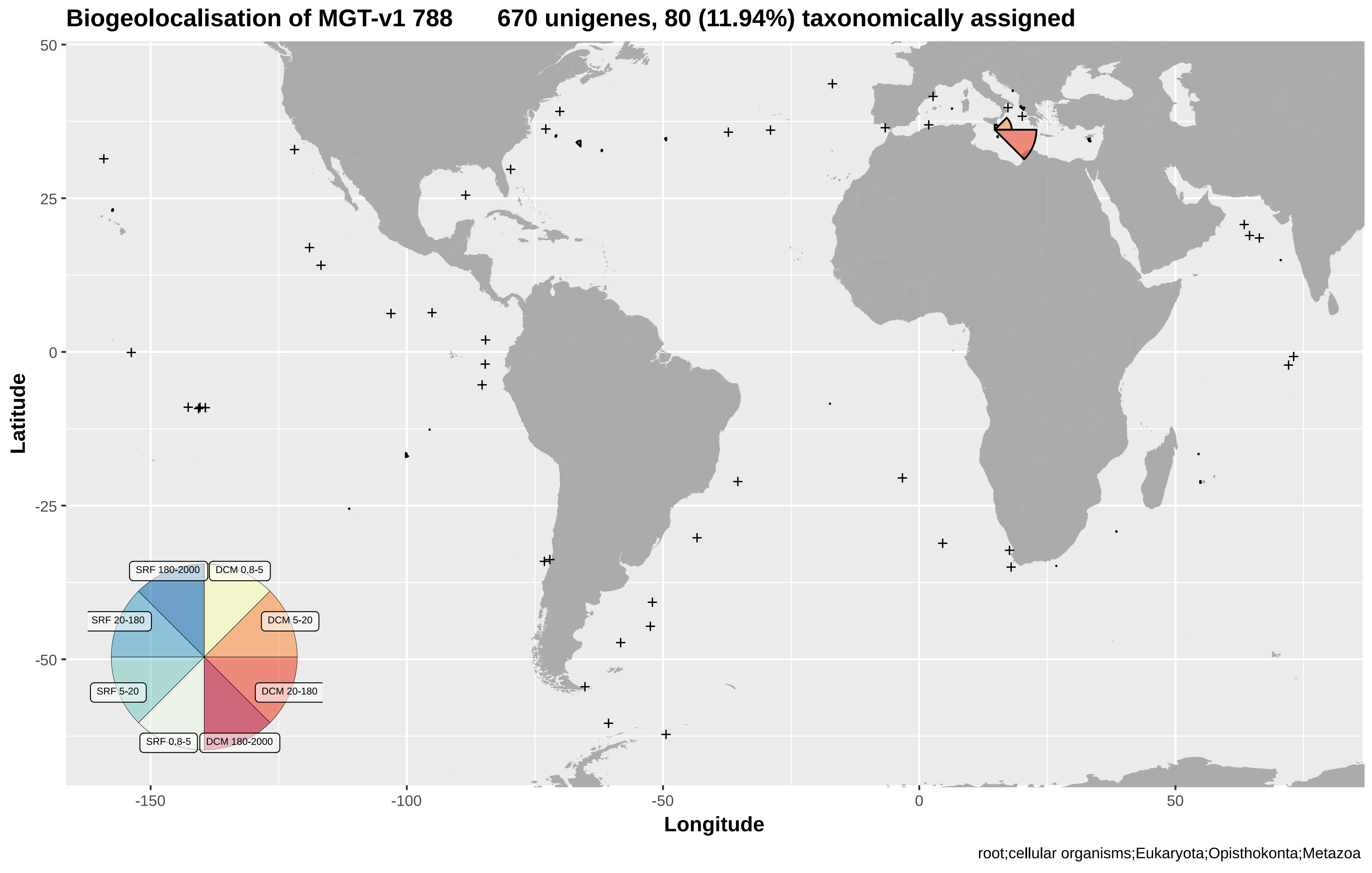

Biogeolocalisation of MGT-v1 788      670 unigenes, 80 (11.94%) taxonomically assigned
Latitude
Longitude
50
25
0
-25
-50
-150
-100
-50
0
50
SRF 180-2000
DCM 0.8-5
SRF 20-180
DCM 5-20
SRF 5-20
DCM 20-180
SRF 0.8-5
DCM 180-2000
root;cellular organisms;Eukaryota;Opisthokonta;Metazoa

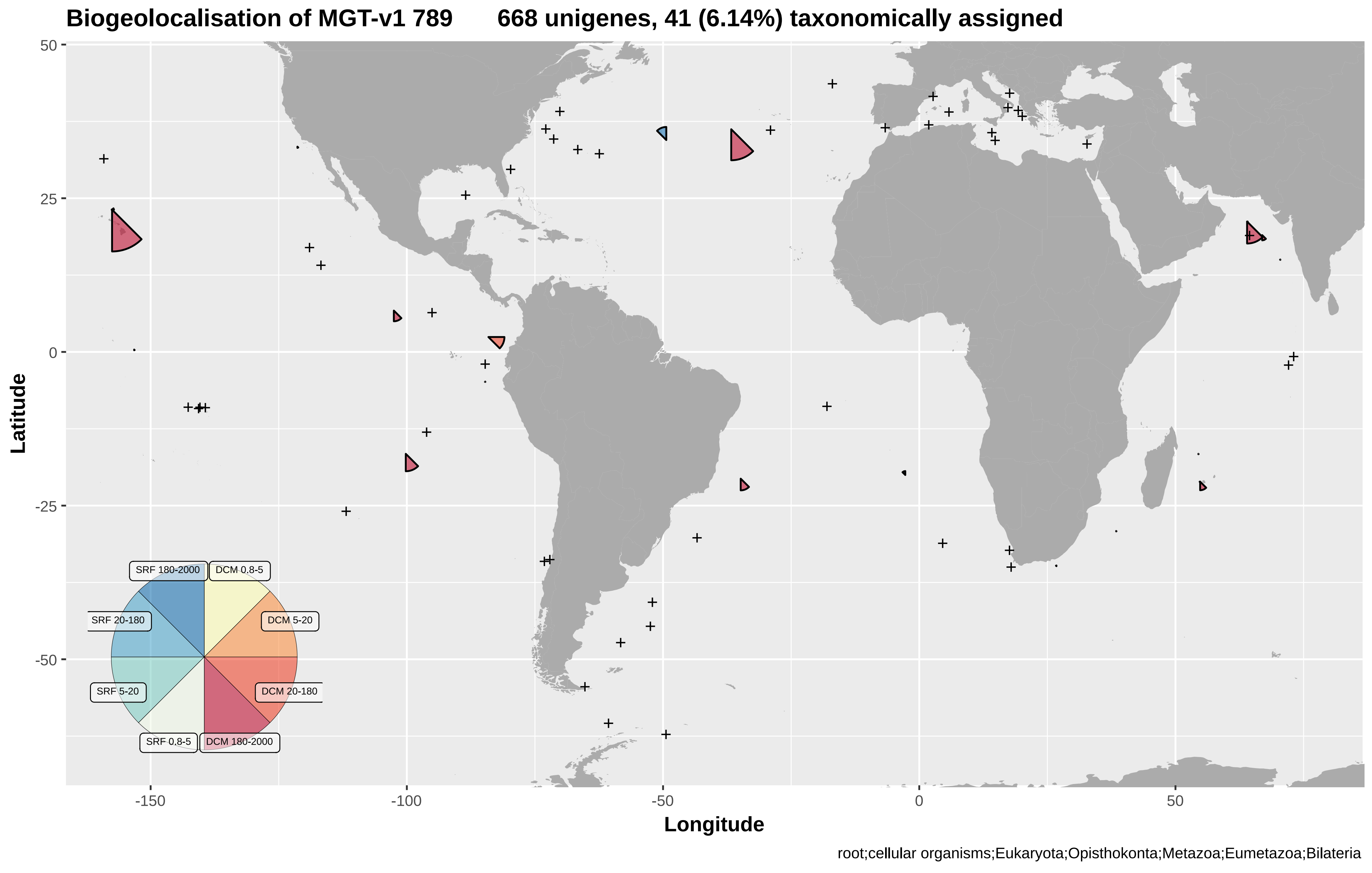
Biogeolocalisation of MGT-v1 789 668 unigenes, 41 (6.14%) taxonomically assigned
50
25
0
-25
-50
Latitude
-150
-100
-50
0
50
Longitude
SRF 180-2000
DCM 0.8-5
SRF 20-180
DCM 5-20
SRF 5-20
DCM 20-180
SRF 0.8-5
DCM 180-2000
root;cellular organisms;Eukaryota;Opisthokonta;Metazoa;Eumetazoa;Bilateria

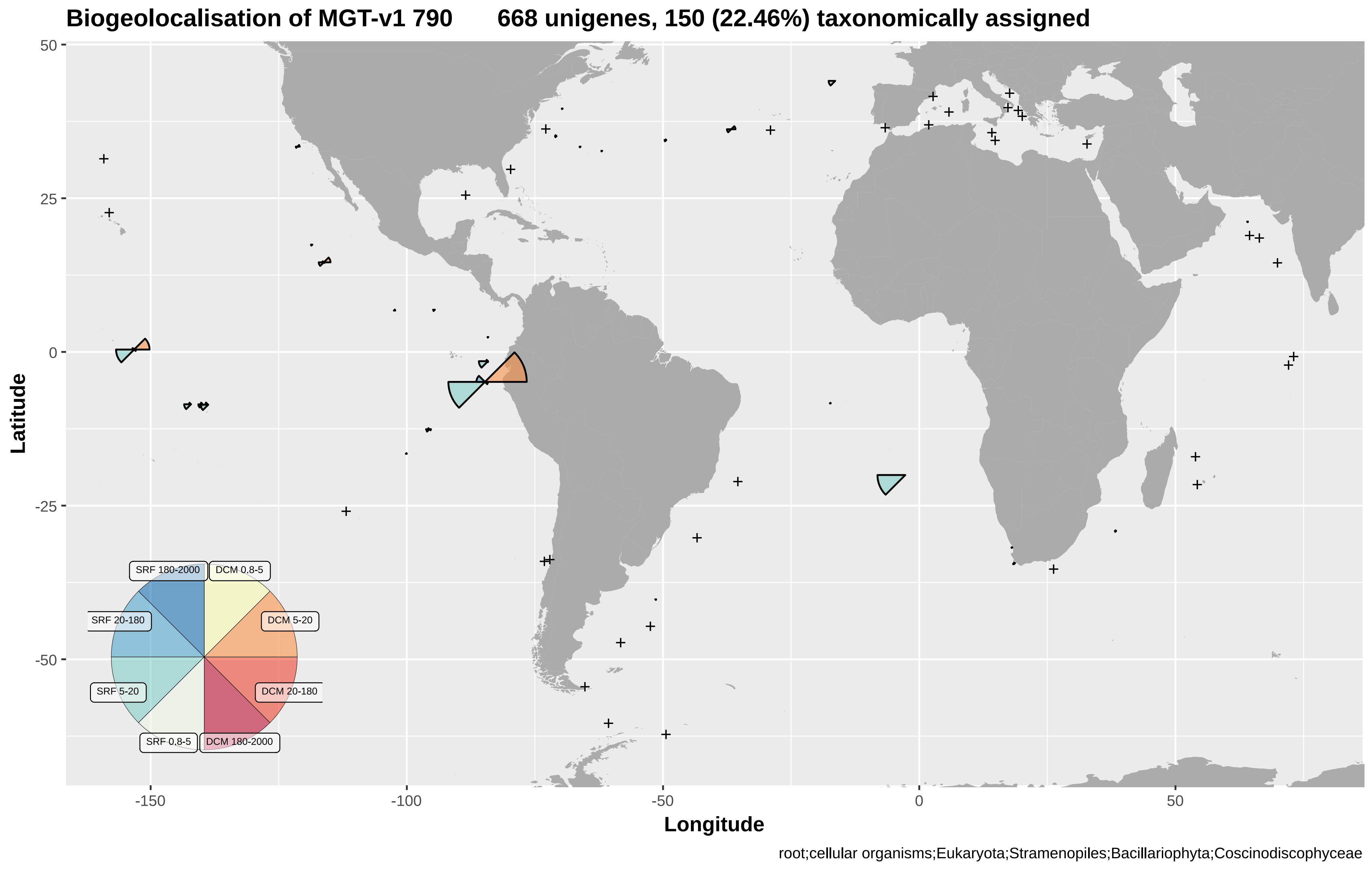

Biogeolocalisation of MGT-v1 790 668 unigenes, 150 (22.46%) taxonomically assigned
50
25
0
-25
-50
Latitude
-150
-100
-50
0
50
Longitude
SRF 180-2000
DCM 0.8-5
SRF 20-180
DCM 5-20
SRF 5-20
DCM 20-180
SRF 0.8-5
DCM 180-2000
root;cellular organisms;Eukaryota;Stramenopiles;Bacillariophyta;Coscinodiscophyceae

## Biogeolocalisation of MGT-v1 791      664 unigenes, 26 (3.92%) taxonomically assigned

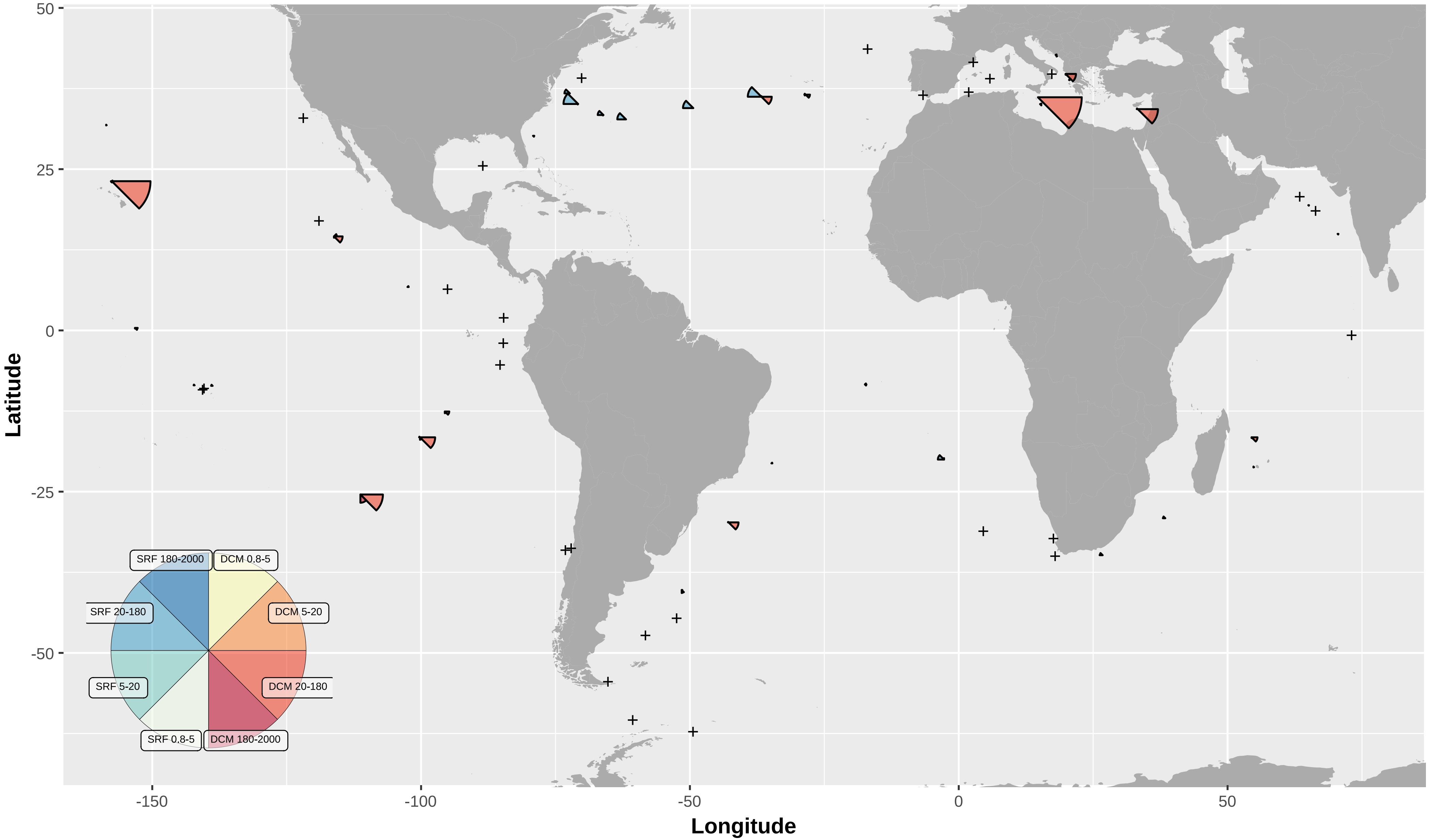


root;cellular organisms;Eukaryota

### Biogeolocalisation of MGT-v1 792 663 unigenes, 565 (85.22%) taxonomically assigned

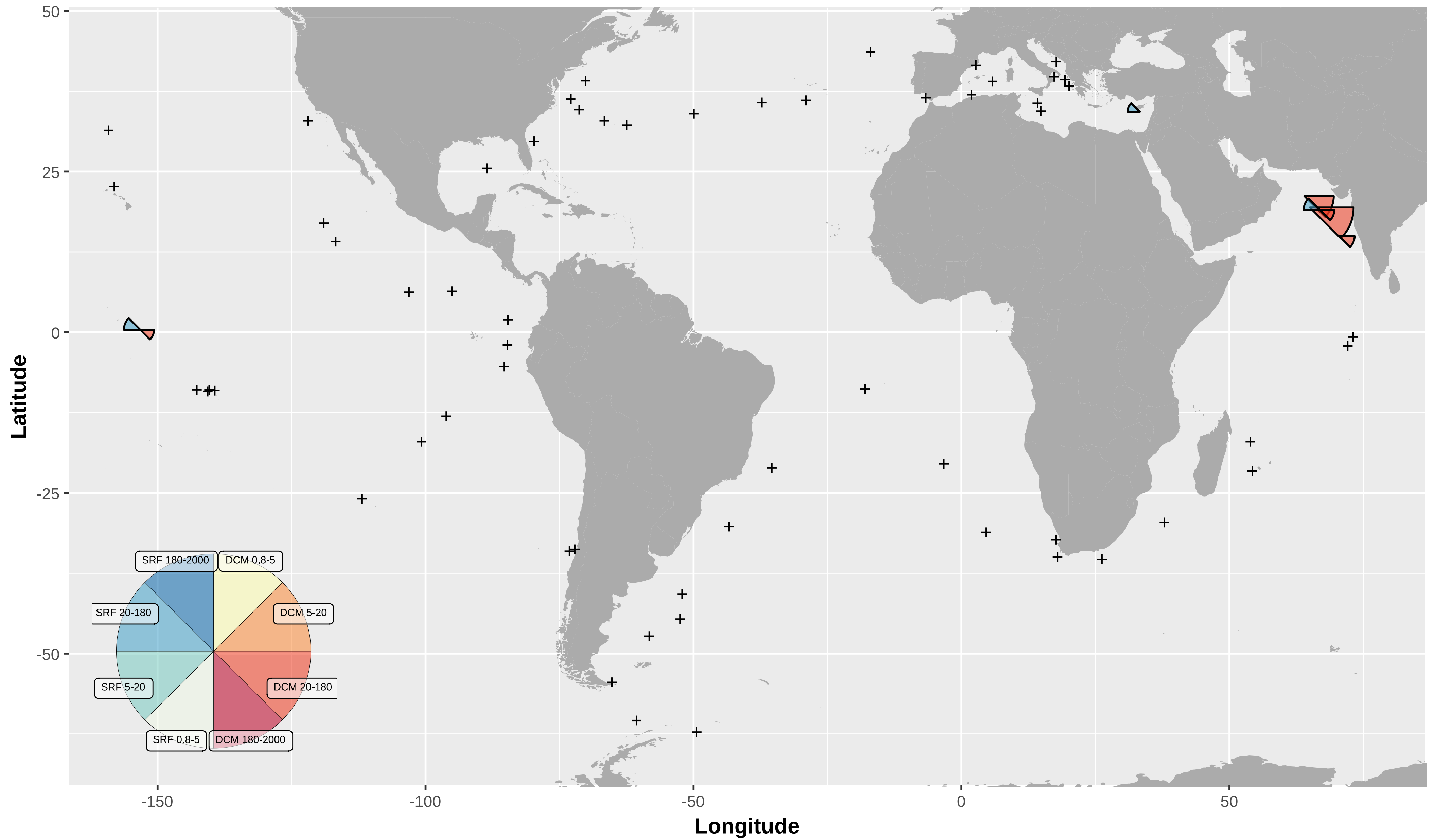


karyota;Opisthokonta;Metazoa;Eumetazoa;Bilateria;Protostomia;Ecdysozoa;Panarthropoda;Arthropoda;Mandibulata;Pancrustacea;Crustacea;Maxillopoda;Copepoda;Neocopepoda;Podoplea;Cyclopoida

## Biogeolocalisation of MGT-v1 793      663 unigenes, 411 (61.99%) taxonomically assigned

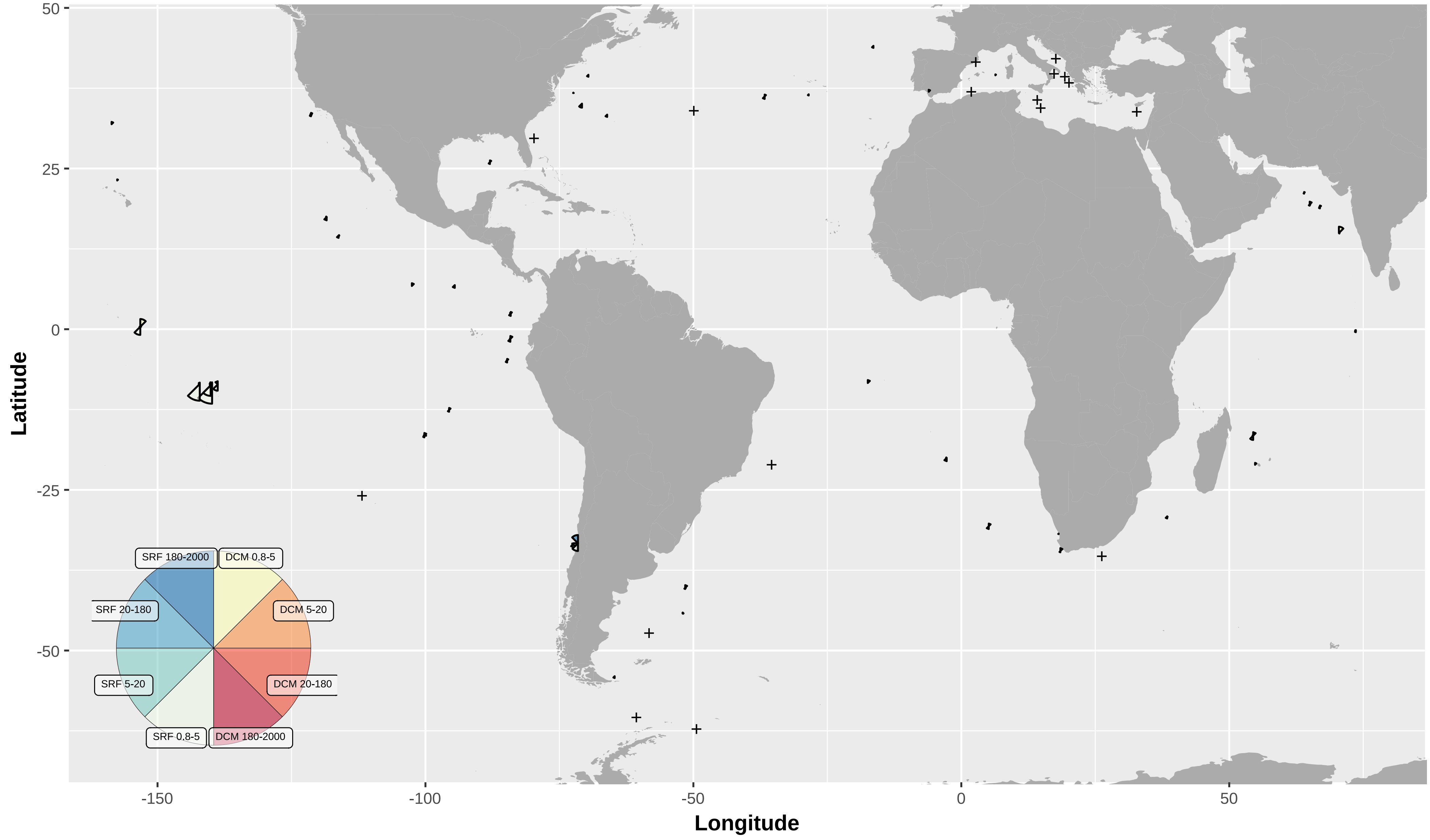


root;cellular organisms;Eukaryota;Haptophyceae;Phaeocystales

## Biogeolocalisation of MGT-v1 794      663 unigenes, 380 (57.32%) taxonomically assigned

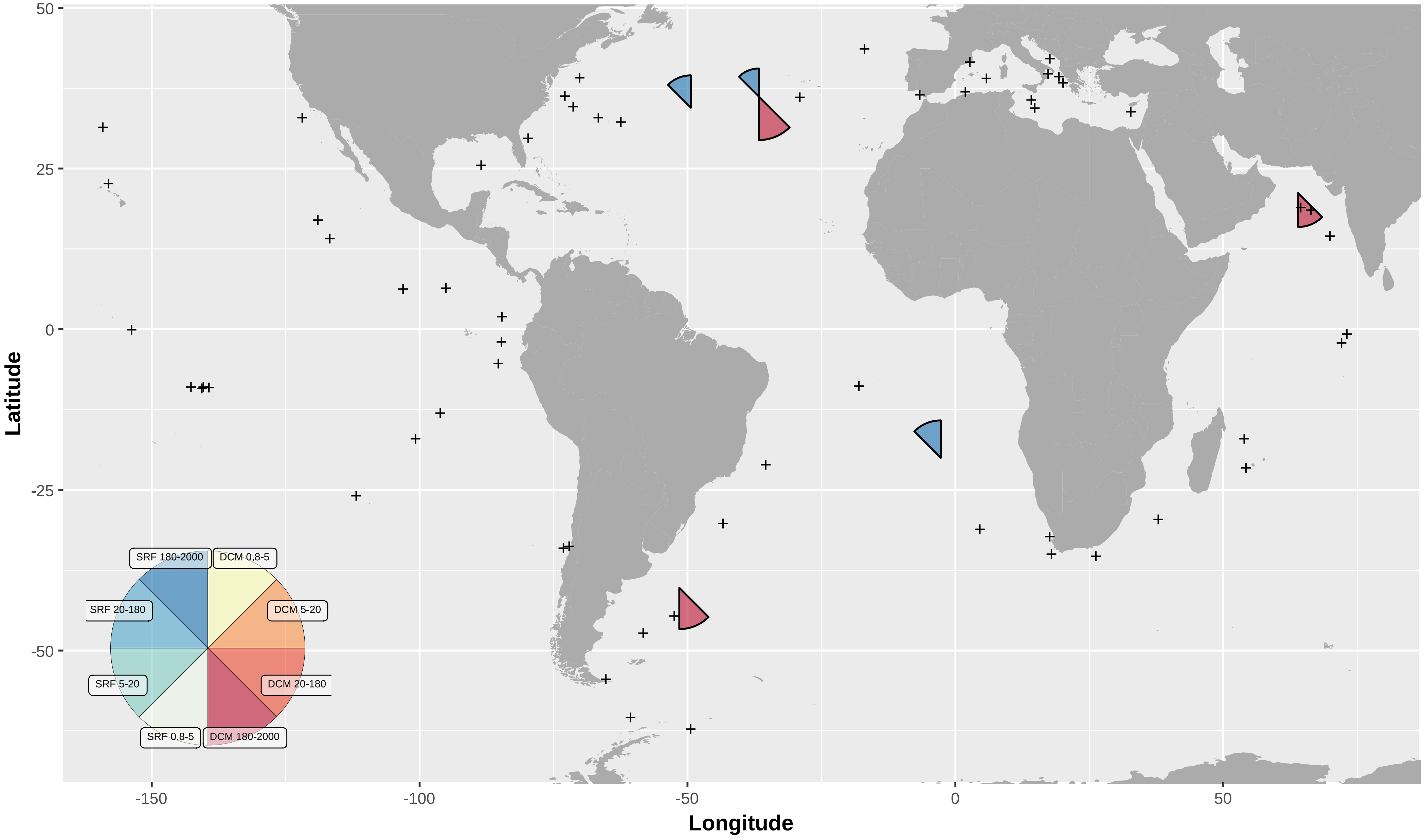


root;cellular organisms;Eukaryota;Opisthokonta;Metazoa;Eumetazoa;Bilateria;Deuterostomia;Chordata

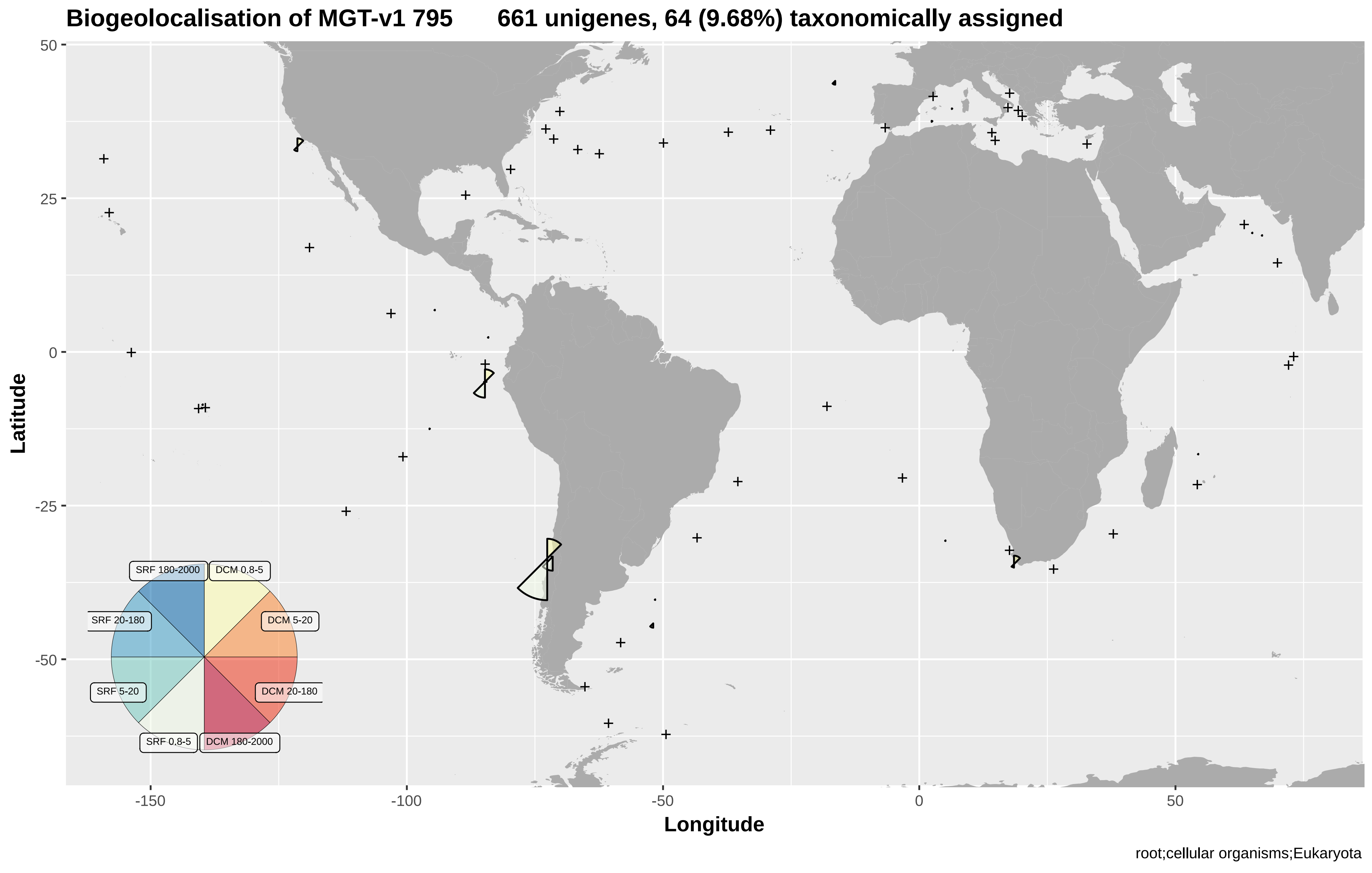

Biogeolocalisation of MGT-v1 795 661 unigenes, 64 (9.68%) taxonomically assigned
50
25
0
-25
-50
Latitude
-150
-100
-50
0
50
Longitude
SRF 180-2000
DCM 0.8-5
SRF 20-180
DCM 5-20
SRF 5-20
DCM 20-180
SRF 0.8-5
DCM 180-2000
root;cellular organisms;Eukaryota

# Biogeolocalisation of MGT-v1 796      660 unigenes, 62 (9.39%) taxonomically assigned

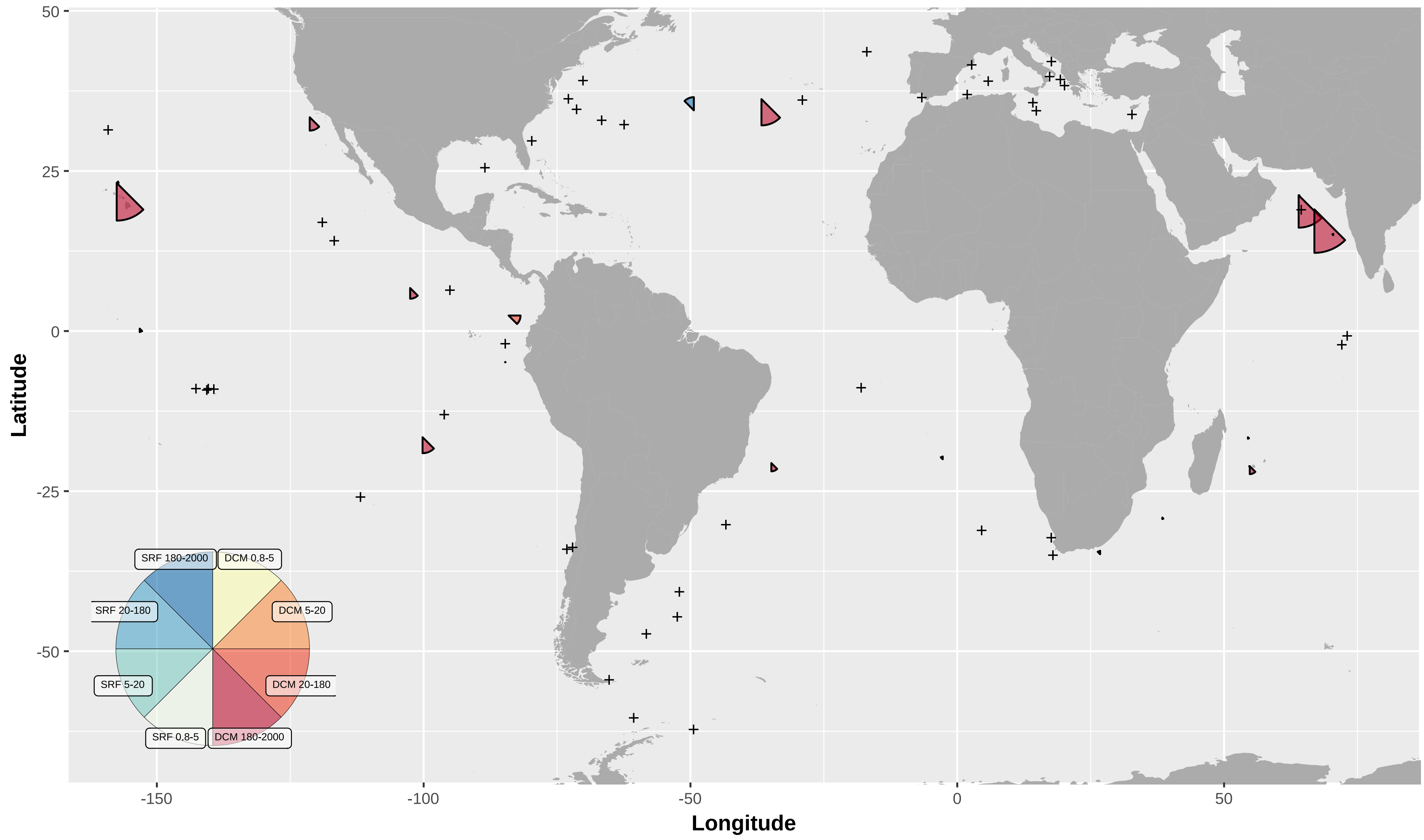


root;cellular organisms;Eukaryota;Opisthokonta;Metazoa;Eumetazoa

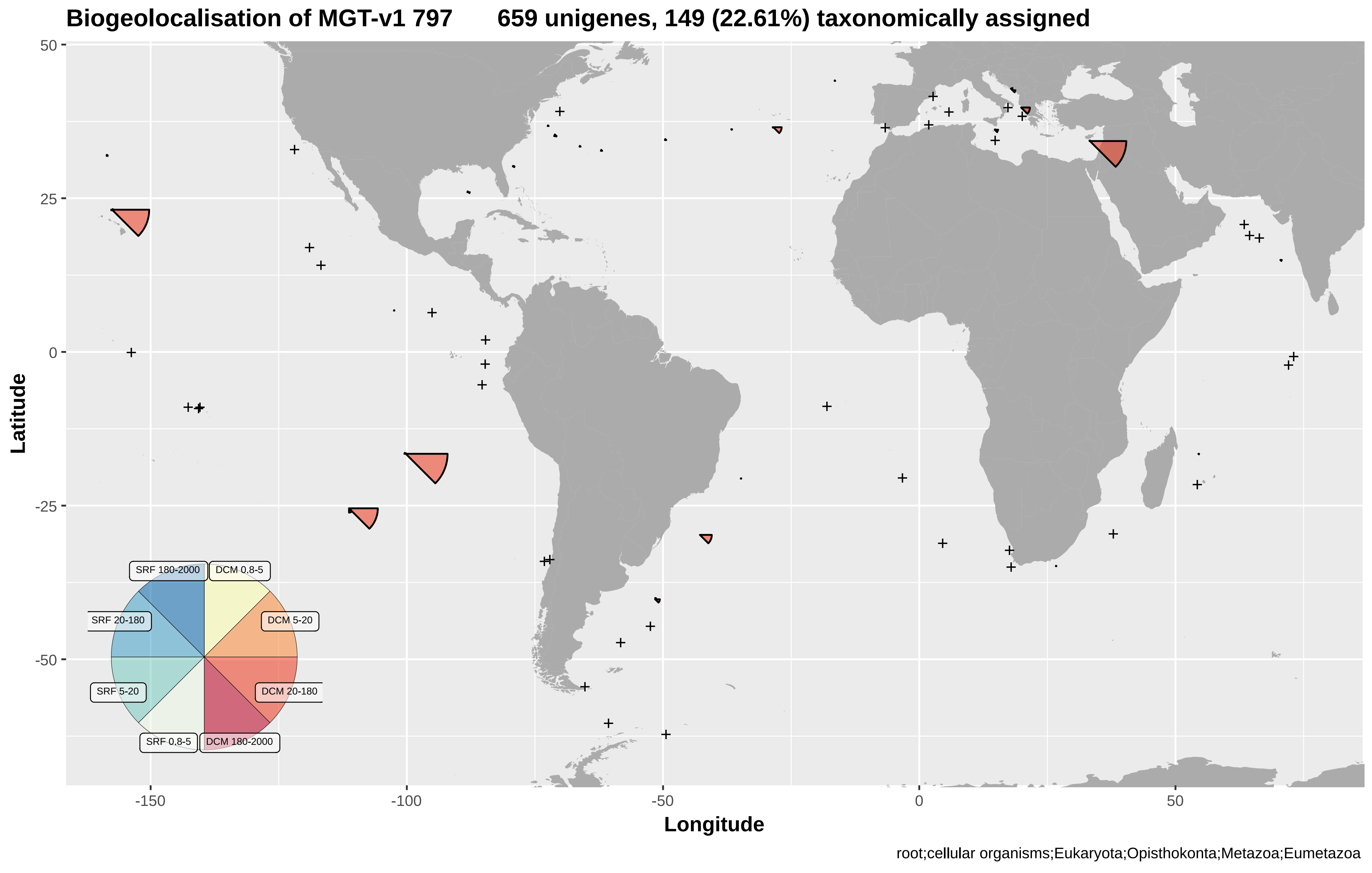
Biogeolocalisation of MGT-v1 797 659 unigenes, 149 (22.61%) taxonomically assigned
Latitude
Longitude
50
25
0
-25
-50
-150
-100
-50
0
50
SRF 180-2000
DCM 0.8-5
SRF 20-180
DCM 5-20
SRF 5-20
DCM 20-180
SRF 0.8-5
DCM 180-2000
root;cellular organisms;Eukaryota;Opisthokonta;Metazoa;Eumetazoa

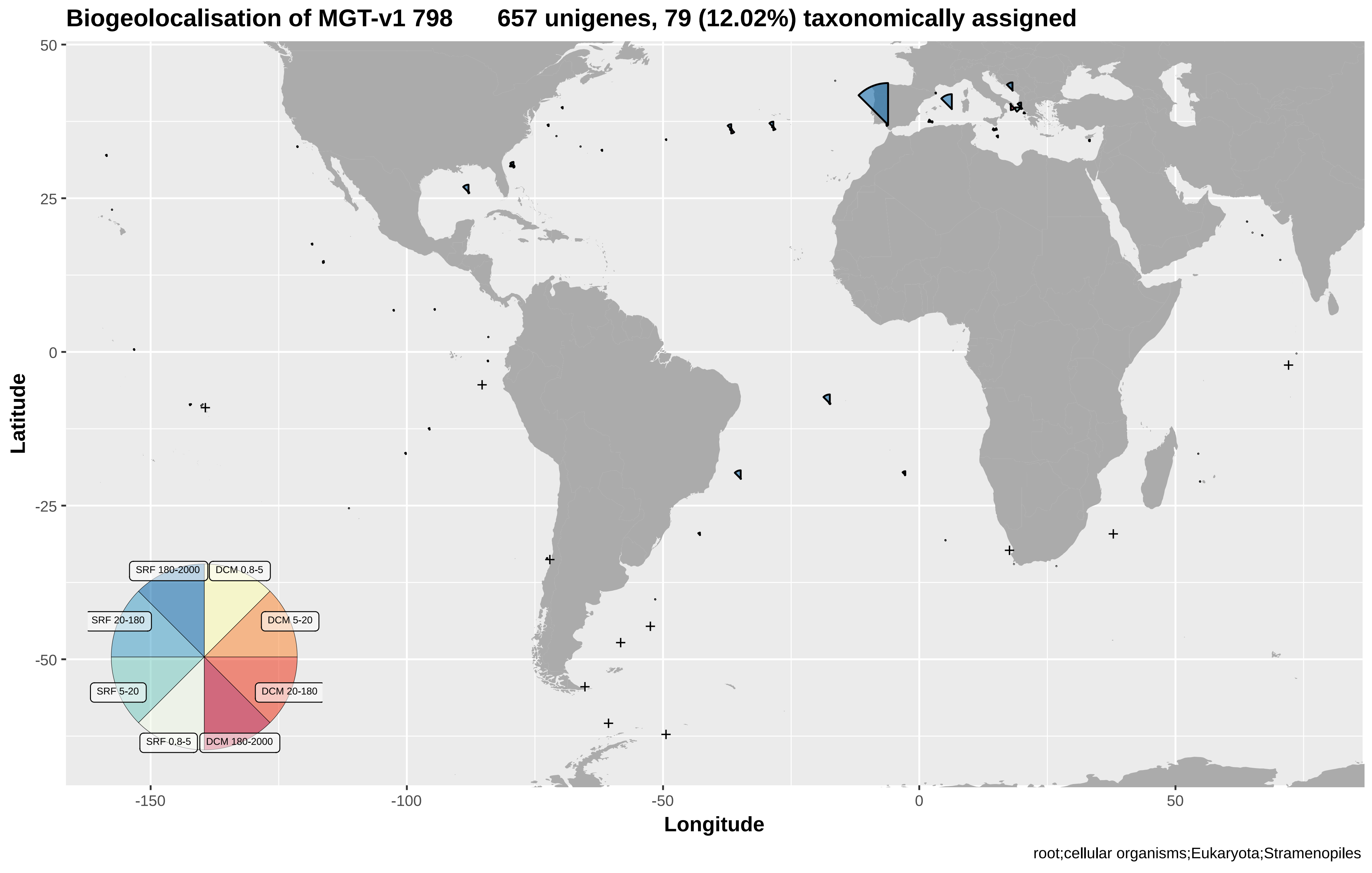

Biogeolocalisation of MGT-v1 798 657 unigenes, 79 (12.02%) taxonomically assigned
50
25
0
-25
-50
Latitude
-150
-100
-50
0
50
Longitude
SRF 180-2000
DCM 0.8-5
SRF 20-180
DCM 5-20
SRF 5-20
DCM 20-180
SRF 0.8-5
DCM 180-2000
root;cellular organisms;Eukaryota;Stramenopiles

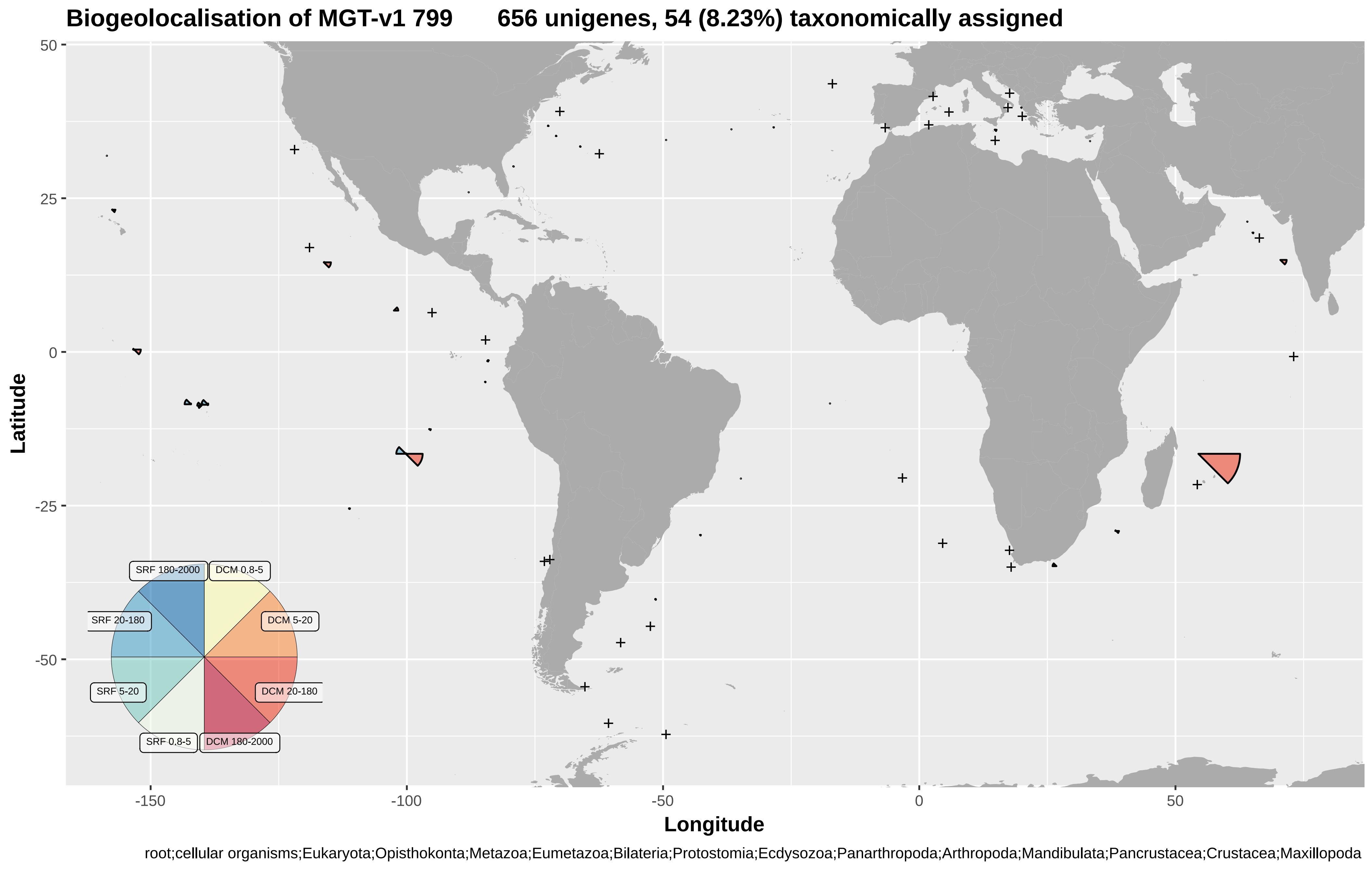

Biogeolocalisation of MGT-v1 799
656 unigenes, 54 (8.23%) taxonomically assigned
Latitude
Longitude
50
25
0
-25
-50
-150
-100
-50
0
50
SRF 180-2000
DCM 0.8-5
SRF 20-180
DCM 5-20
SRF 5-20
DCM 20-180
SRF 0.8-5
DCM 180-2000
root;cellular organisms;Eukaryota;Opisthokonta;Metazoa;Eumetazoa;Bilateria;Protostomia;Ecdysozoa;Panarthropoda;Arthropoda;Mandibulata;Pancrustacea;Crustacea;Maxillopoda

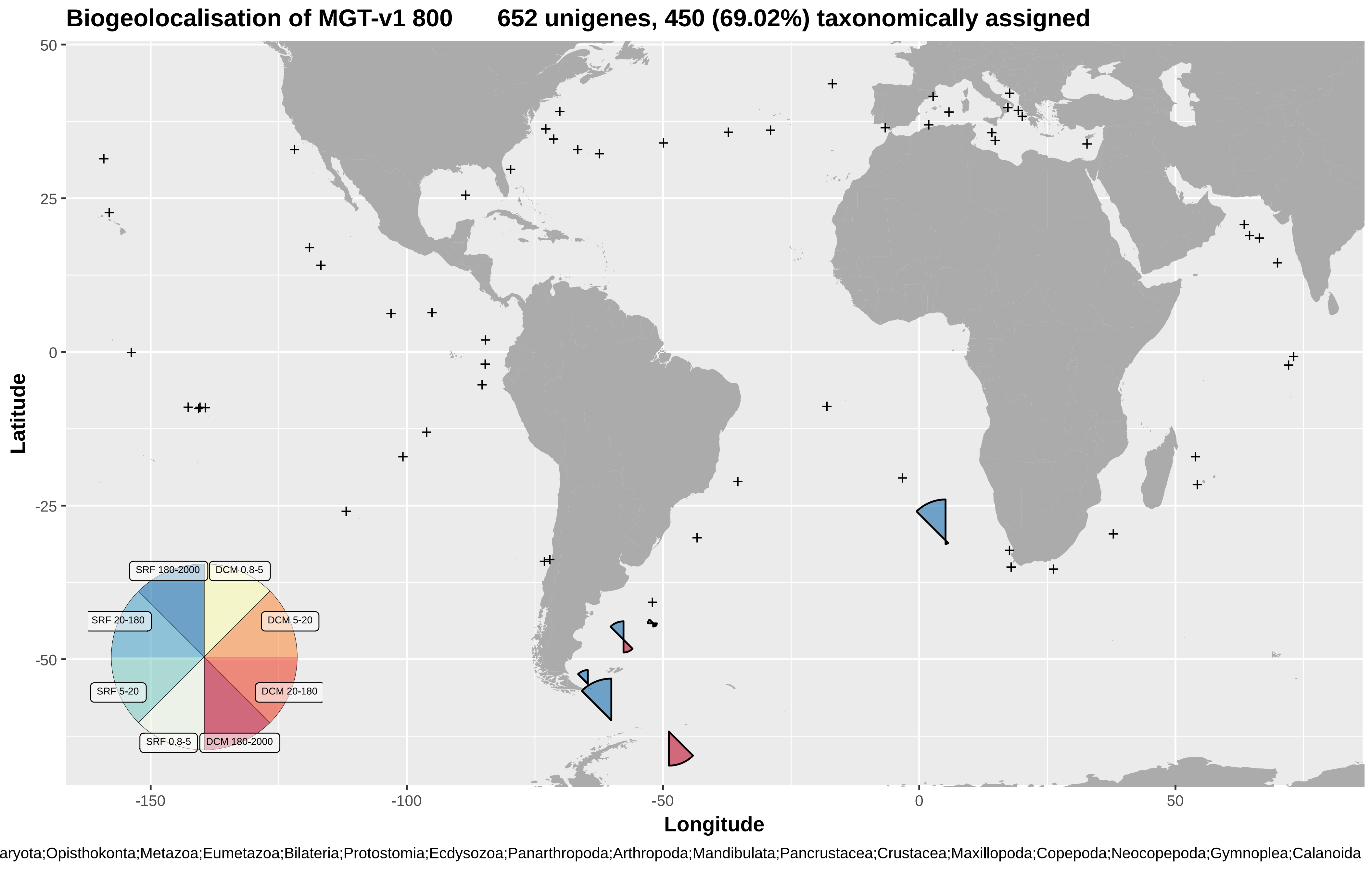
Biogeolocalisation of MGT-v1 800 652 unigenes, 450 (69.02%) taxonomically assigned
50
25
0
-25
-50
Latitude
-150
-100
-50
0
50
Longitude
SRF 180-2000
DCM 0.8-5
SRF 20-180
DCM 5-20
SRF 5-20
DCM 20-180
SRF 0.8-5
DCM 180-2000
aryota;Opisthokonta;Metazoa;Eumetazoa;Bilateria;Protostomia;Ecdysozoa;Panarthropoda;Arthropoda;Mandibulata;Pancrustacea;Crustacea;Maxillopoda;Copepoda;Neocopepoda;Gymnoplea;Calanoida

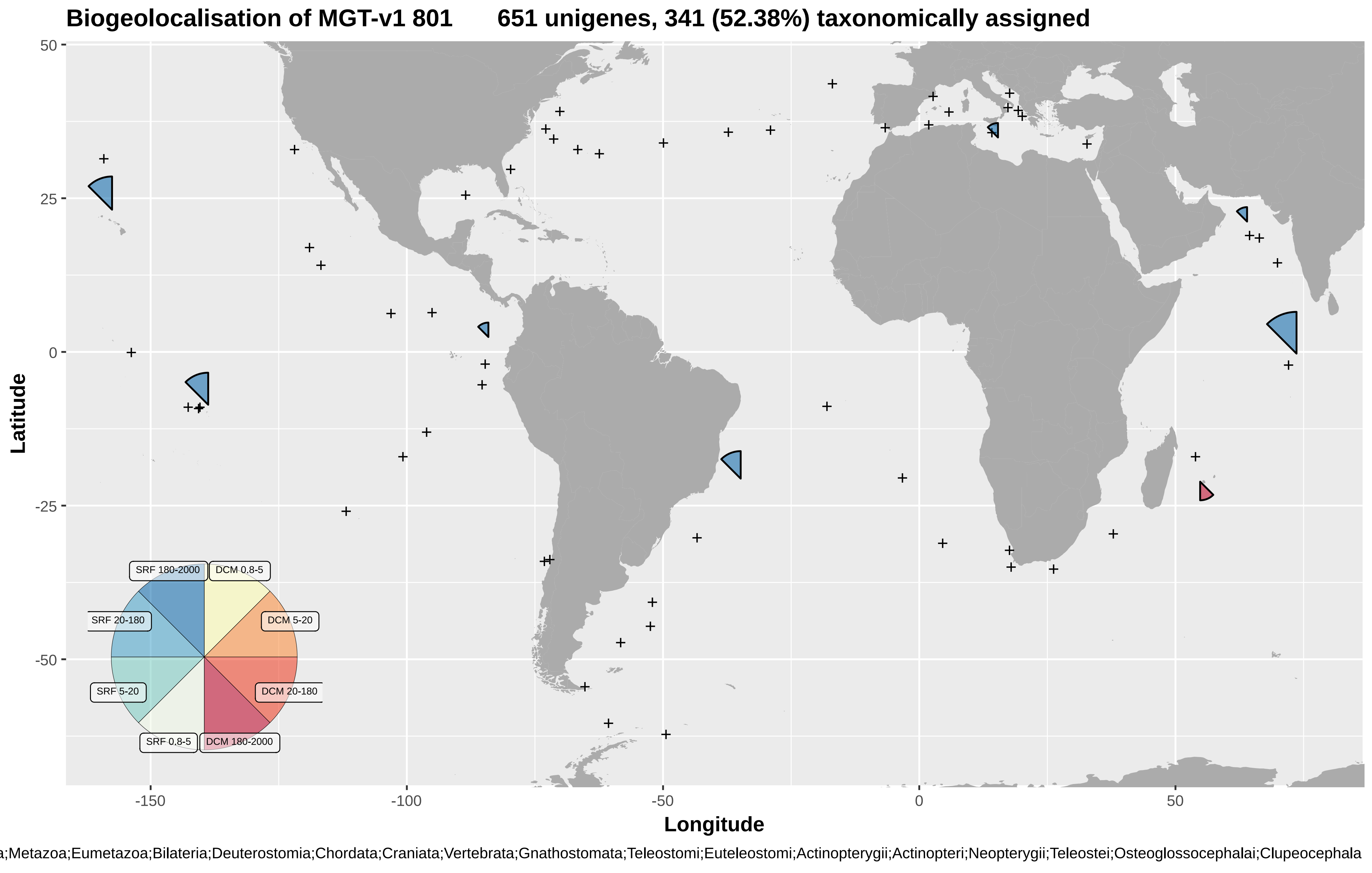

Biogeolocalisation of MGT-v1 801      651 unigenes, 341 (52.38%) taxonomically assigned
50
25
0
-25
-50
Latitude
-150
-100
-50
0
50
Longitude
SRF 180-2000
DCM 0.8-5
SRF 20-180
DCM 5-20
SRF 5-20
DCM 20-180
SRF 0.8-5
DCM 180-2000
a;Metazoa;Eumetazoa;Bilateria;Deuterostomia;Chordata;Craniata;Vertebrata;Gnathostomata;Teleostomi;Euteleostomi;Actinopterygii;Actinopteri;Neopterygii;Teleostei;Osteoglossocephalai;Clupeocephala

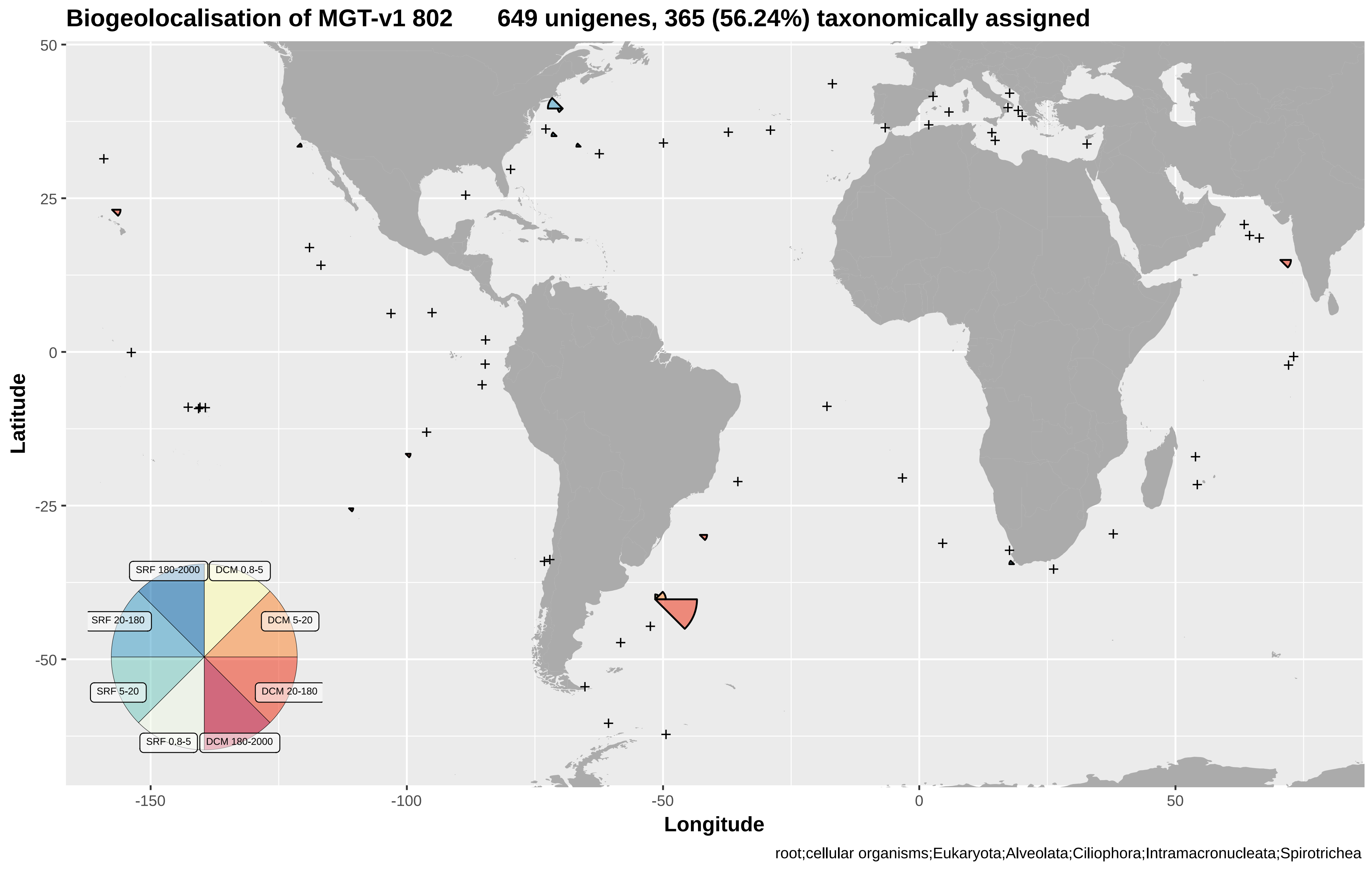

Biogeolocalisation of MGT-v1 802 649 unigenes, 365 (56.24%) taxonomically assigned
Latitude
50
25
0
-25
-50
Longitude
-150
-100
-50
0
50
SRF 180-2000
DCM 0.8-5
SRF 20-180
DCM 5-20
SRF 5-20
DCM 20-180
SRF 0.8-5
DCM 180-2000
root;cellular organisms;Eukaryota;Alveolata;Ciliophora;Intramacronucleata;Spirotrichea

### Biogeolocalisation of MGT-v1 803 647 unigenes, 596 (92.12%) taxonomically assigned

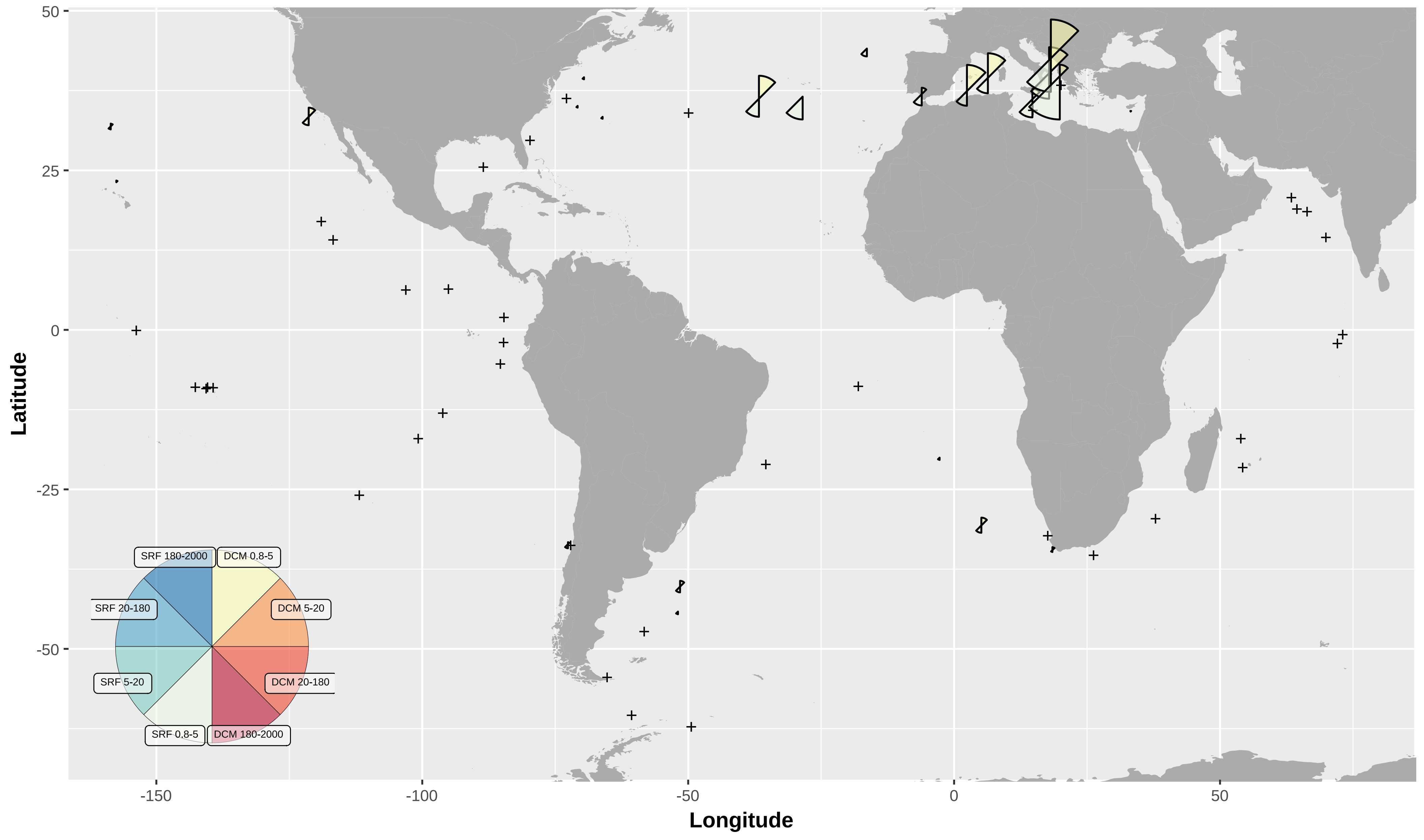


root;cellular organisms;Bacteria;Proteobacteria

## Biogeolocalisation of MGT-v1 804      644 unigenes, 76 (11.8%) taxonomically assigned

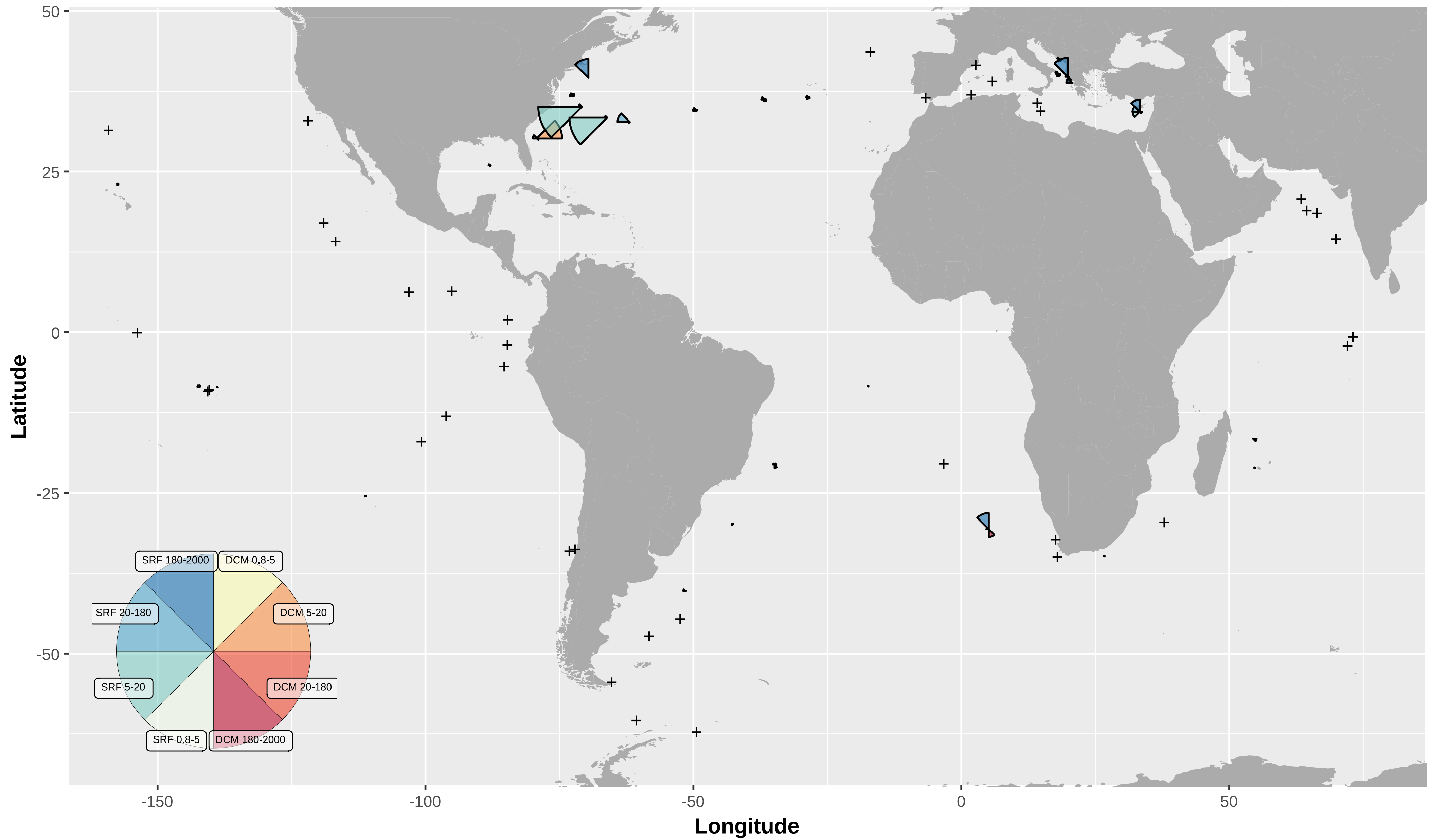


root;cellular organisms;Eukaryota;Opisthokonta;Metazoa;Eumetazoa

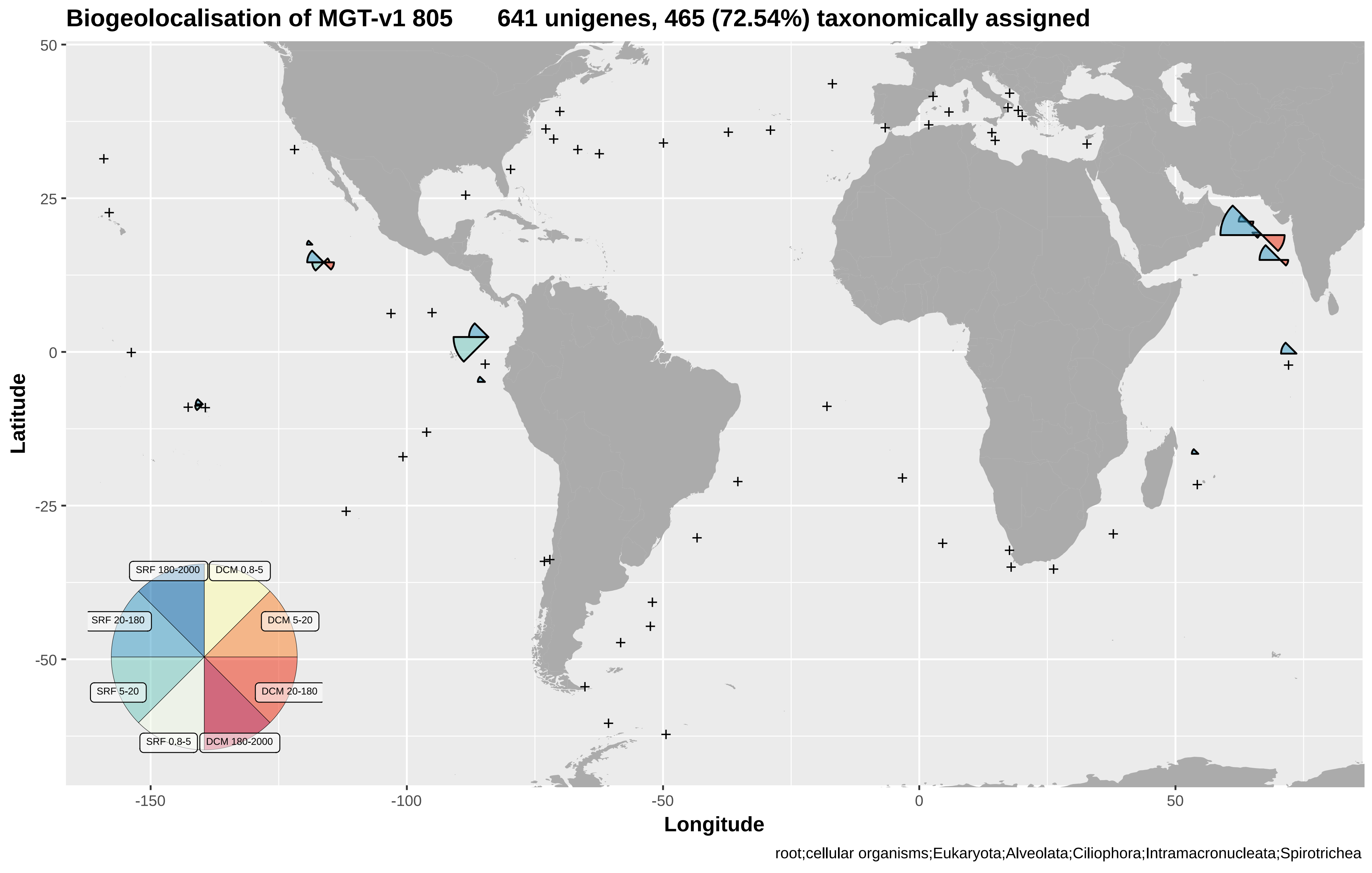

Biogeolocalisation of MGT-v1 805 641 unigenes, 465 (72.54%) taxonomically assigned
50
25
0
-25
-50
Latitude
-150
-100
-50
0
50
Longitude
SRF 180-2000
DCM 0.8-5
SRF 20-180
DCM 5-20
SRF 5-20
DCM 20-180
SRF 0.8-5
DCM 180-2000
root;cellular organisms;Eukaryota;Alveolata;Ciliophora;Intramacronucleata;Spirotrichea

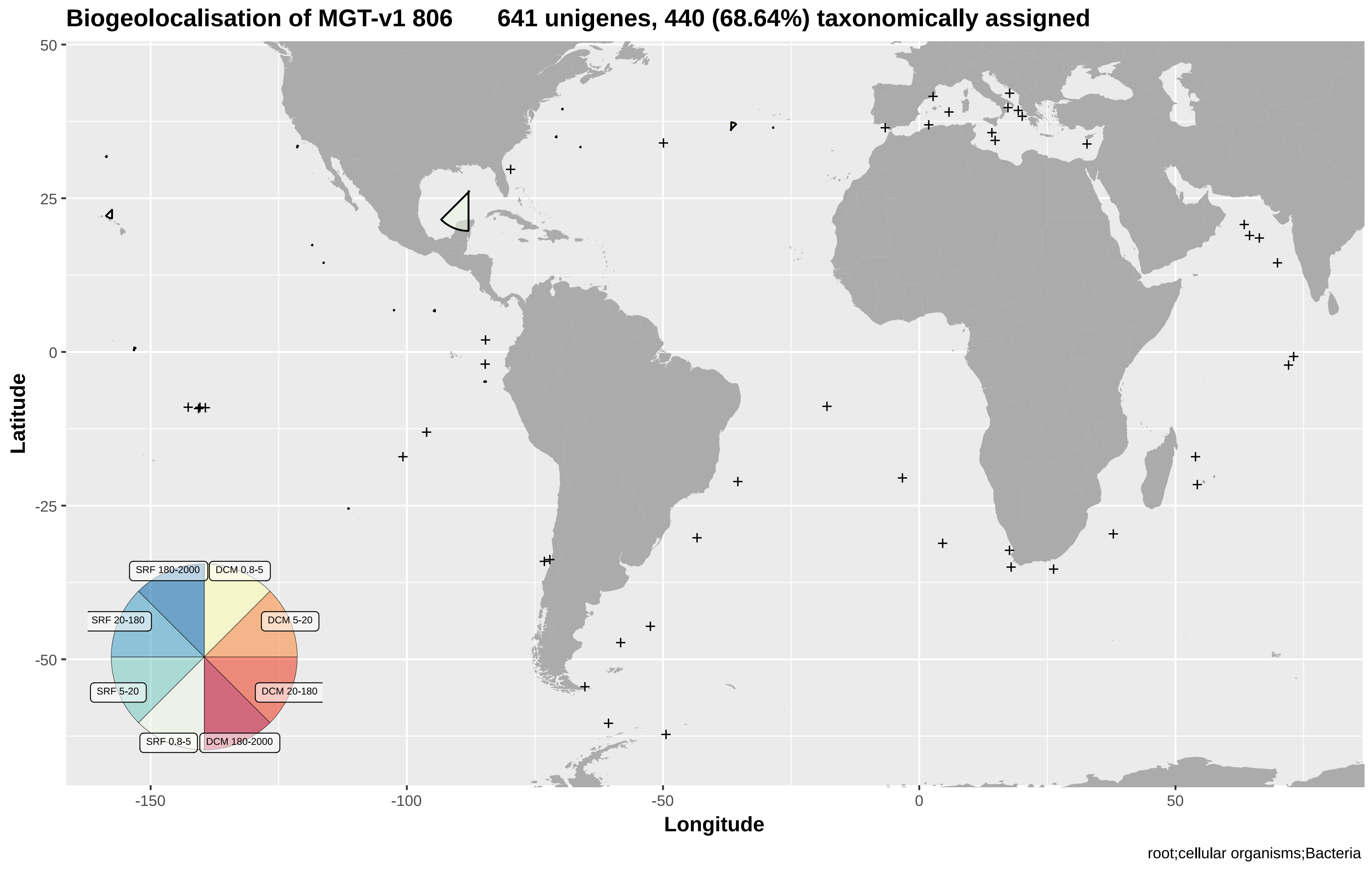
Biogeolocalisation of MGT-v1 806
641 unigenes, 440 (68.64%) taxonomically assigned
Latitude
Longitude
50
25
0
-25
-50
-150
-100
-50
0
50
SRF 180-2000
DCM 0.8-5
SRF 20-180
DCM 5-20
SRF 5-20
DCM 20-180
SRF 0.8-5
DCM 180-2000
root;cellular organisms;Bacteria

## Biogeolocalisation of MGT-v1 807      640 unigenes, 85 (13.28%) taxonomically assigned

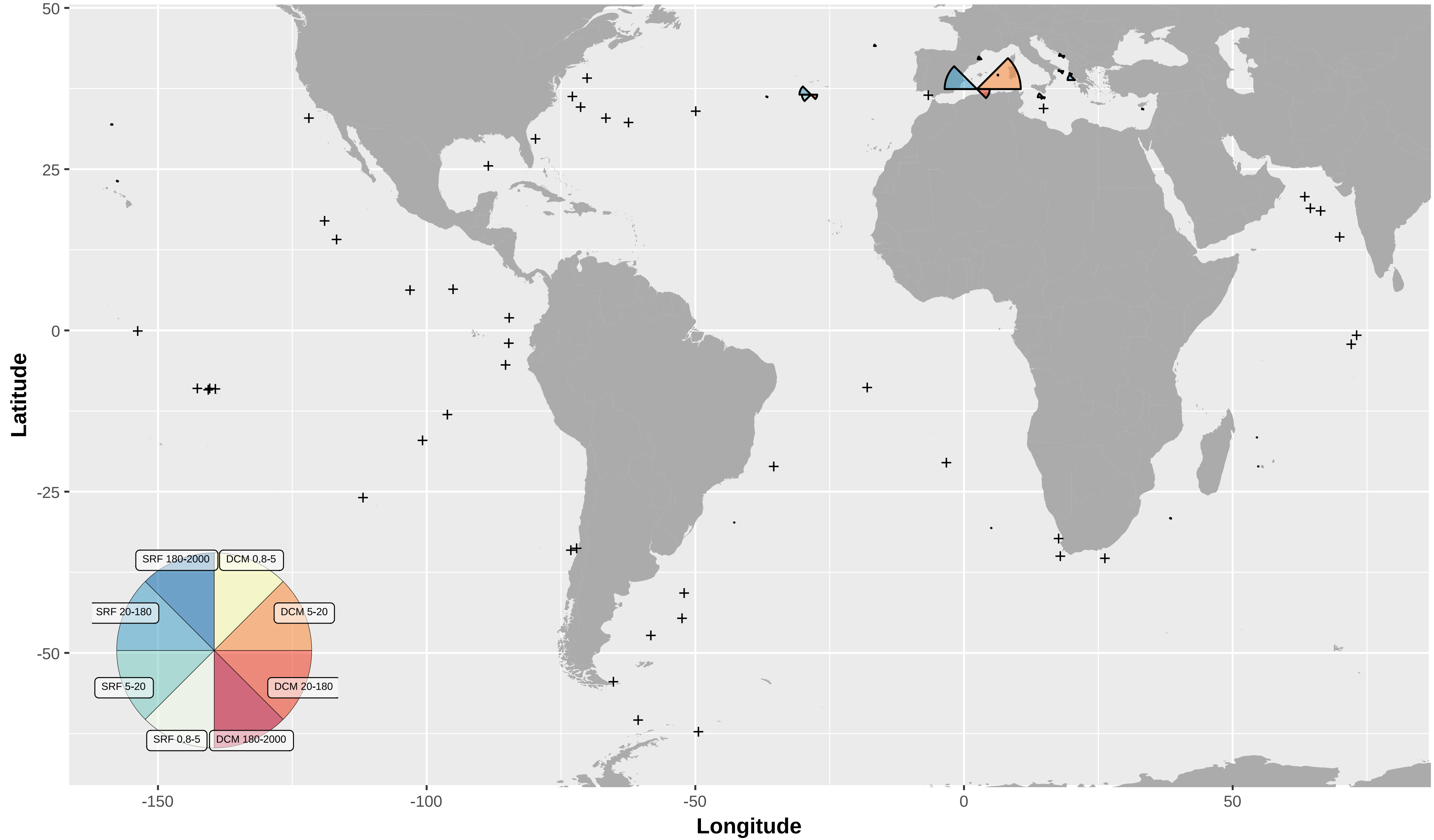


root;cellular organisms;Eukaryota;Opisthokonta;Metazoa;Eumetazoa;Bilateria

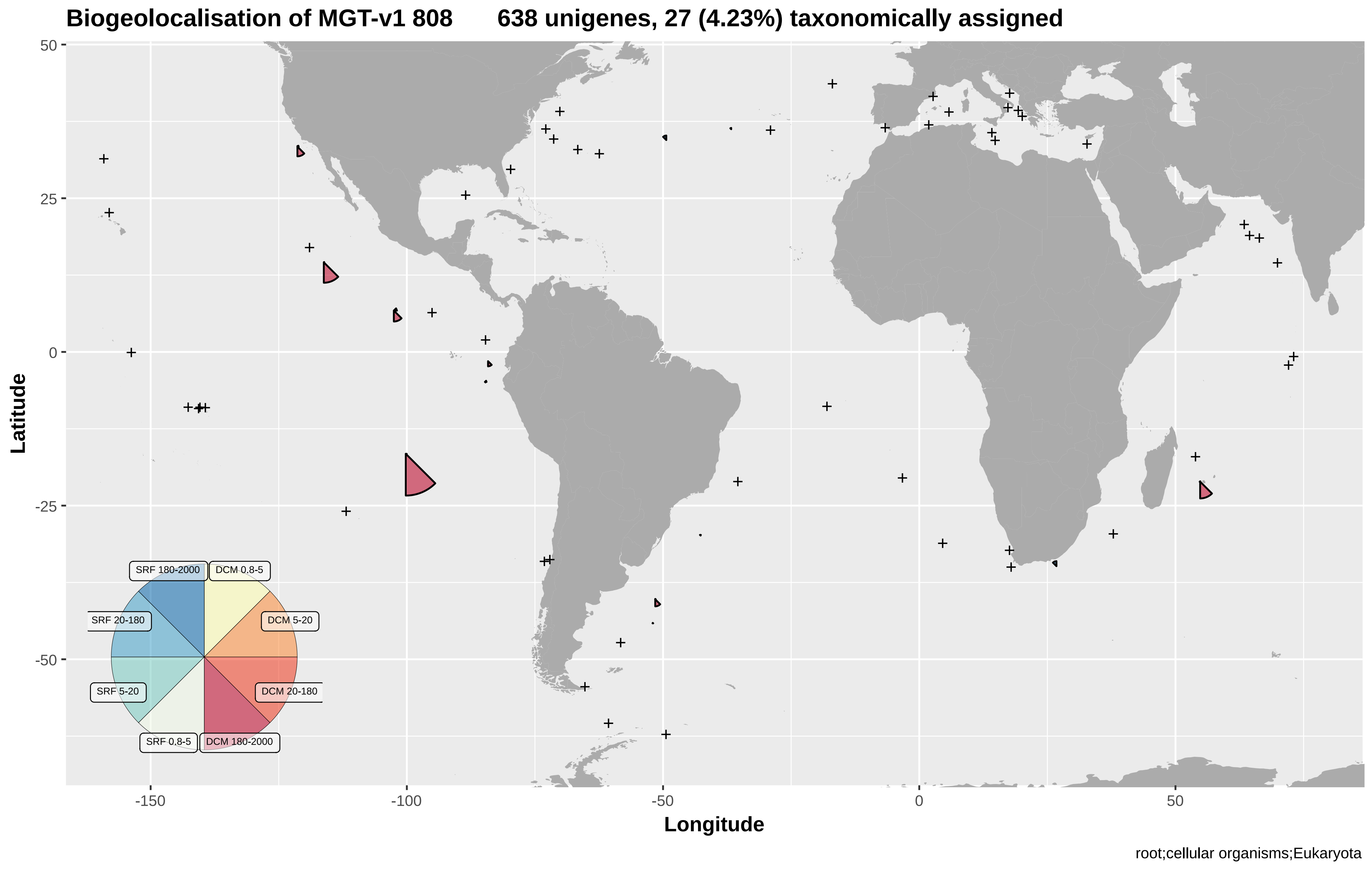
Biogeolocalisation of MGT-v1 808 638 unigenes, 27 (4.23%) taxonomically assigned
Latitude
50
25
0
-25
-50
-150
-100
-50
0
50
Longitude
SRF 180-2000
DCM 0.8-5
SRF 20-180
DCM 5-20
SRF 5-20
DCM 20-180
SRF 0.8-5
DCM 180-2000
root;cellular organisms;Eukaryota

## Biogeolocalisation of MGT-v1 809      636 unigenes, 69 (10.85%) taxonomically assigned

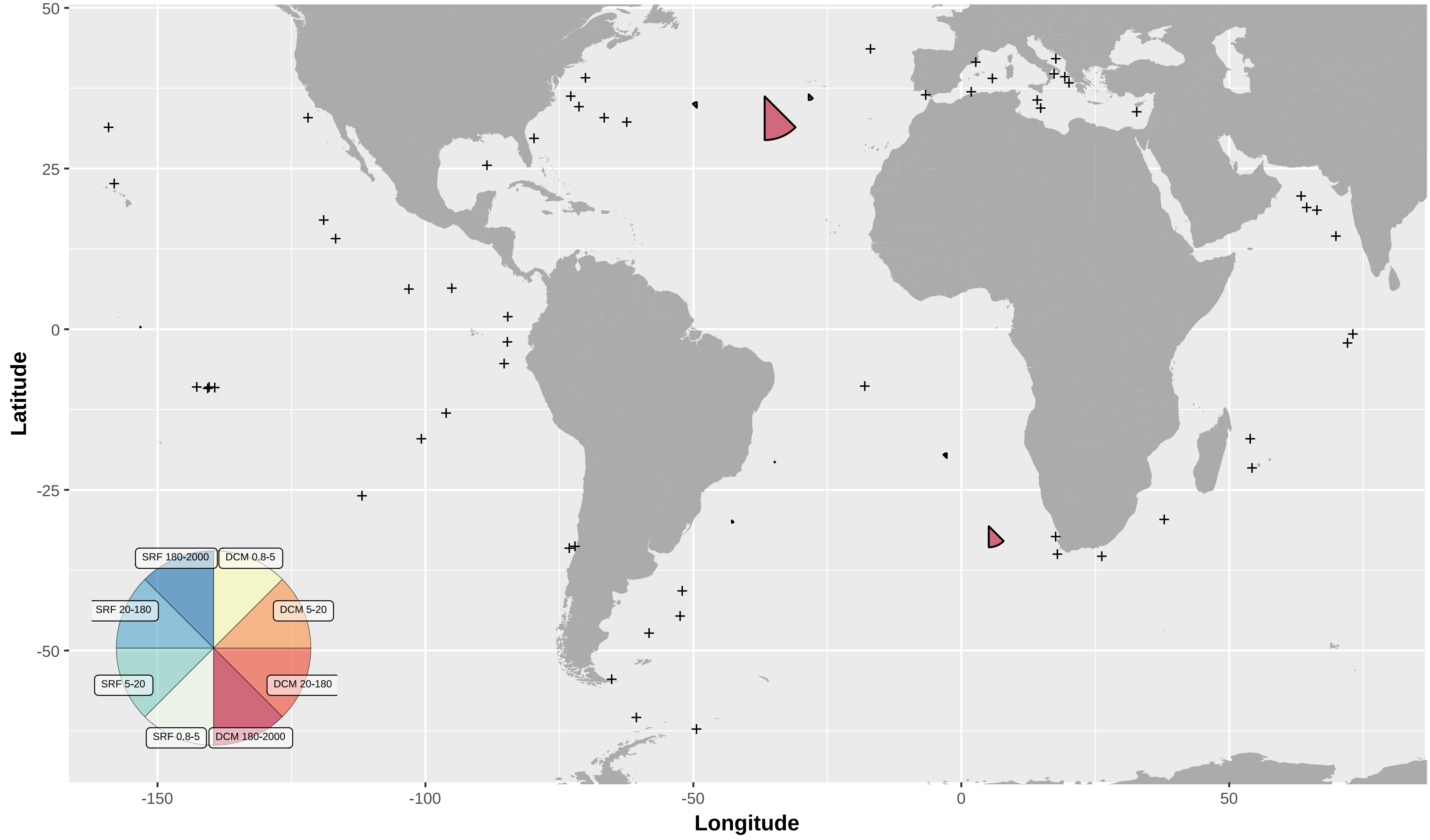


root;cellular organisms;Eukaryota;Opisthokonta;Metazoa;Eumetazoa;Bilateria

Biogeolocalisation of MGT-v1 810 636 unigenes, 515 (80.97%) taxonomically assigned

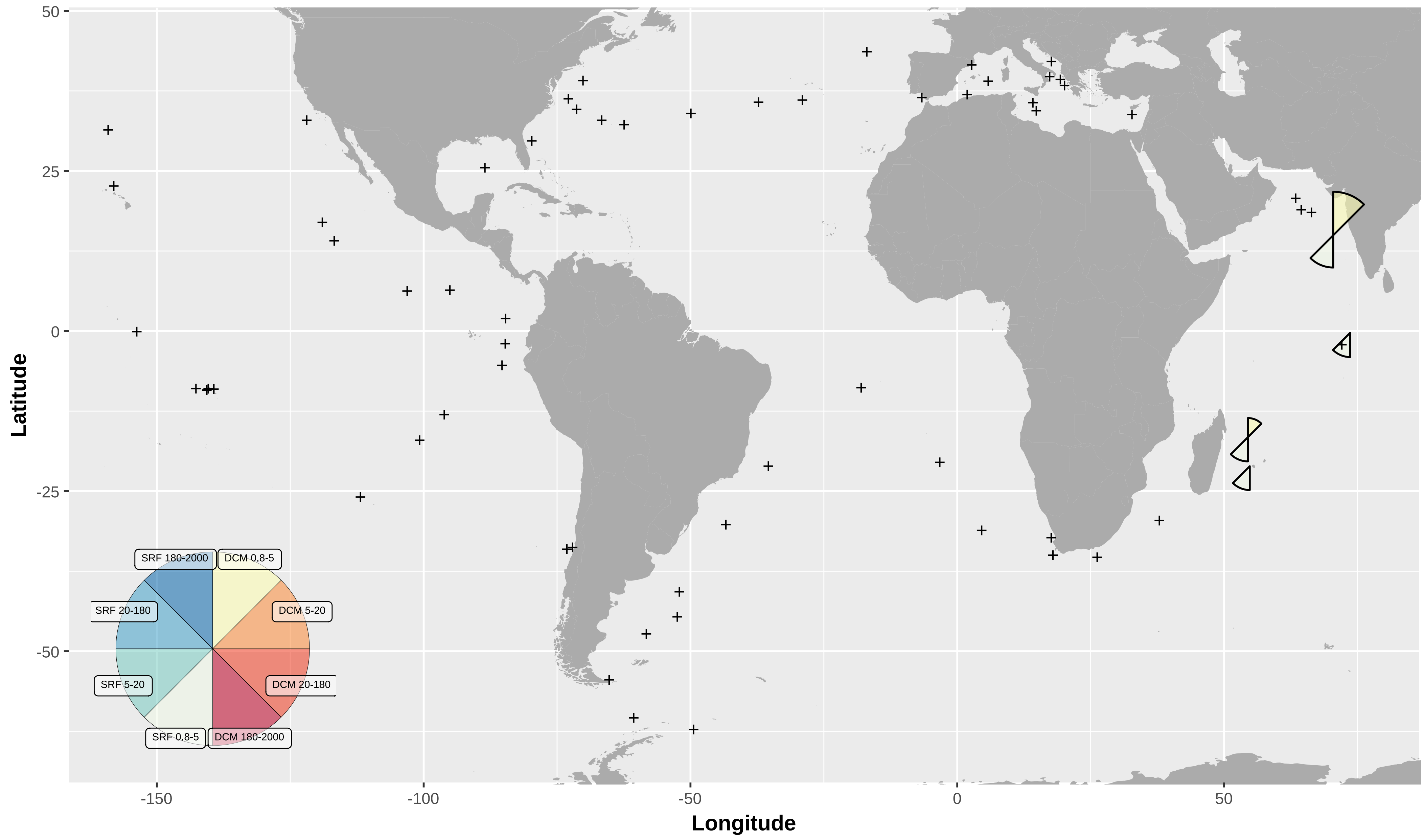


root;cellular organisms;Bacteria

#### Biogeolocalisation of MGT-v1 811 635 unigenes, 106 (16.69%) taxonomically assigned

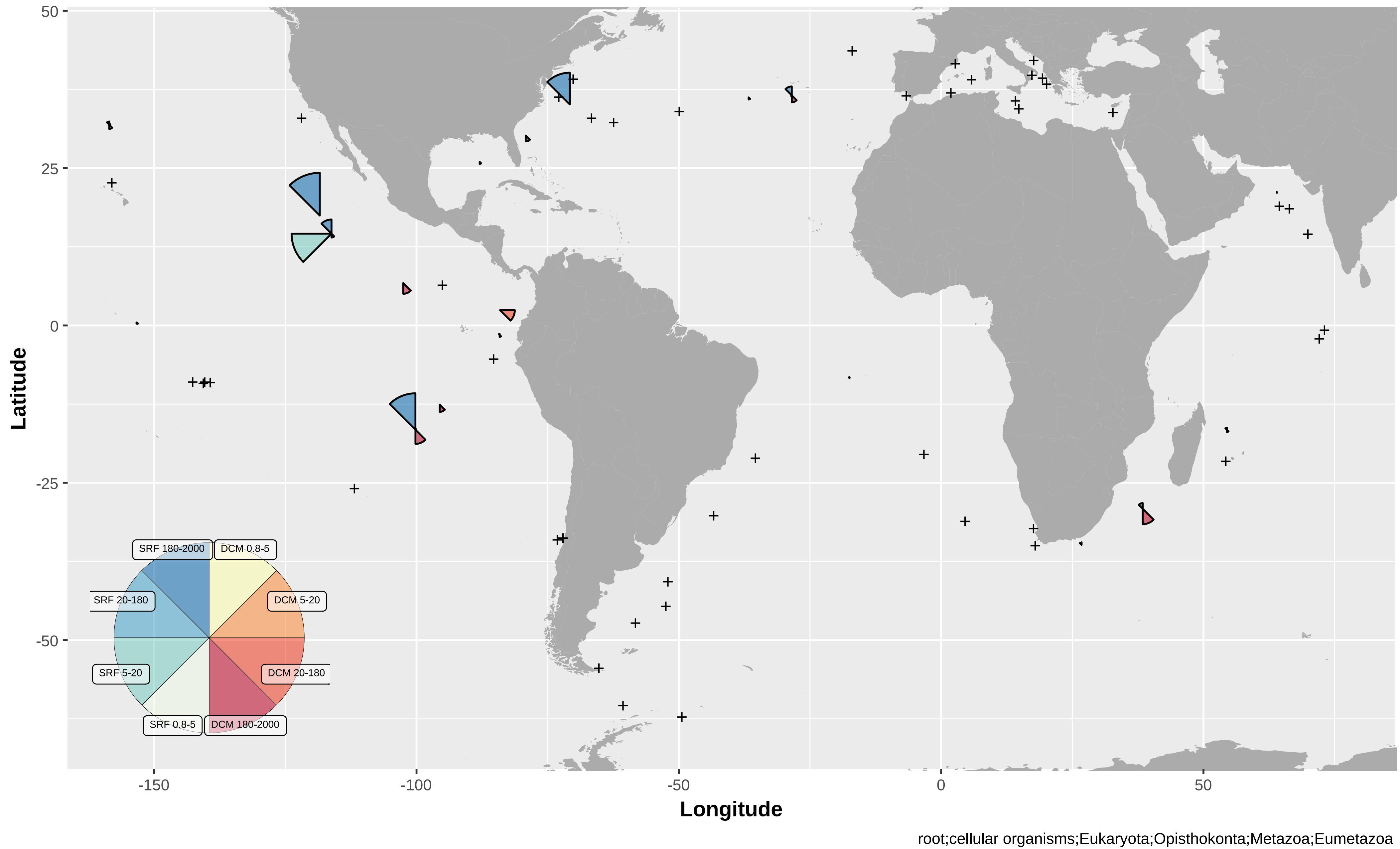

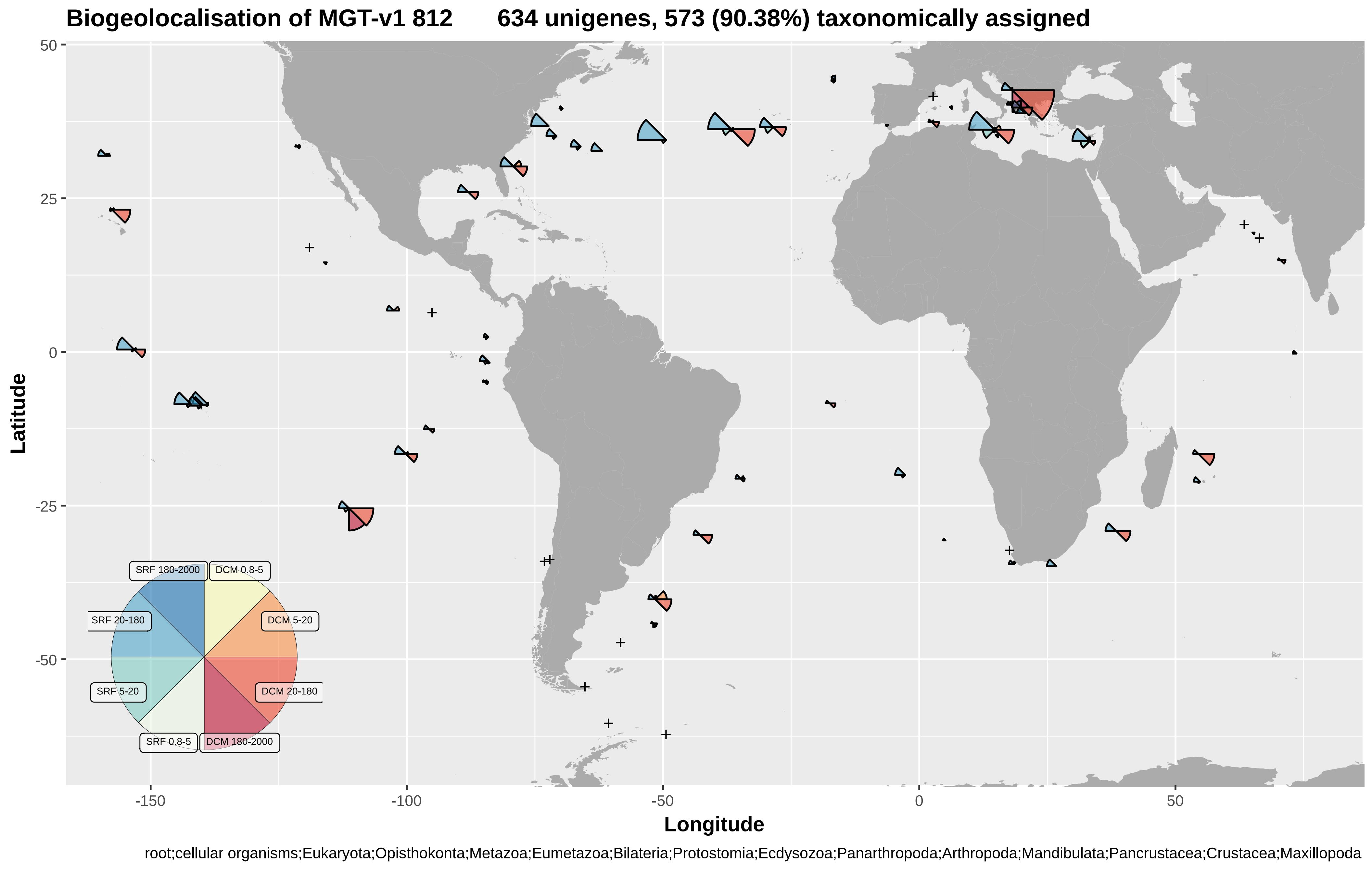

Biogeolocalisation of MGT-v1 812
634 unigenes, 573 (90.38%) taxonomically assigned
50
25
0
-25
-50
Latitude
-150
-100
-50
0
50
Longitude
SRF 180-2000
DCM 0.8-5
SRF 20-180
DCM 5-20
SRF 5-20
DCM 20-180
SRF 0.8-5
DCM 180-2000
root;cellular organisms;Eukaryota;Opisthokonta;Metazoa;Eumetazoa;Bilateria;Protostomia;Ecdysozoa;Panarthropoda;Arthropoda;Mandibulata;Pancrustacea;Crustacea;Maxillopoda

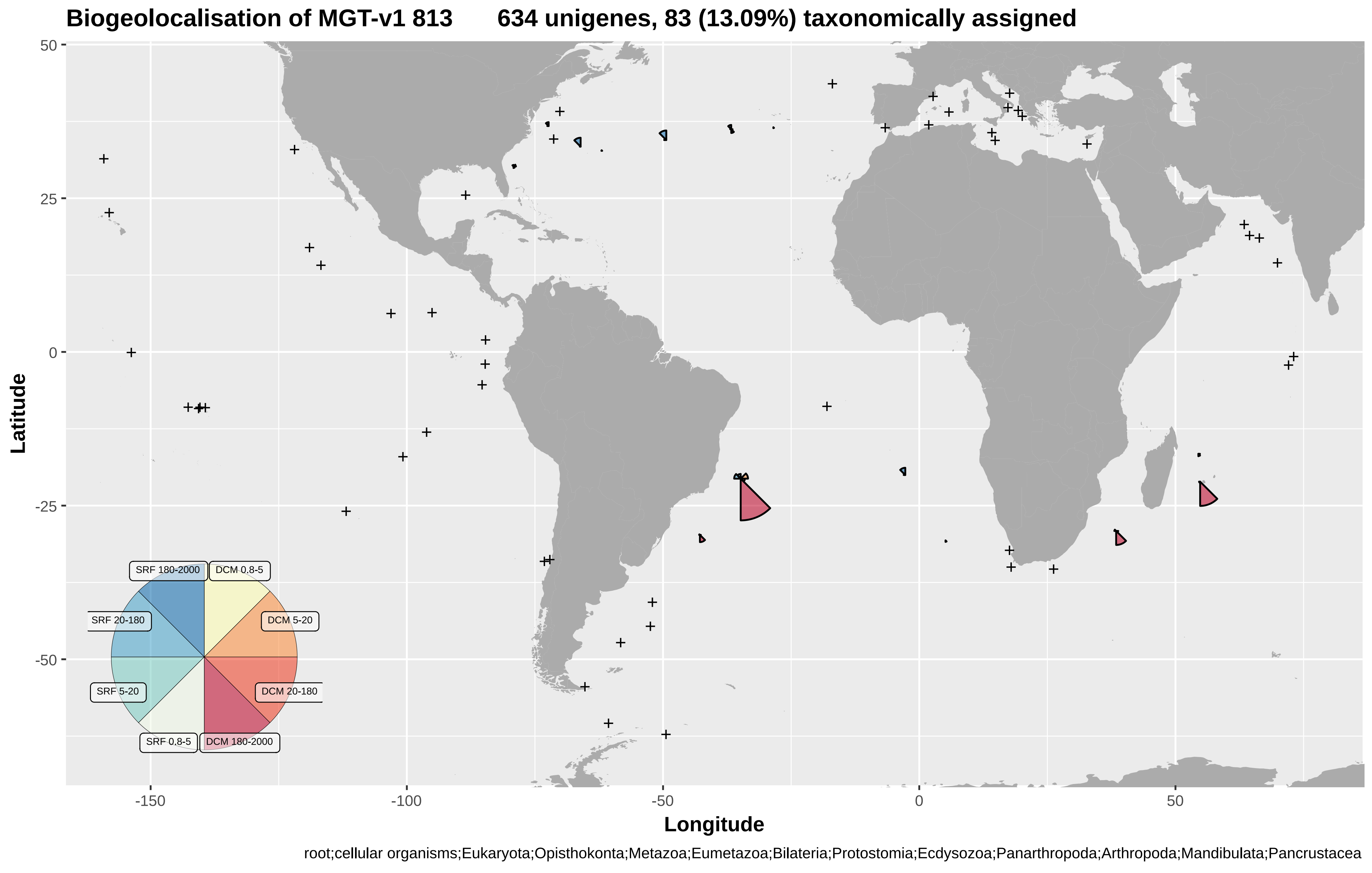
Biogeolocalisation of MGT-v1 813 634 unigenes, 83 (13.09%) taxonomically assigned
Latitude
Longitude
50
25
0
-25
-50
-150
-100
-50
0
50
SRF 180-2000
DCM 0.8-5
SRF 20-180
DCM 5-20
SRF 5-20
DCM 20-180
SRF 0.8-5
DCM 180-2000
root;cellular organisms;Eukaryota;Opisthokonta;Metazoa;Eumetazoa;Bilateria;Protostomia;Ecdysozoa;Panarthropoda;Arthropoda;Mandibulata;Pancrustacea

# Biogeolocalisation of MGT-v1 814      630 unigenes, 145 (23.02%) taxonomically assigned

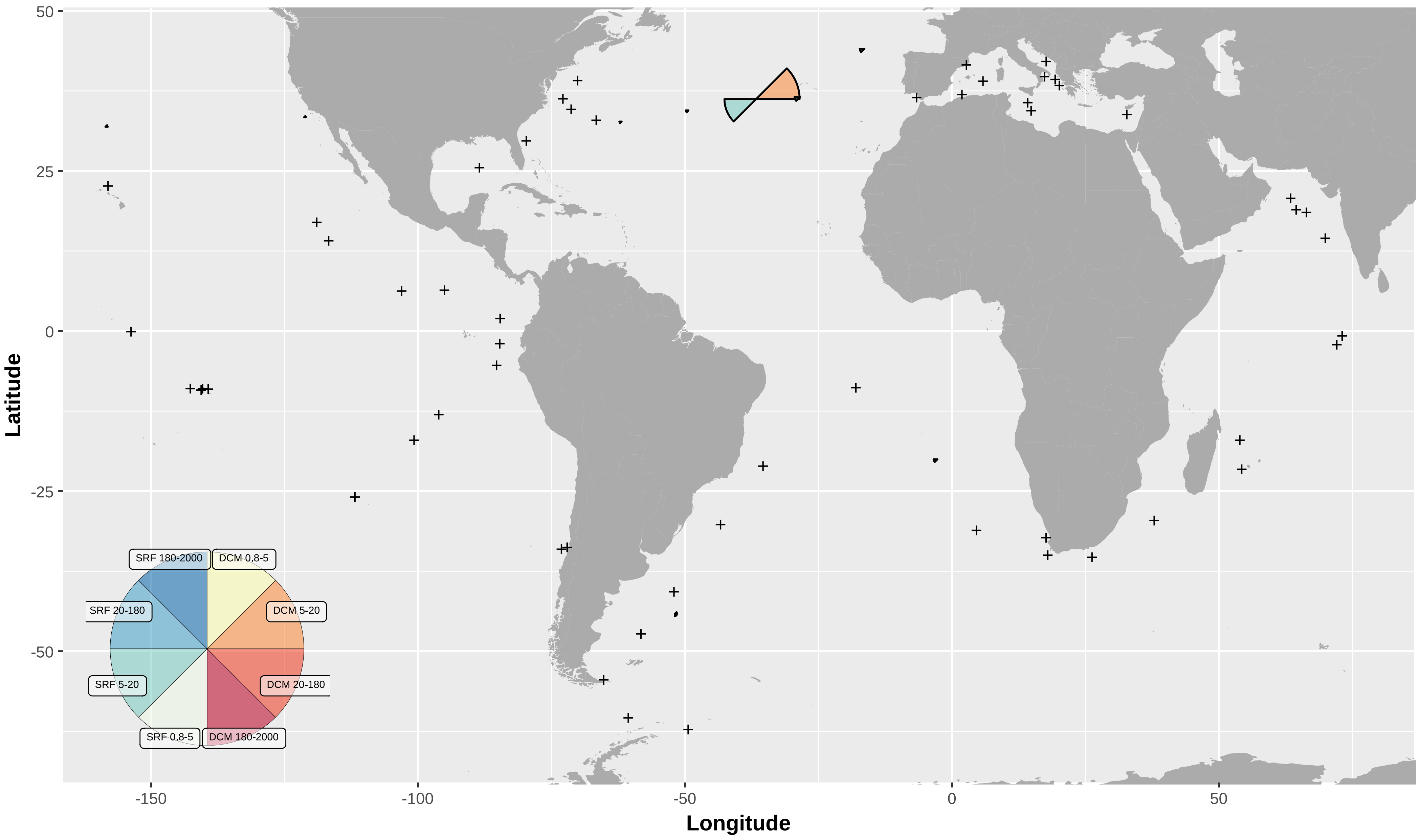


root;cellular organisms;Eukaryota

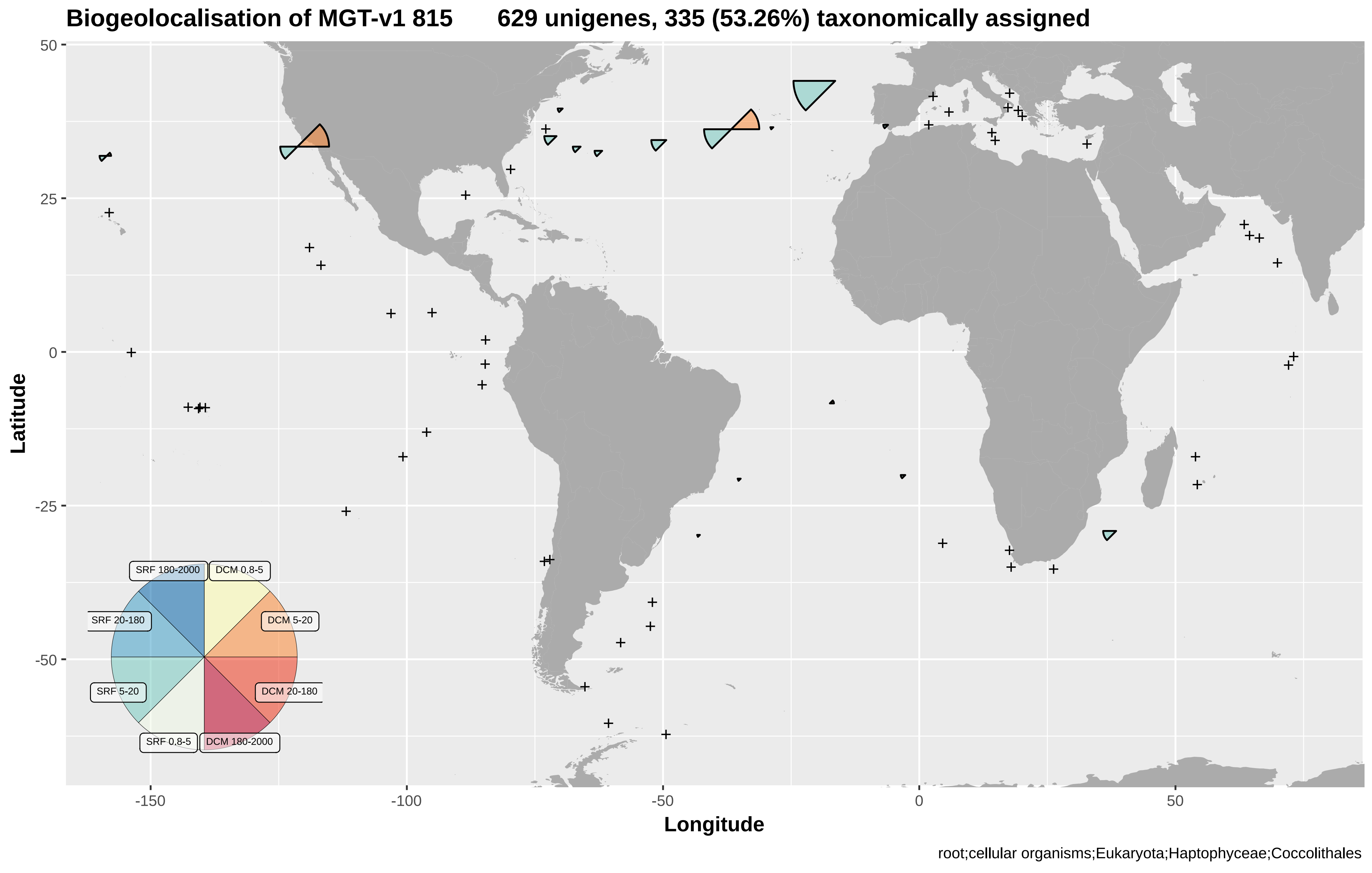
Biogeolocalisation of MGT-v1 815 629 unigenes, 335 (53.26%) taxonomically assigned
SRF 180-2000
DCM 0.8-5
SRF 20-180
DCM 5-20
SRF 5-20
DCM 20-180
SRF 0.8-5
DCM 180-2000
Latitude
50
25
0
-25
-50
Longitude
-150
-100
-50
0
50
root;cellular organisms;Eukaryota;Haptophyceae;Coccolithales

#### Biogeolocalisation of MGT-v1 816 628 unigenes, 593 (94.43%) taxonomically assigned

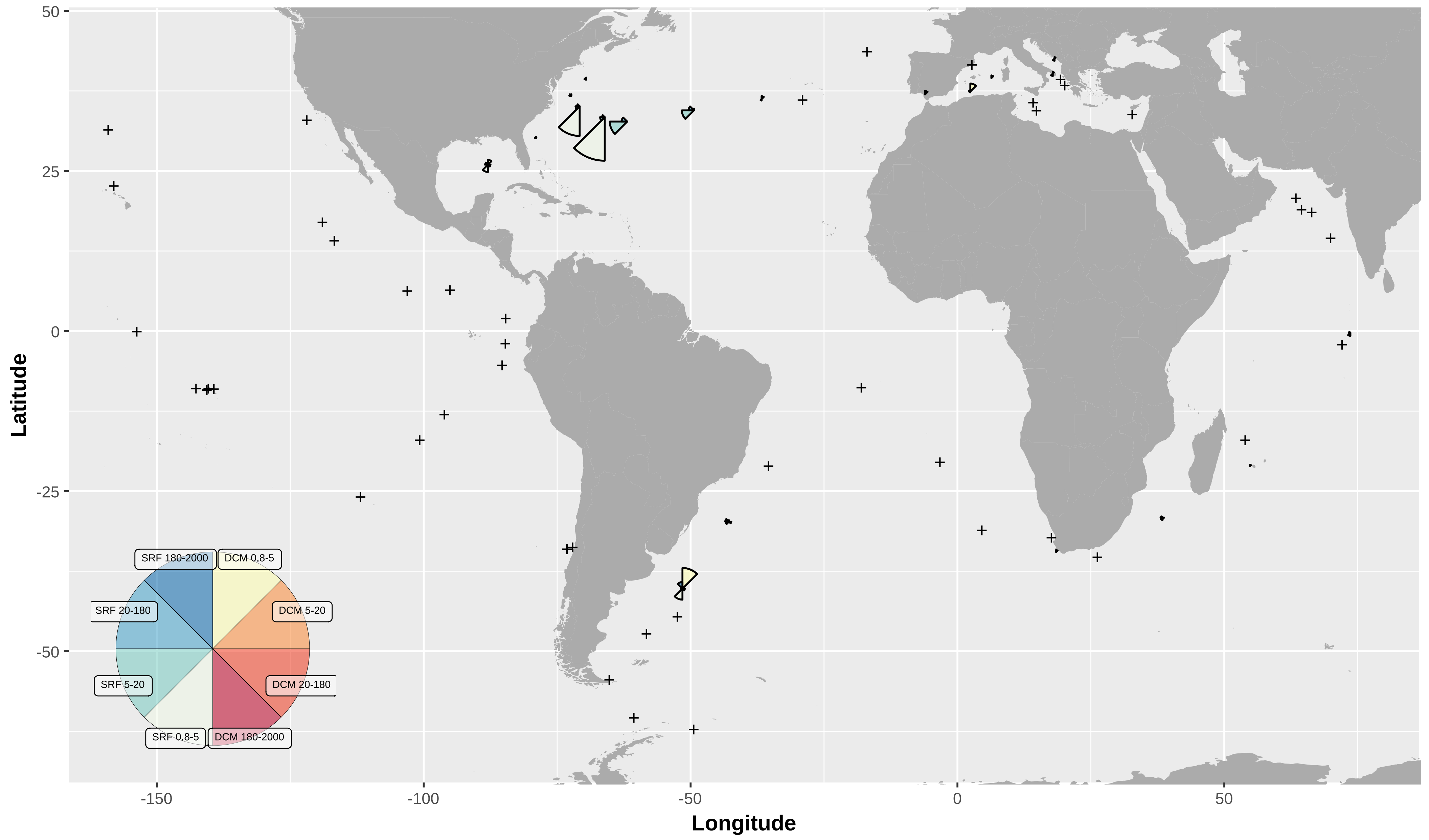


root;cellular organisms;Bacteria;Cyanobacteria;Oscillatoriophycideae;Chroococcales;Synechococcus

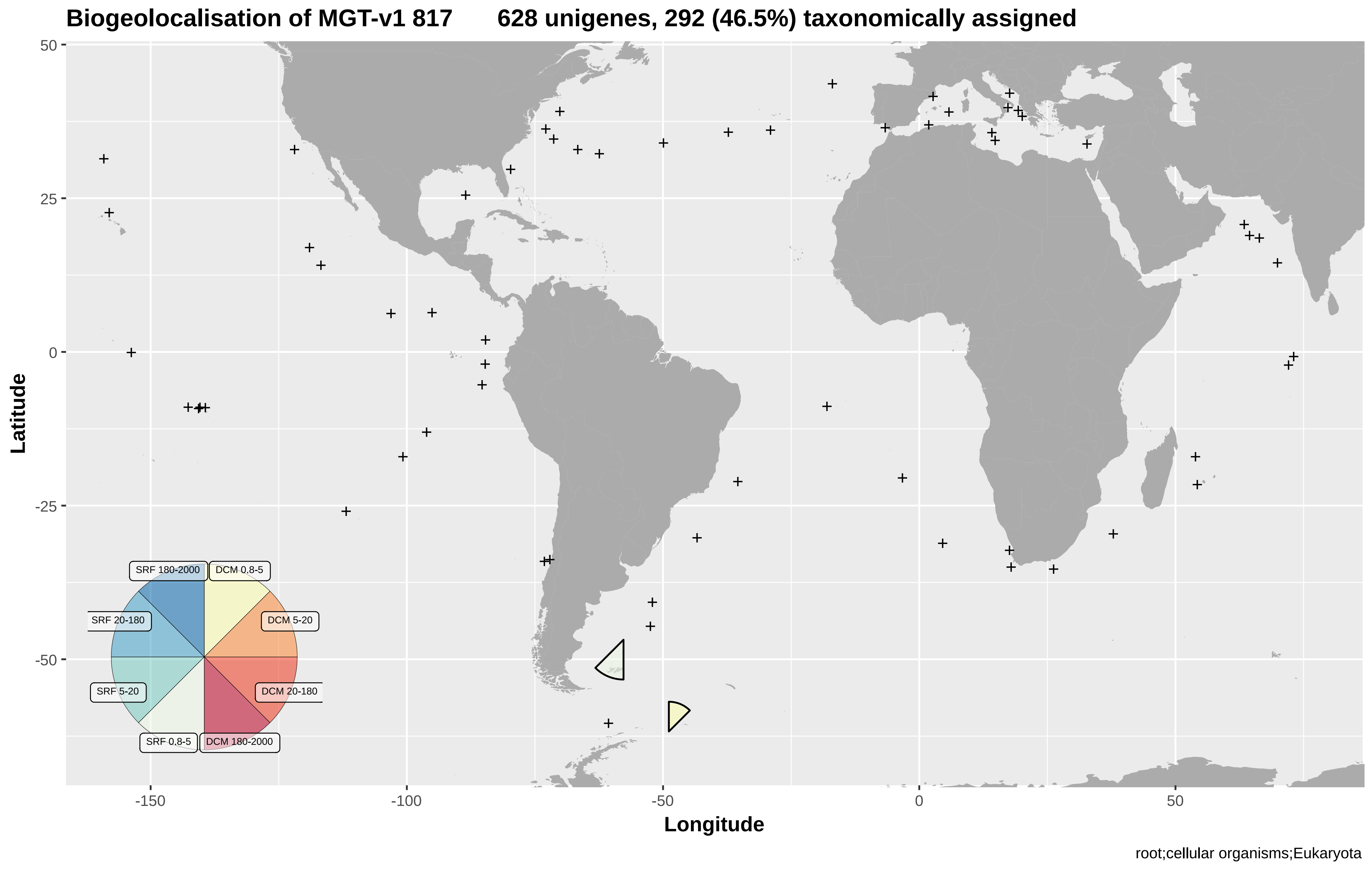

Biogeolocalisation of MGT-v1 817 628 unigenes, 292 (46.5%) taxonomically assigned
50
25
0
-25
-50
Latitude
-150
-100
-50
0
50
Longitude
SRF 180-2000
DCM 0.8-5
SRF 20-180
DCM 5-20
SRF 5-20
DCM 20-180
SRF 0.8-5
DCM 180-2000
root;cellular organisms;Eukaryota

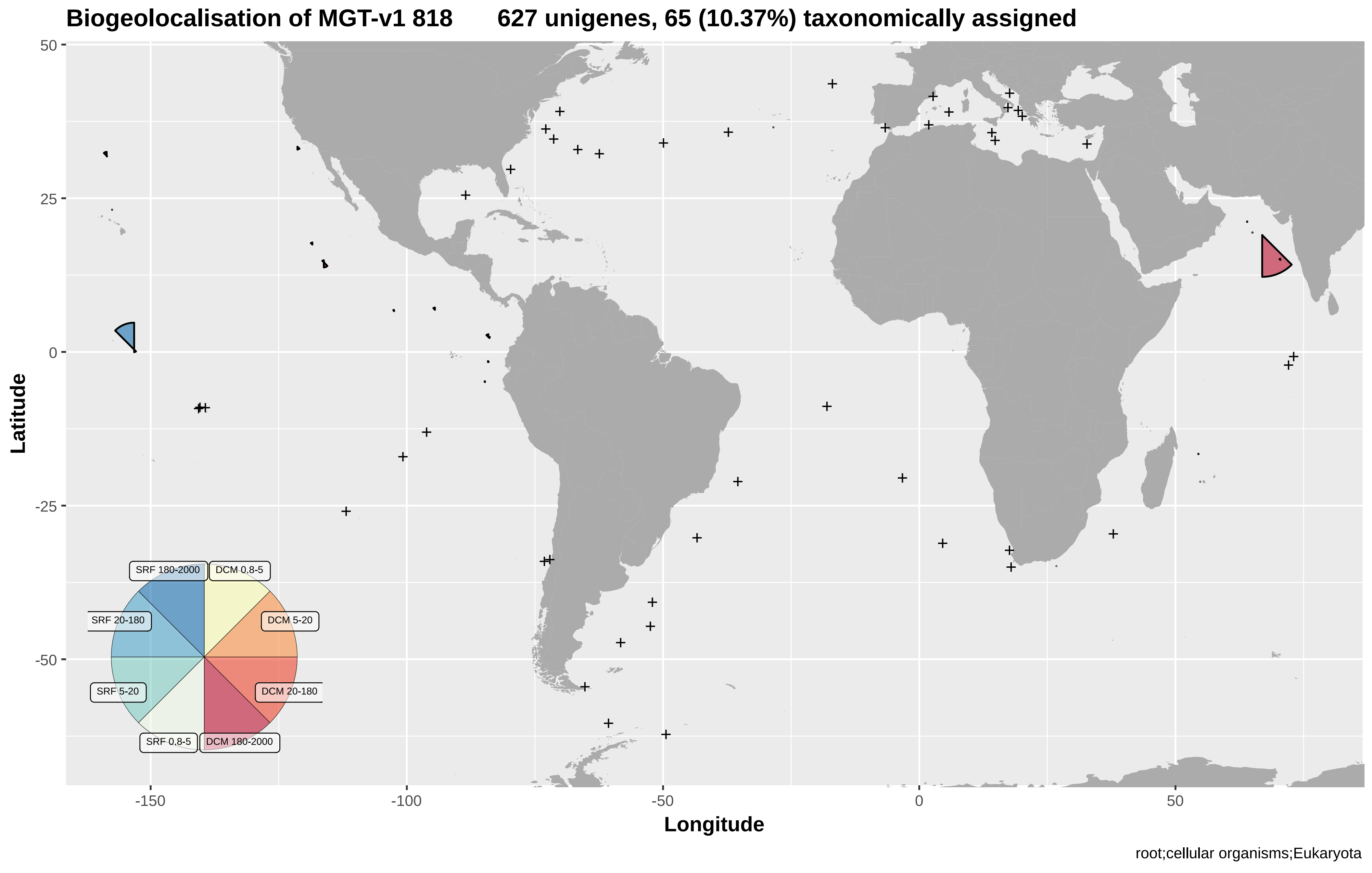

Biogeolocalisation of MGT-v1 818 627 unigenes, 65 (10.37%) taxonomically assigned
50
25
0
-25
-50
Latitude
-150
-100
-50
0
50
Longitude
SRF 180-2000
DCM 0.8-5
SRF 20-180
DCM 5-20
SRF 5-20
DCM 20-180
SRF 0.8-5
DCM 180-2000
root;cellular organisms;Eukaryota

## Biogeolocalisation of MGT-v1 819      627 unigenes, 143 (22.81%) taxonomically assigned

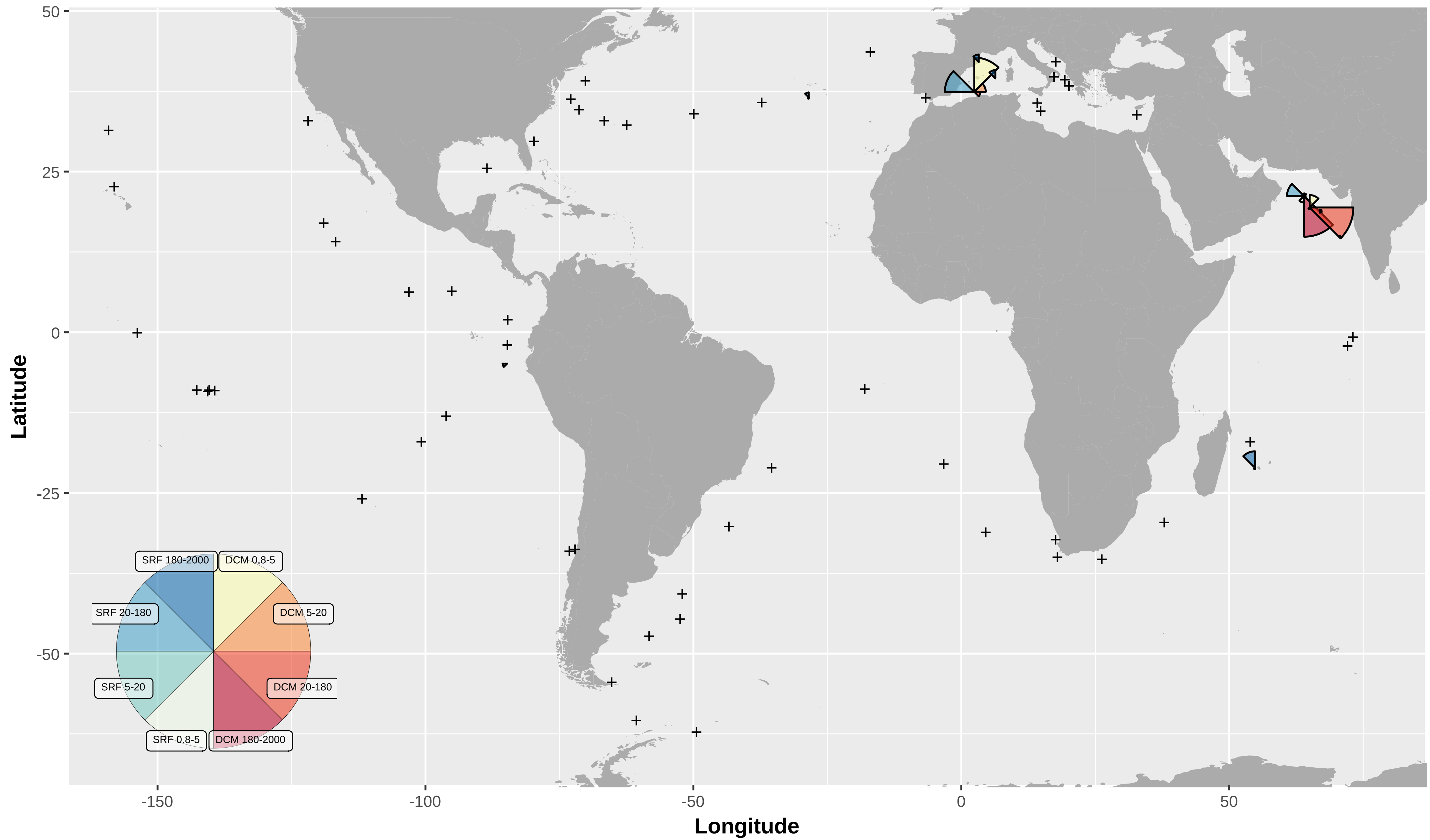


root;cellular organisms;Eukaryota;Opisthokonta;Metazoa;Eumetazoa

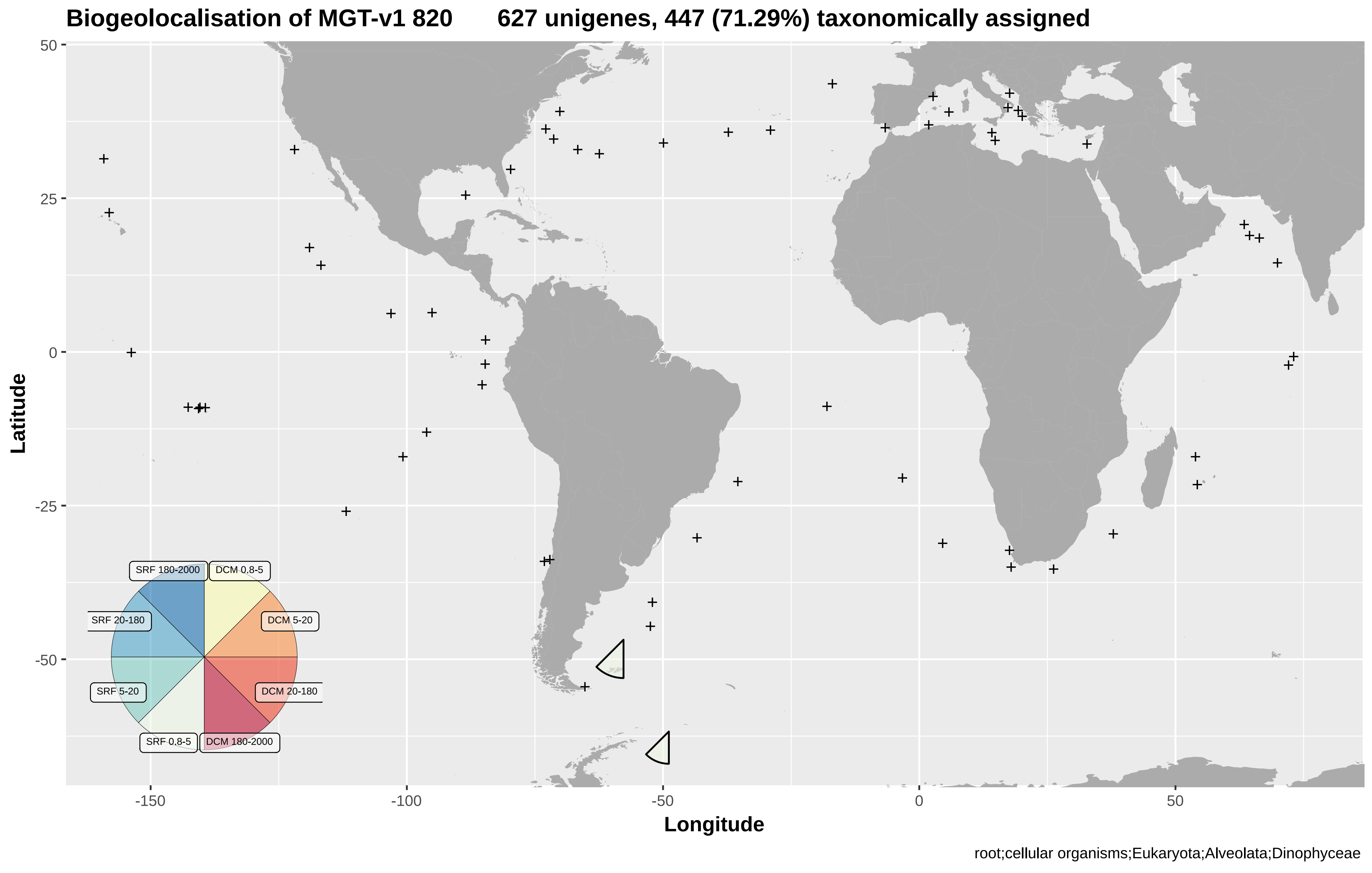
Biogeolocalisation of MGT-v1 820 627 unigenes, 447 (71.29%) taxonomically assigned
50
25
0
-25
-50
Latitude
-150
-100
-50
0
50
Longitude
SRF 180-2000
DCM 0.8-5
SRF 20-180
DCM 5-20
SRF 5-20
DCM 20-180
SRF 0.8-5
DCM 180-2000
root;cellular organisms;Eukaryota;Alveolata;Dinophyceae

**Biogeolocalisation of MGT-v1 821 626 unigenes, 419 (66.93%) taxonomically assigned**

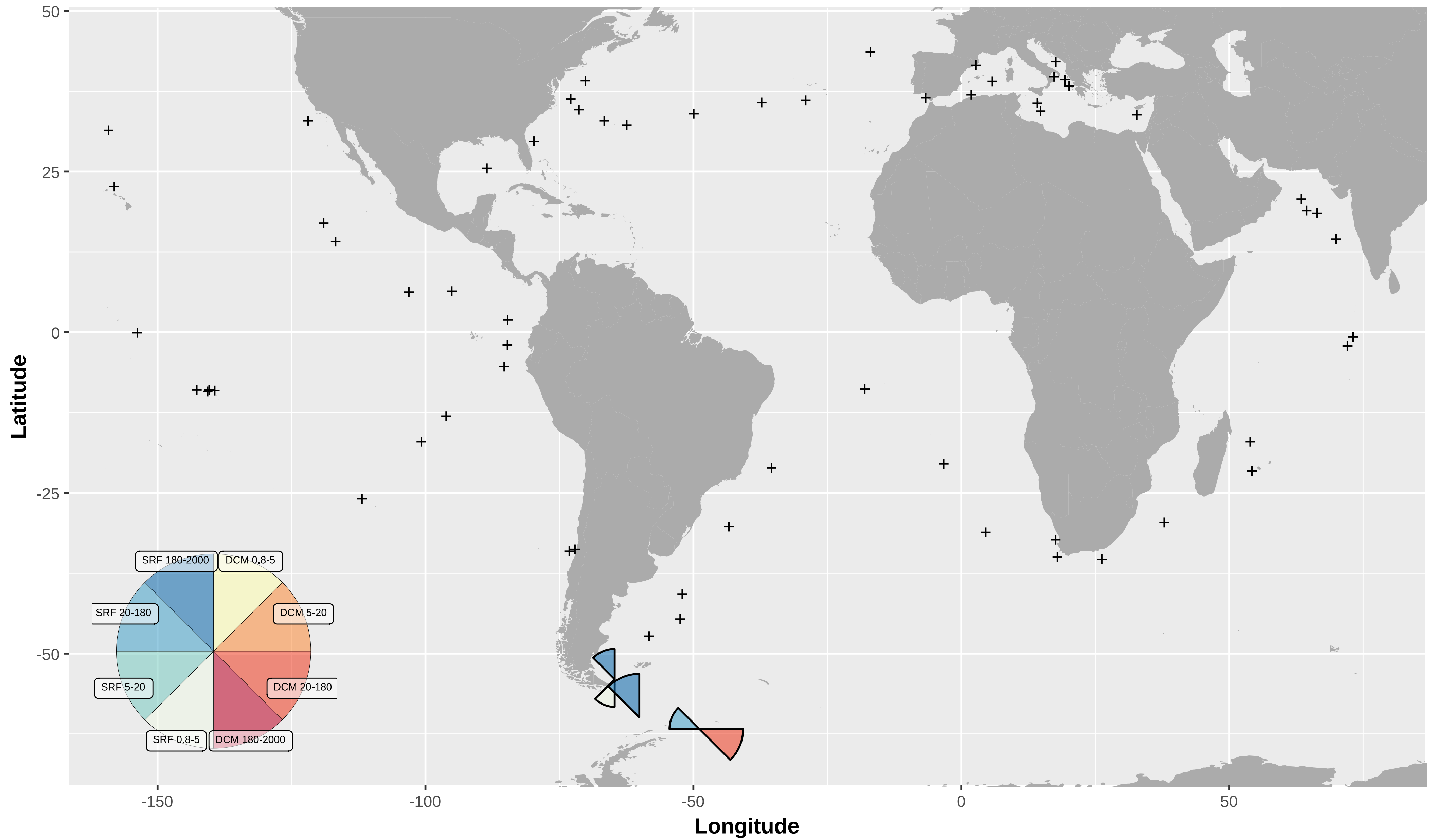


root;cellular organisms;Eukaryota;Opisthokonta;Metazoa;Eumetazoa;Bilateria

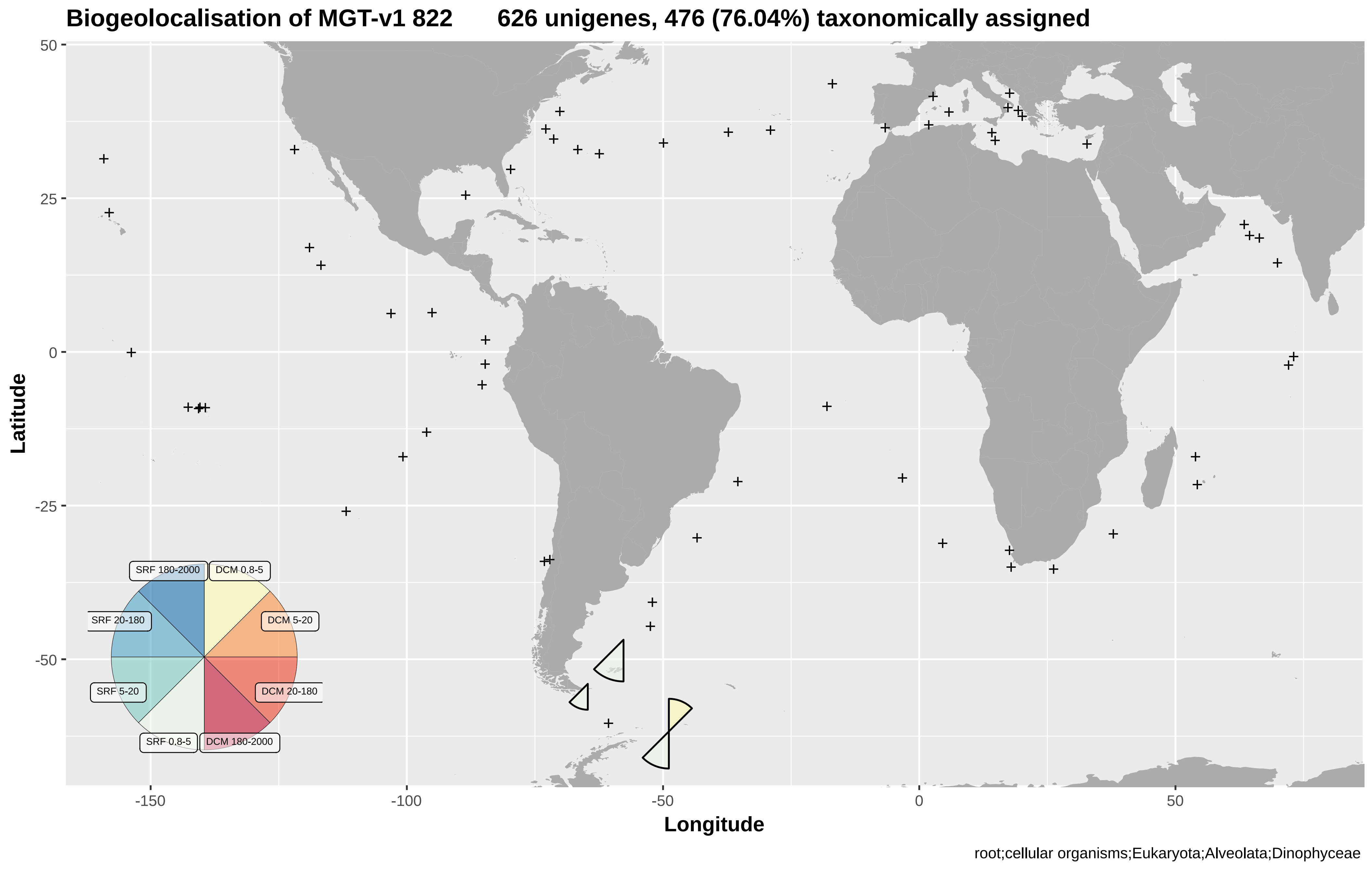
Biogeolocalisation of MGT-v1 822 626 unigenes, 476 (76.04%) taxonomically assigned
50
25
0
-25
-50
Latitude
-150
-100
-50
0
50
Longitude
SRF 180-2000
DCM 0.8-5
SRF 20-180
DCM 5-20
SRF 5-20
DCM 20-180
SRF 0.8-5
DCM 180-2000
root;cellular organisms;Eukaryota;Alveolata;Dinophyceae

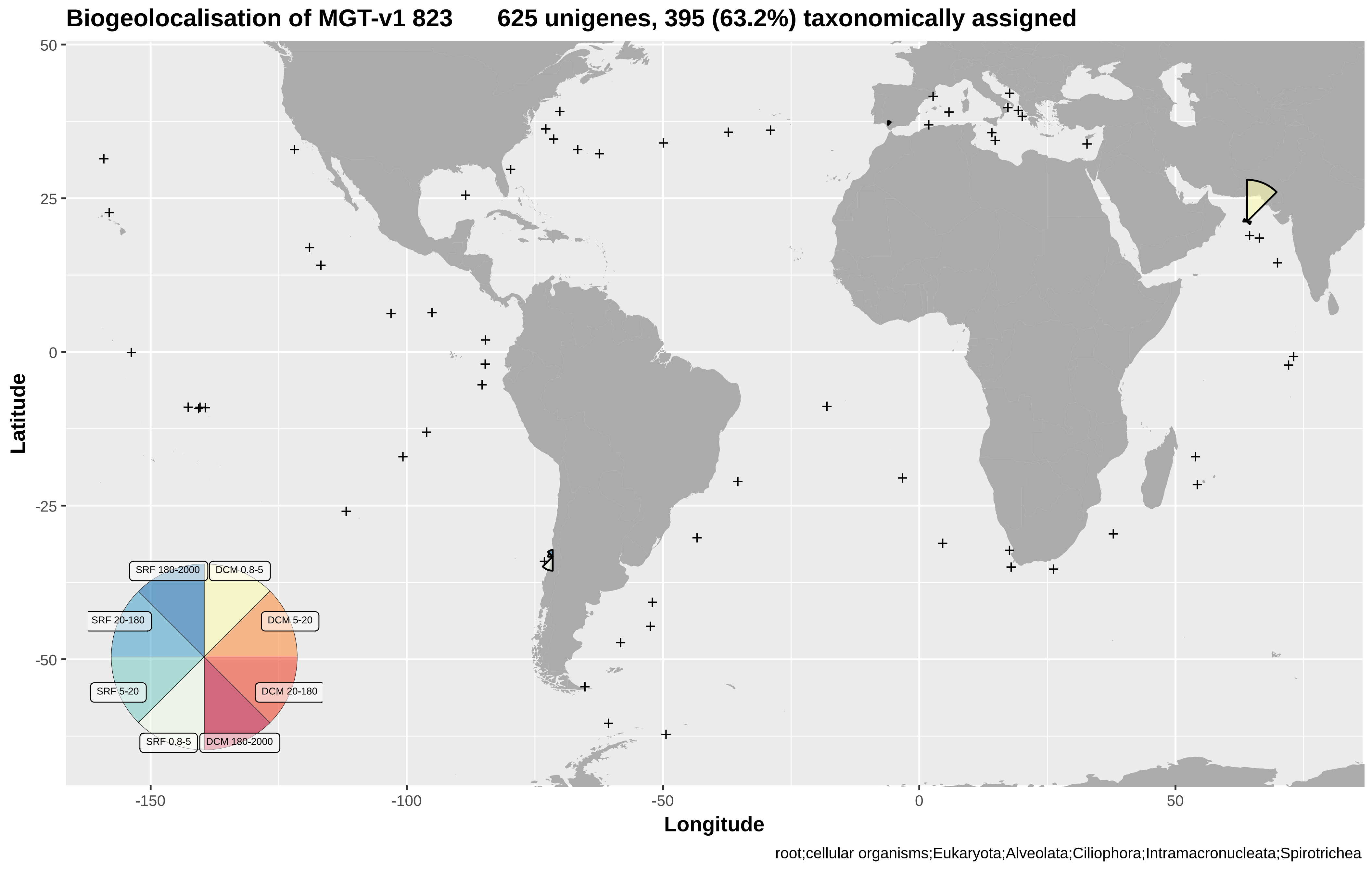

Biogeolocalisation of MGT-v1 823 625 unigenes, 395 (63.2%) taxonomically assigned
50
25
0
-25
-50
Latitude
-150
-100
-50
0
50
Longitude
SRF 180-2000
DCM 0.8-5
SRF 20-180
DCM 5-20
SRF 5-20
DCM 20-180
SRF 0.8-5
DCM 180-2000
root;cellular organisms;Eukaryota;Alveolata;Ciliophora;Intramacronucleata;Spirotrichea

## Biogeolocalisation of MGT-v1 824      625 unigenes, 170 (27.2%) taxonomically assigned

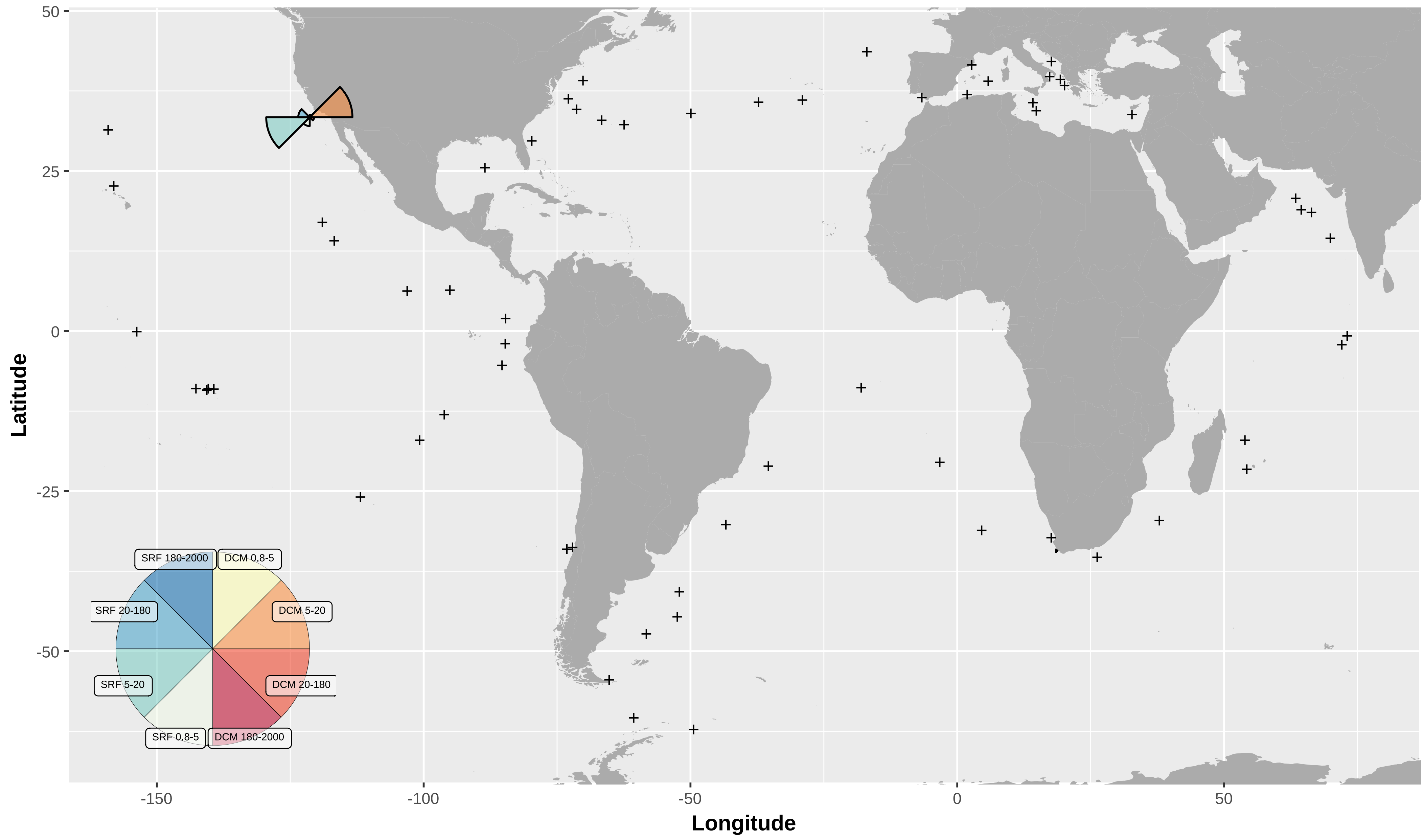


root;cellular organisms

## Biogeolocalisation of MGT-v1 825      625 unigenes, 32 (5.12%) taxonomically assigned

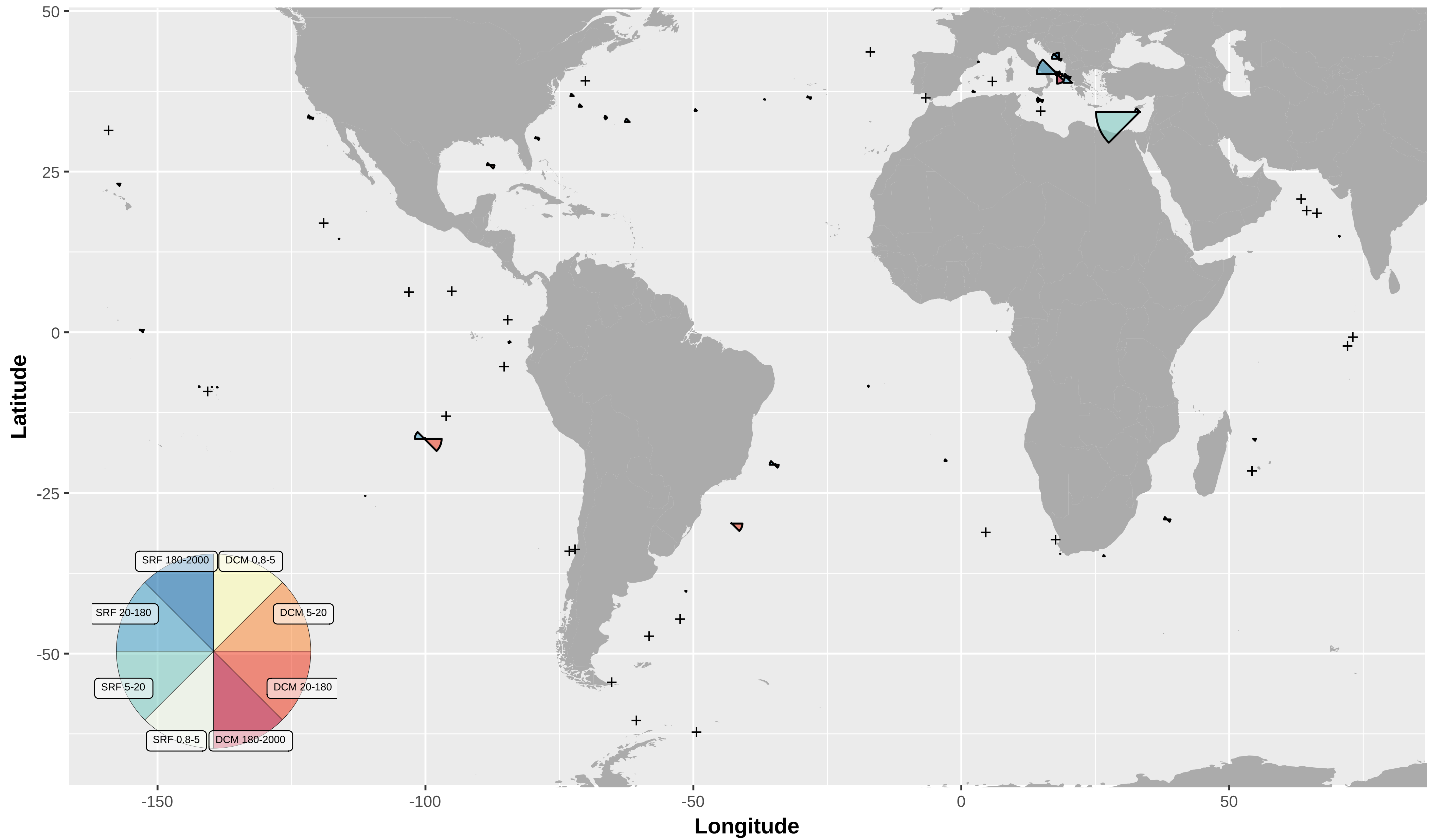


root;cellular organisms;Eukaryota;Opisthokonta;Metazoa;Eumetazoa;Bilateria;Protostomia

### Biogeolocalisation of MGT-v1 826 624 unigenes, 396 (63.46%) taxonomically assigned

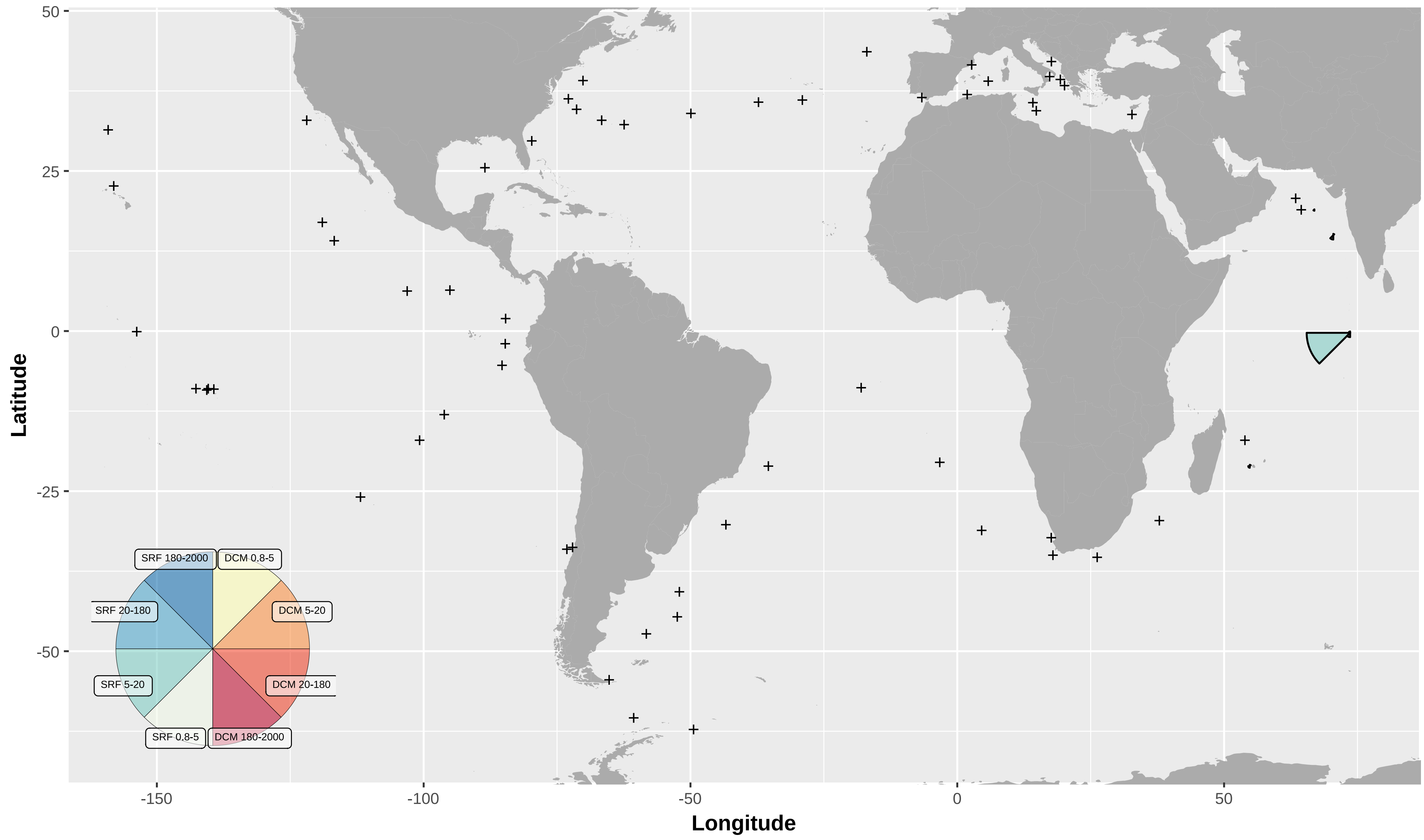


root;cellular organisms;Bacteria

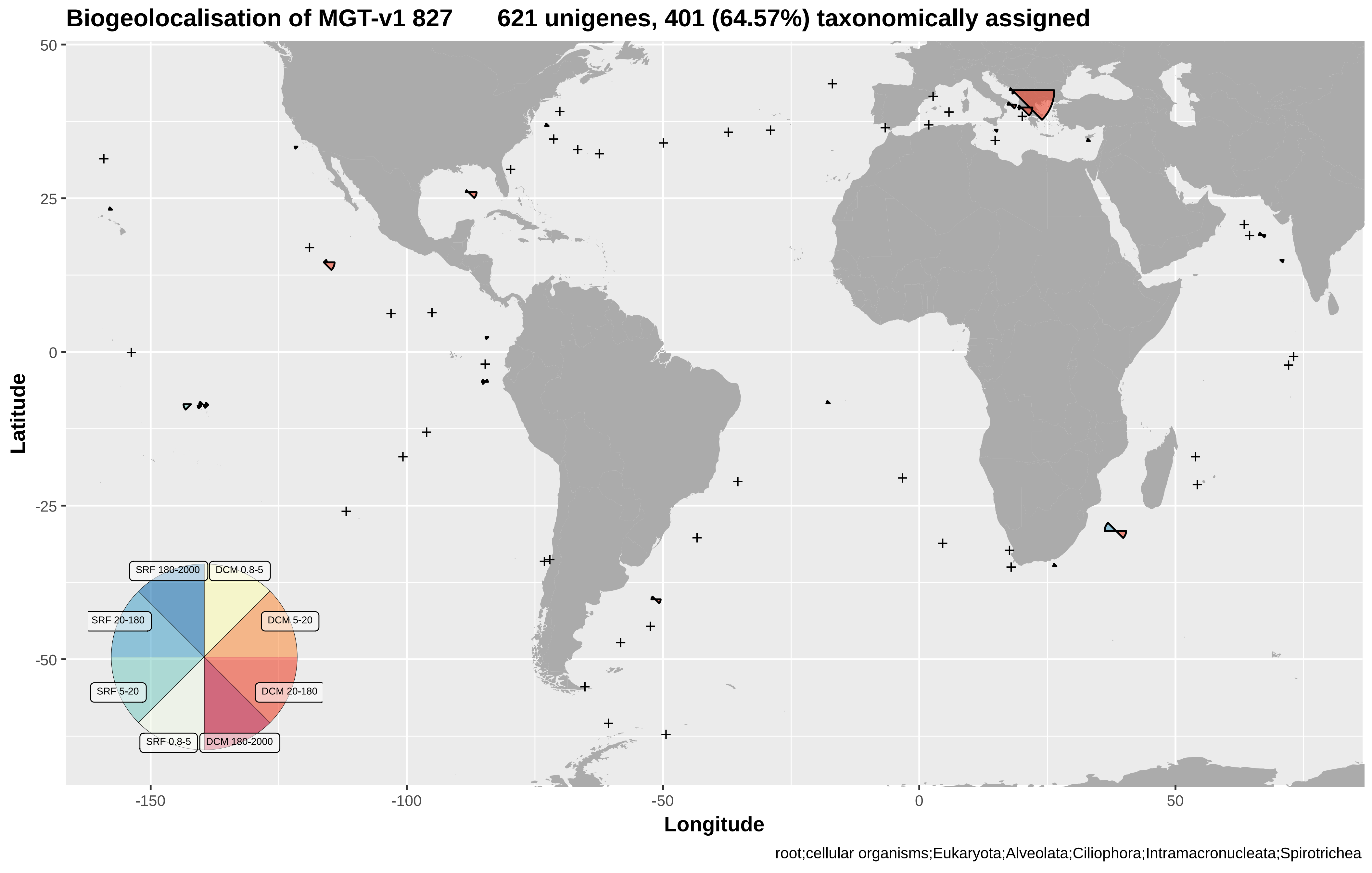
Biogeolocalisation of MGT-v1 827 621 unigenes, 401 (64.57%) taxonomically assigned
Latitude
Longitude
50
25
0
-25
-50
-150
-100
-50
0
50
SRF 180-2000
DCM 0.8-5
SRF 20-180
DCM 5-20
SRF 5-20
DCM 20-180
SRF 0.8-5
DCM 180-2000
root;cellular organisms;Eukaryota;Alveolata;Ciliophora;Intramacronucleata;Spirotrichea

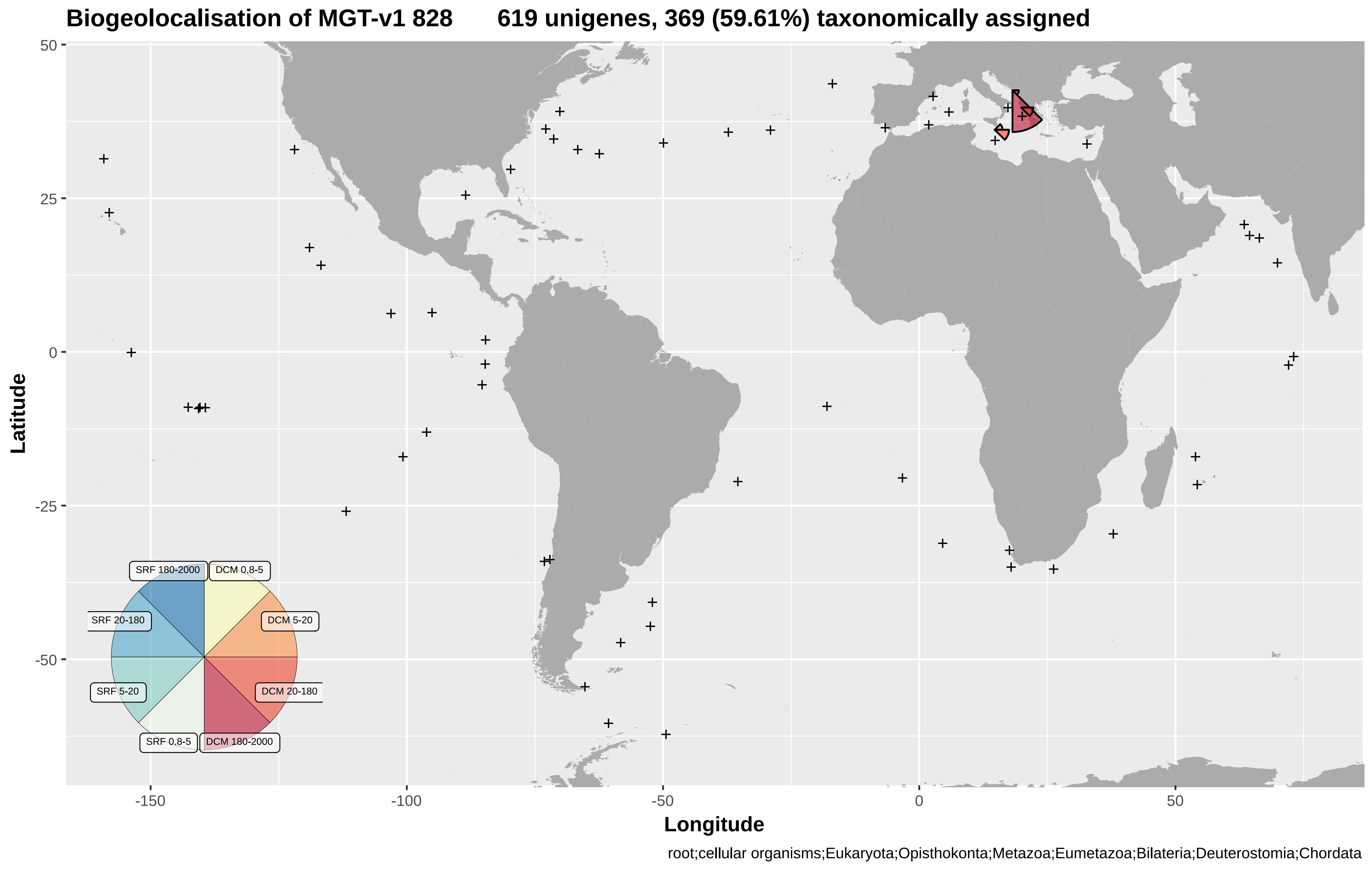
Biogeolocalisation of MGT-v1 828 619 unigenes, 369 (59.61%) taxonomically assigned
50
25
0
-25
-50
Latitude
-150
-100
-50
0
50
Longitude
SRF 180-2000
DCM 0.8-5
SRF 20-180
DCM 5-20
SRF 5-20
DCM 20-180
SRF 0.8-5
DCM 180-2000
root;cellular organisms;Eukaryota;Opisthokonta;Metazoa;Eumetazoa;Bilateria;Deuterostomia;Chordata

# Biogeolocalisation of MGT-v1 829      619 unigenes, 556 (89.82%) taxonomically assigned

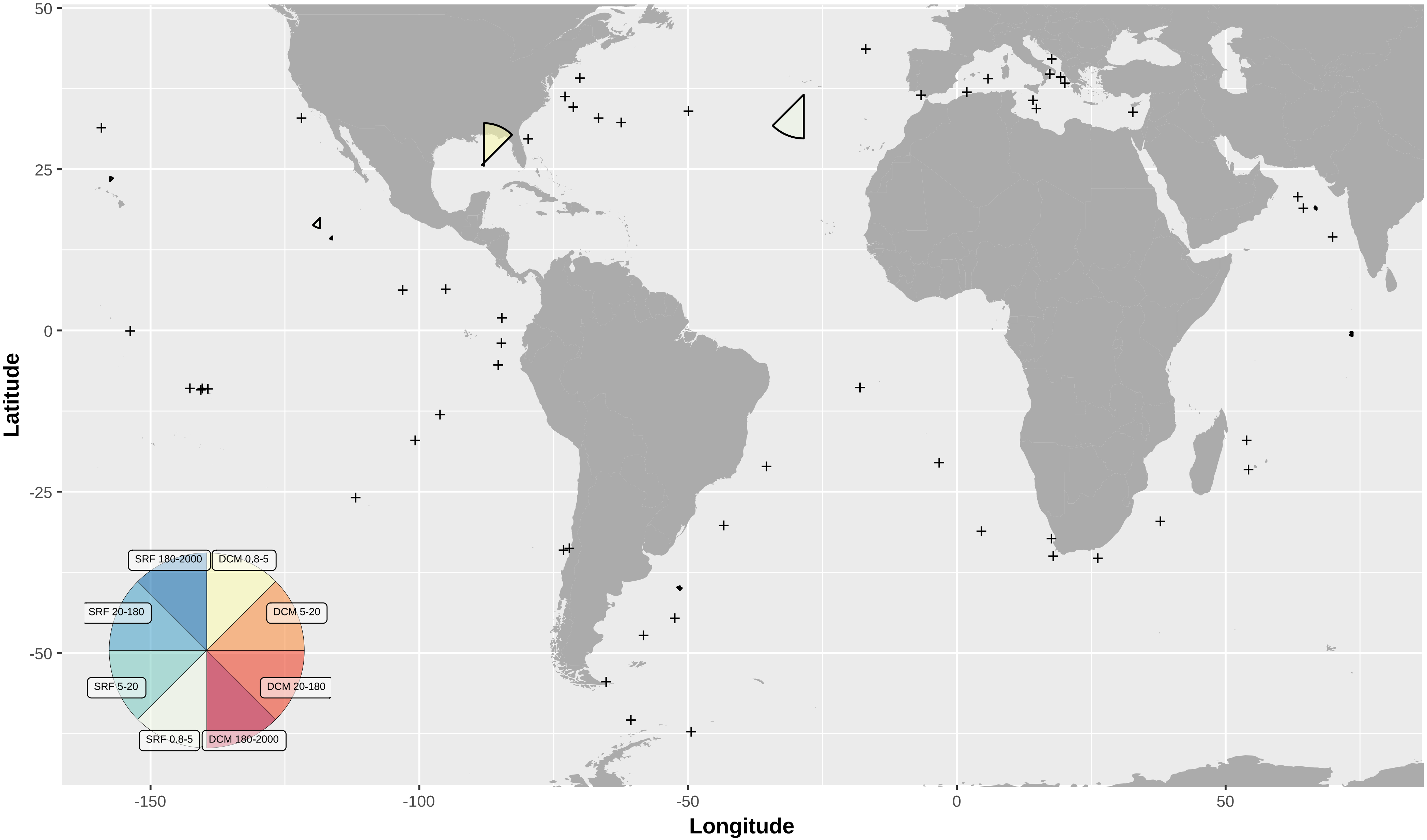


root;cellular organisms;Bacteria;Proteobacteria;Betaproteobacteria;Burkholderiales

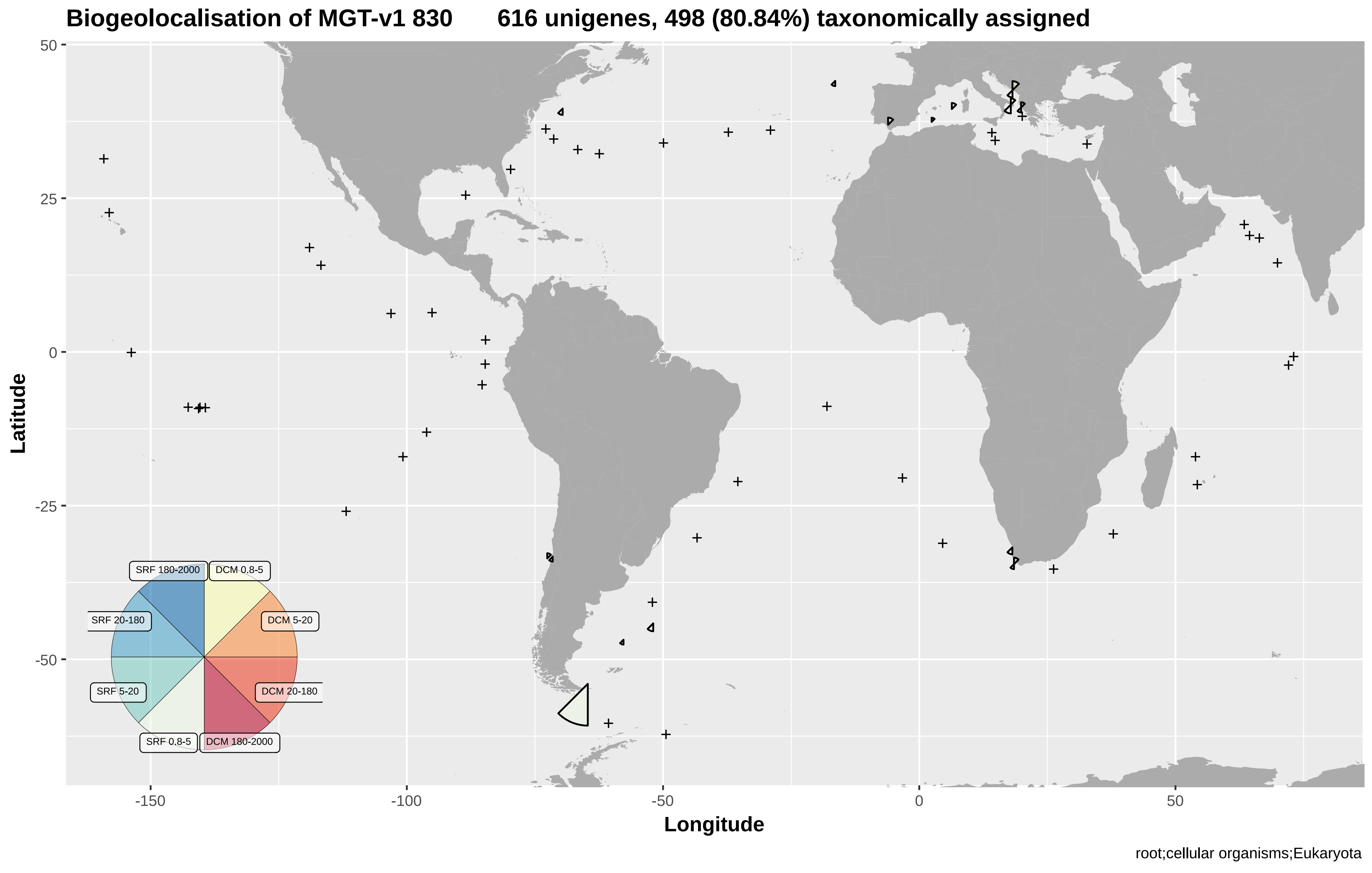
Biogeolocalisation of MGT-v1 830 616 unigenes, 498 (80.84%) taxonomically assigned
50
25
0
-25
-50
Latitude
-150
-100
-50
0
50
Longitude
SRF 180-2000
DCM 0.8-5
SRF 20-180
DCM 5-20
SRF 5-20
DCM 20-180
SRF 0.8-5
DCM 180-2000
root;cellular organisms;Eukaryota

Biogeolocalisation of MGT-v1 831 615 unigenes, 46 (7.48%) taxonomically assigned

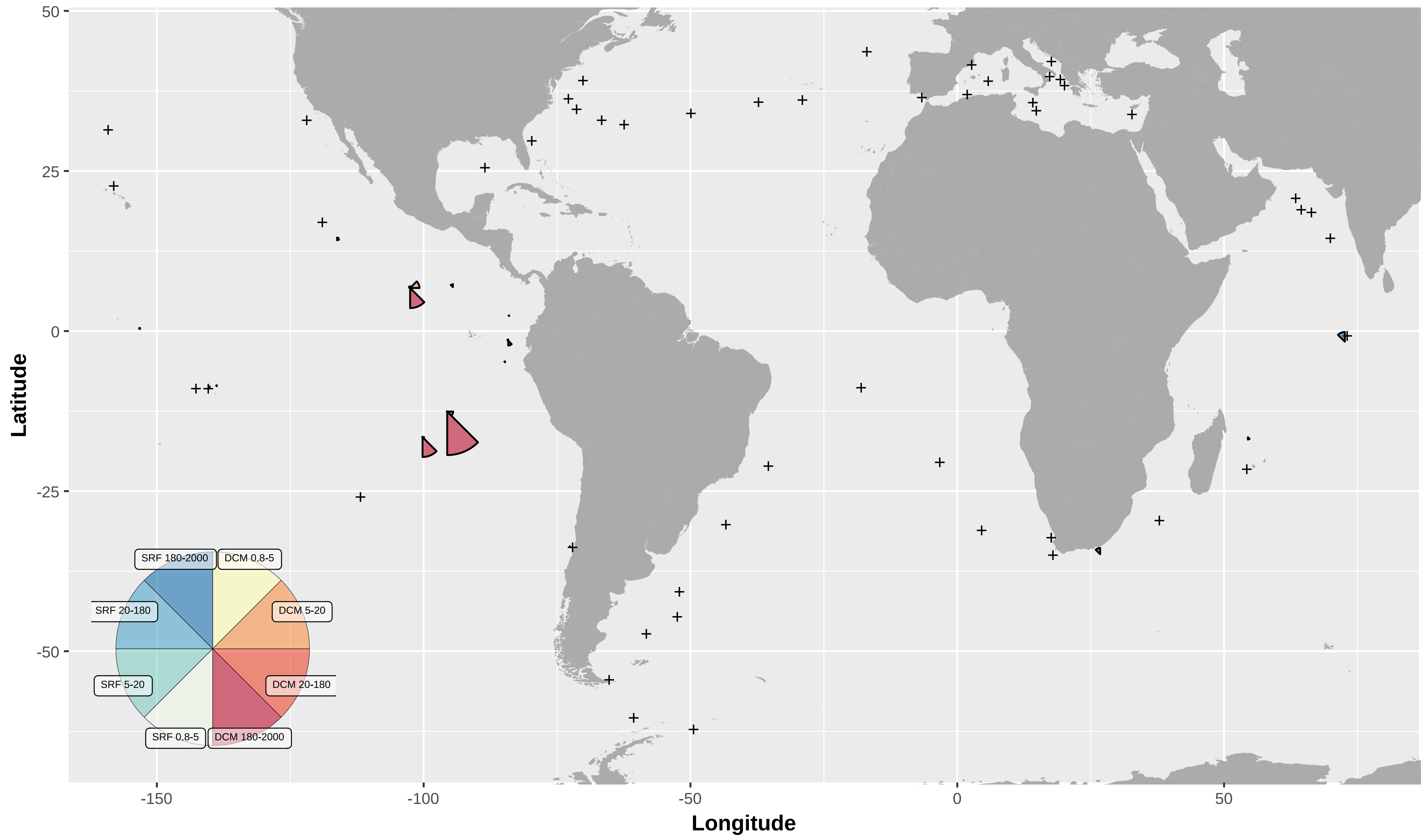


root;cellular organisms;Eukaryota;Opisthokonta;Metazoa;Eumetazoa;Bilateria

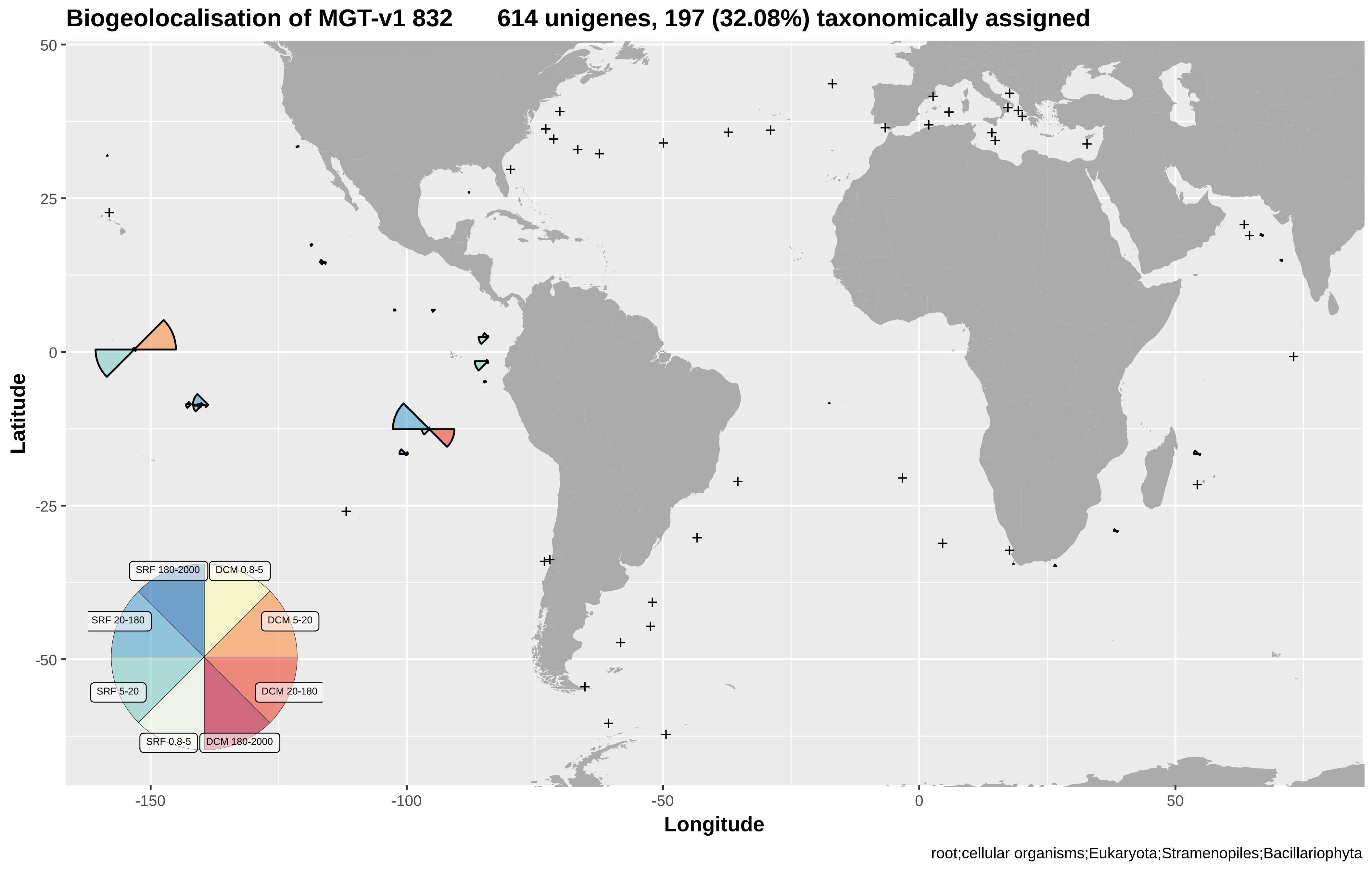
Biogeolocalisation of MGT-v1 832 614 unigenes, 197 (32.08%) taxonomically assigned
50
25
0
-25
-50
Latitude
-150
-100
-50
0
50
Longitude
SRF 180-2000
DCM 0.8-5
SRF 20-180
DCM 5-20
SRF 5-20
DCM 20-180
SRF 0.8-5
DCM 180-2000
root;cellular organisms;Eukaryota;Stramenopiles;Bacillariophyta

## Biogeolocalisation of MGT-v1 833      614 unigenes, 141 (22.96%) taxonomically assigned

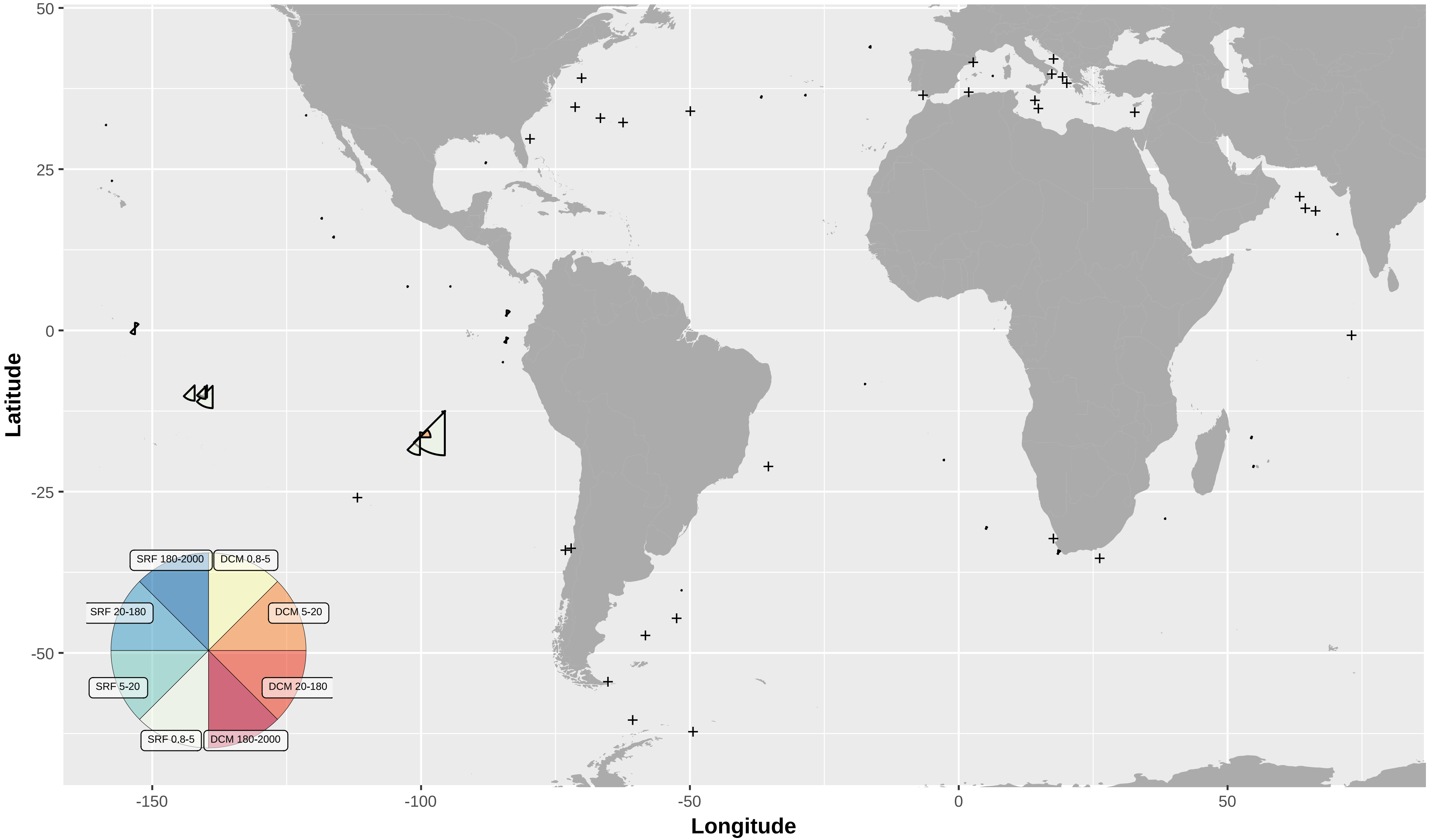


root;cellular organisms;Eukaryota

### Biogeolocalisation of MGT-v1 834 614 unigenes, 371 (60.42%) taxonomically assigned

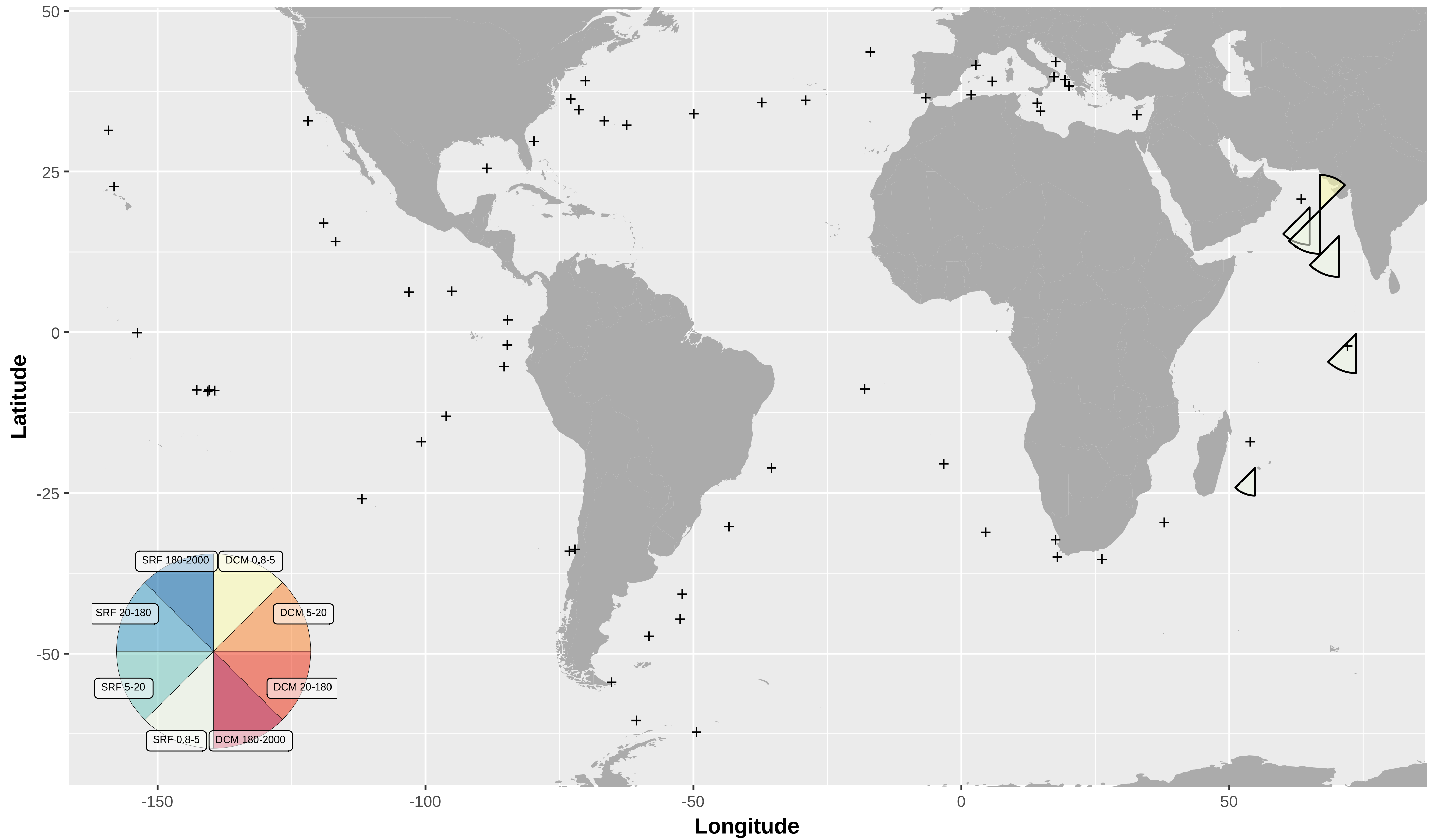


root;cellular organisms;Eukaryota

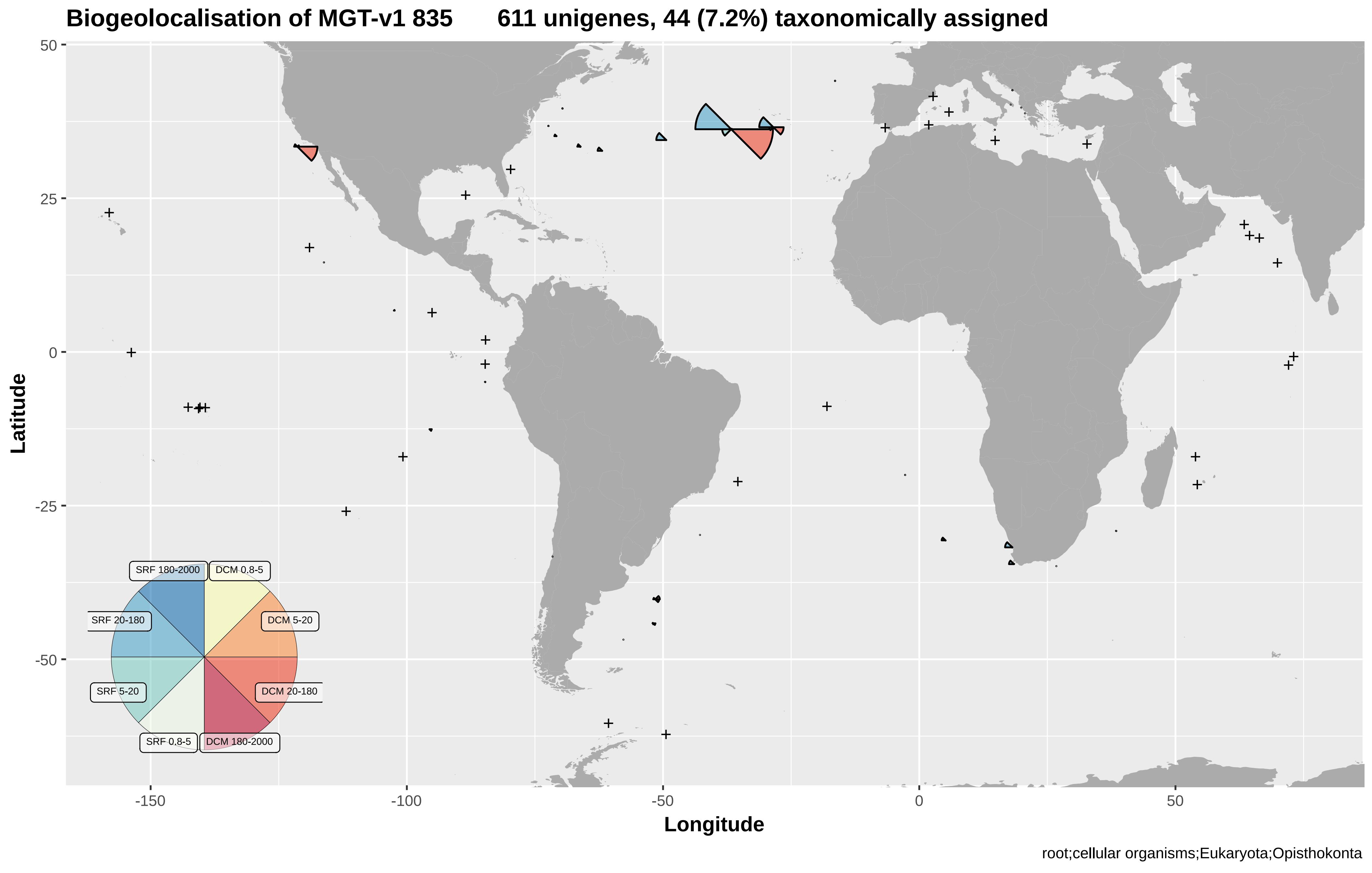

Biogeolocalisation of MGT-v1 835 611 unigenes, 44 (7.2%) taxonomically assigned
50
25
0
-25
-50
Latitude
-150
-100
-50
0
50
Longitude
SRF 180-2000
DCM 0.8-5
SRF 20-180
DCM 5-20
SRF 5-20
DCM 20-180
SRF 0.8-5
DCM 180-2000
root;cellular organisms;Eukaryota;Opisthokonta

#### Biogeolocalisation of MGT-v1 836 610 unigenes, 404 (66.23%) taxonomically assigned

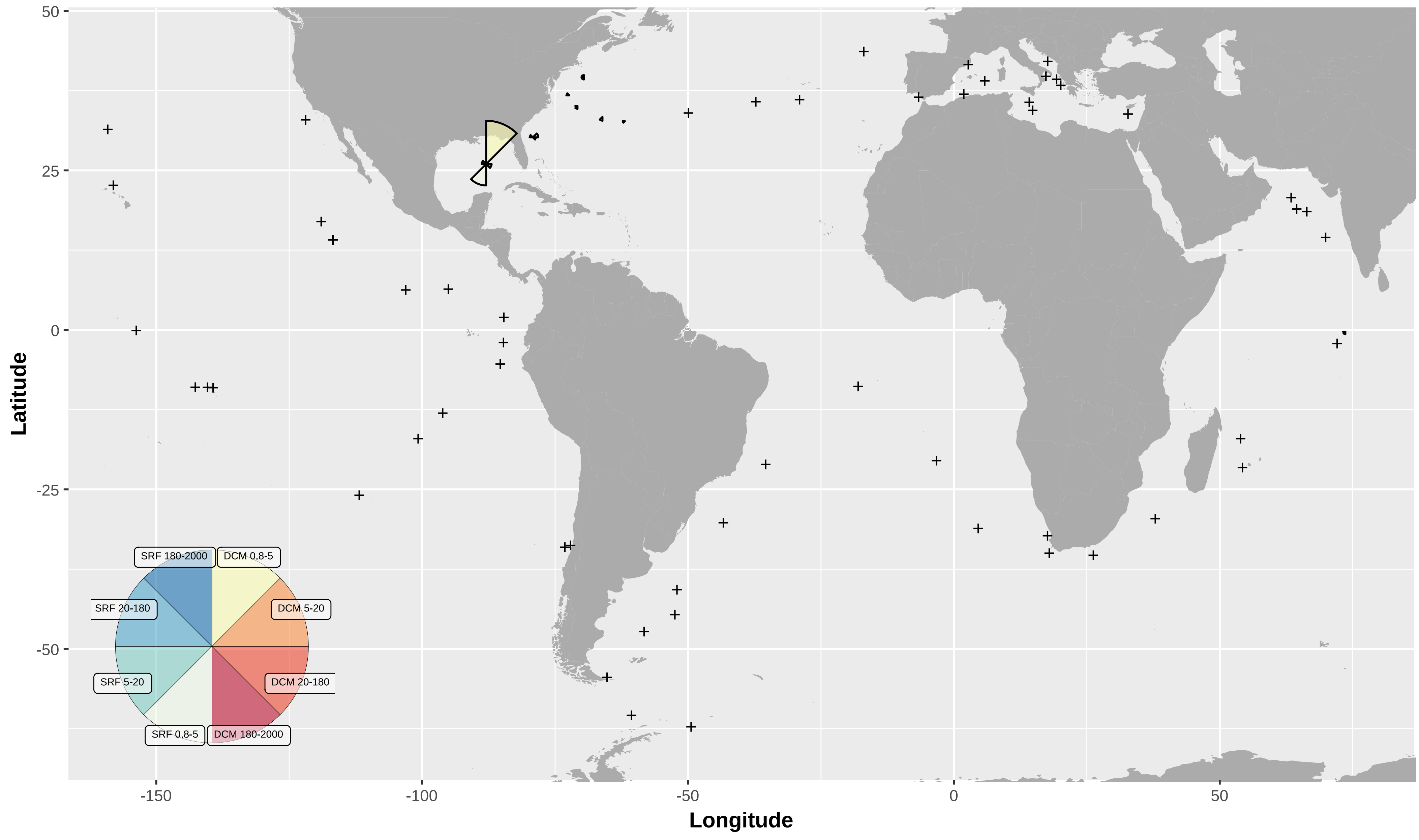


root;cellular organisms;Bacteria

## Biogeolocalisation of MGT-v1 837      608 unigenes, 366 (60.2%) taxonomically assigned

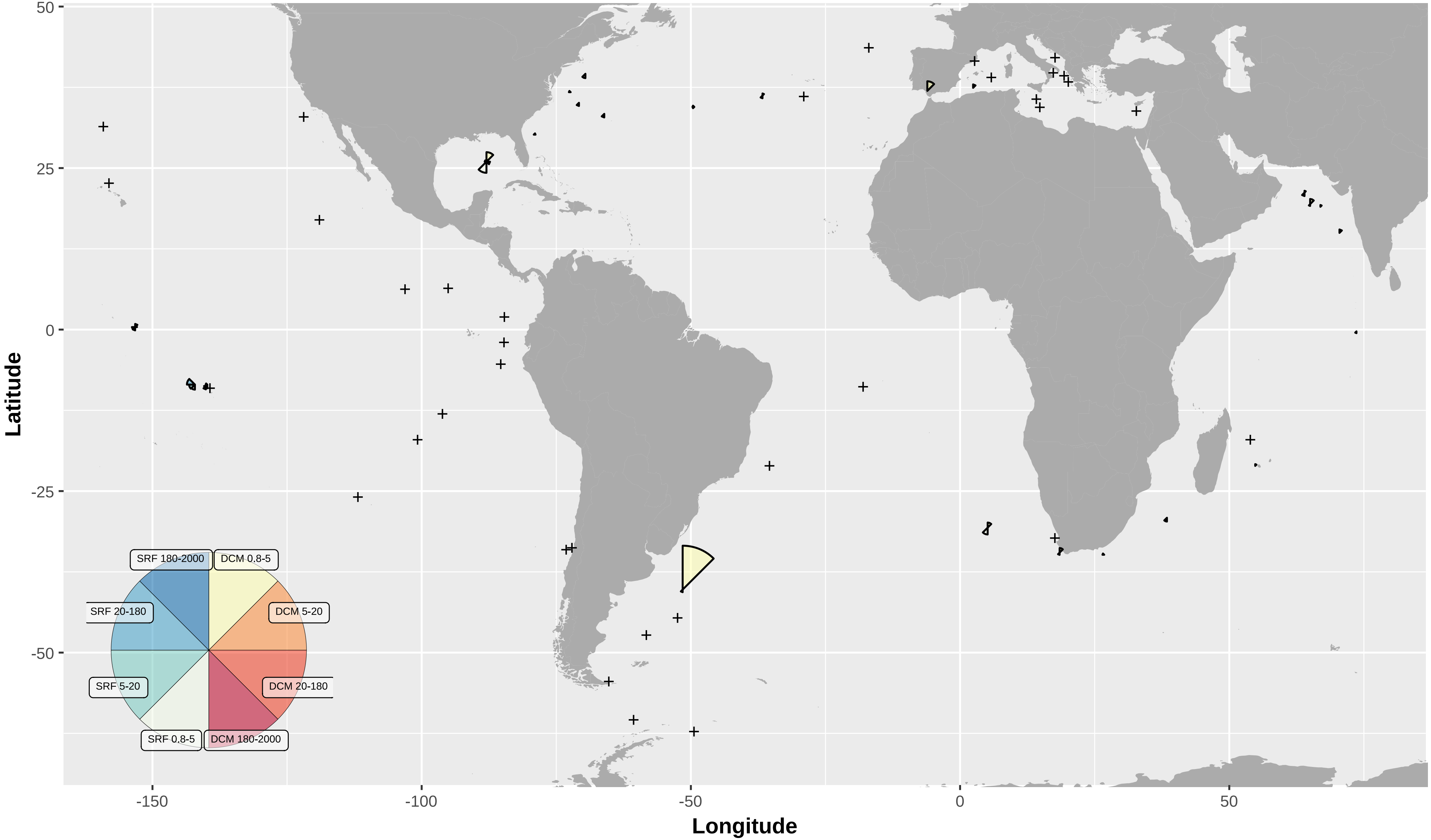


root

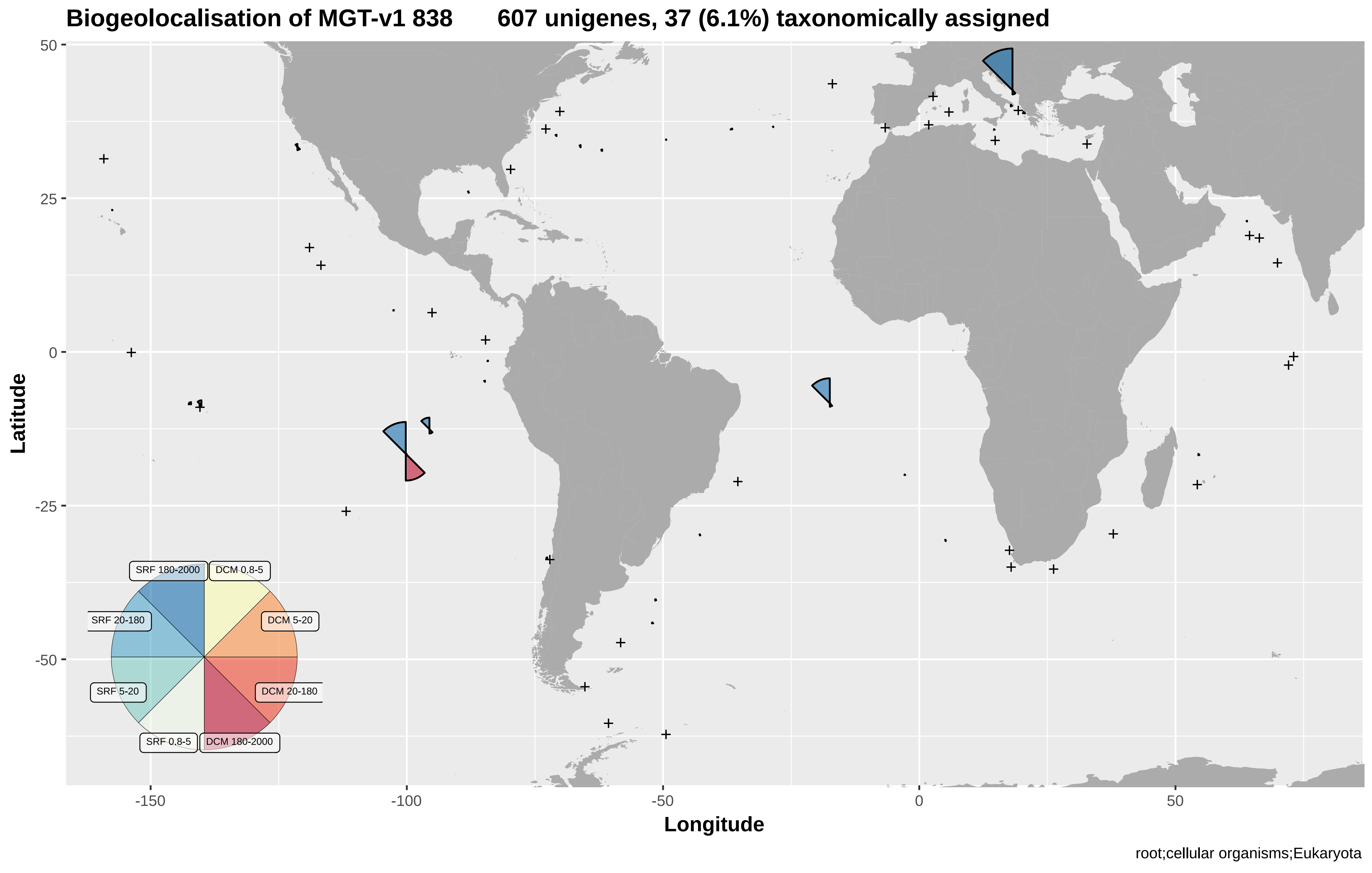
Biogeolocalisation of MGT-v1 838 607 unigenes, 37 (6.1%) taxonomically assigned
50
25
0
-25
-50
Latitude
-150
-100
-50
0
50
Longitude
SRF 180-2000
DCM 0.8-5
SRF 20-180
DCM 5-20
SRF 5-20
DCM 20-180
SRF 0.8-5
DCM 180-2000
root;cellular organisms;Eukaryota

## Biogeolocalisation of MGT-v1 839      607 unigenes, 85 (14%) taxonomically assigned

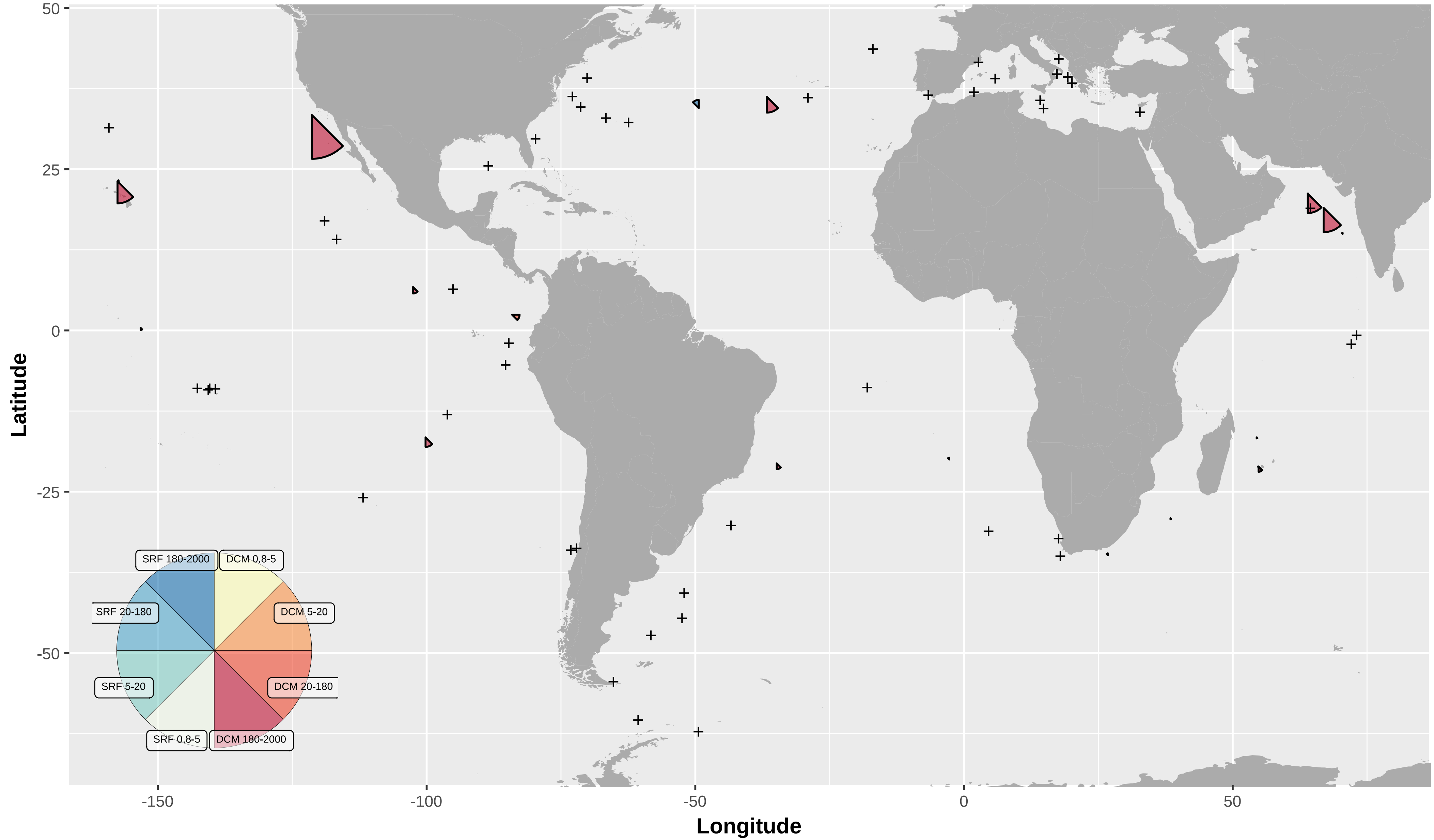


root;cellular organisms;Eukaryota;Opisthokonta;Metazoa;Eumetazoa

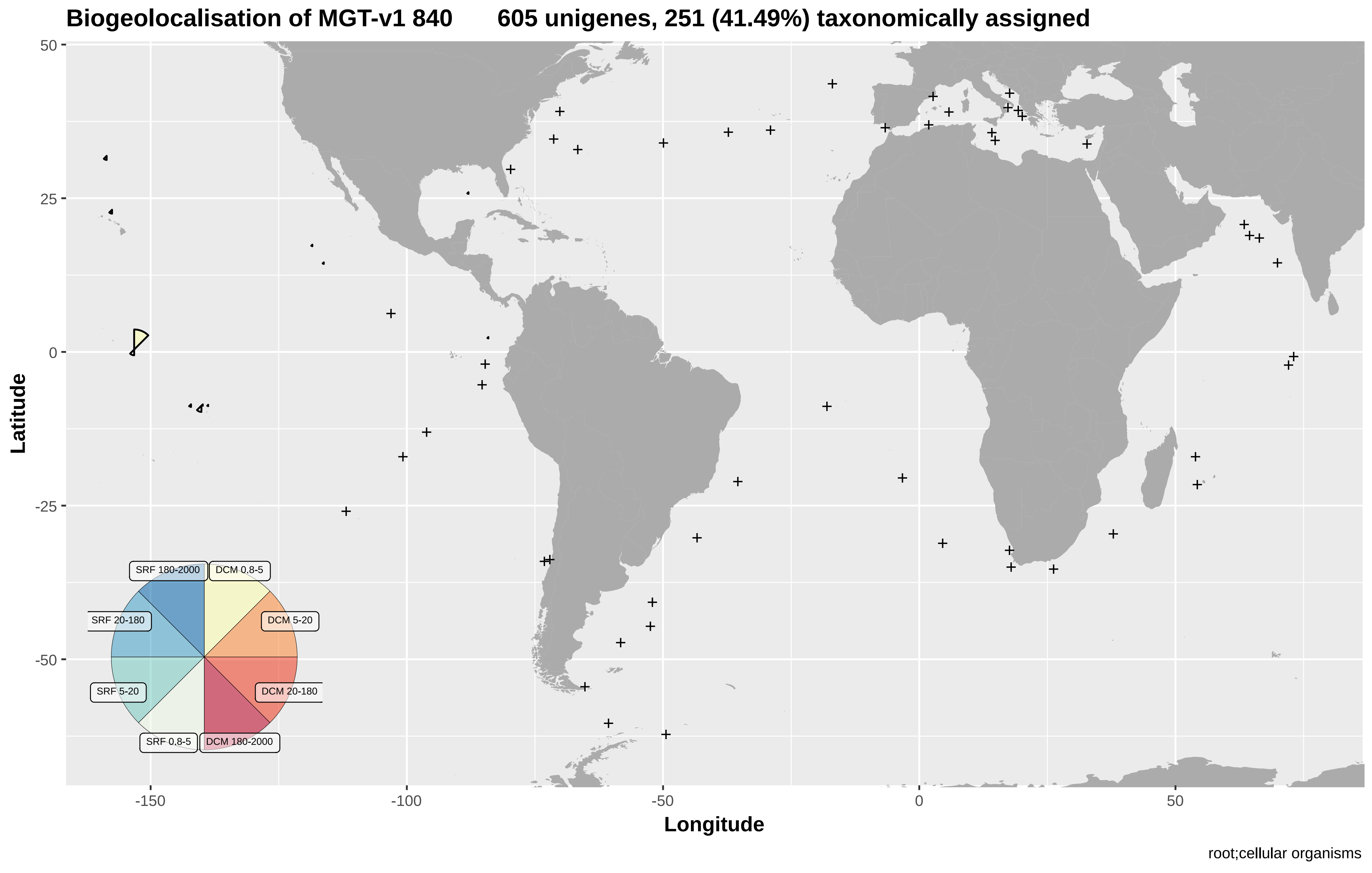

Biogeolocalisation of MGT-v1 840 605 unigenes, 251 (41.49%) taxonomically assigned
50
25
0
-25
-50
Latitude
-150
-100
-50
0
50
Longitude
SRF 180-2000
DCM 0.8-5
SRF 20-180
DCM 5-20
SRF 5-20
DCM 20-180
SRF 0.8-5
DCM 180-2000
root;cellular organisms

# Biogeolocalisation of MGT-v1 841      604 unigenes, 77 (12.75%) taxonomically assigned

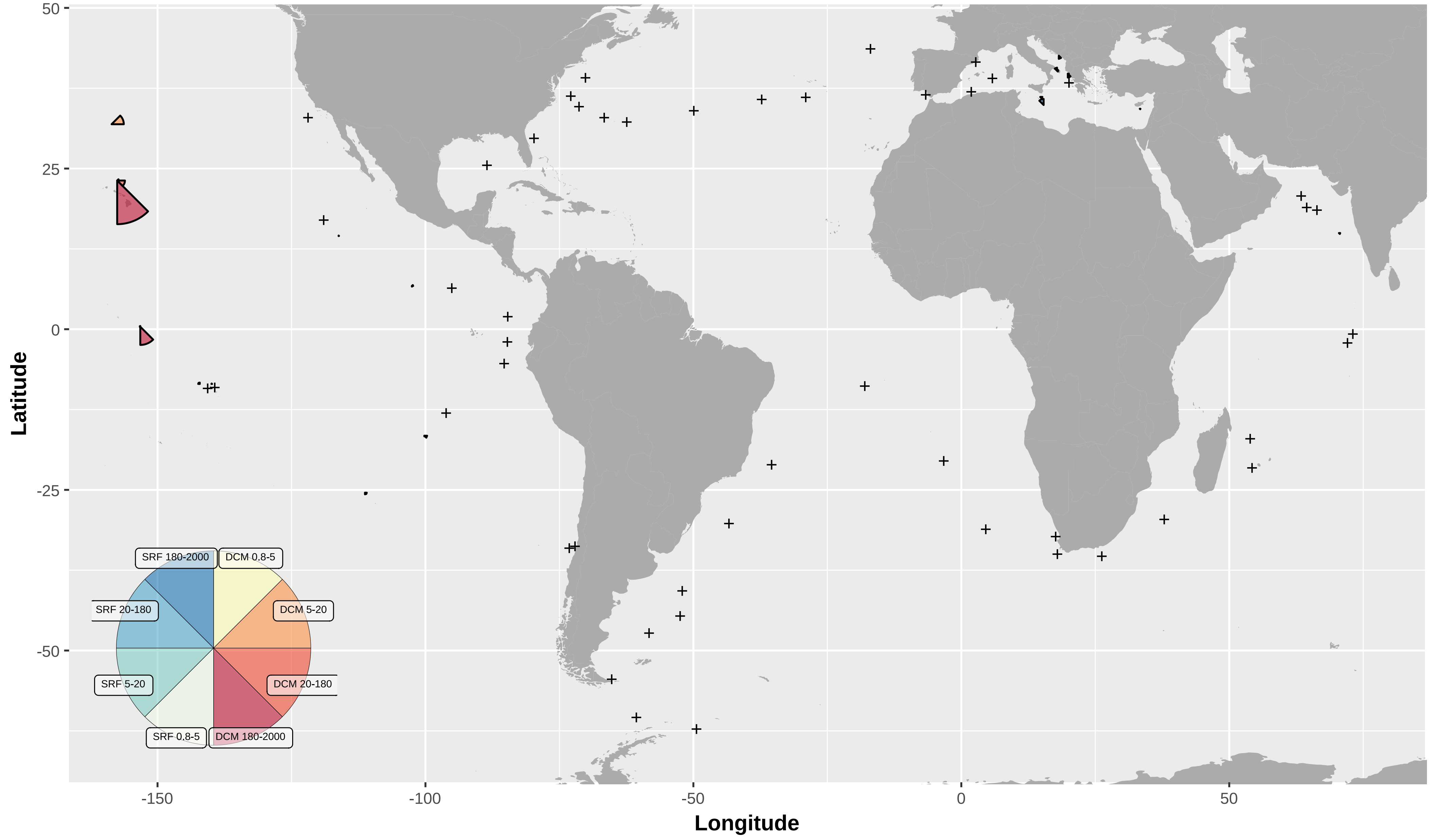


root;cellular organisms;Eukaryota;Opisthokonta;Metazoa;Eumetazoa;Bilateria

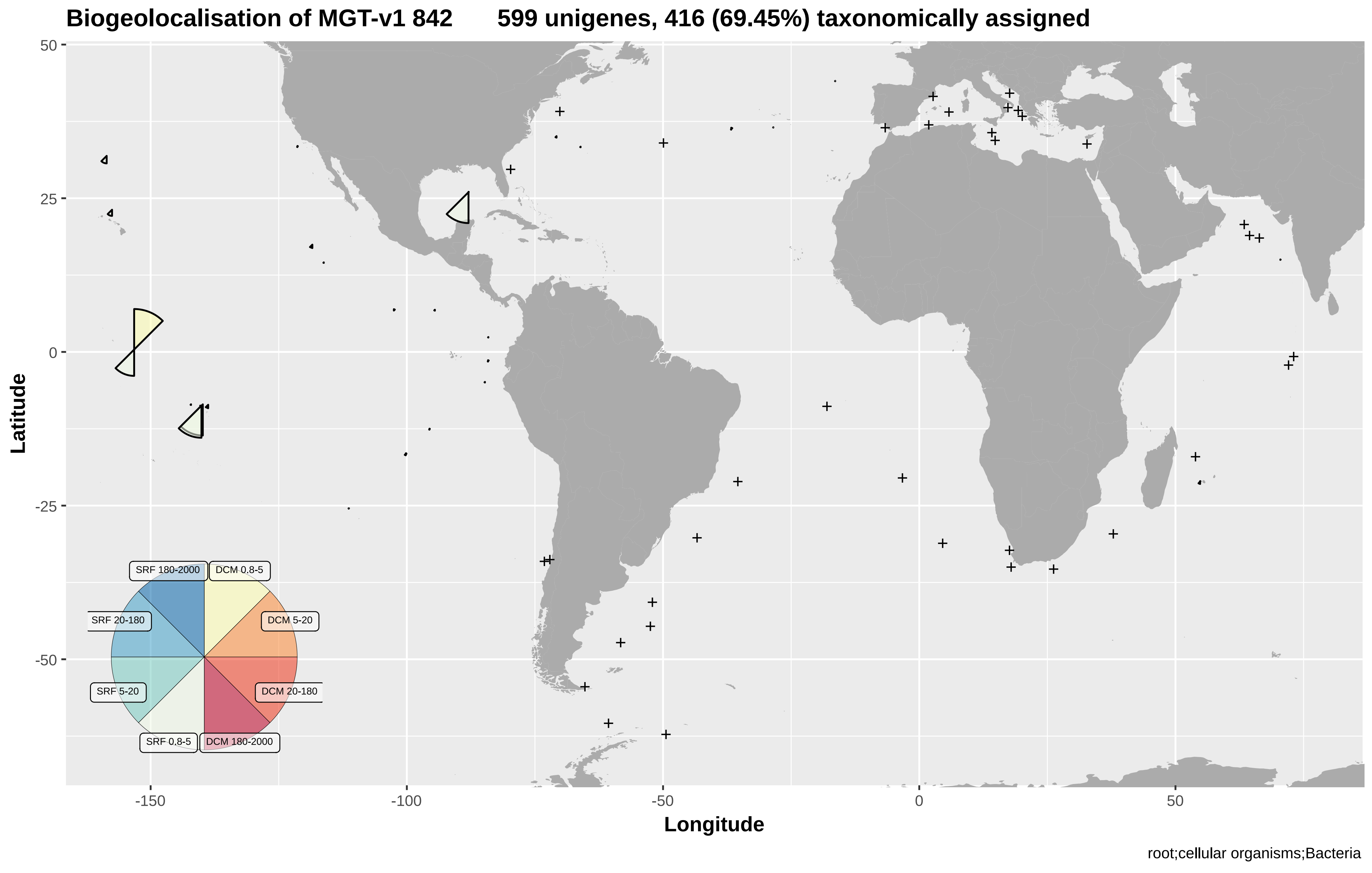
Biogeolocalisation of MGT-v1 842 599 unigenes, 416 (69.45%) taxonomically assigned
50
25
0
-25
-50
Latitude
-150
-100
-50
0
50
Longitude
SRF 180-2000
DCM 0.8-5
SRF 20-180
DCM 5-20
SRF 5-20
DCM 20-180
SRF 0.8-5
DCM 180-2000
root;cellular organisms;Bacteria

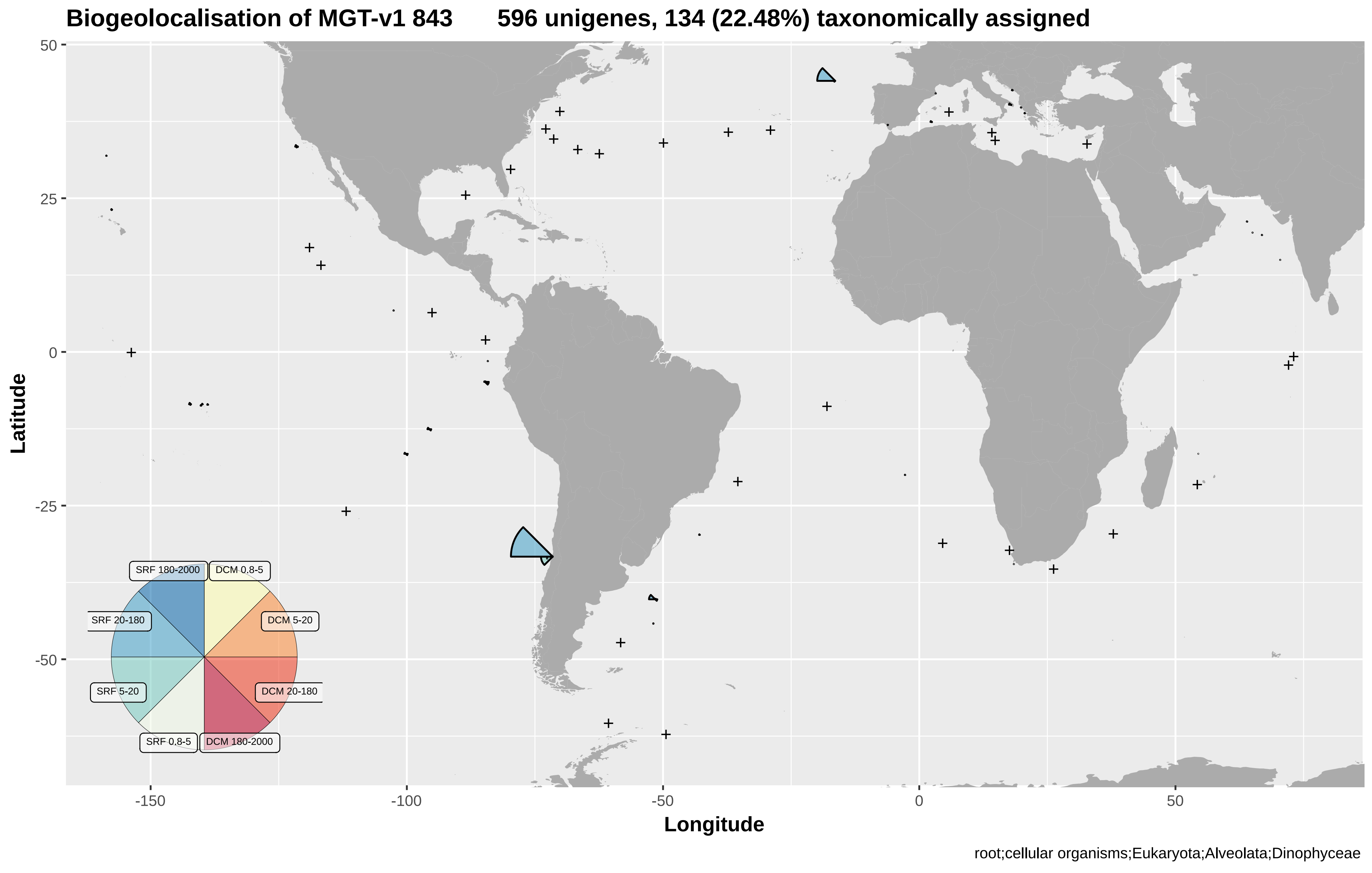
Biogeolocalisation of MGT-v1 843 596 unigenes, 134 (22.48%) taxonomically assigned
Latitude
Longitude
50
25
0
-25
-50
-150
-100
-50
0
50
SRF 180-2000
DCM 0.8-5
SRF 20-180
DCM 5-20
SRF 5-20
DCM 20-180
SRF 0.8-5
DCM 180-2000
root;cellular organisms;Eukaryota;Alveolata;Dinophyceae

Biogeolocalisation of MGT-v1 844 595 unigenes, 25 (4.2%) taxonomically assigned

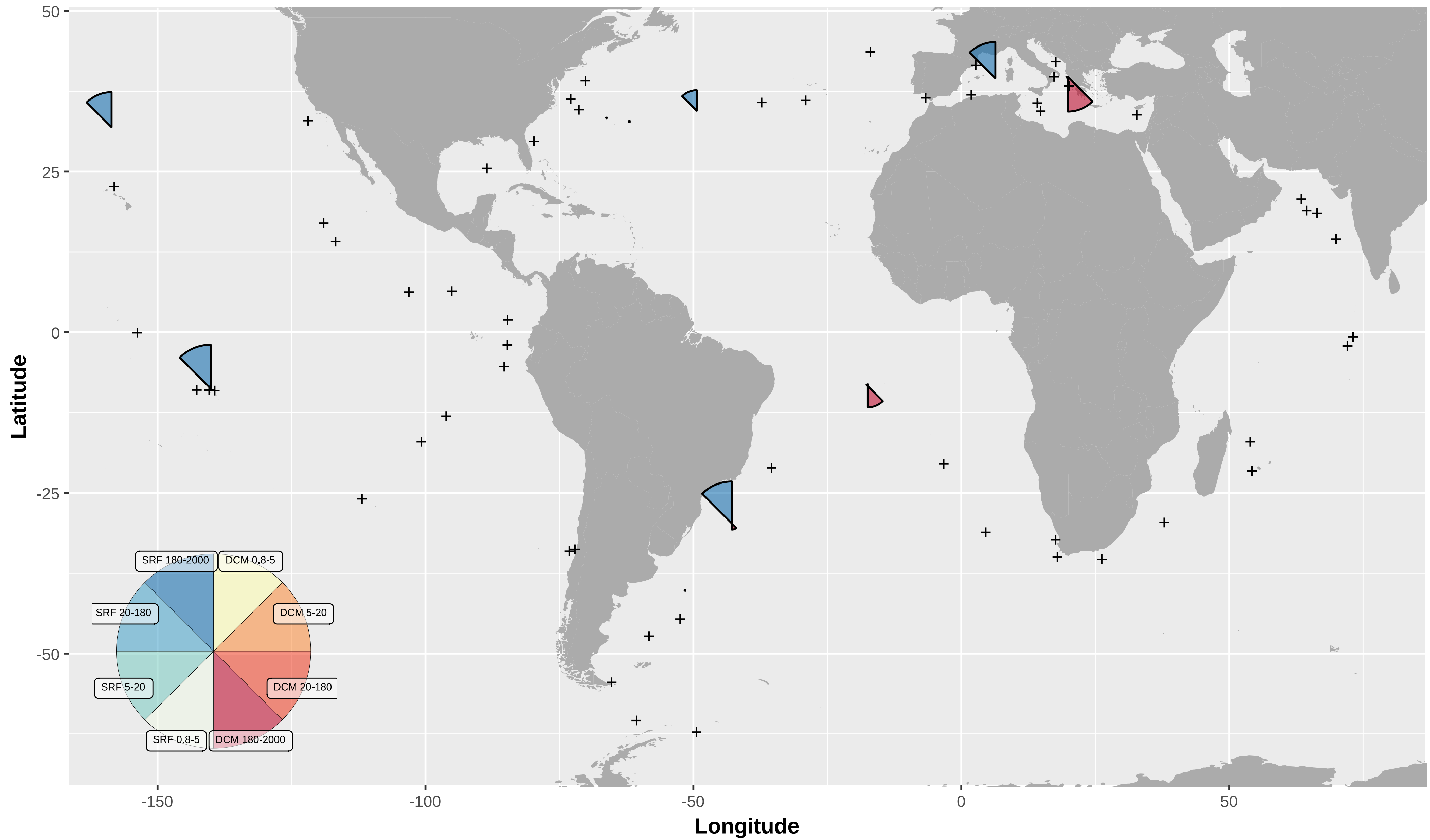


root;cellular organisms;Eukaryota;Opisthokonta;Metazoa;Eumetazoa

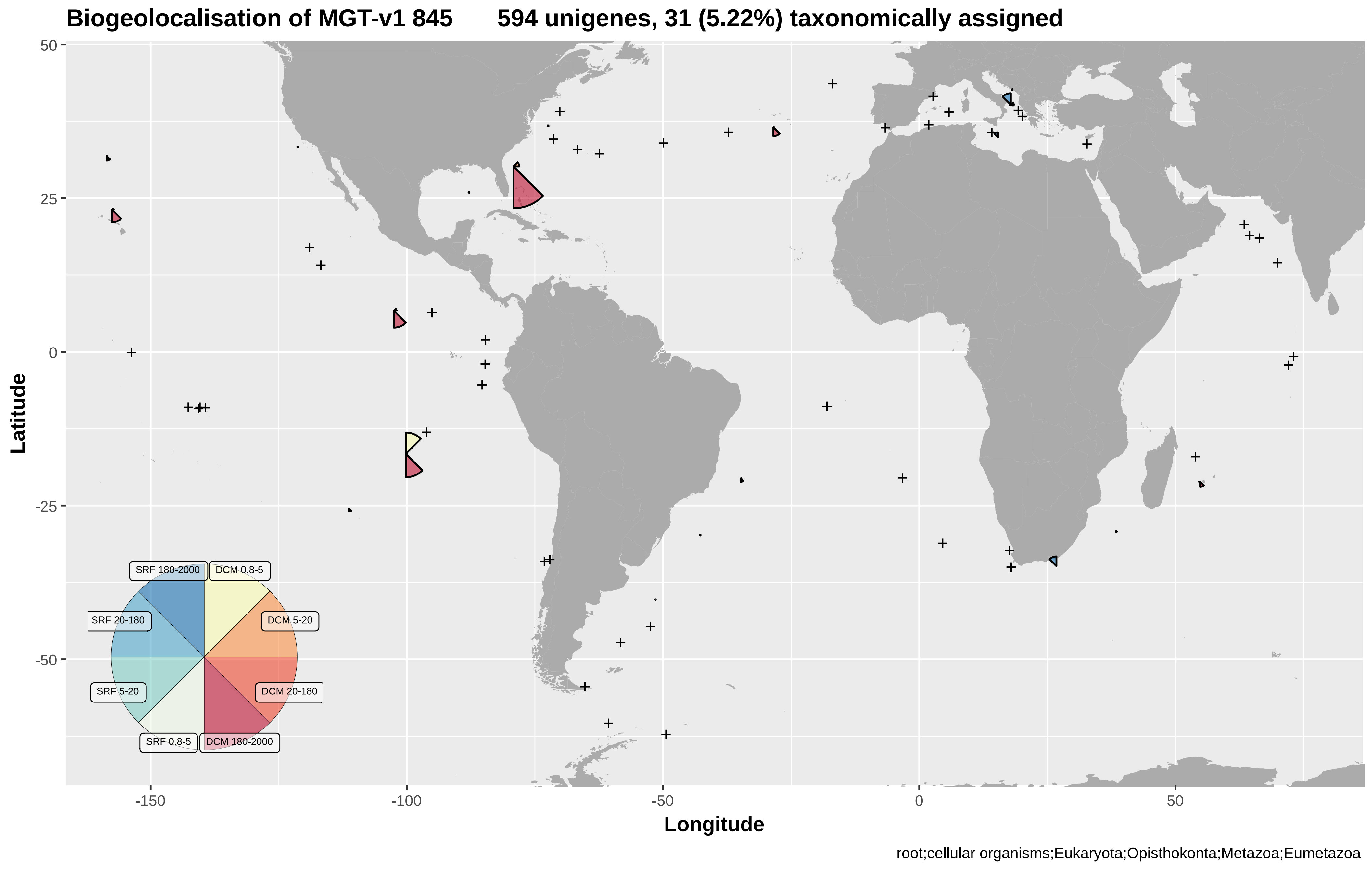
Biogeolocalisation of MGT-v1 845 594 unigenes, 31 (5.22%) taxonomically assigned
Latitude
Longitude
50
25
0
-25
-50
-150
-100
-50
0
50
SRF 180-2000
DCM 0.8-5
SRF 20-180
DCM 5-20
SRF 5-20
DCM 20-180
SRF 0.8-5
DCM 180-2000
root;cellular organisms;Eukaryota;Opisthokonta;Metazoa;Eumetazoa

## Biogeolocalisation of MGT-v1 846      591 unigenes, 295 (49.92%) taxonomically assigned

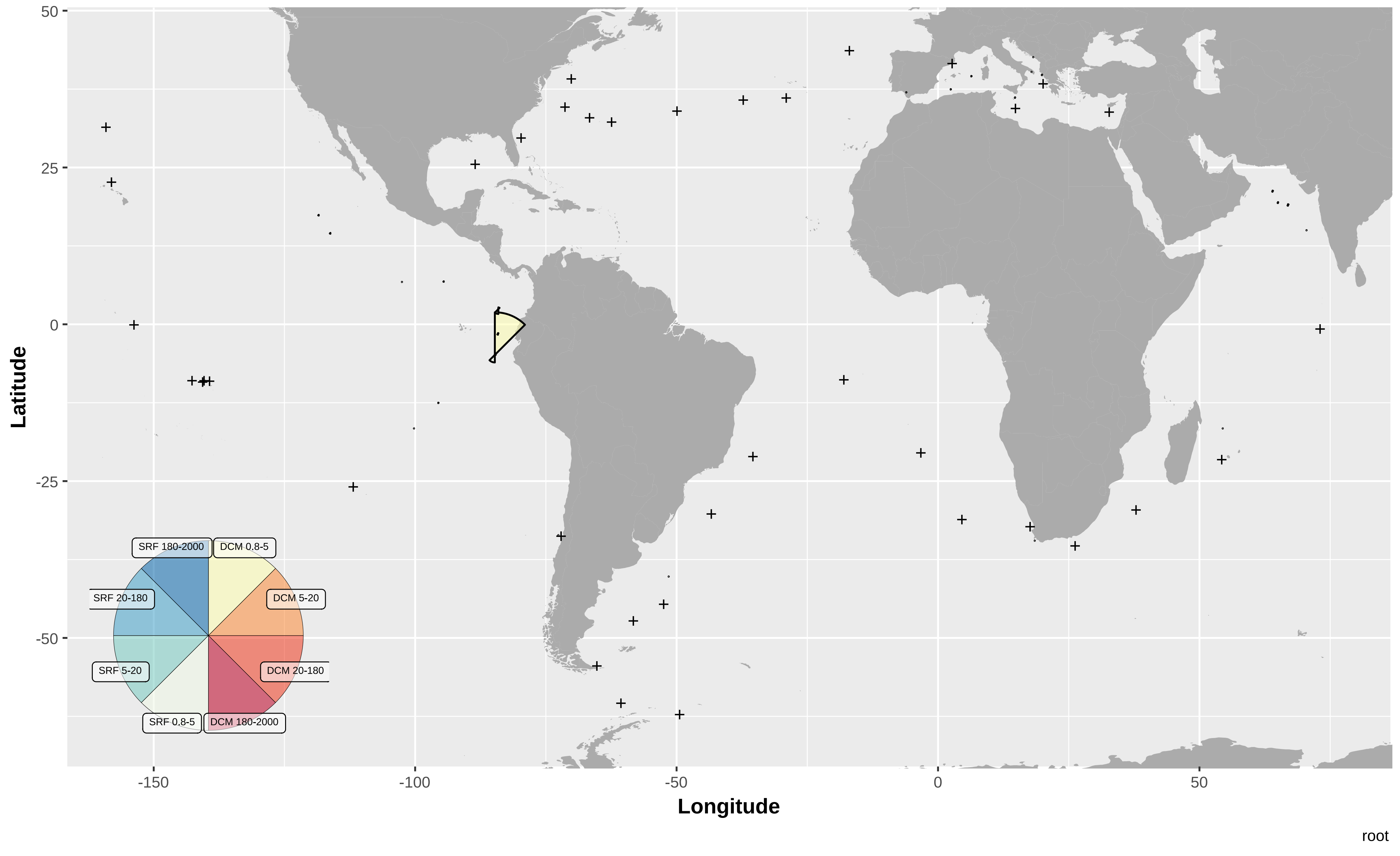

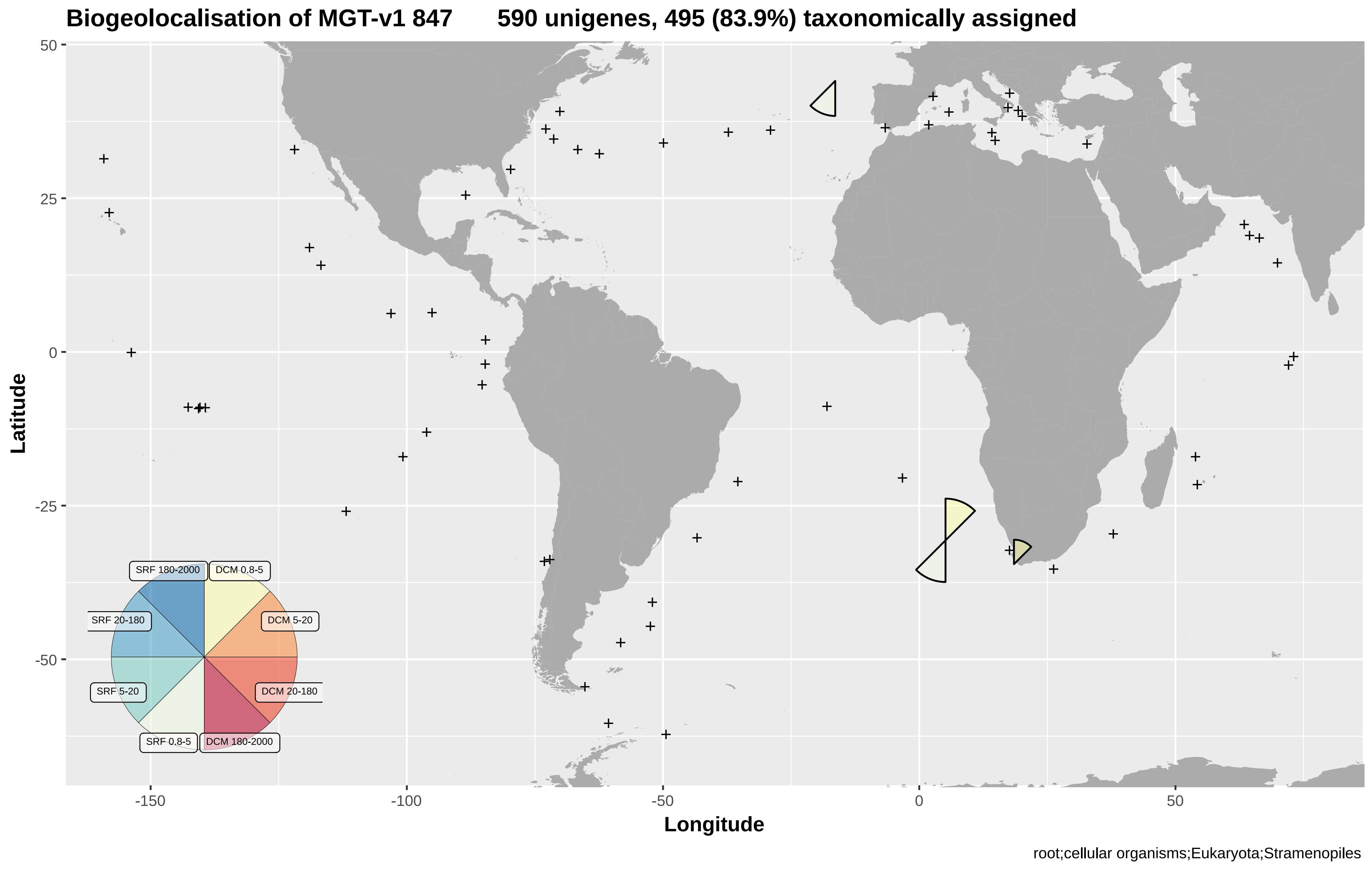

Biogeolocalisation of MGT-v1 847 590 unigenes, 495 (83.9%) taxonomically assigned
Latitude
Longitude
50
25
0
-25
-50
-150
-100
-50
0
50
SRF 180-2000
DCM 0.8-5
SRF 20-180
DCM 5-20
SRF 5-20
DCM 20-180
SRF 0.8-5
DCM 180-2000
root;cellular organisms;Eukaryota;Stramenopiles

Biogeolocalisation of MGT-v1 848 589 unigenes, 35 (5.94%) taxonomically assigned

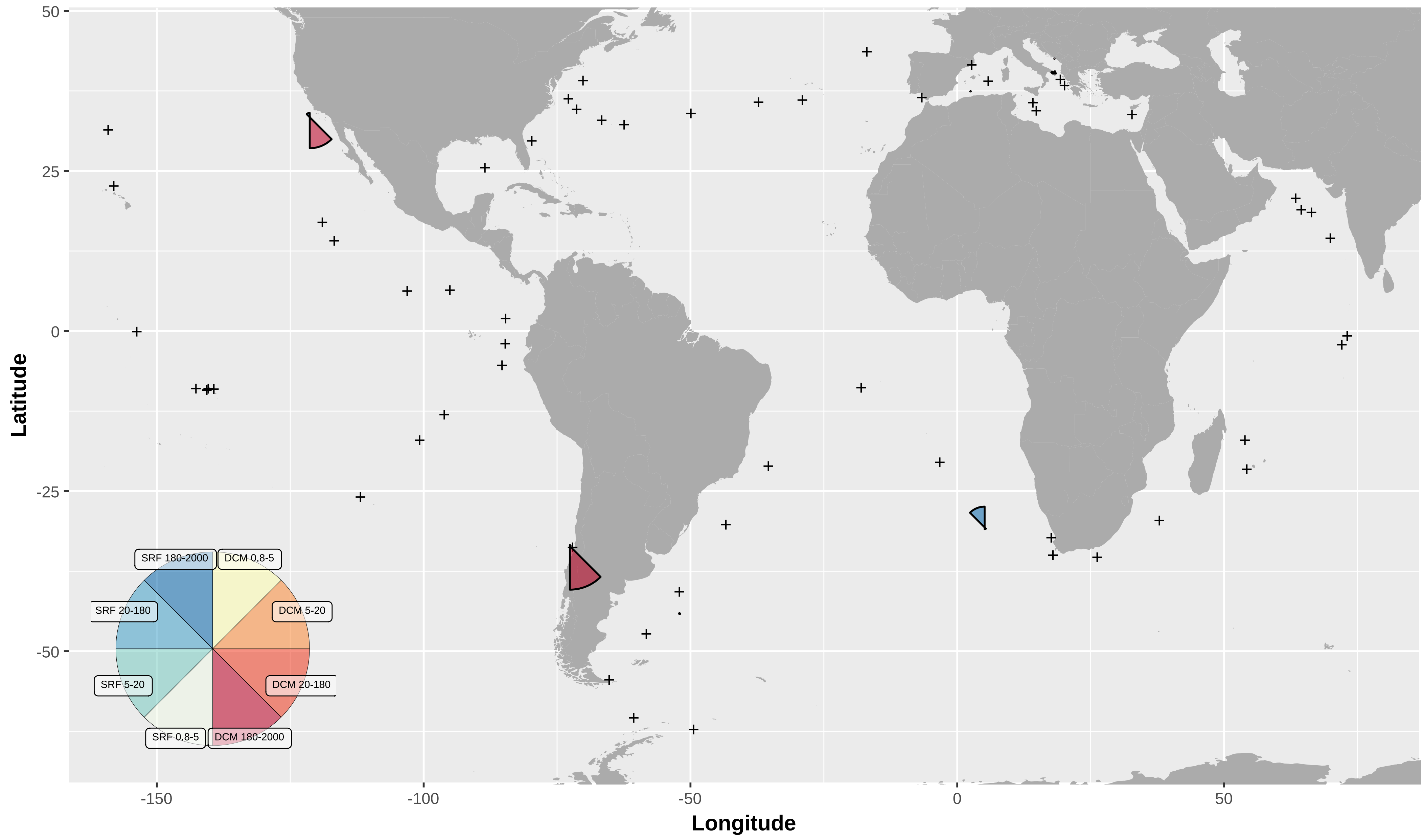


root;cellular organisms;Eukaryota;Opisthokonta;Metazoa;Eumetazoa

# Biogeolocalisation of MGT-v1 849      587 unigenes, 529 (90.12%) taxonomically assigned

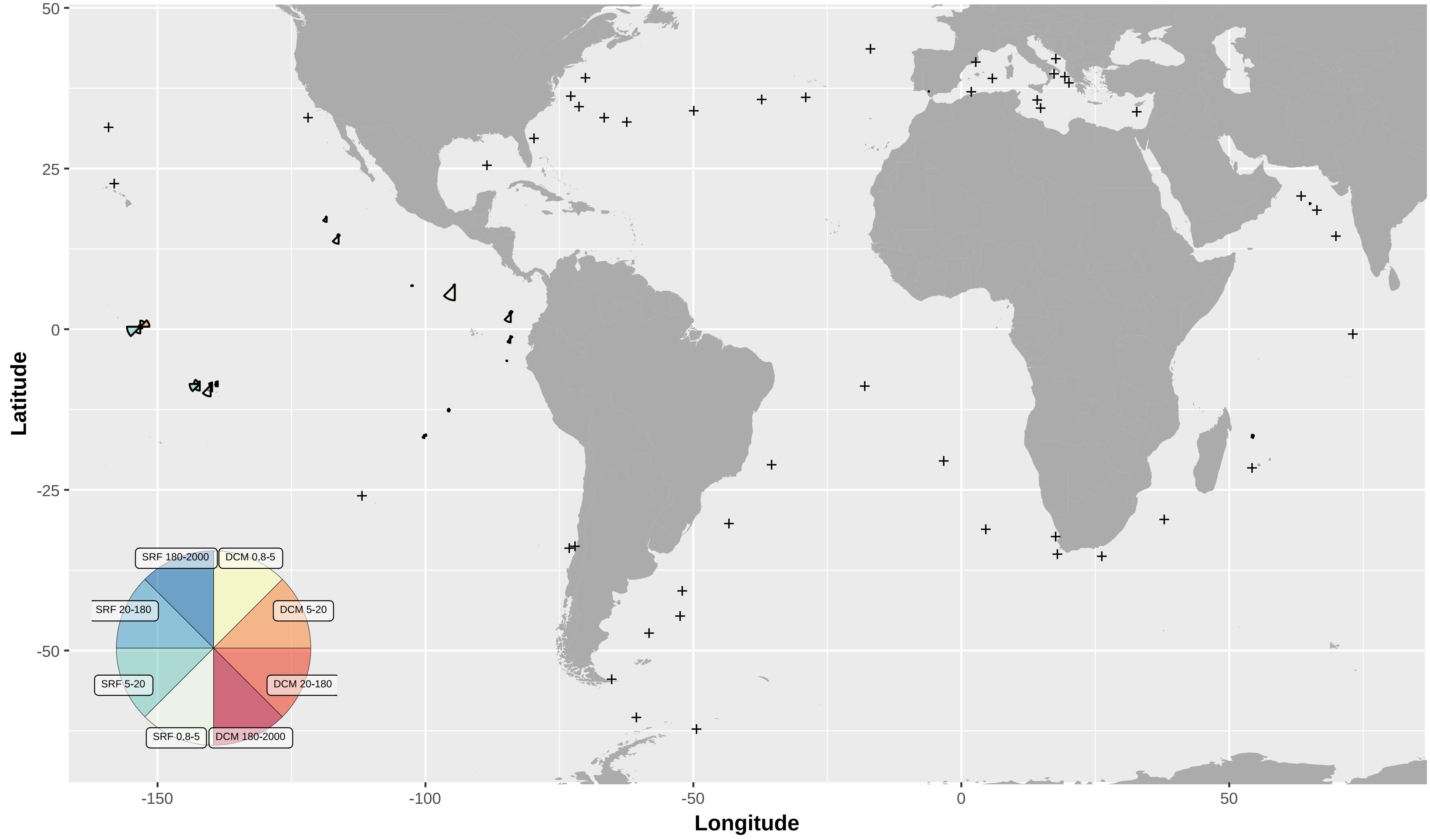


root;cellular organisms;Bacteria;Cyanobacteria

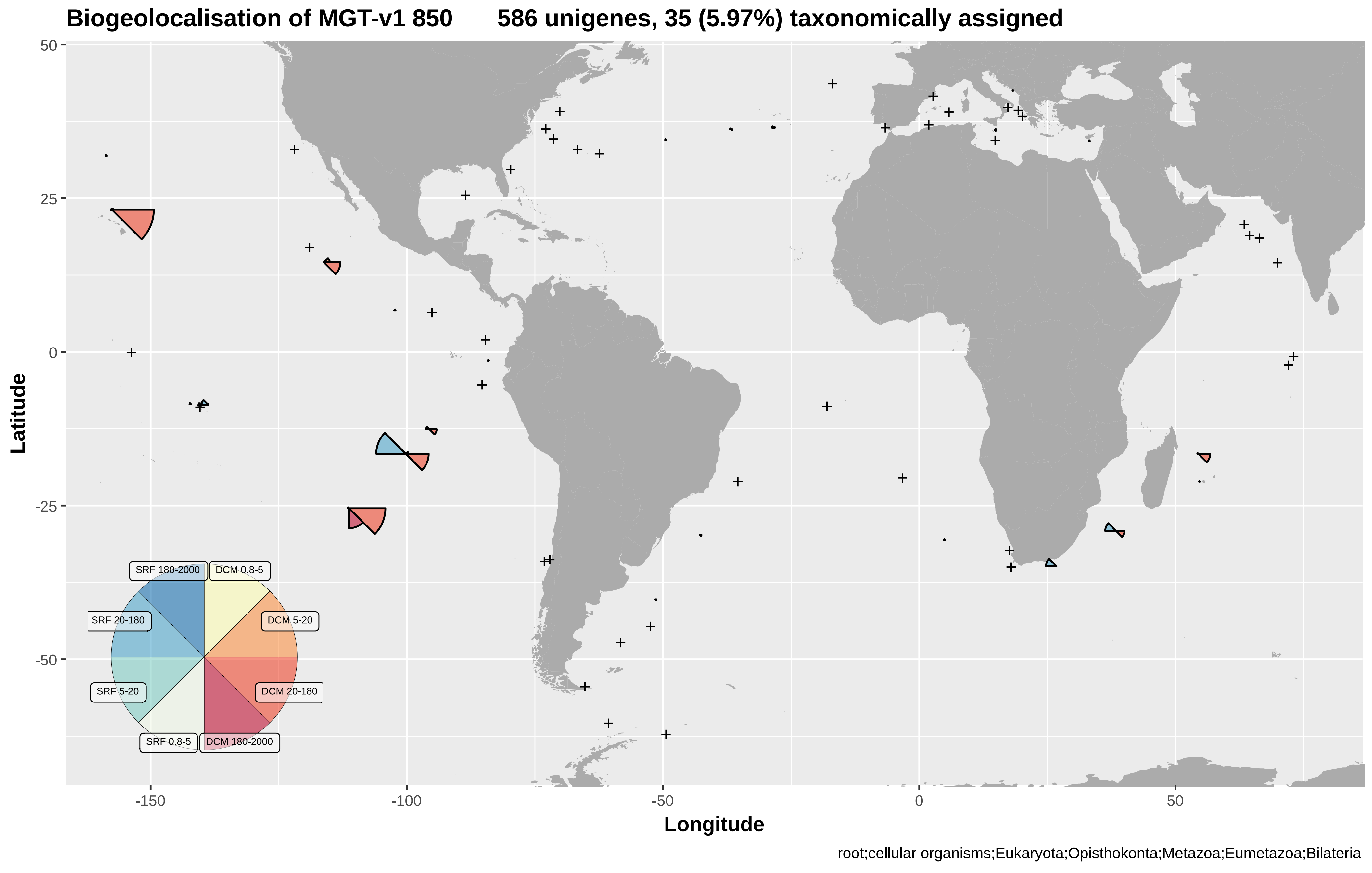

Biogeolocalisation of MGT-v1 850 586 unigenes, 35 (5.97%) taxonomically assigned
50
25
0
-25
-50
Latitude
-150
-100
-50
0
50
Longitude
SRF 180-2000
DCM 0.8-5
SRF 20-180
DCM 5-20
SRF 5-20
DCM 20-180
SRF 0.8-5
DCM 180-2000
root;cellular organisms;Eukaryota;Opisthokonta;Metazoa;Eumetazoa;Bilateria

# Biogeolocalisation of MGT-v1 851      586 unigenes, 376 (64.16%) taxonomically assigned

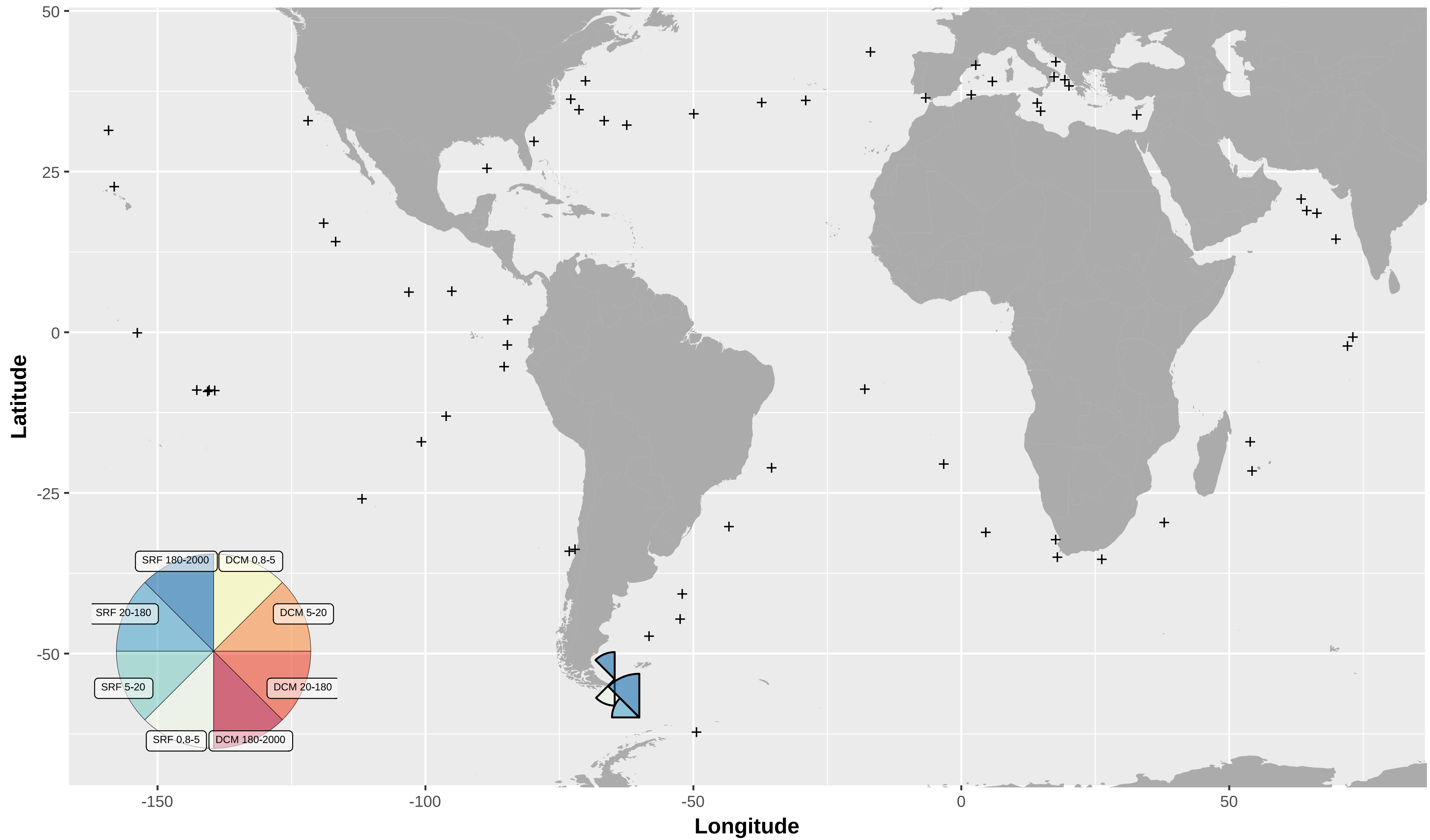


root;cellular organisms;Eukaryota;Opisthokonta;Metazoa;Eumetazoa;Bilateria

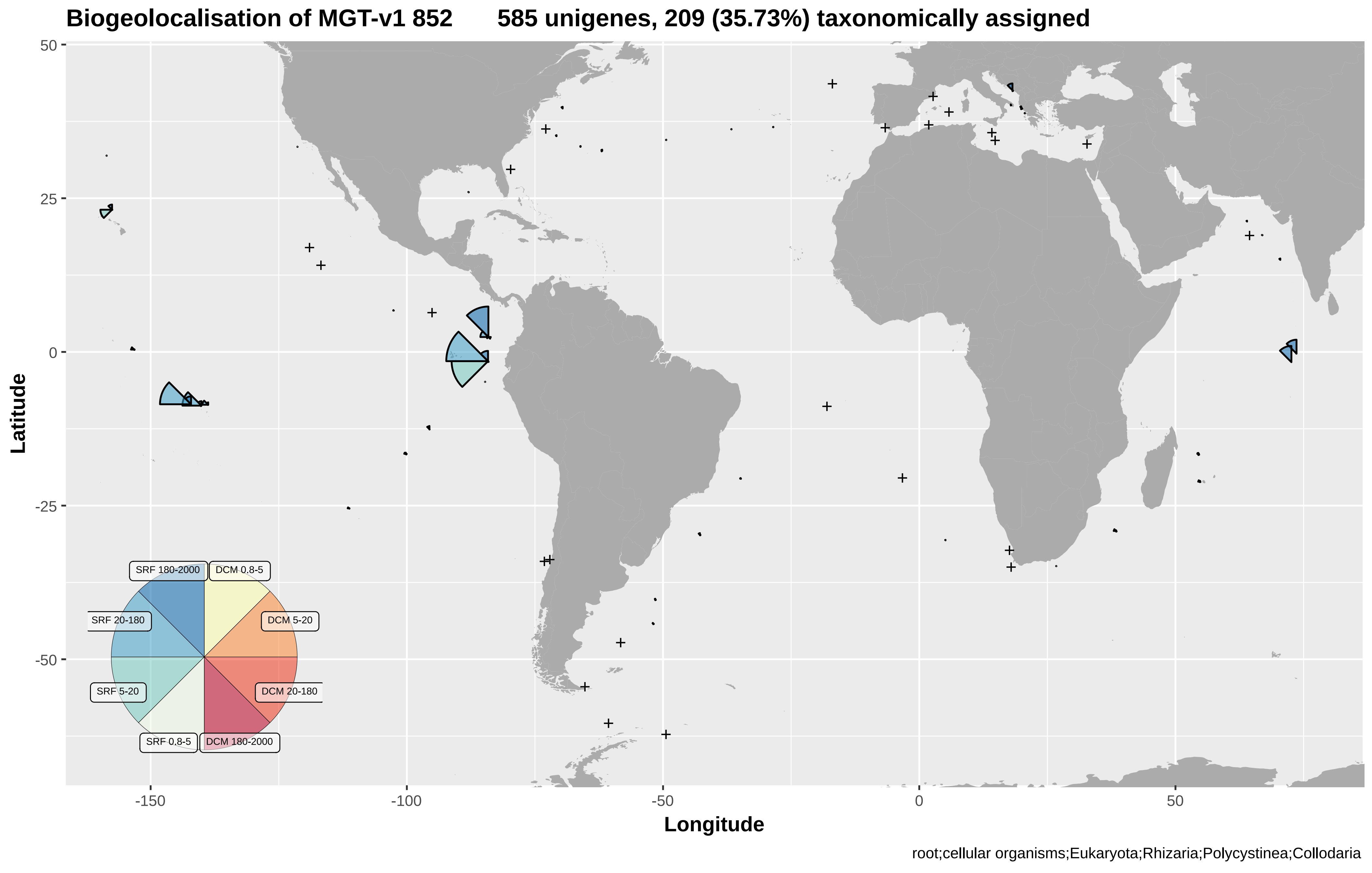

Biogeolocalisation of MGT-v1 852 585 unigenes, 209 (35.73%) taxonomically assigned
50
25
0
-25
-50
Latitude
-150
-100
-50
0
50
Longitude
SRF 180-2000
DCM 0.8-5
SRF 20-180
DCM 5-20
SRF 5-20
DCM 20-180
SRF 0.8-5
DCM 180-2000
root;cellular organisms;Eukaryota;Rhizaria;Polycystinea;Collodaria

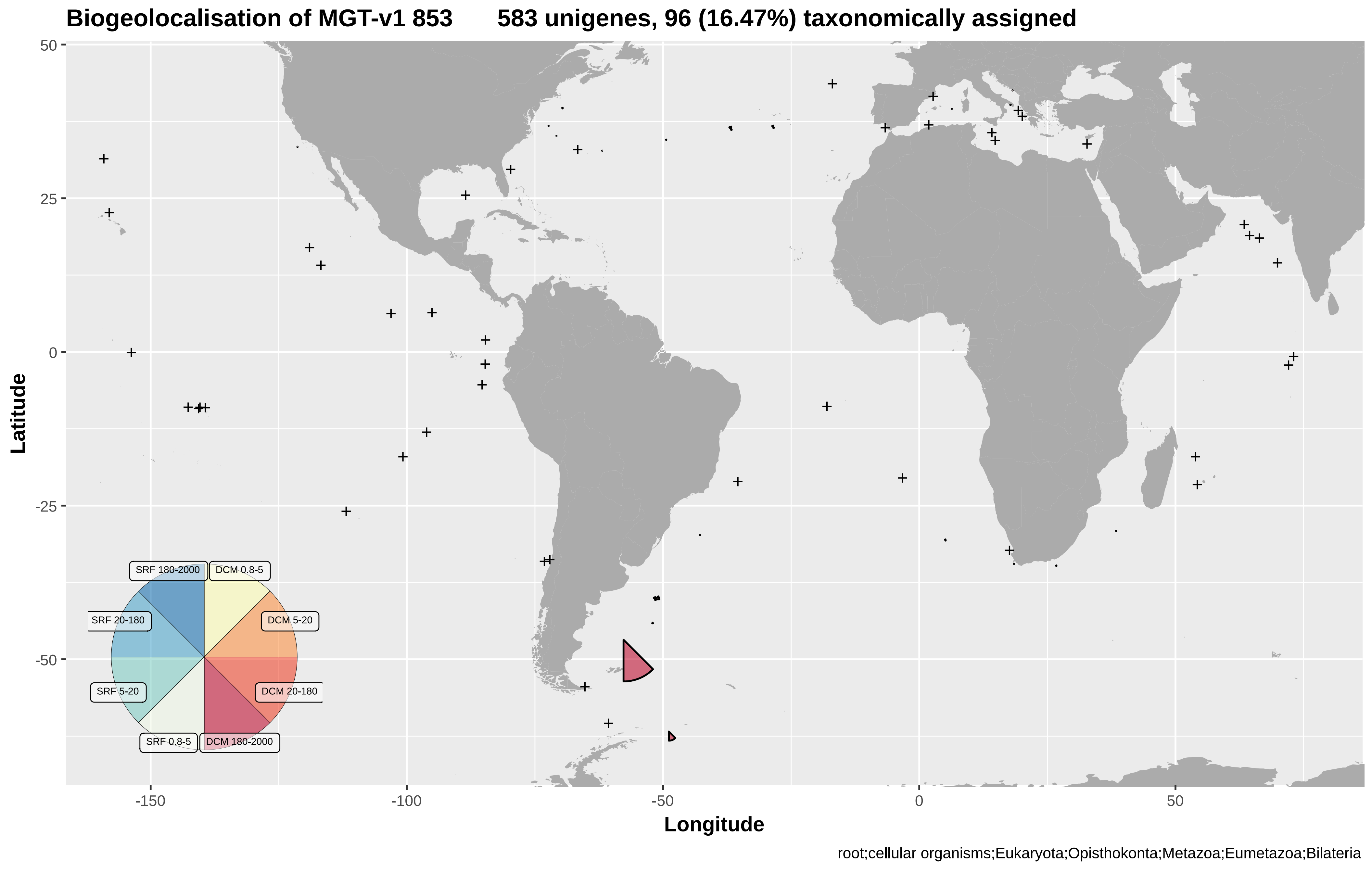

Biogeolocalisation of MGT-v1 853 583 unigenes, 96 (16.47%) taxonomically assigned
Latitude
50
25
0
-25
-50
-150
-100
-50
0
50
Longitude
SRF 180-2000
DCM 0.8-5
SRF 20-180
DCM 5-20
SRF 5-20
DCM 20-180
SRF 0.8-5
DCM 180-2000
root;cellular organisms;Eukaryota;Opisthokonta;Metazoa;Eumetazoa;Bilateria

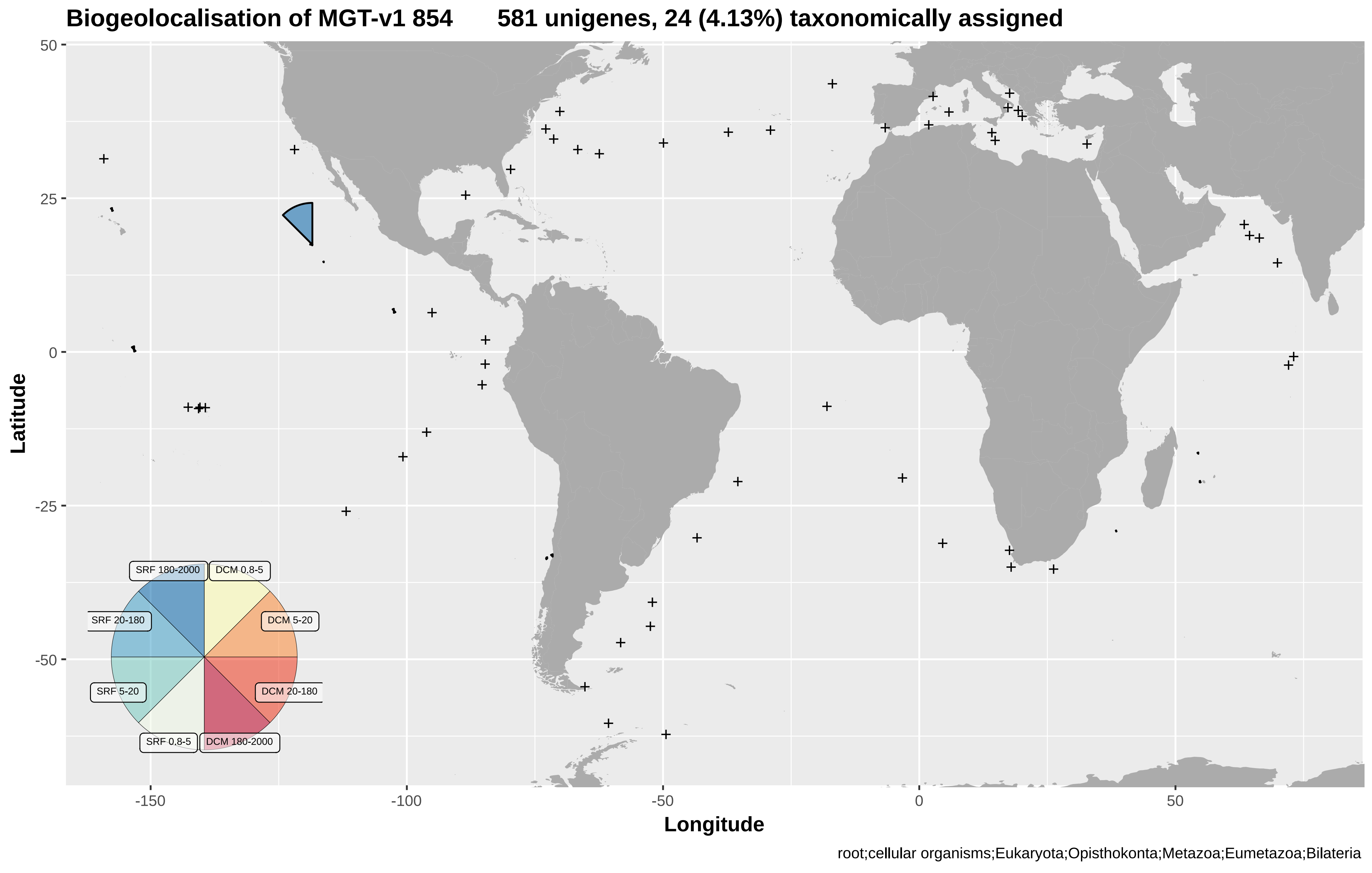
Biogeolocalisation of MGT-v1 854 581 unigenes, 24 (4.13%) taxonomically assigned
Latitude
Longitude
50
25
0
-25
-50
-150
-100
-50
0
50
SRF 180-2000
DCM 0.8-5
SRF 20-180
DCM 5-20
SRF 5-20
DCM 20-180
SRF 0.8-5
DCM 180-2000
root;cellular organisms;Eukaryota;Opisthokonta;Metazoa;Eumetazoa;Bilateria

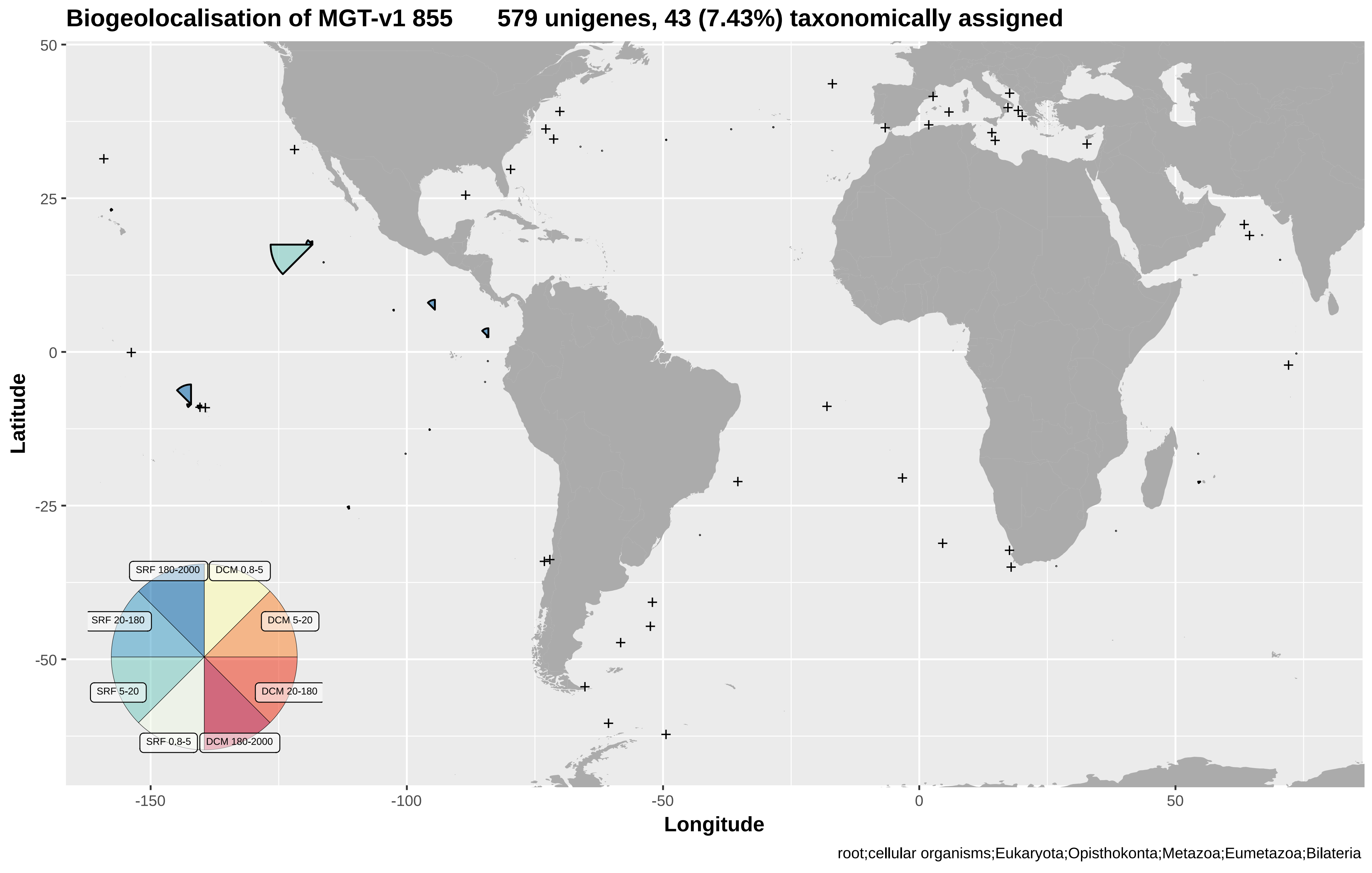
Biogeolocalisation of MGT-v1 855 579 unigenes, 43 (7.43%) taxonomically assigned
Latitude
50
25
0
-25
-50
Longitude
-150
-100
-50
0
50
SRF 180-2000
DCM 0.8-5
SRF 20-180
DCM 5-20
SRF 5-20
DCM 20-180
SRF 0.8-5
DCM 180-2000
root;cellular organisms;Eukaryota;Opisthokonta;Metazoa;Eumetazoa;Bilateria

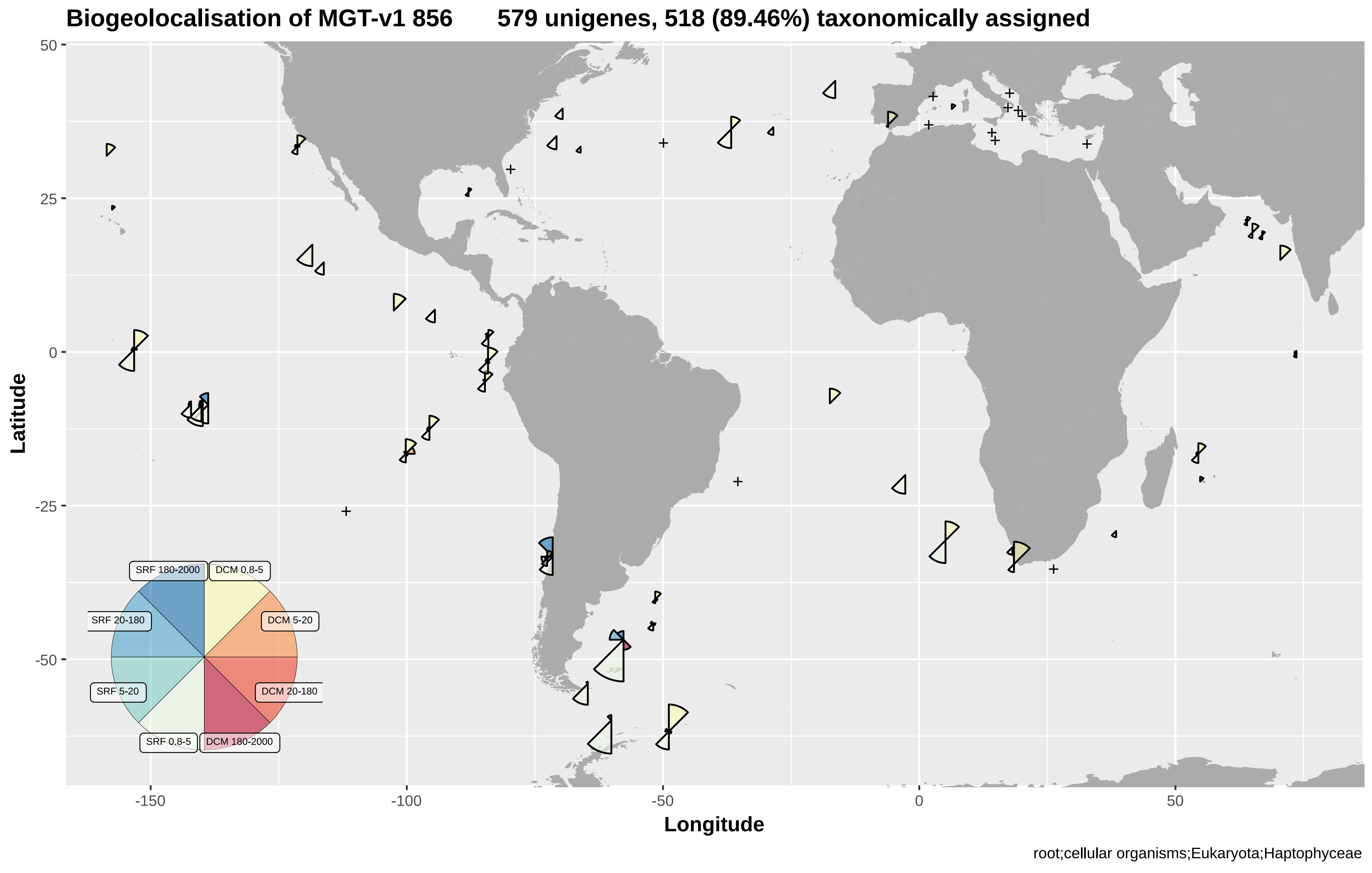
Biogeolocalisation of MGT-v1 856
579 unigenes, 518 (89.46%) taxonomically assigned
50
25
0
-25
-50
Latitude
-150
-100
-50
0
50
Longitude
SRF 180-2000
DCM 0.8-5
SRF 20-180
DCM 5-20
SRF 5-20
DCM 20-180
SRF 0.8-5
DCM 180-2000
root;cellular organisms;Eukaryota;Haptophyceae

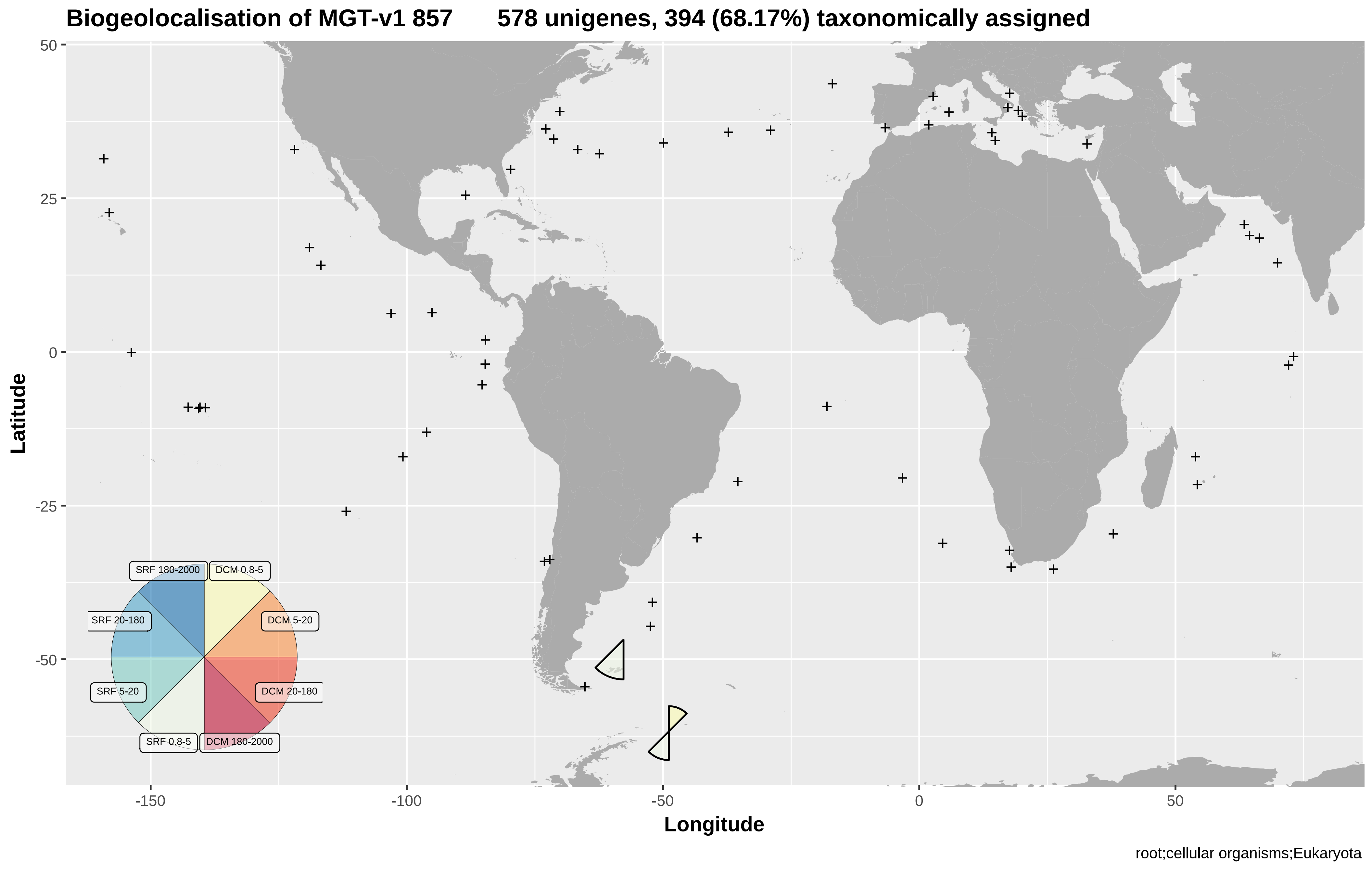

Biogeolocalisation of MGT-v1 857 578 unigenes, 394 (68.17%) taxonomically assigned
50
25
0
-25
-50
Latitude
-150
-100
-50
0
50
Longitude
SRF 180-2000
DCM 0.8-5
SRF 20-180
DCM 5-20
SRF 5-20
DCM 20-180
SRF 0.8-5
DCM 180-2000
root;cellular organisms;Eukaryota

## Biogeolocalisation of MGT-v1 858      578 unigenes, 25 (4.33%) taxonomically assigned

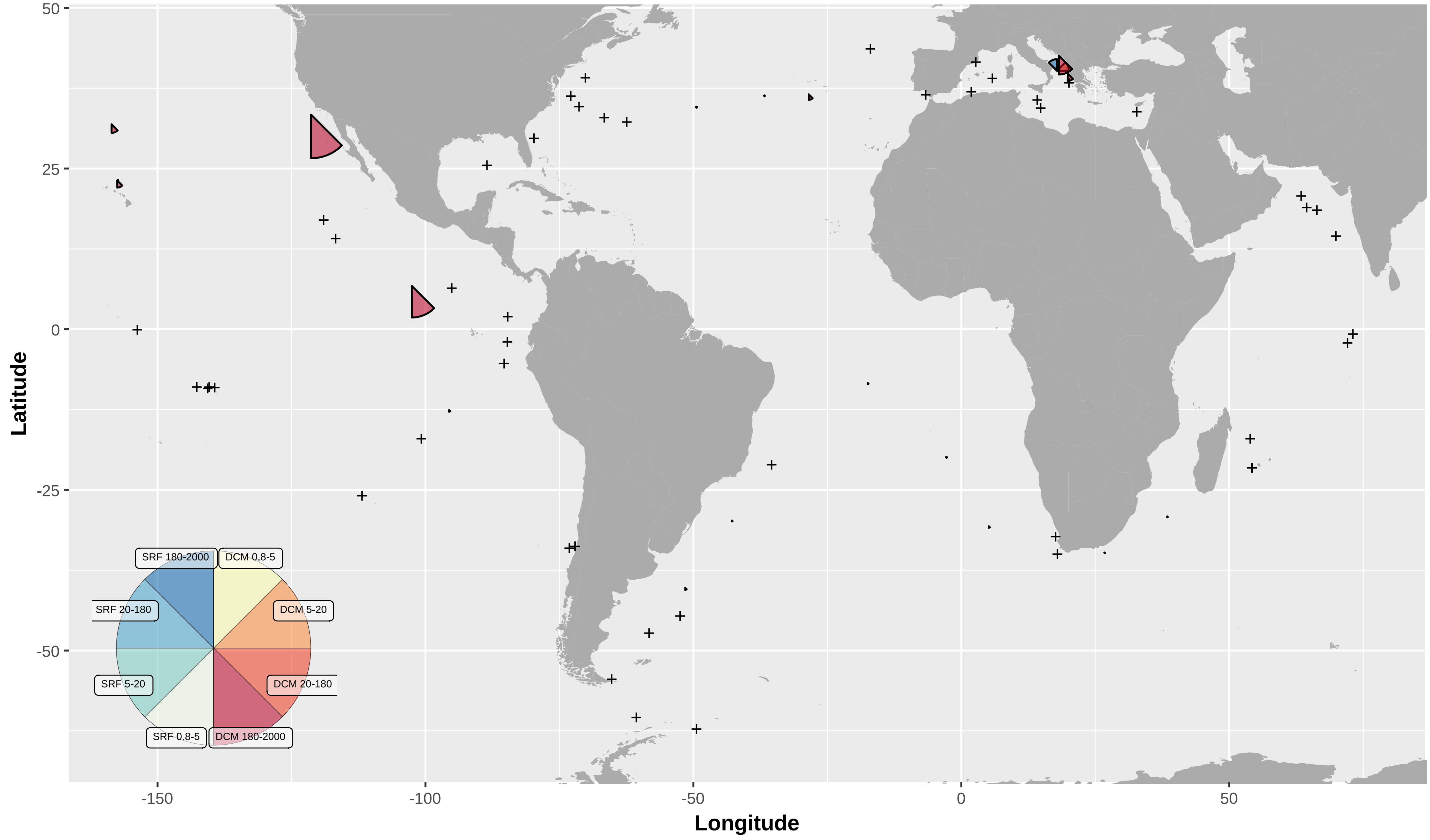


root;cellular organisms;Eukaryota;Opisthokonta;Metazoa;Eumetazoa

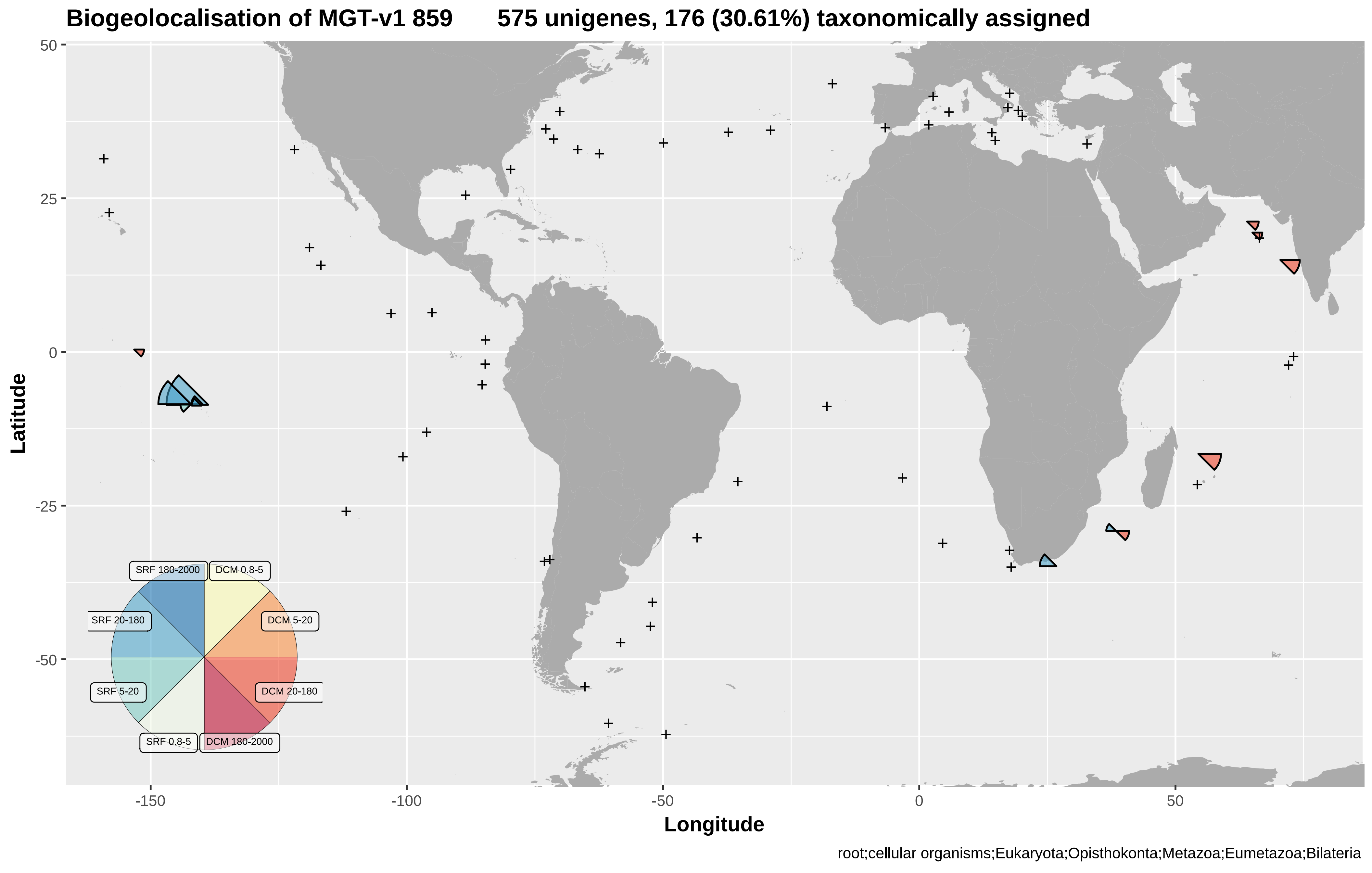
Biogeolocalisation of MGT-v1 859 575 unigenes, 176 (30.61%) taxonomically assigned
Latitude
Longitude
50
25
0
-25
-50
-150
-100
-50
0
50
SRF 180-2000
DCM 0.8-5
SRF 20-180
DCM 5-20
SRF 5-20
DCM 20-180
SRF 0.8-5
DCM 180-2000
root;cellular organisms;Eukaryota;Opisthokonta;Metazoa;Eumetazoa;Bilateria

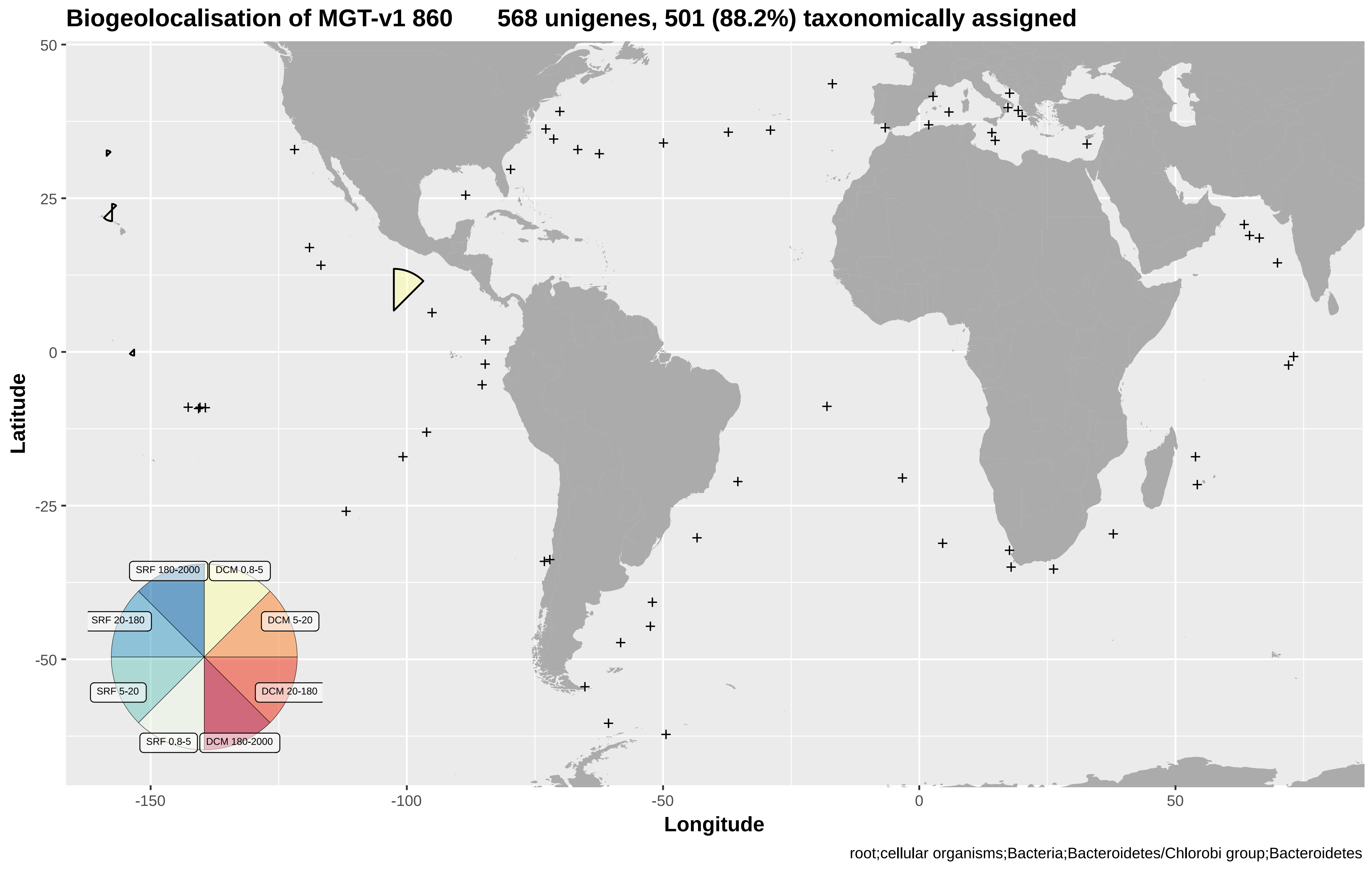

Biogeolocalisation of MGT-v1 860
568 unigenes, 501 (88.2%) taxonomically assigned
Latitude
Longitude
50
25
0
-25
-50
-150
-100
-50
0
50
SRF 180-2000
DCM 0.8-5
SRF 20-180
DCM 5-20
SRF 5-20
DCM 20-180
SRF 0.8-5
DCM 180-2000
root;cellular organisms;Bacteria;Bacteroidetes/Chlorobi group;Bacteroidetes

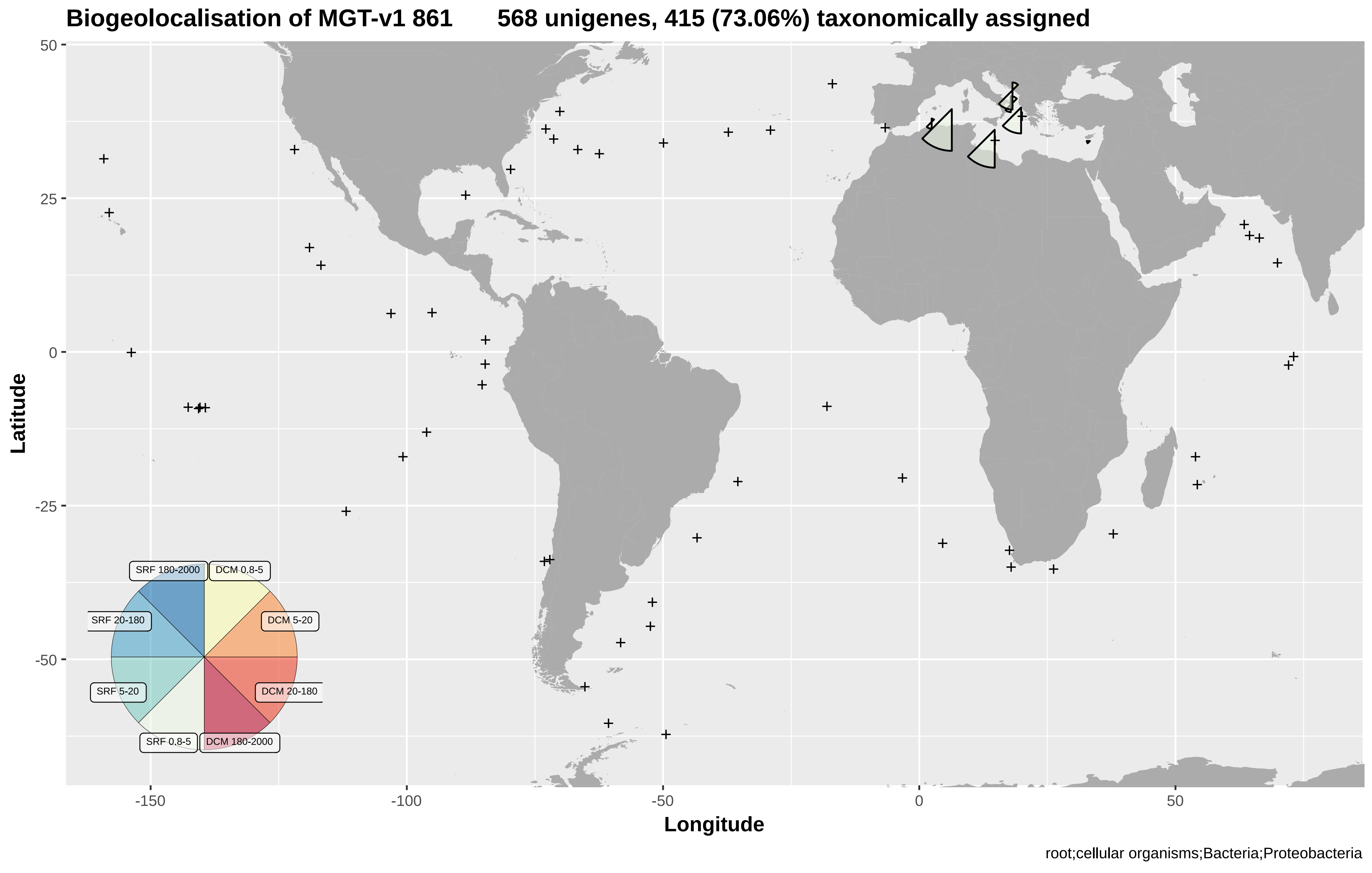
Biogeolocalisation of MGT-v1 861 568 unigenes, 415 (73.06%) taxonomically assigned
Latitude
Longitude
50
25
0
-25
-50
-150
-100
-50
0
50
SRF 180-2000
DCM 0.8-5
SRF 20-180
DCM 5-20
SRF 5-20
DCM 20-180
SRF 0.8-5
DCM 180-2000
root;cellular organisms;Bacteria;Proteobacteria

## Biogeolocalisation of MGT-v1 862      562 unigenes, 58 (10.32%) taxonomically assigned

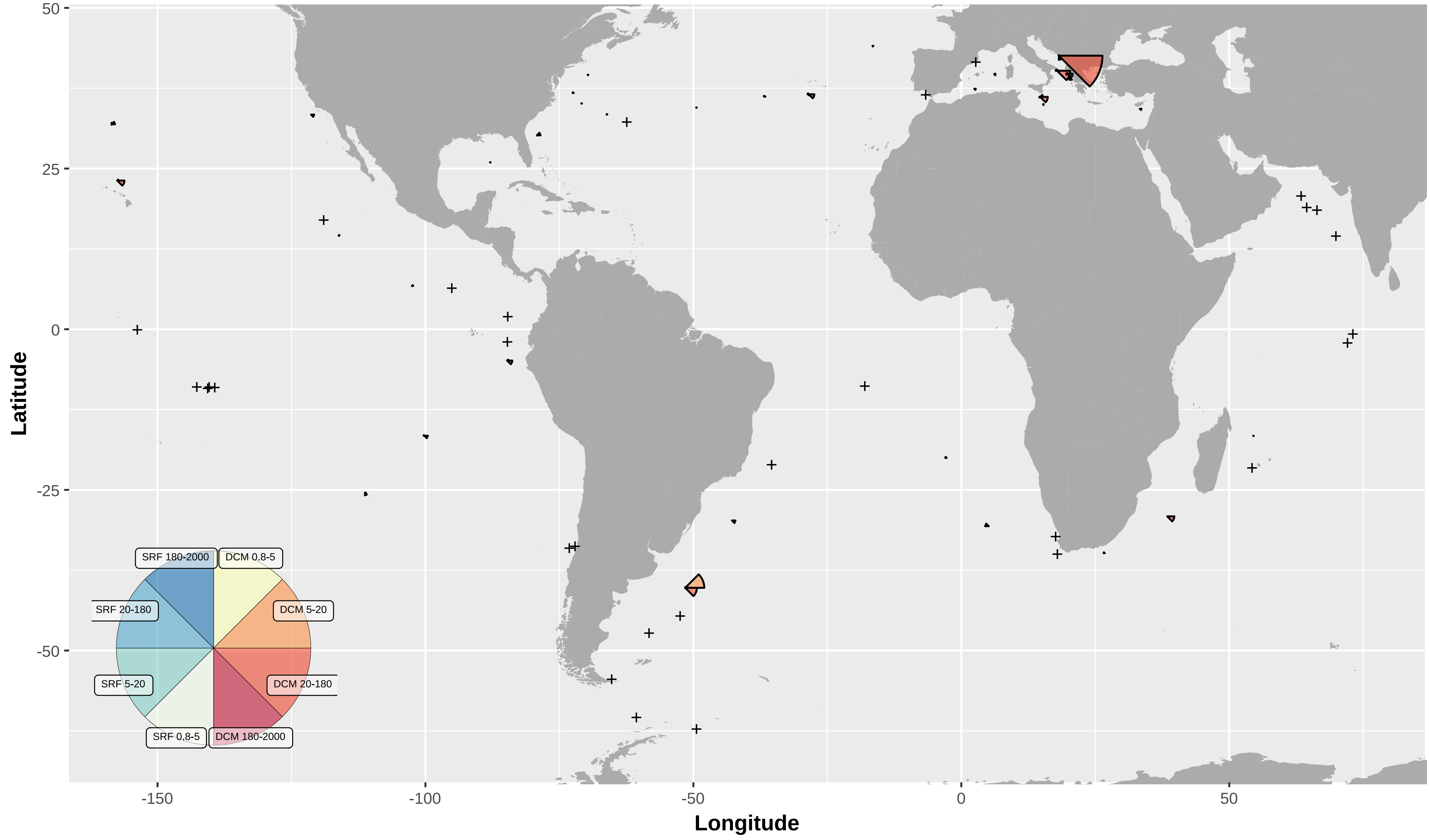


root;cellular organisms;Eukaryota

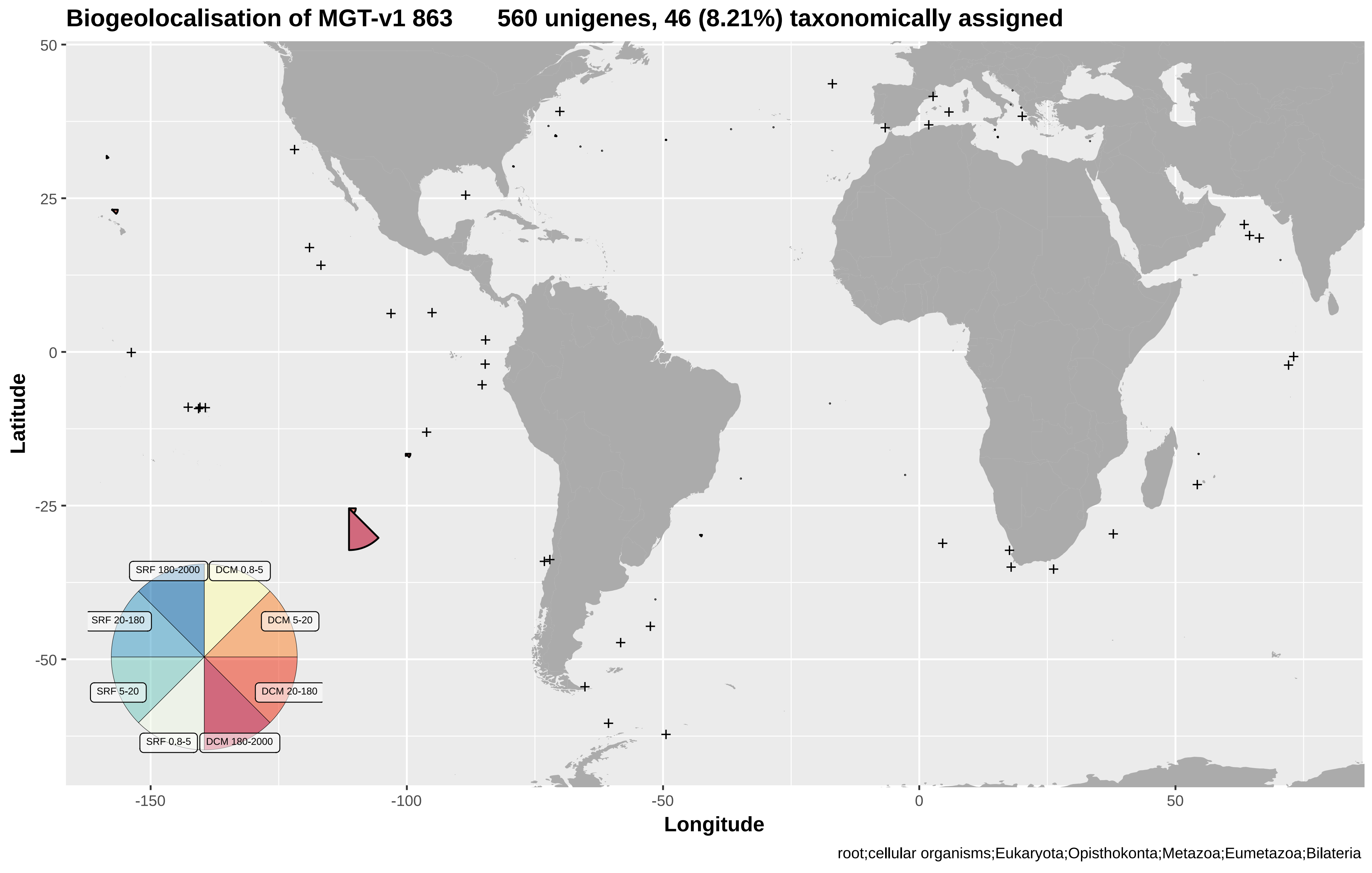
Biogeolocalisation of MGT-v1 863
560 unigenes, 46 (8.21%) taxonomically assigned
Latitude
Longitude
50
25
0
-25
-50
-150
-100
-50
0
50
SRF 180-2000
DCM 0.8-5
SRF 20-180
DCM 5-20
SRF 5-20
DCM 20-180
SRF 0.8-5
DCM 180-2000
root;cellular organisms;Eukaryota;Opisthokonta;Metazoa;Eumetazoa;Bilateria

# Biogeolocalisation of MGT-v1 864      560 unigenes, 17 (3.04%) taxonomically assigned

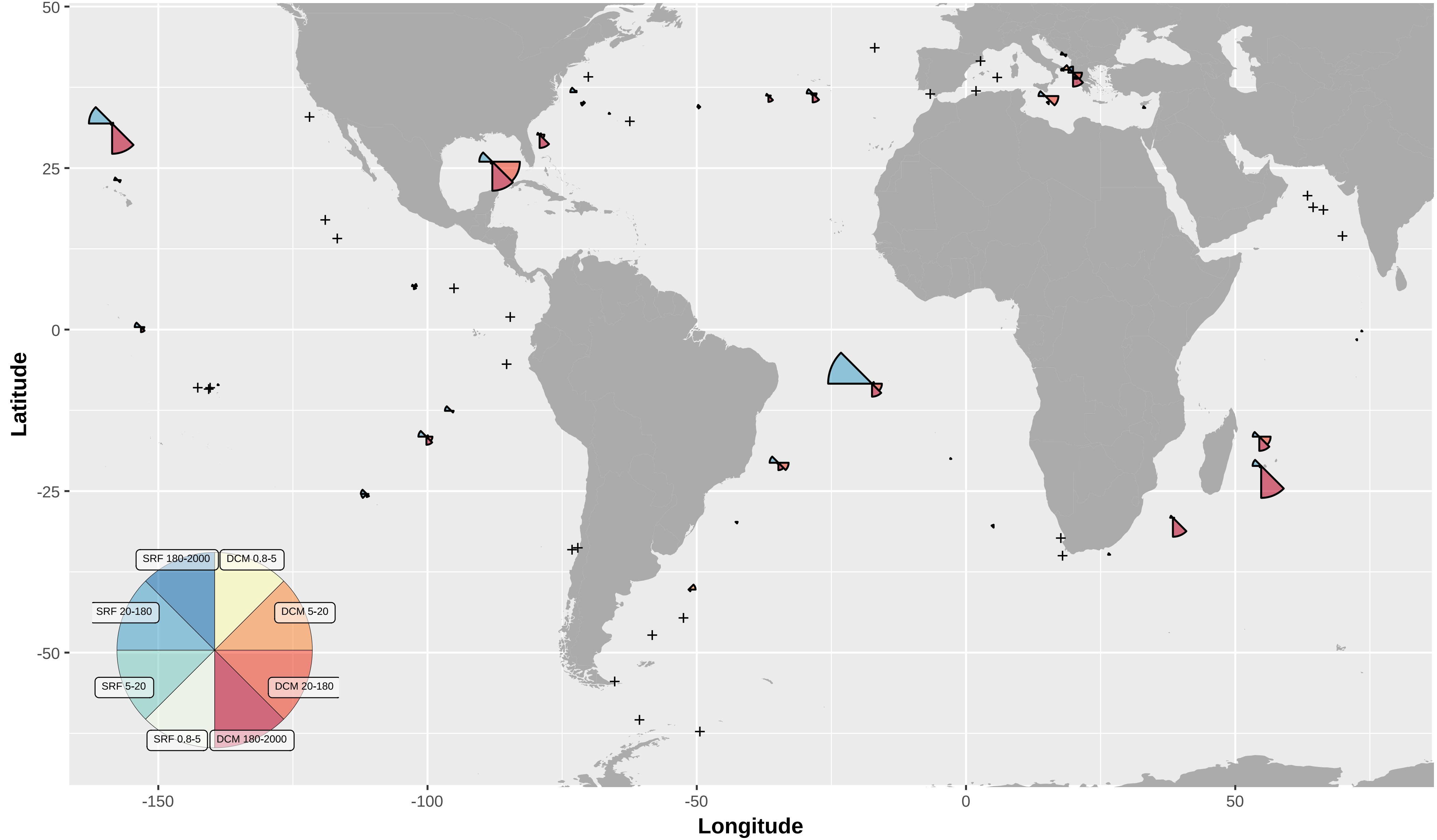


root;cellular organisms;Eukaryota;Opisthokonta;Metazoa;Eumetazoa;Bilateria

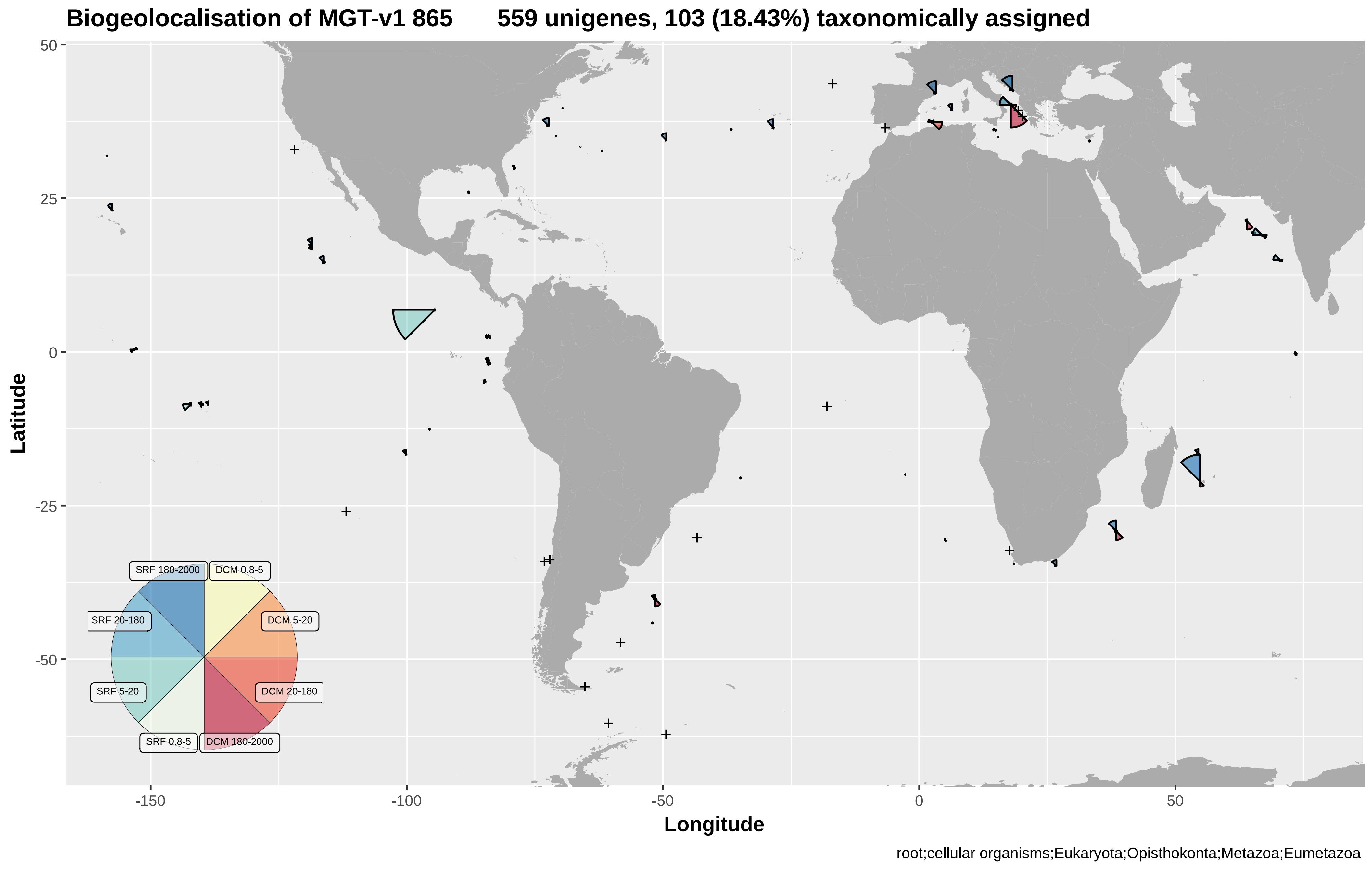

Biogeolocalisation of MGT-v1 865 559 unigenes, 103 (18.43%) taxonomically assigned
50
25
0
-25
-50
Latitude
-150
-100
-50
0
50
Longitude
SRF 180-2000
DCM 0.8-5
SRF 20-180
DCM 5-20
SRF 5-20
DCM 20-180
SRF 0.8-5
DCM 180-2000
root;cellular organisms;Eukaryota;Opisthokonta;Metazoa;Eumetazoa

## Biogeolocalisation of MGT-v1 866      556 unigenes, 433 (77.88%) taxonomically assigned

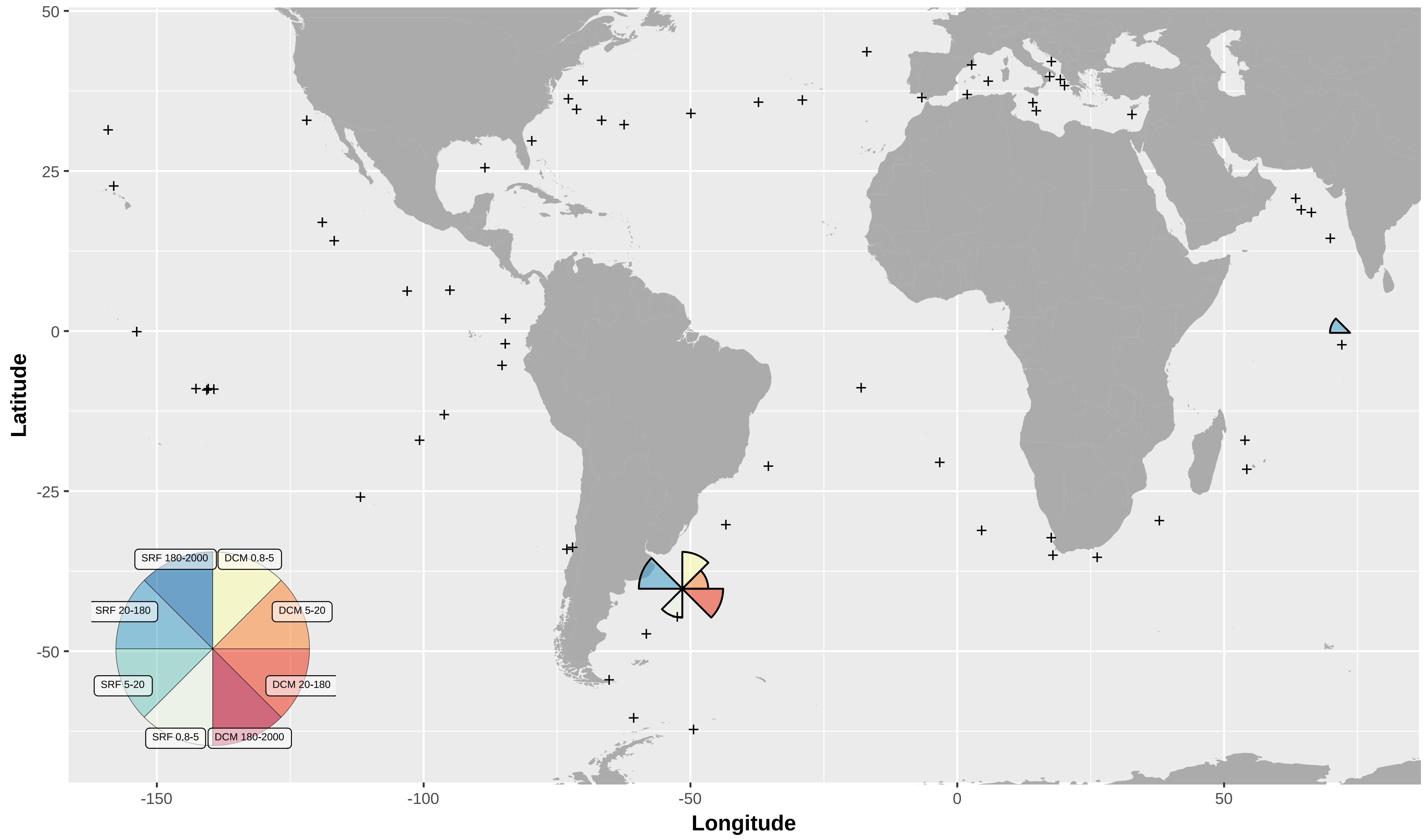


root;cellular organisms;Eukaryota;Stramenopiles;Bacillariophyta;Bacillariophyceae

#### Biogeolocalisation of MGT-v1 867 555 unigenes, 26 (4.68%) taxonomically assigned

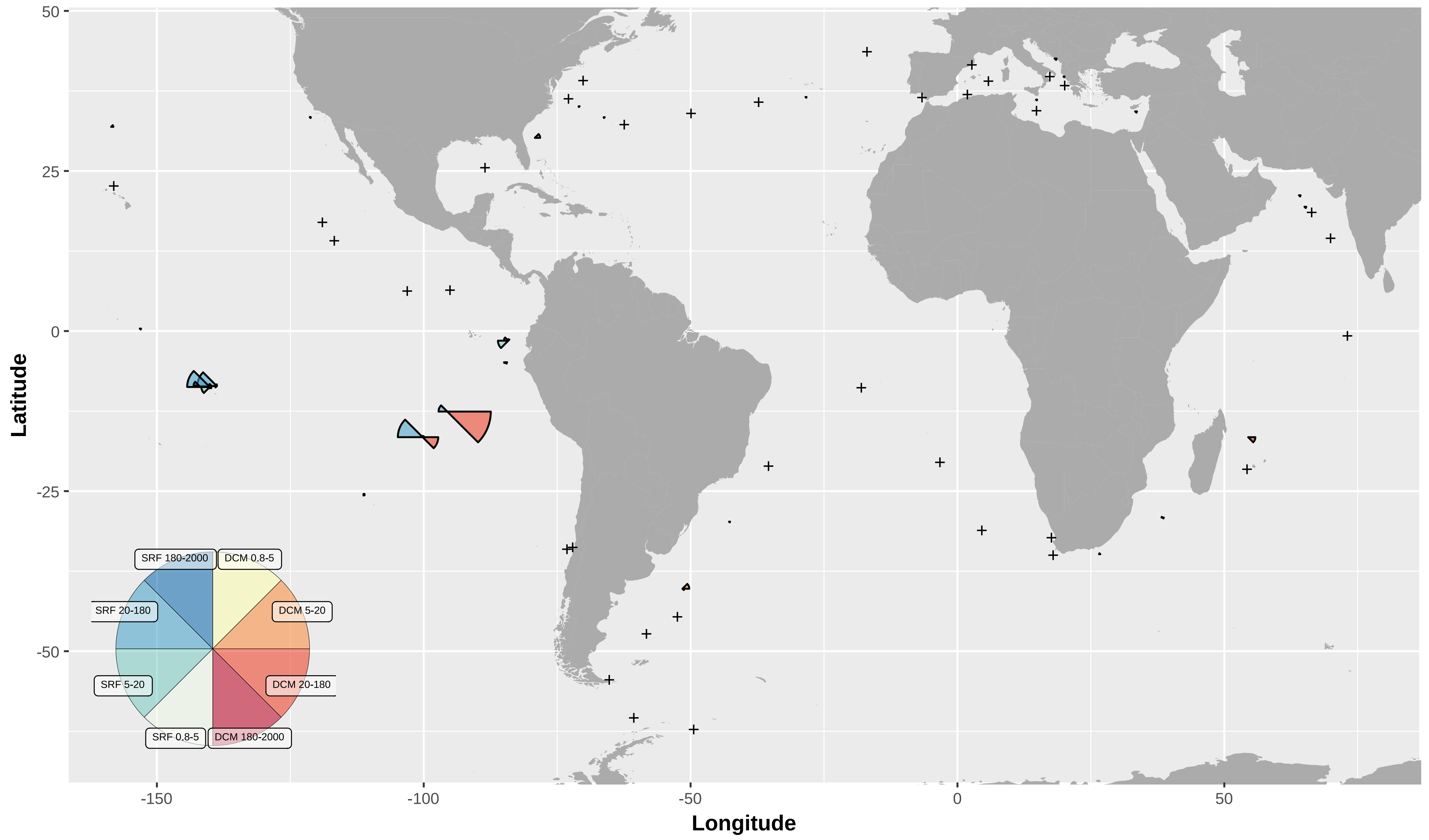


root;cellular organisms;Eukaryota

# Biogeolocalisation of MGT-v1 868      554 unigenes, 44 (7.94%) taxonomically assigned

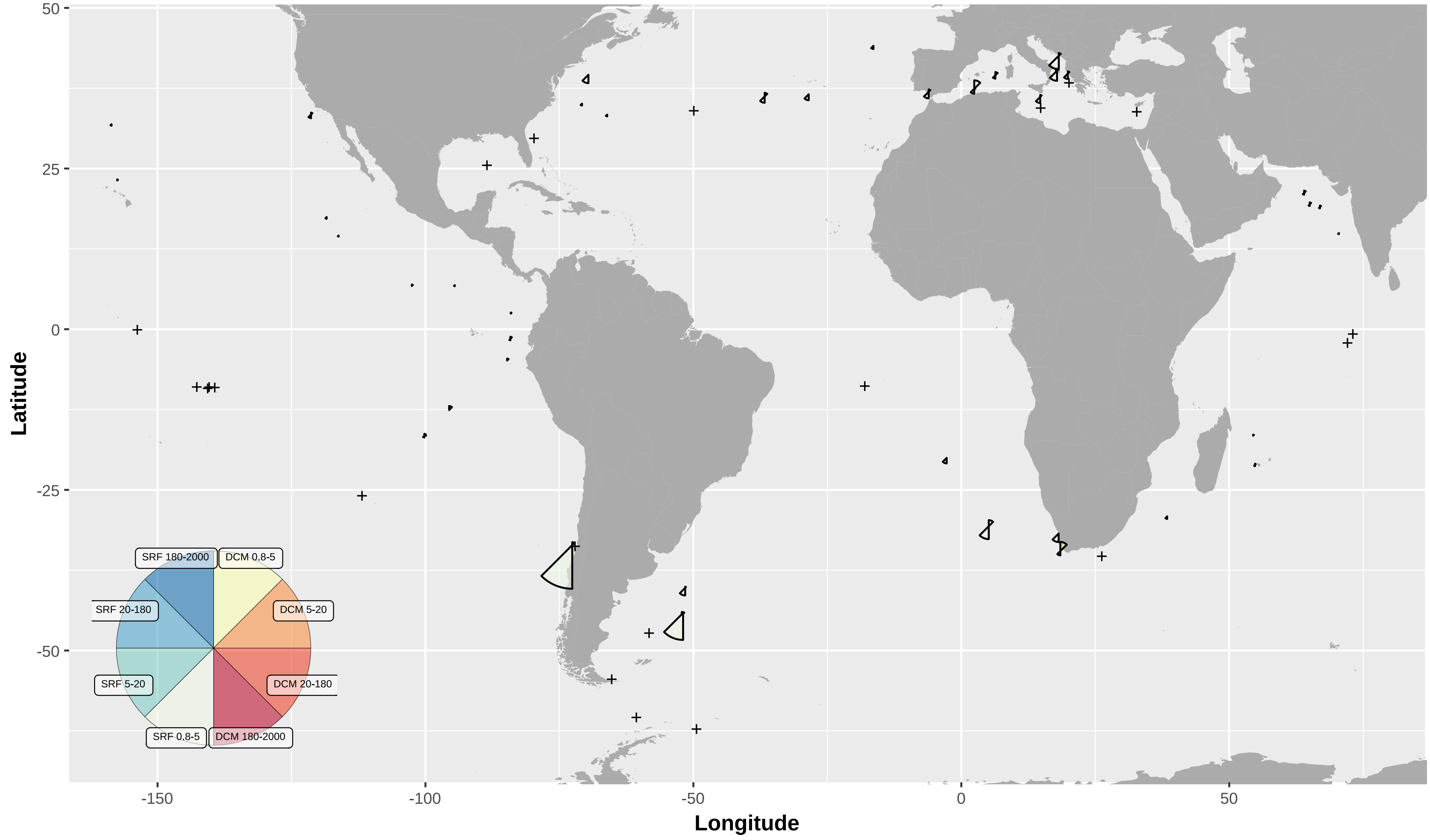


root;cellular organisms;Eukaryota

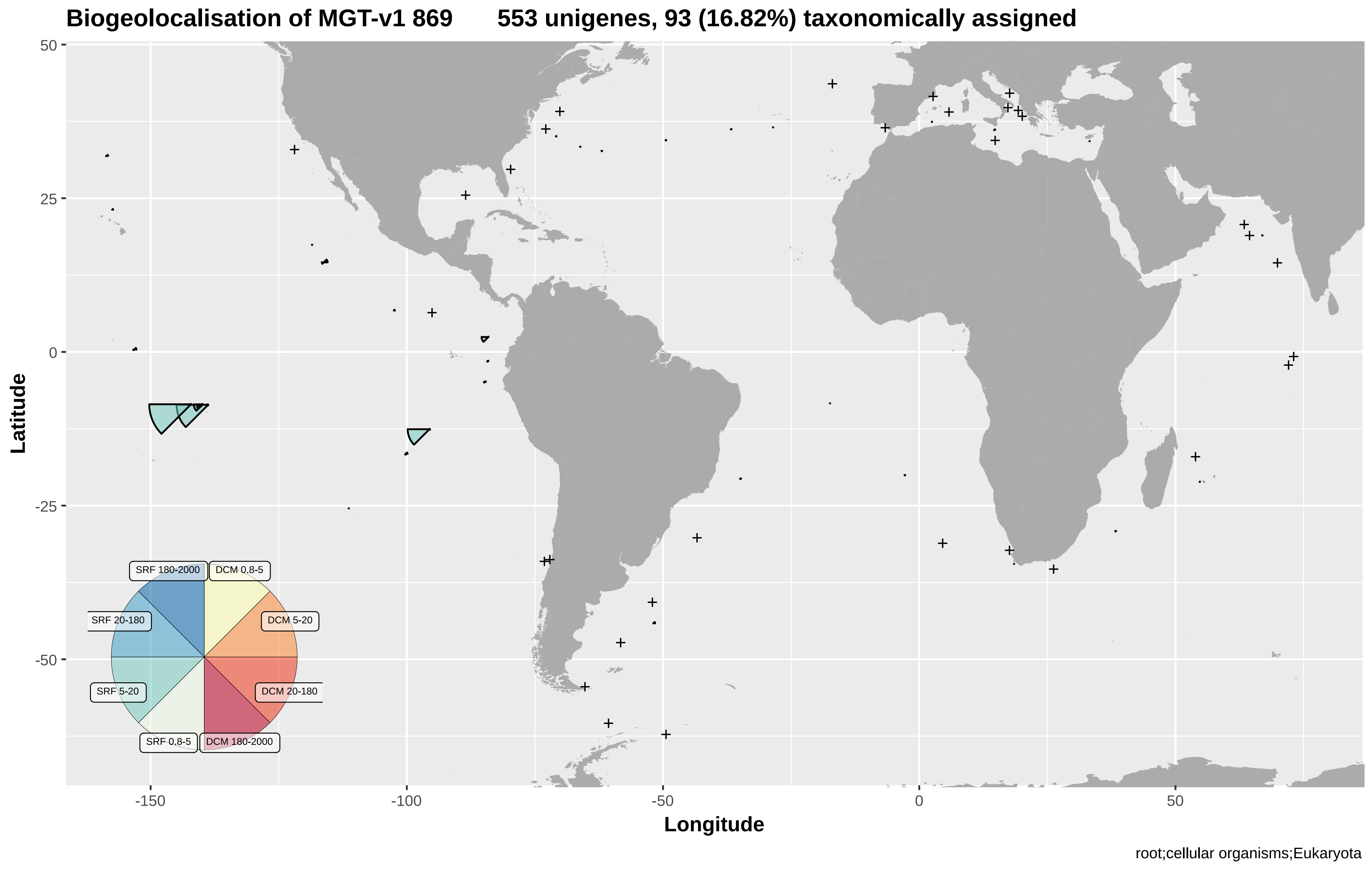

Biogeolocalisation of MGT-v1 869 553 unigenes, 93 (16.82%) taxonomically assigned
50
25
0
-25
-50
Latitude
-150
-100
-50
0
50
Longitude
SRF 180-2000
DCM 0.8-5
SRF 20-180
DCM 5-20
SRF 5-20
DCM 20-180
SRF 0.8-5
DCM 180-2000
root;cellular organisms;Eukaryota

## Biogeolocalisation of MGT-v1 870      552 unigenes, 542 (98.19%) taxonomically assigned

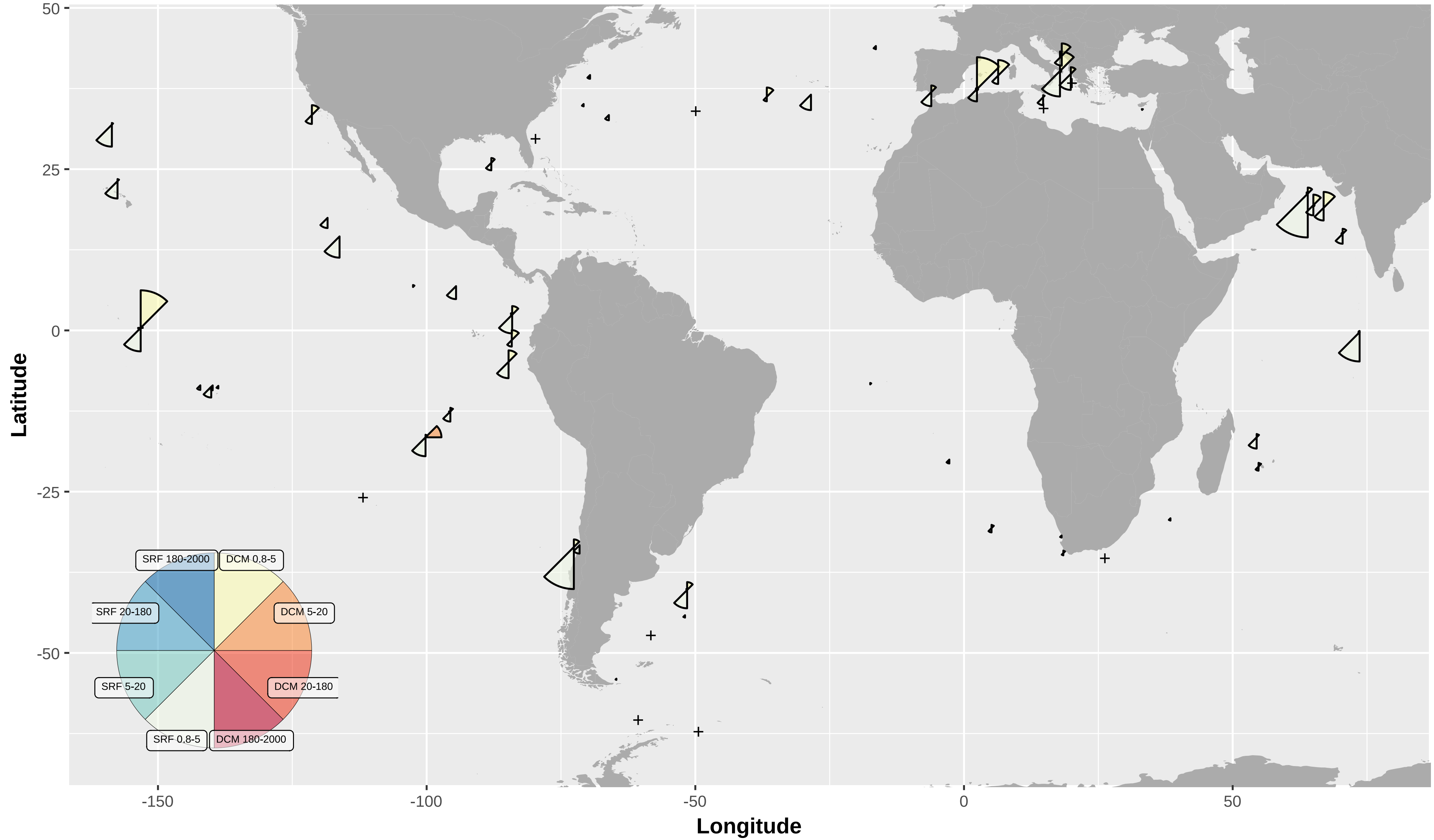


root;cellular organisms;Bacteria;Proteobacteria;Alphaproteobacteria;Pelagibacterales

#### Biogeolocalisation of MGT-v1 871 551 unigenes, 37 (6.72%) taxonomically assigned

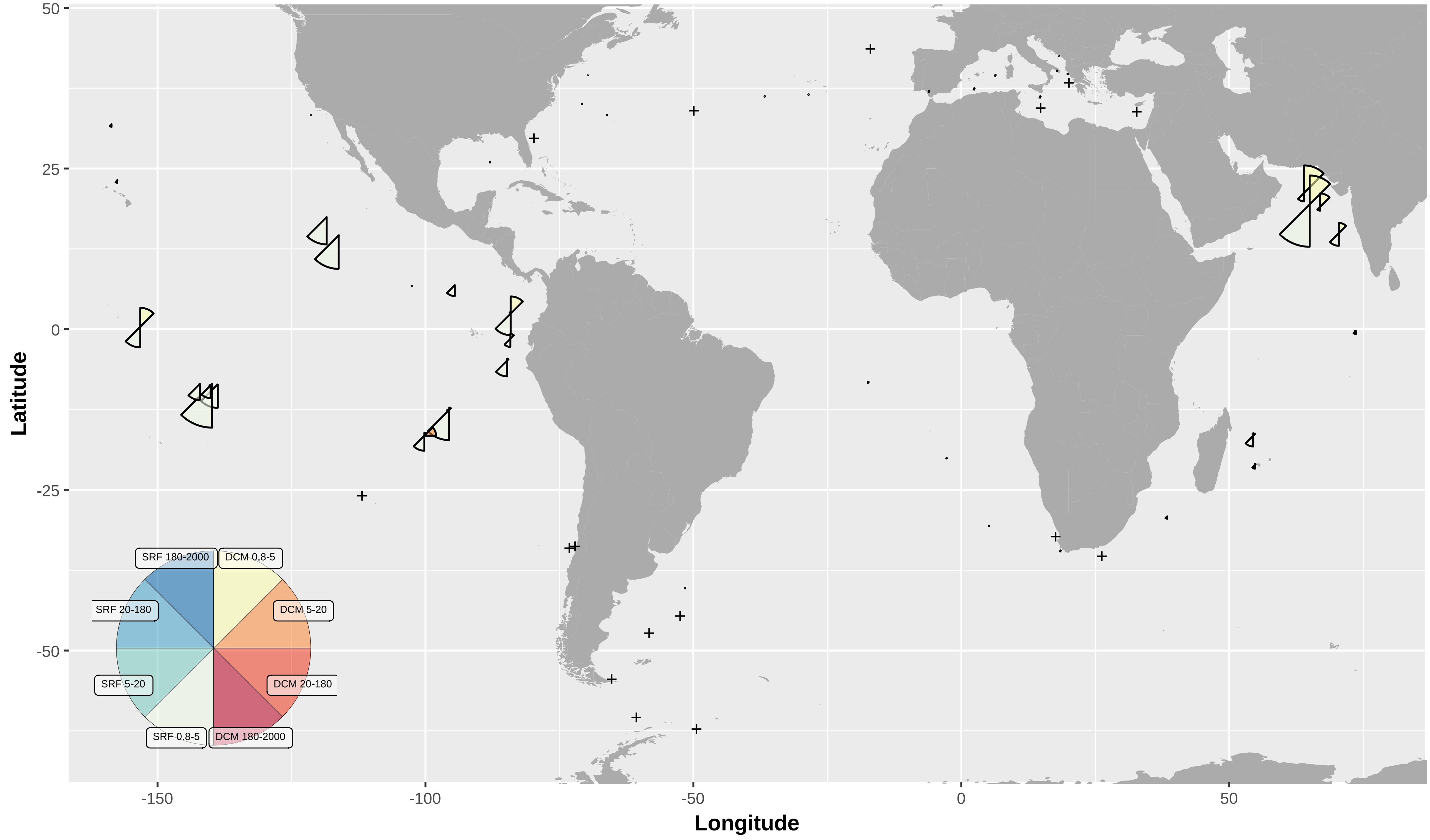


root;cellular organisms;Eukaryota

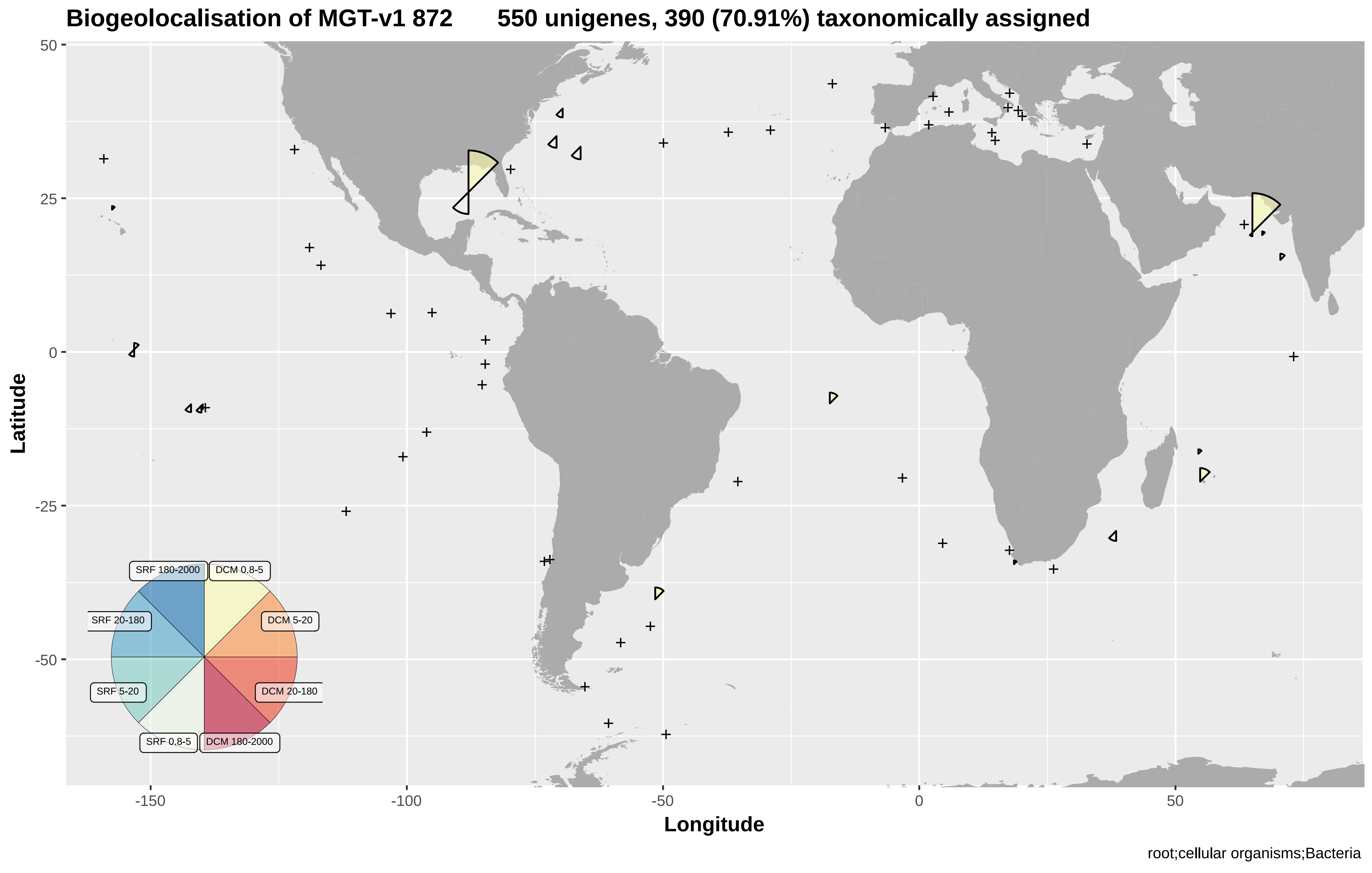

Biogeolocalisation of MGT-v1 872
550 unigenes, 390 (70.91%) taxonomically assigned
50
25
0
-25
-50
Latitude
-150
-100
-50
0
50
Longitude
SRF 180-2000
DCM 0.8-5
SRF 20-180
DCM 5-20
SRF 5-20
DCM 20-180
SRF 0.8-5
DCM 180-2000
root;cellular organisms;Bacteria

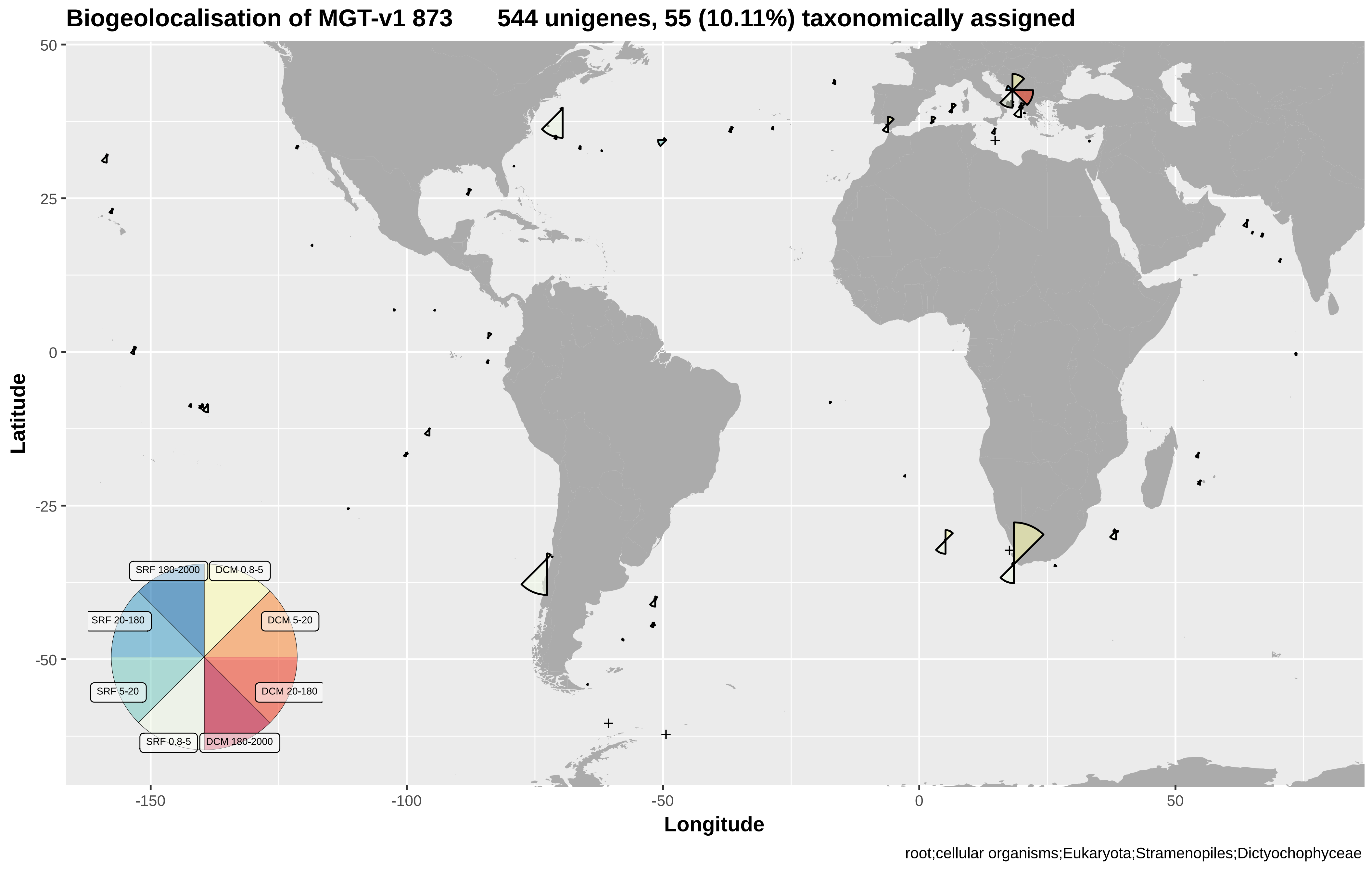

Biogeolocalisation of MGT-v1 873      544 unigenes, 55 (10.11%) taxonomically assigned
50
25
0
-25
-50
Latitude
-150
-100
-50
0
50
Longitude
SRF 180-2000
DCM 0.8-5
SRF 20-180
DCM 5-20
SRF 5-20
DCM 20-180
SRF 0.8-5
DCM 180-2000
root;cellular organisms;Eukaryota;Stramenopiles;Dictyochophyceae

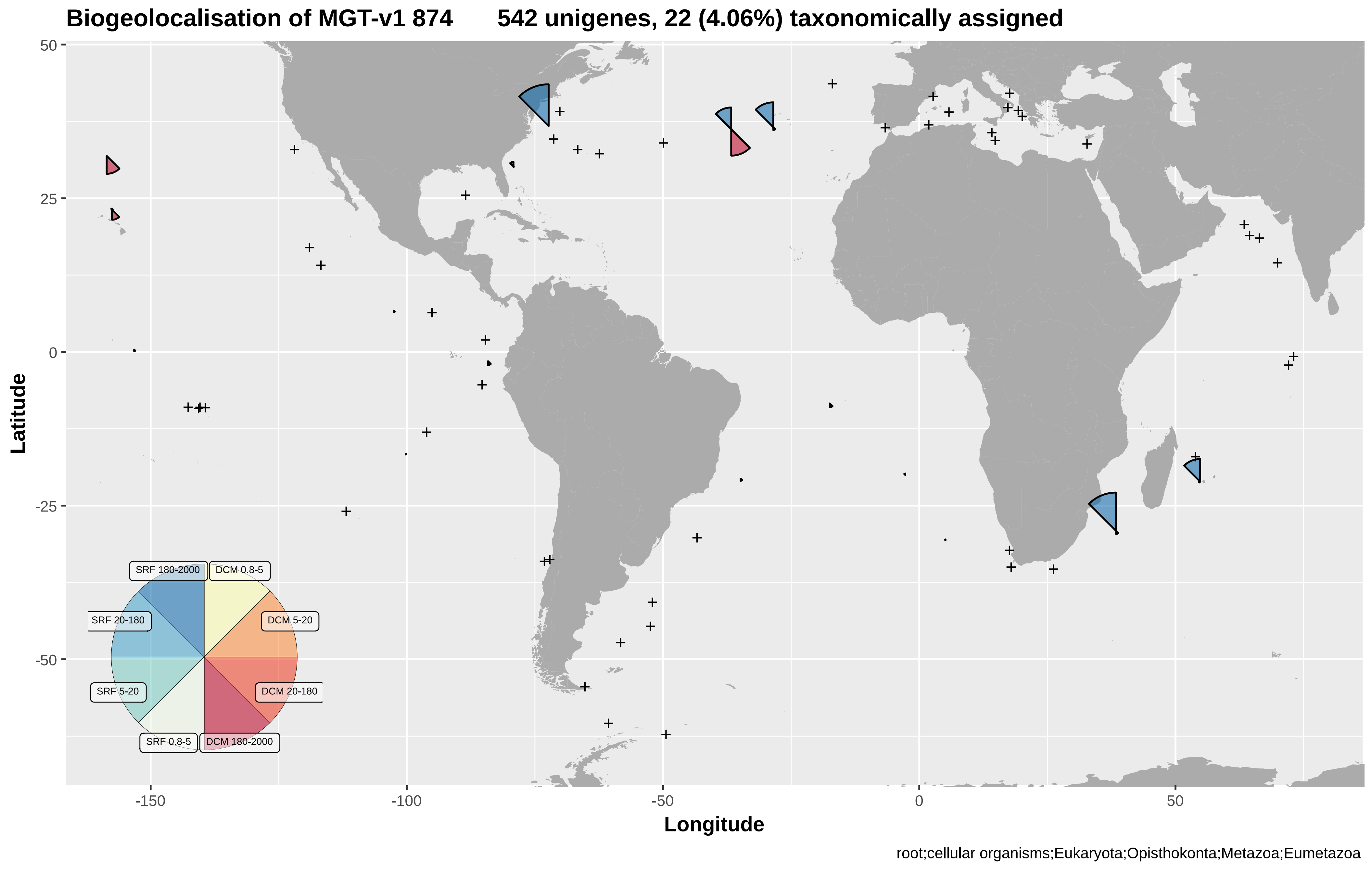
Biogeolocalisation of MGT-v1 874 542 unigenes, 22 (4.06%) taxonomically assigned
50
25
0
-25
-50
Latitude
-150
-100
-50
0
50
Longitude
SRF 180-2000
DCM 0.8-5
SRF 20-180
DCM 5-20
SRF 5-20
DCM 20-180
SRF 0.8-5
DCM 180-2000
root;cellular organisms;Eukaryota;Opisthokonta;Metazoa;Eumetazoa

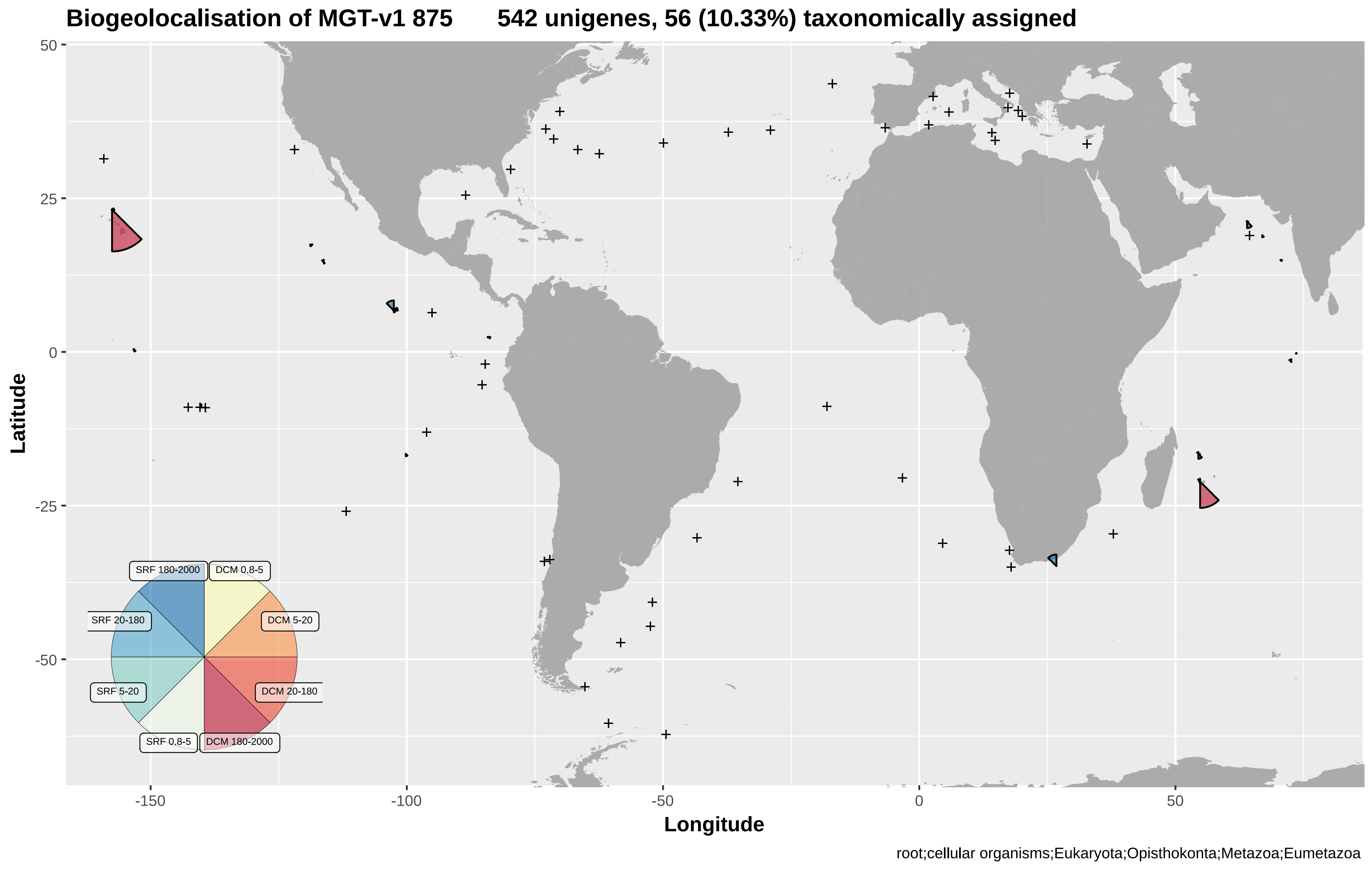
Biogeolocalisation of MGT-v1 875 542 unigenes, 56 (10.33%) taxonomically assigned
Latitude
Longitude
50
25
0
-25
-50
-150
-100
-50
0
50
SRF 180-2000
DCM 0.8-5
SRF 20-180
DCM 5-20
SRF 5-20
DCM 20-180
SRF 0.8-5
DCM 180-2000
root;cellular organisms;Eukaryota;Opisthokonta;Metazoa;Eumetazoa

# Biogeolocalisation of MGT-v1 876      540 unigenes, 416 (77.04%) taxonomically assigned

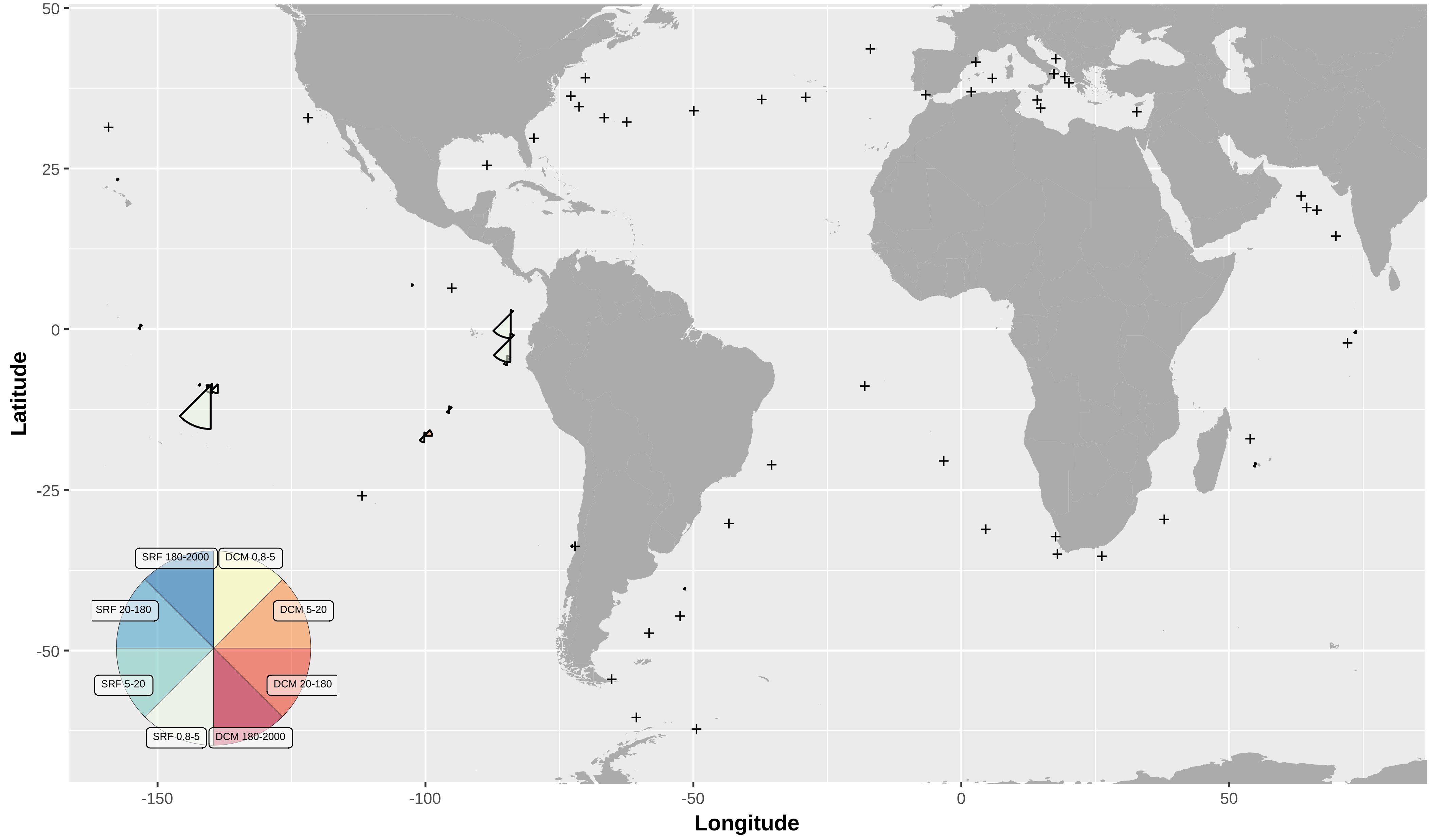


root;cellular organisms;Bacteria;Proteobacteria;Alphaproteobacteria

## Biogeolocalisation of MGT-v1 877      539 unigenes, 463 (85.9%) taxonomically assigned

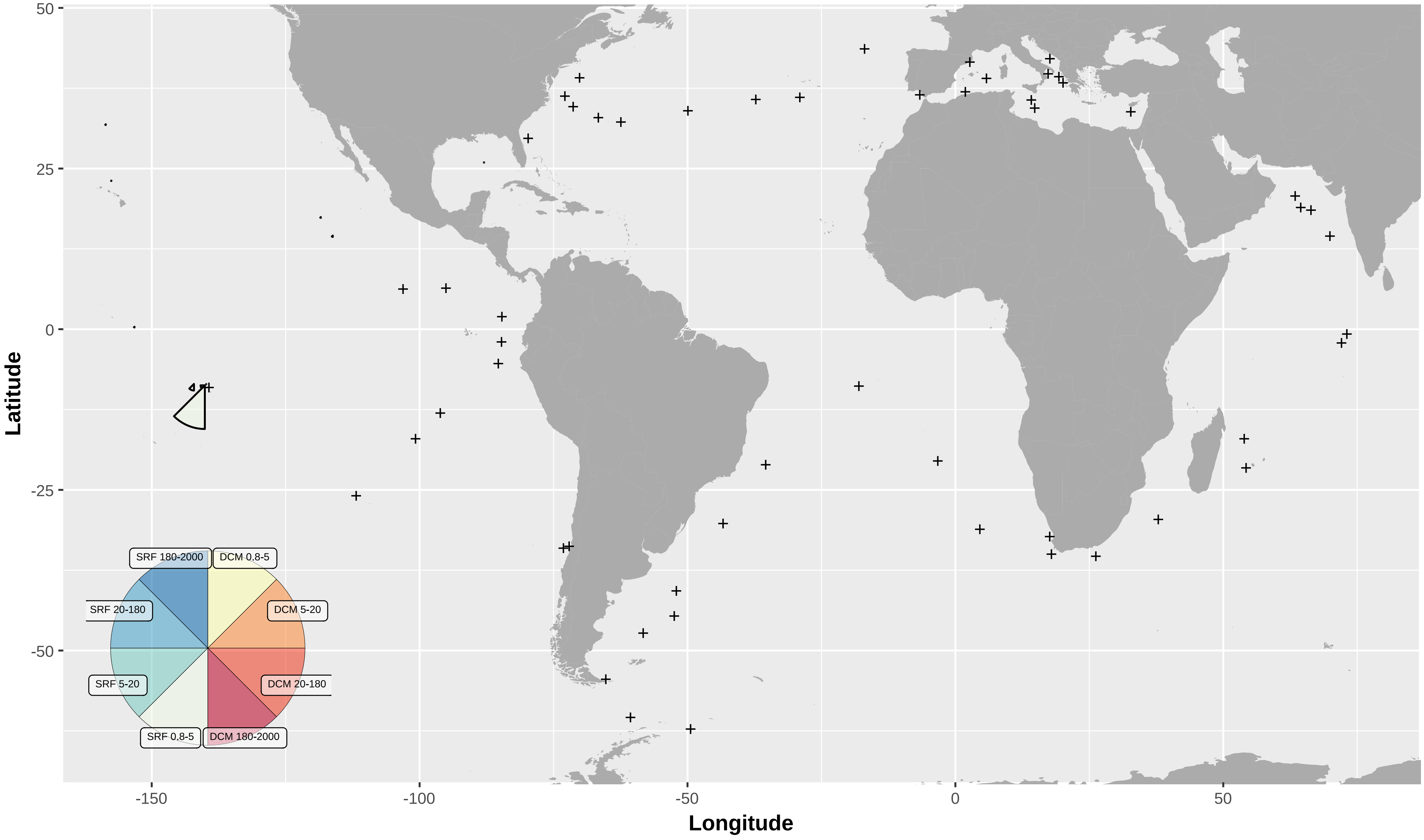


root

Biogeolocalisation of MGT-v1 878 537 unigenes, 309 (57.54%) taxonomically assigned

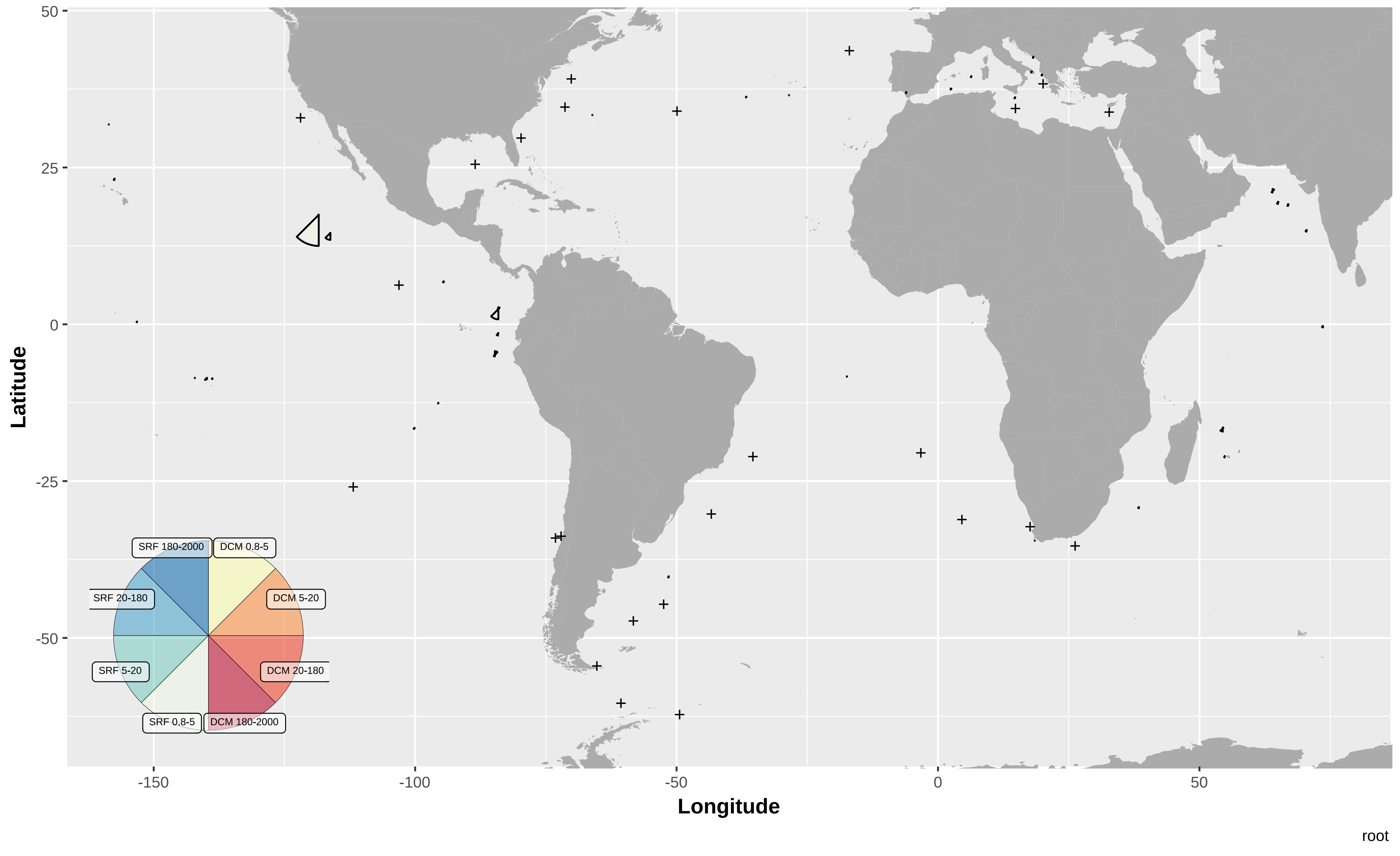

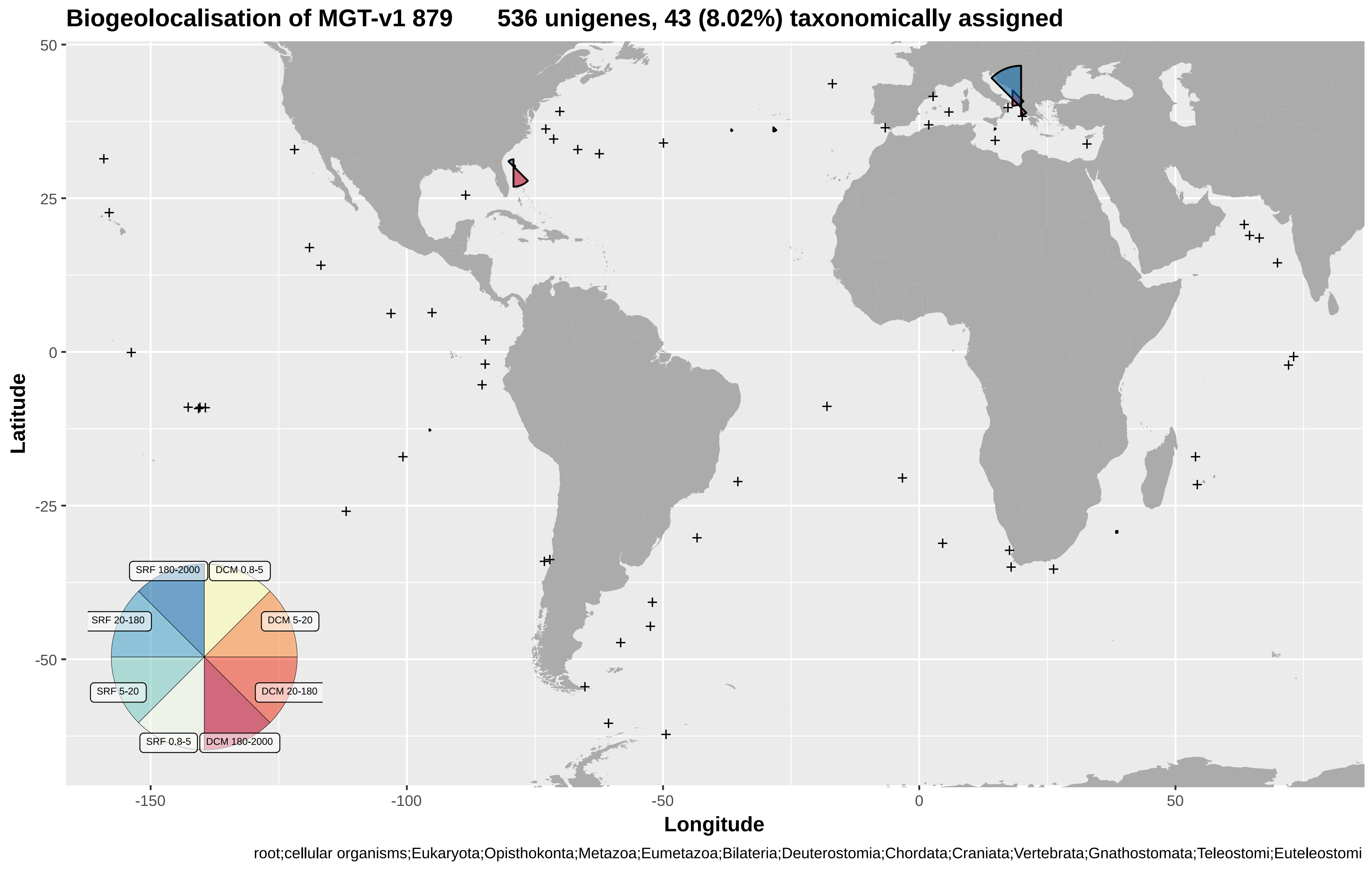

Biogeolocalisation of MGT-v1 879 536 unigenes, 43 (8.02%) taxonomically assigned
Latitude
Longitude
50
25
0
-25
-50
-150
-100
-50
0
50
SRF 180-2000
DCM 0.8-5
SRF 20-180
DCM 5-20
SRF 5-20
DCM 20-180
SRF 0.8-5
DCM 180-2000
root;cellular organisms;Eukaryota;Opisthokonta;Metazoa;Eumetazoa;Bilateria;Deuterostomia;Chordata;Craniata;Vertebrata;Gnathostomata;Teleostomi;Euteleostomi

## Biogeolocalisation of MGT-v1 880      532 unigenes, 71 (13.35%) taxonomically assigned

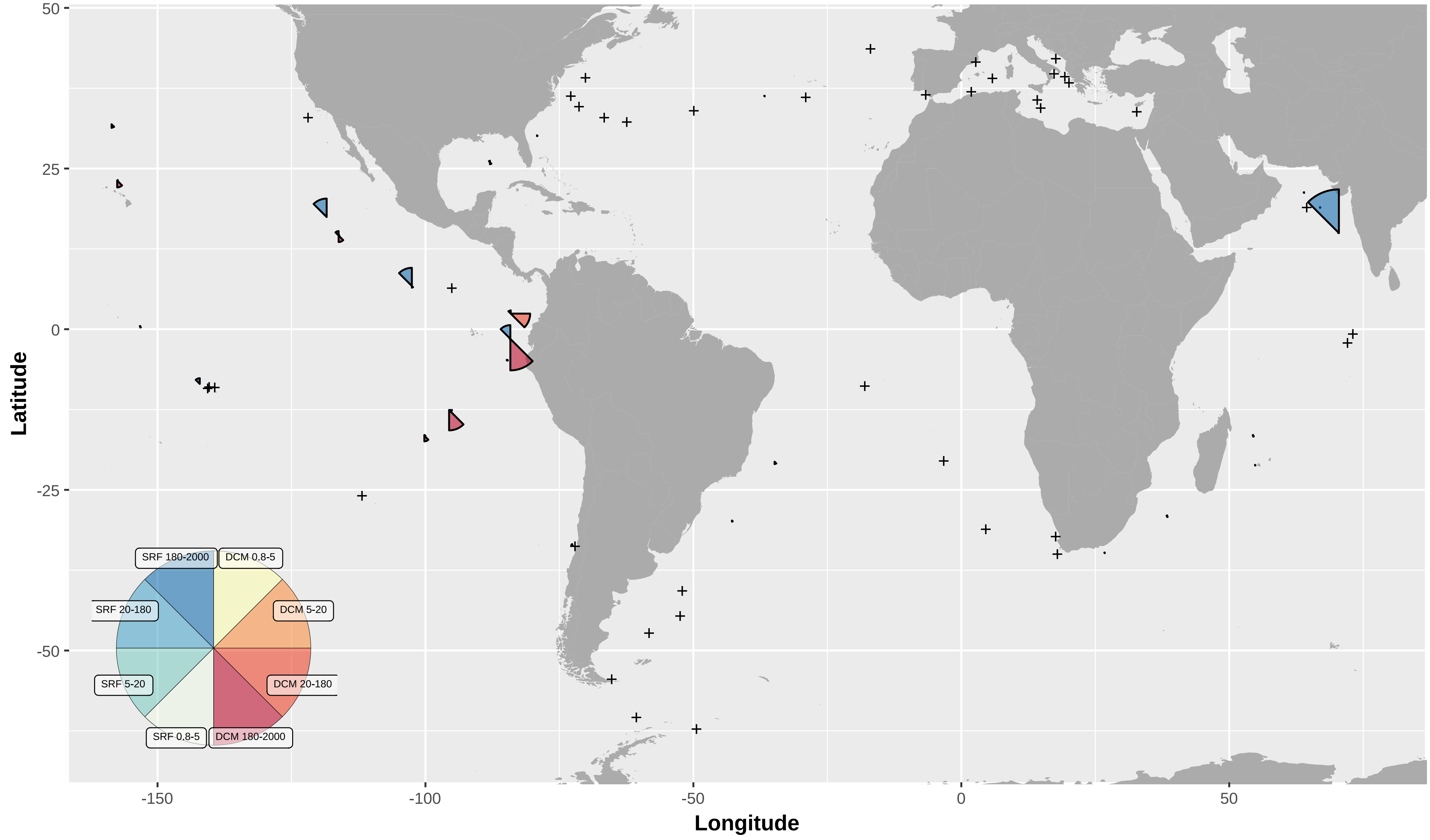


root;cellular organisms;Eukaryota;Opisthokonta;Metazoa;Eumetazoa;Bilateria;Deuterostomia;Chordata;Craniata;Vertebrata;Gnathostomata;Teleostomi;Euteleostomi

### Biogeolocalisation of MGT-v1 881 531 unigenes, 47 (8.85%) taxonomically assigned

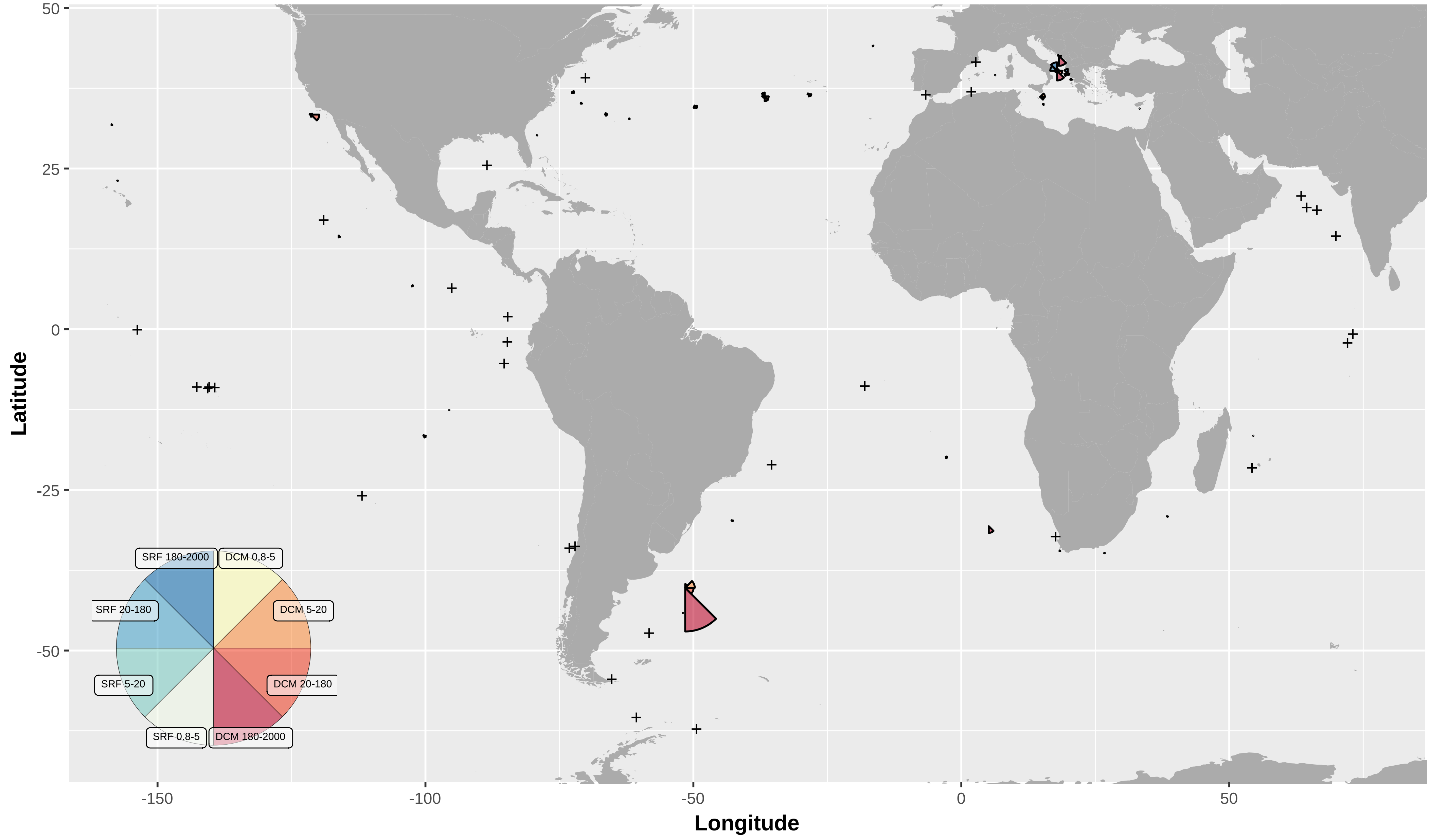


root;cellular organisms;Eukaryota;Opisthokonta;Metazoa;Eumetazoa

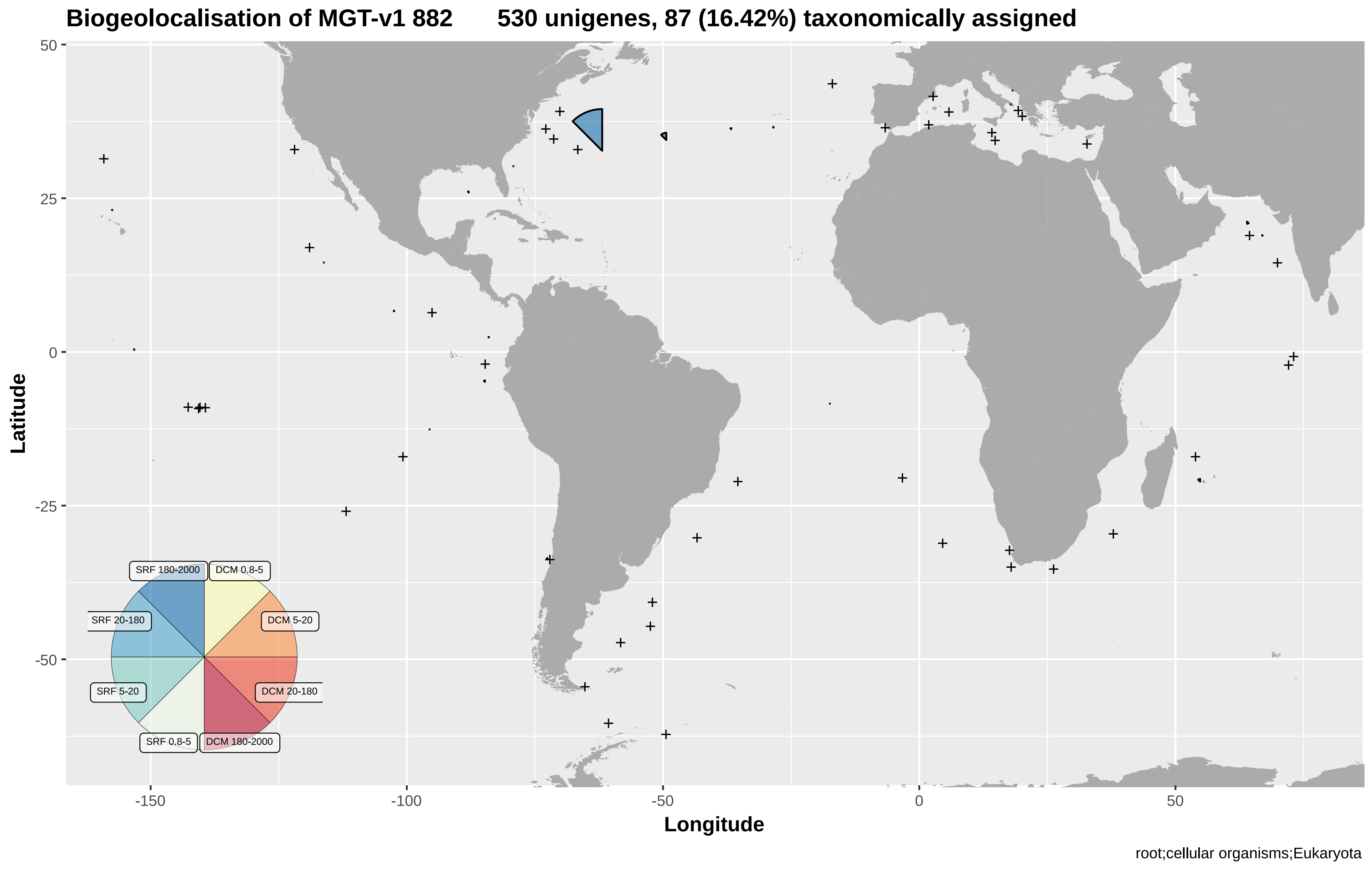

Biogeolocalisation of MGT-v1 882 530 unigenes, 87 (16.42%) taxonomically assigned
50
25
0
-25
-50
Latitude
-150
-100
-50
0
50
Longitude
SRF 180-2000
DCM 0.8-5
SRF 20-180
DCM 5-20
SRF 5-20
DCM 20-180
SRF 0.8-5
DCM 180-2000
root;cellular organisms;Eukaryota

# Biogeolocalisation of MGT-v1 883      529 unigenes, 62 (11.72%) taxonomically assigned

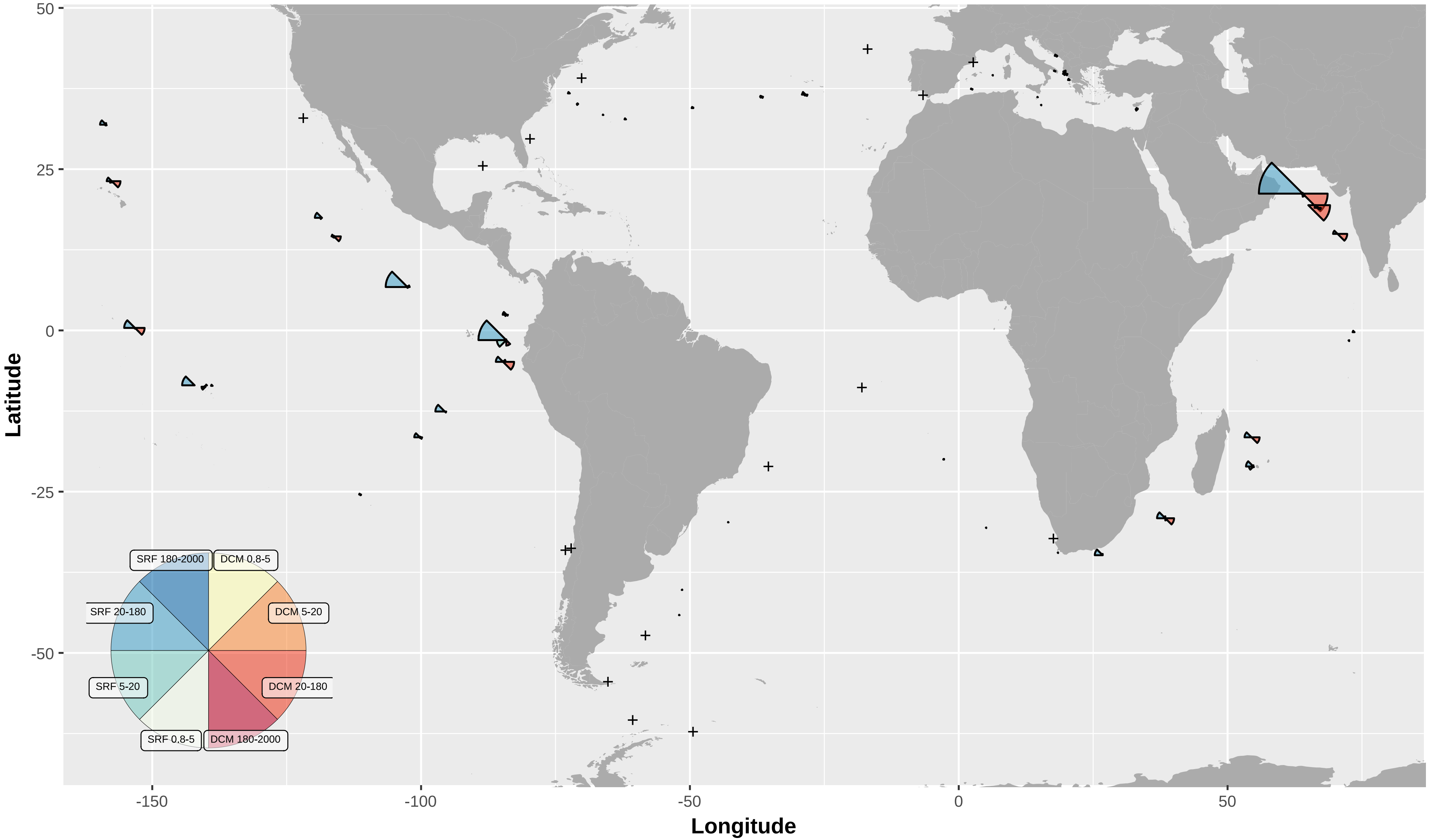


root;cellular organisms;Eukaryota;Opisthokonta;Metazoa;Eumetazoa;Bilateria

#### Biogeolocalisation of MGT-v1 884 529 unigenes, 15 (2.84%) taxonomically assigned

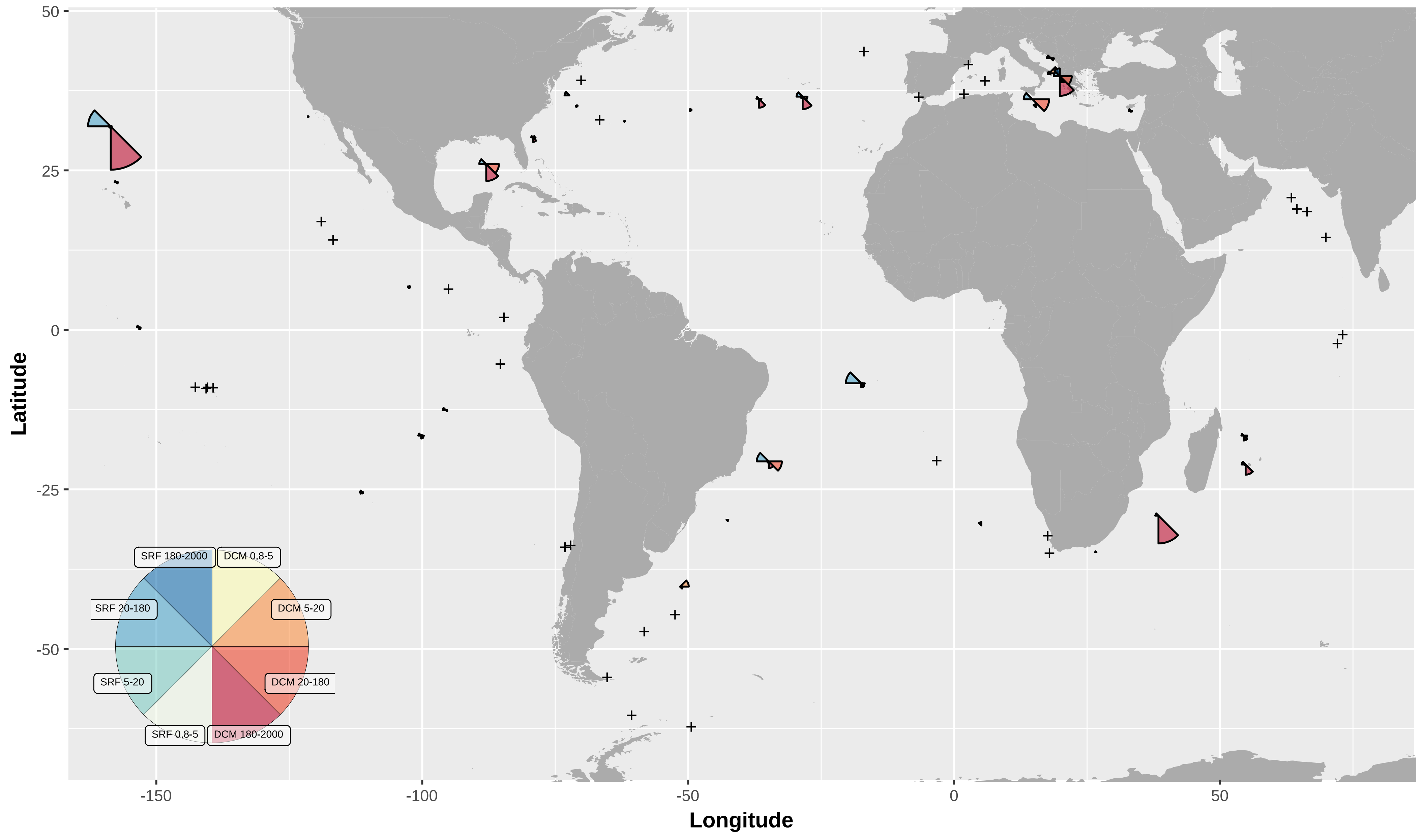


root;cellular organisms;Eukaryota;Opisthokonta

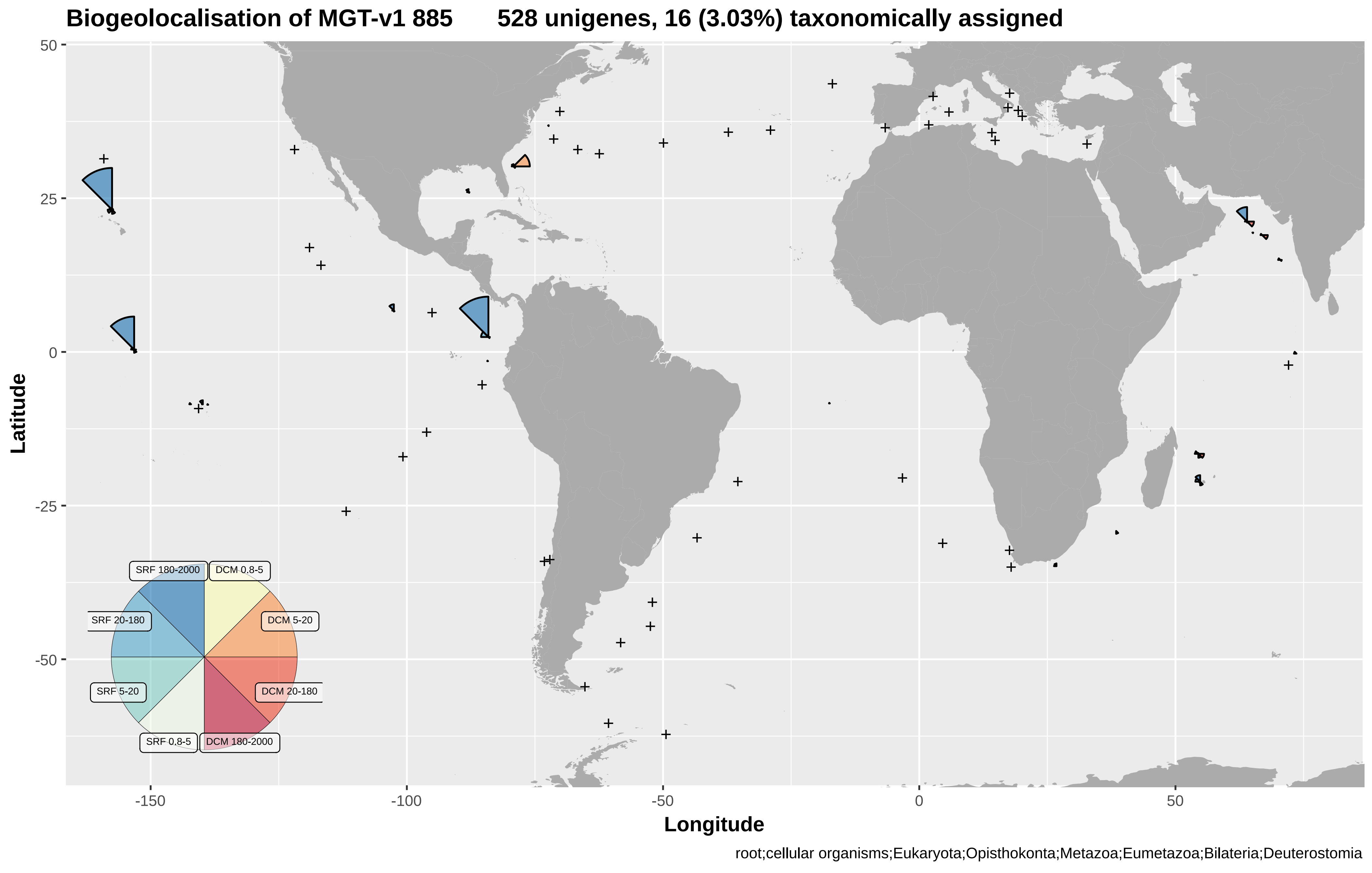

Biogeolocalisation of MGT-v1 885 528 unigenes, 16 (3.03%) taxonomically assigned
50
25
0
-25
-50
Latitude
-150
-100
-50
0
50
Longitude
SRF 180-2000
DCM 0.8-5
SRF 20-180
DCM 5-20
SRF 5-20
DCM 20-180
SRF 0.8-5
DCM 180-2000
root;cellular organisms;Eukaryota;Opisthokonta;Metazoa;Eumetazoa;Bilateria;Deuterostomia

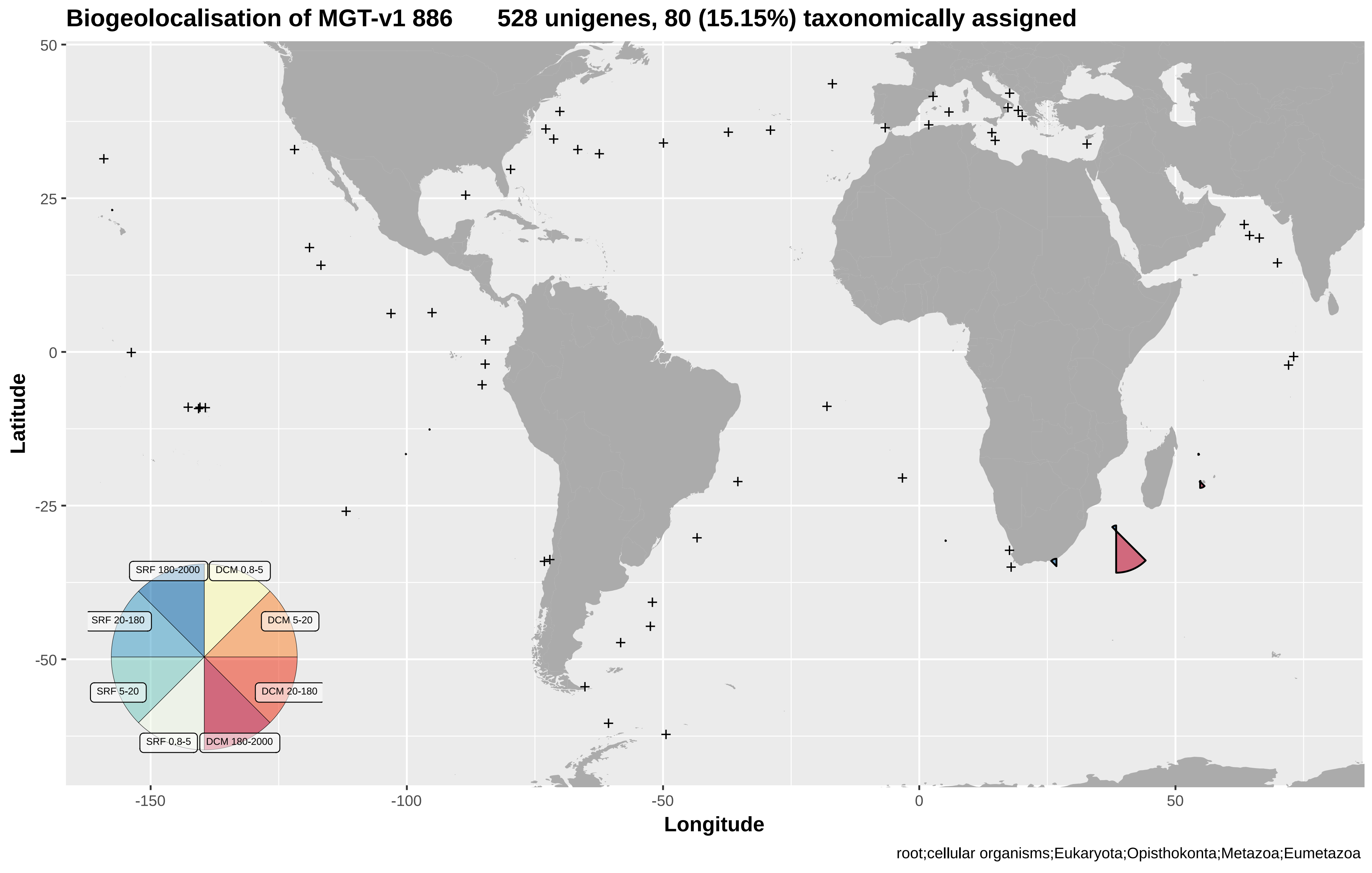
Biogeolocalisation of MGT-v1 886 528 unigenes, 80 (15.15%) taxonomically assigned
50
25
0
-25
-50
Latitude
-150
-100
-50
0
50
Longitude
SRF 180-2000
DCM 0.8-5
SRF 20-180
DCM 5-20
SRF 5-20
DCM 20-180
SRF 0.8-5
DCM 180-2000
root;cellular organisms;Eukaryota;Opisthokonta;Metazoa;Eumetazoa

### Biogeolocalisation of MGT-v1 887 526 unigenes, 323 (61.41%) taxonomically assigned

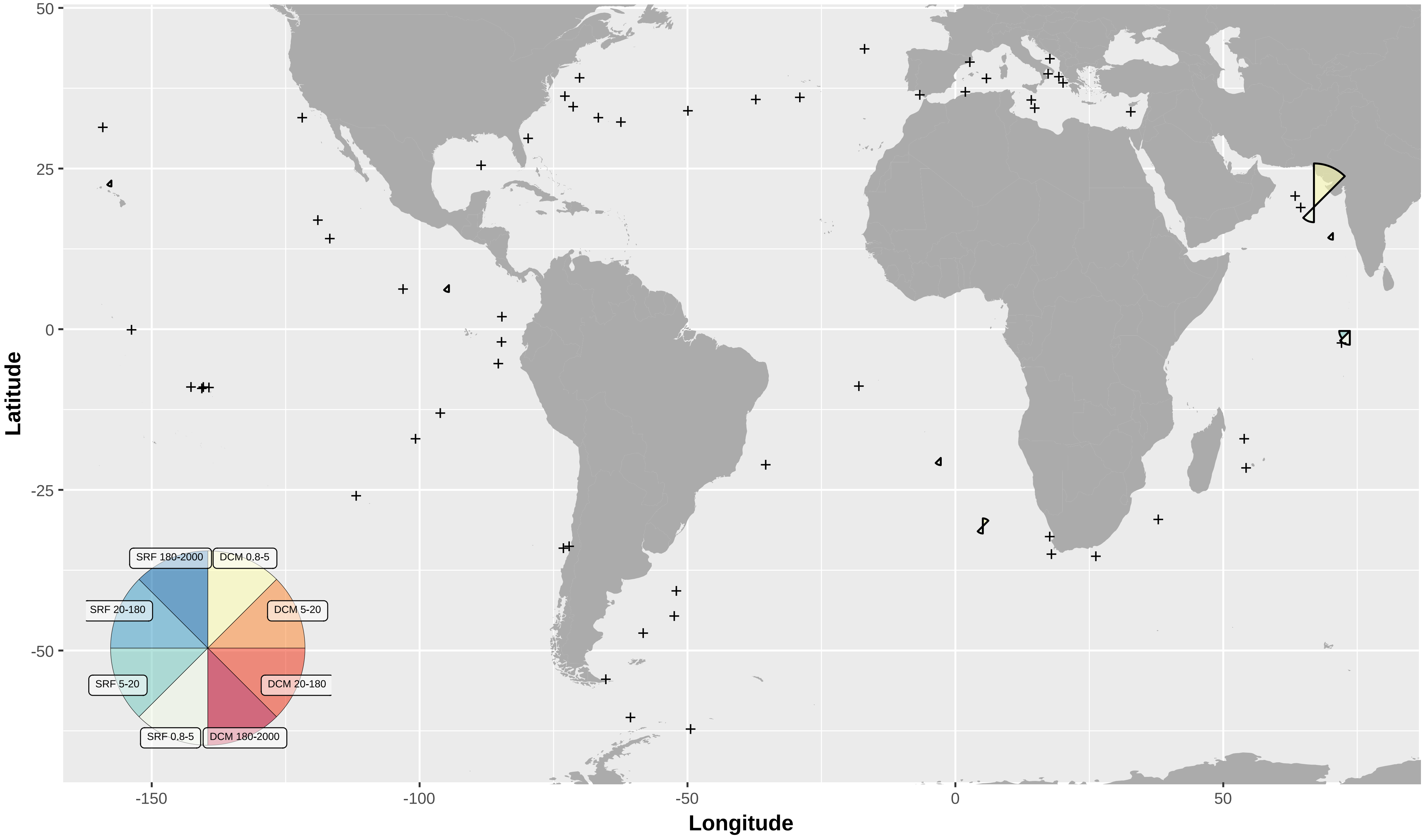


root;cellular organisms;Bacteria

#### Biogeolocalisation of MGT-v1 888 526 unigenes, 225 (42.78%) taxonomically assigned

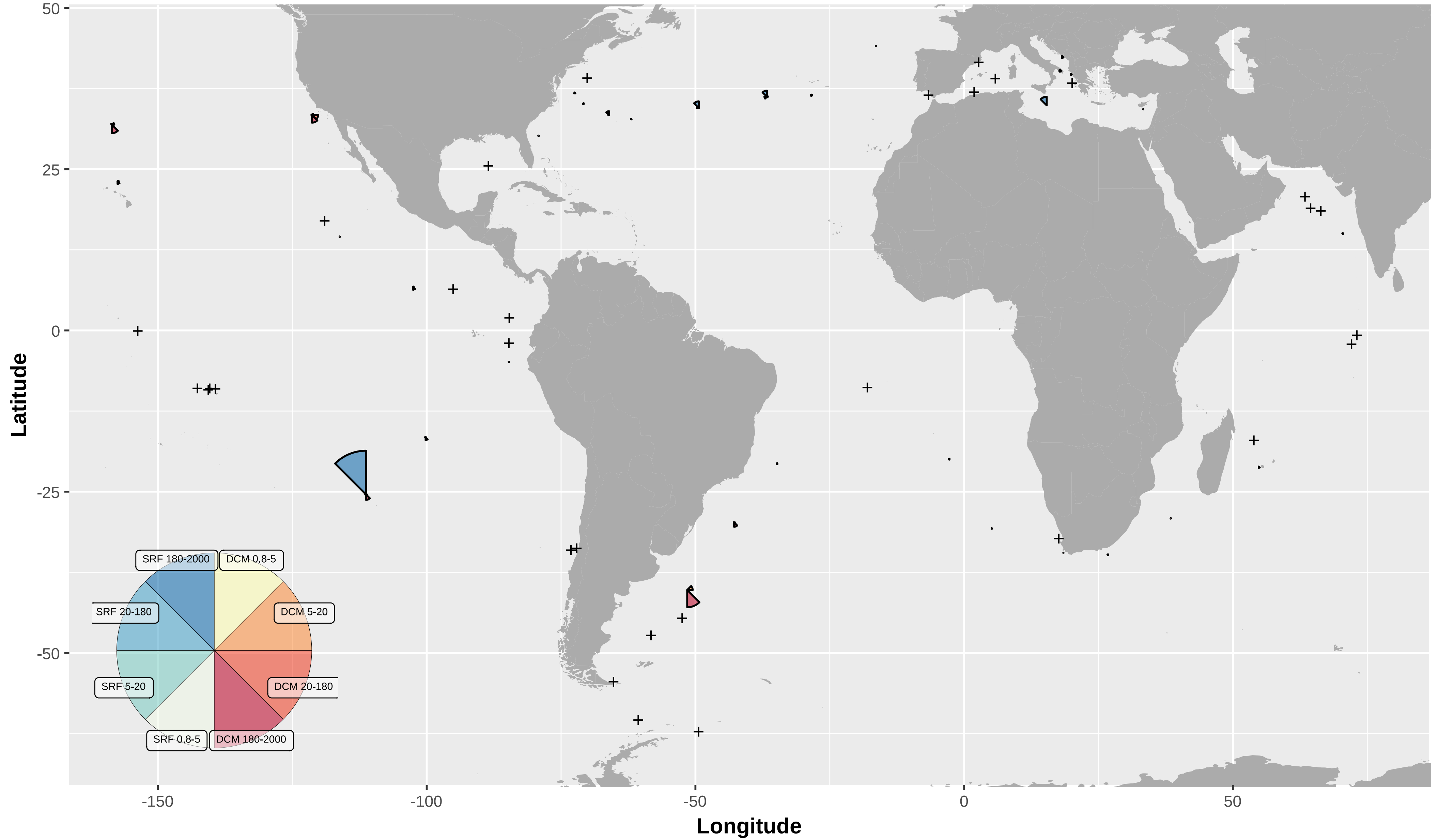


cellular organisms;Eukaryota;Opisthokonta;Metazoa;Eumetazoa;Bilateria;Protostomia;Ecdysozoa;Panarthropoda;Arthropoda;Mandibulata;Pancrustacea;Crustacea;Maxillopoda;Copepoda;Neocopepoda

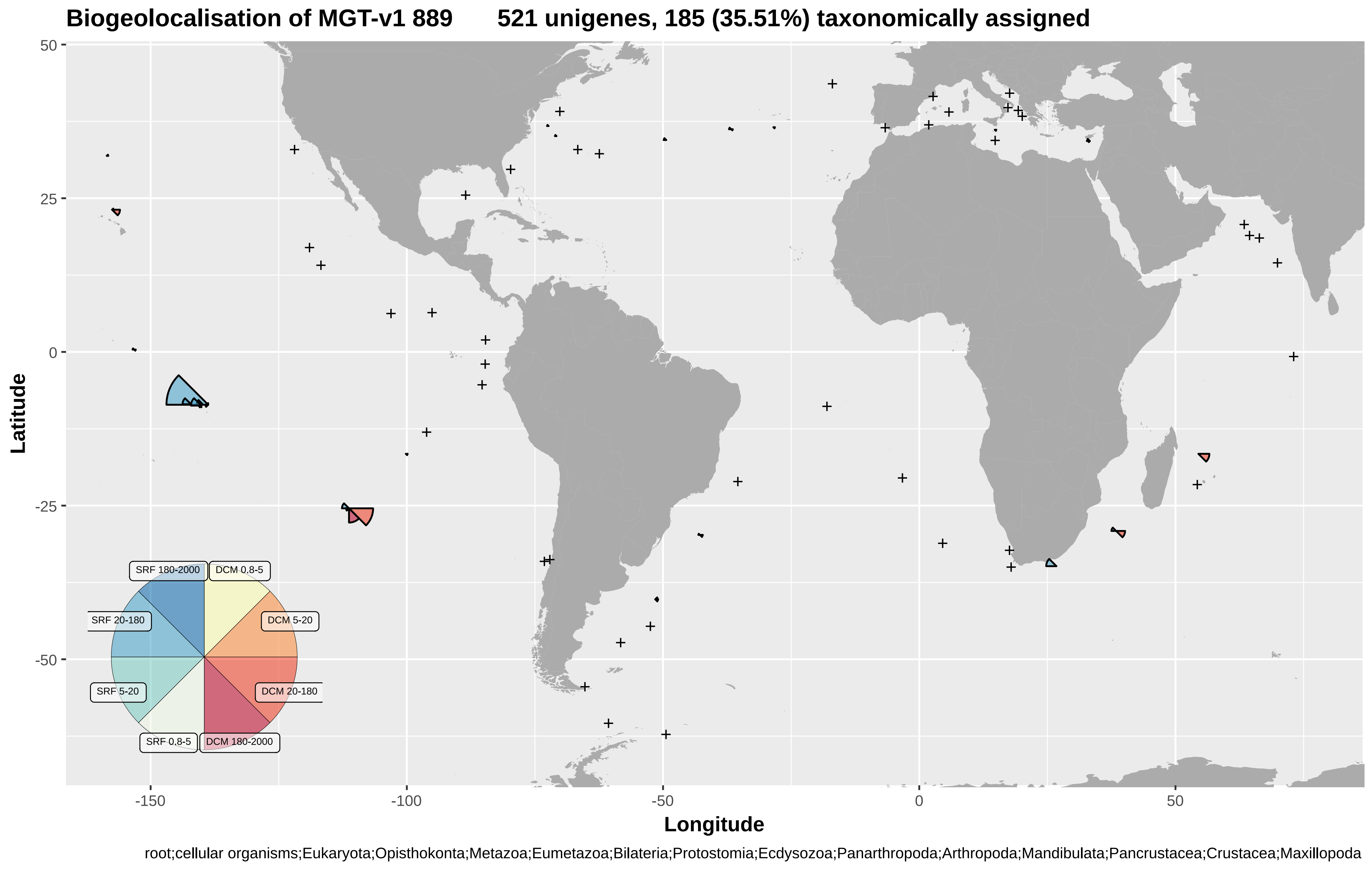
Biogeolocalisation of MGT-v1 889
521 unigenes, 185 (35.51%) taxonomically assigned
Latitude
Longitude
50
25
0
-25
-50
-150
-100
-50
0
50
SRF 180-2000
DCM 0.8-5
SRF 20-180
DCM 5-20
SRF 5-20
DCM 20-180
SRF 0.8-5
DCM 180-2000
root;cellular organisms;Eukaryota;Opisthokonta;Metazoa;Eumetazoa;Bilateria;Protostomia;Ecdysozoa;Panarthropoda;Arthropoda;Mandibulata;Pancrustacea;Crustacea;Maxillopoda

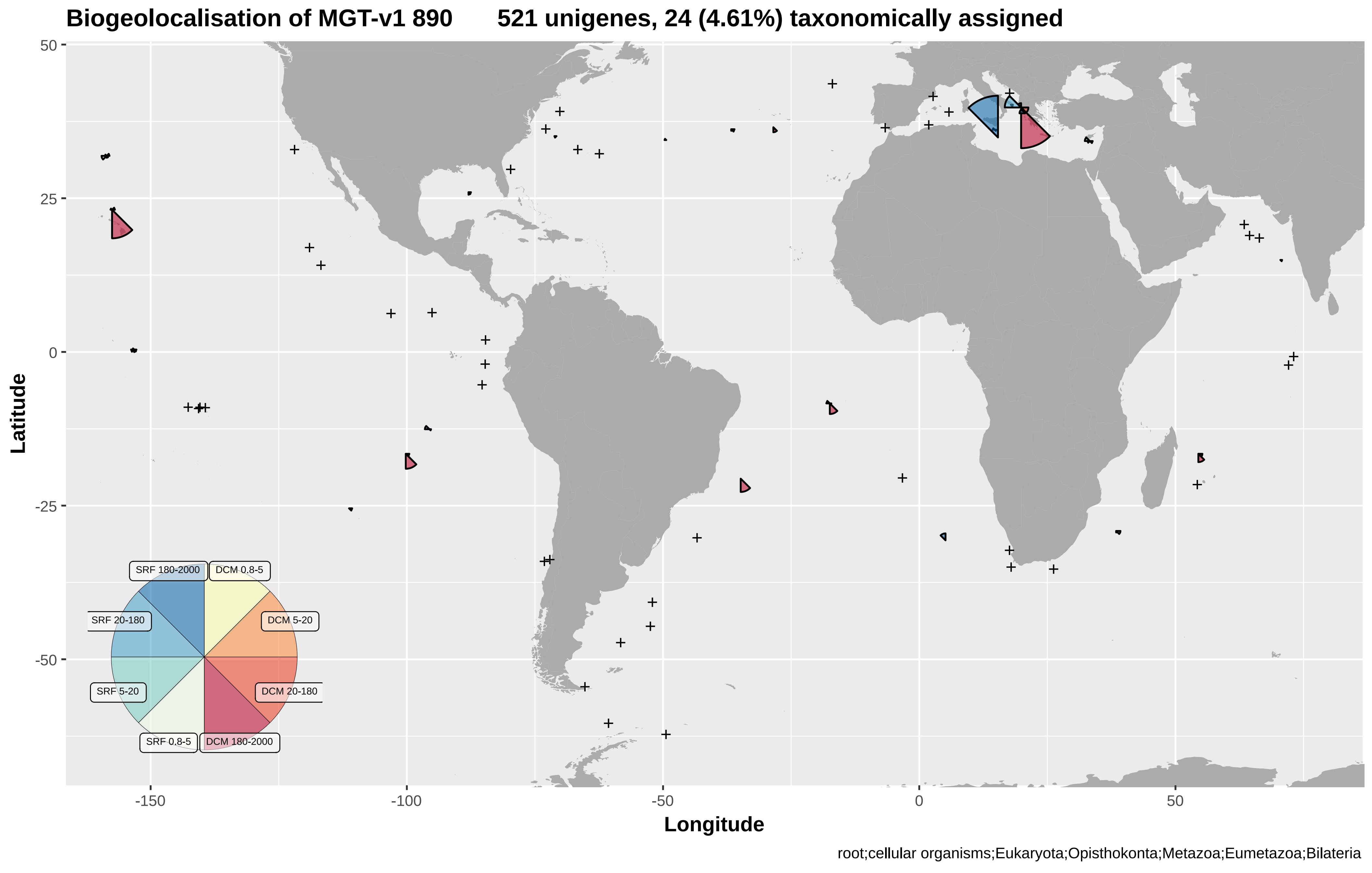
Biogeolocalisation of MGT-v1 890 521 unigenes, 24 (4.61%) taxonomically assigned
50
25
0
-25
-50
Latitude
-150
-100
-50
0
50
Longitude
SRF 180-2000
DCM 0.8-5
SRF 20-180
DCM 5-20
SRF 5-20
DCM 20-180
SRF 0.8-5
DCM 180-2000
root;cellular organisms;Eukaryota;Opisthokonta;Metazoa;Eumetazoa;Bilateria

#### Biogeolocalisation of MGT-v1 891 521 unigenes, 59 (11.32%) taxonomically assigned

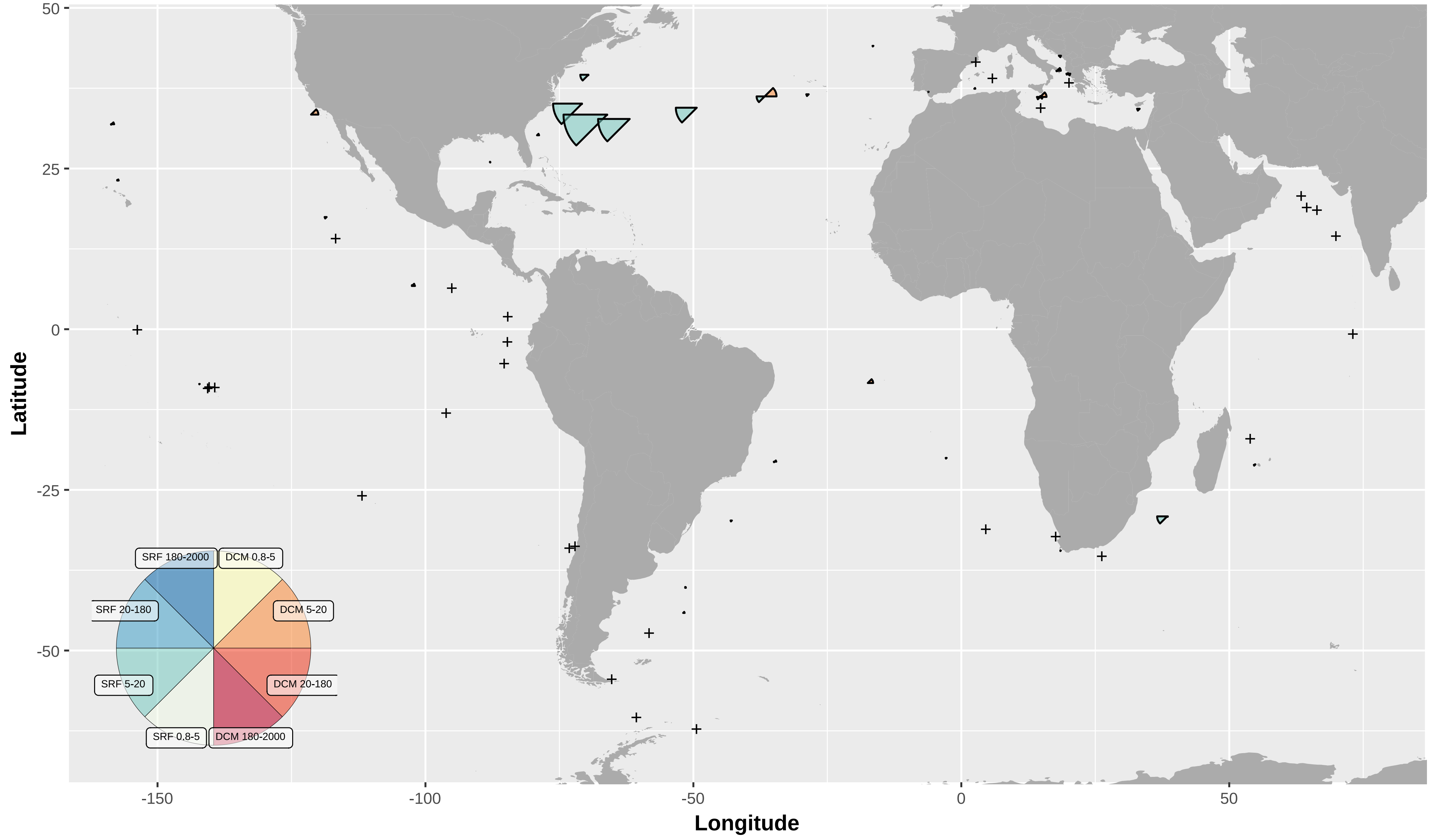


root;cellular organisms;Eukaryota

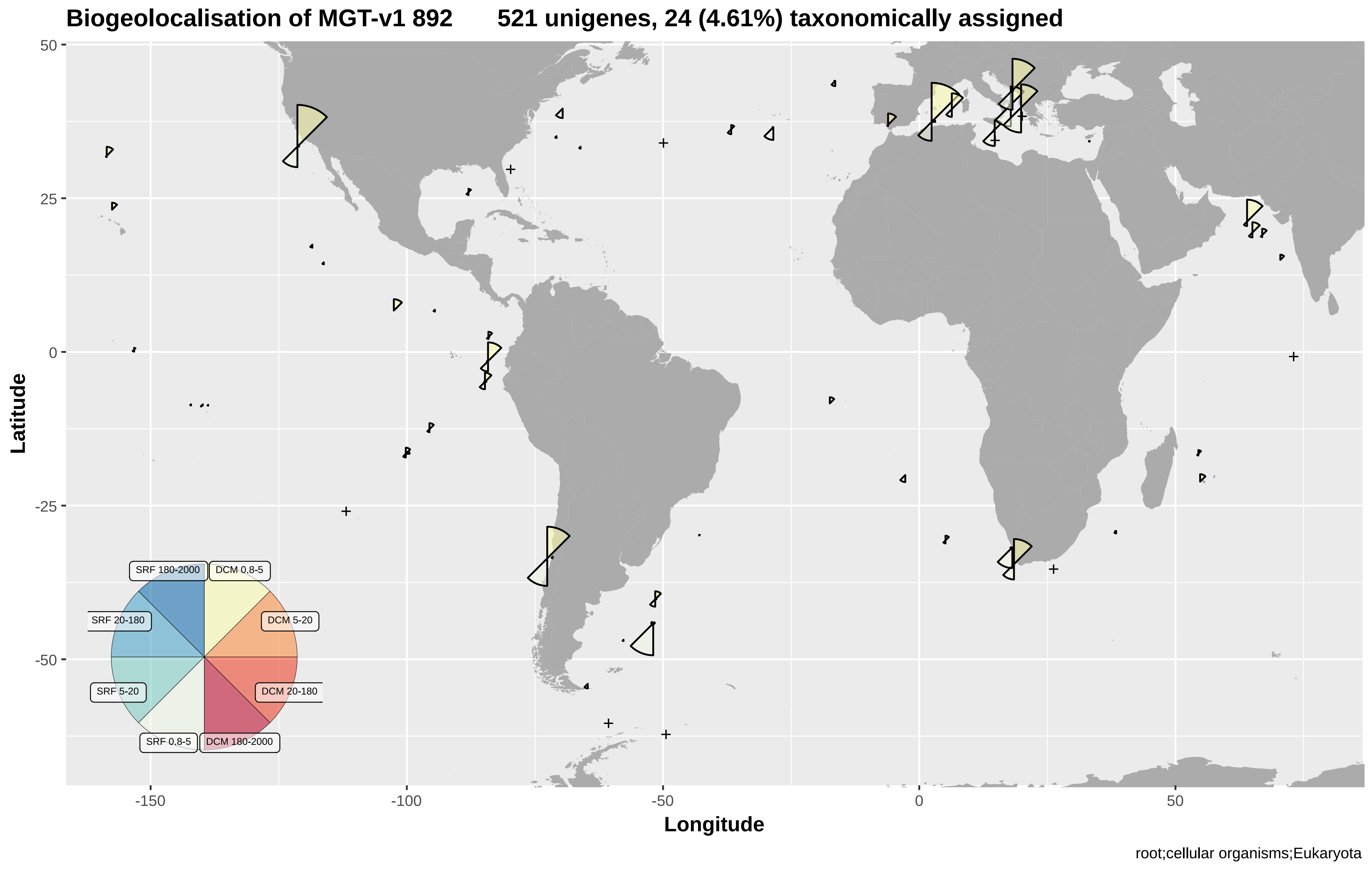

Biogeolocalisation of MGT-v1 892 521 unigenes, 24 (4.61%) taxonomically assigned
50
25
0
-25
-50
Latitude
-150
-100
-50
0
50
Longitude
SRF 180-2000
DCM 0.8-5
SRF 20-180
DCM 5-20
SRF 5-20
DCM 20-180
SRF 0.8-5
DCM 180-2000
root;cellular organisms;Eukaryota

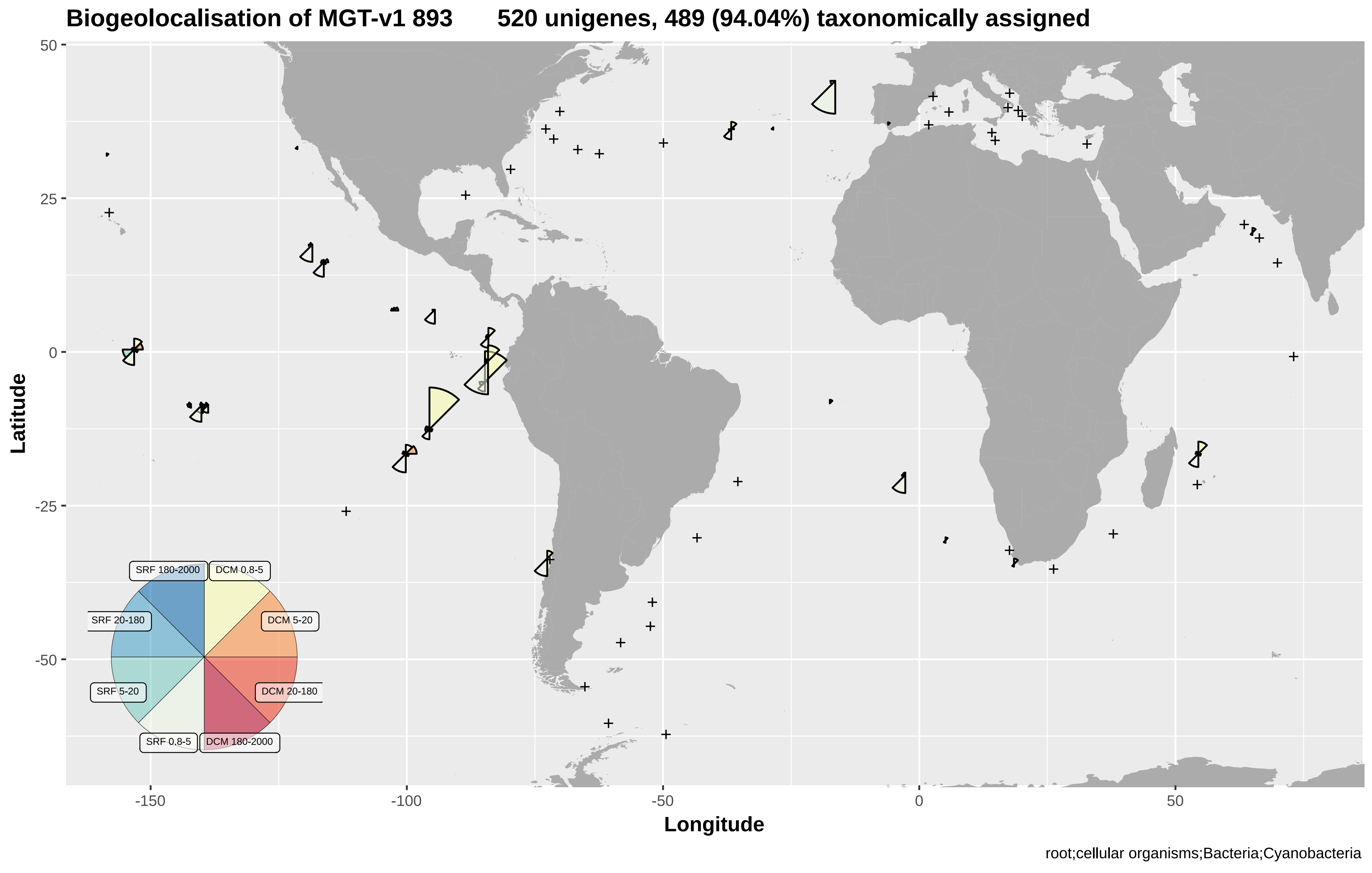

Biogeolocalisation of MGT-v1 893 520 unigenes, 489 (94.04%) taxonomically assigned
Latitude
Longitude
50
25
0
-25
-50
-150
-100
-50
0
50
SRF 180-2000
DCM 0.8-5
SRF 20-180
DCM 5-20
SRF 5-20
DCM 20-180
SRF 0.8-5
DCM 180-2000
root;cellular organisms;Bacteria;Cyanobacteria

## Biogeolocalisation of MGT-v1 894      519 unigenes, 32 (6.17%) taxonomically assigned

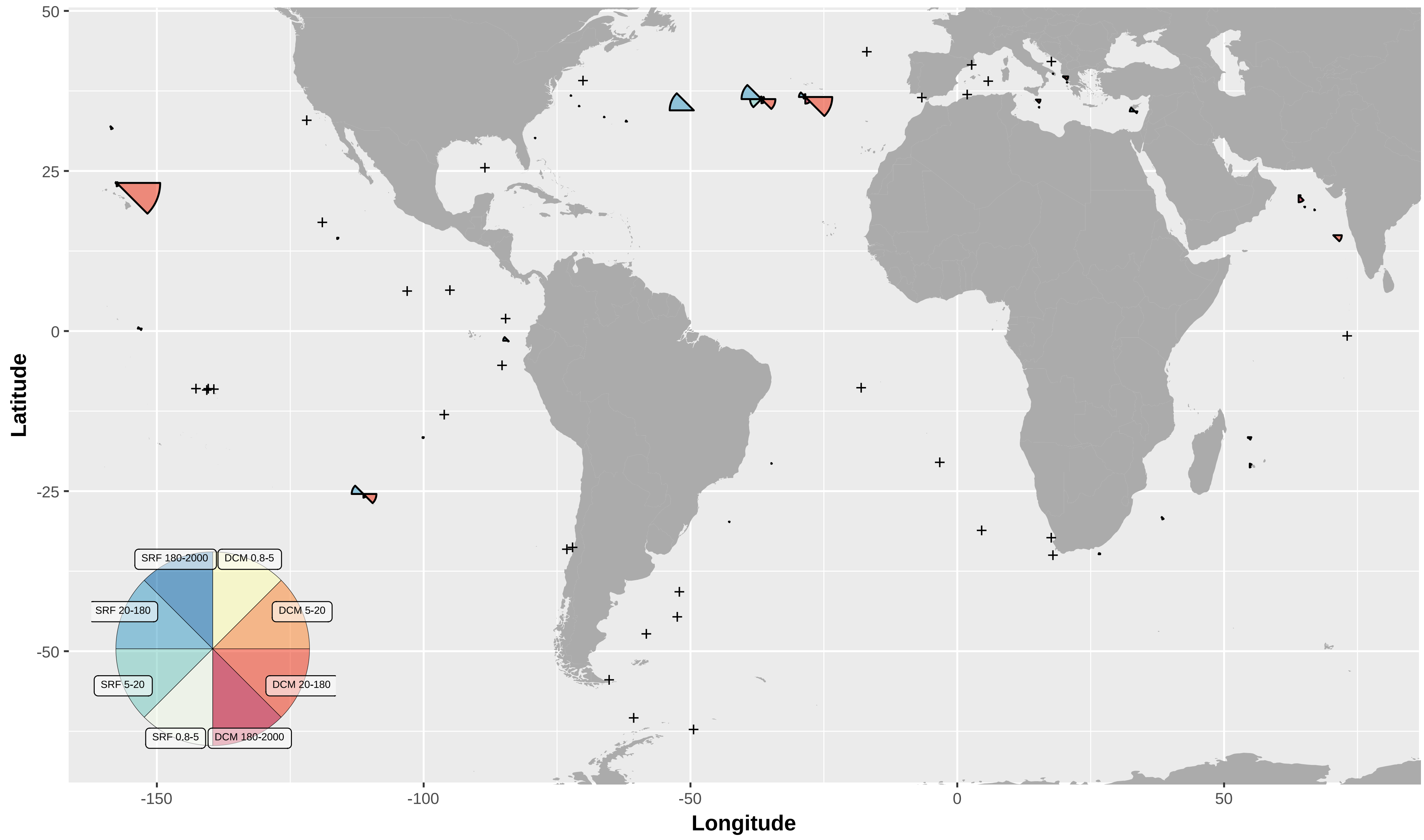


root;cellular organisms;Eukaryota;Opisthokonta;Metazoa;Eumetazoa;Bilateria

### Biogeolocalisation of MGT-v1 895 518 unigenes, 149 (28.76%) taxonomically assigned

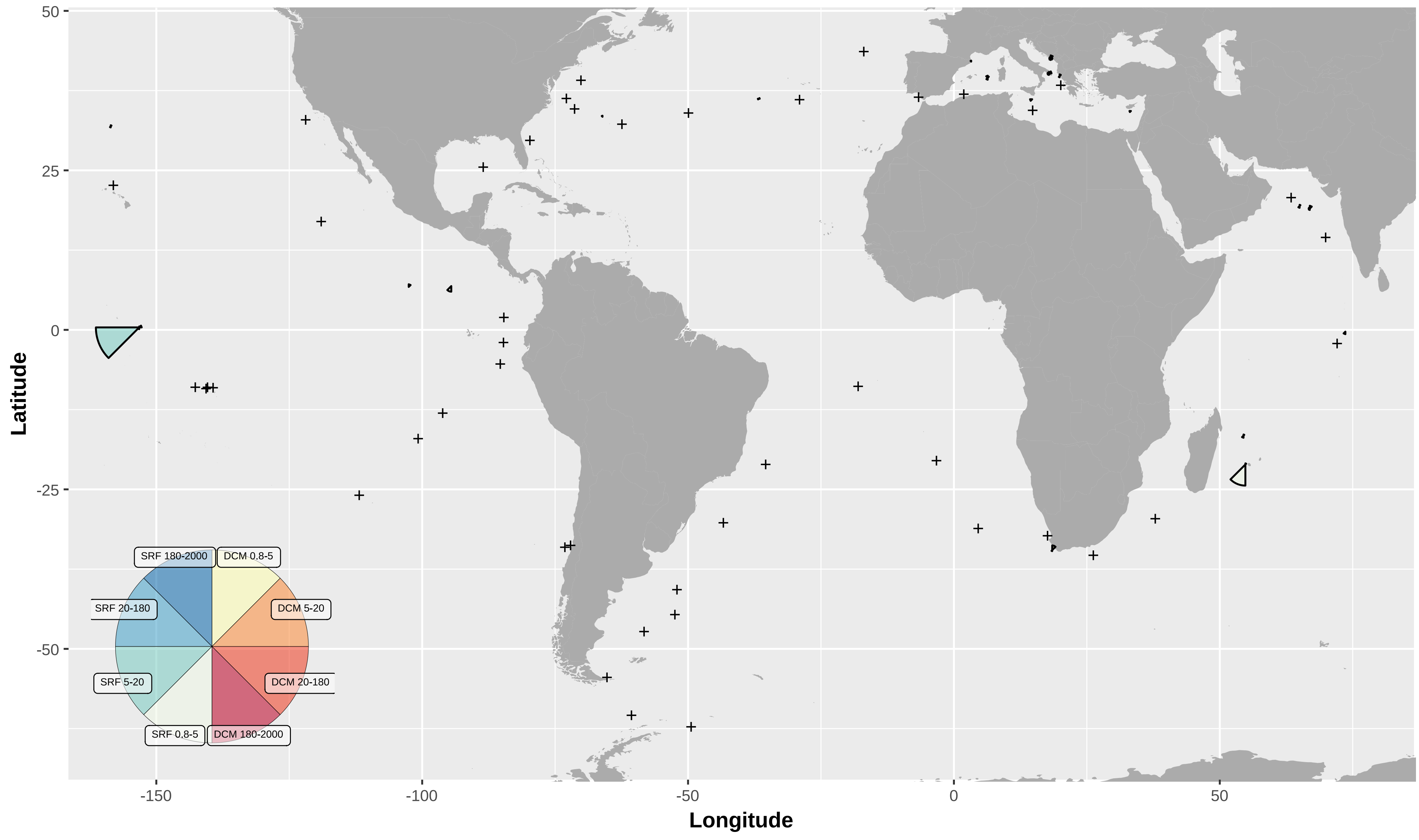


root;cellular organisms;Bacteria

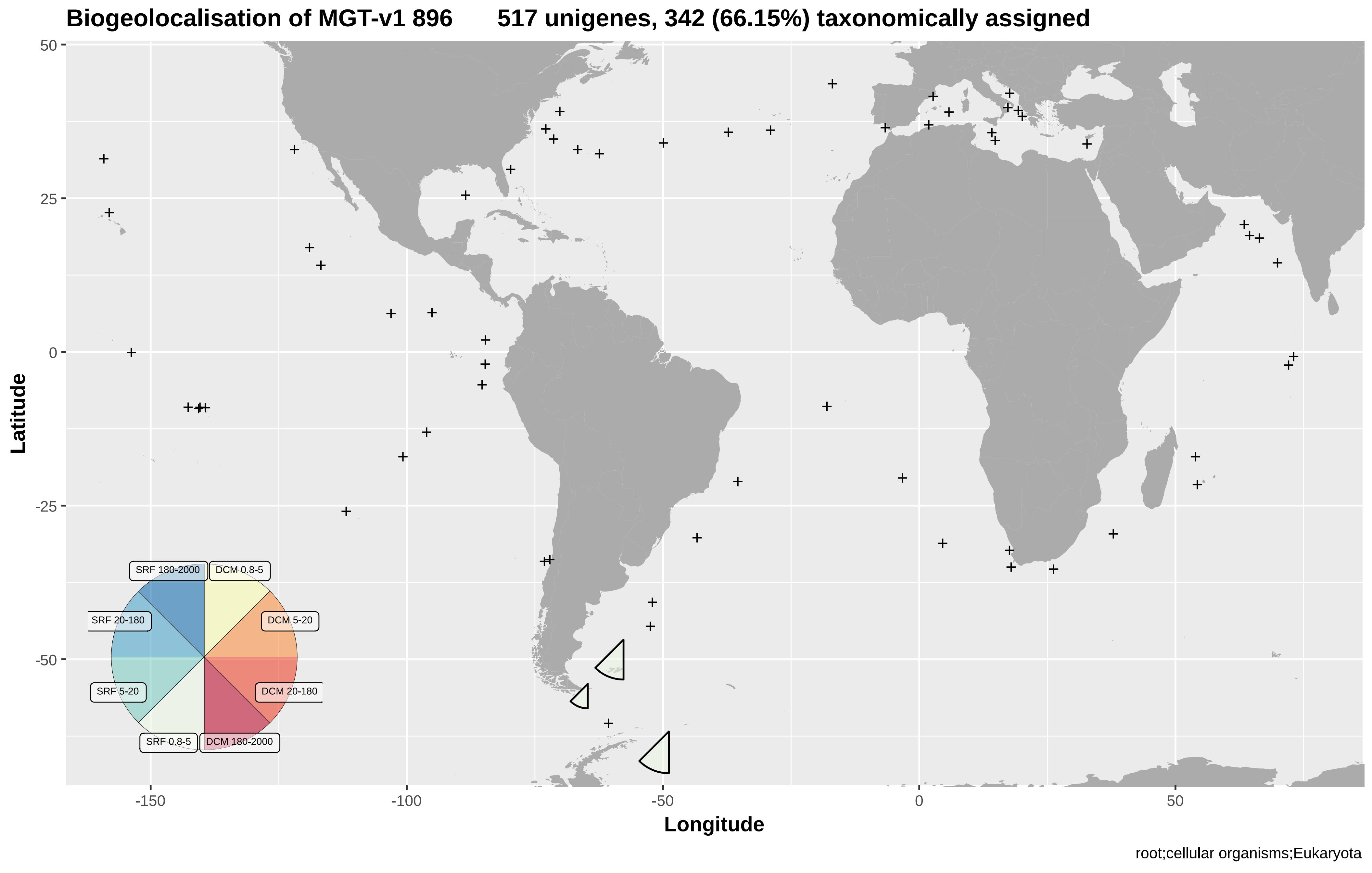
Biogeolocalisation of MGT-v1 896
517 unigenes, 342 (66.15%) taxonomically assigned
Latitude
Longitude
50
25
0
-25
-50
-150
-100
-50
0
50
SRF 180-2000
DCM 0.8-5
SRF 20-180
DCM 5-20
SRF 5-20
DCM 20-180
SRF 0.8-5
DCM 180-2000
root;cellular organisms;Eukaryota

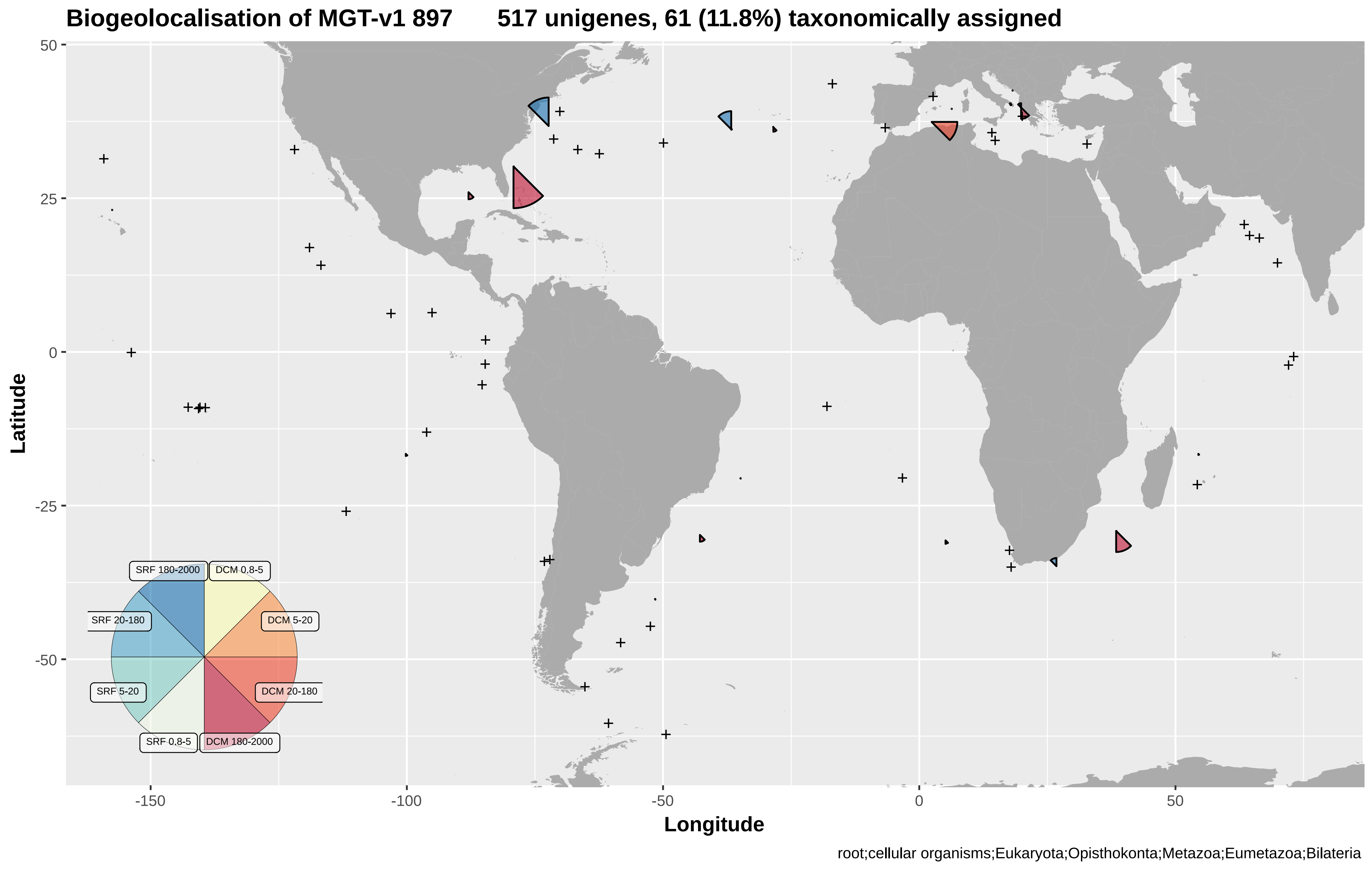
Biogeolocalisation of MGT-v1 897 517 unigenes, 61 (11.8%) taxonomically assigned
50
25
0
-25
-50
Latitude
-150
-100
-50
0
50
Longitude
SRF 180-2000
DCM 0.8-5
SRF 20-180
DCM 5-20
SRF 5-20
DCM 20-180
SRF 0.8-5
DCM 180-2000
root;cellular organisms;Eukaryota;Opisthokonta;Metazoa;Eumetazoa;Bilateria

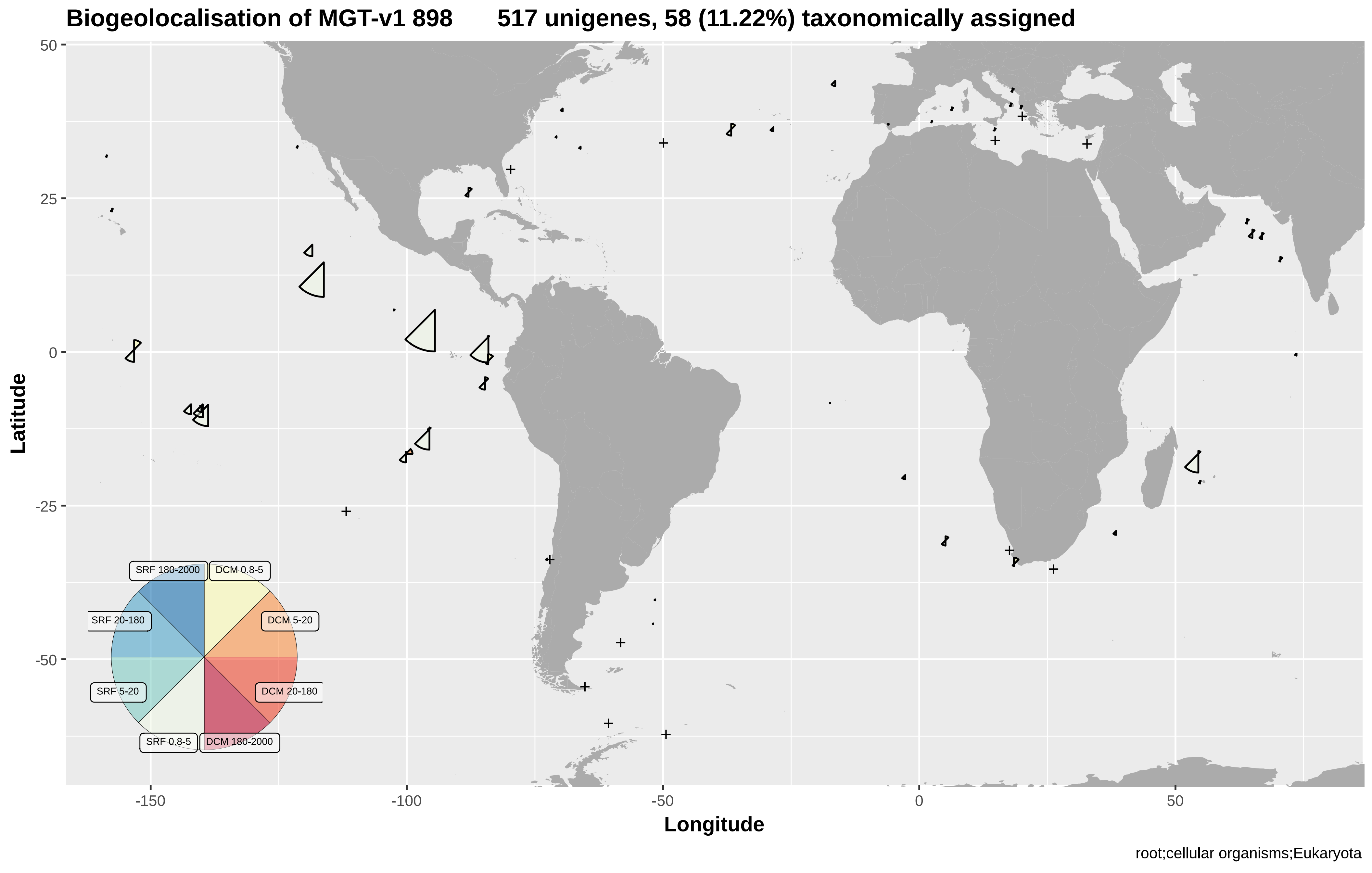
Biogeolocalisation of MGT-v1 898
517 unigenes, 58 (11.22%) taxonomically assigned
Latitude
50
25
0
-25
-50
Longitude
-150
-100
-50
0
50
SRF 180-2000
DCM 0.8-5
SRF 20-180
DCM 5-20
SRF 5-20
DCM 20-180
SRF 0.8-5
DCM 180-2000
root;cellular organisms;Eukaryota

Biogeolocalisation of MGT-v1 899      516 unigenes, 153 (29.65%) taxonomically assigned

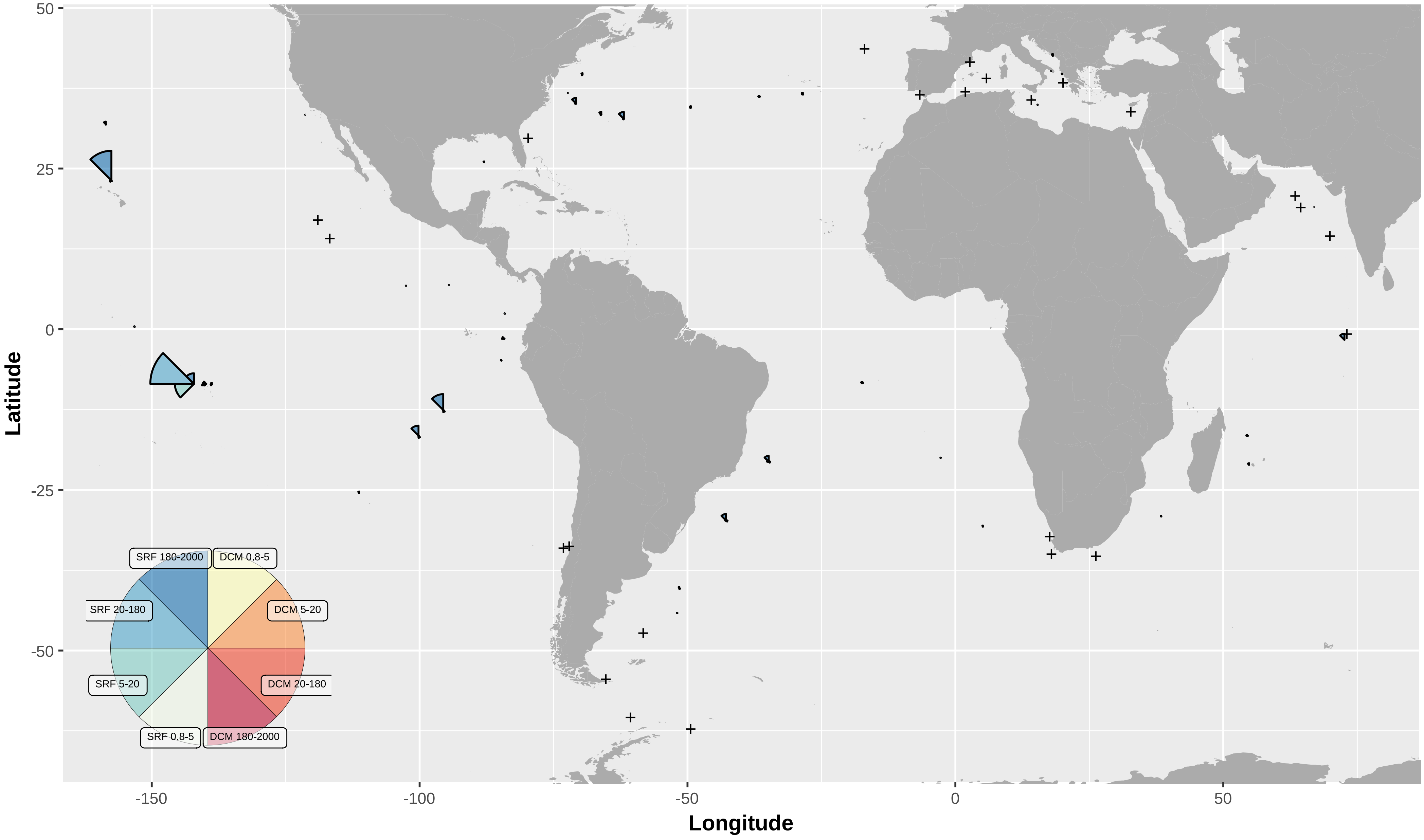


root;cellular organisms;Eukaryota;Rhizaria;Polycystinea;Collodaria

Biogeolocalisation of MGT-v1 900 515 unigenes, 481 (93.4%) taxonomically assigned

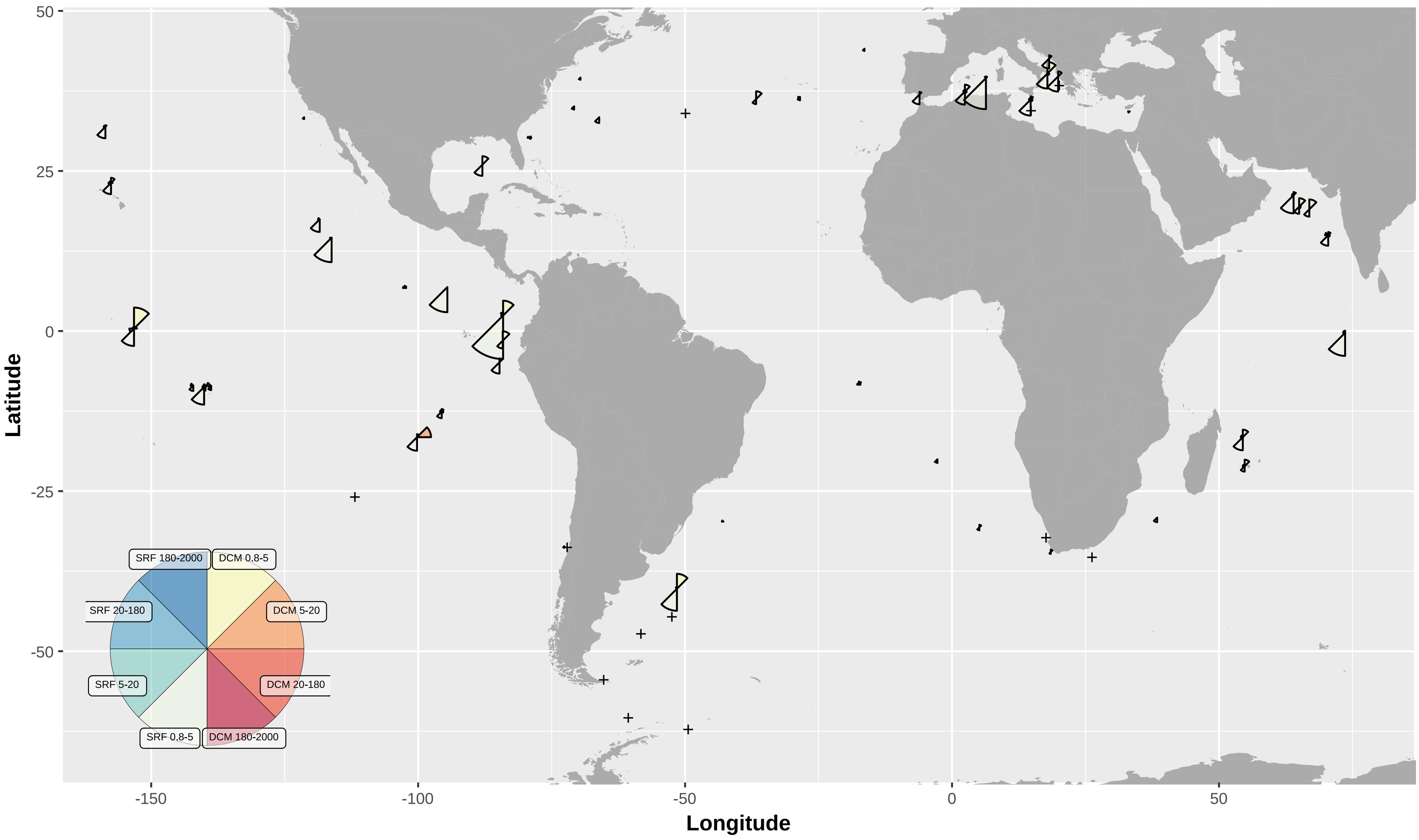


root;cellular organisms;Bacteria;Proteobacteria

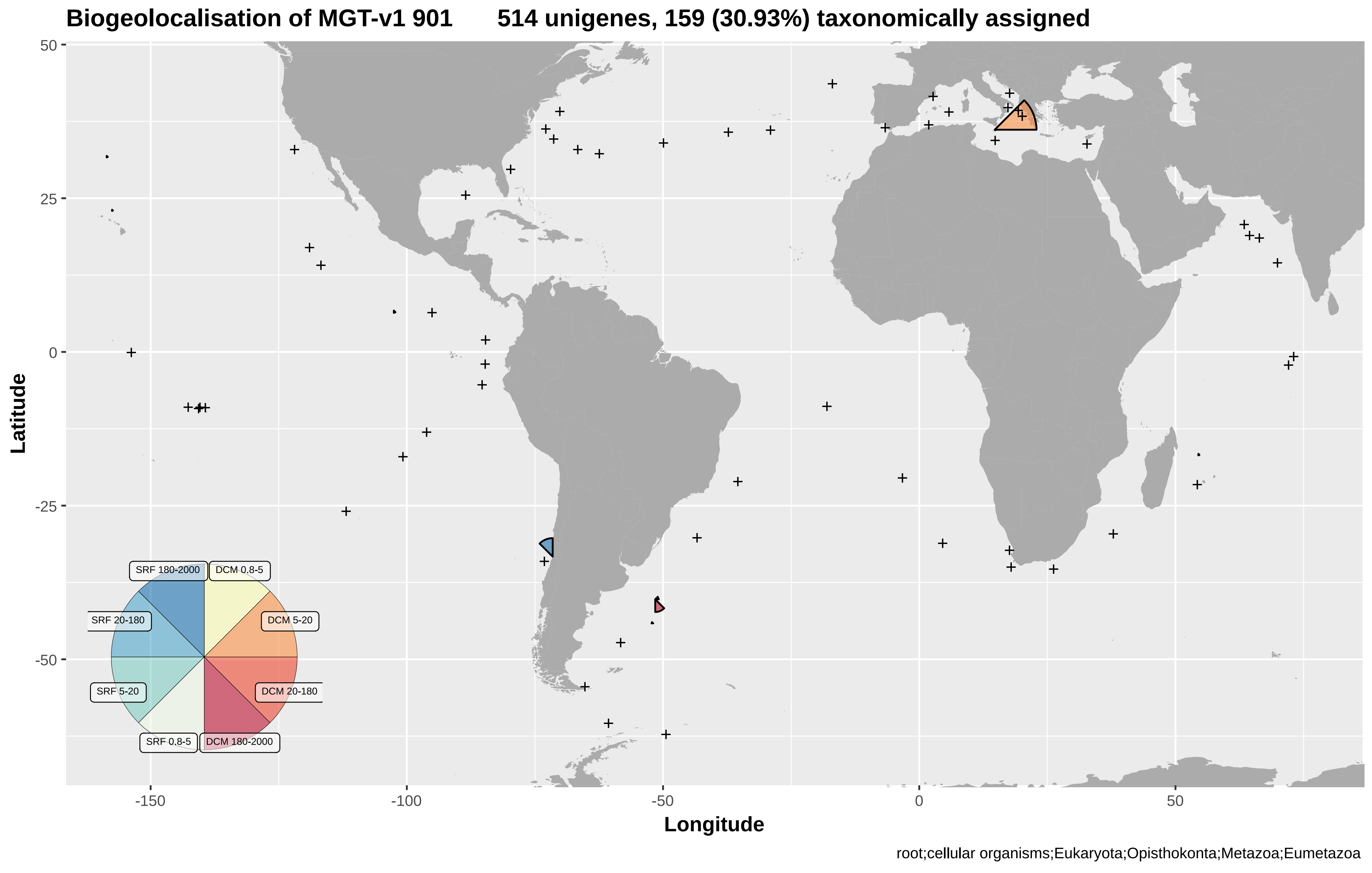

Biogeolocalisation of MGT-v1 901 514 unigenes, 159 (30.93%) taxonomically assigned
50
25
0
-25
-50
Latitude
-150
-100
-50
0
50
Longitude
SRF 180-2000
DCM 0.8-5
SRF 20-180
DCM 5-20
SRF 5-20
DCM 20-180
SRF 0.8-5
DCM 180-2000
root;cellular organisms;Eukaryota;Opisthokonta;Metazoa;Eumetazoa

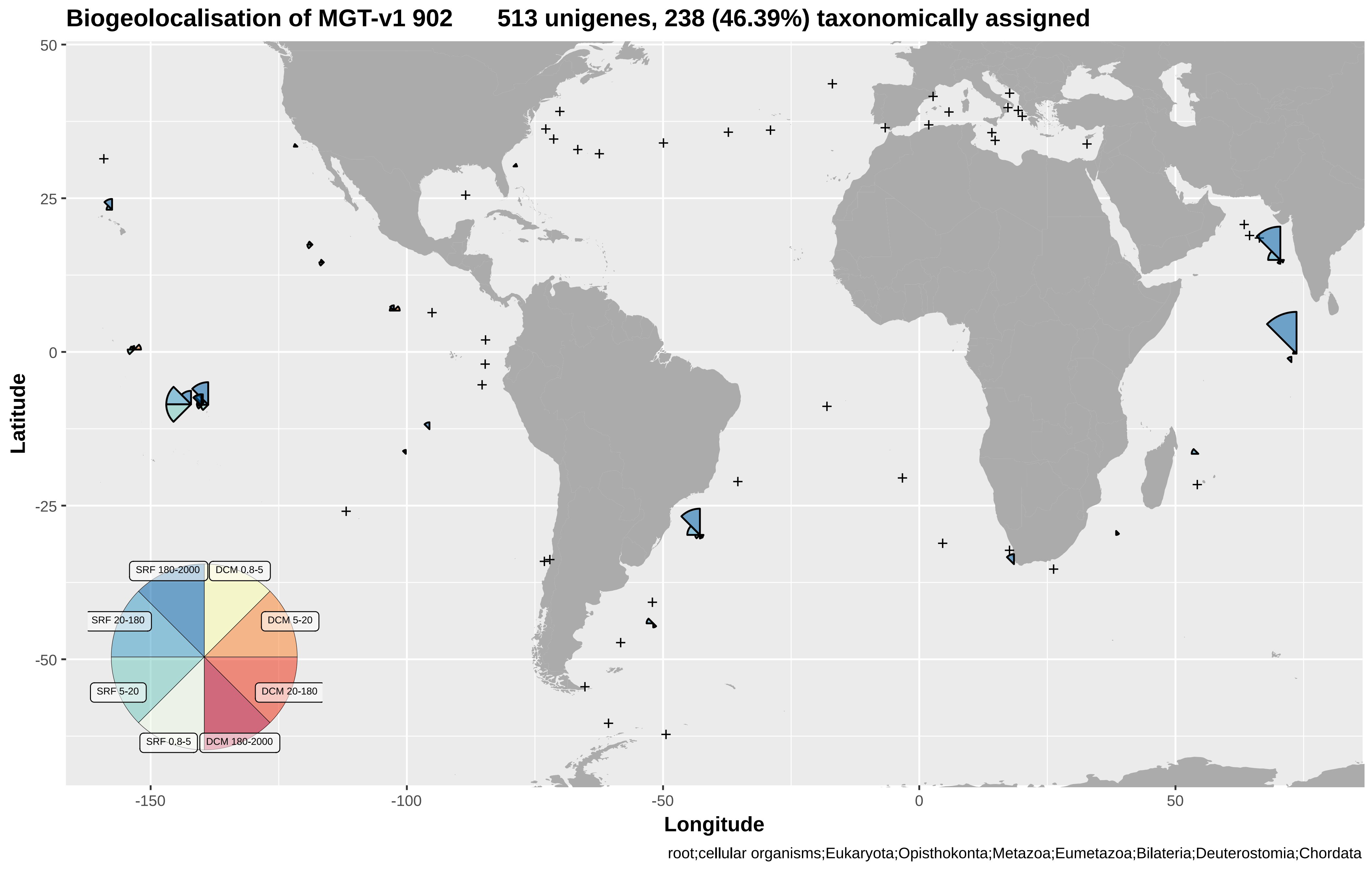

Biogeolocalisation of MGT-v1 902 513 unigenes, 238 (46.39%) taxonomically assigned
50
25
0
-25
-50
Latitude
-150
-100
-50
0
50
Longitude
SRF 180-2000
DCM 0.8-5
SRF 20-180
DCM 5-20
SRF 5-20
DCM 20-180
SRF 0.8-5
DCM 180-2000
root;cellular organisms;Eukaryota;Opisthokonta;Metazoa;Eumetazoa;Bilateria;Deuterostomia;Chordata

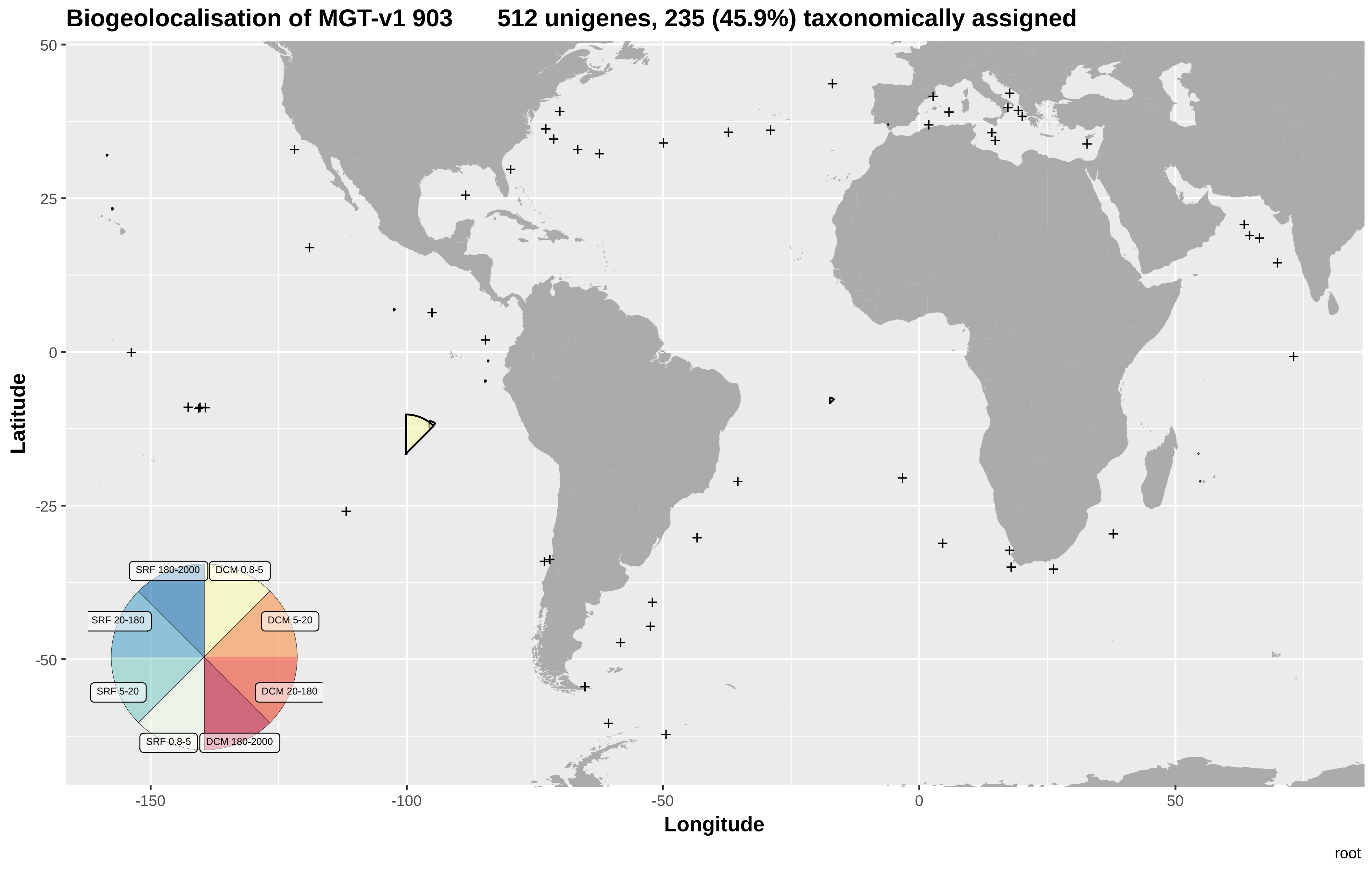

Biogeolocalisation of MGT-v1 903 512 unigenes, 235 (45.9%) taxonomically assigned
50
25
0
-25
-50
Latitude
-150
-100
-50
0
50
Longitude
SRF 180-2000
DCM 0.8-5
SRF 20-180
DCM 5-20
SRF 5-20
DCM 20-180
SRF 0.8-5
DCM 180-2000
root

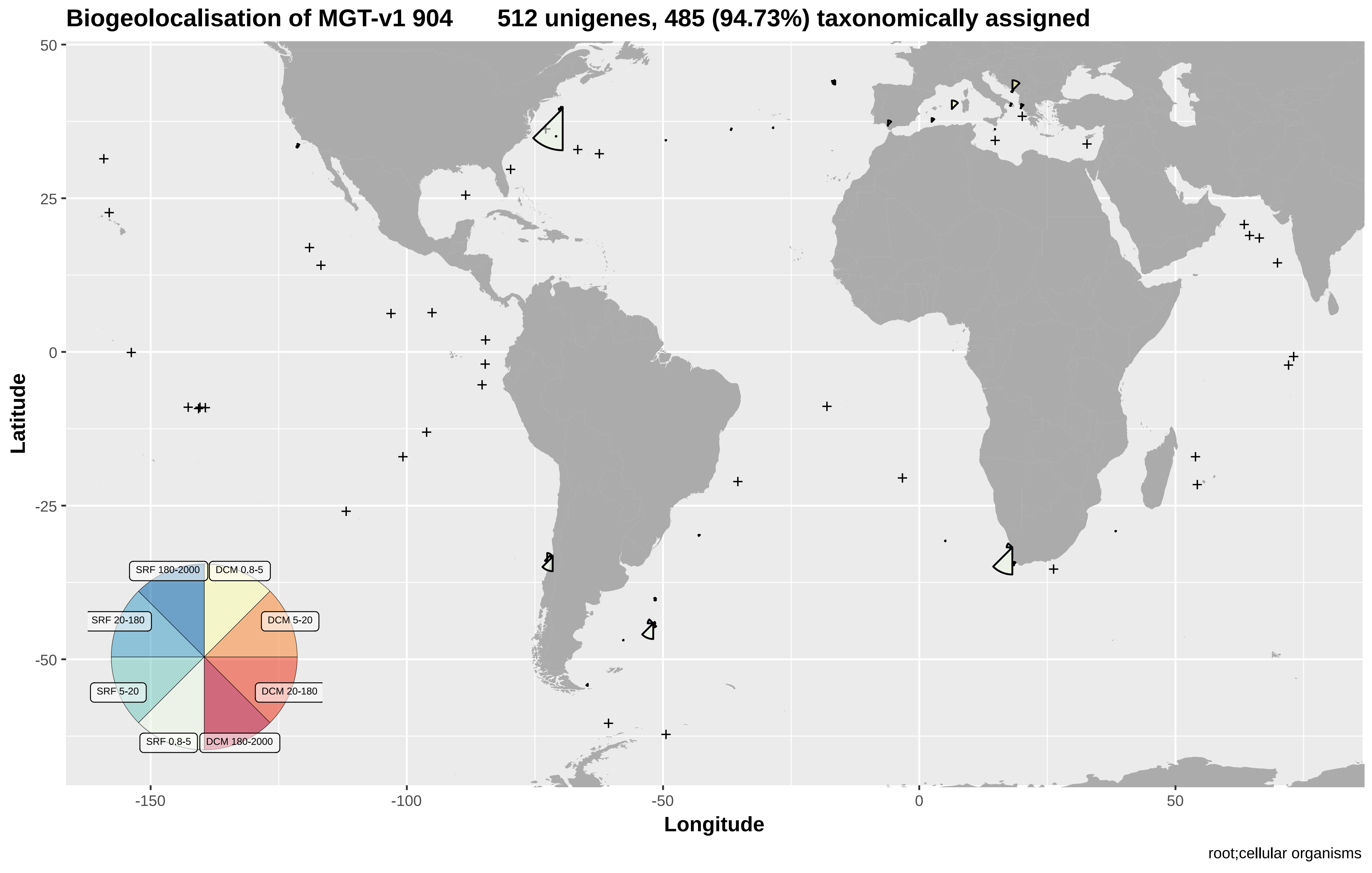
Biogeolocalisation of MGT-v1 904
512 unigenes, 485 (94.73%) taxonomically assigned
Latitude
Longitude
50
25
0
-25
-50
-150
-100
-50
0
50
SRF 180-2000
DCM 0.8-5
SRF 20-180
DCM 5-20
SRF 5-20
DCM 20-180
SRF 0.8-5
DCM 180-2000
root;cellular organisms

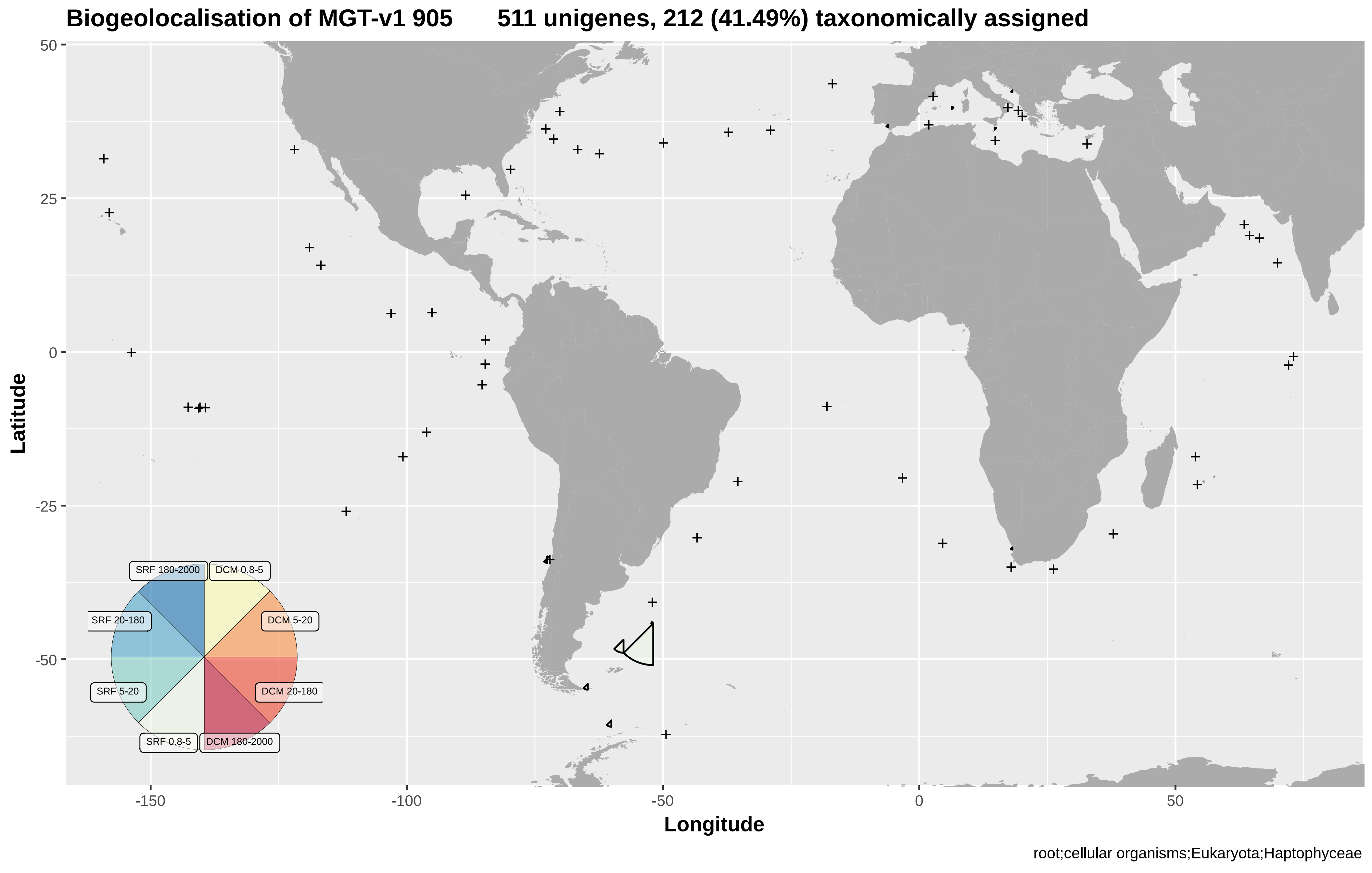
Biogeolocalisation of MGT-v1 905 511 unigenes, 212 (41.49%) taxonomically assigned
Latitude
50
25
0
-25
-50
Longitude
-150
-100
-50
0
50
SRF 180-2000
DCM 0.8-5
SRF 20-180
DCM 5-20
SRF 5-20
DCM 20-180
SRF 0.8-5
DCM 180-2000
root;cellular organisms;Eukaryota;Haptophyceae

## Biogeolocalisation of MGT-v1 906      510 unigenes, 260 (50.98%) taxonomically assigned

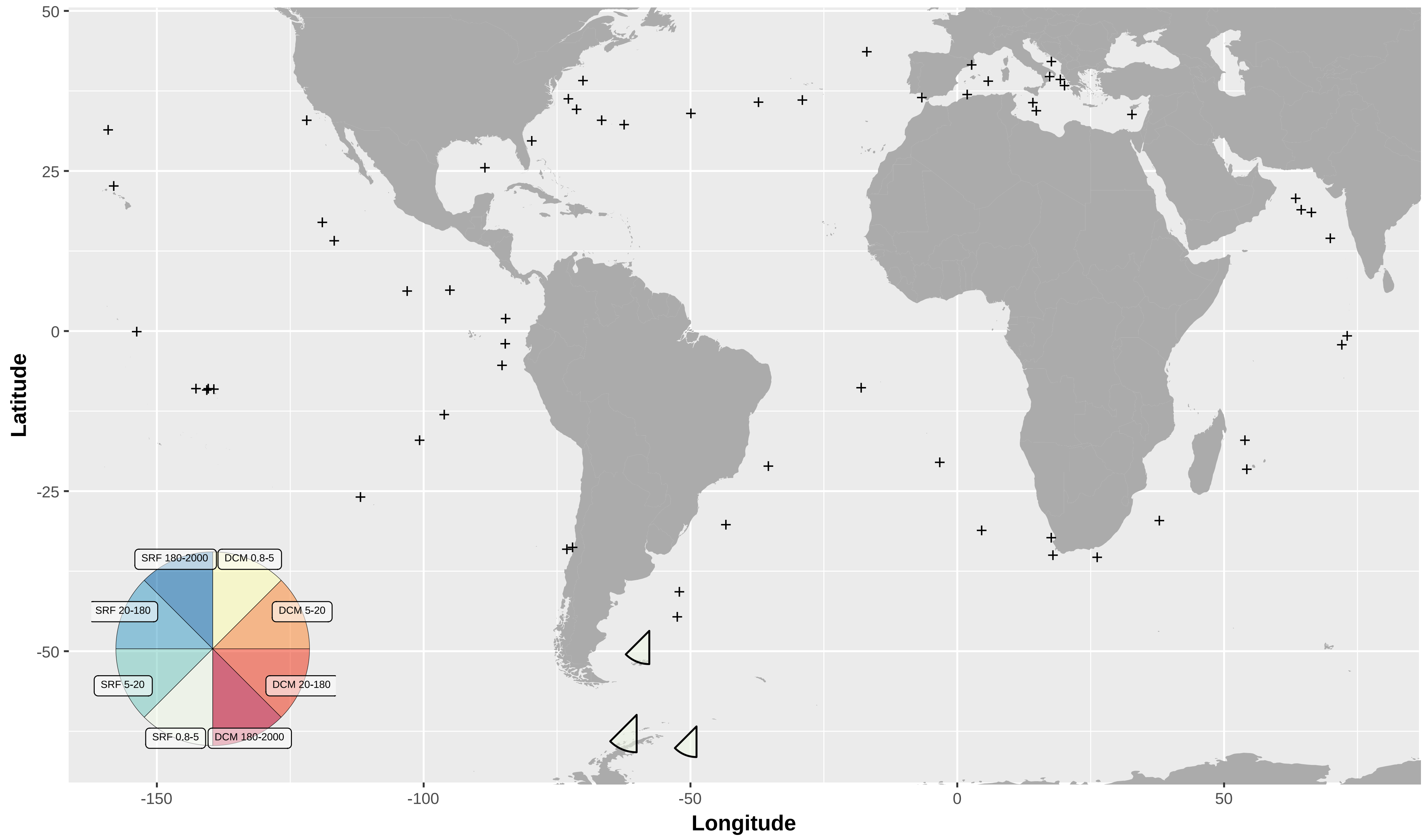


root;cellular organisms;Eukaryota

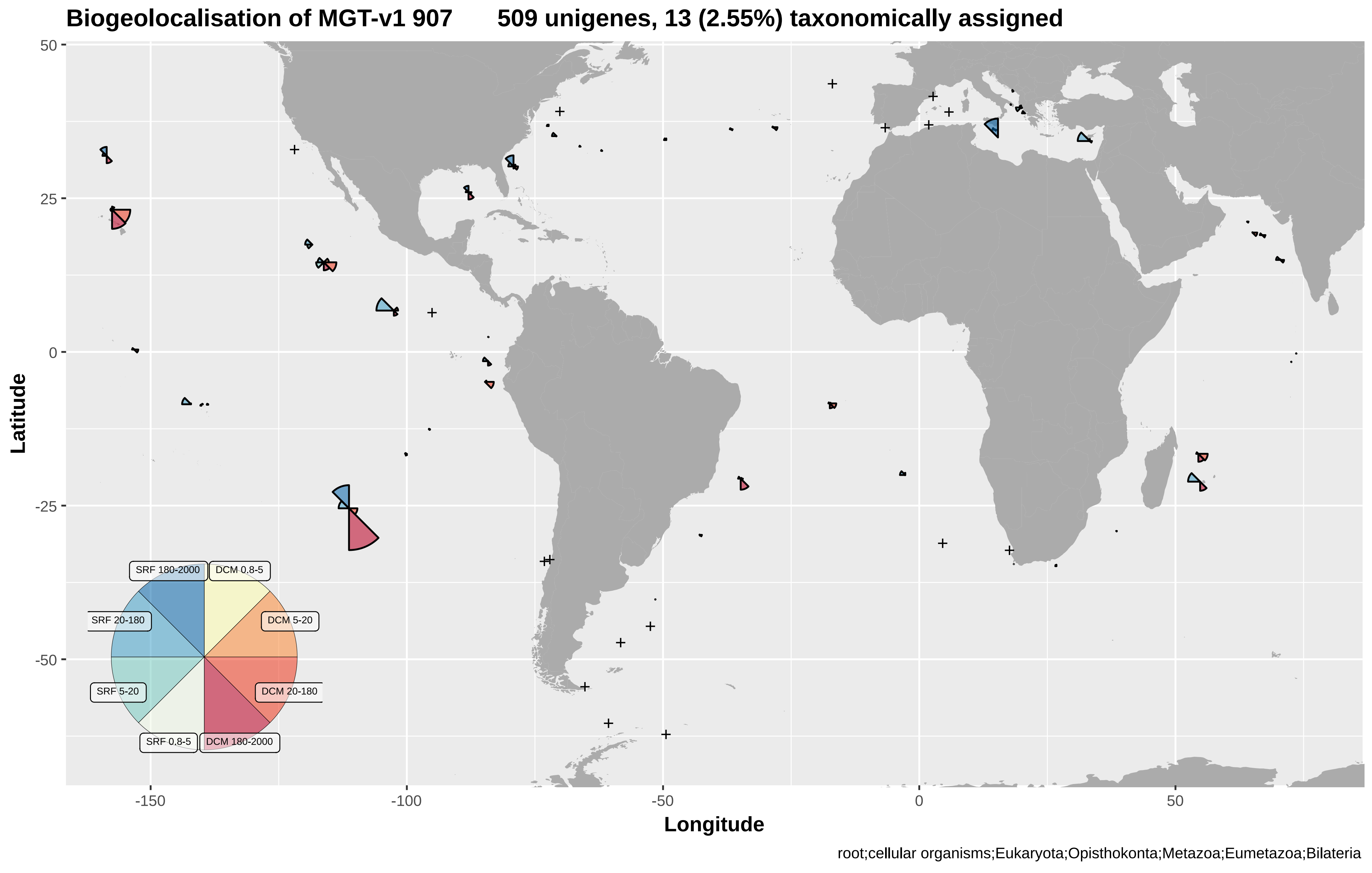

Biogeolocalisation of MGT-v1 907      509 unigenes, 13 (2.55%) taxonomically assigned
50
25
0
-25
-50
Latitude
-150
-100
-50
0
50
Longitude
SRF 180-2000
DCM 0.8-5
SRF 20-180
DCM 5-20
SRF 5-20
DCM 20-180
SRF 0.8-5
DCM 180-2000
root;cellular organisms;Eukaryota;Opisthokonta;Metazoa;Eumetazoa;Bilateria

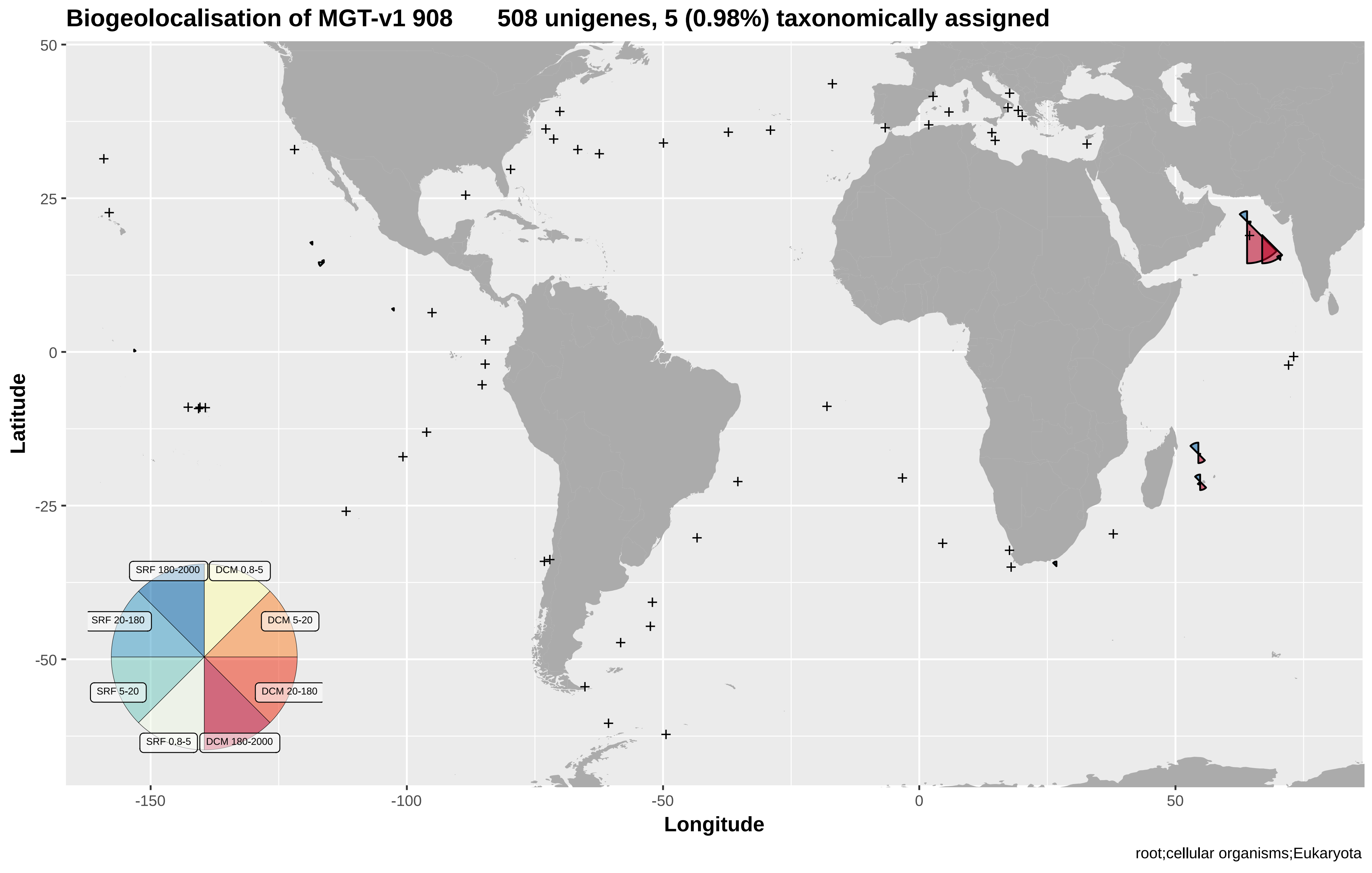

Biogeolocalisation of MGT-v1 908 508 unigenes, 5 (0.98%) taxonomically assigned
50
25
0
-25
-50
Latitude
-150
-100
-50
0
50
Longitude
SRF 180-2000
DCM 0.8-5
SRF 20-180
DCM 5-20
SRF 5-20
DCM 20-180
SRF 0.8-5
DCM 180-2000
root;cellular organisms;Eukaryota

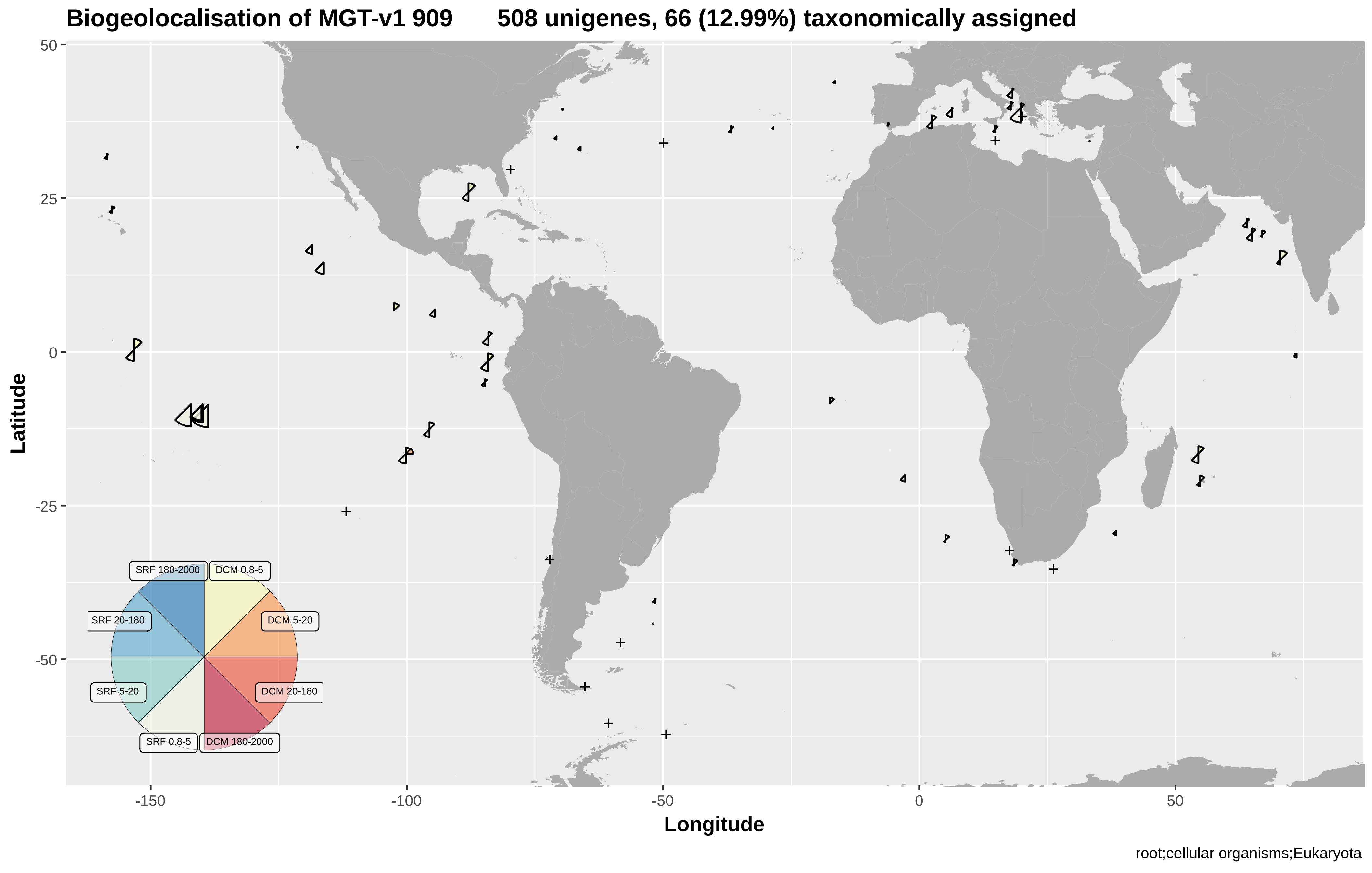
Biogeolocalisation of MGT-v1 909 508 unigenes, 66 (12.99%) taxonomically assigned
50
25
0
-25
-50
Latitude
-150
-100
-50
0
50
Longitude
SRF 180-2000
DCM 0.8-5
SRF 20-180
DCM 5-20
SRF 5-20
DCM 20-180
SRF 0.8-5
DCM 180-2000
root;cellular organisms;Eukaryota

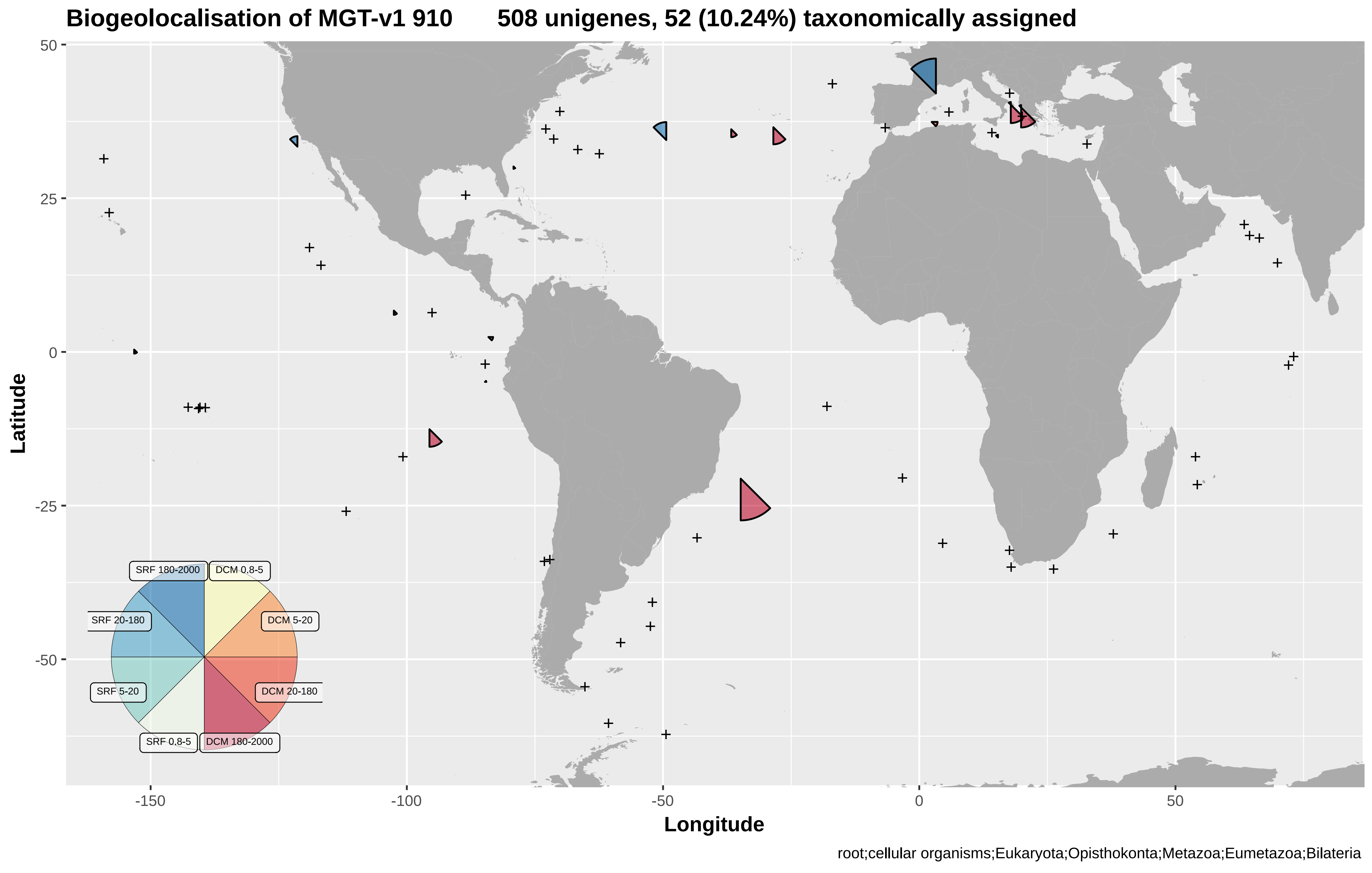

Biogeolocalisation of MGT-v1 910 508 unigenes, 52 (10.24%) taxonomically assigned
50
25
0
-25
-50
Latitude
-150
-100
-50
0
50
Longitude
SRF 180-2000
DCM 0.8-5
SRF 20-180
DCM 5-20
SRF 5-20
DCM 20-180
SRF 0.8-5
DCM 180-2000
root;cellular organisms;Eukaryota;Opisthokonta;Metazoa;Eumetazoa;Bilateria

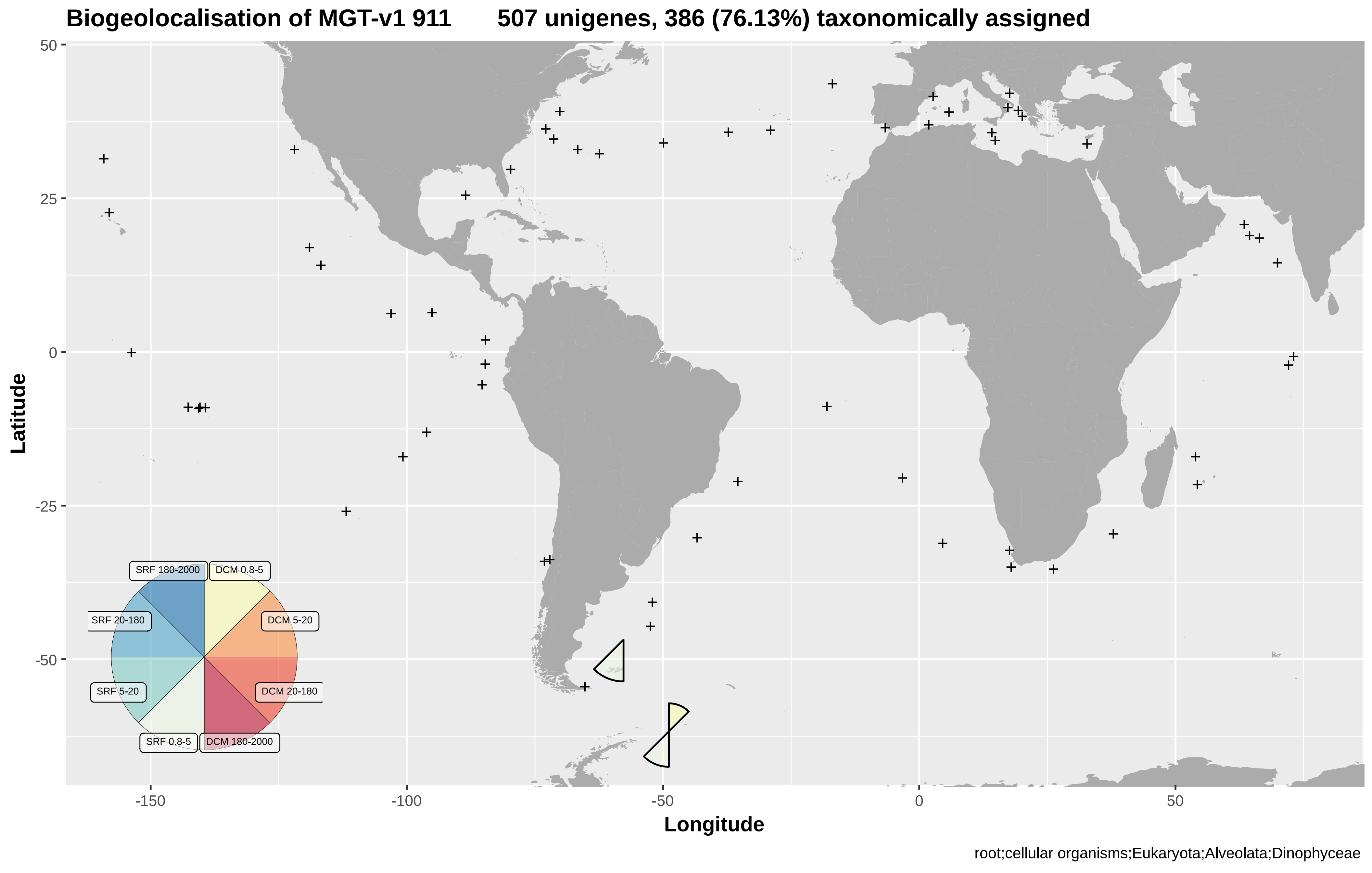

Biogeolocalisation of MGT-v1 911
507 unigenes, 386 (76.13%) taxonomically assigned
50
25
0
-25
-50
Latitude
-150
-100
-50
0
50
Longitude
SRF 180-2000
DCM 0.8-5
SRF 20-180
DCM 5-20
SRF 5-20
DCM 20-180
SRF 0.8-5
DCM 180-2000
root;cellular organisms;Eukaryota;Alveolata;Dinophyceae

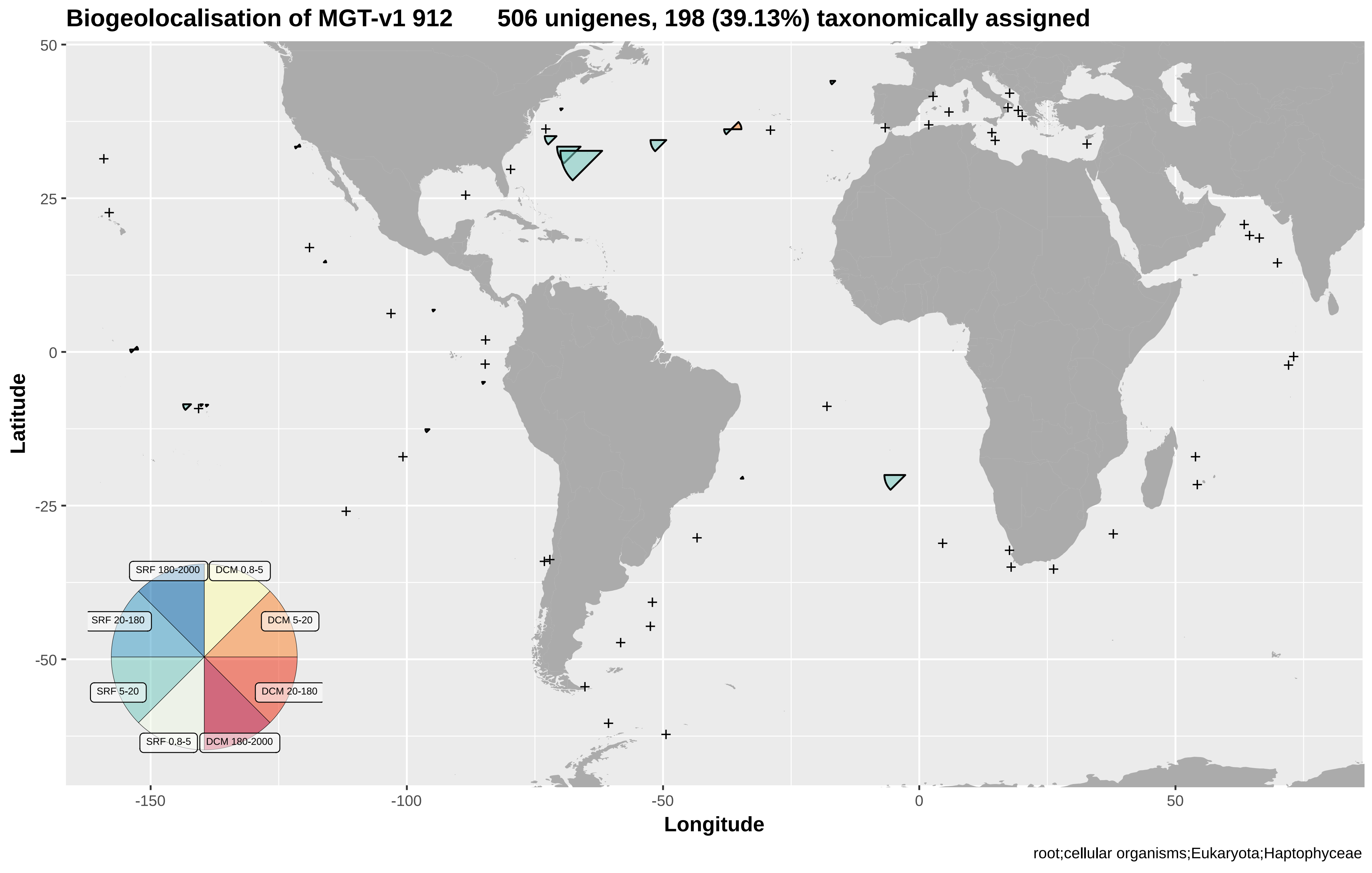

Biogeolocalisation of MGT-v1 912 506 unigenes, 198 (39.13%) taxonomically assigned
50
25
0
-25
-50
Latitude
-150
-100
-50
0
50
Longitude
SRF 180-2000
DCM 0.8-5
SRF 20-180
DCM 5-20
SRF 5-20
DCM 20-180
SRF 0.8-5
DCM 180-2000
root;cellular organisms;Eukaryota;Haptophyceae

# Biogeolocalisation of MGT-v1 913      505 unigenes, 489 (96.83%) taxonomically assigned

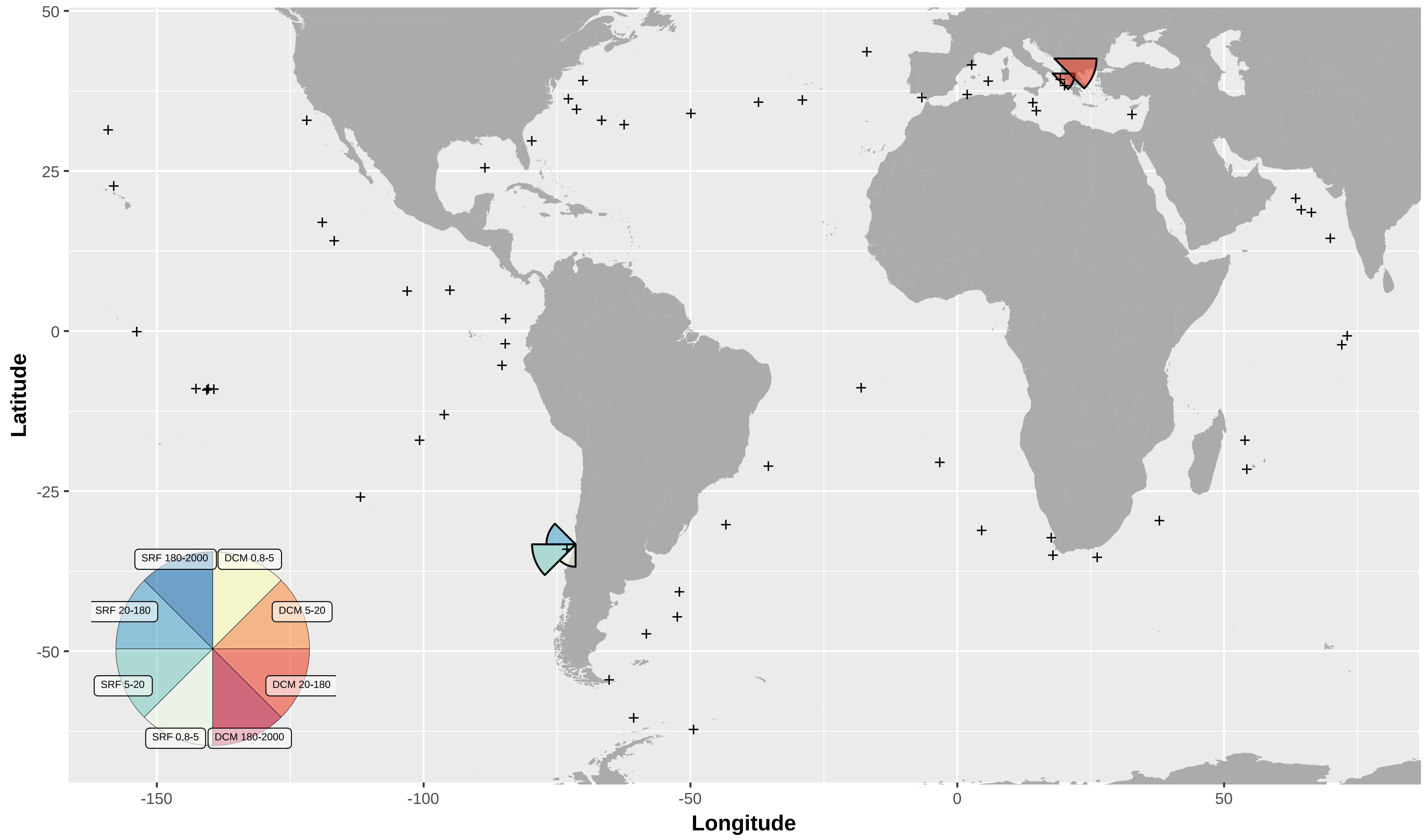


root;cellular organisms;Eukaryota;Stramenopiles;Bacillariophyta;Coscinodiscophyceae;Chaetocerotophycidae;Leptocylindrales;Leptocylindraceae;Leptocylindrus

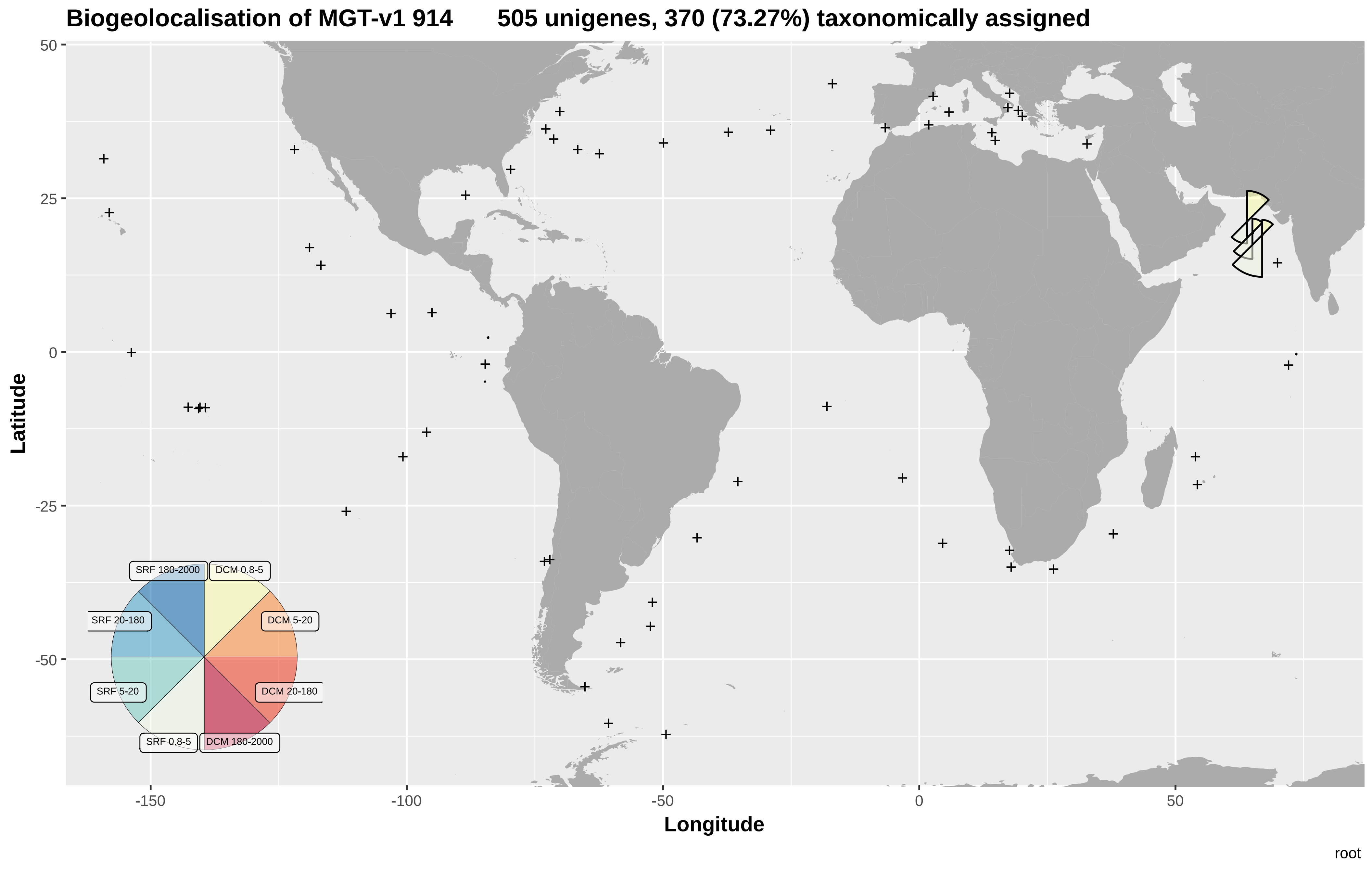

Biogeolocalisation of MGT-v1 914 505 unigenes, 370 (73.27%) taxonomically assigned
50
25
0
-25
-50
Latitude
-150
-100
-50
0
50
Longitude
SRF 180-2000
DCM 0.8-5
SRF 20-180
DCM 5-20
SRF 5-20
DCM 20-180
SRF 0.8-5
DCM 180-2000
root

# Biogeolocalisation of MGT-v1 915      505 unigenes, 294 (58.22%) taxonomically assigned

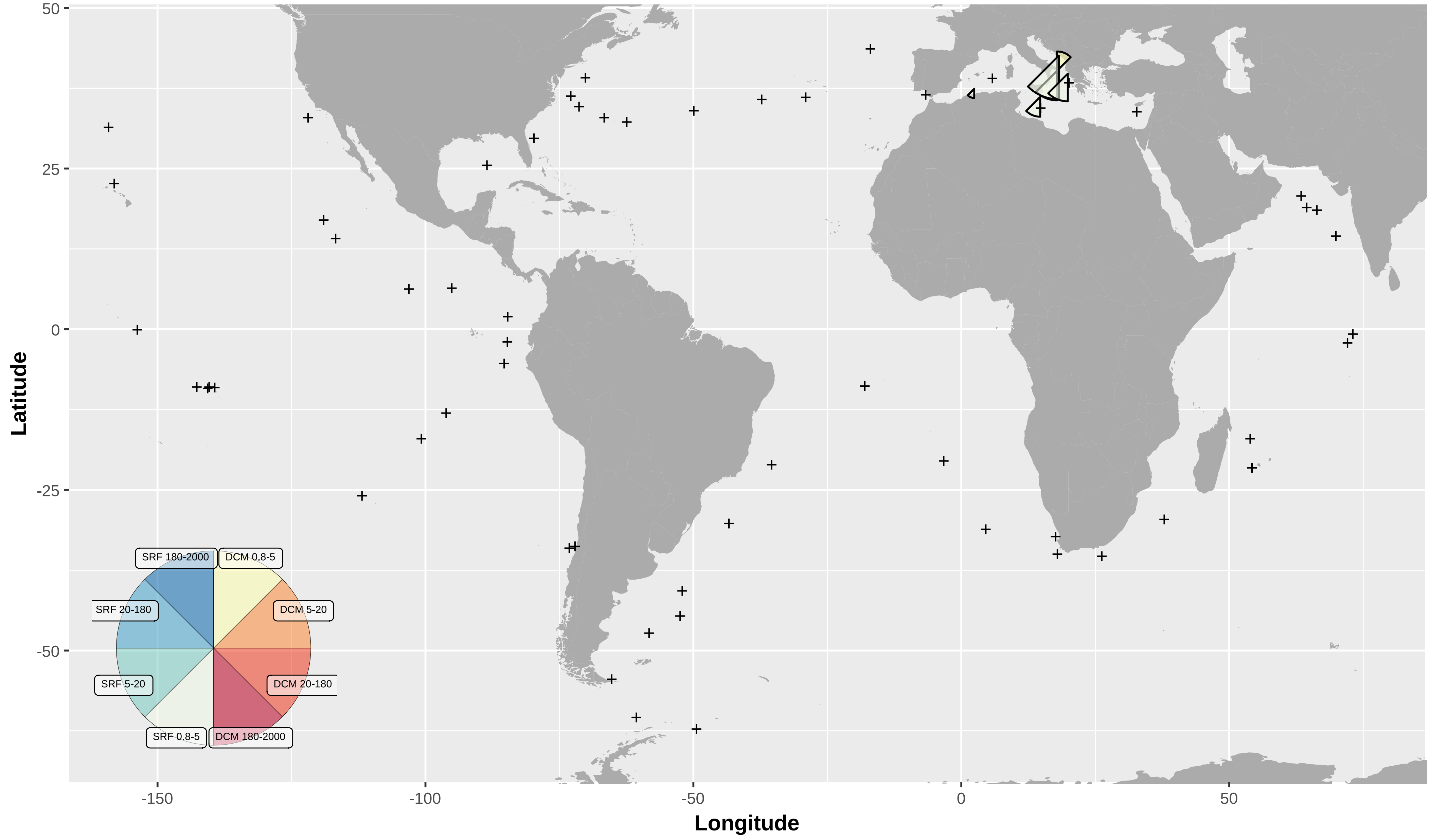


root;cellular organisms;Eukaryota

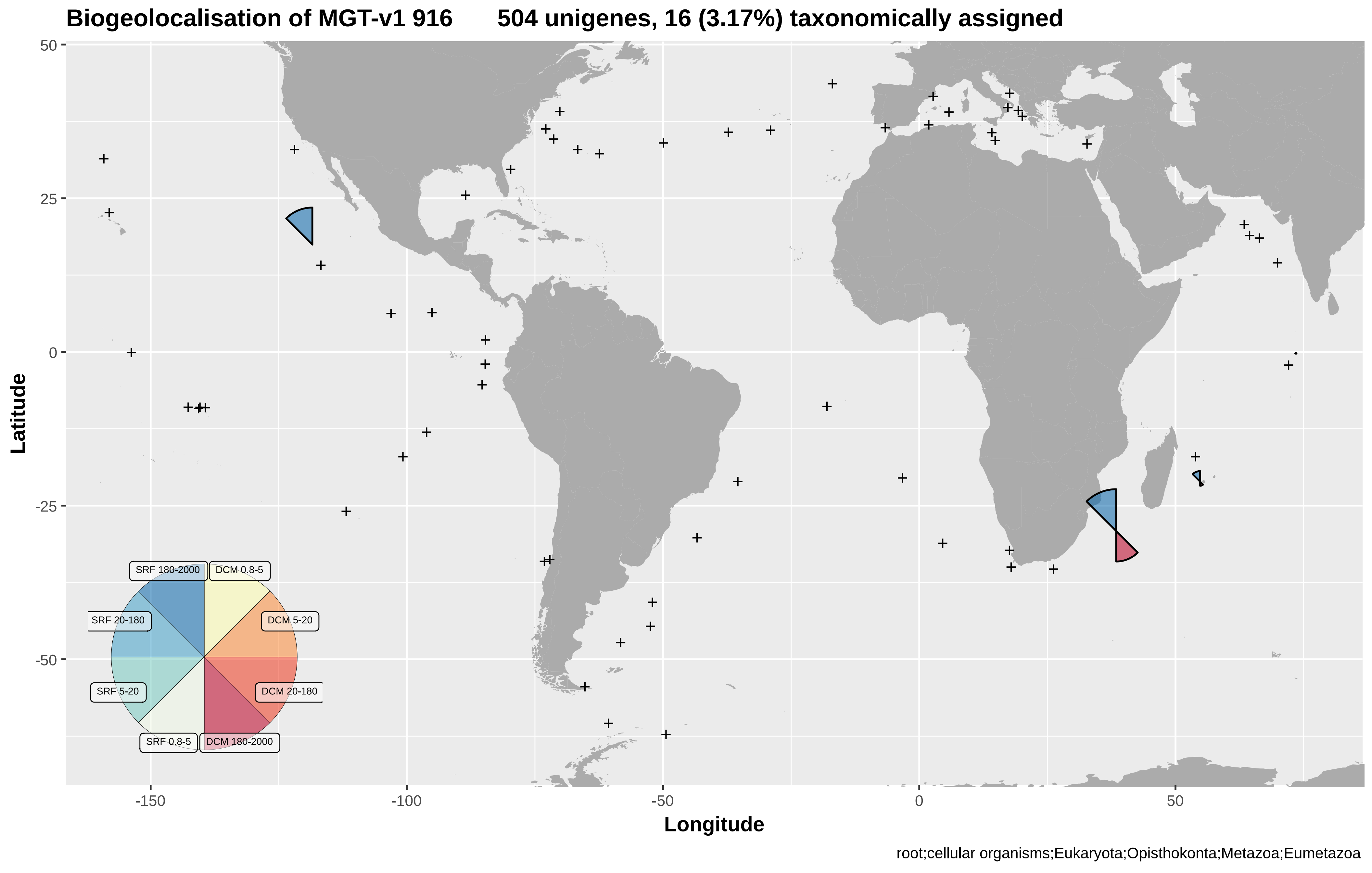
Biogeolocalisation of MGT-v1 916 504 unigenes, 16 (3.17%) taxonomically assigned
50
25
0
-25
-50
Latitude
-150
-100
-50
0
50
Longitude
SRF 180-2000
DCM 0.8-5
SRF 20-180
DCM 5-20
SRF 5-20
DCM 20-180
SRF 0.8-5
DCM 180-2000
root;cellular organisms;Eukaryota;Opisthokonta;Metazoa;Eumetazoa

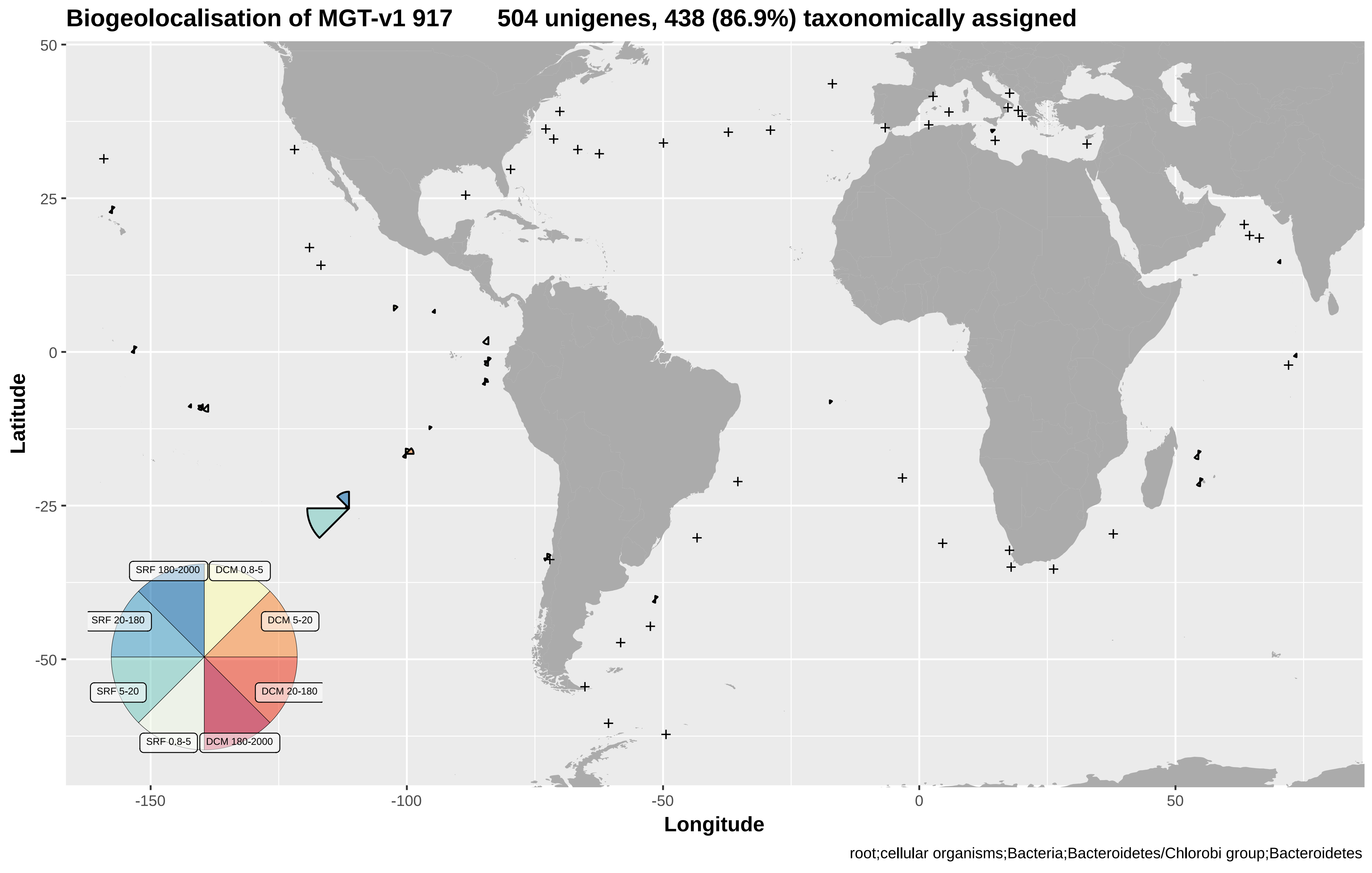

Biogeolocalisation of MGT-v1 917 504 unigenes, 438 (86.9%) taxonomically assigned
Latitude
50
25
0
-25
-50
Longitude
-150
-100
-50
0
50
SRF 180-2000
DCM 0.8-5
SRF 20-180
DCM 5-20
SRF 5-20
DCM 20-180
SRF 0.8-5
DCM 180-2000
root;cellular organisms;Bacteria;Bacteroidetes/Chlorobi group;Bacteroidetes

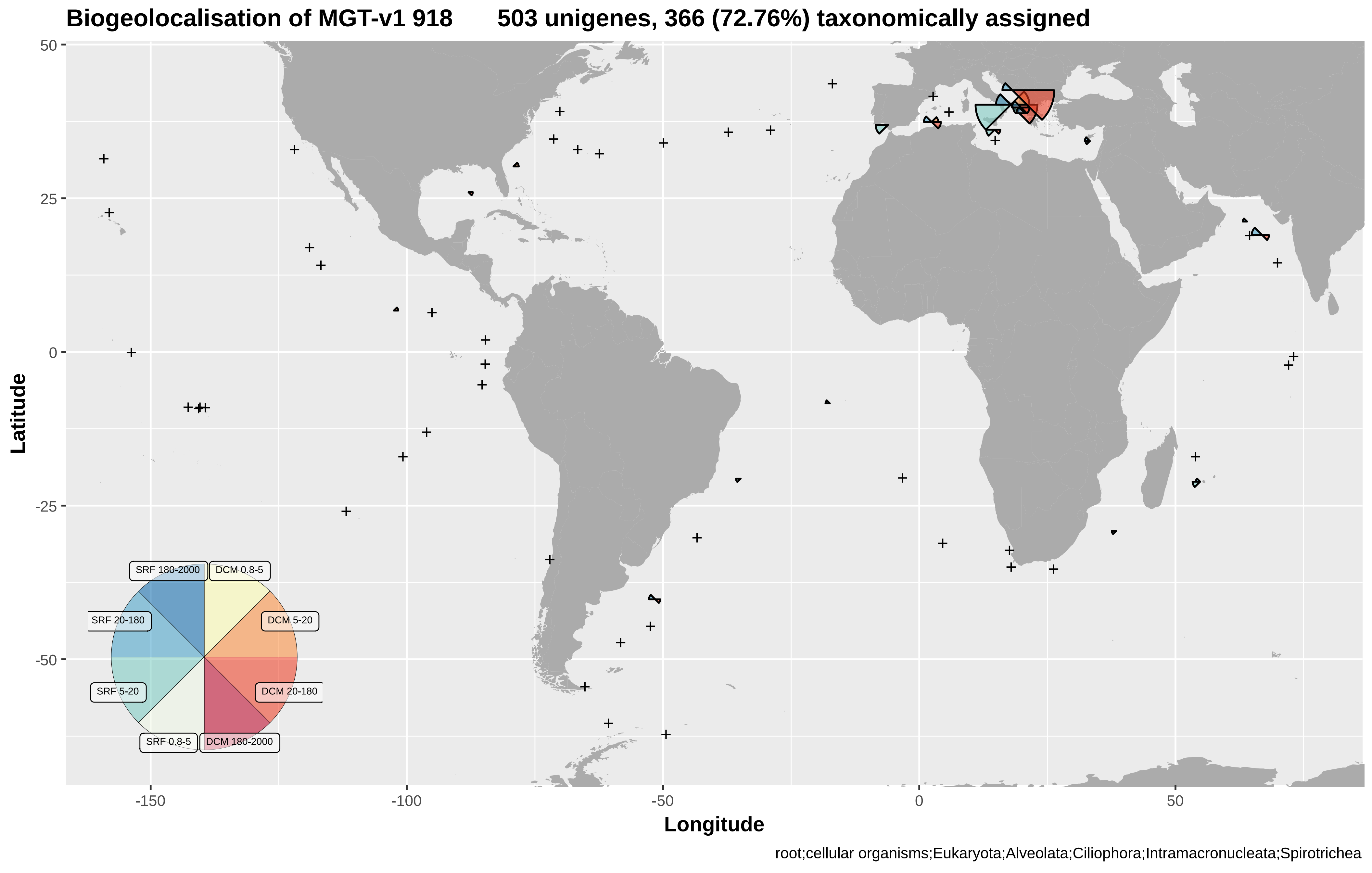
Biogeolocalisation of MGT-v1 918 503 unigenes, 366 (72.76%) taxonomically assigned
50
25
0
-25
-50
Latitude
-150
-100
-50
0
50
Longitude
SRF 180-2000
DCM 0.8-5
SRF 20-180
DCM 5-20
SRF 5-20
DCM 20-180
SRF 0.8-5
DCM 180-2000
root;cellular organisms;Eukaryota;Alveolata;Ciliophora;Intramacronucleata;Spirotrichea

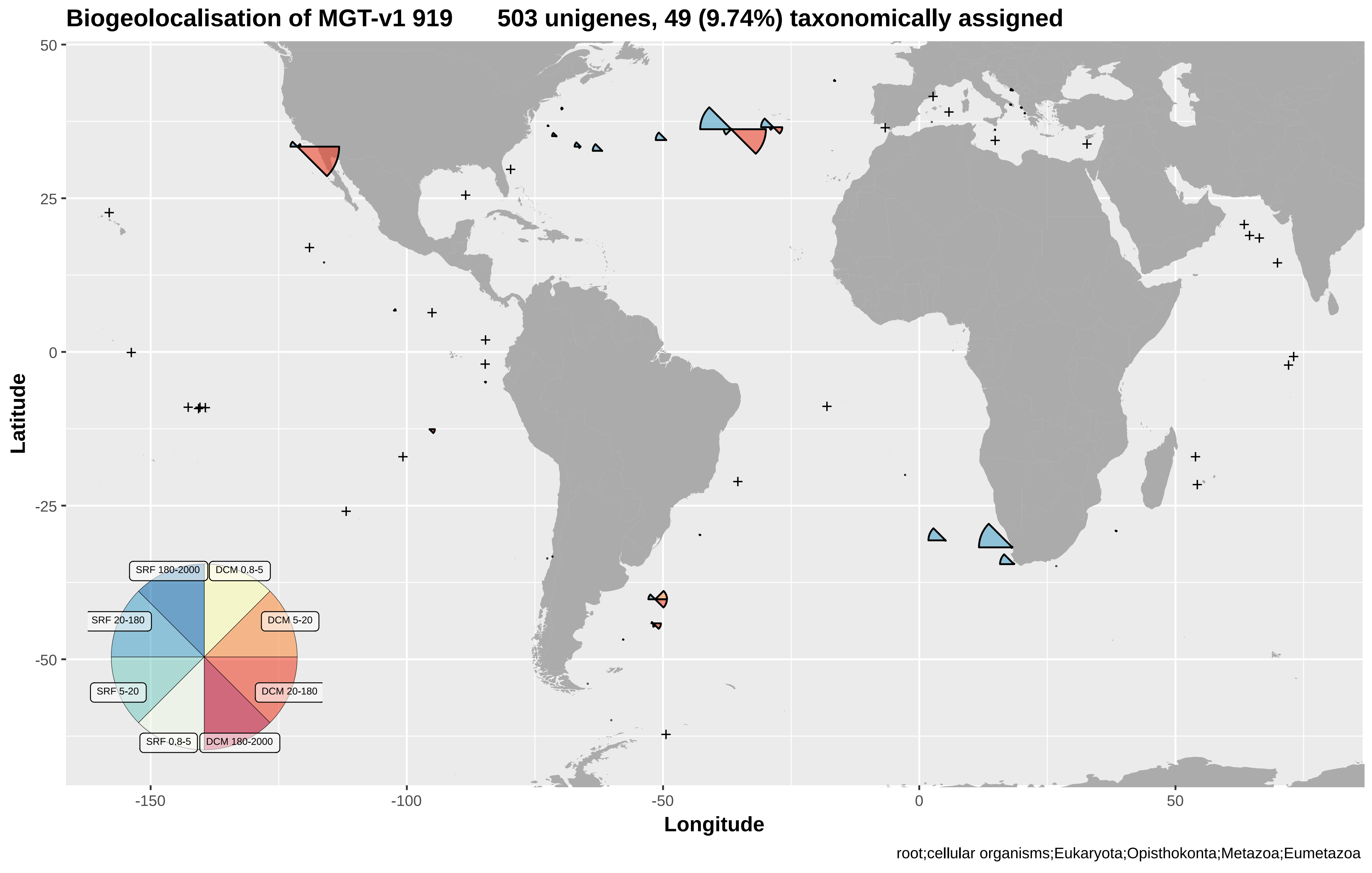

Biogeolocalisation of MGT-v1 919 503 unigenes, 49 (9.74%) taxonomically assigned
50
25
0
-25
-50
Latitude
-150
-100
-50
0
50
Longitude
SRF 180-2000
DCM 0.8-5
SRF 20-180
DCM 5-20
SRF 5-20
DCM 20-180
SRF 0.8-5
DCM 180-2000
root;cellular organisms;Eukaryota;Opisthokonta;Metazoa;Eumetazoa

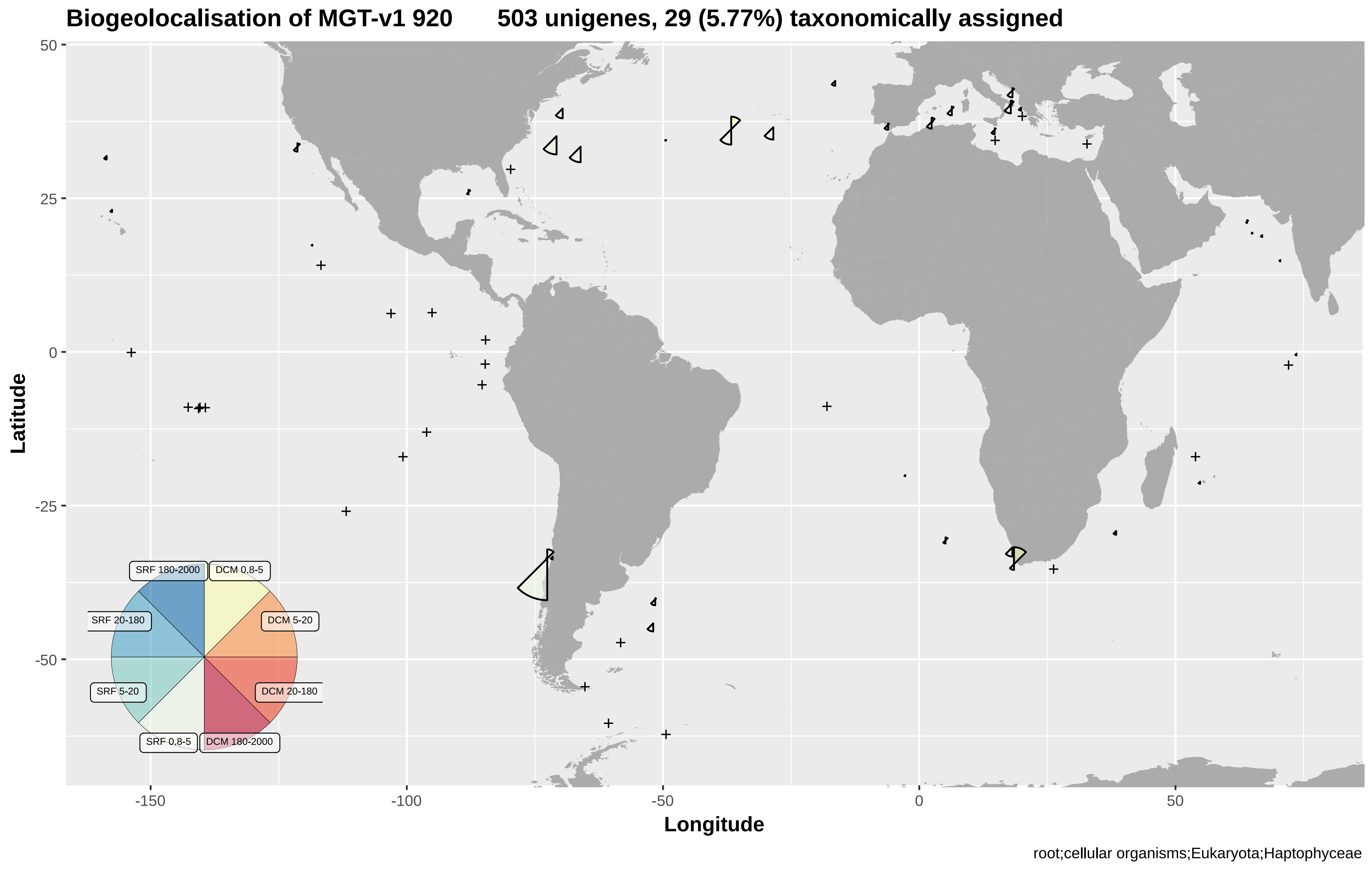

Biogeolocalisation of MGT-v1 920      503 unigenes, 29 (5.77%) taxonomically assigned
50
25
0
-25
-50
Latitude
-150
-100
-50
0
50
Longitude
SRF 180-2000
DCM 0.8-5
SRF 20-180
DCM 5-20
SRF 5-20
DCM 20-180
SRF 0.8-5
DCM 180-2000
root;cellular organisms;Eukaryota;Haptophyceae

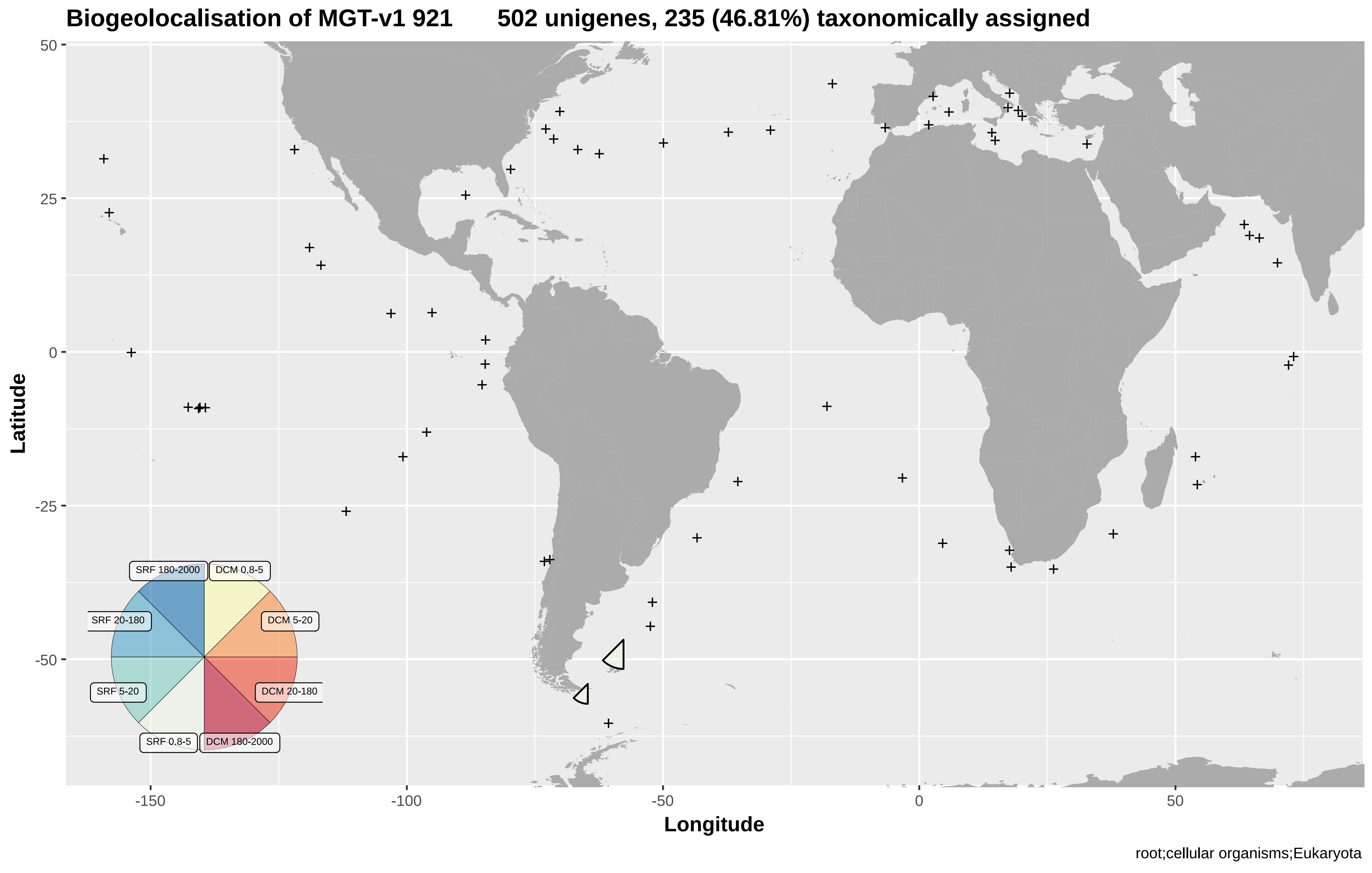

Biogeolocalisation of MGT-v1 921 502 unigenes, 235 (46.81%) taxonomically assigned
50
25
0
-25
-50
Latitude
-150
-100
-50
0
50
Longitude
SRF 180-2000
DCM 0.8-5
SRF 20-180
DCM 5-20
SRF 5-20
DCM 20-180
SRF 0.8-5
DCM 180-2000
root;cellular organisms;Eukaryota

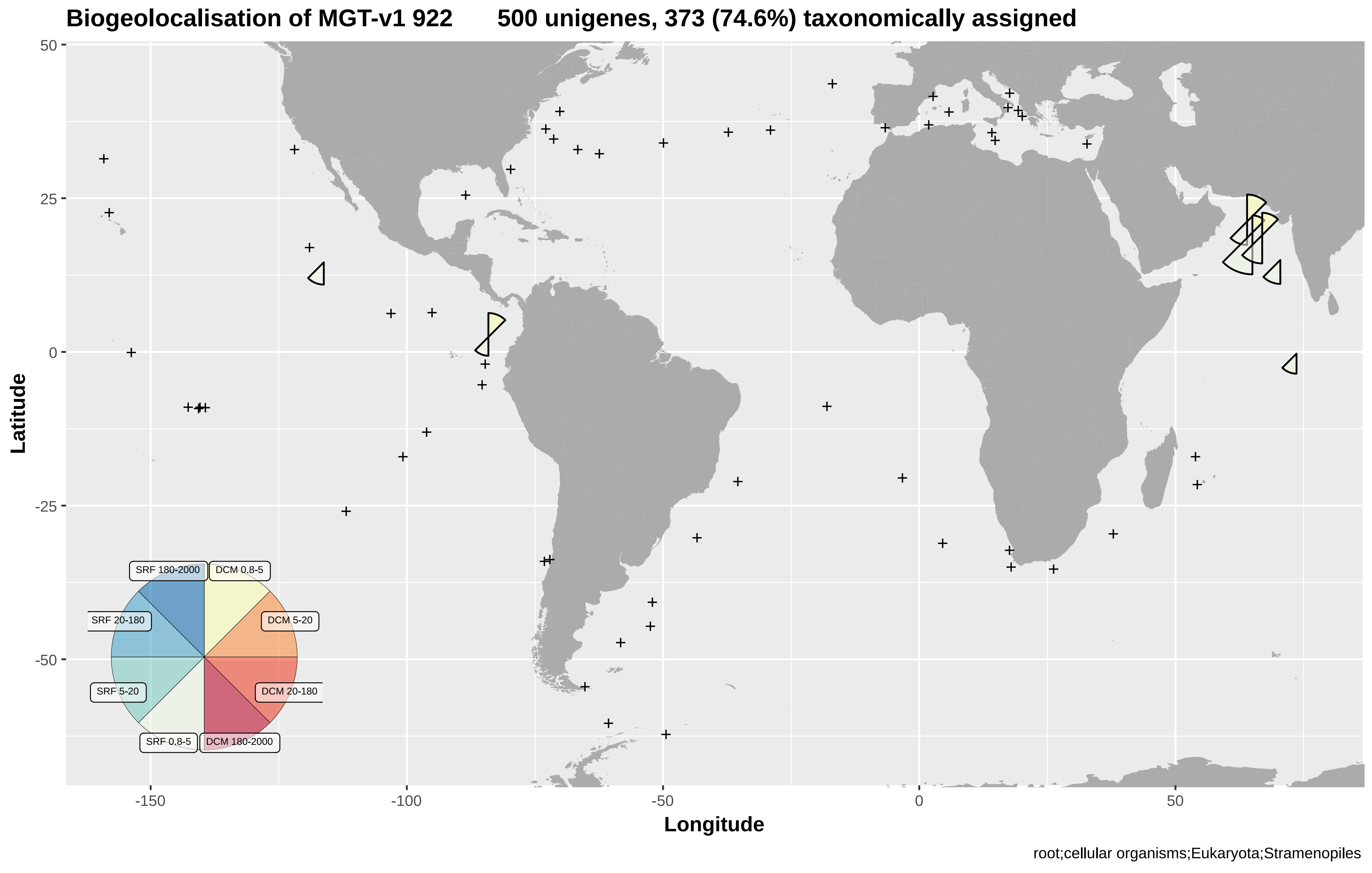

Biogeolocalisation of MGT-v1 922 500 unigenes, 373 (74.6%) taxonomically assigned
50
25
0
-25
-50
Latitude
-150
-100
-50
0
50
Longitude
SRF 180-2000
DCM 0.8-5
SRF 20-180
DCM 5-20
SRF 5-20
DCM 20-180
SRF 0.8-5
DCM 180-2000
root;cellular organisms;Eukaryota;Stramenopiles

#### Biogeolocalisation of MGT-v1 923 500 unigenes, 475 (95%) taxonomically assigned

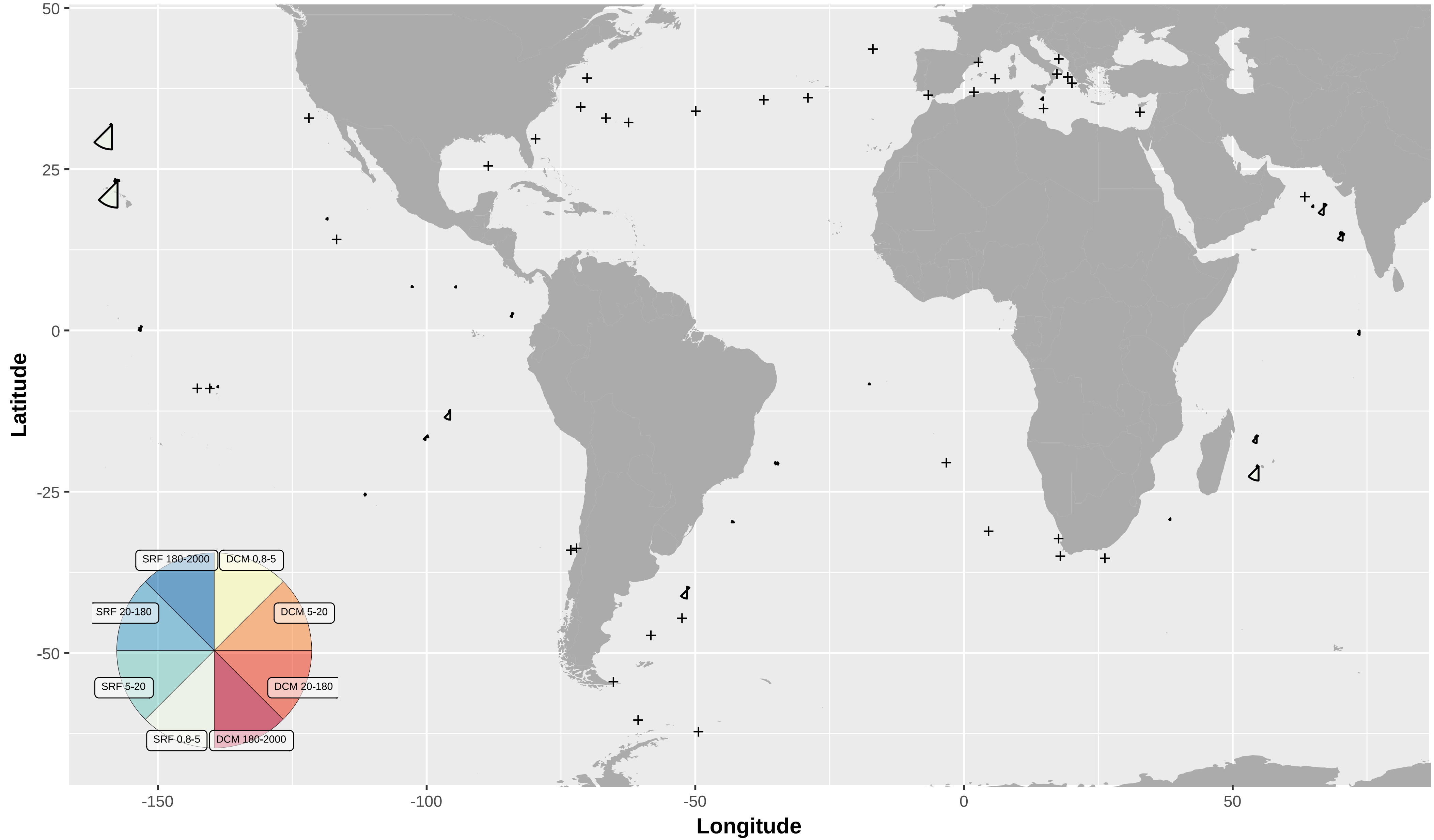


root;cellular organisms;Bacteria;Proteobacteria;Gammaproteobacteria

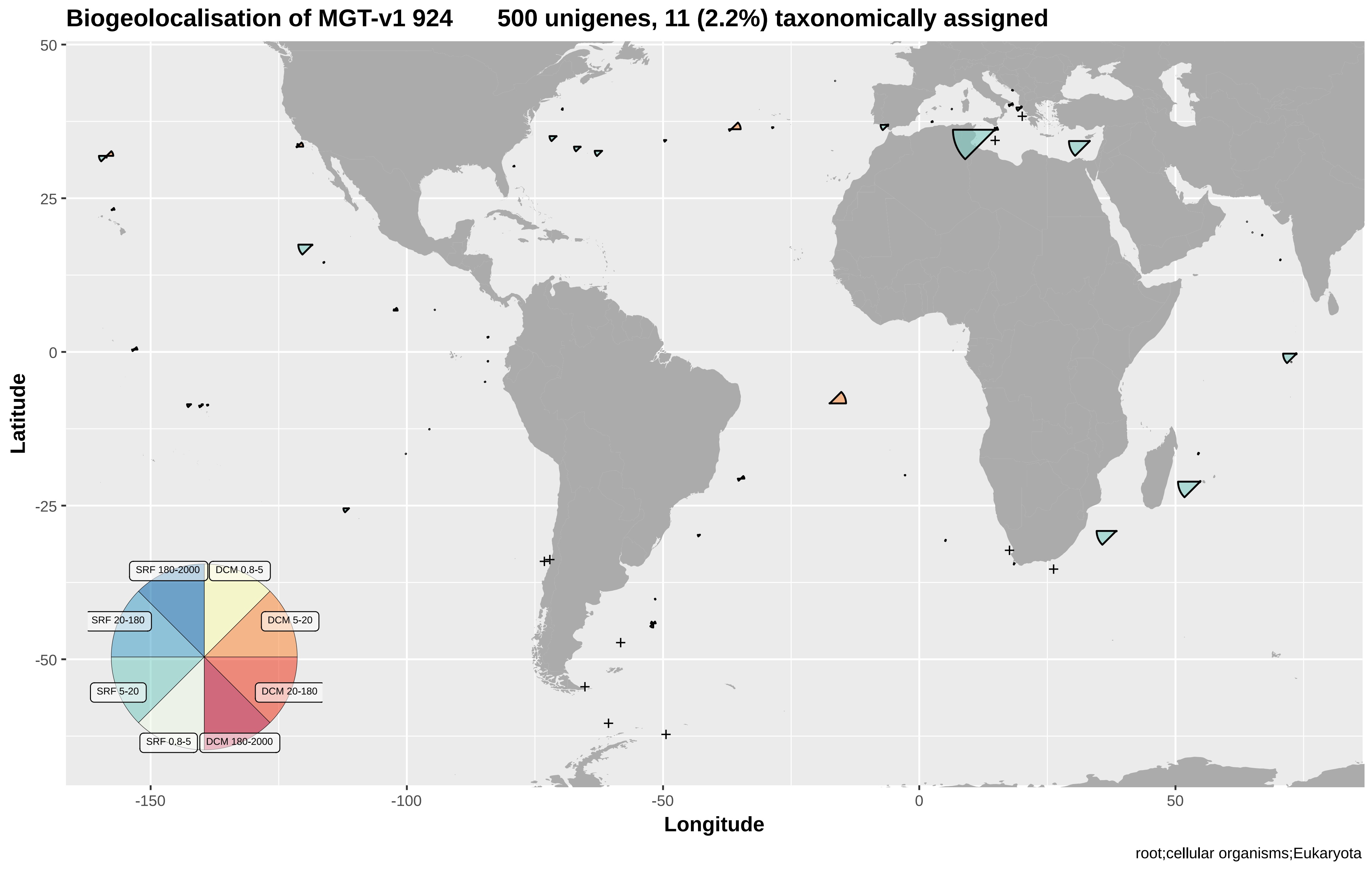

Biogeolocalisation of MGT-v1 924 500 unigenes, 11 (2.2%) taxonomically assigned
50
25
0
-25
-50
Latitude
-150
-100
-50
0
50
Longitude
SRF 180-2000
DCM 0.8-5
SRF 20-180
DCM 5-20
SRF 5-20
DCM 20-180
SRF 0.8-5
DCM 180-2000
root;cellular organisms;Eukaryota
